# Discovery of a Linked Constellation of Gene Expression Revealed by Local Editing of Fibroblasts in Tumors

**DOI:** 10.1101/2024.07.11.603114

**Authors:** Nicholas F. Kuhn, Itzia Zaleta-Linares, Kenneth H. Hu, Tristan Courau, Brittany Davidson, Tammie Tam, Alexis J. Combes, William A. Nyberg, Justin Eyquem, Matthew F. Krummel

## Abstract

Fibroblasts play critical roles in regulating cellular relationships during tissue homeostasis, immunity, and tumor biology at multiple sites. However, tools to perturb fibroblasts at just one site in vivo are limited, restricting our understanding of how these cellular relationships develop on a local level. We optimized local gene editing of fibroblasts in multiple mouse tumor models to investigate how locally restricted fibroblast perturbations affect the cellular tumor microenvironment (TME). By knocking out surface receptors *Osmr, Tgfbr2,* or *Il1r1* on cancer-associated fibroblasts (CAFs), we uncover that TGFBR2 signaling loss uniquely induces the emergence of a *Col18a1*^hi^ CAF cell state that is distinct from previously described fibroblast states and is associated with worse survival in human PDAC patients. Further application of a local as well as combinatorial gene knockout technology in CAFs reveals a circuit in which these *Col18a1*^hi^ CAFs reshape the TME by recruiting Siglec-F^hi^ neutrophils via *Cxcl5* expression; and that the *Col18a1*^hi^ CAF cell state is further dependent on TNFR1 and canonical Wnt signaling. Together, a fast, affordable, and modular engineering method is demonstrated, allowing discovery of a modified fibroblast identify, as well as the network details of a local inter-cellular circuitry in a tumor.

**Graphical Abstract:** 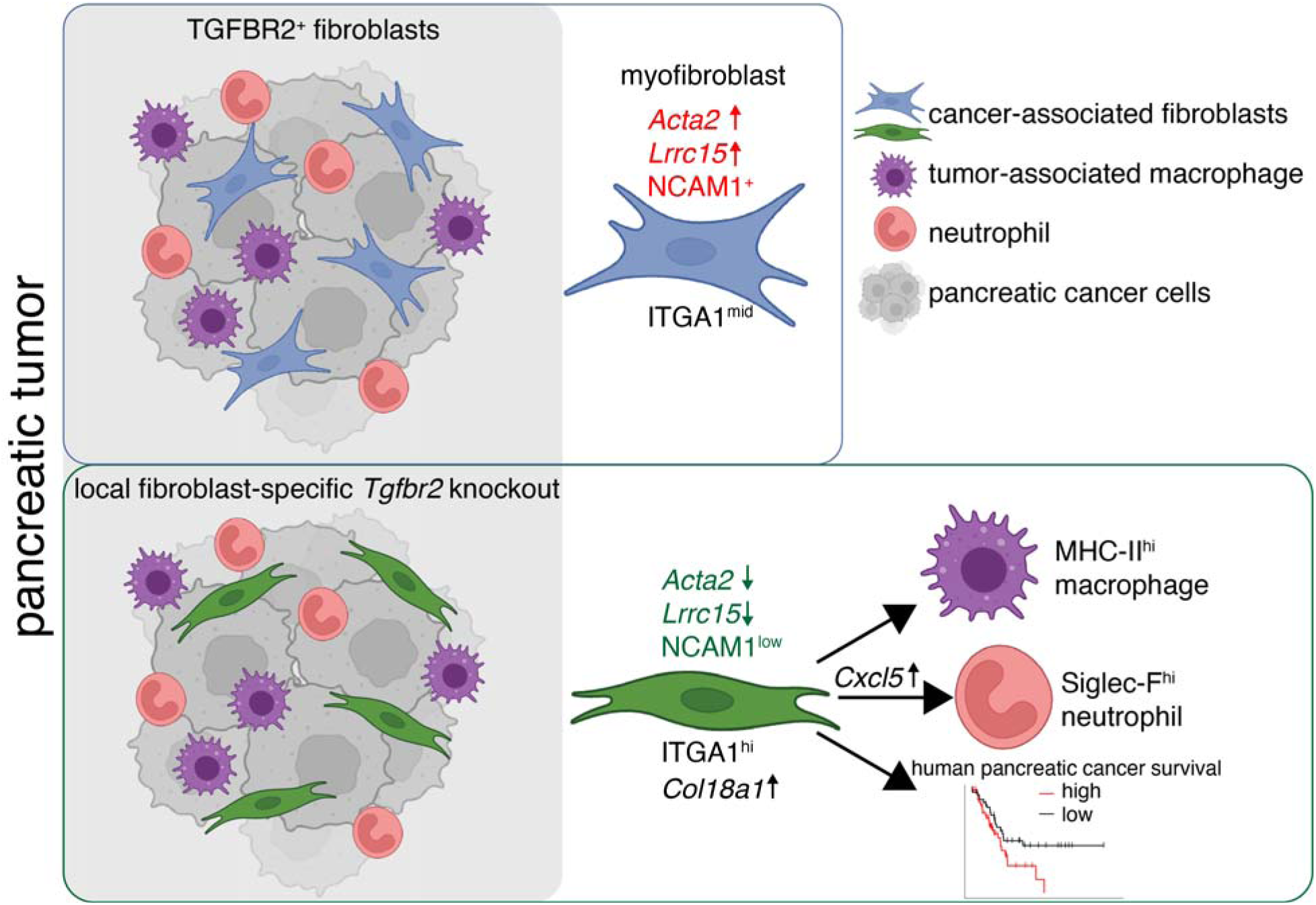

## Introduction

Fibroblasts play essential roles as both mediators of homeostasis and in driving transitions in tissues, for example during repair.^1–6^ However, fibroblast dysregulation has been associated with many chronic diseases, including fibrosis, autoimmunity and cancer.^7–11^ In cancer specifically, the term ‘cancer-associated fibroblast’ (CAF) refers to a cell that is defined by absence of epithelial, endothelial, and immune cell lineage markers and lack of cancer cell mutations^9^. Recent work has greatly expanded our understanding of the heterogenous CAF cell states in human cancer and mouse tumor models,^12–15^ where CAF subset identity is informed by their transcriptional state and ability to influence tumor, endothelial, and immune cell function *ex vivo* in experimental co-culture conditions.^16–18^

However, studying the interaction of CAFs with other cells in replete settings has been challenging, as *in vitro* models do not fully model complex three-dimensional TMEs and testing gene functions in mice has suffered from a lack of methods to modify fibroblast gene expression locally at scale.^9^ In some cases and to circumvent these limitations, tumor cells have been co-injected with defined CAF subsets, which revealed different CAF subset functions in these tumor models.^19^ However, the injected CAFs are often outcompeted by host fibroblasts, leading to poor engraftment of the co-injected CAFs.^9^ Mouse models using Cre-loxP provide a highly controllable method to perturb individual genes in fibroblasts by using Cre-driver lines such as *Pdgfra*-Cre, *Col1a2*-Cre or others that have been developed recently.^1^ These models require mouse breeding of Cre-driver lines with floxed genes-of-interest, are time- and cost-intensive, and typically deplete large populations of fibroblasts globally, which limits the ability to draw conclusions about their specific roles in a defined environment. Thus, we set out to develop an *in vivo* fibroblast perturbation platform that is localized, fast, affordable, and modular.

Viral vectors have been extensively used to modify cells *in vivo*, with recombinant adeno-associated viruses (AAVs) being a popular choice due to their low immunogenicity and cytotoxicity.^20^ We hypothesized that local delivery of a guide RNA (gRNA) via AAV in combination with conditional expression of Cas9 in fibroblasts could generate local KOs of target genes in a defined area of the skin, allowing subsequent study of tumor growth within that tissue area of fibroblast perturbation.

To test our approach, we targeted the following three receptors known to shape CAF biology: (1) *Tgfbr2* (transforming growth factor beta type II receptor) encoding a surface receptor that upon binding to its ligand TGFβ1 together with TGFBR1 has been demonstrated to drive profound phenotypic changes in fibroblasts.^21^ In tumors, TGFβ1-mediated activation of fibroblasts leads to a contractile phenotype that is associated with increased extracellular matrix (ECM) production, which leads to an overall remodeling of the ECM in the growing tumor.^22–24^ As TGFBR2 is expressed on a multitude of epithelial, stromal and immune cells, deciphering how TGFBR2-signaling affects tumor growth requires targeted deletion.^14^ (2) *Osmr* (Oncostatin M receptor) encoding a surface receptor that has been implicated in stimulating a pro-inflammatory gene program in fibroblasts.^6,25,26^ (3) *Il1r1* (interleukin 1 receptor type 1), which—in colorectal cancer lesions—stimulates CAFs to adopt an inflammatory cell state that is associated with increased tumor infiltration of immunosuppressive macrophages^27^ and results in therapy-induced senescence of CAFs,^28^ both contributing to increased tumor growth.

Here, we sought to develop tools that are adaptable for knocking out genes, singly or in combination. Focusing on these three genes that have produced data indicating intratumoral roles for them in fibroblasts, we sought to develop both the method for localized fibroblast engineering and to answer questions about the localized role of these genes and pathways in intratumoral populations of fibroblasts.

## Results

### Targeted local knockout of gene-of-interest in fibroblasts

We hypothesized that two elements were necessary for an approach to locally genetically engineer fibroblasts. First is a vector system which, together with pre-existing fibroblast-specific alleles that regulate expression of Cas9, provide modular flexibility for gene expression and editing. Second is identification of a highly efficient AAV serotype that delivers necessary elements to fibroblasts locally *in vivo*. Toward the first, we designed a self-complementary AAV (scAAV)^29–32^ cargo cassette comprised of the human U6 promoter (hU6) driving RNA polymerase III-mediated expression of a gRNA and a hybrid form of the CBA promoter (CBh)^33^ driving mCherry expression as a transduction readout (**Figure 1A**). Cargo plasmids encoding gRNAs against *Thy1* (gene encoding Thy1/CD90) or *Pdpn* (gene encoding podoplanin, PDPN) were packaged into scAAVs with the AAV serotype 1 (AAV1) capsid—from here on referred to as AAV-gRNA—and used to transduce the mouse fibroblast cell line NIH/3T3 expressing Cas9 (NIH/3T3.Cas9) (**Figure 1B**). The scAAV transduced at an efficiency of >75% and induced knockout (KO) noticeable by loss of surface expression of Thy1 or PDPN (**Figures 1C** and **1D**). A comparison of the AAV1 capsid to recently developed fibroblast-tropic synthetic capsids 7m8^34^ and DJ^35^ showed similar transduction and KO efficiency *in vitro* (**Figure S1A**).

**Figure 1.**
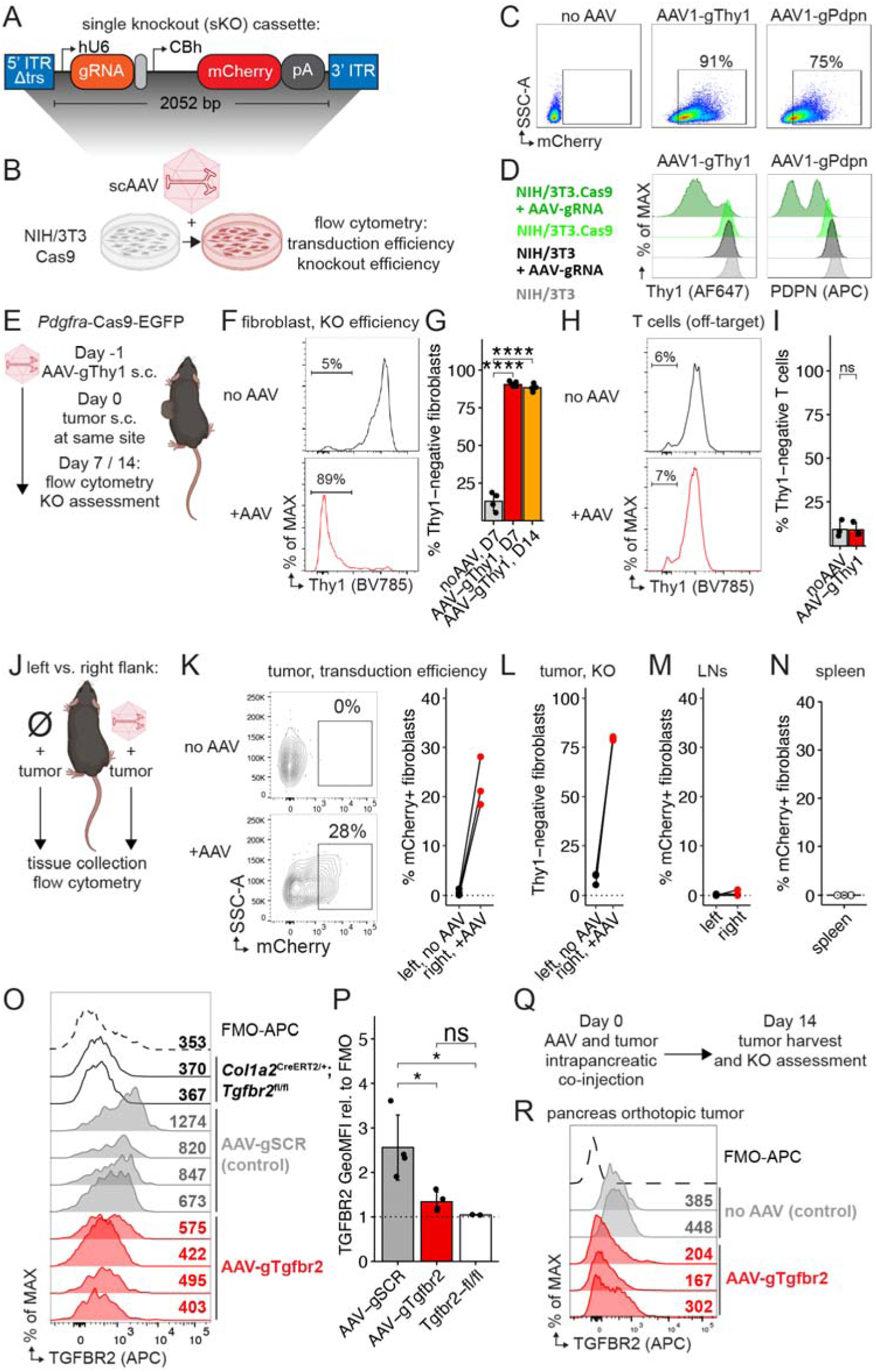
Local knockout of *Thy1* or *Tgfbr2* in fibroblasts. (A) Self-complementary adeno-associated virus (scAAV) cassette for single gene knockout (sKO). ITR, inverted terminal repeat. Δtrs, no terminal resolution site. hU6, human U6 promoter. CBh, chicken beta-actin hybrid promoter. pA, SV40 late polyadenylation signal. bp, base pairs. (B) Experimental scheme for testing transduction and knockout efficiency of scAAV in vitro. NIH/3T3.Cas9 cells were transduced with scAAV serotype 1 (AAV1) at an MOI of 1e5. 72 h later, cells are analyzed via flow cytometry. (C) Flow cytometry plots showing mCherry expression in NIH/3T3.Cas9 cells after transduction with AAV as outlined in (B). One of two representative experiments shown. (D) Flow cytometry overlay plots of (left) Thy1 or (right) of PDPN protein surface expression in NIH/3T3 cells expressing Cas9 (=green) or not (=black) after transduction with AAV as outlined in (B). Only Cas9-expressing NIH/3T3 cells exposed to AAV-gRNA lose surface expression of target protein. One of two representative experiments is shown. (E) Experimental scheme for gene knockout (KO) in tumor fibroblasts using scAAV. (F) Representative histograms showing Thy1 surface expression analyzed by flow cytometry of all EGFP+ tumor fibroblasts from mice treated as outlined in (E) and analyzed on day 7. (G) Quantification of KO efficiency in all EGFP+ tumor fibroblasts on day 7 or 14 after tumor challenge (n=4-6 mice/group). Data are mean ± s.d. and significance was tested using one-way ANOVA with Tukey’s multiple comparisons test. (H) Same as (F) in CD45+ CD3e+ T cells. (I) Quantification of (H) on day 7 (n=3/group). Data are mean ± s.d. ns, non-significant. (J) Experimental scheme for testing local KO in tumor fibroblasts. (K) Left, representative flow cytometry plots showing mCherry expression as a readout for AAV transduction. Cells gated on EGFP+ tumor fibroblasts. Right, quantification of mCherry+ EGFP+ tumor fibroblasts in paired flank tumors from same mouse (n=3 mice). (L) Quantification of Thy1 KO efficiency in all EGFP+ fibroblasts from paired flank tumors on same mouse (n=3 mice). (M) Quantification of mCherry expression in fibroblasts of brachial and inguinal lymph nodes (LNs) pooled from left or right flanks (n=3 mice). (N) Quantification of mCherry expression in splenic fibroblasts after flank AAV injection (n=3 mice). (O) Overlay histograms of TGFBR2 surface expression by flow cytometry on all tumor fibroblasts from mice treated with 3e10 viral genomes (vg) AAV-gTgfbr2 or AAV-gSCR (control). Tumor fibroblasts of *Col1a2*^CreERT2/+^;*Tgfbr2*^fl/fl^ included as negative staining control. GeoMFI value listed for each sample. FMO, fluorescence minus one. (P) Quantification of TGFBR2 (GeoMFI) surface expression on all tumor fibroblasts from (M) relative to FMO (n=2-4 mice/group). Data are mean ± s.d. and significance was tested using one-way ANOVA with Tukey’s multiple comparisons test. (Q) Experimental scheme for KO assessment in orthotopically transplanted pancreatic tumors. (R) Overlay histograms of TGFBR2 surface expression by flow cytometry on all tumor fibroblasts isolated from orthotopically grown HY19636 pancreatic tumors from *Pdgfra*-Cas9-EGFP mice after intrapancreatic co-injection of 3e10 vg AAV-gTgfbr2 and 2.5e5 HY19636 cells. GeoMFI values listed for each sample. fl, floxed. GeoMFI, geometric mean fluorescence intensity. SCR, scramble. ns, non-significant. *p<0.05, **p<0.01, ***p<0.001, ****p<0.0001.

This prompted us to find a solution to the second requirement to locally deliver gene editing elements to fibroblasts *in vivo*. For this, Cas9 expression in host fibroblasts was achieved by combining a *Pdgfra*-driven Cre allele^36,37^ with the *Rosa26*-LSL-Cas9-EGFP allele^38^ (**Figures S1B and S1C**, and Methods for details), hereafter referred to as *Pdgfra*-Cas9-EGFP mice. Testing local delivery of AAV-gRNA was performed by injecting 1e10 viral genomes (vg) of AAV-gThy1-CBh-mCherry subcutaneously on the dorsal flank of mice and marking the injection site to allow subsequent injection of YUMM5.2 mouse melanoma tumor cells at the same site on the next day (**Figure S1D**). Measuring mCherry expression in the cancer-associated fibroblasts (CAFs) that populate the growing tumor mass was used as a readout for transduction efficiency and surface expression of Thy1 was used as a readout for knockout efficiency of the AAV. *In vivo* comparison of the transduction mediated by AAV1, DJ, and 7m8 capsids led to 68%, 52%, and 29% respectively, and 90%, 82%, and 56% Thy1-negative CAFs, respectively (**Figure S1E**). On this basis, our future experiments used AAV1.

Next, we tested efficiency of local gene KO in these CAFs by targeting *Thy1* for deletion (**Figure 1E**). More than 85% of all CAFs were Thy1-negative 7 days after tumor challenge, compared to ∼10% without AAV (**Figures 1F** and **1G**). Targeted *Thy1* KO was stable in CAFs, as the same percentage of about 85% of all CAFs were still Thy1-negative when sampled 14 days after tumor challenge (**Figure 1G**); and *Thy1* KO was specific to CAFs, as Thy1+ T cells in the same tumors did not lose their Thy1 surface expression (**Figures 1H** and **1I**). This efficient and persistent KO of a target gene in CAFs was consistent at higher doses of AAV (**Figure S1F**). To test if the KO was truly local, the same *Pdgfra*-Cas9-EGFP mouse was injected on only one flank s.c. with AAV-gThy1 but challenged with YUMM5.2 tumor on both flanks on the following day (**Figure 1J**). CAF transduction and *Thy1* KO were only detected in the tumor that was grown at the site of prior AAV-gThy1 injection (**Figures 1K** and **1L**), not in tumor-draining lymph nodes (**Figure 1M**), nor in the spleen (**Figure 1N**). This provided evidence that s.c. injection of AAV-gRNA allows local KO and does not lead to systemic spread of AAV, preventing KO at unintended locations.

As part of a goal to understand localized roles for specific genes in fibroblasts, we next generated AAV-gTgfbr2 to target the *Tgfbr2* gene in fibroblasts for KO (**Figure S1G**). AAV-gTgfbr2 decreased TGFBR2 levels on all CAFs in *Pdgfra*-Cas9-EGFP mice to similar levels as conventional *Col1a2*-CreERT2-mediated knockout of the floxed *Tgfbr2* gene in *Col1a2*^CreERT^^2/wt^;*Tgfbr2*^fl/fl^ mice (**Figures 1O** and **1P**), demonstrating that AAV-delivered gRNAs have comparable *in vivo* knockout efficiency as conventional Cre/loxP-based approaches, without having to rely on lengthy mouse breeding to get to the desired genotype for experimental analysis. Instead, AAV-delivered gRNAs required only 2 weeks of cloning and AAV generation until injection of AAV-gRNA for targeted, local knockout of any gene-of-interest. This allows subsequent functional studies of any gene-of-interest expressed by CAFs via local gene knockout.

To assess the KO efficiency of our AAV-based system at a different site, we co-injected the pancreatic cancer cell line HY19636, which is derived from a mouse pancreatic ductal adenocarcinoma (PDAC) KPC mouse (*Kras*^LSL-G12D/+^; *Trp53*^lox/+^*; p48/Ptf1a*-Cre)^39^, together with AAV-gTgfbr2 orthotopically in the pancreas of *Pdgfra*-Cas9-EGFP mice (**Figure 1Q**). Fibroblasts isolated from orthotopically grown pancreatic tumors also lost TGFBR2 surface expression when AAV-gTgfbr2 was co-injected (**Figure 1R**), demonstrating the adaptability of this approach to different tissue sites.

### Local *Osmr* or *Tgfbr2* knockout in CAFs repolarizes them towards homeostatic state but differentially affects TREM2+ tumor-associated macrophage phenotype

As a proof-of-concept and test of the specificity achievable with selective gene knockouts, we set out to compare how blocking of a putatively pro-inflammatory gene program in CAFs via *Osmr* KO (**Figures S2A-S2C**) or blocking of a putatively pro-fibrotic gene program via *Tgfbr2* KO (**Figure 1P**) can affect the fibroblast cell states in growing YUMM5.2 melanoma or HY19636 pancreatic cancer tumors (**Figure 2A**). The HY19636 PDAC model represents a more desmoplastic tumor with high CAF involvement (**Figure S2D**). To assay changes in fibroblast identity, we initially measured surface markers that are typically indicative of different activity states: expression of SCA-I+ Ly6C+ are associated with homeostatic fibroblasts in skin^2^, skin repair^6^ (**Figures S2E and S2F**), and tumor growth^14^ (**Figures S2H and 2I**), whereas loss of SCA-I and Ly6C with concomitant upregulation of *Ncam1* marks activated fibroblasts in these settings (**Figures S2G and S2J**). We thus distinguished double-positive (DP) SCA-I+ Ly6C+, intermediate (Int), double-negative (DN) SCA-I- Ly6C-, and DN NCAM1^-/+^ CAF subsets (**Figures S2K and S2L**). We corroborated these cell states as defined by the aforementioned markers by measuring increased expression of homeostatic marker *Dpt* in the DP CAFs and, inversely, increased expression of *Acta2* and *Postn* in DN NCAM1+ CAFs (**Figure S2M**). We also noted that DN CAFs had the highest expression of *Cthrc1*, a recently described marker of a pro-fibrotic fibroblast subset (**Figures S2N and S2O)**.^11,40^

**Fig. 2:**
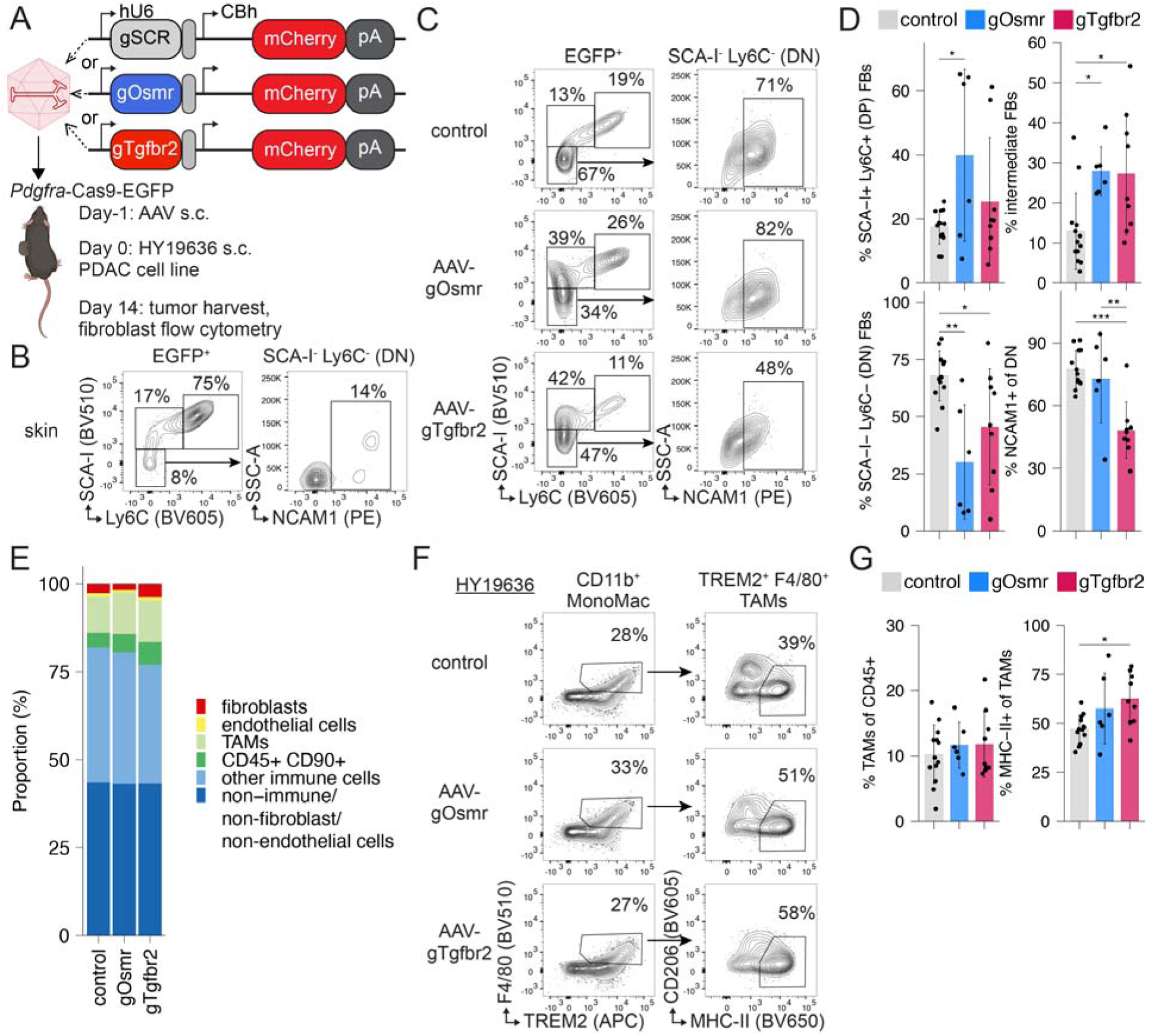
Local *Osmr* or *Tgfbr2* knockout in CAFs repolarizes them towards homeostatic state but differentially affects TREM2+ tumor-associated macrophage phenotype. (A) Different AAV-gRNA cassettes used in outlined experiment for local KO in tumor fibroblasts. Each AAV-gRNA was injected s.c. in HY19636 pancreatic cancer model at a dose of 1-3e10 vg AAV. (B) Flow cytometry plots of (left) EGFP^+^ skin fibroblasts from *Pdgfra*-Cas9-EGFP mice and (right) the double-negative (DN) SCA-I^-^ Ly6C^-^ subset. (C) Representative flow cytometry plots of (left) EGFP^+^ tumor fibroblasts and (right) DN tumor fibroblasts from control-, AAV-gOsmr, or AAV-gTgfbr2-injected *Pdgfra*-Cas9-EGFP mice on day 14 after HY19636 tumor challenge. (D) Quantification of (C). Data are mean ± s.d. and significance was tested using one-way ANOVA with Tukey’s multiple comparisons test (n=6-13 tumors/group, pooled from four independent experiments). (E) Stacked bar chart of main cell populations identified in tumors from (C) via flow cytometry plotted as proportion of all live cells. (F) Representative flow cytometry plots of (left) CD11b^+^ monocyte/macrophages and (right) F4/80+ TREM2+ tumor-associated macrophages (TAMs) from HY19636 tumors with *Osmr* or *Tgfbr2* KO in fibroblasts. (G) Quantification of (F). Data are mean ± s.d. and significance was tested using one-way ANOVA with Tukey’s multiple comparison test (n=6-13 tumors/group, pooled from four independent experiments). AAV, adeno-associated virus. hU6, human U6 promoter. CBh, chicken beta-actin hybrid promoter. pA, polyadenylation signal. s.c., subcutaneous. SCR, scramble. ns, non-significant. *p<0.05, **p<0.01, ***p<0.001, ****p<0.0001.

While homeostatic skin fibroblasts expectedly had mostly DP fibroblast and low NCAM1 activation-marker expression (**Figure 2B**), CAFs exhibited a more DN phenotype (**Figure 2C**), which was decreased in both AAV-gOsmr and AAV-Tgfbr2 groups when compared to controls (**Figure 2D**), suggesting that *Osmr* or *Tgfbr2* KO in CAFs is sufficient to shift the subset balance away from the activated DN CAF population. *Tgfbr2* KO also decreased the frequency of the NCAM1+ DN population (**Figure 2D**). We did not observe a significant effect on tumor weight or fibroblast, endothelial, and immune cell numbers (**Figures 2E and S3A**). These findings were replicated in the YUMM5.2 melanoma model (**Figures S3B-S3D**), demonstrating robust adaptability of this local fibroblast gene editing tool to other settings.

For the control group, we decided to pool tumor samples that had received AAV with a scrambled gRNA sequence (AAV-gSCR) that does not recognize any sequence in the mouse,^41^ AAV with a gRNA targeting the T cell receptor α constant (*Trac*) locus—which is not expressed in fibroblasts and, thus, serves as a genome cutting control (**Figure S3E**)—and samples that had received no AAV. These three control groups all showed no significant differences on measured fibroblast phenotypes when compared to each other (**Figure S3F**) and so are presented as one ‘control’ group for the rest of the study.

A benefit of engineering fibroblasts specifically in the TME is the ability to study the effect of their modification upon other local cells. Given the relationship between macrophages and fibroblasts in a multitude of tissue functions and pathology,^42,43^ we sought to apply our method to understand modulation in tumor-associated macrophages (TAMs) (**Figure S3G**). While TAMs did not change in their frequency within all immune cells, both *Osmr* and *Tgfbr2* KO in CAFs led to a slight but statistically significant increase in the frequency of those that expressed MHCII^+^ (**Figures 2F** and **2G**)—a phenotype associated with immunostimulatory myeloid cells.^44,45^ This increase in MHCII+ TAMs reproduced in the YUMM5.2 model again without an increase in total intratumoral CD90+ lymphocytes or monocyte/macrophages (**Figures S3H-S3J**). These findings highlight the utility of modulating genes-of-interest in CAFs to induce changes in their phenotype, with secondary effects on other cells of the tumor microenvironment, such as TAMs.

Extending the analysis of CAF pathway perturbations beyond OSM and TGFβ signaling, we tested how *Il1r1* KO in CAFs (**Figures S3K and S3L**) in the PDAC model affects their subset polarization because IL-1 pathway activity in CAFs has been shown to promote disease progression. ^27,28^ Here, we found no changes compared to control (**Figure S3M**), as well as no changes when measuring TAM frequencies in the tumor, although MHCII^+^ TAMs were again more abundant in CAF *Il1r1* KO tumors (**Figure S3N**). Thus, *Osmr*, *Tgfbr2*, or *Il1r1* KO in CAFs repolarizes CAFs differently, prompting us to investigate those differences and their effect on the networks of gene expression in the TME in more detail.

### Local *Tgfbr2* knockout in cancer-associated fibroblasts reveals emergence of unique *Col18a1*^hi^ CAF cell state

To study the effect of *Osmr*, *Tgfbr2*, or *Il1r1* knockout in CAFs on a transcriptional level and to capture its effect on other cells of the TME, immune, stromal, and tumor cells of subcutaneously grown KPC tumors were collected for single cell RNA sequencing (scRNA-Seq) analysis after targeted, local knockout by AAV in *Pdgfra*-Cas9-EGFP mice (**Figures 3A and S4A**). A total of 115,637 high-quality cells were captured from a total of 16 tumors (n=4 per AAV-gRNA group), covering all major cell populations in the TME: neutrophils, monocyte/macrophages (MonoMacs), dendritic cells (DCs), mast cells, T cells, natural killer (NK) cells, endothelial cells, pericytes, and fibroblasts (**Figures S4B-S4D).**

**Figure 3.**
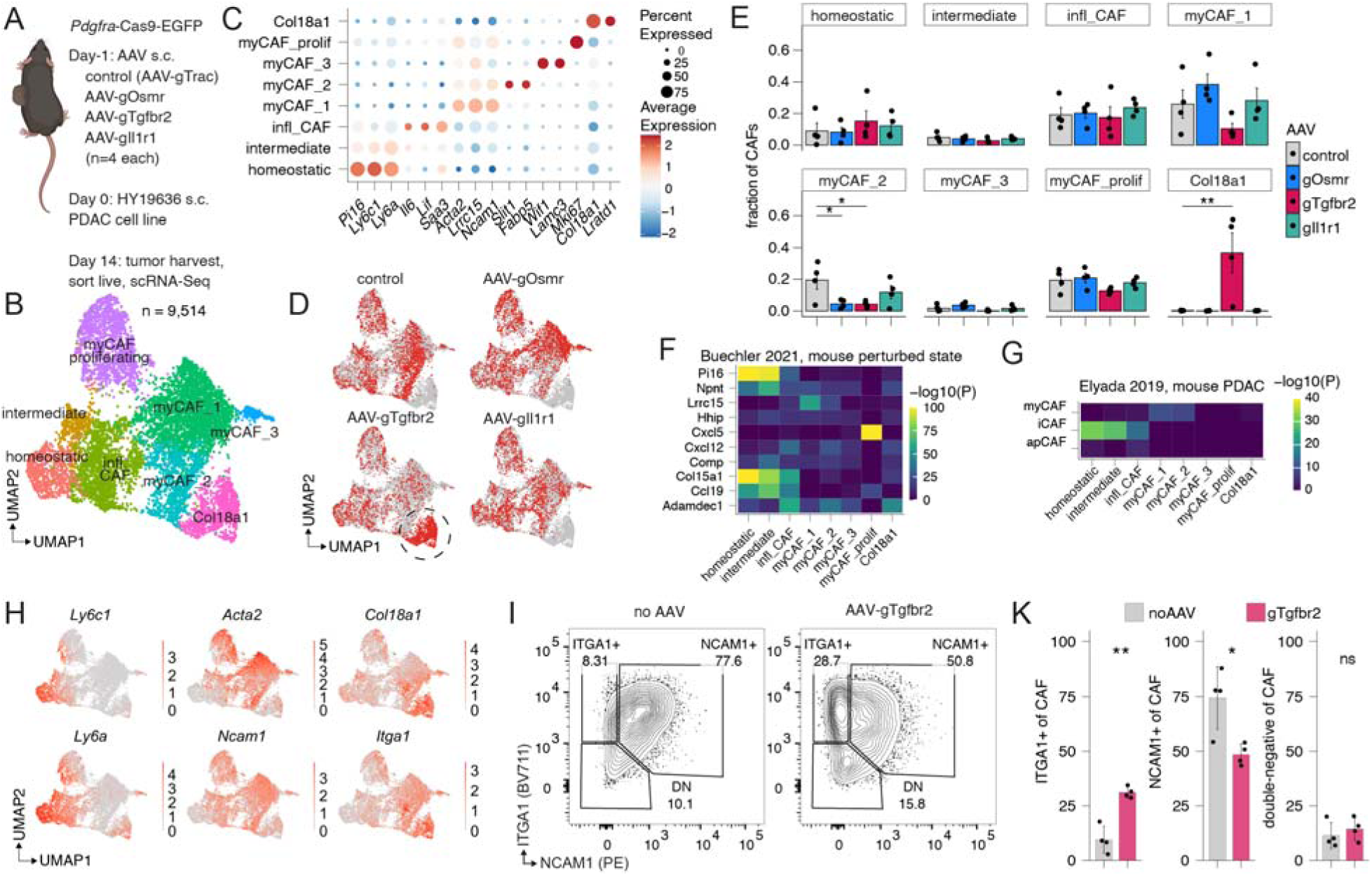
Local *Tgfbr2* knockout in cancer-associated fibroblasts reveals emergence of unique *Col18a1*^hi^ CAF cell state. (A) Experimental layout. (B) Uniform manifold approximation and projection (UMAP) of cancer-associated fibroblast (CAF) subsets. (C) Marker gene expression of identified CAF subsets. (D) UMAP of CAFs highlighted by AAV-gRNA condition. (E) Bar charts of frequency of CAF subsets (scRNA-Seq) as a fraction of all fibroblasts by AAV-gRNA condition plotted as mean±SEM (n=4 per group). *p<0.05 one-way ANOVA with Dunnett’s multiple comparison test. (F and G) Similarity of DEGs between CAF subsets (columns) and (F) mouse fibroblasts in perturbed states from Buechler et al., 2021^2^ (rows) or (G) mouse PDAC CAF subsets from Elyada et al, 2019^18^. Statistical significance of DEG overlap (-log10(P value), hypergeometric test, *P* value adjusted using Bonferroni correction) for each pairwise comparison plotted as heatmap. Yellow indicates high statistically significant overlap of DEG lists. (H) UMAP visualization of select genes (homeostatic *Ly6c1* and *Ly6a*; myofibroblast-associated *Acta2* and *Ncam1*; AAV-gTgfbr2-associated *Col18a1* and *Itga1*) in CAFs. (I) Flow cytometry contour plots of tumor fibroblasts on day 14 after 2.5e5 KPC s.c. cell injection in *Pdgfra*-Cas9-EGFP mice with or without 1e10 vg AAV-gRNA injection on day −1. ITGA1+ NCAM1- gate (top left) highlights AAV-gTgfbr2-specific CAF population. (K) Quantification of (I). Data are mean ± s.d. and significance was tested using Welch’s t-test (n=4/group). One of two representative experiments shown. *p<0.05, **p<0.01, ns, non-significant.

Sub-clustering of the CAF compartment revealed eight different subsets (**Figures 3B** and **3C**). Homeostatic fibroblasts expressed *Pi16*, *Ly6c1*, and *Ly6a*, genes associated with ‘universal’ fibroblasts (**Figure S2K**); inflammatory CAFs expressed *Il6*, *Lif*, and *Saa3*, genes previously described in pro-inflammatory settings^16,46^; myofibroblastic CAFs (myCAFs) were characterized by myofibroblast-associated genes, such as *Acta2*, *Lrrc15*, and *Ncam1* ^47–49^ and could be further subdivided into 4 groups based on proliferative markers (*Mki67*, myCAF_prolif), *Slit1* and *Fabp5* (myCAF_2), or *Wif1* and *Lamc3* (myCAF_3). The intermediate cluster harbored genes of both homeostatic and inflammatory fibroblasts, whereas the Col18a1 cluster was named after one of its most differentially expressed gene, *Col18a1*, and noticeably has low expression of the other homeostatic, inflammatory, or myofibroblast-defining genes.

The Col18a1 cluster was only present in the AAV-gTgfbr2 group (**Figure 3D**) and represented the most abundant CAF subset within those samples at about one half of all AAV-gTgfbr2 CAFs (**Figure 3E**). Conversely, *Tgfbr2* knockout in CAFs lead to a decrease in all myCAF subsets (**Figure 3E**). This could be explained by lack of a recently published fibroblast TGFβ1 response signature specifically in AAV-gTgfbr2 CAFs (**Figure S4E**), a prominent driver of myofibroblast differentiation.^50^ To assess if the emergence of the *Col18a1* expressing CAFs in mouse PDAC model was unique to our AAV-based *Tgfbr2* knockout, we re-analyzed a published mouse PDAC scRNAseq data set using conditional *Tgfbr2* knockout in universal fibroblasts^14^ and found that a population with the Col18a1 CAF cluster signature can also be identified specifically when *Tgfbr2* was knocked out using Cre/LoxP-based mouse genetics (**Figure S4F**), demonstrating the methodological robustness of using AAV-gRNA or Cre/LoxP-based gene KOs.

To gain a better understanding of how *Osmr*, *Tgfbr2*, or *Il1r1* knockout affects CAF transcriptional polarization, we compared the overlap of DEG lists from each knockout group and noticed that the majority of DEGs were unique to their respective knockout group (**Figures S4G and S4H and Supplementary Table 3**). For example, AAV-gOsmr and AAV-gTgfbr2 only shared 20 and 24 genes that were up- or downregulated, respectively, when compared to the unperturbed control CAFs (**Figure S4H**). Gene set enrichment analysis (GSEA) revealed that AAV-gOsmr CAFs were enriched for inflammatory response genes (*Ifitm1*, *Il1a*, *Irf1*, *Lyn*, *P2rx4*), but depleted of hypoxia-related genes that are involved in carbohydrate metabolic processes (*Slc2a1*, *Pfkl*, *Pdk1*, *Gys1*), whereas AAV-gTgfbr2 CAFs were enriched for hypoxia-related (*Ndrg1*, *S100a3*) and glycolytic genes (*Pgk1*, *Ldha*, *Ugp2*), but depleted of epithelial-to-mesenchymal transition (EMT) genes (*Acta2*, *Tagln*, *Lrrc15*, *Ccn1*) (**Figures S4I and S4J**), a known consequence of absent TGFR2 signaling in fibroblasts.^2,18,48^ Together, these results demonstrate that the AAV-mediated knockout of individual receptors can induce distinct transcriptional changes in CAFs.

With the Col18a1 cluster uniquely emerging in the AAV-gTgfbr2 condition and it not being characterized by homeostatic, inflammatory, or myofibroblastic gene signatures, we also compared its transcriptional profile to published mouse scRNAseq fibroblast data sets.^2,18^ As expected, the DEGs of the homeostatic and intermediate clusters had the most overlap with the *Pi16*+ and *Col15a1*+ homeostatic/universal state, the inflammatory CAF cluster had the most overlap with the colitis-associated *Adamdec1*+ state, and the myCAF_1 and myCAF_2 clusters were most similar to *Lrrc15*+ myofibroblasts (**Figure 3F**) and PDAC myCAFs (**Figure 3G**). However, the Col18a1 cluster did not uniquely map to any previously identified mouse fibroblast states (**Figures 3F** and **3G**). Even when comparing the Col18a1 gene signature to published human fibroblast profiles,^51,52^ the Col18a1 cluster did not have the highest overlap with any identified human fibroblast subset (**Figures S4K and S4L**), further suggesting that *Tgfbr2* KO CAFs adopt a unique transcriptional cell state.

To identify the Col18a1 CAFs orthogonally via flow cytometry, we identified surface expressed proteins that were transcriptionally expressed at high levels within the Col18a1 cluster. While *Itga1* (encoding integrin alpha 1 / ITGA1) was not exclusively expressed in the Col18a1 cluster (**Figure 3H**), when combining ITGA1 staining with SCA-I, Ly6C, and NCAM1, we identified a ITGA1+ NCAM1- CAF population that specifically emerged in AAV-gTgfbr2 tumors (**Figures 3I** and **3K**) and was low for the homeostatic markers SCA-I and Ly6C (**Figure S4M**).

*Tgfbr2* knockout-induced *Col18a1*^hi^ CAFs are distinct from iCAFs and myCAFs and correlate with worse survival in human PDAC patients.

The emergence of the *Col18a1*^hi^ CAF state prompted us to investigate the possibility that local *Tgfbr2* knockout in fibroblasts during tumor growth led to the polarization of a distinct CAF cell state. *In silico* pseudotime-based trajectory analysis using Monocle3^53^ of all captured CAFs, sans proliferating myCAFs, placed Col18a1 CAFs at the end of the trajectory (**Figure 4A**). The analysis suggested that CAFs, in the setting of *Tgfbr2* knockout, progressed from homeostatic to inflammatory through the myCAF cell state until they eventually arrive at the Col18a1 cell state (**Figure 4B**). This progression along the pseudotime axis was characterized by a loss of the universal marker *Pi16*, sequential emergence of inflammatory markers *Saa3* and *Cthrc1*, prior to upregulation of myofibroblast markers *Lrrc15*, *Acta*, and *Ncam1*, which then were lost when the Col18a1 state was reached (**Figure S5A**).

**Figure 4.**
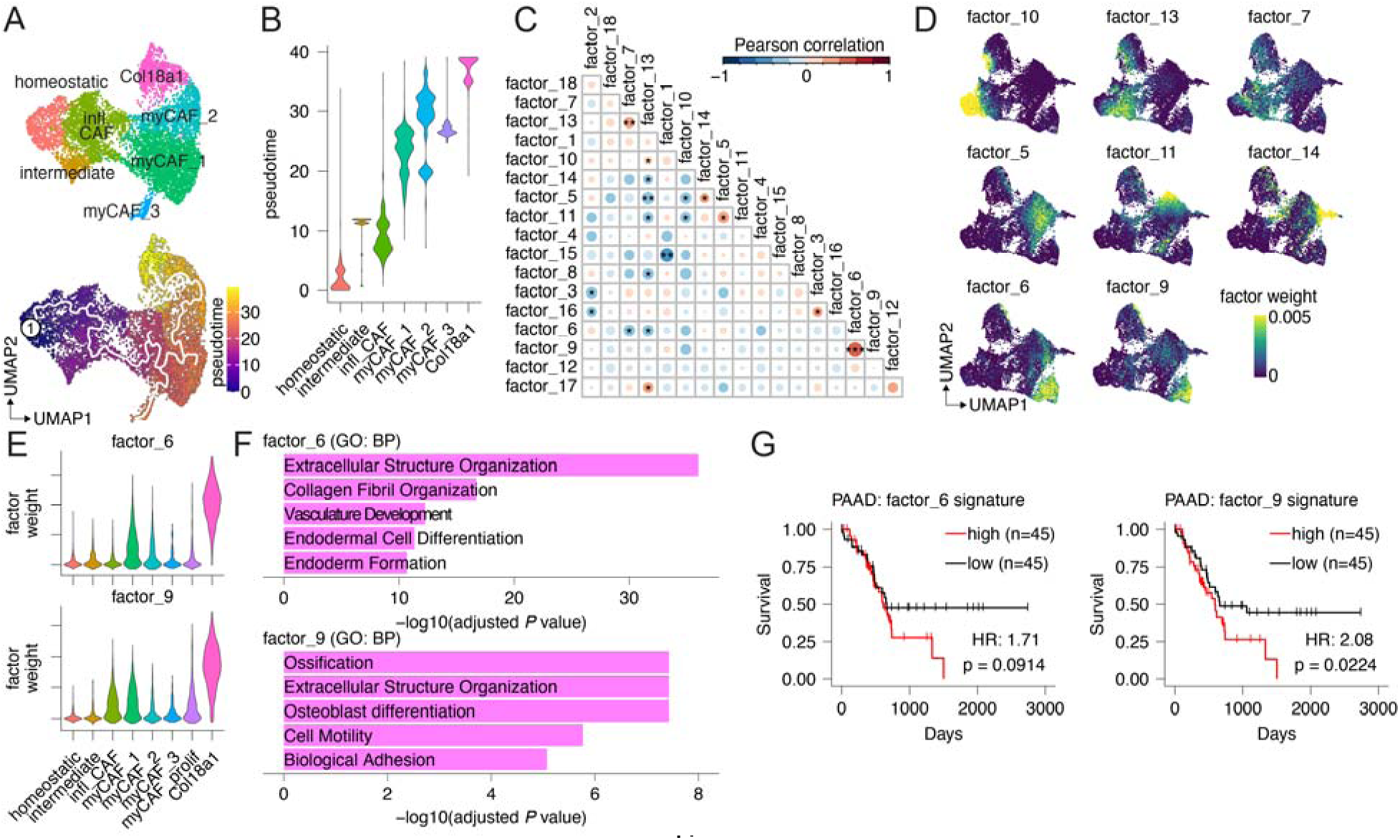
*Tgfbr2* knockout-induced *Col18a1*^hi^ CAFs are distinct from iCAFs and myCAFs and correlate with worse survival in human PDAC patients. (A) UMAP of (top) CAF subsets without myCAF_prolif subset and (bottom) pseudotime trajectory starting at the homeostatic fibroblast subset and ending at cluster myCAF_3 or Col18a1. (B) Violin plot of CAF subsets plotted according to their distribution in pseudotime. (C) Pairwise Pearson correlation coefficient of CAF NMF factor expression across all CAFs. Rows and columns are ordered by hierarchical clustering. (D) UMAP visualization of select NMF factor expression in CAFs. (E) Violin plots of (top) NMF factor_6 and (bottom) NMF factor_9 expression in the CAF subsets. (F) Enriched gene sets (Gene Ontology biological process) based on top contributing genes in (top) NMF factor_6 and (bottom) NMF factor_9. (G) Kaplan-Meier survival plots of TCGA PAAD patients categorized into upper and lower quartiles by (left) the NMF factor_6 signature gene expression and (right) the NMF factor_9 signature gene expression. Hazard ratio (HR) and p value of Cox regression fit are shown.

With the Col18a1 cluster not previously classified, we therefore identified de novo signatures across all identified CAF states using non-negative matrix factorization (NMF).^54^ This approach allows non-cluster-based analysis of transcriptional cell states by decomposing the cell-by-gene scRNA-Seq count matrix into two smaller matrices, where one matrix represents the factors, i.e. groupings of co-expressed genes, and the other matrix represents the expression of those individual factors across each cell. Especially for cell populations with mixed cell states or without clearly defined subsets, such as fibroblasts, this approach can uncover heterogeneous gene expression patterns that are masked by clustering analysis.

We identified a total of 18 different factors within the CAF population (**Figure S5B and Supplementary Table 1**). While most averaged factor weights did not differ between the AAV-gRNA groups, factor-6 and factor-9 were elevated in AAV-gTgfbr2 CAFs and factor-14 was elevated in the AAV-gOsmr CAFs (**Figure S5C**). Correlation of factor weights across all CAFs further revealed that there were 3 major groupings of gene factors: (1) factors 7 and 13 that were enriched in inflammatory CAFs; (2) the factors 5, 11, and 14 that were enriched in myCAFs; (3) and the AAV-gTgfbr2-enriched factors 6 and 9 (**Figures 4C** and **4D**). Noticeably, the inflammatory factors 7 and 13 were anti-correlated with both other groupings, suggesting that the inflammatory cell state is mutually exclusive with other CAF cell states detected here. CAFs within the Col18a1 cluster were enriched for factors-6 and 9 (**Figure 4E**), which were identified as gene signatures of various developmental states (factor_6) and ossification/osteoblast differentiation (*Fzd1*, *Pth1r*, *Spp1*) (factor_9) (**Figure 4 and Supplementary Table 1**). With osteoblasts representing a unique mesenchymal lineage within the bone tissue,^55^ we speculated whether *Tgfbr2* knockout in CAFs reprograms them to an osteoblast-like cell. However, comparison of Col18a1 DEGs to a recently published mouse bone marrow stromal cell atlas^56^ did not suggest that the Col18a1 cluster adopted a specific bone marrow stroma program (**Figure S5D**). This suggested that the AAV-gTgfbr2 induced CAFs harbored a cell state that is distinct of myofibroblasts, homeostatic, and inflammatory CAFs, instead presenting as a mix of ECM-producing CAFs with an osteoblast-like gene signature.

While, both, a high stromal gene signature,^57^ as well as a high fibroblast TGFβ1 response signature^50^ in human cancers is associated with worse survival (**Figures S5E and S5F**), we noticed that a signature derived of the DEGs from AAV-gTgfbr2 CAFs—which lack the TGFβ1 response signature (**Figure S4E**)—does not associate with better survival (**Figure S5G**). In fact, a high AAV-gTgfbr2 CAF signature was associated with a hazard ratio of >1.5 in bladder carcinoma, glioblastoma multiforme, low-grade gliomas, stomach adenocarcinoma, and pancreatic ductal adenocarcinoma (PAAD) (**Figure S5G**). Additionally, the AAV-gTgfbr2 CAF-specific gene programs factor_6 and factor_9 (top25 weighted genes from each) were associated with worse survival in PAAD, with factor_9 reaching a significant hazard ratio of 2.08 (p = 0.0224) (**Figure 4G**). Together, this suggests that the AAV-gTgfbr2-induced CAF cell state, characterized by an osteoblast-like gene signature, is pro-tumorigenic.^58^

### *Tgfbr2* knockout-induced *Col18a1*^hi^ CAFs recruit Siglec-F^hi^ neutrophils via high *Cxcl5* expression

To gain a better understanding how the AAV-gTgfbr2-induced *Col18a1*^hi^ CAFs interact with their TME, the percentage of all identified major immune cell subsets (**Figures S6A-S6I**) were correlated across all samples (**Figure 5A**). This revealed several cell networks, with Col18a1 CAFs positively correlating with two populations; the TAM.Ndrg1 subset and the neutrophil.4 subset. The neutrophil.4 subset was most abundant in AAV-gTgfbr2 tumors (**Figures 5B**) and characterized by high *Siglecf* expression (**Figure 5C**). The TAM.Ndrg1 subset had elevated MHC-II gene (*H2-Ab1*, *H2-Aa*, *H2-Eb1*) expression in AAV-gTgfbr2 tumors (**Figure S6J**), mirroring our previous findings of increased MHCII+ TAMs, as analyzed by flow cytometry (**Figure 3F**).

**Figure 5.**
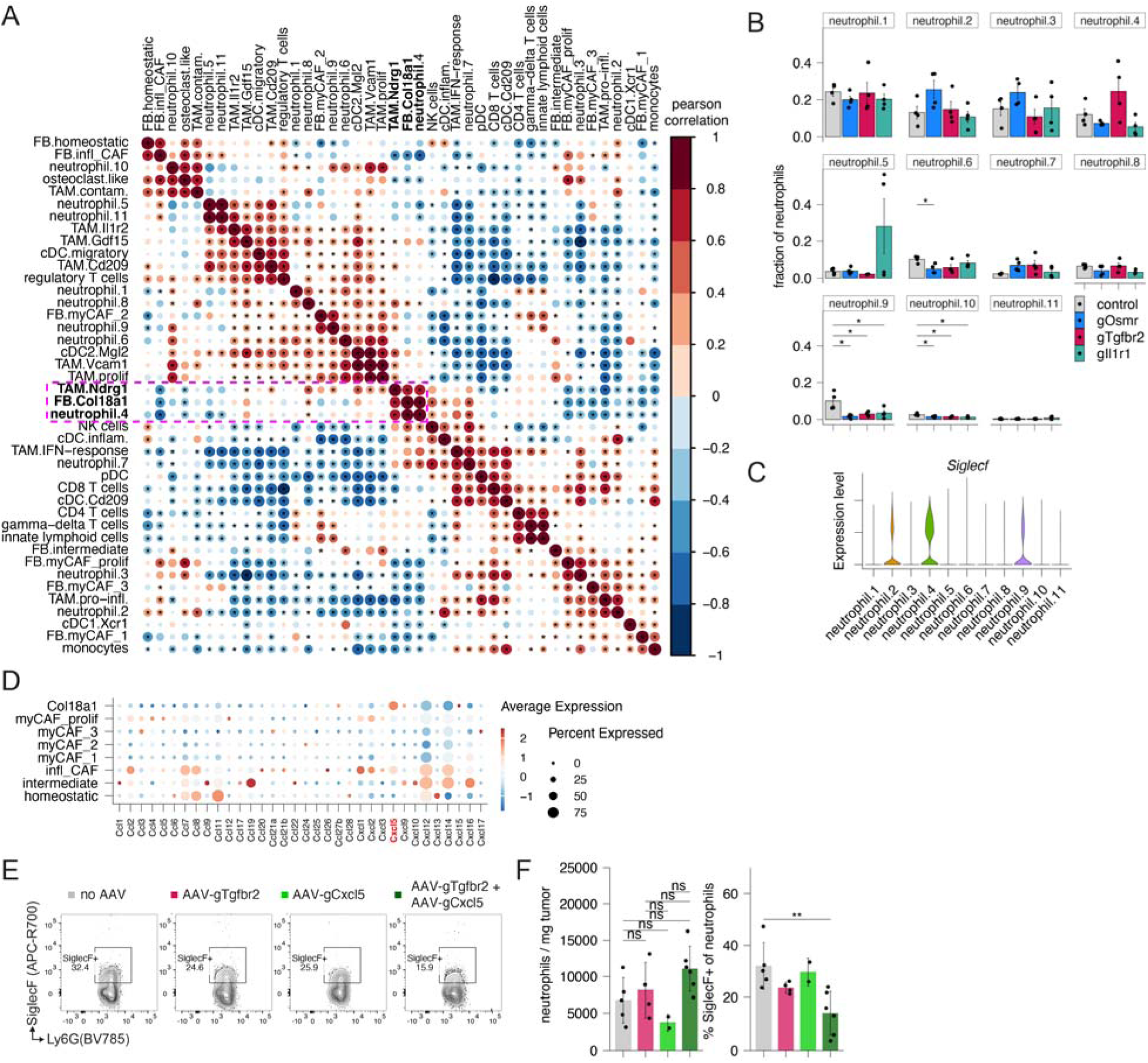
*Tgfbr2* knockout-induced *Col18a1*^hi^ CAFs recruit Siglec-F^hi^ neutrophils via high *Cxcl5* expression. (A) Heatmap showing the pairwise Pearson correlation of cell type frequencies among AAV-gRNA samples. Box highlights grouping of Col18a1 CAF-associated cell types. (B) Bar charts of frequency of neutrophil subsets (scRNA-Seq) as a fraction of all neutrophils by AAV-gRNA condition plotted as mean±SEM (n=4 per group). *p<0.05 one-way ANOVA with Dunnett’s multiple comparison test. (C) Violin plot of *Siglecf* expression among all neutrophil subsets. (D) Bubble plot of scaled expression of all detected *Ccl* and *Cxcl* genes in all CAF subsets. *Cxcl5* highlighted in red as most highly expressed in Col18a1 cluster. (E) Flow cytometry contour plots of DAPI- CD45+ CD11b+ Ly6G+ neutrophils on day 14 after 2.5e5 KPC s.c. cell injection in *Pdgfra*-Cas9-EGFP mice with single or combinatorial 1e10 vg AAV-gRNA injection on day −1. (F) Quantification of (E): (left) total neutrophil numbers normalized by tumor weight and (right) percentage of Siglec-F+ neutrophils among all neutrophils.

To discover whether *Col18a1*^hi^ CAF-mediated recruitment of the neutrophil.4 subset, we analyzed transcriptional expression of all detected CCL/CXCL chemokine family members in the CAF subsets and noted the elevated expression of *Cxcl5* (**Figure 5D**), a chemokine binding to the constitutively expressed chemokine receptor CXCR2 on neutrophils.^59^ Cancer cell-derived CXCL5 has been described to promote neutrophil infiltration into tumors,^60^ with *Tgfbr2* deletion in breast cancer cells resulting in increased CXCL5 production.^61^ To test if *Cxcl5* expression in AAV-gTgfbr2-induced *Col18a1*^hi^ CAFs is recruiting the neutrophil.4 subset to the PDAC TME, we used Siglec-F staining as a marker for the neutrophil.4 cluster (**Figure 5C**) and employed combinatorial AAV-gRNA administration (AAV-gCxcl5 plus AAV-gTgfbr2) to knock out *Cxcl5* (**Figure S6K**) together with *Tgfbr2* in CAFs. While Siglec-F staining and analysis by flow cytometry did not show a significant increase of Siglec-F+ neutrophils in AAV-gTgfbr2 tumors, dual *Cxcl5* and *Tgfbr2* knockout locally in CAFs via combinatorial AAV-Tgfbr2 + AAV-gCxcl5 delivery did reduce Siglec-F+ neutrophil accumulation in the TME (**Figures 5E** and **5F**), implying that CAF-derived CXCL5 contributes to Siglec-F+ neutrophil accumulation in PDAC tumors.

Combinatorial knockout shows that emergence of *Tgfbr2* knockout-induced *Col18a1*^hi^ CAFs is dependent on TNFR1 and canonical Wnt signaling.

Finally, to nominate pathways that could potentially positively amplify the generation of the osteoblast-like AAV-gTgfbr2-inhibited *Col18a1*^hi^ CAF subset, we used two computational approaches: ligand-receptor analysis using CellChat^62^ and transcription factor activity inference using decoupleR.^63^

Ligand-receptor based analysis using CellChat^62^ to identify potential signaling pathways between the Col18a1 CAF cluster and the TAM.Ndrg1 cluster, as well as the neutrophil.4 cluster, nominated TNF-TNFR1 (*Tnf*-to-*Tnfrsf1a*) signaling as a major contributor to the Col18a1 CAF cell state (**Figure 6A**). Both, the TAM.Ndrg1 and the neutrophil.4 clusters were most abundant in AAV-gTgfbr2 tumors (**Figures S7A and S7C**) and were transcriptionally enriched for the GO hallmark term “TNFa signaling via NFkB” (**Figures S7B and S7D**), suggesting that they together with the Col18a1 CAFs exist in a TME with high TNF⍺ signaling. Thus, we hypothesized if TNFR1-mediated signaling in AAV-gTgfbr2 CAFs mediates the emergence of the Col18a1 / factor-9 osteoblast-like cell state.

**Figure 6.**
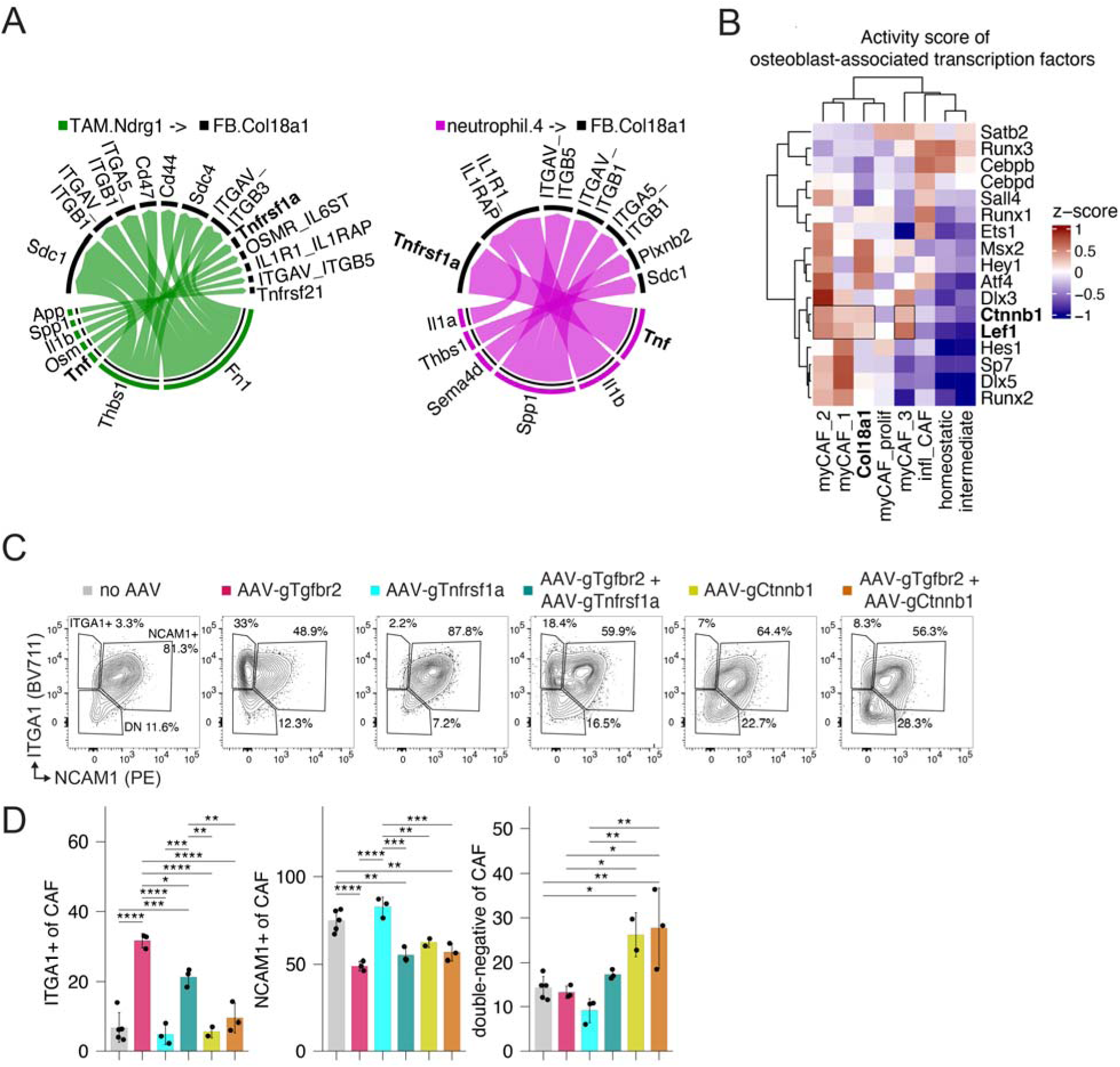
Combinatorial knockout shows that emergence of *Tgfbr2* knockout-induced *Col18a1*^hi^ CAFs is dependent on TNFR1 and canonical Wnt signaling. (A) Chord diagrams of ligand-receptor pairs between (left) TAM.Ndrg1 cells (senders) and Col18a1 CAFs (receivers) and (right) neutrophil.4 subset (senders) and Col18a1 CAFs (receivers). Tnf/Tnfrsf1a ligand-receptor pairing highlighted as bold. (B) Heatmap of average activity score calculated by PROGENy^63^ of osteoblast-associated transcription factors^55^ in CAF subsets. (C) Flow cytometry contour plots of tumor fibroblasts on day 14 after 2.5e5 KPC s.c. cell injection in *Pdgfra*-Cas9-EGFP mice with single or combinatorial 1e10 vg AAV-gRNA injection on day −1. (D) Quantification of (C). Data are mean ± s.d. and significance was tested using one-way ANOVA with Tukey’s multiple comparison test (n=2-5 tumors/group, pooled from two independent experiments).

For transcription factor activity inference, we focused on osteoblast-associated transcription factors,^55^ because NMF gene program analysis revealed that the Col18a1 cluster was enriched for osteoblast-related genes (**Figure 4F**). *Ctnnb1* and *Lef1*—two regulators of the Wnt pathway—were nominated as highly active in Col18a1 and all myCAF subsets (**Figure 6B**). This prompted us to hypothesize that Wnt pathway activity—a known contributor to the myofibroblast phenotype in cancer^64^ and skin fibrosis^65^—in the context of TGFBR2 signaling loss produces the Col18a1 osteoblast-like CAF phenotype. To test the contributions of TNFR1 or Wnt pathway signaling in CAFs, we generated AAV-gTnfrsf1a and AAV-gCtnnb1 (**Figures S7E and S7F**) and found that combinatorial knockout of *Tgfbr2* with *Tnfrsf1a* or *Ctnnb1* in CAFs decreased the ITGA1+ NCAM1- CAF population compared to AAV-gTgfbr2 alone (**Figures 6C** and **6D**), indicating that both pathways are contributing to ITGA1+ Col18a1 CAF cluster generation. Furthermore, AAV-gCtnnb1 alone decreased the myofibroblast NCAM1+ population to similar levels as AAV-gTgfbr2, while simultaneously maintaining the ITGA1- NCAM1- double-negative, homeostatic-like CAF population (**Figures 6C** and **6D**).

Using AAV-gRNA-mediated knockouts in fibroblasts, we present an experimental toolbox that allows local *in vivo* gene editing of fibroblasts to study gene functions—both on an individual, as well as on a combinatorial level—in the context of mouse tumor models (**Graphical Abstract**).

## Discussion

The modularity of the localized *in vivo* CAF gene editing approach presented here—relying only on delivering a gRNA packaged in an AAV and conditional Cas9 expression in PDGFRA+ fibroblasts—allows customizable, user-defined investigation of individual or combinatorial gene perturbations, while the local application allows spatially confined perturbations that limit potentially confounding effects induced by systemic gene perturbations, such as conventional Cre-loxP-based mouse models. We believe this approach will be beneficial to the advancement of understanding the complexity of fibroblast biology within different disease settings, as gene candidates can be more quickly and cost-effectively evaluated by not fully relying on complex mouse breeding to generate genetically engineered single- or multiple-knockout mouse strains.

Previous to our study, AAV-mediated gene delivery in fibroblasts has been demonstrated *in vivo*, resulting in transient target gene expression from a classically introduced promoter.^66–68^ However, our study goes further in achieving true genome editing as well as improving selectivity for fibroblast lineages as well as to the local environment. As an alternative approach to our CRISPR/Cas9 method, another recent study reported the knockdown of a target in fibroblasts *in vivo* using shRNA.^69^ While this will be beneficial in situations when Cas9 expression cannot be achieved in the target cell population, downregulation of mRNA in this method is transient, whereas CRISPR/Cas9-mediated KO allows sustained gene KO and more long-term analysis of gene perturbations—we observed stable KO of up to 14 days in the tumor fibroblast population (**Figure 1**). Generating a sustained gene KO is especially valuable when studying long-term processes, excluding confounding effects via residual gene expression in knockdown scenarios. Additionally, the AAV-mediated gene delivery in fibroblasts allows editing in adult animals of genes that can cause embryonic lethality when knocked out constitutively.

Importantly, transduction of fibroblasts with a non-cutting gRNA (gSCR) or a gRNA cutting an irrelevant, non-expressed gene (gTrac) *in vivo* did not affect their polarization noticeably compared to non-transduced fibroblasts (**Figure S3F**). This indicated that at least in the context of tumor growth, the use of AAV1-serotyped scAAV does not affect fibroblast biology on its own and can be considered a fairly innocuous transduction method^70^ that does not mask effects of targeted gene KO.

One clear opportunity afforded here is the possibility of sculpting fibroblasts through a series of edits. Fibroblasts receive a multitude of signaling input during tumor growth, as demonstrated by their expression of different receptor families,^2,62^ as well as their transcriptional and functional polarization when receiving these signals.^1,17,47,71^ Sequential gene edits via subsequent injection of multiple sKO-AAVs or simultaneous edits via co-injection of multiple sKO-AAVs allows uncovering of redundant, synergistic, compensatory, or agonistic effects of gene functions in CAF biology. For example, inflammatory and myofibroblast CAF states have been described as two distinct fates fibroblasts can adopt in PDAC.^14,17^ While inhibition of either the JAK/STAT or the TGFBR signaling pathway via systemic drug application polarized CAFs to opposite cell states,^17^ we noticed adoption of a separate osteoblast-like transcriptional cell state when TGFBR2 signaling was specifically inhibited in CAFs only, as opposed to pathway inhibition in any cell that a systemically administered drug could reach. Furthermore, combinatorial knockout of the TNF pathway or the canonical Wnt pathway (via *Ctnnb1* knockout) in combination with the TGFBR2 pathway diminished CAF polarization towards both myofibroblast and osteoblast-like phenotypes, demonstrating how multiple gene edits via AAV-gRNA delivery can uncover hierarchies of pathway activities in CAFs (**Figure 6**).

Lack of TGFBR2 signaling in FSP1+ fibroblasts with a resulting increase in secreted hepatocyte growth factor (HGF) has been described as a transforming factor in epithelial cells, specifically contributing to squamous cell carcinoma of the forestomach, intraepithelial neoplasia in the prostate of mice, and developmental abnormalities in mammary duct development.^58,72,73^ Other tissues appeared histologically normal, emphasizing the importance of how organ-specific and temporally controlled fibroblast perturbations (e.g. constitutively throughout development or acute perturbation in the adult animal) can have distinct effects on their local tissue environments. The local fibroblast-specific gene editing afforded us the possibility of temporally controlled perturbation, where *Tgfbr2* knockout induced a unique CAF cell state that led to recruitment of distinct neutrophil and TAM subsets. This observation was similar to recent findings of conditional fibroblast-specific knockout of *Tgfbr2* in either alveolar fibroblasts in the setting of lung fibrosis or in brain border fibroblasts in stroke, where an increase in monocytes or neutrophils, respectively, were observed when fibroblast-specific TGFBR2 signaling was disrupted.^46,74^ Intriguingly, mice fared worse in both settings when *Tgfbr2* was knocked out in fibroblasts, mirroring the worse survival of PAAD in patients with a high *Tgfbr2* knockout-associated, osteoblast-like gene signature (**Figure 4G**). While the higher morbidity in the lung fibrosis and stroke models could be attributed to a disrupted resolution of inflammation, further work is required to determine how a lack of *Tgfbr2* signaling in CAFs contributes to worse survival.

Single cell studies of tissue ecosystems have highlighted the relationship of immune, stroma, and parenchymal cells in settings of disease and homeostasis.^57,75–77^ Observing these processes over space and time allows generation of experimentally and computationally inferred cell state trajectories within individual cells, as defined by transcriptional changes occurring throughout the process.^6,78^ These type of changes of transcriptional states by individual cells are linked through paracrine signaling with their neighboring cells and has been formulated in two-cell circuits between heterotypic cells, such as macrophages and fibroblasts.^79^ By manipulating fibroblasts locally with high genetic precision, we were able to extend these findings to test how perturbations in fibroblasts alter the relationships between different cell types and which cell states are linked during multicellular tissue processes. Crucially, this goes beyond computationally inferred cell-cell interactions and affords rapid validation of which mediators orchestrate the observed multicellular changes, which are not necessarily observed on a compositional level of changed major cell type proportions but instead hidden within linked transcriptional cell state changes. One previously underappreciated linkage discovered here is between TGFBR2 signaling inhibited CAFs recruiting Siglec-F^hi^ neutrophils via increased *Cxcl5* expression (**Figure 5**). Siglec-F^hi^ neutrophils in the TME have been described as CD8 T cell inhibitory,^80^ inversely correlated with immunotherapy response,^81^ and being recruited by cancer cell-derived *Cxcl5*.^60^ Here we extend that cell-cell relationship to include *Cxcl5*+ CAFs, demonstrating how their altered state can negatively affect the cellular TME by recruiting these immune inhibitory neutrophils. This highlights how fibroblasts occupy an important niche within the cellular hierarchy in the TME^82^ and how their local perturbation allows discovery of linked constellations of cell states at site.

The method defined here thus provides a rapid and site-selective means to modulate fibroblasts and record the resulting changes within them and the pathways that are modified via single or combinatorial knockouts of target genes. We envisage that with this quicker turnaround time from gene target identification to combinatorial gene perturbations, we can learn more, discover faster, and describe better fibroblast functions in their native tissue context.

## Limitations

The outlined approach requires use of a transgenic mouse line expressing Cas9 driven by Cre expression in fibroblasts. While choice of Cre-driver can be further modified to control conditional Cas9 expression in a desired cell population to potentially investigate more defined fibroblast subpopulations or other cell types,^1^ it does prevent this technology from being directly adopted for use in genetically engineered mouse models of cancer that rely on Cre expression in the cancer-initiating cell. Genetically engineered mouse models (GEMMs) of cancer generally recapitulate aspects of tumor initiation, progression, metastasis, and outgrowth more faithfully than transplanted tumor models, as used in this study. Fibroblasts have been described to be involved all of these processes,^9^ emphasizing the need to develop more sophisticated approaches. It is conceivable to combine the Cre-driven Cas9 expression in the fibroblast with alternative recombinase systems, such as the Flp/Frt or Dre/rox systems,^83–85^ or non-recombinase-dependent tumor models such as the MMTV-PyMT mammary carcinoma model,^86^ which would allow studying these gene perturbations in GEMMs of cancer.

This study investigated cancer-associated fibroblast cell states as collected from subcutaneously transplanted pancreatic cancer cells. The heterotopic growth of the tumor in the skin does not fully recapitulate the tumor microenvironment in pancreatic cancer. Thus, aspects of pancreas-specific fibroblasts, such as pancreatic stellate cells, which contribute unique aspects of CAF biology,^87^ are not considered here. Orthotopic injection of AAVs into the organ of interest, such as the pancreas, is conceivable, albeit requiring invasive or ultrasound-guided surgical procedures.^88^

## Supporting information

Supplementary Table Plasmids

Supplementary Table Reagents

Supplementary Table gRNA sequences

## Acknowledgements

We would like to thank members of the Krummel and Ari Molofsky labs for scientific discussions and comments on the manuscript. We would like to thank Bushra Samad in the DataScience CoLab at UCSF for guidance on computational analysis. This work was supported by funds from NIH R01CA197363 and the Parker Institute for Cancer Immunotherapy. N.F.K. is the recipient of a CRI / Merck Postdoctoral Fellowship (CRI4546). Flow cytometry was performed at the UCSF Parnassus Flow CoLab (RRID:SCR_018206) and supported in part by the DRC Center Grant NIH P30 DK063720. Sequencing was performed at the UCSF CAT, supported by UCSF PBBR, RRP IMIA, and NIH 1S10OD028511-01 grants.

## Author Contributions

Conceptualization, N.F.K. and M.F.K.; Methodology, N.F.K., W.A.N., J.E., and M.F.K.; Investigation, N.F.K. and I.Z.-L.; Formal Analysis, N.F.K.; Supportive Analysis, K.H., T.C., B.D, T.T., A.J.C.; Visualization, N.F.K.; Writing – Original Draft, N.F.K.; Writing – Review & Editing, N.F.K., K.H., A.J.C., W.A.N., J.E., and M.F.K.; Funding Acquisition, N.F.K, J.E., and M.F.K.; Resources, M.F.K.; Supervision, N.F.K, and M.F.K.

## MATERIALS & METHODS

### Animals

#### Mouse strains and breeding

All mice were housed in an American Association for the Accreditation of Laboratory Animal Care (AALAC)-accredited animal facility and maintained in specific pathogen-free conditions. All animal experiments were approved and performed in accordance with the Institutional Animal Care and Use (IACUC) Program protocol number AN200424. Wild-type C57BL/6J (#000664), *Pdgfra*-Cre (#013148), and *Rosa26*-LSL-Cas9-EGFP (#026175) mice were purchased from The Jackson Laboratory. *Pdgfra*-CreERT2 (stock #032770) mice were a kind gift from Dr. Tien Peng (UCSF). *Cthrc1*-CreERT2;Ai14 mice were a kind gift from Drs. Tatsuya Tsukui and Dean Sheppard (UCSF). Transgenic *Pdgfra*-Cre mice were crossed with *Rosa26*-LSL-Cas9-EGFP^+/+^ mice to generate F1 *Pdgfra*-Cre;*Rosa26*-LSL-Cas9-EGFP^+/wt^ mice. F1 *Pdgfra*-Cre;*Rosa26*-LSL-Cas9-EGFP^+/wt^ mice were crossed with *Rosa26*-LSL-Cas9-EGFP mice to generate F2 *Pdgfra*-Cre;*Rosa26*-LSL-Cas9-EGFP^+/+^ mice, which were used for experiments. Knockin Pdgfra-CreERT2 mice were bred in the same manner to generate *Pdgfra*-CreERT2^+/wt^;*Rosa26*-LSL-Cas9-EGFP^+/+ or +/wt^ mice and used in experiments after tamoxifen treatment (see below). *Col1a2*-CreERT2 (MGI 6721050 from Bin Zhou)^89^ were crossed with *Tgfbr2*^Exon^^2-fl/fl^ mice (MGI 2384513, kind gift from Dr. Ari Molofsky (UCSF))^90^ to generate *Col1a2*-CreERT2^+/wt^;*Tgfbr2*^Exon^^2-fl/fl^ mice. All mice were housed at the University of California, San Francisco (UCSF) animal facility with typical light/dark cycles and standard chow.

#### Tamoxifen treatment

For tamoxifen-induced Cre recombination in *Cthrc1*-CreERT2, *Pdgfra*-CreERT2, and *Col1a2*-CreERT2 mice, adult mice age 6-12 weeks were injected i.p with 100 µl of tamoxifen (Sigma-Aldrich #) dissolved in corn oil or ChremophorEL:EtOH:PBS (ratio 1:1:2) at 20 mg/ml, yielding a dose of 2 mg per mouse, on 5 consecutive days one week prior to tumor engraftment.

#### Tumor models

For tumor growth studies and engraftment, YUMM5.2 mouse melanoma cancer cells derived from male *Tyr*-Cre;*Braf*^V600E/wt^;*Trp53*^-/-^ mice^91^ (5×10^5^ cells / 50 ul PBS) or HY19636 mouse pancreatic cancer cells derived from female KPC mice (*Kras*^LSL-G12D/+^;*Trp53*^lox/+^*;p48/Ptf1a*-Cre)^92^ (2.5×10^5^ cells / 50 ul PBS) were transplanted into the subcutaneous mouse flank of male or female mice, respectively. For tumor growth studies, tumor size was monitored twice a week using the formula 0.5 x (length x width^2^). Once reaching endpoint of size >1000 mm^3^, mice were euthanized. For indicated analyses, mice were sacrificed at indicated timepoints, tumors were excised and processed for downstream analysis.

For intrapancreatic orthotopic growth of HY19636 cells, mice were anesthetized with 3% isoflurane and subcutaneously injected with 50 ul of 0.25% bupivacaine and 50 ul of 50 ug/ml buprenorphine for analgesia. The hair was removed on the left side flank and cleaned with a ChloraPrep swab stick (2% w/v chlorhexidine gluconate and 70% v/v isopropyl alcohol). A small incision was made in proximity to the spleen, both in the skin and the peritoneum, and the pancreas was carefully exposed by gently pulling on the spleen with a forceps. For injection, 2.5e5 HY19636 cells and 3e10 vg of AAV were co-injected in a total volume of 50 ul of PBS into the tail of the pancreas using a 31G insulin syringe. After injection, the peritoneal incision was closed using 4-0 PERMA-HAND SILK sutures (Ethicon, #663) and then the skin incision was closed using the BD AutoClip Wound Closing System. Mice were monitored for 48 h post-surgery for any signs of distress.

### Cell lines

The YUMM5.2 mouse melanoma cell line (ATCC, CRL-3367), the HEK293T cell line (ATCC, CRL-3216), and the mouse fibroblast cell line NIH/3T3 (ATCC, CRL-1658) were purchased from ATCC. The HY19636 pancreatic tumor cell line was a kind gift from Dr. Hiaoqing Ying (MD Anderson Cancer Center, Houston, TX). The Lenti-X 293T cell line was purchased from Takara (#632180). All cell lines were cultured at 37°C in 5% CO_2_ in cell growth media: DMEM (Gibco, 11995065) supplemented with 10% FBS (Foundation C, Gemini Bio, 900-308), 2 mM L-glutamine, 100 U/ml penicillin, 100 µg/ml streptomycin, and 50 µM beta-mercaptoethanol (Gibco, 21985-023). For all tumor injections, cells were used within the first two passages after thawing. The HY19636 cell line was kept below passage #10 and the YUMM5.2 cell line was kept below passage #20 for experiments.

To generate NIH/3T3 cells expressing Cas9 (=NIH/3T3.Cas9.EGFP), NIH/3T3 cells were lentivirally transduced with the viral supernatant produced in Lenti-X 293T cells after transfection with the following plasmids: cargo plasmid containing Cas9 and EGFP (pL-CRISPR.EFS-GFP, Addgene #57818), VSV-G envelope containing plasmid (pMD2.G, Addgene #12259), and envelope plasmid (pCMV-dR8.91, Creative Biogene #OVT2971) at a molar ratio of 1:0.44:0.8, respectively, and added polyethylenimine (PEI), linear, MW 25000 (Polysciences #23966) at a PEI-to-DNA ratio of 4-to-1. Two days after Lenti-X 293T transfection, viral supernatant was filtered through a 0.45 µm filter (Whatman) and added to NIH/3T3 cells with supplemented polybrene (Sigma-Aldrich, #TR-1003-G) at final concentration of 8 µg/ml. Cas9-expressing NIH/3T3 cells were enriched by sorting EGFP^+^ cells.

### AAV production

The helper plasmid (pHelper, Takara, #6234) was used in conjunction with one of three different RepCap plasmids (7m8,^34^ Addgene, #64839; pAAV-DJ,^35^ Cell Biolabs, VPK-420-DJ; pAAV2/1, Addgene, #112862), and a self-complementary AAV cargo vector (see design below) to package vector genomes into double-stranded AAV particles. For AAV production, on day 1, three 15 cm dishes were seeded each with 9e6 HEK293T cells (ATCC, CRL-3216) in 23 ml of cell growth media DMEM (Gibco, 11995065) supplemented with 10% FBS (Foundation C, Gemini Bio, 900-308), 2 mM L-glutamine, 100 U/ml penicillin, 100 µg/ml streptomycin, and 50 µM beta-mercaptoethanol (Gibco, 21985-023). On day 2, the HEK293T cells were transfected using PEI at a PEI-to-DNA ratio of 8-to-1 (660 µg total PEI), with the DNA comprising 20.25 µg cargo plasmid, 26 µg RepCap plasmid, and 36.36 µg pHelper plasmid, all in a total volume of 7.5 ml of a 150 mM NaCl solution. Each plate of HEK293T received 2.5 ml of the PEI-DNA transfection reagent by dropwise adding it on top. Transfected HEK293T cells were cultured for 3 more days and then AAV was collected using the AAVpro Purification Kit Midi (Takara, 6675) following the manufacturer’s instructions. After purification, aliquots were taken to determine titers by qPCR after DNase I (NEB #B0303S) treatment and proteinase K (Qiagen #1114886) digestion. For qPCR, primers amplifying a 138 bp region of the human U6 promoter on the cargo plasmid within the ITRs were used (titer_U6_Forward, gagggcctatttcccatgattcc; titer_U6_Reverse, cccaagaaattattactttctacgtcacg) with the SsoAdvanced Universal SYBR Green Supermix (Bio-Rad #1725270) on CFX384 BioRad Real-Time PCR System with the following cycling protocol: 95C for 30s, 95C for 5s, 58C for 20s, repeat steps 2 and 3 for 40 times. Relative quantity was determined by comparison to a serial dilution of the cargo plasmid standard of known concentration. AAVs were stored in autoclaved protein low-binding tubes (Costar, 3207) and handled with protein low-binding pipets (Corning, 4151).

### AAV cargo plasmid design

To generate the pscAAV-hU6-gSCR-CBh-mCherry plasmid for generation of self-complementary (sc) AAVs carrying a gRNA cassette plus mCherry as a reporter gene for transduction efficiency, the pscAAV-CAG-EGFP plasmid (Addgene, 83279) was cut using AvrII and NotI and then Gibson assembled^93^ with a PCR-amplified fragment from pAAV-U6-gRNA-CBh-mCherry (forward primer, gttcctggaggggtggagtcgtgacctagggagggcctatttcccatgattcctt; reverse primer, aaagcatcgagatcgcaggtgaggcctagcggccgcttacttgtacagctcgtccatgcc). This yielded the intermediate plasmid pscAAV-hU6-gRNA-CBh-mCherry which was never used for scAAV production because it had a 77 bp sequence between the RNA polymerase III termination sequence at the 3’ end of the gRNA scaffold sequence and the KpnI cut site at the 5’ end of the CBh promoter sequence. The pscAAV-hU6-gRNA-CBh-mCherry plasmid was then cut with AvrII and KpnI and Gibson assembled with a 402 bp long double-stranded DNA (dsDNA) oligo synthesized by Twist Biosciences containing the scrambled gSCR sequence (GCTTAGTTACGCGTGGACGA),^41^ which does not exist in the murine genome, gRNA scaffold, and shortened sequence between gRNA scaffold and CBh promoter, yielding the pscAAV-hU6-gSCR-CBh-mCherry plasmid with a 2053 bp insert from 5’ ITR to 3’ ITR sequence. This plasmid was then used to clone all other plasmids containing a single gRNA and used for scAAV packaging. All sequences are found in Supplementary Table and plasmids are in the process of being deposited on Addgene.

### AAV transduction in vitro

For validation of AAV-mediated knock-out (KO), NIH/3T3.Cas9 cells were used as target cells and analyzed by flow cytometry to assess KO efficiencies. On day 1, 1e4 NIH/3T3.Cas9 cells were seeded in 500 µl culture media per well in a 24-well cell culture plate. On day 2, 16 h later, AAV was added at specified multiplicity of infection (MOI). After 3-5 days of incubation, culture media was aspirated, cells washed 1x with PBS, and collected using trypsin (0.05% Trypsin-EDTA, Gibco, #25300054) to detach the cells. Cells were further processed for flow cytometry analysis of surface protein expression of target gene knockout and mCherry reporter gene expression transduction efficiency. See below for details on flow cytometry analysis.

### AAV application in vivo

All AAV injections in this study were done subcutaneously (s.c.) on the dorsal lateral flank of mice 24 h prior to tumor engraftment at the same site. The hair at the site of injection was removed using clippers and cleaned using alcohol wipe pads. The doses used were either 1e10-3e10 vg AAV per injection in 50 ul PBS using a 31G insulin syringe (Sol-M, 163311564B). The actual dose used for each experiment can be found in the figure legends. AAV injection sites were demarcated using green tattoo paste (Fisher Scientific, 2420101) allowing engraftment of tumor cells at the same site the next day (see **Figure S1**).

### Indel frequency analysis

Indel frequencies within target cell populations at gRNA target sites were estimated using genomic DNA as input for amplicon PCR surrounding the gRNA target sites followed by Sanger sequencing and ICE analysis (Synthego Performance Analysis, ICE Analysis. 2019. v3.0. Synthego).

### Tissue collection and processing for flow cytometry

#### Tumor and skin samples

At time of analysis, mice were euthanized and tumor mass was carefully excised from the subcutaneous region without collecting the adjacent skin and surrounding adipose tissue. For skin collection, a 3×3 cm^2^ region on the dorsal flank cleared from hair using clippers was excised using scissors and excess fat was removed. On ice, samples were finely minced with scissors and then placed in a 2 ml tube containing 1.5 ml of digestion medium (2 mg/ml collagenase XI, 0.5 mg/ml hyaluronidase, 0.1 mg/ml DNase in DMEM with 10% DMEM (Gibco, 11995065) supplemented with 10% FBS (Foundation C, Gemini Bio, 900-308), 2 mM L-glutamine, 100 U/ml penicillin, 100 µg/ml streptomycin, and 50 µM beta-mercaptoethanol (Gibco, 21985-023). The tube was placed horizontally in a bacterial shaker for 45 min at 37°C and 225 rpm. The tumor samples were then filtered through a 40 µm filter and skin samples were filtered through a 100 µm filter. Samples were then washed with 10 ml cold PBS. The generated single cell suspension was then processed further depending on analysis method.

#### Spleen and lymph nodes

Inguinal and brachial lymph nodes were collected and the capsule was mechanically torn open using two 25 G needles prior to digestion. LNs and spleens were then separately digested in 1 ml digestion media (2 mg/ml collagenase D, collagenase IV (50 µg/ml) in DMEM with 2% FBS, 10 mM HEPES (Sigma, H0887)) for 30 min at 37°C without agitation. Every 10 min samples were pipetted up and down 5-times using a P1000 pipette. Samples were filtered through a 30 µm filter and washed with 3 ml of cold PBS. After pelleting the spleen sample at 500 g for 5 min in a centrifuge, the sample was resuspended in 1 ml of RBC lysis buffer (Roche, 11814389001) and incubated for 10 min on ice to lyse red blood cells. The lysis was quenched by adding 4 ml of fluorescence-activated cell sorting (FACS) buffer (staining buffer (PBS, 2% FBS, 2 mM EDTA)). Cells were pelleted again at 500 g for 5 min in a centrifuge and resuspended in FACS buffer and used for further analysis.

### Flow Cytometry

#### AAV validation in vitro

NIH/3T3.Cas9 cells transduced with AAVs in vitro were collected, washed and pelleted at 500 g for 5’ centrifugation in FACS staining buffer (PBS, 2% FBS, 2 mM EDTA) and resuspended in 50 µl of staining buffer plus 0.5 µg purified anti-mouse CD16/32 antibody (clone 2.4G2, Tonbo Biosciences). Samples were incubated for 10’ on ice to block FC receptors. Following this incubation, the cells were stained with antibodies against surface receptors that were targeted by AAV. Antibodies were diluted in 50 µl staining buffer prior to addition of cells to yield a final volume of 100 µl staining volume. Antibodies used for staining are listed in the Materials section. The cells were incubated for further 10’ on ice in the dark. After this staining incubation, cells were washed with 1 ml of staining buffer, pelleted again at 500 g for 5’ and resuspended in staining buffer containing DAPI (1 µg/ml) as a live/dead stain. Cells were analyzed on a BD LSRFortessa Cell Analyzer (BD Biosciences) at the Parnassus Flow Cytometry core at UCSF and gated on live (= DAPI-negative) singlets. Loss of surface receptor expression compared to naïve (= not exposed to AAV) NIH/3T3.Cas9 cells was used as a readout of KO efficiency and expression of mCherry was used as a readout for transient AAV expression in the KO constructs.

#### Staining of tumor and tissue samples

Single cell suspensions from tumors and other tissues were generated as described above, pelleted by centrifugation at 500 g for 5 mins, resuspended in 5e6 cells / 50 ul FACS staining buffer containing 1 µg purified anti-mouse CD16/32 antibody (clone 2.4G2, Tonbo Biosciences). Samples were incubated for 10’ on ice to block FC receptors. Following this incubation, 50 µl FACS staining buffer was added containing antibodies for surface staining and incubated for 20 min at room temperature. Antibodies used for staining are listed in the Materials section. Cells were washed with 1 ml of FACS staining buffer, pelleted and resuspended in FACS staining buffer containing DAPI (1 µg/ml) for live cell discrimination prior to analysis on a BD LSRFortessa Cell Analyzer (BD Biosciences).

#### Fluorescence-activated cell sorting of fibroblast subsets

For sorting of tumor fibroblast subsets, tumors were processed as described above to generate a single cell suspension and stained as described in the previous section. After staining, cells were filtered on a 40 µm filter right before being sorted on a BD FACSAria II cell sorter (BD Biosciences) Parnassus Flow Cytometry core at UCSF. Cells were collected based on the gating scheme outlined in **Figure S1B** to yield live singlet fibroblasts as defined by DAPI^-^ CD31^-^ CD45^-^ E-Cadherin^-^ MCAM^-^ Thy1^+^ PDPN^+^ EGFP^+^ and then further subsetted as outlined in **Figure S2J** to sort four fibroblast subsets: double-positive (DP) SCA-I^+^ Ly6C^+^, intermediate (Int) SCA-I^mid^ Ly6C^mid^, double-negative (DN) SCA-I^-^ Ly6C^-^ NCAM1^-^, and DN SCA-I^-^ Ly6C^-^ NCAM1^+^. Sorted cells were collected in 4°C cold cell culture media (DMEM with 10% FCS, 2 mM L-glutamine, 100 U/ml penicillin, 100 µg/ml streptomycin, 50 µM beta-mercaptoethanol). For each sample, 2,000 to 24,000 cells were collected per fibroblast subset. Individual samples were immediately pelleted by centrifugation at 500 g for 5 min at 4°C and processed for qPCR analysis as described below.

### Fluorescence and second-harmonic generation imaging using 2-photon microscopy

Fluorescence and second-harmonic generation (SHG) imaging was performed on 200 µm thick cryosections. The cryosections were cut using a cryostat and placed into PBS to wash OCT residue away. Samples were mounted using VECTASHIELD PLUS (Vector Laboratories, Inc., H-1900). For imaging, a custom-built 2-photon setup equipped with two infrafred lasers (8W Mai Tai Ti:Sapphire, Spectra Physics; 18W Chameleon Vision II, Coherent) was used. The Mai Tai laser was tuned to 920 nm for excitation of EGFP. The Chameleon laser was tuned to 780 nm for simultaneous excitation of mCherry and detection of SHG. The two lasers were exciting in alternating sequences. Emitted light was detected using a 25x 1.05-NA water lens (Olympus, XLPlan N) coupled to a 6-color detector array (custom; utilizing Hamamatsu H9433MOD detectors). Emission filters used were: violet 417/50, green 510/42, red 607/70 to detect SHG, EGFP, and mCherry emission, respectively. The microscope was controlled using the MicroManager software suite,^94^ 15 z-stacks at z-depth of 10 µm were acquired to cover 150 µm total z-distance of the 200 µm thick samples. Data analysis was performed using the Imaris sotware suite (Bitplane).

### Reverse transcription real-time quantitative PCR of sorted fibroblast subsets

Fibroblasts from tumors were isolated and sorted as described using the gating strategy outlined in **Figures S1 and S2J**. Cells were washed and pelleted in 1X cold PBS and RNA was isolated using the Qiagen RNeasy Micro Kit (#74004) according to the manufacturer’s instructions. All isolated RNA was used as input for cDNA generation using the iScript Reverse Transcription Supermix (Bio-Rad, #1708840) following the manufacturer’s instructions. The reaction mix was used for qPCR analysis using the SsoAdvanced Universal SYBR Green Supermix (Bio-Rad #1725270) on a Bio-Rad thermocycler with the following cycling conditions: 95C for 30s, 95C for 5s, 60C for 10s, repeat of step 2+3 39-times, and then melt-curve analysis 65C to 95C with 0.5C increments at 5s each. Each sample was run in technical triplicates for each assayed gene. 18s expression was used as the reference gene normalization. Primer sequences used can be found in the Supplementary Table.

### scRNAseq reference data sets

The previously published data sets from Hu et al.^6^ (mouse wound fibroblasts) and Krishnamurty et al.^14^ (mouse pancreatic tumor fibroblasts) were used as reference data sets for single cell RNA sequencing (scRNAseq) analysis. Deposited data was downloaded from GEO (Hu et al., GSE204777) or ArrayExpress (Krishnamurty et al., E-MTAB-12028). To generate the wound fibroblast Seurat object from Hu et al., the data was processed as described.^6^ To generate the tumor fibroblast Seurat object, the ReadMtx function from the Seurat R package (v5.0.1)^95^ was used to read in matrix.mtx, genes.tsv, and barcodes.tsv files. The CreateSeuratObject function was then used to generate the tumor fibroblast Seurat object. Cells were annotated using the meta.txt file and fibroblasts were extracted based on rows matching ‘fibroblast’ in the ‘CellType’ column using the subset function. Cells were removed according to the feature count (<300) and mitochondrial count (>5%) parameters outlined in the original paper. The remaining fibroblasts were then further processed using the following Seurat functions in the listed order: NormalizeData (log-normalize RNA transcript counts), FindVariableFeatures (selection.method ‘vst’, nfeatures = 2000), ScaleData (default parameters), RunPCA (default parameters), FindNeighbors (15 PCs), and RunUMAP (13 PCs) was used for 2D visualization. Clustering was performed using the FindClusters function (resolution of 0.3) and cluster 5 was removed as it was identified as a contaminating immune cell cluster (∼1% of total object, 66 of 6589 cells). The remaining tumor fibroblasts and wound fibroblasts were used to plot gene expression of homeostatic and activation marker genes using the DotPlot, FeaturePlot, and VlnPlot functions in the Seurat package.

### Tumor collection and processing for single cell RNA sequencing

For tumor collection and subsequent analysis by single cell RNA sequencing, live cells from day 14 HY19636 tumors (after subcutaneous injection of 2.5e5 HY19636 on day 0 into *Pdgfra*-Cas9-EGFP mice) that had received 3e10 vg scAAV1-gTrac, scAAV1-gOsmr, scAAV1-gTgfbr2, or scAAV1-gIl1r1 to generate local KOs on day-1, were isolated and processed for sorting as described above on a BD FACSAria II cell sorter (BD Biosciences) Parnassus Flow Cytometry core at UCSF with minor modifications: live cells were identified as negative for Zombie-NIR (BioLegend, 1:200) and positive for Calcein blue AM (Thermo Fisher Scientific, C1429, 10 µM working concentration). Samples were additionally stained with anti-mouse CD45-BUV395 (clone 30-F11, BD Biosciences, 1:500), anti-Thy1.2/CD90.2-BV786 (clone 53-2.1, BD Biosciences, 1:500, for Thy1 detection), and anti-E-Cadherin/CD324-PE-Cy7 (clone DECMA-1, BioLegend, 1:200). Four live cell populations were sorted: (1) CD45+ (immune cells), (2) CD45- EGFP+ Thy1+ (fibroblasts), (3) CD45- EGFP- E-Cadherin- (miscellaneous), (4) CD45- EGFP- E-Cadherin+ (tumor cells) (see **Figure S4A** for gating scheme). The four populations from each sample were then pooled in a 15 ml Falcon tube and pelleted for 7 minutes at 500 g. The supernatant was removed, and cells were further processed using the Chromium Fixed RNA Profiling (Gene Expression Flex) pipeline. In brief, each sample was first fixed using the Single Cell Fixed RNA Sample Preparation Kit (10x Genomics, PN-1000414) following the manufacturer’s instructions. Samples were fixed for 18 h at 4 °C and, after addition of quenching buffer and storage buffer, stored at −80 °C until further processing. After thawing of samples, samples were processed using the Chromium Fixed RNA Kit, Mouse Transcriptome (10x Genomics, PN-1000496) following the manufacturer’s instructions to generate a fixed RNA gene expression library. For multiplexing, each sample (4 replicates * 4 AAV groups = 16 total) was hybridized with a separate Probe Barcode library including added custom probes (*Egfp*, 8casd09, GGTAGTGGTCGGCGAGCTGCACGCTGCCGTCCTCGATGTTGTGGCGGATC; and *Egfp*, 8casd10, AGGGTGTCGCCCTCGAACTTCACCTCGGCGCGGGTCTTGTAGTTGCCGTC). Approximately 2.5e5 total cells were encapsulated and sequenced on two lanes of a NovaSeq X flow cell (1.25 B reads/lane) to reach about 10.000 reads/cell. Sequencing output was processed using cellranger-7.2.0 and the mouse transcriptome probe set v1.0.1_mm10-2020-A with added custom *Egfp* probes from above. After demultiplexing the 16 samples using the provided Probe Barcodes, the samples were integrated in Seurat v5.1.0^95^ and low-quality cells were filtered out if they had <150 genes or >20% mitochondrial genes. Features with counts in <3 cells were also filtered out. After merging all samples using the Seurat merge() function, the gene expression matrix was normalized using Seurat NormalizeData() function, with default parameters. The top 2000 variable genes were identified, and the gene expression matrix was scaled prior to principal component analysis (PCA). The first 23 PCs were used for UMAP visualization based on elbow plot cut off and the FindClusters() function at a resolution of 0.2 was used to identify individual clusters. The following markers were used to identify major cell types: fibroblasts (*Pdgfra*, *Col1a2*, *Egfp*), pericytes (*Rgs5*), epidermal basal cells (*Krt14, Krt5*), endothelial cells (*Pecam1*), tumor cells (*Cdh1, Msln*), T cells and NK cells (*Trac*, *Ncr1*), neutrophils (*S100a8*), mast cells (*Mcpt4*), and myeloid cells (*H2-Ab1, Lyz2*). Major cell types were subsetted and further processed to remove doublets. Dendritic cells were further subsetted from the ‘myeloid cells’ object. This generated 9 separate objects: fibroblasts (n=9,514), pericytes (n=745), endothelial cells (n=3,895), tumor cells (n=81,588), T and NK cells (n=5,209), neutrophils (n=43,824), mast cells (n=1,581), monocyte/macrophages (n=35,810), and dendritic cells (n=5,059). Tumor cells were subsampled to a total of 10,000 cells and then all objects were remerged for **Figure S4B**. Fibroblasts, monocyte/macrophages (MonoMac), dendritic cells, neutrophils, and T/NK cell objects were further sub-clustered and the DEGs of the individual sub-clusters can be found in **Supplementary Table 2**.

### Pseudobulk differential gene expression analysis

For identification of differentially expressed genes (DEGs) between CAFs of different AAV-gRNA groups, we used a pairwise pseudobulk approach comparing the control group (=AAV-gTrac) to the three different experimental groups (AAV-gOsmr, AAV-gTgfbr2, or AAV-gIl1r1) individually. Each group was represented by 4 replicates and DEGs were identified using MAST with random effects.^96,97^ The number of DEGs are grouped by cell type with a threshold of p-value < 0.05 after false discovery rate correction using the Benjamini-Hochberg procedure. The list of DEGs for each experimental group can be found in **Supplementary Table 3**.

### Pseudotime analysis of the fibroblast single cell RNA sequencing object

Prior to pseudotime analysis of the fibroblast object, the ‘myCAF proliferating’ cluster was removed. After re-running PCA, a UMAP representation was generated using the first 18 PCs (**Figure 4A**) in Seurat. The Seurat object was converted to a cell_data_set object using Monocle3.^98^ The pseudotime trajectory was calculated using Monocle3 with the homeostatic fibroblast cluster used as the root cell state. To plot the distribution of cells in pseudotime as a violin plot in Seurat, the position of each cell within the pseudotime graph was extracted from the cell_data_set object and then added as meta data to the Seurat object.

### Comparison of gene expression similarity between cancer-associated fibroblasts and existing data sets

To measure the similarity of DEGs between cancer-associated fibroblast clusters in this study to published data sets, we performed the hypergeometric test to find overlapping genes using the phyper() function from the stats package in R. When comparing to published human data sets, the human genes from those data sets were label-transferred to mouse genes using the homologene (v1.4.68.19.3.27)^99^ R package. DEGs from each individual data set were filtered based on p-value < 0.05 and average log2 fold change > 0.25.

### Non-negative matrix factorization (NMF) decomposition

NMF was performed on the normalized gene-by-cell count matrix of the fibroblast object using the nmf function from the RcppML (v0.3.7) R package.^100^ All genes expressed in >2% of all cells were used as input. Mitochondrial (starting with ‘mt-‘) and ribosomal (starting with ‘Rps’ or ‘Rpl’) genes were excluded. The factor-weight-by-cell matrix (= h matrix output) was added to the fibroblast Seurat object as a new ‘NMF’ assay using the Seurat CreateAssayObject function. The factor weight values from the ‘NMF’ assay were used to visualize NMF factor weights per cell in UMAP representation (**Figure 4D**). The AverageExpression function in Seurat was used to calculate the average NMF factor weights per sample.

### NMF factor weight correlation across cancer-associated fibroblasts

The NMF factor weights of each cell were correlated in pairwise fashion using the cor() function and ‘method=pearson’ from the stats package. Statistical significance was measured using cor.mtest() function from the corrplot (v0.95) R package.^101^

### Gene set enrichment analysis

Gene set enrichment analysis (GSEA) for fibroblast NMF factors was done using the hypeR (v2.0.0) package.^102^ The top 50 genes of each NMF factor were inputted for GSEA and statistically significant enrichment among the gene ontology term ‘Biological Process’ from the Molecular Sigantures Database (MSigDB v7.2.1) was computed using the hypergeometric test.

### Survival analysis of TCGA data based on gene signatures

#### Data download and processing

The TCGA transcriptomic TPM data was downloaded from the Toil recompute^103^ in the TCGA Pan-Cancer cohort on the UCSC Xena browser (http://xena.ucsc.edu) and imported into R. The tumor samples were filtered down to only include primary solid tumors (sample type code = 01) and tumor types with <100 samples were excluded from further analysis. Prior to calculation of gene signature scores derived from mouse genes, the mouse gene names were transferred to their human orthologs using the gprofiler2 (v0.2.3) R package.^104^ Mouse genes without human orthologs or human orthologs not present in gene-by-sample matrix of TCGA data set were dropped from further analysis. From each NMF factor, the top 25 genes ranked by gene contribution in descending order were used to calculate NMF factor-specific gene signatures. For the ‘stromal signature’, the identified genes in Combes et al.^57^ were used. For the ‘fibroblast TGFβ response signature’, the genes from Mariathasan et al.^50^ were used. For the ‘AAV-gTgfbr2 up DEG signature’, the top 25 upregulated genes ranked in descending order were selected.

#### Gene signature score calculation

Gene signature scores were calculated on log10(TPM+1) normalized data and the expression of each gene was converted to percentile ranks across the samples. The final gene signature score of each sample was calculated as the average percentile rank from all genes belonging to one gene signature.

#### Survival analysis

For survival analysis, the Cox proportional hazards model from the survival (v3.8-3) R package was used to test overall survival differences between patients in the top and bottom quartiles based on gene signature scores, with age and sex as covariates for each cancer type separately. Kaplan-Meier survival curves were plotted using the survminer (v0.5.0) R package. Forest plots of all analyzed cancer types were generated after meta-analysis using a random effect model using the meta (v8.1-0) R package.

### Identification of cell frequency correlation between fibroblasts and immune cells

To identify potential co-occurring cell populations of CAF subsets and captured immune cells in the scRNAseq data set, the proportion of each cell subset within its corresponding major cell compartment (fibroblasts, monocyte/macrophages, dendritic cells, neutrophils, T and NK cells) was calculated. The proportion of each cell subset was correlated in pairwise fashion with all other cell subsets across all 16 samples (= 4 replicates * 4 AAV groups) using the cor() function and ‘method=pearson’ from the stats package. Statistical significance of correlation as measured using cor.mtest() function from the corrplot package^101^ and results were plotted as a heatmap with samples arranged by hierarchical clustering.

### Candidate identification of ligand-receptor interactions

The CellChat^62^ (v2.1.2) package was used to identify putative ligand-receptor interactions between two target cell populations. First, all cell subsets belonging to one experimental AAV group were subsetted from the full Seurat object. Then a CellChat object was created and processed using default parameters. Source and target cells were defined prior to plotting the top 6 inferred ligand-receptor interactions.

### Transcription factor activity prediction with decoupleR

To infer osteoblast-related transcription factor activity in the CAF subsets, the decoupleR (v2.10.0) package was used to run a univariate linear model (function ‘ulm’) with default parameters. All genes were used as input and all mouse transcription factors were initially considered. For plotting osteoblast-related transcription factor activity,^55^ the mean activity of each transcription factor per CAF subset was used.

### Quantification and statistical analysis

Statistical analyses were done using R version 4.3.2. Details of individual experiments can be found in their respective figure legends.

## SUPPLEMENTARY FIGURES

**Supplementary Figure 1.**
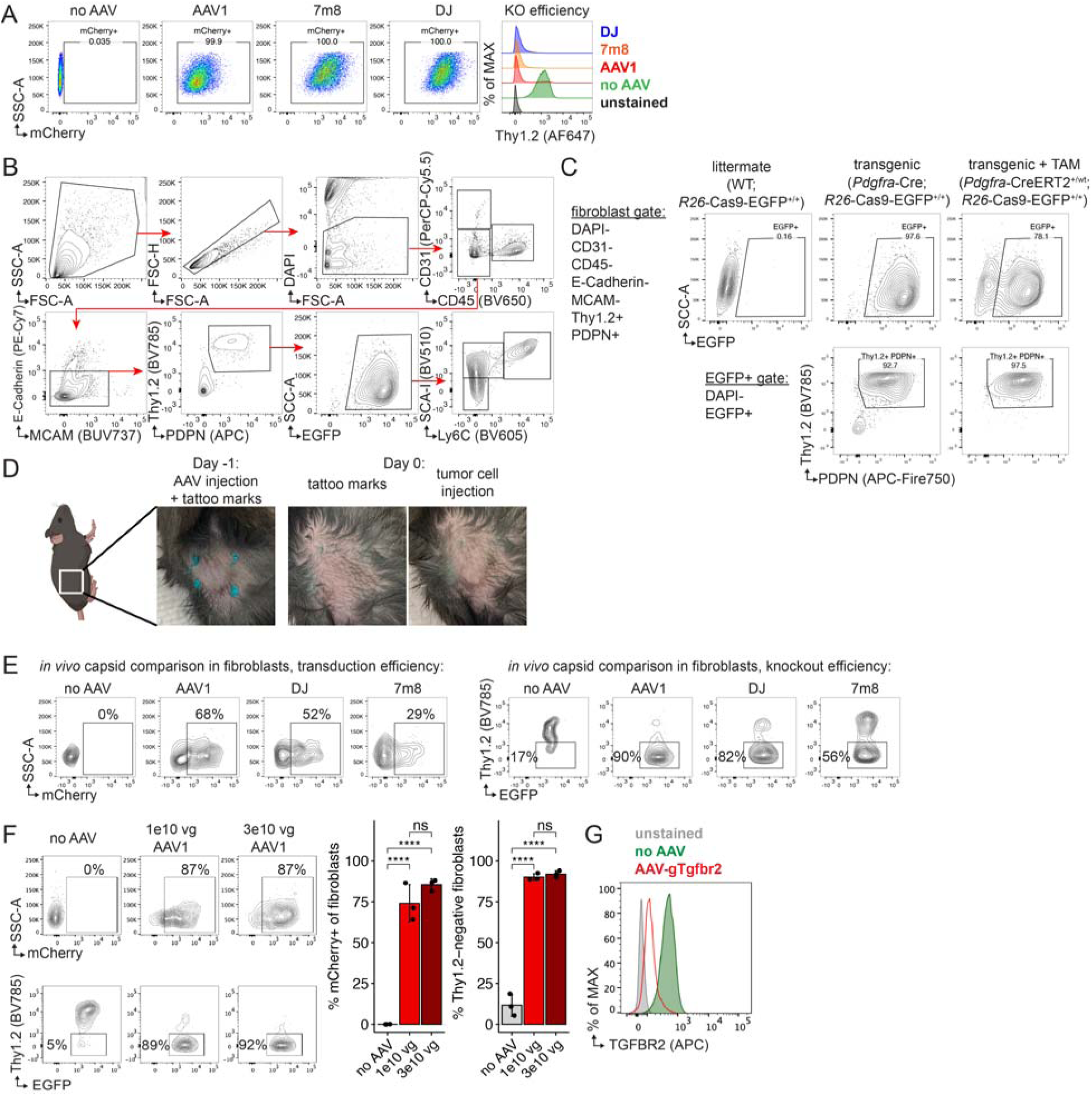
(A) NIH/3T3.Cas9 cells were transduced with three different AAVs as outlined in Figure 1B all carrying the same hU6-gThy1-mCherry cassette but generated with three different capsids (AAV1, 7m8, DJ). Left, flow cytometry plots depicting mCherry expression in NIH/3T3.Cas9 cells as a readout for transduction efficiency of AAVs after 96 h at an MOI of 1e5. Right, flow cytometry overlay plots of those NIH/3T3.Cas9 cells showing KO efficiency of the AAVs with different capsids at day 6 post AAV transduction. One of two representative experiments is shown. (B) *In vivo* gating scheme to identify tumor fibroblasts: DAPI^-^ CD31^-^ CD45^-^ E-Cadherin^-^ MCAM^-^ THY1^+^ PDPN^+^ EGFP^+^ in *Pdgfra*-Cas9-EGFP mice. For *Col1a2*^CreERT2/+^;*Tgfbr2*^fl/fl^ mice in Figures 1O and 1P, tumor fibroblasts were gated as DAPI^-^ CD31^-^ CD45^-^ E-Cadherin^-^ MCAM^-^ THY1^+^ PDPN^+^. (C) (Top) Flow cytometry contour plots of EGFP expression in tumor fibroblasts as gated in (B) on day 14 after 2.5e5 HY19636 pancreatic tumor cell s.c. injection in *Pdgfra*-Cre;*R26*-Cas9-EGFP^+/+^ or *Pdgfra*-CreERT2^+/wt^;*R26*-Cas9-EGFP^+/+^ mice (both referred to as *Pdgfra*-Cas9-EGFP mice). *Pdgfra*-CreERT2^+/wt^;*R26*-Cas9-EGFP^+/+^ mice were injected with 2 mg of tamoxifen (TAM) on 5 consecutive days one week prior to tumor engraftment to induce Cas9-EGFP expression. (Bottom) Contour plots of fibroblast markers THY1 and PDPN expression in DAPI^-^ EGFP^+^ cells. (D) Pictures of subsequent injections of AAV and tumor cells on consecutive days to generate local gene KO in tumor fibroblasts. Local AAV injection is marked by tattoo paste using a 31 gauge needle on day −1 around wheal from AAV injection. Tattoo pricks are visible on day 0 and used as a guide for tumor cell injection that day. (E) Flow cytometry contour plots of EGFP+ tumor fibroblasts from *Pdgfra-*Cas9-EGFP mice injected s.c. with 1e10 vg AAV-gThy1 on day −1 and 5e5 YUMM5.2 tumor cells on day 0. Tumors were excised on day 7 and (left) mCherry expression in tumor fibroblasts (gated as outlined in (B)) was measured as a readout for transduction efficiency and (right) surface level of THY1 was measured as a readout for knockout efficiency by the different AAV capsids. (F) Test of different scAAV1 doses for transduction and KO in tumor fibroblasts. *Pdgfra*-Cas9-EGFP mice were injected s.c. on day −1 with 1e10 or 3e10 vg of scAAV1-gThy1-mCherry (self-complementary AAV carrying gRNA targeting *Thy1* and an mCherry ORF). On day 0, 5e5 YUMM5.2 cells were injected s.c. at site of prior AAV injection. Tumors were harvested on day 7 and transduction and *Thy1* knockout efficiency was assessed. Top, flow cytometry plots of all tumor fibroblasts (gated as outlined in (B)) showing mCherry expression. Bottom, flow cytometry plots of all tumor fibroblasts for THY1 surface expression. Right, quantification of transduction and KO efficiency at two different doses of scAAV1-gThy1. Data are mean ± s.d. and significance was tested using one-way ANOVA with Tukey’s multiple comparison test. ns, non-significant. ****p<0.0001. (G) Flow cytometry overlay plot of surface TGFBR2 expression on NIH/3T3.Cas9 cells transduced with or without scAAV1-gTgfbr2 at an MOI of 1e6 and analyzed 72h later. MOI, multiplicity of infection. scAAV, self-complementary adeno-associated virus. s.c., subcutaneously. vg, viral genome.

**Supplementary Figure 2.**
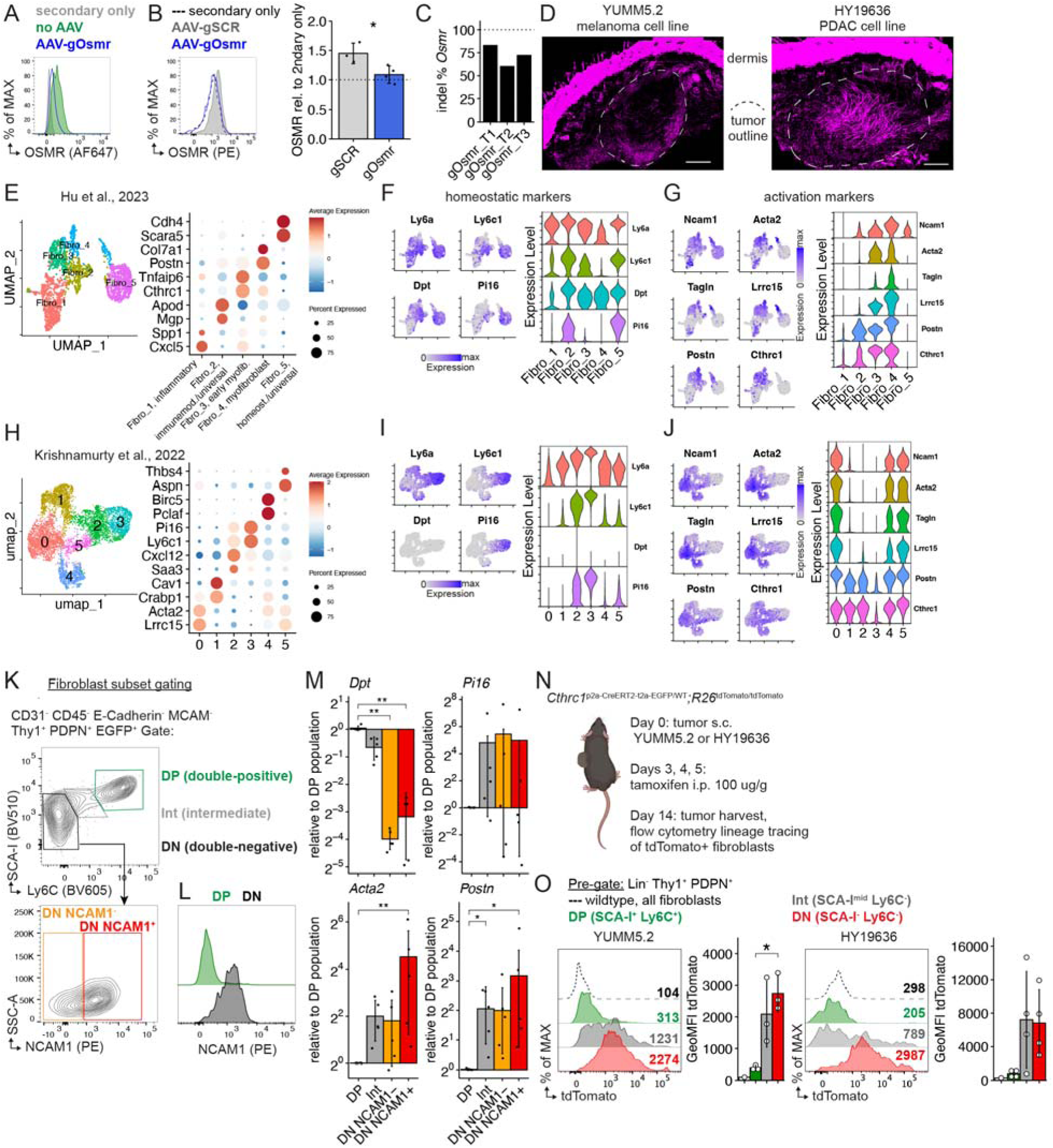
(A) Flow cytometry overlay plot of surface OSMR expression on NIH/3T3.Cas9 cells treated with or without scAAV1-gOsmr at an MOI of 1e6 and analyzed 6 days later. (B) (Left) Representative flow cytometry overlay plot of EGFP+ tumor fibroblasts from *Pdgfra*-Cas9-EGFP mice injected s.c. with 3e10 vg AAV-gSCR (control) or AAV-gOsmr and YUMM5.2 tumor cells on subsequent days. Flow analysis done on day 14 after tumor challenge. (Right) Quantification of OSMR (GeoMFI) surface expression on EGFP+ tumor fibroblasts treated with control AAV-gSCR (control) or *Osmr* targeting AAV-gOsmr (n=3-4 tumors/group) normalized to secondary only stain control. Data are mean ± s.d. and significance was tested using Welch’s t-test. GeoMFI, geometric mean fluorescence intensity. (C) Indel percentages as measured by ICE analysis of the *Osmr* KO locus from sorted CAFs from three different HY19636 tumors on day 14 after tumor injection s.c. and prior injection of 3e10 vg AAV-gOsmr on day −1. (D) Second-harmonic generation (SHG) imaging of subcutaneously injected (left) HY19636 mouse pancreatic cancer and (right) YUMM5.2 mouse melanoma cell lines to visualize fibrillar collagen deposition in tumor mass on day 10 in 150 µm thick sections via maximum projection. Dermal skin layer at top of images is thresholded to visualize collagen deposition in tumor mass. White dotted line demarcates outline of tumor mass. (E) (Left) UMAP of mouse wound fibroblasts from Hu, Kuhn et al. analyzed by scRNAseq and colored by cluster. (Right) Dot plot showing relative average expression of marker genes across all clusters. (F) Fibroblast homeostatic marker gene expression in wound fibroblasts is highlighted (left) on the UMAP and (right) by cluster annotation. (G) Fibroblast activation marker gene expression in wound fibroblasts is highlighted (left) on the UMAP and (right) by cluster annotation. (H) (Left) UMAP of mouse pancreatic tumor fibroblasts from Krishnamurty et al. analyzed by scRNAseq and colored by cluster. (Right) Dot plot showing relative average expression of marker genes across all clusters. (I) Fibroblast homeostatic marker gene expression in tumor fibroblasts is highlighted (left) on the UMAP and (right) by cluster annotation. (J) Fibroblast activation marker gene expression in tumor fibroblasts is highlighted (left) on the UMAP and (right) by cluster annotation. (K) Gating strategy to identify three different tumor fibroblast subsets: DP (double-positive) SCA-I^+^ Ly6C^+^, Int (intermediate) SCA-I^mid^ Ly6C^mid^, and DN (double-negative) SCA-I^-^ Ly6C^-^. The fibroblast DN subset is further divided by NCAM1 low and high expression. (L) NCAM1 expression in DP (SCA-I+ Ly6C+) and DN (SCA-I- Ly6C+) tumor fibroblasts. NCAM1 expression is absent in DP subset. (M) Tumor fibroblast subsets were isolated and sorted based on gating strategy in (J) from WT C57BL/6 mice challenged with 2.5e5 HY19636 s.c. on day 8-10 after tumor challenge. Gene expression of homeostatic genes *Dpt* and *Pi16* and activation genes *Acta2* and *Postn* were analyzed by quantitative real-time PCR. Data are mean ± s.d. and significance was tested using one-way ANOVA with Tukey’s multiple comparison test (n=5 tumors, pooled from 2 experiments). (N) Experimental scheme to lineage trace *Cthrc1-*expressing cells in tumor-(YUMM5.2 or HY19636) challenged mice. (O) Representative overlay histograms of tdTomato expression in (Lin- [CD31 CD45 E-Cadherin MCAM] CD90.2+ PDPN+) tumor fibroblast cell subsets harvested from (left) YUMM5.2 or (right) HY19636 tumors. Quantification of tdTomato mean fluorescence intensity is plotted as mean ± s.d. (n=3 YUMM5.2 tumors; n=4 HY19636 tumors) and significance was tested using one-way ANOVA with Tukey’s multiple comparison test.

**Supplementary Figure 3.**
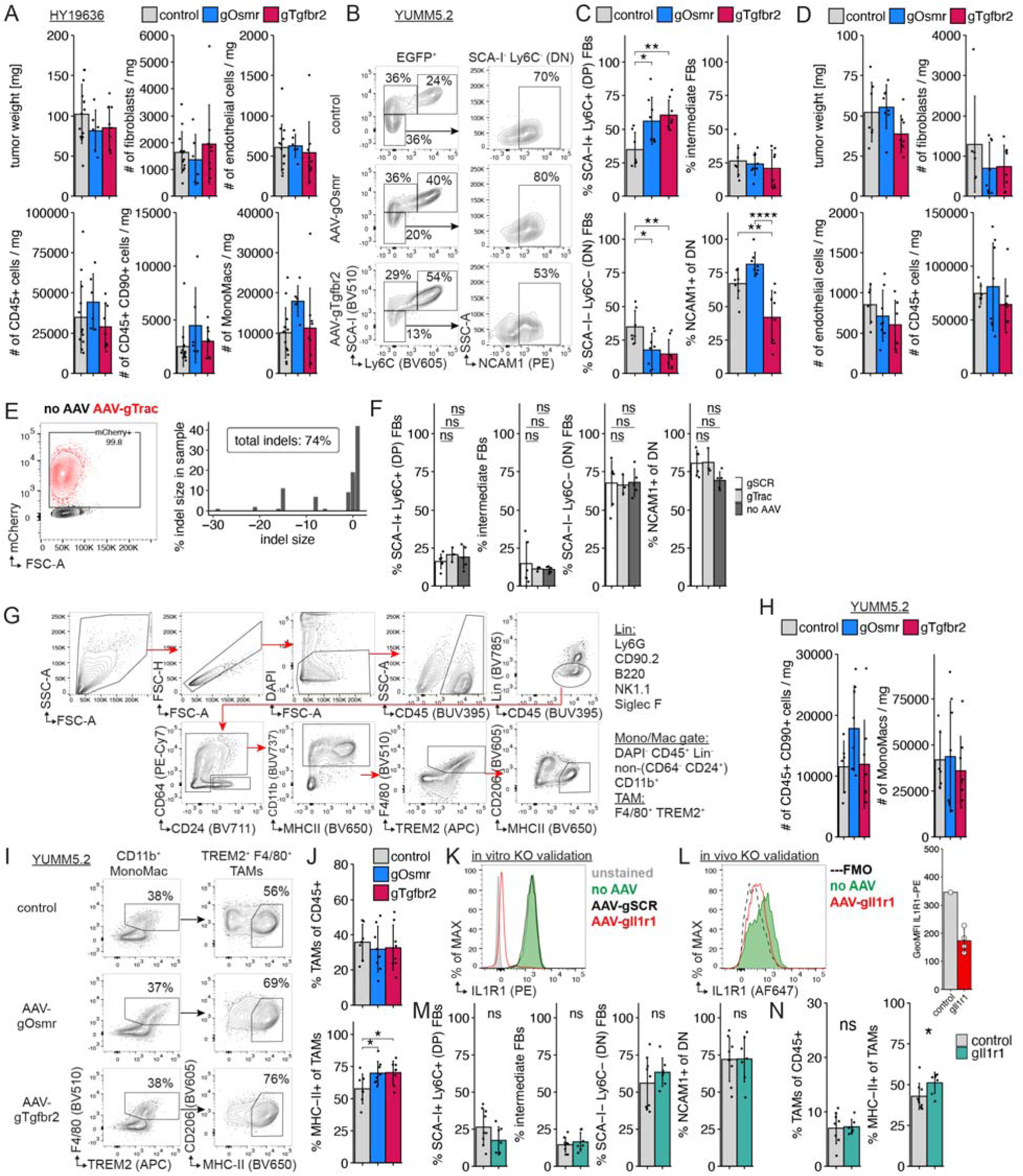
(A) Quantification of tumor weight, fibroblasts (Lin- CD90.2+ PDPN+ EGFP+), endothelial cells (CD45- CD31+), immune cells (CD45+ CD31-), and CD45+ CD90+ lymphocytes, and monocyte/macrophages (MonoMacs, CD45+ Lin- CD11b+) normalized by tumor weight in HY19636 pancreatic tumors on D14 after tumor challenge in mice treated with AAV outlined in Figure 2A. Data are mean ± s.d (n=6-13 tumors/group, pooled from four independent experiments). (B) Representative flow cytometry plots of (left) EGFP^+^ tumor fibroblasts and (right) DN tumor fibroblasts from control-, AAV-gOsmr, or AAV-gTgfbr2-injected *Pdgfra*-Cas9-EGFP mice on day 14 after YUMM5.2 tumor challenge. (C) Quantification of EGFP^+^ tumor fibroblast subsets from tumors with *Osmr* or *Tgfbr2* KO in fibroblasts. Data are mean ± s.d. and significance was tested using one-way ANOVA with Tukey’s multiple comparisons test (n=7-8 tumors/group, pooled from two independent experiments). (D) Quantification of YUMM5.2 tumor weight and fibroblasts (Lin- CD90.2+ PDPN+ EGFP+), endothelial cells (CD45- CD31+) and immune cells (CD45+ CD31-) normalized by tumor weight. Data are mean ± s.d (n=6-8 tumors/group, pooled from two independent experiments). (E) NIH/3T3.Cas9 cells were transduced with genome cutting control AAV-gTrac at an MOI of 1e6 and three days later analyzed for (top) transduction efficiency by mCherry expression readout via flow cytometry and for (bottom) indel frequency as a readout of cutting at the *Trac* locus via ICE analysis. (F) Quantification of EGFP^+^ tumor fibroblast subsets from tumors of different control groups— 1-3e10 vg AAV-gSCR (scrambled gRNA) or AAV-gTrac (genome cutting control), and ‘no AAV’—in (left) YUMM5.2 mouse melanoma and (right) HY19636 mouse pancreatic cancer models on day 14 after tumor challenge. Data are mean ± s.d. and significance was tested using (left, YUMM5.2, two groups) Welch’s t-test; and (right, HY19636, 3 groups) one-way ANOVA with Tukey’s multiple comparison test (n=3-6 tumors/group). (G) Gating strategy for monocyte/macrophage (Mono/Mac) and tumor-associated macrophage (TAM) identification in mouse tumors. (H) Quantification of CD45+ CD90+ lymphocytes, and monocyte/macrophages (MonoMacs, CD45+ Lin- CD11b+) normalized by tumor weight in YUMM5.2 melanoma on D14 after tumor challenge in mice treated with AAV outlined in Figure 2A. Data are mean ± s.d (n=6-8 tumors/group, pooled from two independent experiments). (I) Representative flow cytometry plots of (left) CD11b^+^ monocyte/macrophages and (right) F4/80+ TREM2+ tumor-associated macrophages (TAMs) from YUMM5.2 tumors with *Osmr* or *Tgfbr2* KO in fibroblasts as treated in Figure 2A. (J) Quantification of (I). Data are mean ± s.d. and significance was tested using one-way ANOVA with Dunnett’s multiple comparisons test (n=7-8 tumors/group, pooled from two independent experiments). (K) In vitro validation of scAAV1-gIl1r1. Flow cytometry overlay plot of surface expression on NIH/3T3.Cas9 cells infected with control AAV (scAAV1-gSCR) or scAAV1-gIl1r1 at a MOI of 1e6 and analyzed 6 days later. One of two representative experiments is shown. (L) In vivo validation of AAV-mediated *Il1r1* knockout. Representative flow cytometry overlay plot of EGFP+ tumor fibroblasts from *Pdgfra*-Cas9-EGFP mice injected s.c. with 3e10 vg AAV-gIl1r1 and HY19636 tumor cells on subsequent days. Flow analysis done on day 14 after tumor challenge. Quantification of IL1R1 surface expression (GeoMFI) on EGFP+ tumor fibroblasts treated with AAV-gIl1r1 are plotted as mean ± s.d. GeoMFI, geometric mean fluorescence intensity. (M) Quantification of EGFP^+^ tumor fibroblast subsets in *Pdgfra*-Cas9-EGFP mice 14 days after s.c. challenge with HY19636 tumor cells and prior local KO of *Il1r1* in fibroblasts via 3e10 vg AAV-gIl1r1 s.c. injection. Data are mean ± s.d. and significance was tested using Welch’s t-test (n=7-9 tumors/group, pooled from two independent experiments). (N) Quantification of TAMs and MHCII-hi TAMs in tumors from (M).

**Supplementary Figure 4.**
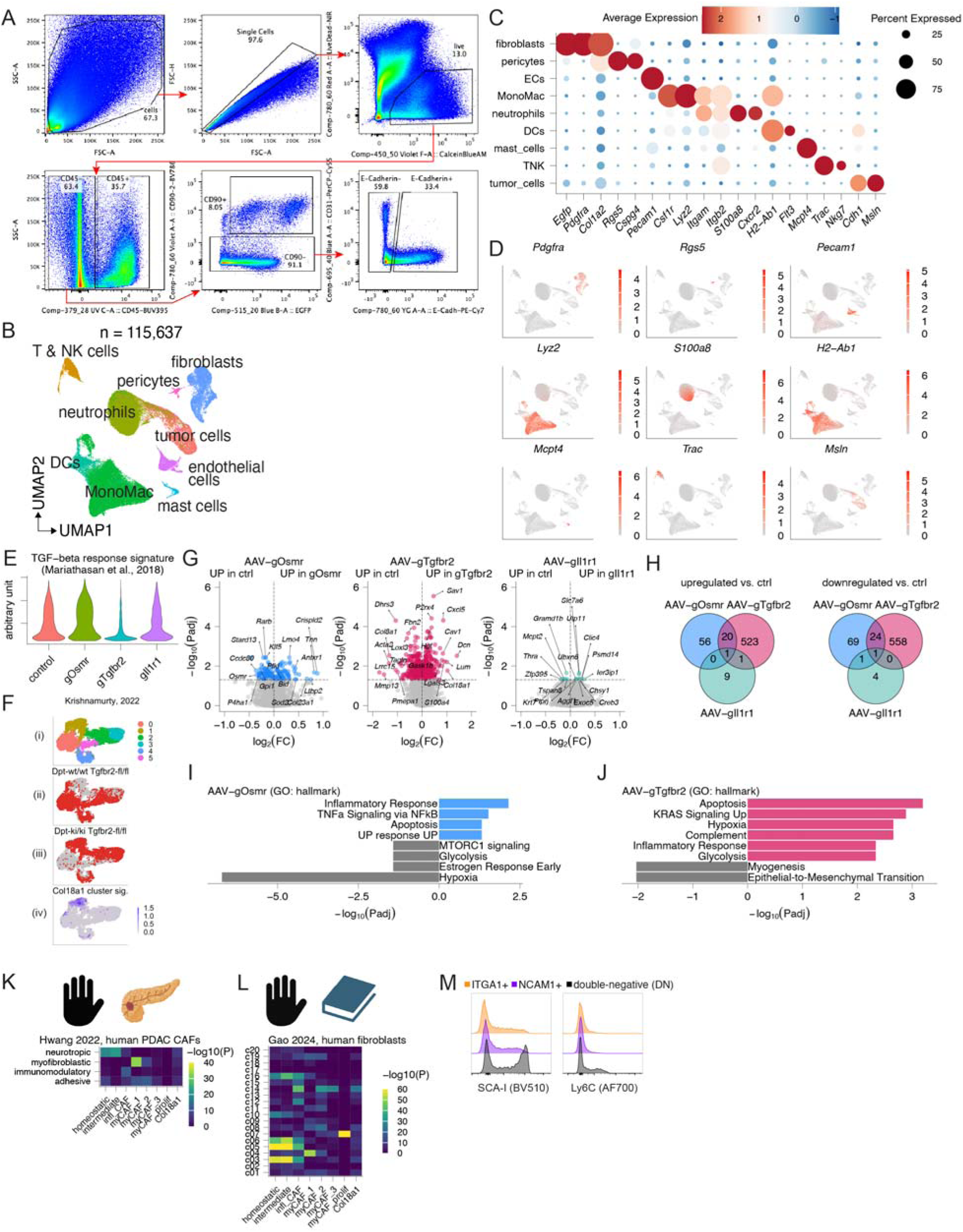
(A) Gating scheme for sorting cells in Figure 3A. (B) Uniform manifold approximation and projection (UMAP) of all collected cells by scRNA-seq. (C) Bubble plot of scaled marker gene expression in main cell types identified in Figure 7B. EC, endothelial cells. DCs, dendritic cells. TNK, T and NK cells. (D) UMAP visualization of marker gene expression in all captured cells. (E) Violin plot depicting mean expression of TGFbeta response signature from Mariathasan et al., 2018^50^ in CAFs grouped by AAV-gRNA condition. (F) UMAP visualization of CAFs from Krishnamurthy et al. 2022 (re-analyzed mouse PDAC scRNA-Seq data set) colored by (i) cluster, CAFs identified in (ii) *Dpt^wt/wt^;Tgfbr2^fl/fl^* control or (iii) *Dpt^ki/ki^;Tgfbr2^fl/fl^* mice with conditional *Tgfbr2* knockout in fibroblasts, and (iv) Col18a1 CAF cluster signature expression. AAV-gTgfbr2-induced Col18a1 CAFs also appear in *Tgfbr2* knockout mice challenged with PDAC tumor. (G) Volcano plots of differentially expressed genes (DEG, pseudobulk scRNA-Seq) in CAFs between control (AAV-gTrac) and (left) AAV-gOsmr, (middle) AAV-gTgfbr2, and (right) AAV-gIl1r1. Selected genes are highlighted. FC, fold change. (H) Venn diagram of (top) upregulated and (bottom) downregulated DEGs between control (ctrl) and AAV-gOsmr, AAV-gTgfbr2, or AAV-gIl1r1. (I and J) Enriched gene sets (Gene Ontology hallmark) based on the up- and downregulated DEGs between control and (I) AAV-gOsmr and (J) AAV-gTgfbr2. (K and L) Similarity of DEGs between mouse CAF subsets (columns) and (K) human PDAC CAF from Hwang et al., 2022^51^ (rows) or (L) cross-tissue human fibroblast atlas subtypes from Gao et al, 2024.^52^ Statistical significance of DEG overlap (-log10(P value), hypergeometric test, *P* value adjusted using Bonferroni correction) for each pairwise comparison plotted as heatmap. Yellow indicates high statistically significant overlap of DEG lists. Human genes from gene lists in Hwang et al. and Gao et al. were re-labelled as their mouse orthologs prior to comparison. (M) Flow cytometry histograms of homeostatic fibroblast markers (left) SCA-I and (right) Ly6C in CAF subpopulations as gated in Figure 3K. Double-negative CAFs have the highest expression of homeostatic markers.

**Supplementary Figure 5.**
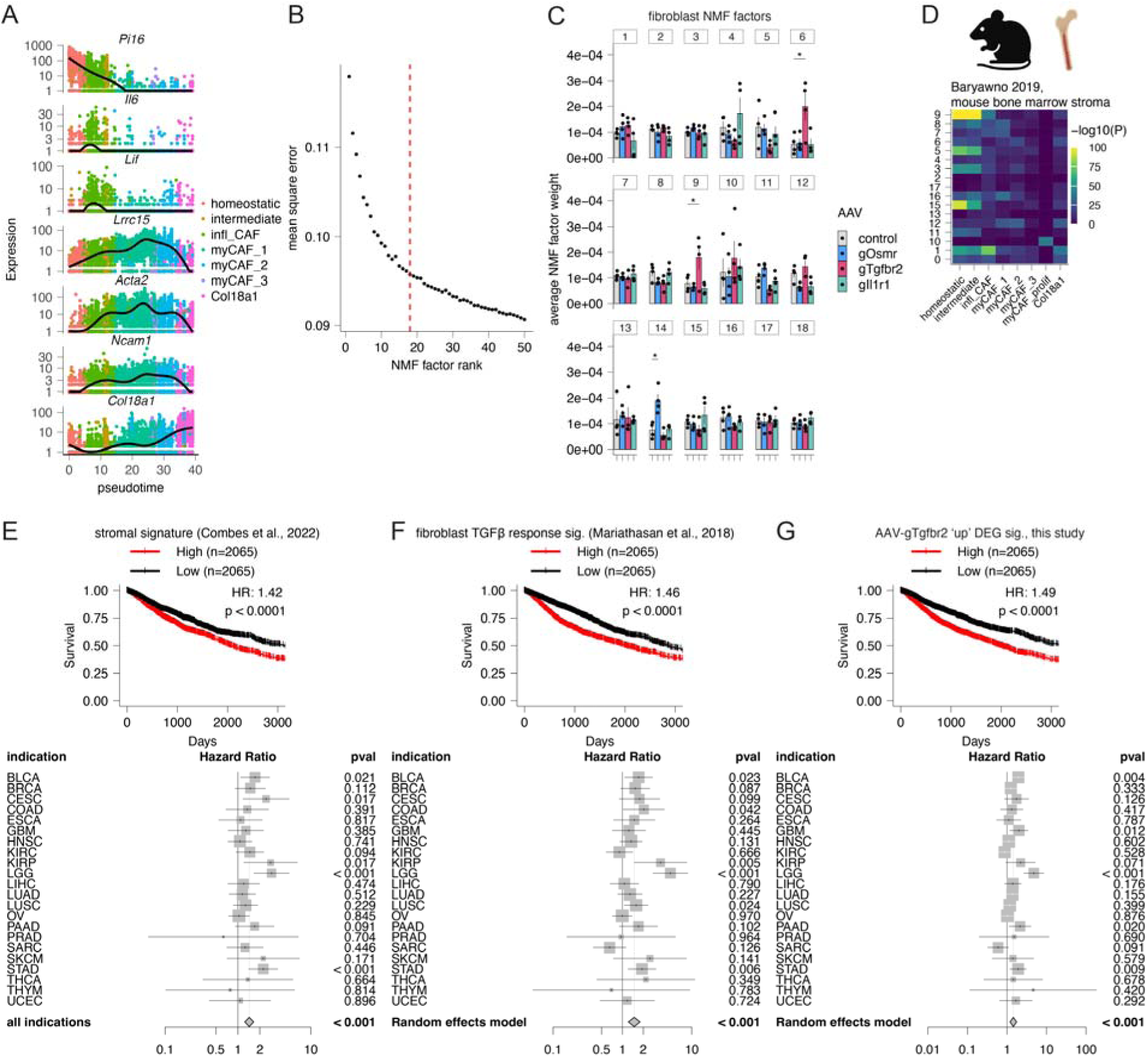
(A) Gene expression of select genes (homeostatic *Pi16*; inflammatory *Saa3* and *Cthrc1*, myofibroblast-associated *Lrrc15*, *Acta2*, and *Ncam1*; AAV-gTgfbr2-associated *Col18a1*) along pseudotime in CAFs from Figure 4A. (B) Mean squared error of an NMF model vs. model factorization rank (k = 1-50) in the fibroblast object. Vertical dotted line at k = 18 highlights model selected for analysis. (C) Bar charts of average expression of NMF factors in CAFs grouped by AAV-gRNA condition plotted as mean±SEM (n=4 per group). *p<0.05 one-way ANOVA with Dunnett’s multiple comparison test. (D) Similarity of DEGs between mouse CAF subsets (columns) and mouse bone marrow stroma cells from Baryawno et al., 2019^56^ (rows) Statistical significance of DEG overlap (-log10(P value), hypergeometric test, *P* value adjusted using Bonferroni correction) for each pairwise comparison plotted as heatmap. Yellow indicates high statistically significant overlap of DEG lists. (E) (Top) Kaplan-Meier survival plots of TCGA patients categorized into upper and lower quartiles by the stroma score identified in Combes et al., 2022.^50,57^ Hazard ratio (HR) and p value of Cox regression fit are shown. (Bottom) Forest plot of hazard ratios ± 95% confidence interval and p-values in Cox regression model for overall survival of TCGA patients separated by indication and categorized into upper and lower quartiles by stroma score. (F) (Top) Kaplan-Meier survival plots of TCGA patients categorized into upper and lower quartiles by the fibroblast TGFβ response signature from Mariathasan et al., 2018.^50^ Hazard ratio (HR) and p value of Cox regression fit are shown. (Bottom) Forest plot of hazard ratios ± 95% confidence interval and p-values in Cox regression model for overall survival of TCGA patients separated by indication and categorized into upper and lower quartiles by fibroblast TGFβ response signature. (G) (Top) Kaplan-Meier survival plots of TCGA patients categorized into upper and lower quartiles by the AAV-gTgfbr2 upregulated DEG signature. Hazard ratio (HR) and p value of Cox regression fit are shown. (Bottom) Forest plot of hazard ratios ± 95% confidence interval and p-values in Cox regression model for overall survival of TCGA patients separated by indication and categorized into upper and lower quartiles by AAV-gTgfbr2 upregulated DEG signature.

**Supplementary Figure 6.**
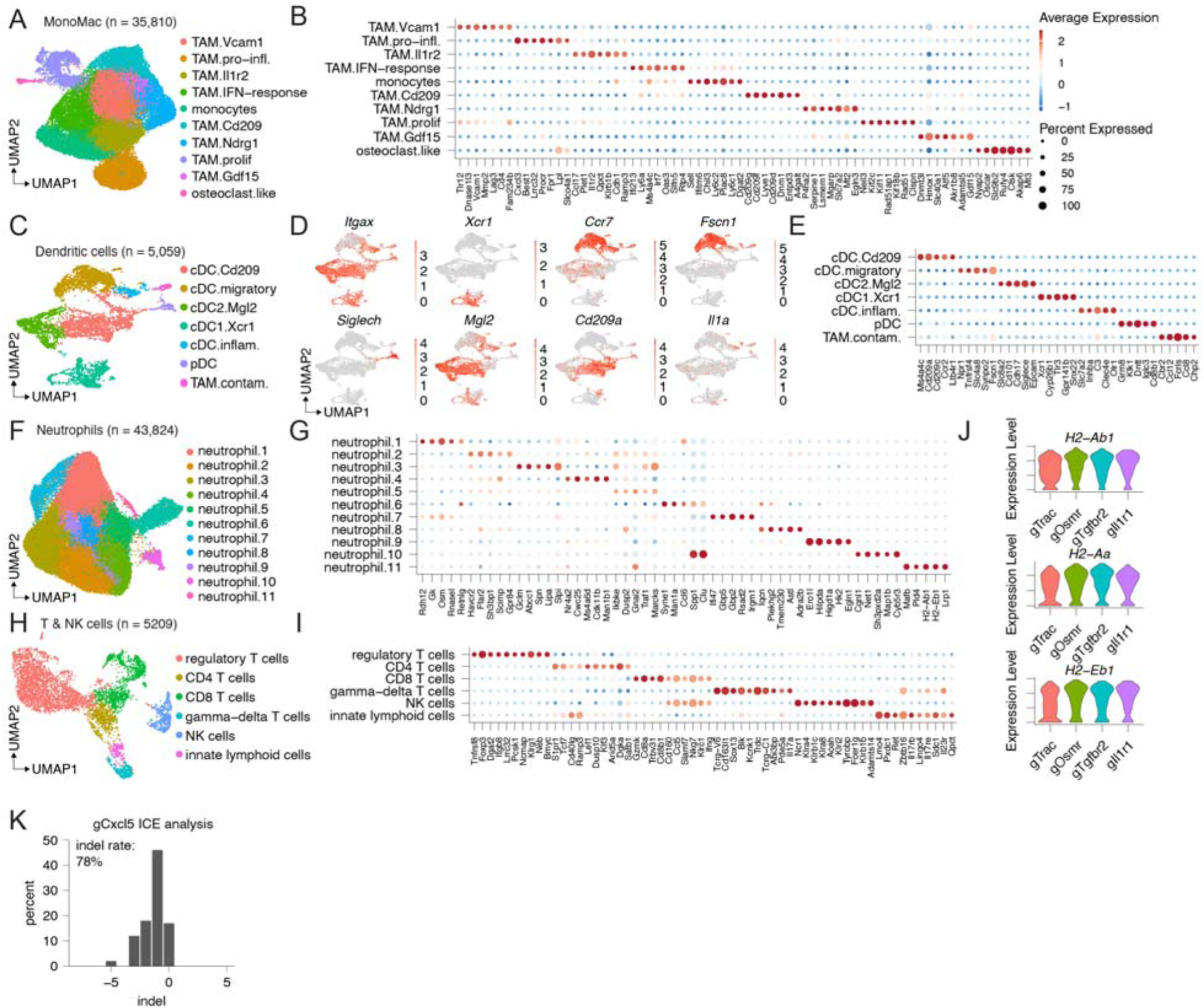
(A) UMAP of monocyte/macrophage (MonoMac) cell subsets from all AAV-gRNA tumors. (B) Bubble plot of scaled differentially expressed genes (DEGs) in MonoMac cell subsets. (C) UMAP of dendritic cell (DC) subsets from all AAV-gRNA tumors. (D) UMAP visualization of select DC marker genes in the DC object. (E) Bubble plot of scaled DEGs in DC subsets. (F) UMAP of neutrophil subsets from all AAV-gRNA tumors. (G) Bubble plot of scaled DEGs in neutrophil subsets. (H) UMAP of lymphoid cell subsets from all AAV-gRNA tumors. (I) Bubble plot of scaled DEGs in lymphoid cell subsets. (J) Violin plots of *H2-Ab1, H2-Aa, H2-Eb1* expression in the TAM.Ndrg1 subset, grouped by AAV-gRNA tumors. (K) Histogram depicting insertion/deletion (indel) distribution in NIH/3T3.Cas9 cells at gCxcl5 cut site 3 days after transfection with gCxcl5, as analyzed by ICE analysis.

**Supplementary Figure 7.**
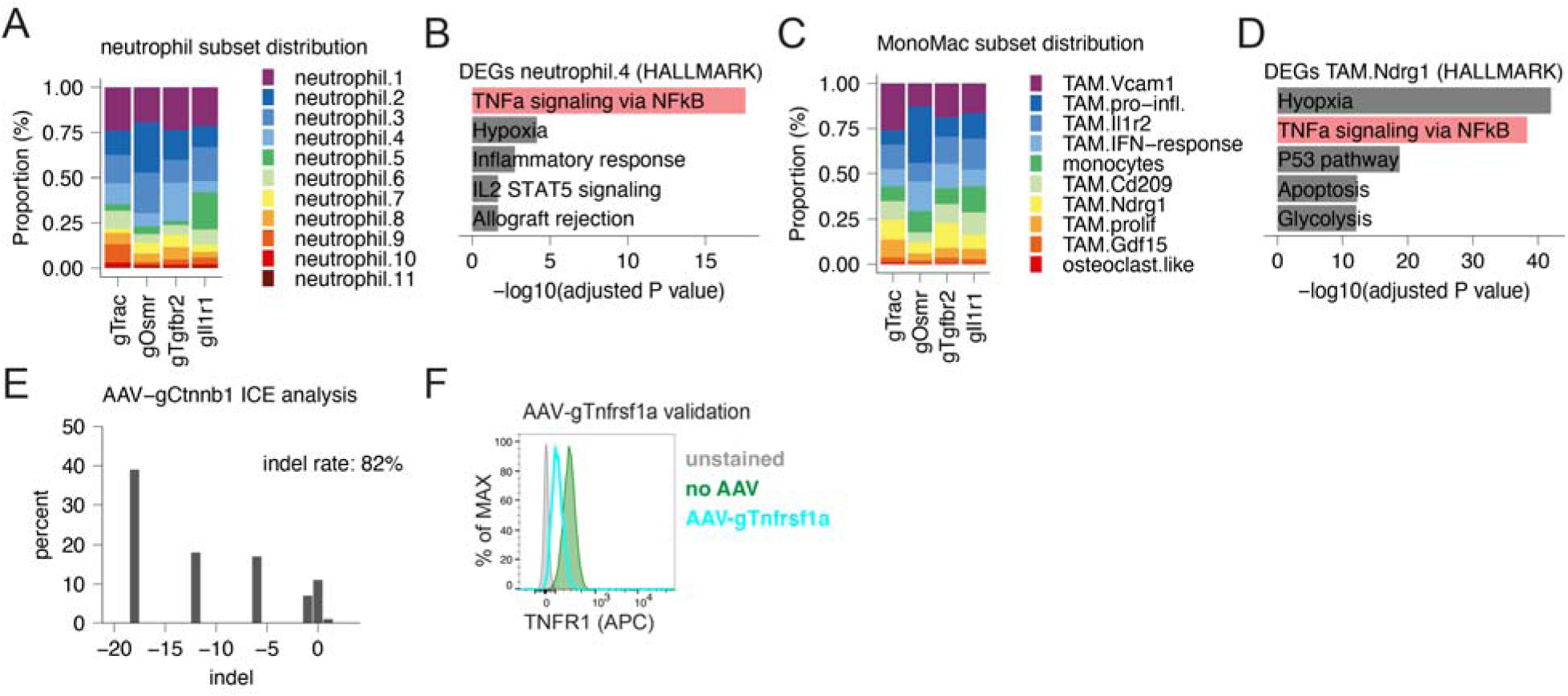
(A) Proportion of neutrophil subsets (scRNA-seq) in each AAV-gRNA group. (B) Enriched gene sets (Gene Ontology hallmark) based on DEGs from neutrophil.4 cluster. (C) Proportion of MonoMac subsets (scRNA-seq) in each AAV-gRNA group. (D) Enriched gene sets (Gene Ontology hallmark) based on DEGs from TAM.Ndrg1 cluster. (E) Histogram depicting insertion/deletion (indel) distribution in NIH/3T3.Cas9 cells at gCtnnb1 cut site 3 days after transduction with AAV-gCtnnb1, as analyzed by ICE analysis. (F) Flow cytometry histogram overlay plot of surface tumor necrosis factor receptor 1 (TNFR1) expression on NIH/3T3.Cas9 cells transduced with or without scAAV1-gTnfrsf1a at an MOI of 1e6 and analyzed 72 h later.

