## Supplementary Table 1 for "Discovery of a Linked Constellation of Gene Expression Revealed by Local Editing of Fibroblasts in Tumors"

weighted gene output ordered by non-negative matrix factorization (NMF) factors from fibroblast object

| | V1 | V2 | V3 | V4 | V5 | V6 | V7 | V8 | V9 | V10 | V11 | V12 | V13 | V14 | V15 | V16 | V17 | V18 |
|---|---|---|---|---|---|---|---|---|---|---|---|---|---|---|---|---|---|---|
| Crabp1 | 0 | 0 | 0.00054 | 0 | 0.00177 | 0.00085 | 0 | 0.00019 | 0.0043 | 0 | 0 | 0.00058 | 0 | 0.0021 | 0.00027 | 0.00054 | 0.00023 | 0 |
| Spp1 | 0.00013 | 0 | 0.00087 | 0.00043 | 0.00169 | 0.00114 | 0 | 0.00436 | 0.00215 | 0 | 0.00039 | 0 | 0.00047 | 0.00107 | 0.00154 | 0.00036 | 0 | 0 |
| Thbs2 | 0.00027 | 0.00056 | 0.00072 | 0 | 0.0015 | 0.00073 | 0.00268 | 0.00096 | 0.00179 | 0.00075 | 0.00121 | 0.00037 | 0.0013 | 0.0014 | 0.00057 | 0.0003 | 0.00056 | 0 |
| Col12a1 | 2.11E-05 | 0 | 0.00053 | 0.00013 | 0.00187 | 0.00163 | 0.00232 | 0.00061 | 0.00174 | 0 | 0.00218 | 0.00095 | 0.00115 | 0.00229 | 0.00172 | 0.00036 | 8.12E-05 | 0 |
| Cxcl5 | 0 | 0 | 0 | 0 | 0 | 0.00021 | 0 | 0 | 0.00169 | 0 | 0 | 0 | 0.00139 | 0.00037 | 0.00015 | 0 | 0 | 0 |
| Cdh11 | 0.00037 | 0.0003 | 0.00016 | 0 | 0.00109 | 0.0006 | 0.00088 | 0.00097 | 0.00156 | 0 | 0.00105 | 2.10E-05 | 0.00013 | 0.00085 | 6.53E-05 | 0.00025 | 0.0004 | 0.00065 |
| Col3a1 | 0.00023 | 0 | 0.00059 | 0.00161 | 0.00214 | 0.00227 | 0.00449 | 0.00296 | 0.00153 | 0.00259 | 0.00129 | 0.00108 | 0.00347 | 0.00166 | 0.00356 | 0.0007 | 0.00097 | 0.00226 |
| Tnc | 0 | 0 | 0.00215 | 0 | 0.00139 | 0.00114 | 0.00201 | 0.00283 | 0.00151 | 0 | 0.00192 | 0.00055 | 0.00132 | 0.00153 | 0.00252 | 0.0004 | 0 | 0 |
| Gpc1 | 0 | 0.00029 | 0.00047 | 0.00072 | 0.00061 | 0.00053 | 0.00098 | 0.00136 | 0.00141 | 9.45E-05 | 0.00037 | 0.00041 | 0.00084 | 0.00115 | 0.00115 | 0.0002 | 0 | 0.00048 |
| Ccn1 | 0 | 0 | 0.00017 | 0.00025 | 0.00078 | 2.39E-05 | 0.00034 | 0 | 0.0013 | 0.00052 | 0.00255 | 0.00047 | 0.00232 | 0.00104 | 0.00109 | 0 | 0.00184 | 7.46E-05 |
| Itm2a | 1.17E-05 | 0.00015 | 0 | 0 | 0.00065 | 0 | 0.00093 | 0.00052 | 0.00123 | 0.00052 | 0.00027 | 0 | 0.00105 | 0 | 0 | 3.47E-05 | 0 | 0.00012 |
| Slc16a1 | 0.00013 | 0 | 0.0003 | 0 | 0 | 0 | 0.00032 | 0.00013 | 0.00117 | 0.00015 | 0 | 0.00015 | 0.0014 | 0.00039 | 0 | 9.12E-05 | 0 | 0 |
| Sema7a | 0 | 0 | 0.00074 | 0 | 0.00122 | 0.00064 | 0.00034 | 0.00256 | 0.00117 | 0 | 6.85E-05 | 0.00021 | 3.43E-05 | 0.00059 | 0.0003 | 3.33E-05 | 0 | 0 |
| Cpxm1 | 0 | 9.45E-05 | 0.00011 | 0 | 0.00193 | 0.00219 | 0.00185 | 0.00068 | 0.00114 | 0.00069 | 0.00018 | 0 | 0.00432 | 0.00095 | 0.00025 | 0.0002 | 0 | 0.00016 |
| Timp1 | 0.0005 | 0 | 0.00164 | 0.00062 | 1.80E-05 | 0.00058 | 0.00177 | 0.002 | 0.00113 | 0.00016 | 8.86E-05 | 0.00043 | 0.00237 | 0.00021 | 0.00015 | 0.0004 | 0 | 0.00059 |
| Lum | 0.00047 | 0 | 0 | 0 | 0.00041 | 0.00111 | 0.00084 | 0.00062 | 0.00112 | 0.00089 | 0 | 0.00059 | 0.00056 | 0 | 0 | 0.00015 | 0 | 0.00112 |
| Ctsb | 0.00019 | 0.00069 | 0.00019 | 0.00032 | 0.00134 | 0.00115 | 0.00125 | 0.00162 | 0.00112 | 0.00103 | 0.00052 | 0.00071 | 0.00167 | 0.00091 | 0.00024 | 0.00013 | 0.00026 | 0.00062 |
| Tpbg | 0.00019 | 0.00019 | 0.00052 | 0.00017 | 0.00026 | 0.0004 | 0.00018 | 0.0003 | 0.00111 | 0.00032 | 0.00013 | 0.00012 | 0.00073 | 0.00083 | 0 | 0.0003 | 0.00029 | 0.00016 |
| Thy1 | 0.00033 | 0.00034 | 0.00055 | 0.00071 | 0.00106 | 0.00171 | 0.00068 | 0.00232 | 0.00109 | 0.00114 | 0.00103 | 0.00041 | 0.00115 | 0.00144 | 0.00021 | 0.00027 | 0.0005 | 0.00107 |
| Fstl1 | 0.00026 | 0.00012 | 0.00042 | 0.00096 | 0.00101 | 0.00127 | 0.00222 | 0.00151 | 0.00109 | 0.00187 | 0.00129 | 0.00057 | 0.00186 | 0.00127 | 0.0009 | 0.00031 | 0.00025 | 0.00153 |
| Ptk7 | 0 | 4.33E-05 | 0.00027 | 0 | 0.00113 | 0.0007 | 0.00028 | 0.00075 | 0.00107 | 0 | 0.00078 | 0.00027 | 0 | 0.00125 | 0.00077 | 3.69E-05 | 4.19E-05 | 0 |
| S100a6 | 0.00047 | 0.00018 | 0.00027 | 0.00096 | 5.62E-05 | 0.00099 | 0.00059 | 0.00122 | 0.00107 | 0.00108 | 0.00103 | 0.0011 | 0.00047 | 0.00094 | 0.00048 | 0.00032 | 0.00021 | 0.00111 |
| Gas1 | 0 | 0.00027 | 0 | 0.00041 | 0.00032 | 0.00042 | 0.00034 | 0.00099 | 0.00106 | 0 | 0 | 0 | 0.00071 | 0.00088 | 0.0009 | 9.34E-05 | 3.34E-05 | 0 |
| Spats2l | 5.10E-05 | 0 | 3.87E-05 | 0 | 0.00069 | 0.00052 | 0.00037 | 0 | 0.00106 | 0 | 0.00027 | 0.00016 | 0.00038 | 0.00051 | 9.04E-05 | 1.64E-06 | 3.24E-06 | 0 |
| Rarres2 | 0 | 0 | 0.00068 | 6.72E-05 | 0 | 0.00031 | 0.00084 | 0 | 0.00105 | 0.0012 | 3.66E-05 | 0.00095 | 0.00445 | 0.00051 | 0 | 0.00039 | 0 | 0 |
| Tcf4 | 6.77E-05 | 0.00063 | 0 | 0 | 0.00098 | 0.00047 | 6.32E-05 | 0.00051 | 0.00104 | 0.00012 | 0.00025 | 0.00022 | 0 | 0.00011 | 0.00125 | 0.00013 | 0.00097 | 0.00015 |
| Col6a2 | 0.00029 | 0 | 0.00046 | 0.0008 | 0.00166 | 0.00239 | 0.00234 | 0.00271 | 0.00103 | 0.00102 | 0.00091 | 0.00058 | 0 | 0.00178 | 0.00182 | 0.00039 | 0.00067 | 0.00144 |
| Fzd1 | 0 | 0.00031 | 0.00011 | 0.00018 | 0.00092 | 0.00037 | 0.00026 | 0.00081 | 0.00102 | 0 | 0.00033 | 0 | 0 | 0.00081 | 0.00067 | 0.00019 | 0.00014 | 0.00073 |
| Sfrp1 | 0 | 0 | 0 | 0 | 0.0004 | 0.00024 | 0.00093 | 0 | 0.00102 | 0 | 0.00097 | 0 | 0.00149 | 0 | 0.00026 | 5.27E-05 | 0 | 0.00138 |

| | | | | | | | | | | | | | | | | | | |
|---|---|---|---|---|---|---|---|---|---|---|---|---|---|---|---|---|---|---|
| Cnn2 | 0.00023 | 2.27E-05 | 0.00026 | 0.00021 | 0.00038 | 0 | 1.65E-05 | 5.98E-05 | 0.00101 | 0 | 0.00255 | 0.00032 | 0 | 0.00107 | 0 | 0.00013 | 7.87E-05 | 0.00042 |
| Lgi2 | 0 | 0 | 0 | 0 | 0 | 0 | 0.00103 | 0 | 0.00101 | 0.00054 | 0 | 0 | 0.0023 | 0 | 0 | 0 | 0 | 0.00022 |
| Twist1 | 0 | 0.00049 | 0.00013 | 0.00063 | 0.00013 | 0 | 0.00064 | 8.09E-05 | 0.00099 | 0.00044 | 0 | 0 | 0.00029 | 0.0002 | 0.0008 | 0.00038 | 0.00114 | 0.00024 |
| Spon1 | 0 | 0.0002 | 0 | 0 | 0.00196 | 0.00143 | 0.00109 | 0.00167 | 0.00098 | 4.91E-05 | 0.00091 | 0.00054 | 0.00048 | 0.00087 | 0.00108 | 0 | 4.58E-05 | 0 |
| Fbln1 | 0.00013 | 9.50E-05 | 3.73E-05 | 0 | 0.00037 | 0.00036 | 0.00104 | 0.00093 | 0.00098 | 0.00091 | 0 | 4.82E-05 | 0.00174 | 0.00117 | 6.10E-06 | 0.00028 | 2.45E-05 | 0.00064 |
| Col6a1 | 0.00049 | 5.01E-05 | 0.00036 | 0.00042 | 0.00155 | 0.00227 | 0.00201 | 0.00243 | 0.00097 | 0.00087 | 0.00088 | 0.00062 | 0 | 0.00153 | 0.00181 | 0.00036 | 0.00059 | 0.00112 |
| Pth1r | 3.10E-05 | 0.00026 | 0 | 0.0002 | 0.00013 | 0 | 0.00038 | 0.00022 | 0.00096 | 0.00016 | 0.00026 | 0 | 0.00044 | 0.00031 | 0.00018 | 0.00015 | 0 | 0.00019 |
| Ldlr | 2.87E-05 | 9.82E-05 | 0 | 0 | 0.00013 | 0 | 0 | 0.00023 | 0.00096 | 4.89E-05 | 0.00045 | 8.59E-05 | 0 | 0.00022 | 0.00021 | 7.63E-05 | 0.00048 | 0.00012 |
| Hmgcs1 | 0.0003 | 8.88E-05 | 0 | 5.26E-05 | 0.00012 | 1.24E-05 | 0 | 0.00021 | 0.00095 | 3.83E-05 | 0.00031 | 5.04E-05 | 0 | 0.00018 | 0 | 0 | 0.00124 | 0.00017 |
| Sdc1 | 0 | 4.18E-05 | 0.00067 | 0.00042 | 0.00069 | 0.00152 | 0.00053 | 0.00225 | 0.00095 | 0 | 0.00045 | 0.00065 | 0 | 0.00101 | 0.0006 | 0.00025 | 0 | 0.00022 |
| Dcakd | 8.64E-05 | 9.77E-05 | 0.00025 | 0.00038 | 0.00095 | 0.00017 | 0.00034 | 0.00067 | 0.00094 | 0 | 0 | 0 | 0.00035 | 0.00075 | 0 | 0.00022 | 0 | 0 |
| Pros1 | 0.00022 | 0.0002 | 0.00014 | 0.00025 | 0.00063 | 0.00107 | 0.00043 | 0.00104 | 0.00093 | 0.00053 | 0.00021 | 0.00031 | 0.0016 | 0.00088 | 0 | 0.00017 | 0.00014 | 0.00041 |
| Mgst1 | 0.00025 | 0.00014 | 0 | 0.0001 | 0.00036 | 0.00016 | 0.00082 | 0 | 0.00093 | 0.00064 | 8.70E-05 | 0 | 0.00098 | 0.00017 | 0 | 5.98E-05 | 0 | 0.00038 |
| Col1a1 | 0.00042 | 3.56E-05 | 0.00061 | 0.00182 | 0.00248 | 0.00261 | 0.00472 | 0.00266 | 0.00092 | 0.00209 | 0.00202 | 0.00151 | 0.00284 | 0.00268 | 0.00426 | 0.00081 | 0.00122 | 0.00224 |
| Svep1 | 1.07E-05 | 2.30E-05 | 0 | 0 | 0.0002 | 0.00056 | 0.00043 | 0 | 0.00091 | 0.00084 | 0.00031 | 0 | 0.00263 | 0.00073 | 0.00021 | 1.04E-05 | 0 | 0.0007 |
| Plat | 0 | 1.12E-06 | 0.00046 | 0.00028 | 0.00115 | 0.00092 | 0.00183 | 0.00181 | 0.0009 | 0.00101 | 0.00045 | 0.0002 | 0.00057 | 0.0006 | 0 | 0.0002 | 0.00082 | 0.00018 |
| Slit2 | 0.00022 | 2.03E-06 | 5.47E-05 | 0 | 0.00015 | 0 | 0 | 0.00101 | 0.0009 | 0 | 0.00025 | 0 | 0 | 0 | 0 | 0.00014 | 9.72E-05 | 0.00053 |
| Col1a2 | 0.00011 | 4.09E-05 | 0.00044 | 0.00199 | 0.00233 | 0.00218 | 0.00425 | 0.00256 | 0.00089 | 0.00209 | 0.00188 | 0.00112 | 0.00284 | 0.00231 | 0.00376 | 0.0007 | 0.00099 | 0.00222 |
| Tnfrsf19 | 0 | 0 | 0 | 0 | 0.00026 | 0 | 0 | 0.00042 | 0.00089 | 0 | 0 | 0 | 0 | 0 | 0 | 0 | 0 | 5.42E-05 |
| Abi3bp | 0 | 0 | 0 | 0 | 0.00059 | 0 | 0.00175 | 0 | 0.00088 | 0.00022 | 1.21E-05 | 0 | 0.00206 | 0 | 0 | 0 | 0 | 0.00063 |
| Igfbp3 | 0.00016 | 0 | 0 | 0 | 0.00304 | 0 | 0.00034 | 0.00081 | 0.00088 | 0 | 0.00055 | 0.00065 | 0 | 0.00147 | 0 | 0.00033 | 0.00024 | 0.00016 |
| Nxn | 0.00016 | 0 | 0.00051 | 0.00051 | 0.00062 | 0.00043 | 0.00081 | 0.00087 | 0.00087 | 1.47E-05 | 0.00046 | 0.00016 | 0.00053 | 0.00021 | 0.00035 | 0.0002 | 0.00023 | 0 |
| Tgfbr3 | 0 | 0.00022 | 0 | 0.0003 | 0.00022 | 0.00066 | 0.00045 | 0 | 0.00086 | 0.00089 | 0 | 0 | 0.00152 | 0.0005 | 0.00018 | 0 | 0.00028 | 0.00019 |
| Insig1 | 0.00018 | 0.00014 | 0 | 0.00017 | 0.00015 | 0 | 0 | 0.00028 | 0.00086 | 1.33E-05 | 0.00018 | 6.92E-05 | 0.00012 | 0.00035 | 0 | 0 | 0.00177 | 9.21E-05 |
| Cfh | 0 | 0.00017 | 0 | 0 | 0.00167 | 0.00129 | 0.00049 | 0.00104 | 0.00084 | 0.00027 | 9.45E-05 | 0.00017 | 0.00049 | 0 | 3.24E-05 | 0 | 0 | 0.00069 |
| Oaf | 0 | 0 | 0.00092 | 0.00027 | 0.00049 | 0.00028 | 0.00077 | 0.00142 | 0.00083 | 0.00039 | 0.00051 | 0.00026 | 0 | 0.00068 | 0.00029 | 0.00016 | 4.81E-05 | 0.00037 |
| Ddah1 | 0.00023 | 0 | 0.00033 | 0.00013 | 0.00051 | 0.00028 | 0.00068 | 0 | 0.00081 | 0 | 0.00121 | 0.00018 | 0.0011 | 0.00025 | 0 | 0.00021 | 0 | 0 |
| Plec | 0 | 0.00044 | 0.00059 | 0.00076 | 0.00057 | 0.00046 | 0.00059 | 2.90E-05 | 0.00081 | 0.00044 | 0.0006 | 0.00045 | 0.00013 | 0.00061 | 0.00157 | 0.00014 | 0.00043 | 0.00092 |
| Cyp51 | 0.00017 | 7.65E-06 | 0 | 0 | 0.00032 | 0 | 0 | 0.00023 | 0.0008 | 0 | 0.00025 | 0 | 0 | 0.0003 | 0 | 4.37E-05 | 0.00075 | 0.00013 |
| Spred1 | 0.00021 | 0.00034 | 0.0003 | 0.00016 | 0.00042 | 0.00038 | 0.00019 | 0.0012 | 0.0008 | 0.00019 | 0 | 0.00032 | 0.00013 | 0.00041 | 0.00043 | 0.00025 | 0.00067 | 0.0002 |
| Tgfbr2 | 0 | 0.00043 | 0.00011 | 0.00027 | 0.00114 | 0.00035 | 0.00158 | 0.00095 | 0.0008 | 0.0016 | 0.00042 | 0.00043 | 0.00084 | 0 | 0.00043 | 0.00016 | 0.00012 | 0.00106 |
| Ecrg4 | 8.35E-05 | 0 | 0 | 0 | 0.00031 | 0 | 0.00033 | 0 | 0.0008 | 0 | 0.00016 | 0 | 0 | 0.00014 | 0 | 0.00024 | 0 | 0.00013 |
| Lhfp | 0 | 0.00021 | 0.00022 | 0.00066 | 0.00088 | 0.001 | 0.00048 | 0.00112 | 0.0008 | 0.00073 | 0.00032 | 7.69E-05 | 0.00092 | 0.00079 | 0.00083 | 9.28E-05 | 0.00021 | 0.00093 |

| | | | | | | | | | | | | | | | | | | |
|---|---|---|---|---|---|---|---|---|---|---|---|---|---|---|---|---|---|---|
| Rnf149 | 0 | 5.64E-05 | 0.00066 | 5.52E-05 | 0.00092 | 0.00056 | 0.00031 | 0.00156 | 0.0008 | 0 | 0.00015 | 0.00041 | 0.00062 | 0.00073 | 0.00102 | 0 | 4.24E-05 | 0 |
| Bgn | 0.00033 | 0.00031 | 0.00057 | 0.00102 | 0.00192 | 0.00173 | 0.00271 | 0.00242 | 0.0008 | 0.00071 | 0.00141 | 0.0012 | 0.00287 | 0.00198 | 0.00119 | 0.0005 | 0.00037 | 0.00235 |
| Sh3pxd2b | 0 | 0.00034 | 0.00071 | 0.00017 | 0.00068 | 0.00056 | 0.00116 | 0.0007 | 0.0008 | 0.0005 | 0 | 0.00014 | 0.00095 | 0.00102 | 0.0004 | 0.00016 | 0.00024 | 4.62E-05 |
| Emp1 | 0.0007 | 0.0006 | 0.0003 | 0.00012 | 0.00116 | 0.00081 | 0.00077 | 0.00111 | 0.00079 | 0.00084 | 0.00021 | 0.00078 | 0.00039 | 0.0004 | 0 | 0.00047 | 0.00084 | 0.00089 |
| Prrx1 | 0.00034 | 0.0005 | 0.0004 | 0.00039 | 0.00041 | 0.0005 | 0.00022 | 0.00026 | 0.00079 | 0.00051 | 0.0002 | 0.00064 | 0.00034 | 0.00089 | 0.00069 | 0.00036 | 0.00075 | 0.00026 |
| Lrp1 | 0 | 0.00057 | 0.00052 | 0.00078 | 0.00118 | 0.00129 | 0.00125 | 0.00128 | 0.00078 | 0.00134 | 0.00063 | 0.00082 | 0.00195 | 0.0011 | 0.00164 | 0.0001 | 0.00035 | 0.00116 |
| Jak2 | 0.00022 | 0.00018 | 0.00026 | 7.58E-05 | 0 | 0.00013 | 0.00021 | 1.67E-05 | 0.00078 | 0 | 0 | 0.00014 | 0.00064 | 0.00058 | 0.00035 | 0.00013 | 0.00035 | 0.00012 |
| Slit3 | 0 | 3.44E-05 | 0.00029 | 0.00033 | 0.00065 | 0.00099 | 0.00068 | 0.00073 | 0.00078 | 0.00092 | 0.00057 | 0.00023 | 0.00152 | 0.00137 | 0.00056 | 5.99E-05 | 0.00025 | 0.0011 |
| Sparc | 0.00045 | 0 | 0.00057 | 0.00145 | 0.00189 | 0.00213 | 0.00388 | 0.00244 | 0.00078 | 0.00178 | 0.00168 | 0.00115 | 0.0025 | 0.00212 | 0.00217 | 0.0006 | 0.00048 | 0.00194 |
| Fdps | 0.00025 | 3.95E-05 | 0 | 0.00014 | 4.72E-05 | 0 | 0 | 0.00012 | 0.00078 | 0 | 0.00041 | 0 | 0 | 0.00018 | 0 | 0 | 3.66E-05 | 3.72E-05 |
| Loxl1 | 0.00035 | 0 | 0.00049 | 0.00023 | 0.00117 | 0.00121 | 0.00113 | 0.00126 | 0.00077 | 0.00135 | 0.00112 | 0.0002 | 0.00187 | 0.00114 | 0.00018 | 0.00028 | 6.71E-05 | 0.00058 |
| Fblim1 | 0 | 0 | 0.00013 | 0.0003 | 0.00017 | 0 | 0 | 0.00067 | 0.00077 | 0 | 0.00056 | 0.00011 | 0.0005 | 0 | 0.00032 | 8.28E-06 | 4.17E-05 | 0.00014 |
| Zyx | 0 | 8.87E-05 | 0.00079 | 0.00045 | 0.00015 | 0 | 0 | 0.00028 | 0.00077 | 0.00084 | 0.00137 | 0.00046 | 0.00023 | 0.0006 | 1.71E-05 | 0.00019 | 0.00033 | 0 |
| Masp1 | 0 | 0 | 0 | 0 | 0.00035 | 0.00023 | 0 | 0 | 0.00077 | 0.00012 | 0 | 0 | 0 | 0.00042 | 0 | 0 | 0 | 0.00047 |
| Slc39a1 | 0 | 0.00031 | 0.00042 | 0.00093 | 0.00086 | 0.00062 | 0.00106 | 0.00113 | 0.00077 | 0.00075 | 0.00045 | 0.00046 | 0.00094 | 0.00073 | 0.00013 | 0.0002 | 0.00014 | 0.00082 |
| Calm3 | 0.00036 | 0.0003 | 0.00029 | 0.00105 | 0.00042 | 0.00048 | 0.00032 | 0.00082 | 0.00076 | 0.00032 | 0.00037 | 0.00019 | 0.00023 | 0.00047 | 0 | 0.00043 | 0.00029 | 0.00048 |
| Srebf2 | 6.25E-05 | 0.00022 | 3.19E-05 | 0.00024 | 0.00026 | 3.23E-05 | 0 | 0.00034 | 0.00076 | 0.00011 | 0.00036 | 0.00026 | 0 | 0.0002 | 0.00037 | 8.10E-05 | 0.00029 | 0.00021 |
| Stra6 | 0 | 0 | 0 | 7.21E-06 | 0.00174 | 0.00051 | 0 | 0.00083 | 0.00076 | 0 | 0.00064 | 0.00043 | 0 | 0 | 0.00047 | 0 | 0 | 0 |
| Igf1 | 0 | 0.0001 | 0 | 0 | 0.00044 | 0.00058 | 0.00071 | 0 | 0.00076 | 0.0002 | 0 | 0 | 0.00041 | 0 | 0 | 0 | 0 | 0.00087 |
| Ybx3 | 0.00032 | 0.00031 | 0.0004 | 0.00037 | 0.00076 | 0.0009 | 0 | 0.00149 | 0.00076 | 0.00029 | 6.78E-06 | 0.00034 | 0 | 0.0004 | 0.00016 | 0.0003 | 0 | 0.00021 |
| Svil | 0 | 0.00023 | 4.07E-05 | 0 | 0.00034 | 0.0002 | 0.00037 | 0.0005 | 0.00076 | 0.00025 | 0 | 0 | 0.00025 | 0.00019 | 0.00052 | 0.00011 | 0.00036 | 0.00015 |
| Sema5a | 0 | 0.00027 | 0 | 0 | 0.00021 | 0.00026 | 0 | 0 | 0.00075 | 0 | 0 | 0 | 0 | 0.0003 | 0 | 1.21E-05 | 0.00014 | 0 |
| Dcn | 0.00055 | 0.00039 | 0.00037 | 0.00014 | 0.00035 | 0.0027 | 0.00105 | 0.00067 | 0.00075 | 0.00151 | 0 | 0.00043 | 0.00277 | 0.00065 | 0 | 0.00015 | 9.25E-05 | 0.00091 |
| S100a11 | 0.00092 | 0.00014 | 0.00042 | 0.00074 | 0.00053 | 0.00062 | 0.00089 | 0.0008 | 0.00074 | 0.00088 | 0.001 | 0.00057 | 0.00057 | 0.00075 | 0.00064 | 0.00038 | 0.00042 | 0.00072 |
| Dio3 | 0 | 0 | 0.00049 | 8.19E-05 | 0 | 0 | 0.00017 | 0 | 0.00074 | 0 | 0.00031 | 0 | 0.00038 | 0 | 0 | 0.00023 | 0 | 9.01E-05 |
| Tmem176a | 0.00016 | 0 | 0 | 0 | 0 | 0 | 0.00027 | 0.00029 | 0.00073 | 0 | 0.00039 | 0 | 0.00104 | 0.00079 | 0 | 8.22E-05 | 0 | 0.00055 |
| Ugdh | 0.00026 | 9.00E-05 | 0.00064 | 0.00041 | 0.00038 | 0.00037 | 0.00049 | 0.0001 | 0.00073 | 0.00076 | 0 | 0.00085 | 0.00056 | 0.00011 | 0.00034 | 0.0005 | 0.00039 | 3.69E-05 |
| Fndc1 | 0 | 0 | 0 | 0 | 0.00141 | 0 | 0.00085 | 0 | 0.00073 | 0.0008 | 0.00055 | 0.00047 | 0 | 0.00039 | 0.00026 | 0 | 0 | 0.00051 |
| Nlgn2 | 0.00023 | 0.00043 | 6.24E-05 | 0 | 0.00036 | 0.00035 | 0.00058 | 0.00018 | 0.00073 | 0.0002 | 7.62E-05 | 0.00021 | 0.00046 | 0.00024 | 0.00089 | 0 | 0.0004 | 0.0003 |
| Actg1 | 0.00053 | 0.00034 | 0.00099 | 0.00055 | 0.00029 | 0 | 0.00051 | 7.33E-05 | 0.00072 | 0.00052 | 0.0021 | 0.00087 | 2.77E-05 | 0.00047 | 0 | 0.00058 | 0.00075 | 0.00071 |
| Mfap2 | 5.56E-05 | 0 | 0 | 0.00048 | 0.00073 | 0.00068 | 0.001 | 0.00091 | 0.00072 | 0.00051 | 0.00039 | 0 | 0.00084 | 0.00118 | 4.97E-05 | 1.95E-05 | 0 | 0.00047 |
| Efemp2 | 0.00041 | 0 | 0.00036 | 0.00014 | 0.0005 | 0.00101 | 0.00126 | 0.00089 | 0.00072 | 0.00073 | 0.00053 | 0.00067 | 0.00179 | 0.00065 | 0.00021 | 0.00022 | 0 | 0.00043 |
| Cavin1 | 1.34E-05 | 0 | 0.00025 | 0.00086 | 0 | 0.0002 | 0.00053 | 0 | 0.00072 | 0.00128 | 0.00039 | 0.00037 | 0.00032 | 0.00053 | 7.06E-05 | 0.00029 | 0.00037 | 0.00098 |

| | | | | | | | | | | | | | | | | | | |
|---|---|---|---|---|---|---|---|---|---|---|---|---|---|---|---|---|---|---|
| Lbh | 0 | 4.36E-05 | 0 | 0.0002 | 0.00133 | 0.00079 | 0 | 0.00034 | 0.00072 | 0.00012 | 0.001 | 0.00019 | 0.00046 | 0.0006 | 0.00029 | 0 | 0 | 0.00025 |
| Prickle1 | 4.54E-05 | 0.00012 | 0.00012 | 0 | 0.00066 | 0.00023 | 0.00055 | 4.06E-05 | 0.00071 | 2.07E-05 | 0.00028 | 0.00027 | 4.20E-05 | 0.00011 | 4.59E-05 | 0 | 0.00051 | 0.00017 |
| Col6a3 | 0.00042 | 0 | 0.00054 | 0.00056 | 0.00136 | 0.00237 | 0.00179 | 0.00262 | 0.0007 | 0.00111 | 0.00096 | 0.0006 | 1.41E-07 | 0.00223 | 0.00241 | 0.00042 | 0.00046 | 0.00093 |
| Angpt4 | 0 | 0 | 0 | 0 | 0.00022 | 0 | 0.00019 | 0.00017 | 0.0007 | 0 | 0.00021 | 0 | 0.00049 | 0 | 0 | 2.00E-05 | 0 | 7.43E-05 |
| Nid1 | 4.98E-05 | 0.00021 | 2.01E-05 | 0.00059 | 0.00103 | 0.00126 | 0.00082 | 0.00136 | 0.0007 | 0.00157 | 0.00094 | 0.00077 | 0.00143 | 0.00072 | 0.00062 | 0.00011 | 0.00048 | 0.00048 |
| Ctsl | 4.16E-05 | 0.0003 | 0.00031 | 0.00025 | 0.00097 | 0.00145 | 0.00066 | 0.00073 | 0.00069 | 0.00075 | 0.00015 | 0.00057 | 0.00159 | 0.00096 | 0 | 5.31E-05 | 9.76E-06 | 0.00032 |
| Arhgef7 | 7.48E-05 | 0.0002 | 0.00026 | 0 | 0.0004 | 0.0003 | 0.00026 | 0.00029 | 0.00069 | 0.00014 | 0.00021 | 5.27E-06 | 0.00025 | 0.00055 | 0 | 0.00019 | 0.00026 | 1.85E-06 |
| 9-Sep | 0.00033 | 0.00035 | 0.00048 | 0.00027 | 0.00035 | 0.00053 | 0.00028 | 0.00097 | 0.00069 | 0 | 0.00025 | 0.00049 | 0 | 0.00022 | 0.00035 | 0.0002 | 0.00042 | 0.00024 |
| Hsd11b1 | 0 | 4.89E-05 | 0 | 0 | 0.00016 | 0.00042 | 8.56E-05 | 8.87E-07 | 0.00068 | 4.65E-05 | 0 | 0 | 0.00068 | 0 | 0 | 3.93E-05 | 0 | 0.00039 |
| Scd2 | 0.00016 | 9.12E-05 | 0 | 0.00042 | 0.00029 | 0 | 0.00015 | 0.0004 | 0.00068 | 0.00011 | 0.00011 | 0.00056 | 0 | 0.00015 | 6.00E-05 | 0 | 0 | 0.00011 |
| Pla1a | 0 | 0 | 4.09E-05 | 0 | 0.00093 | 0.00153 | 0 | 0.00133 | 0.00068 | 0.00097 | 0.00072 | 0.00031 | 0.0005 | 0.00149 | 0.00043 | 0 | 0 | 0 |
| Lpar1 | 0 | 6.57E-05 | 2.15E-05 | 0.00049 | 0.00047 | 0.00016 | 0.00039 | 0.00025 | 0.00068 | 0.00055 | 0.00036 | 0 | 0.0004 | 0 | 2.91E-05 | 0.00016 | 0.00013 | 0.00024 |
| Fgl2 | 0 | 3.94E-05 | 0.00024 | 0.00017 | 1.13E-05 | 6.65E-05 | 3.32E-05 | 0 | 0.00067 | 0.00051 | 0 | 0 | 0.00065 | 0 | 6.73E-05 | 8.11E-05 | 0.00031 | 0.00023 |
| Rgma | 6.79E-05 | 0.00033 | 0 | 0.00014 | 0.00053 | 0.00014 | 8.82E-05 | 0 | 0.00067 | 0.00016 | 0.00014 | 0 | 0 | 1.87E-05 | 5.30E-05 | 0.00014 | 3.33E-05 | 0.00014 |
| Nucb1 | 0.00013 | 0.00021 | 0.00012 | 0.00048 | 0.00065 | 0.00064 | 0.00111 | 0.00092 | 0.00067 | 0.00077 | 0.00046 | 0.00046 | 0.00104 | 0.00091 | 0.00015 | 0.00014 | 0 | 0.00078 |
| Gpm6b | 0 | 0 | 0.00033 | 0.0002 | 0.0006 | 0.00072 | 5.43E-06 | 0.00011 | 0.00066 | 5.93E-05 | 1.87E-05 | 0 | 0.00086 | 0.00075 | 0 | 4.30E-05 | 0 | 0.00043 |
| Man2a1 | 6.71E-05 | 7.39E-06 | 0.00038 | 0.00035 | 0.00023 | 0.00051 | 0.00091 | 0.00054 | 0.00066 | 0.00074 | 0.00025 | 0.00019 | 0.00143 | 0.00044 | 9.20E-06 | 0.0002 | 0.00022 | 0.00053 |
| Itgb5 | 0.0002 | 0.00043 | 7.35E-05 | 9.89E-05 | 0.00112 | 0.00086 | 0.00071 | 0.00142 | 0.00066 | 0.00052 | 0.00073 | 8.56E-05 | 0.00073 | 0.00102 | 0.00043 | 8.82E-05 | 0.00022 | 0.00039 |
| Tpm1 | 0.0004 | 0.00017 | 0.00025 | 0.00057 | 0.0008 | 0.00023 | 0 | 0.00033 | 0.00066 | 5.16E-05 | 0.00285 | 0.00027 | 0 | 0.00074 | 0.00118 | 0.00027 | 0.00033 | 0.00044 |
| Ldha | 0.00036 | 0 | 0.00033 | 0.0009 | 0.00021 | 0.00035 | 0.00013 | 0.00043 | 0.00066 | 0.00039 | 0.00054 | 0.00165 | 0.00072 | 0.00036 | 0.0002 | 0.0002 | 0.00042 | 0.00012 |
| Egfp | 0.00017 | 0.00039 | 0.00027 | 0 | 0.00082 | 0.00082 | 0.00058 | 0.00075 | 0.00066 | 7.48E-05 | 0.0005 | 0.00036 | 0.00079 | 0.00066 | 0 | 7.87E-05 | 0.00035 | 0.00031 |
| Ermp1 | 1.06E-05 | 0.00011 | 0 | 0.00013 | 0.00107 | 0 | 0.00018 | 0.00081 | 0.00066 | 6.61E-05 | 9.65E-05 | 4.69E-05 | 0 | 0 | 0 | 0.00017 | 0 | 5.04E-05 |
| Il1r1 | 0.00043 | 0 | 3.21E-05 | 0.00021 | 0.00032 | 0.00083 | 9.22E-05 | 0.0004 | 0.00065 | 0.00021 | 0 | 3.64E-05 | 0.00177 | 0.00093 | 0.00025 | 0.00014 | 0.00028 | 0.00042 |
| Il4ra | 0.00028 | 6.66E-05 | 0.00025 | 0.00061 | 0.00032 | 9.98E-05 | 0.00075 | 0.00027 | 0.00065 | 0.00028 | 0.00051 | 0.00043 | 0.00217 | 0.00037 | 0.00088 | 0.00021 | 0 | 0 |
| Rgcc | 0 | 0 | 0.00029 | 2.40E-05 | 0.00013 | 0.00031 | 0.00012 | 0 | 0.00065 | 0 | 0 | 0.00079 | 0.00034 | 0.00065 | 0 | 0 | 0.00095 | 0 |
| Prkar1a | 0.00015 | 0.00032 | 0.00034 | 0.00063 | 0.00061 | 0.00063 | 0 | 0.00104 | 0.00065 | 0.00017 | 0.00047 | 0.00013 | 0.00016 | 0.00056 | 0.00025 | 8.81E-05 | 0.00016 | 0.00028 |
| 11-Sep | 0.00055 | 0.00016 | 0.0001 | 1.09E-05 | 2.22E-05 | 0.00019 | 0 | 0.00065 | 0.00064 | 7.88E-05 | 0.00015 | 0.00039 | 0 | 0.00016 | 0.00028 | 0.00035 | 0.00046 | 0.00035 |
| Txndc5 | 0.00035 | 1.67E-05 | 0.00049 | 0.00087 | 0.00052 | 0.00058 | 0.0015 | 0.00087 | 0.00064 | 0.00069 | 0.00055 | 0.0002 | 0.00053 | 0.0009 | 0.00063 | 0.00022 | 0.0002 | 0.00065 |
| Tmem176b | 5.43E-05 | 4.55E-05 | 0 | 0.00017 | 2.24E-05 | 0 | 0.00029 | 0.00027 | 0.00064 | 0 | 0.00041 | 0 | 0.00127 | 0.00094 | 0 | 8.62E-05 | 0 | 0.0007 |
| Mgst3 | 0 | 0 | 0 | 7.86E-06 | 5.84E-05 | 0 | 0 | 0 | 0.00064 | 3.43E-05 | 0 | 4.29E-05 | 0 | 0.00068 | 0 | 2.12E-05 | 0 | 0.00039 |
| Ak3 | 0.00019 | 0.00019 | 6.50E-05 | 9.59E-05 | 0.00015 | 0.00037 | 0.00052 | 0 | 0.00064 | 0.00022 | 3.18E-05 | 0 | 0.00014 | 0.00055 | 0 | 0 | 0 | 0.00031 |
| Ly6e | 0.00043 | 5.01E-05 | 0.00023 | 0.00057 | 0.00036 | 0.00039 | 0.00042 | 0.00084 | 0.00064 | 0.00112 | 0.00039 | 0.00026 | 0.00118 | 0.00055 | 0 | 0.00034 | 7.21E-05 | 0.00084 |
| Eif2s2 | 0.00037 | 0 | 5.44E-05 | 0 | 6.30E-05 | 0.00049 | 4.56E-05 | 0 | 0.00064 | 0 | 9.00E-05 | 7.63E-05 | 0 | 0.00013 | 3.28E-05 | 6.48E-05 | 0 | 0 |

| | | | | | | | | | | | | | | | | | | |
|---|---|---|---|---|---|---|---|---|---|---|---|---|---|---|---|---|---|---|
| Ctsk | 0 | 0.00033 | 5.97E-05 | 0.00078 | 0.00109 | 0.00131 | 0.00147 | 0.00155 | 0.00064 | 0.00057 | 0.00038 | 0.00012 | 0.00134 | 0.00169 | 0 | 9.67E-05 | 0 | 0.00093 |
| Cnn3 | 0.00047 | 0.00023 | 0.0005 | 7.44E-05 | 0.00033 | 0.00034 | 0.00079 | 0.00066 | 0.00063 | 0.00017 | 0.00108 | 0.00018 | 0.00037 | 0.00026 | 0 | 0.00013 | 0.00027 | 0.00046 |
| Palld | 4.97E-05 | 5.84E-05 | 0.00043 | 2.77E-05 | 0.00118 | 0.00021 | 0.00102 | 0.0005 | 0.00063 | 0.00025 | 0.00202 | 0.00024 | 0.00011 | 0.00025 | 0.00025 | 0.00018 | 0.0005 | 0.0004 |
| Ehd2 | 0.00027 | 7.13E-05 | 0.00032 | 0.00059 | 0.00051 | 0.0006 | 0.00043 | 0.00043 | 0.00063 | 0.00071 | 0.00055 | 0.00012 | 0.00015 | 0.00039 | 4.99E-05 | 0.00015 | 0.00016 | 0.00045 |
| Eef1a1 | 0.00101 | 0.00078 | 0.00062 | 0.00093 | 0.00073 | 0.0013 | 0.00146 | 0.00101 | 0.00062 | 0.00084 | 0.00067 | 0.00109 | 0.001 | 0.00099 | 0.00047 | 0.00038 | 0.00101 | 0.00064 |
| Parva | 0.00031 | 0.00015 | 0.00019 | 0.00019 | 0.00049 | 0.00044 | 0.00031 | 0.00034 | 0.00062 | 0.00023 | 0.00102 | 0.0002 | 0.00033 | 0.00047 | 0 | 0.00019 | 3.20E-06 | 0.00039 |
| Ctdsp2 | 5.63E-05 | 0.00076 | 0.00014 | 0.00086 | 0.00061 | 0.00056 | 0.00049 | 0.00052 | 0.00062 | 0.00076 | 0.00051 | 0.00058 | 0.00058 | 0.0007 | 0.00049 | 0.00031 | 0.00043 | 0.00065 |
| Marcks | 0.0002 | 0.00048 | 0 | 0.00124 | 0.00062 | 0.00019 | 0.00015 | 0.00075 | 0.00062 | 0.00104 | 0.00063 | 0 | 0 | 0.00046 | 0.00131 | 0.00056 | 0.0004 | 0.0007 |
| Tpm4 | 0.0008 | 9.28E-05 | 0.00059 | 0.00044 | 0.00045 | 0.00058 | 0.00055 | 0.00068 | 0.00062 | 0.00022 | 0.00146 | 0.00065 | 0.00037 | 0.0007 | 0 | 0.00042 | 0.00026 | 0.00027 |
| Ltbp1 | 0 | 5.68E-05 | 0 | 0 | 0 | 0.00035 | 0 | 0 | 0.00061 | 0.00067 | 0 | 0 | 2.82E-05 | 0 | 0 | 0 | 0 | 0.00011 |
| Igfbp4 | 0 | 0.00026 | 0 | 0.00021 | 0.00104 | 0.0005 | 0.00029 | 0.00167 | 0.00061 | 0.00089 | 0 | 0 | 0.0014 | 0.00106 | 0.001 | 0 | 0 | 0.00303 |
| Timp2 | 6.39E-05 | 0.00044 | 3.16E-05 | 0 | 0.0009 | 0.00102 | 0.00083 | 0.00061 | 0.00061 | 0.00117 | 0.00078 | 0 | 0.0009 | 0.00092 | 0.00032 | 0 | 0 | 0.00082 |
| Arhgdib | 0.00012 | 6.09E-05 | 0 | 0 | 0.00013 | 0.00066 | 0 | 0.0002 | 0.00061 | 0 | 0 | 0.00038 | 0 | 0.0009 | 0 | 0 | 0 | 0.00043 |
| Rnase4 | 0.00032 | 0.00042 | 0.00019 | 0 | 0.00028 | 0.00096 | 0.00128 | 0.00066 | 0.0006 | 0.00116 | 8.90E-05 | 0.0001 | 0.00195 | 0.00086 | 0 | 0 | 0 | 0.00066 |
| Ugcg | 0.00029 | 0.0001 | 0 | 0 | 0 | 5.31E-05 | 0.00032 | 4.37E-05 | 0.0006 | 0.00031 | 0.00013 | 0.00101 | 0.00174 | 0.00023 | 1.62E-05 | 2.10E-05 | 0 | 0.00032 |
| Medag | 0.00028 | 0 | 0.00018 | 0 | 0 | 0 | 0.00104 | 0.00044 | 0.0006 | 0.00057 | 0.00012 | 0.00044 | 0.00061 | 0.00035 | 6.02E-05 | 0.00017 | 0 | 0.00028 |
| Pde5a | 0 | 0.00012 | 0 | 0 | 0.00017 | 0.00023 | 0 | 0.00031 | 0.0006 | 0 | 0 | 0 | 0 | 0.00029 | 2.41E-05 | 6.15E-05 | 0 | 0 |
| Inhbb | 0 | 0.00055 | 0 | 0 | 0.00029 | 0.00109 | 0 | 0 | 0.0006 | 0.0001 | 0.00023 | 0.0004 | 0.00013 | 0.00056 | 0 | 0 | 0.00016 | 0 |
| C1qtnf2 | 0 | 8.82E-05 | 0 | 5.39E-05 | 0.00051 | 9.97E-05 | 0.00058 | 0 | 0.0006 | 0.00022 | 0 | 0 | 0.00014 | 0.00058 | 0 | 4.55E-05 | 0 | 0.00027 |
| Cfl1 | 0.00034 | 0.00027 | 0.00057 | 0.00154 | 0.00031 | 0.00028 | 0.00043 | 0.00073 | 0.0006 | 0.00037 | 0.00102 | 0.00045 | 0.00031 | 0.00065 | 6.09E-06 | 0.0004 | 0.00078 | 0.00042 |
| Ptms | 0.00029 | 0.00043 | 0.00016 | 0.00152 | 0.00037 | 0.00028 | 0.00054 | 0.00085 | 0.0006 | 0.00048 | 0.00078 | 0.00026 | 0 | 0.0006 | 0.00182 | 0.0007 | 0.001 | 0.00087 |
| Flna | 0.00063 | 0.00039 | 0.00063 | 0 | 0.00041 | 0.00018 | 0.00018 | 0.00031 | 0.00059 | 0.0002 | 0.00196 | 0.00057 | 0 | 0.00056 | 0.00072 | 0.00025 | 0.00032 | 0.00034 |
| Fosl2 | 0.00015 | 0.00034 | 2.49E-05 | 0.00018 | 0.00037 | 0.00032 | 0.00032 | 0 | 0.00059 | 0.00012 | 0.00036 | 0.00037 | 0.0004 | 0.00026 | 0.00068 | 0.00012 | 0.00043 | 5.87E-05 |
| Csrp1 | 0 | 7.48E-05 | 0.00042 | 0.00055 | 0.00069 | 0 | 0.00035 | 0.00021 | 0.00059 | 0 | 0.00248 | 0.00027 | 0 | 0.00079 | 0.00014 | 0.00026 | 0 | 0.00023 |
| S100a16 | 0.00021 | 0 | 0.00046 | 0.00049 | 0.00016 | 0.00024 | 0.00075 | 0.00114 | 0.00059 | 0.00112 | 0.0003 | 0.00028 | 0.00031 | 5.23E-06 | 0 | 0 | 0 | 0.00052 |
| Myo1c | 5.38E-05 | 0.0002 | 0.0004 | 0.00057 | 0.00026 | 0.00025 | 0.00032 | 0.00038 | 0.00059 | 0.00041 | 0.0009 | 0.00054 | 0.00027 | 0.0004 | 0.00038 | 0.00015 | 7.48E-05 | 0.00061 |
| Pkm | 0.00037 | 0.00019 | 0.00035 | 0.00052 | 0.00031 | 0.00031 | 0.00032 | 0.00046 | 0.00059 | 0.00012 | 0.0003 | 0.00162 | 0.0006 | 0.00034 | 0.00078 | 0.00025 | 0.00058 | 0 |
| Nfe2l2 | 0.00026 | 0.0004 | 0.00019 | 2.10E-05 | 0.00021 | 0.00044 | 0 | 0.00043 | 0.00059 | 0.00031 | 0 | 0.00034 | 0.00029 | 0.00036 | 0.00011 | 0.00013 | 0.00037 | 4.48E-05 |
| Sc5d | 0.0001 | 0 | 2.62E-05 | 0 | 0.00018 | 9.57E-05 | 0 | 0 | 0.00058 | 0 | 0.00022 | 2.82E-05 | 2.67E-07 | 2.10E-05 | 0 | 0 | 0.00013 | 0 |
| Ston1 | 9.19E-06 | 0.00021 | 0 | 0.00016 | 0.00064 | 0.00022 | 0 | 0.00028 | 0.00058 | 0 | 0.00038 | 4.97E-05 | 0 | 7.71E-05 | 6.59E-05 | 4.07E-05 | 7.38E-05 | 0.00012 |
| Lgals1 | 0.00047 | 0 | 0.0003 | 0.00031 | 0.00039 | 0.00039 | 0.00068 | 0 | 0.00058 | 0 | 0.00064 | 0.00039 | 2.38E-05 | 3.71E-06 | 0 | 0.00024 | 0.00012 | 0.00016 |
| Igsf10 | 0.00036 | 0.00011 | 0 | 0 | 0.00028 | 0.00024 | 0.00038 | 0 | 0.00058 | 0.00051 | 0 | 0 | 0.00021 | 0.00047 | 0 | 0.00011 | 0 | 0.00013 |
| Sqle | 6.14E-05 | 3.66E-05 | 0 | 0 | 5.77E-05 | 0 | 0 | 1.29E-05 | 0.00058 | 0 | 0.00017 | 0.0001 | 0 | 8.62E-05 | 0 | 0 | 0.00039 | 6.09E-05 |

| | | | | | | | | | | | | | | | | | | |
|---|---|---|---|---|---|---|---|---|---|---|---|---|---|---|---|---|---|---|
| Fbln2 | 4.02E-07 | 0 | 0.00071 | 0.00057 | 0.00122 | 0.00114 | 0.00212 | 0.00092 | 0.00058 | 0.00097 | 0.00084 | 0.00093 | 0.00179 | 0.00186 | 0.0015 | 0.00036 | 7.82E-05 | 0.00089 |
| Aebp1 | 0 | 0.00035 | 0.00039 | 0.00015 | 0.00128 | 0.00141 | 0.00182 | 0.00091 | 0.00058 | 0.00119 | 0.00124 | 0.00091 | 0.00242 | 0.00134 | 0.00092 | 0.00022 | 0.00011 | 0.00067 |
| Tuba1a | 0.00037 | 0.00015 | 0.00046 | 0.00111 | 5.23E-05 | 4.73E-05 | 0.00031 | 0.00039 | 0.00058 | 0.00063 | 0.00113 | 0.00035 | 7.20E-05 | 0.00032 | 0 | 0.00083 | 0.00091 | 0.00073 |
| Plscr3 | 0.00017 | 0.00032 | 0.0002 | 0.0005 | 0.00023 | 0.00021 | 0.0005 | 0.00023 | 0.00058 | 0.0003 | 0.00052 | 0.00023 | 0.00031 | 0.00038 | 7.17E-05 | 0 | 2.42E-05 | 0.00055 |
| Mxra8 | 0.00011 | 0.00054 | 0.0002 | 0 | 0.00133 | 0.00082 | 0.00164 | 0.00062 | 0.00058 | 0.00021 | 0.00037 | 0 | 0.00173 | 0.0013 | 0.00035 | 0 | 0.00012 | 0.00066 |
| Sptbn1 | 0.00016 | 0.00037 | 0.00026 | 5.71E-05 | 0.00049 | 0.0004 | 0.00058 | 0.00086 | 0.00057 | 0.00095 | 0.00055 | 0.0004 | 0.00015 | 0.00025 | 0.00036 | 0.00015 | 0.00042 | 0.00106 |
| Anxa2 | 0.00059 | 0 | 0.00091 | 0.00082 | 3.23E-05 | 0.0003 | 0.00103 | 0.0004 | 0.00057 | 0.00126 | 0.00077 | 0.00126 | 0 | 0.00024 | 0.00032 | 0.00036 | 0.00061 | 0.00136 |
| Sh3gl1 | 0.00029 | 0.00014 | 0.00032 | 0.00069 | 0.00044 | 0.00025 | 0.00064 | 0.00042 | 0.00057 | 0.00033 | 0.00049 | 0.00039 | 0.00033 | 0.00047 | 0.00069 | 0.00015 | 0.00017 | 0.00025 |
| Pdpn | 0 | 0.00018 | 0.00056 | 0.00026 | 0.00078 | 0.00038 | 0.00057 | 0 | 0.00057 | 0.00105 | 7.78E-05 | 0.00048 | 0.00163 | 0.00061 | 0 | 4.20E-05 | 0.00027 | 0.00045 |
| Hs3st1 | 0 | 2.69E-05 | 9.86E-05 | 0.00013 | 0.00099 | 0.00038 | 0.00011 | 0.00084 | 0.00057 | 0 | 0 | 0 | 0 | 9.37E-05 | 0.00016 | 0 | 0 | 0 |
| Enc1 | 0 | 0.00027 | 0.00068 | 6.50E-06 | 0.00028 | 0.00011 | 0.00033 | 0 | 0.00056 | 0.00016 | 9.08E-05 | 0 | 0.0007 | 0.00068 | 0.00019 | 9.06E-06 | 0.00053 | 0.00017 |
| Gja1 | 0 | 0.00048 | 0.00042 | 0.00013 | 0.00074 | 0 | 0.00106 | 0.00072 | 0.00056 | 0.00085 | 0 | 0.00114 | 0.00043 | 0.00031 | 9.53E-05 | 0.00039 | 0 | 0 |
| Vim | 0.00072 | 0.00056 | 0.00079 | 0.00152 | 0.00053 | 0.00066 | 0.00139 | 0.00113 | 0.00056 | 0.00139 | 0.00119 | 0.00185 | 0.00047 | 0.00139 | 0.00189 | 0.00074 | 0.00207 | 0.00164 |
| Fat1 | 0.00027 | 0.00029 | 0.0002 | 0 | 0.00038 | 0.00021 | 0 | 0.0002 | 0.00056 | 9.63E-05 | 0.00065 | 6.53E-05 | 0.00021 | 0.00067 | 8.65E-05 | 0.00014 | 0.0005 | 0.00071 |
| Map4k4 | 0.00023 | 0.00059 | 0.00059 | 0.00014 | 5.95E-05 | 0.00017 | 8.87E-05 | 0.00088 | 0.00056 | 6.76E-06 | 5.16E-05 | 0.00028 | 0.00029 | 5.21E-05 | 4.17E-05 | 0.00032 | 0.00051 | 0.00021 |
| Pdgfrb | 0.0001 | 0.00042 | 0.0003 | 0.00032 | 0.00088 | 0.00111 | 0.00013 | 0.00179 | 0.00056 | 0 | 0.0003 | 0.00024 | 0.00044 | 0.00118 | 0.00115 | 0.00031 | 0.00042 | 0.00028 |
| Vcl | 0 | 0.0002 | 0.00033 | 0.00017 | 4.41E-06 | 0 | 0 | 0.00032 | 0.00055 | 0.00031 | 0.00196 | 0.00046 | 0 | 0.00042 | 0.00018 | 0.00014 | 0.00029 | 0.00055 |
| Flnc | 0 | 0.00016 | 0.00041 | 0.00028 | 0.0002 | 0 | 0.00044 | 0.00011 | 0.00055 | 0.00045 | 0.00053 | 0.00017 | 0 | 0 | 9.53E-05 | 0 | 0.00035 | 5.92E-06 |
| Npc2 | 0.00013 | 0.00035 | 0.00026 | 0.00063 | 0.00041 | 0.0004 | 0.00049 | 0.00078 | 0.00055 | 0.00084 | 0.0002 | 0.00057 | 0.00159 | 0.00079 | 0 | 0.00016 | 0.00018 | 0.00051 |
| Tln1 | 0.00038 | 0.00029 | 0.00038 | 0.00026 | 0.00028 | 0.00014 | 0.00012 | 0.00047 | 0.00055 | 0.00024 | 0.00093 | 0.00029 | 4.30E-05 | 0.00034 | 0.00049 | 0.00017 | 0 | 0.00025 |
| Fhl3 | 4.44E-05 | 0.00019 | 0.00025 | 0.00037 | 2.17E-05 | 1.85E-05 | 4.12E-05 | 3.67E-05 | 0.00055 | 0 | 0.00055 | 0.00017 | 1.88E-05 | 0 | 8.57E-05 | 0.00019 | 0.00017 | 0 |
| Rhoj | 0.00014 | 0.00013 | 0.00044 | 0.00016 | 0.00083 | 0.00054 | 0.00066 | 0.00093 | 0.00055 | 0.00094 | 0.00029 | 3.81E-05 | 0.00011 | 0.00052 | 0 | 9.27E-05 | 0.00019 | 0.00056 |
| DEPRECATED | 0 | 0 | 0 | 0 | 0.00025 | 0.00013 | 0 | 0 | 0.00055 | 8.28E-05 | 0 | 0 | 0 | 0.00029 | 0 | 0 | 0 | 0.00034 |
| Dap | 0.00044 | 7.58E-05 | 0.0002 | 0.00026 | 0.00037 | 0.0005 | 0.00087 | 0.00033 | 0.00055 | 0.00017 | 8.42E-05 | 0.00022 | 0 | 0.00019 | 6.00E-07 | 9.02E-05 | 0 | 0.00048 |
| Ctnnb1 | 0.00043 | 0.00058 | 0.0003 | 0.0005 | 0.00058 | 0.00046 | 0.00032 | 0.00033 | 0.00055 | 0.00012 | 0.00024 | 0.00046 | 0.00033 | 0.00039 | 0.00026 | 0.00033 | 0.00053 | 0.00026 |
| Tecr | 0.00023 | 0.00016 | 5.03E-05 | 0.00044 | 0.00024 | 9.16E-05 | 0.0001 | 0.00026 | 0.00054 | 0.00016 | 0.00021 | 9.08E-05 | 0.00017 | 0.0003 | 0 | 0.00019 | 0.00019 | 0.00026 |
| Cpxm2 | 0 | 0 | 0 | 0 | 0.00146 | 0.00101 | 0.00018 | 0.00015 | 0.00054 | 0 | 0.00123 | 0.00032 | 0 | 0.00015 | 0.00022 | 0 | 0 | 0.00031 |
| S100a4 | 3.42E-05 | 0 | 0.0002 | 0 | 0 | 0.0002 | 0 | 0 | 0.00054 | 0.00064 | 0 | 0.00037 | 0 | 0.00075 | 0 | 0.00031 | 0 | 0.0012 |
| Extl3 | 5.90E-05 | 0.00035 | 0.00021 | 0.00012 | 0.00038 | 7.41E-05 | 0.00016 | 0.00041 | 0.00054 | 0.0002 | 0.00023 | 0 | 0.00061 | 0.00042 | 9.11E-05 | 0.0002 | 1.90E-05 | 0.00033 |
| Arap1 | 0 | 0.00021 | 0.00011 | 0.00017 | 0.0003 | 0.00014 | 0.00019 | 0.00047 | 0.00054 | 0.00035 | 0.00015 | 0 | 0 | 0.00023 | 0.00027 | 2.39E-05 | 0.0002 | 0.00084 |
| Adgra2 | 0 | 0.00014 | 0.00042 | 0.00048 | 0.00031 | 0 | 0.00048 | 0.00013 | 0.00054 | 0.00054 | 0.0003 | 2.41E-05 | 0.00101 | 0.00037 | 0.00048 | 0.00019 | 1.25E-05 | 0.00011 |
| Cd81 | 0.00014 | 0.00051 | 0.00021 | 0.00031 | 0.00054 | 0.00071 | 0.00078 | 0.00091 | 0.00054 | 0.00073 | 0.00055 | 0.00087 | 0.0005 | 0.00019 | 0 | 0.00011 | 0.00013 | 0.00089 |
| Plpp3 | 0 | 0.00019 | 1.74E-05 | 0.00027 | 0 | 0.00055 | 0 | 0.00069 | 0.00054 | 0.00113 | 0 | 0 | 0.00126 | 0.00069 | 0.00019 | 0 | 0.00011 | 0.00103 |

| | | | | | | | | | | | | | | | | | | |
|---|---|---|---|---|---|---|---|---|---|---|---|---|---|---|---|---|---|---|
| Zmiz1 | 0.0001 | 0.00036 | 0.00024 | 0 | 0.00025 | 0.00032 | 0 | 0.00048 | 0.00054 | 0.00023 | 0.00026 | 0.00047 | 0 | 9.75E-05 | 9.50E-05 | 0.00011 | 0.00044 | 0.00042 |
| Osmr | 0.00031 | 0.0002 | 0.00023 | 0 | 0.00017 | 0.00076 | 0.00022 | 0.00032 | 0.00053 | 0.0002 | 0 | 0.00057 | 0.00073 | 0.00013 | 0.00048 | 3.86E-05 | 0 | 0.00013 |
| Cyp26b1 | 0 | 8.87E-06 | 0 | 0 | 0.00016 | 0.00093 | 0 | 0 | 0.00053 | 0 | 0 | 0.00045 | 0.00021 | 0.00189 | 0.00033 | 0 | 7.31E-05 | 2.13E-06 |
| Hmgcr | 8.54E-05 | 0 | 0.00011 | 0.00017 | 6.41E-05 | 0 | 0 | 0.00029 | 0.00053 | 2.89E-05 | 0.00038 | 6.51E-07 | 0 | 0.00016 | 0.00013 | 4.15E-05 | 0.00062 | 5.03E-05 |
| Cald1 | 0.00039 | 0.00034 | 0.00068 | 0.00045 | 0.00075 | 0.00037 | 0.00069 | 0.00096 | 0.00053 | 0.00027 | 0.00159 | 0.00054 | 0.0006 | 0.00048 | 0.00103 | 0.00025 | 0.00052 | 0.00075 |
| Slc16a2 | 0.00011 | 0.00015 | 0 | 0 | 0.00021 | 0.00022 | 0.00036 | 0 | 0.00053 | 0.00013 | 0.00012 | 9.73E-06 | 0.00099 | 0.00016 | 0 | 2.23E-05 | 0 | 2.68E-05 |
| Nectin2 | 0 | 2.55E-05 | 8.71E-06 | 0.00017 | 0.00026 | 0.00016 | 0 | 0 | 0.00053 | 3.02E-05 | 0.00076 | 0.00028 | 0.00112 | 0.00043 | 0.00016 | 0 | 9.18E-05 | 0 |
| Glul | 0.00011 | 8.60E-05 | 3.96E-05 | 0 | 9.07E-05 | 0.00028 | 0 | 0 | 0.00053 | 0.0003 | 0 | 0.00045 | 0.00028 | 0 | 0 | 1.90E-05 | 0.00068 | 2.32E-05 |
| Arpc1b | 0.00018 | 0.00025 | 0.0004 | 0.00094 | 0.00031 | 0.00038 | 0.00052 | 0.00092 | 0.00053 | 0.00055 | 0.0003 | 0.00021 | 0.00061 | 0.00046 | 0.0001 | 0.00015 | 0.00043 | 0.00049 |
| App | 0.00024 | 0.00029 | 0.00018 | 0.00024 | 0.00099 | 0.00098 | 0.00107 | 0.00129 | 0.00052 | 0.00082 | 0.00104 | 0.00064 | 0.00102 | 0.0007 | 0.00112 | 0.00021 | 0.00032 | 0.0009 |
| Adprh | 5.22E-05 | 0.00021 | 0.00021 | 0.00054 | 0.00053 | 0.00056 | 0.00034 | 0.00095 | 0.00052 | 0.0004 | 0.00041 | 0.00025 | 0.00021 | 0.00036 | 0.00014 | 0.00042 | 9.18E-05 | 0.0003 |
| Draxin | 0 | 0 | 0 | 5.83E-06 | 0.00013 | 0.00037 | 0 | 0 | 0.00052 | 0 | 0 | 0 | 0.00042 | 0.00049 | 8.56E-07 | 0 | 0 | 0 |
| Itm2b | 0.00031 | 0.00044 | 6.19E-05 | 0.00043 | 0.00097 | 0.0009 | 0.00062 | 0.00106 | 0.00052 | 0.00088 | 0.00053 | 0.00045 | 0.00135 | 0.00083 | 0.00065 | 0.00015 | 0.00029 | 0.00099 |
| Rbms1 | 0.00058 | 0.00031 | 0.00025 | 0 | 0.0004 | 0.00047 | 8.16E-05 | 0.00032 | 0.00052 | 0.00016 | 0.0002 | 0.0004 | 0.00016 | 0.00034 | 0.00027 | 0.00025 | 0.00045 | 0.00018 |
| Raly | 0.00044 | 0.00036 | 0.00022 | 0.00034 | 0.00029 | 0.00035 | 2.19E-05 | 0.00011 | 0.00052 | 0.00011 | 0.00026 | 0.00026 | 8.34E-05 | 0.00028 | 0.00022 | 0.00051 | 0.00062 | 6.45E-05 |
| Zeb2 | 0.00029 | 0.00037 | 0.0003 | 0.0002 | 0.00026 | 0.00049 | 0.00011 | 0.0006 | 0.00052 | 0.00028 | 0.0001 | 0.00017 | 0.0004 | 0.00022 | 0.00088 | 0.00018 | 3.58E-05 | 0.0003 |
| Uck2 | 0 | 0 | 0.00081 | 0.00039 | 0.00142 | 0.00046 | 0.00021 | 0.00093 | 0.00052 | 0 | 0.00106 | 0.00026 | 2.44E-05 | 0.00113 | 0.00077 | 0.00025 | 6.32E-05 | 0 |
| Vcp | 0.00031 | 0.00033 | 0.0006 | 0.00111 | 0.00021 | 0.00011 | 0.00036 | 0.00044 | 0.00052 | 0.0004 | 0.00041 | 0.00014 | 0.00025 | 0.00037 | 0.00046 | 0.0003 | 0.00059 | 0.00044 |
| Cxcl14 | 0 | 0 | 0.00069 | 0.00035 | 0.00107 | 0 | 0 | 0 | 0.00052 | 0 | 0.00174 | 1.28E-05 | 0.00436 | 0 | 0.00107 | 0 | 0 | 0.00242 |
| Akr1a1 | 0.00035 | 0.00031 | 0.00035 | 0.00123 | 0.00049 | 0.00046 | 0.0008 | 0.0007 | 0.00052 | 0.00049 | 0.00057 | 0.00041 | 0.00044 | 0.0006 | 0.00019 | 0.00025 | 0.00019 | 0.00062 |
| Plekho2 | 0 | 0.00037 | 0.00031 | 9.48E-05 | 0.00023 | 7.48E-05 | 0 | 0.00011 | 0.00052 | 0.00042 | 0.00033 | 0.0004 | 0.00024 | 0.00025 | 0.0005 | 5.12E-05 | 0.00017 | 0.00017 |
| Tmsb10 | 0.00055 | 5.09E-05 | 0.00019 | 0.00102 | 0.00054 | 0.00068 | 0.00087 | 0.00021 | 0.00051 | 0.00051 | 0.00137 | 0.00093 | 0.00057 | 0.00058 | 0.00065 | 0.00031 | 0.00064 | 0.00048 |
| Rassf8 | 5.98E-05 | 0.00029 | 0.00053 | 8.67E-05 | 0.00042 | 0.00096 | 0.00015 | 0.00038 | 0.00051 | 0.00032 | 0.00029 | 0.00036 | 3.34E-05 | 0.00053 | 0.0003 | 0.00011 | 0.00046 | 0.00036 |
| Serpinh1 | 0.00068 | 0.00015 | 0.00065 | 0.00177 | 0.00115 | 0.00139 | 0.00249 | 0.00169 | 0.00051 | 0.00119 | 0.00117 | 0.00079 | 0.0016 | 0.00147 | 0.00133 | 0.0005 | 0.00092 | 0.00113 |
| Nisch | 0.00024 | 0.00042 | 0.00018 | 0.00063 | 0.00053 | 0.00042 | 0.00056 | 0.00045 | 0.00051 | 0.00064 | 0.00045 | 0.00053 | 0.00041 | 0.0005 | 0.00133 | 0.00013 | 0.00046 | 0.00044 |
| Fdft1 | 2.08E-06 | 0.00016 | 0 | 0.00029 | 8.12E-05 | 0 | 0 | 6.94E-05 | 0.00051 | 0 | 0.0002 | 2.87E-05 | 0 | 9.86E-05 | 0 | 7.02E-05 | 0.00018 | 7.05E-05 |
| Tbc1d2b | 0.00019 | 0.00042 | 0 | 0 | 0.00061 | 0.00087 | 0.00046 | 0.00021 | 0.00051 | 0.00016 | 0 | 0 | 0.00046 | 0.00076 | 9.73E-05 | 0.00025 | 0 | 8.30E-05 |
| Fads1 | 0.00019 | 0.00017 | 1.45E-05 | 3.15E-06 | 0.00012 | 0 | 7.30E-05 | 0.00019 | 0.00051 | 4.35E-05 | 9.82E-05 | 0 | 0.00017 | 0.00012 | 0 | 0.00016 | 0 | 5.00E-05 |
| Anxa7 | 0.00035 | 0.00036 | 0.00019 | 2.27E-05 | 8.93E-05 | 0.00014 | 6.36E-05 | 8.43E-05 | 0.00051 | 0.00027 | 6.86E-05 | 0.00041 | 0 | 0 | 2.66E-05 | 4.28E-05 | 0.00014 | 0.00026 |
| Hic1 | 0.0001 | 0.00021 | 0.00024 | 0.00047 | 0.00051 | 3.52E-05 | 0.00027 | 0.00111 | 0.00051 | 0.00022 | 0.00021 | 5.29E-05 | 0.00018 | 0 | 0.00037 | 9.36E-05 | 0.00058 | 0.00021 |
| Steap2 | 8.32E-05 | 0 | 9.04E-05 | 4.57E-05 | 0.00012 | 0.00019 | 2.88E-05 | 6.32E-05 | 0.00051 | 8.78E-05 | 0.00016 | 0.00027 | 0.00051 | 0.00016 | 0 | 0.00023 | 5.21E-05 | 0 |
| Cd63 | 0.00035 | 0.00034 | 0.00053 | 0.0015 | 0.00108 | 0.0013 | 0.0014 | 0.00114 | 0.00051 | 0.00047 | 0.0007 | 0.00076 | 0.00163 | 0.00122 | 0.00057 | 0.0004 | 0.00041 | 0.00049 |
| Mt2 | 0 | 0 | 6.78E-06 | 0.00093 | 0 | 0.00122 | 0.00068 | 0.00017 | 0.0005 | 0.00111 | 0 | 0.00248 | 0.00407 | 0 | 0 | 0 | 0.00107 | 0 |

| | | | | | | | | | | | | | | | | | | |
|---|---|---|---|---|---|---|---|---|---|---|---|---|---|---|---|---|---|---|
| Ccdc80 | 0.00033 | 0.00014 | 0 | 0.00047 | 0.00059 | 0.00038 | 0.00099 | 0.00092 | 0.0005 | 0.00098 | 0.00111 | 0.00015 | 0.00103 | 0.00041 | 0.00041 | 0.00022 | 0.0002 | 0.00086 |
| Hspg2 | 0 | 0.00025 | 0.00021 | 0.00056 | 0.00091 | 0.00098 | 0.00095 | 0.0011 | 0.0005 | 0.00106 | 0.00097 | 0.00064 | 0.00116 | 0.00047 | 0.00086 | 0.00011 | 8.40E-05 | 0.00081 |
| Pear1 | 0 | 6.03E-05 | 0 | 0 | 0.0001 | 0.0002 | 0 | 0.00106 | 0.0005 | 0 | 0 | 0 | 0.00015 | 0.00038 | 0.00027 | 0 | 0 | 0.00038 |
| Cthrc1 | 0.00031 | 0 | 0.00046 | 0.00025 | 0 | 0 | 0.0032 | 0.00063 | 0.0005 | 0.0004 | 0 | 0.00042 | 5.35E-05 | 0 | 0 | 0.00037 | 0 | 0 |
| Mrc2 | 6.84E-05 | 1.55E-05 | 0.00024 | 0 | 0.0012 | 0.00084 | 0.00158 | 0.00145 | 0.0005 | 0.00059 | 0.00074 | 0.00032 | 0.00125 | 0.00099 | 0.00084 | 0.00018 | 0 | 0.00077 |
| Lima1 | 0.00033 | 0.00016 | 0.00011 | 0 | 0.00032 | 0.00023 | 0.00029 | 0 | 0.0005 | 0.00037 | 0.00037 | 0.00017 | 0.00025 | 0.00017 | 0.00016 | 6.00E-05 | 0.0002 | 0.00019 |
| Pmepa1 | 0 | 0.00039 | 0.00057 | 0.00052 | 0.00181 | 0.00014 | 0.00126 | 0.00127 | 0.0005 | 0.0002 | 0.0006 | 0.00035 | 0.00024 | 0.00101 | 0.00082 | 3.91E-05 | 0.00046 | 0.0008 |
| Farp1 | 8.71E-05 | 0.00014 | 0.00016 | 1.46E-05 | 0.00025 | 0.00015 | 4.59E-05 | 0.00031 | 0.0005 | 0.00023 | 0.00036 | 0.00028 | 0 | 0.00028 | 0.00012 | 0.00017 | 0.00019 | 0.00016 |
| Wdr1 | 0.00031 | 0.00023 | 0.00054 | 0.00023 | 9.65E-05 | 0 | 0.00031 | 0.00016 | 0.0005 | 0.00025 | 0.00105 | 0.00025 | 4.70E-05 | 0.00012 | 6.67E-05 | 0.00028 | 0.0001 | 0.00027 |
| Tead1 | 8.98E-05 | 0.00015 | 0.00034 | 1.29E-05 | 0.00017 | 0.0003 | 0 | 0.00031 | 0.0005 | 9.77E-05 | 0.00077 | 1.98E-05 | 0.0001 | 0.00027 | 0.00031 | 0.00016 | 0.00043 | 0.00019 |
| Pfn1 | 0.00038 | 0.0002 | 0.00043 | 0.00116 | 0.00019 | 0.0001 | 0.0003 | 0.00044 | 0.0005 | 0.00028 | 0.00077 | 0.00031 | 0.00023 | 0.00048 | 0.00046 | 0.00033 | 0.00023 | 0.00027 |
| Ywhab | 0.00037 | 0.00034 | 0.00053 | 0.00042 | 0.00029 | 0.00032 | 0.00026 | 0.0004 | 0.00049 | 0.00032 | 0.00035 | 0.00014 | 0.00029 | 0.00023 | 3.83E-05 | 0.00021 | 1.81E-06 | 0.00024 |
| Flot1 | 0.0002 | 0.00019 | 8.06E-05 | 0.00021 | 0.00023 | 0.00012 | 0.00021 | 3.63E-05 | 0.00049 | 0.00028 | 0.00024 | 0 | 0.00034 | 0.00019 | 0.00034 | 0 | 0 | 0.00025 |
| Cdon | 0 | 9.49E-05 | 0 | 4.07E-05 | 0.00031 | 0 | 0.00066 | 0 | 0.00049 | 0.00032 | 7.96E-05 | 0 | 0.00018 | 0 | 7.24E-05 | 0.00018 | 6.03E-05 | 0.00045 |
| Palm | 3.57E-05 | 0.00026 | 2.72E-05 | 0.00023 | 0.00023 | 4.75E-05 | 1.36E-05 | 0.0002 | 0.00049 | 0.00051 | 0.00017 | 0.00016 | 0 | 0.00011 | 0.00034 | 0.00011 | 5.18E-05 | 0.00042 |
| Cst3 | 0.00013 | 0.00032 | 0.00022 | 0.00081 | 0.0006 | 0.00092 | 0.00058 | 0.00061 | 0.00049 | 0.00055 | 0.0007 | 0.00053 | 0.00132 | 0.00127 | 0 | 0.00031 | 0.0001 | 0.00136 |
| Pdlim2 | 0 | 0.00015 | 0 | 0.00019 | 0.00042 | 0.00017 | 0 | 8.45E-05 | 0.00049 | 2.59E-05 | 0.0003 | 0 | 0 | 0.00048 | 0 | 0 | 0 | 0.00039 |
| Tspan11 | 0 | 0.00027 | 3.11E-05 | 0.00015 | 0.00036 | 0.00049 | 7.16E-05 | 0 | 0.00049 | 6.47E-05 | 0 | 0 | 0.00058 | 0.00054 | 0.00012 | 2.09E-05 | 0 | 0.00096 |
| Ahr | 0 | 3.41E-05 | 0 | 0 | 0 | 0.00011 | 7.62E-05 | 0 | 0.00049 | 7.25E-05 | 0 | 0 | 0.00037 | 0 | 0 | 5.36E-05 | 0 | 0.00055 |
| Ahnak | 8.31E-05 | 0.00043 | 0.00029 | 0.00039 | 0.00018 | 0.00042 | 0.00035 | 9.38E-05 | 0.00048 | 0.00125 | 0.00082 | 0.00076 | 0 | 0.00039 | 0.00055 | 7.11E-05 | 0.00129 | 0.00136 |
| Psap | 9.48E-05 | 0.00051 | 7.66E-05 | 0 | 0.00069 | 0.00093 | 0.0007 | 0.00054 | 0.00048 | 0.00061 | 0.00031 | 0.00045 | 0.00097 | 0.00088 | 0 | 7.26E-05 | 0 | 0.0006 |
| Map1lc3a | 0.00018 | 0.0003 | 0 | 0.00017 | 0.00051 | 0.00055 | 0.00015 | 0.00017 | 0.00048 | 0.00017 | 0.00062 | 0.00025 | 0.00074 | 0.00041 | 0 | 0 | 0 | 0.00029 |
| Atp2a2 | 0.00025 | 0.00045 | 0.00049 | 0.00021 | 0.00049 | 0.00032 | 0.00056 | 0.00028 | 0.00048 | 0.00041 | 0.00063 | 0.00014 | 0.00075 | 0.00032 | 0.00059 | 0.00025 | 0.00058 | 0.00028 |
| Synpo | 7.23E-05 | 9.78E-05 | 0.00027 | 1.38E-05 | 0.00055 | 0.00011 | 0.00054 | 0.00035 | 0.00048 | 1.38E-05 | 0.00022 | 0.00039 | 0.00053 | 0.00047 | 0.00022 | 0.00014 | 7.09E-05 | 0.00067 |
| Rtn4 | 0.0005 | 0.00037 | 0.00054 | 0.00014 | 0.00025 | 0.0003 | 0.00023 | 0.00029 | 0.00048 | 0.0005 | 0.00083 | 0.00076 | 0.00057 | 0.00057 | 0 | 0.00021 | 0.00018 | 0.0003 |
| Cttn | 0.00023 | 0.00039 | 0.00045 | 0.00036 | 0.0004 | 0.00024 | 0.00055 | 0.00037 | 0.00048 | 0.0003 | 0.00053 | 0.0004 | 0.00044 | 0.00033 | 0.0004 | 0.00015 | 0.00017 | 0.00041 |
| Fscn1 | 0.00015 | 5.67E-05 | 0.00066 | 0.00067 | 0.00041 | 0 | 0.00033 | 0.00039 | 0.00048 | 0.00018 | 0.0007 | 0.00028 | 0.00031 | 0.00013 | 0.00022 | 6.36E-05 | 0.00036 | 0.00024 |
| Vmp1 | 0.00028 | 0.0005 | 0.00031 | 0.00035 | 0.00059 | 0.00018 | 0.00015 | 0.00056 | 0.00048 | 7.26E-05 | 0.00028 | 0.00017 | 0.00028 | 0.00051 | 0.00183 | 0.00015 | 9.28E-05 | 0.0003 |
| Dclk1 | 2.44E-05 | 8.98E-05 | 0.00047 | 0.00023 | 0.00013 | 0 | 0.00076 | 8.90E-05 | 0.00048 | 0.00012 | 0 | 0.00011 | 0.00046 | 0.0007 | 0.0001 | 0.00012 | 6.06E-05 | 0.00066 |
| Actn4 | 5.78E-05 | 0.00029 | 0.00023 | 0.00071 | 0.00027 | 0 | 0 | 0.00031 | 0.00048 | 0.00035 | 0.00162 | 0.00025 | 0.00027 | 0.00043 | 0.0006 | 0.00059 | 0.00033 | 0.00059 |
| Abhd2 | 0 | 0.0002 | 0.00025 | 3.22E-05 | 0.00066 | 0.00058 | 0.00081 | 0.00162 | 0.00048 | 0.00032 | 0 | 0.00035 | 0 | 0.00052 | 0.00018 | 0.00018 | 0 | 9.97E-06 |
| Nudt4 | 0.00021 | 0.00017 | 0.00015 | 0.0003 | 0.0007 | 0.0002 | 0.0001 | 0.00096 | 0.00048 | 0 | 0.00041 | 5.08E-05 | 0.00013 | 0.00026 | 0.00031 | 0.00027 | 0.00043 | 0.00027 |
| Tnfrsf1a | 0.00038 | 0.00046 | 0.00014 | 0.00069 | 0.00057 | 0.00066 | 0.00045 | 0.0006 | 0.00048 | 0.00064 | 0.00014 | 0.00029 | 0.00086 | 0.00029 | 0.00023 | 0.00038 | 0.00021 | 0.00037 |

| | | | | | | | | | | | | | | | | | | |
|---|---|---|---|---|---|---|---|---|---|---|---|---|---|---|---|---|---|---|
| Uba1 | 0.00037 | 0.00035 | 0.00051 | 0.00065 | 0.00026 | 0.00018 | 0.00028 | 0.00041 | 0.00048 | 0.00039 | 0.00037 | 0.00021 | 0.00019 | 0.00037 | 0.00025 | 0.0002 | 0.0002 | 0.00039 |
| Nedd4 | 0.00043 | 0.00045 | 5.76E-05 | 0.0007 | 0.0004 | 0.00043 | 0.0006 | 0.0003 | 0.00048 | 0.0004 | 0.00052 | 0.00031 | 0.00052 | 0.00036 | 6.39E-05 | 0.00028 | 0.00036 | 0.0004 |
| Litaf | 0.00016 | 0.00045 | 0.00021 | 0.00027 | 0.00041 | 0.00059 | 0 | 0.00057 | 0.00048 | 0.00034 | 0.00027 | 0.00038 | 0.00061 | 0.00045 | 5.27E-05 | 0.0002 | 0.00031 | 0.0002 |
| Twist2 | 0 | 8.94E-05 | 0.00052 | 0 | 0.00065 | 0.00109 | 9.21E-05 | 0 | 0.00048 | 0.00011 | 0 | 0.00027 | 0 | 0.00111 | 0.00055 | 2.25E-05 | 0.00035 | 0 |
| Acly | 0.00025 | 0.00025 | 0.00013 | 0.00029 | 0.00013 | 0 | 0 | 7.94E-05 | 0.00048 | 0.00038 | 0.00025 | 0.00015 | 0.00015 | 6.88E-05 | 4.95E-05 | 7.02E-06 | 0.00084 | 4.36E-05 |
| Itgb1 | 0.00026 | 0.00028 | 0.00059 | 0.00044 | 0.00081 | 0.00061 | 0.00096 | 0.00085 | 0.00048 | 0.00056 | 0.00144 | 0.00076 | 0.00058 | 0.00084 | 0.00031 | 0.00029 | 0.0002 | 0.00049 |
| Chsy1 | 0.00015 | 5.33E-05 | 0.00023 | 0.00015 | 0.00031 | 0.00012 | 0.00064 | 0.00043 | 0.00048 | 0 | 0.00024 | 5.98E-06 | 0.00041 | 0.00033 | 0.0001 | 0.00015 | 0.00013 | 0.00015 |
| Igfbp7 | 0.00011 | 0.00033 | 0.0002 | 0.0008 | 0.00137 | 0.00121 | 0.00099 | 0.00219 | 0.00047 | 0.00103 | 0.00162 | 0.00061 | 0.00164 | 0.00212 | 0.00171 | 0.00028 | 0.00028 | 0.00262 |
| Zfp36l2 | 1.92E-05 | 0.00045 | 3.40E-05 | 0.00059 | 0 | 0.00039 | 0 | 0.0001 | 0.00047 | 0.00027 | 0 | 6.13E-05 | 0.00061 | 0.00023 | 0.00032 | 0.00031 | 0.00105 | 0.00068 |
| Bcat1 | 0.00019 | 0 | 0.00082 | 0.0006 | 0.0001 | 8.26E-07 | 0.00095 | 0.00014 | 0.00047 | 0 | 0 | 0.00027 | 0.00013 | 1.45E-05 | 0.00022 | 0.00028 | 0 | 0 |
| Pdia4 | 0.00054 | 5.08E-05 | 0.00029 | 0.00069 | 0.00023 | 0.00032 | 0.00061 | 0.00077 | 0.00047 | 0.00035 | 0.00031 | 0.00032 | 0.00066 | 0.00036 | 0 | 0.00025 | 6.82E-05 | 0.0004 |
| Ywhag | 0.00016 | 0.00042 | 0.00069 | 0.00085 | 0.00019 | 5.78E-05 | 3.94E-05 | 0.00041 | 0.00047 | 0.00042 | 0.00046 | 0.00068 | 6.39E-05 | 0.00011 | 0.00075 | 0.00029 | 0.00048 | 0.00032 |
| Nkd2 | 0 | 0.00027 | 0 | 0 | 0.00074 | 0.00011 | 0 | 0.00011 | 0.00047 | 0 | 0.00019 | 0 | 0 | 0.00037 | 0.00026 | 0 | 0 | 5.96E-05 |
| Postn | 0.00066 | 2.96E-05 | 0.00024 | 0 | 0.00157 | 0.00189 | 0.00297 | 0.00225 | 0.00047 | 0 | 0.00133 | 0.00025 | 0 | 0.00163 | 0.00091 | 0.00034 | 0 | 0.00098 |
| Adamts2 | 0 | 0 | 0 | 0.00065 | 0.00116 | 0.00198 | 0.00101 | 0.0009 | 0.00047 | 0.00111 | 0.00104 | 0.00035 | 0.00132 | 0.00166 | 0.00159 | 0.00014 | 9.37E-05 | 0.00083 |
| Gpx1 | 0.00043 | 0.00046 | 0.00023 | 0.00092 | 0 | 6.19E-05 | 0.00055 | 6.59E-05 | 0.00047 | 0.00035 | 0.00051 | 0 | 0.00011 | 0.00022 | 7.39E-05 | 0.0002 | 0.00056 | 0.00059 |
| Colec12 | 0.00032 | 0.00028 | 2.47E-05 | 0 | 0.00045 | 0.001 | 0.00032 | 0.00052 | 0.00047 | 4.29E-06 | 0.0007 | 0.00075 | 0.00074 | 0.00071 | 0 | 0.00012 | 6.02E-05 | 6.32E-05 |
| Ptgfrn | 0 | 0.00011 | 0.00051 | 0.00044 | 0.00089 | 0.00075 | 0.00107 | 0.00067 | 0.00047 | 0.00046 | 0.00029 | 0 | 0.00073 | 0.00104 | 0.00032 | 0.00016 | 0 | 0.00091 |
| Id2 | 5.52E-05 | 9.87E-05 | 0 | 0 | 0.00031 | 0.00033 | 0 | 0.00016 | 0.00046 | 0.00021 | 0.00011 | 0.00049 | 7.24E-05 | 0 | 4.14E-06 | 9.77E-05 | 0.00012 | 0.00022 |
| Arf6 | 0.00019 | 0.00021 | 0.00045 | 9.26E-05 | 0.00033 | 0.00013 | 0.00024 | 0.00032 | 0.00046 | 0.0002 | 0.00023 | 0.00046 | 0 | 6.16E-05 | 0.00018 | 0.00041 | 0.0003 | 0.00014 |
| Socs3 | 0.00014 | 0.00019 | 0.0001 | 0.00034 | 0.0002 | 0.00013 | 9.14E-05 | 7.39E-05 | 0.00046 | 0.00031 | 0 | 2.50E-05 | 0.00103 | 0.00044 | 0.00021 | 0.00013 | 0.0008 | 0 |
| Mmp3 | 0 | 0 | 0.00014 | 0 | 0 | 0 | 0.00077 | 0.00103 | 0.00046 | 0.00031 | 0 | 0 | 0.00367 | 0.00118 | 0 | 0.00016 | 0 | 0.00033 |
| Sh3bgrl3 | 0.00033 | 9.49E-05 | 0.00047 | 0.0005 | 0.00019 | 0.00038 | 0.00052 | 0.00076 | 0.00046 | 0.00023 | 0.00016 | 0.00033 | 0 | 0 | 0 | 0.0002 | 6.01E-05 | 0.0004 |
| Itga1 | 8.37E-06 | 9.52E-05 | 0 | 0 | 0.00017 | 0.00072 | 0 | 0.00067 | 0.00046 | 0 | 0.00012 | 0.00027 | 3.33E-05 | 0.00038 | 0 | 0 | 0 | 0.00032 |
| Adcy3 | 0 | 9.12E-05 | 3.42E-05 | 0.00011 | 0.00042 | 0.0004 | 7.64E-05 | 0.00031 | 0.00046 | 0.00015 | 3.59E-05 | 0.00014 | 0.00014 | 0.00036 | 0 | 4.00E-05 | 5.71E-05 | 0 |
| Gng12 | 0.00051 | 0.00019 | 0.00013 | 0 | 0.00025 | 0.00053 | 0 | 9.33E-05 | 0.00046 | 0.00048 | 0.00026 | 0.00022 | 0.00063 | 0.00017 | 3.39E-05 | 0.00013 | 0.00024 | 0.00012 |
| Vstm4 | 0 | 0.00018 | 0 | 9.41E-05 | 0.00011 | 0.00024 | 1.41E-05 | 0.00026 | 0.00046 | 1.86E-05 | 0 | 0 | 0.0001 | 2.48E-05 | 0 | 0 | 0 | 0.0003 |
| Xbp1 | 0.00051 | 0.00018 | 0.00038 | 0.00047 | 0.00021 | 0.00054 | 0.00044 | 0.00056 | 0.00046 | 0.00035 | 0.00025 | 0.00041 | 0.00046 | 0.00029 | 0.00073 | 4.59E-05 | 0 | 0.00041 |
| Ociad2 | 9.51E-06 | 0.00011 | 0 | 0 | 0.00059 | 0.00063 | 0 | 0.00015 | 0.00046 | 0 | 0 | 0.00014 | 0 | 0 | 0 | 0 | 0 | 0 |
| F2r | 0 | 7.54E-05 | 0.00031 | 0.00011 | 0.00039 | 0 | 0.00016 | 1.02E-05 | 0.00046 | 0 | 0.0002 | 0.00014 | 0 | 0.00137 | 0 | 0.00016 | 0.00015 | 0.00093 |
| Adgrl2 | 8.93E-05 | 0.00013 | 0 | 0 | 4.41E-05 | 1.52E-05 | 7.38E-05 | 7.90E-06 | 0.00046 | 0 | 9.64E-05 | 0 | 0 | 0 | 0 | 3.84E-05 | 0.00024 | 8.11E-05 |
| Rack1 | 0.00041 | 0.00045 | 0.00054 | 0.0011 | 0.00043 | 0.00066 | 0.00066 | 0.00064 | 0.00045 | 0.00036 | 0.00026 | 0.0007 | 0.0005 | 0.00059 | 0.0002 | 0.00028 | 0.0006 | 0.00029 |
| Cped1 | 0 | 0.00037 | 2.27E-05 | 4.32E-05 | 0.00042 | 0.00027 | 0.00022 | 0.00024 | 0.00045 | 0.00025 | 0 | 0 | 0.00037 | 0.00036 | 0.00037 | 2.71E-05 | 4.50E-05 | 0.00033 |

| | | | | | | | | | | | | | | | | | | |
|---|---|---|---|---|---|---|---|---|---|---|---|---|---|---|---|---|---|---|
| Bmp1 | 0 | 0.00015 | 0.00041 | 0.0006 | 0.00102 | 0.00101 | 0.00185 | 0.00162 | 0.00045 | 0.00105 | 0.00082 | 0.00034 | 0.00194 | 0.00158 | 0.00145 | 0.00021 | 0.00026 | 0.00034 |
| Aldh7a1 | 0 | 0.00011 | 0.00013 | 0.00054 | 0.0003 | 7.30E-05 | 0.00033 | 0.00044 | 0.00045 | 0.00035 | 4.98E-05 | 3.00E-05 | 0.00024 | 0.0002 | 0 | 8.32E-06 | 0.00026 | 0.00014 |
| Gnas | 0.00029 | 0.00033 | 0.00023 | 0.00135 | 0.00049 | 0.00048 | 0.00086 | 0.00058 | 0.00045 | 0.00063 | 0.00058 | 0.00045 | 0.00044 | 0.00075 | 0.00031 | 0.00033 | 0.00069 | 0.00073 |
| Fam20a | 9.44E-05 | 0.00015 | 0 | 0.00011 | 0.00051 | 0.00026 | 0.00019 | 0.00046 | 0.00045 | 0.00023 | 0.00039 | 4.70E-05 | 0 | 0.00081 | 0.00015 | 0 | 0 | 0.00014 |
| Rflna | 0 | 2.37E-05 | 0 | 0 | 0.00074 | 0 | 0 | 0.00054 | 0.00045 | 0 | 0 | 0 | 0 | 0.00011 | 0 | 2.80E-05 | 0 | 0 |
| C1s1 | 0 | 0.00033 | 0 | 0 | 0.00058 | 0.00081 | 0.00032 | 0.00027 | 0.00045 | 0.00124 | 0.00022 | 0.00046 | 0.00188 | 0.0003 | 0.00013 | 0 | 0.0002 | 0.00135 |
| Lxn | 0.00023 | 0 | 2.89E-05 | 0 | 0 | 6.63E-05 | 1.15E-05 | 0 | 0.00045 | 0 | 3.60E-05 | 9.81E-05 | 0 | 0.00019 | 0 | 6.25E-06 | 1.37E-05 | 0.00014 |
| Lamb1 | 4.53E-05 | 0.00016 | 6.35E-05 | 0.00027 | 0.00063 | 0.00076 | 0.0006 | 0.00132 | 0.00045 | 0.00056 | 0.00032 | 0.0003 | 0.00086 | 0.00096 | 0.00047 | 0.00014 | 0.00029 | 0 |
| Mid1ip1 | 0 | 0.00016 | 7.53E-05 | 0.00018 | 0.00012 | 0.00011 | 0 | 9.00E-06 | 0.00044 | 0 | 0.00044 | 0 | 0.00013 | 0.00025 | 0.00018 | 6.24E-05 | 0 | 0.00029 |
| Lamc1 | 0 | 0.0002 | 0.00054 | 9.54E-05 | 0.00058 | 0.00113 | 0.0008 | 0.00118 | 0.00044 | 0.00127 | 0.00028 | 0.00026 | 0.00059 | 0.00078 | 0.00064 | 0.00012 | 0.00036 | 0.00062 |
| Kremen1 | 3.29E-05 | 0.00012 | 0.00028 | 0 | 0.00073 | 0.00079 | 0.00036 | 0.00116 | 0.00044 | 0.00015 | 0.00052 | 0.00087 | 0 | 0.00085 | 0 | 8.33E-05 | 0.00015 | 0.00016 |
| Slc25a51 | 0.00022 | 0.0003 | 0.00012 | 9.60E-05 | 0.00045 | 0.00015 | 0.0003 | 0.00036 | 0.00044 | 9.99E-05 | 0.00012 | 0.00034 | 0.00012 | 2.96E-05 | 8.40E-05 | 0.00018 | 0 | 7.58E-05 |
| Adra2a | 0 | 1.94E-05 | 0 | 0 | 0.00038 | 8.05E-05 | 0 | 0.00134 | 0.00044 | 0 | 0 | 0 | 0 | 0.00034 | 0 | 0 | 0 | 0 |
| Tkt | 0.00033 | 0.00017 | 0.00025 | 0.00033 | 6.42E-06 | 9.47E-05 | 0 | 0.00019 | 0.00044 | 0.00011 | 0.00019 | 0.00031 | 3.58E-05 | 0.0002 | 4.81E-06 | 0.00024 | 0.00011 | 0.00015 |
| Adam9 | 0.00036 | 0.00018 | 0.00018 | 0.00013 | 0.00044 | 0.00054 | 0.00035 | 0.00048 | 0.00044 | 0.00058 | 0.0004 | 0.00048 | 0.00106 | 0.00052 | 0.00014 | 0.00016 | 6.92E-05 | 0.00029 |
| Myl12a | 0.00032 | 0.00012 | 0.00038 | 0.00078 | 0.00042 | 0.00018 | 0.00031 | 0.00056 | 0.00044 | 0.00023 | 0.00137 | 0.00048 | 0.00011 | 0.00059 | 6.61E-06 | 0.00022 | 0.00035 | 0.00053 |
| Pik3r1 | 0.00016 | 0.00017 | 0 | 5.09E-05 | 0.00027 | 0.00032 | 0 | 0.00069 | 0.00044 | 0.00014 | 0 | 0 | 0.00019 | 0.00031 | 4.28E-05 | 3.15E-05 | 0 | 0.0002 |
| Sod2 | 3.45E-05 | 0.00012 | 0.00055 | 0 | 0.00018 | 0.00081 | 0 | 2.77E-05 | 0.00044 | 2.25E-05 | 0 | 1.34E-06 | 0.0018 | 0.00094 | 0 | 0 | 0 | 0 |
| Sdk1 | 0 | 2.57E-05 | 1.29E-05 | 0 | 0.00018 | 9.21E-05 | 0.00019 | 0 | 0.00044 | 0.00033 | 0 | 8.04E-05 | 0.00039 | 6.00E-05 | 0 | 6.81E-05 | 0 | 0 |
| Pcsk9 | 0 | 0 | 0 | 5.88E-05 | 4.63E-05 | 0 | 8.47E-05 | 0 | 0.00044 | 0 | 0.00018 | 0 | 5.68E-05 | 0.00032 | 8.62E-05 | 2.38E-05 | 0 | 0 |
| Ap2m1 | 0.00035 | 0.0003 | 0.00042 | 0.00083 | 0.00026 | 0.00036 | 0.00039 | 0.00045 | 0.00043 | 0.00029 | 0.00028 | 0.00024 | 0.00028 | 6.76E-06 | 0 | 0.00033 | 8.66E-05 | 0.00036 |
| Smim3 | 0 | 0 | 0.00032 | 1.39E-06 | 0.00035 | 0 | 0.00032 | 0.00036 | 0.00043 | 0 | 8.84E-05 | 0.00013 | 0.00026 | 0.00019 | 0 | 5.53E-05 | 0 | 2.59E-05 |
| Selenow | 0.00023 | 0.00026 | 0.00015 | 0.00105 | 0.00053 | 0.00049 | 0.00069 | 0.0003 | 0.00043 | 0.00066 | 0.00047 | 0.00042 | 0.00086 | 0.00052 | 9.04E-05 | 9.17E-05 | 0.00017 | 0.00051 |
| Gda | 0 | 0.00011 | 0 | 0 | 0 | 0.00025 | 0 | 0.00011 | 0.00043 | 0.00086 | 0 | 1.09E-05 | 0.00074 | 0 | 0 | 0 | 0 | 0.00025 |
| Fau | 0.00043 | 0.00053 | 0.00034 | 0.00076 | 0.00038 | 0.00061 | 0.00049 | 0.00059 | 0.00043 | 0.00035 | 0.00029 | 0.00116 | 0.00042 | 0.0005 | 0.00037 | 0.00025 | 0.00065 | 0.00031 |
| Ckap4 | 0.00056 | 0 | 0.00076 | 0.00099 | 0.00059 | 0.0006 | 0.00149 | 0.00063 | 0.00043 | 0.00037 | 0.0008 | 0.00032 | 0.00093 | 0.00068 | 0.00083 | 0.0009 | 0.00036 | 0.00027 |
| Irf2bp2 | 1.89E-05 | 0.00059 | 0.00061 | 0.00046 | 0.00064 | 0.00036 | 0.00015 | 0.0004 | 0.00043 | 0.0001 | 0.00027 | 0.00048 | 0.00012 | 0.0002 | 0.00112 | 0.00015 | 0.00064 | 0.00016 |
| Tspan3 | 0.00011 | 0.00017 | 0.00025 | 0.00047 | 0.00078 | 0.00055 | 0.00074 | 0.00075 | 0.00043 | 0.00047 | 0.00085 | 0.00027 | 0.00065 | 0.00061 | 0 | 0.00032 | 3.39E-05 | 0.00042 |
| Akt1 | 0.00036 | 0.00039 | 0.00053 | 0.00056 | 0.00038 | 0.00022 | 0.00035 | 0.00035 | 0.00043 | 0.0003 | 0.00051 | 0.00013 | 0.00011 | 0.00035 | 0.00061 | 0.00035 | 0.00025 | 0.00033 |
| Cav1 | 0.00013 | 0 | 0.00058 | 3.18E-05 | 0 | 0.00061 | 0.0002 | 0 | 0.00043 | 0.00043 | 0 | 0.00063 | 0 | 0.00019 | 0 | 0.00028 | 0 | 0.00035 |
| Tpt1 | 0.00098 | 0.00059 | 0.00031 | 0.00048 | 0.00079 | 0.00142 | 0.00113 | 0.00094 | 0.00043 | 0.00062 | 0.00047 | 0.00107 | 0.00071 | 0.00071 | 1.17E-05 | 0.00025 | 0.00118 | 0.00044 |
| Mcoln2 | 0 | 0 | 6.74E-05 | 0 | 0.00021 | 0.00022 | 0 | 0.00051 | 0.00043 | 0.00011 | 0 | 0 | 0.0004 | 0.00025 | 8.23E-05 | 1.90E-05 | 0 | 0 |
| Rab20 | 0 | 3.53E-05 | 9.19E-05 | 0 | 0.0002 | 7.89E-05 | 0 | 0.00069 | 0.00043 | 0 | 0 | 0 | 0.00022 | 0.00032 | 1.95E-05 | 4.73E-05 | 0 | 3.83E-05 |

| | | | | | | | | | | | | | | | | | | |
|---|---|---|---|---|---|---|---|---|---|---|---|---|---|---|---|---|---|---|
| Calr | 0.00061 | 5.55E-05 | 0.00051 | 0.0007 | 0.00043 | 0.00044 | 0.00064 | 0.00059 | 0.00043 | 0.00045 | 0.00048 | 0.00035 | 0.00089 | 0.00051 | 0 | 0.00012 | 0.00023 | 0.00051 |
| Myadm | 6.05E-05 | 6.00E-05 | 0.00039 | 0.0007 | 0.00017 | 0.00046 | 0.00032 | 0.00053 | 0.00043 | 0.00049 | 0.00152 | 0.00092 | 0 | 0.00032 | 0 | 0.00032 | 0.0003 | 0.00082 |
| Arf5 | 0.0003 | 0.00017 | 0.00031 | 0.00096 | 0.00049 | 0.00044 | 0.00073 | 0.00061 | 0.00043 | 0.00035 | 0.00039 | 0.00019 | 0.00029 | 0.00044 | 0.00029 | 0.00018 | 0.00015 | 0.00044 |
| Lpp | 0.00012 | 0.00022 | 0.00017 | 0.0003 | 0.00024 | 0.00027 | 0.00018 | 0.0004 | 0.00042 | 0.00032 | 0.00149 | 0.00036 | 0.00027 | 0.00049 | 0.00026 | 0.00011 | 0.0005 | 0.00032 |
| Eid1 | 0.00016 | 0.00037 | 0 | 0.00125 | 0.0007 | 0.00066 | 0.00063 | 0.00028 | 0.00042 | 0.00021 | 0.00032 | 0.00044 | 0.00058 | 0.0007 | 0.00019 | 0 | 0.00025 | 0.00035 |
| Pcolce | 0.00028 | 0 | 9.24E-05 | 0.00127 | 0.00062 | 0.0011 | 0.00174 | 0.0011 | 0.00042 | 0.00102 | 0.00079 | 0.00044 | 0.00177 | 0.00105 | 0.00087 | 0.00035 | 0.00031 | 0.00127 |
| Stat3 | 0.00034 | 0.00036 | 0.0003 | 0.00014 | 0.00021 | 0.00035 | 0.00026 | 0.00018 | 0.00042 | 0.0002 | 5.12E-05 | 0.00056 | 0.00053 | 0.00033 | 0.00041 | 2.40E-05 | 0.00031 | 0.00025 |
| Uqcrc1 | 0.00038 | 0.00024 | 0.00044 | 0.00061 | 0.0003 | 0.00024 | 0.0002 | 0.00018 | 0.00042 | 0.0002 | 0.00046 | 3.59E-05 | 2.86E-05 | 0.00058 | 0.00045 | 0.00027 | 0.00016 | 0.00017 |
| Man2a2 | 0 | 0.00021 | 0 | 0.0002 | 0.0004 | 0.00055 | 0.00015 | 0.00041 | 0.00042 | 0.00018 | 0.00032 | 0 | 0.00024 | 0.00055 | 0.00012 | 8.45E-05 | 0.00011 | 0.00027 |
| S100a10 | 0.00025 | 2.27E-05 | 0.00033 | 5.32E-05 | 0 | 0.0002 | 0.00077 | 0 | 0.00042 | 0.00125 | 0.00073 | 0.00056 | 0 | 0.00042 | 0 | 0.00026 | 0.00035 | 0.00108 |
| Tmem173 | 0.0003 | 0 | 0.00025 | 0.00012 | 0.00012 | 0.00013 | 0.00018 | 0.0005 | 0.00042 | 7.83E-05 | 3.03E-05 | 0 | 0.00027 | 0.00019 | 0 | 4.32E-05 | 0 | 0.00016 |
| Pdgfra | 0.00034 | 0.00035 | 0 | 0.00041 | 0.00117 | 0.00083 | 0.00053 | 0 | 0.00042 | 0.00049 | 0 | 1.60E-06 | 0.00124 | 0.00111 | 0.00084 | 0.00021 | 0.00022 | 0.00012 |
| Prdx5 | 0.00014 | 0.00021 | 0.00035 | 0.00112 | 0.00028 | 0.00035 | 0.00054 | 0.00054 | 0.00042 | 0.00077 | 0.00078 | 0.00063 | 0.00107 | 0.00067 | 6.69E-05 | 5.17E-05 | 0.00022 | 0.00017 |
| Sbno2 | 0.00016 | 0.00047 | 3.77E-05 | 0.00029 | 0.0004 | 0.00029 | 0.00018 | 3.16E-05 | 0.00042 | 0.00022 | 0.00018 | 0.00027 | 0.0006 | 0.00046 | 0.00077 | 0.00014 | 9.63E-05 | 0 |
| Tnfaip2 | 3.11E-05 | 0.00026 | 0.0004 | 0.00028 | 0.00024 | 0.00087 | 0.00017 | 0.00077 | 0.00042 | 0.00034 | 0.00017 | 0.00037 | 0.00061 | 0.00052 | 0.00057 | 0 | 0 | 4.94E-06 |
| Mbnl2 | 0.00028 | 0.00025 | 0.00044 | 0.00033 | 0.0004 | 0.00035 | 0.00054 | 0.00041 | 0.00042 | 0.00027 | 0.00022 | 0.00062 | 0.00015 | 0.0001 | 0.00028 | 0.00017 | 0.00043 | 0.00031 |
| Rnd3 | 0.00028 | 0.00021 | 0 | 0.00021 | 0.00013 | 0.00045 | 0 | 0.00019 | 0.00042 | 0.0002 | 0.00036 | 0.00043 | 0 | 0.00055 | 0.00039 | 5.68E-05 | 0.00025 | 0.00022 |
| Kirrel | 0 | 0.00034 | 0.00018 | 0.00043 | 0.0004 | 0.00032 | 0.00029 | 0.00024 | 0.00042 | 0.00032 | 0.00054 | 9.86E-05 | 0.00049 | 0.00069 | 0.0005 | 0.00035 | 0.00025 | 0.00053 |
| Il11ra1 | 0 | 0 | 0.0002 | 0 | 0 | 9.99E-05 | 0.00022 | 7.01E-05 | 0.00042 | 0.00032 | 0 | 5.21E-05 | 0.00078 | 2.62E-05 | 5.30E-05 | 8.62E-06 | 0 | 0.00012 |
| Mtch1 | 7.69E-05 | 0.0003 | 0.00025 | 0.00097 | 0.00114 | 0.00055 | 0.00096 | 0.00107 | 0.00042 | 0.0005 | 0.00078 | 0 | 0.00078 | 0.00076 | 0.00018 | 0.00026 | 0.00015 | 0.00076 |
| Ergic1 | 0.00037 | 9.83E-05 | 0.00025 | 0.00018 | 0.00021 | 0.00031 | 0.00036 | 0.00033 | 0.00042 | 0.00058 | 0.0003 | 0.00018 | 0.0002 | 0.00024 | 0 | 0.0002 | 0 | 0.00018 |
| Mbd3 | 0.00038 | 0.00019 | 0.00031 | 0.00062 | 0.00025 | 0.00018 | 0.0003 | 0.00028 | 0.00042 | 0.00014 | 0.00032 | 0.00021 | 0.00012 | 0.00023 | 0.00021 | 0.00031 | 8.89E-05 | 0.00026 |
| Itm2c | 0 | 0.00026 | 1.49E-05 | 0.00111 | 0.00092 | 0.00099 | 0.00086 | 0.00108 | 0.00042 | 0.00055 | 0.00049 | 0.00042 | 0.0011 | 0.0011 | 0 | 0 | 0.00014 | 0.00061 |
| Aplp2 | 0.00017 | 0.0002 | 9.48E-05 | 7.99E-05 | 0.00019 | 0.00016 | 0 | 0.00024 | 0.00042 | 0.00036 | 0.00024 | 0.00017 | 0.00032 | 0.00048 | 7.77E-05 | 0.00012 | 0.00019 | 0 |
| Asph | 0.00026 | 0.00013 | 9.88E-05 | 0.00046 | 0.00047 | 0.00037 | 0.00049 | 0.00018 | 0.00041 | 0.00024 | 0.00019 | 0.00071 | 0.00042 | 0.0001 | 0.00046 | 0.00015 | 0.00034 | 8.16E-05 |
| Islr | 0 | 0.00021 | 0 | 0.0003 | 0.00127 | 0.00049 | 0 | 0.00114 | 0.00041 | 0.00103 | 5.50E-05 | 0 | 0.0001 | 0 | 0 | 0 | 0 | 0.00033 |
| Cd9 | 2.33E-05 | 0.00069 | 0.00023 | 0.00012 | 0.00041 | 0.00079 | 0.00053 | 0.00035 | 0.00041 | 6.30E-05 | 0.00035 | 0.00045 | 0 | 0.00057 | 9.88E-05 | 0.0002 | 0.00017 | 0.00105 |
| Fasn | 7.71E-05 | 0.00013 | 0.00026 | 0.00045 | 7.93E-05 | 0 | 0 | 0.00011 | 0.00041 | 6.58E-05 | 0.00013 | 2.10E-05 | 6.96E-05 | 0.00024 | 0.00011 | 0 | 0.00031 | 6.36E-05 |
| Lrig1 | 0 | 0.00023 | 0.00024 | 0 | 0 | 0.00018 | 0.00011 | 0.00021 | 0.00041 | 0 | 0 | 3.88E-05 | 0.00011 | 0 | 0 | 0.00017 | 0.00025 | 9.47E-05 |
| Fbln5 | 3.33E-05 | 0 | 0 | 0 | 0.00071 | 0.00018 | 5.07E-05 | 0 | 0.00041 | 0.00019 | 0.00054 | 0 | 5.99E-05 | 0.00087 | 0 | 0 | 0 | 0.00035 |
| Emilin2 | 0.00015 | 9.04E-05 | 0.0002 | 0.00024 | 0 | 0.00048 | 0 | 0.00033 | 0.00041 | 0.00118 | 0 | 0.00031 | 0.00012 | 8.56E-05 | 9.24E-05 | 0.00017 | 0 | 0.00016 |
| Sdc2 | 0.00034 | 0.00026 | 0.00011 | 0 | 9.70E-05 | 0.00046 | 0.00012 | 0.00073 | 0.00041 | 0.00034 | 0.00049 | 0.0003 | 0 | 0.00022 | 0 | 2.80E-05 | 0 | 0.00023 |
| Rcn2 | 0.00041 | 8.01E-05 | 0.00017 | 6.40E-05 | 0.00042 | 0.00042 | 0.00032 | 0.00034 | 0.00041 | 0.00014 | 0.00059 | 0.0002 | 0.00035 | 0.00036 | 0 | 0.00023 | 6.68E-05 | 8.28E-05 |

| | | | | | | | | | | | | | | | | | | |
|---|---|---|---|---|---|---|---|---|---|---|---|---|---|---|---|---|---|---|
| Tnrc18 | 1.78E-06 | 0.00035 | 0 | 0.00054 | 0.00027 | 5.31E-05 | 0.00013 | 5.89E-05 | 0.00041 | 0.00016 | 0.00046 | 6.75E-05 | 0.00015 | 0.00022 | 0.00062 | 0.0003 | 0.00032 | 0.00019 |
| Ddost | 0.00062 | 5.59E-05 | 0.00037 | 0.00051 | 0.0002 | 0.00042 | 0.00098 | 0.00064 | 0.00041 | 0.00042 | 0.00037 | 0.00045 | 0.00077 | 0.00067 | 2.44E-05 | 0.00033 | 0 | 0.00055 |
| Msmo1 | 3.34E-05 | 5.02E-05 | 0 | 0 | 8.38E-05 | 0 | 0 | 6.35E-05 | 0.00041 | 1.64E-05 | 8.82E-05 | 2.83E-05 | 0 | 4.70E-05 | 0 | 0 | 0.00029 | 5.53E-05 |
| P4ha3 | 0.00024 | 0 | 0.00024 | 0.0003 | 0.0006 | 0.00052 | 0.00135 | 0.00046 | 0.0004 | 0.0003 | 0.00067 | 0 | 0.00066 | 0.00011 | 0.00033 | 0.00034 | 0.00015 | 0.00035 |
| Akr1b8 | 0.00017 | 2.35E-06 | 0.00044 | 6.73E-05 | 0.00102 | 0.00042 | 0.00045 | 0.00017 | 0.0004 | 0.00011 | 0.00014 | 0.00061 | 0.00036 | 0.0005 | 0.00022 | 0.00011 | 0 | 0.00022 |
| Mgrn1 | 9.40E-05 | 0.00017 | 0.0002 | 0.00019 | 0.00022 | 0.00016 | 0.00012 | 0.00041 | 0.0004 | 0.0002 | 0.00032 | 0.0002 | 0.00021 | 0.00029 | 5.37E-05 | 5.97E-06 | 2.54E-05 | 0.00018 |
| Il6st | 0.00075 | 0.00024 | 0.00018 | 0 | 0 | 0.00038 | 0.00035 | 0.00033 | 0.0004 | 0.0005 | 0 | 0.00016 | 0.00137 | 0 | 0.0002 | 0.00021 | 3.96E-05 | 0.0011 |
| Scn1b | 0 | 0.00017 | 0 | 0.00012 | 9.84E-05 | 0.00012 | 0.00032 | 0.00022 | 0.0004 | 0.00043 | 0 | 0 | 0.00038 | 0.00049 | 0 | 0 | 0 | 0.00079 |
| Ryk | 0.00039 | 0.00016 | 0.00038 | 8.55E-05 | 0.00014 | 0.00054 | 0.00064 | 0.00062 | 0.0004 | 0.00012 | 0.00029 | 0.00025 | 0.00034 | 0.0001 | 3.47E-05 | 0.00021 | 7.52E-05 | 9.81E-05 |
| Sigmar1 | 0 | 0 | 0.00048 | 0.00043 | 0.00011 | 2.02E-05 | 0.00018 | 0.00022 | 0.0004 | 0.00061 | 0.00015 | 0.00033 | 0.00057 | 0.00014 | 0 | 0.0001 | 0 | 0.00044 |
| Tle3 | 5.56E-05 | 0.00027 | 0.00027 | 0.00012 | 0.00028 | 0.00024 | 0.00031 | 0.00019 | 0.0004 | 9.50E-05 | 2.12E-05 | 0.00022 | 0 | 0.00025 | 0.00023 | 0.00013 | 0.00013 | 0.00019 |
| Lamp2 | 0.00062 | 0.0004 | 0 | 0 | 0.00052 | 0.00102 | 0.00055 | 0.00047 | 0.0004 | 0.00044 | 0.00012 | 0.00047 | 0.00078 | 0.0004 | 7.72E-05 | 4.44E-05 | 0.00021 | 0.00039 |
| Bhlhe40 | 0 | 0.00034 | 0.00055 | 8.91E-05 | 0.00025 | 0.00013 | 0.00042 | 0.00043 | 0.0004 | 0 | 0.00011 | 0.00109 | 5.59E-05 | 0.00021 | 0.00022 | 0.00017 | 0.00151 | 0.00047 |
| Erg28 | 0.00018 | 3.21E-05 | 1.23E-05 | 7.05E-05 | 0.00014 | 2.41E-05 | 3.67E-05 | 1.94E-05 | 0.0004 | 1.66E-05 | 0.00012 | 1.49E-06 | 0.00013 | 0.00018 | 0 | 0.00019 | 0 | 4.42E-05 |
| Fus | 0.00032 | 0.00023 | 0.00036 | 0.0003 | 9.21E-05 | 8.08E-05 | 6.44E-05 | 0.00029 | 0.0004 | 0.00017 | 0.00025 | 0.00028 | 7.29E-05 | 9.95E-05 | 0.00037 | 0.00034 | 0.00113 | 0.00014 |
| Tspan4 | 0.00017 | 0.00017 | 0.00029 | 0.00069 | 0.00061 | 0.00103 | 3.70E-05 | 0.00035 | 0.0004 | 0 | 0.00048 | 0.00038 | 0.0014 | 0.00099 | 0 | 0.00035 | 0 | 0.0002 |
| Ms4a4d | 0.00038 | 5.44E-06 | 0 | 0 | 9.81E-05 | 0.00028 | 1.94E-05 | 0.00012 | 0.0004 | 0.00052 | 4.86E-07 | 0 | 0.00049 | 0.00014 | 0 | 3.43E-05 | 0 | 3.48E-05 |
| Irx1 | 0 | 0.00011 | 0 | 0.00017 | 0 | 4.76E-05 | 0 | 0 | 0.0004 | 0.00036 | 6.69E-05 | 0.00013 | 0.00092 | 0.00039 | 5.70E-05 | 0.00018 | 0.00028 | 0 |
| Hnrnpa2b1 | 0.00071 | 0.00023 | 0.0005 | 0.00027 | 0.00017 | 0.00013 | 0.00012 | 8.74E-05 | 0.00039 | 6.26E-05 | 0.00029 | 9.16E-05 | 0.00016 | 0.00017 | 0.0002 | 0.00055 | 0.00029 | 0 |
| Capn2 | 0.00012 | 0.00017 | 0.0005 | 0.00018 | 0 | 7.82E-05 | 0.00056 | 0 | 0.00039 | 0.0008 | 0.00098 | 0.00035 | 6.23E-05 | 0 | 3.74E-05 | 0.00028 | 0.00012 | 0.00078 |
| Mrgprf | 0 | 0 | 0 | 0.00014 | 8.98E-05 | 0.00017 | 0.00029 | 0 | 0.00039 | 0.0004 | 7.66E-05 | 0 | 0 | 0.00021 | 0 | 0 | 0 | 0.00013 |
| Mmp14 | 6.36E-05 | 0.0009 | 0.00032 | 0.00052 | 0.00115 | 0.00111 | 0.00187 | 0.00187 | 0.00039 | 0.00071 | 0.0009 | 0.00035 | 0.00144 | 0.00116 | 0.00031 | 0.0002 | 0.00035 | 0.00089 |
| Dnm1 | 0 | 0.00034 | 0 | 0.00012 | 0.00039 | 0.00024 | 0.00061 | 0.00024 | 0.00039 | 0.00053 | 0 | 8.39E-05 | 7.23E-05 | 0.00026 | 0.0006 | 0 | 0.00034 | 0.00036 |
| Chst15 | 3.84E-05 | 0.00018 | 0 | 0 | 5.66E-05 | 0.00041 | 0 | 9.20E-05 | 0.00039 | 0 | 0 | 3.59E-06 | 0 | 0.00048 | 0 | 0 | 8.17E-06 | 3.12E-05 |
| Nsg1 | 5.56E-06 | 5.53E-05 | 0 | 0.00013 | 0 | 8.91E-07 | 0.00011 | 0 | 0.00039 | 0.00061 | 1.29E-05 | 0 | 0.00107 | 0.00017 | 0 | 0.00051 | 0 | 0.00014 |
| Emc10 | 7.58E-06 | 8.61E-05 | 4.70E-05 | 0.00073 | 0.00035 | 0.0003 | 0.00041 | 0.00058 | 0.00039 | 0.00031 | 0.0003 | 0.00022 | 0.00041 | 0.0005 | 0.00022 | 0.00011 | 0.00014 | 0.00042 |
| Apod | 0 | 0.00022 | 0 | 0 | 0 | 0 | 0 | 0 | 0.00039 | 0.00121 | 0 | 0 | 0.00182 | 0.00106 | 0 | 0 | 0 | 0.00181 |
| Kank2 | 9.31E-06 | 0.0003 | 0.00014 | 0.00035 | 0.00018 | 0.00011 | 0.00011 | 0.00025 | 0.00039 | 0.00031 | 0.00058 | 0.00022 | 7.13E-05 | 0.00027 | 0.00016 | 0.00015 | 0.00013 | 0.00047 |
| Emp3 | 0.00023 | 0.00024 | 0.00034 | 0.00036 | 9.11E-06 | 0.00023 | 0.0004 | 0 | 0.00039 | 0.00066 | 0.00023 | 0.00025 | 8.03E-06 | 0.00037 | 0 | 0.00025 | 0 | 0.00069 |
| Dhcr24 | 2.93E-05 | 1.88E-05 | 0 | 0.00011 | 7.17E-06 | 0 | 0 | 9.25E-05 | 0.00039 | 2.39E-05 | 0.00018 | 0 | 0 | 8.54E-05 | 0 | 0 | 0 | 6.65E-05 |
| Pebp1 | 0.00022 | 0.00018 | 0.00023 | 0.00051 | 0.00012 | 0.00011 | 4.26E-06 | 5.96E-05 | 0.00039 | 3.12E-05 | 7.89E-05 | 0.0003 | 0.00016 | 0.00032 | 0 | 0.00011 | 0.00029 | 0.00012 |
| Pten | 0.00072 | 0.00034 | 0 | 0 | 0.00018 | 0.0006 | 7.81E-05 | 0.00017 | 0.00039 | 0 | 0.0002 | 0.00039 | 0.0002 | 0.00064 | 0.00013 | 0.0001 | 0.00037 | 0.00029 |
| Marcksl1 | 4.34E-05 | 0.00025 | 0.00015 | 0.00028 | 0.00018 | 0 | 0 | 0.00028 | 0.00039 | 0 | 0.00041 | 0.00025 | 0.00032 | 0.00045 | 0.00023 | 0.00043 | 0.0011 | 0 |

| | | | | | | | | | | | | | | | | | | |
|---|---|---|---|---|---|---|---|---|---|---|---|---|---|---|---|---|---|---|
| Serpina3g | 0.00012 | 0 | 5.66E-05 | 0 | 1.35E-05 | 0.0008 | 0 | 0.00037 | 0.00039 | 0 | 0 | 0 | 0.00067 | 0.00108 | 0 | 3.87E-05 | 0 | 1.46E-05 |
| Mrfap1 | 0.00013 | 0.00023 | 0.00028 | 0.00109 | 0.00033 | 0.0002 | 0.0003 | 0.00046 | 0.00039 | 0.00023 | 0.00031 | 0.00017 | 0.00045 | 0.0003 | 0.00024 | 0.0003 | 0.00031 | 0.00031 |
| Ptov1 | 0.00039 | 0.00031 | 0.00026 | 0.0006 | 0.00027 | 0.00034 | 0.00045 | 0.00032 | 0.00039 | 0.0002 | 0.00032 | 0.00011 | 0.00015 | 0.00029 | 0.00038 | 0.00018 | 0.00038 | 0.00024 |
| Smo | 6.09E-05 | 0.00022 | 0.00011 | 0.00048 | 0.00034 | 0.00015 | 0.00026 | 0.00018 | 0.00039 | 9.85E-05 | 0.00025 | 0.00015 | 0.00022 | 0.00029 | 3.54E-05 | 0.00015 | 0 | 0.00021 |
| Angptl2 | 0 | 0.00024 | 0.00011 | 0.00067 | 0.00172 | 0.00129 | 0.00091 | 0.00089 | 0.00039 | 0.00068 | 0.00105 | 0.00085 | 0.00138 | 0.00159 | 0.00054 | 0.00015 | 0.00038 | 0 |
| Fkbp8 | 0.00023 | 0.00031 | 0.00016 | 0.00041 | 0.00025 | 0.00028 | 0.00023 | 0.00028 | 0.00039 | 0.00027 | 0.00027 | 0.00034 | 0.00022 | 0.00026 | 0.0003 | 0.00015 | 0.00014 | 0.00023 |
| Irs1 | 0 | 0.00018 | 0 | 0.00018 | 0.00029 | 0 | 0.00039 | 0.00058 | 0.00038 | 0.00026 | 0.00016 | 0 | 0 | 0 | 0.00033 | 4.90E-05 | 3.52E-05 | 0.00027 |
| a | 0 | 8.30E-05 | 0 | 0 | 7.54E-05 | 0.00021 | 0 | 0 | 0.00038 | 0 | 0 | 0 | 3.37E-05 | 0.00022 | 6.01E-05 | 0 | 0 | 0 |
| Slc25a1 | 5.41E-05 | 0.00019 | 6.31E-08 | 0.0002 | 0.00025 | 0.0001 | 0.00012 | 0.0002 | 0.00038 | 0.00015 | 0.00025 | 0.00033 | 0.00016 | 0.00022 | 0 | 0 | 0 | 0.00025 |
| Ephb2 | 1.78E-05 | 0 | 9.31E-05 | 0 | 0.00032 | 6.05E-05 | 7.09E-05 | 0 | 0.00038 | 0 | 0.00022 | 0.00019 | 6.34E-05 | 0.0004 | 3.29E-06 | 0.00012 | 0.00014 | 0.00064 |
| Zfp36l1 | 4.08E-05 | 0.00042 | 0 | 6.69E-05 | 0.00056 | 0.00029 | 0.00012 | 0 | 0.00038 | 0.00046 | 7.17E-05 | 0.00046 | 0.00065 | 0.00039 | 0 | 8.18E-05 | 0.00114 | 0.00029 |
| Selenon | 8.60E-05 | 0.00013 | 0.00015 | 0.00042 | 0.00026 | 0.00024 | 0.00032 | 0.00042 | 0.00038 | 0.00047 | 0.00027 | 0.0002 | 0.00041 | 0.00031 | 0 | 4.96E-05 | 0.00013 | 0.00032 |
| Ifngr1 | 5.09E-05 | 0.00018 | 9.03E-05 | 0.00026 | 0.00037 | 0.00016 | 0 | 0.00022 | 0.00038 | 0.00039 | 0.00027 | 0.00012 | 0.00038 | 0.00013 | 0.00012 | 0.0002 | 3.27E-05 | 0.00041 |
| Cd164 | 0.00042 | 0.00034 | 0.00033 | 0.00067 | 0.0005 | 0.0004 | 0.00069 | 0.00038 | 0.00038 | 0.00076 | 0.00011 | 0.00035 | 0.0008 | 0.00026 | 2.56E-05 | 0.00035 | 0.00025 | 9.61E-05 |
| Midn | 0.00017 | 0.00032 | 0.0003 | 0.0005 | 0.00016 | 1.78E-05 | 0 | 0.00014 | 0.00038 | 0.00012 | 0.00026 | 0.00022 | 0.00016 | 0.00026 | 0.00056 | 0.00012 | 0.00097 | 9.11E-05 |
| Srsf5 | 6.81E-05 | 0.00043 | 0.00018 | 0.00059 | 0.00035 | 0.00024 | 8.53E-05 | 0.00024 | 0.00038 | 0.00022 | 0.00029 | 0.00033 | 0.00024 | 0.00031 | 0.00068 | 1.27E-05 | 0.00017 | 0.00019 |
| Ilk | 0.00043 | 0.00031 | 0.00027 | 0.00034 | 0.00022 | 0.0002 | 0.00022 | 0.00035 | 0.00038 | 0.00018 | 0.00105 | 0.00028 | 0.00012 | 0.00038 | 9.56E-05 | 0.00022 | 0 | 0.00011 |
| Tmbim6 | 0.00034 | 0.00027 | 0.00021 | 0.00037 | 0.00028 | 0.00039 | 0.00022 | 0.00036 | 0.00038 | 0.00028 | 0.00028 | 0.00037 | 0.00029 | 0.00029 | 0 | 0.0001 | 0.00015 | 0.00035 |
| Ptpn9 | 7.17E-05 | 0.00023 | 0.00012 | 7.40E-05 | 0.00012 | 0.00018 | 0 | 0.00028 | 0.00038 | 0.00014 | 0.00024 | 0.00013 | 0.0001 | 0.00015 | 7.42E-05 | 6.59E-05 | 5.51E-05 | 0.00023 |
| Tmsb4x | 0.00069 | 0.00024 | 0.00056 | 0.00056 | 0.00105 | 8.38E-05 | 0.00113 | 0.00108 | 0.00038 | 0.00103 | 0.00209 | 0.00038 | 0 | 0.00068 | 0.00105 | 0.00039 | 0.00051 | 0.00124 |
| Fads2 | 9.34E-05 | 7.73E-05 | 5.69E-05 | 2.93E-05 | 9.92E-05 | 0 | 8.35E-05 | 0.00045 | 0.00038 | 0.00011 | 0.00011 | 0 | 6.72E-05 | 0 | 0 | 0.00021 | 0 | 0.00012 |
| Pcca | 7.97E-05 | 4.98E-05 | 8.70E-05 | 2.85E-05 | 7.10E-05 | 8.48E-05 | 7.88E-05 | 0.00022 | 0.00038 | 7.27E-05 | 0 | 0 | 6.64E-05 | 0 | 0.00026 | 6.73E-05 | 0 | 0 |
| Il13ra1 | 0.0004 | 5.13E-05 | 0.00032 | 0 | 0 | 0.00036 | 0.00036 | 0.00051 | 0.00038 | 0.00026 | 0 | 0.00051 | 0.00069 | 8.09E-05 | 0 | 5.80E-05 | 0 | 0 |
| Sema3a | 1.16E-06 | 0 | 0 | 0 | 0.00016 | 0.00036 | 0 | 0 | 0.00037 | 0 | 0 | 6.01E-05 | 0 | 0 | 0 | 0 | 0 | 0 |
| Ctnna1 | 0.00031 | 0.00036 | 0.00027 | 5.91E-05 | 0.00013 | 0.00025 | 3.51E-05 | 0.00034 | 0.00037 | 0.0002 | 0.00048 | 0.00035 | 0 | 0.00023 | 0.00027 | 0.00025 | 0.00035 | 0.00031 |
| Nrg1 | 0 | 0 | 0 | 0 | 0.00011 | 0 | 0.00034 | 0 | 0.00037 | 0 | 0.00035 | 0 | 0 | 0 | 4.61E-05 | 9.45E-05 | 8.07E-05 | 0 |
| Arf4 | 0.00105 | 8.93E-06 | 0.00043 | 0 | 0.00026 | 0.00053 | 0.00093 | 0.00042 | 0.00037 | 0.00024 | 0.00033 | 0.00044 | 0.00041 | 0.00022 | 5.23E-05 | 4.66E-05 | 0.00057 | 0.00012 |
| Myo1b | 0.00012 | 0.00015 | 0.00013 | 4.93E-05 | 0.00016 | 0.0003 | 4.53E-05 | 0.00048 | 0.00037 | 0 | 0.00016 | 9.06E-06 | 0 | 0.00043 | 0.00017 | 0.00015 | 0.00027 | 1.43E-05 |
| Il15ra | 1.55E-05 | 0 | 0.00015 | 0 | 0.00022 | 0.00016 | 0.00049 | 0 | 0.00037 | 0.00021 | 0 | 0 | 0.001 | 0.0006 | 0.00015 | 8.12E-05 | 0 | 9.35E-05 |
| Rab1b | 0.00025 | 0.00014 | 0.00026 | 0.00055 | 0.00027 | 0.00025 | 0.00023 | 0.00044 | 0.00037 | 0.00026 | 0.00027 | 9.96E-05 | 9.52E-05 | 0.00033 | 0 | 0.0002 | 8.64E-05 | 0.00027 |
| Ptar1 | 0.0002 | 0.00024 | 0.00026 | 0 | 0.00035 | 0.00011 | 0.00023 | 0.00012 | 0.00037 | 7.13E-05 | 0.00039 | 9.30E-05 | 0 | 0.00015 | 0.00015 | 0.00019 | 0.0004 | 5.85E-05 |
| Fbn2 | 0.00014 | 2.27E-05 | 0 | 7.77E-05 | 0.00073 | 0 | 0.00038 | 0.00029 | 0.00037 | 0.0002 | 0.00058 | 0 | 7.28E-05 | 0 | 0.0001 | 9.23E-05 | 0 | 7.57E-05 |
| Atn1 | 0.00012 | 0.00035 | 0.00031 | 0.00043 | 0.00021 | 0.0002 | 8.32E-05 | 0.00019 | 0.00037 | 0.0002 | 0.00037 | 0.00025 | 0.00019 | 0.00031 | 0.00034 | 0.00041 | 0.00031 | 0.00027 |

| | | | | | | | | | | | | | | | | | | |
|---|---|---|---|---|---|---|---|---|---|---|---|---|---|---|---|---|---|---|
| Prrc2b | 0.00018 | 0.00038 | 0.00019 | 0.00014 | 0.00025 | 0.00019 | 8.30E-05 | 7.10E-05 | 0.00037 | 0.00016 | 0.00019 | 0.00019 | 7.26E-05 | 0.00017 | 0.00041 | 0.0002 | 0.00012 | 0.00017 |
| Pttg1ip | 7.04E-05 | 0.00042 | 0.00026 | 0.00038 | 0.00026 | 0.00032 | 0.00015 | 0.00048 | 0.00037 | 0.00041 | 0.00031 | 0.0006 | 0.00021 | 0.0002 | 0.0001 | 9.40E-05 | 0.00012 | 0.00042 |
| Wipi1 | 0.00021 | 0.00016 | 2.95E-06 | 0.00039 | 0.00044 | 0.0006 | 0.00069 | 0.00032 | 0.00037 | 0.00023 | 0.00025 | 0.0002 | 0.00047 | 0.0007 | 0.00011 | 2.29E-05 | 0 | 0.00021 |
| Soat1 | 9.27E-05 | 7.89E-05 | 0.00025 | 5.72E-05 | 0.00034 | 0.00042 | 0.00025 | 0.00072 | 0.00037 | 0 | 0.00044 | 0.00072 | 0.00031 | 0.00034 | 0 | 0.00017 | 0 | 0 |
| Ptprs | 2.95E-05 | 0.00013 | 4.48E-05 | 0.00033 | 0.00033 | 0.00037 | 0.00058 | 0.00064 | 0.00037 | 0.00024 | 0.00034 | 0.00025 | 0 | 0.00023 | 0.00049 | 0.00035 | 7.49E-05 | 0.00054 |
| Bace1 | 0 | 6.98E-05 | 0.00038 | 0.00012 | 0.00073 | 3.02E-05 | 0.0006 | 0.00064 | 0.00037 | 0 | 0.00029 | 0 | 0.00036 | 0.00036 | 4.75E-05 | 7.41E-05 | 0 | 0.00054 |
| Vwa5a | 0.00019 | 0.00031 | 0 | 0 | 0.0002 | 0.00024 | 0.0002 | 0 | 0.00037 | 0.00037 | 7.74E-05 | 0.00012 | 0.00022 | 0 | 1.85E-05 | 0 | 6.79E-05 | 0.00031 |
| Pdia6 | 0.00078 | 0 | 0.00043 | 0.00052 | 0.0002 | 0.00052 | 0.00059 | 0.00053 | 0.00037 | 0.00039 | 0.00035 | 0.00051 | 0.00079 | 0.00046 | 0.00026 | 0.00023 | 0.00019 | 0.00046 |
| Mvp | 0 | 0.0002 | 0.00036 | 0.00038 | 0.00016 | 7.60E-05 | 0.00023 | 0.00019 | 0.00037 | 0.00028 | 0.00012 | 0.00014 | 0.00031 | 0.00017 | 0.00026 | 0 | 0.00014 | 0.00028 |
| Ttyh2 | 0 | 0.00029 | 0.00042 | 0.00017 | 0.0003 | 0.00011 | 0.00024 | 0.00153 | 0.00037 | 0.00046 | 0 | 9.53E-06 | 0 | 3.61E-05 | 0 | 4.46E-05 | 2.73E-05 | 6.22E-05 |
| Hspa8 | 0.00046 | 0.00039 | 0.00053 | 0.00044 | 0.00024 | 0.00047 | 0.0003 | 0.00036 | 0.00037 | 0.00033 | 0.00031 | 0.00029 | 0.00031 | 0.00037 | 0.00011 | 0.00054 | 0.00287 | 0.00032 |
| P4hb | 0.00032 | 7.30E-05 | 0.00034 | 0.00123 | 0.00037 | 0.00055 | 0.00133 | 0.00097 | 0.00037 | 0.0006 | 0.00044 | 0.00106 | 0.00078 | 0.00068 | 0.00058 | 0.00029 | 0.00048 | 0.0007 |
| Mfge8 | 0 | 0.00025 | 0 | 0.00028 | 0.00092 | 0.0006 | 0.00066 | 0.00095 | 0.00037 | 0.0003 | 0.00076 | 0.00058 | 0.00034 | 0.00119 | 0.00022 | 0 | 0 | 0.00038 |
| Tuba1b | 9.93E-05 | 0 | 0.00033 | 0.0007 | 0 | 4.78E-05 | 0 | 6.26E-05 | 0.00037 | 0.0002 | 0.00028 | 0.0004 | 2.69E-05 | 0 | 0 | 0.00148 | 0.00046 | 0.00026 |
| Prr16 | 0 | 0.00014 | 0 | 0 | 4.84E-05 | 0.00045 | 0 | 0 | 0.00037 | 0 | 0 | 8.82E-06 | 0 | 0.00021 | 0.00011 | 0 | 2.41E-05 | 0.00012 |
| Ctbp1 | 0.00011 | 0.00028 | 0.00037 | 0.00049 | 0.00029 | 0.00014 | 0.00056 | 0.00033 | 0.00037 | 0.00034 | 0.00031 | 0.00018 | 0.00018 | 0.00026 | 2.31E-05 | 0.00034 | 0.00014 | 0.00042 |
| Lims1 | 0.00032 | 0.0002 | 0.00025 | 6.52E-05 | 0.00014 | 0.00018 | 3.73E-05 | 8.32E-05 | 0.00037 | 0.00012 | 0.001 | 0.0003 | 0.00015 | 0.00023 | 0 | 0.00034 | 0.00017 | 0.00011 |
| Esyt1 | 0.00019 | 0.00011 | 0.00018 | 0.00015 | 2.61E-05 | 7.27E-05 | 0.00039 | 0.00025 | 0.00037 | 0.00029 | 0.00046 | 0.0004 | 0.00023 | 0.00013 | 0.00018 | 7.46E-05 | 8.15E-05 | 0.00045 |
| Errfi1 | 0 | 0 | 0.00036 | 0 | 0 | 0.00043 | 0 | 0.0004 | 0.00036 | 0.00061 | 0.00081 | 0.00083 | 0.00024 | 0.00075 | 0.00028 | 0 | 0.00123 | 0 |
| Ptbp1 | 0.00046 | 0.00029 | 0.00052 | 0.00062 | 0.00019 | 0.00018 | 0.00023 | 0.0003 | 0.00036 | 0.0001 | 0.0002 | 0.00028 | 7.88E-05 | 0.00022 | 0.00039 | 0.00045 | 0.00121 | 7.99E-05 |
| Rarg | 0.00015 | 0.00022 | 0.00014 | 0.00029 | 0.00017 | 0.00022 | 0.00028 | 8.25E-05 | 0.00036 | 7.81E-05 | 0.00013 | 0.00027 | 0 | 0.00011 | 0.0002 | 0 | 0.00014 | 0.00013 |
| Tmem86a | 0 | 0.00015 | 0 | 0.00039 | 0.00079 | 0.00037 | 9.15E-06 | 0.00077 | 0.00036 | 0 | 0.00033 | 0 | 0.00016 | 0.00023 | 0 | 0 | 0 | 0 |
| Nfix | 0.00024 | 0.00038 | 0 | 0.00058 | 0.0003 | 0.00017 | 0.00088 | 5.85E-05 | 0.00036 | 0.00071 | 0.00034 | 0.0002 | 0.00064 | 0 | 7.38E-06 | 0.00035 | 0.00046 | 0.00096 |
| Avpr1a | 0 | 0 | 0 | 0.00016 | 0.00058 | 0 | 0 | 0.00024 | 0.00036 | 0 | 0.00014 | 1.50E-06 | 0.00092 | 0.00052 | 0 | 0.00013 | 0 | 0 |
| St3gal1 | 1.72E-05 | 5.38E-05 | 0.00021 | 0 | 0 | 5.09E-05 | 6.31E-05 | 0 | 0.00036 | 0.00057 | 0 | 0.00013 | 0.0004 | 0.00011 | 0 | 6.56E-05 | 0.00017 | 0.00017 |
| Scpep1 | 0.00015 | 0.00049 | 0.00014 | 0.00043 | 0.00042 | 0.00071 | 0.00061 | 0.00034 | 0.00036 | 0.00032 | 5.14E-05 | 0.0006 | 0.00059 | 0.00044 | 0 | 0.00027 | 0.00024 | 0.00083 |
| Gsr | 0.00015 | 0.00015 | 0.0003 | 0.00011 | 0 | 0.00026 | 0.00024 | 0.00021 | 0.00036 | 2.68E-05 | 0 | 0.00023 | 4.11E-05 | 0.00043 | 0 | 0.00019 | 0.00019 | 9.91E-05 |
| Vegfd | 2.82E-05 | 0 | 0 | 0 | 0 | 0.00017 | 0 | 0 | 0.00036 | 0.00111 | 0 | 8.39E-05 | 0.00117 | 0 | 0 | 0 | 0 | 0 |
| Ppib | 0.00047 | 3.45E-05 | 0.0003 | 0.00103 | 0.00045 | 0.00072 | 0.00115 | 0.00099 | 0.00036 | 0.00055 | 0.00052 | 0.00044 | 0.00097 | 0.00091 | 0.00011 | 0.00026 | 2.83E-05 | 0.00061 |
| Kctd10 | 0 | 0.00034 | 0.00026 | 0.00034 | 0.00039 | 9.13E-05 | 0.00016 | 0.00016 | 0.00036 | 0.00015 | 0.00084 | 0.00046 | 0.00031 | 0.0003 | 1.90E-05 | 0.00019 | 4.86E-05 | 0.00033 |
| C1ra | 4.38E-05 | 0.00012 | 0 | 0 | 0.00041 | 0.00068 | 0.00043 | 0.00013 | 0.00036 | 0.0011 | 0.0001 | 0.00023 | 0.00211 | 0.00047 | 0 | 0 | 9.55E-06 | 0.00077 |
| Pgk1 | 0.00035 | 0 | 0.00024 | 7.45E-05 | 6.38E-05 | 0.00033 | 0.00018 | 6.89E-05 | 0.00036 | 2.16E-05 | 0.00023 | 0.00156 | 0.00021 | 0.00012 | 0 | 0.00015 | 0.00015 | 0 |
| Fzd2 | 7.21E-05 | 5.88E-05 | 0 | 0.00053 | 0.00062 | 0.00026 | 0.00066 | 0 | 0.00036 | 0.00014 | 0.00089 | 0.00022 | 0.00035 | 0.00025 | 0 | 0.00016 | 0 | 0.00027 |

| | | | | | | | | | | | | | | | | | | |
|---|---|---|---|---|---|---|---|---|---|---|---|---|---|---|---|---|---|---|
| Rnf10 | 0.00031 | 0.00042 | 0.00035 | 0.00054 | 0.00033 | 0.00033 | 0.00023 | 0.00035 | 0.00035 | 0.0003 | 0.00036 | 0.00031 | 0.00023 | 0.00025 | 0.00031 | 0.00016 | 0.00027 | 0.00029 |
| Axl | 0 | 0.0004 | 0.00047 | 0.00041 | 0.00043 | 0.00062 | 0.00013 | 0.00043 | 0.00035 | 0.00136 | 0.00098 | 0.00033 | 0.00035 | 0.00082 | 0.00082 | 0.00017 | 0.00036 | 0.00057 |
| Sesn3 | 1.00E-05 | 0.00021 | 0 | 0 | 0.00013 | 0.00016 | 0.00014 | 0.00014 | 0.00035 | 8.58E-05 | 0 | 0 | 0 | 0.00034 | 0 | 6.75E-05 | 0 | 0.00017 |
| Sdcbp | 0.00043 | 0.00038 | 0.00034 | 0.00012 | 0.00033 | 0.00051 | 0.00014 | 0.00046 | 0.00035 | 0.0004 | 6.50E-05 | 0.00024 | 0.00015 | 0.00034 | 0 | 0.00016 | 0.00029 | 0.0002 |
| Mlf2 | 0.00037 | 0.00024 | 0.00056 | 0.00055 | 0.00026 | 0.00022 | 0.00033 | 0.00049 | 0.00035 | 0.0002 | 0.00046 | 0.00016 | 5.10E-05 | 0.00034 | 0.00029 | 0.00032 | 0.00012 | 0.00024 |
| Pbx1 | 2.97E-05 | 0.00035 | 4.37E-05 | 0.0003 | 0.00027 | 0.00027 | 0 | 0 | 0.00035 | 0.00019 | 0.00017 | 4.96E-05 | 0.0001 | 0.0003 | 0.0005 | 9.06E-05 | 0.00024 | 0.0002 |
| Myl6 | 3.95E-05 | 0 | 0.00016 | 0.0007 | 0.00019 | 2.58E-05 | 0.00014 | 0.00047 | 0.00035 | 0.00018 | 0.00099 | 0.0002 | 0 | 0.00029 | 0.0001 | 1.88E-05 | 9.44E-05 | 0.00033 |
| Capzb | 0.00046 | 0.00047 | 0.00035 | 0.0003 | 0.0004 | 0.00026 | 0.00047 | 0.00017 | 0.00035 | 0.00025 | 0.00064 | 0.00029 | 0.0004 | 0.00034 | 0 | 0.00013 | 0 | 0.00018 |
| Qsox1 | 2.37E-06 | 0.00018 | 0.00012 | 0.00048 | 0.00054 | 0.00043 | 0.00083 | 0.00027 | 0.00035 | 0.00027 | 0.00076 | 0.0003 | 0.00055 | 0.0007 | 0.00013 | 9.45E-05 | 0.00011 | 0.00033 |
| Mlec | 0.0006 | 1.91E-05 | 0.00043 | 0.00077 | 0.00037 | 0.00027 | 0.00081 | 0.00064 | 0.00035 | 0.00035 | 0.00026 | 2.05E-05 | 0.00035 | 0.00041 | 0.00031 | 0.00022 | 0.00016 | 0.00028 |
| Glrx | 8.36E-05 | 2.78E-05 | 8.58E-05 | 0 | 6.47E-05 | 0.00033 | 0 | 0 | 0.00035 | 0 | 2.01E-07 | 8.14E-05 | 0.00037 | 0.0004 | 0 | 0 | 0 | 0 |
| Igf1r | 0 | 0.00036 | 0.00014 | 0 | 0.00011 | 0.00024 | 0 | 0 | 0.00035 | 8.51E-05 | 0.00049 | 0.00021 | 0.00023 | 0.00044 | 0.00011 | 8.72E-05 | 0.00026 | 5.53E-05 |
| Cyb5r3 | 0.00036 | 0.00032 | 0.00035 | 0.00055 | 0.00042 | 0.00029 | 0.00073 | 0.00029 | 0.00035 | 0.00043 | 0.00041 | 0.00056 | 3.75E-05 | 0.00042 | 0 | 0.00023 | 0 | 0.00092 |
| Fndc3b | 0.00029 | 0.00037 | 0.00038 | 0.00029 | 0.00071 | 0.00077 | 0.00081 | 0.00064 | 0.00035 | 0.00044 | 0.00038 | 0.00037 | 0.00074 | 0.00059 | 0.00083 | 0 | 0.00056 | 0.0002 |
| Dbi | 0.0003 | 3.60E-05 | 1.27E-05 | 0.0001 | 6.90E-05 | 3.37E-05 | 0 | 7.69E-05 | 0.00035 | 7.45E-05 | 0.00011 | 0 | 6.93E-05 | 4.34E-05 | 0 | 0.00013 | 1.11E-05 | 7.09E-05 |
| Slco3a1 | 0 | 4.79E-05 | 0.00012 | 6.01E-05 | 9.25E-05 | 0 | 0.00015 | 2.44E-05 | 0.00035 | 0.00054 | 0.00017 | 0 | 7.57E-05 | 0.00014 | 0 | 0.00014 | 0.00012 | 0.00037 |
| Tceal9 | 0.00061 | 0.0002 | 7.00E-07 | 0.00056 | 0.00047 | 0.00044 | 0.00079 | 0.00046 | 0.00035 | 0 | 0.00039 | 0.00014 | 0.00041 | 0.00059 | 0.0004 | 0.00049 | 0.00031 | 0.00014 |
| Psme1 | 0.00035 | 0.00014 | 2.85E-05 | 0.00019 | 7.64E-05 | 5.78E-05 | 0 | 9.43E-05 | 0.00035 | 0.00021 | 5.77E-05 | 0.00011 | 0.00016 | 0.00013 | 4.29E-05 | 7.35E-05 | 0.00015 | 0.00027 |
| Nuak1 | 0 | 3.39E-05 | 0 | 0 | 0.00069 | 0.00013 | 0 | 0 | 0.00035 | 2.52E-05 | 0.00112 | 0 | 7.61E-05 | 0.00044 | 0.00032 | 0 | 0.0002 | 0 |
| Dag1 | 0.00015 | 0.00019 | 0 | 4.05E-05 | 0.00022 | 0.00027 | 0.00037 | 0.00017 | 0.00035 | 0.00053 | 0.0002 | 0.00057 | 0.00032 | 0.00023 | 0 | 0.0001 | 0.00012 | 0.00057 |
| Vegfc | 0 | 0 | 0.00015 | 8.36E-05 | 8.77E-05 | 9.36E-05 | 0 | 0.00018 | 0.00035 | 0 | 0.0002 | 2.86E-05 | 0.00036 | 0.00022 | 9.00E-05 | 8.82E-05 | 0 | 0.00012 |
| Prss23 | 0.00015 | 0 | 0.00048 | 0.0001 | 0 | 0.00015 | 0 | 0 | 0.00035 | 0.0015 | 0.00053 | 0 | 7.20E-05 | 0 | 0 | 0.00028 | 0 | 0.00082 |
| Tbl1x | 0.00024 | 0.00019 | 6.86E-05 | 0 | 3.55E-05 | 0.00017 | 4.94E-05 | 6.41E-05 | 0.00034 | 9.86E-05 | 0.00011 | 0.00017 | 7.78E-05 | 0 | 0 | 0.00011 | 0.00011 | 0.00013 |
| Ccnd3 | 0.00015 | 9.46E-05 | 0.00039 | 0.00034 | 0.00025 | 0.00016 | 3.52E-05 | 0.00046 | 0.00034 | 0.00028 | 0.00014 | 3.26E-05 | 0.00012 | 0.00017 | 0 | 0.00023 | 0 | 0.00014 |
| Gna12 | 0.00018 | 0.00021 | 0.00024 | 0.00013 | 0.0003 | 0.00049 | 0.00011 | 0.00057 | 0.00034 | 0.00014 | 0.00017 | 3.39E-06 | 0 | 0.00035 | 0.00011 | 0.00014 | 2.60E-05 | 0.00024 |
| Ptger3 | 0 | 1.36E-05 | 0 | 0 | 0 | 0 | 0.00011 | 0 | 0.00034 | 0 | 0 | 0.00012 | 7.63E-05 | 0 | 0 | 0 | 0 | 0.00025 |
| Arf3 | 0 | 0.00022 | 0.00017 | 0.00042 | 0.00032 | 0.00023 | 0.00017 | 0.0004 | 0.00034 | 0.00036 | 0.00031 | 0.00021 | 0.00013 | 0.00027 | 7.18E-05 | 1.90E-05 | 5.71E-05 | 0.0003 |
| Myh10 | 0.00011 | 0.00015 | 8.32E-05 | 0 | 0.00054 | 5.36E-05 | 0 | 6.13E-05 | 0.00034 | 0.00014 | 0.00085 | 0.00012 | 0 | 0 | 0 | 0.00053 | 8.08E-05 | 0.00013 |
| Pgd | 0.00023 | 0.00017 | 0.00033 | 0.00023 | 9.64E-08 | 2.86E-05 | 0 | 2.11E-05 | 0.00034 | 0.00025 | 8.38E-05 | 0.00027 | 0 | 9.73E-05 | 0.00011 | 5.03E-05 | 1.88E-05 | 7.18E-05 |
| Ndufa13 | 0.00029 | 0.00017 | 0.00018 | 0.00056 | 0.00036 | 0.00027 | 0.00042 | 0.00039 | 0.00034 | 0.00029 | 0.00032 | 0.00013 | 0.00036 | 0.00031 | 9.09E-05 | 5.79E-05 | 0 | 0.00033 |
| Actn1 | 0.00025 | 0.00015 | 0.00065 | 3.81E-05 | 0.00035 | 0 | 0.00041 | 0.00011 | 0.00034 | 0 | 0.00222 | 0.00044 | 0.00038 | 0.00047 | 0.00052 | 0.00034 | 0.00048 | 0.00084 |
| Cpz | 0 | 0 | 0 | 0 | 0.00017 | 0.00063 | 0.0006 | 0 | 0.00034 | 3.37E-05 | 0 | 0 | 0.00048 | 0.00057 | 0.00019 | 0 | 0 | 0.00011 |
| Laptm4a | 0.00071 | 0.00027 | 0.00017 | 0 | 0.00052 | 0.00078 | 0.00059 | 0.00065 | 0.00034 | 0.00032 | 0.00035 | 0.00019 | 0.00043 | 0.00038 | 0 | 7.74E-05 | 0 | 0.00037 |

| | | | | | | | | | | | | | | | | | | |
|---|---|---|---|---|---|---|---|---|---|---|---|---|---|---|---|---|---|---|
| Ifitm2 | 0.00071 | 0.00035 | 0.00028 | 0.00099 | 0.00048 | 0.0012 | 0.00072 | 0.00083 | 0.00034 | 0.00074 | 0.00037 | 0.00018 | 0.00098 | 0.00105 | 0 | 0.00022 | 0.00035 | 0.00061 |
| Ppp2cb | 0.00015 | 0.00027 | 0.0004 | 0.00013 | 0.00022 | 0.00017 | 0.00018 | 0.00028 | 0.00034 | 0.00016 | 0.0003 | 0.00032 | 0 | 0.00017 | 0.00014 | 0.0002 | 0.00025 | 0.00019 |
| Srsf6 | 0.00038 | 0.00035 | 0.00038 | 0.00043 | 0.00019 | 0.00015 | 0.00018 | 0.0002 | 0.00034 | 0.00014 | 0.00019 | 0.00018 | 5.34E-05 | 0.00014 | 0.00127 | 0.0003 | 0.00047 | 0.00018 |
| Tmem9 | 6.19E-05 | 0.00023 | 0 | 0.00022 | 0.00031 | 0.00014 | 0.00028 | 8.42E-05 | 0.00034 | 0.00013 | 0.00014 | 1.42E-05 | 0.00016 | 0.00031 | 0 | 4.81E-05 | 0 | 0.00018 |
| Rnf187 | 7.08E-05 | 0.00023 | 0.00032 | 0.00082 | 0.00032 | 8.69E-05 | 0.00023 | 0.00028 | 0.00034 | 0.00022 | 0.00025 | 6.33E-05 | 0.00013 | 0.00029 | 0.00025 | 0.00026 | 0.00014 | 0.00036 |
| Lhfpl2 | 6.09E-06 | 0.00035 | 0.00033 | 6.27E-05 | 0.00034 | 0.00072 | 0.00042 | 0.00034 | 0.00034 | 5.65E-05 | 0 | 2.75E-06 | 0.00029 | 0.00014 | 0 | 0.00026 | 0.0004 | 0.00026 |
| Tns3 | 0 | 0.00013 | 0 | 0 | 0.00024 | 0.00045 | 0 | 0.00034 | 0.00034 | 0 | 0.00037 | 0.00039 | 0 | 0.00067 | 0.00015 | 7.61E-05 | 0 | 0.00042 |
| Gfpt2 | 0.00012 | 7.44E-05 | 0.00045 | 0 | 0.00033 | 0 | 0.00023 | 0.00031 | 0.00034 | 0.001 | 7.24E-05 | 0.00025 | 0.00074 | 9.75E-06 | 0.00013 | 0 | 0.00124 | 0.00025 |
| Eva1b | 8.96E-05 | 0.00028 | 3.84E-05 | 0.00045 | 0.00043 | 0.00046 | 0 | 0.00042 | 0.00034 | 2.02E-05 | 0.00026 | 0 | 6.44E-05 | 0.00056 | 0.00036 | 0 | 0.00021 | 0.00031 |
| Sptan1 | 0.00019 | 0.00029 | 0.00027 | 0.00011 | 0.00015 | 0.00013 | 0.0003 | 0.00038 | 0.00034 | 0.00045 | 0.00039 | 0.00035 | 0 | 0.00018 | 0.00025 | 7.88E-05 | 0.00057 | 0.0006 |
| Tcf25 | 9.44E-05 | 0.00025 | 0.00027 | 0.00077 | 0.00026 | 0.0002 | 0.00027 | 0.00048 | 0.00033 | 0.00039 | 0.00045 | 0.00067 | 0.00011 | 0.00029 | 0.00065 | 0.0001 | 0.00042 | 0.00037 |
| Twsg1 | 0 | 0.00033 | 8.69E-05 | 0.00022 | 0.00045 | 0.00029 | 0.00038 | 4.08E-05 | 0.00033 | 0.00054 | 0.00017 | 6.81E-05 | 0.00068 | 0.00031 | 0 | 0.0003 | 2.76E-05 | 0.00032 |
| Itpripl2 | 1.09E-05 | 0.00026 | 0.00045 | 0.00047 | 0.00027 | 0.0002 | 0.00038 | 0.00046 | 0.00033 | 0.00014 | 0.00037 | 0.00029 | 0.0004 | 0.00038 | 0.00049 | 0.00014 | 6.76E-05 | 0.0002 |
| Ssr1 | 0.00075 | 0 | 0.00058 | 0.00083 | 0.00019 | 0.00037 | 0.00103 | 0.0005 | 0.00033 | 0.00024 | 0.00028 | 0.00033 | 0.00038 | 0.00036 | 0.00023 | 0.00048 | 1.74E-05 | 0.00015 |
| Aldoa | 0.00026 | 0.00017 | 0.00032 | 0.00069 | 0.00016 | 0.00034 | 0.00016 | 0.00036 | 0.00033 | 0.00029 | 0.00038 | 0.0014 | 0.00037 | 0.00024 | 0 | 1.70E-05 | 0.00027 | 0 |
| Tle5 | 0.00018 | 0.00025 | 0 | 0.00025 | 0.00035 | 0 | 0.00013 | 0 | 0.00033 | 0.0004 | 0.00015 | 0.00015 | 0.00018 | 5.45E-05 | 0 | 9.91E-05 | 1.24E-05 | 0.00022 |
| Slc25a24 | 0.00025 | 0.00017 | 0.00019 | 2.34E-05 | 0.00016 | 0.00013 | 0.00018 | 1.62E-05 | 0.00033 | 0.00017 | 0.00028 | 0.0002 | 0 | 0.0001 | 0 | 0.00013 | 0.00014 | 1.00E-04 |
| Ptpa | 0.00021 | 0.00021 | 0.0002 | 0.00016 | 0.00017 | 9.95E-05 | 0 | 0.00011 | 0.00033 | 6.68E-05 | 0.0001 | 0.00015 | 0 | 3.50E-05 | 0.0001 | 0.00023 | 0 | 0.00014 |
| Mafb | 0 | 0.00047 | 0 | 4.01E-05 | 0.00033 | 0 | 0.00066 | 0.00098 | 0.00033 | 0 | 0 | 0 | 0 | 0 | 0.00035 | 1.53E-05 | 0.0006 | 0.00047 |
| Aamp | 0.00017 | 0.00031 | 0.00024 | 0.0006 | 0.00027 | 0.00014 | 0.00015 | 0.00025 | 0.00033 | 0.00016 | 0.00016 | 0.00026 | 0.00012 | 0.00024 | 0.00041 | 0 | 0.00026 | 0.00016 |
| Prelid1 | 0.00011 | 8.01E-05 | 0.00027 | 0.001 | 0.00015 | 0.00019 | 0.00033 | 0.00046 | 0.00033 | 0.00024 | 0.00031 | 0.0008 | 9.76E-05 | 0.00035 | 0.00011 | 0.00023 | 0.00018 | 0.00021 |
| Dusp7 | 0.00026 | 0.00025 | 0.00023 | 0.00021 | 0.00012 | 0.00017 | 0.0002 | 0 | 0.00033 | 0.00016 | 0 | 0 | 4.81E-05 | 0.00041 | 4.17E-06 | 3.91E-05 | 0 | 2.29E-05 |
| Ctdsp1 | 6.66E-05 | 0.00035 | 5.82E-05 | 0.00069 | 0.00033 | 0.00024 | 0.00012 | 0.00018 | 0.00033 | 0.0003 | 0.00021 | 0.00019 | 0.00015 | 0.00027 | 0.00019 | 0.00026 | 9.48E-05 | 0.00033 |
| Tnfrsf12a | 0 | 0 | 0.00075 | 0.00029 | 0.00021 | 0 | 0 | 0 | 0.00033 | 0.0001 | 0.00088 | 0.00014 | 0.00022 | 0.00033 | 0.00033 | 1.94E-05 | 0.00145 | 0.00019 |
| Cd44 | 0.00028 | 8.78E-05 | 0.00094 | 0 | 7.45E-05 | 0.00011 | 0.00018 | 0.00013 | 0.00033 | 0.00032 | 0.00061 | 0.00021 | 0.00032 | 0.00049 | 0.0002 | 0.00035 | 0.00058 | 0.00043 |
| Pabpc1 | 0.00051 | 0.00022 | 0.00035 | 0.00011 | 0.00013 | 0.00036 | 0.00043 | 0.0003 | 0.00033 | 6.17E-05 | 0 | 0.00049 | 0.0003 | 0.00014 | 0 | 0.00016 | 0.00031 | 2.62E-06 |
| Smco4 | 0.00013 | 9.65E-06 | 0.00012 | 4.17E-05 | 0.00017 | 2.89E-05 | 0.00015 | 7.57E-05 | 0.00033 | 0 | 0.00011 | 1.82E-05 | 3.54E-06 | 0 | 0 | 6.70E-05 | 4.86E-05 | 0 |
| Tcf12 | 0.0003 | 0.00034 | 9.61E-05 | 0 | 0.00031 | 0.0003 | 0.00015 | 0.00026 | 0.00033 | 0 | 6.60E-05 | 0.00011 | 0 | 0.00014 | 3.83E-05 | 0.00029 | 0.00036 | 0.00011 |
| Ada | 8.00E-05 | 0 | 0.0002 | 0.00011 | 1.29E-05 | 0 | 0.00024 | 0.00025 | 0.00033 | 0 | 0 | 0 | 0 | 0 | 5.96E-05 | 0.0001 | 0 | 0 |
| Yap1 | 0.00018 | 0.00027 | 0.00018 | 0.00017 | 0.00038 | 0.0001 | 0.00021 | 0.00011 | 0.00033 | 8.07E-05 | 0.00045 | 0.00015 | 0.00024 | 0.00016 | 0.00021 | 0.00012 | 0.00028 | 1.54E-05 |
| Eps8 | 0 | 3.23E-05 | 2.96E-07 | 0 | 0 | 6.00E-05 | 7.41E-05 | 3.82E-05 | 0.00033 | 0.00018 | 0 | 5.86E-05 | 0.00021 | 0 | 0 | 5.83E-05 | 6.07E-05 | 0.00023 |
| Rarres1 | 0 | 0 | 0 | 0 | 0 | 0.00014 | 0 | 0 | 0.00033 | 0 | 0 | 0 | 0 | 0.00021 | 0 | 1.26E-06 | 0 | 0.00012 |
| Tceal8 | 0.00039 | 0.00013 | 0.00036 | 0.00089 | 0.00035 | 0.00025 | 0.00068 | 0.0004 | 0.00033 | 0.00019 | 0.00057 | 0.00016 | 0.00021 | 0.00037 | 0.00023 | 0.00025 | 0.00023 | 0.00019 |

| | | | | | | | | | | | | | | | | | | |
|---|---|---|---|---|---|---|---|---|---|---|---|---|---|---|---|---|---|---|
| Fkbp1a | 0.00018 | 9.69E-05 | 0.00044 | 0.00065 | 0.00014 | 6.98E-05 | 0.00026 | 0.00035 | 0.00033 | 0.00033 | 0.0005 | 0.0002 | 0.00012 | 0.00024 | 0.00016 | 0.00015 | 1.42E-05 | 0.00025 |
| Ahcyl1 | 0.00028 | 0.00022 | 0.00032 | 0.0003 | 0.00027 | 0.00012 | 0.00012 | 0.00028 | 0.00033 | 0.00031 | 0.00015 | 0.00024 | 0.00021 | 0.00011 | 0.0003 | 0.00026 | 0.00043 | 0.0002 |
| Rhoc | 0.0003 | 0.00029 | 0.00047 | 0.00028 | 0.00037 | 0.00019 | 0.00041 | 0.00033 | 0.00033 | 0.00035 | 0.0006 | 0.00068 | 0.00022 | 0.00028 | 6.82E-05 | 0.0002 | 0.00019 | 0.0002 |
| Epn1 | 0.00019 | 0.00028 | 0.00018 | 0.0005 | 0.00026 | 0.00016 | 0.0002 | 0.0002 | 0.00033 | 0.00024 | 0.0002 | 0.00016 | 4.76E-05 | 0.00024 | 0.00037 | 9.29E-05 | 0.00024 | 0.00016 |
| Ifitm3 | 0.00103 | 0.00024 | 0.00038 | 0.00016 | 0.00015 | 0.00061 | 4.08E-05 | 0.00056 | 0.00033 | 0.00073 | 0.00029 | 6.00E-05 | 0.00097 | 0.00053 | 0 | 0.00011 | 0.00026 | 0.00028 |
| Prdx2 | 0.0004 | 0.00016 | 0.00015 | 0.00068 | 0.00021 | 0.00017 | 0.00034 | 0.00019 | 0.00033 | 0.00016 | 0.00025 | 0.00015 | 0.00022 | 0.00024 | 0 | 0.00014 | 2.02E-05 | 0.00022 |
| Wnt2b | 0 | 0 | 0.00011 | 0 | 0.00121 | 0.0002 | 0.00038 | 0 | 0.00033 | 0 | 0.0004 | 7.71E-05 | 0 | 0 | 0 | 0 | 0 | 0 |
| Rab34 | 0.00012 | 0.00024 | 0.00016 | 0.00039 | 0.00026 | 0.00023 | 0.0002 | 0 | 0.00033 | 9.42E-05 | 0.00011 | 0.00021 | 0.00027 | 0.00033 | 0.00014 | 6.82E-06 | 0 | 0.00027 |
| Csnk1g2 | 0.00018 | 0.00024 | 0.00026 | 0.0002 | 0.00044 | 0.00033 | 0.00019 | 0.00067 | 0.00033 | 0.00018 | 0.00026 | 0.00017 | 8.21E-05 | 0.00039 | 0.00028 | 7.50E-05 | 0.00013 | 0.00016 |
| Rufy1 | 2.83E-05 | 9.36E-05 | 9.96E-05 | 5.93E-05 | 0.00016 | 2.79E-05 | 8.95E-05 | 0 | 0.00033 | 7.54E-05 | 0.00017 | 0 | 5.29E-05 | 0.00012 | 5.24E-05 | 4.41E-05 | 0 | 8.67E-05 |
| Inf2 | 0 | 0.00023 | 8.48E-06 | 0.00021 | 0.00011 | 0.00018 | 4.51E-05 | 0 | 0.00033 | 0 | 0.00021 | 0.00043 | 2.24E-05 | 0.00026 | 0.00082 | 0.00012 | 6.09E-05 | 0.00028 |
| Scarb1 | 5.90E-05 | 5.57E-05 | 0 | 1.42E-05 | 0.00017 | 0.00022 | 0 | 0 | 0.00032 | 0 | 0.0002 | 0.00013 | 0 | 0.0003 | 0 | 0.00012 | 0 | 1.45E-05 |
| Zcchc24 | 3.50E-06 | 0.00035 | 0.00024 | 0.00069 | 0.00059 | 0.00024 | 0.00038 | 0.00079 | 0.00032 | 0.0003 | 0 | 3.91E-06 | 0.00031 | 0.00051 | 0.00059 | 0.00023 | 0.00039 | 0.00024 |
| Cebpd | 2.18E-05 | 0.0003 | 0 | 0.00056 | 0 | 0.00027 | 0 | 3.03E-05 | 0.00032 | 0.00089 | 8.42E-05 | 0.0002 | 0.00072 | 0.00028 | 0.00032 | 0 | 0.00071 | 0.00052 |
| Gnb1 | 0.00022 | 0.0003 | 0.00043 | 0.00074 | 0.00028 | 0.00027 | 0.00019 | 0.00037 | 0.00032 | 0.00036 | 0.0005 | 0.00049 | 5.54E-05 | 0.00032 | 5.82E-05 | 0.00026 | 0.00028 | 0.00038 |
| Mmp2 | 0 | 0.0007 | 0.0006 | 3.24E-05 | 0.00124 | 0.00173 | 0.00217 | 0.00063 | 0.00032 | 0.00124 | 0 | 0 | 0.00258 | 0.00097 | 0.00066 | 0 | 0.00016 | 0.00093 |
| Lifr | 0.00015 | 4.96E-05 | 1.24E-05 | 0 | 0.00017 | 0.00012 | 4.37E-05 | 0 | 0.00032 | 0 | 4.91E-06 | 0.00011 | 0.00067 | 0.0002 | 7.90E-05 | 0.00014 | 8.52E-05 | 0.0001 |
| Lasp1 | 0.00024 | 0.00024 | 0.00018 | 0.00046 | 6.71E-05 | 3.41E-05 | 3.05E-05 | 0.0002 | 0.00032 | 0.0003 | 0.00051 | 0.00022 | 2.64E-05 | 0.00021 | 0.0002 | 0.00022 | 0.00026 | 0.00018 |
| Khsrp | 0.00027 | 0.00029 | 0.00028 | 0.00033 | 0.00015 | 0.00011 | 4.10E-05 | 0.00015 | 0.00032 | 8.05E-05 | 0.00011 | 0.00029 | 1.37E-05 | 0.00016 | 0.00037 | 0.00039 | 0.00013 | 0.00013 |
| Actr1b | 0.00012 | 0.00018 | 0.00051 | 0.00032 | 0.00045 | 0.00021 | 0.0004 | 0.0004 | 0.00032 | 0.00016 | 0.00037 | 0.00015 | 0.00017 | 0.00037 | 0.00015 | 0.00017 | 0 | 0.00022 |
| Rcn1 | 0.00046 | 0 | 0.00046 | 0.00088 | 0.0006 | 0.00075 | 0.00167 | 0.00103 | 0.00032 | 0.00027 | 0.00041 | 0.00025 | 0.00068 | 0.00054 | 0.00045 | 0.00019 | 0.00023 | 0.00031 |
| Enah | 0.00017 | 3.72E-05 | 0.00047 | 8.87E-05 | 0.0002 | 0.00031 | 0.00034 | 0.00026 | 0.00032 | 0.00018 | 0.00046 | 0.00011 | 0 | 0.00025 | 0.00033 | 0.00011 | 0 | 0 |
| Ap1s1 | 0.00043 | 5.63E-05 | 0.00028 | 0.00032 | 5.50E-05 | 0.00012 | 9.68E-05 | 0.00012 | 0.00032 | 0.00014 | 0.00027 | 0.00018 | 0.00025 | 0.00013 | 0 | 0.00012 | 0.00015 | 6.28E-05 |
| B4galt5 | 0 | 0.00012 | 0.00079 | 0 | 8.68E-05 | 0.00032 | 0 | 0.00054 | 0.00032 | 0.00035 | 5.23E-05 | 0.0005 | 0.0009 | 0.00019 | 0 | 8.91E-05 | 0.00017 | 0.00019 |
| Arhgdia | 0.00052 | 0.00032 | 0.00054 | 0.00044 | 0.00024 | 0.00022 | 0.00031 | 0.0003 | 0.00032 | 0.00032 | 0.00047 | 0.00052 | 0 | 0.00021 | 0.00037 | 0.00037 | 2.79E-06 | 0.00029 |
| Plxna1 | 0.00011 | 0.00026 | 0.00024 | 4.76E-05 | 0.00023 | 0.00029 | 0.00013 | 7.71E-05 | 0.00032 | 0.00028 | 0.00034 | 0 | 0.00012 | 0.00022 | 0.00018 | 7.97E-05 | 0 | 0.00017 |
| Trim8 | 0 | 0.00029 | 0.00054 | 0.00027 | 0.00016 | 0 | 7.33E-05 | 0.00052 | 0.00032 | 0.00029 | 8.56E-05 | 0.0002 | 8.24E-05 | 0 | 0.00019 | 8.12E-05 | 0.00022 | 0.00019 |
| Lmna | 0.00038 | 0.00043 | 0.00075 | 0.00098 | 0.00014 | 0.00051 | 0.00089 | 0 | 0.00032 | 0.00051 | 0.00044 | 0.00036 | 0 | 0.0005 | 0.00104 | 0.00053 | 0.00179 | 0.00108 |
| Fez1 | 4.64E-07 | 5.89E-05 | 4.67E-05 | 0 | 0.00017 | 0.00016 | 0 | 0 | 0.00032 | 0.00027 | 6.56E-05 | 6.28E-06 | 3.35E-05 | 0 | 0 | 4.08E-05 | 0 | 0 |
| Asah1 | 0.00034 | 0.00022 | 0 | 0 | 0.00023 | 0.00041 | 0 | 5.83E-05 | 0.00032 | 6.00E-05 | 0.00012 | 0.00012 | 7.22E-05 | 0.00011 | 0 | 0 | 8.03E-06 | 0.0002 |
| Ano6 | 9.66E-05 | 0.0002 | 0.0002 | 0 | 0.00022 | 0.00016 | 0.00018 | 8.27E-05 | 0.00032 | 0.00032 | 0.00037 | 0.0002 | 0.00016 | 0.00011 | 2.33E-05 | 0.00016 | 0.0002 | 0.00027 |
| Adamts12 | 0 | 0 | 0 | 0.00036 | 0.00153 | 0.00099 | 0.00074 | 0.00089 | 0.00032 | 0.00014 | 0.00064 | 5.41E-06 | 0.00044 | 0.00097 | 0.00053 | 0 | 0 | 0.00052 |
| Sat1 | 0 | 0.00033 | 0.00015 | 0 | 0.00021 | 0.00033 | 4.90E-05 | 0.00028 | 0.00032 | 0.00021 | 0.0002 | 0.00048 | 5.55E-06 | 0.00044 | 0.00026 | 0 | 0.0032 | 4.67E-05 |

| | | | | | | | | | | | | | | | | | | |
|---|---|---|---|---|---|---|---|---|---|---|---|---|---|---|---|---|---|---|
| 2-Sep | 0.00061 | 0.0002 | 0.0003 | 0.0004 | 0.0002 | 0.00044 | 0.00043 | 0.0004 | 0.00032 | 0.0003 | 0.00039 | 0.00027 | 0.00013 | 0.00022 | 0.00026 | 0.00022 | 0.00022 | 0.00021 |
| Glipr2 | 0.00036 | 0 | 0.00041 | 0.00018 | 0 | 0.00017 | 0.00029 | 0.00014 | 0.00032 | 0.00052 | 0.00048 | 0.00042 | 0.00039 | 0 | 0 | 0.00013 | 0 | 0 |
| Tapbp | 0 | 0.00019 | 0 | 0.00037 | 0.00035 | 0 | 6.66E-05 | 0.00029 | 0.00032 | 0.00037 | 0.00018 | 4.64E-05 | 0.00069 | 0.0003 | 0.00025 | 0 | 0.00014 | 0.00071 |
| Crtap | 0.00023 | 0 | 0.00025 | 0.0006 | 0.00054 | 0.00094 | 0.00132 | 0.00096 | 0.00032 | 0.00043 | 0.0005 | 0.00041 | 0.00121 | 0.00081 | 0 | 0.00021 | 0 | 0.00075 |
| Txnip | 0.00016 | 0.00065 | 0 | 0.0004 | 0.00029 | 0.00049 | 0.00017 | 0.0002 | 0.00032 | 0.00061 | 0.00021 | 0 | 0.00016 | 0.0002 | 9.60E-05 | 9.02E-05 | 0 | 0.0006 |
| Surf4 | 0.0008 | 6.66E-05 | 0.00053 | 0.00043 | 0.00041 | 0.0006 | 0.0011 | 0.00053 | 0.00032 | 0.00036 | 0.00046 | 0.00039 | 0.00063 | 0.00052 | 0 | 0.00012 | 0 | 0.00022 |
| Peak1 | 6.86E-05 | 0.00017 | 6.01E-05 | 0 | 0.00014 | 0.00034 | 0 | 0.00023 | 0.00031 | 0.00015 | 1.21E-05 | 0.00082 | 1.40E-05 | 6.55E-05 | 0 | 0 | 0.00024 | 0.00026 |
| Eif4a1 | 0.00048 | 0.00019 | 0.00044 | 9.35E-05 | 2.43E-05 | 5.90E-05 | 0 | 0 | 0.00031 | 0 | 0.00019 | 0.0002 | 5.76E-05 | 5.40E-05 | 0.00023 | 0.00023 | 0.00058 | 0 |
| Lats2 | 8.97E-05 | 0.00025 | 3.53E-05 | 4.56E-05 | 0.00028 | 0.00022 | 1.75E-05 | 0 | 0.00031 | 0.00018 | 0.00055 | 8.73E-05 | 0.00044 | 0.00052 | 0.00026 | 2.67E-05 | 2.97E-05 | 0.00011 |
| Jak1 | 0.00039 | 0.00027 | 0.00024 | 7.71E-05 | 0.00012 | 0.00041 | 0.00021 | 0.00052 | 0.00031 | 0.00055 | 1.07E-05 | 0.00051 | 0.00037 | 0.00011 | 0 | 0.0001 | 0.00044 | 0.0002 |
| Add3 | 0.00011 | 0.00013 | 0 | 0 | 1.01E-06 | 0.00011 | 0 | 0 | 0.00031 | 0.00068 | 9.42E-06 | 0.00023 | 0 | 0 | 0 | 0.00015 | 0 | 0.0006 |
| Eif5a | 0.00055 | 0.00017 | 0.00076 | 0.00065 | 2.57E-05 | 0.00011 | 0.0002 | 9.78E-05 | 0.00031 | 4.41E-05 | 0.00022 | 0.00017 | 8.04E-05 | 5.44E-05 | 0 | 0.00028 | 0.00021 | 3.90E-05 |
| Mvk | 1.63E-05 | 6.78E-05 | 0 | 0.00024 | 6.82E-05 | 0 | 0 | 3.53E-05 | 0.00031 | 5.83E-05 | 0.00012 | 0 | 0 | 8.44E-05 | 1.44E-05 | 8.32E-05 | 0 | 9.81E-05 |
| Efemp1 | 2.17E-05 | 0 | 0 | 0 | 0 | 0.00012 | 0 | 0 | 0.00031 | 0.00141 | 0 | 0 | 0.00042 | 0 | 0 | 0 | 0 | 0.00016 |
| Ddx3x | 0.00046 | 0.00042 | 0.0004 | 0.00037 | 0.00028 | 0.00033 | 0.00019 | 0.00011 | 0.00031 | 0.00013 | 0.00022 | 0.00014 | 0.00032 | 0.0003 | 0.00035 | 0.00024 | 0.00115 | 0.00017 |
| Mcl1 | 5.21E-05 | 0.00038 | 0.00033 | 0.00043 | 0.00026 | 0.00018 | 9.90E-05 | 0.00034 | 0.00031 | 0.00031 | 0.00021 | 0.00052 | 0.00032 | 0.00043 | 0.00074 | 0.0002 | 0.00206 | 0.00031 |
| Cemip2 | 3.74E-05 | 0.00014 | 0 | 0 | 0.0005 | 0 | 0 | 0.00051 | 0.00031 | 0 | 0.00043 | 0.00016 | 0 | 0.00013 | 0.00025 | 6.04E-05 | 0.00027 | 0 |
| Dynlrb1 | 0.00052 | 0.00027 | 0.00016 | 0.00046 | 0.00056 | 0.0005 | 0.00047 | 0.00017 | 0.00031 | 0.00031 | 0.00068 | 0.00029 | 0.00066 | 0.00041 | 0 | 0.00015 | 0 | 0.00015 |
| Col15a1 | 0 | 0 | 0.00043 | 5.78E-06 | 0.00012 | 0.0015 | 0.00057 | 0.00358 | 0.00031 | 0.0002 | 0.0003 | 0.00067 | 0 | 0 | 0.00056 | 0.00018 | 0 | 0.00116 |
| Amotl2 | 0 | 0 | 6.45E-05 | 0.0003 | 0.00025 | 0 | 0 | 0 | 0.00031 | 0.00048 | 0.00098 | 0.00016 | 0.00099 | 0.00037 | 0.00036 | 3.55E-05 | 0.00013 | 0.00017 |
| Flot2 | 0.00013 | 0.00021 | 5.70E-05 | 9.44E-05 | 9.29E-05 | 0.00016 | 4.58E-05 | 7.50E-05 | 0.00031 | 0.00031 | 8.69E-05 | 0.00017 | 0.00025 | 0.00012 | 0 | 0.00017 | 0 | 0.00035 |
| R3hdm4 | 0.00012 | 0.0002 | 0.00015 | 0.00039 | 0.00027 | 0.00019 | 0.00015 | 0.0004 | 0.00031 | 0.00019 | 0.00014 | 0.00011 | 7.49E-05 | 0.0002 | 0.00012 | 7.22E-05 | 2.95E-05 | 0.00024 |
| Tspan31 | 0.00037 | 0.0001 | 0.00018 | 0.00024 | 0.00032 | 0.00025 | 0.00029 | 0.00046 | 0.00031 | 0.00019 | 0.00018 | 0.00025 | 0.00019 | 0.00025 | 0 | 2.58E-05 | 2.65E-05 | 0.00022 |
| Scarf2 | 0 | 0.00013 | 5.48E-05 | 0 | 0.0005 | 9.81E-05 | 0.00025 | 5.90E-05 | 0.00031 | 0 | 0.00025 | 0 | 0.00017 | 0.00056 | 0.00036 | 3.53E-05 | 0 | 0.00031 |
| Atl3 | 0.00031 | 0.00034 | 0.00015 | 0 | 0.00032 | 0.00039 | 0.00023 | 0.00013 | 0.00031 | 0.00042 | 0.00046 | 0.00028 | 0.00028 | 0.0003 | 2.48E-05 | 0.00026 | 2.48E-05 | 0.00044 |
| Capns1 | 0.00056 | 0.00018 | 0.00016 | 0 | 0.0002 | 0.00032 | 0.00023 | 0.00024 | 0.00031 | 0.0003 | 0.00013 | 0.00021 | 6.45E-05 | 0.00016 | 0 | 5.93E-05 | 2.58E-05 | 0.00027 |
| P3h3 | 4.04E-05 | 0 | 1.50E-05 | 0.00048 | 0.00035 | 0.00052 | 0.00098 | 0.00081 | 0.00031 | 0.00034 | 0.00035 | 0.00042 | 0.00055 | 0.00057 | 0.00055 | 0.00015 | 6.17E-05 | 0.00049 |
| Sdhc | 0.0005 | 0.00017 | 0.00025 | 0.00044 | 0.00014 | 0.00022 | 0.00012 | 0.00017 | 0.00031 | 0.00011 | 0.00017 | 0.00018 | 0.0002 | 0.00019 | 0 | 0.00015 | 0 | 0.00019 |
| Atp6v0c | 0.00025 | 0.00035 | 0.00035 | 0.00019 | 0.00026 | 0.00017 | 0.0001 | 0.00018 | 0.00031 | 8.41E-05 | 0.00027 | 0.00021 | 0.00014 | 0.00024 | 0 | 7.46E-05 | 0.00043 | 0.00013 |
| Bcl3 | 0.00015 | 0.00024 | 0.00021 | 0.00031 | 0.00022 | 0.00011 | 0.00011 | 0.00031 | 0.00031 | 0.00022 | 0 | 4.69E-05 | 0.0005 | 0.00027 | 0.00019 | 0.00022 | 0.0004 | 0 |
| Ccnd1 | 0 | 0 | 5.82E-05 | 7.13E-06 | 0 | 0 | 0 | 0 | 0.00031 | 0.0001 | 0 | 2.54E-05 | 0 | 0 | 1.51E-05 | 4.23E-05 | 5.93E-06 | 0.00018 |
| Gpr153 | 0 | 1.73E-05 | 0.00029 | 0.00084 | 0.0007 | 0.00075 | 0.00118 | 0.00076 | 0.00031 | 0.00047 | 0.00014 | 0.00021 | 0.00065 | 0.00074 | 0.00049 | 0 | 0.00034 | 0.00017 |
| Ier5l | 0 | 0.00024 | 7.15E-05 | 0.00037 | 0.00022 | 1.87E-05 | 0 | 0.00035 | 0.00031 | 0 | 0.00039 | 2.09E-05 | 0.0003 | 0.00018 | 0.0004 | 0.00017 | 0.00017 | 0.00015 |

| | | | | | | | | | | | | | | | | | | |
|---|---|---|---|---|---|---|---|---|---|---|---|---|---|---|---|---|---|---|
| Gnb5 | 3.02E-05 | 6.80E-05 | 2.58E-05 | 7.81E-05 | 0.00015 | 0.00013 | 0.00035 | 0.00023 | 0.00031 | 0.00015 | 0.00013 | 4.01E-05 | 0.00017 | 0 | 0 | 0 | 0 | 0 |
| Gns | 0.00012 | 0.00032 | 0.00016 | 0.00018 | 0.00026 | 0.00044 | 0.00074 | 5.96E-05 | 0.00031 | 0.00048 | 4.09E-05 | 5.67E-05 | 0.00114 | 0.00043 | 0 | 0 | 0 | 0.00031 |
| Ap2a2 | 0.00018 | 0.00018 | 9.84E-05 | 0.00046 | 0.00012 | 0 | 0.00021 | 0.00022 | 0.00031 | 0.00048 | 0.00029 | 0.00033 | 0.00028 | 0 | 0.00017 | 0.00018 | 0.00013 | 0.00037 |
| Gpat4 | 0.00016 | 0.00024 | 0.00024 | 0.00032 | 0.00019 | 0.00011 | 0.00025 | 0.00015 | 0.00031 | 7.04E-05 | 0.00012 | 6.18E-05 | 3.80E-05 | 0.00019 | 9.83E-05 | 0.00019 | 0 | 7.75E-05 |
| Dpysl2 | 0.0002 | 0.00019 | 1.72E-05 | 0 | 0 | 0.00016 | 7.07E-05 | 0.00038 | 0.00031 | 0.00056 | 1.54E-05 | 0.00077 | 0 | 0.00024 | 0 | 0.00037 | 5.77E-05 | 0.00052 |
| Sdha | 0.00029 | 0.00031 | 0.00031 | 0.0004 | 0.00019 | 0.00019 | 0.00022 | 0.00023 | 0.00031 | 0.00021 | 0.00021 | 8.97E-05 | 0.00016 | 0.0002 | 9.23E-05 | 0.00036 | 3.89E-05 | 0.00024 |
| Adpgk | 0.00017 | 0.00012 | 0.00026 | 6.12E-05 | 0.00026 | 0.00017 | 6.38E-05 | 0.00026 | 0.00031 | 3.21E-06 | 0.00014 | 6.99E-05 | 0.00015 | 0 | 3.02E-05 | 0.00018 | 8.09E-05 | 1.34E-05 |
| Aldh2 | 4.84E-05 | 0.00028 | 5.40E-05 | 0.00066 | 0.00032 | 1.31E-05 | 0.00023 | 0.00028 | 0.00031 | 0.00061 | 0.00016 | 0.00015 | 0.00048 | 0.00036 | 0 | 5.52E-05 | 0.00026 | 0.00039 |
| Lman2 | 0.00032 | 0.00017 | 0.00045 | 0.0003 | 0.00031 | 0.00029 | 0.00025 | 0.00025 | 0.00031 | 0.00024 | 0.00028 | 0.00031 | 0.00039 | 0.00023 | 0 | 0.00018 | 0 | 0.00023 |
| Atp5d | 0.00029 | 0.00025 | 0.00026 | 0.00058 | 0.00015 | 0.00018 | 0.00016 | 0.00019 | 0.00031 | 0.00015 | 0.00025 | 0.00029 | 2.91E-05 | 0.00018 | 9.98E-05 | 0.00017 | 0.00026 | 0.0002 |
| Prrc2a | 0.00016 | 0.00034 | 0.00033 | 0.00041 | 0.00023 | 9.73E-05 | 0 | 0.00011 | 0.00031 | 0.00016 | 0.00021 | 0.00018 | 5.38E-05 | 6.30E-05 | 0.00051 | 0.00044 | 0.00044 | 0.00021 |
| Apcdd1 | 0 | 0.00014 | 0 | 0 | 0.00041 | 0.00035 | 0 | 0 | 0.00031 | 8.58E-05 | 8.06E-05 | 0 | 0 | 0.0004 | 0 | 4.01E-05 | 0 | 0 |
| Eif1a | 0.00022 | 0.00024 | 0.00034 | 0.00029 | 7.53E-05 | 5.62E-05 | 0.0001 | 2.17E-05 | 0.00031 | 7.23E-05 | 0.00013 | 0.00029 | 7.94E-05 | 0.00017 | 9.30E-05 | 8.30E-05 | 9.86E-05 | 0.0001 |
| Limd2 | 0.0002 | 5.34E-05 | 6.00E-05 | 0 | 0.0001 | 2.49E-05 | 0 | 0.00027 | 0.00031 | 0 | 6.87E-05 | 0 | 0.00011 | 0.00042 | 0 | 9.02E-05 | 0 | 4.09E-06 |
| Hnrnpm | 0.0005 | 0.00027 | 0.0004 | 0.00049 | 7.90E-05 | 5.15E-05 | 0.00012 | 6.89E-05 | 0.00031 | 7.59E-05 | 9.32E-05 | 8.67E-05 | 8.45E-05 | 0.00011 | 0.00064 | 0.00038 | 0.00019 | 7.32E-05 |
| Oaz1 | 0.00018 | 0.00015 | 0.00016 | 0.00058 | 0.00013 | 5.54E-05 | 0.00011 | 0.00013 | 0.00031 | 0.00013 | 0.00019 | 8.62E-05 | 5.29E-05 | 0.00013 | 0.00024 | 0.00019 | 0.00034 | 0.00016 |
| Prdx6 | 0.00013 | 2.43E-05 | 0.0003 | 9.37E-05 | 0.00015 | 0 | 0.00027 | 0 | 0.0003 | 4.10E-05 | 0 | 0.00035 | 0.00073 | 4.35E-05 | 0 | 0 | 0.00015 | 0 |
| Tsen34 | 0.00013 | 0.00014 | 0.00017 | 0.00041 | 0.00017 | 4.04E-05 | 9.65E-05 | 8.87E-05 | 0.0003 | 0.00023 | 0.00015 | 7.89E-05 | 0.00019 | 0.00016 | 0.00011 | 0.00048 | 4.71E-05 | 0.00029 |
| Ndufa3 | 0.0003 | 0.00012 | 0.00012 | 0.00078 | 0.00034 | 0.00029 | 0.0003 | 0.0003 | 0.0003 | 0.00023 | 0.00036 | 5.65E-05 | 0.00034 | 0.00033 | 0 | 0.00027 | 4.15E-05 | 0.00023 |
| Fnbp1 | 0.00018 | 0.0003 | 0 | 3.18E-06 | 0.00047 | 0.00034 | 3.91E-05 | 0.00032 | 0.0003 | 0 | 0.00028 | 0 | 0 | 0.0003 | 0.00062 | 0.00012 | 0.00037 | 0.00017 |
| Copa | 0.00048 | 0.00025 | 0.00042 | 0.00063 | 0.00034 | 0.00042 | 0.00073 | 0.00049 | 0.0003 | 0.00036 | 0.00039 | 0.00034 | 0.00034 | 0.00038 | 0.00042 | 0.00013 | 0.00011 | 0.00036 |
| Camk2d | 0.00014 | 0.00019 | 0 | 0 | 0.00012 | 0.00013 | 0.00014 | 2.83E-06 | 0.0003 | 0.00026 | 0.00011 | 0.00012 | 4.10E-05 | 0 | 0.00037 | 6.68E-05 | 0.00035 | 0.00021 |
| Slc39a14 | 0 | 0 | 0.00047 | 0.00061 | 0.00081 | 0.00062 | 0.00104 | 0.00048 | 0.0003 | 0.0004 | 0 | 0.00074 | 0.00213 | 0.00082 | 0.00082 | 0 | 6.54E-05 | 0.00018 |
| Esr1 | 1.21E-05 | 0 | 0 | 2.51E-06 | 4.62E-05 | 0 | 0.00016 | 4.71E-05 | 0.0003 | 0.00035 | 8.70E-05 | 0.00021 | 3.20E-05 | 0 | 5.05E-05 | 5.97E-05 | 2.51E-05 | 3.87E-06 |
| Sh3d19 | 0 | 0.00018 | 5.83E-05 | 0.00023 | 0.00019 | 0.00019 | 0.00061 | 3.22E-05 | 0.0003 | 0.00039 | 0.00012 | 2.88E-05 | 0.00023 | 0.00019 | 4.83E-05 | 8.47E-05 | 0.00028 | 0.00024 |
| Shc4 | 0 | 1.10E-05 | 4.39E-05 | 0 | 0.00041 | 0.00015 | 0 | 0.00025 | 0.0003 | 0 | 3.96E-05 | 0.00013 | 0.00034 | 0.0002 | 0 | 0 | 0 | 2.10E-05 |
| Eef1b2 | 0.00064 | 0.00029 | 0.00034 | 0.00057 | 0.0002 | 0.00063 | 0.00073 | 0.00051 | 0.0003 | 0.00035 | 9.85E-05 | 0.0005 | 0.00045 | 0.00044 | 0.00014 | 0.00026 | 0.00052 | 0.00019 |
| Wsb1 | 0.00013 | 0.00021 | 0.00016 | 5.88E-05 | 0.00011 | 0 | 3.78E-05 | 0.00013 | 0.0003 | 0.00016 | 0.00034 | 0.00061 | 0.00011 | 0.00032 | 0.00039 | 0.00026 | 0.00128 | 0.00021 |
| Nptn | 0.00035 | 0.00022 | 0.00023 | 4.99E-05 | 0.0002 | 0.00062 | 0.00024 | 0.00054 | 0.0003 | 0.00042 | 0.00037 | 0.0006 | 0.00014 | 0.00042 | 1.23E-05 | 0.00024 | 0.0002 | 0.00016 |
| S1pr2 | 4.74E-05 | 0.00018 | 0.00032 | 0.00011 | 0.00024 | 0.00022 | 9.80E-05 | 0.00041 | 0.0003 | 0.00013 | 0.00013 | 5.01E-05 | 0.00018 | 0.00018 | 0 | 4.17E-06 | 0 | 0.00022 |
| Fstl3 | 0 | 0 | 0.00021 | 0.00016 | 0.00026 | 0.00019 | 0 | 0.00012 | 0.0003 | 0 | 0.00049 | 0.00087 | 0.00029 | 0.00022 | 4.81E-05 | 0 | 0 | 0.00019 |
| Flrt2 | 0.00017 | 0.00019 | 7.71E-05 | 0 | 0 | 0 | 0.00042 | 0 | 0.0003 | 0.00082 | 0 | 0 | 0.0006 | 0 | 0 | 0.00015 | 0.0002 | 0.00036 |
| Tmem259 | 0.0001 | 0.00034 | 0.00027 | 0.00035 | 0.00028 | 0.00026 | 0.00017 | 0.00024 | 0.0003 | 0.00021 | 0.00027 | 0.00014 | 0.00014 | 0.00032 | 0.00059 | 0.00017 | 0.00029 | 0.00027 |

| | | | | | | | | | | | | | | | | | | |
|---|---|---|---|---|---|---|---|---|---|---|---|---|---|---|---|---|---|---|
| Cdk16 | 0.0002 | 0.00022 | 0.00032 | 0.00033 | 0.00025 | 0.00015 | 0.00071 | 0.00022 | 0.0003 | 0.00027 | 0.0002 | 0.00021 | 0.00017 | 0.0002 | 0.00016 | 0.00024 | 0 | 0.00023 |
| Adamts4 | 1.94E-05 | 8.79E-05 | 0.00069 | 9.76E-05 | 0.00095 | 0.00061 | 0.00101 | 0.00141 | 0.0003 | 9.34E-05 | 2.31E-05 | 0.00026 | 0.00035 | 0.00111 | 0.00015 | 0.00022 | 0.00018 | 0 |
| Map4 | 0.00017 | 0.00027 | 0.00038 | 0.00044 | 0.00018 | 0.00013 | 0.00031 | 0.00032 | 0.0003 | 0.00031 | 0.00047 | 0.00032 | 6.93E-05 | 0.00021 | 0 | 0.00036 | 0.00029 | 0.0004 |
| Trim35 | 0.00014 | 0.00029 | 0.00028 | 0.0004 | 0.00027 | 0.00013 | 0.00021 | 0.00033 | 0.0003 | 0.00032 | 0.00034 | 0.00024 | 2.48E-05 | 0.00025 | 0.00036 | 0.00015 | 0 | 0.00027 |
| Szrd1 | 0.00022 | 0.00024 | 0.00026 | 0.00033 | 0.00043 | 0.00019 | 0.00024 | 0.00016 | 0.0003 | 0.00013 | 0.00017 | 0.00033 | 0.00012 | 0.0004 | 0.00026 | 0.00022 | 6.46E-05 | 0.00018 |
| Col23a1 | 0 | 0 | 0 | 0.00012 | 0.00065 | 0.00024 | 0 | 0.00109 | 0.0003 | 0 | 2.09E-05 | 0 | 0.00025 | 0.00179 | 0.00074 | 4.38E-05 | 0 | 0.00011 |
| Nsdhl | 7.53E-05 | 2.44E-05 | 0 | 0 | 5.51E-05 | 0 | 0 | 1.86E-05 | 0.0003 | 2.49E-05 | 8.65E-05 | 0 | 0 | 6.51E-05 | 0 | 3.82E-06 | 0 | 6.50E-05 |
| Il16 | 0 | 5.64E-06 | 0 | 0 | 0 | 5.70E-05 | 0 | 0 | 0.0003 | 6.42E-05 | 0 | 0 | 0.00025 | 0 | 0 | 0 | 0 | 5.53E-05 |
| Lgals3bp | 9.90E-05 | 9.62E-05 | 0 | 0.00037 | 0.00018 | 0.00026 | 0.00039 | 0.00083 | 0.0003 | 0.00098 | 0.00013 | 0.00022 | 0.00138 | 0.00072 | 8.66E-05 | 0.00011 | 4.77E-05 | 0.00095 |
| Klf3 | 7.36E-05 | 0.00031 | 0.00016 | 6.59E-06 | 0.00017 | 0.00012 | 8.17E-05 | 5.94E-05 | 0.0003 | 0.00042 | 0.0002 | 0.0001 | 0 | 0.00021 | 5.07E-05 | 7.30E-05 | 5.53E-06 | 0.00016 |
| Ndufs2 | 0.00026 | 0.00031 | 0.00023 | 0.00059 | 0.00023 | 0.00019 | 0.00025 | 0.00026 | 0.0003 | 0.00014 | 0.00016 | 7.47E-05 | 0.00011 | 0.00027 | 0.00018 | 0.00025 | 0.00012 | 0.00019 |
| Fth1 | 0.00041 | 0.00057 | 0.00023 | 0.00086 | 0.00033 | 0.00046 | 0.00014 | 0.00038 | 0.0003 | 0.00108 | 2.45E-05 | 0.00031 | 0.00164 | 0.00046 | 0.00105 | 0.00021 | 0.00099 | 0.00124 |
| Hivep3 | 0 | 0 | 0.00011 | 0 | 0.00016 | 0.00021 | 0 | 0.00029 | 0.0003 | 2.11E-05 | 2.30E-05 | 7.64E-05 | 0.00041 | 6.29E-05 | 0.00028 | 0 | 0 | 2.11E-05 |
| Rab4b | 0.00021 | 0.00014 | 0.00012 | 0.00031 | 0.00022 | 0.00015 | 0.00038 | 0.00029 | 0.0003 | 0.00015 | 0.00019 | 4.82E-05 | 0.00025 | 0.00023 | 0.00032 | 3.49E-05 | 4.22E-05 | 8.56E-05 |
| Psmb8 | 0.00017 | 4.98E-05 | 1.44E-05 | 9.11E-05 | 0 | 0 | 0 | 1.25E-05 | 0.0003 | 0.00014 | 1.17E-06 | 3.41E-05 | 0.00018 | 0.00015 | 0 | 0.00024 | 0.00012 | 0.00021 |
| Ctnnd1 | 0.00022 | 0.00026 | 5.49E-05 | 0.0003 | 0.00027 | 0.00041 | 8.23E-05 | 0.00053 | 0.0003 | 0.00016 | 0.00062 | 0.00036 | 0.00011 | 0.00033 | 0.00015 | 0.00022 | 4.68E-05 | 0.00026 |
| Rab1a | 0.00067 | 0.00026 | 0.00046 | 0.0004 | 0.00025 | 0.00044 | 0.00062 | 0.00025 | 0.0003 | 0.00035 | 0.00035 | 0.0003 | 0.00019 | 0.00026 | 0.00037 | 0.00018 | 0.00043 | 0.00029 |
| Tubb5 | 0.00021 | 0.0003 | 0.00048 | 0.0008 | 0.00025 | 0.00044 | 0.00024 | 0.00034 | 0.0003 | 0.00035 | 0.00086 | 0.00073 | 0.00029 | 0.00047 | 0 | 0.00156 | 0.00015 | 0.00043 |
| Tpi1 | 0.00025 | 0 | 0.00014 | 0.00038 | 7.98E-05 | 0.00017 | 0.00029 | 0.00016 | 0.0003 | 0.00017 | 0.00024 | 0.00165 | 0.00041 | 0.00012 | 0 | 0.00024 | 0.00018 | 0 |
| Ddx17 | 0.00028 | 0.00051 | 0.00027 | 0.0003 | 0.00043 | 0.00025 | 0.00022 | 0.00024 | 0.0003 | 0.00025 | 0.0003 | 0.00043 | 0.00021 | 0.0003 | 0.00155 | 7.57E-05 | 6.73E-05 | 0.00031 |
| Pea15a | 6.93E-05 | 0.00013 | 0.00013 | 0.00014 | 0.00018 | 0 | 3.04E-05 | 0.00034 | 0.0003 | 0.00019 | 0.0004 | 0.00013 | 0.0002 | 5.15E-05 | 0 | 0.00027 | 0 | 0.00036 |
| Mgat1 | 0.00013 | 0.00023 | 0.00025 | 0.00022 | 0.00037 | 0.00018 | 0.00027 | 7.19E-05 | 0.0003 | 0.00024 | 0.00035 | 0.00023 | 0.00029 | 0.00021 | 9.44E-05 | 0.00019 | 3.53E-05 | 0.00024 |
| Rhoa | 0.00024 | 0.00028 | 0.00033 | 0.00069 | 0.00022 | 0.00024 | 0.0003 | 0.0003 | 0.0003 | 0.00042 | 0.00043 | 0.00025 | 0.00013 | 0.00032 | 0 | 0.00025 | 0.00026 | 0.00038 |
| Maz | 0.00028 | 0.00029 | 0.00015 | 0.00017 | 0.00021 | 0.00021 | 9.19E-05 | 0.00011 | 0.0003 | 8.93E-05 | 0.00013 | 0.0002 | 0 | 0.00012 | 0.00045 | 0.00049 | 8.80E-05 | 0.00011 |
| Sgms1 | 4.41E-05 | 7.71E-05 | 7.36E-05 | 0 | 3.42E-05 | 4.71E-05 | 0.00017 | 2.89E-05 | 0.0003 | 6.38E-05 | 0.00013 | 6.94E-05 | 0 | 0 | 0 | 2.69E-05 | 0.00014 | 4.82E-06 |
| Zic1 | 1.78E-05 | 5.39E-05 | 0 | 0 | 8.60E-05 | 1.64E-05 | 0 | 0 | 0.00029 | 1.46E-06 | 2.79E-05 | 0.0001 | 0 | 0.0002 | 3.63E-05 | 6.05E-05 | 1.70E-05 | 1.04E-05 |
| Gli2 | 9.57E-06 | 0.0002 | 0 | 0 | 2.99E-05 | 7.67E-05 | 0 | 0.00017 | 0.00029 | 0 | 0 | 4.92E-05 | 0 | 5.64E-05 | 0 | 3.57E-05 | 8.92E-05 | 0.00016 |
| Rcn3 | 0.00039 | 0 | 0.00034 | 0.0013 | 0.00048 | 0.00084 | 0.00222 | 0.00112 | 0.00029 | 0.00051 | 0.00066 | 0.0004 | 0.00114 | 0.00094 | 0.00109 | 0.00031 | 0.00014 | 0.00079 |
| Cbx6 | 0 | 0.00014 | 0 | 0.00012 | 0.00022 | 6.96E-05 | 9.91E-06 | 0 | 0.00029 | 0.00014 | 0 | 0 | 6.77E-05 | 9.00E-05 | 0.00017 | 0.00019 | 0 | 0.00027 |
| Sumo3 | 0.00023 | 0.00012 | 0.00016 | 0.00034 | 0.00016 | 0.00015 | 0.0001 | 0.00012 | 0.00029 | 0.00016 | 0.00015 | 6.40E-06 | 0.00016 | 0.00017 | 5.43E-05 | 0.00022 | 3.38E-05 | 0.00017 |
| Unc93b1 | 6.49E-05 | 0.00013 | 0 | 9.93E-05 | 0.00014 | 0.00015 | 9.50E-05 | 0.00017 | 0.00029 | 0.00014 | 3.10E-05 | 4.63E-05 | 0.00024 | 0 | 0 | 0 | 8.52E-05 | 0.00013 |
| Hsbp1 | 0.00028 | 0.00032 | 0.00019 | 0.00044 | 0.00042 | 0.00025 | 0.00026 | 0.00032 | 0.00029 | 0.00026 | 0.00035 | 0.0002 | 0.00019 | 0.00016 | 1.16E-05 | 0.00014 | 0 | 0.00029 |
| Nnmt | 0.00017 | 9.63E-05 | 4.44E-05 | 0.0002 | 0 | 0.00022 | 0 | 0 | 0.00029 | 0.0003 | 4.58E-05 | 4.98E-05 | 0.00031 | 0.00018 | 0 | 1.28E-05 | 0 | 0.0002 |

| | | | | | | | | | | | | | | | | | | |
|---|---|---|---|---|---|---|---|---|---|---|---|---|---|---|---|---|---|---|
| Sipa1l1 | 8.91E-06 | 4.92E-05 | 0 | 0 | 1.76E-05 | 0.00024 | 3.38E-05 | 0 | 0.00029 | 5.54E-05 | 8.53E-05 | 0.00022 | 5.22E-05 | 0.00016 | 8.21E-05 | 2.52E-05 | 8.35E-05 | 9.66E-05 |
| Stambpl1 | 0.00011 | 3.69E-05 | 5.30E-05 | 2.29E-05 | 0.0001 | 0.0002 | 0 | 0.00023 | 0.00029 | 0.00015 | 0.00019 | 5.54E-05 | 0.00015 | 0.00028 | 0.0002 | 0.00015 | 0 | 0 |
| Ide | 0.00034 | 0.00017 | 0.00019 | 0 | 0.0002 | 0 | 0.00026 | 0 | 0.00029 | 0.00012 | 9.83E-05 | 1.30E-05 | 2.22E-05 | 6.98E-05 | 6.68E-05 | 5.23E-05 | 0 | 8.87E-05 |
| Lgmn | 0.00019 | 0.00028 | 8.00E-06 | 7.42E-05 | 0.00048 | 0.00038 | 0.00071 | 0.00039 | 0.00029 | 0.00043 | 0 | 0 | 0.00125 | 0.00024 | 0 | 0.00014 | 0 | 0.0003 |
| Olfm1 | 2.21E-05 | 8.58E-05 | 0 | 0 | 8.04E-05 | 0.0004 | 0 | 0.00035 | 0.00029 | 0.00012 | 0 | 8.50E-05 | 0 | 0.00053 | 0 | 7.55E-05 | 0 | 0.00019 |
| Ntrk2 | 0 | 0.00012 | 0 | 0 | 0 | 0.00015 | 0 | 0 | 0.00029 | 0.00039 | 0 | 0 | 0.00024 | 0 | 0 | 0 | 0 | 0.00059 |
| Pdcd6ip | 0.00023 | 0.00027 | 0.00045 | 0.00014 | 0.00025 | 0.00024 | 0.0002 | 0.00018 | 0.00029 | 0.00023 | 0.00044 | 0.00031 | 0.00014 | 0.00021 | 0.00015 | 9.94E-05 | 0.00018 | 0.00019 |
| Gtf2i | 0.00025 | 0.00029 | 0.00011 | 0.00024 | 0.00026 | 0.00017 | 7.29E-05 | 0.00025 | 0.00029 | 0.00013 | 9.47E-05 | 5.86E-05 | 0.00011 | 0.00024 | 0.00018 | 7.42E-05 | 0.00024 | 0.00014 |
| Arxes2 | 9.05E-05 | 0 | 0 | 0 | 0.00034 | 4.30E-05 | 0 | 4.51E-05 | 0.00029 | 0 | 7.06E-05 | 0 | 0 | 0.00017 | 0 | 3.07E-05 | 0 | 0 |
| Clec2d | 9.32E-05 | 0.00058 | 0 | 0.00026 | 0.0008 | 0.00046 | 0 | 0 | 0.00029 | 0.00081 | 0.00034 | 0 | 0.00046 | 0.00035 | 0.00064 | 0 | 0 | 0.00038 |
| Wnk1 | 0.0004 | 0.00033 | 0.00023 | 0 | 0.00017 | 0.00029 | 0.00013 | 3.30E-05 | 0.00029 | 0.00023 | 0.00025 | 0.0003 | 0 | 8.76E-05 | 0.00011 | 0.0003 | 0.00027 | 0.00017 |
| Runx2 | 0 | 0.0001 | 0.00026 | 0.00014 | 0 | 0.00012 | 0.00013 | 0.00071 | 0.00029 | 0 | 0 | 2.78E-06 | 0 | 5.68E-05 | 0.00087 | 8.18E-05 | 0.00023 | 3.75E-05 |
| Clip2 | 1.56E-05 | 0.00021 | 0.00024 | 0.00019 | 0 | 0.00011 | 0 | 0.0004 | 0.00029 | 0.00015 | 0 | 0.00029 | 0 | 0.00015 | 0.00043 | 3.06E-05 | 7.61E-05 | 4.49E-05 |
| Serpinf1 | 0.00035 | 5.62E-05 | 0.00028 | 0.00012 | 0.00074 | 0.00097 | 0.00153 | 0.00126 | 0.00029 | 0.0007 | 0.00084 | 0.00069 | 0.00131 | 0.00121 | 0.00021 | 0.00013 | 0 | 0.00115 |
| Irx3 | 0 | 3.22E-05 | 3.36E-05 | 0.00015 | 7.71E-05 | 0 | 6.01E-05 | 0.00027 | 0.00029 | 0 | 3.36E-05 | 8.89E-05 | 0 | 0 | 4.60E-05 | 2.78E-05 | 0.00016 | 0 |
| Akr1b3 | 0.00038 | 0 | 0.00021 | 0.00021 | 8.35E-05 | 7.25E-05 | 0.00027 | 0 | 0.00029 | 6.65E-05 | 0 | 0 | 0.0001 | 0.00023 | 0 | 0.00022 | 0.00014 | 6.26E-05 |
| Puf60 | 0.00026 | 0.00027 | 0.00035 | 0.00058 | 0.00017 | 0.00012 | 0.00025 | 0.00022 | 0.00029 | 0.00021 | 0.00029 | 0.00014 | 5.10E-05 | 0.00014 | 0.0003 | 0.00035 | 0.00042 | 0.00021 |
| Eif4e | 0.00034 | 0.00013 | 0.00041 | 9.59E-05 | 0.00013 | 6.91E-05 | 0.00014 | 2.17E-05 | 0.00029 | 4.75E-05 | 0.00011 | 0.00017 | 9.22E-05 | 1.06E-05 | 0 | 0.00032 | 7.80E-05 | 0 |
| Hnrnpk | 0.00054 | 0.00026 | 0.00033 | 0.00027 | 0.00016 | 0.00033 | 9.34E-05 | 0.00021 | 0.00029 | 9.01E-05 | 0.00026 | 0.00026 | 3.96E-05 | 0.00022 | 0 | 0.00034 | 0.0003 | 7.29E-05 |
| Myo10 | 0 | 0.0002 | 6.69E-05 | 9.83E-06 | 0.00018 | 5.78E-05 | 1.91E-05 | 2.91E-05 | 0.00029 | 0 | 0.00022 | 0.00012 | 0 | 0.00044 | 0 | 0.00011 | 6.44E-05 | 0.00016 |
| Add1 | 0.00026 | 0.00036 | 0.00012 | 0.00022 | 0.00019 | 0.0003 | 0.00031 | 0.0003 | 0.00029 | 0.00042 | 0.00031 | 0.00035 | 0.00016 | 0.0002 | 0 | 0.00024 | 0.00011 | 0.00047 |
| Pdzrn3 | 0.00011 | 0.00012 | 0.00029 | 2.32E-05 | 0.00056 | 0.0004 | 0.00068 | 0.00016 | 0.00029 | 0 | 0.00078 | 0 | 0.00022 | 0.00017 | 0.00047 | 0.00011 | 0.00061 | 0.00051 |
| Axin2 | 0 | 0.00029 | 0 | 0 | 9.28E-05 | 0.00013 | 0.00017 | 0 | 0.00029 | 3.40E-05 | 0.00013 | 8.37E-05 | 0 | 0.00024 | 0.00018 | 3.74E-05 | 0 | 0 |
| Runx1 | 0 | 0.00021 | 0.00033 | 8.90E-05 | 0.0007 | 5.75E-05 | 0.00072 | 0.00066 | 0.00029 | 0 | 0.00054 | 0.00043 | 0.00072 | 0.00048 | 0.0008 | 0.00014 | 0.00094 | 0.0002 |
| Gstm2 | 0.0003 | 2.30E-05 | 2.20E-06 | 0.00032 | 0.00033 | 0.00021 | 0.00049 | 0.00018 | 0.00029 | 0.00058 | 0.0002 | 0.00026 | 0.00057 | 0.0001 | 0 | 0 | 0 | 0.00048 |
| Eif4g3 | 7.82E-05 | 0.00026 | 0.00031 | 0.0002 | 0.00035 | 0.00031 | 0.0001 | 0.00042 | 0.00029 | 0.00032 | 0.00022 | 6.52E-05 | 0.00019 | 0.00024 | 0.00036 | 0.00014 | 0.00031 | 0.00026 |
| Rgs16 | 0 | 0 | 5.21E-05 | 0 | 0.00014 | 0.00032 | 0 | 0.00014 | 0.00029 | 0 | 0.00042 | 0.00059 | 0 | 2.76E-05 | 3.94E-05 | 0 | 0.00124 | 0.00013 |
| Hsp90ab1 | 0.0006 | 0.00041 | 0.00058 | 0.00043 | 7.14E-05 | 0.00032 | 5.52E-06 | 0.00023 | 0.00029 | 8.91E-05 | 0.00015 | 0.00054 | 0 | 8.45E-05 | 9.22E-05 | 0.00024 | 0.00073 | 0.00016 |
| Lpcat3 | 0.00016 | 8.88E-05 | 0.00016 | 0.00012 | 5.49E-05 | 0 | 0.00022 | 0 | 0.00029 | 0.00017 | 0.00011 | 0 | 0.00034 | 0.00011 | 0 | 0 | 0 | 0.00013 |
| Ptpra | 0.00018 | 0.00029 | 0.00015 | 9.44E-05 | 0.00028 | 0.00037 | 0.00019 | 0.0002 | 0.00029 | 0.00028 | 0.00023 | 0.00017 | 0.00027 | 0.00022 | 0 | 0.00019 | 0.00024 | 0.0002 |
| Cenpb | 6.25E-05 | 0.00023 | 0.00024 | 0.00068 | 0.00027 | 6.41E-05 | 0.00026 | 0.00021 | 0.00029 | 0.00017 | 0.00021 | 0.0003 | 5.93E-05 | 0.00017 | 0.00035 | 0.00035 | 0.0001 | 0.00035 |
| Hspa5 | 0.00056 | 0.00019 | 0.00071 | 0.00066 | 0.00014 | 5.56E-05 | 0.00014 | 0.00023 | 0.00029 | 0.00032 | 0.00039 | 0.00056 | 0.00042 | 0.00016 | 7.60E-05 | 0 | 0.00057 | 0.00022 |
| Nedd8 | 0.00038 | 0.00018 | 0.00036 | 0.00062 | 0.00028 | 0.00024 | 0.0002 | 0.00029 | 0.00029 | 0.00019 | 0.00031 | 0.00011 | 0.00021 | 0.0003 | 8.51E-05 | 0.00027 | 9.39E-05 | 0.00022 |

| | | | | | | | | | | | | | | | | | | |
|---|---|---|---|---|---|---|---|---|---|---|---|---|---|---|---|---|---|---|
| Ecscr | 0.0002 | 0 | 0.00018 | 0 | 0 | 0.00019 | 0 | 0.00015 | 0.00029 | 0 | 0.00021 | 0.0001 | 7.90E-06 | 9.28E-05 | 0 | 2.94E-05 | 0 | 0.00027 |
| Atp5b | 0.00045 | 0.00025 | 0.00056 | 0.00055 | 9.20E-05 | 0.00018 | 0.00013 | 0.0002 | 0.00029 | 8.66E-05 | 0.00037 | 0.00015 | 4.96E-05 | 0.00014 | 0 | 0.00031 | 0 | 8.59E-05 |
| Srf | 0 | 0.00012 | 0.00042 | 1.10E-05 | 9.39E-05 | 6.33E-05 | 0 | 7.05E-05 | 0.00029 | 0.0001 | 0.00087 | 0.00016 | 0 | 0.00022 | 0.00046 | 0.00021 | 0.0007 | 0 |
| Myof | 0.00016 | 0.00028 | 0.00018 | 0.00038 | 0.00039 | 0.00036 | 0.00031 | 0.00011 | 0.00029 | 0.00049 | 0.00068 | 0.00047 | 0.00012 | 0.00033 | 0.00038 | 0.0003 | 0.00024 | 0.00023 |
| Cyfip1 | 0.00025 | 0.00015 | 0.00014 | 0.00013 | 8.47E-05 | 0.00013 | 5.87E-05 | 9.86E-05 | 0.00029 | 0.0002 | 0.00014 | 7.61E-05 | 0 | 0.00015 | 4.63E-05 | 0.00013 | 0 | 0.00021 |
| Col5a2 | 0.00071 | 6.85E-05 | 0.00019 | 5.17E-05 | 0.00132 | 0.00173 | 0.00202 | 0.00156 | 0.00029 | 0.00067 | 0.00126 | 0.00092 | 0.0012 | 0.0013 | 0.00096 | 0.00015 | 0 | 0.00075 |
| Lama4 | 0.0004 | 0.00013 | 0.00019 | 0 | 0.00035 | 0.00108 | 0.0008 | 0.00083 | 0.00029 | 0.00077 | 8.38E-05 | 0.00025 | 0.00068 | 0.00023 | 0.00038 | 0.00012 | 4.50E-05 | 4.64E-07 |
| Tmem64 | 0.00017 | 0.00011 | 0 | 0 | 9.99E-05 | 0 | 0.00023 | 0.00015 | 0.00028 | 0 | 8.39E-05 | 0 | 0 | 0 | 0 | 0.0001 | 0 | 0.00012 |
| Cebpb | 0 | 0.00042 | 0.00045 | 0.00096 | 0.00055 | 0.00084 | 0 | 0.00017 | 0.00028 | 0.00023 | 6.59E-05 | 0.00045 | 0.00129 | 0.00103 | 0.00145 | 2.95E-05 | 0.00187 | 0.00014 |
| Dlg5 | 0.00013 | 0.00023 | 2.93E-05 | 4.31E-05 | 0.00027 | 9.01E-05 | 1.63E-05 | 6.01E-05 | 0.00028 | 4.94E-05 | 0.00011 | 0 | 4.77E-05 | 7.26E-05 | 0.00029 | 0.00019 | 0.00021 | 7.39E-05 |
| B4galt1 | 0.00026 | 0.00028 | 0.00034 | 0.00037 | 0.00058 | 0.00081 | 0.00115 | 0.00055 | 0.00028 | 0.00061 | 0.00022 | 0.00062 | 0.00089 | 0.0003 | 0 | 0.00028 | 9.37E-05 | 0.00049 |
| Eln | 0 | 0 | 0.00021 | 1.62E-05 | 0.00103 | 0 | 2.06E-06 | 7.29E-05 | 0.00028 | 0.00083 | 0 | 0 | 0 | 0 | 0.0004 | 2.11E-05 | 0 | 0.00159 |
| Cap1 | 0.00051 | 0.00035 | 0.00037 | 0.00043 | 0.00022 | 0.0003 | 0.00023 | 0.00021 | 0.00028 | 0.00036 | 0.00089 | 0.00037 | 3.93E-05 | 0.00038 | 0 | 0.00016 | 0.00012 | 0.00048 |
| Grb10 | 0.00046 | 6.57E-05 | 0 | 8.99E-05 | 0 | 9.43E-05 | 0.00061 | 0 | 0.00028 | 0.00032 | 2.26E-05 | 0.00023 | 0 | 0 | 0 | 0.00019 | 0.00023 | 0.00039 |
| Sav1 | 0.00021 | 0.00027 | 4.43E-05 | 2.75E-05 | 8.49E-05 | 0.00047 | 0.00012 | 0 | 0.00028 | 0.0002 | 5.69E-05 | 0.00022 | 0.00021 | 0.00019 | 0.00023 | 0.00018 | 0 | 0.00016 |
| Vasp | 0.00024 | 0.00022 | 0.00037 | 0.00042 | 8.23E-05 | 1.92E-05 | 4.40E-05 | 0.00034 | 0.00028 | 0.00011 | 0.00077 | 0.00018 | 1.77E-05 | 0.00027 | 0.00028 | 3.27E-05 | 0.00033 | 0.00017 |
| Idh1 | 0.00037 | 0.00015 | 0 | 0.00022 | 1.98E-05 | 8.85E-05 | 0.00019 | 8.50E-05 | 0.00028 | 0.00034 | 1.45E-05 | 0 | 2.49E-05 | 0.00011 | 0 | 0 | 0 | 0.0002 |
| Anxa6 | 0.00045 | 0.00022 | 0.00031 | 0.00052 | 0.00024 | 0.00036 | 0.00056 | 0.00037 | 0.00028 | 0.00038 | 0.00072 | 0.00023 | 0 | 0.00036 | 0.00015 | 0 | 0.00018 | 8.56E-05 |
| Sar1a | 0.00079 | 0.00013 | 0.00039 | 0.00034 | 0.00034 | 0.00053 | 0.00079 | 0.00036 | 0.00028 | 0.00032 | 0.0003 | 0.00028 | 0.00036 | 0.00028 | 0.00027 | 0.00013 | 0.00026 | 0.00028 |
| Xdh | 0 | 0.00049 | 0 | 0 | 2.56E-05 | 0.00076 | 0 | 0 | 0.00028 | 0.00072 | 0 | 0.00021 | 0.0009 | 0.00019 | 0.00017 | 0 | 0 | 2.35E-05 |
| Son | 0.00028 | 0.00047 | 0.00028 | 0.00022 | 0.00024 | 0.00022 | 0.00014 | 0.00019 | 0.00028 | 0.00023 | 0.00021 | 0.00025 | 8.93E-05 | 0.0002 | 0.00101 | 0.00017 | 0.00046 | 0.00017 |
| Aph1a | 0.00013 | 0.00016 | 0.00029 | 0.00052 | 0.00019 | 0.00016 | 0.00019 | 0.00029 | 0.00028 | 0.00023 | 0.00016 | 0.00017 | 0.00012 | 0.00018 | 0 | 5.84E-05 | 4.09E-05 | 0.00025 |
| Ubtd2 | 1.06E-05 | 6.16E-05 | 0.00026 | 1.25E-05 | 0.00015 | 0.00022 | 0.00024 | 0.00013 | 0.00028 | 0.00015 | 1.46E-05 | 9.17E-05 | 0.00016 | 0.00023 | 0.00023 | 2.64E-05 | 0 | 0.00014 |
| Mapk6 | 0.00024 | 0.00011 | 0.00043 | 0 | 0.00019 | 5.91E-05 | 0.00033 | 0 | 0.00028 | 5.97E-05 | 0.00042 | 0.00019 | 0.0003 | 0.00028 | 4.39E-05 | 6.90E-05 | 0.00041 | 0 |
| Efnb2 | 0 | 6.34E-05 | 0.00015 | 0.00014 | 6.40E-05 | 0 | 0.00017 | 8.09E-05 | 0.00028 | 4.21E-05 | 0.00012 | 0.00018 | 0 | 0 | 6.31E-05 | 0.00013 | 0.00059 | 0 |
| Tnfsf12 | 3.55E-05 | 0.00019 | 0 | 0.00016 | 0.00026 | 0.00018 | 0.00021 | 0.00022 | 0.00028 | 0.00028 | 0.00015 | 3.76E-05 | 0.00036 | 0.00016 | 0 | 0 | 0 | 0.00031 |
| Dynll2 | 0.00016 | 0.0003 | 0.0003 | 0.00046 | 0.00024 | 0.0001 | 0.00015 | 4.81E-05 | 0.00028 | 0.0001 | 0.00021 | 0.00013 | 0.00011 | 0.00011 | 3.96E-05 | 0.00037 | 0.00022 | 6.20E-05 |
| Sdc4 | 0 | 0 | 0.00011 | 0.00025 | 6.09E-05 | 0.00047 | 0 | 0 | 0.00028 | 0.00031 | 0.00016 | 0.00078 | 0.00139 | 0.00036 | 6.57E-05 | 3.64E-05 | 0.00023 | 0 |
| Hnrnpu | 0.00038 | 0.0003 | 0.00042 | 0.00025 | 0.00011 | 0.00014 | 0 | 0.0001 | 0.00028 | 8.54E-05 | 0.00023 | 0.00018 | 3.28E-05 | 0.00016 | 0.00018 | 0.00041 | 0.00024 | 5.30E-05 |
| Phc2 | 8.69E-05 | 0.00034 | 0.00024 | 0.00026 | 0.00018 | 2.26E-05 | 0 | 0.00048 | 0.00028 | 9.64E-06 | 0.00015 | 7.21E-05 | 0 | 0.00013 | 0.00029 | 0.0002 | 5.82E-05 | 0.00016 |
| Cltb | 3.24E-05 | 0.0002 | 0.00021 | 0.00036 | 0.00026 | 9.41E-05 | 0 | 0.00017 | 0.00028 | 0.00052 | 0.00037 | 0.0001 | 4.34E-06 | 1.32E-05 | 5.17E-05 | 9.47E-05 | 9.57E-05 | 0.00073 |
| Plxnb2 | 0.00014 | 0.00023 | 0.00016 | 0.00037 | 0.00029 | 0.00036 | 0.00032 | 0.00046 | 0.00028 | 0.00023 | 0.0002 | 0.00024 | 0.00027 | 0.00023 | 7.56E-05 | 0.00026 | 0.00021 | 0.00048 |
| Trim44 | 0.00023 | 0.00023 | 0.00024 | 8.58E-05 | 0.0002 | 4.93E-05 | 6.71E-05 | 0.00023 | 0.00028 | 7.10E-05 | 0.00018 | 0.00018 | 7.66E-05 | 6.75E-05 | 0.00031 | 8.41E-05 | 4.87E-06 | 0.00014 |

| | | | | | | | | | | | | | | | | | | |
|---|---|---|---|---|---|---|---|---|---|---|---|---|---|---|---|---|---|---|
| Cops6 | 0.00044 | 0.00021 | 0.00034 | 0.00043 | 0.0002 | 0.00027 | 0.00015 | 0.0002 | 0.00028 | 0.00013 | 0.00023 | 9.41E-05 | 0.00015 | 0.00022 | 0 | 0.00025 | 0 | 0.0001 |
| Rab2a | 0.00051 | 0.00021 | 0.00036 | 0.00042 | 0.00046 | 0.00054 | 0.0007 | 0.00043 | 0.00028 | 0.00028 | 0.00032 | 0.00038 | 0.00027 | 0.00041 | 0 | 0.00016 | 0.00036 | 0.00034 |
| Arsb | 5.69E-05 | 6.24E-05 | 0 | 0.0001 | 0.00017 | 0.00017 | 0.00023 | 7.56E-05 | 0.00028 | 0.00023 | 0 | 0 | 0.00019 | 0.00019 | 0 | 0.00016 | 0 | 0.00032 |
| Fryl | 3.50E-05 | 0.00012 | 0 | 5.29E-05 | 0.00019 | 0.00021 | 4.55E-06 | 0.00019 | 0.00028 | 3.20E-05 | 2.27E-05 | 3.66E-05 | 0 | 1.12E-05 | 4.73E-05 | 0.00015 | 0.00013 | 1.56E-05 |
| Slk | 0.00019 | 0.00019 | 0.00011 | 0 | 8.92E-05 | 0.00024 | 0.00012 | 0 | 0.00028 | 9.58E-05 | 0 | 9.26E-05 | 0 | 4.88E-05 | 0.00025 | 0.00017 | 0.00017 | 0 |
| Mbd2 | 0.00019 | 0.00032 | 0.00016 | 0.00052 | 0.00023 | 0.00024 | 0.00027 | 0.00022 | 0.00028 | 0.00044 | 0.00015 | 0.00051 | 0.00028 | 0.00019 | 0.00073 | 0.00028 | 0.0004 | 0.00029 |
| Tshz3 | 0.00012 | 0.00022 | 5.24E-05 | 0 | 0.0002 | 0.00037 | 0 | 0.00018 | 0.00028 | 0 | 0 | 0.00021 | 0 | 0.00018 | 1.73E-05 | 4.22E-05 | 0.0003 | 3.18E-05 |
| Ywhah | 0.00046 | 0.0002 | 0.00031 | 0 | 0.00024 | 2.44E-05 | 0.00012 | 0 | 0.00028 | 0.00014 | 0.00059 | 0.00036 | 0.00014 | 5.32E-05 | 0 | 0.00058 | 0.00012 | 0.00014 |
| B4galnt1 | 2.44E-05 | 4.12E-05 | 2.05E-06 | 0 | 0.00014 | 0.00022 | 0 | 0.0005 | 0.00028 | 0 | 0.00015 | 7.26E-05 | 0 | 0.00039 | 0 | 0 | 0 | 7.44E-05 |
| Spin1 | 0.00033 | 0.00024 | 7.37E-05 | 0 | 0.00019 | 0.0003 | 3.97E-05 | 0.00021 | 0.00028 | 9.29E-05 | 0.00012 | 8.06E-05 | 0.00015 | 7.12E-05 | 0 | 0.00014 | 0 | 3.94E-05 |
| Zc3h7b | 6.28E-05 | 0.00018 | 8.83E-05 | 0.00023 | 0.00019 | 7.94E-05 | 2.25E-05 | 0.00016 | 0.00028 | 0.00013 | 0.00014 | 1.56E-05 | 7.92E-05 | 0.00017 | 0 | 4.62E-05 | 0.0002 | 0.00024 |
| Tax1bp3 | 0.00035 | 0.00033 | 0.00016 | 0.00023 | 0.00028 | 9.15E-05 | 0.00023 | 0.00017 | 0.00028 | 0.00017 | 0.00039 | 0.00015 | 0.00016 | 0.00029 | 0 | 0.00017 | 1.13E-05 | 0.00022 |
| Tspan9 | 5.23E-05 | 7.48E-05 | 0 | 0 | 0.00027 | 0.00032 | 0 | 0.00021 | 0.00028 | 0 | 0.00014 | 0.00056 | 0 | 0.00032 | 0 | 1.54E-05 | 6.86E-05 | 0.00027 |
| Serpine2 | 0.00012 | 0.00034 | 0 | 0 | 0.00206 | 0.00265 | 0 | 0.00266 | 0.00028 | 0.00023 | 0.00032 | 0 | 0 | 0.00099 | 0 | 0 | 0 | 0 |
| Mark1 | 0 | 0.00026 | 0 | 2.03E-05 | 0.00042 | 0.00049 | 0.00017 | 0.00016 | 0.00028 | 8.30E-05 | 0.0002 | 0.00011 | 0.00089 | 0 | 0.00022 | 0 | 0.00017 | 0 |
| Tram2 | 2.40E-05 | 9.96E-05 | 0.00024 | 0.00015 | 0.00023 | 0.00015 | 7.41E-05 | 0.00021 | 0.00028 | 0.00044 | 0.00049 | 0.00033 | 9.99E-05 | 0.00016 | 0 | 0.0002 | 0 | 0.00033 |
| C1qtnf6 | 0.00013 | 0 | 0 | 0.00088 | 0.00074 | 0.00091 | 0.00123 | 0.00145 | 0.00028 | 0.0005 | 0.00036 | 0 | 0.00052 | 0.00082 | 0 | 3.95E-05 | 0 | 0.00052 |
| Ptpn11 | 0.00022 | 0.0002 | 0.00037 | 4.60E-05 | 0.00021 | 0.00017 | 0.00011 | 0.0001 | 0.00028 | 0.00018 | 0.00039 | 9.99E-05 | 0.00012 | 0.00016 | 9.59E-05 | 0.00024 | 8.41E-05 | 0.00012 |
| Usp5 | 0.00029 | 0.00017 | 0.00031 | 0.00015 | 8.03E-05 | 1.90E-05 | 3.17E-05 | 2.23E-05 | 0.00028 | 7.08E-05 | 0.00014 | 5.65E-05 | 0 | 3.21E-05 | 0.00028 | 0.00019 | 0 | 0.00014 |
| Cavin3 | 0.00025 | 2.90E-05 | 5.33E-05 | 0.00016 | 0 | 0 | 0.00022 | 0.00016 | 0.00028 | 0.00068 | 0.00054 | 0.00047 | 0 | 0.00033 | 0 | 0.00033 | 0 | 0.00076 |
| Lix1l | 0.00016 | 0.00035 | 7.94E-05 | 0.00046 | 0.00047 | 0.0005 | 0.00042 | 0.00042 | 0.00028 | 0.00025 | 0.00025 | 0.00022 | 0.00029 | 0.00062 | 0.0002 | 0.00014 | 0.00032 | 0.00046 |
| Grn | 0 | 0.00042 | 0 | 0.00072 | 0.00045 | 0.00059 | 0.00052 | 0.00058 | 0.00028 | 0.0007 | 0.00037 | 0.00035 | 0.00107 | 0.00084 | 0.00034 | 0.00012 | 0 | 0.00099 |
| Mtap | 0.00023 | 0.00012 | 0.00032 | 0.00024 | 2.51E-06 | 2.17E-05 | 3.34E-05 | 7.87E-05 | 0.00028 | 1.71E-05 | 0.00017 | 0 | 0 | 6.56E-05 | 2.71E-05 | 0.0002 | 0 | 4.09E-05 |
| Map7d1 | 0.00017 | 0.00033 | 0.00032 | 0.00038 | 0.00016 | 9.16E-05 | 0.00032 | 0.00019 | 0.00028 | 0.00032 | 0.00032 | 0.00013 | 0.00022 | 0.00028 | 0.00092 | 0.00018 | 0.0005 | 0.00031 |
| Cmtm3 | 7.35E-05 | 0.00048 | 0.00013 | 0.00057 | 0.0005 | 0.00041 | 0.00054 | 0.00026 | 0.00028 | 0.00034 | 0.00017 | 1.73E-05 | 0.0007 | 0.00065 | 0 | 6.29E-05 | 7.44E-05 | 0.00032 |
| Junb | 3.25E-05 | 0.00028 | 0.0002 | 0.00053 | 0.0002 | 0.00025 | 6.98E-05 | 0.00058 | 0.00027 | 0.00026 | 6.57E-05 | 0.00051 | 0.00056 | 0.00019 | 0.00059 | 8.17E-05 | 0.00616 | 0.00023 |
| Mmp23 | 0.00017 | 0 | 0 | 0.00027 | 0.00025 | 0.00035 | 0.00095 | 0 | 0.00027 | 0.00045 | 0.00043 | 0 | 0.00058 | 0.00041 | 0.00015 | 0.00013 | 0 | 0.00036 |
| Ggta1 | 8.63E-05 | 0.0002 | 6.07E-06 | 0 | 0 | 0.00014 | 0 | 0 | 0.00027 | 0 | 0 | 0.00017 | 0.00013 | 0.00023 | 0 | 0.00011 | 0.00018 | 3.42E-05 |
| Ndufs7 | 9.74E-05 | 0.00013 | 0.0002 | 0.00057 | 0.00013 | 3.38E-05 | 4.11E-05 | 0.00024 | 0.00027 | 0.00015 | 0.00015 | 8.77E-05 | 9.86E-05 | 0.00016 | 9.50E-06 | 0.00015 | 0.00011 | 0.00016 |
| Tmem14c | 0.00034 | 9.29E-05 | 0.00021 | 0.00028 | 0.0002 | 0.00018 | 0.00015 | 8.69E-05 | 0.00027 | 9.07E-05 | 0.00019 | 0 | 0.00023 | 0.00021 | 0 | 0.0002 | 0 | 0.00011 |
| Cdc42 | 0.00057 | 0.00022 | 0.00021 | 0.00035 | 0.00018 | 0.00039 | 0.00022 | 0.00018 | 0.00027 | 0.00016 | 0.00023 | 0.00033 | 5.20E-05 | 0.00023 | 0 | 0.00015 | 0.0003 | 0.0002 |
| Gdnf | 0 | 0 | 1.92E-05 | 0 | 9.07E-05 | 0 | 4.75E-05 | 0 | 0.00027 | 0 | 1.44E-05 | 8.08E-06 | 7.62E-05 | 0 | 3.56E-05 | 0 | 0.00014 | 0 |
| Cyb5b | 0.00012 | 0.0001 | 0.00022 | 0.00039 | 7.70E-05 | 1.43E-05 | 0 | 9.62E-05 | 0.00027 | 5.41E-05 | 7.02E-05 | 0 | 6.47E-05 | 0.00015 | 0 | 0.00015 | 5.87E-05 | 0.0001 |

| | | | | | | | | | | | | | | | | | | |
|---|---|---|---|---|---|---|---|---|---|---|---|---|---|---|---|---|---|---|
| Bex3 | 8.07E-05 | 0.00013 | 0 | 0.00015 | 0.00025 | 0.00014 | 0 | 0 | 0.00027 | 0 | 0.00034 | 0 | 0 | 0.0004 | 0 | 0.00028 | 5.22E-05 | 3.03E-05 |
| Nono | 0.00026 | 0.0003 | 0.00028 | 0.00061 | 0.00015 | 0.00015 | 5.03E-05 | 0.00023 | 0.00027 | 0.00015 | 0.00021 | 0.00024 | 9.23E-05 | 0.0003 | 0 | 0.00063 | 0.00012 | 0.00028 |
| Aacs | 7.56E-05 | 0.0001 | 9.51E-05 | 0.00019 | 1.67E-05 | 0 | 0 | 1.48E-05 | 0.00027 | 6.43E-05 | 0.00012 | 5.69E-05 | 0 | 6.37E-05 | 0.00014 | 0 | 0.00019 | 3.65E-05 |
| Lgalsl | 0.00017 | 0 | 4.53E-05 | 0 | 0.00052 | 8.28E-05 | 0.00021 | 0 | 0.00027 | 0.0001 | 0.00072 | 0 | 0.00032 | 0.00041 | 0 | 4.15E-05 | 0 | 6.17E-05 |
| Col5a1 | 0.00049 | 0 | 0.00056 | 0.0008 | 0.0013 | 0.00181 | 0.003 | 0.00122 | 0.00027 | 0.0011 | 0.0014 | 0.00062 | 0.00115 | 0.00158 | 0.00237 | 0.00046 | 0.00033 | 0.00109 |
| Man1c1 | 0 | 4.75E-06 | 1.25E-05 | 0.00028 | 0.00014 | 3.50E-05 | 0 | 0.00031 | 0.00027 | 0.00015 | 2.50E-05 | 4.28E-05 | 9.61E-05 | 5.05E-05 | 0.00034 | 4.93E-05 | 6.26E-05 | 0.00032 |
| Ccdc102a | 7.09E-05 | 0.00012 | 7.32E-05 | 0.00024 | 0.00027 | 0 | 0.0002 | 7.28E-06 | 0.00027 | 5.73E-05 | 0.00037 | 0 | 0.00019 | 0.00024 | 0.00038 | 6.71E-05 | 0.00011 | 6.15E-05 |
| Ywhae | 0.00043 | 0.00023 | 0.00038 | 0.00051 | 9.03E-05 | 9.40E-05 | 0.00012 | 0.00017 | 0.00027 | 0.0002 | 0.00027 | 0.00031 | 1.72E-05 | 0.00014 | 0.00017 | 0.00033 | 0.00018 | 0.00015 |
| Smim7 | 0.00027 | 0.00016 | 0.00022 | 0.00018 | 0.00016 | 0.00012 | 0.00025 | 0.00023 | 0.00027 | 0.00017 | 0.00016 | 0.00018 | 0.00014 | 0.00014 | 0 | 0.00011 | 0 | 0.00018 |
| Glg1 | 0.00045 | 0.00016 | 0.00011 | 1.07E-05 | 0.00034 | 0.00058 | 0.00055 | 0.00047 | 0.00027 | 0.00022 | 0.00025 | 0.0004 | 0.00036 | 0.00035 | 9.82E-05 | 0.00018 | 0.00026 | 0.00033 |
| Ncor2 | 2.55E-05 | 0.00028 | 0.00056 | 0.00021 | 0.00016 | 5.08E-05 | 0.00014 | 0.00055 | 0.00027 | 0.00014 | 0.00028 | 6.97E-05 | 8.88E-05 | 0.00039 | 0.00081 | 0.00021 | 0.00058 | 0.0004 |
| Pde7b | 0 | 7.67E-05 | 0 | 0 | 2.94E-05 | 0.00036 | 0 | 1.88E-05 | 0.00027 | 0 | 0 | 6.47E-05 | 0 | 0 | 0 | 0 | 0 | 2.96E-05 |
| Phlda1 | 2.63E-06 | 2.72E-05 | 0.00017 | 0 | 0 | 0.00021 | 0 | 0.00027 | 0.00027 | 0.00015 | 0.00012 | 9.18E-05 | 0.00025 | 0.00034 | 0 | 0 | 0.00201 | 0 |
| Lratd1 | 0 | 1.54E-05 | 0 | 0 | 0 | 0.00036 | 0 | 1.13E-05 | 0.00027 | 4.94E-07 | 0 | 0 | 0 | 0 | 3.80E-06 | 0 | 2.73E-05 | 0 |
| Nod1 | 0 | 4.07E-05 | 0.00014 | 0 | 0 | 5.81E-05 | 0.00018 | 0.00017 | 0.00027 | 0.00035 | 0 | 2.80E-07 | 0.00058 | 0 | 5.73E-05 | 0 | 0 | 0.00046 |
| Klhl9 | 0.00026 | 0.00022 | 0.00012 | 0.00035 | 0.00019 | 0.00021 | 0.00031 | 0.00026 | 0.00027 | 0.00021 | 0.00021 | 0.00018 | 9.28E-05 | 0.00024 | 7.70E-05 | 0.00012 | 9.48E-05 | 0.00023 |
| Ddr2 | 0.00014 | 0.00036 | 0.00013 | 0.00015 | 6.93E-05 | 0.00018 | 0.00012 | 0.00022 | 0.00027 | 0.00078 | 0.0002 | 0.00031 | 0.00016 | 0.00038 | 0.00017 | 0.00022 | 0.00024 | 0.00026 |
| Ehmt2 | 0.00018 | 0.00043 | 0.00011 | 0.00049 | 0.00027 | 7.93E-05 | 0.00016 | 7.65E-05 | 0.00027 | 0.00017 | 0.00012 | 0.00015 | 0.00018 | 8.14E-05 | 0.00047 | 0.00031 | 0.00017 | 0.00026 |
| Nfil3 | 9.69E-05 | 8.17E-05 | 4.36E-05 | 4.67E-06 | 0 | 0.00024 | 0.0002 | 0 | 0.00027 | 0.00021 | 2.85E-05 | 0.00037 | 0.00049 | 0.00022 | 0.00032 | 0 | 6.93E-05 | 0 |
| Aldh1l1 | 0 | 0.0001 | 0 | 3.79E-05 | 0.00013 | 0 | 0 | 0 | 0.00027 | 6.21E-05 | 0 | 4.64E-05 | 0.0002 | 1.62E-05 | 7.41E-06 | 0 | 0 | 2.32E-06 |
| Maged1 | 0.00066 | 0.0001 | 0.00047 | 0.00067 | 0.0005 | 0.00065 | 0.00138 | 0.00073 | 0.00027 | 0.00037 | 0.00043 | 0.00034 | 0.00077 | 0.00054 | 0.00069 | 0.00016 | 0.00048 | 0.00049 |
| Reep5 | 0.00062 | 0.00011 | 0.00051 | 0.00037 | 0.00028 | 0.0003 | 0.00103 | 0.00027 | 0.00027 | 0.0002 | 0.00014 | 0.00013 | 0.00026 | 0.00022 | 0.00015 | 0.00041 | 0 | 0.00055 |
| Mt1 | 0 | 0.00017 | 0 | 0.00096 | 0.00015 | 0.00096 | 0.00085 | 9.36E-05 | 0.00027 | 0.00067 | 0 | 0.0022 | 0.0034 | 0.00012 | 0.00033 | 0 | 0.00069 | 0 |
| Rsu1 | 0.00021 | 0.00022 | 0.00011 | 0.00031 | 0.00011 | 4.91E-05 | 3.42E-05 | 0.00017 | 0.00027 | 0.00043 | 0.00065 | 0.00018 | 0.00014 | 0.00029 | 0 | 6.19E-06 | 8.65E-05 | 0.00036 |
| Snx18 | 9.85E-05 | 0.0002 | 8.17E-05 | 0 | 9.96E-05 | 3.53E-05 | 0.0002 | 0.0002 | 0.00027 | 0.00036 | 0.00019 | 0.00014 | 0.00012 | 0.00027 | 3.21E-05 | 0.00012 | 0.0001 | 0.0005 |
| Scd1 | 6.12E-05 | 0.00014 | 5.06E-05 | 9.66E-05 | 8.97E-06 | 0 | 0 | 0.00023 | 0.00027 | 0.00015 | 0 | 5.31E-05 | 1.61E-05 | 0 | 0 | 0 | 1.11E-06 | 0.00015 |
| Hyou1 | 0.00033 | 8.99E-05 | 0.00052 | 0.00028 | 0.00012 | 0 | 5.82E-05 | 0.00026 | 0.00027 | 0.00017 | 0.00018 | 0.00012 | 0.00016 | 7.99E-05 | 0.00023 | 0 | 0 | 0.00013 |
| Snai1 | 0 | 5.79E-05 | 0.00081 | 0.0005 | 0.00016 | 0.00052 | 0.00022 | 0.00022 | 0.00027 | 0 | 0.00011 | 1.62E-05 | 0.00064 | 0.00048 | 0.00043 | 0 | 0.00019 | 0.0003 |
| Mmp19 | 1.19E-05 | 4.72E-05 | 0.00042 | 0.00037 | 0.00043 | 0.00115 | 0.00085 | 0.00085 | 0.00027 | 0.00082 | 0 | 0 | 0.00059 | 0.00099 | 0 | 5.57E-05 | 0.00054 | 0.00064 |
| Impad1 | 0.00031 | 0.00019 | 0.00028 | 0.00014 | 0.00022 | 0.00024 | 0.00017 | 0.00027 | 0.00027 | 0.00011 | 0.00021 | 0.00011 | 0.00013 | 0.00015 | 7.17E-06 | 0.00025 | 6.46E-05 | 0.00016 |
| Traf7 | 0.00027 | 0.00019 | 0.0002 | 9.63E-05 | 0.00019 | 0.00015 | 0.00015 | 7.01E-05 | 0.00027 | 0.0001 | 0.00027 | 8.43E-05 | 0.00015 | 0.00016 | 0.00011 | 0.0002 | 0 | 4.12E-05 |
| Lamb2 | 0 | 0.00021 | 0.00015 | 1.73E-05 | 2.88E-05 | 0.00035 | 0.0002 | 0.00042 | 0.00027 | 0.00071 | 0.00023 | 0.00037 | 0 | 0 | 0.00045 | 9.20E-05 | 0 | 0.00077 |
| Fam126a | 0.00015 | 3.86E-05 | 0.00015 | 0 | 9.74E-05 | 0.00016 | 0.00011 | 0 | 0.00027 | 2.80E-05 | 0 | 8.59E-05 | 1.40E-05 | 0.00019 | 0 | 3.63E-05 | 6.25E-05 | 0 |

| | | | | | | | | | | | | | | | | | | |
|---|---|---|---|---|---|---|---|---|---|---|---|---|---|---|---|---|---|---|
| Glud1 | 0.00034 | 0.00029 | 0.00021 | 0.00021 | 9.59E-05 | 0.00027 | 0.00019 | 1.35E-05 | 0.00027 | 0.00042 | 0.0003 | 0.0001 | 0 | 3.96E-05 | 0 | 0.00023 | 0.00057 | 0.00042 |
| Nmt1 | 0.00031 | 0.00028 | 0.0003 | 0.00023 | 0.00034 | 0.00011 | 0.00012 | 0.00014 | 0.00027 | 0 | 0.00024 | 4.41E-05 | 0.00018 | 0.00029 | 0.00016 | 0.0002 | 0 | 5.17E-05 |
| Cldnd1 | 7.32E-05 | 0.00019 | 0.00036 | 0.00018 | 0.00024 | 0.00019 | 8.75E-05 | 5.81E-05 | 0.00027 | 0.00015 | 0.00032 | 0.00019 | 0.00022 | 0.00026 | 0 | 0.0002 | 0.00025 | 0.00016 |
| Agfg1 | 0.00028 | 0.00018 | 0.00033 | 0 | 0.00014 | 0 | 0.00034 | 0.00023 | 0.00027 | 0.00033 | 6.56E-05 | 0.00027 | 0.00058 | 0 | 0.00011 | 0.00019 | 0.00015 | 0.00017 |
| Gpi1 | 0.00017 | 0.00019 | 0.00012 | 0.00056 | 6.42E-05 | 0.00021 | 0.00012 | 0.0002 | 0.00027 | 0.00017 | 5.37E-05 | 0.00151 | 0.00014 | 8.89E-05 | 0 | 0.00011 | 0.00045 | 0 |
| Snai2 | 0.00014 | 0 | 0 | 6.60E-05 | 0.00014 | 0.00011 | 0 | 0.00011 | 0.00027 | 0.00021 | 0.00028 | 0 | 0 | 0.00035 | 4.76E-05 | 0.00018 | 0 | 0 |
| Arrdc4 | 5.44E-05 | 0.00019 | 0.0001 | 0.00023 | 9.02E-05 | 0.00011 | 0 | 1.06E-05 | 0.00027 | 8.49E-05 | 0 | 0 | 3.80E-05 | 2.50E-06 | 0.00012 | 0 | 0 | 5.87E-05 |
| Prpf8 | 0.00047 | 0.00028 | 0.00022 | 4.55E-06 | 0.0001 | 0.0002 | 4.49E-05 | 7.47E-06 | 0.00027 | 4.79E-05 | 8.03E-05 | 0.0002 | 0 | 0 | 0.00018 | 0.00024 | 1.00E-05 | 3.43E-05 |
| DEPRECATED | 0 | 3.92E-05 | 0 | 0.00043 | 0.00032 | 0 | 0.00011 | 0.00014 | 0.00027 | 0 | 9.15E-06 | 0 | 2.95E-05 | 0.0003 | 0.00045 | 3.21E-05 | 9.69E-05 | 0.00036 |
| Gsn | 0 | 0.00048 | 6.48E-05 | 0.00067 | 0.00069 | 0.00055 | 0.00031 | 0.00023 | 0.00027 | 0.00184 | 0.00077 | 0.0004 | 0.00161 | 0.00062 | 0.00042 | 3.26E-05 | 0.00036 | 0.00139 |
| Ednra | 0 | 0.00011 | 0 | 0 | 0.00012 | 0.00048 | 0 | 0.00185 | 0.00026 | 0 | 0 | 0 | 0 | 0.00039 | 0 | 4.92E-06 | 0 | 0.00014 |
| Chtop | 0.00023 | 0.00029 | 0.00031 | 0.00038 | 0.00019 | 0.00017 | 9.17E-05 | 0.00021 | 0.00026 | 0.00019 | 0.00023 | 8.35E-05 | 0.00014 | 0.0002 | 0.00021 | 0.00029 | 0.00021 | 0.00014 |
| Atox1 | 0.00039 | 0.00014 | 0.00023 | 0.00044 | 0.00028 | 0.00022 | 0.00027 | 0.00025 | 0.00026 | 0.00023 | 0.00017 | 9.10E-05 | 0.00017 | 0.00023 | 0 | 7.15E-05 | 0 | 0.00022 |
| Copg1 | 0.00066 | 0.00013 | 0.00044 | 0.00024 | 0.00021 | 0.00031 | 0.00082 | 0.00031 | 0.00026 | 0.00024 | 0.00028 | 0.0002 | 0.00015 | 0.00028 | 0.00022 | 0.00019 | 0 | 9.95E-05 |
| Lrrc59 | 0.00058 | 3.50E-05 | 0.00089 | 0.00112 | 5.05E-05 | 2.69E-05 | 0.00071 | 0.00027 | 0.00026 | 0.00023 | 0.00041 | 0 | 0.00012 | 0.00017 | 0.0003 | 0.00034 | 0.0002 | 5.39E-05 |
| Tm9sf3 | 0.00045 | 0.00028 | 0.00047 | 0.00038 | 0.00038 | 0.00051 | 0.00062 | 0.00059 | 0.00026 | 0.00029 | 0.00045 | 0.00018 | 0.00039 | 0.00043 | 0 | 0.00015 | 4.78E-05 | 0.00025 |
| Calm1 | 0.00064 | 8.78E-05 | 0.00036 | 0.0005 | 8.03E-05 | 0.00013 | 0.00031 | 0.00021 | 0.00026 | 0.00019 | 0.00057 | 0.00035 | 0 | 0.00013 | 9.20E-05 | 0.00019 | 0.00029 | 0.00024 |
| Otub1 | 0.00028 | 0.00019 | 0.00026 | 0.00018 | 0.00016 | 0.00012 | 8.49E-05 | 0.00014 | 0.00026 | 0.00011 | 0.00024 | 0.0001 | 6.17E-05 | 0.00017 | 0 | 4.27E-05 | 0 | 0.00013 |
| Gli3 | 6.19E-05 | 0.00012 | 5.68E-05 | 0 | 0.00024 | 6.13E-05 | 6.04E-05 | 8.67E-05 | 0.00026 | 1.42E-05 | 7.80E-05 | 4.50E-05 | 1.14E-05 | 0.00021 | 8.86E-05 | 0.00019 | 0.00026 | 4.76E-05 |
| Cacna1g | 0 | 0 | 0 | 0 | 0.00029 | 0.00047 | 0.00049 | 0 | 0.00026 | 0 | 0.00014 | 0 | 0.00024 | 0.0005 | 0.00022 | 0 | 2.64E-05 | 0 |
| Dse | 0.00038 | 0.00014 | 0.00015 | 8.66E-05 | 0.0001 | 0.00033 | 0.0003 | 3.02E-05 | 0.00026 | 0.00021 | 0.0003 | 8.85E-05 | 0.0005 | 0 | 0.00015 | 0.0001 | 3.46E-05 | 0.00015 |
| Snx17 | 0.00019 | 0.00019 | 0.00023 | 0.00041 | 0.00014 | 0.00011 | 7.85E-05 | 0.00015 | 0.00026 | 0.00014 | 0.00018 | 0.00013 | 7.07E-05 | 0.00017 | 4.56E-05 | 6.89E-05 | 0 | 0.00016 |
| Oat | 0.00031 | 5.78E-05 | 0.00015 | 0.00053 | 0.00024 | 0.00028 | 0.00056 | 0.00021 | 0.00026 | 0.00035 | 0.00033 | 0.00012 | 0.00057 | 0.00022 | 0 | 0.00023 | 0.00032 | 0.00022 |
| Sptlc2 | 0.00014 | 0.00014 | 0.00021 | 0 | 0.0002 | 0.00016 | 0.0002 | 4.91E-05 | 0.00026 | 0.00018 | 0.00011 | 9.93E-05 | 0.00016 | 0.00016 | 4.84E-05 | 5.06E-05 | 0 | 0 |
| Eif3k | 0.00039 | 0.00035 | 0.00024 | 0.00055 | 0.00021 | 0.00039 | 0.00029 | 0.00022 | 0.00026 | 0.00014 | 0.00019 | 0.00034 | 0.00019 | 0.00035 | 0 | 0.00021 | 0.00014 | 0.0002 |
| Purb | 2.08E-05 | 0.00019 | 0.00028 | 0.00035 | 8.43E-05 | 6.52E-05 | 0 | 0.00035 | 0.00026 | 0.0001 | 0.00027 | 0.00015 | 0 | 0.00017 | 0.00055 | 0.00029 | 0.00026 | 0.00015 |
| Wdfy3 | 0 | 0.00029 | 0.00018 | 0.00018 | 0.00024 | 0.00018 | 6.60E-05 | 0.00019 | 0.00026 | 0.0002 | 0.00013 | 0.00013 | 1.21E-05 | 0.0002 | 0.00033 | 0.00015 | 0.00038 | 0.00021 |
| Mdh1 | 0.00029 | 9.44E-05 | 0.00014 | 0.00039 | 0.00013 | 0.00014 | 9.86E-05 | 0.00017 | 0.00026 | 0.00014 | 0.00012 | 2.29E-05 | 0.00014 | 0.00015 | 0 | 0.00028 | 7.73E-05 | 0.00017 |
| Cebpa | 6.24E-06 | 0.00021 | 0 | 0.00019 | 5.40E-05 | 0.00025 | 0 | 0.00029 | 0.00026 | 6.12E-05 | 0 | 0 | 0.00014 | 0.00039 | 0.00022 | 4.76E-06 | 0.00054 | 0.00014 |
| Hacd3 | 7.75E-05 | 0.00014 | 0.00012 | 2.41E-05 | 0.00013 | 5.75E-05 | 2.59E-05 | 0.00015 | 0.00026 | 3.26E-05 | 6.80E-05 | 0 | 0.00021 | 7.77E-05 | 5.54E-06 | 5.82E-06 | 0 | 0.00013 |
| Pgpep1 | 0 | 0.00012 | 2.72E-05 | 6.68E-05 | 0.00018 | 7.60E-05 | 2.20E-05 | 0 | 0.00026 | 0.00011 | 8.56E-05 | 8.16E-05 | 7.48E-05 | 0 | 1.15E-05 | 0 | 0 | 5.02E-05 |
| Rbbp7 | 0.00033 | 0.0002 | 0.00027 | 0.00048 | 0.00014 | 0.00014 | 8.15E-05 | 0.00012 | 0.00026 | 0.00016 | 0.00028 | 8.48E-05 | 0.00016 | 9.30E-05 | 3.96E-05 | 0.00068 | 0.00024 | 0.00021 |
| Papss1 | 0.00023 | 9.57E-05 | 2.94E-05 | 0.00022 | 0.00011 | 9.97E-05 | 0.00021 | 0.00028 | 0.00026 | 0.00011 | 9.39E-05 | 0.00016 | 0 | 9.12E-05 | 0.00014 | 0.00021 | 4.02E-06 | 0.00013 |

| | | | | | | | | | | | | | | | | | | |
|---|---|---|---|---|---|---|---|---|---|---|---|---|---|---|---|---|---|---|
| Shisa5 | 3.04E-05 | 0.00033 | 0.00023 | 0.00044 | 0.00028 | 0.00027 | 0.00056 | 0.00053 | 0.00026 | 0.00062 | 0.00017 | 0.00019 | 0.00072 | 0.0003 | 0 | 0.00016 | 5.75E-05 | 0.00062 |
| Trir | 8.27E-05 | 0.00034 | 0.00032 | 0.00043 | 0.0003 | 0.00021 | 0.00026 | 0.00039 | 0.00026 | 0.00028 | 0.00036 | 0.00022 | 0.0002 | 0.0003 | 0.00038 | 0.0004 | 7.32E-05 | 0.00029 |
| Tgif1 | 7.41E-05 | 0.00023 | 0.00012 | 0 | 5.38E-05 | 0 | 9.83E-05 | 5.00E-05 | 0.00026 | 0 | 5.02E-05 | 0.00015 | 7.86E-05 | 0.00022 | 0.00011 | 9.25E-05 | 0.00066 | 0 |
| Ndufa2 | 0.00035 | 0.00018 | 0.00022 | 0.00064 | 0.00029 | 0.00024 | 0.00026 | 0.00023 | 0.00026 | 0.00017 | 0.00032 | 0.00019 | 0.00024 | 0.00035 | 1.66E-05 | 0.0003 | 4.17E-05 | 0.00018 |
| Akt2 | 0.00025 | 0.0002 | 0.00013 | 2.57E-05 | 0.00015 | 0.00017 | 0.00019 | 0.00023 | 0.00026 | 0.00013 | 6.10E-05 | 0.00016 | 0.00013 | 2.21E-05 | 4.85E-05 | 6.25E-05 | 2.53E-05 | 8.57E-05 |
| Casp6 | 0.00012 | 5.58E-05 | 3.68E-05 | 0.00021 | 0.00017 | 0.00021 | 0.00019 | 7.63E-05 | 0.00026 | 0 | 8.32E-05 | 0.00035 | 0.00017 | 0.00024 | 4.11E-05 | 7.92E-05 | 0 | 0 |
| Ifitm1 | 1.28E-05 | 5.01E-05 | 0 | 2.38E-05 | 0 | 0.00044 | 0 | 0.00011 | 0.00026 | 3.88E-05 | 0 | 0 | 0 | 0.00109 | 0 | 3.73E-05 | 0 | 4.11E-05 |
| Scaf1 | 6.55E-05 | 0.00023 | 0.00015 | 0.00033 | 0.00012 | 6.74E-05 | 5.18E-05 | 0.00012 | 0.00026 | 0.00015 | 0.00025 | 0.0002 | 6.34E-05 | 0.00014 | 0.00036 | 0.00022 | 0.00016 | 0.00016 |
| Cstb | 0.00033 | 0.00038 | 0.00052 | 0.00039 | 0.00023 | 0.00031 | 0.00011 | 2.36E-05 | 0.00026 | 2.49E-05 | 0.00014 | 0.00033 | 0.00072 | 0.00015 | 0 | 0 | 0.00048 | 0.00028 |
| Atp8b1 | 2.03E-05 | 0.00013 | 0.00015 | 0 | 0 | 0 | 4.15E-05 | 0.00017 | 0.00026 | 0.00033 | 0.00036 | 2.56E-05 | 0.00057 | 0 | 0 | 0 | 0.00016 | 0.00018 |
| Rrbp1 | 0.00074 | 0.00022 | 0.00065 | 0.00053 | 0.00038 | 0.00053 | 0.00102 | 0.00053 | 0.00026 | 0.00034 | 0.00039 | 0.00044 | 0.00048 | 0.00044 | 0.00147 | 7.10E-05 | 0.00026 | 0.00023 |
| Atxn10 | 0.00052 | 0.00013 | 0.00015 | 0 | 0.00012 | 0.00023 | 0.00022 | 0.00013 | 0.00026 | 0.00016 | 0.0001 | 0.0002 | 4.99E-05 | 0.00013 | 0.00021 | 6.90E-05 | 0.00018 | 0.00016 |
| Tmem165 | 0.00026 | 0.00014 | 0.00021 | 7.88E-05 | 0.00015 | 0.00018 | 0.00029 | 0.00013 | 0.00026 | 0.00021 | 0.00017 | 4.43E-06 | 0.00013 | 0.00016 | 0.00012 | 0.00021 | 0 | 8.10E-05 |
| Rgmb | 0 | 0.00034 | 9.74E-05 | 0.00014 | 0.00014 | 8.91E-05 | 2.02E-05 | 0.00037 | 0.00026 | 0.00029 | 9.30E-05 | 6.41E-06 | 0 | 0 | 3.42E-05 | 0.0001 | 0.00025 | 0.00026 |
| Ifngr2 | 0.00018 | 0.00012 | 0.00037 | 0 | 0.00012 | 0.00035 | 0.0001 | 0.00043 | 0.00026 | 0.00021 | 7.16E-05 | 0.0002 | 0.00018 | 0 | 0 | 0 | 0 | 0.00015 |
| H2afy2 | 7.78E-05 | 1.18E-05 | 0 | 7.91E-05 | 5.49E-05 | 4.65E-05 | 5.35E-05 | 0 | 0.00026 | 0 | 7.85E-05 | 0 | 9.75E-06 | 8.18E-05 | 9.78E-07 | 0.00018 | 0 | 1.77E-05 |
| Lman1 | 0.00053 | 0 | 0.00055 | 0.00069 | 0.00028 | 0.00045 | 0.00116 | 0.00058 | 0.00026 | 0.00026 | 0.00025 | 0.00025 | 0.00052 | 0.00037 | 0.00045 | 0.0002 | 8.19E-05 | 0.00017 |
| Hnrnph1 | 0.00029 | 0.00013 | 0.00019 | 0.00014 | 4.81E-05 | 8.17E-05 | 2.35E-05 | 5.39E-05 | 0.00026 | 5.96E-05 | 9.35E-05 | 0.0001 | 0 | 7.51E-05 | 0.00031 | 0.00027 | 0.00025 | 1.27E-05 |
| Tead3 | 0 | 0.00014 | 5.11E-05 | 0.00013 | 0.00017 | 9.33E-05 | 2.92E-05 | 4.92E-05 | 0.00026 | 0.0001 | 0.00043 | 8.32E-05 | 0.00021 | 0.00021 | 0.00031 | 7.83E-05 | 7.12E-06 | 3.08E-05 |
| Dennd5a | 0.00012 | 0.0003 | 0.00013 | 0.00015 | 0.00027 | 0.00024 | 0.00013 | 0.00017 | 0.00026 | 0.00026 | 0.00034 | 0.00016 | 0.00018 | 0.00016 | 0.0001 | 0.00028 | 0.0002 | 0.00029 |
| Psmc3 | 0.00053 | 0.00027 | 0.00047 | 0.00033 | 0.00016 | 0.00026 | 0.00023 | 8.90E-05 | 0.00026 | 8.66E-05 | 0.00031 | 6.42E-05 | 6.83E-05 | 0.0002 | 9.32E-05 | 0.00041 | 0 | 7.93E-05 |
| Bin1 | 0.00017 | 0.00021 | 0.00025 | 0.0002 | 0 | 4.52E-05 | 2.74E-05 | 0 | 0.00026 | 0.00021 | 0.00022 | 0.00018 | 0.00031 | 0.00012 | 0.00035 | 0 | 0.00015 | 0 |
| Dlst | 0.0003 | 0.00021 | 0.00038 | 4.54E-05 | 8.12E-05 | 0.00015 | 3.61E-05 | 9.05E-05 | 0.00026 | 9.25E-05 | 0.00019 | 8.87E-05 | 0 | 0.00014 | 0.0004 | 0.00025 | 1.54E-06 | 3.59E-05 |
| Arf1 | 0.00015 | 0.00022 | 0.00045 | 0.00099 | 0.00028 | 0.00018 | 0.00053 | 0.00035 | 0.00026 | 0.00027 | 0.0004 | 0.00027 | 0.00024 | 0.0004 | 0.00019 | 0.00031 | 0.00033 | 0.00023 |
| Cpd | 0.00028 | 0.00018 | 0 | 0 | 0.00024 | 0.00029 | 0.00013 | 0.00012 | 0.00026 | 0.0002 | 0.00027 | 4.01E-05 | 0.00039 | 6.42E-05 | 0 | 0.00024 | 7.35E-05 | 2.17E-05 |
| Ociad1 | 0.00025 | 0.00016 | 0.00017 | 0.00016 | 0.00019 | 0.00016 | 7.58E-05 | 0.00021 | 0.00026 | 4.21E-05 | 8.12E-05 | 0.00013 | 0 | 6.97E-05 | 0.00012 | 0.00013 | 0 | 5.68E-05 |
| Adamtsl5 | 0 | 0.00015 | 2.24E-05 | 0 | 6.96E-05 | 0.00019 | 0 | 6.20E-05 | 0.00026 | 0.00019 | 5.74E-05 | 7.49E-05 | 0 | 0.00016 | 0.00024 | 0 | 0 | 0.00025 |
| Rap1b | 0.00065 | 0.0003 | 0.00028 | 0 | 0.0002 | 0.00033 | 0.00019 | 0.00031 | 0.00026 | 0.00038 | 0.00028 | 0.00032 | 0 | 0.0001 | 0.00023 | 0.00011 | 0.00048 | 0.00021 |
| Fis1 | 0.00024 | 0.00024 | 0.00023 | 0.0006 | 0.00023 | 0.00026 | 0.00021 | 0.00032 | 0.00026 | 0.00025 | 0.0003 | 5.77E-05 | 0.00021 | 0.00031 | 0 | 8.80E-05 | 0 | 0.00025 |
| Mat2a | 0.00013 | 0.00023 | 0.00039 | 0.00018 | 4.41E-05 | 0 | 0 | 0.00012 | 0.00026 | 0.00017 | 0.00015 | 0.00021 | 0.00015 | 6.31E-05 | 0.00026 | 0.00027 | 0.00189 | 0 |
| Nap1l1 | 0.00032 | 2.00E-05 | 0.00023 | 0 | 0 | 0.00015 | 0 | 0 | 0.00026 | 1.06E-05 | 1.60E-05 | 9.00E-05 | 9.91E-05 | 0 | 0 | 0.00035 | 0.00015 | 6.86E-05 |
| Gsdmd | 0.00013 | 0.0001 | 5.99E-05 | 0.00012 | 6.15E-05 | 6.86E-05 | 9.59E-05 | 4.57E-05 | 0.00026 | 0.00015 | 0.00011 | 3.07E-06 | 8.69E-05 | 0.00011 | 0.00011 | 5.02E-05 | 6.24E-05 | 0.00017 |
| Map2k2 | 0.00026 | 0.00026 | 0.00034 | 0.00038 | 0.00019 | 9.26E-05 | 0.00022 | 0.00015 | 0.00026 | 0.00012 | 0.00019 | 6.60E-05 | 0.0001 | 0.00013 | 0.00046 | 7.03E-05 | 8.04E-06 | 7.60E-05 |

| | | | | | | | | | | | | | | | | | | |
|---|---|---|---|---|---|---|---|---|---|---|---|---|---|---|---|---|---|---|
| Tmem256 | 0.00032 | 0.00017 | 2.58E-05 | 0.00051 | 9.09E-05 | 0.00015 | 0.00023 | 6.01E-05 | 0.00025 | 0.00011 | 7.51E-05 | 2.95E-05 | 0.00023 | 0.00024 | 0 | 0.0002 | 7.30E-05 | 0.00023 |
| Mesd | 0.00012 | 9.46E-05 | 0.00027 | 0.00029 | 0.00015 | 6.08E-05 | 5.98E-05 | 0.00025 | 0.00025 | 0.00012 | 0.00018 | 5.86E-05 | 0.00022 | 0.00015 | 6.80E-05 | 0.00015 | 2.31E-05 | 7.90E-05 |
| Rere | 9.24E-05 | 0.00041 | 0 | 0.00012 | 0.00012 | 0.00013 | 4.81E-05 | 3.94E-06 | 0.00025 | 0.00015 | 0 | 0.00025 | 6.17E-05 | 4.08E-05 | 0.0004 | 0.00014 | 0.00022 | 0.0002 |
| Myh9 | 9.44E-06 | 0.00019 | 0.00044 | 0.00058 | 0.00035 | 0.00031 | 0.00047 | 0.00053 | 0.00025 | 9.58E-05 | 0.00142 | 0.00056 | 0.00032 | 0.00052 | 0.00038 | 0.00016 | 4.36E-06 | 0.00035 |
| Fam78b | 0 | 2.41E-06 | 0.00025 | 0 | 0 | 0.00017 | 0 | 0.00025 | 0.00025 | 0 | 0 | 0.00048 | 0.00026 | 0 | 0 | 0 | 4.94E-05 | 0.00027 |
| Pafah1b2 | 0.00038 | 0.00023 | 0.00045 | 0.00021 | 0.00017 | 0.00019 | 0.00015 | 0.00017 | 0.00025 | 0.00015 | 0.0002 | 0.0002 | 0.00011 | 0.00017 | 6.28E-05 | 0.00023 | 3.29E-05 | 0.00013 |
| Dtnbp1 | 0.00017 | 0.0003 | 0.00035 | 0.00021 | 8.90E-05 | 0.00012 | 0.00021 | 7.51E-05 | 0.00025 | 0.00051 | 0.00013 | 9.30E-05 | 0.0002 | 0.00018 | 0.00031 | 0.00023 | 2.17E-05 | 0.00018 |
| Spg20 | 0.00014 | 0.00013 | 1.50E-05 | 0.00012 | 0.00023 | 0.00027 | 0.00018 | 0.00018 | 0.00025 | 1.87E-05 | 0 | 1.56E-05 | 6.15E-05 | 0.00021 | 8.46E-05 | 3.66E-05 | 0 | 1.16E-05 |
| Rbm3 | 0.00034 | 0.00014 | 0.00034 | 0.00072 | 5.22E-05 | 0.00014 | 4.57E-05 | 0.00013 | 0.00025 | 0 | 0.00032 | 6.92E-05 | 3.35E-05 | 0.00036 | 0.00012 | 0.00046 | 1.50E-05 | 2.30E-05 |
| Ndst1 | 0.00012 | 0.00029 | 0.00031 | 0.00018 | 0.00024 | 0.0002 | 3.07E-05 | 8.90E-05 | 0.00025 | 9.30E-05 | 0.00017 | 8.89E-05 | 0 | 0.00034 | 0.00028 | 7.24E-05 | 0.00024 | 0.00019 |
| M6pr | 0.0002 | 0.00017 | 0.00026 | 0.00014 | 0.00018 | 0.00027 | 0 | 4.05E-05 | 0.00025 | 0.00015 | 0.00061 | 0.00022 | 0.00017 | 0.00033 | 0 | 0.00011 | 0 | 1.96E-05 |
| Ror2 | 9.57E-06 | 0.0002 | 0.00011 | 0.00016 | 0.00041 | 8.60E-05 | 0.0003 | 0.0003 | 0.00025 | 6.11E-05 | 0.00036 | 0.0003 | 0 | 6.53E-05 | 0.00014 | 0.00016 | 0.00043 | 4.81E-06 |
| Cltc | 0.00057 | 0.00019 | 0.00018 | 0 | 8.43E-05 | 0.00025 | 0.00023 | 0 | 0.00025 | 4.10E-05 | 9.64E-05 | 0.00013 | 5.18E-07 | 3.99E-05 | 4.37E-05 | 6.48E-05 | 0.00042 | 5.93E-05 |
| Msn | 0.00031 | 0.00033 | 0.00083 | 0.00044 | 0.00018 | 0.00019 | 0.00043 | 0.00043 | 0.00025 | 0.0005 | 0.00137 | 0.00058 | 0.00033 | 0.00053 | 0.00055 | 0.00035 | 0.00045 | 0.00059 |
| Sf3b3 | 0.00021 | 0.00025 | 0.00046 | 0.00019 | 0.0002 | 0.0001 | 0.00011 | 0.00016 | 0.00025 | 0.00011 | 9.94E-05 | 6.06E-05 | 9.97E-05 | 0.00017 | 0.00031 | 0.00031 | 0 | 0.00013 |
| Ctsz | 0.00012 | 0.00027 | 0.00018 | 0.00058 | 0.00046 | 0.00049 | 0.00055 | 0.00069 | 0.00025 | 0.00052 | 0.00037 | 0.00044 | 0.00101 | 0.00068 | 2.50E-05 | 0.00012 | 0.00023 | 0.00075 |
| Cacna1c | 0 | 0.00019 | 6.78E-05 | 0 | 0.00048 | 0.00036 | 3.00E-05 | 0.00045 | 0.00025 | 0 | 0.00028 | 3.32E-05 | 2.34E-07 | 0.00038 | 0.00053 | 1.40E-05 | 0.00051 | 4.12E-06 |
| Mta2 | 0.00029 | 0.00029 | 0.00031 | 0.00015 | 0.00018 | 8.45E-05 | 0 | 7.95E-05 | 0.00025 | 8.89E-05 | 0.00013 | 8.32E-05 | 0.00011 | 5.69E-05 | 0.00019 | 0.00017 | 0.0002 | 6.08E-05 |
| Susd6 | 0.00011 | 0.00033 | 0.00018 | 0.0002 | 0.00026 | 0.00018 | 0.00033 | 0.00074 | 0.00025 | 0.00015 | 9.39E-05 | 5.81E-06 | 0.00094 | 0.00025 | 0.00019 | 0.00017 | 0.00031 | 0.00039 |
| Ganab | 0.00014 | 0.0002 | 0.00027 | 0.0004 | 0.00019 | 0.00026 | 0.00048 | 0.00051 | 0.00025 | 0.00024 | 0.00018 | 0.00012 | 0.00031 | 0.00025 | 0.00025 | 0.00027 | 0.00013 | 0.00017 |
| Ppp2r1a | 0.00032 | 0.00029 | 0.0003 | 0.00038 | 0.00014 | 0.00014 | 2.43E-05 | 5.85E-05 | 0.00025 | 0.0001 | 0.00021 | 0.00023 | 1.57E-06 | 0.00015 | 1.96E-05 | 0.00015 | 0.0001 | 0.00017 |
| Arpc5 | 0.00038 | 0.00023 | 0.00033 | 0.00019 | 5.01E-05 | 0.00012 | 0.00011 | 0.00014 | 0.00025 | 0.00022 | 0.0007 | 0.00021 | 3.24E-05 | 0.00013 | 0 | 0.00017 | 3.17E-05 | 0.00019 |
| Rab14 | 0.00056 | 0.00033 | 0.00027 | 0 | 0.00016 | 0.00033 | 2.40E-05 | 4.06E-05 | 0.00025 | 0.00012 | 0.00024 | 0.00024 | 1.08E-05 | 0.0001 | 0.00033 | 0.00013 | 0.00017 | 0.00011 |
| Tspan17 | 0 | 0.00014 | 0 | 5.37E-05 | 0.00013 | 0.00018 | 0 | 1.38E-05 | 0.00025 | 4.92E-05 | 3.67E-05 | 7.43E-05 | 0.00011 | 0.00014 | 0.00014 | 0 | 0 | 0.00019 |
| Bsg | 0.00052 | 1.08E-05 | 9.76E-05 | 5.39E-05 | 0.00012 | 0.00031 | 0.00052 | 0.00041 | 0.00025 | 0.00018 | 0.00027 | 0.0013 | 0.00047 | 0.00029 | 0 | 5.75E-05 | 0 | 0.0002 |
| Pakap.1 | 2.38E-05 | 0.00017 | 0.0004 | 0 | 0.00012 | 0.00061 | 0 | 0 | 0.00025 | 0 | 0.00034 | 0.00039 | 0 | 0.00011 | 0 | 7.22E-05 | 0.0002 | 5.17E-05 |
| Nt5e | 0 | 0 | 0 | 1.72E-06 | 4.04E-05 | 8.00E-05 | 6.71E-05 | 1.63E-05 | 0.00025 | 0 | 5.19E-05 | 0.00033 | 6.99E-05 | 0.00017 | 0 | 0 | 0 | 0 |
| Hnrnpa1 | 0.00057 | 7.28E-05 | 0.00043 | 0.00079 | 2.59E-05 | 7.20E-05 | 0.00018 | 5.81E-05 | 0.00025 | 2.95E-05 | 0.00016 | 0.0002 | 0 | 0.00025 | 0 | 0.00059 | 0.00036 | 0 |
| Osbpl5 | 2.66E-05 | 0.00021 | 0.00016 | 0 | 0.00026 | 0.00012 | 4.74E-05 | 8.65E-05 | 0.00025 | 0.00015 | 0.0003 | 0.00022 | 4.29E-05 | 0.00014 | 0.00017 | 0 | 0 | 0.00025 |
| Neo1 | 9.14E-05 | 0.00023 | 0 | 0 | 0.00017 | 5.97E-07 | 0.00057 | 0.00024 | 0.00025 | 0 | 6.63E-05 | 0 | 7.40E-05 | 4.70E-05 | 0 | 4.71E-05 | 0.00023 | 0.00021 |
| Tmod3 | 0.00015 | 0.00032 | 0.0002 | 0.00047 | 0.00024 | 9.94E-05 | 0.00018 | 0.0003 | 0.00025 | 0.00027 | 0.00018 | 0.00035 | 0.00014 | 0.00011 | 0.0002 | 0.00025 | 0.00031 | 0.0005 |
| Chst12 | 8.72E-05 | 0.00018 | 9.16E-05 | 0.0003 | 0.00039 | 0.00037 | 0.00072 | 0.0002 | 0.00025 | 0.00028 | 0.00048 | 0.00019 | 0.00022 | 0.00033 | 9.12E-05 | 1.28E-05 | 0 | 0.00026 |
| Ctsa | 7.91E-05 | 0.00046 | 0 | 0.00012 | 0.0005 | 0.00064 | 0.00046 | 0.00045 | 0.00025 | 0.00026 | 0.0002 | 0.00028 | 0.0008 | 0.00076 | 0 | 0 | 0 | 0.00037 |

| | | | | | | | | | | | | | | | | | | |
|---|---|---|---|---|---|---|---|---|---|---|---|---|---|---|---|---|---|---|
| Huwe1 | 0.00023 | 0.00033 | 0.00023 | 0.00022 | 0.00024 | 0.00017 | 0.00014 | 0.00012 | 0.00025 | 0.0002 | 0.00015 | 8.74E-05 | 8.76E-05 | 9.71E-05 | 0.00028 | 0.00021 | 0.00023 | 0.00013 |
| Pigt | 0.00011 | 6.96E-05 | 0.00012 | 0.00021 | 0.00022 | 0.00026 | 0.00018 | 0.00026 | 0.00025 | 0.00032 | 7.46E-05 | 0.00014 | 0.00032 | 0.00018 | 0 | 3.99E-05 | 0 | 0.00043 |
| Ddx21 | 0.00029 | 0.00017 | 0.00077 | 0.00018 | 0 | 3.98E-06 | 0 | 0 | 0.00025 | 0.00011 | 8.92E-05 | 0 | 2.51E-05 | 8.36E-05 | 0.00035 | 0.00015 | 0.00034 | 4.90E-05 |
| Lamtor2 | 0.00037 | 6.03E-05 | 6.90E-05 | 0.00022 | 0.00017 | 0.00021 | 0.00022 | 0.00018 | 0.00025 | 0.00013 | 8.28E-05 | 0.00015 | 4.02E-05 | 0.00014 | 0 | 0 | 8.66E-06 | 0.00014 |
| Nnt | 0.00013 | 0.00011 | 0 | 9.75E-05 | 0.00011 | 4.15E-05 | 0.0001 | 4.43E-05 | 0.00025 | 0.00015 | 0.00013 | 0 | 7.13E-05 | 0.00012 | 0 | 8.10E-05 | 0 | 0.00015 |
| Ncstn | 0.00017 | 0.00017 | 0.00018 | 0.00022 | 0.00017 | 0.00035 | 0.00023 | 0.00038 | 0.00025 | 0.0003 | 0.0003 | 0.00027 | 0.00033 | 0.00032 | 0 | 1.69E-05 | 0 | 0.00031 |
| Mapkapk2 | 0.00018 | 0.00044 | 0.00043 | 0.00023 | 0.0003 | 0.00036 | 7.80E-05 | 9.34E-05 | 0.00025 | 8.82E-05 | 0.00026 | 0.00038 | 0.00016 | 0.00022 | 0.00048 | 0.00019 | 0.00062 | 0.0002 |
| Adamts9 | 3.86E-05 | 1.61E-05 | 0 | 1.25E-05 | 8.89E-05 | 0 | 1.73E-05 | 0 | 0.00025 | 1.02E-06 | 0.00022 | 0 | 0 | 2.57E-05 | 2.95E-05 | 3.43E-05 | 0.00011 | 0 |
| Acss3 | 0 | 4.45E-05 | 0 | 0 | 0 | 7.42E-06 | 0 | 0.0001 | 0.00025 | 0.00012 | 0 | 0 | 0 | 0 | 0 | 0 | 0 | 0 |
| B3gnt9 | 5.75E-05 | 5.42E-05 | 0.00013 | 0.0004 | 0.00045 | 0.00053 | 0.00077 | 0.0004 | 0.00025 | 0.00018 | 0.00016 | 0.00014 | 0.00042 | 0.00072 | 0 | 0 | 0 | 0.00023 |
| Lbr | 0.0001 | 0.00014 | 0.00017 | 2.06E-05 | 0 | 4.25E-05 | 0 | 4.05E-05 | 0.00025 | 1.44E-05 | 5.63E-06 | 5.38E-05 | 0 | 0 | 3.47E-05 | 0.00063 | 0.00012 | 0 |
| Qk | 0.00039 | 0.00038 | 0.00036 | 1.42E-05 | 0.00022 | 0.00028 | 0.00017 | 0.00022 | 0.00025 | 0.00022 | 0.00029 | 0.0003 | 0.00018 | 0.00021 | 0.00024 | 0.00042 | 0.00035 | 0.00015 |
| Nfasc | 0 | 0 | 0 | 0 | 0 | 0.00015 | 1.29E-05 | 2.40E-06 | 0.00025 | 0.00013 | 0 | 3.34E-05 | 0.00043 | 6.00E-05 | 5.49E-05 | 2.65E-05 | 0 | 0.00052 |
| Ptprg | 0.00015 | 0.00014 | 0 | 0 | 5.30E-05 | 0.0002 | 1.20E-05 | 0.00016 | 0.00025 | 0 | 0 | 4.63E-05 | 0 | 1.34E-05 | 0 | 4.52E-05 | 5.51E-05 | 0 |
| Rab5c | 0.00046 | 0.00027 | 0.00024 | 6.61E-05 | 0.00033 | 0.00032 | 0.00018 | 0.00042 | 0.00025 | 0.00024 | 0.00021 | 9.36E-05 | 0.0002 | 0.00026 | 0 | 0.00012 | 0.00026 | 0.00016 |
| Serping1 | 0 | 0.00023 | 0.00011 | 0 | 0.00066 | 0.00113 | 0.00086 | 0.00104 | 0.00025 | 0.0015 | 0.00049 | 0.00091 | 0.00285 | 0.00133 | 0 | 0 | 0 | 0.00185 |
| Snx9 | 0.00012 | 0.00037 | 0.0004 | 0.00014 | 0.00057 | 0.00055 | 0.00023 | 0.0003 | 0.00025 | 0.00018 | 0.00028 | 0.0004 | 0.00032 | 0.00061 | 0.00017 | 0.00019 | 0.00015 | 0.00033 |
| Sdf4 | 0.00024 | 0.00023 | 0.00018 | 0.00022 | 0.00039 | 0.00052 | 0.00039 | 0.00038 | 0.00025 | 0.00034 | 0.00027 | 0.00037 | 0.00054 | 0.00032 | 0.00024 | 8.37E-05 | 0 | 0.0005 |
| Acvr1 | 0.00013 | 0.0003 | 0.00023 | 0 | 0.00024 | 0.00018 | 0.0003 | 0.00034 | 0.00025 | 5.27E-05 | 0.00016 | 0.00014 | 2.01E-05 | 0.00013 | 7.66E-05 | 8.30E-05 | 0.00027 | 0.0002 |
| Ppp2ca | 0.00046 | 0.0003 | 0.00044 | 0.00011 | 0.00022 | 0.00022 | 0.00032 | 3.35E-05 | 0.00025 | 0.00014 | 0.00016 | 0.00022 | 0.00014 | 0.00013 | 0.0002 | 0.00023 | 0.0003 | 6.29E-05 |
| Capn1 | 0.0001 | 0.00014 | 0.00013 | 0.00012 | 4.17E-05 | 0.00022 | 6.93E-05 | 4.89E-05 | 0.00025 | 0.00027 | 7.79E-05 | 0.00036 | 6.22E-05 | 0 | 6.26E-05 | 0 | 0 | 0.00029 |
| Sfxn3 | 5.69E-05 | 0.0004 | 8.30E-05 | 7.17E-05 | 0.0004 | 0.0003 | 0.00015 | 3.46E-05 | 0.00025 | 0.00029 | 0.00035 | 0.00027 | 0.00012 | 0.0005 | 8.17E-05 | 0 | 0 | 0.00041 |
| Atp2b1 | 0.00014 | 0.00021 | 0.0002 | 0 | 3.66E-05 | 0.00018 | 9.31E-05 | 0.00014 | 0.00025 | 0.00016 | 0.00019 | 0.00016 | 0 | 9.37E-05 | 3.71E-05 | 0.00013 | 0.0002 | 5.92E-05 |
| Tpd52l2 | 0.00014 | 0.0003 | 0.0002 | 0.00039 | 0.00025 | 0.00022 | 0.00011 | 0.00036 | 0.00025 | 0.00026 | 0.00014 | 0.00015 | 0.00017 | 0.00025 | 4.22E-05 | 0.00014 | 1.81E-05 | 0.0002 |
| Tmem50a | 0.0002 | 0.00039 | 0.00016 | 0.00026 | 0.00034 | 0.00054 | 0.00025 | 0.00049 | 0.00025 | 0.0004 | 0.00025 | 0.00046 | 0.0004 | 0.00049 | 0 | 4.58E-05 | 0 | 0.00059 |
| Sf3b2 | 0.00025 | 0.00033 | 0.00021 | 0.00035 | 0.00014 | 9.16E-05 | 9.16E-05 | 0.00012 | 0.00025 | 0.00018 | 0.00016 | 0.00014 | 8.69E-05 | 0.0001 | 0.00036 | 0.00037 | 0.00029 | 0.0002 |
| Cyp7b1 | 0.0003 | 0 | 0 | 7.70E-07 | 0 | 0.00017 | 0.00018 | 0 | 0.00025 | 0.00013 | 0 | 0 | 0.0014 | 0 | 0.00011 | 5.22E-05 | 0 | 0 |
| Hnrnpl | 0.00026 | 0.00015 | 0.00024 | 0.00059 | 0.0001 | 0.00019 | 0.0002 | 0.00027 | 0.00025 | 0.00016 | 0.00024 | 6.30E-05 | 0.0001 | 0.00026 | 5.25E-06 | 0.00049 | 0.00041 | 0.00017 |
| Ube2m | 0.00021 | 0.00022 | 0.00038 | 0.00049 | 0.0001 | 4.64E-05 | 0.00012 | 0.00027 | 0.00025 | 8.86E-05 | 0.00027 | 0.00017 | 0 | 0.0001 | 0.00013 | 0.0002 | 0.00013 | 0.00011 |
| Clta | 0.00029 | 0.00043 | 0.00032 | 0.001 | 0.00037 | 0.00028 | 0.00046 | 0.00028 | 0.00025 | 0.0003 | 0.00044 | 0.00037 | 0.00026 | 0.00052 | 0.0004 | 0.0002 | 0.00046 | 0.00048 |
| Slc9a1 | 0 | 0.00016 | 0.00021 | 0.00023 | 0.00029 | 0.00016 | 0.00017 | 0.0004 | 0.00025 | 0.00026 | 0 | 6.60E-05 | 0.00022 | 0.00018 | 4.60E-05 | 0.00015 | 0.00038 | 0.00019 |
| Cnksr3 | 0 | 6.48E-05 | 0 | 1.57E-08 | 8.87E-05 | 0.00025 | 0 | 0 | 0.00025 | 1.06E-05 | 4.90E-05 | 3.19E-05 | 1.13E-06 | 9.03E-05 | 8.62E-05 | 0 | 6.50E-05 | 3.69E-05 |
| Hectd1 | 0.0001 | 0.00024 | 9.87E-05 | 0.00027 | 0.0002 | 6.83E-05 | 7.03E-05 | 3.91E-05 | 0.00025 | 6.34E-05 | 0.00016 | 9.73E-05 | 9.39E-05 | 0.00024 | 0.00026 | 9.54E-05 | 0.00016 | 6.35E-05 |

| | | | | | | | | | | | | | | | | | | |
|---|---|---|---|---|---|---|---|---|---|---|---|---|---|---|---|---|---|---|
| Hs6st1 | 6.19E-05 | 0.00011 | 0.0002 | 0 | 0.00021 | 0.0002 | 0 | 0.00027 | 0.00025 | 5.25E-05 | 0.00073 | 0 | 0 | 0.00063 | 0 | 0.00012 | 0 | 4.15E-05 |
| Acsl4 | 0.00025 | 3.39E-05 | 0.00032 | 0 | 0.00017 | 1.51E-05 | 0.00036 | 0.00049 | 0.00025 | 0.00013 | 0.00029 | 0.00023 | 0.00039 | 0 | 2.31E-05 | 3.59E-06 | 2.38E-05 | 0 |
| Shc1 | 0.00023 | 0.00039 | 6.68E-05 | 0.00024 | 0.00041 | 0.00047 | 0.00013 | 0.0004 | 0.00025 | 0.00012 | 7.26E-05 | 0.00027 | 0.00033 | 0.00033 | 0.00021 | 2.17E-05 | 3.18E-05 | 0.00017 |
| Irak1 | 0.00015 | 0.00023 | 0.00021 | 0.00016 | 0.00024 | 0.00021 | 0.00018 | 0.00012 | 0.00025 | 0.00026 | 0.00012 | 8.72E-05 | 0.00014 | 0.0001 | 1.22E-05 | 0.00014 | 4.49E-05 | 0.00022 |
| Flnb | 0 | 0.00012 | 0.00059 | 0 | 0.00025 | 0 | 4.46E-05 | 0.00038 | 0.00025 | 0.00049 | 0.00135 | 0.00076 | 0 | 0 | 0.00031 | 0.00017 | 0 | 0.00018 |
| Cul4a | 0.00015 | 0.0002 | 0.00021 | 9.09E-05 | 0.00017 | 0.00013 | 0.00017 | 8.11E-05 | 0.00025 | 8.39E-05 | 0.00017 | 0.00019 | 0.0001 | 0.00011 | 0.00021 | 0.00014 | 0 | 4.18E-05 |
| Pomp | 0.00029 | 0.00011 | 0.00026 | 0.00023 | 0.00012 | 6.35E-05 | 1.07E-05 | 9.97E-05 | 0.00025 | 5.78E-05 | 0.00013 | 6.46E-05 | 5.41E-05 | 7.04E-05 | 4.36E-05 | 0.00014 | 3.46E-05 | 8.26E-05 |
| Ndufa8 | 0.00024 | 0.00017 | 0.00037 | 0.00063 | 0.00021 | 0.00013 | 0.00021 | 0.00042 | 0.00025 | 0.0002 | 0.00026 | 0.00017 | 0.00024 | 0.00029 | 1.87E-05 | 0.00018 | 0 | 0.00022 |
| Ppp3ca | 0.00012 | 0.00019 | 5.64E-05 | 0.00027 | 8.54E-05 | 0 | 0.00043 | 0.00021 | 0.00024 | 7.82E-05 | 0 | 3.69E-05 | 0.00016 | 0 | 0.00022 | 0 | 0.00032 | 0.00016 |
| Galnt2 | 0.00014 | 0.00019 | 0.00032 | 0.00055 | 0.0004 | 0.00019 | 0.00044 | 0.00023 | 0.00024 | 0.00033 | 0.00041 | 0.00015 | 0.00075 | 0.00031 | 0.00017 | 0.0002 | 0.00013 | 0.00058 |
| Arcn1 | 0.00075 | 0.00021 | 0.00047 | 0.0003 | 0.00024 | 0.00038 | 0.00065 | 0.00024 | 0.00024 | 0.00023 | 0.00024 | 0.00032 | 0.00016 | 0.00027 | 0.00028 | 0.00019 | 0 | 0.00017 |
| Dync1h1 | 0.00028 | 0.00041 | 0.00036 | 0.00012 | 0.00024 | 0.00033 | 0.00041 | 0.00034 | 0.00024 | 0.00036 | 0.00033 | 0.00028 | 0.00016 | 0.00027 | 0.00023 | 0.00014 | 0.00024 | 0.00031 |
| Khdrbs1 | 0.00037 | 0.00025 | 0.00026 | 0.00034 | 0.00012 | 0.00016 | 9.91E-05 | 0.00012 | 0.00024 | 6.20E-05 | 0.00015 | 0.00014 | 6.56E-05 | 0.00015 | 0.00014 | 0.00027 | 0.00014 | 7.50E-05 |
| Cox5b | 0.00027 | 0.00012 | 0.0003 | 0.00082 | 0.00012 | 5.28E-05 | 0.00013 | 0.00017 | 0.00024 | 8.70E-05 | 0.00022 | 0.00013 | 3.48E-05 | 0.00019 | 0.00012 | 0.00022 | 2.72E-05 | 0.00015 |
| Srsf7 | 0.00022 | 0.00017 | 0.00039 | 0.00034 | 0 | 0 | 0 | 0 | 0.00024 | 3.26E-05 | 0.00015 | 0.00017 | 9.60E-05 | 8.77E-05 | 0.00033 | 0.00057 | 0.00086 | 0 |
| Lin7a | 0 | 2.54E-05 | 5.16E-05 | 0 | 0 | 2.80E-05 | 0 | 0.00011 | 0.00024 | 0 | 0 | 0 | 0 | 0 | 0 | 7.56E-06 | 0 | 2.81E-05 |
| Stip1 | 0.00038 | 0.00012 | 0.00036 | 3.83E-05 | 7.93E-05 | 9.71E-06 | 0 | 4.91E-06 | 0.00024 | 1.29E-05 | 0.00013 | 0.00015 | 6.55E-05 | 0.00013 | 0 | 0.00035 | 5.46E-05 | 0 |
| Atf1 | 7.19E-05 | 0.00018 | 0.00016 | 0.0005 | 0.00011 | 8.01E-05 | 5.02E-05 | 7.98E-05 | 0.00024 | 0.00012 | 6.62E-05 | 0.00017 | 8.62E-05 | 0.00014 | 0.0002 | 0.00033 | 0.00041 | 0.00015 |
| Slc25a3 | 0.00032 | 0.00034 | 0.00045 | 0.00059 | 0.00013 | 0.0003 | 0.00017 | 0.00021 | 0.00024 | 0.00026 | 0.00027 | 0.00014 | 0.00012 | 0.00039 | 0.00011 | 0.00025 | 0.00073 | 0.00034 |
| Gnb2 | 0.00037 | 0.00028 | 0.0003 | 0.00029 | 0.00021 | 0.00019 | 0.00017 | 0.00021 | 0.00024 | 0.0002 | 0.00035 | 0.00029 | 6.46E-05 | 8.28E-05 | 0 | 0.00021 | 7.96E-05 | 0.0002 |
| Ch25h | 0 | 0 | 9.18E-05 | 0 | 0 | 0 | 0 | 9.80E-05 | 0.00024 | 0 | 0 | 1.82E-05 | 0.00035 | 0 | 0 | 4.06E-05 | 3.50E-05 | 0 |
| Mitf | 0 | 0.00026 | 0 | 0 | 4.52E-05 | 0.0004 | 0 | 4.50E-06 | 0.00024 | 0.00012 | 0 | 0.00038 | 0.00026 | 0 | 0 | 0 | 0.00016 | 0 |
| Pi15 | 3.12E-05 | 0 | 3.13E-05 | 0 | 2.56E-05 | 0.00019 | 0 | 0 | 0.00024 | 7.41E-05 | 0 | 0 | 0.00017 | 0.00015 | 0 | 2.46E-05 | 0 | 8.44E-06 |
| Gcnt4 | 0 | 0 | 0 | 0 | 0.00015 | 0.00072 | 0 | 0.00014 | 0.00024 | 0 | 5.22E-05 | 0 | 0 | 0.00021 | 0 | 0 | 0 | 0 |
| Mapk1ip1l | 0.00017 | 0.00018 | 0.00036 | 0.00038 | 0.00011 | 4.42E-05 | 4.60E-06 | 0.0002 | 0.00024 | 0.00015 | 0.00016 | 0.00016 | 0.0001 | 0.00016 | 7.33E-05 | 0.00017 | 0.00016 | 0.00014 |
| 2-Mar | 0.00019 | 0.0002 | 0 | 0.00026 | 0.00012 | 0.00013 | 9.19E-05 | 0.00011 | 0.00024 | 0.00019 | 0.00011 | 0.00021 | 0.00031 | 0.00011 | 2.10E-05 | 0 | 1.22E-05 | 0.00014 |
| Arhgef25 | 0.00022 | 0.00017 | 0 | 0.0003 | 0.00023 | 0.00032 | 0.0004 | 0.00029 | 0.00024 | 0.00016 | 0.00028 | 0.00019 | 0.00016 | 0.00018 | 0.0006 | 0 | 6.88E-05 | 0.00044 |
| Pcbp1 | 0.00046 | 0.00029 | 0.00047 | 0.00033 | 0.0001 | 0.00013 | 0.00017 | 0.0001 | 0.00024 | 0.0001 | 0.00015 | 0.0002 | 6.33E-05 | 0.00017 | 0.00018 | 0.00025 | 0.00021 | 0.00012 |
| Dlgap4 | 7.46E-05 | 0.00035 | 0.00046 | 0.00035 | 0.00016 | 0.00012 | 0.00014 | 0.0003 | 0.00024 | 0.00039 | 0.0004 | 0.00026 | 9.62E-05 | 0.00023 | 0.00043 | 0 | 0.00056 | 0.00023 |
| Prrc2c | 0.00048 | 0.0003 | 0.00026 | 0 | 0.00016 | 0.00025 | 0.00013 | 5.03E-05 | 0.00024 | 7.30E-05 | 7.99E-05 | 0.00025 | 8.51E-05 | 6.39E-05 | 0.00039 | 0.00023 | 0.00041 | 2.84E-05 |
| Zfand3 | 0.00019 | 0.00033 | 0.00035 | 0.00016 | 0.00029 | 0.00035 | 0.00025 | 0.00033 | 0.00024 | 0.0004 | 0.00024 | 0.00035 | 0.00014 | 0.00025 | 0.00023 | 6.61E-05 | 0.00036 | 0.00033 |
| Bzw1 | 0.0007 | 0.00024 | 0.00048 | 1.18E-05 | 4.39E-05 | 0.00029 | 0.00015 | 5.51E-05 | 0.00024 | 0.00011 | 0.00018 | 0.00018 | 0 | 6.50E-05 | 9.40E-05 | 0.00026 | 0.0003 | 6.23E-05 |
| Atp5a1 | 0.00022 | 0.00011 | 0.00029 | 0.00051 | 5.79E-05 | 6.09E-05 | 0.00016 | 0.00015 | 0.00024 | 2.06E-05 | 9.44E-05 | 7.05E-05 | 6.56E-05 | 6.50E-05 | 0 | 0.00017 | 0.00013 | 7.27E-05 |

| | | | | | | | | | | | | | | | | | | |
|---|---|---|---|---|---|---|---|---|---|---|---|---|---|---|---|---|---|---|
| Pknox2 | 5.32E-05 | 0 | 0.00019 | 0 | 0 | 0 | 0.00028 | 2.80E-05 | 0.00024 | 0 | 0.00012 | 0 | 0.00048 | 0 | 0 | 6.99E-05 | 0 | 0 |
| Cdc42se1 | 6.54E-05 | 0.00015 | 0.00049 | 0.00033 | 0.00025 | 7.58E-05 | 6.55E-05 | 0.00031 | 0.00024 | 0.00018 | 0.00025 | 0.00017 | 0.00012 | 0.00021 | 7.46E-05 | 0.00012 | 0.00012 | 5.33E-05 |
| Klhl5 | 0.00012 | 9.89E-05 | 0.00011 | 6.42E-06 | 0.00021 | 0.00024 | 0.00015 | 0 | 0.00024 | 9.51E-05 | 9.16E-05 | 0.00011 | 0.00017 | 0.00021 | 0 | 0.00013 | 3.82E-05 | 0.00012 |
| Tmed7 | 0.0004 | 0.00022 | 0.00033 | 0.00026 | 0.00023 | 0.00035 | 0.00017 | 0.00029 | 0.00024 | 0.00016 | 0.00025 | 0.00019 | 0.00021 | 0.00025 | 6.38E-05 | 0.00016 | 0.00033 | 0.00019 |
| Mfap5 | 0 | 0 | 0.00081 | 1.12E-05 | 0 | 0 | 0.00207 | 0.00112 | 0.00024 | 0.00122 | 0.00077 | 0.00036 | 0.00118 | 0 | 0 | 0.00015 | 0 | 0.00125 |
| Etv5 | 0 | 0.00018 | 6.07E-05 | 2.25E-05 | 0.00025 | 0 | 0 | 0.00032 | 0.00024 | 8.91E-05 | 0 | 0.00021 | 0.00018 | 0.00016 | 0 | 0.0002 | 0.00014 | 0.00011 |
| Snrpb | 0.00019 | 0.00013 | 0.00031 | 0.00019 | 4.79E-05 | 3.51E-05 | 0 | 3.30E-05 | 0.00024 | 5.51E-05 | 0.00015 | 6.60E-05 | 1.68E-05 | 9.00E-05 | 0 | 0.0003 | 8.62E-05 | 4.05E-05 |
| Setd7 | 3.40E-05 | 0.0002 | 7.68E-05 | 0.00027 | 0.00017 | 2.46E-05 | 0.00032 | 0.0001 | 0.00024 | 0.00024 | 0.0002 | 0 | 0.00024 | 0.00014 | 3.38E-06 | 3.07E-05 | 2.83E-05 | 0.0001 |
| Klf5 | 0 | 5.74E-05 | 3.77E-05 | 0 | 0 | 9.08E-06 | 0 | 0.0002 | 0.00024 | 0 | 0 | 0.00027 | 0 | 0 | 0 | 6.39E-05 | 4.82E-05 | 0.00021 |
| Serbp1 | 0.00036 | 0.00024 | 0.00049 | 0.00037 | 3.67E-05 | 0.00012 | 0 | 6.52E-05 | 0.00024 | 3.99E-05 | 0.00016 | 0.00019 | 0 | 6.95E-05 | 1.90E-05 | 0.0004 | 0.00015 | 8.60E-05 |
| Pdha1 | 0.00039 | 0.00016 | 0.00019 | 0.00025 | 8.54E-05 | 0.00014 | 0.00011 | 0.00013 | 0.00024 | 0.00013 | 0.00025 | 0.00011 | 9.60E-05 | 0.00021 | 0 | 0.00011 | 0.00012 | 0.00018 |
| Penk | 0 | 0 | 1.05E-05 | 0 | 0 | 2.08E-05 | 0 | 0 | 0.00024 | 6.90E-05 | 0 | 0 | 0 | 6.47E-05 | 0 | 1.08E-05 | 0 | 0.00032 |
| Nsd1 | 0.00028 | 0.00033 | 0.00015 | 0.00024 | 0.0002 | 0.00022 | 0.0002 | 0.00012 | 0.00024 | 0.00011 | 0.00015 | 0.00012 | 7.06E-05 | 0.00015 | 0.00059 | 0.00012 | 7.00E-05 | 0.00013 |
| Ext2 | 0.00026 | 9.22E-05 | 0.00017 | 0.00031 | 0.00014 | 0.00023 | 0.00027 | 0.00025 | 0.00024 | 0.00027 | 0.0002 | 8.92E-05 | 0.00033 | 0.00026 | 5.59E-05 | 0.00013 | 6.07E-06 | 0.00025 |
| Clstn1 | 5.65E-05 | 0.00023 | 0.00017 | 0.00012 | 0.00021 | 0.00014 | 3.32E-05 | 0.00028 | 0.00024 | 0.00032 | 4.83E-05 | 0.0001 | 8.33E-05 | 0.00014 | 0.00017 | 5.70E-05 | 0.00011 | 0.00029 |
| Hnrnpa0 | 7.13E-05 | 0.00021 | 0.00029 | 0.00041 | 0.0001 | 0 | 0 | 0.0002 | 0.00024 | 7.17E-05 | 0.00012 | 0.00044 | 0 | 2.23E-05 | 0.00061 | 0.00026 | 9.52E-05 | 0.00014 |
| Srsf1 | 0.00017 | 0.00014 | 0.00044 | 0.00078 | 5.56E-05 | 0 | 0 | 5.88E-05 | 0.00024 | 6.54E-05 | 0.00021 | 0 | 0.00011 | 0.0001 | 0.00063 | 0.0006 | 0 | 6.44E-05 |
| Snrpd3 | 0.00041 | 0.00012 | 0.00044 | 0.00035 | 3.75E-05 | 2.21E-06 | 1.39E-05 | 6.16E-06 | 0.00024 | 0 | 0.00011 | 0.00011 | 7.07E-05 | 5.60E-05 | 0 | 0.00031 | 0 | 0 |
| Cast | 0.00019 | 0.00028 | 0.00017 | 0.00015 | 0.00024 | 0.00023 | 0.00027 | 8.20E-05 | 0.00024 | 0.0004 | 0.00015 | 0.00043 | 0.00023 | 0.00016 | 0.00015 | 0.00012 | 0.00019 | 0.00037 |
| Apeh | 6.01E-05 | 0.0001 | 3.46E-05 | 0.00012 | 8.17E-05 | 1.02E-05 | 3.63E-05 | 7.42E-05 | 0.00024 | 8.41E-05 | 0.00011 | 0.00011 | 3.30E-05 | 0.0001 | 0.00023 | 0 | 0 | 0.00012 |
| Tmed9 | 0.00067 | 0.00012 | 0.00052 | 0.00036 | 0.00038 | 0.00044 | 0.00079 | 0.00042 | 0.00024 | 0.00021 | 0.00031 | 0.0003 | 0.00037 | 0.00035 | 0.00026 | 0.00013 | 0 | 0.00022 |
| Spcs1 | 0.00055 | 0.00014 | 0.00032 | 0.00048 | 0.00033 | 0.00055 | 0.00055 | 0.00049 | 0.00024 | 0.00015 | 0.00036 | 0.00019 | 0.00032 | 0.0004 | 0 | 7.83E-05 | 0 | 0.00024 |
| Elob | 0.00068 | 0.00019 | 0.00031 | 0.00038 | 0.00037 | 0.0005 | 0.0005 | 0.00022 | 0.00024 | 0.00017 | 0.00052 | 0.00042 | 0.00024 | 0.00026 | 0 | 0.00023 | 1.48E-05 | 0.00021 |
| Grcc10 | 0.00024 | 0.00026 | 2.70E-05 | 0.00053 | 0.00039 | 0.00023 | 0.00029 | 0.00053 | 0.00024 | 0.00019 | 0.00011 | 0.00018 | 0.00015 | 0.00025 | 0 | 2.74E-05 | 0.00018 | 0.00032 |
| Mib1 | 0.00023 | 0.00036 | 0.00032 | 7.11E-06 | 0.00018 | 0.00021 | 0.00023 | 0.00017 | 0.00024 | 8.73E-05 | 0.00019 | 0.00015 | 0 | 0.00014 | 0.00017 | 0.0001 | 0.00024 | 0.00015 |
| Ap2a1 | 0.00019 | 0.0002 | 0.00027 | 0.00012 | 0.00015 | 0.00012 | 0.0001 | 0.00014 | 0.00024 | 0.00011 | 0.00021 | 6.70E-05 | 4.31E-05 | 0.00017 | 0.00017 | 0.00017 | 0.00013 | 0.00014 |
| Dhx9 | 0.00042 | 0.00022 | 0.00027 | 5.66E-05 | 0.00022 | 0.00025 | 7.41E-05 | 0.00016 | 0.00024 | 2.87E-05 | 7.64E-05 | 4.94E-06 | 0.00011 | 0.00012 | 0.0002 | 0.0004 | 0.00017 | 0 |
| Fto | 0.00019 | 0.00019 | 8.81E-06 | 0.00024 | 0.00018 | 0.0001 | 0.00029 | 0.00021 | 0.00024 | 0.00013 | 0.00014 | 7.01E-05 | 0.00017 | 6.80E-05 | 0.00038 | 0.00017 | 0.00012 | 6.01E-05 |
| Etnk1 | 0.00023 | 0.00024 | 0.00015 | 4.88E-05 | 8.16E-05 | 0.00015 | 5.60E-05 | 8.12E-05 | 0.00024 | 6.49E-05 | 0.00018 | 0.00012 | 1.65E-05 | 9.67E-05 | 0.00032 | 0.00017 | 0.00027 | 1.31E-05 |
| Phldb2 | 0 | 7.14E-05 | 0 | 0.00025 | 0.00022 | 0 | 0.0004 | 0 | 0.00024 | 0.00025 | 0.00126 | 0 | 0.00054 | 0.00019 | 0.00045 | 9.03E-05 | 8.80E-05 | 0.0003 |
| Nudt3 | 7.62E-05 | 0.00024 | 0.00031 | 0.0004 | 0.00018 | 0.00011 | 6.42E-06 | 0.00019 | 0.00024 | 0.00013 | 0.00022 | 0.00013 | 0.00013 | 0.00012 | 0.00036 | 0.00022 | 2.86E-05 | 0.00016 |
| Pls3 | 0.00035 | 0.0002 | 0.00013 | 0.00011 | 0.00017 | 0.00018 | 0.00016 | 0 | 0.00024 | 0.00026 | 0.001 | 0 | 0.00016 | 0.0003 | 0 | 0.00018 | 0 | 0.00026 |
| Dhcr7 | 3.55E-05 | 1.82E-05 | 0 | 1.14E-05 | 8.11E-05 | 0 | 0 | 9.90E-05 | 0.00024 | 4.99E-05 | 0.00011 | 0 | 0 | 4.76E-05 | 0 | 1.58E-05 | 0 | 7.23E-05 |

| | | | | | | | | | | | | | | | | | | |
|---|---|---|---|---|---|---|---|---|---|---|---|---|---|---|---|---|---|---|
| Cyb5a | 0.00032 | 0.00018 | 0 | 0 | 8.49E-06 | 0.00034 | 0 | 1.57E-05 | 0.00024 | 0.00044 | 0 | 9.27E-05 | 0.00029 | 0 | 0 | 0 | 0 | 0.0002 |
| Klf9 | 0 | 0.00049 | 0.00011 | 0.00017 | 0.00018 | 0.00045 | 0.00014 | 0.00038 | 0.00023 | 0.00019 | 9.18E-05 | 0.00011 | 0.00031 | 0.00045 | 0.00057 | 0 | 0.00069 | 7.70E-05 |
| Cblb | 0.00015 | 0.00024 | 0.00014 | 0 | 0.00014 | 0.00033 | 0 | 0.00051 | 0.00023 | 0.00015 | 0 | 0.00045 | 0.00029 | 0.00019 | 0.0005 | 0 | 0.00048 | 0 |
| Wasf2 | 0.00011 | 0.00033 | 0.00014 | 0.00011 | 6.04E-05 | 4.16E-05 | 5.11E-05 | 0.00032 | 0.00023 | 0.00024 | 0.00018 | 0.00035 | 0 | 7.02E-05 | 0.00017 | 0.00022 | 0.00014 | 0.00023 |
| Mboat7 | 2.57E-05 | 0.00012 | 0.00023 | 0.00034 | 0.00023 | 0.00021 | 0.0002 | 0.00038 | 0.00023 | 0.00012 | 0.00014 | 0.00025 | 0.00032 | 0.00016 | 9.49E-05 | 0.00019 | 0 | 0.00012 |
| Tacc1 | 7.18E-05 | 0.00026 | 7.92E-05 | 0.00019 | 0.0002 | 0.00055 | 2.01E-05 | 0.00021 | 0.00023 | 0.00027 | 0.00019 | 0.00033 | 0.00045 | 8.25E-05 | 0.00031 | 0.00013 | 8.85E-05 | 0.00017 |
| Pde11a | 3.65E-05 | 4.74E-06 | 0 | 5.64E-07 | 0.00015 | 6.45E-05 | 2.76E-05 | 4.08E-05 | 0.00023 | 0 | 0 | 0 | 0 | 3.80E-05 | 9.21E-05 | 8.05E-06 | 8.93E-05 | 0 |
| Mapk1 | 0.00039 | 0.00025 | 0.00022 | 1.25E-05 | 7.60E-05 | 0.00022 | 7.32E-05 | 0.00014 | 0.00023 | 0.00013 | 0.0001 | 0.00025 | 0 | 0.00013 | 0.00022 | 0.00027 | 0.00016 | 0.00012 |
| Ncln | 0.00018 | 0.00017 | 0.00039 | 0.00034 | 0.00019 | 0.00013 | 1.68E-05 | 0.00019 | 0.00023 | 0.00016 | 0.00021 | 0.00022 | 0.0001 | 0.00019 | 0.00017 | 0.00022 | 0 | 0.00013 |
| Nradd | 9.23E-05 | 0.00011 | 0.00014 | 0 | 1.18E-05 | 0 | 0 | 7.93E-05 | 0.00023 | 0 | 0.0002 | 5.15E-05 | 1.56E-05 | 0.00015 | 0 | 0 | 1.63E-05 | 0.00016 |
| Tma7 | 0.00047 | 8.07E-05 | 0.00013 | 0.00017 | 0.00011 | 0.0002 | 8.74E-05 | 0.00014 | 0.00023 | 5.36E-05 | 0.00011 | 0 | 7.31E-05 | 8.53E-05 | 0 | 0.00014 | 0 | 4.68E-05 |
| Anapc11 | 0.00036 | 0.0002 | 0.00019 | 0.00018 | 8.55E-05 | 0.00023 | 0.00029 | 0.00027 | 0.00023 | 0.00013 | 0.00013 | 0.00023 | 3.11E-06 | 9.19E-05 | 0 | 0.00072 | 0 | 0.00024 |
| Tnks1bp1 | 3.94E-05 | 0.0002 | 0.00034 | 0.00014 | 0.00014 | 9.61E-05 | 0.00014 | 0.00029 | 0.00023 | 0.00043 | 0.00062 | 0.00012 | 0.0001 | 0.00022 | 0.00073 | 0 | 8.32E-05 | 0.00031 |
| Wbp1l | 0 | 0.00037 | 6.39E-05 | 0.00053 | 0.00029 | 8.91E-05 | 0.00049 | 0.00023 | 0.00023 | 0.00031 | 9.93E-05 | 0.00049 | 0.00034 | 0.00019 | 0.0003 | 2.00E-05 | 0.00022 | 0.0003 |
| Ifi205 | 0.00025 | 0 | 0.0001 | 0 | 0 | 0.00015 | 0 | 0.00026 | 0.00023 | 0.00144 | 0 | 0 | 0.00138 | 0 | 0 | 0 | 0.00024 | 0.00018 |
| Sdhd | 0.00019 | 0.00017 | 0.00026 | 0.00045 | 6.52E-05 | 0.00014 | 0.00018 | 0.00013 | 0.00023 | 0.00014 | 0.00014 | 8.09E-05 | 9.43E-05 | 0.0002 | 0 | 0.00024 | 0 | 8.67E-05 |
| Hdac1 | 0.00032 | 9.28E-05 | 0.00014 | 5.60E-05 | 7.63E-05 | 0.00014 | 0 | 0.00012 | 0.00023 | 0.0001 | 0.0001 | 3.01E-05 | 0.0001 | 6.61E-05 | 6.04E-05 | 0.00023 | 0.00017 | 9.58E-06 |
| Tpst1 | 0.00019 | 2.68E-05 | 0.00012 | 1.31E-05 | 0.0002 | 0.00069 | 0.00027 | 0.00027 | 0.00023 | 0.00024 | 0 | 0 | 0.00038 | 0.00038 | 4.04E-05 | 0.00021 | 0.0001 | 0.0002 |
| Stk3 | 0.00013 | 0.00014 | 1.53E-05 | 4.20E-05 | 0.00017 | 0.00017 | 0.00022 | 0.00019 | 0.00023 | 0.00023 | 0 | 0.00014 | 0.00014 | 0 | 0.00043 | 0.00017 | 0.0002 | 7.22E-05 |
| Cdc42ep4 | 0.00013 | 0.00026 | 0.00011 | 5.49E-05 | 0.0001 | 0.00013 | 4.66E-05 | 0.0002 | 0.00023 | 7.00E-05 | 7.68E-05 | 0.00012 | 0.00013 | 0.00013 | 4.51E-05 | 0.00016 | 0.00021 | 0.00015 |
| Dpp9 | 0.00017 | 0.00014 | 0.00027 | 0.00028 | 0.0001 | 0.00013 | 3.53E-05 | 0.00019 | 0.00023 | 0.00014 | 0.00015 | 8.00E-05 | 8.75E-05 | 9.79E-05 | 0.0002 | 0.00016 | 0.00015 | 5.88E-05 |
| Lgals9 | 0.00035 | 2.56E-05 | 4.19E-05 | 3.40E-06 | 8.15E-05 | 0 | 4.49E-06 | 0.00014 | 0.00023 | 0.00049 | 0.00013 | 2.15E-05 | 0.00071 | 3.76E-05 | 0 | 0 | 0 | 0.00024 |
| Cp | 0 | 0.00013 | 0 | 0 | 5.08E-05 | 0.00012 | 0 | 0 | 0.00023 | 2.12E-05 | 0 | 5.61E-06 | 0.00086 | 0.0002 | 0 | 0 | 0 | 0.00027 |
| Fxyd1 | 0 | 0.00019 | 0 | 3.60E-05 | 0.0001 | 9.53E-05 | 0.00016 | 0 | 0.00023 | 0.00049 | 6.94E-06 | 0.00012 | 0.00085 | 0.00017 | 0 | 0 | 0 | 0.00045 |
| Chmp4b | 0.00034 | 0.00021 | 0.00018 | 0.00015 | 0.00026 | 0.0003 | 0.00013 | 0.00019 | 0.00023 | 0.00013 | 0.00014 | 0.00019 | 0.00019 | 0.00014 | 3.20E-05 | 1.43E-05 | 7.77E-05 | 9.86E-05 |
| Ripor3 | 0 | 1.82E-05 | 0 | 6.55E-05 | 6.75E-05 | 0.00023 | 0 | 4.60E-05 | 0.00023 | 0 | 0 | 0 | 0.00048 | 0.00026 | 0.00044 | 0 | 0 | 0 |
| Plekhf1 | 3.54E-05 | 0.00011 | 0 | 7.62E-05 | 0.00022 | 0.00011 | 2.79E-06 | 0 | 0.00023 | 5.24E-05 | 0.00022 | 7.33E-05 | 6.29E-05 | 0.00015 | 6.51E-05 | 5.34E-05 | 0 | 0.00013 |
| Ncs1 | 1.57E-05 | 5.88E-05 | 0.00036 | 0 | 5.03E-05 | 5.25E-05 | 0 | 0.00017 | 0.00023 | 1.60E-05 | 0.00032 | 0.00015 | 0 | 7.60E-05 | 0 | 0 | 0 | 0.00019 |
| Galk1 | 0.00022 | 0 | 0.00031 | 0.00053 | 3.41E-05 | 5.07E-05 | 0.00015 | 0.00013 | 0.00023 | 0 | 0.00015 | 0.0005 | 0.00014 | 0.00025 | 0 | 0.00011 | 6.41E-05 | 1.34E-05 |
| Chl1 | 9.89E-05 | 0 | 0 | 0 | 0 | 6.06E-05 | 0.00012 | 0 | 0.00023 | 0.00038 | 0 | 0 | 0.0015 | 0 | 0 | 5.00E-05 | 0 | 0.00019 |
| Tpm2 | 7.52E-05 | 0 | 0.00027 | 0.00055 | 0.00014 | 0 | 0.00023 | 0.00031 | 0.00023 | 0 | 0.00324 | 0.00047 | 0 | 0.00066 | 0.00035 | 0.00041 | 0 | 0 |
| Metrnl | 8.88E-05 | 0.00014 | 0.00025 | 5.96E-05 | 1.89E-06 | 1.07E-05 | 7.95E-05 | 0.00019 | 0.00023 | 0.00052 | 0.00058 | 0.00014 | 0 | 0 | 0 | 6.07E-05 | 0.00021 | 9.65E-05 |
| Dbnl | 0.00021 | 0.00022 | 0.00021 | 0.00024 | 0.00017 | 0.00011 | 0.00013 | 0 | 0.00023 | 0.00013 | 0.0003 | 0.0003 | 0.0001 | 0.00027 | 0.00016 | 7.64E-05 | 0 | 0.00015 |

| | | | | | | | | | | | | | | | | | | |
|---|---|---|---|---|---|---|---|---|---|---|---|---|---|---|---|---|---|---|
| Tap2 | 1.83E-05 | 0.00018 | 4.93E-05 | 0.0002 | 9.44E-05 | 6.58E-05 | 0 | 9.83E-05 | 0.00023 | 0.00019 | 4.72E-05 | 9.02E-06 | 9.86E-05 | 0.00012 | 0.00018 | 8.10E-06 | 0.00013 | 0.00028 |
| Pde4b | 5.19E-05 | 0 | 0 | 0 | 0.00017 | 0.00046 | 0 | 0 | 0.00023 | 0 | 0 | 0.00012 | 2.15E-05 | 0.00052 | 0.00012 | 9.17E-05 | 0 | 0 |
| Zfp809 | 0.00014 | 5.51E-05 | 1.79E-05 | 0 | 0.00018 | 1.75E-05 | 8.19E-05 | 1.54E-05 | 0.00023 | 1.74E-05 | 0 | 0 | 4.84E-05 | 0 | 5.56E-05 | 0 | 0 | 0 |
| Cadm2 | 0 | 0 | 0 | 0 | 0 | 8.01E-05 | 0 | 0 | 0.00023 | 0 | 0 | 1.99E-06 | 0.00033 | 0 | 0 | 1.35E-05 | 0 | 0 |
| Cyth3 | 9.09E-05 | 0.00027 | 0.00026 | 2.93E-05 | 7.74E-05 | 0.0008 | 0 | 0.00072 | 0.00023 | 0.00036 | 8.13E-05 | 0.00021 | 0.00041 | 0.00056 | 7.83E-05 | 0.00014 | 0.00042 | 0.00049 |
| Rab18 | 0.00041 | 0.00034 | 0.00028 | 0.00041 | 0.00037 | 0.00046 | 0.0004 | 0.00035 | 0.00023 | 0.00024 | 0.0003 | 0.00028 | 0.00033 | 0.00037 | 0.0001 | 0.0001 | 0.00013 | 0.00025 |
| Med13l | 5.07E-05 | 0.00032 | 0.00011 | 0.00038 | 0.00017 | 8.20E-05 | 0.00015 | 0 | 0.00023 | 3.95E-05 | 0.00045 | 0.00019 | 0.0004 | 0.00023 | 0.00072 | 0.00012 | 0.0006 | 0.00022 |
| Actr2 | 0.00035 | 0.0002 | 0.00029 | 6.15E-05 | 6.89E-05 | 5.42E-05 | 6.97E-05 | 0.00021 | 0.00023 | 7.93E-05 | 0.00023 | 0.0001 | 2.41E-05 | 7.20E-05 | 5.28E-05 | 0.00012 | 0 | 0.00013 |
| Tram1 | 0.00056 | 7.30E-05 | 0.00047 | 0.00048 | 0.00027 | 0.00053 | 0.001 | 0.0005 | 0.00023 | 0.00038 | 0.00032 | 0.0002 | 0.00043 | 0.00037 | 4.64E-06 | 0.00015 | 0 | 0.00035 |
| Plp2 | 0.00014 | 0.00014 | 0.00028 | 0.0008 | 0.0002 | 0.00013 | 0.00059 | 0.0005 | 0.00023 | 0.00023 | 0.00044 | 0.00053 | 0.00023 | 0.00039 | 6.39E-05 | 0.00049 | 0.00014 | 0.00047 |
| Hsd17b10 | 0.0002 | 0.00016 | 0.00012 | 0.00023 | 8.43E-05 | 0.00015 | 3.73E-05 | 9.14E-05 | 0.00023 | 4.37E-05 | 9.37E-05 | 0.00016 | 0.00013 | 4.57E-05 | 0 | 1.51E-05 | 0 | 0.00012 |
| Ube2v1 | 0.00043 | 0.00032 | 0.0003 | 0.0003 | 0.0002 | 0.00031 | 0.00015 | 0.00028 | 0.00023 | 0.00014 | 0.00023 | 0.00019 | 1.05E-05 | 0.00019 | 0 | 0.00016 | 8.97E-05 | 6.98E-05 |
| Sidt2 | 0 | 0.0002 | 0.00014 | 0.00025 | 0.00028 | 0.00024 | 0.00025 | 0.00031 | 0.00023 | 0.00036 | 0.00037 | 8.53E-05 | 0.00025 | 0.00029 | 0.00016 | 0 | 0 | 0.00027 |
| Ikbip | 0.00039 | 0 | 0.0002 | 0.00052 | 0.00025 | 0.00035 | 0.00051 | 0.00058 | 0.00023 | 9.79E-05 | 0.00019 | 0.00022 | 0.00016 | 0.0003 | 0.00054 | 8.76E-05 | 6.92E-05 | 0.00017 |
| Pld3 | 0 | 5.34E-05 | 2.15E-05 | 0.00017 | 0.0002 | 0.00053 | 0.00033 | 0.00061 | 0.00023 | 0.00017 | 0.00027 | 0.00019 | 0.00082 | 0.00059 | 0 | 0 | 0 | 0.00057 |
| Ppfibp1 | 0.00019 | 0.00024 | 0.00028 | 4.02E-05 | 0.00046 | 0.00011 | 0.00049 | 0 | 0.00023 | 5.93E-05 | 0.00059 | 0.0004 | 0 | 0.00041 | 0.00019 | 0.00028 | 0.0003 | 0.00016 |
| Dgkz | 1.89E-05 | 0.00014 | 0.00018 | 0.00017 | 8.08E-05 | 0.00021 | 1.36E-06 | 4.19E-05 | 0.00023 | 6.29E-05 | 6.91E-05 | 0 | 0 | 7.77E-05 | 7.76E-05 | 0.0002 | 0 | 0.00019 |
| Tnfsf11 | 0 | 0 | 0.00012 | 0 | 0 | 0.00017 | 0 | 0.00015 | 0.00023 | 0 | 0 | 0 | 0.00041 | 0.0005 | 0 | 0.00011 | 0 | 0 |
| Stub1 | 0.0002 | 0.00026 | 0.00033 | 0.00067 | 0.00027 | 0.00023 | 0.00028 | 0.00028 | 0.00023 | 0.0002 | 0.00025 | 0.00024 | 0.00019 | 0.00023 | 0.00028 | 0.00056 | 0.00016 | 0.0002 |
| Ptma | 0.00024 | 0.00032 | 0.00028 | 0.00062 | 0.00011 | 6.98E-05 | 0 | 7.73E-05 | 0.00023 | 0 | 0.00021 | 0.0002 | 5.91E-05 | 0.00018 | 0.00021 | 0.00078 | 0.00082 | 0.00017 |
| Tsku | 3.05E-05 | 6.59E-06 | 0.00064 | 0.00029 | 0.00026 | 0 | 0.00083 | 0.00036 | 0.00023 | 0.00025 | 0.00025 | 0.00043 | 0.00059 | 0.00029 | 4.83E-05 | 1.44E-05 | 0 | 0.00023 |
| Ppp1r2 | 1.69E-05 | 0.0002 | 0.0001 | 0.00011 | 7.76E-05 | 9.06E-05 | 0 | 4.97E-05 | 0.00023 | 0.00026 | 4.49E-05 | 1.86E-05 | 0.00021 | 0.00016 | 0 | 0.00021 | 0.00022 | 0.00034 |
| Scap | 0 | 0.0002 | 0.00012 | 0.00025 | 0.00026 | 7.27E-05 | 9.27E-05 | 5.66E-05 | 0.00023 | 4.44E-05 | 7.52E-05 | 4.19E-05 | 0.00023 | 1.39E-05 | 0.0002 | 4.51E-07 | 5.09E-05 | 0.00017 |
| Hsp90b1 | 0.00073 | 2.95E-06 | 0.00028 | 0.0007 | 0.00012 | 0.00028 | 0.00035 | 0.00039 | 0.00023 | 0.0003 | 0.00026 | 0.0007 | 0.00051 | 0.00027 | 0.0006 | 0.00023 | 0.0004 | 0.00031 |
| Canx | 0.00059 | 0.00022 | 0.00035 | 0.00014 | 0.00022 | 0.00024 | 0.00019 | 0.00023 | 0.00023 | 0.00021 | 0.00023 | 0.00035 | 0.00031 | 0.00017 | 0 | 0.00012 | 0 | 0.0002 |
| Ipo4 | 0.00021 | 0.00012 | 0.00044 | 0.00047 | 7.58E-05 | 4.54E-05 | 0.00021 | 8.16E-05 | 0.00023 | 0.0002 | 6.40E-05 | 7.97E-05 | 0.00018 | 3.66E-05 | 0.00032 | 0 | 0.00025 | 0.00012 |
| Tiparp | 9.57E-05 | 4.94E-05 | 0 | 0 | 0 | 0.00023 | 0 | 0.00017 | 0.00023 | 1.31E-05 | 0 | 0.00063 | 0 | 0 | 0 | 1.15E-05 | 0.00052 | 2.84E-05 |
| Cct3 | 0.00038 | 7.66E-05 | 0.00046 | 0.00017 | 0 | 0 | 3.21E-05 | 0 | 0.00023 | 4.74E-05 | 0.0001 | 6.06E-06 | 3.26E-05 | 0 | 9.62E-05 | 0.00012 | 9.02E-05 | 1.76E-05 |
| Tgfb1i1 | 9.53E-05 | 0.00017 | 0.00031 | 0.00058 | 0.00013 | 0 | 0.00011 | 0.00015 | 0.00023 | 0.00029 | 0.00105 | 0.00043 | 0 | 0.00031 | 0.00017 | 2.28E-05 | 4.73E-05 | 0.00035 |
| Strn4 | 0.00011 | 0.00027 | 0.00026 | 0.00017 | 0.00014 | 0.00014 | 1.36E-06 | 0.00014 | 0.00023 | 0.00012 | 0.00016 | 0.00017 | 6.06E-05 | 0.00013 | 0.00024 | 0.00023 | 0.00018 | 8.18E-05 |
| Sel1l | 0.00017 | 0.00025 | 0.00019 | 0.00021 | 0.00032 | 0.00027 | 0.00018 | 0.00026 | 0.00023 | 0.00032 | 0.00022 | 0.00037 | 0.00021 | 0.00024 | 0.00016 | 7.97E-05 | 0 | 0.0003 |
| Dcaf7 | 0.00011 | 0.0002 | 0.00016 | 0.00026 | 0.00021 | 0.0002 | 8.05E-05 | 0.00022 | 0.00023 | 0.00018 | 0.00018 | 9.79E-05 | 0.00014 | 0.00023 | 2.20E-06 | 0.00027 | 0 | 0.00018 |
| Adgrl1 | 1.77E-05 | 3.23E-05 | 0.00011 | 0 | 0.00026 | 0.00024 | 3.95E-05 | 0.00025 | 0.00023 | 2.49E-05 | 0.0003 | 0 | 0 | 0.00021 | 7.40E-05 | 0.00018 | 0 | 0.00028 |

| | | | | | | | | | | | | | | | | | | |
|---|---|---|---|---|---|---|---|---|---|---|---|---|---|---|---|---|---|---|
| Pgls | 0.00014 | 0.00016 | 0.00017 | 0.00033 | 0.00019 | 0.0002 | 0.00022 | 0.00022 | 0.00023 | 0.00015 | 0.00025 | 0.00019 | 8.73E-05 | 0.00017 | 0 | 0.00014 | 0 | 9.28E-05 |
| Prpf19 | 0.00021 | 0.00022 | 0.00038 | 0.00022 | 8.61E-05 | 1.97E-06 | 1.58E-05 | 0 | 0.00023 | 4.83E-05 | 7.26E-05 | 1.96E-05 | 2.11E-05 | 3.75E-05 | 0 | 0.00028 | 4.01E-05 | 0 |
| Mapk14 | 0.00019 | 0.0002 | 9.73E-05 | 3.88E-05 | 0.00016 | 0.00018 | 0.00019 | 4.91E-05 | 0.00023 | 0.00017 | 7.34E-05 | 6.50E-05 | 0.00011 | 0.00021 | 0.00021 | 0.00014 | 0 | 4.13E-05 |
| Tusc3 | 0.00022 | 9.70E-05 | 0.0002 | 9.54E-05 | 0.00023 | 0.00042 | 0.00029 | 0.00042 | 0.00023 | 0.00014 | 0.00012 | 0.00017 | 0.00013 | 0.0002 | 0.00012 | 0.00016 | 7.72E-05 | 0.00015 |
| Adamtsl1 | 9.59E-06 | 0 | 0.00011 | 0 | 6.89E-05 | 0 | 0.00071 | 0.00043 | 0.00023 | 0.00082 | 7.26E-05 | 6.14E-05 | 0.00068 | 0 | 0.00018 | 7.88E-05 | 0 | 0.0004 |
| Bdkrb1 | 0 | 0 | 0.00034 | 0 | 0 | 0 | 4.53E-05 | 8.55E-05 | 0.00023 | 0 | 0 | 0 | 1.15E-05 | 0 | 0 | 0 | 0.0002 | 8.33E-05 |
| Alad | 8.04E-05 | 5.70E-05 | 0.00012 | 0.00023 | 3.60E-05 | 2.85E-05 | 0 | 0.00031 | 0.00023 | 0 | 3.40E-05 | 0 | 0 | 9.66E-05 | 0 | 0.00026 | 0 | 0 |
| Plekhg2 | 0 | 8.39E-05 | 9.54E-05 | 0.00015 | 0.00016 | 0.00014 | 8.12E-06 | 0.00062 | 0.00023 | 8.11E-05 | 0.0003 | 0.00015 | 0.00025 | 0.00042 | 0.00047 | 0.00014 | 0.00102 | 9.89E-05 |
| Col18a1 | 0 | 0.00028 | 0 | 0 | 0 | 0.00167 | 0 | 0.00063 | 0.00023 | 0.00026 | 0 | 0.00014 | 0 | 0.00025 | 0.00012 | 0 | 2.24E-05 | 0.00048 |
| Etv4 | 0 | 6.00E-05 | 0.00011 | 0 | 0.00024 | 0 | 0 | 0.00045 | 0.00023 | 0 | 0.0001 | 0.00012 | 0 | 0.00027 | 0.00018 | 5.24E-05 | 6.13E-05 | 0 |
| Gmfb | 0.00017 | 0.00014 | 0.00029 | 0.00038 | 0.00023 | 8.82E-05 | 0.00026 | 0.00016 | 0.00023 | 0.00016 | 0.00027 | 0.00018 | 0.00037 | 0.00014 | 3.97E-05 | 2.02E-05 | 0 | 9.15E-05 |
| Inpp5a | 8.69E-05 | 0.00015 | 0.00022 | 0 | 0.00041 | 0.00028 | 0.00021 | 8.26E-05 | 0.00023 | 9.14E-05 | 0.00046 | 0.00016 | 0.00014 | 4.30E-05 | 0 | 2.91E-05 | 0.00012 | 0 |
| Elmo2 | 3.27E-05 | 0.00011 | 7.79E-05 | 0 | 8.86E-05 | 0.00016 | 0 | 0 | 0.00023 | 8.76E-05 | 0.00011 | 3.15E-05 | 8.09E-05 | 0.00017 | 2.56E-05 | 0.00019 | 8.25E-05 | 6.55E-05 |
| Tbx15 | 0.00024 | 0.00012 | 0 | 0 | 5.37E-05 | 0.00031 | 0.00025 | 0 | 0.00023 | 0.00013 | 0.00016 | 0 | 0.00022 | 0.0002 | 0 | 0.00014 | 8.12E-05 | 0.00022 |
| Cope | 0.00051 | 9.02E-05 | 0.00033 | 0.0004 | 0.00018 | 0.00032 | 0.00065 | 0.00033 | 0.00023 | 0.00017 | 0.0002 | 0.00031 | 0.00022 | 0.00035 | 0.00013 | 5.00E-06 | 0 | 0.00016 |
| Cdo1 | 0 | 0 | 0 | 0 | 0 | 0 | 8.38E-05 | 0 | 0.00023 | 0 | 2.51E-05 | 0.00017 | 0 | 0.00023 | 0 | 4.61E-05 | 1.43E-06 | 0 |
| Cox4i1 | 0.00024 | 0.00022 | 0.0002 | 0.00043 | 0.00013 | 0.00016 | 0.00013 | 0.00017 | 0.00023 | 7.55E-05 | 0.00019 | 0.00019 | 0.00011 | 0.00022 | 0 | 0.00014 | 3.09E-05 | 0.00011 |
| Ccdc85b | 0 | 6.12E-05 | 0 | 0.00025 | 1.87E-06 | 0.00014 | 0 | 0.00023 | 0.00023 | 3.13E-05 | 0.00014 | 0.00034 | 0 | 0 | 0.00022 | 0 | 0 | 0 |
| Clmp | 0 | 0.00033 | 0.00025 | 0.0006 | 0.00052 | 0.00041 | 0.00054 | 0 | 0.00023 | 0.00067 | 0 | 0.00062 | 0.0007 | 0.00063 | 0.0002 | 0.00019 | 0 | 0.00036 |
| Rab11b | 0 | 0.00027 | 0.00011 | 0.00044 | 0.0002 | 0.00014 | 2.99E-05 | 0.00025 | 0.00023 | 0.0002 | 0.00021 | 0.00015 | 7.09E-05 | 0.00014 | 0 | 0.00022 | 0.00019 | 0.00021 |
| Rac1 | 0.00024 | 0.00014 | 0.00026 | 9.78E-05 | 5.22E-05 | 8.94E-05 | 0.0001 | 0.00017 | 0.00023 | 0.00016 | 0.0002 | 0.00024 | 1.01E-05 | 0.00014 | 0 | 0.00023 | 0 | 0.00012 |
| Plppr3 | 0 | 0.0002 | 0 | 7.68E-05 | 0.00014 | 0 | 0 | 7.28E-05 | 0.00022 | 0 | 0 | 9.69E-05 | 1.84E-05 | 0 | 0.00017 | 1.71E-05 | 0 | 5.42E-05 |
| Cers2 | 0.00024 | 0.00024 | 0.00028 | 0.00013 | 0.00014 | 0.00014 | 6.95E-05 | 0.00015 | 0.00022 | 0.00015 | 0.00023 | 7.39E-05 | 2.96E-05 | 8.75E-05 | 9.35E-05 | 0.00012 | 0 | 0.00012 |
| Wtip | 1.65E-05 | 0.00018 | 0.00014 | 0.00016 | 0.00024 | 0.00018 | 0.00013 | 9.73E-05 | 0.00022 | 7.58E-05 | 0.00026 | 0.00016 | 0.00015 | 0.0003 | 5.42E-05 | 4.51E-05 | 0.0002 | 9.38E-05 |
| Bcap31 | 0.00022 | 0.0002 | 0.00031 | 0.00043 | 0.00021 | 0.00027 | 0.00029 | 0.00027 | 0.00022 | 0.00023 | 0.0002 | 0.00022 | 0.00031 | 0.00021 | 3.70E-05 | 0.00016 | 0 | 0.00033 |
| Fibin | 0 | 0 | 0.00043 | 0.00057 | 0.00027 | 2.92E-05 | 0.00092 | 0.00197 | 0.00022 | 0.00021 | 8.86E-05 | 0 | 0.00044 | 0 | 0.00056 | 0.00014 | 0 | 0 |
| Stard4 | 4.28E-05 | 0 | 0 | 0.00021 | 6.28E-05 | 0 | 0 | 9.78E-05 | 0.00022 | 6.51E-06 | 0.00015 | 0.00011 | 0 | 0.0001 | 8.42E-06 | 0.00022 | 0.0003 | 2.45E-05 |
| Tlnrd1 | 0 | 0.00015 | 0.0002 | 0.00034 | 0 | 0 | 0 | 0.0002 | 0.00022 | 0.00017 | 0.00017 | 0 | 5.93E-06 | 0.00018 | 0.00032 | 0.00038 | 0.00035 | 8.85E-05 |
| Pkd2 | 5.90E-05 | 0.00031 | 0.0004 | 0 | 0.00071 | 0.00041 | 0.0005 | 0.00051 | 0.00022 | 0.00033 | 0.00031 | 0.00011 | 0.00062 | 0.00034 | 0.00022 | 0.00017 | 6.56E-05 | 0.0002 |
| S100a13 | 0.00018 | 5.70E-05 | 9.78E-06 | 0.00046 | 4.34E-05 | 0.00018 | 0.00036 | 0.00057 | 0.00022 | 0.00071 | 0.00012 | 0.00019 | 0.00053 | 0.00019 | 0 | 1.77E-05 | 6.92E-05 | 0.00039 |
| Src | 2.86E-05 | 0.00024 | 0.00019 | 5.34E-05 | 0.0002 | 8.14E-05 | 0.00014 | 9.80E-05 | 0.00022 | 0.00012 | 0.00056 | 0.00018 | 0.00021 | 0.00019 | 7.65E-05 | 0 | 9.12E-05 | 0.00013 |
| Polr2e | 0.00015 | 0.00015 | 0.00034 | 0.00054 | 7.09E-05 | 6.22E-05 | 0.00011 | 0.00014 | 0.00022 | 6.87E-05 | 4.64E-05 | 2.73E-05 | 9.57E-06 | 6.26E-05 | 9.20E-05 | 0.00014 | 0.0001 | 0.00011 |
| Dpp3 | 0.00012 | 0.00013 | 0.00016 | 0.00024 | 8.78E-05 | 0 | 7.32E-05 | 8.39E-05 | 0.00022 | 0.00012 | 0.00025 | 0.00011 | 2.32E-05 | 0.00011 | 1.88E-05 | 0 | 0 | 0.0001 |

| | | | | | | | | | | | | | | | | | | |
|---|---|---|---|---|---|---|---|---|---|---|---|---|---|---|---|---|---|---|
| 7-Sep | 0.00066 | 0.0002 | 0.00031 | 3.95E-05 | 0.00023 | 0.0004 | 0.00032 | 0.00029 | 0.00022 | 0.00026 | 0.0002 | 0.00031 | 0 | 9.81E-05 | 0.00019 | 0.00034 | 0.00016 | 0.00034 |
| Selenom | 0.00027 | 1.61E-05 | 0.00013 | 0.00126 | 0.00038 | 0.00028 | 0.0012 | 0.0008 | 0.00022 | 0.00047 | 0.00046 | 0.00015 | 0.00047 | 0.00052 | 6.09E-05 | 3.42E-05 | 0.00017 | 0.00055 |
| Fam168b | 0.00025 | 0.00033 | 0.00024 | 0.00029 | 0.00027 | 0.00015 | 0.00016 | 0.00012 | 0.00022 | 9.31E-05 | 0.00016 | 0.00027 | 0.00011 | 0.00021 | 0 | 0.00029 | 6.35E-05 | 0.00015 |
| Epb41l1 | 0 | 0.00013 | 6.24E-06 | 4.86E-05 | 0.00014 | 0.00025 | 4.11E-05 | 3.70E-05 | 0.00022 | 0.00011 | 0.00038 | 0.00021 | 5.75E-05 | 0 | 2.52E-05 | 3.82E-05 | 0 | 1.26E-05 |
| Trip12 | 0.00052 | 0.0003 | 0.00028 | 0 | 0.00027 | 0.00029 | 0.00029 | 0.00016 | 0.00022 | 0.00015 | 0.00011 | 0.00036 | 0 | 0.00011 | 0.0004 | 0.00011 | 0.00036 | 0.00018 |
| Lnpep | 0.00025 | 0.00034 | 0.00011 | 0.00011 | 9.12E-05 | 0.00022 | 0.00013 | 0.0002 | 0.00022 | 0.00024 | 0.00018 | 0.00033 | 0.00012 | 9.73E-05 | 0.00019 | 9.45E-05 | 0.00036 | 0.00021 |
| Elovl1 | 0.00031 | 9.73E-05 | 0.00028 | 4.54E-05 | 0.00014 | 0.0002 | 0.00017 | 0.00018 | 0.00022 | 0.00011 | 0.00018 | 2.58E-05 | 0 | 0.00013 | 0 | 4.58E-05 | 0 | 9.43E-05 |
| H1fx | 0 | 5.35E-05 | 0 | 0.00018 | 0.00035 | 7.45E-06 | 0 | 0.00018 | 0.00022 | 0 | 9.85E-05 | 0 | 0 | 4.44E-05 | 3.90E-05 | 0.00044 | 0 | 2.49E-05 |
| Copz1 | 0.00035 | 8.36E-05 | 0.00027 | 0.00077 | 0.00029 | 0.00027 | 0.00049 | 0.00037 | 0.00022 | 0.00021 | 0.00017 | 9.94E-05 | 0.0003 | 0.00032 | 0 | 5.75E-05 | 0 | 0.00017 |
| Abhd17b | 0.00014 | 0.00014 | 8.43E-05 | 5.83E-06 | 0.00021 | 0.00011 | 6.52E-05 | 0.00014 | 0.00022 | 6.98E-05 | 2.71E-05 | 3.31E-05 | 1.45E-05 | 9.85E-05 | 7.19E-05 | 0.00013 | 0.00014 | 0 |
| Frmd4a | 3.22E-05 | 0.00015 | 0.00013 | 8.51E-06 | 4.10E-05 | 8.18E-05 | 0 | 0.00036 | 0.00022 | 3.19E-05 | 0.00014 | 0.00015 | 0 | 0.00016 | 0.00011 | 0.00015 | 0.0002 | 6.78E-05 |
| Comt | 0.00016 | 4.41E-05 | 0.00018 | 0.00026 | 0 | 0 | 0.00043 | 0.00017 | 0.00022 | 0.00051 | 0.00018 | 0.00015 | 0.00055 | 0 | 0 | 1.92E-05 | 0 | 0.00045 |
| Ssr3 | 0.00062 | 0 | 0.00037 | 0.0005 | 0.00023 | 0.00054 | 0.00072 | 0.0006 | 0.00022 | 0.00023 | 0.00024 | 0.00031 | 0.00015 | 0.00032 | 0.00016 | 0.00027 | 0.00025 | 0.00017 |
| Larp1 | 0.00016 | 0.00022 | 0.00044 | 0.00021 | 0.00011 | 7.57E-05 | 0 | 3.35E-05 | 0.00022 | 4.43E-05 | 0.0002 | 0.00019 | 6.33E-05 | 0.0001 | 0.00019 | 0.00012 | 0.0001 | 9.32E-05 |
| Wdr82 | 0.00021 | 0.00023 | 0.00021 | 0.00019 | 0.00013 | 0.00015 | 3.17E-05 | 7.25E-05 | 0.00022 | 0.00017 | 0.00014 | 9.57E-06 | 8.35E-05 | 0.00014 | 5.52E-05 | 0.00035 | 0.00018 | 0.00014 |
| Slc25a5 | 0.00048 | 0 | 0.0003 | 0 | 0 | 0.0002 | 0 | 1.69E-05 | 0.00022 | 0 | 0 | 0 | 1.72E-05 | 4.74E-05 | 0 | 0.00017 | 0 | 0 |
| Dkk3 | 1.19E-05 | 0.00034 | 8.32E-05 | 0 | 0.00153 | 0.00047 | 0.00104 | 0.00032 | 0.00022 | 0 | 0.00056 | 0.00014 | 0 | 0.00122 | 0 | 0 | 0 | 0.00072 |
| H3f3b | 0.00043 | 0.00038 | 0.00032 | 0 | 0.00032 | 0.00031 | 6.61E-06 | 0.00018 | 0.00022 | 0 | 0.00028 | 0.00049 | 0.00027 | 0.00026 | 0 | 0.00042 | 0.00129 | 5.79E-05 |
| Hsd17b7 | 1.21E-05 | 2.80E-05 | 3.71E-06 | 0 | 4.60E-05 | 0 | 0 | 0 | 0.00022 | 2.35E-05 | 0.00014 | 1.14E-05 | 0 | 4.91E-05 | 0 | 0 | 0.00012 | 7.45E-06 |
| Hp1bp3 | 0.00027 | 0.00044 | 3.26E-05 | 0.00039 | 0.0002 | 0.00024 | 0.00012 | 0 | 0.00022 | 0.00019 | 3.52E-05 | 0.00017 | 0.00024 | 0.00015 | 0.00053 | 0.00061 | 0.00023 | 0.00025 |
| Rbfox2 | 0.00032 | 0.00029 | 0.00017 | 0.00013 | 0.00031 | 0.00027 | 0.00021 | 0.00016 | 0.00022 | 0.00019 | 0.00036 | 0.00028 | 1.13E-05 | 0.00019 | 0.00068 | 0.00016 | 0.00054 | 0.00012 |
| Gstp1 | 0.00023 | 0.0002 | 0.00029 | 0.00067 | 1.44E-05 | 0.00012 | 6.63E-05 | 0.0002 | 0.00022 | 0.00015 | 3.36E-05 | 3.37E-05 | 9.92E-06 | 0.0002 | 0 | 3.20E-05 | 0.00011 | 0.00013 |
| Trappc4 | 0.00014 | 0.00011 | 0.00033 | 0.0003 | 9.70E-05 | 5.97E-05 | 6.80E-05 | 0.00025 | 0.00022 | 0.00012 | 0.00017 | 0.00019 | 0 | 0.00011 | 1.57E-05 | 3.95E-06 | 5.38E-05 | 0.00016 |
| Cpne1 | 9.52E-05 | 0.00019 | 0.00013 | 0.00031 | 0.00015 | 0.00011 | 0 | 0.0001 | 0.00022 | 8.01E-05 | 0.00015 | 0.0001 | 4.07E-05 | 0.00023 | 3.76E-05 | 0.00017 | 6.32E-05 | 0.00015 |
| Vgll3 | 0.00011 | 0 | 0 | 5.68E-05 | 0.0001 | 0 | 0 | 0 | 0.00022 | 0.00104 | 0.00124 | 1.67E-05 | 0.00098 | 0.00024 | 0 | 0.00018 | 0 | 0 |
| Mbtps1 | 0.00025 | 0.00024 | 0.00012 | 0 | 0.00023 | 0.0003 | 0.00018 | 8.07E-05 | 0.00022 | 0.00018 | 0.00014 | 4.86E-05 | 0.00016 | 0.00013 | 4.58E-05 | 9.79E-05 | 3.51E-05 | 0.0002 |
| Dctn2 | 0.00032 | 0.00035 | 0.00026 | 0.00034 | 0.00023 | 0.00026 | 0.00023 | 0.00021 | 0.00022 | 0.00031 | 0.00023 | 0.00024 | 4.54E-05 | 0.00022 | 0.00012 | 0.0001 | 0.00012 | 0.00039 |
| Zfp207 | 0.0004 | 0.0002 | 0.00027 | 0.00012 | 0.00015 | 0.00011 | 8.08E-05 | 8.80E-05 | 0.00022 | 0.00011 | 0.00011 | 8.68E-05 | 5.77E-05 | 0.0002 | 0.00061 | 0.00019 | 1.91E-06 | 8.92E-05 |
| Rab12 | 0.00031 | 0.00031 | 0.00018 | 0 | 0.0004 | 0.00043 | 0.00016 | 0.0002 | 0.00022 | 0.00014 | 7.37E-05 | 0.00016 | 0.00013 | 0.00014 | 0 | 0.00014 | 9.71E-05 | 0.00019 |
| Man1a2 | 0.00012 | 0.00012 | 0.00012 | 9.48E-05 | 0.00014 | 0.00017 | 0.00013 | 0.00011 | 0.00022 | 9.08E-05 | 0.00024 | 1.81E-05 | 9.82E-05 | 0.0002 | 0 | 0.00016 | 7.64E-05 | 0.00013 |
| Pik3r2 | 3.93E-05 | 0.00019 | 0 | 0.00024 | 0.00021 | 0.00011 | 0.00013 | 0.00026 | 0.00022 | 0.00024 | 8.05E-05 | 1.69E-05 | 0 | 0.00016 | 0.0002 | 0 | 4.68E-05 | 0.00017 |
| Cachd1 | 8.91E-06 | 0.00016 | 1.43E-05 | 4.30E-06 | 9.54E-05 | 7.22E-05 | 6.81E-05 | 0.0001 | 0.00022 | 8.08E-05 | 2.56E-05 | 0 | 0 | 6.43E-05 | 3.56E-05 | 0.00012 | 0.0002 | 0.00011 |
| Prkaca | 0.0002 | 0.0001 | 0.00013 | 0.00031 | 0.00019 | 0.00017 | 9.34E-05 | 0.00021 | 0.00022 | 0.00016 | 0.00012 | 0.00013 | 8.64E-05 | 0.00014 | 0 | 0.00025 | 0 | 0.00022 |

| | | | | | | | | | | | | | | | | | | |
|---|---|---|---|---|---|---|---|---|---|---|---|---|---|---|---|---|---|---|
| Echs1 | 0.00028 | 0.00016 | 8.65E-05 | 0.00016 | 0.00019 | 0.00023 | 7.28E-05 | 2.30E-05 | 0.00022 | 4.24E-05 | 2.50E-05 | 5.29E-05 | 0.00035 | 0.00013 | 0 | 7.21E-05 | 0 | 2.25E-05 |
| Eef1g | 0.00058 | 0.00026 | 0.0005 | 0.00036 | 9.40E-05 | 0.00031 | 0.0002 | 0.00016 | 0.00022 | 0.00012 | 6.87E-05 | 0.00026 | 0.00015 | 0 | 0 | 0.00012 | 0 | 0 |
| Crk | 8.45E-06 | 0.00019 | 0.00023 | 0.00047 | 0.00011 | 2.21E-05 | 3.11E-06 | 0.00024 | 0.00022 | 0.00016 | 0.00026 | 0.00017 | 0 | 0.00015 | 0.00041 | 0.00018 | 0.00034 | 0.00016 |
| Jund | 0 | 0.00027 | 0.00021 | 0.00064 | 0.00021 | 0.00023 | 0.00045 | 0.00045 | 0.00022 | 0.00076 | 0.00042 | 0.00066 | 0.00015 | 5.90E-05 | 0.00106 | 0 | 0.00622 | 0.00062 |
| Pgam5 | 2.25E-05 | 0.00011 | 0.00026 | 2.21E-05 | 0.00011 | 0 | 7.79E-05 | 0 | 0.00022 | 5.62E-05 | 0.00016 | 2.17E-05 | 6.11E-05 | 4.62E-05 | 0 | 0.00011 | 0 | 0 |
| Sfrp2 | 0 | 0 | 0 | 0.00073 | 0.00147 | 0.0008 | 0.00216 | 0 | 0.00022 | 0.00082 | 0 | 0 | 0.00226 | 0.00033 | 0.00016 | 0.00032 | 0 | 9.10E-05 |
| Kif1c | 6.83E-05 | 0.00022 | 0.00015 | 0.00013 | 0.00026 | 7.76E-05 | 6.55E-05 | 2.81E-05 | 0.00022 | 0.00016 | 0.00036 | 0.00012 | 5.38E-05 | 0.00014 | 0.00016 | 0.00014 | 0 | 0.00025 |
| Drap1 | 0.00035 | 0.0003 | 0.0003 | 0.00036 | 0.00027 | 0.00032 | 0.0003 | 0.00019 | 0.00022 | 0.00018 | 0.00023 | 0.00024 | 0.00012 | 0.00023 | 4.04E-05 | 0.00012 | 0 | 0.00022 |
| Kif26b | 0 | 0 | 0.0001 | 9.54E-05 | 0.00092 | 0.00035 | 0.00061 | 0.00097 | 0.00022 | 0 | 0.00052 | 2.06E-05 | 0.00022 | 0.00064 | 0.0006 | 0 | 0.00043 | 0.00016 |
| Babam1 | 0.00019 | 0.00016 | 0.00011 | 0.00021 | 0.00012 | 0.00015 | 0.00011 | 0.00016 | 0.00022 | 9.40E-05 | 0.00011 | 8.81E-05 | 8.51E-05 | 0.00014 | 0 | 0.00018 | 0 | 0.00013 |
| Ddx6 | 0.00034 | 0.00036 | 0.00014 | 9.39E-05 | 0.0002 | 0.00029 | 0.00017 | 0.00014 | 0.00022 | 8.22E-05 | 0.00017 | 0.0002 | 0.00012 | 0.00016 | 0.00013 | 0.00025 | 0.00012 | 5.99E-05 |
| Gk | 0 | 0 | 4.06E-05 | 0 | 0.0001 | 0.00022 | 0 | 1.03E-05 | 0.00022 | 0 | 0 | 0.00011 | 5.47E-05 | 0.00012 | 1.54E-05 | 8.48E-05 | 1.70E-05 | 0 |
| Tmem250-ps | 6.06E-05 | 0.00018 | 0.00012 | 0.00061 | 0.00028 | 0.00017 | 0.00014 | 0.0003 | 0.00022 | 0.0002 | 9.14E-05 | 0 | 0.00018 | 0.00024 | 0 | 0.00011 | 0.00013 | 0.00017 |
| Fibp | 5.54E-05 | 0.00017 | 4.65E-05 | 0.00023 | 0.00012 | 0.00017 | 6.69E-05 | 0.00012 | 0.00022 | 8.90E-05 | 0.00013 | 0.00014 | 0.00016 | 0.00016 | 4.31E-05 | 3.52E-05 | 0 | 0.00018 |
| Rbbp4 | 0.00033 | 0.00026 | 0.00018 | 0.00012 | 0.00015 | 0.00023 | 1.57E-05 | 8.04E-05 | 0.00022 | 8.71E-05 | 0.00014 | 0.00015 | 8.80E-05 | 0.00014 | 5.96E-06 | 0.00056 | 0.00019 | 5.57E-05 |
| Ndufb8 | 0.00038 | 0.00012 | 0.00026 | 0.00038 | 5.08E-05 | 4.99E-05 | 4.06E-05 | 0.00015 | 0.00022 | 6.31E-05 | 0.00017 | 1.59E-05 | 4.08E-05 | 0.00018 | 0 | 0.00019 | 0 | 2.15E-05 |
| Gps1 | 0.00037 | 0.00014 | 0.00034 | 9.30E-05 | 0.00015 | 0.00017 | 0.00018 | 1.97E-05 | 0.00022 | 7.04E-05 | 8.65E-05 | 0 | 6.95E-05 | 4.36E-05 | 0.00013 | 0.00021 | 6.39E-05 | 4.60E-05 |
| Adk | 5.92E-05 | 6.44E-05 | 3.04E-05 | 0 | 6.35E-05 | 0 | 0.00016 | 0 | 0.00022 | 0.0002 | 0.00023 | 0 | 0.00017 | 0 | 0 | 0.00014 | 0 | 0.00017 |
| Ctso | 2.35E-05 | 0.00013 | 0 | 0 | 9.85E-05 | 0.00011 | 2.26E-05 | 9.56E-05 | 0.00022 | 0.00022 | 8.16E-06 | 2.94E-06 | 0.00023 | 0 | 0 | 0 | 0 | 0.00027 |
| Rheb | 0.00034 | 0.00028 | 0.00028 | 0 | 0.00015 | 0.0002 | 0.00018 | 0.00013 | 0.00022 | 0.00016 | 0.00044 | 0.00031 | 0.00022 | 6.88E-05 | 0 | 0.00031 | 9.10E-05 | 0.00015 |
| Syncrip | 0.00035 | 0.00031 | 0.00051 | 0.00048 | 0.00013 | 0.00017 | 0.0002 | 0.00011 | 0.00022 | 0.0001 | 0.00021 | 8.07E-05 | 0.0002 | 0.00014 | 0.00054 | 0.00049 | 0.00041 | 5.63E-05 |
| Isyna1 | 8.13E-05 | 8.75E-05 | 7.77E-05 | 0.00017 | 0 | 0.0001 | 0 | 7.66E-05 | 0.00022 | 6.80E-06 | 0 | 0 | 0.00021 | 3.23E-05 | 2.99E-05 | 4.26E-05 | 7.36E-05 | 0.00054 |
| Calu | 0.00047 | 0 | 0.00018 | 0 | 0.0001 | 0.00033 | 0.00033 | 0.00024 | 0.00022 | 0.00016 | 0.00017 | 0.00025 | 6.34E-05 | 0.00014 | 4.28E-05 | 6.65E-05 | 0.00011 | 7.04E-05 |
| Snx3 | 0.00041 | 0.00015 | 0.00034 | 0.00058 | 0.00019 | 0.00025 | 0.00027 | 0.00035 | 0.00022 | 0.00019 | 0.00018 | 0.00035 | 9.61E-06 | 9.86E-05 | 0 | 0.00029 | 0.00022 | 0.00025 |
| Lpgat1 | 4.16E-05 | 0.00021 | 0.00016 | 0.00011 | 0.00012 | 0.00018 | 0.00018 | 0.00015 | 0.00022 | 0.00038 | 0.00021 | 0.00017 | 0.00078 | 0.00026 | 6.84E-05 | 0.00016 | 0.00013 | 0.00011 |
| Wwtr1 | 0.00024 | 0.0004 | 0.0002 | 5.94E-05 | 0.00053 | 0.00044 | 0.00032 | 0.0003 | 0.00022 | 7.52E-05 | 0.00022 | 0.00021 | 0.00014 | 0.00048 | 0 | 0.00018 | 0.0005 | 0.00049 |
| Scara3 | 0 | 0.00025 | 4.22E-05 | 0 | 3.98E-05 | 0 | 0.00035 | 1.73E-05 | 0.00022 | 0.00105 | 0 | 0 | 0.00111 | 0.00021 | 0 | 0.0002 | 0 | 0.00015 |
| Srebf1 | 8.08E-05 | 0.00022 | 7.26E-05 | 0.00016 | 8.85E-05 | 9.68E-05 | 0 | 7.25E-05 | 0.00022 | 0.00011 | 0.0001 | 0.00021 | 4.05E-05 | 0.00011 | 0.00011 | 9.12E-05 | 0 | 0.00011 |
| Pxylp1 | 6.27E-05 | 0 | 0.00019 | 1.06E-05 | 0.00031 | 2.89E-05 | 0.00062 | 0.0001 | 0.00022 | 0 | 0 | 0.0002 | 0.00016 | 0.00025 | 0 | 0.00031 | 6.39E-05 | 0 |
| Pign | 9.68E-05 | 0.00012 | 0.00016 | 0 | 4.99E-05 | 0.00015 | 0 | 0.00014 | 0.00022 | 4.45E-05 | 0 | 8.22E-05 | 0.00018 | 2.09E-05 | 0.00011 | 0.00013 | 2.68E-05 | 1.75E-05 |
| Maea | 0.00013 | 0.00029 | 0.00028 | 0.00029 | 0.00018 | 4.55E-05 | 0.00027 | 7.50E-05 | 0.00022 | 0.00017 | 0.0003 | 0.00016 | 8.56E-05 | 5.93E-05 | 9.82E-05 | 0.00014 | 8.02E-05 | 0.00028 |
| Tenm3 | 1.65E-05 | 2.73E-05 | 0 | 0 | 0.00061 | 0.00011 | 2.94E-05 | 6.94E-05 | 0.00022 | 0 | 0.00024 | 0 | 0 | 0.00022 | 8.33E-05 | 0.00014 | 0 | 0.00026 |
| Derl1 | 0.00033 | 0.00019 | 0.00045 | 0.00027 | 0.00044 | 0.00029 | 0.00066 | 0.00041 | 0.00022 | 0.0002 | 0.00015 | 0.00033 | 0.00029 | 0.00019 | 7.07E-05 | 6.03E-05 | 6.08E-06 | 0.0002 |

| | | | | | | | | | | | | | | | | | | |
|---|---|---|---|---|---|---|---|---|---|---|---|---|---|---|---|---|---|---|
| Lgr5 | 0 | 0 | 0 | 0 | 0.00024 | 0 | 0.00066 | 0 | 0.00022 | 0 | 6.07E-05 | 0 | 0.00022 | 0 | 0 | 7.02E-05 | 0 | 0 |
| Wdr83os | 0.00026 | 0.00015 | 0.00034 | 0.00048 | 9.53E-05 | 0.00018 | 9.19E-05 | 0.0002 | 0.00022 | 0.00016 | 0.00016 | 9.85E-05 | 0.00012 | 0.00014 | 0 | 0.00015 | 7.71E-05 | 0.00021 |
| Gcn1 | 0.00014 | 0.00015 | 0.00026 | 0.00025 | 0.00016 | 1.06E-05 | 0.00012 | 6.88E-05 | 0.00022 | 7.62E-05 | 0.00012 | 1.72E-05 | 4.38E-05 | 7.50E-05 | 0.00042 | 4.04E-05 | 0 | 6.32E-05 |
| Fgfr1 | 0.00026 | 0.00066 | 0.00037 | 0.00028 | 0.00124 | 0.00076 | 0.00069 | 4.09E-05 | 0.00022 | 0.00052 | 0.0005 | 0.00062 | 0.00046 | 0.00118 | 0.00137 | 0.00017 | 0.00058 | 0.00017 |
| Megf8 | 0 | 0.00013 | 7.66E-05 | 0.00024 | 0.00018 | 0.00013 | 0.00015 | 0.00021 | 0.00022 | 0.00018 | 0.00014 | 8.05E-05 | 6.96E-05 | 0.0001 | 0.0004 | 5.18E-05 | 0 | 0.00016 |
| Rapgef1 | 0.00014 | 0.00021 | 0.00022 | 0 | 0.00023 | 0.00016 | 4.26E-05 | 4.38E-05 | 0.00022 | 0.00027 | 0.00027 | 0.00031 | 0 | 0.00013 | 3.90E-05 | 0.00015 | 2.80E-05 | 6.88E-05 |
| Edf1 | 0.00027 | 0.00011 | 0.00028 | 0.00033 | 0.00012 | 0.00011 | 8.28E-05 | 0.00013 | 0.00022 | 0.00012 | 0.00014 | 8.74E-05 | 8.77E-05 | 8.28E-05 | 2.76E-05 | 8.99E-05 | 3.43E-05 | 0.00026 |
| Ppp1ca | 0.00033 | 0.00018 | 0.00028 | 0.00041 | 0.00012 | 0.00018 | 0.00012 | 0.00022 | 0.00022 | 7.11E-05 | 0.00025 | 8.95E-05 | 0 | 0.00015 | 0 | 0.0002 | 0 | 5.98E-05 |
| Yy1 | 5.19E-05 | 0.00032 | 0.00021 | 0.00039 | 0.00011 | 1.12E-05 | 0 | 8.12E-05 | 0.00021 | 0.00017 | 0.00014 | 0.00031 | 0 | 9.28E-05 | 0.00101 | 0.00023 | 0.00055 | 0.00016 |
| Mycbp2 | 0.00019 | 0.00018 | 1.44E-05 | 4.48E-05 | 8.11E-05 | 0.00034 | 8.28E-05 | 1.52E-05 | 0.00021 | 6.73E-05 | 9.83E-05 | 2.79E-06 | 0.00069 | 0.00033 | 0.00044 | 1.37E-05 | 0.00028 | 0.00034 |
| Rbm42 | 0.00013 | 0.00026 | 0.00027 | 0.00046 | 0.00012 | 0.00013 | 7.10E-05 | 0.00019 | 0.00021 | 0.00017 | 0.00022 | 9.55E-05 | 4.03E-05 | 0.00017 | 5.00E-06 | 0.00016 | 0.00028 | 0.00024 |
| E2f4 | 0.00011 | 0.00016 | 0.00035 | 0.00021 | 0.00016 | 6.08E-05 | 7.80E-05 | 0 | 0.00021 | 0.00015 | 0.00021 | 8.31E-05 | 0.00019 | 0.00012 | 0.0002 | 0.00016 | 8.61E-05 | 8.21E-05 |
| Mpdu1 | 0.00012 | 0.00013 | 0.00011 | 0.00036 | 0.00016 | 0.00014 | 0.00021 | 0.00022 | 0.00021 | 3.64E-05 | 0.00013 | 1.27E-05 | 0.00019 | 0.00023 | 5.64E-05 | 0.00021 | 4.65E-05 | 4.43E-05 |
| Gaa | 0 | 0.00032 | 5.11E-06 | 0.00017 | 0.00057 | 0.00064 | 0.00048 | 0.00049 | 0.00021 | 0.00053 | 0.00016 | 0.00034 | 0.00098 | 0.00052 | 0.00013 | 0 | 4.41E-05 | 0.00055 |
| Idh2 | 0.00021 | 6.43E-05 | 7.03E-05 | 0.00045 | 0 | 0.00012 | 0.00029 | 0.00015 | 0.00021 | 0 | 0.00015 | 0 | 0 | 0.00049 | 0.00015 | 0.00066 | 0.00016 | 3.09E-05 |
| Tmed2 | 0.00059 | 0.00011 | 0.00043 | 0.00025 | 0.00027 | 0.00041 | 0.00063 | 0.00031 | 0.00021 | 0.00016 | 0.00023 | 0.00013 | 0.00028 | 0.00022 | 0 | 0.00018 | 0 | 0.00012 |
| St3gal2 | 0 | 0.0002 | 0.00013 | 7.92E-05 | 0.00023 | 5.90E-05 | 0.00018 | 0.00012 | 0.00021 | 0.00032 | 0.00019 | 5.06E-05 | 0.0002 | 6.19E-05 | 0.00012 | 0.00011 | 2.41E-05 | 0 |
| Gpc4 | 6.59E-05 | 9.86E-05 | 0 | 0 | 0.00015 | 0 | 0.00028 | 7.85E-06 | 0.00021 | 0.00014 | 0.00032 | 0 | 0.00045 | 0.00014 | 0 | 8.02E-05 | 0.00013 | 0 |
| Trim28 | 0.00021 | 8.99E-05 | 0.00019 | 0.00013 | 7.27E-05 | 0 | 3.87E-05 | 8.06E-05 | 0.00021 | 2.05E-05 | 0.00011 | 4.94E-05 | 6.17E-06 | 5.20E-05 | 2.25E-05 | 0.00027 | 8.10E-05 | 0 |
| Tmem45a | 0.00039 | 0 | 0.0001 | 0 | 0 | 0.00022 | 0.00053 | 6.28E-05 | 0.00021 | 0.00028 | 0 | 0.00136 | 0.0003 | 0 | 0 | 8.05E-05 | 0 | 2.50E-05 |
| S1pr1 | 0 | 0 | 7.73E-05 | 2.35E-05 | 0 | 0 | 1.29E-05 | 5.71E-05 | 0.00021 | 0 | 0 | 0 | 0 | 0 | 3.51E-05 | 4.35E-05 | 0 | 0 |
| Ttyh3 | 4.78E-05 | 0.00023 | 0 | 0.00065 | 0.00043 | 0.00018 | 0.0003 | 0.00013 | 0.00021 | 0.00013 | 0.0005 | 0 | 0.00023 | 0.0002 | 0.0003 | 0.00017 | 8.26E-05 | 9.55E-05 |
| Psmc5 | 0.0003 | 0.00015 | 0.0005 | 0.00045 | 9.35E-05 | 2.23E-05 | 4.97E-05 | 9.58E-05 | 0.00021 | 0.00019 | 0.00019 | 0 | 6.87E-05 | 0.00015 | 0.00013 | 0.00022 | 0 | 0.00019 |
| Efna5 | 0 | 3.22E-05 | 0 | 8.91E-05 | 7.62E-06 | 1.38E-05 | 4.78E-06 | 0 | 0.00021 | 0.00023 | 0 | 0 | 0.00011 | 3.72E-05 | 0.0001 | 0.00012 | 9.63E-05 | 0 |
| C1qtnf12 | 8.38E-06 | 5.81E-05 | 8.67E-05 | 0.00027 | 3.62E-05 | 9.22E-05 | 5.81E-06 | 0.00011 | 0.00021 | 0 | 0.00011 | 0.00018 | 0.00012 | 6.68E-05 | 0 | 4.06E-05 | 0.0001 | 0.00026 |
| Pex19 | 0.00019 | 0.00016 | 0 | 0.00013 | 0.00026 | 1.47E-05 | 0 | 0.00019 | 0.00021 | 9.95E-06 | 0.00029 | 1.89E-05 | 4.99E-05 | 9.08E-05 | 7.33E-05 | 3.63E-05 | 3.29E-06 | 5.84E-05 |
| Ipo9 | 0.00011 | 0.00021 | 0.00016 | 0.00012 | 0.0002 | 9.52E-05 | 7.18E-05 | 0.00013 | 0.00021 | 0.00011 | 0.00013 | 4.14E-05 | 8.63E-05 | 7.95E-05 | 0 | 0.0006 | 7.40E-05 | 0.00014 |
| Matn2 | 0.00016 | 0 | 7.47E-06 | 0.00015 | 0.00033 | 0.00024 | 0.0004 | 2.95E-05 | 0.00021 | 2.13E-05 | 7.80E-05 | 0 | 0 | 0.00044 | 5.09E-05 | 0.00022 | 0 | 0.00038 |
| Gnai3 | 0.00035 | 0.00027 | 0.00042 | 0.00041 | 0.00014 | 0.00027 | 0.00017 | 0.00042 | 0.00021 | 0.00027 | 0.00026 | 0.00033 | 6.23E-05 | 0.00019 | 0 | 0.0003 | 0.00023 | 0.00022 |
| Sec13 | 0.00052 | 1.62E-05 | 0.00053 | 0.00079 | 0.00018 | 0.00024 | 0.00053 | 0.0003 | 0.00021 | 9.85E-05 | 0.00025 | 0 | 0.00018 | 0.00027 | 0.00023 | 0.00013 | 0 | 0.00014 |
| Ppm1a | 0.00025 | 0.00039 | 0.00026 | 0 | 0.00026 | 0.00032 | 0.00026 | 0.00016 | 0.00021 | 0.0003 | 0.00029 | 0.00032 | 0.00011 | 0.00022 | 0.00015 | 0.00017 | 0.00032 | 0.00011 |
| S100a1 | 3.99E-05 | 7.50E-05 | 1.38E-05 | 0.0002 | 0.0003 | 0.00041 | 8.09E-05 | 0.00024 | 0.00021 | 0.00017 | 0.00014 | 0.0002 | 0.00043 | 0 | 0 | 0 | 0 | 8.07E-05 |
| Ccdc12 | 0.00014 | 0.00012 | 0.00012 | 0.00028 | 7.51E-05 | 0 | 0.00017 | 0.00024 | 0.00021 | 0.00011 | 9.42E-05 | 0 | 0.00012 | 8.16E-05 | 9.04E-05 | 4.60E-05 | 5.74E-05 | 0.00016 |

| | | | | | | | | | | | | | | | | | | |
|---|---|---|---|---|---|---|---|---|---|---|---|---|---|---|---|---|---|---|
| Cul7 | 0.00013 | 0.00011 | 0.00013 | 0.00028 | 4.88E-05 | 1.29E-05 | 0.00025 | 0.00014 | 0.00021 | 0.00013 | 1.99E-05 | 3.30E-05 | 6.97E-05 | 3.10E-06 | 0.00014 | 0 | 0 | 0.00011 |
| Eng | 0.00023 | 3.42E-05 | 0.00011 | 0 | 0.00016 | 0 | 0 | 0.00042 | 0.00021 | 0.00015 | 0.0003 | 9.20E-05 | 0.0002 | 0.00067 | 2.86E-05 | 5.55E-05 | 0 | 0.00056 |
| Amotl1 | 5.31E-06 | 0.00022 | 0.00019 | 5.81E-05 | 0.00043 | 0.00014 | 0.0002 | 0.00109 | 0.00021 | 0.00018 | 0.00011 | 0.00046 | 0 | 0 | 0.00013 | 8.15E-05 | 7.98E-05 | 0.00011 |
| Hgsnat | 0.00012 | 0.00015 | 0.00012 | 0 | 0.00024 | 0.00031 | 0.00023 | 0.00019 | 0.00021 | 0.00016 | 0.00012 | 7.13E-05 | 0.00022 | 0.00011 | 0 | 0 | 0 | 0.00022 |
| Il1rap | 0 | 1.08E-05 | 0.00011 | 0 | 0 | 0.00067 | 0 | 0 | 0.00021 | 0 | 0 | 0 | 0.00063 | 0.00038 | 0 | 2.55E-05 | 0 | 0 |
| Fbn1 | 0.00041 | 8.04E-05 | 0.00041 | 0.00011 | 0.00074 | 0.00092 | 0.00139 | 0.00139 | 0.00021 | 0.00154 | 0.00091 | 0.00037 | 0.00081 | 0.00094 | 0.00102 | 0.0002 | 0.00032 | 0.0011 |
| Pcyt1a | 8.37E-05 | 6.40E-05 | 0.00035 | 0.00015 | 0.00023 | 0.00018 | 0.0002 | 0.00028 | 0.00021 | 0.00017 | 0.00012 | 0 | 0.00012 | 0.00015 | 5.16E-06 | 0.00011 | 1.25E-05 | 0.00014 |
| Trio | 0.00012 | 0.00024 | 0.00029 | 0 | 0.00023 | 3.08E-05 | 0.0001 | 2.85E-05 | 0.00021 | 2.66E-05 | 0.0003 | 0.00023 | 0 | 8.65E-05 | 0.00053 | 7.22E-05 | 0.00052 | 0 |
| Got2 | 0.00016 | 0.00021 | 0.00025 | 0.00074 | 0.00022 | 8.69E-05 | 6.65E-05 | 6.27E-05 | 0.00021 | 8.93E-05 | 0.00018 | 8.00E-05 | 0.0002 | 0.00022 | 0 | 0.00033 | 0.00015 | 0.00013 |
| Bcl10 | 0.00019 | 9.24E-05 | 0.0002 | 0 | 8.29E-05 | 9.07E-05 | 2.76E-05 | 4.01E-05 | 0.00021 | 3.97E-06 | 0.00012 | 0.00015 | 7.38E-05 | 6.78E-05 | 1.64E-05 | 4.92E-05 | 0.00041 | 0 |
| Mob1a | 0.00029 | 0.00022 | 0.00029 | 2.68E-05 | 0.00013 | 0.00015 | 0.00013 | 8.94E-05 | 0.00021 | 0.00013 | 0.00023 | 0.00019 | 4.47E-05 | 2.95E-05 | 0 | 0.00017 | 6.23E-05 | 0.00014 |
| Nadk | 0.00034 | 0.00019 | 0.00032 | 0 | 6.23E-05 | 0.00012 | 0 | 7.01E-05 | 0.00021 | 0.00021 | 0.0001 | 0.00012 | 0.00012 | 0.00013 | 0.00017 | 2.54E-05 | 6.38E-05 | 0.00011 |
| Selenok | 0.00034 | 0.00013 | 0.00012 | 0.00026 | 0.00019 | 0.00026 | 0.00016 | 0.00022 | 0.00021 | 0.0001 | 0.00017 | 0.00023 | 8.96E-05 | 0.00018 | 5.63E-05 | 0 | 0.00027 | 0.00013 |
| Psma4 | 0.00063 | 0.00021 | 0.00041 | 0.00033 | 0.0001 | 0.00019 | 0.00017 | 2.11E-05 | 0.00021 | 9.81E-05 | 0.00011 | 4.02E-05 | 0.0001 | 6.24E-05 | 0 | 0.00027 | 0 | 7.45E-05 |
| Slc44a1 | 0.00029 | 0.00018 | 2.38E-05 | 0 | 0.00029 | 0.00062 | 0.00017 | 0.00028 | 0.00021 | 0.00034 | 0 | 3.57E-05 | 0.0003 | 0.00016 | 0 | 0.00011 | 5.58E-05 | 0.00022 |
| Tm9sf1 | 0.00019 | 0.00012 | 0.00024 | 0.0002 | 0.00022 | 0.00025 | 0.00034 | 0.00023 | 0.00021 | 0.00021 | 0.00017 | 0 | 0.00025 | 0.00023 | 0 | 0.00035 | 4.38E-07 | 0.00021 |
| Psip1 | 0.00014 | 0.00015 | 0.00012 | 0.00029 | 0 | 5.15E-05 | 5.28E-05 | 0.00026 | 0.00021 | 9.67E-05 | 2.44E-05 | 0 | 0 | 9.88E-05 | 7.58E-05 | 0.00095 | 0.00013 | 0.00018 |
| Rasa1 | 0.00023 | 0.00021 | 1.93E-05 | 0 | 9.74E-05 | 0.0004 | 0 | 0.00018 | 0.00021 | 0 | 3.03E-06 | 8.53E-05 | 0 | 0.00016 | 9.17E-05 | 6.92E-05 | 0.00014 | 8.55E-05 |
| Use1 | 0.00016 | 0.00028 | 0 | 0.00025 | 0.00031 | 0.00033 | 0.00028 | 3.31E-05 | 0.00021 | 0.00014 | 0 | 0.00039 | 0.00012 | 0.00019 | 7.79E-06 | 0 | 0.00018 | 0.00012 |
| Cox8a | 0.00018 | 0.00014 | 0.00025 | 0.0007 | 0.00017 | 0.0002 | 7.40E-05 | 0.00036 | 0.00021 | 5.65E-05 | 0.00021 | 0.00018 | 0.00016 | 0.00013 | 9.56E-05 | 0.00023 | 0.00027 | 0.00014 |
| Sh3bgrl | 0.00055 | 0.00015 | 4.27E-05 | 0.00013 | 0.00019 | 0.00024 | 0 | 0.00025 | 0.00021 | 8.59E-06 | 0.00048 | 7.34E-05 | 0.00023 | 0.00017 | 0 | 0.00017 | 4.33E-05 | 0.00028 |
| Lrp3 | 0 | 0.00011 | 2.86E-05 | 1.33E-05 | 6.19E-05 | 9.57E-05 | 2.80E-05 | 7.81E-05 | 0.00021 | 0.00016 | 0 | 0 | 0.00021 | 0.00018 | 6.45E-05 | 0 | 0 | 5.12E-05 |
| Nkd1 | 0 | 0.00011 | 0 | 0 | 0.00036 | 7.58E-05 | 0 | 0 | 0.00021 | 0 | 0.00028 | 1.53E-05 | 0 | 3.39E-06 | 0 | 0 | 3.89E-05 | 3.43E-05 |
| Tmem127 | 0.00014 | 0.00022 | 0.00013 | 8.61E-05 | 0.00029 | 0.00017 | 0.00026 | 0.00022 | 0.00021 | 0.00018 | 0.00013 | 0.00018 | 0.00013 | 0.00016 | 0 | 3.08E-05 | 0 | 0.00018 |
| Hipk3 | 0.00027 | 0.00028 | 0.00018 | 0 | 0.00023 | 0.00032 | 0.00019 | 0.00017 | 0.00021 | 0.00018 | 0.0003 | 0.0002 | 0.00028 | 0.00015 | 6.92E-05 | 0.0002 | 0.00039 | 0.00016 |
| Chmp3 | 0.00018 | 0.00019 | 0.00013 | 0.00011 | 0.00018 | 0.00019 | 0.00014 | 0.00017 | 0.00021 | 0.00025 | 0.00022 | 0.00018 | 9.75E-05 | 0.00014 | 0.00016 | 9.04E-05 | 0 | 0.00022 |
| Gatad1 | 0.00042 | 0.00018 | 0.00025 | 0 | 0.00012 | 0.00031 | 0.00012 | 0.00024 | 0.00021 | 3.22E-05 | 0.00017 | 0.00013 | 6.83E-05 | 5.06E-05 | 0 | 0.00031 | 8.23E-05 | 3.50E-05 |
| Tomm6 | 0.0002 | 4.80E-05 | 0.0002 | 0.00043 | 7.60E-05 | 8.43E-05 | 6.38E-05 | 0.00013 | 0.00021 | 4.34E-05 | 6.52E-05 | 0.00021 | 0.0002 | 0.00019 | 0 | 6.86E-05 | 0.00011 | 5.45E-05 |
| Sipa1 | 0 | 6.51E-05 | 0.00012 | 4.08E-05 | 0.00015 | 4.05E-05 | 5.44E-06 | 0.00026 | 0.00021 | 0.00011 | 8.88E-06 | 9.43E-05 | 0 | 6.51E-05 | 8.76E-05 | 8.00E-05 | 1.85E-05 | 0 |
| Tmem100 | 0 | 8.42E-06 | 6.43E-05 | 0 | 0 | 0.00015 | 0.00037 | 0.00059 | 0.00021 | 0.00054 | 0 | 0 | 0 | 0 | 0 | 6.29E-05 | 0 | 0.00012 |
| Ogdh | 0.00014 | 0.0002 | 0.00029 | 0.00052 | 0.00026 | 0.00021 | 0.00034 | 0.00022 | 0.00021 | 0.00022 | 0.00034 | 0 | 0.00026 | 0.00031 | 2.42E-05 | 0.00033 | 0 | 0.00032 |
| Snu13 | 0.00028 | 0.00013 | 0.00044 | 0.00026 | 0.00018 | 0.00024 | 0 | 0.00013 | 0.00021 | 0.00012 | 0.00013 | 0.00016 | 6.30E-05 | 0.00016 | 0 | 0.00035 | 0.00034 | 0.00018 |
| Hcfc1 | 0.00014 | 0.00018 | 0.00023 | 0.00018 | 8.73E-05 | 0 | 7.24E-05 | 8.58E-05 | 0.00021 | 6.54E-05 | 0.00014 | 3.55E-05 | 0 | 4.34E-05 | 5.09E-05 | 0.0003 | 0.00015 | 0.0001 |

| | | | | | | | | | | | | | | | | | | |
|---|---|---|---|---|---|---|---|---|---|---|---|---|---|---|---|---|---|---|
| Megf6 | 0 | 0 | 0 | 0 | 0.00036 | 0.00021 | 0 | 0 | 0.00021 | 0 | 0 | 0 | 0 | 0.00028 | 0 | 0 | 0 | 2.75E-06 |
| DEPRECATED | 9.88E-05 | 0.00019 | 0.00033 | 0.00043 | 6.80E-05 | 1.34E-07 | 5.59E-05 | 8.74E-05 | 0.00021 | 7.85E-05 | 7.23E-05 | 9.45E-05 | 2.98E-05 | 1.51E-05 | 0.0003 | 0.00015 | 0 | 4.46E-05 |
| Arid1b | 0.00016 | 0.00022 | 5.65E-05 | 0 | 0.00034 | 0.00018 | 7.32E-05 | 5.82E-05 | 0.00021 | 6.14E-05 | 9.83E-05 | 0.00011 | 0.0001 | 3.07E-05 | 3.80E-05 | 0.00012 | 0.00032 | 9.49E-05 |
| Dynll1 | 0.00064 | 7.65E-05 | 0.00027 | 2.86E-05 | 1.16E-05 | 0.00022 | 0.00012 | 6.52E-05 | 0.00021 | 9.52E-05 | 0.0004 | 0.00014 | 0 | 0.00015 | 0 | 0.00019 | 0.00048 | 0.00012 |
| Tbc1d9b | 0.00017 | 0.00022 | 7.96E-05 | 0 | 0.00022 | 0.00021 | 0.0002 | 0.00028 | 0.00021 | 0.00019 | 0.00025 | 0.00018 | 0.0002 | 8.57E-05 | 9.54E-05 | 5.25E-05 | 6.82E-05 | 0.00024 |
| G3bp2 | 0.00023 | 0.00014 | 0.00016 | 5.05E-05 | 7.86E-05 | 3.62E-05 | 0 | 7.03E-05 | 0.00021 | 0.00012 | 0.00013 | 0.00012 | 4.89E-05 | 0.00012 | 0 | 9.89E-05 | 8.36E-05 | 9.60E-05 |
| Stt3b | 0.00051 | 0.00015 | 0.00031 | 0.00014 | 0.00018 | 0.00026 | 0.0003 | 0.00013 | 0.00021 | 0.00027 | 0.00018 | 0.00035 | 0.00026 | 0.00015 | 0 | 0.00039 | 0.00014 | 0.00028 |
| Vps28 | 0.00024 | 0.00011 | 0.00016 | 0.0001 | 0.00014 | 0.00017 | 7.28E-05 | 0.00014 | 0.00021 | 0.00015 | 0.00011 | 6.03E-05 | 3.39E-05 | 0.00011 | 0 | 0.00011 | 0 | 0.00018 |
| Ube2z | 0.00014 | 0.00024 | 0.00037 | 0.00032 | 0.00019 | 0.00013 | 0.0003 | 0.0002 | 0.00021 | 0.00018 | 0.0002 | 0.00039 | 0.00016 | 0.0002 | 0 | 0.00032 | 0 | 0.00016 |
| Emg1 | 0.00019 | 0.00017 | 0.00034 | 0.00043 | 8.64E-05 | 0 | 0 | 6.54E-05 | 0.00021 | 7.68E-05 | 7.86E-05 | 0.00014 | 9.43E-05 | 0.00013 | 3.86E-05 | 0.00029 | 8.80E-05 | 7.10E-05 |
| Smad2 | 0.00021 | 0.00014 | 0.00014 | 0 | 5.24E-05 | 0.00014 | 4.35E-05 | 6.81E-05 | 0.00021 | 3.48E-05 | 0.00013 | 8.69E-05 | 0 | 0 | 0 | 0.00011 | 0.00017 | 6.78E-05 |
| Slc25a4 | 0.00014 | 0.00017 | 0.00025 | 0.00086 | 0.00034 | 0.00022 | 0.00025 | 0.0002 | 0.00021 | 0.0003 | 0.00026 | 0.00011 | 0.00022 | 0.00036 | 8.04E-05 | 0.00024 | 0.00039 | 0.00042 |
| Atp6ap1 | 0.00017 | 0.00018 | 0.00022 | 0.00026 | 0.00024 | 0.00028 | 0.00018 | 0.00035 | 0.00021 | 0.00024 | 0.00028 | 0.00028 | 0.00032 | 0.00039 | 0 | 0.00013 | 3.87E-05 | 0.00038 |
| Klf6 | 3.33E-05 | 0.00016 | 2.38E-05 | 0.0003 | 0.00019 | 0.00013 | 0.00028 | 0.00017 | 0.00021 | 0.00049 | 0.00058 | 0.00013 | 7.75E-05 | 0.00031 | 0.00035 | 0.00026 | 0.00279 | 0.00041 |
| Pgs1 | 0 | 0.00017 | 0.00023 | 0.00029 | 0.00016 | 6.18E-05 | 0 | 3.43E-05 | 0.00021 | 0.00011 | 0.00013 | 0 | 0.00035 | 0.00027 | 0.00037 | 0.00012 | 2.13E-05 | 2.40E-05 |
| Nfkb2 | 0 | 0.00027 | 0.00049 | 0.00034 | 0.00011 | 4.75E-05 | 0 | 0 | 0.00021 | 3.81E-05 | 5.91E-05 | 0.00017 | 0.00018 | 0.00027 | 0.00014 | 8.53E-05 | 0.00078 | 0.00039 |
| Sap30l | 0.00023 | 0.00019 | 0.00013 | 3.84E-05 | 0.00017 | 0.00015 | 0.00029 | 5.88E-05 | 0.00021 | 6.47E-05 | 4.07E-05 | 8.45E-05 | 0 | 0.00014 | 3.99E-05 | 0.00014 | 0.00011 | 4.96E-05 |
| Hmgn3 | 0.00012 | 8.36E-05 | 0 | 9.39E-05 | 0.00013 | 0 | 8.21E-05 | 0.00018 | 0.00021 | 0.00015 | 0 | 0 | 0 | 3.37E-05 | 3.76E-05 | 5.91E-05 | 0 | 7.42E-05 |
| Peli1 | 3.82E-05 | 8.37E-05 | 0.0002 | 2.17E-05 | 0.00017 | 4.44E-05 | 8.94E-05 | 0.00025 | 0.00021 | 5.41E-05 | 0.00018 | 0.00014 | 0 | 0.00013 | 1.91E-05 | 8.31E-05 | 0.00071 | 9.87E-05 |
| Hcfc1r1 | 0.00022 | 0.00016 | 0 | 0.00021 | 0.00034 | 0.00017 | 0.00039 | 0 | 0.00021 | 0.0002 | 0.00039 | 0.00049 | 0 | 0 | 0 | 0 | 1.66E-05 | 0.00057 |
| Nfat5 | 0.00015 | 0.00029 | 0.00011 | 0 | 0.00017 | 0.00022 | 0.00017 | 0 | 0.00021 | 7.49E-05 | 2.01E-05 | 0.00014 | 4.52E-05 | 0.00022 | 0.00057 | 0 | 0.00053 | 0.00013 |
| Tanc1 | 4.04E-05 | 0.00017 | 8.37E-05 | 0 | 0.00046 | 0.0003 | 0.00011 | 5.15E-05 | 0.00021 | 0 | 9.96E-05 | 0 | 0 | 0.0005 | 0.00022 | 0 | 4.54E-05 | 0 |
| Ppp3r1 | 0.00035 | 0.00021 | 0.00026 | 0 | 0.00018 | 0.00019 | 8.82E-05 | 0.00012 | 0.00021 | 2.29E-07 | 0.00025 | 0.00018 | 0 | 8.79E-06 | 0 | 0.00012 | 2.98E-06 | 6.32E-05 |
| Zdhhc8 | 0 | 0.00014 | 0.00016 | 0.00024 | 0.00015 | 4.34E-05 | 0.00015 | 6.52E-05 | 0.00021 | 0.0003 | 0.00016 | 0.00011 | 7.21E-05 | 0.00019 | 2.78E-05 | 6.01E-05 | 0.0002 | 0.00031 |
| Eef2 | 0.00011 | 0.00063 | 0.0003 | 0.00103 | 0.00029 | 0.00045 | 0.00052 | 0.00051 | 0.00021 | 0.0003 | 0.00017 | 0.00075 | 0.00029 | 0.00046 | 0.00033 | 0.00018 | 0.00061 | 0.00031 |
| Myl12b | 0.00017 | 0.00017 | 0 | 5.79E-05 | 0.00011 | 0.00012 | 7.62E-05 | 9.33E-05 | 0.00021 | 0.00028 | 0 | 0 | 0.00011 | 0 | 0 | 3.58E-05 | 0 | 0.00025 |
| Psmb10 | 0.00014 | 4.64E-05 | 0 | 0.00016 | 0 | 0 | 0 | 8.82E-05 | 0.00021 | 0.00011 | 0 | 0 | 7.80E-05 | 0 | 0 | 0 | 0 | 0.00019 |
| Spen | 7.08E-05 | 0.0002 | 0.00012 | 5.60E-05 | 5.73E-05 | 9.32E-05 | 2.30E-06 | 7.73E-05 | 0.00021 | 2.96E-05 | 9.51E-05 | 1.39E-05 | 8.64E-05 | 0.00014 | 0.00016 | 7.81E-05 | 0.00018 | 0.00011 |
| Gpx8 | 0.00052 | 0 | 0.00018 | 0.00018 | 0.00033 | 0.00044 | 0.00067 | 0.00033 | 0.00021 | 0.0002 | 0.00019 | 0.00029 | 0.00048 | 0.00029 | 0 | 0.00012 | 0 | 0.00039 |
| Ap2b1 | 0.0003 | 0.00012 | 0.00021 | 0.00031 | 0.0001 | 0.00014 | 0.00012 | 8.51E-05 | 0.00021 | 0.00023 | 0.00022 | 0.00021 | 4.95E-05 | 6.99E-05 | 0 | 0.00011 | 9.68E-05 | 0.00022 |
| Adam17 | 0.0001 | 0.00019 | 4.60E-05 | 1.64E-05 | 0.00015 | 0.00015 | 0.00014 | 0.00029 | 0.00021 | 0.00033 | 0 | 0.00022 | 0.00027 | 0.00022 | 0 | 0.00019 | 0.00025 | 0.00022 |
| Ttpal | 0.00012 | 7.72E-05 | 1.29E-05 | 4.11E-05 | 0.00014 | 8.55E-05 | 3.69E-05 | 0.00021 | 0.00021 | 0 | 0 | 0 | 4.61E-05 | 7.64E-05 | 2.87E-05 | 9.19E-05 | 0 | 0 |
| Rnh1 | 0.00018 | 0.00032 | 0.0003 | 0.00044 | 6.88E-05 | 9.34E-05 | 0.00026 | 5.75E-06 | 0.00021 | 0.00021 | 0.00012 | 0.00018 | 6.01E-05 | 4.69E-05 | 5.99E-05 | 6.65E-05 | 6.76E-05 | 0.00021 |

| | | | | | | | | | | | | | | | | | | |
|---|---|---|---|---|---|---|---|---|---|---|---|---|---|---|---|---|---|---|
| Irf1 | 0.00017 | 0.00014 | 3.07E-05 | 7.98E-05 | 0.00022 | 0 | 0 | 7.82E-05 | 0.00021 | 0 | 4.36E-05 | 0 | 0.00029 | 0.00037 | 0.00017 | 0.00012 | 0.001 | 0.00012 |
| Ppp2r5a | 0.00019 | 0.00022 | 8.09E-05 | 3.27E-05 | 2.99E-05 | 8.83E-05 | 9.92E-05 | 5.94E-05 | 0.00021 | 0.00013 | 4.61E-05 | 8.26E-05 | 0 | 4.92E-05 | 1.70E-06 | 5.10E-05 | 0.00014 | 4.10E-05 |
| Id3 | 0 | 0.00013 | 0 | 4.30E-05 | 0.00031 | 0.00038 | 0 | 0 | 0.00021 | 0.00031 | 0.00095 | 2.48E-06 | 0.00043 | 0 | 8.25E-05 | 0 | 0.00198 | 0.00089 |
| Slc20a2 | 0 | 7.35E-05 | 0.00017 | 0.0001 | 0.00011 | 2.81E-05 | 7.08E-06 | 0 | 0.00021 | 3.63E-06 | 5.72E-05 | 7.81E-05 | 0.00014 | 0.00022 | 3.42E-05 | 0.0002 | 3.14E-05 | 0.0001 |
| Rap2c | 0.00023 | 7.32E-05 | 7.49E-05 | 0 | 3.61E-05 | 0.00016 | 0 | 1.99E-05 | 0.00021 | 3.37E-05 | 3.90E-05 | 4.77E-05 | 0 | 7.19E-05 | 6.87E-05 | 1.17E-05 | 5.60E-05 | 0 |
| Srsf2 | 0.00015 | 0.00016 | 0.00019 | 0.00023 | 0 | 1.89E-05 | 0 | 8.96E-05 | 0.00021 | 3.96E-05 | 9.76E-05 | 0.00022 | 0 | 4.22E-05 | 0.00026 | 0.00037 | 0.0009 | 4.71E-05 |
| Upf1 | 9.55E-05 | 0.0002 | 0.0003 | 0.00017 | 0.00012 | 2.92E-05 | 0 | 8.23E-05 | 0.00021 | 0.00011 | 0.00016 | 1.67E-06 | 0 | 0.00014 | 0.00023 | 0.0002 | 0.00023 | 0.00018 |
| Manf | 0.00049 | 7.21E-05 | 0.00083 | 0.00081 | 0.00031 | 0.00014 | 0.00048 | 0.00042 | 0.00021 | 0.00039 | 0.00048 | 0.00034 | 0.00068 | 0.00041 | 0 | 0.00028 | 0 | 0.00037 |
| Bnip2 | 0.00037 | 0.00017 | 0.00025 | 0 | 0.00017 | 0.00025 | 2.67E-05 | 0.00011 | 0.00021 | 5.30E-05 | 0.00021 | 0.00017 | 7.87E-05 | 0.0001 | 0.00023 | 0.00017 | 0 | 6.93E-05 |
| Atp11a | 0.0001 | 0.0003 | 1.32E-05 | 0 | 0.00039 | 0.00023 | 0.00021 | 0.00069 | 0.00021 | 0.00026 | 0.00026 | 0.0002 | 5.97E-05 | 0.00012 | 0.00011 | 0.00017 | 0.00034 | 0.00014 |
| Arhgef17 | 1.46E-05 | 0.00023 | 4.06E-05 | 0.00012 | 0.00015 | 0.00026 | 0 | 0.00026 | 0.00021 | 1.15E-05 | 0.00036 | 0.00012 | 4.35E-05 | 0.00024 | 0.00012 | 8.22E-05 | 6.46E-05 | 5.53E-05 |
| Gpc6 | 3.72E-05 | 6.17E-05 | 0 | 0 | 0.00014 | 0.00022 | 0.00022 | 0 | 0.00021 | 0.00022 | 0.00019 | 0.00023 | 0 | 0 | 0 | 6.46E-05 | 0 | 0.00039 |
| Morf4l1 | 0.00052 | 0.00033 | 0.00023 | 0.00052 | 0.00028 | 0.00045 | 0.00039 | 0.00027 | 0.00021 | 0.00027 | 0.0003 | 0.0003 | 0.00022 | 0.00035 | 0.00023 | 0.00028 | 0.00039 | 0.00027 |
| Pofut1 | 0 | 0.00012 | 9.66E-05 | 0.00017 | 0.00016 | 7.48E-05 | 0.00039 | 0.00012 | 0.00021 | 0.00024 | 4.37E-05 | 3.15E-05 | 0.00015 | 6.25E-05 | 0.00015 | 0.00022 | 4.83E-05 | 0.00017 |
| Rnpepl1 | 2.83E-05 | 0.00018 | 8.52E-05 | 0.00024 | 6.91E-05 | 8.82E-05 | 0.00012 | 0.00012 | 0.00021 | 0.00013 | 0.00012 | 0.00012 | 0 | 0.00018 | 4.35E-05 | 4.68E-05 | 5.46E-05 | 0.00016 |
| Frzb | 0 | 0 | 0 | 0 | 0.00011 | 0.00015 | 0 | 0.0002 | 0.00021 | 0 | 0 | 0 | 0 | 0.00022 | 0 | 0 | 0 | 1.96E-05 |
| Smdt1 | 0.00022 | 0.00016 | 0.0002 | 0.00036 | 0.00028 | 0.00016 | 0.00031 | 0.00025 | 0.0002 | 0.00022 | 0.00027 | 0 | 4.43E-06 | 9.32E-05 | 0 | 0.00017 | 0.00016 | 0.00029 |
| Plpp1 | 0.00019 | 0 | 5.55E-05 | 0 | 8.72E-05 | 0.00024 | 0.00058 | 0 | 0.0002 | 2.80E-05 | 7.91E-05 | 0.00023 | 0.00016 | 0 | 0 | 6.30E-05 | 0 | 0.00038 |
| Reep3 | 0.00016 | 0.00025 | 0.00018 | 0 | 0.00014 | 0.00027 | 0.00013 | 0.00019 | 0.0002 | 0.00031 | 4.97E-05 | 0.00023 | 0 | 0.0003 | 0 | 7.71E-05 | 6.46E-05 | 0.00039 |
| Ndufb9 | 0.00036 | 0.00016 | 0.0003 | 0.00042 | 0.00014 | 0.00017 | 9.16E-05 | 0.00015 | 0.0002 | 0.00011 | 0.00027 | 7.11E-05 | 5.08E-05 | 0.00018 | 0 | 0.00015 | 0 | 0.00019 |
| Ank | 3.43E-05 | 8.90E-05 | 6.54E-05 | 0 | 5.72E-05 | 0 | 1.83E-05 | 0.00138 | 0.0002 | 0 | 0 | 0 | 0 | 0.00035 | 0 | 0.00011 | 0 | 0.0004 |
| Clptm1l | 0.00025 | 0.0001 | 0.00038 | 0.00054 | 0.00034 | 0.00028 | 0.00034 | 0.00031 | 0.0002 | 0.00027 | 0.00024 | 6.01E-05 | 0.00034 | 0.00018 | 0.00012 | 6.19E-06 | 0 | 0.00028 |
| Atp1a1 | 0.00018 | 0.0002 | 0.00051 | 0.00063 | 0.00028 | 0.00041 | 0.00067 | 0.00037 | 0.0002 | 0.00017 | 0.00034 | 4.72E-05 | 0.00039 | 0.00041 | 0.00023 | 0.00026 | 0.0005 | 0.00019 |
| Nr2f2 | 0 | 5.75E-05 | 0 | 1.12E-05 | 0 | 0.00033 | 0 | 0.00034 | 0.0002 | 0 | 0 | 2.19E-06 | 0 | 3.61E-06 | 4.92E-05 | 3.39E-05 | 0 | 0.00061 |
| Ifnar1 | 0.00026 | 0.00018 | 0.00013 | 0 | 8.61E-05 | 0.00017 | 9.25E-05 | 9.14E-05 | 0.0002 | 0.00011 | 2.91E-06 | 0.00018 | 0.00019 | 7.86E-05 | 6.03E-05 | 3.73E-05 | 4.91E-06 | 9.12E-05 |
| Rela | 7.97E-05 | 0.00032 | 0.00028 | 8.42E-05 | 0.00015 | 1.62E-05 | 2.31E-05 | 0.00021 | 0.0002 | 0.00019 | 0.00015 | 0.00027 | 5.12E-05 | 0.00021 | 0.00021 | 8.56E-05 | 0.00036 | 0.0002 |
| Odc1 | 0.00029 | 0 | 0.00049 | 0.00042 | 0.00061 | 0.00011 | 0.00032 | 0 | 0.0002 | 0 | 0.00092 | 3.89E-05 | 0.00063 | 0.00087 | 0 | 0.00023 | 0.001 | 4.87E-05 |
| Usp4 | 0.00021 | 0.0002 | 0.00024 | 9.50E-05 | 9.58E-05 | 5.93E-05 | 7.55E-05 | 4.07E-05 | 0.0002 | 0.00017 | 0.00014 | 4.68E-05 | 7.07E-06 | 5.48E-05 | 0.00012 | 0.00011 | 0 | 0.00016 |
| Ssbp4 | 3.51E-05 | 0.00014 | 7.24E-05 | 0.00039 | 7.00E-05 | 8.32E-05 | 0 | 0.00044 | 0.0002 | 0.00017 | 0 | 0 | 0.00012 | 0.00016 | 8.03E-05 | 0.00016 | 5.81E-05 | 0.00026 |
| Ankrd52 | 8.89E-05 | 0.00012 | 0.00029 | 0.00026 | 9.56E-05 | 4.45E-06 | 6.14E-05 | 0.00019 | 0.0002 | 8.38E-05 | 0.00019 | 6.60E-05 | 0 | 0.00014 | 0.00022 | 0.00016 | 0 | 4.69E-05 |
| Mtpn | 0.00018 | 0.00017 | 0.00044 | 0.00027 | 9.41E-05 | 0.00014 | 0 | 8.15E-05 | 0.0002 | 0.00014 | 0.00036 | 0.00021 | 0.00015 | 0.00017 | 0 | 0.00018 | 6.31E-05 | 0.00011 |
| Pitpnc1 | 0 | 8.64E-05 | 0.00011 | 0 | 0.00029 | 0.00039 | 9.71E-05 | 7.43E-05 | 0.0002 | 0 | 0.00014 | 0 | 2.69E-05 | 0.00033 | 0.00016 | 0 | 0.00011 | 0 |
| Atxn7l3b | 3.06E-05 | 0.00028 | 8.26E-05 | 0.00042 | 0.00028 | 0.00021 | 4.21E-05 | 0.00031 | 0.0002 | 0.00019 | 0.00027 | 0.0002 | 0.00017 | 0.00025 | 0.00015 | 3.49E-05 | 0 | 0.00023 |

| | | | | | | | | | | | | | | | | | | |
|---|---|---|---|---|---|---|---|---|---|---|---|---|---|---|---|---|---|---|
| Ddah2 | 0 | 0.00017 | 0 | 0.00031 | 0.00015 | 0 | 0 | 0.00022 | 0.0002 | 0.00045 | 0.00028 | 0.0002 | 0 | 6.91E-05 | 5.27E-07 | 1.00E-04 | 8.33E-05 | 0.00041 |
| Clic4 | 0.00018 | 0.00038 | 0.0004 | 0.00018 | 0.00026 | 0.00018 | 0.00028 | 1.13E-05 | 0.0002 | 0.0007 | 0.00059 | 0.00048 | 0.00032 | 0.00041 | 0 | 0.00034 | 0.00065 | 0.00047 |
| Gpr27 | 0 | 0 | 2.53E-05 | 5.78E-05 | 0 | 0 | 0 | 0.0001 | 0.0002 | 3.34E-05 | 0 | 0 | 0.00017 | 0.00011 | 6.31E-05 | 0 | 0 | 0.00016 |
| Get4 | 0.00021 | 0.00013 | 0.00026 | 0.00015 | 8.83E-05 | 9.01E-05 | 0.00014 | 5.29E-05 | 0.0002 | 7.80E-05 | 0.00016 | 3.41E-05 | 3.75E-05 | 7.39E-05 | 3.87E-05 | 0.0002 | 0 | 3.14E-05 |
| Cmtm6 | 3.03E-05 | 0.00017 | 0.00018 | 2.40E-05 | 0.00017 | 0.00015 | 0 | 0.00037 | 0.0002 | 0.00022 | 0.00014 | 0.00014 | 7.47E-05 | 0.00012 | 2.22E-05 | 7.59E-05 | 0 | 0.0002 |
| Hdac3 | 0.0002 | 0.00015 | 0.00019 | 0.00012 | 8.94E-05 | 5.36E-05 | 0.0001 | 9.54E-05 | 0.0002 | 6.79E-05 | 0.00012 | 0.0002 | 5.17E-05 | 0.00012 | 5.42E-05 | 0.00023 | 0 | 4.74E-05 |
| Arl6ip5 | 0.00022 | 0.00017 | 0.00016 | 4.27E-05 | 0.00014 | 0.00022 | 0.00015 | 2.75E-05 | 0.0002 | 0.00025 | 0.00016 | 0.00025 | 0.00016 | 0.00011 | 0 | 0 | 0.0002 | 0.00036 |
| Ranbp1 | 0.00048 | 4.63E-05 | 0.00052 | 0.0003 | 3.11E-05 | 0.00013 | 0.00014 | 8.19E-05 | 0.0002 | 4.38E-05 | 0.00017 | 0 | 0.00012 | 9.79E-05 | 8.31E-05 | 0.00076 | 9.38E-05 | 1.00E-05 |
| Bri3bp | 0 | 0.00013 | 0.00013 | 0.00032 | 0.00016 | 0.00013 | 0 | 0.00018 | 0.0002 | 7.38E-06 | 8.14E-05 | 0.0002 | 0.00013 | 0.00012 | 2.61E-05 | 8.30E-05 | 0 | 5.93E-05 |
| Igdcc4 | 0 | 6.01E-05 | 0 | 0.00015 | 0.00027 | 0.00012 | 0.00032 | 0 | 0.0002 | 8.69E-05 | 0.00012 | 3.70E-05 | 7.11E-05 | 0.00012 | 0.00019 | 0 | 0 | 4.44E-05 |
| Sirpa | 9.73E-05 | 0.00018 | 0.00014 | 2.25E-05 | 2.62E-05 | 0.00023 | 0.00031 | 0 | 0.0002 | 0.00048 | 1.48E-05 | 0 | 0.00047 | 0.00026 | 9.50E-05 | 0.00016 | 0 | 0.00017 |
| U2af2 | 0.00026 | 0.00029 | 0.00041 | 0.00037 | 6.31E-05 | 0 | 1.79E-05 | 3.31E-05 | 0.0002 | 9.48E-05 | 0.00012 | 0.00012 | 4.54E-05 | 9.89E-05 | 0.00053 | 0.00038 | 0.00048 | 9.89E-05 |
| Sfpq | 0.00021 | 0.00022 | 0.00027 | 0.00048 | 9.13E-05 | 4.36E-05 | 0 | 0.00015 | 0.0002 | 7.50E-05 | 9.76E-05 | 0.00017 | 0 | 0.0002 | 0.00052 | 0.00037 | 0.00091 | 7.86E-05 |
| Parp3 | 6.98E-05 | 0.00023 | 0 | 0.00013 | 0.00014 | 0.00022 | 0.00013 | 0 | 0.0002 | 0.00034 | 1.40E-05 | 0 | 0.00021 | 0.00017 | 0.00015 | 0 | 0 | 0.00047 |
| Iffo2 | 4.33E-06 | 8.51E-05 | 0.00013 | 0 | 8.96E-05 | 8.10E-05 | 0 | 0.00022 | 0.0002 | 4.58E-05 | 0.00015 | 7.29E-05 | 0 | 0.00018 | 0.00011 | 2.64E-06 | 0.00033 | 0 |
| Por | 6.30E-05 | 0.00025 | 0.00015 | 0.00014 | 0.00022 | 0 | 0 | 0.00014 | 0.0002 | 3.57E-05 | 5.11E-05 | 0.00023 | 4.16E-05 | 0.00014 | 6.30E-05 | 8.41E-06 | 5.55E-05 | 3.49E-05 |
| D8Ertd738e | 0.00022 | 0.00018 | 0.00015 | 0.00046 | 0.00013 | 7.78E-05 | 7.99E-05 | 0.00011 | 0.0002 | 9.68E-05 | 0.00016 | 0.00011 | 0 | 0.00012 | 0.00017 | 0.00021 | 0.00011 | 0.00013 |
| Ifi27 | 0.00031 | 0.00019 | 0.00014 | 0.0002 | 0.00015 | 1.97E-05 | 0.00029 | 0.00015 | 0.0002 | 0 | 0.00019 | 0.0002 | 0.00068 | 1.66E-05 | 0 | 0.0001 | 0.00029 | 0.00057 |
| Dram1 | 0 | 0.00012 | 0.0003 | 0.00027 | 0.00029 | 0.00069 | 0.00051 | 0 | 0.0002 | 0.00037 | 0 | 0 | 0.00104 | 0.00044 | 0.00014 | 5.46E-05 | 0 | 0.00021 |
| Gatad2a | 0.00017 | 0.00021 | 0.0004 | 0.00011 | 9.68E-05 | 5.32E-05 | 2.75E-05 | 7.13E-05 | 0.0002 | 0.0001 | 0.00016 | 1.75E-05 | 0 | 5.46E-05 | 3.18E-05 | 0.00027 | 0.0002 | 7.74E-05 |
| Hoxc8 | 0.0001 | 0.0001 | 2.81E-05 | 0.00029 | 0.00021 | 0.00018 | 0.00026 | 0 | 0.0002 | 7.17E-05 | 0.00026 | 0 | 0.00015 | 0 | 7.55E-05 | 7.86E-05 | 0 | 0.00027 |
| Thbd | 0 | 0.00015 | 4.73E-05 | 0 | 0 | 0 | 0 | 0.00029 | 0.0002 | 0.00072 | 0 | 0 | 6.98E-05 | 0 | 0 | 5.50E-05 | 5.23E-05 | 0.00038 |
| Cdr2l | 0.00019 | 0.00013 | 0.00042 | 0.00038 | 9.94E-05 | 0 | 0.00077 | 0.00048 | 0.0002 | 0.00028 | 0 | 0.00028 | 0.00017 | 0 | 0.00055 | 0 | 7.93E-05 | 0 |
| Timm13 | 0.00032 | 0.00013 | 0.00043 | 0.00074 | 0.00021 | 0.00022 | 0.00017 | 0.00022 | 0.0002 | 6.97E-05 | 0.00021 | 5.91E-06 | 7.26E-05 | 0.00026 | 0 | 0.00021 | 0.00018 | 0.00014 |
| Dgcr2 | 2.31E-05 | 0.00017 | 8.33E-05 | 0.00018 | 0.00018 | 0.00016 | 8.05E-05 | 0.00016 | 0.0002 | 0.0002 | 0.00012 | 4.98E-05 | 0.00013 | 0.0002 | 0.00016 | 0.00019 | 0 | 0.00011 |
| Chp1 | 9.59E-05 | 0.00029 | 0.00023 | 0.0003 | 0.00022 | 0.00027 | 0.00013 | 0.00021 | 0.0002 | 0.00037 | 0.0002 | 0.00011 | 0.00039 | 0.00031 | 0 | 8.33E-05 | 0 | 0.00032 |
| Ppp1cc | 0.00012 | 0.00021 | 0.00019 | 8.34E-05 | 0.00021 | 0.00011 | 6.04E-05 | 0.00011 | 0.0002 | 0.00014 | 2.50E-05 | 0.00019 | 0.00019 | 8.35E-05 | 0 | 0.00027 | 1.56E-05 | 0 |
| Pcyt2 | 2.59E-05 | 7.15E-05 | 0 | 2.05E-05 | 5.32E-05 | 0 | 0 | 0 | 0.0002 | 3.02E-05 | 3.23E-05 | 1.67E-05 | 0 | 8.14E-05 | 0.00019 | 0 | 0.00015 | 1.42E-05 |
| Papola | 0.00047 | 0.00029 | 0.00036 | 0.00023 | 0.00021 | 0.00027 | 0.0002 | 0.00023 | 0.0002 | 0.00014 | 0.00022 | 0.00024 | 0.00014 | 0.00017 | 0.00037 | 0.00027 | 2.03E-05 | 8.41E-05 |
| P2rx4 | 4.35E-05 | 0.00019 | 4.81E-07 | 0 | 1.38E-05 | 0.00033 | 0.00012 | 3.56E-05 | 0.0002 | 0.0002 | 7.71E-05 | 0.00011 | 0.00048 | 0.0003 | 0 | 0 | 0 | 0.00013 |
| Cbl | 0 | 0.00016 | 0.00022 | 0.00012 | 0.00016 | 8.54E-05 | 0 | 0.00018 | 0.0002 | 8.10E-05 | 0.00017 | 0.0001 | 1.90E-05 | 0.00018 | 6.92E-05 | 7.43E-05 | 0.00017 | 3.77E-05 |
| Glo1 | 0.00017 | 0.00015 | 0.00014 | 0.00029 | 0.00018 | 0.00011 | 0.00012 | 0.00011 | 0.0002 | 0.00013 | 0.00014 | 7.53E-05 | 8.45E-05 | 5.85E-05 | 0 | 1.58E-05 | 0 | 0.00019 |
| Tomm40 | 0.00015 | 7.67E-05 | 0.00046 | 0.00032 | 3.22E-05 | 0 | 0 | 9.31E-05 | 0.0002 | 2.87E-05 | 0.00012 | 9.16E-05 | 2.32E-06 | 6.96E-05 | 3.70E-05 | 0.00016 | 3.39E-05 | 1.54E-05 |

| | | | | | | | | | | | | | | | | | | |
|---|---|---|---|---|---|---|---|---|---|---|---|---|---|---|---|---|---|---|
| Snrnp70 | 0.00031 | 0.00023 | 0.00035 | 0.00034 | 0.00013 | 5.69E-05 | 3.24E-05 | 4.51E-05 | 0.0002 | 9.02E-05 | 0.00012 | 0.00011 | 7.25E-05 | 4.38E-05 | 0.00082 | 0.00044 | 0.0004 | 6.89E-05 |
| Hsd17b4 | 0.00012 | 0.00019 | 7.18E-06 | 0.00013 | 0.00015 | 8.11E-05 | 6.87E-05 | 0 | 0.0002 | 0.00017 | 0.0001 | 0 | 0.00019 | 1.54E-05 | 0 | 5.47E-05 | 0 | 0.00023 |
| Man1b1 | 0.00013 | 0.00016 | 0.00014 | 0.00016 | 0.00018 | 0.00014 | 9.55E-05 | 3.84E-05 | 0.0002 | 0.00012 | 0.00014 | 0.00011 | 6.93E-05 | 7.31E-05 | 0.00014 | 0.00017 | 0 | 0.00013 |
| Zfp503 | 0 | 0.00021 | 0 | 0.00022 | 1.88E-05 | 0 | 0 | 0 | 0.0002 | 0 | 5.16E-05 | 5.04E-05 | 7.72E-05 | 7.82E-05 | 0.00028 | 0.00011 | 0.0003 | 0.00016 |
| Osr2 | 0 | 2.05E-05 | 0 | 0 | 0.00016 | 0.00017 | 0.00059 | 0 | 0.0002 | 0.00053 | 0 | 7.52E-05 | 3.56E-05 | 0 | 2.54E-05 | 3.70E-05 | 0.00051 | 0 |
| Ptbp3 | 0.00025 | 0.00031 | 0.00028 | 0.00015 | 0.00011 | 0.00023 | 0.00016 | 0.00032 | 0.0002 | 0.00013 | 0.00025 | 0.00034 | 0.00012 | 0.0003 | 0.00017 | 0.00038 | 0.00031 | 0.00015 |
| Stx4a | 0.00013 | 0.00024 | 0.00014 | 0.00012 | 0.00011 | 5.46E-05 | 0.00015 | 0.00016 | 0.0002 | 0.00027 | 8.75E-05 | 0.00022 | 0 | 3.89E-05 | 0.00025 | 0 | 4.98E-05 | 0.00024 |
| Man2b2 | 0 | 8.96E-05 | 0 | 0 | 0.00035 | 0.00029 | 0.00022 | 0.0002 | 0.0002 | 0.00036 | 0.0002 | 0.00019 | 0.00074 | 0.00044 | 0 | 0 | 0 | 0.00046 |
| Tmem164 | 0.00016 | 9.63E-05 | 4.80E-05 | 0.00018 | 0.00017 | 0.00017 | 3.54E-05 | 0.00014 | 0.0002 | 3.67E-05 | 4.00E-05 | 2.28E-05 | 9.70E-05 | 0.00021 | 7.93E-05 | 9.36E-05 | 8.91E-05 | 5.23E-05 |
| Rbms2 | 0.00024 | 0.00026 | 9.38E-05 | 0.00021 | 0.00023 | 0.0003 | 0.00032 | 6.82E-05 | 0.0002 | 0.00021 | 0.00017 | 0.00085 | 0.00015 | 5.51E-05 | 2.60E-05 | 0.00018 | 8.50E-05 | 0.00015 |
| B4galt2 | 5.38E-05 | 5.74E-06 | 0.00013 | 0.00013 | 8.81E-05 | 4.86E-05 | 0.00033 | 0.00016 | 0.0002 | 7.45E-05 | 0.00011 | 0 | 0.00017 | 7.92E-05 | 0 | 5.22E-05 | 4.60E-05 | 5.52E-05 |
| Iah1 | 0.0001 | 0.00028 | 0 | 0.00023 | 0.00013 | 0.00022 | 8.53E-05 | 0.00023 | 0.0002 | 0.00034 | 5.80E-05 | 3.20E-06 | 0.00013 | 0.00027 | 0 | 0 | 0 | 0.00025 |
| Degs1 | 0.00033 | 0.00017 | 0.00031 | 1.28E-05 | 0.00017 | 0.00017 | 0 | 0 | 0.0002 | 0.00015 | 0.00017 | 0.00023 | 0.00021 | 0.00025 | 0 | 0.00016 | 0.00016 | 0.00011 |
| Mcfd2 | 0.00028 | 5.79E-05 | 0.00053 | 0.00049 | 0.00039 | 0.00049 | 0.00037 | 0.00035 | 0.0002 | 0.00024 | 0.00045 | 0.00051 | 0.00093 | 0.00025 | 0 | 0 | 0 | 0.00033 |
| Tfdp1 | 0.00014 | 8.20E-05 | 0.00033 | 0.0003 | 0.00016 | 0.00017 | 0.00018 | 2.49E-05 | 0.0002 | 0.00015 | 0.00012 | 0 | 0.00025 | 0.00016 | 7.68E-05 | 0.00085 | 0.00016 | 0.00015 |
| Gnai2 | 3.03E-05 | 0.00016 | 6.07E-05 | 0.001 | 0.00017 | 9.11E-05 | 0.00019 | 0.00047 | 0.0002 | 0.00027 | 0.00026 | 0.00023 | 0.00014 | 0.00034 | 0.00037 | 0.0002 | 0.00055 | 0.00039 |
| Anxa5 | 0.00044 | 0.00029 | 0.00063 | 0.00094 | 0.0003 | 0.00064 | 0.00114 | 0.00076 | 0.0002 | 0.00074 | 0.00046 | 0.00079 | 0.00018 | 0.00061 | 0.00017 | 0.00017 | 0.00057 | 0.00067 |
| Adamts1 | 0.0002 | 4.09E-05 | 0 | 0.0001 | 0.00012 | 0 | 0 | 0.00014 | 0.0002 | 0.0007 | 0 | 0.00011 | 0.00043 | 0.00035 | 0 | 5.27E-05 | 0.0014 | 0.00013 |
| Ltbr | 0.00022 | 0.00029 | 7.88E-05 | 0.00043 | 0.00014 | 0.00016 | 0.00012 | 0.00012 | 0.0002 | 0.00025 | 5.98E-05 | 0 | 0.00032 | 5.11E-06 | 6.77E-05 | 0.00012 | 5.59E-05 | 0.00028 |
| Auts2 | 3.91E-05 | 0.00021 | 3.40E-05 | 1.45E-05 | 2.14E-05 | 3.46E-06 | 0.00017 | 8.19E-07 | 0.0002 | 0 | 0 | 0 | 0.00012 | 0.00029 | 0.00051 | 0.00017 | 0.00028 | 0 |
| Cpq | 6.69E-05 | 6.88E-05 | 0 | 0 | 5.96E-06 | 0.00032 | 0 | 1.10E-05 | 0.0002 | 0.00031 | 0 | 9.29E-05 | 0.00028 | 2.24E-05 | 0 | 0 | 0 | 0.00012 |
| Ppp1r18 | 0.00016 | 0.00022 | 0.00022 | 0.00013 | 0.00019 | 6.35E-05 | 6.96E-05 | 0.00033 | 0.0002 | 0.00014 | 0.00053 | 0.00036 | 6.68E-05 | 0.00018 | 0 | 0.00012 | 0.00022 | 7.64E-05 |
| Ndufc1 | 0.00028 | 6.01E-05 | 0.00018 | 0.00034 | 8.18E-05 | 5.27E-05 | 0.0001 | 6.73E-05 | 0.0002 | 2.48E-05 | 0.00023 | 1.92E-06 | 7.83E-05 | 7.29E-05 | 0 | 0.00016 | 0 | 4.46E-05 |
| Slc30a4 | 0 | 0.00016 | 0.00017 | 0.00019 | 0.00011 | 3.47E-05 | 0 | 0.00011 | 0.0002 | 0.00018 | 7.73E-05 | 6.58E-05 | 4.59E-05 | 0.00022 | 0.0001 | 6.40E-05 | 0.00014 | 0.00011 |
| Mcrip1 | 0.00019 | 0.00027 | 0.00013 | 0.00033 | 0.00051 | 0.00036 | 0.00054 | 0.00038 | 0.0002 | 0.00023 | 0.00032 | 0.00019 | 0.0003 | 0.0003 | 9.06E-05 | 0.00012 | 0 | 0.00022 |
| Bak1 | 0.00025 | 0.00016 | 0.00019 | 0.00044 | 1.00E-04 | 6.32E-05 | 0.00013 | 0.00011 | 0.0002 | 0.00014 | 0.00014 | 2.13E-05 | 0 | 9.88E-05 | 0 | 0.00017 | 0 | 0.00012 |
| Atxn2l | 0.00015 | 0.00036 | 0.00037 | 0.00033 | 0.0002 | 0.0001 | 0.00015 | 0.00019 | 0.0002 | 0.00013 | 0.00013 | 0.00032 | 9.30E-05 | 0.00014 | 0.0005 | 0.00023 | 0.0003 | 0.0001 |
| Mllt1 | 2.36E-05 | 8.73E-05 | 0.00011 | 0.00015 | 0.00012 | 0 | 0 | 0.00011 | 0.0002 | 9.40E-05 | 7.90E-05 | 4.67E-05 | 6.55E-05 | 9.51E-05 | 4.50E-05 | 0.00023 | 7.35E-05 | 0.00011 |
| Ephb4 | 2.23E-06 | 0.00018 | 7.22E-06 | 0.00018 | 8.53E-05 | 7.38E-06 | 0 | 4.45E-05 | 0.0002 | 3.63E-05 | 0.00028 | 0.00012 | 0.00014 | 0.0002 | 0.0002 | 7.86E-05 | 0 | 0.00018 |
| Eif4g2 | 0.00071 | 0.00033 | 0.00051 | 0.00018 | 0.0003 | 0.00051 | 0.00043 | 0.00033 | 0.0002 | 0.00027 | 0.00037 | 0.00027 | 7.89E-05 | 0.0003 | 2.35E-06 | 0.00033 | 0.00062 | 9.84E-05 |
| Hnrnpd | 0.00023 | 0.00011 | 0.00028 | 0.00026 | 5.03E-05 | 5.86E-05 | 0 | 1.94E-05 | 0.0002 | 3.06E-05 | 0.0001 | 9.42E-05 | 2.75E-05 | 9.52E-05 | 0.00025 | 0.00045 | 0.00023 | 9.95E-06 |
| Cyba | 1.40E-05 | 0.00014 | 0.00018 | 0.0003 | 0.00032 | 0.00027 | 0 | 0.00028 | 0.0002 | 0.00018 | 0.00036 | 0.00018 | 0.00033 | 0.00023 | 1.34E-05 | 8.06E-05 | 0 | 0.00018 |
| Ipmk | 2.03E-05 | 6.39E-05 | 3.61E-05 | 3.16E-05 | 7.91E-05 | 0.00025 | 0 | 0 | 0.0002 | 3.47E-05 | 6.26E-06 | 0.00013 | 9.28E-05 | 0.00011 | 1.04E-05 | 5.04E-05 | 0.00013 | 7.54E-06 |

| | | | | | | | | | | | | | | | | | | |
|---|---|---|---|---|---|---|---|---|---|---|---|---|---|---|---|---|---|---|
| Il3ra | 0 | 8.95E-05 | 0 | 0.00056 | 0.00021 | 0.00021 | 0.00028 | 0.00032 | 0.0002 | 0.00041 | 8.93E-05 | 0.00019 | 0.00082 | 0.00015 | 0.00037 | 0 | 0.00016 | 0.00034 |
| Tmem234 | 0.00026 | 0.00024 | 0.00022 | 0.00031 | 0.00021 | 0.00025 | 0.00015 | 0.00013 | 0.0002 | 0.00015 | 9.43E-05 | 0.0002 | 0.00011 | 0.00018 | 0 | 0.00019 | 0 | 0.00012 |
| Ppp4r2 | 0.0003 | 0.00024 | 0.00026 | 0.00012 | 7.18E-05 | 0.00015 | 0.00015 | 0 | 0.0002 | 0.00013 | 0.0001 | 0.00018 | 8.96E-05 | 0.00013 | 0.00015 | 0.00017 | 0.00023 | 7.05E-05 |
| Numbl | 0.00011 | 0.00018 | 0.00017 | 0.0002 | 0.00015 | 0.0002 | 0.00028 | 5.50E-05 | 0.0002 | 0.00018 | 0.0003 | 0.0002 | 0.00014 | 0.00029 | 0.00035 | 8.43E-06 | 0 | 0.00019 |
| Ndufv1 | 0.00017 | 0.0002 | 0.00026 | 0.00041 | 0.00014 | 0.00013 | 6.42E-05 | 0.0001 | 0.0002 | 8.62E-05 | 0.00015 | 1.66E-05 | 0.00012 | 0.00017 | 5.53E-05 | 0.00029 | 0.00011 | 0.00015 |
| Cnih1 | 0.00048 | 8.38E-05 | 0.00036 | 3.46E-05 | 0.00018 | 0.00027 | 0.00042 | 0.00012 | 0.0002 | 0.00027 | 0.00015 | 0.00015 | 0.00015 | 0.00016 | 0 | 0.00023 | 0 | 3.89E-05 |
| Snrnp200 | 0.00021 | 0.0002 | 0.00018 | 0.00014 | 6.45E-05 | 3.69E-05 | 4.67E-05 | 5.79E-07 | 0.0002 | 3.86E-05 | 4.81E-05 | 0.00013 | 2.06E-05 | 9.78E-05 | 6.33E-05 | 0.00016 | 1.61E-05 | 6.98E-05 |
| Cic | 6.54E-05 | 0.00027 | 5.17E-05 | 0.00015 | 0.00019 | 8.70E-05 | 0.0001 | 1.98E-05 | 0.0002 | 9.62E-05 | 0.00011 | 0.00025 | 7.87E-05 | 2.84E-05 | 2.21E-05 | 0.00016 | 0.00013 | 7.58E-05 |
| Hnrnpul1 | 0.0002 | 0.00039 | 0.00023 | 0.00064 | 0.00027 | 0.00025 | 0.00031 | 0.00035 | 0.0002 | 0.00031 | 0.0002 | 0.00025 | 7.52E-05 | 0.00021 | 0.00046 | 0.00068 | 0.00035 | 0.00028 |
| Msrb3 | 0.00013 | 0.00012 | 9.64E-05 | 0 | 0.00035 | 0.00033 | 0.00026 | 4.77E-05 | 0.00019 | 7.34E-05 | 0.00079 | 4.98E-06 | 3.31E-05 | 0.00025 | 0 | 8.24E-05 | 0 | 0.00043 |
| Smarca2 | 0.00023 | 0.00017 | 0 | 0 | 0.00011 | 0.00014 | 0.00023 | 0 | 0.00019 | 0.00016 | 0 | 4.12E-05 | 0 | 0 | 0 | 3.03E-05 | 0.00011 | 0.00022 |
| Clec11a | 0 | 0 | 0 | 0.00016 | 1.28E-05 | 0 | 0.0009 | 0.00057 | 0.00019 | 0 | 0.00023 | 0 | 0.00013 | 0 | 0 | 0.00016 | 0 | 0.00168 |
| Samd4 | 0 | 7.72E-05 | 0.00033 | 0 | 0.00017 | 0.0001 | 0.00014 | 0 | 0.00019 | 0.00012 | 0.00061 | 0.00028 | 0.00012 | 0.00033 | 0 | 5.25E-05 | 0.00015 | 0 |
| Clic1 | 0.00026 | 0.00017 | 0.00038 | 0.0003 | 6.12E-05 | 3.20E-05 | 0.00017 | 0.00023 | 0.00019 | 0.00019 | 0.00033 | 0.00041 | 0 | 0.00011 | 0 | 0.00035 | 1.60E-05 | 0.00012 |
| Mcpt8 | 0 | 0 | 0 | 0 | 9.32E-05 | 0 | 0 | 0.00014 | 0.00019 | 0 | 0 | 0 | 0 | 0.00033 | 0 | 1.07E-05 | 0 | 0 |
| Ppp4c | 0.00021 | 0.00017 | 0.0002 | 0.0002 | 9.86E-05 | 1.81E-05 | 2.43E-05 | 0.0001 | 0.00019 | 6.05E-05 | 9.06E-05 | 5.69E-05 | 0 | 7.83E-05 | 0 | 0.00024 | 8.83E-05 | 4.80E-05 |
| Ptpn12 | 0.00016 | 0.00026 | 0.00031 | 6.11E-05 | 0.00016 | 0.00017 | 0 | 0.00026 | 0.00019 | 0.00013 | 8.50E-05 | 0.00015 | 0.00012 | 0.0002 | 0.0002 | 0.00023 | 0.00035 | 8.46E-05 |
| Chmp1a | 5.00E-05 | 0.00022 | 0.00014 | 0.00047 | 0.00023 | 0.00013 | 0.00012 | 0.00023 | 0.00019 | 0.00014 | 0.00017 | 0.00015 | 0.00011 | 0.00017 | 1.85E-05 | 2.52E-05 | 3.86E-06 | 0.00017 |
| Tns2 | 0 | 0.00029 | 0 | 0.00034 | 4.86E-05 | 0.00019 | 0 | 0.00055 | 0.00019 | 0.00032 | 0 | 0 | 0.00011 | 0.00015 | 0.00026 | 0.00019 | 0.00019 | 0.00022 |
| Tmem26 | 0 | 0 | 0 | 0 | 9.62E-05 | 0.0002 | 0 | 0 | 0.00019 | 0 | 0 | 1.61E-05 | 0 | 2.98E-05 | 0 | 1.88E-05 | 0 | 0 |
| Med13 | 0.00016 | 0.00025 | 7.18E-05 | 8.22E-05 | 0.00018 | 0.00019 | 9.97E-05 | 0.00011 | 0.00019 | 8.66E-05 | 8.82E-05 | 0.00015 | 0.00017 | 0.00014 | 0.00016 | 9.90E-05 | 0.00069 | 8.12E-05 |
| Pdcd4 | 9.14E-05 | 0.00016 | 0 | 0.00014 | 0 | 0.00011 | 0.00017 | 0.00029 | 0.00019 | 0.00027 | 0 | 0.00015 | 0.00016 | 4.79E-05 | 0.00022 | 1.70E-05 | 0.00022 | 0.00037 |
| AW551984 | 3.24E-05 | 0 | 0 | 0 | 3.42E-05 | 0.00011 | 0.00032 | 0 | 0.00019 | 0.0006 | 0 | 0 | 0.00011 | 0 | 3.31E-05 | 0.0001 | 0 | 0 |
| Eftud2 | 0.00019 | 0.00012 | 0.0003 | 0.00031 | 8.45E-05 | 6.63E-05 | 5.87E-05 | 7.28E-05 | 0.00019 | 7.53E-05 | 8.16E-05 | 0 | 4.71E-05 | 0.00014 | 0 | 0.00044 | 5.75E-05 | 8.58E-05 |
| Ifi207 | 0.00015 | 0.00021 | 0 | 0 | 0 | 0.00039 | 0 | 0.00019 | 0.00019 | 0.00038 | 0 | 4.08E-05 | 0.00054 | 2.76E-05 | 2.18E-05 | 0 | 0.00013 | 0 |
| Nrp2 | 6.48E-05 | 0.00014 | 5.66E-06 | 0 | 4.54E-05 | 1.17E-05 | 1.03E-05 | 7.55E-05 | 0.00019 | 0 | 0.00028 | 7.66E-05 | 0 | 0.00033 | 0 | 4.20E-05 | 3.12E-05 | 5.62E-05 |
| Calm2 | 0.00031 | 0.0003 | 0.00018 | 5.19E-05 | 0.00036 | 0.00026 | 0.00041 | 7.33E-06 | 0.00019 | 9.61E-05 | 0.00035 | 0.00031 | 3.33E-05 | 0 | 0 | 0.00049 | 0.00026 | 0.00021 |
| Lrrc32 | 0 | 0.00021 | 0.00031 | 0.00038 | 0 | 0 | 0.0009 | 2.85E-05 | 0.00019 | 0.00013 | 0 | 0.00047 | 0.00168 | 0 | 0 | 0 | 0.00025 | 0.00088 |
| Ehd1 | 7.02E-05 | 4.73E-05 | 0.00028 | 0.00015 | 1.96E-05 | 0 | 7.99E-05 | 0.00011 | 0.00019 | 0.00031 | 0.00036 | 0.00021 | 0 | 0 | 1.99E-05 | 7.24E-05 | 0.00024 | 8.88E-05 |
| Zmynd11 | 0.00037 | 0.00033 | 0.00011 | 1.75E-05 | 0.00021 | 0.00026 | 9.98E-05 | 0.00012 | 0.00019 | 0.00016 | 0.00016 | 0.00018 | 0.0001 | 0.00014 | 0.00027 | 0.00019 | 0.00026 | 0.00015 |
| Igfbp6 | 0 | 0 | 0.00015 | 0.0002 | 0.0003 | 7.00E-05 | 0.0011 | 3.10E-05 | 0.00019 | 0.00144 | 2.52E-05 | 0.00043 | 0 | 0 | 0 | 0 | 0 | 0.00122 |
| Chd4 | 5.90E-05 | 0.00026 | 0.00023 | 0.0004 | 0.00013 | 6.01E-05 | 4.01E-05 | 0.00022 | 0.00019 | 8.39E-05 | 0.00016 | 6.46E-05 | 0.00013 | 0.00011 | 0.00029 | 0.00027 | 0.0011 | 8.49E-05 |
| Nfatc4 | 0 | 0.00025 | 5.76E-05 | 0.00025 | 0.00033 | 0.00018 | 0.0005 | 0 | 0.00019 | 9.54E-05 | 0.00032 | 5.11E-05 | 0.00041 | 0.0005 | 0.00083 | 7.70E-05 | 0 | 2.12E-05 |

| | | | | | | | | | | | | | | | | | | |
|---|---|---|---|---|---|---|---|---|---|---|---|---|---|---|---|---|---|---|
| Idi1 | 1.39E-05 | 1.07E-05 | 0 | 0 | 0 | 0 | 0 | 0 | 0.00019 | 0 | 3.74E-05 | 4.53E-06 | 0 | 9.67E-06 | 0 | 0 | 0.00027 | 7.74E-06 |
| Eml4 | 0.00011 | 7.98E-05 | 0.00014 | 4.40E-05 | 0.0002 | 0 | 0.00022 | 0.00021 | 0.00019 | 0 | 0.0001 | 0 | 6.45E-05 | 1.33E-06 | 0.00018 | 0.00011 | 0.00018 | 0 |
| Ndufv3 | 0.00018 | 1.22E-05 | 1.73E-05 | 0.00054 | 0.00017 | 9.04E-05 | 0.00026 | 0.00024 | 0.00019 | 0.00019 | 0.00014 | 0.00062 | 0.00021 | 0.00022 | 0.00012 | 0 | 0 | 0.00026 |
| Twf1 | 0.00033 | 0.00018 | 0.00037 | 0.00034 | 0.00017 | 0.00016 | 0.00054 | 0.00018 | 0.00019 | 0.00022 | 0.00019 | 1.21E-06 | 0.00016 | 0.00014 | 0 | 0.00017 | 6.94E-05 | 0.00012 |
| Samhd1 | 0.00017 | 4.30E-05 | 0 | 0 | 2.48E-05 | 0 | 1.18E-05 | 0 | 0.00019 | 0.00046 | 0.00026 | 0 | 8.33E-05 | 0.00018 | 0.0001 | 0.00011 | 0 | 0.00032 |
| Abl1 | 5.06E-05 | 0.00032 | 0.00014 | 0.00017 | 0.00013 | 0.00013 | 4.95E-05 | 3.96E-05 | 0.00019 | 0.00035 | 0.00022 | 7.26E-05 | 5.52E-05 | 0.00024 | 0.00025 | 0.0002 | 0.00029 | 4.66E-05 |
| Tra2a | 0.00022 | 0.00023 | 0.00023 | 0.00011 | 6.93E-05 | 1.73E-05 | 6.89E-06 | 4.79E-05 | 0.00019 | 7.85E-05 | 0.00016 | 0.00026 | 5.48E-05 | 0.00011 | 0.00033 | 0.00025 | 0.00069 | 8.96E-05 |
| Aco1 | 0.00017 | 0.00012 | 0.00017 | 0.00013 | 8.23E-05 | 4.36E-05 | 0 | 4.16E-05 | 0.00019 | 0.00013 | 0.00012 | 2.16E-05 | 5.73E-05 | 5.27E-05 | 8.34E-05 | 6.10E-05 | 5.59E-05 | 0.0001 |
| Rhobtb3 | 0.00023 | 9.79E-05 | 5.38E-05 | 0 | 0 | 0.00016 | 6.17E-05 | 0.00013 | 0.00019 | 0.00016 | 2.81E-05 | 1.07E-06 | 8.29E-05 | 0.00012 | 0 | 5.97E-05 | 3.03E-05 | 0 |
| Gorasp2 | 0.00062 | 0.0001 | 0.0005 | 0.00017 | 0.00019 | 0.0004 | 0.00063 | 0.00024 | 0.00019 | 0.00012 | 0.00026 | 0.0002 | 0.00017 | 0.00024 | 6.27E-05 | 0.00016 | 0 | 7.29E-05 |
| Tnfsf9 | 0 | 9.60E-05 | 0 | 0 | 1.85E-05 | 9.94E-05 | 0 | 0 | 0.00019 | 1.05E-05 | 0 | 0.0003 | 0 | 0 | 0 | 2.50E-05 | 0.00029 | 0 |
| Anapc2 | 0.00015 | 0.0002 | 0.00012 | 0.00027 | 0.00012 | 0.00012 | 0.00014 | 4.72E-05 | 0.00019 | 0.0002 | 0.00018 | 0.00016 | 2.51E-05 | 0.0001 | 0.00012 | 0.00022 | 6.54E-05 | 0.00023 |
| Mfsd1 | 0.00016 | 0.00026 | 5.66E-05 | 0.0002 | 0.00021 | 0.00024 | 0.00016 | 3.97E-05 | 0.00019 | 0.00013 | 0.00017 | 3.85E-06 | 0.00015 | 0.00026 | 0 | 9.99E-05 | 4.62E-05 | 6.20E-05 |
| Snrpf | 0.00034 | 0.00013 | 0.00046 | 0.00058 | 3.40E-06 | 8.60E-05 | 6.76E-05 | 3.00E-05 | 0.00019 | 3.24E-05 | 0.00021 | 8.74E-05 | 3.11E-05 | 5.59E-05 | 0 | 0.00047 | 0.00014 | 2.60E-05 |
| Rhbdf1 | 0 | 0.00019 | 0.00019 | 0.00018 | 0.0001 | 0.00019 | 0.0001 | 0.00044 | 0.00019 | 0.00029 | 0.00012 | 4.40E-05 | 4.65E-05 | 0.0002 | 0.00034 | 0 | 0.00018 | 0.00027 |
| Ubap2l | 0.00047 | 0.00029 | 0.00021 | 5.92E-05 | 0.00019 | 0.00025 | 0.00033 | 1.31E-05 | 0.00019 | 0.0001 | 8.49E-05 | 0.00019 | 0.00013 | 0.00015 | 0.0002 | 0.00013 | 6.93E-05 | 1.63E-05 |
| H1f0 | 0.00022 | 0.00046 | 0 | 0.00032 | 0.0006 | 0.00043 | 0.00037 | 0.00036 | 0.00019 | 0.00019 | 0.00027 | 0.00052 | 6.30E-05 | 0.00029 | 0.00025 | 0.00098 | 0.0001 | 0.00043 |
| Cops9 | 0.00033 | 0.00014 | 0.00022 | 0.00076 | 0.00029 | 0.00037 | 0.00027 | 0.00032 | 0.00019 | 0.00018 | 0.00029 | 0.00022 | 0.00023 | 0.00026 | 0 | 0.00013 | 9.94E-05 | 0.00024 |
| Arl3 | 0.00022 | 0.00012 | 0.00018 | 0.00034 | 0.00014 | 0.00012 | 0.00015 | 0.00025 | 0.00019 | 0.00014 | 9.71E-05 | 0.00025 | 8.01E-05 | 7.60E-05 | 3.15E-07 | 2.12E-05 | 5.23E-05 | 0.00015 |
| Pisd | 0.00011 | 0.00011 | 0.00026 | 0 | 5.13E-05 | 9.19E-05 | 2.07E-05 | 0.00033 | 0.00019 | 6.48E-05 | 9.22E-05 | 0.00018 | 6.46E-05 | 5.73E-05 | 2.71E-05 | 0 | 1.95E-05 | 4.03E-05 |
| Hmg20b | 0.00021 | 0.00015 | 0.00011 | 4.54E-05 | 0.00019 | 0.00011 | 0.00021 | 0.00017 | 0.00019 | 4.77E-05 | 0.0002 | 0 | 0 | 0.0002 | 0.00031 | 0.00012 | 8.26E-05 | 0.00013 |
| Sdhb | 0.00029 | 0.00012 | 0.00032 | 0.0004 | 0.00011 | 5.11E-05 | 0.0002 | 0.00012 | 0.00019 | 0.00013 | 0.00022 | 0 | 3.81E-06 | 0.00016 | 0 | 0.00015 | 0 | 0.0001 |
| Dab2ip | 6.03E-05 | 0.0002 | 0.00015 | 0 | 0.00013 | 9.70E-05 | 0 | 0.00013 | 0.00019 | 0.00014 | 0.00035 | 0.00012 | 0 | 0.00023 | 0.00021 | 0.00011 | 7.04E-05 | 0.00022 |
| Stk16 | 0.00018 | 9.89E-05 | 0.00015 | 0.00016 | 0.00012 | 7.07E-05 | 6.55E-05 | 7.60E-05 | 0.00019 | 6.39E-05 | 0.0001 | 0 | 0 | 9.09E-05 | 0.00014 | 0.00015 | 0 | 9.44E-05 |
| Pgr | 0 | 0 | 0 | 0 | 0 | 6.42E-05 | 0 | 0 | 0.00019 | 0.00024 | 0 | 3.31E-05 | 0 | 0 | 0 | 0 | 0 | 0 |
| 8-Sep | 0.00021 | 0.00033 | 0.00024 | 0.00032 | 0.00026 | 0.00018 | 0.00033 | 0.00032 | 0.00019 | 0.00055 | 0.00014 | 0.00022 | 0.00012 | 0.00027 | 0.00018 | 0.00027 | 0.00013 | 0.00061 |
| Cdc37 | 0.00027 | 0.00024 | 0.00034 | 0.00059 | 0.00021 | 0.00016 | 0.00017 | 0.00022 | 0.00019 | 0.00013 | 0.00018 | 0.00029 | 5.45E-05 | 0.00024 | 3.34E-05 | 0.00022 | 0.00019 | 0.00018 |
| Kank1 | 0 | 8.61E-05 | 0 | 0 | 0.00017 | 0 | 0.00011 | 0 | 0.00019 | 9.35E-05 | 0.00015 | 0.00011 | 1.98E-05 | 2.54E-05 | 0 | 8.71E-06 | 0 | 3.88E-05 |
| Mtdh | 0.00022 | 0.00021 | 0.00022 | 0.00011 | 6.20E-05 | 9.52E-05 | 0 | 0.00018 | 0.00019 | 4.42E-05 | 6.37E-05 | 0.00011 | 0 | 7.73E-05 | 0.00044 | 0.00012 | 0.00028 | 3.83E-05 |
| Ube2d2a | 0.00039 | 0.00023 | 0.00024 | 6.33E-05 | 0.00016 | 0.00027 | 9.10E-05 | 0.00017 | 0.00019 | 0.00017 | 0.00025 | 0.00022 | 4.52E-05 | 0.00017 | 0.00012 | 0.00026 | 0.00043 | 0.00021 |
| Atp5o | 0.00052 | 6.19E-05 | 0.0002 | 0.00028 | 7.76E-05 | 0.00021 | 0.00014 | 8.90E-05 | 0.00019 | 3.63E-05 | 0.00018 | 0.0001 | 5.18E-05 | 0.00012 | 0 | 0.00029 | 0 | 3.10E-05 |
| Tead2 | 0 | 0.00014 | 8.33E-05 | 8.40E-05 | 3.24E-05 | 9.27E-05 | 0 | 0.00024 | 0.00019 | 0 | 2.32E-05 | 0 | 8.55E-05 | 7.15E-05 | 0 | 0.00019 | 4.68E-05 | 8.18E-05 |
| Rtcb | 0.00024 | 0.00025 | 0.00032 | 0.00045 | 0.0001 | 0.00013 | 0.00019 | 0.00011 | 0.00019 | 0.00015 | 0.00013 | 0.00018 | 0.0001 | 0.00015 | 0 | 0.00017 | 7.29E-05 | 0.00019 |

| | | | | | | | | | | | | | | | | | | |
|---|---|---|---|---|---|---|---|---|---|---|---|---|---|---|---|---|---|---|
| Nus1 | 0.00021 | 0.00013 | 0.00041 | 0.00028 | 0.00019 | 0.00018 | 0.0002 | 0.00012 | 0.00019 | 1.79E-05 | 0.00025 | 0.0002 | 0.00029 | 0.00036 | 0.00024 | 0 | 3.74E-05 | 1.41E-05 |
| Ash1l | 0.00015 | 0.00022 | 7.98E-05 | 1.35E-05 | 0.00016 | 0.00015 | 0.00012 | 3.67E-05 | 0.00019 | 9.07E-05 | 6.09E-05 | 0.00015 | 6.52E-05 | 0.00012 | 0.00024 | 0.0001 | 0.00032 | 5.70E-05 |
| Lsm7 | 0.00025 | 0.00013 | 0.00033 | 0.00033 | 4.11E-05 | 0 | 0 | 5.23E-06 | 0.00019 | 2.37E-05 | 9.56E-05 | 0 | 7.74E-05 | 0.0001 | 0.00012 | 0.00012 | 0 | 0 |
| Arhgap1 | 0.00034 | 0.00011 | 0.00016 | 0.00023 | 0.00012 | 0.00026 | 0.0002 | 0.00024 | 0.00019 | 0.00021 | 0.00032 | 4.90E-05 | 0.00017 | 0.00013 | 0 | 5.42E-05 | 0.00012 | 0.00022 |
| Pank3 | 0.00018 | 0.00012 | 8.87E-05 | 0 | 0.00014 | 9.08E-05 | 0 | 8.90E-05 | 0.00019 | 1.43E-05 | 0.00017 | 7.38E-05 | 0 | 0.0001 | 0 | 0.00011 | 0.00065 | 3.64E-05 |
| Irx5 | 1.02E-05 | 5.55E-05 | 4.09E-05 | 6.03E-05 | 0.00011 | 1.47E-05 | 5.32E-05 | 2.55E-05 | 0.00019 | 6.50E-06 | 8.93E-05 | 4.02E-05 | 2.45E-05 | 0 | 1.67E-05 | 7.40E-05 | 0.00012 | 0 |
| Tsc2 | 0.00013 | 0.00016 | 0.00011 | 4.66E-05 | 0.00011 | 0.00017 | 3.87E-05 | 0.00018 | 0.00019 | 8.71E-05 | 8.08E-05 | 3.92E-05 | 5.27E-05 | 0.0001 | 5.96E-05 | 5.32E-05 | 2.57E-05 | 9.79E-05 |
| Tmem63a | 0 | 0.00015 | 0.00012 | 0.00012 | 4.27E-05 | 0.00018 | 0 | 3.01E-05 | 0.00019 | 0.0003 | 0.00027 | 0.0002 | 0 | 0.0003 | 7.57E-05 | 3.88E-05 | 6.82E-05 | 0.00018 |
| Sfr1 | 0.00061 | 0.00034 | 0.00045 | 0.00074 | 0.00016 | 0.00028 | 0.00049 | 0.00012 | 0.00019 | 0.00022 | 0.00024 | 0.00036 | 7.42E-05 | 0.00019 | 0.00024 | 0.00028 | 0.00052 | 0.00015 |
| Msrb1 | 0.0002 | 0.00012 | 9.91E-05 | 0.00026 | 9.22E-05 | 6.77E-05 | 9.78E-05 | 0.00013 | 0.00019 | 0.00018 | 0.00016 | 0.00015 | 0.00014 | 0.00019 | 0 | 5.60E-05 | 0 | 0.00011 |
| Gnb4 | 1.50E-05 | 0.00017 | 2.25E-06 | 3.31E-05 | 0.00032 | 0.00011 | 0 | 7.68E-05 | 0.00019 | 0 | 0.00012 | 4.06E-05 | 0.00011 | 0.0001 | 0 | 3.59E-05 | 0.00012 | 0.00019 |
| Csnk1d | 9.29E-05 | 0.00032 | 0.0003 | 0.00039 | 0.00017 | 0.00011 | 0.00018 | 0.00025 | 0.00019 | 0.00019 | 0.00015 | 0.00026 | 0.00017 | 0.00021 | 0.00031 | 0.00012 | 0.0004 | 0.00021 |
| Frmd6 | 0.00013 | 3.56E-05 | 9.09E-05 | 0 | 0.00014 | 0.00025 | 0 | 0 | 0.00019 | 0.00017 | 0.00031 | 0.00038 | 0 | 0.00021 | 0 | 0 | 0 | 0.00011 |
| Adam12 | 0.00013 | 0 | 0.00046 | 0.00015 | 0.00095 | 0.00033 | 0.00165 | 0 | 0.00019 | 0 | 0.00096 | 0 | 0.00088 | 0.00085 | 0.00033 | 0.00042 | 0 | 0.00026 |
| Txnrd1 | 0.00018 | 0.00019 | 0.00049 | 0.00014 | 9.09E-05 | 5.88E-05 | 6.87E-05 | 6.33E-05 | 0.00019 | 4.21E-05 | 0.00012 | 2.71E-05 | 0.00023 | 0.00017 | 9.89E-05 | 0.00045 | 0.00025 | 6.99E-07 |
| Lss | 2.96E-07 | 3.88E-05 | 0 | 4.82E-05 | 0 | 0 | 0 | 0 | 0.00019 | 1.95E-05 | 2.33E-05 | 2.05E-05 | 0 | 0.00014 | 3.64E-05 | 0 | 1.01E-05 | 1.13E-05 |
| Phb2 | 0.00016 | 0.00014 | 0.00034 | 0.0004 | 2.26E-05 | 4.02E-05 | 7.71E-07 | 6.36E-05 | 0.00019 | 6.87E-05 | 0.00011 | 0.00011 | 0 | 8.86E-05 | 0 | 0.00018 | 0.00013 | 6.58E-05 |
| Med10 | 0.00021 | 0.00021 | 0.00012 | 0.0003 | 0.00024 | 0.00016 | 6.73E-05 | 0.00017 | 0.00019 | 0 | 0.0002 | 0.00013 | 0 | 0.00013 | 4.20E-05 | 0.00016 | 6.09E-05 | 4.38E-05 |
| Adamts14 | 0 | 0 | 0.00023 | 5.17E-05 | 0.00012 | 0.00013 | 6.76E-05 | 6.93E-05 | 0.00019 | 0.00021 | 9.92E-05 | 0.00033 | 0 | 0.00016 | 0.00025 | 0 | 0 | 0 |
| Trp53inp2 | 0 | 0.00019 | 0 | 0.00027 | 0.00045 | 0.00013 | 0 | 0.00014 | 0.00019 | 7.83E-05 | 0.00032 | 0 | 0.00056 | 0.0003 | 2.41E-05 | 0 | 0 | 0.00016 |
| Naa38 | 0.00034 | 5.41E-05 | 6.45E-05 | 0.0002 | 2.54E-05 | 5.50E-05 | 5.39E-05 | 0.00014 | 0.00019 | 3.31E-05 | 9.64E-05 | 1.97E-05 | 3.04E-05 | 5.52E-07 | 0 | 9.49E-05 | 0 | 9.96E-05 |
| Epn2 | 0 | 0.00012 | 9.59E-05 | 0 | 9.19E-05 | 5.68E-05 | 0 | 0.00016 | 0.00019 | 5.15E-05 | 0.00018 | 4.70E-05 | 0 | 8.16E-05 | 8.80E-05 | 3.35E-05 | 0 | 0.00012 |
| Casc4 | 2.46E-05 | 8.44E-05 | 5.12E-05 | 0.00015 | 0.00021 | 5.86E-05 | 0 | 0.00025 | 0.00019 | 3.51E-05 | 5.69E-05 | 0 | 0.00015 | 0.00035 | 4.02E-05 | 0.00013 | 1.65E-05 | 0.00024 |
| Pmvk | 3.35E-05 | 2.61E-05 | 0 | 6.73E-05 | 4.52E-05 | 0 | 0 | 7.96E-05 | 0.00019 | 1.60E-05 | 5.40E-05 | 2.07E-05 | 0 | 5.17E-05 | 3.39E-05 | 0 | 6.87E-05 | 2.00E-05 |
| Samd4b | 0.00019 | 0.00021 | 0.0003 | 3.48E-05 | 0.00016 | 5.64E-05 | 0.00015 | 8.96E-05 | 0.00019 | 0.00032 | 0.00026 | 0.0002 | 0 | 0.00013 | 9.20E-05 | 0.00017 | 0.00021 | 0.00011 |
| Pnrc1 | 0.00014 | 0.00041 | 0 | 0.00014 | 0.00021 | 0.00026 | 0.0001 | 0.00011 | 0.00019 | 0.00028 | 0 | 0.00059 | 0.00041 | 0.00026 | 0.0002 | 0 | 0.00209 | 0.00012 |
| Snx12 | 8.88E-05 | 0.00021 | 0.00018 | 0.00046 | 0.00011 | 0.00011 | 7.01E-05 | 0.00012 | 0.00019 | 0.00017 | 0.00016 | 7.90E-05 | 8.71E-05 | 0.00013 | 0 | 0.00019 | 0.00015 | 0.0002 |
| Acadsb | 0.00024 | 0.00015 | 0 | 0.00011 | 0.00015 | 0.00023 | 7.73E-05 | 2.65E-05 | 0.00019 | 0.00027 | 4.05E-05 | 0.0002 | 0.00014 | 2.35E-05 | 0 | 0 | 7.85E-05 | 5.09E-05 |
| Ssb | 0.00045 | 0.00019 | 0.00036 | 0.00035 | 6.99E-05 | 0.00017 | 0.00013 | 7.84E-05 | 0.00019 | 7.84E-05 | 0.00012 | 0.00012 | 9.55E-05 | 9.08E-05 | 0.00043 | 0.00036 | 0.00016 | 4.00E-05 |
| Mybbp1a | 0.00019 | 0.00012 | 0.00049 | 0.00026 | 0 | 0 | 0 | 0 | 0.00019 | 2.32E-05 | 7.06E-05 | 4.25E-05 | 1.94E-05 | 4.12E-05 | 0.00017 | 2.24E-05 | 6.56E-05 | 0 |
| Morf4l2 | 0.0008 | 0.00015 | 0.00036 | 0 | 0.00013 | 0.00039 | 0.00034 | 0.00036 | 0.00019 | 0.00011 | 0.00011 | 0.00027 | 2.30E-05 | 9.73E-05 | 0 | 0.00014 | 0.00042 | 0 |
| Atp5j2 | 0.00036 | 0.00011 | 0.00021 | 0.00048 | 0.00018 | 0.00017 | 0.00032 | 0.00013 | 0.00019 | 0.00013 | 0.0002 | 4.69E-05 | 6.79E-05 | 0.0002 | 0 | 0.0002 | 2.54E-05 | 0.00014 |
| Kdelr2 | 0.00067 | 0 | 0.00055 | 0.00098 | 0.00041 | 0.00071 | 0.00137 | 0.00073 | 0.00019 | 0.00038 | 0.00052 | 0.00042 | 0.00059 | 0.00057 | 0 | 0.0003 | 0.00011 | 0.00026 |

| | | | | | | | | | | | | | | | | | | |
|---|---|---|---|---|---|---|---|---|---|---|---|---|---|---|---|---|---|---|
| Gls | 0.00023 | 0.00016 | 0.00026 | 0 | 7.05E-05 | 0.00017 | 7.64E-05 | 0.00016 | 0.00019 | 0.00011 | 0.00013 | 0.00021 | 7.81E-05 | 9.11E-05 | 5.86E-05 | 0.00014 | 0.00077 | 0.00016 |
| Sgce | 4.32E-05 | 9.17E-05 | 0 | 0.00013 | 0.00018 | 0.00022 | 0.00013 | 0.00017 | 0.00019 | 0.0004 | 0.00011 | 0.0002 | 0.00028 | 0.00012 | 0 | 7.59E-05 | 7.82E-05 | 0.00032 |
| Tubb6 | 0 | 9.42E-05 | 0.00062 | 0.00043 | 0.00013 | 0 | 0 | 0.00013 | 0.00019 | 0.00021 | 0.00078 | 0.00016 | 0.00019 | 0.00019 | 0.00021 | 0.00125 | 0.00037 | 0 |
| Dennd2a | 0 | 0.00018 | 0.00014 | 4.72E-05 | 0.00014 | 8.37E-05 | 0 | 0.0008 | 0.00019 | 0.00028 | 0 | 2.56E-05 | 0 | 0 | 0 | 9.26E-05 | 0.00015 | 7.81E-05 |
| Bmp7 | 0 | 0.00013 | 0 | 0 | 0 | 0.00042 | 0 | 0 | 0.00019 | 4.07E-06 | 0 | 0 | 0 | 0 | 0 | 0 | 0 | 0 |
| Farp2 | 0.00015 | 6.21E-06 | 0.00018 | 1.60E-06 | 5.41E-05 | 1.21E-05 | 0.00022 | 0 | 0.00019 | 7.69E-05 | 0.00017 | 0 | 0.00011 | 6.63E-05 | 9.25E-05 | 2.60E-05 | 0 | 5.83E-05 |
| Rab22a | 0.00014 | 0.00023 | 0.0001 | 0.00013 | 0.00016 | 0.00013 | 0.00011 | 6.20E-05 | 0.00019 | 0.00017 | 0.00013 | 0.00015 | 0 | 0.00014 | 0.00013 | 8.41E-05 | 0.00024 | 0.00018 |
| Tspan5 | 1.97E-05 | 7.35E-05 | 8.46E-05 | 0 | 0 | 0.00013 | 7.90E-05 | 0.00012 | 0.00019 | 0.00024 | 0 | 0.00026 | 2.87E-06 | 0 | 0 | 5.99E-05 | 9.48E-05 | 0.0002 |
| C1qtnf1 | 4.78E-05 | 0.00021 | 0 | 0.00011 | 0.00015 | 0.00039 | 0.00058 | 8.24E-05 | 0.00019 | 0.00051 | 0 | 0 | 0 | 0.00121 | 0 | 1.01E-05 | 0 | 0.00037 |
| Glmp | 9.91E-05 | 0.00031 | 4.45E-05 | 0.00039 | 0.00032 | 0.00038 | 0.00034 | 0.00033 | 0.00019 | 0.00025 | 0.00015 | 0.0003 | 0.00036 | 0.00057 | 5.12E-05 | 1.24E-05 | 0.00012 | 0.00042 |
| Anxa1 | 0.00054 | 0.00013 | 0.00011 | 0.00016 | 0.00012 | 0.00045 | 6.45E-05 | 9.15E-05 | 0.00019 | 0.00099 | 0.00131 | 0.00028 | 7.25E-05 | 0.00054 | 0 | 0.00025 | 0.00098 | 0.00041 |
| Blmh | 0.00039 | 0.00011 | 0.00018 | 9.00E-05 | 9.17E-05 | 0.00014 | 8.38E-05 | 0.00012 | 0.00019 | 1.79E-05 | 7.12E-05 | 4.65E-05 | 0 | 7.84E-05 | 0 | 0.0002 | 2.56E-06 | 8.92E-05 |
| Rap1gds1 | 0.00024 | 0.00013 | 0.00021 | 0 | 0.00014 | 0.00013 | 6.49E-05 | 0.00015 | 0.00019 | 7.23E-05 | 0.00024 | 0.00024 | 5.40E-05 | 6.16E-05 | 1.33E-05 | 0.00015 | 0.00016 | 5.59E-05 |
| Man1a | 0 | 0.0003 | 1.08E-05 | 0.00018 | 0.00018 | 0.00024 | 0.00044 | 8.67E-05 | 0.00019 | 0.00092 | 0 | 0.00033 | 0.00055 | 0.00028 | 0.00031 | 0.0002 | 0.0004 | 0.00049 |
| Arpc4 | 0.00038 | 0.00015 | 0.00026 | 8.53E-05 | 8.83E-05 | 0.00013 | 0.00013 | 0.00015 | 0.00019 | 6.59E-05 | 0.00026 | 0.00014 | 0 | 0.00011 | 0 | 8.92E-05 | 0 | 5.80E-05 |
| Ror1 | 3.47E-05 | 0 | 0 | 0 | 3.48E-05 | 0.00021 | 6.06E-05 | 0.00028 | 0.00019 | 0 | 0 | 9.40E-05 | 6.09E-05 | 1.30E-06 | 0 | 0 | 0.00011 | 0.00021 |
| Rbms3 | 0.00031 | 0.00029 | 0 | 0 | 0.00032 | 0.00031 | 0 | 0.00025 | 0.00019 | 0.00021 | 0.00022 | 0.00033 | 0.00014 | 0.00028 | 0.00097 | 0 | 0.00051 | 0.00029 |
| Ptgir | 0 | 0.00014 | 3.67E-05 | 0 | 0 | 0.00031 | 0 | 0 | 0.00019 | 9.73E-06 | 0 | 0 | 0.00012 | 0.00019 | 0 | 0 | 7.46E-05 | 0.00018 |
| Qpct | 1.07E-05 | 8.99E-05 | 0 | 0 | 0 | 0.00014 | 0 | 3.34E-05 | 0.00019 | 0.00048 | 0 | 1.62E-05 | 7.69E-05 | 0.00024 | 0 | 0 | 0 | 0 |
| Dnaja1 | 0.00036 | 0.00014 | 5.60E-05 | 0 | 7.55E-05 | 0.00018 | 0 | 0.00028 | 0.00019 | 0.00019 | 4.49E-05 | 0.00012 | 5.17E-05 | 0 | 0 | 4.65E-05 | 0.00176 | 4.02E-05 |
| Rfng | 4.08E-05 | 5.48E-05 | 5.71E-05 | 8.60E-05 | 8.25E-05 | 4.17E-05 | 5.88E-05 | 6.32E-05 | 0.00019 | 0.0001 | 3.11E-05 | 2.40E-05 | 0.0001 | 3.81E-05 | 0 | 3.84E-05 | 8.09E-05 | 7.18E-05 |
| Cdc42ep2 | 5.19E-05 | 1.30E-05 | 1.83E-05 | 0 | 0 | 0 | 9.88E-05 | 5.81E-05 | 0.00019 | 0.00035 | 0 | 0.00027 | 0 | 0 | 0 | 0 | 0 | 0.00017 |
| Slc41a1 | 4.56E-05 | 0.00014 | 0.00016 | 2.87E-05 | 0.00016 | 0.00012 | 0.00011 | 0.0002 | 0.00019 | 7.83E-05 | 7.13E-05 | 0.00033 | 0.00018 | 0.00011 | 2.62E-05 | 2.16E-05 | 0 | 0.00013 |
| Pak2 | 0.00022 | 0.00024 | 0.00035 | 0.00014 | 0.00015 | 0.00013 | 5.12E-05 | 0.00014 | 0.00019 | 0.00014 | 0.00027 | 0.00022 | 5.00E-05 | 0.00019 | 5.31E-05 | 0.0002 | 0.00014 | 0.00014 |
| Ddx24 | 0.00021 | 0.00024 | 0.00027 | 0.00022 | 7.84E-05 | 2.15E-05 | 5.85E-05 | 0 | 0.00019 | 0.00014 | 0.00019 | 6.35E-05 | 0.00011 | 9.40E-05 | 0.00015 | 0.00013 | 0 | 9.74E-05 |
| Pdxk | 9.53E-05 | 0.00012 | 9.59E-05 | 0 | 5.95E-05 | 3.67E-05 | 4.20E-05 | 0 | 0.00019 | 9.70E-05 | 0 | 0 | 0.00011 | 5.97E-05 | 0 | 0 | 1.20E-05 | 7.47E-06 |
| Nacc1 | 9.54E-05 | 7.47E-05 | 0.00019 | 0.00012 | 7.34E-05 | 3.90E-05 | 0 | 0.00011 | 0.00019 | 7.97E-05 | 0.00012 | 2.43E-05 | 6.17E-05 | 6.61E-05 | 0 | 0.00021 | 4.34E-05 | 4.25E-05 |
| Amn | 0 | 0 | 0 | 0 | 2.85E-05 | 0.0003 | 0 | 0.00023 | 0.00019 | 0 | 0 | 0 | 0 | 3.74E-05 | 5.46E-05 | 0 | 0 | 3.75E-05 |
| Mob3a | 9.59E-05 | 6.18E-05 | 0.00018 | 1.87E-05 | 0 | 0 | 0 | 6.81E-05 | 0.00019 | 3.48E-05 | 0.00015 | 0.00011 | 3.76E-05 | 7.33E-05 | 6.40E-05 | 0 | 3.95E-05 | 7.63E-05 |
| Eci2 | 0.00021 | 5.38E-05 | 6.56E-06 | 2.35E-05 | 1.61E-05 | 6.34E-05 | 4.76E-06 | 0.00011 | 0.00019 | 9.86E-05 | 0 | 3.71E-05 | 4.71E-05 | 8.72E-05 | 0 | 2.98E-05 | 1.92E-05 | 9.94E-05 |
| Hspa14 | 4.97E-05 | 5.03E-05 | 0.00023 | 5.39E-05 | 8.02E-05 | 9.40E-05 | 0 | 5.92E-05 | 0.00019 | 8.27E-05 | 6.27E-05 | 6.48E-05 | 4.02E-05 | 4.87E-05 | 3.07E-05 | 0.00041 | 3.62E-05 | 1.51E-05 |
| Zdhhc3 | 5.93E-05 | 0.00015 | 0.00022 | 0.00038 | 0.00016 | 0.00013 | 0.00014 | 0.00017 | 0.00019 | 0.00029 | 0.0002 | 4.17E-05 | 0.00013 | 0.00021 | 0 | 0.00025 | 0 | 0.00017 |
| Etv6 | 9.22E-05 | 9.77E-05 | 0.00017 | 8.61E-05 | 0.00016 | 0.0001 | 7.59E-05 | 0.00026 | 0.00019 | 0 | 0.00017 | 0 | 0.00012 | 0.00029 | 0.00016 | 8.82E-05 | 0.00029 | 9.97E-05 |

| | | | | | | | | | | | | | | | | | | |
|---|---|---|---|---|---|---|---|---|---|---|---|---|---|---|---|---|---|---|
| Dnaja2 | 0.00034 | 0.00027 | 0.00041 | 0.00028 | 0.00013 | 0.00022 | 0 | 0.00016 | 0.00018 | 6.28E-05 | 0.00016 | 3.75E-05 | 0.00014 | 0.00013 | 7.57E-05 | 0.00018 | 7.16E-05 | 0.00011 |
| Qrich1 | 0.00032 | 0.00021 | 0.00019 | 2.46E-05 | 9.86E-05 | 0.00018 | 9.48E-05 | 8.53E-05 | 0.00018 | 0.00011 | 9.42E-05 | 0.00014 | 0 | 5.82E-05 | 0.00026 | 0.00014 | 0.00023 | 0.00012 |
| Gspt1 | 0.00019 | 7.50E-05 | 0.00042 | 0.00035 | 8.26E-05 | 1.92E-05 | 5.68E-05 | 5.28E-05 | 0.00018 | 9.57E-05 | 0.00013 | 1.17E-05 | 4.60E-05 | 5.06E-05 | 0.00027 | 0.00028 | 0.00014 | 9.70E-05 |
| Cox7c | 0.00032 | 0.00018 | 0.00027 | 0.00048 | 0.00019 | 0.00029 | 0.00015 | 0.00033 | 0.00018 | 6.82E-05 | 0.00026 | 0.0003 | 0.00015 | 0.0002 | 5.84E-05 | 0.00027 | 0 | 0.00014 |
| Leprot | 0.00025 | 0.00015 | 8.79E-05 | 0.00017 | 0.00032 | 0.00046 | 0.00028 | 0.00038 | 0.00018 | 0.00038 | 0.00021 | 0.00014 | 0.0002 | 0.00021 | 0 | 0.0002 | 0 | 0.00029 |
| Inhba | 0 | 0 | 0.00044 | 0 | 0.0007 | 0.00121 | 0 | 0.00051 | 0.00018 | 0 | 0.00143 | 0.00108 | 0 | 0.00104 | 0.00068 | 0 | 0 | 0 |
| Ndufa1 | 0.00036 | 0 | 0.00011 | 0.00031 | 0.00022 | 0.00041 | 0.00038 | 0.00017 | 0.00018 | 6.08E-05 | 0.00015 | 0 | 0.00028 | 0.0004 | 0 | 7.11E-05 | 0 | 0.0002 |
| Fbxw8 | 0.00015 | 0.00013 | 0.00033 | 0.00053 | 0.00012 | 6.71E-05 | 0.00033 | 0.0002 | 0.00018 | 0.00024 | 0.0002 | 0.0001 | 0.00012 | 0.00013 | 0.00039 | 1.79E-05 | 1.94E-05 | 0.00023 |
| Smarca4 | 0.00027 | 0.00018 | 0.00027 | 0.00017 | 4.62E-05 | 2.09E-05 | 3.76E-05 | 9.86E-05 | 0.00018 | 8.55E-05 | 0.00016 | 3.86E-05 | 0 | 6.87E-05 | 0.00042 | 0.00037 | 0.00023 | 0.00013 |
| Cnbp | 0.00011 | 0.00015 | 0.00042 | 0.00048 | 8.10E-05 | 8.28E-05 | 0 | 8.17E-05 | 0.00018 | 8.57E-05 | 7.47E-05 | 5.76E-05 | 0.00012 | 0.00011 | 3.71E-05 | 0.00023 | 0.00016 | 1.97E-05 |
| Gtf2h5 | 0.00026 | 0.00017 | 0.00023 | 0.00044 | 0.00018 | 0.00013 | 0.00014 | 7.69E-05 | 0.00018 | 0.00012 | 0.00016 | 1.71E-05 | 0.0002 | 0.00017 | 0 | 0.00022 | 0 | 9.97E-05 |
| Smarcb1 | 0.00029 | 0.00018 | 0.00012 | 0.00024 | 0.00016 | 0.00012 | 7.82E-05 | 5.32E-05 | 0.00018 | 6.69E-05 | 5.29E-05 | 6.27E-05 | 5.27E-05 | 0.00014 | 5.76E-05 | 0.00015 | 0 | 1.77E-05 |
| Peli2 | 0 | 0.00016 | 0 | 4.49E-05 | 0.00012 | 0.00017 | 3.77E-05 | 0.00011 | 0.00018 | 8.50E-05 | 0 | 0 | 9.47E-05 | 0.00019 | 0.0002 | 1.61E-06 | 0 | 9.00E-05 |
| Bcat2 | 0.00011 | 0.00015 | 6.72E-05 | 0.00038 | 0.00011 | 0 | 4.19E-05 | 8.40E-05 | 0.00018 | 0.0001 | 6.68E-05 | 0.00013 | 8.76E-05 | 1.14E-05 | 4.32E-05 | 2.16E-05 | 2.37E-05 | 9.12E-05 |
| Srsf9 | 0.00019 | 0.00017 | 0.00038 | 0.00069 | 0.00029 | 0.00021 | 0.00021 | 0.00038 | 0.00018 | 0.00012 | 0.00029 | 0.00013 | 0.00018 | 0.00029 | 0.00031 | 0.00028 | 0.00022 | 0.00019 |
| Tmem263 | 0.00058 | 3.74E-05 | 0.00031 | 0.00048 | 0.00032 | 0.00028 | 0.0008 | 0.00033 | 0.00018 | 0.00016 | 0.00019 | 0.00034 | 0.00022 | 0.00021 | 0.00068 | 0.00015 | 0.00018 | 2.27E-06 |
| Foxn3 | 4.70E-05 | 0.00018 | 4.33E-05 | 0 | 0.00051 | 0.00012 | 0.00022 | 8.69E-05 | 0.00018 | 0.00012 | 0.00026 | 0.00026 | 2.78E-07 | 2.80E-05 | 0.00019 | 5.94E-05 | 0.00019 | 9.09E-05 |
| Ndufc2 | 0.00023 | 0.00014 | 0.00026 | 0.00063 | 0.00013 | 0.00013 | 0.00013 | 0.00022 | 0.00018 | 0.00017 | 0.00021 | 0.00015 | 0.00013 | 0.00025 | 0 | 0.00026 | 0 | 0.00022 |
| Fndc4 | 3.44E-05 | 4.56E-05 | 9.59E-05 | 2.90E-05 | 0.00018 | 4.74E-05 | 0.0001 | 0 | 0.00018 | 1.52E-05 | 7.01E-05 | 0 | 0.00012 | 7.50E-05 | 0 | 7.97E-05 | 4.04E-05 | 0 |
| Mcmbp | 0.00015 | 0.00019 | 0.00024 | 0.00015 | 1.43E-05 | 8.94E-05 | 0 | 0.00011 | 0.00018 | 6.79E-05 | 9.85E-05 | 0.00011 | 5.30E-06 | 6.40E-05 | 0.00021 | 0.00024 | 6.32E-05 | 6.64E-05 |
| Ifrd1 | 0.00012 | 0.00013 | 0.00028 | 0 | 1.79E-05 | 7.13E-05 | 0 | 0 | 0.00018 | 0 | 0.00019 | 0.00027 | 8.68E-05 | 0.00012 | 0 | 0.00011 | 0.00372 | 0 |
| Apbb1ip | 9.79E-06 | 0 | 0.00026 | 0.00027 | 8.76E-05 | 0.00012 | 0 | 0.00051 | 0.00018 | 0.00069 | 0.00016 | 0.00036 | 0.00055 | 0.00034 | 0.00012 | 7.94E-05 | 4.09E-05 | 0.00033 |
| Sike1 | 0.00015 | 0.00018 | 5.79E-05 | 0.00018 | 0.00019 | 0.00018 | 9.17E-05 | 0.00012 | 0.00018 | 8.68E-05 | 0.00018 | 6.00E-05 | 0 | 0.00019 | 0.00011 | 3.64E-05 | 0 | 9.12E-05 |
| Eif3f | 0.00049 | 0.00035 | 0.00018 | 0.00015 | 0.0002 | 0.0005 | 0.00032 | 0.0002 | 0.00018 | 0.00011 | 0 | 0.0003 | 0.00015 | 0.00028 | 0 | 4.67E-05 | 0.00012 | 0.00013 |
| Sumf1 | 0.00011 | 0.00015 | 5.78E-05 | 5.35E-05 | 0.00026 | 0.00027 | 0.00019 | 0.00017 | 0.00018 | 0.00015 | 0.00014 | 0 | 0.0003 | 0.00023 | 0 | 4.11E-05 | 0 | 0.00028 |
| Htr1b | 0 | 0 | 8.17E-05 | 0 | 0.00024 | 0.00027 | 0 | 0.00016 | 0.00018 | 0 | 0.00026 | 4.50E-05 | 0 | 0.00019 | 0 | 4.28E-05 | 0 | 0 |
| Txndc17 | 0.00028 | 0.00018 | 0.00028 | 0.00052 | 0.00015 | 0.00026 | 0.0003 | 0.00023 | 0.00018 | 0.0003 | 0.0003 | 0.00041 | 0.00015 | 0.0002 | 0.00011 | 3.00E-05 | 7.35E-05 | 0.00028 |
| Bag6 | 0.00014 | 0.00023 | 0.00014 | 0.00045 | 0.00013 | 0.00012 | 0.00012 | 0.0002 | 0.00018 | 9.74E-05 | 0.00013 | 0.00019 | 5.13E-06 | 0.00019 | 0.00028 | 0.0002 | 0.00013 | 0.00016 |
| Marveld1 | 0 | 0.00012 | 0.00023 | 0.00068 | 0.00032 | 0.00033 | 0.00049 | 0.00048 | 0.00018 | 0.00029 | 0.00031 | 6.02E-05 | 4.95E-05 | 0.00039 | 0.00034 | 0 | 0.00014 | 0.00029 |
| Dync1i2 | 0.00025 | 0.00021 | 0.00019 | 4.10E-05 | 9.47E-05 | 0.00014 | 0.0001 | 6.61E-05 | 0.00018 | 0.00014 | 0.00021 | 0.00023 | 0 | 0.00014 | 2.92E-05 | 0.00014 | 5.96E-05 | 0.00018 |
| Hsp90aa1 | 0.0003 | 0.00015 | 0.00036 | 0.00024 | 0.00011 | 9.10E-05 | 0 | 6.50E-05 | 0.00018 | 0.00014 | 0.00031 | 0.00042 | 0.00011 | 5.78E-05 | 0 | 0.00038 | 0.00193 | 8.29E-05 |
| Fgfr1op2 | 0.00028 | 0.00018 | 0.00024 | 0.00016 | 0.00017 | 0.00024 | 8.24E-05 | 0.00027 | 0.00018 | 0.00015 | 0.00011 | 0.00013 | 0 | 0.00016 | 0.00032 | 0.00014 | 0 | 9.52E-05 |
| Ppme1 | 6.04E-05 | 6.57E-05 | 0.0002 | 7.80E-05 | 0.00014 | 4.98E-06 | 0.00011 | 0.00012 | 0.00018 | 6.50E-05 | 0.00023 | 0.00058 | 7.01E-05 | 0 | 0.00022 | 3.73E-05 | 7.99E-05 | 6.21E-05 |

| | | | | | | | | | | | | | | | | | | |
|---|---|---|---|---|---|---|---|---|---|---|---|---|---|---|---|---|---|---|
| Xpo6 | 5.13E-05 | 0.00022 | 0.00025 | 0.00031 | 0.0001 | 3.89E-05 | 4.22E-05 | 2.25E-05 | 0.00018 | 0.0001 | 0.00013 | 9.96E-05 | 3.27E-05 | 0.00012 | 6.74E-05 | 0.00013 | 0.00033 | 9.94E-05 |
| Nfe2l1 | 0.00013 | 0.00036 | 0.00042 | 0.00031 | 0.00031 | 0.0002 | 9.34E-05 | 0.00031 | 0.00018 | 0.0002 | 0.00034 | 0.00033 | 0.00019 | 0.0001 | 5.95E-05 | 0.00012 | 0.00034 | 0.00042 |
| Mindy1 | 8.43E-05 | 0.00016 | 7.92E-05 | 0 | 5.50E-05 | 3.75E-05 | 7.13E-05 | 0.0001 | 0.00018 | 0.00017 | 0 | 6.50E-05 | 0 | 0.00013 | 0.0001 | 9.94E-06 | 0 | 0.00014 |
| Vat1 | 0 | 0.00055 | 0.00036 | 0.00085 | 0.00028 | 0.00047 | 0.00073 | 0.00031 | 0.00018 | 0.00066 | 0.0004 | 0.00061 | 0.00023 | 0.00029 | 7.05E-05 | 0 | 0.00021 | 0.00047 |
| Mllt6 | 2.02E-05 | 0.00027 | 7.87E-05 | 0.00023 | 0.00022 | 0.00019 | 9.04E-05 | 0.00039 | 0.00018 | 0.00011 | 2.54E-05 | 0.0002 | 0.00014 | 0.00014 | 0.00034 | 7.11E-05 | 8.80E-05 | 2.51E-05 |
| Pde4d | 4.43E-05 | 2.65E-05 | 0.00018 | 0 | 0.0002 | 0.00024 | 0.00019 | 0 | 0.00018 | 0 | 0 | 0 | 6.50E-06 | 0.00036 | 7.63E-05 | 2.05E-05 | 3.72E-05 | 8.02E-05 |
| Skp1a | 0.00027 | 0.0002 | 0.00012 | 0.00037 | 0.00015 | 0.00015 | 2.46E-05 | 8.26E-05 | 0.00018 | 0.00016 | 0.00013 | 6.73E-05 | 0.00023 | 8.02E-05 | 0 | 0.00015 | 0.00023 | 0.00014 |
| Ssr2 | 0.00042 | 0 | 0.00057 | 0.00112 | 0.00026 | 0.00035 | 0.00098 | 0.00065 | 0.00018 | 0.00021 | 0.0003 | 0.00024 | 0.00065 | 0.0005 | 0.00016 | 0.00023 | 0 | 0.00023 |
| Hdgf | 0.00012 | 0.00017 | 0.00028 | 0.00068 | 6.52E-05 | 2.70E-05 | 0 | 2.61E-05 | 0.00018 | 0.00017 | 0.00022 | 0.00011 | 0 | 0.00014 | 0.00036 | 0.00041 | 0.00025 | 0.00013 |
| Tpr | 0.00035 | 0.0002 | 8.99E-05 | 0 | 8.55E-05 | 0.00015 | 4.38E-05 | 5.92E-05 | 0.00018 | 3.96E-05 | 2.73E-05 | 0.00015 | 0 | 0 | 7.91E-05 | 0.00026 | 0.00016 | 3.46E-05 |
| Pabpn1 | 0.00036 | 0.00027 | 0.00044 | 0.00035 | 0.00018 | 0.0001 | 4.33E-05 | 8.98E-05 | 0.00018 | 0.00011 | 0.00018 | 0.00019 | 0.00012 | 0.0001 | 0.00099 | 0.00031 | 0.00033 | 9.23E-05 |
| Adamts15 | 0 | 0.00024 | 0 | 0 | 0.00024 | 0.00015 | 0.0004 | 0 | 0.00018 | 0.00086 | 0 | 0 | 0.0006 | 0 | 0 | 0 | 0 | 2.74E-05 |
| Brd4 | 0.00025 | 0.00043 | 0.00026 | 0.00019 | 0.0002 | 0.00019 | 0.00016 | 0.00013 | 0.00018 | 0.00013 | 0.00015 | 0.00017 | 5.11E-05 | 0.00022 | 0.00059 | 0.0003 | 0.00031 | 0.00011 |
| Tmed1 | 0.00023 | 0.0001 | 0.00017 | 0.00026 | 5.49E-05 | 0.00012 | 0.0002 | 3.55E-05 | 0.00018 | 2.20E-05 | 8.06E-05 | 3.10E-06 | 0.00016 | 0.00017 | 0 | 0.00017 | 5.04E-05 | 9.11E-06 |
| Suv39h1 | 5.29E-06 | 2.41E-05 | 5.35E-05 | 2.23E-05 | 6.51E-05 | 1.67E-05 | 0 | 3.50E-05 | 0.00018 | 0 | 6.51E-05 | 1.27E-05 | 6.65E-06 | 0.00012 | 0.00012 | 0.00053 | 9.62E-05 | 0 |
| Commd10 | 0.00014 | 6.12E-05 | 7.69E-05 | 5.47E-05 | 2.70E-05 | 1.88E-05 | 4.02E-05 | 5.23E-05 | 0.00018 | 7.13E-05 | 7.34E-05 | 0.00012 | 0 | 3.85E-05 | 7.25E-05 | 0.0002 | 0 | 7.01E-05 |
| Fam120a | 0.00045 | 0.00016 | 0.00022 | 0 | 8.65E-05 | 0.00024 | 0.00017 | 5.62E-05 | 0.00018 | 0.00011 | 0.00012 | 0.00018 | 0 | 0.0001 | 6.53E-05 | 0.00016 | 0.00018 | 0.00013 |
| Trim56 | 8.00E-05 | 0.00018 | 4.80E-05 | 9.04E-05 | 9.23E-05 | 1.46E-05 | 4.49E-05 | 8.27E-05 | 0.00018 | 0.00014 | 0.00014 | 0.00011 | 0.00017 | 9.79E-05 | 0.00019 | 3.58E-05 | 0 | 3.07E-05 |
| Snd1 | 0.00051 | 0.00019 | 0.00045 | 0.00047 | 0.0001 | 0.00021 | 0.00047 | 0.0004 | 0.00018 | 0.00018 | 0.00015 | 0.0003 | 7.97E-05 | 0.00025 | 0.00098 | 0.00019 | 0.0004 | 0.00015 |
| Arfgap1 | 0.00041 | 7.23E-05 | 0.00033 | 0.00027 | 0.00013 | 0.0002 | 0.00064 | 0.00019 | 0.00018 | 8.04E-05 | 0.00014 | 9.71E-05 | 0.00021 | 0.00022 | 0.00099 | 0 | 0.00019 | 1.49E-05 |
| Tpst2 | 5.89E-05 | 0.00016 | 0.00019 | 2.39E-05 | 8.32E-05 | 2.01E-05 | 6.11E-05 | 5.89E-05 | 0.00018 | 0.0001 | 0.00039 | 0.00016 | 0.00012 | 0.00017 | 2.71E-05 | 9.84E-05 | 0 | 8.58E-05 |
| Egr2 | 0 | 0 | 0.00015 | 0 | 0 | 0.0002 | 0 | 0.00049 | 0.00018 | 0 | 0.00023 | 0.00049 | 0 | 0 | 0 | 0 | 0.00115 | 1.66E-05 |
| Sec31a | 0.00045 | 0.00018 | 0.0003 | 0.0005 | 0.00023 | 0.00036 | 0.00068 | 0.00034 | 0.00018 | 0.00023 | 0.0002 | 0.00015 | 0.00022 | 0.00029 | 0.00031 | 5.42E-05 | 0.00033 | 0.00018 |
| Asap1 | 4.83E-05 | 0.00013 | 4.76E-05 | 0 | 7.81E-05 | 0.00016 | 0.00012 | 0 | 0.00018 | 0.00015 | 0.00013 | 0.00014 | 0 | 0.00018 | 0 | 0.00014 | 0.00022 | 0.00014 |
| Ppp1r9b | 8.93E-05 | 0.00031 | 0.00024 | 0.00042 | 0.0002 | 0.00015 | 0.00019 | 0.00024 | 0.00018 | 0.00022 | 0.00015 | 8.55E-06 | 0.00011 | 0.00017 | 0.00037 | 8.52E-05 | 0.00023 | 0.00024 |
| Ankrd40 | 0.00019 | 0.0002 | 0.0001 | 0.0003 | 0.00028 | 0.00021 | 6.29E-05 | 6.13E-05 | 0.00018 | 0.00032 | 0.00026 | 5.61E-05 | 0.00041 | 8.98E-05 | 0 | 0.00019 | 0.00012 | 0.00012 |
| Kazn | 0 | 7.81E-05 | 0 | 0 | 0.00014 | 0.00027 | 0 | 0.00014 | 0.00018 | 3.84E-05 | 4.53E-05 | 0 | 0 | 0.00066 | 0.00015 | 0 | 0 | 8.33E-05 |
| Armcx4 | 0.00011 | 5.03E-06 | 7.92E-05 | 0.00023 | 0.00013 | 0.0001 | 9.76E-05 | 0.00033 | 0.00018 | 0 | 1.34E-05 | 0 | 0.00011 | 0.00024 | 0.00029 | 6.86E-05 | 0 | 0 |
| Ube2e3 | 9.68E-05 | 0.00011 | 4.62E-05 | 4.22E-05 | 0.00013 | 0.00015 | 0 | 8.26E-05 | 0.00018 | 5.15E-05 | 0.00013 | 2.06E-05 | 6.74E-05 | 0.00017 | 4.83E-05 | 0.00022 | 1.79E-05 | 7.18E-05 |
| Tcaf1 | 0.00022 | 0.00015 | 0.00013 | 8.06E-05 | 0.00013 | 0.00015 | 0.00016 | 8.76E-05 | 0.00018 | 0.00013 | 0.00022 | 8.69E-05 | 9.23E-05 | 0.00012 | 0 | 0.00011 | 3.19E-06 | 0.00012 |
| Pla2r1 | 0 | 0.00012 | 0 | 0 | 1.69E-05 | 0.00019 | 0 | 0 | 0.00018 | 0.00023 | 1.23E-05 | 0 | 0.00012 | 0.00017 | 5.15E-05 | 0 | 0 | 0.0001 |
| Vars | 7.28E-05 | 8.71E-05 | 0.0004 | 0.00045 | 2.83E-05 | 0 | 1.26E-06 | 0.00016 | 0.00018 | 9.53E-05 | 7.84E-05 | 0.00014 | 9.13E-05 | 4.08E-05 | 0.00025 | 0.00053 | 0 | 7.25E-05 |
| Ar | 3.55E-05 | 8.19E-05 | 0 | 2.57E-06 | 7.76E-05 | 7.76E-06 | 0.00011 | 3.96E-05 | 0.00018 | 0.00025 | 9.98E-06 | 0 | 5.22E-05 | 0 | 4.31E-07 | 6.69E-05 | 6.88E-05 | 1.66E-05 |

| | | | | | | | | | | | | | | | | | | |
|---|---|---|---|---|---|---|---|---|---|---|---|---|---|---|---|---|---|---|
| Desi1 | 6.44E-05 | 4.86E-05 | 0.00015 | 0.00027 | 0.00029 | 0.00014 | 7.52E-05 | 0.00015 | 0.00018 | 4.34E-05 | 0.00013 | 0.00011 | 3.19E-05 | 0.00015 | 0 | 9.83E-05 | 9.03E-05 | 0 |
| Osbpl9 | 0.00012 | 0.00024 | 4.79E-05 | 0 | 1.40E-05 | 0.00016 | 0.00031 | 0 | 0.00018 | 0.00024 | 6.85E-05 | 0.00011 | 0.00025 | 0.00012 | 0 | 0.00015 | 6.20E-05 | 8.65E-05 |
| Atp5j | 0.00039 | 1.65E-05 | 6.85E-05 | 0 | 5.56E-05 | 0.00018 | 0.00012 | 7.92E-05 | 0.00018 | 0 | 7.79E-05 | 5.11E-05 | 6.14E-05 | 2.65E-05 | 3.02E-05 | 0.0001 | 2.65E-05 | 2.28E-05 |
| Txn2 | 0.0004 | 7.72E-05 | 0.0004 | 0.00013 | 6.42E-05 | 0.00016 | 8.65E-05 | 0.00016 | 0.00018 | 5.50E-05 | 4.61E-05 | 0 | 7.16E-06 | 0 | 0 | 3.02E-05 | 8.72E-05 | 3.54E-05 |
| Rad23a | 0.00016 | 0.00016 | 0.00019 | 0.00017 | 0.00011 | 7.26E-05 | 0.00017 | 0 | 0.00018 | 9.60E-05 | 6.41E-05 | 9.09E-05 | 0.00011 | 4.71E-05 | 0 | 0.00021 | 0.0003 | 8.52E-05 |
| Nr2f6 | 1.99E-05 | 8.93E-05 | 0.00014 | 0.00021 | 0.00015 | 0.0001 | 3.25E-06 | 0.0002 | 0.00018 | 3.02E-05 | 0.00015 | 0 | 0 | 7.79E-05 | 0.00014 | 5.10E-05 | 1.88E-05 | 0.0001 |
| Nenf | 0.00024 | 7.68E-05 | 0.00012 | 0.00062 | 0.00024 | 0.0004 | 0.00069 | 0.00029 | 0.00018 | 0.00046 | 0.00031 | 0.00013 | 0.00059 | 0.00036 | 0 | 0.00017 | 0 | 0.00052 |
| Slc39a7 | 0.00036 | 0.00024 | 0.00047 | 0.00068 | 0.00022 | 0.00027 | 0.00062 | 0.00029 | 0.00018 | 0.00023 | 0.00016 | 0.00021 | 0.00022 | 0.00028 | 0.00066 | 3.80E-05 | 0.00024 | 0.00018 |
| En1 | 2.05E-05 | 0.00011 | 0 | 0.00022 | 0.0003 | 0 | 3.77E-05 | 4.91E-05 | 0.00018 | 0.00013 | 0.00016 | 6.45E-05 | 0.00034 | 0.00015 | 7.32E-05 | 6.36E-05 | 9.45E-06 | 0 |
| Dad1 | 0.00029 | 7.69E-05 | 0.0004 | 0.00105 | 0.00034 | 0.00031 | 0.00066 | 0.00084 | 0.00018 | 0.00026 | 0.00048 | 0.00028 | 0.0004 | 0.00039 | 0 | 0.00011 | 0.00027 | 0.00039 |
| Oxct1 | 0.00018 | 3.89E-06 | 0 | 4.35E-05 | 0.00013 | 5.87E-05 | 0.00022 | 0 | 0.00018 | 0.00017 | 0.00016 | 0 | 6.85E-06 | 3.69E-05 | 0 | 0.00022 | 0 | 0.00016 |
| Stx7 | 0.00023 | 0.00017 | 0.00011 | 0 | 0.00015 | 4.60E-05 | 8.74E-05 | 0.00038 | 0.00018 | 8.62E-05 | 8.59E-05 | 0.00024 | 0 | 0 | 0.0001 | 0.00012 | 2.80E-05 | 0.00012 |
| Samm50 | 0.00031 | 9.61E-05 | 0.00025 | 0.00014 | 3.06E-05 | 9.28E-05 | 6.60E-05 | 3.28E-05 | 0.00018 | 4.63E-05 | 0.00014 | 7.65E-05 | 0 | 6.77E-05 | 8.65E-06 | 0.0001 | 3.51E-05 | 2.47E-05 |
| Chd3 | 0.00011 | 0.00026 | 0 | 0.00032 | 0.00023 | 0 | 0.00028 | 0 | 0.00018 | 0.00022 | 0.00017 | 0 | 0.00024 | 0.0001 | 0.00035 | 4.77E-05 | 0.00013 | 0.00012 |
| Casp12 | 0.00017 | 8.91E-05 | 0 | 0 | 0.00011 | 7.11E-05 | 0 | 0 | 0.00018 | 0.0001 | 0 | 0 | 2.17E-05 | 1.18E-05 | 9.11E-05 | 5.31E-05 | 0 | 6.30E-05 |
| Pdia3 | 0.00067 | 0 | 0.00033 | 0.00071 | 0.00035 | 0.00055 | 0.00077 | 0.00052 | 0.00018 | 0.00038 | 0.00044 | 0.0003 | 0.00085 | 0.00056 | 0 | 0.0002 | 0.00018 | 0.00053 |
| Atp6v1a | 0.00011 | 0.0003 | 4.53E-05 | 0.00028 | 0.00017 | 0.00014 | 0 | 6.58E-05 | 0.00018 | 0.00011 | 3.49E-05 | 0.0002 | 9.87E-05 | 0.00014 | 2.03E-05 | 7.61E-05 | 0.00011 | 0.00012 |
| Btf3 | 0.00052 | 0.00024 | 0.00039 | 0.00037 | 1.52E-05 | 0.00017 | 5.51E-05 | 0 | 0.00018 | 4.48E-05 | 0.00013 | 0.00035 | 1.03E-05 | 0.00014 | 0 | 0.00015 | 0.00028 | 1.74E-05 |
| Tmem184b | 6.04E-05 | 1.49E-05 | 0.00036 | 0.00034 | 0.00043 | 0.00033 | 0.00064 | 0.0005 | 0.00018 | 0.00032 | 0.00036 | 6.99E-05 | 0.00014 | 0.00033 | 7.29E-05 | 5.38E-05 | 0.0002 | 0.00031 |
| Crip1 | 0.00029 | 0 | 0.00041 | 0.00082 | 0.00013 | 0.00034 | 0.00112 | 0.00093 | 0.00018 | 0.001 | 0.00073 | 0.00027 | 0 | 0.00013 | 0 | 0.00055 | 0.00022 | 0.001 |
| Maged2 | 0.00058 | 0 | 0.00026 | 0.0004 | 0.00015 | 0.00032 | 0.0007 | 0.00036 | 0.00018 | 0.00022 | 0.00014 | 0.00013 | 0.00012 | 0.00024 | 7.36E-05 | 0 | 6.55E-06 | 0.00018 |
| Pitpna | 0.00024 | 0.00015 | 0.00041 | 0.00029 | 9.17E-05 | 0.00014 | 0.00017 | 0.00025 | 0.00018 | 0.00025 | 0.00023 | 0.00013 | 1.95E-05 | 0.00015 | 9.75E-05 | 0.00011 | 1.98E-05 | 0.00031 |
| Col16a1 | 4.17E-05 | 1.40E-06 | 0.00014 | 0.00027 | 0.00054 | 0.00061 | 0.00119 | 0.00059 | 0.00018 | 0.00042 | 0.0004 | 0.00078 | 0.00097 | 0.00094 | 0.00091 | 0.00016 | 0.0002 | 0.00094 |
| Pofut2 | 0.0001 | 0 | 0.00035 | 0.00068 | 0.00044 | 0.00047 | 0.00085 | 0.00064 | 0.00018 | 0.00025 | 0.0003 | 0.00014 | 0.00039 | 0.00055 | 0.00046 | 7.12E-05 | 0.00012 | 0.00033 |
| Zswim8 | 1.55E-05 | 0.00026 | 0.00014 | 0.00031 | 0.00016 | 9.18E-05 | 8.80E-05 | 0.00015 | 0.00018 | 0.00017 | 0.00019 | 7.03E-05 | 7.16E-05 | 0.00022 | 0.00048 | 5.16E-05 | 0.00042 | 0.00018 |
| Socs5 | 0.00015 | 0.00011 | 0.00025 | 7.69E-05 | 2.41E-05 | 0.00021 | 0 | 0.00015 | 0.00018 | 7.75E-05 | 0.00019 | 6.99E-05 | 0.00033 | 5.06E-05 | 4.26E-05 | 7.21E-05 | 6.45E-05 | 0.00016 |
| Hes1 | 2.10E-05 | 6.47E-05 | 1.81E-05 | 0 | 0.00016 | 0.00024 | 0.0002 | 0.00076 | 0.00018 | 0.00016 | 9.14E-05 | 8.52E-05 | 0 | 0.00018 | 9.52E-05 | 0.00019 | 0.00263 | 0.00031 |
| Pds5a | 0.00016 | 0.00016 | 0.00015 | 7.66E-06 | 5.89E-05 | 0.00011 | 4.60E-05 | 4.96E-05 | 0.00018 | 5.29E-05 | 0.00012 | 0.00011 | 0 | 7.81E-05 | 0.0003 | 0.00028 | 0.00016 | 5.90E-05 |
| Tex261 | 0.00029 | 0.00021 | 0.00022 | 0.00017 | 0.00017 | 0.00011 | 0.00016 | 0.00016 | 0.00018 | 0.00017 | 0.00014 | 0.0002 | 7.42E-05 | 5.65E-05 | 0 | 0.00018 | 0 | 8.93E-05 |
| Vapb | 6.91E-05 | 0.00012 | 0.0002 | 0.00044 | 0.00017 | 0.00016 | 0.00011 | 8.68E-05 | 0.00018 | 0.00027 | 0.0002 | 0.00015 | 9.45E-05 | 7.81E-05 | 5.78E-06 | 7.54E-05 | 6.85E-05 | 0.00019 |
| Hibadh | 0.00022 | 0.0002 | 0.00014 | 0.00015 | 0.00012 | 9.22E-05 | 0 | 8.97E-05 | 0.00018 | 0.00012 | 3.45E-05 | 0.00014 | 0.00015 | 5.93E-05 | 5.90E-05 | 5.96E-05 | 9.45E-06 | 0.00011 |
| Lrp6 | 0.00023 | 0.00028 | 7.12E-05 | 4.02E-05 | 0.00021 | 0.0002 | 0.00016 | 0.00018 | 0.00018 | 0.00029 | 0.00011 | 0.00016 | 0.0003 | 0.00014 | 0.00037 | 8.80E-05 | 0.00026 | 0.00066 |
| Rassf1 | 0.00017 | 0.00024 | 0.0004 | 1.67E-06 | 8.74E-05 | 8.53E-05 | 0.00012 | 6.20E-05 | 0.00018 | 1.58E-05 | 0.00019 | 0.00049 | 0 | 0.00021 | 0.00026 | 0.00024 | 0.00018 | 0 |

| | | | | | | | | | | | | | | | | | | |
|---|---|---|---|---|---|---|---|---|---|---|---|---|---|---|---|---|---|---|
| Sash1 | 8.20E-05 | 0.00022 | 0.00015 | 0.0001 | 0.00021 | 0.0002 | 0.00026 | 7.30E-05 | 0.00018 | 0.00047 | 0 | 0.00025 | 4.30E-05 | 4.00E-05 | 0.0002 | 0.00016 | 0.00026 | 0.00019 |
| Steap1 | 0.00012 | 0 | 0.00018 | 8.74E-05 | 0 | 0 | 5.42E-05 | 0 | 0.00018 | 1.58E-05 | 0 | 6.86E-05 | 0.00028 | 0 | 0 | 0.00014 | 0 | 0 |
| Sra1 | 0.00042 | 7.86E-05 | 0.00033 | 0.0004 | 0.00019 | 0.00018 | 0.00048 | 0.00017 | 0.00018 | 0.00014 | 0.00024 | 0.00032 | 0.00027 | 0.00013 | 0.00019 | 0 | 0 | 0.00013 |
| Atxn7l3 | 0.00012 | 0.00021 | 0.00024 | 0.00012 | 0.00017 | 0.00012 | 0.00013 | 0.00012 | 0.00018 | 9.24E-05 | 0.00026 | 9.92E-05 | 9.22E-05 | 0.00014 | 0.00016 | 0.0001 | 0.00016 | 0 |
| Map6 | 1.00E-06 | 2.00E-05 | 0.00022 | 0 | 0 | 0 | 0.00014 | 0.00011 | 0.00018 | 7.63E-05 | 0.00032 | 0.0002 | 0 | 6.57E-05 | 9.99E-05 | 2.44E-06 | 0 | 0.00054 |
| Ifi211 | 0.00028 | 1.33E-05 | 0 | 0 | 0 | 0.00011 | 0 | 0.00013 | 0.00018 | 0.0006 | 0 | 0.00025 | 0.00049 | 0.00013 | 4.93E-05 | 7.16E-05 | 0 | 6.64E-05 |
| Jam3 | 0.00011 | 0.00011 | 8.95E-05 | 8.00E-05 | 0.00013 | 0.00016 | 1.97E-05 | 0.00012 | 0.00018 | 8.72E-05 | 0.00012 | 0 | 0.00018 | 0.00018 | 0 | 2.66E-05 | 0 | 0.0001 |
| Rras2 | 0.00017 | 4.55E-05 | 0.00012 | 0 | 4.94E-05 | 3.08E-05 | 0 | 0 | 0.00018 | 0.00067 | 0.0004 | 8.44E-05 | 1.53E-05 | 5.02E-05 | 0 | 6.11E-05 | 8.20E-05 | 0 |
| Rassf5 | 0 | 1.07E-05 | 8.68E-05 | 0 | 0 | 8.84E-06 | 0.00034 | 0.0003 | 0.00018 | 0.00023 | 0 | 0.00011 | 0 | 0 | 2.10E-05 | 0 | 6.92E-05 | 0 |
| Nrd1 | 0.00029 | 0.00017 | 0.00019 | 0 | 3.52E-05 | 7.43E-05 | 4.88E-05 | 9.08E-05 | 0.00018 | 0.00013 | 8.01E-05 | 0.00013 | 0 | 7.95E-05 | 0.00026 | 0.0001 | 5.39E-05 | 4.75E-05 |
| Nhp2 | 0.00025 | 3.77E-05 | 0.00049 | 0.00036 | 2.10E-05 | 5.99E-06 | 5.64E-05 | 2.00E-05 | 0.00018 | 0 | 4.51E-05 | 0 | 3.38E-05 | 0.0001 | 0 | 0.00031 | 0 | 2.39E-05 |
| Bcl7c | 0.00013 | 0.00011 | 0.00018 | 0.00039 | 8.46E-05 | 4.66E-05 | 0.00011 | 0.00018 | 0.00018 | 0.0001 | 3.40E-05 | 8.01E-06 | 1.67E-05 | 6.88E-05 | 0.00025 | 0.0002 | 0.00015 | 0.00017 |
| Slc12a4 | 0.00021 | 0.00017 | 0.00024 | 0 | 0.00016 | 0.00021 | 0.00013 | 0.00014 | 0.00018 | 8.45E-05 | 0.00029 | 0.00013 | 4.14E-05 | 5.68E-05 | 0.00014 | 1.22E-05 | 4.73E-05 | 0.00014 |
| Carhsp1 | 6.33E-05 | 0.00026 | 0.00024 | 0.0002 | 2.19E-05 | 0 | 0 | 0.0002 | 0.00018 | 0.00069 | 0.00018 | 0.00043 | 0.00021 | 0 | 0 | 0.00032 | 0.00021 | 0.00019 |
| Adam15 | 8.72E-06 | 9.79E-05 | 0.0001 | 0.00018 | 9.76E-05 | 7.76E-05 | 0.00023 | 0.00016 | 0.00018 | 0.00044 | 0.00016 | 0.00025 | 0.00016 | 0.00016 | 4.52E-05 | 8.10E-06 | 0 | 0.00049 |
| Mta1 | 0.00016 | 0.00024 | 0.00032 | 0.00028 | 0.00019 | 0.00018 | 3.16E-05 | 0.00018 | 0.00018 | 7.67E-05 | 0.00015 | 0.00034 | 6.29E-05 | 0.00015 | 0.00018 | 0.00029 | 0.0002 | 9.57E-05 |
| Scarb2 | 0.0003 | 0.00027 | 0.00015 | 0 | 0.00027 | 0.00046 | 0.00027 | 0.00028 | 0.00018 | 0.0002 | 0 | 6.83E-05 | 0.00034 | 0.00017 | 0 | 7.33E-05 | 0 | 0.00035 |
| Cab39 | 0.00018 | 0.00027 | 0.00031 | 9.75E-05 | 0.00021 | 0.0002 | 4.58E-05 | 0.00013 | 0.00018 | 0.00011 | 0.00017 | 0.0002 | 0 | 0.00018 | 0.00027 | 0.00018 | 0.0001 | 0.00013 |
| Ptk2 | 0.00018 | 0.0001 | 0.00013 | 0 | 5.93E-05 | 0.0001 | 0.00012 | 0.00017 | 0.00018 | 9.37E-05 | 0.00014 | 0.00012 | 1.72E-06 | 6.87E-05 | 0.00024 | 6.67E-05 | 0.00012 | 0.00013 |
| Cant1 | 0 | 0.00022 | 8.00E-05 | 0.00025 | 0.00018 | 0.00012 | 0.0002 | 0.00021 | 0.00018 | 0.00024 | 0.00012 | 0.00011 | 0.00012 | 0.00015 | 4.51E-05 | 0.00016 | 7.91E-05 | 0.00015 |
| Ndufb7 | 0.00024 | 8.60E-05 | 0.00022 | 0.00045 | 9.33E-05 | 5.30E-05 | 0.0001 | 8.74E-05 | 0.00018 | 7.00E-05 | 0.00021 | 4.29E-06 | 6.16E-05 | 0.00013 | 1.61E-05 | 0.00019 | 0 | 0.00013 |
| Atic | 0.00016 | 0.00014 | 0.00044 | 0.0004 | 4.98E-05 | 4.57E-05 | 0 | 7.18E-05 | 0.00018 | 8.55E-05 | 0.00012 | 3.98E-05 | 0 | 3.69E-06 | 0 | 0.00019 | 4.16E-05 | 9.27E-05 |
| Psmd3 | 0.00026 | 0.00015 | 0.00037 | 0.0002 | 7.27E-05 | 4.28E-05 | 2.78E-05 | 7.49E-05 | 0.00018 | 2.00E-05 | 9.52E-05 | 6.77E-05 | 9.24E-05 | 5.82E-05 | 4.78E-06 | 0.00012 | 0 | 3.48E-05 |
| Atp5k | 0.00027 | 3.57E-05 | 0.00017 | 0.0004 | 0.00011 | 7.99E-05 | 9.40E-05 | 8.48E-05 | 0.00018 | 9.99E-06 | 0.00021 | 0 | 8.90E-06 | 0.00011 | 0 | 0.00017 | 0 | 2.86E-05 |
| Ptgr1 | 0.00018 | 0 | 3.88E-05 | 0.00012 | 0 | 0 | 3.66E-05 | 1.79E-06 | 0.00018 | 6.03E-05 | 5.46E-05 | 8.22E-05 | 0.00012 | 1.43E-05 | 0 | 0.00016 | 0 | 0.00024 |
| Csnk1a1 | 0.00029 | 0.00024 | 0.00038 | 0.00018 | 9.93E-05 | 0.00022 | 0.00011 | 0.00018 | 0.00018 | 0.00015 | 0.0002 | 0.00027 | 0.00025 | 0.00024 | 0.00022 | 0.00012 | 7.98E-05 | 0.00011 |
| Nudc | 0.00016 | 0.00024 | 0.0004 | 0.00043 | 0.00011 | 3.49E-05 | 0 | 8.07E-05 | 0.00017 | 0.00011 | 0.00021 | 0.00014 | 8.11E-05 | 0.0001 | 0.00023 | 0.00066 | 0.0001 | 0.00011 |
| Acap2 | 0.00013 | 0.00022 | 4.82E-05 | 5.01E-05 | 4.24E-05 | 0.00015 | 8.75E-05 | 0.00018 | 0.00017 | 0.00024 | 0 | 2.82E-05 | 9.85E-05 | 5.54E-05 | 0.00015 | 0.00017 | 0.00029 | 0.00019 |
| Sp3 | 0.00043 | 0.00027 | 2.95E-05 | 0 | 7.50E-05 | 0.00034 | 3.00E-06 | 0 | 0.00017 | 6.79E-05 | 0 | 0.00029 | 0.00011 | 2.16E-05 | 0.00016 | 0.0002 | 0.00018 | 0 |
| Cdk9 | 1.65E-05 | 0.00029 | 0.00024 | 0.00038 | 6.63E-05 | 2.62E-05 | 0 | 0.00027 | 0.00017 | 9.45E-05 | 0.00023 | 0.00012 | 2.55E-05 | 0.00018 | 0.0003 | 8.03E-05 | 0.00021 | 8.31E-05 |
| Stoml2 | 0.00018 | 9.66E-05 | 0.0002 | 0.00023 | 2.42E-05 | 1.32E-05 | 0 | 0 | 0.00017 | 3.59E-05 | 3.02E-05 | 5.46E-05 | 0 | 7.63E-05 | 0 | 0.00017 | 0.00018 | 1.66E-05 |
| Hnrnpf | 0.00032 | 0.00022 | 0.00039 | 0.00028 | 4.54E-05 | 9.95E-05 | 5.33E-06 | 4.00E-05 | 0.00017 | 7.00E-05 | 0.00017 | 0.00021 | 7.31E-05 | 9.00E-05 | 1.70E-05 | 0.00039 | 0.0003 | 5.30E-05 |
| Asna1 | 0.00027 | 0.00023 | 0.00018 | 0.00028 | 0.0001 | 2.48E-05 | 8.92E-06 | 3.19E-05 | 0.00017 | 3.28E-05 | 0.0001 | 4.73E-05 | 2.97E-06 | 0.00014 | 0 | 9.71E-05 | 3.52E-05 | 4.00E-05 |

| | | | | | | | | | | | | | | | | | | |
|---|---|---|---|---|---|---|---|---|---|---|---|---|---|---|---|---|---|---|
| Rap1a | 0.00026 | 0.00016 | 0.00026 | 0.00016 | 0.00018 | 0.00025 | 0.00027 | 8.83E-05 | 0.00017 | 0.00028 | 4.63E-05 | 0.00015 | 5.60E-05 | 0.00015 | 4.80E-05 | 0.00018 | 2.40E-05 | 0.0001 |
| Tmx3 | 0.00027 | 9.75E-05 | 0.00013 | 3.96E-06 | 4.04E-05 | 0.00015 | 1.21E-05 | 0.00023 | 0.00017 | 0.00011 | 9.93E-05 | 9.76E-05 | 5.90E-05 | 6.02E-05 | 9.46E-05 | 7.28E-05 | 1.73E-05 | 0.00014 |
| Ndfip1 | 0.0003 | 0.00031 | 0.00028 | 0.00051 | 0.00038 | 0.00056 | 0.00044 | 0.0008 | 0.00017 | 0.0005 | 0.0004 | 0.00067 | 0.00018 | 0.00065 | 0 | 0 | 0.00011 | 0.00053 |
| Prr5 | 0.0001 | 0 | 0.00028 | 0.00024 | 7.47E-05 | 3.66E-05 | 0.00019 | 0.00033 | 0.00017 | 0 | 0.00017 | 0 | 0 | 0.00013 | 3.06E-05 | 0 | 0 | 7.60E-05 |
| Nudt16l1 | 4.55E-05 | 0.00012 | 4.83E-05 | 0.00025 | 6.74E-05 | 0 | 0 | 8.18E-05 | 0.00017 | 0.00011 | 5.90E-05 | 5.13E-05 | 3.02E-05 | 6.64E-05 | 0.00012 | 4.69E-05 | 7.04E-06 | 0.00015 |
| Hadh | 0.00018 | 0.0001 | 0 | 9.07E-05 | 5.42E-05 | 6.33E-05 | 0 | 9.32E-05 | 0.00017 | 4.89E-05 | 1.15E-05 | 9.82E-05 | 0 | 0.00012 | 0 | 3.32E-06 | 3.13E-05 | 0.0002 |
| Lipo3 | 6.37E-05 | 0 | 0 | 0 | 8.95E-05 | 0.00013 | 0 | 4.81E-05 | 0.00017 | 0 | 9.27E-05 | 6.11E-05 | 0 | 2.84E-05 | 0 | 0 | 0 | 0 |
| Lrrc41 | 0.00012 | 0.0002 | 0.00025 | 0.00033 | 0.00012 | 2.25E-05 | 0.00013 | 5.04E-05 | 0.00017 | 0.00015 | 0.00016 | 7.88E-05 | 0.00015 | 7.31E-05 | 0.00012 | 0 | 0 | 0.00011 |
| Hnrnph2 | 0.00026 | 0.00012 | 0.00013 | 0.00029 | 8.59E-05 | 0.00012 | 9.41E-05 | 9.11E-05 | 0.00017 | 5.69E-05 | 4.93E-05 | 6.44E-05 | 0.00011 | 0.00017 | 4.78E-05 | 0.00022 | 0 | 3.12E-05 |
| Snx6 | 0.0002 | 0.00013 | 2.51E-05 | 1.48E-05 | 1.17E-05 | 3.34E-05 | 0.00013 | 4.74E-05 | 0.00017 | 0.00014 | 9.57E-05 | 0.00021 | 0 | 5.15E-05 | 5.17E-05 | 7.92E-05 | 0 | 9.70E-05 |
| Mapre1 | 0.00028 | 0.00018 | 0.00041 | 0.00021 | 0.00012 | 0.00014 | 0.00016 | 0.00019 | 0.00017 | 0.00018 | 0.00037 | 0.00018 | 0.00015 | 0.0001 | 0 | 0.00037 | 0.00061 | 0.00012 |
| Xxylt1 | 9.04E-05 | 0.00011 | 0.00013 | 4.12E-05 | 0.00021 | 0.00018 | 0 | 5.73E-05 | 0.00017 | 6.57E-05 | 4.85E-05 | 5.24E-05 | 4.45E-05 | 0.00028 | 0 | 9.07E-05 | 6.50E-05 | 0.00014 |
| Smarcc1 | 0.00028 | 0.00022 | 0.00024 | 0.00011 | 0.00012 | 5.70E-05 | 8.44E-05 | 1.19E-05 | 0.00017 | 0 | 0.0001 | 5.78E-05 | 9.03E-05 | 4.24E-05 | 0.00018 | 0.00034 | 6.67E-05 | 1.32E-06 |
| Npepps | 0.00023 | 0.00026 | 8.62E-05 | 9.07E-05 | 8.53E-05 | 7.87E-05 | 0.00016 | 1.64E-05 | 0.00017 | 9.78E-05 | 7.16E-05 | 0.00057 | 0.00016 | 0 | 5.11E-05 | 0.00013 | 0.00011 | 9.86E-05 |
| Fnta | 0.00019 | 0.00015 | 0.00019 | 0.00029 | 0.00013 | 0.00015 | 0.00014 | 0.0001 | 0.00017 | 0.00012 | 6.80E-05 | 0.00013 | 0 | 5.03E-05 | 7.14E-05 | 9.18E-05 | 7.15E-05 | 0.00017 |
| Park7 | 0.00033 | 0.00012 | 2.00E-05 | 0.00041 | 0.00011 | 8.53E-05 | 0.00026 | 0 | 0.00017 | 0.00021 | 0.00016 | 0.00014 | 0.0003 | 0.00017 | 0 | 0.00017 | 7.17E-05 | 0.00013 |
| Ski | 0.0001 | 0.00043 | 0.00017 | 0.00055 | 0.00046 | 0.00024 | 0.00026 | 0.00048 | 0.00017 | 0.00049 | 0.00028 | 2.45E-05 | 0.00039 | 0.00052 | 0.00067 | 0.00032 | 0.00068 | 0.00072 |
| Cggbp1 | 0.00027 | 0.00025 | 0.00029 | 0.00019 | 0.00019 | 0.00017 | 0.00024 | 0.00023 | 0.00017 | 0.00019 | 0.00024 | 0.00019 | 4.67E-05 | 0.00014 | 7.59E-05 | 0.00039 | 0.00044 | 0.00023 |
| Rab6a | 0.00058 | 0.00015 | 0.00033 | 0 | 7.40E-05 | 0.00029 | 0.00024 | 0.00014 | 0.00017 | 4.90E-05 | 0.00017 | 0.00041 | 1.17E-05 | 0 | 0 | 9.03E-06 | 0.0002 | 1.94E-05 |
| Slc38a2 | 0.00026 | 0.00046 | 7.00E-05 | 0.00062 | 0.00021 | 0.00021 | 0.00054 | 0.00013 | 0.00017 | 0.00011 | 0.00036 | 7.09E-05 | 0.0003 | 0.00032 | 0.00024 | 0.00027 | 0.00167 | 0.00012 |
| Kars | 0.0004 | 0.00014 | 0.00042 | 0.00036 | 9.51E-06 | 6.53E-05 | 5.85E-05 | 3.31E-05 | 0.00017 | 5.44E-05 | 0.0001 | 9.75E-05 | 0 | 0 | 0 | 0.00025 | 0.00011 | 3.15E-05 |
| Tfe3 | 0.00013 | 0.00022 | 0.00021 | 0.00012 | 0.0001 | 4.42E-05 | 5.24E-05 | 9.39E-05 | 0.00017 | 0.00011 | 5.31E-05 | 8.51E-05 | 2.28E-05 | 4.57E-05 | 0.00012 | 4.64E-05 | 1.69E-05 | 4.98E-05 |
| Trrap | 0.00015 | 0.00018 | 0.00015 | 0.00014 | 9.84E-05 | 0.00012 | 8.68E-05 | 0.00012 | 0.00017 | 4.17E-05 | 5.75E-05 | 0.0001 | 1.43E-05 | 5.25E-05 | 0.0003 | 9.46E-05 | 0.0003 | 4.41E-05 |
| Selenoi | 0.00015 | 5.69E-05 | 0.00027 | 0.00011 | 7.64E-05 | 0.00011 | 0.00013 | 1.77E-05 | 0.00017 | 8.21E-05 | 0.00021 | 0.0002 | 4.88E-05 | 3.40E-05 | 3.22E-05 | 6.28E-05 | 4.81E-05 | 0 |
| Nr3c1 | 0.00011 | 0.00023 | 0 | 7.26E-05 | 7.63E-05 | 0.00038 | 0 | 7.28E-05 | 0.00017 | 0.00018 | 0 | 0.00029 | 0.00013 | 0.00012 | 3.67E-05 | 0.00014 | 0.00026 | 0.00018 |
| Ubr4 | 0.0002 | 0.00027 | 0.00015 | 4.41E-05 | 0.00014 | 0.00014 | 0.0002 | 0.00013 | 0.00017 | 0.00013 | 0.00011 | 0.0002 | 7.36E-05 | 9.35E-05 | 0.00024 | 9.60E-05 | 0.00018 | 0.00014 |
| Traf3ip2 | 0.0001 | 7.76E-05 | 3.15E-05 | 7.17E-05 | 7.41E-06 | 8.11E-05 | 0 | 0 | 0.00017 | 9.79E-05 | 0 | 1.85E-05 | 0.00028 | 0.00013 | 0.00017 | 4.21E-05 | 0 | 0 |
| Irf2bpl | 2.35E-05 | 0.00025 | 8.27E-05 | 0.00047 | 0.00035 | 9.73E-05 | 0.00015 | 0.0001 | 0.00017 | 7.52E-05 | 0.0002 | 3.34E-05 | 0.00012 | 0.00027 | 0.0004 | 0.00016 | 0 | 0.0002 |
| Hmgn1 | 0.0002 | 8.72E-05 | 0.00011 | 0.00024 | 9.14E-05 | 3.35E-05 | 0 | 6.77E-05 | 0.00017 | 0 | 0.00017 | 2.61E-05 | 0 | 1.35E-05 | 0.00011 | 0.00023 | 4.98E-05 | 5.74E-05 |
| Rin1 | 0 | 6.38E-05 | 3.45E-05 | 2.35E-05 | 5.09E-05 | 8.18E-05 | 0 | 0.0002 | 0.00017 | 0 | 0.00011 | 0.00024 | 0 | 0.00026 | 0.00017 | 0 | 0 | 3.20E-05 |
| Setd5 | 0.00026 | 0.00026 | 0.00021 | 0.00014 | 0.00017 | 0.00018 | 0.00019 | 8.06E-05 | 0.00017 | 9.67E-05 | 0.00015 | 0.00033 | 0.0001 | 0.00014 | 0.00035 | 5.85E-05 | 0.00026 | 3.10E-05 |
| Qdpr | 0.00011 | 0.00014 | 7.01E-05 | 0.0002 | 8.59E-05 | 4.42E-05 | 0 | 0.00017 | 0.00017 | 0.0001 | 1.41E-05 | 0.00013 | 9.36E-05 | 0 | 0 | 0.00012 | 0 | 0.00013 |
| Rhou | 1.43E-05 | 0.00018 | 0.00018 | 0.00013 | 4.08E-05 | 0.00012 | 0.00043 | 4.37E-05 | 0.00017 | 0.00053 | 0.00022 | 2.13E-05 | 8.67E-05 | 5.03E-05 | 7.20E-05 | 4.48E-05 | 0.0001 | 0.00022 |

| | | | | | | | | | | | | | | | | | | |
|---|---|---|---|---|---|---|---|---|---|---|---|---|---|---|---|---|---|---|
| Hoxd8 | 0.00011 | 7.12E-06 | 0 | 0 | 9.09E-05 | 8.62E-05 | 6.55E-05 | 0 | 0.00017 | 3.12E-05 | 0.00016 | 0 | 0 | 0.00024 | 0 | 4.51E-07 | 0 | 0.00027 |
| Eri3 | 0.00021 | 0.00013 | 0.0001 | 0.0001 | 0.00012 | 0.00012 | 0.00027 | 0.00011 | 0.00017 | 9.43E-05 | 7.77E-05 | 4.07E-05 | 8.59E-05 | 6.38E-05 | 0.00022 | 4.96E-05 | 4.41E-05 | 0.0001 |
| Ago1 | 3.73E-05 | 0.00022 | 4.69E-05 | 4.87E-05 | 0.00012 | 0.00012 | 0 | 7.09E-05 | 0.00017 | 9.65E-05 | 0.0001 | 8.97E-05 | 0 | 4.63E-05 | 0 | 9.70E-06 | 0 | 0.0001 |
| Epb41l3 | 0.00015 | 6.18E-05 | 0.00014 | 5.09E-05 | 0.00017 | 0.00046 | 0.00044 | 0 | 0.00017 | 0.00014 | 7.23E-05 | 0.00019 | 0 | 0.00041 | 0.00015 | 0.00013 | 0.00019 | 3.72E-05 |
| Srcap | 8.64E-05 | 0.00027 | 0.00011 | 0.00033 | 0.00013 | 7.19E-05 | 5.79E-05 | 0.00017 | 0.00017 | 0.0001 | 0.00012 | 0.0001 | 7.05E-05 | 9.12E-05 | 0.00055 | 0.00024 | 0.00034 | 0.00012 |
| Mme | 0 | 1.72E-05 | 0 | 0 | 0 | 9.97E-05 | 0 | 0 | 0.00017 | 0 | 0 | 0 | 3.26E-06 | 7.91E-05 | 0 | 0 | 0 | 0.00014 |
| Atp6v1g1 | 0.00023 | 0.00039 | 0.00024 | 0.00023 | 0.00027 | 0.00027 | 0.00017 | 0.00025 | 0.00017 | 5.72E-05 | 0.0003 | 0.00031 | 0.00012 | 0.00023 | 1.08E-05 | 0.00014 | 0 | 0.00016 |
| Eif4a2 | 0.00043 | 0.00045 | 0.00014 | 0 | 0.00022 | 0.0003 | 7.67E-05 | 1.20E-05 | 0.00017 | 0.00015 | 4.73E-05 | 0.00045 | 1.43E-05 | 0.00011 | 0.00073 | 0.00012 | 0.00058 | 0.00023 |
| Rbck1 | 0.00011 | 0.00015 | 6.87E-05 | 0.00015 | 7.49E-05 | 2.15E-05 | 0 | 4.31E-05 | 0.00017 | 8.66E-05 | 4.15E-05 | 8.56E-05 | 0 | 2.82E-05 | 0.00014 | 8.53E-05 | 1.33E-05 | 0.00016 |
| Uqcr10 | 0.0002 | 4.12E-05 | 0.00016 | 0.00017 | 7.87E-05 | 9.66E-05 | 9.98E-05 | 2.88E-05 | 0.00017 | 0 | 6.94E-05 | 0 | 0 | 3.84E-05 | 0 | 0.00017 | 0 | 3.90E-06 |
| Gsto1 | 0.00018 | 0.00013 | 0.00047 | 0.00021 | 0.00036 | 0.00054 | 0 | 0.00019 | 0.00017 | 0 | 0.0004 | 4.30E-05 | 0.00062 | 0.00078 | 0 | 0 | 0.00083 | 0.00015 |
| Sppl3 | 8.87E-05 | 0.00027 | 3.73E-05 | 8.55E-05 | 0.00022 | 0.00022 | 0.00018 | 0.00019 | 0.00017 | 0.00016 | 0.00015 | 0.00015 | 7.70E-05 | 0.0002 | 0 | 0.00017 | 0.00026 | 0.00012 |
| Gatd3a | 8.56E-05 | 0.0001 | 0 | 0.00019 | 0.0001 | 2.18E-05 | 1.37E-05 | 7.53E-05 | 0.00017 | 7.10E-05 | 4.45E-05 | 0.00014 | 6.26E-05 | 6.60E-05 | 2.35E-05 | 0 | 0 | 9.37E-05 |
| Lrfn4 | 0 | 5.37E-05 | 0.00013 | 0.00019 | 8.52E-05 | 2.08E-05 | 1.38E-05 | 3.14E-05 | 0.00017 | 0 | 9.54E-05 | 0 | 7.33E-05 | 0.00016 | 0.0001 | 7.96E-05 | 1.52E-05 | 1.99E-05 |
| Sec24c | 0.00018 | 0.00022 | 0.00018 | 0.00025 | 0.00011 | 0.00013 | 0 | 0.00018 | 0.00017 | 0.00015 | 5.14E-05 | 0.00018 | 0 | 8.98E-05 | 0.00021 | 0.00015 | 0.00015 | 4.16E-05 |
| Kctd11 | 0 | 7.03E-05 | 0.0006 | 0.00041 | 0.00027 | 1.34E-05 | 0.00018 | 0.00048 | 0.00017 | 6.43E-05 | 0.0002 | 0.0009 | 2.30E-05 | 0.00042 | 0.00048 | 0 | 0.00034 | 0.00023 |
| Myl9 | 0 | 0 | 0 | 0 | 0.00042 | 0 | 0 | 0 | 0.00017 | 3.46E-05 | 0.00274 | 0 | 0 | 0.00065 | 0 | 0 | 0 | 0 |
| Adnp | 0.00021 | 0.00021 | 0.00016 | 4.13E-05 | 0.00016 | 0.00018 | 9.81E-05 | 9.86E-05 | 0.00017 | 0.00011 | 0.00017 | 0.00011 | 5.32E-05 | 9.83E-05 | 0.00014 | 0.00021 | 0.00022 | 0.00012 |
| Dctn4 | 0.00022 | 0.00022 | 0.00017 | 0.00016 | 0.00014 | 8.61E-05 | 0.00013 | 8.28E-05 | 0.00017 | 8.22E-05 | 9.10E-05 | 0.00015 | 3.28E-05 | 5.85E-05 | 0.00024 | 8.64E-05 | 0 | 0.00018 |
| Pam | 0.00016 | 0.0001 | 0.0001 | 0 | 0.00011 | 9.53E-05 | 0.00012 | 3.04E-05 | 0.00017 | 0.00077 | 0 | 0.0001 | 0.0011 | 8.89E-05 | 0 | 7.51E-05 | 0 | 0.00052 |
| Bcar1 | 0 | 0.00016 | 6.51E-05 | 0.00016 | 0.00019 | 0.00012 | 0 | 1.67E-05 | 0.00017 | 0 | 0.0006 | 0.00061 | 9.66E-05 | 0.00048 | 0.00024 | 9.22E-05 | 0 | 0.00018 |
| Arhgap42 | 8.29E-05 | 0.00011 | 9.55E-05 | 7.51E-05 | 0.00016 | 0.00051 | 0 | 0.00044 | 0.00017 | 0.00032 | 0.00018 | 0 | 0 | 0.00053 | 0.00075 | 0 | 0.00015 | 1.17E-05 |
| Crlf1 | 0 | 0 | 0.00044 | 0 | 0.00028 | 0 | 0 | 0.00046 | 0.00017 | 0 | 0.00023 | 0 | 0 | 0 | 0.00024 | 0 | 0 | 0.00061 |
| Cand1 | 0.0004 | 0.0002 | 0.0003 | 6.42E-05 | 0.00011 | 0.00014 | 0.00018 | 0 | 0.00017 | 0.0001 | 0.00012 | 0.00015 | 8.09E-05 | 5.37E-05 | 5.73E-05 | 0.00026 | 0.00013 | 8.67E-05 |
| Pdlim7 | 2.13E-05 | 0.00018 | 0.00039 | 0.00037 | 8.77E-05 | 0 | 0.00018 | 0.00029 | 0.00017 | 0.00021 | 0.00089 | 0.00034 | 0 | 0.00012 | 0.0008 | 0.00021 | 0.00038 | 0.00013 |
| Kifap3 | 0.00018 | 0.00013 | 1.86E-05 | 1.15E-05 | 9.72E-05 | 7.36E-05 | 8.20E-05 | 0.00014 | 0.00017 | 0.00011 | 5.17E-05 | 3.28E-05 | 0 | 8.72E-05 | 9.96E-05 | 7.67E-05 | 0.0001 | 0.00017 |
| Naca | 0.0006 | 0.00026 | 0.00032 | 0.00018 | 0 | 0.00031 | 9.12E-05 | 0 | 0.00017 | 7.50E-06 | 9.09E-05 | 0.00036 | 0 | 4.09E-05 | 0.00011 | 0.0002 | 0.00021 | 0 |
| Ubqln2 | 0.00037 | 0.00019 | 0.00019 | 0.00019 | 0.0001 | 0.00016 | 0.00011 | 4.55E-05 | 0.00017 | 0 | 0.00014 | 8.56E-05 | 4.74E-05 | 0.00018 | 0 | 0.00021 | 0.00029 | 0.00016 |
| Tm7sf3 | 0.00014 | 0.00011 | 0.00012 | 0.00034 | 0.00022 | 0.00034 | 0.00062 | 0.00033 | 0.00017 | 8.50E-05 | 8.91E-05 | 0 | 0.00016 | 0.00036 | 1.92E-05 | 0 | 0 | 0.00019 |
| Slc35a4 | 0.00017 | 0.00013 | 0.00032 | 0.0003 | 0.00016 | 0.0001 | 9.65E-05 | 8.57E-05 | 0.00017 | 8.17E-05 | 0.00018 | 2.59E-05 | 9.35E-05 | 9.47E-05 | 5.37E-05 | 0.00024 | 0 | 4.95E-05 |
| Ube2g2 | 9.95E-05 | 0.00017 | 0.00044 | 0.00032 | 0.00019 | 0.00014 | 0.00029 | 0.0001 | 0.00017 | 0.00024 | 0.00027 | 0.00012 | 0 | 0.00011 | 3.01E-05 | 0.00014 | 4.52E-05 | 0.00013 |
| Hk1 | 2.47E-05 | 0.00014 | 5.75E-05 | 0.0003 | 0.00012 | 0 | 4.25E-05 | 0.00028 | 0.00017 | 0.00023 | 0.00021 | 0.00109 | 0.00016 | 0.00015 | 0.00012 | 1.60E-05 | 3.36E-05 | 0.00026 |
| Itgav | 0.00037 | 0.00016 | 0.00027 | 0 | 0.00026 | 0.00039 | 1.42E-05 | 0.0006 | 0.00017 | 0.00017 | 0.00055 | 0.00051 | 3.60E-05 | 0.00058 | 7.27E-05 | 0.00014 | 0.00011 | 0.00012 |

| | | | | | | | | | | | | | | | | | | |
|---|---|---|---|---|---|---|---|---|---|---|---|---|---|---|---|---|---|---|
| Dctn1 | 0.00011 | 0.00023 | 0.00012 | 0.00017 | 9.85E-05 | 9.02E-05 | 0.00013 | 0.00013 | 0.00017 | 0.00013 | 0.00019 | 0.00019 | 0 | 0.00012 | 0.00028 | 0.00018 | 2.33E-05 | 0.00019 |
| Limk1 | 9.26E-05 | 4.80E-05 | 0.0003 | 0.00024 | 3.61E-06 | 0.00013 | 0.00012 | 0.00018 | 0.00017 | 0.00028 | 8.17E-06 | 0.00025 | 6.33E-05 | 8.09E-06 | 0 | 0 | 0 | 0.00027 |
| Ctps2 | 0.00017 | 0.0002 | 0 | 0 | 9.17E-05 | 0.00019 | 0.00013 | 0 | 0.00017 | 0.00011 | 6.87E-05 | 3.43E-06 | 0.00031 | 0.00011 | 4.42E-05 | 0.00016 | 0 | 0.00011 |
| Rab7 | 0.00016 | 0.00023 | 0.00021 | 0.00018 | 0.00026 | 0.00024 | 0.00011 | 0.00014 | 0.00017 | 0.00013 | 0.00034 | 0.00012 | 0.00013 | 0.00023 | 0 | 2.86E-05 | 0.00025 | 0.00012 |
| Kdelr1 | 0.00073 | 0.00016 | 0.00029 | 0.00041 | 0.00032 | 0.00067 | 0.00102 | 0.00055 | 0.00017 | 0.0003 | 0.00023 | 0.0004 | 0.00033 | 0.00038 | 0 | 0.0002 | 0.00024 | 0.00032 |
| Serinc3 | 0.00027 | 0.00055 | 0 | 0.00041 | 0.00071 | 0.00073 | 0.00065 | 0.00072 | 0.00017 | 0.00019 | 0.00031 | 0.00045 | 0.00038 | 0.00057 | 8.87E-05 | 4.92E-05 | 0.00058 | 0.00032 |
| Rcor1 | 0.00016 | 0.00012 | 0.00015 | 1.03E-05 | 6.36E-05 | 7.61E-05 | 4.98E-05 | 0.00011 | 0.00017 | 1.70E-05 | 8.26E-05 | 0.00024 | 4.33E-06 | 3.74E-05 | 0.00027 | 9.03E-05 | 0.00017 | 1.68E-05 |
| Spats2 | 6.90E-05 | 0.00011 | 0.00013 | 0.00026 | 0.0003 | 6.72E-05 | 7.05E-05 | 0.00022 | 0.00017 | 4.37E-05 | 0.00021 | 0.00026 | 0 | 5.36E-05 | 0.00013 | 0.00016 | 0.00012 | 0 |
| Arfip1 | 0.00025 | 0.00013 | 0.00018 | 0.00018 | 0.00013 | 0.00011 | 0.00023 | 0.00018 | 0.00017 | 5.20E-05 | 0.00019 | 0.00022 | 2.06E-05 | 0.00012 | 0.00031 | 0 | 0.00017 | 0.00015 |
| Ech1 | 0.00029 | 0.00018 | 0 | 0.00014 | 6.32E-05 | 0.00013 | 7.34E-05 | 9.77E-05 | 0.00017 | 0.00023 | 0 | 8.81E-05 | 0.00012 | 5.09E-05 | 1.92E-05 | 0 | 0.00015 | 0.00033 |
| Dynlt3 | 0.00023 | 6.93E-05 | 7.67E-05 | 6.85E-05 | 0.00016 | 0.0002 | 0.00019 | 0.00025 | 0.00017 | 2.67E-05 | 0.00028 | 0.00012 | 0 | 0 | 0 | 0 | 0 | 9.77E-05 |
| Cuedc2 | 9.26E-05 | 0.00017 | 2.27E-05 | 0.00018 | 0.00017 | 0.00012 | 7.91E-05 | 0.00013 | 0.00017 | 7.42E-05 | 0.00016 | 0.00011 | 8.35E-05 | 0.0002 | 7.60E-05 | 6.55E-05 | 0 | 0.00014 |
| Ppic | 0 | 0.00012 | 0.00013 | 0.00062 | 0.00144 | 0.00113 | 0.0012 | 0.00117 | 0.00017 | 0.00079 | 0.00088 | 0.00033 | 0.00113 | 0.00113 | 0.00016 | 0.00019 | 0 | 0.00068 |
| Eif4e2 | 0.00038 | 0.00018 | 0.00032 | 0.00024 | 0.00017 | 0.0002 | 0.00036 | 0.00011 | 0.00017 | 0.00014 | 0.00026 | 0.00016 | 3.53E-06 | 0.00013 | 0.0002 | 0.00011 | 0 | 6.07E-05 |
| Elk3 | 0.00017 | 0.00016 | 0.00023 | 0 | 0.00052 | 0.00018 | 0.00027 | 8.31E-05 | 0.00017 | 6.49E-05 | 0.0003 | 0.00018 | 0.00026 | 0.00034 | 0.00016 | 3.45E-05 | 0 | 0.00011 |
| Senp3 | 0.00013 | 0.00011 | 0.00016 | 0.00011 | 5.29E-05 | 2.00E-05 | 1.16E-05 | 2.79E-05 | 0.00017 | 3.29E-05 | 0.0001 | 0 | 0 | 7.51E-05 | 0.00016 | 7.19E-05 | 7.30E-05 | 2.47E-05 |
| Scamp2 | 0.0001 | 0.00011 | 9.44E-05 | 0.00013 | 0.00011 | 7.29E-05 | 0.00014 | 0.00015 | 0.00017 | 6.58E-05 | 6.96E-05 | 4.58E-06 | 0.00011 | 7.02E-05 | 0 | 0 | 0 | 7.84E-05 |
| Ddb1 | 0.00043 | 0.0001 | 0.00017 | 6.81E-05 | 0.00011 | 0.00024 | 0.00027 | 6.94E-05 | 0.00017 | 2.63E-05 | 0.00028 | 0.00014 | 4.31E-05 | 7.70E-05 | 2.60E-05 | 3.93E-05 | 4.30E-05 | 4.00E-05 |
| Myo5a | 0.00012 | 0.00018 | 0.00021 | 0 | 0.00012 | 0 | 0.00027 | 0.00018 | 0.00017 | 0 | 7.27E-05 | 8.05E-05 | 0.00012 | 0.00012 | 0.00016 | 0.00019 | 7.87E-05 | 0.00017 |
| Drg1 | 0.00016 | 0.00021 | 0.00021 | 0.00035 | 8.96E-05 | 5.95E-05 | 8.91E-06 | 0.00013 | 0.00017 | 0.00012 | 0.00018 | 0.00017 | 3.64E-05 | 0.00013 | 5.73E-05 | 0.00034 | 0 | 0.00018 |
| Srp14 | 0.00011 | 0.00013 | 0.0001 | 0.00029 | 0.00014 | 0.00014 | 7.74E-05 | 0.00015 | 0.00017 | 0.00011 | 0.00015 | 8.52E-05 | 0.00011 | 3.95E-05 | 7.80E-06 | 1.56E-05 | 6.07E-05 | 0.00012 |
| Npepl1 | 6.52E-05 | 0.00019 | 0.00011 | 0.00014 | 0.00011 | 8.23E-05 | 0.00012 | 0 | 0.00017 | 0.00028 | 0.00012 | 5.04E-05 | 0 | 2.95E-05 | 0.00026 | 7.39E-05 | 9.21E-05 | 7.21E-05 |
| Ap2s1 | 0.0002 | 9.31E-05 | 0.00016 | 0.00044 | 0.00022 | 0.00012 | 0.00012 | 0.00028 | 0.00017 | 6.46E-05 | 0.00014 | 2.61E-05 | 5.01E-05 | 0.00031 | 6.81E-05 | 0.00015 | 1.48E-05 | 0.00012 |
| H3f3a | 3.40E-05 | 0.00034 | 4.60E-05 | 0.00101 | 0.00026 | 0.00021 | 0.00019 | 0.0002 | 0.00017 | 0.0002 | 0.00025 | 0.00013 | 0.00012 | 0.0004 | 8.13E-05 | 0.00034 | 0.00058 | 0.00021 |
| Hs6st2 | 4.28E-06 | 0 | 9.01E-05 | 0 | 0.00017 | 2.86E-05 | 3.18E-06 | 1.11E-05 | 0.00017 | 0 | 0.0016 | 0.00035 | 5.56E-05 | 0.00012 | 1.29E-05 | 0.00026 | 0 | 0 |
| Cxxc5 | 6.17E-05 | 0.00022 | 0.00032 | 0.00014 | 0.00024 | 0.00019 | 0.00015 | 0.00039 | 0.00017 | 0.0002 | 6.87E-05 | 0.00049 | 0.00011 | 5.37E-05 | 0.00024 | 0.00012 | 0 | 0.00019 |
| Sppl2b | 9.07E-07 | 8.41E-05 | 0.00012 | 0.00022 | 0.00011 | 0 | 5.24E-06 | 8.61E-05 | 0.00017 | 6.15E-05 | 1.04E-05 | 0 | 1.23E-05 | 9.23E-05 | 6.51E-05 | 1.82E-05 | 0 | 0.00011 |
| Eif4e3 | 9.86E-05 | 0.00011 | 2.08E-05 | 0 | 0 | 0.00018 | 0 | 0.0002 | 0.00017 | 6.00E-05 | 0 | 7.26E-05 | 0.00026 | 0.0002 | 0.00012 | 2.75E-06 | 4.26E-06 | 0.00024 |
| Dlg1 | 9.58E-05 | 0.00015 | 0.0002 | 3.35E-05 | 0.00014 | 0.00021 | 0.00016 | 6.40E-05 | 0.00017 | 7.16E-05 | 0.00017 | 8.83E-05 | 8.65E-05 | 0.0002 | 0.00018 | 5.51E-05 | 0.00024 | 0.00013 |
| Ppard | 0.00012 | 0.00017 | 0.00013 | 0 | 0.0001 | 0.00015 | 0 | 0.00019 | 0.00017 | 1.76E-05 | 4.71E-05 | 0.00021 | 5.47E-05 | 0.00024 | 3.42E-05 | 3.31E-05 | 9.27E-06 | 0 |
| Brk1 | 0.00029 | 0.00028 | 0.00029 | 0.00043 | 0.00027 | 0.00029 | 0.00033 | 0.00017 | 0.00017 | 0.00026 | 0.00028 | 0.00029 | 0.00019 | 0.00019 | 0 | 0.00026 | 0 | 0.00023 |
| Map2k1 | 9.23E-05 | 7.66E-05 | 0.00032 | 0.00011 | 0.00024 | 9.08E-05 | 0 | 0.00012 | 0.00017 | 9.85E-05 | 0.00021 | 0.00096 | 0.0004 | 0.00016 | 1.61E-05 | 4.22E-05 | 8.05E-05 | 0 |
| Supt20 | 0.00011 | 0.0002 | 9.96E-05 | 0.00022 | 1.02E-05 | 2.92E-05 | 9.47E-05 | 3.93E-05 | 0.00017 | 5.32E-05 | 9.92E-05 | 0.00012 | 0 | 4.25E-05 | 0.00047 | 0.00017 | 0.00011 | 6.64E-05 |

| | | | | | | | | | | | | | | | | | | |
|---|---|---|---|---|---|---|---|---|---|---|---|---|---|---|---|---|---|---|
| Naxe | 0.00026 | 9.55E-05 | 3.03E-05 | 6.48E-05 | 0.00012 | 0.00014 | 0.00014 | 3.38E-05 | 0.00017 | 8.33E-05 | 3.41E-05 | 0.0001 | 5.46E-05 | 6.52E-05 | 0 | 7.06E-05 | 0 | 0.00012 |
| Ppp1r15b | 0.00013 | 0.00015 | 0.00016 | 0.00011 | 6.69E-06 | 7.17E-05 | 1.75E-05 | 8.25E-05 | 0.00017 | 0.00015 | 9.38E-05 | 0.00017 | 0.00014 | 8.08E-05 | 0.0001 | 7.76E-05 | 0.00039 | 0 |
| Zfyve21 | 7.64E-05 | 7.51E-05 | 4.60E-05 | 0.00023 | 9.08E-05 | 2.80E-05 | 0.00019 | 0.00011 | 0.00017 | 0.00016 | 0.00011 | 0.0003 | 0 | 6.29E-05 | 6.22E-05 | 0 | 0 | 0.00019 |
| Naa50 | 0.00036 | 0.00012 | 0.00051 | 0.00032 | 7.91E-05 | 7.49E-05 | 0.00013 | 5.37E-05 | 0.00017 | 0 | 3.80E-05 | 0.00013 | 0 | 5.46E-05 | 7.46E-05 | 0.0005 | 0.00025 | 6.53E-05 |
| Arpc1a | 0.0002 | 9.08E-05 | 0.00011 | 5.95E-06 | 8.36E-05 | 0.00016 | 0.00012 | 0.00019 | 0.00017 | 9.82E-05 | 0.00012 | 0.00014 | 0.00012 | 9.68E-05 | 0 | 0.0001 | 0 | 2.73E-05 |
| Erf | 0.00021 | 0.00026 | 0.00018 | 8.79E-05 | 4.90E-05 | 0 | 0 | 7.46E-05 | 0.00017 | 0.0001 | 0.00013 | 4.18E-05 | 0.00012 | 3.62E-05 | 0 | 0.00022 | 0.00123 | 8.11E-05 |
| Psme2 | 0.00027 | 9.18E-05 | 8.66E-05 | 6.86E-05 | 5.50E-05 | 0 | 1.52E-05 | 2.47E-05 | 0.00017 | 5.20E-05 | 7.29E-05 | 0 | 2.45E-05 | 5.87E-06 | 0 | 5.66E-05 | 0 | 4.27E-05 |
| Abcc9 | 0 | 2.37E-05 | 0 | 0 | 0 | 1.81E-05 | 0.00022 | 0 | 0.00017 | 1.28E-05 | 0 | 0.00013 | 0.00028 | 0 | 0 | 0 | 0 | 0.00042 |
| Mfsd5 | 0.00012 | 0.00018 | 0.00026 | 0.00017 | 0.00016 | 0.00012 | 0.00018 | 0.00017 | 0.00017 | 0.00015 | 0.00012 | 6.79E-05 | 7.87E-05 | 8.17E-05 | 0 | 2.91E-07 | 0 | 0.00015 |
| Cbx5 | 6.48E-05 | 3.43E-05 | 9.76E-05 | 0 | 6.45E-05 | 7.26E-05 | 5.36E-05 | 0.00013 | 0.00017 | 0.00015 | 5.59E-05 | 2.99E-05 | 0 | 6.39E-05 | 6.30E-05 | 0.00077 | 3.36E-05 | 0.00025 |
| Stk25 | 0.00018 | 0.00022 | 0.00011 | 0.0003 | 0.0002 | 6.30E-05 | 0.00032 | 0.00035 | 0.00017 | 0.0002 | 0.00017 | 0.00023 | 4.75E-06 | 0.00018 | 0.00055 | 4.64E-05 | 0.00018 | 0.00016 |
| Poglut3 | 0.00024 | 0.0001 | 0 | 0 | 0.00011 | 0.0001 | 0.0002 | 7.24E-05 | 0.00017 | 0.00032 | 2.26E-07 | 0 | 2.81E-05 | 0 | 0 | 2.41E-05 | 1.70E-05 | 0.00018 |
| Rnasek | 0.00028 | 0.00026 | 0.00036 | 0.00058 | 0.00029 | 0.00033 | 0.00037 | 0.00037 | 0.00017 | 0.00025 | 0.00031 | 9.49E-05 | 0.00012 | 0.00024 | 0 | 7.78E-05 | 8.70E-05 | 0.00015 |
| Birc3 | 0.00013 | 0.00012 | 0.00018 | 3.08E-05 | 2.94E-05 | 7.95E-05 | 1.12E-05 | 0 | 0.00017 | 5.26E-05 | 0 | 9.05E-05 | 0.00037 | 0.00017 | 1.78E-05 | 3.04E-05 | 0.00049 | 0.00014 |
| Far1 | 0.00024 | 0.00014 | 0.00035 | 0.00011 | 6.52E-05 | 0.00015 | 0.00013 | 0.00024 | 0.00017 | 0.00017 | 0.00026 | 0.00015 | 0 | 0.00021 | 0.00037 | 0.00011 | 0.00016 | 0.00018 |
| Sephs1 | 0.00015 | 9.95E-05 | 8.96E-05 | 7.10E-05 | 9.91E-05 | 6.50E-05 | 0.00011 | 5.27E-05 | 0.00017 | 5.39E-05 | 3.64E-05 | 0 | 2.83E-05 | 6.66E-05 | 0 | 0.00047 | 5.25E-05 | 0.0001 |
| Cyc1 | 0.00023 | 0.00018 | 0.0004 | 0.00033 | 7.83E-05 | 8.57E-05 | 2.29E-05 | 5.67E-05 | 0.00017 | 3.43E-05 | 0.00021 | 2.70E-06 | 0 | 0.00017 | 3.95E-05 | 0.00027 | 0 | 5.27E-05 |
| Ndufb11 | 0.0004 | 0.00015 | 0.00011 | 0.00036 | 0.00015 | 0.00027 | 0.00024 | 0.00016 | 0.00017 | 0.0001 | 5.44E-05 | 9.95E-05 | 0.00013 | 0.00017 | 0 | 0.00018 | 0.00011 | 9.41E-05 |
| Dennd1a | 5.44E-05 | 0.00016 | 0 | 0 | 0.0001 | 0.00012 | 0 | 2.99E-05 | 0.00017 | 0.00012 | 4.16E-05 | 6.71E-05 | 0.00013 | 0.0001 | 0 | 5.88E-05 | 0.00013 | 2.72E-06 |
| Atp5e | 0.0003 | 4.82E-05 | 0.00011 | 0.00065 | 0.00019 | 0.00027 | 0.00037 | 0.00021 | 0.00017 | 8.91E-05 | 0.0002 | 0 | 0.00024 | 0.00025 | 0 | 0.0001 | 0 | 0.00014 |
| Ampd3 | 0 | 9.22E-05 | 0 | 0 | 0.00016 | 7.02E-05 | 7.53E-05 | 0 | 0.00017 | 0.00044 | 0 | 0.00035 | 0.00068 | 0 | 3.49E-05 | 0 | 0 | 0 |
| Ubqln4 | 0.00016 | 0.00012 | 0.0002 | 0.00013 | 4.21E-05 | 4.43E-05 | 5.84E-05 | 0 | 0.00017 | 0 | 7.28E-05 | 3.34E-05 | 0 | 4.29E-05 | 0.00022 | 0.00015 | 0.00018 | 0 |
| Ncl | 0.00042 | 0.00016 | 0.00046 | 0.00017 | 4.24E-05 | 4.67E-05 | 0 | 0 | 0.00017 | 0 | 4.42E-05 | 0.00026 | 0 | 1.70E-05 | 0.00023 | 0.00032 | 0.00035 | 1.13E-05 |
| Patz1 | 2.01E-05 | 0.00018 | 3.34E-05 | 0.00018 | 0.00013 | 3.45E-05 | 0 | 5.08E-05 | 0.00017 | 3.92E-06 | 7.14E-07 | 6.93E-05 | 8.27E-06 | 0.00011 | 0.00013 | 3.20E-05 | 0 | 3.81E-05 |
| Usp22 | 0.00015 | 0.00024 | 0.00018 | 0.00019 | 9.65E-05 | 1.81E-05 | 9.13E-05 | 0.00011 | 0.00017 | 7.11E-05 | 0.00031 | 0.00031 | 2.81E-05 | 0.00015 | 2.53E-05 | 6.90E-05 | 0 | 0.00027 |
| Rarb | 9.67E-06 | 1.27E-05 | 0 | 1.04E-05 | 7.97E-05 | 0 | 1.68E-05 | 0.00017 | 0.00017 | 1.87E-05 | 0 | 8.82E-05 | 0 | 0 | 0 | 2.10E-05 | 0 | 1.97E-06 |
| Psmc4 | 0.00025 | 0.00016 | 0.00035 | 0.00015 | 0.0001 | 4.79E-05 | 0 | 2.52E-05 | 0.00017 | 6.38E-05 | 0.00013 | 0 | 0 | 5.82E-05 | 7.14E-05 | 0.00023 | 0 | 5.12E-05 |
| Stk24 | 0.00017 | 0.00022 | 0.00011 | 8.03E-05 | 0.00015 | 0.00023 | 2.92E-05 | 0 | 0.00017 | 0.00017 | 0.00011 | 0.00062 | 0.00017 | 0.00015 | 0.00027 | 0.00019 | 0.00023 | 7.24E-05 |
| Btg3 | 0.00028 | 0 | 0.00031 | 1.85E-06 | 0.00012 | 0.00033 | 0.00021 | 0.00015 | 0.00017 | 0 | 0.00025 | 8.31E-05 | 4.92E-05 | 0.00018 | 0 | 4.18E-05 | 0.00041 | 0 |
| Flywch1 | 1.12E-05 | 0.00016 | 5.74E-05 | 0.00014 | 6.86E-05 | 0 | 1.06E-05 | 5.34E-05 | 0.00017 | 0.00011 | 0.0001 | 0.00019 | 4.32E-05 | 9.96E-05 | 0.00014 | 0.00011 | 1.03E-05 | 0.00012 |
| Meis1 | 7.29E-05 | 0.0002 | 0 | 9.31E-05 | 0.00012 | 0.00026 | 0 | 0 | 0.00017 | 3.29E-05 | 0.00012 | 6.76E-05 | 1.62E-05 | 0.00018 | 0.00043 | 0.00012 | 7.76E-05 | 0 |
| Slc35b2 | 0.00013 | 6.81E-05 | 0.00015 | 0.00049 | 0.00019 | 0.00018 | 0.00033 | 0.00019 | 0.00017 | 0.00011 | 0.00012 | 2.11E-06 | 0.00014 | 0.00024 | 6.97E-05 | 0.00022 | 0.0003 | 0.0002 |
| Ubc | 0.00019 | 0.00035 | 0.00011 | 0.00037 | 0.00017 | 0.00032 | 0.00033 | 0.00029 | 0.00017 | 0.00024 | 0.00033 | 0.00041 | 0.00025 | 0.00026 | 0.00023 | 0.00019 | 0.00441 | 0.00042 |

| | | | | | | | | | | | | | | | | | | |
|---|---|---|---|---|---|---|---|---|---|---|---|---|---|---|---|---|---|---|
| Xpo1 | 0.00035 | 8.44E-05 | 0.00014 | 0 | 1.51E-05 | 0.00014 | 0 | 0 | 0.00017 | 1.26E-05 | 1.74E-05 | 0.00018 | 0 | 0 | 0.00014 | 0.00055 | 6.01E-05 | 1.67E-05 |
| Ubxn4 | 0.00034 | 0.00026 | 0.0003 | 9.14E-05 | 0.00014 | 7.57E-05 | 0.00024 | 6.09E-06 | 0.00017 | 0.00011 | 0.0002 | 0.00013 | 8.77E-05 | 8.41E-05 | 0.00031 | 2.73E-05 | 2.93E-05 | 4.69E-05 |
| Psmb5 | 0.00025 | 0.00014 | 0.00042 | 0.00063 | 0.00017 | 0.00017 | 0.0001 | 0.00017 | 0.00017 | 8.64E-05 | 0.00021 | 4.16E-05 | 7.58E-05 | 0.00016 | 6.59E-05 | 0.00011 | 3.34E-05 | 7.94E-05 |
| Dpp8 | 0.00011 | 0.00017 | 0.00015 | 0.0002 | 0.00016 | 0.00016 | 8.54E-05 | 0.00016 | 0.00017 | 0.00015 | 0.00015 | 9.03E-05 | 0 | 0.00011 | 0.00023 | 7.55E-05 | 8.65E-05 | 0.00013 |
| BC031181 | 0.00019 | 0.0001 | 0.00019 | 7.72E-05 | 8.15E-05 | 7.68E-05 | 3.42E-05 | 0.00014 | 0.00017 | 7.73E-05 | 0.00022 | 8.48E-05 | 6.95E-05 | 1.16E-05 | 0 | 0 | 4.05E-05 | 4.40E-05 |
| Mfsd10 | 4.38E-05 | 9.36E-05 | 0.00013 | 0.00027 | 2.86E-05 | 9.50E-05 | 1.84E-05 | 3.46E-05 | 0.00017 | 8.36E-05 | 0.00013 | 0.00015 | 0.00023 | 0.0001 | 9.96E-06 | 0.00023 | 0 | 0.00022 |
| Rbm17 | 0.00021 | 0.00019 | 0.0004 | 0.00038 | 0.00019 | 0.00015 | 0.00025 | 2.05E-05 | 0.00017 | 6.17E-05 | 0.0001 | 0.00018 | 0.00014 | 0.0002 | 0.00034 | 0.00035 | 0.00013 | 4.23E-05 |
| Hmgb1 | 9.13E-05 | 0.00018 | 0 | 0.00019 | 3.00E-05 | 6.86E-05 | 1.93E-05 | 0 | 0.00017 | 4.08E-05 | 8.54E-05 | 0 | 1.55E-05 | 0.00012 | 0.00036 | 0.00015 | 0 | 7.92E-05 |
| Casp1 | 8.58E-05 | 8.02E-05 | 0 | 0 | 0 | 0 | 0 | 1.83E-05 | 0.00017 | 2.96E-05 | 0 | 0 | 0 | 1.01E-06 | 5.62E-06 | 1.45E-05 | 0 | 4.48E-05 |
| Csde1 | 0.00043 | 0.00024 | 0.0003 | 0.00013 | 0.00017 | 0.00032 | 0.00016 | 0.00013 | 0.00017 | 7.64E-05 | 0.0002 | 0.00024 | 0.0001 | 0.00011 | 0 | 0.00018 | 0.00011 | 6.32E-05 |
| Zfp281 | 0.00015 | 0.00011 | 0 | 0 | 0.00018 | 0.00018 | 0 | 3.67E-05 | 0.00017 | 0 | 0.00012 | 3.33E-05 | 0.00028 | 0.00025 | 0.00031 | 2.10E-05 | 0 | 0 |
| Ccnd2 | 2.92E-05 | 0.00016 | 0.00027 | 0.0001 | 0.00019 | 0.00048 | 0.00025 | 6.84E-05 | 0.00017 | 7.73E-05 | 0.00059 | 0.00046 | 0.00023 | 0.00011 | 0 | 0 | 0 | 0 |
| Polr2a | 0.00033 | 0.00032 | 0.0001 | 9.14E-05 | 0.00014 | 0.00019 | 0.00018 | 5.53E-05 | 0.00017 | 1.83E-05 | 3.71E-05 | 0.0002 | 0 | 0.00014 | 0.00012 | 0.00037 | 0.00039 | 3.19E-05 |
| Set | 0.0002 | 0.00014 | 0.00025 | 0.00029 | 2.39E-05 | 9.13E-05 | 0 | 4.81E-05 | 0.00016 | 0 | 4.47E-05 | 1.87E-05 | 0 | 1.06E-05 | 5.80E-05 | 0.00029 | 0.00011 | 0 |
| Ube2r2 | 0.00034 | 0.00038 | 0.00024 | 0.00026 | 0.00032 | 0.00035 | 0.00038 | 0.00037 | 0.00016 | 0.00037 | 0.00027 | 0.00032 | 0.00031 | 0.00035 | 0.00044 | 0.00011 | 0.00034 | 0.00026 |
| Cfl2 | 0.00017 | 8.51E-05 | 9.69E-05 | 1.25E-05 | 3.10E-05 | 0 | 0 | 8.44E-05 | 0.00016 | 0.00016 | 0.00033 | 0.00019 | 3.06E-05 | 0 | 4.21E-05 | 0.00016 | 0 | 7.32E-06 |
| Npm1 | 0.00036 | 0.00024 | 0.00062 | 0.00102 | 1.28E-05 | 9.60E-05 | 0.00017 | 0.00015 | 0.00016 | 6.05E-05 | 4.62E-05 | 0.00047 | 0.00025 | 0.0002 | 0.00021 | 0.00044 | 0.00071 | 0.00013 |
| Atp7a | 0.00011 | 9.04E-05 | 4.32E-05 | 0 | 7.63E-05 | 0.00011 | 0 | 8.48E-05 | 0.00016 | 4.51E-05 | 0 | 0.00016 | 6.49E-06 | 0.00014 | 0 | 4.78E-05 | 0.0001 | 6.71E-05 |
| Acin1 | 0.00025 | 0.0003 | 0.00026 | 0.00011 | 6.73E-05 | 0 | 5.17E-05 | 0 | 0.00016 | 7.14E-05 | 3.45E-05 | 4.91E-05 | 2.47E-05 | 8.41E-05 | 0.00064 | 0.00024 | 0.00022 | 5.13E-05 |
| H6pd | 0.00015 | 0.00029 | 8.67E-05 | 0.0002 | 0.00062 | 0.0007 | 4.44E-05 | 0.00032 | 0.00016 | 0.00036 | 0.00013 | 0.0003 | 0.00156 | 0.00035 | 0.00014 | 1.56E-05 | 0 | 0.0004 |
| Gnai1 | 6.50E-05 | 0 | 2.54E-05 | 0 | 0 | 6.86E-05 | 4.43E-05 | 0 | 0.00016 | 2.15E-05 | 1.38E-05 | 0 | 0.00018 | 0.0001 | 0 | 5.45E-06 | 0 | 0 |
| Fbxo6 | 0.00013 | 0.00022 | 0 | 0.0001 | 0.00015 | 5.07E-05 | 0 | 1.80E-05 | 0.00016 | 5.53E-05 | 0.0001 | 5.72E-05 | 4.38E-05 | 2.59E-06 | 0 | 1.87E-05 | 0 | 7.22E-05 |
| Ptprm | 2.52E-05 | 0.00012 | 7.54E-05 | 0 | 0.00033 | 0.00015 | 9.92E-05 | 0.0002 | 0.00016 | 0 | 0.00032 | 0.00013 | 0 | 0.00027 | 0.00016 | 4.41E-05 | 0.0002 | 0.00014 |
| Psmb2 | 0.00043 | 0.00014 | 0.00039 | 0.00037 | 7.38E-05 | 0.00011 | 0.00012 | 4.78E-05 | 0.00016 | 5.82E-05 | 0.00014 | 7.92E-05 | 0.00015 | 6.78E-05 | 0 | 0.00022 | 0 | 9.21E-05 |
| Tardbp | 0.00034 | 0.00023 | 0.00037 | 0.00013 | 9.87E-05 | 6.48E-05 | 1.07E-05 | 0 | 0.00016 | 0 | 0.00012 | 0.00011 | 0 | 8.90E-05 | 0.00058 | 0.00029 | 0 | 0 |
| Bzw2 | 0.00022 | 2.21E-05 | 0.0002 | 9.54E-05 | 0 | 3.54E-05 | 3.87E-05 | 0 | 0.00016 | 0 | 0 | 0 | 3.29E-05 | 0 | 5.70E-05 | 0.00011 | 2.84E-05 | 4.34E-05 |
| Ccny | 0.0003 | 0.00022 | 0.0002 | 7.98E-05 | 0.00019 | 0.00025 | 9.67E-05 | 0.0002 | 0.00016 | 8.20E-05 | 0.00017 | 0.00017 | 4.84E-05 | 0.00014 | 0 | 7.04E-05 | 7.56E-05 | 2.43E-05 |
| Spr | 0.00016 | 0.00014 | 0.00024 | 0.00024 | 3.73E-05 | 3.31E-06 | 3.69E-06 | 8.09E-05 | 0.00016 | 7.95E-05 | 0.00012 | 0.00011 | 0 | 0.0001 | 0 | 8.65E-05 | 0 | 8.92E-05 |
| Cops7a | 0.00016 | 0.00014 | 0.0003 | 0.00057 | 7.69E-05 | 0.00011 | 0.00027 | 0.0002 | 0.00016 | 0.00012 | 0.00012 | 9.58E-05 | 0.00017 | 0.00016 | 0 | 0.00013 | 1.48E-05 | 0.00022 |
| Rap2b | 5.71E-05 | 0.00012 | 4.69E-05 | 0.0002 | 0.00011 | 7.25E-06 | 0 | 9.12E-05 | 0.00016 | 0.00021 | 7.13E-05 | 5.94E-05 | 9.19E-05 | 0.0001 | 1.32E-05 | 7.47E-05 | 0.00012 | 0.00011 |
| Dbn1 | 4.63E-05 | 0.00021 | 0.00061 | 0.0002 | 6.27E-05 | 0.00011 | 0.00027 | 0.00054 | 0.00016 | 0.00054 | 0.00091 | 0.00026 | 0 | 0.00012 | 0.00031 | 4.81E-05 | 0 | 0.00029 |
| Dpt | 0 | 0 | 0.00032 | 0 | 0 | 0.00015 | 0.00107 | 0 | 0.00016 | 0.00216 | 0 | 0 | 0.00168 | 0.00024 | 0 | 0.00016 | 0 | 0.00122 |
| Rnf220 | 8.21E-05 | 0.00028 | 0.00017 | 0.00015 | 0.00018 | 0.0001 | 6.37E-05 | 7.58E-05 | 0.00016 | 0.00013 | 0.00013 | 1.84E-05 | 0.00014 | 0.00012 | 0.00029 | 0.0001 | 0 | 4.87E-05 |

| | | | | | | | | | | | | | | | | | | |
|---|---|---|---|---|---|---|---|---|---|---|---|---|---|---|---|---|---|---|
| Rab8b | 0.00016 | 0.00019 | 0.00015 | 0 | 4.70E-06 | 0 | 0.00035 | 0 | 0.00016 | 0.00019 | 1.03E-05 | 0.00013 | 0.0002 | 5.05E-05 | 0 | 4.44E-05 | 0.00023 | 5.34E-05 |
| Cdc42bpb | 0.00017 | 0.00031 | 0.00024 | 6.84E-05 | 0.00014 | 0.00012 | 0.00012 | 8.89E-05 | 0.00016 | 0.0002 | 0.0002 | 0.00017 | 0 | 9.12E-05 | 0.00053 | 0.00011 | 0.0001 | 0.00016 |
| Mmadhc | 0.00021 | 8.36E-05 | 0.00023 | 0.00014 | 6.64E-05 | 8.67E-05 | 5.07E-05 | 6.36E-05 | 0.00016 | 5.32E-05 | 0.00018 | 6.99E-05 | 0 | 8.28E-05 | 0 | 0.00015 | 0 | 0 |
| Fes | 4.23E-05 | 1.82E-05 | 0 | 0.00013 | 0 | 0 | 6.74E-05 | 7.29E-05 | 0.00016 | 0.0001 | 0 | 0.00013 | 2.63E-05 | 0 | 0.00015 | 0 | 0 | 0.00019 |
| Ppia | 0.00038 | 0.00013 | 0.00031 | 0.00053 | 5.66E-05 | 8.56E-05 | 8.62E-05 | 0.00011 | 0.00016 | 2.01E-05 | 0.00044 | 0.0002 | 0.00015 | 0.00024 | 0 | 0.00041 | 4.58E-05 | 9.96E-05 |
| Zfand5 | 0.00028 | 0.00043 | 0.00019 | 5.44E-05 | 0.00013 | 0.00015 | 0 | 0.00016 | 0.00016 | 3.28E-05 | 9.98E-05 | 0.00018 | 0 | 0.00028 | 0.00015 | 0.00015 | 0.00176 | 1.45E-05 |
| Lrrc17 | 0.00034 | 0 | 0 | 0.0002 | 2.93E-05 | 0 | 0.00194 | 0.00029 | 0.00016 | 0.00055 | 0.00043 | 0 | 0 | 0.00027 | 0 | 0.00027 | 0 | 0 |
| Adora2b | 1.34E-06 | 0 | 4.48E-05 | 0 | 4.08E-05 | 1.48E-05 | 0 | 6.06E-05 | 0.00016 | 0.0003 | 0 | 9.63E-05 | 0.00013 | 0 | 0 | 0 | 0 | 0 |
| Cct6a | 0.00025 | 9.83E-05 | 0.00046 | 0.00036 | 0 | 3.08E-05 | 0 | 1.01E-05 | 0.00016 | 2.11E-05 | 0.00016 | 0.0001 | 0 | 3.70E-05 | 1.99E-05 | 0.00018 | 0 | 7.04E-06 |
| Galnt1 | 0.00025 | 0.00011 | 0.00032 | 0 | 0.00019 | 0.00033 | 0.00035 | 0.00018 | 0.00016 | 0.00042 | 0.00016 | 6.98E-05 | 0.0002 | 0.00034 | 9.99E-05 | 0.00026 | 4.69E-05 | 0.00031 |
| Ythdf3 | 0.0002 | 0.00028 | 0.00022 | 9.60E-05 | 0.00021 | 0.00023 | 0.00016 | 0.00023 | 0.00016 | 0.00014 | 0.00019 | 0.00022 | 9.17E-05 | 0.00021 | 0.00018 | 0.00011 | 0.00029 | 0.00013 |
| Snx1 | 0.00028 | 0.0002 | 8.82E-05 | 9.56E-05 | 7.38E-05 | 9.34E-05 | 0.00021 | 5.99E-05 | 0.00016 | 0.00025 | 1.19E-05 | 0.00028 | 2.47E-05 | 0 | 5.09E-05 | 0.00016 | 2.37E-05 | 7.86E-05 |
| Zbp1 | 4.86E-05 | 6.34E-05 | 0 | 5.24E-05 | 9.73E-05 | 0 | 0.00013 | 5.04E-05 | 0.00016 | 0.00061 | 0 | 0 | 0.00047 | 1.36E-06 | 8.37E-05 | 6.09E-05 | 0.00025 | 0.00013 |
| Pkdcc | 7.35E-05 | 3.54E-06 | 8.86E-05 | 0.00023 | 3.07E-05 | 0 | 0.00057 | 7.85E-05 | 0.00016 | 0.00036 | 0.00015 | 0 | 0.00068 | 3.18E-05 | 0.00029 | 0.00016 | 0.00015 | 0.00021 |
| Wdr41 | 4.12E-05 | 6.27E-05 | 1.58E-05 | 7.34E-05 | 6.01E-05 | 2.86E-05 | 6.25E-05 | 3.19E-05 | 0.00016 | 9.33E-05 | 3.12E-05 | 0 | 6.93E-05 | 5.48E-05 | 4.54E-05 | 2.87E-05 | 0 | 8.27E-05 |
| Mtmr9 | 0.00014 | 0.00017 | 7.69E-05 | 1.68E-05 | 0.00014 | 7.88E-05 | 6.22E-05 | 6.35E-05 | 0.00016 | 0.00013 | 0.00013 | 8.69E-05 | 0 | 0 | 0 | 0.00013 | 0 | 0.00011 |
| Sntb2 | 1.24E-05 | 0.0002 | 9.69E-05 | 3.38E-05 | 9.48E-05 | 0.00016 | 8.04E-05 | 0.00013 | 0.00016 | 0.00032 | 0 | 1.32E-05 | 0.00031 | 0.00014 | 0 | 0.00012 | 8.80E-05 | 0.00016 |
| Plcb1 | 0 | 0 | 3.36E-05 | 0 | 0 | 0.00015 | 0 | 0 | 0.00016 | 1.47E-05 | 0 | 0 | 0.00015 | 0.00018 | 4.67E-05 | 8.24E-07 | 0 | 1.71E-05 |
| Fbln7 | 0 | 8.71E-05 | 0 | 0 | 0.00061 | 0 | 4.82E-05 | 0.00037 | 0.00016 | 3.99E-06 | 0 | 0.00018 | 0 | 0 | 0 | 0 | 0 | 0.00017 |
| Il17ra | 4.62E-05 | 0.00013 | 3.37E-05 | 0.00011 | 0 | 0.0001 | 6.95E-06 | 0.0002 | 0.00016 | 0.00046 | 0 | 0.00033 | 0.00078 | 0.00014 | 0.00018 | 0.0001 | 1.01E-05 | 0.00011 |
| Rpn1 | 0.00056 | 4.20E-06 | 0.00059 | 0.00083 | 0.00026 | 0.00035 | 0.00087 | 0.00052 | 0.00016 | 0.00037 | 0.00029 | 7.62E-05 | 0.00057 | 0.00037 | 0.00014 | 0.00023 | 9.76E-05 | 0.00034 |
| Pgrmc1 | 0.00018 | 0.00027 | 0.00015 | 0.00022 | 0.00028 | 0.0003 | 0.00018 | 0 | 0.00016 | 0.00011 | 0.00036 | 0.00047 | 0.00013 | 0.00023 | 0 | 0.00011 | 0.00024 | 6.97E-05 |
| Pim1 | 0 | 0.00026 | 2.75E-05 | 0.0002 | 0.0001 | 0.00022 | 0.00033 | 0 | 0.00016 | 0.00039 | 0.00062 | 2.50E-05 | 0.00052 | 0.00016 | 0.00023 | 0.00018 | 0.00109 | 3.30E-05 |
| Seh1l | 0.00016 | 0.00018 | 0.00018 | 0.00014 | 9.78E-05 | 0.00011 | 0.00011 | 0 | 0.00016 | 0.00012 | 5.05E-05 | 0.0003 | 0.00012 | 6.76E-06 | 8.75E-05 | 0.00026 | 3.52E-05 | 7.71E-05 |
| Arl8a | 0 | 0.00034 | 0.00025 | 0.00025 | 0.00023 | 0.00018 | 0 | 0.00029 | 0.00016 | 0.00018 | 0.00041 | 0.00031 | 0.00012 | 0.0002 | 0.00016 | 0 | 0.0002 | 0.00017 |
| Nploc4 | 0.0001 | 0.00025 | 0.00033 | 0.00015 | 6.64E-05 | 3.50E-05 | 1.10E-05 | 7.66E-05 | 0.00016 | 8.90E-05 | 0.00019 | 8.28E-05 | 6.40E-05 | 9.40E-05 | 7.75E-05 | 0.00017 | 9.33E-05 | 4.14E-05 |
| Raf1 | 0.00013 | 0.00025 | 0.00013 | 0.0002 | 9.18E-05 | 6.60E-05 | 4.57E-05 | 8.16E-05 | 0.00016 | 0.00017 | 0.00019 | 3.19E-05 | 3.97E-05 | 0.00012 | 9.95E-05 | 0.00022 | 0.00031 | 0.00011 |
| Taldo1 | 0.00021 | 0.00014 | 0.00011 | 0.00023 | 0.00012 | 9.91E-07 | 9.48E-05 | 0 | 0.00016 | 0.00015 | 0.00017 | 0 | 0 | 7.36E-05 | 0 | 8.75E-05 | 0 | 0.00013 |
| Tmem214 | 0.0002 | 0.00011 | 0.00028 | 0.00028 | 6.39E-05 | 9.86E-05 | 0.00026 | 5.84E-05 | 0.00016 | 7.67E-05 | 4.87E-05 | 9.32E-05 | 7.79E-05 | 0.00013 | 0.00036 | 5.27E-05 | 0 | 1.88E-05 |
| Gga1 | 0.00012 | 0.00017 | 0.00017 | 0.00018 | 8.91E-05 | 0.00011 | 6.02E-05 | 0.00015 | 0.00016 | 0.00011 | 7.47E-05 | 5.93E-05 | 3.34E-05 | 8.19E-05 | 0.0002 | 0.0001 | 0.00016 | 8.46E-05 |
| Acsl3 | 7.44E-05 | 6.89E-05 | 2.01E-06 | 0 | 2.27E-05 | 0 | 0 | 0 | 0.00016 | 0 | 7.29E-05 | 3.32E-05 | 0 | 5.59E-05 | 0 | 9.09E-05 | 0.00027 | 0 |
| Pon2 | 8.37E-05 | 0.0001 | 2.02E-05 | 4.37E-05 | 7.23E-05 | 0.00013 | 3.60E-05 | 0.00025 | 0.00016 | 7.44E-05 | 8.00E-05 | 6.69E-05 | 4.14E-05 | 0.00015 | 0 | 7.85E-05 | 0 | 0.00014 |
| Fubp3 | 0.00014 | 0.00011 | 0.00015 | 9.26E-05 | 5.21E-05 | 7.29E-05 | 1.15E-05 | 3.52E-05 | 0.00016 | 2.29E-05 | 0.00012 | 7.20E-05 | 1.21E-05 | 5.77E-05 | 2.52E-05 | 6.38E-05 | 6.13E-05 | 5.01E-05 |

| | | | | | | | | | | | | | | | | | | |
|---|---|---|---|---|---|---|---|---|---|---|---|---|---|---|---|---|---|---|
| Flii | 0.00014 | 0.00017 | 0.00015 | 9.64E-05 | 3.59E-05 | 2.92E-05 | 5.77E-05 | 2.52E-05 | 0.00016 | 0.00021 | 0.00014 | 0.00015 | 0 | 7.04E-05 | 0.00016 | 4.36E-05 | 3.75E-05 | 0.00018 |
| Smarcc2 | 0.00028 | 0.00036 | 9.14E-05 | 0.00015 | 0.00023 | 0.00027 | 0.00029 | 0.00017 | 0.00016 | 0.00022 | 8.06E-05 | 0.00017 | 0.00025 | 0.00017 | 0.00053 | 0.00027 | 0 | 0.00023 |
| Anapc1 | 0.00022 | 0.00017 | 0.00021 | 0.00018 | 0.00013 | 6.50E-05 | 7.30E-05 | 7.65E-05 | 0.00016 | 7.72E-05 | 0.00013 | 7.44E-05 | 0 | 5.49E-05 | 0.00017 | 0.00032 | 0 | 7.73E-05 |
| Alas1 | 7.19E-05 | 3.42E-05 | 1.13E-05 | 8.37E-05 | 1.07E-05 | 2.69E-05 | 0 | 0 | 0.00016 | 0.00011 | 8.29E-05 | 2.62E-05 | 0.00023 | 0.00017 | 3.69E-05 | 0 | 0.00018 | 4.89E-05 |
| Mia3 | 0.00025 | 0.00025 | 0.00026 | 0.00033 | 0.00021 | 0.00032 | 0.00049 | 0.00025 | 0.00016 | 0.00041 | 0.00011 | 0.00023 | 0.00037 | 0.00025 | 0.00057 | 0.00012 | 8.89E-06 | 0.00018 |
| Commd1 | 0.00039 | 0.00015 | 0.00021 | 0.00024 | 0.0001 | 8.67E-05 | 0.00012 | 4.89E-06 | 0.00016 | 8.26E-05 | 0.00014 | 8.72E-05 | 2.06E-05 | 2.05E-05 | 0 | 0.00036 | 3.85E-06 | 7.30E-05 |
| Nf2 | 6.77E-05 | 0.0002 | 0.00016 | 0.00024 | 0.00016 | 8.74E-05 | 0 | 0.00012 | 0.00016 | 9.28E-05 | 0.00013 | 4.33E-05 | 5.11E-05 | 4.73E-05 | 0.00032 | 0.00013 | 5.76E-05 | 7.56E-05 |
| Plod1 | 0 | 0 | 0 | 0.00039 | 0.00036 | 0.00076 | 0.0007 | 0.0007 | 0.00016 | 0.00026 | 9.91E-05 | 0.0013 | 0.00048 | 0.00026 | 2.78E-05 | 4.97E-05 | 0 | 0.00028 |
| Sf3b1 | 0.00042 | 0.00031 | 0.00027 | 0.00016 | 0.00019 | 0.00022 | 0.00011 | 0.00011 | 0.00016 | 9.80E-05 | 0.00014 | 0.00052 | 5.31E-05 | 0.00014 | 0.00057 | 0.00021 | 0.00082 | 0.00013 |
| Atf6b | 3.34E-05 | 9.66E-05 | 0.00013 | 0.00025 | 6.95E-05 | 0 | 0 | 6.72E-05 | 0.00016 | 6.07E-05 | 3.41E-05 | 0 | 1.49E-05 | 3.06E-05 | 4.68E-05 | 7.67E-05 | 5.85E-05 | 4.87E-05 |
| Afg3l1 | 9.54E-05 | 7.04E-05 | 0.00018 | 0.00043 | 0.0001 | 3.43E-05 | 0.00015 | 9.00E-05 | 0.00016 | 0.00013 | 0.00014 | 1.67E-05 | 0.00014 | 0.00014 | 0.0001 | 0.0001 | 9.34E-05 | 0.00011 |
| Galnt17 | 0 | 0 | 0 | 0.00012 | 4.07E-06 | 6.39E-06 | 0.00023 | 0 | 0.00016 | 0.00043 | 0.00045 | 0 | 0.00055 | 0 | 0 | 5.58E-05 | 0 | 0.00015 |
| Prkci | 0.0001 | 8.83E-05 | 6.62E-05 | 1.67E-05 | 0.00011 | 4.06E-05 | 0 | 1.85E-05 | 0.00016 | 6.54E-05 | 0.00013 | 6.85E-05 | 0 | 5.40E-06 | 2.28E-05 | 0.00011 | 0.00019 | 1.33E-05 |
| Nova1 | 7.21E-05 | 1.65E-05 | 0 | 0 | 1.29E-05 | 2.42E-05 | 2.58E-05 | 0 | 0.00016 | 0.00049 | 8.10E-06 | 0 | 6.80E-05 | 0 | 1.00E-05 | 2.29E-05 | 7.59E-05 | 0 |
| Btg1 | 0.00019 | 0.00027 | 0 | 0 | 3.88E-05 | 0.00041 | 0 | 5.52E-05 | 0.00016 | 0 | 0 | 0.00047 | 0 | 3.43E-05 | 2.83E-05 | 2.97E-05 | 0.00167 | 1.65E-05 |
| Arih1 | 0.00041 | 0.00026 | 0.00019 | 0 | 0.00012 | 0.00012 | 8.30E-06 | 0 | 0.00016 | 0 | 0.00012 | 0.0002 | 0 | 0 | 0.00034 | 7.07E-05 | 0.00052 | 7.36E-05 |
| Rnf122 | 9.44E-05 | 1.87E-05 | 0.00014 | 2.35E-05 | 0.00036 | 0.00017 | 0.00017 | 5.06E-05 | 0.00016 | 1.92E-05 | 0 | 0 | 6.46E-05 | 0.00033 | 0 | 0.0001 | 0 | 0 |
| Arnt | 0.00016 | 0.00027 | 6.41E-05 | 0.00019 | 0.00024 | 0.0002 | 0.00025 | 0.00016 | 0.00016 | 0.00024 | 3.77E-05 | 0.00018 | 0.00024 | 0.00015 | 0.00029 | 0.00011 | 9.61E-05 | 0.00022 |
| Arhgef12 | 0.00026 | 0.00035 | 0.00013 | 0 | 0.00029 | 0.00029 | 1.49E-05 | 0.00014 | 0.00016 | 0.00015 | 4.77E-05 | 0.00023 | 2.90E-05 | 0.00013 | 0.00015 | 0.00015 | 0.00033 | 0.00016 |
| Foxk2 | 0.00018 | 9.10E-05 | 0.0003 | 0 | 8.99E-05 | 4.01E-05 | 6.36E-05 | 7.34E-05 | 0.00016 | 3.13E-05 | 0.00017 | 0 | 0 | 4.74E-05 | 1.31E-05 | 0.00021 | 0.00015 | 3.08E-06 |
| Ptpn13 | 2.55E-05 | 0.00013 | 5.18E-05 | 1.71E-05 | 0 | 0 | 6.29E-05 | 0.00015 | 0.00016 | 0.00015 | 0 | 4.05E-05 | 6.98E-05 | 7.07E-05 | 4.20E-05 | 0 | 0.00015 | 0.0002 |
| Rab11fip5 | 1.90E-05 | 0.00029 | 0.00017 | 7.08E-05 | 0.00013 | 0.00011 | 0.00014 | 6.72E-05 | 0.00016 | 0.00023 | 0.00013 | 0.00022 | 0 | 0.00013 | 0.00019 | 8.24E-05 | 2.07E-05 | 0.00021 |
| Ifi204 | 0.00021 | 3.85E-05 | 0 | 0 | 0 | 5.20E-05 | 0 | 0.00013 | 0.00016 | 0.00058 | 0 | 0.00014 | 0.00048 | 0 | 0 | 5.70E-05 | 0 | 2.24E-05 |
| Slco2a1 | 0 | 0 | 0.00032 | 0 | 0 | 0.00025 | 0 | 0.00047 | 0.00016 | 0.00031 | 0 | 0.00031 | 0 | 0 | 0 | 4.06E-05 | 0 | 9.53E-06 |
| Impa1 | 2.75E-06 | 0.00016 | 0.00022 | 0.00014 | 0.00016 | 6.43E-05 | 0.00018 | 8.96E-05 | 0.00016 | 0.00011 | 0.00016 | 3.88E-06 | 0.0001 | 5.60E-05 | 0 | 0.00015 | 0 | 0.00013 |
| Tmem50b | 1.44E-06 | 8.18E-05 | 0.00014 | 4.05E-05 | 7.57E-05 | 1.60E-05 | 0 | 6.70E-05 | 0.00016 | 0.00013 | 8.24E-05 | 5.46E-06 | 9.61E-05 | 9.56E-05 | 0 | 0 | 0 | 0.00013 |
| Acadl | 0.00022 | 0.00017 | 1.59E-05 | 1.30E-05 | 0.00018 | 0.00019 | 8.07E-05 | 0 | 0.00016 | 0.0001 | 0.00024 | 0.00011 | 0.00016 | 0.00019 | 4.38E-05 | 0.00024 | 0 | 0.00019 |
| Wdfy2 | 5.03E-05 | 0.00011 | 2.63E-05 | 0.00012 | 1.97E-05 | 5.18E-05 | 0 | 0.00012 | 0.00016 | 0.00016 | 0 | 5.38E-05 | 7.20E-05 | 0 | 0.00016 | 0 | 0.00021 | 7.84E-05 |
| B4gat1 | 1.16E-05 | 0.00012 | 0 | 0.00026 | 0.00017 | 0.00021 | 3.86E-05 | 0.00025 | 0.00016 | 0.00019 | 9.67E-05 | 0.00011 | 3.45E-05 | 0.00028 | 0 | 2.18E-05 | 2.85E-05 | 0.00014 |
| Pip4k2b | 0.00012 | 0.00016 | 0.00019 | 0.00017 | 5.81E-05 | 5.86E-05 | 0 | 7.60E-05 | 0.00016 | 8.22E-05 | 5.41E-05 | 6.46E-05 | 2.51E-05 | 2.91E-05 | 5.79E-05 | 5.60E-05 | 7.37E-05 | 0.00011 |
| Pex5 | 6.57E-05 | 0.00021 | 0.00015 | 0.00019 | 9.72E-05 | 4.30E-05 | 2.49E-05 | 0 | 0.00016 | 7.76E-05 | 7.74E-05 | 0.00016 | 9.24E-05 | 0.0001 | 0.00018 | 2.64E-05 | 0.0003 | 2.45E-05 |
| Dazap1 | 0.00034 | 0.00029 | 0.00035 | 0.00067 | 0.0001 | 0.00013 | 0.00019 | 0.00022 | 0.00016 | 7.33E-05 | 0.00016 | 0.00022 | 0 | 0.00019 | 0.00054 | 0.00041 | 0.00025 | 0.00014 |
| Rnps1 | 7.05E-05 | 0.00023 | 0.00036 | 0.00048 | 9.00E-05 | 5.71E-05 | 0 | 0.00017 | 0.00016 | 0.00011 | 0.00018 | 0.00019 | 0.00011 | 0.00017 | 9.35E-05 | 0.0005 | 0.00023 | 0.00012 |

| | | | | | | | | | | | | | | | | | | |
|---|---|---|---|---|---|---|---|---|---|---|---|---|---|---|---|---|---|---|
| Ypel5 | 0.00023 | 0.00024 | 4.16E-05 | 5.62E-06 | 0.00019 | 0.00014 | 0.00025 | 4.69E-05 | 0.00016 | 0.0002 | 5.98E-05 | 0.0002 | 2.71E-05 | 0 | 7.05E-05 | 0 | 0.00016 | 1.60E-05 |
| Fkbp4 | 0.0001 | 5.98E-05 | 0.0002 | 0.00012 | 7.33E-05 | 4.10E-05 | 0 | 7.19E-05 | 0.00016 | 1.35E-05 | 0.00011 | 0 | 0 | 0 | 0 | 0.00019 | 0 | 4.40E-05 |
| Clptm1 | 0.0002 | 0.00027 | 0.00036 | 0.00033 | 0.00024 | 0.00028 | 9.07E-05 | 0.00024 | 0.00016 | 0.00016 | 0.00025 | 0.00011 | 0.00019 | 0.00018 | 0 | 0.00019 | 3.45E-05 | 0.00016 |
| Cstf2t | 9.68E-05 | 0.00014 | 8.11E-05 | 0.0002 | 0.00014 | 0 | 1.34E-05 | 0 | 0.00016 | 3.32E-05 | 4.19E-05 | 2.08E-05 | 0 | 3.23E-05 | 0.0001 | 9.43E-05 | 0 | 5.35E-05 |
| Cox6b1 | 0.00015 | 4.41E-05 | 0.00022 | 0.00094 | 0.00011 | 0.0001 | 0.00018 | 0.00025 | 0.00016 | 0.00017 | 0.00021 | 0.00011 | 0.00013 | 0.00025 | 0 | 0.00015 | 0.00015 | 0.0002 |
| Arl5a | 0.00018 | 0.00015 | 0.00013 | 8.03E-05 | 6.34E-05 | 0.00016 | 0.00022 | 4.15E-05 | 0.00016 | 0.00012 | 0.00025 | 7.03E-05 | 0.00018 | 9.11E-05 | 0 | 0.00011 | 2.61E-05 | 1.99E-05 |
| Ppdpf | 0.00016 | 0.00014 | 0.00011 | 0.00026 | 0.00014 | 0.0001 | 7.40E-05 | 0.00015 | 0.00016 | 5.20E-05 | 5.42E-05 | 2.35E-05 | 7.43E-05 | 0.00011 | 0 | 1.96E-05 | 6.07E-05 | 3.89E-05 |
| Unc119b | 7.13E-05 | 0.00012 | 0.00013 | 0.00012 | 0.0001 | 7.36E-06 | 2.03E-06 | 7.62E-05 | 0.00016 | 0.00013 | 8.67E-05 | 0.00012 | 0 | 5.84E-05 | 8.22E-05 | 4.58E-05 | 0 | 6.07E-05 |
| Sub1 | 0.00026 | 5.45E-05 | 0.00013 | 4.89E-05 | 3.59E-05 | 8.58E-05 | 0 | 0 | 0.00016 | 7.81E-06 | 5.04E-05 | 1.00E-04 | 3.63E-05 | 6.03E-05 | 0 | 8.18E-06 | 3.11E-05 | 0 |
| Hnrnpr | 0.00029 | 0.00015 | 0.00012 | 0 | 6.60E-05 | 0.00015 | 0 | 2.11E-05 | 0.00016 | 0 | 1.23E-05 | 9.00E-05 | 0 | 0 | 0.00011 | 0.00026 | 0 | 5.14E-05 |
| Arhgef40 | 6.42E-05 | 0.00025 | 0.00046 | 9.35E-06 | 0.00035 | 6.40E-05 | 0.00028 | 0.00042 | 0.00016 | 0.00014 | 0.00025 | 8.22E-05 | 0.00017 | 0.00026 | 0.00073 | 0.00023 | 0.00024 | 0.00042 |
| Eaf1 | 6.39E-05 | 0.00017 | 0.00038 | 5.10E-05 | 0.00011 | 0.00011 | 0 | 6.08E-05 | 0.00016 | 1.34E-05 | 9.92E-05 | 0.00049 | 4.31E-06 | 0.00018 | 0.00023 | 1.86E-05 | 5.57E-05 | 1.55E-05 |
| Rgl1 | 9.84E-05 | 0.0003 | 0 | 7.65E-05 | 0.00027 | 0 | 7.38E-05 | 0 | 0.00016 | 0.00028 | 0.00012 | 0.00018 | 0.00032 | 0 | 0 | 0.00012 | 9.89E-05 | 0.00027 |
| Dnm2 | 0.00012 | 0.00023 | 0.00017 | 0.0001 | 0.00013 | 0.00011 | 5.86E-06 | 0.00013 | 0.00016 | 0.0001 | 0.00017 | 7.50E-05 | 3.83E-05 | 7.69E-05 | 0.0003 | 0.00013 | 9.27E-05 | 4.69E-05 |
| Rpn2 | 0.00039 | 0 | 0.00032 | 0.00048 | 6.19E-05 | 0.00032 | 0.0007 | 0.00041 | 0.00016 | 0.00033 | 0.00021 | 0.00025 | 0.0006 | 0.00043 | 0 | 0.00029 | 1.77E-05 | 0.00038 |
| H2afy | 0.00031 | 0.00023 | 0.0004 | 0.00031 | 4.85E-05 | 9.65E-05 | 8.47E-05 | 0.00013 | 0.00016 | 0.00018 | 0.00015 | 5.27E-05 | 0.0001 | 0.00011 | 0.00042 | 0.00049 | 0.00017 | 9.95E-05 |
| Rnf126 | 6.94E-05 | 9.35E-05 | 0.00033 | 0.00027 | 6.83E-05 | 0 | 2.67E-05 | 0.00011 | 0.00016 | 4.73E-05 | 0.00015 | 0.0005 | 0 | 0.00012 | 0.00027 | 0.0003 | 3.76E-06 | 5.34E-05 |
| Adipor1 | 0.00019 | 0.00041 | 0.0003 | 0.00054 | 0.00031 | 0.00037 | 0.00042 | 0.00037 | 0.00016 | 0.00018 | 0.00046 | 0.00042 | 0.00019 | 0.00048 | 0 | 0.00019 | 7.21E-05 | 0.0003 |
| Ciao2a | 0.00022 | 0.00013 | 0.00017 | 0.00036 | 5.76E-05 | 4.26E-05 | 0.00012 | 0.00014 | 0.00016 | 0.0001 | 4.90E-05 | 5.06E-05 | 6.48E-05 | 0.00011 | 6.75E-05 | 0.00028 | 0 | 0 |
| Tcp1 | 0.00026 | 0.00014 | 0.00041 | 0.00049 | 3.42E-05 | 4.98E-05 | 2.74E-06 | 0.00011 | 0.00016 | 7.86E-05 | 0.00017 | 0.00028 | 0 | 0.00011 | 0.00014 | 0.00041 | 2.52E-05 | 6.75E-05 |
| Ola1 | 0.00029 | 8.63E-05 | 0.0002 | 6.26E-05 | 2.32E-05 | 6.07E-05 | 0 | 1.66E-05 | 0.00016 | 2.11E-05 | 3.08E-05 | 6.55E-05 | 3.46E-05 | 0 | 0 | 9.08E-05 | 4.23E-07 | 9.02E-06 |
| Pigs | 1.35E-05 | 0.00012 | 0.00014 | 9.10E-05 | 0.00012 | 8.89E-05 | 0.00039 | 0.00017 | 0.00016 | 0.00022 | 8.44E-05 | 9.90E-05 | 0.00024 | 0.0002 | 0 | 1.42E-05 | 0 | 0.00026 |
| Vdac2 | 0.00029 | 8.11E-05 | 0.00011 | 0 | 3.55E-05 | 0.00024 | 0 | 9.12E-05 | 0.00016 | 0 | 0.00011 | 0.00028 | 0 | 8.35E-05 | 0 | 0.00012 | 0 | 0 |
| Trmt112 | 0.00012 | 0.00014 | 0.00017 | 0.00063 | 9.24E-05 | 5.96E-05 | 0.00013 | 0.0001 | 0.00016 | 8.15E-05 | 5.04E-05 | 0.00022 | 0.00011 | 0.00011 | 0 | 0.00021 | 0.00018 | 0.00016 |
| Hdlbp | 0.00082 | 0.00014 | 0.00058 | 0.00048 | 0.00041 | 0.00068 | 0.00121 | 0.0004 | 0.00016 | 0.00038 | 0.00049 | 0.00056 | 0.00047 | 0.00044 | 0.00035 | 0.00017 | 0.00052 | 0.00033 |
| Cers6 | 0.00011 | 4.13E-05 | 0.00012 | 0 | 0 | 0 | 0.00014 | 0 | 0.00016 | 0.00014 | 0.00013 | 0 | 0.00018 | 6.60E-05 | 9.99E-05 | 0.00012 | 9.91E-05 | 0.00011 |
| Jdp2 | 0 | 0.00019 | 0 | 0.00046 | 0.00033 | 7.02E-05 | 0.00038 | 0 | 0.00016 | 0.0003 | 0.00037 | 7.68E-05 | 0 | 0.00031 | 0.0004 | 0.0002 | 0.00087 | 0.0002 |
| Nubp1 | 0.00014 | 0.0001 | 0.0002 | 0.00015 | 4.95E-05 | 0 | 0 | 5.64E-05 | 0.00016 | 2.92E-05 | 9.01E-05 | 0 | 5.23E-05 | 0.0001 | 0.00011 | 0.00027 | 0 | 3.71E-05 |
| Gxylt1 | 8.33E-05 | 7.41E-05 | 0.00015 | 0.00015 | 6.66E-05 | 1.54E-05 | 7.04E-05 | 3.92E-05 | 0.00016 | 5.91E-05 | 9.56E-05 | 8.09E-05 | 9.62E-05 | 0.00013 | 8.74E-05 | 0.00026 | 0.00015 | 5.38E-05 |
| Necap2 | 6.28E-05 | 0.00017 | 0.00017 | 0.00029 | 0.00012 | 5.03E-05 | 5.53E-05 | 0.00012 | 0.00016 | 5.55E-05 | 0.00011 | 0.00017 | 4.59E-05 | 0.00011 | 0 | 2.58E-05 | 0 | 0.00011 |
| Rai1 | 4.24E-05 | 0.00019 | 0.00013 | 0.0001 | 0.00014 | 3.17E-05 | 8.82E-06 | 0.00011 | 0.00016 | 3.57E-05 | 0.00012 | 0.00022 | 0 | 0.00014 | 0.00061 | 7.46E-05 | 0.00024 | 6.83E-05 |
| Sh2b3 | 7.22E-06 | 0.00018 | 9.05E-05 | 0 | 1.21E-05 | 0.0003 | 0 | 0.00065 | 0.00016 | 4.42E-06 | 1.08E-05 | 0.00014 | 0 | 0.00019 | 0.00012 | 3.64E-05 | 1.81E-05 | 2.58E-05 |
| Pdzd11 | 0.00024 | 4.24E-05 | 0.00012 | 0.00013 | 8.90E-05 | 7.70E-05 | 0.00012 | 9.37E-05 | 0.00016 | 0 | 0.00014 | 2.30E-05 | 1.84E-05 | 1.53E-05 | 8.21E-05 | 7.28E-05 | 0 | 2.10E-06 |

| | | | | | | | | | | | | | | | | | | |
|---|---|---|---|---|---|---|---|---|---|---|---|---|---|---|---|---|---|---|
| Dtnb | 1.69E-05 | 8.57E-05 | 0 | 0 | 0.00014 | 0.00012 | 0 | 0 | 0.00016 | 0 | 9.37E-06 | 0.00017 | 8.38E-05 | 0.0001 | 0.0001 | 5.99E-05 | 1.23E-05 | 0 |
| Hnrnpab | 0.00014 | 8.88E-05 | 0.00043 | 0.00041 | 5.46E-05 | 1.10E-05 | 7.45E-05 | 0.00013 | 0.00016 | 2.59E-05 | 0.00018 | 0 | 2.06E-05 | 0.00012 | 0.00012 | 0.0004 | 0.0005 | 4.93E-05 |
| Cdk20 | 0 | 6.20E-05 | 8.32E-06 | 5.86E-05 | 3.59E-05 | 0.00011 | 6.77E-06 | 5.66E-05 | 0.00016 | 0.0001 | 0 | 7.05E-06 | 0.00011 | 3.93E-05 | 5.14E-05 | 4.59E-05 | 0 | 0 |
| Ikbke | 0 | 8.86E-05 | 0 | 8.07E-05 | 2.93E-05 | 7.13E-05 | 0 | 0.00017 | 0.00016 | 7.54E-05 | 0 | 8.00E-06 | 0.00013 | 0 | 0.00021 | 0 | 7.21E-05 | 2.93E-05 |
| Mbnl1 | 7.35E-05 | 0.00014 | 5.76E-05 | 1.37E-05 | 5.95E-05 | 0.00011 | 0 | 4.96E-05 | 0.00016 | 0.00013 | 0.00018 | 0 | 0 | 5.87E-05 | 3.32E-05 | 0.00012 | 4.49E-05 | 0.00042 |
| Alkbh5 | 0.00032 | 0.0003 | 0.00019 | 0.00015 | 0.00021 | 0.0003 | 0.00024 | 0.00015 | 0.00016 | 0.00018 | 0.00017 | 0.00069 | 0.00018 | 0.00019 | 0.0002 | 8.21E-05 | 0.00057 | 0 |
| Nelfb | 6.33E-05 | 0.00015 | 0.0001 | 0.00028 | 5.58E-05 | 0 | 4.25E-05 | 0.00013 | 0.00016 | 0.00014 | 3.27E-05 | 0.0002 | 0.00012 | 6.94E-05 | 7.56E-05 | 9.24E-05 | 0.00028 | 8.72E-05 |
| Ppp3cb | 0.0003 | 0.00021 | 0.00015 | 2.66E-05 | 8.04E-05 | 0.0002 | 8.84E-05 | 0.00014 | 0.00016 | 0.00014 | 0.00018 | 0.00017 | 7.82E-05 | 0.00014 | 0.00018 | 0.00029 | 1.40E-05 | 0.00011 |
| Zadh2 | 0 | 0.00012 | 2.28E-05 | 0.00026 | 0.00011 | 0.00011 | 4.44E-05 | 4.80E-05 | 0.00016 | 0.00016 | 0 | 5.24E-05 | 9.70E-05 | 2.23E-05 | 2.93E-05 | 0.00017 | 0 | 0.00013 |
| Tmem120a | 5.50E-05 | 8.72E-05 | 0.00015 | 0.00015 | 8.73E-05 | 0 | 3.67E-05 | 7.17E-05 | 0.00016 | 2.44E-05 | 5.82E-06 | 4.25E-05 | 0 | 4.98E-05 | 3.29E-05 | 9.74E-06 | 0 | 5.38E-06 |
| Cuta | 0.00021 | 0.00017 | 8.83E-05 | 0.00044 | 0.00018 | 0.00014 | 0.00011 | 0.00012 | 0.00016 | 0.00014 | 8.41E-05 | 2.63E-05 | 0.00011 | 0.00024 | 0 | 0.00017 | 0 | 0.00018 |
| H2afz | 0.00037 | 2.26E-05 | 0.00014 | 0 | 0 | 3.66E-05 | 0.00016 | 5.34E-05 | 0.00016 | 0 | 0 | 3.86E-05 | 0 | 0 | 0 | 0.00113 | 0.00059 | 0 |
| Pcdh7 | 0.00024 | 0.00015 | 0 | 0 | 6.07E-05 | 0.00019 | 0.00011 | 0 | 0.00016 | 7.98E-05 | 0.00011 | 5.70E-05 | 0.00026 | 1.18E-05 | 4.68E-05 | 8.68E-05 | 5.33E-05 | 0.00019 |
| Slc27a4 | 2.42E-05 | 8.10E-05 | 0.00011 | 0.00013 | 8.25E-05 | 6.70E-05 | 6.43E-05 | 8.44E-05 | 0.00016 | 0.00016 | 0.00014 | 0.00022 | 4.53E-05 | 0.00012 | 0.00011 | 7.46E-05 | 0 | 0.00015 |
| Hexim1 | 0.00019 | 0.00027 | 0 | 8.39E-05 | 0.00011 | 0.00011 | 0 | 2.79E-06 | 0.00016 | 3.10E-05 | 5.77E-05 | 0.00013 | 0 | 0.00013 | 5.74E-05 | 4.93E-06 | 0.00126 | 0.00012 |
| Smim10l1 | 6.47E-05 | 0.00013 | 0.00011 | 0.00042 | 0.00025 | 0.00018 | 9.16E-05 | 0.00023 | 0.00016 | 0.00012 | 0.00012 | 6.17E-05 | 3.65E-05 | 0.00013 | 1.23E-05 | 0 | 0 | 0.00022 |
| Apex1 | 0.00012 | 9.24E-05 | 0.00022 | 0.00017 | 4.02E-05 | 4.23E-05 | 0 | 5.52E-05 | 0.00016 | 0 | 4.89E-05 | 0 | 1.03E-06 | 7.73E-05 | 2.17E-05 | 4.98E-05 | 0 | 2.82E-05 |
| Lamp1 | 0.00028 | 0.00029 | 0.00019 | 0.00013 | 0.00036 | 0.00049 | 0.00036 | 0.00022 | 0.00016 | 0.00037 | 0.0003 | 0.00015 | 0.00044 | 0.00032 | 0 | 0.0001 | 0 | 0.00043 |
| Apbb2 | 6.86E-05 | 0.00016 | 0 | 5.29E-05 | 0.00032 | 0.00037 | 0 | 0.00053 | 0.00016 | 8.49E-05 | 0.00014 | 0.00022 | 0 | 0.00032 | 6.85E-05 | 7.43E-05 | 0.00025 | 0.00023 |
| Abcc1 | 2.12E-05 | 8.63E-05 | 0.00025 | 0 | 5.93E-05 | 7.19E-05 | 0 | 8.46E-05 | 0.00016 | 5.93E-05 | 0.00029 | 0.00027 | 7.59E-06 | 0.00011 | 0.00022 | 0 | 9.12E-05 | 0 |
| Gna14 | 0 | 6.57E-05 | 0 | 0 | 0.00041 | 0.00013 | 0 | 0 | 0.00016 | 2.77E-05 | 0.00016 | 0 | 0 | 0 | 0 | 0 | 0 | 0.0001 |
| Adh5 | 0.00022 | 0.00017 | 8.72E-05 | 0.00034 | 7.94E-05 | 0.00013 | 0 | 4.40E-05 | 0.00016 | 7.54E-05 | 9.41E-05 | 0.00017 | 8.91E-05 | 0.00012 | 0 | 0.00013 | 2.16E-05 | 0.00014 |
| Mea1 | 0.00014 | 0.00012 | 0.00014 | 0.00021 | 7.19E-05 | 4.05E-05 | 6.81E-05 | 1.70E-05 | 0.00016 | 7.37E-05 | 0.0001 | 8.61E-05 | 0 | 1.21E-05 | 0 | 0 | 1.88E-05 | 8.67E-05 |
| Cox7b | 0.00045 | 4.75E-05 | 0.00016 | 0.00032 | 5.67E-05 | 0.0002 | 9.22E-05 | 7.24E-05 | 0.00015 | 5.25E-05 | 7.58E-05 | 7.90E-05 | 5.19E-05 | 6.17E-05 | 1.88E-05 | 0.00012 | 3.51E-05 | 2.85E-05 |
| Etfb | 0.00026 | 0.00011 | 0.00017 | 0.00061 | 5.31E-05 | 0.00014 | 7.01E-05 | 7.84E-05 | 0.00015 | 0.00024 | 0.00019 | 0 | 0.00014 | 0.00014 | 6.95E-05 | 0.00017 | 6.98E-06 | 0.00017 |
| Nynrin | 8.34E-05 | 0.00018 | 0 | 0.00025 | 0.00026 | 8.87E-05 | 0.0002 | 8.86E-05 | 0.00015 | 7.81E-05 | 0.00015 | 3.73E-05 | 0 | 0.00024 | 0.00024 | 2.06E-05 | 0 | 7.58E-05 |
| Epm2aip1 | 8.67E-05 | 0.00013 | 3.32E-05 | 0.00012 | 0.00012 | 7.26E-05 | 4.32E-05 | 9.19E-05 | 0.00015 | 2.62E-05 | 2.21E-05 | 7.54E-05 | 0 | 0.0001 | 0.00026 | 0 | 4.54E-05 | 8.72E-06 |
| Eif2b5 | 0.00014 | 0.00012 | 0.00017 | 0.00028 | 5.02E-05 | 9.10E-05 | 4.28E-05 | 3.01E-05 | 0.00015 | 3.80E-05 | 0.00011 | 6.50E-05 | 0 | 0.00012 | 7.47E-05 | 9.45E-05 | 2.06E-05 | 2.59E-05 |
| Txnl1 | 0.00036 | 0.00016 | 0.0004 | 0.00026 | 3.96E-05 | 5.21E-05 | 4.63E-05 | 2.29E-05 | 0.00015 | 2.55E-05 | 0.00019 | 0.00019 | 0 | 6.96E-05 | 6.89E-05 | 0.00027 | 0 | 1.42E-05 |
| Sorbs3 | 5.21E-05 | 0.00011 | 0 | 6.27E-05 | 7.34E-05 | 5.84E-05 | 0.00012 | 4.59E-05 | 0.00015 | 5.49E-05 | 4.14E-05 | 9.70E-05 | 5.40E-05 | 0.00013 | 9.64E-05 | 3.64E-05 | 2.80E-05 | 5.30E-05 |
| Ilrun | 0.0002 | 0.00033 | 0.00022 | 8.77E-05 | 0.00025 | 0.00028 | 0 | 0 | 0.00015 | 0.0001 | 2.24E-05 | 0.00033 | 0.00012 | 0.00017 | 9.09E-05 | 0.00016 | 0.00012 | 6.75E-05 |
| Acads | 0.00012 | 0.00013 | 7.92E-05 | 0.00017 | 6.24E-05 | 6.72E-05 | 4.78E-05 | 7.59E-05 | 0.00015 | 4.98E-05 | 0.00011 | 1.22E-05 | 9.17E-06 | 0.00011 | 0.0001 | 0 | 0 | 5.67E-05 |
| Gypc | 7.92E-05 | 0.00011 | 5.83E-05 | 7.35E-05 | 3.55E-05 | 2.93E-05 | 7.59E-05 | 0 | 0.00015 | 0.00017 | 6.58E-05 | 3.54E-05 | 6.28E-05 | 4.24E-05 | 3.29E-05 | 8.93E-05 | 1.88E-05 | 6.03E-05 |

| | | | | | | | | | | | | | | | | | | |
|---|---|---|---|---|---|---|---|---|---|---|---|---|---|---|---|---|---|---|
| Pygb | 5.35E-05 | 0.00013 | 0 | 0.00028 | 0.00012 | 1.98E-05 | 0.00021 | 9.27E-05 | 0.00015 | 0.00025 | 0.00022 | 0 | 1.35E-05 | 0.0001 | 1.25E-05 | 0.00021 | 0.00012 | 0.00031 |
| Scaf11 | 0.00029 | 0.00022 | 0.0002 | 1.87E-05 | 0.00011 | 0.00014 | 9.59E-05 | 1.42E-06 | 0.00015 | 7.29E-05 | 0.00013 | 0.00025 | 4.34E-05 | 0.00011 | 0.0002 | 0.00033 | 7.71E-06 | 5.46E-05 |
| Ubl3 | 0.00014 | 0.00021 | 0.00012 | 7.22E-05 | 0.00013 | 0.00011 | 8.47E-05 | 0.00023 | 0.00015 | 0.00015 | 0.00012 | 0 | 2.37E-05 | 6.43E-05 | 0 | 5.37E-05 | 0.00011 | 0.00024 |
| Elavl1 | 0.00037 | 0.00022 | 0.00045 | 0.0001 | 0.00016 | 0.00016 | 6.17E-05 | 0.00018 | 0.00015 | 6.25E-05 | 0.00018 | 0.00014 | 2.34E-05 | 0.00011 | 0.00017 | 0.00031 | 0.00011 | 7.46E-05 |
| Ppp1r12a | 8.73E-05 | 0.00024 | 0.00012 | 2.16E-05 | 0.00011 | 7.42E-05 | 3.31E-06 | 0.00015 | 0.00015 | 0.00014 | 0.00033 | 0.00015 | 0 | 6.88E-05 | 0.00023 | 0.0002 | 0.00042 | 0.00014 |
| Men1 | 1.08E-05 | 0.0001 | 4.92E-05 | 0.00015 | 8.87E-05 | 6.68E-05 | 1.09E-05 | 4.54E-05 | 0.00015 | 6.78E-05 | 0.0001 | 8.65E-05 | 6.73E-05 | 9.09E-05 | 8.13E-05 | 0.00011 | 2.61E-05 | 9.80E-05 |
| Xpnpep1 | 0.00018 | 0.0001 | 0.00029 | 0.00036 | 0.00012 | 2.82E-05 | 0.00029 | 0.0002 | 0.00015 | 0.00015 | 0.00022 | 0.00026 | 4.10E-06 | 8.25E-05 | 5.11E-05 | 9.19E-05 | 0 | 0.00029 |
| Pdap1 | 0.00041 | 0.00016 | 0.00039 | 0.00058 | 0.00015 | 0.00017 | 0.00016 | 8.09E-05 | 0.00015 | 2.98E-05 | 0.00017 | 0.00012 | 2.93E-05 | 7.55E-05 | 0.00031 | 0.00033 | 9.81E-05 | 8.20E-05 |
| Ube2d1 | 9.43E-05 | 9.62E-05 | 0.0001 | 3.80E-05 | 7.05E-05 | 8.28E-05 | 5.32E-05 | 4.93E-05 | 0.00015 | 7.89E-05 | 0.00014 | 5.49E-05 | 1.12E-05 | 9.23E-05 | 0 | 3.46E-05 | 0 | 0.00011 |
| Pfkl | 2.81E-05 | 0 | 4.82E-05 | 0.00034 | 8.11E-05 | 0.00015 | 4.57E-05 | 0.0002 | 0.00015 | 0.00015 | 0.00011 | 0.00141 | 0.00041 | 0.00015 | 4.62E-05 | 0 | 0.0002 | 0 |
| Tgm1 | 0 | 3.32E-06 | 1.62E-05 | 2.47E-05 | 4.79E-05 | 2.01E-05 | 0 | 3.14E-05 | 0.00015 | 3.13E-05 | 0 | 0.0001 | 0.00015 | 9.84E-05 | 2.89E-05 | 3.43E-05 | 0 | 0 |
| Mtx2 | 0.00025 | 0.0001 | 0.00014 | 0.00014 | 0.00011 | 0.00017 | 9.13E-05 | 4.17E-05 | 0.00015 | 4.79E-05 | 8.15E-05 | 4.81E-05 | 0 | 0.00014 | 0 | 0.00012 | 0 | 6.23E-05 |
| Atp1b3 | 0.0002 | 0.00018 | 0.00024 | 9.96E-06 | 0.00013 | 0.00013 | 0 | 0.00022 | 0.00015 | 0.00021 | 0.0002 | 4.71E-05 | 6.28E-05 | 0.00011 | 0 | 0.00031 | 0.00028 | 0.00015 |
| Cs | 0.00025 | 7.91E-05 | 0.00021 | 0.00014 | 0 | 3.46E-05 | 2.65E-05 | 0 | 0.00015 | 2.14E-05 | 0.00011 | 0 | 0 | 2.06E-05 | 0 | 0.00014 | 0 | 5.59E-05 |
| Aldh9a1 | 0.00018 | 0.00011 | 0.00017 | 0.00019 | 0.00014 | 4.06E-05 | 4.39E-05 | 9.64E-05 | 0.00015 | 8.50E-05 | 0 | 0.00014 | 1.78E-05 | 0.00041 | 0 | 5.77E-05 | 2.26E-05 | 0 |
| Limd1 | 2.35E-05 | 0.00024 | 0.00014 | 0.00023 | 0.00025 | 0.00029 | 1.14E-05 | 0.00014 | 0.00015 | 0.00012 | 0.00021 | 0.00057 | 0.00013 | 0.00026 | 0.00018 | 0.00011 | 1.75E-05 | 0.00018 |
| Lrig2 | 5.96E-05 | 0.00012 | 5.75E-06 | 0 | 2.28E-05 | 5.45E-05 | 4.11E-05 | 1.49E-05 | 0.00015 | 4.59E-05 | 0 | 4.56E-05 | 9.67E-05 | 4.84E-05 | 4.77E-05 | 0 | 6.75E-05 | 4.80E-05 |
| Rlim | 0.00035 | 0.00017 | 0.00034 | 0.00046 | 6.95E-05 | 0.00015 | 0.00037 | 0.00017 | 0.00015 | 8.23E-05 | 0.00021 | 0.00016 | 0 | 0.00021 | 0.00031 | 0.00025 | 0.00042 | 4.82E-05 |
| Eif4b | 0.00011 | 0.00036 | 0.0001 | 0.00081 | 3.88E-05 | 0.0002 | 7.92E-05 | 0.00017 | 0.00015 | 0.00012 | 0 | 0.00046 | 2.44E-05 | 0.00018 | 0.00011 | 0.0002 | 0.00041 | 0.00021 |
| Eif3b | 3.37E-05 | 0.00012 | 0.0004 | 0.00066 | 3.96E-05 | 9.08E-06 | 0 | 0.00016 | 0.00015 | 9.17E-05 | 0.00014 | 4.39E-05 | 5.47E-05 | 9.07E-05 | 0.00035 | 0.00016 | 0.00013 | 8.62E-05 |
| Dda1 | 0.00014 | 8.62E-05 | 0.00023 | 0.00023 | 9.00E-05 | 6.87E-05 | 6.32E-05 | 0.00019 | 0.00015 | 9.28E-05 | 0.00018 | 8.70E-05 | 0 | 2.60E-05 | 1.31E-05 | 9.40E-05 | 9.13E-05 | 4.92E-05 |
| Fah | 7.22E-05 | 5.98E-05 | 0 | 4.52E-05 | 6.44E-05 | 0.00015 | 5.32E-05 | 6.66E-05 | 0.00015 | 9.39E-06 | 7.10E-05 | 2.61E-05 | 0.00018 | 0.00046 | 2.61E-05 | 0 | 0 | 9.37E-05 |
| Tmem167 | 0.00071 | 7.80E-05 | 0.00027 | 0 | 0.00018 | 0.00037 | 0.00044 | 0.00018 | 0.00015 | 9.72E-05 | 0.00016 | 4.88E-05 | 0.00021 | 2.04E-05 | 0 | 0.00018 | 0 | 5.72E-05 |
| Fyttd1 | 0.00028 | 0.00017 | 0.00019 | 1.61E-05 | 7.90E-05 | 0.00011 | 4.58E-05 | 0.00011 | 0.00015 | 4.28E-05 | 0.0001 | 0.00024 | 5.78E-06 | 3.81E-05 | 0.00013 | 6.11E-05 | 0 | 2.16E-05 |
| Ankrd12 | 6.12E-05 | 0.00017 | 0 | 0 | 0.00015 | 7.87E-05 | 7.15E-05 | 0 | 0.00015 | 2.19E-06 | 0 | 0 | 0 | 0.00014 | 3.53E-05 | 0 | 4.60E-05 | 0 |
| Metap2 | 0.00037 | 0.00016 | 0.00041 | 0.00015 | 3.21E-05 | 0.00021 | 0 | 0.00015 | 0.00015 | 5.00E-05 | 0.00016 | 0.00015 | 0 | 3.43E-05 | 0.00013 | 0.00018 | 0 | 0.00011 |
| Poldip3 | 0.00022 | 0.00033 | 0.00023 | 0.00018 | 0.00013 | 0.00014 | 7.94E-05 | 0 | 0.00015 | 6.13E-05 | 0.00014 | 0.00029 | 0 | 8.67E-05 | 0.00017 | 0.00026 | 0 | 0.00012 |
| Dnajc3 | 0.00035 | 0.00015 | 0.00039 | 0.00031 | 0.00027 | 0.00022 | 0.00046 | 0.00034 | 0.00015 | 0.00037 | 0.00028 | 0.0001 | 0.00054 | 0.00028 | 5.86E-05 | 0.00026 | 0 | 0.00025 |
| Ado | 1.13E-05 | 0.00011 | 0.00014 | 6.84E-05 | 3.93E-05 | 6.47E-05 | 0 | 0.00011 | 0.00015 | 2.67E-05 | 0.00013 | 7.61E-05 | 8.76E-08 | 4.37E-05 | 2.96E-05 | 0.00013 | 0 | 6.26E-05 |
| Dars | 0.00029 | 0.00015 | 0.00033 | 0.00028 | 0.00011 | 4.12E-05 | 6.06E-05 | 7.15E-05 | 0.00015 | 0.00013 | 0.00014 | 0.00034 | 6.67E-05 | 7.65E-05 | 3.55E-05 | 0.00017 | 0 | 0.00012 |
| Zwint | 0.00013 | 0.00025 | 0.0003 | 1.54E-05 | 4.62E-05 | 6.63E-05 | 0 | 5.23E-05 | 0.00015 | 3.39E-05 | 0.00014 | 0.00021 | 4.60E-05 | 4.79E-05 | 0.00046 | 0.00031 | 3.86E-07 | 0 |
| Hars | 0.00021 | 8.49E-05 | 0.00036 | 0.0002 | 4.51E-05 | 2.02E-05 | 8.27E-06 | 1.28E-05 | 0.00015 | 7.18E-05 | 9.45E-05 | 1.19E-06 | 8.38E-05 | 3.53E-05 | 4.10E-05 | 0.00019 | 1.71E-05 | 7.24E-05 |
| Bscl2 | 0.00017 | 0.00014 | 0.00015 | 0.00022 | 9.08E-05 | 0.0002 | 8.66E-07 | 9.90E-05 | 0.00015 | 0.00021 | 0 | 0 | 0.00025 | 0.00011 | 0 | 1.27E-05 | 0 | 0.00019 |

| | | | | | | | | | | | | | | | | | | |
|---|---|---|---|---|---|---|---|---|---|---|---|---|---|---|---|---|---|---|
| Cbx3 | 0.00045 | 6.85E-05 | 7.34E-05 | 0 | 0 | 0.00018 | 0.00013 | 1.14E-06 | 0.00015 | 0 | 6.58E-06 | 5.38E-05 | 3.03E-05 | 0 | 3.01E-06 | 0.00036 | 0.00011 | 0 |
| Nsun4 | 9.17E-05 | 0.0001 | 0.0001 | 8.24E-05 | 7.41E-05 | 1.06E-05 | 2.92E-05 | 0.00013 | 0.00015 | 2.27E-05 | 0 | 2.26E-05 | 3.27E-05 | 0.00011 | 0.00025 | 0 | 0 | 5.17E-05 |
| Uqcrfs1 | 0.0004 | 0.00024 | 0.00028 | 0.0001 | 9.65E-05 | 0.00016 | 4.03E-05 | 2.65E-05 | 0.00015 | 3.70E-05 | 0.00012 | 0.00019 | 8.86E-05 | 5.80E-05 | 0 | 0.00028 | 0 | 1.58E-05 |
| Med8 | 0.00018 | 9.46E-05 | 0.00018 | 5.68E-05 | 5.48E-05 | 2.20E-05 | 3.01E-05 | 0 | 0.00015 | 7.08E-05 | 5.24E-05 | 0 | 1.98E-05 | 6.05E-05 | 7.48E-05 | 0.0001 | 0 | 2.09E-05 |
| Polr2f | 0.00027 | 8.90E-05 | 0.00025 | 0.00024 | 8.75E-05 | 7.41E-05 | 2.58E-05 | 5.30E-05 | 0.00015 | 6.18E-05 | 9.20E-05 | 2.86E-05 | 6.20E-05 | 9.64E-05 | 6.74E-05 | 0.00024 | 6.61E-05 | 9.79E-05 |
| Nsd2 | 7.31E-05 | 7.00E-05 | 0.00024 | 0 | 0.00015 | 3.78E-06 | 6.22E-05 | 0.0001 | 0.00015 | 0 | 3.88E-05 | 0.00013 | 0 | 0.00014 | 0.00029 | 0.00075 | 8.77E-05 | 6.90E-05 |
| Tcf3 | 7.56E-06 | 0.00014 | 0.00013 | 0.00022 | 4.45E-05 | 1.47E-05 | 5.64E-06 | 7.55E-05 | 0.00015 | 5.51E-05 | 0.00011 | 5.22E-05 | 0 | 8.03E-05 | 0.00016 | 0.00018 | 3.60E-05 | 7.88E-05 |
| Ccm2 | 8.34E-05 | 0.00016 | 0.0002 | 0.00018 | 0.00012 | 7.83E-05 | 2.82E-06 | 0.00021 | 0.00015 | 0.0001 | 0.00018 | 6.60E-05 | 0.00011 | 0.00012 | 0 | 0.00021 | 0 | 7.18E-05 |
| Stxbp5 | 7.48E-05 | 0.00015 | 3.29E-05 | 0 | 8.33E-05 | 4.83E-05 | 3.46E-05 | 0 | 0.00015 | 0.00013 | 1.69E-05 | 0.00015 | 0.00012 | 2.21E-05 | 8.79E-05 | 5.62E-05 | 0.00021 | 0 |
| Chpf | 0 | 0.00023 | 0.00044 | 0.00044 | 0.00039 | 0.00021 | 0.0013 | 0.00064 | 0.00015 | 0.00027 | 0 | 0 | 0.0005 | 4.15E-05 | 0.00028 | 0.0001 | 0.00017 | 0.00058 |
| Lsm14a | 0.0003 | 0.00037 | 0.00023 | 0.00018 | 0.0002 | 0.00022 | 4.96E-05 | 4.65E-05 | 0.00015 | 9.32E-05 | 0.00021 | 0.00033 | 0.00011 | 0.00012 | 0.0004 | 0.00031 | 0.00017 | 0.0001 |
| Fam98b | 0.00013 | 6.40E-05 | 0.00018 | 0.00011 | 0.00012 | 3.11E-05 | 3.66E-05 | 0.00025 | 0.00015 | 1.30E-05 | 0.00011 | 0 | 0 | 5.43E-05 | 3.12E-05 | 0.00019 | 3.74E-05 | 2.06E-05 |
| Sae1 | 0.00025 | 0.0001 | 0.00027 | 0.00028 | 6.21E-05 | 6.76E-05 | 7.79E-05 | 0.0001 | 0.00015 | 0.00013 | 8.03E-05 | 0 | 4.36E-05 | 8.63E-05 | 0.00012 | 0.00074 | 0 | 8.78E-05 |
| Nek6 | 0.00035 | 0.00013 | 0.00016 | 0.00034 | 0.00027 | 0.00039 | 0.00011 | 1.40E-05 | 0.00015 | 0 | 0.0003 | 0 | 0.0009 | 0.00037 | 0.00041 | 8.37E-05 | 9.57E-05 | 0.00013 |
| Ipo13 | 5.73E-05 | 0.00011 | 0.00012 | 5.80E-05 | 2.77E-05 | 0 | 1.15E-05 | 2.65E-06 | 0.00015 | 8.73E-05 | 8.35E-05 | 1.07E-07 | 9.35E-05 | 2.20E-05 | 9.71E-05 | 4.00E-05 | 8.32E-06 | 5.46E-05 |
| Glrx5 | 0.0001 | 0.00011 | 0.00019 | 0.00033 | 8.18E-05 | 6.38E-05 | 0 | 8.12E-05 | 0.00015 | 3.41E-05 | 0.00012 | 0 | 2.44E-05 | 9.89E-05 | 8.90E-05 | 0.00012 | 7.06E-05 | 6.69E-05 |
| Cndp2 | 7.89E-05 | 0.00011 | 0.00028 | 9.23E-05 | 4.83E-05 | 0.00014 | 6.28E-05 | 0.00032 | 0.00015 | 0.00022 | 0 | 7.51E-05 | 0.00019 | 8.61E-05 | 0.00011 | 5.16E-07 | 3.98E-06 | 0 |
| Ptpn23 | 0.00012 | 0.00028 | 0.00017 | 8.77E-05 | 0.00011 | 2.21E-05 | 3.12E-05 | 8.68E-06 | 0.00015 | 7.68E-05 | 9.25E-05 | 0.00014 | 0.00015 | 9.33E-05 | 0.00016 | 0.0001 | 0.00019 | 7.22E-05 |
| Slbp | 2.18E-05 | 7.57E-05 | 0.00011 | 0.00017 | 3.42E-05 | 0 | 0 | 0.00012 | 0.00015 | 7.87E-05 | 7.05E-05 | 6.24E-05 | 8.48E-05 | 9.60E-05 | 0.00013 | 0.00055 | 0.00071 | 2.06E-07 |
| Icosl | 0 | 0 | 0 | 0 | 0 | 0.00011 | 0 | 0 | 0.00015 | 5.09E-05 | 0 | 3.94E-05 | 0 | 0 | 0 | 8.45E-06 | 0 | 0.00012 |
| Zfp703 | 0 | 9.73E-05 | 0.00028 | 0.00027 | 0 | 0 | 0 | 0.00019 | 0.00015 | 0.00018 | 2.66E-05 | 0.00026 | 0 | 0.00013 | 0.00032 | 7.94E-05 | 0.00085 | 0.00012 |
| Trip6 | 5.24E-06 | 8.56E-05 | 0.0001 | 0.00019 | 9.65E-06 | 1.28E-05 | 0 | 0 | 0.00015 | 7.14E-05 | 0.00017 | 6.04E-05 | 0.00011 | 0.00013 | 9.61E-05 | 2.55E-05 | 5.69E-05 | 5.15E-05 |
| Micu2 | 0.00023 | 0.0001 | 7.31E-05 | 5.96E-05 | 7.33E-05 | 6.70E-05 | 0.00014 | 9.90E-05 | 0.00015 | 7.26E-05 | 0.00015 | 0.00014 | 1.51E-05 | 1.59E-05 | 4.97E-05 | 7.08E-05 | 0 | 0.00014 |
| Nek9 | 0.00018 | 0.00018 | 8.75E-05 | 2.95E-05 | 0.00019 | 0.00012 | 0.00019 | 7.95E-05 | 0.00015 | 6.90E-05 | 0.00011 | 9.45E-05 | 8.37E-05 | 9.06E-05 | 6.04E-06 | 5.89E-05 | 4.25E-05 | 5.50E-05 |
| Timm22 | 0.00014 | 7.87E-05 | 0.00013 | 0.00014 | 0.00012 | 2.71E-05 | 7.01E-05 | 5.53E-05 | 0.00015 | 3.46E-05 | 9.65E-05 | 4.16E-06 | 3.55E-06 | 4.23E-05 | 0 | 0.00014 | 0 | 4.29E-05 |
| Fcgr2b | 0 | 0 | 0 | 0 | 0.00048 | 0.00145 | 0 | 0 | 0.00015 | 0 | 0 | 0 | 0 | 0.00055 | 0 | 0 | 0 | 0 |
| Tmem97 | 0.00028 | 0 | 0.00033 | 0.00017 | 0 | 2.96E-05 | 0.0003 | 0 | 0.00015 | 0 | 8.49E-05 | 5.37E-05 | 0 | 3.86E-05 | 0 | 0.00029 | 0 | 0 |
| Tulp3 | 4.16E-05 | 9.30E-05 | 3.65E-05 | 0.00015 | 3.54E-05 | 2.88E-05 | 0 | 7.20E-05 | 0.00015 | 5.74E-05 | 7.75E-05 | 1.77E-05 | 3.54E-05 | 5.37E-05 | 3.02E-05 | 2.93E-05 | 0 | 2.65E-05 |
| Rab21 | 0.0001 | 0.00034 | 0.00028 | 0.00032 | 0.00033 | 0.00021 | 0.00024 | 0.00023 | 0.00015 | 0.00016 | 0.00036 | 0.00034 | 5.97E-05 | 0.00024 | 0.00034 | 0.00011 | 0.00053 | 0.00035 |
| Evc2 | 3.38E-06 | 0.00012 | 0.00012 | 5.73E-05 | 7.88E-05 | 7.63E-05 | 0.00014 | 8.41E-05 | 0.00015 | 0.00011 | 4.59E-05 | 1.89E-05 | 0.00013 | 5.68E-05 | 0.00024 | 2.40E-05 | 3.71E-05 | 0.00018 |
| Carm1 | 0.00035 | 0.00011 | 0.00034 | 0.00026 | 0.00011 | 0.00024 | 0.00032 | 0.00013 | 0.00015 | 9.41E-05 | 0.00019 | 0.00012 | 0.00018 | 0.00011 | 0.00016 | 0.00013 | 0.00015 | 0.00011 |
| Fmnl3 | 0 | 0.00015 | 0.00032 | 6.88E-05 | 0.00029 | 0.00022 | 0.00021 | 0.00038 | 0.00015 | 0 | 0.00021 | 0.00014 | 4.82E-06 | 0.00036 | 0.00018 | 0.00016 | 0.00025 | 0.00016 |
| Mdh2 | 0.00021 | 9.83E-05 | 0.00041 | 0.00028 | 7.24E-05 | 4.72E-05 | 3.25E-05 | 0.00013 | 0.00015 | 1.80E-05 | 0.00021 | 6.05E-05 | 0 | 3.67E-05 | 0 | 0.00016 | 0 | 2.20E-05 |

| | | | | | | | | | | | | | | | | | | |
|---|---|---|---|---|---|---|---|---|---|---|---|---|---|---|---|---|---|---|
| Srrt | 0.0002 | 0.00014 | 0.00019 | 0.00019 | 2.21E-05 | 0 | 0 | 0 | 0.00015 | 3.02E-05 | 3.53E-05 | 0 | 0 | 4.42E-05 | 0.00036 | 0.0004 | 0.00011 | 4.22E-05 |
| Pum3 | 0.00021 | 0.00012 | 0.0003 | 4.69E-05 | 0 | 2.02E-05 | 0 | 0 | 0.00015 | 2.69E-05 | 7.70E-05 | 5.33E-05 | 1.02E-05 | 5.78E-06 | 0.00014 | 2.00E-05 | 3.74E-05 | 6.65E-05 |
| Pgap1 | 3.36E-05 | 0 | 0 | 0 | 0 | 0.00025 | 0 | 8.38E-05 | 0.00015 | 8.92E-06 | 1.42E-05 | 9.88E-05 | 0 | 1.09E-05 | 0 | 1.86E-05 | 0 | 1.84E-05 |
| Pip5k1a | 5.51E-05 | 0.00011 | 0.00017 | 6.18E-05 | 0.00013 | 0.00019 | 3.37E-05 | 0.00015 | 0.00015 | 4.31E-05 | 0.00014 | 9.37E-05 | 3.45E-05 | 6.96E-05 | 7.74E-05 | 0 | 0.00026 | 0 |
| Nup188 | 5.33E-05 | 0.0001 | 0.00019 | 0.0001 | 6.10E-05 | 1.31E-05 | 0 | 1.50E-06 | 0.00015 | 6.37E-05 | 4.57E-05 | 7.62E-05 | 2.33E-05 | 6.02E-05 | 8.14E-05 | 0.00024 | 3.64E-05 | 2.10E-05 |
| Pi4k2a | 9.40E-05 | 0.00021 | 0.00016 | 1.24E-05 | 7.44E-05 | 1.87E-05 | 1.44E-05 | 9.35E-05 | 0.00015 | 6.47E-05 | 3.48E-05 | 0.00011 | 2.58E-05 | 6.99E-05 | 5.88E-05 | 3.68E-05 | 0.00018 | 2.01E-06 |
| Sema6b | 0 | 8.17E-06 | 0 | 8.68E-05 | 0 | 0 | 0 | 0.00013 | 0.00015 | 8.61E-05 | 0 | 0.00017 | 0.00016 | 0 | 0 | 0.00017 | 0 | 3.99E-06 |
| Snrpd1 | 0.00029 | 3.51E-05 | 0.00032 | 0.00013 | 4.93E-06 | 2.22E-05 | 0 | 0 | 0.00015 | 8.37E-06 | 0.00011 | 3.64E-05 | 3.64E-05 | 1.52E-05 | 1.50E-05 | 0.00064 | 1.98E-05 | 0 |
| Bcl2l2 | 4.51E-05 | 0.00016 | 0.00023 | 0.00027 | 0.00026 | 0.00016 | 0.00015 | 0.00023 | 0.00015 | 0.00013 | 0.00015 | 0.0003 | 0.00015 | 0.00018 | 0.00039 | 0 | 0 | 8.16E-05 |
| Rtn3 | 0.00029 | 0.00026 | 0.00024 | 0.00016 | 9.78E-05 | 0.00013 | 0.0002 | 0.00023 | 0.00015 | 0.00031 | 0.00028 | 0.00033 | 0.00016 | 0.00011 | 0 | 0.00036 | 3.71E-05 | 0.00024 |
| Rnf44 | 4.16E-05 | 0.00026 | 0.0002 | 0.00021 | 0.00019 | 0.00018 | 0 | 0.00019 | 0.00015 | 0.00014 | 0.0002 | 0.00015 | 0.00017 | 0.0002 | 0.00011 | 7.37E-05 | 0.00019 | 0.00012 |
| Glod4 | 0.00023 | 4.52E-05 | 9.78E-05 | 7.80E-05 | 4.87E-05 | 9.24E-05 | 8.72E-05 | 4.68E-05 | 0.00015 | 0.00011 | 2.20E-05 | 0.00011 | 0.00011 | 0 | 0 | 0.00011 | 0.00013 | 4.57E-05 |
| Myd88 | 0.00019 | 0.00021 | 0.00034 | 0.00044 | 0.00015 | 0.0001 | 0.00012 | 0.00044 | 0.00015 | 0.00023 | 0 | 2.90E-05 | 0.00041 | 0.00016 | 0 | 8.25E-05 | 0.00019 | 8.71E-05 |
| Kdm5c | 6.66E-05 | 0.00017 | 0.00015 | 0.00034 | 9.44E-05 | 1.89E-05 | 0.00016 | 0.00022 | 0.00015 | 0.00014 | 0.00014 | 0.00024 | 7.89E-06 | 7.70E-05 | 0.00025 | 9.28E-06 | 3.26E-05 | 0.00016 |
| Wwc2 | 0.00014 | 0.00019 | 0.00014 | 0 | 0.00022 | 2.77E-05 | 0.00015 | 4.96E-05 | 0.00015 | 4.23E-05 | 0.0005 | 0.00012 | 8.44E-05 | 0.00012 | 0.00012 | 9.39E-05 | 0.00032 | 0.00017 |
| Cdyl | 4.20E-05 | 0.00014 | 9.10E-05 | 2.63E-06 | 4.34E-05 | 3.44E-05 | 4.83E-05 | 0.00013 | 0.00015 | 0.00012 | 0.00011 | 1.84E-05 | 3.26E-05 | 0.00013 | 0.00024 | 8.76E-05 | 5.12E-05 | 6.17E-05 |
| Smg6 | 0 | 0.00014 | 0.0002 | 0.00014 | 0.00021 | 0.00012 | 0.00025 | 0.00042 | 0.00015 | 9.74E-05 | 0 | 0.00033 | 0 | 2.10E-05 | 0.00029 | 0.00011 | 0.00028 | 8.16E-05 |
| Zc3h14 | 0.00024 | 0.00016 | 0.00014 | 9.43E-06 | 0.00015 | 0.00024 | 1.35E-05 | 0.00012 | 0.00015 | 0.00012 | 0.00016 | 0.00016 | 2.56E-05 | 7.14E-05 | 0.0001 | 0.0002 | 4.09E-06 | 7.07E-05 |
| Gbp2 | 4.21E-05 | 8.03E-05 | 0 | 0 | 0 | 0 | 0 | 0 | 0.00015 | 0.00027 | 0 | 0 | 9.63E-05 | 0 | 0 | 2.84E-05 | 1.01E-05 | 0.00034 |
| Fxyd6 | 0 | 0 | 0 | 0 | 0.00096 | 0 | 0 | 0 | 0.00015 | 0.00038 | 0.00044 | 0 | 0.00158 | 0 | 0 | 8.37E-05 | 0 | 0.00093 |
| Hnrnpul2 | 0.00034 | 0.00034 | 0.00029 | 0.00034 | 0.00021 | 0.0002 | 0.0004 | 0.00019 | 0.00015 | 0.00018 | 0.00017 | 0.00024 | 3.79E-05 | 0.00017 | 0.00022 | 0.00034 | 0.00011 | 0.00017 |
| Stat5a | 6.20E-05 | 9.09E-05 | 0.00022 | 9.70E-06 | 0 | 1.27E-05 | 3.18E-05 | 1.91E-05 | 0.00015 | 6.04E-05 | 0 | 8.43E-05 | 0.0002 | 0 | 0.00011 | 8.28E-05 | 4.17E-05 | 0 |
| Atrx | 0.00028 | 0.00026 | 2.83E-05 | 0 | 7.02E-05 | 0.00016 | 4.43E-05 | 5.20E-05 | 0.00015 | 5.53E-05 | 8.51E-05 | 6.94E-05 | 3.06E-05 | 4.77E-05 | 0.00032 | 0.00017 | 0.0002 | 0 |
| Ppfia1 | 9.69E-05 | 0.00021 | 0.00017 | 4.64E-05 | 0.00018 | 0.00012 | 8.05E-05 | 4.28E-07 | 0.00015 | 8.10E-05 | 0.00026 | 5.96E-05 | 5.78E-05 | 0.00024 | 0.00029 | 0.00015 | 0.00024 | 0.00011 |
| Pitpnb | 0.00016 | 0.00016 | 0.00022 | 0.00017 | 8.85E-05 | 7.33E-05 | 3.92E-05 | 5.56E-05 | 0.00015 | 9.47E-05 | 0.00015 | 7.71E-05 | 3.85E-05 | 8.74E-05 | 0.00011 | 0.00018 | 0 | 8.30E-05 |
| Tsn | 0.00036 | 0.00012 | 0.00022 | 0.00028 | 0.00013 | 8.67E-05 | 0.00021 | 6.08E-05 | 0.00015 | 7.04E-05 | 0.00012 | 4.81E-05 | 0.00011 | 0.00012 | 4.84E-05 | 0.00031 | 0 | 8.74E-05 |
| Slc39a13 | 9.71E-05 | 4.13E-05 | 0.00017 | 0.0002 | 0.00034 | 0.00023 | 0.00052 | 9.30E-06 | 0.00015 | 0.00011 | 0.00028 | 0.0003 | 0.00042 | 0.0004 | 0.00035 | 0 | 0 | 0.00025 |
| Agpat2 | 0.00013 | 0.0001 | 0.00011 | 1.28E-05 | 7.55E-05 | 0.00016 | 0 | 0 | 0.00015 | 0 | 0.00011 | 9.64E-05 | 3.82E-05 | 0.00012 | 0 | 0 | 0 | 4.85E-05 |
| Gusb | 0.00017 | 8.29E-05 | 8.87E-05 | 0.0001 | 5.71E-05 | 0.00013 | 0.0002 | 0.00018 | 0.00015 | 0.00013 | 2.79E-05 | 0 | 0.00018 | 0.00031 | 0 | 9.39E-05 | 0 | 0.00022 |
| Vps29 | 0.00022 | 0.00014 | 0.00021 | 0.00018 | 9.48E-05 | 0.00013 | 6.21E-05 | 0.00014 | 0.00015 | 0.00013 | 0.00014 | 0.00012 | 1.06E-05 | 0.00013 | 0 | 0.00013 | 0 | 0.00015 |
| Minpp1 | 0.00022 | 5.87E-05 | 0.00011 | 0 | 5.34E-05 | 0.00012 | 0 | 3.79E-05 | 0.00015 | 1.17E-06 | 4.34E-05 | 2.02E-05 | 2.72E-05 | 0 | 0 | 0.0001 | 6.61E-05 | 2.82E-05 |
| Cul1 | 0.00037 | 0.00035 | 0.00022 | 0 | 0.00032 | 0.00016 | 0.00022 | 0.00016 | 0.00015 | 4.05E-05 | 0 | 3.78E-05 | 0 | 0.00012 | 0.00018 | 0.00019 | 9.53E-05 | 5.20E-05 |
| Tbc1d13 | 5.07E-05 | 0.00011 | 8.86E-05 | 0.00011 | 0.0001 | 5.11E-05 | 7.14E-05 | 8.88E-05 | 0.00015 | 9.51E-05 | 2.84E-05 | 6.89E-06 | 5.17E-05 | 8.36E-05 | 6.76E-05 | 0 | 0 | 8.78E-05 |

| | | | | | | | | | | | | | | | | | | |
|---|---|---|---|---|---|---|---|---|---|---|---|---|---|---|---|---|---|---|
| Ralb | 0.00025 | 0.00014 | 0.0002 | 8.58E-05 | 6.94E-05 | 9.15E-05 | 0.0001 | 0.00012 | 0.00015 | 5.38E-05 | 0.0002 | 0.00017 | 1.53E-05 | 8.43E-05 | 9.30E-05 | 4.18E-05 | 1.23E-05 | 5.95E-08 |
| H2-K1 | 0 | 0.00017 | 0.00022 | 0.00034 | 2.18E-05 | 1.14E-05 | 5.03E-05 | 3.74E-06 | 0.00015 | 0.00054 | 0.00013 | 0.00042 | 0.00099 | 0.00055 | 0.0002 | 2.71E-05 | 0.00044 | 0.00061 |
| Eif4g1 | 0.0003 | 0.00017 | 0.00042 | 0.00025 | 0.00012 | 9.90E-05 | 0.00023 | 7.69E-05 | 0.00015 | 0.00012 | 0.0002 | 6.23E-05 | 2.94E-05 | 0.00018 | 2.71E-05 | 0.00021 | 0 | 8.42E-05 |
| Iars2 | 5.95E-05 | 0.00018 | 0.00018 | 0.00012 | 4.36E-05 | 2.91E-05 | 9.47E-05 | 1.91E-05 | 0.00015 | 0.00012 | 0.00011 | 6.18E-05 | 4.28E-05 | 7.57E-05 | 6.06E-05 | 0.00011 | 0 | 7.76E-05 |
| Pcdh19 | 0.0001 | 2.03E-05 | 0 | 8.64E-05 | 0 | 2.92E-05 | 2.03E-05 | 0.00017 | 0.00015 | 0.00022 | 2.50E-05 | 0 | 0.00014 | 0 | 0 | 0.00014 | 0 | 0.00025 |
| Golim4 | 0.00023 | 8.85E-05 | 0.00018 | 0.00024 | 0.00026 | 0.00025 | 0.0005 | 0.0003 | 0.00015 | 0.00025 | 0.00025 | 0 | 5.55E-05 | 0.00021 | 0.00027 | 0.00016 | 0.00011 | 0.0003 |
| Lynx1 | 0 | 5.19E-05 | 0 | 7.56E-05 | 0.00012 | 0.00011 | 6.93E-05 | 7.71E-05 | 0.00015 | 0.00023 | 0.00011 | 6.08E-05 | 0.00015 | 7.27E-05 | 0 | 0 | 0 | 0.00025 |
| Gdi1 | 7.64E-05 | 0.00028 | 8.05E-05 | 5.18E-05 | 0.00018 | 0.00011 | 0 | 0.00021 | 0.00015 | 0.00024 | 0.00025 | 0.00024 | 3.28E-05 | 3.39E-05 | 0.00038 | 3.52E-05 | 0 | 0.00025 |
| Epb41l2 | 0.0003 | 0.00015 | 2.75E-05 | 3.64E-05 | 0 | 0.00021 | 0.00015 | 0.00022 | 0.00015 | 0.00048 | 0.00031 | 0.00023 | 0.00031 | 3.04E-05 | 0.00026 | 0.00029 | 0.00042 | 0.00066 |
| Becn1 | 0.00011 | 0.00017 | 0.00017 | 0.00022 | 0.0002 | 0.00014 | 7.82E-05 | 0.00018 | 0.00015 | 0.00015 | 0.00017 | 0.00011 | 7.47E-05 | 0.00013 | 0.00016 | 9.94E-05 | 0 | 0.00014 |
| Snf8 | 0.0002 | 7.88E-05 | 0.00018 | 0.00047 | 0.00015 | 0.00015 | 0.00018 | 8.24E-05 | 0.00015 | 5.22E-05 | 0.00017 | 0 | 4.49E-05 | 0.00019 | 9.56E-05 | 0.0001 | 0 | 5.30E-05 |
| Homer3 | 8.00E-05 | 0.00026 | 0.00016 | 0 | 0.00016 | 7.16E-05 | 3.44E-05 | 0.00022 | 0.00015 | 6.49E-05 | 3.55E-05 | 0.00014 | 1.49E-06 | 0.00013 | 0.00011 | 0.00012 | 0 | 0.00011 |
| Bckdk | 0.00018 | 0.00011 | 0.00011 | 8.54E-05 | 9.57E-05 | 0.00014 | 0.00014 | 9.67E-06 | 0.00015 | 3.64E-05 | 7.13E-05 | 6.05E-05 | 7.59E-05 | 9.48E-05 | 0 | 0.00025 | 0 | 7.07E-05 |
| Ephb3 | 7.01E-07 | 0.00012 | 0 | 0.00011 | 0.00016 | 5.98E-05 | 0.00021 | 0 | 0.00015 | 9.35E-05 | 0.00015 | 0 | 0.00013 | 6.44E-05 | 6.75E-06 | 2.91E-05 | 0 | 0.00033 |
| Fut8 | 9.69E-05 | 0.00017 | 0 | 0 | 0.00018 | 0.0001 | 0 | 0.00019 | 0.00015 | 6.62E-05 | 0.00039 | 0.00012 | 0 | 0.00022 | 0 | 9.55E-05 | 0 | 4.54E-05 |
| Grk5 | 5.34E-05 | 3.43E-05 | 2.75E-05 | 0 | 6.63E-05 | 0 | 0 | 0 | 0.00015 | 0.00033 | 6.88E-05 | 4.99E-05 | 0.00024 | 5.23E-05 | 0.00015 | 0 | 2.21E-05 | 0.00025 |
| Ddx1 | 0.00044 | 0.00012 | 0.00038 | 0.00015 | 3.62E-05 | 0.00011 | 0.00019 | 4.97E-05 | 0.00015 | 4.74E-05 | 7.68E-05 | 6.51E-05 | 0 | 7.38E-05 | 7.74E-05 | 0.00022 | 0 | 5.10E-05 |
| Cnot2 | 0.00027 | 0.00021 | 0.00013 | 0 | 9.59E-05 | 0.0001 | 7.34E-05 | 1.72E-05 | 0.00015 | 8.21E-05 | 6.42E-05 | 0.00012 | 0 | 2.14E-05 | 0.00023 | 0.00014 | 0 | 0 |
| Faf2 | 0.00012 | 0.00014 | 0.00017 | 0.0001 | 0.00011 | 6.05E-05 | 0.00011 | 8.44E-05 | 0.00015 | 7.14E-05 | 0.0001 | 0.00011 | 7.85E-05 | 9.44E-05 | 4.82E-05 | 0.00012 | 0 | 5.35E-05 |
| Efr3a | 0.00022 | 0.00011 | 8.95E-05 | 0 | 8.56E-05 | 0.00017 | 2.39E-05 | 3.67E-05 | 0.00015 | 5.36E-05 | 9.25E-05 | 0.0001 | 0 | 8.00E-05 | 0 | 5.44E-05 | 2.54E-05 | 7.29E-05 |
| Immt | 0.00023 | 0.00014 | 0.0002 | 8.91E-05 | 1.54E-05 | 4.97E-05 | 0 | 2.14E-05 | 0.00015 | 4.42E-05 | 0.00012 | 0.00014 | 0 | 9.18E-05 | 3.64E-05 | 0.00025 | 0 | 3.66E-05 |
| Snrpa | 0.00027 | 0.0001 | 0.00023 | 0.00031 | 1.89E-05 | 0 | 5.34E-06 | 0 | 0.00015 | 6.67E-06 | 9.79E-05 | 4.45E-05 | 3.76E-06 | 7.58E-05 | 0 | 0.00018 | 0 | 3.18E-05 |
| Zfp512b | 7.59E-05 | 0.00018 | 1.97E-05 | 4.40E-05 | 0.00014 | 0.00014 | 3.26E-06 | 0 | 0.00015 | 0.00014 | 0.00011 | 0 | 8.94E-05 | 0.00011 | 8.12E-05 | 7.42E-06 | 2.02E-05 | 7.82E-05 |
| Fance | 0 | 8.11E-05 | 0.00014 | 0.00023 | 0.00015 | 0.00022 | 0 | 0.00016 | 0.00015 | 1.66E-05 | 0.00023 | 0 | 0 | 9.78E-05 | 0.00034 | 8.08E-05 | 0 | 0 |
| Vps16 | 1.13E-05 | 0.00012 | 0.00012 | 0.00012 | 0.00018 | 0.00018 | 0.00019 | 0 | 0.00015 | 8.12E-05 | 0.00013 | 6.67E-05 | 0.00022 | 0.00017 | 0.00022 | 0.00015 | 0 | 3.11E-05 |
| Napa | 0.00022 | 0.00024 | 0.00023 | 0.00028 | 0.00021 | 0.0002 | 0.00028 | 0.00013 | 0.00015 | 5.27E-05 | 0.00015 | 0.00016 | 0.0001 | 0.00022 | 0.00019 | 8.73E-05 | 0.0002 | 7.83E-05 |
| Heg1 | 0 | 0 | 0.00037 | 5.59E-05 | 0.00021 | 5.02E-05 | 0.00065 | 0.00063 | 0.00015 | 0.00166 | 9.66E-05 | 0 | 0.00021 | 0 | 0 | 0.0002 | 0.00011 | 0.00084 |
| Smc6 | 0.00022 | 9.12E-05 | 8.76E-05 | 0 | 0.00015 | 0.00011 | 6.24E-05 | 0 | 0.00015 | 7.70E-05 | 0.00015 | 3.77E-05 | 0 | 5.49E-05 | 8.06E-05 | 0.00038 | 0 | 8.70E-05 |
| Prmt1 | 0.00013 | 5.87E-05 | 0.00033 | 0.00035 | 0 | 0 | 0 | 5.79E-05 | 0.00015 | 0 | 9.58E-05 | 4.74E-05 | 0 | 3.08E-05 | 9.32E-06 | 0.00019 | 0.00018 | 2.66E-05 |
| Cmpk1 | 0.00026 | 6.64E-05 | 0.00015 | 0.00016 | 0.00019 | 0.00022 | 0.00023 | 0.00011 | 0.00015 | 9.78E-05 | 0.00022 | 3.31E-05 | 1.14E-05 | 0.00015 | 0 | 0.00012 | 0 | 0.00012 |
| Sfxn1 | 0.00024 | 0.00011 | 0.00017 | 0.00041 | 0 | 0.00033 | 2.87E-05 | 8.29E-05 | 0.00015 | 0.00017 | 0 | 5.66E-05 | 0.00034 | 9.30E-05 | 0 | 6.28E-05 | 6.42E-05 | 2.62E-05 |
| Sh3glb2 | 7.54E-05 | 8.54E-05 | 2.43E-05 | 0.00014 | 7.85E-05 | 0 | 1.94E-05 | 0.00024 | 0.00015 | 8.19E-05 | 5.73E-05 | 7.55E-05 | 2.03E-05 | 1.50E-05 | 6.32E-05 | 5.61E-06 | 5.70E-05 | 0.00012 |
| Mef2d | 0.00018 | 0.00033 | 0.00028 | 0.0001 | 0.00016 | 0.0002 | 8.05E-05 | 3.35E-05 | 0.00015 | 0.00012 | 0.0001 | 0.00014 | 1.61E-05 | 0.00013 | 0.00033 | 7.63E-06 | 0.00028 | 5.29E-05 |

| | | | | | | | | | | | | | | | | | | |
|---|---|---|---|---|---|---|---|---|---|---|---|---|---|---|---|---|---|---|
| Zdhhc18 | 0 | 0.00014 | 0.00018 | 0.00018 | 0.00014 | 3.15E-05 | 6.28E-05 | 0.00016 | 0.00015 | 0.00017 | 0.00016 | 1.88E-05 | 0 | 0.00012 | 9.17E-05 | 0.00018 | 0.00014 | 0.00016 |
| Glyr1 | 0.0002 | 0.00022 | 0.00017 | 0.00017 | 0.00011 | 0.00011 | 8.81E-05 | 0.00014 | 0.00015 | 7.65E-05 | 8.27E-05 | 7.62E-05 | 6.77E-05 | 0.00011 | 0.00035 | 0.00014 | 0.00023 | 5.20E-05 |
| Stau1 | 0.00025 | 0.00023 | 0.00017 | 9.01E-05 | 0.00015 | 0.00014 | 5.68E-05 | 0.00013 | 0.00015 | 4.05E-05 | 4.39E-05 | 0.00016 | 9.95E-05 | 0.00011 | 8.85E-05 | 6.20E-05 | 0.0002 | 0.00014 |
| Top2b | 0.00026 | 0.0002 | 7.69E-05 | 6.79E-05 | 0.00013 | 0.00022 | 8.91E-05 | 7.90E-05 | 0.00015 | 0.00011 | 0.00012 | 1.28E-05 | 0 | 0.00012 | 0.00015 | 0.00014 | 0.00014 | 7.68E-05 |
| Ube2q1 | 0.00023 | 0.00021 | 0.00019 | 2.07E-05 | 0.00013 | 0.00021 | 0.00014 | 0.00019 | 0.00015 | 6.79E-05 | 0.00014 | 0.00012 | 6.31E-05 | 0.00013 | 6.57E-05 | 5.04E-05 | 0.00013 | 3.93E-06 |
| Nab2 | 7.55E-05 | 0.00016 | 3.74E-05 | 0.00029 | 0.00013 | 0.00022 | 9.52E-05 | 0 | 0.00015 | 0.00014 | 0.00019 | 0.00032 | 0.00023 | 9.47E-05 | 0.00034 | 0.00023 | 0.00011 | 0 |
| Psmd2 | 0.00032 | 0.00017 | 0.00033 | 0.00021 | 8.87E-05 | 3.29E-05 | 9.31E-05 | 8.85E-05 | 0.00015 | 8.10E-05 | 0.00013 | 6.78E-05 | 4.09E-05 | 6.23E-05 | 0.0003 | 0.00023 | 0 | 6.51E-05 |
| Dusp3 | 5.23E-05 | 0.00033 | 0.00016 | 9.80E-05 | 0.00024 | 9.71E-05 | 0.00012 | 0.00024 | 0.00015 | 0.00024 | 0.0005 | 0.00033 | 2.85E-05 | 0.00013 | 0 | 0 | 8.98E-05 | 0.00022 |
| Gskip | 0.00014 | 0.0001 | 0.00019 | 7.31E-05 | 0.00012 | 0.00017 | 4.90E-05 | 7.55E-05 | 0.00015 | 8.68E-05 | 0.00014 | 2.86E-05 | 5.99E-05 | 7.17E-05 | 0 | 0 | 0 | 9.51E-05 |
| Abcf1 | 0.00012 | 0.00012 | 0.00019 | 0.00013 | 6.14E-05 | 1.22E-05 | 0 | 8.43E-06 | 0.00015 | 2.98E-05 | 5.43E-05 | 5.39E-05 | 1.83E-05 | 3.70E-05 | 8.77E-05 | 0.00011 | 0 | 6.52E-05 |
| Dnajc14 | 0.00011 | 0.00022 | 0.00016 | 0.00034 | 0.00018 | 0.00016 | 0.00016 | 0.0002 | 0.00015 | 0.00016 | 0.00021 | 0.00014 | 7.25E-05 | 0.00018 | 0.00024 | 0.00015 | 0.00025 | 0.00018 |
| Zfp651 | 2.78E-05 | 0.0001 | 6.25E-05 | 4.00E-05 | 0.00024 | 6.81E-05 | 0.00031 | 6.44E-05 | 0.00015 | 0.00015 | 0.00013 | 7.31E-05 | 0 | 0.0001 | 0.00013 | 0 | 8.81E-05 | 9.48E-05 |
| Ncdn | 4.02E-05 | 7.88E-05 | 0.00019 | 0.0002 | 7.54E-05 | 1.21E-05 | 0 | 0.0001 | 0.00015 | 6.03E-05 | 5.39E-05 | 7.61E-06 | 0 | 7.08E-05 | 7.58E-05 | 0 | 0 | 8.07E-05 |
| Prr13 | 9.44E-05 | 0.00012 | 9.48E-05 | 0.00026 | 0 | 2.83E-05 | 0.0002 | 7.85E-05 | 0.00015 | 0.00047 | 0.00013 | 0.00027 | 0 | 0 | 0 | 0 | 7.04E-05 | 0.00036 |
| Abhd17a | 0.00012 | 0.00012 | 0.00015 | 0.00036 | 0.00016 | 8.12E-05 | 0.00015 | 0.00017 | 0.00015 | 8.76E-05 | 0.00014 | 8.80E-05 | 6.98E-05 | 0.00014 | 0 | 5.27E-06 | 1.88E-05 | 0.00014 |
| Fam114a1 | 0.00033 | 2.98E-05 | 0.00018 | 0.00049 | 0.00025 | 0.00023 | 0.00087 | 0.00041 | 0.00015 | 0.00038 | 0.00024 | 0.00034 | 0.00036 | 0.00027 | 0.00032 | 8.58E-05 | 0.00016 | 0.00036 |
| Eif3l | 0.00038 | 0.00018 | 0.00035 | 0.00041 | 0.00011 | 0.00026 | 0.00014 | 0.00016 | 0.00015 | 8.58E-05 | 8.19E-05 | 0.00011 | 2.54E-06 | 0.00017 | 0 | 0.00034 | 0.00023 | 0.00011 |
| Ptpn1 | 1.02E-05 | 8.49E-05 | 0.00026 | 0 | 0.00014 | 0.00044 | 0 | 0.0003 | 0.00015 | 5.75E-05 | 3.49E-05 | 0.0004 | 0.00021 | 0.00026 | 9.74E-05 | 1.58E-05 | 0 | 0.00019 |
| Ndel1 | 0.00038 | 0.00017 | 0.00032 | 0 | 5.17E-05 | 0.00015 | 0.00041 | 0.00016 | 0.00015 | 0.00014 | 0.00019 | 0.00021 | 8.40E-05 | 0.00011 | 0.00026 | 5.53E-05 | 0.00056 | 1.73E-05 |
| Rae1 | 0.00017 | 0.00013 | 0.00028 | 9.58E-05 | 4.56E-05 | 5.56E-06 | 5.14E-05 | 0 | 0.00015 | 4.74E-05 | 5.12E-05 | 1.31E-05 | 1.45E-05 | 6.75E-05 | 0.00014 | 4.15E-05 | 6.21E-05 | 5.67E-05 |
| Srpk2 | 0.0001 | 0.00014 | 0 | 3.80E-05 | 0.00021 | 9.93E-05 | 1.50E-05 | 5.42E-05 | 0.00015 | 6.29E-05 | 1.61E-05 | 4.63E-05 | 6.00E-05 | 7.31E-05 | 0.00021 | 7.74E-05 | 0.00012 | 5.34E-05 |
| Txn1 | 0.0006 | 0.00014 | 0.00051 | 0.00035 | 0 | 0.00024 | 8.47E-05 | 0 | 0.00015 | 0.00024 | 3.94E-05 | 0.00028 | 0 | 3.93E-05 | 0 | 0.0004 | 0.00046 | 0.00022 |
| Pum1 | 0.00021 | 0.0002 | 0.00015 | 0 | 0.00012 | 0.00013 | 3.03E-05 | 6.53E-05 | 0.00015 | 8.79E-05 | 0.00011 | 0.00015 | 4.64E-05 | 4.43E-05 | 0.00025 | 7.37E-05 | 0.00035 | 4.91E-05 |
| Lig3 | 8.96E-05 | 7.71E-05 | 0.00011 | 0.00016 | 7.28E-05 | 2.72E-05 | 7.68E-05 | 9.71E-05 | 0.00015 | 3.98E-06 | 9.32E-05 | 2.61E-05 | 0 | 5.69E-05 | 0.00011 | 9.89E-05 | 0 | 3.76E-05 |
| Pcnp | 0.00023 | 0.00017 | 0.0002 | 6.44E-05 | 0.0001 | 0.0001 | 4.34E-05 | 1.37E-05 | 0.00014 | 4.75E-05 | 0.00013 | 0.00013 | 6.19E-05 | 3.54E-05 | 0.00019 | 0.00015 | 7.01E-05 | 0.00013 |
| Arpc2 | 0.00042 | 0.00015 | 0.00035 | 0 | 8.97E-05 | 0.00017 | 0.00012 | 0.00016 | 0.00014 | 0.00014 | 0.00033 | 0.00034 | 0 | 0.00011 | 0 | 9.33E-05 | 0 | 9.30E-05 |
| Rfk | 0.00018 | 0.00015 | 9.46E-05 | 5.82E-05 | 0.00017 | 0.00011 | 8.00E-05 | 9.48E-05 | 0.00014 | 3.97E-05 | 1.44E-05 | 5.77E-05 | 0 | 4.64E-06 | 0 | 0.00012 | 1.36E-05 | 0 |
| Leng8 | 0.00011 | 0.00035 | 0.0001 | 0.00012 | 0.00027 | 0.00011 | 0 | 0.00017 | 0.00014 | 0.00017 | 0.00014 | 0.00019 | 0 | 0.00014 | 0.00099 | 0 | 0 | 0.00013 |
| Ptp4a2 | 0.00027 | 0.00013 | 0.00015 | 2.28E-05 | 4.93E-05 | 0.00011 | 0.00012 | 0.00015 | 0.00014 | 0.00011 | 4.30E-05 | 0.00014 | 0 | 3.29E-05 | 4.84E-05 | 0.00013 | 0.00018 | 7.74E-05 |
| Ddx39b | 0.00027 | 0.00018 | 0.00026 | 6.87E-05 | 2.49E-05 | 4.27E-05 | 0 | 4.30E-06 | 0.00014 | 1.47E-05 | 9.34E-05 | 0.00011 | 0 | 3.44E-05 | 0.00047 | 0.00036 | 0.00019 | 4.98E-05 |
| Aak1 | 0.00014 | 5.16E-05 | 8.33E-05 | 3.57E-05 | 8.13E-05 | 9.89E-05 | 0.00024 | 2.32E-05 | 0.00014 | 9.79E-05 | 8.15E-05 | 0.00019 | 0.00011 | 0 | 0.00021 | 0.00014 | 0.0002 | 8.98E-05 |
| Atp6ap2 | 0.00011 | 0.00025 | 0.00014 | 8.49E-06 | 0.00021 | 0.00016 | 0.0002 | 0.00014 | 0.00014 | 8.35E-05 | 9.91E-05 | 8.65E-05 | 0.00011 | 0.00013 | 0 | 0.00011 | 9.77E-05 | 0.0003 |
| Sec11a | 0.00036 | 9.77E-05 | 0.0003 | 0.0003 | 0.00022 | 0.00027 | 0.00027 | 0.00024 | 0.00014 | 0.00017 | 0.00019 | 0.00023 | 0.00029 | 0.00021 | 9.51E-05 | 0.00017 | 0 | 0.00027 |

| | | | | | | | | | | | | | | | | | | |
|---|---|---|---|---|---|---|---|---|---|---|---|---|---|---|---|---|---|---|
| Ptpmt1 | 0.00021 | 5.70E-05 | 0.00013 | 0 | 1.46E-05 | 7.04E-05 | 0 | 8.16E-06 | 0.00014 | 4.42E-05 | 2.38E-05 | 7.29E-05 | 3.35E-05 | 3.50E-06 | 5.21E-05 | 3.80E-05 | 0 | 0 |
| Ik | 0.00026 | 0.00024 | 0.00015 | 0.00028 | 0.00014 | 5.65E-05 | 0.00011 | 1.75E-05 | 0.00014 | 0.0001 | 2.37E-05 | 0.00013 | 4.77E-05 | 8.51E-05 | 0.00026 | 0.0002 | 6.06E-05 | 0.00015 |
| Scamp3 | 0.00015 | 9.35E-05 | 0.00017 | 0.00037 | 9.24E-05 | 2.46E-05 | 6.97E-05 | 8.95E-05 | 0.00014 | 0.0001 | 9.37E-05 | 0 | 0.00012 | 1.11E-05 | 0.00011 | 5.75E-05 | 9.67E-05 | 9.91E-05 |
| Kitl | 0 | 9.31E-06 | 0 | 0 | 4.59E-05 | 0.00016 | 0 | 0 | 0.00014 | 9.61E-05 | 0 | 2.52E-05 | 7.22E-05 | 0 | 0 | 0 | 0.00012 | 5.97E-05 |
| Evc | 0 | 5.75E-05 | 0 | 5.48E-05 | 7.33E-05 | 8.86E-06 | 4.00E-06 | 8.27E-05 | 0.00014 | 0.00015 | 1.79E-05 | 4.37E-06 | 0 | 5.98E-05 | 0.00014 | 0.00014 | 0 | 0.00017 |
| Antxr2 | 0.00026 | 0.00027 | 0 | 0 | 0.00028 | 0.00025 | 8.03E-05 | 0.00033 | 0.00014 | 0.00053 | 0 | 0 | 4.26E-05 | 0.00026 | 0.00019 | 6.54E-05 | 0.00012 | 0.00012 |
| Capg | 0.00035 | 0.00029 | 0.00053 | 0.00023 | 0.00026 | 0.00036 | 0.00014 | 0.00082 | 0.00014 | 9.14E-05 | 0.00029 | 0.00055 | 0.00012 | 0.00015 | 0 | 1.30E-05 | 0.00015 | 0.00032 |
| Gid8 | 0.00022 | 0.00015 | 0.00014 | 0.00011 | 0.00012 | 0.00016 | 8.52E-05 | 0.00012 | 0.00014 | 9.39E-05 | 0.00011 | 4.91E-05 | 5.50E-05 | 8.98E-05 | 4.63E-05 | 0.00017 | 0 | 8.60E-05 |
| Lrpap1 | 0.00019 | 0.0001 | 0.00013 | 9.18E-05 | 0.00033 | 0.00041 | 0.00023 | 0.00029 | 0.00014 | 0.00025 | 0.00017 | 0.00026 | 0.00052 | 0.0002 | 6.24E-05 | 3.54E-05 | 7.37E-05 | 0.00029 |
| Psme3 | 0.00028 | 0.0002 | 0.00052 | 0.00023 | 9.15E-05 | 0.0001 | 5.19E-05 | 3.44E-05 | 0.00014 | 9.34E-05 | 0.00016 | 2.62E-05 | 9.30E-05 | 4.11E-05 | 0.00028 | 0.00025 | 8.26E-05 | 4.12E-05 |
| Prdx3 | 0.00028 | 0.00013 | 0.00029 | 0.00021 | 4.17E-05 | 9.27E-05 | 0 | 1.92E-05 | 0.00014 | 2.85E-05 | 2.49E-05 | 7.37E-05 | 1.65E-06 | 0 | 0 | 3.74E-05 | 0 | 2.02E-05 |
| Hnrnpdl | 0.00018 | 6.58E-05 | 0.00016 | 6.35E-05 | 4.49E-07 | 1.84E-05 | 0 | 0 | 0.00014 | 2.13E-05 | 3.90E-05 | 5.13E-05 | 1.60E-05 | 2.20E-06 | 0.00026 | 0.0002 | 0.00027 | 0 |
| Rnf19b | 0.00024 | 0.0002 | 0.00049 | 0 | 0.00056 | 0.00037 | 0.00049 | 0.00017 | 0.00014 | 0 | 0.00077 | 0.0003 | 0.00076 | 0.00036 | 0.0003 | 0.00012 | 0.0009 | 4.15E-05 |
| Mvd | 2.68E-05 | 6.53E-05 | 0 | 8.25E-05 | 4.94E-05 | 0 | 0 | 1.29E-05 | 0.00014 | 0 | 3.84E-05 | 0 | 0 | 2.58E-05 | 0 | 0 | 7.92E-06 | 1.10E-05 |
| Eif3h | 0.00035 | 0.00014 | 8.39E-05 | 0 | 7.78E-05 | 0.00023 | 6.34E-05 | 3.69E-05 | 0.00014 | 0 | 0 | 0.00021 | 2.75E-06 | 5.94E-05 | 0 | 4.52E-05 | 0.00015 | 2.00E-05 |
| Mad2l2 | 5.12E-05 | 2.93E-05 | 0 | 6.94E-05 | 6.24E-05 | 6.82E-05 | 0 | 0 | 0.00014 | 6.58E-06 | 3.66E-05 | 0 | 5.84E-05 | 8.50E-05 | 0 | 5.53E-05 | 0 | 0 |
| Cmtm4 | 7.03E-06 | 8.52E-05 | 1.37E-05 | 0 | 0.00012 | 0.0001 | 3.93E-05 | 0 | 0.00014 | 0 | 0.00024 | 3.34E-05 | 6.97E-05 | 0.00026 | 0 | 0.00015 | 0 | 8.92E-05 |
| Preb | 0.00019 | 0.00013 | 0.00027 | 0.00032 | 4.18E-05 | 2.56E-05 | 0.00011 | 9.02E-05 | 0.00014 | 7.57E-05 | 4.73E-05 | 8.14E-05 | 5.77E-05 | 8.89E-05 | 0.00042 | 4.69E-05 | 0.00021 | 4.85E-05 |
| Osbpl3 | 2.86E-06 | 2.10E-05 | 5.85E-05 | 0 | 0.00016 | 0 | 9.18E-05 | 0.0001 | 0.00014 | 7.74E-05 | 2.01E-06 | 9.05E-05 | 0 | 0.00014 | 6.92E-05 | 9.15E-05 | 2.43E-05 | 0 |
| Nudt21 | 0.00037 | 0.00013 | 0.00015 | 1.32E-05 | 4.54E-05 | 0.00015 | 3.13E-05 | 5.24E-05 | 0.00014 | 3.31E-05 | 3.37E-05 | 0.00019 | 3.56E-05 | 8.34E-05 | 0 | 0.00039 | 9.74E-05 | 0 |
| Arl2 | 4.93E-05 | 0.00013 | 2.98E-05 | 9.88E-05 | 0.00019 | 0.00014 | 2.96E-05 | 4.82E-05 | 0.00014 | 1.49E-05 | 0.00016 | 0.00011 | 2.10E-05 | 0.00011 | 9.34E-05 | 0.00011 | 0 | 8.93E-05 |
| Tpgs1 | 2.23E-05 | 0.00012 | 4.30E-05 | 0.00035 | 0.00013 | 3.15E-05 | 5.93E-05 | 0.00014 | 0.00014 | 9.83E-05 | 5.67E-05 | 0.00011 | 4.11E-05 | 0.00012 | 3.59E-05 | 7.00E-05 | 6.95E-05 | 0.00015 |
| Rnf24 | 7.79E-05 | 8.15E-05 | 7.51E-05 | 0 | 0.0001 | 0.00019 | 2.05E-05 | 0 | 0.00014 | 1.06E-05 | 0 | 9.75E-05 | 0 | 4.82E-06 | 0 | 1.88E-05 | 8.03E-05 | 2.60E-05 |
| Hsd17b12 | 0.00029 | 4.34E-05 | 0.0002 | 0 | 8.29E-05 | 7.11E-05 | 0 | 1.12E-06 | 0.00014 | 1.76E-05 | 6.44E-05 | 0.00012 | 7.27E-06 | 3.27E-05 | 0 | 0.00014 | 0 | 0 |
| Gng10 | 0.00038 | 7.34E-05 | 0.00019 | 0.00028 | 3.93E-05 | 0.00011 | 0.00013 | 3.97E-05 | 0.00014 | 0.00011 | 0.00013 | 0.0001 | 0.00018 | 0 | 0 | 0.00011 | 0.0002 | 0.0002 |
| Nudcd3 | 7.50E-05 | 0.00022 | 0.00016 | 0.00022 | 0.00015 | 5.30E-05 | 0 | 9.84E-05 | 0.00014 | 0.00011 | 0.00015 | 9.93E-05 | 5.47E-06 | 0.00011 | 0.00026 | 0.0002 | 0 | 9.51E-05 |
| Ift43 | 3.69E-05 | 0.00011 | 0.0001 | 0.00025 | 6.72E-05 | 0.00013 | 9.40E-05 | 0.00022 | 0.00014 | 5.70E-05 | 0.00025 | 4.09E-05 | 6.36E-05 | 0.00016 | 2.74E-05 | 1.34E-05 | 0.00012 | 7.95E-05 |
| Srrm1 | 0.00024 | 0.00018 | 0.0002 | 1.37E-05 | 7.00E-05 | 7.34E-05 | 0 | 0 | 0.00014 | 6.96E-05 | 4.43E-05 | 9.17E-05 | 5.06E-07 | 1.50E-05 | 0.00046 | 0.00025 | 0.00018 | 3.69E-05 |
| Arl6ip4 | 0.00012 | 9.07E-05 | 8.49E-05 | 0.00012 | 9.16E-05 | 4.52E-05 | 3.39E-05 | 6.28E-05 | 0.00014 | 5.06E-05 | 6.47E-05 | 3.34E-05 | 1.83E-05 | 0.0001 | 0.00015 | 0.00017 | 7.91E-06 | 9.36E-05 |
| Coa5 | 0.00017 | 0.00012 | 0.00015 | 0.00029 | 0.00013 | 0.00011 | 0.00014 | 0.00012 | 0.00014 | 0.00013 | 7.85E-05 | 0.00014 | 0.0001 | 7.03E-05 | 3.64E-06 | 0 | 0 | 0.00012 |
| Steap3 | 0 | 6.15E-05 | 0 | 8.92E-06 | 0.00023 | 7.72E-05 | 2.96E-05 | 0 | 0.00014 | 0.00052 | 0.0001 | 0 | 0.00027 | 0.00015 | 0 | 4.79E-06 | 0 | 0.00044 |
| Prr14 | 0.00013 | 0.00017 | 0.00013 | 0 | 0.00011 | 0.00019 | 3.42E-05 | 7.77E-05 | 0.00014 | 4.18E-05 | 0.00011 | 2.46E-05 | 1.18E-05 | 0.00012 | 1.48E-05 | 0 | 2.36E-05 | 1.12E-05 |
| Alg10b | 0.00012 | 0.00014 | 0.00018 | 0.00015 | 0.00013 | 7.00E-05 | 0.00017 | 0.00015 | 0.00014 | 9.08E-05 | 0.0001 | 3.58E-05 | 3.10E-05 | 3.50E-05 | 0 | 0.0002 | 7.77E-05 | 0.00019 |

| | | | | | | | | | | | | | | | | | | |
|---|---|---|---|---|---|---|---|---|---|---|---|---|---|---|---|---|---|---|
| Chchd2 | 0.00037 | 0.00017 | 0.00026 | 0.00118 | 0.00024 | 0.00034 | 0.00015 | 0.00035 | 0.00014 | 0.00032 | 0.00039 | 0.00023 | 0.00048 | 0.0004 | 4.37E-05 | 0.00014 | 0.00109 | 0.00021 |
| Gtpbp4 | 0.00015 | 0.00014 | 0.00035 | 5.11E-05 | 1.74E-05 | 0 | 0 | 0 | 0.00014 | 2.73E-05 | 9.70E-05 | 2.99E-05 | 3.70E-05 | 7.93E-07 | 8.94E-05 | 2.26E-05 | 0 | 0 |
| Rab43 | 0.00013 | 0.00024 | 0.00016 | 6.74E-05 | 0.00014 | 0.00013 | 1.19E-05 | 0.0003 | 0.00014 | 0.00012 | 0 | 0.00016 | 0 | 0.00017 | 0.00021 | 0 | 0.00039 | 0.0001 |
| Mocs2 | 0.00022 | 0.00019 | 0.00018 | 0.00024 | 0.00027 | 0.00028 | 0.00014 | 7.43E-05 | 0.00014 | 0.00018 | 7.04E-06 | 8.55E-05 | 0.00017 | 7.29E-05 | 0.00012 | 5.03E-05 | 0 | 8.05E-05 |
| Slc4a2 | 3.26E-06 | 0.00021 | 0.00014 | 0.00032 | 3.92E-05 | 5.62E-06 | 8.36E-05 | 0.00017 | 0.00014 | 0.00017 | 0.00017 | 0.00018 | 0 | 4.72E-05 | 0.00048 | 0.0002 | 4.67E-05 | 0.00019 |
| Ipo5 | 0.00034 | 0.00014 | 0.00037 | 0.00013 | 0 | 8.73E-05 | 6.48E-05 | 9.75E-06 | 0.00014 | 9.78E-05 | 0.00012 | 0.00023 | 0.0002 | 0 | 0.00012 | 0.0004 | 0.00019 | 0 |
| Vrk3 | 4.91E-05 | 0.00011 | 3.32E-05 | 0.00021 | 9.73E-05 | 1.58E-05 | 6.72E-05 | 0.00013 | 0.00014 | 3.63E-05 | 5.17E-06 | 5.13E-05 | 0.00013 | 6.21E-05 | 0.00021 | 0.00013 | 9.84E-05 | 5.59E-05 |
| Zfp385a | 1.06E-05 | 9.89E-05 | 0.00017 | 3.74E-05 | 0.00021 | 8.09E-05 | 0 | 0.00026 | 0.00014 | 0.0007 | 6.88E-05 | 0 | 5.81E-06 | 0.00028 | 2.58E-05 | 0 | 2.02E-05 | 7.22E-05 |
| Ier2 | 1.86E-06 | 0.00012 | 6.68E-05 | 0 | 0.00025 | 0.00018 | 3.23E-05 | 0.00046 | 0.00014 | 0.00014 | 0 | 0.00012 | 0.00022 | 6.48E-05 | 0.00011 | 0 | 0.00553 | 0.00018 |
| Prcc | 9.36E-05 | 0.00024 | 0.00015 | 0.00032 | 9.43E-05 | 3.65E-05 | 7.92E-05 | 0.00013 | 0.00014 | 8.64E-05 | 7.83E-05 | 8.91E-05 | 0 | 9.28E-05 | 0.00011 | 0.00017 | 0.00019 | 2.54E-05 |
| Fam20b | 0.00011 | 8.81E-05 | 0.00016 | 0.00013 | 0.00015 | 0.00018 | 0.00013 | 0.00019 | 0.00014 | 0.00014 | 0.00011 | 2.87E-05 | 7.04E-05 | 6.31E-06 | 0 | 0.00013 | 0 | 0.00012 |
| Serinc5 | 6.36E-05 | 0.00013 | 8.60E-05 | 9.53E-05 | 0.00025 | 0.00025 | 0.00031 | 5.42E-05 | 0.00014 | 6.78E-05 | 0.00011 | 7.29E-06 | 0 | 0.00022 | 3.84E-05 | 0.00012 | 4.02E-06 | 6.18E-05 |
| Myo9b | 0.00015 | 9.65E-05 | 0.00012 | 0 | 1.22E-05 | 9.85E-05 | 7.95E-05 | 8.07E-05 | 0.00014 | 2.12E-05 | 8.28E-05 | 0.00012 | 2.56E-05 | 5.20E-05 | 0.00015 | 6.26E-05 | 0.00015 | 5.03E-06 |
| Sf3b4 | 0.00014 | 0.0001 | 0.00029 | 0.00013 | 1.03E-05 | 1.27E-05 | 0 | 5.62E-05 | 0.00014 | 1.90E-05 | 8.34E-05 | 2.02E-05 | 0 | 9.26E-05 | 4.47E-05 | 0.00019 | 0.00011 | 3.15E-05 |
| Ppp2r5c | 0.00028 | 0.00021 | 0.00015 | 0.00014 | 0.00019 | 0.00026 | 0.00022 | 7.74E-05 | 0.00014 | 0.00012 | 0.00011 | 0.00019 | 0.00011 | 7.94E-05 | 0.00013 | 0.00016 | 7.98E-05 | 9.44E-05 |
| Rnf4 | 0.0004 | 0.00027 | 0.00049 | 0.0002 | 0.00014 | 0.00018 | 0.00021 | 8.64E-05 | 0.00014 | 0.0001 | 0.00023 | 0.00019 | 0.00011 | 0.00015 | 0.00028 | 0.00035 | 6.79E-05 | 5.40E-05 |
| Apba3 | 0.00013 | 0.00015 | 0.00018 | 0.00021 | 0.00016 | 5.42E-05 | 0.00012 | 6.43E-05 | 0.00014 | 0.00012 | 0.00014 | 0.00025 | 4.03E-07 | 4.66E-05 | 9.60E-05 | 0 | 0 | 4.58E-05 |
| Sardh | 0 | 5.62E-06 | 2.43E-05 | 9.35E-05 | 7.65E-05 | 9.55E-05 | 2.00E-05 | 0.00019 | 0.00014 | 7.85E-05 | 5.59E-05 | 0 | 4.31E-05 | 0 | 3.55E-05 | 0 | 0 | 9.76E-05 |
| Sec23ip | 0.00026 | 0.00013 | 0.00031 | 0.0003 | 9.16E-05 | 0.00015 | 0.00033 | 4.68E-05 | 0.00014 | 0.00012 | 0.00011 | 3.24E-05 | 8.99E-05 | 0.00012 | 0.00016 | 0 | 5.61E-05 | 4.39E-05 |
| Acat2 | 2.70E-05 | 1.01E-05 | 0 | 3.07E-05 | 3.29E-05 | 0 | 0 | 0 | 0.00014 | 1.24E-05 | 3.78E-05 | 0 | 7.21E-05 | 6.84E-05 | 0 | 5.78E-05 | 0 | 2.93E-05 |
| Ercc3 | 0.00013 | 0.00015 | 0.00011 | 0.00011 | 3.28E-05 | 0 | 0 | 0 | 0.00014 | 6.74E-05 | 8.26E-05 | 0 | 1.61E-05 | 4.05E-05 | 0.00016 | 6.08E-05 | 0 | 5.96E-05 |
| Birc6 | 0.0003 | 0.00022 | 0.0002 | 3.88E-05 | 0.00011 | 0.00018 | 0.00014 | 7.37E-05 | 0.00014 | 9.06E-05 | 0.00013 | 9.92E-05 | 1.71E-05 | 7.97E-05 | 0.00021 | 0.00014 | 0.00026 | 4.14E-05 |
| Ube2d3 | 0.00038 | 0.00017 | 0.00022 | 0 | 5.49E-05 | 0.00018 | 7.27E-05 | 0 | 0.00014 | 7.02E-05 | 0.00011 | 0.00022 | 7.96E-05 | 0.00012 | 0 | 8.78E-05 | 0.00038 | 0 |
| Stx12 | 0.00031 | 0.00024 | 0.00021 | 0 | 0.00013 | 0.00021 | 0.00012 | 0.00013 | 0.00014 | 0.00017 | 0.00018 | 0.00018 | 0.00011 | 8.83E-05 | 5.94E-05 | 0.00026 | 9.10E-05 | 0.00012 |
| Psma5 | 0.00039 | 0.00018 | 0.00026 | 0.00022 | 4.97E-05 | 0.00014 | 2.07E-05 | 0 | 0.00014 | 2.01E-05 | 6.05E-05 | 0.00013 | 4.33E-05 | 0.0001 | 1.47E-05 | 0.00021 | 6.81E-05 | 0.0001 |
| Sin3b | 0.00033 | 0.00025 | 0.00025 | 0.00035 | 0.00016 | 0.00022 | 0.00015 | 0.00024 | 0.00014 | 0.00011 | 0.00025 | 0.00027 | 7.59E-05 | 0.00022 | 0.00024 | 6.96E-05 | 0.00024 | 9.20E-05 |
| Abracl | 0.00025 | 4.32E-06 | 0.00026 | 0.00011 | 0.00012 | 1.43E-05 | 7.33E-05 | 0.0001 | 0.00014 | 0.00012 | 0.00077 | 8.87E-05 | 1.75E-06 | 0.00016 | 0 | 0.00012 | 0.00011 | 0 |
| Actr3 | 0.00039 | 0.00011 | 0.00025 | 0 | 4.84E-06 | 7.99E-05 | 1.66E-05 | 0 | 0.00014 | 4.92E-05 | 0.00039 | 0.00016 | 0 | 4.69E-05 | 0 | 0.00014 | 3.82E-05 | 0 |
| Cdkn2a | 0 | 2.43E-05 | 0 | 4.02E-05 | 0 | 0 | 0 | 2.33E-05 | 0.00014 | 6.50E-06 | 3.75E-06 | 5.57E-05 | 0 | 6.21E-05 | 9.05E-05 | 8.50E-05 | 0 | 6.05E-05 |
| Zfp362 | 3.92E-06 | 0.0002 | 1.48E-05 | 0.00015 | 0.00012 | 1.02E-05 | 5.59E-06 | 0.00017 | 0.00014 | 1.35E-05 | 7.83E-05 | 0 | 0 | 0.0001 | 4.63E-05 | 0.0001 | 6.33E-05 | 0.00022 |
| Bloc1s1 | 0.00013 | 7.99E-05 | 0 | 0.00013 | 0.00013 | 5.97E-05 | 0.0001 | 8.67E-05 | 0.00014 | 3.83E-05 | 7.96E-05 | 0 | 6.86E-05 | 3.47E-05 | 0 | 3.72E-05 | 0 | 4.75E-05 |
| Gnaq | 0.0002 | 0.00018 | 8.31E-05 | 0 | 0.00017 | 0.00012 | 7.49E-05 | 0.00015 | 0.00014 | 8.05E-05 | 0.00019 | 7.96E-05 | 0 | 2.67E-05 | 7.20E-05 | 9.35E-05 | 0.00035 | 0.00023 |
| Nckap1 | 0.00032 | 0.00019 | 0.00017 | 0 | 6.46E-05 | 0.00022 | 5.73E-05 | 3.26E-05 | 0.00014 | 0.00021 | 0.00011 | 0.00012 | 0 | 0 | 0 | 0.00014 | 8.47E-05 | 8.23E-05 |

| | | | | | | | | | | | | | | | | | | |
|---|---|---|---|---|---|---|---|---|---|---|---|---|---|---|---|---|---|---|
| Sacs | 3.81E-06 | 9.38E-06 | 7.81E-05 | 0 | 1.90E-05 | 0 | 3.24E-05 | 1.54E-05 | 0.00014 | 0 | 0.00017 | 0.00017 | 0 | 5.16E-05 | 1.42E-05 | 0.00011 | 0 | 0 |
| Nap1l4 | 0.00025 | 0.00011 | 0.00022 | 4.04E-05 | 8.11E-05 | 3.69E-05 | 0 | 5.47E-05 | 0.00014 | 1.97E-05 | 0.00011 | 0.00028 | 0 | 0 | 0.00016 | 0.00022 | 0 | 2.36E-05 |
| Ap1b1 | 0.00011 | 0.00011 | 8.90E-05 | 7.26E-05 | 9.64E-05 | 6.25E-05 | 9.32E-05 | 5.26E-05 | 0.00014 | 4.23E-05 | 1.05E-05 | 2.32E-05 | 2.91E-05 | 2.74E-05 | 6.57E-05 | 2.33E-05 | 7.40E-05 | 8.46E-05 |
| Krtcap2 | 0.00034 | 2.79E-05 | 5.55E-05 | 0.00032 | 0.00015 | 0.00023 | 0.00027 | 0.00015 | 0.00014 | 8.63E-05 | 4.81E-05 | 0.00014 | 0.00021 | 5.85E-05 | 5.22E-05 | 0.00015 | 5.21E-05 | 5.57E-05 |
| Galnt11 | 2.84E-05 | 8.74E-05 | 3.63E-06 | 1.96E-05 | 7.64E-05 | 7.48E-05 | 5.37E-05 | 0 | 0.00014 | 0.0001 | 9.50E-05 | 8.96E-05 | 0.00011 | 1.72E-05 | 4.64E-05 | 1.61E-05 | 0 | 5.07E-05 |
| Tmem242 | 0.00011 | 0.00011 | 3.82E-05 | 0.0003 | 0.00011 | 0.00011 | 9.92E-05 | 8.18E-05 | 0.00014 | 9.30E-05 | 4.83E-05 | 7.09E-05 | 4.73E-05 | 0.00012 | 0 | 0.00011 | 9.06E-06 | 9.37E-05 |
| Hexa | 0.00014 | 0.00026 | 0 | 6.61E-06 | 0.00055 | 0.00066 | 0.00045 | 0.0005 | 0.00014 | 0.00027 | 0.00021 | 0.00017 | 0.00073 | 0.00045 | 0 | 8.85E-06 | 0 | 0.00057 |
| Arhgap23 | 9.13E-07 | 0.0001 | 6.74E-05 | 0.00018 | 8.96E-05 | 4.21E-05 | 0 | 1.76E-05 | 0.00014 | 2.83E-05 | 8.04E-05 | 5.65E-05 | 7.54E-06 | 0.00019 | 0.00027 | 0 | 0.00024 | 8.89E-05 |
| Dgcr6 | 8.05E-05 | 0.00014 | 3.10E-05 | 0.00021 | 0.00014 | 0.00014 | 1.17E-05 | 7.05E-05 | 0.00014 | 6.21E-05 | 0.00015 | 1.77E-05 | 5.71E-05 | 0.00014 | 0 | 4.06E-05 | 0 | 7.61E-05 |
| Hprt | 0.00024 | 0.0001 | 0.00031 | 4.21E-05 | 2.74E-05 | 7.52E-05 | 0 | 0 | 0.00014 | 0.00011 | 0.00014 | 0.00015 | 0 | 0 | 0 | 0.00024 | 1.68E-05 | 0 |
| Spcs3 | 0.00042 | 1.34E-05 | 0.00032 | 4.67E-05 | 5.54E-05 | 0.00016 | 7.53E-05 | 0.0002 | 0.00014 | 5.01E-05 | 4.94E-05 | 0.00028 | 6.80E-05 | 1.50E-05 | 0 | 0.00023 | 5.02E-05 | 7.21E-05 |
| Anp32e | 0.00011 | 6.22E-05 | 0.00028 | 0.00038 | 0 | 0 | 2.91E-05 | 3.42E-05 | 0.00014 | 0.00011 | 0 | 0.00025 | 3.58E-05 | 0 | 6.18E-05 | 0.00098 | 0.00027 | 7.95E-05 |
| Ndufa10 | 0.0004 | 0.00017 | 0.00023 | 0.00021 | 8.61E-05 | 0.0002 | 0.00011 | 3.29E-05 | 0.00014 | 0 | 4.21E-05 | 4.94E-05 | 0 | 0.00012 | 2.08E-05 | 0.00015 | 3.31E-06 | 6.21E-05 |
| Eif3d | 0.0004 | 0.00018 | 0.00046 | 0.00023 | 3.80E-05 | 9.17E-05 | 2.92E-05 | 9.55E-06 | 0.00014 | 4.55E-05 | 0.00011 | 0.00012 | 0 | 3.39E-05 | 5.04E-05 | 0.00022 | 2.29E-05 | 6.57E-05 |
| Cnrip1 | 0.00013 | 0.0001 | 2.78E-05 | 2.15E-05 | 0.00014 | 0.00019 | 0.00015 | 0.00019 | 0.00014 | 0.00011 | 0 | 0.00016 | 9.33E-05 | 0.00014 | 0 | 0 | 7.67E-05 | 4.54E-05 |
| Pi4ka | 0.00012 | 0.00019 | 0.00015 | 9.71E-05 | 8.55E-05 | 0.00013 | 4.44E-05 | 0.00016 | 0.00014 | 0.00013 | 7.73E-05 | 0.00011 | 4.98E-05 | 4.78E-05 | 0.00013 | 9.98E-05 | 9.88E-05 | 0.00017 |
| Bicc1 | 3.60E-05 | 0.00024 | 0 | 0.00046 | 0.00068 | 0.00049 | 0.00051 | 0.00046 | 0.00014 | 0.00037 | 0.00014 | 0.00042 | 0.00089 | 0.00035 | 0.00022 | 0.00015 | 0.00032 | 0.00065 |
| Zmpste24 | 0.00012 | 0.00014 | 0.00028 | 0.00043 | 0.00015 | 9.65E-05 | 0.00021 | 0.00013 | 0.00014 | 0.00014 | 0.0002 | 3.51E-05 | 0.00012 | 0.0002 | 0 | 0.00017 | 6.15E-05 | 0.00024 |
| Ctps | 0.00014 | 0 | 0.00012 | 0 | 0 | 4.40E-05 | 0 | 0 | 0.00014 | 0.00033 | 0.00032 | 0 | 0.00057 | 9.74E-05 | 0 | 9.22E-05 | 0 | 0 |
| Tmem59 | 0.00045 | 0.00029 | 4.80E-05 | 4.78E-05 | 0.00045 | 0.00073 | 0.00041 | 0.00047 | 0.00014 | 0.00019 | 0.00022 | 0.0004 | 0.00047 | 0.00049 | 0 | 1.44E-05 | 5.60E-06 | 0.00031 |
| Lmf2 | 4.85E-05 | 0 | 6.80E-05 | 0.00029 | 5.06E-05 | 7.10E-05 | 0.00015 | 0.00015 | 0.00014 | 0.00011 | 0.00016 | 0.00023 | 0.00013 | 0.00012 | 2.89E-05 | 0.00038 | 7.17E-05 | 0.00016 |
| Pigk | 0.0003 | 6.14E-05 | 9.82E-05 | 0 | 0.00016 | 0.00017 | 7.84E-05 | 0 | 0.00014 | 9.77E-05 | 0.00015 | 0 | 0.00013 | 9.75E-05 | 0 | 2.37E-05 | 0 | 0.00013 |
| Tmem132c | 0 | 7.24E-05 | 0 | 0 | 0.00014 | 2.54E-05 | 0 | 0 | 0.00014 | 1.89E-05 | 0 | 0 | 0 | 2.51E-05 | 0 | 0 | 0 | 0 |
| Tspo | 5.41E-05 | 0.00011 | 0.00029 | 0.0009 | 0 | 7.77E-05 | 0.00028 | 0.00029 | 0.00014 | 0.00041 | 0.0001 | 0.00039 | 0.00025 | 0.0002 | 0 | 0.00025 | 0.00023 | 0.00082 |
| Manbal | 9.28E-05 | 0.00017 | 0.00016 | 0.00017 | 0.00012 | 5.33E-05 | 1.61E-05 | 4.46E-06 | 0.00014 | 9.91E-05 | 0.00014 | 0.00012 | 5.19E-05 | 9.56E-05 | 0.0001 | 0.00011 | 0 | 0.0001 |
| Psmc2 | 0.00031 | 0.00016 | 0.00036 | 0.00022 | 1.06E-05 | 0 | 3.73E-05 | 3.42E-05 | 0.00014 | 7.14E-05 | 0.00011 | 0 | 2.00E-06 | 0.0001 | 1.42E-05 | 0.00021 | 0 | 6.80E-05 |
| Rnpep | 6.38E-05 | 7.34E-05 | 8.94E-06 | 3.97E-05 | 5.23E-05 | 5.25E-05 | 0 | 0 | 0.00014 | 7.94E-05 | 0.00024 | 3.66E-05 | 5.48E-05 | 0.00019 | 3.65E-06 | 1.88E-05 | 2.10E-06 | 9.50E-05 |
| Prpsap1 | 0.00015 | 0.00018 | 0.00014 | 0.0002 | 0.00011 | 4.84E-05 | 0 | 0.00016 | 0.00014 | 6.38E-05 | 4.41E-05 | 6.54E-05 | 0 | 6.30E-05 | 2.65E-05 | 0.00015 | 2.35E-05 | 6.07E-05 |
| Serpina3i | 0 | 0 | 0 | 0 | 4.77E-05 | 0.00012 | 0 | 0 | 0.00014 | 4.24E-05 | 0 | 3.67E-05 | 0.00035 | 0.00015 | 6.36E-05 | 2.96E-05 | 0 | 0 |
| Srsf4 | 3.24E-05 | 0.00017 | 0.00019 | 0.00039 | 9.71E-05 | 1.56E-05 | 0 | 7.49E-05 | 0.00014 | 8.20E-05 | 0.00011 | 9.30E-05 | 0 | 6.96E-05 | 0.00032 | 0.00037 | 3.88E-05 | 7.48E-05 |
| Fkrp | 0 | 0.00012 | 0.00013 | 0.00026 | 0.00011 | 2.61E-05 | 0 | 0.00017 | 0.00014 | 4.77E-05 | 0.00011 | 4.63E-05 | 1.99E-05 | 0.00016 | 0.00013 | 8.83E-05 | 3.05E-05 | 5.90E-05 |
| Pura | 0.00013 | 0.00025 | 0.00017 | 0.00014 | 0.0002 | 0.00018 | 0.00013 | 0.00017 | 0.00014 | 0.00012 | 0.00017 | 0.00029 | 1.94E-06 | 0.00011 | 0.00055 | 3.72E-05 | 0.0002 | 0.00025 |
| Mlkl | 0.00013 | 0 | 1.62E-05 | 0 | 0 | 4.14E-05 | 0 | 0.00011 | 0.00014 | 5.64E-05 | 0 | 0.00013 | 6.47E-05 | 0 | 0 | 7.49E-05 | 0 | 8.68E-05 |

| | | | | | | | | | | | | | | | | | | |
|---|---|---|---|---|---|---|---|---|---|---|---|---|---|---|---|---|---|---|
| Ppp1cb | 0.0002 | 0.00023 | 0.00023 | 0.00033 | 6.87E-05 | 4.21E-05 | 0.0002 | 0.00018 | 0.00014 | 0.00017 | 0.00016 | 0.00023 | 5.30E-05 | 5.81E-05 | 0.00022 | 0.00025 | 0.00013 | 9.01E-05 |
| Nagk | 8.20E-05 | 9.60E-05 | 2.43E-05 | 0.00034 | 1.17E-05 | 6.32E-05 | 8.31E-05 | 0.00015 | 0.00014 | 0.0002 | 0.00029 | 0.00011 | 0 | 0.0001 | 4.57E-05 | 7.97E-05 | 0 | 0.0002 |
| Smyd2 | 0.00019 | 0.00011 | 0.00023 | 0.00011 | 6.08E-05 | 0.00021 | 0 | 8.41E-05 | 0.00014 | 1.52E-05 | 0.00014 | 0.00018 | 5.06E-05 | 7.33E-05 | 0.00014 | 9.62E-05 | 0 | 0.00012 |
| Med25 | 0.00015 | 0.0002 | 0.00017 | 0.00026 | 0.00011 | 6.96E-05 | 6.14E-05 | 7.33E-05 | 0.00014 | 0.00012 | 0.00012 | 2.77E-05 | 1.16E-05 | 0.00017 | 0.0002 | 8.59E-05 | 8.84E-05 | 0.00012 |
| Coa3 | 7.77E-05 | 0.0001 | 9.25E-05 | 0.00033 | 0.00011 | 4.60E-05 | 4.60E-05 | 8.34E-05 | 0.00014 | 0.0001 | 7.88E-05 | 4.93E-05 | 0 | 5.33E-05 | 0 | 9.61E-05 | 0 | 0.00012 |
| Galt | 4.13E-06 | 0 | 6.66E-05 | 3.65E-05 | 0 | 2.83E-05 | 0.00012 | 3.60E-05 | 0.00014 | 0.00011 | 4.26E-06 | 2.38E-05 | 0.00017 | 6.41E-05 | 1.18E-05 | 0 | 0 | 8.01E-05 |
| Tra2b | 0.00028 | 0.00011 | 0.00019 | 3.85E-05 | 2.33E-05 | 0.00011 | 0 | 5.80E-05 | 0.00014 | 1.39E-05 | 0.00012 | 8.18E-05 | 0 | 8.00E-05 | 4.41E-05 | 0.00024 | 0.00057 | 2.81E-05 |
| Vti1a | 5.30E-05 | 0.00013 | 9.89E-05 | 0.0001 | 3.49E-05 | 1.49E-05 | 0 | 0 | 0.00014 | 9.11E-05 | 9.42E-05 | 2.98E-05 | 3.18E-05 | 9.89E-05 | 0.00034 | 8.97E-05 | 5.50E-05 | 0.00013 |
| Kansl2 | 0.00012 | 0.00022 | 0.00025 | 0.0003 | 0.0003 | 0.00022 | 0.00011 | 0.00017 | 0.00014 | 5.11E-05 | 0.00013 | 0.00011 | 5.93E-05 | 0.00034 | 0.00041 | 0.00011 | 8.54E-05 | 4.97E-05 |
| Phb | 0.00011 | 8.83E-05 | 0.00018 | 0.00019 | 4.96E-05 | 0 | 0 | 6.65E-06 | 0.00014 | 1.93E-05 | 8.85E-05 | 0 | 5.34E-05 | 2.91E-05 | 0 | 3.73E-05 | 7.44E-06 | 0 |
| Mpg | 8.92E-05 | 1.00E-04 | 3.06E-05 | 0.00011 | 6.42E-05 | 9.04E-05 | 1.57E-05 | 0.00022 | 0.00014 | 5.44E-05 | 0 | 5.43E-05 | 5.31E-05 | 6.59E-05 | 0.00012 | 3.99E-05 | 0 | 5.24E-05 |
| Spag7 | 0.00017 | 9.27E-05 | 0.00014 | 0.00022 | 1.02E-05 | 2.04E-06 | 3.29E-05 | 5.97E-05 | 0.00014 | 5.55E-05 | 0.00012 | 2.40E-05 | 0 | 5.98E-05 | 9.25E-05 | 0.0001 | 0 | 8.24E-05 |
| Ier3 | 0 | 3.79E-05 | 0.00022 | 0.00014 | 3.54E-05 | 0 | 0.00031 | 0 | 0.00014 | 0.00017 | 1.65E-05 | 0.00128 | 0.0008 | 0.00052 | 0.00023 | 7.47E-05 | 0.00425 | 4.26E-05 |
| Spata13 | 0 | 0 | 0.0001 | 0 | 9.35E-06 | 6.40E-05 | 0 | 0 | 0.00014 | 0 | 0.00017 | 0 | 0 | 0.00023 | 3.56E-05 | 6.19E-05 | 2.15E-05 | 3.59E-06 |
| Golm1 | 0.00014 | 2.84E-05 | 9.16E-05 | 0.00018 | 0.00033 | 9.83E-05 | 8.15E-05 | 0.00048 | 0.00014 | 2.65E-05 | 0.00031 | 0.00018 | 0.00014 | 0.00041 | 5.48E-05 | 0.00016 | 6.49E-06 | 0.00012 |
| Zfp523 | 4.21E-05 | 7.90E-05 | 3.45E-05 | 9.14E-05 | 9.62E-05 | 3.94E-05 | 4.55E-05 | 6.73E-05 | 0.00014 | 3.58E-05 | 6.79E-05 | 0 | 2.23E-05 | 0.00013 | 0.00011 | 4.44E-05 | 1.14E-05 | 4.50E-05 |
| Acaca | 0.00014 | 0.00015 | 0.00017 | 5.47E-05 | 0.00013 | 0.00014 | 3.24E-05 | 2.10E-05 | 0.00014 | 7.54E-05 | 4.94E-05 | 0 | 8.86E-06 | 7.18E-05 | 0.00024 | 7.19E-05 | 8.18E-05 | 1.12E-05 |
| Tnk2 | 0 | 8.13E-05 | 0 | 4.16E-05 | 0.0001 | 0 | 0 | 0 | 0.00014 | 5.02E-05 | 0.00047 | 4.66E-05 | 7.62E-05 | 0.00014 | 0.00027 | 8.67E-05 | 5.51E-05 | 6.28E-05 |
| Dmwd | 5.04E-05 | 0.00014 | 0.00017 | 0 | 5.27E-05 | 6.44E-05 | 0 | 0.00018 | 0.00014 | 4.76E-05 | 0.00014 | 0.00018 | 1.13E-05 | 9.00E-05 | 0.00021 | 7.23E-05 | 0.00017 | 7.89E-05 |
| Fkbp14 | 0.00038 | 0 | 0.0002 | 0.00016 | 6.21E-05 | 0.00016 | 0.00062 | 0.00035 | 0.00014 | 0.00016 | 7.76E-05 | 0 | 0.00018 | 0.00014 | 5.76E-05 | 2.76E-05 | 0 | 0 |
| Zhx1 | 0.00014 | 6.87E-05 | 3.38E-05 | 2.26E-06 | 4.56E-05 | 8.65E-05 | 1.69E-06 | 6.69E-05 | 0.00014 | 4.86E-05 | 0 | 4.28E-05 | 4.63E-05 | 2.29E-05 | 5.22E-05 | 6.27E-06 | 4.25E-05 | 8.57E-05 |
| Pkn1 | 3.37E-05 | 0.00019 | 0 | 9.95E-05 | 0.00021 | 3.55E-05 | 2.96E-05 | 2.95E-05 | 0.00014 | 8.66E-05 | 8.76E-05 | 7.83E-05 | 4.95E-05 | 0.00015 | 0.00018 | 5.84E-05 | 0 | 0.00014 |
| Gsk3a | 8.97E-05 | 0.00021 | 0.00018 | 0.00023 | 0.00014 | 8.81E-05 | 3.30E-05 | 0.00011 | 0.00014 | 0.00012 | 0.0001 | 0.00016 | 0.00011 | 7.87E-05 | 0.00011 | 0.00026 | 0.0004 | 5.42E-05 |
| Lsm4 | 0.00023 | 9.79E-05 | 0.00023 | 0.00043 | 2.13E-05 | 0 | 2.62E-05 | 1.94E-05 | 0.00014 | 0 | 2.53E-05 | 2.13E-05 | 2.77E-05 | 8.32E-05 | 0.00013 | 0.00033 | 8.87E-05 | 0 |
| Rsrc2 | 0.00021 | 0.00018 | 0.00012 | 2.16E-05 | 8.44E-05 | 5.94E-05 | 0 | 6.92E-05 | 0.00014 | 5.04E-05 | 0.00011 | 0.00017 | 0 | 3.02E-05 | 0.00047 | 0.00018 | 0.00026 | 6.41E-05 |
| Elk4 | 0.00016 | 0.00022 | 0.00015 | 0 | 0.00023 | 0.00021 | 0.00022 | 0.0003 | 0.00014 | 0.00013 | 0.00016 | 0.00015 | 8.47E-05 | 0.0001 | 0.00017 | 0.00017 | 0.00012 | 0.00016 |
| Cetn3 | 0.00018 | 0.00014 | 9.31E-05 | 0.00023 | 3.77E-05 | 2.14E-05 | 9.28E-05 | 0 | 0.00014 | 0.00013 | 0.0001 | 0.00011 | 7.95E-05 | 0 | 5.07E-05 | 0.00023 | 2.58E-05 | 0.00017 |
| Ykt6 | 0.00036 | 0.00012 | 0.0004 | 0.00033 | 0.0002 | 0.0001 | 0.00035 | 3.28E-05 | 0.00014 | 0.00011 | 0.00014 | 0 | 0.00011 | 0.00016 | 7.01E-05 | 5.50E-05 | 3.02E-05 | 0.00018 |
| Wdr26 | 0.00034 | 0.00041 | 0.00021 | 8.69E-05 | 0.00031 | 0.0004 | 0.00029 | 0.00018 | 0.00014 | 0.0002 | 0.00022 | 0.00029 | 0.00014 | 0.00026 | 0.00066 | 0.00012 | 0.00024 | 0.00017 |
| Atxn7l1 | 6.77E-05 | 5.82E-05 | 0 | 3.39E-05 | 6.95E-07 | 0.00017 | 0 | 0 | 0.00014 | 0.00013 | 7.85E-05 | 0.00049 | 0.0004 | 9.35E-06 | 0.00028 | 0 | 9.70E-05 | 0 |
| Pmpca | 0.00016 | 8.19E-05 | 0.00017 | 4.91E-05 | 2.32E-06 | 1.35E-05 | 3.44E-05 | 9.43E-06 | 0.00014 | 3.26E-05 | 0.00012 | 8.67E-05 | 0 | 1.97E-05 | 7.95E-05 | 0.00013 | 3.73E-06 | 1.49E-05 |
| Os9 | 3.93E-06 | 0.00016 | 2.96E-05 | 0.00034 | 0.00022 | 0.00028 | 0.00027 | 0.00032 | 0.00014 | 0.00024 | 0.00021 | 0.00025 | 0.00057 | 0.00032 | 0.00023 | 0 | 0.00014 | 0.00031 |
| Mtfr1l | 1.35E-05 | 0.00013 | 9.94E-05 | 0.0002 | 0.00012 | 3.81E-05 | 6.45E-05 | 7.39E-05 | 0.00014 | 0.0001 | 7.13E-05 | 0 | 5.90E-05 | 5.71E-05 | 1.94E-05 | 0 | 3.31E-05 | 8.58E-05 |

| | | | | | | | | | | | | | | | | | | |
|---|---|---|---|---|---|---|---|---|---|---|---|---|---|---|---|---|---|---|
| Gemin7 | 0.00017 | 0.00016 | 0.00013 | 0.00031 | 0.00013 | 7.78E-05 | 0.00012 | 0.00011 | 0.00014 | 0.00013 | 0.00019 | 4.48E-05 | 5.57E-05 | 0.00015 | 0 | 0.00025 | 0 | 0.00013 |
| Fam91a1 | 0.00037 | 0.00014 | 0.00025 | 0 | 0.00013 | 0.00019 | 8.61E-05 | 3.41E-05 | 0.00014 | 8.98E-06 | 0.00024 | 0.00021 | 2.25E-05 | 7.69E-05 | 0.00024 | 8.22E-05 | 0.00027 | 2.85E-05 |
| Zfp106 | 0.0003 | 0.0002 | 0.00029 | 5.60E-05 | 0.00012 | 0.00011 | 0.00019 | 0 | 0.00014 | 2.72E-05 | 7.53E-05 | 0.00028 | 7.42E-05 | 4.36E-05 | 7.22E-05 | 0.00013 | 0 | 3.58E-05 |
| Rnf11 | 0.00023 | 0.00024 | 0.00024 | 0 | 0.00027 | 7.12E-05 | 9.91E-05 | 0.00014 | 0.00014 | 0.00015 | 0.00034 | 0.0002 | 0.00013 | 0.00018 | 2.84E-05 | 0 | 0.00017 | 0.00014 |
| Mpnd | 0.00014 | 0.00012 | 6.28E-05 | 0.00029 | 3.14E-05 | 5.38E-05 | 0.00014 | 5.11E-06 | 0.00014 | 4.60E-05 | 7.26E-05 | 7.88E-05 | 4.34E-05 | 0.00011 | 2.68E-05 | 4.34E-05 | 5.93E-05 | 0.00022 |
| Pnrc2 | 0.00019 | 0.00024 | 1.08E-05 | 3.53E-05 | 0.00011 | 0.00018 | 0.00011 | 0.00016 | 0.00014 | 0.00017 | 7.33E-07 | 7.95E-05 | 4.87E-05 | 0.00011 | 0.00016 | 0.00032 | 0 | 0.00017 |
| Keap1 | 0.00012 | 0.00016 | 7.14E-05 | 0.00033 | 0.00015 | 8.81E-05 | 0.00023 | 5.34E-05 | 0.00014 | 0.00013 | 6.51E-05 | 0.00013 | 0.00011 | 0.00011 | 3.43E-05 | 0.00013 | 0 | 9.24E-05 |
| Egln2 | 1.35E-05 | 0.00023 | 0.0001 | 0.00029 | 0.00011 | 0.00012 | 7.95E-05 | 5.94E-05 | 0.00014 | 0.00015 | 0.00012 | 3.87E-05 | 7.89E-05 | 0.00018 | 0.00013 | 0.00012 | 0 | 0.0001 |
| Tada1 | 0.00013 | 0.00011 | 7.02E-05 | 1.74E-05 | 8.87E-05 | 2.78E-05 | 2.19E-05 | 4.22E-05 | 0.00014 | 2.18E-05 | 0 | 0 | 0 | 4.06E-06 | 0.00013 | 6.23E-05 | 0 | 8.64E-06 |
| Borcs8 | 0.00017 | 0.00015 | 0 | 0.00015 | 0.00015 | 0.00013 | 8.14E-05 | 2.73E-05 | 0.00014 | 5.17E-05 | 6.64E-05 | 8.61E-05 | 6.75E-05 | 6.00E-05 | 0.00011 | 0 | 0 | 0.00011 |
| Lbp | 0 | 8.67E-05 | 0 | 0 | 0 | 0.00022 | 0 | 0.00044 | 0.00014 | 0.00163 | 0 | 4.96E-06 | 0.00183 | 0.00044 | 3.49E-05 | 0 | 0 | 0.0013 |
| Sh3bp5 | 6.93E-05 | 6.85E-05 | 3.88E-05 | 0.00018 | 4.22E-05 | 6.92E-05 | 1.48E-05 | 0 | 0.00014 | 0.00018 | 1.25E-05 | 0 | 6.84E-05 | 0 | 7.41E-05 | 0 | 7.91E-05 | 9.29E-05 |
| Nqo2 | 0.00018 | 3.65E-05 | 5.51E-05 | 0.00011 | 0 | 2.62E-05 | 0 | 9.60E-05 | 0.00014 | 0.00026 | 0 | 0 | 4.42E-05 | 0 | 0 | 0.00012 | 0 | 6.02E-05 |
| Hook3 | 0.00024 | 0.00014 | 0.00011 | 0 | 0.00014 | 0.00022 | 0.00026 | 0.00016 | 0.00014 | 0.00019 | 0.00011 | 9.96E-05 | 7.27E-05 | 0.00011 | 0.00023 | 2.25E-05 | 5.91E-05 | 0.00018 |
| Uba2 | 0.00031 | 0.0001 | 0.0003 | 6.84E-05 | 0.00012 | 0.00017 | 1.94E-05 | 3.96E-05 | 0.00014 | 7.95E-05 | 0.00014 | 0.00015 | 4.48E-05 | 5.91E-05 | 5.41E-06 | 0.00039 | 5.41E-05 | 0 |
| Nsmf | 0 | 0.00014 | 0.00018 | 0.0002 | 0.00019 | 0.00017 | 0.00012 | 0.0002 | 0.00014 | 6.34E-05 | 0.0001 | 4.80E-05 | 0.00011 | 0.00024 | 0.00027 | 3.02E-05 | 5.19E-05 | 9.45E-05 |
| Mtor | 9.71E-05 | 9.06E-05 | 7.16E-05 | 2.27E-05 | 8.44E-05 | 4.32E-05 | 3.41E-06 | 8.15E-05 | 0.00014 | 4.77E-05 | 1.55E-05 | 1.31E-05 | 0 | 0 | 0.00026 | 3.94E-05 | 2.27E-05 | 2.13E-05 |
| Kctd17 | 0.0002 | 0.00016 | 0.00026 | 0.00032 | 4.08E-05 | 0.00014 | 0.00024 | 0.00044 | 0.00014 | 0.00028 | 0.00014 | 0.00011 | 0 | 0 | 0.00011 | 6.70E-05 | 0 | 0.00035 |
| Cul3 | 0.00033 | 0.00032 | 0.00029 | 0.00011 | 0.00021 | 0.00021 | 0.00012 | 0.00013 | 0.00014 | 9.30E-05 | 0.00015 | 0.00023 | 0 | 0.00013 | 0.00023 | 0.00036 | 0.00028 | 0.00011 |
| Plbd2 | 0 | 0.00022 | 0.00016 | 0.00018 | 0.0003 | 0.00021 | 0.00025 | 0.00013 | 0.00014 | 0.00024 | 0.00016 | 0.00017 | 0.00041 | 0.00026 | 6.35E-05 | 1.66E-05 | 0 | 0.00038 |
| Slc48a1 | 0 | 0.00024 | 8.91E-06 | 7.93E-05 | 2.41E-05 | 5.87E-05 | 9.99E-05 | 0 | 0.00014 | 9.75E-06 | 1.47E-05 | 8.37E-05 | 9.43E-05 | 0.00026 | 0 | 0 | 0 | 0.00023 |
| Ahsa2 | 0.00022 | 9.90E-05 | 9.97E-05 | 3.14E-05 | 6.73E-05 | 2.95E-05 | 2.92E-05 | 9.45E-05 | 0.00014 | 3.09E-05 | 5.69E-05 | 5.38E-05 | 0 | 6.72E-05 | 0.00022 | 2.72E-05 | 0.00014 | 1.76E-05 |
| Slc7a8 | 0 | 2.61E-05 | 0 | 1.02E-05 | 0.00014 | 0 | 0 | 2.32E-06 | 0.00014 | 6.05E-05 | 0 | 6.69E-05 | 2.99E-05 | 0 | 0 | 0 | 6.67E-05 | 0 |
| Sypl | 0.00029 | 0.00026 | 0.00037 | 4.85E-05 | 0.00012 | 0.00019 | 0.00015 | 0.00013 | 0.00014 | 0.00021 | 0.00017 | 0.00035 | 0.0001 | 8.97E-05 | 0 | 0.00023 | 0.00011 | 0.00011 |
| Ttll5 | 1.35E-05 | 1.35E-05 | 0 | 0 | 0.00014 | 5.59E-05 | 2.77E-05 | 0 | 0.00014 | 6.32E-06 | 1.61E-05 | 5.37E-05 | 3.79E-05 | 0.00021 | 1.12E-05 | 8.44E-05 | 8.17E-06 | 9.48E-06 |
| Skap2 | 8.71E-05 | 9.42E-05 | 9.40E-06 | 3.43E-05 | 0.00012 | 7.06E-05 | 0.00019 | 0.00018 | 0.00014 | 0.00012 | 7.49E-05 | 9.25E-05 | 4.30E-05 | 0 | 0.00017 | 0.00012 | 0 | 6.00E-05 |
| Kdm2a | 0.00018 | 0.0003 | 7.88E-05 | 0.00024 | 0.00021 | 0.00015 | 0.00017 | 9.44E-05 | 0.00014 | 0.00021 | 6.76E-05 | 0.00066 | 0.00013 | 0.00017 | 0.00075 | 0.00019 | 0.00015 | 0.0001 |
| Zfp608 | 9.45E-05 | 0.00013 | 0 | 0 | 3.62E-05 | 0.00014 | 0.00016 | 0.00013 | 0.00014 | 9.13E-05 | 0 | 5.16E-05 | 0.00016 | 0 | 0 | 0.0001 | 0.00016 | 0 |
| Tut7 | 0.00024 | 0.00021 | 6.18E-05 | 0 | 0.00012 | 0.00017 | 0.00013 | 0.0002 | 0.00014 | 0.0001 | 3.74E-05 | 0.00018 | 0 | 0.00019 | 0.00038 | 4.61E-05 | 0.00038 | 0.00014 |
| Zfp516 | 7.02E-05 | 0.00021 | 0.00012 | 0.00022 | 0.00021 | 0.00014 | 0.00037 | 0.00018 | 0.00014 | 8.91E-05 | 7.93E-06 | 0.00027 | 0 | 3.97E-05 | 0.0002 | 2.78E-05 | 0.00012 | 7.25E-05 |
| Pygo2 | 4.63E-05 | 0.00013 | 0.0001 | 0.00016 | 9.00E-05 | 1.30E-05 | 3.98E-05 | 0.0001 | 0.00014 | 5.35E-05 | 7.27E-05 | 4.73E-05 | 5.45E-05 | 0.00012 | 0.00021 | 9.35E-05 | 6.69E-06 | 5.92E-05 |
| Chmp5 | 0.00024 | 0.00019 | 0.00013 | 0.00013 | 0.00014 | 0.00017 | 0.0001 | 0.0002 | 0.00014 | 0.00013 | 0.00018 | 0.00024 | 7.99E-05 | 0.00014 | 3.27E-05 | 6.93E-05 | 9.14E-05 | 0.00015 |
| Selenos | 0.00037 | 6.48E-05 | 0.00031 | 0.00033 | 0.00021 | 5.70E-05 | 0.00028 | 0.0002 | 0.00014 | 0.00013 | 0.00011 | 0.00022 | 0.00018 | 0.00016 | 0.00021 | 1.50E-05 | 0 | 5.57E-05 |

| | | | | | | | | | | | | | | | | | | |
|---|---|---|---|---|---|---|---|---|---|---|---|---|---|---|---|---|---|---|
| Sec24b | 9.57E-05 | 0.00019 | 0.00019 | 6.70E-05 | 5.31E-05 | 6.56E-05 | 7.23E-05 | 7.16E-05 | 0.00014 | 0.00013 | 7.84E-05 | 0.00028 | 3.25E-06 | 2.31E-05 | 0.00013 | 0.00018 | 0.00026 | 0.00011 |
| Sf3b6 | 0.00045 | 0.00012 | 0.00023 | 7.31E-05 | 7.49E-05 | 0.00012 | 9.84E-05 | 0 | 0.00014 | 8.78E-05 | 0.0001 | 5.91E-05 | 7.15E-05 | 1.08E-05 | 0 | 0.00016 | 8.50E-06 | 2.91E-05 |
| Tomm40l | 8.29E-05 | 0.00011 | 0.00012 | 0.00018 | 6.40E-05 | 2.48E-05 | 0 | 1.20E-05 | 0.00014 | 5.75E-05 | 4.38E-05 | 0 | 0 | 0 | 0 | 4.87E-05 | 0 | 0.0001 |
| Rftn1 | 3.22E-05 | 0.00013 | 0.00018 | 0.00015 | 0.00041 | 0.00021 | 0.00029 | 0.00017 | 0.00014 | 0.00015 | 0.00028 | 5.38E-05 | 0.00036 | 0.00041 | 0.00029 | 0 | 5.25E-05 | 0.00028 |
| Ndufa7 | 0.00026 | 0.00013 | 0.00013 | 0.00029 | 0.00011 | 0.00013 | 0.00012 | 5.27E-05 | 0.00014 | 6.76E-05 | 0.00013 | 0.00015 | 0.00011 | 0.00011 | 0 | 8.24E-05 | 5.39E-05 | 0.00012 |
| Polr1a | 5.14E-05 | 5.19E-05 | 0.00022 | 5.69E-05 | 0 | 7.41E-06 | 0 | 0 | 0.00014 | 3.29E-05 | 0 | 0 | 1.74E-05 | 7.25E-07 | 5.77E-05 | 2.15E-05 | 7.02E-05 | 1.02E-05 |
| Map4k3 | 0.00014 | 0.00022 | 0 | 0 | 3.64E-06 | 0.00012 | 0 | 4.14E-05 | 0.00014 | 0 | 0.0002 | 4.25E-05 | 0.0001 | 5.62E-05 | 0.00025 | 2.01E-05 | 0.00018 | 0.00014 |
| Zfp217 | 8.70E-06 | 0.00021 | 8.38E-05 | 0.00012 | 6.96E-05 | 5.61E-05 | 0 | 0.00015 | 0.00014 | 0.00021 | 0.00011 | 0.00012 | 5.71E-05 | 0.00021 | 0.00019 | 0.00012 | 0.00016 | 0.0001 |
| Micos10 | 0.00037 | 0.00012 | 0.00024 | 0.00044 | 0.00026 | 0.00018 | 0.00029 | 0.00022 | 0.00014 | 9.56E-05 | 0.00015 | 4.94E-05 | 0.00012 | 2.14E-05 | 0 | 0.0002 | 7.00E-05 | 0.00012 |
| Commd4 | 0.00011 | 9.61E-05 | 4.87E-05 | 0.00015 | 5.88E-05 | 3.96E-05 | 0 | 5.01E-05 | 0.00014 | 7.31E-05 | 6.63E-05 | 4.84E-05 | 4.77E-05 | 0.00011 | 0.00011 | 3.52E-05 | 5.19E-05 | 5.80E-05 |
| Irx2 | 0 | 0.00017 | 4.42E-05 | 0.00016 | 6.16E-05 | 0.00012 | 0 | 0 | 0.00013 | 8.20E-07 | 0.00012 | 7.02E-05 | 0.00017 | 0.00033 | 0.0002 | 8.29E-05 | 0.0004 | 0 |
| Nek7 | 0.00031 | 0.00013 | 4.65E-06 | 0 | 0.00023 | 0.00026 | 0.00015 | 4.91E-05 | 0.00013 | 5.25E-05 | 4.59E-05 | 0.00022 | 0.00033 | 1.81E-05 | 0 | 0.00025 | 0.00017 | 8.59E-05 |
| Pced1a | 1.21E-06 | 8.17E-05 | 0 | 8.84E-05 | 0.00013 | 0.00011 | 0.00011 | 0 | 0.00013 | 6.69E-05 | 1.01E-05 | 4.21E-05 | 0.00017 | 7.22E-05 | 0.00025 | 1.04E-05 | 0 | 7.91E-06 |
| Arhgap32 | 4.89E-05 | 3.75E-05 | 5.30E-05 | 0 | 0 | 2.29E-05 | 7.27E-05 | 0.00014 | 0.00013 | 8.17E-06 | 0 | 5.31E-05 | 0 | 0 | 0 | 1.57E-05 | 8.21E-05 | 6.42E-05 |
| Usf2 | 7.92E-05 | 0.00029 | 0.00012 | 0.00044 | 0.0002 | 0.0002 | 0.00026 | 0.00032 | 0.00013 | 0.00016 | 0.00014 | 0.00023 | 0.00012 | 0.00025 | 0.0004 | 0.00021 | 0.00022 | 0.00018 |
| Zdhhc9 | 0.00018 | 4.39E-06 | 9.86E-05 | 8.50E-05 | 6.34E-05 | 0.00011 | 0.0004 | 4.18E-05 | 0.00013 | 0.00018 | 0.00014 | 0.00025 | 0.00016 | 7.20E-05 | 0 | 0 | 0 | 0.00015 |
| R3hdm2 | 0.00017 | 0.00024 | 0.00015 | 0.00017 | 0.00017 | 0.00023 | 0.00014 | 0.00024 | 0.00013 | 0.00017 | 0.00016 | 0.00014 | 5.75E-05 | 0.00012 | 0.00012 | 0.00016 | 0.00024 | 0.00015 |
| Ptprd | 4.48E-05 | 9.12E-06 | 0 | 0 | 0.00012 | 0 | 5.19E-05 | 7.81E-05 | 0.00013 | 0 | 0 | 0 | 0 | 8.28E-07 | 0 | 3.77E-05 | 0 | 0 |
| Nfkb1 | 2.18E-05 | 0.00015 | 0.00037 | 3.09E-05 | 0.0001 | 0.00014 | 0 | 0.00017 | 0.00013 | 0.00017 | 3.88E-05 | 0.00022 | 0.00015 | 0.00017 | 0.00014 | 7.11E-05 | 0.00027 | 0.00019 |
| Kmt2d | 9.86E-05 | 0.00027 | 0.00016 | 0.00018 | 0.00015 | 0.00011 | 2.45E-05 | 0.0001 | 0.00013 | 9.07E-05 | 0.00014 | 9.18E-05 | 0 | 4.70E-05 | 0.00036 | 8.94E-05 | 0.00031 | 5.52E-05 |
| Dlg3 | 5.09E-05 | 4.89E-05 | 3.12E-05 | 0 | 8.02E-05 | 2.33E-05 | 3.52E-05 | 4.74E-06 | 0.00013 | 0 | 0.00011 | 4.53E-05 | 1.35E-05 | 7.73E-05 | 4.16E-05 | 3.42E-05 | 2.93E-05 | 6.71E-05 |
| Dbt | 6.08E-05 | 2.12E-05 | 5.27E-05 | 5.19E-06 | 9.87E-05 | 1.86E-06 | 4.86E-06 | 0.0001 | 0.00013 | 4.02E-05 | 0 | 1.21E-05 | 2.01E-05 | 0 | 0 | 0 | 0 | 1.18E-05 |
| Sf3a1 | 0.00018 | 0.00013 | 0.00027 | 0.00025 | 4.88E-05 | 0 | 4.57E-05 | 8.16E-06 | 0.00013 | 4.87E-05 | 7.84E-05 | 0 | 1.35E-05 | 0 | 0.0002 | 0.00025 | 4.59E-05 | 1.14E-05 |
| Vdac1 | 0.0003 | 7.36E-05 | 0.00017 | 6.41E-05 | 5.17E-05 | 4.01E-05 | 0 | 5.26E-05 | 0.00013 | 2.45E-05 | 8.27E-05 | 0.00068 | 0 | 5.94E-05 | 0 | 0.00023 | 0 | 7.15E-05 |
| Ly6a | 0.0002 | 0 | 1.49E-05 | 0 | 0 | 0 | 0.00019 | 0 | 0.00013 | 0.00213 | 0 | 3.88E-05 | 0.00133 | 0 | 0 | 0.00016 | 0 | 0.00134 |
| Cdc42ep1 | 2.01E-05 | 0.00012 | 0.00017 | 0.00012 | 0.00014 | 2.91E-05 | 0 | 0.00011 | 0.00013 | 2.97E-05 | 0.00026 | 0 | 0.0002 | 0.00013 | 9.93E-05 | 5.68E-05 | 3.03E-05 | 6.97E-05 |
| Dusp11 | 0.00037 | 0.00018 | 0.00012 | 0 | 1.89E-05 | 0.00014 | 9.86E-05 | 0 | 0.00013 | 4.93E-05 | 2.55E-05 | 0.00023 | 0 | 6.44E-05 | 0.00033 | 7.34E-05 | 9.34E-05 | 1.24E-05 |
| Pcbp2 | 0.00053 | 0.00016 | 0.0002 | 0 | 0.00015 | 0.00042 | 9.89E-05 | 2.75E-05 | 0.00013 | 0 | 2.44E-05 | 0.00018 | 0 | 0.00012 | 0 | 0.00026 | 0.00045 | 8.38E-06 |
| Cdipt | 0.00016 | 9.97E-05 | 0.00018 | 9.69E-05 | 0.00013 | 6.46E-05 | 4.29E-05 | 8.34E-05 | 0.00013 | 7.98E-05 | 0.00016 | 0 | 2.77E-05 | 0.00011 | 0 | 6.52E-05 | 3.87E-05 | 7.77E-05 |
| Eif4h | 0.0002 | 0.00013 | 0.00022 | 0.00019 | 7.90E-05 | 8.08E-05 | 3.32E-05 | 4.18E-05 | 0.00013 | 3.99E-05 | 0.00014 | 0.00015 | 6.55E-05 | 5.63E-05 | 0.00018 | 0.00022 | 0 | 5.11E-05 |
| Tbca | 0.00049 | 0.00022 | 0.00028 | 0.00031 | 4.28E-05 | 0.00024 | 7.63E-06 | 0.00015 | 0.00013 | 1.14E-05 | 6.52E-05 | 0.0003 | 0.00026 | 0.0001 | 0 | 0.00015 | 0.00015 | 7.69E-05 |
| Rbm14 | 0.00022 | 0.00018 | 0.0002 | 0.00027 | 6.78E-05 | 2.60E-05 | 3.69E-05 | 7.86E-05 | 0.00013 | 4.87E-05 | 5.53E-05 | 9.90E-05 | 0 | 0.0001 | 0.00061 | 0.00022 | 0 | 5.89E-05 |
| Scand1 | 4.40E-05 | 9.83E-05 | 6.75E-05 | 0.00049 | 0.00013 | 0.0001 | 8.51E-05 | 0.00034 | 0.00013 | 9.41E-05 | 0.0001 | 7.74E-05 | 9.99E-05 | 0.0001 | 0.00021 | 6.96E-05 | 1.34E-05 | 0.00013 |

| | | | | | | | | | | | | | | | | | | |
|---|---|---|---|---|---|---|---|---|---|---|---|---|---|---|---|---|---|---|
| Rock2 | 0.00034 | 0.00019 | 0.00017 | 0 | 9.85E-06 | 0.00019 | 0 | 0 | 0.00013 | 0.00026 | 0.00033 | 0.00025 | 6.47E-05 | 8.93E-05 | 0.00032 | 0.00022 | 0.00031 | 0.00012 |
| Pgrmc2 | 0.0002 | 0.00024 | 0.00025 | 0.00026 | 0.00024 | 0.00012 | 0.00029 | 7.12E-05 | 0.00013 | 8.58E-05 | 0.00029 | 0.00039 | 0.00015 | 5.05E-05 | 0.00017 | 0.00014 | 0 | 0.00017 |
| Hdgfl2 | 9.62E-05 | 0.00014 | 0.00015 | 0.0001 | 6.18E-05 | 0 | 0 | 1.01E-05 | 0.00013 | 7.31E-05 | 8.92E-05 | 2.70E-05 | 5.60E-05 | 4.40E-05 | 0.00036 | 0.00021 | 0.00036 | 7.87E-05 |
| Il34 | 0 | 0.00014 | 0 | 0 | 0.00014 | 1.53E-05 | 0 | 0.00021 | 0.00013 | 2.40E-05 | 0 | 3.88E-06 | 0 | 0.00017 | 0 | 0 | 0 | 8.44E-05 |
| Nrbp1 | 0.00021 | 0.00016 | 0.00026 | 0.00035 | 5.73E-05 | 3.98E-05 | 0.00013 | 9.03E-05 | 0.00013 | 6.18E-05 | 0.00014 | 3.09E-05 | 6.34E-06 | 6.14E-05 | 0.00017 | 4.86E-05 | 0 | 5.63E-05 |
| Fip1l1 | 0.00018 | 0.00012 | 8.72E-05 | 2.38E-05 | 0.00011 | 2.90E-05 | 1.37E-05 | 0 | 0.00013 | 5.13E-05 | 2.97E-05 | 0.0001 | 4.54E-07 | 6.85E-05 | 0.0002 | 0.00019 | 2.59E-05 | 6.80E-05 |
| Smyd5 | 0.00013 | 6.24E-06 | 0.00038 | 0.00013 | 0 | 0 | 1.77E-05 | 1.18E-06 | 0.00013 | 1.32E-05 | 6.85E-05 | 1.77E-05 | 2.44E-05 | 0 | 4.85E-05 | 0 | 0 | 0 |
| Renbp | 0 | 0.0002 | 2.70E-05 | 9.24E-05 | 0.00015 | 7.56E-05 | 3.71E-05 | 0.00012 | 0.00013 | 6.19E-05 | 1.88E-05 | 7.09E-05 | 6.54E-05 | 0.00013 | 0.00011 | 0 | 0.00012 | 0.00014 |
| Wdr13 | 5.32E-05 | 0.00012 | 5.38E-05 | 5.64E-05 | 9.27E-05 | 0.00014 | 3.32E-06 | 0.00013 | 0.00013 | 7.69E-05 | 0.00013 | 5.15E-05 | 3.67E-05 | 3.74E-05 | 0.00012 | 0 | 0.00013 | 7.62E-05 |
| Exoc7 | 0.00019 | 0.00017 | 5.48E-05 | 9.29E-05 | 7.24E-05 | 5.98E-05 | 2.73E-05 | 1.57E-05 | 0.00013 | 7.20E-05 | 5.41E-05 | 8.58E-05 | 4.39E-05 | 1.99E-05 | 7.14E-05 | 5.60E-05 | 2.62E-05 | 7.65E-05 |
| Nhlrc2 | 3.88E-05 | 8.62E-05 | 0.00014 | 4.35E-05 | 6.59E-05 | 8.93E-06 | 0 | 5.51E-05 | 0.00013 | 0.00015 | 6.47E-05 | 7.35E-05 | 7.77E-05 | 4.62E-05 | 8.32E-05 | 9.24E-05 | 0 | 9.25E-05 |
| Rexo2 | 0.00025 | 0.00012 | 0.00033 | 0.00075 | 0.00019 | 8.61E-05 | 0.0004 | 9.69E-05 | 0.00013 | 0.00029 | 0.00046 | 0 | 0.00024 | 0.00014 | 0.00015 | 5.62E-05 | 0 | 0.0003 |
| Rbm22 | 0.00019 | 0.00012 | 0.00015 | 4.09E-06 | 7.85E-05 | 8.66E-05 | 4.60E-05 | 3.85E-05 | 0.00013 | 0 | 1.58E-05 | 8.21E-05 | 2.84E-05 | 4.74E-05 | 7.19E-05 | 0.00015 | 0.00012 | 3.81E-05 |
| Scyl1 | 0.00023 | 0.00022 | 0.00028 | 0.0005 | 0.00024 | 0.00018 | 0.00036 | 0.00024 | 0.00013 | 0.00016 | 0.00012 | 0.00019 | 0.00019 | 0.00024 | 0.00036 | 0.0001 | 0.00021 | 0.00011 |
| Arsj | 1.26E-05 | 0 | 0 | 0 | 1.91E-05 | 6.35E-05 | 0 | 0 | 0.00013 | 2.03E-06 | 0 | 0 | 0 | 0 | 0 | 2.88E-06 | 0 | 3.97E-05 |
| Vamp2 | 6.37E-05 | 0.00012 | 1.99E-05 | 9.89E-05 | 0.00011 | 5.05E-05 | 0 | 0.00011 | 0.00013 | 7.43E-05 | 3.27E-05 | 0.00014 | 0 | 5.96E-05 | 0.0004 | 0 | 0 | 0.00012 |
| Dpy19l1 | 0.00017 | 6.06E-05 | 0.00011 | 0 | 0.00012 | 0.00017 | 0.00023 | 0.00012 | 0.00013 | 0.00021 | 7.80E-05 | 0 | 0.00021 | 0.00012 | 0 | 0.00023 | 5.50E-05 | 0.00018 |
| Thrap3 | 3.08E-05 | 0.00015 | 0.00018 | 0.00029 | 6.76E-05 | 2.81E-05 | 1.38E-05 | 0.0001 | 0.00013 | 5.89E-05 | 4.70E-05 | 5.89E-05 | 3.64E-05 | 7.10E-05 | 0.00023 | 0.00025 | 9.54E-05 | 9.90E-05 |
| Tnpo1 | 0.00026 | 0.0002 | 0.00022 | 9.87E-05 | 4.91E-05 | 0.00016 | 8.14E-05 | 5.68E-05 | 0.00013 | 3.11E-05 | 3.86E-05 | 0.00014 | 3.05E-06 | 0.00012 | 0.00013 | 0.00025 | 0.00024 | 3.48E-05 |
| Chst14 | 0.00014 | 0.00013 | 8.39E-05 | 0.00018 | 0.00015 | 0.00018 | 0.00014 | 0.00013 | 0.00013 | 0.00012 | 0.00014 | 0.00012 | 2.17E-05 | 0.00024 | 0 | 7.77E-05 | 0.00013 | 0.00011 |
| Wars | 0.00015 | 9.18E-05 | 0.00017 | 3.83E-05 | 1.18E-05 | 0 | 0 | 7.28E-05 | 0.00013 | 9.28E-05 | 3.62E-05 | 7.17E-05 | 0 | 0 | 5.09E-05 | 8.00E-05 | 0 | 7.46E-05 |
| Wipf1 | 6.43E-05 | 0.00036 | 9.91E-05 | 0.00034 | 0.00037 | 0.00041 | 8.80E-05 | 0.00013 | 0.00013 | 0.00022 | 0.00014 | 0.00047 | 6.17E-06 | 0.00031 | 7.56E-05 | 0.00021 | 0.00032 | 0.00022 |
| Bub3 | 2.58E-05 | 0.00013 | 0.00013 | 0.0002 | 6.77E-05 | 0 | 7.92E-06 | 6.41E-05 | 0.00013 | 6.87E-05 | 0.00012 | 1.08E-06 | 5.08E-06 | 3.67E-05 | 6.05E-06 | 0.00053 | 0 | 4.67E-05 |
| Usp19 | 8.74E-05 | 0.00026 | 0.00019 | 0.0002 | 9.02E-05 | 6.56E-05 | 0.00012 | 0.00013 | 0.00013 | 8.18E-05 | 6.98E-05 | 4.42E-05 | 1.53E-05 | 7.40E-05 | 0.00037 | 7.51E-05 | 4.81E-05 | 0.00014 |
| Arhgap35 | 0.00033 | 0.00023 | 7.86E-05 | 0 | 1.79E-05 | 0.00023 | 0 | 3.55E-05 | 0.00013 | 6.01E-05 | 1.77E-06 | 0.00011 | 0 | 5.28E-05 | 7.83E-05 | 0.0001 | 0.00018 | 0 |
| Sec63 | 0.00041 | 0.00011 | 0.00029 | 0 | 8.57E-05 | 0.00022 | 0.00011 | 9.39E-05 | 0.00013 | 4.64E-05 | 0.00019 | 0.00027 | 0.00019 | 0.00014 | 6.41E-05 | 6.81E-05 | 8.42E-05 | 6.33E-05 |
| Pias2 | 1.04E-05 | 0.00015 | 6.05E-05 | 7.57E-05 | 9.00E-05 | 7.23E-05 | 2.11E-05 | 0.00013 | 0.00013 | 7.13E-05 | 0.00021 | 5.07E-05 | 0 | 0.00015 | 0.00023 | 0.00011 | 0.0002 | 2.06E-05 |
| Fhdc1 | 7.96E-05 | 6.05E-06 | 2.63E-05 | 0 | 1.84E-05 | 0 | 0.00019 | 0.00016 | 0.00013 | 0 | 4.11E-05 | 0.00011 | 0 | 0 | 0 | 1.18E-05 | 6.79E-05 | 0 |
| Lclat1 | 2.29E-05 | 5.54E-05 | 0.00014 | 0.00014 | 0.00023 | 0.00028 | 8.37E-05 | 0.00011 | 0.00013 | 0 | 0.00016 | 3.33E-05 | 4.17E-05 | 4.99E-05 | 0 | 0.00028 | 8.03E-05 | 0 |
| Lrp10 | 0 | 0.00036 | 0.00011 | 0.00046 | 0.00037 | 0.00018 | 0.00019 | 0.00045 | 0.00013 | 0.00041 | 0.00029 | 0.00037 | 0.00015 | 0.00032 | 0.00021 | 0 | 4.95E-05 | 0.00038 |
| Sf3a2 | 0.0002 | 0.00016 | 0.00023 | 1.93E-05 | 0 | 3.18E-05 | 0 | 0 | 0.00013 | 3.57E-05 | 8.02E-05 | 5.54E-05 | 0 | 6.76E-05 | 7.76E-05 | 0.00024 | 0 | 1.53E-05 |
| Prkag2 | 1.70E-05 | 4.81E-05 | 0 | 0 | 0 | 0 | 6.63E-05 | 4.38E-06 | 0.00013 | 9.03E-05 | 0.00014 | 2.98E-05 | 0.00012 | 0 | 3.78E-06 | 3.32E-05 | 2.06E-05 | 0.00011 |
| Plcb3 | 0.00012 | 0.00014 | 0.00016 | 0.00014 | 7.59E-06 | 1.80E-05 | 5.17E-05 | 0.00012 | 0.00013 | 0.00016 | 0.00028 | 0.0001 | 0.00015 | 7.10E-05 | 0.00018 | 1.17E-05 | 1.95E-05 | 0.00011 |

| | | | | | | | | | | | | | | | | | | |
|---|---|---|---|---|---|---|---|---|---|---|---|---|---|---|---|---|---|---|
| Idh3b | 0.00033 | 0.00025 | 0.00021 | 0.00015 | 0.00012 | 0.00016 | 5.12E-05 | 8.35E-05 | 0.00013 | 5.44E-05 | 5.33E-05 | 0.00013 | 3.67E-05 | 0.00013 | 0.00014 | 8.32E-05 | 1.41E-05 | 5.07E-05 |
| Pigq | 3.92E-05 | 9.34E-05 | 7.59E-05 | 0.00015 | 6.41E-05 | 5.58E-05 | 0 | 0.00011 | 0.00013 | 9.74E-05 | 8.58E-05 | 0 | 4.20E-05 | 8.23E-05 | 7.20E-05 | 0.0001 | 0 | 7.96E-05 |
| Ddx54 | 7.19E-05 | 0.00015 | 0.00022 | 0.00029 | 3.77E-05 | 0 | 0 | 2.23E-05 | 0.00013 | 5.15E-05 | 0.00013 | 4.72E-05 | 0 | 8.86E-05 | 0.00013 | 0 | 0 | 5.25E-05 |
| Ywhaq | 0.00036 | 9.75E-05 | 0.00012 | 0 | 3.29E-05 | 0.00013 | 0 | 1.23E-05 | 0.00013 | 2.58E-05 | 3.56E-05 | 6.14E-05 | 0 | 1.52E-05 | 0 | 0.0002 | 0.00026 | 2.84E-05 |
| Cfdp1 | 0.00045 | 0.00016 | 0.00023 | 0.0002 | 0.0001 | 0.00019 | 0.00015 | 8.13E-05 | 0.00013 | 1.79E-05 | 0.0001 | 0.00015 | 0.00011 | 0.0001 | 0.00012 | 0.00025 | 0.0001 | 8.93E-06 |
| Hfe | 0 | 4.37E-05 | 0 | 1.48E-05 | 0 | 2.28E-05 | 0 | 5.01E-05 | 0.00013 | 7.67E-05 | 0 | 0 | 0.00013 | 1.24E-05 | 0 | 8.45E-06 | 0 | 0.0001 |
| Fbxl14 | 0 | 0.00011 | 9.97E-05 | 0.00011 | 8.23E-05 | 2.62E-05 | 0 | 0.00017 | 0.00013 | 9.16E-05 | 9.62E-05 | 0.00019 | 0 | 6.45E-05 | 8.80E-05 | 0.00017 | 0.00019 | 0.00014 |
| Ccdc3 | 0 | 1.71E-05 | 0 | 0 | 0 | 2.70E-05 | 0 | 0 | 0.00013 | 0 | 0 | 2.01E-06 | 0 | 0 | 0 | 0 | 0 | 0.001 |
| Cbfa2t2 | 0.00011 | 0.00011 | 2.53E-05 | 0 | 8.50E-05 | 5.57E-05 | 0.00015 | 0 | 0.00013 | 3.86E-06 | 1.46E-05 | 6.17E-05 | 1.84E-05 | 4.93E-05 | 5.78E-05 | 6.02E-05 | 5.58E-05 | 6.22E-05 |
| Poldip2 | 0.00014 | 8.04E-05 | 0.0001 | 0.00016 | 6.59E-05 | 3.66E-05 | 0.00011 | 0 | 0.00013 | 6.64E-06 | 4.20E-05 | 1.50E-05 | 8.78E-05 | 3.22E-05 | 3.52E-05 | 4.98E-05 | 0 | 2.83E-05 |
| Rnf6 | 0.00013 | 0.00013 | 0.00018 | 0.00013 | 7.61E-05 | 2.35E-05 | 6.40E-05 | 0 | 0.00013 | 8.25E-05 | 0.00011 | 0.00024 | 7.57E-06 | 9.08E-05 | 0.0002 | 0.00016 | 0 | 0.00011 |
| Supt16 | 0.00014 | 0.00018 | 0.00025 | 0.00018 | 7.90E-05 | 3.53E-05 | 0 | 5.70E-07 | 0.00013 | 6.71E-05 | 4.14E-05 | 2.19E-05 | 1.31E-05 | 4.18E-05 | 8.31E-05 | 0.00047 | 7.95E-05 | 0.00011 |
| Srrm2 | 0.00014 | 0.00039 | 0.00018 | 0.00029 | 0.00024 | 0.00013 | 8.41E-05 | 0.00014 | 0.00013 | 9.94E-05 | 8.55E-05 | 0.00034 | 6.46E-05 | 0.00016 | 0.00078 | 0.0002 | 0.00029 | 0.00011 |
| Gosr2 | 0.00041 | 0.00016 | 0.00026 | 0.00013 | 0.00025 | 0.00022 | 0.00036 | 0.00015 | 0.00013 | 0.00011 | 0.00016 | 0.00012 | 0.00016 | 5.39E-05 | 2.25E-05 | 4.58E-05 | 0 | 0.00012 |
| Pja1 | 0.00018 | 8.80E-05 | 0.00014 | 0.00022 | 0 | 5.08E-05 | 7.65E-05 | 8.36E-05 | 0.00013 | 5.79E-05 | 7.39E-05 | 5.73E-05 | 0 | 0.00013 | 0.00014 | 2.59E-05 | 0 | 8.34E-05 |
| Crebbp | 0.00026 | 0.00026 | 7.42E-05 | 0 | 0.00012 | 0.00016 | 0.00011 | 4.25E-05 | 0.00013 | 3.83E-05 | 5.28E-05 | 0.00013 | 2.93E-06 | 7.34E-05 | 0.00018 | 9.21E-05 | 0.00018 | 4.67E-05 |
| Cdc27 | 0.00021 | 0.00012 | 0.00014 | 0 | 6.13E-05 | 6.77E-05 | 6.56E-05 | 5.60E-05 | 0.00013 | 2.45E-05 | 3.72E-05 | 3.76E-05 | 0 | 3.30E-05 | 5.34E-05 | 0.00028 | 0.00012 | 1.78E-05 |
| Amfr | 0.00014 | 0.00015 | 0.00028 | 0.00025 | 0.00017 | 0.00018 | 0.00011 | 0.00022 | 0.00013 | 0.00017 | 0.00018 | 0.00014 | 6.26E-05 | 7.16E-05 | 5.20E-05 | 0.00018 | 0 | 0.00014 |
| Pcdh18 | 0.00012 | 9.71E-05 | 0 | 1.79E-05 | 3.92E-05 | 0.00013 | 0 | 7.77E-06 | 0.00013 | 1.62E-05 | 3.68E-05 | 0 | 0 | 0.00019 | 6.73E-06 | 6.09E-05 | 0 | 6.68E-05 |
| Edem3 | 0.00022 | 7.39E-05 | 0.00012 | 4.69E-05 | 0.00015 | 0.00015 | 0.00017 | 0.00013 | 0.00013 | 9.95E-05 | 0.00011 | 4.74E-05 | 0.0001 | 9.00E-05 | 0.00011 | 0 | 9.05E-05 | 8.16E-05 |
| Sec61b | 0.00054 | 0 | 0.00034 | 0.00031 | 6.54E-05 | 0.00012 | 0.0005 | 1.59E-05 | 0.00013 | 0 | 0.00012 | 0 | 4.24E-05 | 0.0001 | 0 | 1.98E-05 | 0 | 0 |
| Brd1 | 0 | 0.00024 | 0.00018 | 0.00015 | 0.0001 | 2.13E-05 | 8.89E-05 | 0.00012 | 0.00013 | 8.92E-05 | 0.00018 | 0.00014 | 6.05E-06 | 0.00013 | 0.00025 | 6.57E-05 | 0.0003 | 3.97E-05 |
| Tubb4b | 0.00026 | 0.00014 | 0.0002 | 0.00021 | 0 | 4.56E-05 | 0 | 4.05E-07 | 0.00013 | 0.00013 | 0.00015 | 0.00028 | 0.00012 | 0 | 3.76E-05 | 0.00112 | 0.00148 | 0.0001 |
| Fundc2 | 0.00032 | 0.00012 | 4.06E-05 | 0.00022 | 0.00011 | 0.00018 | 1.88E-05 | 9.44E-05 | 0.00013 | 0.00012 | 1.65E-05 | 0.00014 | 0.00011 | 0.00012 | 0 | 0.00023 | 7.00E-05 | 0.00013 |
| Dnm1l | 0.00016 | 8.03E-05 | 0.00013 | 4.98E-05 | 6.72E-05 | 8.85E-05 | 4.42E-05 | 0.00012 | 0.00013 | 9.60E-05 | 0.00012 | 9.98E-05 | 1.43E-05 | 5.80E-05 | 9.51E-05 | 0.00015 | 5.71E-05 | 0.00012 |
| Hdac4 | 6.35E-06 | 0.00023 | 0.00014 | 1.37E-05 | 8.86E-05 | 5.55E-05 | 4.03E-05 | 0.0001 | 0.00013 | 0.00016 | 7.95E-05 | 8.89E-05 | 7.77E-05 | 5.12E-05 | 0.00016 | 6.32E-05 | 0.00024 | 0.00012 |
| Bahcc1 | 7.78E-07 | 8.22E-05 | 7.04E-05 | 8.59E-05 | 0 | 0 | 0 | 0 | 0.00013 | 3.54E-05 | 0 | 0 | 3.79E-05 | 4.26E-06 | 7.15E-05 | 0.00013 | 0.00011 | 0.00023 |
| Sertad1 | 8.69E-05 | 0.00011 | 3.29E-05 | 9.68E-05 | 4.61E-06 | 0 | 0 | 0 | 0.00013 | 3.57E-05 | 0.00013 | 0.00038 | 0 | 6.83E-05 | 0 | 0 | 0.00252 | 1.93E-05 |
| Vps52 | 8.69E-05 | 6.45E-05 | 8.83E-05 | 0.00014 | 0.0001 | 5.70E-05 | 5.31E-05 | 0.0001 | 0.00013 | 0.0001 | 6.91E-05 | 6.33E-05 | 3.72E-05 | 6.99E-05 | 7.36E-05 | 2.39E-05 | 0 | 0.00012 |
| Isca2 | 5.83E-05 | 8.66E-05 | 5.13E-05 | 8.12E-05 | 7.10E-05 | 4.46E-05 | 0 | 6.01E-05 | 0.00013 | 7.59E-05 | 5.85E-05 | 5.69E-05 | 2.05E-06 | 7.71E-05 | 5.69E-05 | 9.28E-05 | 0 | 3.31E-05 |
| Il17rc | 3.11E-05 | 5.49E-05 | 7.53E-05 | 6.91E-05 | 3.40E-05 | 0 | 3.73E-05 | 1.23E-05 | 0.00013 | 6.67E-05 | 5.17E-05 | 2.81E-05 | 5.67E-05 | 4.37E-05 | 3.69E-05 | 3.31E-05 | 3.62E-05 | 5.49E-05 |
| Nop53 | 0.00011 | 0.00031 | 5.22E-05 | 0.00033 | 5.62E-05 | 0.00012 | 0 | 0.00012 | 0.00013 | 8.95E-05 | 0 | 0.00037 | 4.87E-05 | 7.44E-05 | 0.00026 | 0.0001 | 0.00011 | 0.00011 |
| Fam53c | 2.59E-06 | 0.00013 | 7.18E-05 | 0.00017 | 8.44E-05 | 5.30E-06 | 1.38E-05 | 4.51E-05 | 0.00013 | 0.00013 | 0.00013 | 9.70E-05 | 5.64E-05 | 9.95E-05 | 5.38E-05 | 5.06E-05 | 0.00017 | 7.86E-05 |

| | | | | | | | | | | | | | | | | | | |
|---|---|---|---|---|---|---|---|---|---|---|---|---|---|---|---|---|---|---|
| Smad4 | 0.00024 | 0.00042 | 0.00028 | 0.00018 | 0.00023 | 0.00028 | 0.00036 | 0.00023 | 0.00013 | 0.00018 | 0.00035 | 0.00044 | 0.00013 | 0.00021 | 6.33E-05 | 0.00022 | 0.00012 | 0.00018 |
| Wdr35 | 3.45E-05 | 5.02E-05 | 3.50E-06 | 0 | 0.00015 | 0 | 0 | 7.98E-06 | 0.00013 | 3.67E-05 | 5.11E-05 | 0 | 4.20E-06 | 0 | 2.13E-05 | 0.00012 | 4.70E-05 | 6.62E-05 |
| Zfp652 | 9.89E-05 | 0.00016 | 4.04E-05 | 9.25E-05 | 0.00021 | 0.00015 | 0.00027 | 9.16E-05 | 0.00013 | 0.0001 | 1.39E-05 | 9.75E-05 | 0.00012 | 0.00011 | 0.00026 | 6.75E-05 | 0.00014 | 0.00016 |
| Tmtc3 | 0.00016 | 6.63E-05 | 0.00011 | 3.47E-05 | 7.22E-05 | 0.00013 | 3.87E-05 | 9.37E-05 | 0.00013 | 5.09E-05 | 0.00011 | 4.09E-05 | 1.65E-05 | 7.45E-05 | 0 | 0.00018 | 1.29E-05 | 4.57E-05 |
| Zbtb38 | 0.00012 | 0.00021 | 0.00012 | 0 | 0.00016 | 0 | 0.00023 | 6.93E-05 | 0.00013 | 4.29E-05 | 0.00018 | 0.00016 | 0.00044 | 8.12E-05 | 7.61E-05 | 0.0001 | 0.00032 | 5.83E-05 |
| Pank2 | 5.93E-05 | 0.00014 | 8.34E-05 | 0.0001 | 8.65E-05 | 4.85E-05 | 3.29E-05 | 0.00016 | 0.00013 | 9.05E-05 | 0.00011 | 2.08E-06 | 4.49E-05 | 7.80E-05 | 0.00015 | 0.00021 | 0 | 9.56E-05 |
| Fdx2 | 0.00015 | 0.00014 | 0.00027 | 0.00028 | 0 | 0 | 5.09E-05 | 5.53E-05 | 0.00013 | 2.03E-05 | 0.0001 | 2.46E-05 | 1.66E-05 | 0 | 0.0001 | 0 | 0.00019 | 3.74E-05 |
| Cdca4 | 3.09E-05 | 0.00012 | 0.00017 | 4.11E-05 | 8.03E-05 | 5.49E-05 | 0 | 8.81E-05 | 0.00013 | 3.53E-05 | 0.00012 | 0.00014 | 2.82E-05 | 9.68E-05 | 0.00015 | 0.00049 | 0 | 5.61E-05 |
| Fbxo31 | 5.78E-05 | 0.00011 | 6.77E-05 | 8.88E-05 | 3.36E-05 | 1.19E-05 | 0 | 6.45E-05 | 0.00013 | 4.25E-05 | 0 | 0 | 0 | 3.25E-05 | 4.56E-05 | 1.28E-05 | 0 | 5.25E-05 |
| Ago2 | 0.00012 | 0.00024 | 0.0003 | 4.71E-05 | 0.00012 | 0.00017 | 6.85E-05 | 0.00014 | 0.00013 | 0.00012 | 0.00019 | 0.00053 | 0 | 0.00015 | 0.00013 | 0.00023 | 0.00017 | 0.00011 |
| Snapin | 9.86E-05 | 0.00014 | 0 | 0 | 0.00011 | 9.01E-05 | 7.59E-05 | 4.56E-05 | 0.00013 | 7.81E-05 | 0.00011 | 0.00012 | 2.53E-05 | 5.36E-05 | 0 | 0 | 0 | 0.00017 |
| Tomm22 | 0.00027 | 0.00013 | 0.00023 | 6.48E-05 | 7.37E-05 | 0.0001 | 0 | 5.32E-05 | 0.00013 | 3.70E-05 | 9.78E-05 | 7.95E-05 | 0 | 5.69E-05 | 0 | 0.00013 | 1.50E-05 | 5.11E-05 |
| Atp9b | 8.07E-05 | 0.00016 | 9.77E-05 | 0 | 0.00011 | 8.54E-05 | 8.78E-06 | 0.00012 | 0.00013 | 0.00011 | 8.99E-05 | 0.00018 | 6.31E-05 | 5.34E-05 | 0.00034 | 5.70E-05 | 0.00016 | 0.00015 |
| Srsf3 | 0.00032 | 3.76E-05 | 7.28E-05 | 0 | 0 | 4.10E-05 | 0 | 0 | 0.00013 | 6.73E-06 | 0 | 0.0003 | 8.56E-06 | 0 | 0 | 0.00024 | 0.00075 | 0 |
| Hs2st1 | 0.0002 | 0.00017 | 8.14E-05 | 0 | 0.00021 | 0.00018 | 0.0002 | 0.00016 | 0.00013 | 3.32E-05 | 0.00016 | 0.00022 | 0.00017 | 7.07E-05 | 0 | 0.00018 | 0.00011 | 0.00027 |
| Sik3 | 0.00011 | 0.00014 | 0.0001 | 1.38E-05 | 0.00015 | 0.00012 | 5.73E-05 | 9.83E-05 | 0.00013 | 0.00013 | 6.88E-05 | 0.00018 | 0 | 0.00014 | 0.0002 | 8.14E-05 | 0.00021 | 0 |
| Cflar | 2.34E-05 | 0.00013 | 4.84E-05 | 6.46E-05 | 0.00019 | 8.84E-05 | 4.50E-05 | 0.00012 | 0.00013 | 0.0001 | 0.00016 | 0.00014 | 0.00017 | 0.0002 | 9.25E-05 | 3.39E-05 | 4.73E-05 | 0.00018 |
| Arglu1 | 0.00018 | 0.00024 | 0.00025 | 0.00014 | 0.00012 | 0 | 0 | 9.27E-05 | 0.00013 | 8.29E-05 | 9.97E-06 | 0 | 0 | 7.76E-05 | 0.002 | 0.00026 | 0 | 7.01E-05 |
| Mmab | 5.41E-05 | 3.10E-05 | 0 | 2.37E-05 | 4.53E-05 | 0 | 1.20E-05 | 3.21E-05 | 0.00013 | 6.27E-06 | 2.91E-05 | 0 | 0 | 0 | 8.92E-05 | 1.70E-06 | 0 | 1.96E-05 |
| Dnajc15 | 0.00016 | 4.80E-06 | 7.08E-05 | 0.00012 | 3.48E-05 | 5.98E-05 | 0 | 0.0001 | 0.00013 | 0 | 0 | 0 | 6.50E-05 | 0.00012 | 0 | 5.35E-05 | 0 | 0 |
| Trpc4ap | 0.0004 | 0.00025 | 0.0002 | 0.00021 | 0.00014 | 0.00022 | 0.00036 | 0.00011 | 0.00013 | 0.00014 | 0.0001 | 0.00018 | 0.00016 | 8.01E-05 | 0.00032 | 5.08E-05 | 7.95E-05 | 0.00012 |
| G6pdx | 0.00013 | 5.69E-05 | 0.00011 | 0.00013 | 0 | 6.58E-06 | 0 | 0 | 0.00013 | 0.00014 | 0 | 5.38E-05 | 4.25E-05 | 2.88E-05 | 0 | 3.31E-05 | 0 | 2.50E-05 |
| Bola2 | 0.00018 | 0.00012 | 0.00033 | 0.00041 | 6.21E-05 | 7.54E-05 | 0.00011 | 0.00015 | 0.00013 | 5.76E-05 | 0.00016 | 0.00012 | 0.00011 | 0.00017 | 0 | 0.00013 | 9.57E-05 | 6.06E-05 |
| Icam1 | 0 | 8.25E-05 | 0.00024 | 0 | 0 | 0 | 0 | 3.18E-05 | 0.00013 | 0.0001 | 0 | 0 | 0.00083 | 0.00026 | 0 | 0 | 0.00059 | 0.00042 |
| Golph3 | 0.00026 | 0.00026 | 0.00036 | 0.00021 | 0.00023 | 0.00019 | 0.0004 | 0.00014 | 0.00013 | 0.00013 | 0.00018 | 0.00021 | 5.07E-06 | 0.00025 | 0.00057 | 8.13E-05 | 0.00062 | 0.00017 |
| Luzp1 | 0.00012 | 0.00021 | 0.00012 | 1.39E-05 | 0.00018 | 0.00012 | 4.45E-05 | 6.12E-05 | 0.00013 | 2.12E-05 | 0.0002 | 1.01E-05 | 0.0003 | 0.00031 | 0.0002 | 9.93E-05 | 9.17E-05 | 0.0002 |
| Abhd16a | 6.33E-05 | 7.16E-05 | 0.0001 | 7.81E-05 | 7.78E-05 | 9.37E-05 | 0.00012 | 7.82E-05 | 0.00013 | 0.00017 | 0.00011 | 4.66E-05 | 8.55E-05 | 7.00E-05 | 8.29E-05 | 2.69E-05 | 6.32E-05 | 0.00018 |
| Rnf166 | 5.92E-05 | 0.00012 | 7.40E-05 | 0.00013 | 0.00017 | 0.00012 | 0.00014 | 8.51E-05 | 0.00013 | 6.51E-05 | 8.91E-05 | 2.35E-05 | 0.00014 | 6.54E-05 | 0.00012 | 1.87E-05 | 0 | 0.00011 |
| Rbm25 | 0.0002 | 0.00024 | 0.00023 | 9.91E-05 | 8.50E-05 | 5.35E-05 | 4.70E-05 | 0 | 0.00013 | 5.10E-05 | 0.00011 | 0.00013 | 2.79E-05 | 7.12E-05 | 0.00044 | 0.0002 | 0.00015 | 4.25E-05 |
| Pfdn6 | 0.00011 | 0.0001 | 0.00022 | 0.00037 | 4.60E-05 | 2.06E-05 | 0 | 0 | 0.00013 | 2.43E-05 | 0.00013 | 3.63E-05 | 0 | 6.30E-05 | 0 | 0.00025 | 0.00011 | 1.78E-05 |
| Shisa4 | 0 | 2.12E-05 | 6.73E-06 | 7.26E-05 | 0.00019 | 7.63E-05 | 0 | 7.77E-05 | 0.00013 | 1.94E-05 | 0.00025 | 0 | 0.00012 | 0 | 0 | 4.92E-05 | 0 | 8.66E-05 |
| Rab5if | 0.00041 | 0.00015 | 0.00039 | 0.00026 | 0.00017 | 0.00028 | 0.00021 | 7.16E-05 | 0.00013 | 7.35E-05 | 0.00024 | 4.71E-05 | 0.00018 | 0.00021 | 0 | 9.15E-05 | 3.81E-07 | 0 |
| F2rl1 | 0 | 0 | 0.00098 | 0 | 0.00035 | 5.90E-05 | 0.0004 | 0.00035 | 0.00013 | 0 | 0.00017 | 0.00018 | 0 | 2.35E-05 | 0 | 0.00017 | 0 | 0 |

| | | | | | | | | | | | | | | | | | | |
|---|---|---|---|---|---|---|---|---|---|---|---|---|---|---|---|---|---|---|
| Ubac2 | 9.84E-05 | 7.87E-05 | 0.00011 | 7.14E-05 | 7.33E-05 | 2.73E-05 | 0.00015 | 3.25E-05 | 0.00013 | 7.22E-05 | 5.96E-05 | 0 | 2.51E-05 | 6.57E-05 | 3.25E-05 | 3.70E-05 | 0 | 7.16E-05 |
| Rhog | 0.00011 | 0.00013 | 0.00023 | 0.00015 | 8.72E-05 | 4.03E-06 | 0 | 0.00024 | 0.00013 | 8.43E-05 | 8.80E-05 | 0.00026 | 7.43E-06 | 0.00015 | 0 | 0.00013 | 5.15E-05 | 0.00013 |
| Dek | 0.00016 | 0.00015 | 6.25E-05 | 0 | 0.00012 | 0.00018 | 5.26E-05 | 0 | 0.00013 | 6.99E-05 | 2.65E-05 | 1.31E-05 | 0.00011 | 8.55E-05 | 0.00017 | 0.00099 | 0.00012 | 7.73E-05 |
| Isoc1 | 8.53E-05 | 1.64E-05 | 0 | 5.77E-05 | 3.87E-05 | 2.74E-05 | 1.63E-05 | 4.70E-06 | 0.00013 | 2.56E-05 | 2.86E-05 | 0 | 0 | 0 | 1.06E-05 | 5.61E-05 | 8.96E-06 | 3.45E-05 |
| Rraga | 0.00038 | 0.0001 | 0.00014 | 0 | 5.59E-05 | 0.00015 | 4.10E-05 | 0 | 0.00013 | 4.28E-05 | 5.83E-05 | 0.00014 | 1.53E-06 | 2.87E-05 | 0 | 0 | 0 | 5.89E-05 |
| Psmb6 | 0.00031 | 6.48E-05 | 0.00023 | 0.00024 | 4.01E-05 | 2.33E-05 | 4.56E-05 | 0 | 0.00013 | 0 | 8.79E-05 | 2.20E-06 | 0 | 4.03E-05 | 3.89E-05 | 0.00013 | 0.00021 | 4.47E-05 |
| Dcbld2 | 8.60E-05 | 2.10E-05 | 0.00022 | 0 | 3.50E-05 | 0 | 7.22E-05 | 0.00079 | 0.00013 | 0.00046 | 0.00024 | 0.00042 | 0 | 6.23E-05 | 0.00011 | 0.00029 | 9.25E-05 | 0 |
| Myef2 | 4.87E-05 | 0.00016 | 0.00013 | 0 | 6.79E-05 | 0 | 0 | 0.00027 | 0.00013 | 4.04E-05 | 0.00013 | 5.29E-05 | 0 | 0 | 0.00026 | 0.00048 | 5.45E-05 | 0 |
| H2afv | 0.00014 | 0.00033 | 0 | 0.00026 | 0.00014 | 0.00018 | 0.00011 | 5.85E-05 | 0.00013 | 9.10E-05 | 3.85E-05 | 7.14E-06 | 7.62E-05 | 0.00017 | 0.00013 | 0.00099 | 0 | 0.00012 |
| Tbc1d14 | 3.73E-05 | 0.00013 | 4.91E-05 | 7.91E-05 | 8.90E-05 | 2.75E-05 | 7.79E-05 | 2.92E-05 | 0.00013 | 0.00015 | 8.56E-05 | 8.45E-05 | 0.00016 | 4.12E-05 | 7.18E-05 | 6.04E-05 | 0 | 0.00012 |
| Lama2 | 0 | 0.00041 | 0 | 0 | 0.00025 | 0.00038 | 0 | 0.0006 | 0.00013 | 0.00053 | 0 | 0 | 0 | 0.00053 | 0 | 0 | 0 | 0.00059 |
| Hgs | 9.45E-05 | 0.00022 | 0.00021 | 0.00023 | 6.92E-05 | 4.54E-06 | 0 | 6.91E-05 | 0.00013 | 0.00012 | 0.00011 | 0.00025 | 3.12E-05 | 5.06E-05 | 0.00019 | 5.94E-05 | 6.64E-05 | 0.00013 |
| Tm2d2 | 0.00011 | 0.00018 | 8.30E-05 | 0.00033 | 0.00023 | 0.00017 | 0.00014 | 0.00021 | 0.00013 | 0.00028 | 0.00019 | 0.00017 | 0.00025 | 0.00014 | 0 | 7.20E-05 | 0 | 0.0003 |
| Hpse | 9.00E-06 | 0 | 1.62E-06 | 0 | 0 | 0.00011 | 4.53E-05 | 2.73E-05 | 0.00013 | 0.0002 | 0 | 0.00026 | 0.00011 | 0 | 0 | 8.74E-05 | 6.41E-06 | 0.0001 |
| Psmd7 | 0.00044 | 0.00027 | 0.00043 | 0.00039 | 0.00014 | 0.00014 | 0.00012 | 1.52E-05 | 0.00013 | 0.00011 | 0.00014 | 7.73E-05 | 6.41E-05 | 2.55E-05 | 0.00025 | 0.00028 | 0.00016 | 7.05E-05 |
| Dlg2 | 2.98E-05 | 1.05E-05 | 9.57E-06 | 0 | 0 | 0.0001 | 3.78E-05 | 3.30E-05 | 0.00013 | 2.14E-05 | 0 | 0 | 0 | 0 | 0 | 2.12E-05 | 0 | 0 |
| Zfp445 | 8.16E-05 | 0.00017 | 0.00012 | 9.88E-05 | 0.00011 | 1.70E-05 | 2.74E-05 | 3.20E-05 | 0.00013 | 8.12E-05 | 5.52E-05 | 6.19E-05 | 5.26E-05 | 6.43E-05 | 0.00034 | 4.00E-05 | 0 | 4.54E-05 |
| Ubtf | 0.0003 | 0.0002 | 0.00027 | 4.95E-05 | 6.66E-05 | 0.00018 | 0.00016 | 0 | 0.00013 | 0.00014 | 3.66E-05 | 7.89E-05 | 0.00012 | 3.54E-05 | 0.00019 | 0.00029 | 0.00019 | 7.92E-05 |
| Kdm4a | 6.88E-05 | 0.00017 | 3.48E-05 | 0.00017 | 9.35E-05 | 5.76E-05 | 5.22E-05 | 4.07E-05 | 0.00013 | 9.44E-05 | 7.82E-05 | 9.95E-05 | 6.87E-05 | 4.54E-05 | 0.0002 | 7.06E-05 | 4.55E-05 | 0.00011 |
| Med15 | 8.55E-05 | 0.00021 | 0.00029 | 0.00028 | 0.00013 | 0.00012 | 0.00013 | 3.57E-05 | 0.00013 | 0.00017 | 0.00028 | 0.00013 | 0.0001 | 0.00016 | 6.02E-05 | 0.00012 | 0.00015 | 0.00011 |
| Fut10 | 0 | 0.00011 | 0 | 0.00011 | 8.54E-05 | 4.17E-05 | 2.65E-05 | 4.66E-05 | 0.00013 | 7.52E-05 | 0 | 0 | 5.69E-05 | 2.99E-05 | 1.78E-05 | 0.00028 | 0 | 9.06E-05 |
| Isg20 | 4.48E-05 | 1.44E-05 | 0 | 0 | 0.0001 | 6.38E-05 | 0.00025 | 0 | 0.00013 | 3.68E-05 | 0.00012 | 0.0004 | 0.00011 | 0.00011 | 0 | 0 | 0 | 0.00013 |
| Map3k6 | 0 | 4.87E-05 | 0.0001 | 0 | 0 | 5.20E-05 | 0 | 0.00021 | 0.00013 | 9.02E-05 | 0 | 3.52E-05 | 0 | 3.09E-05 | 0.00012 | 0 | 0.00011 | 7.27E-05 |
| Agpat3 | 0 | 0.0001 | 2.99E-05 | 0.0003 | 0 | 0.00016 | 0 | 8.25E-05 | 0.00013 | 0.00027 | 0 | 1.71E-05 | 0.00024 | 0 | 0 | 6.96E-05 | 5.29E-06 | 0.00031 |
| Acsl5 | 0.00029 | 9.40E-05 | 0.0002 | 5.35E-05 | 4.68E-05 | 0.00011 | 1.76E-05 | 4.11E-05 | 0.00013 | 0.00016 | 0 | 1.00E-05 | 0 | 2.80E-05 | 8.68E-07 | 0.00014 | 4.49E-05 | 0.00014 |
| Pole4 | 7.83E-05 | 4.45E-05 | 0.00025 | 0.00022 | 6.18E-05 | 0 | 5.61E-05 | 8.12E-06 | 0.00013 | 4.73E-05 | 0.00034 | 0.00011 | 5.68E-05 | 0.00011 | 0.00016 | 0.00023 | 0.00022 | 2.65E-05 |
| Pacsin2 | 6.37E-05 | 0.00011 | 0.0002 | 8.34E-05 | 2.34E-05 | 0 | 0 | 4.81E-05 | 0.00013 | 0.00016 | 0.00017 | 0.00011 | 9.26E-06 | 5.92E-05 | 7.02E-05 | 0.00027 | 0.00019 | 0.00014 |
| Pitrm1 | 0.00024 | 0.00011 | 0.0003 | 7.17E-05 | 6.28E-05 | 7.64E-05 | 0.00025 | 0 | 0.00013 | 1.62E-05 | 0.00015 | 3.30E-05 | 0.00012 | 9.14E-05 | 0 | 0.00027 | 0.00028 | 0 |
| Tor1aip1 | 0.00023 | 0.00019 | 0.00019 | 0.00015 | 3.64E-05 | 0.00022 | 7.80E-05 | 5.85E-05 | 0.00013 | 0.0004 | 0.00011 | 0.00025 | 0.00027 | 7.52E-05 | 8.82E-05 | 0.00028 | 0.00011 | 0.00023 |
| Arfgap2 | 1.97E-05 | 0.0002 | 0.00013 | 0.00023 | 0.00012 | 5.00E-05 | 1.59E-05 | 0.00011 | 0.00013 | 6.74E-05 | 9.14E-05 | 6.34E-05 | 5.60E-05 | 7.03E-05 | 0.00023 | 0.00016 | 0 | 0.00015 |
| Gna13 | 3.93E-05 | 0.00033 | 0.00023 | 0.00033 | 0.00017 | 6.92E-05 | 0 | 0.00015 | 0.00013 | 0.00015 | 8.88E-05 | 0.00054 | 0 | 0.00022 | 0.00039 | 0.00025 | 7.50E-05 | 0.00023 |
| Tagln2 | 0.00018 | 0.00023 | 0.00108 | 0.00089 | 5.31E-06 | 0.00049 | 0.0002 | 0.00077 | 0.00013 | 0.00068 | 0.00138 | 0.00083 | 0.00019 | 0.00065 | 0.00023 | 0.00087 | 0.00053 | 0.00082 |
| Slc9a3r2 | 0 | 6.12E-05 | 0 | 2.11E-05 | 9.54E-05 | 0.00011 | 0 | 0.00024 | 0.00013 | 0 | 0.00021 | 0 | 0 | 0.00016 | 4.08E-05 | 0 | 0 | 8.47E-05 |

| | | | | | | | | | | | | | | | | | | |
|---|---|---|---|---|---|---|---|---|---|---|---|---|---|---|---|---|---|---|
| Nr4a1 | 0 | 0 | 0.00019 | 6.85E-05 | 4.61E-05 | 0.00028 | 0 | 8.26E-06 | 0.00013 | 0 | 2.98E-05 | 0 | 0 | 0 | 3.58E-07 | 0 | 0.00749 | 0.00028 |
| Cmtr1 | 5.58E-05 | 0.00013 | 7.55E-05 | 9.84E-05 | 7.47E-05 | 1.05E-05 | 1.86E-05 | 5.56E-05 | 0.00013 | 0.00014 | 4.90E-05 | 2.86E-05 | 0.00011 | 2.29E-05 | 0.00013 | 0.00014 | 6.95E-05 | 4.95E-05 |
| Cers5 | 5.52E-05 | 0.00015 | 0.00014 | 0.00015 | 0.00015 | 0.00012 | 0.00029 | 0.00018 | 0.00013 | 0.00011 | 0.0002 | 0.00018 | 0.00013 | 0.00015 | 1.15E-05 | 7.71E-05 | 7.73E-06 | 0.00019 |
| Qars | 0.00019 | 0.00016 | 0.00018 | 0.00024 | 6.80E-05 | 6.79E-05 | 6.96E-05 | 9.12E-05 | 0.00013 | 6.49E-05 | 0 | 0.00011 | 0 | 0.00013 | 6.28E-05 | 1.00E-04 | 0.00014 | 5.50E-05 |
| Pom121 | 8.88E-05 | 0.0001 | 0.00015 | 3.63E-05 | 4.76E-05 | 0 | 5.77E-06 | 3.14E-05 | 0.00013 | 3.42E-05 | 8.23E-05 | 8.86E-05 | 1.38E-05 | 2.86E-05 | 0.00015 | 0.00022 | 0.00026 | 0 |
| Gba | 0.00013 | 9.61E-05 | 4.56E-05 | 5.97E-05 | 8.86E-05 | 0.00013 | 0.00015 | 0.00017 | 0.00013 | 0.00013 | 9.48E-05 | 0.00015 | 0.00026 | 0.00021 | 0 | 0 | 0 | 0.00024 |
| Gzme | 0 | 0 | 0 | 0 | 8.28E-05 | 0 | 0 | 0.00012 | 0.00013 | 0 | 5.74E-05 | 0 | 0 | 0.00049 | 0 | 7.39E-06 | 0 | 0 |
| Plekhj1 | 8.59E-05 | 0.00017 | 0.00021 | 0.00029 | 6.82E-05 | 0.00012 | 0.00011 | 9.11E-05 | 0.00013 | 0.0002 | 0.00017 | 9.91E-05 | 9.64E-05 | 0.00011 | 0.00017 | 0.00012 | 0 | 0.00014 |
| Smap1 | 0.00021 | 0.00012 | 0.00012 | 0 | 0.00013 | 0.00016 | 0.00012 | 9.58E-05 | 0.00013 | 0.0001 | 0.0001 | 0.00014 | 0 | 6.35E-05 | 4.52E-05 | 1.39E-05 | 7.28E-05 | 4.28E-05 |
| Dock1 | 0.00015 | 0.00029 | 1.39E-05 | 0.0001 | 0.00023 | 0.00024 | 6.29E-05 | 0.00023 | 0.00013 | 8.68E-05 | 6.37E-05 | 0.00011 | 0.00022 | 0.00011 | 0.00065 | 0.00025 | 0.00041 | 0.00014 |
| Babam2 | 0.00017 | 8.37E-05 | 1.05E-06 | 0.00012 | 2.26E-05 | 1.47E-05 | 0.00017 | 3.39E-05 | 0.00013 | 2.24E-05 | 1.86E-05 | 0.00011 | 0.00017 | 0 | 3.79E-05 | 9.21E-05 | 3.67E-05 | 3.63E-05 |
| Trap1 | 0.0002 | 6.89E-05 | 0.00011 | 5.61E-05 | 3.22E-05 | 7.03E-05 | 0 | 0 | 0.00013 | 3.21E-05 | 1.64E-05 | 1.18E-05 | 1.74E-05 | 9.44E-06 | 4.96E-05 | 0.00011 | 6.67E-05 | 5.87E-05 |
| Plekhm2 | 1.75E-05 | 0.00019 | 9.65E-05 | 5.45E-05 | 5.14E-05 | 2.81E-05 | 2.78E-05 | 2.94E-05 | 0.00013 | 6.64E-05 | 5.04E-05 | 3.97E-05 | 6.62E-05 | 6.10E-05 | 0.0001 | 3.84E-05 | 1.77E-05 | 8.12E-05 |
| Leo1 | 9.92E-05 | 4.04E-05 | 0.00015 | 5.18E-05 | 5.02E-05 | 0 | 1.34E-05 | 9.59E-06 | 0.00013 | 6.79E-05 | 5.79E-05 | 5.20E-05 | 4.04E-05 | 0 | 0.0001 | 0.00014 | 6.84E-06 | 6.46E-07 |
| Taok2 | 0.00016 | 0.0002 | 0.00012 | 0.0001 | 0.00015 | 8.77E-05 | 8.76E-05 | 6.75E-05 | 0.00013 | 0.00017 | 5.64E-05 | 4.67E-05 | 0 | 5.53E-05 | 0.00043 | 0.00014 | 0 | 0.00011 |
| Syde1 | 5.35E-05 | 0.00012 | 0.0001 | 0 | 4.15E-05 | 4.33E-05 | 0 | 6.73E-05 | 0.00013 | 4.52E-05 | 0.00017 | 5.41E-05 | 0.00012 | 8.48E-05 | 5.32E-05 | 6.10E-05 | 0 | 5.86E-05 |
| Capza2 | 0.00032 | 8.51E-05 | 9.34E-05 | 0 | 5.24E-05 | 0.00017 | 1.34E-05 | 3.69E-06 | 0.00013 | 8.19E-05 | 0.00012 | 0.00017 | 0 | 1.78E-05 | 0 | 0.00014 | 0 | 0.00011 |
| Cebpzos | 5.77E-05 | 5.30E-05 | 0 | 0.00013 | 7.91E-05 | 4.49E-05 | 5.46E-05 | 0 | 0.00013 | 4.71E-05 | 4.29E-06 | 3.59E-05 | 4.59E-05 | 2.61E-05 | 0 | 5.34E-05 | 0 | 4.34E-05 |
| Drg2 | 0.00022 | 0.0001 | 0.00022 | 0.00014 | 3.67E-05 | 4.70E-05 | 2.20E-05 | 5.13E-05 | 0.00013 | 4.66E-05 | 6.45E-05 | 7.27E-05 | 1.90E-05 | 8.42E-05 | 0 | 8.21E-05 | 0 | 4.59E-05 |
| Ylpm1 | 0.00011 | 0.00013 | 0.00011 | 3.84E-05 | 8.85E-05 | 2.09E-05 | 5.14E-06 | 4.68E-05 | 0.00013 | 2.28E-05 | 3.17E-05 | 6.06E-05 | 4.73E-06 | 1.96E-05 | 0.00024 | 0.00016 | 0.00012 | 4.02E-05 |
| Rabif | 0.00014 | 7.88E-05 | 0.00013 | 0.00013 | 2.88E-05 | 5.08E-06 | 9.11E-05 | 0.00011 | 0.00013 | 6.75E-05 | 7.38E-05 | 4.53E-05 | 3.89E-05 | 5.73E-05 | 0.00012 | 0 | 0 | 5.70E-05 |
| Smarce1 | 0.00018 | 0.00017 | 0.00011 | 0.00025 | 0.00013 | 0.0001 | 0 | 8.98E-05 | 0.00013 | 7.81E-05 | 0.00015 | 3.94E-06 | 8.79E-06 | 0.00015 | 0 | 0.00019 | 0.00016 | 0.00013 |
| Azin1 | 0.00015 | 4.90E-06 | 8.12E-05 | 0 | 0 | 0.00014 | 0 | 0 | 0.00013 | 1.67E-05 | 4.75E-05 | 8.30E-05 | 0 | 0 | 0 | 0.00013 | 0.00102 | 1.14E-05 |
| Fbf1 | 2.72E-05 | 0.0001 | 1.97E-05 | 3.46E-05 | 5.17E-05 | 5.30E-05 | 0 | 0.00011 | 0.00013 | 1.99E-05 | 0 | 5.96E-06 | 0 | 7.51E-05 | 0.00029 | 0.00011 | 9.10E-05 | 5.45E-05 |
| Ube4b | 0.00028 | 0.00022 | 0.00011 | 0.00018 | 0.00016 | 0.00022 | 0.00023 | 7.34E-05 | 0.00013 | 9.99E-05 | 7.83E-05 | 0.0001 | 1.00E-04 | 0.00014 | 0.00034 | 9.95E-05 | 0.00026 | 8.24E-06 |
| Atp5g3 | 0.00033 | 5.03E-05 | 0.00023 | 9.76E-05 | 1.48E-05 | 0.0001 | 1.36E-05 | 9.20E-05 | 0.00013 | 0 | 0.00013 | 4.74E-05 | 6.36E-05 | 6.18E-05 | 0 | 0.00017 | 0 | 0 |
| Nacc2 | 0 | 5.06E-05 | 6.87E-06 | 5.90E-06 | 2.70E-05 | 0 | 4.51E-05 | 0 | 0.00013 | 7.28E-05 | 0.00015 | 1.88E-05 | 3.33E-05 | 0.00011 | 1.59E-05 | 0 | 3.20E-05 | 0.00011 |
| Swi5 | 0.00043 | 1.96E-05 | 0.00024 | 0.0005 | 0.00011 | 0.00016 | 0.00035 | 0.00019 | 0.00013 | 8.52E-05 | 0.00019 | 0.00014 | 0.00012 | 9.69E-05 | 0 | 0.00011 | 1.26E-05 | 0.0001 |
| Dhrs7 | 0.00019 | 0.00011 | 0 | 5.96E-05 | 0.00012 | 0.00013 | 0.00024 | 2.46E-05 | 0.00013 | 0.00024 | 0 | 0.00039 | 0.00016 | 5.49E-05 | 0 | 0 | 0 | 1.42E-05 |
| Fubp1 | 0.00028 | 0.00021 | 0.00036 | 0.00013 | 1.87E-05 | 0 | 0 | 1.68E-05 | 0.00013 | 4.83E-05 | 0.00016 | 0.0001 | 4.53E-05 | 5.27E-05 | 0.00058 | 0.00026 | 2.40E-05 | 0 |
| H13 | 0.00045 | 0.00014 | 0.00052 | 0.00024 | 0.00023 | 0.00031 | 0.00061 | 0.00042 | 0.00013 | 5.39E-05 | 0.00029 | 0.00032 | 0.00023 | 0.0002 | 0.00016 | 0 | 0 | 0.00013 |
| Spag9 | 0.00017 | 0.00025 | 0.00014 | 0 | 0.00011 | 0.00014 | 0.00025 | 0.00016 | 0.00013 | 0.00022 | 0.00012 | 0.00013 | 8.05E-07 | 0.00013 | 0.00029 | 5.67E-05 | 0.00049 | 0.00012 |
| Lsm12 | 0.00029 | 0.00011 | 0.00027 | 0.00021 | 7.24E-05 | 9.94E-05 | 8.16E-05 | 8.23E-05 | 0.00013 | 1.50E-05 | 0.00011 | 0.00011 | 0 | 1.88E-05 | 6.63E-05 | 0.00013 | 4.25E-05 | 1.23E-05 |

| | | | | | | | | | | | | | | | | | | |
|---|---|---|---|---|---|---|---|---|---|---|---|---|---|---|---|---|---|---|
| Tmem65 | 0.00025 | 0.00021 | 0.00014 | 0 | 3.75E-05 | 0.00028 | 5.69E-05 | 0 | 0.00013 | 0.00016 | 0.00022 | 0.00031 | 0 | 0.00013 | 0 | 0.00018 | 0.00013 | 0.00021 |
| Ecpas | 0.00025 | 0.00017 | 9.22E-05 | 0 | 5.03E-05 | 0.0001 | 1.02E-05 | 1.31E-05 | 0.00013 | 6.42E-05 | 5.71E-06 | 0.00014 | 8.35E-05 | 0 | 0.00028 | 0.00012 | 0.0001 | 2.72E-05 |
| Slc39a6 | 0.00012 | 6.53E-05 | 0.00019 | 0 | 7.04E-05 | 0.00013 | 6.46E-05 | 0 | 0.00013 | 0.00014 | 9.41E-05 | 0.00059 | 0.00018 | 6.25E-05 | 0 | 5.83E-05 | 3.22E-05 | 0 |
| Ptges | 6.28E-05 | 0.0002 | 0.00026 | 0.00011 | 0.00028 | 0.0002 | 4.61E-05 | 0 | 0.00013 | 0.0002 | 3.02E-05 | 0.0004 | 0.00029 | 0.00049 | 0 | 0 | 0.00017 | 0.00014 |
| Slc44a2 | 3.92E-05 | 0.00017 | 0 | 0.00015 | 0.00035 | 0.0002 | 1.35E-05 | 0.00031 | 0.00013 | 0.0001 | 0.00047 | 0.00029 | 0.00038 | 0.00023 | 0.00024 | 9.85E-05 | 0 | 0.00064 |
| Psmd9 | 7.63E-05 | 0.00011 | 0.00013 | 0.00017 | 9.64E-05 | 7.04E-05 | 1.53E-05 | 0.00012 | 0.00013 | 4.45E-05 | 5.05E-05 | 1.71E-05 | 7.88E-05 | 0.00012 | 0 | 0.00018 | 4.42E-06 | 4.41E-05 |
| Psenen | 0.00033 | 0.0003 | 0.00012 | 0.00039 | 0.00013 | 0.00023 | 0.00028 | 0.00016 | 0.00013 | 0.00018 | 5.10E-05 | 0.00014 | 0.00016 | 0.00017 | 0 | 0.00016 | 0.0001 | 0.00013 |
| Ranbp3 | 0.00019 | 0.00018 | 0.00021 | 2.58E-05 | 6.67E-05 | 3.47E-05 | 2.31E-05 | 6.63E-05 | 0.00013 | 4.86E-05 | 0.00011 | 6.50E-05 | 1.53E-05 | 2.91E-05 | 0.00028 | 0.00027 | 2.46E-05 | 4.81E-05 |
| Polr1d | 0.00035 | 0.0002 | 0.00027 | 0.00021 | 0.00015 | 0.00018 | 3.90E-05 | 0.00012 | 0.00013 | 6.93E-05 | 8.21E-05 | 0.00027 | 2.84E-05 | 6.76E-05 | 0.00019 | 0.00017 | 0.00014 | 3.96E-05 |
| Jmjd8 | 1.60E-05 | 4.45E-05 | 4.87E-08 | 0.0001 | 5.55E-05 | 5.36E-05 | 1.49E-06 | 9.85E-05 | 0.00013 | 0.00013 | 1.97E-05 | 4.33E-05 | 0.00013 | 7.23E-05 | 0 | 0 | 6.02E-05 | 0.00012 |
| Syne3 | 0 | 9.45E-05 | 1.06E-05 | 1.60E-05 | 1.40E-05 | 0.00014 | 0 | 0 | 0.00013 | 2.85E-05 | 2.64E-05 | 3.63E-05 | 1.07E-05 | 0.00017 | 6.93E-05 | 0.00024 | 8.93E-05 | 0.00022 |
| Uqcr11 | 0.00023 | 4.39E-05 | 0.00014 | 0.00033 | 7.09E-05 | 3.21E-05 | 7.86E-05 | 8.04E-05 | 0.00013 | 5.00E-05 | 0.00012 | 6.44E-05 | 3.25E-05 | 8.42E-05 | 7.97E-05 | 8.84E-05 | 0 | 8.15E-05 |
| Limk2 | 0 | 0.00011 | 0.00014 | 5.09E-05 | 0.0002 | 7.69E-05 | 1.19E-05 | 0.00011 | 0.00013 | 0 | 0.00036 | 2.80E-05 | 4.51E-05 | 0.00015 | 0.00011 | 3.58E-07 | 4.52E-05 | 3.76E-05 |
| Dnajb1 | 7.16E-05 | 2.46E-05 | 0 | 4.70E-06 | 1.44E-05 | 0 | 0.00011 | 7.41E-05 | 0.00013 | 4.89E-05 | 0 | 0.00029 | 0.00039 | 0 | 0 | 1.95E-05 | 0.00368 | 3.68E-05 |
| Nt5c | 0.00016 | 0.00012 | 0.00014 | 0.00021 | 4.22E-05 | 8.10E-05 | 7.27E-05 | 3.52E-05 | 0.00013 | 0.00013 | 0.00016 | 1.01E-05 | 0 | 0.00012 | 0 | 0 | 0 | 0.00016 |
| Azi2 | 0.00016 | 0.0001 | 7.56E-05 | 1.15E-05 | 6.96E-05 | 0.00013 | 1.79E-05 | 0.00013 | 0.00013 | 3.14E-05 | 5.94E-05 | 8.54E-06 | 0 | 9.82E-05 | 0.00019 | 5.06E-05 | 0 | 1.30E-05 |
| C1d | 0.00024 | 7.85E-05 | 7.47E-05 | 6.75E-05 | 6.98E-05 | 0.00014 | 7.14E-05 | 7.96E-05 | 0.00013 | 2.93E-05 | 8.17E-05 | 3.15E-05 | 4.37E-05 | 7.02E-05 | 3.90E-06 | 0.0001 | 0 | 4.69E-05 |
| Chst11 | 0 | 0 | 0.001 | 4.07E-05 | 0.00039 | 3.82E-05 | 0.00034 | 0.00063 | 0.00013 | 0 | 0.00076 | 0.00066 | 0.00074 | 0 | 0.00018 | 0 | 0 | 0 |
| Bud23 | 7.69E-05 | 7.38E-05 | 0.00019 | 0.00023 | 2.38E-05 | 0 | 0 | 2.94E-05 | 0.00013 | 2.51E-05 | 6.03E-05 | 0 | 1.89E-07 | 3.87E-05 | 1.32E-05 | 9.50E-05 | 0 | 2.56E-05 |
| Gna11 | 0.00014 | 0.00021 | 0.0003 | 0.00026 | 0.00026 | 0.00021 | 0.00016 | 0.00026 | 0.00013 | 0.00017 | 0.00019 | 5.75E-05 | 8.79E-05 | 0.00015 | 0 | 0.0003 | 0.00013 | 0.00019 |
| Ctif | 7.81E-06 | 0.00015 | 0.00011 | 0 | 9.22E-05 | 7.68E-06 | 0.00011 | 0.00022 | 0.00013 | 7.84E-05 | 0.00029 | 0.00022 | 0 | 4.67E-05 | 0.00023 | 1.98E-05 | 0.00013 | 7.92E-05 |
| Brms1l | 8.86E-05 | 9.61E-05 | 6.35E-05 | 7.01E-05 | 0.00011 | 3.22E-05 | 2.43E-05 | 0 | 0.00013 | 6.61E-05 | 5.12E-05 | 0 | 4.21E-05 | 5.81E-05 | 0.0002 | 0.00025 | 1.73E-05 | 5.93E-05 |
| Prpf40a | 0.00021 | 0.00011 | 0.00017 | 7.32E-05 | 2.86E-05 | 0.0001 | 0 | 7.57E-05 | 0.00013 | 4.73E-05 | 0.00014 | 4.95E-05 | 4.18E-05 | 4.65E-05 | 3.53E-05 | 0.00015 | 7.80E-05 | 4.41E-05 |
| Akt1s1 | 0.00011 | 0.00019 | 0.00018 | 0.00076 | 0.00011 | 5.23E-05 | 0.00019 | 0.00015 | 0.00013 | 5.52E-05 | 5.16E-05 | 8.74E-05 | 7.61E-06 | 0.00011 | 0.00034 | 0 | 0.00025 | 0.00017 |
| Rad23b | 0.00024 | 0.0002 | 0.00041 | 0.00038 | 2.28E-05 | 3.94E-05 | 0.00018 | 2.93E-05 | 0.00013 | 0.00014 | 0.0004 | 0.00023 | 0.0001 | 8.46E-05 | 4.77E-05 | 0.00019 | 0.00022 | 0 |
| Fut11 | 0.00015 | 7.33E-05 | 9.75E-06 | 0 | 6.08E-05 | 0.00016 | 0.00015 | 3.08E-05 | 0.00013 | 0.00014 | 0.00014 | 8.79E-05 | 8.05E-05 | 0.00013 | 0 | 3.05E-05 | 0 | 0.00011 |
| Armcx2 | 0.00026 | 6.93E-05 | 9.17E-05 | 0.00013 | 1.64E-05 | 3.24E-05 | 0.00012 | 7.42E-05 | 0.00013 | 3.74E-05 | 3.86E-05 | 8.66E-05 | 4.63E-06 | 0 | 1.73E-05 | 6.12E-05 | 0.00015 | 7.19E-05 |
| Trnau1ap | 9.91E-05 | 6.88E-05 | 0.00014 | 0.00013 | 4.43E-05 | 4.64E-06 | 0 | 2.33E-05 | 0.00013 | 7.04E-05 | 5.45E-05 | 5.15E-05 | 4.48E-05 | 2.76E-05 | 2.09E-05 | 1.01E-05 | 0.00017 | 1.72E-05 |
| Slc50a1 | 0.00037 | 6.25E-05 | 4.77E-05 | 8.96E-05 | 0.00011 | 0.00047 | 0.00029 | 0.00031 | 0.00013 | 5.46E-05 | 3.86E-05 | 0 | 0.00016 | 0.00017 | 0.00012 | 0 | 0 | 0.00016 |
| Uba7 | 2.96E-05 | 7.15E-05 | 0 | 8.76E-06 | 0.00014 | 0 | 0 | 9.34E-05 | 0.00013 | 0.00017 | 0 | 2.71E-05 | 1.16E-05 | 0.00014 | 0.00022 | 0 | 4.71E-06 | 0.00016 |
| Anapc13 | 0.00023 | 9.49E-05 | 5.10E-05 | 0.00034 | 0.0001 | 0.0001 | 0.00016 | 5.00E-05 | 0.00013 | 8.27E-05 | 4.70E-05 | 8.39E-05 | 0.00013 | 5.96E-05 | 0 | 0.00016 | 3.75E-05 | 0.00012 |
| Notch2 | 0 | 0.00021 | 0.00019 | 0 | 5.30E-05 | 0 | 0.00021 | 0 | 0.00013 | 0.00024 | 0.00013 | 2.12E-05 | 0.00011 | 8.50E-05 | 0.00012 | 0.0001 | 0.0001 | 0.00013 |
| Tmem106a | 1.82E-05 | 6.94E-05 | 0 | 6.09E-05 | 0 | 3.70E-06 | 0 | 0 | 0.00013 | 0.00028 | 1.77E-05 | 0 | 0.00015 | 6.83E-05 | 1.65E-05 | 1.20E-05 | 0 | 0.00014 |

| | | | | | | | | | | | | | | | | | | |
|---|---|---|---|---|---|---|---|---|---|---|---|---|---|---|---|---|---|---|
| Mdfi | 0 | 0 | 7.99E-05 | 0.0003 | 0.00011 | 0 | 0.00013 | 0.00016 | 0.00013 | 0 | 4.51E-05 | 0 | 0.00028 | 0.00018 | 0.00011 | 0 | 0 | 7.54E-05 |
| Imp3 | 5.10E-05 | 0.00019 | 0.00011 | 0.00027 | 3.43E-05 | 7.32E-05 | 0 | 0.0002 | 0.00013 | 2.22E-05 | 0 | 6.41E-05 | 6.58E-05 | 0.0003 | 1.01E-05 | 1.86E-05 | 1.45E-05 | 4.48E-05 |
| Zfhx4 | 0.00015 | 0.00012 | 0 | 7.52E-06 | 0.00022 | 8.34E-05 | 6.19E-05 | 2.90E-05 | 0.00013 | 8.86E-05 | 9.05E-05 | 0 | 0 | 0 | 5.60E-05 | 2.90E-05 | 0.00024 | 0.00014 |
| Vegfb | 0 | 0.00027 | 0 | 7.16E-05 | 7.63E-05 | 1.03E-05 | 4.57E-05 | 0 | 0.00013 | 7.29E-05 | 0 | 0.00011 | 0.0002 | 7.74E-05 | 0 | 0.00013 | 9.36E-05 | 7.59E-05 |
| Mtmr6 | 0.0002 | 0.00015 | 0.00011 | 5.60E-05 | 0.00013 | 0.00021 | 0.00016 | 0.00011 | 0.00013 | 0.00014 | 0.00017 | 0.00014 | 1.89E-05 | 7.11E-05 | 0 | 0.0001 | 4.58E-05 | 0.00014 |
| Atrnl1 | 0.00025 | 0.00034 | 0 | 0 | 0.00046 | 0.00036 | 0.00016 | 0 | 0.00013 | 0.00018 | 0 | 0.00014 | 0.00037 | 0 | 3.61E-05 | 5.06E-06 | 0.0003 | 0 |
| Ppp6r1 | 0.00011 | 0.00019 | 0.00015 | 0.00013 | 0.00015 | 0.00011 | 6.34E-05 | 9.03E-05 | 0.00013 | 0.00011 | 0.00011 | 0.00011 | 2.09E-05 | 0.00012 | 4.84E-05 | 3.97E-05 | 9.06E-06 | 8.87E-05 |
| Gltp | 7.96E-05 | 7.53E-06 | 5.38E-05 | 8.09E-05 | 0.00014 | 0.00023 | 8.43E-05 | 0.00015 | 0.00012 | 0.00015 | 7.93E-05 | 0 | 3.00E-05 | 0.00014 | 0 | 0.00017 | 0 | 4.78E-05 |
| Pcyox1 | 8.71E-06 | 0.00015 | 0 | 0.00037 | 0.00026 | 0.00019 | 0.00017 | 0.00031 | 0.00012 | 0.00021 | 0.00021 | 0.00019 | 0.00031 | 0.00019 | 5.78E-06 | 4.00E-05 | 3.91E-05 | 0.00039 |
| Ctbp2 | 0.00022 | 0.00025 | 0.00038 | 0.00014 | 3.32E-05 | 8.46E-05 | 5.17E-05 | 1.76E-06 | 0.00012 | 5.08E-05 | 0.00021 | 0.00026 | 0.00013 | 0.0001 | 0.00011 | 9.10E-05 | 0.00021 | 0 |
| Ndnf | 0 | 0 | 0 | 0 | 0 | 3.62E-05 | 0 | 0 | 0.00012 | 0 | 0 | 0 | 0.00024 | 0 | 0 | 0 | 0 | 0.00025 |
| Ndufv2 | 0.00045 | 0.00011 | 0.00022 | 0.00018 | 8.16E-05 | 0.00016 | 5.84E-05 | 3.28E-05 | 0.00012 | 4.00E-05 | 8.85E-05 | 3.37E-05 | 0 | 4.07E-05 | 0 | 0.00018 | 0 | 4.10E-05 |
| Copb1 | 0.00073 | 0.00012 | 0.00039 | 0.00018 | 9.65E-05 | 0.0003 | 0.00045 | 8.92E-05 | 0.00012 | 7.74E-05 | 0.00011 | 0.00018 | 7.67E-05 | 0.00012 | 0.00012 | 9.85E-05 | 0.00011 | 0 |
| Rtca | 0.00021 | 5.49E-05 | 0.00019 | 0.00014 | 4.57E-05 | 2.03E-05 | 3.13E-05 | 6.74E-05 | 0.00012 | 5.30E-05 | 1.32E-05 | 0 | 3.10E-05 | 8.20E-05 | 0 | 0.00012 | 3.99E-06 | 1.72E-05 |
| Cnpy2 | 0.00048 | 0.00011 | 8.85E-05 | 0.00052 | 0.00027 | 0.0004 | 0.00065 | 0.00039 | 0.00012 | 0.00015 | 0.00016 | 5.92E-05 | 0.00053 | 0.0004 | 0 | 9.70E-05 | 0 | 0.00022 |
| Zfp740 | 0.00021 | 0.0002 | 3.87E-05 | 0 | 7.84E-05 | 0.00015 | 7.90E-05 | 1.18E-05 | 0.00012 | 1.38E-05 | 2.39E-05 | 0.00011 | 0 | 4.89E-05 | 0.00023 | 8.77E-05 | 0 | 6.60E-05 |
| Lin7c | 0.00013 | 0.00015 | 0.00019 | 4.80E-05 | 7.65E-05 | 4.42E-05 | 0 | 9.43E-05 | 0.00012 | 7.41E-05 | 0.00018 | 0.00013 | 1.78E-05 | 6.49E-05 | 0.0002 | 0.00012 | 0.00013 | 7.05E-05 |
| Cct7 | 0.00025 | 0.00014 | 0.00045 | 0.00037 | 2.80E-05 | 4.48E-05 | 0 | 2.46E-05 | 0.00012 | 9.93E-06 | 0.00015 | 6.00E-05 | 0 | 9.61E-06 | 0 | 0.0002 | 0.00015 | 9.34E-06 |
| Timm17a | 0.00018 | 0.00013 | 0.00039 | 0.00023 | 2.70E-05 | 7.08E-05 | 0 | 1.62E-05 | 0.00012 | 3.59E-05 | 0.0004 | 0.00011 | 2.45E-05 | 8.32E-05 | 0 | 0.00014 | 0 | 0 |
| Tmem134 | 2.47E-05 | 0.00014 | 2.22E-05 | 0.00045 | 0.0002 | 0.00014 | 0.00014 | 0.00023 | 0.00012 | 0.0001 | 0.00017 | 0.00012 | 1.21E-05 | 0.00015 | 0.0001 | 0.00013 | 0.00014 | 0.00014 |
| Dstn | 0.00031 | 0.00032 | 0.00021 | 0.00033 | 0.00014 | 0.00022 | 0.00031 | 2.69E-05 | 0.00012 | 0.00031 | 0.00133 | 0.00021 | 0 | 0.00046 | 0 | 0.00012 | 5.24E-05 | 0.00032 |
| Znhit1 | 0.00017 | 7.92E-05 | 0.00014 | 0.00033 | 0.00013 | 4.83E-05 | 7.77E-05 | 3.12E-05 | 0.00012 | 4.85E-05 | 0.00016 | 0.00023 | 0.00012 | 8.38E-05 | 8.04E-05 | 8.81E-05 | 0 | 8.61E-05 |
| Sh3bp1 | 3.38E-05 | 7.17E-05 | 6.11E-05 | 9.15E-05 | 6.52E-05 | 0 | 0 | 3.69E-05 | 0.00012 | 4.59E-05 | 7.26E-05 | 4.44E-05 | 0 | 4.06E-05 | 0.00018 | 1.45E-05 | 5.93E-05 | 4.20E-05 |
| Tcea1 | 0.00041 | 0.00012 | 0.00018 | 5.78E-05 | 4.04E-05 | 0.00018 | 0 | 3.19E-05 | 0.00012 | 6.63E-06 | 5.55E-06 | 0.00013 | 0 | 1.51E-05 | 0 | 0.00013 | 4.06E-05 | 6.27E-05 |
| Cnep1r1 | 0.00021 | 0.00012 | 0.0001 | 7.92E-05 | 0.0001 | 0.00017 | 0.00012 | 0.00011 | 0.00012 | 0.00013 | 0.00014 | 0.0001 | 4.22E-05 | 7.70E-05 | 2.40E-05 | 0.00027 | 3.25E-05 | 7.27E-05 |
| Efnb1 | 2.94E-05 | 3.19E-05 | 0.00014 | 0.00021 | 0.00016 | 3.03E-05 | 0 | 0.00032 | 0.00012 | 1.29E-05 | 0.0005 | 0 | 0 | 0.00017 | 0.00015 | 6.97E-05 | 0 | 0.00034 |
| Wdtc1 | 8.52E-05 | 0.00016 | 3.13E-05 | 0 | 0.00013 | 8.82E-05 | 0.00011 | 3.01E-05 | 0.00012 | 7.28E-05 | 5.45E-05 | 9.37E-05 | 2.36E-05 | 4.66E-05 | 1.48E-05 | 3.69E-05 | 5.31E-05 | 4.88E-05 |
| Atp5l | 0.00019 | 0.00011 | 0.00014 | 0.00034 | 0.00014 | 0.00013 | 9.65E-05 | 0.00011 | 0.00012 | 5.26E-05 | 0.00013 | 4.48E-05 | 9.00E-05 | 9.90E-05 | 0 | 8.83E-05 | 0 | 9.03E-05 |
| L3hypdh | 0.00014 | 2.52E-05 | 0 | 0.00012 | 6.08E-05 | 8.16E-05 | 6.79E-05 | 4.36E-05 | 0.00012 | 4.67E-05 | 3.20E-05 | 1.70E-05 | 1.85E-05 | 2.26E-05 | 8.47E-05 | 1.81E-06 | 0 | 0.0002 |
| Klhdc3 | 0.00016 | 0.00015 | 0.00016 | 0.0003 | 0.00015 | 0.00011 | 0.0002 | 0.00019 | 0.00012 | 5.15E-05 | 0.00015 | 4.00E-05 | 0 | 9.57E-05 | 6.94E-05 | 0.00011 | 0 | 2.52E-05 |
| Npdc1 | 0 | 0.00018 | 0.00013 | 0.00051 | 0.00014 | 5.85E-05 | 0.00038 | 0.00037 | 0.00012 | 0.00033 | 6.83E-05 | 8.05E-05 | 0.00043 | 6.00E-05 | 0.00031 | 0 | 7.87E-05 | 0.00047 |
| Nras | 0.00025 | 0.00014 | 0.0002 | 0 | 0 | 6.51E-05 | 7.23E-05 | 0.00013 | 0.00012 | 0.00012 | 0.00015 | 0.00021 | 0 | 1.90E-05 | 0.0002 | 0.00011 | 0.00015 | 8.00E-05 |
| Psmc1 | 0.00037 | 0.00015 | 0.00027 | 0.00013 | 2.77E-05 | 8.32E-05 | 0 | 0 | 0.00012 | 4.70E-06 | 8.78E-05 | 6.70E-05 | 4.31E-06 | 3.97E-05 | 0 | 0.0002 | 0 | 2.39E-05 |

| | | | | | | | | | | | | | | | | | | |
|---|---|---|---|---|---|---|---|---|---|---|---|---|---|---|---|---|---|---|
| Ppm1b | 4.19E-05 | 0.00023 | 0.00013 | 6.44E-05 | 0.0001 | 6.44E-05 | 0 | 0.00012 | 0.00012 | 7.24E-05 | 0.0002 | 0.00019 | 2.44E-06 | 0.0001 | 0.00021 | 0.00022 | 6.43E-05 | 7.77E-05 |
| Ddx23 | 0.0002 | 0.0002 | 0.0003 | 0.00034 | 9.67E-05 | 2.37E-05 | 5.49E-05 | 5.72E-05 | 0.00012 | 8.08E-05 | 6.10E-05 | 0.00014 | 0 | 0.00011 | 0.00038 | 0.00027 | 0.00022 | 7.64E-05 |
| Apobec3 | 9.34E-05 | 6.62E-05 | 0 | 0.00013 | 8.98E-05 | 0 | 1.40E-05 | 9.57E-05 | 0.00012 | 4.83E-06 | 1.24E-05 | 0 | 9.16E-05 | 5.74E-05 | 8.60E-05 | 7.22E-05 | 0 | 0.00022 |
| Klhl26 | 9.45E-06 | 7.40E-05 | 0.00012 | 3.36E-05 | 0.00013 | 2.96E-05 | 2.71E-05 | 9.83E-05 | 0.00012 | 5.03E-05 | 5.52E-05 | 5.78E-05 | 5.12E-05 | 2.16E-05 | 8.86E-05 | 0 | 0 | 0.00017 |
| Lrch3 | 0.00013 | 0.00018 | 8.43E-05 | 0 | 6.13E-05 | 0.0001 | 2.16E-05 | 4.43E-05 | 0.00012 | 0.00015 | 1.56E-05 | 0.0001 | 4.41E-06 | 0.00011 | 0.00026 | 4.79E-05 | 0.0001 | 3.60E-05 |
| Ppt1 | 0.00016 | 0.00013 | 0 | 0 | 0.00027 | 0.00029 | 5.34E-05 | 0.00014 | 0.00012 | 0.00015 | 0.00012 | 6.64E-05 | 0.00026 | 0.00012 | 0 | 1.25E-05 | 0 | 0.00017 |
| Col27a1 | 0 | 0 | 0.00036 | 0.00017 | 0.00075 | 0.00036 | 0.00044 | 0.0007 | 0.00012 | 0 | 8.35E-05 | 0 | 0 | 0.00041 | 0.00107 | 0 | 0.00014 | 0 |
| Slc31a1 | 0.00028 | 4.43E-05 | 0.00011 | 0 | 8.55E-05 | 0.0001 | 0.00012 | 7.34E-05 | 0.00012 | 0 | 5.26E-06 | 9.00E-05 | 6.39E-05 | 5.52E-05 | 0 | 4.11E-05 | 8.91E-05 | 0 |
| Tcf20 | 0.00015 | 0.00016 | 4.22E-05 | 0 | 0.00015 | 0.00023 | 0 | 3.82E-05 | 0.00012 | 1.03E-05 | 0.00014 | 0.00012 | 0 | 4.21E-06 | 5.61E-05 | 0.0001 | 0.00023 | 0 |
| Naxd | 0.00016 | 0.00011 | 0.00011 | 5.70E-05 | 0.00015 | 0.00022 | 0.00021 | 0.00017 | 0.00012 | 0.00012 | 6.03E-05 | 5.47E-05 | 0.00021 | 0.00012 | 6.63E-05 | 9.48E-06 | 0 | 0.00016 |
| Foxp4 | 3.11E-05 | 0.00023 | 0.00022 | 0.00023 | 0.00013 | 3.53E-05 | 0.00015 | 0.00012 | 0.00012 | 0 | 0.00012 | 6.29E-05 | 0.0001 | 5.16E-06 | 0.0001 | 9.43E-05 | 0.00013 | 0.00017 |
| Spry2 | 4.53E-05 | 0.00014 | 0 | 0.00019 | 3.78E-05 | 3.05E-05 | 0 | 0.00017 | 0.00012 | 0.00045 | 0 | 4.71E-05 | 0.00034 | 0.00011 | 5.62E-06 | 0.00013 | 0.00048 | 3.55E-05 |
| Creld1 | 0 | 0.00014 | 0 | 0.00031 | 0.00014 | 0 | 0.00029 | 6.82E-05 | 0.00012 | 0.00025 | 7.14E-05 | 0.0002 | 0.00029 | 5.07E-05 | 4.12E-05 | 0 | 7.06E-05 | 0.00012 |
| Ptpn21 | 0 | 0.00012 | 7.37E-05 | 0 | 9.65E-05 | 2.97E-05 | 0 | 2.94E-05 | 0.00012 | 8.01E-05 | 0.00022 | 8.91E-05 | 3.64E-06 | 0 | 2.94E-05 | 0.00014 | 0.00012 | 6.60E-05 |
| Chm | 0.00013 | 8.78E-05 | 1.33E-05 | 0 | 7.04E-05 | 0.00011 | 0 | 3.89E-05 | 0.00012 | 1.72E-05 | 0 | 8.92E-05 | 0 | 9.39E-06 | 0.00015 | 1.44E-05 | 3.43E-05 | 2.62E-05 |
| Sod1 | 0.00027 | 0.00019 | 0.00011 | 0.00029 | 6.29E-05 | 0.00021 | 0.00011 | 4.76E-05 | 0.00012 | 0.00013 | 4.07E-05 | 0.0002 | 9.89E-05 | 0.00013 | 0 | 0.00024 | 0.00023 | 0.00019 |
| Amdhd2 | 2.28E-05 | 0.00012 | 0 | 0.00015 | 1.94E-05 | 4.15E-05 | 3.81E-05 | 4.61E-05 | 0.00012 | 5.50E-06 | 8.05E-05 | 1.89E-05 | 4.25E-05 | 0.00018 | 2.37E-05 | 0 | 0 | 6.37E-05 |
| Higd2a | 0.00012 | 0.00015 | 5.38E-05 | 0.00019 | 0.00016 | 0.00015 | 0.00011 | 9.10E-05 | 0.00012 | 0.00011 | 0.00016 | 0.00011 | 0.00014 | 7.39E-05 | 0 | 0 | 0 | 0.00022 |
| Card19 | 0 | 0.00015 | 5.15E-05 | 7.28E-05 | 5.82E-05 | 6.59E-06 | 6.57E-06 | 0 | 0.00012 | 6.92E-05 | 0.00015 | 0.0004 | 0 | 0.00011 | 1.28E-05 | 2.30E-05 | 4.79E-05 | 9.64E-05 |
| Actr1a | 0.0002 | 0.00025 | 0.00032 | 0.00044 | 0.0002 | 0.00019 | 0.00013 | 0.00017 | 0.00012 | 0.00016 | 0.00031 | 9.47E-05 | 6.35E-05 | 0.00027 | 0 | 0.00031 | 0 | 0.00025 |
| Rprd2 | 0.00016 | 0.00018 | 7.84E-05 | 9.61E-05 | 6.13E-05 | 5.89E-05 | 6.24E-05 | 1.23E-05 | 0.00012 | 2.30E-05 | 4.59E-06 | 0.00016 | 0 | 5.48E-05 | 0.00023 | 0.00011 | 0 | 5.72E-05 |
| Glb1 | 0 | 6.40E-05 | 0 | 0.00013 | 0.00011 | 0.00021 | 0.00027 | 0.00036 | 0.00012 | 0.00016 | 5.56E-05 | 0.00023 | 0.00032 | 0.00021 | 0 | 2.82E-05 | 0 | 0.00025 |
| Tmem168 | 0.00013 | 6.60E-05 | 3.59E-05 | 0 | 4.80E-05 | 8.02E-05 | 2.10E-05 | 0 | 0.00012 | 3.42E-06 | 8.11E-05 | 3.20E-05 | 1.07E-05 | 2.70E-06 | 0 | 4.05E-05 | 1.75E-05 | 8.09E-05 |
| Gtf2ird1 | 7.02E-05 | 0.00014 | 0.00013 | 0 | 9.73E-05 | 7.14E-05 | 0.00014 | 0.00017 | 0.00012 | 0 | 3.56E-05 | 0.00012 | 2.41E-05 | 0 | 0.00019 | 7.60E-05 | 6.50E-05 | 0 |
| Gjc1 | 8.24E-05 | 9.63E-05 | 4.50E-05 | 0 | 5.47E-05 | 0.00015 | 0 | 0.00013 | 0.00012 | 0 | 8.66E-05 | 1.44E-05 | 0 | 0.00019 | 0 | 0.00032 | 9.49E-05 | 9.07E-06 |
| Chic2 | 0.00016 | 0.00017 | 0.00012 | 0 | 0.00026 | 0.00012 | 1.81E-05 | 5.72E-06 | 0.00012 | 3.76E-05 | 8.54E-05 | 0.00018 | 0.00019 | 0.00011 | 4.05E-05 | 4.74E-05 | 7.65E-05 | 0 |
| Zdhhc5 | 0.00026 | 0.00022 | 0.0003 | 3.31E-05 | 0.00025 | 0.00023 | 0.00015 | 0.00019 | 0.00012 | 0.00015 | 0.00023 | 0.00024 | 0.00017 | 8.77E-05 | 0 | 0.00024 | 8.49E-06 | 9.12E-05 |
| Hspa4 | 0.0002 | 9.13E-05 | 0.00034 | 8.74E-05 | 9.37E-05 | 2.06E-05 | 1.93E-05 | 0 | 0.00012 | 4.38E-05 | 7.92E-05 | 5.41E-05 | 9.29E-05 | 0 | 0 | 0.00011 | 6.64E-05 | 4.13E-05 |
| Rai14 | 0.00027 | 0.00027 | 0.00037 | 0 | 0.00062 | 9.84E-05 | 0 | 0.00028 | 0.00012 | 0 | 0.00043 | 0.00032 | 9.28E-10 | 0.00033 | 0.00056 | 0.00013 | 0.00059 | 0.00017 |
| Prmt5 | 0.00011 | 8.22E-05 | 0.00023 | 9.79E-05 | 1.32E-05 | 0 | 0 | 2.83E-05 | 0.00012 | 2.77E-05 | 7.37E-05 | 3.82E-05 | 0 | 0 | 0 | 8.82E-05 | 0 | 9.95E-06 |
| Camk1 | 0.00016 | 0.00014 | 8.90E-05 | 6.46E-05 | 0.0001 | 0.00014 | 0.0001 | 4.82E-06 | 0.00012 | 0.00014 | 0.00011 | 5.77E-05 | 0.00011 | 0.00012 | 6.19E-05 | 2.72E-05 | 9.39E-05 | 0.00019 |
| Rsrp1 | 0.00019 | 0.00059 | 3.68E-05 | 5.67E-05 | 0.00039 | 6.71E-05 | 8.97E-05 | 5.05E-05 | 0.00012 | 6.57E-05 | 5.17E-05 | 0.00049 | 0.00017 | 0.00031 | 0.00157 | 0 | 0.00137 | 7.89E-05 |
| Vps41 | 0.00012 | 0.0001 | 0 | 1.74E-05 | 0.00012 | 9.56E-05 | 7.79E-05 | 1.77E-05 | 0.00012 | 5.98E-05 | 3.91E-07 | 5.08E-05 | 1.27E-05 | 2.92E-05 | 3.82E-05 | 0 | 3.96E-05 | 9.16E-05 |

| | | | | | | | | | | | | | | | | | | |
|---|---|---|---|---|---|---|---|---|---|---|---|---|---|---|---|---|---|---|
| Ggcx | 0.00015 | 1.04E-06 | 0.00014 | 0.00025 | 0.00018 | 0.00022 | 0.00058 | 0.00041 | 0.00012 | 0.00014 | 9.34E-05 | 7.89E-05 | 0.00029 | 0.00012 | 0.0002 | 0 | 0 | 0.00025 |
| Tgfbr1 | 0.0003 | 0.00019 | 0.00011 | 0 | 0.00017 | 0.00049 | 1.84E-05 | 0.00036 | 0.00012 | 2.81E-05 | 0.00011 | 0.00023 | 0.00036 | 9.98E-06 | 0 | 9.14E-05 | 0.00025 | 0 |
| Dnmt1 | 6.56E-05 | 5.54E-05 | 0.00022 | 4.26E-05 | 6.84E-05 | 2.84E-05 | 2.84E-05 | 0 | 0.00012 | 7.24E-05 | 3.51E-05 | 0 | 9.73E-05 | 7.91E-05 | 0.00019 | 0.00075 | 0 | 5.11E-05 |
| Wdr36 | 0.00015 | 5.32E-05 | 0.00025 | 2.44E-05 | 0 | 3.49E-05 | 0 | 0 | 0.00012 | 7.29E-05 | 6.12E-05 | 4.38E-05 | 2.75E-05 | 0 | 1.86E-05 | 7.92E-05 | 0 | 6.66E-06 |
| Prr12 | 7.86E-05 | 0.00023 | 6.35E-05 | 0.00021 | 0.00014 | 0.00013 | 5.79E-05 | 9.19E-06 | 0.00012 | 9.09E-05 | 9.99E-05 | 0.00011 | 8.27E-05 | 8.66E-05 | 0.00029 | 0.00016 | 7.54E-05 | 8.35E-05 |
| Slc35f6 | 9.56E-05 | 0.00011 | 0.00014 | 4.23E-05 | 6.34E-05 | 6.63E-05 | 6.02E-05 | 6.44E-05 | 0.00012 | 1.64E-05 | 2.09E-05 | 5.13E-05 | 4.26E-05 | 7.10E-05 | 0 | 6.71E-06 | 0 | 1.09E-05 |
| Cherp | 0.00015 | 0.00018 | 0.00017 | 0.00011 | 7.08E-05 | 3.88E-05 | 3.16E-05 | 1.80E-05 | 0.00012 | 3.44E-05 | 1.93E-05 | 6.12E-05 | 3.29E-05 | 6.13E-05 | 0.00033 | 0.00019 | 0.00014 | 3.61E-05 |
| Akap8 | 7.91E-05 | 0.00021 | 0.00016 | 0.00013 | 1.49E-05 | 9.29E-06 | 0 | 7.57E-06 | 0.00012 | 5.06E-05 | 2.92E-05 | 1.05E-05 | 0 | 2.80E-05 | 0.00041 | 0.00013 | 0.00016 | 1.94E-05 |
| Ndufs3 | 0.00029 | 6.36E-05 | 0.00014 | 0.00022 | 7.62E-05 | 0.00014 | 0.00011 | 0.00011 | 0.00012 | 3.89E-05 | 0.0001 | 7.12E-05 | 6.99E-05 | 0.00011 | 6.49E-05 | 8.94E-05 | 0 | 6.97E-05 |
| Chmp6 | 7.95E-05 | 0.0001 | 8.82E-05 | 0.00011 | 6.82E-05 | 2.76E-05 | 2.65E-05 | 2.58E-05 | 0.00012 | 4.91E-05 | 5.38E-05 | 0 | 0 | 6.61E-05 | 0.00012 | 9.39E-05 | 1.61E-05 | 6.92E-05 |
| Setdb1 | 6.75E-05 | 8.92E-05 | 8.35E-05 | 3.88E-05 | 2.34E-05 | 7.05E-06 | 2.94E-05 | 5.89E-05 | 0.00012 | 2.26E-05 | 3.00E-05 | 5.12E-05 | 0 | 5.79E-05 | 0.00014 | 0.00011 | 4.03E-05 | 4.73E-05 |
| Zzef1 | 4.27E-05 | 7.72E-05 | 6.91E-05 | 2.07E-05 | 6.23E-05 | 3.58E-05 | 2.98E-05 | 3.75E-05 | 0.00012 | 5.20E-05 | 0.00015 | 0.0001 | 6.70E-05 | 9.13E-05 | 0.00015 | 0 | 6.18E-05 | 1.80E-05 |
| Retreg2 | 0 | 0.0003 | 1.99E-05 | 0.00034 | 0.00017 | 0.00012 | 0.0001 | 5.54E-05 | 0.00012 | 0.00015 | 7.89E-05 | 0.00015 | 7.16E-05 | 0.00014 | 0.00024 | 0.00015 | 6.96E-05 | 0.00012 |
| Api5 | 0.00043 | 0.00012 | 0.00018 | 0 | 7.64E-05 | 0.00024 | 6.51E-05 | 2.77E-05 | 0.00012 | 0 | 0.00011 | 4.66E-05 | 0 | 3.66E-05 | 0.00014 | 0.00013 | 0.00014 | 1.28E-05 |
| Spred2 | 5.84E-05 | 0.00015 | 9.00E-05 | 0 | 0.00017 | 0.0001 | 3.66E-05 | 0.0001 | 0.00012 | 5.67E-05 | 6.42E-05 | 0.0003 | 4.95E-05 | 0.0001 | 1.41E-05 | 8.08E-05 | 5.90E-05 | 0.00011 |
| Taok1 | 0.00029 | 0.00021 | 0.00011 | 0 | 2.35E-05 | 0.0002 | 7.27E-06 | 9.91E-06 | 0.00012 | 9.24E-05 | 8.31E-05 | 0.00019 | 5.67E-06 | 1.94E-05 | 0.00023 | 0.00011 | 0.00031 | 7.33E-05 |
| Psmd13 | 0.00026 | 0.00014 | 0.00031 | 0.00029 | 0.00012 | 8.73E-05 | 8.33E-05 | 4.06E-05 | 0.00012 | 5.65E-05 | 0.00014 | 0 | 2.31E-05 | 0.00013 | 0 | 0.00026 | 0 | 9.50E-05 |
| Ssrp1 | 0.00017 | 9.63E-05 | 0.0002 | 4.11E-05 | 4.52E-05 | 5.60E-05 | 4.88E-06 | 4.94E-06 | 0.00012 | 3.07E-05 | 4.06E-05 | 0.00011 | 0 | 0 | 0 | 0.00053 | 1.31E-05 | 1.78E-05 |
| Camk2g | 0.00012 | 7.16E-05 | 6.44E-05 | 0 | 1.08E-06 | 0.00013 | 0.00012 | 0 | 0.00012 | 8.08E-05 | 0.00023 | 4.24E-05 | 0.00016 | 8.66E-05 | 9.88E-05 | 1.95E-05 | 4.21E-05 | 0 |
| Dync1li2 | 0.00016 | 0.00026 | 0.00022 | 0.00029 | 0.0002 | 0.00012 | 0.0003 | 0.0001 | 0.00012 | 0.00013 | 0.00025 | 0.00012 | 0.00011 | 0.00016 | 0.00039 | 9.80E-05 | 0 | 0.00023 |
| Runx1t1 | 3.33E-05 | 5.49E-05 | 0 | 2.64E-05 | 7.42E-05 | 1.25E-05 | 7.67E-06 | 0 | 0.00012 | 3.64E-05 | 7.60E-05 | 0 | 0.0001 | 8.17E-05 | 6.18E-06 | 4.60E-07 | 0 | 0.00011 |
| Tapbpl | 8.34E-05 | 4.31E-05 | 0 | 2.77E-05 | 3.87E-05 | 0 | 0 | 0 | 0.00012 | 5.24E-05 | 1.69E-05 | 0 | 9.55E-05 | 9.22E-06 | 0 | 0 | 0 | 7.81E-05 |
| Hspa9 | 0.00019 | 0.00019 | 0.00038 | 0.00011 | 0 | 5.83E-05 | 0 | 0 | 0.00012 | 0 | 2.44E-05 | 0.00038 | 5.61E-06 | 4.33E-05 | 2.02E-06 | 0.00012 | 0 | 0 |
| Cyp20a1 | 0.00031 | 1.67E-05 | 0.00015 | 3.09E-07 | 1.49E-05 | 0.00013 | 0.00016 | 0 | 0.00012 | 5.98E-05 | 3.54E-05 | 0.00017 | 7.19E-05 | 0 | 1.16E-05 | 5.96E-05 | 4.38E-05 | 8.05E-05 |
| Hoxc6 | 0.00011 | 0.00027 | 0 | 0.00025 | 0.00027 | 0.00018 | 0.00029 | 0 | 0.00012 | 3.33E-05 | 5.13E-05 | 9.40E-05 | 1.19E-05 | 0.00016 | 0.00051 | 0.0001 | 0 | 0.00019 |
| Tm2d3 | 0.0001 | 7.26E-05 | 4.17E-05 | 6.46E-05 | 6.70E-05 | 5.82E-05 | 3.74E-05 | 2.55E-05 | 0.00012 | 9.39E-05 | 3.01E-05 | 1.74E-05 | 0.0001 | 6.39E-05 | 0 | 1.33E-05 | 0 | 5.08E-05 |
| Lzts2 | 1.05E-06 | 0.00034 | 8.66E-05 | 0.00044 | 0.00027 | 7.29E-05 | 0.00014 | 1.55E-05 | 0.00012 | 0.00017 | 0.00036 | 0.00014 | 0.00015 | 0.00012 | 0.00064 | 8.53E-05 | 6.56E-05 | 4.36E-05 |
| Sympk | 0.00017 | 0.00025 | 0.00017 | 0.00017 | 9.26E-05 | 9.66E-05 | 5.05E-05 | 5.85E-05 | 0.00012 | 3.27E-05 | 7.59E-05 | 0.00018 | 0 | 5.48E-05 | 0.00023 | 0.0001 | 0 | 0 |
| Bag3 | 0.00022 | 0.0002 | 0.00033 | 0.00035 | 0.0001 | 5.75E-05 | 0.00019 | 4.55E-05 | 0.00012 | 0.00015 | 2.71E-05 | 8.26E-05 | 0.00023 | 6.72E-05 | 7.60E-05 | 8.21E-05 | 0.00119 | 0.00029 |
| Erbin | 0.00028 | 0.0003 | 0.0001 | 1.94E-05 | 0.00018 | 0.00022 | 0.00018 | 0.0001 | 0.00012 | 6.45E-05 | 0.00011 | 6.12E-05 | 0.00011 | 0.00021 | 7.06E-05 | 8.22E-05 | 0.0002 | 1.70E-05 |
| Wdr61 | 0.00017 | 3.89E-05 | 0.00012 | 0.0001 | 4.52E-05 | 6.78E-05 | 3.54E-05 | 3.44E-05 | 0.00012 | 7.35E-06 | 4.06E-05 | 4.57E-05 | 4.59E-05 | 5.19E-05 | 0 | 4.69E-05 | 0 | 0 |
| Mapk3 | 0.00016 | 0.00025 | 0 | 0.00012 | 0.00012 | 0.00028 | 2.89E-05 | 0 | 0.00012 | 0.00011 | 0.00016 | 0.0003 | 5.25E-05 | 2.81E-05 | 0 | 0.00015 | 0.00018 | 0.00021 |
| Slc7a6 | 0.00024 | 5.04E-06 | 0.00017 | 0.00014 | 0.0001 | 0.00018 | 0.00024 | 6.65E-05 | 0.00012 | 7.79E-05 | 0.00022 | 0 | 0.00024 | 0.00022 | 0.00011 | 5.11E-05 | 0 | 0 |

| | | | | | | | | | | | | | | | | | | |
|---|---|---|---|---|---|---|---|---|---|---|---|---|---|---|---|---|---|---|
| Coq2 | 5.56E-05 | 5.96E-05 | 0.00016 | 0.00011 | 9.42E-05 | 7.81E-05 | 5.39E-05 | 7.84E-05 | 0.00012 | 9.68E-05 | 9.67E-05 | 4.01E-05 | 9.70E-05 | 3.72E-05 | 5.80E-05 | 0.00019 | 0 | 9.33E-05 |
| Naa20 | 0.00031 | 0.00023 | 0.00021 | 0 | 2.73E-05 | 0.00015 | 6.84E-05 | 0 | 0.00012 | 9.90E-05 | 0.0001 | 3.15E-05 | 0.00013 | 0.00014 | 2.49E-05 | 0.00023 | 0 | 0 |
| Chka | 2.73E-05 | 3.95E-05 | 2.66E-06 | 0 | 0 | 2.32E-05 | 0 | 0 | 0.00012 | 0 | 1.99E-05 | 0.00016 | 4.66E-06 | 7.97E-06 | 2.73E-05 | 1.76E-05 | 0.00086 | 0 |
| Prxl2a | 0 | 2.65E-05 | 0 | 0 | 0.00014 | 0.00013 | 0 | 0 | 0.00012 | 1.43E-05 | 5.74E-05 | 0 | 2.18E-05 | 8.53E-05 | 0 | 0 | 0 | 0 |
| Gadd45gip1 | 0.00021 | 9.79E-05 | 0.00024 | 0.00027 | 2.33E-05 | 0 | 2.55E-05 | 3.17E-05 | 0.00012 | 9.79E-06 | 5.53E-05 | 0.00013 | 0 | 5.86E-05 | 7.44E-05 | 0.00014 | 0 | 4.57E-05 |
| Otud7b | 0 | 0.00016 | 0.00012 | 0.00016 | 0.00014 | 5.66E-05 | 0.00011 | 0.00012 | 0.00012 | 0.00015 | 0.00016 | 6.74E-05 | 0 | 0.00013 | 9.17E-05 | 5.64E-05 | 2.87E-05 | 0.00017 |
| Vps35l | 0.00011 | 0.00018 | 9.08E-05 | 0.00015 | 0.00015 | 7.82E-05 | 0.00012 | 5.48E-05 | 0.00012 | 0.00012 | 6.80E-05 | 9.29E-05 | 2.04E-05 | 8.83E-05 | 0 | 5.34E-05 | 8.52E-05 | 0.00012 |
| Phf2 | 3.71E-05 | 0.00023 | 0.00016 | 4.63E-05 | 0.00014 | 0.0001 | 1.71E-05 | 4.63E-05 | 0.00012 | 6.88E-05 | 9.35E-05 | 0.0001 | 4.86E-05 | 0.00013 | 0.00017 | 0.00016 | 0.00021 | 7.49E-05 |
| Borcs6 | 0 | 8.89E-05 | 5.66E-05 | 0.00015 | 5.96E-05 | 2.31E-05 | 2.44E-05 | 0.00011 | 0.00012 | 4.93E-05 | 2.56E-05 | 0.00012 | 0 | 4.41E-05 | 7.08E-05 | 0 | 5.65E-05 | 6.65E-05 |
| Fam53a | 3.27E-05 | 7.90E-05 | 3.78E-05 | 8.19E-05 | 3.04E-05 | 0 | 0 | 0 | 0.00012 | 5.23E-05 | 0.00012 | 0 | 2.66E-05 | 3.51E-05 | 6.77E-05 | 0 | 0.0001 | 2.56E-05 |
| Slc25a11 | 0.00013 | 9.42E-05 | 6.55E-05 | 0.0002 | 5.28E-05 | 5.44E-05 | 0.00014 | 0.00014 | 0.00012 | 4.95E-05 | 7.93E-05 | 8.66E-05 | 4.32E-06 | 2.27E-05 | 0 | 0.00012 | 0.0001 | 0.00013 |
| Dvl2 | 3.76E-05 | 0.00017 | 0 | 0.00015 | 7.21E-05 | 3.15E-05 | 4.06E-05 | 0 | 0.00012 | 1.63E-05 | 0 | 6.20E-05 | 2.19E-05 | 7.81E-05 | 0.00014 | 0.00011 | 0.00019 | 3.31E-05 |
| Ndufs8 | 0.00031 | 5.61E-05 | 0.00016 | 0.00023 | 0.00011 | 0.0001 | 0.00012 | 0.00011 | 0.00012 | 0 | 0.00016 | 0 | 0 | 8.53E-05 | 0 | 0.00011 | 0 | 9.78E-05 |
| Vangl1 | 5.69E-05 | 0.00014 | 2.30E-05 | 5.18E-06 | 8.77E-05 | 8.46E-05 | 7.17E-05 | 0.00013 | 0.00012 | 9.80E-05 | 2.19E-06 | 6.38E-05 | 0 | 4.44E-05 | 4.34E-05 | 0.00016 | 0 | 7.03E-05 |
| Ddx5 | 0.00058 | 0.00028 | 0.00012 | 0 | 8.47E-05 | 0.00039 | 6.48E-05 | 2.68E-05 | 0.00012 | 3.93E-05 | 3.71E-05 | 0.0003 | 0 | 0 | 0 | 0.00014 | 0.0013 | 0 |
| Stk40 | 0 | 0.0002 | 0.00035 | 4.59E-05 | 9.06E-05 | 0.00019 | 0 | 0.00028 | 0.00012 | 9.63E-05 | 0.00012 | 0.00018 | 0 | 0.00011 | 9.52E-05 | 0 | 0.00023 | 8.94E-05 |
| Slirp | 0.0003 | 2.96E-05 | 0.00014 | 0.00012 | 1.42E-05 | 8.05E-05 | 3.87E-05 | 1.31E-05 | 0.00012 | 4.16E-05 | 6.25E-05 | 4.72E-05 | 0 | 1.37E-05 | 0 | 0.00011 | 3.71E-05 | 1.36E-05 |
| Dcp2 | 0.0001 | 8.45E-05 | 7.67E-05 | 2.55E-06 | 8.52E-05 | 4.39E-05 | 0 | 6.71E-05 | 0.00012 | 6.21E-05 | 4.71E-05 | 2.73E-05 | 4.82E-05 | 7.83E-05 | 2.96E-05 | 0.00023 | 0 | 4.41E-05 |
| Slc3a2 | 0.00013 | 0.00039 | 0.00045 | 0.0003 | 0.00018 | 0.0002 | 7.53E-05 | 0.0002 | 0.00012 | 1.83E-05 | 7.82E-05 | 0.00037 | 0 | 0.00017 | 0.00027 | 0.00024 | 0.00058 | 0.0002 |
| Tmed4 | 2.04E-06 | 0.00021 | 0 | 0.00013 | 0.00024 | 5.87E-05 | 4.98E-05 | 0.00018 | 0.00012 | 0.00018 | 0.00016 | 0.00023 | 0.00018 | 6.15E-05 | 0 | 1.88E-05 | 0 | 0.00012 |
| Slc25a22 | 3.69E-05 | 4.49E-05 | 0.00019 | 0.00016 | 3.48E-05 | 0 | 1.88E-05 | 6.72E-05 | 0.00012 | 7.70E-05 | 8.10E-05 | 3.29E-05 | 7.23E-05 | 4.15E-05 | 8.43E-05 | 0 | 6.29E-05 | 4.28E-05 |
| Cep170 | 0.0002 | 0.00022 | 0.0002 | 3.92E-05 | 3.19E-05 | 0 | 6.46E-05 | 0.00023 | 0.00012 | 0.0003 | 4.16E-05 | 0.00048 | 0.00017 | 2.42E-05 | 0.00046 | 0.00017 | 5.19E-05 | 0.00012 |
| Exoc6b | 0.00013 | 0.0002 | 2.14E-05 | 0 | 7.48E-05 | 4.99E-05 | 2.04E-05 | 0 | 0.00012 | 4.88E-05 | 3.48E-05 | 0.00015 | 6.00E-05 | 3.78E-05 | 0.00022 | 0 | 0.00017 | 8.05E-05 |
| Aars | 0.0003 | 0.00027 | 0.00043 | 0.00015 | 0.00012 | 8.80E-05 | 0.00013 | 8.44E-05 | 0.00012 | 6.15E-05 | 0.00011 | 0.00018 | 0 | 7.33E-05 | 0.00011 | 0.00025 | 0 | 3.35E-05 |
| Wdr77 | 0.00018 | 0.00013 | 0.00038 | 5.63E-05 | 2.69E-05 | 0 | 1.65E-05 | 0 | 0.00012 | 2.64E-05 | 1.83E-05 | 0 | 0.0001 | 1.18E-05 | 0.00015 | 9.15E-05 | 0 | 2.28E-05 |
| Ubl7 | 0.0002 | 0.00021 | 0.0001 | 0.0001 | 0.00015 | 0.00013 | 0.00013 | 4.91E-05 | 0.00012 | 8.39E-05 | 8.64E-05 | 4.87E-05 | 4.04E-05 | 0.00021 | 0 | 5.08E-05 | 0 | 6.36E-05 |
| Ints7 | 5.83E-05 | 6.42E-05 | 8.14E-05 | 9.45E-05 | 3.23E-05 | 1.06E-05 | 1.38E-05 | 3.99E-05 | 0.00012 | 7.17E-05 | 6.24E-07 | 4.83E-06 | 2.60E-05 | 4.86E-05 | 5.71E-05 | 0.00014 | 0 | 8.12E-06 |
| Ptbp2 | 0.00015 | 0.00017 | 3.56E-05 | 0.00012 | 0.0001 | 8.96E-05 | 5.10E-05 | 4.20E-05 | 0.00012 | 1.38E-06 | 0.00018 | 6.47E-05 | 0 | 6.72E-05 | 0.00072 | 0.00011 | 6.58E-05 | 0 |
| Cog1 | 0.00011 | 0.0001 | 0.00015 | 2.90E-05 | 9.30E-05 | 5.80E-05 | 7.93E-05 | 9.73E-05 | 0.00012 | 5.32E-05 | 0.0001 | 3.83E-06 | 0 | 4.81E-05 | 7.97E-05 | 4.73E-05 | 5.03E-05 | 8.55E-05 |
| Yipf1 | 0.00015 | 0.00011 | 7.04E-05 | 0.00014 | 0.00012 | 9.93E-05 | 0.00016 | 8.34E-05 | 0.00012 | 7.25E-05 | 4.23E-05 | 1.16E-06 | 0.00012 | 0.00014 | 0 | 5.88E-05 | 0 | 0.00013 |
| Btbd2 | 6.48E-05 | 0.00012 | 0 | 0.00014 | 0.00011 | 4.35E-05 | 1.56E-05 | 7.77E-05 | 0.00012 | 8.41E-05 | 2.01E-05 | 0.00012 | 0.0001 | 5.37E-05 | 0 | 3.77E-05 | 0.00031 | 5.79E-05 |
| Syngr2 | 1.32E-05 | 8.54E-05 | 8.89E-05 | 0.00016 | 4.58E-05 | 4.02E-05 | 5.23E-05 | 8.15E-05 | 0.00012 | 0.00012 | 0.00013 | 2.48E-05 | 0.00015 | 6.14E-05 | 0.00011 | 0 | 0 | 0.00013 |
| Hmbs | 0.00019 | 6.19E-05 | 9.02E-05 | 0.00011 | 3.52E-05 | 8.92E-07 | 3.93E-05 | 1.34E-05 | 0.00012 | 2.93E-05 | 3.78E-05 | 0 | 1.95E-05 | 3.14E-05 | 0 | 0.00014 | 0 | 1.63E-05 |

| | | | | | | | | | | | | | | | | | | |
|---|---|---|---|---|---|---|---|---|---|---|---|---|---|---|---|---|---|---|
| Snx19 | 9.36E-05 | 0.00012 | 0.00011 | 4.56E-05 | 0.00012 | 8.07E-05 | 5.56E-05 | 0.0001 | 0.00012 | 9.58E-05 | 7.57E-05 | 0 | 9.06E-05 | 5.09E-05 | 0.00011 | 7.94E-05 | 0 | 9.48E-05 |
| Sdf2 | 0.00019 | 0.00025 | 0.00016 | 0.00026 | 0.00021 | 0.00031 | 0.00021 | 0.00029 | 0.00012 | 0.00013 | 0.00013 | 0.00031 | 0.00013 | 0.00022 | 2.05E-05 | 5.13E-05 | 0 | 0.00024 |
| Selenot | 0.00052 | 5.32E-05 | 9.09E-05 | 0 | 1.99E-05 | 0.00024 | 0 | 4.35E-05 | 0.00012 | 0 | 4.78E-05 | 8.20E-05 | 0 | 0 | 0 | 0.0001 | 4.32E-05 | 0 |
| Stox2 | 4.76E-05 | 5.69E-05 | 3.27E-05 | 0 | 0.00013 | 7.69E-05 | 7.00E-06 | 5.51E-05 | 0.00012 | 0 | 0.00012 | 0 | 0 | 3.68E-05 | 0 | 0 | 6.13E-05 | 2.30E-05 |
| Anxa4 | 0.0002 | 0.0002 | 8.98E-05 | 0.00017 | 0 | 5.81E-05 | 6.99E-05 | 2.60E-05 | 0.00012 | 0.00018 | 0.00015 | 0.00035 | 1.00E-04 | 3.91E-05 | 4.05E-05 | 4.29E-05 | 9.62E-05 | 0.00044 |
| Pum2 | 0.00036 | 0.00035 | 0.00023 | 0 | 0.00019 | 0.00034 | 0.0001 | 0.00016 | 0.00012 | 8.61E-05 | 0.00018 | 0.00031 | 2.33E-05 | 8.42E-05 | 1.46E-05 | 0.00037 | 0.00045 | 9.29E-05 |
| Fyn | 0.00013 | 0.00018 | 0.00012 | 0 | 0 | 3.29E-05 | 6.73E-05 | 5.30E-05 | 0.00012 | 0.00048 | 0.00011 | 0.00014 | 0 | 0 | 0 | 0 | 8.30E-05 | 4.96E-05 |
| Tmx2 | 8.04E-05 | 7.77E-05 | 8.30E-05 | 6.13E-05 | 5.46E-05 | 2.70E-05 | 5.10E-06 | 4.63E-05 | 0.00012 | 9.35E-05 | 4.66E-05 | 7.82E-05 | 1.75E-05 | 0 | 0 | 0.0003 | 0 | 6.31E-05 |
| Bptf | 6.91E-05 | 0.00025 | 9.76E-05 | 0.00017 | 8.68E-05 | 3.57E-05 | 0.0001 | 7.58E-05 | 0.00012 | 6.10E-05 | 0.0001 | 5.08E-05 | 8.55E-06 | 0.00015 | 0.00034 | 9.14E-05 | 0.00032 | 5.41E-05 |
| Pxn | 0 | 8.26E-05 | 8.45E-05 | 9.49E-05 | 5.58E-05 | 0 | 1.11E-05 | 6.04E-05 | 0.00012 | 0.00032 | 4.61E-05 | 5.49E-05 | 8.85E-06 | 0 | 7.65E-05 | 4.97E-05 | 2.98E-06 | 9.33E-05 |
| Inppl1 | 0.00012 | 0.00027 | 0.00023 | 0.00025 | 0.00015 | 0.00015 | 5.28E-05 | 0.00015 | 0.00012 | 5.70E-05 | 0.0004 | 0.00018 | 6.28E-05 | 0.00012 | 0.00028 | 0.00022 | 0.00017 | 5.38E-05 |
| Tcirg1 | 1.92E-05 | 0.00015 | 0.00012 | 0.0002 | 6.93E-05 | 0 | 7.08E-05 | 0.00025 | 0.00012 | 0.00012 | 4.24E-05 | 0 | 0.00012 | 7.76E-05 | 0.00039 | 0.00024 | 0.00011 | 9.29E-05 |
| Def8 | 0.00012 | 5.68E-05 | 5.13E-05 | 3.70E-05 | 7.22E-05 | 6.47E-05 | 4.80E-05 | 2.83E-05 | 0.00012 | 3.92E-05 | 2.74E-05 | 0 | 2.18E-05 | 5.06E-05 | 5.65E-05 | 5.54E-05 | 0 | 4.11E-05 |
| Zfp810 | 5.89E-05 | 9.88E-06 | 0 | 0 | 9.80E-05 | 9.00E-05 | 4.39E-05 | 3.51E-05 | 0.00012 | 0 | 0 | 0 | 5.16E-06 | 0.00011 | 0.00011 | 0 | 0 | 0 |
| Magt1 | 0.00027 | 0.00011 | 0.00029 | 0.00038 | 0 | 0 | 0.00013 | 0.00021 | 0.00012 | 0.00015 | 3.50E-05 | 0.00028 | 9.19E-05 | 6.84E-05 | 5.31E-05 | 0.00023 | 0 | 0.00011 |
| Kat2a | 8.60E-05 | 8.75E-05 | 0.00015 | 8.73E-05 | 2.23E-05 | 7.11E-06 | 0 | 5.93E-05 | 0.00012 | 2.96E-05 | 0 | 0 | 0 | 3.01E-05 | 0.0001 | 7.69E-05 | 1.78E-05 | 2.85E-05 |
| Akr7a5 | 2.58E-05 | 0.00011 | 3.09E-05 | 0.0002 | 0.0001 | 1.80E-05 | 5.21E-05 | 0.00011 | 0.00012 | 6.57E-05 | 1.65E-05 | 6.91E-05 | 0.0001 | 3.09E-05 | 0 | 1.02E-05 | 6.28E-05 | 9.99E-05 |
| Zfp91 | 0.00026 | 0.00026 | 0.00018 | 0.00011 | 0.00016 | 0.00021 | 8.24E-05 | 0.00013 | 0.00012 | 4.63E-05 | 9.14E-05 | 0.00013 | 0 | 0.00014 | 6.72E-05 | 0.00033 | 8.49E-05 | 3.48E-05 |
| Wtap | 0.00022 | 0.00013 | 0.00011 | 0 | 3.07E-05 | 0.00011 | 1.09E-05 | 0 | 0.00012 | 4.63E-05 | 7.80E-05 | 0 | 8.33E-05 | 3.10E-05 | 0.00014 | 0.00011 | 6.71E-05 | 0 |
| Pdcl3 | 0.00019 | 0.00017 | 0.00015 | 0.00018 | 8.09E-05 | 0.00011 | 3.57E-05 | 3.50E-05 | 0.00012 | 7.16E-05 | 0.00034 | 0.00015 | 0 | 7.16E-05 | 0.00013 | 0.00011 | 1.47E-05 | 9.45E-05 |
| Gatd1 | 0.00016 | 0.00021 | 2.99E-06 | 0.00017 | 0.00018 | 0.00017 | 0.00023 | 0.00014 | 0.00012 | 5.32E-05 | 2.97E-05 | 0.00015 | 3.11E-05 | 6.72E-05 | 0 | 1.51E-05 | 0 | 5.04E-05 |
| Psmd12 | 0.00034 | 0.00013 | 0.00031 | 0.0001 | 2.57E-05 | 3.97E-05 | 2.85E-05 | 3.20E-06 | 0.00012 | 2.81E-05 | 8.46E-05 | 5.47E-05 | 0 | 6.15E-05 | 7.33E-05 | 0.00022 | 0 | 6.51E-05 |
| Lamtor3 | 0.00018 | 0.00024 | 0.00028 | 0.00017 | 0.00024 | 0.00029 | 0.00012 | 0.00016 | 0.00012 | 0.00012 | 9.30E-05 | 0.00019 | 0.00011 | 0.00013 | 0 | 0.00014 | 0.00011 | 7.89E-05 |
| Ist1 | 0.00018 | 0.00033 | 0.00022 | 0.0002 | 0.00016 | 0.00011 | 0.00012 | 7.50E-05 | 0.00012 | 0.00011 | 0.00012 | 0.00018 | 0 | 0.00017 | 0.00038 | 0.00014 | 0.00029 | 7.01E-05 |
| Tbc1d20 | 0.00015 | 0.00022 | 0.00022 | 0.00028 | 0.00024 | 0.00019 | 0.00033 | 0.00014 | 0.00012 | 0.00017 | 0.00016 | 0.00015 | 0.0002 | 0.00018 | 4.20E-07 | 2.69E-06 | 0 | 7.84E-05 |
| Utp20 | 8.84E-05 | 6.23E-05 | 0.00018 | 6.63E-06 | 1.20E-05 | 2.93E-06 | 0 | 0 | 0.00012 | 1.27E-05 | 3.65E-05 | 1.41E-06 | 0 | 2.76E-05 | 3.35E-05 | 4.32E-05 | 4.96E-05 | 0 |
| Sec23a | 0.00047 | 4.14E-05 | 0.00031 | 0.00021 | 6.78E-05 | 0.00015 | 0.00057 | 0.00017 | 0.00012 | 0.00012 | 0.00019 | 0.00015 | 0.00014 | 6.49E-05 | 0.00024 | 3.38E-05 | 0.00032 | 4.07E-05 |
| Auh | 4.49E-05 | 0.00019 | 0 | 0.00016 | 6.34E-05 | 0.0002 | 0.00016 | 0 | 0.00012 | 0.00011 | 6.14E-05 | 8.15E-05 | 0.00029 | 0.00013 | 0.00011 | 7.41E-06 | 7.80E-05 | 0 |
| Spryd3 | 4.76E-05 | 7.20E-05 | 0.00021 | 0.00039 | 8.84E-05 | 3.91E-06 | 0.00027 | 0 | 0.00012 | 0.00012 | 0.00016 | 0.00012 | 0 | 0.00016 | 0.00033 | 7.80E-05 | 0 | 0.00011 |
| Dctn3 | 0.0001 | 7.87E-05 | 0.00013 | 0.00018 | 6.72E-05 | 4.41E-05 | 9.00E-05 | 9.16E-05 | 0.00012 | 9.54E-05 | 0.00013 | 0.00014 | 7.01E-05 | 3.84E-05 | 5.85E-05 | 5.31E-05 | 0 | 0.00012 |
| Gdi2 | 0.00069 | 0.00013 | 0.00024 | 2.15E-06 | 7.27E-05 | 0.00031 | 0.00033 | 0.00011 | 0.00012 | 8.70E-05 | 6.03E-05 | 0.00022 | 1.84E-05 | 1.87E-05 | 0 | 0.00012 | 3.46E-05 | 3.92E-05 |
| Dtymk | 0.00021 | 6.48E-05 | 0.00025 | 0.00017 | 6.64E-05 | 7.26E-05 | 6.19E-06 | 5.84E-05 | 0.00012 | 4.80E-05 | 2.63E-05 | 0 | 3.10E-05 | 7.20E-05 | 9.33E-06 | 0.00071 | 0 | 8.68E-05 |
| Dhx15 | 0.00039 | 0.00021 | 0.00039 | 4.40E-05 | 3.64E-05 | 0.00017 | 0 | 6.80E-05 | 0.00012 | 3.51E-05 | 0.00011 | 0.0001 | 2.95E-05 | 9.10E-05 | 0.00037 | 0.00043 | 0.0003 | 1.20E-05 |

| | | | | | | | | | | | | | | | | | | |
|---|---|---|---|---|---|---|---|---|---|---|---|---|---|---|---|---|---|---|
| Pja2 | 0.00012 | 0.00022 | 0.00012 | 0.00028 | 0.00013 | 0.00013 | 9.84E-05 | 0.0001 | 0.00012 | 0.00015 | 0.00019 | 0.00018 | 2.82E-05 | 4.70E-05 | 0.00023 | 8.56E-05 | 7.20E-05 | 0.00014 |
| Plpbp | 0.00012 | 9.03E-05 | 0.00015 | 9.68E-05 | 1.63E-05 | 2.40E-05 | 9.77E-05 | 0 | 0.00012 | 0.00012 | 0 | 7.25E-05 | 0.00017 | 5.46E-06 | 0.0002 | 4.38E-05 | 0.00025 | 0 |
| Rhob | 0 | 0.00023 | 0.00025 | 0.00019 | 0.00012 | 7.94E-06 | 0.00022 | 0.00054 | 0.00012 | 0.00039 | 0.00025 | 0.00047 | 0 | 7.71E-05 | 0.00027 | 0 | 0.00339 | 0.00046 |
| Gsk3b | 0.00034 | 0.0003 | 0.00011 | 0 | 0.0001 | 0.00027 | 6.90E-05 | 0.00016 | 0.00012 | 8.07E-05 | 0.00018 | 0.00034 | 0 | 0.0001 | 8.92E-05 | 0.00019 | 0.0004 | 0.00015 |
| Prkd3 | 0.0002 | 0.00015 | 5.22E-05 | 0 | 8.18E-05 | 0.00013 | 8.91E-05 | 5.41E-05 | 0.00012 | 6.10E-05 | 3.88E-05 | 9.69E-05 | 7.80E-07 | 6.68E-05 | 0.0001 | 0.00014 | 9.98E-05 | 4.90E-05 |
| Smim20 | 0.00016 | 7.35E-05 | 1.27E-05 | 0.00018 | 0.0002 | 0.00014 | 0.00015 | 0.00011 | 0.00012 | 0.0002 | 9.87E-05 | 0 | 4.15E-05 | 0.0001 | 0 | 7.27E-05 | 0 | 5.87E-05 |
| Cox6c | 0.00014 | 1.55E-05 | 5.92E-05 | 0.00044 | 5.03E-05 | 0.00012 | 6.66E-05 | 0.00016 | 0.00012 | 4.23E-05 | 0.00015 | 1.70E-05 | 8.94E-05 | 0.00013 | 0 | 0.00015 | 0.00012 | 0.00013 |
| Pcdhgc3 | 0.00015 | 0.00039 | 5.75E-05 | 0.00042 | 0.00017 | 0.00017 | 0.00011 | 0.00024 | 0.00012 | 0.00037 | 3.85E-05 | 0 | 0.00063 | 0.00016 | 0 | 0.00025 | 0.00021 | 0.00044 |
| AI837181 | 0.00012 | 0.00019 | 7.99E-05 | 0.0002 | 9.55E-05 | 6.75E-05 | 0.00011 | 5.21E-05 | 0.00012 | 9.41E-05 | 0.00011 | 0.00017 | 8.69E-05 | 0.00011 | 0.00011 | 1.80E-05 | 0 | 6.28E-05 |
| Ank2 | 5.42E-05 | 0.00014 | 3.94E-06 | 2.49E-05 | 0 | 0.00016 | 0 | 0.00023 | 0.00012 | 6.67E-05 | 0 | 6.68E-05 | 8.41E-06 | 0.00011 | 2.38E-05 | 2.79E-05 | 0.00012 | 0.00026 |
| Cyp39a1 | 2.93E-05 | 3.11E-05 | 0 | 0 | 0 | 0.00012 | 0 | 0.00013 | 0.00012 | 5.53E-05 | 0 | 0 | 0 | 3.35E-05 | 0 | 0.00019 | 0 | 0 |
| Snx4 | 5.56E-05 | 9.27E-05 | 0.00011 | 0.00023 | 3.84E-05 | 0 | 2.89E-05 | 9.25E-05 | 0.00012 | 9.39E-05 | 0.0001 | 6.10E-05 | 0 | 7.78E-05 | 0.00015 | 1.70E-05 | 0 | 7.83E-05 |
| Slc25a39 | 0.00024 | 0.00019 | 0.00028 | 0.00063 | 0.00025 | 0.00029 | 0.00034 | 0.00025 | 0.00012 | 0.00023 | 0.00014 | 0.0001 | 0.00027 | 0.00016 | 0 | 0.00018 | 2.03E-05 | 0.00012 |
| Pxmp2 | 0 | 0 | 0 | 0 | 3.19E-05 | 0 | 1.28E-05 | 0 | 0.00012 | 1.36E-06 | 5.81E-05 | 1.21E-05 | 0 | 0 | 0 | 9.80E-05 | 0 | 0 |
| Ino80e | 8.41E-05 | 0.00011 | 0.00022 | 0.00022 | 4.71E-05 | 0 | 0 | 3.57E-05 | 0.00012 | 4.81E-05 | 6.84E-05 | 3.94E-05 | 0 | 5.58E-05 | 0.00011 | 0.00021 | 0 | 5.41E-05 |
| Fam110a | 0 | 9.57E-05 | 0 | 0.00013 | 8.60E-05 | 0 | 0.00026 | 0 | 0.00012 | 0 | 1.27E-05 | 0 | 0.00018 | 2.63E-06 | 4.24E-05 | 0.00024 | 0 | 0 |
| Srgap2 | 0.00014 | 0.00013 | 0 | 0 | 1.42E-05 | 7.55E-05 | 1.39E-05 | 3.30E-05 | 0.00012 | 9.14E-05 | 0 | 3.45E-05 | 0 | 0 | 0.00029 | 8.57E-05 | 0.0002 | 5.42E-05 |
| Magee1 | 5.13E-05 | 6.90E-05 | 6.01E-05 | 2.73E-05 | 7.25E-05 | 3.53E-05 | 4.97E-05 | 6.62E-05 | 0.00012 | 9.97E-05 | 4.60E-05 | 5.71E-05 | 1.58E-05 | 2.71E-05 | 0 | 2.99E-05 | 0 | 8.71E-05 |
| Ubxn1 | 8.33E-05 | 0.00027 | 0.0002 | 0.00053 | 0.00016 | 6.93E-05 | 2.43E-05 | 7.48E-05 | 0.00012 | 0.00017 | 0.00013 | 0.00024 | 7.20E-05 | 0.00016 | 0.00023 | 0.00013 | 0.00017 | 0.00012 |
| Mgat4b | 0.00012 | 0.00016 | 8.27E-05 | 5.70E-05 | 6.27E-05 | 0.00012 | 0 | 4.81E-05 | 0.00012 | 4.52E-05 | 6.89E-05 | 0.00013 | 0 | 8.51E-06 | 0 | 7.86E-05 | 6.28E-06 | 0.00017 |
| Nfic | 0.00026 | 0.00025 | 0.00032 | 0.00034 | 0.00029 | 8.17E-05 | 0.00043 | 8.84E-06 | 0.00012 | 0.00025 | 0.00012 | 1.91E-05 | 0.00013 | 8.59E-05 | 0.00022 | 0.00036 | 7.84E-05 | 0.00032 |
| Rcl1 | 8.95E-05 | 5.48E-05 | 0.00022 | 0 | 0 | 0 | 0 | 0 | 0.00012 | 0 | 0 | 0 | 0.00018 | 5.55E-05 | 0 | 1.28E-05 | 9.18E-07 | 0 |
| Cct2 | 0.00012 | 0.00016 | 0.00038 | 0.00069 | 1.18E-05 | 3.23E-05 | 0 | 0.00016 | 0.00012 | 3.77E-05 | 0.00014 | 7.67E-05 | 0 | 0.00013 | 3.11E-05 | 0.00018 | 0.00024 | 6.89E-05 |
| Exoc3 | 0.00017 | 7.90E-05 | 8.00E-05 | 3.94E-05 | 3.37E-05 | 5.16E-05 | 1.63E-05 | 2.43E-05 | 0.00012 | 7.22E-05 | 4.50E-05 | 3.58E-06 | 0 | 0 | 3.99E-05 | 6.72E-05 | 0 | 8.46E-05 |
| Zmat2 | 0.00019 | 0.00014 | 7.58E-05 | 0.00022 | 6.48E-05 | 4.49E-05 | 0 | 3.60E-05 | 0.00012 | 9.54E-05 | 9.16E-05 | 9.40E-05 | 0.0001 | 3.85E-05 | 0.00012 | 0.00015 | 0 | 0.00012 |
| Ndufa4 | 0.0004 | 0 | 9.37E-05 | 0.00012 | 0 | 1.48E-05 | 8.95E-05 | 0 | 0.00012 | 8.52E-06 | 8.04E-05 | 0 | 0.0001 | 8.33E-05 | 0 | 0.00012 | 0 | 0.0001 |
| Pgp | 7.65E-05 | 1.20E-05 | 0.0002 | 0.00011 | 0.00011 | 2.34E-05 | 0 | 8.68E-05 | 0.00012 | 5.55E-05 | 0.00012 | 0.00044 | 0 | 4.90E-05 | 0 | 0.00026 | 0 | 7.79E-05 |
| Zbtb20 | 0.00012 | 0.00019 | 0 | 0 | 0.0001 | 2.57E-05 | 2.08E-05 | 0 | 0.00012 | 0.00032 | 2.08E-05 | 0 | 9.80E-05 | 0 | 0 | 2.38E-05 | 0.00016 | 0.00026 |
| Edc3 | 8.50E-05 | 8.62E-05 | 0.00012 | 4.51E-05 | 5.21E-05 | 2.52E-05 | 0.0001 | 8.65E-05 | 0.00012 | 2.97E-05 | 8.31E-05 | 8.81E-05 | 5.30E-06 | 8.21E-05 | 8.43E-05 | 2.90E-05 | 0 | 3.63E-05 |
| Evl | 1.82E-05 | 0.0001 | 0.0001 | 0.00023 | 0.00017 | 0.00021 | 3.17E-05 | 0.00013 | 0.00012 | 0 | 0.00029 | 0.00035 | 9.28E-05 | 0.00026 | 0.00017 | 0.00044 | 0.00018 | 0.00013 |
| Cnot6 | 0.00028 | 0.0001 | 7.61E-05 | 0 | 6.57E-05 | 0.0001 | 9.62E-05 | 9.88E-05 | 0.00012 | 7.35E-05 | 9.73E-05 | 9.83E-05 | 3.45E-05 | 6.19E-05 | 0 | 0.00031 | 7.74E-05 | 6.42E-05 |
| Larp7 | 0.00013 | 4.88E-05 | 0.00012 | 7.20E-05 | 0 | 1.20E-05 | 9.20E-06 | 2.54E-05 | 0.00012 | 5.80E-06 | 1.82E-05 | 1.16E-05 | 0 | 1.58E-05 | 3.86E-05 | 0.00032 | 7.59E-06 | 5.38E-05 |
| Cpsf3 | 0.00024 | 4.97E-05 | 5.87E-05 | 0 | 2.92E-05 | 7.06E-05 | 2.69E-06 | 1.63E-05 | 0.00012 | 1.74E-05 | 0 | 6.57E-05 | 8.44E-05 | 0 | 7.23E-05 | 6.84E-05 | 7.01E-06 | 4.58E-05 |

| | | | | | | | | | | | | | | | | | | |
|---|---|---|---|---|---|---|---|---|---|---|---|---|---|---|---|---|---|---|
| Atp6v1f | 0.00011 | 0.00028 | 7.64E-06 | 0.00018 | 0.0002 | 0.00018 | 6.71E-05 | 0.00012 | 0.00012 | 8.97E-05 | 0.0001 | 0.00018 | 0.00015 | 0.00011 | 0 | 0 | 0.00012 | 0.00014 |
| Mapkbp1 | 1.63E-05 | 4.63E-05 | 6.55E-05 | 2.51E-06 | 8.75E-07 | 0 | 2.51E-05 | 1.64E-05 | 0.00012 | 2.67E-05 | 2.97E-05 | 0 | 3.05E-05 | 0 | 5.15E-05 | 2.39E-05 | 0.00012 | 2.99E-05 |
| Clns1a | 0.00014 | 8.13E-05 | 0.00013 | 0.00017 | 2.32E-05 | 1.93E-05 | 7.55E-06 | 1.19E-05 | 0.00012 | 2.75E-06 | 3.76E-05 | 4.81E-05 | 2.24E-05 | 4.19E-05 | 2.67E-06 | 0.00011 | 0 | 0 |
| Pml | 1.48E-05 | 0.00011 | 0 | 0.00013 | 0.00025 | 0 | 0 | 0.00017 | 0.00012 | 0.00024 | 0 | 1.40E-05 | 0.00018 | 4.17E-05 | 0.00017 | 0.00034 | 0 | 0.00013 |
| Znhit6 | 0.00011 | 3.82E-05 | 6.66E-05 | 9.78E-06 | 6.61E-06 | 0 | 0 | 0 | 0.00012 | 6.07E-05 | 1.44E-05 | 6.03E-06 | 0 | 5.02E-06 | 8.11E-05 | 1.92E-05 | 0 | 0 |
| Ndufaf8 | 3.12E-05 | 6.42E-05 | 8.47E-05 | 0.00022 | 3.95E-05 | 0 | 0 | 7.74E-05 | 0.00012 | 7.60E-05 | 6.68E-05 | 4.03E-05 | 4.98E-05 | 3.17E-05 | 0 | 6.96E-05 | 7.72E-05 | 0.0001 |
| Fam189b | 3.24E-05 | 0.00011 | 0 | 9.42E-05 | 7.90E-05 | 7.17E-05 | 0.00014 | 3.76E-05 | 0.00012 | 0.00014 | 9.70E-05 | 1.98E-05 | 0 | 7.24E-05 | 0 | 0 | 6.70E-06 | 9.44E-05 |
| Lpcat1 | 6.02E-05 | 3.71E-05 | 0.00011 | 1.80E-05 | 0.00022 | 1.93E-06 | 2.46E-05 | 0.00014 | 0.00012 | 5.23E-05 | 0.00012 | 9.29E-05 | 1.23E-05 | 0.00014 | 4.99E-05 | 0.00017 | 0 | 1.66E-05 |
| Psmb7 | 0.00022 | 0.00013 | 0.00026 | 0.00029 | 6.76E-05 | 4.49E-06 | 7.67E-05 | 3.23E-05 | 0.00012 | 2.66E-05 | 0.00013 | 9.56E-06 | 4.78E-06 | 9.63E-05 | 0 | 0.0002 | 0.00023 | 4.01E-05 |
| Tia1 | 8.30E-05 | 9.00E-05 | 4.58E-05 | 5.26E-05 | 4.20E-05 | 2.65E-05 | 0 | 4.41E-05 | 0.00012 | 2.67E-05 | 6.36E-05 | 0 | 3.09E-05 | 6.70E-05 | 0.00024 | 0.00011 | 0 | 2.35E-05 |
| Ano10 | 0.0001 | 0.00013 | 3.37E-05 | 0.00018 | 0.00015 | 0.00013 | 0.00014 | 0.00013 | 0.00012 | 0.00017 | 0.00016 | 4.75E-05 | 0.00013 | 0.00017 | 0 | 0 | 5.19E-06 | 0.00016 |
| U2af1 | 0.0001 | 6.22E-05 | 0.00018 | 0.00016 | 0 | 0 | 0 | 3.13E-05 | 0.00012 | 5.82E-05 | 8.27E-05 | 6.37E-05 | 0 | 3.79E-05 | 0.0002 | 0.00023 | 0.00015 | 8.46E-06 |
| Spg21 | 0.0003 | 0.00023 | 4.63E-05 | 0 | 7.80E-05 | 0.00011 | 0.00011 | 8.36E-05 | 0.00012 | 6.96E-05 | 8.22E-06 | 0.00038 | 4.86E-05 | 0 | 0 | 9.65E-06 | 0 | 4.32E-05 |
| Timm10b | 0.00022 | 2.05E-05 | 0.00015 | 0.00031 | 8.42E-05 | 5.13E-05 | 0.00016 | 0.00017 | 0.00012 | 5.37E-05 | 8.97E-05 | 9.70E-06 | 8.09E-05 | 0.0001 | 6.60E-05 | 4.04E-05 | 0 | 3.72E-05 |
| Thra | 8.37E-05 | 0.00016 | 0 | 0.0002 | 0.00017 | 0.00026 | 0 | 0 | 0.00012 | 0.00015 | 3.96E-06 | 0.00019 | 0.00016 | 2.92E-05 | 4.04E-05 | 1.42E-05 | 0 | 0.00015 |
| Man2c1 | 2.29E-05 | 0.00014 | 6.39E-05 | 0.00012 | 4.60E-05 | 3.33E-05 | 0 | 2.75E-05 | 0.00012 | 0.00011 | 3.35E-05 | 6.40E-05 | 6.41E-05 | 2.58E-05 | 0.00029 | 4.91E-05 | 4.42E-05 | 0.00012 |
| Plgrkt | 0.00014 | 0.00014 | 0 | 0 | 5.52E-05 | 1.18E-05 | 7.53E-06 | 0 | 0.00012 | 0.00014 | 4.78E-05 | 1.88E-05 | 2.39E-05 | 9.27E-05 | 8.07E-05 | 1.01E-05 | 4.17E-05 | 0.00011 |
| Rras | 8.84E-05 | 0.00027 | 1.55E-05 | 0.00028 | 0.00022 | 0.00025 | 0.00012 | 0.00022 | 0.00012 | 0.00035 | 0.00057 | 0.00051 | 0.00031 | 0.00033 | 0 | 0 | 9.04E-05 | 0.00035 |
| Snrnp27 | 0.00015 | 5.77E-05 | 9.49E-05 | 3.00E-05 | 6.88E-05 | 5.47E-05 | 8.12E-07 | 4.31E-05 | 0.00012 | 4.02E-05 | 8.87E-05 | 9.97E-05 | 2.52E-05 | 2.63E-05 | 1.77E-05 | 7.98E-05 | 0 | 4.69E-05 |
| Usp25 | 0.0002 | 0.00018 | 2.72E-05 | 0.0001 | 0.00014 | 0.00014 | 0 | 8.53E-05 | 0.00012 | 0.00014 | 8.33E-05 | 0.00017 | 6.66E-05 | 0.00012 | 0.00027 | 0.00016 | 0.00025 | 0.00027 |
| Zeb1 | 1.00E-05 | 0.00015 | 5.91E-05 | 0 | 0.00017 | 8.87E-05 | 0.00023 | 0.00027 | 0.00012 | 0.00024 | 6.09E-05 | 7.33E-05 | 0 | 0 | 2.18E-05 | 6.08E-05 | 0.00033 | 0.00011 |
| Hif1a | 0.00053 | 0 | 0.00024 | 0 | 7.37E-05 | 0.00022 | 0.00043 | 0.00012 | 0.00012 | 0 | 4.00E-05 | 0.00028 | 0.00047 | 0.00016 | 9.18E-06 | 0.00022 | 0 | 0 |
| Med1 | 0.00024 | 0.00013 | 0.00016 | 0 | 5.83E-05 | 0.0001 | 2.53E-06 | 0 | 0.00012 | 2.06E-05 | 5.56E-05 | 7.23E-05 | 0 | 8.31E-06 | 9.51E-05 | 0.00011 | 5.68E-05 | 0 |
| Washc1 | 0.00011 | 0.00022 | 1.12E-05 | 0.00031 | 7.21E-05 | 4.85E-05 | 7.18E-05 | 2.79E-05 | 0.00012 | 7.37E-05 | 0 | 0 | 0 | 8.38E-05 | 0.00012 | 0 | 9.92E-05 | 0.00011 |
| Slc35b4 | 0.00029 | 5.21E-05 | 7.90E-05 | 3.15E-05 | 0.00016 | 0.00018 | 0.0002 | 3.51E-05 | 0.00012 | 9.48E-05 | 0.00016 | 0 | 6.69E-05 | 0.0002 | 0 | 0.00022 | 0 | 9.49E-06 |
| Mettl9 | 0.00048 | 0.00011 | 0.00023 | 9.61E-05 | 0.00011 | 0.00047 | 0.00041 | 0.0003 | 0.00012 | 8.84E-05 | 0.0001 | 0.00054 | 0.0002 | 9.06E-05 | 0 | 0.00034 | 0 | 0.00018 |
| Tomm70a | 0.00029 | 1.51E-05 | 0.0002 | 0 | 0 | 0.00013 | 0 | 0 | 0.00012 | 0 | 6.55E-06 | 4.47E-05 | 0 | 0 | 0 | 6.98E-05 | 0 | 0 |
| Tfpi | 0.00026 | 1.46E-06 | 2.49E-05 | 0.00012 | 0.00025 | 0.00057 | 2.42E-05 | 0.00021 | 0.00012 | 0.00017 | 0.00016 | 0.00052 | 0.00074 | 0.00011 | 0 | 7.75E-05 | 2.27E-05 | 0.00013 |
| Atf7 | 0.00018 | 0.00026 | 7.21E-05 | 0 | 0.00019 | 0.00023 | 0.00017 | 8.55E-05 | 0.00012 | 0.00012 | 7.63E-05 | 0.00038 | 7.24E-05 | 0.00013 | 8.93E-05 | 2.27E-05 | 1.50E-05 | 0.00022 |
| Kpnb1 | 0.00026 | 9.37E-05 | 0.00036 | 4.93E-05 | 0 | 4.88E-05 | 0.00014 | 2.14E-05 | 0.00012 | 4.45E-05 | 0.0001 | 0.00015 | 0 | 0 | 0 | 0.00043 | 8.76E-05 | 2.10E-05 |
| Msl1 | 0.0001 | 0.00025 | 0.00015 | 0.00026 | 0.00014 | 0.00016 | 6.85E-05 | 0.00015 | 0.00012 | 0.00014 | 0.00013 | 0.00014 | 1.83E-05 | 0.00014 | 0.00022 | 0.00011 | 0.00012 | 0.00014 |
| Sec23b | 0.00019 | 7.67E-05 | 0.0002 | 6.57E-05 | 2.86E-05 | 1.66E-05 | 3.39E-05 | 2.70E-06 | 0.00012 | 4.54E-05 | 0.0001 | 1.18E-05 | 4.35E-05 | 2.28E-05 | 5.16E-05 | 4.84E-05 | 0 | 2.18E-06 |
| Ppm1f | 3.52E-05 | 9.56E-05 | 6.23E-05 | 7.51E-05 | 6.99E-05 | 1.82E-05 | 0 | 2.44E-05 | 0.00012 | 3.23E-05 | 1.21E-05 | 2.07E-05 | 6.71E-05 | 3.21E-05 | 2.35E-05 | 1.43E-05 | 4.10E-06 | 4.61E-05 |

| | | | | | | | | | | | | | | | | | | |
|---|---|---|---|---|---|---|---|---|---|---|---|---|---|---|---|---|---|---|
| Oaz2 | 1.02E-05 | 0.00024 | 0 | 0.00051 | 0.00045 | 0.00023 | 0.00016 | 0.00035 | 0.00012 | 0.00022 | 0.00027 | 6.59E-05 | 0.00023 | 0.00036 | 0 | 3.03E-05 | 0.00012 | 0.00025 |
| Srp68 | 0.00022 | 0.00022 | 0.00035 | 0.00034 | 0.00011 | 5.92E-05 | 0.00016 | 0.00015 | 0.00012 | 3.24E-05 | 9.15E-05 | 0.0002 | 0 | 5.46E-05 | 0.00028 | 6.44E-05 | 0 | 5.90E-05 |
| Rab35 | 0.00018 | 0.00024 | 0.00022 | 0.00016 | 0.00012 | 7.43E-05 | 0 | 2.68E-05 | 0.00012 | 9.74E-05 | 0.00018 | 0.0001 | 0 | 6.68E-05 | 0.00011 | 0.00025 | 0.00011 | 0.00011 |
| Upf3a | 8.84E-05 | 0.00011 | 0.00014 | 8.28E-05 | 6.56E-05 | 3.89E-05 | 3.86E-05 | 3.20E-05 | 0.00012 | 7.54E-05 | 0.00017 | 1.15E-06 | 4.77E-05 | 0.00011 | 0.00013 | 0.00011 | 9.62E-05 | 8.24E-05 |
| Uxs1 | 0.00016 | 5.00E-05 | 0.00018 | 9.77E-05 | 3.75E-05 | 8.83E-05 | 0.00015 | 6.10E-05 | 0.00012 | 0.00012 | 8.86E-05 | 1.79E-07 | 0.0002 | 0 | 0 | 0.00025 | 0 | 8.10E-05 |
| Lrrc42 | 3.52E-05 | 5.76E-05 | 5.80E-05 | 0.00011 | 2.20E-05 | 0 | 0.00015 | 0 | 0.00012 | 3.69E-05 | 0.00011 | 3.97E-05 | 1.28E-05 | 7.05E-05 | 3.05E-06 | 3.74E-05 | 0 | 0.00021 |
| Emc1 | 8.35E-05 | 0.00012 | 0.00017 | 0.0001 | 4.61E-05 | 7.52E-05 | 9.76E-06 | 9.77E-05 | 0.00012 | 0.00013 | 5.78E-05 | 7.79E-05 | 0.00011 | 5.61E-05 | 0.00024 | 0.00017 | 1.40E-05 | 0.0001 |
| Mcm4 | 9.33E-05 | 0 | 0.00015 | 0 | 0 | 0 | 0 | 2.61E-05 | 0.00012 | 4.00E-05 | 2.44E-05 | 0 | 2.31E-05 | 5.12E-05 | 5.17E-06 | 0.00053 | 0 | 0 |
| Tor4a | 0 | 5.88E-05 | 3.10E-06 | 2.34E-05 | 0.00013 | 8.91E-05 | 0 | 0.0001 | 0.00012 | 0 | 1.97E-05 | 0.00012 | 5.57E-05 | 3.12E-05 | 0 | 2.37E-05 | 0.00012 | 4.86E-05 |
| Tmem41b | 0.00017 | 0.00014 | 7.14E-05 | 0 | 6.62E-05 | 8.06E-05 | 5.84E-06 | 0 | 0.00012 | 3.07E-05 | 4.66E-05 | 0.00015 | 3.03E-05 | 1.68E-05 | 0 | 8.11E-05 | 5.06E-05 | 8.43E-05 |
| Colgalt1 | 0.0004 | 2.93E-05 | 0.00024 | 0.00037 | 0.00037 | 0.00067 | 0.00086 | 0.00067 | 0.00012 | 0.00031 | 0.00054 | 0.00046 | 0.0005 | 0.00068 | 1.17E-05 | 0.00019 | 6.32E-05 | 0.00037 |
| Mcts1 | 0.0003 | 4.18E-05 | 0.00012 | 1.92E-05 | 7.02E-05 | 0.0001 | 4.65E-05 | 4.12E-05 | 0.00012 | 4.48E-05 | 5.55E-06 | 1.56E-05 | 3.56E-05 | 0 | 8.93E-05 | 8.39E-06 | 0 | 0 |
| Dhrs4 | 0.00017 | 0.00011 | 5.91E-05 | 0.00022 | 0.00012 | 0.00011 | 0.00011 | 0.0001 | 0.00012 | 7.50E-05 | 2.05E-05 | 0 | 8.05E-05 | 0.00013 | 2.31E-05 | 0.00021 | 0 | 0.0001 |
| Ctdspl | 0 | 7.67E-05 | 7.64E-06 | 1.26E-05 | 0 | 4.42E-06 | 0 | 7.96E-05 | 0.00012 | 2.52E-05 | 4.54E-05 | 4.44E-05 | 0 | 9.40E-05 | 6.44E-06 | 6.59E-05 | 8.17E-05 | 9.79E-05 |
| Ankrd46 | 9.25E-05 | 8.97E-05 | 1.35E-05 | 4.12E-05 | 7.48E-05 | 1.25E-05 | 0.00013 | 1.07E-05 | 0.00012 | 0.00025 | 3.58E-05 | 9.49E-05 | 0 | 2.46E-05 | 8.45E-05 | 6.86E-05 | 0 | 7.42E-05 |
| Ripk1 | 0.00021 | 0.00015 | 0.00011 | 3.36E-05 | 8.94E-05 | 0.00011 | 0.00014 | 0 | 0.00012 | 0.00011 | 7.39E-05 | 0.0001 | 0.00017 | 2.29E-05 | 0.00012 | 2.95E-05 | 0.00013 | 5.93E-05 |
| Hspd1 | 0.00022 | 7.38E-05 | 0.00052 | 0.0002 | 0 | 2.05E-05 | 0 | 5.60E-05 | 0.00012 | 1.80E-06 | 7.46E-05 | 5.47E-05 | 0 | 0 | 3.03E-05 | 0.00026 | 0.00023 | 0 |
| N4bp1 | 0.00013 | 0.00014 | 8.00E-05 | 3.24E-05 | 0.0001 | 0.00015 | 0.00011 | 0 | 0.00012 | 6.77E-05 | 7.68E-05 | 9.44E-05 | 0.00021 | 0.00013 | 0.00017 | 7.12E-05 | 9.58E-05 | 2.20E-05 |
| Nucks1 | 0.00029 | 6.34E-05 | 0.0002 | 0 | 3.21E-05 | 5.85E-05 | 0 | 2.06E-05 | 0.00012 | 2.99E-05 | 3.42E-05 | 3.55E-05 | 0 | 0 | 7.99E-05 | 0.00075 | 4.63E-05 | 1.76E-05 |
| Tbcb | 0.00029 | 0.00021 | 0.0003 | 0.00049 | 9.84E-05 | 5.69E-05 | 8.17E-05 | 9.97E-06 | 0.00012 | 0.00012 | 0.00022 | 4.65E-05 | 9.96E-05 | 6.01E-05 | 0 | 0.00019 | 0 | 0.00021 |
| P3h4 | 0.00025 | 0 | 0.00019 | 0.00027 | 0.00015 | 0.00022 | 0.00066 | 0.0003 | 0.00012 | 8.88E-05 | 0.00027 | 7.05E-05 | 0.0001 | 0.00023 | 0 | 2.48E-05 | 8.85E-05 | 6.52E-05 |
| Asb6 | 2.37E-06 | 0.00012 | 0.00017 | 0.00016 | 5.72E-05 | 0 | 0 | 3.18E-05 | 0.00011 | 3.57E-05 | 0.00011 | 4.67E-06 | 2.67E-05 | 9.51E-05 | 9.62E-05 | 6.39E-05 | 0 | 6.74E-05 |
| Vopp1 | 4.13E-05 | 8.88E-05 | 0.0001 | 0 | 0.00015 | 0.00012 | 4.33E-05 | 9.00E-05 | 0.00011 | 9.98E-05 | 8.96E-05 | 0.0002 | 0.0001 | 4.65E-05 | 0 | 0.00017 | 0 | 8.64E-05 |
| Mydgf | 0.00052 | 1.46E-05 | 0.0004 | 0.0006 | 0.0002 | 0.00028 | 0.00064 | 0.00025 | 0.00011 | 0.00014 | 0.00015 | 0.00014 | 0.00038 | 0.00031 | 0 | 0.00014 | 0 | 0.00016 |
| Sgcd | 1.94E-05 | 0 | 2.41E-05 | 0 | 5.30E-06 | 0 | 0.00019 | 0 | 0.00011 | 0 | 0 | 0.00039 | 0 | 0 | 0 | 0.00013 | 0 | 0.00013 |
| Slc30a7 | 0.00026 | 1.45E-06 | 0.00011 | 2.11E-05 | 9.83E-05 | 0.00017 | 0.00023 | 0.00016 | 0.00011 | 5.22E-05 | 1.44E-05 | 3.22E-05 | 0.00013 | 7.66E-05 | 8.84E-05 | 0 | 0.00011 | 4.43E-05 |
| Mtmr4 | 0.00011 | 7.61E-05 | 3.16E-05 | 4.58E-05 | 7.81E-05 | 1.80E-05 | 0 | 3.71E-05 | 0.00011 | 1.81E-05 | 0.00011 | 4.12E-05 | 5.65E-06 | 5.35E-05 | 1.47E-05 | 0.00013 | 2.47E-05 | 3.94E-05 |
| Scfd1 | 0.00047 | 2.49E-05 | 0.00014 | 2.11E-05 | 4.26E-05 | 0.00013 | 0.00026 | 8.05E-05 | 0.00011 | 7.74E-05 | 7.89E-06 | 2.36E-05 | 0 | 2.01E-05 | 0.00027 | 0 | 0 | 4.70E-05 |
| Pcif1 | 3.33E-05 | 0.00016 | 2.53E-05 | 0.00014 | 5.50E-05 | 6.78E-06 | 3.43E-07 | 0 | 0.00011 | 4.88E-05 | 7.57E-05 | 6.69E-05 | 0 | 6.74E-05 | 0.00013 | 0.00016 | 0 | 6.88E-05 |
| Tmed10 | 0.00034 | 0.00013 | 0.00028 | 0.00019 | 0.00011 | 0.00021 | 0.00029 | 0.00018 | 0.00011 | 0.00013 | 0.00015 | 0.00018 | 0.00024 | 0.0002 | 0 | 9.93E-05 | 0 | 0.00016 |
| Nab1 | 0.00022 | 0.00017 | 0.00012 | 3.92E-05 | 9.49E-05 | 0.00011 | 0.00018 | 0.00033 | 0.00011 | 1.29E-05 | 0 | 0 | 0.00019 | 0.00019 | 5.71E-05 | 0.0002 | 0.00018 | 0.00022 |
| Ube3b | 8.29E-05 | 0.0001 | 3.69E-05 | 7.52E-05 | 5.32E-05 | 3.70E-05 | 2.53E-05 | 4.37E-06 | 0.00011 | 2.98E-05 | 6.87E-05 | 0.00012 | 2.02E-05 | 2.65E-05 | 0.00014 | 5.64E-05 | 7.35E-07 | 3.25E-05 |
| Ehbp1l1 | 0 | 0.00011 | 0.00029 | 0.00024 | 4.66E-05 | 0 | 4.96E-05 | 0.00016 | 0.00011 | 0.00022 | 0.00057 | 9.32E-05 | 0 | 6.67E-05 | 0.00031 | 0.0001 | 0 | 0.00017 |

| | | | | | | | | | | | | | | | | | | |
|---|---|---|---|---|---|---|---|---|---|---|---|---|---|---|---|---|---|---|
| Farsa | 0.00013 | 0.0001 | 0.00025 | 0.00016 | 2.12E-05 | 2.77E-06 | 0 | 0 | 0.00011 | 5.13E-05 | 8.28E-05 | 1.19E-05 | 1.73E-05 | 3.93E-05 | 3.84E-05 | 8.04E-05 | 1.41E-05 | 0 |
| Mta3 | 0.00012 | 0.0001 | 4.73E-05 | 5.72E-05 | 3.79E-05 | 2.89E-05 | 5.75E-05 | 4.79E-05 | 0.00011 | 9.44E-05 | 7.77E-05 | 6.42E-05 | 1.25E-05 | 5.54E-05 | 0.00025 | 1.94E-05 | 3.79E-05 | 0.00012 |
| Dvl1 | 1.99E-05 | 0.00018 | 0.00019 | 0 | 7.76E-05 | 0 | 7.36E-05 | 0 | 0.00011 | 0.0001 | 0.00012 | 0.00012 | 1.65E-05 | 0 | 0.00044 | 0.00011 | 0.00017 | 0.00016 |
| Ndufa12 | 0.00026 | 6.32E-05 | 0.00016 | 6.97E-05 | 5.17E-05 | 5.21E-05 | 0 | 2.35E-05 | 0.00011 | 0 | 8.01E-05 | 2.52E-05 | 0 | 1.69E-05 | 0 | 8.41E-05 | 0.00012 | 0 |
| Drosha | 0.00013 | 7.06E-05 | 9.93E-05 | 8.59E-05 | 4.75E-05 | 7.54E-06 | 3.80E-05 | 1.22E-05 | 0.00011 | 4.23E-05 | 4.31E-05 | 0 | 0 | 5.62E-05 | 0 | 0.0001 | 9.96E-05 | 3.74E-05 |
| Selenof | 0.00049 | 5.30E-06 | 3.37E-05 | 0 | 0.00012 | 0.00032 | 0.00031 | 0.00018 | 0.00011 | 9.32E-05 | 0.00013 | 1.32E-05 | 3.92E-05 | 0.00014 | 0 | 5.43E-05 | 0.00015 | 0.0001 |
| Fchsd2 | 0.00011 | 9.23E-05 | 4.24E-05 | 0 | 0.00013 | 0.00016 | 0 | 1.90E-05 | 0.00011 | 7.19E-06 | 1.13E-05 | 0.00013 | 0.00015 | 0.00019 | 0.00016 | 3.98E-05 | 0.00025 | 7.47E-05 |
| Cnot3 | 0.0002 | 0.00014 | 0.00017 | 0.00013 | 0.00012 | 6.83E-05 | 5.58E-05 | 9.97E-05 | 0.00011 | 6.34E-05 | 4.37E-05 | 7.09E-05 | 3.40E-05 | 8.55E-05 | 0.00029 | 0.00021 | 1.97E-05 | 2.78E-05 |
| Ptges3 | 0.00044 | 0.00012 | 0.00027 | 0 | 1.66E-05 | 0.00016 | 6.71E-06 | 0 | 0.00011 | 2.03E-05 | 6.74E-05 | 0.00012 | 8.27E-07 | 1.94E-05 | 0.00012 | 0.00034 | 9.05E-05 | 3.15E-05 |
| Kif5b | 0.00031 | 0.00026 | 0.00036 | 0 | 0.00017 | 0.00011 | 0 | 3.35E-05 | 0.00011 | 6.48E-05 | 0.00035 | 9.90E-05 | 0 | 3.45E-05 | 0.0003 | 0.00024 | 1.66E-05 | 5.21E-05 |
| Tpp2 | 0.00026 | 0.00014 | 8.49E-05 | 0 | 7.06E-05 | 9.13E-05 | 0 | 5.31E-05 | 0.00011 | 2.46E-05 | 5.89E-05 | 0.00011 | 0 | 8.66E-05 | 0.00022 | 0.00011 | 7.15E-05 | 7.00E-06 |
| Aldh3a2 | 0.0001 | 9.85E-05 | 3.74E-05 | 0 | 6.07E-05 | 8.48E-05 | 3.25E-05 | 3.47E-05 | 0.00011 | 9.71E-05 | 3.17E-05 | 3.57E-05 | 2.06E-05 | 0 | 4.70E-05 | 2.22E-05 | 0 | 5.14E-05 |
| Jpt1 | 0 | 0.00022 | 6.70E-05 | 0.00051 | 0 | 0 | 0 | 0.00027 | 0.00011 | 0.00033 | 0.00022 | 0.00025 | 0 | 3.30E-05 | 0 | 0.00101 | 0.00052 | 0.00029 |
| Wdr91 | 0 | 0.00017 | 1.11E-05 | 3.69E-05 | 0.00011 | 2.68E-05 | 2.95E-06 | 0.00011 | 0.00011 | 4.39E-05 | 0 | 8.22E-05 | 4.45E-07 | 0 | 0.00014 | 4.95E-05 | 4.41E-05 | 0 |
| Ncaph2 | 2.80E-05 | 9.48E-05 | 7.99E-05 | 0.00018 | 3.54E-05 | 9.65E-06 | 0 | 5.70E-05 | 0.00011 | 5.50E-05 | 6.77E-05 | 0.00019 | 0 | 2.57E-05 | 0.00016 | 0.00042 | 7.64E-06 | 2.62E-05 |
| Ivd | 8.22E-05 | 9.89E-05 | 0 | 5.91E-05 | 8.90E-05 | 8.08E-05 | 0 | 1.70E-05 | 0.00011 | 4.74E-05 | 0 | 3.61E-06 | 4.54E-05 | 2.55E-05 | 3.94E-05 | 0.00011 | 6.39E-06 | 0 |
| Ogfr | 0.00013 | 0.00021 | 0.00011 | 0.0004 | 0.00016 | 9.40E-05 | 2.95E-05 | 9.06E-05 | 0.00011 | 0.00015 | 7.59E-05 | 0.00011 | 4.22E-05 | 0.0002 | 0.00016 | 0.00015 | 0.00016 | 0.00011 |
| Hip1 | 1.54E-05 | 0.00016 | 0 | 1.51E-05 | 6.49E-05 | 5.35E-05 | 7.02E-05 | 0 | 0.00011 | 4.75E-06 | 0.00027 | 0.00012 | 0 | 9.49E-05 | 8.48E-05 | 5.34E-05 | 0 | 0.0002 |
| BC029722 | 0 | 0.00012 | 4.53E-05 | 0.00029 | 0.00012 | 6.24E-05 | 1.89E-05 | 0.00017 | 0.00011 | 0.00015 | 6.15E-06 | 5.82E-05 | 0.0001 | 2.97E-05 | 0 | 0 | 4.37E-05 | 0.00013 |
| Rex1bd | 0.00014 | 8.50E-05 | 7.37E-05 | 0.00024 | 3.15E-05 | 4.36E-05 | 0.00011 | 3.18E-06 | 0.00011 | 5.87E-05 | 2.21E-05 | 6.34E-06 | 1.12E-05 | 9.13E-05 | 1.81E-06 | 9.11E-05 | 1.85E-05 | 8.73E-05 |
| Cwc15 | 0.00029 | 0.00025 | 0.00029 | 0.00034 | 0.00012 | 0.00015 | 0.00019 | 0.00018 | 0.00011 | 9.92E-05 | 0.00014 | 0.00012 | 1.08E-05 | 0.00013 | 4.15E-05 | 0.00025 | 1.03E-05 | 6.41E-05 |
| Ammecr1l | 7.05E-05 | 0.00012 | 8.79E-05 | 3.54E-05 | 7.32E-05 | 4.23E-05 | 4.93E-05 | 6.91E-05 | 0.00011 | 6.51E-05 | 0.00014 | 9.50E-05 | 5.51E-06 | 0.0001 | 6.40E-05 | 1.74E-05 | 0.00016 | 3.53E-05 |
| Chrac1 | 0.00013 | 8.86E-05 | 6.15E-05 | 5.69E-05 | 4.34E-05 | 6.89E-05 | 7.76E-05 | 2.63E-05 | 0.00011 | 3.63E-05 | 4.23E-05 | 5.29E-05 | 2.74E-05 | 6.12E-05 | 1.51E-05 | 4.12E-05 | 0 | 5.16E-05 |
| Sema4c | 0 | 0.00015 | 0.00016 | 0.00028 | 8.72E-05 | 0.00012 | 0 | 0.00061 | 0.00011 | 9.38E-05 | 0 | 0 | 0.00012 | 0.00028 | 0.00032 | 5.01E-05 | 7.63E-05 | 8.60E-05 |
| Arl1 | 0.00067 | 1.98E-05 | 0.00033 | 7.47E-05 | 8.95E-05 | 0.00041 | 0.00051 | 0.00012 | 0.00011 | 6.57E-05 | 0.00016 | 0.00017 | 0.00017 | 4.36E-05 | 0 | 0 | 0.00018 | 6.06E-05 |
| Golga7 | 0.00021 | 0.00023 | 0.00026 | 0.00013 | 0.00031 | 0.00032 | 0.00033 | 0.00028 | 0.00011 | 5.87E-05 | 0.00038 | 0.00025 | 0.0001 | 1.18E-05 | 0.00019 | 7.66E-05 | 4.05E-05 | 0.00044 |
| Ric1 | 0.00022 | 0.00016 | 5.61E-05 | 0 | 0.00019 | 0.00013 | 0.00012 | 5.16E-05 | 0.00011 | 0 | 3.97E-05 | 9.37E-05 | 0 | 0.00016 | 0.00013 | 2.07E-05 | 0.0002 | 0 |
| Ncbp2 | 0.00021 | 0.0001 | 0.00019 | 0.00031 | 0.00012 | 1.09E-05 | 3.16E-05 | 9.35E-05 | 0.00011 | 7.68E-05 | 2.54E-05 | 0 | 2.95E-05 | 5.88E-05 | 8.09E-05 | 0.00011 | 0 | 0.00012 |
| Prdx4 | 0.00057 | 0 | 6.92E-05 | 0.00034 | 0 | 0.00016 | 0.00054 | 0 | 0.00011 | 0 | 0 | 0.00039 | 0.00039 | 0.0002 | 0 | 0.00048 | 0 | 0.00032 |
| Dhrs1 | 0.00021 | 5.99E-05 | 0.00012 | 0.00026 | 5.49E-05 | 4.52E-05 | 0.00024 | 2.56E-05 | 0.00011 | 9.14E-05 | 8.31E-05 | 8.38E-05 | 0 | 9.99E-05 | 0 | 4.25E-05 | 0 | 6.62E-05 |
| Nop58 | 0.00023 | 6.17E-06 | 0.00041 | 3.95E-06 | 0 | 1.88E-05 | 0 | 0 | 0.00011 | 2.05E-05 | 4.16E-06 | 5.62E-06 | 5.75E-05 | 0 | 0.00014 | 0.00028 | 0.0002 | 0 |
| Pink1 | 1.59E-05 | 0.0005 | 0 | 2.88E-05 | 0.00036 | 0.00049 | 8.01E-05 | 5.15E-05 | 0.00011 | 0.00017 | 3.11E-05 | 0.00022 | 0.00012 | 0.00029 | 0 | 0 | 0 | 0.00022 |
| Bud31 | 0.00042 | 0.00014 | 0.00027 | 0.00015 | 8.08E-05 | 7.41E-05 | 0.00018 | 6.34E-05 | 0.00011 | 0 | 0.00011 | 6.25E-05 | 4.86E-05 | 7.97E-05 | 7.30E-05 | 0.00015 | 6.51E-05 | 9.00E-06 |

| | | | | | | | | | | | | | | | | | | |
|---|---|---|---|---|---|---|---|---|---|---|---|---|---|---|---|---|---|---|
| Fam171a1 | 4.37E-05 | 4.86E-05 | 0.00012 | 0.0001 | 5.80E-05 | 0 | 0.0001 | 0.00011 | 0.00011 | 0.00027 | 0.00011 | 1.11E-05 | 8.09E-06 | 2.66E-05 | 3.24E-05 | 0.00011 | 8.45E-05 | 0.0002 |
| Anapc4 | 0.00027 | 0.00012 | 0.0002 | 0.00017 | 2.52E-05 | 7.35E-05 | 0.00022 | 4.18E-05 | 0.00011 | 8.23E-05 | 7.34E-05 | 9.24E-05 | 2.90E-05 | 7.66E-05 | 0.00021 | 0.00019 | 0.0001 | 0.00014 |
| Nipa2 | 0.00013 | 6.45E-05 | 0.00012 | 0.00013 | 9.52E-05 | 0.00011 | 7.64E-05 | 7.82E-05 | 0.00011 | 4.78E-05 | 0.00016 | 2.27E-06 | 6.79E-05 | 0.00011 | 0 | 7.42E-05 | 1.36E-05 | 8.32E-06 |
| Pard3 | 4.35E-05 | 0.00019 | 8.20E-05 | 7.03E-05 | 4.15E-05 | 6.96E-05 | 4.90E-05 | 0.00011 | 0.00011 | 8.70E-05 | 5.98E-05 | 9.25E-05 | 5.55E-05 | 3.65E-05 | 0.0002 | 4.03E-05 | 0.0003 | 9.71E-05 |
| Ncoa2 | 9.64E-05 | 0.00014 | 0 | 0 | 2.39E-05 | 3.47E-05 | 1.46E-05 | 0 | 0.00011 | 8.48E-05 | 4.23E-05 | 3.54E-05 | 4.19E-05 | 3.78E-05 | 2.02E-05 | 5.54E-05 | 0.00017 | 3.48E-05 |
| Ppp2r5b | 1.77E-06 | 0.00011 | 0.00015 | 7.89E-05 | 7.86E-05 | 5.79E-05 | 5.16E-05 | 0.00012 | 0.00011 | 0.00018 | 0.00029 | 3.62E-05 | 0 | 8.85E-05 | 3.58E-05 | 8.27E-06 | 1.74E-06 | 0.00011 |
| Zhx2 | 9.40E-05 | 7.22E-05 | 8.20E-05 | 0 | 0.00013 | 2.23E-06 | 8.33E-05 | 9.53E-05 | 0.00011 | 8.16E-05 | 0 | 3.23E-05 | 3.36E-05 | 0 | 0 | 8.00E-06 | 6.15E-05 | 4.84E-05 |
| Ostc | 0.00055 | 0 | 0.0004 | 0.00115 | 0.0002 | 0.0004 | 0.00092 | 0.0005 | 0.00011 | 0.00029 | 0.00027 | 0.00016 | 0.00028 | 0.00038 | 0 | 0.00014 | 0 | 0.00019 |
| Erlin2 | 6.47E-05 | 0.00012 | 9.12E-05 | 7.24E-05 | 9.91E-05 | 0.00012 | 0.0001 | 0.00016 | 0.00011 | 0.00018 | 1.83E-05 | 0.00017 | 9.57E-05 | 6.21E-05 | 0 | 0.0001 | 0 | 0.0001 |
| Sntb1 | 0 | 1.45E-05 | 0 | 3.01E-07 | 4.03E-05 | 6.21E-05 | 0 | 0 | 0.00011 | 3.94E-05 | 0 | 0 | 3.29E-05 | 0 | 2.03E-05 | 0 | 6.04E-06 | 0 |
| Numa1 | 0.00019 | 0.00033 | 0 | 0 | 0.00013 | 0.00019 | 0.00011 | 0 | 0.00011 | 8.13E-05 | 7.06E-05 | 0.00017 | 0.00011 | 6.97E-05 | 0.00031 | 0.00033 | 0.00017 | 0.00017 |
| Rnf213 | 0.00016 | 0.00014 | 0 | 2.93E-07 | 0 | 3.27E-05 | 0 | 0 | 0.00011 | 0.00039 | 0 | 0 | 0.00027 | 1.42E-05 | 7.29E-06 | 6.97E-05 | 0.00013 | 0.00022 |
| Gpsm1 | 0 | 0.00013 | 9.65E-05 | 0.00021 | 3.18E-05 | 1.49E-05 | 0 | 6.29E-05 | 0.00011 | 8.70E-05 | 0.00012 | 0 | 0 | 0.00014 | 0.00025 | 0 | 0.00017 | 0.00014 |
| Ism1 | 0 | 0 | 5.65E-05 | 0 | 0 | 5.65E-05 | 0 | 4.21E-05 | 0.00011 | 6.76E-05 | 0 | 4.08E-05 | 0 | 0 | 0 | 0 | 0 | 0.00013 |
| Cdk6 | 6.00E-05 | 7.71E-06 | 2.91E-05 | 0 | 0 | 9.13E-05 | 0 | 6.32E-05 | 0.00011 | 1.26E-05 | 8.41E-05 | 0 | 0 | 9.73E-05 | 0 | 6.01E-05 | 1.45E-05 | 3.67E-05 |
| Syn1 | 9.91E-06 | 0 | 5.45E-06 | 0 | 0.00017 | 0 | 0 | 5.44E-05 | 0.00011 | 0 | 8.63E-05 | 8.02E-05 | 0 | 3.78E-06 | 0 | 0 | 0 | 0 |
| Itfg1 | 0.00042 | 0.0001 | 6.94E-05 | 0 | 0.00012 | 0.00034 | 0.00012 | 0.00015 | 0.00011 | 9.34E-05 | 8.42E-05 | 9.44E-05 | 6.89E-05 | 4.60E-05 | 0 | 3.99E-05 | 5.44E-05 | 0.0001 |
| Fzd3 | 2.78E-05 | 3.49E-05 | 8.58E-06 | 0 | 0.00011 | 2.95E-05 | 1.85E-05 | 4.24E-05 | 0.00011 | 0 | 0 | 0 | 0 | 6.67E-05 | 0 | 0 | 0 | 0 |
| Snw1 | 0.00023 | 0.00016 | 8.33E-05 | 5.94E-05 | 3.68E-05 | 8.38E-05 | 0 | 0 | 0.00011 | 4.23E-05 | 2.66E-05 | 8.22E-05 | 0 | 0 | 6.50E-05 | 0.00014 | 0 | 5.16E-05 |
| Gnpat | 0.00022 | 0.00012 | 0.00012 | 0.00011 | 0.0001 | 0.00012 | 0.00012 | 0.0001 | 0.00011 | 4.66E-05 | 5.56E-05 | 5.57E-05 | 8.74E-06 | 9.72E-05 | 0.0001 | 0.00012 | 0 | 0 |
| Dnpep | 0.00032 | 0.00012 | 0.00013 | 0.00012 | 1.41E-05 | 4.32E-05 | 0.00021 | 0 | 0.00011 | 0.00015 | 0.00015 | 0.00013 | 7.89E-05 | 0 | 8.11E-05 | 0 | 0 | 9.97E-05 |
| Yaf2 | 0.00016 | 7.84E-05 | 0.00011 | 0 | 7.61E-05 | 0.0001 | 4.81E-05 | 0.00018 | 0.00011 | 7.73E-06 | 7.43E-05 | 9.24E-05 | 7.67E-06 | 5.23E-06 | 0.00012 | 0.00018 | 0.00013 | 4.23E-05 |
| Smarcd2 | 0.00013 | 0.00011 | 8.65E-05 | 0.00011 | 7.20E-05 | 1.14E-05 | 8.58E-05 | 0 | 0.00011 | 0.00014 | 6.17E-06 | 0 | 0 | 4.95E-05 | 0.00011 | 7.51E-05 | 9.50E-05 | 0.00018 |
| Pbx2 | 7.10E-05 | 0.00018 | 9.03E-05 | 0.00019 | 9.54E-05 | 8.17E-05 | 2.91E-05 | 0.00018 | 0.00011 | 5.26E-05 | 7.70E-05 | 3.31E-05 | 0 | 6.50E-05 | 0.00019 | 3.96E-05 | 5.60E-05 | 3.96E-05 |
| Lypla2 | 9.31E-05 | 8.84E-05 | 0.00015 | 0.00014 | 6.19E-05 | 9.07E-05 | 6.61E-05 | 9.10E-05 | 0.00011 | 4.75E-05 | 7.71E-05 | 3.33E-05 | 4.86E-06 | 6.91E-05 | 7.32E-05 | 2.76E-05 | 0 | 4.64E-05 |
| Jkamp | 0.00015 | 9.74E-05 | 5.73E-05 | 4.93E-05 | 8.88E-05 | 0.00014 | 0.00013 | 0.00011 | 0.00011 | 7.04E-05 | 0.00012 | 0.0001 | 1.59E-05 | 5.31E-05 | 0 | 6.12E-05 | 0 | 4.69E-05 |
| Paics | 8.43E-05 | 0.00012 | 6.09E-05 | 0.00014 | 2.65E-06 | 5.44E-05 | 0 | 9.61E-06 | 0.00011 | 6.78E-05 | 5.90E-05 | 7.97E-05 | 6.53E-05 | 4.10E-05 | 0 | 0.00018 | 1.35E-05 | 8.42E-05 |
| Rpp25l | 1.65E-05 | 5.98E-05 | 8.78E-05 | 0.00015 | 3.64E-05 | 0 | 1.95E-05 | 5.11E-05 | 0.00011 | 7.35E-05 | 2.94E-05 | 0 | 4.29E-05 | 5.48E-05 | 4.36E-06 | 7.71E-05 | 0 | 7.35E-05 |
| Prdx1 | 0.00051 | 8.36E-05 | 0.0002 | 0.00034 | 5.08E-06 | 0.00022 | 0.00025 | 0.00014 | 0.00011 | 7.32E-05 | 4.21E-05 | 6.55E-05 | 0.00011 | 0.00015 | 0 | 0.00021 | 0.00037 | 3.80E-05 |
| Eef1d | 0.00027 | 7.50E-05 | 0.00031 | 0.00028 | 0 | 3.49E-05 | 7.09E-05 | 0 | 0.00011 | 1.89E-05 | 7.50E-05 | 0.00012 | 2.08E-06 | 3.40E-05 | 1.50E-05 | 0.00023 | 0.00012 | 5.08E-05 |
| Sfxn5 | 1.78E-05 | 4.98E-05 | 0 | 0 | 3.16E-05 | 0 | 8.27E-05 | 2.35E-05 | 0.00011 | 2.56E-05 | 0 | 2.91E-06 | 6.06E-05 | 0 | 0.00014 | 6.01E-05 | 2.17E-05 | 0 |
| Camsap2 | 0.0001 | 0.00018 | 0.00017 | 0 | 0.00016 | 6.65E-05 | 4.80E-05 | 5.68E-05 | 0.00011 | 6.26E-06 | 0.00011 | 0.00016 | 0 | 5.14E-05 | 8.99E-05 | 0.00011 | 7.04E-05 | 3.60E-05 |
| Lacc1 | 2.93E-05 | 6.64E-05 | 0 | 0 | 0 | 3.56E-05 | 0 | 7.10E-05 | 0.00011 | 3.30E-05 | 0 | 0 | 3.31E-06 | 2.70E-05 | 6.59E-05 | 5.58E-05 | 3.29E-05 | 0 |

| | | | | | | | | | | | | | | | | | | |
|---|---|---|---|---|---|---|---|---|---|---|---|---|---|---|---|---|---|---|
| Gnl1 | 0.00013 | 0.00014 | 0.00031 | 0.00041 | 0.00016 | 0.00017 | 0.0001 | 0.00012 | 0.00011 | 4.64E-05 | 0.00011 | 0.00025 | 8.63E-05 | 0.00014 | 0.00012 | 0.00011 | 6.03E-05 | 0.00013 |
| Irak4 | 8.82E-05 | 7.01E-05 | 8.67E-06 | 0.00011 | 3.79E-05 | 2.59E-05 | 5.18E-05 | 0.0001 | 0.00011 | 0.00012 | 1.10E-05 | 4.70E-05 | 0 | 1.55E-05 | 3.91E-05 | 0 | 0 | 6.00E-05 |
| Gart | 0.00015 | 5.39E-05 | 0.00026 | 0.00018 | 7.47E-06 | 0 | 0 | 0 | 0.00011 | 2.95E-05 | 3.08E-05 | 0 | 4.98E-05 | 0 | 0 | 0.00017 | 0 | 2.12E-05 |
| Basp1 | 1.27E-05 | 6.56E-06 | 0.00027 | 0.00051 | 3.31E-05 | 0 | 0 | 0.00014 | 0.00011 | 0.00068 | 0.00077 | 0.00036 | 0.00022 | 0 | 0.00031 | 0.00027 | 0.0001 | 0.00017 |
| Fam220a.1 | 0.00013 | 0.00012 | 0.00012 | 0 | 3.13E-05 | 0.00012 | 0 | 2.18E-05 | 0.00011 | 8.91E-05 | 9.65E-05 | 4.37E-05 | 2.08E-05 | 2.45E-05 | 0 | 3.63E-05 | 8.88E-05 | 5.48E-05 |
| Ppp1r12b | 4.40E-06 | 3.32E-05 | 0 | 0 | 5.53E-05 | 9.33E-05 | 2.59E-05 | 0 | 0.00011 | 4.89E-05 | 8.79E-05 | 0 | 1.16E-05 | 1.85E-05 | 0.00016 | 1.90E-05 | 0.00015 | 7.70E-05 |
| Hmox2 | 0.0002 | 0.00011 | 6.51E-05 | 9.79E-05 | 9.14E-05 | 7.13E-05 | 9.74E-05 | 9.51E-05 | 0.00011 | 3.51E-05 | 2.62E-05 | 0.00017 | 5.14E-05 | 9.40E-05 | 0 | 0.00013 | 0 | 2.99E-05 |
| Supt6 | 0.00015 | 0.00024 | 0.00017 | 0.00017 | 8.34E-05 | 7.13E-05 | 0.00011 | 5.58E-05 | 0.00011 | 7.07E-05 | 0.00011 | 0.0001 | 9.57E-06 | 7.38E-05 | 0.00026 | 4.70E-05 | 0.00013 | 0.00012 |
| Chmp2b | 0.00013 | 0.00015 | 4.10E-05 | 0 | 0.0001 | 0.00014 | 0.00018 | 5.49E-05 | 0.00011 | 8.52E-05 | 7.48E-05 | 0.00013 | 9.86E-05 | 7.91E-05 | 0 | 1.14E-05 | 0 | 0.00018 |
| Sgpp1 | 1.65E-05 | 0.00012 | 0.00021 | 0.00041 | 0.0001 | 0.00033 | 0 | 0.00013 | 0.00011 | 0.0004 | 0.00016 | 0.00036 | 0.0004 | 0.00027 | 8.18E-05 | 0.00017 | 4.93E-05 | 0.00025 |
| Appbp2 | 0.00025 | 0.00013 | 0.00022 | 6.13E-05 | 0.00011 | 0.0001 | 0.00015 | 0.00013 | 0.00011 | 6.62E-05 | 0.00013 | 6.74E-05 | 8.25E-05 | 7.59E-05 | 0.0002 | 3.95E-05 | 0.00018 | 0 |
| Ankrd13a | 0.00018 | 0.00014 | 0.00018 | 0.00036 | 0.00018 | 0.0002 | 0.00026 | 0.00019 | 0.00011 | 0.00022 | 0.00026 | 6.58E-06 | 0 | 0.00013 | 0 | 0.00026 | 0.0001 | 0.00022 |
| Csnk2a1 | 0.00021 | 0.00022 | 0.00023 | 7.56E-05 | 0.00011 | 0.00014 | 9.63E-05 | 0.00012 | 0.00011 | 0.00011 | 0.00013 | 0.00016 | 3.93E-05 | 8.28E-05 | 9.16E-05 | 9.95E-05 | 0 | 6.82E-05 |
| Ireb2 | 0.00015 | 0.00015 | 0.00011 | 7.00E-05 | 0.00011 | 9.69E-05 | 3.18E-05 | 7.42E-05 | 0.00011 | 5.16E-05 | 9.33E-05 | 0.00012 | 2.48E-05 | 9.13E-05 | 0.00013 | 9.86E-05 | 0 | 5.21E-05 |
| Dcun1d5 | 0.00043 | 9.93E-05 | 0.00022 | 8.63E-05 | 0.00015 | 0.00019 | 0.00015 | 0.00012 | 0.00011 | 2.76E-05 | 3.03E-05 | 5.63E-05 | 5.96E-05 | 0.00014 | 0.00014 | 0.00017 | 0 | 0 |
| Ptdss2 | 7.81E-05 | 8.56E-05 | 4.20E-05 | 1.92E-05 | 9.05E-05 | 6.09E-05 | 2.95E-05 | 1.38E-05 | 0.00011 | 3.83E-05 | 3.02E-05 | 2.25E-05 | 6.15E-05 | 5.61E-05 | 0 | 4.43E-05 | 0 | 5.95E-05 |
| Fam49b | 0.00011 | 1.89E-05 | 9.83E-05 | 0 | 0 | 2.54E-05 | 0 | 0 | 0.00011 | 2.91E-05 | 9.73E-05 | 3.60E-05 | 0 | 2.01E-05 | 3.36E-05 | 4.74E-05 | 7.68E-05 | 1.16E-05 |
| Mif4gd | 1.55E-05 | 7.69E-05 | 1.92E-05 | 1.78E-05 | 6.59E-05 | 2.99E-05 | 0.00011 | 5.41E-05 | 0.00011 | 0.00015 | 0.00011 | 3.67E-05 | 4.41E-05 | 8.29E-05 | 4.80E-05 | 3.28E-05 | 0 | 0.00012 |
| Fam117a | 3.79E-06 | 3.39E-05 | 6.28E-06 | 4.86E-05 | 8.64E-06 | 5.48E-05 | 2.91E-05 | 4.43E-05 | 0.00011 | 3.67E-05 | 8.44E-08 | 0 | 0 | 0 | 0 | 0 | 9.16E-05 | 0 |
| Tmcc3 | 5.63E-05 | 1.66E-05 | 0 | 0 | 0.00019 | 7.29E-05 | 0 | 0.00027 | 0.00011 | 8.05E-05 | 0.00019 | 3.78E-05 | 0 | 0.00035 | 0 | 6.25E-05 | 0 | 1.90E-05 |
| Sipa1l3 | 0 | 4.41E-05 | 0.00028 | 2.87E-05 | 0.00014 | 0 | 5.22E-05 | 0.00011 | 0.00011 | 7.95E-05 | 3.69E-05 | 0.00038 | 0.0001 | 3.95E-05 | 0.00021 | 0.00011 | 0.00024 | 0.00013 |
| Vti1b | 0.00033 | 9.80E-05 | 0.00014 | 1.02E-05 | 0.00011 | 0.00016 | 6.32E-05 | 4.01E-05 | 0.00011 | 9.82E-05 | 0.00023 | 0.00016 | 3.62E-05 | 1.66E-05 | 5.98E-06 | 0 | 0 | 6.32E-05 |
| Fbxw5 | 9.58E-05 | 0.00014 | 7.03E-05 | 6.76E-05 | 9.16E-05 | 8.75E-05 | 6.17E-05 | 6.76E-05 | 0.00011 | 7.09E-05 | 8.46E-05 | 0 | 4.78E-05 | 7.20E-05 | 0.00015 | 2.47E-05 | 0 | 5.62E-05 |
| Dip2a | 1.01E-05 | 0.00012 | 3.67E-05 | 0.00012 | 0.00014 | 6.58E-05 | 0 | 0.00033 | 0.00011 | 0.00012 | 0.00039 | 0.00021 | 0 | 0.00023 | 8.27E-05 | 9.03E-05 | 0 | 0.00011 |
| Mdp1 | 0.00015 | 7.51E-05 | 5.85E-05 | 4.83E-05 | 6.15E-05 | 6.18E-05 | 9.09E-05 | 0.00011 | 0.00011 | 0.00011 | 8.80E-05 | 0 | 0 | 5.39E-05 | 6.84E-05 | 5.08E-05 | 0 | 6.85E-05 |
| Gtf3c2 | 0.00016 | 0.00023 | 0.00012 | 8.91E-05 | 0.00012 | 0.00015 | 0.00011 | 7.84E-05 | 0.00011 | 6.23E-05 | 6.10E-05 | 0.00012 | 3.79E-05 | 9.62E-05 | 0.00042 | 0.00035 | 5.78E-05 | 4.63E-05 |
| Eif3a | 0.00024 | 0.00015 | 0.00023 | 0.00014 | 5.14E-05 | 5.40E-05 | 1.77E-06 | 2.71E-05 | 0.00011 | 2.20E-05 | 4.31E-05 | 6.71E-05 | 0 | 6.21E-05 | 0.00016 | 9.95E-05 | 0.00018 | 0 |
| Ppp2r2a | 0.00019 | 0.00015 | 0.00014 | 1.55E-05 | 7.06E-05 | 3.72E-05 | 0 | 1.35E-05 | 0.00011 | 3.95E-05 | 8.59E-05 | 8.03E-05 | 0 | 2.67E-05 | 0 | 0.00012 | 0.00018 | 2.58E-05 |
| Kif1b | 0.00012 | 0.00021 | 0.00011 | 1.32E-06 | 0 | 2.18E-05 | 4.37E-05 | 2.93E-05 | 0.00011 | 0.00012 | 0.00017 | 8.21E-05 | 0 | 5.00E-05 | 4.50E-05 | 5.69E-05 | 2.50E-05 | 0.00014 |
| Czib | 0.00015 | 5.41E-05 | 7.89E-05 | 8.13E-05 | 9.13E-05 | 4.72E-05 | 0.00013 | 6.99E-05 | 0.00011 | 8.92E-05 | 9.14E-05 | 1.61E-06 | 6.18E-05 | 2.24E-05 | 0.00011 | 6.14E-05 | 0 | 0.00012 |
| Psd3 | 3.24E-05 | 9.40E-05 | 0 | 0 | 6.17E-05 | 0 | 0.00018 | 0.00011 | 0.00011 | 0.00055 | 6.98E-05 | 0.00013 | 0.00023 | 5.97E-05 | 4.88E-05 | 0.00013 | 0.00019 | 0 |
| Nop56 | 0.00012 | 6.57E-05 | 0.00042 | 0.00012 | 0 | 0 | 0 | 0 | 0.00011 | 2.92E-06 | 6.50E-05 | 0 | 1.53E-05 | 2.65E-05 | 0.00029 | 0.00026 | 0.00039 | 0 |
| Bhlhe22 | 0 | 8.00E-06 | 0 | 6.75E-05 | 2.52E-05 | 8.85E-06 | 0 | 4.85E-05 | 0.00011 | 0 | 5.00E-05 | 0 | 3.63E-05 | 3.82E-06 | 8.71E-05 | 5.21E-05 | 0 | 2.95E-06 |

| | | | | | | | | | | | | | | | | | | |
|---|---|---|---|---|---|---|---|---|---|---|---|---|---|---|---|---|---|---|
| Bag5 | 0.00015 | 9.71E-05 | 0.00013 | 1.38E-05 | 0.00014 | 7.27E-05 | 0.00011 | 0 | 0.00011 | 0.00013 | 0.00013 | 3.81E-05 | 1.41E-05 | 0.0001 | 7.85E-05 | 3.52E-05 | 0 | 0.00011 |
| Pcgf3 | 0.00016 | 0.00015 | 0.00014 | 0 | 0.00012 | 5.14E-05 | 3.39E-05 | 9.27E-05 | 0.00011 | 5.64E-05 | 0.00015 | 0.00016 | 2.61E-05 | 5.12E-05 | 1.87E-05 | 8.13E-05 | 0 | 6.21E-05 |
| Grasp | 5.12E-06 | 0.00024 | 6.39E-05 | 0.00011 | 7.40E-05 | 2.01E-05 | 0 | 0 | 0.00011 | 0 | 3.28E-06 | 1.29E-05 | 3.62E-05 | 0.00014 | 0.00014 | 1.26E-05 | 0.00051 | 0.00012 |
| Rnf115 | 0.00015 | 0.00014 | 1.92E-05 | 0 | 3.52E-05 | 0.00013 | 4.86E-05 | 0 | 0.00011 | 2.60E-05 | 0 | 0 | 8.39E-06 | 3.23E-05 | 4.19E-05 | 7.24E-05 | 6.36E-05 | 5.54E-05 |
| Atrn | 8.01E-05 | 0.00011 | 0.00014 | 0 | 0.00011 | 9.71E-05 | 8.51E-05 | 1.22E-05 | 0.00011 | 6.26E-05 | 7.67E-05 | 2.98E-05 | 3.16E-05 | 0.00011 | 8.66E-05 | 2.65E-05 | 0.00012 | 9.30E-05 |
| N6amt1 | 3.28E-05 | 4.02E-05 | 7.54E-05 | 7.01E-05 | 6.75E-05 | 4.74E-05 | 2.12E-06 | 4.70E-05 | 0.00011 | 3.55E-05 | 5.71E-05 | 8.04E-06 | 9.30E-05 | 4.93E-05 | 5.24E-06 | 0 | 0 | 4.51E-05 |
| Arhgef1 | 0.00013 | 0.00021 | 0.00022 | 0.00051 | 8.48E-05 | 6.41E-05 | 0.00016 | 9.13E-05 | 0.00011 | 0.00013 | 9.03E-05 | 3.43E-05 | 6.81E-05 | 0.0001 | 0.00067 | 0.00012 | 0.00037 | 0.0002 |
| Irf2bp1 | 1.49E-05 | 0.00014 | 0.00012 | 0.00014 | 9.74E-05 | 2.04E-05 | 0 | 6.48E-05 | 0.00011 | 2.01E-05 | 3.53E-05 | 2.00E-05 | 1.86E-05 | 9.62E-05 | 0.00012 | 0.00017 | 0 | 3.74E-05 |
| Anks1 | 0 | 0.0001 | 4.11E-05 | 4.44E-05 | 0 | 8.37E-05 | 0 | 9.96E-05 | 0.00011 | 0.00014 | 1.46E-05 | 9.02E-06 | 0 | 0.00014 | 6.59E-05 | 0 | 9.08E-05 | 4.67E-05 |
| Layn | 4.49E-05 | 6.33E-05 | 4.80E-05 | 4.97E-05 | 2.15E-06 | 9.58E-05 | 0 | 0.00018 | 0.00011 | 0.00011 | 0 | 0.00042 | 0.00033 | 4.95E-05 | 1.38E-05 | 6.35E-05 | 0 | 0.00022 |
| Dpf2 | 5.75E-05 | 0.00025 | 0.00016 | 0.00032 | 5.90E-05 | 2.61E-05 | 0 | 0 | 0.00011 | 0.00013 | 2.70E-05 | 0.00015 | 0 | 2.38E-05 | 0.00018 | 0.00013 | 0 | 3.69E-05 |
| Tmem131 | 0.0002 | 0.00019 | 0.00016 | 0 | 0.00011 | 0.00015 | 5.68E-05 | 6.02E-05 | 0.00011 | 0.00012 | 4.29E-05 | 0.00013 | 5.19E-05 | 1.55E-06 | 0.00025 | 9.71E-05 | 0.00021 | 0.00013 |
| Prkacb | 0.00016 | 0.00025 | 0.00028 | 4.95E-05 | 0.00022 | 0.00016 | 7.54E-05 | 0.00055 | 0.00011 | 7.47E-05 | 0.00012 | 0.0002 | 9.75E-05 | 9.96E-05 | 0 | 0.00021 | 0.00033 | 0.00024 |
| Irf3 | 4.19E-05 | 0.00011 | 9.46E-05 | 0.00017 | 7.56E-05 | 2.12E-05 | 1.07E-05 | 0.00011 | 0.00011 | 5.03E-05 | 5.55E-05 | 0 | 0 | 9.89E-05 | 0.00011 | 3.26E-05 | 2.04E-05 | 7.57E-05 |
| Csk | 0.00013 | 0.00015 | 0.00018 | 5.79E-05 | 8.05E-05 | 3.64E-05 | 4.68E-05 | 0.00014 | 0.00011 | 7.93E-05 | 0.00011 | 8.16E-05 | 0 | 5.00E-05 | 6.16E-05 | 8.21E-05 | 0 | 6.74E-05 |
| Klhl22 | 8.96E-05 | 0.00013 | 8.29E-05 | 0.0001 | 0.0001 | 0.00011 | 6.30E-05 | 6.68E-05 | 0.00011 | 5.66E-05 | 0.00022 | 1.44E-05 | 2.99E-05 | 0.0001 | 9.35E-05 | 5.76E-05 | 0 | 0.00012 |
| Wdr33 | 0.00022 | 0.00023 | 0.00012 | 5.98E-05 | 9.48E-05 | 0.00011 | 3.12E-05 | 8.82E-06 | 0.00011 | 4.12E-05 | 4.57E-05 | 0.00011 | 1.10E-06 | 3.34E-05 | 0.00023 | 0.00013 | 5.06E-05 | 3.23E-05 |
| Erap1 | 0.00017 | 2.56E-05 | 1.08E-06 | 0 | 0 | 1.50E-05 | 0 | 0 | 0.00011 | 1.20E-05 | 0 | 0 | 0.00018 | 7.32E-06 | 0 | 3.73E-05 | 0 | 0.00014 |
| Cirbp | 0 | 0.00026 | 0 | 0.00026 | 0.00013 | 0.00016 | 0 | 5.46E-05 | 0.00011 | 7.81E-05 | 9.45E-05 | 0.00011 | 1.35E-05 | 0.00017 | 0.00013 | 9.56E-05 | 4.81E-05 | 6.45E-05 |
| Chst1 | 0 | 1.53E-05 | 1.76E-05 | 3.36E-05 | 0 | 3.22E-05 | 6.68E-05 | 0 | 0.00011 | 0.00012 | 6.43E-06 | 1.19E-05 | 0 | 0 | 0 | 5.70E-05 | 3.98E-05 | 0 |
| Spry4 | 0 | 8.91E-05 | 0 | 0 | 0.00012 | 0.0002 | 0 | 0.0003 | 0.00011 | 5.80E-07 | 0.00011 | 9.37E-05 | 9.55E-06 | 0.00026 | 9.04E-06 | 0 | 0.00023 | 5.39E-05 |
| Nf1 | 0.00014 | 0.00021 | 0.00016 | 0 | 0.00013 | 0.00017 | 4.39E-05 | 0.00014 | 0.00011 | 5.01E-05 | 6.99E-05 | 0.00016 | 1.01E-05 | 6.12E-05 | 2.04E-05 | 0.00016 | 0.00028 | 0.00012 |
| Dok6 | 2.72E-05 | 1.49E-05 | 0 | 0 | 3.68E-05 | 6.31E-05 | 2.66E-05 | 7.51E-06 | 0.00011 | 0 | 6.29E-05 | 6.02E-05 | 0 | 8.07E-06 | 6.40E-05 | 0 | 5.46E-05 | 0 |
| Adi1 | 6.85E-05 | 9.36E-05 | 2.42E-05 | 0.00017 | 3.64E-05 | 3.48E-05 | 0 | 0 | 0.00011 | 1.01E-05 | 0 | 3.09E-05 | 8.01E-05 | 1.58E-05 | 3.09E-06 | 7.81E-05 | 0 | 5.23E-05 |
| Plin3 | 0.0002 | 0.00028 | 6.66E-06 | 0.00034 | 6.46E-06 | 0.00011 | 0.00018 | 9.16E-05 | 0.00011 | 0.00022 | 0 | 0.0002 | 0.0001 | 9.65E-06 | 6.10E-05 | 2.39E-05 | 0 | 0.00023 |
| Fam107b | 0 | 2.23E-05 | 4.85E-05 | 3.03E-05 | 3.33E-05 | 2.80E-06 | 0 | 0.00013 | 0.00011 | 5.15E-05 | 0.0001 | 5.31E-05 | 0 | 6.27E-05 | 0 | 9.43E-05 | 0.00013 | 3.79E-05 |
| Nr2c2 | 9.23E-05 | 0.00013 | 2.16E-05 | 0 | 9.02E-05 | 9.12E-05 | 9.76E-05 | 5.73E-05 | 0.00011 | 3.06E-05 | 9.56E-05 | 6.35E-05 | 6.28E-05 | 9.80E-05 | 6.20E-05 | 0.00011 | 2.66E-05 | 3.58E-05 |
| Ndufs1 | 0.00015 | 7.01E-05 | 0.00017 | 0.00013 | 3.46E-05 | 2.89E-05 | 3.98E-05 | 4.28E-05 | 0.00011 | 4.43E-05 | 6.19E-05 | 0 | 0 | 4.98E-05 | 6.21E-05 | 9.53E-05 | 3.58E-05 | 2.92E-05 |
| Nfia | 0.00014 | 0.00017 | 0 | 4.74E-05 | 0 | 0 | 0.00022 | 0.00015 | 0.00011 | 0.00046 | 0 | 0 | 0.00035 | 0 | 0 | 0.00013 | 0.0001 | 0.00027 |
| Mocs1 | 2.68E-06 | 0.00012 | 1.37E-06 | 8.06E-05 | 6.69E-05 | 0.00011 | 0 | 3.92E-05 | 0.00011 | 0.00017 | 0 | 2.05E-05 | 0.00011 | 8.86E-05 | 8.94E-05 | 0 | 0 | 2.17E-05 |
| Vamp3 | 0.00033 | 0.00014 | 0.00016 | 0 | 8.10E-05 | 0.00019 | 0 | 0.00011 | 0.00011 | 8.94E-05 | 9.78E-05 | 0.00019 | 0 | 1.61E-06 | 0 | 0.00013 | 0 | 6.00E-05 |
| Sys1 | 9.21E-05 | 0.00012 | 0.00019 | 0.00033 | 0.00011 | 5.09E-05 | 2.68E-05 | 0.00019 | 0.00011 | 9.31E-05 | 0.00011 | 5.77E-05 | 8.49E-05 | 0.00015 | 1.17E-05 | 7.58E-05 | 7.19E-05 | 0.00014 |
| Oga | 0.0002 | 0.00025 | 0.00012 | 0 | 6.34E-05 | 0.0001 | 3.39E-06 | 7.52E-06 | 0.00011 | 9.68E-05 | 8.44E-05 | 0.00025 | 0 | 3.49E-05 | 0.00032 | 4.41E-05 | 0.00024 | 6.90E-05 |

| | | | | | | | | | | | | | | | | | | |
|---|---|---|---|---|---|---|---|---|---|---|---|---|---|---|---|---|---|---|
| Dnajc13 | 0.00015 | 0.00022 | 0.00015 | 0.00015 | 0.00012 | 8.81E-05 | 7.49E-05 | 9.15E-06 | 0.00011 | 0.00012 | 5.48E-05 | 8.51E-05 | 3.07E-05 | 0.00013 | 6.74E-05 | 0.00015 | 5.48E-05 | 9.07E-05 |
| Nol6 | 0.00012 | 9.97E-05 | 0.00021 | 0.00014 | 4.14E-05 | 0 | 3.55E-05 | 6.85E-06 | 0.00011 | 9.29E-06 | 4.35E-05 | 0 | 6.08E-05 | 0 | 6.16E-05 | 0 | 5.21E-05 | 4.60E-05 |
| Tfpt | 6.21E-05 | 0.00015 | 9.77E-05 | 0.00026 | 4.79E-05 | 1.32E-05 | 4.42E-05 | 0 | 0.00011 | 4.71E-05 | 0.0001 | 4.23E-05 | 9.44E-05 | 8.26E-05 | 9.97E-05 | 6.15E-05 | 5.88E-06 | 0 |
| Oxa1l | 0.00011 | 0.00017 | 7.54E-05 | 0.00016 | 0.00011 | 8.21E-05 | 7.93E-05 | 0.0001 | 0.00011 | 6.55E-05 | 3.84E-06 | 4.26E-05 | 3.78E-05 | 0.00011 | 2.61E-05 | 0.00013 | 0 | 6.57E-05 |
| Etfa | 0.00033 | 0.00022 | 0.00025 | 0.00033 | 6.66E-05 | 0.0001 | 0.0001 | 2.86E-05 | 0.00011 | 0.00032 | 0.0001 | 0.0001 | 0.00015 | 9.30E-05 | 0 | 0.00033 | 0 | 7.19E-05 |
| Smad1 | 3.29E-05 | 0.00012 | 0.00011 | 2.26E-05 | 8.15E-05 | 3.82E-05 | 4.62E-05 | 2.07E-05 | 0.00011 | 0.00023 | 0.00019 | 0.00011 | 7.22E-05 | 0.00011 | 3.96E-05 | 9.94E-05 | 7.61E-05 | 2.23E-06 |
| Prkcsh | 0.00017 | 9.38E-05 | 0.00038 | 0.00047 | 0.00019 | 0.00019 | 0.00042 | 0.00052 | 0.00011 | 0.00015 | 0.00026 | 0.0004 | 0.00025 | 0.00031 | 0.00031 | 0.00015 | 0 | 0.00018 |
| Cybc1 | 9.32E-05 | 0.00012 | 8.50E-05 | 8.29E-05 | 9.08E-05 | 2.32E-05 | 0.00011 | 0.00017 | 0.00011 | 8.69E-05 | 2.70E-05 | 0.00016 | 1.76E-06 | 2.58E-05 | 8.89E-05 | 4.57E-05 | 0 | 0.00011 |
| Ldlrap1 | 4.53E-05 | 0.00013 | 9.44E-05 | 0.00014 | 0.00015 | 4.98E-05 | 8.31E-05 | 5.65E-05 | 0.00011 | 2.59E-06 | 8.65E-05 | 0.00015 | 4.84E-05 | 0.00018 | 0.00013 | 9.44E-07 | 0 | 4.58E-05 |
| Carmil1 | 1.07E-06 | 6.21E-05 | 2.41E-05 | 2.83E-05 | 2.60E-05 | 1.98E-05 | 0 | 5.18E-05 | 0.00011 | 1.13E-05 | 9.51E-05 | 2.66E-06 | 2.65E-05 | 4.51E-05 | 5.63E-05 | 4.21E-05 | 0.00011 | 0.00012 |
| Phf20 | 8.43E-05 | 0.00012 | 2.36E-05 | 3.87E-06 | 4.59E-05 | 1.93E-05 | 2.11E-05 | 1.92E-05 | 0.00011 | 2.55E-05 | 0 | 6.85E-05 | 2.95E-05 | 2.98E-05 | 0.00015 | 6.76E-05 | 6.55E-05 | 1.36E-05 |
| Fbxw11 | 0.00013 | 0.00028 | 0.00023 | 8.60E-05 | 0.00014 | 6.96E-05 | 5.10E-05 | 0.00011 | 0.00011 | 0.00011 | 0.00018 | 0.00017 | 0 | 3.87E-05 | 0.00011 | 0.00011 | 0.00021 | 3.99E-05 |
| Usp10 | 7.72E-05 | 7.75E-05 | 0.00015 | 0 | 2.79E-05 | 0 | 0 | 0 | 0.00011 | 4.34E-05 | 5.09E-05 | 0 | 1.01E-05 | 0 | 0.00016 | 0.00012 | 0 | 3.69E-05 |
| Cox14 | 0.00029 | 0.0002 | 0.00013 | 0.00029 | 0.00018 | 0.00023 | 0.00016 | 0.00013 | 0.00011 | 9.53E-05 | 0.0001 | 8.95E-05 | 0 | 9.54E-05 | 0 | 0.00011 | 1.51E-06 | 0.00022 |
| Pex14 | 9.93E-05 | 0.00016 | 0.00016 | 0.0001 | 7.40E-05 | 2.25E-05 | 7.62E-05 | 3.29E-05 | 0.00011 | 2.79E-05 | 6.72E-05 | 0.00012 | 3.49E-05 | 7.14E-05 | 3.96E-05 | 5.96E-05 | 6.33E-05 | 3.72E-05 |
| Ankrd50 | 4.86E-05 | 0.00013 | 9.90E-05 | 4.67E-06 | 0.0001 | 2.24E-05 | 0 | 0.00027 | 0.00011 | 0 | 0.0002 | 0.0003 | 3.95E-05 | 0.0001 | 2.29E-05 | 5.03E-05 | 3.24E-05 | 7.40E-05 |
| Dhrs3 | 0 | 0.00027 | 0 | 0.00037 | 0.00095 | 0 | 0.00048 | 0.00032 | 0.00011 | 0.00027 | 0.00027 | 0.00032 | 0.00017 | 0 | 0.00023 | 0 | 0 | 0.00028 |
| Txndc15 | 0.00014 | 0.00011 | 7.10E-05 | 0.00021 | 0.00019 | 0.00014 | 0.00022 | 0.00015 | 0.00011 | 0.00017 | 6.09E-05 | 8.22E-05 | 0.00017 | 0.00017 | 0 | 2.11E-05 | 0 | 0.0002 |
| Rcc2 | 0.0001 | 0.00015 | 0.00014 | 0.00029 | 0.00013 | 0.0002 | 0.00011 | 0 | 0.00011 | 0 | 0 | 0.00012 | 0.00012 | 0.00022 | 0.00013 | 0.0002 | 6.36E-05 | 6.05E-06 |
| Ndufb6 | 0.00015 | 0.00014 | 0.00026 | 0.00077 | 5.76E-05 | 6.34E-06 | 0 | 6.67E-05 | 0.00011 | 7.13E-05 | 0.00022 | 2.71E-05 | 4.80E-05 | 9.70E-05 | 3.72E-05 | 0.00019 | 0.00026 | 0.00014 |
| Tmem123 | 0.00048 | 2.39E-05 | 0 | 0 | 5.50E-05 | 0.0002 | 3.39E-05 | 0 | 0.00011 | 0 | 0.00042 | 4.10E-05 | 6.75E-05 | 0.00016 | 0 | 4.34E-05 | 0.00011 | 7.49E-05 |
| Dab2 | 0.00013 | 0.00017 | 3.50E-06 | 0 | 9.52E-05 | 0.00011 | 4.86E-05 | 0 | 0.00011 | 0.0001 | 0 | 0.00013 | 0.00035 | 0.00023 | 1.67E-05 | 2.29E-05 | 0.00019 | 0.00011 |
| Rab11a | 0.00036 | 0.00013 | 0.00019 | 0 | 0.00011 | 0.00012 | 0.0001 | 0.00011 | 0.00011 | 5.70E-05 | 0.00013 | 0.00014 | 3.97E-05 | 0.00015 | 2.59E-06 | 1.82E-05 | 1.12E-05 | 3.93E-05 |
| Pcbp4 | 5.97E-05 | 0.00012 | 0.00026 | 0.00025 | 0 | 0 | 0 | 9.48E-06 | 0.00011 | 5.16E-05 | 0.00021 | 4.05E-05 | 6.35E-05 | 0 | 3.41E-05 | 6.30E-05 | 3.64E-05 | 0.00019 |
| Pelp1 | 1.53E-05 | 8.63E-05 | 0.00016 | 0.0002 | 2.39E-05 | 0 | 0 | 4.22E-05 | 0.00011 | 4.37E-05 | 9.16E-05 | 1.97E-05 | 3.85E-05 | 2.27E-05 | 1.84E-05 | 0.00025 | 0 | 5.72E-05 |
| Eif4ebp2 | 0.00022 | 0.00026 | 0.00021 | 0.00023 | 0.00018 | 0.00035 | 0.00013 | 0.00015 | 0.00011 | 9.30E-05 | 0.00011 | 2.31E-05 | 3.78E-05 | 0.00021 | 3.29E-05 | 0.0002 | 0.00011 | 7.21E-05 |
| Nbea | 4.27E-05 | 0.00015 | 2.19E-05 | 0 | 4.96E-05 | 0.00024 | 0 | 2.22E-05 | 0.00011 | 7.76E-06 | 0 | 0 | 0 | 5.88E-05 | 0.0001 | 0 | 8.06E-05 | 0 |
| Gnl3l | 5.95E-05 | 8.81E-05 | 0.00016 | 0.00014 | 4.48E-05 | 9.51E-06 | 4.64E-05 | 0.00012 | 0.00011 | 3.83E-05 | 9.99E-05 | 3.73E-05 | 9.67E-06 | 5.01E-05 | 4.19E-05 | 4.47E-05 | 0 | 7.11E-05 |
| Egr1 | 0.00033 | 0.00021 | 9.80E-05 | 0.00012 | 0.00045 | 0.00054 | 0.00073 | 0.00055 | 0.00011 | 0.00019 | 0 | 0.00065 | 0.0008 | 0.00024 | 0.00054 | 0.00013 | 0.0101 | 0.00023 |
| Eprs | 0.00048 | 9.56E-05 | 0.00037 | 0.00012 | 6.64E-05 | 0.00017 | 0.0003 | 0 | 0.00011 | 4.47E-05 | 7.32E-05 | 0.00025 | 4.54E-05 | 1.60E-05 | 0.00022 | 7.53E-05 | 9.50E-05 | 0 |
| Srpx | 0.00028 | 0 | 0 | 0.00022 | 0.00021 | 0.00028 | 0.00026 | 2.36E-05 | 0.00011 | 0.00048 | 0.00028 | 0 | 0.0004 | 0.00011 | 0 | 0.00014 | 0 | 0.00079 |
| Gria3 | 0.00011 | 5.54E-05 | 0 | 0 | 0.00017 | 0.00012 | 2.81E-05 | 0 | 0.00011 | 0 | 0.00011 | 0 | 0.00011 | 0.00015 | 7.81E-05 | 0 | 3.23E-05 | 9.88E-05 |
| Tor1aip2 | 0.00046 | 0.00028 | 0.00017 | 0.00016 | 0.0003 | 0.00031 | 0.00018 | 0.00014 | 0.00011 | 0.00026 | 0.00033 | 0.00065 | 0.00011 | 0.00012 | 0.0003 | 0.0002 | 0.00034 | 0.00019 |

| | | | | | | | | | | | | | | | | | | |
|---|---|---|---|---|---|---|---|---|---|---|---|---|---|---|---|---|---|---|
| Hacd4 | 0.0001 | 1.53E-05 | 0 | 0 | 8.08E-05 | 1.36E-05 | 0.00012 | 0 | 0.00011 | 0.00021 | 8.49E-05 | 0 | 0 | 0 | 0 | 7.60E-06 | 0 | 7.80E-05 |
| Med16 | 5.74E-05 | 9.53E-05 | 5.52E-05 | 0.00013 | 4.12E-05 | 4.77E-05 | 0 | 4.21E-05 | 0.00011 | 5.40E-05 | 1.47E-05 | 4.59E-05 | 2.59E-05 | 3.87E-05 | 6.96E-05 | 7.78E-05 | 0 | 8.71E-05 |
| Zbed3 | 3.65E-05 | 0.00012 | 3.53E-05 | 0.0002 | 8.41E-05 | 0 | 0 | 0.00013 | 0.00011 | 2.65E-05 | 0 | 2.10E-05 | 5.00E-05 | 5.15E-05 | 0.00012 | 0.00023 | 0 | 8.10E-05 |
| Nt5c2 | 5.51E-05 | 9.00E-05 | 3.32E-05 | 4.39E-05 | 0.00011 | 8.07E-05 | 9.85E-05 | 0 | 0.00011 | 5.80E-05 | 0.00013 | 0.00019 | 0.00024 | 8.15E-05 | 6.56E-05 | 4.62E-05 | 0 | 0 |
| Mast2 | 5.47E-05 | 0.00016 | 0.00038 | 7.64E-05 | 0.0002 | 8.15E-05 | 0.00024 | 0.00016 | 0.00011 | 0.00013 | 0.0004 | 0.0003 | 0.00011 | 0 | 0.00017 | 0.00034 | 4.63E-05 | 6.03E-05 |
| Denr | 0.00027 | 9.01E-05 | 0.00033 | 0.00016 | 4.19E-05 | 6.35E-05 | 9.13E-05 | 5.49E-05 | 0.00011 | 9.31E-05 | 0.0002 | 0.00019 | 4.76E-05 | 0.00012 | 5.44E-05 | 0.00015 | 0 | 6.26E-05 |
| Osbp | 0.00026 | 0.00013 | 0.00017 | 1.43E-05 | 8.24E-05 | 0.00011 | 0.00016 | 8.82E-05 | 0.00011 | 4.10E-05 | 6.62E-05 | 0.0001 | 0 | 8.47E-05 | 0.00017 | 0.00012 | 0.00011 | 3.69E-05 |
| Fkbp3 | 0.0002 | 6.89E-05 | 6.97E-05 | 5.62E-05 | 1.79E-05 | 0.00011 | 0 | 0 | 0.00011 | 1.48E-05 | 2.56E-05 | 3.59E-05 | 1.99E-06 | 0 | 0 | 0.00021 | 4.26E-05 | 0 |
| Ptgr2 | 0.00011 | 0.0001 | 3.59E-05 | 0.00018 | 0.0001 | 0.00011 | 0.00011 | 9.07E-05 | 0.00011 | 0.00012 | 6.91E-05 | 4.78E-05 | 0.00011 | 2.91E-05 | 0 | 5.65E-05 | 0 | 0.00011 |
| Ppm1m | 0 | 4.88E-05 | 7.02E-05 | 0.00029 | 0.00013 | 9.66E-05 | 0.00012 | 0.00018 | 0.00011 | 0.00013 | 9.14E-05 | 0 | 6.93E-05 | 0.00017 | 0.00012 | 0.00019 | 0.00013 | 0.00011 |
| Fkbp10 | 0.00051 | 0 | 0.00047 | 0.00086 | 0.00044 | 0.00062 | 0.00159 | 0.00085 | 0.00011 | 0.00034 | 0.0006 | 0.0002 | 0.00069 | 0.0008 | 0.00048 | 0.00012 | 0.00013 | 0.00041 |
| Dock7 | 0.00016 | 0.00019 | 3.41E-05 | 0.00016 | 0.00013 | 0.00013 | 4.56E-05 | 7.14E-05 | 0.00011 | 0.0002 | 0.00019 | 0.00054 | 5.46E-05 | 6.86E-05 | 0.00019 | 0.00022 | 0.00013 | 0.0002 |
| G6pc3 | 4.79E-05 | 2.36E-05 | 0.00012 | 0.00032 | 5.60E-05 | 5.23E-05 | 0.00014 | 0.00013 | 0.00011 | 4.15E-05 | 7.89E-05 | 3.85E-05 | 6.08E-05 | 4.63E-05 | 4.54E-05 | 5.34E-05 | 0 | 4.23E-05 |
| Zbtb7a | 1.37E-05 | 0.00026 | 0.00026 | 0.00022 | 6.96E-05 | 9.69E-05 | 3.22E-05 | 9.29E-05 | 0.00011 | 0.00019 | 0.00011 | 0.00022 | 5.71E-05 | 0.00016 | 0.00043 | 0 | 0.00019 | 0.00012 |
| Aip | 0.00026 | 0.00017 | 0.00013 | 0.00016 | 8.83E-05 | 0.00013 | 0.00011 | 0.00011 | 0.00011 | 0.0001 | 6.46E-05 | 5.42E-05 | 3.79E-05 | 6.92E-05 | 0 | 0.00014 | 5.43E-05 | 0.00014 |
| Arhgap17 | 6.73E-05 | 0.00012 | 5.86E-05 | 2.56E-05 | 7.74E-05 | 9.13E-05 | 2.35E-05 | 0 | 0.00011 | 8.40E-05 | 1.31E-05 | 6.44E-05 | 0 | 5.35E-05 | 0.00011 | 6.27E-05 | 0.00014 | 5.49E-05 |
| Ptpn5 | 0 | 0 | 0 | 0 | 0 | 3.97E-05 | 0 | 0 | 0.00011 | 0.00018 | 0 | 5.49E-05 | 4.01E-05 | 0 | 0 | 0 | 0 | 0 |
| Usp40 | 0.00012 | 0.0001 | 3.70E-05 | 0 | 6.15E-05 | 3.13E-05 | 5.64E-05 | 2.86E-05 | 0.00011 | 1.68E-05 | 1.65E-05 | 2.33E-05 | 7.06E-05 | 5.07E-06 | 0.00015 | 3.98E-05 | 0 | 2.23E-05 |
| Map4k5 | 0.0002 | 0.00018 | 0.00017 | 0 | 0.00018 | 0.00011 | 0.00026 | 3.73E-06 | 0.00011 | 1.22E-06 | 0.00018 | 0.00022 | 4.64E-05 | 0.00011 | 8.65E-05 | 0.00014 | 0.00015 | 2.39E-05 |
| Tomm20 | 0.0002 | 9.95E-05 | 0.0002 | 3.05E-05 | 6.84E-05 | 0.00011 | 0 | 6.02E-05 | 0.00011 | 0 | 5.96E-05 | 8.75E-05 | 0 | 4.95E-05 | 5.62E-06 | 4.97E-05 | 0.00014 | 0 |
| Reps1 | 0.00012 | 0.00012 | 0.00012 | 0 | 5.71E-05 | 2.38E-05 | 1.34E-05 | 2.28E-05 | 0.00011 | 5.80E-05 | 8.67E-05 | 9.13E-05 | 0 | 7.59E-06 | 4.08E-05 | 0.00016 | 0.00012 | 3.03E-05 |
| Alg9 | 0.00014 | 4.12E-05 | 0.00015 | 0.0001 | 0 | 0 | 8.19E-05 | 1.49E-05 | 0.00011 | 5.55E-05 | 9.78E-05 | 0.00013 | 0 | 0 | 4.90E-05 | 2.68E-05 | 0 | 2.62E-05 |
| Extl2 | 0.00011 | 5.99E-05 | 3.20E-05 | 2.20E-05 | 2.53E-05 | 3.96E-05 | 6.39E-06 | 4.70E-05 | 0.00011 | 8.00E-05 | 0 | 2.29E-05 | 0 | 1.20E-07 | 0 | 3.77E-05 | 0 | 6.46E-05 |
| Bhlhb9 | 2.99E-05 | 4.94E-05 | 3.83E-06 | 9.51E-05 | 8.21E-05 | 3.42E-06 | 0 | 0 | 0.00011 | 1.11E-05 | 2.69E-05 | 0 | 5.80E-05 | 9.15E-05 | 6.55E-06 | 0.00012 | 0 | 8.97E-05 |
| Eif6 | 6.77E-05 | 9.50E-05 | 0.00041 | 0.0004 | 2.44E-05 | 0 | 3.18E-05 | 7.95E-05 | 0.00011 | 3.10E-05 | 7.44E-05 | 0.00011 | 0.0001 | 0.00011 | 0.00011 | 9.22E-05 | 0 | 6.26E-05 |
| Cdkn2b | 0 | 2.72E-05 | 0 | 0 | 0 | 0 | 0 | 6.06E-05 | 0.00011 | 0 | 5.76E-05 | 1.26E-05 | 0 | 4.13E-05 | 8.04E-06 | 0 | 0 | 4.54E-05 |
| Ube2l3 | 0.00021 | 6.91E-05 | 0.00015 | 0.00011 | 7.11E-05 | 5.82E-05 | 3.99E-05 | 2.60E-05 | 0.00011 | 3.77E-05 | 5.02E-05 | 3.61E-05 | 3.58E-05 | 9.39E-06 | 0 | 0.00014 | 5.45E-05 | 4.55E-05 |
| Sin3a | 0.0001 | 0.00015 | 5.91E-05 | 4.30E-05 | 5.93E-05 | 0.00011 | 6.08E-05 | 3.86E-05 | 0.00011 | 5.94E-05 | 3.45E-06 | 8.73E-05 | 8.87E-05 | 6.63E-05 | 9.68E-05 | 0.00023 | 0.00012 | 9.28E-06 |
| Dcaf8 | 0.00022 | 0.00026 | 0 | 8.85E-05 | 0.00022 | 0.00016 | 0.0001 | 8.52E-05 | 0.00011 | 0.0001 | 9.15E-05 | 2.62E-05 | 0.00011 | 8.67E-05 | 0.00021 | 2.17E-05 | 0 | 9.03E-05 |
| Dnajb12 | 9.86E-05 | 0.00024 | 0.00022 | 0.00029 | 0.00021 | 0.00022 | 0.00019 | 0.0002 | 0.00011 | 9.10E-05 | 0.00016 | 0.00016 | 0.00013 | 0.00018 | 0.00018 | 0.00017 | 0.00015 | 9.69E-05 |
| Arid1a | 5.34E-05 | 0.00016 | 0.00012 | 5.98E-05 | 6.84E-05 | 0.00011 | 5.52E-06 | 0.00013 | 0.00011 | 5.63E-05 | 6.37E-05 | 8.28E-05 | 0 | 4.86E-05 | 6.58E-05 | 0.00019 | 0.0003 | 5.47E-05 |
| Pmpcb | 0.00024 | 0.00011 | 0.00021 | 9.51E-05 | 0.00025 | 2.12E-05 | 0.00012 | 2.05E-05 | 0.00011 | 1.33E-05 | 0.00021 | 5.10E-05 | 5.04E-05 | 0 | 7.03E-05 | 0.0001 | 1.20E-05 | 0 |
| Zswim6 | 5.79E-05 | 2.95E-05 | 0.00012 | 0 | 0.00015 | 0.00019 | 0 | 9.64E-05 | 0.00011 | 0 | 0.0001 | 0.00018 | 7.24E-05 | 0.00018 | 2.21E-05 | 0 | 0.00039 | 0 |

| | | | | | | | | | | | | | | | | | | |
|---|---|---|---|---|---|---|---|---|---|---|---|---|---|---|---|---|---|---|
| Ino80 | 9.92E-05 | 0.00011 | 9.49E-05 | 1.21E-05 | 1.94E-05 | 5.89E-05 | 0 | 1.37E-05 | 0.00011 | 2.78E-05 | 3.31E-06 | 8.07E-05 | 5.73E-06 | 0 | 0.00017 | 2.04E-05 | 0.00015 | 5.37E-05 |
| Micos13 | 0.00017 | 0.00011 | 0.00018 | 0.0004 | 0.00012 | 0.00013 | 0.00016 | 0.00018 | 0.00011 | 0.00011 | 0.00011 | 0.00014 | 9.69E-05 | 0.00026 | 0 | 3.21E-05 | 0 | 0.00015 |
| Bri3 | 0.0001 | 0.00046 | 0.00011 | 0.00035 | 0.00027 | 0.00032 | 5.70E-05 | 0.00011 | 0.00011 | 0.00016 | 0.00026 | 0.00036 | 0.00021 | 0.00026 | 8.94E-05 | 0.00011 | 0.00015 | 0.00026 |
| Usp7 | 0.00023 | 0.00022 | 0.00019 | 7.09E-05 | 9.98E-05 | 0.00011 | 6.71E-05 | 6.07E-05 | 0.00011 | 3.89E-05 | 7.53E-05 | 0.00015 | 1.51E-05 | 5.80E-05 | 0.00029 | 0.00014 | 0.00021 | 4.89E-05 |
| Naa60 | 0.00012 | 0.00014 | 0.00013 | 0.0001 | 0.00017 | 7.36E-05 | 0.00013 | 0.00016 | 0.00011 | 8.87E-05 | 4.45E-05 | 0 | 5.76E-05 | 6.13E-05 | 1.56E-06 | 9.71E-05 | 0 | 0.00017 |
| Ppp6r3 | 0.00026 | 0.00018 | 9.68E-05 | 0 | 7.42E-05 | 0.00016 | 3.25E-05 | 4.26E-05 | 0.00011 | 3.33E-05 | 3.34E-05 | 0.00011 | 8.92E-06 | 6.84E-06 | 0.00028 | 0.00022 | 0.00023 | 2.20E-05 |
| Prrx2 | 6.20E-05 | 0.00012 | 7.06E-05 | 0.00021 | 4.75E-05 | 0 | 0.00036 | 0.00021 | 0.00011 | 0.00021 | 0.00038 | 0 | 0.00028 | 0.00023 | 0 | 8.40E-05 | 0 | 0.00057 |
| Plxnd1 | 7.62E-05 | 0.00015 | 5.10E-05 | 3.80E-05 | 5.83E-05 | 0.00016 | 0 | 0.00038 | 0.00011 | 0 | 0 | 0 | 9.05E-05 | 0.00041 | 0.00015 | 6.56E-05 | 0 | 0.00015 |
| Tmem147 | 0.00018 | 9.93E-05 | 0.00014 | 0.00022 | 0.00015 | 0.00011 | 0.00018 | 5.88E-05 | 0.00011 | 7.65E-05 | 0.00011 | 0 | 9.36E-05 | 0.00012 | 0 | 4.88E-05 | 0 | 0.0001 |
| Pcnx | 8.51E-05 | 0.00019 | 0.00021 | 0 | 0.00012 | 0.00017 | 6.55E-05 | 0.00014 | 0.00011 | 3.45E-06 | 6.23E-05 | 0.00011 | 1.08E-06 | 0.00014 | 9.43E-05 | 5.11E-05 | 0.00018 | 4.48E-05 |
| Ctdnep1 | 4.39E-05 | 9.68E-05 | 0.00014 | 0.00045 | 0.00011 | 0.00014 | 0.0002 | 0.00024 | 0.00011 | 0.00015 | 0.0001 | 9.16E-05 | 5.49E-05 | 0.00016 | 1.66E-05 | 0.0002 | 0.00011 | 0.00016 |
| Blcap | 9.17E-05 | 0.00021 | 0.00022 | 9.19E-05 | 0.00018 | 4.20E-05 | 5.48E-06 | 6.82E-05 | 0.00011 | 3.70E-05 | 0.00017 | 0.00031 | 6.62E-06 | 2.01E-05 | 0.00011 | 0 | 0 | 2.21E-05 |
| Usp47 | 0.00033 | 0.00018 | 0.0002 | 0 | 0.00011 | 0.00016 | 0.00013 | 1.41E-05 | 0.00011 | 1.91E-05 | 0.0002 | 9.53E-05 | 5.46E-05 | 0.00019 | 0.0003 | 0.00021 | 0.00014 | 1.81E-05 |
| Ppp1r7 | 0.00017 | 9.27E-05 | 0.00013 | 8.96E-05 | 1.44E-05 | 7.84E-05 | 0 | 3.27E-05 | 0.00011 | 6.39E-05 | 8.57E-05 | 5.75E-05 | 1.98E-06 | 1.96E-05 | 0 | 0.00026 | 0 | 2.93E-05 |
| Tmem222 | 0.00012 | 0.00011 | 9.62E-05 | 0.00011 | 5.43E-05 | 5.43E-05 | 4.27E-06 | 5.93E-05 | 0.00011 | 2.48E-05 | 9.03E-05 | 8.33E-05 | 6.48E-05 | 7.91E-05 | 0 | 9.25E-05 | 0 | 5.21E-05 |
| Ints6 | 0.00014 | 6.36E-05 | 4.94E-05 | 0 | 2.00E-05 | 0 | 3.69E-05 | 5.59E-06 | 0.00011 | 2.43E-05 | 0 | 0.00013 | 9.68E-05 | 1.93E-05 | 0.00011 | 5.60E-05 | 0.00102 | 0 |
| Pwwp3a | 5.80E-05 | 0.00011 | 5.37E-05 | 3.98E-05 | 4.87E-05 | 8.40E-06 | 0 | 3.86E-05 | 0.00011 | 1.50E-05 | 1.91E-05 | 1.67E-06 | 0 | 1.09E-05 | 0.00032 | 0.00013 | 5.10E-05 | 6.57E-05 |
| Phip | 8.60E-05 | 0.00019 | 0 | 5.30E-05 | 8.98E-05 | 1.63E-05 | 6.12E-05 | 7.00E-05 | 0.00011 | 4.83E-05 | 0 | 0 | 2.91E-05 | 8.81E-06 | 0.00019 | 0.00037 | 0.00027 | 9.39E-05 |
| Sema4b | 5.49E-05 | 0.00015 | 1.22E-05 | 0.00013 | 0.00012 | 0.00016 | 8.50E-05 | 6.40E-05 | 0.00011 | 9.85E-05 | 8.60E-05 | 4.12E-05 | 0.00015 | 0 | 9.11E-05 | 0 | 0 | 2.25E-05 |
| Tmbim1 | 0.00013 | 0.00024 | 0.00033 | 0 | 0.00011 | 0.00021 | 9.14E-06 | 0 | 0.00011 | 0.00028 | 0.00014 | 0.00016 | 9.05E-06 | 0.00015 | 0 | 1.11E-05 | 4.05E-05 | 0.00026 |
| Dcaf5 | 3.72E-05 | 0.00012 | 5.19E-05 | 4.51E-05 | 3.83E-05 | 2.88E-06 | 9.22E-05 | 2.61E-05 | 0.00011 | 5.45E-05 | 0 | 2.54E-05 | 7.33E-05 | 0 | 2.47E-05 | 3.09E-05 | 6.04E-05 | 2.37E-05 |
| Htra2 | 7.13E-05 | 0.00012 | 0.00019 | 0.00031 | 5.45E-05 | 6.29E-05 | 0.00018 | 0 | 0.00011 | 6.62E-05 | 9.99E-05 | 0 | 0.00027 | 3.31E-05 | 0.0002 | 0.00023 | 0 | 9.47E-05 |
| Grb2 | 0.00014 | 0.00011 | 0.00011 | 6.78E-05 | 7.26E-05 | 5.34E-05 | 9.57E-05 | 5.09E-05 | 0.00011 | 5.44E-05 | 0.00012 | 0.00012 | 4.93E-05 | 6.85E-05 | 4.66E-05 | 4.70E-05 | 1.87E-05 | 4.80E-06 |
| Rnf168 | 7.30E-05 | 5.13E-05 | 8.23E-05 | 0 | 3.48E-05 | 4.19E-05 | 1.63E-05 | 9.24E-06 | 0.00011 | 3.05E-05 | 3.99E-05 | 1.09E-05 | 0 | 4.16E-05 | 3.13E-05 | 0.00021 | 0 | 1.19E-05 |
| Psma6 | 0.00041 | 0.0002 | 0.00035 | 0.00033 | 8.41E-05 | 8.48E-05 | 0 | 0 | 0.00011 | 5.73E-05 | 0.00016 | 8.55E-05 | 4.29E-05 | 0.00011 | 0 | 0.00021 | 0 | 3.07E-05 |
| Dnaja3 | 0.0002 | 0.00011 | 0.00021 | 6.14E-05 | 5.59E-05 | 1.49E-05 | 1.27E-05 | 0 | 0.00011 | 3.07E-05 | 4.02E-05 | 0.00012 | 9.98E-05 | 0 | 0.00011 | 5.35E-05 | 1.68E-05 | 3.34E-05 |
| Ibtk | 0.0002 | 9.12E-05 | 0.0002 | 0 | 3.35E-05 | 4.51E-05 | 9.88E-05 | 0 | 0.00011 | 0.0001 | 7.44E-05 | 4.61E-05 | 0.00027 | 1.77E-05 | 0.00017 | 0.00011 | 2.63E-05 | 0 |
| Dnajb2 | 7.89E-06 | 9.04E-05 | 3.95E-05 | 0.00011 | 6.01E-05 | 1.19E-05 | 0 | 0 | 0.00011 | 3.27E-05 | 9.02E-05 | 7.74E-05 | 3.28E-05 | 6.05E-05 | 2.12E-05 | 0 | 6.55E-05 | 6.58E-05 |
| Grem1 | 0 | 0.00012 | 3.50E-06 | 0 | 0.00095 | 0.00012 | 0 | 0 | 0.00011 | 0 | 0 | 0.00011 | 0 | 0.0014 | 0 | 0 | 0 | 0 |
| Prep | 0.00023 | 0.00012 | 0.00039 | 0.0002 | 4.73E-05 | 5.74E-05 | 1.21E-05 | 0 | 0.00011 | 4.44E-05 | 0.00021 | 0.0001 | 0 | 6.34E-05 | 3.31E-05 | 0.00015 | 0 | 1.16E-05 |
| Plekha2 | 0.00011 | 9.32E-05 | 0 | 1.88E-05 | 5.03E-05 | 4.12E-05 | 0 | 0.00014 | 0.00011 | 0.00017 | 7.54E-05 | 0.00057 | 0 | 0 | 1.60E-05 | 0.00013 | 0 | 5.83E-05 |
| Ostf1 | 0.00023 | 4.91E-05 | 0.00023 | 0.00029 | 0.0003 | 0.00011 | 0.00038 | 0.00025 | 0.00011 | 0.00022 | 0.00046 | 0.00061 | 0 | 0.00027 | 3.55E-05 | 0.0001 | 0 | 0.00013 |
| Cdip1 | 7.53E-05 | 0.00019 | 8.10E-05 | 2.86E-05 | 0.00011 | 0.00012 | 6.09E-05 | 5.04E-05 | 0.00011 | 2.02E-05 | 7.86E-05 | 2.57E-05 | 1.16E-05 | 8.45E-05 | 0.00018 | 0 | 2.25E-05 | 2.05E-05 |

| | | | | | | | | | | | | | | | | | | |
|---|---|---|---|---|---|---|---|---|---|---|---|---|---|---|---|---|---|---|
| Ncor1 | 0.00017 | 0.00025 | 0.00014 | 0 | 0.00016 | 0.00016 | 7.32E-05 | 0.00013 | 0.00011 | 0.00011 | 0.00015 | 9.66E-05 | 4.15E-05 | 0.00019 | 0.00036 | 6.57E-05 | 0.00015 | 0.00021 |
| Luc7l2 | 0.00028 | 0.00034 | 0.0002 | 0.00014 | 0.00018 | 0.00016 | 0.00027 | 0.0001 | 0.00011 | 0.00014 | 7.47E-05 | 0.00032 | 0 | 0.00011 | 0.00125 | 0.00028 | 0.00064 | 0.00017 |
| Sh3bp4 | 0 | 0.00013 | 0 | 1.25E-05 | 8.14E-05 | 0 | 7.73E-05 | 0 | 0.00011 | 3.87E-05 | 0.00017 | 0 | 2.13E-05 | 2.26E-05 | 1.48E-05 | 2.53E-05 | 0.00013 | 4.76E-05 |
| Atp5md | 0.00032 | 0 | 0.00015 | 0.00011 | 0.00012 | 0.00018 | 8.78E-05 | 8.08E-05 | 0.00011 | 0 | 0.00022 | 5.04E-05 | 1.32E-05 | 8.26E-05 | 0 | 2.72E-05 | 4.74E-05 | 0 |
| Ube2e2 | 0 | 9.93E-05 | 1.27E-05 | 0.00011 | 0.00011 | 2.47E-05 | 9.57E-05 | 0.00014 | 0.00011 | 0.0002 | 0.00014 | 0.00013 | 0.00024 | 1.46E-05 | 0.0003 | 2.48E-05 | 9.50E-05 | 0.00015 |
| Bax | 0.00018 | 0.00052 | 0.00027 | 0.0006 | 0.00021 | 9.35E-05 | 0 | 0 | 0.00011 | 0.00018 | 0.00018 | 0.00016 | 0.00011 | 0.00014 | 0.00017 | 5.83E-05 | 0.0002 | 0.00013 |
| Tbrg1 | 0.00059 | 0.00021 | 0.00042 | 0.00041 | 0 | 0.00011 | 0.00046 | 8.14E-05 | 0.00011 | 0.00021 | 6.87E-05 | 0.00018 | 0.00011 | 1.83E-05 | 0.00017 | 0.00013 | 0.00011 | 0.00025 |
| Hgf | 7.98E-05 | 7.47E-05 | 0 | 3.36E-05 | 0 | 0.00034 | 0 | 5.25E-06 | 0.00011 | 5.40E-05 | 0 | 0 | 0.00015 | 2.39E-05 | 0 | 2.15E-05 | 0 | 0.00012 |
| Abcd2 | 0 | 1.73E-05 | 0 | 0 | 0 | 0 | 0 | 1.81E-05 | 0.00011 | 0.00014 | 0 | 0 | 0.00013 | 0 | 0 | 1.51E-05 | 0 | 0 |
| Rbm26 | 0.00013 | 0.00016 | 9.29E-05 | 0.00016 | 6.73E-05 | 5.40E-05 | 2.49E-05 | 5.84E-05 | 0.00011 | 5.22E-05 | 3.58E-05 | 8.16E-05 | 0 | 1.48E-05 | 0.00034 | 6.04E-05 | 0.00036 | 3.42E-05 |
| Paip1 | 0.00044 | 0.0002 | 0.00019 | 0 | 0.00013 | 0.00027 | 3.13E-06 | 9.17E-05 | 0.00011 | 0 | 0.00011 | 0.0002 | 0 | 4.69E-05 | 8.39E-05 | 0.00012 | 0.00015 | 0 |
| Cdk13 | 0.00017 | 0.00017 | 0.00013 | 0 | 0.00011 | 0.00016 | 8.67E-05 | 0.0001 | 0.00011 | 4.05E-05 | 0.00012 | 0.00015 | 4.79E-05 | 5.15E-05 | 0.00014 | 6.25E-05 | 0.0001 | 2.83E-05 |
| Dock5 | 3.03E-05 | 3.75E-05 | 3.26E-05 | 0 | 4.99E-05 | 0 | 1.88E-05 | 6.07E-05 | 0.00011 | 0 | 0.00019 | 0.00017 | 0 | 0 | 0 | 3.49E-05 | 0 | 2.45E-05 |
| Med22 | 0.0001 | 0.00011 | 0.00013 | 3.29E-05 | 9.20E-05 | 5.34E-05 | 0 | 0.00011 | 0.00011 | 3.02E-05 | 3.32E-05 | 5.46E-05 | 0 | 9.61E-05 | 3.78E-05 | 7.18E-05 | 2.33E-05 | 4.66E-05 |
| Lsm2 | 0.00015 | 5.20E-05 | 0.00022 | 8.12E-05 | 3.39E-05 | 2.11E-05 | 4.97E-05 | 4.80E-05 | 0.00011 | 4.45E-05 | 3.27E-05 | 0 | 4.04E-06 | 8.15E-05 | 2.65E-05 | 0.00071 | 0 | 5.05E-05 |
| Wdr18 | 0.00012 | 7.59E-05 | 0.00021 | 0.00017 | 1.22E-05 | 0 | 0 | 2.37E-06 | 0.00011 | 1.87E-05 | 6.16E-05 | 9.76E-08 | 2.51E-05 | 1.14E-05 | 3.72E-05 | 8.37E-05 | 0 | 0 |
| Arhgap24 | 0.0001 | 7.92E-05 | 0 | 0 | 6.37E-05 | 9.95E-05 | 0.00012 | 6.07E-05 | 0.00011 | 6.39E-05 | 0 | 0 | 7.43E-07 | 0.00012 | 0.00014 | 1.82E-05 | 0.00014 | 1.83E-05 |
| Elf1 | 0.00012 | 0.00022 | 1.03E-05 | 0 | 4.41E-05 | 0.00011 | 8.72E-05 | 0 | 0.00011 | 0.0002 | 0.0001 | 0.00014 | 6.69E-05 | 0 | 0 | 8.61E-05 | 0.00027 | 0.00012 |
| Slc25a17 | 0.00016 | 7.01E-05 | 0.00016 | 8.58E-06 | 8.62E-05 | 0.00014 | 8.92E-05 | 6.30E-05 | 0.00011 | 9.63E-05 | 9.52E-05 | 5.94E-05 | 6.61E-05 | 5.83E-05 | 2.40E-05 | 0.00012 | 9.15E-05 | 0.00011 |
| Tmem243 | 5.66E-05 | 4.12E-05 | 0 | 0 | 1.87E-05 | 5.21E-06 | 1.32E-05 | 0 | 0.00011 | 9.97E-07 | 0 | 1.65E-06 | 0.00017 | 1.58E-05 | 0 | 1.36E-05 | 2.89E-05 | 0 |
| Fam104a | 0.0001 | 0.00013 | 0.00022 | 0.0002 | 9.07E-05 | 9.42E-05 | 4.81E-06 | 6.68E-05 | 0.00011 | 4.00E-05 | 0.00011 | 0.0001 | 3.82E-05 | 7.38E-05 | 0 | 0.00012 | 0 | 8.16E-05 |
| Fech | 0.00025 | 0.00011 | 4.58E-05 | 0.00016 | 0.00015 | 0.00023 | 0.00019 | 4.02E-05 | 0.00011 | 0.00018 | 0.00012 | 0 | 0.00015 | 9.40E-05 | 4.96E-05 | 0.00011 | 0 | 0.00015 |
| Mon2 | 0.00016 | 0.00018 | 0.00015 | 0.00013 | 0.00016 | 9.88E-05 | 0.00029 | 8.34E-05 | 0.00011 | 9.72E-05 | 5.11E-05 | 0.00011 | 6.26E-05 | 6.89E-05 | 0.00027 | 2.53E-06 | 0.00015 | 6.86E-05 |
| Phf6 | 8.51E-05 | 1.11E-05 | 3.84E-06 | 0 | 6.00E-05 | 3.27E-06 | 4.07E-05 | 0 | 0.00011 | 2.15E-06 | 2.72E-05 | 3.54E-06 | 0 | 0 | 0 | 0.00013 | 3.21E-05 | 1.67E-05 |
| Jade2 | 0 | 0.00011 | 0 | 7.37E-06 | 1.33E-05 | 3.46E-05 | 0 | 0 | 0.00011 | 3.98E-05 | 8.55E-07 | 0 | 9.33E-05 | 4.54E-05 | 1.39E-05 | 4.84E-05 | 0 | 0.00012 |
| Txndc12 | 0.00025 | 8.26E-05 | 6.43E-05 | 5.96E-05 | 0.00013 | 0.00011 | 7.15E-05 | 5.14E-05 | 0.00011 | 4.77E-05 | 0 | 0.00014 | 0.00016 | 1.86E-05 | 0 | 0.00028 | 0 | 9.48E-05 |
| Sde2 | 8.36E-05 | 0.00012 | 7.24E-05 | 3.31E-05 | 4.94E-05 | 4.10E-05 | 0 | 5.20E-06 | 0.00011 | 1.52E-05 | 0 | 0.00012 | 0 | 5.28E-05 | 0.00021 | 2.55E-06 | 0.00046 | 5.70E-05 |
| Thoc7 | 0.00025 | 0.00013 | 0.00014 | 0.00023 | 0.00011 | 0.00015 | 3.97E-05 | 3.27E-05 | 0.00011 | 7.01E-05 | 0.00014 | 0.00018 | 9.88E-05 | 0.00013 | 0.00016 | 0.00031 | 2.49E-05 | 9.95E-05 |
| Rce1 | 5.90E-05 | 0.00011 | 0.00018 | 0.00023 | 4.32E-05 | 0.00011 | 0.0001 | 0.00012 | 0.00011 | 2.23E-05 | 0.00014 | 4.74E-05 | 8.45E-06 | 0.00014 | 1.63E-05 | 0.00013 | 1.16E-05 | 2.84E-05 |
| Usp45 | 9.39E-05 | 8.38E-05 | 5.07E-05 | 5.30E-05 | 7.19E-05 | 5.43E-05 | 4.39E-05 | 4.21E-05 | 0.00011 | 1.13E-05 | 4.22E-05 | 8.56E-08 | 2.28E-05 | 2.26E-05 | 7.65E-05 | 7.31E-05 | 9.32E-05 | 1.46E-05 |
| Ptger4 | 6.40E-05 | 0 | 8.44E-06 | 2.18E-05 | 8.00E-05 | 4.84E-05 | 2.75E-05 | 0.00011 | 0.00011 | 0 | 3.82E-05 | 0 | 0 | 0.00014 | 3.00E-05 | 7.76E-05 | 8.10E-06 | 5.09E-05 |
| Fam32a | 0.00018 | 0.00029 | 9.45E-05 | 0.00025 | 0.00023 | 0.0002 | 0.00013 | 0.00018 | 0.00011 | 0.00019 | 0.00017 | 0.0001 | 0.00013 | 0.00016 | 0 | 6.36E-05 | 0 | 0.00011 |
| Wdr11 | 6.88E-05 | 8.67E-05 | 4.89E-05 | 3.67E-06 | 5.07E-05 | 3.36E-05 | 4.62E-05 | 5.08E-05 | 0.00011 | 4.68E-05 | 1.28E-05 | 3.68E-05 | 7.97E-06 | 3.34E-05 | 4.10E-05 | 3.61E-05 | 2.43E-05 | 5.64E-05 |

| | | | | | | | | | | | | | | | | | | |
|---|---|---|---|---|---|---|---|---|---|---|---|---|---|---|---|---|---|---|
| Sap30bp | 0.00017 | 0.00012 | 0.00011 | 4.98E-05 | 7.82E-05 | 2.03E-05 | 1.86E-05 | 1.15E-05 | 0.00011 | 1.42E-05 | 4.84E-05 | 7.60E-05 | 9.76E-06 | 5.34E-06 | 8.92E-05 | 9.58E-05 | 0 | 4.90E-06 |
| Ubfd1 | 0.00033 | 9.96E-05 | 0.00023 | 0.00014 | 0.00011 | 8.48E-05 | 0.0003 | 0.00011 | 0.00011 | 4.74E-05 | 3.38E-05 | 0 | 9.11E-05 | 8.08E-06 | 0.0001 | 1.19E-05 | 3.15E-05 | 8.45E-06 |
| Ubxn6 | 8.13E-05 | 0.00017 | 0 | 0.00017 | 4.87E-05 | 0.00019 | 1.72E-05 | 0.00012 | 0.00011 | 4.99E-05 | 5.40E-05 | 0.00014 | 4.55E-05 | 0.0001 | 2.65E-05 | 0 | 0 | 3.89E-05 |
| Map2k7 | 0.00011 | 0.00017 | 8.23E-05 | 7.50E-05 | 7.04E-05 | 3.50E-05 | 0.0001 | 6.92E-05 | 0.00011 | 3.82E-05 | 7.70E-05 | 0.00015 | 4.01E-05 | 8.03E-05 | 0.00031 | 2.10E-05 | 8.74E-05 | 2.49E-05 |
| Epas1 | 0 | 0.00013 | 0 | 0.00021 | 0.00025 | 0.00116 | 0 | 0.00014 | 0.00011 | 0 | 0 | 0.00021 | 0.0008 | 0.00069 | 6.07E-05 | 0 | 0 | 4.18E-06 |
| Wipi2 | 8.89E-05 | 0.00019 | 6.03E-05 | 8.25E-05 | 8.03E-05 | 0.00012 | 5.65E-05 | 0 | 0.00011 | 4.96E-05 | 7.31E-05 | 0.00013 | 0.00012 | 0.00014 | 0.00015 | 7.10E-05 | 0.00019 | 4.49E-05 |
| Sec24d | 0.00037 | 4.63E-05 | 0.00026 | 0.00027 | 0.00015 | 0.00024 | 0.00056 | 0.00018 | 0.00011 | 0.00012 | 0.00022 | 0.00016 | 0.0001 | 0.00024 | 0.00012 | 5.43E-05 | 8.84E-06 | 0.00012 |
| Kdm3b | 3.01E-05 | 0.00025 | 2.10E-05 | 0.00024 | 9.89E-05 | 7.26E-05 | 1.42E-05 | 5.59E-05 | 0.00011 | 0.00014 | 3.20E-05 | 0.00021 | 1.98E-05 | 9.53E-05 | 0.00029 | 0.00017 | 2.93E-05 | 9.68E-05 |
| Ints1 | 6.41E-05 | 9.14E-05 | 0.00012 | 0.00017 | 3.98E-05 | 1.02E-06 | 2.39E-05 | 5.13E-05 | 0.00011 | 6.96E-05 | 6.11E-05 | 1.96E-05 | 0 | 3.22E-05 | 0.00017 | 9.86E-05 | 2.32E-06 | 6.31E-05 |
| Lsm14b | 0.00022 | 0.00023 | 0.00024 | 0.00016 | 0.00015 | 0.00018 | 4.96E-07 | 0.0001 | 0.00011 | 8.90E-05 | 9.67E-05 | 0.00032 | 3.67E-05 | 0.00015 | 0.00022 | 0.00024 | 7.56E-05 | 1.21E-05 |
| Nelfe | 6.50E-05 | 8.00E-05 | 0.0001 | 0.00013 | 3.12E-05 | 8.18E-05 | 2.80E-05 | 6.48E-05 | 0.0001 | 5.63E-06 | 6.73E-05 | 2.53E-05 | 2.59E-05 | 4.56E-05 | 2.27E-05 | 0.00027 | 0 | 5.79E-05 |
| Kmt5a | 0.00021 | 0.00012 | 0.00033 | 2.76E-05 | 0 | 3.12E-05 | 5.25E-06 | 0 | 0.0001 | 8.32E-05 | 0.00022 | 0 | 8.56E-05 | 1.06E-05 | 0.00036 | 0.0003 | 3.95E-05 | 0 |
| Tgif2 | 0 | 1.85E-05 | 5.02E-05 | 0 | 3.75E-05 | 0 | 0 | 7.79E-05 | 0.0001 | 0 | 6.13E-05 | 0 | 0 | 5.57E-05 | 1.07E-05 | 1.67E-05 | 0.00017 | 3.37E-05 |
| Zer1 | 7.55E-05 | 0.00015 | 0 | 2.60E-05 | 8.13E-05 | 0.00012 | 5.15E-05 | 4.74E-05 | 0.0001 | 3.76E-05 | 1.87E-05 | 3.40E-05 | 3.62E-06 | 9.27E-05 | 7.67E-05 | 1.08E-05 | 0 | 5.97E-05 |
| Hspa12b | 0 | 5.30E-05 | 0 | 2.18E-05 | 5.72E-06 | 0 | 0 | 2.57E-05 | 0.0001 | 0.00023 | 0 | 0 | 0 | 4.11E-05 | 4.65E-05 | 0 | 0 | 5.39E-05 |
| Spty2d1 | 0.00018 | 9.79E-05 | 9.46E-05 | 6.90E-05 | 4.92E-05 | 0 | 4.74E-05 | 0.00013 | 0.0001 | 8.39E-05 | 4.34E-05 | 0.00015 | 1.05E-05 | 0 | 0.00034 | 0 | 0.00099 | 2.55E-06 |
| Hadha | 0.00021 | 0.00015 | 0.00012 | 0.00027 | 7.99E-05 | 0.00012 | 0.00019 | 0.00014 | 0.0001 | 8.38E-05 | 6.35E-05 | 3.08E-05 | 8.11E-05 | 0.00015 | 0 | 0.0001 | 0 | 0.00014 |
| Erp29 | 0.00041 | 4.83E-05 | 0.00024 | 0.00042 | 0.00021 | 0.00028 | 0.0004 | 0.00031 | 0.0001 | 0.0002 | 0.00012 | 0.00031 | 0.00031 | 0.00019 | 9.70E-05 | 0.00024 | 9.75E-05 | 0.00034 |
| Asxl1 | 0.00012 | 0.00015 | 1.27E-06 | 0 | 7.36E-05 | 0.00016 | 9.73E-05 | 0 | 0.0001 | 1.95E-05 | 0.00012 | 0.00014 | 5.28E-05 | 6.66E-05 | 0.00023 | 0.00019 | 0.0001 | 3.52E-05 |
| Phf23 | 0.00012 | 0.00027 | 0.00027 | 0.00017 | 7.03E-05 | 7.65E-05 | 5.14E-05 | 7.64E-05 | 0.0001 | 6.79E-05 | 0.00013 | 3.62E-05 | 7.34E-05 | 0.00014 | 0.00026 | 0.0001 | 0.00011 | 6.39E-05 |
| Cd47 | 0.00047 | 0.00023 | 0.00016 | 6.65E-07 | 0.00043 | 0.0009 | 0.0005 | 0 | 0.0001 | 0.00066 | 0 | 0.00071 | 0.00061 | 0.00071 | 0 | 9.88E-05 | 2.79E-05 | 0.0005 |
| Pdhb | 0.00021 | 0.00014 | 6.72E-05 | 5.24E-05 | 2.80E-05 | 5.18E-05 | 0 | 4.71E-05 | 0.0001 | 3.39E-05 | 2.67E-05 | 7.91E-05 | 0 | 3.70E-05 | 0.00012 | 5.90E-05 | 0.00014 | 3.62E-05 |
| Hspb8 | 0.0001 | 0.0002 | 0.00029 | 0 | 0.00012 | 7.27E-06 | 0.00021 | 0 | 0.0001 | 0.00038 | 9.52E-05 | 0.00022 | 5.40E-05 | 0 | 0 | 1.53E-05 | 0.00058 | 0.00019 |
| Tnfrsf11b | 0 | 0 | 3.73E-05 | 0 | 0 | 0.00025 | 0 | 0 | 0.0001 | 0.00022 | 0.00021 | 0.00018 | 0 | 0.00088 | 0 | 0 | 0 | 4.36E-05 |
| Aprt | 6.21E-05 | 5.87E-05 | 9.19E-05 | 0.00019 | 4.03E-05 | 7.02E-05 | 0 | 5.57E-05 | 0.0001 | 0 | 5.28E-05 | 7.37E-05 | 3.70E-05 | 6.50E-05 | 3.09E-05 | 0 | 0.00017 | 6.34E-05 |
| Echdc1 | 4.49E-05 | 4.45E-05 | 1.11E-06 | 5.37E-05 | 1.26E-05 | 0 | 1.60E-06 | 2.70E-05 | 0.0001 | 2.17E-05 | 2.28E-05 | 1.81E-05 | 4.02E-05 | 2.01E-07 | 0 | 8.05E-06 | 0 | 3.90E-05 |
| Fbrs | 5.20E-05 | 0.00025 | 0.00016 | 0.00037 | 8.11E-05 | 2.13E-05 | 1.66E-05 | 0.0001 | 0.0001 | 8.95E-05 | 7.94E-05 | 9.28E-05 | 2.29E-05 | 0.00011 | 0.00036 | 4.51E-05 | 0.00048 | 9.29E-05 |
| Hnrnpll | 0.00034 | 0.00013 | 0.00016 | 0 | 7.06E-05 | 0.00021 | 0 | 0 | 0.0001 | 5.62E-07 | 4.33E-05 | 0.00015 | 0 | 6.74E-05 | 4.00E-05 | 0.00019 | 0.00027 | 0 |
| Myc | 0.00013 | 0 | 0.00036 | 9.29E-05 | 0 | 0 | 0 | 0 | 0.0001 | 9.30E-05 | 0 | 0.00026 | 0.00036 | 0 | 9.53E-05 | 6.23E-05 | 0.00113 | 0 |
| Emd | 0.00024 | 0.00014 | 0.0002 | 5.25E-05 | 0 | 1.70E-05 | 0 | 0 | 0.0001 | 2.57E-05 | 0.00026 | 6.53E-05 | 0 | 0 | 3.04E-05 | 0.00024 | 0.00014 | 4.45E-05 |
| Zfp771 | 2.75E-05 | 7.31E-05 | 4.76E-05 | 0.00014 | 8.30E-05 | 5.09E-05 | 5.06E-05 | 0.00013 | 0.0001 | 4.91E-05 | 6.25E-05 | 6.30E-05 | 7.29E-05 | 6.32E-05 | 0.00012 | 5.37E-05 | 0 | 4.13E-05 |
| Vps13c | 9.81E-05 | 0.00022 | 9.03E-06 | 0 | 9.40E-05 | 0.00023 | 6.57E-05 | 3.27E-05 | 0.0001 | 5.70E-05 | 0 | 0 | 0.0002 | 3.06E-05 | 2.37E-05 | 0 | 0 | 1.97E-05 |
| Pip5k1c | 2.19E-05 | 0.00023 | 6.67E-05 | 0.00023 | 0.00014 | 8.48E-05 | 0 | 8.19E-05 | 0.0001 | 9.85E-05 | 0.00017 | 5.15E-05 | 0 | 0.00023 | 0.00017 | 0.00011 | 0.00014 | 0.00014 |

| | | | | | | | | | | | | | | | | | | |
|---|---|---|---|---|---|---|---|---|---|---|---|---|---|---|---|---|---|---|
| Larp6 | 0 | 0.00012 | 0 | 9.20E-06 | 8.37E-05 | 8.78E-05 | 0 | 0 | 0.0001 | 0.00014 | 0 | 7.97E-05 | 0.00014 | 9.77E-05 | 8.89E-05 | 0 | 0 | 9.59E-06 |
| Mccc1 | 5.37E-05 | 5.62E-05 | 9.03E-06 | 3.15E-05 | 0 | 1.39E-05 | 5.12E-05 | 0 | 0.0001 | 3.29E-06 | 4.03E-05 | 1.03E-05 | 5.89E-05 | 0 | 3.18E-05 | 0 | 0 | 2.81E-05 |
| Nfyc | 0.00015 | 6.07E-05 | 9.87E-05 | 9.91E-05 | 8.34E-06 | 5.20E-06 | 1.92E-05 | 6.02E-05 | 0.0001 | 2.15E-05 | 4.80E-05 | 1.82E-05 | 0 | 3.93E-05 | 2.36E-05 | 0.00013 | 4.31E-05 | 3.33E-05 |
| Asb8 | 8.31E-05 | 7.81E-05 | 0.00012 | 0.00013 | 0.0001 | 6.79E-05 | 4.72E-05 | 5.80E-05 | 0.0001 | 7.28E-05 | 0.00011 | 0 | 4.07E-05 | 9.88E-05 | 5.11E-06 | 3.48E-05 | 0 | 6.90E-05 |
| Elof1 | 8.11E-05 | 6.44E-05 | 9.04E-05 | 0.00011 | 6.00E-05 | 2.95E-05 | 3.44E-05 | 8.26E-05 | 0.0001 | 6.36E-05 | 7.19E-05 | 1.92E-05 | 0 | 6.18E-05 | 0 | 0.00017 | 0 | 7.34E-06 |
| Dld | 0.00036 | 0.00013 | 0.0002 | 1.38E-05 | 5.81E-05 | 0.00014 | 1.65E-06 | 0 | 0.0001 | 2.98E-05 | 4.62E-05 | 3.49E-05 | 3.85E-05 | 1.63E-05 | 0.00011 | 0.00015 | 1.53E-05 | 0 |
| Sacm1l | 0.00026 | 0.00012 | 0.00011 | 0 | 8.03E-05 | 0.00014 | 1.40E-05 | 0 | 0.0001 | 4.47E-05 | 8.62E-05 | 0.00018 | 7.50E-06 | 1.02E-05 | 6.56E-05 | 9.87E-05 | 0.00012 | 7.30E-05 |
| Nup62 | 8.10E-05 | 0.00015 | 0.00024 | 6.02E-05 | 3.62E-05 | 5.00E-05 | 0 | 0 | 0.0001 | 8.45E-05 | 0.00013 | 0.00014 | 0 | 4.38E-05 | 9.94E-05 | 0.00033 | 0 | 4.06E-05 |
| Crtc3 | 5.03E-05 | 0.0002 | 3.23E-05 | 8.89E-05 | 4.10E-05 | 0.00011 | 0 | 0 | 0.0001 | 9.00E-05 | 0.00011 | 0.00013 | 1.38E-05 | 4.03E-05 | 0.00011 | 6.87E-05 | 0.00022 | 0.0001 |
| Mark4 | 9.76E-05 | 0.00015 | 0.00014 | 9.94E-05 | 0.00011 | 5.64E-05 | 5.97E-05 | 7.12E-05 | 0.0001 | 5.60E-05 | 0.00016 | 0.00012 | 0.0001 | 0.00011 | 0.00015 | 0 | 0.00014 | 2.59E-05 |
| Atp5g1 | 0.00019 | 0 | 0.00023 | 0.0004 | 1.93E-05 | 0 | 7.29E-05 | 0.00012 | 0.0001 | 4.31E-06 | 9.90E-05 | 0 | 9.82E-06 | 6.58E-05 | 0 | 7.26E-05 | 0 | 8.04E-05 |
| Gngt2 | 2.63E-06 | 8.82E-05 | 0 | 1.31E-05 | 9.93E-05 | 0.00012 | 0 | 9.69E-05 | 0.0001 | 0.00026 | 2.19E-05 | 0.00011 | 5.11E-05 | 0.00014 | 4.91E-05 | 1.26E-05 | 0 | 4.69E-05 |
| Rbpj | 0.00027 | 0.00017 | 0.00012 | 0 | 5.90E-05 | 0.00018 | 9.80E-06 | 0 | 0.0001 | 0.00026 | 0.00015 | 0.00059 | 0.00013 | 7.77E-06 | 0 | 0.0001 | 5.53E-05 | 0 |
| Ints3 | 8.74E-05 | 0.00013 | 0.0001 | 0.00015 | 5.97E-05 | 9.07E-05 | 4.91E-05 | 3.10E-05 | 0.0001 | 0.00015 | 6.35E-05 | 5.35E-05 | 0 | 6.55E-06 | 1.83E-05 | 0.00018 | 8.63E-05 | 3.62E-05 |
| Isg20l2 | 0.00012 | 9.29E-05 | 0.00013 | 7.39E-06 | 4.05E-05 | 1.34E-05 | 1.03E-05 | 0 | 0.0001 | 0 | 3.46E-05 | 6.19E-06 | 6.55E-06 | 2.07E-05 | 8.62E-05 | 4.61E-05 | 0 | 0 |
| Ndufa11 | 0.00025 | 9.14E-05 | 0.00014 | 0.00024 | 0.0001 | 0.00011 | 1.83E-05 | 5.30E-05 | 0.0001 | 5.22E-05 | 0.00015 | 0 | 6.92E-05 | 8.21E-05 | 0 | 0.00011 | 0 | 0.00012 |
| Prkx | 1.55E-05 | 8.35E-05 | 2.38E-05 | 0 | 0 | 0.00012 | 0 | 0.00012 | 0.0001 | 3.02E-05 | 3.07E-05 | 5.08E-05 | 0 | 9.45E-06 | 1.48E-05 | 0 | 0.00011 | 7.44E-05 |
| Plrg1 | 0.00014 | 0.00013 | 0.00013 | 8.62E-05 | 0.00012 | 3.92E-05 | 0 | 5.94E-05 | 0.0001 | 2.17E-05 | 6.75E-05 | 8.37E-05 | 0 | 2.38E-06 | 0 | 0.00018 | 0 | 1.91E-05 |
| Tnfrsf26 | 0.00014 | 6.69E-05 | 0.00015 | 0 | 0.00045 | 0.00065 | 0 | 0.00049 | 0.0001 | 0 | 0.00016 | 0.00025 | 0 | 0.00041 | 0 | 1.49E-05 | 0 | 0 |
| Pigc | 4.77E-05 | 0.0002 | 2.61E-05 | 0.00011 | 0.00011 | 0.0001 | 0.00024 | 8.25E-05 | 0.0001 | 3.73E-05 | 2.29E-05 | 1.84E-05 | 0 | 6.19E-05 | 0 | 8.21E-05 | 0 | 9.18E-05 |
| Fkbp5 | 8.28E-05 | 2.47E-05 | 0.00012 | 2.96E-05 | 5.31E-05 | 0 | 6.90E-05 | 1.04E-05 | 0.0001 | 0 | 0.00019 | 3.04E-05 | 0.00023 | 0.00014 | 4.80E-05 | 0.00028 | 4.53E-05 | 0.00026 |
| Zfpl1 | 0.00019 | 7.22E-05 | 0.00014 | 0.00015 | 5.26E-05 | 4.70E-05 | 0.00018 | 0.00011 | 0.0001 | 4.37E-05 | 9.88E-05 | 6.81E-05 | 5.11E-05 | 9.37E-05 | 0.00011 | 0 | 0 | 2.75E-05 |
| Ccdc124 | 0.0002 | 0.00015 | 0.00027 | 0.00052 | 0.00012 | 0.00011 | 2.88E-05 | 0.00017 | 0.0001 | 5.78E-05 | 0.00024 | 0 | 0 | 0.00014 | 0.00022 | 0.00019 | 0 | 0.00011 |
| Slc35a2 | 0.00013 | 0 | 0.00015 | 0.00029 | 5.60E-05 | 9.09E-05 | 0.00028 | 0.00012 | 0.0001 | 3.92E-05 | 7.09E-05 | 0.00012 | 7.52E-05 | 7.61E-05 | 0.00011 | 0 | 0.00013 | 7.05E-05 |
| Aebp2 | 0.00016 | 0.00016 | 0.00023 | 0 | 5.49E-05 | 9.40E-05 | 7.73E-05 | 0 | 0.0001 | 0.00012 | 0.00021 | 0.00053 | 0.00024 | 6.00E-05 | 0.00018 | 0.00019 | 0.00024 | 0 |
| Ubxn8 | 0.00014 | 7.54E-05 | 0.00011 | 0.00013 | 4.01E-05 | 0.00014 | 9.49E-05 | 0.00018 | 0.0001 | 1.03E-05 | 0 | 1.50E-05 | 6.78E-05 | 0.00016 | 0 | 6.16E-05 | 0 | 4.82E-05 |
| Ulk2 | 0.00016 | 0.00011 | 3.50E-05 | 3.37E-05 | 0.00012 | 0.00018 | 0.00017 | 7.43E-05 | 0.0001 | 8.05E-05 | 0.00011 | 0.00013 | 2.94E-05 | 0.0002 | 0 | 7.23E-05 | 4.74E-05 | 8.71E-05 |
| Vav3 | 0 | 4.19E-05 | 1.89E-05 | 0 | 0 | 2.29E-05 | 4.56E-05 | 0 | 0.0001 | 0 | 0 | 2.21E-05 | 5.03E-06 | 0 | 8.99E-06 | 0 | 7.17E-06 | 0 |
| Ralgapb | 0.00011 | 0.00012 | 9.44E-05 | 9.18E-06 | 7.76E-05 | 9.12E-05 | 0.00011 | 4.93E-05 | 0.0001 | 5.38E-05 | 0.00014 | 1.79E-05 | 3.34E-06 | 8.46E-05 | 4.60E-05 | 9.82E-05 | 8.47E-06 | 7.79E-05 |
| Ebna1bp2 | 0.00022 | 0.00011 | 0.00051 | 0.00028 | 2.20E-05 | 0 | 0 | 0 | 0.0001 | 1.75E-05 | 0.00011 | 4.90E-05 | 2.05E-05 | 5.89E-05 | 5.57E-05 | 0.00013 | 4.87E-05 | 0 |
| Pcgf2 | 6.72E-05 | 0.00014 | 5.36E-05 | 9.66E-05 | 0.00011 | 2.44E-05 | 6.18E-05 | 8.77E-06 | 0.0001 | 1.43E-05 | 5.38E-05 | 0.00017 | 4.38E-05 | 2.70E-05 | 0.00023 | 4.06E-05 | 6.40E-05 | 6.01E-05 |
| Cenpx | 5.95E-05 | 2.64E-05 | 2.54E-05 | 0.00022 | 3.56E-05 | 2.83E-05 | 7.31E-05 | 7.88E-05 | 0.0001 | 8.58E-05 | 6.22E-05 | 4.25E-05 | 3.54E-05 | 8.39E-05 | 0 | 0.00025 | 7.97E-05 | 7.94E-05 |
| Kcnip3 | 5.51E-06 | 0 | 2.30E-05 | 0 | 0 | 3.80E-05 | 8.23E-06 | 5.39E-05 | 0.0001 | 0 | 0 | 0 | 0 | 0 | 0 | 2.31E-05 | 0 | 2.61E-05 |

| | | | | | | | | | | | | | | | | | | |
|---|---|---|---|---|---|---|---|---|---|---|---|---|---|---|---|---|---|---|
| Tbx2 | 0 | 8.63E-05 | 0 | 0 | 3.02E-05 | 0.00014 | 0 | 0.00024 | 0.0001 | 0 | 0 | 5.12E-05 | 0 | 0 | 0 | 0 | 0 | 0.00011 |
| Gucd1 | 8.43E-05 | 9.06E-05 | 6.89E-05 | 9.90E-05 | 4.25E-05 | 5.61E-05 | 7.01E-05 | 6.60E-05 | 0.0001 | 6.78E-05 | 5.07E-05 | 1.93E-05 | 2.89E-05 | 1.47E-05 | 0 | 7.70E-05 | 0 | 2.25E-05 |
| Cluh | 0.00015 | 7.98E-05 | 0.0003 | 6.58E-05 | 1.14E-05 | 0 | 1.32E-05 | 0 | 0.0001 | 0 | 5.25E-05 | 0 | 8.55E-05 | 1.08E-05 | 0.00013 | 4.24E-05 | 7.82E-05 | 0 |
| Ddx42 | 0.00017 | 0.00018 | 0.00014 | 4.46E-05 | 7.51E-05 | 5.68E-05 | 0 | 2.99E-05 | 0.0001 | 3.75E-05 | 5.70E-05 | 0.00013 | 0 | 0 | 0.00025 | 0.0001 | 0 | 2.93E-05 |
| Tmem185b | 6.93E-05 | 7.32E-05 | 0.00018 | 1.17E-05 | 7.40E-05 | 3.70E-05 | 0 | 0.00012 | 0.0001 | 2.01E-05 | 8.73E-05 | 8.20E-05 | 3.82E-05 | 6.76E-05 | 7.24E-05 | 0 | 1.39E-05 | 3.91E-05 |
| Rxra | 5.60E-05 | 0.00024 | 6.13E-05 | 0.00022 | 0.0001 | 0.00022 | 2.46E-05 | 2.07E-05 | 0.0001 | 0.00014 | 3.28E-05 | 2.48E-05 | 0.00014 | 0.00018 | 6.77E-05 | 9.97E-05 | 4.48E-05 | 0.00021 |
| Ergic3 | 0 | 0.00014 | 7.36E-05 | 0.00056 | 0.00022 | 0.00027 | 0.00015 | 0.00031 | 0.0001 | 0.00021 | 0.00014 | 0.00026 | 0.00038 | 0.00028 | 0 | 3.50E-06 | 0 | 0.0004 |
| Mfsd14a | 9.69E-05 | 6.36E-05 | 8.25E-05 | 7.34E-06 | 6.25E-05 | 5.45E-05 | 0 | 6.21E-05 | 0.0001 | 1.53E-05 | 5.31E-05 | 5.67E-05 | 0 | 5.42E-05 | 4.82E-05 | 1.84E-05 | 0.00015 | 2.01E-05 |
| Gtf3c5 | 8.72E-05 | 5.69E-05 | 5.82E-05 | 2.34E-05 | 3.78E-05 | 1.47E-05 | 1.70E-05 | 0 | 0.0001 | 4.38E-05 | 4.42E-06 | 5.69E-05 | 6.56E-06 | 7.20E-05 | 6.37E-05 | 0.00019 | 6.91E-05 | 2.24E-05 |
| Ip6k1 | 0.00012 | 0.00048 | 0 | 0.00031 | 0.00024 | 0.00023 | 0.00015 | 0.00015 | 0.0001 | 0.00017 | 0.00013 | 0.00022 | 7.97E-05 | 0.00024 | 0.00051 | 4.87E-05 | 0.00021 | 0.00021 |
| Tmem160 | 6.53E-05 | 5.22E-05 | 9.57E-05 | 0.00034 | 0.00011 | 4.27E-05 | 5.51E-05 | 3.71E-05 | 0.0001 | 0.0001 | 0.0001 | 1.11E-05 | 4.65E-05 | 0.00016 | 0 | 3.21E-05 | 1.46E-05 | 0.00021 |
| Sem1 | 0.00014 | 0.00012 | 0.00018 | 0.00081 | 0.0001 | 0.00014 | 0.00013 | 0.0003 | 0.0001 | 0.00021 | 0.00021 | 9.09E-05 | 6.10E-05 | 0.00026 | 0.00031 | 0.00013 | 0.00017 | 0.0002 |
| Psme2b | 6.85E-05 | 6.05E-05 | 4.46E-05 | 0.00015 | 5.34E-05 | 0 | 0 | 8.50E-05 | 0.0001 | 8.39E-05 | 8.78E-05 | 0 | 1.90E-05 | 4.91E-05 | 0 | 4.94E-05 | 0 | 7.81E-05 |
| Arhgap21 | 0.00011 | 8.33E-05 | 0.00012 | 0 | 3.25E-05 | 1.72E-05 | 4.55E-05 | 0.0002 | 0.0001 | 0.0003 | 0 | 5.56E-05 | 0 | 0 | 0.00017 | 0.00013 | 0.00026 | 0.00026 |
| Pfn2 | 5.40E-05 | 8.76E-05 | 0 | 2.16E-05 | 0.00033 | 0 | 0 | 0 | 0.0001 | 6.20E-05 | 6.95E-05 | 3.33E-05 | 0 | 0 | 0 | 4.80E-07 | 0 | 6.79E-05 |
| Tufm | 0.00014 | 8.23E-05 | 0.00023 | 0.00032 | 3.11E-05 | 3.44E-06 | 2.39E-05 | 4.60E-05 | 0.0001 | 4.70E-05 | 5.61E-05 | 3.62E-06 | 0 | 8.08E-05 | 8.20E-06 | 8.81E-05 | 2.39E-05 | 6.14E-05 |
| Slc30a5 | 0.00025 | 7.55E-05 | 0.0002 | 0.00022 | 0.00012 | 0.00015 | 0.00025 | 0.00013 | 0.0001 | 4.30E-05 | 7.95E-05 | 0 | 5.92E-05 | 0.00017 | 0 | 0.00018 | 0 | 7.54E-05 |
| Icmt | 6.19E-05 | 0.00012 | 0.0002 | 0.00017 | 0.00013 | 0.00011 | 0.00014 | 0.00016 | 0.0001 | 5.95E-05 | 0.00017 | 3.25E-05 | 6.95E-05 | 8.08E-05 | 3.02E-05 | 0.00013 | 8.92E-05 | 9.25E-05 |
| Gss | 4.80E-05 | 3.82E-05 | 0.00017 | 8.59E-05 | 0 | 0.00015 | 0 | 0.00024 | 0.0001 | 9.13E-06 | 8.37E-05 | 0 | 0 | 3.14E-05 | 2.25E-05 | 5.20E-06 | 0 | 0 |
| Rgs19 | 0.00014 | 4.36E-05 | 0.00021 | 0.00015 | 3.29E-06 | 1.99E-05 | 8.09E-05 | 0 | 0.0001 | 4.63E-05 | 9.08E-05 | 1.57E-05 | 7.04E-05 | 0.00013 | 9.71E-05 | 0 | 0 | 0.00012 |
| Ten1 | 8.38E-05 | 5.98E-05 | 2.24E-05 | 7.29E-05 | 6.04E-05 | 2.23E-05 | 6.83E-06 | 4.16E-05 | 0.0001 | 1.37E-05 | 1.38E-05 | 3.06E-05 | 0 | 2.62E-05 | 0 | 2.20E-05 | 0 | 2.43E-05 |
| Sppl2a | 0.00022 | 9.76E-05 | 9.07E-05 | 3.86E-05 | 6.91E-05 | 0.00013 | 0.00019 | 0.00013 | 0.0001 | 0.00026 | 0.00012 | 4.65E-05 | 0.00011 | 9.16E-05 | 0 | 0.00011 | 2.77E-05 | 0.00019 |
| Ric8b | 0.00011 | 0.00015 | 5.37E-05 | 0 | 0.00015 | 3.26E-05 | 3.11E-05 | 6.19E-05 | 0.0001 | 3.47E-05 | 7.22E-05 | 4.47E-05 | 0 | 3.40E-05 | 0.00013 | 4.54E-05 | 0 | 3.42E-05 |
| Dnttip1 | 4.21E-05 | 0.00013 | 0.00012 | 0.00021 | 6.47E-05 | 2.26E-05 | 5.82E-05 | 3.87E-05 | 0.0001 | 5.61E-05 | 3.36E-05 | 4.03E-05 | 1.61E-05 | 6.48E-05 | 0.00014 | 9.27E-05 | 4.22E-05 | 5.66E-05 |
| Sccpdh | 0.00019 | 7.67E-05 | 7.37E-05 | 0 | 6.29E-05 | 4.94E-05 | 2.54E-06 | 8.13E-06 | 0.0001 | 0 | 5.83E-05 | 0.00027 | 2.52E-05 | 3.83E-05 | 3.81E-05 | 3.83E-05 | 0 | 7.03E-05 |
| BC005537 | 0.00039 | 0.00032 | 0.00024 | 0.00028 | 0.00019 | 0.00037 | 0.00032 | 0.00022 | 0.0001 | 0.00032 | 0.00029 | 0.00054 | 0.00015 | 0.00021 | 0 | 0.00027 | 0.001 | 0.00023 |
| Srpr | 0.00025 | 8.70E-05 | 0.00037 | 0.00036 | 0.00019 | 0.0002 | 0.00041 | 0.00033 | 0.0001 | 6.82E-05 | 9.95E-05 | 0.0001 | 0.00018 | 0.00023 | 0.00013 | 2.57E-05 | 8.54E-05 | 3.98E-05 |
| Cyp4b1 | 0 | 0 | 0 | 0 | 0 | 6.29E-05 | 0 | 0 | 0.0001 | 0.00048 | 0 | 0 | 0 | 0.0007 | 0 | 0 | 0 | 0 |
| Fbxw2 | 6.68E-05 | 0.00016 | 5.67E-05 | 0.00013 | 0.0001 | 7.28E-05 | 9.12E-05 | 6.33E-05 | 0.0001 | 0.00011 | 0.00013 | 6.46E-05 | 5.57E-05 | 0.00012 | 0 | 0.00016 | 0 | 0.00012 |
| Ap1g1 | 0.00027 | 0.00018 | 0.00023 | 7.67E-05 | 0.00014 | 0.00017 | 0.00018 | 0.00012 | 0.0001 | 9.69E-05 | 0.00017 | 0.00012 | 0.00011 | 3.77E-05 | 0.00014 | 8.57E-05 | 0.00011 | 5.52E-05 |
| Ddx19a | 0.00014 | 0.00013 | 0.00015 | 8.37E-05 | 5.82E-05 | 2.32E-05 | 4.78E-05 | 4.90E-05 | 0.0001 | 2.25E-05 | 1.55E-05 | 8.46E-05 | 0 | 8.19E-05 | 0.00013 | 0.00019 | 0 | 4.16E-05 |
| Ythdf1 | 0.00019 | 0.00013 | 0.00019 | 2.91E-06 | 8.25E-05 | 0.00017 | 0.00011 | 4.24E-05 | 0.0001 | 6.21E-05 | 7.42E-05 | 7.60E-05 | 3.06E-05 | 5.83E-05 | 9.30E-05 | 0.00013 | 5.14E-05 | 4.28E-05 |
| Tyro3 | 9.11E-07 | 3.74E-05 | 1.47E-05 | 3.25E-05 | 4.07E-05 | 3.31E-05 | 0 | 4.80E-05 | 0.0001 | 1.06E-05 | 6.60E-05 | 0 | 0 | 3.43E-05 | 1.60E-06 | 0.00012 | 0 | 3.17E-05 |

| | | | | | | | | | | | | | | | | | | |
|---|---|---|---|---|---|---|---|---|---|---|---|---|---|---|---|---|---|---|
| Atp6v0a2 | 0.00017 | 0.00012 | 0.00024 | 0.00017 | 0.00019 | 8.33E-05 | 0.00028 | 9.58E-05 | 0.0001 | 6.28E-05 | 8.52E-05 | 0 | 0.0001 | 9.48E-05 | 9.68E-05 | 0 | 0 | 0.00014 |
| Eif2s1 | 0.00033 | 0.00011 | 0.0004 | 6.45E-05 | 1.10E-05 | 0 | 0 | 0 | 0.0001 | 1.25E-05 | 0.0001 | 5.28E-05 | 2.29E-05 | 0 | 0 | 0.00019 | 0 | 0 |
| Erc1 | 0.0002 | 0.00011 | 9.68E-05 | 0 | 5.95E-05 | 3.26E-05 | 8.68E-05 | 6.63E-06 | 0.0001 | 0 | 0.00018 | 0.00013 | 6.36E-06 | 7.08E-05 | 0.00043 | 0.00014 | 0.00032 | 5.28E-05 |
| Vamp8 | 0.00029 | 0.00017 | 0.00014 | 0.00044 | 0.0001 | 0.00017 | 0.00018 | 0.0002 | 0.0001 | 6.49E-05 | 0.00026 | 0.00022 | 1.92E-05 | 0.00019 | 2.11E-06 | 9.31E-05 | 8.22E-05 | 4.23E-05 |
| Casd1 | 7.42E-05 | 8.93E-05 | 4.63E-05 | 7.27E-05 | 9.01E-05 | 0.00012 | 1.40E-05 | 3.59E-05 | 0.0001 | 4.72E-05 | 4.11E-05 | 2.80E-05 | 0.00011 | 0.00013 | 8.85E-05 | 2.63E-05 | 2.57E-05 | 4.37E-05 |
| Scmh1 | 2.99E-05 | 9.99E-05 | 4.39E-05 | 0 | 7.84E-05 | 5.04E-05 | 0.00015 | 8.60E-05 | 0.0001 | 5.04E-05 | 2.13E-05 | 1.13E-05 | 0 | 1.15E-06 | 5.91E-05 | 8.50E-05 | 0.00013 | 6.45E-05 |
| Aspscr1 | 5.79E-05 | 0.00014 | 4.24E-05 | 0.00014 | 7.73E-05 | 5.18E-05 | 5.33E-05 | 5.63E-06 | 0.0001 | 6.21E-05 | 7.14E-06 | 0 | 5.43E-05 | 1.95E-05 | 0.0001 | 6.23E-05 | 0 | 3.69E-05 |
| Nubp2 | 1.20E-05 | 0.00012 | 0.00013 | 0.00022 | 4.91E-05 | 0 | 1.20E-05 | 5.28E-05 | 0.0001 | 6.61E-05 | 8.99E-05 | 0 | 8.13E-05 | 5.84E-05 | 0 | 9.34E-05 | 1.42E-05 | 5.98E-05 |
| Chpt1 | 0.00012 | 0.00013 | 4.75E-05 | 8.86E-05 | 0.0001 | 0.00014 | 5.33E-05 | 0 | 0.0001 | 7.94E-05 | 8.27E-05 | 0 | 0.00015 | 0.00012 | 0 | 9.59E-05 | 4.79E-06 | 0.00011 |
| Ift27 | 0.00017 | 7.52E-05 | 3.63E-05 | 0.00015 | 6.56E-05 | 2.23E-05 | 3.28E-05 | 1.59E-05 | 0.0001 | 0.00014 | 0 | 0 | 0.0001 | 0 | 0 | 0.00015 | 1.32E-05 | 0.00013 |
| Rnf14 | 0.00014 | 0.00015 | 0.00011 | 0.00012 | 0.00012 | 0.00011 | 0 | 8.49E-05 | 0.0001 | 5.27E-05 | 0.00015 | 0.00011 | 5.08E-05 | 9.14E-05 | 0.00011 | 0.0001 | 0 | 0.00011 |
| Ppa2 | 9.26E-05 | 4.32E-05 | 5.11E-05 | 4.31E-05 | 7.34E-05 | 4.58E-05 | 9.17E-05 | 5.97E-06 | 0.0001 | 2.61E-05 | 5.71E-05 | 2.72E-05 | 8.19E-06 | 2.65E-05 | 0 | 4.81E-05 | 8.38E-06 | 1.38E-05 |
| Nbdy | 6.47E-05 | 5.80E-05 | 1.03E-05 | 7.90E-05 | 7.64E-05 | 6.51E-05 | 1.60E-05 | 5.64E-05 | 0.0001 | 2.12E-05 | 3.73E-05 | 4.14E-05 | 2.03E-05 | 0 | 0 | 0 | 0 | 6.11E-05 |
| St6galnac6 | 0.00012 | 0.00013 | 8.99E-05 | 0.00027 | 0.00014 | 0.00025 | 0.00019 | 4.31E-05 | 0.0001 | 0.00014 | 0.0001 | 3.10E-05 | 0 | 8.75E-05 | 5.61E-05 | 0 | 4.87E-06 | 6.12E-05 |
| Gar1 | 0.00013 | 5.24E-05 | 0.0004 | 0.00027 | 0 | 0 | 0 | 0 | 0.0001 | 5.75E-06 | 2.71E-05 | 3.43E-05 | 6.79E-05 | 0 | 0.00014 | 0.00017 | 0.00014 | 2.35E-05 |
| Rpe | 0.00018 | 0.00011 | 9.86E-05 | 0 | 6.58E-05 | 8.20E-05 | 0 | 1.96E-06 | 0.0001 | 6.92E-05 | 8.22E-05 | 2.85E-05 | 7.59E-05 | 0 | 1.97E-05 | 6.13E-05 | 0 | 5.46E-05 |
| Acsf2 | 6.59E-05 | 8.52E-05 | 0 | 0 | 6.53E-05 | 9.57E-05 | 1.56E-05 | 0.0001 | 0.0001 | 3.43E-06 | 0 | 0 | 0 | 5.96E-05 | 1.52E-05 | 2.81E-05 | 0 | 6.08E-05 |
| Psmd5 | 0.00013 | 8.28E-05 | 0.00013 | 7.10E-06 | 7.01E-05 | 8.43E-05 | 4.48E-05 | 0 | 0.0001 | 3.90E-05 | 0.00013 | 0 | 5.48E-06 | 0 | 0 | 0.00013 | 0 | 1.80E-05 |
| Ddx56 | 0.0001 | 0.00011 | 0.00024 | 0.00016 | 7.89E-06 | 1.67E-06 | 0 | 2.39E-06 | 0.0001 | 2.84E-05 | 5.65E-05 | 1.32E-05 | 1.63E-06 | 5.94E-05 | 6.61E-05 | 9.26E-05 | 0 | 3.61E-05 |
| Setd2 | 0.00017 | 0.00017 | 7.33E-05 | 0 | 0.0001 | 9.45E-05 | 6.42E-05 | 9.61E-06 | 0.0001 | 3.76E-05 | 4.04E-05 | 8.91E-05 | 4.34E-05 | 5.74E-05 | 0.00017 | 0.00013 | 0.0001 | 6.55E-05 |
| Eif3c | 0.0003 | 0.00016 | 0.00026 | 0.00022 | 5.15E-05 | 7.21E-05 | 4.27E-05 | 1.16E-06 | 0.0001 | 3.18E-05 | 5.83E-05 | 0.00028 | 0 | 2.67E-05 | 7.51E-05 | 0.0002 | 7.76E-05 | 5.94E-05 |
| Elp2 | 0.00014 | 5.62E-05 | 6.52E-05 | 0 | 2.92E-05 | 4.61E-05 | 1.32E-05 | 1.53E-05 | 0.0001 | 2.41E-05 | 2.11E-05 | 5.17E-05 | 0 | 6.36E-06 | 4.59E-05 | 3.59E-05 | 0 | 1.93E-05 |
| Pdgfd | 2.14E-05 | 7.95E-07 | 0 | 0 | 6.13E-05 | 0 | 4.71E-06 | 0 | 0.0001 | 0.00032 | 0.00013 | 0 | 6.23E-05 | 0 | 0 | 0 | 0 | 4.01E-05 |
| Kat6a | 6.53E-05 | 0.00019 | 0.00012 | 0.00011 | 8.73E-05 | 5.79E-05 | 8.94E-05 | 7.21E-05 | 0.0001 | 8.95E-05 | 7.51E-05 | 7.70E-05 | 7.50E-05 | 5.55E-05 | 0.00016 | 0.00013 | 0.00024 | 8.49E-05 |
| Kat7 | 0.00015 | 0.00012 | 0.00011 | 0 | 4.57E-05 | 9.37E-05 | 0 | 2.11E-05 | 0.0001 | 2.75E-05 | 5.09E-05 | 7.21E-05 | 0 | 1.63E-05 | 0.00013 | 0.00016 | 0 | 4.66E-05 |
| Angpt1 | 6.77E-05 | 9.74E-06 | 0 | 0 | 1.35E-05 | 0 | 0 | 0.00014 | 0.0001 | 0 | 0 | 0 | 0 | 0 | 0 | 3.46E-05 | 0 | 0.00015 |
| Adamts5 | 6.53E-05 | 0.00013 | 0 | 0 | 0.00012 | 6.82E-07 | 0 | 0 | 0.0001 | 0.00104 | 0 | 0 | 0.00092 | 0.00031 | 0 | 8.28E-05 | 0 | 0.00049 |
| Nr1h2 | 4.81E-05 | 0.00018 | 8.14E-05 | 0.00019 | 0.00012 | 5.26E-05 | 5.14E-05 | 9.49E-05 | 0.0001 | 6.19E-05 | 6.65E-05 | 0.00018 | 0 | 8.37E-05 | 0.00023 | 3.27E-06 | 4.87E-05 | 8.42E-05 |
| Secisbp2l | 3.44E-05 | 0.00017 | 0.00011 | 0.00013 | 0.00013 | 6.05E-05 | 4.18E-05 | 8.44E-05 | 0.0001 | 0.00013 | 0.00012 | 2.43E-05 | 0.00013 | 0.00014 | 0.00012 | 7.01E-05 | 0.00011 | 0.00011 |
| Gpalpp1 | 0.00012 | 0.00011 | 6.81E-05 | 1.68E-05 | 5.26E-05 | 0.00012 | 7.32E-05 | 0 | 0.0001 | 2.41E-05 | 5.74E-05 | 6.07E-05 | 5.04E-05 | 1.20E-05 | 7.19E-05 | 8.97E-05 | 0 | 8.07E-06 |
| Snx8 | 3.09E-05 | 0.00011 | 3.25E-05 | 0.00015 | 0.00017 | 4.18E-05 | 0.00012 | 0 | 0.0001 | 1.56E-06 | 5.60E-05 | 0 | 3.28E-05 | 8.74E-05 | 0.00012 | 0 | 0 | 6.62E-05 |
| Gab1 | 7.15E-05 | 0.00015 | 0 | 0 | 0 | 0.00015 | 0 | 0.00021 | 0.0001 | 1.04E-05 | 0 | 1.68E-05 | 0 | 1.39E-06 | 2.27E-05 | 9.02E-05 | 0.00013 | 0.00011 |
| Pdcd6 | 0.00044 | 0.00015 | 0.00023 | 0 | 0.00014 | 0.00023 | 0.00018 | 5.95E-05 | 0.0001 | 7.99E-05 | 9.16E-05 | 0.0002 | 0.00014 | 0.00011 | 0 | 5.18E-05 | 0.00011 | 3.80E-05 |

| | | | | | | | | | | | | | | | | | | |
|---|---|---|---|---|---|---|---|---|---|---|---|---|---|---|---|---|---|---|
| Mlxip | 0.00012 | 0.00021 | 0.00011 | 0.00016 | 0.00015 | 0.00022 | 6.90E-05 | 0.0002 | 0.0001 | 1.67E-05 | 0.00015 | 6.28E-05 | 0.00017 | 0.00025 | 0.00016 | 6.12E-05 | 0.00022 | 7.48E-07 |
| Clcn4 | 0.00012 | 0.00014 | 0.00016 | 5.54E-05 | 0.0002 | 0.00017 | 0.00021 | 0.00023 | 0.0001 | 0.00012 | 0.00027 | 2.03E-05 | 0.00015 | 0.00018 | 6.52E-05 | 0 | 3.81E-05 | 0.00014 |
| Cox7a2 | 0.00023 | 4.49E-05 | 0.00016 | 0.0002 | 4.05E-05 | 4.70E-05 | 1.50E-05 | 0.00011 | 0.0001 | 0 | 9.33E-05 | 8.98E-05 | 0 | 4.97E-05 | 0 | 0.00027 | 4.43E-05 | 0 |
| Tm4sf1 | 1.79E-05 | 3.76E-05 | 0.00066 | 0.00022 | 0 | 0 | 0.00041 | 0.00025 | 0.0001 | 0.00033 | 0.0002 | 0.00013 | 0.00046 | 0 | 0.00021 | 0.00039 | 2.03E-05 | 0.0006 |
| Exosc3 | 0.0001 | 7.72E-05 | 0.00018 | 7.42E-05 | 3.09E-05 | 1.97E-05 | 0 | 2.52E-05 | 0.0001 | 4.71E-05 | 1.44E-05 | 2.30E-05 | 4.12E-05 | 1.03E-05 | 9.43E-05 | 8.91E-05 | 0 | 6.76E-05 |
| Tmem87a | 0.0002 | 0.00012 | 9.09E-05 | 0 | 6.69E-05 | 0.00011 | 6.02E-05 | 7.15E-05 | 0.0001 | 1.10E-05 | 0 | 6.67E-05 | 6.23E-05 | 7.46E-05 | 6.43E-05 | 2.48E-05 | 2.05E-05 | 4.81E-05 |
| Eml3 | 6.90E-05 | 0.00012 | 0.00011 | 0.00013 | 8.19E-05 | 3.61E-05 | 7.90E-05 | 0.00011 | 0.0001 | 6.80E-05 | 2.78E-06 | 3.60E-05 | 2.19E-05 | 4.91E-05 | 0.00025 | 3.84E-05 | 1.81E-05 | 7.44E-05 |
| Hoxa9 | 0 | 3.60E-05 | 0 | 6.08E-06 | 2.64E-05 | 0 | 9.43E-05 | 2.12E-05 | 0.0001 | 2.63E-05 | 2.81E-05 | 3.75E-05 | 0 | 1.78E-06 | 0.00012 | 3.19E-05 | 0 | 1.83E-05 |
| Pphln1 | 0.00012 | 0.0001 | 0.00014 | 4.24E-05 | 7.47E-05 | 9.07E-05 | 1.13E-05 | 1.31E-05 | 0.0001 | 2.81E-05 | 0.00013 | 5.45E-05 | 0 | 0.00011 | 0.00011 | 0.00022 | 3.30E-05 | 7.06E-05 |
| Epdr1 | 5.02E-05 | 5.09E-05 | 6.19E-05 | 0 | 0 | 2.36E-05 | 0.00024 | 2.76E-05 | 0.0001 | 0.00035 | 0 | 0 | 6.21E-05 | 1.84E-05 | 0 | 3.77E-05 | 0 | 0.00037 |
| Plcl1 | 0 | 1.94E-05 | 0 | 0 | 6.40E-05 | 0 | 2.48E-05 | 5.79E-05 | 0.0001 | 3.46E-05 | 2.04E-05 | 0 | 0 | 0 | 0 | 0 | 0 | 9.45E-05 |
| Adsl | 0.00013 | 4.41E-05 | 0.00019 | 7.86E-05 | 0 | 1.88E-05 | 0 | 0 | 0.0001 | 3.31E-05 | 9.12E-06 | 1.30E-06 | 2.43E-05 | 3.45E-05 | 2.84E-05 | 0.00012 | 0 | 4.77E-05 |
| Arl2bp | 0.00013 | 0.00013 | 7.17E-05 | 0.00023 | 9.23E-05 | 0.00011 | 7.41E-05 | 9.32E-05 | 0.0001 | 0.00013 | 9.61E-05 | 0 | 3.70E-05 | 7.42E-05 | 0 | 0.00013 | 6.10E-05 | 0.00012 |
| Phf5a | 0.00016 | 8.18E-05 | 0.00018 | 0.0002 | 7.07E-05 | 3.27E-05 | 4.89E-05 | 7.94E-06 | 0.0001 | 0 | 4.57E-05 | 0 | 0 | 8.03E-05 | 0 | 0.00032 | 0 | 5.32E-06 |
| Adss | 0.00019 | 0.00017 | 0.00024 | 8.60E-05 | 4.03E-05 | 2.70E-05 | 1.84E-05 | 3.62E-05 | 0.0001 | 0.0001 | 7.44E-05 | 0.00015 | 0 | 1.44E-05 | 0.00014 | 0.00018 | 0.00015 | 0.00013 |
| Atxn1 | 5.06E-06 | 0.00014 | 0 | 0.00011 | 0.0002 | 0.00016 | 3.73E-05 | 1.43E-05 | 0.0001 | 3.45E-05 | 0.00044 | 0.00013 | 0.00018 | 0.00025 | 5.89E-05 | 0 | 0 | 0.00017 |
| Rab24 | 0.00012 | 0.00019 | 4.99E-05 | 1.31E-05 | 9.57E-05 | 0.00018 | 0.00011 | 0.00011 | 0.0001 | 8.15E-05 | 6.37E-05 | 0.00032 | 8.45E-06 | 0.00019 | 0.00039 | 0.0001 | 0 | 0.00014 |
| Srpk1 | 0.00018 | 0.00019 | 0.00022 | 7.67E-05 | 4.70E-05 | 1.61E-05 | 0 | 0 | 0.0001 | 3.84E-05 | 8.35E-05 | 1.91E-05 | 1.19E-05 | 0.00013 | 0.00051 | 0.0003 | 0.00023 | 2.69E-05 |
| Aldh6a1 | 8.16E-05 | 9.93E-05 | 0 | 0 | 5.71E-05 | 0.00012 | 8.21E-06 | 0 | 0.0001 | 6.95E-05 | 0 | 4.98E-05 | 6.32E-05 | 0 | 0 | 0 | 0 | 1.79E-05 |
| Rrn3 | 0.00017 | 0.0001 | 0.00025 | 2.86E-05 | 5.55E-05 | 4.50E-05 | 0 | 8.07E-06 | 0.0001 | 3.59E-05 | 7.52E-05 | 7.59E-05 | 1.85E-05 | 1.50E-05 | 5.48E-05 | 0.00018 | 4.17E-05 | 0 |
| Dapk3 | 0.00014 | 0.00015 | 0.00011 | 0.00024 | 4.92E-05 | 2.33E-05 | 0.00014 | 9.20E-05 | 0.0001 | 5.12E-05 | 0.00016 | 0.00014 | 0 | 5.34E-05 | 0.00026 | 3.49E-05 | 0.00026 | 0.0001 |
| Nt5dc2 | 0.00025 | 0 | 0.00021 | 0.00055 | 0.00014 | 0 | 0.00053 | 1.92E-05 | 0.0001 | 0 | 0.0001 | 0 | 0 | 0.00012 | 0 | 0.0007 | 0 | 0 |
| Dnajc7 | 0.00028 | 0.00012 | 0.00018 | 2.60E-05 | 5.31E-05 | 0.00012 | 1.04E-05 | 4.98E-05 | 0.0001 | 6.95E-06 | 1.38E-05 | 9.86E-05 | 0 | 4.48E-05 | 0.00024 | 9.55E-05 | 0 | 0 |
| Pfdn5 | 0.00026 | 0.00017 | 3.93E-05 | 0.00024 | 0.00017 | 0.00034 | 0.00014 | 0.00022 | 0.0001 | 7.63E-05 | 6.04E-05 | 0.00022 | 4.73E-05 | 0.0002 | 0 | 5.83E-05 | 0.0002 | 0.00017 |
| Cmtm7 | 4.50E-05 | 0.00013 | 0.00014 | 0.00027 | 0.00011 | 0.00011 | 0.00011 | 6.59E-05 | 0.0001 | 5.59E-05 | 0.00027 | 4.51E-05 | 0.0001 | 0.00015 | 1.64E-05 | 0.00011 | 2.12E-05 | 5.26E-05 |
| Irf2 | 4.14E-05 | 0.00011 | 1.43E-05 | 3.82E-05 | 3.24E-05 | 0.00014 | 0 | 0 | 0.0001 | 8.70E-05 | 0 | 3.83E-05 | 0.0001 | 7.63E-05 | 4.44E-05 | 3.56E-05 | 0.00012 | 2.87E-05 |
| Vkorc1l1 | 0.00027 | 7.41E-05 | 0.00017 | 0 | 4.58E-05 | 0.00015 | 5.73E-05 | 4.60E-05 | 0.0001 | 2.27E-05 | 9.38E-05 | 8.53E-05 | 4.90E-05 | 0 | 0 | 0.00015 | 1.51E-05 | 2.28E-05 |
| Fst | 0 | 0 | 0.00054 | 6.38E-06 | 0 | 0 | 0.00012 | 0 | 0.0001 | 0.0003 | 0.00048 | 0 | 0.00065 | 0.00014 | 0.00012 | 0.00025 | 0 | 0.0002 |
| Memo1 | 0.00022 | 0.00013 | 0.00016 | 0 | 6.83E-05 | 0.00013 | 2.72E-05 | 1.64E-05 | 0.0001 | 1.83E-05 | 6.95E-05 | 6.84E-05 | 0 | 6.69E-05 | 7.44E-05 | 0.00014 | 6.33E-05 | 0 |
| Brd7 | 0.00014 | 0.00013 | 0.00011 | 4.51E-05 | 8.23E-05 | 3.02E-05 | 2.75E-05 | 0 | 0.0001 | 1.67E-05 | 9.15E-05 | 0 | 0 | 4.71E-05 | 8.69E-05 | 0.00012 | 0 | 7.91E-05 |
| Ndufs4 | 0.00027 | 8.11E-05 | 0.00012 | 7.86E-05 | 4.13E-05 | 0.00012 | 4.35E-05 | 2.57E-06 | 0.0001 | 3.44E-05 | 8.64E-05 | 3.83E-05 | 6.79E-05 | 3.57E-05 | 0.00011 | 9.70E-05 | 0 | 2.77E-05 |
| Dhx38 | 4.00E-05 | 0.00014 | 0.00012 | 0.00026 | 5.71E-05 | 1.78E-05 | 0.0001 | 9.22E-05 | 0.0001 | 4.83E-05 | 8.07E-05 | 2.75E-05 | 4.31E-05 | 6.07E-05 | 0.00022 | 0.00015 | 0.00023 | 9.27E-05 |
| Rhod | 2.52E-05 | 0.0001 | 3.31E-05 | 4.87E-05 | 0.00014 | 2.14E-05 | 0 | 0 | 0.0001 | 0 | 0.00012 | 0 | 7.28E-05 | 0.00016 | 0 | 0 | 0 | 9.53E-05 |

| | | | | | | | | | | | | | | | | | | |
|---|---|---|---|---|---|---|---|---|---|---|---|---|---|---|---|---|---|---|
| Capza1 | 0.0003 | 6.83E-05 | 8.52E-05 | 0 | 0 | 0.00015 | 0.0001 | 5.65E-05 | 0.0001 | 3.90E-05 | 1.86E-05 | 8.53E-05 | 0 | 0 | 0 | 6.21E-05 | 4.58E-05 | 2.61E-05 |
| Thbs1 | 0.00026 | 0 | 0.00068 | 0 | 1.46E-05 | 0 | 0.00016 | 0 | 0.0001 | 1.77E-05 | 0.00415 | 0 | 0.00163 | 0.00057 | 0.00098 | 0.00023 | 0.00128 | 0 |
| Gtpbp6 | 2.12E-05 | 0.00011 | 4.38E-05 | 5.86E-05 | 1.10E-05 | 1.97E-05 | 0 | 6.89E-05 | 0.0001 | 2.72E-05 | 4.66E-05 | 7.62E-05 | 0 | 4.16E-05 | 0.0002 | 1.44E-05 | 0 | 6.10E-05 |
| Baz1b | 0.00011 | 0.00017 | 0.00019 | 6.59E-05 | 0.0001 | 3.17E-05 | 4.03E-05 | 3.49E-05 | 1.00E-04 | 4.07E-05 | 4.14E-05 | 3.67E-05 | 5.44E-05 | 5.85E-05 | 0.00033 | 0.00039 | 6.90E-05 | 2.63E-05 |
| Ufc1 | 0.00041 | 7.65E-05 | 0.0002 | 0.00048 | 0.00019 | 0.00027 | 0.00056 | 0.00021 | 9.99E-05 | 0.00012 | 8.84E-05 | 8.43E-05 | 0.00017 | 0.00016 | 0 | 0.0001 | 0 | 0.00014 |
| Ap3d1 | 0.00012 | 0.00026 | 0.0002 | 0.00039 | 0.00011 | 7.12E-05 | 0.0002 | 7.50E-05 | 9.99E-05 | 0.00014 | 0.00013 | 0.00015 | 7.45E-05 | 0.00013 | 0.00033 | 1.98E-05 | 0.00014 | 0.00017 |
| Zbtb4 | 0 | 0.00024 | 0 | 0.00031 | 0.00015 | 7.07E-05 | 1.54E-05 | 0 | 9.99E-05 | 0.00011 | 5.27E-05 | 0.00024 | 3.76E-05 | 0.00016 | 0.00032 | 0 | 2.74E-05 | 0.00017 |
| Acyp2 | 2.93E-05 | 6.38E-06 | 0 | 0 | 3.36E-05 | 3.21E-05 | 0 | 4.14E-05 | 9.99E-05 | 8.48E-05 | 0 | 0 | 0 | 0 | 2.75E-06 | 2.43E-05 | 0 | 7.91E-05 |
| Nipsnap3b | 0.00015 | 5.07E-05 | 6.99E-05 | 0 | 5.10E-06 | 1.25E-05 | 3.25E-05 | 6.53E-07 | 9.99E-05 | 5.14E-05 | 6.92E-05 | 6.20E-05 | 0 | 0 | 0 | 0.00011 | 4.82E-06 | 3.76E-05 |
| Slc35b1 | 0.00025 | 5.46E-05 | 0.00024 | 0.00023 | 5.82E-05 | 8.17E-06 | 2.33E-05 | 4.39E-05 | 9.99E-05 | 9.46E-06 | 0.00013 | 0 | 6.19E-05 | 4.37E-05 | 0 | 4.63E-05 | 0 | 6.30E-05 |
| Afap1 | 0 | 0.00023 | 0.00025 | 0.00014 | 0.0002 | 6.64E-06 | 0.00016 | 0.0002 | 9.99E-05 | 0.00023 | 0.00036 | 0.00058 | 0 | 2.86E-05 | 0.00056 | 6.52E-05 | 0.00012 | 0.00023 |
| Map1s | 3.60E-05 | 9.75E-05 | 0.00011 | 0.00019 | 1.02E-05 | 0 | 2.18E-05 | 0 | 9.99E-05 | 5.88E-05 | 0.00013 | 8.93E-05 | 5.53E-05 | 5.00E-06 | 0.00024 | 4.78E-05 | 0 | 6.96E-05 |
| Zzz3 | 0.00021 | 0.00019 | 0.00012 | 1.74E-05 | 0.00012 | 0.00021 | 0.00015 | 8.09E-05 | 9.98E-05 | 1.96E-05 | 0.00015 | 3.84E-05 | 0.00014 | 0.00014 | 9.55E-05 | 6.42E-05 | 6.31E-05 | 1.81E-05 |
| Pde1a | 2.90E-05 | 1.64E-05 | 1.34E-05 | 0 | 0 | 9.43E-05 | 0 | 1.25E-05 | 9.98E-05 | 1.55E-05 | 0 | 0 | 0.00021 | 0 | 0 | 2.93E-05 | 0 | 3.20E-05 |
| Nt5c3b | 0.00017 | 6.34E-05 | 6.14E-05 | 0 | 1.24E-05 | 7.63E-05 | 4.27E-05 | 0 | 9.98E-05 | 0 | 1.90E-05 | 2.98E-05 | 2.22E-05 | 4.24E-05 | 4.04E-06 | 9.63E-05 | 6.53E-05 | 3.27E-05 |
| Fut4 | 1.54E-06 | 0 | 2.47E-05 | 9.86E-05 | 7.85E-05 | 0.00011 | 0 | 0.00057 | 9.97E-05 | 0 | 0 | 0 | 2.03E-05 | 0.00026 | 0 | 0 | 0 | 0 |
| Sgpl1 | 0.00019 | 0.00033 | 0.00021 | 0 | 0.0003 | 0.00031 | 6.06E-05 | 0 | 9.96E-05 | 2.13E-06 | 0.00034 | 0.0001 | 6.61E-05 | 5.36E-05 | 0 | 9.10E-05 | 0.00018 | 8.97E-05 |
| Slc39a10 | 0.00015 | 3.84E-05 | 4.51E-06 | 0 | 0 | 6.69E-05 | 0 | 4.69E-05 | 9.96E-05 | 0 | 0 | 4.20E-05 | 3.87E-05 | 0 | 0 | 0.00011 | 8.42E-05 | 0 |
| Tsta3 | 0.00012 | 8.22E-05 | 7.70E-05 | 0.00013 | 6.11E-05 | 4.05E-05 | 3.00E-05 | 5.05E-05 | 9.96E-05 | 1.13E-05 | 6.81E-05 | 5.12E-05 | 0 | 6.85E-05 | 2.21E-05 | 9.29E-05 | 0 | 4.50E-05 |
| Pnp | 0.00019 | 0.00014 | 0 | 0.00012 | 2.12E-05 | 0 | 0.00011 | 0 | 9.96E-05 | 0.00046 | 0 | 0.00017 | 0.00058 | 6.79E-05 | 0 | 6.83E-05 | 0.00134 | 0.00023 |
| Hdac7 | 4.37E-05 | 0.00018 | 1.18E-05 | 0.00015 | 3.21E-06 | 0.00011 | 0 | 0 | 9.95E-05 | 0.00055 | 0.00027 | 8.01E-05 | 0.00018 | 0.00026 | 0.00023 | 0.00014 | 0.0002 | 0.00013 |
| Ppp1r11 | 0.00022 | 8.56E-05 | 0.00021 | 0.00018 | 6.60E-05 | 2.62E-05 | 3.95E-05 | 6.62E-05 | 9.95E-05 | 0 | 0.00012 | 7.94E-05 | 0 | 4.93E-05 | 0 | 0.00014 | 0 | 3.21E-05 |
| Frg1 | 0.00011 | 0.00015 | 1.45E-05 | 3.73E-05 | 4.32E-05 | 3.52E-05 | 0 | 2.92E-05 | 9.94E-05 | 5.99E-05 | 6.08E-05 | 0.0001 | 2.82E-05 | 6.29E-05 | 6.06E-05 | 8.78E-05 | 0 | 3.81E-05 |
| Mospd2 | 0.00011 | 7.45E-05 | 0 | 0 | 3.14E-05 | 2.60E-05 | 1.68E-05 | 0 | 9.94E-05 | 7.58E-05 | 1.22E-05 | 2.17E-05 | 2.11E-05 | 3.02E-05 | 7.48E-05 | 3.53E-05 | 4.69E-05 | 1.23E-05 |
| Nup98 | 0.00029 | 0.0001 | 9.88E-05 | 0 | 2.78E-05 | 0.00018 | 1.27E-05 | 0 | 9.93E-05 | 2.32E-05 | 7.35E-06 | 7.92E-05 | 3.40E-05 | 2.67E-05 | 2.54E-05 | 0.00018 | 0.00021 | 0 |
| Utp11 | 0.00023 | 0.00012 | 0.00023 | 0.00019 | 3.62E-05 | 6.77E-05 | 9.46E-05 | 0 | 9.93E-05 | 5.34E-05 | 7.38E-05 | 2.73E-05 | 5.08E-05 | 0 | 0.00016 | 0.00014 | 5.40E-05 | 3.45E-05 |
| Exoc8 | 2.01E-05 | 8.35E-05 | 7.77E-05 | 9.44E-05 | 5.16E-05 | 1.19E-05 | 4.87E-06 | 4.25E-05 | 9.93E-05 | 6.99E-05 | 8.53E-05 | 2.96E-05 | 0 | 5.45E-05 | 8.23E-05 | 2.49E-05 | 0 | 7.69E-05 |
| Irgm2 | 9.69E-05 | 4.85E-05 | 0 | 0 | 0 | 0 | 0 | 0 | 9.93E-05 | 0.00036 | 0 | 0 | 4.74E-05 | 0 | 5.03E-05 | 0 | 0 | 0.0002 |
| Plscr4 | 4.08E-06 | 0.00012 | 0 | 0 | 2.46E-05 | 0 | 6.41E-05 | 0.00013 | 9.93E-05 | 0.00014 | 0 | 0 | 0 | 0 | 0 | 1.01E-05 | 0 | 0.0002 |
| Unc50 | 0.00024 | 9.37E-05 | 0.00012 | 5.23E-05 | 0.00012 | 0.00016 | 0.00023 | 7.59E-05 | 9.92E-05 | 8.91E-05 | 0.00013 | 3.69E-05 | 5.26E-05 | 8.70E-05 | 0.0001 | 5.02E-05 | 0 | 3.68E-05 |
| Sestd1 | 6.28E-05 | 0.00016 | 0.00013 | 9.00E-06 | 4.55E-05 | 7.83E-05 | 0 | 2.08E-05 | 9.92E-05 | 0.00013 | 4.07E-05 | 8.39E-05 | 4.90E-06 | 5.98E-05 | 3.85E-05 | 0.00014 | 0.00017 | 1.73E-05 |
| Ubl4a | 0.0002 | 4.44E-05 | 0.00023 | 0.00027 | 3.18E-05 | 3.00E-05 | 0.00017 | 0.00013 | 9.92E-05 | 6.41E-05 | 8.44E-05 | 0 | 1.14E-05 | 6.37E-05 | 0 | 0.00012 | 0 | 6.67E-05 |
| Tm7sf2 | 0 | 1.67E-05 | 0 | 4.49E-06 | 4.20E-05 | 1.31E-05 | 0 | 2.39E-05 | 9.92E-05 | 2.16E-05 | 1.17E-05 | 1.05E-05 | 0 | 0 | 2.73E-05 | 1.81E-06 | 0 | 3.59E-05 |

| | | | | | | | | | | | | | | | | | | |
|---|---|---|---|---|---|---|---|---|---|---|---|---|---|---|---|---|---|---|
| Ddx27 | 4.59E-05 | 0.00012 | 0.00025 | 8.66E-05 | 1.40E-05 | 6.70E-06 | 0 | 0 | 9.92E-05 | 5.40E-06 | 3.48E-05 | 3.02E-05 | 0 | 3.57E-05 | 0.00021 | 0.00013 | 3.34E-05 | 2.06E-05 |
| Acvrl1 | 0 | 5.52E-05 | 0.00012 | 0.0003 | 9.15E-05 | 0 | 6.19E-05 | 0.00027 | 9.92E-05 | 0.00024 | 0.00024 | 0.00031 | 0.00039 | 0.00014 | 0.00012 | 1.72E-05 | 0 | 0.00032 |
| Atp6v0e | 0.00035 | 0.00015 | 0.00017 | 7.73E-05 | 0.00014 | 0.00022 | 0.00025 | 0.00018 | 9.91E-05 | 0.0001 | 0.00028 | 0.00019 | 0.00012 | 0.00014 | 0 | 0 | 3.87E-05 | 0.00017 |
| Slc39a9 | 0.00016 | 0.0001 | 0.00022 | 0.00023 | 0.00011 | 0.00011 | 0.00029 | 0.00012 | 9.91E-05 | 0.0001 | 7.27E-05 | 3.19E-05 | 8.70E-05 | 0.0001 | 2.30E-05 | 0.00012 | 0 | 6.70E-05 |
| Gys1 | 4.60E-05 | 0 | 1.77E-05 | 0.0003 | 9.37E-06 | 0 | 4.49E-05 | 0.00011 | 9.91E-05 | 9.20E-05 | 8.83E-05 | 0.00105 | 0.00016 | 3.87E-05 | 0.00018 | 8.53E-06 | 0 | 0 |
| Fundc1 | 7.34E-05 | 0.00013 | 4.79E-05 | 0.00027 | 9.06E-05 | 5.80E-05 | 8.30E-05 | 0.00013 | 9.90E-05 | 0.00013 | 6.23E-05 | 0.00011 | 4.66E-05 | 4.67E-05 | 6.35E-05 | 9.53E-05 | 4.36E-05 | 0.00017 |
| Exosc4 | 0.00016 | 8.68E-05 | 0.00015 | 0.00014 | 5.26E-05 | 9.40E-06 | 2.47E-06 | 2.83E-05 | 9.90E-05 | 1.87E-05 | 1.11E-05 | 0 | 2.34E-05 | 4.54E-05 | 5.88E-05 | 2.69E-05 | 0 | 1.23E-05 |
| Cav2 | 1.06E-05 | 2.47E-05 | 5.44E-05 | 0 | 0 | 0.00014 | 8.73E-06 | 0 | 9.89E-05 | 3.15E-05 | 5.71E-05 | 0.0001 | 8.61E-06 | 5.45E-05 | 0 | 9.88E-05 | 0 | 7.39E-05 |
| Srsf10 | 0.00022 | 0.00013 | 0.00025 | 0.00018 | 6.68E-05 | 8.59E-05 | 0 | 1.46E-05 | 9.89E-05 | 5.48E-05 | 8.06E-05 | 3.00E-05 | 2.58E-05 | 5.85E-06 | 0.00034 | 0.0003 | 4.86E-05 | 5.60E-05 |
| Ube2h | 0.00013 | 0.00027 | 2.75E-05 | 0 | 0.00021 | 0.00023 | 9.73E-05 | 0.00015 | 9.88E-05 | 0.00019 | 0.00017 | 0.00019 | 3.35E-05 | 0.00023 | 0 | 0 | 0.00017 | 0.00015 |
| Atp6v1c1 | 0.00013 | 0.00016 | 4.57E-05 | 3.88E-05 | 5.95E-05 | 0.00017 | 3.62E-05 | 0 | 9.88E-05 | 2.45E-05 | 1.18E-05 | 0.00015 | 2.06E-05 | 0.00013 | 0 | 9.48E-05 | 8.12E-05 | 3.68E-05 |
| Atp8b2 | 0.00027 | 0.0001 | 0.00021 | 8.64E-05 | 0.00016 | 0.00031 | 0.00035 | 0.00017 | 9.87E-05 | 0.00018 | 0.00018 | 0.00021 | 0.00012 | 0.00018 | 0.00017 | 0 | 0.00013 | 0.00016 |
| Ahsa1 | 0.00038 | 0.00025 | 0.00046 | 0.00038 | 0.00016 | 0.00015 | 3.35E-05 | 2.92E-05 | 9.87E-05 | 9.45E-05 | 8.04E-05 | 2.27E-05 | 0.00011 | 6.77E-05 | 0 | 0.00046 | 0.00015 | 0.00015 |
| Psma3 | 0.00041 | 0.00012 | 0.00017 | 0 | 0 | 0.00015 | 0 | 0 | 9.87E-05 | 8.30E-06 | 2.90E-05 | 8.50E-05 | 0 | 0 | 5.14E-05 | 9.09E-05 | 2.52E-05 | 1.28E-05 |
| Krcc1 | 0.00021 | 7.15E-05 | 3.55E-05 | 5.98E-05 | 2.79E-05 | 9.61E-05 | 0.00011 | 8.95E-05 | 9.87E-05 | 0.0001 | 9.66E-05 | 4.65E-05 | 4.65E-05 | 4.94E-05 | 6.02E-05 | 8.86E-05 | 0 | 8.08E-05 |
| Abi1 | 8.01E-05 | 0.00032 | 0.00025 | 0.00023 | 0.00018 | 7.51E-05 | 0.00013 | 0.00025 | 9.87E-05 | 0.00026 | 0.00019 | 0.00032 | 0.00018 | 8.49E-05 | 0.00016 | 0.00014 | 0.00039 | 0.00014 |
| Sall2 | 2.44E-05 | 5.14E-05 | 2.81E-05 | 3.89E-05 | 7.76E-05 | 0 | 3.57E-05 | 9.08E-05 | 9.87E-05 | 0 | 1.44E-05 | 0 | 1.47E-05 | 6.15E-05 | 2.40E-06 | 2.11E-05 | 0 | 6.04E-05 |
| Rdh10 | 3.44E-05 | 0.00012 | 0.00031 | 0.00016 | 0 | 0.00025 | 0.00015 | 0.00056 | 9.86E-05 | 7.11E-05 | 0 | 0 | 0.0007 | 0.00014 | 0.00042 | 0.00011 | 0.00022 | 0.00054 |
| Caprin1 | 0.00038 | 0.00019 | 0.00029 | 2.29E-05 | 9.85E-05 | 9.28E-05 | 5.04E-05 | 0 | 9.86E-05 | 6.91E-05 | 0.00015 | 0.00013 | 1.65E-05 | 0 | 6.76E-05 | 0.00022 | 0.0001 | 2.35E-05 |
| Il15 | 0 | 2.64E-05 | 0 | 0 | 0 | 8.31E-05 | 0 | 0.00013 | 9.86E-05 | 1.92E-05 | 0 | 0 | 0 | 0 | 0 | 0 | 0 | 0 |
| Fmnl2 | 0 | 6.81E-05 | 0.00012 | 0 | 0 | 0 | 0.00027 | 0 | 9.86E-05 | 0.00021 | 0 | 8.32E-05 | 0.00033 | 0 | 0 | 9.82E-05 | 6.33E-05 | 0.00019 |
| Rab8a | 0.00031 | 0.00014 | 9.24E-05 | 0.00013 | 0.00011 | 0.00019 | 0.00011 | 8.49E-05 | 9.85E-05 | 0.00013 | 0.0002 | 0.00023 | 8.37E-05 | 0.00012 | 3.33E-05 | 0.00017 | 0 | 0.00012 |
| Ccdc8 | 0.00019 | 8.60E-05 | 7.99E-05 | 0.00019 | 6.55E-05 | 0.0001 | 0.00018 | 0.00017 | 9.85E-05 | 7.19E-05 | 6.32E-05 | 9.55E-05 | 0 | 8.10E-05 | 0.00022 | 0 | 0 | 0.00018 |
| Tmco3 | 6.08E-05 | 8.58E-05 | 6.74E-05 | 3.42E-05 | 0.0001 | 8.62E-05 | 0.00012 | 0.00012 | 9.85E-05 | 0.00012 | 0.0001 | 9.87E-05 | 0.0001 | 6.65E-05 | 0 | 4.87E-05 | 0 | 0.00012 |
| Wbp4 | 0.00015 | 0.00012 | 9.14E-05 | 5.47E-05 | 5.35E-05 | 6.12E-05 | 2.20E-05 | 2.30E-05 | 9.85E-05 | 1.01E-05 | 7.14E-05 | 4.26E-05 | 2.17E-05 | 6.12E-06 | 0.00017 | 0.00011 | 3.48E-05 | 6.53E-05 |
| Nid2 | 0.00019 | 0 | 0 | 0.00015 | 0 | 1.42E-05 | 0 | 0.00011 | 9.85E-05 | 0 | 0 | 0 | 0.00038 | 0.00027 | 8.80E-05 | 0.00019 | 0 | 0.0003 |
| Rwdd4a | 0.00012 | 3.83E-05 | 0.00017 | 4.83E-05 | 5.67E-05 | 1.02E-05 | 2.51E-06 | 0 | 9.85E-05 | 2.91E-06 | 5.38E-05 | 6.72E-05 | 1.13E-05 | 2.02E-05 | 1.66E-05 | 4.63E-05 | 0 | 0 |
| Tll1 | 0 | 3.85E-05 | 0 | 0 | 5.05E-06 | 0 | 0.00017 | 0 | 9.85E-05 | 1.19E-05 | 7.22E-05 | 0 | 3.83E-05 | 0 | 0 | 2.18E-05 | 3.81E-05 | 0.0001 |
| Cd248 | 0.00018 | 0 | 0.00032 | 0.0007 | 0 | 0.0008 | 0.00041 | 0.00177 | 9.84E-05 | 0.00151 | 0.00055 | 0.00024 | 0.00021 | 0.00048 | 0 | 0.00014 | 0.00013 | 0.00096 |
| Pigp | 0.00021 | 4.54E-05 | 0 | 2.96E-05 | 4.93E-05 | 0.00017 | 6.88E-05 | 7.50E-05 | 9.84E-05 | 6.95E-05 | 8.54E-05 | 2.86E-05 | 0 | 4.02E-05 | 0 | 0 | 0 | 0.0001 |
| Lrrc47 | 0.00011 | 0.00018 | 0.00026 | 0.00016 | 9.30E-05 | 4.51E-05 | 0 | 2.13E-05 | 9.84E-05 | 2.60E-05 | 5.72E-05 | 8.40E-05 | 0 | 9.70E-05 | 0.00015 | 6.27E-05 | 2.17E-05 | 0 |
| Exosc10 | 0.00016 | 9.96E-05 | 0.00024 | 6.86E-05 | 3.91E-05 | 2.54E-05 | 3.32E-07 | 0 | 9.84E-05 | 3.44E-05 | 3.13E-05 | 7.78E-05 | 2.89E-05 | 2.02E-05 | 8.92E-05 | 0.00021 | 1.91E-05 | 3.71E-05 |
| Ddx41 | 0.00016 | 0.00015 | 0.00012 | 0.0001 | 8.24E-05 | 1.51E-05 | 5.72E-05 | 2.95E-05 | 9.84E-05 | 0 | 0 | 0 | 5.78E-05 | 0.00013 | 0.00011 | 7.35E-05 | 9.60E-05 | 1.27E-05 |

| | | | | | | | | | | | | | | | | | | |
|---|---|---|---|---|---|---|---|---|---|---|---|---|---|---|---|---|---|---|
| Ralbp1 | 0.00014 | 0.00028 | 0.00013 | 0.00015 | 9.91E-05 | 0.00022 | 1.98E-05 | 3.95E-05 | 9.83E-05 | 0.00025 | 0.0001 | 0.00012 | 7.21E-05 | 9.67E-05 | 8.19E-05 | 0.00016 | 0.00014 | 0.00012 |
| Ocrl | 0.00014 | 9.38E-05 | 2.99E-05 | 0 | 7.37E-05 | 9.24E-05 | 6.86E-05 | 0.00012 | 9.83E-05 | 8.60E-05 | 4.65E-05 | 2.97E-05 | 1.44E-05 | 3.34E-05 | 5.51E-05 | 6.35E-05 | 4.97E-05 | 5.39E-05 |
| Trappc1 | 0.00029 | 5.54E-05 | 3.96E-05 | 7.23E-05 | 2.25E-06 | 7.53E-05 | 0.0001 | 3.97E-05 | 9.83E-05 | 7.05E-05 | 8.06E-05 | 0 | 3.34E-05 | 4.48E-05 | 0 | 0.00017 | 0 | 3.40E-05 |
| Pef1 | 9.49E-05 | 0.00015 | 0.00015 | 0.00023 | 0.00012 | 5.01E-05 | 0.00022 | 5.31E-05 | 9.83E-05 | 6.99E-05 | 0.00011 | 7.24E-05 | 2.39E-05 | 8.43E-05 | 0 | 9.38E-05 | 0 | 0.00015 |
| Rmnd5b | 5.23E-05 | 9.66E-05 | 7.65E-05 | 7.03E-05 | 7.09E-05 | 2.34E-05 | 0 | 3.09E-05 | 9.83E-05 | 7.29E-05 | 5.92E-05 | 1.91E-06 | 0 | 4.17E-05 | 3.06E-05 | 6.63E-06 | 0 | 0.00013 |
| Mb21d2 | 1.65E-05 | 7.83E-06 | 2.19E-05 | 0 | 0 | 1.60E-05 | 7.37E-05 | 6.67E-05 | 9.83E-05 | 5.62E-06 | 0 | 8.48E-05 | 1.65E-05 | 0 | 0 | 8.50E-06 | 0 | 3.40E-05 |
| Rhbdd2 | 0 | 4.45E-05 | 7.70E-05 | 7.77E-05 | 5.16E-05 | 0 | 1.07E-05 | 7.80E-05 | 9.83E-05 | 4.44E-05 | 3.65E-05 | 5.55E-05 | 6.98E-05 | 1.04E-05 | 5.42E-05 | 1.97E-05 | 7.77E-05 | 7.32E-06 |
| Rbm39 | 0.00021 | 0.00025 | 0.00013 | 0.00015 | 0.00015 | 0.00014 | 7.46E-05 | 2.83E-05 | 9.82E-05 | 6.81E-05 | 5.50E-05 | 0.00023 | 0 | 7.41E-05 | 0.00063 | 0.00015 | 0.00017 | 6.44E-05 |
| Slc37a3 | 0.00022 | 5.69E-05 | 0 | 0 | 3.94E-05 | 9.84E-05 | 0.00025 | 0.00011 | 9.82E-05 | 5.61E-05 | 0 | 1.57E-05 | 0.00013 | 5.87E-05 | 6.32E-05 | 0 | 0 | 0 |
| Tiprl | 0.00016 | 0.00016 | 0.00021 | 0.00024 | 8.44E-05 | 5.42E-05 | 0.00011 | 5.32E-05 | 9.82E-05 | 0.0001 | 0.00013 | 0.00018 | 4.90E-05 | 7.15E-05 | 9.17E-05 | 0.00021 | 0 | 9.92E-05 |
| Prickle2 | 0 | 0.00014 | 0 | 5.82E-05 | 6.06E-05 | 4.25E-05 | 3.97E-05 | 0 | 9.82E-05 | 8.77E-05 | 0 | 0 | 0.00019 | 0.00013 | 6.38E-05 | 0 | 0 | 0.00012 |
| Tbcd | 0.00012 | 6.12E-05 | 0.00012 | 0.00017 | 3.20E-05 | 4.24E-05 | 0.0001 | 9.18E-06 | 9.82E-05 | 7.66E-05 | 0.00011 | 0 | 9.97E-05 | 7.48E-05 | 8.38E-05 | 4.25E-05 | 0 | 2.95E-05 |
| Usf1 | 0.00012 | 0.00016 | 5.04E-05 | 0.00012 | 0.00012 | 3.81E-05 | 4.37E-05 | 7.00E-05 | 9.82E-05 | 5.04E-05 | 5.71E-05 | 7.08E-05 | 3.23E-05 | 0.0001 | 0.00013 | 8.70E-05 | 0 | 5.72E-05 |
| Gabpa | 0.00017 | 0.00011 | 6.34E-05 | 5.38E-05 | 2.37E-05 | 5.47E-05 | 2.16E-05 | 4.25E-05 | 9.81E-05 | 3.78E-05 | 4.31E-05 | 0.00011 | 2.27E-05 | 8.27E-05 | 0.00013 | 0.0002 | 6.60E-05 | 3.95E-05 |
| Nutf2-ps1 | 0.00015 | 5.48E-05 | 0.00018 | 7.04E-05 | 5.87E-06 | 2.72E-05 | 2.43E-05 | 0 | 9.81E-05 | 2.33E-05 | 8.09E-05 | 1.90E-05 | 0 | 0 | 0 | 0.00016 | 3.22E-06 | 0 |
| Opa3 | 4.92E-05 | 8.40E-05 | 0.00012 | 9.89E-05 | 5.85E-05 | 5.92E-06 | 0 | 9.88E-06 | 9.81E-05 | 5.69E-05 | 6.32E-05 | 9.74E-06 | 1.17E-06 | 2.16E-05 | 4.38E-05 | 0.00018 | 0.00012 | 3.79E-05 |
| Zmym3 | 2.65E-05 | 9.10E-05 | 0 | 7.86E-05 | 5.92E-05 | 5.10E-06 | 2.26E-05 | 4.14E-05 | 9.81E-05 | 4.46E-05 | 2.34E-05 | 2.86E-05 | 0 | 5.08E-05 | 0.00015 | 5.02E-05 | 0 | 6.05E-05 |
| Plod3 | 2.17E-05 | 0 | 0.00051 | 0.00072 | 0.00016 | 0.00026 | 0.00086 | 0.00069 | 9.80E-05 | 0.00034 | 0.00042 | 0.00057 | 0.00092 | 0.0005 | 0.00027 | 0.00018 | 0.0001 | 0.00048 |
| Stard3 | 2.69E-06 | 0.00011 | 5.72E-05 | 0.00012 | 0.00011 | 3.58E-05 | 3.21E-05 | 0.00023 | 9.80E-05 | 5.10E-05 | 2.05E-06 | 8.81E-05 | 0 | 0.00015 | 0.00011 | 1.79E-05 | 4.62E-05 | 3.78E-05 |
| Dhx36 | 0.00021 | 0.00014 | 0.0001 | 0 | 9.32E-05 | 0.00013 | 1.04E-05 | 1.60E-05 | 9.79E-05 | 2.49E-05 | 3.59E-06 | 5.97E-05 | 0 | 8.27E-05 | 0.00017 | 0.00011 | 0.00011 | 5.66E-05 |
| Polr2b | 0.00016 | 0.00012 | 0.00011 | 3.73E-05 | 8.19E-05 | 4.31E-05 | 2.71E-05 | 0 | 9.79E-05 | 3.64E-05 | 2.62E-05 | 6.88E-05 | 0 | 3.39E-05 | 3.58E-05 | 0.00018 | 0 | 4.03E-06 |
| Trim2 | 5.67E-05 | 5.47E-05 | 0 | 3.39E-05 | 4.88E-05 | 1.85E-05 | 9.23E-05 | 0 | 9.79E-05 | 0.00012 | 0 | 0 | 2.31E-06 | 7.36E-05 | 6.89E-06 | 3.93E-05 | 0 | 0.00014 |
| Rassf2 | 0 | 5.48E-05 | 0 | 0 | 1.47E-05 | 3.26E-07 | 5.69E-05 | 6.15E-05 | 9.79E-05 | 0.00028 | 0 | 0 | 0 | 0 | 0 | 0 | 0 | 0.00034 |
| Large1 | 8.64E-05 | 0.00012 | 1.51E-05 | 0 | 0.00013 | 8.87E-05 | 6.18E-05 | 4.72E-05 | 9.78E-05 | 7.83E-05 | 0.00026 | 0.00014 | 8.30E-05 | 2.61E-05 | 0.00014 | 5.29E-05 | 2.59E-05 | 8.05E-05 |
| Art3 | 3.34E-05 | 2.49E-05 | 0 | 0 | 0 | 0.0001 | 0 | 6.01E-05 | 9.77E-05 | 3.04E-05 | 0 | 4.19E-05 | 3.82E-05 | 0 | 0 | 0 | 2.04E-05 | 0.00016 |
| Itga9 | 3.60E-05 | 0 | 0.00015 | 0 | 0 | 0.00023 | 0 | 0.0004 | 9.76E-05 | 3.21E-05 | 0.00025 | 5.45E-05 | 0.00013 | 0.00011 | 2.67E-05 | 0 | 3.78E-05 | 0.00017 |
| Map2k3 | 0.00013 | 0.00014 | 0.00037 | 0.00012 | 5.10E-05 | 0.00017 | 1.44E-05 | 0.00018 | 9.76E-05 | 0.00013 | 6.67E-05 | 7.16E-05 | 4.70E-05 | 0.00026 | 0.00022 | 0.00014 | 8.84E-05 | 0.0001 |
| Pbrm1 | 0.00017 | 0.00018 | 9.80E-05 | 6.43E-05 | 7.33E-05 | 6.20E-05 | 6.80E-05 | 4.44E-05 | 9.76E-05 | 5.44E-05 | 0.0001 | 8.52E-05 | 7.25E-05 | 9.74E-05 | 0.00014 | 0.00024 | 0.00014 | 6.79E-05 |
| Ufd1 | 0.00024 | 0.00016 | 0.0002 | 4.74E-05 | 7.36E-05 | 5.57E-05 | 3.38E-05 | 8.21E-08 | 9.76E-05 | 2.87E-05 | 3.24E-05 | 0.00014 | 4.87E-06 | 4.77E-05 | 0.00019 | 0.00015 | 1.56E-05 | 1.65E-05 |
| Smad5 | 0.00015 | 0.00019 | 0.00016 | 6.05E-05 | 0.0001 | 2.59E-05 | 3.70E-05 | 4.31E-05 | 9.76E-05 | 9.28E-05 | 5.84E-05 | 5.13E-05 | 6.70E-06 | 0.00014 | 0.00019 | 0.00016 | 0.00015 | 0.00012 |
| Aldh3b1 | 1.74E-05 | 2.63E-05 | 0 | 4.80E-05 | 2.39E-05 | 1.49E-05 | 0 | 0 | 9.76E-05 | 0.00011 | 3.17E-05 | 9.71E-06 | 9.65E-05 | 3.18E-06 | 2.96E-05 | 0 | 0 | 2.46E-05 |
| Enoph1 | 9.63E-05 | 4.87E-05 | 0.00016 | 7.78E-05 | 1.97E-05 | 0 | 0 | 4.57E-05 | 9.76E-05 | 4.53E-06 | 7.19E-05 | 0 | 0 | 1.28E-05 | 6.93E-05 | 8.12E-05 | 0 | 2.87E-05 |
| Dvl3 | 8.41E-05 | 0.00015 | 0.00012 | 9.60E-05 | 0.00014 | 0.00012 | 0.00019 | 0.00012 | 9.75E-05 | 6.34E-05 | 0.00014 | 1.75E-05 | 0 | 0.00017 | 3.75E-05 | 1.30E-05 | 0.00016 | 3.51E-05 |

| | | | | | | | | | | | | | | | | | | |
|---|---|---|---|---|---|---|---|---|---|---|---|---|---|---|---|---|---|---|
| Plekha6 | 0 | 0.00014 | 0 | 0 | 0 | 7.38E-05 | 0 | 0.00014 | 9.74E-05 | 0 | 0 | 4.63E-05 | 0 | 0 | 0 | 0 | 0 | 3.45E-06 |
| Dap3 | 0.00028 | 9.27E-05 | 0.00011 | 0.00017 | 7.77E-05 | 0.00015 | 0.00012 | 0 | 9.73E-05 | 6.56E-05 | 0.00013 | 0.00013 | 0.00012 | 0.00012 | 4.92E-05 | 7.22E-05 | 0 | 3.36E-05 |
| Arf2 | 9.94E-05 | 0.00011 | 0.00012 | 0 | 0.00014 | 9.16E-05 | 9.58E-05 | 0 | 9.73E-05 | 0.00015 | 0.00012 | 3.63E-06 | 0 | 6.43E-05 | 0 | 0.00021 | 0 | 0.00018 |
| Actr10 | 0.00019 | 0.00012 | 0.00022 | 0.00017 | 6.67E-05 | 6.42E-05 | 1.19E-05 | 4.28E-05 | 9.73E-05 | 6.60E-05 | 0.00016 | 0.00013 | 0 | 1.89E-05 | 0 | 0.00014 | 0 | 5.71E-05 |
| Zfp326 | 0.0001 | 0.00011 | 7.53E-05 | 8.21E-05 | 6.77E-05 | 4.53E-05 | 1.95E-06 | 1.01E-05 | 9.73E-05 | 1.22E-05 | 5.60E-05 | 1.09E-05 | 2.34E-06 | 4.37E-05 | 8.94E-05 | 0.0001 | 0.0001 | 2.69E-05 |
| Haghl | 2.58E-05 | 5.90E-05 | 0 | 5.79E-05 | 7.78E-05 | 1.84E-05 | 0 | 2.59E-05 | 9.73E-05 | 3.62E-05 | 2.93E-05 | 4.79E-05 | 0 | 4.34E-05 | 4.45E-05 | 0 | 0 | 8.82E-05 |
| Txndc11 | 0.00017 | 9.48E-05 | 0.00025 | 0.00047 | 5.92E-05 | 6.62E-05 | 0.0002 | 0.00012 | 9.72E-05 | 0.00012 | 7.50E-05 | 8.28E-05 | 9.64E-05 | 0.00019 | 0.00014 | 1.04E-06 | 0 | 8.17E-05 |
| Pgghg | 3.89E-05 | 0.00014 | 0 | 0.00012 | 0.0001 | 3.98E-05 | 0.00011 | 5.30E-05 | 9.71E-05 | 0.00012 | 3.83E-05 | 0 | 7.68E-05 | 3.16E-05 | 0.00016 | 0.0001 | 0 | 9.66E-05 |
| Zfp518b | 3.92E-05 | 5.88E-05 | 3.89E-05 | 1.61E-05 | 6.29E-05 | 1.75E-05 | 2.50E-05 | 0 | 9.71E-05 | 3.27E-05 | 7.15E-05 | 0 | 9.48E-06 | 6.25E-05 | 3.62E-05 | 0.00012 | 0 | 2.85E-05 |
| Tnks2 | 0.0004 | 0.00023 | 0.00014 | 0 | 0.00011 | 0.00024 | 0.00013 | 0.00021 | 9.70E-05 | 4.64E-05 | 6.21E-05 | 0.00031 | 5.74E-06 | 4.67E-05 | 3.66E-05 | 2.81E-05 | 0 | 0 |
| Lrrc8a | 0 | 0.00026 | 0.0002 | 6.85E-06 | 0.00015 | 0.00012 | 0 | 0.00022 | 9.70E-05 | 0.00012 | 0.00037 | 0.00023 | 0.00018 | 0.00022 | 1.17E-05 | 6.24E-05 | 0.00019 | 0.00016 |
| Ehmt1 | 9.77E-05 | 0.00019 | 8.07E-05 | 8.75E-05 | 9.90E-05 | 4.63E-05 | 1.52E-05 | 3.74E-05 | 9.70E-05 | 4.14E-05 | 8.28E-05 | 6.02E-05 | 6.22E-05 | 7.03E-05 | 0.00039 | 0.00013 | 0.00013 | 6.15E-05 |
| P3h1 | 0.00037 | 0 | 0.00037 | 0.00061 | 0.00018 | 0.00042 | 0.00084 | 0.00062 | 9.69E-05 | 0.0001 | 0.00019 | 0.00012 | 0.00026 | 0.0004 | 0.0002 | 0 | 7.92E-05 | 0.00014 |
| Glt8d1 | 0.00037 | 2.15E-05 | 0 | 0 | 5.74E-05 | 0.00028 | 0.0004 | 0.00012 | 9.69E-05 | 6.63E-05 | 1.51E-05 | 6.42E-05 | 9.37E-05 | 9.20E-05 | 0.00016 | 0 | 0 | 6.85E-05 |
| Pycr2 | 5.55E-05 | 6.89E-05 | 0.00021 | 0.00015 | 3.04E-05 | 4.44E-05 | 0 | 7.80E-05 | 9.69E-05 | 5.49E-05 | 4.23E-05 | 1.39E-05 | 0 | 9.78E-05 | 4.15E-05 | 7.21E-05 | 0 | 4.46E-05 |
| Slc39a11 | 3.14E-05 | 1.70E-05 | 7.81E-05 | 0.00027 | 5.91E-05 | 6.19E-06 | 0.00027 | 0.00022 | 9.69E-05 | 7.91E-05 | 4.05E-05 | 0 | 4.78E-05 | 5.38E-06 | 9.10E-05 | 5.37E-05 | 7.43E-05 | 0.00014 |
| Rrs1 | 4.97E-05 | 5.13E-05 | 0.00024 | 0.00013 | 4.38E-06 | 0 | 0 | 9.72E-06 | 9.69E-05 | 4.92E-05 | 3.34E-05 | 0 | 5.68E-05 | 1.94E-05 | 0 | 1.51E-05 | 0 | 5.28E-05 |
| Dact3 | 0 | 0.00014 | 0 | 0.00021 | 0.00013 | 0 | 0 | 0 | 9.69E-05 | 1.20E-05 | 0.00011 | 0 | 0 | 0.00016 | 0.00058 | 4.87E-05 | 6.52E-05 | 0.00035 |
| Mia2 | 0.00019 | 0.0002 | 0.00024 | 0.00021 | 1.00E-04 | 0.00021 | 0.00028 | 9.28E-05 | 9.68E-05 | 5.09E-05 | 5.57E-05 | 0.00011 | 0.00016 | 0.00015 | 0.0001 | 9.32E-05 | 4.82E-05 | 0 |
| Mpdz | 0.00013 | 6.72E-05 | 9.17E-05 | 0 | 0 | 3.04E-05 | 5.33E-05 | 2.57E-05 | 9.68E-05 | 0.00015 | 4.22E-05 | 6.35E-06 | 7.12E-05 | 0 | 6.66E-05 | 0.00017 | 9.57E-07 | 8.68E-05 |
| Ccdc88a | 6.86E-05 | 0.00011 | 0.00011 | 7.18E-05 | 2.67E-05 | 4.77E-05 | 2.36E-05 | 0.0001 | 9.67E-05 | 9.00E-05 | 0.00011 | 5.25E-05 | 0 | 7.34E-05 | 9.35E-05 | 0.0003 | 7.15E-05 | 8.98E-05 |
| Cdkn2d | 1.96E-06 | 7.62E-05 | 0 | 0 | 5.86E-05 | 3.21E-05 | 1.19E-05 | 0.00016 | 9.67E-05 | 6.37E-05 | 1.25E-05 | 0.00016 | 2.05E-05 | 0 | 0 | 0.00071 | 0 | 4.15E-05 |
| Rab28 | 0.00017 | 0.00014 | 9.93E-05 | 0.00019 | 9.06E-05 | 0.0002 | 7.36E-05 | 0.0001 | 9.66E-05 | 0.0002 | 7.24E-06 | 8.06E-05 | 9.51E-05 | 0.00011 | 4.07E-05 | 8.02E-05 | 0.00014 | 0.00017 |
| Btbd1 | 0.0002 | 0.00013 | 8.78E-05 | 5.54E-07 | 7.91E-05 | 0.0002 | 6.35E-05 | 0.00012 | 9.66E-05 | 6.16E-05 | 3.75E-05 | 0.00012 | 4.54E-05 | 2.78E-05 | 0 | 7.24E-05 | 1.40E-05 | 6.27E-05 |
| Tfg | 0.00058 | 0.00016 | 0.00042 | 0.00021 | 0.00019 | 0.0002 | 0.00026 | 4.64E-05 | 9.66E-05 | 9.62E-05 | 0.00015 | 0.00025 | 0 | 7.25E-05 | 0 | 0.00016 | 0 | 5.68E-05 |
| Romo1 | 0.00017 | 2.55E-05 | 0.00014 | 0.00015 | 6.73E-05 | 8.66E-05 | 7.65E-05 | 3.83E-05 | 9.66E-05 | 5.58E-05 | 0.00013 | 8.60E-06 | 7.57E-05 | 9.15E-05 | 0 | 1.34E-05 | 0 | 7.96E-05 |
| Dlat | 0.00019 | 0.00014 | 0.00015 | 5.40E-05 | 7.24E-05 | 8.44E-05 | 9.74E-06 | 1.10E-05 | 9.66E-05 | 1.74E-05 | 0.00015 | 4.55E-05 | 3.52E-05 | 1.55E-05 | 0 | 0.00012 | 7.43E-05 | 9.20E-06 |
| Trappc3 | 0.00014 | 0.00017 | 0.00015 | 0.00022 | 9.33E-05 | 7.94E-05 | 9.53E-05 | 6.49E-05 | 9.66E-05 | 9.14E-05 | 0.00016 | 8.74E-05 | 3.13E-05 | 0.00014 | 0 | 0.00015 | 0 | 0.00013 |
| Cdk14 | 0.00018 | 0.00014 | 0 | 0 | 5.43E-05 | 0.00015 | 1.71E-05 | 0 | 9.65E-05 | 0.00013 | 4.39E-05 | 0 | 0.00024 | 0.00033 | 9.56E-05 | 9.23E-05 | 5.84E-05 | 0.0003 |
| Rwdd1 | 0.00019 | 5.41E-05 | 0.00013 | 0 | 2.64E-05 | 0.00013 | 4.13E-05 | 4.10E-05 | 9.65E-05 | 0 | 4.91E-05 | 4.28E-05 | 0 | 4.16E-06 | 4.11E-05 | 6.74E-05 | 2.01E-05 | 1.02E-05 |
| Herc2 | 8.38E-05 | 0.00014 | 0.00011 | 0 | 0.00011 | 7.58E-05 | 4.19E-05 | 4.54E-05 | 9.65E-05 | 0.0001 | 8.06E-05 | 0 | 4.77E-05 | 8.18E-05 | 5.63E-05 | 0.0001 | 0 | 8.72E-05 |
| Cdk5 | 6.83E-05 | 0.00016 | 2.15E-05 | 0.00015 | 9.15E-05 | 6.66E-05 | 8.49E-05 | 0.0001 | 9.65E-05 | 8.39E-05 | 0.00012 | 1.62E-05 | 5.44E-05 | 9.61E-05 | 0.00014 | 5.35E-05 | 0 | 5.60E-05 |
| Fez2 | 0.00013 | 0.00011 | 0.00012 | 0.00011 | 9.77E-05 | 4.68E-05 | 0 | 1.97E-05 | 9.65E-05 | 0.00018 | 0.00023 | 0.00018 | 0 | 3.69E-05 | 4.31E-05 | 4.10E-05 | 0.00012 | 5.73E-05 |

| | | | | | | | | | | | | | | | | | | |
|---|---|---|---|---|---|---|---|---|---|---|---|---|---|---|---|---|---|---|
| Gpaa1 | 3.79E-05 | 4.86E-05 | 0.00012 | 0.00023 | 6.48E-05 | 4.02E-05 | 1.22E-05 | 0.00012 | 9.65E-05 | 6.34E-05 | 6.96E-05 | 0 | 9.95E-05 | 6.27E-05 | 3.00E-05 | 0.00021 | 0 | 6.79E-05 |
| Itpr1 | 0 | 3.17E-05 | 0 | 0 | 7.25E-05 | 1.89E-05 | 0 | 3.23E-06 | 9.65E-05 | 0 | 7.18E-05 | 0 | 0 | 0.00012 | 5.99E-05 | 7.34E-05 | 0 | 0.00019 |
| Zfp866 | 5.18E-05 | 0.00012 | 0 | 5.02E-05 | 0.0001 | 4.44E-05 | 0 | 4.02E-05 | 9.64E-05 | 6.03E-05 | 2.46E-05 | 1.24E-05 | 0 | 6.26E-05 | 0.00015 | 4.28E-05 | 0 | 5.77E-05 |
| Farsb | 0.00022 | 1.64E-05 | 0.00023 | 0 | 0 | 3.51E-05 | 0 | 0 | 9.64E-05 | 1.16E-05 | 3.41E-05 | 2.29E-05 | 4.51E-05 | 4.91E-06 | 2.51E-05 | 5.85E-05 | 9.48E-06 | 0 |
| Cops5 | 0.00024 | 0.00015 | 0.00023 | 0.0002 | 5.64E-05 | 5.99E-05 | 4.11E-05 | 5.93E-06 | 9.63E-05 | 9.15E-05 | 9.40E-05 | 0.00011 | 1.89E-05 | 3.16E-06 | 0.00015 | 0.00015 | 0 | 6.59E-05 |
| Nabp2 | 0.00018 | 9.30E-05 | 6.09E-05 | 8.55E-05 | 4.42E-05 | 5.83E-05 | 8.00E-06 | 4.36E-06 | 9.63E-05 | 2.93E-05 | 0 | 2.65E-05 | 0 | 1.67E-05 | 0 | 6.76E-05 | 5.85E-05 | 4.93E-05 |
| Adamts10 | 5.86E-05 | 0.00015 | 1.64E-05 | 0.00012 | 5.57E-05 | 5.11E-05 | 0.00014 | 0.00023 | 9.63E-05 | 0.00019 | 0 | 0 | 8.69E-05 | 0.0002 | 0.00062 | 3.92E-05 | 0.00015 | 0.00047 |
| Taf1 | 9.55E-05 | 8.61E-05 | 8.13E-05 | 0 | 6.80E-05 | 6.16E-05 | 7.75E-05 | 3.67E-05 | 9.62E-05 | 5.92E-05 | 3.63E-05 | 8.96E-05 | 0 | 5.34E-05 | 0.00013 | 0.00021 | 0.00014 | 6.26E-05 |
| Fam50a | 0.00017 | 0.00015 | 7.41E-05 | 8.27E-05 | 9.52E-05 | 4.32E-05 | 4.93E-05 | 0 | 9.62E-05 | 2.40E-05 | 3.10E-05 | 0.00015 | 0 | 8.65E-05 | 6.21E-05 | 7.21E-05 | 2.92E-05 | 7.16E-05 |
| Psmb9 | 5.87E-05 | 2.51E-05 | 0 | 2.15E-07 | 0 | 0 | 0 | 0 | 9.61E-05 | 0.00011 | 0 | 0 | 7.12E-05 | 2.19E-05 | 0 | 7.12E-05 | 0 | 0.00013 |
| Slc41a2 | 0.00013 | 3.37E-05 | 0.00033 | 0 | 0.00012 | 6.74E-05 | 0.00044 | 0.00015 | 9.60E-05 | 0.00012 | 0.00019 | 0.00045 | 7.97E-05 | 0 | 2.18E-05 | 0 | 4.03E-05 | 7.32E-05 |
| Taf2 | 0.00013 | 5.42E-05 | 0.00015 | 1.66E-05 | 8.87E-05 | 6.30E-05 | 5.46E-05 | 5.28E-05 | 9.60E-05 | 3.42E-05 | 8.24E-05 | 0 | 2.87E-05 | 6.90E-05 | 4.38E-05 | 0.00011 | 0 | 4.34E-05 |
| Npm3 | 0.00016 | 1.91E-05 | 0.0002 | 0.00018 | 0 | 0 | 0 | 0 | 9.60E-05 | 0 | 0 | 0 | 7.11E-05 | 9.11E-06 | 0 | 0.00013 | 9.01E-05 | 1.29E-05 |
| Syf2 | 9.36E-05 | 0.00034 | 8.19E-05 | 0.00035 | 7.92E-05 | 0.00014 | 1.75E-05 | 9.36E-05 | 9.59E-05 | 0.0001 | 0.00014 | 0.0002 | 0.00015 | 0.00016 | 0.00024 | 7.51E-05 | 0.00072 | 7.74E-05 |
| Cops8 | 0.00022 | 0.00018 | 0.00035 | 0.00021 | 9.56E-05 | 4.92E-05 | 0 | 6.62E-05 | 9.59E-05 | 3.71E-05 | 0.00013 | 9.37E-05 | 0 | 9.18E-05 | 8.12E-05 | 0.00024 | 9.97E-05 | 4.11E-05 |
| Tmem29 | 0.00011 | 2.20E-05 | 3.76E-05 | 1.95E-05 | 0 | 4.87E-07 | 3.29E-05 | 0 | 9.59E-05 | 2.00E-05 | 3.39E-06 | 1.90E-05 | 2.98E-05 | 6.28E-07 | 0 | 3.42E-05 | 2.72E-05 | 7.74E-06 |
| Tm9sf2 | 0.00037 | 0.00018 | 0.00032 | 0.00026 | 0.00028 | 0.00045 | 0.00044 | 0.00029 | 9.58E-05 | 0.0002 | 0.00019 | 0.00016 | 0.00027 | 0.00027 | 0 | 0.0002 | 0.00018 | 0.00027 |
| Kpna6 | 0.00015 | 0.00013 | 0.00019 | 0.00015 | 8.89E-05 | 4.36E-05 | 5.47E-06 | 4.63E-05 | 9.57E-05 | 6.19E-05 | 9.77E-05 | 4.41E-05 | 1.13E-05 | 3.92E-05 | 0 | 0.00013 | 0 | 6.69E-05 |
| Ehd4 | 6.75E-06 | 6.73E-05 | 6.01E-05 | 2.26E-06 | 3.09E-05 | 3.66E-05 | 0 | 0 | 9.57E-05 | 8.14E-05 | 0.00016 | 0.00011 | 0 | 8.08E-05 | 0 | 7.63E-05 | 1.83E-05 | 5.52E-05 |
| Tbx18 | 8.81E-05 | 4.40E-05 | 0 | 0 | 0 | 9.83E-05 | 0 | 6.01E-05 | 9.56E-05 | 2.90E-05 | 0.00015 | 0 | 3.18E-05 | 0.00016 | 2.63E-05 | 1.00E-05 | 0 | 6.23E-05 |
| Clcc1 | 0.00011 | 4.05E-05 | 6.78E-05 | 3.38E-06 | 0.00011 | 7.65E-05 | 2.65E-06 | 5.14E-05 | 9.56E-05 | 6.84E-05 | 5.91E-07 | 2.46E-05 | 0.0001 | 5.82E-05 | 4.68E-05 | 0.00026 | 1.44E-05 | 1.16E-05 |
| Usp48 | 0.00013 | 0.00013 | 9.96E-05 | 0.00015 | 6.24E-05 | 7.01E-05 | 9.98E-05 | 3.36E-05 | 9.56E-05 | 6.01E-05 | 7.83E-05 | 9.72E-05 | 7.72E-06 | 3.14E-05 | 0.00023 | 0.00013 | 0.00014 | 9.16E-05 |
| Larp4b | 9.41E-05 | 0.00014 | 0.00021 | 6.60E-05 | 0.00013 | 5.75E-05 | 4.31E-05 | 6.02E-05 | 9.56E-05 | 4.21E-05 | 0.00013 | 0.00015 | 4.36E-06 | 7.39E-05 | 0.0002 | 0.00018 | 0.00014 | 9.30E-05 |
| Chmp7 | 7.75E-05 | 0.00016 | 0.00011 | 5.89E-05 | 8.26E-05 | 2.38E-05 | 0 | 2.12E-05 | 9.56E-05 | 6.42E-05 | 0.00011 | 7.54E-05 | 1.90E-05 | 4.69E-05 | 8.93E-05 | 8.69E-05 | 0 | 5.62E-05 |
| Dnajb11 | 0.00027 | 0.00011 | 0.00048 | 0.00042 | 0.00029 | 0.0002 | 0.00033 | 0.00027 | 9.55E-05 | 0.00022 | 0.00028 | 2.75E-06 | 0.00035 | 0.00024 | 0 | 0.00021 | 0 | 0.00027 |
| Med9 | 6.49E-05 | 6.83E-05 | 7.08E-05 | 0.00022 | 3.06E-05 | 3.77E-05 | 1.60E-05 | 3.18E-05 | 9.55E-05 | 3.09E-05 | 0.0001 | 7.78E-05 | 7.86E-05 | 4.56E-05 | 0 | 0.00025 | 0 | 2.02E-05 |
| Deaf1 | 3.97E-05 | 0.00013 | 3.33E-05 | 8.99E-05 | 9.72E-05 | 6.38E-05 | 7.01E-05 | 5.10E-05 | 9.54E-05 | 6.15E-05 | 6.51E-05 | 1.53E-05 | 0 | 8.85E-05 | 0.00026 | 0.00011 | 5.89E-05 | 8.77E-05 |
| Prkaa1 | 9.19E-05 | 0.00014 | 9.02E-05 | 7.76E-05 | 5.02E-05 | 4.04E-05 | 4.15E-05 | 3.45E-05 | 9.54E-05 | 5.15E-05 | 0.00012 | 4.52E-05 | 0 | 0 | 7.62E-06 | 9.86E-05 | 0.00011 | 9.91E-05 |
| Tbcc | 1.39E-05 | 0.00011 | 0.00011 | 0.00011 | 4.49E-05 | 8.53E-05 | 0 | 5.49E-05 | 9.53E-05 | 6.30E-05 | 7.55E-05 | 2.89E-05 | 0 | 9.40E-05 | 1.23E-05 | 2.42E-05 | 0 | 0.00012 |
| Pthlh | 0 | 1.84E-05 | 1.93E-06 | 0 | 0.00017 | 5.68E-05 | 9.17E-05 | 0 | 9.53E-05 | 0 | 0 | 2.91E-05 | 0 | 0.00023 | 9.25E-05 | 0 | 2.75E-05 | 0 |
| Wbp11 | 7.66E-05 | 0.00011 | 0.00015 | 0.00011 | 6.63E-05 | 4.98E-05 | 0 | 3.49E-05 | 9.53E-05 | 2.09E-05 | 3.20E-05 | 9.41E-05 | 9.44E-06 | 1.90E-05 | 0.00015 | 0.00029 | 4.76E-05 | 5.86E-05 |
| Tcf7 | 0 | 6.30E-05 | 6.40E-06 | 0 | 6.74E-05 | 3.15E-05 | 0 | 6.88E-05 | 9.53E-05 | 0 | 2.65E-05 | 1.99E-05 | 0 | 3.38E-05 | 0 | 0 | 0 | 0 |
| Fam162a | 0.00027 | 0 | 9.93E-05 | 0.00022 | 6.24E-05 | 9.37E-05 | 0.00041 | 2.23E-05 | 9.52E-05 | 0.00014 | 4.96E-05 | 0.00171 | 0.00052 | 0 | 0 | 0.00013 | 0.0002 | 0 |

| | | | | | | | | | | | | | | | | | | |
|---|---|---|---|---|---|---|---|---|---|---|---|---|---|---|---|---|---|---|
| Gnpnat1 | 0.00024 | 0 | 0.00025 | 3.81E-05 | 2.39E-05 | 0.0001 | 0.00011 | 0.00015 | 9.51E-05 | 0.00033 | 7.88E-05 | 7.22E-05 | 0 | 9.35E-05 | 9.05E-05 | 0 | 0 | 0 |
| Hyal2 | 5.91E-05 | 8.72E-05 | 6.71E-05 | 9.58E-05 | 3.59E-05 | 2.70E-05 | 2.09E-05 | 3.67E-06 | 9.51E-05 | 1.88E-05 | 9.60E-05 | 0 | 4.14E-05 | 4.43E-05 | 0 | 0 | 0 | 4.89E-05 |
| Csad | 0 | 0.00012 | 0 | 0.00015 | 2.58E-05 | 8.91E-06 | 9.75E-05 | 0.0001 | 9.51E-05 | 6.84E-05 | 1.10E-06 | 0 | 2.49E-05 | 4.80E-05 | 0.0002 | 2.18E-05 | 8.09E-06 | 0.00014 |
| Sart1 | 0.00011 | 0.00016 | 0.00019 | 0.00024 | 2.56E-05 | 0 | 0 | 5.64E-05 | 9.51E-05 | 3.76E-05 | 5.66E-05 | 5.21E-05 | 5.41E-06 | 5.27E-05 | 0.00018 | 0.00011 | 6.03E-06 | 2.26E-05 |
| Tmem30a | 0.00043 | 7.02E-05 | 7.44E-05 | 0 | 8.22E-05 | 0.00047 | 6.89E-05 | 0.00018 | 9.50E-05 | 8.83E-05 | 0.0001 | 0.00021 | 6.48E-05 | 5.48E-05 | 0 | 0.00011 | 0.00014 | 0.00015 |
| Oxr1 | 8.96E-05 | 0.00015 | 0 | 1.30E-05 | 2.00E-06 | 0.00013 | 0 | 0.00013 | 9.50E-05 | 8.11E-05 | 0 | 0.00011 | 0 | 3.57E-05 | 4.09E-05 | 0.00011 | 0.00013 | 9.14E-05 |
| Mri1 | 0.00014 | 9.18E-05 | 0.00011 | 8.56E-05 | 0 | 5.75E-06 | 1.06E-05 | 7.57E-05 | 9.50E-05 | 0 | 4.26E-07 | 2.19E-05 | 0 | 2.45E-05 | 6.85E-05 | 0 | 0 | 2.82E-05 |
| Ppp2r1b | 5.25E-05 | 0.00011 | 0.00014 | 0.0001 | 2.27E-05 | 0 | 3.30E-05 | 3.51E-05 | 9.50E-05 | 6.02E-05 | 2.76E-05 | 1.68E-05 | 0 | 6.00E-06 | 1.99E-05 | 0.00013 | 0.00011 | 1.59E-07 |
| Zfp592 | 0.0001 | 0.00017 | 8.22E-05 | 4.67E-06 | 9.11E-05 | 0.00016 | 0 | 8.70E-05 | 9.49E-05 | 5.01E-05 | 7.22E-05 | 0.00011 | 0 | 2.22E-05 | 7.82E-05 | 0.0001 | 6.88E-06 | 7.22E-05 |
| Aurkaip1 | 0.00026 | 0.00014 | 0.00025 | 0.00015 | 0.00011 | 0.00011 | 0.0001 | 3.59E-05 | 9.49E-05 | 0 | 0.00011 | 0.0001 | 1.36E-06 | 6.24E-05 | 5.37E-05 | 0.00012 | 7.78E-05 | 7.19E-05 |
| Sdhaf2 | 0.00018 | 8.60E-05 | 9.36E-05 | 0 | 6.28E-05 | 5.99E-05 | 5.23E-07 | 3.31E-05 | 9.49E-05 | 2.01E-05 | 3.06E-05 | 5.95E-05 | 0 | 2.97E-05 | 0 | 4.32E-05 | 0 | 4.52E-05 |
| Bin3 | 0.00013 | 0.00013 | 8.93E-05 | 0.0001 | 6.93E-05 | 5.79E-05 | 2.57E-05 | 6.30E-05 | 9.49E-05 | 7.62E-05 | 4.06E-05 | 6.97E-05 | 2.33E-05 | 5.87E-05 | 0.00011 | 2.07E-05 | 9.45E-05 | 3.34E-05 |
| Dok1 | 4.81E-05 | 4.74E-05 | 0.00019 | 0.00038 | 0.00025 | 0 | 0.00013 | 0.00053 | 9.49E-05 | 0 | 0.00037 | 0 | 0.00011 | 0.0002 | 0.00016 | 0.00026 | 0 | 0.00012 |
| Csf1 | 0 | 0 | 0.00025 | 0.00016 | 0 | 0 | 0 | 0.0001 | 9.49E-05 | 0.00073 | 0.00018 | 0 | 0.0012 | 0 | 0 | 0.00014 | 4.13E-05 | 0.00046 |
| Timm8b | 0.00022 | 9.53E-05 | 0.00011 | 0.00013 | 7.17E-05 | 0.00019 | 3.26E-05 | 9.23E-05 | 9.48E-05 | 1.82E-05 | 4.98E-05 | 6.71E-05 | 0 | 7.14E-05 | 0 | 4.76E-05 | 0 | 2.73E-05 |
| Cct8 | 0.00041 | 8.24E-05 | 0.0003 | 6.16E-06 | 0 | 0.00012 | 0 | 0 | 9.48E-05 | 2.22E-05 | 8.12E-05 | 0.00017 | 1.68E-05 | 0 | 2.01E-05 | 0.00011 | 0 | 0 |
| Cdc23 | 0.00019 | 0.00013 | 0.00016 | 0.00011 | 0.00011 | 0.00011 | 0.00011 | 2.00E-05 | 9.48E-05 | 6.07E-05 | 9.90E-05 | 2.09E-05 | 2.48E-05 | 1.10E-05 | 4.63E-05 | 0.00028 | 0 | 5.21E-05 |
| Galnt4 | 7.99E-05 | 2.13E-05 | 6.23E-05 | 6.22E-05 | 5.00E-05 | 2.41E-05 | 2.21E-05 | 0.00015 | 9.48E-05 | 4.23E-05 | 6.79E-05 | 0 | 9.63E-05 | 4.92E-05 | 0 | 3.12E-05 | 0 | 0.0001 |
| Isy1 | 0.00014 | 8.01E-05 | 0.00017 | 0.00018 | 5.67E-05 | 1.63E-05 | 0 | 4.68E-05 | 9.48E-05 | 7.61E-05 | 3.24E-05 | 5.44E-05 | 1.58E-05 | 2.15E-05 | 0.00014 | 0.00022 | 0.00023 | 7.79E-05 |
| Ggh | 8.03E-05 | 2.00E-05 | 2.98E-05 | 1.95E-06 | 0.00012 | 0.00014 | 0.00027 | 0.00012 | 9.47E-05 | 0.00014 | 0 | 0.00011 | 0.00015 | 0 | 0 | 3.81E-05 | 0 | 0.00014 |
| Pde10a | 5.92E-06 | 5.59E-05 | 6.10E-05 | 4.36E-06 | 6.03E-05 | 0.00012 | 0 | 7.23E-05 | 9.47E-05 | 9.79E-05 | 7.66E-05 | 2.18E-05 | 2.58E-05 | 2.35E-05 | 1.80E-05 | 0 | 8.45E-05 | 0 |
| Ttc37 | 0.00022 | 0 | 0.00024 | 8.97E-05 | 5.25E-05 | 0.00011 | 0.00023 | 4.09E-05 | 9.47E-05 | 0.0001 | 1.29E-05 | 3.72E-05 | 0.00015 | 2.11E-05 | 0.00018 | 0 | 5.18E-05 | 0 |
| Tfip11 | 9.68E-05 | 0.00015 | 0.00011 | 6.97E-05 | 6.86E-05 | 9.87E-07 | 1.91E-05 | 1.31E-05 | 9.47E-05 | 2.83E-05 | 6.23E-05 | 7.11E-05 | 0 | 6.65E-05 | 0.00012 | 9.05E-05 | 9.04E-06 | 4.90E-05 |
| Cldn12 | 4.93E-06 | 4.09E-05 | 2.28E-05 | 0.00015 | 5.22E-05 | 0 | 0 | 0 | 9.47E-05 | 6.61E-05 | 0.00018 | 8.01E-05 | 0.00013 | 7.31E-05 | 5.57E-05 | 2.88E-05 | 0 | 0 |
| Golga2 | 0.0002 | 0.00016 | 0.00024 | 0.00024 | 0.00016 | 0.00011 | 0.00039 | 0.00017 | 9.46E-05 | 0.00011 | 0.00013 | 0.00015 | 1.13E-05 | 0.00012 | 0.00029 | 7.90E-05 | 0.00017 | 0.00012 |
| Taf9b | 7.43E-05 | 0 | 2.93E-05 | 0 | 3.80E-05 | 8.47E-05 | 0 | 0 | 9.46E-05 | 2.27E-05 | 4.85E-05 | 0 | 0 | 0.0001 | 0 | 3.31E-05 | 0 | 0 |
| Uqcrc2 | 0.00046 | 0.00011 | 0.00024 | 5.99E-05 | 5.97E-05 | 0.00018 | 5.96E-05 | 3.91E-05 | 9.45E-05 | 0 | 8.41E-05 | 0.00013 | 0 | 4.67E-05 | 0 | 8.31E-05 | 4.09E-05 | 0 |
| Bcl6 | 6.90E-05 | 0.00021 | 2.44E-06 | 1.29E-05 | 0.0002 | 0.00011 | 4.82E-05 | 0.0001 | 9.45E-05 | 9.65E-05 | 5.18E-05 | 0.00026 | 0.00029 | 9.80E-05 | 0.00011 | 9.28E-05 | 0.00019 | 8.13E-05 |
| Ngly1 | 0.00011 | 8.97E-05 | 4.96E-05 | 0 | 6.32E-05 | 9.03E-05 | 8.88E-05 | 4.16E-05 | 9.45E-05 | 8.45E-05 | 5.10E-06 | 7.21E-05 | 4.21E-05 | 4.46E-05 | 3.88E-07 | 1.57E-05 | 0 | 8.33E-05 |
| Ash2l | 0.00012 | 0.00016 | 0.00018 | 0.00015 | 7.66E-05 | 8.17E-05 | 0 | 1.77E-05 | 9.45E-05 | 5.30E-05 | 5.38E-05 | 7.08E-05 | 3.31E-05 | 2.75E-05 | 0.00011 | 0.00019 | 0 | 5.31E-05 |
| G3bp1 | 4.83E-06 | 7.63E-05 | 0.00026 | 0.00042 | 3.08E-06 | 0 | 0 | 2.55E-05 | 9.45E-05 | 3.47E-05 | 0.0001 | 1.85E-05 | 3.76E-05 | 3.03E-05 | 6.71E-06 | 0.00031 | 0.00017 | 4.87E-05 |
| Kctd5 | 0.00011 | 6.95E-05 | 0.00013 | 6.56E-05 | 0.00019 | 3.94E-05 | 0.00013 | 0 | 9.43E-05 | 4.58E-05 | 0.00053 | 6.57E-05 | 0.00019 | 0.00017 | 0 | 7.36E-05 | 2.24E-05 | 0 |
| Cdk4 | 0.00013 | 0.00013 | 0.00017 | 0.00047 | 7.47E-05 | 6.95E-05 | 7.87E-05 | 8.36E-05 | 9.42E-05 | 1.59E-06 | 5.47E-05 | 9.70E-05 | 9.47E-05 | 0.00014 | 0 | 0.00033 | 5.19E-05 | 1.18E-05 |

| | | | | | | | | | | | | | | | | | | |
|---|---|---|---|---|---|---|---|---|---|---|---|---|---|---|---|---|---|---|
| Nudcd2 | 0.00029 | 1.64E-05 | 0.00023 | 0.00017 | 2.88E-05 | 4.39E-05 | 0.00019 | 7.88E-05 | 9.42E-05 | 1.33E-05 | 6.02E-05 | 0 | 4.96E-05 | 8.93E-05 | 4.21E-05 | 0.00021 | 0 | 0 |
| Rab11fip3 | 5.32E-05 | 0.00015 | 6.23E-05 | 0 | 6.83E-05 | 3.87E-05 | 8.65E-05 | 0 | 9.42E-05 | 4.89E-05 | 8.33E-05 | 3.57E-06 | 0 | 0.00015 | 0.00018 | 6.16E-05 | 5.23E-05 | 0.00011 |
| Klhdc2 | 0.00034 | 0.00024 | 0.00021 | 0.00012 | 0.00014 | 0.00036 | 3.18E-05 | 0.0001 | 9.41E-05 | 5.46E-05 | 9.14E-05 | 0.00016 | 7.28E-05 | 7.86E-05 | 0.00012 | 0.0002 | 1.49E-05 | 3.60E-05 |
| Zdhhc6 | 0.00012 | 7.49E-05 | 0.00011 | 7.08E-05 | 8.92E-05 | 8.03E-05 | 2.56E-05 | 3.00E-05 | 9.41E-05 | 4.13E-05 | 5.55E-05 | 2.16E-05 | 1.52E-05 | 6.85E-05 | 6.16E-06 | 0.00016 | 0 | 7.21E-05 |
| Larp4 | 0.00013 | 0.00016 | 0.00041 | 9.10E-05 | 9.11E-05 | 6.88E-05 | 3.60E-05 | 5.95E-06 | 9.41E-05 | 2.02E-05 | 9.71E-05 | 0.00012 | 9.65E-06 | 1.66E-05 | 0.00011 | 8.99E-05 | 0.00025 | 1.56E-05 |
| Ppp1r8 | 0.0001 | 9.98E-05 | 0.00016 | 9.85E-05 | 3.51E-05 | 7.02E-06 | 0 | 0 | 9.41E-05 | 3.69E-05 | 3.58E-05 | 0 | 2.87E-05 | 2.45E-05 | 1.62E-05 | 0.00023 | 0 | 1.91E-05 |
| Atad3a | 4.21E-05 | 7.54E-05 | 0.00021 | 6.34E-05 | 0 | 0 | 0 | 0 | 9.41E-05 | 2.49E-05 | 3.47E-05 | 1.36E-05 | 2.30E-05 | 7.02E-06 | 0.00013 | 6.27E-05 | 4.63E-05 | 1.92E-05 |
| Celf1 | 9.64E-05 | 0.00013 | 0.00018 | 2.03E-05 | 6.75E-05 | 0.0001 | 0 | 6.32E-05 | 9.40E-05 | 5.23E-06 | 0.00013 | 6.41E-05 | 0 | 4.79E-05 | 2.28E-05 | 0.0001 | 1.89E-05 | 0 |
| Agps | 0.00014 | 6.06E-05 | 0.00011 | 0 | 3.59E-05 | 8.93E-05 | 0 | 0 | 9.40E-05 | 9.63E-06 | 8.20E-05 | 6.14E-05 | 2.08E-06 | 3.70E-05 | 4.03E-05 | 0.00011 | 0.00012 | 5.47E-05 |
| Cdk2ap2 | 0.00012 | 0.00018 | 7.88E-05 | 0.00012 | 1.96E-05 | 0 | 0 | 0 | 9.40E-05 | 7.83E-05 | 3.01E-05 | 7.30E-05 | 2.23E-05 | 3.09E-05 | 5.03E-05 | 0 | 0.00012 | 7.77E-05 |
| Entpd7 | 7.96E-05 | 5.13E-06 | 0.00025 | 0.00035 | 0.0003 | 0.00017 | 0.00025 | 0.00026 | 9.39E-05 | 8.36E-08 | 0.00024 | 0 | 3.01E-05 | 0.00023 | 0.00014 | 0 | 4.11E-05 | 0 |
| Agtrap | 0 | 5.71E-05 | 0 | 7.74E-05 | 2.82E-05 | 7.78E-05 | 0 | 0.00012 | 9.39E-05 | 0.00019 | 6.34E-05 | 0 | 4.72E-05 | 3.19E-05 | 0 | 6.02E-06 | 0 | 0.00011 |
| Fbxl5 | 0.0002 | 0.00015 | 6.41E-05 | 0 | 2.95E-05 | 0.00019 | 0 | 0 | 9.38E-05 | 2.46E-05 | 6.38E-09 | 6.43E-05 | 1.01E-05 | 3.91E-05 | 0.00012 | 0.00012 | 0 | 2.28E-05 |
| Txndc9 | 0.00026 | 0.0002 | 0.00018 | 0.00016 | 5.01E-05 | 6.89E-05 | 2.28E-05 | 0 | 9.38E-05 | 0.00011 | 0.00015 | 0.0001 | 0 | 9.64E-05 | 0.00019 | 8.60E-05 | 0 | 0.00013 |
| Vps11 | 8.09E-05 | 0.00019 | 4.61E-05 | 2.71E-05 | 6.99E-05 | 5.51E-05 | 1.35E-05 | 3.92E-05 | 9.38E-05 | 4.03E-05 | 0 | 7.62E-05 | 1.26E-06 | 9.02E-05 | 0.00016 | 1.53E-05 | 3.73E-05 | 7.07E-05 |
| Atp6v0a1 | 1.62E-05 | 7.07E-05 | 0 | 4.09E-05 | 0 | 3.19E-05 | 0 | 8.55E-06 | 9.38E-05 | 0.00011 | 4.66E-05 | 1.78E-05 | 2.32E-05 | 5.24E-05 | 4.91E-05 | 0 | 6.83E-05 | 0.00011 |
| Ppp1r14b | 0.00029 | 8.92E-05 | 0.00044 | 0.00044 | 0 | 0 | 0.00015 | 0 | 9.38E-05 | 0.00023 | 0.00016 | 3.76E-05 | 9.29E-05 | 1.74E-05 | 9.36E-05 | 9.52E-05 | 4.67E-05 | 3.52E-05 |
| Zfr | 0.00025 | 0.00022 | 0.00021 | 2.79E-05 | 0.00016 | 0.00024 | 0.00021 | 6.99E-05 | 9.37E-05 | 9.59E-05 | 0.00012 | 0.00013 | 0 | 6.05E-05 | 0.0001 | 7.53E-05 | 6.66E-05 | 8.65E-05 |
| Ergic2 | 0.00017 | 6.90E-05 | 0.00011 | 2.19E-06 | 8.05E-05 | 6.63E-05 | 6.15E-05 | 4.62E-05 | 9.37E-05 | 1.68E-05 | 7.96E-05 | 3.87E-05 | 0 | 7.58E-05 | 1.99E-05 | 0.00017 | 0 | 2.95E-05 |
| Rcbtb2 | 0.0002 | 5.74E-05 | 5.68E-05 | 0.00013 | 8.41E-05 | 3.67E-05 | 0.00013 | 2.72E-05 | 9.37E-05 | 7.64E-05 | 1.29E-05 | 0 | 0.00015 | 0 | 7.60E-05 | 0.00021 | 0 | 7.99E-05 |
| Pin1 | 7.64E-05 | 5.01E-05 | 0.00014 | 3.10E-05 | 3.61E-05 | 0 | 0 | 4.66E-05 | 9.37E-05 | 2.01E-05 | 5.17E-05 | 5.09E-05 | 1.09E-05 | 1.07E-06 | 5.15E-05 | 0.00014 | 3.36E-05 | 3.49E-06 |
| Cacna2d1 | 6.74E-05 | 0.00014 | 0 | 0.00031 | 8.98E-05 | 0.00038 | 0.00013 | 0.00044 | 9.36E-05 | 0.0003 | 0.00015 | 0.00015 | 0.00055 | 0.00014 | 0.00034 | 3.49E-05 | 0.00021 | 0.00026 |
| Ttc3 | 0.00016 | 0.00014 | 1.40E-05 | 0.00011 | 0.00011 | 0.00015 | 0.00015 | 0.00016 | 9.36E-05 | 9.10E-05 | 8.85E-05 | 7.05E-05 | 0 | 3.70E-05 | 0.00013 | 3.38E-05 | 3.03E-05 | 9.75E-05 |
| Smim13 | 7.24E-05 | 4.20E-05 | 5.14E-05 | 1.44E-05 | 7.78E-05 | 0.00011 | 3.30E-05 | 8.03E-05 | 9.36E-05 | 1.38E-05 | 5.35E-05 | 1.44E-05 | 2.62E-05 | 4.54E-05 | 0 | 0 | 0 | 2.61E-05 |
| Zmym4 | 8.09E-06 | 8.69E-05 | 9.23E-05 | 5.57E-05 | 8.52E-05 | 5.40E-05 | 3.79E-06 | 8.55E-05 | 9.36E-05 | 6.06E-05 | 3.81E-05 | 5.14E-05 | 1.81E-05 | 3.00E-05 | 4.10E-05 | 7.67E-05 | 1.91E-05 | 7.09E-05 |
| Ccnk | 0.00013 | 0.00012 | 8.76E-05 | 8.36E-05 | 6.40E-05 | 5.26E-05 | 2.85E-05 | 1.98E-05 | 9.36E-05 | 2.83E-05 | 4.15E-05 | 0.00012 | 1.92E-05 | 5.02E-05 | 5.40E-05 | 0.00013 | 0.00013 | 0 |
| Adipor2 | 3.81E-05 | 0.00013 | 6.04E-05 | 9.87E-05 | 8.10E-05 | 3.49E-05 | 6.62E-06 | 5.17E-06 | 9.36E-05 | 2.85E-05 | 0.00012 | 0.00019 | 3.06E-05 | 7.50E-05 | 0 | 5.02E-05 | 9.57E-05 | 3.49E-05 |
| Ctnnbip1 | 3.83E-05 | 6.27E-05 | 4.25E-05 | 0.0001 | 7.71E-05 | 2.64E-05 | 6.83E-05 | 3.04E-05 | 9.36E-05 | 6.31E-05 | 0.00014 | 3.71E-05 | 0.0001 | 3.77E-05 | 0 | 6.55E-05 | 0 | 0.0002 |
| Fmr1 | 0.00023 | 0.00019 | 9.25E-05 | 0.00015 | 2.07E-05 | 0.00026 | 0 | 0 | 9.35E-05 | 3.28E-05 | 7.75E-05 | 7.56E-06 | 0.00068 | 0 | 0.0003 | 0.0004 | 0.00022 | 5.13E-05 |
| Rreb1 | 8.44E-05 | 0.00013 | 4.81E-05 | 5.20E-05 | 5.69E-05 | 7.69E-05 | 7.80E-05 | 8.29E-05 | 9.35E-05 | 1.89E-05 | 0 | 4.56E-05 | 0 | 8.09E-05 | 0.00011 | 0 | 0.00025 | 7.85E-05 |
| Slc2a3 | 0 | 2.20E-05 | 0 | 0 | 0 | 5.13E-05 | 0 | 0.00035 | 9.35E-05 | 0 | 0 | 0.00109 | 0 | 8.85E-06 | 0 | 0 | 0 | 3.46E-06 |
| Rab5b | 0.0002 | 0.00018 | 1.09E-05 | 0 | 0.00012 | 0.00014 | 5.08E-05 | 5.23E-05 | 9.34E-05 | 8.14E-05 | 7.70E-05 | 0.00018 | 1.24E-05 | 9.24E-06 | 0 | 8.90E-07 | 0 | 5.61E-05 |
| Casp4 | 0.00014 | 7.54E-05 | 0 | 0 | 2.42E-05 | 0.00013 | 2.66E-06 | 2.21E-05 | 9.34E-05 | 9.14E-05 | 0 | 0 | 0.00028 | 0 | 0.00015 | 5.11E-05 | 0.00018 | 3.34E-06 |

| | | | | | | | | | | | | | | | | | | |
|---|---|---|---|---|---|---|---|---|---|---|---|---|---|---|---|---|---|---|
| Aasdhppt | 0.00011 | 5.37E-05 | 8.30E-05 | 5.57E-05 | 5.30E-05 | 4.84E-05 | 2.83E-05 | 3.39E-05 | 9.34E-05 | 5.93E-06 | 1.51E-06 | 9.23E-06 | 1.52E-05 | 2.61E-05 | 9.04E-05 | 9.31E-06 | 0 | 4.02E-05 |
| Itpa | 0.00014 | 8.11E-05 | 0.00014 | 0.00017 | 6.48E-05 | 3.51E-05 | 0 | 1.21E-07 | 9.34E-05 | 1.39E-05 | 3.29E-05 | 0 | 2.09E-05 | 2.45E-05 | 1.76E-05 | 0.00018 | 0 | 2.68E-05 |
| Smap2 | 0.00012 | 0.00011 | 0.00018 | 0.00017 | 0.00012 | 6.71E-05 | 1.97E-05 | 0.00012 | 9.33E-05 | 4.37E-05 | 0.00013 | 0.00012 | 0 | 1.08E-06 | 2.96E-05 | 0.00011 | 9.81E-05 | 0.00013 |
| Meis2 | 0.00011 | 9.38E-05 | 0 | 4.22E-05 | 7.95E-05 | 5.30E-05 | 4.92E-05 | 1.23E-05 | 9.33E-05 | 3.17E-05 | 0 | 0.00017 | 0 | 0 | 0.00032 | 7.94E-05 | 0.00012 | 0 |
| Arhgap6 | 2.48E-05 | 6.28E-05 | 0 | 0 | 0 | 3.84E-05 | 0 | 6.15E-05 | 9.33E-05 | 6.44E-05 | 0 | 2.65E-05 | 1.44E-05 | 0 | 0 | 2.17E-05 | 8.02E-05 | 4.37E-05 |
| Wls | 0.0004 | 0.00041 | 1.21E-05 | 0 | 0.00046 | 0.00054 | 0.00052 | 0.00042 | 9.32E-05 | 0.00026 | 0.0005 | 7.08E-05 | 0 | 0.00026 | 1.92E-05 | 0.00018 | 0.00016 | 0.00032 |
| Robo1 | 0 | 3.64E-05 | 0 | 0 | 0.00023 | 0.00027 | 0 | 0.00102 | 9.32E-05 | 8.84E-06 | 0 | 0 | 0 | 0.00049 | 0 | 2.34E-05 | 0 | 0 |
| Gpr107 | 0.00016 | 0.00015 | 0.00012 | 0.00013 | 0.0002 | 0.00014 | 0.00022 | 0.00013 | 9.32E-05 | 6.60E-05 | 0.00015 | 9.20E-05 | 9.03E-05 | 0.00014 | 7.59E-05 | 0 | 1.42E-05 | 9.98E-05 |
| Pacs1 | 7.74E-05 | 8.86E-05 | 8.47E-05 | 0 | 6.16E-05 | 8.34E-05 | 2.71E-05 | 4.10E-05 | 9.32E-05 | 4.72E-05 | 5.46E-05 | 0.00025 | 5.65E-05 | 0.00012 | 0 | 7.60E-05 | 5.96E-05 | 5.44E-05 |
| B3galnt1 | 3.19E-05 | 1.69E-05 | 5.98E-05 | 4.20E-05 | 7.24E-05 | 8.15E-05 | 0.00012 | 6.82E-05 | 9.32E-05 | 0.00023 | 0.00012 | 7.16E-05 | 5.66E-05 | 2.52E-05 | 0 | 0.00023 | 2.33E-05 | 5.92E-05 |
| Psma1 | 0.00033 | 0.00014 | 0.00025 | 0.00011 | 1.22E-05 | 4.34E-05 | 1.27E-05 | 0 | 9.32E-05 | 6.64E-05 | 5.60E-05 | 8.10E-05 | 1.46E-05 | 2.93E-05 | 1.38E-05 | 0.00018 | 0 | 3.82E-05 |
| Cox5a | 0.00026 | 8.24E-05 | 0.00033 | 0.00042 | 8.33E-05 | 7.20E-05 | 4.90E-05 | 4.79E-05 | 9.31E-05 | 4.01E-06 | 0.00019 | 0 | 1.38E-05 | 0.00011 | 0 | 0.00027 | 0 | 6.69E-06 |
| Decr2 | 7.94E-05 | 6.42E-05 | 0 | 1.72E-05 | 4.36E-05 | 6.64E-05 | 4.84E-05 | 3.24E-05 | 9.31E-05 | 3.65E-05 | 1.06E-05 | 7.14E-05 | 0 | 4.14E-05 | 0 | 0 | 0 | 9.01E-05 |
| Cisd3 | 0.00016 | 5.87E-05 | 4.38E-05 | 0.00013 | 5.20E-05 | 4.77E-05 | 1.39E-05 | 0.0001 | 9.31E-05 | 4.57E-05 | 3.64E-05 | 0.00016 | 5.47E-05 | 2.10E-05 | 0 | 4.57E-05 | 0 | 1.87E-05 |
| Pdk3 | 0.00015 | 0 | 2.44E-05 | 2.68E-05 | 4.76E-05 | 4.73E-05 | 0.0001 | 0 | 9.31E-05 | 0 | 0.0001 | 7.62E-05 | 9.43E-05 | 0.00016 | 0 | 9.20E-05 | 0 | 1.60E-05 |
| Ube2s | 0.00017 | 9.15E-05 | 0.00033 | 0.00029 | 5.79E-05 | 2.73E-05 | 4.94E-05 | 0.00012 | 9.31E-05 | 1.84E-05 | 0.00015 | 0.00013 | 0 | 1.69E-05 | 0 | 0.00103 | 0.00045 | 0 |
| Zfp451 | 6.59E-05 | 0.00016 | 0.00013 | 7.72E-05 | 2.14E-05 | 4.29E-06 | 0.00012 | 4.88E-05 | 9.31E-05 | 8.20E-05 | 0 | 4.61E-05 | 6.79E-05 | 1.65E-05 | 0.00021 | 6.61E-05 | 0.00011 | 0.00015 |
| Washc5 | 3.87E-05 | 0.00011 | 5.19E-05 | 0.0001 | 5.27E-05 | 9.66E-05 | 3.85E-05 | 3.78E-05 | 9.30E-05 | 7.13E-05 | 5.38E-05 | 8.37E-06 | 0 | 9.35E-05 | 0 | 2.14E-05 | 1.92E-05 | 8.01E-05 |
| Lsm1 | 9.74E-05 | 5.44E-05 | 6.05E-05 | 5.25E-05 | 9.92E-05 | 7.06E-05 | 7.41E-05 | 0.0001 | 9.30E-05 | 5.45E-05 | 2.34E-05 | 5.36E-05 | 6.03E-05 | 5.02E-05 | 0 | 7.16E-05 | 0 | 2.26E-05 |
| Ilf3 | 0.00019 | 0.00018 | 0.00013 | 5.55E-05 | 6.10E-05 | 6.59E-05 | 0 | 0 | 9.30E-05 | 0 | 1.95E-05 | 2.27E-05 | 2.18E-05 | 3.40E-05 | 0.00014 | 0.00034 | 0.00026 | 0 |
| Ggnbp2 | 0.00027 | 0.00018 | 0.00011 | 0 | 5.15E-05 | 0.00016 | 6.94E-05 | 0 | 9.29E-05 | 5.13E-05 | 2.11E-05 | 0.00013 | 7.03E-05 | 1.48E-05 | 0.00029 | 4.97E-05 | 7.20E-05 | 3.24E-05 |
| Mosmo | 3.86E-05 | 0.00013 | 0.00012 | 2.85E-06 | 0.0001 | 0.00014 | 0 | 0.00012 | 9.29E-05 | 4.18E-05 | 0.0001 | 7.17E-05 | 3.99E-05 | 0.00018 | 0.00014 | 6.93E-05 | 8.11E-05 | 7.95E-05 |
| Rhno1 | 6.48E-05 | 5.87E-05 | 4.34E-05 | 5.99E-05 | 4.67E-05 | 5.14E-05 | 6.04E-05 | 4.19E-05 | 9.29E-05 | 7.50E-05 | 8.48E-05 | 6.16E-06 | 3.49E-05 | 0.00011 | 6.64E-05 | 0.00035 | 0 | 5.07E-05 |
| Emc6 | 8.54E-05 | 9.91E-05 | 0.00018 | 0.00016 | 2.51E-05 | 0 | 7.40E-07 | 4.02E-05 | 9.29E-05 | 3.77E-05 | 6.54E-05 | 4.69E-05 | 0 | 3.88E-05 | 0 | 7.47E-05 | 2.17E-05 | 0 |
| Fhl2 | 3.51E-05 | 5.14E-05 | 0 | 0 | 0.00018 | 0 | 0 | 9.44E-05 | 9.29E-05 | 0 | 0.00065 | 9.85E-05 | 0 | 0.00015 | 0 | 4.14E-06 | 0 | 0.00036 |
| Agpat1 | 0.00011 | 0.0001 | 0.00016 | 0.0002 | 0.0001 | 0.00011 | 9.12E-05 | 0.00016 | 9.28E-05 | 5.49E-05 | 0.00016 | 5.18E-05 | 2.83E-05 | 5.64E-05 | 3.96E-07 | 4.56E-05 | 0 | 7.92E-05 |
| Wdr45b | 0.00017 | 0.00011 | 0.00014 | 0.00011 | 8.04E-05 | 8.10E-05 | 7.34E-05 | 1.63E-05 | 9.28E-05 | 6.08E-05 | 7.50E-05 | 8.03E-05 | 6.90E-05 | 0.00011 | 4.28E-05 | 7.04E-05 | 1.83E-05 | 8.37E-05 |
| Zdhhc7 | 4.52E-05 | 8.08E-05 | 9.57E-05 | 0.00022 | 7.15E-05 | 5.35E-05 | 6.50E-05 | 8.95E-05 | 9.28E-05 | 0.00011 | 7.93E-05 | 0 | 2.69E-05 | 8.78E-05 | 5.76E-05 | 5.85E-05 | 1.18E-06 | 9.52E-05 |
| Nufip1 | 5.93E-05 | 3.84E-05 | 0.00011 | 4.03E-05 | 8.15E-06 | 1.56E-05 | 2.02E-05 | 0 | 9.28E-05 | 1.28E-05 | 4.45E-05 | 7.31E-05 | 2.32E-05 | 3.90E-06 | 0.00015 | 7.40E-05 | 0 | 0 |
| Ascc3 | 0.0003 | 9.64E-05 | 0.00012 | 0 | 3.17E-05 | 0.00013 | 8.74E-05 | 1.68E-05 | 9.27E-05 | 2.68E-05 | 5.67E-05 | 0.00015 | 0 | 7.14E-05 | 0.00042 | 2.19E-05 | 0.00031 | 1.04E-05 |
| Il10rb | 8.68E-05 | 0.00014 | 0.00012 | 1.07E-05 | 6.66E-05 | 9.25E-05 | 9.16E-05 | 1.00E-05 | 9.27E-05 | 0.00029 | 4.18E-05 | 0.00024 | 0.00014 | 9.48E-05 | 1.83E-05 | 2.56E-05 | 0 | 3.14E-06 |
| Akr1b10 | 6.66E-05 | 6.88E-05 | 0 | 3.28E-05 | 5.50E-05 | 6.80E-05 | 1.69E-05 | 0 | 9.27E-05 | 6.10E-06 | 4.86E-05 | 1.39E-05 | 2.88E-05 | 0.00017 | 3.95E-05 | 2.65E-05 | 0 | 0.00012 |
| Robo2 | 0 | 3.73E-05 | 0 | 0 | 0 | 3.43E-05 | 0 | 0 | 9.27E-05 | 0 | 0 | 0 | 0 | 0.00012 | 0 | 0 | 0 | 3.62E-06 |

| | | | | | | | | | | | | | | | | | | |
|---|---|---|---|---|---|---|---|---|---|---|---|---|---|---|---|---|---|---|
| Caml | 7.52E-05 | 8.44E-05 | 0.00012 | 8.97E-05 | 5.40E-05 | 2.04E-05 | 0 | 8.11E-05 | 9.27E-05 | 8.98E-05 | 4.75E-05 | 3.93E-05 | 0 | 1.90E-05 | 0.00021 | 7.28E-05 | 0 | 6.32E-05 |
| Dhx30 | 6.64E-05 | 0.0001 | 0.00012 | 4.00E-05 | 4.03E-05 | 6.16E-06 | 0 | 4.17E-06 | 9.27E-05 | 2.63E-05 | 2.39E-05 | 1.67E-05 | 5.64E-06 | 3.76E-05 | 0.00012 | 5.29E-05 | 5.70E-05 | 2.99E-05 |
| Phf13 | 0 | 0.0002 | 8.84E-05 | 0.00018 | 0.00012 | 0 | 7.78E-06 | 3.11E-05 | 9.27E-05 | 8.57E-05 | 8.39E-05 | 0.00013 | 3.52E-05 | 0.00011 | 0.00014 | 0.00026 | 0.00017 | 1.49E-05 |
| Ctu2 | 3.07E-05 | 4.10E-05 | 0.00012 | 0.0001 | 3.11E-05 | 0 | 0 | 0 | 9.27E-05 | 3.02E-05 | 4.76E-05 | 0 | 3.75E-05 | 2.70E-05 | 4.17E-05 | 7.45E-05 | 0 | 3.37E-05 |
| Abcc5 | 5.35E-05 | 0.00026 | 0 | 0.00012 | 0.00012 | 0.00014 | 0.00012 | 4.00E-05 | 9.26E-05 | 0.00016 | 0.00011 | 0 | 6.36E-05 | 0.0001 | 0.00034 | 0 | 0 | 0.00013 |
| Klc1 | 0.00011 | 0.00019 | 0.00013 | 8.30E-06 | 3.69E-05 | 0.00011 | 0 | 1.21E-05 | 9.26E-05 | 0.00016 | 0.00028 | 0.00016 | 0 | 0.00016 | 0.00059 | 4.30E-05 | 8.91E-05 | 0.00015 |
| Ccdc47 | 0.00026 | 0.00012 | 0.00015 | 6.84E-05 | 4.85E-05 | 9.32E-05 | 7.86E-05 | 5.60E-05 | 9.26E-05 | 3.94E-05 | 6.53E-05 | 0.00019 | 2.36E-05 | 4.81E-05 | 4.91E-05 | 8.20E-05 | 0 | 5.53E-05 |
| Ankrd17 | 0.00021 | 0.00028 | 0.00021 | 0.00017 | 0.00014 | 0.00013 | 0.00018 | 0.00011 | 9.25E-05 | 9.96E-05 | 9.30E-05 | 0.00014 | 0.00013 | 0.00013 | 0.00026 | 7.22E-05 | 0.00024 | 8.65E-05 |
| Socs6 | 0.00012 | 0.00011 | 7.82E-05 | 1.36E-05 | 8.46E-05 | 1.71E-05 | 3.25E-05 | 7.14E-05 | 9.25E-05 | 8.17E-06 | 9.38E-05 | 3.58E-05 | 3.56E-05 | 9.62E-05 | 0 | 5.18E-05 | 7.19E-05 | 0 |
| Sri | 0.00048 | 0.00024 | 3.34E-05 | 0.00017 | 0.0002 | 0.00036 | 0.00019 | 7.75E-05 | 9.24E-05 | 0.00027 | 0.00027 | 0.00024 | 0.00011 | 0.00017 | 2.65E-05 | 8.87E-05 | 1.67E-06 | 0.00026 |
| Atf7ip | 0.00021 | 0.00024 | 0 | 8.72E-05 | 0.00023 | 0.00029 | 0.00012 | 0.00025 | 9.24E-05 | 6.47E-05 | 2.69E-05 | 0 | 0.00014 | 6.72E-05 | 0.00022 | 0.00017 | 9.54E-05 | 3.88E-05 |
| Ccar2 | 0.00012 | 0.00012 | 0.00011 | 0.00028 | 1.26E-05 | 6.04E-06 | 4.91E-05 | 5.13E-05 | 9.24E-05 | 6.98E-05 | 2.27E-05 | 4.82E-06 | 1.38E-05 | 0.0001 | 0.00016 | 5.76E-05 | 0.00013 | 7.45E-05 |
| Zfp422 | 2.04E-05 | 6.25E-05 | 0 | 3.12E-05 | 2.00E-05 | 1.99E-06 | 0 | 5.23E-05 | 9.24E-05 | 2.67E-05 | 3.19E-05 | 0 | 3.03E-05 | 3.94E-05 | 0 | 5.93E-05 | 0 | 0 |
| Trim26 | 0.0001 | 9.21E-05 | 5.61E-05 | 2.85E-05 | 1.84E-05 | 1.49E-06 | 9.31E-06 | 6.26E-05 | 9.24E-05 | 5.72E-05 | 2.27E-05 | 1.01E-05 | 3.90E-05 | 0 | 0.00022 | 2.67E-05 | 4.37E-05 | 6.77E-05 |
| Maf | 0 | 0.00026 | 0 | 0.00035 | 0.00116 | 0.00046 | 0.00027 | 0.00045 | 9.23E-05 | 0.00032 | 0 | 9.85E-05 | 0 | 0.00106 | 0.00111 | 9.10E-05 | 4.03E-05 | 0 |
| Gpr176 | 1.03E-07 | 6.84E-05 | 0.0002 | 0 | 0.00031 | 0.00031 | 0 | 0 | 9.23E-05 | 0 | 0.00108 | 0.00021 | 0.00076 | 0.00051 | 0 | 0 | 0 | 0 |
| Gtf3c1 | 3.35E-05 | 0.00013 | 8.83E-05 | 7.81E-05 | 6.68E-05 | 1.97E-05 | 1.60E-05 | 6.92E-05 | 9.23E-05 | 3.75E-05 | 8.42E-05 | 4.31E-05 | 0 | 4.15E-05 | 0.00012 | 7.63E-05 | 8.51E-05 | 6.10E-05 |
| Rnf41 | 2.98E-05 | 0.00016 | 0.00011 | 6.49E-05 | 0.00013 | 0 | 7.36E-05 | 8.84E-05 | 9.23E-05 | 1.35E-05 | 7.65E-05 | 7.48E-05 | 2.80E-06 | 1.87E-05 | 0.00013 | 0 | 8.33E-05 | 7.21E-05 |
| Pfdn2 | 0.00014 | 0.00015 | 0.00022 | 0.0001 | 7.85E-05 | 0 | 0 | 4.10E-05 | 9.23E-05 | 4.14E-05 | 6.90E-05 | 7.76E-05 | 5.08E-06 | 0 | 4.78E-05 | 8.06E-05 | 8.91E-05 | 1.31E-05 |
| Stard3nl | 0.00027 | 9.84E-05 | 9.98E-05 | 0 | 6.73E-05 | 0.00017 | 6.03E-05 | 0.00013 | 9.22E-05 | 0.0001 | 0.00012 | 6.50E-05 | 5.74E-05 | 2.90E-05 | 0 | 0.00014 | 0 | 6.17E-05 |
| Phf12 | 0.00013 | 0.00021 | 0.00013 | 0.00012 | 0.00014 | 0.00013 | 0.0001 | 1.78E-05 | 9.22E-05 | 5.74E-05 | 0.00014 | 3.26E-05 | 9.39E-05 | 9.87E-05 | 0.00043 | 0.00014 | 6.06E-05 | 2.64E-05 |
| Cnot1 | 0.0002 | 0.00017 | 0.00012 | 5.10E-05 | 0.00011 | 7.85E-05 | 0.00011 | 1.19E-05 | 9.22E-05 | 7.14E-05 | 6.99E-05 | 0.00016 | 2.02E-05 | 3.97E-05 | 0.00014 | 0.00034 | 0.00014 | 3.30E-05 |
| Helb | 6.27E-05 | 8.26E-05 | 0 | 1.33E-05 | 0.00011 | 5.39E-05 | 2.19E-05 | 0 | 9.22E-05 | 7.35E-06 | 6.84E-05 | 8.36E-05 | 1.69E-05 | 5.74E-05 | 6.83E-05 | 9.49E-05 | 0 | 0 |
| Rgp1 | 4.58E-05 | 8.87E-05 | 7.73E-05 | 6.29E-05 | 8.32E-05 | 4.14E-05 | 9.81E-05 | 7.57E-05 | 9.22E-05 | 3.71E-05 | 5.19E-05 | 6.73E-05 | 2.59E-05 | 6.19E-05 | 6.67E-05 | 0 | 8.81E-05 | 2.71E-05 |
| Kif2a | 7.60E-05 | 8.02E-05 | 0.00012 | 0 | 4.16E-05 | 3.33E-05 | 8.20E-07 | 9.16E-06 | 9.22E-05 | 0.00012 | 4.18E-05 | 0.00011 | 0 | 0 | 9.30E-05 | 0.00013 | 0 | 0 |
| Zfp704 | 0 | 0.00015 | 3.15E-05 | 2.77E-05 | 0.00013 | 1.54E-05 | 4.18E-05 | 1.90E-05 | 9.22E-05 | 4.37E-05 | 2.35E-05 | 0 | 0 | 0.0002 | 0.00028 | 8.43E-05 | 0.00014 | 0.00015 |
| D16Ertd472e | 9.81E-05 | 6.64E-05 | 7.19E-05 | 0 | 7.06E-05 | 3.23E-06 | 0 | 0 | 9.22E-05 | 0 | 9.33E-05 | 0 | 6.02E-06 | 3.21E-05 | 2.50E-05 | 1.84E-05 | 0.0001 | 0 |
| Gtf2a1 | 0.00012 | 0.00019 | 9.08E-05 | 5.15E-05 | 5.90E-05 | 7.66E-05 | 5.93E-05 | 8.82E-05 | 9.20E-05 | 6.67E-05 | 6.40E-05 | 0.00016 | 0 | 1.78E-05 | 2.20E-05 | 0.00019 | 0.00013 | 3.83E-05 |
| Tcf7l1 | 1.70E-05 | 9.01E-05 | 5.55E-05 | 7.62E-05 | 0 | 6.54E-05 | 0 | 0 | 9.20E-05 | 0.00015 | 3.57E-05 | 4.85E-05 | 4.55E-05 | 7.28E-05 | 4.76E-05 | 0.00012 | 8.87E-05 | 0.00012 |
| Nmt2 | 0.00018 | 8.61E-05 | 0.00018 | 0 | 9.85E-05 | 2.42E-05 | 0.00016 | 8.97E-05 | 9.20E-05 | 0.0003 | 8.03E-05 | 4.35E-05 | 0.0001 | 8.17E-05 | 0.00015 | 0.0002 | 0.00012 | 8.82E-05 |
| Tle2 | 0 | 7.47E-05 | 0 | 2.06E-05 | 1.27E-06 | 1.29E-05 | 0 | 0 | 9.20E-05 | 5.80E-05 | 2.93E-05 | 0 | 8.91E-05 | 0.00011 | 0.00016 | 0 | 1.11E-05 | 6.15E-05 |
| Aifm2 | 1.51E-05 | 0.0001 | 2.59E-05 | 6.18E-05 | 9.32E-05 | 5.48E-06 | 3.99E-05 | 2.59E-05 | 9.20E-05 | 7.00E-05 | 0 | 7.35E-05 | 1.82E-05 | 9.30E-07 | 0 | 0 | 2.24E-07 | 7.25E-05 |
| Btbd7 | 0.00016 | 0.00015 | 8.41E-05 | 0 | 0.00011 | 0.00017 | 6.92E-05 | 7.75E-05 | 9.19E-05 | 9.69E-05 | 0.00011 | 9.88E-05 | 3.07E-05 | 7.30E-05 | 0 | 4.25E-05 | 0.00021 | 0.00012 |

| | | | | | | | | | | | | | | | | | | |
|---|---|---|---|---|---|---|---|---|---|---|---|---|---|---|---|---|---|---|
| Trappc5 | 7.23E-05 | 0.00012 | 0.00015 | 0.00018 | 8.59E-05 | 7.83E-05 | 5.54E-06 | 9.05E-05 | 9.19E-05 | 6.16E-05 | 0.00012 | 0.00018 | 7.61E-05 | 0.00012 | 0 | 0.00015 | 0 | 6.24E-05 |
| Ankfy1 | 0.00019 | 0.00011 | 6.63E-05 | 0 | 3.59E-05 | 5.62E-05 | 3.14E-05 | 3.25E-05 | 9.19E-05 | 8.16E-05 | 3.47E-05 | 6.46E-05 | 4.15E-05 | 0 | 6.02E-05 | 4.84E-05 | 8.75E-05 | 9.79E-05 |
| Zkscan3 | 1.92E-05 | 0.00013 | 7.60E-05 | 0.00012 | 7.23E-05 | 3.76E-05 | 5.62E-05 | 8.43E-05 | 9.19E-05 | 7.05E-05 | 4.22E-05 | 2.19E-05 | 2.18E-05 | 6.17E-05 | 0.00021 | 0 | 0 | 5.18E-05 |
| Sh3kbp1 | 0.00017 | 4.46E-05 | 8.14E-05 | 0 | 2.51E-05 | 0 | 0.00021 | 0.00011 | 9.19E-05 | 5.08E-06 | 0.00034 | 0.00025 | 0 | 0 | 3.56E-06 | 0.0001 | 4.48E-05 | 5.98E-05 |
| Lars | 0.0002 | 0.0001 | 0.00016 | 0.00012 | 4.47E-05 | 5.56E-05 | 3.69E-05 | 2.09E-05 | 9.18E-05 | 7.93E-05 | 1.81E-05 | 0.00013 | 0 | 3.00E-05 | 0.0001 | 0.00021 | 4.00E-05 | 4.79E-05 |
| Ewsr1 | 0.00012 | 4.22E-05 | 5.28E-05 | 7.68E-05 | 2.15E-07 | 4.49E-05 | 1.26E-06 | 0 | 9.18E-05 | 0 | 0 | 5.27E-05 | 0 | 2.62E-05 | 9.62E-05 | 9.81E-05 | 4.06E-05 | 0 |
| Epha3 | 0.00011 | 0 | 0 | 0 | 0.00015 | 0 | 0 | 0.00051 | 9.18E-05 | 0 | 0.00036 | 0 | 4.66E-05 | 0 | 0 | 0.00013 | 0 | 0.00087 |
| Hps1 | 2.05E-05 | 9.70E-05 | 9.00E-06 | 0.00013 | 5.05E-05 | 0 | 0.0001 | 0 | 9.18E-05 | 5.18E-05 | 1.94E-05 | 6.81E-05 | 1.91E-05 | 5.79E-05 | 0.00017 | 1.55E-05 | 7.48E-05 | 8.71E-05 |
| Senp5 | 0.00024 | 0.00012 | 0.00016 | 0 | 6.70E-05 | 0.00013 | 6.68E-05 | 9.53E-06 | 9.17E-05 | 3.65E-06 | 0.00011 | 5.46E-05 | 6.69E-06 | 3.64E-05 | 0.00017 | 5.20E-06 | 1.53E-05 | 0 |
| Rala | 0.00019 | 0.0001 | 0.00021 | 5.22E-05 | 4.87E-05 | 6.00E-05 | 2.06E-05 | 3.25E-05 | 9.17E-05 | 0.0001 | 0.00015 | 0.00023 | 0 | 0 | 0 | 8.06E-05 | 3.64E-05 | 6.48E-05 |
| Fap | 0.00016 | 8.26E-05 | 0 | 0 | 7.18E-05 | 0 | 0.00011 | 0.00088 | 9.17E-05 | 0.00026 | 0 | 0 | 8.23E-05 | 0 | 0 | 0 | 0 | 0.00013 |
| D5Ertd579e | 9.14E-05 | 0.00016 | 0.00011 | 6.59E-06 | 9.77E-05 | 5.90E-05 | 0 | 9.35E-05 | 9.16E-05 | 0.00011 | 0.00012 | 4.86E-05 | 4.17E-05 | 6.85E-05 | 0.00015 | 6.25E-05 | 0 | 5.40E-05 |
| Mtmr11 | 6.58E-06 | 0.00011 | 0 | 6.10E-05 | 8.80E-05 | 1.56E-05 | 0.00013 | 0 | 9.16E-05 | 0.00017 | 7.66E-05 | 8.88E-05 | 0 | 5.34E-05 | 8.88E-05 | 4.72E-06 | 5.69E-06 | 0.00022 |
| Stxbp1 | 4.18E-05 | 8.92E-05 | 0.00018 | 3.25E-05 | 4.22E-05 | 0.00017 | 0 | 0.00014 | 9.15E-05 | 8.08E-05 | 0.00012 | 0.00021 | 0 | 3.94E-05 | 8.85E-05 | 3.36E-05 | 4.54E-05 | 5.92E-05 |
| Tnfsf13 | 1.79E-05 | 7.79E-05 | 0 | 0 | 8.94E-05 | 8.38E-05 | 1.96E-06 | 0 | 9.15E-05 | 5.12E-05 | 7.13E-06 | 1.26E-05 | 9.89E-06 | 5.92E-05 | 5.17E-06 | 0 | 4.24E-05 | 4.04E-05 |
| Ubr7 | 5.62E-05 | 6.60E-05 | 6.77E-05 | 0.00011 | 7.54E-05 | 4.43E-05 | 4.15E-06 | 4.44E-05 | 9.15E-05 | 5.67E-05 | 5.89E-05 | 3.46E-05 | 4.79E-05 | 3.26E-05 | 8.59E-06 | 0.00029 | 0 | 5.89E-05 |
| Rbm28 | 7.16E-05 | 0.00011 | 0.00018 | 8.92E-05 | 3.76E-05 | 0 | 0 | 0 | 9.15E-05 | 7.42E-05 | 3.25E-05 | 0 | 0 | 0 | 0.0002 | 0.00012 | 0.00011 | 6.70E-05 |
| Lmbrd1 | 0.00017 | 0.00014 | 8.79E-06 | 0 | 4.74E-05 | 0.00021 | 1.69E-05 | 4.43E-05 | 9.14E-05 | 3.76E-05 | 1.24E-05 | 9.22E-05 | 0 | 4.22E-05 | 1.21E-05 | 0 | 5.34E-05 | 0.00011 |
| Ost4 | 0.00031 | 0 | 0.00015 | 0.00028 | 0.00011 | 0.00019 | 0.00036 | 0.00022 | 9.14E-05 | 5.86E-05 | 0.00021 | 0.00014 | 0.00014 | 0.00011 | 0 | 4.67E-05 | 0 | 0 |
| Tsnax | 0.00014 | 4.97E-05 | 7.04E-05 | 7.13E-05 | 5.01E-05 | 4.02E-05 | 0 | 4.55E-05 | 9.14E-05 | 3.99E-05 | 4.00E-05 | 0.00011 | 6.32E-06 | 5.04E-05 | 8.13E-05 | 3.15E-05 | 0 | 5.46E-05 |
| E2f1 | 0 | 0 | 6.10E-05 | 0 | 5.83E-05 | 0 | 2.87E-05 | 7.67E-05 | 9.14E-05 | 4.92E-05 | 6.92E-06 | 0.00014 | 2.47E-05 | 7.19E-06 | 0 | 0.00027 | 5.70E-05 | 1.58E-05 |
| Tap1 | 2.73E-05 | 5.35E-05 | 1.29E-05 | 0.00012 | 0 | 0 | 0 | 2.49E-05 | 9.14E-05 | 0.00023 | 0 | 6.74E-05 | 0.00015 | 5.12E-05 | 3.99E-05 | 8.36E-05 | 7.96E-05 | 0.00018 |
| Ube2q2 | 0.0002 | 0.00012 | 6.10E-05 | 0 | 4.24E-05 | 0.00018 | 1.11E-05 | 8.82E-06 | 9.13E-05 | 0 | 0 | 0.00011 | 3.37E-05 | 3.62E-05 | 1.03E-05 | 3.21E-05 | 0.00015 | 0 |
| Metap1 | 0.00014 | 9.54E-05 | 0.00014 | 3.98E-05 | 3.55E-05 | 5.39E-05 | 1.99E-05 | 0 | 9.13E-05 | 5.70E-05 | 9.20E-05 | 3.42E-05 | 6.01E-05 | 6.18E-05 | 5.88E-05 | 9.06E-05 | 0 | 1.05E-05 |
| Fzr1 | 3.01E-05 | 0.00014 | 4.36E-05 | 9.21E-05 | 4.93E-05 | 3.75E-05 | 6.95E-05 | 6.72E-05 | 9.13E-05 | 8.46E-05 | 9.62E-05 | 9.47E-05 | 7.05E-06 | 4.10E-05 | 0.00013 | 0.00059 | 1.56E-05 | 8.63E-05 |
| Dnajc10 | 0.00031 | 0.00012 | 0.00031 | 0.00032 | 0.00015 | 0.00031 | 0.00039 | 0.00033 | 9.11E-05 | 0.0004 | 9.17E-05 | 3.13E-05 | 0.00026 | 0.00028 | 0 | 0.00042 | 7.01E-05 | 0.00043 |
| Cxcl12 | 0.00042 | 0 | 6.07E-05 | 0.00069 | 3.20E-05 | 0.00027 | 0 | 0.0002 | 9.11E-05 | 0.00049 | 0 | 0 | 0.00416 | 0.00021 | 2.58E-05 | 0.00017 | 0 | 0.00137 |
| Socs1 | 0 | 3.08E-05 | 3.36E-05 | 0.00031 | 3.01E-05 | 2.84E-05 | 0 | 0.00014 | 9.11E-05 | 0.0001 | 0 | 7.95E-05 | 0.00014 | 0 | 4.71E-05 | 4.08E-05 | 0.00115 | 0.00011 |
| Top1 | 0.00036 | 0.00013 | 0.00018 | 0 | 9.88E-05 | 0.00016 | 0 | 0.00012 | 9.10E-05 | 0 | 6.43E-05 | 3.90E-06 | 1.74E-05 | 5.71E-05 | 0.00018 | 0.0002 | 0.0004 | 0 |
| Ube2k | 0.00032 | 0.00014 | 0.00027 | 1.88E-05 | 0.0001 | 0.00014 | 4.95E-05 | 3.53E-05 | 9.10E-05 | 4.18E-05 | 8.28E-05 | 5.66E-05 | 2.29E-05 | 7.60E-05 | 0 | 0.00022 | 9.54E-05 | 0 |
| Tjp1 | 0.00018 | 0.00025 | 0 | 0 | 0.0001 | 6.71E-05 | 0 | 0.00016 | 9.10E-05 | 5.89E-05 | 9.16E-05 | 0.0003 | 0 | 8.78E-05 | 6.92E-06 | 5.04E-05 | 0.00019 | 5.04E-05 |
| Cisd1 | 0.00012 | 2.01E-05 | 0.00019 | 4.25E-05 | 5.15E-05 | 6.00E-05 | 0 | 5.72E-06 | 9.10E-05 | 2.91E-05 | 5.23E-05 | 0 | 1.87E-05 | 3.92E-05 | 0 | 0.00014 | 0 | 5.34E-05 |
| Srp19 | 0.00043 | 5.40E-05 | 0.00025 | 0.0002 | 8.71E-05 | 0.00015 | 0.00021 | 0.00011 | 9.09E-05 | 5.64E-05 | 0.00011 | 0.00011 | 6.25E-05 | 0.00011 | 0.00016 | 6.06E-05 | 0 | 7.18E-06 |

| | | | | | | | | | | | | | | | | | | |
|---|---|---|---|---|---|---|---|---|---|---|---|---|---|---|---|---|---|---|
| Tmem159 | 1.84E-05 | 0.00016 | 0 | 0.0001 | 0.0001 | 0.00013 | 5.45E-05 | 0.0001 | 9.09E-05 | 0.00012 | 8.18E-05 | 0.00016 | 7.46E-05 | 0.00015 | 0 | 0 | 0 | 0.00017 |
| Gulp1 | 0.00015 | 5.06E-05 | 0 | 0 | 2.18E-05 | 0.0001 | 1.71E-05 | 0 | 9.09E-05 | 0.00016 | 8.56E-05 | 0 | 0.0001 | 4.07E-05 | 2.51E-05 | 2.16E-05 | 2.29E-05 | 6.59E-05 |
| Ccn4 | 9.65E-05 | 0 | 0.00048 | 0 | 0.00057 | 0.00058 | 0.00028 | 0.0002 | 9.08E-05 | 0 | 0.00068 | 0 | 0.00021 | 0.00078 | 0.00015 | 0.00011 | 0 | 0 |
| Ube3a | 0.00029 | 0.00023 | 0.0002 | 9.11E-06 | 0.00014 | 0.00015 | 4.76E-05 | 3.02E-05 | 9.08E-05 | 6.36E-05 | 7.38E-05 | 0.00014 | 1.37E-05 | 2.80E-05 | 0.00034 | 8.91E-05 | 2.20E-05 | 2.78E-05 |
| Tmx1 | 3.23E-05 | 9.36E-05 | 0.00029 | 0.00037 | 8.17E-05 | 8.68E-05 | 0.00025 | 0.00028 | 9.08E-05 | 0.00016 | 8.29E-05 | 0.00022 | 0.0001 | 0.00016 | 3.53E-05 | 0.00049 | 0.00016 | 0.00018 |
| Psmb1 | 0.00034 | 0.00015 | 0.00016 | 0.00012 | 5.01E-05 | 0.00023 | 9.39E-05 | 6.64E-05 | 9.07E-05 | 3.36E-05 | 0.00011 | 0.0001 | 8.25E-05 | 0.00012 | 0 | 0.00011 | 0 | 4.58E-05 |
| Idh3a | 0.00023 | 9.52E-05 | 0.00037 | 4.59E-05 | 0.00011 | 0.00013 | 0 | 4.13E-05 | 9.07E-05 | 5.25E-05 | 0.00039 | 0 | 0 | 0 | 0 | 0.00026 | 0 | 6.12E-05 |
| Ankmy2 | 0.00011 | 5.02E-05 | 0 | 4.16E-07 | 3.34E-05 | 4.50E-05 | 4.49E-05 | 5.31E-05 | 9.07E-05 | 3.11E-05 | 0 | 1.57E-05 | 6.91E-06 | 0 | 7.50E-05 | 2.31E-05 | 0 | 5.15E-05 |
| Ppp1r16a | 0 | 7.31E-05 | 8.31E-05 | 0.00014 | 4.61E-05 | 0 | 4.25E-05 | 5.69E-05 | 9.07E-05 | 9.05E-05 | 2.55E-05 | 0 | 3.90E-05 | 9.42E-05 | 0.00015 | 9.19E-05 | 4.45E-05 | 9.05E-05 |
| Eif3g | 0.00044 | 0.00023 | 0.0005 | 0.0003 | 7.57E-05 | 0.00014 | 8.40E-05 | 0 | 9.06E-05 | 2.79E-07 | 0.00016 | 4.69E-05 | 1.05E-05 | 0.00011 | 0.00016 | 0.00012 | 0 | 0 |
| Nsmaf | 5.53E-05 | 0.00014 | 0 | 7.81E-05 | 7.13E-05 | 7.96E-05 | 9.63E-05 | 0 | 9.06E-05 | 7.46E-05 | 4.14E-06 | 2.10E-05 | 8.44E-06 | 5.70E-05 | 9.18E-05 | 3.36E-05 | 1.59E-05 | 6.38E-05 |
| Tmco4 | 3.78E-07 | 1.54E-05 | 2.69E-05 | 6.18E-05 | 5.69E-05 | 5.65E-05 | 3.24E-05 | 3.43E-05 | 9.06E-05 | 5.63E-05 | 0 | 0.00011 | 0.00011 | 4.70E-05 | 0 | 0.00016 | 0 | 7.69E-05 |
| Rab10 | 0.00025 | 0.00015 | 0.00019 | 0 | 0.00011 | 0.00015 | 3.70E-05 | 0.00013 | 9.05E-05 | 0.00011 | 0.00011 | 0.00017 | 2.15E-05 | 5.93E-05 | 0.00011 | 6.95E-05 | 0.00013 | 7.74E-05 |
| Psmd6 | 0.00041 | 0.00014 | 0.00032 | 5.51E-05 | 5.18E-05 | 9.69E-05 | 2.22E-05 | 0 | 9.05E-05 | 4.19E-05 | 0.00014 | 2.77E-05 | 3.61E-06 | 4.79E-05 | 7.49E-05 | 9.71E-05 | 0 | 0 |
| Dnajc5 | 0.00019 | 0.00018 | 0.00023 | 6.78E-05 | 0.00013 | 8.89E-05 | 4.76E-05 | 0.00015 | 9.05E-05 | 0.00013 | 0.00015 | 0.00043 | 0.00011 | 9.45E-05 | 0.00013 | 8.36E-05 | 0 | 6.08E-05 |
| Casp7 | 1.91E-05 | 2.41E-05 | 6.24E-05 | 1.65E-05 | 8.35E-06 | 3.13E-05 | 0 | 9.72E-05 | 9.05E-05 | 3.27E-05 | 3.87E-06 | 0 | 7.46E-05 | 2.85E-05 | 0 | 0.00021 | 0 | 6.38E-05 |
| Rdh11 | 5.05E-05 | 9.21E-05 | 2.88E-05 | 5.32E-05 | 1.72E-05 | 1.14E-06 | 0 | 0 | 9.05E-05 | 2.03E-06 | 6.81E-05 | 7.28E-05 | 0 | 1.05E-05 | 0 | 0.00021 | 3.94E-05 | 4.95E-05 |
| Htt | 6.81E-05 | 6.81E-05 | 0.00015 | 0 | 5.26E-05 | 0 | 6.73E-05 | 6.40E-05 | 9.05E-05 | 8.74E-05 | 8.32E-05 | 0.0001 | 0 | 0 | 0.00012 | 7.64E-05 | 0.00011 | 3.29E-05 |
| Sumo1 | 0.00035 | 0.00013 | 0.00011 | 0.00022 | 0.00013 | 0.00023 | 9.75E-05 | 9.10E-05 | 9.04E-05 | 3.04E-05 | 6.04E-05 | 9.58E-05 | 0 | 0.00011 | 0 | 0.00016 | 6.64E-05 | 5.52E-05 |
| Kctd9 | 6.67E-05 | 5.35E-05 | 3.48E-05 | 0 | 2.01E-05 | 4.00E-05 | 5.53E-05 | 0 | 9.04E-05 | 0.00022 | 0.00015 | 0.00032 | 0 | 0 | 5.98E-05 | 0.00035 | 0 | 0.00016 |
| Dpm2 | 0.0001 | 0.0001 | 0.00013 | 0.00015 | 4.53E-05 | 3.36E-05 | 0.00014 | 6.06E-05 | 9.04E-05 | 7.54E-05 | 6.58E-05 | 1.07E-05 | 7.90E-05 | 4.33E-05 | 0 | 2.19E-05 | 0 | 5.52E-05 |
| Hmgn5 | 6.26E-05 | 0 | 4.85E-05 | 0.00012 | 6.35E-06 | 1.29E-06 | 0 | 0 | 9.03E-05 | 0 | 6.72E-06 | 0 | 0 | 3.63E-05 | 2.13E-05 | 0.00019 | 2.16E-06 | 5.95E-06 |
| Nkain1 | 0.00011 | 7.14E-05 | 0.00017 | 7.72E-05 | 3.54E-06 | 0 | 0.0002 | 0 | 9.03E-05 | 0 | 4.74E-05 | 5.03E-05 | 2.78E-05 | 0 | 0 | 0 | 0 | 0 |
| Rbm5 | 0.00017 | 0.00032 | 3.89E-05 | 0.00015 | 0.00016 | 9.74E-05 | 7.54E-05 | 5.31E-05 | 9.02E-05 | 0.00013 | 3.58E-05 | 0.00016 | 0 | 0.00014 | 0.00045 | 8.55E-05 | 0 | 0.0001 |
| Tspan14 | 5.57E-05 | 9.88E-05 | 3.69E-06 | 0 | 8.12E-05 | 7.02E-05 | 0 | 0.00013 | 9.02E-05 | 2.37E-05 | 8.06E-05 | 7.11E-05 | 0 | 8.41E-05 | 0 | 9.68E-05 | 0 | 7.93E-05 |
| Magi3 | 1.57E-05 | 4.26E-05 | 0 | 0 | 1.46E-05 | 1.83E-05 | 5.76E-05 | 0 | 9.02E-05 | 9.69E-05 | 2.93E-05 | 4.89E-05 | 3.98E-05 | 5.37E-06 | 0 | 5.92E-05 | 5.39E-07 | 3.41E-05 |
| Acbd6 | 7.34E-05 | 0.00012 | 8.70E-05 | 0.00018 | 8.17E-05 | 5.53E-05 | 5.00E-06 | 0 | 9.00E-05 | 1.61E-05 | 5.02E-05 | 0.00019 | 2.33E-05 | 5.80E-05 | 0.0002 | 8.29E-05 | 4.75E-05 | 6.98E-05 |
| Ap1m1 | 0.00023 | 0.00014 | 0.00015 | 0.0001 | 4.01E-05 | 4.76E-05 | 9.02E-05 | 2.81E-05 | 9.00E-05 | 0.00014 | 9.17E-05 | 3.97E-05 | 0 | 4.71E-05 | 9.80E-05 | 0.00011 | 0 | 9.69E-05 |
| Nsun2 | 0.00029 | 7.16E-05 | 0.0004 | 7.80E-05 | 0 | 0 | 8.39E-05 | 1.66E-05 | 9.00E-05 | 0 | 2.33E-05 | 0.00024 | 0.00013 | 0 | 7.17E-05 | 5.14E-05 | 1.29E-05 | 0 |
| Tmem258 | 0.00046 | 3.29E-05 | 0.00036 | 0.00059 | 0.00012 | 0.00021 | 0.0006 | 0.00023 | 8.99E-05 | 6.04E-05 | 0.00015 | 0.00015 | 0.00032 | 0.00019 | 0 | 0.00014 | 0 | 5.58E-06 |
| Sox12 | 1.04E-05 | 2.42E-05 | 2.74E-05 | 8.41E-05 | 2.27E-05 | 2.62E-05 | 0 | 7.91E-05 | 8.99E-05 | 0 | 2.64E-05 | 1.59E-05 | 7.21E-05 | 0 | 0.00021 | 4.57E-05 | 0 | 0 |
| Gxylt2 | 3.09E-05 | 0.00011 | 0 | 0.00016 | 0.00061 | 1.44E-05 | 0.00046 | 0 | 8.99E-05 | 0.00014 | 0.00018 | 0 | 0.00022 | 0.00031 | 0 | 2.17E-05 | 0 | 0.00059 |
| Sucla2 | 0.00024 | 0.00013 | 0.00015 | 5.27E-05 | 5.80E-05 | 0.00011 | 3.16E-07 | 0 | 8.98E-05 | 7.75E-05 | 0.00011 | 3.84E-05 | 5.09E-05 | 6.33E-05 | 0.0001 | 0.00013 | 3.25E-05 | 3.92E-05 |

| | | | | | | | | | | | | | | | | | | |
|---|---|---|---|---|---|---|---|---|---|---|---|---|---|---|---|---|---|---|
| Zfp384 | 0.00014 | 0.00017 | 0.00011 | 0.00017 | 4.79E-05 | 8.24E-05 | 7.11E-05 | 6.89E-05 | 8.98E-05 | 4.80E-05 | 1.71E-05 | 9.53E-05 | 3.85E-05 | 8.19E-05 | 0.00011 | 0.00014 | 7.53E-06 | 7.65E-05 |
| Cpsf2 | 0.00015 | 6.20E-05 | 0.00018 | 4.69E-05 | 3.66E-05 | 2.17E-05 | 1.57E-05 | 0 | 8.98E-05 | 7.13E-05 | 7.72E-05 | 0 | 3.31E-05 | 4.20E-05 | 2.79E-05 | 0.00016 | 0 | 1.89E-05 |
| Ascc2 | 7.14E-05 | 0.00011 | 9.07E-05 | 5.79E-05 | 8.19E-06 | 1.01E-05 | 4.32E-06 | 7.37E-06 | 8.97E-05 | 3.31E-05 | 4.83E-05 | 5.48E-05 | 0 | 1.21E-05 | 0.00016 | 3.12E-05 | 8.13E-05 | 4.84E-05 |
| Slc6a8 | 0 | 0.0004 | 0.00039 | 0.00049 | 0.00062 | 0.00042 | 6.29E-05 | 4.69E-06 | 8.96E-05 | 0 | 0 | 0.00043 | 0.00026 | 0.00027 | 0.0004 | 0 | 0.00081 | 0 |
| P2ry2 | 0 | 8.59E-05 | 8.73E-05 | 0 | 2.96E-06 | 0.00016 | 0 | 0 | 8.96E-05 | 0 | 0 | 0.00035 | 2.25E-05 | 8.76E-05 | 0 | 0 | 0 | 0 |
| Zdhhc16 | 0.0001 | 7.04E-05 | 0.00016 | 0.00017 | 0.00016 | 1.85E-06 | 7.70E-05 | 0.0001 | 8.96E-05 | 1.88E-05 | 0.00029 | 0 | 0 | 9.70E-05 | 9.81E-05 | 0.00019 | 0 | 6.33E-05 |
| Snap47 | 0.00013 | 7.69E-05 | 1.87E-05 | 0.00017 | 0.00011 | 0 | 0.00019 | 1.42E-05 | 8.96E-05 | 5.15E-05 | 9.86E-05 | 3.88E-05 | 6.73E-05 | 4.59E-05 | 4.05E-05 | 2.88E-05 | 0 | 0.00016 |
| Cad | 0.00017 | 1.46E-05 | 0.00031 | 0.00044 | 1.22E-05 | 0 | 0.00014 | 4.73E-05 | 8.95E-05 | 2.99E-05 | 3.77E-05 | 0 | 0 | 2.74E-05 | 0.00015 | 0.00021 | 0 | 2.44E-05 |
| Sec61a1 | 0.00071 | 2.64E-05 | 0.00047 | 0.00016 | 0.00016 | 0.00037 | 0.00077 | 0.00029 | 8.94E-05 | 4.99E-05 | 0.0002 | 0.00013 | 0.00013 | 0.00026 | 8.10E-05 | 0.00012 | 0 | 2.15E-05 |
| Nsa2 | 0.00024 | 0.00012 | 2.31E-06 | 0 | 0 | 0.0002 | 6.80E-05 | 0 | 8.94E-05 | 4.09E-07 | 0 | 0.00017 | 0 | 0 | 0 | 3.92E-05 | 4.78E-05 | 9.31E-06 |
| Uqcc2 | 0.00034 | 3.88E-05 | 0.00026 | 0.00019 | 0.00012 | 8.50E-05 | 0.00015 | 7.26E-05 | 8.94E-05 | 0.0001 | 7.31E-05 | 0 | 0 | 0.00019 | 0 | 0.00013 | 0 | 0.0001 |
| Mapkap1 | 0.00015 | 0.0001 | 0.00012 | 4.13E-05 | 9.22E-05 | 8.10E-05 | 0.0001 | 2.69E-05 | 8.94E-05 | 6.64E-05 | 7.57E-05 | 0 | 1.34E-05 | 1.49E-05 | 0 | 0.0001 | 8.88E-05 | 0.00011 |
| Smad3 | 0 | 0.00014 | 0.00032 | 9.47E-05 | 0 | 5.24E-05 | 0.00023 | 0.00043 | 8.94E-05 | 6.19E-05 | 0 | 0.0001 | 0.00022 | 0 | 5.05E-05 | 5.51E-05 | 0.00023 | 0.00028 |
| Erlec1 | 0.00034 | 0 | 0 | 0 | 7.21E-05 | 0.00037 | 0.00028 | 0.00025 | 8.93E-05 | 5.07E-05 | 5.17E-05 | 9.00E-05 | 0.00018 | 0.00019 | 5.74E-05 | 1.70E-05 | 0 | 0.00013 |
| Ube4a | 0.0002 | 0.00015 | 7.03E-05 | 4.82E-05 | 9.41E-05 | 0.00013 | 9.74E-05 | 6.83E-05 | 8.93E-05 | 5.11E-05 | 4.12E-05 | 8.59E-05 | 0 | 0.00011 | 9.06E-05 | 2.37E-05 | 3.33E-05 | 3.25E-05 |
| Pigu | 6.59E-05 | 4.82E-05 | 7.31E-05 | 8.05E-05 | 3.65E-05 | 6.06E-05 | 1.79E-05 | 5.13E-05 | 8.93E-05 | 4.18E-05 | 4.64E-05 | 1.92E-05 | 0.00011 | 7.11E-05 | 3.67E-05 | 9.36E-05 | 0 | 5.96E-05 |
| Rfx5 | 2.55E-05 | 3.08E-05 | 1.10E-05 | 0 | 2.36E-05 | 2.56E-05 | 1.98E-05 | 0 | 8.93E-05 | 2.62E-05 | 2.77E-06 | 1.44E-05 | 5.92E-05 | 4.45E-05 | 4.22E-05 | 1.11E-05 | 4.01E-06 | 8.17E-05 |
| Ripor1 | 2.19E-05 | 0.00017 | 0.00014 | 0.00025 | 4.81E-05 | 1.88E-06 | 0 | 0.00016 | 8.93E-05 | 0.0001 | 2.40E-05 | 0.00022 | 0 | 1.95E-05 | 7.69E-05 | 5.14E-05 | 7.14E-05 | 0.00013 |
| Ube2w | 0.00023 | 0.0001 | 6.49E-05 | 9.59E-06 | 0.00014 | 0.00017 | 0.0001 | 9.46E-05 | 8.92E-05 | 0.00014 | 0.00024 | 0.00011 | 0.00014 | 1.37E-05 | 0.00019 | 0 | 0 | 7.75E-05 |
| Ube2v2 | 0.00024 | 1.53E-05 | 0.00018 | 0 | 4.18E-05 | 0.00012 | 7.57E-06 | 0 | 8.92E-05 | 0 | 0 | 0.00023 | 0 | 0 | 0 | 0 | 0.00102 | 0 |
| Abcd3 | 0.00015 | 5.94E-05 | 4.56E-05 | 0 | 4.89E-06 | 4.74E-05 | 0 | 0 | 8.92E-05 | 5.38E-05 | 2.84E-05 | 4.09E-05 | 4.49E-05 | 4.77E-05 | 1.29E-06 | 5.78E-05 | 0 | 0.0001 |
| Cox17 | 0.00012 | 9.32E-05 | 0.00034 | 0.0002 | 1.61E-05 | 0 | 2.44E-05 | 0.00016 | 8.92E-05 | 3.06E-05 | 0.00038 | 0.00012 | 6.09E-05 | 5.72E-05 | 8.11E-05 | 0.0001 | 0 | 0 |
| Rab5a | 0.00042 | 0.0003 | 0.00024 | 1.98E-05 | 0.00024 | 0.00038 | 0.00025 | 0.00018 | 8.91E-05 | 9.80E-05 | 0.00024 | 0.0003 | 2.89E-05 | 0.00018 | 0.00012 | 9.14E-05 | 0.00045 | 8.83E-05 |
| Slmap | 0.00018 | 0.00017 | 7.26E-05 | 0 | 9.37E-05 | 0.00015 | 4.89E-05 | 3.85E-05 | 8.91E-05 | 4.68E-05 | 5.58E-05 | 3.99E-05 | 0 | 3.47E-05 | 9.49E-05 | 7.76E-05 | 0.00017 | 6.44E-05 |
| Psmc6 | 0.00043 | 0.00026 | 0.00031 | 0.00019 | 4.45E-05 | 0.00015 | 0 | 1.14E-05 | 8.91E-05 | 3.29E-05 | 1.95E-05 | 0.00019 | 0 | 8.57E-05 | 0.00026 | 0.00027 | 0.00014 | 0 |
| Ric8a | 7.72E-05 | 0.0002 | 0.00016 | 0.00022 | 8.34E-05 | 8.09E-05 | 0.00017 | 0.00012 | 8.91E-05 | 0.00016 | 0.00015 | 0.00023 | 0 | 0.00012 | 0.00014 | 0 | 0 | 0.00017 |
| Aamdc | 5.29E-05 | 7.17E-05 | 0 | 6.29E-05 | 9.28E-05 | 3.12E-06 | 9.55E-05 | 0 | 8.91E-05 | 7.53E-05 | 5.21E-05 | 0 | 0 | 4.93E-05 | 7.50E-06 | 0 | 0 | 0.00011 |
| Ctcf | 0.00014 | 0.00014 | 7.22E-05 | 0 | 5.36E-05 | 5.33E-05 | 3.52E-05 | 3.58E-05 | 8.90E-05 | 1.98E-05 | 1.85E-05 | 2.67E-05 | 2.72E-05 | 0 | 0.00022 | 0.00038 | 3.83E-05 | 2.27E-05 |
| Ppig | 0.00012 | 0.00012 | 0.00011 | 8.87E-05 | 4.43E-05 | 5.28E-05 | 2.78E-05 | 4.38E-05 | 8.90E-05 | 5.18E-05 | 4.34E-05 | 7.02E-05 | 6.19E-06 | 9.67E-05 | 3.33E-05 | 0.00016 | 0 | 1.89E-05 |
| Lgi4 | 0 | 1.39E-06 | 0 | 6.71E-06 | 0 | 2.28E-05 | 0 | 0.00011 | 8.90E-05 | 0.00032 | 0 | 1.14E-05 | 6.11E-05 | 0 | 0 | 0 | 0 | 0.00027 |
| Slc38a10 | 6.28E-05 | 0 | 0.00019 | 0.00078 | 0.00027 | 0.00037 | 0.00096 | 0.00049 | 8.89E-05 | 0.00033 | 0.00018 | 0.00024 | 0.00054 | 0.00046 | 0.0004 | 8.10E-05 | 3.16E-05 | 0.00034 |
| Atpif1 | 0.00025 | 5.96E-05 | 0.00018 | 0.00039 | 6.41E-05 | 0.00014 | 0.00018 | 0.00012 | 8.89E-05 | 2.73E-05 | 0.00014 | 2.47E-05 | 0.0001 | 0.00017 | 7.44E-05 | 0.00021 | 4.16E-05 | 0.00017 |
| Agap3 | 0 | 0.00031 | 0.00015 | 0.00027 | 0.0002 | 0.00012 | 0.00021 | 7.93E-05 | 8.89E-05 | 0.0001 | 6.82E-05 | 8.98E-05 | 4.91E-05 | 0.00014 | 0.00015 | 0.00013 | 0.00024 | 0.00018 |

| | | | | | | | | | | | | | | | | | | |
|---|---|---|---|---|---|---|---|---|---|---|---|---|---|---|---|---|---|---|
| Fastk | 0.00017 | 0.00011 | 0.00017 | 0.00041 | 5.64E-05 | 6.69E-05 | 0.00022 | 0.00018 | 8.88E-05 | 0.00012 | 8.65E-05 | 0.00018 | 2.00E-05 | 7.14E-05 | 0.00046 | 4.54E-05 | 4.30E-05 | 0.0001 |
| Timm50 | 9.00E-05 | 4.38E-05 | 0.00023 | 0.00016 | 4.22E-05 | 1.91E-05 | 0 | 5.44E-05 | 8.88E-05 | 4.44E-05 | 9.78E-05 | 2.91E-05 | 2.38E-05 | 2.33E-06 | 0 | 0.00039 | 6.16E-05 | 3.40E-05 |
| Vps18 | 1.95E-05 | 0.0001 | 4.83E-05 | 2.85E-05 | 5.08E-05 | 8.78E-06 | 0 | 0 | 8.88E-05 | 4.36E-05 | 1.45E-05 | 5.09E-05 | 0 | 5.58E-05 | 9.40E-05 | 2.75E-05 | 0 | 4.62E-05 |
| Fxr2 | 4.52E-05 | 0.00016 | 0.00019 | 0.00012 | 5.23E-05 | 4.03E-06 | 4.60E-05 | 9.17E-05 | 8.88E-05 | 4.43E-05 | 9.36E-05 | 7.63E-05 | 1.59E-05 | 5.36E-05 | 0.0003 | 0.00014 | 0.0001 | 7.36E-05 |
| Dnm3 | 0 | 1.15E-05 | 0 | 1.30E-05 | 0.00032 | 0 | 0.00032 | 0 | 8.88E-05 | 1.06E-05 | 0 | 0 | 0.0003 | 7.54E-05 | 3.96E-05 | 5.53E-05 | 5.81E-05 | 0 |
| Ssna1 | 0.00016 | 0.00015 | 0.00016 | 0.00024 | 0.00014 | 0.00015 | 0.00011 | 0.0001 | 8.87E-05 | 0.00013 | 0.00014 | 3.78E-05 | 2.19E-05 | 7.01E-05 | 3.91E-05 | 0.00046 | 0 | 0.00014 |
| Wsb2 | 5.97E-05 | 9.78E-05 | 0.00016 | 0.00014 | 0.00011 | 6.97E-05 | 2.40E-05 | 9.39E-05 | 8.87E-05 | 5.67E-05 | 0.00019 | 4.33E-05 | 6.01E-05 | 2.02E-05 | 9.82E-05 | 3.21E-05 | 0 | 5.01E-05 |
| Gse1 | 1.30E-05 | 8.68E-05 | 8.56E-05 | 2.72E-05 | 1.67E-05 | 1.83E-05 | 0 | 0 | 8.87E-05 | 4.88E-05 | 1.70E-05 | 2.22E-05 | 2.82E-05 | 2.53E-05 | 3.84E-05 | 4.78E-05 | 0.00012 | 2.35E-05 |
| Elmsan1 | 3.77E-06 | 9.70E-05 | 4.74E-05 | 7.27E-05 | 4.11E-05 | 1.35E-05 | 4.60E-05 | 2.21E-05 | 8.87E-05 | 9.52E-05 | 7.69E-05 | 0.00015 | 1.58E-06 | 3.55E-05 | 0.00014 | 7.16E-05 | 0.00028 | 5.15E-05 |
| Arl5b | 7.12E-05 | 7.57E-05 | 4.66E-05 | 4.48E-06 | 2.43E-05 | 4.06E-06 | 1.20E-05 | 0.00012 | 8.87E-05 | 4.63E-05 | 3.96E-05 | 0 | 0 | 4.88E-05 | 0 | 0 | 0.00109 | 0.00011 |
| Chpf2 | 0.00017 | 0.0001 | 0.00021 | 0.00039 | 0.0003 | 0.0004 | 0.00061 | 0.00043 | 8.86E-05 | 0.00019 | 0.00028 | 9.39E-05 | 0.00041 | 0.00036 | 0.00038 | 0 | 0 | 0.00016 |
| Gtf2b | 0.00017 | 0.00013 | 0.00014 | 0.00016 | 0.00012 | 0.00018 | 6.99E-05 | 7.06E-05 | 8.86E-05 | 1.77E-05 | 4.51E-05 | 3.41E-05 | 6.16E-05 | 0.00015 | 0 | 0.00013 | 0 | 1.33E-05 |
| Abi2 | 0.00011 | 9.12E-05 | 4.61E-05 | 0 | 4.30E-05 | 0.00012 | 0 | 4.20E-05 | 8.86E-05 | 3.78E-05 | 1.44E-05 | 8.28E-05 | 4.99E-05 | 5.40E-05 | 5.83E-05 | 8.61E-05 | 4.85E-05 | 6.14E-07 |
| Psma7 | 0.00037 | 6.66E-05 | 0.00018 | 0 | 0 | 9.27E-05 | 1.86E-06 | 2.99E-05 | 8.86E-05 | 0 | 1.65E-05 | 6.99E-05 | 0 | 0 | 0 | 4.92E-05 | 0 | 1.98E-05 |
| Mcm2 | 8.43E-05 | 0 | 0.00017 | 0 | 0 | 0 | 6.77E-06 | 0 | 8.85E-05 | 4.09E-06 | 2.82E-05 | 0 | 0 | 3.52E-05 | 0 | 0.00061 | 0 | 0 |
| Hadhb | 0.00031 | 0.00017 | 0.00018 | 0.00019 | 0.00014 | 0.00022 | 0.00017 | 0.00013 | 8.84E-05 | 0.00015 | 7.83E-05 | 5.93E-05 | 0.00014 | 0.0001 | 0 | 0.00013 | 0 | 7.89E-05 |
| Zbtb22 | 1.94E-05 | 0.00012 | 4.70E-05 | 9.63E-05 | 5.43E-05 | 1.76E-05 | 2.27E-05 | 0 | 8.84E-05 | 4.98E-05 | 6.02E-05 | 7.30E-05 | 0 | 6.43E-05 | 8.01E-05 | 8.39E-05 | 5.00E-05 | 7.15E-05 |
| Mms19 | 3.09E-05 | 7.43E-05 | 4.70E-05 | 3.88E-05 | 6.09E-05 | 1.72E-05 | 3.52E-06 | 6.86E-05 | 8.84E-05 | 1.85E-05 | 2.12E-06 | 9.42E-05 | 0 | 3.82E-05 | 4.92E-05 | 4.75E-05 | 0 | 3.85E-05 |
| Wdr6 | 2.90E-05 | 5.24E-05 | 0 | 0.00015 | 8.08E-05 | 0 | 7.62E-05 | 0 | 8.84E-05 | 0.0001 | 7.38E-05 | 0 | 0 | 0.00015 | 0 | 4.08E-05 | 3.14E-05 | 8.38E-05 |
| Ap1ar | 0.00012 | 6.69E-05 | 0.00011 | 9.20E-05 | 5.59E-05 | 6.30E-05 | 0 | 9.04E-05 | 8.83E-05 | 2.98E-05 | 2.82E-05 | 0 | 7.74E-05 | 6.19E-05 | 0.00017 | 0.00011 | 0.00015 | 0.00012 |
| Cdc123 | 0.00027 | 7.62E-05 | 0.0002 | 0.00016 | 6.79E-05 | 5.86E-05 | 0.0002 | 3.57E-05 | 8.83E-05 | 7.38E-05 | 3.98E-05 | 4.32E-05 | 9.06E-05 | 6.15E-05 | 7.84E-05 | 0.00031 | 0 | 3.61E-05 |
| Yipf5 | 0.00042 | 5.82E-05 | 0.00033 | 0.00036 | 0.0002 | 0.00024 | 0.00056 | 0.00032 | 8.82E-05 | 0.00017 | 0.00012 | 0.00019 | 0.00021 | 0.00011 | 0 | 0.00011 | 0 | 0.00012 |
| Ang | 2.55E-05 | 9.35E-05 | 3.35E-05 | 3.91E-05 | 0 | 5.35E-05 | 0.00013 | 5.56E-05 | 8.82E-05 | 0.00019 | 0 | 1.60E-05 | 0.00027 | 9.03E-05 | 9.59E-06 | 0 | 0 | 7.67E-05 |
| Tmem18 | 4.94E-06 | 5.65E-05 | 6.95E-05 | 6.85E-05 | 7.31E-05 | 3.68E-05 | 0 | 6.07E-05 | 8.82E-05 | 8.73E-05 | 6.01E-05 | 0 | 4.22E-05 | 5.11E-05 | 6.59E-06 | 0.00015 | 0 | 5.24E-05 |
| Rida | 1.56E-05 | 4.81E-05 | 0 | 4.31E-05 | 0.00012 | 0 | 2.96E-07 | 7.68E-05 | 8.82E-05 | 1.07E-05 | 7.43E-05 | 1.47E-05 | 0 | 0 | 0 | 0 | 0 | 2.94E-05 |
| Nfkbia | 0 | 4.56E-05 | 0.00032 | 4.95E-05 | 5.78E-05 | 0 | 0.00012 | 0 | 8.82E-05 | 0.00028 | 7.11E-05 | 0.00018 | 0.00071 | 0.00049 | 0.00023 | 1.43E-05 | 0.0055 | 0 |
| Eif3i | 0.00032 | 0.00013 | 0.00028 | 0.00033 | 4.98E-05 | 0.00023 | 9.41E-05 | 5.09E-05 | 8.81E-05 | 2.82E-05 | 4.30E-05 | 0.00013 | 4.15E-05 | 0.00011 | 0 | 9.62E-05 | 5.04E-05 | 2.35E-05 |
| Hdac2 | 0.00023 | 8.54E-05 | 0.00017 | 5.66E-05 | 4.60E-05 | 0.00011 | 0 | 9.10E-05 | 8.81E-05 | 5.67E-05 | 0.0001 | 3.23E-05 | 0 | 4.40E-06 | 6.04E-06 | 0.00022 | 0 | 0 |
| Pelo | 9.76E-05 | 6.81E-05 | 0.0001 | 0.00014 | 5.09E-05 | 6.79E-05 | 8.29E-05 | 3.58E-05 | 8.81E-05 | 1.36E-05 | 0 | 2.74E-05 | 0 | 4.22E-05 | 0.00022 | 1.86E-05 | 0 | 4.64E-05 |
| AI413582 | 2.64E-06 | 0.00011 | 2.84E-05 | 0 | 5.59E-05 | 4.58E-05 | 0 | 1.92E-06 | 8.81E-05 | 3.75E-05 | 0 | 5.92E-05 | 3.52E-05 | 3.57E-05 | 0 | 0 | 0 | 6.70E-05 |
| Snrnp40 | 0.00017 | 6.34E-05 | 0.00018 | 7.31E-05 | 2.68E-05 | 3.73E-05 | 3.60E-06 | 0 | 8.81E-05 | 2.35E-05 | 2.72E-05 | 5.28E-06 | 1.80E-05 | 6.25E-06 | 6.12E-05 | 0.00031 | 0 | 9.26E-06 |
| Txlna | 0.00015 | 0.00014 | 0.00026 | 0.00032 | 8.43E-05 | 2.79E-05 | 0.00023 | 0.00011 | 8.81E-05 | 8.09E-05 | 4.68E-05 | 0 | 8.21E-05 | 8.84E-05 | 0.00049 | 0.00024 | 0.0001 | 0.00014 |
| Dedd | 8.21E-05 | 0.00011 | 9.50E-05 | 0.0001 | 3.14E-05 | 1.73E-05 | 8.28E-05 | 5.92E-05 | 8.81E-05 | 5.15E-05 | 8.35E-05 | 9.08E-05 | 0 | 2.20E-05 | 0.00017 | 5.36E-05 | 0 | 5.16E-05 |

| | | | | | | | | | | | | | | | | | | |
|---|---|---|---|---|---|---|---|---|---|---|---|---|---|---|---|---|---|---|
| Adrm1 | 3.62E-05 | 9.28E-05 | 0.00021 | 0.00016 | 2.11E-05 | 0 | 0 | 1.97E-05 | 8.81E-05 | 3.67E-05 | 5.57E-05 | 0 | 0 | 4.18E-05 | 4.47E-05 | 0.0001 | 0.00011 | 5.26E-05 |
| Mfn2 | 5.20E-05 | 0.00012 | 0.00014 | 0.00012 | 6.60E-05 | 5.64E-05 | 3.33E-05 | 5.76E-05 | 8.80E-05 | 0.0001 | 8.77E-05 | 2.49E-05 | 2.58E-05 | 4.96E-05 | 0 | 7.58E-05 | 4.38E-05 | 9.86E-05 |
| Samd5 | 0 | 0 | 3.70E-05 | 5.30E-06 | 0 | 0 | 0 | 0 | 8.80E-05 | 3.34E-05 | 6.00E-06 | 2.12E-05 | 6.96E-06 | 0 | 6.03E-05 | 4.27E-05 | 6.17E-05 | 0.00026 |
| Col13a1 | 0 | 1.69E-05 | 0 | 0 | 0 | 8.17E-05 | 0 | 4.71E-06 | 8.79E-05 | 0 | 0 | 0 | 0 | 0 | 0 | 0 | 0 | 0 |
| Stoml1 | 0 | 0.00011 | 0 | 0.00015 | 7.46E-05 | 1.69E-05 | 2.34E-05 | 0.00011 | 8.79E-05 | 8.82E-05 | 5.52E-05 | 4.28E-05 | 4.79E-05 | 4.57E-05 | 0.0001 | 0 | 1.11E-05 | 0.00014 |
| Wdr55 | 8.15E-05 | 4.96E-05 | 0.00015 | 0.0001 | 0 | 1.20E-05 | 0 | 0 | 8.79E-05 | 1.03E-05 | 4.57E-06 | 3.64E-05 | 2.74E-05 | 3.84E-05 | 3.38E-05 | 2.98E-05 | 0 | 1.23E-06 |
| Gatad2b | 0.00023 | 0.0003 | 0.0001 | 4.28E-05 | 0.0002 | 0.00015 | 8.94E-05 | 4.60E-06 | 8.78E-05 | 7.15E-05 | 0.00011 | 0.0002 | 4.60E-05 | 0.00013 | 0.00034 | 0.00015 | 0.00013 | 0.00015 |
| Xrn2 | 0.00023 | 0.00017 | 0.0002 | 5.07E-05 | 2.56E-05 | 8.90E-05 | 5.51E-05 | 3.47E-05 | 8.78E-05 | 3.08E-05 | 3.39E-05 | 6.15E-05 | 3.12E-05 | 7.35E-05 | 0.0002 | 0.00013 | 3.77E-05 | 2.09E-05 |
| Bap1 | 9.16E-05 | 0.00012 | 6.55E-05 | 7.59E-05 | 6.54E-05 | 7.77E-05 | 4.31E-05 | 3.87E-05 | 8.78E-05 | 3.16E-05 | 7.05E-05 | 5.96E-05 | 1.80E-05 | 0.00011 | 1.44E-05 | 8.82E-05 | 0 | 1.53E-05 |
| Map1a | 0.00011 | 3.91E-05 | 0.00012 | 5.72E-05 | 4.47E-05 | 5.80E-05 | 0.00013 | 5.03E-05 | 8.78E-05 | 2.66E-05 | 3.35E-05 | 7.27E-06 | 0 | 0 | 2.87E-05 | 0 | 0 | 0.00012 |
| Fcf1 | 0.0002 | 0.00011 | 0.00017 | 1.73E-05 | 0 | 5.66E-05 | 0 | 0 | 8.78E-05 | 3.61E-05 | 1.83E-05 | 4.98E-05 | 3.04E-05 | 2.14E-05 | 7.16E-05 | 3.04E-05 | 8.89E-05 | 3.33E-05 |
| BC028528 | 9.38E-05 | 8.38E-05 | 0 | 0 | 0 | 0.00025 | 1.16E-05 | 0.00026 | 8.77E-05 | 3.00E-05 | 0 | 6.00E-06 | 0 | 6.18E-05 | 0 | 0 | 1.83E-06 | 0 |
| Ick | 8.22E-05 | 0.00016 | 3.46E-05 | 0.00011 | 0.00011 | 0.00012 | 8.86E-05 | 4.48E-05 | 8.77E-05 | 9.69E-05 | 4.80E-06 | 0 | 6.11E-05 | 0.00011 | 9.82E-05 | 5.67E-05 | 0 | 9.97E-05 |
| Fam172a | 3.81E-05 | 7.37E-05 | 4.82E-05 | 2.94E-05 | 6.99E-05 | 6.56E-05 | 3.33E-05 | 1.11E-05 | 8.77E-05 | 0.00012 | 4.99E-05 | 4.72E-05 | 2.64E-05 | 4.33E-05 | 6.26E-05 | 8.35E-05 | 0 | 6.70E-05 |
| Stk11 | 0.00015 | 0.00028 | 0.00018 | 0.00035 | 7.92E-05 | 5.74E-05 | 1.26E-05 | 0 | 8.77E-05 | 0.00012 | 0.00011 | 0.00021 | 0 | 4.62E-05 | 0.00024 | 0.00031 | 0.0003 | 0.00011 |
| Cinp | 0.00015 | 8.04E-05 | 0.00022 | 7.62E-05 | 6.11E-05 | 4.87E-05 | 0 | 8.11E-05 | 8.77E-05 | 4.92E-05 | 7.37E-05 | 0.00012 | 5.45E-05 | 4.10E-05 | 7.86E-06 | 0.00032 | 0 | 2.73E-05 |
| H2-T22 | 5.03E-05 | 2.50E-05 | 1.33E-05 | 7.66E-06 | 8.13E-05 | 4.31E-05 | 0 | 2.26E-05 | 8.77E-05 | 0.0001 | 5.59E-05 | 4.95E-05 | 6.60E-05 | 4.77E-05 | 0 | 2.34E-05 | 2.16E-05 | 0.00011 |
| Trip10 | 9.02E-05 | 0.00023 | 0.00016 | 0.00016 | 2.15E-05 | 3.64E-05 | 5.10E-05 | 7.79E-05 | 8.77E-05 | 0.00029 | 0.00012 | 0.00036 | 7.72E-05 | 1.61E-05 | 0.00037 | 4.38E-05 | 6.25E-05 | 0.00017 |
| Dnajb4 | 8.58E-05 | 6.07E-05 | 1.53E-05 | 1.79E-05 | 0 | 0 | 0 | 1.70E-05 | 8.77E-05 | 0.00013 | 0.00033 | 3.06E-05 | 5.46E-06 | 0 | 4.36E-05 | 3.40E-05 | 0.00033 | 7.49E-05 |
| Fcgrt | 0 | 0.00034 | 0 | 0.00031 | 0.00037 | 0.00047 | 0.00046 | 0.00048 | 8.76E-05 | 0.0005 | 3.92E-05 | 0.00026 | 0.00039 | 0.00037 | 0.00016 | 0 | 0.0001 | 0.0005 |
| Dstyk | 4.01E-05 | 0.00017 | 7.27E-05 | 1.04E-05 | 0.00014 | 0.00011 | 0.00013 | 3.12E-05 | 8.76E-05 | 2.36E-05 | 6.36E-05 | 9.04E-05 | 5.26E-05 | 0.00014 | 0.0001 | 2.42E-05 | 0 | 8.51E-05 |
| Eif2ak1 | 0.0001 | 0.00011 | 0.0001 | 9.75E-06 | 0.00012 | 0.0001 | 4.24E-05 | 1.53E-05 | 8.76E-05 | 5.99E-05 | 9.28E-05 | 3.03E-06 | 4.60E-05 | 8.48E-06 | 0 | 3.27E-05 | 1.11E-05 | 3.66E-05 |
| Chchd3 | 0.00019 | 9.87E-05 | 0.00017 | 0.00026 | 0.00013 | 9.36E-05 | 2.81E-05 | 0.0001 | 8.76E-05 | 8.53E-05 | 7.94E-05 | 2.81E-05 | 1.18E-06 | 4.84E-05 | 0.00017 | 0.00026 | 0.00012 | 4.54E-05 |
| Csnk2b | 0.00025 | 0.00012 | 0.00022 | 9.66E-05 | 6.35E-05 | 9.07E-05 | 3.68E-05 | 0.00012 | 8.76E-05 | 1.32E-05 | 9.98E-05 | 4.97E-05 | 0 | 6.36E-05 | 0 | 0.00012 | 0 | 0 |
| Alg14 | 0.00011 | 5.80E-05 | 5.28E-05 | 0.00014 | 8.08E-05 | 8.28E-05 | 5.58E-05 | 6.38E-05 | 8.76E-05 | 6.71E-05 | 7.61E-05 | 1.90E-06 | 1.94E-05 | 3.09E-05 | 0 | 0.00013 | 0 | 9.78E-05 |
| Rbx1 | 0.0002 | 7.48E-05 | 0.00013 | 8.95E-05 | 3.30E-05 | 6.12E-05 | 8.52E-05 | 1.80E-05 | 8.76E-05 | 1.99E-05 | 0.0001 | 1.74E-05 | 5.02E-05 | 3.61E-05 | 0 | 0.00011 | 3.84E-06 | 4.38E-06 |
| Gosr1 | 0.00014 | 0.00012 | 0.00012 | 3.42E-05 | 7.45E-05 | 6.02E-05 | 3.57E-05 | 4.25E-05 | 8.76E-05 | 4.86E-05 | 4.84E-05 | 5.92E-05 | 5.90E-05 | 6.94E-05 | 7.80E-05 | 7.26E-05 | 1.62E-05 | 2.12E-05 |
| Utp25 | 5.67E-05 | 4.14E-05 | 4.60E-05 | 2.55E-05 | 9.45E-05 | 3.66E-05 | 0 | 3.01E-05 | 8.76E-05 | 9.30E-06 | 0 | 0 | 0 | 5.03E-06 | 7.81E-05 | 6.22E-06 | 0 | 2.10E-05 |
| Usp12 | 6.88E-05 | 8.28E-05 | 8.96E-05 | 3.02E-05 | 0.00014 | 3.41E-06 | 0 | 2.56E-05 | 8.76E-05 | 3.15E-05 | 0.00012 | 0.00011 | 0.00017 | 0.00011 | 0.00023 | 4.16E-05 | 9.42E-05 | 0 |
| Sbds | 0.00026 | 0.00022 | 0.00016 | 0 | 4.56E-05 | 0.00017 | 0 | 0 | 8.75E-05 | 7.39E-05 | 9.45E-05 | 0.00027 | 1.31E-05 | 2.74E-05 | 6.07E-05 | 5.65E-05 | 9.32E-05 | 0.00014 |
| Dsel | 0.00032 | 8.55E-05 | 5.18E-05 | 0 | 0.00014 | 0.00015 | 0.00039 | 0.00013 | 8.75E-05 | 0.00023 | 0 | 1.46E-05 | 0.00012 | 5.91E-05 | 2.70E-05 | 4.58E-05 | 0 | 8.60E-06 |
| Snx33 | 7.70E-05 | 0.00021 | 0 | 2.85E-06 | 0.00013 | 0.00014 | 5.90E-05 | 1.53E-05 | 8.75E-05 | 0.0002 | 0 | 0 | 9.47E-05 | 0.00019 | 0.00011 | 3.14E-05 | 0 | 5.52E-05 |
| Ppp2r5e | 0.0002 | 0.00018 | 9.86E-05 | 0 | 9.18E-05 | 0.00012 | 2.05E-05 | 0 | 8.75E-05 | 4.17E-05 | 3.97E-05 | 9.67E-05 | 3.93E-06 | 0 | 0.00013 | 0.00014 | 0.0001 | 7.33E-06 |

| | | | | | | | | | | | | | | | | | | |
|---|---|---|---|---|---|---|---|---|---|---|---|---|---|---|---|---|---|---|
| Dnmbp | 4.09E-05 | 6.90E-05 | 2.71E-05 | 3.04E-05 | 3.54E-05 | 1.13E-05 | 4.66E-05 | 1.23E-05 | 8.75E-05 | 0.0001 | 9.53E-06 | 1.38E-05 | 2.24E-05 | 4.48E-05 | 6.98E-05 | 6.90E-05 | 8.81E-05 | 9.16E-05 |
| Pgm1 | 0.00011 | 0 | 0 | 0.00028 | 0.00011 | 0 | 0.00023 | 6.97E-05 | 8.75E-05 | 0.00016 | 5.26E-05 | 0.001 | 0.0003 | 1.06E-05 | 2.25E-05 | 2.18E-05 | 0 | 0 |
| Celf2 | 0 | 2.05E-05 | 2.66E-05 | 0 | 1.68E-05 | 0 | 0.00019 | 0.00013 | 8.75E-05 | 0.00043 | 3.75E-05 | 0.00012 | 0.00025 | 0 | 0.00012 | 0.00012 | 5.22E-05 | 0.00036 |
| Pfdn1 | 0.00017 | 4.81E-05 | 8.82E-05 | 9.53E-05 | 3.46E-05 | 0 | 3.16E-05 | 8.54E-05 | 8.75E-05 | 3.19E-05 | 9.97E-05 | 6.52E-05 | 0 | 1.43E-05 | 9.84E-05 | 8.53E-05 | 4.49E-05 | 4.41E-05 |
| Yars | 0.00018 | 8.60E-05 | 0.00029 | 0.00014 | 3.10E-05 | 9.16E-06 | 7.11E-05 | 5.39E-05 | 8.74E-05 | 4.36E-05 | 6.63E-05 | 0.00018 | 0 | 8.79E-06 | 0.00016 | 6.81E-05 | 0 | 5.13E-05 |
| Atp5f1 | 0.00038 | 9.96E-05 | 0.00019 | 0.00016 | 8.21E-05 | 0.00024 | 0.00015 | 8.85E-05 | 8.73E-05 | 4.18E-05 | 0.00015 | 6.39E-05 | 0.00011 | 6.93E-05 | 0 | 0.00019 | 2.16E-06 | 0 |
| Ndrg3 | 0.0002 | 6.94E-05 | 0 | 0 | 9.24E-05 | 0.00012 | 1.22E-05 | 0 | 8.73E-05 | 2.28E-05 | 8.12E-06 | 4.70E-05 | 0 | 0 | 0 | 3.11E-05 | 2.75E-05 | 7.69E-05 |
| Mapk9 | 0.00015 | 9.60E-05 | 6.07E-05 | 2.84E-05 | 9.50E-05 | 6.42E-05 | 8.47E-05 | 9.38E-05 | 8.73E-05 | 0.00015 | 5.90E-05 | 9.65E-05 | 0.00011 | 3.50E-05 | 2.70E-05 | 9.44E-05 | 0.00022 | 0.00011 |
| Tmem63b | 9.18E-05 | 0.00013 | 0.00017 | 7.29E-05 | 5.97E-05 | 5.47E-05 | 8.50E-05 | 5.99E-05 | 8.73E-05 | 7.08E-05 | 0.00012 | 0.00012 | 1.40E-05 | 7.93E-05 | 0.00018 | 0 | 1.84E-05 | 4.81E-05 |
| Axin1 | 4.57E-05 | 0.00014 | 0.00011 | 2.90E-05 | 5.33E-05 | 3.15E-05 | 1.55E-05 | 8.58E-06 | 8.73E-05 | 4.93E-05 | 0.00012 | 7.22E-05 | 0 | 8.43E-05 | 7.32E-05 | 0.0001 | 0.00011 | 5.34E-05 |
| Tmem47 | 2.69E-05 | 2.89E-05 | 2.91E-05 | 0 | 7.71E-06 | 0 | 0 | 0.00021 | 8.73E-05 | 0 | 0.00031 | 7.96E-05 | 0 | 5.25E-05 | 0 | 6.17E-05 | 0 | 4.69E-05 |
| Cdyl2 | 0 | 6.82E-05 | 8.27E-05 | 4.03E-05 | 9.51E-05 | 0.00011 | 0 | 0.0002 | 8.72E-05 | 0 | 8.83E-05 | 0.00017 | 0 | 8.31E-05 | 1.43E-05 | 1.02E-05 | 4.64E-05 | 0 |
| Herc4 | 9.61E-05 | 0.00014 | 5.22E-05 | 0 | 1.98E-05 | 9.78E-05 | 0 | 1.12E-05 | 8.72E-05 | 6.06E-05 | 4.06E-06 | 8.80E-05 | 2.00E-05 | 7.44E-05 | 0.00012 | 3.29E-05 | 6.93E-05 | 3.44E-05 |
| Topors | 0.00013 | 0.00013 | 6.27E-05 | 1.76E-05 | 4.73E-05 | 5.41E-05 | 1.05E-06 | 2.18E-06 | 8.72E-05 | 0 | 3.38E-05 | 4.42E-05 | 0 | 5.20E-05 | 8.79E-05 | 0.00014 | 0.00017 | 4.58E-05 |
| Grk6 | 0.00012 | 0.00014 | 0.00012 | 0 | 2.87E-05 | 2.32E-05 | 0 | 0.00011 | 8.72E-05 | 1.97E-05 | 0.0002 | 4.99E-05 | 0 | 4.91E-05 | 0.00022 | 0.00032 | 1.83E-05 | 0 |
| Slc25a25 | 0 | 8.56E-05 | 0.00012 | 0 | 2.74E-05 | 4.97E-06 | 0 | 2.70E-05 | 8.72E-05 | 0 | 7.29E-05 | 0 | 0 | 3.41E-05 | 1.74E-05 | 9.49E-05 | 0.00045 | 0 |
| Sumf2 | 4.87E-05 | 8.45E-05 | 4.70E-05 | 0.00016 | 4.27E-05 | 0 | 8.60E-06 | 3.45E-05 | 8.72E-05 | 8.75E-05 | 1.63E-05 | 1.21E-06 | 7.25E-05 | 3.45E-05 | 2.47E-07 | 5.85E-05 | 4.87E-05 | 0.00011 |
| Lgals3 | 0 | 0.00019 | 0 | 0 | 0 | 0.00029 | 0 | 0 | 8.71E-05 | 0.0006 | 0 | 0.00127 | 0.00057 | 0.00023 | 0 | 0 | 0 | 0.0006 |
| Slc35e1 | 0.00022 | 6.60E-05 | 0.00028 | 0.00019 | 0.00016 | 0.00013 | 0.00044 | 0.00024 | 8.71E-05 | 9.19E-05 | 0.00015 | 0.00018 | 5.99E-05 | 0.00012 | 0 | 0.00031 | 7.72E-05 | 7.40E-05 |
| Utp6 | 0.00013 | 9.73E-05 | 0.00021 | 0.00011 | 5.59E-05 | 3.39E-05 | 0 | 0 | 8.71E-05 | 7.18E-05 | 4.46E-05 | 0 | 4.91E-05 | 0 | 1.63E-06 | 0.00017 | 0 | 4.20E-05 |
| AW549877 | 5.72E-05 | 0.00018 | 0 | 0 | 0.00017 | 0.00015 | 0.00012 | 6.62E-05 | 8.70E-05 | 9.29E-05 | 1.62E-05 | 7.95E-05 | 2.23E-05 | 5.04E-05 | 0.00026 | 2.82E-06 | 0 | 0.00012 |
| Ctr9 | 0.00017 | 0.00012 | 6.83E-05 | 1.94E-05 | 3.79E-05 | 3.29E-05 | 4.73E-05 | 0 | 8.70E-05 | 3.04E-05 | 7.65E-05 | 8.86E-06 | 1.53E-05 | 4.29E-05 | 3.40E-05 | 8.76E-05 | 0 | 2.57E-05 |
| Zfp414 | 0.00012 | 7.10E-05 | 2.64E-05 | 0.00012 | 3.19E-05 | 3.15E-05 | 0.00018 | 0.00013 | 8.70E-05 | 0.00011 | 2.42E-05 | 7.92E-05 | 0 | 0 | 0.00011 | 0.00014 | 0 | 0.00014 |
| Mier2 | 2.81E-05 | 7.60E-05 | 1.00E-04 | 9.38E-06 | 3.87E-05 | 6.81E-06 | 0 | 4.99E-05 | 8.70E-05 | 3.52E-05 | 7.24E-05 | 8.35E-05 | 4.17E-06 | 6.63E-05 | 0.00013 | 1.67E-05 | 6.58E-05 | 0 |
| Snrnp35 | 2.82E-05 | 5.86E-05 | 2.66E-05 | 0.0001 | 3.82E-05 | 0 | 0 | 8.57E-05 | 8.70E-05 | 1.28E-05 | 2.51E-05 | 0 | 2.73E-06 | 3.91E-05 | 0.00011 | 0.00013 | 0 | 2.23E-05 |
| Akirin2 | 0.00027 | 0.00029 | 8.33E-05 | 0 | 9.71E-05 | 0.00015 | 0.00012 | 3.96E-05 | 8.69E-05 | 9.97E-05 | 9.67E-05 | 0.00014 | 0 | 5.17E-06 | 0.0002 | 0.00041 | 7.84E-05 | 1.95E-05 |
| Tmx4 | 5.69E-05 | 0.0001 | 7.03E-05 | 1.80E-05 | 5.21E-05 | 8.39E-05 | 4.22E-05 | 0.00015 | 8.69E-05 | 8.48E-05 | 0 | 0.00016 | 6.89E-05 | 5.12E-05 | 4.60E-05 | 8.38E-05 | 0.00016 | 0.00011 |
| Tor1a | 0.00012 | 0.00016 | 0.00011 | 6.96E-05 | 8.66E-05 | 7.20E-05 | 8.38E-05 | 6.60E-05 | 8.69E-05 | 9.20E-05 | 8.19E-05 | 0.00013 | 3.01E-05 | 5.23E-05 | 2.22E-05 | 0.00018 | 7.26E-05 | 3.61E-05 |
| Cc2d1a | 3.23E-05 | 0.00011 | 6.05E-05 | 0.00015 | 0 | 4.62E-05 | 3.29E-05 | 1.58E-05 | 8.69E-05 | 7.41E-05 | 3.43E-05 | 6.02E-05 | 7.81E-06 | 0 | 0.00011 | 5.56E-05 | 0 | 0.00012 |
| Ccdc71l | 0 | 4.48E-05 | 3.42E-05 | 0.00011 | 4.63E-05 | 9.31E-06 | 0 | 0.00012 | 8.69E-05 | 9.73E-05 | 8.64E-05 | 0.00016 | 2.38E-05 | 0.00013 | 0.00034 | 1.51E-05 | 0.00011 | 5.23E-06 |
| Brix1 | 0.00018 | 8.52E-05 | 0.00027 | 0.00012 | 2.25E-05 | 0 | 7.19E-06 | 0 | 8.69E-05 | 2.63E-05 | 2.92E-05 | 8.23E-06 | 1.71E-05 | 2.06E-05 | 0.00014 | 0.00011 | 0 | 0 |
| Rbm18 | 0.00016 | 6.76E-05 | 5.28E-05 | 0 | 5.55E-05 | 0.00012 | 0 | 3.80E-05 | 8.68E-05 | 0 | 3.88E-05 | 7.17E-05 | 0 | 1.63E-05 | 4.03E-05 | 4.30E-05 | 5.96E-05 | 0 |
| Xpo5 | 7.34E-05 | 0.0001 | 0.00022 | 7.65E-05 | 7.02E-05 | 4.54E-05 | 0 | 4.26E-06 | 8.68E-05 | 3.05E-05 | 4.63E-05 | 4.58E-06 | 7.07E-05 | 4.76E-05 | 4.04E-05 | 0.00012 | 1.52E-05 | 2.58E-05 |

| | | | | | | | | | | | | | | | | | | |
|---|---|---|---|---|---|---|---|---|---|---|---|---|---|---|---|---|---|---|
| Banf1 | 0.00014 | 4.55E-05 | 0.00016 | 0.00015 | 5.58E-06 | 0 | 0 | 2.53E-05 | 8.68E-05 | 0 | 7.92E-05 | 2.39E-05 | 0 | 0 | 3.94E-05 | 0.00022 | 4.34E-05 | 0 |
| Sarnp | 0.00024 | 9.64E-05 | 0.0001 | 5.63E-05 | 2.77E-05 | 8.79E-05 | 3.84E-06 | 7.48E-06 | 8.67E-05 | 2.92E-05 | 5.85E-05 | 5.32E-05 | 1.84E-05 | 2.76E-05 | 0 | 0.00011 | 0.00015 | 2.21E-05 |
| Tsc22d4 | 8.62E-06 | 0.00013 | 3.43E-06 | 0.00029 | 9.99E-05 | 4.10E-05 | 6.74E-05 | 0.00017 | 8.67E-05 | 9.40E-05 | 8.84E-05 | 0.00018 | 5.30E-05 | 9.40E-05 | 0.00017 | 5.94E-05 | 2.28E-05 | 8.64E-05 |
| Msi1 | 0 | 0 | 8.63E-05 | 7.93E-05 | 6.13E-05 | 2.84E-05 | 4.31E-05 | 8.50E-05 | 8.67E-05 | 0 | 4.66E-05 | 7.85E-07 | 4.30E-05 | 0 | 7.74E-07 | 0 | 2.55E-05 | 0 |
| Stx16 | 0.00011 | 0.00016 | 3.03E-05 | 8.62E-05 | 0.00012 | 0.00011 | 7.00E-05 | 2.67E-05 | 8.66E-05 | 0.00022 | 5.87E-05 | 6.33E-05 | 4.00E-06 | 1.55E-05 | 0.00036 | 9.33E-05 | 6.75E-05 | 9.31E-05 |
| Gng2 | 0 | 3.48E-05 | 0 | 0 | 0.00019 | 5.49E-05 | 0 | 0.00012 | 8.66E-05 | 0 | 0.00014 | 0.00017 | 0 | 0.00031 | 0 | 0 | 0 | 0 |
| Parp1 | 0.00014 | 3.95E-05 | 0.0001 | 7.99E-05 | 0 | 0 | 5.22E-05 | 0 | 8.66E-05 | 4.03E-05 | 0 | 0 | 1.98E-05 | 3.76E-05 | 7.83E-06 | 0.0003 | 0 | 2.64E-05 |
| Fhod3 | 0 | 0 | 0 | 0 | 0.00062 | 0 | 1.03E-05 | 0 | 8.66E-05 | 0 | 0.00037 | 0 | 0 | 0 | 0.00015 | 1.52E-05 | 0 | 0 |
| Slc11a2 | 0.00017 | 7.27E-05 | 0.00023 | 1.51E-05 | 0.00012 | 0.00011 | 0.00013 | 5.77E-07 | 8.65E-05 | 2.01E-05 | 7.38E-06 | 0 | 0.00021 | 0.00015 | 7.83E-05 | 1.52E-05 | 6.43E-05 | 0 |
| Ppm1g | 0.00022 | 0.00019 | 0.00036 | 0.00019 | 0.00012 | 8.87E-05 | 2.31E-05 | 2.60E-05 | 8.65E-05 | 5.45E-05 | 0.00011 | 0 | 2.56E-05 | 1.83E-05 | 2.76E-05 | 0.00043 | 6.65E-06 | 5.00E-05 |
| Ppp2r5d | 0.00013 | 0.00011 | 9.60E-05 | 7.72E-05 | 2.86E-05 | 8.80E-06 | 4.46E-05 | 8.22E-05 | 8.65E-05 | 6.52E-05 | 3.68E-05 | 2.60E-06 | 0 | 3.38E-05 | 0 | 0.00036 | 3.72E-05 | 8.53E-05 |
| Rhoq | 0.00031 | 0.00021 | 0.00029 | 0 | 0.00025 | 0.00042 | 0.00052 | 0.00034 | 8.64E-05 | 0.00031 | 0.00021 | 0.00043 | 0.00052 | 0.00038 | 0.00026 | 4.74E-05 | 0.00021 | 7.26E-05 |
| As3mt | 7.08E-05 | 2.68E-05 | 2.30E-05 | 8.63E-05 | 9.36E-05 | 7.99E-05 | 3.73E-05 | 6.09E-05 | 8.63E-05 | 3.32E-05 | 1.42E-05 | 0 | 0 | 9.82E-05 | 0 | 0 | 0 | 0 |
| Rbfa | 0.00011 | 0.00012 | 8.61E-05 | 0.00012 | 8.09E-05 | 7.11E-05 | 6.15E-06 | 3.72E-05 | 8.63E-05 | 6.45E-05 | 8.88E-05 | 2.88E-05 | 2.06E-05 | 5.18E-05 | 8.07E-05 | 1.37E-05 | 0 | 0.00011 |
| Npr2 | 0 | 0.00014 | 0 | 0 | 0.00012 | 3.57E-05 | 2.73E-05 | 0 | 8.63E-05 | 0.00011 | 4.84E-05 | 9.33E-05 | 9.81E-05 | 0.00022 | 0.00014 | 0 | 0.00012 | 1.09E-05 |
| Ftsj1 | 0.0001 | 7.56E-05 | 8.80E-05 | 0.00016 | 4.30E-05 | 3.57E-06 | 2.70E-05 | 0 | 8.63E-05 | 0 | 8.28E-05 | 0 | 7.58E-05 | 9.98E-05 | 2.39E-05 | 8.20E-05 | 0 | 3.13E-05 |
| Sap18b | 0.00017 | 5.29E-05 | 0.0001 | 0.00018 | 2.59E-05 | 2.07E-05 | 2.30E-05 | 0 | 8.62E-05 | 3.78E-05 | 7.97E-05 | 0.00011 | 7.29E-05 | 5.09E-05 | 0 | 0.00013 | 0.00039 | 2.41E-05 |
| Pid1 | 0 | 0 | 0 | 0 | 0.00018 | 0 | 1.07E-05 | 0 | 8.62E-05 | 0.00014 | 0 | 6.18E-06 | 0 | 9.51E-05 | 0 | 0 | 0 | 0.00032 |
| Ppan | 0.00011 | 5.73E-05 | 0.0003 | 0.00017 | 0 | 0 | 0 | 0 | 8.62E-05 | 4.09E-06 | 0 | 0 | 8.52E-06 | 0 | 0.00016 | 0 | 0 | 2.78E-06 |
| Usp9x | 0.00045 | 0.00024 | 0.00024 | 0 | 0.00015 | 0.00024 | 0.00029 | 0.00011 | 8.61E-05 | 3.08E-05 | 0.00017 | 0.00022 | 0.00015 | 3.68E-05 | 9.30E-05 | 0.0001 | 0.00041 | 0.00016 |
| Bad | 0.00012 | 0.00012 | 7.12E-05 | 0.00035 | 7.85E-05 | 7.89E-05 | 7.01E-05 | 8.17E-05 | 8.60E-05 | 0.00017 | 2.47E-05 | 0 | 6.76E-05 | 8.86E-05 | 0 | 2.39E-05 | 0 | 0.00015 |
| Nck2 | 1.40E-05 | 0.00011 | 0 | 0 | 0.00013 | 7.16E-05 | 0 | 0.00025 | 8.60E-05 | 0 | 0.00025 | 4.33E-05 | 0 | 0.00015 | 0 | 4.88E-05 | 5.43E-05 | 1.71E-05 |
| Vav2 | 1.48E-05 | 3.80E-05 | 5.24E-05 | 0 | 9.46E-06 | 5.55E-05 | 0 | 0.00014 | 8.60E-05 | 4.90E-05 | 3.84E-05 | 2.65E-05 | 0 | 0 | 9.72E-05 | 1.03E-05 | 2.67E-05 | 0.00027 |
| Sptlc1 | 8.84E-05 | 9.75E-05 | 0.0001 | 5.62E-06 | 5.28E-05 | 4.98E-05 | 0 | 3.72E-05 | 8.60E-05 | 5.21E-05 | 7.90E-05 | 3.15E-05 | 1.88E-05 | 4.11E-05 | 1.05E-05 | 4.44E-05 | 0 | 5.35E-05 |
| Rbm4 | 3.56E-05 | 0.00022 | 0.00011 | 0.0003 | 0.00011 | 4.12E-05 | 1.61E-05 | 0.00013 | 8.60E-05 | 9.35E-05 | 6.12E-05 | 8.15E-05 | 3.46E-05 | 9.25E-05 | 0.00033 | 8.02E-05 | 0 | 7.27E-05 |
| Wbp2 | 8.63E-05 | 0.00018 | 0 | 4.80E-05 | 5.31E-05 | 3.82E-05 | 0 | 3.39E-05 | 8.60E-05 | 3.29E-05 | 6.41E-05 | 0.00015 | 0 | 1.50E-05 | 4.01E-05 | 9.73E-05 | 8.12E-05 | 5.97E-05 |
| Dpy19l3 | 1.51E-05 | 4.31E-05 | 2.47E-06 | 0 | 0.0001 | 3.53E-05 | 7.61E-05 | 0 | 8.60E-05 | 4.77E-05 | 5.21E-05 | 6.35E-06 | 7.43E-05 | 0 | 0 | 0 | 1.19E-07 | 0 |
| Fam43a | 0 | 0.00012 | 0 | 0.00014 | 9.25E-05 | 0.00026 | 0 | 0.0002 | 8.59E-05 | 1.65E-05 | 4.09E-05 | 0.00011 | 0.00028 | 0.00036 | 0.00014 | 0 | 9.77E-05 | 0.00013 |
| Pdcd10 | 0.00016 | 0.00012 | 0.0001 | 7.48E-05 | 3.01E-05 | 0.00014 | 0 | 3.31E-05 | 8.59E-05 | 2.53E-05 | 7.51E-05 | 0.00011 | 0 | 5.15E-05 | 6.88E-05 | 3.51E-05 | 7.05E-06 | 1.39E-05 |
| Tmem198b | 0 | 0.00012 | 6.59E-05 | 0.00016 | 8.53E-05 | 7.38E-05 | 3.17E-06 | 0.00029 | 8.59E-05 | 0.00016 | 0.00016 | 5.67E-05 | 0 | 9.00E-05 | 0 | 3.92E-05 | 0 | 0.00014 |
| Rbm45 | 0.00013 | 7.96E-05 | 9.01E-05 | 3.74E-05 | 3.71E-05 | 5.17E-05 | 5.06E-05 | 3.28E-05 | 8.59E-05 | 3.11E-05 | 4.97E-05 | 3.97E-05 | 4.19E-06 | 2.73E-05 | 1.99E-05 | 5.74E-05 | 0 | 5.81E-05 |
| Dnlz | 9.26E-05 | 3.54E-05 | 9.87E-05 | 0.00028 | 2.87E-05 | 4.30E-05 | 0.00016 | 8.28E-05 | 8.59E-05 | 2.05E-05 | 8.23E-05 | 1.16E-05 | 5.58E-05 | 0.00011 | 9.09E-06 | 7.51E-05 | 3.79E-05 | 7.64E-05 |
| Usp34 | 0.00022 | 0.00014 | 9.86E-05 | 0 | 6.29E-05 | 0.00012 | 0.00011 | 1.67E-05 | 8.58E-05 | 5.54E-06 | 5.32E-05 | 6.57E-05 | 7.71E-06 | 4.51E-05 | 0.00013 | 7.48E-05 | 0.00018 | 2.66E-05 |

| | | | | | | | | | | | | | | | | | | |
|---|---|---|---|---|---|---|---|---|---|---|---|---|---|---|---|---|---|---|
| Trim25 | 0.00016 | 0.00015 | 6.10E-05 | 5.90E-05 | 7.51E-05 | 0.00011 | 0 | 4.25E-05 | 8.58E-05 | 0.0002 | 0 | 8.72E-05 | 6.40E-05 | 5.18E-05 | 0.00012 | 1.26E-05 | 2.95E-05 | 0.00011 |
| Tent4a | 4.41E-05 | 0.00016 | 0.00015 | 0 | 6.72E-05 | 4.26E-05 | 1.81E-05 | 4.02E-05 | 8.58E-05 | 5.88E-05 | 0.00014 | 6.64E-05 | 1.44E-05 | 2.36E-05 | 0.00016 | 0.00031 | 9.74E-05 | 7.50E-05 |
| Slc16a3 | 0 | 0 | 0.00025 | 0.0003 | 5.29E-05 | 3.54E-05 | 0.00041 | 0.00021 | 8.58E-05 | 6.13E-05 | 0.00024 | 0.00211 | 0.00019 | 4.54E-06 | 0 | 8.84E-05 | 0.00032 | 0 |
| Pnkp | 3.99E-05 | 6.50E-05 | 5.73E-05 | 3.86E-06 | 3.28E-05 | 2.77E-05 | 0 | 7.55E-05 | 8.58E-05 | 3.07E-05 | 4.42E-05 | 0 | 0 | 3.99E-05 | 7.65E-05 | 5.50E-05 | 0 | 6.64E-05 |
| Ndufab1 | 0.00018 | 4.57E-05 | 0.00017 | 0.00013 | 3.45E-05 | 2.23E-05 | 1.90E-05 | 4.68E-05 | 8.58E-05 | 0 | 5.56E-05 | 3.61E-05 | 5.39E-05 | 0 | 0 | 0.00015 | 8.49E-07 | 0 |
| Tlcd2 | 1.85E-05 | 5.86E-06 | 1.66E-06 | 3.51E-05 | 0 | 0 | 0 | 1.90E-05 | 8.58E-05 | 6.77E-05 | 0 | 1.32E-05 | 4.30E-05 | 2.51E-05 | 2.87E-06 | 0 | 8.46E-05 | 2.76E-05 |
| Mark3 | 0.00022 | 0.00017 | 0.00015 | 0 | 0.0001 | 0.0001 | 6.79E-06 | 0 | 8.57E-05 | 3.65E-05 | 0.00011 | 3.01E-05 | 0 | 6.69E-05 | 0.00025 | 7.85E-05 | 0.00012 | 2.05E-05 |
| Abce1 | 0.00036 | 5.05E-05 | 0.00029 | 0 | 7.63E-07 | 9.49E-05 | 1.71E-05 | 0 | 8.57E-05 | 0 | 5.45E-05 | 0 | 1.58E-05 | 0 | 6.15E-06 | 0.00012 | 0 | 0 |
| Cpsf1 | 8.06E-05 | 9.57E-05 | 0.00011 | 8.29E-05 | 3.58E-05 | 1.69E-05 | 0 | 8.60E-06 | 8.57E-05 | 4.78E-05 | 3.84E-05 | 2.64E-05 | 0 | 1.76E-05 | 6.85E-05 | 9.39E-05 | 2.48E-05 | 5.05E-05 |
| Cdv3 | 0.00041 | 0.00018 | 0.00042 | 0 | 2.12E-05 | 0.00018 | 0.00012 | 0 | 8.56E-05 | 2.59E-05 | 0.00017 | 0.00026 | 0 | 4.17E-05 | 0 | 0.00018 | 0.00012 | 8.46E-06 |
| Suclg1 | 0.00016 | 4.74E-05 | 0.00014 | 0.00016 | 2.48E-05 | 3.66E-05 | 5.56E-05 | 0 | 8.56E-05 | 1.15E-05 | 5.04E-05 | 1.99E-06 | 9.88E-06 | 1.84E-05 | 5.01E-05 | 1.13E-05 | 0 | 1.53E-05 |
| Phrf1 | 4.16E-05 | 0.00019 | 0.00011 | 0.00021 | 5.01E-05 | 7.02E-06 | 0 | 1.02E-05 | 8.56E-05 | 3.58E-05 | 9.14E-05 | 0.0001 | 3.47E-05 | 4.68E-05 | 0.0003 | 0.00012 | 0 | 3.95E-05 |
| Psmg4 | 7.34E-05 | 6.46E-05 | 9.57E-05 | 0.00019 | 6.02E-05 | 0 | 1.79E-06 | 2.67E-05 | 8.56E-05 | 2.00E-05 | 1.46E-05 | 4.86E-05 | 5.40E-05 | 6.23E-06 | 0 | 8.21E-05 | 5.80E-05 | 7.79E-05 |
| Lonp2 | 0.00012 | 8.38E-05 | 5.45E-05 | 3.60E-05 | 3.21E-05 | 7.14E-05 | 1.24E-05 | 3.75E-05 | 8.55E-05 | 7.25E-05 | 2.65E-05 | 6.81E-05 | 4.82E-05 | 5.23E-06 | 1.77E-05 | 1.54E-05 | 3.05E-05 | 7.07E-05 |
| Ttc19 | 0.00014 | 5.92E-05 | 0.00011 | 2.02E-05 | 9.68E-05 | 3.52E-05 | 9.39E-05 | 7.86E-05 | 8.55E-05 | 3.36E-05 | 9.97E-05 | 3.93E-05 | 6.21E-05 | 0 | 0.00032 | 0 | 6.03E-05 | 1.44E-05 |
| Rerg | 0 | 0.00011 | 0 | 0 | 0 | 0 | 9.20E-06 | 0 | 8.55E-05 | 0 | 0 | 1.60E-05 | 5.73E-05 | 0 | 0 | 0 | 0 | 8.69E-05 |
| Kcmf1 | 0.00023 | 0.00026 | 0.00029 | 3.09E-05 | 0.00016 | 0.0002 | 2.94E-05 | 8.35E-05 | 8.54E-05 | 8.30E-05 | 0.00017 | 0.00027 | 0 | 4.41E-05 | 8.48E-05 | 0.00019 | 0.00019 | 9.62E-05 |
| Creb1 | 0.00016 | 0.00015 | 9.03E-05 | 5.47E-05 | 9.58E-05 | 9.81E-05 | 0 | 9.53E-05 | 8.53E-05 | 3.57E-05 | 4.60E-05 | 8.59E-05 | 2.77E-05 | 3.97E-05 | 7.82E-05 | 0.00013 | 0.0001 | 6.02E-05 |
| Grpel1 | 0.00031 | 0.00011 | 0.00029 | 0.00023 | 9.12E-05 | 7.50E-05 | 3.83E-05 | 4.75E-05 | 8.53E-05 | 0 | 0.00015 | 9.18E-05 | 0 | 3.66E-05 | 1.99E-07 | 9.61E-05 | 0 | 0 |
| Rnf215 | 3.26E-05 | 0.00014 | 5.86E-05 | 0.00015 | 0.00012 | 9.88E-05 | 3.06E-05 | 0.00012 | 8.52E-05 | 6.96E-05 | 3.38E-05 | 1.95E-06 | 9.52E-05 | 7.45E-05 | 0.00016 | 2.76E-05 | 0 | 0.00016 |
| Tor1b | 0.00015 | 8.93E-05 | 0.00012 | 6.85E-06 | 8.87E-05 | 8.59E-05 | 7.00E-05 | 3.55E-05 | 8.52E-05 | 6.15E-05 | 8.16E-05 | 5.98E-05 | 9.87E-05 | 1.08E-05 | 3.92E-05 | 6.37E-05 | 0 | 3.75E-05 |
| Psma2 | 0.00032 | 0.00011 | 0.00016 | 7.15E-05 | 1.85E-05 | 8.24E-05 | 8.51E-06 | 0 | 8.52E-05 | 1.31E-05 | 6.75E-05 | 6.19E-05 | 0 | 1.52E-05 | 8.46E-07 | 0.00011 | 5.50E-06 | 9.55E-06 |
| Slc41a3 | 1.04E-05 | 5.40E-05 | 0 | 0 | 5.52E-05 | 3.33E-05 | 0.00012 | 2.34E-05 | 8.52E-05 | 7.38E-05 | 4.36E-05 | 6.14E-05 | 3.85E-05 | 2.81E-05 | 0 | 3.73E-05 | 0 | 0.00012 |
| Llgl1 | 2.63E-05 | 8.80E-05 | 2.15E-05 | 4.28E-05 | 5.32E-05 | 0 | 1.24E-05 | 3.32E-05 | 8.52E-05 | 5.14E-05 | 7.26E-05 | 0.00011 | 2.57E-06 | 4.81E-05 | 8.26E-05 | 7.90E-05 | 0 | 8.77E-05 |
| Ggps1 | 0.00013 | 0.00012 | 9.36E-05 | 2.24E-05 | 9.61E-05 | 0.00011 | 2.25E-05 | 9.30E-05 | 8.51E-05 | 4.64E-05 | 5.26E-05 | 4.88E-05 | 3.58E-05 | 4.57E-05 | 6.50E-05 | 1.54E-05 | 0 | 1.82E-05 |
| Afg3l2 | 8.32E-05 | 0.00015 | 0.0002 | 0.00024 | 8.12E-05 | 7.37E-05 | 0 | 7.36E-05 | 8.51E-05 | 5.28E-05 | 9.20E-05 | 0.00015 | 9.92E-05 | 8.58E-05 | 0.00026 | 8.08E-06 | 8.40E-05 | 3.43E-05 |
| Rom1 | 4.00E-05 | 4.25E-05 | 0 | 6.21E-05 | 3.20E-05 | 2.93E-05 | 2.78E-05 | 7.16E-05 | 8.51E-05 | 6.87E-05 | 0 | 0 | 0 | 1.76E-05 | 0 | 0 | 4.25E-05 | 3.77E-05 |
| Dph6 | 7.54E-05 | 4.82E-05 | 6.53E-05 | 1.99E-05 | 1.19E-05 | 1.91E-05 | 0 | 3.13E-05 | 8.51E-05 | 2.69E-05 | 3.79E-05 | 2.85E-05 | 1.08E-05 | 2.07E-05 | 0.00013 | 4.29E-06 | 2.14E-05 | 1.03E-05 |
| Kcns3 | 0 | 1.42E-06 | 0 | 0 | 7.26E-05 | 5.34E-05 | 0 | 0 | 8.50E-05 | 0 | 1.87E-05 | 0 | 0 | 6.59E-05 | 0 | 2.28E-05 | 0 | 3.43E-05 |
| Tlr2 | 0 | 5.18E-05 | 0 | 1.12E-05 | 3.33E-05 | 3.58E-05 | 1.82E-05 | 6.65E-05 | 8.50E-05 | 3.94E-05 | 0 | 0 | 0 | 7.44E-05 | 2.42E-05 | 0 | 0 | 4.27E-05 |
| Tubgcp4 | 8.40E-05 | 0.0001 | 9.23E-05 | 4.02E-05 | 7.30E-05 | 2.34E-05 | 3.91E-05 | 1.14E-05 | 8.50E-05 | 5.19E-05 | 2.49E-06 | 2.09E-05 | 0 | 3.46E-05 | 0.00014 | 0.00017 | 2.53E-05 | 4.40E-05 |
| Mief1 | 5.78E-05 | 9.26E-05 | 0.00014 | 4.84E-05 | 5.85E-05 | 1.89E-05 | 8.88E-06 | 6.66E-05 | 8.50E-05 | 3.01E-05 | 3.52E-05 | 0.0001 | 3.35E-05 | 3.54E-05 | 2.08E-05 | 6.59E-05 | 0 | 0 |
| Ttll12 | 9.36E-05 | 7.23E-05 | 9.02E-05 | 1.61E-05 | 6.50E-05 | 1.67E-05 | 0 | 2.96E-05 | 8.50E-05 | 3.64E-05 | 5.62E-05 | 5.33E-05 | 0 | 1.74E-06 | 0 | 0.00014 | 0 | 1.98E-05 |

| | | | | | | | | | | | | | | | | | | |
|---|---|---|---|---|---|---|---|---|---|---|---|---|---|---|---|---|---|---|
| Rbm19 | 6.16E-05 | 7.39E-05 | 0.00015 | 5.87E-05 | 8.77E-06 | 0 | 0 | 0 | 8.50E-05 | 1.28E-05 | 3.78E-05 | 0 | 1.59E-05 | 2.17E-05 | 8.69E-05 | 4.24E-05 | 0 | 0 |
| Ifnar2 | 0.00023 | 0.00014 | 0.00013 | 0.0002 | 0.00014 | 0.00033 | 0.00017 | 0.00024 | 8.49E-05 | 0.0003 | 7.67E-05 | 0.00037 | 0.00088 | 0.00017 | 0 | 0.00011 | 0 | 0.00013 |
| BC005624 | 5.98E-05 | 0.0001 | 4.98E-05 | 0.00016 | 5.26E-05 | 1.74E-05 | 2.03E-05 | 6.62E-05 | 8.49E-05 | 5.80E-05 | 5.92E-05 | 6.31E-05 | 7.85E-05 | 5.45E-05 | 4.46E-05 | 9.86E-06 | 1.46E-06 | 8.06E-05 |
| Lrrfip1 | 0.0001 | 0.00023 | 0.00038 | 6.37E-05 | 5.67E-05 | 0 | 0.00013 | 7.10E-05 | 8.49E-05 | 0 | 0.00034 | 0.00034 | 4.46E-05 | 0.00011 | 0.00025 | 5.18E-05 | 3.51E-05 | 0.00011 |
| Slc25a36 | 0.00015 | 0.00015 | 1.64E-06 | 0 | 0.00011 | 0.00012 | 3.10E-05 | 6.73E-05 | 8.48E-05 | 4.74E-05 | 0 | 0.00024 | 4.16E-05 | 2.39E-05 | 0.00016 | 0 | 0.00011 | 4.99E-05 |
| Rnf214 | 7.85E-05 | 0.00014 | 3.60E-05 | 5.92E-05 | 6.72E-05 | 7.70E-05 | 1.74E-05 | 3.28E-05 | 8.48E-05 | 6.03E-05 | 7.36E-05 | 0.00015 | 7.52E-06 | 8.72E-05 | 0.00012 | 7.91E-05 | 9.35E-05 | 6.66E-05 |
| Rilpl1 | 8.23E-06 | 0.00011 | 1.34E-05 | 0.00017 | 7.73E-05 | 5.29E-05 | 5.62E-05 | 1.33E-05 | 8.48E-05 | 6.92E-05 | 0.00011 | 3.14E-05 | 5.32E-05 | 5.96E-05 | 8.98E-05 | 1.12E-05 | 0.00011 | 0.00011 |
| Mcu | 8.56E-05 | 9.70E-05 | 0.0001 | 0 | 0.00011 | 4.10E-05 | 4.11E-05 | 1.91E-05 | 8.48E-05 | 0.00014 | 6.57E-06 | 0.0001 | 0 | 0 | 8.28E-06 | 0.00011 | 4.42E-05 | 8.29E-05 |
| Bcl2l1 | 0 | 0.0002 | 6.19E-05 | 6.55E-05 | 0 | 0 | 0 | 0 | 8.48E-05 | 8.20E-05 | 0 | 6.42E-05 | 7.47E-05 | 0 | 3.79E-05 | 4.31E-05 | 1.06E-05 | 7.82E-05 |
| Klf13 | 0 | 0.00017 | 0.00011 | 0 | 6.63E-05 | 0.00015 | 0 | 0.00013 | 8.47E-05 | 0.00014 | 0.00039 | 0.00017 | 0 | 0.00039 | 0.00011 | 7.78E-05 | 0.00017 | 0.00013 |
| Ensa | 0.00014 | 0.0001 | 9.66E-05 | 2.76E-05 | 0.00013 | 0.00011 | 3.62E-05 | 4.03E-05 | 8.47E-05 | 5.24E-05 | 6.05E-05 | 8.95E-05 | 9.36E-06 | 0 | 2.72E-05 | 0.00011 | 5.19E-05 | 8.88E-05 |
| Dhx16 | 9.33E-05 | 0.00011 | 6.38E-05 | 6.92E-05 | 5.88E-05 | 5.02E-05 | 3.55E-05 | 9.53E-06 | 8.47E-05 | 5.32E-05 | 0 | 9.13E-05 | 1.11E-05 | 8.63E-05 | 0.00013 | 9.36E-05 | 8.30E-05 | 4.92E-05 |
| Csrnp1 | 0 | 0 | 9.51E-05 | 0.00016 | 9.07E-05 | 0 | 0 | 0.0002 | 8.47E-05 | 0 | 6.65E-05 | 1.32E-05 | 0 | 0.00013 | 0.00019 | 9.48E-06 | 0.00136 | 0 |
| Abl2 | 7.59E-05 | 0.00019 | 0.00013 | 1.53E-05 | 5.69E-05 | 0 | 6.23E-05 | 3.29E-05 | 8.47E-05 | 0.0001 | 8.10E-05 | 8.29E-05 | 2.93E-06 | 5.19E-05 | 0.00016 | 3.52E-05 | 0.00042 | 9.62E-05 |
| Ssu72 | 0.00027 | 0.00018 | 0.00026 | 0.00029 | 0.00011 | 0.00016 | 0.00012 | 6.74E-05 | 8.46E-05 | 6.05E-05 | 0.00019 | 0 | 4.82E-05 | 0.00011 | 3.81E-05 | 0.00011 | 0 | 4.52E-05 |
| Hras | 0.00021 | 7.64E-05 | 0.0002 | 3.47E-05 | 6.24E-05 | 9.66E-05 | 3.00E-05 | 3.92E-05 | 8.46E-05 | 5.15E-05 | 0.0002 | 8.30E-05 | 0 | 1.87E-05 | 0 | 0.00016 | 0 | 6.33E-06 |
| Hoxc9 | 1.45E-05 | 8.57E-05 | 0 | 0.00015 | 3.74E-05 | 0 | 3.12E-05 | 1.76E-05 | 8.46E-05 | 4.85E-05 | 6.37E-05 | 0 | 0 | 3.76E-05 | 9.90E-05 | 8.12E-05 | 0 | 0.00012 |
| Epc1 | 0.00011 | 0.00023 | 6.07E-05 | 4.28E-05 | 8.22E-05 | 7.81E-05 | 4.30E-05 | 0.00013 | 8.45E-05 | 6.85E-05 | 5.31E-05 | 8.13E-05 | 0 | 7.52E-05 | 0.00041 | 0.00011 | 0.00053 | 5.34E-05 |
| Mroh1 | 1.25E-05 | 0.00018 | 0 | 9.39E-05 | 4.17E-05 | 2.84E-05 | 4.07E-06 | 0 | 8.45E-05 | 3.83E-05 | 1.47E-06 | 0.00015 | 1.28E-05 | 0.00017 | 0.00017 | 0 | 0 | 4.67E-05 |
| Spg7 | 2.38E-05 | 0.00012 | 0.00012 | 0.0002 | 0.00011 | 9.29E-05 | 0.00011 | 0.00014 | 8.44E-05 | 0.00016 | 0.00016 | 8.66E-05 | 3.53E-05 | 0.00013 | 0.0002 | 0.00027 | 9.86E-05 | 0.00016 |
| Kmt5b | 6.81E-05 | 0.00018 | 8.89E-05 | 6.44E-05 | 8.53E-05 | 8.07E-05 | 5.13E-05 | 0.00014 | 8.44E-05 | 9.06E-05 | 7.85E-05 | 0.00019 | 2.11E-05 | 0.00014 | 0.00025 | 0 | 2.36E-06 | 6.58E-05 |
| Svbp | 0.00011 | 4.57E-06 | 3.38E-05 | 0.00011 | 0.00012 | 0.00015 | 0.00011 | 0.00021 | 8.43E-05 | 9.10E-06 | 0.00011 | 0 | 4.59E-05 | 0.00015 | 0 | 0 | 0 | 3.04E-06 |
| Sbf2 | 0.00014 | 0.00014 | 1.35E-05 | 0 | 7.57E-05 | 0.00014 | 0 | 4.81E-05 | 8.43E-05 | 8.36E-05 | 5.47E-05 | 0.0002 | 4.13E-06 | 1.48E-05 | 0.00014 | 5.63E-05 | 0.00022 | 4.88E-05 |
| Tmem183a | 0.0002 | 0.00016 | 0.0002 | 0.0001 | 3.19E-05 | 8.02E-05 | 1.16E-05 | 2.11E-06 | 8.43E-05 | 2.08E-05 | 0.00013 | 7.16E-05 | 0 | 2.07E-05 | 0.00015 | 0.00024 | 7.85E-05 | 1.13E-05 |
| Prpf6 | 0.0002 | 0.00017 | 0.00013 | 7.09E-05 | 5.66E-05 | 7.62E-05 | 8.75E-05 | 0 | 8.43E-05 | 1.94E-05 | 8.03E-06 | 7.92E-05 | 5.41E-05 | 4.10E-05 | 0.00019 | 7.36E-05 | 0 | 6.78E-05 |
| Uap1l1 | 0 | 0.00034 | 0 | 9.54E-05 | 0.00011 | 0.00017 | 1.41E-05 | 0 | 8.42E-05 | 1.83E-06 | 0 | 0 | 9.77E-05 | 0.00021 | 3.02E-05 | 0 | 0 | 0.00018 |
| Stc1 | 0 | 0 | 4.05E-05 | 0 | 0 | 0.00012 | 0 | 0 | 8.42E-05 | 4.53E-06 | 0 | 0.0002 | 1.72E-05 | 0.00018 | 0 | 0 | 0 | 0 |
| Rabl6 | 0.00024 | 0.0001 | 0.00026 | 0.00021 | 0.00013 | 0.00011 | 0.00019 | 0.00011 | 8.42E-05 | 3.23E-05 | 6.67E-05 | 0.00013 | 0 | 0.00013 | 0.00023 | 0.00015 | 0.00015 | 4.74E-05 |
| Begain | 0 | 2.07E-05 | 3.50E-05 | 0 | 5.81E-05 | 7.92E-05 | 0 | 4.99E-05 | 8.42E-05 | 3.79E-05 | 1.93E-05 | 4.94E-06 | 2.69E-05 | 4.74E-05 | 0.00017 | 5.17E-05 | 0 | 5.46E-05 |
| Nit2 | 9.94E-05 | 5.64E-05 | 1.68E-05 | 2.81E-05 | 2.72E-05 | 3.97E-05 | 0 | 4.32E-06 | 8.42E-05 | 3.10E-05 | 1.17E-05 | 0 | 2.12E-06 | 3.02E-05 | 0 | 2.31E-05 | 5.72E-05 | 6.34E-05 |
| Polr3e | 5.59E-05 | 8.02E-05 | 0.00015 | 9.53E-05 | 2.97E-05 | 0 | 3.99E-05 | 0 | 8.42E-05 | 9.19E-06 | 4.85E-05 | 0 | 2.29E-05 | 9.89E-06 | 0.00013 | 0 | 0 | 2.51E-05 |
| Ncoa4 | 0.00028 | 0.00016 | 0.00019 | 2.63E-05 | 0.00015 | 0.00016 | 1.63E-05 | 5.56E-05 | 8.40E-05 | 0.00014 | 5.28E-05 | 0.00012 | 1.16E-05 | 2.88E-05 | 0 | 2.51E-05 | 0 | 0.00011 |
| Cdk7 | 0.00025 | 7.11E-05 | 0.00012 | 0.00019 | 7.65E-05 | 0.00013 | 0.00015 | 0 | 8.40E-05 | 1.86E-05 | 0.00013 | 1.15E-05 | 9.15E-05 | 6.24E-05 | 0.00013 | 0 | 0 | 7.67E-05 |

| | | | | | | | | | | | | | | | | | | |
|---|---|---|---|---|---|---|---|---|---|---|---|---|---|---|---|---|---|---|
| Lrrk2 | 5.52E-05 | 4.55E-05 | 7.06E-05 | 0 | 5.91E-06 | 4.95E-05 | 4.61E-05 | 3.08E-05 | 8.40E-05 | 3.54E-05 | 5.52E-05 | 3.10E-05 | 0.00015 | 3.55E-05 | 0 | 6.86E-05 | 9.50E-05 | 0.00015 |
| Rbmxl1 | 0.00012 | 3.81E-05 | 0.00012 | 0.00012 | 1.68E-05 | 5.37E-07 | 0 | 1.43E-05 | 8.40E-05 | 0 | 3.05E-05 | 1.11E-05 | 1.49E-05 | 3.35E-05 | 9.07E-06 | 0.00012 | 1.19E-05 | 1.10E-05 |
| Ank3 | 3.10E-05 | 0 | 0 | 0 | 3.03E-06 | 0 | 0.00014 | 9.03E-07 | 8.40E-05 | 6.41E-05 | 0 | 0.00031 | 0.0005 | 0.00011 | 3.42E-05 | 3.50E-05 | 4.27E-05 | 0 |
| Ptch1 | 1.02E-05 | 8.98E-05 | 3.44E-05 | 0 | 0 | 0 | 0 | 2.07E-05 | 8.40E-05 | 0 | 6.92E-05 | 8.75E-05 | 0 | 0 | 0 | 5.09E-06 | 0 | 0.00032 |
| Selenop | 0.00015 | 0.00046 | 0 | 0 | 0.00017 | 0.0004 | 0 | 0.00028 | 8.39E-05 | 0.00072 | 0 | 0 | 0.00064 | 0 | 0 | 0 | 0 | 0.00107 |
| Uba3 | 0.00025 | 0.00014 | 0.00019 | 6.45E-05 | 4.13E-05 | 0.00017 | 5.26E-05 | 5.43E-05 | 8.39E-05 | 3.58E-05 | 0.00013 | 0.0002 | 0 | 2.69E-05 | 4.23E-05 | 0.00013 | 0 | 0.0001 |
| Sltm | 0.00011 | 0.00012 | 0.00015 | 0.00014 | 2.29E-06 | 0 | 3.23E-05 | 0 | 8.39E-05 | 5.49E-05 | 0 | 8.51E-05 | 0 | 3.03E-05 | 0.00034 | 9.72E-05 | 0.00026 | 3.09E-05 |
| Copz2 | 0.00045 | 0 | 0.00021 | 0.00061 | 0.00027 | 0.0004 | 0.00115 | 0.00027 | 8.38E-05 | 0.00018 | 0.00019 | 0.00015 | 8.47E-05 | 0.00023 | 0.00011 | 8.52E-05 | 0 | 0.0003 |
| Tmed8 | 2.60E-05 | 7.65E-05 | 7.86E-05 | 7.13E-05 | 7.03E-05 | 6.88E-05 | 2.15E-06 | 9.19E-05 | 8.38E-05 | 2.99E-05 | 4.96E-05 | 0 | 4.13E-05 | 9.57E-05 | 4.58E-05 | 7.73E-05 | 1.57E-05 | 4.85E-05 |
| Glrx3 | 0.00019 | 0.00012 | 0.00019 | 0.00023 | 3.50E-05 | 4.73E-05 | 1.64E-05 | 0.00011 | 8.38E-05 | 4.51E-05 | 7.77E-05 | 5.07E-05 | 0 | 9.39E-06 | 0 | 7.03E-05 | 0 | 0 |
| Spout1 | 0.00011 | 7.74E-05 | 0.00016 | 0.00021 | 3.02E-05 | 2.50E-05 | 0 | 1.81E-05 | 8.38E-05 | 0 | 1.61E-05 | 4.71E-05 | 3.25E-05 | 0 | 8.25E-05 | 0.00013 | 0 | 1.75E-07 |
| Sh3rf1 | 0 | 5.54E-05 | 6.03E-05 | 0.0001 | 0.0001 | 1.14E-05 | 0 | 0.00023 | 8.38E-05 | 1.78E-05 | 0.00015 | 9.67E-05 | 0 | 1.41E-05 | 0.00018 | 1.77E-05 | 0.00023 | 0 |
| Dmac1 | 6.53E-05 | 1.87E-05 | 4.32E-05 | 0.00018 | 6.47E-05 | 9.40E-06 | 1.92E-05 | 3.95E-05 | 8.38E-05 | 5.30E-05 | 2.49E-05 | 2.34E-05 | 4.48E-05 | 3.18E-05 | 0 | 7.03E-05 | 0 | 5.25E-05 |
| Dcaf12 | 0.00017 | 0.00016 | 0.00013 | 7.21E-05 | 0.0001 | 8.95E-05 | 7.26E-05 | 0.0001 | 8.37E-05 | 8.46E-05 | 9.05E-05 | 0.00022 | 6.18E-06 | 8.70E-05 | 9.24E-05 | 6.09E-05 | 0 | 4.78E-05 |
| Pdcd11 | 8.41E-05 | 8.68E-05 | 0.00018 | 0.0001 | 3.76E-05 | 2.63E-05 | 0 | 0 | 8.37E-05 | 2.96E-05 | 5.57E-06 | 1.46E-05 | 4.04E-06 | 3.36E-05 | 0.00023 | 3.65E-05 | 0 | 3.47E-05 |
| Has2 | 0.00027 | 0 | 0.00081 | 0 | 0 | 0 | 0.00017 | 0 | 8.37E-05 | 0 | 8.33E-05 | 0 | 0.00011 | 8.71E-05 | 0.00013 | 0.00016 | 4.28E-06 | 0 |
| Cpt1c | 5.43E-06 | 0.0002 | 0 | 0.0001 | 0.00012 | 6.11E-05 | 1.98E-05 | 0.00014 | 8.36E-05 | 0.00012 | 3.51E-05 | 4.22E-05 | 0.00011 | 5.02E-05 | 0.00022 | 0 | 2.22E-05 | 6.10E-05 |
| Polr2g | 0.00017 | 0.00012 | 0.00014 | 0 | 0.00011 | 9.87E-05 | 6.52E-05 | 7.92E-05 | 8.35E-05 | 0 | 5.62E-05 | 7.13E-05 | 0 | 6.73E-05 | 0 | 4.89E-05 | 1.54E-05 | 2.83E-05 |
| Nars | 0.00023 | 0.00017 | 0.00031 | 0.00011 | 3.03E-05 | 2.65E-05 | 0 | 0 | 8.35E-05 | 1.30E-05 | 0.00029 | 0.00024 | 1.18E-05 | 4.38E-05 | 0 | 7.48E-05 | 1.65E-06 | 0 |
| Ltbp3 | 0 | 0.00027 | 2.97E-05 | 0.00012 | 0.00029 | 0.00034 | 0.00031 | 0.00031 | 8.34E-05 | 0.00038 | 0.00011 | 0.00022 | 0.00047 | 0.00015 | 0.00029 | 4.28E-06 | 6.78E-05 | 0.00088 |
| Zbtb1 | 0.00013 | 0.00014 | 0.00012 | 2.27E-05 | 9.19E-05 | 0.00015 | 0.00011 | 8.77E-05 | 8.34E-05 | 8.67E-05 | 7.97E-05 | 0.00012 | 3.68E-05 | 0.00016 | 0.0001 | 5.16E-05 | 0 | 0.00012 |
| Ino80c | 0.00011 | 6.36E-05 | 0.00013 | 2.93E-06 | 1.23E-05 | 3.76E-05 | 2.38E-05 | 1.36E-05 | 8.34E-05 | 0.00013 | 0 | 4.72E-05 | 2.60E-05 | 2.52E-06 | 0 | 8.05E-05 | 0 | 6.22E-05 |
| Smg5 | 1.49E-05 | 0.00016 | 0.00023 | 0.00021 | 8.74E-05 | 1.75E-05 | 0 | 4.54E-05 | 8.34E-05 | 4.27E-05 | 0.00018 | 2.42E-05 | 0 | 3.78E-05 | 0.00028 | 0.00025 | 2.70E-05 | 0 |
| Smarcd3 | 5.43E-05 | 1.60E-05 | 3.50E-05 | 4.91E-05 | 2.64E-05 | 1.58E-05 | 3.65E-05 | 0 | 8.34E-05 | 3.34E-05 | 0.00012 | 3.25E-05 | 0 | 4.52E-05 | 3.91E-05 | 4.39E-07 | 0 | 8.08E-05 |
| Cacybp | 0.00017 | 7.34E-05 | 0.00021 | 0.00013 | 1.01E-05 | 9.24E-06 | 0 | 0 | 8.34E-05 | 5.61E-05 | 7.50E-05 | 0 | 0 | 2.60E-05 | 0 | 0.00024 | 3.45E-05 | 6.47E-05 |
| St6galnac5 | 1.16E-05 | 6.36E-05 | 0 | 0 | 0 | 0 | 0 | 0 | 8.34E-05 | 6.56E-06 | 0 | 0 | 3.38E-07 | 0.00018 | 0 | 2.91E-05 | 2.24E-05 | 3.64E-05 |
| Leprotl1 | 5.28E-05 | 0.00015 | 0.00025 | 0.00026 | 0.00018 | 0.00011 | 0.0001 | 0.00031 | 8.33E-05 | 0.00012 | 0.0003 | 0.00017 | 5.57E-05 | 0.00023 | 1.37E-05 | 0.00011 | 0 | 0.0002 |
| Slc25a20 | 0.00018 | 0.00014 | 8.55E-05 | 1.51E-05 | 7.31E-05 | 9.56E-05 | 0.00013 | 3.61E-05 | 8.33E-05 | 8.72E-05 | 4.29E-05 | 0 | 7.92E-05 | 8.94E-05 | 0 | 0 | 0 | 7.20E-05 |
| Washc2 | 0.00015 | 0.00025 | 2.96E-05 | 0.00018 | 5.34E-05 | 8.00E-05 | 3.11E-05 | 3.09E-05 | 8.33E-05 | 6.31E-05 | 5.43E-05 | 7.23E-05 | 5.56E-06 | 0.0001 | 0.00023 | 1.12E-05 | 0.00013 | 0.00013 |
| Myo9a | 6.72E-05 | 7.51E-05 | 6.49E-05 | 2.59E-05 | 5.07E-05 | 4.24E-05 | 7.68E-05 | 2.35E-05 | 8.33E-05 | 8.95E-05 | 6.81E-05 | 7.45E-05 | 3.11E-06 | 0 | 0.00011 | 6.19E-05 | 0.0002 | 6.08E-05 |
| Amz2 | 0.00014 | 4.49E-05 | 1.81E-05 | 0.00011 | 8.01E-05 | 2.43E-05 | 0.00012 | 0.00013 | 8.33E-05 | 3.36E-05 | 0 | 9.29E-05 | 3.29E-06 | 0 | 0 | 7.20E-05 | 0 | 3.16E-05 |
| Sqor | 7.82E-07 | 6.49E-05 | 6.95E-06 | 1.55E-06 | 3.67E-05 | 2.24E-05 | 0 | 1.69E-05 | 8.33E-05 | 0.00038 | 0 | 1.89E-05 | 0.00018 | 0 | 0 | 0 | 0 | 8.40E-05 |
| Clasp1 | 5.67E-05 | 0.00019 | 0.00012 | 6.35E-05 | 0.0001 | 2.03E-05 | 6.74E-05 | 9.72E-05 | 8.33E-05 | 0.00016 | 0.00017 | 0.00018 | 7.19E-05 | 4.88E-05 | 0.00016 | 0.00026 | 0.00014 | 0.00014 |

| | | | | | | | | | | | | | | | | | | |
|---|---|---|---|---|---|---|---|---|---|---|---|---|---|---|---|---|---|---|
| Yipf3 | 0.00018 | 0.00019 | 5.42E-05 | 0.00032 | 0.00023 | 0.0003 | 0.00044 | 0.00025 | 8.32E-05 | 0.00024 | 0.00017 | 0.00022 | 0.00016 | 0.00019 | 0.00019 | 0 | 8.86E-05 | 0.00029 |
| Usp14 | 0.00015 | 9.32E-05 | 0.00019 | 0 | 1.63E-05 | 4.44E-05 | 0 | 2.38E-05 | 8.32E-05 | 1.59E-05 | 0.00013 | 4.61E-05 | 0 | 2.00E-05 | 5.16E-05 | 0.00019 | 0 | 2.49E-05 |
| Wdr86 | 0 | 1.39E-05 | 0 | 0 | 0 | 9.22E-06 | 0 | 0 | 8.32E-05 | 0 | 0 | 0 | 0 | 9.15E-05 | 1.27E-05 | 0 | 0 | 1.26E-05 |
| Synj2bp | 0.0001 | 0.00017 | 0.00012 | 0.00013 | 0.00015 | 7.53E-05 | 9.85E-05 | 9.17E-05 | 8.31E-05 | 0.00011 | 0.0001 | 4.32E-05 | 0.00011 | 6.81E-05 | 0.00012 | 0.0001 | 0 | 0.00014 |
| Mpc2 | 0.00026 | 9.41E-05 | 5.93E-05 | 0.00012 | 0.00012 | 0.00019 | 0.0001 | 8.53E-05 | 8.30E-05 | 0.00012 | 8.10E-05 | 4.72E-05 | 0.00014 | 4.34E-05 | 0 | 2.91E-05 | 0 | 0.00016 |
| Fbxl6 | 2.76E-05 | 9.81E-05 | 9.60E-05 | 0.00012 | 7.78E-05 | 7.19E-05 | 0 | 0.0001 | 8.30E-05 | 5.58E-05 | 6.34E-05 | 2.36E-05 | 0 | 9.63E-05 | 6.43E-05 | 7.29E-06 | 0 | 5.91E-05 |
| Cdk17 | 7.06E-05 | 5.65E-05 | 0.00014 | 0 | 0 | 6.27E-05 | 4.49E-05 | 6.62E-05 | 8.30E-05 | 2.07E-05 | 9.78E-05 | 0.00015 | 7.41E-05 | 0 | 0.00012 | 3.37E-05 | 0.00014 | 3.25E-05 |
| Aarsd1 | 0.00012 | 9.47E-05 | 0.00011 | 0.00017 | 3.00E-05 | 4.67E-05 | 0 | 1.52E-06 | 8.30E-05 | 1.35E-05 | 0 | 0 | 0 | 4.70E-05 | 3.70E-05 | 0.00036 | 7.23E-06 | 4.88E-05 |
| Eid2 | 2.24E-05 | 7.77E-05 | 9.57E-05 | 0.0001 | 7.42E-05 | 0.00011 | 0 | 6.98E-05 | 8.28E-05 | 5.22E-05 | 6.26E-05 | 5.21E-05 | 3.86E-05 | 8.41E-05 | 9.88E-05 | 0 | 0 | 2.98E-05 |
| Unc45a | 2.20E-05 | 0.00012 | 8.26E-05 | 0.00018 | 2.97E-05 | 6.07E-05 | 1.96E-05 | 1.70E-05 | 8.28E-05 | 3.11E-05 | 0.00019 | 0 | 0 | 9.49E-05 | 0.00012 | 0 | 8.84E-05 | 6.84E-05 |
| Wdr46 | 4.67E-05 | 8.95E-05 | 0.00029 | 0.00019 | 0 | 0 | 0 | 0 | 8.28E-05 | 2.66E-05 | 1.39E-05 | 0 | 4.83E-05 | 3.58E-05 | 7.08E-05 | 9.15E-05 | 8.56E-06 | 3.39E-05 |
| Fam207a | 9.26E-05 | 0.00014 | 0.0002 | 0.0003 | 6.85E-05 | 0 | 5.10E-05 | 8.06E-05 | 8.28E-05 | 2.86E-05 | 6.24E-05 | 5.31E-05 | 4.71E-05 | 3.26E-05 | 0.00018 | 6.56E-05 | 0 | 7.08E-05 |
| Mgat2 | 0.00038 | 0.00012 | 0.00043 | 0 | 0.00012 | 0.00012 | 0.00033 | 0.00011 | 8.27E-05 | 0.00019 | 0.00014 | 0.00023 | 0.00013 | 4.69E-05 | 0 | 0.00015 | 0 | 0.00012 |
| Cerk | 4.03E-07 | 0.00011 | 5.30E-06 | 6.97E-07 | 3.62E-05 | 7.97E-05 | 0 | 0.00012 | 8.27E-05 | 4.43E-05 | 0 | 3.85E-05 | 5.28E-05 | 8.38E-05 | 1.58E-05 | 4.81E-05 | 3.55E-05 | 3.65E-05 |
| Ndufb1-ps | 0.00037 | 0.00012 | 9.75E-05 | 0.00021 | 0.00011 | 0.00021 | 0.00012 | 5.79E-05 | 8.26E-05 | 5.81E-05 | 0.00016 | 5.96E-05 | 0.00014 | 0.00012 | 0 | 0.00011 | 5.55E-05 | 4.02E-06 |
| Heatr5b | 9.73E-05 | 6.67E-05 | 5.24E-05 | 8.47E-06 | 4.46E-05 | 6.90E-05 | 5.27E-05 | 2.50E-05 | 8.26E-05 | 6.96E-05 | 5.15E-05 | 1.78E-05 | 4.86E-05 | 3.25E-05 | 2.76E-05 | 2.25E-05 | 8.13E-05 | 3.26E-05 |
| Rpp21 | 7.20E-05 | 7.85E-05 | 2.48E-05 | 0.0001 | 7.00E-05 | 5.86E-05 | 1.84E-05 | 4.36E-05 | 8.26E-05 | 2.97E-05 | 6.87E-05 | 2.71E-05 | 3.46E-05 | 2.89E-06 | 0.0001 | 0.00013 | 4.38E-05 | 1.73E-05 |
| Ttc14 | 0.00012 | 0.00015 | 4.07E-05 | 0 | 9.19E-05 | 5.60E-05 | 0 | 5.71E-05 | 8.26E-05 | 4.18E-05 | 8.93E-05 | 9.51E-05 | 0 | 2.44E-05 | 0.00042 | 2.70E-05 | 0 | 4.07E-05 |
| Adam33 | 0 | 1.30E-05 | 0 | 0 | 0 | 5.40E-05 | 0 | 0 | 8.26E-05 | 0.00017 | 0 | 0 | 3.63E-05 | 0.00015 | 0.00019 | 0 | 0 | 7.20E-05 |
| Tnpo3 | 0.00018 | 0.00014 | 0.00017 | 0.00013 | 5.65E-05 | 4.76E-05 | 7.85E-05 | 0 | 8.26E-05 | 5.93E-05 | 6.82E-05 | 0.00014 | 0 | 3.12E-05 | 8.52E-05 | 0.00021 | 9.99E-05 | 7.93E-05 |
| Ccdc71 | 9.18E-05 | 0.00014 | 6.05E-05 | 8.44E-05 | 4.70E-05 | 1.30E-05 | 6.24E-06 | 3.74E-06 | 8.26E-05 | 4.90E-05 | 3.45E-05 | 5.43E-05 | 0 | 2.62E-05 | 0.00011 | 9.36E-05 | 0 | 5.57E-05 |
| Urgcp | 2.13E-05 | 9.89E-05 | 5.48E-05 | 6.78E-05 | 5.37E-05 | 7.49E-06 | 7.34E-05 | 0 | 8.26E-05 | 6.58E-05 | 5.68E-05 | 5.06E-05 | 4.17E-05 | 5.08E-05 | 0.00021 | 3.26E-05 | 0 | 6.55E-05 |
| Uhmk1 | 0.00012 | 0.00011 | 0.00013 | 1.35E-05 | 5.94E-05 | 7.27E-05 | 0 | 1.55E-05 | 8.25E-05 | 3.14E-05 | 0.0001 | 9.70E-05 | 0 | 6.11E-05 | 3.44E-05 | 6.71E-05 | 0 | 0 |
| Sptssa | 0.00014 | 0.00015 | 3.65E-05 | 6.02E-05 | 9.39E-05 | 7.08E-05 | 7.98E-05 | 6.84E-05 | 8.25E-05 | 0.00012 | 0 | 8.00E-05 | 2.09E-05 | 4.19E-05 | 0 | 0.0002 | 3.77E-05 | 0.00015 |
| Tssc4 | 4.22E-05 | 8.88E-05 | 0.00015 | 0.00018 | 0 | 0 | 7.56E-05 | 5.38E-05 | 8.25E-05 | 4.01E-05 | 2.76E-05 | 7.05E-05 | 8.57E-05 | 0 | 0.0001 | 0.00026 | 0.00054 | 8.94E-05 |
| Tox4 | 0.00014 | 0.00018 | 0.00015 | 0.00013 | 6.98E-05 | 5.20E-05 | 2.66E-05 | 5.30E-05 | 8.24E-05 | 0 | 4.96E-05 | 0.00013 | 3.27E-06 | 6.05E-05 | 1.83E-05 | 3.79E-05 | 8.72E-05 | 3.60E-05 |
| Rnf7 | 0.00016 | 7.40E-05 | 0.0001 | 0.00011 | 4.41E-05 | 4.47E-05 | 0.0001 | 4.90E-05 | 8.24E-05 | 2.70E-05 | 0.00011 | 8.27E-06 | 7.16E-05 | 5.01E-05 | 1.12E-05 | 7.00E-05 | 0 | 1.60E-05 |
| D10Wsu102e | 6.65E-05 | 7.14E-05 | 9.92E-05 | 5.72E-05 | 3.66E-05 | 1.02E-06 | 0 | 1.72E-05 | 8.24E-05 | 0 | 2.27E-05 | 0 | 0 | 8.78E-06 | 6.88E-05 | 0.00011 | 0.00045 | 0 |
| Gtf3c6 | 0.00017 | 6.99E-05 | 0.00018 | 0.00027 | 3.84E-05 | 0 | 0 | 0 | 8.24E-05 | 2.33E-05 | 6.16E-05 | 0 | 1.65E-05 | 0 | 3.76E-05 | 5.56E-05 | 0 | 5.50E-05 |
| Nufip2 | 0.00025 | 0.00018 | 0.00016 | 8.61E-05 | 0.00011 | 0.00017 | 9.24E-05 | 3.62E-05 | 8.23E-05 | 8.21E-05 | 0.00014 | 5.87E-05 | 0.00012 | 0.00012 | 0.00013 | 5.14E-05 | 0.00074 | 1.19E-05 |
| Ndufb5 | 0.00019 | 7.88E-05 | 8.01E-05 | 0.00023 | 5.10E-05 | 0.00013 | 0.00012 | 9.13E-05 | 8.23E-05 | 2.70E-05 | 6.06E-05 | 4.41E-05 | 0 | 0.00011 | 0 | 8.53E-05 | 7.21E-05 | 3.74E-05 |
| Hspe1 | 0.00031 | 3.55E-05 | 0.00024 | 0.00012 | 2.57E-05 | 0.00011 | 3.47E-06 | 3.43E-05 | 8.23E-05 | 1.17E-05 | 7.50E-06 | 4.56E-05 | 5.74E-07 | 0 | 6.97E-05 | 0.00011 | 0.00022 | 3.22E-05 |
| Klhl42 | 7.25E-05 | 0.00011 | 3.88E-05 | 8.20E-06 | 8.60E-05 | 2.80E-05 | 1.36E-05 | 6.35E-05 | 8.23E-05 | 1.03E-05 | 0 | 2.07E-05 | 0 | 6.32E-05 | 0 | 1.80E-05 | 0 | 6.72E-05 |

| | | | | | | | | | | | | | | | | | | |
|---|---|---|---|---|---|---|---|---|---|---|---|---|---|---|---|---|---|---|
| Xylt2 | 0 | 0.0001 | 1.11E-05 | 6.66E-05 | 7.43E-05 | 2.40E-05 | 3.43E-05 | 8.72E-05 | 8.23E-05 | 6.54E-05 | 2.35E-05 | 9.95E-05 | 2.83E-05 | 6.09E-05 | 9.79E-05 | 3.44E-05 | 1.40E-05 | 5.84E-05 |
| Smarcd1 | 4.85E-05 | 7.53E-05 | 9.79E-05 | 4.96E-05 | 4.75E-05 | 5.57E-05 | 2.88E-05 | 2.79E-05 | 8.22E-05 | 1.91E-05 | 7.67E-05 | 1.66E-05 | 0 | 8.42E-05 | 8.69E-05 | 9.43E-05 | 0 | 4.97E-05 |
| Tmem51 | 0 | 2.70E-05 | 4.67E-06 | 0 | 0 | 5.27E-05 | 0 | 2.96E-05 | 8.22E-05 | 5.37E-06 | 0 | 6.21E-05 | 0 | 5.21E-05 | 0 | 1.77E-06 | 0 | 0 |
| Zfp513 | 4.96E-05 | 9.30E-05 | 6.84E-05 | 2.90E-05 | 4.11E-05 | 2.56E-05 | 3.61E-05 | 1.76E-05 | 8.22E-05 | 3.77E-05 | 6.67E-05 | 1.11E-05 | 1.91E-06 | 4.57E-05 | 9.24E-05 | 1.67E-05 | 0 | 2.65E-05 |
| 6-Mar | 3.27E-05 | 0.00019 | 0.00015 | 0.00018 | 0.00019 | 0.00015 | 0.00011 | 0.00026 | 8.21E-05 | 8.80E-05 | 0.00016 | 0.00016 | 4.90E-05 | 7.48E-05 | 0.00015 | 0.00014 | 0.00012 | 0.00012 |
| Atg9a | 8.81E-05 | 0.00012 | 4.18E-05 | 0.00012 | 9.81E-05 | 0.00012 | 0.00017 | 8.88E-05 | 8.21E-05 | 0.00011 | 6.73E-05 | 0.00012 | 8.75E-05 | 0.00012 | 0.00013 | 0 | 0 | 7.01E-05 |
| Ctdspl2 | 0.00013 | 7.73E-05 | 3.93E-05 | 0 | 5.78E-05 | 7.10E-05 | 0 | 1.81E-05 | 8.21E-05 | 2.43E-05 | 2.21E-05 | 0.00016 | 0 | 1.29E-05 | 0.00015 | 0.00022 | 0.00013 | 6.77E-06 |
| Sh2b1 | 3.45E-06 | 0.00012 | 5.45E-05 | 0.00018 | 0.00012 | 3.16E-05 | 8.31E-05 | 0.00017 | 8.21E-05 | 8.33E-05 | 2.45E-05 | 8.85E-05 | 0 | 9.99E-05 | 0.0002 | 3.68E-05 | 0.00011 | 6.02E-05 |
| Prpf31 | 0.00011 | 7.03E-05 | 0.00019 | 0.00017 | 1.64E-05 | 0 | 0 | 0 | 8.21E-05 | 7.97E-06 | 3.10E-05 | 0 | 9.90E-06 | 3.31E-05 | 3.29E-05 | 0.00022 | 0 | 1.16E-06 |
| Mpzl1 | 0.00019 | 7.14E-05 | 7.26E-05 | 0 | 0.00011 | 0.00011 | 0.00018 | 0.00014 | 8.19E-05 | 0 | 0.00011 | 0 | 0 | 0.00012 | 0 | 0.00017 | 4.22E-05 | 8.03E-05 |
| Mrtfb | 8.28E-05 | 0.00016 | 4.13E-05 | 0 | 4.69E-05 | 6.37E-05 | 7.03E-05 | 0.00016 | 8.19E-05 | 9.18E-05 | 5.96E-05 | 7.73E-05 | 0 | 3.65E-05 | 0.00024 | 1.80E-05 | 0 | 2.26E-05 |
| Dzip1 | 3.83E-05 | 9.51E-05 | 0 | 4.67E-05 | 1.70E-05 | 1.35E-07 | 6.99E-05 | 0 | 8.19E-05 | 0.00022 | 0 | 0 | 9.87E-05 | 1.77E-05 | 2.81E-05 | 5.47E-05 | 3.68E-05 | 8.65E-05 |
| Tm9sf4 | 0.00025 | 5.21E-05 | 0.00016 | 1.99E-06 | 0.00012 | 0.00017 | 0.0003 | 0.00014 | 8.18E-05 | 0.00012 | 8.51E-05 | 6.12E-05 | 0.00019 | 4.16E-05 | 5.88E-05 | 4.17E-05 | 0 | 6.60E-05 |
| Mecp2 | 6.59E-05 | 0.00018 | 5.90E-05 | 3.75E-05 | 0.00011 | 9.62E-05 | 6.43E-05 | 2.42E-05 | 8.18E-05 | 4.85E-05 | 2.56E-05 | 0.00013 | 1.47E-05 | 2.95E-05 | 9.02E-05 | 3.71E-05 | 2.05E-05 | 3.94E-05 |
| Ttc21b | 2.69E-05 | 4.90E-05 | 0 | 3.69E-05 | 4.30E-05 | 3.82E-06 | 8.52E-06 | 0 | 8.18E-05 | 2.24E-05 | 2.31E-06 | 5.49E-05 | 2.53E-05 | 1.15E-05 | 8.98E-06 | 5.47E-05 | 6.89E-06 | 4.81E-05 |
| Matr3 | 0.00024 | 0.00019 | 0.00015 | 6.19E-05 | 7.91E-05 | 0.00016 | 0 | 0 | 8.17E-05 | 2.37E-05 | 9.11E-06 | 0.00014 | 0 | 2.29E-05 | 0.00026 | 0.00026 | 0.00013 | 1.51E-05 |
| Arpc5l | 0.00019 | 0.00011 | 0.00017 | 9.44E-05 | 8.96E-05 | 0.00011 | 5.86E-06 | 0.00013 | 8.17E-05 | 4.93E-05 | 2.33E-05 | 0 | 6.00E-06 | 4.00E-05 | 0 | 0.00019 | 0 | 4.23E-05 |
| Pdcl | 7.47E-05 | 7.32E-05 | 5.58E-05 | 7.25E-05 | 3.80E-05 | 5.58E-05 | 5.35E-05 | 7.22E-05 | 8.17E-05 | 7.53E-05 | 9.81E-05 | 4.93E-05 | 2.01E-05 | 7.65E-05 | 8.07E-05 | 2.90E-05 | 6.81E-05 | 6.67E-05 |
| Tesk1 | 0 | 0.00013 | 9.55E-05 | 0.00044 | 0.00015 | 5.65E-05 | 0.00011 | 0.00011 | 8.16E-05 | 2.98E-05 | 0.00014 | 0 | 3.37E-05 | 7.25E-05 | 0.00013 | 1.34E-05 | 8.80E-05 | 0.00013 |
| Ccdc127 | 0.00013 | 4.61E-05 | 9.31E-05 | 0 | 6.23E-05 | 4.56E-05 | 9.34E-05 | 5.52E-05 | 8.16E-05 | 4.78E-05 | 5.56E-05 | 6.15E-05 | 2.91E-05 | 3.50E-05 | 6.21E-06 | 7.70E-05 | 0 | 2.94E-05 |
| Tspyl1 | 4.26E-05 | 0.0002 | 0 | 0.00037 | 0.00013 | 2.72E-05 | 2.31E-05 | 7.61E-05 | 8.16E-05 | 0.00012 | 0.00011 | 0.00013 | 4.91E-06 | 6.61E-05 | 0.00039 | 1.68E-07 | 0 | 0.00013 |
| Nol12 | 9.64E-05 | 6.26E-05 | 8.61E-05 | 0.00017 | 2.80E-05 | 5.48E-06 | 3.36E-05 | 1.49E-05 | 8.16E-05 | 5.93E-05 | 5.49E-05 | 2.48E-05 | 0 | 0 | 9.22E-05 | 9.79E-05 | 0 | 3.64E-05 |
| Stat1 | 0.00023 | 6.43E-05 | 1.94E-06 | 0.00013 | 4.61E-05 | 2.39E-05 | 3.75E-05 | 7.21E-05 | 8.15E-05 | 0.00025 | 0 | 0 | 0.0002 | 2.33E-05 | 1.52E-05 | 0.00024 | 0 | 0.00032 |
| Kat5 | 9.01E-05 | 0.00012 | 7.16E-05 | 0.00011 | 2.36E-05 | 1.17E-05 | 5.33E-05 | 2.58E-05 | 8.15E-05 | 4.14E-05 | 6.49E-05 | 2.83E-05 | 3.71E-06 | 5.21E-05 | 0.00015 | 8.07E-05 | 4.47E-05 | 3.15E-05 |
| 7-Mar | 0.00024 | 0.00024 | 6.90E-05 | 0 | 5.80E-05 | 0.00018 | 5.47E-05 | 1.05E-05 | 8.14E-05 | 6.04E-06 | 7.82E-05 | 0.00023 | 0 | 6.35E-05 | 0.00037 | 8.93E-05 | 3.75E-05 | 1.37E-06 |
| Psen1 | 0.00026 | 0.00018 | 0.00018 | 0 | 7.02E-05 | 0.00013 | 9.26E-05 | 6.42E-05 | 8.14E-05 | 0.00014 | 3.95E-05 | 0.00011 | 9.35E-05 | 7.05E-05 | 2.25E-05 | 9.12E-05 | 3.91E-05 | 8.30E-05 |
| Stat5b | 3.18E-05 | 0.00011 | 7.69E-05 | 6.76E-05 | 3.89E-05 | 3.21E-05 | 4.18E-05 | 8.52E-06 | 8.14E-05 | 0.00017 | 9.45E-05 | 8.92E-05 | 0.00011 | 1.26E-05 | 4.37E-05 | 9.90E-05 | 0.00017 | 0.00012 |
| Arxes1 | 2.08E-05 | 0 | 0 | 0 | 0.00012 | 1.30E-05 | 0 | 0 | 8.14E-05 | 0 | 6.67E-05 | 0 | 0 | 6.59E-05 | 0 | 5.13E-06 | 0 | 0 |
| Rybp | 4.10E-05 | 0.00012 | 5.24E-05 | 4.56E-05 | 3.74E-05 | 0 | 0 | 1.12E-05 | 8.14E-05 | 3.67E-05 | 4.84E-05 | 0.00011 | 1.27E-05 | 4.32E-07 | 0 | 7.10E-05 | 0.00039 | 5.92E-05 |
| Ube2b | 0.00036 | 0.00034 | 7.61E-05 | 0.00013 | 0.00023 | 0.00038 | 0.00027 | 0.00027 | 8.13E-05 | 0.00012 | 0.00022 | 0.00034 | 0.00013 | 0.00019 | 0.00028 | 7.73E-05 | 0.0003 | 0.00018 |
| Wdr5 | 0.00017 | 0.00011 | 0.0002 | 3.36E-05 | 2.64E-05 | 4.07E-05 | 0 | 0 | 8.13E-05 | 2.36E-06 | 6.76E-05 | 4.42E-05 | 1.28E-05 | 4.44E-05 | 6.22E-05 | 0.00018 | 0 | 5.79E-06 |
| Ndufs5 | 0.0002 | 5.73E-05 | 0.00013 | 0.00021 | 3.86E-05 | 3.63E-05 | 5.04E-05 | 7.05E-05 | 8.13E-05 | 1.53E-05 | 0.0001 | 0 | 1.94E-05 | 3.81E-05 | 0 | 5.78E-05 | 0 | 7.38E-05 |
| Champ1 | 0.0001 | 0.00011 | 9.51E-05 | 8.30E-05 | 2.92E-05 | 2.83E-05 | 2.06E-05 | 0 | 8.13E-05 | 3.00E-05 | 4.90E-05 | 4.70E-05 | 2.57E-05 | 8.68E-06 | 0.0001 | 5.29E-05 | 0 | 6.07E-05 |

| | | | | | | | | | | | | | | | | | | |
|---|---|---|---|---|---|---|---|---|---|---|---|---|---|---|---|---|---|---|
| Vps72 | 5.86E-05 | 0.00014 | 7.30E-05 | 0.0001 | 5.45E-05 | 1.71E-05 | 0 | 2.59E-05 | 8.12E-05 | 3.20E-05 | 1.91E-05 | 5.20E-05 | 0 | 5.70E-05 | 6.37E-05 | 6.92E-05 | 3.06E-05 | 1.58E-05 |
| Erh | 0.00012 | 2.45E-06 | 0.0001 | 0.0001 | 8.52E-07 | 0 | 0 | 0 | 8.12E-05 | 0 | 3.42E-05 | 1.63E-06 | 2.33E-05 | 0 | 1.52E-05 | 0.00015 | 0 | 3.15E-08 |
| Pdlim5 | 0.00019 | 6.70E-05 | 0.00015 | 0 | 0 | 0 | 8.57E-05 | 4.55E-05 | 8.12E-05 | 0.0002 | 0.00068 | 0.00015 | 0.00025 | 0 | 0.00012 | 6.95E-06 | 9.38E-05 | 0.00033 |
| Tax1bp1 | 0.00031 | 0.00027 | 0.00016 | 0.00011 | 8.26E-05 | 0.00019 | 4.10E-05 | 1.37E-05 | 8.11E-05 | 0.00014 | 6.86E-05 | 4.48E-05 | 0.00015 | 0.00013 | 0.00011 | 0.00019 | 0.00019 | 0.00017 |
| Sec22b | 0.00047 | 2.00E-05 | 0.00017 | 0.00039 | 0.00019 | 0.00016 | 0.00047 | 0.00026 | 8.11E-05 | 8.78E-05 | 7.67E-05 | 0.00014 | 0.00019 | 3.71E-05 | 0.00026 | 4.17E-05 | 0.00011 | 0.00013 |
| Polr2m | 0.00012 | 7.21E-05 | 0.0001 | 1.97E-05 | 0.00014 | 0.00014 | 1.77E-05 | 6.74E-05 | 8.11E-05 | 4.12E-05 | 4.50E-05 | 0.00012 | 4.49E-05 | 1.51E-05 | 3.03E-05 | 0.00017 | 1.94E-05 | 2.76E-05 |
| Chd1 | 0.00013 | 0.00014 | 0.00023 | 2.57E-05 | 2.86E-05 | 6.21E-05 | 0 | 2.60E-05 | 8.11E-05 | 0 | 2.79E-05 | 3.19E-05 | 0 | 6.90E-05 | 0.00027 | 0.00016 | 0.00039 | 5.67E-05 |
| Timm10 | 6.22E-05 | 9.34E-05 | 0.00028 | 0.00015 | 1.10E-05 | 0 | 0 | 0 | 8.11E-05 | 0 | 3.91E-05 | 0 | 1.35E-05 | 3.52E-05 | 0 | 2.40E-06 | 0 | 2.15E-06 |
| Lrrfip2 | 7.52E-05 | 0.00015 | 0.00021 | 0 | 9.68E-05 | 8.74E-05 | 4.90E-05 | 7.21E-06 | 8.10E-05 | 6.52E-05 | 4.73E-05 | 0.00011 | 0 | 2.49E-05 | 5.01E-05 | 8.61E-05 | 5.24E-05 | 2.75E-05 |
| Rp9 | 0.00017 | 0.00017 | 5.60E-05 | 0.00011 | 4.66E-05 | 0.00011 | 3.99E-05 | 8.73E-05 | 8.09E-05 | 5.57E-05 | 0 | 0.00022 | 0 | 9.16E-05 | 0 | 0.0001 | 1.82E-05 | 6.93E-05 |
| Lamtor1 | 6.83E-05 | 8.39E-05 | 2.79E-05 | 0.00014 | 7.91E-05 | 6.03E-05 | 6.01E-05 | 6.50E-05 | 8.09E-05 | 1.45E-05 | 2.56E-05 | 5.20E-05 | 2.21E-05 | 2.88E-05 | 0 | 8.99E-05 | 5.49E-06 | 5.53E-05 |
| Fbl | 6.92E-05 | 6.04E-05 | 0.00031 | 0.00041 | 0 | 0 | 0 | 0 | 8.09E-05 | 3.25E-05 | 3.99E-05 | 9.81E-07 | 2.95E-05 | 1.65E-06 | 6.25E-05 | 0.00013 | 0.00011 | 2.36E-05 |
| Csgalnact1 | 0.00013 | 0.00016 | 0.00019 | 0 | 0.00042 | 0.0002 | 0.00059 | 0.00021 | 8.08E-05 | 0 | 6.82E-05 | 0 | 0.00072 | 0.00037 | 0 | 4.94E-05 | 0 | 8.75E-05 |
| Smc1a | 0.00021 | 0.00011 | 0.00013 | 8.82E-06 | 7.05E-05 | 0.00015 | 7.67E-05 | 0.00015 | 8.08E-05 | 8.35E-05 | 0 | 4.56E-05 | 6.88E-05 | 2.07E-05 | 0.00016 | 0.00049 | 0.00013 | 0.00016 |
| Ei24 | 0.00021 | 0.00018 | 3.77E-05 | 0 | 0.0001 | 0.00013 | 0 | 6.31E-05 | 8.08E-05 | 8.37E-05 | 4.64E-05 | 1.13E-05 | 0 | 0 | 0 | 7.02E-05 | 2.83E-07 | 4.31E-05 |
| Tbce | 9.48E-05 | 6.35E-05 | 2.48E-05 | 4.32E-05 | 1.61E-05 | 5.82E-05 | 0 | 2.58E-05 | 8.08E-05 | 2.07E-05 | 0 | 3.66E-05 | 2.24E-06 | 1.16E-05 | 0.00019 | 1.82E-05 | 0 | 7.48E-05 |
| Krit1 | 8.90E-05 | 6.59E-05 | 2.49E-05 | 0 | 3.34E-05 | 4.55E-05 | 1.41E-05 | 2.08E-05 | 8.08E-05 | 3.90E-05 | 2.95E-05 | 0 | 0 | 3.15E-05 | 0.00016 | 4.46E-05 | 6.02E-05 | 5.22E-05 |
| Casp3 | 0.00014 | 7.23E-05 | 9.83E-05 | 5.13E-05 | 0.00016 | 0 | 7.47E-05 | 0.00024 | 8.08E-05 | 3.31E-05 | 1.54E-05 | 2.29E-05 | 6.92E-05 | 0.00013 | 3.63E-05 | 0.00021 | 0.00015 | 0 |
| Sphk1 | 0 | 0.00012 | 0.00032 | 0.00022 | 0.00041 | 0.00027 | 0 | 0.00024 | 8.07E-05 | 0 | 5.81E-05 | 0.00025 | 0.00026 | 0.00054 | 0.00024 | 0 | 0.00016 | 0 |
| Setd3 | 0.00013 | 0.00021 | 0.00014 | 0.00011 | 0.0001 | 0.00018 | 3.30E-05 | 9.49E-05 | 8.06E-05 | 5.71E-05 | 0.0002 | 0.00026 | 1.46E-05 | 5.60E-05 | 0.00019 | 0.00019 | 7.49E-06 | 0.00014 |
| Dmxl1 | 0.00013 | 0.00012 | 4.63E-05 | 0 | 4.77E-05 | 6.36E-05 | 2.00E-05 | 0 | 8.06E-05 | 2.82E-05 | 3.85E-05 | 9.49E-05 | 1.62E-05 | 0 | 4.69E-05 | 4.27E-05 | 0.00012 | 1.21E-05 |
| Mvb12b | 5.86E-05 | 9.19E-05 | 4.76E-05 | 0 | 5.06E-05 | 2.21E-05 | 0 | 0.0001 | 8.06E-05 | 0 | 4.72E-05 | 5.31E-05 | 3.43E-05 | 0 | 6.51E-05 | 2.88E-05 | 0.00012 | 7.16E-05 |
| Kng2 | 0 | 7.15E-06 | 0 | 0 | 1.05E-05 | 0.00013 | 0 | 0 | 8.05E-05 | 2.84E-06 | 0 | 1.32E-05 | 0.00066 | 0 | 0 | 0 | 0 | 0.00018 |
| Gramd4 | 4.38E-05 | 0.00014 | 5.80E-05 | 0 | 0.0001 | 0.00011 | 0 | 0.00034 | 8.05E-05 | 1.92E-05 | 5.43E-05 | 0.0003 | 0 | 9.36E-05 | 3.54E-05 | 0 | 0.00014 | 1.03E-05 |
| Rnf34 | 3.11E-05 | 0.00012 | 4.65E-05 | 0.00013 | 6.21E-05 | 1.45E-05 | 5.88E-05 | 9.14E-05 | 8.05E-05 | 3.46E-05 | 0.00012 | 0.00011 | 3.28E-05 | 5.72E-05 | 5.53E-05 | 4.69E-05 | 2.98E-05 | 4.52E-05 |
| Cbr3 | 4.93E-05 | 6.16E-05 | 2.98E-05 | 4.03E-06 | 0 | 4.50E-06 | 0 | 0 | 8.05E-05 | 5.59E-05 | 0 | 0 | 0 | 2.48E-05 | 0 | 0 | 0 | 0.0001 |
| Pprc1 | 0.00011 | 6.51E-05 | 0.00026 | 0.00025 | 0 | 0 | 5.32E-05 | 2.58E-05 | 8.05E-05 | 6.92E-05 | 0.0001 | 0 | 4.68E-05 | 6.71E-05 | 0.00017 | 1.37E-05 | 0.00012 | 3.61E-05 |
| Hint1 | 0.00038 | 0.00011 | 0.00017 | 2.13E-05 | 4.61E-05 | 0.00018 | 4.78E-05 | 0 | 8.04E-05 | 1.42E-05 | 2.45E-05 | 7.45E-05 | 1.67E-05 | 5.67E-05 | 0 | 0.0003 | 0 | 0 |
| Top3b | 0.0001 | 0.00012 | 5.03E-05 | 2.84E-05 | 7.06E-05 | 2.87E-05 | 7.78E-05 | 3.04E-05 | 8.04E-05 | 4.84E-05 | 1.89E-05 | 2.66E-05 | 4.52E-06 | 0 | 0.00025 | 3.71E-05 | 6.74E-07 | 1.97E-05 |
| Tbc1d10a | 1.77E-05 | 0.00017 | 0.00022 | 0.00014 | 2.88E-05 | 1.21E-05 | 8.14E-05 | 5.63E-05 | 8.04E-05 | 9.35E-05 | 0.00024 | 4.45E-05 | 0 | 9.39E-05 | 0.00029 | 9.65E-05 | 0.00013 | 0.00018 |
| Rabggta | 2.38E-05 | 8.04E-05 | 8.91E-05 | 9.50E-05 | 6.14E-06 | 0 | 1.92E-05 | 7.77E-06 | 8.04E-05 | 1.72E-05 | 3.00E-05 | 0.00011 | 1.11E-05 | 6.81E-05 | 0.0002 | 2.66E-05 | 5.09E-06 | 3.44E-05 |
| Msto1 | 8.13E-05 | 3.53E-05 | 8.35E-05 | 8.75E-05 | 0 | 0 | 0 | 1.88E-05 | 8.04E-05 | 4.58E-05 | 1.91E-05 | 0 | 0 | 0 | 6.90E-05 | 2.35E-05 | 0 | 8.03E-05 |
| Ankrd28 | 6.29E-05 | 0.00018 | 2.93E-05 | 0 | 1.12E-05 | 0.0002 | 3.89E-05 | 0 | 8.03E-05 | 1.36E-05 | 6.02E-05 | 7.83E-05 | 0 | 0.00028 | 3.56E-05 | 9.92E-05 | 0.00027 | 0.00012 |

| | | | | | | | | | | | | | | | | | | |
|---|---|---|---|---|---|---|---|---|---|---|---|---|---|---|---|---|---|---|
| Fam168a | 0.00013 | 0.00015 | 4.14E-05 | 0.00013 | 0.00014 | 0.00015 | 0.00018 | 8.46E-05 | 8.03E-05 | 0.00016 | 0.00014 | 0.00023 | 0.0001 | 8.11E-05 | 0 | 0.00011 | 8.27E-05 | 6.45E-05 |
| Commd7 | 0.00014 | 0.00014 | 0.00014 | 0.00023 | 0.00014 | 6.73E-05 | 7.28E-05 | 0.00014 | 8.03E-05 | 9.10E-05 | 5.14E-05 | 9.11E-05 | 5.62E-05 | 9.24E-05 | 5.13E-05 | 4.52E-05 | 0 | 0.00015 |
| Derl2 | 0.00017 | 0.00012 | 0.00015 | 0.0002 | 0.0001 | 5.52E-05 | 5.52E-05 | 0.00011 | 8.03E-05 | 0.00012 | 5.60E-05 | 0.00014 | 3.12E-05 | 5.48E-05 | 2.59E-05 | 0.00016 | 0 | 8.67E-05 |
| Gak | 8.07E-05 | 0.00015 | 0.00012 | 0.00012 | 6.61E-05 | 1.75E-05 | 4.25E-05 | 4.49E-05 | 8.03E-05 | 6.83E-05 | 8.08E-05 | 0.00012 | 1.78E-05 | 8.26E-05 | 0.00014 | 4.88E-05 | 3.66E-05 | 8.51E-05 |
| Tmem200a | 0 | 0 | 8.15E-05 | 0 | 0 | 0 | 0 | 5.30E-05 | 8.03E-05 | 0 | 8.04E-05 | 3.26E-05 | 0 | 0 | 0 | 0 | 0 | 6.53E-05 |
| Srek1 | 0.00017 | 0.00022 | 9.36E-05 | 0.00011 | 7.94E-05 | 9.42E-05 | 9.64E-05 | 2.76E-05 | 8.02E-05 | 1.68E-05 | 7.32E-07 | 0 | 4.98E-05 | 0.00011 | 0.0007 | 0.00018 | 0.00029 | 4.35E-05 |
| Glrx2 | 0.00024 | 0.00011 | 8.52E-05 | 0 | 9.66E-05 | 8.65E-05 | 9.84E-05 | 0 | 8.02E-05 | 4.22E-05 | 7.11E-05 | 0 | 2.08E-05 | 5.85E-05 | 6.33E-05 | 0 | 0 | 8.01E-05 |
| Nkiras2 | 6.41E-05 | 0.00017 | 6.21E-05 | 0.00024 | 0.00011 | 7.87E-05 | 9.99E-06 | 0.00013 | 8.02E-05 | 2.36E-05 | 9.51E-05 | 0.00011 | 0 | 6.71E-05 | 1.18E-06 | 6.86E-05 | 0 | 5.63E-05 |
| Ube3c | 0.00022 | 0.00015 | 0.00024 | 0 | 6.53E-05 | 7.57E-05 | 0.00012 | 5.71E-05 | 8.02E-05 | 8.25E-05 | 7.74E-05 | 3.12E-05 | 1.99E-05 | 8.70E-06 | 0.0001 | 0.00017 | 0.00011 | 6.76E-05 |
| Lrrc45 | 6.08E-05 | 7.93E-05 | 2.88E-05 | 4.32E-06 | 4.76E-05 | 1.52E-05 | 7.68E-05 | 0 | 8.02E-05 | 6.46E-05 | 8.28E-08 | 3.18E-06 | 0 | 3.18E-05 | 0.00019 | 6.00E-05 | 5.23E-05 | 2.70E-05 |
| Soga1 | 0 | 0.00018 | 0.00012 | 2.50E-05 | 7.20E-05 | 5.82E-06 | 4.22E-06 | 0.0002 | 8.02E-05 | 0.00012 | 0.00016 | 0.00029 | 0 | 0 | 0.00049 | 0.00031 | 0.00025 | 7.28E-05 |
| Nfatc3 | 0.00025 | 0.00013 | 0.00011 | 0 | 7.19E-05 | 0.00011 | 0.00014 | 0 | 8.01E-05 | 2.88E-05 | 5.05E-05 | 2.41E-05 | 8.82E-06 | 1.14E-05 | 3.49E-05 | 0.00013 | 4.96E-05 | 2.76E-05 |
| Abhd11 | 0.0001 | 8.16E-05 | 0.00014 | 0.00014 | 9.85E-06 | 1.63E-05 | 4.08E-05 | 0 | 8.01E-05 | 9.73E-05 | 3.19E-05 | 1.47E-05 | 9.47E-05 | 0 | 8.66E-05 | 0 | 0 | 1.75E-05 |
| Letm1 | 0 | 4.54E-05 | 6.53E-05 | 0.00017 | 2.81E-05 | 1.51E-05 | 0 | 4.53E-05 | 8.01E-05 | 2.64E-05 | 2.66E-05 | 0 | 3.40E-05 | 0.0001 | 0.00014 | 1.77E-05 | 0 | 8.88E-05 |
| Ccdc9 | 9.73E-06 | 0.00025 | 0.00014 | 0.00025 | 1.77E-05 | 4.09E-06 | 7.31E-05 | 5.11E-05 | 8.01E-05 | 5.54E-05 | 2.26E-05 | 0.00011 | 0 | 9.00E-05 | 0.00034 | 5.11E-05 | 0.0004 | 9.01E-05 |
| Cc2d2a | 1.43E-05 | 7.03E-05 | 0 | 7.12E-05 | 0 | 0 | 1.45E-05 | 7.74E-06 | 8.01E-05 | 9.59E-05 | 0 | 0 | 0 | 1.20E-05 | 9.37E-05 | 0.00011 | 0 | 0.00011 |
| Unc5c | 0 | 0 | 1.38E-06 | 0 | 2.45E-05 | 0 | 8.60E-05 | 0.00014 | 8.01E-05 | 0 | 6.52E-06 | 0 | 0 | 0 | 0 | 3.68E-05 | 9.02E-06 | 3.90E-05 |
| Slfn5 | 5.94E-05 | 0.00018 | 0 | 5.49E-06 | 2.35E-05 | 0 | 0.00027 | 6.68E-05 | 8.01E-05 | 0.00108 | 0 | 2.41E-05 | 0.00082 | 0 | 0 | 0 | 0 | 0.00053 |
| Aco2 | 0.00016 | 6.53E-05 | 7.71E-05 | 5.11E-05 | 6.68E-05 | 8.81E-05 | 2.07E-05 | 3.73E-05 | 8.00E-05 | 5.81E-06 | 6.01E-05 | 2.39E-05 | 1.74E-05 | 2.83E-05 | 3.74E-05 | 9.07E-05 | 0 | 1.56E-05 |
| Snx13 | 0.0001 | 8.97E-05 | 1.79E-05 | 1.40E-05 | 4.08E-05 | 9.68E-05 | 8.55E-05 | 2.24E-05 | 7.99E-05 | 3.42E-05 | 2.95E-05 | 4.99E-05 | 5.31E-07 | 6.42E-05 | 0.00016 | 2.30E-05 | 8.03E-05 | 3.98E-05 |
| Ro60 | 0.00013 | 0.0001 | 1.60E-05 | 0 | 5.81E-05 | 7.59E-05 | 4.89E-05 | 0 | 7.99E-05 | 2.46E-05 | 1.77E-05 | 8.10E-05 | 4.02E-05 | 2.31E-05 | 8.83E-06 | 0 | 3.99E-05 | 4.76E-05 |
| Ddx49 | 0.00014 | 6.38E-05 | 0.0002 | 0.00016 | 3.83E-05 | 4.86E-05 | 1.61E-05 | 1.97E-05 | 7.99E-05 | 1.37E-05 | 7.29E-05 | 1.96E-05 | 2.23E-05 | 1.85E-05 | 7.00E-05 | 4.87E-05 | 0 | 3.09E-05 |
| Rnf25 | 8.61E-05 | 9.34E-05 | 0.0001 | 1.64E-05 | 3.98E-05 | 1.77E-05 | 5.92E-05 | 0 | 7.99E-05 | 5.49E-05 | 1.64E-06 | 4.28E-05 | 3.02E-06 | 2.32E-06 | 0.00012 | 0 | 0 | 9.90E-06 |
| Med24 | 0.00016 | 6.86E-05 | 4.60E-05 | 4.16E-05 | 4.65E-05 | 3.54E-05 | 0 | 0 | 7.98E-05 | 2.70E-05 | 1.44E-06 | 0 | 5.71E-05 | 0 | 8.60E-05 | 9.18E-05 | 0 | 1.66E-05 |
| Snrpc | 0.00022 | 0.00013 | 0.00024 | 0.00053 | 2.00E-05 | 2.56E-05 | 4.88E-05 | 2.18E-05 | 7.98E-05 | 2.01E-05 | 4.96E-05 | 1.09E-05 | 0 | 7.08E-05 | 0 | 0.00032 | 0.00014 | 6.32E-05 |
| Ddx46 | 0.00023 | 0.00017 | 0.0002 | 0.0001 | 3.52E-05 | 1.76E-05 | 3.09E-05 | 0 | 7.98E-05 | 3.10E-05 | 6.09E-05 | 6.80E-05 | 0 | 4.26E-05 | 0.00024 | 0.00019 | 0 | 0 |
| Repin1 | 1.30E-05 | 0.00016 | 3.43E-05 | 2.77E-05 | 3.47E-05 | 1.69E-05 | 0 | 0 | 7.98E-05 | 6.17E-05 | 1.80E-05 | 9.37E-05 | 0.00016 | 1.20E-05 | 2.17E-05 | 1.81E-05 | 0 | 0 |
| Paqr8 | 3.27E-06 | 6.65E-06 | 0 | 2.25E-06 | 6.86E-07 | 1.40E-05 | 0 | 4.45E-05 | 7.98E-05 | 5.28E-06 | 2.79E-05 | 2.90E-05 | 0 | 1.70E-05 | 0 | 0 | 2.19E-05 | 3.92E-05 |
| Lcmt1 | 2.21E-05 | 7.29E-05 | 0 | 7.49E-05 | 5.85E-05 | 2.38E-06 | 1.25E-05 | 3.58E-05 | 7.98E-05 | 6.44E-05 | 3.36E-05 | 4.02E-05 | 3.87E-05 | 3.05E-05 | 8.36E-06 | 4.37E-05 | 0 | 3.96E-05 |
| Efna4 | 8.30E-06 | 7.11E-05 | 3.26E-06 | 7.59E-05 | 3.93E-05 | 0 | 2.10E-06 | 1.48E-05 | 7.98E-05 | 0 | 4.03E-05 | 0 | 3.84E-05 | 0.00011 | 0 | 0 | 0 | 3.49E-05 |
| Tpp1 | 9.28E-05 | 0.00038 | 0 | 0.00013 | 0.00038 | 0.00045 | 8.34E-05 | 0.00014 | 7.97E-05 | 0.00017 | 0.00013 | 0.00023 | 0.00024 | 0.00021 | 0 | 0 | 0 | 4.40E-05 |
| Zdhhc4 | 7.32E-05 | 9.96E-05 | 5.87E-05 | 0.00016 | 0.0001 | 0.00011 | 3.95E-05 | 2.98E-05 | 7.97E-05 | 7.63E-05 | 6.59E-05 | 2.05E-05 | 3.81E-05 | 9.64E-05 | 2.09E-07 | 4.18E-05 | 0 | 0.00013 |
| Tbcel | 7.37E-05 | 0.00011 | 1.56E-05 | 3.58E-05 | 0.00011 | 9.59E-05 | 5.24E-05 | 4.21E-05 | 7.97E-05 | 8.48E-05 | 2.38E-05 | 5.47E-05 | 9.70E-05 | 0.00012 | 4.72E-05 | 0 | 5.28E-05 | 1.98E-05 |

| | | | | | | | | | | | | | | | | | | |
|---|---|---|---|---|---|---|---|---|---|---|---|---|---|---|---|---|---|---|
| Bicd2 | 5.97E-05 | 0.00019 | 0.00024 | 0 | 7.08E-05 | 4.48E-05 | 5.33E-05 | 0.00028 | 7.97E-05 | 0.00014 | 0.00031 | 0.0002 | 0 | 8.22E-05 | 0.00013 | 0.00011 | 0.00015 | 0.00013 |
| Il19 | 0 | 0 | 0 | 0 | 0 | 3.85E-05 | 0 | 0 | 7.97E-05 | 2.33E-07 | 0 | 0 | 0 | 0 | 0 | 8.12E-06 | 0 | 0 |
| Mthfd1 | 6.28E-05 | 0 | 0.00011 | 1.24E-06 | 0 | 2.56E-05 | 0 | 0 | 7.97E-05 | 4.18E-05 | 4.34E-05 | 0 | 3.55E-05 | 3.93E-05 | 0 | 0.00016 | 0 | 3.07E-05 |
| Hspbp1 | 6.99E-05 | 8.96E-05 | 0.00011 | 0.0001 | 3.05E-05 | 0 | 0 | 0 | 7.96E-05 | 2.61E-05 | 5.27E-05 | 0 | 0 | 3.23E-05 | 4.17E-05 | 7.44E-05 | 2.84E-05 | 3.35E-05 |
| Il33 | 2.23E-06 | 0 | 0.00014 | 0 | 0 | 0 | 0.00031 | 0 | 7.96E-05 | 0.0007 | 0 | 0.00076 | 0.00085 | 0 | 0 | 0 | 0 | 0.00101 |
| Gtf2f1 | 0.00015 | 7.02E-05 | 0.00014 | 9.13E-05 | 0 | 0 | 1.96E-06 | 0 | 7.96E-05 | 5.00E-05 | 2.72E-05 | 8.01E-05 | 0 | 0 | 0.00012 | 0.00012 | 0.00033 | 2.15E-05 |
| St3gal3 | 2.20E-05 | 3.57E-05 | 3.02E-05 | 0 | 0.00013 | 0.00011 | 0 | 0.0001 | 7.95E-05 | 4.09E-05 | 0.00018 | 6.04E-05 | 6.32E-05 | 9.43E-06 | 4.22E-06 | 3.64E-06 | 0 | 1.09E-05 |
| Dnajb5 | 2.87E-05 | 4.23E-05 | 3.32E-05 | 0 | 8.14E-05 | 3.34E-05 | 2.51E-05 | 6.76E-05 | 7.95E-05 | 3.47E-05 | 7.78E-05 | 0 | 5.38E-05 | 6.38E-05 | 3.45E-06 | 3.67E-05 | 0 | 1.27E-06 |
| Pex11b | 4.39E-05 | 8.89E-05 | 7.29E-05 | 4.39E-05 | 5.06E-05 | 2.63E-05 | 0 | 4.37E-05 | 7.95E-05 | 3.49E-05 | 3.80E-05 | 3.76E-05 | 2.44E-05 | 4.13E-05 | 2.08E-05 | 5.90E-05 | 3.95E-05 | 1.66E-05 |
| Rasgrp2 | 0 | 1.05E-05 | 0 | 0 | 0 | 0.0001 | 0 | 2.63E-05 | 7.94E-05 | 4.26E-06 | 0 | 0 | 0 | 4.34E-05 | 0 | 0 | 1.26E-05 | 6.63E-05 |
| Phax | 0.00019 | 0.00019 | 9.54E-05 | 1.99E-05 | 7.72E-05 | 9.47E-05 | 0 | 2.58E-05 | 7.94E-05 | 3.86E-05 | 2.24E-05 | 0.00013 | 0 | 2.20E-05 | 0.00015 | 0.00013 | 0 | 6.43E-05 |
| Uspl1 | 7.22E-05 | 5.29E-05 | 5.16E-05 | 7.53E-05 | 1.12E-05 | 3.97E-05 | 3.56E-06 | 1.57E-06 | 7.94E-05 | 1.94E-05 | 4.02E-05 | 2.15E-05 | 0 | 7.38E-05 | 0.0001 | 4.46E-05 | 0 | 5.01E-06 |
| Cdk2 | 3.44E-05 | 5.23E-05 | 9.70E-05 | 4.25E-06 | 3.00E-05 | 1.27E-05 | 5.88E-05 | 6.81E-05 | 7.94E-05 | 3.07E-05 | 8.04E-05 | 3.92E-05 | 8.34E-05 | 8.20E-05 | 0 | 0.00054 | 8.82E-05 | 1.89E-05 |
| Arl15 | 1.08E-05 | 0.00013 | 2.21E-05 | 6.22E-05 | 0.00012 | 2.76E-07 | 0.00012 | 6.08E-05 | 7.94E-05 | 0.00024 | 4.88E-05 | 0 | 0.00029 | 0.00012 | 0.00014 | 1.41E-05 | 0.00017 | 2.90E-05 |
| Mageh1 | 0.00013 | 0 | 0 | 7.69E-05 | 4.43E-05 | 8.12E-05 | 0.00029 | 0.00023 | 7.93E-05 | 0.00012 | 0 | 0 | 0.0001 | 0 | 0 | 0 | 0 | 1.34E-05 |
| Klhdc10 | 0.00014 | 0.00014 | 0.00017 | 0.00019 | 0.00011 | 7.01E-05 | 7.67E-06 | 0.00011 | 7.93E-05 | 3.52E-05 | 8.72E-05 | 0 | 2.70E-05 | 0.00011 | 4.92E-05 | 0.00014 | 1.38E-05 | 4.98E-05 |
| Strap | 0.00027 | 7.41E-05 | 0.00025 | 4.96E-05 | 4.95E-05 | 0.00012 | 5.42E-05 | 3.95E-06 | 7.92E-05 | 7.19E-06 | 7.11E-05 | 3.67E-05 | 0 | 5.53E-05 | 0 | 0.00015 | 3.97E-06 | 2.00E-05 |
| Clk2 | 6.63E-05 | 0.00017 | 0.0001 | 6.95E-05 | 0.00011 | 9.68E-05 | 6.18E-05 | 6.89E-05 | 7.92E-05 | 5.42E-05 | 4.80E-05 | 8.61E-05 | 7.45E-05 | 0.00012 | 0.00026 | 6.54E-05 | 0.00012 | 5.54E-05 |
| Prpf38b | 0.00014 | 0.00018 | 0.00011 | 3.00E-05 | 3.82E-05 | 2.93E-05 | 4.12E-05 | 4.25E-05 | 7.92E-05 | 3.97E-05 | 6.99E-05 | 7.18E-05 | 0 | 5.64E-05 | 0.00062 | 0.00014 | 0.00012 | 1.35E-05 |
| Mgat3 | 0 | 6.96E-05 | 0 | 4.86E-05 | 0.00017 | 0.00016 | 0 | 0.00021 | 7.91E-05 | 0 | 0.00015 | 0.0001 | 0 | 0.00016 | 9.41E-05 | 0 | 0 | 0.00012 |
| Ncbp3 | 0.00013 | 9.97E-05 | 8.69E-05 | 4.25E-05 | 7.08E-05 | 3.16E-05 | 6.36E-05 | 0 | 7.91E-05 | 3.48E-05 | 2.91E-05 | 3.51E-05 | 0 | 9.18E-05 | 8.76E-05 | 8.72E-05 | 5.13E-06 | 3.43E-05 |
| Kri1 | 2.06E-05 | 0.0001 | 0.00016 | 5.49E-05 | 1.97E-05 | 0 | 0 | 4.93E-05 | 7.91E-05 | 6.30E-05 | 2.55E-05 | 3.82E-05 | 6.81E-05 | 5.09E-05 | 0.0003 | 6.63E-05 | 0 | 2.95E-05 |
| Evi5 | 8.09E-05 | 9.25E-05 | 5.59E-05 | 1.20E-05 | 5.98E-05 | 7.30E-05 | 6.87E-05 | 3.63E-05 | 7.90E-05 | 4.64E-05 | 2.80E-05 | 4.19E-05 | 2.75E-05 | 6.34E-05 | 3.29E-05 | 4.53E-05 | 4.09E-05 | 4.46E-05 |
| Ccnh | 8.24E-05 | 5.02E-05 | 3.30E-05 | 1.02E-05 | 6.12E-05 | 6.94E-05 | 0 | 0 | 7.90E-05 | 3.43E-05 | 0 | 1.12E-05 | 6.91E-05 | 5.71E-05 | 0 | 4.49E-05 | 1.16E-05 | 7.37E-05 |
| Chd8 | 0.00014 | 0.00021 | 3.78E-05 | 3.11E-05 | 6.76E-05 | 5.51E-05 | 6.43E-05 | 2.49E-05 | 7.90E-05 | 1.39E-05 | 4.99E-05 | 8.34E-05 | 0 | 6.77E-05 | 0.00025 | 9.30E-05 | 3.49E-05 | 4.51E-05 |
| Lats1 | 0.00014 | 0.00011 | 0.00018 | 0 | 0.00011 | 0.00013 | 0.00014 | 0.0001 | 7.89E-05 | 5.04E-05 | 6.67E-05 | 9.71E-05 | 4.83E-05 | 6.14E-05 | 2.57E-06 | 4.55E-05 | 6.18E-05 | 5.37E-05 |
| Sumo2 | 0.00039 | 8.77E-05 | 0.00012 | 0 | 3.26E-05 | 0.00012 | 0 | 2.46E-06 | 7.89E-05 | 0 | 3.78E-05 | 8.10E-05 | 1.28E-06 | 0 | 0 | 0.00023 | 0 | 0 |
| Itgb1bp1 | 0.00015 | 0.00012 | 6.07E-05 | 4.07E-05 | 0.00011 | 6.68E-05 | 8.01E-05 | 0 | 7.89E-05 | 0.00013 | 1.81E-05 | 1.08E-06 | 2.98E-06 | 4.78E-05 | 0 | 1.58E-05 | 0 | 2.67E-05 |
| Spsb3 | 8.12E-05 | 0.00015 | 0.00011 | 3.62E-05 | 9.43E-05 | 5.51E-05 | 8.89E-05 | 4.96E-05 | 7.89E-05 | 7.99E-05 | 0.00012 | 6.69E-06 | 1.70E-05 | 0.0001 | 0.00011 | 0.00011 | 0 | 8.66E-05 |
| Eif1b | 0.00025 | 8.49E-05 | 9.43E-05 | 0 | 0.00011 | 4.45E-05 | 6.67E-05 | 4.44E-05 | 7.89E-05 | 2.53E-05 | 0 | 0.00038 | 5.51E-05 | 4.30E-05 | 0 | 0.0001 | 1.93E-05 | 4.63E-05 |
| Cdk10 | 1.20E-05 | 5.44E-05 | 1.26E-05 | 1.01E-05 | 4.19E-05 | 3.35E-05 | 1.52E-05 | 4.29E-05 | 7.88E-05 | 4.14E-05 | 2.85E-05 | 3.00E-05 | 3.04E-05 | 1.61E-05 | 7.77E-05 | 2.86E-05 | 0 | 4.91E-05 |
| Mak16 | 0.0001 | 7.48E-05 | 0.00023 | 0 | 3.06E-06 | 2.84E-05 | 0 | 4.61E-06 | 7.88E-05 | 2.63E-06 | 4.99E-05 | 6.06E-06 | 0 | 0 | 8.32E-05 | 3.22E-05 | 0 | 0 |
| Agfg2 | 0 | 6.58E-05 | 0 | 3.50E-05 | 6.50E-05 | 0 | 0 | 3.91E-05 | 7.88E-05 | 5.81E-05 | 1.87E-05 | 0 | 0 | 0 | 0 | 0.00011 | 3.04E-05 | 4.10E-05 |

| | | | | | | | | | | | | | | | | | | |
|---|---|---|---|---|---|---|---|---|---|---|---|---|---|---|---|---|---|---|
| Tmem39a | 0.00031 | 2.31E-05 | 0.00017 | 7.59E-06 | 0.00012 | 0.00029 | 0.00023 | 0.00017 | 7.87E-05 | 6.23E-05 | 1.52E-06 | 0.00016 | 0.00011 | 7.25E-05 | 0.00011 | 2.90E-05 | 0.00022 | 1.98E-05 |
| Gpr137 | 1.15E-05 | 9.55E-05 | 4.52E-05 | 0.00014 | 0.00012 | 3.64E-05 | 3.60E-05 | 0.00011 | 7.87E-05 | 6.26E-05 | 4.65E-05 | 4.36E-06 | 4.29E-05 | 1.59E-05 | 5.22E-05 | 0 | 1.15E-05 | 8.96E-05 |
| Guca1a | 5.26E-05 | 0 | 5.39E-05 | 2.39E-05 | 2.68E-05 | 2.66E-05 | 0.00016 | 0 | 7.87E-05 | 0 | 2.05E-05 | 4.44E-05 | 0 | 1.21E-05 | 0 | 0 | 0 | 0 |
| Gpn1 | 6.70E-05 | 5.21E-05 | 0.0001 | 5.41E-05 | 3.64E-05 | 1.20E-06 | 0 | 0 | 7.87E-05 | 2.00E-05 | 1.51E-05 | 3.75E-05 | 1.11E-05 | 0 | 5.68E-05 | 0.0001 | 0 | 1.62E-05 |
| Mex3d | 0 | 0.00026 | 0.00014 | 0.0002 | 0.00011 | 0 | 0 | 0 | 7.87E-05 | 3.22E-05 | 0.00027 | 0.00045 | 5.17E-05 | 0.00016 | 0.00024 | 7.95E-05 | 0.00018 | 0.00012 |
| Magoh | 0.00038 | 9.24E-05 | 0.00021 | 0.00015 | 1.22E-05 | 0.00014 | 1.91E-05 | 0 | 7.86E-05 | 0 | 6.00E-05 | 9.28E-05 | 0 | 2.59E-05 | 0 | 0.00026 | 0.00019 | 9.26E-06 |
| Elp3 | 0.00023 | 0.0001 | 0.00017 | 0.00033 | 3.69E-05 | 7.92E-05 | 0.00022 | 6.48E-05 | 7.86E-05 | 3.16E-05 | 0 | 7.41E-05 | 8.84E-05 | 1.74E-05 | 0.00017 | 0.00011 | 0 | 2.93E-05 |
| Uqcc3 | 5.79E-05 | 0.00014 | 0 | 0.00025 | 9.70E-05 | 7.90E-05 | 0.00012 | 0.00012 | 7.86E-05 | 8.54E-05 | 4.26E-06 | 6.46E-05 | 3.85E-05 | 0.00011 | 3.83E-05 | 0 | 0 | 7.71E-05 |
| Phtf1 | 0.00015 | 7.23E-05 | 8.66E-05 | 0 | 7.82E-05 | 4.39E-05 | 6.62E-05 | 5.05E-05 | 7.86E-05 | 6.38E-05 | 6.27E-05 | 9.17E-05 | 2.84E-05 | 6.99E-05 | 6.79E-05 | 0.00014 | 9.45E-05 | 7.72E-05 |
| Gtpbp1 | 0.00011 | 6.77E-05 | 0.00013 | 4.57E-05 | 4.48E-05 | 4.30E-05 | 7.22E-05 | 1.47E-05 | 7.86E-05 | 3.90E-05 | 6.14E-05 | 0 | 1.71E-05 | 6.25E-05 | 0.00011 | 0.00014 | 8.18E-05 | 3.04E-05 |
| Tgfb3 | 1.93E-05 | 7.63E-05 | 7.78E-05 | 0 | 0.00041 | 2.67E-05 | 0.00035 | 0.00068 | 7.86E-05 | 0 | 0.00081 | 0 | 0 | 0.00064 | 0 | 2.27E-05 | 0 | 0.0007 |
| Heatr3 | 8.95E-05 | 3.80E-05 | 0.00019 | 4.00E-05 | 2.03E-05 | 2.82E-06 | 0 | 0 | 7.86E-05 | 4.34E-05 | 1.90E-05 | 0 | 1.08E-05 | 8.64E-06 | 6.78E-05 | 2.42E-05 | 1.16E-05 | 5.11E-05 |
| Washc3 | 0.0002 | 6.83E-05 | 0 | 0.00012 | 6.82E-05 | 0.00012 | 0.00017 | 5.92E-05 | 7.85E-05 | 7.41E-05 | 3.71E-06 | 6.36E-05 | 3.18E-05 | 4.72E-05 | 8.73E-05 | 5.59E-05 | 0 | 8.59E-05 |
| Cnpy4 | 0.00013 | 3.86E-05 | 3.14E-05 | 0.00024 | 0.0001 | 0.00011 | 8.52E-05 | 8.03E-05 | 7.85E-05 | 9.28E-05 | 5.53E-05 | 6.14E-05 | 0.0001 | 6.93E-05 | 0 | 0.00023 | 3.52E-05 | 9.94E-05 |
| Pafah1b1 | 0.00018 | 0.00024 | 0.00026 | 0.00022 | 0.00012 | 0.00011 | 0.00013 | 5.76E-05 | 7.85E-05 | 0.00012 | 0.00022 | 0.00018 | 0 | 6.70E-05 | 0.00011 | 0.00016 | 0.00015 | 0.00012 |
| Anxa11 | 6.72E-05 | 0.00017 | 9.99E-05 | 0.00019 | 9.28E-05 | 0.00011 | 2.68E-05 | 0 | 7.85E-05 | 0.00026 | 0.00013 | 0.00014 | 0 | 0.00028 | 0 | 5.82E-05 | 0 | 0.00042 |
| Stard7 | 0.00016 | 7.84E-05 | 0.00016 | 0.00012 | 5.46E-05 | 7.81E-05 | 8.78E-05 | 0 | 7.85E-05 | 7.58E-05 | 4.33E-05 | 1.63E-06 | 5.39E-05 | 5.78E-05 | 0.00014 | 0.00021 | 0 | 2.54E-05 |
| Thada | 9.20E-05 | 7.84E-05 | 7.61E-05 | 0 | 2.40E-05 | 5.16E-05 | 2.33E-05 | 1.41E-05 | 7.85E-05 | 2.35E-05 | 0 | 0 | 8.90E-06 | 0 | 9.02E-05 | 9.39E-05 | 3.93E-05 | 4.66E-05 |
| Pafah1b3 | 0.00014 | 3.31E-05 | 0 | 0.00018 | 3.89E-05 | 2.18E-05 | 0.00021 | 1.75E-05 | 7.85E-05 | 0 | 0.00011 | 0.00055 | 3.30E-05 | 6.20E-05 | 0 | 7.91E-05 | 1.13E-05 | 0 |
| Eapp | 0.00019 | 0.0002 | 5.78E-05 | 0.0001 | 0.0001 | 0.00012 | 6.76E-05 | 3.53E-05 | 7.84E-05 | 6.84E-05 | 0.00011 | 0.00013 | 5.25E-05 | 4.71E-05 | 8.63E-05 | 2.99E-05 | 0 | 0.00011 |
| Pla2g4a | 3.16E-05 | 4.45E-05 | 5.11E-05 | 0 | 0.00019 | 0.00011 | 0 | 0.00061 | 7.84E-05 | 0.00012 | 7.73E-05 | 0.0002 | 5.72E-05 | 0 | 0 | 0.00012 | 2.95E-05 | 0.00019 |
| Sugp2 | 6.22E-05 | 6.37E-05 | 7.49E-05 | 4.85E-05 | 5.37E-05 | 3.02E-05 | 3.91E-05 | 2.41E-05 | 7.84E-05 | 5.82E-05 | 4.89E-05 | 2.67E-05 | 5.54E-05 | 4.23E-05 | 0.00023 | 8.44E-05 | 0 | 5.99E-05 |
| Ubp1 | 0.00014 | 0.00025 | 0.00025 | 1.55E-05 | 0.00012 | 8.71E-05 | 4.15E-05 | 7.06E-05 | 7.83E-05 | 7.11E-05 | 7.66E-05 | 0.00012 | 4.30E-05 | 0.0002 | 0.00059 | 0.00014 | 0.00023 | 7.62E-05 |
| Rgs10 | 6.20E-05 | 8.14E-05 | 6.14E-06 | 0 | 0.00021 | 4.97E-05 | 0.00012 | 0 | 7.83E-05 | 2.42E-05 | 2.96E-05 | 7.38E-06 | 0 | 0 | 0 | 7.76E-05 | 3.32E-05 | 4.68E-05 |
| Ell | 2.64E-05 | 0.00016 | 2.50E-05 | 2.79E-05 | 7.48E-05 | 4.04E-05 | 3.65E-05 | 0 | 7.83E-05 | 2.88E-05 | 3.82E-05 | 0.00021 | 0.00015 | 9.95E-05 | 0.00025 | 0 | 0.00015 | 1.53E-05 |
| Lap3 | 0.00031 | 0.00011 | 0.00024 | 0.00014 | 0 | 9.90E-05 | 0.00013 | 3.32E-05 | 7.82E-05 | 2.73E-05 | 0.00017 | 1.10E-05 | 5.47E-05 | 0 | 0 | 3.87E-05 | 8.67E-07 | 0 |
| Smu1 | 0.00019 | 0.0002 | 0.00021 | 0.00022 | 1.63E-05 | 9.72E-06 | 0 | 0 | 7.82E-05 | 4.97E-05 | 9.00E-05 | 8.70E-05 | 0 | 1.54E-05 | 9.94E-05 | 0.00019 | 8.35E-05 | 3.39E-05 |
| Tnfaip1 | 0.0002 | 8.48E-05 | 0.00018 | 0 | 3.86E-05 | 0 | 0.00015 | 0 | 7.82E-05 | 0.0001 | 0.00022 | 0.00013 | 0 | 2.13E-07 | 0 | 0.00013 | 0 | 7.01E-05 |
| Per1 | 4.09E-05 | 0.00016 | 4.16E-05 | 0.00014 | 6.88E-05 | 0.00012 | 0 | 8.76E-05 | 7.81E-05 | 3.97E-05 | 3.22E-05 | 0.00018 | 0 | 5.09E-05 | 0.00035 | 0 | 0.00047 | 6.68E-05 |
| Pqbp1 | 0.00021 | 0.0001 | 0.00015 | 0.00013 | 6.86E-05 | 6.68E-05 | 7.26E-05 | 6.02E-05 | 7.81E-05 | 6.06E-05 | 8.59E-06 | 4.46E-05 | 0 | 7.03E-05 | 0.00016 | 9.36E-05 | 0 | 5.31E-05 |
| Snx11 | 7.73E-06 | 6.13E-05 | 0.00015 | 5.09E-05 | 9.38E-05 | 0 | 1.60E-05 | 0.0001 | 7.81E-05 | 4.44E-05 | 7.19E-05 | 0 | 3.63E-05 | 8.81E-05 | 0.00014 | 0.00012 | 0 | 6.16E-05 |
| Hikeshi | 0.00016 | 8.06E-05 | 7.97E-05 | 0.00016 | 1.90E-05 | 0.00011 | 3.28E-05 | 4.11E-05 | 7.80E-05 | 7.63E-05 | 7.30E-05 | 9.08E-05 | 9.22E-05 | 0 | 5.34E-06 | 0.00017 | 0 | 3.80E-05 |
| Prune1 | 3.36E-05 | 7.82E-05 | 5.11E-05 | 1.94E-05 | 4.26E-05 | 3.31E-05 | 1.72E-05 | 2.34E-06 | 7.80E-05 | 1.63E-05 | 4.47E-05 | 6.05E-05 | 1.37E-05 | 2.52E-05 | 0 | 5.84E-05 | 3.19E-05 | 3.33E-05 |

| | | | | | | | | | | | | | | | | | | |
|---|---|---|---|---|---|---|---|---|---|---|---|---|---|---|---|---|---|---|
| Fkbp2 | 0.00034 | 0.00013 | 0.00026 | 0.00051 | 4.51E-05 | 2.00E-05 | 0.00019 | 3.43E-05 | 7.80E-05 | 0.00017 | 0.00012 | 0.00018 | 0.00015 | 7.12E-05 | 2.95E-05 | 0.00046 | 0 | 0.00016 |
| Tnks | 0.00012 | 0.00019 | 0.0001 | 9.00E-05 | 0.00014 | 0.00011 | 0.00016 | 0.00012 | 7.79E-05 | 0.00012 | 0.00016 | 0.00022 | 0.0001 | 9.03E-05 | 6.22E-05 | 0.00013 | 0.00012 | 0.0001 |
| Arid3a | 0 | 0.00011 | 2.93E-05 | 7.41E-05 | 0.00014 | 9.04E-05 | 0 | 0 | 7.79E-05 | 0 | 6.47E-05 | 0.0001 | 4.40E-05 | 0.00012 | 0.00028 | 0 | 0 | 0 |
| Gcc1 | 9.05E-05 | 0.00012 | 5.76E-05 | 0.00014 | 0.0001 | 6.94E-05 | 7.35E-05 | 5.46E-05 | 7.79E-05 | 1.03E-05 | 0 | 0 | 0 | 6.52E-05 | 0.00016 | 3.66E-06 | 0 | 4.09E-05 |
| Vps45 | 7.24E-05 | 3.51E-05 | 8.70E-06 | 6.34E-06 | 1.64E-05 | 2.52E-05 | 1.09E-05 | 2.38E-05 | 7.79E-05 | 5.16E-05 | 3.53E-05 | 7.99E-06 | 4.77E-06 | 0 | 3.23E-05 | 3.91E-05 | 0 | 2.51E-05 |
| Tbrg4 | 0.00011 | 9.32E-05 | 0.00028 | 0.00013 | 0 | 0 | 0 | 0 | 7.79E-05 | 1.88E-05 | 4.16E-05 | 1.64E-05 | 0 | 1.95E-05 | 0.00018 | 0.0001 | 5.14E-05 | 0 |
| Tmem128 | 0.00022 | 4.42E-05 | 9.19E-05 | 4.58E-05 | 2.53E-05 | 0.00015 | 6.23E-05 | 5.83E-05 | 7.78E-05 | 6.71E-05 | 0.00012 | 5.12E-06 | 3.90E-05 | 8.29E-05 | 2.27E-05 | 8.58E-05 | 0 | 1.99E-07 |
| Sugt1 | 0.00029 | 0.00012 | 9.38E-05 | 0.0001 | 0 | 0.00015 | 4.70E-05 | 4.62E-05 | 7.78E-05 | 6.90E-06 | 6.90E-05 | 0.00015 | 0 | 1.98E-05 | 3.90E-05 | 0.00016 | 0.00013 | 9.04E-05 |
| Il6ra | 0 | 5.19E-05 | 0 | 1.38E-05 | 0 | 3.53E-05 | 0 | 0 | 7.78E-05 | 0.00014 | 0 | 6.59E-05 | 3.77E-05 | 0 | 0 | 2.83E-07 | 4.46E-05 | 0 |
| Fam102a | 0 | 0.00019 | 0 | 0.00017 | 6.39E-05 | 1.64E-05 | 0.00036 | 0 | 7.78E-05 | 0.00025 | 0 | 0 | 0.00035 | 0.0001 | 0 | 4.46E-05 | 0 | 6.73E-05 |
| Srp9 | 0.00042 | 0.00013 | 0.00034 | 0.00049 | 0.00018 | 0.00031 | 0.00049 | 0.00029 | 7.77E-05 | 9.15E-05 | 0.00012 | 0.00021 | 0.00013 | 0.00023 | 0 | 0.00018 | 7.82E-05 | 8.94E-05 |
| Aup1 | 0.00026 | 0.0003 | 0.00031 | 0.00025 | 0.0002 | 0.00023 | 0.00019 | 0.0002 | 7.77E-05 | 8.34E-05 | 0.00018 | 0.00011 | 9.42E-05 | 0.0002 | 0.00022 | 0.00017 | 3.29E-05 | 3.88E-05 |
| Ttc28 | 0 | 0.00018 | 0 | 9.92E-06 | 0.00022 | 9.78E-05 | 2.98E-05 | 0 | 7.77E-05 | 0.00015 | 0.0004 | 5.89E-05 | 1.99E-05 | 0.00022 | 0.00021 | 5.68E-05 | 6.78E-05 | 0.0002 |
| Chordc1 | 0.00011 | 8.19E-05 | 9.60E-05 | 3.58E-05 | 5.08E-05 | 3.24E-05 | 0 | 3.71E-05 | 7.77E-05 | 1.56E-05 | 3.98E-05 | 8.47E-06 | 0 | 4.59E-05 | 6.95E-05 | 4.97E-05 | 0 | 1.99E-05 |
| Rnaseh1 | 7.76E-05 | 3.18E-05 | 6.66E-05 | 6.52E-05 | 3.73E-05 | 2.81E-05 | 3.75E-05 | 0 | 7.77E-05 | 3.02E-05 | 4.26E-05 | 0 | 3.56E-05 | 8.92E-06 | 6.80E-05 | 6.15E-05 | 0 | 1.01E-05 |
| Ilf2 | 0.00016 | 0.00011 | 0.00019 | 0.00024 | 3.11E-05 | 2.70E-05 | 0 | 4.23E-05 | 7.77E-05 | 3.28E-05 | 3.31E-05 | 8.28E-06 | 0 | 4.77E-05 | 6.72E-05 | 0.00042 | 7.65E-05 | 0 |
| Topbp1 | 3.55E-05 | 0 | 8.51E-05 | 0 | 0 | 1.52E-05 | 2.52E-05 | 2.38E-05 | 7.77E-05 | 5.14E-05 | 3.35E-05 | 0 | 2.71E-05 | 4.62E-05 | 2.18E-05 | 0.00059 | 0 | 2.45E-05 |
| Anp32b | 5.64E-05 | 3.37E-05 | 0.00019 | 0.00023 | 0 | 0 | 0 | 1.66E-05 | 7.77E-05 | 2.62E-06 | 2.46E-05 | 8.26E-05 | 4.11E-08 | 0 | 8.74E-05 | 0.00029 | 0 | 0 |
| Fosl1 | 0 | 0 | 4.38E-05 | 0 | 0 | 0.00033 | 0 | 0 | 7.76E-05 | 0 | 0 | 0.00093 | 0.00024 | 8.23E-05 | 0 | 4.51E-05 | 0.00015 | 0 |
| Slc35a1 | 0.0002 | 5.56E-05 | 7.13E-05 | 0 | 7.46E-05 | 0.00015 | 0 | 9.10E-05 | 7.76E-05 | 6.30E-05 | 4.86E-05 | 6.06E-05 | 4.07E-05 | 2.25E-05 | 1.16E-05 | 5.72E-05 | 0 | 7.80E-05 |
| Wbp1 | 3.63E-05 | 0.00015 | 0 | 0.00017 | 0.00017 | 6.43E-06 | 2.83E-05 | 7.22E-05 | 7.76E-05 | 0.0001 | 9.57E-05 | 7.83E-06 | 0 | 7.16E-05 | 2.18E-05 | 4.53E-05 | 0 | 0.00015 |
| Slc4a3 | 0 | 4.72E-05 | 0 | 2.76E-05 | 3.45E-05 | 0 | 1.83E-05 | 0 | 7.76E-05 | 8.65E-05 | 1.66E-05 | 2.41E-05 | 5.33E-05 | 0 | 0.0001 | 0 | 6.80E-05 | 8.73E-05 |
| Nipsnap2 | 0.00019 | 9.47E-05 | 5.85E-05 | 0.00014 | 9.05E-05 | 0.00015 | 0.00013 | 7.90E-05 | 7.75E-05 | 8.25E-05 | 0.00012 | 0.00011 | 0 | 7.67E-05 | 1.34E-07 | 7.23E-05 | 0 | 5.68E-05 |
| Csnk1e | 3.72E-05 | 0.00033 | 0.00032 | 0 | 0.0002 | 0.00013 | 0 | 0.00024 | 7.75E-05 | 0.00021 | 0.00034 | 0.00017 | 0 | 0.00023 | 0.00023 | 0.00011 | 3.93E-05 | 0.00018 |
| Slc1a5 | 6.54E-05 | 8.60E-05 | 0.00035 | 0.00018 | 0 | 0 | 9.18E-05 | 6.51E-05 | 7.75E-05 | 0.00043 | 0 | 4.51E-05 | 0.00012 | 0 | 0.0002 | 9.04E-05 | 0.00041 | 0.00033 |
| Ppil1 | 7.82E-05 | 1.64E-05 | 0.00012 | 0.00013 | 2.63E-05 | 0 | 1.99E-05 | 2.79E-05 | 7.75E-05 | 1.79E-05 | 3.09E-05 | 0 | 2.96E-05 | 4.56E-05 | 0 | 0.00043 | 0 | 1.60E-06 |
| Mvb12a | 0.00013 | 8.67E-05 | 5.61E-05 | 0.00029 | 9.07E-05 | 7.12E-05 | 0.0002 | 8.66E-05 | 7.74E-05 | 5.12E-05 | 2.97E-05 | 4.16E-05 | 7.76E-05 | 8.73E-05 | 0 | 0 | 0 | 9.78E-05 |
| Jazf1 | 3.31E-05 | 4.01E-06 | 0 | 0 | 5.52E-05 | 8.93E-06 | 0 | 0 | 7.74E-05 | 1.14E-05 | 7.06E-06 | 0.0001 | 4.47E-05 | 0 | 0 | 3.51E-05 | 0 | 2.02E-05 |
| Cpne2 | 1.09E-05 | 9.31E-05 | 0 | 7.20E-06 | 0 | 0 | 3.36E-05 | 0 | 7.74E-05 | 0.00017 | 0 | 1.71E-05 | 0.00026 | 0 | 0 | 8.26E-05 | 0 | 3.10E-05 |
| Nop2 | 3.03E-05 | 2.55E-05 | 0.00033 | 0.00011 | 0 | 0 | 0 | 0 | 7.74E-05 | 3.85E-05 | 3.49E-05 | 2.01E-05 | 0.00028 | 5.39E-05 | 7.34E-06 | 2.94E-05 | 0.00069 | 0 |
| Smim41 | 0 | 0 | 0 | 0 | 0 | 7.11E-05 | 0 | 0.00017 | 7.73E-05 | 0 | 0 | 8.48E-05 | 2.04E-05 | 0 | 0 | 0 | 0 | 9.73E-05 |
| Tor2a | 5.16E-05 | 1.83E-05 | 0.00014 | 0.00011 | 0.00011 | 6.41E-05 | 0.00015 | 0.00011 | 7.73E-05 | 6.10E-05 | 2.16E-05 | 0 | 9.85E-05 | 7.31E-05 | 0 | 5.49E-05 | 0 | 6.96E-05 |
| Pcx | 0 | 2.56E-05 | 0 | 4.90E-05 | 5.48E-05 | 2.69E-05 | 0 | 0 | 7.73E-05 | 5.45E-06 | 0 | 5.40E-05 | 0.00013 | 0 | 0 | 7.23E-05 | 0 | 0 |

| | | | | | | | | | | | | | | | | | | |
|---|---|---|---|---|---|---|---|---|---|---|---|---|---|---|---|---|---|---|
| Mff | 0.00038 | 0.00014 | 2.89E-05 | 0 | 0.00011 | 0.0002 | 7.33E-05 | 1.90E-05 | 7.72E-05 | 1.13E-05 | 0.00011 | 0.00019 | 3.23E-05 | 3.62E-05 | 7.10E-05 | 9.66E-05 | 0 | 1.46E-05 |
| Etf1 | 0.0003 | 6.88E-05 | 0.00027 | 0 | 0 | 0.00013 | 6.66E-06 | 1.46E-05 | 7.72E-05 | 0 | 0.0001 | 5.38E-05 | 2.78E-05 | 1.99E-05 | 0 | 0.00012 | 0.00026 | 0 |
| Rnf111 | 0.00011 | 0.00016 | 0.00011 | 0 | 6.07E-05 | 7.60E-05 | 6.91E-05 | 2.47E-05 | 7.72E-05 | 6.80E-05 | 4.91E-05 | 9.37E-05 | 2.94E-05 | 5.12E-05 | 0.00016 | 7.37E-05 | 0.0002 | 3.50E-05 |
| Tcf7l2 | 0 | 0.00015 | 0 | 0 | 0 | 0 | 0.00015 | 0 | 7.72E-05 | 0.00014 | 5.20E-06 | 1.05E-05 | 0.00021 | 6.49E-05 | 1.72E-05 | 5.59E-05 | 0.00019 | 0.00019 |
| Pccb | 0.0001 | 3.55E-05 | 0 | 8.43E-05 | 2.15E-05 | 0 | 0.00016 | 3.90E-06 | 7.72E-05 | 5.16E-05 | 3.84E-05 | 0 | 1.57E-05 | 3.44E-05 | 1.43E-05 | 0 | 0 | 8.57E-05 |
| Rubcn | 2.40E-05 | 8.84E-05 | 4.17E-05 | 7.30E-05 | 6.05E-05 | 0 | 5.79E-05 | 0 | 7.72E-05 | 0.00011 | 6.44E-05 | 0.00014 | 0 | 4.86E-05 | 0.00011 | 5.73E-05 | 0 | 0.00011 |
| Pfas | 1.44E-05 | 2.24E-05 | 0.00013 | 2.45E-05 | 2.05E-05 | 0 | 0 | 1.71E-06 | 7.72E-05 | 3.59E-05 | 2.69E-05 | 6.46E-06 | 8.12E-07 | 0 | 1.27E-05 | 0.0002 | 5.16E-05 | 2.58E-05 |
| Rest | 0.00011 | 0.00012 | 0.0001 | 4.47E-05 | 5.64E-05 | 8.63E-05 | 2.30E-05 | 6.28E-05 | 7.71E-05 | 0 | 9.05E-05 | 0.00018 | 0 | 7.58E-05 | 0.00012 | 4.61E-05 | 0.00011 | 5.30E-06 |
| Map3k14 | 1.33E-05 | 7.75E-05 | 3.92E-05 | 0.00012 | 1.21E-05 | 0 | 0.00014 | 0 | 7.71E-05 | 7.72E-05 | 0 | 0 | 0.00016 | 4.33E-05 | 0.00015 | 1.05E-05 | 8.68E-05 | 0.00015 |
| Rptor | 0 | 0.00012 | 0.00011 | 0.00016 | 5.61E-05 | 0 | 0 | 5.40E-05 | 7.71E-05 | 5.54E-05 | 2.91E-05 | 2.92E-05 | 7.74E-06 | 1.64E-05 | 0.00036 | 7.33E-05 | 0.0002 | 5.95E-05 |
| Ablim1 | 0 | 4.40E-05 | 0 | 0 | 3.67E-05 | 0 | 0 | 0 | 7.71E-05 | 0.00043 | 0 | 7.11E-06 | 0 | 0 | 0 | 0 | 0 | 0.00024 |
| Fam149a | 0.00013 | 5.57E-05 | 0.00011 | 3.94E-05 | 7.79E-05 | 0.0001 | 0.00015 | 3.11E-05 | 7.70E-05 | 6.60E-05 | 0 | 0 | 2.97E-05 | 0 | 0 | 8.48E-05 | 0 | 0.0001 |
| Snx21 | 6.85E-06 | 0.00014 | 7.80E-06 | 0.00018 | 9.46E-05 | 7.51E-05 | 0.0001 | 6.58E-05 | 7.70E-05 | 0.00017 | 3.31E-05 | 0.00022 | 0 | 4.00E-05 | 0.00012 | 5.58E-05 | 0 | 8.60E-05 |
| Tipin | 0.0001 | 0 | 0.00014 | 2.47E-05 | 2.20E-05 | 4.32E-05 | 2.13E-05 | 3.01E-05 | 7.70E-05 | 5.37E-05 | 2.90E-05 | 4.94E-05 | 4.30E-05 | 4.47E-05 | 6.14E-06 | 0.00059 | 0 | 1.86E-05 |
| Nelfcd | 3.70E-05 | 6.07E-05 | 2.88E-05 | 9.50E-05 | 4.15E-05 | 2.93E-05 | 0 | 0 | 7.70E-05 | 2.17E-05 | 1.91E-05 | 0 | 4.76E-05 | 5.30E-05 | 3.28E-05 | 0.00018 | 0 | 3.62E-05 |
| Pag1 | 0 | 2.86E-05 | 0 | 0 | 9.15E-05 | 2.02E-05 | 0 | 0.00013 | 7.70E-05 | 0 | 0 | 5.04E-05 | 0 | 0 | 0 | 0 | 0 | 0 |
| Pde8a | 0 | 6.30E-05 | 5.86E-05 | 3.91E-05 | 0 | 2.23E-05 | 7.33E-05 | 0 | 7.69E-05 | 0.00048 | 5.42E-06 | 6.21E-06 | 6.40E-05 | 0 | 0 | 5.75E-05 | 1.77E-05 | 4.70E-05 |
| Usp39 | 9.64E-05 | 5.84E-05 | 0.00011 | 6.49E-06 | 2.99E-05 | 0.00014 | 0 | 0 | 7.68E-05 | 1.48E-05 | 7.48E-05 | 0.00011 | 1.67E-05 | 1.85E-05 | 0 | 0.00019 | 0 | 1.29E-05 |
| Ndn | 8.91E-06 | 4.75E-05 | 0.00012 | 0.00034 | 0.00016 | 0 | 0.00025 | 0.00025 | 7.68E-05 | 0.00057 | 0.00011 | 9.77E-05 | 0.00037 | 0 | 0 | 7.79E-05 | 0 | 0.00053 |
| Nectin3 | 0.0001 | 0.00012 | 0.00014 | 7.40E-05 | 8.63E-05 | 0.00017 | 9.83E-05 | 0.00024 | 7.67E-05 | 0.0002 | 0.00011 | 7.14E-05 | 0 | 0.0001 | 4.31E-05 | 9.76E-05 | 0.00011 | 5.68E-05 |
| Akap7 | 1.30E-05 | 1.84E-05 | 0 | 0.0001 | 0.00017 | 0.00015 | 0 | 8.21E-05 | 7.67E-05 | 0 | 0 | 8.67E-05 | 3.20E-05 | 6.04E-06 | 0.00014 | 0 | 0 | 0 |
| Ube2a | 0.0002 | 8.16E-05 | 8.34E-05 | 8.29E-05 | 2.26E-05 | 8.53E-05 | 1.09E-05 | 7.74E-06 | 7.67E-05 | 2.06E-05 | 7.19E-05 | 2.48E-05 | 0 | 4.00E-05 | 7.57E-05 | 6.93E-05 | 8.18E-05 | 0 |
| Dhdh | 1.08E-05 | 3.91E-05 | 0 | 4.83E-05 | 3.43E-06 | 0 | 0 | 3.02E-06 | 7.67E-05 | 5.15E-05 | 0 | 7.76E-05 | 0.00011 | 0 | 1.76E-06 | 5.88E-06 | 1.12E-05 | 0 |
| Ppil2 | 0.00011 | 0.00017 | 3.72E-05 | 0.00016 | 6.95E-05 | 7.34E-05 | 5.38E-05 | 4.91E-05 | 7.66E-05 | 7.68E-05 | 3.32E-05 | 5.87E-05 | 5.38E-05 | 0.0001 | 1.14E-05 | 0.0001 | 0 | 8.03E-05 |
| Arpin | 2.85E-05 | 0.00011 | 4.78E-05 | 3.85E-05 | 7.29E-05 | 7.07E-05 | 0.0001 | 6.98E-05 | 7.66E-05 | 0.00015 | 1.86E-05 | 3.78E-05 | 0 | 3.84E-05 | 0 | 0.00049 | 0 | 0.00013 |
| Nipa1 | 2.63E-05 | 5.30E-05 | 5.50E-05 | 0 | 4.22E-05 | 6.93E-05 | 5.21E-05 | 1.79E-05 | 7.66E-05 | 5.30E-05 | 7.64E-05 | 2.63E-05 | 4.20E-05 | 0 | 0 | 3.46E-05 | 0 | 7.71E-05 |
| Eef2kmt | 0.00013 | 5.02E-05 | 3.51E-05 | 7.40E-05 | 1.94E-05 | 3.56E-05 | 6.38E-05 | 3.66E-05 | 7.66E-05 | 5.11E-05 | 0 | 7.73E-05 | 4.77E-05 | 0 | 0 | 0 | 0 | 0 |
| Fam222b | 2.17E-05 | 8.70E-05 | 7.41E-05 | 3.53E-05 | 3.46E-05 | 0 | 1.52E-05 | 3.63E-05 | 7.66E-05 | 3.40E-05 | 5.33E-05 | 4.95E-06 | 0 | 1.15E-05 | 0.0001 | 2.76E-05 | 0 | 3.04E-05 |
| Srd5a3 | 9.10E-05 | 6.19E-05 | 4.80E-05 | 4.30E-05 | 8.22E-05 | 7.48E-05 | 7.09E-05 | 2.88E-05 | 7.65E-05 | 6.61E-05 | 2.40E-05 | 2.85E-05 | 5.32E-05 | 5.37E-05 | 0 | 2.19E-05 | 0 | 1.95E-05 |
| Arl14ep | 0.00011 | 0.00013 | 8.28E-05 | 0.00012 | 8.45E-05 | 6.23E-05 | 8.50E-05 | 6.06E-05 | 7.65E-05 | 0 | 4.68E-05 | 4.94E-05 | 5.63E-05 | 3.29E-05 | 0.00016 | 7.10E-05 | 0 | 2.13E-05 |
| Gdf11 | 2.57E-05 | 4.87E-05 | 5.79E-05 | 4.59E-05 | 9.23E-05 | 2.81E-05 | 1.18E-05 | 7.82E-05 | 7.65E-05 | 0 | 4.67E-05 | 0 | 5.69E-06 | 3.04E-05 | 0 | 9.71E-06 | 0.00013 | 0 |
| Bop1 | 9.72E-05 | 3.22E-05 | 0.00018 | 6.56E-05 | 2.33E-05 | 0 | 0 | 3.27E-05 | 7.65E-05 | 2.50E-05 | 2.87E-05 | 0 | 0 | 1.62E-05 | 5.61E-05 | 5.84E-05 | 0 | 1.51E-05 |
| Tnrc6b | 0.00011 | 0.00022 | 0 | 6.57E-05 | 7.35E-05 | 0.00015 | 5.49E-05 | 0.0001 | 7.64E-05 | 8.96E-05 | 0 | 6.48E-05 | 0 | 4.20E-05 | 0.00014 | 9.36E-05 | 0.00025 | 6.54E-05 |

| | | | | | | | | | | | | | | | | | | |
|---|---|---|---|---|---|---|---|---|---|---|---|---|---|---|---|---|---|---|
| B4galt6 | 8.25E-05 | 3.05E-05 | 0.00014 | 0 | 0.00014 | 8.16E-05 | 6.19E-05 | 3.27E-05 | 7.64E-05 | 2.61E-06 | 0.00012 | 2.90E-05 | 0 | 0 | 2.55E-06 | 9.57E-05 | 0.00011 | 0 |
| BC034090 | 5.87E-06 | 0.0001 | 0.0001 | 6.62E-05 | 0.00013 | 1.55E-05 | 3.76E-05 | 8.20E-05 | 7.64E-05 | 3.08E-05 | 0.00014 | 1.20E-05 | 9.70E-05 | 3.52E-05 | 0.00021 | 0 | 9.94E-05 | 3.08E-06 |
| Mad1l1 | 3.63E-05 | 4.20E-05 | 6.35E-05 | 4.56E-05 | 1.01E-05 | 0 | 0 | 2.93E-05 | 7.64E-05 | 2.12E-05 | 2.32E-05 | 0 | 0 | 2.69E-05 | 5.78E-05 | 0.00029 | 2.40E-05 | 1.36E-05 |
| Ndufb10 | 0.00037 | 0.0002 | 0.00022 | 0.0005 | 0.00018 | 0.00025 | 0.00023 | 8.29E-05 | 7.63E-05 | 0.0001 | 0.00011 | 8.09E-05 | 4.03E-05 | 0.00012 | 0 | 0.00022 | 2.28E-05 | 0.00015 |
| Atp6v1h | 0.00026 | 0.00015 | 0.00012 | 0 | 0.0001 | 0.00022 | 2.66E-05 | 0 | 7.63E-05 | 3.96E-05 | 0.00011 | 6.59E-05 | 9.06E-05 | 8.31E-05 | 3.04E-05 | 0 | 0.00014 | 4.15E-05 |
| Zmynd8 | 4.95E-05 | 8.13E-05 | 1.61E-05 | 0.00013 | 9.33E-05 | 3.96E-05 | 0.0001 | 4.02E-05 | 7.63E-05 | 6.77E-05 | 1.41E-06 | 8.27E-05 | 7.25E-05 | 0 | 0.00012 | 7.13E-05 | 0.00011 | 3.98E-05 |
| Usp42 | 1.91E-05 | 9.38E-05 | 9.90E-05 | 3.18E-05 | 5.54E-05 | 5.75E-05 | 0 | 9.56E-05 | 7.62E-05 | 4.96E-05 | 6.80E-05 | 2.22E-06 | 9.14E-05 | 6.57E-05 | 0.00017 | 8.53E-05 | 0 | 2.70E-05 |
| Polr3k | 0.00013 | 8.15E-05 | 0.00015 | 0.00017 | 4.19E-05 | 3.69E-05 | 9.86E-05 | 0 | 7.62E-05 | 6.23E-05 | 9.84E-05 | 1.43E-06 | 6.14E-05 | 6.55E-05 | 0 | 0.00019 | 0 | 9.98E-05 |
| Elp5 | 0.00014 | 0.00012 | 0.00024 | 0.00025 | 2.88E-05 | 1.28E-05 | 8.32E-05 | 3.37E-05 | 7.62E-05 | 0.00011 | 0.00015 | 8.52E-05 | 3.34E-05 | 4.86E-05 | 2.59E-05 | 0.00012 | 2.74E-05 | 6.36E-05 |
| B4galt3 | 4.53E-05 | 7.90E-05 | 0.0001 | 0.00011 | 5.65E-05 | 1.29E-06 | 4.50E-05 | 5.20E-05 | 7.62E-05 | 0 | 0.00013 | 1.03E-05 | 9.22E-05 | 7.13E-05 | 8.12E-05 | 5.05E-05 | 0.00011 | 0 |
| Man2b1 | 0 | 0.0001 | 0 | 4.60E-05 | 0.00016 | 0.00014 | 0.00032 | 0.00024 | 7.61E-05 | 0.00025 | 5.73E-05 | 1.54E-05 | 0.00041 | 0.00029 | 4.76E-05 | 0 | 0 | 0.00034 |
| Emc4 | 0.00025 | 0.00012 | 0.00014 | 0 | 5.71E-05 | 0.00011 | 0 | 0 | 7.61E-05 | 7.28E-05 | 9.40E-05 | 7.61E-05 | 8.17E-06 | 1.44E-05 | 0 | 6.28E-05 | 0 | 5.76E-06 |
| Ubald1 | 7.80E-05 | 0.00013 | 8.56E-05 | 5.38E-05 | 4.28E-05 | 4.22E-05 | 2.44E-06 | 9.75E-05 | 7.61E-05 | 3.33E-05 | 0.00027 | 0.00028 | 2.26E-05 | 5.85E-05 | 0.00013 | 9.93E-07 | 4.74E-05 | 3.09E-05 |
| Zbtb21 | 7.58E-05 | 0.00011 | 5.72E-05 | 0 | 4.69E-05 | 3.94E-05 | 1.82E-05 | 0 | 7.61E-05 | 2.73E-05 | 5.14E-05 | 0.00011 | 1.85E-05 | 1.72E-05 | 0.00012 | 2.52E-05 | 0.00012 | 1.58E-05 |
| Atg101 | 6.15E-05 | 0.0001 | 8.28E-05 | 8.36E-05 | 1.53E-05 | 2.27E-05 | 0 | 0.00014 | 7.61E-05 | 4.69E-05 | 4.24E-06 | 6.99E-05 | 0 | 0.00011 | 9.29E-05 | 7.36E-05 | 0.00043 | 0.00011 |
| Etfbkmt | 9.37E-06 | 0.0001 | 0 | 1.45E-05 | 3.76E-05 | 7.78E-06 | 7.94E-05 | 0 | 7.61E-05 | 0.00014 | 1.17E-05 | 2.73E-05 | 1.08E-05 | 0 | 1.75E-05 | 0 | 1.77E-05 | 7.02E-05 |
| Exosc9 | 0.00011 | 5.14E-05 | 0.00011 | 5.07E-05 | 1.13E-05 | 0 | 0 | 0 | 7.61E-05 | 3.78E-05 | 4.50E-05 | 0 | 2.89E-05 | 6.30E-06 | 8.51E-05 | 0.00011 | 0 | 1.68E-05 |
| Hdgfl3 | 6.43E-05 | 9.07E-05 | 6.29E-05 | 0 | 1.14E-05 | 0 | 4.32E-05 | 3.56E-05 | 7.61E-05 | 6.31E-05 | 0 | 6.55E-06 | 5.21E-05 | 0 | 2.95E-05 | 0.00011 | 5.96E-05 | 8.99E-05 |
| Rabggtb | 0.00015 | 0.00017 | 0.0003 | 0.00024 | 4.87E-05 | 0.00013 | 0 | 7.46E-05 | 7.60E-05 | 2.66E-05 | 0.00011 | 0.00015 | 0 | 1.40E-05 | 0.00019 | 0.0002 | 0.00011 | 2.03E-05 |
| Mtmr3 | 0.00011 | 0.00017 | 1.70E-05 | 1.42E-05 | 5.56E-05 | 7.07E-05 | 6.55E-05 | 0 | 7.60E-05 | 5.00E-05 | 3.78E-05 | 8.41E-05 | 4.34E-05 | 4.78E-05 | 8.78E-05 | 5.92E-05 | 0.0001 | 4.00E-05 |
| Zfand6 | 0.00016 | 0.00011 | 0 | 0 | 3.82E-05 | 4.27E-05 | 6.99E-05 | 4.42E-05 | 7.60E-05 | 3.83E-05 | 4.24E-05 | 0.00012 | 0 | 7.95E-05 | 0.00014 | 7.91E-05 | 0 | 5.48E-05 |
| Hsd3b7 | 8.55E-06 | 0.00019 | 0 | 0 | 0.00029 | 0 | 0.00027 | 2.19E-05 | 7.60E-05 | 0.00028 | 0 | 0.00013 | 8.85E-06 | 2.27E-05 | 3.81E-05 | 0 | 0 | 0.00018 |
| Fam76b | 9.70E-05 | 0.00011 | 3.04E-06 | 0 | 5.45E-05 | 6.49E-05 | 1.94E-05 | 1.20E-05 | 7.59E-05 | 4.51E-05 | 4.05E-05 | 9.77E-05 | 3.43E-06 | 6.47E-05 | 0.00037 | 0.0002 | 0 | 2.86E-05 |
| Wnt4 | 0 | 0 | 0 | 0 | 0 | 2.80E-05 | 0 | 0 | 7.59E-05 | 7.33E-07 | 2.05E-08 | 1.18E-05 | 0 | 1.22E-05 | 0 | 0 | 9.34E-06 | 3.92E-06 |
| Ccs | 3.94E-05 | 5.96E-05 | 2.38E-05 | 0.00015 | 6.36E-05 | 2.53E-05 | 2.24E-05 | 4.89E-05 | 7.59E-05 | 4.02E-05 | 7.29E-05 | 0 | 5.00E-05 | 4.65E-05 | 0.00012 | 0 | 0 | 3.02E-05 |
| Dcun1d3 | 1.13E-05 | 0.00011 | 0.0001 | 8.47E-05 | 1.89E-05 | 1.13E-05 | 1.80E-05 | 1.01E-05 | 7.59E-05 | 7.90E-05 | 6.90E-05 | 1.44E-05 | 2.53E-05 | 8.24E-05 | 8.19E-05 | 1.11E-06 | 0.00013 | 2.02E-05 |
| Ddrgk1 | 8.76E-05 | 0.00013 | 0.0001 | 0.00031 | 0.00012 | 0.00014 | 0.00021 | 0.00013 | 7.58E-05 | 0.00015 | 9.32E-05 | 8.34E-05 | 0.00011 | 0.00017 | 9.49E-05 | 0.00011 | 4.73E-05 | 0.00019 |
| Maml1 | 3.75E-05 | 0.0002 | 0.00012 | 0.0002 | 0.00015 | 0.00011 | 6.01E-05 | 0.00012 | 7.58E-05 | 2.04E-05 | 0.00015 | 0.00027 | 9.69E-05 | 0.00013 | 0.00013 | 0.0001 | 5.44E-05 | 5.69E-06 |
| Atp13a1 | 0.00012 | 0.00015 | 0.00022 | 0.00012 | 7.10E-05 | 8.15E-05 | 5.61E-05 | 1.88E-05 | 7.58E-05 | 5.95E-05 | 5.61E-05 | 1.50E-05 | 5.63E-06 | 8.37E-05 | 0.00024 | 6.36E-05 | 0 | 4.68E-05 |
| Pias3 | 0.00011 | 0.00013 | 0.00011 | 9.86E-05 | 0.00012 | 6.61E-05 | 0.00012 | 8.78E-05 | 7.58E-05 | 5.46E-05 | 5.24E-05 | 0 | 0 | 4.86E-05 | 0.00027 | 0.00015 | 0 | 0 |
| Zfp334 | 7.10E-05 | 5.89E-05 | 5.06E-05 | 3.38E-05 | 4.74E-05 | 3.31E-05 | 4.98E-05 | 3.35E-05 | 7.58E-05 | 9.01E-07 | 1.37E-05 | 3.54E-05 | 1.17E-05 | 6.34E-05 | 0.00012 | 5.65E-05 | 0 | 0 |
| Msl3l2 | 0 | 2.89E-05 | 2.55E-06 | 0.00012 | 4.76E-05 | 0 | 5.94E-05 | 1.93E-05 | 7.58E-05 | 4.75E-05 | 6.86E-06 | 2.85E-06 | 1.75E-06 | 2.72E-05 | 2.38E-05 | 4.20E-05 | 0 | 4.68E-05 |
| Shoc2 | 0.0002 | 0.00011 | 7.11E-05 | 0 | 3.93E-05 | 0.00012 | 6.59E-05 | 0.00011 | 7.57E-05 | 2.54E-05 | 2.66E-05 | 0.00013 | 0 | 3.66E-05 | 4.19E-05 | 8.58E-05 | 5.25E-05 | 0 |

| | | | | | | | | | | | | | | | | | | |
|---|---|---|---|---|---|---|---|---|---|---|---|---|---|---|---|---|---|---|
| Zbtb44 | 0.00012 | 0.00018 | 5.75E-05 | 0 | 2.40E-05 | 0.0001 | 4.58E-05 | 2.43E-05 | 7.57E-05 | 9.55E-05 | 2.48E-05 | 0.00013 | 4.42E-05 | 2.76E-05 | 7.70E-06 | 6.53E-05 | 9.51E-05 | 6.01E-05 |
| Pcgf5 | 5.95E-05 | 1.32E-05 | 0 | 0 | 0 | 7.02E-05 | 0 | 0 | 7.57E-05 | 1.18E-05 | 5.82E-05 | 0.00046 | 6.81E-05 | 0 | 0 | 5.90E-05 | 0 | 7.82E-05 |
| Trappc13 | 0.00014 | 6.55E-05 | 5.54E-05 | 1.93E-05 | 5.85E-05 | 5.77E-05 | 4.65E-05 | 2.65E-05 | 7.57E-05 | 2.18E-05 | 6.87E-05 | 0 | 3.23E-05 | 1.84E-05 | 1.03E-05 | 0.00018 | 0 | 3.98E-05 |
| Aifm1 | 0.0001 | 3.00E-05 | 9.16E-05 | 8.43E-05 | 3.72E-05 | 1.08E-05 | 2.28E-05 | 0 | 7.57E-05 | 1.01E-05 | 2.24E-05 | 0 | 1.49E-05 | 1.96E-05 | 5.34E-05 | 0.00013 | 0 | 7.80E-06 |
| BC003965 | 3.90E-05 | 6.11E-05 | 0.0001 | 9.87E-05 | 5.10E-05 | 0 | 0 | 0 | 7.57E-05 | 3.39E-05 | 1.81E-05 | 0 | 0 | 3.64E-05 | 0.00018 | 4.37E-05 | 0 | 2.34E-05 |
| Dennd4c | 0.00014 | 0.00013 | 3.09E-05 | 0 | 5.38E-05 | 0.00015 | 2.78E-05 | 0 | 7.56E-05 | 5.42E-05 | 7.00E-06 | 4.89E-05 | 0 | 1.40E-05 | 0.00011 | 8.35E-05 | 4.90E-05 | 1.30E-05 |
| Bloc1s2 | 0.00013 | 0.00012 | 0.00018 | 0.00013 | 5.23E-05 | 5.82E-05 | 4.10E-05 | 4.65E-05 | 7.56E-05 | 3.78E-05 | 2.16E-05 | 9.32E-05 | 0 | 2.89E-05 | 1.93E-05 | 5.28E-05 | 0 | 0 |
| Osbpl1a | 4.84E-05 | 0.00012 | 2.25E-05 | 0 | 6.92E-06 | 4.45E-05 | 0 | 0 | 7.56E-05 | 0.00016 | 2.18E-05 | 4.13E-05 | 8.36E-05 | 2.80E-05 | 5.21E-05 | 0.00018 | 2.82E-05 | 0.00012 |
| Igtp | 6.44E-05 | 0 | 0 | 0 | 0 | 0 | 0 | 0 | 7.56E-05 | 0.00015 | 0 | 0 | 4.16E-05 | 0 | 0 | 1.14E-05 | 0 | 9.55E-05 |
| Erbb2 | 6.38E-06 | 5.51E-05 | 5.18E-05 | 4.29E-05 | 1.30E-05 | 0 | 1.60E-05 | 6.32E-06 | 7.56E-05 | 3.43E-05 | 3.54E-05 | 0 | 0 | 3.49E-05 | 6.94E-05 | 3.93E-06 | 0 | 6.13E-05 |
| Impdh2 | 0.00021 | 8.19E-05 | 0.00026 | 0.00027 | 0 | 8.86E-05 | 0 | 6.51E-05 | 7.55E-05 | 5.01E-05 | 0 | 0.00013 | 9.29E-05 | 9.18E-05 | 0 | 0.00044 | 0.0005 | 9.52E-05 |
| Ints14 | 0.0001 | 8.44E-05 | 0.00014 | 0.00012 | 6.05E-05 | 7.67E-05 | 2.57E-05 | 3.82E-05 | 7.55E-05 | 4.72E-05 | 5.97E-05 | 6.03E-05 | 5.23E-05 | 5.10E-05 | 0 | 0.00015 | 0 | 0 |
| Ndufa5 | 0.00025 | 5.11E-05 | 0.00016 | 0.00025 | 7.23E-05 | 6.82E-05 | 2.04E-05 | 0 | 7.55E-05 | 1.84E-05 | 6.28E-05 | 5.58E-06 | 5.27E-05 | 4.67E-05 | 4.06E-06 | 0.00027 | 0 | 3.69E-05 |
| Dpm3 | 0.00014 | 6.77E-05 | 0.00015 | 0.00066 | 0.00014 | 0.00011 | 0.00032 | 0.00022 | 7.54E-05 | 0.00011 | 3.99E-05 | 3.91E-05 | 0.00014 | 7.54E-05 | 0 | 0.00013 | 2.34E-05 | 0.00014 |
| Lypla1 | 0.00016 | 5.70E-05 | 7.49E-05 | 1.14E-05 | 4.53E-05 | 3.95E-05 | 5.41E-05 | 3.32E-05 | 7.54E-05 | 5.38E-05 | 4.50E-05 | 0 | 0 | 4.80E-05 | 0 | 4.44E-05 | 2.01E-05 | 4.99E-05 |
| Zfp687 | 5.88E-05 | 0.00012 | 2.86E-05 | 6.64E-05 | 3.17E-05 | 1.76E-05 | 4.09E-06 | 1.72E-05 | 7.54E-05 | 1.77E-05 | 7.16E-05 | 7.22E-05 | 0 | 1.68E-05 | 0.0001 | 7.95E-05 | 0 | 3.84E-06 |
| Nme2 | 0.00067 | 1.83E-05 | 0.00032 | 0.00022 | 0.00016 | 0.00027 | 0.00042 | 0.00017 | 7.53E-05 | 4.17E-05 | 0.00019 | 4.22E-05 | 0.00013 | 0.00012 | 0 | 0.00014 | 0 | 6.98E-06 |
| Lrrc14 | 3.65E-06 | 9.42E-05 | 4.48E-05 | 8.48E-05 | 5.03E-05 | 2.19E-05 | 2.48E-05 | 4.21E-05 | 7.53E-05 | 5.87E-05 | 6.38E-05 | 1.64E-05 | 3.52E-06 | 6.70E-05 | 9.35E-05 | 5.23E-05 | 0 | 3.52E-05 |
| Ncoa3 | 0.00013 | 0.00018 | 1.64E-05 | 1.42E-05 | 0.00012 | 1.53E-05 | 1.93E-05 | 2.47E-05 | 7.53E-05 | 5.67E-05 | 0.00011 | 8.10E-05 | 0.00011 | 5.21E-05 | 5.17E-05 | 0.0001 | 0.0002 | 0.00015 |
| Xrcc6 | 9.14E-05 | 2.38E-05 | 4.93E-05 | 5.42E-05 | 2.06E-05 | 1.33E-05 | 2.09E-06 | 0 | 7.53E-05 | 7.97E-06 | 5.25E-06 | 2.04E-05 | 2.43E-06 | 0 | 1.90E-05 | 0.00012 | 0 | 0 |
| Dele1 | 7.67E-05 | 6.46E-05 | 4.08E-05 | 4.04E-05 | 4.27E-05 | 5.96E-05 | 2.41E-05 | 7.19E-05 | 7.51E-05 | 4.54E-05 | 2.97E-05 | 7.49E-05 | 1.84E-05 | 6.39E-05 | 0.00013 | 0.00019 | 0 | 6.60E-05 |
| Naprt | 2.09E-05 | 2.90E-05 | 0 | 0.0001 | 0 | 2.36E-05 | 6.94E-05 | 5.68E-05 | 7.51E-05 | 0.0001 | 0 | 1.31E-06 | 0 | 6.86E-06 | 4.52E-05 | 1.28E-05 | 0 | 9.07E-05 |
| Cstf1 | 5.18E-05 | 9.44E-05 | 7.18E-05 | 3.89E-05 | 3.09E-05 | 1.49E-05 | 0 | 0 | 7.51E-05 | 2.86E-05 | 3.13E-05 | 3.19E-05 | 0 | 3.08E-05 | 2.24E-05 | 0.00015 | 0 | 9.34E-06 |
| Zfp462 | 3.14E-05 | 3.88E-05 | 3.07E-05 | 1.02E-05 | 8.93E-05 | 5.99E-06 | 0.00017 | 0 | 7.51E-05 | 5.52E-05 | 0.00013 | 1.56E-05 | 0 | 0 | 0.00011 | 2.18E-05 | 8.47E-05 | 1.69E-05 |
| Thoc5 | 8.44E-05 | 6.52E-05 | 9.56E-05 | 2.23E-05 | 1.02E-05 | 5.23E-06 | 0 | 0 | 7.51E-05 | 3.10E-05 | 2.00E-05 | 1.59E-05 | 0 | 0 | 7.35E-05 | 2.90E-05 | 2.54E-07 | 5.24E-05 |
| Dcbld1 | 2.00E-05 | 0 | 0 | 0 | 0.00017 | 0.00025 | 0 | 0.00037 | 7.50E-05 | 0 | 5.46E-05 | 0 | 0 | 0.00035 | 0 | 7.15E-05 | 0 | 0 |
| Cisd2 | 0.00029 | 0.00012 | 0.00026 | 0.0001 | 8.97E-05 | 0.00023 | 0.00015 | 0.00013 | 7.50E-05 | 4.56E-05 | 4.27E-05 | 7.13E-05 | 6.68E-05 | 0.00015 | 0.00011 | 0.00027 | 0 | 3.83E-05 |
| Cop1 | 0.00026 | 0.00019 | 0.00014 | 0 | 0.0001 | 0.00022 | 1.95E-05 | 5.57E-05 | 7.50E-05 | 4.40E-05 | 0.00012 | 0.00013 | 0 | 5.04E-05 | 0 | 9.87E-05 | 0.00018 | 4.45E-05 |
| Rab7b | 2.54E-05 | 5.75E-05 | 1.53E-05 | 0 | 7.19E-05 | 0.00012 | 4.64E-05 | 0 | 7.50E-05 | 0.00019 | 2.45E-05 | 1.12E-05 | 6.52E-06 | 1.47E-05 | 0 | 0 | 0 | 7.73E-05 |
| Cdkn2c | 0 | 6.39E-05 | 0 | 0 | 5.18E-05 | 8.10E-05 | 4.38E-05 | 2.32E-05 | 7.50E-05 | 5.30E-05 | 1.94E-05 | 0 | 0 | 0 | 0 | 0.00096 | 0 | 4.40E-05 |
| Fopnl | 0.00013 | 3.99E-05 | 2.05E-05 | 2.80E-05 | 5.02E-05 | 2.96E-05 | 4.12E-05 | 0 | 7.50E-05 | 2.91E-05 | 2.22E-05 | 2.11E-05 | 0 | 9.60E-06 | 1.16E-05 | 0.00016 | 0 | 1.90E-05 |
| Casp9 | 2.22E-05 | 8.00E-05 | 7.56E-06 | 4.02E-05 | 1.48E-05 | 2.71E-05 | 0 | 6.23E-05 | 7.50E-05 | 2.37E-05 | 0 | 1.14E-05 | 0 | 7.19E-06 | 1.62E-05 | 0 | 0 | 4.80E-05 |
| Trim16 | 0 | 5.94E-05 | 9.33E-06 | 0 | 0 | 0 | 0 | 0 | 7.50E-05 | 9.95E-05 | 0.00012 | 0 | 0 | 6.59E-05 | 4.51E-05 | 4.56E-06 | 0 | 4.25E-05 |

| | | | | | | | | | | | | | | | | | | |
|---|---|---|---|---|---|---|---|---|---|---|---|---|---|---|---|---|---|---|
| Nme1 | 0.00029 | 0 | 0.00046 | 0.00044 | 0 | 0 | 1.42E-05 | 3.84E-05 | 7.50E-05 | 0 | 0.0001 | 0 | 0 | 8.24E-05 | 0 | 0.00025 | 0 | 1.20E-05 |
| Trps1 | 0 | 0.00012 | 0 | 0 | 0 | 0.0001 | 0 | 0 | 7.49E-05 | 0 | 0 | 0 | 0 | 0.0004 | 9.08E-05 | 0 | 2.40E-05 | 0.00017 |
| Polr2j | 0.00021 | 5.06E-05 | 4.83E-05 | 0 | 6.17E-05 | 8.97E-05 | 1.95E-05 | 0 | 7.49E-05 | 0 | 2.07E-05 | 7.78E-05 | 2.30E-05 | 1.74E-05 | 0 | 7.65E-05 | 0 | 0 |
| Sun1 | 3.56E-05 | 0.00013 | 7.92E-05 | 3.42E-05 | 6.65E-05 | 3.66E-05 | 2.00E-05 | 4.96E-05 | 7.49E-05 | 7.33E-05 | 0.00012 | 0.00043 | 1.90E-05 | 5.62E-05 | 0.0001 | 7.01E-05 | 6.07E-05 | 0.00011 |
| Zranb2 | 9.45E-05 | 7.74E-05 | 0.00013 | 9.11E-05 | 7.48E-05 | 0 | 3.11E-05 | 7.91E-06 | 7.49E-05 | 3.53E-05 | 3.95E-05 | 5.67E-05 | 0 | 3.59E-05 | 0.00022 | 0.00012 | 0 | 6.17E-05 |
| Gtf2h4 | 8.94E-05 | 4.46E-05 | 0.00011 | 0.00013 | 3.58E-05 | 0 | 0 | 0 | 7.49E-05 | 2.67E-07 | 1.41E-05 | 1.57E-05 | 0 | 4.88E-06 | 0 | 8.87E-05 | 0 | 3.56E-05 |
| Strn3 | 0.00037 | 0.00019 | 0.00012 | 0 | 5.89E-05 | 0.0002 | 4.75E-05 | 2.05E-05 | 7.48E-05 | 4.97E-05 | 7.94E-05 | 0.00021 | 0 | 0 | 0.00015 | 3.35E-05 | 0.00022 | 5.73E-05 |
| Zfp871 | 0.00014 | 0.00018 | 2.77E-05 | 0 | 0.00011 | 0.00015 | 1.76E-05 | 4.74E-05 | 7.48E-05 | 4.46E-05 | 1.84E-05 | 0.00013 | 0 | 4.73E-05 | 0.00012 | 0.00011 | 2.22E-05 | 6.15E-05 |
| Vps26b | 0.00019 | 0.00016 | 0.00018 | 0.00028 | 0.00017 | 0.00012 | 0.00021 | 0.00017 | 7.48E-05 | 0.00013 | 0.00012 | 0.00018 | 0.0001 | 0.00015 | 0 | 0.00011 | 5.45E-06 | 0.00012 |
| Ciz1 | 7.14E-05 | 0.00018 | 6.02E-05 | 0.00015 | 3.31E-05 | 4.28E-05 | 4.35E-05 | 0 | 7.48E-05 | 8.17E-05 | 2.64E-05 | 4.87E-05 | 0 | 4.93E-05 | 0.00038 | 0.00013 | 0.00013 | 7.41E-05 |
| Meaf6 | 7.02E-05 | 8.89E-05 | 7.84E-05 | 6.86E-05 | 6.75E-05 | 3.69E-05 | 2.37E-05 | 1.23E-05 | 7.48E-05 | 2.30E-05 | 6.35E-05 | 5.91E-06 | 4.34E-05 | 1.18E-05 | 0.00027 | 0.00017 | 1.80E-05 | 5.15E-05 |
| Pa2g4 | 0.0002 | 4.70E-05 | 0.00027 | 6.68E-05 | 1.32E-05 | 3.32E-05 | 0 | 0 | 7.48E-05 | 1.13E-05 | 1.57E-05 | 1.31E-05 | 1.19E-05 | 0 | 0 | 0.00027 | 4.87E-05 | 0 |
| Dst | 0.00017 | 0.00025 | 0.00017 | 0 | 0.00011 | 2.87E-05 | 0.00044 | 0 | 7.48E-05 | 0.00036 | 0.00021 | 0.00013 | 0.00032 | 0.00016 | 0.00034 | 7.37E-05 | 0.00025 | 0.00027 |
| Traf4 | 2.96E-05 | 0.00012 | 5.11E-05 | 7.90E-05 | 2.43E-05 | 0 | 0 | 0.00037 | 7.48E-05 | 0 | 4.32E-05 | 0 | 0 | 0.00017 | 0 | 5.11E-05 | 0.00015 | 0.00011 |
| Orc3 | 0.00011 | 6.63E-05 | 7.55E-05 | 1.39E-05 | 5.13E-05 | 3.57E-05 | 5.39E-05 | 1.81E-05 | 7.47E-05 | 4.61E-05 | 1.55E-06 | 3.95E-05 | 9.74E-06 | 2.53E-06 | 5.37E-05 | 6.62E-05 | 0 | 4.91E-05 |
| Hoxa10 | 1.88E-06 | 4.13E-05 | 0 | 1.66E-06 | 1.71E-05 | 2.20E-06 | 8.62E-05 | 0 | 7.47E-05 | 2.53E-05 | 1.58E-05 | 2.06E-05 | 3.50E-05 | 0 | 2.43E-05 | 1.20E-05 | 0 | 1.44E-06 |
| Naa35 | 9.18E-05 | 6.65E-05 | 5.81E-05 | 2.55E-05 | 2.31E-05 | 0 | 1.81E-05 | 0 | 7.47E-05 | 2.79E-05 | 1.24E-06 | 5.04E-05 | 8.20E-06 | 2.64E-05 | 4.88E-05 | 7.16E-05 | 0 | 5.64E-05 |
| Gipc1 | 3.41E-05 | 6.24E-05 | 7.06E-05 | 0.00025 | 5.34E-05 | 0 | 7.58E-05 | 0.00023 | 7.47E-05 | 3.79E-05 | 0.00011 | 7.71E-05 | 9.57E-07 | 3.22E-05 | 7.59E-05 | 7.59E-05 | 3.47E-05 | 0.00012 |
| Copb2 | 0.00055 | 0.0001 | 0.00045 | 0.00057 | 0.00018 | 0.00029 | 0.00069 | 0.00025 | 7.46E-05 | 0.00012 | 0.00026 | 9.84E-05 | 0.00026 | 0.00018 | 5.87E-05 | 0.00011 | 0.00015 | 0.00018 |
| Sp1 | 0.00012 | 0.00015 | 5.40E-05 | 0 | 6.80E-05 | 0.00013 | 1.74E-05 | 9.13E-05 | 7.46E-05 | 4.04E-05 | 3.22E-05 | 0.00014 | 0 | 4.75E-05 | 1.99E-05 | 0.00015 | 0.00013 | 7.19E-05 |
| Jagn1 | 0.00015 | 7.82E-05 | 0.00022 | 7.33E-05 | 7.22E-05 | 6.12E-05 | 0 | 4.41E-05 | 7.46E-05 | 1.12E-05 | 6.48E-05 | 0 | 5.56E-08 | 4.01E-05 | 0 | 3.13E-05 | 0 | 0 |
| Actl6a | 0.00018 | 9.53E-05 | 0.00017 | 0 | 5.58E-05 | 4.42E-05 | 0 | 0 | 7.46E-05 | 5.45E-06 | 5.48E-05 | 0 | 0 | 4.17E-05 | 4.94E-05 | 0.00025 | 0 | 0 |
| Cyld | 4.79E-05 | 8.40E-05 | 4.02E-06 | 0 | 2.28E-05 | 3.79E-05 | 1.99E-05 | 0 | 7.46E-05 | 0.0001 | 6.01E-05 | 7.05E-05 | 6.50E-05 | 5.95E-05 | 9.76E-05 | 6.20E-05 | 0 | 9.18E-05 |
| Ilvbl | 2.24E-06 | 9.12E-05 | 5.01E-06 | 6.90E-05 | 5.59E-05 | 6.02E-06 | 6.69E-06 | 0 | 7.46E-05 | 4.98E-05 | 9.42E-05 | 5.26E-05 | 1.86E-05 | 4.78E-05 | 4.96E-05 | 5.71E-06 | 0 | 6.77E-05 |
| Gapvd1 | 0.00014 | 0.00017 | 0.00013 | 0 | 9.33E-05 | 7.10E-05 | 7.49E-05 | 1.97E-05 | 7.45E-05 | 6.88E-05 | 6.68E-05 | 0.00014 | 8.25E-06 | 6.11E-05 | 0.00015 | 9.89E-05 | 4.02E-05 | 4.31E-05 |
| Tpcn2 | 2.91E-05 | 8.31E-05 | 3.81E-05 | 6.57E-05 | 5.14E-05 | 5.19E-05 | 4.94E-05 | 0 | 7.45E-05 | 8.86E-05 | 0 | 0 | 5.04E-05 | 6.09E-05 | 0.0001 | 1.04E-05 | 0 | 6.55E-05 |
| Kctd15 | 5.54E-05 | 4.79E-05 | 9.49E-05 | 0 | 4.14E-05 | 8.48E-05 | 3.44E-05 | 9.32E-05 | 7.44E-05 | 0 | 0.00021 | 0.0001 | 0 | 0.00023 | 0 | 0 | 4.87E-05 | 0.00011 |
| Acox1 | 6.21E-05 | 0.00021 | 6.61E-06 | 0.00012 | 0.0001 | 7.50E-05 | 3.02E-05 | 7.66E-05 | 7.44E-05 | 8.41E-05 | 4.57E-05 | 0.00011 | 6.43E-05 | 6.98E-05 | 3.03E-05 | 1.44E-05 | 0.00019 | 0.00012 |
| Plppr2 | 0 | 1.92E-05 | 4.80E-05 | 0.00024 | 9.94E-05 | 6.07E-05 | 0.00015 | 0.00013 | 7.44E-05 | 0.0001 | 5.27E-05 | 2.72E-05 | 3.84E-05 | 7.52E-05 | 7.56E-05 | 0 | 0 | 0.00013 |
| Pld2 | 1.95E-05 | 5.98E-05 | 0 | 4.66E-06 | 6.34E-05 | 0 | 0.00012 | 0 | 7.44E-05 | 7.77E-05 | 0.0001 | 9.87E-05 | 4.53E-05 | 6.67E-05 | 5.00E-05 | 1.53E-06 | 0 | 6.54E-05 |
| Fuca1 | 0.00019 | 0.0001 | 1.74E-05 | 0 | 0.00022 | 0.00031 | 0.00033 | 0.00022 | 7.43E-05 | 0.00021 | 5.52E-05 | 0.00015 | 0.00043 | 0.00016 | 0 | 8.77E-05 | 0 | 0.00038 |
| Cyp2f2 | 0 | 0 | 0 | 0 | 0 | 5.44E-05 | 0 | 0 | 7.43E-05 | 1.73E-05 | 0 | 3.94E-07 | 0.00013 | 0 | 0 | 0 | 0 | 7.10E-05 |
| Mpv17 | 0.0001 | 0.00017 | 0 | 0.00019 | 0.00016 | 0.00015 | 0.00015 | 3.36E-05 | 7.42E-05 | 0.00012 | 2.96E-05 | 7.13E-05 | 0.00011 | 0.00016 | 0 | 0 | 0 | 0.00015 |

| | | | | | | | | | | | | | | | | | | |
|---|---|---|---|---|---|---|---|---|---|---|---|---|---|---|---|---|---|---|
| Fer | 0.00019 | 8.85E-05 | 3.00E-05 | 0 | 0.0001 | 6.60E-05 | 6.89E-05 | 0 | 7.42E-05 | 0.00021 | 0.00023 | 0.00018 | 5.13E-05 | 0 | 7.74E-05 | 0.0001 | 0.00017 | 0 |
| Lonp1 | 9.30E-05 | 0.00014 | 0.00021 | 0.00026 | 7.23E-05 | 0 | 0.00013 | 0 | 7.42E-05 | 2.16E-05 | 5.91E-05 | 0.0006 | 5.10E-05 | 2.86E-05 | 0.00014 | 0.00018 | 1.54E-05 | 0 |
| Mxd4 | 0.00029 | 0.00031 | 0 | 0.00036 | 0.00039 | 0.00054 | 0.0007 | 0.00023 | 7.41E-05 | 0.00033 | 0.00016 | 9.33E-05 | 0.00022 | 0.00045 | 0.00012 | 0 | 0 | 0.00041 |
| Tvp23b | 0.00021 | 4.50E-05 | 8.19E-05 | 0.00023 | 0.00016 | 0.00026 | 0.00036 | 0.0002 | 7.41E-05 | 9.56E-05 | 0.00013 | 0.00012 | 0.00021 | 0.00013 | 1.42E-05 | 5.08E-06 | 1.53E-05 | 8.88E-05 |
| Paqr7 | 0 | 0.00011 | 0 | 8.32E-05 | 0.00013 | 0.00011 | 0 | 0.00013 | 7.41E-05 | 2.74E-05 | 0 | 0 | 0 | 0.00016 | 0 | 0 | 0 | 8.12E-05 |
| Tank | 0.00012 | 7.08E-05 | 7.83E-05 | 0 | 5.02E-05 | 0.00011 | 0 | 7.47E-06 | 7.41E-05 | 5.16E-05 | 0 | 0 | 0.00011 | 8.59E-05 | 0.00018 | 5.34E-05 | 0 | 5.66E-06 |
| Dtd2 | 2.77E-05 | 3.96E-05 | 1.95E-05 | 8.19E-05 | 5.84E-05 | 2.15E-05 | 0 | 6.16E-05 | 7.41E-05 | 4.51E-05 | 3.58E-05 | 2.94E-05 | 1.51E-05 | 2.07E-05 | 6.19E-06 | 1.72E-05 | 0 | 4.68E-05 |
| Flvcr1 | 2.19E-05 | 8.05E-05 | 8.21E-05 | 2.20E-05 | 0 | 0 | 0 | 0 | 7.41E-05 | 1.97E-05 | 5.48E-05 | 3.04E-06 | 5.69E-06 | 6.68E-05 | 5.02E-05 | 2.82E-05 | 7.84E-05 | 0 |
| Eif2b1 | 0.0002 | 5.98E-05 | 0.00018 | 7.97E-05 | 5.52E-05 | 6.97E-05 | 2.47E-05 | 1.01E-06 | 7.40E-05 | 1.24E-05 | 1.77E-05 | 4.56E-06 | 6.73E-06 | 3.47E-05 | 2.03E-05 | 0.00015 | 0 | 4.46E-06 |
| Suclg2 | 9.48E-05 | 5.01E-05 | 0 | 2.34E-06 | 3.53E-05 | 6.82E-05 | 5.50E-05 | 0 | 7.40E-05 | 4.46E-05 | 0 | 5.10E-05 | 2.69E-05 | 0 | 0 | 5.40E-05 | 8.25E-05 | 0 |
| Fdx1 | 9.77E-05 | 4.81E-05 | 2.69E-05 | 0 | 0 | 6.51E-05 | 4.58E-05 | 0 | 7.40E-05 | 9.03E-05 | 0 | 0.0001 | 5.19E-05 | 0 | 6.67E-05 | 1.73E-05 | 0.00013 | 5.67E-05 |
| Htatsf1 | 0.00019 | 5.55E-05 | 0.00011 | 8.76E-05 | 3.72E-05 | 3.65E-05 | 4.12E-05 | 0 | 7.40E-05 | 1.15E-05 | 1.76E-05 | 3.83E-05 | 0 | 0 | 5.25E-05 | 7.77E-05 | 0 | 2.82E-05 |
| Brd9 | 6.92E-05 | 8.49E-05 | 3.36E-05 | 0 | 1.78E-05 | 1.88E-05 | 1.27E-05 | 0 | 7.40E-05 | 6.20E-05 | 4.06E-05 | 1.20E-05 | 4.11E-05 | 3.74E-07 | 0.00013 | 0.00018 | 8.20E-06 | 4.15E-05 |
| Snx15 | 1.86E-05 | 5.22E-05 | 1.88E-05 | 9.15E-05 | 4.64E-05 | 7.24E-05 | 0 | 0 | 7.39E-05 | 3.71E-05 | 8.21E-05 | 0 | 3.90E-05 | 5.73E-05 | 2.68E-05 | 1.92E-05 | 2.76E-05 | 7.62E-05 |
| Vps26a | 0.00016 | 0.00019 | 3.09E-05 | 0 | 7.62E-05 | 6.31E-05 | 3.55E-05 | 0 | 7.39E-05 | 6.29E-05 | 1.07E-05 | 0.00013 | 8.43E-05 | 6.44E-05 | 0 | 6.26E-05 | 0 | 0.00011 |
| Mbip | 3.85E-05 | 3.27E-05 | 4.47E-07 | 0 | 3.20E-05 | 3.17E-05 | 7.57E-06 | 2.22E-06 | 7.39E-05 | 3.77E-05 | 1.62E-06 | 2.89E-05 | 5.28E-06 | 1.89E-05 | 0.00011 | 1.61E-05 | 0 | 3.92E-05 |
| Efcab14 | 0.00018 | 4.70E-05 | 4.00E-05 | 0 | 0.0002 | 0.00027 | 1.95E-05 | 2.50E-06 | 7.38E-05 | 0.00012 | 9.37E-05 | 2.31E-05 | 7.82E-05 | 0.00013 | 0 | 5.26E-05 | 0.00012 | 0 |
| Esrra | 0 | 0.00012 | 0.00019 | 0.00017 | 4.78E-05 | 5.65E-05 | 0 | 0.00012 | 7.38E-05 | 7.74E-05 | 0.00012 | 0.00012 | 1.25E-05 | 7.16E-05 | 0.00011 | 0 | 0 | 2.10E-05 |
| Mcrs1 | 7.63E-05 | 6.40E-05 | 6.21E-05 | 0.00015 | 3.54E-05 | 5.20E-05 | 3.70E-05 | 6.51E-05 | 7.38E-05 | 2.92E-05 | 1.23E-05 | 7.13E-06 | 0 | 2.99E-05 | 7.47E-05 | 7.61E-05 | 0 | 5.82E-05 |
| Stat6 | 0.00016 | 0.00021 | 0.00013 | 0.00013 | 4.02E-05 | 0.00017 | 5.59E-05 | 6.58E-05 | 7.37E-05 | 0.00018 | 2.02E-05 | 0.00038 | 0 | 5.79E-05 | 0.00018 | 0 | 0 | 0.00024 |
| Senp6 | 0.00023 | 0.00018 | 0.00015 | 4.88E-05 | 5.92E-05 | 0.00015 | 8.02E-05 | 7.44E-05 | 7.37E-05 | 5.80E-05 | 0.00011 | 0.00011 | 0 | 4.91E-05 | 0.00015 | 6.41E-05 | 0.00023 | 0 |
| Micu1 | 0.00015 | 0.00017 | 5.41E-05 | 3.92E-05 | 0.00016 | 8.24E-05 | 2.19E-05 | 0 | 7.37E-05 | 0.00012 | 2.89E-05 | 0.00015 | 0 | 0 | 0 | 0.0001 | 5.58E-05 | 2.42E-05 |
| Snx14 | 0.00012 | 4.81E-05 | 0 | 0 | 0 | 7.52E-05 | 5.20E-05 | 0 | 7.37E-05 | 4.53E-05 | 4.58E-05 | 0.00011 | 2.81E-05 | 6.79E-05 | 3.93E-05 | 6.62E-06 | 5.47E-06 | 4.51E-05 |
| Slx4 | 2.36E-05 | 2.72E-05 | 5.20E-05 | 0 | 9.29E-06 | 1.63E-05 | 2.38E-05 | 4.35E-05 | 7.37E-05 | 4.05E-05 | 3.67E-05 | 0 | 3.57E-05 | 3.96E-05 | 9.97E-05 | 8.06E-05 | 4.16E-06 | 1.26E-05 |
| Aida | 0.00043 | 8.85E-05 | 7.82E-05 | 0 | 0 | 0.00027 | 0 | 0 | 7.36E-05 | 0.00022 | 0 | 0.00025 | 0 | 0 | 0 | 0.00023 | 0 | 0.00015 |
| Ubqln1 | 0.00043 | 0.00023 | 0.00046 | 0.00012 | 0.00012 | 0.00024 | 0.00013 | 5.92E-05 | 7.36E-05 | 4.35E-05 | 0.00024 | 0.00017 | 2.59E-05 | 8.01E-05 | 0 | 0.0002 | 0.00021 | 1.50E-05 |
| Hmg20a | 0.00013 | 0.00017 | 0 | 5.80E-05 | 0.00013 | 9.53E-05 | 9.19E-05 | 5.46E-05 | 7.36E-05 | 8.04E-05 | 3.10E-05 | 9.27E-05 | 2.14E-05 | 1.94E-05 | 0.00037 | 8.61E-05 | 0.00012 | 5.37E-05 |
| Exoc2 | 9.45E-05 | 9.99E-05 | 4.84E-05 | 1.17E-06 | 2.93E-05 | 4.29E-05 | 2.90E-05 | 8.84E-05 | 7.36E-05 | 7.40E-05 | 3.85E-05 | 2.57E-05 | 6.09E-05 | 5.51E-05 | 9.13E-05 | 1.32E-05 | 1.62E-05 | 5.05E-05 |
| Crat | 0.00011 | 0.00013 | 0 | 1.18E-05 | 4.78E-05 | 6.69E-05 | 1.37E-06 | 0 | 7.35E-05 | 4.58E-05 | 6.74E-05 | 2.06E-05 | 9.56E-06 | 2.63E-05 | 0 | 3.28E-05 | 0 | 4.16E-05 |
| Sgsm2 | 0 | 6.76E-05 | 1.10E-05 | 3.90E-05 | 7.22E-05 | 3.45E-05 | 1.94E-05 | 0 | 7.35E-05 | 0.00016 | 0 | 1.67E-05 | 0 | 0 | 8.37E-05 | 2.26E-07 | 1.70E-05 | 0.00014 |
| Selenoo | 4.22E-05 | 6.55E-05 | 4.11E-05 | 4.74E-05 | 4.51E-05 | 2.67E-05 | 0 | 3.02E-05 | 7.35E-05 | 3.58E-05 | 3.78E-05 | 4.93E-05 | 0 | 5.86E-05 | 4.78E-05 | 0 | 0 | 4.67E-05 |
| Kras | 1.54E-05 | 9.74E-05 | 0.00021 | 6.38E-05 | 4.15E-05 | 2.54E-05 | 0.00013 | 0.00012 | 7.35E-05 | 0.00018 | 0.00022 | 5.57E-05 | 0 | 0.00011 | 0.00011 | 0.00021 | 9.81E-05 | 0.00023 |
| Etv1 | 0 | 5.13E-05 | 0 | 0 | 3.69E-05 | 2.17E-05 | 0 | 4.05E-05 | 7.35E-05 | 0 | 0 | 0 | 5.54E-06 | 0.00027 | 1.51E-06 | 3.52E-06 | 0 | 9.46E-05 |

| | | | | | | | | | | | | | | | | | | |
|---|---|---|---|---|---|---|---|---|---|---|---|---|---|---|---|---|---|---|
| Ltv1 | 9.44E-05 | 4.82E-05 | 0.00022 | 0.00012 | 6.34E-06 | 0 | 5.61E-05 | 0 | 7.35E-05 | 5.31E-05 | 4.56E-05 | 0 | 6.12E-05 | 0 | 0 | 1.62E-05 | 5.26E-05 | 1.85E-05 |
| Sh3rf3 | 0 | 5.71E-06 | 0 | 8.70E-05 | 0.00012 | 0 | 5.75E-05 | 0 | 7.35E-05 | 4.37E-05 | 0 | 0 | 1.25E-05 | 0.00016 | 0.00018 | 0 | 5.78E-06 | 0.00015 |
| Prps1l3 | 0.00011 | 8.99E-05 | 0.00013 | 9.21E-05 | 2.17E-05 | 1.94E-05 | 2.19E-05 | 3.33E-05 | 7.34E-05 | 3.13E-05 | 0 | 3.93E-05 | 1.49E-05 | 4.44E-05 | 8.09E-05 | 0.00013 | 0 | 4.22E-05 |
| Egln1 | 0.00012 | 6.74E-05 | 0.00012 | 0.00016 | 0.00035 | 0.00024 | 0.00031 | 0.00035 | 7.33E-05 | 0.00024 | 0.00033 | 0.00181 | 0.00021 | 0.00023 | 0 | 0.00022 | 0.00021 | 0.00014 |
| Rspo3 | 0 | 0 | 0 | 0 | 0 | 8.79E-05 | 0 | 0 | 7.33E-05 | 4.92E-06 | 0 | 0 | 3.64E-06 | 4.00E-05 | 0 | 2.80E-06 | 0 | 0 |
| Xrcc5 | 2.17E-05 | 5.15E-05 | 0 | 2.98E-05 | 6.88E-06 | 1.66E-06 | 1.92E-05 | 9.40E-07 | 7.33E-05 | 6.12E-05 | 2.90E-06 | 1.20E-05 | 2.21E-05 | 0 | 1.84E-07 | 2.73E-05 | 0 | 3.80E-05 |
| Cox6a1 | 0.00014 | 4.03E-05 | 0.0002 | 0.00045 | 2.10E-05 | 0 | 0.00017 | 0.0001 | 7.33E-05 | 5.34E-05 | 0.00019 | 4.46E-05 | 4.06E-05 | 5.60E-05 | 0.00017 | 0.00011 | 6.94E-05 | 6.67E-05 |
| Alg2 | 4.99E-05 | 6.82E-05 | 9.10E-05 | 0.00014 | 5.46E-05 | 0 | 2.86E-05 | 6.00E-05 | 7.33E-05 | 2.29E-05 | 5.66E-05 | 6.12E-05 | 0 | 4.62E-05 | 0 | 0 | 0 | 0 |
| Lrp5 | 8.50E-06 | 0.00024 | 1.77E-05 | 7.63E-05 | 5.61E-05 | 9.74E-05 | 0 | 0.00015 | 7.32E-05 | 0.00011 | 0.00015 | 0.0004 | 0.00013 | 0.00044 | 0.00022 | 0.00022 | 0.00024 | 0.00028 |
| Eps15 | 0.00017 | 0.00018 | 0.00011 | 0 | 4.09E-05 | 9.72E-05 | 4.09E-05 | 7.41E-05 | 7.32E-05 | 3.54E-05 | 4.67E-05 | 0.00017 | 1.40E-05 | 4.10E-05 | 0.0001 | 0.00012 | 7.92E-05 | 9.62E-05 |
| Idnk | 5.83E-05 | 0.00011 | 2.28E-05 | 8.94E-05 | 9.68E-05 | 4.53E-05 | 0 | 4.16E-05 | 7.32E-05 | 3.64E-05 | 1.67E-06 | 4.60E-05 | 0 | 4.44E-05 | 0.00011 | 0 | 0 | 7.31E-05 |
| Thap7 | 8.64E-05 | 0.00011 | 0.00011 | 0.00011 | 0.00011 | 3.31E-05 | 6.04E-05 | 0 | 7.32E-05 | 7.33E-05 | 7.71E-05 | 3.90E-05 | 1.64E-05 | 5.55E-05 | 9.42E-05 | 4.77E-05 | 0 | 7.60E-05 |
| Cox15 | 3.23E-05 | 7.46E-05 | 4.08E-05 | 0.00013 | 2.58E-05 | 2.00E-05 | 1.64E-05 | 3.28E-05 | 7.32E-05 | 5.25E-05 | 2.51E-05 | 1.60E-05 | 1.15E-05 | 1.16E-05 | 3.19E-05 | 7.27E-05 | 8.31E-05 | 6.97E-05 |
| Rbpms | 0.00028 | 0.00028 | 8.79E-05 | 6.64E-06 | 0 | 0.00018 | 0 | 0.0001 | 7.31E-05 | 0.00026 | 0.00032 | 0.00021 | 0.00046 | 7.64E-05 | 0.00017 | 9.26E-05 | 0.00019 | 0 |
| Klhl11 | 9.37E-05 | 0.00015 | 7.09E-05 | 8.10E-07 | 5.53E-05 | 6.97E-05 | 0 | 1.59E-06 | 7.31E-05 | 2.20E-05 | 4.54E-05 | 7.41E-05 | 1.99E-05 | 6.79E-05 | 0 | 0.00012 | 0.00021 | 4.97E-05 |
| Arel1 | 5.55E-05 | 7.56E-05 | 9.26E-05 | 4.21E-05 | 5.76E-05 | 5.79E-05 | 1.63E-05 | 2.96E-05 | 7.31E-05 | 5.20E-05 | 0.0001 | 4.08E-06 | 0.00011 | 4.77E-05 | 1.35E-05 | 0 | 0 | 5.10E-05 |
| Jtb | 0.00027 | 9.11E-05 | 8.49E-05 | 7.26E-05 | 0.00011 | 0.0002 | 0.0003 | 0.0001 | 7.30E-05 | 9.61E-05 | 5.32E-05 | 0 | 9.09E-05 | 8.41E-05 | 0.00014 | 0 | 0 | 9.35E-05 |
| Sec16a | 0.00025 | 0.00012 | 0.00015 | 0.00026 | 0.00014 | 0.00016 | 0.00032 | 0.00014 | 7.30E-05 | 7.59E-05 | 3.53E-05 | 0.00018 | 0.00015 | 0.00013 | 0.00023 | 0 | 0.00019 | 1.26E-06 |
| Smarca5 | 0.00022 | 5.20E-05 | 0.00017 | 0 | 4.36E-05 | 0.00013 | 0 | 5.96E-05 | 7.30E-05 | 0 | 3.22E-05 | 8.56E-05 | 0 | 0 | 0 | 0.00021 | 0.0001 | 1.59E-05 |
| Anks3 | 4.57E-05 | 0.00015 | 6.87E-05 | 6.68E-05 | 7.79E-05 | 5.13E-05 | 0.00011 | 1.35E-05 | 7.30E-05 | 6.26E-05 | 3.56E-05 | 2.59E-05 | 6.72E-05 | 3.00E-05 | 0.0001 | 2.19E-05 | 1.64E-05 | 0.00012 |
| Ice1 | 0.00013 | 0.00011 | 0.00016 | 2.38E-06 | 4.89E-05 | 4.69E-05 | 1.78E-05 | 8.80E-05 | 7.30E-05 | 3.52E-05 | 0.00028 | 5.20E-05 | 3.96E-05 | 0.00019 | 0.00035 | 7.90E-05 | 4.84E-06 | 0 |
| Usp46 | 7.39E-05 | 3.90E-05 | 4.72E-05 | 6.19E-05 | 8.64E-05 | 3.58E-05 | 4.37E-05 | 0 | 7.30E-05 | 1.45E-05 | 7.78E-05 | 0 | 0 | 6.40E-07 | 0 | 4.81E-05 | 3.14E-05 | 6.46E-05 |
| Arhgap29 | 0.00013 | 5.06E-06 | 3.52E-05 | 0 | 0 | 0 | 0.0002 | 0 | 7.30E-05 | 0.00045 | 0.00011 | 0.00029 | 0.00059 | 0.00028 | 0 | 8.60E-06 | 0 | 0.00024 |
| Rab29 | 4.87E-05 | 9.87E-06 | 0 | 1.99E-05 | 5.84E-05 | 0 | 7.28E-05 | 0 | 7.30E-05 | 0.0001 | 5.41E-05 | 0 | 0.00032 | 5.82E-05 | 7.10E-06 | 0 | 0 | 0.0002 |
| Fgf10 | 0 | 0 | 0 | 0 | 0 | 0.00012 | 0 | 0 | 7.29E-05 | 3.19E-05 | 0 | 0 | 0.00012 | 0 | 0 | 0 | 0 | 0.00019 |
| Ndufs6 | 0.00024 | 2.67E-05 | 8.11E-05 | 5.75E-05 | 5.38E-05 | 8.85E-05 | 4.01E-05 | 0 | 7.29E-05 | 1.16E-05 | 3.94E-05 | 0 | 2.74E-05 | 3.34E-05 | 0 | 4.87E-05 | 0 | 0 |
| Pcnx3 | 1.49E-05 | 0.00016 | 9.44E-05 | 0.00021 | 7.77E-05 | 5.05E-05 | 8.04E-05 | 2.79E-05 | 7.29E-05 | 6.67E-05 | 3.45E-05 | 0 | 3.51E-05 | 0.00012 | 0.00039 | 4.24E-05 | 2.47E-05 | 8.04E-05 |
| Cd2bp2 | 0.00011 | 0.00021 | 9.74E-05 | 0.00027 | 0.00016 | 7.56E-05 | 1.21E-05 | 0.00014 | 7.28E-05 | 6.27E-05 | 0.00013 | 2.31E-05 | 5.45E-05 | 0.00011 | 0.00037 | 0.00024 | 3.16E-05 | 6.50E-05 |
| Vash2 | 4.71E-05 | 5.43E-05 | 4.98E-05 | 0 | 7.25E-05 | 6.69E-05 | 0.00012 | 0 | 7.28E-05 | 0 | 0.00014 | 0 | 0 | 0.00016 | 0 | 5.34E-05 | 0 | 0 |
| Ttc7b | 3.58E-05 | 8.07E-05 | 0.00012 | 0 | 6.14E-05 | 5.39E-05 | 5.85E-05 | 0.00011 | 7.28E-05 | 7.93E-05 | 9.90E-05 | 7.89E-05 | 5.97E-05 | 5.10E-05 | 6.37E-05 | 8.14E-05 | 7.49E-05 | 2.60E-05 |
| B2m | 0.00062 | 0.0003 | 6.95E-05 | 0 | 3.93E-05 | 0.00037 | 0 | 7.70E-05 | 7.27E-05 | 0.00052 | 3.30E-06 | 0.00048 | 0.00064 | 0.00041 | 0 | 3.35E-05 | 0.00033 | 0.00066 |
| Lpin3 | 0 | 0 | 6.38E-05 | 9.34E-05 | 0.00029 | 5.40E-05 | 0.00031 | 0 | 7.27E-05 | 5.42E-05 | 0 | 7.29E-05 | 4.11E-05 | 2.83E-05 | 0.00016 | 1.70E-05 | 0 | 0 |
| Trub1 | 2.86E-05 | 5.03E-05 | 8.49E-05 | 2.96E-05 | 3.93E-05 | 7.44E-06 | 0 | 0 | 7.27E-05 | 1.39E-05 | 8.39E-06 | 0 | 2.15E-06 | 0 | 0.00018 | 2.38E-05 | 0 | 3.77E-05 |

| | | | | | | | | | | | | | | | | | | |
|---|---|---|---|---|---|---|---|---|---|---|---|---|---|---|---|---|---|---|
| Mul1 | 6.23E-05 | 9.77E-05 | 4.74E-05 | 7.06E-05 | 5.01E-05 | 5.04E-05 | 3.44E-05 | 1.58E-05 | 7.26E-05 | 1.94E-05 | 1.15E-05 | 5.22E-05 | 6.50E-06 | 4.83E-05 | 5.62E-05 | 4.49E-05 | 0 | 3.46E-05 |
| Pih1d1 | 0.00011 | 9.01E-05 | 4.07E-05 | 0.00011 | 1.88E-05 | 3.27E-05 | 6.25E-05 | 6.01E-05 | 7.26E-05 | 5.72E-06 | 1.85E-05 | 8.71E-05 | 2.68E-07 | 0 | 0 | 0.00028 | 0 | 7.69E-05 |
| Eef1akmt1 | 0.00018 | 1.48E-05 | 0.00011 | 0.00011 | 0 | 0 | 5.55E-05 | 0 | 7.26E-05 | 1.29E-05 | 4.07E-05 | 0 | 0 | 1.37E-05 | 0 | 8.36E-05 | 0 | 2.26E-05 |
| Shkbp1 | 5.62E-05 | 4.86E-05 | 3.88E-05 | 0.00012 | 3.34E-05 | 0 | 3.13E-07 | 3.10E-05 | 7.26E-05 | 2.01E-05 | 0.0001 | 0 | 0 | 1.79E-05 | 7.15E-05 | 9.36E-05 | 0 | 1.80E-05 |
| Smpd1 | 0.00012 | 9.11E-05 | 2.69E-06 | 0.00019 | 0.00021 | 0.00018 | 9.53E-05 | 5.71E-05 | 7.25E-05 | 0.00016 | 0.00013 | 9.47E-05 | 0.00014 | 0.00014 | 0 | 3.15E-05 | 2.43E-05 | 0.00025 |
| Atxn7 | 4.13E-05 | 8.80E-05 | 8.12E-05 | 0 | 4.86E-05 | 5.49E-05 | 5.38E-05 | 7.37E-05 | 7.25E-05 | 3.24E-05 | 8.39E-05 | 5.08E-05 | 7.04E-05 | 0.00011 | 0.00014 | 2.10E-05 | 0.00012 | 2.15E-05 |
| Sgta | 8.25E-05 | 9.53E-05 | 6.36E-05 | 7.82E-05 | 1.91E-05 | 3.83E-06 | 0 | 2.95E-05 | 7.25E-05 | 3.18E-05 | 3.25E-05 | 4.97E-05 | 0 | 3.95E-05 | 4.70E-05 | 1.30E-05 | 3.57E-05 | 2.18E-05 |
| Bmpr1a | 0.00037 | 0.0002 | 9.85E-05 | 0 | 0.0002 | 0.00035 | 0.00017 | 0.00016 | 7.24E-05 | 5.98E-05 | 0.00019 | 0.00014 | 0.00015 | 2.40E-05 | 5.05E-05 | 0.00011 | 0 | 0.00019 |
| Zfp865 | 6.47E-05 | 0.00015 | 7.25E-05 | 9.11E-05 | 7.98E-05 | 6.74E-05 | 7.30E-05 | 6.53E-05 | 7.24E-05 | 4.71E-05 | 2.29E-05 | 2.54E-05 | 2.12E-06 | 3.27E-05 | 0.00023 | 3.86E-05 | 0 | 6.04E-05 |
| Polg | 3.17E-05 | 8.50E-05 | 6.65E-05 | 8.17E-05 | 4.89E-05 | 3.71E-05 | 4.14E-05 | 7.09E-05 | 7.24E-05 | 4.34E-05 | 5.58E-05 | 4.21E-05 | 0 | 8.65E-05 | 0.00015 | 0.00013 | 2.70E-05 | 6.68E-05 |
| Thop1 | 5.64E-05 | 4.29E-05 | 0.00029 | 0.0002 | 1.28E-05 | 0 | 0 | 2.04E-05 | 7.24E-05 | 5.54E-05 | 8.62E-07 | 0 | 2.19E-06 | 4.35E-05 | 2.59E-05 | 0.00017 | 6.87E-05 | 2.24E-05 |
| Nectin1 | 0 | 0 | 0.00036 | 8.61E-06 | 0.00028 | 7.65E-05 | 0.00025 | 0.00044 | 7.23E-05 | 8.65E-05 | 0.00028 | 0.00049 | 0 | 0 | 0.00029 | 5.27E-05 | 0 | 5.23E-05 |
| Pheta2 | 4.32E-05 | 2.03E-05 | 2.44E-05 | 0.00014 | 7.76E-05 | 2.16E-05 | 0.00019 | 8.20E-05 | 7.23E-05 | 1.18E-05 | 5.14E-05 | 0 | 5.38E-05 | 0.00011 | 6.75E-05 | 0 | 0 | 8.32E-05 |
| Rer1 | 0.00063 | 8.68E-05 | 0.0003 | 3.82E-05 | 0.00013 | 0.00045 | 0.00039 | 0.00023 | 7.22E-05 | 0.00014 | 0.00022 | 0.00029 | 0.00019 | 0.00015 | 3.53E-05 | 0.00026 | 0 | 0.00011 |
| Fam169b | 0 | 5.01E-07 | 1.47E-06 | 1.49E-05 | 1.63E-05 | 3.53E-05 | 0 | 0 | 7.22E-05 | 0 | 1.37E-05 | 0 | 7.96E-07 | 5.69E-05 | 1.19E-05 | 1.04E-05 | 0 | 0 |
| Hsf1 | 7.15E-05 | 0.0001 | 0.00015 | 3.77E-05 | 5.80E-05 | 2.78E-05 | 3.72E-07 | 4.05E-05 | 7.22E-05 | 4.67E-05 | 6.02E-05 | 0 | 4.73E-06 | 0 | 0.0002 | 6.12E-05 | 0.00013 | 4.95E-05 |
| Sufu | 2.46E-05 | 3.94E-05 | 5.83E-05 | 0 | 3.04E-05 | 1.08E-05 | 0 | 2.02E-05 | 7.22E-05 | 1.82E-05 | 2.60E-05 | 1.09E-05 | 4.48E-05 | 0 | 6.85E-05 | 1.71E-05 | 2.58E-05 | 1.98E-05 |
| Dhodh | 4.78E-05 | 4.22E-05 | 6.98E-05 | 6.05E-05 | 1.52E-05 | 0 | 0 | 0 | 7.22E-05 | 1.81E-05 | 1.14E-05 | 1.30E-07 | 2.08E-05 | 1.34E-06 | 0 | 7.30E-05 | 0 | 0 |
| Notch1 | 0 | 1.12E-05 | 0.00025 | 0 | 5.02E-05 | 0.00028 | 0 | 0.00088 | 7.21E-05 | 0 | 0 | 6.05E-05 | 0 | 0.00025 | 0.00036 | 0.00011 | 0 | 0 |
| Iigp1 | 3.92E-05 | 0 | 0 | 0 | 0 | 0 | 0 | 0 | 7.21E-05 | 0.00026 | 0 | 0 | 0.00026 | 0 | 0 | 0 | 0 | 0.0002 |
| Eif5b | 0.00023 | 9.83E-05 | 0.00018 | 1.84E-05 | 2.29E-05 | 8.64E-05 | 1.85E-05 | 0 | 7.20E-05 | 2.60E-05 | 6.88E-05 | 3.60E-05 | 1.64E-05 | 2.42E-05 | 0.00012 | 4.82E-05 | 0 | 3.66E-05 |
| Tradd | 9.48E-05 | 9.08E-05 | 7.07E-05 | 0.00014 | 4.92E-05 | 6.20E-05 | 0.00012 | 3.06E-05 | 7.20E-05 | 7.78E-05 | 5.42E-05 | 8.28E-05 | 0 | 3.76E-05 | 7.71E-06 | 1.15E-05 | 0 | 0.00013 |
| Gamt | 7.30E-05 | 2.78E-05 | 0 | 7.31E-05 | 0 | 4.80E-05 | 0 | 0.00026 | 7.20E-05 | 5.86E-05 | 1.03E-05 | 0 | 0 | 0 | 0 | 0 | 0 | 4.32E-05 |
| Zc3h18 | 8.48E-05 | 0.00011 | 0.00017 | 0.00013 | 3.66E-05 | 3.01E-05 | 2.55E-05 | 9.06E-07 | 7.20E-05 | 2.09E-05 | 3.00E-05 | 3.69E-05 | 1.03E-05 | 3.41E-05 | 0.00019 | 0.0002 | 0.00013 | 2.61E-05 |
| Dcun1d1 | 9.57E-05 | 6.55E-05 | 7.62E-05 | 5.74E-06 | 3.05E-05 | 1.62E-05 | 2.57E-05 | 2.22E-05 | 7.20E-05 | 2.81E-05 | 5.40E-05 | 2.66E-05 | 7.26E-06 | 2.03E-05 | 0 | 4.55E-05 | 2.27E-05 | 8.08E-06 |
| Stard5 | 0 | 7.19E-05 | 0 | 2.42E-05 | 0 | 3.42E-05 | 0 | 1.18E-05 | 7.19E-05 | 5.68E-05 | 0 | 0 | 5.56E-05 | 0.00025 | 0 | 5.57E-05 | 0 | 5.56E-05 |
| Uck1 | 7.76E-05 | 0.00013 | 6.81E-05 | 0 | 0.00012 | 2.47E-05 | 3.57E-05 | 4.18E-05 | 7.19E-05 | 6.03E-05 | 3.17E-05 | 6.93E-05 | 1.07E-05 | 4.13E-06 | 0 | 9.14E-05 | 0 | 6.45E-05 |
| Cds2 | 0.00012 | 0.00022 | 1.37E-08 | 0 | 3.73E-05 | 0.00018 | 0 | 0 | 7.18E-05 | 0.00012 | 0.00011 | 0.00023 | 0.00015 | 9.23E-05 | 9.65E-05 | 4.13E-05 | 9.09E-05 | 2.93E-05 |
| Sgsh | 9.75E-06 | 9.82E-05 | 0 | 3.33E-05 | 0.00012 | 0.00011 | 0 | 3.55E-05 | 7.18E-05 | 9.80E-05 | 0 | 0 | 4.08E-05 | 8.31E-05 | 0 | 7.46E-06 | 0 | 5.59E-05 |
| Hdac5 | 5.17E-07 | 0.0003 | 0 | 0.0002 | 0.00016 | 0.00034 | 4.16E-05 | 0 | 7.17E-05 | 6.03E-05 | 5.55E-05 | 0.00049 | 0.00022 | 0.00018 | 0.00014 | 0 | 0.00017 | 7.65E-06 |
| Sirt2 | 8.71E-05 | 0.0003 | 2.05E-05 | 0.0003 | 0.00026 | 0.00024 | 0.00023 | 0.00016 | 7.17E-05 | 0.00012 | 8.84E-05 | 0.00017 | 0.00011 | 0.00027 | 5.39E-05 | 1.40E-05 | 0.00013 | 0.00018 |
| Nbr1 | 0.00017 | 0.00017 | 0 | 0 | 9.59E-05 | 0.00013 | 4.94E-05 | 8.71E-05 | 7.17E-05 | 3.03E-05 | 0 | 8.05E-05 | 4.26E-05 | 2.79E-05 | 0.00017 | 3.23E-05 | 0 | 3.05E-05 |
| Prdm5 | 6.30E-05 | 4.96E-05 | 3.84E-05 | 4.66E-05 | 1.91E-05 | 7.20E-05 | 3.49E-05 | 3.52E-05 | 7.17E-05 | 1.74E-05 | 0 | 8.14E-06 | 8.73E-05 | 5.98E-05 | 0.00023 | 0 | 0 | 7.90E-05 |

| | | | | | | | | | | | | | | | | | | |
|---|---|---|---|---|---|---|---|---|---|---|---|---|---|---|---|---|---|---|
| Spire1 | 4.76E-05 | 6.14E-05 | 0.00013 | 1.08E-05 | 3.39E-05 | 5.66E-05 | 1.47E-05 | 1.70E-05 | 7.17E-05 | 8.92E-06 | 9.64E-05 | 0.00012 | 5.31E-05 | 4.18E-05 | 0 | 1.00E-05 | 0 | 8.48E-08 |
| Slc25a12 | 6.53E-05 | 7.36E-05 | 8.50E-05 | 8.45E-05 | 3.92E-05 | 5.49E-05 | 7.36E-05 | 2.48E-05 | 7.16E-05 | 0.00018 | 0 | 4.94E-06 | 1.97E-05 | 5.28E-05 | 7.15E-06 | 6.10E-05 | 3.29E-05 | 9.18E-05 |
| DEPRECATEI | 3.73E-05 | 6.83E-05 | 7.51E-05 | 2.62E-05 | 4.44E-05 | 0 | 3.75E-05 | 0.00011 | 7.16E-05 | 2.43E-05 | 0 | 5.41E-05 | 3.13E-06 | 8.89E-05 | 3.57E-05 | 2.78E-05 | 0.00017 | 1.10E-05 |
| Dctn6 | 0.00029 | 0.00011 | 0.00011 | 0 | 0.00011 | 0.00019 | 7.99E-05 | 6.96E-06 | 7.15E-05 | 7.65E-05 | 0.00024 | 9.04E-05 | 3.67E-05 | 4.87E-05 | 0 | 2.99E-05 | 0 | 6.47E-05 |
| Ppp6r2 | 9.08E-05 | 8.80E-05 | 2.30E-05 | 1.69E-05 | 8.66E-05 | 6.42E-05 | 6.81E-05 | 7.64E-05 | 7.15E-05 | 6.73E-05 | 9.27E-06 | 3.93E-05 | 0 | 8.86E-05 | 0.0002 | 4.51E-05 | 3.45E-05 | 7.20E-05 |
| Otud6b | 0.00014 | 3.88E-05 | 0.00014 | 0 | 1.12E-05 | 1.73E-05 | 0 | 0 | 7.15E-05 | 3.60E-05 | 2.08E-05 | 3.72E-05 | 0 | 3.21E-05 | 0 | 0.00011 | 0 | 3.16E-05 |
| Tmem245 | 4.19E-05 | 9.23E-05 | 9.20E-05 | 1.73E-05 | 4.23E-05 | 1.71E-05 | 1.80E-06 | 7.16E-05 | 7.15E-05 | 2.44E-05 | 9.09E-05 | 4.95E-05 | 0 | 4.51E-05 | 8.50E-05 | 0.00012 | 6.21E-05 | 7.53E-05 |
| Nr2c2ap | 6.45E-05 | 6.41E-05 | 0.00011 | 3.72E-05 | 2.96E-05 | 8.68E-06 | 0 | 0 | 7.15E-05 | 2.75E-05 | 3.01E-05 | 1.02E-05 | 0 | 2.97E-05 | 2.32E-05 | 0 | 0 | 1.22E-05 |
| Capn10 | 2.99E-05 | 5.86E-05 | 4.10E-05 | 0.00014 | 2.80E-05 | 0 | 3.42E-05 | 3.31E-05 | 7.15E-05 | 3.92E-05 | 2.81E-05 | 3.11E-06 | 0 | 5.62E-05 | 0.00017 | 3.18E-05 | 0 | 3.54E-05 |
| Pex6 | 5.42E-05 | 0.00012 | 7.53E-05 | 0.00014 | 8.10E-05 | 6.63E-05 | 4.39E-05 | 0.0001 | 7.14E-05 | 6.19E-05 | 2.58E-06 | 8.15E-05 | 0 | 3.18E-05 | 0.00014 | 0 | 6.34E-06 | 6.55E-05 |
| Chmp2a | 6.74E-05 | 6.02E-05 | 7.93E-05 | 0.00016 | 3.76E-05 | 5.62E-05 | 3.54E-05 | 4.01E-05 | 7.14E-05 | 7.37E-05 | 0.00014 | 9.34E-05 | 2.62E-05 | 4.36E-05 | 0 | 3.06E-06 | 0 | 6.99E-05 |
| Cog2 | 0.0001 | 8.22E-05 | 7.54E-05 | 0.00011 | 4.60E-05 | 4.70E-05 | 0.00011 | 1.83E-05 | 7.14E-05 | 2.97E-05 | 5.61E-05 | 2.27E-05 | 6.87E-06 | 7.86E-05 | 0.0001 | 0 | 0 | 2.28E-05 |
| Capn5 | 0 | 6.59E-05 | 0 | 9.61E-05 | 0 | 3.83E-06 | 1.58E-05 | 2.90E-05 | 7.14E-05 | 0.00019 | 0 | 8.91E-05 | 6.77E-05 | 0 | 0 | 0 | 6.91E-05 | 0.00015 |
| Hnrnph3 | 9.46E-05 | 9.43E-05 | 3.54E-05 | 0.00011 | 5.57E-05 | 0 | 0 | 0 | 7.14E-05 | 2.09E-05 | 7.62E-05 | 6.89E-05 | 0 | 9.02E-06 | 0.00026 | 0.00013 | 9.68E-05 | 4.97E-05 |
| Mov10 | 3.69E-05 | 4.89E-05 | 0 | 0.00016 | 2.37E-06 | 0 | 0 | 7.11E-05 | 7.14E-05 | 6.44E-05 | 1.01E-05 | 0 | 4.45E-05 | 5.47E-06 | 9.89E-05 | 5.99E-05 | 0 | 5.13E-05 |
| Hdac6 | 1.70E-05 | 5.16E-05 | 0 | 5.86E-05 | 4.66E-05 | 0 | 1.88E-05 | 2.16E-05 | 7.14E-05 | 1.29E-05 | 0 | 0 | 6.86E-05 | 2.20E-05 | 7.85E-05 | 6.42E-05 | 5.63E-05 | 1.76E-05 |
| Cpsf7 | 0.00016 | 0.00016 | 0.00012 | 7.37E-05 | 7.37E-05 | 7.27E-05 | 2.26E-05 | 0 | 7.13E-05 | 2.94E-05 | 3.78E-05 | 7.61E-05 | 0 | 5.46E-05 | 0.00019 | 0.00012 | 3.54E-05 | 0 |
| Rbm6 | 9.96E-05 | 0.00021 | 9.06E-05 | 9.85E-05 | 5.33E-05 | 5.57E-05 | 4.44E-05 | 1.92E-05 | 7.13E-05 | 5.96E-05 | 7.30E-05 | 7.09E-05 | 3.77E-05 | 6.98E-05 | 0.0004 | 0.00013 | 3.76E-05 | 6.01E-05 |
| Dnajc9 | 2.33E-05 | 1.21E-05 | 3.73E-05 | 0.00011 | 0 | 2.98E-05 | 4.12E-05 | 0 | 7.13E-05 | 3.93E-05 | 0 | 0 | 3.20E-06 | 8.07E-05 | 0 | 0.00096 | 0 | 7.19E-05 |
| Fbxl19 | 3.70E-06 | 6.68E-05 | 5.16E-05 | 0.00017 | 8.16E-06 | 0 | 5.55E-06 | 7.36E-05 | 7.13E-05 | 2.27E-05 | 4.77E-05 | 5.03E-05 | 0 | 2.05E-05 | 0.00022 | 1.23E-05 | 5.70E-05 | 6.48E-05 |
| Arpc3 | 0.00036 | 0.00014 | 0.0002 | 0 | 0.00011 | 0.00022 | 0.00011 | 0.0001 | 7.12E-05 | 6.83E-05 | 0.00012 | 0.00013 | 0.00012 | 8.22E-05 | 0 | 1.64E-05 | 3.04E-05 | 7.89E-05 |
| Gmppa | 0.00025 | 6.88E-05 | 8.51E-05 | 0.00016 | 6.42E-05 | 8.02E-05 | 0.00032 | 0.00011 | 7.12E-05 | 9.19E-05 | 0 | 9.21E-05 | 5.69E-06 | 0 | 0.00011 | 3.74E-05 | 1.16E-05 | 7.59E-05 |
| Pnkd | 9.52E-05 | 7.56E-05 | 7.45E-06 | 6.14E-05 | 4.40E-05 | 5.16E-05 | 1.64E-05 | 5.22E-06 | 7.12E-05 | 0 | 2.57E-05 | 1.75E-05 | 0 | 5.68E-05 | 3.09E-06 | 1.29E-05 | 4.78E-05 | 2.62E-05 |
| Nom1 | 9.52E-05 | 7.64E-05 | 0.00011 | 4.06E-05 | 3.19E-05 | 2.30E-05 | 0 | 0 | 7.12E-05 | 2.25E-05 | 3.05E-05 | 2.52E-05 | 4.99E-05 | 2.10E-05 | 0.0001 | 6.18E-05 | 7.80E-06 | 8.00E-06 |
| Dyrk2 | 0 | 0.00018 | 0.0002 | 0.00011 | 0.00013 | 0 | 8.97E-05 | 1.20E-05 | 7.12E-05 | 0.00013 | 0.00035 | 0.00014 | 4.68E-05 | 3.18E-05 | 9.24E-05 | 6.33E-05 | 0.00014 | 6.97E-05 |
| Clint1 | 0.00042 | 0.00018 | 0.00029 | 8.61E-05 | 6.35E-05 | 0.00012 | 0.00037 | 0.00023 | 7.11E-05 | 0.0001 | 0.00015 | 0.00014 | 0.00012 | 4.41E-05 | 0.00022 | 6.52E-05 | 0.00017 | 0 |
| Fam210a | 0.00015 | 5.77E-05 | 0.00011 | 0 | 1.50E-05 | 3.48E-05 | 5.69E-06 | 0 | 7.11E-05 | 2.48E-05 | 0 | 0.00018 | 2.08E-05 | 4.90E-05 | 5.89E-05 | 6.26E-05 | 0 | 0 |
| Spata2 | 2.20E-06 | 6.40E-05 | 0.00011 | 5.88E-05 | 1.76E-05 | 1.57E-05 | 3.96E-05 | 4.83E-05 | 7.11E-05 | 3.83E-05 | 4.83E-05 | 0 | 0 | 6.13E-05 | 4.86E-05 | 1.90E-05 | 6.82E-05 | 4.14E-05 |
| Zc3h4 | 0.00012 | 0.00015 | 7.92E-05 | 0.00024 | 5.82E-05 | 5.57E-05 | 0.00011 | 7.05E-05 | 7.10E-05 | 5.25E-05 | 7.58E-05 | 5.79E-05 | 0 | 6.31E-05 | 8.19E-05 | 0.00027 | 0.00022 | 5.74E-05 |
| Thap4 | 0.00012 | 0.00011 | 0.00012 | 1.98E-05 | 1.34E-05 | 5.06E-05 | 6.32E-06 | 0 | 7.10E-05 | 1.66E-05 | 4.14E-05 | 9.06E-05 | 0 | 1.80E-05 | 3.54E-05 | 3.74E-05 | 5.35E-05 | 4.11E-05 |
| Wiz | 5.10E-05 | 0.00012 | 0.00011 | 3.15E-05 | 5.97E-05 | 3.29E-05 | 0 | 3.43E-05 | 7.10E-05 | 3.34E-05 | 4.51E-05 | 7.85E-05 | 2.04E-06 | 2.38E-05 | 0.00011 | 4.25E-05 | 0 | 3.07E-05 |
| Atg2a | 4.14E-06 | 0.00018 | 5.33E-05 | 0.00012 | 4.39E-05 | 1.07E-05 | 0 | 2.78E-05 | 7.10E-05 | 0.00011 | 7.34E-05 | 6.08E-05 | 0 | 9.03E-05 | 0.00024 | 2.29E-06 | 0.0002 | 6.63E-05 |
| C2 | 0 | 5.08E-05 | 0 | 0 | 0.00015 | 0 | 0.00019 | 0 | 7.10E-05 | 0.00021 | 6.38E-05 | 0 | 0.00103 | 0 | 0 | 0 | 0 | 0.00081 |

| | | | | | | | | | | | | | | | | | | |
|---|---|---|---|---|---|---|---|---|---|---|---|---|---|---|---|---|---|---|
| Il1rl2 | 0.00026 | 0.00011 | 6.22E-05 | 0 | 0.00013 | 0.0003 | 0.0001 | 8.04E-05 | 7.09E-05 | 0.00028 | 0 | 0 | 0.00024 | 7.23E-05 | 1.43E-05 | 1.53E-05 | 8.19E-05 | 0 |
| Ak2 | 0.0002 | 0.00021 | 0.0005 | 0.00069 | 0.00036 | 0.00022 | 9.35E-05 | 0.00014 | 7.09E-05 | 0 | 0.00043 | 0.00018 | 0.00026 | 0.00025 | 0.00022 | 0.00042 | 0.00018 | 0 |
| Alkbh3 | 3.21E-05 | 5.07E-05 | 6.37E-05 | 0.00011 | 1.30E-05 | 5.53E-06 | 1.26E-05 | 3.16E-05 | 7.09E-05 | 3.47E-05 | 4.52E-05 | 3.37E-05 | 0 | 1.62E-05 | 1.98E-05 | 2.98E-05 | 5.59E-05 | 7.94E-05 |
| Smc3 | 0.00018 | 7.58E-05 | 7.35E-05 | 0 | 5.08E-05 | 8.09E-05 | 3.87E-05 | 0 | 7.08E-05 | 2.59E-05 | 0 | 9.01E-05 | 2.64E-05 | 4.64E-05 | 0.00023 | 0.00034 | 9.86E-06 | 7.35E-05 |
| Arl5c | 6.71E-07 | 2.04E-05 | 0 | 0 | 5.01E-05 | 3.27E-05 | 0 | 4.77E-05 | 7.08E-05 | 2.69E-05 | 1.87E-05 | 0.00011 | 0 | 0 | 0 | 0 | 0 | 4.14E-06 |
| Aqp5 | 0 | 0 | 0 | 5.58E-05 | 0.00045 | 0.00018 | 0 | 0 | 7.07E-05 | 0 | 0.00015 | 0 | 0 | 0.0003 | 2.61E-05 | 0 | 0 | 1.92E-05 |
| Tnfaip8 | 4.28E-05 | 6.11E-05 | 0 | 0 | 1.01E-05 | 8.51E-05 | 0 | 0 | 7.07E-05 | 0 | 0 | 5.65E-05 | 0 | 0 | 1.04E-05 | 6.89E-05 | 1.37E-05 | 0.0001 |
| Il1r2 | 2.80E-05 | 0 | 0 | 1.66E-05 | 5.60E-05 | 4.53E-05 | 0 | 2.81E-05 | 7.07E-05 | 0.00028 | 0 | 0 | 0.00011 | 3.75E-05 | 2.80E-05 | 6.70E-05 | 6.44E-06 | 0 |
| Pgpep1l | 0 | 1.31E-05 | 5.33E-06 | 0 | 2.93E-05 | 2.25E-05 | 0 | 8.09E-06 | 7.07E-05 | 0 | 0 | 0 | 1.03E-06 | 1.61E-05 | 1.18E-05 | 0 | 0 | 0 |
| Trafd1 | 7.52E-05 | 0.00016 | 0 | 0.00017 | 6.69E-05 | 3.19E-05 | 8.68E-05 | 0 | 7.06E-05 | 0.00019 | 3.05E-05 | 2.75E-05 | 5.43E-05 | 0.0001 | 0.00014 | 0.0001 | 8.39E-05 | 0.00019 |
| Traf3 | 6.60E-05 | 3.65E-05 | 0.00016 | 3.15E-05 | 3.97E-05 | 1.86E-05 | 4.76E-05 | 0 | 7.06E-05 | 3.21E-05 | 1.23E-05 | 1.99E-05 | 7.66E-05 | 2.18E-05 | 6.61E-05 | 5.26E-05 | 0.00012 | 4.13E-05 |
| Arhgap31 | 3.85E-05 | 0.00012 | 0.00023 | 0 | 0.0003 | 0.00021 | 9.29E-05 | 0.00021 | 7.05E-05 | 3.57E-05 | 0.00039 | 0.00011 | 0 | 0.00014 | 0 | 9.85E-05 | 0.00012 | 0 |
| Rcbtb1 | 0.00015 | 0.00011 | 6.48E-05 | 3.53E-05 | 5.53E-05 | 8.49E-05 | 0.00018 | 0 | 7.05E-05 | 1.65E-05 | 5.25E-05 | 1.96E-05 | 5.58E-05 | 9.88E-05 | 7.30E-05 | 0.00015 | 0 | 4.99E-05 |
| Cetn2 | 0.00012 | 3.17E-05 | 1.02E-05 | 7.44E-05 | 3.88E-05 | 3.33E-05 | 8.18E-06 | 1.74E-05 | 7.05E-05 | 1.74E-05 | 1.62E-05 | 3.10E-06 | 0 | 8.43E-06 | 0 | 7.49E-06 | 4.53E-05 | 5.95E-05 |
| Polr2d | 0.0001 | 2.71E-05 | 8.39E-05 | 6.08E-05 | 4.50E-05 | 7.81E-06 | 2.31E-05 | 4.57E-05 | 7.05E-05 | 4.63E-05 | 3.00E-05 | 0.00017 | 0 | 1.41E-05 | 3.67E-05 | 9.55E-05 | 0 | 2.25E-05 |
| Fam98c | 0 | 9.50E-05 | 5.94E-05 | 8.99E-05 | 1.64E-05 | 3.77E-05 | 2.12E-06 | 0.00013 | 7.04E-05 | 6.24E-05 | 1.90E-05 | 0.00015 | 0 | 7.96E-05 | 0.00013 | 0 | 0.0001 | 8.82E-05 |
| C1qbp | 0.00018 | 2.08E-05 | 0.00021 | 2.13E-05 | 0 | 3.70E-05 | 0 | 0 | 7.04E-05 | 0 | 5.14E-07 | 1.06E-05 | 1.76E-05 | 0 | 0 | 5.88E-05 | 4.94E-05 | 0 |
| Slc26a2 | 0.00015 | 1.18E-05 | 6.58E-05 | 0 | 1.57E-05 | 3.55E-05 | 0.00013 | 0.00014 | 7.04E-05 | 1.33E-05 | 3.64E-05 | 3.78E-05 | 0 | 0 | 2.78E-05 | 9.10E-06 | 6.40E-05 | 0 |
| Prpf38a | 0.00015 | 8.12E-05 | 0.00011 | 7.49E-05 | 2.45E-05 | 1.77E-05 | 0 | 2.68E-05 | 7.04E-05 | 3.59E-05 | 3.65E-05 | 2.61E-05 | 0 | 1.65E-05 | 2.74E-05 | 0.00019 | 0 | 1.19E-05 |
| Amd1 | 9.63E-05 | 2.89E-05 | 0.00016 | 6.29E-06 | 7.42E-07 | 1.68E-05 | 0 | 0 | 7.04E-05 | 0 | 3.33E-05 | 3.90E-06 | 1.44E-05 | 5.48E-06 | 0 | 0.00011 | 4.38E-05 | 1.10E-05 |
| Vangl2 | 6.78E-06 | 9.57E-05 | 1.95E-05 | 0.00012 | 0.00012 | 0 | 0 | 0.00011 | 7.04E-05 | 2.69E-05 | 5.37E-05 | 0 | 8.32E-06 | 9.77E-05 | 1.85E-05 | 9.89E-05 | 1.10E-05 | 7.48E-05 |
| Pym1 | 4.96E-05 | 7.30E-05 | 6.87E-05 | 8.39E-05 | 0 | 0 | 0 | 7.98E-06 | 7.04E-05 | 1.51E-05 | 3.44E-05 | 1.02E-05 | 0 | 4.16E-06 | 9.96E-05 | 8.51E-05 | 0 | 0 |
| Cept1 | 0.00035 | 0.00011 | 0.0001 | 0 | 0.00017 | 0.00055 | 7.03E-05 | 0.00033 | 7.03E-05 | 2.30E-05 | 8.42E-05 | 0.0001 | 4.13E-05 | 0.00011 | 0.0002 | 1.82E-05 | 0 | 0.0001 |
| Emc3 | 0.00037 | 0.00012 | 0.00012 | 0 | 0.00013 | 0.00021 | 0.00014 | 1.63E-05 | 7.03E-05 | 3.56E-05 | 6.16E-05 | 0.00014 | 4.61E-05 | 3.38E-05 | 0 | 0.00016 | 0 | 3.94E-05 |
| Tjap1 | 3.14E-05 | 0.00016 | 0.00012 | 0.00016 | 7.75E-05 | 9.68E-05 | 3.92E-05 | 0 | 7.03E-05 | 8.40E-05 | 0.00038 | 7.43E-05 | 3.94E-05 | 0.00022 | 0.00028 | 0.00014 | 6.30E-05 | 7.56E-05 |
| Bicra | 4.77E-06 | 5.25E-05 | 9.86E-05 | 3.34E-05 | 0.00012 | 9.51E-05 | 5.23E-05 | 0 | 7.03E-05 | 2.97E-05 | 2.99E-05 | 1.99E-05 | 9.92E-05 | 0.0001 | 7.85E-05 | 3.28E-05 | 3.45E-05 | 0 |
| Ankrd11 | 0.0001 | 0.00025 | 0.00017 | 0.00012 | 7.46E-05 | 0.00015 | 7.87E-05 | 0.00012 | 7.02E-05 | 6.50E-05 | 5.38E-05 | 0.00019 | 8.82E-05 | 0.00017 | 0.00053 | 0.00013 | 0.00033 | 5.42E-05 |
| Sgcb | 0.00015 | 6.48E-05 | 6.26E-05 | 1.47E-05 | 8.68E-05 | 9.42E-05 | 0.00012 | 4.86E-05 | 7.02E-05 | 3.95E-05 | 5.20E-05 | 2.70E-05 | 0.00026 | 5.79E-05 | 6.31E-05 | 3.89E-05 | 0 | 0.00017 |
| Dhdds | 0.00015 | 6.13E-05 | 8.53E-05 | 2.04E-05 | 5.43E-05 | 7.96E-05 | 9.91E-05 | 0 | 7.02E-05 | 6.92E-05 | 3.11E-05 | 5.70E-05 | 6.55E-05 | 4.39E-05 | 0.00012 | 0.00013 | 0 | 3.17E-05 |
| Nol10 | 6.37E-05 | 3.82E-05 | 0.00014 | 0 | 3.86E-05 | 0 | 0 | 0 | 7.02E-05 | 1.26E-05 | 2.77E-05 | 0 | 2.06E-05 | 0 | 9.35E-05 | 5.55E-05 | 0 | 1.39E-05 |
| Grwd1 | 7.75E-06 | 2.83E-05 | 0.00022 | 0.00011 | 0 | 0 | 0 | 0 | 7.02E-05 | 4.10E-05 | 2.70E-05 | 0 | 1.38E-06 | 1.82E-05 | 1.77E-05 | 7.68E-05 | 0 | 1.21E-05 |
| Cox7a2l | 0.00021 | 0.00039 | 2.27E-05 | 0.00021 | 0.00017 | 0.00031 | 0.00013 | 0.0002 | 7.01E-05 | 0.00011 | 0 | 0.0004 | 0.00014 | 0.00014 | 0 | 8.43E-05 | 0 | 9.05E-05 |
| Crtc2 | 2.98E-05 | 0.00017 | 0.0001 | 0.00013 | 3.78E-05 | 4.48E-05 | 0 | 8.88E-05 | 7.01E-05 | 3.57E-05 | 5.58E-06 | 8.95E-05 | 0 | 8.50E-05 | 0.00015 | 7.64E-05 | 0.00022 | 4.75E-05 |

| | | | | | | | | | | | | | | | | | | |
|---|---|---|---|---|---|---|---|---|---|---|---|---|---|---|---|---|---|---|
| Zcchc2 | 4.99E-05 | 8.44E-05 | 4.54E-05 | 0 | 4.54E-05 | 1.74E-05 | 0 | 0 | 7.01E-05 | 1.66E-05 | 0 | 4.05E-05 | 0 | 2.74E-05 | 5.58E-05 | 2.59E-05 | 4.13E-05 | 1.56E-05 |
| Dnttip2 | 0.00018 | 9.82E-05 | 0.00017 | 8.86E-05 | 3.11E-05 | 1.71E-05 | 4.26E-05 | 0 | 7.01E-05 | 0 | 1.39E-05 | 3.91E-05 | 5.53E-06 | 1.88E-05 | 4.24E-05 | 9.38E-05 | 0 | 0 |
| Traf2 | 4.30E-05 | 0.00013 | 0.00014 | 9.60E-05 | 3.36E-05 | 6.54E-06 | 9.30E-05 | 0 | 7.01E-05 | 4.97E-05 | 0 | 3.93E-05 | 5.49E-05 | 6.29E-05 | 6.14E-05 | 0.00014 | 0 | 0.00012 |
| Dkc1 | 0.00012 | 0 | 0.0002 | 3.95E-06 | 0 | 0 | 0 | 0 | 7.01E-05 | 2.81E-05 | 0 | 0 | 2.61E-05 | 0 | 0 | 0.00017 | 0 | 9.03E-06 |
| Trim21 | 3.09E-05 | 4.11E-05 | 0 | 2.07E-05 | 1.27E-05 | 0 | 0 | 2.56E-05 | 7.01E-05 | 7.81E-05 | 0 | 0 | 4.20E-05 | 0 | 2.68E-05 | 0 | 0 | 5.11E-05 |
| Snrpb2 | 0.00023 | 8.92E-05 | 0.00014 | 4.25E-05 | 7.68E-06 | 8.71E-05 | 3.17E-08 | 0 | 7.00E-05 | 0 | 6.15E-05 | 4.00E-05 | 1.51E-05 | 7.15E-05 | 0 | 0.00012 | 8.23E-05 | 0 |
| Scaf4 | 0.00013 | 0.0001 | 0.00012 | 0 | 5.66E-05 | 7.02E-05 | 4.28E-05 | 0 | 7.00E-05 | 2.09E-05 | 3.72E-06 | 3.91E-05 | 5.72E-06 | 3.26E-06 | 0.00019 | 9.00E-05 | 4.94E-06 | 1.14E-05 |
| Rab11fip2 | 8.16E-05 | 0.00012 | 0 | 3.21E-05 | 8.94E-05 | 6.10E-05 | 2.51E-05 | 0 | 7.00E-05 | 9.97E-05 | 6.76E-06 | 0 | 9.32E-05 | 4.14E-05 | 0.00024 | 6.15E-05 | 7.23E-05 | 7.97E-05 |
| Eif3m | 0.00011 | 4.75E-05 | 6.08E-05 | 5.53E-05 | 1.08E-05 | 5.45E-05 | 0 | 1.30E-05 | 7.00E-05 | 0 | 0 | 5.83E-05 | 0 | 1.56E-05 | 0 | 3.03E-05 | 1.29E-05 | 1.17E-05 |
| Ncoa5 | 0.00012 | 0.00015 | 0.00012 | 3.19E-05 | 2.48E-05 | 2.91E-05 | 0 | 2.71E-06 | 7.00E-05 | 4.76E-05 | 4.45E-05 | 4.73E-05 | 3.14E-05 | 2.80E-05 | 0.00015 | 9.83E-05 | 0 | 8.77E-06 |
| Gpsm2 | 5.89E-05 | 8.08E-05 | 0 | 2.81E-05 | 1.11E-05 | 0 | 0.00019 | 2.61E-05 | 7.00E-05 | 0.00015 | 0 | 2.13E-05 | 0 | 5.21E-05 | 0 | 0.00041 | 0 | 7.70E-05 |
| Srxn1 | 2.17E-05 | 9.20E-05 | 0.00018 | 9.21E-05 | 3.11E-05 | 0 | 0.00011 | 3.73E-05 | 7.00E-05 | 9.03E-05 | 2.11E-05 | 8.81E-05 | 1.07E-05 | 0 | 0 | 7.94E-05 | 0.00047 | 5.61E-05 |
| Tbc1d10b | 6.35E-05 | 0.0002 | 0.00012 | 0.00019 | 0.00012 | 7.53E-05 | 0 | 0.00013 | 6.99E-05 | 7.59E-05 | 0.00019 | 0.00016 | 1.02E-05 | 0.00012 | 0.00013 | 0.0002 | 0 | 5.89E-05 |
| DEPRECATEI | 2.60E-05 | 6.31E-05 | 5.00E-05 | 2.20E-05 | 3.31E-06 | 7.44E-05 | 8.78E-06 | 3.29E-05 | 6.99E-05 | 2.57E-05 | 2.70E-05 | 0 | 0 | 0 | 3.65E-05 | 0 | 6.07E-07 | 3.66E-05 |
| Itfg2 | 3.03E-05 | 8.55E-05 | 7.49E-05 | 5.14E-05 | 3.22E-05 | 6.82E-05 | 2.64E-05 | 4.96E-05 | 6.99E-05 | 1.13E-05 | 1.49E-05 | 1.92E-05 | 5.20E-05 | 4.78E-05 | 3.73E-05 | 4.20E-05 | 3.21E-05 | 3.69E-05 |
| Parp4 | 6.51E-05 | 0.0001 | 0 | 3.83E-05 | 2.49E-05 | 5.56E-05 | 2.20E-05 | 3.91E-05 | 6.99E-05 | 5.56E-05 | 3.48E-05 | 9.00E-05 | 4.59E-05 | 1.18E-06 | 1.92E-06 | 1.69E-05 | 4.01E-05 | 0.0001 |
| Alox5ap | 1.34E-05 | 2.32E-06 | 2.40E-06 | 5.37E-05 | 0 | 2.11E-05 | 6.83E-06 | 1.16E-06 | 6.99E-05 | 1.37E-06 | 1.74E-06 | 0 | 0 | 2.01E-05 | 4.02E-05 | 3.54E-06 | 0 | 2.64E-05 |
| Dolk | 8.38E-05 | 7.50E-05 | 0.00011 | 8.13E-05 | 8.38E-05 | 6.03E-05 | 4.78E-05 | 4.86E-05 | 6.98E-05 | 2.96E-05 | 4.57E-05 | 1.74E-05 | 2.09E-05 | 3.70E-05 | 0 | 4.78E-06 | 0 | 1.02E-05 |
| Iqsec2 | 4.53E-05 | 8.23E-05 | 3.01E-05 | 3.50E-05 | 6.09E-05 | 1.64E-05 | 4.10E-05 | 8.51E-05 | 6.98E-05 | 0.00016 | 1.21E-05 | 5.16E-05 | 4.50E-05 | 1.99E-05 | 7.16E-05 | 0 | 8.42E-05 | 6.15E-05 |
| Arfgap3 | 0.00022 | 9.22E-05 | 0.00016 | 0.00031 | 9.12E-05 | 0.0002 | 0.00043 | 0.00011 | 6.97E-05 | 0.00017 | 6.49E-05 | 0.00037 | 0.00011 | 0.00012 | 0.00024 | 0 | 5.58E-06 | 0.00017 |
| Mtch2 | 8.87E-05 | 9.38E-05 | 6.02E-05 | 5.02E-05 | 4.25E-05 | 4.45E-05 | 8.47E-06 | 0 | 6.97E-05 | 6.22E-05 | 2.91E-05 | 0 | 2.15E-05 | 5.28E-05 | 0 | 6.29E-05 | 0 | 4.87E-05 |
| Trappc12 | 8.64E-05 | 0.00011 | 3.62E-05 | 1.90E-05 | 4.84E-05 | 3.12E-05 | 5.00E-05 | 1.47E-05 | 6.97E-05 | 7.76E-05 | 2.87E-05 | 6.24E-05 | 2.51E-05 | 2.40E-05 | 0.00016 | 0.00011 | 2.32E-05 | 4.25E-05 |
| Zfp746 | 3.46E-05 | 0.0001 | 6.41E-05 | 4.35E-05 | 4.90E-05 | 0 | 0 | 0 | 6.97E-05 | 3.12E-05 | 4.97E-05 | 0 | 0 | 4.50E-05 | 0.00013 | 5.23E-05 | 0 | 3.47E-05 |
| Ethe1 | 8.37E-05 | 8.55E-05 | 2.13E-05 | 3.01E-05 | 5.36E-05 | 6.93E-05 | 3.51E-05 | 2.36E-05 | 6.96E-05 | 2.72E-05 | 0.00018 | 3.74E-05 | 2.49E-05 | 0.00016 | 0 | 0.00013 | 1.90E-05 | 0 |
| Ube2j1 | 0.00018 | 8.21E-05 | 0.00015 | 6.57E-05 | 3.39E-05 | 4.74E-05 | 7.83E-05 | 2.25E-06 | 6.96E-05 | 1.67E-05 | 0.00011 | 0.00014 | 0 | 0 | 0 | 5.15E-05 | 0 | 5.63E-05 |
| S100pbp | 9.05E-05 | 8.10E-05 | 3.85E-05 | 0 | 3.03E-05 | 3.86E-05 | 4.00E-05 | 1.29E-05 | 6.96E-05 | 0 | 3.52E-06 | 9.04E-05 | 0 | 1.51E-05 | 0.00013 | 3.07E-05 | 0 | 3.04E-06 |
| Aqr | 0.0001 | 9.30E-05 | 0.00014 | 3.71E-05 | 4.26E-05 | 3.83E-05 | 0 | 0 | 6.96E-05 | 3.64E-05 | 2.37E-05 | 1.39E-05 | 4.05E-05 | 3.38E-05 | 9.18E-05 | 9.33E-05 | 0 | 1.74E-05 |
| Dera | 0.00011 | 2.86E-05 | 2.99E-05 | 7.52E-05 | 9.00E-06 | 3.71E-05 | 2.20E-05 | 3.74E-05 | 6.96E-05 | 2.82E-05 | 3.31E-05 | 4.78E-05 | 1.50E-06 | 0 | 1.14E-05 | 3.16E-05 | 0 | 8.96E-06 |
| Ntpcr | 5.23E-05 | 5.29E-05 | 1.03E-05 | 5.55E-05 | 5.83E-05 | 3.32E-05 | 3.69E-05 | 0 | 6.96E-05 | 5.13E-05 | 3.86E-05 | 3.80E-05 | 3.81E-05 | 1.80E-05 | 2.56E-05 | 0 | 2.98E-05 | 4.45E-05 |
| Resf1 | 0.00012 | 0.00011 | 0 | 0 | 0 | 2.22E-05 | 7.26E-05 | 2.01E-05 | 6.96E-05 | 2.95E-05 | 2.58E-05 | 4.69E-05 | 0 | 0 | 7.88E-05 | 7.66E-05 | 6.01E-05 | 4.95E-05 |
| Nsmce1 | 0.00011 | 1.57E-05 | 5.36E-07 | 1.20E-05 | 0 | 1.70E-05 | 1.69E-05 | 4.64E-05 | 6.96E-05 | 2.61E-05 | 0 | 2.95E-05 | 1.53E-05 | 0 | 0 | 0.00017 | 0 | 3.40E-05 |
| Lrpprc | 8.11E-05 | 7.79E-05 | 9.85E-05 | 4.83E-05 | 6.97E-06 | 1.38E-05 | 0 | 1.06E-05 | 6.96E-05 | 0 | 2.39E-05 | 1.18E-05 | 0 | 7.17E-06 | 3.11E-05 | 3.82E-05 | 0 | 2.12E-05 |
| Enpp2 | 0 | 0.00014 | 3.84E-06 | 0 | 0 | 0 | 0 | 0 | 6.96E-05 | 0 | 0 | 0 | 0 | 2.11E-07 | 0 | 2.22E-05 | 0 | 6.72E-05 |

| | | | | | | | | | | | | | | | | | | |
|---|---|---|---|---|---|---|---|---|---|---|---|---|---|---|---|---|---|---|
| Dynlt1f | 0.00022 | 0.00012 | 1.56E-05 | 5.45E-05 | 2.58E-05 | 0.00014 | 4.69E-05 | 6.07E-05 | 6.95E-05 | 7.24E-05 | 0 | 0 | 0 | 6.85E-05 | 0 | 0.00027 | 0 | 0 |
| Ssh2 | 5.15E-05 | 0.00012 | 0 | 0 | 1.54E-05 | 4.43E-05 | 0 | 0.00012 | 6.95E-05 | 8.62E-05 | 0 | 0 | 8.91E-05 | 1.39E-05 | 0 | 1.29E-05 | 1.08E-05 | 4.84E-05 |
| Casc3 | 0.0001 | 0.00021 | 0.00012 | 0.0001 | 4.56E-05 | 3.02E-05 | 4.47E-05 | 2.85E-05 | 6.95E-05 | 5.77E-05 | 8.51E-05 | 0.0001 | 2.16E-07 | 4.72E-05 | 0.00029 | 0.00014 | 4.79E-05 | 5.21E-05 |
| Stx8 | 0.00014 | 0.00016 | 7.19E-05 | 9.70E-05 | 9.86E-05 | 0.0001 | 6.96E-05 | 4.40E-05 | 6.94E-05 | 7.69E-05 | 8.74E-05 | 3.14E-05 | 0 | 7.08E-05 | 0.00024 | 0 | 4.25E-05 | 9.09E-05 |
| Fat4 | 0 | 4.81E-05 | 0 | 0 | 0.00019 | 1.48E-06 | 0.00011 | 0 | 6.94E-05 | 0 | 3.71E-05 | 0 | 0 | 0.00011 | 0 | 0 | 0 | 7.98E-05 |
| Dnajc21 | 8.29E-05 | 0.00011 | 0.00015 | 3.16E-05 | 7.69E-05 | 0 | 0 | 0 | 6.94E-05 | 4.76E-06 | 3.19E-05 | 0.00024 | 0 | 4.40E-05 | 0.00012 | 7.01E-05 | 0.0001 | 0 |
| Ppp4r3a | 9.75E-05 | 0.00012 | 0.00012 | 8.72E-05 | 2.71E-05 | 6.17E-05 | 0 | 2.45E-05 | 6.93E-05 | 1.85E-05 | 4.56E-05 | 7.34E-05 | 0 | 2.77E-05 | 0.00011 | 0.00017 | 0.00017 | 1.30E-05 |
| Iws1 | 0.00014 | 0.00011 | 0.00011 | 0 | 6.01E-05 | 1.80E-05 | 1.17E-05 | 2.29E-05 | 6.93E-05 | 3.86E-05 | 3.70E-05 | 6.48E-05 | 0 | 2.60E-05 | 0.00014 | 0.00016 | 0.00011 | 0 |
| Knop1 | 6.37E-05 | 5.49E-05 | 9.48E-05 | 5.91E-06 | 3.21E-05 | 9.84E-06 | 0 | 8.41E-06 | 6.93E-05 | 3.81E-05 | 3.71E-05 | 5.71E-06 | 0 | 1.89E-05 | 6.01E-05 | 8.05E-05 | 0 | 1.71E-05 |
| Sirt6 | 3.66E-05 | 0.00013 | 7.70E-05 | 0.00013 | 7.89E-05 | 0 | 0.00016 | 0.00013 | 6.93E-05 | 3.69E-05 | 6.05E-05 | 0.00012 | 5.27E-05 | 9.41E-05 | 0.00018 | 0 | 0 | 6.26E-05 |
| Lipa | 0.00019 | 0.0002 | 1.54E-05 | 0 | 0.00017 | 0.00028 | 0.00023 | 0.0002 | 6.92E-05 | 0.00016 | 0 | 2.26E-05 | 0.00016 | 9.48E-05 | 0 | 0 | 0 | 0.00025 |
| Mospd3 | 0.0002 | 0.00011 | 3.78E-05 | 0.00028 | 0.00012 | 0.00015 | 0.00026 | 0.00029 | 6.92E-05 | 8.45E-05 | 8.45E-05 | 7.44E-05 | 2.57E-05 | 5.51E-05 | 0 | 7.77E-05 | 0 | 0.00013 |
| Carnmt1 | 0.00013 | 2.09E-05 | 4.28E-05 | 0 | 2.41E-05 | 7.52E-05 | 3.34E-06 | 0 | 6.92E-05 | 0 | 2.46E-05 | 4.06E-05 | 0 | 0 | 1.99E-05 | 9.22E-05 | 2.72E-05 | 0 |
| Plxna3 | 1.72E-05 | 3.93E-06 | 0 | 1.07E-05 | 0 | 1.47E-05 | 3.83E-05 | 0 | 6.92E-05 | 2.08E-05 | 0 | 2.15E-05 | 4.83E-06 | 5.84E-06 | 3.83E-06 | 1.21E-05 | 2.69E-05 | 4.44E-05 |
| Psmg3 | 5.82E-05 | 6.00E-05 | 8.75E-05 | 0.00011 | 4.29E-05 | 1.11E-05 | 0 | 1.25E-05 | 6.92E-05 | 1.89E-05 | 3.90E-05 | 0 | 2.16E-05 | 1.63E-05 | 2.59E-05 | 7.03E-05 | 0 | 4.67E-06 |
| Cacnb3 | 0.00012 | 0.00022 | 1.37E-05 | 0.00013 | 0.0004 | 0.00023 | 0.0003 | 1.40E-05 | 6.91E-05 | 2.74E-05 | 0.00036 | 0.00016 | 0.00017 | 0.00053 | 0.00021 | 2.36E-06 | 9.94E-05 | 0.00028 |
| Akap11 | 0.00015 | 0.00015 | 0.00013 | 0 | 7.75E-05 | 0.00016 | 5.79E-05 | 9.78E-05 | 6.91E-05 | 0 | 4.08E-06 | 0.00011 | 0.00011 | 0.00014 | 3.03E-05 | 0.00011 | 4.76E-05 | 3.59E-06 |
| Acad8 | 7.77E-05 | 7.72E-05 | 4.14E-05 | 4.62E-05 | 5.77E-05 | 9.26E-05 | 3.20E-05 | 2.76E-05 | 6.91E-05 | 6.64E-05 | 5.81E-06 | 9.68E-06 | 5.72E-05 | 3.56E-05 | 2.98E-05 | 7.16E-05 | 0 | 9.01E-05 |
| Skiv2l | 1.72E-05 | 7.16E-05 | 2.47E-05 | 0.00011 | 3.60E-05 | 0 | 0 | 2.75E-05 | 6.91E-05 | 3.69E-05 | 8.11E-06 | 1.66E-05 | 3.27E-05 | 2.27E-05 | 7.98E-05 | 1.81E-05 | 0 | 1.99E-05 |
| Parp12 | 0.00015 | 7.59E-05 | 0 | 8.34E-05 | 2.58E-05 | 0 | 2.34E-05 | 1.79E-05 | 6.91E-05 | 0.00015 | 2.43E-05 | 0 | 2.55E-05 | 4.23E-05 | 8.38E-05 | 8.71E-05 | 3.39E-05 | 0.00013 |
| Pds5b | 0.00013 | 0.00011 | 1.40E-05 | 0 | 5.71E-05 | 0.00014 | 0 | 3.97E-05 | 6.90E-05 | 2.39E-05 | 2.54E-05 | 4.45E-05 | 0 | 9.03E-06 | 8.90E-05 | 0.00025 | 9.70E-05 | 4.86E-05 |
| Iqsec1 | 8.42E-05 | 0.00025 | 5.44E-05 | 0 | 0.00015 | 0.00029 | 7.40E-05 | 0 | 6.89E-05 | 0.0002 | 0.00022 | 0.00052 | 5.72E-05 | 9.28E-05 | 5.02E-05 | 9.64E-05 | 3.41E-05 | 7.13E-05 |
| Znrf1 | 0 | 0.0002 | 0.00022 | 9.73E-06 | 0.00016 | 0.00022 | 0 | 0.00014 | 6.89E-05 | 6.31E-05 | 0.00019 | 0.0004 | 0.00022 | 0.00019 | 0.00017 | 5.96E-05 | 0.00019 | 7.45E-05 |
| Dnajc8 | 0.00025 | 0.00014 | 0.00011 | 4.05E-05 | 0.00012 | 0.00018 | 0 | 2.24E-05 | 6.89E-05 | 3.73E-05 | 7.91E-05 | 8.48E-05 | 9.17E-06 | 7.42E-05 | 5.35E-05 | 0.00027 | 0 | 1.78E-05 |
| Mef2c | 2.32E-05 | 8.15E-05 | 0 | 4.46E-05 | 0 | 8.73E-05 | 5.89E-05 | 2.12E-05 | 6.89E-05 | 8.00E-05 | 0 | 0 | 3.52E-06 | 0 | 0 | 9.46E-05 | 5.86E-05 | 0.00015 |
| Armc1 | 0.00013 | 9.19E-05 | 0.00012 | 6.64E-05 | 9.51E-05 | 4.79E-05 | 0.00013 | 5.99E-05 | 6.89E-05 | 3.55E-05 | 6.49E-05 | 8.68E-06 | 1.40E-06 | 0 | 0 | 0.00011 | 2.46E-06 | 2.50E-05 |
| Samd1 | 4.98E-05 | 3.09E-05 | 2.98E-05 | 1.93E-05 | 1.91E-05 | 2.32E-05 | 7.03E-06 | 2.56E-05 | 6.89E-05 | 7.81E-06 | 1.66E-05 | 4.31E-05 | 0 | 2.59E-05 | 7.73E-05 | 8.56E-05 | 0 | 1.42E-06 |
| Vps4b | 8.50E-05 | 9.65E-05 | 5.69E-05 | 4.27E-05 | 5.76E-05 | 7.91E-05 | 6.77E-05 | 3.49E-05 | 6.88E-05 | 7.93E-05 | 5.09E-05 | 0.00012 | 1.22E-05 | 2.53E-05 | 9.39E-05 | 6.40E-05 | 5.62E-05 | 4.67E-05 |
| Tlk2 | 0.00017 | 0.00015 | 8.01E-05 | 0 | 5.65E-05 | 6.56E-05 | 1.93E-05 | 0 | 6.87E-05 | 3.48E-05 | 2.14E-05 | 0.00013 | 0 | 0 | 0.00021 | 5.05E-05 | 0.00021 | 5.04E-06 |
| Gmpr2 | 0.00013 | 3.83E-05 | 7.37E-05 | 9.52E-05 | 1.85E-05 | 3.27E-05 | 1.57E-05 | 5.60E-05 | 6.87E-05 | 2.91E-05 | 4.22E-05 | 3.58E-05 | 0 | 4.81E-05 | 0 | 6.33E-05 | 0 | 1.91E-05 |
| Bphl | 7.64E-06 | 6.01E-05 | 0 | 3.83E-05 | 3.62E-05 | 2.98E-05 | 3.20E-05 | 0 | 6.87E-05 | 8.65E-05 | 5.04E-05 | 6.00E-05 | 8.12E-05 | 6.74E-05 | 0 | 0 | 0 | 0.00011 |
| Ebp | 0.00013 | 5.72E-05 | 2.37E-06 | 0.00029 | 0 | 0 | 0.00021 | 0 | 6.87E-05 | 1.55E-05 | 0.00025 | 0 | 0.00011 | 0.0002 | 0 | 0.00017 | 0 | 0.00026 |
| Rit1 | 7.68E-05 | 0.00018 | 4.58E-05 | 0.00015 | 0.00012 | 0.00012 | 2.98E-05 | 7.61E-05 | 6.86E-05 | 0.00016 | 0.00014 | 6.15E-05 | 1.33E-05 | 0.00011 | 7.40E-05 | 4.34E-05 | 0 | 0.00018 |

| | | | | | | | | | | | | | | | | | | |
|---|---|---|---|---|---|---|---|---|---|---|---|---|---|---|---|---|---|---|
| Mcur1 | 5.38E-05 | 6.56E-05 | 4.37E-05 | 6.44E-05 | 2.54E-05 | 8.20E-05 | 1.87E-05 | 8.14E-05 | 6.86E-05 | 7.08E-05 | 5.54E-05 | 4.60E-06 | 2.50E-05 | 3.56E-05 | 0 | 0 | 7.43E-05 | 0.00016 |
| Apip | 0.00011 | 4.76E-05 | 5.11E-05 | 0 | 4.00E-05 | 6.21E-05 | 0 | 0 | 6.86E-05 | 2.43E-05 | 3.13E-05 | 3.94E-06 | 6.82E-07 | 5.38E-05 | 6.55E-06 | 0.00012 | 0 | 2.96E-05 |
| Tbc1d19 | 9.80E-05 | 6.43E-05 | 2.35E-05 | 0 | 7.48E-05 | 4.22E-05 | 0 | 1.47E-05 | 6.86E-05 | 0.00011 | 2.96E-05 | 6.95E-05 | 2.59E-05 | 9.55E-06 | 9.22E-05 | 9.43E-05 | 2.88E-05 | 4.08E-05 |
| Cog4 | 6.13E-05 | 9.03E-05 | 2.26E-05 | 5.59E-05 | 8.87E-05 | 1.84E-05 | 3.55E-05 | 7.46E-05 | 6.86E-05 | 2.06E-05 | 7.76E-06 | 1.52E-05 | 0 | 6.94E-05 | 1.20E-05 | 4.52E-05 | 0 | 6.36E-05 |
| Noc2l | 0.00023 | 0.00015 | 0.0005 | 0.00011 | 7.71E-06 | 7.57E-06 | 0 | 4.36E-07 | 6.86E-05 | 0 | 5.20E-05 | 0.00014 | 0 | 0 | 0.00014 | 4.18E-05 | 9.17E-05 | 0 |
| Ift20 | 0.00045 | 0 | 0.00029 | 0.00044 | 0.00016 | 0.00021 | 0.0007 | 0.00016 | 6.85E-05 | 8.47E-05 | 0.00017 | 8.47E-06 | 0.00013 | 0.00014 | 0.00011 | 0 | 0 | 2.37E-05 |
| Btg2 | 2.23E-05 | 0 | 1.01E-05 | 0 | 0 | 0.00016 | 6.17E-05 | 7.67E-05 | 6.85E-05 | 1.97E-06 | 0 | 8.31E-05 | 7.32E-05 | 0 | 0.0001 | 3.91E-05 | 0.00564 | 7.49E-05 |
| Pnpt1 | 0.00012 | 2.76E-05 | 5.79E-05 | 1.52E-05 | 6.13E-06 | 8.21E-06 | 0 | 0 | 6.85E-05 | 4.67E-05 | 3.83E-07 | 0 | 2.47E-05 | 8.42E-06 | 4.34E-05 | 4.82E-05 | 0 | 6.21E-06 |
| Mpp1 | 0.00018 | 6.46E-05 | 1.54E-05 | 4.17E-05 | 5.12E-06 | 6.33E-06 | 0 | 0 | 6.85E-05 | 2.76E-05 | 1.46E-05 | 1.25E-05 | 0 | 4.98E-05 | 3.37E-05 | 0.00012 | 0 | 0.00011 |
| Daxx | 6.38E-05 | 8.01E-05 | 0.00011 | 0.00014 | 3.83E-05 | 3.42E-06 | 7.34E-05 | 4.21E-05 | 6.85E-05 | 0.00019 | 7.46E-05 | 1.72E-05 | 0.00012 | 6.66E-05 | 9.61E-05 | 5.45E-05 | 4.29E-05 | 5.75E-05 |
| Tmem106c | 5.61E-06 | 4.29E-05 | 0 | 9.91E-06 | 3.40E-05 | 0 | 8.34E-05 | 3.41E-05 | 6.85E-05 | 0.00011 | 0 | 6.99E-05 | 0.0001 | 1.44E-05 | 0 | 9.11E-05 | 0 | 7.95E-05 |
| Ubxn7 | 0.00017 | 0.00025 | 1.34E-05 | 0 | 0.00015 | 0.00012 | 0.00011 | 2.20E-05 | 6.84E-05 | 1.90E-05 | 7.55E-05 | 0.00019 | 3.75E-05 | 0.00011 | 0.00027 | 6.11E-05 | 4.85E-05 | 4.74E-05 |
| Arl6ip6 | 7.88E-05 | 4.96E-05 | 0.00013 | 8.69E-05 | 5.87E-05 | 0.00011 | 2.19E-05 | 0.00012 | 6.84E-05 | 4.86E-05 | 5.91E-05 | 0.00015 | 2.16E-05 | 7.26E-05 | 0 | 0.00024 | 0.00019 | 0 |
| Filip1l | 8.24E-05 | 2.43E-06 | 0 | 0 | 5.91E-05 | 0 | 5.53E-06 | 0 | 6.84E-05 | 0.00011 | 0.00056 | 0 | 0 | 2.16E-05 | 6.82E-05 | 2.98E-05 | 0 | 0 |
| Psd | 0 | 2.03E-05 | 0 | 0 | 0.00017 | 8.33E-05 | 0 | 0 | 6.83E-05 | 1.10E-05 | 1.84E-05 | 1.50E-06 | 7.84E-06 | 5.67E-05 | 7.55E-06 | 0 | 3.04E-05 | 3.12E-05 |
| Slc25a10 | 0 | 4.71E-05 | 3.43E-05 | 0.00014 | 3.82E-05 | 9.46E-06 | 0 | 0.00016 | 6.83E-05 | 1.45E-05 | 1.22E-05 | 0 | 1.60E-05 | 8.05E-05 | 1.72E-05 | 0.00047 | 0 | 5.72E-05 |
| Pgm2 | 6.55E-05 | 0.00011 | 0.00018 | 8.34E-05 | 6.27E-05 | 0 | 0 | 7.27E-05 | 6.83E-05 | 7.86E-05 | 0.00029 | 0.00018 | 0 | 2.19E-05 | 3.82E-05 | 6.99E-05 | 8.68E-05 | 8.35E-05 |
| Pgam1 | 0 | 8.63E-06 | 0.00021 | 0.00106 | 0.00012 | 0 | 0.00019 | 0.0003 | 6.83E-05 | 0.00019 | 0.00028 | 0.00123 | 0.00035 | 0.00013 | 9.54E-05 | 0.00025 | 0.00036 | 0 |
| Bclaf1 | 0.00024 | 0.00024 | 0.00019 | 0.00019 | 0.00012 | 0.00014 | 4.97E-05 | 2.29E-05 | 6.82E-05 | 5.75E-05 | 8.07E-05 | 0.00016 | 0 | 6.54E-05 | 0.00069 | 0.00027 | 0.00032 | 3.31E-05 |
| Pgap2 | 6.77E-05 | 7.60E-05 | 1.41E-05 | 0.00012 | 3.65E-05 | 4.63E-05 | 6.70E-05 | 5.58E-05 | 6.82E-05 | 5.92E-05 | 7.77E-06 | 6.04E-05 | 2.85E-05 | 0 | 0 | 5.81E-06 | 3.78E-05 | 7.21E-05 |
| Fyco1 | 4.90E-05 | 0.00013 | 1.00E-05 | 0 | 0.00017 | 3.87E-05 | 0.00017 | 4.27E-06 | 6.82E-05 | 7.28E-05 | 2.83E-05 | 4.26E-05 | 9.41E-05 | 0.00014 | 7.83E-05 | 4.18E-05 | 0 | 4.79E-05 |
| Fnbp4 | 8.00E-05 | 0.00014 | 6.60E-05 | 4.99E-05 | 3.65E-05 | 2.73E-05 | 2.07E-05 | 5.69E-06 | 6.82E-05 | 3.03E-05 | 8.10E-06 | 5.04E-05 | 0 | 1.30E-05 | 0.00022 | 0.00015 | 0.00025 | 2.90E-05 |
| Unc5a | 0 | 6.07E-06 | 0 | 3.49E-05 | 0.00019 | 0.0002 | 0 | 0.00023 | 6.81E-05 | 6.44E-06 | 3.41E-05 | 0 | 0 | 0 | 3.78E-06 | 0 | 0 | 2.89E-05 |
| Id1 | 0 | 4.07E-05 | 0 | 0 | 0 | 5.34E-05 | 0 | 8.35E-05 | 6.81E-05 | 0 | 0 | 0 | 0 | 0 | 9.57E-06 | 7.53E-05 | 0.0006 | 5.95E-05 |
| Plin2 | 1.40E-05 | 0.0004 | 0.00013 | 0.00036 | 0 | 3.72E-05 | 0 | 0.00019 | 6.81E-05 | 0.00011 | 0.00013 | 0.00028 | 0 | 0.00079 | 0.00014 | 0.00014 | 8.79E-05 | 8.37E-05 |
| H2afx | 0 | 3.50E-05 | 1.55E-05 | 0.00026 | 0 | 0 | 0 | 5.58E-05 | 6.81E-05 | 2.40E-05 | 3.78E-05 | 0.00011 | 0 | 0 | 0 | 0.00147 | 0.00068 | 3.72E-05 |
| Atg4b | 0.00015 | 0.00019 | 0.00014 | 3.89E-05 | 9.07E-05 | 9.31E-05 | 3.72E-05 | 0 | 6.80E-05 | 5.58E-05 | 4.03E-05 | 9.53E-05 | 1.14E-05 | 5.32E-05 | 0.00013 | 8.74E-05 | 0 | 7.95E-05 |
| Tas2r119 | 0 | 4.71E-06 | 0 | 0 | 0.00011 | 8.62E-05 | 0 | 7.02E-05 | 6.80E-05 | 0 | 0 | 0 | 0 | 3.60E-05 | 0 | 0 | 0 | 0 |
| Med6 | 0.00014 | 6.42E-05 | 5.38E-05 | 0 | 2.50E-06 | 6.49E-05 | 6.01E-06 | 2.60E-06 | 6.80E-05 | 1.56E-05 | 1.81E-05 | 2.35E-05 | 1.27E-05 | 3.88E-05 | 2.47E-05 | 8.77E-05 | 7.84E-07 | 4.71E-06 |
| Phactr4 | 5.72E-05 | 7.12E-05 | 3.78E-05 | 2.06E-05 | 0.00011 | 7.05E-06 | 0 | 0 | 6.80E-05 | 1.57E-05 | 7.65E-05 | 0.00024 | 6.09E-05 | 6.92E-05 | 0.00016 | 9.21E-05 | 2.40E-05 | 5.75E-05 |
| Ccdc97 | 2.94E-05 | 0.00016 | 7.47E-05 | 0.00018 | 5.37E-05 | 0 | 3.73E-05 | 0.00021 | 6.80E-05 | 8.56E-05 | 4.79E-05 | 3.23E-05 | 0 | 4.42E-05 | 0.00011 | 3.15E-05 | 0.00024 | 5.48E-05 |
| Ddx50 | 0.00032 | 0.00013 | 8.03E-05 | 4.17E-05 | 0.00012 | 0.00023 | 0.00019 | 1.09E-05 | 6.79E-05 | 6.58E-05 | 6.17E-05 | 0.00016 | 0.00011 | 9.28E-05 | 0.0003 | 1.36E-05 | 5.36E-05 | 0.0001 |
| Hnrnpc | 0.0003 | 8.41E-05 | 0.00014 | 0 | 2.06E-05 | 8.59E-05 | 0 | 0 | 6.79E-05 | 0 | 1.89E-05 | 0.0001 | 0 | 0 | 0 | 6.70E-05 | 0.00018 | 0 |

| | | | | | | | | | | | | | | | | | | |
|---|---|---|---|---|---|---|---|---|---|---|---|---|---|---|---|---|---|---|
| Fam173a | 0 | 0.0001 | 0 | 0.00022 | 0.00013 | 5.17E-05 | 0 | 0.00014 | 6.79E-05 | 7.64E-05 | 4.81E-05 | 7.77E-05 | 1.54E-05 | 7.22E-05 | 0.0001 | 5.91E-05 | 0 | 0.00018 |
| Nav1 | 5.11E-05 | 0.00022 | 0.00052 | 0 | 0.00026 | 0.00028 | 0.00051 | 0.00033 | 6.78E-05 | 0.00047 | 0 | 0.00019 | 5.58E-05 | 5.50E-05 | 0.00051 | 0.00015 | 0.00054 | 0.0005 |
| Ifi203 | 9.09E-05 | 0.00011 | 0 | 0 | 0 | 0.00019 | 0 | 9.19E-05 | 6.78E-05 | 0.0005 | 0 | 0.00011 | 0.00045 | 0 | 0 | 0 | 0 | 0.00013 |
| Wasl | 0.00019 | 0.00022 | 9.53E-05 | 1.22E-05 | 0.00012 | 0.00014 | 6.59E-05 | 7.34E-05 | 6.78E-05 | 8.48E-05 | 4.07E-05 | 0.00012 | 0 | 0.00013 | 0.00018 | 6.26E-05 | 0.00021 | 2.34E-05 |
| Gtf2h3 | 8.24E-05 | 2.51E-05 | 5.10E-05 | 3.10E-05 | 5.23E-06 | 4.82E-05 | 2.12E-05 | 0 | 6.78E-05 | 4.14E-05 | 0 | 2.36E-05 | 4.90E-05 | 4.27E-05 | 2.10E-05 | 2.86E-05 | 0 | 0 |
| Pcdhga11 | 4.02E-05 | 7.29E-05 | 2.30E-05 | 3.76E-06 | 4.12E-05 | 4.17E-05 | 3.93E-05 | 2.84E-05 | 6.78E-05 | 2.50E-05 | 1.70E-05 | 1.67E-05 | 2.98E-05 | 2.60E-05 | 7.65E-05 | 1.93E-06 | 2.06E-05 | 3.61E-05 |
| Ulbp1 | 0 | 0 | 3.70E-05 | 0 | 2.20E-05 | 3.11E-06 | 6.22E-05 | 2.96E-05 | 6.78E-05 | 4.97E-06 | 4.79E-05 | 0 | 0 | 0 | 0 | 0.00046 | 0 | 0 |
| B3galnt2 | 9.59E-05 | 7.94E-05 | 9.16E-05 | 5.01E-05 | 9.62E-05 | 6.96E-05 | 0.00012 | 2.56E-05 | 6.77E-05 | 4.56E-05 | 2.24E-05 | 8.21E-05 | 4.39E-05 | 3.17E-05 | 8.81E-05 | 2.61E-05 | 0 | 9.18E-06 |
| Kif3b | 8.15E-05 | 0.00011 | 6.44E-05 | 2.18E-05 | 7.15E-05 | 4.77E-05 | 0 | 5.21E-05 | 6.77E-05 | 9.67E-05 | 5.36E-05 | 5.05E-05 | 3.57E-05 | 6.45E-05 | 6.73E-05 | 5.00E-05 | 0 | 9.12E-05 |
| Gpkow | 6.11E-05 | 4.03E-05 | 6.97E-05 | 9.88E-06 | 4.83E-05 | 2.15E-05 | 1.31E-05 | 4.37E-05 | 6.77E-05 | 3.62E-05 | 3.54E-05 | 1.11E-05 | 0 | 2.47E-05 | 2.10E-05 | 3.35E-05 | 0 | 3.00E-05 |
| Ywhaz | 9.43E-05 | 8.48E-05 | 0.00014 | 4.00E-05 | 4.39E-05 | 9.88E-06 | 0 | 9.71E-05 | 6.77E-05 | 5.68E-05 | 9.01E-05 | 6.94E-05 | 0 | 2.39E-05 | 0 | 8.56E-05 | 7.55E-05 | 5.06E-05 |
| Commd2 | 0.00015 | 6.46E-05 | 0.00015 | 9.06E-05 | 9.23E-06 | 2.52E-06 | 4.76E-05 | 0 | 6.77E-05 | 4.13E-05 | 4.94E-05 | 0 | 3.41E-06 | 5.05E-05 | 1.17E-05 | 8.37E-05 | 0 | 1.77E-05 |
| Nol7 | 0.00016 | 7.91E-05 | 0.00011 | 5.63E-05 | 0 | 0 | 4.39E-06 | 0 | 6.77E-05 | 5.42E-05 | 6.75E-05 | 6.59E-05 | 0 | 1.37E-05 | 0 | 0.00011 | 0 | 0 |
| Nfkbiz | 2.20E-05 | 0 | 4.38E-05 | 0 | 4.25E-05 | 0 | 4.85E-05 | 0.00018 | 6.77E-05 | 0.0002 | 0 | 8.68E-05 | 0.00053 | 0.00015 | 0.00035 | 1.49E-05 | 0.0032 | 0 |
| Rbl2 | 8.96E-05 | 0.00015 | 0 | 0 | 0.00013 | 0.00012 | 0.00014 | 4.27E-05 | 6.76E-05 | 6.02E-05 | 6.27E-06 | 5.99E-05 | 7.12E-05 | 4.39E-05 | 2.46E-05 | 0.0001 | 5.26E-05 | 7.75E-05 |
| Rbbp9 | 8.66E-05 | 3.49E-05 | 0 | 2.35E-05 | 7.49E-05 | 8.44E-05 | 1.98E-05 | 5.70E-05 | 6.76E-05 | 1.89E-05 | 1.38E-05 | 1.96E-05 | 3.31E-05 | 4.29E-05 | 0 | 2.83E-06 | 0 | 7.13E-05 |
| Mccc2 | 5.47E-05 | 5.84E-05 | 0 | 7.64E-05 | 2.02E-05 | 0 | 3.23E-05 | 0 | 6.76E-05 | 1.31E-05 | 6.75E-06 | 2.24E-06 | 7.81E-06 | 2.80E-05 | 1.04E-05 | 0 | 5.85E-05 | 3.61E-07 |
| Nifk | 0.00019 | 7.68E-05 | 0.00023 | 0.00011 | 0 | 0 | 0 | 0 | 6.76E-05 | 2.09E-05 | 2.82E-06 | 0 | 9.60E-07 | 7.76E-06 | 1.84E-05 | 5.22E-06 | 0 | 2.05E-05 |
| Calcoco1 | 0 | 0.00038 | 0 | 0.00015 | 0.00022 | 0.00014 | 3.34E-05 | 0 | 6.75E-05 | 5.27E-05 | 6.09E-05 | 0.0002 | 3.62E-05 | 0.00022 | 0.00024 | 0 | 4.36E-05 | 0.00011 |
| Virma | 0.00016 | 8.50E-05 | 8.27E-05 | 5.45E-06 | 3.08E-05 | 5.16E-05 | 7.21E-05 | 5.32E-06 | 6.75E-05 | 1.46E-05 | 5.14E-05 | 4.39E-05 | 0 | 1.25E-06 | 7.58E-05 | 8.34E-05 | 3.02E-05 | 7.84E-06 |
| Senp7 | 5.77E-05 | 0.00012 | 0 | 1.43E-05 | 5.62E-05 | 3.99E-06 | 7.10E-05 | 8.50E-06 | 6.75E-05 | 2.40E-05 | 0 | 1.90E-05 | 0 | 2.91E-05 | 0.00012 | 5.18E-06 | 7.89E-05 | 3.73E-05 |
| Ndufa6 | 0.00026 | 0.00031 | 7.66E-05 | 0.00024 | 0.00021 | 0.00029 | 0.00018 | 0.00015 | 6.74E-05 | 6.66E-05 | 0.00013 | 0.00032 | 3.70E-05 | 0 | 1.54E-05 | 7.96E-05 | 0 | 0.00013 |
| Foxo4 | 5.68E-05 | 0.00015 | 0 | 0.00017 | 8.56E-05 | 8.66E-05 | 0.00013 | 5.86E-05 | 6.74E-05 | 3.92E-05 | 3.38E-05 | 3.08E-05 | 2.79E-05 | 9.66E-05 | 2.98E-05 | 2.53E-05 | 1.81E-06 | 5.12E-05 |
| Araf | 7.98E-05 | 0.00013 | 6.12E-05 | 2.08E-05 | 9.71E-05 | 5.89E-05 | 6.17E-05 | 4.93E-05 | 6.74E-05 | 9.48E-05 | 4.94E-05 | 7.23E-05 | 2.57E-05 | 0 | 0.00016 | 4.50E-05 | 0 | 7.44E-05 |
| Arhgap18 | 1.84E-05 | 3.80E-05 | 0 | 0 | 2.47E-05 | 4.91E-06 | 0 | 0 | 6.74E-05 | 8.88E-05 | 0 | 3.68E-05 | 1.12E-05 | 0 | 4.11E-06 | 6.57E-05 | 9.94E-06 | 0 |
| Lztr1 | 0.00016 | 0.00015 | 5.92E-05 | 9.93E-05 | 0.00013 | 0.00015 | 0.00021 | 9.55E-05 | 6.73E-05 | 5.34E-05 | 9.03E-05 | 6.94E-05 | 5.74E-05 | 0.00012 | 8.79E-05 | 9.25E-05 | 7.20E-06 | 0.00011 |
| Usp21 | 5.45E-05 | 0.00011 | 2.17E-05 | 0.00012 | 6.90E-05 | 7.21E-05 | 1.77E-05 | 2.49E-05 | 6.73E-05 | 5.84E-05 | 2.69E-05 | 0 | 2.77E-05 | 4.09E-05 | 2.18E-05 | 0.00013 | 0 | 5.96E-05 |
| Smurf2 | 0.00018 | 0.00013 | 9.04E-05 | 0 | 4.22E-05 | 6.82E-05 | 3.52E-05 | 5.16E-05 | 6.73E-05 | 0.00019 | 9.87E-05 | 0.00013 | 0 | 5.44E-06 | 8.75E-05 | 3.84E-05 | 0.00018 | 0 |
| Nol9 | 0.00011 | 7.06E-05 | 0.00018 | 2.13E-05 | 6.10E-06 | 1.19E-05 | 0 | 0 | 6.73E-05 | 4.74E-06 | 3.04E-05 | 0 | 7.39E-05 | 0 | 6.34E-05 | 8.97E-06 | 9.32E-05 | 0 |
| Sphk2 | 0 | 8.18E-05 | 3.65E-05 | 0.00012 | 7.89E-05 | 7.99E-06 | 0 | 5.78E-05 | 6.73E-05 | 3.94E-05 | 4.57E-05 | 1.66E-05 | 3.35E-05 | 5.27E-05 | 7.43E-05 | 4.35E-05 | 0 | 1.41E-05 |
| Slc25a28 | 5.19E-05 | 0.00013 | 0.00011 | 7.19E-06 | 2.46E-05 | 5.22E-07 | 3.25E-06 | 2.58E-06 | 6.73E-05 | 0.00018 | 3.78E-05 | 2.79E-05 | 0 | 0 | 0.0003 | 0.00017 | 0 | 5.46E-05 |
| Wfs1 | 4.38E-05 | 5.01E-05 | 9.76E-05 | 6.14E-05 | 6.26E-05 | 0 | 4.59E-05 | 9.11E-05 | 6.73E-05 | 7.34E-05 | 7.68E-05 | 0.00013 | 9.51E-05 | 6.84E-05 | 0 | 4.52E-07 | 0 | 1.55E-05 |
| Wdsub1 | 2.17E-05 | 8.16E-05 | 0 | 2.68E-05 | 1.84E-05 | 4.75E-05 | 1.65E-05 | 0 | 6.72E-05 | 5.76E-05 | 4.95E-05 | 1.63E-05 | 1.69E-05 | 4.30E-05 | 0 | 0 | 0 | 8.56E-05 |

| | | | | | | | | | | | | | | | | | | |
|---|---|---|---|---|---|---|---|---|---|---|---|---|---|---|---|---|---|---|
| Ttc13 | 8.27E-05 | 7.45E-05 | 6.39E-05 | 2.03E-05 | 3.42E-05 | 3.35E-05 | 7.28E-05 | 8.15E-05 | 6.72E-05 | 4.14E-05 | 5.57E-05 | 3.05E-05 | 4.58E-05 | 1.24E-06 | 0 | 0.00014 | 0 | 2.04E-05 |
| Dis3l | 8.44E-05 | 4.40E-05 | 3.12E-05 | 3.21E-05 | 3.41E-05 | 1.89E-05 | 3.31E-05 | 2.82E-05 | 6.72E-05 | 4.56E-05 | 3.73E-05 | 0 | 0 | 4.83E-06 | 3.70E-05 | 9.40E-05 | 2.56E-05 | 4.62E-05 |
| Ecd | 0.00013 | 8.09E-05 | 7.67E-05 | 7.26E-05 | 1.12E-05 | 6.75E-06 | 0 | 0 | 6.72E-05 | 3.06E-05 | 2.11E-05 | 3.75E-05 | 0 | 0 | 6.21E-05 | 0.00011 | 0 | 0 |
| Tom1l2 | 6.80E-05 | 0.00015 | 3.74E-05 | 1.67E-05 | 0.00011 | 8.88E-05 | 3.17E-05 | 6.77E-05 | 6.71E-05 | 9.89E-05 | 7.88E-05 | 0.00011 | 3.91E-05 | 5.82E-05 | 6.11E-05 | 4.40E-05 | 9.40E-05 | 0.00012 |
| Lrch4 | 0 | 0.00019 | 0 | 0.00021 | 7.36E-05 | 8.21E-05 | 0 | 0.0002 | 6.71E-05 | 7.44E-05 | 5.52E-05 | 0 | 0 | 0.00012 | 0.00025 | 0.00016 | 0 | 0.00015 |
| Rpia | 4.74E-05 | 4.79E-05 | 7.30E-05 | 0 | 0 | 7.91E-05 | 0 | 2.04E-05 | 6.71E-05 | 7.30E-05 | 9.13E-06 | 3.15E-05 | 4.61E-05 | 0 | 1.80E-05 | 0.00012 | 2.18E-05 | 0.00011 |
| Acot11 | 2.34E-06 | 4.53E-07 | 0 | 5.28E-05 | 3.99E-05 | 3.63E-05 | 5.93E-05 | 7.12E-05 | 6.71E-05 | 6.37E-05 | 4.25E-06 | 0 | 0 | 9.13E-05 | 6.02E-05 | 4.46E-05 | 0 | 0.00015 |
| Pspc1 | 0.00014 | 0.0001 | 0.00013 | 0 | 5.20E-05 | 2.77E-05 | 1.50E-05 | 5.38E-05 | 6.71E-05 | 5.61E-07 | 0 | 2.30E-05 | 0 | 2.13E-05 | 0.00016 | 0.00023 | 0.00036 | 1.32E-05 |
| Rnf31 | 1.12E-05 | 8.23E-05 | 3.58E-05 | 5.40E-05 | 3.64E-05 | 1.35E-05 | 0 | 1.02E-05 | 6.71E-05 | 8.22E-05 | 2.38E-05 | 4.92E-05 | 0 | 2.87E-05 | 6.70E-05 | 4.43E-05 | 0.00014 | 7.73E-05 |
| Dpy30 | 0.00013 | 8.46E-05 | 0.0001 | 5.30E-05 | 3.45E-05 | 6.70E-06 | 6.33E-05 | 0 | 6.71E-05 | 4.98E-05 | 3.51E-05 | 1.44E-05 | 3.34E-06 | 2.49E-05 | 0 | 0.00026 | 3.54E-06 | 3.54E-05 |
| Atad1 | 0.00016 | 7.62E-05 | 8.02E-05 | 0 | 3.72E-05 | 0.00012 | 0 | 5.84E-05 | 6.70E-05 | 1.77E-05 | 3.52E-05 | 3.78E-05 | 4.98E-05 | 4.96E-05 | 4.57E-05 | 5.32E-05 | 1.94E-05 | 1.84E-05 |
| Grk2 | 0.00013 | 0.00015 | 0.00023 | 0 | 6.47E-05 | 6.16E-05 | 0 | 5.16E-05 | 6.70E-05 | 4.31E-05 | 0.00011 | 6.50E-05 | 0 | 2.09E-05 | 0.00016 | 0.0002 | 0.00018 | 2.92E-05 |
| Ankrd27 | 0.00015 | 7.15E-05 | 0.0002 | 8.76E-05 | 1.14E-05 | 4.56E-05 | 0.00011 | 9.23E-06 | 6.70E-05 | 3.18E-05 | 6.33E-05 | 8.44E-05 | 3.45E-05 | 2.78E-05 | 6.92E-05 | 3.02E-05 | 2.95E-05 | 2.11E-05 |
| Ikbkg | 4.86E-05 | 0.00011 | 4.01E-05 | 4.57E-05 | 5.60E-05 | 2.17E-05 | 2.50E-05 | 4.37E-05 | 6.70E-05 | 7.82E-05 | 2.10E-05 | 4.53E-05 | 3.32E-05 | 4.60E-05 | 6.64E-05 | 1.41E-05 | 0 | 8.28E-05 |
| Ptges3l | 2.77E-05 | 0 | 2.39E-05 | 0.00012 | 1.43E-05 | 0 | 6.87E-05 | 6.51E-05 | 6.70E-05 | 2.24E-05 | 0.00011 | 1.14E-05 | 0 | 1.38E-05 | 6.21E-05 | 0 | 0 | 3.36E-05 |
| Ntmt1 | 6.61E-05 | 6.70E-05 | 0.00018 | 0.00015 | 1.64E-05 | 0 | 4.85E-05 | 4.73E-05 | 6.70E-05 | 1.45E-05 | 9.28E-05 | 0.00013 | 7.10E-05 | 3.45E-05 | 2.02E-05 | 1.48E-05 | 0 | 3.23E-05 |
| Pnpla8 | 0.0002 | 0.00022 | 4.35E-05 | 0 | 4.04E-05 | 0.00017 | 4.56E-05 | 0 | 6.69E-05 | 7.33E-05 | 2.63E-05 | 0.00016 | 1.13E-05 | 6.98E-05 | 8.80E-06 | 4.77E-05 | 3.59E-05 | 4.15E-05 |
| Psmd11 | 0.00029 | 0.00025 | 0.00032 | 0.00016 | 6.71E-05 | 0.0001 | 2.24E-06 | 0 | 6.69E-05 | 4.22E-05 | 0.00012 | 0.00012 | 6.09E-05 | 7.92E-05 | 1.71E-05 | 0.00014 | 1.89E-05 | 7.07E-05 |
| Nfya | 5.06E-05 | 0.00015 | 0.00012 | 1.54E-05 | 7.94E-05 | 8.56E-05 | 2.50E-05 | 0.00011 | 6.69E-05 | 6.08E-05 | 0.00012 | 9.50E-05 | 4.81E-05 | 5.36E-05 | 6.82E-05 | 0.00023 | 0.00019 | 9.52E-05 |
| Fam136a | 0.00017 | 2.28E-05 | 0.00014 | 6.54E-05 | 1.01E-05 | 4.28E-05 | 0 | 1.56E-05 | 6.69E-05 | 0 | 7.24E-06 | 6.61E-05 | 0 | 0 | 0 | 0 | 0 | 0 |
| Dnajb14 | 0.00011 | 2.25E-05 | 9.21E-05 | 0 | 5.67E-05 | 4.08E-05 | 5.49E-05 | 6.77E-05 | 6.69E-05 | 1.80E-05 | 4.31E-05 | 5.60E-05 | 9.22E-05 | 0 | 0 | 3.99E-05 | 0.00025 | 0 |
| Trappc10 | 6.71E-05 | 9.60E-05 | 0.00012 | 0 | 5.60E-05 | 2.61E-05 | 2.47E-06 | 0 | 6.69E-05 | 4.47E-05 | 4.66E-05 | 4.19E-05 | 1.98E-05 | 2.41E-05 | 7.88E-05 | 8.45E-05 | 7.23E-05 | 6.94E-05 |
| Polr2h | 0.00012 | 6.80E-05 | 0.00017 | 7.96E-05 | 1.35E-05 | 2.01E-05 | 0 | 0 | 6.69E-05 | 4.21E-05 | 3.99E-05 | 5.98E-05 | 0 | 2.07E-05 | 5.74E-05 | 0 | 1.71E-05 | 5.01E-05 |
| Six5 | 5.66E-05 | 9.24E-05 | 0 | 0.00018 | 9.25E-05 | 0.00011 | 0.00015 | 5.30E-05 | 6.68E-05 | 2.83E-05 | 7.43E-05 | 8.22E-05 | 6.52E-05 | 1.55E-05 | 0.00033 | 0 | 0 | 3.49E-05 |
| Zfp638 | 0.00013 | 0.00011 | 2.71E-05 | 0 | 5.92E-05 | 9.96E-05 | 0 | 2.47E-05 | 6.68E-05 | 5.90E-05 | 2.09E-05 | 5.64E-05 | 0 | 4.30E-05 | 0.00012 | 7.38E-05 | 0 | 1.59E-05 |
| Pskh1 | 1.78E-05 | 9.97E-05 | 3.79E-06 | 7.51E-05 | 9.57E-05 | 8.13E-05 | 2.96E-05 | 8.17E-05 | 6.68E-05 | 3.45E-05 | 8.54E-05 | 5.54E-05 | 3.57E-05 | 5.44E-05 | 3.47E-05 | 5.09E-05 | 0 | 3.95E-05 |
| Grin2a | 0 | 0 | 0 | 0 | 0 | 7.39E-05 | 0 | 0 | 6.68E-05 | 0 | 0 | 3.13E-05 | 0 | 0 | 0 | 1.50E-05 | 0 | 0 |
| Twistnb | 7.45E-05 | 5.96E-05 | 6.61E-05 | 0 | 1.71E-05 | 3.57E-05 | 0 | 0 | 6.68E-05 | 5.08E-07 | 2.82E-05 | 3.54E-05 | 3.05E-06 | 3.05E-06 | 5.76E-06 | 6.63E-05 | 2.00E-05 | 0 |
| Dcaf13 | 0.00021 | 8.85E-05 | 0.0002 | 6.12E-05 | 2.53E-05 | 1.92E-05 | 0 | 0 | 6.68E-05 | 3.49E-05 | 1.51E-05 | 3.77E-05 | 7.91E-06 | 1.92E-05 | 7.00E-05 | 0.00011 | 0 | 3.86E-05 |
| Ppie | 9.20E-05 | 4.38E-05 | 0.00016 | 0.00014 | 4.28E-05 | 5.23E-06 | 0 | 3.66E-05 | 6.68E-05 | 3.82E-05 | 2.16E-05 | 4.12E-05 | 0 | 1.70E-05 | 3.45E-05 | 0.00013 | 7.96E-05 | 3.06E-05 |
| Brwd1 | 0.00023 | 0.00021 | 0.00014 | 0 | 0.00011 | 0.00013 | 0.00016 | 0.00011 | 6.67E-05 | 0 | 0.00012 | 0.00014 | 3.14E-05 | 5.90E-05 | 0.00048 | 0 | 0.00013 | 0 |
| Zfc3h1 | 0.00016 | 0.00012 | 4.39E-05 | 0 | 0.0001 | 8.57E-05 | 2.46E-05 | 0 | 6.67E-05 | 1.72E-05 | 6.26E-05 | 0.00012 | 0 | 6.06E-05 | 0.00031 | 0 | 0 | 4.51E-05 |
| Nin | 0.00012 | 7.58E-05 | 0.00011 | 0 | 1.88E-05 | 6.41E-05 | 6.24E-05 | 7.93E-05 | 6.67E-05 | 9.46E-05 | 0.00013 | 0 | 2.78E-05 | 0.0001 | 0.00015 | 0.00025 | 9.08E-05 | 5.06E-05 |

| | | | | | | | | | | | | | | | | | | |
|---|---|---|---|---|---|---|---|---|---|---|---|---|---|---|---|---|---|---|
| Apc | 0.00014 | 0.00014 | 9.64E-05 | 1.99E-05 | 6.02E-05 | 6.30E-05 | 6.24E-06 | 1.63E-05 | 6.67E-05 | 2.82E-05 | 6.62E-05 | 8.66E-05 | 2.25E-06 | 3.72E-05 | 0.00011 | 7.05E-05 | 7.20E-05 | 6.05E-05 |
| Ctnnbl1 | 4.96E-05 | 8.34E-05 | 9.79E-05 | 0.00012 | 3.05E-05 | 2.78E-05 | 0 | 1.18E-05 | 6.67E-05 | 3.31E-05 | 4.51E-05 | 3.13E-05 | 2.26E-05 | 1.93E-05 | 5.07E-05 | 3.97E-05 | 6.22E-05 | 5.02E-05 |
| Faim | 0.00011 | 6.47E-05 | 6.84E-05 | 9.40E-05 | 8.25E-06 | 2.26E-05 | 1.32E-05 | 2.90E-05 | 6.67E-05 | 6.95E-05 | 3.60E-05 | 3.54E-05 | 5.51E-05 | 1.65E-06 | 5.34E-05 | 4.99E-05 | 0 | 2.69E-06 |
| Dus1l | 9.04E-05 | 4.16E-05 | 9.98E-05 | 6.42E-05 | 1.05E-06 | 1.80E-05 | 3.46E-05 | 0 | 6.67E-05 | 1.49E-05 | 3.43E-05 | 0 | 0 | 6.40E-05 | 7.75E-05 | 5.64E-05 | 0 | 0 |
| Cep68 | 7.69E-06 | 6.39E-05 | 2.39E-05 | 7.63E-06 | 3.20E-05 | 2.64E-06 | 0 | 0 | 6.67E-05 | 3.35E-05 | 4.02E-05 | 4.18E-05 | 2.83E-05 | 2.27E-05 | 5.92E-05 | 7.22E-05 | 0 | 6.82E-06 |
| Sectm1b | 0 | 0 | 0 | 0 | 0 | 0 | 0 | 0 | 6.67E-05 | 1.73E-06 | 0 | 1.60E-05 | 0 | 0 | 0 | 0 | 0 | 0 |
| Clip3 | 8.27E-05 | 0.00034 | 0 | 0.00016 | 0.00024 | 0.0002 | 0.00026 | 0.0001 | 6.66E-05 | 9.06E-05 | 0.00014 | 0.00026 | 0.00012 | 0.00017 | 0 | 0 | 6.84E-05 | 0.00029 |
| Slc36a4 | 0.00014 | 6.63E-05 | 8.42E-05 | 0 | 9.85E-05 | 0.00019 | 0.00012 | 0.00011 | 6.66E-05 | 4.16E-05 | 7.55E-05 | 7.15E-05 | 3.07E-05 | 1.43E-05 | 6.28E-05 | 3.70E-05 | 0 | 3.56E-05 |
| Lancl1 | 4.43E-05 | 0.0001 | 0 | 3.12E-05 | 5.24E-05 | 9.22E-05 | 0 | 2.65E-05 | 6.66E-05 | 6.31E-05 | 3.38E-05 | 6.55E-05 | 4.12E-05 | 1.55E-05 | 1.09E-05 | 8.25E-05 | 4.42E-06 | 7.95E-05 |
| Rfx7 | 6.18E-05 | 9.47E-05 | 2.60E-05 | 2.01E-05 | 3.08E-05 | 5.65E-05 | 0 | 0 | 6.66E-05 | 1.52E-05 | 4.12E-05 | 6.37E-05 | 0 | 3.37E-05 | 2.89E-05 | 0 | 3.06E-06 | 2.58E-05 |
| Smpd4 | 2.39E-05 | 7.05E-05 | 4.27E-05 | 4.23E-05 | 6.24E-05 | 4.72E-05 | 0 | 6.92E-05 | 6.66E-05 | 4.10E-05 | 3.29E-05 | 9.06E-07 | 4.32E-05 | 3.71E-05 | 4.49E-05 | 0.00018 | 5.49E-05 | 3.53E-05 |
| Mex3c | 0.0002 | 0.0002 | 0.0003 | 0 | 5.23E-05 | 0.00021 | 0 | 9.47E-05 | 6.65E-05 | 5.90E-05 | 0.00021 | 0.00017 | 4.67E-05 | 0.00014 | 2.64E-05 | 0.00018 | 0.00066 | 4.48E-05 |
| Rbbp6 | 0.00018 | 0.00022 | 5.33E-05 | 2.66E-05 | 8.62E-05 | 8.01E-05 | 0 | 3.44E-05 | 6.65E-05 | 8.70E-06 | 0 | 0.00016 | 0 | 5.56E-05 | 0.0002 | 0.00012 | 0.00097 | 3.10E-05 |
| C1galt1 | 0.00024 | 3.02E-05 | 3.14E-05 | 0 | 0 | 6.82E-05 | 0 | 8.36E-05 | 6.65E-05 | 0.00027 | 0 | 2.75E-05 | 0.00017 | 0 | 4.59E-05 | 4.93E-05 | 3.78E-05 | 0 |
| Hoxc10 | 6.73E-05 | 0.00019 | 0 | 0.00022 | 0.00024 | 0.00013 | 0.00015 | 2.49E-05 | 6.64E-05 | 9.21E-05 | 6.44E-05 | 0 | 0 | 8.80E-06 | 0.00013 | 0.00019 | 0 | 0.00024 |
| Gstm5 | 0.00012 | 3.34E-05 | 0 | 0 | 4.11E-05 | 5.51E-05 | 9.33E-06 | 0 | 6.64E-05 | 4.27E-05 | 0 | 1.91E-05 | 8.76E-05 | 2.22E-05 | 0 | 0 | 0 | 7.22E-05 |
| Gbp7 | 8.62E-05 | 5.27E-05 | 0 | 0 | 2.92E-05 | 6.37E-06 | 0 | 1.49E-05 | 6.64E-05 | 6.82E-05 | 0 | 6.82E-06 | 0 | 2.17E-06 | 0 | 8.72E-06 | 2.50E-07 | 8.18E-05 |
| Acad9 | 9.61E-05 | 4.41E-05 | 4.23E-05 | 6.17E-06 | 1.58E-06 | 0 | 6.27E-05 | 0 | 6.64E-05 | 2.99E-05 | 1.94E-05 | 1.98E-05 | 1.73E-05 | 0 | 0.00015 | 5.45E-05 | 0 | 5.51E-05 |
| Ssbp3 | 0.00024 | 0.0002 | 8.59E-05 | 0.00022 | 0.00021 | 0.00011 | 4.11E-05 | 5.34E-05 | 6.63E-05 | 5.84E-05 | 0.00014 | 0.00012 | 2.95E-05 | 0.00014 | 8.57E-06 | 0.00018 | 0.00022 | 9.70E-05 |
| Tmem115 | 7.83E-05 | 0.00015 | 0.00018 | 0.00031 | 0.00011 | 5.19E-05 | 0.00011 | 0.0001 | 6.63E-05 | 0.0001 | 8.40E-05 | 0.00023 | 3.94E-05 | 6.81E-05 | 0.00017 | 5.40E-05 | 4.16E-06 | 0.0001 |
| Slc9a3r1 | 0 | 0 | 0 | 0.00028 | 8.14E-05 | 0 | 0.00031 | 0 | 6.63E-05 | 3.09E-06 | 5.58E-05 | 0 | 0.00033 | 0.00017 | 0.00019 | 1.94E-05 | 0.00023 | 0 |
| Blvrb | 7.58E-05 | 9.57E-05 | 2.95E-05 | 0.00014 | 0 | 7.04E-05 | 0 | 8.89E-06 | 6.62E-05 | 0.00027 | 0 | 0.00011 | 1.66E-05 | 1.53E-05 | 7.52E-06 | 0.00013 | 7.17E-05 | 0.00025 |
| Parl | 0.00011 | 2.64E-05 | 0.00012 | 2.05E-05 | 2.77E-05 | 4.28E-05 | 1.17E-05 | 2.66E-05 | 6.62E-05 | 6.40E-06 | 1.22E-05 | 0 | 3.95E-05 | 2.78E-05 | 1.11E-06 | 4.33E-05 | 6.33E-06 | 6.40E-05 |
| Ahdc1 | 1.55E-05 | 0.00022 | 7.75E-05 | 0.00019 | 6.06E-05 | 0 | 2.81E-05 | 0.00041 | 6.62E-05 | 8.93E-05 | 0.00022 | 0.00012 | 0 | 0.00011 | 0.00043 | 7.62E-05 | 0.00022 | 0.00028 |
| Fadd | 0 | 0.0001 | 0 | 0.00022 | 5.18E-05 | 0 | 0 | 1.22E-05 | 6.62E-05 | 5.25E-05 | 3.48E-05 | 0 | 3.85E-05 | 2.17E-05 | 0.00018 | 0.00016 | 0 | 0.00012 |
| Strada | 7.02E-05 | 6.14E-05 | 3.20E-05 | 4.76E-05 | 1.61E-05 | 9.97E-06 | 5.41E-06 | 6.45E-06 | 6.61E-05 | 4.98E-06 | 1.16E-05 | 1.91E-05 | 0 | 2.47E-05 | 6.61E-05 | 6.01E-05 | 0 | 2.45E-05 |
| Nfx1 | 9.76E-05 | 0.00013 | 0.00012 | 0.00011 | 2.38E-05 | 9.85E-06 | 0 | 0 | 6.61E-05 | 1.46E-05 | 8.92E-07 | 0.0001 | 7.32E-06 | 2.77E-05 | 0.00032 | 6.37E-05 | 0 | 1.16E-05 |
| Dip2c | 3.34E-05 | 0.00015 | 0.00013 | 0 | 3.47E-05 | 0 | 7.77E-05 | 3.13E-05 | 6.61E-05 | 2.98E-05 | 4.04E-05 | 0.00015 | 1.43E-05 | 0 | 0.0001 | 3.99E-05 | 0.00012 | 5.00E-05 |
| Ruvbl1 | 0.00014 | 3.57E-05 | 0.00017 | 0.00013 | 0 | 2.33E-05 | 0 | 3.77E-05 | 6.60E-05 | 0 | 0 | 8.83E-06 | 0 | 1.81E-05 | 1.69E-05 | 7.80E-05 | 3.08E-05 | 0 |
| Pigyl | 3.03E-05 | 2.49E-05 | 2.82E-05 | 0.00011 | 7.85E-05 | 7.56E-06 | 2.48E-05 | 2.58E-05 | 6.60E-05 | 3.07E-05 | 4.08E-05 | 0 | 5.22E-06 | 5.59E-05 | 0 | 1.82E-05 | 0 | 2.75E-05 |
| Wdr74 | 7.07E-05 | 7.75E-05 | 0.00024 | 0.00013 | 0 | 0 | 0 | 0 | 6.60E-05 | 2.40E-05 | 4.89E-05 | 6.73E-06 | 0 | 3.75E-05 | 0.0001 | 4.65E-05 | 0 | 0 |
| Ndrg1 | 0 | 8.52E-05 | 4.01E-05 | 6.72E-05 | 3.75E-05 | 0.00025 | 0.0003 | 0 | 6.59E-05 | 0.00015 | 0 | 0.00249 | 0 | 0.00036 | 4.77E-05 | 0 | 0.00118 | 0.00025 |
| Tnrc6c | 4.22E-05 | 0.00022 | 5.89E-05 | 0.0002 | 0.00011 | 6.23E-05 | 8.21E-05 | 6.15E-05 | 6.59E-05 | 8.48E-05 | 0.0001 | 9.04E-05 | 5.23E-05 | 8.34E-05 | 0.00027 | 0.00014 | 0.00019 | 0.00012 |

| | | | | | | | | | | | | | | | | | | |
|---|---|---|---|---|---|---|---|---|---|---|---|---|---|---|---|---|---|---|
| Zfp318 | 1.55E-05 | 0.00011 | 1.04E-05 | 5.58E-05 | 7.49E-05 | 3.02E-05 | 1.86E-05 | 5.57E-06 | 6.59E-05 | 5.27E-05 | 3.01E-05 | 3.59E-05 | 1.91E-05 | 7.49E-05 | 0.00013 | 4.52E-05 | 8.84E-05 | 6.23E-05 |
| Dgcr8 | 1.98E-05 | 5.53E-05 | 5.55E-05 | 1.75E-05 | 1.23E-05 | 2.24E-05 | 5.76E-05 | 8.21E-05 | 6.59E-05 | 4.05E-05 | 8.86E-05 | 9.52E-06 | 1.56E-05 | 2.17E-05 | 0.00017 | 0.00016 | 0 | 6.62E-05 |
| Eif4enif1 | 0.00013 | 9.44E-05 | 0.00011 | 1.20E-05 | 2.92E-05 | 3.53E-06 | 4.51E-05 | 0 | 6.59E-05 | 2.52E-05 | 3.61E-05 | 9.12E-06 | 5.09E-05 | 0 | 3.56E-06 | 0.00012 | 3.95E-05 | 2.01E-05 |
| Fam53b | 4.46E-05 | 8.33E-05 | 0 | 7.28E-05 | 6.76E-05 | 0 | 2.58E-05 | 2.95E-05 | 6.59E-05 | 3.89E-05 | 3.45E-06 | 0 | 0 | 0.00013 | 0.00013 | 0 | 5.67E-05 | 0.00016 |
| Sfxn2 | 6.85E-05 | 2.53E-05 | 8.52E-05 | 8.30E-05 | 1.80E-05 | 0 | 5.49E-05 | 1.52E-05 | 6.59E-05 | 2.37E-05 | 0 | 0 | 6.55E-05 | 0 | 0 | 6.73E-05 | 0 | 1.10E-05 |
| Uhrf1bp1l | 0.00013 | 0.00011 | 0.00014 | 0 | 8.64E-05 | 9.44E-05 | 1.05E-05 | 0 | 6.58E-05 | 0 | 0.0001 | 0.00013 | 2.23E-05 | 4.46E-05 | 3.25E-05 | 0 | 9.95E-05 | 0 |
| Brpf3 | 1.48E-05 | 9.74E-05 | 5.42E-05 | 2.88E-05 | 5.21E-05 | 6.80E-05 | 5.11E-05 | 0 | 6.58E-05 | 4.26E-05 | 3.84E-05 | 5.90E-05 | 6.13E-05 | 8.67E-05 | 0.00015 | 7.05E-05 | 7.86E-05 | 4.21E-07 |
| Evi5l | 2.26E-05 | 5.96E-05 | 3.52E-05 | 5.13E-05 | 7.82E-05 | 2.74E-05 | 3.55E-05 | 8.22E-05 | 6.58E-05 | 0.00018 | 4.14E-05 | 0.00014 | 0 | 2.07E-05 | 0.00016 | 4.56E-05 | 6.16E-05 | 0.00012 |
| Borcs7 | 8.04E-05 | 3.68E-05 | 1.32E-05 | 5.63E-05 | 1.90E-05 | 2.57E-05 | 6.33E-05 | 3.06E-05 | 6.58E-05 | 5.83E-06 | 3.80E-05 | 2.73E-05 | 3.59E-05 | 1.84E-05 | 3.65E-05 | 6.85E-06 | 0 | 0 |
| Ccdc107 | 6.72E-05 | 8.12E-05 | 3.55E-05 | 8.84E-05 | 9.72E-05 | 2.54E-05 | 4.19E-05 | 5.60E-05 | 6.58E-05 | 4.89E-05 | 0.00018 | 1.82E-05 | 7.08E-05 | 9.63E-05 | 6.09E-06 | 8.42E-05 | 0 | 1.96E-05 |
| Pcf11 | 5.06E-05 | 0.0002 | 6.40E-05 | 0.00022 | 3.96E-05 | 0 | 0 | 3.00E-05 | 6.58E-05 | 6.64E-05 | 0 | 0.00021 | 0 | 6.11E-05 | 0.00052 | 0.00019 | 0.00123 | 3.98E-05 |
| Ppp1r37 | 8.91E-05 | 0.00019 | 0.00012 | 7.68E-05 | 0.00012 | 0.00011 | 5.74E-05 | 4.50E-05 | 6.57E-05 | 7.29E-05 | 8.71E-05 | 7.70E-05 | 3.05E-05 | 5.61E-05 | 0.0002 | 9.30E-05 | 6.29E-05 | 4.45E-05 |
| Glis3 | 7.69E-05 | 0.00025 | 0.00025 | 0 | 0.00016 | 0.00035 | 0.00094 | 0.00034 | 6.56E-05 | 0.00031 | 0 | 0.00053 | 0.00114 | 0.00065 | 0.00046 | 0 | 0.00014 | 1.56E-05 |
| Usp38 | 0.0001 | 8.29E-05 | 7.11E-05 | 0 | 4.11E-05 | 6.76E-05 | 1.83E-05 | 3.89E-05 | 6.56E-05 | 1.86E-05 | 4.44E-05 | 2.93E-05 | 8.52E-06 | 5.52E-05 | 1.46E-05 | 2.00E-05 | 1.87E-05 | 1.30E-05 |
| Phf20l1 | 0.00011 | 0.00017 | 8.28E-05 | 2.79E-05 | 0.00011 | 2.56E-05 | 1.16E-05 | 1.85E-05 | 6.56E-05 | 1.31E-05 | 1.52E-05 | 0 | 0 | 7.14E-05 | 0.00047 | 0.00011 | 0.0002 | 4.87E-05 |
| Slc52a2 | 0 | 1.79E-05 | 3.40E-05 | 0.00011 | 4.36E-05 | 9.82E-06 | 0 | 8.70E-05 | 6.56E-05 | 1.80E-05 | 5.24E-05 | 2.09E-06 | 7.52E-06 | 2.12E-05 | 3.75E-05 | 1.13E-05 | 1.18E-05 | 3.78E-05 |
| Zrsr2 | 5.98E-05 | 7.00E-05 | 0.00013 | 0.00017 | 1.14E-05 | 0 | 0 | 0.0001 | 6.56E-05 | 0 | 4.16E-05 | 6.69E-05 | 2.28E-05 | 6.87E-05 | 0.00024 | 9.03E-05 | 0.00014 | 4.15E-05 |
| Ptdss1 | 0.00013 | 0.00013 | 0.00011 | 0.00021 | 0.00019 | 0.00017 | 0.00022 | 0.00014 | 6.55E-05 | 0.0002 | 0.00013 | 0.00013 | 4.44E-05 | 0.00015 | 0 | 0.00011 | 0 | 0.00022 |
| Mdm4 | 0.00013 | 0.00019 | 1.58E-05 | 0 | 5.99E-05 | 9.17E-05 | 2.19E-05 | 0 | 6.55E-05 | 3.90E-05 | 3.85E-05 | 6.19E-05 | 2.96E-06 | 6.25E-05 | 0.00031 | 0.0001 | 8.61E-05 | 7.48E-05 |
| Agk | 4.43E-05 | 6.03E-05 | 5.60E-05 | 4.66E-05 | 3.23E-05 | 2.63E-05 | 2.08E-05 | 2.00E-05 | 6.55E-05 | 3.89E-05 | 1.62E-05 | 1.46E-05 | 4.89E-05 | 2.52E-05 | 0 | 5.81E-05 | 3.94E-06 | 6.05E-05 |
| Ttc7 | 0 | 5.17E-05 | 0.00012 | 0.00014 | 8.29E-05 | 7.26E-05 | 5.62E-05 | 0 | 6.54E-05 | 5.60E-05 | 0.00011 | 9.48E-05 | 0.00018 | 8.67E-05 | 0.00011 | 6.92E-05 | 3.06E-05 | 0.00019 |
| Ccnt1 | 6.44E-05 | 9.93E-05 | 8.36E-05 | 1.97E-05 | 4.29E-05 | 4.69E-05 | 2.40E-05 | 5.60E-05 | 6.54E-05 | 1.36E-05 | 3.07E-05 | 0.00011 | 0 | 7.48E-05 | 0.00017 | 3.38E-05 | 0.00014 | 1.03E-05 |
| Napg | 4.60E-05 | 8.33E-05 | 0.00012 | 8.34E-05 | 3.85E-05 | 3.59E-05 | 5.77E-06 | 7.08E-05 | 6.54E-05 | 5.41E-05 | 9.08E-05 | 3.00E-05 | 6.97E-06 | 4.86E-05 | 0.00018 | 4.94E-05 | 5.81E-05 | 3.45E-05 |
| Pwp1 | 0.00013 | 3.29E-05 | 0.00015 | 2.53E-05 | 2.78E-06 | 1.98E-05 | 0 | 0 | 6.54E-05 | 8.93E-06 | 2.12E-05 | 2.79E-06 | 2.75E-05 | 0 | 1.39E-05 | 6.96E-05 | 0 | 0 |
| Ube2f | 0.00017 | 0.00015 | 0.00025 | 0.00016 | 0.00014 | 0 | 2.18E-05 | 0.00011 | 6.54E-05 | 8.12E-05 | 0.0004 | 1.39E-05 | 9.66E-05 | 0.00016 | 0.00025 | 1.76E-05 | 0.00017 | 9.96E-06 |
| Lancl2 | 8.30E-05 | 2.94E-05 | 6.27E-06 | 3.29E-05 | 3.29E-06 | 0 | 5.23E-05 | 0 | 6.54E-05 | 0 | 5.91E-05 | 0 | 0 | 3.83E-05 | 0 | 0.00014 | 0 | 5.28E-05 |
| Mafg | 0 | 0.00025 | 0.00024 | 8.44E-05 | 0.00014 | 3.03E-05 | 2.86E-05 | 0.0001 | 6.53E-05 | 3.20E-05 | 0.0001 | 0.00019 | 0 | 0.00012 | 0.00054 | 5.39E-05 | 0.00018 | 4.21E-05 |
| Rbsn | 0.0001 | 9.59E-05 | 0.00013 | 6.97E-05 | 4.95E-05 | 2.68E-05 | 7.70E-05 | 2.10E-05 | 6.53E-05 | 5.22E-05 | 1.84E-05 | 3.12E-05 | 5.48E-05 | 5.98E-05 | 6.47E-05 | 6.92E-05 | 0 | 2.70E-05 |
| Timm23 | 0.00014 | 5.44E-05 | 0.00029 | 0.00023 | 5.91E-05 | 2.29E-05 | 0.0001 | 3.22E-05 | 6.53E-05 | 4.72E-05 | 8.66E-05 | 3.95E-05 | 1.52E-05 | 2.18E-05 | 5.81E-05 | 0.00012 | 0 | 1.59E-05 |
| Vac14 | 1.11E-05 | 9.94E-05 | 7.47E-05 | 6.74E-05 | 4.44E-05 | 1.66E-05 | 0 | 4.40E-05 | 6.53E-05 | 4.46E-05 | 3.62E-05 | 0 | 0 | 8.44E-05 | 9.20E-05 | 9.02E-05 | 6.70E-05 | 4.01E-05 |
| Pdcd7 | 2.34E-05 | 7.45E-05 | 6.86E-05 | 5.41E-05 | 2.05E-05 | 0 | 0 | 2.88E-05 | 6.53E-05 | 3.18E-05 | 2.86E-06 | 6.64E-05 | 0 | 2.92E-05 | 0.00017 | 7.13E-05 | 2.61E-05 | 4.17E-05 |
| Tomm5 | 0.0004 | 0.00011 | 0.00036 | 0.00024 | 7.57E-05 | 0.00016 | 6.99E-05 | 3.81E-06 | 6.52E-05 | 5.94E-05 | 6.70E-05 | 6.53E-05 | 0.00012 | 2.00E-05 | 0 | 0 | 0 | 1.04E-05 |
| Max | 0.00018 | 0.00018 | 0.00017 | 0.00012 | 5.29E-05 | 9.76E-05 | 4.35E-05 | 8.20E-06 | 6.52E-05 | 6.30E-05 | 9.28E-05 | 0.00018 | 6.18E-05 | 4.48E-05 | 9.72E-05 | 0 | 5.25E-05 | 0.00013 |

| | | | | | | | | | | | | | | | | | | |
|---|---|---|---|---|---|---|---|---|---|---|---|---|---|---|---|---|---|---|
| Dpagt1 | 0.00016 | 8.91E-05 | 0.00011 | 0.00014 | 1.17E-05 | 2.79E-05 | 0.00015 | 2.60E-05 | 6.52E-05 | 8.81E-05 | 4.85E-05 | 0.00014 | 2.87E-05 | 4.51E-05 | 3.91E-05 | 0.00018 | 0 | 0.00011 |
| Gabarapl2 | 0.00011 | 0.00018 | 4.88E-05 | 0.0002 | 0.00015 | 0.00018 | 0 | 0.00019 | 6.51E-05 | 9.43E-05 | 0.00015 | 0.00012 | 6.34E-05 | 0.0001 | 6.60E-05 | 3.93E-05 | 0.00013 | 8.19E-05 |
| Cd99l2 | 0.00012 | 0.0001 | 7.75E-06 | 7.00E-05 | 7.31E-05 | 0.00012 | 3.05E-05 | 6.33E-05 | 6.51E-05 | 8.76E-05 | 6.44E-05 | 6.05E-05 | 1.87E-05 | 7.73E-05 | 0 | 1.69E-05 | 3.48E-06 | 0.00013 |
| Smg1 | 0.00019 | 0.00021 | 0.00015 | 5.32E-05 | 6.16E-05 | 7.45E-05 | 7.28E-05 | 3.67E-05 | 6.51E-05 | 7.79E-05 | 6.05E-05 | 0.00017 | 7.81E-06 | 1.72E-07 | 0.00022 | 9.37E-05 | 0.00031 | 3.82E-05 |
| Slc25a44 | 7.27E-05 | 5.63E-05 | 5.42E-05 | 2.59E-05 | 6.31E-05 | 2.40E-05 | 1.06E-06 | 1.04E-05 | 6.51E-05 | 2.36E-05 | 2.49E-05 | 1.51E-05 | 5.06E-05 | 3.35E-05 | 2.59E-05 | 1.46E-05 | 9.36E-05 | 0 |
| Rbm10 | 5.63E-05 | 4.52E-05 | 2.14E-05 | 2.81E-05 | 3.25E-05 | 1.36E-05 | 0 | 2.04E-05 | 6.51E-05 | 1.10E-05 | 1.48E-05 | 0 | 0 | 1.90E-05 | 4.58E-05 | 0.00011 | 0 | 2.30E-05 |
| Ankra2 | 3.86E-05 | 5.40E-05 | 0 | 4.77E-05 | 6.48E-05 | 1.07E-05 | 4.60E-06 | 0 | 6.51E-05 | 6.03E-05 | 3.64E-05 | 5.81E-05 | 2.20E-05 | 3.68E-05 | 3.18E-05 | 9.63E-06 | 0 | 3.28E-05 |
| Pop4 | 9.47E-05 | 6.53E-05 | 7.05E-05 | 0 | 4.98E-05 | 1.18E-05 | 0 | 3.55E-05 | 6.50E-05 | 3.57E-05 | 7.85E-05 | 5.65E-05 | 0 | 4.62E-05 | 4.27E-05 | 0.00026 | 0 | 2.25E-05 |
| Disp1 | 0 | 2.91E-05 | 1.77E-05 | 0 | 2.22E-05 | 2.74E-06 | 2.00E-05 | 2.05E-05 | 6.50E-05 | 0.00011 | 1.56E-05 | 1.23E-05 | 0 | 0 | 5.48E-06 | 3.53E-05 | 0 | 3.20E-06 |
| Mfsd14b | 0.00026 | 0.00014 | 8.04E-05 | 0 | 0.00016 | 0.00029 | 5.70E-05 | 6.04E-05 | 6.49E-05 | 6.04E-05 | 4.26E-05 | 0.00011 | 1.91E-05 | 2.79E-05 | 0 | 0.00028 | 0.00013 | 0.00013 |
| Fam193a | 9.40E-05 | 0.00018 | 7.42E-05 | 5.19E-05 | 7.86E-05 | 8.29E-05 | 5.33E-05 | 4.47E-05 | 6.49E-05 | 6.11E-05 | 3.43E-05 | 9.77E-05 | 0 | 8.04E-05 | 0.00032 | 7.17E-05 | 0.00025 | 0.00012 |
| Glb1l | 0 | 5.29E-05 | 0 | 1.66E-05 | 6.97E-05 | 4.98E-05 | 4.42E-05 | 1.17E-05 | 6.49E-05 | 8.88E-05 | 2.53E-05 | 0 | 0.00014 | 9.16E-05 | 0.0001 | 1.37E-05 | 2.07E-05 | 0.00014 |
| Nadk2 | 3.32E-05 | 3.80E-05 | 1.15E-05 | 2.59E-05 | 2.16E-05 | 1.11E-05 | 2.09E-05 | 0 | 6.49E-05 | 5.41E-06 | 0 | 0.00018 | 5.01E-05 | 3.61E-05 | 0 | 9.05E-05 | 0 | 1.34E-05 |
| Psme4 | 0.00028 | 0.00016 | 0.00015 | 9.31E-05 | 8.41E-05 | 0.00011 | 0.00016 | 2.50E-05 | 6.48E-05 | 7.07E-06 | 6.83E-05 | 1.05E-05 | 0 | 0.00013 | 0.00018 | 8.61E-05 | 0.00014 | 3.19E-06 |
| Tmem161a | 9.47E-06 | 0.0001 | 9.99E-05 | 0.00016 | 0.0001 | 8.78E-05 | 2.47E-05 | 0.0001 | 6.48E-05 | 5.31E-05 | 6.31E-05 | 8.71E-05 | 8.87E-05 | 6.72E-05 | 8.12E-05 | 8.73E-05 | 0 | 9.99E-05 |
| Necap1 | 3.10E-05 | 8.49E-05 | 0.00011 | 4.60E-05 | 5.02E-05 | 1.79E-05 | 8.69E-05 | 5.00E-05 | 6.48E-05 | 3.80E-05 | 4.76E-05 | 0.00011 | 9.60E-05 | 3.87E-05 | 8.26E-05 | 5.30E-05 | 9.01E-06 | 1.93E-05 |
| Tgm2 | 0 | 0 | 0.00016 | 6.93E-05 | 0 | 0 | 0 | 0.00058 | 6.48E-05 | 0.0005 | 0 | 0.00014 | 0.00067 | 0 | 0 | 2.33E-05 | 7.94E-05 | 0.0004 |
| Fam216a | 9.97E-05 | 5.17E-05 | 3.58E-05 | 9.23E-06 | 1.64E-05 | 0 | 1.82E-05 | 0 | 6.48E-05 | 1.87E-05 | 2.74E-05 | 1.77E-05 | 1.06E-05 | 7.16E-05 | 8.20E-05 | 0.00016 | 0 | 1.31E-05 |
| Rspo4 | 0 | 0 | 0 | 0 | 0 | 0.00025 | 0 | 0 | 6.47E-05 | 0 | 0 | 0 | 0 | 0 | 0 | 0 | 0 | 0 |
| Prxl2b | 1.15E-05 | 4.03E-05 | 0 | 5.41E-05 | 2.89E-05 | 4.60E-05 | 0 | 0 | 6.47E-05 | 2.41E-05 | 3.82E-05 | 2.04E-05 | 2.75E-05 | 6.78E-05 | 0 | 0 | 0 | 5.99E-06 |
| Iars | 0.00019 | 0.00011 | 0.00029 | 7.50E-05 | 2.49E-05 | 0 | 4.85E-05 | 0 | 6.47E-05 | 4.31E-05 | 0.00014 | 0.0002 | 0 | 0 | 0.0001 | 4.18E-05 | 0 | 4.24E-05 |
| Desi2 | 0.0002 | 0.00016 | 0.00015 | 7.61E-05 | 0.00012 | 0.00017 | 0.0003 | 0.00015 | 6.46E-05 | 9.71E-05 | 8.96E-05 | 0.00018 | 7.05E-05 | 6.78E-05 | 2.73E-05 | 0 | 0.00011 | 5.61E-05 |
| Sspn | 1.87E-05 | 3.74E-06 | 0 | 3.20E-06 | 6.96E-05 | 6.31E-05 | 6.70E-05 | 0.00011 | 6.46E-05 | 0.00011 | 6.23E-05 | 0 | 0 | 9.44E-05 | 0 | 3.19E-05 | 0 | 0.00019 |
| Nup155 | 7.62E-05 | 5.53E-05 | 6.91E-05 | 0 | 2.70E-05 | 4.03E-05 | 0 | 1.21E-05 | 6.46E-05 | 1.60E-05 | 1.11E-05 | 2.53E-05 | 2.84E-05 | 1.51E-05 | 0.00011 | 0.00041 | 0 | 2.54E-05 |
| Cops3 | 0.00019 | 2.26E-05 | 0.0001 | 2.62E-05 | 0 | 3.46E-05 | 7.03E-05 | 0 | 6.46E-05 | 4.32E-05 | 5.26E-06 | 1.51E-05 | 3.90E-05 | 1.38E-05 | 0.00011 | 9.64E-05 | 0 | 3.54E-05 |
| Zyg11b | 6.13E-05 | 9.17E-05 | 5.69E-05 | 3.88E-05 | 4.04E-05 | 3.34E-05 | 0 | 0 | 6.46E-05 | 4.91E-05 | 3.18E-05 | 5.42E-05 | 2.19E-05 | 4.57E-05 | 4.57E-05 | 3.81E-05 | 3.29E-05 | 5.42E-05 |
| Polr3d | 0.00013 | 4.35E-05 | 0.0003 | 0.00017 | 4.94E-05 | 7.64E-06 | 0.0002 | 0.00011 | 6.46E-05 | 5.82E-05 | 0.00011 | 2.27E-05 | 8.79E-05 | 6.47E-05 | 0.00023 | 0 | 0.00014 | 0 |
| Bms1 | 9.86E-05 | 5.60E-05 | 9.45E-05 | 0 | 1.74E-05 | 0 | 0 | 0 | 6.46E-05 | 7.88E-06 | 0 | 3.53E-05 | 9.43E-06 | 0 | 8.27E-05 | 6.73E-05 | 2.41E-05 | 0 |
| Josd1 | 7.75E-05 | 0.00017 | 0.00013 | 5.48E-05 | 0.00011 | 0.00013 | 9.49E-05 | 5.37E-05 | 6.45E-05 | 5.15E-05 | 7.48E-05 | 9.49E-05 | 7.83E-05 | 0.0001 | 0 | 9.82E-05 | 7.66E-06 | 0.00012 |
| Rock1 | 0.00022 | 0.00017 | 9.13E-05 | 0 | 2.63E-05 | 9.93E-05 | 6.72E-06 | 9.27E-06 | 6.45E-05 | 9.51E-05 | 9.49E-05 | 0.00013 | 0 | 1.13E-05 | 0.00032 | 0.00014 | 0.00035 | 2.27E-05 |
| Psmd1 | 0.00026 | 0.00027 | 0.00042 | 0.00029 | 0.00022 | 7.41E-05 | 2.02E-05 | 3.27E-05 | 6.45E-05 | 8.10E-05 | 0.00014 | 7.33E-05 | 4.23E-05 | 3.58E-05 | 5.01E-05 | 0.00026 | 3.91E-06 | 6.31E-05 |
| Uqcrb | 0.00022 | 7.49E-05 | 0.0002 | 0.00014 | 2.01E-05 | 5.57E-05 | 2.28E-07 | 1.10E-05 | 6.45E-05 | 4.80E-05 | 0.0001 | 1.92E-05 | 1.71E-05 | 3.07E-05 | 0 | 0.0001 | 7.04E-05 | 4.53E-05 |
| Crppa | 7.21E-05 | 3.91E-05 | 0 | 0 | 3.81E-05 | 3.73E-05 | 0 | 3.18E-05 | 6.45E-05 | 7.96E-06 | 0 | 0 | 0 | 3.14E-05 | 0.0001 | 8.00E-06 | 0 | 7.68E-06 |

| | | | | | | | | | | | | | | | | | | |
|---|---|---|---|---|---|---|---|---|---|---|---|---|---|---|---|---|---|---|
| Pmm2 | 0.00015 | 8.27E-05 | 0.00012 | 9.02E-05 | 0.0001 | 6.27E-05 | 3.33E-05 | 0 | 6.44E-05 | 4.45E-05 | 3.01E-05 | 4.35E-05 | 0 | 5.54E-05 | 6.35E-05 | 9.76E-05 | 0 | 5.81E-05 |
| Bbip1 | 0.00013 | 5.14E-05 | 1.68E-06 | 1.26E-05 | 4.95E-05 | 5.20E-05 | 5.96E-05 | 7.00E-05 | 6.44E-05 | 0 | 5.25E-05 | 2.93E-05 | 8.58E-06 | 0 | 0 | 0.00015 | 0 | 4.37E-05 |
| Mettl1 | 0.00013 | 2.66E-05 | 7.33E-05 | 0.00016 | 5.01E-05 | 4.05E-05 | 7.16E-05 | 0 | 6.44E-05 | 1.52E-05 | 5.82E-05 | 9.10E-06 | 6.45E-05 | 3.14E-05 | 6.86E-05 | 4.68E-05 | 1.01E-05 | 2.27E-06 |
| Nol11 | 0.0001 | 6.54E-05 | 0.00012 | 4.58E-05 | 9.24E-06 | 7.92E-06 | 0 | 0 | 6.44E-05 | 3.68E-05 | 2.19E-05 | 3.55E-06 | 3.05E-05 | 0 | 1.88E-06 | 7.27E-05 | 0 | 0 |
| Sntg2 | 6.46E-06 | 1.13E-05 | 0 | 0 | 5.65E-05 | 5.86E-06 | 3.86E-05 | 0 | 6.44E-05 | 5.26E-06 | 0 | 0 | 4.55E-06 | 6.06E-06 | 1.29E-05 | 9.12E-06 | 0 | 3.12E-07 |
| Cbx4 | 4.83E-05 | 5.94E-05 | 1.56E-05 | 0.00012 | 3.92E-05 | 0 | 0 | 1.89E-05 | 6.44E-05 | 3.11E-05 | 0 | 1.19E-05 | 7.92E-05 | 1.27E-05 | 7.53E-05 | 0 | 0.00084 | 4.45E-05 |
| Mdfic | 0.00019 | 0.00032 | 9.92E-05 | 0 | 0.00033 | 0.00027 | 6.03E-05 | 0.00028 | 6.43E-05 | 2.25E-05 | 0.00016 | 6.53E-05 | 0.00021 | 6.81E-05 | 0.00037 | 0.00016 | 3.63E-05 | 0.00013 |
| Tmem9b | 0.00021 | 5.34E-05 | 0.00016 | 2.19E-05 | 0.00013 | 0.00024 | 7.12E-05 | 0.00011 | 6.43E-05 | 0.00012 | 0.0001 | 3.94E-05 | 0.00017 | 9.98E-05 | 0 | 0 | 0 | 0.00019 |
| Hmcn1 | 0 | 3.44E-05 | 0 | 0 | 0.00019 | 0.00021 | 5.99E-05 | 0 | 6.43E-05 | 2.17E-05 | 0 | 0 | 0 | 6.45E-05 | 1.89E-05 | 0 | 0 | 0.00017 |
| Plekha3 | 8.60E-05 | 0.00013 | 0.00028 | 4.63E-05 | 0.00014 | 0.00013 | 0 | 0.00014 | 6.43E-05 | 5.00E-05 | 0.00022 | 6.12E-05 | 0.00014 | 3.44E-05 | 3.50E-05 | 0.00018 | 0 | 2.15E-05 |
| Ep300 | 0.00017 | 0.00024 | 5.81E-05 | 1.28E-05 | 7.26E-05 | 0.00013 | 3.46E-05 | 2.11E-05 | 6.43E-05 | 2.60E-05 | 4.74E-05 | 0.00012 | 1.80E-05 | 9.14E-05 | 0.00021 | 9.00E-05 | 0.00026 | 3.64E-05 |
| Csnk2a2 | 0.00019 | 0.00017 | 0.00028 | 2.31E-05 | 9.88E-05 | 0.00011 | 2.82E-05 | 9.23E-06 | 6.43E-05 | 3.06E-05 | 8.35E-05 | 0.00016 | 0 | 5.16E-05 | 4.53E-05 | 0.00014 | 0.00014 | 1.93E-05 |
| Med29 | 0.00018 | 8.62E-05 | 0.00014 | 4.50E-05 | 3.56E-05 | 6.16E-05 | 2.62E-05 | 0 | 6.43E-05 | 0 | 7.60E-05 | 0 | 1.39E-05 | 1.51E-05 | 0 | 6.54E-05 | 0 | 1.37E-06 |
| Rnf121 | 0.00016 | 3.39E-05 | 0.00021 | 2.61E-06 | 4.83E-05 | 5.39E-05 | 0.00012 | 4.73E-05 | 6.43E-05 | 6.16E-05 | 8.17E-05 | 0 | 8.08E-05 | 5.46E-05 | 0.00011 | 7.42E-05 | 0 | 2.31E-05 |
| Ptp4a3 | 0.00012 | 7.41E-05 | 0.00011 | 0.00018 | 9.42E-05 | 3.49E-05 | 9.07E-05 | 4.42E-05 | 6.43E-05 | 4.04E-05 | 3.23E-05 | 0 | 0 | 2.53E-05 | 5.43E-05 | 0 | 0 | 3.73E-05 |
| Tuba1c | 7.99E-05 | 4.50E-05 | 0.00067 | 0.00027 | 0.0002 | 0 | 0.00017 | 3.20E-05 | 6.43E-05 | 1.88E-05 | 0.00056 | 0.00028 | 0.00019 | 0.0002 | 0 | 0.00101 | 0.00069 | 0 |
| Rab3gap1 | 4.60E-05 | 7.45E-05 | 7.40E-05 | 9.89E-05 | 0.0001 | 7.18E-05 | 0.00013 | 7.23E-05 | 6.42E-05 | 9.49E-05 | 0.0001 | 7.61E-05 | 7.98E-05 | 4.92E-05 | 0.00014 | 3.58E-05 | 1.19E-05 | 7.05E-05 |
| Lnpk | 5.90E-05 | 7.09E-05 | 0.00017 | 0 | 3.01E-05 | 6.52E-05 | 1.57E-05 | 6.28E-05 | 6.42E-05 | 6.00E-05 | 7.25E-05 | 5.96E-05 | 0 | 5.71E-05 | 2.28E-05 | 0.00036 | 7.52E-05 | 6.21E-05 |
| Ccndbp1 | 7.27E-05 | 0.00015 | 2.53E-05 | 8.42E-05 | 6.88E-05 | 7.81E-05 | 7.61E-05 | 6.57E-05 | 6.41E-05 | 5.07E-05 | 6.05E-05 | 9.77E-05 | 2.61E-05 | 9.18E-05 | 6.36E-05 | 0 | 0 | 8.17E-05 |
| Nedd1 | 3.04E-05 | 6.88E-05 | 4.98E-05 | 2.63E-05 | 6.90E-05 | 7.24E-05 | 8.10E-07 | 2.77E-05 | 6.41E-05 | 4.23E-05 | 7.14E-05 | 9.49E-05 | 2.30E-05 | 1.98E-05 | 0 | 0.00039 | 9.93E-06 | 4.22E-05 |
| Trappc11 | 0.00013 | 9.39E-05 | 7.16E-05 | 3.14E-05 | 7.09E-05 | 6.15E-05 | 4.21E-05 | 2.26E-05 | 6.41E-05 | 2.22E-05 | 1.88E-05 | 3.54E-05 | 1.84E-05 | 2.38E-05 | 0.00011 | 4.35E-05 | 1.04E-05 | 0 |
| Kmt2b | 2.86E-05 | 0.00017 | 5.75E-05 | 0.0001 | 5.08E-05 | 3.11E-05 | 4.68E-06 | 1.14E-05 | 6.41E-05 | 5.11E-05 | 3.83E-05 | 3.88E-05 | 0 | 2.50E-05 | 0.00033 | 9.71E-05 | 7.18E-05 | 1.44E-05 |
| Scrib | 0 | 9.35E-05 | 4.42E-05 | 0.00012 | 3.48E-05 | 6.99E-06 | 0 | 2.40E-06 | 6.41E-05 | 5.22E-05 | 0 | 1.93E-05 | 0 | 2.27E-05 | 0.00018 | 0.00023 | 8.79E-05 | 4.86E-05 |
| Irgm1 | 0.00012 | 9.97E-05 | 0 | 2.63E-05 | 2.43E-05 | 0 | 0 | 0 | 6.41E-05 | 0.00027 | 0.00011 | 0 | 0 | 3.42E-05 | 0 | 5.09E-05 | 0 | 0.00019 |
| Sgk1 | 0.00012 | 9.11E-05 | 0.0003 | 0.0002 | 0 | 8.75E-05 | 0.00041 | 9.01E-05 | 6.40E-05 | 0 | 6.47E-05 | 0 | 0.00038 | 0.00084 | 1.14E-05 | 0.0002 | 0.00109 | 0.00018 |
| Nprl3 | 3.43E-05 | 0.00011 | 3.68E-05 | 8.87E-05 | 7.56E-05 | 5.77E-05 | 1.70E-05 | 4.48E-05 | 6.40E-05 | 2.19E-05 | 0 | 0 | 3.65E-05 | 1.89E-05 | 9.42E-05 | 1.27E-05 | 3.67E-05 | 4.33E-05 |
| Satb1 | 0 | 2.86E-05 | 0 | 0 | 1.98E-05 | 4.81E-05 | 0 | 0 | 6.40E-05 | 2.67E-06 | 0 | 0 | 4.04E-05 | 5.89E-05 | 0 | 0 | 0 | 1.12E-05 |
| Rab3gap2 | 0.00019 | 0.00015 | 7.39E-05 | 3.39E-05 | 1.07E-05 | 4.74E-05 | 0.00014 | 0 | 6.40E-05 | 6.63E-05 | 7.38E-05 | 0.00024 | 0.00015 | 2.50E-06 | 0.00014 | 0 | 8.79E-05 | 1.42E-06 |
| Pygo1 | 5.05E-05 | 3.57E-05 | 1.83E-05 | 2.16E-05 | 7.83E-05 | 3.63E-05 | 6.38E-05 | 5.08E-05 | 6.40E-05 | 1.64E-05 | 2.89E-05 | 1.62E-05 | 7.08E-05 | 2.53E-05 | 1.25E-05 | 9.71E-06 | 0 | 3.83E-05 |
| Tcerg1 | 9.41E-05 | 0.00011 | 0.00019 | 5.85E-05 | 4.24E-05 | 3.36E-05 | 0 | 0 | 6.40E-05 | 4.52E-05 | 5.77E-05 | 0.00011 | 8.10E-06 | 0 | 0.0002 | 0.00023 | 2.66E-05 | 2.76E-05 |
| Ap3b1 | 0.00019 | 0.00022 | 0.00011 | 0.00014 | 8.49E-05 | 0.00013 | 0.00016 | 0.00011 | 6.39E-05 | 9.92E-05 | 5.25E-05 | 0.0002 | 5.46E-05 | 0.00012 | 0.00029 | 5.22E-05 | 1.29E-05 | 0.00011 |
| Wdr37 | 0.00012 | 9.68E-05 | 3.17E-05 | 0 | 4.99E-05 | 4.80E-05 | 0 | 0 | 6.39E-05 | 3.90E-05 | 4.03E-06 | 1.96E-05 | 1.25E-05 | 0 | 0.00021 | 8.56E-05 | 0 | 9.89E-06 |
| Otulinl | 9.99E-06 | 3.16E-05 | 0 | 0 | 0.00012 | 2.22E-05 | 0 | 2.11E-05 | 6.39E-05 | 0 | 3.38E-05 | 0 | 0 | 0.00013 | 4.63E-05 | 0 | 0 | 2.00E-05 |

| | | | | | | | | | | | | | | | | | | |
|---|---|---|---|---|---|---|---|---|---|---|---|---|---|---|---|---|---|---|
| Ndufa9 | 9.63E-05 | 3.99E-05 | 7.10E-05 | 6.30E-05 | 2.23E-05 | 2.00E-05 | 0 | 0 | 6.39E-05 | 7.68E-06 | 4.83E-05 | 1.75E-05 | 0 | 3.05E-06 | 0 | 3.70E-05 | 0 | 2.49E-05 |
| Cep164 | 0 | 4.13E-05 | 4.37E-05 | 9.52E-05 | 3.97E-05 | 1.12E-05 | 4.56E-05 | 2.24E-05 | 6.39E-05 | 2.82E-05 | 3.47E-05 | 0 | 0 | 6.89E-05 | 9.08E-05 | 8.82E-05 | 2.95E-05 | 6.60E-05 |
| Foxk1 | 1.04E-05 | 0.00015 | 8.20E-05 | 0.00018 | 3.91E-05 | 0 | 6.74E-05 | 0.00022 | 6.39E-05 | 3.08E-05 | 0 | 0.00016 | 7.00E-07 | 0.00014 | 0.00023 | 6.85E-05 | 7.03E-05 | 8.65E-05 |
| Tmem260 | 7.47E-05 | 4.22E-05 | 1.55E-05 | 1.08E-05 | 8.23E-05 | 9.08E-05 | 4.72E-05 | 4.96E-05 | 6.38E-05 | 4.60E-05 | 0 | 2.40E-05 | 3.92E-05 | 2.14E-05 | 0.00011 | 5.85E-05 | 0 | 4.45E-05 |
| Phf14 | 0.00016 | 0.00013 | 5.41E-05 | 7.04E-06 | 7.20E-05 | 2.96E-05 | 6.26E-05 | 0 | 6.38E-05 | 4.46E-05 | 3.19E-05 | 5.72E-05 | 2.35E-06 | 4.86E-05 | 0.00036 | 7.73E-05 | 1.91E-05 | 3.53E-05 |
| Taok3 | 6.95E-05 | 0.00011 | 3.89E-05 | 0 | 5.00E-05 | 2.86E-05 | 3.43E-05 | 2.37E-06 | 6.38E-05 | 0.00017 | 7.89E-05 | 6.62E-06 | 4.01E-05 | 1.88E-05 | 9.93E-05 | 1.29E-05 | 6.71E-05 | 0.00011 |
| Ano8 | 0 | 8.85E-05 | 4.72E-05 | 9.69E-05 | 1.51E-05 | 2.36E-06 | 0 | 6.59E-05 | 6.38E-05 | 6.92E-05 | 3.50E-05 | 2.83E-05 | 1.16E-05 | 3.58E-05 | 9.77E-05 | 9.32E-07 | 6.24E-05 | 9.40E-05 |
| Zdhhc20 | 0.0003 | 0.00012 | 8.19E-05 | 0 | 9.55E-05 | 0.00022 | 0.00015 | 0.00018 | 6.37E-05 | 0 | 1.22E-05 | 0.00015 | 0 | 3.94E-05 | 7.86E-05 | 1.65E-05 | 9.28E-05 | 4.35E-05 |
| Usb1 | 8.96E-05 | 0.00011 | 9.70E-05 | 8.98E-05 | 6.72E-05 | 7.62E-05 | 3.05E-05 | 6.00E-05 | 6.37E-05 | 2.92E-05 | 5.71E-05 | 4.79E-05 | 2.08E-05 | 2.30E-06 | 5.55E-05 | 2.07E-05 | 0 | 9.57E-07 |
| Cdc40 | 0.00011 | 7.42E-05 | 4.07E-05 | 0 | 2.09E-05 | 6.25E-05 | 0 | 0 | 6.37E-05 | 2.26E-05 | 3.98E-05 | 7.62E-05 | 7.29E-06 | 2.64E-05 | 2.31E-05 | 5.34E-05 | 2.85E-05 | 0 |
| Retsat | 6.59E-05 | 2.89E-05 | 4.56E-05 | 1.33E-05 | 0 | 4.24E-05 | 0 | 5.38E-05 | 6.37E-05 | 3.06E-05 | 0 | 0 | 3.04E-05 | 8.40E-06 | 0 | 4.19E-05 | 0 | 2.81E-05 |
| Ncoa7 | 4.55E-05 | 4.14E-05 | 0 | 3.27E-06 | 6.89E-06 | 3.74E-05 | 0 | 1.29E-06 | 6.37E-05 | 0.00013 | 1.37E-05 | 3.37E-05 | 4.62E-05 | 1.92E-05 | 4.61E-05 | 9.71E-05 | 0.00012 | 5.89E-05 |
| Zfp628 | 2.34E-05 | 7.21E-05 | 5.19E-05 | 8.19E-05 | 6.35E-05 | 3.35E-05 | 4.18E-05 | 2.55E-05 | 6.37E-05 | 4.92E-05 | 2.85E-05 | 3.77E-05 | 0 | 4.48E-05 | 3.92E-05 | 4.11E-05 | 1.76E-05 | 4.91E-05 |
| Ccdc58 | 0.00011 | 4.66E-05 | 7.78E-05 | 6.60E-05 | 2.78E-05 | 1.83E-05 | 1.54E-05 | 0 | 6.37E-05 | 7.04E-06 | 0 | 0.00019 | 3.09E-05 | 0 | 4.95E-05 | 5.60E-05 | 0 | 0 |
| Emc7 | 0.00028 | 0.00016 | 0.00031 | 0.0004 | 0.00015 | 0.00037 | 4.14E-05 | 0.00028 | 6.36E-05 | 0.00017 | 0.00017 | 0.00019 | 0.00026 | 0.00019 | 0 | 0.00019 | 5.59E-05 | 0.00011 |
| Gpsm3 | 0 | 0.00016 | 0 | 0.0004 | 0.00015 | 0.00013 | 0 | 0.00041 | 6.36E-05 | 0.00013 | 3.20E-05 | 1.77E-05 | 6.75E-05 | 3.54E-05 | 7.59E-05 | 0 | 0.00013 | 0.00017 |
| Anapc16 | 0.00019 | 0.00018 | 0.00016 | 0.00028 | 0.00015 | 0.0001 | 9.28E-05 | 4.99E-05 | 6.36E-05 | 0 | 1.99E-05 | 0.0002 | 3.31E-05 | 8.31E-05 | 5.47E-05 | 0.00015 | 3.46E-05 | 4.33E-05 |
| Rab4a | 5.27E-05 | 4.34E-05 | 5.60E-06 | 0 | 6.38E-05 | 3.86E-05 | 1.35E-05 | 2.10E-05 | 6.36E-05 | 5.83E-05 | 5.64E-05 | 5.42E-07 | 1.16E-05 | 1.79E-05 | 6.40E-06 | 0 | 0 | 9.38E-06 |
| Ndufaf7 | 9.13E-05 | 4.26E-05 | 4.12E-05 | 5.05E-05 | 6.09E-06 | 8.54E-06 | 6.09E-06 | 0 | 6.36E-05 | 2.72E-05 | 2.07E-06 | 1.00E-05 | 3.07E-05 | 0 | 1.77E-05 | 3.53E-05 | 0 | 2.42E-05 |
| Adprhl2 | 3.47E-05 | 9.83E-05 | 9.15E-05 | 9.46E-05 | 3.65E-05 | 5.70E-06 | 3.49E-06 | 1.86E-05 | 6.36E-05 | 4.08E-05 | 2.14E-05 | 1.57E-05 | 1.08E-05 | 2.34E-05 | 4.30E-05 | 5.84E-05 | 0 | 2.35E-05 |
| Nsmce2 | 0.00012 | 0.00015 | 8.11E-06 | 3.35E-05 | 6.89E-05 | 0.00011 | 0 | 2.97E-05 | 6.35E-05 | 3.40E-05 | 3.05E-05 | 0.00011 | 2.17E-05 | 0.00011 | 0.00022 | 0.00026 | 5.77E-06 | 6.28E-05 |
| Ninj1 | 2.67E-05 | 0.00015 | 2.61E-05 | 9.45E-05 | 0.00043 | 8.42E-05 | 0.0001 | 0.00022 | 6.35E-05 | 0 | 5.31E-05 | 0.00056 | 5.03E-05 | 0.0002 | 0 | 0 | 5.82E-05 | 1.06E-05 |
| Tbc1d5 | 6.97E-05 | 7.83E-05 | 1.94E-05 | 0 | 5.41E-05 | 4.81E-05 | 2.58E-05 | 4.12E-05 | 6.35E-05 | 8.71E-05 | 1.62E-05 | 3.26E-05 | 4.69E-05 | 5.45E-05 | 2.61E-05 | 7.14E-05 | 8.84E-05 | 9.01E-05 |
| Hira | 7.10E-05 | 8.64E-05 | 9.27E-05 | 5.41E-05 | 1.98E-05 | 4.24E-05 | 2.95E-05 | 0 | 6.35E-05 | 2.41E-05 | 6.26E-05 | 3.77E-05 | 3.32E-05 | 2.67E-05 | 3.00E-05 | 9.39E-05 | 4.20E-05 | 3.11E-05 |
| Ddx28 | 4.27E-05 | 3.40E-05 | 3.06E-05 | 1.22E-05 | 2.43E-05 | 3.52E-05 | 0 | 4.82E-07 | 6.35E-05 | 1.32E-06 | 2.50E-05 | 5.39E-06 | 0 | 2.55E-05 | 9.50E-06 | 1.02E-05 | 0 | 1.53E-05 |
| Slc30a1 | 2.15E-05 | 0.00011 | 0.00018 | 9.00E-05 | 8.56E-06 | 1.09E-05 | 3.35E-05 | 1.48E-05 | 6.35E-05 | 4.54E-05 | 6.46E-05 | 2.41E-05 | 0.00024 | 4.64E-05 | 0.00011 | 0 | 1.22E-05 | 8.33E-05 |
| Crlf2 | 0 | 5.10E-05 | 5.62E-05 | 0.00017 | 5.20E-05 | 0 | 0 | 0.00032 | 6.35E-05 | 2.44E-05 | 3.91E-05 | 0.00011 | 7.98E-05 | 6.01E-05 | 0.00024 | 2.34E-05 | 0 | 0.00016 |
| Rexo4 | 7.79E-05 | 7.21E-05 | 0.00012 | 0.00013 | 1.54E-05 | 0 | 4.00E-05 | 2.62E-05 | 6.35E-05 | 1.39E-05 | 1.61E-05 | 3.54E-06 | 1.59E-07 | 4.24E-05 | 0.00023 | 9.93E-05 | 0 | 7.97E-06 |
| Adcy2 | 5.38E-05 | 9.04E-05 | 0 | 0 | 6.41E-05 | 0.00023 | 0 | 0.00012 | 6.34E-05 | 8.57E-05 | 0 | 0 | 0.00011 | 5.23E-05 | 4.71E-05 | 4.53E-05 | 0.00016 | 1.17E-05 |
| Retreg3 | 3.33E-05 | 0.00024 | 8.18E-05 | 0.00013 | 0.00015 | 0.00016 | 0.00017 | 0.0002 | 6.34E-05 | 0.00012 | 9.10E-05 | 8.86E-05 | 0.00013 | 0.00012 | 4.95E-05 | 0.00015 | 0 | 0.00013 |
| Pak1ip1 | 0.00017 | 0.00013 | 0.00029 | 2.79E-05 | 1.68E-05 | 4.44E-05 | 1.16E-05 | 0 | 6.34E-05 | 2.94E-05 | 7.79E-05 | 4.33E-05 | 6.64E-05 | 0 | 3.80E-05 | 0.00014 | 3.75E-05 | 0 |
| Rabgef1 | 9.64E-05 | 7.57E-05 | 0.00015 | 0 | 1.11E-05 | 4.21E-05 | 4.26E-05 | 0 | 6.34E-05 | 3.81E-05 | 6.13E-05 | 0.00014 | 2.00E-05 | 5.52E-05 | 0.0001 | 4.10E-05 | 0.00026 | 1.59E-05 |
| Tmem265 | 7.90E-05 | 8.26E-05 | 2.77E-05 | 2.59E-05 | 1.69E-05 | 3.21E-05 | 5.73E-05 | 1.23E-05 | 6.34E-05 | 1.07E-05 | 3.02E-05 | 6.82E-05 | 1.95E-06 | 2.20E-05 | 8.75E-05 | 0.00011 | 9.88E-05 | 0 |

| | | | | | | | | | | | | | | | | | | |
|---|---|---|---|---|---|---|---|---|---|---|---|---|---|---|---|---|---|---|
| Dip2b | 5.84E-05 | 9.35E-05 | 5.84E-05 | 4.63E-05 | 4.52E-05 | 8.19E-06 | 0 | 1.58E-05 | 6.34E-05 | 3.35E-05 | 1.00E-05 | 5.89E-05 | 0 | 1.69E-05 | 5.91E-05 | 4.22E-05 | 5.12E-05 | 4.91E-05 |
| 5-Mar | 2.64E-05 | 5.32E-05 | 5.74E-05 | 0.00013 | 3.45E-05 | 1.82E-08 | 0 | 2.90E-05 | 6.34E-05 | 4.51E-05 | 9.18E-06 | 1.72E-05 | 8.27E-05 | 4.95E-05 | 7.19E-05 | 4.50E-05 | 1.61E-05 | 4.86E-05 |
| Wdr4 | 2.28E-05 | 5.00E-05 | 0.00013 | 0.0001 | 1.08E-05 | 0 | 0 | 0 | 6.34E-05 | 1.88E-05 | 3.15E-05 | 5.36E-05 | 0 | 1.02E-05 | 6.05E-05 | 1.78E-05 | 0 | 1.88E-05 |
| Decr1 | 0.00021 | 6.52E-05 | 4.85E-05 | 4.92E-05 | 3.03E-05 | 0.00012 | 7.49E-05 | 1.98E-05 | 6.33E-05 | 6.56E-05 | 6.12E-06 | 2.71E-05 | 2.18E-05 | 0 | 0 | 5.48E-05 | 4.60E-06 | 0.00011 |
| Blvra | 7.74E-05 | 4.49E-05 | 5.09E-05 | 6.75E-05 | 0 | 0 | 1.35E-05 | 1.77E-05 | 6.33E-05 | 6.26E-05 | 2.68E-05 | 1.35E-05 | 9.12E-06 | 6.21E-05 | 0 | 0.00018 | 0 | 8.19E-05 |
| Yeats2 | 6.85E-05 | 3.22E-05 | 2.15E-05 | 0 | 5.09E-05 | 4.61E-05 | 2.37E-05 | 1.53E-05 | 6.32E-05 | 9.06E-06 | 2.30E-05 | 3.72E-05 | 0 | 2.64E-05 | 0.00013 | 4.46E-05 | 0.0001 | 8.76E-06 |
| Bcl9 | 3.20E-05 | 9.18E-05 | 2.28E-05 | 0.00011 | 2.10E-05 | 3.61E-05 | 4.31E-05 | 3.63E-05 | 6.32E-05 | 3.68E-05 | 1.85E-05 | 2.00E-05 | 9.42E-05 | 1.34E-05 | 6.67E-05 | 1.30E-05 | 2.20E-05 | 4.78E-06 |
| Eif3e | 0.00026 | 3.67E-05 | 5.50E-05 | 0 | 0 | 0.00018 | 0 | 4.94E-07 | 6.31E-05 | 3.15E-05 | 0 | 9.78E-05 | 0 | 8.96E-06 | 0 | 3.97E-05 | 0.00011 | 5.87E-06 |
| Ddx47 | 0.00013 | 9.29E-05 | 0.0001 | 6.56E-05 | 6.41E-05 | 5.17E-05 | 1.02E-05 | 0 | 6.31E-05 | 2.23E-05 | 2.59E-05 | 6.26E-05 | 5.06E-05 | 3.91E-05 | 4.57E-05 | 7.34E-05 | 2.52E-05 | 4.73E-05 |
| Ggt7 | 3.68E-06 | 9.60E-06 | 0 | 4.52E-05 | 2.92E-07 | 3.26E-05 | 0 | 6.85E-05 | 6.31E-05 | 3.92E-06 | 3.51E-05 | 4.00E-05 | 0 | 6.93E-05 | 5.05E-05 | 0 | 0 | 0 |
| Sp140 | 1.89E-05 | 2.93E-06 | 0 | 0 | 0 | 6.35E-06 | 3.37E-05 | 0 | 6.31E-05 | 9.70E-05 | 0 | 2.18E-05 | 7.60E-05 | 0 | 0 | 0 | 0 | 5.62E-05 |
| Arhgef10 | 3.12E-05 | 1.37E-05 | 0 | 1.84E-05 | 2.92E-05 | 0 | 0.00013 | 0 | 6.31E-05 | 0.00016 | 8.79E-05 | 4.19E-05 | 1.83E-05 | 9.08E-05 | 1.11E-05 | 2.93E-05 | 1.77E-05 | 0.00024 |
| Akap12 | 2.95E-05 | 7.20E-05 | 3.80E-05 | 0 | 3.21E-05 | 7.67E-05 | 0.00017 | 1.34E-05 | 6.30E-05 | 0.00018 | 0.00016 | 1.64E-06 | 0 | 0 | 0.0001 | 7.74E-05 | 0 | 0.00029 |
| Gripap1 | 6.01E-05 | 7.86E-05 | 3.99E-05 | 5.00E-05 | 2.04E-05 | 1.53E-05 | 8.23E-07 | 0 | 6.30E-05 | 4.33E-05 | 2.47E-06 | 2.06E-05 | 1.90E-05 | 0 | 0.00015 | 9.12E-06 | 0.00011 | 2.46E-05 |
| Fgf11 | 0 | 1.84E-05 | 0 | 1.84E-05 | 5.75E-05 | 7.64E-06 | 6.79E-06 | 0 | 6.30E-05 | 7.82E-06 | 0 | 0.00028 | 0 | 0 | 5.74E-05 | 0 | 0 | 5.12E-05 |
| Nbl1 | 0 | 0.00098 | 6.10E-05 | 0 | 0.00115 | 0.00026 | 0.00135 | 0.00012 | 6.29E-05 | 0.00079 | 0.00017 | 0.00024 | 0.00125 | 5.27E-05 | 9.42E-05 | 5.78E-05 | 0.00018 | 0.00096 |
| Gnptab | 4.65E-05 | 0.00013 | 0.00029 | 2.15E-05 | 9.31E-05 | 0.00017 | 0.00016 | 8.66E-05 | 6.29E-05 | 9.29E-05 | 3.55E-05 | 0 | 0.0002 | 8.44E-05 | 0.00011 | 9.18E-05 | 4.11E-06 | 0.00021 |
| Mfn1 | 0.00011 | 0.00015 | 9.05E-05 | 0.00011 | 6.58E-05 | 5.90E-05 | 2.32E-05 | 0 | 6.29E-05 | 6.36E-05 | 6.29E-05 | 7.64E-05 | 2.15E-05 | 7.78E-05 | 0.00018 | 6.72E-05 | 0.00012 | 0.00011 |
| Ift80 | 4.63E-05 | 6.82E-05 | 0 | 0 | 6.03E-05 | 4.59E-05 | 1.55E-05 | 9.77E-06 | 6.29E-05 | 4.26E-05 | 0 | 0 | 4.15E-05 | 0 | 2.47E-05 | 0.00019 | 4.56E-05 | 6.53E-05 |
| Trabd | 7.36E-05 | 0.00012 | 0.00015 | 0.00018 | 5.31E-05 | 3.08E-05 | 1.14E-05 | 1.73E-05 | 6.29E-05 | 2.65E-05 | 7.91E-05 | 0.00015 | 0 | 4.29E-05 | 0.00021 | 8.76E-05 | 0 | 3.19E-05 |
| Lzic | 9.63E-05 | 0.00013 | 0.00016 | 7.37E-05 | 5.84E-05 | 2.27E-05 | 0 | 2.01E-05 | 6.29E-05 | 4.59E-05 | 5.31E-05 | 4.01E-08 | 1.24E-05 | 0 | 1.35E-05 | 0.00012 | 1.47E-05 | 2.93E-05 |
| Tmem101 | 2.29E-05 | 3.65E-05 | 3.32E-05 | 0.0001 | 1.33E-05 | 3.49E-06 | 3.19E-05 | 3.17E-05 | 6.29E-05 | 2.18E-05 | 1.41E-05 | 5.15E-06 | 0 | 2.83E-05 | 1.11E-05 | 1.32E-05 | 0 | 1.48E-05 |
| Sart3 | 1.78E-05 | 7.78E-05 | 5.39E-05 | 7.61E-05 | 2.31E-05 | 0 | 0 | 5.29E-05 | 6.29E-05 | 1.32E-05 | 2.60E-05 | 4.78E-05 | 1.35E-05 | 2.90E-05 | 5.69E-05 | 0.00016 | 0 | 5.82E-06 |
| Tmem109 | 0.00027 | 0.0001 | 0.00019 | 0.00018 | 9.06E-05 | 0.00013 | 1.82E-05 | 4.09E-05 | 6.28E-05 | 7.59E-05 | 0.00011 | 0.00016 | 4.72E-05 | 0 | 0 | 0.00042 | 6.50E-05 | 0.00021 |
| Idh3g | 0.00031 | 0.00018 | 0.00013 | 0.00023 | 2.44E-05 | 0.00011 | 0.00014 | 5.55E-05 | 6.28E-05 | 4.81E-05 | 1.44E-05 | 5.39E-05 | 3.56E-08 | 8.13E-05 | 9.97E-05 | 0.00017 | 6.99E-05 | 9.05E-05 |
| Cpsf6 | 0.00025 | 0.00024 | 0.0002 | 0.00014 | 9.27E-05 | 7.93E-05 | 0 | 6.73E-05 | 6.28E-05 | 2.49E-05 | 8.58E-05 | 0.00011 | 1.22E-05 | 9.93E-05 | 0.00046 | 0.00032 | 0 | 0 |
| Zmat5 | 7.75E-05 | 4.99E-05 | 1.55E-05 | 6.92E-05 | 3.30E-05 | 9.96E-06 | 1.44E-05 | 0 | 6.28E-05 | 4.67E-05 | 7.97E-06 | 2.03E-05 | 0 | 0 | 0 | 5.02E-06 | 0 | 1.48E-05 |
| Gtpbp3 | 7.18E-06 | 9.59E-05 | 5.93E-05 | 0.00011 | 2.70E-05 | 5.10E-06 | 0 | 3.41E-05 | 6.28E-05 | 3.86E-05 | 1.00E-05 | 0 | 1.77E-05 | 4.09E-05 | 0.0001 | 6.14E-05 | 1.01E-05 | 8.78E-05 |
| Sbf1 | 0 | 7.37E-05 | 6.13E-05 | 5.80E-05 | 2.54E-05 | 0 | 0 | 2.58E-05 | 6.28E-05 | 2.74E-05 | 5.90E-05 | 3.75E-05 | 0 | 1.70E-05 | 0.0001 | 3.97E-05 | 3.35E-05 | 3.78E-05 |
| Cnot8 | 0.00023 | 0.00011 | 0.00013 | 6.86E-05 | 0.00011 | 0.00016 | 0.0002 | 0.00011 | 6.27E-05 | 2.15E-05 | 7.62E-05 | 9.03E-05 | 9.09E-05 | 6.58E-05 | 8.41E-05 | 7.06E-05 | 0 | 3.30E-05 |
| Fem1b | 0.00019 | 0.00015 | 0.00022 | 0 | 6.95E-05 | 9.57E-05 | 0 | 2.34E-05 | 6.27E-05 | 0 | 0.00013 | 0.00053 | 0 | 0 | 0 | 0.00013 | 0.00024 | 8.92E-05 |
| Psmd4 | 0.00025 | 0.00018 | 0.00022 | 0.00012 | 4.93E-05 | 8.04E-05 | 1.05E-05 | 0 | 6.27E-05 | 2.48E-05 | 5.30E-05 | 7.71E-05 | 0 | 0.0001 | 4.74E-05 | 0.0001 | 8.15E-05 | 1.70E-05 |
| Kmt2a | 6.04E-05 | 0.0002 | 9.26E-05 | 1.24E-05 | 6.91E-05 | 4.03E-05 | 4.69E-05 | 5.18E-05 | 6.27E-05 | 6.10E-05 | 3.81E-05 | 0.00012 | 1.74E-05 | 1.99E-05 | 0.00035 | 9.75E-05 | 0.00021 | 6.19E-05 |

| | | | | | | | | | | | | | | | | | | |
|---|---|---|---|---|---|---|---|---|---|---|---|---|---|---|---|---|---|---|
| Aox1 | 0 | 4.17E-05 | 0 | 0 | 5.27E-05 | 3.65E-05 | 1.08E-05 | 0 | 6.27E-05 | 4.28E-05 | 0 | 0 | 4.64E-05 | 2.98E-05 | 5.64E-05 | 0 | 4.50E-06 | 3.83E-05 |
| Ddx10 | 0.00014 | 0.00013 | 0.00011 | 1.72E-05 | 7.63E-06 | 3.94E-06 | 0 | 1.57E-05 | 6.27E-05 | 1.79E-05 | 1.59E-05 | 0.00012 | 0 | 1.30E-06 | 0.00014 | 3.31E-05 | 3.23E-06 | 0 |
| Rdx | 0.00042 | 0.00014 | 0.00019 | 5.90E-05 | 5.26E-05 | 0.00014 | 0.00015 | 3.71E-05 | 6.26E-05 | 7.65E-05 | 6.72E-05 | 8.90E-05 | 6.05E-06 | 2.82E-05 | 9.59E-05 | 0.00029 | 0.00015 | 6.22E-05 |
| Rnft1 | 0.00024 | 0.0001 | 0 | 0 | 2.63E-05 | 0.00012 | 0.00015 | 1.54E-05 | 6.26E-05 | 5.33E-05 | 3.01E-05 | 0.00028 | 2.61E-05 | 3.64E-05 | 1.21E-05 | 9.30E-05 | 0 | 5.76E-05 |
| Ddhd2 | 8.29E-05 | 6.46E-05 | 3.54E-05 | 8.16E-05 | 0.00016 | 9.25E-05 | 6.66E-05 | 6.92E-05 | 6.26E-05 | 6.87E-05 | 3.13E-05 | 1.07E-05 | 1.02E-06 | 8.00E-05 | 0.00014 | 1.52E-05 | 0 | 4.97E-05 |
| Rem1 | 0 | 0 | 1.57E-05 | 2.17E-05 | 0.00033 | 0.00042 | 0 | 0.0003 | 6.25E-05 | 0 | 0.00019 | 0 | 0 | 0 | 0.00056 | 0 | 0 | 5.57E-05 |
| Ints13 | 0.00013 | 5.73E-05 | 7.74E-05 | 1.26E-05 | 1.27E-05 | 5.86E-05 | 0 | 3.23E-05 | 6.25E-05 | 1.48E-06 | 4.11E-05 | 5.07E-05 | 0 | 2.75E-05 | 5.25E-05 | 9.43E-05 | 0.0001 | 1.56E-05 |
| Arid2 | 0.00017 | 0.00022 | 6.35E-05 | 3.84E-08 | 0.00014 | 4.49E-05 | 1.00E-05 | 5.62E-05 | 6.25E-05 | 1.19E-05 | 8.02E-05 | 0.00025 | 0 | 7.43E-05 | 0.00031 | 0.00012 | 0.00017 | 3.15E-05 |
| Zfp319 | 2.13E-05 | 0.0001 | 3.25E-05 | 3.76E-05 | 3.78E-05 | 2.84E-05 | 3.05E-05 | 5.68E-05 | 6.25E-05 | 3.96E-05 | 3.00E-05 | 4.01E-05 | 0 | 3.19E-05 | 6.61E-05 | 3.04E-05 | 0 | 4.77E-05 |
| Mex3b | 0 | 0.00015 | 1.36E-05 | 9.90E-05 | 9.52E-05 | 0 | 5.88E-05 | 8.38E-05 | 6.25E-05 | 0.00015 | 9.93E-05 | 0 | 0 | 0.0002 | 1.53E-05 | 0.00012 | 0.0003 | 0 |
| Gng8 | 0 | 8.92E-05 | 0 | 0 | 0 | 5.92E-05 | 0 | 0 | 6.24E-05 | 0 | 0 | 5.12E-05 | 0 | 9.02E-05 | 6.37E-05 | 0 | 4.28E-06 | 9.90E-05 |
| Ino80b | 3.83E-05 | 0.00016 | 4.02E-05 | 0.00019 | 6.28E-05 | 4.16E-05 | 0 | 2.35E-05 | 6.24E-05 | 6.69E-05 | 2.28E-05 | 2.38E-05 | 1.20E-05 | 5.35E-05 | 0.00011 | 5.26E-05 | 0 | 8.77E-05 |
| Elovl6 | 3.73E-05 | 0 | 5.69E-06 | 4.14E-05 | 0 | 0 | 0 | 1.33E-05 | 6.24E-05 | 5.12E-05 | 0 | 0 | 0 | 3.34E-05 | 0 | 3.62E-05 | 0 | 0 |
| Tatdn2 | 8.29E-05 | 0.00019 | 0.00024 | 9.42E-05 | 9.06E-05 | 0 | 3.82E-05 | 0 | 6.24E-05 | 4.22E-05 | 0.00022 | 6.12E-05 | 5.02E-05 | 0.00014 | 0.0002 | 8.44E-05 | 0.00022 | 3.39E-05 |
| Bcas2 | 0.00027 | 0.00017 | 6.05E-05 | 6.14E-05 | 6.61E-05 | 0.00011 | 6.96E-05 | 0 | 6.23E-05 | 3.66E-05 | 4.04E-05 | 0.00012 | 2.17E-05 | 0.00012 | 0.00013 | 0.00023 | 0 | 2.70E-05 |
| Trmt61a | 3.99E-05 | 2.66E-05 | 0.00019 | 5.97E-05 | 1.47E-05 | 0 | 0 | 0 | 6.23E-05 | 1.57E-05 | 7.25E-05 | 0 | 9.13E-05 | 1.54E-05 | 3.42E-05 | 0 | 0 | 0 |
| Nop16 | 9.70E-05 | 6.31E-05 | 0.00027 | 0.00016 | 5.35E-06 | 0 | 0 | 0 | 6.23E-05 | 1.47E-05 | 4.65E-05 | 1.82E-06 | 3.97E-05 | 3.86E-06 | 0 | 7.47E-05 | 0 | 9.22E-06 |
| Vapa | 0.00022 | 9.89E-05 | 0.00013 | 4.14E-05 | 2.55E-05 | 8.68E-05 | 2.56E-05 | 4.01E-05 | 6.22E-05 | 0 | 7.50E-05 | 0.00021 | 0 | 4.47E-05 | 0 | 3.98E-05 | 0 | 0 |
| Eea1 | 9.73E-05 | 0.00017 | 0.00016 | 0 | 0.00011 | 7.54E-05 | 5.91E-05 | 3.84E-05 | 6.22E-05 | 7.20E-05 | 0 | 0.00016 | 1.12E-05 | 0 | 0.00012 | 9.80E-05 | 0.00019 | 5.39E-05 |
| Cript | 0.00017 | 7.60E-05 | 6.72E-05 | 7.19E-05 | 5.30E-05 | 6.08E-05 | 2.65E-05 | 2.65E-05 | 6.22E-05 | 2.76E-05 | 0.0001 | 9.42E-05 | 4.82E-05 | 2.46E-05 | 5.79E-05 | 7.17E-05 | 1.80E-05 | 3.99E-05 |
| Glis2 | 0 | 0.00018 | 0.0002 | 0.0002 | 0.00029 | 5.47E-05 | 0.00045 | 0.00019 | 6.22E-05 | 0.00023 | 0.0001 | 7.73E-06 | 0.00031 | 0 | 0.00022 | 0.00011 | 0.00032 | 0.00066 |
| Pts | 7.28E-05 | 1.97E-05 | 0 | 1.42E-05 | 1.92E-05 | 2.90E-05 | 0 | 1.65E-05 | 6.22E-05 | 7.85E-05 | 5.08E-06 | 1.33E-05 | 0 | 7.04E-06 | 1.66E-05 | 1.93E-07 | 0 | 3.62E-05 |
| Pdzd4 | 3.27E-05 | 2.78E-05 | 1.17E-05 | 8.89E-05 | 0.0001 | 2.60E-05 | 0.0001 | 0 | 6.22E-05 | 8.85E-06 | 9.18E-06 | 0 | 5.54E-05 | 7.38E-05 | 3.17E-05 | 6.20E-05 | 1.45E-05 | 5.47E-05 |
| Mien1 | 8.38E-05 | 8.72E-05 | 5.94E-05 | 0.00028 | 6.65E-05 | 4.29E-06 | 0.00014 | 0 | 6.22E-05 | 8.56E-05 | 7.78E-05 | 7.67E-05 | 2.88E-05 | 3.51E-05 | 0 | 0 | 3.47E-05 | 7.54E-05 |
| Ep400 | 0.0002 | 0.00018 | 0.00012 | 0 | 3.41E-05 | 6.58E-05 | 4.32E-05 | 0 | 6.21E-05 | 4.04E-05 | 4.35E-05 | 0.00013 | 5.11E-05 | 4.49E-05 | 0.0002 | 0.00013 | 0.00075 | 0 |
| Smim15 | 0.00015 | 0.0001 | 0.00014 | 0 | 5.32E-05 | 5.64E-05 | 0.00012 | 6.87E-05 | 6.21E-05 | 5.19E-05 | 9.22E-05 | 4.21E-05 | 0 | 0 | 2.95E-05 | 7.71E-05 | 8.16E-05 | 0.00012 |
| Hif1an | 8.75E-05 | 0.00013 | 7.88E-05 | 4.51E-05 | 3.67E-05 | 2.72E-05 | 0 | 1.63E-05 | 6.21E-05 | 1.45E-05 | 3.21E-05 | 4.69E-05 | 0 | 3.27E-05 | 0.00016 | 2.16E-05 | 0 | 0 |
| Khk | 0 | 2.88E-05 | 0 | 1.41E-05 | 0 | 2.36E-05 | 0 | 0 | 6.21E-05 | 6.50E-05 | 0 | 1.97E-05 | 3.48E-05 | 0 | 6.19E-06 | 1.35E-05 | 1.12E-05 | 1.92E-05 |
| Zfp768 | 3.15E-05 | 7.77E-05 | 4.92E-05 | 0.00011 | 4.74E-05 | 4.86E-06 | 2.71E-05 | 1.56E-05 | 6.21E-05 | 3.46E-05 | 1.63E-05 | 3.22E-05 | 0 | 6.00E-05 | 5.83E-05 | 1.50E-05 | 0 | 4.30E-05 |
| Wif1 | 0 | 4.23E-05 | 0 | 0 | 0 | 0 | 0 | 0 | 6.21E-05 | 0 | 0 | 0 | 0 | 0.0004 | 0 | 0 | 0 | 0.00014 |
| Mob1b | 0.00015 | 0.00014 | 0.00015 | 1.30E-05 | 6.93E-05 | 9.81E-05 | 4.33E-05 | 6.41E-05 | 6.20E-05 | 2.53E-05 | 4.85E-05 | 7.97E-05 | 4.39E-05 | 4.49E-05 | 1.53E-05 | 0.0001 | 6.77E-05 | 1.44E-05 |
| Acadm | 6.47E-05 | 8.76E-05 | 2.23E-05 | 6.32E-05 | 0 | 7.25E-06 | 7.71E-06 | 2.10E-05 | 6.20E-05 | 2.60E-05 | 2.16E-05 | 6.06E-05 | 0 | 1.21E-05 | 2.80E-05 | 2.04E-05 | 3.49E-05 | 4.45E-05 |
| Acss2 | 8.33E-06 | 3.25E-05 | 0 | 0 | 1.39E-05 | 5.50E-07 | 0 | 0 | 6.20E-05 | 4.53E-05 | 0 | 2.33E-05 | 0 | 8.78E-06 | 1.56E-06 | 0 | 9.35E-06 | 4.02E-05 |

| | | | | | | | | | | | | | | | | | | |
|---|---|---|---|---|---|---|---|---|---|---|---|---|---|---|---|---|---|---|
| Aldh4a1 | 0 | 4.74E-05 | 1.47E-05 | 1.18E-05 | 3.34E-05 | 0 | 1.90E-05 | 6.07E-05 | 6.20E-05 | 4.88E-05 | 4.39E-05 | 0 | 4.05E-05 | 0 | 0 | 0 | 9.59E-05 | 6.09E-05 |
| Fgfr2 | 8.27E-07 | 4.03E-05 | 0 | 0 | 7.53E-06 | 0 | 0 | 0 | 6.20E-05 | 0 | 2.16E-05 | 0 | 0 | 2.06E-05 | 0 | 3.34E-06 | 0 | 3.25E-05 |
| Uggt1 | 0.00028 | 6.43E-05 | 0.00027 | 0.00022 | 0.0001 | 0.00023 | 0.00028 | 0.00015 | 6.19E-05 | 0.00022 | 0.00012 | 5.84E-05 | 0.00034 | 0.00012 | 4.20E-05 | 0.00022 | 9.09E-05 | 0.00018 |
| Snx25 | 7.92E-05 | 4.87E-05 | 6.74E-05 | 0 | 9.71E-05 | 0.00011 | 0 | 9.89E-05 | 6.19E-05 | 0 | 3.37E-05 | 0.0002 | 0 | 5.93E-05 | 1.49E-05 | 0.00011 | 5.94E-05 | 0 |
| Cdc26 | 0.0002 | 3.84E-05 | 8.97E-05 | 0.00011 | 5.23E-05 | 0.0001 | 0.00014 | 4.03E-05 | 6.19E-05 | 1.38E-05 | 8.60E-06 | 0 | 3.97E-05 | 5.54E-05 | 0.00012 | 6.73E-05 | 0 | 0 |
| Usp24 | 0.0001 | 0.00014 | 0.00019 | 0 | 4.72E-05 | 4.54E-05 | 4.69E-06 | 0 | 6.19E-05 | 0.00015 | 4.01E-05 | 2.41E-05 | 3.84E-05 | 1.83E-05 | 0.00015 | 7.18E-05 | 0.00013 | 6.15E-05 |
| Epha7 | 0 | 0 | 0 | 0 | 0 | 3.28E-05 | 0 | 0 | 6.19E-05 | 0 | 0 | 0 | 0 | 2.20E-05 | 0 | 3.44E-06 | 0 | 1.34E-05 |
| Lta4h | 0.00029 | 0.00015 | 0.00015 | 7.75E-05 | 5.38E-05 | 0.00012 | 0.0002 | 0 | 6.18E-05 | 7.64E-05 | 0.00015 | 0.00018 | 0 | 0.0001 | 0 | 0.00018 | 0 | 1.75E-05 |
| Urod | 0.00011 | 0.00014 | 2.23E-05 | 0.00011 | 8.84E-05 | 0.0001 | 6.75E-05 | 4.84E-05 | 6.18E-05 | 4.78E-05 | 4.83E-05 | 2.49E-05 | 5.79E-05 | 9.70E-05 | 1.21E-05 | 0.00012 | 1.40E-05 | 5.00E-05 |
| Fkbp15 | 0.00015 | 0.00016 | 9.40E-05 | 5.65E-05 | 6.24E-05 | 9.25E-05 | 4.67E-05 | 8.64E-05 | 6.18E-05 | 7.98E-05 | 7.07E-05 | 7.09E-05 | 2.77E-05 | 7.14E-05 | 0.00013 | 3.32E-05 | 0 | 7.22E-05 |
| Rad54l2 | 6.53E-05 | 0.00011 | 0.00014 | 5.21E-05 | 9.29E-05 | 7.10E-05 | 0 | 0.00011 | 6.18E-05 | 4.62E-06 | 4.13E-05 | 1.74E-05 | 5.34E-06 | 0.00011 | 0.00021 | 6.31E-05 | 0.00011 | 0 |
| Clpp | 0.00017 | 0.00011 | 0.00022 | 6.39E-05 | 5.70E-05 | 2.03E-05 | 0 | 1.04E-05 | 6.18E-05 | 0 | 2.78E-05 | 0 | 2.70E-05 | 0 | 7.34E-05 | 6.47E-05 | 0 | 0 |
| Ranbp10 | 3.24E-05 | 7.27E-05 | 4.73E-05 | 3.23E-05 | 4.40E-05 | 2.02E-05 | 5.43E-05 | 3.75E-05 | 6.18E-05 | 2.58E-05 | 3.70E-05 | 4.12E-05 | 2.09E-05 | 4.96E-05 | 7.38E-05 | 3.85E-05 | 5.38E-05 | 4.32E-05 |
| Ern1 | 0 | 0.00014 | 0.00021 | 0.00014 | 0 | 0 | 2.23E-05 | 3.95E-05 | 6.18E-05 | 2.01E-05 | 0 | 0.0002 | 0.00013 | 4.54E-05 | 0.00023 | 3.00E-06 | 0.00051 | 3.73E-05 |
| Jarid2 | 2.89E-05 | 0.00017 | 1.24E-05 | 0 | 8.09E-05 | 0.00011 | 0 | 0.00017 | 6.17E-05 | 0 | 0 | 0.00045 | 0 | 0 | 0.00016 | 0 | 0.00016 | 0 |
| Trp53bp2 | 1.44E-05 | 0.00011 | 6.02E-05 | 6.84E-05 | 6.31E-06 | 2.28E-05 | 0 | 4.97E-05 | 6.17E-05 | 2.96E-05 | 2.32E-05 | 1.13E-05 | 0 | 5.12E-05 | 4.64E-05 | 4.41E-05 | 0.00015 | 1.64E-06 |
| Prdm11 | 6.08E-06 | 2.75E-05 | 1.56E-05 | 2.71E-05 | 2.30E-05 | 9.86E-06 | 1.63E-08 | 8.79E-06 | 6.17E-05 | 0 | 0 | 0 | 1.18E-05 | 4.52E-05 | 5.58E-05 | 1.46E-06 | 0 | 2.68E-05 |
| Nsfl1c | 0.00018 | 0.00013 | 0.00017 | 0.00022 | 0 | 0 | 3.66E-06 | 5.36E-06 | 6.17E-05 | 2.52E-05 | 6.48E-06 | 9.13E-05 | 0 | 0 | 0.00035 | 7.28E-05 | 3.80E-05 | 3.71E-05 |
| Wdr70 | 6.32E-05 | 5.15E-05 | 6.66E-05 | 5.97E-05 | 1.78E-05 | 0 | 1.86E-05 | 9.52E-06 | 6.17E-05 | 1.54E-05 | 2.84E-05 | 1.04E-05 | 0 | 3.48E-06 | 0.00019 | 6.02E-05 | 4.25E-05 | 2.64E-05 |
| Prkag1 | 0.00026 | 6.78E-05 | 0.00014 | 2.42E-05 | 5.21E-05 | 0.00014 | 0.00012 | 7.97E-05 | 6.16E-05 | 4.75E-05 | 4.19E-05 | 5.50E-05 | 0 | 8.02E-05 | 6.68E-05 | 0.00017 | 0 | 4.18E-05 |
| Scaf8 | 4.35E-05 | 0.00015 | 0.00011 | 0.00013 | 6.96E-05 | 2.22E-05 | 3.53E-05 | 6.43E-07 | 6.16E-05 | 5.31E-05 | 4.05E-06 | 6.36E-05 | 0 | 2.44E-05 | 0.00015 | 6.80E-05 | 6.59E-05 | 5.95E-05 |
| Abr | 7.16E-05 | 0.00018 | 8.35E-05 | 0.00012 | 7.32E-05 | 7.98E-05 | 0 | 5.16E-05 | 6.15E-05 | 0 | 2.60E-05 | 6.23E-05 | 1.86E-05 | 0.00017 | 0.00015 | 8.36E-05 | 3.89E-05 | 7.95E-05 |
| Crmp1 | 0 | 0 | 0 | 0 | 0.00014 | 6.94E-05 | 0 | 0 | 6.15E-05 | 0 | 5.84E-05 | 3.99E-05 | 0 | 0 | 0 | 0 | 0 | 0 |
| Slc18b1 | 3.90E-05 | 2.20E-05 | 4.78E-06 | 2.60E-05 | 4.42E-05 | 3.50E-05 | 9.69E-06 | 0 | 6.15E-05 | 5.36E-05 | 4.02E-05 | 0 | 2.66E-05 | 4.47E-05 | 7.79E-07 | 1.30E-05 | 0 | 4.85E-05 |
| Adnp2 | 7.49E-05 | 0.00011 | 4.71E-05 | 8.86E-05 | 8.03E-05 | 3.38E-05 | 7.74E-05 | 8.23E-06 | 6.15E-05 | 2.54E-05 | 5.41E-05 | 0.00012 | 0 | 6.54E-05 | 6.55E-05 | 0.00011 | 0.00025 | 5.30E-05 |
| Ubiad1 | 7.26E-05 | 2.15E-05 | 0.00012 | 8.06E-05 | 3.26E-05 | 1.38E-05 | 1.89E-05 | 1.74E-05 | 6.15E-05 | 6.25E-06 | 2.28E-05 | 7.89E-06 | 2.42E-05 | 0 | 0 | 4.60E-05 | 0 | 6.43E-06 |
| Diaph2 | 0.00014 | 0.00014 | 2.36E-06 | 0 | 3.04E-05 | 0.00017 | 0 | 6.58E-05 | 6.14E-05 | 5.76E-05 | 4.46E-05 | 3.81E-05 | 9.53E-05 | 0 | 0.00026 | 5.24E-06 | 0.00015 | 6.83E-05 |
| Slc35a5 | 8.30E-05 | 9.87E-05 | 0 | 0 | 5.65E-05 | 7.36E-05 | 3.33E-05 | 8.36E-05 | 6.14E-05 | 6.00E-05 | 2.62E-05 | 6.75E-05 | 2.20E-05 | 6.13E-05 | 0 | 1.88E-05 | 9.69E-06 | 6.96E-05 |
| Vipas39 | 7.15E-05 | 0.00011 | 2.62E-05 | 0.00011 | 5.97E-05 | 5.79E-05 | 5.20E-05 | 5.84E-05 | 6.14E-05 | 5.32E-05 | 5.34E-05 | 1.42E-05 | 5.41E-05 | 4.87E-05 | 0 | 2.62E-05 | 0 | 7.16E-05 |
| Thg1l | 8.93E-05 | 2.50E-05 | 0.0001 | 5.35E-05 | 3.80E-06 | 4.60E-05 | 4.18E-06 | 0 | 6.14E-05 | 1.54E-05 | 7.64E-05 | 6.84E-06 | 6.73E-05 | 3.77E-05 | 0 | 1.84E-05 | 0 | 0 |
| Nup50 | 0.00011 | 0.0001 | 0.00018 | 5.11E-05 | 7.42E-05 | 2.85E-05 | 3.43E-05 | 2.65E-06 | 6.14E-05 | 6.25E-06 | 8.72E-05 | 0.00011 | 3.43E-05 | 2.86E-05 | 8.00E-05 | 0.00025 | 9.26E-05 | 0 |
| Nfrkb | 3.07E-05 | 7.88E-05 | 3.50E-05 | 1.20E-05 | 4.48E-05 | 3.95E-06 | 1.63E-05 | 3.94E-06 | 6.14E-05 | 2.25E-05 | 2.24E-05 | 2.86E-05 | 1.96E-05 | 3.35E-05 | 0.00011 | 7.05E-07 | 7.38E-05 | 1.87E-05 |
| Atp1a3 | 0 | 0 | 0 | 2.40E-05 | 0.00026 | 5.96E-05 | 0 | 0.00018 | 6.13E-05 | 1.08E-05 | 3.02E-05 | 6.68E-05 | 0 | 0 | 2.66E-06 | 0 | 0 | 0 |

| | | | | | | | | | | | | | | | | | | |
|---|---|---|---|---|---|---|---|---|---|---|---|---|---|---|---|---|---|---|
| Chkb | 3.15E-05 | 0.00017 | 7.51E-06 | 0.00021 | 6.75E-05 | 3.03E-05 | 0.00014 | 6.42E-05 | 6.13E-05 | 7.54E-05 | 4.43E-05 | 6.77E-05 | 4.11E-05 | 0.00011 | 0.00059 | 7.52E-05 | 9.02E-05 | 9.92E-05 |
| Crot | 0.0001 | 5.83E-05 | 0 | 0 | 4.58E-05 | 2.76E-05 | 4.42E-05 | 0 | 6.13E-05 | 7.09E-05 | 0 | 0 | 0 | 4.40E-05 | 5.75E-05 | 3.62E-05 | 0 | 8.21E-05 |
| Hipk2 | 1.66E-05 | 7.30E-05 | 0.00013 | 0 | 3.42E-05 | 1.82E-05 | 0 | 4.37E-05 | 6.13E-05 | 5.96E-05 | 0.0002 | 7.03E-05 | 0.00012 | 2.48E-05 | 6.08E-05 | 2.91E-05 | 0.00014 | 0 |
| Zfp395 | 0 | 6.72E-05 | 0 | 6.44E-05 | 3.93E-05 | 1.30E-05 | 1.82E-05 | 2.76E-05 | 6.13E-05 | 5.12E-05 | 0 | 0.00064 | 0 | 2.99E-06 | 7.26E-08 | 5.50E-05 | 0 | 8.70E-05 |
| Mtr | 4.77E-05 | 5.54E-05 | 7.44E-05 | 4.14E-05 | 2.51E-05 | 0 | 5.73E-05 | 0 | 6.13E-05 | 7.50E-06 | 4.79E-05 | 5.75E-06 | 1.35E-05 | 0 | 0.00011 | 0.00011 | 3.32E-05 | 2.65E-05 |
| Gcnt2 | 2.63E-05 | 2.29E-05 | 8.31E-05 | 0 | 0 | 0.00022 | 0 | 0.00016 | 6.12E-05 | 0 | 0.00015 | 0 | 1.72E-05 | 0.00044 | 3.84E-05 | 0 | 6.35E-05 | 0 |
| Yrdc | 0.00013 | 3.22E-05 | 0.00033 | 0 | 4.84E-05 | 8.34E-05 | 0 | 6.09E-05 | 6.12E-05 | 0 | 0.00012 | 7.52E-05 | 0 | 6.14E-05 | 1.72E-05 | 0 | 9.48E-05 | 0 |
| Nav3 | 7.36E-05 | 7.46E-05 | 9.04E-05 | 0 | 0.00018 | 7.96E-05 | 0.00013 | 0.0002 | 6.12E-05 | 6.53E-05 | 0 | 3.05E-05 | 0.00011 | 0 | 0.00031 | 0.00013 | 0.00035 | 0 |
| Tmbim4 | 0.00015 | 0.00014 | 9.35E-05 | 0.00017 | 0.00011 | 7.60E-05 | 9.93E-05 | 7.11E-05 | 6.12E-05 | 7.25E-05 | 6.79E-05 | 0.00011 | 7.86E-05 | 6.09E-05 | 0 | 3.22E-05 | 0 | 7.06E-05 |
| Lmbrd2 | 0.0001 | 4.71E-05 | 4.38E-05 | 0 | 4.61E-05 | 4.01E-05 | 2.04E-05 | 7.17E-06 | 6.12E-05 | 1.75E-05 | 1.15E-05 | 5.91E-05 | 0 | 3.31E-05 | 0 | 0.00011 | 6.50E-05 | 2.48E-05 |
| Slc25a30 | 5.61E-05 | 0.00013 | 7.88E-05 | 0 | 4.80E-05 | 3.10E-05 | 0.00011 | 7.43E-05 | 6.12E-05 | 0.00011 | 2.23E-05 | 0.00016 | 5.82E-05 | 3.06E-05 | 0 | 0.00013 | 0.00015 | 0 |
| Srp72 | 0.00039 | 0.00017 | 0.00025 | 0.00025 | 7.55E-05 | 0.00015 | 0.00029 | 9.65E-05 | 6.11E-05 | 4.84E-05 | 0.00011 | 0.00021 | 7.47E-05 | 8.54E-05 | 0.00017 | 3.48E-05 | 0 | 2.11E-05 |
| Zfp260 | 0.00023 | 9.48E-05 | 0 | 3.21E-05 | 7.59E-05 | 0.00014 | 0.00025 | 0.00012 | 6.11E-05 | 8.01E-05 | 4.08E-05 | 9.89E-05 | 3.74E-05 | 0.00013 | 0.00025 | 0 | 0 | 0.00011 |
| Togaram1 | 6.12E-05 | 0.00013 | 9.77E-05 | 6.43E-05 | 4.65E-05 | 2.89E-05 | 0 | 2.95E-05 | 6.11E-05 | 3.37E-05 | 3.14E-05 | 4.07E-05 | 0 | 4.17E-05 | 0.00014 | 3.45E-05 | 0.00019 | 4.41E-05 |
| Pet100 | 5.76E-05 | 4.49E-05 | 1.09E-05 | 0.00016 | 2.11E-05 | 1.80E-05 | 3.95E-05 | 9.39E-06 | 6.11E-05 | 4.69E-05 | 0.00011 | 3.41E-05 | 4.70E-05 | 5.69E-05 | 2.97E-05 | 1.31E-05 | 2.11E-05 | 6.95E-05 |
| Ciapin1 | 1.08E-05 | 5.24E-05 | 0.00013 | 0.00014 | 5.22E-05 | 2.23E-06 | 0 | 1.65E-05 | 6.11E-05 | 3.31E-05 | 9.28E-05 | 0 | 4.04E-05 | 0.00013 | 7.19E-06 | 0.00013 | 0 | 4.05E-05 |
| Tbl3 | 6.28E-05 | 4.22E-05 | 0.00017 | 0.00013 | 0 | 0 | 0 | 0 | 6.11E-05 | 8.64E-05 | 5.48E-05 | 0 | 8.63E-06 | 0 | 0.0001 | 0.00012 | 0 | 1.68E-05 |
| Bcl2l12 | 4.03E-05 | 1.85E-05 | 3.08E-05 | 9.59E-05 | 0 | 0 | 0 | 6.56E-05 | 6.11E-05 | 3.16E-05 | 6.06E-05 | 1.77E-06 | 0 | 5.01E-05 | 8.77E-05 | 0.00013 | 0 | 3.91E-05 |
| Ccne1 | 2.01E-05 | 0 | 6.90E-05 | 0 | 0 | 0 | 1.13E-05 | 0 | 6.11E-05 | 0 | 2.60E-05 | 0 | 3.66E-05 | 1.06E-05 | 0 | 0.00038 | 1.75E-05 | 0 |
| Tprkb | 0.0001 | 9.27E-05 | 8.18E-05 | 5.41E-05 | 2.81E-05 | 7.17E-05 | 6.11E-05 | 0 | 6.10E-05 | 4.28E-05 | 0 | 2.84E-05 | 3.72E-05 | 0 | 3.71E-05 | 2.16E-05 | 0 | 9.58E-06 |
| Naa15 | 0.00026 | 0.00011 | 0.00024 | 0 | 0 | 7.05E-05 | 0 | 0 | 6.10E-05 | 1.12E-05 | 7.35E-05 | 0 | 0 | 0 | 0.00022 | 9.19E-05 | 0.00018 | 0 |
| Ccdc134 | 5.72E-05 | 1.98E-05 | 9.21E-05 | 6.52E-05 | 3.90E-05 | 0 | 2.85E-05 | 1.19E-06 | 6.10E-05 | 2.67E-05 | 2.68E-05 | 3.18E-05 | 2.13E-05 | 0 | 3.62E-05 | 4.04E-05 | 4.14E-05 | 0 |
| Atraid | 0.00032 | 0.0002 | 0 | 6.09E-06 | 0.00027 | 0.00039 | 0.00016 | 0.00014 | 6.09E-05 | 0.00017 | 2.75E-05 | 0.0002 | 0.00018 | 0.00012 | 0 | 2.65E-05 | 0 | 0.00021 |
| Rtf2 | 5.73E-05 | 0.00016 | 0.0001 | 0.00021 | 6.84E-05 | 4.85E-05 | 2.26E-05 | 2.58E-05 | 6.09E-05 | 7.45E-05 | 0.0001 | 0.00013 | 4.53E-05 | 5.22E-05 | 6.88E-05 | 4.43E-05 | 0 | 8.71E-05 |
| Trmt2a | 8.34E-05 | 7.03E-05 | 0.00013 | 5.34E-05 | 3.52E-05 | 6.59E-06 | 4.59E-05 | 3.10E-05 | 6.09E-05 | 5.52E-05 | 4.57E-05 | 5.61E-05 | 7.68E-06 | 2.22E-05 | 0.00013 | 6.42E-05 | 0 | 6.23E-05 |
| Tmem201 | 1.14E-05 | 4.20E-05 | 0.00011 | 1.59E-05 | 4.74E-05 | 3.41E-06 | 0 | 6.43E-05 | 6.09E-05 | 2.40E-05 | 2.37E-05 | 0.00011 | 4.18E-05 | 1.43E-06 | 7.84E-05 | 9.66E-05 | 0 | 3.36E-05 |
| Prr14l | 9.01E-05 | 8.96E-05 | 5.50E-05 | 1.01E-05 | 6.03E-05 | 0.0001 | 9.61E-05 | 0.00011 | 6.08E-05 | 5.22E-05 | 6.69E-05 | 4.08E-05 | 7.06E-05 | 4.35E-05 | 0.00017 | 7.70E-05 | 0 | 0 |
| Setd1a | 0 | 0.00019 | 0.00019 | 0.00026 | 7.99E-05 | 8.13E-05 | 0 | 8.73E-05 | 6.08E-05 | 8.25E-05 | 6.20E-05 | 3.64E-05 | 5.11E-05 | 0.00011 | 0.00035 | 0.00023 | 0.00033 | 4.15E-05 |
| Nmd3 | 0.00015 | 8.27E-05 | 0.00014 | 8.04E-07 | 0 | 7.53E-05 | 0 | 0 | 6.08E-05 | 7.42E-06 | 2.43E-05 | 9.83E-06 | 0 | 0 | 6.19E-05 | 5.14E-05 | 0 | 7.11E-06 |
| Narf | 5.10E-05 | 0.00016 | 0 | 1.26E-06 | 0.00012 | 8.70E-05 | 6.60E-05 | 8.53E-05 | 6.07E-05 | 0.00011 | 4.14E-05 | 0.00026 | 2.93E-05 | 2.10E-05 | 1.21E-05 | 0 | 0 | 0 |
| Nfyb | 0.00013 | 7.16E-05 | 7.70E-05 | 0 | 6.67E-05 | 7.18E-05 | 5.06E-05 | 2.62E-05 | 6.07E-05 | 1.71E-05 | 6.98E-05 | 5.39E-05 | 5.47E-05 | 3.80E-05 | 6.46E-05 | 0.00022 | 1.63E-05 | 7.78E-06 |
| Pomk | 2.75E-05 | 3.03E-05 | 0 | 4.61E-06 | 1.57E-05 | 8.26E-06 | 3.59E-05 | 3.31E-05 | 6.07E-05 | 4.16E-05 | 2.42E-06 | 7.41E-06 | 6.52E-06 | 2.33E-05 | 0 | 2.60E-07 | 3.46E-05 | 6.47E-05 |
| Mthfsd | 2.19E-05 | 6.27E-05 | 0 | 0.00014 | 5.17E-05 | 0 | 5.03E-05 | 2.64E-05 | 6.07E-05 | 1.48E-05 | 8.44E-05 | 7.88E-06 | 1.75E-05 | 5.00E-05 | 0.00017 | 0 | 0 | 0.0001 |

| | | | | | | | | | | | | | | | | | | |
|---|---|---|---|---|---|---|---|---|---|---|---|---|---|---|---|---|---|---|
| Gtf2a2 | 0.00023 | 6.53E-05 | 8.38E-05 | 9.11E-06 | 5.68E-05 | 0.00017 | 5.01E-05 | 3.38E-05 | 6.06E-05 | 2.68E-05 | 3.93E-06 | 2.19E-05 | 8.07E-05 | 1.83E-05 | 2.89E-05 | 0.00026 | 2.49E-05 | 1.71E-07 |
| Sbspon | 0 | 0 | 0 | 0 | 0.00017 | 0.00016 | 0 | 0 | 6.06E-05 | 0 | 0.00034 | 0 | 0 | 0 | 0 | 0 | 0 | 0 |
| Gmcl1 | 8.46E-05 | 7.82E-05 | 3.25E-05 | 2.68E-05 | 5.84E-05 | 9.36E-05 | 4.60E-05 | 6.78E-05 | 6.06E-05 | 2.78E-05 | 3.87E-05 | 3.10E-05 | 1.99E-05 | 1.22E-05 | 2.74E-05 | 0.0001 | 6.25E-06 | 2.01E-05 |
| B3galt6 | 4.64E-05 | 3.05E-05 | 0.00011 | 4.40E-06 | 3.68E-05 | 3.90E-05 | 2.63E-05 | 7.26E-05 | 6.06E-05 | 1.34E-05 | 7.36E-05 | 1.51E-05 | 3.65E-05 | 0 | 0 | 5.39E-05 | 0 | 9.08E-06 |
| Pomt2 | 6.60E-05 | 5.39E-05 | 7.81E-05 | 0.00011 | 7.52E-05 | 3.39E-05 | 5.94E-05 | 2.64E-05 | 6.06E-05 | 2.27E-05 | 0 | 1.48E-05 | 3.08E-05 | 3.38E-05 | 6.25E-05 | 3.50E-05 | 1.59E-06 | 3.42E-05 |
| Nosip | 7.74E-05 | 0.00017 | 6.46E-05 | 0.00029 | 3.46E-05 | 1.79E-05 | 3.54E-05 | 3.44E-05 | 6.06E-05 | 3.22E-05 | 6.08E-05 | 6.30E-05 | 0 | 5.57E-05 | 0.0001 | 0.00015 | 0 | 5.39E-05 |
| Gabpb2 | 5.13E-05 | 5.69E-05 | 4.48E-05 | 5.86E-05 | 4.86E-05 | 1.68E-05 | 4.14E-05 | 0 | 6.06E-05 | 3.41E-05 | 1.32E-05 | 2.56E-05 | 2.45E-05 | 5.36E-05 | 0.00017 | 0.00014 | 0 | 6.12E-05 |
| Prmt2 | 3.64E-05 | 8.10E-05 | 4.80E-06 | 1.40E-06 | 2.16E-05 | 4.22E-06 | 0 | 0 | 6.06E-05 | 4.77E-05 | 4.86E-05 | 0.00012 | 0 | 0 | 5.09E-05 | 0 | 6.45E-05 | 9.26E-05 |
| Hsph1 | 6.49E-05 | 6.00E-05 | 0.00015 | 1.07E-05 | 2.74E-05 | 4.22E-06 | 0 | 0 | 6.06E-05 | 4.95E-05 | 3.07E-05 | 0 | 5.38E-05 | 1.20E-05 | 2.47E-05 | 8.69E-05 | 0.00061 | 2.06E-05 |
| Gli1 | 0 | 3.49E-05 | 0 | 0 | 0 | 0 | 0 | 0 | 6.06E-05 | 0 | 0 | 0 | 0 | 0 | 3.21E-05 | 0 | 0 | 0.00039 |
| Aggf1 | 0.00011 | 0.0002 | 0.00013 | 6.71E-05 | 0.00015 | 0.00012 | 0.0002 | 6.82E-05 | 6.05E-05 | 5.55E-05 | 4.74E-05 | 0 | 0.00016 | 0.00014 | 3.83E-05 | 0.00012 | 0 | 0.00013 |
| Gmps | 0.00014 | 0.00012 | 0.0002 | 0.00011 | 3.97E-05 | 2.97E-05 | 0 | 0 | 6.05E-05 | 3.15E-05 | 9.61E-05 | 2.30E-05 | 1.44E-05 | 5.83E-05 | 9.70E-05 | 0.00019 | 1.57E-05 | 0 |
| Pus1 | 7.68E-05 | 8.54E-05 | 0.00024 | 0.00012 | 0 | 0 | 2.88E-05 | 0 | 6.05E-05 | 1.90E-05 | 8.84E-06 | 6.64E-05 | 1.92E-05 | 2.99E-05 | 7.07E-05 | 3.04E-05 | 3.20E-05 | 3.53E-06 |
| Slc39a3 | 0 | 5.22E-05 | 1.24E-05 | 9.75E-05 | 5.46E-05 | 0 | 0 | 4.71E-05 | 6.05E-05 | 6.65E-05 | 1.64E-05 | 4.28E-05 | 1.64E-05 | 3.43E-05 | 5.62E-05 | 3.40E-05 | 0 | 5.56E-05 |
| Mindy4 | 0 | 4.12E-05 | 0 | 4.06E-05 | 0.00012 | 0.00013 | 2.98E-05 | 7.15E-05 | 6.04E-05 | 3.22E-05 | 0 | 9.65E-05 | 2.30E-05 | 0 | 0.00013 | 6.61E-05 | 8.25E-05 | 7.23E-05 |
| Nlk | 5.43E-05 | 6.25E-05 | 5.59E-05 | 0 | 0.00012 | 0.00012 | 0 | 0.00015 | 6.04E-05 | 9.88E-06 | 4.53E-05 | 1.97E-05 | 0 | 9.54E-05 | 3.73E-05 | 0 | 7.30E-05 | 0 |
| Cryzl1 | 0.00016 | 0.00017 | 1.75E-05 | 4.78E-05 | 5.99E-05 | 0.0001 | 7.17E-05 | 5.13E-05 | 6.04E-05 | 7.89E-05 | 0 | 0.00011 | 8.80E-06 | 7.17E-05 | 6.31E-05 | 6.83E-06 | 0 | 8.98E-05 |
| Foxj3 | 0.00012 | 0.00011 | 4.60E-05 | 0 | 3.00E-05 | 9.34E-05 | 0 | 0 | 6.04E-05 | 0 | 0 | 0.00013 | 0 | 3.10E-06 | 0.00021 | 3.42E-05 | 6.16E-05 | 1.96E-05 |
| Rp2 | 0.00015 | 5.27E-05 | 0 | 0 | 5.79E-05 | 7.26E-05 | 0 | 2.27E-05 | 6.04E-05 | 6.80E-05 | 1.98E-05 | 7.43E-05 | 0 | 2.79E-05 | 0 | 7.49E-05 | 0 | 4.19E-05 |
| Atg7 | 0 | 7.72E-05 | 5.98E-05 | 6.08E-05 | 2.76E-05 | 2.24E-05 | 2.85E-05 | 1.42E-05 | 6.04E-05 | 5.91E-05 | 3.18E-05 | 2.89E-05 | 2.04E-05 | 2.27E-05 | 7.53E-05 | 2.94E-05 | 0 | 5.79E-05 |
| Ppp5c | 0.00016 | 9.15E-05 | 0.00018 | 0.00012 | 6.91E-05 | 3.25E-05 | 1.52E-05 | 4.57E-06 | 6.03E-05 | 3.42E-05 | 3.03E-05 | 3.39E-05 | 6.90E-07 | 0 | 0.00011 | 0.00024 | 0.00012 | 5.46E-05 |
| Nln | 7.70E-05 | 0.0001 | 7.45E-05 | 1.62E-05 | 3.17E-06 | 1.56E-05 | 2.43E-05 | 0 | 6.03E-05 | 5.73E-05 | 8.51E-05 | 2.23E-05 | 3.63E-05 | 3.72E-05 | 2.86E-05 | 8.75E-05 | 4.34E-05 | 4.23E-05 |
| Cxcl9 | 3.21E-05 | 0 | 1.10E-05 | 0 | 0 | 0 | 0 | 0 | 6.03E-05 | 3.67E-05 | 0 | 0 | 0 | 0 | 0 | 9.62E-06 | 0 | 0.00013 |
| Plcd3 | 0 | 4.86E-05 | 2.94E-06 | 1.57E-05 | 1.11E-05 | 0 | 0 | 0 | 6.03E-05 | 8.13E-06 | 1.72E-05 | 2.75E-05 | 0 | 6.43E-05 | 4.45E-05 | 0 | 4.35E-05 | 1.77E-05 |
| Rnf123 | 1.32E-05 | 0.00011 | 9.84E-06 | 3.02E-05 | 7.16E-05 | 4.93E-05 | 0 | 0 | 6.02E-05 | 3.96E-05 | 5.76E-05 | 1.27E-05 | 2.92E-05 | 7.42E-05 | 0.00014 | 1.18E-05 | 2.42E-05 | 5.62E-05 |
| Stimate | 2.17E-05 | 3.79E-05 | 6.19E-05 | 7.98E-05 | 4.89E-05 | 2.91E-05 | 4.81E-05 | 3.56E-05 | 6.02E-05 | 5.48E-05 | 5.73E-05 | 0 | 4.30E-06 | 7.55E-05 | 1.43E-05 | 5.32E-05 | 5.20E-05 | 0.00015 |
| Uprt | 6.27E-05 | 0.00013 | 0.00012 | 1.43E-05 | 7.37E-05 | 2.49E-05 | 0.00012 | 6.73E-05 | 6.02E-05 | 6.94E-05 | 8.74E-06 | 1.05E-07 | 0.0001 | 0 | 0 | 3.23E-05 | 2.16E-05 | 0.00015 |
| Tysnd1 | 3.47E-05 | 5.54E-05 | 9.84E-05 | 0.00022 | 3.03E-05 | 0 | 9.90E-05 | 5.74E-05 | 6.02E-05 | 2.86E-05 | 3.44E-05 | 2.27E-06 | 0 | 2.19E-05 | 0.00013 | 6.40E-05 | 6.53E-06 | 4.65E-05 |
| Dnajc11 | 4.19E-05 | 6.66E-05 | 0.00013 | 0.00012 | 2.04E-05 | 0 | 0 | 0 | 6.02E-05 | 2.19E-05 | 4.31E-05 | 1.02E-05 | 2.25E-05 | 3.41E-05 | 8.03E-05 | 4.65E-05 | 0 | 0 |
| Psmg1 | 0.00021 | 1.08E-05 | 0.00016 | 8.75E-05 | 0 | 3.80E-05 | 4.14E-05 | 0 | 6.01E-05 | 1.05E-05 | 0 | 2.90E-05 | 2.94E-05 | 2.67E-05 | 3.36E-05 | 7.67E-05 | 0.00017 | 0 |
| Fam111a | 6.18E-05 | 2.37E-05 | 0 | 0 | 0 | 2.83E-05 | 0 | 5.47E-05 | 6.01E-05 | 0.00011 | 0 | 6.66E-06 | 0 | 0 | 0 | 0.00043 | 0 | 5.06E-06 |
| Pdhx | 5.84E-05 | 4.81E-05 | 7.01E-05 | 6.32E-05 | 2.74E-05 | 1.47E-05 | 0 | 4.94E-06 | 6.01E-05 | 1.36E-05 | 0 | 0 | 0 | 2.18E-05 | 0.00014 | 7.47E-05 | 0 | 0 |
| Cpped1 | 3.95E-05 | 6.40E-05 | 3.31E-05 | 3.38E-05 | 2.67E-05 | 1.17E-05 | 0 | 2.28E-06 | 6.01E-05 | 1.37E-05 | 1.18E-05 | 1.90E-05 | 2.37E-05 | 0 | 1.60E-05 | 0 | 0 | 3.30E-05 |

| | | | | | | | | | | | | | | | | | | |
|---|---|---|---|---|---|---|---|---|---|---|---|---|---|---|---|---|---|---|
| Arl8b | 0.00022 | 0.00021 | 0.00013 | 4.66E-06 | 0.00014 | 0.00013 | 6.97E-05 | 0.0001 | 6.00E-05 | 5.68E-05 | 3.63E-05 | 0.00015 | 6.05E-05 | 0.00011 | 0 | 8.87E-05 | 9.42E-05 | 7.05E-05 |
| Habp4 | 3.40E-05 | 0.00011 | 9.09E-05 | 0.00015 | 0.00011 | 8.67E-05 | 9.12E-05 | 6.66E-05 | 6.00E-05 | 8.50E-05 | 9.78E-05 | 0 | 5.32E-05 | 0.0001 | 0 | 7.12E-05 | 1.06E-05 | 8.18E-05 |
| Ring1 | 4.17E-05 | 8.90E-05 | 5.22E-05 | 0.00017 | 5.30E-05 | 5.28E-05 | 4.27E-05 | 4.24E-05 | 5.99E-05 | 8.20E-05 | 3.18E-05 | 3.00E-05 | 2.56E-05 | 5.99E-05 | 0.00016 | 0 | 9.83E-05 | 9.70E-05 |
| Cdc42ep5 | 7.77E-05 | 1.69E-05 | 5.24E-05 | 0.00014 | 5.42E-05 | 3.53E-05 | 5.53E-05 | 4.86E-05 | 5.99E-05 | 0 | 5.42E-05 | 0.00011 | 0 | 4.80E-05 | 2.19E-05 | 0 | 0 | 4.93E-05 |
| Mavs | 5.34E-05 | 8.34E-05 | 2.07E-05 | 2.08E-05 | 3.07E-05 | 2.41E-05 | 0 | 0 | 5.99E-05 | 3.30E-05 | 2.97E-05 | 3.90E-05 | 3.17E-05 | 0 | 0 | 3.19E-05 | 0 | 3.65E-05 |
| Msantd3 | 4.82E-05 | 4.57E-05 | 0.0001 | 2.76E-05 | 0.00012 | 9.00E-06 | 4.34E-05 | 7.07E-06 | 5.99E-05 | 8.74E-06 | 0.00012 | 1.84E-05 | 0 | 5.27E-05 | 2.47E-05 | 7.19E-05 | 0 | 1.94E-05 |
| Tiam2 | 1.14E-06 | 3.12E-05 | 0 | 8.56E-06 | 2.94E-05 | 0 | 2.28E-07 | 2.70E-05 | 5.99E-05 | 0 | 6.09E-05 | 9.10E-06 | 0 | 7.49E-05 | 2.25E-05 | 3.56E-06 | 0.00012 | 0.00011 |
| DEPRECATED | 7.23E-06 | 4.57E-05 | 1.13E-05 | 9.94E-05 | 5.60E-05 | 7.29E-05 | 2.46E-05 | 0.00016 | 5.98E-05 | 4.92E-05 | 5.20E-05 | 4.29E-05 | 3.81E-05 | 0.00011 | 9.38E-06 | 4.93E-05 | 0 | 7.70E-05 |
| Dus2 | 4.17E-05 | 1.03E-05 | 4.16E-05 | 3.78E-05 | 3.61E-05 | 3.30E-06 | 6.24E-06 | 2.31E-05 | 5.98E-05 | 3.00E-05 | 4.01E-05 | 9.40E-06 | 1.32E-05 | 0 | 4.99E-05 | 1.20E-05 | 0 | 1.41E-05 |
| Znhit2 | 5.52E-05 | 5.51E-05 | 9.33E-05 | 0.00012 | 5.11E-05 | 0 | 2.04E-05 | 5.01E-05 | 5.98E-05 | 1.87E-05 | 3.55E-05 | 0 | 0 | 1.54E-05 | 1.89E-05 | 3.90E-05 | 0 | 6.03E-05 |
| Utp23 | 2.22E-05 | 4.42E-05 | 3.07E-05 | 2.58E-05 | 1.19E-05 | 0 | 7.76E-06 | 0 | 5.98E-05 | 3.03E-05 | 2.71E-05 | 0 | 1.57E-05 | 6.55E-06 | 7.04E-05 | 4.46E-05 | 0 | 2.08E-05 |
| Wapl | 0.00022 | 0.00011 | 7.71E-05 | 0 | 5.62E-05 | 0.00018 | 0 | 9.54E-06 | 5.97E-05 | 7.22E-06 | 2.08E-05 | 0.00011 | 0 | 0 | 0.00011 | 0.00015 | 0.00016 | 2.84E-05 |
| Tmcc1 | 0.00012 | 0.00018 | 0.00015 | 1.77E-05 | 8.73E-05 | 9.56E-05 | 0.00012 | 4.50E-05 | 5.97E-05 | 6.31E-05 | 8.05E-05 | 5.16E-05 | 3.12E-05 | 7.52E-05 | 0.00029 | 3.91E-05 | 0.00029 | 6.07E-05 |
| Galk2 | 0.00013 | 0.00012 | 0 | 8.37E-05 | 7.04E-05 | 9.46E-05 | 1.90E-05 | 1.53E-05 | 5.97E-05 | 6.73E-05 | 0 | 1.54E-05 | 0.00035 | 4.86E-05 | 0.00012 | 0 | 0 | 0.00011 |
| Cabin1 | 8.25E-05 | 0.00014 | 6.10E-05 | 6.54E-05 | 5.72E-05 | 5.87E-05 | 0 | 2.23E-05 | 5.97E-05 | 7.75E-05 | 2.35E-05 | 0 | 6.26E-05 | 7.19E-05 | 0.00016 | 4.18E-05 | 0.00015 | 4.78E-05 |
| Nkapd1 | 7.09E-05 | 7.99E-05 | 6.31E-05 | 4.91E-05 | 5.64E-05 | 5.49E-05 | 3.35E-05 | 2.74E-05 | 5.97E-05 | 4.35E-05 | 0 | 2.21E-05 | 2.54E-05 | 4.63E-05 | 8.30E-05 | 2.11E-05 | 0 | 2.49E-05 |
| Ppm1d | 2.69E-05 | 0.00014 | 6.20E-06 | 4.71E-05 | 2.99E-05 | 5.27E-05 | 0 | 3.51E-05 | 5.97E-05 | 4.54E-06 | 1.33E-05 | 7.66E-07 | 0 | 3.44E-05 | 4.34E-05 | 0.00015 | 0 | 2.89E-05 |
| Psmd10 | 0.00015 | 6.57E-05 | 0.00013 | 7.01E-05 | 9.79E-06 | 4.47E-05 | 0 | 4.86E-05 | 5.97E-05 | 2.65E-06 | 1.43E-05 | 4.22E-05 | 8.00E-05 | 5.97E-05 | 0 | 2.12E-05 | 0 | 1.71E-06 |
| Cd33 | 0 | 2.18E-05 | 0 | 0 | 0 | 3.10E-05 | 0 | 0 | 5.97E-05 | 0 | 0 | 0 | 0 | 0.00011 | 0 | 0 | 0 | 0 |
| Nptxr | 2.59E-05 | 7.64E-06 | 5.31E-06 | 6.01E-05 | 2.91E-05 | 1.52E-05 | 1.54E-05 | 0 | 5.97E-05 | 0 | 0 | 1.12E-05 | 0 | 0.00016 | 0 | 0.00026 | 7.79E-05 | 1.52E-05 |
| Stc2 | 0 | 0 | 0 | 0 | 4.13E-06 | 0 | 0 | 0 | 5.97E-05 | 4.34E-06 | 1.10E-05 | 0.00062 | 0.00014 | 2.22E-05 | 0 | 0 | 0 | 0 |
| Hipk1 | 0.00026 | 0.00031 | 0.00023 | 0 | 0.00017 | 0.00025 | 9.05E-05 | 0 | 5.96E-05 | 6.75E-05 | 0.00015 | 0.00019 | 0.00012 | 0.00012 | 1.41E-05 | 0.00026 | 0.00026 | 8.27E-05 |
| Dennd6a | 9.38E-05 | 9.51E-05 | 8.48E-06 | 0 | 6.23E-05 | 0.00012 | 6.28E-05 | 4.31E-05 | 5.96E-05 | 3.59E-05 | 4.09E-05 | 3.21E-05 | 2.33E-06 | 7.69E-05 | 0 | 3.64E-05 | 2.12E-05 | 5.06E-05 |
| AU022252 | 8.36E-05 | 5.86E-05 | 5.42E-05 | 3.00E-05 | 4.45E-05 | 3.60E-05 | 5.66E-05 | 3.06E-05 | 5.96E-05 | 2.72E-05 | 4.72E-05 | 5.41E-05 | 6.00E-06 | 5.16E-05 | 3.87E-05 | 3.37E-06 | 0 | 1.92E-05 |
| Supt7l | 7.45E-05 | 3.19E-05 | 2.46E-05 | 6.41E-05 | 2.30E-05 | 3.05E-05 | 1.83E-05 | 3.24E-05 | 5.96E-05 | 2.14E-05 | 1.44E-05 | 6.15E-06 | 3.20E-05 | 2.69E-05 | 1.51E-05 | 2.27E-05 | 0 | 2.38E-05 |
| Fam160b2 | 3.58E-05 | 7.82E-05 | 4.70E-05 | 4.92E-05 | 4.46E-05 | 1.66E-05 | 1.32E-05 | 4.30E-05 | 5.96E-05 | 2.68E-05 | 4.42E-05 | 7.06E-06 | 0 | 3.50E-05 | 0.00017 | 2.77E-06 | 0 | 3.59E-05 |
| Tmem186 | 8.32E-06 | 8.18E-05 | 5.01E-05 | 8.81E-05 | 3.82E-05 | 1.61E-06 | 0 | 2.20E-05 | 5.96E-05 | 3.32E-05 | 2.53E-05 | 2.66E-05 | 2.70E-05 | 1.97E-05 | 1.75E-05 | 0.00013 | 0 | 3.21E-05 |
| Fam76a | 1.73E-05 | 0.00015 | 6.51E-05 | 9.88E-05 | 9.84E-05 | 0.00012 | 6.37E-05 | 6.67E-05 | 5.95E-05 | 6.97E-05 | 8.97E-05 | 4.19E-05 | 6.51E-05 | 7.85E-05 | 9.72E-06 | 9.63E-05 | 0 | 8.28E-05 |
| Lyst | 7.67E-05 | 7.82E-05 | 0 | 0 | 5.33E-05 | 0.00011 | 1.03E-05 | 1.03E-05 | 5.95E-05 | 3.80E-05 | 7.87E-05 | 0.00011 | 8.23E-05 | 0 | 0.00011 | 3.06E-05 | 0.00012 | 4.63E-06 |
| Gdf6 | 0 | 0 | 0 | 0 | 1.32E-05 | 6.01E-05 | 0 | 0 | 5.95E-05 | 1.28E-05 | 0.00017 | 0 | 8.43E-05 | 0 | 1.39E-05 | 0 | 0 | 2.20E-05 |
| Gins4 | 4.52E-05 | 4.70E-05 | 5.76E-06 | 6.85E-05 | 2.57E-05 | 5.61E-05 | 0.0001 | 4.67E-05 | 5.95E-05 | 2.09E-05 | 6.20E-05 | 0 | 5.19E-05 | 2.24E-05 | 0 | 0.00039 | 0 | 0.00011 |
| Wdr24 | 4.36E-05 | 4.85E-05 | 1.15E-05 | 4.56E-05 | 4.42E-05 | 3.04E-05 | 3.06E-05 | 2.04E-05 | 5.95E-05 | 3.19E-05 | 9.38E-06 | 2.53E-05 | 0 | 4.38E-05 | 6.38E-05 | 1.82E-05 | 0 | 2.86E-05 |
| Ndst2 | 2.09E-05 | 6.61E-05 | 2.70E-05 | 2.36E-05 | 2.78E-05 | 2.92E-05 | 3.37E-05 | 0 | 5.95E-05 | 7.10E-05 | 2.22E-05 | 0 | 4.69E-05 | 5.45E-05 | 5.79E-05 | 4.44E-06 | 0 | 2.48E-05 |

| | | | | | | | | | | | | | | | | | | |
|---|---|---|---|---|---|---|---|---|---|---|---|---|---|---|---|---|---|---|
| Zfp275 | 6.81E-05 | 5.73E-05 | 0.0001 | 6.15E-05 | 7.29E-05 | 8.52E-06 | 4.07E-05 | 5.14E-05 | 5.95E-05 | 1.49E-05 | 0 | 1.21E-05 | 0 | 6.14E-05 | 0.00015 | 4.87E-05 | 0.00012 | 3.85E-05 |
| Dtx3l | 9.39E-05 | 8.52E-05 | 0 | 1.83E-05 | 6.71E-05 | 4.83E-06 | 5.43E-05 | 0 | 5.95E-05 | 0.00016 | 2.86E-05 | 7.09E-05 | 0.00011 | 0.0001 | 0 | 2.40E-05 | 0 | 0.00013 |
| Ctsd | 8.12E-05 | 0.00028 | 6.87E-05 | 0 | 0.00026 | 0.00032 | 0.00024 | 0.00019 | 5.94E-05 | 0.00021 | 0.00012 | 0.00018 | 0.00045 | 0.00023 | 0 | 0 | 0 | 0.00043 |
| Suds3 | 0.00016 | 0.0003 | 0.00019 | 0.00023 | 0.00016 | 0.00017 | 0.00017 | 7.47E-05 | 5.94E-05 | 9.85E-05 | 0.00012 | 3.76E-05 | 5.76E-05 | 3.33E-05 | 0.00021 | 0.00028 | 0.00014 | 0.00013 |
| Tulp4 | 5.83E-05 | 0.00022 | 0.00011 | 5.18E-05 | 0.00018 | 0.00011 | 9.47E-05 | 0.00012 | 5.94E-05 | 9.38E-05 | 0.00023 | 1.61E-05 | 0.00018 | 0 | 8.90E-05 | 9.44E-06 | 0.00013 | 0.00018 |
| Acp1 | 0.00027 | 9.11E-05 | 0.00017 | 0.00012 | 5.40E-05 | 6.21E-05 | 9.16E-06 | 1.80E-06 | 5.94E-05 | 2.62E-05 | 1.80E-05 | 0.00013 | 0 | 0 | 0 | 0.00026 | 0 | 7.83E-06 |
| Taf6 | 6.64E-05 | 9.31E-05 | 5.50E-05 | 5.14E-05 | 5.06E-05 | 3.58E-05 | 3.57E-05 | 2.45E-05 | 5.94E-05 | 3.82E-05 | 6.94E-05 | 1.58E-05 | 3.83E-05 | 3.98E-05 | 1.90E-05 | 0.00019 | 0 | 2.85E-05 |
| Qpctl | 0 | 6.32E-05 | 0.00011 | 0.00016 | 4.90E-05 | 3.27E-05 | 2.59E-05 | 2.35E-05 | 5.94E-05 | 0.00011 | 6.89E-05 | 3.34E-05 | 0.00021 | 9.64E-05 | 0 | 2.27E-05 | 0 | 7.74E-05 |
| Ppp1r10 | 0.00014 | 0.00011 | 6.53E-05 | 3.20E-05 | 1.58E-05 | 1.88E-05 | 8.89E-05 | 0 | 5.94E-05 | 1.45E-05 | 0 | 0.00011 | 0.00011 | 5.38E-06 | 5.78E-05 | 0.00013 | 0.00137 | 0 |
| Tmem223 | 1.79E-05 | 7.12E-05 | 2.81E-06 | 0.00025 | 0.00012 | 5.10E-05 | 0 | 4.79E-05 | 5.93E-05 | 5.12E-05 | 3.97E-05 | 2.84E-06 | 5.61E-05 | 4.96E-05 | 4.02E-05 | 0 | 0.00014 | 0.00011 |
| Rilpl2 | 5.69E-05 | 7.41E-05 | 1.56E-05 | 0.00017 | 6.63E-05 | 3.87E-05 | 3.67E-05 | 9.28E-05 | 5.93E-05 | 1.41E-05 | 9.96E-06 | 0 | 4.64E-05 | 9.82E-05 | 9.92E-05 | 5.31E-06 | 2.31E-05 | 3.84E-05 |
| Pigm | 7.14E-05 | 0.0001 | 0.00012 | 4.49E-05 | 4.06E-05 | 3.83E-05 | 4.62E-06 | 4.67E-05 | 5.93E-05 | 4.40E-05 | 7.22E-05 | 5.90E-05 | 0 | 4.56E-05 | 0.00012 | 4.77E-05 | 0 | 3.33E-05 |
| Specc1 | 3.47E-05 | 8.34E-05 | 0 | 0 | 8.25E-05 | 2.10E-05 | 0 | 1.61E-05 | 5.93E-05 | 0 | 0.00024 | 1.08E-05 | 0 | 6.51E-05 | 8.79E-05 | 9.55E-05 | 1.80E-05 | 4.90E-05 |
| Fam192a | 0.00018 | 0.00013 | 6.10E-05 | 7.86E-05 | 1.76E-05 | 1.72E-05 | 2.53E-05 | 2.72E-05 | 5.93E-05 | 5.92E-06 | 4.16E-05 | 6.00E-05 | 2.99E-05 | 8.12E-06 | 0.00015 | 6.13E-05 | 0 | 1.24E-05 |
| Klc2 | 1.57E-07 | 9.86E-05 | 9.67E-05 | 9.07E-05 | 3.32E-05 | 6.85E-06 | 0 | 6.00E-05 | 5.93E-05 | 0 | 0.00016 | 0 | 0 | 7.37E-05 | 0.00016 | 0 | 0.00016 | 2.04E-05 |
| Rpusd1 | 3.52E-05 | 4.26E-05 | 1.69E-05 | 1.23E-05 | 6.99E-06 | 1.68E-06 | 4.32E-06 | 2.05E-05 | 5.93E-05 | 1.53E-05 | 2.80E-05 | 7.08E-06 | 6.47E-06 | 3.12E-06 | 3.47E-05 | 6.28E-05 | 0 | 3.24E-05 |
| Slc7a5 | 4.42E-05 | 7.08E-05 | 0.00042 | 5.40E-05 | 0 | 0 | 0 | 0.00038 | 5.93E-05 | 1.94E-05 | 4.00E-05 | 0.00025 | 0 | 0 | 0 | 0.00026 | 0.00043 | 0 |
| Zmynd19 | 0.00012 | 1.17E-05 | 0.00021 | 1.06E-05 | 3.98E-05 | 0 | 7.96E-05 | 2.57E-05 | 5.93E-05 | 1.89E-05 | 4.41E-05 | 2.01E-05 | 2.31E-05 | 2.78E-06 | 3.64E-05 | 8.93E-05 | 5.63E-05 | 1.19E-06 |
| Pcdhgc5 | 1.25E-05 | 7.80E-05 | 0 | 4.47E-05 | 0.00017 | 0.00011 | 8.23E-06 | 0.00018 | 5.92E-05 | 2.35E-05 | 0 | 0 | 0.00011 | 1.62E-05 | 9.45E-05 | 0 | 1.15E-05 | 4.12E-05 |
| Zfp777 | 6.86E-05 | 9.38E-05 | 0.0001 | 3.93E-05 | 3.65E-05 | 3.10E-05 | 2.29E-06 | 2.79E-05 | 5.92E-05 | 3.15E-05 | 1.82E-05 | 7.23E-05 | 0 | 2.31E-05 | 0.00014 | 1.44E-05 | 0 | 1.77E-05 |
| Tecpr1 | 0 | 9.93E-05 | 9.51E-07 | 4.76E-05 | 3.70E-05 | 9.25E-06 | 0 | 3.47E-05 | 5.92E-05 | 6.38E-05 | 3.62E-05 | 2.34E-05 | 0 | 3.60E-05 | 0.00012 | 0 | 0 | 4.99E-05 |
| Oxsr1 | 9.21E-05 | 9.13E-05 | 5.78E-05 | 2.75E-05 | 2.42E-05 | 0 | 8.07E-06 | 0 | 5.92E-05 | 6.02E-05 | 0.00011 | 0.00023 | 0 | 0 | 3.23E-05 | 6.67E-05 | 8.21E-05 | 2.69E-05 |
| Polr1c | 0.00016 | 0.00016 | 0.00015 | 0.00022 | 4.83E-06 | 4.55E-05 | 5.73E-05 | 6.60E-06 | 5.91E-05 | 2.85E-05 | 0 | 6.31E-05 | 1.19E-05 | 4.90E-05 | 0.0001 | 0.00013 | 0 | 9.34E-06 |
| Irs2 | 9.23E-06 | 0.00016 | 0 | 9.65E-05 | 4.95E-05 | 0.00014 | 0 | 0.00022 | 5.90E-05 | 2.65E-06 | 0.00013 | 5.85E-05 | 0.00015 | 8.12E-05 | 0 | 1.00E-06 | 0.00185 | 0.00012 |
| Zkscan17 | 7.23E-05 | 0.00014 | 7.11E-05 | 5.98E-05 | 5.75E-05 | 5.41E-05 | 1.29E-05 | 0 | 5.90E-05 | 2.18E-05 | 0 | 8.98E-05 | 0 | 2.29E-05 | 2.44E-05 | 9.99E-05 | 6.16E-05 | 3.78E-05 |
| Sf3a3 | 0.00019 | 8.29E-05 | 0.00015 | 0.00012 | 0 | 3.94E-05 | 2.19E-05 | 0 | 5.90E-05 | 2.90E-05 | 4.66E-05 | 0 | 3.33E-05 | 3.32E-05 | 6.42E-05 | 0.00025 | 7.51E-05 | 0 |
| Exoc3l4 | 0 | 5.60E-06 | 0 | 2.04E-06 | 0 | 3.25E-05 | 0 | 0 | 5.90E-05 | 2.15E-06 | 0 | 0 | 2.30E-05 | 0 | 0 | 0 | 0 | 7.27E-05 |
| Trappc2l | 0.00018 | 0.00012 | 0.00011 | 0.00014 | 0.00016 | 0.00021 | 8.53E-05 | 0.00019 | 5.89E-05 | 8.30E-05 | 7.50E-05 | 3.72E-05 | 0.00011 | 0.0001 | 0 | 1.24E-05 | 0 | 2.59E-05 |
| Alg5 | 0.00019 | 1.53E-05 | 0.00014 | 0.00012 | 2.22E-05 | 7.79E-05 | 0.00025 | 5.32E-05 | 5.89E-05 | 3.18E-05 | 6.54E-05 | 9.78E-06 | 6.81E-05 | 2.80E-05 | 6.05E-05 | 8.02E-05 | 0 | 1.75E-05 |
| Cul2 | 0.00018 | 3.75E-05 | 6.38E-05 | 0 | 2.58E-05 | 7.63E-05 | 0 | 0 | 5.89E-05 | 1.75E-05 | 1.79E-05 | 5.13E-05 | 0 | 0 | 0 | 5.96E-05 | 6.22E-06 | 1.23E-05 |
| Hmgxb3 | 6.86E-05 | 0.00011 | 0.00011 | 6.37E-05 | 3.54E-05 | 7.51E-05 | 2.07E-05 | 1.67E-05 | 5.89E-05 | 1.32E-05 | 3.23E-05 | 2.03E-05 | 1.60E-05 | 7.31E-05 | 0.00013 | 3.46E-06 | 3.08E-05 | 6.70E-06 |
| Timm17b | 0.00016 | 0 | 6.90E-05 | 0.00017 | 3.47E-05 | 5.16E-05 | 0.00014 | 6.48E-05 | 5.89E-05 | 3.53E-06 | 1.97E-05 | 0 | 0 | 4.64E-05 | 0 | 4.66E-05 | 7.55E-06 | 0 |
| Zfp410 | 6.69E-05 | 0.00012 | 9.56E-05 | 4.84E-05 | 9.95E-05 | 4.47E-05 | 6.31E-05 | 8.64E-05 | 5.89E-05 | 2.70E-05 | 5.29E-05 | 7.05E-05 | 0 | 9.06E-05 | 9.31E-05 | 0.0001 | 0 | 3.63E-05 |

| | | | | | | | | | | | | | | | | | | |
|---|---|---|---|---|---|---|---|---|---|---|---|---|---|---|---|---|---|---|
| Rmdn2 | 6.59E-05 | 2.26E-05 | 7.68E-06 | 0 | 5.29E-05 | 1.78E-05 | 3.24E-05 | 3.80E-05 | 5.89E-05 | 1.52E-05 | 4.03E-05 | 3.06E-05 | 2.83E-05 | 0 | 4.63E-06 | 3.69E-05 | 9.25E-06 | 4.05E-05 |
| Mcm6 | 8.79E-05 | 0 | 0.00022 | 0 | 0 | 0 | 8.23E-06 | 0 | 5.89E-05 | 5.15E-05 | 3.17E-05 | 0 | 3.83E-05 | 1.06E-05 | 0 | 0.00074 | 2.36E-05 | 5.05E-05 |
| Exoc4 | 0.00016 | 0.00032 | 0 | 0 | 0.00012 | 8.29E-05 | 3.70E-06 | 6.61E-05 | 5.88E-05 | 0.00017 | 0 | 2.19E-05 | 9.94E-05 | 0 | 0.00044 | 0.00019 | 0.00015 | 0.00013 |
| Eml2 | 0 | 7.21E-05 | 1.76E-05 | 8.48E-05 | 8.08E-06 | 7.23E-05 | 0 | 8.80E-05 | 5.88E-05 | 1.42E-05 | 7.71E-05 | 0.0001 | 1.24E-05 | 3.23E-05 | 3.59E-05 | 0 | 0 | 0.00011 |
| Tbck | 8.72E-05 | 6.34E-05 | 3.05E-06 | 0 | 3.93E-05 | 4.22E-05 | 6.51E-06 | 2.58E-06 | 5.88E-05 | 3.36E-05 | 2.81E-05 | 5.31E-05 | 0 | 0 | 8.31E-05 | 2.33E-05 | 2.50E-05 | 3.91E-05 |
| Ehbp1 | 3.60E-05 | 7.35E-05 | 0 | 0 | 2.08E-05 | 1.21E-05 | 0 | 0 | 5.88E-05 | 0.00017 | 2.33E-05 | 7.81E-06 | 1.14E-06 | 5.18E-05 | 0.00015 | 8.32E-06 | 0.00012 | 8.59E-05 |
| Ppox | 0 | 0.00012 | 0 | 6.71E-05 | 2.77E-05 | 0 | 2.36E-05 | 6.18E-05 | 5.88E-05 | 8.44E-05 | 1.57E-05 | 7.50E-05 | 4.43E-05 | 0 | 0.00023 | 6.90E-05 | 0 | 9.64E-05 |
| Ftsj3 | 5.48E-05 | 2.59E-05 | 0.00015 | 3.61E-05 | 0 | 0 | 0 | 0 | 5.88E-05 | 2.19E-05 | 3.57E-05 | 0 | 6.31E-05 | 0 | 2.90E-05 | 3.24E-05 | 0 | 1.33E-05 |
| Ube2o | 5.65E-05 | 0.00014 | 0.00015 | 3.65E-05 | 7.79E-05 | 0 | 0 | 8.28E-05 | 5.88E-05 | 1.71E-05 | 2.78E-05 | 0.0004 | 0 | 3.87E-06 | 0.00013 | 4.02E-05 | 6.23E-05 | 5.22E-05 |
| Dync1li1 | 0.0002 | 0.00026 | 0.0003 | 5.19E-05 | 0.00014 | 0.00014 | 0 | 7.72E-05 | 5.87E-05 | 9.42E-05 | 0.00011 | 0.00036 | 0 | 0.00012 | 0.00012 | 8.64E-05 | 0.00012 | 7.75E-05 |
| Miga2 | 0 | 2.79E-05 | 0.00011 | 3.83E-05 | 4.06E-05 | 3.28E-05 | 2.20E-05 | 4.17E-05 | 5.87E-05 | 2.98E-05 | 5.03E-05 | 0 | 3.49E-05 | 4.82E-05 | 0.00013 | 4.83E-05 | 4.32E-07 | 1.82E-05 |
| Katna1 | 5.69E-05 | 5.74E-05 | 3.62E-05 | 1.56E-06 | 0 | 0 | 4.36E-06 | 0 | 5.87E-05 | 1.51E-05 | 1.01E-05 | 6.37E-05 | 7.43E-06 | 0 | 9.26E-05 | 1.95E-05 | 0.00015 | 2.01E-05 |
| Reg1 | 0 | 0 | 0 | 0 | 0 | 0 | 0 | 3.76E-05 | 5.87E-05 | 0 | 0 | 0 | 0.00026 | 0 | 3.32E-06 | 1.43E-05 | 0 | 0.00018 |
| Afdn | 7.05E-05 | 0.00014 | 8.96E-06 | 0 | 2.11E-05 | 0 | 9.40E-06 | 4.54E-05 | 5.86E-05 | 1.23E-06 | 7.18E-06 | 8.67E-05 | 0 | 4.93E-05 | 0.00011 | 8.71E-05 | 0.00014 | 9.85E-05 |
| Aff1 | 8.59E-05 | 0.00029 | 0.00025 | 0 | 0.00025 | 8.66E-05 | 0.00019 | 8.37E-05 | 5.85E-05 | 0.00013 | 0.00021 | 6.93E-05 | 0.00027 | 0.00028 | 0.00036 | 0.00011 | 0.00043 | 6.36E-05 |
| Cstf3 | 8.10E-05 | 5.10E-05 | 8.56E-05 | 7.69E-05 | 1.70E-05 | 1.10E-05 | 2.84E-05 | 5.25E-06 | 5.85E-05 | 2.35E-05 | 5.74E-05 | 2.22E-05 | 1.73E-06 | 0 | 3.66E-05 | 0.00012 | 1.56E-05 | 1.03E-05 |
| Ubr5 | 0.00017 | 0.0003 | 0.00022 | 0.0002 | 0.0003 | 0.0003 | 0.00019 | 0.00015 | 5.84E-05 | 0.00013 | 0.00023 | 0.0002 | 8.63E-05 | 0.00018 | 0.00024 | 0.00018 | 0.00037 | 0.00016 |
| Setbp1 | 5.39E-05 | 0.0002 | 0 | 6.59E-05 | 0.00012 | 0.00017 | 7.29E-06 | 0 | 5.84E-05 | 6.77E-05 | 5.43E-05 | 6.63E-05 | 0.00041 | 0 | 0.00028 | 0 | 0.00026 | 0.00019 |
| Coq7 | 7.27E-05 | 2.38E-05 | 7.44E-05 | 7.89E-05 | 2.29E-05 | 4.02E-05 | 0 | 3.51E-05 | 5.84E-05 | 0 | 4.28E-05 | 1.47E-05 | 3.51E-05 | 0 | 0 | 0.00019 | 0 | 2.83E-05 |
| Nudt19 | 0.00013 | 2.11E-05 | 9.87E-05 | 3.52E-05 | 2.22E-06 | 2.45E-05 | 1.75E-06 | 1.04E-05 | 5.84E-05 | 1.90E-05 | 2.92E-05 | 1.10E-05 | 4.16E-05 | 0 | 0 | 4.46E-05 | 0 | 7.24E-05 |
| Znfx1 | 4.31E-05 | 6.49E-05 | 0 | 1.83E-06 | 3.17E-05 | 2.33E-05 | 0 | 5.96E-07 | 5.84E-05 | 0.0001 | 2.54E-05 | 4.36E-05 | 2.12E-05 | 1.25E-05 | 3.30E-05 | 4.12E-05 | 0 | 0.00011 |
| Slc35c1 | 4.20E-05 | 7.11E-05 | 0.00013 | 0.00017 | 4.96E-06 | 1.45E-05 | 7.90E-05 | 5.81E-05 | 5.84E-05 | 3.03E-05 | 5.91E-05 | 1.51E-05 | 1.12E-05 | 3.59E-05 | 0 | 0 | 0 | 1.08E-05 |
| Cask | 0.00026 | 0.00012 | 5.84E-05 | 0 | 0.00029 | 0.00051 | 0.00013 | 3.56E-05 | 5.83E-05 | 1.13E-05 | 0.00011 | 7.15E-05 | 8.84E-05 | 0.00016 | 0 | 8.97E-05 | 0.00023 | 0 |
| Rnf130 | 0.00018 | 0.00018 | 1.54E-05 | 0 | 0.00013 | 0.00021 | 0 | 0 | 5.83E-05 | 7.79E-05 | 5.82E-05 | 0.00019 | 0.00012 | 5.11E-05 | 0 | 8.04E-06 | 0 | 0.00014 |
| Chchd7 | 0.00018 | 4.53E-05 | 5.15E-05 | 4.57E-05 | 7.15E-05 | 8.36E-05 | 7.50E-05 | 6.46E-05 | 5.83E-05 | 0 | 7.61E-05 | 5.43E-06 | 9.92E-05 | 6.39E-05 | 0 | 4.48E-05 | 0 | 0 |
| Ube2j2 | 9.71E-05 | 9.69E-05 | 0.00022 | 0.00017 | 4.07E-05 | 5.30E-05 | 5.21E-05 | 3.28E-05 | 5.83E-05 | 2.92E-05 | 9.17E-05 | 6.82E-05 | 0 | 4.48E-05 | 2.50E-05 | 7.99E-05 | 4.08E-05 | 3.54E-05 |
| Ssbp1 | 0.00016 | 6.11E-05 | 8.70E-05 | 0.00012 | 1.58E-05 | 4.58E-05 | 1.50E-05 | 7.82E-06 | 5.83E-05 | 9.82E-06 | 0 | 0 | 1.73E-05 | 0 | 0.00014 | 8.33E-05 | 0 | 3.40E-05 |
| Prkra | 9.14E-05 | 4.43E-05 | 0 | 0 | 2.94E-05 | 4.26E-05 | 3.02E-05 | 1.90E-05 | 5.83E-05 | 1.97E-05 | 5.67E-06 | 3.70E-05 | 2.24E-06 | 2.67E-06 | 0 | 1.38E-05 | 2.25E-05 | 2.07E-05 |
| Senp1 | 7.46E-05 | 4.80E-05 | 7.22E-05 | 4.08E-06 | 3.98E-05 | 1.59E-05 | 1.10E-05 | 3.76E-05 | 5.83E-05 | 1.48E-05 | 1.37E-05 | 1.08E-05 | 1.90E-05 | 3.05E-07 | 0.00012 | 0.00019 | 7.53E-05 | 1.59E-05 |
| Trmt1l | 9.99E-05 | 8.24E-05 | 2.56E-05 | 9.58E-06 | 3.24E-05 | 1.29E-05 | 9.18E-06 | 0 | 5.83E-05 | 1.65E-05 | 1.38E-05 | 1.06E-05 | 1.71E-06 | 3.61E-05 | 9.81E-05 | 3.66E-05 | 0 | 4.17E-05 |
| Sac3d1 | 8.65E-06 | 7.06E-05 | 0.00017 | 7.97E-05 | 2.55E-05 | 5.64E-07 | 3.42E-05 | 3.39E-05 | 5.83E-05 | 2.85E-05 | 4.94E-05 | 0 | 0 | 3.84E-05 | 0 | 0.00038 | 0 | 6.22E-05 |
| Spopl | 0.0001 | 0.0001 | 6.63E-05 | 3.28E-06 | 6.57E-05 | 7.75E-05 | 0 | 3.96E-05 | 5.82E-05 | 5.50E-05 | 4.87E-05 | 8.47E-05 | 1.34E-05 | 2.82E-05 | 0.00013 | 3.24E-05 | 5.99E-05 | 5.81E-05 |
| Eif2ak2 | 0.00016 | 5.33E-05 | 8.38E-06 | 3.76E-06 | 3.02E-05 | 3.45E-05 | 2.55E-05 | 2.28E-05 | 5.82E-05 | 0.00021 | 0 | 3.24E-05 | 0.00014 | 1.18E-05 | 4.35E-05 | 7.09E-05 | 0 | 6.94E-05 |

| | | | | | | | | | | | | | | | | | | |
|---|---|---|---|---|---|---|---|---|---|---|---|---|---|---|---|---|---|---|
| Gpatch2l | 1.96E-05 | 6.49E-05 | 2.88E-05 | 5.51E-05 | 5.16E-05 | 3.05E-05 | 9.28E-06 | 4.33E-05 | 5.82E-05 | 5.02E-05 | 5.94E-05 | 3.64E-05 | 0 | 5.45E-05 | 0.00013 | 1.58E-05 | 2.00E-05 | 4.84E-05 |
| Nup88 | 9.66E-05 | 7.20E-05 | 0.0001 | 0 | 2.50E-05 | 1.79E-05 | 0 | 3.73E-06 | 5.82E-05 | 4.58E-05 | 8.08E-05 | 5.47E-05 | 0 | 9.11E-06 | 0.00016 | 0.00011 | 1.31E-05 | 0 |
| Brinp3 | 0 | 0 | 0 | 0.00015 | 0.00075 | 0.0011 | 0 | 0 | 5.81E-05 | 0 | 0.00041 | 0 | 0 | 0 | 0.00014 | 0 | 0 | 0 |
| Alg1 | 7.66E-05 | 9.69E-05 | 0.00012 | 0.00018 | 0.00011 | 5.92E-05 | 0.00011 | 6.83E-05 | 5.81E-05 | 6.50E-05 | 8.12E-05 | 0 | 0.00014 | 7.70E-05 | 0.00016 | 0 | 0 | 7.04E-05 |
| Zcchc9 | 4.44E-05 | 5.01E-05 | 4.50E-05 | 1.57E-05 | 2.06E-05 | 2.25E-05 | 1.59E-05 | 2.53E-05 | 5.81E-05 | 8.96E-06 | 1.25E-05 | 1.47E-05 | 0 | 3.96E-05 | 6.15E-05 | 2.24E-05 | 0 | 3.92E-06 |
| Sugp1 | 8.98E-05 | 8.71E-05 | 2.65E-05 | 4.72E-05 | 5.59E-05 | 1.64E-05 | 3.30E-05 | 1.23E-05 | 5.81E-05 | 3.37E-05 | 1.93E-05 | 1.79E-05 | 9.25E-06 | 6.08E-06 | 0.00015 | 2.09E-05 | 0 | 5.20E-06 |
| Abhd4 | 0.00019 | 0.00035 | 0 | 3.94E-05 | 0.00025 | 0.00029 | 0.00024 | 0.00039 | 5.80E-05 | 0.00012 | 4.19E-05 | 6.93E-05 | 0.00011 | 0.00014 | 0 | 0 | 0 | 0.00016 |
| Tmem33 | 0.00031 | 0.00014 | 0.0002 | 2.28E-05 | 8.91E-05 | 0.00015 | 2.85E-05 | 3.14E-06 | 5.80E-05 | 2.49E-05 | 9.01E-05 | 8.47E-05 | 7.57E-05 | 5.75E-05 | 1.02E-05 | 6.86E-05 | 1.54E-05 | 0 |
| Acvr1b | 0 | 3.35E-05 | 4.12E-05 | 2.52E-05 | 7.70E-05 | 0.00013 | 0 | 7.36E-05 | 5.80E-05 | 9.93E-05 | 9.54E-05 | 0.00016 | 0.00045 | 0.00018 | 9.49E-05 | 0 | 0 | 0.00013 |
| Galc | 8.51E-06 | 5.93E-05 | 6.73E-06 | 2.05E-05 | 3.61E-05 | 4.60E-05 | 1.60E-05 | 5.67E-05 | 5.80E-05 | 2.70E-06 | 1.08E-05 | 1.53E-05 | 1.29E-05 | 5.00E-05 | 0 | 3.80E-05 | 5.25E-06 | 1.79E-05 |
| Spry1 | 0 | 0.00036 | 0 | 4.08E-06 | 0.0005 | 0 | 0 | 0.00067 | 5.80E-05 | 0 | 0.00013 | 5.23E-05 | 0 | 0.00037 | 0.00015 | 0.00011 | 0.00024 | 0.00059 |
| Gyg | 5.43E-05 | 0.00014 | 1.41E-05 | 2.73E-05 | 5.64E-05 | 0 | 6.21E-05 | 0 | 5.80E-05 | 7.52E-05 | 1.28E-05 | 0 | 0.00013 | 2.99E-05 | 0 | 9.48E-05 | 0 | 0.00015 |
| Kif21a | 5.12E-06 | 0 | 0 | 0 | 0 | 0 | 4.07E-05 | 0 | 5.80E-05 | 0.00017 | 0 | 1.25E-05 | 0.00019 | 0 | 0 | 6.39E-06 | 3.12E-05 | 0 |
| Myl3 | 0 | 0 | 0 | 0 | 1.01E-06 | 0 | 1.46E-05 | 4.53E-06 | 5.80E-05 | 0 | 0 | 0 | 2.34E-05 | 0 | 4.81E-06 | 2.20E-06 | 0 | 0 |
| Kctd2 | 4.92E-05 | 0.00011 | 2.86E-05 | 0.0001 | 0.00012 | 6.59E-05 | 5.36E-05 | 5.48E-05 | 5.79E-05 | 8.09E-05 | 4.02E-05 | 4.00E-05 | 0 | 5.73E-05 | 2.19E-05 | 7.16E-05 | 6.66E-05 | 8.65E-05 |
| Phc3 | 7.70E-05 | 0.0002 | 3.31E-05 | 2.65E-05 | 9.42E-05 | 6.37E-05 | 2.50E-05 | 2.72E-06 | 5.79E-05 | 8.65E-05 | 6.18E-05 | 0.00012 | 2.87E-05 | 5.67E-05 | 0.00029 | 3.96E-05 | 0.00016 | 8.78E-05 |
| Ppil3 | 0.00014 | 2.81E-05 | 4.58E-05 | 1.93E-06 | 1.65E-05 | 3.03E-05 | 6.58E-06 | 0 | 5.79E-05 | 3.27E-05 | 2.27E-05 | 7.43E-06 | 3.47E-06 | 0 | 0 | 2.20E-05 | 0 | 3.10E-05 |
| Golga3 | 0.00016 | 0.00011 | 0.0002 | 0.00021 | 0.00012 | 7.25E-05 | 0.00027 | 0.00015 | 5.78E-05 | 7.36E-05 | 0.00013 | 0.00011 | 9.49E-05 | 7.06E-05 | 0.00035 | 1.27E-05 | 0.00019 | 3.62E-05 |
| St5 | 8.16E-05 | 4.30E-05 | 5.28E-05 | 0.00022 | 2.93E-05 | 3.01E-05 | 8.43E-05 | 0 | 5.78E-05 | 0.00014 | 0.00017 | 0 | 0.0003 | 6.82E-05 | 0.00021 | 0.00015 | 0.00015 | 0.00013 |
| Eci1 | 7.97E-05 | 5.79E-05 | 4.00E-05 | 8.85E-05 | 2.82E-05 | 7.16E-06 | 1.25E-05 | 0 | 5.78E-05 | 3.25E-05 | 3.13E-05 | 0 | 2.25E-05 | 5.23E-05 | 1.07E-05 | 0 | 3.38E-05 | 5.04E-05 |
| Relch | 0.00011 | 5.75E-05 | 4.63E-05 | 0 | 4.06E-05 | 0.00012 | 0 | 5.48E-05 | 5.77E-05 | 3.08E-05 | 3.25E-05 | 3.13E-05 | 4.04E-05 | 4.06E-05 | 6.65E-05 | 0 | 7.77E-05 | 4.81E-06 |
| Cct5 | 0.0002 | 4.85E-05 | 0.0002 | 5.89E-05 | 0 | 7.39E-05 | 0 | 2.74E-05 | 5.77E-05 | 2.06E-05 | 4.82E-05 | 6.01E-05 | 0 | 0 | 0 | 9.05E-05 | 0 | 0 |
| Fam122a | 6.55E-05 | 6.44E-05 | 6.32E-05 | 2.65E-05 | 6.43E-05 | 5.83E-05 | 1.98E-05 | 5.60E-05 | 5.77E-05 | 2.64E-05 | 5.41E-05 | 5.50E-05 | 2.00E-05 | 4.93E-05 | 1.90E-05 | 4.00E-05 | 1.35E-05 | 7.15E-05 |
| Rspry1 | 0.0002 | 0.00012 | 0.00011 | 5.18E-05 | 3.43E-05 | 5.27E-05 | 0.00016 | 0.00011 | 5.77E-05 | 6.95E-05 | 0 | 9.39E-05 | 0 | 1.63E-05 | 0.00014 | 4.35E-05 | 0 | 8.09E-05 |
| Mllt3 | 9.40E-06 | 3.16E-05 | 0 | 0 | 8.56E-06 | 2.24E-05 | 0 | 0 | 5.77E-05 | 1.00E-07 | 0 | 5.58E-05 | 0 | 1.96E-05 | 0 | 1.47E-05 | 8.62E-06 | 3.97E-05 |
| Bcl9l | 8.14E-05 | 0.0002 | 0 | 0.00038 | 0.00021 | 0.00023 | 0.00019 | 0 | 5.76E-05 | 0.00014 | 0.00023 | 0.0003 | 0.00029 | 0.00029 | 0.00039 | 8.37E-05 | 0 | 0.00019 |
| AU040320 | 9.09E-05 | 7.56E-05 | 2.78E-05 | 1.93E-05 | 5.81E-05 | 0.00013 | 8.47E-05 | 8.44E-05 | 5.76E-05 | 0.0001 | 7.39E-05 | 0.00011 | 1.18E-05 | 5.38E-05 | 5.22E-05 | 8.82E-06 | 0.00011 | 4.39E-05 |
| Mnt | 4.57E-05 | 0.00012 | 3.01E-05 | 2.36E-05 | 3.60E-05 | 2.41E-05 | 8.14E-06 | 2.45E-05 | 5.76E-05 | 3.29E-05 | 2.21E-05 | 0.00014 | 0 | 3.34E-05 | 0.00014 | 1.67E-05 | 9.90E-05 | 1.57E-05 |
| Tada2b | 6.72E-05 | 0.00011 | 0.00016 | 1.40E-05 | 3.18E-05 | 1.30E-05 | 5.97E-05 | 1.82E-05 | 5.76E-05 | 4.89E-05 | 6.16E-05 | 0 | 6.52E-05 | 3.13E-05 | 4.05E-05 | 6.21E-05 | 0 | 5.08E-05 |
| Abcf2 | 0.00015 | 0.00012 | 0.00023 | 4.94E-05 | 5.08E-05 | 3.73E-06 | 0 | 0 | 5.76E-05 | 1.24E-05 | 5.20E-05 | 0 | 3.58E-05 | 5.60E-05 | 1.15E-05 | 0.00016 | 0 | 0 |
| Mcat | 3.64E-05 | 4.21E-05 | 5.76E-05 | 8.57E-05 | 2.78E-05 | 0 | 7.68E-06 | 1.49E-05 | 5.76E-05 | 0 | 4.82E-05 | 1.15E-06 | 1.00E-05 | 2.69E-05 | 2.66E-06 | 3.51E-05 | 0 | 0 |
| Med11 | 6.71E-05 | 3.97E-05 | 4.22E-05 | 0.00017 | 2.48E-05 | 0 | 6.37E-05 | 4.50E-06 | 5.76E-05 | 1.24E-05 | 5.42E-05 | 0 | 5.46E-05 | 4.82E-05 | 0 | 5.86E-05 | 1.79E-05 | 7.75E-05 |
| Ruvbl2 | 0.00014 | 3.42E-05 | 0.00025 | 8.54E-05 | 0 | 0 | 0 | 0 | 5.76E-05 | 3.40E-05 | 0 | 9.00E-05 | 0 | 0 | 7.11E-05 | 0.00026 | 0 | 0 |

| | | | | | | | | | | | | | | | | | | |
|---|---|---|---|---|---|---|---|---|---|---|---|---|---|---|---|---|---|---|
| Noc4l | 4.82E-05 | 3.44E-05 | 0.00013 | 5.70E-05 | 2.32E-06 | 0 | 0 | 0 | 5.76E-05 | 1.64E-05 | 1.29E-05 | 0 | 7.00E-06 | 2.14E-05 | 6.65E-06 | 0.00012 | 3.70E-05 | 1.54E-05 |
| Mcoln1 | 8.81E-05 | 0.00016 | 0 | 0 | 5.40E-05 | 0.00011 | 8.20E-05 | 1.71E-05 | 5.75E-05 | 6.33E-05 | 4.21E-05 | 8.27E-05 | 5.14E-05 | 9.21E-05 | 6.19E-05 | 6.51E-05 | 2.10E-05 | 3.89E-05 |
| Smg7 | 0.00014 | 0.00013 | 0.00018 | 0 | 4.51E-05 | 9.90E-05 | 0 | 5.58E-05 | 5.75E-05 | 4.80E-05 | 3.66E-05 | 0.00014 | 0 | 7.01E-05 | 0.00013 | 0.00014 | 0.00014 | 0 |
| Atxn2 | 0.00014 | 0.00017 | 0.00012 | 0 | 0.00011 | 9.72E-05 | 9.33E-05 | 0 | 5.75E-05 | 2.17E-05 | 9.68E-05 | 6.64E-05 | 9.93E-05 | 5.49E-05 | 0.00019 | 4.10E-05 | 0.00014 | 0 |
| Cul5 | 0.0002 | 9.75E-05 | 0.0001 | 3.90E-06 | 2.84E-05 | 6.22E-05 | 5.18E-05 | 4.01E-05 | 5.75E-05 | 4.35E-05 | 4.57E-05 | 4.57E-05 | 0 | 1.39E-05 | 8.75E-05 | 4.91E-05 | 5.32E-05 | 0 |
| Ubtd1 | 4.05E-05 | 0.00023 | 0.00021 | 0.0001 | 0.00013 | 6.13E-05 | 9.11E-05 | 0.00025 | 5.75E-05 | 9.24E-05 | 8.38E-05 | 6.75E-05 | 0.00012 | 0.00016 | 0.00012 | 0.00011 | 5.10E-05 | 8.50E-05 |
| Tsfm | 0.00011 | 6.60E-05 | 0.00013 | 0.00012 | 2.21E-06 | 0 | 7.91E-05 | 0 | 5.75E-05 | 1.48E-05 | 3.57E-05 | 1.26E-05 | 2.71E-05 | 3.34E-06 | 0.00013 | 4.41E-05 | 0 | 2.05E-05 |
| Telo2 | 6.00E-06 | 3.14E-05 | 7.59E-05 | 8.47E-05 | 1.56E-05 | 0 | 0 | 4.08E-05 | 5.75E-05 | 4.09E-05 | 1.29E-05 | 2.43E-05 | 0 | 1.64E-05 | 9.73E-05 | 5.83E-06 | 0 | 5.90E-05 |
| Lrrk1 | 3.74E-05 | 0.00012 | 0 | 4.38E-05 | 0.00014 | 0 | 0.00011 | 0 | 5.75E-05 | 0.00014 | 0.00023 | 0 | 0.00013 | 0.00013 | 0.0002 | 5.95E-05 | 0 | 0.00031 |
| Armcx1 | 0.00012 | 0.00014 | 0 | 6.44E-05 | 8.55E-05 | 0.00016 | 0.00013 | 3.57E-05 | 5.74E-05 | 4.14E-05 | 0 | 0.00015 | 9.97E-06 | 0 | 8.57E-05 | 0 | 0 | 2.75E-05 |
| Commd8 | 0.00016 | 0.00018 | 8.61E-05 | 9.71E-05 | 0.00016 | 9.77E-05 | 0.00014 | 5.70E-05 | 5.74E-05 | 0.00011 | 4.32E-05 | 7.24E-05 | 9.12E-05 | 7.78E-05 | 1.51E-05 | 0.00013 | 0 | 9.57E-05 |
| Rnaseh2a | 3.59E-05 | 5.68E-05 | 4.82E-05 | 0.00012 | 0 | 4.88E-05 | 2.65E-05 | 7.70E-05 | 5.74E-05 | 4.90E-05 | 7.84E-05 | 6.45E-05 | 8.77E-05 | 2.74E-05 | 8.75E-05 | 0.0002 | 1.49E-05 | 8.94E-07 |
| Sertad3 | 3.57E-05 | 5.49E-05 | 0 | 9.47E-05 | 5.30E-05 | 3.71E-06 | 3.75E-05 | 1.75E-05 | 5.74E-05 | 9.06E-05 | 3.35E-05 | 0 | 0 | 0 | 7.88E-05 | 0 | 0 | 9.95E-05 |
| Spaca6 | 1.29E-05 | 0.00015 | 1.49E-06 | 0.00016 | 6.53E-05 | 2.30E-06 | 0 | 7.94E-05 | 5.74E-05 | 1.29E-05 | 6.42E-05 | 4.95E-05 | 2.86E-05 | 1.53E-05 | 0.00039 | 0 | 0 | 5.80E-05 |
| Aff4 | 0.00037 | 0.00031 | 0.00014 | 0 | 0.0003 | 0.00031 | 0.00014 | 3.98E-05 | 5.73E-05 | 4.31E-05 | 0.00024 | 0.00057 | 5.54E-05 | 0.00018 | 0.00012 | 3.84E-05 | 0.00035 | 5.96E-05 |
| Atp6v1e1 | 0.00021 | 0.00027 | 0.00015 | 8.51E-05 | 0.00017 | 0.00017 | 9.00E-05 | 9.87E-05 | 5.73E-05 | 0.00015 | 0.00018 | 0.00011 | 7.99E-05 | 0.00016 | 2.96E-05 | 0 | 0.00024 | 2.83E-05 |
| Lamtor5 | 0.00015 | 7.57E-05 | 1.69E-05 | 6.30E-05 | 2.46E-05 | 0.00011 | 4.02E-05 | 3.84E-05 | 5.73E-05 | 3.19E-05 | 6.04E-05 | 3.32E-05 | 0 | 6.44E-05 | 3.82E-05 | 3.17E-05 | 0 | 3.05E-05 |
| Agpat4 | 6.47E-05 | 0.00012 | 0.00011 | 0.00015 | 3.65E-05 | 6.87E-05 | 8.66E-05 | 1.43E-05 | 5.73E-05 | 0.00022 | 0.00012 | 0.00011 | 0 | 3.86E-05 | 7.69E-06 | 4.99E-05 | 3.64E-05 | 2.61E-05 |
| Tdrd7 | 0.0001 | 8.66E-05 | 5.00E-05 | 0 | 0 | 3.63E-05 | 6.88E-05 | 0 | 5.73E-05 | 0.00011 | 1.52E-06 | 6.42E-05 | 6.03E-05 | 0 | 5.90E-05 | 0 | 1.43E-05 | 8.69E-05 |
| Ercc6l2 | 5.97E-05 | 4.39E-05 | 3.61E-05 | 1.41E-05 | 3.39E-05 | 3.57E-05 | 2.37E-05 | 8.56E-06 | 5.73E-05 | 2.15E-05 | 3.23E-05 | 1.81E-05 | 1.10E-05 | 7.82E-06 | 4.72E-06 | 8.33E-05 | 0 | 2.25E-05 |
| Pogz | 5.97E-05 | 0.00012 | 3.52E-05 | 8.14E-05 | 4.10E-05 | 2.03E-05 | 6.69E-06 | 6.75E-05 | 5.73E-05 | 3.21E-05 | 4.29E-05 | 5.06E-05 | 1.34E-05 | 5.25E-05 | 9.09E-05 | 6.58E-05 | 0.00018 | 2.42E-05 |
| Imp4 | 0.00015 | 8.71E-05 | 0.00024 | 0.00014 | 2.37E-05 | 9.74E-06 | 0 | 9.98E-06 | 5.73E-05 | 3.83E-06 | 4.56E-05 | 4.59E-05 | 0 | 2.68E-05 | 3.18E-06 | 2.28E-05 | 0 | 4.67E-06 |
| Yes1 | 0.00012 | 0.0001 | 6.07E-05 | 0 | 2.13E-05 | 0.00013 | 4.30E-05 | 8.41E-06 | 5.72E-05 | 0 | 4.88E-05 | 3.91E-05 | 0 | 7.36E-05 | 0 | 6.66E-05 | 5.49E-05 | 3.58E-05 |
| Fam174a | 3.03E-05 | 9.75E-05 | 1.61E-05 | 8.37E-05 | 0.00011 | 0.00012 | 1.30E-05 | 7.13E-05 | 5.72E-05 | 9.95E-05 | 8.78E-05 | 2.85E-05 | 5.72E-05 | 6.93E-05 | 8.63E-05 | 1.50E-05 | 1.79E-05 | 0.00011 |
| Prelid3b | 0.00019 | 4.54E-05 | 0.00018 | 2.39E-06 | 9.31E-06 | 6.91E-05 | 7.12E-06 | 9.31E-06 | 5.72E-05 | 1.56E-05 | 6.15E-05 | 0 | 1.31E-05 | 0 | 0 | 0.00018 | 0 | 5.97E-06 |
| Unk | 1.72E-05 | 0.0001 | 2.00E-05 | 8.06E-05 | 6.28E-05 | 4.31E-05 | 0 | 7.04E-05 | 5.72E-05 | 3.28E-05 | 0 | 5.18E-05 | 2.10E-05 | 3.99E-05 | 0.00016 | 7.68E-05 | 3.56E-06 | 7.78E-05 |
| Ccdc32 | 4.50E-05 | 4.72E-05 | 3.33E-05 | 3.29E-05 | 4.70E-05 | 4.15E-05 | 9.17E-06 | 0 | 5.72E-05 | 2.79E-05 | 3.02E-05 | 1.08E-05 | 0 | 2.03E-05 | 8.65E-06 | 1.20E-05 | 0 | 2.45E-05 |
| 6-Sep | 3.67E-05 | 2.10E-05 | 2.27E-05 | 9.93E-06 | 0 | 0 | 3.93E-05 | 1.22E-06 | 5.72E-05 | 0 | 0 | 9.03E-06 | 3.36E-05 | 0 | 0 | 5.21E-05 | 9.87E-06 | 2.50E-05 |
| Coro1c | 0.00012 | 0.00011 | 0.00016 | 0 | 0.0001 | 0.0001 | 0 | 0.00011 | 5.71E-05 | 0.00014 | 0.00034 | 8.60E-05 | 0 | 2.14E-05 | 0 | 0.00019 | 0 | 7.58E-05 |
| Rnf216 | 0.00014 | 0.00015 | 9.74E-05 | 4.04E-05 | 8.52E-05 | 8.89E-05 | 5.32E-05 | 9.59E-06 | 5.71E-05 | 4.08E-05 | 7.66E-05 | 8.74E-05 | 8.41E-05 | 7.53E-05 | 0.00021 | 5.36E-05 | 2.29E-07 | 7.39E-06 |
| Numb | 2.82E-05 | 0.00012 | 5.25E-05 | 0 | 7.63E-05 | 7.20E-05 | 0.0001 | 8.33E-05 | 5.71E-05 | 0.00011 | 8.59E-05 | 0.00013 | 0.00011 | 9.07E-05 | 5.00E-05 | 1.07E-05 | 7.08E-05 | 0.0001 |
| Slc36a1 | 2.13E-05 | 0.0001 | 3.19E-05 | 0.00015 | 1.40E-05 | 3.70E-05 | 7.24E-05 | 8.89E-05 | 5.71E-05 | 6.17E-05 | 2.46E-05 | 1.33E-05 | 7.59E-06 | 9.64E-05 | 0.0001 | 0 | 0 | 8.58E-05 |
| Vps51 | 8.93E-05 | 0.00014 | 2.55E-05 | 0.00015 | 4.11E-05 | 2.80E-05 | 3.58E-05 | 1.56E-05 | 5.71E-05 | 2.32E-05 | 2.61E-05 | 5.63E-05 | 0 | 3.78E-05 | 0.00021 | 6.72E-05 | 0 | 5.60E-05 |

| | | | | | | | | | | | | | | | | | | |
|---|---|---|---|---|---|---|---|---|---|---|---|---|---|---|---|---|---|---|
| Umps | 6.28E-05 | 4.56E-05 | 0.00015 | 7.46E-05 | 5.26E-05 | 2.12E-05 | 0 | 0 | 5.71E-05 | 4.65E-05 | 6.46E-05 | 0 | 2.72E-05 | 3.90E-05 | 0 | 0.00029 | 0 | 1.50E-05 |
| Baalc | 0 | 0 | 4.67E-06 | 0 | 0 | 0 | 0 | 0 | 5.71E-05 | 4.70E-06 | 0 | 1.06E-05 | 0 | 7.70E-06 | 0 | 4.14E-06 | 0 | 4.97E-05 |
| Hebp1 | 0 | 0.00011 | 0 | 0 | 0.00016 | 0.00011 | 0 | 3.63E-05 | 5.70E-05 | 0.00015 | 0 | 0.00011 | 7.46E-06 | 0.00013 | 3.65E-05 | 0 | 0 | 0.00013 |
| Agl | 2.95E-05 | 3.76E-05 | 8.21E-06 | 3.17E-05 | 5.22E-05 | 2.96E-05 | 2.19E-05 | 5.85E-05 | 5.70E-05 | 2.29E-05 | 4.63E-05 | 0.00017 | 0 | 1.82E-05 | 4.08E-05 | 9.25E-06 | 0 | 4.36E-05 |
| Atp10a | 3.36E-05 | 0 | 0.00016 | 0 | 0 | 2.42E-05 | 0.00015 | 0 | 5.70E-05 | 0.00024 | 9.32E-05 | 0 | 4.08E-05 | 0 | 0 | 0.00019 | 3.21E-05 | 0.00018 |
| R3hcc1 | 7.52E-06 | 7.37E-05 | 6.84E-05 | 0.00017 | 5.75E-05 | 0 | 1.40E-05 | 4.05E-05 | 5.70E-05 | 2.65E-05 | 8.72E-05 | 7.12E-05 | 0 | 6.38E-05 | 0 | 8.24E-05 | 0 | 3.33E-05 |
| Tlk1 | 0.00011 | 9.22E-05 | 1.32E-05 | 0 | 4.23E-05 | 8.46E-05 | 7.65E-05 | 0 | 5.69E-05 | 1.18E-05 | 4.33E-05 | 4.81E-05 | 3.00E-06 | 9.15E-06 | 4.30E-05 | 3.35E-05 | 6.76E-05 | 2.46E-05 |
| Daglb | 0 | 6.47E-05 | 1.15E-05 | 1.97E-05 | 0 | 6.23E-05 | 0 | 0 | 5.69E-05 | 0.00026 | 0 | 0 | 0 | 5.30E-05 | 1.57E-05 | 2.83E-05 | 0 | 5.48E-05 |
| Mdn1 | 5.44E-05 | 5.03E-05 | 0.00011 | 0 | 1.77E-05 | 8.71E-06 | 0 | 0 | 5.69E-05 | 0 | 2.30E-06 | 0 | 1.31E-05 | 8.03E-06 | 3.19E-05 | 0 | 3.74E-06 | 6.64E-06 |
| Cox16 | 0.0002 | 3.86E-05 | 0.0001 | 0.00015 | 8.14E-05 | 0.00011 | 8.28E-05 | 6.24E-05 | 5.68E-05 | 3.06E-05 | 4.99E-08 | 2.03E-05 | 3.33E-06 | 2.78E-06 | 7.05E-05 | 2.67E-05 | 0 | 6.09E-05 |
| Abcd1 | 7.25E-05 | 0.00013 | 6.31E-07 | 0.00011 | 3.59E-05 | 2.80E-05 | 1.91E-05 | 2.66E-05 | 5.68E-05 | 7.69E-05 | 3.76E-05 | 2.27E-05 | 3.19E-05 | 6.43E-05 | 3.51E-05 | 3.67E-05 | 0.00012 | 6.04E-06 |
| Gbp9 | 0 | 1.38E-05 | 0 | 0 | 0 | 5.97E-07 | 0 | 0 | 5.68E-05 | 6.56E-05 | 0 | 4.66E-07 | 1.05E-05 | 0 | 0 | 0 | 0 | 4.00E-05 |
| Cdk5rap3 | 0.00014 | 9.82E-05 | 5.57E-05 | 0.00047 | 8.33E-05 | 9.57E-05 | 0.00031 | 0.00016 | 5.67E-05 | 0.00012 | 2.92E-05 | 8.71E-05 | 0.00012 | 0.00012 | 0.0004 | 1.65E-05 | 3.78E-05 | 0.00011 |
| Tmem268 | 2.67E-05 | 1.44E-05 | 5.06E-05 | 4.11E-05 | 9.96E-05 | 3.92E-05 | 0 | 5.92E-05 | 5.67E-05 | 0 | 1.91E-05 | 0 | 5.24E-06 | 6.90E-05 | 9.60E-05 | 0 | 0 | 0 |
| Utp15 | 3.69E-05 | 7.91E-05 | 0.00011 | 5.22E-06 | 7.08E-06 | 3.90E-05 | 0 | 4.18E-07 | 5.67E-05 | 1.70E-05 | 4.18E-05 | 4.72E-05 | 1.48E-05 | 5.51E-05 | 1.40E-05 | 7.07E-05 | 4.29E-06 | 7.69E-06 |
| Ncoa1 | 4.66E-05 | 8.84E-05 | 0 | 0 | 4.58E-06 | 3.64E-05 | 1.77E-05 | 0 | 5.67E-05 | 4.15E-05 | 1.50E-05 | 3.68E-05 | 9.87E-06 | 0 | 7.52E-06 | 0 | 8.90E-05 | 3.99E-05 |
| Cnpy3 | 0 | 6.34E-05 | 0.0001 | 0.00041 | 9.98E-05 | 2.51E-05 | 0 | 7.89E-05 | 5.67E-05 | 0.00017 | 0 | 1.72E-05 | 0.00016 | 4.27E-05 | 0.0001 | 0.00014 | 0.00014 | 0.00022 |
| Depdc5 | 4.77E-06 | 7.87E-05 | 0 | 6.31E-05 | 4.54E-05 | 1.85E-05 | 2.94E-05 | 0 | 5.67E-05 | 4.23E-05 | 2.67E-05 | 2.90E-05 | 4.23E-05 | 5.33E-05 | 0.00014 | 0.00013 | 8.20E-05 | 1.34E-05 |
| Meis3 | 3.87E-05 | 0.00014 | 2.31E-05 | 2.14E-05 | 5.31E-05 | 7.46E-05 | 3.91E-05 | 8.95E-05 | 5.66E-05 | 7.54E-05 | 2.96E-05 | 3.10E-05 | 3.10E-05 | 0.00015 | 7.49E-05 | 0 | 2.37E-05 | 6.96E-05 |
| Fam204a | 8.75E-05 | 7.99E-05 | 8.03E-05 | 6.47E-05 | 2.17E-05 | 4.59E-05 | 0 | 1.96E-05 | 5.66E-05 | 4.82E-05 | 3.38E-05 | 3.05E-05 | 0 | 0 | 0.00012 | 0.0001 | 3.65E-05 | 1.55E-06 |
| Strip1 | 8.70E-05 | 7.43E-05 | 7.52E-05 | 1.01E-05 | 3.21E-05 | 2.94E-05 | 0 | 2.55E-05 | 5.66E-05 | 3.46E-05 | 3.28E-05 | 0.00013 | 0 | 2.76E-05 | 4.50E-05 | 2.54E-05 | 0 | 3.85E-05 |
| Asb7 | 7.54E-05 | 4.63E-05 | 1.49E-05 | 3.48E-06 | 4.93E-05 | 2.02E-05 | 1.02E-05 | 1.60E-05 | 5.66E-05 | 1.76E-05 | 4.14E-05 | 4.32E-05 | 0 | 4.63E-05 | 3.10E-05 | 9.09E-06 | 0 | 4.29E-06 |
| Sergef | 2.62E-05 | 2.89E-05 | 4.31E-05 | 3.39E-05 | 1.29E-05 | 1.14E-05 | 3.12E-06 | 1.17E-05 | 5.66E-05 | 3.27E-05 | 3.99E-05 | 3.22E-05 | 5.45E-06 | 2.33E-05 | 3.66E-05 | 0 | 0 | 1.08E-05 |
| Arhgef5 | 0 | 2.46E-05 | 6.44E-06 | 6.58E-06 | 8.92E-05 | 3.00E-05 | 0 | 8.78E-05 | 5.65E-05 | 0 | 0.00011 | 2.03E-05 | 0 | 1.24E-05 | 2.92E-05 | 0 | 1.03E-05 | 8.11E-05 |
| Mphosph8 | 6.63E-05 | 7.72E-05 | 3.51E-05 | 1.96E-05 | 1.74E-05 | 2.02E-05 | 2.08E-05 | 4.96E-06 | 5.65E-05 | 7.71E-06 | 1.74E-05 | 3.35E-06 | 0 | 3.55E-05 | 0.00015 | 8.59E-05 | 0 | 2.24E-05 |
| Lmf1 | 8.17E-05 | 6.32E-05 | 4.79E-05 | 0.00013 | 0.00011 | 0.00011 | 0.00014 | 0.00013 | 5.64E-05 | 6.63E-05 | 6.29E-05 | 0.00024 | 0.00015 | 0.00011 | 0 | 0 | 2.72E-05 | 7.97E-05 |
| Setd1b | 0.00011 | 0.00021 | 4.80E-05 | 0.00022 | 3.76E-05 | 9.08E-05 | 9.42E-05 | 1.15E-05 | 5.64E-05 | 9.06E-05 | 6.39E-05 | 4.31E-05 | 1.04E-05 | 4.75E-05 | 0.00027 | 0.00012 | 7.90E-05 | 3.57E-05 |
| Rab13 | 0.00018 | 0.00011 | 0 | 0.0001 | 5.65E-06 | 9.07E-05 | 0.00018 | 0.00014 | 5.64E-05 | 5.99E-05 | 0 | 9.01E-05 | 0 | 0 | 2.17E-05 | 0.00015 | 0 | 1.52E-06 |
| Ttl | 7.14E-05 | 9.45E-05 | 7.53E-05 | 9.20E-05 | 6.54E-05 | 6.01E-05 | 0.00014 | 4.75E-05 | 5.64E-05 | 1.32E-05 | 0.0001 | 5.98E-06 | 2.14E-05 | 4.40E-05 | 0 | 7.15E-05 | 0 | 5.44E-05 |
| Prkd2 | 9.04E-06 | 6.95E-05 | 1.37E-05 | 2.80E-05 | 5.09E-05 | 1.14E-05 | 0 | 9.38E-06 | 5.64E-05 | 4.30E-05 | 1.89E-05 | 6.13E-06 | 0 | 2.14E-05 | 5.11E-05 | 6.15E-05 | 0.00016 | 7.15E-05 |
| Mterf4 | 3.45E-05 | 3.21E-05 | 3.58E-06 | 1.37E-05 | 1.88E-05 | 1.12E-05 | 1.91E-05 | 1.55E-05 | 5.64E-05 | 0 | 1.13E-05 | 2.71E-05 | 0 | 4.07E-05 | 2.07E-05 | 2.19E-05 | 0 | 6.09E-06 |
| Rnf139 | 0.00014 | 0.00013 | 3.69E-05 | 0 | 7.66E-05 | 0.00011 | 2.35E-05 | 1.83E-05 | 5.63E-05 | 5.58E-05 | 8.89E-05 | 4.00E-05 | 2.04E-05 | 0 | 7.04E-08 | 5.00E-05 | 8.38E-06 | 0.00011 |
| Cse1l | 0.00017 | 4.71E-05 | 0.00017 | 0 | 1.83E-05 | 5.59E-05 | 0 | 0 | 5.63E-05 | 3.25E-05 | 5.28E-05 | 0 | 0 | 0 | 1.68E-05 | 0.00055 | 0 | 2.96E-05 |

| | | | | | | | | | | | | | | | | | | |
|---|---|---|---|---|---|---|---|---|---|---|---|---|---|---|---|---|---|---|
| Dnajc16 | 2.85E-05 | 6.45E-05 | 5.83E-05 | 7.13E-05 | 3.22E-05 | 1.43E-05 | 0 | 3.16E-05 | 5.63E-05 | 2.91E-05 | 7.76E-06 | 4.60E-05 | 2.44E-05 | 6.01E-05 | 0.00011 | 0 | 7.47E-06 | 4.27E-06 |
| Orai3 | 5.80E-05 | 0.00016 | 7.31E-05 | 0.00017 | 7.60E-05 | 7.85E-05 | 8.29E-05 | 0.00012 | 5.62E-05 | 6.85E-05 | 1.05E-05 | 3.08E-05 | 9.93E-05 | 8.03E-05 | 0 | 0 | 0 | 9.96E-05 |
| Zbtb14 | 6.02E-05 | 5.83E-05 | 3.89E-05 | 1.14E-05 | 3.16E-05 | 4.21E-05 | 4.29E-06 | 7.65E-05 | 5.62E-05 | 1.34E-05 | 2.14E-05 | 6.48E-06 | 0 | 2.50E-05 | 2.00E-05 | 7.11E-05 | 0 | 5.31E-05 |
| Riok3 | 0.00026 | 0.00023 | 0.00039 | 2.68E-05 | 9.69E-05 | 0.00022 | 0.00012 | 0 | 5.61E-05 | 9.44E-05 | 0.00011 | 0.00027 | 0.00011 | 6.66E-05 | 0.0002 | 5.13E-05 | 0.00022 | 5.28E-05 |
| Gnpda2 | 0.00012 | 5.99E-05 | 0 | 0 | 7.97E-05 | 0.00016 | 7.14E-05 | 3.86E-05 | 5.61E-05 | 2.98E-05 | 0 | 5.44E-05 | 3.04E-05 | 3.00E-05 | 8.38E-06 | 7.14E-06 | 0 | 1.90E-05 |
| Homer1 | 9.44E-05 | 0 | 0.00018 | 0 | 0 | 0 | 0 | 0.00012 | 5.61E-05 | 0 | 0.00013 | 0.0003 | 0 | 0 | 0 | 5.19E-06 | 0.00139 | 0 |
| Supt5 | 8.17E-05 | 0.00012 | 9.24E-05 | 0.00012 | 4.42E-05 | 0 | 1.92E-05 | 1.26E-05 | 5.61E-05 | 1.29E-05 | 3.61E-05 | 8.72E-05 | 3.73E-05 | 1.01E-05 | 0.00014 | 3.99E-05 | 0.00025 | 3.20E-05 |
| Dph5 | 3.40E-05 | 7.38E-06 | 0.00011 | 2.82E-05 | 0 | 0 | 0 | 4.79E-06 | 5.61E-05 | 1.62E-05 | 0 | 8.35E-06 | 2.37E-05 | 0 | 9.34E-05 | 3.86E-05 | 8.60E-06 | 1.94E-05 |
| Appl1 | 0.00025 | 0.00014 | 0.00014 | 0 | 7.10E-05 | 0.00015 | 0.0001 | 6.30E-05 | 5.60E-05 | 8.20E-05 | 0.00013 | 0.00019 | 0 | 0 | 0.00019 | 0.0001 | 0.00014 | 9.01E-05 |
| Pip4p2 | 0.00016 | 0.00015 | 8.48E-05 | 8.78E-05 | 7.05E-05 | 8.16E-05 | 3.53E-05 | 2.34E-05 | 5.60E-05 | 5.22E-07 | 7.63E-05 | 9.90E-06 | 0 | 2.65E-05 | 0 | 0.00012 | 4.48E-05 | 2.37E-05 |
| Nup133 | 1.41E-05 | 5.14E-05 | 6.13E-05 | 5.56E-05 | 1.29E-05 | 1.13E-05 | 0 | 9.52E-06 | 5.60E-05 | 3.33E-05 | 4.48E-05 | 3.65E-05 | 1.15E-05 | 3.15E-05 | 0 | 0.00027 | 2.24E-06 | 4.35E-05 |
| Arid5b | 0.00018 | 0.00026 | 0 | 0 | 0.00014 | 0.0004 | 0 | 0 | 5.59E-05 | 6.62E-05 | 0.00049 | 0.00018 | 0.00034 | 0.00032 | 0.00017 | 0 | 0.00031 | 0 |
| Zfyve1 | 4.87E-06 | 0.00022 | 0 | 6.20E-05 | 0.00017 | 0.00015 | 0.00013 | 5.30E-05 | 5.59E-05 | 0.0001 | 5.71E-05 | 0.00012 | 0.00011 | 0.00013 | 0.00011 | 4.24E-06 | 0 | 7.89E-05 |
| Psmf1 | 7.13E-05 | 8.43E-05 | 6.56E-05 | 0.00013 | 7.23E-05 | 9.20E-05 | 0 | 1.91E-05 | 5.59E-05 | 4.75E-05 | 4.02E-05 | 6.59E-05 | 5.29E-05 | 3.89E-05 | 0 | 5.53E-05 | 0 | 8.44E-05 |
| Usf3 | 0.0001 | 6.69E-05 | 1.85E-05 | 0 | 5.07E-05 | 9.15E-05 | 3.46E-05 | 6.87E-05 | 5.59E-05 | 1.95E-05 | 2.53E-05 | 7.87E-05 | 2.15E-06 | 1.77E-05 | 0.00019 | 6.56E-05 | 6.40E-05 | 1.80E-05 |
| Ogfrl1 | 1.59E-05 | 1.30E-05 | 5.04E-06 | 1.66E-05 | 4.72E-05 | 4.01E-06 | 0 | 3.73E-06 | 5.59E-05 | 0 | 2.13E-05 | 0 | 3.87E-05 | 3.83E-05 | 2.18E-05 | 3.06E-05 | 1.04E-05 | 4.49E-05 |
| Aldh16a1 | 2.76E-05 | 4.39E-05 | 1.40E-05 | 9.67E-05 | 1.69E-05 | 0 | 0 | 1.94E-05 | 5.59E-05 | 5.55E-06 | 8.77E-06 | 1.23E-05 | 0 | 1.63E-05 | 6.73E-05 | 0.00012 | 0 | 3.72E-05 |
| Ipo8 | 0.00019 | 0.00011 | 5.49E-05 | 5.28E-06 | 2.67E-05 | 9.64E-05 | 7.34E-05 | 5.18E-05 | 5.58E-05 | 4.58E-05 | 3.16E-05 | 5.01E-05 | 0 | 0 | 6.72E-05 | 0.00014 | 1.65E-05 | 8.92E-05 |
| Dicer1 | 9.64E-05 | 0.00013 | 0.00013 | 4.47E-05 | 4.93E-05 | 6.62E-05 | 4.61E-05 | 1.23E-05 | 5.58E-05 | 3.35E-05 | 6.85E-05 | 0 | 0 | 4.05E-05 | 0.00015 | 7.15E-05 | 0 | 4.89E-05 |
| Hs1bp3 | 2.33E-05 | 0.00015 | 0 | 8.34E-05 | 8.30E-05 | 6.40E-05 | 5.18E-05 | 6.40E-05 | 5.58E-05 | 6.29E-05 | 0 | 2.37E-05 | 0 | 0.00012 | 2.75E-05 | 0 | 0 | 0.0001 |
| Tmem230 | 0.00016 | 0.00011 | 8.53E-05 | 0.00011 | 7.31E-05 | 5.41E-05 | 5.82E-05 | 6.79E-05 | 5.58E-05 | 2.11E-05 | 3.28E-05 | 0.00017 | 8.48E-05 | 2.46E-05 | 0 | 3.60E-05 | 0.00021 | 1.30E-05 |
| Ndufaf2 | 6.54E-05 | 0.00011 | 0.00013 | 0.00028 | 5.38E-05 | 4.07E-05 | 0 | 5.87E-05 | 5.58E-05 | 5.98E-05 | 2.58E-05 | 0 | 1.38E-05 | 8.17E-05 | 4.78E-05 | 0.0003 | 4.87E-05 | 6.98E-05 |
| Nae1 | 7.90E-05 | 9.64E-05 | 5.87E-05 | 0 | 3.55E-05 | 2.86E-05 | 0 | 0 | 5.58E-05 | 5.66E-05 | 8.57E-05 | 3.27E-05 | 0 | 9.79E-06 | 0.00016 | 0.00012 | 0 | 5.62E-05 |
| Mapkapk3 | 0 | 1.30E-05 | 8.03E-05 | 1.15E-05 | 0 | 2.50E-05 | 0 | 0 | 5.58E-05 | 1.89E-06 | 0 | 0 | 0.0001 | 4.44E-05 | 1.57E-05 | 1.46E-06 | 0 | 7.21E-05 |
| Smg8 | 1.49E-05 | 6.37E-05 | 4.73E-05 | 4.76E-05 | 4.04E-05 | 7.90E-06 | 0 | 2.29E-06 | 5.58E-05 | 1.86E-05 | 0 | 4.71E-05 | 1.76E-05 | 1.71E-05 | 5.07E-05 | 4.15E-05 | 0 | 1.48E-05 |
| Pik3r4 | 3.96E-05 | 6.82E-05 | 0 | 8.04E-05 | 5.53E-05 | 4.90E-05 | 2.19E-05 | 4.59E-05 | 5.57E-05 | 5.93E-05 | 2.47E-05 | 3.39E-05 | 4.34E-05 | 8.40E-05 | 4.38E-05 | 0 | 0 | 4.40E-05 |
| Twf2 | 7.19E-05 | 2.65E-05 | 3.59E-05 | 0.00012 | 0 | 1.37E-05 | 0 | 4.16E-05 | 5.57E-05 | 3.67E-05 | 8.98E-06 | 2.44E-05 | 0 | 2.42E-05 | 0 | 1.51E-05 | 2.23E-05 | 3.94E-05 |
| Pfdn4 | 0.00012 | 4.78E-05 | 0.00011 | 4.96E-05 | 1.16E-05 | 1.26E-05 | 0 | 1.85E-05 | 5.57E-05 | 0 | 6.22E-05 | 4.59E-05 | 0 | 3.30E-05 | 6.37E-05 | 8.61E-05 | 1.63E-05 | 0 |
| Creld2 | 0.0002 | 5.85E-05 | 0.00034 | 0.00027 | 5.59E-05 | 0 | 9.44E-05 | 6.97E-05 | 5.57E-05 | 0.00013 | 9.14E-05 | 0.0001 | 0.00014 | 2.84E-05 | 0.00018 | 0.00015 | 0 | 0.00015 |
| Zdhhc12 | 1.96E-05 | 2.15E-05 | 3.62E-05 | 9.60E-05 | 2.49E-06 | 0 | 1.24E-05 | 2.81E-05 | 5.57E-05 | 1.23E-05 | 0.00011 | 7.10E-06 | 2.56E-05 | 5.61E-05 | 2.19E-05 | 1.83E-05 | 0 | 0 |
| Rnf152 | 0 | 8.37E-06 | 0 | 7.82E-06 | 4.07E-05 | 0 | 1.34E-05 | 2.26E-05 | 5.57E-05 | 0 | 2.22E-06 | 9.66E-05 | 0 | 0 | 4.03E-06 | 0 | 0 | 1.65E-05 |
| Klhdc4 | 9.59E-05 | 7.30E-05 | 0.00016 | 0.00013 | 0 | 0 | 0 | 0 | 5.57E-05 | 0 | 5.09E-05 | 0 | 0 | 5.25E-06 | 8.25E-05 | 0.00012 | 0 | 0 |
| Syap1 | 0.00017 | 0.00015 | 0.00011 | 2.45E-05 | 0.00011 | 0.00019 | 0.00021 | 8.17E-05 | 5.56E-05 | 0.00013 | 3.75E-05 | 0.00013 | 7.49E-05 | 9.43E-05 | 0 | 9.39E-05 | 0.00015 | 0.00014 |

| | | | | | | | | | | | | | | | | | | |
|---|---|---|---|---|---|---|---|---|---|---|---|---|---|---|---|---|---|---|
| Mpc1 | 0.00029 | 5.89E-05 | 8.76E-05 | 5.04E-05 | 0.00011 | 0.00011 | 7.27E-05 | 2.93E-05 | 5.56E-05 | 3.50E-05 | 0.00012 | 5.79E-05 | 9.13E-06 | 8.59E-05 | 0 | 8.08E-05 | 8.55E-05 | 9.58E-05 |
| Igsf8 | 0 | 0.00011 | 0.00018 | 0.00041 | 2.10E-05 | 5.92E-05 | 2.40E-05 | 0.00019 | 5.56E-05 | 0.00034 | 0.00025 | 0.00032 | 0.00027 | 0.00013 | 0.00025 | 0.00027 | 0.00014 | 0.00027 |
| Rabepk | 6.99E-05 | 9.94E-06 | 8.76E-05 | 4.00E-05 | 9.04E-06 | 2.90E-05 | 0 | 5.54E-06 | 5.56E-05 | 2.33E-05 | 8.60E-06 | 4.79E-06 | 3.46E-05 | 3.07E-05 | 0 | 4.52E-05 | 0 | 4.18E-05 |
| Vps37b | 1.39E-05 | 7.87E-05 | 7.81E-05 | 5.14E-05 | 1.90E-05 | 1.24E-05 | 0 | 0.00014 | 5.56E-05 | 7.77E-07 | 5.33E-05 | 2.22E-05 | 0 | 5.07E-05 | 6.35E-05 | 6.46E-05 | 8.63E-05 | 1.35E-05 |
| Katnal1 | 1.75E-05 | 3.01E-05 | 5.54E-05 | 1.07E-05 | 2.96E-05 | 6.86E-06 | 2.58E-05 | 1.30E-05 | 5.56E-05 | 3.76E-05 | 5.90E-05 | 3.20E-05 | 2.18E-05 | 1.79E-05 | 2.61E-05 | 1.95E-05 | 0 | 1.64E-05 |
| Prpsap2 | 5.92E-05 | 2.58E-05 | 2.63E-05 | 9.16E-06 | 0 | 0 | 0 | 2.62E-05 | 5.56E-05 | 8.60E-06 | 0 | 2.01E-05 | 0 | 0 | 6.10E-06 | 2.87E-05 | 9.96E-05 | 7.75E-06 |
| Rtraf | 0.00036 | 5.24E-05 | 9.31E-05 | 1.26E-05 | 4.75E-05 | 0.0002 | 0.00022 | 8.72E-05 | 5.55E-05 | 0 | 0 | 8.18E-05 | 0 | 3.44E-05 | 3.62E-05 | 5.93E-05 | 4.88E-05 | 1.27E-05 |
| Ranbp2 | 0.00035 | 0.00023 | 0.00022 | 0 | 5.27E-05 | 0.00018 | 8.95E-05 | 1.62E-05 | 5.55E-05 | 8.99E-06 | 8.66E-05 | 0.0002 | 0 | 6.32E-05 | 0.00024 | 0.0002 | 0.00032 | 0 |
| Abhd8 | 9.30E-05 | 6.80E-05 | 7.94E-05 | 0.00013 | 0.00023 | 0.00011 | 0.00014 | 0.00017 | 5.55E-05 | 2.11E-05 | 5.90E-05 | 0 | 0 | 0.00018 | 0 | 9.81E-05 | 0 | 7.98E-05 |
| Pggt1b | 8.82E-05 | 0.00011 | 9.50E-05 | 0.0001 | 8.41E-05 | 9.95E-05 | 6.51E-05 | 8.62E-05 | 5.55E-05 | 4.78E-05 | 7.86E-05 | 1.31E-05 | 2.62E-05 | 4.20E-05 | 4.62E-05 | 2.38E-05 | 1.19E-06 | 7.25E-05 |
| Hacd1 | 0.00019 | 3.12E-05 | 0.00011 | 7.46E-06 | 6.17E-05 | 8.29E-05 | 6.65E-05 | 0 | 5.55E-05 | 0.0001 | 0.00025 | 0.00019 | 3.88E-05 | 0.00014 | 0 | 6.90E-05 | 0 | 0 |
| Ankib1 | 0.00017 | 7.12E-05 | 0.0001 | 0 | 3.23E-05 | 6.47E-05 | 1.55E-05 | 1.21E-05 | 5.55E-05 | 5.53E-05 | 8.06E-05 | 1.99E-05 | 2.65E-05 | 4.87E-06 | 0 | 5.73E-05 | 7.04E-05 | 2.01E-05 |
| Ripk3 | 0.0002 | 0 | 0.00033 | 0.00019 | 0 | 3.81E-05 | 8.62E-05 | 0.00013 | 5.55E-05 | 1.97E-05 | 0.00033 | 0.00015 | 0 | 0.00012 | 0.00011 | 0.00034 | 2.09E-05 | 0 |
| Capn15 | 3.19E-05 | 9.74E-05 | 8.66E-05 | 4.90E-05 | 4.12E-05 | 2.03E-05 | 6.07E-05 | 1.62E-05 | 5.55E-05 | 2.54E-05 | 4.27E-05 | 0 | 1.38E-05 | 3.43E-05 | 0.00021 | 4.61E-05 | 0 | 2.89E-05 |
| Nemf | 0.00011 | 0.00012 | 0.0001 | 2.09E-05 | 4.25E-05 | 1.75E-05 | 5.23E-06 | 0 | 5.55E-05 | 2.79E-05 | 2.43E-05 | 4.32E-05 | 1.31E-06 | 2.20E-05 | 0.00043 | 4.27E-05 | 7.60E-06 | 7.34E-06 |
| Prtn3 | 0 | 9.54E-06 | 0 | 5.32E-05 | 2.15E-05 | 0 | 1.45E-05 | 4.02E-05 | 5.55E-05 | 8.36E-06 | 0 | 8.11E-05 | 6.24E-05 | 0 | 0 | 0 | 0.00012 | 0 |
| Uri1 | 0.00013 | 0.00014 | 8.14E-05 | 0.00013 | 4.80E-05 | 9.48E-05 | 6.18E-05 | 4.21E-05 | 5.54E-05 | 9.41E-06 | 4.22E-05 | 3.27E-05 | 3.80E-05 | 6.27E-05 | 5.53E-05 | 2.57E-05 | 0 | 9.62E-05 |
| Fam171a2 | 0 | 5.99E-05 | 0 | 0.00014 | 2.77E-05 | 4.44E-05 | 6.24E-05 | 1.40E-05 | 5.54E-05 | 0.00017 | 6.36E-05 | 0 | 1.03E-05 | 5.60E-05 | 8.26E-06 | 1.20E-05 | 7.62E-05 | 0.00014 |
| Nck1 | 0.0001 | 9.84E-05 | 7.31E-05 | 1.48E-05 | 4.84E-05 | 4.04E-05 | 5.98E-05 | 4.00E-05 | 5.54E-05 | 0.00011 | 4.28E-05 | 2.76E-05 | 9.39E-05 | 0.0001 | 0.00012 | 1.18E-05 | 0 | 0.00016 |
| Crtc1 | 1.26E-05 | 4.23E-05 | 2.65E-05 | 6.03E-05 | 4.12E-05 | 2.91E-05 | 3.59E-05 | 5.33E-05 | 5.54E-05 | 3.15E-05 | 9.84E-06 | 1.63E-05 | 2.53E-05 | 8.55E-06 | 8.61E-06 | 0 | 1.42E-05 | 6.05E-05 |
| LTO1 | 3.33E-05 | 6.02E-05 | 4.48E-05 | 3.80E-05 | 5.92E-05 | 2.63E-05 | 6.64E-06 | 1.46E-05 | 5.54E-05 | 6.76E-05 | 3.06E-05 | 4.62E-05 | 1.72E-05 | 3.93E-05 | 3.95E-05 | 5.92E-06 | 0 | 3.98E-05 |
| Ndufaf1 | 7.97E-05 | 1.87E-05 | 3.57E-05 | 2.34E-05 | 3.46E-05 | 2.07E-05 | 5.70E-06 | 4.56E-06 | 5.54E-05 | 2.74E-05 | 1.62E-05 | 4.70E-06 | 0 | 9.61E-06 | 4.06E-05 | 1.35E-05 | 0 | 1.93E-05 |
| Smpd2 | 1.26E-05 | 6.47E-05 | 8.09E-06 | 6.24E-05 | 5.96E-05 | 1.27E-05 | 2.50E-05 | 3.67E-05 | 5.54E-05 | 7.70E-05 | 5.59E-05 | 2.44E-05 | 3.08E-05 | 4.32E-05 | 5.33E-05 | 0 | 5.02E-05 | 8.05E-05 |
| Ubap1 | 6.89E-05 | 0.00013 | 9.86E-05 | 5.56E-06 | 4.59E-05 | 4.44E-06 | 2.14E-05 | 2.26E-05 | 5.54E-05 | 4.80E-05 | 1.77E-05 | 0.00018 | 0 | 1.03E-05 | 6.89E-05 | 4.34E-05 | 0.00014 | 2.50E-05 |
| Spsb2 | 4.03E-05 | 0.0001 | 7.40E-06 | 0.00015 | 4.47E-05 | 0 | 2.48E-05 | 9.20E-06 | 5.54E-05 | 5.38E-05 | 3.87E-05 | 0 | 0 | 2.77E-05 | 4.60E-05 | 2.87E-05 | 0 | 0.00015 |
| Ivns1abp | 0.00029 | 0.00014 | 0.00035 | 0 | 7.15E-05 | 0.00025 | 0.00011 | 3.01E-05 | 5.53E-05 | 2.18E-05 | 0 | 5.26E-05 | 4.78E-05 | 0.00016 | 0.00031 | 0.00013 | 0.00035 | 0 |
| D17H6S53E | 2.53E-05 | 9.52E-05 | 7.75E-05 | 0.00014 | 3.51E-05 | 4.72E-05 | 6.47E-05 | 9.41E-05 | 5.53E-05 | 6.30E-05 | 6.84E-05 | 1.49E-05 | 7.63E-06 | 9.17E-05 | 5.55E-05 | 2.28E-05 | 0 | 4.55E-05 |
| Slc45a4 | 3.02E-05 | 4.18E-05 | 8.39E-05 | 0.0001 | 3.39E-05 | 6.37E-07 | 8.91E-05 | 5.22E-05 | 5.53E-05 | 5.50E-05 | 7.45E-05 | 8.30E-06 | 3.58E-05 | 6.48E-05 | 4.53E-05 | 0 | 0.0001 | 3.87E-05 |
| Cebpz | 0.0001 | 6.31E-05 | 8.27E-05 | 0 | 6.42E-06 | 5.50E-05 | 7.26E-06 | 0 | 5.52E-05 | 3.59E-06 | 0 | 2.98E-05 | 5.97E-06 | 0 | 3.79E-05 | 3.11E-05 | 0 | 2.12E-05 |
| Sertad2 | 7.77E-05 | 0.00013 | 8.30E-05 | 0 | 5.05E-05 | 5.07E-05 | 0 | 2.75E-05 | 5.52E-05 | 0 | 0.00014 | 0.00017 | 0 | 5.08E-05 | 1.81E-05 | 4.30E-05 | 0.0002 | 3.79E-05 |
| Higd1a | 0.00039 | 0 | 0.00023 | 0.00014 | 0.00012 | 1.98E-05 | 0.00034 | 4.05E-05 | 5.52E-05 | 0.00017 | 0.00018 | 0.00133 | 0.00012 | 0.00011 | 0 | 0.00019 | 6.50E-05 | 0 |
| Brpf1 | 3.57E-05 | 0.0001 | 6.69E-05 | 3.64E-05 | 2.97E-05 | 4.20E-06 | 0 | 2.16E-05 | 5.52E-05 | 9.05E-06 | 6.37E-05 | 1.21E-05 | 1.34E-05 | 5.21E-05 | 0.00017 | 7.35E-05 | 5.27E-05 | 7.65E-06 |
| Ppp4r1 | 9.64E-05 | 0.00012 | 5.96E-05 | 5.54E-06 | 7.07E-05 | 9.52E-07 | 2.28E-05 | 0 | 5.52E-05 | 9.02E-06 | 7.51E-05 | 8.98E-05 | 0 | 1.16E-05 | 7.05E-05 | 8.60E-05 | 4.28E-05 | 1.98E-05 |

| | | | | | | | | | | | | | | | | | | |
|---|---|---|---|---|---|---|---|---|---|---|---|---|---|---|---|---|---|---|
| Zfp593 | 5.09E-05 | 3.47E-05 | 0.00014 | 5.51E-05 | 2.95E-05 | 0 | 0 | 0 | 5.52E-05 | 1.09E-05 | 5.06E-05 | 1.32E-05 | 1.12E-05 | 3.78E-05 | 0 | 0 | 2.83E-05 | 7.22E-06 |
| Mknk2 | 5.22E-05 | 0.00013 | 3.60E-05 | 0.0002 | 3.07E-05 | 0 | 6.14E-05 | 5.63E-05 | 5.52E-05 | 9.42E-05 | 9.12E-05 | 0.00016 | 0.00014 | 0.00011 | 9.14E-05 | 0 | 0.00162 | 5.98E-05 |
| Zcchc8 | 9.28E-05 | 8.26E-05 | 6.80E-05 | 6.11E-07 | 3.49E-05 | 7.10E-05 | 0 | 0 | 5.51E-05 | 2.83E-05 | 2.81E-05 | 9.46E-07 | 0 | 1.91E-05 | 0.00017 | 0.00022 | 9.37E-05 | 1.24E-05 |
| Mmp27 | 2.70E-07 | 4.36E-06 | 0 | 0 | 1.26E-05 | 1.29E-05 | 5.18E-05 | 0 | 5.51E-05 | 9.52E-05 | 0 | 0 | 8.14E-05 | 5.81E-05 | 1.97E-05 | 2.58E-05 | 0 | 0 |
| Dhx33 | 8.58E-05 | 9.36E-05 | 0.00018 | 0.00012 | 5.74E-05 | 0 | 2.71E-05 | 3.30E-05 | 5.51E-05 | 8.10E-06 | 3.70E-05 | 0 | 1.63E-05 | 5.80E-06 | 0.00013 | 2.53E-05 | 7.81E-05 | 5.16E-05 |
| Laptm4b | 0.00019 | 0.00015 | 0 | 8.79E-05 | 0.00029 | 0.00029 | 0.00019 | 0.0002 | 5.50E-05 | 0 | 0.00015 | 0 | 0 | 0.00025 | 0 | 1.87E-05 | 0 | 0.00015 |
| Rpp14 | 0.00012 | 5.42E-05 | 9.42E-05 | 5.82E-05 | 4.16E-06 | 2.62E-05 | 0 | 0 | 5.50E-05 | 7.73E-06 | 3.31E-05 | 0 | 0 | 9.06E-06 | 2.98E-05 | 5.54E-05 | 1.42E-05 | 4.33E-06 |
| Pno1 | 0.00019 | 3.09E-05 | 0.00024 | 1.55E-05 | 0 | 2.12E-05 | 0 | 0 | 5.50E-05 | 0 | 2.72E-05 | 0 | 4.99E-06 | 0 | 0 | 6.14E-05 | 2.76E-05 | 0 |
| Gpd1l | 3.83E-05 | 3.31E-05 | 2.92E-05 | 3.87E-05 | 2.33E-05 | 5.97E-06 | 2.98E-06 | 3.22E-06 | 5.50E-05 | 6.20E-05 | 0 | 0 | 0 | 0 | 6.99E-06 | 2.06E-05 | 0 | 7.46E-05 |
| Erlin1 | 5.43E-05 | 1.68E-05 | 4.70E-05 | 5.42E-05 | 1.23E-06 | 0 | 0 | 0 | 5.50E-05 | 9.56E-05 | 1.41E-05 | 2.32E-05 | 9.86E-05 | 1.75E-05 | 0 | 7.98E-05 | 1.90E-06 | 4.95E-05 |
| Ccz1 | 0.0002 | 7.20E-05 | 6.10E-05 | 2.31E-05 | 1.18E-05 | 8.73E-05 | 5.08E-05 | 0 | 5.49E-05 | 1.36E-05 | 1.37E-05 | 3.95E-05 | 2.12E-05 | 3.19E-05 | 0.00012 | 5.09E-05 | 5.30E-06 | 0 |
| Ids | 0.00015 | 0.00012 | 0 | 3.22E-05 | 9.89E-05 | 7.67E-05 | 2.96E-05 | 7.38E-05 | 5.49E-05 | 0.00015 | 5.21E-05 | 0.00012 | 0 | 0 | 0 | 0 | 9.16E-06 | 0.00022 |
| Naa30 | 7.66E-05 | 0.00013 | 0.0001 | 7.50E-05 | 4.55E-05 | 6.97E-05 | 1.96E-05 | 4.30E-05 | 5.49E-05 | 2.26E-05 | 5.28E-05 | 2.91E-05 | 9.38E-06 | 7.25E-05 | 6.40E-05 | 7.64E-05 | 0.00016 | 2.72E-05 |
| Rbm15b | 4.78E-05 | 0.00012 | 0.00011 | 0.00012 | 6.26E-05 | 2.70E-05 | 0 | 5.07E-05 | 5.49E-05 | 1.54E-05 | 0.00011 | 2.75E-05 | 2.16E-05 | 2.98E-05 | 9.83E-05 | 7.04E-05 | 0.00018 | 4.20E-05 |
| Chfr | 4.78E-05 | 0.0001 | 3.96E-05 | 0.00011 | 9.11E-05 | 1.62E-05 | 0 | 3.12E-05 | 5.49E-05 | 8.60E-05 | 2.12E-05 | 0 | 0 | 6.62E-05 | 0.00044 | 6.03E-06 | 0 | 9.31E-05 |
| Smim19 | 7.04E-05 | 7.77E-05 | 0 | 3.16E-05 | 8.32E-05 | 7.92E-05 | 8.10E-05 | 5.13E-05 | 5.48E-05 | 7.86E-05 | 3.46E-05 | 8.78E-05 | 2.37E-05 | 9.48E-05 | 0 | 5.81E-05 | 9.09E-06 | 0.0001 |
| Dr1 | 0.0002 | 0.00014 | 0.00016 | 1.49E-05 | 5.57E-05 | 7.19E-05 | 5.62E-05 | 2.58E-05 | 5.48E-05 | 3.84E-05 | 5.68E-05 | 5.26E-05 | 1.78E-05 | 3.72E-05 | 2.04E-07 | 7.53E-05 | 2.09E-05 | 1.35E-05 |
| Gkap1 | 9.72E-05 | 0.00012 | 3.24E-05 | 0 | 5.74E-05 | 6.47E-05 | 0 | 0 | 5.48E-05 | 1.80E-05 | 2.56E-05 | 1.06E-05 | 5.31E-05 | 0 | 0.00015 | 0.00012 | 4.23E-05 | 5.43E-05 |
| Cdk2ap1 | 0.00018 | 8.38E-05 | 0.00027 | 0.00032 | 1.93E-05 | 6.25E-05 | 0 | 3.31E-05 | 5.48E-05 | 4.85E-05 | 0.0001 | 0 | 0 | 9.15E-05 | 0 | 0.00028 | 0 | 0.00013 |
| Cbx1 | 0.00011 | 7.26E-05 | 6.25E-05 | 2.00E-05 | 4.52E-05 | 1.68E-05 | 2.25E-05 | 4.24E-05 | 5.48E-05 | 6.01E-06 | 1.68E-05 | 1.85E-05 | 0 | 3.51E-05 | 0.00027 | 0.00022 | 0 | 6.27E-06 |
| Arhgef10l | 0 | 0.00013 | 0 | 1.84E-05 | 6.75E-05 | 0 | 0 | 0 | 5.48E-05 | 0.00018 | 0.0001 | 6.02E-05 | 0.00013 | 1.63E-05 | 2.84E-05 | 0 | 3.55E-05 | 2.04E-05 |
| Mepce | 1.03E-05 | 0.00013 | 3.30E-05 | 9.85E-05 | 3.33E-05 | 0 | 0 | 2.73E-05 | 5.48E-05 | 3.16E-05 | 3.22E-05 | 0.00013 | 2.39E-05 | 6.57E-05 | 6.76E-05 | 0.00015 | 0.00036 | 5.01E-05 |
| Dzip3 | 6.47E-05 | 1.93E-05 | 2.27E-05 | 0 | 3.66E-05 | 0 | 1.18E-05 | 2.91E-05 | 5.48E-05 | 9.98E-06 | 3.32E-05 | 2.03E-05 | 1.59E-05 | 3.25E-05 | 3.15E-05 | 8.70E-05 | 0 | 2.63E-05 |
| Ginm1 | 0.00022 | 0.00033 | 5.48E-05 | 6.86E-06 | 0.00031 | 0.00039 | 9.25E-05 | 6.80E-05 | 5.47E-05 | 0.00016 | 0.00017 | 0.00022 | 0.00029 | 0.00018 | 0 | 0 | 0 | 0.00016 |
| Snrpg | 0.00031 | 4.55E-05 | 0.00011 | 0.00013 | 3.14E-05 | 0.00014 | 5.05E-05 | 0 | 5.47E-05 | 3.04E-06 | 0 | 2.41E-05 | 1.02E-05 | 6.77E-05 | 0 | 0.00028 | 0.00017 | 0 |
| Prdm2 | 0.00019 | 0.00017 | 4.43E-05 | 0 | 9.64E-05 | 0.00014 | 4.78E-05 | 2.01E-05 | 5.47E-05 | 4.91E-05 | 3.05E-05 | 0.00013 | 0 | 3.60E-05 | 9.61E-06 | 4.91E-05 | 0.0003 | 4.85E-05 |
| Rfc1 | 0.00011 | 5.16E-05 | 8.27E-05 | 2.65E-05 | 3.57E-05 | 6.32E-05 | 6.45E-05 | 2.27E-05 | 5.47E-05 | 2.80E-05 | 1.66E-05 | 2.96E-05 | 3.81E-05 | 1.40E-05 | 1.24E-05 | 0.00026 | 6.15E-05 | 5.79E-05 |
| Stx2 | 0.00012 | 8.39E-05 | 6.68E-05 | 9.09E-05 | 3.66E-05 | 6.22E-05 | 0.00018 | 7.45E-05 | 5.47E-05 | 0.0003 | 9.30E-05 | 0.00013 | 0 | 5.84E-06 | 0.00017 | 0.00012 | 0 | 0.00028 |
| Pcna | 0.00013 | 2.14E-05 | 0.00027 | 0.00012 | 1.77E-05 | 4.93E-05 | 0 | 5.07E-05 | 5.47E-05 | 3.25E-05 | 2.67E-05 | 8.90E-05 | 3.06E-05 | 6.52E-05 | 0 | 0.00098 | 0.00095 | 3.29E-05 |
| DEPRECATED | 6.53E-05 | 3.32E-05 | 8.21E-05 | 7.68E-06 | 2.31E-05 | 1.15E-05 | 4.44E-05 | 0 | 5.47E-05 | 5.23E-05 | 7.72E-05 | 3.26E-05 | 0 | 0 | 0 | 4.82E-05 | 4.51E-06 | 4.52E-05 |
| Rnasel | 4.11E-06 | 3.75E-05 | 0 | 0 | 0 | 4.99E-06 | 0 | 8.35E-06 | 5.47E-05 | 6.06E-05 | 0 | 0 | 0 | 0 | 1.99E-05 | 0 | 0 | 5.42E-05 |
| Ammecr1 | 0.00013 | 5.13E-05 | 0.00012 | 8.56E-05 | 5.53E-06 | 0 | 0.00016 | 0.00025 | 5.47E-05 | 6.90E-05 | 9.71E-05 | 0.00019 | 0 | 7.59E-05 | 9.95E-05 | 0.00014 | 0 | 0.00022 |
| Urm1 | 0.0001 | 0.00013 | 0.00019 | 0.00026 | 8.23E-05 | 5.42E-05 | 0.00013 | 1.70E-05 | 5.46E-05 | 5.37E-06 | 3.06E-05 | 5.55E-05 | 5.14E-05 | 6.58E-05 | 8.03E-05 | 7.38E-05 | 0 | 2.16E-05 |

| | | | | | | | | | | | | | | | | | | |
|---|---|---|---|---|---|---|---|---|---|---|---|---|---|---|---|---|---|---|
| Dhx8 | 9.61E-05 | 8.77E-05 | 0.00011 | 0.00014 | 4.62E-05 | 3.80E-05 | 9.09E-05 | 2.57E-06 | 5.46E-05 | 2.43E-05 | 5.28E-05 | 2.65E-05 | 5.08E-05 | 7.16E-05 | 0.00014 | 2.38E-05 | 0 | 1.66E-06 |
| Exosc5 | 4.54E-05 | 6.72E-05 | 0.00012 | 0.00025 | 2.61E-05 | 3.75E-05 | 0 | 5.74E-05 | 5.46E-05 | 5.72E-05 | 1.52E-05 | 1.15E-05 | 1.77E-05 | 3.52E-05 | 6.40E-05 | 0.00011 | 8.37E-06 | 2.94E-05 |
| Slc19a1 | 8.09E-06 | 5.60E-06 | 8.06E-05 | 4.91E-05 | 2.69E-05 | 1.31E-05 | 0 | 0 | 5.46E-05 | 0 | 2.79E-05 | 8.60E-05 | 4.49E-05 | 4.41E-05 | 0 | 2.27E-05 | 0 | 7.24E-05 |
| Mpst | 6.23E-05 | 9.98E-05 | 0 | 0.0001 | 9.72E-06 | 5.80E-05 | 5.37E-05 | 2.03E-05 | 5.45E-05 | 9.54E-05 | 0 | 0.00011 | 0 | 0 | 0 | 6.87E-05 | 3.58E-05 | 9.88E-05 |
| Tubg1 | 9.34E-05 | 4.38E-06 | 0.00016 | 1.72E-05 | 3.85E-05 | 4.13E-05 | 7.05E-06 | 8.38E-05 | 5.45E-05 | 3.39E-05 | 9.57E-05 | 0 | 7.92E-06 | 0 | 0 | 0.00047 | 0 | 2.77E-05 |
| Smim11 | 0.00013 | 7.72E-05 | 2.33E-05 | 0.00018 | 8.96E-05 | 3.77E-05 | 4.90E-05 | 1.99E-05 | 5.45E-05 | 7.54E-05 | 1.72E-05 | 1.43E-05 | 5.41E-05 | 4.58E-06 | 0 | 5.99E-05 | 0 | 5.93E-05 |
| D11Wsu47e | 9.97E-06 | 4.58E-05 | 4.59E-05 | 6.48E-05 | 5.30E-05 | 3.08E-05 | 1.64E-05 | 6.89E-05 | 5.45E-05 | 2.95E-05 | 2.91E-05 | 0 | 4.30E-05 | 5.26E-05 | 6.85E-05 | 7.21E-06 | 0 | 1.76E-05 |
| Gtf3c3 | 8.94E-05 | 0.0001 | 5.54E-05 | 8.55E-05 | 2.17E-05 | 2.47E-05 | 1.27E-05 | 3.14E-05 | 5.45E-05 | 2.55E-05 | 5.76E-05 | 4.32E-05 | 1.00E-05 | 3.69E-05 | 1.12E-05 | 5.28E-05 | 3.38E-05 | 3.73E-05 |
| Ldb1 | 0.00029 | 0.00045 | 8.45E-05 | 0.00015 | 9.27E-05 | 0.00021 | 0 | 0 | 5.44E-05 | 8.22E-05 | 9.19E-05 | 0.00024 | 9.36E-05 | 0.00022 | 0.00056 | 7.01E-05 | 0.00013 | 3.58E-05 |
| Osbpl8 | 0.00011 | 0.00015 | 0.00012 | 2.34E-05 | 9.21E-05 | 0.00011 | 0.00013 | 0 | 5.44E-05 | 7.38E-05 | 3.40E-05 | 0.00012 | 0 | 0.00013 | 0.00011 | 7.65E-05 | 0.00016 | 5.48E-05 |
| Katnb1 | 3.67E-05 | 5.41E-05 | 2.72E-05 | 6.97E-05 | 4.03E-05 | 1.21E-05 | 1.96E-05 | 2.52E-05 | 5.44E-05 | 2.76E-05 | 2.89E-05 | 1.19E-05 | 5.06E-06 | 3.66E-05 | 2.07E-05 | 0.00011 | 0 | 0 |
| Prmt7 | 7.98E-05 | 3.20E-05 | 0.00013 | 9.46E-05 | 0 | 0 | 0 | 0 | 5.44E-05 | 3.89E-05 | 3.53E-05 | 0 | 0 | 0 | 7.23E-05 | 7.18E-05 | 5.14E-05 | 2.22E-05 |
| Znrf3 | 3.42E-06 | 6.24E-05 | 5.63E-05 | 2.17E-05 | 2.34E-05 | 0 | 0 | 0 | 5.44E-05 | 2.91E-06 | 5.87E-05 | 3.16E-05 | 0 | 4.12E-05 | 2.39E-05 | 3.44E-05 | 3.81E-05 | 2.38E-05 |
| Cnih4 | 0.00021 | 0.00012 | 0.00015 | 0 | 4.64E-05 | 0.00017 | 0 | 0 | 5.43E-05 | 6.01E-05 | 9.92E-05 | 7.28E-05 | 3.02E-05 | 7.28E-05 | 0 | 0.00031 | 0 | 1.96E-05 |
| Fbh1 | 0.00017 | 0.00011 | 7.35E-05 | 9.94E-05 | 0.00011 | 0.00012 | 0.00019 | 8.73E-05 | 5.43E-05 | 0.00014 | 5.48E-05 | 5.08E-05 | 0.00011 | 0.0001 | 0.00015 | 0 | 0 | 9.12E-05 |
| Rlf | 0.00011 | 0.00011 | 8.56E-06 | 0 | 0.00011 | 9.11E-05 | 2.30E-05 | 4.31E-05 | 5.43E-05 | 3.03E-05 | 2.70E-05 | 0.00033 | 3.11E-05 | 0 | 5.85E-05 | 0 | 0.0002 | 1.39E-05 |
| Pex2 | 0.00023 | 6.08E-05 | 4.82E-05 | 0 | 4.27E-05 | 5.62E-05 | 4.41E-05 | 0 | 5.43E-05 | 7.18E-05 | 5.84E-05 | 1.76E-05 | 0 | 1.96E-06 | 3.47E-05 | 9.85E-05 | 0 | 7.87E-06 |
| Fgfr1op | 5.43E-05 | 5.12E-05 | 4.23E-05 | 0 | 3.05E-05 | 2.24E-05 | 0 | 0 | 5.43E-05 | 7.48E-06 | 9.14E-06 | 0 | 5.39E-05 | 5.77E-05 | 0.00012 | 9.91E-05 | 3.33E-05 | 0 |
| Nudt22 | 3.01E-05 | 3.30E-05 | 3.70E-05 | 5.14E-05 | 5.38E-05 | 0 | 2.12E-05 | 0 | 5.43E-05 | 2.52E-05 | 3.46E-05 | 0 | 1.27E-05 | 7.07E-06 | 4.25E-05 | 2.30E-05 | 0 | 2.44E-05 |
| Arhgef11 | 9.79E-05 | 0.00011 | 0 | 0 | 5.70E-05 | 0.0001 | 0 | 5.89E-05 | 5.42E-05 | 2.79E-05 | 3.74E-05 | 0.0002 | 9.59E-07 | 2.76E-05 | 5.77E-05 | 9.44E-05 | 8.84E-05 | 6.09E-05 |
| Stn1 | 4.45E-05 | 7.68E-05 | 4.17E-05 | 8.79E-05 | 5.03E-05 | 4.43E-05 | 3.55E-05 | 0 | 5.42E-05 | 7.25E-05 | 0 | 3.77E-05 | 1.33E-05 | 0 | 7.22E-05 | 9.63E-05 | 1.69E-05 | 5.17E-05 |
| Ppm1h | 6.34E-06 | 0 | 2.49E-05 | 0 | 5.51E-05 | 1.08E-05 | 5.24E-05 | 0 | 5.42E-05 | 8.00E-06 | 0 | 5.97E-05 | 0.00011 | 2.25E-05 | 5.05E-05 | 2.14E-05 | 0 | 0 |
| Col6a6 | 7.92E-06 | 0 | 0 | 0 | 0 | 0 | 0 | 0 | 5.42E-05 | 4.98E-05 | 0 | 0 | 4.09E-05 | 4.91E-05 | 0 | 6.45E-06 | 0 | 0 |
| Aga | 0.00013 | 2.56E-05 | 0 | 6.96E-05 | 0.0001 | 0.00017 | 0.00012 | 0.00018 | 5.41E-05 | 7.92E-05 | 1.32E-05 | 8.45E-05 | 0.00018 | 3.48E-05 | 0 | 4.43E-05 | 0 | 0.00013 |
| Mfsd11 | 0.00014 | 0.00016 | 4.20E-06 | 4.44E-05 | 0.0001 | 0.00013 | 0.00016 | 6.00E-05 | 5.41E-05 | 1.79E-05 | 0 | 4.06E-05 | 5.16E-05 | 0.00019 | 3.46E-06 | 0.00015 | 0 | 5.16E-05 |
| Grpel2 | 4.51E-05 | 4.43E-05 | 6.77E-05 | 0 | 4.47E-05 | 1.42E-05 | 5.06E-06 | 0 | 5.41E-05 | 8.16E-06 | 1.67E-05 | 3.06E-05 | 2.77E-06 | 2.23E-05 | 0 | 2.00E-05 | 0 | 1.56E-09 |
| Gtf2h2 | 9.67E-05 | 5.97E-05 | 2.92E-05 | 3.74E-05 | 0 | 3.54E-06 | 0 | 7.30E-06 | 5.41E-05 | 0 | 1.49E-06 | 8.12E-06 | 0 | 0 | 9.51E-05 | 4.90E-05 | 0 | 8.89E-06 |
| Tmem132a | 0 | 8.41E-05 | 2.90E-05 | 0.00011 | 8.08E-05 | 0 | 0 | 0.0002 | 5.41E-05 | 3.61E-05 | 0.00022 | 4.70E-05 | 4.98E-05 | 0.00011 | 0.00021 | 1.53E-05 | 0 | 4.00E-05 |
| Fgd1 | 6.13E-05 | 6.84E-05 | 9.39E-05 | 8.34E-05 | 8.68E-05 | 6.35E-05 | 5.58E-05 | 0.00012 | 5.40E-05 | 0 | 0 | 9.80E-06 | 0 | 6.47E-05 | 1.91E-05 | 0.00022 | 1.68E-05 | 6.90E-05 |
| Prkab1 | 8.34E-05 | 6.77E-05 | 8.81E-05 | 9.75E-06 | 3.48E-05 | 6.20E-05 | 8.09E-06 | 3.09E-05 | 5.40E-05 | 8.89E-06 | 0 | 2.57E-05 | 2.29E-05 | 0 | 0 | 6.10E-05 | 0 | 0 |
| Mlx | 0.00012 | 5.85E-05 | 9.68E-05 | 1.27E-05 | 6.45E-05 | 5.41E-05 | 3.19E-05 | 1.23E-05 | 5.40E-05 | 9.84E-07 | 1.03E-05 | 1.30E-05 | 1.06E-05 | 7.21E-06 | 4.97E-05 | 8.61E-06 | 0 | 0 |
| Pgap3 | 1.80E-05 | 5.10E-05 | 4.89E-06 | 6.33E-05 | 5.02E-05 | 3.78E-05 | 9.92E-06 | 7.92E-06 | 5.40E-05 | 5.92E-06 | 2.84E-05 | 2.97E-05 | 2.82E-05 | 2.78E-05 | 3.61E-05 | 0 | 2.28E-05 | 1.78E-05 |
| Snrpa1 | 0.00021 | 2.91E-05 | 0.00026 | 0.00014 | 0 | 1.69E-05 | 0 | 0 | 5.40E-05 | 1.56E-05 | 5.68E-05 | 2.92E-05 | 0 | 0 | 2.77E-05 | 0.00031 | 0 | 1.34E-06 |

| | | | | | | | | | | | | | | | | | | |
|---|---|---|---|---|---|---|---|---|---|---|---|---|---|---|---|---|---|---|
| Elf2 | 0.00014 | 0.00022 | 6.12E-05 | 3.79E-05 | 8.20E-05 | 0.00012 | 0.00015 | 8.21E-05 | 5.39E-05 | 7.25E-05 | 4.99E-05 | 5.45E-05 | 2.53E-05 | 0.00011 | 0.00016 | 0.00018 | 0.0001 | 0.0001 |
| Atxn3 | 5.30E-05 | 7.04E-05 | 3.37E-05 | 1.38E-05 | 2.99E-05 | 7.59E-05 | 0 | 4.32E-05 | 5.39E-05 | 4.12E-05 | 3.95E-05 | 5.07E-05 | 2.84E-05 | 3.75E-05 | 6.69E-05 | 0 | 0 | 2.95E-05 |
| Irak3 | 6.33E-05 | 9.86E-05 | 0 | 3.40E-05 | 0 | 7.07E-05 | 0 | 2.62E-06 | 5.39E-05 | 0.00041 | 0 | 4.27E-05 | 0.00023 | 5.95E-05 | 0 | 0 | 0 | 6.60E-05 |
| Tmem138 | 0.0001 | 5.35E-05 | 3.66E-05 | 1.72E-05 | 4.90E-05 | 1.95E-05 | 4.64E-06 | 1.29E-05 | 5.39E-05 | 8.34E-06 | 6.25E-06 | 3.67E-05 | 0 | 0 | 0 | 0.00012 | 0 | 0 |
| Gclc | 4.46E-05 | 4.54E-05 | 8.70E-05 | 0 | 0 | 1.22E-05 | 0 | 9.37E-06 | 5.39E-05 | 1.83E-05 | 0 | 5.46E-05 | 0 | 6.36E-05 | 0 | 5.31E-05 | 0.00035 | 0 |
| Tmf1 | 0.00035 | 0.00016 | 0.00024 | 9.69E-06 | 0.00012 | 0.00025 | 0.00033 | 0.00014 | 5.38E-05 | 1.45E-05 | 0.00016 | 7.99E-05 | 7.38E-05 | 0.00017 | 0.00025 | 2.61E-05 | 0.00021 | 8.13E-06 |
| Rad21 | 0.00015 | 0.00019 | 8.38E-05 | 3.14E-05 | 7.01E-05 | 0.00011 | 3.53E-05 | 7.92E-05 | 5.38E-05 | 5.00E-05 | 5.21E-05 | 0.00021 | 0 | 7.89E-05 | 0 | 0.00103 | 0 | 5.32E-05 |
| Wdr7 | 6.21E-05 | 9.71E-05 | 2.02E-05 | 2.99E-05 | 9.48E-05 | 6.61E-05 | 8.14E-05 | 3.29E-05 | 5.38E-05 | 8.48E-05 | 2.76E-05 | 0 | 7.61E-06 | 5.75E-05 | 0.00013 | 1.10E-05 | 2.58E-05 | 6.32E-05 |
| Wdr59 | 1.50E-05 | 2.82E-05 | 6.56E-06 | 3.17E-05 | 0 | 6.58E-05 | 0 | 1.48E-06 | 5.38E-05 | 1.34E-05 | 0 | 0.00023 | 0 | 0 | 3.70E-05 | 8.75E-06 | 0 | 5.37E-05 |
| Zfp655 | 7.92E-05 | 0.0002 | 4.71E-05 | 1.43E-05 | 6.40E-05 | 4.58E-05 | 1.62E-05 | 5.88E-06 | 5.38E-05 | 8.18E-05 | 0 | 0.00015 | 7.91E-05 | 7.36E-05 | 0.00036 | 0 | 0.00033 | 0 |
| Tgs1 | 3.68E-05 | 3.67E-05 | 4.88E-05 | 1.81E-06 | 1.51E-05 | 2.39E-05 | 0 | 0 | 5.38E-05 | 3.16E-05 | 8.95E-06 | 5.97E-06 | 0 | 2.89E-05 | 0.0001 | 2.85E-05 | 0 | 1.61E-05 |
| Nfu1 | 2.45E-05 | 3.36E-05 | 4.47E-05 | 0.00013 | 2.20E-05 | 1.56E-05 | 0 | 1.07E-05 | 5.38E-05 | 4.56E-05 | 8.91E-05 | 1.40E-07 | 2.13E-05 | 2.27E-05 | 0 | 2.40E-05 | 6.80E-05 | 4.62E-05 |
| Ramp2 | 2.66E-05 | 1.63E-05 | 2.89E-05 | 0 | 0 | 0 | 0 | 1.69E-05 | 5.38E-05 | 0.00038 | 0 | 0 | 0 | 0 | 0 | 0 | 0 | 0.00011 |
| Tprgl | 0.00011 | 0.00025 | 3.84E-06 | 6.17E-05 | 0.00017 | 0.00019 | 0.00012 | 3.68E-05 | 5.37E-05 | 0.00012 | 0.0001 | 2.00E-05 | 0 | 5.61E-05 | 1.54E-05 | 0 | 0.0001 | 0.00016 |
| Elk1 | 7.83E-05 | 8.18E-05 | 4.34E-05 | 4.02E-05 | 8.81E-05 | 6.19E-05 | 4.08E-05 | 6.05E-05 | 5.37E-05 | 3.39E-05 | 2.57E-05 | 1.23E-05 | 2.55E-05 | 5.44E-05 | 4.89E-05 | 9.45E-06 | 9.04E-05 | 0 |
| Smarcad1 | 7.37E-05 | 0.00013 | 0.00011 | 0.00018 | 5.94E-05 | 5.95E-05 | 3.54E-05 | 9.46E-05 | 5.37E-05 | 3.64E-05 | 5.10E-05 | 3.14E-05 | 2.54E-05 | 0.00013 | 0.00025 | 7.66E-05 | 0 | 4.09E-05 |
| Hccs | 0.0001 | 3.55E-05 | 8.82E-05 | 2.41E-05 | 4.59E-05 | 5.16E-05 | 0 | 1.01E-05 | 5.37E-05 | 1.19E-05 | 2.73E-05 | 0 | 3.85E-05 | 1.75E-05 | 2.63E-05 | 0.0001 | 9.79E-05 | 0 |
| Ghdc | 2.08E-05 | 6.29E-05 | 2.62E-05 | 4.75E-05 | 4.56E-05 | 2.43E-05 | 0 | 2.88E-05 | 5.37E-05 | 6.95E-05 | 3.85E-05 | 8.31E-05 | 5.10E-05 | 2.20E-05 | 0 | 0 | 0 | 7.16E-05 |
| Hunk | 0 | 0 | 0 | 0 | 0 | 1.29E-05 | 0 | 3.67E-05 | 5.37E-05 | 1.74E-06 | 0 | 4.46E-06 | 0 | 0 | 0 | 1.06E-05 | 0 | 9.54E-06 |
| Thyn1 | 9.48E-05 | 2.14E-05 | 7.60E-05 | 2.53E-05 | 0 | 1.03E-05 | 0 | 1.27E-05 | 5.37E-05 | 1.77E-06 | 2.81E-05 | 3.71E-06 | 0 | 0 | 3.29E-05 | 7.93E-05 | 0 | 2.22E-06 |
| Tnip1 | 5.86E-05 | 0.00011 | 9.58E-05 | 9.71E-06 | 6.77E-05 | 1.14E-06 | 0 | 7.15E-05 | 5.37E-05 | 6.99E-05 | 8.18E-06 | 4.76E-05 | 0.00021 | 0.00023 | 0.0001 | 1.90E-05 | 1.45E-05 | 0.00016 |
| Eva1a | 0 | 0 | 2.13E-05 | 0 | 5.21E-05 | 0 | 0 | 8.29E-05 | 5.37E-05 | 0 | 0.00017 | 0 | 0 | 0.0003 | 1.27E-05 | 6.49E-05 | 0 | 0 |
| Kti12 | 6.86E-05 | 6.12E-05 | 0.00018 | 0.00012 | 0 | 0 | 0 | 2.79E-05 | 5.37E-05 | 2.11E-05 | 1.58E-05 | 0 | 1.55E-05 | 6.45E-06 | 9.03E-05 | 2.26E-05 | 2.70E-06 | 1.35E-05 |
| Fbxo4 | 7.14E-05 | 4.06E-05 | 3.33E-05 | 2.37E-05 | 5.71E-05 | 8.50E-05 | 9.18E-05 | 0 | 5.36E-05 | 0.00013 | 1.57E-05 | 0 | 8.24E-05 | 4.31E-05 | 0 | 2.65E-06 | 5.60E-05 | 7.70E-05 |
| Fbxw4 | 2.35E-05 | 4.27E-05 | 1.18E-05 | 1.81E-05 | 3.31E-05 | 2.91E-05 | 2.56E-05 | 3.32E-05 | 5.36E-05 | 4.51E-06 | 2.63E-05 | 3.81E-05 | 0 | 2.20E-05 | 1.85E-05 | 1.05E-06 | 3.07E-05 | 5.44E-05 |
| Acp2 | 5.31E-05 | 0.00017 | 1.03E-06 | 0 | 2.95E-05 | 0.0002 | 5.28E-05 | 7.76E-05 | 5.35E-05 | 5.01E-05 | 0 | 1.65E-05 | 9.77E-05 | 8.98E-05 | 0 | 0 | 0 | 6.34E-05 |
| Ube2g1 | 0.00022 | 0.0002 | 0.0001 | 1.03E-06 | 7.46E-05 | 0.00018 | 4.16E-05 | 2.69E-05 | 5.35E-05 | 4.93E-05 | 0.00014 | 0.00025 | 3.64E-05 | 3.36E-05 | 6.28E-05 | 7.39E-05 | 7.68E-05 | 7.00E-05 |
| Vcpip1 | 0.00013 | 0.0001 | 2.70E-05 | 0 | 2.36E-05 | 8.82E-05 | 1.33E-05 | 0 | 5.35E-05 | 2.29E-05 | 7.49E-06 | 8.55E-05 | 2.22E-06 | 5.91E-06 | 7.88E-05 | 5.03E-05 | 0 | 1.45E-05 |
| Plekhh3 | 0 | 9.17E-05 | 1.86E-05 | 4.53E-05 | 0 | 1.49E-05 | 0 | 0.00017 | 5.35E-05 | 2.92E-05 | 1.64E-05 | 7.86E-06 | 0 | 0 | 0.00012 | 3.87E-05 | 1.93E-05 | 5.72E-05 |
| Rnf217 | 9.21E-05 | 0.0001 | 5.35E-05 | 0 | 1.42E-05 | 1.45E-05 | 0 | 0 | 5.35E-05 | 3.26E-05 | 5.42E-05 | 0.00041 | 0 | 4.36E-05 | 0.0001 | 3.41E-05 | 5.79E-05 | 2.04E-05 |
| Vps37c | 1.65E-05 | 6.85E-05 | 6.17E-05 | 0.00011 | 4.10E-05 | 1.43E-06 | 0 | 2.45E-05 | 5.35E-05 | 3.78E-05 | 5.21E-05 | 5.76E-05 | 5.14E-05 | 3.74E-05 | 0.00014 | 1.34E-05 | 8.50E-05 | 8.32E-05 |
| Anln | 1.79E-05 | 0 | 0 | 0 | 0 | 0 | 0 | 0 | 5.35E-05 | 0 | 2.67E-06 | 0 | 0 | 0 | 0.00013 | 0.00163 | 0 | 0 |
| Zmiz2 | 6.42E-06 | 0.00012 | 2.25E-05 | 0.0001 | 5.72E-05 | 0 | 0 | 3.19E-05 | 5.35E-05 | 4.96E-05 | 5.51E-05 | 8.66E-05 | 8.31E-06 | 1.11E-05 | 7.85E-05 | 5.56E-05 | 2.94E-05 | 6.04E-05 |

| | | | | | | | | | | | | | | | | | | |
|---|---|---|---|---|---|---|---|---|---|---|---|---|---|---|---|---|---|---|
| Fam234a | 0 | 0.00017 | 2.47E-05 | 0.00016 | 0.00019 | 0.00012 | 0.00019 | 5.78E-05 | 5.34E-05 | 0.00019 | 0.00016 | 0.0001 | 0.00014 | 7.06E-05 | 0 | 0 | 0 | 0.00028 |
| Smim1 | 1.67E-05 | 2.53E-05 | 0 | 2.86E-05 | 0.00013 | 0.0001 | 0 | 6.68E-05 | 5.34E-05 | 4.77E-05 | 1.59E-05 | 1.95E-05 | 2.80E-05 | 0 | 1.86E-05 | 0 | 0 | 6.46E-05 |
| Entr1 | 0.00012 | 0.00012 | 0.00019 | 0 | 9.17E-05 | 9.02E-05 | 3.88E-05 | 5.01E-05 | 5.34E-05 | 5.80E-05 | 0.00013 | 6.57E-06 | 6.30E-06 | 9.26E-05 | 0.00017 | 6.81E-05 | 9.83E-05 | 3.02E-05 |
| Nhsl2 | 2.34E-05 | 6.24E-05 | 0 | 1.48E-05 | 7.49E-05 | 8.39E-05 | 0 | 0.00017 | 5.34E-05 | 1.01E-05 | 0.00013 | 0 | 0 | 0.00011 | 2.46E-05 | 3.07E-05 | 4.68E-05 | 6.21E-05 |
| Fbxo33 | 7.43E-05 | 7.54E-05 | 2.08E-05 | 0 | 4.83E-05 | 7.87E-05 | 5.03E-06 | 0 | 5.34E-05 | 6.30E-06 | 0 | 2.58E-05 | 0 | 4.06E-05 | 0 | 0 | 0.00032 | 1.71E-05 |
| Mtrex | 0.00013 | 9.09E-05 | 0.00013 | 6.43E-06 | 3.49E-05 | 6.23E-05 | 0 | 0 | 5.34E-05 | 2.75E-05 | 1.76E-05 | 3.25E-06 | 0 | 2.31E-06 | 7.08E-05 | 0.00019 | 0 | 0 |
| Tbc1d22a | 3.96E-05 | 0.00012 | 1.83E-05 | 3.87E-05 | 6.41E-05 | 3.38E-05 | 6.61E-05 | 7.05E-05 | 5.34E-05 | 5.46E-05 | 0 | 0.00012 | 0 | 0 | 0.00016 | 0 | 0.0001 | 0.00011 |
| Supt3 | 6.78E-05 | 2.22E-05 | 2.13E-05 | 8.86E-06 | 1.76E-05 | 3.32E-05 | 3.85E-05 | 1.55E-05 | 5.34E-05 | 0 | 0 | 3.10E-05 | 4.28E-06 | 8.81E-06 | 5.62E-05 | 4.94E-05 | 0.00011 | 0 |
| Iqce | 3.51E-05 | 6.70E-05 | 7.85E-06 | 4.04E-05 | 1.79E-05 | 2.82E-05 | 2.76E-05 | 2.84E-05 | 5.34E-05 | 4.65E-05 | 0 | 8.67E-06 | 0 | 1.15E-05 | 0.00019 | 2.78E-05 | 6.39E-07 | 2.40E-05 |
| Ddx51 | 4.92E-05 | 4.39E-05 | 7.53E-05 | 1.11E-05 | 1.78E-05 | 6.96E-07 | 0 | 0 | 5.34E-05 | 0 | 1.30E-05 | 2.55E-06 | 8.22E-07 | 8.85E-06 | 3.25E-05 | 2.71E-05 | 0 | 0 |
| Egfr | 0.00019 | 0.00025 | 0.00017 | 9.17E-05 | 3.91E-05 | 0.00039 | 0.00022 | 0.0006 | 5.33E-05 | 0.00065 | 6.90E-05 | 0.00024 | 0.00127 | 0.00018 | 0.00026 | 4.60E-05 | 0.00022 | 0.00018 |
| Speg | 1.69E-05 | 5.83E-05 | 1.31E-06 | 3.80E-05 | 4.71E-05 | 7.52E-05 | 3.42E-05 | 4.19E-05 | 5.33E-05 | 0 | 5.53E-05 | 3.25E-06 | 2.28E-05 | 0 | 0.0001 | 0 | 0 | 0 |
| Wdr20 | 4.41E-05 | 7.72E-05 | 3.67E-05 | 1.06E-05 | 5.69E-05 | 1.56E-05 | 0 | 2.43E-05 | 5.33E-05 | 3.17E-05 | 9.19E-06 | 5.46E-05 | 2.29E-05 | 1.18E-05 | 0.00012 | 3.19E-05 | 0 | 7.35E-06 |
| Eif2b2 | 8.56E-05 | 9.25E-05 | 0.0002 | 9.55E-05 | 1.58E-05 | 0 | 3.47E-05 | 2.61E-05 | 5.33E-05 | 4.79E-05 | 0.00016 | 0.00011 | 0 | 0.00014 | 2.50E-05 | 6.34E-05 | 0 | 0 |
| Dnmt3a | 0.00011 | 0.00012 | 3.21E-05 | 1.19E-05 | 0.00015 | 0.00015 | 0.00011 | 0.00018 | 5.32E-05 | 2.35E-05 | 8.20E-05 | 8.28E-05 | 0 | 8.23E-05 | 0 | 2.02E-05 | 0.00023 | 5.25E-05 |
| Snrpe | 0.00023 | 7.70E-05 | 0.00016 | 0.00013 | 9.85E-06 | 8.75E-05 | 2.98E-05 | 0 | 5.32E-05 | 0 | 2.35E-05 | 7.67E-05 | 2.11E-05 | 4.82E-05 | 0 | 0.00033 | 0.00018 | 0 |
| Zc3h15 | 0.00018 | 0.00012 | 0.00024 | 0.00015 | 3.77E-05 | 4.21E-05 | 5.00E-05 | 0.00013 | 5.32E-05 | 4.27E-05 | 7.36E-05 | 9.63E-05 | 3.95E-05 | 6.84E-05 | 5.04E-05 | 0.00014 | 0.00018 | 5.90E-06 |
| Wrn | 7.38E-05 | 8.29E-05 | 0 | 0 | 1.53E-05 | 2.47E-05 | 0 | 0 | 5.32E-05 | 2.18E-05 | 1.41E-06 | 5.67E-05 | 0 | 0 | 0.00012 | 8.90E-05 | 2.83E-05 | 3.23E-05 |
| Sec22a | 0.00012 | 2.27E-05 | 2.68E-05 | 5.03E-05 | 3.15E-05 | 2.46E-05 | 0.00015 | 4.62E-06 | 5.32E-05 | 3.47E-05 | 4.68E-05 | 3.31E-05 | 5.97E-05 | 5.58E-05 | 0 | 0 | 0 | 5.52E-05 |
| Slc9a8 | 6.61E-05 | 4.13E-05 | 2.45E-05 | 5.65E-06 | 2.99E-05 | 2.37E-05 | 2.85E-06 | 3.71E-05 | 5.32E-05 | 1.45E-05 | 0 | 2.61E-05 | 2.81E-05 | 0 | 0.0001 | 2.79E-05 | 4.29E-05 | 1.11E-05 |
| 2-Mar | 2.17E-05 | 0.00013 | 0 | 5.57E-05 | 7.57E-05 | 5.06E-06 | 0 | 2.64E-05 | 5.32E-05 | 0.00015 | 9.28E-05 | 4.63E-05 | 2.86E-06 | 7.69E-06 | 0 | 0 | 0 | 0.00015 |
| Golt1b | 0.00029 | 0 | 0.0005 | 0.00051 | 0.00012 | 0.00028 | 0.00055 | 0.00034 | 5.31E-05 | 0.00012 | 0.00028 | 9.92E-05 | 0.00014 | 0.00027 | 2.68E-05 | 0.00025 | 0 | 4.03E-05 |
| Cdc16 | 0.00031 | 5.75E-05 | 0.00018 | 0.00011 | 8.28E-05 | 0.00014 | 0.00034 | 0.0001 | 5.31E-05 | 2.00E-05 | 0.00015 | 3.55E-05 | 6.73E-05 | 9.03E-05 | 3.15E-05 | 0.00013 | 0 | 0.0001 |
| Gzf1 | 4.88E-05 | 7.93E-05 | 1.21E-06 | 5.42E-05 | 4.80E-05 | 5.75E-05 | 1.20E-05 | 6.06E-05 | 5.31E-05 | 4.91E-05 | 8.19E-06 | 0.00013 | 5.46E-05 | 7.04E-07 | 0.00013 | 7.37E-05 | 0 | 8.52E-06 |
| Zbtb8a | 7.15E-05 | 2.94E-05 | 0 | 5.86E-05 | 6.69E-06 | 4.37E-05 | 3.41E-05 | 0 | 5.31E-05 | 1.95E-05 | 7.22E-06 | 0 | 7.45E-06 | 6.52E-05 | 0 | 7.24E-05 | 0 | 0.0001 |
| Senp2 | 6.77E-05 | 7.21E-05 | 0.00011 | 0 | 5.34E-05 | 2.48E-05 | 8.39E-06 | 0 | 5.31E-05 | 3.29E-05 | 5.60E-05 | 0.00011 | 2.83E-05 | 2.54E-05 | 0.00011 | 0.00013 | 9.34E-05 | 2.87E-05 |
| Kat14 | 5.47E-05 | 8.54E-05 | 0.00011 | 0.00019 | 3.70E-05 | 1.92E-05 | 0.00014 | 4.00E-06 | 5.31E-05 | 6.16E-05 | 4.75E-05 | 0 | 1.20E-05 | 0.0001 | 5.45E-05 | 9.97E-05 | 0 | 6.11E-05 |
| Pus7 | 9.66E-05 | 2.84E-05 | 0.0001 | 0 | 0 | 2.32E-06 | 0 | 0 | 5.31E-05 | 0 | 4.27E-06 | 3.43E-05 | 0 | 1.43E-05 | 0 | 2.14E-05 | 3.20E-05 | 2.10E-07 |
| Vta1 | 6.59E-05 | 3.89E-05 | 7.18E-05 | 7.32E-05 | 1.48E-05 | 0 | 3.48E-05 | 2.89E-05 | 5.31E-05 | 1.16E-05 | 1.39E-05 | 3.50E-05 | 2.79E-06 | 0 | 2.71E-05 | 2.70E-05 | 1.26E-05 | 1.93E-05 |
| Cyb5r4 | 0.00011 | 0.00011 | 7.24E-05 | 0.00016 | 0.00013 | 9.66E-05 | 9.62E-05 | 8.79E-05 | 5.30E-05 | 0.00011 | 6.49E-05 | 7.76E-05 | 3.11E-05 | 8.71E-05 | 1.74E-05 | 4.37E-05 | 6.67E-06 | 2.33E-05 |
| E2f3 | 0.00011 | 3.10E-05 | 0.00013 | 0 | 6.56E-05 | 7.59E-05 | 0 | 6.28E-05 | 5.30E-05 | 0 | 3.44E-05 | 4.14E-05 | 7.72E-05 | 7.87E-05 | 1.27E-05 | 4.04E-05 | 9.05E-05 | 0 |
| Tbk1 | 4.40E-05 | 0.00012 | 4.42E-05 | 1.09E-05 | 4.08E-05 | 6.95E-05 | 0 | 0 | 5.30E-05 | 9.86E-05 | 4.35E-05 | 4.08E-05 | 4.93E-05 | 8.97E-05 | 0.00016 | 5.50E-05 | 4.91E-05 | 1.14E-05 |
| Zfp664 | 7.34E-05 | 0.0001 | 6.81E-05 | 1.01E-05 | 6.69E-05 | 6.32E-05 | 1.17E-06 | 5.24E-05 | 5.30E-05 | 2.33E-05 | 8.05E-05 | 0.00011 | 2.47E-05 | 8.40E-05 | 1.17E-05 | 0.0001 | 0 | 1.82E-05 |

| | | | | | | | | | | | | | | | | | | |
|---|---|---|---|---|---|---|---|---|---|---|---|---|---|---|---|---|---|---|
| Usp32 | 7.83E-05 | 0.00016 | 2.92E-05 | 5.70E-06 | 3.84E-05 | 3.38E-05 | 2.12E-05 | 1.61E-06 | 5.30E-05 | 5.52E-05 | 5.19E-05 | 6.68E-05 | 4.37E-05 | 5.87E-05 | 0.00029 | 0 | 0.00021 | 4.44E-05 |
| Bcl7b | 3.37E-05 | 5.42E-05 | 2.53E-05 | 5.27E-05 | 1.07E-05 | 2.15E-05 | 1.12E-05 | 4.68E-05 | 5.30E-05 | 2.82E-05 | 3.44E-05 | 4.16E-06 | 7.94E-07 | 3.32E-05 | 6.37E-05 | 3.07E-05 | 6.22E-05 | 2.73E-05 |
| Fam160a2 | 5.78E-06 | 5.74E-05 | 3.68E-05 | 6.02E-05 | 2.20E-05 | 1.92E-05 | 6.61E-06 | 3.38E-05 | 5.30E-05 | 5.43E-05 | 6.08E-05 | 4.71E-05 | 0 | 2.62E-05 | 6.83E-05 | 6.79E-06 | 2.90E-05 | 4.98E-05 |
| Apopt1 | 7.64E-05 | 5.75E-05 | 3.42E-05 | 3.03E-05 | 4.09E-05 | 1.60E-05 | 1.24E-05 | 7.84E-06 | 5.30E-05 | 3.73E-05 | 2.58E-05 | 1.36E-05 | 2.54E-05 | 2.66E-05 | 9.86E-06 | 2.10E-05 | 0 | 1.04E-06 |
| Srrd | 1.11E-05 | 3.64E-05 | 3.89E-05 | 5.97E-05 | 3.30E-05 | 7.98E-06 | 2.82E-06 | 1.46E-05 | 5.30E-05 | 2.96E-05 | 3.51E-05 | 1.77E-05 | 4.13E-06 | 8.57E-06 | 1.06E-05 | 2.80E-05 | 0 | 1.48E-05 |
| Mpi | 6.10E-05 | 3.22E-05 | 6.41E-05 | 9.97E-05 | 3.99E-05 | 0 | 0 | 1.31E-05 | 5.30E-05 | 3.92E-05 | 4.90E-05 | 0 | 2.21E-06 | 3.05E-05 | 7.93E-05 | 6.86E-05 | 0 | 6.35E-05 |
| Fbxo11 | 0.00025 | 0.00018 | 9.52E-05 | 0 | 0.00015 | 0.00015 | 0.0001 | 4.44E-05 | 5.29E-05 | 2.63E-05 | 0.00014 | 9.41E-05 | 7.56E-06 | 3.33E-05 | 6.32E-05 | 4.26E-05 | 0.00034 | 9.58E-05 |
| Prkcd | 7.73E-05 | 0.00012 | 0.00016 | 1.04E-06 | 3.51E-05 | 6.93E-05 | 6.93E-05 | 9.09E-05 | 5.29E-05 | 2.27E-06 | 0.00012 | 0.00016 | 0 | 4.79E-05 | 0.00019 | 0 | 7.21E-05 | 0.00011 |
| Stambp | 9.56E-05 | 5.74E-05 | 4.69E-05 | 1.80E-05 | 2.93E-05 | 4.99E-05 | 4.10E-05 | 9.38E-06 | 5.29E-05 | 4.14E-05 | 3.40E-05 | 5.38E-05 | 0 | 0 | 2.70E-05 | 0 | 0 | 4.55E-05 |
| Eif4a3 | 0.00018 | 5.38E-05 | 0.00019 | 6.63E-05 | 2.29E-05 | 4.26E-05 | 3.84E-05 | 2.48E-05 | 5.29E-05 | 0 | 5.95E-05 | 0 | 0 | 5.48E-05 | 0 | 0.00013 | 5.15E-05 | 0 |
| Dctn5 | 0.00011 | 9.22E-05 | 6.72E-05 | 0.00011 | 5.94E-05 | 4.07E-05 | 0.0001 | 8.56E-05 | 5.29E-05 | 6.72E-05 | 3.49E-05 | 5.34E-05 | 4.89E-06 | 2.11E-05 | 0.00025 | 4.24E-05 | 3.28E-05 | 5.98E-05 |
| Mapk11 | 1.53E-05 | 1.05E-06 | 5.08E-05 | 0 | 0 | 2.94E-05 | 0 | 8.30E-05 | 5.29E-05 | 1.51E-05 | 2.27E-05 | 3.48E-05 | 0 | 0 | 0 | 3.83E-05 | 1.88E-05 | 1.33E-05 |
| Nop10 | 0.00028 | 0.00011 | 0.00026 | 0.00032 | 0.00014 | 0.00016 | 6.78E-05 | 4.92E-05 | 5.28E-05 | 3.94E-05 | 1.43E-05 | 0 | 3.72E-05 | 3.33E-05 | 0 | 0.00021 | 7.88E-05 | 3.49E-05 |
| Cyhr1 | 7.81E-05 | 0.00018 | 4.12E-05 | 0.00014 | 0.00014 | 0.00011 | 5.70E-05 | 0 | 5.28E-05 | 1.45E-05 | 3.45E-06 | 0 | 0 | 8.00E-05 | 0.00019 | 4.10E-05 | 4.87E-05 | 6.76E-05 |
| Cdk11b | 0.00016 | 0.00012 | 0.00018 | 0.00014 | 3.17E-05 | 4.58E-05 | 7.16E-05 | 3.08E-05 | 5.28E-05 | 2.67E-05 | 0.00012 | 3.07E-05 | 0 | 3.09E-05 | 0.00039 | 0.00015 | 0.00012 | 4.81E-05 |
| Map3k7 | 8.75E-05 | 8.06E-05 | 7.16E-05 | 0 | 7.93E-05 | 3.45E-05 | 7.79E-06 | 1.31E-05 | 5.28E-05 | 2.48E-05 | 3.96E-05 | 4.01E-05 | 3.67E-05 | 1.75E-08 | 2.29E-05 | 6.80E-05 | 1.25E-05 | 3.38E-05 |
| Atp13a2 | 7.51E-06 | 7.57E-05 | 0 | 3.93E-05 | 0.0001 | 3.08E-05 | 2.04E-06 | 1.81E-05 | 5.28E-05 | 7.67E-05 | 4.27E-05 | 2.71E-05 | 8.09E-07 | 3.05E-05 | 8.47E-05 | 2.80E-06 | 0 | 3.32E-05 |
| Dcaf11 | 0.00011 | 0.0001 | 1.74E-05 | 3.54E-06 | 5.50E-05 | 0.00012 | 1.61E-05 | 3.49E-05 | 5.27E-05 | 6.77E-05 | 4.89E-05 | 9.76E-05 | 1.20E-05 | 3.19E-05 | 1.41E-05 | 0 | 0 | 5.18E-05 |
| Naf1 | 9.92E-05 | 5.92E-05 | 5.85E-05 | 0 | 2.57E-05 | 6.72E-05 | 0 | 0 | 5.27E-05 | 9.83E-07 | 9.29E-07 | 3.11E-05 | 4.87E-06 | 1.55E-05 | 0 | 2.17E-05 | 5.84E-05 | 0 |
| Nup214 | 6.59E-05 | 9.41E-05 | 8.31E-05 | 4.27E-05 | 5.80E-05 | 5.85E-05 | 4.35E-06 | 2.93E-05 | 5.27E-05 | 1.80E-05 | 4.20E-05 | 1.06E-05 | 3.48E-05 | 4.34E-05 | 5.85E-05 | 0.0001 | 4.34E-06 | 2.87E-05 |
| Emc8 | 2.98E-05 | 3.89E-05 | 7.47E-05 | 0.00013 | 2.44E-05 | 1.40E-05 | 0 | 3.02E-05 | 5.27E-05 | 7.86E-06 | 3.86E-05 | 2.47E-06 | 0 | 2.52E-05 | 0.00011 | 0.00012 | 3.17E-05 | 4.00E-05 |
| Erp44 | 0.00057 | 1.18E-05 | 0.00012 | 0 | 5.93E-05 | 0.00029 | 0.00015 | 5.84E-05 | 5.26E-05 | 3.53E-05 | 0 | 0.00023 | 0.00015 | 0 | 6.00E-07 | 7.15E-05 | 0.00012 | 3.94E-05 |
| Cfap36 | 0.00013 | 3.83E-05 | 2.82E-05 | 5.00E-06 | 2.46E-05 | 5.75E-05 | 0 | 0 | 5.26E-05 | 1.23E-05 | 5.08E-06 | 3.10E-05 | 0 | 8.90E-06 | 1.11E-05 | 2.89E-05 | 0 | 0 |
| Gorab | 0.00012 | 4.25E-05 | 5.65E-05 | 0 | 2.60E-05 | 4.99E-05 | 1.73E-05 | 0 | 5.26E-05 | 0 | 9.95E-06 | 4.58E-05 | 0 | 4.04E-05 | 8.63E-05 | 1.00E-05 | 0 | 1.04E-05 |
| Foxred1 | 8.27E-05 | 4.81E-05 | 6.03E-05 | 8.06E-05 | 3.41E-05 | 1.18E-05 | 1.64E-05 | 5.29E-05 | 5.26E-05 | 7.45E-06 | 9.90E-06 | 0 | 6.27E-05 | 2.51E-05 | 0.00014 | 7.31E-05 | 2.82E-05 | 8.10E-06 |
| Mpv17l2 | 2.15E-05 | 6.76E-05 | 7.05E-05 | 0.00024 | 4.24E-05 | 0 | 7.23E-05 | 1.59E-05 | 5.26E-05 | 0.0001 | 7.76E-05 | 0 | 3.27E-05 | 6.96E-05 | 0 | 0.00014 | 0 | 9.60E-05 |
| Ccnq | 4.74E-05 | 4.89E-05 | 3.50E-05 | 3.51E-05 | 2.71E-05 | 2.40E-05 | 0 | 2.40E-05 | 5.25E-05 | 1.21E-05 | 8.18E-06 | 3.13E-06 | 4.86E-06 | 0 | 0 | 5.01E-05 | 0 | 1.77E-05 |
| Slc29a3 | 0 | 8.73E-05 | 4.67E-05 | 0.00017 | 5.44E-05 | 1.65E-05 | 4.40E-05 | 4.92E-05 | 5.25E-05 | 6.29E-05 | 2.42E-05 | 3.46E-05 | 3.90E-05 | 4.22E-05 | 2.76E-05 | 0 | 1.61E-05 | 9.06E-05 |
| Rdh13 | 1.84E-05 | 4.42E-05 | 9.42E-05 | 4.78E-06 | 7.39E-06 | 0 | 0 | 0 | 5.25E-05 | 2.54E-05 | 5.27E-05 | 1.45E-05 | 1.58E-05 | 4.69E-05 | 7.48E-05 | 1.92E-05 | 1.58E-05 | 5.61E-06 |
| Snrpd2 | 0.0002 | 1.78E-05 | 7.93E-05 | 0.00013 | 0 | 8.58E-05 | 0 | 4.48E-05 | 5.24E-05 | 0 | 5.86E-05 | 8.06E-05 | 0 | 7.11E-05 | 0 | 0.00015 | 4.91E-05 | 0 |
| Zfp639 | 0.00017 | 8.94E-05 | 2.60E-05 | 0 | 1.24E-05 | 8.39E-05 | 2.35E-05 | 0 | 5.24E-05 | 9.84E-07 | 2.27E-05 | 8.89E-05 | 1.88E-06 | 1.62E-05 | 9.20E-05 | 6.93E-05 | 0 | 1.37E-05 |
| Gtf3a | 0.00016 | 8.31E-05 | 6.89E-05 | 0.0001 | 5.54E-05 | 5.50E-05 | 8.30E-05 | 8.09E-06 | 5.24E-05 | 6.68E-05 | 2.74E-05 | 5.21E-05 | 3.81E-05 | 6.52E-06 | 6.73E-05 | 0.00018 | 0 | 0.00012 |
| Sdad1 | 0.00013 | 4.90E-05 | 0.00018 | 0 | 1.46E-05 | 1.19E-05 | 9.57E-06 | 0 | 5.24E-05 | 0 | 1.03E-05 | 8.81E-05 | 1.12E-05 | 4.92E-05 | 0.00012 | 7.99E-06 | 5.06E-07 | 0 |

| | | | | | | | | | | | | | | | | | | |
|---|---|---|---|---|---|---|---|---|---|---|---|---|---|---|---|---|---|---|
| Wdr3 | 7.79E-05 | 6.98E-05 | 0.00011 | 1.33E-05 | 1.03E-05 | 1.51E-05 | 0 | 0 | 5.23E-05 | 3.65E-06 | 2.61E-05 | 0 | 5.54E-06 | 2.55E-05 | 3.24E-05 | 1.56E-05 | 0 | 2.58E-06 |
| Asb1 | 0 | 6.09E-05 | 6.18E-05 | 8.30E-05 | 4.63E-05 | 5.44E-07 | 3.51E-06 | 7.07E-05 | 5.23E-05 | 3.22E-05 | 5.25E-05 | 5.79E-05 | 0 | 1.21E-05 | 1.45E-05 | 0 | 0 | 1.75E-05 |
| Coro7 | 3.09E-05 | 8.01E-05 | 0 | 2.29E-05 | 6.70E-05 | 0 | 5.45E-05 | 9.07E-05 | 5.23E-05 | 8.89E-05 | 1.94E-05 | 1.75E-05 | 0 | 6.15E-05 | 8.51E-05 | 3.44E-06 | 0 | 9.06E-05 |
| Kcnd1 | 5.33E-05 | 1.56E-05 | 2.15E-05 | 5.86E-05 | 0 | 0 | 6.59E-05 | 4.16E-05 | 5.23E-05 | 5.49E-05 | 7.30E-07 | 0 | 5.56E-05 | 0 | 3.94E-05 | 0 | 3.41E-05 | 1.74E-05 |
| Pi4k2b | 0.0002 | 1.89E-05 | 0.0002 | 8.68E-05 | 5.39E-05 | 0.00014 | 0.0003 | 0.00016 | 5.22E-05 | 0 | 5.50E-05 | 8.02E-05 | 0.00018 | 0.00011 | 0 | 0.00018 | 0.00011 | 0 |
| Camta2 | 2.14E-06 | 0.00011 | 8.66E-06 | 0.00015 | 8.55E-05 | 8.76E-05 | 3.16E-05 | 0.00014 | 5.22E-05 | 8.46E-05 | 0.0001 | 9.51E-06 | 0 | 4.94E-05 | 0.00015 | 0 | 0.00012 | 8.33E-05 |
| Fam135a | 6.05E-05 | 8.15E-05 | 1.30E-05 | 0 | 4.88E-05 | 7.59E-05 | 1.32E-05 | 0 | 5.22E-05 | 1.25E-05 | 1.90E-06 | 5.57E-05 | 2.39E-05 | 4.01E-05 | 0.00012 | 4.85E-05 | 7.45E-05 | 8.85E-06 |
| Zfp26 | 6.89E-05 | 5.03E-05 | 2.64E-05 | 0 | 7.04E-05 | 6.55E-05 | 2.59E-05 | 2.05E-05 | 5.22E-05 | 2.63E-05 | 0 | 3.27E-05 | 5.70E-06 | 1.87E-05 | 9.08E-05 | 4.17E-05 | 0 | 6.39E-05 |
| Naa25 | 6.93E-05 | 8.57E-05 | 0.00019 | 2.24E-05 | 0 | 2.57E-05 | 0 | 2.28E-05 | 5.22E-05 | 2.89E-05 | 2.12E-05 | 1.99E-05 | 3.46E-05 | 0 | 9.88E-05 | 2.41E-05 | 0 | 0 |
| Ppfibp2 | 7.24E-06 | 7.22E-05 | 0 | 0 | 5.09E-06 | 0 | 0 | 0.00013 | 5.22E-05 | 2.05E-06 | 0 | 0 | 0 | 1.67E-06 | 2.99E-05 | 0 | 0 | 2.36E-05 |
| Map3k2 | 7.55E-05 | 0.00015 | 0.00012 | 0 | 0.00012 | 0.00013 | 7.39E-05 | 0.00014 | 5.21E-05 | 7.07E-05 | 2.23E-05 | 0.00016 | 1.91E-05 | 0.00012 | 0.0001 | 9.06E-05 | 0.0003 | 9.03E-05 |
| Znrd1 | 0.0002 | 5.95E-05 | 7.53E-05 | 7.12E-05 | 6.75E-05 | 8.99E-05 | 0.00012 | 0 | 5.21E-05 | 3.33E-05 | 0 | 0 | 3.09E-05 | 1.71E-05 | 3.32E-05 | 8.48E-05 | 6.66E-05 | 3.91E-05 |
| Rsf1 | 8.38E-05 | 0.0001 | 1.71E-05 | 1.76E-06 | 6.22E-05 | 8.94E-05 | 7.42E-05 | 6.45E-05 | 5.21E-05 | 3.34E-05 | 4.14E-05 | 9.60E-05 | 9.93E-06 | 4.99E-05 | 0.00015 | 2.39E-05 | 0.00015 | 2.73E-05 |
| Cep131 | 4.79E-06 | 3.91E-05 | 0 | 3.26E-05 | 1.54E-05 | 2.62E-06 | 0 | 0 | 5.21E-05 | 1.63E-05 | 0 | 1.59E-05 | 2.77E-05 | 5.32E-05 | 5.21E-05 | 3.33E-05 | 8.33E-06 | 1.97E-05 |
| Psmd8 | 0.00025 | 0.0002 | 0.00025 | 0.00014 | 0.00013 | 0.00017 | 8.70E-05 | 7.33E-05 | 5.20E-05 | 0.00012 | 0.00041 | 0.00017 | 1.81E-05 | 0.00019 | 0 | 2.32E-05 | 0.00012 | 0 |
| Ccdc90b | 0.00019 | 7.61E-05 | 2.24E-05 | 4.08E-05 | 2.37E-05 | 0.00012 | 5.21E-05 | 3.12E-05 | 5.20E-05 | 5.12E-05 | 2.33E-05 | 6.55E-05 | 3.73E-05 | 1.30E-05 | 0 | 4.15E-05 | 0 | 7.57E-05 |
| Vma21 | 0.00017 | 1.36E-05 | 2.20E-05 | 0 | 7.82E-06 | 0.00012 | 0 | 0 | 5.20E-05 | 7.54E-06 | 0 | 5.79E-05 | 0 | 0 | 0 | 3.27E-05 | 4.48E-05 | 1.64E-05 |
| Mndal | 7.76E-05 | 5.34E-05 | 0 | 0 | 0 | 7.76E-05 | 0 | 7.38E-06 | 5.20E-05 | 0.00021 | 0 | 3.68E-05 | 4.75E-05 | 0 | 0 | 0 | 0 | 9.70E-05 |
| Gramd1a | 7.98E-05 | 0.0002 | 7.25E-05 | 0 | 2.84E-05 | 4.83E-05 | 6.89E-05 | 0.00022 | 5.20E-05 | 1.49E-05 | 0 | 0.00019 | 3.83E-05 | 0.00017 | 0.00033 | 5.28E-05 | 0.00033 | 9.32E-05 |
| Hhat | 4.57E-05 | 4.73E-05 | 3.35E-05 | 9.83E-05 | 1.56E-05 | 1.59E-05 | 4.11E-05 | 0.00015 | 5.20E-05 | 0 | 5.95E-05 | 2.79E-05 | 0 | 1.81E-05 | 9.55E-05 | 2.38E-05 | 0 | 0.00011 |
| Kpna2 | 0 | 4.69E-05 | 3.81E-06 | 0 | 1.44E-05 | 0 | 0 | 0 | 5.20E-05 | 5.40E-05 | 4.96E-05 | 0 | 1.21E-05 | 7.96E-06 | 1.66E-05 | 0.00087 | 0 | 1.08E-05 |
| Slc12a2 | 2.87E-05 | 7.67E-05 | 3.34E-05 | 0 | 0 | 0 | 5.84E-06 | 0 | 5.20E-05 | 1.01E-05 | 5.22E-05 | 8.33E-06 | 0 | 6.75E-05 | 2.26E-05 | 3.77E-05 | 7.32E-05 | 1.15E-05 |
| Pnpla6 | 1.33E-05 | 9.27E-05 | 0 | 4.78E-05 | 3.57E-05 | 0 | 3.28E-05 | 1.60E-05 | 5.20E-05 | 4.22E-05 | 2.89E-05 | 4.50E-05 | 8.53E-06 | 1.28E-05 | 0.0002 | 0 | 0 | 6.37E-05 |
| Maf1 | 0.00013 | 0.00021 | 6.38E-06 | 2.13E-05 | 6.26E-05 | 9.01E-05 | 3.66E-05 | 0 | 5.19E-05 | 2.63E-05 | 0 | 9.66E-05 | 0 | 6.19E-05 | 7.09E-05 | 3.90E-05 | 0 | 2.31E-05 |
| Gnl2 | 0.00012 | 0.0001 | 0.00013 | 0 | 0 | 5.39E-05 | 0 | 0 | 5.19E-05 | 0 | 0 | 3.60E-05 | 3.49E-05 | 1.70E-06 | 0.00012 | 4.45E-05 | 9.74E-05 | 0 |
| Sec61a2 | 0.00011 | 7.63E-05 | 9.25E-05 | 3.94E-05 | 5.38E-05 | 4.70E-05 | 0 | 3.65E-05 | 5.19E-05 | 1.34E-05 | 0 | 6.64E-05 | 8.87E-06 | 3.32E-05 | 0.00015 | 4.36E-05 | 0 | 2.87E-05 |
| Lrrc49 | 3.64E-05 | 6.36E-05 | 1.11E-06 | 0 | 6.73E-06 | 1.16E-05 | 3.55E-05 | 0 | 5.19E-05 | 0.00011 | 0 | 8.75E-05 | 0 | 4.53E-05 | 2.58E-05 | 6.25E-05 | 3.68E-05 | 2.72E-05 |
| Dph7 | 4.00E-05 | 2.52E-05 | 2.54E-05 | 1.15E-05 | 2.44E-05 | 7.02E-06 | 0 | 1.94E-05 | 5.19E-05 | 1.07E-05 | 0 | 2.70E-05 | 1.09E-05 | 7.69E-06 | 8.33E-06 | 1.59E-05 | 0 | 1.69E-05 |
| Mfap3 | 0.00015 | 8.24E-05 | 8.30E-05 | 0.00012 | 8.91E-05 | 9.83E-05 | 7.15E-05 | 6.62E-05 | 5.18E-05 | 5.10E-05 | 5.25E-05 | 0.00011 | 4.40E-05 | 9.87E-05 | 0 | 6.24E-05 | 0 | 6.86E-05 |
| Ambra1 | 0.00012 | 0.00017 | 9.07E-05 | 2.07E-05 | 6.36E-05 | 6.44E-05 | 4.57E-05 | 4.14E-05 | 5.18E-05 | 2.46E-05 | 3.69E-05 | 4.29E-05 | 0 | 3.95E-05 | 0.00017 | 4.77E-06 | 0.00011 | 1.81E-05 |
| Arsk | 6.15E-05 | 7.04E-05 | 0 | 0 | 1.96E-05 | 5.80E-05 | 0 | 4.29E-05 | 5.18E-05 | 1.30E-05 | 0 | 2.28E-05 | 0 | 3.36E-07 | 0 | 0 | 0 | 5.31E-05 |
| Nsf | 0.00015 | 4.76E-05 | 0.00016 | 6.54E-05 | 6.99E-05 | 4.49E-05 | 0.0001 | 0 | 5.18E-05 | 4.11E-05 | 9.01E-05 | 6.43E-06 | 2.46E-05 | 6.13E-05 | 8.17E-05 | 6.20E-05 | 0 | 0 |
| Fignl2 | 0 | 0.00011 | 0 | 0.00013 | 3.20E-05 | 4.15E-05 | 0 | 3.18E-05 | 5.18E-05 | 3.07E-05 | 0 | 0.00011 | 6.69E-06 | 0 | 0.00024 | 0 | 0.00012 | 8.93E-05 |

| | | | | | | | | | | | | | | | | | | |
|---|---|---|---|---|---|---|---|---|---|---|---|---|---|---|---|---|---|---|
| Dido1 | 7.89E-05 | 0.0001 | 3.78E-05 | 4.96E-05 | 4.08E-05 | 8.06E-06 | 1.22E-05 | 0 | 5.18E-05 | 1.88E-05 | 2.51E-05 | 2.38E-05 | 0 | 3.39E-05 | 0.00031 | 4.14E-05 | 0 | 0 |
| Esf1 | 7.40E-05 | 4.56E-05 | 9.95E-05 | 6.04E-06 | 2.04E-05 | 4.48E-05 | 0 | 0 | 5.17E-05 | 0 | 4.83E-05 | 4.81E-05 | 0 | 3.01E-05 | 9.43E-05 | 7.33E-05 | 0 | 0 |
| Atg16l1 | 3.49E-05 | 0.0001 | 0.00014 | 9.01E-05 | 3.84E-05 | 0 | 3.43E-05 | 0 | 5.17E-05 | 2.45E-05 | 4.31E-05 | 5.30E-05 | 3.15E-06 | 3.69E-05 | 0.00014 | 9.09E-06 | 6.60E-05 | 3.48E-05 |
| Rnf40 | 7.31E-05 | 0.00015 | 8.85E-05 | 0.0002 | 6.92E-05 | 4.78E-05 | 3.82E-05 | 8.24E-05 | 5.16E-05 | 4.74E-05 | 2.10E-05 | 2.54E-05 | 2.68E-05 | 8.10E-05 | 0.00022 | 0.00012 | 0.00014 | 6.04E-05 |
| Brd3 | 6.66E-05 | 0.00017 | 9.87E-05 | 4.91E-05 | 7.63E-05 | 3.11E-05 | 3.17E-05 | 8.20E-05 | 5.16E-05 | 3.07E-05 | 7.48E-05 | 1.06E-05 | 4.78E-05 | 5.68E-05 | 0.00011 | 0.0002 | 0 | 2.41E-05 |
| Pptc7 | 2.13E-05 | 7.01E-05 | 5.75E-05 | 7.40E-07 | 5.56E-05 | 4.44E-05 | 1.38E-05 | 2.78E-05 | 5.15E-05 | 4.70E-05 | 4.57E-05 | 2.25E-05 | 2.30E-05 | 4.12E-05 | 2.43E-05 | 3.58E-06 | 0 | 3.67E-06 |
| Hoxc5 | 1.21E-05 | 4.58E-05 | 0 | 4.54E-05 | 7.01E-05 | 1.30E-05 | 9.20E-06 | 0 | 5.15E-05 | 3.19E-05 | 7.83E-05 | 0 | 0 | 2.27E-05 | 7.88E-05 | 1.69E-05 | 0 | 5.75E-05 |
| Dazap2 | 0.00019 | 0.00017 | 3.65E-05 | 9.25E-05 | 9.80E-05 | 0.00017 | 8.47E-05 | 5.71E-05 | 5.14E-05 | 7.02E-05 | 0.0001 | 0.00017 | 6.20E-05 | 7.99E-05 | 0 | 3.27E-05 | 0 | 8.21E-05 |
| Taz | 0.00011 | 0.00016 | 4.01E-05 | 1.91E-05 | 9.58E-05 | 6.97E-05 | 4.49E-05 | 2.29E-05 | 5.14E-05 | 7.62E-05 | 4.00E-06 | 6.82E-05 | 0 | 1.57E-05 | 0.00025 | 7.82E-08 | 0 | 0.0001 |
| Sik2 | 5.90E-05 | 0.00012 | 4.37E-05 | 1.81E-05 | 5.17E-05 | 5.76E-05 | 1.57E-05 | 4.18E-05 | 5.14E-05 | 9.69E-05 | 0 | 0.00012 | 5.72E-05 | 4.17E-05 | 2.87E-05 | 0.00011 | 0.00012 | 6.68E-05 |
| Btaf1 | 8.11E-05 | 7.09E-05 | 0.00012 | 8.03E-06 | 1.10E-05 | 5.39E-05 | 0 | 1.08E-05 | 5.14E-05 | 2.79E-05 | 3.97E-05 | 3.37E-05 | 5.47E-05 | 2.91E-05 | 0.00011 | 5.71E-05 | 9.37E-05 | 0 |
| Gdap2 | 8.05E-05 | 8.89E-05 | 7.22E-05 | 0 | 8.86E-05 | 4.31E-05 | 0.00011 | 0 | 5.14E-05 | 6.49E-05 | 9.53E-05 | 9.59E-06 | 1.15E-05 | 8.33E-06 | 5.50E-05 | 6.75E-07 | 4.09E-05 | 5.71E-05 |
| Ttc4 | 9.71E-05 | 9.10E-05 | 6.40E-05 | 6.09E-05 | 4.07E-05 | 3.32E-05 | 0 | 0 | 5.14E-05 | 1.46E-05 | 6.26E-06 | 2.85E-05 | 1.92E-06 | 9.84E-06 | 0.00017 | 0.00011 | 0 | 2.72E-05 |
| Nup37 | 8.20E-05 | 3.97E-05 | 3.83E-05 | 5.15E-05 | 1.69E-05 | 1.13E-05 | 2.73E-05 | 0 | 5.14E-05 | 1.48E-05 | 0 | 1.36E-05 | 5.50E-05 | 0 | 2.86E-05 | 0.00042 | 0 | 2.45E-05 |
| Tmem203 | 3.22E-05 | 9.65E-05 | 1.60E-05 | 0.00015 | 6.59E-05 | 9.50E-05 | 4.17E-05 | 8.70E-05 | 5.13E-05 | 5.26E-05 | 8.08E-05 | 4.00E-05 | 5.79E-05 | 7.46E-05 | 4.32E-05 | 1.10E-06 | 0 | 7.91E-05 |
| Polr3h | 9.52E-05 | 5.72E-05 | 0.00019 | 8.98E-05 | 2.92E-05 | 2.48E-05 | 0 | 7.39E-06 | 5.13E-05 | 1.47E-05 | 2.51E-05 | 8.33E-06 | 7.14E-06 | 3.66E-05 | 4.38E-05 | 5.50E-05 | 0 | 5.02E-07 |
| Aste1 | 3.47E-05 | 3.14E-05 | 2.49E-05 | 0 | 6.81E-06 | 2.11E-05 | 0 | 1.53E-05 | 5.13E-05 | 2.28E-05 | 2.71E-05 | 1.07E-06 | 0 | 2.02E-05 | 5.55E-05 | 3.46E-05 | 0 | 0 |
| Gps2 | 0.00015 | 0.0001 | 5.69E-05 | 0 | 0 | 1.57E-05 | 0 | 0 | 5.13E-05 | 0 | 0 | 2.66E-05 | 0 | 2.17E-05 | 0.00016 | 0.00011 | 3.00E-06 | 1.12E-05 |
| Cwc22 | 6.96E-05 | 7.69E-05 | 5.70E-05 | 7.91E-05 | 1.13E-05 | 1.23E-05 | 5.09E-06 | 0 | 5.13E-05 | 1.14E-05 | 2.31E-05 | 7.31E-06 | 2.09E-07 | 0 | 0.00016 | 5.42E-05 | 0 | 1.69E-05 |
| Aen | 3.26E-05 | 0.0002 | 0.00029 | 0.00015 | 8.85E-05 | 0 | 0 | 0 | 5.13E-05 | 2.50E-05 | 5.75E-05 | 0 | 5.89E-05 | 2.47E-05 | 0.00014 | 2.83E-05 | 0 | 2.38E-06 |
| Eogt | 3.29E-05 | 1.81E-05 | 3.76E-05 | 4.49E-06 | 3.62E-06 | 0 | 4.00E-06 | 0 | 5.13E-05 | 3.10E-05 | 0.0002 | 0 | 0 | 4.36E-05 | 0 | 5.80E-05 | 0 | 5.53E-05 |
| Mipep | 4.76E-05 | 3.50E-05 | 4.71E-05 | 3.50E-05 | 1.59E-05 | 0 | 1.05E-05 | 1.81E-05 | 5.13E-05 | 2.78E-05 | 2.89E-06 | 1.95E-05 | 0 | 0 | 0.00012 | 8.30E-05 | 3.05E-05 | 1.03E-05 |
| Tomm34 | 0.00015 | 0.00014 | 0.00016 | 6.38E-05 | 5.83E-05 | 6.13E-05 | 1.75E-05 | 3.67E-05 | 5.12E-05 | 5.27E-05 | 2.55E-05 | 6.75E-05 | 0 | 0.0001 | 6.21E-05 | 8.18E-05 | 8.32E-05 | 9.54E-05 |
| Map3k5 | 1.16E-05 | 9.69E-06 | 4.20E-05 | 0 | 0 | 4.07E-05 | 0 | 4.48E-05 | 5.12E-05 | 5.52E-05 | 0 | 0 | 7.46E-05 | 3.61E-05 | 1.11E-05 | 8.45E-05 | 4.96E-05 | 4.89E-05 |
| Bckdhb | 1.82E-05 | 3.44E-05 | 0 | 4.94E-05 | 2.09E-05 | 2.47E-05 | 8.91E-05 | 2.28E-05 | 5.12E-05 | 9.14E-05 | 0 | 0 | 6.55E-06 | 0 | 2.33E-05 | 1.67E-05 | 5.01E-05 | 0.00019 |
| Bola3 | 0.00017 | 5.92E-06 | 0.00014 | 0.0001 | 1.21E-05 | 2.21E-05 | 1.04E-06 | 0 | 5.12E-05 | 3.79E-05 | 5.10E-05 | 0 | 6.25E-05 | 0 | 2.81E-05 | 0.0001 | 3.75E-05 | 3.45E-05 |
| Cyth2 | 6.32E-05 | 0.00028 | 0.0002 | 0.00024 | 0.00012 | 4.88E-05 | 0 | 0.00015 | 5.11E-05 | 5.20E-05 | 0.00016 | 0.00014 | 0 | 0.00024 | 0.0003 | 0.00025 | 0.00026 | 8.90E-06 |
| Ddt | 0.00011 | 7.62E-05 | 0 | 8.99E-05 | 3.80E-05 | 3.96E-05 | 3.86E-05 | 0 | 5.11E-05 | 9.24E-05 | 0 | 3.00E-05 | 0.0001 | 0 | 0 | 0 | 4.10E-05 | 0 |
| Paf1 | 0.00015 | 9.98E-05 | 4.87E-05 | 7.32E-06 | 6.42E-05 | 3.72E-05 | 0 | 1.72E-06 | 5.11E-05 | 3.41E-05 | 7.17E-05 | 6.78E-05 | 7.43E-07 | 2.93E-05 | 0.00012 | 0.00011 | 4.27E-05 | 3.78E-05 |
| Fbrsl1 | 2.12E-05 | 9.68E-05 | 5.28E-05 | 3.74E-05 | 5.19E-05 | 3.11E-05 | 0 | 1.31E-05 | 5.11E-05 | 6.36E-06 | 2.85E-05 | 2.52E-05 | 7.04E-06 | 0.0001 | 0.00018 | 1.38E-05 | 0.00018 | 4.38E-05 |
| Fbxo10 | 2.18E-05 | 7.78E-05 | 0 | 4.52E-05 | 5.63E-05 | 2.92E-05 | 0.00012 | 4.00E-05 | 5.11E-05 | 4.40E-05 | 0 | 0.00016 | 1.68E-05 | 6.62E-05 | 7.55E-05 | 1.58E-05 | 0 | 4.72E-05 |
| Cryzl2 | 3.56E-06 | 1.28E-05 | 0 | 5.67E-05 | 7.01E-05 | 2.13E-05 | 0.00011 | 6.63E-05 | 5.11E-05 | 3.47E-05 | 3.02E-06 | 1.97E-06 | 5.87E-05 | 1.33E-05 | 0 | 0 | 0 | 7.69E-05 |
| Armc5 | 2.50E-06 | 6.52E-05 | 3.45E-05 | 0.00013 | 3.55E-05 | 0 | 3.09E-06 | 3.43E-05 | 5.11E-05 | 2.11E-05 | 3.23E-05 | 1.86E-05 | 3.95E-08 | 1.41E-05 | 8.82E-05 | 1.75E-05 | 0 | 4.98E-05 |

| | | | | | | | | | | | | | | | | | | |
|---|---|---|---|---|---|---|---|---|---|---|---|---|---|---|---|---|---|---|
| Spindoc | 1.48E-05 | 9.79E-05 | 9.71E-05 | 0.0002 | 4.20E-05 | 0 | 0 | 7.05E-05 | 5.11E-05 | 2.46E-05 | 0.00012 | 6.71E-05 | 0 | 6.42E-05 | 0.00036 | 5.39E-05 | 0.00034 | 1.97E-05 |
| Cct4 | 0.00034 | 0.00013 | 0.0003 | 3.18E-05 | 2.46E-05 | 0.00016 | 0 | 0 | 5.10E-05 | 4.40E-05 | 6.61E-05 | 0.00015 | 4.30E-06 | 1.89E-05 | 4.27E-06 | 0.00014 | 0.00014 | 5.18E-05 |
| DEPRECATED | 0.00013 | 6.52E-05 | 1.91E-05 | 2.75E-05 | 5.75E-05 | 8.39E-05 | 7.93E-05 | 6.63E-05 | 5.10E-05 | 5.09E-05 | 2.95E-05 | 3.85E-05 | 0 | 3.35E-05 | 0 | 1.55E-05 | 8.35E-05 | 2.20E-05 |
| Mterf3 | 0.00012 | 9.32E-05 | 7.41E-05 | 0 | 3.26E-05 | 6.27E-05 | 0 | 0 | 5.10E-05 | 1.29E-05 | 2.14E-05 | 1.04E-05 | 2.65E-05 | 0 | 0.00012 | 4.05E-05 | 0 | 0 |
| Lrfn3 | 3.33E-06 | 2.29E-05 | 2.14E-05 | 3.42E-05 | 6.52E-05 | 2.97E-05 | 2.70E-05 | 4.61E-05 | 5.10E-05 | 2.23E-05 | 3.85E-06 | 1.57E-05 | 3.22E-05 | 6.34E-05 | 4.19E-06 | 0 | 0 | 2.29E-05 |
| Mettl3 | 5.70E-05 | 8.90E-05 | 7.06E-05 | 0.00011 | 1.04E-05 | 2.75E-05 | 0 | 3.16E-05 | 5.10E-05 | 3.18E-05 | 3.73E-05 | 2.88E-05 | 1.33E-05 | 1.47E-06 | 0.00013 | 4.24E-05 | 0 | 3.54E-05 |
| Fars2 | 3.94E-05 | 9.05E-05 | 1.63E-05 | 6.06E-05 | 3.27E-05 | 2.49E-05 | 4.12E-05 | 3.26E-05 | 5.10E-05 | 6.17E-05 | 5.40E-06 | 0 | 6.12E-05 | 2.12E-05 | 0.00017 | 4.50E-05 | 6.23E-05 | 4.80E-05 |
| Lyrm2 | 0.00012 | 4.82E-05 | 9.03E-06 | 2.03E-05 | 6.99E-06 | 1.98E-05 | 1.64E-05 | 0 | 5.10E-05 | 2.70E-05 | 2.57E-05 | 4.32E-05 | 1.61E-05 | 4.00E-06 | 0 | 2.15E-05 | 0 | 4.29E-05 |
| Rpain | 1.86E-05 | 9.36E-05 | 0 | 4.81E-05 | 3.10E-05 | 1.06E-05 | 3.84E-05 | 1.50E-05 | 5.10E-05 | 8.16E-05 | 2.05E-05 | 1.50E-05 | 1.22E-05 | 4.16E-06 | 4.45E-05 | 4.64E-05 | 6.23E-07 | 2.43E-05 |
| Uhrf1bp1 | 2.45E-05 | 4.47E-05 | 5.25E-05 | 5.32E-05 | 2.00E-05 | 9.92E-06 | 1.55E-05 | 0 | 5.10E-05 | 2.52E-05 | 0 | 2.82E-05 | 3.67E-05 | 8.98E-06 | 0.00011 | 0 | 0 | 3.32E-05 |
| Pus7l | 5.22E-05 | 1.67E-05 | 6.64E-05 | 8.38E-06 | 2.71E-06 | 6.65E-06 | 1.02E-05 | 1.19E-05 | 5.10E-05 | 1.89E-05 | 0 | 0 | 0 | 0 | 2.06E-05 | 4.91E-06 | 0 | 0 |
| Fbxo28 | 8.98E-05 | 0.00011 | 0.00011 | 6.77E-05 | 4.01E-05 | 0 | 5.21E-05 | 1.17E-05 | 5.10E-05 | 6.08E-05 | 4.09E-05 | 6.87E-05 | 0 | 6.15E-05 | 0.00012 | 7.03E-05 | 0 | 2.17E-05 |
| Creb3 | 0.00017 | 6.04E-05 | 0.00019 | 0.00022 | 0.00013 | 0.00013 | 0.00038 | 0.00013 | 5.09E-05 | 0.00017 | 0.00028 | 8.94E-05 | 6.93E-05 | 0.00016 | 0.00011 | 0 | 0 | 0.00013 |
| Dtx4 | 4.52E-08 | 0.00031 | 0 | 0 | 0.00013 | 0.00012 | 0 | 0.00063 | 5.09E-05 | 0.00019 | 0 | 0 | 0 | 0.00014 | 8.39E-06 | 1.55E-05 | 0 | 0.00034 |
| Nmnat3 | 2.01E-06 | 1.80E-05 | 0 | 1.49E-05 | 0.00015 | 9.13E-05 | 0 | 1.63E-05 | 5.09E-05 | 1.80E-05 | 9.44E-05 | 1.58E-05 | 8.30E-05 | 0.0001 | 9.98E-05 | 0 | 0 | 1.25E-05 |
| Fam110b | 3.56E-05 | 3.88E-05 | 7.26E-06 | 0 | 2.67E-05 | 7.07E-05 | 0 | 0.00025 | 5.09E-05 | 0 | 0 | 0 | 0 | 0 | 0 | 2.30E-05 | 0.00012 | 8.74E-05 |
| Stx17 | 0.00011 | 0.00011 | 1.76E-05 | 3.62E-05 | 7.45E-05 | 7.02E-05 | 3.52E-05 | 1.97E-05 | 5.09E-05 | 5.10E-05 | 8.02E-07 | 1.59E-05 | 3.97E-05 | 7.32E-05 | 8.53E-05 | 7.06E-05 | 0 | 3.91E-05 |
| Rbbp5 | 8.55E-05 | 0.00011 | 5.49E-05 | 2.52E-05 | 4.75E-05 | 3.83E-05 | 0 | 0 | 5.09E-05 | 2.17E-05 | 3.42E-05 | 5.72E-05 | 0 | 1.38E-05 | 0.00013 | 0.00012 | 0 | 3.76E-05 |
| Crlf3 | 9.77E-06 | 7.21E-05 | 9.09E-05 | 2.48E-05 | 3.79E-05 | 1.61E-05 | 0 | 7.46E-05 | 5.09E-05 | 4.33E-05 | 8.10E-05 | 5.12E-05 | 0 | 3.55E-05 | 0 | 0.00011 | 4.80E-05 | 5.96E-05 |
| Nampt | 0.00012 | 6.95E-05 | 0 | 0 | 0 | 0 | 0 | 0 | 5.09E-05 | 9.42E-05 | 0 | 0.00041 | 6.86E-05 | 0 | 0 | 9.22E-05 | 9.46E-05 | 2.01E-06 |
| Rchy1 | 0.00026 | 0.00014 | 4.68E-05 | 0 | 3.77E-05 | 0.00017 | 8.20E-05 | 7.16E-05 | 5.08E-05 | 1.59E-05 | 0 | 8.15E-05 | 1.82E-06 | 0 | 4.34E-05 | 2.60E-05 | 0 | 2.44E-05 |
| Map2k4 | 0.00012 | 3.91E-05 | 4.36E-05 | 0 | 3.58E-05 | 0.00014 | 6.43E-06 | 7.65E-05 | 5.08E-05 | 0 | 8.60E-05 | 0.00012 | 8.02E-07 | 9.88E-06 | 1.18E-05 | 1.98E-05 | 1.94E-05 | 1.84E-05 |
| Traf5 | 0 | 3.38E-05 | 1.66E-05 | 9.66E-06 | 3.83E-05 | 2.38E-05 | 0 | 4.17E-05 | 5.08E-05 | 7.84E-05 | 8.05E-05 | 5.55E-06 | 0 | 4.39E-05 | 6.04E-05 | 0 | 5.65E-05 | 4.09E-05 |
| Heatr1 | 9.06E-05 | 3.53E-05 | 0.00019 | 1.06E-06 | 0 | 2.14E-05 | 0 | 0 | 5.08E-05 | 1.06E-05 | 8.67E-07 | 0 | 1.52E-05 | 3.24E-06 | 1.23E-05 | 0 | 6.97E-06 | 3.69E-07 |
| Vcpkmt | 5.37E-05 | 3.99E-05 | 9.87E-05 | 5.10E-05 | 2.91E-05 | 1.88E-05 | 0 | 2.99E-05 | 5.08E-05 | 1.45E-05 | 1.74E-05 | 4.12E-05 | 0 | 0 | 2.12E-05 | 4.30E-05 | 2.99E-05 | 1.47E-07 |
| Gatc | 4.62E-05 | 3.78E-05 | 2.79E-05 | 5.27E-05 | 1.30E-05 | 1.23E-05 | 0 | 0 | 5.08E-05 | 4.14E-05 | 0 | 0 | 4.81E-06 | 2.87E-05 | 7.57E-06 | 9.96E-05 | 0 | 2.62E-05 |
| Nme4 | 2.48E-05 | 2.78E-05 | 0 | 2.83E-05 | 2.55E-06 | 0 | 0 | 0 | 5.08E-05 | 0 | 0 | 2.08E-05 | 1.32E-05 | 3.51E-05 | 8.01E-06 | 4.93E-06 | 3.52E-05 | 6.07E-06 |
| Dgkd | 8.07E-06 | 0.00015 | 6.72E-05 | 0.00011 | 5.09E-05 | 0 | 0 | 0 | 5.08E-05 | 0 | 3.86E-05 | 1.15E-05 | 0 | 4.34E-05 | 9.12E-05 | 4.56E-05 | 0.00013 | 0 |
| Ntan1 | 0.00018 | 7.26E-05 | 2.82E-05 | 0 | 1.35E-05 | 6.92E-05 | 3.24E-05 | 4.76E-05 | 5.07E-05 | 2.98E-05 | 7.32E-05 | 0.00017 | 0 | 7.19E-06 | 0 | 4.29E-05 | 0 | 0.00011 |
| Ankle2 | 9.86E-05 | 0.00012 | 7.66E-05 | 1.15E-05 | 3.98E-05 | 4.35E-05 | 3.93E-05 | 4.56E-06 | 5.07E-05 | 2.76E-05 | 0 | 6.86E-05 | 0 | 3.74E-05 | 5.32E-05 | 0.00011 | 0.00016 | 3.19E-05 |
| Kctd3 | 9.55E-05 | 8.66E-05 | 0.00011 | 4.91E-05 | 3.04E-05 | 3.01E-05 | 3.64E-05 | 0 | 5.07E-05 | 1.18E-05 | 8.09E-05 | 3.68E-05 | 1.15E-05 | 2.22E-05 | 8.75E-05 | 5.40E-05 | 1.42E-05 | 1.03E-05 |
| Tm2d1 | 8.75E-05 | 4.79E-05 | 2.57E-05 | 1.98E-05 | 5.04E-05 | 8.11E-05 | 1.34E-05 | 4.43E-05 | 5.06E-05 | 3.31E-05 | 2.34E-05 | 6.26E-05 | 3.93E-05 | 3.75E-05 | 0 | 3.45E-05 | 0 | 7.26E-06 |
| Ckap5 | 0.00016 | 0.00011 | 1.82E-05 | 0 | 3.14E-05 | 7.87E-05 | 3.95E-05 | 2.14E-05 | 5.06E-05 | 3.19E-05 | 2.04E-05 | 0.00027 | 1.57E-05 | 0 | 8.70E-05 | 0.00078 | 2.14E-05 | 2.27E-05 |

| | | | | | | | | | | | | | | | | | | |
|---|---|---|---|---|---|---|---|---|---|---|---|---|---|---|---|---|---|---|
| Lactb | 3.91E-05 | 3.12E-05 | 1.42E-06 | 3.47E-05 | 0.0001 | 7.04E-05 | 9.05E-06 | 0 | 5.06E-05 | 1.40E-05 | 0.00013 | 0 | 2.71E-05 | 3.01E-06 | 1.87E-05 | 9.66E-06 | 0 | 4.51E-05 |
| Def6 | 6.93E-06 | 7.43E-05 | 3.36E-05 | 0.00011 | 9.42E-05 | 1.79E-05 | 1.39E-05 | 3.39E-06 | 5.06E-05 | 3.38E-05 | 1.81E-06 | 3.31E-05 | 0 | 0.00011 | 3.09E-05 | 0 | 4.66E-06 | 3.44E-05 |
| Ints10 | 6.46E-05 | 3.96E-05 | 7.94E-05 | 7.29E-05 | 2.17E-05 | 3.09E-06 | 0 | 1.62E-05 | 5.06E-05 | 3.31E-05 | 1.56E-05 | 0 | 1.00E-06 | 3.11E-05 | 1.62E-05 | 0.00012 | 2.91E-05 | 4.26E-05 |
| Rab23 | 0.00012 | 5.16E-05 | 7.57E-05 | 0 | 1.75E-05 | 0 | 0.0002 | 0 | 5.06E-05 | 0.00027 | 0.00012 | 0 | 0.00024 | 0 | 0 | 0.0002 | 0 | 7.40E-05 |
| Prepl | 5.56E-05 | 4.80E-05 | 2.67E-05 | 5.81E-05 | 1.00E-05 | 0 | 1.86E-05 | 2.58E-05 | 5.06E-05 | 1.38E-05 | 7.77E-05 | 7.32E-06 | 1.46E-05 | 0 | 5.88E-06 | 9.22E-06 | 2.33E-06 | 1.10E-05 |
| Diaph1 | 5.58E-05 | 0.00012 | 0.00011 | 2.81E-05 | 9.97E-06 | 0 | 0 | 2.40E-05 | 5.06E-05 | 0.00021 | 9.04E-05 | 3.79E-05 | 0 | 3.47E-05 | 4.65E-05 | 0.00014 | 7.84E-05 | 0.0001 |
| Ccar1 | 0.00027 | 0.0002 | 0.00017 | 4.50E-06 | 4.74E-05 | 9.25E-05 | 2.09E-05 | 0 | 5.05E-05 | 2.08E-05 | 5.22E-05 | 2.95E-05 | 0 | 4.44E-05 | 0.00049 | 0.00019 | 3.57E-05 | 5.19E-05 |
| Mef2a | 4.77E-05 | 0.00024 | 9.29E-05 | 0 | 0.00027 | 9.07E-05 | 4.34E-06 | 0.00016 | 5.05E-05 | 8.27E-05 | 0.00017 | 5.78E-05 | 0 | 0.00021 | 1.06E-06 | 6.00E-05 | 0.00033 | 9.73E-05 |
| Lsm10 | 8.67E-05 | 0.00012 | 3.84E-05 | 0.00011 | 5.27E-05 | 6.20E-05 | 2.80E-05 | 0 | 5.05E-05 | 6.34E-05 | 2.52E-05 | 5.10E-05 | 0 | 4.36E-05 | 0 | 1.27E-06 | 0 | 4.97E-05 |
| Iscu | 7.20E-05 | 7.81E-05 | 4.33E-05 | 7.68E-05 | 7.76E-05 | 5.36E-05 | 0 | 3.28E-05 | 5.05E-05 | 4.87E-05 | 7.81E-05 | 0 | 5.91E-05 | 4.24E-05 | 0 | 3.01E-05 | 0 | 5.00E-05 |
| Gtpbp8 | 4.99E-05 | 2.87E-05 | 0 | 1.22E-05 | 2.42E-05 | 2.82E-05 | 0 | 3.41E-05 | 5.05E-05 | 4.92E-05 | 7.68E-06 | 8.17E-06 | 0 | 2.90E-05 | 5.64E-05 | 5.60E-06 | 0 | 4.58E-05 |
| Ptrh1 | 1.28E-05 | 7.68E-05 | 0.0002 | 0.00019 | 6.46E-05 | 2.78E-05 | 0 | 0 | 5.05E-05 | 3.77E-05 | 0.00014 | 0.00042 | 2.68E-06 | 0.00014 | 0.00012 | 0 | 0 | 0.00011 |
| Ppp1r35 | 2.66E-06 | 7.08E-05 | 2.30E-05 | 0.00013 | 5.36E-05 | 0 | 0 | 4.07E-05 | 5.05E-05 | 4.77E-05 | 4.08E-05 | 1.60E-05 | 0 | 3.58E-05 | 0.00011 | 5.39E-05 | 0 | 4.96E-05 |
| Vps9d1 | 3.30E-05 | 3.93E-05 | 5.07E-05 | 4.80E-05 | 2.83E-05 | 3.25E-05 | 2.85E-05 | 1.67E-05 | 5.04E-05 | 3.39E-05 | 1.04E-05 | 1.72E-05 | 0 | 2.68E-05 | 7.71E-05 | 1.99E-05 | 1.21E-05 | 2.15E-05 |
| Elac2 | 8.24E-05 | 4.54E-05 | 0.00014 | 3.50E-05 | 1.05E-05 | 2.68E-05 | 0 | 0 | 5.04E-05 | 2.10E-05 | 2.83E-05 | 0 | 1.56E-05 | 0 | 1.45E-05 | 1.58E-05 | 0 | 2.31E-05 |
| B3gat3 | 9.29E-06 | 7.72E-05 | 7.74E-05 | 0.00023 | 5.26E-05 | 0 | 8.08E-05 | 0.00011 | 5.04E-05 | 5.94E-05 | 1.70E-05 | 2.08E-05 | 5.01E-06 | 4.25E-05 | 0.00017 | 1.18E-05 | 3.35E-05 | 2.08E-05 |
| Gpx7 | 0.00039 | 0 | 0.0003 | 0.00101 | 0 | 0.00013 | 0.00088 | 0.00012 | 5.03E-05 | 0.00015 | 9.95E-05 | 0 | 0.00046 | 0.00042 | 0 | 0.00036 | 5.33E-05 | 0.00043 |
| Zfp644 | 0.00012 | 9.14E-05 | 0 | 0 | 3.35E-05 | 0.00013 | 0 | 2.05E-05 | 5.03E-05 | 2.01E-05 | 2.04E-05 | 6.01E-05 | 0 | 3.23E-05 | 4.87E-05 | 5.59E-05 | 7.15E-05 | 0 |
| Lemd3 | 5.37E-05 | 0.00012 | 0.00011 | 6.01E-05 | 5.35E-05 | 0.00012 | 3.25E-05 | 4.16E-05 | 5.03E-05 | 3.02E-05 | 6.02E-05 | 8.52E-05 | 0 | 0.00016 | 0.00011 | 8.32E-05 | 0.00017 | 8.33E-05 |
| DEPRECATED | 7.27E-05 | 0.00016 | 5.37E-05 | 4.10E-05 | 6.29E-05 | 0.00012 | 7.08E-05 | 4.24E-05 | 5.03E-05 | 4.89E-05 | 3.98E-05 | 4.28E-05 | 2.25E-05 | 4.19E-05 | 0 | 6.35E-05 | 0.00018 | 2.82E-05 |
| Ercc4 | 4.25E-05 | 3.48E-05 | 8.01E-06 | 3.62E-06 | 2.15E-05 | 2.23E-05 | 0 | 3.69E-05 | 5.03E-05 | 2.79E-05 | 2.46E-05 | 6.33E-06 | 2.39E-05 | 1.01E-05 | 6.62E-05 | 0 | 0 | 1.46E-06 |
| Rnf5 | 2.74E-05 | 9.42E-05 | 3.62E-05 | 0.00014 | 3.63E-05 | 1.82E-05 | 5.15E-05 | 7.82E-05 | 5.03E-05 | 3.10E-05 | 7.34E-05 | 7.39E-05 | 1.85E-06 | 1.39E-05 | 4.79E-05 | 0.00014 | 1.88E-05 | 1.75E-05 |
| Aven | 9.45E-05 | 3.30E-05 | 0.00012 | 3.13E-05 | 2.67E-05 | 2.32E-06 | 7.71E-06 | 7.65E-05 | 5.03E-05 | 2.95E-05 | 2.51E-05 | 0 | 0 | 3.77E-05 | 2.55E-05 | 8.03E-05 | 1.08E-05 | 6.67E-05 |
| Ankrd10 | 7.59E-05 | 0.00019 | 9.60E-05 | 0 | 7.58E-05 | 0 | 0 | 7.27E-05 | 5.03E-05 | 1.76E-05 | 4.42E-06 | 1.15E-05 | 3.02E-05 | 9.28E-05 | 0.00034 | 7.08E-05 | 9.03E-05 | 3.18E-05 |
| Rpa1 | 6.18E-05 | 2.21E-05 | 7.43E-05 | 2.49E-05 | 2.13E-05 | 4.06E-05 | 2.20E-06 | 1.09E-05 | 5.02E-05 | 5.58E-05 | 4.14E-05 | 0 | 2.59E-05 | 3.29E-05 | 0 | 0.0006 | 0 | 0.00012 |
| Slc35e3 | 3.85E-05 | 9.73E-05 | 3.83E-05 | 0 | 5.30E-05 | 2.05E-05 | 1.79E-05 | 0 | 5.02E-05 | 5.19E-05 | 1.49E-05 | 0 | 3.53E-06 | 3.51E-05 | 5.54E-05 | 6.02E-05 | 3.82E-05 | 4.49E-05 |
| Cox10 | 5.02E-05 | 4.13E-05 | 8.91E-05 | 3.11E-05 | 4.92E-05 | 0 | 0 | 6.65E-06 | 5.02E-05 | 2.35E-06 | 2.99E-05 | 0 | 4.76E-06 | 0 | 4.34E-05 | 8.20E-05 | 0 | 6.74E-06 |
| Thoc6 | 3.04E-05 | 0 | 0 | 0.00011 | 1.91E-05 | 0 | 0 | 2.62E-05 | 5.02E-05 | 2.44E-05 | 9.57E-05 | 0.00013 | 0 | 2.86E-05 | 0.0001 | 4.52E-05 | 0.00129 | 4.91E-05 |
| Pcmt1 | 0.00019 | 0.00014 | 0.00023 | 0 | 0.0001 | 0.00014 | 9.12E-05 | 6.52E-05 | 5.01E-05 | 7.02E-05 | 0.00011 | 0.00013 | 0 | 5.72E-06 | 0 | 0.00021 | 7.80E-05 | 9.14E-05 |
| Pomgnt1 | 5.52E-05 | 3.98E-05 | 0.00018 | 1.15E-05 | 5.66E-05 | 0.00011 | 9.68E-05 | 6.89E-05 | 5.01E-05 | 4.12E-05 | 0.00014 | 3.34E-05 | 6.34E-06 | 1.93E-05 | 4.30E-05 | 1.64E-05 | 0 | 6.64E-06 |
| Phactr2 | 8.35E-05 | 0.00013 | 0 | 0 | 5.50E-05 | 0.00011 | 4.45E-05 | 0 | 5.01E-05 | 0.00014 | 0 | 0 | 6.79E-05 | 2.23E-05 | 0 | 4.15E-05 | 0 | 4.48E-05 |
| H2afj | 0 | 7.63E-05 | 3.28E-05 | 0.00013 | 3.05E-05 | 5.81E-05 | 0 | 9.73E-05 | 5.01E-05 | 0 | 0 | 2.26E-06 | 0 | 4.08E-05 | 0.00012 | 0 | 0 | 1.23E-05 |
| Ahctf1 | 0.00012 | 0.00012 | 0.00015 | 5.17E-06 | 5.18E-05 | 5.26E-05 | 1.03E-05 | 1.95E-05 | 5.01E-05 | 4.23E-05 | 8.41E-05 | 6.57E-05 | 0 | 8.55E-05 | 0.00016 | 0.00025 | 9.64E-05 | 4.17E-06 |

| | | | | | | | | | | | | | | | | | | |
|---|---|---|---|---|---|---|---|---|---|---|---|---|---|---|---|---|---|---|
| Pim3 | 0 | 7.83E-05 | 0 | 5.56E-05 | 0 | 2.04E-05 | 0 | 0 | 5.01E-05 | 2.42E-05 | 8.60E-05 | 0.00023 | 0.00011 | 4.26E-05 | 0.00018 | 2.20E-06 | 0.00092 | 0 |
| Cpsf4 | 5.03E-05 | 6.61E-05 | 0.00012 | 3.28E-05 | 3.06E-05 | 1.83E-05 | 1.17E-05 | 1.66E-05 | 5.01E-05 | 1.72E-05 | 6.45E-05 | 2.39E-05 | 2.77E-05 | 4.88E-05 | 7.65E-05 | 0.00017 | 0 | 0 |
| Pdxdc1 | 0.0002 | 3.37E-05 | 8.68E-05 | 3.22E-05 | 3.52E-05 | 0.00011 | 0.00018 | 4.06E-05 | 5.00E-05 | 2.27E-05 | 2.67E-05 | 1.54E-05 | 2.41E-05 | 8.20E-05 | 1.70E-05 | 2.12E-06 | 0 | 3.74E-05 |
| Dnajb6 | 0.0002 | 0.00024 | 0.00014 | 0.00011 | 7.01E-05 | 9.69E-05 | 6.56E-05 | 4.98E-05 | 5.00E-05 | 2.71E-05 | 0.00012 | 0.00013 | 0.0001 | 7.72E-05 | 0.00013 | 0.00015 | 0.00012 | 1.61E-05 |
| Coq10b | 0.00016 | 8.95E-05 | 0.00019 | 0 | 8.47E-05 | 7.75E-05 | 0 | 5.41E-05 | 5.00E-05 | 0 | 0.00021 | 9.43E-06 | 0 | 2.42E-05 | 9.49E-05 | 4.78E-05 | 0.00052 | 0 |
| Gopc | 9.74E-05 | 6.50E-05 | 0.00014 | 9.85E-05 | 9.22E-05 | 5.54E-05 | 1.44E-05 | 3.29E-05 | 5.00E-05 | 3.31E-05 | 9.91E-05 | 0.00035 | 2.67E-05 | 0.00017 | 0.00013 | 3.56E-05 | 2.19E-05 | 7.84E-05 |
| Mindy2 | 0.00013 | 0.0001 | 7.87E-07 | 0 | 1.58E-05 | 3.78E-05 | 4.68E-05 | 2.55E-05 | 5.00E-05 | 0.00011 | 1.66E-05 | 0.00022 | 8.12E-05 | 0 | 2.13E-05 | 1.14E-06 | 0.00015 | 4.69E-06 |
| Sectm1a | 0 | 0 | 0 | 0 | 0 | 0 | 0 | 0 | 5.00E-05 | 3.62E-05 | 0 | 0 | 8.06E-05 | 0 | 0 | 0 | 0 | 8.74E-05 |
| Lgr4 | 0.00013 | 7.69E-05 | 1.27E-05 | 0 | 3.30E-07 | 0 | 1.96E-05 | 0 | 5.00E-05 | 5.31E-05 | 1.63E-05 | 0 | 0 | 6.78E-05 | 0 | 7.24E-05 | 2.61E-05 | 0.00024 |
| Ubl5 | 0.00025 | 0.00011 | 9.08E-05 | 0.00025 | 0.00012 | 0.00017 | 0.00017 | 6.47E-05 | 4.99E-05 | 7.35E-05 | 0.00015 | 4.39E-05 | 0.00011 | 0.00012 | 0 | 9.98E-05 | 1.20E-05 | 9.02E-05 |
| Gnptg | 0 | 0.00011 | 3.37E-05 | 8.80E-05 | 0.00011 | 9.93E-05 | 2.10E-05 | 7.59E-05 | 4.99E-05 | 0.00012 | 0 | 5.17E-05 | 0.00015 | 0.00014 | 0 | 3.68E-05 | 3.28E-06 | 0.00021 |
| Bdnf | 0 | 1.41E-06 | 0 | 0 | 8.97E-06 | 2.07E-05 | 0 | 0 | 4.99E-05 | 6.44E-06 | 2.23E-05 | 0 | 0.00012 | 3.59E-07 | 3.08E-06 | 0 | 6.50E-06 | 0.00012 |
| Utrn | 5.38E-05 | 0.00015 | 3.85E-05 | 8.67E-05 | 2.56E-05 | 0 | 0.00011 | 0 | 4.99E-05 | 0.00012 | 0.00024 | 0.00035 | 0 | 6.01E-06 | 0.00025 | 8.43E-05 | 0.00013 | 0.00013 |
| Tubgcp3 | 8.16E-05 | 9.76E-05 | 8.98E-05 | 0 | 3.14E-05 | 5.57E-05 | 0 | 0 | 4.98E-05 | 5.11E-05 | 1.64E-05 | 1.09E-05 | 2.97E-05 | 1.82E-05 | 0.00018 | 0.00026 | 6.79E-05 | 4.55E-05 |
| Ap3s1 | 5.50E-05 | 1.05E-05 | 0 | 0 | 0 | 4.65E-05 | 3.14E-06 | 3.27E-06 | 4.98E-05 | 5.27E-06 | 0 | 6.63E-05 | 4.28E-05 | 0 | 1.39E-05 | 1.03E-05 | 1.37E-05 | 0 |
| Casp2 | 3.51E-05 | 4.85E-05 | 4.24E-06 | 2.86E-05 | 3.65E-05 | 2.87E-05 | 1.10E-05 | 1.45E-05 | 4.98E-05 | 2.03E-05 | 2.65E-05 | 4.57E-05 | 4.52E-06 | 3.83E-05 | 0 | 7.05E-05 | 0 | 0 |
| Hmces | 3.77E-05 | 0.0001 | 7.51E-05 | 7.49E-05 | 8.61E-05 | 2.69E-05 | 1.57E-05 | 0 | 4.98E-05 | 1.29E-05 | 1.54E-05 | 2.10E-05 | 0 | 3.87E-05 | 0.00014 | 8.00E-05 | 3.85E-05 | 2.30E-05 |
| Prkrip1 | 3.00E-05 | 3.13E-05 | 7.20E-05 | 3.92E-05 | 2.24E-05 | 1.32E-05 | 0 | 5.82E-06 | 4.98E-05 | 2.25E-05 | 1.60E-05 | 5.02E-06 | 0 | 3.23E-05 | 6.89E-05 | 7.05E-05 | 5.58E-05 | 1.92E-05 |
| Toe1 | 5.52E-05 | 6.12E-05 | 4.73E-05 | 5.11E-05 | 3.32E-08 | 5.90E-06 | 2.84E-05 | 0 | 4.98E-05 | 1.77E-05 | 4.28E-05 | 1.79E-06 | 0 | 2.25E-05 | 3.91E-05 | 0.00011 | 0 | 1.17E-05 |
| Apex2 | 3.40E-05 | 3.52E-05 | 6.88E-05 | 0.00011 | 2.09E-05 | 0 | 7.10E-06 | 2.06E-05 | 4.98E-05 | 1.62E-05 | 8.79E-06 | 0 | 1.27E-05 | 4.82E-05 | 0.00023 | 7.97E-05 | 0 | 3.07E-05 |
| Mprip | 0.00023 | 0.00046 | 0.00035 | 0.00049 | 0.00051 | 0.00033 | 0.0003 | 0.00059 | 4.97E-05 | 0.00029 | 0.00089 | 0.00021 | 0.00055 | 0.00057 | 0.00077 | 0.00042 | 0.00054 | 0.00048 |
| Exoc5 | 0.00017 | 0.00018 | 0.00033 | 0.00016 | 0.00012 | 0.00014 | 9.25E-05 | 0.00013 | 4.97E-05 | 0.00013 | 0.00021 | 9.78E-05 | 5.03E-06 | 0.00014 | 7.23E-05 | 0.00014 | 0.00013 | 9.71E-05 |
| Eif5 | 0.00022 | 9.46E-05 | 0.00014 | 0 | 2.09E-05 | 7.23E-05 | 1.39E-05 | 0 | 4.97E-05 | 0 | 6.69E-05 | 0.00015 | 1.77E-05 | 0 | 0 | 8.18E-05 | 0.00012 | 0 |
| Ctbs | 9.01E-05 | 3.02E-05 | 0 | 0 | 1.34E-05 | 5.13E-05 | 0 | 1.07E-05 | 4.97E-05 | 7.82E-05 | 7.36E-06 | 2.38E-05 | 1.12E-05 | 2.75E-05 | 0 | 1.28E-05 | 0 | 1.31E-05 |
| Rtl8b | 6.35E-05 | 3.38E-05 | 3.95E-05 | 3.22E-05 | 5.34E-05 | 4.66E-05 | 2.06E-05 | 3.26E-05 | 4.97E-05 | 2.29E-05 | 2.68E-05 | 2.26E-05 | 3.88E-06 | 2.17E-06 | 0 | 0 | 2.19E-05 | 3.91E-05 |
| Cnot11 | 8.47E-05 | 0.00011 | 0.00012 | 4.56E-05 | 5.26E-05 | 4.38E-06 | 7.06E-05 | 5.00E-05 | 4.97E-05 | 4.92E-05 | 9.91E-05 | 3.19E-06 | 1.53E-05 | 4.35E-05 | 9.97E-05 | 0.00017 | 0 | 8.19E-05 |
| Rft1 | 4.92E-05 | 6.83E-05 | 3.16E-05 | 5.74E-05 | 7.69E-05 | 6.92E-05 | 4.04E-05 | 5.34E-05 | 4.96E-05 | 2.96E-05 | 4.61E-05 | 2.20E-05 | 2.76E-05 | 2.70E-05 | 0 | 5.00E-05 | 0 | 5.94E-05 |
| Dcaf17 | 4.95E-05 | 4.09E-05 | 1.01E-05 | 7.34E-06 | 2.24E-05 | 2.44E-05 | 2.41E-05 | 1.08E-05 | 4.96E-05 | 3.29E-05 | 1.11E-05 | 1.91E-05 | 0 | 0 | 4.76E-05 | 3.04E-06 | 0 | 3.10E-05 |
| Ercc1 | 8.23E-05 | 0.00011 | 0.00013 | 0.00022 | 6.94E-05 | 0 | 5.90E-05 | 3.58E-05 | 4.96E-05 | 0 | 0.00012 | 0.00012 | 0 | 3.12E-05 | 6.59E-05 | 0.0001 | 0.00013 | 4.00E-05 |
| Mex3a | 0 | 2.84E-05 | 7.81E-05 | 6.28E-05 | 4.34E-05 | 0 | 0 | 6.93E-05 | 4.96E-05 | 0 | 5.90E-05 | 0 | 1.25E-05 | 3.41E-05 | 0.00012 | 4.97E-05 | 2.21E-05 | 0 |
| Zc3h7a | 0.00028 | 0.00016 | 0.00012 | 1.74E-05 | 3.09E-05 | 9.40E-05 | 0.00019 | 0.00011 | 4.95E-05 | 0.00012 | 5.54E-05 | 0.00023 | 3.81E-05 | 9.47E-05 | 0.00067 | 0.00014 | 0.00025 | 4.88E-05 |
| Dnajc30 | 4.67E-05 | 6.22E-05 | 1.57E-05 | 0.0001 | 5.53E-05 | 5.71E-05 | 3.50E-05 | 6.26E-05 | 4.95E-05 | 3.47E-05 | 7.90E-06 | 2.62E-05 | 1.10E-05 | 2.36E-05 | 3.53E-06 | 3.85E-05 | 0 | 4.96E-05 |
| Tob2 | 0 | 0.00011 | 3.95E-05 | 0.00015 | 0 | 1.43E-05 | 0 | 1.06E-05 | 4.95E-05 | 2.82E-05 | 1.12E-05 | 3.15E-05 | 0 | 4.79E-05 | 8.29E-05 | 2.82E-05 | 0.00064 | 3.00E-05 |

| | | | | | | | | | | | | | | | | | | |
|---|---|---|---|---|---|---|---|---|---|---|---|---|---|---|---|---|---|---|
| Mtif2 | 7.04E-05 | 5.39E-05 | 3.40E-05 | 3.13E-05 | 2.35E-05 | 1.21E-05 | 7.51E-06 | 5.18E-06 | 4.95E-05 | 1.16E-05 | 5.20E-06 | 1.88E-05 | 0 | 8.70E-06 | 2.74E-05 | 3.32E-05 | 0 | 7.25E-07 |
| Ccser2 | 0.00018 | 9.48E-05 | 4.10E-05 | 0 | 7.83E-05 | 0.00012 | 8.21E-05 | 3.98E-05 | 4.94E-05 | 4.18E-05 | 7.09E-05 | 5.92E-05 | 0 | 5.81E-06 | 1.60E-05 | 4.71E-05 | 6.26E-05 | 0.00013 |
| Rbm27 | 9.32E-05 | 0.0001 | 9.78E-05 | 3.46E-05 | 2.19E-05 | 3.57E-05 | 8.49E-06 | 0 | 4.94E-05 | 3.71E-05 | 4.37E-05 | 1.34E-05 | 0 | 6.88E-05 | 7.95E-05 | 9.79E-05 | 0.00014 | 1.90E-05 |
| Slc16a14 | 0 | 0 | 0 | 0 | 1.21E-05 | 3.50E-05 | 0 | 0 | 4.94E-05 | 0 | 0 | 0 | 0 | 0 | 0 | 0 | 0 | 0 |
| Cbr4 | 5.40E-05 | 4.56E-05 | 4.66E-06 | 3.05E-05 | 3.69E-05 | 2.98E-05 | 4.27E-05 | 3.04E-05 | 4.94E-05 | 2.94E-05 | 7.51E-06 | 0 | 1.62E-05 | 1.48E-05 | 0 | 3.37E-05 | 0 | 6.01E-05 |
| Znrd2 | 7.28E-05 | 4.00E-05 | 0.00012 | 0.00016 | 3.29E-05 | 0 | 0 | 0 | 4.94E-05 | 0 | 9.85E-06 | 0 | 2.77E-06 | 4.33E-05 | 5.57E-05 | 6.25E-05 | 5.77E-05 | 3.31E-05 |
| Gemin5 | 5.99E-05 | 6.66E-05 | 0.00012 | 0.00011 | 0 | 0 | 0 | 0 | 4.94E-05 | 7.48E-06 | 3.46E-05 | 0 | 1.12E-05 | 1.49E-05 | 3.75E-05 | 3.73E-05 | 5.65E-06 | 1.07E-05 |
| Vps35 | 0.00032 | 0.00018 | 0.00011 | 0 | 2.68E-05 | 0.00017 | 0 | 0 | 4.93E-05 | 2.02E-05 | 3.32E-05 | 0.00017 | 0 | 9.34E-06 | 0 | 0.00016 | 0 | 5.99E-05 |
| Kpna4 | 0.0002 | 0.00014 | 0.00021 | 4.68E-05 | 7.66E-05 | 9.14E-05 | 0 | 1.97E-05 | 4.93E-05 | 7.17E-05 | 9.67E-05 | 4.25E-05 | 1.79E-05 | 2.05E-05 | 0.00012 | 0.00019 | 0.00018 | 2.16E-05 |
| Plaa | 0.00023 | 0.00016 | 0.00024 | 0.00016 | 9.44E-05 | 8.48E-05 | 0 | 3.46E-05 | 4.93E-05 | 1.17E-05 | 6.89E-05 | 5.69E-05 | 1.33E-06 | 1.21E-07 | 6.57E-05 | 0.00018 | 0 | 5.60E-05 |
| Eloc | 0.00025 | 1.87E-05 | 0.00019 | 0 | 1.12E-05 | 5.85E-05 | 4.43E-06 | 5.88E-05 | 4.93E-05 | 0 | 0.00038 | 5.97E-05 | 2.57E-05 | 2.72E-05 | 0 | 0.00014 | 0.00013 | 0 |
| Tsr2 | 6.80E-05 | 5.38E-05 | 0.00021 | 1.00E-04 | 5.43E-05 | 4.02E-05 | 0 | 1.34E-05 | 4.93E-05 | 3.12E-05 | 2.18E-05 | 7.32E-06 | 8.04E-05 | 2.91E-05 | 0 | 0.00015 | 0 | 4.26E-05 |
| Nol3 | 1.85E-06 | 1.99E-05 | 0 | 4.56E-05 | 0 | 9.96E-05 | 9.40E-05 | 1.22E-05 | 4.92E-05 | 3.28E-05 | 8.04E-06 | 0.00012 | 9.15E-05 | 9.33E-05 | 0 | 0 | 0 | 9.96E-05 |
| D2hgdh | 4.28E-05 | 8.06E-05 | 0 | 6.25E-05 | 4.17E-05 | 1.83E-05 | 2.95E-05 | 3.06E-06 | 4.92E-05 | 2.28E-05 | 4.23E-06 | 3.20E-05 | 0 | 1.83E-05 | 0.00011 | 2.24E-05 | 0 | 8.73E-05 |
| Thoc3 | 9.49E-05 | 5.31E-05 | 9.95E-05 | 5.40E-05 | 6.27E-05 | 1.20E-05 | 1.03E-05 | 4.14E-05 | 4.92E-05 | 8.24E-06 | 1.62E-05 | 7.38E-05 | 0 | 5.59E-06 | 0 | 0.00016 | 0 | 7.97E-07 |
| Dcaf15 | 0 | 8.92E-05 | 4.03E-05 | 7.54E-05 | 4.47E-05 | 0 | 8.56E-06 | 8.95E-05 | 4.92E-05 | 3.58E-05 | 5.28E-05 | 8.70E-06 | 0 | 9.71E-05 | 0.00038 | 0.0003 | 8.62E-05 | 4.65E-06 |
| Strn | 5.75E-05 | 0.00013 | 8.37E-05 | 4.66E-05 | 5.12E-05 | 8.94E-05 | 8.99E-05 | 0.00016 | 4.91E-05 | 6.33E-05 | 4.05E-05 | 3.14E-05 | 6.03E-06 | 0.00012 | 4.04E-05 | 0.00011 | 8.65E-05 | 9.97E-05 |
| Psen2 | 0 | 0.00017 | 0 | 0.00014 | 8.65E-05 | 5.34E-05 | 0 | 1.40E-05 | 4.91E-05 | 0.00011 | 1.51E-05 | 2.68E-05 | 0.00019 | 3.90E-05 | 5.31E-05 | 0 | 1.57E-05 | 0.00015 |
| Riok1 | 0.00012 | 4.47E-05 | 7.20E-05 | 5.75E-05 | 2.74E-05 | 4.85E-05 | 0 | 3.47E-06 | 4.91E-05 | 1.11E-05 | 1.35E-05 | 1.87E-05 | 0 | 0 | 0.00014 | 4.29E-05 | 0 | 1.51E-05 |
| Pycrl | 8.01E-05 | 4.55E-05 | 7.76E-05 | 9.77E-05 | 0 | 0 | 0 | 7.86E-06 | 4.91E-05 | 2.35E-06 | 0 | 0 | 0 | 1.42E-05 | 0 | 6.16E-05 | 0 | 2.65E-05 |
| Rangrf | 7.08E-05 | 5.97E-05 | 0.00014 | 0.00011 | 0 | 0 | 0 | 0 | 4.91E-05 | 1.87E-05 | 2.06E-05 | 4.82E-06 | 0 | 0 | 0 | 2.84E-05 | 2.82E-05 | 3.10E-05 |
| Plekhh2 | 1.16E-06 | 5.20E-05 | 0 | 0 | 0 | 0 | 0 | 0 | 4.91E-05 | 4.36E-05 | 0 | 0 | 2.89E-05 | 0 | 2.66E-05 | 3.26E-05 | 2.62E-06 | 7.34E-05 |
| Esyt2 | 0.00016 | 0.00026 | 0.00032 | 0 | 0.00031 | 0.00025 | 0.00017 | 9.61E-05 | 4.90E-05 | 0.00018 | 0.0017 | 0.00044 | 0 | 0.00028 | 0.00017 | 0.00021 | 0.00035 | 0.00036 |
| Reck | 2.02E-05 | 0.00011 | 0 | 0 | 9.70E-05 | 0.00018 | 0 | 0 | 4.90E-05 | 0.00036 | 9.16E-05 | 2.50E-05 | 0.00017 | 0.00017 | 0 | 0 | 2.01E-05 | 0.00014 |
| Dyrk1a | 0.00019 | 0.00016 | 7.37E-05 | 0 | 9.42E-05 | 0.00012 | 1.45E-05 | 4.70E-07 | 4.90E-05 | 8.64E-06 | 1.73E-05 | 0.00011 | 0 | 8.71E-06 | 5.97E-05 | 0.00023 | 8.67E-05 | 1.57E-05 |
| Map11 | 6.47E-06 | 5.66E-05 | 7.70E-05 | 9.20E-05 | 4.01E-05 | 0 | 4.51E-05 | 7.27E-05 | 4.90E-05 | 4.84E-05 | 4.23E-05 | 0 | 0 | 4.10E-05 | 0.00016 | 6.79E-06 | 0 | 3.92E-05 |
| Nudt1 | 1.49E-05 | 0 | 9.08E-06 | 4.54E-05 | 0 | 6.55E-06 | 2.37E-07 | 2.15E-05 | 4.89E-05 | 1.23E-05 | 7.92E-06 | 4.45E-06 | 1.00E-06 | 0 | 0 | 7.47E-05 | 1.64E-05 | 1.88E-05 |
| Jcad | 3.67E-05 | 0.0001 | 0.00015 | 0 | 0 | 0 | 3.76E-05 | 0 | 4.89E-05 | 0.00021 | 0.00013 | 0.00024 | 5.12E-05 | 0 | 3.69E-06 | 8.37E-05 | 0 | 0.00019 |
| Lpin2 | 7.77E-05 | 6.28E-05 | 6.72E-06 | 0 | 6.12E-05 | 0 | 6.64E-05 | 0 | 4.89E-05 | 3.93E-06 | 1.98E-05 | 5.10E-05 | 0 | 6.31E-05 | 0.00011 | 7.56E-05 | 8.68E-05 | 1.12E-05 |
| Cd276 | 7.68E-06 | 7.71E-05 | 0.00027 | 0.00027 | 0.00012 | 0.00019 | 0.00061 | 0.00068 | 4.88E-05 | 0.00027 | 0 | 7.63E-05 | 6.75E-05 | 0.00036 | 0 | 4.38E-05 | 0 | 0 |
| Med28 | 0.00023 | 0.00018 | 0.00018 | 0.00012 | 0.00015 | 0.00018 | 7.69E-05 | 9.77E-05 | 4.88E-05 | 8.11E-05 | 0.00014 | 0.00016 | 3.63E-05 | 5.01E-05 | 0 | 0.00021 | 8.30E-05 | 0.00011 |
| Tirap | 9.10E-06 | 6.75E-05 | 5.24E-05 | 6.04E-05 | 4.15E-05 | 5.33E-05 | 3.16E-05 | 4.62E-05 | 4.88E-05 | 6.01E-05 | 2.23E-05 | 9.84E-06 | 6.01E-05 | 6.90E-05 | 1.23E-05 | 1.64E-05 | 0 | 6.46E-05 |
| Ankrd13b | 6.31E-06 | 6.92E-05 | 4.36E-05 | 8.59E-05 | 2.29E-06 | 5.02E-05 | 1.27E-05 | 3.31E-05 | 4.88E-05 | 2.60E-05 | 1.38E-05 | 0 | 9.19E-06 | 1.62E-05 | 6.63E-05 | 1.99E-06 | 1.43E-05 | 2.61E-05 |

| | | | | | | | | | | | | | | | | | | |
|---|---|---|---|---|---|---|---|---|---|---|---|---|---|---|---|---|---|---|
| Cep85 | 4.80E-05 | 5.80E-05 | 5.35E-05 | 0 | 3.05E-05 | 4.41E-05 | 7.20E-05 | 1.33E-05 | 4.88E-05 | 0 | 0 | 1.54E-05 | 3.22E-05 | 1.81E-05 | 0.00017 | 0.00011 | 1.09E-05 | 0 |
| Prkce | 2.00E-05 | 8.92E-05 | 5.16E-06 | 0 | 9.22E-06 | 2.65E-05 | 1.71E-05 | 1.06E-05 | 4.88E-05 | 7.69E-05 | 6.11E-06 | 0 | 2.19E-05 | 2.04E-05 | 8.93E-05 | 9.78E-05 | 8.88E-05 | 0.00011 |
| Polr2k | 9.66E-05 | 2.52E-05 | 1.81E-05 | 2.93E-05 | 7.15E-06 | 2.30E-05 | 0 | 0 | 4.88E-05 | 3.22E-06 | 7.69E-06 | 2.97E-05 | 5.72E-06 | 2.22E-05 | 8.38E-06 | 5.57E-05 | 1.77E-05 | 0 |
| Trim41 | 5.79E-05 | 0.00015 | 7.22E-05 | 7.78E-05 | 6.85E-05 | 1.71E-05 | 6.86E-05 | 2.68E-05 | 4.88E-05 | 2.66E-05 | 7.37E-05 | 9.74E-05 | 2.42E-05 | 4.54E-05 | 0.00031 | 2.44E-05 | 7.67E-05 | 4.02E-05 |
| Rbmx | 0.0001 | 4.81E-05 | 9.39E-05 | 9.06E-05 | 0 | 0 | 0 | 0 | 4.88E-05 | 1.60E-05 | 0 | 0 | 0 | 1.49E-05 | 0.00025 | 0.00029 | 0 | 2.70E-06 |
| Alg11 | 0.00012 | 4.40E-05 | 4.46E-05 | 0 | 2.88E-05 | 7.10E-05 | 7.19E-07 | 6.04E-06 | 4.87E-05 | 1.26E-05 | 1.90E-05 | 0.00014 | 2.63E-05 | 0 | 6.43E-06 | 0 | 0 | 2.33E-06 |
| Khdc4 | 0.00017 | 0.00026 | 0.00017 | 6.45E-05 | 8.50E-05 | 3.96E-05 | 1.80E-05 | 0 | 4.87E-05 | 1.97E-05 | 3.34E-05 | 0.00016 | 0 | 9.30E-05 | 0.00058 | 5.97E-05 | 5.36E-05 | 1.20E-05 |
| Plcd1 | 5.14E-05 | 8.42E-05 | 7.72E-05 | 4.56E-05 | 0 | 1.97E-05 | 6.39E-05 | 0 | 4.87E-05 | 5.43E-05 | 0.00014 | 7.90E-05 | 5.05E-05 | 9.09E-05 | 0.00013 | 0 | 5.61E-05 | 0.00014 |
| Utp4 | 7.47E-05 | 8.28E-05 | 0.0002 | 2.49E-05 | 1.95E-06 | 1.69E-05 | 0 | 0 | 4.87E-05 | 3.03E-05 | 1.22E-05 | 2.19E-05 | 3.35E-05 | 0 | 7.64E-05 | 5.55E-06 | 0 | 1.22E-05 |
| Qtrt2 | 4.24E-05 | 5.52E-05 | 8.68E-05 | 3.95E-05 | 8.55E-06 | 9.09E-06 | 0 | 0 | 4.87E-05 | 1.41E-05 | 1.38E-05 | 1.79E-05 | 0 | 2.67E-05 | 6.51E-05 | 1.18E-05 | 7.88E-06 | 0 |
| Nsun5 | 1.76E-05 | 5.66E-05 | 0.0001 | 8.58E-05 | 9.05E-06 | 8.38E-06 | 0 | 0 | 4.87E-05 | 3.40E-05 | 1.57E-05 | 0 | 1.90E-05 | 2.06E-05 | 4.25E-05 | 7.89E-05 | 0 | 7.00E-06 |
| Wnt2 | 0 | 0 | 0 | 0 | 4.52E-05 | 0 | 0.00037 | 0 | 4.87E-05 | 0.00033 | 0 | 7.50E-05 | 0 | 0 | 0 | 0 | 0 | 9.25E-05 |
| Pabpc4 | 0.00027 | 0.0001 | 0.0004 | 0.00057 | 5.33E-05 | 0.00017 | 0.00042 | 7.27E-05 | 4.86E-05 | 0 | 3.09E-05 | 9.30E-05 | 7.88E-05 | 4.90E-05 | 0.00021 | 9.49E-05 | 0.00024 | 3.57E-06 |
| H2-M3 | 7.24E-05 | 3.32E-05 | 6.08E-07 | 1.96E-06 | 6.11E-05 | 6.35E-05 | 0 | 9.90E-05 | 4.86E-05 | 0.00018 | 2.01E-05 | 9.06E-05 | 0.00011 | 3.87E-05 | 2.07E-06 | 2.54E-05 | 0 | 7.42E-05 |
| Slc30a6 | 0.0001 | 1.97E-05 | 3.00E-05 | 0 | 2.76E-05 | 5.37E-05 | 4.39E-05 | 3.36E-05 | 4.86E-05 | 2.07E-06 | 2.33E-05 | 3.75E-05 | 0 | 0 | 2.53E-05 | 2.45E-05 | 0 | 0 |
| Hdac10 | 1.34E-05 | 3.45E-05 | 1.68E-05 | 1.95E-05 | 4.83E-06 | 2.43E-05 | 9.53E-06 | 2.29E-05 | 4.86E-05 | 5.97E-05 | 2.61E-05 | 1.04E-05 | 0 | 0 | 0.00011 | 4.23E-05 | 0 | 7.49E-05 |
| Adprm | 4.52E-05 | 5.19E-05 | 6.08E-06 | 1.88E-05 | 9.50E-07 | 2.30E-05 | 1.10E-05 | 5.53E-07 | 4.86E-05 | 2.79E-05 | 1.62E-05 | 5.36E-06 | 6.53E-06 | 2.14E-05 | 0 | 2.66E-05 | 0 | 2.80E-05 |
| Cdpf1 | 4.83E-05 | 1.57E-05 | 5.50E-05 | 9.66E-05 | 2.68E-05 | 1.15E-05 | 5.46E-05 | 5.16E-05 | 4.86E-05 | 6.02E-06 | 3.10E-05 | 2.27E-05 | 0 | 9.03E-06 | 0 | 1.92E-05 | 0 | 1.88E-05 |
| Szt2 | 1.71E-05 | 5.83E-05 | 8.43E-06 | 4.11E-05 | 3.48E-05 | 7.86E-06 | 2.19E-06 | 2.71E-05 | 4.86E-05 | 2.53E-05 | 1.53E-05 | 2.15E-06 | 0 | 2.49E-05 | 0.0001 | 3.16E-05 | 0 | 2.05E-05 |
| Cfap410 | 4.94E-05 | 6.69E-05 | 6.33E-06 | 3.70E-05 | 3.44E-05 | 0 | 0 | 2.47E-05 | 4.86E-05 | 1.63E-05 | 4.43E-06 | 0 | 4.84E-05 | 2.67E-05 | 3.52E-05 | 1.39E-06 | 0 | 5.92E-05 |
| Bysl | 5.37E-05 | 3.03E-05 | 0.00021 | 0.00014 | 0 | 0 | 0 | 0 | 4.86E-05 | 1.69E-05 | 4.30E-05 | 0 | 1.87E-05 | 4.98E-06 | 9.45E-05 | 4.47E-05 | 0 | 2.17E-05 |
| Slc25a23 | 0 | 9.61E-05 | 0 | 0 | 9.68E-05 | 0 | 0 | 0 | 4.86E-05 | 8.65E-05 | 0 | 2.77E-05 | 4.25E-05 | 0 | 3.72E-06 | 0 | 0 | 1.35E-05 |
| Med30 | 0.00017 | 6.24E-05 | 8.34E-05 | 6.96E-06 | 7.40E-05 | 0.00013 | 1.37E-05 | 7.34E-05 | 4.85E-05 | 7.47E-05 | 2.27E-05 | 8.04E-05 | 0 | 3.24E-05 | 0 | 0.00037 | 0 | 0 |
| Thap12 | 0.00018 | 6.14E-05 | 0.00018 | 4.56E-05 | 8.24E-05 | 0.00012 | 3.26E-05 | 0.00015 | 4.85E-05 | 5.64E-05 | 2.08E-05 | 2.11E-05 | 5.48E-05 | 5.31E-05 | 0 | 0.00013 | 2.84E-05 | 8.98E-05 |
| Gpam | 0.00012 | 6.86E-05 | 3.78E-05 | 3.75E-05 | 2.55E-06 | 5.24E-06 | 5.04E-05 | 3.26E-05 | 4.85E-05 | 0.00012 | 0 | 1.63E-05 | 4.28E-05 | 0 | 2.08E-05 | 5.30E-05 | 0 | 6.33E-05 |
| Zfp574 | 0 | 0.00011 | 2.80E-05 | 0.00011 | 5.41E-05 | 0 | 0 | 1.16E-05 | 4.85E-05 | 4.20E-05 | 3.41E-05 | 2.52E-05 | 0 | 3.25E-05 | 8.41E-05 | 5.81E-05 | 0 | 6.89E-05 |
| Atp9a | 0 | 4.20E-05 | 2.24E-05 | 0 | 0 | 0 | 8.48E-05 | 3.67E-05 | 4.85E-05 | 7.09E-05 | 3.37E-05 | 0.00011 | 2.04E-05 | 0.00025 | 0 | 0 | 0 | 0.00022 |
| Nr2f1 | 4.67E-07 | 4.78E-05 | 0 | 0 | 0 | 0.00021 | 0 | 0.0004 | 4.84E-05 | 1.94E-05 | 0 | 0 | 0 | 0 | 0 | 0 | 1.29E-05 | 9.92E-05 |
| Opa1 | 0.00017 | 9.08E-05 | 7.04E-05 | 1.43E-05 | 2.65E-05 | 8.06E-05 | 4.64E-05 | 2.97E-05 | 4.84E-05 | 2.80E-05 | 3.75E-05 | 8.54E-05 | 2.17E-05 | 3.84E-06 | 0.00014 | 8.79E-05 | 3.17E-05 | 0 |
| Vps13b | 5.07E-05 | 8.20E-05 | 0 | 0 | 1.82E-05 | 5.27E-05 | 8.36E-05 | 2.82E-05 | 4.84E-05 | 4.20E-05 | 0 | 4.82E-05 | 9.02E-06 | 0 | 0 | 5.01E-05 | 3.14E-05 | 9.11E-06 |
| Polr3c | 9.45E-05 | 6.36E-05 | 3.31E-05 | 3.45E-05 | 2.51E-05 | 4.93E-05 | 3.39E-05 | 2.01E-05 | 4.84E-05 | 1.48E-05 | 3.40E-05 | 5.58E-05 | 0 | 1.78E-05 | 4.83E-05 | 5.04E-05 | 0 | 1.61E-06 |
| Snapc3 | 7.34E-05 | 0.00011 | 5.04E-05 | 1.84E-05 | 3.84E-05 | 1.05E-05 | 0 | 6.95E-06 | 4.84E-05 | 3.28E-05 | 2.31E-05 | 1.47E-05 | 0 | 3.03E-05 | 0.0004 | 1.28E-05 | 0 | 3.03E-05 |
| Adar | 5.52E-05 | 0.00014 | 6.16E-05 | 9.37E-05 | 4.97E-05 | 9.37E-06 | 0 | 6.21E-06 | 4.84E-05 | 0.00021 | 0 | 5.40E-05 | 8.34E-05 | 6.39E-06 | 0.00016 | 8.73E-05 | 0.00013 | 6.12E-05 |

| | | | | | | | | | | | | | | | | | | |
|---|---|---|---|---|---|---|---|---|---|---|---|---|---|---|---|---|---|---|
| Porcn | 0 | 5.64E-05 | 0 | 0 | 7.94E-05 | 0 | 0.00012 | 0 | 4.84E-05 | 0 | 0.0003 | 0 | 0.00016 | 0.00034 | 3.74E-05 | 0 | 0 | 0.00017 |
| Ttc12 | 0 | 4.14E-05 | 6.28E-06 | 4.17E-06 | 3.83E-05 | 0 | 2.77E-05 | 0 | 4.84E-05 | 0 | 2.85E-05 | 3.94E-05 | 9.34E-05 | 0 | 4.64E-06 | 0 | 4.41E-05 | 1.95E-05 |
| Tsr1 | 0.00018 | 7.92E-05 | 0.0004 | 0.00028 | 4.25E-05 | 0 | 8.94E-05 | 0 | 4.84E-05 | 9.13E-07 | 3.05E-05 | 0 | 0.00011 | 3.78E-05 | 0.00035 | 0 | 0 | 2.34E-05 |
| Pcm1 | 0.00023 | 0.00019 | 4.04E-05 | 0 | 7.46E-05 | 9.24E-05 | 3.08E-05 | 0 | 4.83E-05 | 1.66E-05 | 1.67E-06 | 0.0002 | 1.10E-05 | 0 | 0.00024 | 0.00016 | 8.06E-05 | 6.88E-06 |
| Rhot1 | 0.00013 | 0.00011 | 9.80E-05 | 0 | 6.53E-05 | 7.60E-05 | 0 | 0 | 4.83E-05 | 2.46E-05 | 4.04E-05 | 5.47E-05 | 0 | 7.94E-06 | 6.71E-05 | 4.99E-05 | 5.88E-05 | 2.32E-05 |
| Foxj2 | 8.55E-05 | 0.00014 | 4.02E-05 | 1.29E-05 | 7.12E-05 | 4.79E-05 | 1.65E-05 | 7.70E-06 | 4.83E-05 | 6.57E-05 | 4.54E-05 | 5.79E-05 | 8.18E-06 | 2.69E-05 | 6.90E-05 | 4.77E-05 | 7.95E-05 | 3.95E-05 |
| Mical1 | 3.22E-05 | 6.48E-05 | 0.00021 | 9.48E-05 | 0 | 4.27E-05 | 2.92E-05 | 0.00043 | 4.83E-05 | 0.00016 | 4.32E-05 | 3.07E-05 | 0 | 0.00015 | 0.00018 | 4.83E-05 | 3.01E-05 | 0.00017 |
| Nrf1 | 3.99E-05 | 7.38E-05 | 6.34E-05 | 2.60E-05 | 2.08E-05 | 6.20E-06 | 8.99E-06 | 2.72E-05 | 4.83E-05 | 1.01E-05 | 2.60E-05 | 3.97E-05 | 0 | 2.74E-05 | 8.85E-05 | 0.00011 | 3.19E-06 | 4.98E-06 |
| Bccip | 0.00037 | 0.00011 | 0.00039 | 0.00017 | 0.00022 | 0.00015 | 0.0001 | 5.87E-05 | 4.82E-05 | 7.68E-07 | 5.30E-05 | 0 | 5.26E-05 | 6.56E-05 | 0.00016 | 5.97E-05 | 0 | 0 |
| Dym | 0.00017 | 8.65E-05 | 9.57E-05 | 7.85E-05 | 8.82E-05 | 7.92E-05 | 0.00015 | 5.09E-05 | 4.82E-05 | 4.92E-05 | 1.08E-05 | 0.00016 | 0 | 6.80E-05 | 0.00017 | 7.70E-05 | 0 | 8.40E-06 |
| Cul4b | 0.00022 | 0.00012 | 2.31E-05 | 7.52E-05 | 4.00E-05 | 6.27E-05 | 2.25E-05 | 5.45E-06 | 4.82E-05 | 0 | 4.12E-05 | 7.79E-05 | 1.78E-05 | 2.19E-05 | 8.58E-05 | 0.00011 | 7.72E-05 | 9.48E-06 |
| Eif2b4 | 0.0001 | 6.30E-05 | 0.00015 | 0.00014 | 0 | 1.29E-05 | 4.08E-05 | 0 | 4.82E-05 | 1.73E-05 | 2.78E-05 | 0 | 0 | 4.83E-05 | 5.34E-05 | 7.55E-05 | 2.06E-05 | 0 |
| Siah2 | 1.94E-05 | 3.30E-05 | 0.00018 | 9.43E-06 | 1.93E-06 | 1.12E-05 | 5.41E-06 | 0 | 4.82E-05 | 0 | 4.68E-05 | 0.00016 | 0 | 0.00013 | 0 | 2.32E-05 | 0.00026 | 2.54E-05 |
| Tmem237 | 2.19E-05 | 4.96E-05 | 3.31E-05 | 1.77E-05 | 2.46E-05 | 9.59E-06 | 0 | 4.42E-06 | 4.82E-05 | 6.40E-05 | 2.19E-05 | 4.24E-05 | 1.24E-05 | 4.55E-06 | 1.65E-06 | 9.78E-05 | 0 | 3.14E-05 |
| Usp36 | 5.94E-05 | 8.45E-05 | 0.00016 | 0.00012 | 4.56E-05 | 0 | 2.81E-05 | 2.96E-05 | 4.82E-05 | 4.51E-05 | 3.62E-05 | 1.19E-06 | 5.46E-05 | 3.39E-05 | 0.00026 | 0 | 0.00016 | 8.86E-07 |
| Ppat | 3.67E-05 | 5.86E-05 | 0.00018 | 4.78E-05 | 2.67E-05 | 0 | 0 | 0 | 4.82E-05 | 1.25E-05 | 2.40E-05 | 1.64E-06 | 0 | 2.60E-05 | 2.59E-05 | 9.62E-05 | 3.38E-05 | 0 |
| Cpne5 | 0 | 0 | 0 | 0 | 0 | 5.35E-05 | 0 | 0 | 4.81E-05 | 0 | 0 | 7.15E-05 | 0 | 0 | 0 | 0 | 0 | 6.10E-06 |
| Cops7b | 6.00E-05 | 5.80E-05 | 7.93E-05 | 6.28E-05 | 4.36E-05 | 2.12E-05 | 0 | 2.22E-05 | 4.81E-05 | 1.85E-05 | 2.11E-05 | 0 | 1.65E-05 | 1.40E-05 | 5.10E-05 | 8.14E-05 | 1.84E-05 | 4.63E-06 |
| Bahd1 | 0 | 0.00013 | 0.0001 | 0.00019 | 7.12E-05 | 1.49E-05 | 1.05E-05 | 5.47E-05 | 4.81E-05 | 5.28E-05 | 1.88E-05 | 3.20E-05 | 1.36E-05 | 4.01E-05 | 0.00015 | 0 | 7.91E-05 | 1.50E-05 |
| Myo6 | 5.15E-05 | 6.52E-05 | 0 | 0 | 2.29E-05 | 1.41E-05 | 0 | 0 | 4.81E-05 | 1.34E-05 | 2.48E-05 | 5.40E-05 | 0 | 9.05E-05 | 3.59E-05 | 0 | 0 | 5.77E-05 |
| Tmpo | 1.99E-05 | 9.89E-05 | 5.94E-05 | 1.17E-05 | 0 | 0 | 0 | 0 | 4.81E-05 | 7.95E-06 | 3.02E-06 | 7.25E-05 | 0 | 3.54E-07 | 0 | 0.00095 | 0 | 0 |
| Pwp2 | 3.55E-05 | 5.87E-05 | 0.00013 | 4.61E-05 | 0 | 0 | 0 | 0 | 4.81E-05 | 1.21E-05 | 7.35E-06 | 2.24E-05 | 0 | 1.79E-05 | 4.83E-05 | 1.30E-05 | 0 | 7.31E-06 |
| Sema3d | 5.97E-06 | 8.68E-06 | 0 | 0 | 0 | 0 | 0 | 0 | 4.81E-05 | 0.00018 | 0 | 0 | 0 | 0 | 0 | 0 | 0 | 0.00023 |
| Flad1 | 2.74E-05 | 3.63E-05 | 2.86E-05 | 5.99E-05 | 1.97E-05 | 0 | 1.77E-05 | 3.17E-05 | 4.81E-05 | 2.02E-05 | 3.49E-05 | 2.87E-05 | 1.12E-05 | 2.30E-06 | 4.32E-05 | 0 | 0 | 4.57E-05 |
| Cops2 | 0.00031 | 0.00015 | 0.00012 | 0.00013 | 4.80E-05 | 0.00014 | 3.42E-05 | 0 | 4.80E-05 | 1.70E-05 | 1.85E-05 | 0.00015 | 5.13E-05 | 0 | 4.93E-05 | 0.00018 | 2.78E-05 | 1.61E-05 |
| Zdhhc21 | 6.68E-05 | 2.60E-05 | 4.46E-05 | 0 | 8.09E-06 | 4.03E-05 | 0 | 3.27E-07 | 4.80E-05 | 1.95E-05 | 3.34E-07 | 1.69E-05 | 0 | 0 | 0 | 6.28E-05 | 0 | 2.07E-05 |
| Plpp2 | 0 | 6.45E-05 | 4.92E-05 | 9.15E-05 | 1.91E-05 | 2.33E-05 | 0 | 0 | 4.80E-05 | 0 | 0 | 0.00019 | 6.57E-05 | 1.85E-05 | 8.71E-06 | 1.27E-05 | 7.26E-05 | 9.86E-05 |
| Lefty1 | 1.23E-05 | 1.12E-05 | 0 | 0 | 0 | 2.28E-05 | 0 | 0 | 4.80E-05 | 5.20E-05 | 0 | 2.36E-05 | 0 | 4.59E-05 | 0 | 0 | 0 | 4.16E-05 |
| Ap3s2 | 9.10E-05 | 6.03E-05 | 1.75E-05 | 1.06E-05 | 1.72E-05 | 1.66E-05 | 0 | 1.34E-05 | 4.80E-05 | 2.39E-05 | 2.59E-05 | 2.19E-05 | 2.07E-06 | 0 | 1.33E-05 | 4.89E-05 | 0 | 1.25E-05 |
| Amer1 | 2.18E-05 | 4.22E-05 | 0 | 4.34E-05 | 3.87E-05 | 1.35E-05 | 0 | 3.29E-05 | 4.80E-05 | 1.60E-05 | 1.62E-05 | 9.51E-06 | 0 | 1.19E-05 | 5.91E-05 | 6.01E-05 | 0 | 3.12E-05 |
| Gpatch11 | 7.39E-05 | 7.51E-05 | 3.25E-05 | 0.00016 | 5.34E-05 | 0 | 1.39E-05 | 0 | 4.80E-05 | 3.63E-05 | 2.12E-05 | 0 | 1.86E-05 | 5.26E-06 | 7.44E-05 | 5.78E-05 | 5.50E-06 | 8.14E-06 |
| Gas2l1 | 0 | 0.00014 | 0.00025 | 0.00029 | 7.17E-05 | 7.88E-05 | 9.17E-05 | 0.00025 | 4.79E-05 | 0.00018 | 7.78E-05 | 0.00015 | 0 | 0.0001 | 0.00017 | 3.06E-05 | 0 | 0.00021 |
| Rars2 | 8.53E-05 | 1.87E-05 | 3.37E-05 | 4.12E-06 | 2.06E-06 | 4.77E-05 | 7.36E-07 | 2.40E-06 | 4.79E-05 | 2.18E-05 | 0 | 1.40E-05 | 2.08E-05 | 7.20E-06 | 1.66E-05 | 1.22E-05 | 0 | 4.37E-06 |

| | | | | | | | | | | | | | | | | | | |
|---|---|---|---|---|---|---|---|---|---|---|---|---|---|---|---|---|---|---|
| Tctn2 | 2.44E-05 | 3.51E-05 | 0 | 5.36E-05 | 2.27E-05 | 2.05E-05 | 1.34E-06 | 0 | 4.79E-05 | 3.69E-05 | 3.70E-06 | 6.13E-06 | 4.67E-05 | 2.12E-05 | 0 | 0 | 0 | 2.70E-05 |
| Rab3ip | 7.66E-06 | 5.93E-05 | 1.07E-05 | 5.27E-05 | 3.24E-05 | 2.01E-05 | 0 | 2.24E-05 | 4.79E-05 | 1.43E-05 | 9.98E-05 | 0.00023 | 0 | 7.24E-05 | 8.23E-05 | 8.13E-05 | 0 | 2.27E-05 |
| Coq8b | 4.02E-05 | 0 | 0.00015 | 0.0002 | 6.59E-05 | 1.76E-05 | 0.0001 | 0.0002 | 4.79E-05 | 2.72E-05 | 0 | 0.00013 | 0.00012 | 0.00014 | 0.0002 | 0 | 6.41E-05 | 7.95E-05 |
| Zfp13 | 1.52E-05 | 3.04E-05 | 0 | 2.45E-05 | 2.57E-05 | 3.93E-06 | 0 | 4.76E-06 | 4.79E-05 | 5.16E-06 | 0 | 6.19E-06 | 0 | 1.10E-05 | 6.54E-05 | 3.95E-05 | 0 | 4.60E-05 |
| Srsf11 | 0.00021 | 0.00022 | 0.00014 | 2.93E-05 | 0.0001 | 0.00011 | 8.03E-05 | 5.54E-05 | 4.78E-05 | 4.30E-05 | 1.91E-05 | 0.00015 | 1.14E-05 | 2.84E-05 | 0.00092 | 0.00015 | 0.00018 | 8.81E-05 |
| Zfp799 | 3.14E-05 | 4.31E-05 | 0 | 1.03E-05 | 5.20E-05 | 6.11E-05 | 1.01E-05 | 1.93E-05 | 4.78E-05 | 2.22E-05 | 0 | 1.21E-05 | 0 | 3.37E-05 | 3.23E-05 | 6.13E-06 | 0 | 3.94E-05 |
| Cdc37l1 | 0.00014 | 0.00015 | 8.46E-05 | 2.29E-05 | 0.00011 | 3.94E-05 | 1.64E-05 | 0 | 4.78E-05 | 1.14E-05 | 5.65E-05 | 5.25E-07 | 3.20E-05 | 9.01E-05 | 0.00034 | 6.01E-05 | 0 | 5.51E-05 |
| Zfp710 | 1.76E-05 | 5.49E-05 | 0 | 5.57E-05 | 2.10E-05 | 3.17E-06 | 5.01E-06 | 3.74E-05 | 4.78E-05 | 5.08E-05 | 3.15E-05 | 9.31E-06 | 1.76E-05 | 8.69E-06 | 6.79E-05 | 1.26E-05 | 0.00012 | 4.22E-05 |
| Phf10 | 0.00028 | 0.00011 | 0.00012 | 4.58E-05 | 4.15E-05 | 0.00014 | 0.00018 | 0 | 4.77E-05 | 3.73E-05 | 9.03E-05 | 0.00023 | 3.40E-05 | 3.63E-05 | 3.44E-05 | 0.00016 | 1.07E-05 | 0 |
| Alkbh7 | 1.13E-05 | 4.14E-05 | 3.52E-05 | 6.08E-05 | 2.75E-05 | 3.26E-06 | 2.15E-05 | 2.52E-05 | 4.77E-05 | 3.09E-05 | 2.66E-05 | 2.91E-05 | 0 | 7.91E-06 | 2.78E-05 | 1.02E-05 | 0 | 1.33E-05 |
| Acsf3 | 1.74E-05 | 3.10E-05 | 1.11E-05 | 4.78E-05 | 1.83E-05 | 0 | 5.60E-06 | 1.58E-05 | 4.77E-05 | 1.84E-05 | 1.64E-05 | 1.77E-05 | 1.34E-05 | 2.32E-05 | 3.92E-05 | 1.69E-05 | 1.05E-05 | 1.88E-05 |
| St13 | 0.00024 | 0.00015 | 0.00011 | 7.61E-05 | 4.57E-05 | 0.00013 | 2.45E-05 | 5.79E-06 | 4.76E-05 | 1.41E-05 | 5.31E-05 | 0.00013 | 2.41E-05 | 5.37E-05 | 9.55E-05 | 7.55E-05 | 7.16E-05 | 7.05E-05 |
| Pdpr | 2.71E-05 | 0.0001 | 2.56E-05 | 5.90E-05 | 9.14E-05 | 4.86E-05 | 2.71E-05 | 2.15E-05 | 4.76E-05 | 4.95E-05 | 4.90E-05 | 7.66E-05 | 2.74E-05 | 8.94E-05 | 2.65E-05 | 5.76E-05 | 0 | 0.00011 |
| Zkscan6 | 2.04E-05 | 4.17E-05 | 2.60E-05 | 6.03E-05 | 3.75E-05 | 4.15E-05 | 1.10E-05 | 4.36E-05 | 4.76E-05 | 2.26E-05 | 6.12E-06 | 7.41E-05 | 2.48E-05 | 1.55E-05 | 7.51E-05 | 0 | 0 | 4.06E-05 |
| Cramp1l | 2.83E-05 | 5.42E-05 | 4.56E-05 | 0 | 2.06E-05 | 1.55E-05 | 0 | 0 | 4.76E-05 | 1.76E-05 | 4.45E-05 | 3.46E-05 | 1.62E-05 | 7.94E-06 | 5.53E-05 | 3.18E-05 | 3.83E-05 | 1.59E-05 |
| Elp1 | 0.00011 | 9.77E-05 | 0.00012 | 0.00012 | 2.21E-05 | 1.31E-05 | 2.83E-05 | 0 | 4.76E-05 | 1.19E-05 | 2.55E-05 | 1.03E-05 | 0 | 1.88E-05 | 0.00023 | 4.85E-05 | 0 | 3.10E-05 |
| Eif4ebp3 | 3.36E-05 | 9.96E-05 | 2.56E-05 | 1.85E-05 | 1.31E-06 | 0 | 4.90E-06 | 0 | 4.76E-05 | 3.04E-05 | 1.88E-05 | 4.90E-05 | 2.87E-05 | 1.72E-05 | 8.73E-05 | 8.17E-05 | 0.00011 | 7.66E-06 |
| Gbf1 | 0.00041 | 8.98E-05 | 0.00019 | 0.00011 | 0.00012 | 0.00016 | 0.00032 | 4.91E-05 | 4.75E-05 | 5.15E-05 | 1.25E-05 | 0 | 6.28E-05 | 4.47E-05 | 0.0002 | 9.63E-05 | 7.94E-05 | 3.97E-05 |
| Prkcb | 0 | 2.00E-05 | 0 | 0 | 0 | 6.38E-05 | 0 | 0 | 4.75E-05 | 6.45E-06 | 0 | 7.61E-06 | 0 | 0 | 0 | 1.26E-05 | 0 | 0 |
| E2f6 | 6.98E-05 | 8.31E-05 | 7.08E-05 | 0 | 2.99E-05 | 1.83E-05 | 0 | 7.65E-06 | 4.75E-05 | 3.40E-05 | 1.29E-05 | 1.62E-05 | 5.56E-05 | 2.76E-05 | 0 | 0.00018 | 0 | 8.77E-06 |
| Mob2 | 8.56E-05 | 0.00018 | 5.87E-05 | 6.85E-05 | 8.96E-05 | 0.00011 | 5.68E-05 | 1.63E-05 | 4.74E-05 | 5.86E-05 | 3.87E-05 | 4.94E-05 | 0 | 9.76E-05 | 8.13E-05 | 9.70E-05 | 2.72E-05 | 2.25E-05 |
| Hdhd2 | 0.00011 | 5.20E-05 | 6.83E-05 | 4.10E-05 | 8.96E-05 | 7.47E-05 | 0 | 2.17E-05 | 4.74E-05 | 1.28E-05 | 7.02E-05 | 9.00E-06 | 4.40E-05 | 1.85E-05 | 8.30E-06 | 4.66E-05 | 0 | 1.65E-05 |
| Ube2n | 0.00023 | 3.61E-05 | 0.00015 | 0 | 1.18E-05 | 6.94E-05 | 2.90E-06 | 1.60E-05 | 4.74E-05 | 0 | 5.90E-05 | 4.05E-05 | 0 | 0 | 8.46E-06 | 0.00012 | 4.14E-05 | 0 |
| Zcchc17 | 7.45E-05 | 7.17E-05 | 8.42E-05 | 9.76E-05 | 3.99E-05 | 1.98E-05 | 2.65E-05 | 0 | 4.74E-05 | 2.68E-05 | 4.79E-05 | 1.13E-05 | 3.86E-05 | 1.89E-05 | 2.17E-05 | 0.00011 | 0 | 1.89E-05 |
| Dohh | 1.79E-05 | 6.35E-05 | 0.00016 | 0.00023 | 7.09E-05 | 3.11E-06 | 0 | 3.50E-05 | 4.74E-05 | 2.67E-05 | 9.11E-05 | 6.90E-05 | 3.55E-06 | 4.61E-05 | 4.50E-05 | 5.13E-05 | 1.03E-05 | 5.89E-05 |
| Ccnyl1 | 0.00018 | 0 | 0.00018 | 0 | 0 | 0 | 0 | 3.15E-05 | 4.74E-05 | 0.00011 | 4.19E-05 | 0.0001 | 4.64E-05 | 0 | 0 | 4.46E-05 | 0.00042 | 4.00E-05 |
| Ccdc50 | 0.0002 | 0.00016 | 6.31E-05 | 0 | 0.00011 | 0.00019 | 0 | 2.21E-05 | 4.73E-05 | 0.00012 | 5.83E-05 | 0.00018 | 0 | 4.47E-05 | 0.00013 | 9.84E-05 | 5.38E-05 | 6.20E-05 |
| Aptx | 7.04E-05 | 5.97E-05 | 3.33E-05 | 0 | 3.32E-05 | 4.27E-05 | 0 | 6.19E-06 | 4.73E-05 | 9.45E-06 | 5.46E-05 | 2.86E-05 | 0 | 2.67E-06 | 5.71E-05 | 2.61E-05 | 0 | 1.73E-05 |
| Vdac3 | 0.00019 | 8.85E-05 | 0.00013 | 0.00016 | 2.44E-05 | 1.07E-05 | 2.12E-05 | 0 | 4.73E-05 | 0.0001 | 6.90E-05 | 8.60E-05 | 0 | 2.94E-05 | 0 | 0.00015 | 0 | 3.58E-05 |
| Epb41l4b | 0 | 1.45E-05 | 0 | 0 | 0 | 0 | 0 | 2.64E-05 | 4.73E-05 | 0 | 0 | 0 | 1.80E-05 | 4.41E-05 | 0 | 7.60E-06 | 0 | 4.71E-05 |
| Mzt2 | 3.89E-05 | 7.87E-05 | 0 | 0.00018 | 6.97E-05 | 4.21E-05 | 1.73E-05 | 2.73E-05 | 4.72E-05 | 6.06E-05 | 1.92E-05 | 1.75E-05 | 1.77E-05 | 4.32E-05 | 9.09E-06 | 5.53E-05 | 0 | 4.65E-05 |
| Aimp2 | 0.00014 | 2.62E-05 | 0.00023 | 4.62E-05 | 0 | 0 | 0 | 0 | 4.72E-05 | 4.37E-05 | 0.0001 | 0.0002 | 2.60E-05 | 1.03E-05 | 1.83E-05 | 0.00018 | 0 | 7.22E-05 |
| Polr3gl | 3.91E-05 | 8.14E-05 | 0 | 3.81E-05 | 0.0001 | 0.00013 | 0.0001 | 3.18E-05 | 4.71E-05 | 6.63E-05 | 0 | 0 | 7.18E-05 | 5.04E-05 | 0.00011 | 2.32E-05 | 0 | 5.41E-05 |

| | | | | | | | | | | | | | | | | | | |
|---|---|---|---|---|---|---|---|---|---|---|---|---|---|---|---|---|---|---|
| Ptpn2 | 0.00015 | 5.67E-05 | 0.00014 | 4.62E-05 | 0 | 3.46E-05 | 1.17E-05 | 0 | 4.71E-05 | 5.32E-06 | 0 | 9.45E-05 | 0.00015 | 0 | 4.70E-05 | 4.93E-05 | 6.68E-05 | 3.08E-05 |
| Slc9a9 | 0 | 2.80E-05 | 0 | 2.14E-05 | 1.29E-05 | 0 | 0.00014 | 1.38E-05 | 4.71E-05 | 5.73E-05 | 0 | 0 | 7.85E-05 | 0 | 0 | 0 | 3.28E-05 | 0 |
| Rai2 | 0 | 3.90E-05 | 0 | 0 | 0 | 0 | 0 | 0 | 4.71E-05 | 0.00011 | 0 | 0 | 6.78E-05 | 0 | 0 | 0 | 0 | 7.40E-05 |
| Urb2 | 4.16E-05 | 6.21E-05 | 0.00012 | 6.77E-05 | 7.89E-06 | 0 | 0 | 0 | 4.71E-05 | 2.43E-05 | 4.43E-05 | 2.22E-06 | 2.53E-05 | 3.20E-05 | 2.86E-05 | 6.91E-05 | 0 | 7.17E-06 |
| Yif1a | 0.00043 | 2.05E-05 | 0.00016 | 0.00021 | 4.60E-05 | 0.00024 | 0.00042 | 8.29E-05 | 4.70E-05 | 4.10E-05 | 0.00015 | 9.03E-05 | 6.69E-05 | 7.07E-05 | 0 | 0 | 0 | 2.39E-05 |
| Adam10 | 0.00024 | 0.00013 | 0.00015 | 8.69E-05 | 7.13E-05 | 0.00016 | 0.00013 | 0.00012 | 4.70E-05 | 0.00016 | 0.00012 | 0 | 0.00011 | 0.00016 | 0 | 0.00021 | 8.26E-05 | 9.50E-05 |
| Poglut1 | 0.00012 | 6.19E-05 | 3.95E-05 | 0 | 7.01E-05 | 0.00014 | 3.24E-05 | 3.48E-06 | 4.70E-05 | 5.56E-05 | 4.38E-05 | 0 | 5.79E-05 | 1.71E-05 | 1.89E-05 | 0 | 1.06E-05 | 7.92E-05 |
| Xpo7 | 0.00014 | 0.0001 | 0.00013 | 7.37E-05 | 5.11E-05 | 8.17E-05 | 7.63E-05 | 3.92E-05 | 4.70E-05 | 4.07E-05 | 2.35E-05 | 3.33E-05 | 3.68E-05 | 5.52E-05 | 1.67E-05 | 0.00014 | 0.00011 | 1.85E-05 |
| Terf2 | 8.00E-05 | 0.00013 | 2.60E-05 | 1.83E-05 | 2.53E-05 | 4.39E-05 | 2.57E-05 | 2.78E-05 | 4.70E-05 | 1.54E-05 | 0 | 6.68E-05 | 2.34E-06 | 1.17E-05 | 0.0002 | 0.00016 | 0 | 4.80E-05 |
| Ip6k2 | 8.95E-05 | 0.00016 | 1.30E-05 | 5.13E-05 | 5.62E-05 | 4.06E-05 | 0 | 3.66E-05 | 4.70E-05 | 0 | 3.96E-05 | 7.93E-05 | 0 | 6.35E-05 | 0.00038 | 1.59E-05 | 2.16E-05 | 5.07E-05 |
| Plekhf2 | 2.85E-05 | 6.74E-05 | 3.91E-05 | 1.94E-05 | 3.24E-06 | 1.62E-05 | 0 | 1.61E-05 | 4.70E-05 | 5.03E-05 | 4.99E-05 | 3.68E-05 | 2.71E-06 | 1.93E-05 | 4.13E-06 | 5.60E-05 | 1.31E-05 | 2.30E-05 |
| Prmt9 | 4.53E-05 | 3.92E-05 | 1.51E-05 | 1.31E-05 | 2.57E-05 | 9.62E-06 | 4.68E-06 | 2.35E-06 | 4.70E-05 | 2.33E-05 | 0 | 5.20E-06 | 0 | 0 | 4.75E-05 | 1.50E-05 | 0 | 4.71E-05 |
| Rev3l | 0.00011 | 6.21E-05 | 0 | 7.39E-06 | 9.72E-07 | 6.14E-05 | 5.33E-05 | 0 | 4.69E-05 | 8.92E-05 | 0 | 7.77E-06 | 7.90E-05 | 2.44E-06 | 4.15E-05 | 6.95E-05 | 7.41E-05 | 0 |
| Zfp507 | 7.71E-05 | 9.23E-05 | 4.35E-05 | 6.06E-05 | 5.76E-05 | 5.99E-05 | 5.60E-05 | 9.65E-06 | 4.69E-05 | 3.84E-06 | 4.14E-05 | 4.37E-05 | 2.20E-05 | 3.46E-05 | 8.06E-05 | 1.22E-05 | 0 | 0 |
| Edem2 | 0.00016 | 5.13E-05 | 0.00011 | 0.0001 | 1.85E-05 | 4.76E-05 | 0.00021 | 2.38E-05 | 4.69E-05 | 0.00019 | 2.30E-05 | 7.03E-05 | 7.22E-05 | 6.16E-05 | 7.81E-05 | 0 | 0 | 1.71E-05 |
| Hectd4 | 3.19E-05 | 0.0001 | 4.11E-05 | 1.87E-05 | 4.29E-05 | 4.03E-05 | 3.61E-05 | 2.84E-05 | 4.69E-05 | 5.80E-05 | 8.10E-05 | 0.00011 | 0 | 5.64E-05 | 0.00015 | 3.71E-05 | 2.80E-05 | 6.29E-05 |
| Ate1 | 5.01E-05 | 8.07E-05 | 3.86E-05 | 4.62E-05 | 3.83E-05 | 3.08E-05 | 1.32E-06 | 0 | 4.69E-05 | 2.13E-05 | 8.42E-05 | 2.92E-05 | 1.47E-05 | 5.67E-06 | 0.00013 | 8.88E-05 | 7.31E-05 | 2.22E-05 |
| Shank3 | 0 | 3.38E-05 | 4.19E-05 | 5.91E-05 | 6.41E-05 | 1.76E-05 | 1.20E-05 | 7.90E-05 | 4.69E-05 | 2.55E-05 | 7.35E-05 | 0 | 0.00011 | 0.00013 | 8.38E-05 | 1.40E-05 | 0 | 7.23E-05 |
| Wdr43 | 0.00015 | 5.57E-05 | 0.00016 | 0 | 3.11E-06 | 8.79E-06 | 0 | 0 | 4.69E-05 | 7.52E-06 | 5.34E-06 | 1.48E-05 | 0 | 0 | 0.00023 | 2.46E-05 | 0 | 0 |
| Oxsm | 3.05E-05 | 4.46E-05 | 1.42E-05 | 4.63E-05 | 1.07E-05 | 5.03E-06 | 4.49E-05 | 2.41E-05 | 4.69E-05 | 4.63E-05 | 7.07E-06 | 0 | 3.05E-05 | 0 | 1.87E-05 | 2.56E-05 | 0 | 4.20E-05 |
| Myo18a | 0 | 4.22E-05 | 1.19E-05 | 1.73E-05 | 4.50E-05 | 0 | 1.11E-05 | 0 | 4.69E-05 | 2.13E-05 | 8.94E-05 | 0 | 3.15E-05 | 7.43E-05 | 0.00013 | 2.70E-05 | 7.23E-06 | 9.49E-05 |
| Fiz1 | 8.21E-06 | 8.40E-05 | 6.52E-05 | 0.00013 | 4.24E-05 | 0 | 1.68E-05 | 2.27E-05 | 4.69E-05 | 3.16E-05 | 5.37E-05 | 4.32E-05 | 2.33E-05 | 0 | 7.00E-05 | 4.10E-05 | 0 | 3.21E-05 |
| Med20 | 0.00013 | 0.00017 | 0.00011 | 0.00015 | 8.19E-05 | 3.15E-05 | 7.70E-05 | 2.72E-05 | 4.68E-05 | 2.87E-05 | 0 | 9.57E-05 | 4.10E-05 | 8.39E-05 | 0.00025 | 1.58E-05 | 0 | 1.84E-05 |
| Spata24 | 2.05E-05 | 9.10E-06 | 0 | 3.88E-05 | 0 | 1.25E-05 | 0 | 3.13E-06 | 4.68E-05 | 1.04E-05 | 6.91E-06 | 0 | 6.89E-06 | 3.20E-05 | 2.80E-05 | 7.50E-05 | 9.73E-06 | 0 |
| Lman2l | 4.04E-05 | 7.56E-05 | 0 | 2.08E-05 | 0.0001 | 1.90E-06 | 3.54E-05 | 4.02E-05 | 4.68E-05 | 2.10E-05 | 0.00014 | 4.58E-05 | 0 | 8.01E-05 | 3.75E-05 | 1.08E-05 | 2.79E-06 | 1.26E-05 |
| Hgh1 | 1.88E-05 | 5.17E-05 | 8.61E-05 | 5.11E-05 | 0 | 0 | 0 | 0 | 4.68E-05 | 1.47E-05 | 8.20E-06 | 0 | 0 | 2.26E-05 | 7.58E-06 | 0 | 0 | 0 |
| Dhps | 4.18E-05 | 0.00012 | 0.00015 | 6.43E-05 | 1.25E-05 | 6.55E-05 | 6.17E-06 | 6.17E-05 | 4.67E-05 | 0 | 6.41E-05 | 0.00011 | 0 | 0 | 8.83E-05 | 3.27E-05 | 1.85E-05 | 0 |
| Fbxo22 | 0.00013 | 8.07E-05 | 8.24E-05 | 8.32E-05 | 2.64E-05 | 4.93E-05 | 2.26E-05 | 4.26E-05 | 4.67E-05 | 2.24E-05 | 3.84E-05 | 0 | 3.02E-05 | 3.75E-05 | 1.70E-06 | 6.76E-05 | 7.37E-05 | 5.09E-05 |
| Kif13a | 2.99E-05 | 9.71E-05 | 6.94E-05 | 7.14E-05 | 5.96E-05 | 1.63E-05 | 0.00015 | 3.55E-05 | 4.67E-05 | 9.85E-05 | 6.91E-05 | 3.67E-05 | 0 | 2.56E-05 | 0.00013 | 7.17E-05 | 9.95E-05 | 8.59E-05 |
| Skp2 | 6.36E-06 | 3.63E-06 | 4.22E-05 | 1.29E-05 | 7.54E-06 | 9.39E-07 | 2.00E-05 | 2.04E-05 | 4.67E-05 | 4.81E-06 | 2.13E-05 | 0 | 0 | 2.17E-05 | 4.78E-05 | 0.00029 | 0 | 5.06E-06 |
| Fzd8 | 1.96E-07 | 4.27E-05 | 0 | 0.0001 | 3.26E-05 | 0 | 0 | 7.63E-06 | 4.67E-05 | 1.27E-05 | 8.53E-05 | 1.07E-05 | 5.21E-05 | 0.00011 | 1.26E-05 | 0 | 0.00027 | 9.22E-05 |
| Paip2 | 0.00056 | 0.00025 | 4.76E-05 | 9.21E-05 | 0.00026 | 0.00051 | 0.00048 | 0.0002 | 4.66E-05 | 0.00012 | 8.45E-05 | 0.0004 | 0.00025 | 0.00022 | 0.00023 | 7.56E-05 | 0.00024 | 0.00016 |
| Aig1 | 3.12E-05 | 3.95E-05 | 7.24E-05 | 0 | 0 | 0.00018 | 0 | 0.00022 | 4.66E-05 | 0.00011 | 0 | 0 | 0 | 0.00011 | 0 | 2.36E-05 | 0 | 0.00016 |

| | | | | | | | | | | | | | | | | | | |
|---|---|---|---|---|---|---|---|---|---|---|---|---|---|---|---|---|---|---|
| N4bp2l2 | 0.00022 | 0.00017 | 7.92E-05 | 0 | 7.35E-05 | 8.11E-05 | 9.17E-06 | 0 | 4.66E-05 | 1.23E-05 | 4.72E-05 | 0.00012 | 0 | 0 | 0.00025 | 6.71E-05 | 0 | 3.17E-05 |
| Dcp1a | 5.64E-05 | 6.84E-05 | 8.72E-05 | 5.45E-05 | 3.76E-05 | 3.39E-05 | 1.56E-05 | 5.65E-05 | 4.66E-05 | 1.25E-05 | 5.81E-05 | 4.47E-05 | 4.80E-05 | 5.29E-05 | 9.76E-05 | 1.43E-05 | 0 | 2.73E-05 |
| Ap4e1 | 1.79E-05 | 7.49E-05 | 2.39E-05 | 1.86E-05 | 1.77E-05 | 2.21E-05 | 0 | 0 | 4.66E-05 | 7.87E-05 | 1.32E-05 | 6.11E-05 | 4.52E-05 | 8.53E-06 | 0.0002 | 5.54E-05 | 9.84E-06 | 2.64E-05 |
| Vhl | 4.80E-05 | 7.62E-05 | 4.36E-06 | 7.94E-05 | 0.00018 | 1.98E-05 | 7.28E-05 | 9.78E-05 | 4.66E-05 | 4.39E-05 | 2.67E-05 | 0.00037 | 0.00011 | 0.0001 | 1.65E-05 | 4.13E-05 | 0 | 9.56E-05 |
| Hpf1 | 0.00013 | 2.34E-05 | 0.0001 | 3.90E-05 | 6.51E-06 | 1.79E-05 | 0 | 0 | 4.66E-05 | 3.61E-05 | 5.17E-06 | 1.31E-05 | 3.72E-06 | 0 | 3.02E-05 | 0.00032 | 0 | 3.86E-05 |
| Msh6 | 3.82E-05 | 3.25E-05 | 7.09E-05 | 1.10E-05 | 1.98E-05 | 0 | 1.22E-06 | 0 | 4.66E-05 | 2.14E-06 | 1.43E-05 | 1.21E-05 | 1.46E-05 | 3.98E-06 | 8.25E-06 | 0.00013 | 2.06E-05 | 9.11E-06 |
| Mlst8 | 3.92E-05 | 5.89E-05 | 7.01E-05 | 6.41E-05 | 0 | 0 | 2.50E-06 | 1.06E-05 | 4.66E-05 | 3.40E-05 | 3.15E-05 | 0 | 1.36E-05 | 1.49E-05 | 2.95E-05 | 7.35E-05 | 0 | 1.80E-05 |
| Snapc5 | 8.78E-05 | 7.86E-05 | 0 | 2.29E-05 | 4.09E-05 | 4.53E-05 | 0 | 4.40E-05 | 4.65E-05 | 0.00014 | 4.39E-05 | 0 | 0 | 5.38E-05 | 0 | 3.55E-05 | 0 | 0.00018 |
| Taf11 | 9.05E-05 | 9.44E-05 | 6.26E-05 | 0.00016 | 8.10E-05 | 3.24E-05 | 3.90E-05 | 2.43E-05 | 4.65E-05 | 3.42E-05 | 3.91E-05 | 2.82E-05 | 1.49E-05 | 5.53E-05 | 9.05E-05 | 0.00012 | 0 | 1.83E-05 |
| Hexdc | 8.21E-06 | 3.88E-05 | 0 | 1.10E-05 | 7.86E-06 | 1.37E-05 | 6.08E-05 | 0 | 4.65E-05 | 0 | 1.11E-05 | 2.80E-05 | 1.94E-05 | 2.72E-05 | 2.04E-05 | 4.43E-06 | 0 | 3.79E-05 |
| Pwwp2b | 0 | 3.15E-05 | 5.35E-06 | 1.09E-05 | 0 | 8.26E-06 | 0 | 0 | 4.65E-05 | 3.04E-05 | 1.44E-05 | 1.49E-05 | 1.83E-05 | 5.17E-05 | 4.25E-06 | 0 | 0 | 4.47E-05 |
| Arl13b | 8.93E-05 | 0.00011 | 4.60E-05 | 4.72E-05 | 6.18E-05 | 8.86E-05 | 0 | 6.93E-05 | 4.64E-05 | 5.68E-05 | 2.72E-06 | 0.00022 | 4.71E-05 | 6.26E-05 | 7.55E-05 | 0.00011 | 0.00033 | 4.17E-05 |
| Polr2i | 0.00011 | 5.56E-05 | 8.00E-05 | 8.88E-05 | 2.92E-05 | 2.04E-06 | 3.18E-05 | 6.97E-06 | 4.64E-05 | 0 | 3.30E-05 | 0 | 0 | 3.16E-05 | 4.10E-05 | 6.35E-05 | 0 | 7.80E-06 |
| Atf5 | 0 | 0.00031 | 0.00029 | 0.00055 | 0.00026 | 0.00025 | 3.99E-05 | 0.00017 | 4.63E-05 | 0.00033 | 2.00E-05 | 0.00053 | 0 | 0 | 0.00041 | 1.53E-05 | 0.00039 | 0.00021 |
| Impact | 0.00015 | 0.00021 | 5.87E-05 | 0.0001 | 5.05E-05 | 0.00017 | 3.04E-05 | 3.16E-05 | 4.63E-05 | 7.33E-05 | 0 | 0 | 0.00016 | 0.00013 | 0.00025 | 0 | 0.00022 | 0.00011 |
| Pcdhb17 | 4.06E-05 | 7.61E-05 | 1.62E-05 | 0 | 9.22E-05 | 6.91E-05 | 8.30E-06 | 1.99E-05 | 4.63E-05 | 6.40E-06 | 2.19E-05 | 1.03E-05 | 5.60E-05 | 3.06E-05 | 0 | 0 | 5.02E-05 | 3.71E-05 |
| Ralgapa1 | 0.00012 | 0.00015 | 6.52E-05 | 0 | 4.66E-05 | 4.86E-05 | 4.31E-05 | 8.53E-06 | 4.63E-05 | 6.88E-05 | 2.91E-05 | 8.38E-05 | 0 | 0 | 0.00018 | 7.11E-05 | 0.0002 | 6.37E-05 |
| Slc15a4 | 2.21E-05 | 9.27E-05 | 5.80E-05 | 7.32E-05 | 6.51E-05 | 2.92E-05 | 0 | 4.90E-06 | 4.63E-05 | 4.54E-05 | 3.76E-05 | 0.00015 | 0 | 5.85E-05 | 0.00014 | 4.50E-05 | 0 | 4.38E-05 |
| Ears2 | 3.31E-05 | 3.92E-05 | 5.17E-05 | 4.71E-05 | 8.18E-06 | 9.45E-06 | 0 | 5.90E-06 | 4.63E-05 | 1.52E-05 | 2.34E-05 | 1.70E-05 | 0 | 5.06E-06 | 5.40E-05 | 0 | 0 | 0 |
| Rab9 | 0.00017 | 0.0002 | 0 | 0.00012 | 8.77E-05 | 0.00022 | 2.94E-05 | 0 | 4.62E-05 | 6.62E-05 | 6.31E-05 | 0 | 0.00013 | 0.00018 | 0 | 6.16E-05 | 0 | 3.16E-05 |
| Abhd14b | 4.56E-06 | 7.10E-05 | 0 | 5.19E-05 | 5.86E-05 | 4.37E-05 | 0 | 1.36E-05 | 4.62E-05 | 9.56E-05 | 0 | 2.50E-05 | 1.26E-05 | 4.20E-05 | 3.14E-06 | 0 | 0 | 0.00015 |
| Pole2 | 3.54E-05 | 0 | 6.80E-05 | 2.09E-05 | 0 | 0 | 9.36E-06 | 1.09E-05 | 4.62E-05 | 2.35E-05 | 0 | 1.22E-05 | 8.39E-07 | 0 | 2.17E-05 | 0.00022 | 0 | 8.67E-07 |
| Insr | 9.53E-05 | 0.0002 | 0 | 0 | 0.00013 | 0.0002 | 5.80E-05 | 3.21E-06 | 4.61E-05 | 1.66E-05 | 4.26E-05 | 0.00017 | 0.00013 | 8.53E-05 | 0.00021 | 1.69E-05 | 0.00017 | 0 |
| Usp28 | 2.55E-05 | 4.70E-05 | 8.12E-06 | 0 | 1.62E-05 | 4.14E-05 | 0 | 0 | 4.61E-05 | 1.40E-05 | 0 | 2.22E-05 | 2.03E-05 | 3.41E-05 | 2.81E-05 | 4.30E-05 | 2.11E-05 | 7.58E-06 |
| B3gnt2 | 4.79E-05 | 0.00012 | 5.76E-05 | 0 | 0.0001 | 1.31E-05 | 0 | 3.66E-05 | 4.61E-05 | 0 | 0.00017 | 3.19E-05 | 0 | 8.13E-05 | 0 | 6.88E-05 | 0.00014 | 0.00011 |
| Pla2g6 | 1.11E-05 | 5.66E-05 | 3.98E-05 | 4.20E-05 | 2.97E-05 | 0 | 3.25E-05 | 2.52E-06 | 4.61E-05 | 2.17E-05 | 1.55E-06 | 6.62E-06 | 0 | 3.21E-05 | 4.72E-05 | 0 | 6.29E-05 | 2.42E-05 |
| Gnl3 | 0.00032 | 0.00011 | 0.00042 | 9.85E-05 | 0 | 9.92E-05 | 5.27E-05 | 0 | 4.60E-05 | 0 | 0 | 2.23E-05 | 7.02E-05 | 7.99E-05 | 0.00038 | 2.78E-05 | 4.77E-05 | 0 |
| Spast | 0.00014 | 0.00017 | 7.78E-05 | 8.83E-05 | 0.00014 | 9.65E-05 | 9.27E-05 | 3.74E-05 | 4.60E-05 | 1.35E-05 | 9.61E-05 | 5.74E-05 | 0 | 4.63E-05 | 4.49E-05 | 8.34E-05 | 6.91E-05 | 4.85E-05 |
| Wrnip1 | 6.90E-05 | 0.00011 | 0.0002 | 0.00011 | 6.21E-05 | 7.87E-05 | 0 | 6.31E-05 | 4.60E-05 | 4.37E-05 | 7.59E-05 | 0.00013 | 4.58E-05 | 0 | 2.05E-05 | 0.00022 | 0.00017 | 9.03E-05 |
| Pcdhga9 | 2.34E-06 | 8.14E-05 | 0 | 1.14E-05 | 7.99E-05 | 2.94E-05 | 2.76E-05 | 5.10E-05 | 4.60E-05 | 3.50E-05 | 1.44E-05 | 2.74E-05 | 4.14E-05 | 3.58E-05 | 0.00012 | 0 | 6.56E-05 | 3.71E-05 |
| Txnrd2 | 1.41E-05 | 3.19E-05 | 3.71E-05 | 8.91E-05 | 7.73E-06 | 0 | 7.49E-05 | 2.38E-05 | 4.60E-05 | 1.29E-05 | 3.89E-05 | 4.43E-05 | 3.29E-05 | 7.45E-06 | 6.74E-05 | 2.46E-05 | 0 | 5.57E-05 |
| Mau2 | 0.00012 | 0.00023 | 8.21E-05 | 5.29E-05 | 0.00013 | 0.00012 | 7.22E-05 | 3.68E-05 | 4.59E-05 | 7.01E-05 | 5.86E-05 | 0.00011 | 2.41E-05 | 3.98E-05 | 0.00026 | 2.27E-05 | 5.79E-05 | 7.22E-05 |
| Usp1 | 0.00011 | 3.06E-05 | 0.00011 | 0 | 0 | 8.64E-05 | 3.72E-05 | 0 | 4.59E-05 | 1.12E-05 | 0 | 0 | 3.66E-05 | 1.31E-05 | 0 | 0.00059 | 0 | 0 |

| | | | | | | | | | | | | | | | | | | |
|---|---|---|---|---|---|---|---|---|---|---|---|---|---|---|---|---|---|---|
| Siah1a | 2.40E-05 | 5.28E-05 | 2.66E-05 | 7.12E-06 | 5.66E-05 | 4.83E-05 | 3.82E-05 | 0 | 4.59E-05 | 1.65E-05 | 2.51E-05 | 2.98E-05 | 2.50E-05 | 1.68E-05 | 1.08E-05 | 2.70E-05 | 0 | 6.17E-06 |
| Zfp24 | 6.36E-05 | 8.64E-05 | 4.53E-05 | 5.84E-05 | 1.66E-05 | 4.60E-05 | 7.51E-06 | 4.71E-05 | 4.59E-05 | 1.76E-05 | 3.54E-05 | 4.73E-06 | 0 | 7.57E-05 | 8.65E-05 | 4.98E-05 | 2.74E-05 | 0 |
| Snn | 1.31E-08 | 4.08E-05 | 4.56E-06 | 2.80E-05 | 2.35E-05 | 2.66E-05 | 0 | 2.77E-05 | 4.59E-05 | 9.73E-06 | 2.24E-05 | 0 | 2.09E-05 | 6.23E-05 | 3.50E-05 | 1.31E-05 | 0 | 4.11E-05 |
| Ahcyl2 | 0 | 6.57E-05 | 4.15E-05 | 1.40E-05 | 6.29E-06 | 1.70E-05 | 1.20E-05 | 7.11E-05 | 4.59E-05 | 3.63E-05 | 2.93E-05 | 6.53E-05 | 5.24E-06 | 0 | 4.23E-06 | 0 | 1.69E-05 | 2.81E-05 |
| Rars | 0.00024 | 4.78E-05 | 0.00027 | 0.00012 | 0 | 1.25E-05 | 4.91E-05 | 0 | 4.59E-05 | 5.13E-06 | 4.95E-05 | 6.64E-05 | 2.76E-05 | 3.28E-05 | 5.55E-05 | 0.00011 | 2.40E-05 | 3.70E-05 |
| Armt1 | 8.02E-05 | 4.74E-05 | 2.68E-05 | 2.43E-05 | 5.96E-06 | 7.13E-06 | 4.72E-05 | 2.40E-05 | 4.59E-05 | 3.13E-05 | 8.69E-06 | 0 | 0 | 7.73E-06 | 7.06E-05 | 0.00012 | 0 | 4.30E-05 |
| Helz | 0.00014 | 0.0001 | 4.96E-05 | 3.40E-05 | 8.33E-05 | 9.39E-05 | 7.06E-05 | 2.85E-05 | 4.58E-05 | 3.36E-05 | 4.41E-05 | 9.09E-05 | 8.03E-06 | 5.87E-05 | 0.00019 | 1.84E-05 | 7.43E-05 | 3.52E-05 |
| Pcdhgb6 | 1.79E-05 | 3.69E-05 | 0 | 3.04E-05 | 4.31E-05 | 4.91E-05 | 3.76E-05 | 5.97E-05 | 4.58E-05 | 1.18E-05 | 3.27E-05 | 8.39E-06 | 6.29E-05 | 5.14E-05 | 5.01E-05 | 0 | 6.45E-06 | 2.06E-05 |
| A2m | 0 | 0 | 0 | 0 | 0 | 7.88E-06 | 4.57E-05 | 0 | 4.58E-05 | 0 | 2.53E-05 | 0.00035 | 1.40E-06 | 0 | 0 | 0 | 0 | 0 |
| Srprb | 0.0001 | 4.27E-05 | 0.00032 | 0.00038 | 3.57E-05 | 6.74E-06 | 0.00018 | 0.00011 | 4.58E-05 | 3.07E-05 | 0.00013 | 7.22E-06 | 5.19E-05 | 2.78E-05 | 0.00012 | 0.00021 | 3.25E-06 | 0 |
| Ppif | 2.99E-05 | 1.10E-05 | 8.28E-05 | 6.37E-05 | 7.42E-06 | 7.73E-07 | 0 | 8.06E-06 | 4.58E-05 | 3.75E-06 | 8.17E-06 | 2.55E-05 | 3.99E-05 | 3.84E-06 | 2.84E-05 | 7.76E-05 | 0.00014 | 0 |
| Galns | 4.19E-05 | 3.22E-05 | 2.84E-06 | 6.33E-05 | 7.73E-05 | 0.00012 | 7.49E-05 | 0.00013 | 4.57E-05 | 1.77E-05 | 2.25E-06 | 5.27E-05 | 6.43E-05 | 8.13E-05 | 0 | 0.0001 | 0 | 7.49E-05 |
| Pcmtd2 | 4.86E-05 | 0.00017 | 0 | 0 | 9.70E-05 | 8.04E-05 | 0 | 0 | 4.57E-05 | 5.22E-05 | 0 | 7.37E-05 | 4.11E-05 | 6.09E-05 | 0.00011 | 0 | 0 | 8.83E-05 |
| Ppcs | 5.76E-05 | 3.28E-05 | 1.99E-05 | 1.62E-05 | 4.08E-05 | 3.42E-05 | 1.20E-05 | 4.81E-06 | 4.57E-05 | 1.24E-05 | 0 | 0 | 0 | 6.74E-06 | 0 | 0 | 0 | 2.48E-05 |
| Map3k10 | 6.37E-05 | 7.27E-05 | 7.32E-05 | 3.60E-05 | 5.56E-05 | 3.05E-05 | 8.32E-05 | 2.06E-05 | 4.57E-05 | 4.65E-05 | 7.51E-05 | 1.24E-05 | 3.06E-05 | 4.91E-05 | 2.54E-05 | 7.43E-06 | 6.47E-05 | 4.87E-05 |
| Nckap5l | 1.73E-05 | 4.50E-05 | 2.28E-05 | 8.96E-05 | 3.80E-05 | 1.12E-05 | 6.90E-06 | 3.38E-05 | 4.57E-05 | 0 | 8.56E-06 | 1.82E-06 | 0 | 4.29E-05 | 5.57E-05 | 0 | 0.00025 | 2.36E-05 |
| Ttc27 | 5.37E-05 | 3.33E-05 | 8.28E-05 | 4.06E-05 | 0 | 0 | 0 | 0 | 4.57E-05 | 4.71E-06 | 1.09E-05 | 0 | 1.80E-05 | 0 | 3.91E-05 | 1.36E-05 | 5.29E-05 | 1.21E-05 |
| Mtfr1 | 8.37E-05 | 0.0001 | 0.00012 | 9.11E-05 | 8.57E-05 | 8.06E-05 | 9.33E-05 | 3.90E-05 | 4.56E-05 | 0 | 0.00015 | 4.53E-05 | 1.88E-05 | 0.00011 | 6.51E-05 | 4.73E-05 | 2.16E-05 | 4.70E-06 |
| B3glct | 0.00012 | 2.99E-05 | 2.49E-05 | 0 | 4.90E-05 | 7.96E-05 | 3.83E-05 | 4.46E-05 | 4.56E-05 | 2.47E-05 | 0 | 0 | 7.94E-05 | 0.00011 | 0 | 8.06E-05 | 0 | 6.24E-05 |
| Pex13 | 0.00012 | 8.79E-05 | 4.66E-05 | 0 | 6.15E-05 | 6.71E-05 | 3.45E-05 | 0 | 4.56E-05 | 3.03E-05 | 0 | 6.44E-05 | 1.24E-05 | 5.78E-06 | 2.39E-05 | 4.84E-05 | 9.18E-05 | 2.59E-05 |
| Tmem126b | 9.67E-05 | 2.82E-05 | 0 | 1.46E-05 | 3.92E-05 | 5.52E-05 | 1.21E-05 | 3.45E-05 | 4.56E-05 | 3.74E-05 | 0 | 2.49E-05 | 7.59E-06 | 1.47E-05 | 0 | 3.94E-05 | 0 | 2.44E-05 |
| Ahi1 | 1.03E-05 | 1.37E-05 | 6.31E-05 | 8.77E-06 | 3.05E-05 | 5.09E-05 | 3.65E-05 | 7.17E-05 | 4.56E-05 | 2.68E-05 | 0 | 4.03E-06 | 8.73E-06 | 1.51E-05 | 1.30E-05 | 2.90E-05 | 3.96E-05 | 4.88E-05 |
| Coq9 | 8.00E-05 | 5.78E-05 | 3.25E-05 | 8.32E-05 | 5.66E-05 | 4.12E-05 | 9.39E-05 | 5.04E-05 | 4.56E-05 | 1.85E-05 | 0 | 1.44E-05 | 2.84E-05 | 3.58E-05 | 3.39E-05 | 2.86E-05 | 6.08E-06 | 2.90E-05 |
| Naa40 | 1.03E-05 | 5.68E-05 | 4.92E-05 | 1.05E-05 | 5.86E-05 | 4.11E-05 | 9.09E-06 | 6.42E-05 | 4.56E-05 | 2.31E-05 | 4.56E-05 | 2.37E-05 | 3.68E-05 | 4.35E-05 | 2.76E-05 | 0.00021 | 0 | 2.29E-05 |
| Lrch2 | 3.40E-05 | 2.26E-05 | 0 | 0 | 7.03E-05 | 1.10E-05 | 6.77E-06 | 6.01E-05 | 4.56E-05 | 0 | 0.00017 | 0.0001 | 0 | 0 | 0 | 9.93E-06 | 4.09E-05 | 0 |
| Zbtb45 | 2.13E-05 | 4.49E-05 | 2.08E-05 | 0.0001 | 3.13E-05 | 9.07E-06 | 0 | 6.00E-05 | 4.56E-05 | 2.20E-05 | 2.98E-05 | 0 | 0 | 3.83E-05 | 1.97E-05 | 6.27E-05 | 0 | 5.02E-05 |
| Btbd6 | 1.04E-05 | 1.68E-05 | 7.94E-06 | 6.41E-05 | 2.29E-06 | 5.67E-06 | 5.35E-06 | 0 | 4.56E-05 | 2.16E-05 | 6.86E-06 | 0 | 0 | 5.88E-05 | 0 | 5.24E-05 | 0 | 4.66E-05 |
| Slc30a9 | 0.00018 | 8.47E-05 | 7.61E-05 | 0 | 9.45E-05 | 0.00017 | 3.67E-05 | 8.76E-05 | 4.55E-05 | 2.96E-05 | 7.46E-05 | 0.00029 | 1.47E-05 | 0 | 1.26E-06 | 3.01E-05 | 2.60E-05 | 1.82E-06 |
| Kpna1 | 0.00014 | 0.00012 | 0.00024 | 0 | 3.73E-05 | 5.28E-05 | 0 | 1.16E-05 | 4.55E-05 | 5.00E-05 | 7.94E-05 | 4.91E-05 | 1.49E-06 | 0 | 4.36E-05 | 4.77E-05 | 9.91E-05 | 8.17E-06 |
| Fig4 | 6.01E-05 | 7.76E-05 | 2.87E-05 | 6.66E-06 | 3.56E-05 | 2.83E-05 | 2.72E-05 | 0 | 4.55E-05 | 8.78E-05 | 3.07E-05 | 0 | 1.18E-05 | 2.74E-06 | 0 | 3.21E-05 | 0 | 1.99E-05 |
| Nup54 | 8.31E-05 | 7.48E-05 | 0.00011 | 0 | 1.57E-05 | 2.93E-05 | 0 | 0 | 4.54E-05 | 1.20E-05 | 0 | 9.46E-05 | 1.29E-05 | 0 | 5.64E-05 | 0.00013 | 2.70E-05 | 1.19E-06 |
| Qtrt1 | 8.74E-05 | 7.61E-05 | 0.00014 | 6.34E-05 | 0 | 2.37E-05 | 1.10E-05 | 0 | 4.54E-05 | 0 | 0 | 1.55E-05 | 9.53E-07 | 3.36E-06 | 0.00016 | 0 | 0.00017 | 0 |
| Plekha4 | 0 | 0 | 0 | 9.90E-06 | 0 | 1.28E-05 | 0 | 7.63E-05 | 4.54E-05 | 0.00019 | 0 | 2.91E-05 | 8.99E-05 | 0 | 0.00012 | 0 | 0 | 0.00025 |

| | | | | | | | | | | | | | | | | | | |
|---|---|---|---|---|---|---|---|---|---|---|---|---|---|---|---|---|---|---|
| Sp100 | 5.82E-05 | 1.38E-05 | 0 | 0 | 0 | 8.08E-06 | 0 | 0 | 4.54E-05 | 0.00027 | 0 | 0 | 7.47E-05 | 0 | 2.53E-05 | 1.28E-06 | 0 | 0.00016 |
| Mcm3 | 0 | 0 | 0.00018 | 4.30E-06 | 0 | 0 | 1.10E-06 | 0 | 4.54E-05 | 3.31E-05 | 1.96E-05 | 0 | 4.36E-05 | 1.00E-05 | 0 | 0.00063 | 0 | 1.08E-05 |
| Bdkrb2 | 0 | 3.09E-05 | 0.00021 | 0 | 0 | 0 | 0 | 0.00017 | 4.54E-05 | 0 | 0 | 4.96E-05 | 0 | 0 | 0 | 0 | 6.68E-05 | 6.42E-05 |
| Gpr180 | 0.0003 | 0 | 0.00022 | 0.00031 | 0.00015 | 0.00024 | 0.00052 | 0.00013 | 4.53E-05 | 6.14E-05 | 0.0001 | 0 | 0.00019 | 0.00014 | 0.00011 | 0 | 0 | 3.21E-05 |
| Col6a5 | 0 | 3.53E-06 | 0 | 0 | 0 | 0.00015 | 0 | 4.19E-05 | 4.53E-05 | 2.08E-05 | 0 | 0 | 0 | 0 | 0 | 0 | 0 | 0 |
| Ssbp2 | 6.24E-05 | 0.0001 | 1.09E-05 | 0 | 3.86E-05 | 6.66E-05 | 8.80E-05 | 1.72E-06 | 4.53E-05 | 0.00018 | 0 | 2.86E-05 | 0.00015 | 0 | 9.02E-05 | 1.56E-05 | 0 | 9.18E-05 |
| Bloc1s6 | 7.59E-05 | 5.09E-05 | 2.46E-05 | 6.05E-05 | 2.34E-05 | 5.11E-05 | 1.41E-06 | 2.99E-05 | 4.53E-05 | 4.11E-05 | 2.96E-05 | 2.81E-05 | 2.84E-05 | 0 | 0 | 1.36E-05 | 0 | 5.61E-06 |
| Prpf40b | 2.75E-05 | 9.51E-05 | 2.65E-05 | 2.17E-05 | 2.51E-05 | 1.78E-05 | 0 | 1.12E-05 | 4.53E-05 | 5.57E-06 | 9.75E-06 | 3.69E-05 | 0 | 2.01E-05 | 0.0002 | 0 | 5.25E-05 | 6.73E-06 |
| Atp6v1b2 | 7.43E-05 | 0.00036 | 0.0001 | 0.00014 | 0.00017 | 0.00025 | 2.25E-07 | 0.00033 | 4.52E-05 | 0.00021 | 4.78E-05 | 0.00029 | 0.00013 | 0.00035 | 8.77E-05 | 1.04E-05 | 0.00024 | 0.00011 |
| Srek1ip1 | 6.57E-05 | 3.49E-05 | 2.32E-06 | 4.56E-05 | 3.44E-05 | 6.34E-05 | 1.37E-05 | 2.34E-05 | 4.52E-05 | 8.69E-06 | 7.50E-05 | 3.69E-05 | 1.38E-05 | 3.35E-05 | 4.82E-05 | 3.14E-05 | 0 | 2.19E-05 |
| Nt5dc1 | 6.95E-05 | 0.00011 | 7.10E-05 | 4.84E-05 | 4.16E-05 | 4.05E-05 | 0 | 2.94E-05 | 4.52E-05 | 5.98E-05 | 3.03E-05 | 6.85E-06 | 1.58E-05 | 2.44E-05 | 0.00013 | 2.06E-05 | 0 | 3.05E-05 |
| Proser1 | 5.89E-05 | 6.89E-05 | 2.16E-05 | 0 | 2.33E-05 | 2.89E-05 | 0 | 2.76E-05 | 4.52E-05 | 4.27E-06 | 3.43E-05 | 1.61E-05 | 0 | 6.47E-06 | 6.15E-05 | 2.21E-05 | 6.77E-05 | 0 |
| Zbtb39 | 6.87E-06 | 6.80E-05 | 4.93E-05 | 4.24E-05 | 2.48E-05 | 1.67E-05 | 0 | 2.77E-05 | 4.52E-05 | 2.31E-05 | 3.06E-05 | 2.25E-06 | 1.51E-06 | 3.54E-05 | 5.80E-05 | 1.65E-05 | 6.07E-06 | 0 |
| Lsg1 | 7.91E-05 | 8.94E-05 | 0.00015 | 6.73E-05 | 2.27E-05 | 1.08E-05 | 0 | 0 | 4.52E-05 | 1.35E-05 | 2.03E-05 | 7.34E-06 | 1.24E-05 | 4.10E-05 | 7.41E-05 | 5.03E-05 | 0 | 2.50E-06 |
| Anks6 | 0 | 2.76E-05 | 6.00E-05 | 0 | 3.27E-05 | 7.74E-06 | 1.27E-05 | 9.90E-05 | 4.52E-05 | 6.77E-05 | 4.39E-05 | 0 | 1.68E-05 | 0 | 4.95E-05 | 4.10E-05 | 5.12E-05 | 6.78E-05 |
| Prmt6 | 3.24E-06 | 4.72E-05 | 4.64E-05 | 8.32E-05 | 1.44E-05 | 2.28E-06 | 0 | 4.54E-05 | 4.52E-05 | 7.05E-06 | 2.33E-05 | 0 | 0 | 1.45E-05 | 4.63E-05 | 0 | 0 | 1.18E-05 |
| Tdp1 | 2.83E-05 | 5.51E-05 | 4.83E-05 | 5.79E-05 | 4.54E-05 | 0 | 0 | 2.85E-05 | 4.52E-05 | 1.90E-05 | 2.15E-05 | 0 | 8.84E-07 | 4.29E-06 | 5.39E-05 | 0.00021 | 6.87E-06 | 1.58E-05 |
| Gan | 0 | 0.00012 | 0.00011 | 4.95E-05 | 6.03E-05 | 0 | 8.75E-05 | 8.35E-05 | 4.52E-05 | 0.0002 | 5.91E-05 | 9.17E-05 | 3.04E-05 | 4.26E-05 | 0.00016 | 4.41E-05 | 7.38E-05 | 0 |
| Ak1 | 0 | 0.00012 | 0.00057 | 0.00028 | 0.00071 | 0.00023 | 0.00043 | 0.00059 | 4.51E-05 | 0.00019 | 0.00049 | 0.00022 | 0.00044 | 0.00018 | 0.00024 | 0 | 0 | 0.00054 |
| Dock4 | 1.91E-05 | 3.31E-05 | 5.71E-06 | 0 | 0.0002 | 0.00021 | 0 | 0 | 4.51E-05 | 5.42E-05 | 0 | 0 | 0.00015 | 0.00037 | 5.76E-05 | 0 | 6.74E-05 | 0 |
| Pbdc1 | 0.00018 | 4.14E-05 | 0.00017 | 0.00016 | 0 | 5.21E-05 | 0.00011 | 5.28E-05 | 4.51E-05 | 3.35E-05 | 4.82E-06 | 0 | 1.49E-05 | 9.94E-06 | 1.67E-05 | 0.00029 | 9.72E-05 | 2.15E-05 |
| Zfp330 | 0.00019 | 6.33E-05 | 0.00018 | 9.80E-05 | 2.43E-05 | 4.26E-05 | 5.67E-05 | 1.22E-05 | 4.51E-05 | 0 | 1.45E-05 | 0.00011 | 1.56E-06 | 9.20E-06 | 0.00019 | 6.66E-05 | 0 | 9.17E-06 |
| Lemd2 | 0 | 0.00033 | 0.00029 | 0.00025 | 0.00018 | 0.00014 | 0 | 4.49E-05 | 4.50E-05 | 6.35E-05 | 9.16E-05 | 0.00028 | 0 | 0.00024 | 0.00058 | 7.94E-05 | 0.00016 | 3.49E-05 |
| Fbxl15 | 3.71E-06 | 2.35E-05 | 3.57E-05 | 8.80E-05 | 3.14E-05 | 1.84E-05 | 8.41E-06 | 4.27E-05 | 4.50E-05 | 8.73E-06 | 0 | 6.87E-05 | 2.85E-05 | 3.29E-05 | 3.91E-05 | 1.93E-05 | 7.72E-06 | 9.54E-06 |
| Tti2 | 2.70E-05 | 5.32E-05 | 2.86E-05 | 6.89E-05 | 2.71E-05 | 9.21E-06 | 0 | 7.98E-06 | 4.50E-05 | 1.34E-05 | 2.96E-05 | 3.22E-05 | 1.39E-05 | 5.11E-07 | 8.43E-05 | 2.74E-05 | 0 | 3.71E-05 |
| Tep1 | 8.05E-05 | 7.83E-05 | 0 | 9.84E-05 | 2.52E-05 | 4.76E-06 | 7.30E-05 | 4.91E-05 | 4.50E-05 | 8.91E-05 | 2.62E-05 | 3.34E-05 | 7.49E-06 | 1.16E-05 | 0.00015 | 3.06E-05 | 6.39E-06 | 0.00011 |
| Myg1 | 4.53E-05 | 4.92E-05 | 9.23E-05 | 0.00012 | 2.26E-05 | 0 | 8.50E-06 | 3.00E-05 | 4.50E-05 | 4.41E-05 | 1.99E-05 | 3.76E-05 | 3.60E-05 | 5.54E-05 | 3.00E-08 | 0.0002 | 7.10E-06 | 2.21E-05 |
| Zfp280d | 0.00011 | 0.00014 | 4.74E-05 | 0 | 6.47E-05 | 0.00013 | 7.21E-06 | 3.91E-05 | 4.49E-05 | 0 | 4.59E-05 | 9.09E-05 | 0 | 3.87E-05 | 0.00026 | 0.0001 | 0 | 4.01E-05 |
| Trim3 | 8.17E-05 | 0.00017 | 5.74E-05 | 0 | 8.47E-05 | 0.00011 | 9.73E-05 | 0 | 4.49E-05 | 3.25E-05 | 0.0002 | 0.00017 | 0 | 9.22E-05 | 0.00016 | 4.09E-05 | 0.00014 | 0.00014 |
| Dipk1b | 0 | 2.93E-05 | 0 | 8.07E-06 | 7.03E-05 | 6.71E-05 | 0 | 5.18E-05 | 4.49E-05 | 4.28E-06 | 1.17E-05 | 3.55E-05 | 0 | 5.52E-05 | 0 | 0 | 0 | 2.91E-05 |
| Parn | 0.00011 | 8.48E-05 | 4.27E-05 | 4.80E-05 | 4.40E-05 | 4.67E-05 | 2.93E-05 | 0 | 4.49E-05 | 3.03E-05 | 2.01E-05 | 6.16E-05 | 5.78E-06 | 1.57E-05 | 4.25E-05 | 5.09E-05 | 0 | 9.18E-05 |
| Abi3 | 0 | 1.67E-05 | 0 | 1.90E-05 | 4.41E-05 | 7.02E-05 | 0 | 9.97E-05 | 4.48E-05 | 6.13E-05 | 0 | 5.01E-06 | 0 | 8.94E-05 | 5.21E-05 | 0 | 0 | 3.20E-05 |
| Smim12 | 4.29E-05 | 6.07E-05 | 5.84E-05 | 0.00015 | 5.48E-05 | 3.86E-05 | 7.39E-06 | 4.23E-05 | 4.48E-05 | 3.99E-05 | 5.16E-05 | 3.90E-05 | 7.76E-06 | 3.48E-05 | 2.50E-05 | 5.61E-05 | 0 | 2.00E-05 |

| | | | | | | | | | | | | | | | | | | |
|---|---|---|---|---|---|---|---|---|---|---|---|---|---|---|---|---|---|---|
| Gclm | 8.88E-05 | 9.92E-05 | 4.13E-05 | 0 | 0 | 1.69E-05 | 0 | 0 | 4.48E-05 | 4.81E-05 | 3.96E-05 | 4.76E-05 | 9.19E-05 | 0 | 0 | 7.00E-05 | 0 | 9.08E-05 |
| Mtx1 | 4.02E-05 | 3.05E-05 | 8.38E-05 | 0.00013 | 1.96E-05 | 9.35E-07 | 0 | 1.59E-05 | 4.48E-05 | 9.12E-06 | 4.84E-05 | 2.76E-06 | 3.79E-05 | 2.50E-05 | 3.28E-05 | 6.29E-05 | 1.06E-05 | 1.64E-05 |
| Dnajc24 | 9.30E-05 | 7.18E-05 | 4.28E-05 | 4.55E-05 | 3.42E-05 | 5.03E-05 | 1.03E-05 | 0 | 4.47E-05 | 3.67E-05 | 1.42E-05 | 7.58E-06 | 2.01E-05 | 5.74E-06 | 9.02E-05 | 5.67E-05 | 0 | 2.50E-05 |
| Saysd1 | 1.21E-05 | 2.48E-05 | 0 | 6.62E-05 | 4.45E-05 | 3.55E-05 | 2.73E-05 | 7.10E-05 | 4.47E-05 | 6.65E-05 | 2.70E-05 | 1.91E-05 | 4.83E-05 | 4.18E-05 | 1.76E-05 | 0 | 0 | 3.36E-05 |
| Wdr83 | 4.06E-05 | 4.12E-05 | 0 | 8.45E-05 | 4.14E-05 | 2.29E-05 | 8.04E-06 | 2.54E-05 | 4.47E-05 | 1.66E-05 | 3.01E-05 | 1.09E-05 | 0 | 3.28E-05 | 4.77E-05 | 4.95E-05 | 0 | 5.98E-05 |
| Ptges2 | 8.59E-05 | 6.84E-05 | 0.00015 | 9.43E-05 | 1.87E-05 | 2.10E-05 | 8.28E-06 | 3.35E-05 | 4.47E-05 | 1.67E-05 | 1.85E-05 | 0 | 3.39E-05 | 1.46E-05 | 3.24E-05 | 5.84E-05 | 0 | 2.01E-05 |
| Haus7 | 8.46E-05 | 4.07E-06 | 0.00013 | 7.74E-05 | 3.25E-05 | 0 | 4.21E-05 | 2.19E-05 | 4.47E-05 | 0 | 4.71E-05 | 0 | 1.60E-05 | 1.44E-05 | 7.20E-05 | 0.00028 | 0 | 8.28E-05 |
| Bbx | 0.00012 | 0.00013 | 0 | 8.26E-05 | 9.22E-05 | 0.00014 | 0 | 6.25E-05 | 4.46E-05 | 5.35E-06 | 0 | 0.00018 | 0 | 1.77E-05 | 0.00011 | 0.00011 | 0.00012 | 4.18E-05 |
| Mark2 | 7.30E-05 | 0.00015 | 9.26E-05 | 6.29E-05 | 5.23E-05 | 5.70E-05 | 0 | 3.91E-05 | 4.46E-05 | 4.97E-05 | 3.45E-05 | 3.34E-05 | 0 | 3.49E-05 | 0.00015 | 7.98E-05 | 0.00015 | 4.60E-05 |
| Pik3c3 | 6.18E-05 | 0.00013 | 1.47E-05 | 7.62E-05 | 5.95E-05 | 4.77E-05 | 4.38E-05 | 0 | 4.46E-05 | 4.49E-05 | 1.63E-05 | 7.35E-05 | 1.38E-05 | 4.63E-05 | 0.00011 | 5.90E-05 | 0 | 6.94E-05 |
| Dgkh | 2.86E-05 | 6.69E-05 | 7.63E-05 | 0 | 3.47E-05 | 3.26E-05 | 1.51E-05 | 3.83E-05 | 4.46E-05 | 0 | 9.81E-05 | 7.64E-05 | 0 | 4.53E-05 | 2.00E-05 | 1.53E-05 | 0.00013 | 2.22E-05 |
| Pcdhga12 | 2.07E-05 | 4.21E-05 | 0 | 1.19E-05 | 3.89E-05 | 1.50E-05 | 4.65E-05 | 5.32E-05 | 4.46E-05 | 2.84E-05 | 1.18E-05 | 5.16E-06 | 6.26E-05 | 4.87E-05 | 0.00017 | 0 | 4.39E-05 | 5.19E-05 |
| Tdg | 7.38E-05 | 2.85E-05 | 0.00012 | 7.05E-05 | 1.77E-05 | 1.45E-05 | 0 | 6.37E-05 | 4.46E-05 | 0 | 6.81E-05 | 4.57E-05 | 0 | 3.42E-05 | 0 | 8.45E-05 | 0 | 0 |
| Rhobtb1 | 5.46E-06 | 1.64E-05 | 0 | 0 | 2.38E-05 | 1.12E-05 | 2.01E-05 | 0 | 4.46E-05 | 1.56E-05 | 1.69E-05 | 5.23E-05 | 3.28E-05 | 4.79E-05 | 0 | 0 | 0 | 0 |
| Ikzf2 | 1.75E-05 | 2.73E-05 | 0 | 0 | 4.65E-05 | 0 | 2.01E-05 | 1.50E-05 | 4.46E-05 | 1.38E-05 | 6.90E-05 | 6.12E-06 | 1.97E-05 | 0 | 0.00013 | 0 | 9.54E-05 | 0 |
| Fbxl17 | 0.00019 | 0.00014 | 6.32E-05 | 0 | 9.10E-05 | 0.00015 | 5.96E-06 | 2.63E-05 | 4.45E-05 | 3.17E-05 | 6.20E-05 | 7.79E-05 | 0 | 0 | 0.00017 | 3.15E-05 | 0.0002 | 3.10E-05 |
| Slc33a1 | 0.00019 | 6.51E-05 | 0.00013 | 0 | 4.24E-05 | 9.91E-05 | 6.88E-05 | 6.08E-05 | 4.45E-05 | 4.06E-05 | 6.05E-05 | 1.72E-05 | 1.57E-05 | 9.25E-06 | 2.21E-05 | 1.46E-05 | 0 | 2.84E-05 |
| Wwp1 | 0.00015 | 8.56E-05 | 9.21E-05 | 0 | 5.06E-05 | 7.41E-05 | 5.63E-05 | 1.09E-05 | 4.45E-05 | 0.00012 | 7.92E-05 | 3.13E-05 | 1.82E-05 | 0 | 6.86E-05 | 8.22E-05 | 0.00012 | 0.00015 |
| Rusc2 | 3.56E-05 | 0.00013 | 1.79E-05 | 4.11E-05 | 8.55E-05 | 5.29E-05 | 2.11E-05 | 0 | 4.45E-05 | 8.49E-05 | 0.00021 | 0.00032 | 0 | 1.44E-05 | 0.00014 | 0 | 0.00022 | 6.55E-05 |
| Lpar6 | 4.43E-05 | 2.54E-05 | 0 | 3.80E-05 | 1.78E-06 | 4.39E-06 | 0 | 2.07E-05 | 4.45E-05 | 1.69E-05 | 2.92E-05 | 0 | 0 | 2.50E-05 | 1.07E-05 | 3.29E-05 | 0 | 8.60E-06 |
| Cebpg | 0.00016 | 0.00024 | 0.00021 | 2.84E-05 | 4.86E-06 | 4.83E-05 | 8.78E-05 | 0 | 4.44E-05 | 2.92E-05 | 9.77E-05 | 0.0002 | 2.68E-05 | 0.0001 | 0.00024 | 0.00011 | 0 | 9.89E-05 |
| Snrk | 5.73E-05 | 8.30E-05 | 0 | 0 | 3.62E-05 | 4.22E-05 | 1.02E-05 | 6.75E-05 | 4.44E-05 | 3.68E-05 | 6.28E-05 | 0.00016 | 0.00012 | 2.92E-05 | 5.06E-05 | 0 | 2.07E-05 | 4.70E-05 |
| Ldlrad3 | 8.40E-06 | 9.54E-05 | 7.60E-05 | 5.46E-05 | 5.63E-05 | 3.28E-05 | 1.62E-05 | 0.0001 | 4.44E-05 | 2.41E-05 | 7.11E-05 | 4.23E-05 | 0 | 4.80E-05 | 5.83E-05 | 3.62E-05 | 7.82E-05 | 5.79E-05 |
| Zfp251 | 5.39E-05 | 3.41E-05 | 4.33E-06 | 7.30E-05 | 3.29E-05 | 8.92E-06 | 5.16E-05 | 4.01E-05 | 4.44E-05 | 2.56E-05 | 5.05E-05 | 0 | 0 | 3.96E-05 | 4.95E-05 | 1.31E-05 | 0 | 6.38E-05 |
| Rmnd1 | 5.07E-05 | 3.16E-05 | 0 | 9.65E-06 | 1.13E-06 | 7.46E-06 | 7.61E-06 | 0 | 4.44E-05 | 1.44E-05 | 0 | 1.00E-05 | 1.04E-05 | 0 | 3.66E-05 | 2.34E-05 | 0 | 2.19E-05 |
| Cactin | 1.32E-05 | 5.67E-05 | 6.07E-05 | 8.17E-05 | 1.23E-05 | 0 | 0 | 0 | 4.44E-05 | 1.73E-05 | 3.21E-05 | 1.97E-05 | 1.66E-05 | 2.99E-05 | 8.96E-05 | 8.28E-05 | 0 | 2.99E-05 |
| Fbxo42 | 3.90E-05 | 0.00011 | 0.00012 | 7.40E-05 | 7.77E-05 | 4.42E-05 | 7.19E-05 | 6.12E-06 | 4.43E-05 | 2.55E-05 | 3.90E-05 | 9.36E-05 | 4.05E-05 | 8.27E-05 | 4.47E-05 | 0.0001 | 0 | 5.45E-05 |
| Fam120b | 8.22E-05 | 0.00011 | 1.01E-05 | 3.29E-05 | 4.97E-05 | 3.88E-05 | 4.87E-05 | 1.67E-05 | 4.43E-05 | 5.09E-05 | 1.80E-05 | 1.95E-05 | 6.89E-07 | 5.50E-05 | 0.00016 | 8.79E-06 | 0 | 2.32E-05 |
| Pex7 | 4.57E-05 | 3.20E-05 | 3.75E-05 | 0 | 1.21E-05 | 2.28E-05 | 1.54E-05 | 2.04E-05 | 4.43E-05 | 1.97E-05 | 4.32E-06 | 8.90E-06 | 1.55E-05 | 9.93E-06 | 6.88E-05 | 2.99E-05 | 0 | 1.92E-05 |
| Ifrd2 | 6.06E-05 | 2.97E-05 | 0.00024 | 0.00014 | 0 | 1.98E-05 | 0 | 0 | 4.43E-05 | 1.22E-05 | 4.17E-06 | 1.10E-05 | 5.77E-05 | 0 | 0.00014 | 0.00027 | 0 | 0 |
| Alg12 | 5.74E-05 | 2.23E-05 | 5.44E-05 | 8.95E-05 | 4.18E-05 | 1.56E-05 | 9.33E-07 | 8.38E-06 | 4.43E-05 | 3.83E-05 | 1.67E-06 | 0 | 1.36E-05 | 1.84E-05 | 8.65E-06 | 1.01E-05 | 0 | 1.74E-05 |
| Msh2 | 5.75E-05 | 3.76E-05 | 2.51E-05 | 4.68E-05 | 1.45E-05 | 4.23E-06 | 4.71E-05 | 8.64E-06 | 4.43E-05 | 3.70E-05 | 3.16E-05 | 0 | 0 | 6.10E-05 | 3.44E-06 | 0.00013 | 1.24E-05 | 2.36E-06 |
| DEPRECATEI | 8.54E-06 | 9.49E-05 | 0 | 2.12E-05 | 2.78E-05 | 9.94E-05 | 3.80E-05 | 4.12E-05 | 4.42E-05 | 5.01E-05 | 6.40E-06 | 2.16E-05 | 2.53E-05 | 3.92E-05 | 0 | 0 | 0 | 2.72E-05 |

| | | | | | | | | | | | | | | | | | | |
|---|---|---|---|---|---|---|---|---|---|---|---|---|---|---|---|---|---|---|
| Zfand2b | 1.49E-05 | 9.22E-05 | 2.47E-05 | 0.00016 | 1.63E-05 | 5.43E-05 | 4.28E-05 | 4.11E-05 | 4.42E-05 | 4.03E-05 | 4.20E-05 | 4.76E-05 | 0 | 9.11E-05 | 6.66E-05 | 3.69E-05 | 7.40E-05 | 7.07E-05 |
| Rsrc1 | 9.24E-05 | 3.52E-05 | 0 | 0 | 3.07E-05 | 3.53E-05 | 3.96E-05 | 0 | 4.42E-05 | 6.00E-05 | 2.97E-05 | 3.54E-05 | 5.60E-05 | 1.76E-05 | 6.32E-05 | 7.95E-05 | 2.38E-06 | 7.90E-05 |
| Arhgap12 | 8.30E-05 | 8.19E-05 | 0 | 0 | 3.17E-05 | 1.83E-05 | 7.07E-05 | 4.73E-05 | 4.42E-05 | 6.55E-05 | 1.57E-05 | 0.0001 | 0 | 0 | 0.00017 | 0 | 8.53E-05 | 1.47E-06 |
| Polr3b | 5.13E-05 | 3.60E-05 | 3.04E-05 | 8.03E-06 | 1.48E-05 | 1.59E-05 | 3.93E-05 | 0 | 4.42E-05 | 0 | 5.36E-06 | 2.07E-05 | 1.47E-05 | 1.96E-05 | 0 | 1.83E-05 | 1.25E-05 | 0 |
| Styk1 | 0 | 1.09E-06 | 2.23E-06 | 0 | 0 | 9.21E-06 | 0 | 0 | 4.42E-05 | 7.33E-06 | 6.40E-06 | 0 | 0 | 1.05E-05 | 0 | 2.70E-06 | 0 | 1.42E-06 |
| Pinx1 | 3.45E-05 | 9.79E-06 | 6.77E-05 | 2.12E-05 | 4.51E-07 | 7.70E-06 | 0 | 0 | 4.42E-05 | 5.70E-06 | 1.73E-05 | 0 | 5.41E-07 | 2.59E-05 | 4.95E-05 | 2.16E-05 | 0 | 1.43E-05 |
| Zfp653 | 9.55E-06 | 1.00E-04 | 3.30E-05 | 3.27E-05 | 1.06E-05 | 0 | 0 | 1.27E-05 | 4.42E-05 | 9.59E-06 | 5.17E-05 | 3.33E-05 | 0 | 5.00E-05 | 0.00014 | 2.20E-05 | 0.00016 | 6.73E-06 |
| Tarbp2 | 4.35E-05 | 4.67E-05 | 7.01E-05 | 6.98E-05 | 5.81E-06 | 0 | 1.61E-05 | 2.74E-05 | 4.42E-05 | 2.81E-05 | 1.91E-05 | 8.51E-06 | 0 | 2.47E-05 | 9.27E-05 | 9.49E-05 | 8.50E-05 | 0 |
| Itpr3 | 0 | 4.73E-05 | 0 | 2.55E-05 | 5.92E-06 | 0 | 5.12E-05 | 0 | 4.42E-05 | 3.88E-05 | 2.26E-05 | 2.25E-05 | 4.60E-05 | 6.56E-05 | 1.71E-06 | 0 | 1.56E-05 | 2.88E-05 |
| Shox2 | 8.20E-06 | 4.01E-05 | 0 | 6.64E-05 | 6.72E-05 | 0.00016 | 0.00025 | 0 | 4.41E-05 | 8.46E-05 | 9.32E-05 | 9.31E-05 | 5.50E-05 | 0.00015 | 4.36E-05 | 8.05E-06 | 4.19E-05 | 0.00024 |
| Yif1b | 0.00022 | 4.73E-05 | 0.00018 | 0.00028 | 3.84E-05 | 8.69E-05 | 0.00023 | 0 | 4.41E-05 | 3.44E-05 | 0.00025 | 3.00E-05 | 4.26E-05 | 0.00011 | 0 | 0.00011 | 4.27E-05 | 4.80E-05 |
| Rcor3 | 7.27E-05 | 0.00014 | 3.40E-05 | 3.98E-05 | 4.75E-05 | 7.88E-05 | 4.52E-05 | 0 | 4.41E-05 | 2.40E-05 | 4.25E-05 | 7.57E-05 | 2.15E-05 | 1.29E-06 | 7.35E-05 | 5.46E-05 | 5.58E-05 | 0.00014 |
| Lmbr1l | 3.06E-05 | 0.00016 | 2.98E-05 | 6.70E-05 | 1.70E-05 | 5.34E-05 | 5.47E-05 | 0 | 4.41E-05 | 3.29E-05 | 7.33E-05 | 0.00014 | 2.78E-05 | 0.00014 | 0.00014 | 3.13E-05 | 5.37E-05 | 2.52E-05 |
| Asl | 7.57E-05 | 7.97E-05 | 1.14E-06 | 8.61E-05 | 8.51E-06 | 0 | 1.75E-06 | 0 | 4.41E-05 | 2.54E-05 | 5.15E-05 | 9.65E-05 | 0 | 0 | 2.79E-06 | 0 | 0 | 4.67E-06 |
| Mettl23 | 7.94E-05 | 4.93E-05 | 0 | 1.55E-05 | 4.43E-05 | 5.95E-05 | 5.33E-05 | 1.91E-05 | 4.40E-05 | 3.52E-05 | 0 | 9.57E-05 | 9.31E-06 | 3.80E-05 | 0 | 1.10E-05 | 0 | 1.43E-05 |
| Vps36 | 0.00016 | 0.00011 | 0.00018 | 8.92E-05 | 8.84E-06 | 4.53E-05 | 0 | 0.0001 | 4.40E-05 | 6.41E-05 | 8.57E-05 | 0 | 0 | 3.47E-05 | 3.52E-05 | 0.00019 | 0 | 9.73E-05 |
| Saraf | 0.00013 | 0.00024 | 0.00016 | 0.00032 | 0.00029 | 0.00018 | 0.0002 | 0.00024 | 4.39E-05 | 0.0002 | 0.00028 | 0.00043 | 0.00018 | 0.00024 | 8.75E-05 | 0 | 0.00027 | 0.00039 |
| Nagpa | 3.45E-05 | 8.64E-05 | 6.73E-05 | 3.29E-05 | 7.01E-05 | 6.27E-05 | 4.22E-05 | 1.06E-05 | 4.39E-05 | 7.23E-05 | 3.10E-05 | 0 | 5.12E-05 | 5.04E-05 | 3.76E-06 | 4.82E-06 | 0 | 0.00012 |
| Ccdc28b | 4.85E-05 | 3.90E-05 | 0 | 4.26E-05 | 1.88E-05 | 4.74E-05 | 8.65E-05 | 0.00012 | 4.39E-05 | 5.48E-05 | 0 | 1.84E-05 | 0 | 0 | 0.00012 | 5.21E-06 | 0 | 0.00012 |
| Stim2 | 5.28E-05 | 4.94E-05 | 0.00012 | 0 | 3.02E-05 | 4.55E-05 | 4.74E-05 | 0.00018 | 4.39E-05 | 8.43E-05 | 5.22E-05 | 1.41E-05 | 2.76E-05 | 0 | 3.16E-06 | 7.69E-05 | 8.54E-05 | 5.08E-05 |
| Mrs2 | 4.34E-05 | 9.13E-05 | 0.0001 | 1.46E-05 | 4.58E-05 | 3.05E-05 | 0 | 9.02E-06 | 4.39E-05 | 2.76E-05 | 5.43E-05 | 3.76E-05 | 0 | 3.93E-05 | 0.00013 | 6.98E-05 | 0 | 2.66E-05 |
| Nub1 | 0.0001 | 0.00022 | 3.55E-06 | 0.00014 | 4.55E-05 | 2.86E-05 | 0 | 3.50E-05 | 4.39E-05 | 1.13E-05 | 1.99E-05 | 0.00013 | 0 | 4.37E-05 | 0.0001 | 5.53E-05 | 5.54E-05 | 0.00016 |
| Zfp64 | 5.77E-05 | 5.59E-05 | 7.65E-05 | 3.29E-05 | 2.91E-05 | 2.41E-05 | 4.24E-05 | 1.25E-05 | 4.39E-05 | 3.87E-05 | 9.77E-06 | 2.26E-05 | 2.76E-05 | 3.90E-05 | 0.00011 | 2.76E-05 | 0 | 4.07E-05 |
| Tbc1d22b | 4.14E-05 | 6.14E-05 | 4.21E-05 | 2.38E-05 | 5.36E-05 | 2.30E-05 | 0 | 8.22E-06 | 4.39E-05 | 3.71E-05 | 2.68E-05 | 7.57E-06 | 8.02E-06 | 2.22E-05 | 9.00E-05 | 4.09E-05 | 0 | 5.02E-05 |
| Dennd4b | 0 | 4.79E-05 | 1.77E-05 | 2.45E-05 | 5.36E-06 | 2.03E-06 | 8.16E-06 | 4.58E-05 | 4.39E-05 | 4.68E-05 | 1.39E-05 | 4.71E-05 | 0 | 1.06E-06 | 8.86E-05 | 3.56E-05 | 4.98E-05 | 7.04E-06 |
| Gtf2f2 | 0.00011 | 8.26E-05 | 0.00016 | 3.38E-05 | 3.63E-05 | 6.18E-05 | 0 | 0 | 4.38E-05 | 1.33E-05 | 7.48E-05 | 0 | 6.06E-06 | 1.03E-06 | 7.52E-05 | 8.15E-05 | 0 | 0 |
| Fam89b | 5.38E-05 | 0.00027 | 0.00015 | 0.00035 | 8.47E-05 | 5.09E-05 | 3.65E-05 | 0.00017 | 4.38E-05 | 9.03E-05 | 0.00017 | 2.83E-05 | 1.94E-05 | 0.00014 | 0 | 0.00015 | 0 | 0.00013 |
| Guf1 | 7.65E-05 | 5.82E-05 | 5.99E-06 | 5.48E-06 | 2.68E-05 | 3.93E-05 | 4.82E-06 | 0 | 4.38E-05 | 1.41E-05 | 8.38E-06 | 1.24E-05 | 0 | 0 | 0.00014 | 4.10E-05 | 0 | 0 |
| Tle4 | 0 | 3.68E-05 | 0 | 0 | 0 | 2.20E-05 | 0 | 0 | 4.38E-05 | 6.08E-05 | 0 | 1.20E-06 | 0 | 9.43E-05 | 0 | 1.03E-05 | 0 | 0 |
| Hmbox1 | 6.03E-05 | 9.67E-05 | 2.52E-05 | 2.86E-05 | 3.42E-05 | 1.88E-05 | 0 | 4.46E-05 | 4.38E-05 | 4.80E-05 | 6.09E-05 | 0.0001 | 2.35E-05 | 1.10E-05 | 0.00018 | 4.36E-05 | 6.91E-05 | 4.10E-05 |
| Fbxo34 | 3.80E-05 | 6.63E-05 | 2.26E-05 | 3.09E-05 | 1.88E-05 | 8.73E-07 | 2.75E-05 | 0 | 4.38E-05 | 5.73E-05 | 2.72E-05 | 7.80E-06 | 1.55E-05 | 1.46E-05 | 3.83E-05 | 6.96E-05 | 8.23E-05 | 5.52E-06 |
| Avpi1 | 1.10E-05 | 4.06E-05 | 4.53E-05 | 0.0001 | 7.98E-05 | 0 | 5.63E-05 | 2.47E-05 | 4.38E-05 | 1.88E-05 | 9.05E-05 | 8.70E-05 | 0 | 0 | 1.91E-05 | 1.72E-05 | 0 | 8.23E-05 |
| Trf | 0 | 9.00E-05 | 0 | 0 | 0.00022 | 9.52E-05 | 4.03E-05 | 4.05E-05 | 4.37E-05 | 0.00024 | 6.48E-05 | 5.87E-05 | 0.00052 | 0 | 7.99E-05 | 0 | 0 | 0 |

| | | | | | | | | | | | | | | | | | | |
|---|---|---|---|---|---|---|---|---|---|---|---|---|---|---|---|---|---|---|
| Cbwd1 | 3.39E-05 | 2.08E-05 | 2.40E-05 | 1.23E-05 | 2.97E-05 | 5.11E-05 | 0 | 4.05E-05 | 4.37E-05 | 2.04E-05 | 1.54E-05 | 3.68E-05 | 8.69E-06 | 1.98E-05 | 3.14E-05 | 5.44E-05 | 2.50E-06 | 5.63E-06 |
| Rad52 | 3.14E-05 | 6.80E-05 | 4.36E-05 | 0 | 1.06E-05 | 4.38E-05 | 9.09E-06 | 0 | 4.37E-05 | 2.86E-05 | 7.79E-06 | 4.07E-05 | 1.74E-05 | 3.88E-05 | 3.54E-05 | 2.00E-05 | 7.86E-05 | 1.69E-05 |
| Tubgcp2 | 1.54E-05 | 2.94E-05 | 3.03E-05 | 0.00011 | 1.80E-05 | 1.90E-05 | 3.39E-05 | 1.30E-05 | 4.37E-05 | 4.51E-05 | 2.07E-05 | 5.97E-05 | 0 | 3.97E-05 | 2.38E-05 | 0.00016 | 4.09E-07 | 4.86E-05 |
| Pstk | 4.90E-05 | 2.18E-05 | 1.53E-05 | 2.08E-05 | 0 | 4.22E-06 | 1.54E-05 | 4.75E-06 | 4.37E-05 | 1.20E-05 | 2.73E-05 | 5.56E-06 | 0 | 0 | 5.94E-06 | 1.95E-05 | 0 | 1.94E-05 |
| Trim32 | 9.95E-06 | 5.59E-05 | 2.88E-05 | 6.62E-05 | 4.93E-05 | 2.95E-06 | 2.64E-05 | 3.43E-05 | 4.37E-05 | 3.26E-05 | 1.96E-05 | 1.92E-05 | 0 | 6.02E-05 | 6.61E-05 | 1.87E-05 | 0 | 2.84E-05 |
| Zfp932 | 1.07E-05 | 2.46E-05 | 0 | 3.99E-05 | 4.61E-05 | 2.39E-06 | 0 | 2.95E-05 | 4.37E-05 | 2.19E-05 | 1.72E-07 | 0 | 0 | 4.81E-06 | 2.56E-05 | 8.90E-06 | 0 | 3.15E-05 |
| Itgb8 | 0 | 5.37E-06 | 0 | 0 | 0 | 8.39E-05 | 0 | 2.37E-05 | 4.36E-05 | 0 | 0 | 4.86E-05 | 0 | 0 | 0 | 0 | 1.82E-05 | 3.66E-05 |
| Ppcdc | 3.71E-05 | 4.84E-05 | 0 | 1.16E-05 | 2.65E-05 | 4.61E-05 | 0 | 1.12E-05 | 4.36E-05 | 2.75E-05 | 0 | 1.34E-05 | 1.83E-06 | 0 | 0.00016 | 4.15E-05 | 0 | 3.22E-05 |
| Cdk12 | 0.00016 | 0.00016 | 7.66E-05 | 1.01E-05 | 3.73E-05 | 2.96E-05 | 1.35E-05 | 0 | 4.36E-05 | 1.88E-05 | 2.07E-05 | 9.87E-05 | 0 | 0 | 0.00043 | 9.51E-05 | 0.0002 | 2.52E-06 |
| Pdcd2 | 9.98E-05 | 6.95E-05 | 0.00012 | 5.93E-05 | 9.91E-06 | 8.37E-06 | 6.54E-05 | 1.75E-05 | 4.36E-05 | 3.78E-06 | 0 | 0 | 0 | 0 | 0 | 1.25E-05 | 0 | 1.88E-06 |
| Inpp5f | 4.45E-05 | 7.27E-05 | 6.17E-05 | 0 | 0 | 0 | 0 | 0 | 4.36E-05 | 2.01E-05 | 0 | 2.84E-05 | 2.39E-05 | 2.79E-05 | 1.41E-05 | 1.30E-05 | 0.0001 | 3.48E-05 |
| Rnf19a | 6.63E-05 | 0.00011 | 0.00011 | 0 | 0 | 0 | 7.38E-05 | 0.00011 | 4.36E-05 | 0.00022 | 0.00016 | 0.00045 | 0 | 4.15E-05 | 0.00022 | 4.07E-05 | 0.0004 | 0.00011 |
| Mink1 | 2.81E-05 | 0.00014 | 5.83E-05 | 5.08E-05 | 5.43E-05 | 0 | 0 | 6.16E-05 | 4.36E-05 | 2.10E-05 | 1.78E-05 | 0.0001 | 0 | 5.60E-06 | 0.00016 | 3.35E-06 | 3.73E-05 | 3.87E-05 |
| Bag1 | 0.00058 | 0.00014 | 0.00031 | 0.00023 | 0.00021 | 0.00034 | 0.00039 | 0.00016 | 4.35E-05 | 0.00019 | 0.00016 | 0.0002 | 0 | 0.00013 | 5.73E-05 | 0.0002 | 7.07E-05 | 5.06E-05 |
| Poglut2 | 0.00026 | 6.26E-05 | 0.00013 | 9.18E-05 | 9.31E-05 | 0.0002 | 0.00014 | 0.00013 | 4.35E-05 | 9.93E-06 | 0 | 0 | 0.00011 | 4.16E-05 | 5.73E-05 | 0.00014 | 0 | 3.84E-05 |
| Epha4 | 0 | 5.43E-05 | 0 | 0 | 3.64E-05 | 6.21E-05 | 0 | 0 | 4.35E-05 | 0 | 0.00011 | 5.23E-05 | 4.63E-05 | 6.68E-05 | 1.40E-05 | 0 | 4.61E-05 | 2.04E-05 |
| Endov | 2.26E-05 | 4.19E-05 | 3.13E-05 | 4.07E-05 | 1.68E-05 | 2.02E-05 | 0 | 2.23E-05 | 4.35E-05 | 9.76E-06 | 0 | 7.28E-06 | 2.18E-06 | 2.54E-05 | 6.97E-05 | 4.99E-06 | 0 | 3.69E-05 |
| Pcid2 | 5.71E-05 | 5.64E-05 | 5.26E-05 | 1.11E-05 | 2.88E-05 | 1.49E-05 | 4.99E-06 | 9.24E-06 | 4.35E-05 | 1.64E-05 | 3.42E-05 | 9.44E-05 | 8.88E-06 | 0 | 1.07E-05 | 4.76E-05 | 1.20E-06 | 2.87E-05 |
| Xrcc4 | 2.33E-05 | 2.22E-05 | 7.84E-06 | 1.47E-05 | 3.67E-05 | 1.27E-05 | 2.23E-05 | 2.52E-05 | 4.35E-05 | 3.08E-05 | 1.73E-05 | 2.93E-05 | 6.22E-06 | 7.60E-06 | 4.20E-05 | 4.23E-05 | 5.13E-05 | 3.95E-05 |
| Hoxb3 | 0 | 3.87E-05 | 2.04E-05 | 7.07E-05 | 4.87E-05 | 5.89E-06 | 1.78E-05 | 0.00019 | 4.35E-05 | 4.29E-06 | 2.67E-05 | 0 | 0 | 0 | 9.88E-05 | 2.43E-06 | 0 | 0 |
| 5-Sep | 7.02E-05 | 0.00011 | 0.00018 | 0.00024 | 6.54E-05 | 0 | 0.00029 | 5.18E-05 | 4.35E-05 | 0.00011 | 0.00027 | 0.00021 | 8.23E-06 | 8.09E-05 | 0.00029 | 4.84E-05 | 0.00016 | 0.00024 |
| Fbxw17 | 2.03E-05 | 6.38E-05 | 0 | 8.82E-05 | 1.79E-05 | 0 | 5.75E-05 | 1.20E-05 | 4.35E-05 | 0.00018 | 2.44E-05 | 0 | 0 | 0 | 0.00013 | 4.15E-05 | 0 | 2.44E-05 |
| Grina | 0 | 0.00052 | 0 | 3.61E-05 | 0.00023 | 0.00015 | 0 | 0.00012 | 4.34E-05 | 0.00022 | 0 | 0.00032 | 8.58E-05 | 0.00017 | 0.00012 | 0 | 6.77E-05 | 0.00032 |
| Scyl3 | 3.62E-05 | 7.29E-05 | 3.58E-05 | 1.43E-05 | 3.09E-05 | 4.83E-05 | 1.96E-05 | 1.35E-05 | 4.34E-05 | 3.83E-05 | 2.71E-05 | 6.49E-06 | 2.75E-05 | 4.53E-05 | 5.53E-05 | 2.44E-05 | 0 | 5.15E-05 |
| Cry2 | 0 | 9.70E-05 | 0 | 5.12E-05 | 2.22E-05 | 0 | 0 | 1.84E-05 | 4.34E-05 | 0.00018 | 2.26E-05 | 0.00023 | 4.18E-05 | 7.17E-05 | 8.61E-05 | 0 | 4.38E-05 | 8.26E-05 |
| Stag1 | 8.06E-05 | 8.53E-05 | 4.58E-05 | 0 | 3.25E-05 | 9.26E-05 | 3.47E-05 | 5.01E-05 | 4.33E-05 | 4.13E-05 | 3.73E-05 | 8.01E-05 | 4.12E-05 | 5.75E-05 | 0 | 0.00014 | 2.88E-05 | 1.56E-05 |
| Dpy19l4 | 7.85E-05 | 5.49E-05 | 2.70E-05 | 0 | 4.05E-05 | 9.22E-05 | 1.07E-05 | 3.46E-05 | 4.33E-05 | 2.18E-05 | 4.60E-05 | 4.94E-05 | 0 | 2.78E-05 | 0 | 5.71E-05 | 5.46E-06 | 2.20E-05 |
| Btf3l4 | 8.35E-05 | 7.85E-05 | 1.66E-05 | 1.83E-05 | 1.37E-06 | 6.40E-05 | 9.76E-08 | 1.22E-05 | 4.33E-05 | 1.69E-05 | 2.89E-05 | 2.90E-05 | 1.12E-05 | 3.59E-05 | 6.19E-06 | 8.70E-05 | 3.58E-06 | 4.86E-06 |
| Slc9a6 | 5.10E-05 | 7.50E-05 | 2.76E-05 | 1.36E-05 | 2.45E-05 | 4.52E-05 | 1.44E-05 | 4.89E-05 | 4.33E-05 | 2.60E-05 | 4.77E-05 | 1.64E-05 | 3.74E-05 | 3.54E-05 | 0 | 4.92E-05 | 1.42E-05 | 4.51E-05 |
| Surf2 | 2.41E-05 | 7.49E-05 | 6.99E-05 | 1.53E-05 | 1.50E-05 | 4.02E-05 | 0 | 1.65E-05 | 4.33E-05 | 7.34E-06 | 0 | 2.19E-05 | 9.81E-06 | 2.17E-05 | 4.60E-05 | 0 | 8.15E-05 | 3.19E-05 |
| Xab2 | 4.08E-05 | 8.80E-05 | 5.62E-05 | 5.11E-05 | 2.51E-05 | 3.23E-05 | 5.45E-06 | 0 | 4.33E-05 | 1.84E-05 | 2.78E-05 | 3.45E-05 | 0 | 2.11E-05 | 0 | 3.71E-05 | 0 | 4.59E-05 |
| Abca7 | 0 | 4.07E-05 | 3.36E-05 | 0 | 0 | 1.80E-05 | 0 | 2.84E-05 | 4.33E-05 | 0.00011 | 0 | 2.61E-06 | 0 | 1.50E-05 | 4.82E-05 | 2.40E-05 | 0 | 7.12E-05 |
| Apbb1 | 3.22E-05 | 6.87E-05 | 0 | 3.42E-05 | 3.84E-05 | 1.08E-05 | 4.82E-05 | 5.45E-06 | 4.33E-05 | 5.90E-05 | 8.56E-05 | 5.41E-05 | 0 | 3.89E-05 | 1.64E-05 | 2.74E-06 | 0 | 0.0001 |

| | | | | | | | | | | | | | | | | | | |
|---|---|---|---|---|---|---|---|---|---|---|---|---|---|---|---|---|---|---|
| Nphp1 | 3.29E-05 | 3.43E-05 | 2.58E-06 | 9.87E-06 | 8.28E-05 | 1.06E-05 | 2.06E-05 | 8.83E-06 | 4.33E-05 | 3.46E-05 | 0 | 2.25E-05 | 0 | 1.33E-05 | 2.51E-05 | 6.12E-05 | 1.21E-05 | 6.16E-05 |
| Entpd5 | 4.23E-05 | 1.23E-05 | 1.04E-05 | 7.54E-05 | 7.37E-05 | 1.04E-05 | 0.00031 | 1.85E-05 | 4.33E-05 | 0.00014 | 1.97E-05 | 6.06E-05 | 3.87E-05 | 4.17E-05 | 0 | 6.26E-05 | 0 | 0.00012 |
| Scnm1 | 0.0001 | 6.83E-05 | 6.59E-06 | 0.00011 | 3.04E-05 | 9.55E-06 | 4.45E-05 | 0 | 4.33E-05 | 3.99E-05 | 9.94E-06 | 2.97E-05 | 7.12E-06 | 5.16E-06 | 0.00013 | 5.38E-05 | 0 | 2.12E-05 |
| Yipf2 | 1.37E-05 | 4.35E-05 | 0 | 7.24E-07 | 1.86E-05 | 1.34E-06 | 0 | 0 | 4.33E-05 | 4.30E-06 | 0 | 2.69E-05 | 0 | 3.81E-06 | 0 | 0 | 0 | 4.90E-05 |
| Naaa | 3.39E-06 | 0 | 1.83E-05 | 0 | 0.00014 | 0 | 0.00016 | 8.16E-05 | 4.33E-05 | 0.00034 | 0 | 4.99E-05 | 7.24E-05 | 0.00023 | 4.18E-06 | 1.69E-05 | 0 | 0.00018 |
| Mthfd1l | 4.77E-05 | 0 | 7.25E-05 | 0 | 0 | 0 | 0 | 0 | 4.33E-05 | 0 | 1.24E-05 | 0.00055 | 8.28E-06 | 5.36E-05 | 7.67E-05 | 0.00015 | 0 | 0 |
| Mat2b | 0.00016 | 0.00014 | 0 | 6.87E-05 | 8.43E-05 | 0.00015 | 0.00014 | 4.05E-05 | 4.32E-05 | 0.00011 | 6.97E-05 | 6.51E-05 | 9.88E-05 | 8.45E-05 | 3.19E-05 | 6.28E-05 | 0 | 6.70E-05 |
| Ppp1r12c | 0 | 0.00018 | 0.00015 | 0.00027 | 0.00015 | 4.52E-05 | 1.31E-05 | 7.84E-05 | 4.32E-05 | 0.00012 | 0.00012 | 0.00012 | 0 | 0.00013 | 0.00061 | 0.00018 | 0.00031 | 0.00014 |
| Eif2d | 4.42E-05 | 7.38E-05 | 5.70E-05 | 8.75E-05 | 4.66E-05 | 4.38E-05 | 7.45E-05 | 1.71E-05 | 4.32E-05 | 5.39E-05 | 1.94E-05 | 3.45E-05 | 6.43E-06 | 0 | 4.35E-05 | 8.82E-05 | 0 | 5.43E-05 |
| Ikbkb | 3.13E-05 | 0.00012 | 9.41E-05 | 9.02E-05 | 5.43E-05 | 3.20E-05 | 4.37E-05 | 2.68E-06 | 4.32E-05 | 0.00011 | 6.99E-05 | 2.43E-05 | 4.68E-05 | 4.96E-05 | 0.00018 | 2.59E-05 | 5.54E-05 | 7.02E-05 |
| Nudt2 | 0.0001 | 2.62E-05 | 2.69E-05 | 6.15E-05 | 0 | 3.16E-05 | 5.16E-05 | 0 | 4.32E-05 | 3.37E-05 | 4.84E-06 | 4.24E-05 | 0 | 0 | 0 | 6.79E-05 | 0 | 2.07E-05 |
| Ccdc43 | 5.07E-05 | 7.08E-05 | 6.37E-05 | 6.06E-05 | 3.47E-05 | 2.52E-05 | 1.13E-05 | 0 | 4.32E-05 | 3.02E-05 | 8.98E-06 | 2.35E-05 | 3.36E-05 | 2.47E-05 | 4.63E-05 | 7.35E-05 | 5.60E-05 | 3.56E-05 |
| Nlrx1 | 3.29E-05 | 1.96E-05 | 5.45E-06 | 7.13E-05 | 4.28E-05 | 2.36E-05 | 7.48E-05 | 7.35E-05 | 4.32E-05 | 2.10E-05 | 5.42E-05 | 1.81E-05 | 2.25E-05 | 4.22E-05 | 1.70E-05 | 0 | 0 | 6.49E-05 |
| Manba | 0 | 7.43E-05 | 0 | 7.28E-06 | 0.00014 | 0.00015 | 5.17E-05 | 8.91E-05 | 4.31E-05 | 0.00012 | 1.48E-05 | 1.24E-05 | 0.00012 | 7.24E-05 | 1.90E-06 | 0 | 0 | 0.0001 |
| Igbp1 | 0.00024 | 0.00018 | 6.18E-05 | 0.00013 | 3.81E-05 | 0.00014 | 9.37E-05 | 2.45E-05 | 4.31E-05 | 2.32E-05 | 0 | 0.00012 | 4.63E-05 | 5.35E-05 | 4.89E-05 | 8.08E-05 | 3.67E-05 | 4.76E-05 |
| Zfp706 | 0.00014 | 5.92E-05 | 0.00013 | 2.06E-05 | 4.90E-05 | 4.31E-05 | 1.27E-05 | 4.55E-05 | 4.31E-05 | 3.08E-05 | 5.38E-05 | 1.93E-05 | 2.09E-05 | 1.04E-05 | 0 | 9.72E-05 | 0 | 2.64E-05 |
| Bmpr1b | 8.73E-06 | 0 | 0.00014 | 0 | 4.87E-05 | 1.28E-06 | 1.54E-05 | 0.0001 | 4.31E-05 | 0 | 9.18E-05 | 0 | 0 | 5.64E-05 | 8.48E-06 | 3.17E-05 | 0 | 0 |
| Relb | 0 | 0.00015 | 0.00026 | 4.80E-06 | 6.85E-05 | 8.92E-05 | 0 | 2.95E-05 | 4.30E-05 | 2.00E-05 | 1.36E-06 | 0.00012 | 0.00015 | 0.00022 | 0.00016 | 4.94E-06 | 0.00022 | 0.00015 |
| Cenpc1 | 6.07E-05 | 5.15E-05 | 1.32E-05 | 1.07E-05 | 1.19E-05 | 4.06E-05 | 3.83E-05 | 4.83E-05 | 4.30E-05 | 3.87E-05 | 7.28E-05 | 3.85E-05 | 3.01E-06 | 1.58E-05 | 0.00011 | 0.00032 | 3.74E-05 | 3.54E-05 |
| Parp10 | 4.45E-06 | 5.54E-05 | 0 | 2.67E-05 | 1.75E-05 | 0 | 0 | 4.66E-05 | 4.30E-05 | 7.75E-05 | 2.49E-05 | 0 | 6.31E-06 | 4.90E-05 | 0.00011 | 1.13E-05 | 0 | 8.48E-05 |
| Btd | 4.25E-05 | 9.70E-05 | 0 | 7.20E-05 | 1.76E-05 | 8.87E-05 | 3.18E-05 | 1.76E-05 | 4.29E-05 | 6.88E-05 | 8.61E-06 | 6.75E-05 | 6.19E-05 | 0.00011 | 2.05E-06 | 0.00011 | 5.04E-05 | 0.00012 |
| Cwf19l2 | 5.87E-05 | 6.77E-05 | 3.08E-05 | 2.31E-05 | 4.60E-05 | 8.41E-05 | 2.95E-05 | 3.15E-07 | 4.29E-05 | 3.45E-05 | 0 | 3.63E-05 | 1.53E-05 | 1.40E-05 | 0.00011 | 6.37E-05 | 0 | 3.38E-05 |
| Med23 | 7.94E-05 | 8.47E-05 | 2.60E-05 | 3.07E-06 | 4.88E-05 | 5.56E-05 | 3.10E-05 | 1.12E-05 | 4.29E-05 | 4.43E-06 | 2.44E-05 | 1.89E-05 | 3.22E-05 | 4.95E-06 | 5.58E-05 | 8.47E-05 | 0 | 0 |
| Nicn1 | 7.55E-05 | 7.07E-05 | 0 | 4.20E-06 | 3.82E-05 | 5.24E-05 | 0 | 0 | 4.29E-05 | 3.14E-05 | 0 | 0.0001 | 0 | 0 | 0 | 5.19E-05 | 1.16E-05 | 3.58E-05 |
| Ecsit | 8.29E-05 | 5.55E-05 | 7.16E-05 | 6.98E-05 | 1.62E-05 | 0 | 9.53E-06 | 0 | 4.29E-05 | 2.87E-06 | 4.37E-05 | 2.67E-07 | 1.11E-05 | 1.38E-05 | 4.95E-05 | 0 | 0 | 5.02E-05 |
| Alg3 | 6.90E-05 | 2.94E-06 | 0.00011 | 0.00011 | 2.79E-05 | 0 | 8.68E-06 | 2.12E-05 | 4.29E-05 | 2.34E-05 | 1.91E-05 | 0 | 1.08E-05 | 2.41E-05 | 1.75E-05 | 1.98E-05 | 0 | 4.72E-05 |
| Trpm7 | 0.00016 | 0.0002 | 6.83E-05 | 0 | 8.16E-05 | 0.00013 | 8.19E-05 | 8.33E-06 | 4.28E-05 | 6.03E-05 | 1.80E-05 | 6.17E-05 | 7.87E-05 | 4.45E-05 | 0.00015 | 0.00011 | 1.48E-05 | 4.38E-05 |
| Xpr1 | 8.21E-05 | 7.58E-05 | 1.77E-05 | 0 | 5.25E-05 | 7.96E-05 | 5.73E-05 | 2.10E-05 | 4.28E-05 | 0 | 3.07E-05 | 4.48E-05 | 0 | 2.26E-05 | 0 | 1.86E-05 | 3.42E-05 | 5.60E-05 |
| Ciao1 | 5.33E-05 | 0.00011 | 7.95E-05 | 0.00019 | 7.60E-05 | 6.47E-05 | 4.45E-05 | 8.92E-05 | 4.28E-05 | 5.76E-05 | 1.86E-05 | 2.73E-05 | 2.04E-05 | 6.84E-05 | 1.65E-05 | 7.39E-05 | 0 | 6.05E-05 |
| Brwd3 | 4.52E-05 | 7.76E-05 | 2.11E-05 | 9.18E-05 | 5.24E-05 | 4.90E-05 | 0 | 4.23E-05 | 4.28E-05 | 1.49E-05 | 3.83E-05 | 0.00012 | 2.32E-05 | 3.27E-05 | 9.29E-05 | 1.51E-05 | 2.33E-05 | 0 |
| Dnal1 | 3.18E-05 | 5.24E-05 | 2.91E-05 | 0 | 6.33E-05 | 3.71E-05 | 0 | 4.59E-05 | 4.27E-05 | 3.11E-05 | 1.48E-05 | 1.17E-05 | 1.49E-05 | 4.60E-05 | 9.13E-05 | 0 | 0 | 3.88E-05 |
| Paqr4 | 0 | 1.71E-05 | 1.36E-06 | 2.32E-05 | 3.59E-05 | 3.44E-05 | 0 | 7.52E-05 | 4.27E-05 | 3.85E-05 | 3.86E-05 | 1.15E-05 | 0 | 3.36E-05 | 0 | 7.82E-05 | 0 | 7.16E-05 |
| Nit1 | 4.23E-05 | 7.62E-05 | 5.14E-06 | 3.10E-05 | 4.66E-05 | 2.18E-05 | 1.21E-05 | 7.55E-07 | 4.27E-05 | 5.60E-05 | 2.49E-05 | 4.88E-05 | 2.45E-05 | 0 | 5.70E-06 | 2.24E-05 | 0 | 6.08E-05 |

| | | | | | | | | | | | | | | | | | | |
|---|---|---|---|---|---|---|---|---|---|---|---|---|---|---|---|---|---|---|
| Siae | 4.77E-05 | 5.79E-05 | 0 | 1.23E-05 | 3.89E-05 | 2.17E-05 | 3.84E-05 | 7.60E-06 | 4.27E-05 | 5.50E-05 | 0 | 2.40E-05 | 4.44E-05 | 2.28E-05 | 0 | 5.00E-05 | 0 | 9.15E-05 |
| Micall1 | 0 | 6.83E-05 | 0.00013 | 5.42E-05 | 2.82E-05 | 0 | 1.47E-05 | 0 | 4.27E-05 | 5.19E-05 | 0.00011 | 4.96E-05 | 0 | 0 | 0.00011 | 0 | 8.72E-05 | 2.22E-05 |
| Cdc34 | 9.76E-05 | 0.00015 | 0.00025 | 0.00028 | 2.32E-05 | 0 | 0 | 3.36E-05 | 4.27E-05 | 6.21E-05 | 0.00012 | 0 | 7.23E-05 | 6.32E-05 | 0.00012 | 0.00013 | 0.00026 | 7.03E-05 |
| Msl2 | 0.00015 | 0.00016 | 0.00012 | 1.62E-05 | 8.95E-05 | 0.00012 | 7.09E-05 | 6.09E-05 | 4.26E-05 | 4.31E-05 | 5.65E-05 | 6.69E-05 | 5.91E-05 | 5.90E-05 | 0.00012 | 0.00011 | 0.00023 | 3.38E-05 |
| Map2k5 | 1.01E-05 | 3.53E-05 | 1.36E-05 | 4.77E-05 | 3.26E-05 | 5.77E-05 | 5.90E-05 | 6.85E-05 | 4.26E-05 | 4.41E-05 | 4.01E-05 | 2.30E-05 | 3.54E-05 | 4.94E-05 | 4.01E-07 | 5.10E-05 | 1.14E-05 | 4.22E-05 |
| Urb1 | 1.94E-05 | 5.39E-05 | 8.58E-05 | 2.03E-05 | 0 | 2.39E-05 | 0 | 0 | 4.26E-05 | 5.04E-06 | 0 | 3.75E-07 | 1.83E-05 | 3.07E-06 | 9.11E-05 | 7.59E-07 | 3.85E-05 | 9.77E-06 |
| Zfp553 | 2.57E-05 | 8.64E-05 | 4.60E-05 | 4.08E-05 | 3.02E-05 | 1.17E-05 | 6.96E-06 | 0 | 4.26E-05 | 1.40E-05 | 2.40E-05 | 3.11E-05 | 0 | 5.98E-05 | 6.92E-05 | 1.68E-05 | 0 | 2.45E-05 |
| Armc10 | 4.33E-05 | 6.66E-05 | 0 | 1.66E-05 | 9.63E-06 | 0 | 3.23E-05 | 2.78E-05 | 4.26E-05 | 4.62E-05 | 2.34E-05 | 6.04E-05 | 1.66E-05 | 0 | 0 | 2.78E-05 | 1.54E-05 | 4.76E-05 |
| Orc4 | 0.00014 | 0.00013 | 0.00015 | 1.98E-05 | 9.50E-05 | 7.25E-05 | 0 | 4.41E-05 | 4.25E-05 | 2.51E-05 | 6.80E-05 | 6.35E-05 | 9.82E-06 | 4.53E-05 | 0.00016 | 6.66E-05 | 0 | 5.34E-05 |
| Tmco6 | 8.15E-06 | 4.99E-05 | 0 | 5.32E-05 | 6.41E-06 | 0 | 0 | 0 | 4.25E-05 | 2.36E-05 | 2.54E-05 | 1.44E-05 | 0 | 4.97E-06 | 0.00016 | 3.31E-05 | 0 | 3.57E-05 |
| Ift46 | 0.00015 | 0.00019 | 0 | 0.00017 | 0.00011 | 0.00013 | 0.00021 | 3.15E-05 | 4.24E-05 | 9.49E-05 | 5.29E-05 | 5.68E-05 | 1.82E-05 | 0.00014 | 0 | 0.00012 | 0 | 0.00015 |
| Cnot4 | 9.20E-05 | 0.00012 | 5.70E-05 | 3.77E-05 | 8.37E-05 | 6.25E-05 | 3.91E-06 | 0 | 4.24E-05 | 3.11E-05 | 4.78E-05 | 5.46E-05 | 1.29E-05 | 1.86E-05 | 0.00016 | 6.09E-05 | 0.00016 | 2.57E-05 |
| Gabpb1 | 6.58E-05 | 4.85E-05 | 0.00011 | 0 | 5.73E-05 | 5.82E-05 | 0 | 4.13E-05 | 4.24E-05 | 7.73E-06 | 1.56E-05 | 6.78E-05 | 0 | 2.29E-05 | 3.40E-05 | 0.00014 | 5.25E-05 | 0 |
| Abcb6 | 5.60E-05 | 2.63E-05 | 2.51E-05 | 8.40E-05 | 1.77E-06 | 0 | 0 | 0 | 4.24E-05 | 1.72E-05 | 0 | 5.60E-05 | 0 | 7.13E-06 | 9.90E-06 | 2.03E-05 | 1.98E-05 | 0 |
| Edem1 | 0.00026 | 3.83E-05 | 0.00026 | 0.00033 | 2.17E-05 | 8.29E-05 | 0.00027 | 0.00015 | 4.23E-05 | 0.00026 | 0.00018 | 0.00032 | 0 | 0 | 7.50E-05 | 0.0003 | 0.00039 | 0.00015 |
| Akap8l | 5.11E-05 | 0.00016 | 0 | 1.46E-05 | 5.04E-05 | 4.34E-05 | 1.65E-05 | 1.37E-05 | 4.23E-05 | 3.19E-05 | 0 | 8.15E-05 | 0 | 4.80E-05 | 0.00025 | 0 | 0.0001 | 4.70E-05 |
| Ift22 | 4.84E-05 | 6.33E-05 | 2.77E-05 | 0.00012 | 3.02E-05 | 3.64E-05 | 6.70E-05 | 5.27E-05 | 4.23E-05 | 8.61E-05 | 2.00E-05 | 5.51E-05 | 5.41E-06 | 0.0001 | 0 | 4.39E-05 | 3.28E-05 | 8.73E-05 |
| Tex10 | 9.43E-05 | 0.00012 | 6.76E-05 | 0 | 5.99E-05 | 3.34E-05 | 0 | 1.45E-05 | 4.23E-05 | 0 | 6.17E-05 | 7.69E-05 | 6.06E-07 | 9.17E-06 | 0.00013 | 0.00013 | 0.00011 | 3.21E-05 |
| Thap11 | 4.37E-06 | 0.00011 | 8.22E-05 | 0.00023 | 7.54E-05 | 3.19E-05 | 4.40E-05 | 6.62E-05 | 4.23E-05 | 7.75E-05 | 5.40E-05 | 4.17E-05 | 0 | 5.48E-05 | 6.42E-05 | 7.44E-05 | 2.25E-05 | 8.66E-05 |
| Zfp57 | 1.68E-05 | 1.26E-05 | 6.91E-06 | 0 | 1.53E-05 | 2.59E-05 | 0 | 6.24E-05 | 4.23E-05 | 0 | 8.68E-05 | 3.16E-05 | 0 | 7.03E-05 | 0.00017 | 0 | 4.28E-05 | 2.40E-05 |
| Trim37 | 3.31E-05 | 7.04E-05 | 1.51E-05 | 1.15E-05 | 1.24E-05 | 0 | 3.97E-06 | 0 | 4.23E-05 | 2.34E-05 | 1.51E-05 | 2.09E-05 | 4.33E-05 | 0 | 0 | 0.00019 | 3.23E-06 | 1.68E-05 |
| Zcrb1 | 0.00013 | 7.85E-05 | 0.00015 | 0.00012 | 5.49E-05 | 0.00013 | 5.14E-05 | 8.53E-05 | 4.22E-05 | 3.55E-05 | 7.88E-05 | 7.97E-05 | 1.10E-05 | 5.35E-05 | 2.63E-05 | 0.00019 | 0 | 5.31E-05 |
| Ktn1 | 0.0002 | 0.0002 | 9.00E-05 | 0 | 3.28E-05 | 8.34E-05 | 0 | 2.62E-05 | 4.22E-05 | 0 | 3.25E-06 | 0.0002 | 0 | 1.53E-05 | 0.00039 | 6.02E-05 | 0.00013 | 5.86E-06 |
| Lrrcc1 | 5.56E-05 | 4.69E-05 | 0 | 0 | 2.18E-06 | 2.22E-05 | 1.86E-05 | 0 | 4.22E-05 | 2.84E-05 | 1.05E-05 | 1.12E-05 | 0 | 0 | 0.00013 | 0.0001 | 2.36E-05 | 4.68E-05 |
| Trit1 | 3.74E-05 | 5.19E-05 | 2.91E-05 | 3.12E-06 | 4.57E-05 | 1.40E-05 | 0 | 0 | 4.22E-05 | 9.79E-06 | 3.49E-05 | 2.78E-05 | 0 | 0 | 0.00011 | 1.29E-05 | 0 | 3.61E-05 |
| Ankrd39 | 3.89E-05 | 4.61E-05 | 2.49E-05 | 0.00011 | 1.97E-05 | 9.49E-06 | 2.31E-05 | 6.60E-05 | 4.22E-05 | 3.23E-05 | 2.79E-05 | 0 | 3.06E-05 | 3.87E-05 | 7.42E-05 | 1.28E-05 | 0 | 3.67E-05 |
| Tnfrsf14 | 1.82E-05 | 3.28E-05 | 0 | 2.55E-06 | 3.20E-05 | 3.54E-06 | 0 | 2.17E-05 | 4.22E-05 | 5.30E-07 | 0 | 1.17E-05 | 0 | 2.27E-05 | 1.38E-07 | 0 | 0 | 2.16E-05 |
| Zfp229 | 1.31E-05 | 2.60E-05 | 0 | 1.84E-05 | 1.59E-05 | 0 | 0 | 5.33E-06 | 4.22E-05 | 4.75E-05 | 0 | 0 | 1.71E-05 | 0 | 2.08E-05 | 1.05E-05 | 0 | 2.12E-05 |
| Apmap | 0.0001 | 4.57E-05 | 0.0001 | 8.31E-05 | 5.61E-05 | 0.00013 | 0 | 7.90E-05 | 4.21E-05 | 8.10E-05 | 6.84E-05 | 1.12E-05 | 7.42E-05 | 4.98E-05 | 4.32E-07 | 0.00013 | 0 | 4.16E-05 |
| Faf1 | 0.00015 | 0.00014 | 5.20E-05 | 5.88E-05 | 6.01E-05 | 7.72E-05 | 9.87E-05 | 0 | 4.21E-05 | 7.43E-05 | 6.18E-05 | 5.11E-05 | 4.94E-05 | 4.35E-05 | 0.00028 | 0.00014 | 0.00013 | 5.02E-05 |
| Slc35c2 | 6.00E-05 | 7.21E-05 | 9.40E-05 | 0.00021 | 5.51E-05 | 5.49E-05 | 0.00012 | 4.74E-05 | 4.21E-05 | 4.49E-05 | 1.72E-05 | 4.30E-05 | 3.19E-05 | 1.07E-05 | 6.94E-06 | 0 | 0 | 2.15E-05 |
| Hmgb3 | 4.97E-06 | 4.65E-05 | 5.09E-05 | 0.00017 | 6.37E-05 | 3.02E-05 | 2.78E-06 | 4.80E-05 | 4.21E-05 | 0 | 2.34E-05 | 0 | 0 | 0.00016 | 8.48E-05 | 0.00056 | 0 | 4.30E-05 |
| Tmem131l | 2.45E-05 | 2.22E-05 | 0 | 0 | 4.59E-07 | 2.85E-05 | 0 | 1.25E-05 | 4.21E-05 | 0 | 2.74E-06 | 6.30E-05 | 0 | 4.04E-05 | 0 | 2.37E-05 | 1.94E-05 | 0 |

| | | | | | | | | | | | | | | | | | | |
|---|---|---|---|---|---|---|---|---|---|---|---|---|---|---|---|---|---|---|
| Ddx18 | 0.00015 | 6.46E-05 | 0.00018 | 3.61E-05 | 8.76E-06 | 2.00E-05 | 0 | 0 | 4.21E-05 | 0 | 4.68E-06 | 5.40E-05 | 1.87E-05 | 0 | 5.51E-05 | 4.84E-05 | 0 | 1.96E-07 |
| Xrcc1 | 2.53E-05 | 7.41E-05 | 6.96E-05 | 8.27E-05 | 3.61E-05 | 1.81E-05 | 2.05E-05 | 3.34E-05 | 4.21E-05 | 3.64E-05 | 1.86E-05 | 4.96E-05 | 1.01E-05 | 3.22E-05 | 9.94E-05 | 0.00034 | 8.43E-05 | 3.55E-05 |
| Gabbr1 | 0 | 0.00018 | 0 | 0.00018 | 4.93E-05 | 8.82E-05 | 0 | 6.42E-05 | 4.20E-05 | 7.43E-06 | 5.80E-05 | 0.0001 | 0 | 0.00019 | 0.00048 | 0 | 0.00012 | 3.67E-05 |
| Rfc2 | 0.00015 | 5.79E-05 | 0.00015 | 3.72E-05 | 6.46E-05 | 6.63E-05 | 0.0001 | 2.27E-05 | 4.20E-05 | 1.75E-05 | 2.20E-05 | 3.05E-06 | 2.49E-05 | 2.99E-05 | 2.20E-05 | 0.00064 | 0 | 1.48E-05 |
| Med12 | 2.89E-05 | 3.30E-05 | 2.43E-05 | 1.39E-05 | 6.13E-06 | 8.56E-06 | 0 | 7.83E-06 | 4.20E-05 | 1.99E-05 | 0 | 1.55E-05 | 1.28E-05 | 1.42E-05 | 0 | 1.60E-05 | 6.75E-05 | 1.57E-05 |
| Actr8 | 4.35E-05 | 6.18E-05 | 4.09E-05 | 6.10E-05 | 2.56E-05 | 0 | 1.87E-05 | 3.43E-05 | 4.20E-05 | 2.56E-05 | 4.02E-05 | 1.45E-05 | 2.10E-06 | 2.41E-05 | 9.80E-05 | 3.25E-05 | 1.62E-06 | 1.60E-05 |
| Lipg | 3.80E-06 | 0 | 0 | 4.06E-06 | 3.35E-06 | 0 | 0 | 1.75E-05 | 4.20E-05 | 3.13E-05 | 1.25E-05 | 0 | 0 | 0 | 0 | 1.82E-06 | 0 | 0 |
| Serinc1 | 0.00073 | 0.00033 | 0.00019 | 0 | 0.00019 | 0.00057 | 0.00025 | 0.00016 | 4.19E-05 | 4.86E-05 | 0.00019 | 0.00036 | 0.00014 | 8.41E-05 | 6.21E-05 | 0 | 0 | 6.47E-05 |
| Dnajc1 | 0.00016 | 0.0001 | 9.98E-05 | 0 | 0.00012 | 0.00016 | 0.00015 | 6.99E-05 | 4.19E-05 | 2.22E-05 | 1.93E-05 | 7.72E-05 | 7.07E-05 | 8.40E-05 | 0.00012 | 1.33E-05 | 0 | 5.54E-05 |
| Zfp280b | 3.96E-05 | 4.52E-05 | 3.85E-05 | 4.11E-05 | 4.25E-05 | 3.76E-05 | 1.76E-05 | 2.86E-05 | 4.19E-05 | 1.13E-05 | 0.00011 | 0 | 2.94E-05 | 0 | 2.88E-05 | 8.86E-05 | 0 | 2.59E-05 |
| Rprd1a | 5.63E-05 | 9.32E-05 | 6.88E-05 | 0 | 3.96E-05 | 3.56E-05 | 3.24E-05 | 4.39E-05 | 4.19E-05 | 4.32E-05 | 2.96E-05 | 6.31E-05 | 1.64E-05 | 0 | 0.00015 | 9.77E-06 | 0 | 3.09E-05 |
| Zc3hav1l | 1.91E-05 | 3.75E-05 | 3.47E-05 | 5.07E-05 | 4.71E-05 | 2.47E-05 | 7.44E-05 | 6.10E-05 | 4.19E-05 | 4.03E-05 | 1.11E-05 | 1.89E-05 | 1.69E-05 | 3.64E-05 | 6.69E-05 | 4.40E-05 | 1.10E-06 | 3.67E-05 |
| Arhgef18 | 4.95E-05 | 0.00013 | 1.76E-05 | 8.78E-05 | 5.48E-05 | 7.20E-06 | 1.71E-05 | 0 | 4.19E-05 | 5.73E-05 | 5.58E-05 | 4.10E-06 | 5.14E-06 | 6.42E-05 | 0.00017 | 8.50E-06 | 0.00011 | 0.0001 |
| Mphosph10 | 9.34E-05 | 6.88E-05 | 0.00014 | 2.76E-05 | 0 | 0 | 0 | 0 | 4.19E-05 | 5.21E-06 | 1.94E-05 | 0 | 0 | 1.30E-05 | 8.63E-05 | 6.47E-05 | 0 | 0 |
| Pomt1 | 2.66E-05 | 3.86E-05 | 2.16E-05 | 3.89E-05 | 4.11E-06 | 0 | 0 | 9.83E-06 | 4.19E-05 | 5.07E-05 | 0 | 1.38E-05 | 1.19E-05 | 0 | 4.82E-05 | 0.0001 | 6.21E-05 | 3.01E-05 |
| Ddx39 | 0.00021 | 6.09E-05 | 0.00039 | 0.00019 | 0 | 0 | 0 | 0 | 4.19E-05 | 4.43E-05 | 5.96E-05 | 1.03E-05 | 0 | 1.52E-05 | 0 | 0.00036 | 2.72E-05 | 0 |
| Rara | 0.00016 | 0.00017 | 0.00012 | 4.47E-05 | 7.95E-05 | 0.00023 | 6.57E-05 | 9.64E-05 | 4.18E-05 | 0.00011 | 8.59E-05 | 0.00015 | 0 | 1.01E-05 | 0.0002 | 4.23E-05 | 0.00015 | 0.00017 |
| Pdpk1 | 0.0001 | 0.00014 | 0.00015 | 1.29E-05 | 9.02E-05 | 0.00019 | 8.61E-05 | 0.0001 | 4.18E-05 | 9.66E-05 | 0.00014 | 0.00016 | 5.20E-06 | 8.59E-05 | 0 | 0.00011 | 0.00019 | 4.10E-05 |
| Fxr1 | 0.0003 | 0.00028 | 0.00015 | 0.00012 | 0.0001 | 0.00013 | 5.72E-05 | 0 | 4.18E-05 | 2.91E-05 | 3.78E-05 | 0.00033 | 3.71E-05 | 5.66E-05 | 0.00021 | 0.0002 | 0.00022 | 5.17E-05 |
| Nup153 | 0.00018 | 0.00013 | 0.00016 | 0 | 9.18E-06 | 7.50E-05 | 0 | 0 | 4.18E-05 | 0 | 6.32E-05 | 6.42E-05 | 7.24E-07 | 5.35E-05 | 0 | 0.00026 | 0.00021 | 0 |
| Trp53bp1 | 0.00011 | 9.37E-05 | 5.34E-05 | 4.36E-05 | 1.82E-05 | 4.23E-05 | 4.65E-05 | 8.67E-06 | 4.18E-05 | 4.82E-05 | 0 | 3.01E-05 | 1.32E-05 | 0 | 0.0002 | 2.19E-06 | 0.00013 | 0 |
| Sft2d1 | 1.40E-05 | 2.84E-05 | 0.00011 | 5.35E-05 | 4.97E-05 | 3.92E-05 | 8.23E-06 | 6.74E-05 | 4.18E-05 | 2.91E-05 | 7.22E-05 | 0 | 7.48E-05 | 5.97E-05 | 0 | 5.01E-06 | 0 | 2.95E-05 |
| Caprin2 | 3.08E-05 | 1.28E-05 | 0 | 0 | 0 | 3.85E-05 | 0 | 3.73E-05 | 4.18E-05 | 1.25E-05 | 0 | 7.56E-05 | 0 | 3.30E-06 | 0 | 6.38E-05 | 0.00023 | 5.85E-06 |
| Lyar | 9.85E-05 | 3.29E-05 | 0.00026 | 0.00014 | 5.02E-06 | 3.27E-05 | 0 | 0 | 4.18E-05 | 0 | 2.97E-06 | 0 | 6.21E-05 | 0 | 1.38E-05 | 0.00027 | 3.69E-05 | 0 |
| Mast3 | 4.79E-06 | 6.14E-05 | 3.56E-05 | 7.80E-05 | 1.54E-05 | 5.35E-06 | 0 | 7.78E-05 | 4.18E-05 | 1.46E-05 | 0 | 9.70E-05 | 5.17E-06 | 6.49E-05 | 0.00014 | 4.04E-05 | 3.77E-05 | 4.77E-05 |
| Hes6 | 0 | 8.42E-05 | 0 | 0.00014 | 9.40E-05 | 0 | 3.90E-05 | 1.54E-05 | 4.18E-05 | 4.15E-05 | 4.83E-05 | 0.00011 | 0 | 5.50E-05 | 6.88E-05 | 0 | 0 | 4.98E-05 |
| Lhpp | 3.32E-05 | 6.85E-06 | 3.52E-05 | 6.38E-05 | 0 | 0 | 5.60E-05 | 8.00E-06 | 4.18E-05 | 8.70E-05 | 0 | 0 | 2.85E-05 | 0 | 0 | 7.10E-06 | 0 | 5.31E-05 |
| Mob4 | 0.00022 | 0.00012 | 5.09E-05 | 0 | 6.37E-05 | 0.00015 | 3.10E-06 | 1.30E-05 | 4.17E-05 | 5.25E-06 | 4.70E-05 | 5.91E-05 | 0 | 3.55E-06 | 0.00011 | 0.00011 | 0 | 3.96E-05 |
| Zmym2 | 0.00016 | 0.00012 | 1.28E-05 | 0 | 6.64E-05 | 0.00012 | 5.34E-05 | 0 | 4.17E-05 | 7.24E-07 | 2.97E-05 | 0.00014 | 6.22E-06 | 4.16E-05 | 0.00014 | 0 | 0.00011 | 0 |
| Ift74 | 0.00013 | 6.99E-05 | 0 | 0 | 5.23E-05 | 7.57E-05 | 0 | 6.43E-06 | 4.17E-05 | 0 | 0 | 9.45E-06 | 0 | 1.48E-05 | 9.76E-05 | 6.97E-05 | 4.14E-06 | 3.53E-05 |
| Smarcal1 | 3.98E-05 | 7.32E-05 | 4.05E-06 | 1.23E-05 | 2.29E-05 | 3.42E-05 | 3.33E-05 | 0 | 4.17E-05 | 4.34E-05 | 2.78E-05 | 1.62E-05 | 2.02E-05 | 2.76E-05 | 1.16E-05 | 6.07E-05 | 1.30E-06 | 5.35E-05 |
| Gmeb1 | 3.80E-05 | 6.20E-05 | 6.63E-05 | 3.33E-05 | 3.68E-05 | 2.99E-05 | 0 | 6.14E-05 | 4.17E-05 | 1.74E-05 | 1.54E-05 | 3.91E-05 | 1.43E-05 | 4.01E-05 | 0.00012 | 0.00013 | 3.40E-05 | 6.38E-05 |
| Golph3l | 0.00012 | 8.87E-05 | 2.07E-06 | 4.02E-05 | 4.30E-05 | 1.46E-05 | 5.43E-05 | 0 | 4.17E-05 | 6.63E-05 | 3.02E-06 | 4.80E-05 | 6.53E-06 | 6.71E-06 | 2.07E-06 | 5.61E-05 | 0 | 5.38E-05 |

| | | | | | | | | | | | | | | | | | | |
|---|---|---|---|---|---|---|---|---|---|---|---|---|---|---|---|---|---|---|
| Trim27 | 0.00014 | 0.00012 | 0.00031 | 0.0003 | 2.78E-05 | 1.07E-05 | 0.00015 | 1.72E-05 | 4.17E-05 | 1.96E-05 | 8.98E-05 | 4.90E-05 | 4.41E-05 | 6.66E-05 | 0.00012 | 0.00023 | 0 | 3.68E-06 |
| Dhx37 | 6.43E-05 | 4.06E-05 | 9.82E-05 | 4.81E-05 | 1.14E-05 | 7.60E-06 | 0 | 0 | 4.17E-05 | 9.55E-06 | 1.41E-05 | 0 | 1.36E-05 | 4.03E-06 | 4.19E-05 | 2.34E-05 | 2.15E-05 | 2.45E-05 |
| Pop7 | 2.71E-05 | 1.32E-05 | 5.69E-05 | 4.22E-05 | 8.20E-06 | 4.82E-06 | 1.08E-05 | 2.38E-05 | 4.17E-05 | 1.12E-05 | 3.81E-05 | 1.35E-05 | 0 | 1.34E-06 | 8.99E-06 | 2.51E-05 | 0 | 1.20E-05 |
| Zc3hc1 | 4.84E-05 | 7.48E-05 | 4.85E-05 | 7.03E-05 | 7.98E-06 | 0 | 0 | 1.26E-06 | 4.17E-05 | 1.25E-05 | 1.32E-05 | 2.55E-05 | 0 | 1.06E-05 | 3.33E-05 | 5.95E-05 | 0 | 0 |
| Vps8 | 9.63E-05 | 0.0001 | 8.31E-05 | 1.79E-05 | 4.65E-05 | 7.07E-05 | 9.25E-05 | 7.64E-05 | 4.16E-05 | 4.77E-05 | 5.47E-05 | 0.00013 | 8.15E-05 | 7.12E-05 | 7.23E-05 | 0 | 5.42E-05 | 7.00E-05 |
| Tmem68 | 8.24E-05 | 6.26E-05 | 7.90E-05 | 0 | 6.49E-05 | 3.85E-05 | 0 | 2.16E-05 | 4.16E-05 | 2.40E-06 | 3.16E-05 | 5.19E-06 | 0 | 4.28E-05 | 4.57E-05 | 4.36E-05 | 8.17E-06 | 0 |
| Pxmp4 | 3.29E-05 | 3.85E-05 | 0 | 3.03E-05 | 2.72E-05 | 2.60E-05 | 2.78E-05 | 1.42E-05 | 4.16E-05 | 2.63E-05 | 0 | 1.34E-05 | 2.82E-08 | 1.04E-05 | 0 | 2.01E-05 | 0 | 4.41E-05 |
| Klhl25 | 2.56E-05 | 2.17E-05 | 1.22E-05 | 2.87E-05 | 3.34E-05 | 1.65E-05 | 0 | 8.93E-05 | 4.16E-05 | 5.04E-06 | 0 | 0 | 1.99E-05 | 8.23E-05 | 4.77E-05 | 4.29E-05 | 0 | 0 |
| Stau2 | 5.47E-05 | 5.19E-05 | 3.52E-05 | 0 | 4.06E-05 | 5.64E-05 | 2.01E-05 | 1.19E-05 | 4.15E-05 | 2.43E-05 | 5.09E-05 | 4.27E-05 | 6.46E-06 | 1.07E-06 | 5.24E-05 | 3.27E-05 | 1.36E-05 | 3.44E-05 |
| Cdk1 | 0 | 0 | 0 | 0 | 4.36E-05 | 5.30E-05 | 3.05E-05 | 5.90E-05 | 4.15E-05 | 0 | 1.32E-05 | 0 | 0 | 7.06E-06 | 0 | 0.0017 | 0 | 0 |
| Zfp131 | 8.24E-05 | 0.0001 | 0.00013 | 5.15E-05 | 3.09E-05 | 4.29E-05 | 3.20E-05 | 0 | 4.15E-05 | 5.85E-05 | 0.00011 | 9.43E-05 | 0 | 4.82E-05 | 0.00029 | 0.00015 | 0.00051 | 4.95E-05 |
| Scoc | 8.50E-05 | 2.33E-05 | 4.07E-05 | 0 | 4.39E-05 | 3.08E-05 | 0 | 0 | 4.15E-05 | 3.19E-05 | 3.51E-05 | 1.53E-05 | 6.68E-05 | 2.55E-05 | 0 | 0.00013 | 0 | 1.77E-05 |
| Dcxr | 1.18E-05 | 7.00E-05 | 0 | 9.13E-06 | 4.91E-05 | 2.51E-05 | 1.76E-06 | 1.28E-06 | 4.15E-05 | 3.67E-05 | 0 | 2.90E-05 | 0.00011 | 0 | 1.08E-06 | 0 | 0 | 0 |
| Abhd17c | 8.91E-05 | 7.37E-05 | 0.0002 | 1.48E-05 | 4.16E-05 | 2.18E-05 | 7.03E-06 | 0 | 4.15E-05 | 2.89E-05 | 6.85E-05 | 1.27E-05 | 0.00026 | 5.17E-05 | 0 | 0.00014 | 2.25E-05 | 0 |
| Cdt1 | 3.57E-06 | 7.30E-06 | 0.00015 | 4.21E-05 | 0 | 2.13E-05 | 0 | 5.24E-05 | 4.15E-05 | 0 | 0 | 4.68E-06 | 0 | 9.05E-05 | 3.20E-05 | 0.00029 | 8.14E-05 | 0 |
| Zfyve19 | 4.38E-05 | 2.98E-05 | 1.79E-05 | 4.45E-05 | 1.72E-05 | 1.89E-05 | 1.60E-05 | 0 | 4.15E-05 | 2.50E-05 | 4.55E-06 | 1.47E-05 | 0 | 0 | 6.94E-05 | 1.12E-05 | 0 | 6.77E-06 |
| Ciao3 | 4.84E-05 | 4.50E-05 | 2.71E-05 | 3.45E-05 | 2.69E-05 | 1.26E-05 | 1.59E-05 | 3.16E-06 | 4.15E-05 | 1.50E-05 | 2.87E-05 | 0 | 0 | 2.40E-05 | 3.16E-05 | 1.84E-05 | 0 | 1.62E-05 |
| Tmem167b | 0.00014 | 0.00011 | 0.00013 | 0.00036 | 0.00016 | 0.00019 | 0.0003 | 0.00014 | 4.14E-05 | 0.00013 | 0.00012 | 0.00016 | 0.00012 | 0.00011 | 0 | 3.89E-05 | 5.18E-05 | 0.00018 |
| 8-Mar | 0.00013 | 7.77E-05 | 5.87E-05 | 5.63E-05 | 6.26E-05 | 9.67E-05 | 0.00018 | 4.01E-05 | 4.14E-05 | 0.00015 | 3.48E-05 | 4.13E-05 | 0.00011 | 2.75E-05 | 1.75E-05 | 0 | 0 | 2.89E-05 |
| Rif1 | 5.51E-05 | 5.66E-05 | 4.54E-05 | 0 | 1.37E-06 | 3.77E-05 | 0 | 0 | 4.14E-05 | 1.49E-05 | 6.04E-06 | 0 | 2.90E-05 | 6.66E-06 | 4.63E-05 | 0.00013 | 4.33E-05 | 0 |
| Gch1 | 0 | 5.42E-05 | 9.49E-05 | 0 | 3.38E-05 | 1.97E-05 | 0 | 9.02E-06 | 4.14E-05 | 0 | 0 | 0.00015 | 0.00018 | 2.12E-05 | 0 | 0 | 0.0002 | 0 |
| Swsap1 | 2.10E-05 | 3.12E-05 | 2.64E-05 | 5.53E-05 | 3.30E-05 | 1.08E-05 | 1.49E-05 | 1.78E-05 | 4.14E-05 | 1.05E-05 | 2.24E-06 | 0 | 1.38E-06 | 3.83E-06 | 0 | 1.70E-05 | 0 | 4.00E-05 |
| Prpf3 | 6.29E-05 | 6.37E-05 | 7.92E-05 | 8.85E-05 | 1.14E-05 | 0 | 5.27E-06 | 0 | 4.14E-05 | 3.64E-05 | 5.06E-05 | 1.70E-07 | 0 | 0 | 0.00021 | 9.88E-05 | 4.34E-06 | 1.96E-05 |
| Plaur | 0 | 4.17E-05 | 0.00078 | 0.00023 | 0 | 0.00015 | 0.00031 | 0.00038 | 4.13E-05 | 0 | 0.00032 | 0.00015 | 0.00036 | 0.00025 | 0.00039 | 0.00013 | 0.00022 | 2.01E-05 |
| Fam83h | 0 | 5.38E-05 | 0 | 2.92E-05 | 2.11E-05 | 1.47E-05 | 1.97E-05 | 0 | 4.13E-05 | 0 | 1.72E-05 | 2.99E-05 | 1.12E-06 | 4.36E-05 | 4.28E-05 | 0 | 0 | 3.10E-05 |
| Yod1 | 1.92E-05 | 6.77E-05 | 3.55E-05 | 1.28E-05 | 4.74E-05 | 5.97E-06 | 3.22E-06 | 9.70E-06 | 4.13E-05 | 1.07E-05 | 1.85E-05 | 4.98E-05 | 1.95E-05 | 2.37E-05 | 5.16E-05 | 2.38E-05 | 0.00017 | 3.19E-07 |
| Aatf | 8.66E-05 | 6.18E-05 | 0.00015 | 8.88E-06 | 0 | 1.62E-06 | 0 | 0 | 4.13E-05 | 2.01E-05 | 0 | 1.64E-05 | 1.42E-05 | 1.36E-05 | 0.00014 | 3.60E-05 | 5.77E-05 | 0 |
| Ifi47 | 8.99E-05 | 4.06E-05 | 0 | 0 | 7.16E-05 | 0 | 0 | 0 | 4.13E-05 | 0.00017 | 2.96E-05 | 0 | 0.00011 | 3.03E-05 | 0 | 1.26E-05 | 0 | 0.00018 |
| Tbl2 | 0.00016 | 2.09E-05 | 4.31E-05 | 0.00014 | 3.30E-05 | 0 | 0.00021 | 1.81E-05 | 4.13E-05 | 1.84E-05 | 0 | 0.00038 | 4.86E-05 | 2.21E-05 | 0.00013 | 0.00016 | 0 | 5.71E-05 |
| Col26a1 | 0 | 5.31E-05 | 0 | 0 | 3.93E-06 | 0 | 0 | 2.87E-06 | 4.13E-05 | 0 | 0 | 0 | 0 | 5.10E-05 | 0 | 0 | 0 | 0 |
| Setx | 0.00014 | 9.23E-05 | 1.91E-05 | 0 | 9.94E-06 | 6.62E-05 | 1.78E-05 | 8.81E-06 | 4.12E-05 | 7.89E-06 | 0 | 4.80E-05 | 4.59E-06 | 1.83E-05 | 4.92E-05 | 8.04E-05 | 0 | 4.24E-06 |
| Letmd1 | 4.89E-05 | 7.10E-05 | 3.47E-05 | 0.00012 | 4.70E-05 | 4.92E-05 | 2.31E-05 | 5.69E-05 | 4.12E-05 | 8.67E-05 | 1.39E-05 | 4.64E-05 | 6.04E-06 | 5.94E-05 | 7.77E-06 | 0 | 0 | 6.29E-05 |
| Mapk7 | 5.43E-05 | 4.32E-05 | 1.87E-05 | 0 | 1.14E-05 | 1.18E-05 | 4.22E-05 | 0 | 4.12E-05 | 3.44E-05 | 3.42E-05 | 5.44E-05 | 9.44E-06 | 0 | 1.19E-05 | 5.07E-06 | 0 | 5.03E-05 |

| | | | | | | | | | | | | | | | | | | |
|---|---|---|---|---|---|---|---|---|---|---|---|---|---|---|---|---|---|---|
| Nedd4l | 2.49E-05 | 4.51E-05 | 1.19E-05 | 0 | 0 | 9.89E-06 | 0 | 0 | 4.12E-05 | 5.74E-05 | 5.85E-05 | 0.00019 | 0.00016 | 7.50E-05 | 0 | 6.52E-05 | 6.87E-05 | 2.54E-06 |
| Amot | 0 | 3.82E-05 | 0 | 3.20E-05 | 1.51E-05 | 0 | 5.37E-05 | 2.96E-05 | 4.12E-05 | 0 | 2.88E-05 | 0 | 1.43E-05 | 0 | 0 | 5.41E-05 | 1.54E-05 | 6.85E-05 |
| Rad50 | 5.95E-05 | 6.66E-05 | 2.71E-05 | 3.21E-05 | 2.49E-05 | 4.17E-05 | 2.90E-05 | 2.42E-06 | 4.11E-05 | 4.50E-05 | 4.20E-05 | 5.72E-05 | 1.97E-05 | 4.15E-05 | 6.17E-05 | 0.00023 | 0 | 5.31E-05 |
| Parp14 | 9.56E-05 | 7.44E-05 | 0 | 0 | 1.36E-05 | 2.27E-05 | 0 | 0 | 4.11E-05 | 0.00016 | 0 | 8.47E-06 | 7.69E-05 | 1.30E-05 | 7.17E-05 | 0 | 0 | 9.46E-05 |
| Fem1a | 6.81E-05 | 4.60E-05 | 9.78E-05 | 2.28E-05 | 4.21E-05 | 2.08E-05 | 1.99E-06 | 0 | 4.11E-05 | 9.85E-06 | 1.17E-05 | 0 | 3.70E-06 | 3.20E-05 | 2.44E-05 | 1.79E-05 | 0 | 1.84E-05 |
| Wrap73 | 1.85E-05 | 3.03E-05 | 2.45E-05 | 5.03E-05 | 1.58E-05 | 6.88E-06 | 1.83E-05 | 1.16E-05 | 4.11E-05 | 2.22E-05 | 2.12E-05 | 2.18E-07 | 0 | 8.90E-06 | 1.63E-05 | 3.47E-05 | 0 | 9.35E-06 |
| Tmem135 | 3.20E-05 | 8.30E-05 | 2.53E-05 | 0.0001 | 6.04E-05 | 6.61E-06 | 0 | 8.64E-05 | 4.11E-05 | 8.26E-05 | 1.69E-05 | 6.77E-05 | 8.57E-05 | 3.55E-05 | 5.73E-05 | 9.71E-06 | 5.49E-05 | 8.71E-05 |
| Akr1e1 | 8.38E-05 | 2.74E-05 | 0 | 0 | 9.40E-07 | 6.29E-06 | 2.38E-05 | 3.26E-06 | 4.11E-05 | 4.06E-05 | 1.49E-05 | 3.63E-05 | 3.92E-06 | 2.70E-06 | 2.83E-05 | 3.50E-05 | 0 | 3.74E-05 |
| Rpusd4 | 2.69E-05 | 2.49E-05 | 2.54E-05 | 1.01E-05 | 0 | 0 | 6.29E-06 | 1.60E-05 | 4.11E-05 | 1.32E-05 | 2.30E-05 | 9.92E-06 | 6.14E-06 | 6.69E-06 | 1.59E-05 | 1.01E-05 | 0 | 2.04E-05 |
| Pcbd1 | 3.03E-06 | 0 | 3.35E-06 | 0 | 3.86E-05 | 3.03E-05 | 1.44E-06 | 0 | 4.10E-05 | 1.72E-05 | 3.03E-06 | 3.50E-05 | 0 | 1.23E-05 | 0 | 0 | 4.89E-06 | 8.10E-05 |
| Zfp354a | 1.06E-05 | 3.18E-05 | 0 | 0 | 3.33E-05 | 2.27E-06 | 0 | 0 | 4.10E-05 | 0 | 1.66E-05 | 0 | 2.44E-06 | 1.68E-05 | 4.11E-05 | 0 | 0 | 6.02E-06 |
| Acaa1a | 2.87E-05 | 4.72E-05 | 2.75E-05 | 7.04E-05 | 2.06E-05 | 0 | 8.60E-06 | 1.02E-05 | 4.10E-05 | 1.29E-05 | 2.75E-05 | 1.12E-05 | 0 | 1.17E-05 | 7.51E-05 | 0 | 0 | 2.38E-05 |
| Prkar2a | 0.0001 | 3.78E-05 | 0.00019 | 0 | 4.04E-05 | 4.64E-05 | 0 | 0 | 4.09E-05 | 5.30E-05 | 0.00021 | 4.26E-05 | 0 | 2.95E-05 | 0 | 0.00019 | 0 | 2.71E-05 |
| Spred3 | 0 | 4.63E-05 | 5.87E-06 | 9.04E-05 | 4.97E-05 | 0 | 0 | 0.00012 | 4.09E-05 | 3.55E-06 | 0 | 0.00011 | 3.01E-05 | 0 | 0.00016 | 0 | 0.00021 | 0 |
| Peg12 | 3.08E-05 | 0 | 8.42E-05 | 4.26E-05 | 3.74E-05 | 0 | 4.26E-05 | 5.34E-05 | 4.09E-05 | 0 | 8.80E-06 | 0 | 0 | 3.87E-05 | 2.09E-05 | 5.47E-05 | 0 | 1.43E-06 |
| Anapc5 | 0.00028 | 4.53E-05 | 3.62E-05 | 0 | 1.61E-05 | 0.00015 | 0.00011 | 0 | 4.08E-05 | 4.27E-05 | 0 | 5.51E-05 | 0 | 9.10E-07 | 0.00014 | 0.00022 | 6.95E-05 | 4.50E-05 |
| Arrb1 | 0 | 4.63E-05 | 0 | 0 | 0 | 0.00013 | 0 | 0 | 4.08E-05 | 0.00011 | 0 | 0.00054 | 0 | 0.00012 | 2.99E-05 | 5.49E-05 | 0 | 0.00021 |
| Fbxo21 | 6.21E-05 | 0.00011 | 6.37E-06 | 0.00015 | 0.00011 | 6.29E-05 | 7.45E-05 | 3.76E-05 | 4.08E-05 | 6.12E-05 | 7.61E-05 | 0.00025 | 0.00011 | 0.00012 | 0 | 0 | 0 | 8.61E-05 |
| Recql | 9.22E-05 | 3.24E-05 | 7.85E-05 | 4.67E-06 | 5.43E-05 | 4.45E-05 | 3.79E-05 | 2.13E-05 | 4.08E-05 | 7.55E-05 | 1.95E-05 | 0 | 0 | 4.59E-05 | 0.0002 | 0.00023 | 0 | 8.76E-05 |
| Heatr6 | 8.36E-05 | 7.89E-05 | 5.83E-05 | 2.70E-05 | 8.76E-06 | 2.66E-05 | 2.91E-05 | 2.98E-06 | 4.08E-05 | 2.65E-05 | 3.04E-05 | 2.28E-05 | 3.00E-06 | 6.22E-06 | 7.30E-05 | 7.28E-05 | 7.10E-05 | 1.71E-05 |
| Cox20 | 0.0001 | 8.41E-05 | 0 | 8.51E-05 | 3.81E-05 | 1.39E-05 | 0 | 4.00E-05 | 4.08E-05 | 3.72E-05 | 7.35E-06 | 6.16E-05 | 0 | 1.13E-05 | 2.48E-05 | 6.13E-05 | 0 | 6.91E-05 |
| Fancc | 1.96E-05 | 4.26E-05 | 7.29E-07 | 3.09E-06 | 2.58E-06 | 1.14E-05 | 0 | 0 | 4.08E-05 | 1.37E-05 | 3.41E-05 | 5.05E-05 | 0 | 1.42E-05 | 1.45E-05 | 5.38E-05 | 4.89E-06 | 9.50E-05 |
| Nhs | 7.07E-06 | 1.81E-05 | 0 | 0 | 0 | 2.89E-06 | 0 | 2.11E-05 | 4.08E-05 | 0 | 0 | 0 | 0 | 0 | 5.51E-06 | 1.75E-05 | 7.43E-06 | 2.25E-06 |
| Prmt3 | 8.20E-05 | 5.78E-05 | 0.00014 | 7.57E-06 | 0 | 0 | 0 | 0 | 4.08E-05 | 1.03E-05 | 2.59E-05 | 0 | 1.80E-05 | 1.07E-06 | 4.90E-05 | 5.36E-05 | 6.00E-05 | 1.12E-05 |
| Vill | 1.06E-05 | 4.52E-05 | 0 | 0 | 0 | 3.65E-05 | 0 | 7.48E-05 | 4.07E-05 | 0 | 2.51E-06 | 3.40E-05 | 0 | 4.57E-05 | 1.05E-05 | 7.07E-06 | 1.28E-05 | 0.00026 |
| Irf9 | 9.12E-05 | 9.58E-05 | 8.81E-06 | 6.54E-05 | 1.40E-05 | 0 | 5.82E-06 | 2.97E-06 | 4.07E-05 | 0.00024 | 0 | 4.02E-05 | 9.64E-05 | 0 | 1.30E-05 | 2.67E-05 | 6.29E-05 | 0.00019 |
| Emc2 | 0.00025 | 9.34E-05 | 8.54E-05 | 0.00011 | 5.84E-05 | 9.29E-05 | 2.28E-05 | 0 | 4.06E-05 | 0.00018 | 6.16E-05 | 6.24E-06 | 0.00013 | 1.49E-05 | 0 | 4.88E-05 | 0 | 6.81E-05 |
| Pdzd8 | 0.00014 | 0.00012 | 0.00014 | 0 | 0.0001 | 7.65E-05 | 2.43E-05 | 3.00E-05 | 4.06E-05 | 1.89E-05 | 0.00011 | 6.95E-05 | 6.28E-05 | 6.26E-05 | 4.19E-05 | 0.00016 | 5.51E-05 | 3.36E-05 |
| Rsbn1l | 6.37E-05 | 8.90E-05 | 1.33E-05 | 5.11E-06 | 4.30E-05 | 7.51E-05 | 8.56E-06 | 4.76E-05 | 4.06E-05 | 3.59E-05 | 1.99E-05 | 5.24E-05 | 0 | 1.43E-05 | 5.79E-05 | 6.27E-05 | 0 | 3.31E-05 |
| Crcp | 0.00012 | 6.97E-05 | 0.00011 | 7.46E-05 | 4.14E-05 | 6.06E-05 | 2.07E-05 | 0 | 4.06E-05 | 2.56E-05 | 6.63E-05 | 7.26E-06 | 0 | 1.21E-05 | 3.13E-05 | 1.35E-05 | 0 | 9.32E-06 |
| Atp11c | 4.36E-05 | 9.64E-05 | 1.93E-05 | 0 | 1.39E-05 | 5.11E-05 | 0 | 3.43E-05 | 4.06E-05 | 2.16E-05 | 0 | 2.80E-05 | 0 | 2.74E-05 | 0.00016 | 0.00014 | 6.25E-05 | 4.83E-06 |
| Bag4 | 5.12E-05 | 5.51E-05 | 6.60E-05 | 0 | 8.10E-05 | 3.42E-05 | 9.21E-05 | 3.46E-05 | 4.06E-05 | 3.71E-05 | 0 | 5.08E-05 | 4.32E-05 | 0 | 0 | 0.0001 | 6.54E-06 | 0 |
| Lrp4 | 3.05E-07 | 4.04E-05 | 4.74E-06 | 0 | 0 | 3.46E-06 | 8.07E-06 | 1.10E-05 | 4.06E-05 | 6.12E-06 | 0 | 1.65E-05 | 0 | 2.49E-06 | 0 | 2.52E-05 | 8.63E-06 | 9.36E-05 |

| | | | | | | | | | | | | | | | | | | |
|---|---|---|---|---|---|---|---|---|---|---|---|---|---|---|---|---|---|---|
| Eloa | 0.00028 | 0.00025 | 0.00024 | 0.00014 | 0.00022 | 0.00023 | 0.00013 | 6.89E-05 | 4.05E-05 | 8.77E-05 | 9.71E-05 | 0.00025 | 0 | 6.54E-05 | 0.0001 | 7.40E-05 | 0.00038 | 5.25E-05 |
| Rabgap1 | 0.00016 | 0.00015 | 9.46E-06 | 0 | 8.41E-05 | 0.00017 | 2.62E-05 | 1.64E-05 | 4.05E-05 | 1.85E-05 | 2.51E-05 | 0.0001 | 4.51E-05 | 3.74E-05 | 0.00012 | 8.30E-06 | 3.05E-06 | 5.33E-06 |
| Atp5mpl | 0.00029 | 7.92E-05 | 0.00016 | 0.00032 | 7.69E-05 | 0.00013 | 8.32E-05 | 0.0001 | 4.05E-05 | 4.14E-06 | 0.00016 | 5.09E-05 | 0 | 8.29E-05 | 0 | 7.27E-05 | 0 | 5.25E-05 |
| Pip4k2c | 5.66E-05 | 0.00017 | 5.69E-05 | 0.00014 | 9.24E-05 | 0.00011 | 5.83E-05 | 0.00012 | 4.05E-05 | 3.31E-05 | 6.18E-05 | 0.00012 | 7.30E-06 | 0.00013 | 4.16E-05 | 0 | 8.55E-05 | 7.18E-05 |
| Apobec1 | 0 | 3.26E-05 | 0 | 0 | 0 | 5.96E-05 | 0 | 4.39E-05 | 4.05E-05 | 2.68E-05 | 0 | 0 | 1.07E-07 | 0 | 0 | 1.26E-05 | 0 | 0.00013 |
| Rsbn1 | 7.75E-05 | 9.64E-05 | 7.60E-05 | 0 | 1.82E-05 | 5.82E-05 | 0 | 0 | 4.05E-05 | 6.38E-07 | 0 | 0.00025 | 0 | 7.81E-05 | 0.0001 | 3.04E-05 | 0.00028 | 5.09E-06 |
| Pde3a | 4.30E-06 | 9.01E-05 | 3.62E-05 | 0 | 2.27E-05 | 3.90E-05 | 0 | 0.00012 | 4.05E-05 | 5.89E-05 | 8.81E-05 | 5.07E-05 | 0 | 0.00017 | 4.70E-05 | 0 | 8.40E-05 | 1.37E-05 |
| Inpp1 | 3.41E-05 | 3.20E-05 | 5.47E-05 | 5.15E-06 | 0 | 2.30E-05 | 7.32E-05 | 1.53E-05 | 4.05E-05 | 0.00013 | 0 | 0 | 0 | 8.14E-05 | 7.25E-05 | 2.62E-05 | 0 | 7.09E-05 |
| Tpra1 | 9.40E-06 | 5.35E-05 | 0 | 4.92E-05 | 3.68E-05 | 1.69E-05 | 2.19E-05 | 0 | 4.05E-05 | 2.29E-05 | 8.90E-06 | 1.31E-05 | 1.09E-05 | 4.43E-05 | 3.86E-05 | 0 | 0 | 3.83E-05 |
| Rbm34 | 8.25E-05 | 4.51E-05 | 7.69E-05 | 7.17E-06 | 1.47E-05 | 1.66E-05 | 0 | 0 | 4.05E-05 | 0 | 1.61E-05 | 2.51E-05 | 1.73E-05 | 0 | 6.47E-06 | 3.33E-05 | 0 | 0 |
| Ndufaf3 | 4.65E-05 | 5.37E-05 | 1.73E-05 | 7.85E-05 | 4.66E-05 | 1.39E-05 | 7.07E-06 | 2.26E-05 | 4.05E-05 | 2.24E-05 | 3.49E-05 | 2.64E-05 | 2.53E-05 | 4.61E-07 | 0 | 1.82E-05 | 0 | 3.81E-05 |
| Ing5 | 2.00E-05 | 4.90E-05 | 0.00012 | 8.06E-05 | 1.59E-05 | 1.29E-05 | 0 | 2.31E-05 | 4.05E-05 | 1.77E-05 | 2.46E-05 | 3.29E-05 | 1.99E-05 | 1.04E-06 | 8.12E-05 | 0.00011 | 9.73E-05 | 2.91E-05 |
| Pold1 | 0 | 1.43E-05 | 8.08E-05 | 1.96E-05 | 0 | 0 | 6.99E-06 | 6.14E-06 | 4.05E-05 | 3.82E-05 | 2.84E-05 | 1.22E-05 | 2.78E-05 | 0 | 0.00011 | 0.00048 | 2.31E-05 | 2.92E-05 |
| Gas8 | 5.76E-05 | 6.05E-05 | 7.59E-05 | 5.08E-05 | 0 | 0 | 0 | 0 | 4.05E-05 | 1.88E-05 | 2.41E-05 | 2.40E-05 | 0 | 4.92E-05 | 0.00011 | 4.73E-05 | 4.83E-05 | 5.48E-05 |
| Atp2c1 | 0.0001 | 0.0002 | 0.00019 | 0.00016 | 0.00015 | 0.0002 | 9.77E-06 | 0.00016 | 4.04E-05 | 0.00013 | 0.0003 | 0.00018 | 0.00011 | 0.00021 | 0.00023 | 0.00016 | 5.19E-05 | 0.0001 |
| Rbm7 | 0.00024 | 0.00021 | 0.00013 | 8.96E-05 | 5.64E-05 | 0.00016 | 1.46E-05 | 4.92E-05 | 4.04E-05 | 2.85E-05 | 8.28E-05 | 0.00014 | 2.58E-06 | 5.89E-05 | 2.37E-05 | 0.00013 | 6.23E-05 | 1.37E-05 |
| Paxip1 | 3.86E-05 | 6.68E-05 | 9.97E-05 | 0 | 5.09E-05 | 6.43E-05 | 4.14E-05 | 0 | 4.04E-05 | 3.81E-05 | 3.60E-05 | 1.89E-06 | 6.47E-05 | 8.57E-05 | 0.0001 | 0.00036 | 7.00E-05 | 3.85E-06 |
| Gtpbp10 | 6.60E-05 | 5.55E-05 | 1.08E-05 | 0 | 1.59E-05 | 4.37E-05 | 0 | 0 | 4.04E-05 | 1.12E-05 | 1.71E-05 | 2.41E-05 | 6.74E-06 | 1.85E-05 | 6.23E-05 | 0.00011 | 6.45E-06 | 3.78E-06 |
| Vwa8 | 4.79E-05 | 5.94E-05 | 5.00E-06 | 3.00E-07 | 5.32E-05 | 3.77E-05 | 3.23E-05 | 4.48E-05 | 4.04E-05 | 4.15E-06 | 5.60E-05 | 2.60E-05 | 1.39E-05 | 0 | 0 | 2.34E-05 | 3.52E-05 | 5.56E-05 |
| Zcchc3 | 4.45E-06 | 2.49E-05 | 0 | 4.21E-05 | 2.10E-05 | 4.08E-06 | 0 | 1.64E-05 | 4.04E-05 | 2.29E-05 | 4.11E-06 | 9.73E-06 | 0 | 2.08E-05 | 4.13E-05 | 2.40E-05 | 0 | 1.34E-05 |
| Tmem121 | 0 | 6.69E-07 | 1.86E-05 | 2.46E-05 | 2.46E-05 | 0 | 0 | 1.63E-05 | 4.04E-05 | 2.47E-06 | 2.90E-06 | 0 | 2.42E-05 | 8.63E-05 | 0 | 5.29E-05 | 3.37E-05 | 2.73E-05 |
| Rab36 | 0 | 2.38E-05 | 0 | 1.42E-05 | 1.55E-05 | 0 | 0 | 2.83E-05 | 4.04E-05 | 2.00E-05 | 0 | 1.05E-05 | 6.30E-06 | 6.02E-06 | 2.29E-05 | 0 | 8.25E-07 | 5.61E-05 |
| Focad | 3.94E-05 | 1.95E-05 | 2.36E-05 | 1.23E-05 | 2.04E-05 | 1.84E-05 | 3.18E-05 | 0 | 4.03E-05 | 1.35E-05 | 0 | 8.59E-06 | 1.05E-05 | 1.17E-05 | 2.34E-05 | 0 | 0 | 1.50E-05 |
| Sfi1 | 5.64E-05 | 9.13E-05 | 1.07E-06 | 2.86E-05 | 6.80E-05 | 6.60E-06 | 4.27E-05 | 3.71E-05 | 4.03E-05 | 8.93E-05 | 0 | 3.67E-05 | 2.49E-05 | 0 | 0.00021 | 0 | 0 | 6.73E-05 |
| Krr1 | 8.79E-05 | 7.60E-05 | 9.22E-05 | 3.22E-05 | 2.51E-05 | 5.77E-06 | 0 | 0 | 4.03E-05 | 8.43E-06 | 9.74E-06 | 2.48E-05 | 0 | 1.30E-05 | 5.99E-05 | 7.47E-05 | 0 | 0 |
| Rangap1 | 4.84E-06 | 7.26E-05 | 0.00021 | 0.0002 | 2.05E-05 | 0 | 0 | 3.95E-06 | 4.03E-05 | 1.71E-05 | 5.80E-05 | 8.61E-06 | 0 | 2.68E-05 | 3.18E-05 | 0.00102 | 0 | 3.51E-05 |
| DEPRECATEI | 1.84E-05 | 2.53E-05 | 6.71E-05 | 1.01E-07 | 2.23E-05 | 0 | 1.23E-06 | 3.02E-05 | 4.03E-05 | 2.79E-05 | 6.13E-06 | 2.45E-05 | 1.42E-05 | 0 | 0 | 0.00021 | 0 | 3.83E-05 |
| Khdrbs3 | 6.68E-05 | 5.97E-05 | 0.00015 | 0 | 0 | 0 | 0 | 3.82E-05 | 4.03E-05 | 8.86E-05 | 0.00033 | 3.37E-06 | 0.00013 | 0 | 0 | 0.00029 | 0 | 8.34E-05 |
| Arl4a | 2.59E-05 | 6.18E-05 | 0 | 0 | 0 | 0 | 0 | 4.90E-06 | 4.03E-05 | 5.14E-05 | 0 | 6.42E-05 | 2.56E-05 | 0 | 0 | 2.98E-06 | 0.00018 | 5.97E-05 |
| Raph1 | 4.80E-05 | 5.30E-05 | 1.51E-05 | 0 | 0 | 6.31E-05 | 0 | 0 | 4.02E-05 | 5.47E-05 | 0.00013 | 0.00012 | 0 | 0.00014 | 2.72E-05 | 7.01E-05 | 6.95E-06 | 4.13E-05 |
| Hdac11 | 0 | 4.98E-05 | 0 | 0 | 1.08E-05 | 5.33E-05 | 0 | 0 | 4.02E-05 | 1.96E-05 | 0 | 1.40E-05 | 4.48E-06 | 4.58E-06 | 2.65E-06 | 0 | 0 | 9.12E-06 |
| Abraxas2 | 9.59E-05 | 6.67E-05 | 5.35E-05 | 0 | 4.86E-05 | 5.23E-05 | 5.95E-05 | 0 | 4.02E-05 | 2.16E-05 | 1.91E-05 | 4.43E-05 | 0 | 0 | 8.32E-05 | 6.04E-05 | 0 | 0 |
| Gfm1 | 0.0001 | 7.14E-05 | 0.00018 | 7.94E-05 | 2.39E-05 | 4.92E-05 | 9.88E-06 | 2.93E-05 | 4.02E-05 | 2.05E-05 | 9.75E-05 | 0 | 3.25E-05 | 5.06E-05 | 0 | 0.00016 | 1.78E-05 | 3.63E-05 |

| | | | | | | | | | | | | | | | | | | |
|---|---|---|---|---|---|---|---|---|---|---|---|---|---|---|---|---|---|---|
| Ddhd1 | 6.17E-07 | 0.00015 | 4.09E-05 | 8.99E-06 | 3.40E-05 | 2.24E-05 | 0 | 5.27E-05 | 4.02E-05 | 0.00015 | 1.81E-05 | 2.93E-05 | 0 | 7.51E-06 | 0.0001 | 0.0001 | 0.0001 | 7.85E-05 |
| Tsen15 | 5.30E-05 | 5.01E-05 | 3.61E-05 | 7.71E-05 | 2.77E-05 | 2.15E-05 | 3.09E-06 | 3.96E-06 | 4.02E-05 | 1.56E-05 | 1.56E-05 | 1.98E-05 | 4.15E-06 | 1.02E-05 | 0 | 0.00014 | 0 | 4.58E-06 |
| Gon4l | 2.91E-05 | 0.0001 | 4.99E-05 | 6.93E-05 | 2.08E-05 | 4.30E-06 | 0 | 5.64E-06 | 4.02E-05 | 4.31E-05 | 1.80E-05 | 4.11E-05 | 0 | 3.33E-05 | 0.00021 | 7.68E-05 | 2.29E-05 | 2.52E-05 |
| Usp11 | 4.16E-05 | 3.47E-05 | 0 | 1.82E-05 | 0 | 0 | 0 | 1.53E-05 | 4.02E-05 | 1.75E-05 | 0 | 0 | 3.63E-05 | 0 | 5.32E-05 | 1.91E-05 | 2.64E-05 | 3.15E-05 |
| 10-Sep | 0.00015 | 1.49E-05 | 0 | 0 | 0 | 0.00013 | 0 | 5.22E-05 | 4.01E-05 | 5.40E-05 | 0 | 0.00011 | 9.61E-05 | 3.99E-05 | 2.62E-05 | 0.00023 | 0 | 0.00011 |
| Tmem179b | 0.00012 | 8.63E-05 | 2.87E-05 | 6.81E-05 | 2.42E-05 | 0.0001 | 8.40E-05 | 6.52E-05 | 4.01E-05 | 3.00E-05 | 3.25E-05 | 8.76E-05 | 3.99E-05 | 4.31E-05 | 1.41E-05 | 3.56E-05 | 1.04E-05 | 0 |
| Git1 | 0.00012 | 0.00019 | 0.00017 | 0.00016 | 4.77E-05 | 6.52E-05 | 0.00019 | 6.56E-05 | 4.01E-05 | 0.00014 | 6.74E-05 | 5.60E-05 | 0.00011 | 2.02E-05 | 0.00013 | 0.00012 | 0.00013 | 0.00013 |
| Trappc9 | 6.15E-05 | 4.98E-05 | 0 | 0 | 7.01E-05 | 5.49E-05 | 1.45E-05 | 4.69E-05 | 4.01E-05 | 6.08E-05 | 1.15E-05 | 4.01E-05 | 7.53E-06 | 0 | 0.00012 | 3.66E-05 | 5.24E-05 | 6.00E-05 |
| Camk2n2 | 0 | 0 | 3.74E-05 | 0.00011 | 5.51E-05 | 5.19E-05 | 4.57E-05 | 7.04E-05 | 4.01E-05 | 3.77E-06 | 3.99E-05 | 0 | 0 | 3.03E-05 | 1.72E-05 | 9.51E-05 | 3.70E-05 | 0 |
| Ublcp1 | 5.69E-05 | 6.23E-05 | 3.32E-05 | 3.06E-05 | 4.96E-05 | 3.66E-05 | 0 | 3.61E-05 | 4.01E-05 | 1.96E-05 | 2.68E-05 | 0 | 2.30E-05 | 4.69E-05 | 1.47E-05 | 6.96E-05 | 0 | 3.94E-05 |
| Slc43a3 | 6.88E-05 | 8.79E-05 | 0 | 0 | 1.84E-05 | 3.12E-05 | 1.16E-05 | 0.00019 | 4.01E-05 | 0.00059 | 2.63E-06 | 0 | 0.00048 | 0 | 0 | 0.00056 | 0 | 0.00044 |
| Cep83 | 6.49E-05 | 3.26E-05 | 7.21E-05 | 0 | 9.23E-06 | 8.62E-06 | 0 | 1.80E-05 | 4.01E-05 | 3.54E-06 | 2.89E-05 | 1.38E-05 | 0 | 5.36E-06 | 0.00013 | 0.00014 | 7.57E-05 | 2.80E-05 |
| Exosc8 | 0.00012 | 1.76E-05 | 0.00016 | 1.68E-05 | 0 | 2.04E-06 | 6.83E-06 | 0 | 4.01E-05 | 2.70E-05 | 2.19E-05 | 2.88E-05 | 1.34E-05 | 0 | 0.00018 | 0.00055 | 0 | 3.51E-06 |
| Tbc1d16 | 0 | 9.72E-05 | 0 | 3.36E-05 | 0.0001 | 0 | 1.98E-05 | 0 | 4.01E-05 | 1.75E-05 | 7.18E-05 | 1.90E-05 | 5.27E-05 | 0.00012 | 5.29E-05 | 0 | 4.48E-05 | 5.09E-05 |
| Atg2b | 0.00015 | 0.00011 | 0.00013 | 0 | 9.27E-05 | 7.16E-05 | 8.37E-05 | 4.28E-06 | 4.00E-05 | 1.62E-05 | 0 | 0.00019 | 5.96E-06 | 6.15E-05 | 0.00011 | 2.95E-05 | 0.00011 | 0 |
| Fam45a | 0.00017 | 3.50E-05 | 9.34E-05 | 4.30E-05 | 3.78E-05 | 6.90E-05 | 0.0001 | 2.06E-06 | 4.00E-05 | 5.42E-05 | 3.27E-05 | 4.53E-05 | 9.79E-08 | 5.15E-06 | 5.29E-05 | 9.13E-05 | 0 | 2.70E-05 |
| Jmjd1c | 7.06E-05 | 0.00027 | 2.69E-05 | 4.59E-06 | 3.59E-05 | 5.04E-05 | 2.68E-05 | 4.08E-05 | 4.00E-05 | 0.00013 | 7.65E-07 | 0.00017 | 0 | 0.00013 | 0.0002 | 9.54E-05 | 0.00037 | 0.00015 |
| Ogg1 | 2.23E-05 | 4.95E-05 | 3.68E-05 | 4.32E-05 | 4.69E-06 | 0 | 0 | 1.33E-05 | 4.00E-05 | 3.22E-05 | 0 | 9.39E-06 | 2.43E-06 | 0 | 1.69E-05 | 7.69E-05 | 0 | 8.91E-06 |
| Dram2 | 0.00019 | 0.00013 | 0 | 0 | 6.66E-05 | 0.00029 | 0 | 8.06E-05 | 3.99E-05 | 0 | 4.66E-05 | 0.00016 | 0 | 2.54E-05 | 0 | 0 | 0 | 1.64E-05 |
| Rabep1 | 8.59E-05 | 0.00019 | 0.00013 | 7.34E-05 | 9.07E-05 | 0.00013 | 4.61E-05 | 0.00014 | 3.99E-05 | 0.0001 | 5.79E-05 | 6.64E-05 | 9.97E-05 | 0.00011 | 0.00016 | 5.70E-05 | 6.89E-05 | 9.77E-05 |
| Pik3c2a | 0.00015 | 5.85E-05 | 8.84E-06 | 0 | 3.61E-05 | 8.71E-05 | 2.29E-05 | 2.27E-05 | 3.99E-05 | 0 | 8.50E-05 | 6.65E-05 | 5.80E-05 | 4.16E-05 | 3.26E-05 | 4.58E-05 | 0.0001 | 3.45E-05 |
| Fh1 | 0.00021 | 6.53E-05 | 0.00016 | 6.73E-05 | 1.43E-05 | 6.36E-05 | 0 | 1.66E-05 | 3.99E-05 | 1.81E-05 | 2.29E-05 | 0 | 1.07E-05 | 2.54E-05 | 5.07E-05 | 0.00017 | 5.15E-05 | 5.24E-06 |
| Hdhd5 | 1.64E-05 | 4.78E-05 | 1.60E-05 | 4.05E-05 | 2.82E-05 | 1.68E-05 | 1.50E-05 | 2.97E-05 | 3.99E-05 | 3.60E-05 | 1.37E-05 | 0 | 1.28E-06 | 3.54E-05 | 6.51E-05 | 1.92E-05 | 0 | 2.21E-05 |
| Tars2 | 2.98E-05 | 2.39E-05 | 6.09E-05 | 2.29E-05 | 4.64E-06 | 1.55E-05 | 0 | 8.95E-07 | 3.99E-05 | 1.76E-05 | 1.59E-05 | 0 | 0 | 1.08E-05 | 5.00E-05 | 7.42E-05 | 1.13E-05 | 8.19E-06 |
| Impg2 | 0 | 0 | 0 | 0 | 2.91E-05 | 7.12E-06 | 0 | 0 | 3.99E-05 | 3.48E-07 | 0 | 0 | 0 | 3.08E-05 | 0 | 5.95E-06 | 0 | 1.45E-05 |
| Ccdc61 | 1.20E-06 | 5.61E-05 | 0 | 4.19E-05 | 1.34E-05 | 5.62E-06 | 0 | 3.35E-05 | 3.99E-05 | 4.30E-05 | 9.64E-06 | 3.47E-05 | 0 | 2.13E-05 | 0.0001 | 0.00011 | 0 | 2.23E-05 |
| Atp6v0b | 0.00015 | 0.00034 | 4.55E-05 | 7.04E-05 | 0.00025 | 0.0002 | 0.00016 | 4.03E-05 | 3.98E-05 | 9.76E-05 | 0.00012 | 0.00028 | 7.46E-05 | 7.91E-05 | 0 | 3.91E-05 | 0.00016 | 0.0001 |
| Dpysl3 | 0.00016 | 1.40E-05 | 0.00021 | 0.00013 | 0.00021 | 0 | 0.00072 | 0.00018 | 3.98E-05 | 0.00053 | 0.0004 | 2.93E-05 | 0 | 0 | 0 | 0.00017 | 2.84E-05 | 0.00111 |
| Tada2a | 8.54E-05 | 4.17E-05 | 4.75E-05 | 1.90E-05 | 1.89E-05 | 0 | 4.56E-05 | 0 | 3.98E-05 | 1.37E-05 | 7.55E-06 | 0 | 2.42E-05 | 2.32E-05 | 0.00016 | 6.69E-05 | 0 | 1.46E-05 |
| Tmem43 | 6.76E-05 | 0.00023 | 0.0002 | 0.00023 | 8.13E-05 | 9.54E-05 | 0.00028 | 7.88E-05 | 3.97E-05 | 0.00054 | 0.00012 | 9.88E-05 | 0 | 0.00011 | 0 | 0.00037 | 9.18E-05 | 0.00039 |
| Trip4 | 0.00014 | 7.73E-05 | 4.20E-05 | 1.72E-05 | 4.89E-05 | 3.84E-05 | 5.85E-05 | 5.35E-06 | 3.97E-05 | 4.64E-05 | 7.16E-06 | 2.76E-05 | 3.02E-05 | 1.59E-05 | 0.0002 | 4.20E-05 | 0 | 4.32E-05 |
| Trub2 | 7.38E-05 | 6.89E-05 | 8.38E-05 | 7.19E-05 | 3.79E-05 | 3.57E-05 | 3.74E-06 | 1.29E-07 | 3.97E-05 | 1.50E-05 | 0 | 0 | 6.34E-05 | 2.49E-05 | 9.60E-05 | 3.30E-05 | 0 | 3.25E-05 |
| Ice2 | 5.76E-05 | 3.54E-05 | 1.96E-05 | 0 | 1.89E-05 | 2.06E-05 | 1.43E-05 | 0 | 3.97E-05 | 2.37E-05 | 0 | 1.90E-06 | 4.21E-06 | 1.42E-05 | 4.44E-05 | 6.75E-06 | 0 | 6.39E-06 |

| | | | | | | | | | | | | | | | | | | |
|---|---|---|---|---|---|---|---|---|---|---|---|---|---|---|---|---|---|---|
| Atpaf1 | 4.58E-05 | 2.06E-05 | 4.81E-06 | 3.53E-06 | 1.58E-05 | 5.42E-06 | 0 | 5.51E-06 | 3.97E-05 | 1.41E-05 | 1.08E-05 | 0 | 0 | 2.78E-05 | 1.32E-05 | 2.52E-05 | 0 | 1.15E-05 |
| Thtpa | 3.27E-05 | 4.61E-05 | 8.75E-06 | 3.18E-05 | 3.14E-05 | 0 | 5.16E-05 | 1.43E-05 | 3.97E-05 | 2.56E-05 | 2.23E-05 | 0 | 2.26E-05 | 1.93E-05 | 3.72E-06 | 3.56E-06 | 0 | 4.14E-05 |
| Spop | 0.00033 | 0.00028 | 0.0002 | 9.96E-05 | 0.00027 | 0.00048 | 0.00043 | 0.00011 | 3.96E-05 | 0.00012 | 0.00038 | 0.00027 | 0.00034 | 0.0003 | 0 | 0.00016 | 1.84E-05 | 0.00011 |
| Pip4p1 | 0.00016 | 0.00028 | 1.08E-05 | 2.50E-05 | 0.00017 | 0.00016 | 9.24E-05 | 0 | 3.96E-05 | 4.78E-05 | 9.56E-05 | 0.00016 | 3.71E-05 | 8.70E-05 | 6.64E-05 | 9.41E-05 | 0.00011 | 7.83E-05 |
| Cd151 | 0.00011 | 0.00015 | 0.0002 | 7.42E-05 | 0 | 0.00015 | 0 | 0.00021 | 3.96E-05 | 0.00028 | 0.00016 | 0.00019 | 0 | 0 | 0.00011 | 3.98E-05 | 0 | 0.00038 |
| Jak3 | 0 | 5.58E-05 | 0 | 6.88E-05 | 9.81E-05 | 8.31E-05 | 0 | 0.00024 | 3.96E-05 | 0 | 0 | 0.00026 | 0 | 3.79E-05 | 0.00013 | 0 | 0 | 9.31E-05 |
| Thoc1 | 0.00013 | 8.22E-05 | 7.95E-05 | 0 | 0 | 3.50E-05 | 0 | 0 | 3.96E-05 | 8.12E-06 | 1.21E-05 | 3.59E-05 | 0 | 0 | 0.00016 | 6.72E-05 | 2.72E-05 | 0 |
| Zfp84 | 6.32E-05 | 7.10E-05 | 3.97E-06 | 1.02E-05 | 5.45E-05 | 3.11E-05 | 2.76E-05 | 1.60E-05 | 3.96E-05 | 2.12E-05 | 2.11E-05 | 0 | 0 | 2.10E-05 | 0.00014 | 1.41E-05 | 0 | 2.88E-05 |
| Fbxl18 | 4.52E-06 | 4.06E-05 | 8.14E-06 | 5.20E-05 | 2.89E-05 | 4.47E-06 | 0 | 9.98E-06 | 3.96E-05 | 2.11E-05 | 9.71E-06 | 1.55E-05 | 6.58E-05 | 9.07E-06 | 4.27E-05 | 1.25E-05 | 0.00015 | 0 |
| Dtx2 | 2.14E-05 | 0.00012 | 5.84E-05 | 6.41E-05 | 2.02E-05 | 0 | 0 | 1.75E-05 | 3.96E-05 | 0.00011 | 2.19E-06 | 1.87E-05 | 8.79E-06 | 6.84E-05 | 1.30E-05 | 2.66E-05 | 0.0001 | 3.44E-06 |
| Socs4 | 0.00011 | 9.02E-05 | 6.85E-05 | 0 | 4.12E-05 | 8.14E-05 | 1.50E-05 | 9.38E-06 | 3.95E-05 | 2.91E-05 | 2.87E-05 | 4.81E-05 | 9.37E-06 | 4.55E-05 | 3.51E-05 | 9.83E-05 | 0 | 2.09E-05 |
| Zfp277 | 9.26E-05 | 4.05E-05 | 0 | 0 | 0 | 4.28E-05 | 9.49E-06 | 8.76E-07 | 3.95E-05 | 1.40E-05 | 2.76E-06 | 2.26E-05 | 1.72E-05 | 0 | 1.27E-06 | 3.61E-05 | 7.43E-06 | 1.03E-05 |
| Ppid | 0.00012 | 2.30E-05 | 7.79E-05 | 0 | 8.13E-06 | 3.13E-05 | 0 | 0 | 3.95E-05 | 0 | 0 | 1.18E-05 | 0 | 0 | 4.60E-05 | 3.76E-05 | 0 | 1.05E-05 |
| Btbd9 | 5.63E-05 | 8.06E-05 | 2.45E-05 | 0 | 2.57E-05 | 2.03E-05 | 0 | 3.56E-05 | 3.95E-05 | 3.38E-05 | 4.69E-05 | 6.77E-05 | 0 | 4.91E-05 | 0.00011 | 0 | 7.17E-05 | 3.76E-05 |
| Klhl36 | 1.58E-06 | 3.68E-05 | 2.83E-06 | 3.69E-05 | 3.19E-05 | 1.62E-05 | 2.92E-06 | 4.51E-05 | 3.95E-05 | 5.51E-05 | 0 | 5.84E-05 | 0 | 2.80E-06 | 1.37E-05 | 3.37E-06 | 0 | 3.76E-05 |
| Gars | 0.00022 | 6.70E-06 | 0.00031 | 6.91E-05 | 0 | 4.91E-07 | 0.00025 | 0 | 3.95E-05 | 5.71E-05 | 7.76E-05 | 8.54E-05 | 0 | 0 | 0 | 8.66E-05 | 0 | 2.04E-05 |
| Mrtfa | 0 | 7.43E-05 | 0.00018 | 0.00011 | 9.25E-05 | 0 | 7.36E-05 | 0.00014 | 3.95E-05 | 8.58E-05 | 0.00025 | 5.93E-05 | 0 | 8.52E-05 | 0.00011 | 7.37E-05 | 0.00023 | 0.0002 |
| Gcsh | 0.0002 | 4.51E-05 | 0.00027 | 0.00016 | 4.95E-05 | 0 | 0.0003 | 0 | 3.95E-05 | 8.20E-05 | 7.40E-05 | 0 | 0.00012 | 0 | 0 | 0.00015 | 0 | 0.00011 |
| Fancf | 0 | 3.48E-05 | 5.05E-05 | 4.52E-05 | 1.91E-05 | 0 | 3.39E-06 | 2.83E-05 | 3.95E-05 | 2.44E-05 | 0 | 5.85E-05 | 0 | 4.45E-05 | 7.41E-05 | 0.00011 | 0 | 3.12E-05 |
| Atrip | 2.43E-05 | 3.54E-05 | 4.84E-05 | 0 | 2.96E-05 | 0 | 0 | 0 | 3.95E-05 | 1.46E-05 | 6.54E-06 | 0 | 1.87E-05 | 1.40E-05 | 0.0001 | 6.18E-05 | 0 | 2.06E-05 |
| Snx27 | 0.00013 | 0.0001 | 7.47E-05 | 0 | 6.53E-05 | 3.97E-05 | 0 | 9.78E-06 | 3.94E-05 | 7.18E-05 | 1.81E-05 | 8.43E-05 | 6.33E-06 | 0 | 0.00011 | 0.00016 | 5.46E-05 | 1.27E-05 |
| Mfsd12 | 0 | 7.19E-05 | 2.09E-05 | 2.51E-05 | 6.17E-05 | 1.28E-05 | 9.78E-06 | 0 | 3.94E-05 | 6.13E-05 | 2.92E-05 | 4.16E-05 | 3.76E-05 | 1.55E-05 | 4.09E-05 | 4.55E-05 | 1.33E-05 | 6.01E-05 |
| Exo5 | 6.31E-05 | 5.04E-05 | 3.66E-05 | 2.35E-05 | 1.69E-05 | 7.91E-06 | 1.46E-05 | 0 | 3.94E-05 | 2.67E-05 | 3.89E-05 | 2.41E-05 | 0 | 1.79E-05 | 0 | 2.36E-05 | 0 | 2.20E-05 |
| Plekho1 | 0 | 0.00018 | 4.57E-05 | 0.00012 | 0.00011 | 0 | 0.00013 | 0 | 3.94E-05 | 2.98E-05 | 0.00038 | 8.07E-05 | 0.00016 | 0.00011 | 0 | 9.74E-05 | 1.51E-05 | 0 |
| Mon1b | 3.35E-05 | 6.70E-05 | 2.99E-05 | 0.00011 | 1.37E-05 | 0 | 6.17E-05 | 4.22E-05 | 3.94E-05 | 4.25E-05 | 3.17E-05 | 1.22E-05 | 7.32E-06 | 4.29E-05 | 1.76E-05 | 2.41E-05 | 0 | 5.18E-05 |
| Cgnl1 | 0 | 6.75E-05 | 2.36E-06 | 4.35E-06 | 0 | 0 | 0 | 0 | 3.94E-05 | 6.80E-05 | 8.52E-05 | 0 | 0.00012 | 0 | 1.41E-05 | 1.49E-05 | 1.36E-05 | 0.0002 |
| Kdm6a | 0.00014 | 0.00014 | 2.36E-05 | 0 | 5.37E-06 | 9.13E-05 | 8.48E-06 | 2.06E-05 | 3.93E-05 | 2.59E-05 | 6.29E-06 | 5.83E-05 | 1.37E-05 | 4.56E-05 | 7.49E-05 | 3.44E-05 | 0.0001 | 4.57E-05 |
| Ppp6c | 0.00012 | 0.00013 | 0.00011 | 0.00017 | 4.64E-05 | 6.86E-05 | 2.34E-05 | 2.80E-05 | 3.93E-05 | 4.11E-05 | 6.39E-05 | 9.48E-05 | 0 | 5.13E-05 | 5.55E-05 | 0.00013 | 1.47E-05 | 6.66E-05 |
| Tut4 | 0.0001 | 0.00013 | 0 | 5.01E-05 | 4.95E-06 | 4.69E-05 | 1.34E-05 | 1.75E-05 | 3.93E-05 | 2.60E-05 | 0 | 8.23E-05 | 0 | 0 | 0.00014 | 0.00012 | 7.04E-05 | 3.66E-06 |
| Mitd1 | 6.70E-05 | 2.37E-05 | 1.97E-05 | 1.63E-05 | 1.24E-05 | 7.81E-06 | 0 | 0 | 3.93E-05 | 5.40E-05 | 1.27E-05 | 0 | 3.45E-05 | 2.35E-05 | 2.33E-05 | 2.73E-05 | 0 | 4.45E-05 |
| Tmem158 | 0 | 3.02E-05 | 5.32E-05 | 3.32E-05 | 0 | 4.32E-06 | 0 | 0.0001 | 3.93E-05 | 0.00015 | 0 | 0 | 0 | 0.00022 | 2.84E-05 | 5.34E-05 | 0 | 7.51E-05 |
| Rbm43 | 7.36E-05 | 3.35E-05 | 3.43E-06 | 6.21E-06 | 4.85E-06 | 0 | 1.83E-05 | 0 | 3.93E-05 | 9.42E-05 | 0 | 0 | 0 | 0 | 3.54E-05 | 5.25E-05 | 0 | 1.33E-05 |
| Snx7 | 0.0002 | 0 | 0.00013 | 0 | 0.00012 | 0.00024 | 0.00016 | 0 | 3.92E-05 | 8.88E-05 | 0.00041 | 0 | 0.00015 | 0.00023 | 0 | 9.32E-05 | 0 | 6.82E-05 |

| | | | | | | | | | | | | | | | | | | |
|---|---|---|---|---|---|---|---|---|---|---|---|---|---|---|---|---|---|---|
| Dusp12 | 0.00012 | 5.00E-05 | 0.00013 | 5.80E-05 | 5.62E-05 | 6.52E-05 | 9.93E-05 | 0 | 3.92E-05 | 0 | 4.00E-05 | 0 | 0 | 2.67E-05 | 1.62E-05 | 2.27E-05 | 0 | 0 |
| Dalrd3 | 2.04E-05 | 6.47E-05 | 3.65E-06 | 3.81E-05 | 3.06E-05 | 0 | 0 | 2.19E-05 | 3.92E-05 | 3.67E-05 | 1.33E-06 | 1.11E-05 | 9.34E-06 | 3.62E-05 | 8.19E-05 | 0 | 9.27E-05 | 4.06E-05 |
| Kifc3 | 0 | 5.38E-05 | 0.00013 | 6.39E-05 | 8.83E-05 | 0.00019 | 0 | 8.11E-05 | 3.91E-05 | 0 | 0.00017 | 0 | 2.67E-05 | 0.00028 | 0.00011 | 0 | 1.30E-05 | 2.86E-05 |
| Fam92a | 0.00018 | 5.49E-05 | 3.75E-05 | 0 | 4.28E-05 | 9.90E-05 | 3.82E-05 | 1.03E-05 | 3.91E-05 | 1.11E-05 | 3.36E-05 | 9.96E-06 | 1.60E-05 | 0 | 2.21E-05 | 0.00014 | 0 | 6.63E-05 |
| Ubn2 | 8.89E-05 | 0.00012 | 9.60E-05 | 3.99E-05 | 6.22E-05 | 5.66E-05 | 8.51E-05 | 1.19E-05 | 3.91E-05 | 2.68E-05 | 2.26E-05 | 0.00012 | 2.01E-05 | 0.00011 | 0.00021 | 0 | 0 | 2.97E-05 |
| Lage3 | 9.05E-05 | 3.56E-05 | 4.19E-05 | 3.77E-06 | 2.08E-05 | 3.07E-05 | 1.78E-06 | 0 | 3.91E-05 | 1.62E-05 | 0 | 8.46E-06 | 3.81E-07 | 0 | 0 | 4.49E-05 | 0 | 2.92E-05 |
| Tmem38a | 1.03E-06 | 2.65E-05 | 0 | 2.71E-05 | 1.44E-05 | 3.59E-06 | 1.43E-05 | 0 | 3.91E-05 | 3.27E-05 | 2.30E-05 | 4.78E-05 | 2.17E-06 | 1.72E-05 | 0 | 3.46E-05 | 1.26E-05 | 2.97E-05 |
| Oma1 | 2.64E-05 | 1.19E-05 | 5.20E-06 | 2.24E-05 | 1.95E-05 | 0 | 2.12E-06 | 2.57E-05 | 3.91E-05 | 1.33E-05 | 1.36E-05 | 1.05E-05 | 0 | 2.18E-05 | 1.77E-05 | 2.81E-05 | 0 | 1.93E-05 |
| Ldhb | 1.60E-05 | 1.95E-05 | 0 | 1.20E-05 | 0 | 6.16E-05 | 0 | 0 | 3.90E-05 | 2.24E-06 | 0 | 0 | 4.82E-05 | 9.07E-05 | 0 | 1.82E-05 | 2.33E-05 | 1.16E-06 |
| Cmc1 | 0.00018 | 0.00014 | 1.53E-05 | 0.00014 | 4.05E-05 | 0.00011 | 7.67E-05 | 6.49E-05 | 3.89E-05 | 5.27E-05 | 2.85E-05 | 0.00015 | 1.91E-05 | 2.66E-05 | 0 | 0.0001 | 4.26E-05 | 4.81E-05 |
| Mst1r | 1.35E-05 | 2.11E-05 | 1.07E-05 | 0 | 4.99E-05 | 6.64E-05 | 1.36E-06 | 1.97E-05 | 3.89E-05 | 0.00011 | 4.52E-06 | 3.33E-05 | 5.22E-05 | 1.31E-05 | 7.20E-05 | 0 | 0 | 3.90E-05 |
| Zfp62 | 5.91E-05 | 7.13E-05 | 2.14E-05 | 2.19E-05 | 6.49E-05 | 5.18E-05 | 4.90E-05 | 4.73E-05 | 3.89E-05 | 2.74E-05 | 7.81E-06 | 4.25E-05 | 0 | 3.62E-05 | 7.43E-05 | 3.71E-05 | 0 | 2.42E-05 |
| Bcas3 | 2.91E-05 | 6.93E-05 | 0 | 0 | 2.92E-05 | 3.18E-05 | 1.38E-05 | 2.82E-05 | 3.89E-05 | 1.25E-05 | 0 | 0 | 7.70E-06 | 2.85E-05 | 2.75E-05 | 4.73E-06 | 0 | 3.05E-05 |
| Hars2 | 5.07E-05 | 9.97E-05 | 2.80E-05 | 4.55E-05 | 1.70E-05 | 2.79E-05 | 2.16E-05 | 1.48E-05 | 3.89E-05 | 3.98E-05 | 5.18E-05 | 3.25E-05 | 1.21E-05 | 2.91E-05 | 6.61E-05 | 3.73E-05 | 5.30E-06 | 5.22E-05 |
| Ptgs1 | 0 | 4.10E-05 | 4.97E-05 | 0 | 6.23E-05 | 1.52E-05 | 0 | 7.80E-06 | 3.89E-05 | 0.00039 | 0 | 0 | 0 | 0.00015 | 0 | 1.28E-05 | 0 | 5.17E-05 |
| Naa10 | 8.49E-05 | 2.09E-05 | 8.20E-05 | 0.00012 | 0 | 0 | 0 | 1.11E-05 | 3.89E-05 | 0 | 0 | 2.86E-05 | 0 | 0 | 0 | 8.93E-05 | 3.78E-05 | 1.01E-05 |
| Jmjd6 | 0.00017 | 7.62E-05 | 0.00016 | 1.02E-05 | 3.26E-05 | 0 | 0 | 2.02E-05 | 3.89E-05 | 1.66E-05 | 5.03E-05 | 0.0005 | 0 | 2.17E-05 | 0.00034 | 7.90E-05 | 0.00039 | 9.55E-06 |
| Abcb8 | 1.24E-05 | 1.39E-05 | 9.39E-07 | 2.86E-05 | 1.70E-05 | 0 | 1.49E-05 | 4.26E-05 | 3.89E-05 | 2.27E-05 | 3.34E-06 | 5.90E-06 | 1.71E-05 | 1.07E-05 | 1.00E-05 | 0 | 6.88E-07 | 1.97E-05 |
| Chsy3 | 6.48E-05 | 0 | 0.00013 | 0 | 0 | 0.00021 | 5.65E-05 | 0.00018 | 3.88E-05 | 0 | 0.00013 | 0 | 0 | 1.89E-05 | 3.29E-06 | 1.55E-05 | 7.85E-05 | 0 |
| Cdadc1 | 9.14E-05 | 0.00011 | 4.69E-06 | 0 | 7.43E-05 | 0.00018 | 2.04E-05 | 0 | 3.88E-05 | 4.28E-05 | 1.37E-05 | 2.50E-05 | 2.14E-05 | 6.23E-05 | 0.0001 | 5.72E-05 | 0 | 4.69E-05 |
| Fam114a2 | 0.00026 | 0.00012 | 9.17E-05 | 8.15E-05 | 6.33E-05 | 0.00015 | 0.00017 | 0 | 3.88E-05 | 0 | 2.51E-05 | 0.00028 | 0.00011 | 7.38E-05 | 5.90E-05 | 0 | 0 | 0 |
| Slu7 | 0.00011 | 0.0001 | 2.78E-05 | 7.41E-05 | 6.97E-05 | 8.13E-05 | 4.25E-05 | 2.56E-05 | 3.88E-05 | 5.55E-05 | 2.96E-05 | 5.10E-05 | 0 | 2.76E-05 | 8.03E-06 | 4.55E-05 | 0 | 7.66E-05 |
| Pias1 | 9.28E-05 | 0.00011 | 0 | 0 | 3.36E-05 | 4.69E-05 | 1.20E-05 | 0 | 3.88E-05 | 4.14E-05 | 2.69E-05 | 9.81E-05 | 1.17E-05 | 9.98E-06 | 4.74E-05 | 7.82E-05 | 8.14E-05 | 4.56E-05 |
| Tet3 | 1.35E-05 | 0.00015 | 7.50E-05 | 2.37E-05 | 0.00015 | 4.61E-05 | 6.16E-06 | 0.00022 | 3.88E-05 | 5.29E-05 | 0.00019 | 0.00015 | 0 | 7.50E-05 | 0.00028 | 0 | 0.00013 | 0 |
| Kctd6 | 5.12E-05 | 6.18E-05 | 2.67E-05 | 0 | 5.62E-05 | 1.72E-05 | 1.38E-05 | 0 | 3.88E-05 | 3.37E-05 | 2.14E-05 | 5.34E-05 | 2.79E-05 | 3.35E-05 | 2.92E-05 | 2.47E-05 | 0 | 4.02E-05 |
| Hscb | 3.45E-05 | 5.42E-05 | 0 | 0 | 3.16E-05 | 6.92E-06 | 1.24E-05 | 0 | 3.88E-05 | 0 | 5.26E-06 | 5.10E-05 | 0 | 0 | 0 | 3.17E-05 | 0 | 3.32E-05 |
| Dnmt3l | 3.03E-05 | 0 | 7.95E-05 | 5.02E-05 | 0 | 0 | 0 | 0 | 3.88E-05 | 1.93E-05 | 0 | 3.91E-05 | 6.64E-05 | 0 | 2.52E-05 | 1.72E-05 | 0 | 0 |
| Nudt17 | 0 | 0 | 0 | 1.13E-05 | 3.85E-05 | 2.35E-05 | 0 | 1.58E-05 | 3.87E-05 | 0 | 9.82E-06 | 0 | 0 | 0 | 0 | 0 | 0 | 0 |
| Acbd4 | 6.09E-06 | 5.36E-05 | 0 | 3.10E-05 | 4.39E-05 | 1.58E-05 | 1.05E-05 | 0 | 3.87E-05 | 3.11E-05 | 1.67E-05 | 1.68E-05 | 4.80E-06 | 4.32E-05 | 9.15E-05 | 0 | 0 | 2.28E-05 |
| Cdc42se2 | 5.82E-05 | 7.63E-05 | 9.81E-05 | 2.00E-05 | 3.33E-05 | 7.06E-06 | 0 | 0 | 3.87E-05 | 6.44E-05 | 5.44E-05 | 1.53E-05 | 1.12E-05 | 1.25E-05 | 0 | 6.83E-05 | 0 | 4.21E-05 |
| Cdc42bpg | 0 | 0.00011 | 0 | 5.37E-05 | 7.52E-05 | 0 | 0 | 2.65E-06 | 3.87E-05 | 0 | 1.91E-05 | 7.89E-06 | 1.49E-05 | 3.95E-05 | 8.70E-05 | 0 | 1.13E-05 | 6.80E-05 |
| Mier3 | 5.22E-05 | 6.81E-05 | 5.92E-05 | 3.74E-06 | 2.49E-05 | 3.84E-05 | 0 | 6.91E-06 | 3.86E-05 | 4.13E-05 | 2.99E-05 | 2.26E-05 | 4.28E-05 | 2.43E-05 | 0.00012 | 3.46E-05 | 0.00016 | 7.71E-06 |
| Bcs1l | 8.53E-05 | 2.95E-05 | 1.65E-05 | 4.76E-05 | 1.01E-05 | 2.76E-05 | 3.68E-05 | 8.10E-06 | 3.86E-05 | 1.97E-05 | 0 | 1.71E-05 | 2.66E-05 | 0 | 0 | 5.86E-05 | 0 | 2.97E-05 |

| | | | | | | | | | | | | | | | | | | |
|---|---|---|---|---|---|---|---|---|---|---|---|---|---|---|---|---|---|---|
| Vps33b | 5.95E-05 | 6.79E-05 | 5.76E-05 | 4.19E-05 | 1.19E-05 | 2.28E-05 | 0 | 7.40E-06 | 3.86E-05 | 4.52E-05 | 3.57E-05 | 3.74E-06 | 0 | 2.53E-05 | 8.34E-05 | 0 | 0 | 1.47E-05 |
| Bicdl1 | 0 | 2.67E-05 | 6.29E-06 | 1.09E-05 | 2.45E-05 | 5.03E-06 | 0 | 7.74E-06 | 3.86E-05 | 3.32E-05 | 1.31E-05 | 5.59E-06 | 5.42E-06 | 0 | 4.13E-06 | 0 | 0 | 4.40E-05 |
| Setd6 | 1.75E-05 | 3.78E-05 | 6.42E-05 | 0 | 1.09E-05 | 1.14E-07 | 0 | 0 | 3.86E-05 | 5.45E-05 | 1.81E-05 | 0 | 2.99E-05 | 8.27E-06 | 5.98E-05 | 3.94E-05 | 1.67E-05 | 2.66E-05 |
| Cep250 | 0 | 5.01E-05 | 0 | 5.57E-05 | 1.88E-05 | 0 | 0 | 0 | 3.86E-05 | 3.09E-05 | 1.46E-05 | 0 | 2.94E-06 | 4.25E-05 | 0.00018 | 0.00024 | 4.45E-05 | 4.70E-05 |
| Yme1l1 | 0.00043 | 0.00011 | 0.00015 | 0 | 4.99E-05 | 0.0002 | 0.00013 | 5.07E-05 | 3.85E-05 | 6.26E-05 | 3.13E-05 | 0.00018 | 1.96E-05 | 1.01E-05 | 0.00034 | 0.0001 | 0.00014 | 3.76E-05 |
| Tsc22d1 | 0.00028 | 0.00049 | 0.00013 | 0.00031 | 0.00047 | 0.00013 | 0.00038 | 0.0003 | 3.85E-05 | 0 | 0.0003 | 0.00017 | 0 | 0.00048 | 0.00023 | 8.01E-05 | 0.00119 | 0.00035 |
| Gpr108 | 0.00015 | 0.00012 | 7.81E-05 | 6.40E-05 | 0.00011 | 0.00013 | 0.00015 | 2.20E-05 | 3.85E-05 | 5.15E-05 | 5.85E-05 | 0.0001 | 6.49E-05 | 4.49E-05 | 0.00011 | 1.07E-05 | 0 | 8.81E-05 |
| Rxrb | 9.53E-05 | 0.00019 | 6.33E-05 | 0.00013 | 0.00014 | 7.07E-05 | 0.00014 | 6.47E-05 | 3.85E-05 | 4.67E-05 | 3.75E-05 | 0.00013 | 0 | 5.64E-05 | 0.0003 | 8.35E-05 | 0.00011 | 8.32E-05 |
| Ptcd3 | 0.00014 | 6.78E-05 | 0.00013 | 0 | 1.45E-05 | 6.98E-05 | 0 | 0 | 3.85E-05 | 1.66E-05 | 0 | 2.43E-05 | 0 | 1.39E-05 | 0.00013 | 0.0001 | 0 | 2.88E-06 |
| Mypop | 7.87E-05 | 9.54E-05 | 5.06E-05 | 0.00016 | 7.53E-05 | 5.75E-05 | 0.00011 | 0 | 3.85E-05 | 2.30E-05 | 9.86E-06 | 2.72E-05 | 0 | 1.78E-05 | 2.90E-05 | 4.58E-05 | 1.51E-05 | 8.10E-05 |
| Ints11 | 0.00012 | 8.45E-05 | 0.00013 | 2.01E-05 | 4.72E-05 | 3.73E-05 | 2.91E-06 | 1.44E-05 | 3.85E-05 | 4.00E-05 | 5.12E-05 | 5.59E-05 | 0 | 4.60E-05 | 3.60E-05 | 0.00014 | 2.39E-05 | 3.49E-05 |
| Josd2 | 8.38E-05 | 5.59E-05 | 7.21E-05 | 0.00013 | 2.81E-05 | 3.57E-05 | 9.18E-05 | 1.64E-05 | 3.85E-05 | 1.63E-05 | 2.26E-05 | 3.34E-05 | 1.31E-05 | 4.99E-06 | 0.00012 | 0 | 0 | 2.41E-05 |
| Arl6 | 5.63E-05 | 6.53E-05 | 2.12E-05 | 1.80E-05 | 1.29E-05 | 2.31E-05 | 0 | 0 | 3.85E-05 | 2.77E-05 | 0 | 1.52E-05 | 1.07E-05 | 1.02E-05 | 0 | 6.62E-05 | 0 | 2.83E-05 |
| Traf3ip3 | 0 | 3.45E-06 | 0 | 4.19E-06 | 4.18E-05 | 1.88E-05 | 0 | 3.10E-06 | 3.85E-05 | 0 | 0 | 0 | 0 | 0 | 3.93E-06 | 0 | 0 | 5.53E-06 |
| Zbtb26 | 2.55E-05 | 2.64E-05 | 1.08E-05 | 0 | 1.81E-05 | 1.23E-05 | 0 | 5.67E-06 | 3.85E-05 | 1.51E-06 | 1.20E-05 | 0 | 0 | 7.91E-06 | 3.17E-05 | 0 | 0 | 0 |
| Mtx3 | 1.39E-05 | 4.30E-05 | 4.95E-05 | 3.59E-05 | 4.32E-05 | 8.90E-06 | 1.09E-05 | 1.03E-05 | 3.85E-05 | 2.70E-05 | 2.76E-05 | 1.95E-05 | 4.21E-05 | 0 | 7.30E-05 | 1.17E-05 | 0 | 2.26E-05 |
| Snupn | 2.46E-05 | 2.54E-05 | 6.75E-06 | 4.85E-06 | 1.37E-05 | 2.54E-06 | 0 | 6.76E-06 | 3.85E-05 | 9.12E-06 | 6.20E-06 | 3.44E-05 | 1.76E-07 | 1.62E-05 | 5.61E-06 | 2.41E-05 | 0 | 3.22E-05 |
| Fgfbp3 | 4.66E-06 | 3.45E-05 | 0 | 5.15E-05 | 0 | 1.33E-06 | 0 | 4.78E-07 | 3.85E-05 | 1.81E-05 | 1.24E-05 | 1.33E-05 | 6.45E-06 | 0 | 1.21E-05 | 0 | 0 | 4.74E-05 |
| Fam131b | 0 | 0 | 5.05E-05 | 0 | 0 | 0 | 0 | 0 | 3.85E-05 | 5.30E-05 | 2.90E-05 | 3.29E-05 | 3.27E-05 | 6.83E-06 | 0 | 4.67E-05 | 0 | 0 |
| Mthfr | 5.14E-05 | 9.89E-05 | 6.70E-06 | 5.90E-05 | 7.30E-05 | 0 | 4.22E-05 | 1.08E-05 | 3.85E-05 | 3.35E-05 | 0 | 9.40E-05 | 3.26E-05 | 0 | 0.00024 | 0 | 0.00018 | 7.39E-05 |
| Ghitm | 0.00029 | 0.00013 | 0.00015 | 0 | 7.75E-05 | 0.00012 | 8.89E-05 | 2.58E-05 | 3.84E-05 | 3.97E-05 | 4.55E-05 | 0.00022 | 2.57E-05 | 2.94E-05 | 3.60E-05 | 7.32E-05 | 2.79E-06 | 6.69E-05 |
| Agtpbp1 | 0.00016 | 6.92E-05 | 4.05E-05 | 0 | 7.51E-05 | 0.0001 | 5.96E-05 | 3.49E-06 | 3.84E-05 | 2.01E-05 | 0 | 9.24E-05 | 0 | 0 | 0.00021 | 0.0001 | 4.54E-05 | 9.04E-05 |
| Smndc1 | 0.00017 | 0.00011 | 0.00013 | 0 | 3.62E-05 | 9.86E-05 | 0 | 3.44E-05 | 3.84E-05 | 0 | 6.82E-05 | 4.73E-05 | 0 | 5.89E-05 | 0 | 7.44E-05 | 0.00013 | 0 |
| Cln6 | 3.30E-05 | 7.56E-05 | 0 | 6.69E-05 | 5.33E-05 | 6.67E-05 | 4.89E-05 | 1.70E-05 | 3.84E-05 | 4.93E-05 | 5.35E-05 | 9.18E-05 | 4.84E-05 | 1.44E-05 | 0 | 0.00018 | 0 | 3.61E-05 |
| Sdhaf4 | 9.07E-05 | 2.82E-05 | 0 | 1.49E-05 | 3.50E-05 | 6.34E-05 | 6.99E-05 | 0 | 3.84E-05 | 4.74E-05 | 2.31E-05 | 1.35E-06 | 1.28E-05 | 4.56E-06 | 0 | 4.00E-05 | 0 | 3.38E-05 |
| Psmd14 | 0.00028 | 4.73E-05 | 0.00026 | 0.00012 | 0 | 5.67E-05 | 7.03E-05 | 0 | 3.84E-05 | 5.37E-05 | 3.63E-05 | 1.56E-05 | 1.25E-05 | 0 | 0 | 0.00022 | 5.15E-05 | 0 |
| Ppp1r15a | 6.53E-05 | 0.0002 | 3.21E-05 | 3.35E-05 | 9.02E-06 | 3.47E-05 | 6.09E-06 | 0 | 3.84E-05 | 4.43E-05 | 8.56E-06 | 0.00045 | 0 | 9.26E-05 | 6.04E-05 | 1.37E-05 | 0.00268 | 0.0001 |
| Zscan21 | 4.29E-05 | 7.75E-05 | 0 | 3.01E-05 | 1.23E-05 | 2.34E-05 | 0 | 2.71E-05 | 3.84E-05 | 3.64E-05 | 1.74E-05 | 1.34E-05 | 0 | 3.14E-05 | 2.38E-05 | 4.48E-05 | 0 | 2.88E-05 |
| Las1l | 0.00012 | 8.82E-05 | 0.00013 | 5.15E-05 | 1.33E-05 | 0 | 0 | 0 | 3.84E-05 | 7.19E-06 | 7.84E-05 | 5.73E-05 | 6.31E-06 | 0 | 0.00012 | 6.57E-05 | 2.16E-05 | 1.06E-05 |
| Vasn | 8.19E-05 | 0.00035 | 0.00062 | 0.00019 | 0.00055 | 0.0003 | 0.00052 | 0.00058 | 3.83E-05 | 0.0004 | 0.00061 | 0.00049 | 0.00108 | 0.00068 | 0 | 0.00014 | 0.00117 | 0.00016 |
| Dexi | 3.02E-05 | 8.24E-05 | 2.83E-05 | 0.00015 | 0.00011 | 0.00012 | 0.00014 | 0.00015 | 3.83E-05 | 0.00011 | 4.31E-05 | 4.76E-05 | 0.0001 | 0.00011 | 6.84E-05 | 7.87E-06 | 4.76E-05 | 0.00018 |
| Dpm1 | 0.00022 | 8.64E-05 | 0.00011 | 6.77E-05 | 3.17E-05 | 0.0001 | 6.22E-05 | 0 | 3.83E-05 | 3.83E-05 | 6.53E-05 | 6.40E-05 | 4.84E-05 | 2.32E-05 | 0.00013 | 0 | 3.12E-05 | 0 |
| Slc35a3 | 0.00012 | 4.05E-05 | 0.0001 | 0 | 5.25E-05 | 7.47E-05 | 8.97E-06 | 2.60E-05 | 3.83E-05 | 1.57E-05 | 2.92E-05 | 4.37E-05 | 0 | 2.44E-05 | 2.84E-05 | 0.00013 | 5.09E-05 | 5.11E-05 |

| | | | | | | | | | | | | | | | | | | |
|---|---|---|---|---|---|---|---|---|---|---|---|---|---|---|---|---|---|---|
| Pla2g2d | 0 | 0 | 3.22E-06 | 0 | 0 | 6.66E-05 | 0 | 2.57E-05 | 3.83E-05 | 5.19E-06 | 0 | 0 | 1.36E-05 | 0 | 0 | 1.11E-05 | 0 | 0 |
| Maoa | 0.00012 | 4.41E-05 | 5.18E-05 | 8.82E-06 | 0 | 6.00E-05 | 0 | 0 | 3.83E-05 | 3.43E-05 | 0 | 3.90E-05 | 3.26E-05 | 0 | 0 | 5.32E-05 | 1.39E-05 | 0 |
| Osbpl2 | 0.00015 | 8.23E-05 | 4.14E-05 | 3.49E-05 | 4.75E-05 | 6.00E-05 | 0.00018 | 0 | 3.83E-05 | 2.43E-05 | 2.33E-05 | 4.25E-05 | 0 | 2.84E-05 | 0.0001 | 3.91E-05 | 0 | 4.15E-05 |
| Herc1 | 8.90E-05 | 0.00015 | 7.14E-05 | 6.02E-05 | 4.65E-05 | 5.21E-05 | 2.91E-05 | 7.31E-06 | 3.83E-05 | 7.02E-05 | 3.08E-05 | 2.74E-05 | 5.78E-05 | 4.57E-05 | 0.0001 | 0.00013 | 0.00018 | 4.62E-05 |
| Vegfa | 0 | 4.68E-05 | 0.00014 | 0 | 0.00024 | 0.00032 | 0.00012 | 0 | 3.82E-05 | 0.00012 | 5.58E-05 | 0.00209 | 0.00021 | 0 | 0.00064 | 0 | 0.00108 | 0 |
| Pqlc3 | 0.0001 | 6.66E-05 | 0 | 0 | 0 | 9.60E-05 | 0 | 0 | 3.82E-05 | 4.99E-06 | 0 | 3.51E-05 | 1.88E-05 | 0 | 0 | 0 | 0 | 0.00017 |
| Arntl2 | 3.30E-05 | 6.38E-05 | 1.02E-06 | 0 | 2.52E-05 | 9.17E-05 | 0 | 3.26E-05 | 3.82E-05 | 9.78E-06 | 4.23E-05 | 2.84E-05 | 0 | 5.48E-05 | 0.00014 | 0 | 0 | 5.51E-06 |
| Sfswap | 5.28E-05 | 0.00017 | 0.00011 | 4.52E-05 | 5.23E-05 | 8.66E-05 | 2.81E-05 | 1.32E-05 | 3.82E-05 | 6.75E-05 | 0.00016 | 6.18E-05 | 5.45E-05 | 7.56E-05 | 0.00037 | 0.00016 | 6.70E-06 | 5.02E-05 |
| Acox3 | 3.76E-05 | 0.00013 | 0 | 0 | 6.93E-05 | 7.81E-05 | 1.87E-05 | 2.28E-06 | 3.82E-05 | 6.61E-05 | 5.06E-05 | 2.67E-05 | 4.79E-05 | 7.76E-05 | 6.67E-05 | 7.43E-06 | 0 | 4.17E-05 |
| Taf1d | 0.00017 | 9.85E-05 | 0.00022 | 9.74E-05 | 0 | 5.84E-05 | 0 | 0 | 3.82E-05 | 7.35E-06 | 1.67E-05 | 6.93E-05 | 3.68E-05 | 8.68E-05 | 0.00045 | 0.0002 | 0.00065 | 0 |
| Rab33b | 0.00011 | 7.59E-05 | 0.00013 | 0.00022 | 6.88E-05 | 5.36E-05 | 0.00015 | 0.00013 | 3.82E-05 | 8.45E-05 | 4.13E-05 | 8.28E-05 | 6.51E-05 | 0.0001 | 0.00012 | 0 | 2.40E-05 | 6.96E-05 |
| Il27ra | 0 | 3.94E-06 | 0 | 0 | 0 | 4.89E-05 | 0 | 7.40E-05 | 3.82E-05 | 3.03E-05 | 0 | 3.56E-05 | 0 | 0 | 0 | 0 | 0 | 8.28E-06 |
| Sepsecs | 2.94E-05 | 3.57E-05 | 3.71E-05 | 2.49E-05 | 1.04E-05 | 2.40E-05 | 0 | 1.01E-05 | 3.82E-05 | 3.56E-05 | 6.76E-06 | 1.73E-05 | 1.36E-05 | 0 | 3.94E-05 | 2.03E-05 | 7.96E-06 | 5.62E-06 |
| Tyk2 | 3.05E-05 | 3.63E-05 | 2.50E-06 | 1.59E-05 | 3.92E-05 | 1.96E-05 | 2.42E-05 | 2.41E-05 | 3.82E-05 | 3.66E-05 | 0 | 2.26E-05 | 0 | 2.74E-05 | 7.43E-05 | 0 | 0 | 4.97E-05 |
| Gria4 | 0 | 0 | 0 | 0 | 5.27E-05 | 1.24E-05 | 0 | 0 | 3.82E-05 | 1.30E-05 | 3.33E-07 | 6.45E-07 | 0 | 0 | 1.36E-05 | 1.12E-06 | 0 | 0.00013 |
| Dis3 | 7.09E-05 | 1.81E-05 | 0.0001 | 1.37E-05 | 0 | 6.35E-06 | 0 | 0 | 3.82E-05 | 2.68E-06 | 8.95E-06 | 0 | 2.90E-05 | 0 | 3.16E-05 | 0.00012 | 3.69E-05 | 0 |
| Zbtb33 | 9.93E-05 | 6.64E-05 | 4.74E-05 | 0 | 6.53E-05 | 9.04E-05 | 4.77E-05 | 0 | 3.81E-05 | 2.76E-05 | 4.94E-05 | 2.16E-05 | 2.99E-05 | 3.39E-05 | 0 | 3.16E-05 | 0 | 5.13E-05 |
| Capn7 | 0.00019 | 7.98E-05 | 7.71E-05 | 0 | 4.61E-05 | 6.77E-05 | 0.00013 | 1.53E-05 | 3.81E-05 | 5.76E-05 | 0 | 6.15E-05 | 4.91E-06 | 0 | 5.37E-05 | 4.26E-05 | 4.92E-05 | 2.49E-05 |
| Kif1bp | 0.0001 | 0.00011 | 3.99E-05 | 0.00015 | 5.76E-05 | 3.71E-05 | 4.24E-05 | 5.93E-06 | 3.81E-05 | 6.30E-05 | 1.45E-05 | 7.58E-05 | 2.81E-05 | 6.31E-05 | 8.05E-05 | 5.67E-05 | 0 | 9.02E-05 |
| Zfp606 | 2.57E-05 | 4.73E-05 | 0 | 1.27E-05 | 3.62E-05 | 2.54E-05 | 0 | 4.24E-06 | 3.81E-05 | 5.97E-06 | 0 | 0 | 0 | 2.03E-05 | 5.09E-05 | 6.45E-06 | 0 | 6.54E-06 |
| Eny2 | 9.40E-05 | 2.16E-05 | 7.49E-05 | 0.00011 | 1.70E-05 | 2.13E-05 | 3.82E-05 | 2.02E-05 | 3.81E-05 | 0 | 3.22E-05 | 3.32E-05 | 6.13E-06 | 8.77E-06 | 7.42E-05 | 9.22E-05 | 4.19E-05 | 1.61E-05 |
| Zbtb11 | 5.46E-05 | 0.00013 | 8.60E-05 | 7.54E-05 | 3.55E-05 | 1.62E-05 | 0 | 0 | 3.81E-05 | 1.65E-05 | 0 | 6.77E-06 | 1.45E-05 | 9.99E-06 | 7.60E-05 | 6.68E-05 | 0.00061 | 3.14E-05 |
| Rabl2 | 6.85E-05 | 3.31E-05 | 1.51E-05 | 3.53E-05 | 2.91E-05 | 4.68E-06 | 2.49E-05 | 3.57E-05 | 3.81E-05 | 2.21E-05 | 1.30E-05 | 1.91E-05 | 0 | 3.36E-05 | 0.00012 | 3.55E-05 | 0 | 1.84E-05 |
| Surf6 | 4.56E-05 | 4.77E-05 | 9.01E-05 | 8.46E-05 | 1.43E-05 | 1.64E-06 | 0 | 1.89E-05 | 3.81E-05 | 0 | 0 | 0 | 4.67E-06 | 2.82E-05 | 0.00013 | 3.46E-06 | 4.40E-05 | 5.99E-06 |
| Parp9 | 0.00016 | 7.10E-05 | 0 | 0 | 5.16E-05 | 0 | 5.15E-05 | 0 | 3.81E-05 | 0.00015 | 2.85E-05 | 0.0001 | 0.00013 | 2.36E-05 | 0 | 4.65E-05 | 0 | 0.00014 |
| Leng1 | 2.67E-05 | 5.13E-05 | 3.30E-05 | 7.44E-05 | 3.00E-05 | 0 | 4.60E-06 | 0 | 3.81E-05 | 3.13E-05 | 2.60E-05 | 3.12E-05 | 2.60E-05 | 0 | 0.00011 | 2.63E-05 | 5.06E-05 | 4.39E-05 |
| Fam3a | 7.35E-05 | 8.34E-05 | 0 | 2.53E-05 | 7.92E-05 | 9.48E-05 | 5.11E-05 | 0.00011 | 3.80E-05 | 8.64E-05 | 4.10E-05 | 8.35E-05 | 7.66E-05 | 3.57E-05 | 0 | 1.57E-05 | 0 | 0.00011 |
| Vamp4 | 0.00014 | 0.00011 | 3.43E-05 | 1.33E-05 | 6.64E-05 | 8.10E-05 | 7.19E-05 | 3.46E-05 | 3.80E-05 | 4.71E-05 | 4.38E-05 | 0.0001 | 2.12E-05 | 4.51E-05 | 0.00011 | 1.06E-05 | 0 | 3.46E-05 |
| Trp53 | 0.00019 | 0.00023 | 0.00025 | 0.00021 | 1.41E-05 | 9.82E-06 | 0 | 1.05E-05 | 3.80E-05 | 1.00E-05 | 5.26E-05 | 4.29E-05 | 0.00018 | 0.00014 | 0.0002 | 0.00034 | 7.78E-05 | 0.00012 |
| Sp2 | 6.84E-05 | 5.53E-05 | 4.45E-05 | 8.50E-06 | 2.21E-05 | 5.93E-06 | 1.83E-05 | 2.64E-05 | 3.80E-05 | 5.76E-05 | 5.75E-05 | 3.21E-06 | 0 | 0 | 2.37E-05 | 0.00012 | 4.93E-05 | 3.46E-05 |
| Maff | 0 | 0.00011 | 4.60E-05 | 4.20E-05 | 3.25E-05 | 0 | 0 | 0 | 3.80E-05 | 3.94E-05 | 0.00024 | 0.00064 | 6.78E-06 | 0.00017 | 0.00014 | 0 | 0.00054 | 9.17E-05 |
| Podnl1 | 0 | 0 | 0 | 3.91E-05 | 0.0008 | 0 | 0 | 0.00039 | 3.80E-05 | 0 | 5.27E-05 | 0 | 0 | 0.00064 | 0.00013 | 0 | 0 | 0 |
| Abca9 | 0 | 2.82E-05 | 0 | 7.38E-06 | 0 | 0 | 7.05E-05 | 0.00015 | 3.80E-05 | 0.00015 | 0 | 0 | 2.77E-05 | 0 | 0 | 7.39E-05 | 0 | 0.00023 |

| | | | | | | | | | | | | | | | | | | |
|---|---|---|---|---|---|---|---|---|---|---|---|---|---|---|---|---|---|---|
| Nsmce4a | 0.00016 | 0.00011 | 6.84E-05 | 0 | 0.00012 | 0.00014 | 3.52E-05 | 0 | 3.79E-05 | 6.28E-05 | 2.26E-05 | 0 | 1.55E-05 | 7.80E-05 | 0 | 0.00053 | 1.30E-05 | 1.96E-05 |
| Osgep | 0.00019 | 9.47E-05 | 1.23E-05 | 0 | 0.00012 | 0.00012 | 0.00011 | 2.45E-05 | 3.79E-05 | 1.76E-05 | 4.44E-05 | 0 | 0 | 3.37E-05 | 2.38E-05 | 0.00013 | 9.18E-06 | 8.23E-05 |
| Qsox2 | 8.25E-05 | 2.99E-05 | 0.0001 | 4.67E-06 | 5.08E-05 | 8.46E-05 | 0 | 2.44E-05 | 3.79E-05 | 3.76E-05 | 7.41E-05 | 4.28E-05 | 0 | 4.66E-05 | 2.22E-05 | 3.33E-05 | 5.17E-05 | 0 |
| Kctd12b | 1.26E-05 | 3.54E-05 | 0 | 0 | 1.19E-05 | 5.52E-05 | 0 | 1.10E-05 | 3.79E-05 | 0 | 0 | 0 | 0 | 2.92E-05 | 0 | 2.12E-05 | 0 | 2.64E-05 |
| Timm44 | 0.00011 | 0.00011 | 8.53E-05 | 3.67E-05 | 8.22E-06 | 4.65E-05 | 1.75E-05 | 3.21E-05 | 3.79E-05 | 0 | 7.42E-06 | 7.00E-05 | 4.05E-06 | 2.35E-05 | 0.00016 | 6.79E-05 | 0 | 6.65E-06 |
| Kif13b | 0 | 0.00011 | 0 | 2.62E-06 | 9.83E-05 | 3.80E-05 | 6.57E-05 | 4.55E-05 | 3.79E-05 | 0.00014 | 0 | 8.92E-06 | 4.15E-05 | 8.21E-05 | 2.78E-05 | 0 | 7.75E-05 | 0.00016 |
| Wwp2 | 6.31E-05 | 0.00011 | 0 | 5.18E-05 | 1.74E-05 | 3.23E-05 | 3.14E-05 | 0 | 3.79E-05 | 0 | 0.00011 | 2.28E-05 | 3.96E-05 | 1.89E-05 | 3.82E-05 | 5.83E-05 | 8.24E-05 | 0.00015 |
| Ccdc91 | 6.12E-05 | 9.60E-05 | 7.77E-06 | 9.20E-05 | 0.00018 | 9.91E-05 | 9.25E-05 | 3.82E-05 | 3.78E-05 | 2.54E-05 | 1.42E-05 | 2.90E-05 | 0 | 4.72E-05 | 8.23E-06 | 3.84E-05 | 5.80E-05 | 4.41E-05 |
| Fbxo25 | 3.53E-05 | 6.13E-05 | 7.19E-06 | 0 | 5.77E-05 | 3.17E-05 | 6.63E-05 | 2.44E-05 | 3.78E-05 | 4.62E-05 | 1.06E-05 | 4.84E-05 | 2.15E-06 | 2.06E-05 | 1.32E-05 | 8.90E-06 | 0 | 7.71E-05 |
| Hypk | 0.0002 | 8.08E-05 | 0.0002 | 0.00029 | 1.95E-05 | 2.55E-05 | 0.00015 | 1.73E-05 | 3.78E-05 | 7.38E-05 | 8.88E-05 | 5.39E-05 | 6.59E-05 | 2.31E-05 | 0.00011 | 0.00015 | 7.39E-05 | 5.32E-05 |
| Cdc5l | 8.64E-05 | 0.0001 | 9.98E-05 | 0.00012 | 1.07E-05 | 2.11E-05 | 4.40E-05 | 4.47E-05 | 3.78E-05 | 2.70E-05 | 0 | 6.97E-05 | 0 | 4.05E-05 | 1.02E-05 | 0.00017 | 4.09E-05 | 4.34E-05 |
| Trmt1 | 8.53E-05 | 9.01E-05 | 0.0001 | 4.06E-05 | 9.12E-06 | 2.03E-05 | 0 | 8.27E-07 | 3.78E-05 | 0 | 0 | 7.98E-05 | 0 | 1.55E-05 | 7.43E-05 | 3.82E-05 | 5.49E-05 | 0 |
| Psmg2 | 9.20E-05 | 5.09E-05 | 0.00011 | 4.68E-05 | 1.62E-05 | 1.97E-05 | 5.20E-05 | 2.56E-05 | 3.78E-05 | 4.14E-05 | 2.74E-05 | 0 | 1.49E-06 | 0 | 0 | 0.00018 | 4.60E-06 | 4.15E-05 |
| Sgms2 | 4.01E-05 | 4.03E-06 | 0.00011 | 0 | 4.54E-06 | 3.03E-06 | 0.00022 | 7.50E-05 | 3.78E-05 | 0.00012 | 3.45E-05 | 5.94E-05 | 0 | 0 | 4.14E-05 | 1.87E-05 | 6.00E-05 | 0.00012 |
| Kdm5b | 0.00024 | 0.00027 | 0 | 3.40E-05 | 0.00017 | 0.00021 | 0.00016 | 7.69E-05 | 3.77E-05 | 5.79E-06 | 1.43E-05 | 0.00046 | 3.63E-05 | 5.73E-05 | 0.00033 | 0 | 0.0001 | 0 |
| Ankrd13c | 0.00018 | 0.00012 | 0.00013 | 4.39E-05 | 3.92E-05 | 0.00012 | 0 | 2.59E-05 | 3.77E-05 | 8.04E-05 | 5.07E-05 | 3.27E-05 | 1.86E-06 | 0 | 9.59E-05 | 0.0001 | 5.96E-05 | 4.91E-05 |
| Msi2 | 2.60E-05 | 0.00013 | 2.28E-05 | 2.89E-05 | 0.00029 | 0.00011 | 8.43E-05 | 0.00022 | 3.77E-05 | 8.43E-05 | 8.83E-05 | 0.0002 | 0 | 4.77E-05 | 0.00048 | 0 | 0.00015 | 2.55E-05 |
| Zbtb8os | 0.00013 | 4.88E-05 | 1.20E-05 | 3.47E-05 | 2.32E-05 | 3.49E-05 | 5.98E-05 | 0 | 3.77E-05 | 2.69E-05 | 0 | 3.56E-05 | 8.23E-06 | 0 | 3.60E-06 | 5.65E-05 | 0 | 3.38E-05 |
| Ttc9c | 0.00012 | 0.0001 | 0.00013 | 0.0001 | 4.93E-05 | 2.72E-05 | 5.45E-05 | 2.26E-05 | 3.77E-05 | 2.15E-05 | 3.80E-05 | 1.54E-05 | 3.55E-05 | 4.26E-05 | 0.00011 | 0.00026 | 0 | 8.23E-06 |
| Plagl2 | 1.13E-05 | 6.20E-05 | 3.71E-05 | 0 | 4.51E-05 | 0 | 0 | 1.90E-05 | 3.77E-05 | 2.68E-05 | 4.78E-06 | 3.63E-05 | 8.97E-06 | 2.84E-05 | 3.85E-05 | 5.71E-05 | 1.82E-05 | 2.39E-05 |
| Tarbp1 | 8.35E-06 | 1.63E-05 | 5.35E-05 | 3.11E-05 | 1.64E-05 | 0 | 0 | 2.48E-05 | 3.77E-05 | 0 | 1.62E-05 | 0 | 0 | 2.72E-05 | 2.93E-05 | 5.66E-06 | 0 | 2.42E-06 |
| Cep63 | 9.38E-05 | 8.46E-05 | 2.50E-05 | 7.06E-06 | 2.09E-05 | 4.51E-05 | 0 | 1.54E-05 | 3.76E-05 | 0 | 1.32E-06 | 3.29E-05 | 0 | 1.08E-06 | 0.00012 | 6.44E-05 | 1.74E-06 | 1.38E-05 |
| E2f5 | 4.66E-05 | 4.88E-05 | 5.56E-05 | 4.36E-06 | 5.78E-06 | 1.40E-05 | 0 | 0 | 3.76E-05 | 4.36E-06 | 3.12E-05 | 4.43E-05 | 4.24E-06 | 3.52E-05 | 5.42E-05 | 7.65E-05 | 0 | 1.56E-05 |
| Kxd1 | 5.36E-05 | 6.21E-05 | 5.02E-06 | 1.85E-05 | 3.91E-05 | 5.06E-05 | 1.05E-05 | 2.01E-05 | 3.75E-05 | 2.18E-05 | 4.05E-06 | 7.67E-05 | 1.07E-06 | 0 | 1.32E-05 | 1.36E-05 | 0 | 5.76E-05 |
| Trmt11 | 4.88E-05 | 5.25E-05 | 4.68E-05 | 1.52E-05 | 2.79E-05 | 4.81E-05 | 0 | 4.35E-06 | 3.75E-05 | 2.21E-05 | 0 | 1.19E-05 | 1.25E-05 | 1.63E-05 | 9.86E-05 | 4.87E-05 | 0 | 0 |
| Dis3l2 | 1.46E-05 | 2.44E-05 | 4.46E-05 | 2.74E-05 | 3.41E-05 | 4.03E-05 | 2.71E-05 | 6.76E-05 | 3.75E-05 | 3.71E-05 | 0 | 1.24E-05 | 7.44E-06 | 2.37E-05 | 5.70E-05 | 1.38E-05 | 0 | 5.50E-05 |
| B9d1 | 7.38E-05 | 2.01E-05 | 3.75E-06 | 5.90E-05 | 2.54E-05 | 1.12E-05 | 1.58E-05 | 8.41E-06 | 3.75E-05 | 3.03E-05 | 3.07E-05 | 7.64E-06 | 3.23E-05 | 2.70E-05 | 2.48E-05 | 3.21E-05 | 2.54E-05 | 5.71E-05 |
| Rnf8 | 0 | 6.07E-05 | 2.32E-05 | 8.28E-05 | 2.57E-05 | 4.90E-07 | 0 | 1.03E-05 | 3.75E-05 | 3.50E-05 | 3.63E-05 | 5.92E-06 | 0 | 2.12E-05 | 0.00012 | 7.71E-05 | 0 | 3.96E-05 |
| Klhl18 | 0 | 5.96E-05 | 6.13E-05 | 6.12E-05 | 3.84E-05 | 0 | 1.18E-05 | 0 | 3.75E-05 | 4.13E-05 | 4.39E-05 | 4.06E-05 | 3.38E-05 | 1.92E-05 | 5.41E-05 | 8.83E-06 | 0 | 5.63E-05 |
| Pxdc1 | 2.76E-05 | 2.56E-05 | 0.00019 | 0 | 0.00011 | 0.00015 | 7.57E-05 | 0.00018 | 3.74E-05 | 5.72E-05 | 8.64E-05 | 0.00025 | 0 | 0.00062 | 0 | 1.37E-06 | 5.48E-05 | 4.68E-05 |
| Katnbl1 | 6.43E-05 | 9.96E-05 | 5.34E-05 | 0 | 2.88E-05 | 8.78E-05 | 0 | 3.37E-05 | 3.74E-05 | 8.35E-05 | 6.14E-05 | 7.95E-05 | 0 | 3.58E-05 | 9.14E-05 | 0.00011 | 2.75E-05 | 2.58E-05 |
| Zxdc | 8.02E-05 | 0.00011 | 0 | 0 | 3.75E-05 | 7.68E-05 | 0 | 0 | 3.74E-05 | 5.61E-05 | 6.73E-05 | 4.62E-05 | 0 | 1.17E-05 | 5.27E-05 | 6.49E-06 | 5.40E-05 | 0 |
| Kansl1l | 3.78E-05 | 9.76E-05 | 0 | 0 | 0.00011 | 2.34E-05 | 0 | 5.82E-06 | 3.74E-05 | 0 | 6.67E-05 | 3.65E-05 | 0 | 4.91E-05 | 0 | 6.94E-05 | 0.00012 | 2.68E-06 |

| | | | | | | | | | | | | | | | | | | |
|---|---|---|---|---|---|---|---|---|---|---|---|---|---|---|---|---|---|---|
| Tmem216 | 3.64E-05 | 1.93E-05 | 1.50E-05 | 2.19E-05 | 1.65E-05 | 1.52E-05 | 8.42E-07 | 0 | 3.74E-05 | 1.98E-05 | 0 | 3.29E-06 | 4.07E-05 | 1.51E-05 | 0 | 1.53E-05 | 0 | 3.66E-05 |
| Kdm2b | 3.23E-05 | 4.98E-05 | 7.80E-05 | 4.04E-05 | 3.21E-05 | 1.86E-06 | 7.33E-07 | 2.61E-05 | 3.74E-05 | 6.09E-06 | 4.28E-05 | 4.55E-05 | 1.70E-05 | 2.98E-05 | 6.68E-05 | 6.17E-05 | 0 | 2.00E-05 |
| Luc7l | 6.46E-05 | 0.00012 | 8.06E-05 | 0 | 7.40E-06 | 1.02E-06 | 0 | 0 | 3.74E-05 | 1.24E-05 | 1.50E-05 | 4.68E-05 | 0 | 0 | 0.00041 | 3.56E-05 | 0.00011 | 1.94E-05 |
| Flt3l | 0 | 9.98E-06 | 0 | 2.78E-05 | 0 | 0 | 1.57E-06 | 0 | 3.74E-05 | 9.29E-05 | 0 | 1.42E-05 | 7.17E-06 | 0 | 1.19E-06 | 0 | 1.29E-05 | 0.00011 |
| Pak4 | 3.07E-05 | 4.52E-05 | 4.51E-05 | 5.79E-06 | 1.67E-05 | 0 | 0 | 1.72E-05 | 3.74E-05 | 2.76E-05 | 1.21E-05 | 2.55E-05 | 0 | 0 | 2.83E-05 | 0.00016 | 3.28E-05 | 7.16E-05 |
| Prkd1 | 1.31E-05 | 2.10E-05 | 0 | 0 | 8.63E-05 | 0 | 5.48E-05 | 0 | 3.74E-05 | 0 | 0.00029 | 0 | 0.00011 | 0 | 9.36E-05 | 5.48E-06 | 7.01E-05 | 0 |
| Pik3ca | 0.00024 | 0.00019 | 7.39E-05 | 0 | 9.60E-05 | 0.0002 | 8.44E-05 | 3.82E-05 | 3.73E-05 | 2.16E-06 | 3.27E-05 | 0.00011 | 0 | 7.40E-05 | 8.76E-05 | 0.00011 | 0.00012 | 0 |
| Kansl1 | 0.00014 | 0.00033 | 8.96E-05 | 5.66E-05 | 0.00012 | 0.00016 | 4.91E-05 | 3.20E-05 | 3.73E-05 | 7.82E-05 | 5.06E-05 | 0.00015 | 0 | 0.0001 | 0.00051 | 9.83E-05 | 0.00017 | 0.00012 |
| Pon3 | 0.00018 | 3.89E-05 | 3.22E-05 | 2.26E-05 | 3.30E-05 | 0.00012 | 0.00021 | 5.41E-05 | 3.73E-05 | 6.01E-05 | 0 | 0 | 0.00011 | 9.45E-05 | 0 | 7.90E-05 | 0 | 5.39E-05 |
| Ubash3b | 0 | 1.29E-05 | 4.57E-05 | 0 | 2.66E-05 | 9.61E-05 | 0 | 0.00028 | 3.73E-05 | 0 | 9.78E-05 | 0.00018 | 0 | 0 | 2.67E-06 | 3.13E-05 | 0 | 2.77E-05 |
| Slc10a7 | 0.00014 | 0 | 8.54E-05 | 3.33E-05 | 4.47E-05 | 7.17E-05 | 0.00016 | 8.32E-05 | 3.73E-05 | 2.83E-05 | 9.42E-05 | 1.58E-05 | 2.88E-05 | 4.91E-06 | 0.00011 | 6.19E-05 | 0 | 0 |
| Ndufb4 | 0.00014 | 2.76E-05 | 5.17E-05 | 1.81E-06 | 1.21E-05 | 3.22E-05 | 2.38E-05 | 0 | 3.73E-05 | 1.65E-05 | 8.71E-06 | 0 | 2.73E-05 | 5.57E-06 | 0 | 2.30E-05 | 0 | 6.94E-07 |
| Cog8 | 3.84E-05 | 2.87E-05 | 4.90E-05 | 7.38E-05 | 3.95E-05 | 9.83E-06 | 2.96E-05 | 2.31E-05 | 3.73E-05 | 2.16E-05 | 9.34E-06 | 0 | 2.12E-06 | 0 | 6.76E-05 | 2.36E-05 | 0 | 3.07E-05 |
| Cyb561d2 | 1.87E-05 | 1.73E-05 | 2.07E-05 | 0.00013 | 5.28E-06 | 5.20E-06 | 5.09E-05 | 2.43E-05 | 3.73E-05 | 3.60E-05 | 2.20E-05 | 8.36E-06 | 3.01E-05 | 1.62E-05 | 0 | 0 | 0 | 4.74E-05 |
| Dimt1 | 3.58E-05 | 2.19E-05 | 8.31E-05 | 4.68E-05 | 1.06E-05 | 0 | 0 | 2.35E-06 | 3.73E-05 | 2.36E-05 | 5.80E-06 | 0 | 0 | 0 | 0.00017 | 7.36E-05 | 0 | 2.25E-05 |
| Fn3krp | 1.66E-05 | 3.82E-05 | 0 | 8.27E-05 | 3.03E-05 | 0 | 8.70E-06 | 0 | 3.73E-05 | 6.88E-05 | 1.97E-05 | 1.38E-06 | 3.10E-05 | 2.32E-05 | 1.04E-05 | 4.45E-05 | 0 | 4.57E-05 |
| Med18 | 9.94E-06 | 2.78E-05 | 4.37E-05 | 2.58E-05 | 1.81E-05 | 0 | 1.25E-06 | 5.49E-06 | 3.73E-05 | 1.65E-05 | 1.65E-05 | 9.47E-06 | 0 | 5.24E-06 | 2.65E-05 | 0.00014 | 0 | 2.12E-05 |
| Olfml1 | 8.87E-06 | 7.25E-05 | 0 | 0 | 0 | 0 | 0 | 2.12E-05 | 3.73E-05 | 0.00037 | 0 | 0 | 0.00023 | 0 | 0 | 6.27E-06 | 0 | 0.00024 |
| Commd6 | 0.00016 | 8.60E-05 | 6.28E-05 | 0 | 4.65E-05 | 8.49E-05 | 4.16E-05 | 9.46E-06 | 3.72E-05 | 2.45E-05 | 3.67E-05 | 1.01E-05 | 0 | 0 | 0 | 2.86E-05 | 0 | 5.17E-05 |
| Tbc1d8b | 0.00016 | 0 | 4.52E-05 | 5.30E-05 | 5.18E-05 | 4.88E-05 | 0.0002 | 0 | 3.72E-05 | 3.25E-05 | 0 | 3.29E-05 | 1.21E-05 | 4.45E-05 | 7.38E-05 | 9.12E-05 | 5.50E-05 | 4.87E-05 |
| Clasrp | 3.21E-05 | 0.00015 | 5.74E-05 | 0.00014 | 2.75E-05 | 3.47E-06 | 1.82E-05 | 3.55E-05 | 3.72E-05 | 5.00E-05 | 0 | 4.78E-05 | 1.37E-05 | 0 | 0.0006 | 0.00013 | 0.0001 | 5.50E-05 |
| Pafah2 | 0 | 2.64E-05 | 8.82E-06 | 6.42E-05 | 2.11E-05 | 0 | 1.96E-05 | 2.46E-05 | 3.72E-05 | 2.08E-05 | 2.00E-05 | 8.82E-06 | 2.37E-05 | 6.73E-06 | 5.87E-06 | 9.81E-06 | 0 | 5.10E-05 |
| Snx10 | 4.66E-05 | 4.32E-05 | 7.58E-06 | 0 | 0 | 0 | 2.06E-05 | 3.10E-05 | 3.72E-05 | 4.90E-05 | 0 | 0 | 5.08E-05 | 0 | 0 | 1.01E-05 | 1.68E-05 | 8.17E-05 |
| Rev1 | 5.99E-05 | 8.09E-05 | 8.50E-06 | 2.33E-06 | 3.71E-05 | 3.79E-05 | 4.62E-06 | 0 | 3.71E-05 | 1.07E-05 | 2.47E-05 | 3.92E-05 | 2.79E-05 | 2.23E-05 | 2.53E-05 | 4.47E-05 | 3.38E-05 | 0 |
| Nob1 | 7.02E-05 | 6.12E-05 | 0.00017 | 3.00E-05 | 0 | 2.61E-05 | 0 | 0 | 3.71E-05 | 1.27E-05 | 6.49E-06 | 2.51E-05 | 1.38E-05 | 2.52E-05 | 3.47E-05 | 1.14E-06 | 0 | 0 |
| Kdm1a | 0.00017 | 0.00014 | 0.00012 | 0 | 9.19E-05 | 8.58E-05 | 0.00011 | 1.07E-05 | 3.70E-05 | 2.23E-05 | 6.68E-05 | 0.00011 | 0 | 4.25E-05 | 4.26E-05 | 8.27E-05 | 0 | 7.04E-06 |
| Syvn1 | 0.00029 | 0.00011 | 0.0002 | 5.67E-05 | 3.34E-05 | 6.02E-05 | 0.00018 | 1.24E-05 | 3.70E-05 | 6.55E-06 | 5.49E-05 | 0.00018 | 0 | 2.12E-05 | 0.00025 | 0 | 0 | 2.06E-05 |
| Spryd7 | 8.10E-05 | 4.42E-05 | 8.46E-05 | 0 | 5.56E-05 | 2.06E-05 | 3.08E-05 | 7.04E-07 | 3.70E-05 | 4.99E-05 | 3.01E-05 | 2.50E-05 | 0.00015 | 0 | 0 | 0 | 5.36E-05 | 2.26E-05 |
| Mettl21a | 5.94E-05 | 4.24E-05 | 0 | 0.0001 | 3.48E-05 | 1.56E-05 | 0.00012 | 0.00012 | 3.70E-05 | 2.75E-05 | 0 | 0.00018 | 4.61E-05 | 0 | 1.16E-05 | 3.68E-05 | 0 | 8.90E-05 |
| Tmem94 | 9.20E-06 | 9.39E-05 | 0 | 9.54E-05 | 5.54E-05 | 1.38E-05 | 3.80E-05 | 1.40E-05 | 3.70E-05 | 4.21E-05 | 1.79E-05 | 1.95E-05 | 2.43E-05 | 5.29E-05 | 7.02E-05 | 8.23E-05 | 0 | 2.94E-05 |
| Tamm41 | 3.79E-05 | 1.66E-05 | 3.35E-05 | 3.75E-05 | 0 | 1.03E-05 | 1.65E-05 | 2.32E-06 | 3.70E-05 | 1.42E-05 | 2.32E-05 | 8.70E-06 | 0 | 1.39E-05 | 7.64E-06 | 0.00012 | 0 | 1.06E-05 |
| Ccdc25 | 0.00011 | 8.03E-05 | 0.00013 | 2.75E-05 | 9.06E-05 | 7.85E-06 | 0.00015 | 1.35E-05 | 3.70E-05 | 0.00011 | 1.05E-05 | 0 | 0.00011 | 8.55E-07 | 6.17E-07 | 0.00019 | 0 | 0.00012 |
| Plscr1 | 7.34E-05 | 5.68E-05 | 4.09E-05 | 0 | 1.84E-05 | 0 | 6.52E-05 | 0.0001 | 3.70E-05 | 3.96E-05 | 0 | 2.45E-06 | 1.61E-05 | 0 | 0 | 9.73E-05 | 3.39E-05 | 5.17E-05 |

| | | | | | | | | | | | | | | | | | | |
|---|---|---|---|---|---|---|---|---|---|---|---|---|---|---|---|---|---|---|
| Epb41l4a | 0 | 1.77E-05 | 3.89E-06 | 0 | 0 | 0 | 1.57E-05 | 9.20E-06 | 3.70E-05 | 0 | 0 | 2.20E-05 | 0 | 0 | 0 | 0 | 7.47E-07 | 0 |
| Lrwd1 | 2.18E-05 | 8.66E-05 | 7.34E-05 | 0.0001 | 3.00E-05 | 0 | 5.52E-06 | 2.94E-05 | 3.70E-05 | 4.33E-05 | 8.58E-06 | 3.10E-05 | 0 | 6.48E-05 | 6.48E-05 | 0.00024 | 0 | 3.84E-05 |
| Cbfb | 0.00018 | 0.00041 | 0.00027 | 0.00047 | 0.00039 | 0.00016 | 0.00037 | 0.00027 | 3.69E-05 | 2.60E-05 | 0.00027 | 0.00021 | 0.0004 | 0.00043 | 0.00054 | 0.00065 | 0.00042 | 0.00036 |
| Slc25a32 | 0.00015 | 1.95E-05 | 7.56E-05 | 0 | 0 | 8.91E-05 | 1.44E-05 | 1.28E-05 | 3.69E-05 | 2.82E-06 | 0 | 9.96E-06 | 0 | 0 | 0 | 4.50E-05 | 1.98E-05 | 2.75E-06 |
| Calhm2 | 3.74E-05 | 0.00011 | 6.30E-05 | 9.34E-05 | 0.00011 | 7.54E-05 | 0.00013 | 0.00011 | 3.69E-05 | 0.00019 | 0.00012 | 3.37E-05 | 2.46E-05 | 6.99E-05 | 1.61E-05 | 4.56E-05 | 0 | 0.00024 |
| Nup85 | 4.57E-05 | 4.12E-05 | 0.00015 | 2.19E-05 | 0 | 3.55E-05 | 0 | 0 | 3.69E-05 | 6.04E-05 | 1.67E-05 | 0 | 5.51E-05 | 1.22E-05 | 0.00016 | 0.00057 | 1.73E-05 | 2.52E-05 |
| Hoxa5 | 9.35E-06 | 3.00E-05 | 0 | 2.05E-05 | 2.95E-05 | 4.88E-06 | 2.27E-05 | 1.92E-05 | 3.69E-05 | 2.61E-05 | 0 | 1.27E-05 | 2.47E-06 | 0 | 5.76E-05 | 3.14E-05 | 0 | 9.97E-06 |
| Zmat3 | 0 | 0.00027 | 0.00016 | 4.98E-05 | 0.00013 | 5.83E-05 | 8.32E-05 | 0.00011 | 3.68E-05 | 5.72E-05 | 3.18E-06 | 0 | 0.00014 | 0 | 1.74E-05 | 0 | 0 | 2.58E-05 |
| Clec16a | 1.79E-05 | 5.20E-05 | 2.73E-05 | 0 | 5.33E-05 | 3.24E-05 | 2.36E-05 | 4.04E-05 | 3.68E-05 | 3.53E-05 | 5.89E-06 | 1.79E-05 | 3.08E-05 | 2.06E-05 | 1.16E-05 | 7.37E-06 | 3.29E-07 | 7.70E-05 |
| Alkbh8 | 6.17E-05 | 2.58E-05 | 2.17E-05 | 0 | 2.75E-05 | 2.70E-05 | 0 | 5.85E-06 | 3.68E-05 | 0 | 1.91E-05 | 7.00E-06 | 1.73E-05 | 1.08E-05 | 3.44E-05 | 3.43E-06 | 0 | 1.13E-05 |
| Phkb | 4.38E-05 | 8.13E-05 | 9.76E-06 | 2.57E-05 | 5.05E-05 | 1.83E-05 | 4.76E-05 | 1.16E-06 | 3.68E-05 | 5.53E-05 | 1.41E-05 | 5.40E-06 | 1.39E-05 | 0 | 4.69E-05 | 3.16E-05 | 5.49E-05 | 5.22E-05 |
| Mief2 | 5.78E-06 | 4.00E-05 | 1.58E-05 | 2.71E-05 | 3.20E-05 | 0 | 2.17E-05 | 3.12E-05 | 3.68E-05 | 4.05E-05 | 4.49E-05 | 0 | 2.15E-06 | 3.63E-05 | 5.63E-05 | 5.00E-06 | 0 | 6.81E-05 |
| Nelfa | 0.00012 | 0.00013 | 0.00016 | 0.00021 | 2.57E-05 | 0 | 0.0001 | 0 | 3.68E-05 | 5.00E-05 | 5.20E-05 | 0 | 5.47E-05 | 3.40E-05 | 0.00013 | 0.00012 | 5.01E-07 | 0 |
| Scrn2 | 2.19E-06 | 6.30E-06 | 1.36E-05 | 5.18E-05 | 1.21E-05 | 0 | 0 | 2.84E-05 | 3.68E-05 | 1.43E-05 | 5.01E-07 | 0 | 0 | 1.98E-05 | 4.13E-05 | 2.13E-05 | 0 | 4.09E-05 |
| Fbxw7 | 3.25E-05 | 5.13E-05 | 4.30E-05 | 0 | 3.79E-05 | 4.38E-05 | 2.14E-05 | 0 | 3.67E-05 | 0 | 5.44E-05 | 8.78E-06 | 0 | 0 | 6.37E-05 | 3.17E-05 | 3.41E-05 | 2.30E-06 |
| Plekha5 | 3.72E-05 | 0.00019 | 8.11E-05 | 0 | 0.00015 | 3.96E-05 | 4.86E-06 | 0 | 3.67E-05 | 0 | 0.00025 | 0.00017 | 9.36E-05 | 0.00022 | 0.00021 | 2.28E-05 | 7.89E-05 | 3.45E-06 |
| Rnaseh2c | 5.02E-05 | 0.00011 | 1.84E-05 | 0.00043 | 4.39E-05 | 2.81E-05 | 2.76E-05 | 9.71E-05 | 3.67E-05 | 1.04E-05 | 1.51E-05 | 2.44E-05 | 1.62E-05 | 0.00014 | 8.48E-05 | 0.00047 | 0.0001 | 5.94E-05 |
| Rab3d | 4.96E-05 | 6.79E-05 | 0 | 5.30E-05 | 4.21E-05 | 7.52E-06 | 0.00013 | 0 | 3.67E-05 | 0.0002 | 1.87E-05 | 2.88E-07 | 9.71E-05 | 2.44E-05 | 1.42E-05 | 0 | 0 | 0.00029 |
| Pigv | 0 | 5.10E-05 | 0 | 3.82E-05 | 4.08E-05 | 3.99E-06 | 1.23E-05 | 1.12E-05 | 3.67E-05 | 2.36E-05 | 7.55E-06 | 4.57E-05 | 0 | 4.03E-05 | 7.85E-05 | 8.76E-06 | 0 | 2.48E-05 |
| Sytl4 | 8.23E-06 | 0 | 8.82E-06 | 0 | 0 | 5.41E-07 | 3.84E-05 | 1.87E-05 | 3.67E-05 | 1.58E-05 | 3.30E-06 | 0 | 4.52E-06 | 0 | 2.45E-05 | 0 | 0 | 4.14E-05 |
| Rc3h2 | 0.00024 | 0.00014 | 4.23E-05 | 0 | 0.00011 | 0.00016 | 5.54E-05 | 1.92E-05 | 3.66E-05 | 0 | 7.95E-05 | 0.00014 | 3.79E-06 | 1.85E-05 | 9.34E-05 | 3.01E-05 | 0.00011 | 1.67E-05 |
| Atg5 | 0.00018 | 4.35E-05 | 0.00011 | 0 | 6.30E-05 | 4.93E-05 | 9.02E-05 | 2.70E-05 | 3.66E-05 | 1.91E-05 | 5.01E-05 | 1.44E-05 | 1.45E-05 | 5.96E-05 | 7.32E-05 | 5.94E-05 | 3.00E-05 | 0 |
| Vezt | 3.07E-05 | 7.74E-05 | 4.26E-06 | 3.94E-05 | 1.45E-05 | 3.09E-05 | 0 | 8.15E-05 | 3.66E-05 | 6.24E-06 | 6.47E-05 | 4.81E-05 | 0 | 1.87E-05 | 5.09E-05 | 7.15E-05 | 7.63E-05 | 7.75E-05 |
| Nme7 | 6.88E-05 | 5.73E-05 | 0 | 4.75E-05 | 2.26E-06 | 2.36E-05 | 3.26E-05 | 2.19E-05 | 3.66E-05 | 2.17E-05 | 0 | 2.47E-06 | 0 | 4.88E-06 | 7.64E-05 | 4.14E-05 | 2.81E-05 | 1.97E-05 |
| Adcy4 | 4.40E-06 | 0.00015 | 9.38E-06 | 0 | 2.25E-05 | 1.63E-05 | 1.80E-06 | 5.11E-05 | 3.66E-05 | 0 | 4.79E-05 | 0 | 1.69E-05 | 0.00014 | 0.00013 | 0 | 0.00013 | 2.58E-05 |
| Ube2e1 | 0.00027 | 9.76E-05 | 0.00017 | 0 | 6.54E-05 | 0.00015 | 6.37E-06 | 5.21E-05 | 3.65E-05 | 1.15E-05 | 6.11E-05 | 6.22E-05 | 0 | 1.58E-05 | 0 | 4.95E-05 | 0 | 1.41E-05 |
| Hspa13 | 0.00026 | 5.93E-05 | 0.00018 | 0 | 0.00011 | 0.00014 | 8.51E-05 | 0.00013 | 3.65E-05 | 0 | 3.06E-05 | 6.97E-05 | 6.76E-05 | 4.80E-05 | 1.86E-05 | 0.00014 | 0.00016 | 0 |
| Ccdc9b | 0 | 0.00011 | 1.83E-05 | 6.09E-05 | 9.23E-05 | 0.00013 | 4.70E-06 | 6.85E-05 | 3.65E-05 | 3.04E-05 | 8.97E-05 | 0 | 7.76E-06 | 0.0001 | 0.00025 | 0 | 0 | 0 |
| Vps4a | 0.00014 | 9.35E-05 | 0.00011 | 4.23E-05 | 8.24E-05 | 0.00012 | 5.15E-05 | 3.51E-05 | 3.65E-05 | 5.56E-05 | 0.00011 | 6.49E-05 | 0 | 0 | 4.00E-05 | 6.80E-05 | 0 | 4.06E-05 |
| Tmtc2 | 1.62E-05 | 1.14E-05 | 0 | 0 | 5.17E-06 | 4.53E-05 | 0 | 0 | 3.65E-05 | 1.21E-05 | 7.47E-05 | 0 | 0 | 1.78E-05 | 1.62E-05 | 2.15E-06 | 6.18E-05 | 7.45E-05 |
| Zrsr1 | 1.94E-05 | 6.85E-05 | 0 | 4.15E-06 | 5.52E-05 | 4.02E-05 | 2.50E-05 | 0 | 3.65E-05 | 2.27E-05 | 5.54E-06 | 6.43E-06 | 0 | 3.20E-06 | 5.12E-05 | 0 | 0 | 5.39E-05 |
| Artn | 0 | 0 | 0 | 0 | 0 | 0 | 0 | 0 | 3.65E-05 | 0 | 0 | 0 | 0.00011 | 0 | 5.78E-06 | 0 | 2.27E-05 | 2.20E-05 |
| Mpp5 | 6.46E-05 | 8.78E-05 | 3.38E-05 | 0 | 8.20E-05 | 0 | 8.19E-05 | 0 | 3.65E-05 | 2.47E-05 | 9.93E-05 | 9.63E-05 | 1.25E-05 | 3.14E-05 | 0.00013 | 4.21E-06 | 0.00015 | 0 |

| | | | | | | | | | | | | | | | | | | |
|---|---|---|---|---|---|---|---|---|---|---|---|---|---|---|---|---|---|---|
| Ttf2 | 1.09E-05 | 2.63E-05 | 3.28E-05 | 3.83E-06 | 4.39E-05 | 1.37E-05 | 8.25E-06 | 1.18E-05 | 3.64E-05 | 1.09E-05 | 0 | 0 | 4.24E-05 | 2.20E-05 | 1.13E-05 | 0.00026 | 0 | 1.95E-05 |
| Hsf2 | 5.62E-05 | 4.53E-05 | 1.45E-05 | 1.42E-05 | 1.13E-05 | 9.88E-06 | 0 | 0 | 3.64E-05 | 8.11E-05 | 1.68E-05 | 0 | 1.03E-05 | 6.35E-05 | 3.68E-05 | 3.26E-05 | 5.57E-05 | 0 |
| Rac3 | 0 | 1.67E-06 | 0 | 5.97E-05 | 3.79E-05 | 0 | 0.00022 | 0 | 3.64E-05 | 1.47E-05 | 3.57E-05 | 0.00021 | 0.00026 | 0.00011 | 0 | 0 | 0 | 0 |
| Plekhn1 | 0 | 0.00013 | 0 | 7.73E-05 | 0.0001 | 0.0001 | 1.97E-05 | 4.87E-05 | 3.63E-05 | 1.77E-05 | 4.65E-06 | 0.00019 | 2.23E-05 | 0.0001 | 0.00025 | 0 | 0 | 5.76E-05 |
| Ddx19b | 5.35E-05 | 3.68E-05 | 2.89E-05 | 5.23E-06 | 3.10E-05 | 5.14E-05 | 0 | 1.77E-05 | 3.63E-05 | 2.52E-05 | 2.25E-05 | 4.71E-05 | 5.62E-06 | 1.79E-05 | 2.75E-05 | 9.63E-06 | 0 | 2.98E-06 |
| Arl6ip1 | 2.21E-05 | 0.00012 | 0 | 0 | 6.90E-05 | 3.76E-05 | 0 | 7.92E-06 | 3.63E-05 | 7.62E-05 | 1.43E-05 | 0.0001 | 0 | 0 | 0 | 0.00052 | 0 | 3.78E-05 |
| Ifih1 | 0.0001 | 7.91E-05 | 0 | 0 | 1.79E-05 | 2.94E-05 | 0 | 1.75E-05 | 3.63E-05 | 0.00024 | 0 | 8.42E-06 | 6.93E-05 | 0 | 6.70E-05 | 4.22E-05 | 0 | 9.91E-05 |
| Hibch | 6.42E-05 | 4.34E-05 | 4.00E-06 | 3.17E-05 | 7.73E-06 | 1.20E-05 | 3.01E-06 | 0 | 3.63E-05 | 1.54E-05 | 0 | 0 | 1.91E-05 | 0 | 4.42E-06 | 4.03E-06 | 0 | 1.54E-05 |
| Cep78 | 4.24E-05 | 1.11E-05 | 1.73E-05 | 1.30E-05 | 1.40E-05 | 1.09E-06 | 1.09E-05 | 0 | 3.63E-05 | 1.66E-05 | 0 | 0 | 0 | 2.70E-05 | 0.00012 | 8.41E-05 | 0 | 4.26E-05 |
| Ppp1r21 | 3.18E-05 | 4.61E-05 | 0 | 3.39E-05 | 4.01E-05 | 0 | 0 | 0 | 3.63E-05 | 3.97E-05 | 3.29E-05 | 3.86E-05 | 1.21E-05 | 2.88E-06 | 3.94E-05 | 4.48E-05 | 5.99E-05 | 1.86E-05 |
| Nop14 | 3.39E-05 | 5.05E-05 | 8.41E-05 | 2.77E-05 | 0 | 0 | 0 | 0 | 3.63E-05 | 2.06E-05 | 1.68E-05 | 1.80E-05 | 0 | 0 | 7.88E-05 | 2.90E-05 | 3.67E-05 | 1.68E-05 |
| Tle6 | 0 | 1.61E-05 | 0 | 1.34E-05 | 4.71E-06 | 0 | 2.12E-05 | 0 | 3.63E-05 | 9.57E-05 | 0 | 3.37E-05 | 7.51E-06 | 4.66E-05 | 1.36E-05 | 0 | 0 | 2.63E-05 |
| Fam102b | 7.79E-05 | 0.00013 | 4.31E-05 | 0 | 0 | 0.00012 | 0.00019 | 0 | 3.62E-05 | 0.00061 | 6.13E-05 | 0.00044 | 0.00029 | 0.00025 | 0 | 0.00017 | 0 | 0.00035 |
| Cdc73 | 0.00017 | 0.00011 | 0.0001 | 6.74E-05 | 3.92E-05 | 7.17E-05 | 3.63E-05 | 1.44E-05 | 3.62E-05 | 2.37E-05 | 3.19E-05 | 9.91E-05 | 0 | 1.61E-05 | 0.00025 | 3.60E-05 | 0.00023 | 1.59E-05 |
| Zfp146 | 0.00015 | 7.57E-05 | 0.0001 | 2.29E-07 | 3.44E-05 | 5.43E-05 | 6.43E-05 | 1.05E-05 | 3.62E-05 | 7.91E-06 | 2.33E-05 | 4.28E-05 | 4.30E-05 | 0 | 0.00012 | 9.43E-06 | 0 | 0 |
| Taf10 | 0.00014 | 2.52E-05 | 0.00014 | 0.00012 | 3.78E-05 | 3.21E-05 | 6.55E-05 | 0 | 3.62E-05 | 0 | 5.04E-05 | 0 | 2.48E-05 | 1.59E-05 | 0 | 8.27E-05 | 2.52E-06 | 5.52E-06 |
| Tada3 | 4.58E-05 | 6.03E-05 | 2.33E-05 | 6.51E-05 | 2.34E-05 | 5.11E-06 | 2.31E-05 | 0 | 3.62E-05 | 5.22E-05 | 6.97E-05 | 8.26E-05 | 0 | 1.24E-05 | 0.00011 | 4.52E-05 | 2.48E-05 | 6.55E-05 |
| Tfeb | 2.80E-05 | 8.02E-05 | 3.93E-05 | 1.83E-05 | 6.78E-05 | 0 | 0 | 9.53E-05 | 3.62E-05 | 5.05E-05 | 3.94E-05 | 0 | 1.01E-05 | 5.16E-06 | 4.51E-05 | 0 | 5.82E-05 | 6.48E-05 |
| Rgs2 | 0 | 3.65E-05 | 0 | 0 | 0 | 0.00031 | 0 | 6.90E-05 | 3.61E-05 | 0 | 0 | 3.65E-05 | 0 | 3.26E-05 | 0 | 0 | 0.00056 | 0 |
| Nomo1 | 0.00017 | 3.97E-05 | 0.00037 | 7.88E-05 | 7.40E-06 | 0.00011 | 5.35E-05 | 0.00011 | 3.61E-05 | 8.30E-05 | 0.00013 | 6.54E-05 | 0.00014 | 0 | 0.00012 | 7.99E-05 | 0 | 1.05E-05 |
| Arpp19 | 0.00019 | 7.42E-05 | 0.00013 | 2.07E-05 | 5.79E-05 | 9.38E-05 | 1.45E-05 | 2.94E-05 | 3.61E-05 | 7.71E-08 | 9.42E-05 | 1.40E-05 | 0 | 9.03E-05 | 0 | 0.00019 | 0 | 0 |
| Mios | 0.00014 | 4.94E-05 | 4.41E-05 | 7.08E-07 | 1.21E-05 | 7.72E-05 | 5.36E-07 | 3.63E-05 | 3.61E-05 | 4.34E-05 | 0 | 1.66E-05 | 0.00011 | 0 | 8.94E-05 | 4.66E-05 | 0 | 2.04E-05 |
| Dguok | 6.05E-05 | 8.99E-05 | 1.97E-05 | 8.70E-05 | 7.83E-05 | 7.29E-05 | 6.35E-05 | 5.11E-05 | 3.61E-05 | 6.87E-05 | 9.88E-06 | 5.05E-05 | 3.52E-05 | 0 | 7.44E-05 | 3.46E-05 | 0 | 9.70E-05 |
| Nip7 | 0.0001 | 2.47E-05 | 0.00019 | 0.00012 | 1.40E-05 | 1.86E-05 | 0 | 0 | 3.61E-05 | 7.10E-07 | 3.41E-05 | 0 | 5.11E-05 | 0 | 5.07E-05 | 0 | 0 | 0 |
| Mtif3 | 2.12E-05 | 4.10E-05 | 3.43E-06 | 3.52E-05 | 3.21E-05 | 3.10E-06 | 1.17E-05 | 3.63E-06 | 3.61E-05 | 2.50E-05 | 1.55E-05 | 2.35E-05 | 4.24E-06 | 7.09E-06 | 1.43E-05 | 2.49E-05 | 0 | 2.32E-05 |
| Dusp19 | 4.94E-05 | 2.16E-05 | 1.75E-05 | 1.94E-05 | 1.63E-05 | 0 | 1.95E-05 | 2.52E-05 | 3.61E-05 | 2.50E-05 | 1.77E-05 | 1.61E-05 | 0 | 0 | 0 | 3.41E-05 | 0 | 2.19E-05 |
| Plaat3 | 0.00017 | 7.89E-05 | 0 | 0 | 0.00065 | 0 | 0.00039 | 0 | 3.61E-05 | 0 | 0.00032 | 0 | 0.00012 | 0.00031 | 0 | 6.95E-05 | 0 | 0.00029 |
| Mbnl3 | 1.10E-05 | 0 | 0 | 0 | 8.78E-05 | 7.11E-05 | 0 | 0 | 3.60E-05 | 0 | 4.67E-05 | 6.30E-06 | 0 | 4.97E-05 | 0 | 8.82E-05 | 0 | 0 |
| Apool | 0.00011 | 4.48E-05 | 1.49E-05 | 1.30E-05 | 2.16E-05 | 6.46E-05 | 2.28E-05 | 4.07E-06 | 3.60E-05 | 7.35E-06 | 3.13E-05 | 3.66E-05 | 0 | 3.11E-05 | 0 | 6.13E-05 | 0 | 9.89E-06 |
| Pikfyve | 6.06E-05 | 6.00E-05 | 5.15E-05 | 0 | 5.06E-05 | 4.86E-05 | 4.69E-05 | 2.84E-05 | 3.60E-05 | 9.51E-06 | 1.41E-05 | 8.37E-06 | 4.18E-05 | 1.73E-05 | 5.62E-05 | 2.38E-06 | 2.46E-05 | 0 |
| Pogk | 2.87E-05 | 0.00011 | 7.51E-05 | 6.55E-05 | 3.49E-05 | 4.54E-05 | 1.79E-05 | 0 | 3.60E-05 | 8.07E-07 | 1.33E-05 | 2.99E-05 | 4.03E-06 | 2.13E-05 | 1.90E-05 | 7.22E-06 | 1.18E-05 | 0 |
| Zfp317 | 9.11E-05 | 6.41E-05 | 1.20E-05 | 0 | 1.98E-05 | 2.88E-05 | 1.79E-05 | 0 | 3.60E-05 | 1.95E-05 | 0 | 4.27E-05 | 0 | 0 | 0.00014 | 5.46E-06 | 0 | 1.70E-05 |
| Gigyf2 | 7.31E-05 | 5.18E-05 | 2.26E-05 | 0 | 2.21E-05 | 2.48E-05 | 6.47E-05 | 9.23E-06 | 3.60E-05 | 9.97E-06 | 0 | 3.69E-05 | 1.37E-05 | 0 | 5.42E-05 | 1.23E-05 | 0 | 0 |

| | | | | | | | | | | | | | | | | | | |
|---|---|---|---|---|---|---|---|---|---|---|---|---|---|---|---|---|---|---|
| Uqcc1 | 5.60E-05 | 5.50E-05 | 5.12E-05 | 8.60E-05 | 1.16E-05 | 2.14E-05 | 2.40E-05 | 0 | 3.60E-05 | 9.14E-06 | 9.38E-06 | 2.87E-05 | 1.72E-05 | 9.75E-06 | 8.49E-05 | 6.20E-06 | 1.42E-05 | 2.27E-06 |
| Xpot | 0.00017 | 4.03E-05 | 0.00025 | 4.51E-05 | 1.07E-05 | 1.88E-05 | 0.00018 | 0 | 3.60E-05 | 3.40E-05 | 2.73E-05 | 0.00012 | 5.27E-06 | 4.74E-06 | 7.47E-05 | 5.50E-06 | 0 | 0 |
| Thumpd3 | 4.48E-05 | 1.17E-05 | 2.98E-05 | 0 | 0 | 1.84E-05 | 0 | 2.71E-05 | 3.60E-05 | 8.81E-06 | 5.10E-06 | 1.73E-05 | 0 | 8.71E-07 | 1.53E-05 | 9.66E-06 | 0 | 2.18E-05 |
| Pdlim4 | 0.00013 | 4.27E-05 | 0.00051 | 0 | 0 | 0 | 0.00027 | 0.0002 | 3.60E-05 | 2.55E-05 | 0.0001 | 0.0003 | 0 | 4.20E-06 | 0.00015 | 2.49E-05 | 1.68E-05 | 0.00022 |
| Irgq | 0 | 9.14E-05 | 8.75E-05 | 0.00027 | 7.18E-05 | 0 | 1.66E-05 | 4.05E-05 | 3.60E-05 | 3.96E-05 | 6.62E-05 | 0.00021 | 0 | 0.00011 | 0.00026 | 0 | 5.87E-06 | 0.0001 |
| Ncoa6 | 0.00017 | 0.00016 | 8.12E-05 | 0 | 6.36E-05 | 0.00016 | 7.47E-05 | 0 | 3.59E-05 | 5.54E-05 | 5.51E-05 | 0.00011 | 2.23E-06 | 7.04E-05 | 4.67E-05 | 7.93E-05 | 4.88E-05 | 4.05E-05 |
| Med21 | 0.00023 | 7.35E-05 | 9.89E-05 | 9.24E-05 | 1.15E-05 | 7.66E-05 | 0.00011 | 0 | 3.59E-05 | 7.47E-05 | 0.00012 | 0.0001 | 2.93E-06 | 5.27E-05 | 0 | 0.00012 | 0 | 1.36E-05 |
| Aimp1 | 0.00029 | 0.00013 | 0.00019 | 0.00014 | 2.13E-05 | 5.76E-05 | 0 | 0 | 3.59E-05 | 0 | 2.13E-05 | 9.12E-05 | 0 | 4.20E-05 | 0.00011 | 0.00014 | 6.06E-05 | 1.21E-05 |
| U2surp | 0.00018 | 0.00014 | 9.42E-05 | 0 | 5.16E-05 | 5.14E-05 | 0 | 0 | 3.59E-05 | 2.86E-05 | 2.20E-05 | 0.00014 | 0 | 1.86E-05 | 0.00025 | 0.00016 | 0 | 2.83E-05 |
| Zbtb37 | 5.22E-05 | 6.21E-05 | 0 | 0 | 2.59E-05 | 2.92E-05 | 3.35E-07 | 3.52E-05 | 3.59E-05 | 1.21E-05 | 1.07E-05 | 3.53E-05 | 4.07E-06 | 1.57E-05 | 9.36E-05 | 0 | 1.84E-05 | 1.73E-05 |
| Rdh14 | 5.87E-05 | 3.12E-05 | 0 | 0 | 2.57E-05 | 2.70E-05 | 4.41E-06 | 1.65E-05 | 3.59E-05 | 2.08E-05 | 0 | 3.02E-05 | 7.36E-06 | 1.81E-06 | 6.76E-06 | 1.77E-05 | 0 | 1.42E-05 |
| Orai1 | 1.26E-05 | 5.55E-05 | 5.72E-05 | 4.91E-05 | 3.64E-05 | 0 | 0 | 5.65E-05 | 3.59E-05 | 6.49E-05 | 4.12E-05 | 8.04E-06 | 0 | 4.19E-05 | 3.63E-05 | 1.33E-05 | 0 | 4.94E-05 |
| Pnisr | 0.00015 | 0.00022 | 6.66E-05 | 9.14E-06 | 8.03E-05 | 4.76E-05 | 5.34E-05 | 2.49E-05 | 3.58E-05 | 5.38E-05 | 0 | 5.52E-05 | 1.92E-05 | 4.98E-05 | 0.00083 | 7.83E-05 | 0 | 6.66E-05 |
| Me1 | 7.29E-05 | 7.52E-08 | 2.19E-05 | 0 | 0.00018 | 4.14E-05 | 7.55E-05 | 0 | 3.58E-05 | 9.05E-05 | 0.00024 | 6.61E-05 | 0 | 4.08E-05 | 0 | 6.53E-05 | 2.30E-05 | 3.28E-05 |
| Rfx1 | 8.95E-06 | 7.44E-05 | 8.28E-06 | 0.00014 | 2.22E-05 | 3.10E-05 | 6.70E-06 | 0.00012 | 3.58E-05 | 1.40E-05 | 5.91E-05 | 3.51E-05 | 2.01E-05 | 5.31E-05 | 0.00016 | 9.22E-05 | 0.00017 | 6.57E-05 |
| Acp6 | 1.26E-05 | 5.82E-05 | 0 | 2.23E-05 | 5.17E-05 | 1.65E-05 | 0 | 1.76E-05 | 3.58E-05 | 5.00E-06 | 0 | 4.75E-05 | 0 | 3.37E-05 | 1.44E-05 | 3.57E-06 | 1.98E-05 | 4.46E-06 |
| Rgs12 | 0 | 0.00017 | 0.0001 | 4.80E-05 | 9.58E-05 | 5.87E-06 | 9.04E-05 | 6.11E-05 | 3.58E-05 | 6.75E-05 | 6.05E-05 | 6.59E-05 | 8.16E-05 | 4.19E-05 | 0.0001 | 0.00018 | 7.15E-05 | 0.00013 |
| Cpne8 | 0 | 0.0001 | 3.03E-05 | 0.00012 | 0 | 0 | 9.31E-05 | 0 | 3.58E-05 | 0.00032 | 0.00012 | 0 | 0.00027 | 3.05E-05 | 4.14E-05 | 0 | 0 | 0.00036 |
| Syngap1 | 0 | 0.00019 | 0 | 7.64E-05 | 0.00016 | 0.0002 | 2.93E-05 | 0.0001 | 3.57E-05 | 7.99E-05 | 1.60E-05 | 8.93E-05 | 5.18E-05 | 7.40E-05 | 0.0002 | 0 | 0.00028 | 6.10E-05 |
| Adgra3 | 0.0001 | 0.00011 | 0.00016 | 0 | 0.00011 | 8.23E-05 | 9.64E-05 | 0.00016 | 3.57E-05 | 0 | 3.42E-05 | 0 | 2.12E-05 | 5.15E-05 | 0 | 0.00015 | 4.44E-05 | 3.47E-05 |
| Mynn | 6.64E-05 | 9.66E-05 | 2.61E-05 | 0 | 3.47E-05 | 5.61E-05 | 4.96E-06 | 0 | 3.57E-05 | 1.75E-05 | 0 | 4.39E-05 | 0 | 3.25E-05 | 6.76E-05 | 6.33E-05 | 1.89E-05 | 1.26E-05 |
| Lrif1 | 5.07E-05 | 3.94E-05 | 0 | 0 | 1.21E-05 | 4.60E-05 | 0 | 4.38E-05 | 3.57E-05 | 3.52E-06 | 0 | 3.95E-05 | 0 | 0 | 2.68E-05 | 8.78E-06 | 0 | 1.73E-05 |
| Mbd6 | 1.70E-05 | 6.70E-05 | 1.43E-05 | 6.35E-05 | 4.32E-05 | 3.25E-05 | 1.29E-05 | 1.95E-05 | 3.57E-05 | 2.43E-05 | 2.66E-06 | 4.52E-05 | 1.04E-05 | 4.24E-05 | 9.54E-05 | 2.00E-05 | 1.18E-05 | 3.74E-05 |
| Bcorl1 | 2.80E-05 | 7.17E-05 | 4.57E-05 | 5.55E-05 | 2.30E-05 | 2.17E-05 | 2.49E-05 | 1.60E-05 | 3.57E-05 | 1.24E-05 | 3.77E-05 | 0.0001 | 1.23E-05 | 5.74E-06 | 0.0001 | 7.48E-07 | 7.61E-05 | 3.99E-05 |
| Gfm2 | 5.42E-05 | 4.48E-05 | 1.77E-05 | 3.06E-05 | 2.46E-05 | 1.37E-05 | 2.22E-05 | 8.08E-06 | 3.57E-05 | 8.04E-06 | 4.80E-06 | 3.44E-05 | 0 | 0 | 6.67E-05 | 2.03E-05 | 0 | 1.47E-05 |
| Atp5c1 | 3.38E-05 | 1.68E-05 | 2.28E-05 | 5.25E-05 | 1.16E-05 | 1.29E-05 | 0 | 2.90E-05 | 3.57E-05 | 1.49E-06 | 2.44E-05 | 1.32E-05 | 0 | 2.10E-05 | 1.26E-05 | 1.83E-05 | 0 | 0 |
| Raet1d | 5.38E-06 | 0 | 0 | 0 | 0 | 1.02E-05 | 0 | 2.34E-05 | 3.57E-05 | 0 | 0 | 3.28E-05 | 0 | 0 | 0 | 0.00014 | 0 | 1.97E-05 |
| Gpatch2 | 3.01E-05 | 6.53E-05 | 2.20E-05 | 4.59E-05 | 3.27E-05 | 6.26E-06 | 1.88E-05 | 3.33E-05 | 3.57E-05 | 2.34E-05 | 2.26E-05 | 1.86E-07 | 3.89E-05 | 7.57E-06 | 0.0002 | 2.25E-06 | 9.06E-05 | 4.87E-05 |
| Hinfp | 1.60E-05 | 5.69E-05 | 1.83E-06 | 3.39E-05 | 1.85E-05 | 0 | 1.61E-05 | 3.45E-05 | 3.57E-05 | 5.93E-06 | 7.15E-06 | 3.65E-05 | 0 | 1.68E-05 | 3.05E-05 | 4.51E-05 | 1.55E-05 | 1.52E-05 |
| Sec14l1 | 0.00018 | 0.00015 | 7.93E-05 | 0.00011 | 0 | 5.20E-05 | 0.00021 | 0.00022 | 3.56E-05 | 0.00035 | 4.37E-05 | 5.03E-05 | 0 | 0 | 7.02E-05 | 0.00015 | 7.44E-05 | 0.00016 |
| Slc10a6 | 0 | 1.58E-05 | 0 | 5.55E-06 | 0 | 4.16E-05 | 0 | 0 | 3.56E-05 | 0.00027 | 0 | 0 | 0.00044 | 2.47E-05 | 0 | 0 | 0 | 0 |
| Phka2 | 2.09E-05 | 5.92E-05 | 0 | 0 | 1.94E-05 | 3.27E-05 | 0 | 0 | 3.56E-05 | 3.65E-05 | 1.62E-05 | 2.27E-05 | 7.65E-06 | 1.70E-05 | 9.36E-05 | 0.00011 | 0 | 3.87E-05 |
| Hist3h2a | 5.47E-05 | 5.73E-05 | 0 | 0 | 1.90E-05 | 2.67E-05 | 0 | 2.08E-05 | 3.56E-05 | 7.31E-06 | 0 | 8.61E-05 | 0 | 1.30E-05 | 8.20E-06 | 7.54E-07 | 8.17E-05 | 3.76E-05 |

| | | | | | | | | | | | | | | | | | | |
|---|---|---|---|---|---|---|---|---|---|---|---|---|---|---|---|---|---|---|
| Ptcd2 | 7.54E-05 | 3.40E-05 | 4.63E-05 | 0 | 0 | 2.59E-05 | 0 | 0 | 3.56E-05 | 8.79E-06 | 0 | 0 | 2.05E-06 | 0 | 3.70E-05 | 3.07E-05 | 0 | 5.65E-06 |
| Brsk1 | 3.39E-05 | 6.19E-05 | 5.93E-05 | 3.49E-06 | 4.91E-05 | 2.22E-05 | 2.11E-05 | 2.58E-05 | 3.56E-05 | 3.21E-06 | 4.53E-05 | 4.96E-05 | 0 | 8.74E-06 | 5.92E-05 | 0 | 2.71E-05 | 0 |
| Rad51d | 5.66E-06 | 5.93E-05 | 0 | 6.15E-05 | 3.02E-05 | 1.05E-05 | 3.96E-05 | 2.75E-05 | 3.56E-05 | 3.14E-05 | 1.22E-05 | 2.91E-05 | 0 | 0 | 3.74E-05 | 3.33E-05 | 0 | 6.90E-05 |
| Klf11 | 3.29E-05 | 0.00012 | 9.61E-06 | 1.47E-05 | 0 | 0 | 3.18E-05 | 4.25E-05 | 3.56E-05 | 4.39E-05 | 5.88E-05 | 0 | 2.90E-05 | 6.81E-05 | 3.72E-05 | 7.33E-05 | 8.50E-05 | 8.19E-05 |
| Bloc1s4 | 0 | 4.94E-05 | 6.34E-05 | 0.00015 | 3.41E-05 | 0 | 0 | 3.31E-05 | 3.56E-05 | 2.87E-05 | 4.90E-06 | 2.82E-05 | 1.16E-05 | 0 | 0.00019 | 6.22E-05 | 1.06E-05 | 5.39E-05 |
| Ildr2 | 0 | 0 | 0 | 0 | 1.63E-05 | 0 | 0 | 0 | 3.56E-05 | 0.00016 | 4.22E-05 | 0 | 0 | 0 | 0 | 6.44E-05 | 0 | 0.00025 |
| Taf5l | 5.94E-05 | 9.32E-05 | 0.00013 | 9.76E-05 | 2.51E-05 | 0 | 4.80E-07 | 1.44E-05 | 3.56E-05 | 1.57E-05 | 8.35E-05 | 2.59E-05 | 3.44E-05 | 4.98E-05 | 4.41E-05 | 5.75E-05 | 7.47E-05 | 1.26E-05 |
| Ttbk2 | 5.15E-05 | 5.79E-05 | 1.68E-05 | 0 | 3.13E-05 | 5.14E-05 | 1.29E-05 | 7.86E-07 | 3.55E-05 | 1.97E-05 | 1.63E-05 | 3.41E-05 | 2.85E-05 | 1.50E-05 | 0 | 5.01E-05 | 9.17E-05 | 2.96E-05 |
| Prkar2b | 1.42E-05 | 2.04E-06 | 0 | 0 | 0 | 4.98E-05 | 0 | 0 | 3.55E-05 | 0 | 0 | 6.19E-05 | 0 | 0.0003 | 0 | 3.29E-05 | 0 | 2.02E-05 |
| Pi4kb | 7.49E-05 | 6.56E-05 | 5.42E-05 | 0 | 3.59E-05 | 4.05E-05 | 6.82E-05 | 0 | 3.55E-05 | 3.46E-05 | 0 | 2.09E-05 | 2.88E-05 | 0 | 3.72E-05 | 6.26E-06 | 1.05E-05 | 1.94E-05 |
| Tango2 | 7.64E-05 | 5.56E-05 | 2.75E-05 | 0 | 2.11E-05 | 3.63E-05 | 4.81E-05 | 2.70E-05 | 3.55E-05 | 6.49E-05 | 5.69E-05 | 3.02E-05 | 2.09E-05 | 1.35E-05 | 0 | 4.64E-05 | 0 | 8.06E-05 |
| Eef1e1 | 0.00012 | 0 | 0.00022 | 0 | 0 | 2.79E-05 | 0 | 0 | 3.55E-05 | 0 | 1.90E-05 | 3.15E-05 | 0 | 0 | 8.97E-06 | 5.41E-05 | 0 | 0 |
| Ift140 | 1.97E-05 | 3.27E-05 | 1.08E-05 | 6.51E-06 | 2.89E-05 | 1.92E-05 | 6.58E-05 | 9.27E-06 | 3.55E-05 | 4.77E-05 | 1.47E-05 | 1.16E-05 | 2.84E-05 | 0 | 6.16E-05 | 6.94E-05 | 1.21E-05 | 4.20E-05 |
| Ifi35 | 8.48E-05 | 5.96E-05 | 1.42E-05 | 5.15E-05 | 6.98E-06 | 6.26E-06 | 0 | 0 | 3.55E-05 | 0.00018 | 0 | 1.65E-06 | 2.54E-05 | 2.83E-05 | 5.27E-05 | 4.18E-05 | 8.95E-06 | 0.00015 |
| Taco1 | 6.44E-05 | 3.71E-05 | 3.06E-05 | 2.92E-05 | 1.05E-05 | 4.67E-07 | 0 | 0 | 3.55E-05 | 1.44E-05 | 5.39E-06 | 0 | 0 | 9.17E-06 | 2.43E-05 | 5.70E-05 | 0 | 2.13E-05 |
| Tmco1 | 0.00041 | 1.47E-05 | 0.00022 | 0.00023 | 0.00014 | 0.00024 | 0.00023 | 0.00013 | 3.54E-05 | 0.00011 | 3.56E-05 | 4.37E-05 | 0.00011 | 0.00012 | 0 | 0.00021 | 0 | 2.83E-05 |
| Ttc17 | 8.25E-05 | 0.0001 | 7.92E-05 | 0 | 7.85E-05 | 0.00012 | 8.42E-05 | 6.96E-05 | 3.54E-05 | 4.56E-05 | 3.50E-05 | 6.10E-05 | 1.12E-05 | 5.70E-05 | 4.94E-05 | 1.47E-05 | 3.14E-05 | 5.21E-05 |
| Mtf1 | 8.91E-05 | 4.80E-05 | 3.84E-05 | 0 | 3.17E-05 | 2.92E-05 | 0 | 0 | 3.54E-05 | 1.57E-05 | 2.74E-05 | 2.24E-05 | 1.74E-06 | 0 | 2.96E-05 | 1.51E-05 | 1.07E-05 | 4.05E-06 |
| Zfp263 | 3.16E-05 | 6.71E-05 | 3.11E-05 | 1.58E-05 | 3.19E-05 | 2.24E-05 | 0 | 0 | 3.54E-05 | 4.32E-06 | 0 | 5.28E-05 | 0 | 3.05E-05 | 0.00016 | 1.75E-05 | 0 | 2.21E-05 |
| Faap100 | 0 | 4.09E-05 | 2.20E-05 | 7.50E-05 | 2.29E-05 | 0 | 0 | 0 | 3.54E-05 | 3.57E-05 | 2.43E-05 | 3.93E-05 | 0 | 1.09E-05 | 7.01E-05 | 6.51E-05 | 0.0001 | 2.38E-05 |
| Fam160b1 | 0.0002 | 0.00016 | 0.00017 | 5.60E-05 | 0.00013 | 0.00012 | 0.00023 | 7.09E-05 | 3.53E-05 | 2.08E-05 | 5.32E-05 | 0.00011 | 5.59E-05 | 0.00013 | 0.00031 | 9.49E-05 | 0.00016 | 3.09E-05 |
| Siva1 | 4.42E-05 | 3.79E-05 | 8.34E-06 | 3.45E-05 | 4.82E-05 | 6.37E-05 | 1.97E-05 | 4.10E-05 | 3.53E-05 | 0 | 0.00015 | 5.13E-05 | 3.42E-05 | 2.04E-05 | 0 | 0.00021 | 0 | 1.03E-05 |
| Uba6 | 0.0001 | 4.96E-05 | 9.54E-05 | 0 | 1.85E-05 | 6.17E-05 | 3.55E-05 | 0 | 3.53E-05 | 6.20E-05 | 9.37E-05 | 8.45E-05 | 2.16E-06 | 0 | 0 | 0.0001 | 3.30E-06 | 1.55E-05 |
| Tfam | 0.00011 | 7.75E-05 | 0.00015 | 1.20E-05 | 3.36E-05 | 2.64E-05 | 2.89E-05 | 7.70E-05 | 3.53E-05 | 2.29E-05 | 0 | 2.64E-05 | 3.60E-05 | 4.26E-05 | 0.00012 | 9.70E-05 | 0 | 3.80E-05 |
| Cep162 | 3.08E-05 | 2.87E-05 | 1.16E-05 | 0 | 1.83E-05 | 4.83E-06 | 2.56E-05 | 4.99E-06 | 3.53E-05 | 2.06E-05 | 0 | 6.08E-06 | 1.18E-05 | 2.20E-05 | 5.20E-05 | 5.97E-05 | 0 | 3.74E-05 |
| Fpgs | 0.00011 | 4.72E-05 | 0.00018 | 0.00011 | 3.19E-05 | 0 | 5.26E-05 | 0 | 3.53E-05 | 4.79E-05 | 8.87E-05 | 0 | 9.93E-05 | 0 | 8.10E-05 | 2.53E-05 | 0 | 4.22E-05 |
| Tnfsf8 | 7.23E-06 | 0 | 0 | 0 | 0 | 0 | 5.15E-05 | 0 | 3.53E-05 | 2.65E-05 | 0 | 0 | 4.38E-07 | 0 | 0 | 1.04E-05 | 0 | 0.00018 |
| Obsl1 | 0.00017 | 4.51E-05 | 0.00016 | 0.00036 | 3.16E-05 | 0 | 0.00031 | 0.00015 | 3.53E-05 | 3.63E-05 | 5.75E-05 | 5.03E-05 | 0.00017 | 5.02E-05 | 0.00031 | 8.84E-05 | 8.95E-06 | 0.00015 |
| Cand2 | 0 | 2.87E-05 | 0 | 5.71E-05 | 3.96E-05 | 0 | 0 | 0 | 3.53E-05 | 6.85E-06 | 1.72E-05 | 4.83E-06 | 2.25E-05 | 0 | 2.04E-05 | 0.00011 | 7.31E-06 | 3.20E-05 |
| Parp8 | 0.00014 | 0.00012 | 1.54E-05 | 0 | 0.00026 | 8.45E-05 | 0.00028 | 0.00028 | 3.52E-05 | 0 | 0 | 4.83E-05 | 0 | 0 | 0.00022 | 5.12E-05 | 0.00022 | 2.46E-06 |
| Stam2 | 0.00018 | 0.0002 | 0.00018 | 6.76E-05 | 6.67E-05 | 7.83E-05 | 2.26E-05 | 3.34E-05 | 3.52E-05 | 3.04E-05 | 7.89E-05 | 0.00013 | 4.28E-05 | 5.50E-05 | 3.99E-06 | 5.71E-05 | 8.07E-05 | 2.91E-05 |
| Tfcp2 | 6.62E-05 | 6.65E-05 | 3.38E-05 | 9.39E-06 | 2.88E-05 | 5.13E-05 | 3.54E-05 | 2.44E-05 | 3.52E-05 | 3.08E-05 | 6.17E-06 | 1.41E-05 | 1.43E-06 | 0 | 6.30E-05 | 0 | 3.41E-05 | 3.14E-05 |
| Tmem19 | 8.90E-05 | 0.00011 | 7.14E-06 | 0.00022 | 0.00012 | 0.00013 | 0.00016 | 8.13E-05 | 3.51E-05 | 8.63E-05 | 0.00018 | 5.96E-05 | 7.18E-05 | 7.45E-05 | 3.35E-05 | 7.73E-05 | 0 | 9.45E-05 |

| | | | | | | | | | | | | | | | | | | |
|---|---|---|---|---|---|---|---|---|---|---|---|---|---|---|---|---|---|---|
| Plekha1 | 0.00014 | 7.79E-05 | 0.00011 | 0 | 0.00012 | 7.81E-05 | 9.95E-05 | 1.73E-05 | 3.51E-05 | 4.60E-05 | 0.00011 | 9.49E-05 | 0 | 4.85E-05 | 0.00011 | 2.69E-05 | 4.18E-05 | 0 |
| Rtl8a | 1.90E-06 | 2.18E-05 | 1.11E-05 | 3.09E-05 | 1.81E-05 | 1.61E-05 | 4.57E-07 | 4.64E-05 | 3.51E-05 | 2.24E-05 | 4.00E-05 | 0 | 2.37E-05 | 8.27E-06 | 5.27E-06 | 0 | 4.51E-06 | 4.23E-05 |
| Prtg | 2.29E-06 | 1.98E-05 | 1.40E-05 | 0 | 3.24E-05 | 0 | 3.95E-07 | 3.12E-06 | 3.51E-05 | 0 | 0 | 9.89E-06 | 0 | 0 | 8.02E-06 | 0 | 4.39E-06 | 1.18E-05 |
| Lias | 6.76E-05 | 5.67E-05 | 3.67E-05 | 7.42E-05 | 2.25E-05 | 2.91E-05 | 2.53E-05 | 1.00E-05 | 3.50E-05 | 6.48E-06 | 0 | 1.84E-05 | 0 | 3.81E-05 | 1.64E-05 | 6.05E-05 | 0 | 3.46E-06 |
| Coq10a | 1.72E-05 | 9.60E-05 | 2.20E-06 | 3.46E-05 | 3.97E-05 | 2.70E-05 | 7.29E-05 | 2.60E-05 | 3.50E-05 | 5.61E-05 | 1.44E-05 | 1.93E-05 | 1.22E-05 | 6.03E-05 | 3.02E-05 | 1.30E-05 | 1.79E-05 | 6.44E-05 |
| Fbxl12 | 1.82E-05 | 9.94E-05 | 9.62E-06 | 6.24E-05 | 3.97E-05 | 1.80E-05 | 0 | 3.56E-05 | 3.50E-05 | 0 | 4.27E-05 | 7.31E-05 | 0 | 6.90E-05 | 0.00011 | 2.14E-05 | 0 | 2.16E-05 |
| Nphp3 | 4.29E-05 | 1.69E-05 | 2.65E-06 | 4.08E-05 | 3.32E-05 | 1.67E-05 | 4.86E-05 | 0 | 3.50E-05 | 2.70E-05 | 0 | 3.15E-05 | 7.17E-06 | 1.28E-05 | 7.75E-05 | 2.73E-06 | 5.49E-05 | 2.73E-05 |
| Eif2b3 | 7.94E-05 | 6.37E-05 | 0.00013 | 0 | 0 | 4.75E-06 | 0 | 0 | 3.50E-05 | 1.21E-05 | 9.31E-06 | 1.75E-05 | 0 | 1.04E-05 | 7.75E-05 | 4.78E-05 | 0 | 2.04E-05 |
| Arrdc1 | 6.04E-06 | 4.61E-05 | 0 | 8.71E-06 | 4.88E-06 | 2.35E-06 | 2.87E-05 | 0 | 3.50E-05 | 4.34E-05 | 3.73E-05 | 3.83E-05 | 1.36E-05 | 3.97E-05 | 2.33E-05 | 4.28E-06 | 2.69E-06 | 4.85E-05 |
| Iffo1 | 0 | 0.0001 | 0 | 5.02E-05 | 2.24E-05 | 0 | 3.05E-05 | 0 | 3.50E-05 | 7.16E-05 | 8.75E-06 | 0 | 7.34E-05 | 2.37E-05 | 0.00031 | 9.14E-05 | 0.00021 | 6.47E-05 |
| Fbxo38 | 0.00012 | 0.00018 | 9.50E-05 | 7.94E-06 | 0.00011 | 0.00011 | 7.42E-05 | 2.87E-05 | 3.49E-05 | 4.07E-05 | 8.37E-05 | 9.68E-05 | 4.84E-05 | 6.96E-05 | 7.45E-05 | 4.41E-05 | 0 | 6.21E-05 |
| Sharpin | 3.95E-05 | 6.76E-05 | 3.84E-05 | 0.00011 | 6.92E-05 | 4.59E-05 | 2.54E-05 | 4.19E-05 | 3.49E-05 | 2.89E-05 | 2.81E-05 | 1.70E-05 | 1.89E-05 | 5.44E-05 | 2.75E-05 | 1.99E-05 | 1.36E-05 | 5.69E-05 |
| Iqcc | 3.46E-05 | 4.57E-05 | 7.80E-06 | 3.65E-05 | 1.93E-05 | 2.81E-05 | 0 | 2.47E-05 | 3.49E-05 | 3.98E-05 | 7.22E-06 | 3.48E-05 | 9.20E-06 | 2.20E-05 | 9.29E-05 | 6.92E-06 | 0 | 3.74E-05 |
| Pramef8 | 5.93E-05 | 7.13E-05 | 1.45E-05 | 4.93E-05 | 1.89E-05 | 2.51E-05 | 6.16E-06 | 1.99E-05 | 3.49E-05 | 3.86E-05 | 1.96E-05 | 7.55E-05 | 6.81E-06 | 5.70E-06 | 1.41E-05 | 4.08E-05 | 1.78E-07 | 4.83E-05 |
| Zbtb9 | 3.31E-05 | 5.98E-05 | 1.18E-05 | 3.57E-05 | 2.32E-05 | 1.72E-05 | 1.16E-05 | 1.26E-05 | 3.49E-05 | 8.24E-06 | 1.84E-05 | 0 | 0 | 3.97E-05 | 4.97E-05 | 1.74E-05 | 0 | 2.49E-05 |
| Lrsam1 | 2.45E-07 | 2.56E-05 | 0 | 1.60E-05 | 1.84E-05 | 0 | 2.27E-05 | 1.39E-05 | 3.49E-05 | 6.19E-05 | 1.38E-05 | 1.97E-05 | 0 | 2.94E-06 | 0 | 3.49E-06 | 0 | 5.30E-05 |
| Csrnp2 | 0 | 4.25E-05 | 9.34E-05 | 4.66E-05 | 4.34E-05 | 0 | 0 | 9.18E-05 | 3.49E-05 | 0 | 7.14E-05 | 3.32E-05 | 1.39E-05 | 4.05E-06 | 0.00012 | 8.93E-05 | 0.00029 | 0 |
| Smchd1 | 9.09E-05 | 8.62E-05 | 0.00011 | 1.08E-05 | 4.35E-05 | 8.41E-05 | 4.58E-05 | 0 | 3.48E-05 | 2.25E-05 | 6.45E-05 | 3.33E-05 | 4.56E-05 | 0 | 4.43E-05 | 0.00043 | 8.24E-05 | 6.06E-05 |
| Zfp637 | 0.00012 | 0.0001 | 4.94E-05 | 2.90E-05 | 6.94E-05 | 4.25E-05 | 2.84E-05 | 1.50E-05 | 3.48E-05 | 5.93E-05 | 4.98E-05 | 3.80E-05 | 7.15E-05 | 9.85E-06 | 2.31E-05 | 5.10E-05 | 0 | 2.87E-05 |
| Slain2 | 0.00019 | 0.00012 | 0.00015 | 0 | 5.36E-05 | 3.69E-05 | 0 | 2.58E-05 | 3.48E-05 | 5.83E-05 | 6.29E-05 | 6.07E-05 | 0 | 0 | 6.06E-05 | 0.00012 | 0.0001 | 0 |
| Bbs5 | 3.71E-05 | 2.50E-05 | 2.64E-05 | 1.29E-05 | 3.36E-05 | 1.76E-05 | 0 | 1.83E-06 | 3.48E-05 | 1.97E-06 | 3.56E-05 | 6.64E-05 | 1.53E-05 | 3.44E-05 | 3.00E-05 | 2.49E-05 | 0 | 1.30E-05 |
| Tsen54 | 1.25E-05 | 2.92E-05 | 3.14E-05 | 5.38E-05 | 5.63E-06 | 1.26E-05 | 0 | 0 | 3.48E-05 | 1.50E-05 | 1.65E-05 | 0 | 0 | 3.93E-05 | 2.74E-05 | 9.71E-06 | 0 | 1.39E-05 |
| Kctd13 | 9.33E-07 | 5.37E-05 | 5.76E-05 | 5.83E-05 | 2.98E-05 | 1.14E-05 | 0 | 8.59E-05 | 3.48E-05 | 4.22E-05 | 6.89E-05 | 1.67E-05 | 2.50E-05 | 3.87E-06 | 6.44E-06 | 4.88E-05 | 8.26E-05 | 7.20E-06 |
| Rbl1 | 2.80E-05 | 0 | 1.26E-05 | 0 | 0 | 7.23E-06 | 3.69E-06 | 0 | 3.48E-05 | 2.09E-05 | 9.37E-06 | 0 | 0 | 0 | 0 | 0.00047 | 0 | 0 |
| Gprasp1 | 0 | 6.73E-05 | 0 | 4.85E-05 | 7.59E-05 | 0 | 2.03E-05 | 4.31E-06 | 3.48E-05 | 2.59E-05 | 0 | 8.80E-06 | 1.81E-05 | 1.40E-05 | 3.40E-05 | 7.29E-06 | 6.50E-05 | 7.87E-05 |
| Atf6 | 0.00016 | 0.00019 | 0.00017 | 0 | 0.00017 | 0.00012 | 0 | 0 | 3.47E-05 | 6.33E-06 | 0.00011 | 0.00019 | 4.38E-05 | 0.00012 | 7.06E-05 | 5.92E-05 | 0.00015 | 0 |
| Kctd20 | 1.65E-05 | 0.00014 | 6.98E-05 | 0.00022 | 9.13E-05 | 0.0001 | 8.28E-05 | 8.36E-05 | 3.47E-05 | 9.01E-05 | 8.27E-05 | 3.41E-05 | 0 | 4.41E-05 | 7.61E-05 | 0.00011 | 0.00015 | 7.42E-05 |
| Utp18 | 5.78E-05 | 4.79E-05 | 0.00019 | 3.58E-05 | 1.51E-05 | 9.08E-06 | 0 | 0 | 3.47E-05 | 1.37E-05 | 5.13E-05 | 4.77E-07 | 9.57E-06 | 4.91E-05 | 6.40E-05 | 2.74E-05 | 5.95E-05 | 1.99E-05 |
| Pter | 1.94E-05 | 1.92E-05 | 1.69E-05 | 1.64E-05 | 0 | 0 | 6.36E-05 | 1.13E-06 | 3.47E-05 | 4.21E-05 | 2.80E-05 | 2.21E-05 | 6.14E-05 | 0 | 0 | 1.71E-05 | 1.35E-05 | 3.53E-05 |
| Eif4ebp1 | 0.00036 | 0.00027 | 0.00036 | 0.00028 | 0.0001 | 0.00029 | 0.00041 | 0 | 3.46E-05 | 9.77E-05 | 0.00033 | 0.00092 | 0.00012 | 0.00022 | 0 | 5.48E-05 | 0 | 3.82E-05 |
| Gbe1 | 2.74E-05 | 0 | 0 | 0 | 0 | 3.93E-05 | 0 | 9.25E-07 | 3.46E-05 | 2.04E-05 | 0 | 0.00062 | 6.45E-05 | 2.52E-05 | 1.74E-05 | 8.50E-06 | 0 | 3.18E-05 |
| Cnot9 | 0.00015 | 3.54E-05 | 0.00015 | 0 | 3.38E-05 | 3.86E-05 | 6.66E-05 | 0 | 3.46E-05 | 3.07E-05 | 2.02E-05 | 4.97E-05 | 0 | 2.16E-05 | 0 | 0.00031 | 0 | 0 |
| Hint3 | 6.11E-05 | 3.84E-05 | 7.48E-06 | 8.82E-05 | 3.54E-05 | 2.93E-05 | 6.21E-05 | 2.67E-05 | 3.46E-05 | 2.88E-05 | 4.82E-05 | 2.86E-05 | 3.44E-05 | 1.05E-06 | 3.13E-05 | 3.62E-05 | 0 | 6.19E-05 |

| | | | | | | | | | | | | | | | | | | |
|---|---|---|---|---|---|---|---|---|---|---|---|---|---|---|---|---|---|---|
| Fbxo9 | 7.33E-05 | 8.17E-05 | 0.00013 | 8.88E-05 | 7.87E-05 | 1.62E-05 | 1.98E-05 | 1.27E-05 | 3.46E-05 | 5.59E-05 | 3.14E-05 | 1.74E-05 | 0 | 2.67E-05 | 0.00014 | 1.86E-05 | 0 | 4.94E-05 |
| Vps13a | 3.99E-05 | 4.21E-05 | 0 | 0 | 1.45E-06 | 1.08E-05 | 0 | 1.72E-05 | 3.46E-05 | 4.81E-05 | 2.57E-05 | 1.88E-05 | 6.55E-06 | 0 | 0 | 2.71E-05 | 5.79E-05 | 3.26E-05 |
| Zfp787 | 8.81E-06 | 5.23E-05 | 4.64E-05 | 6.49E-05 | 6.19E-05 | 0 | 0 | 2.62E-05 | 3.46E-05 | 7.98E-06 | 6.18E-06 | 9.50E-05 | 0 | 2.75E-05 | 0.00012 | 1.39E-05 | 8.89E-06 | 3.39E-05 |
| Syde2 | 0 | 3.50E-05 | 0 | 0 | 0.00036 | 0.00013 | 0 | 5.56E-05 | 3.45E-05 | 3.43E-05 | 0.00039 | 0.00011 | 4.16E-05 | 8.08E-05 | 4.04E-05 | 0 | 0 | 0 |
| Ccdc82 | 6.50E-05 | 0.0001 | 0 | 0 | 7.08E-06 | 8.16E-05 | 0 | 6.21E-06 | 3.45E-05 | 0 | 0 | 7.95E-05 | 0 | 0 | 0.00045 | 0.00023 | 5.35E-05 | 2.79E-05 |
| Akap9 | 8.31E-05 | 0.0001 | 6.04E-05 | 0 | 0.00011 | 7.95E-05 | 0.0002 | 0 | 3.45E-05 | 4.99E-05 | 5.01E-05 | 0.00019 | 6.47E-05 | 6.40E-05 | 0.00023 | 0 | 0.00013 | 6.69E-05 |
| Mbd5 | 7.00E-05 | 8.32E-05 | 0 | 0 | 3.75E-05 | 7.17E-05 | 2.22E-05 | 5.19E-05 | 3.45E-05 | 3.51E-05 | 3.92E-05 | 6.41E-05 | 4.87E-05 | 1.13E-05 | 8.85E-05 | 5.84E-06 | 0.0001 | 9.47E-06 |
| Pdp1 | 6.22E-05 | 6.74E-05 | 5.57E-06 | 0 | 0 | 5.87E-05 | 4.30E-05 | 0 | 3.45E-05 | 3.54E-05 | 0 | 3.51E-05 | 0 | 0 | 3.37E-06 | 2.00E-05 | 0 | 6.16E-05 |
| Pcgf6 | 4.20E-05 | 1.97E-05 | 2.23E-05 | 0 | 2.98E-05 | 3.32E-05 | 1.03E-06 | 0 | 3.45E-05 | 0 | 1.33E-05 | 1.25E-05 | 1.87E-05 | 1.39E-05 | 4.51E-05 | 1.14E-05 | 0 | 4.53E-07 |
| Shq1 | 2.34E-05 | 2.68E-05 | 6.69E-05 | 3.28E-05 | 1.22E-05 | 1.56E-05 | 0 | 2.97E-07 | 3.45E-05 | 6.57E-06 | 0 | 2.20E-05 | 2.02E-05 | 2.65E-05 | 5.07E-05 | 1.89E-05 | 0 | 0 |
| Rbm15 | 7.71E-05 | 0.00016 | 8.94E-05 | 0 | 3.25E-05 | 6.17E-05 | 0 | 1.47E-05 | 3.44E-05 | 2.24E-05 | 6.62E-05 | 0.0001 | 0 | 1.28E-05 | 0.00034 | 0.00036 | 5.57E-05 | 2.61E-05 |
| Tpgs2 | 0.00011 | 5.50E-06 | 1.96E-05 | 0 | 1.36E-05 | 2.34E-05 | 4.29E-05 | 9.52E-06 | 3.44E-05 | 5.53E-06 | 4.71E-05 | 0 | 0 | 2.90E-05 | 0 | 8.18E-05 | 0 | 2.31E-06 |
| Inpp4a | 1.17E-05 | 0.00012 | 2.29E-06 | 5.13E-05 | 2.48E-05 | 5.92E-07 | 2.78E-05 | 5.29E-05 | 3.44E-05 | 4.88E-05 | 8.33E-05 | 6.34E-05 | 3.04E-06 | 0 | 0.00013 | 1.16E-05 | 0.0001 | 6.01E-05 |
| Plxna2 | 0 | 4.08E-05 | 3.96E-06 | 0 | 0 | 0 | 1.44E-05 | 0 | 3.44E-05 | 1.06E-05 | 0 | 0 | 2.21E-05 | 3.04E-05 | 2.15E-05 | 2.18E-05 | 1.90E-05 | 5.92E-05 |
| Zfp266 | 0.00026 | 0.00016 | 3.12E-05 | 3.68E-05 | 8.46E-05 | 0.00014 | 0.00016 | 2.55E-05 | 3.43E-05 | 2.71E-05 | 4.44E-05 | 1.00E-05 | 0 | 4.10E-05 | 0.00014 | 2.82E-05 | 8.14E-05 | 5.07E-05 |
| Sos1 | 6.70E-05 | 0.00011 | 5.18E-05 | 4.17E-06 | 6.72E-05 | 2.72E-05 | 6.12E-05 | 3.67E-05 | 3.43E-05 | 1.09E-05 | 4.36E-05 | 5.70E-05 | 2.78E-06 | 2.02E-05 | 5.71E-06 | 5.57E-05 | 4.98E-05 | 2.01E-05 |
| Trim12c | 4.66E-05 | 4.15E-05 | 0 | 0 | 1.72E-05 | 0 | 0 | 0 | 3.43E-05 | 4.05E-05 | 0 | 3.78E-06 | 0 | 0 | 8.95E-06 | 5.35E-06 | 0 | 3.82E-05 |
| Rnf219 | 2.50E-05 | 2.09E-05 | 3.34E-05 | 2.33E-06 | 1.90E-05 | 0 | 0 | 2.32E-05 | 3.43E-05 | 2.51E-05 | 1.42E-05 | 5.53E-06 | 5.16E-06 | 2.91E-05 | 2.01E-05 | 5.02E-05 | 0 | 2.10E-05 |
| Klhl4 | 0 | 1.29E-06 | 0 | 0 | 3.84E-05 | 0 | 0 | 6.14E-05 | 3.43E-05 | 0 | 0 | 0 | 0 | 0 | 0 | 7.82E-07 | 0 | 0 |
| Hmgcl | 8.41E-05 | 0.00013 | 0.00013 | 0.00028 | 0.0001 | 0.00011 | 0.00013 | 8.13E-05 | 3.42E-05 | 6.53E-05 | 2.69E-05 | 0.00034 | 1.93E-05 | 2.86E-05 | 6.71E-05 | 0 | 0 | 2.59E-05 |
| Mapre3 | 8.88E-06 | 0.00012 | 5.29E-05 | 4.00E-05 | 0.00012 | 0.00011 | 2.08E-05 | 9.37E-05 | 3.42E-05 | 7.24E-05 | 8.82E-05 | 6.58E-05 | 8.18E-05 | 8.55E-05 | 0.00011 | 0 | 0.00013 | 0.00011 |
| Rprd1b | 0.00011 | 8.34E-05 | 6.39E-05 | 1.12E-05 | 5.59E-05 | 0.0001 | 0 | 4.27E-06 | 3.42E-05 | 1.94E-05 | 0 | 1.49E-05 | 4.79E-05 | 1.39E-06 | 0.00013 | 0.00019 | 8.22E-06 | 2.79E-05 |
| Vps33a | 0.0001 | 6.48E-05 | 0 | 0 | 4.94E-05 | 6.99E-05 | 0 | 0 | 3.42E-05 | 3.64E-06 | 0 | 2.79E-05 | 0 | 2.71E-06 | 2.73E-05 | 5.64E-06 | 0 | 0 |
| Scamp5 | 1.61E-05 | 7.89E-05 | 3.07E-05 | 0 | 0.00013 | 3.19E-05 | 0 | 6.67E-05 | 3.42E-05 | 0.0001 | 5.69E-05 | 3.34E-05 | 5.60E-05 | 0 | 0 | 0 | 0 | 0 |
| Cox19 | 0.0001 | 4.19E-05 | 6.09E-05 | 8.30E-05 | 4.45E-05 | 2.74E-05 | 3.48E-05 | 1.68E-05 | 3.42E-05 | 1.05E-05 | 3.80E-05 | 2.53E-05 | 0 | 1.86E-05 | 1.98E-05 | 1.32E-05 | 0 | 5.85E-07 |
| Pold2 | 6.12E-05 | 7.59E-07 | 7.47E-05 | 6.31E-05 | 0 | 1.25E-05 | 0 | 0 | 3.42E-05 | 2.17E-05 | 0 | 2.31E-05 | 0 | 0 | 0 | 0.00036 | 0 | 3.74E-05 |
| Clp1 | 4.19E-05 | 4.77E-05 | 2.81E-05 | 1.57E-05 | 3.84E-05 | 5.15E-06 | 1.52E-05 | 4.51E-07 | 3.42E-05 | 2.15E-05 | 5.86E-06 | 1.13E-05 | 0 | 2.08E-05 | 3.09E-05 | 2.81E-05 | 0 | 1.09E-05 |
| Gbp6 | 3.26E-06 | 2.00E-05 | 0 | 0 | 0 | 0 | 0 | 0 | 3.42E-05 | 4.67E-05 | 0 | 0 | 2.10E-05 | 0 | 0 | 2.42E-06 | 0 | 3.21E-05 |
| Zfp467 | 0 | 9.46E-05 | 0 | 5.62E-05 | 5.18E-05 | 0 | 0 | 1.10E-05 | 3.42E-05 | 2.10E-05 | 2.60E-05 | 3.80E-05 | 1.46E-05 | 6.93E-05 | 6.40E-05 | 0 | 0 | 0 |
| Tecpr2 | 1.33E-05 | 2.82E-05 | 5.76E-05 | 1.54E-05 | 7.68E-06 | 0 | 8.00E-06 | 1.87E-06 | 3.42E-05 | 4.52E-05 | 1.33E-05 | 1.86E-05 | 1.50E-06 | 3.78E-05 | 9.61E-05 | 2.90E-05 | 6.80E-05 | 2.13E-05 |
| Ssr4 | 0.0008 | 0 | 0.00027 | 0.00028 | 3.87E-05 | 0.00045 | 0.00078 | 0.00032 | 3.41E-05 | 8.56E-05 | 0.0002 | 0.00018 | 0.00036 | 0.00036 | 0 | 0.00017 | 0 | 0.00014 |
| Copg2 | 6.97E-05 | 6.17E-05 | 0 | 0 | 7.63E-05 | 5.17E-05 | 0 | 0 | 3.41E-05 | 1.45E-05 | 2.63E-05 | 6.08E-05 | 0 | 1.99E-05 | 1.35E-05 | 2.21E-05 | 0 | 2.42E-05 |
| Caap1 | 1.66E-05 | 7.97E-05 | 4.36E-05 | 8.31E-05 | 4.46E-05 | 1.08E-05 | 0 | 1.93E-05 | 3.41E-05 | 1.62E-05 | 4.06E-05 | 9.18E-06 | 3.28E-05 | 0 | 7.38E-05 | 8.33E-05 | 2.19E-06 | 3.08E-05 |

| | | | | | | | | | | | | | | | | | | |
|---|---|---|---|---|---|---|---|---|---|---|---|---|---|---|---|---|---|---|
| Zfp74 | 2.95E-05 | 4.14E-05 | 0 | 2.09E-05 | 1.34E-05 | 5.86E-06 | 1.52E-05 | 0 | 3.41E-05 | 2.98E-05 | 1.88E-05 | 2.24E-05 | 1.68E-05 | 5.10E-05 | 3.17E-05 | 1.77E-05 | 0 | 1.85E-05 |
| Tmem53 | 0 | 1.29E-05 | 0 | 3.48E-05 | 1.09E-06 | 1.70E-06 | 0 | 3.76E-06 | 3.41E-05 | 3.34E-05 | 2.02E-05 | 1.47E-06 | 2.58E-05 | 2.67E-06 | 0 | 6.30E-06 | 0 | 3.47E-05 |
| Abhd10 | 1.76E-05 | 4.22E-05 | 7.79E-06 | 0 | 4.38E-06 | 0 | 2.13E-05 | 0 | 3.41E-05 | 1.72E-05 | 6.39E-05 | 6.82E-06 | 6.32E-05 | 2.04E-05 | 2.49E-05 | 0.00014 | 5.11E-05 | 5.31E-06 |
| Rnf38 | 6.65E-05 | 0.00017 | 4.97E-05 | 0 | 9.87E-05 | 9.44E-05 | 0 | 5.02E-05 | 3.40E-05 | 1.53E-05 | 1.82E-05 | 0.00012 | 2.23E-05 | 0.0001 | 0.00014 | 8.31E-05 | 0.0002 | 4.15E-05 |
| Sirt1 | 3.83E-05 | 6.84E-05 | 0 | 1.46E-06 | 0 | 3.52E-05 | 0 | 0 | 3.40E-05 | 1.00E-06 | 1.94E-05 | 1.40E-05 | 0 | 4.94E-05 | 8.65E-05 | 2.05E-05 | 0.00063 | 0 |
| Zfp87 | 2.45E-05 | 3.01E-05 | 0 | 1.53E-05 | 2.80E-05 | 1.79E-05 | 1.32E-05 | 1.12E-05 | 3.40E-05 | 2.13E-05 | 5.49E-06 | 0 | 0 | 2.71E-05 | 7.05E-07 | 0 | 0 | 3.29E-05 |
| Eya4 | 0 | 9.51E-06 | 0 | 0 | 2.95E-05 | 1.64E-05 | 0.00011 | 0 | 3.40E-05 | 0.00017 | 0 | 7.92E-06 | 9.41E-05 | 0 | 0 | 0 | 6.60E-05 | 0 |
| Lrba | 9.44E-06 | 3.38E-05 | 0 | 2.91E-06 | 2.37E-05 | 6.28E-06 | 4.60E-05 | 0 | 3.40E-05 | 6.70E-05 | 0 | 5.46E-05 | 2.04E-05 | 9.98E-06 | 0 | 3.49E-05 | 1.74E-05 | 6.59E-05 |
| Patl1 | 3.22E-05 | 0.00021 | 0.0001 | 0.00011 | 4.46E-05 | 7.02E-05 | 0 | 2.96E-05 | 3.39E-05 | 3.55E-05 | 5.63E-05 | 0.00021 | 0 | 8.91E-05 | 0.00024 | 0.00013 | 3.08E-05 | 4.54E-05 |
| Plekhm1 | 5.42E-06 | 9.49E-05 | 4.73E-06 | 5.79E-05 | 6.00E-05 | 1.78E-05 | 1.47E-05 | 2.81E-05 | 3.39E-05 | 5.05E-05 | 2.28E-05 | 6.20E-06 | 8.26E-05 | 5.64E-05 | 7.47E-05 | 1.28E-05 | 0 | 3.28E-05 |
| Ap5s1 | 5.33E-06 | 2.75E-05 | 0 | 6.06E-05 | 2.57E-05 | 1.68E-05 | 4.76E-05 | 1.40E-05 | 3.39E-05 | 2.78E-05 | 0 | 1.02E-05 | 1.66E-05 | 8.10E-06 | 2.41E-06 | 5.88E-05 | 0 | 4.32E-05 |
| Efl1 | 9.63E-05 | 6.52E-05 | 6.16E-05 | 0 | 2.87E-05 | 1.47E-05 | 3.68E-06 | 2.88E-05 | 3.39E-05 | 1.80E-05 | 0 | 2.23E-05 | 2.16E-05 | 3.11E-05 | 0.00013 | 1.11E-05 | 0 | 3.14E-05 |
| Zfyve26 | 1.84E-05 | 0.00011 | 0 | 3.68E-05 | 3.47E-05 | 1.06E-06 | 2.16E-05 | 0 | 3.39E-05 | 2.70E-05 | 0 | 3.70E-05 | 1.28E-05 | 5.96E-05 | 0.00015 | 4.24E-05 | 2.89E-05 | 1.10E-05 |
| Cluap1 | 4.97E-05 | 5.95E-05 | 4.75E-05 | 2.32E-05 | 3.96E-05 | 0 | 2.69E-05 | 1.17E-05 | 3.39E-05 | 3.28E-05 | 9.39E-07 | 2.51E-05 | 0 | 0 | 0.0001 | 1.90E-05 | 0 | 6.04E-05 |
| Rab3a | 2.49E-05 | 1.65E-06 | 0 | 2.89E-05 | 2.20E-05 | 0 | 7.13E-06 | 0 | 3.39E-05 | 1.03E-05 | 0 | 0.0001 | 0.00018 | 0 | 0 | 0 | 0.00038 | 0 |
| Cdh13 | 0 | 7.51E-05 | 3.41E-05 | 0 | 0 | 0.00023 | 1.80E-05 | 0 | 3.38E-05 | 0 | 0 | 0 | 0.00011 | 0.00032 | 0 | 0 | 0 | 0.00034 |
| Gcc2 | 0.00012 | 4.78E-05 | 2.96E-05 | 1.63E-05 | 2.97E-05 | 4.46E-05 | 4.31E-05 | 0 | 3.38E-05 | 3.04E-05 | 0 | 4.54E-05 | 3.67E-05 | 1.57E-05 | 8.27E-05 | 3.65E-05 | 1.35E-05 | 3.08E-05 |
| Otud4 | 6.95E-05 | 0.00012 | 0.00014 | 0 | 5.63E-05 | 3.88E-05 | 0 | 1.15E-05 | 3.38E-05 | 5.94E-05 | 4.27E-05 | 0.00011 | 4.11E-06 | 0 | 8.20E-05 | 0.00012 | 8.08E-06 | 6.79E-05 |
| Gpr161 | 1.66E-05 | 8.65E-05 | 4.79E-05 | 1.41E-05 | 3.26E-05 | 3.74E-06 | 3.34E-05 | 1.47E-05 | 3.38E-05 | 4.12E-05 | 7.04E-05 | 2.06E-05 | 1.51E-05 | 5.36E-05 | 6.42E-05 | 5.60E-05 | 1.91E-05 | 4.58E-05 |
| Slc39a8 | 3.17E-05 | 2.50E-05 | 5.28E-06 | 0 | 2.31E-05 | 3.66E-06 | 5.72E-05 | 1.84E-05 | 3.38E-05 | 3.38E-06 | 0 | 3.12E-06 | 3.45E-05 | 1.89E-05 | 0 | 5.60E-05 | 2.18E-05 | 0 |
| Cyren | 2.03E-05 | 0 | 2.77E-05 | 5.78E-05 | 7.67E-07 | 0 | 1.06E-05 | 2.57E-05 | 3.38E-05 | 1.12E-05 | 0 | 0 | 1.10E-06 | 7.07E-06 | 3.84E-05 | 2.79E-05 | 0 | 0 |
| Nat10 | 4.70E-05 | 5.93E-05 | 0.00013 | 8.59E-05 | 1.64E-05 | 0 | 0 | 0 | 3.38E-05 | 2.10E-05 | 2.07E-05 | 1.50E-05 | 1.72E-05 | 1.43E-05 | 8.29E-05 | 4.72E-06 | 4.54E-05 | 3.75E-05 |
| Akap10 | 0.00012 | 9.62E-05 | 3.86E-05 | 0 | 3.85E-05 | 7.31E-05 | 9.75E-06 | 0 | 3.37E-05 | 1.32E-06 | 1.23E-05 | 5.84E-05 | 2.71E-06 | 4.51E-05 | 0.00013 | 3.92E-05 | 1.66E-05 | 2.08E-05 |
| Amacr | 3.89E-05 | 2.22E-06 | 0 | 3.05E-05 | 0.00011 | 0 | 0.00013 | 3.31E-05 | 3.37E-05 | 4.92E-06 | 0 | 0 | 0 | 8.54E-06 | 4.88E-08 | 2.63E-05 | 0 | 4.83E-05 |
| Utp14b | 7.82E-06 | 2.00E-06 | 0 | 9.17E-06 | 0 | 0 | 0 | 1.59E-05 | 3.37E-05 | 0 | 5.52E-06 | 6.16E-07 | 0 | 9.11E-06 | 7.72E-05 | 1.48E-05 | 0.00032 | 1.39E-06 |
| Stx6 | 9.08E-05 | 0.00023 | 0.00019 | 2.23E-05 | 0.00036 | 0.00016 | 1.82E-05 | 0.00023 | 3.36E-05 | 5.27E-05 | 0.00028 | 0 | 0.00017 | 0.00021 | 0.00011 | 2.46E-05 | 6.70E-06 | 5.92E-05 |
| Sema6a | 4.06E-07 | 2.18E-05 | 2.33E-06 | 2.86E-06 | 0 | 0.00014 | 0 | 2.55E-05 | 3.36E-05 | 3.36E-05 | 5.45E-06 | 0 | 0 | 0 | 0 | 1.49E-05 | 3.62E-05 | 3.26E-05 |
| Dnajc18 | 0.00011 | 9.98E-05 | 3.22E-05 | 3.53E-07 | 4.98E-05 | 8.27E-05 | 4.22E-05 | 6.34E-05 | 3.36E-05 | 5.59E-05 | 6.26E-05 | 4.61E-05 | 6.14E-05 | 9.19E-05 | 0 | 4.09E-05 | 0 | 4.45E-05 |
| Slc25a46 | 0.00011 | 9.11E-05 | 4.76E-05 | 2.00E-06 | 3.62E-05 | 6.55E-05 | 2.65E-07 | 1.31E-05 | 3.36E-05 | 2.27E-05 | 2.33E-05 | 6.41E-05 | 0 | 5.51E-06 | 7.09E-05 | 1.78E-05 | 0 | 2.72E-05 |
| Usp2 | 0 | 6.14E-05 | 9.64E-06 | 0 | 2.13E-05 | 5.70E-05 | 0 | 1.68E-05 | 3.36E-05 | 4.12E-05 | 0 | 1.87E-05 | 0 | 4.03E-05 | 1.56E-06 | 0.00013 | 3.55E-05 | 7.36E-05 |
| Zfp658 | 4.74E-06 | 2.25E-05 | 0 | 0 | 6.64E-06 | 3.43E-05 | 0 | 1.67E-05 | 3.36E-05 | 3.29E-06 | 0 | 0 | 0 | 3.04E-05 | 1.03E-05 | 0 | 0 | 1.05E-05 |
| Ing3 | 5.88E-05 | 3.79E-05 | 2.05E-05 | 0 | 5.41E-06 | 2.87E-05 | 1.35E-06 | 7.47E-06 | 3.36E-05 | 0 | 0 | 4.09E-05 | 0 | 1.63E-05 | 3.92E-05 | 4.55E-05 | 4.12E-05 | 0 |
| Poc5 | 2.82E-05 | 5.16E-05 | 2.31E-05 | 0 | 2.70E-05 | 1.88E-05 | 0 | 8.24E-06 | 3.36E-05 | 1.68E-05 | 9.60E-06 | 6.88E-06 | 3.69E-06 | 1.75E-06 | 0.00011 | 0.00015 | 0 | 1.95E-05 |

| | | | | | | | | | | | | | | | | | | |
|---|---|---|---|---|---|---|---|---|---|---|---|---|---|---|---|---|---|---|
| Tex2 | 0 | 0.00013 | 7.80E-05 | 8.77E-05 | 1.56E-05 | 9.81E-06 | 5.20E-07 | 8.59E-05 | 3.36E-05 | 7.52E-05 | 6.34E-07 | 7.93E-05 | 6.17E-05 | 0 | 0.0001 | 0.0001 | 6.00E-05 | 8.44E-05 |
| Tmem143 | 9.71E-06 | 4.80E-05 | 8.68E-06 | 6.19E-05 | 1.95E-05 | 0 | 0 | 1.83E-05 | 3.36E-05 | 1.91E-05 | 1.40E-05 | 3.50E-05 | 2.33E-05 | 2.20E-05 | 1.02E-05 | 7.03E-05 | 2.29E-05 | 4.17E-05 |
| Serpini1 | 0 | 5.03E-06 | 0 | 0 | 0 | 2.17E-05 | 0 | 0 | 3.35E-05 | 0 | 0 | 2.61E-05 | 0 | 0 | 0 | 1.52E-06 | 0 | 0 |
| Snip1 | 2.39E-05 | 4.20E-05 | 3.93E-05 | 5.11E-05 | 2.73E-05 | 8.72E-06 | 0 | 3.11E-06 | 3.35E-05 | 9.67E-06 | 0 | 3.39E-05 | 2.47E-07 | 1.59E-05 | 5.59E-05 | 5.72E-05 | 1.00E-06 | 3.43E-05 |
| Pxk | 0.00029 | 6.04E-05 | 0.00012 | 4.78E-05 | 0.00012 | 0.00015 | 0.00026 | 0 | 3.34E-05 | 0.00015 | 8.49E-05 | 3.94E-05 | 0.00017 | 0.00012 | 0.00029 | 0.00015 | 0 | 0.00013 |
| Bnip3l | 0.00011 | 0.00013 | 0 | 0 | 1.02E-05 | 0.0001 | 2.65E-05 | 0 | 3.34E-05 | 2.16E-05 | 0 | 0.0004 | 1.56E-05 | 2.71E-05 | 0 | 0 | 0 | 5.51E-05 |
| Ppp2r2d | 0.00014 | 0.0001 | 6.37E-05 | 0 | 2.06E-05 | 9.65E-05 | 3.72E-05 | 1.59E-05 | 3.34E-05 | 2.19E-05 | 0 | 3.95E-05 | 0 | 0 | 0 | 0.00013 | 0 | 1.42E-05 |
| Timmdc1 | 7.93E-05 | 4.06E-05 | 0.00012 | 6.46E-05 | 7.03E-05 | 4.38E-05 | 0 | 0.00012 | 3.34E-05 | 3.77E-05 | 6.74E-05 | 2.59E-05 | 6.28E-05 | 1.04E-05 | 2.27E-05 | 5.61E-05 | 1.19E-05 | 2.61E-05 |
| Maip1 | 7.39E-05 | 5.04E-05 | 5.64E-05 | 1.10E-05 | 9.22E-06 | 4.06E-05 | 0 | 0 | 3.34E-05 | 0 | 5.53E-05 | 2.06E-05 | 0 | 2.71E-05 | 4.10E-06 | 4.46E-05 | 0 | 0 |
| Dhx35 | 4.05E-05 | 4.16E-05 | 2.97E-05 | 2.52E-06 | 1.50E-05 | 3.40E-05 | 0 | 2.83E-06 | 3.34E-05 | 0 | 1.14E-05 | 1.33E-05 | 0 | 2.96E-06 | 7.50E-05 | 8.92E-06 | 3.58E-05 | 8.35E-06 |
| Usp37 | 3.83E-05 | 3.90E-05 | 5.97E-05 | 0 | 1.54E-05 | 3.20E-05 | 4.07E-05 | 2.26E-05 | 3.34E-05 | 8.69E-06 | 2.22E-05 | 4.26E-05 | 3.66E-05 | 1.41E-05 | 3.31E-05 | 0.0001 | 1.46E-05 | 0 |
| Mecr | 4.42E-05 | 2.20E-05 | 9.85E-05 | 8.69E-05 | 1.51E-05 | 8.27E-06 | 1.40E-05 | 0 | 3.34E-05 | 3.31E-05 | 5.05E-05 | 0 | 8.57E-06 | 3.09E-05 | 0 | 4.40E-05 | 5.82E-05 | 6.06E-05 |
| Hmox1 | 6.75E-05 | 0.00026 | 4.17E-05 | 0.00021 | 8.51E-05 | 0.00021 | 0 | 0 | 3.33E-05 | 0 | 0 | 0.00085 | 0.00029 | 0.00043 | 0 | 7.59E-06 | 0.00113 | 0 |
| Npat | 5.45E-05 | 4.67E-05 | 9.47E-06 | 0 | 1.91E-05 | 4.29E-05 | 1.46E-05 | 1.50E-05 | 3.33E-05 | 3.41E-06 | 1.74E-05 | 0 | 4.17E-06 | 2.67E-05 | 7.11E-05 | 0.00012 | 5.61E-06 | 1.76E-05 |
| Uvssa | 3.27E-05 | 7.80E-05 | 0 | 7.86E-05 | 3.32E-05 | 4.27E-07 | 4.46E-05 | 3.37E-05 | 3.33E-05 | 4.78E-05 | 0 | 3.04E-05 | 0 | 1.58E-05 | 0.00019 | 4.68E-06 | 0.0001 | 1.78E-05 |
| Dync2li1 | 2.48E-05 | 5.22E-05 | 0 | 4.22E-06 | 1.66E-05 | 0 | 1.56E-06 | 1.85E-05 | 3.33E-05 | 3.48E-05 | 6.39E-06 | 3.35E-06 | 6.72E-06 | 0 | 0 | 0 | 1.68E-05 | 8.45E-05 |
| Acaa2 | 0.00022 | 0.00011 | 4.65E-05 | 0.00022 | 1.74E-05 | 0.00023 | 2.82E-05 | 0.00051 | 3.32E-05 | 0.00023 | 0 | 0 | 0 | 0.0003 | 0 | 5.85E-05 | 0 | 5.38E-05 |
| Tasor2 | 0.00014 | 9.13E-05 | 9.52E-05 | 0 | 4.79E-05 | 5.73E-05 | 1.95E-05 | 2.51E-05 | 3.32E-05 | 1.49E-05 | 1.61E-05 | 5.32E-05 | 6.85E-06 | 1.61E-05 | 0.00011 | 6.66E-05 | 9.67E-05 | 2.11E-05 |
| Tctex1d2 | 5.49E-05 | 1.67E-05 | 3.27E-05 | 0 | 4.14E-05 | 4.20E-05 | 3.41E-05 | 1.39E-05 | 3.32E-05 | 1.51E-05 | 2.06E-05 | 4.98E-07 | 4.10E-05 | 2.86E-05 | 0 | 3.88E-05 | 0 | 0 |
| Sco2 | 8.24E-05 | 7.61E-05 | 0.00013 | 1.70E-05 | 5.38E-06 | 2.95E-05 | 0 | 8.04E-06 | 3.32E-05 | 6.43E-06 | 2.02E-05 | 7.26E-05 | 0 | 4.38E-05 | 0 | 0 | 0 | 3.02E-05 |
| Pde8b | 0 | 7.68E-06 | 0 | 0 | 0 | 2.10E-05 | 0 | 0 | 3.32E-05 | 0 | 2.37E-05 | 4.78E-06 | 0 | 0 | 0 | 0 | 0 | 8.59E-06 |
| Pin4 | 3.27E-05 | 2.22E-05 | 3.59E-05 | 4.70E-05 | 1.23E-05 | 3.24E-06 | 0 | 1.76E-05 | 3.32E-05 | 4.69E-06 | 8.87E-06 | 1.67E-05 | 4.10E-05 | 2.44E-06 | 3.18E-05 | 1.27E-05 | 2.78E-05 | 1.30E-07 |
| Fancl | 1.94E-05 | 1.05E-05 | 7.81E-06 | 1.74E-05 | 2.43E-06 | 0 | 2.10E-05 | 0 | 3.32E-05 | 1.94E-05 | 1.76E-05 | 0 | 0 | 1.10E-05 | 1.17E-05 | 5.36E-05 | 0 | 0 |
| Txndc16 | 1.25E-05 | 0.00017 | 0 | 0 | 0.00017 | 9.30E-05 | 0 | 0 | 3.31E-05 | 3.60E-05 | 3.40E-05 | 4.77E-05 | 0.00014 | 8.88E-05 | 0 | 2.79E-05 | 0 | 0.00011 |
| Stx18 | 0.00017 | 5.17E-05 | 0.00013 | 4.81E-05 | 3.68E-05 | 5.23E-05 | 0.0001 | 7.74E-05 | 3.31E-05 | 0 | 2.41E-05 | 0 | 1.28E-05 | 0 | 0.00021 | 2.85E-05 | 0.00015 | 5.44E-05 |
| Slc7a6os | 7.98E-05 | 8.96E-05 | 9.41E-05 | 0.00011 | 1.49E-05 | 1.42E-05 | 0 | 1.36E-06 | 3.31E-05 | 3.53E-05 | 7.19E-05 | 2.45E-05 | 0 | 2.46E-05 | 0.00018 | 6.55E-05 | 0.00032 | 0 |
| Vars2 | 2.49E-05 | 3.46E-05 | 1.64E-05 | 5.08E-05 | 2.22E-05 | 1.11E-05 | 2.21E-06 | 0 | 3.31E-05 | 2.67E-05 | 0 | 6.37E-06 | 0 | 2.25E-05 | 3.01E-05 | 3.91E-05 | 0 | 3.47E-05 |
| Chn1 | 0 | 6.16E-06 | 0 | 0 | 4.71E-06 | 5.90E-06 | 0 | 2.26E-05 | 3.31E-05 | 0 | 0 | 2.79E-06 | 0 | 0 | 0 | 3.28E-05 | 0 | 1.07E-05 |
| Fam219b | 8.14E-06 | 2.85E-05 | 2.40E-05 | 6.77E-05 | 1.41E-05 | 5.74E-06 | 1.40E-05 | 6.12E-07 | 3.31E-05 | 8.31E-06 | 0 | 9.35E-06 | 0 | 7.91E-06 | 2.92E-05 | 2.14E-05 | 0 | 2.70E-05 |
| Ccdc86 | 5.14E-05 | 6.78E-05 | 0.00019 | 0.00013 | 0 | 0 | 0 | 0 | 3.31E-05 | 0 | 1.43E-05 | 0 | 2.89E-05 | 1.19E-05 | 0.00013 | 3.26E-05 | 0 | 0 |
| Tmtc4 | 2.79E-05 | 0 | 8.12E-05 | 4.71E-05 | 6.23E-05 | 0 | 0.00016 | 0.00015 | 3.31E-05 | 0 | 0.00017 | 0 | 0 | 3.08E-05 | 0 | 0.00017 | 0 | 6.13E-07 |
| Hpcal1 | 1.35E-05 | 3.32E-05 | 6.45E-06 | 2.36E-05 | 3.26E-05 | 0 | 0 | 3.41E-05 | 3.31E-05 | 1.05E-05 | 4.44E-05 | 2.62E-05 | 0 | 3.11E-05 | 0 | 2.31E-05 | 2.14E-05 | 1.99E-05 |
| Eif2a | 0.00028 | 0.0001 | 9.49E-05 | 4.58E-05 | 4.43E-05 | 0.00017 | 3.47E-05 | 0 | 3.30E-05 | 3.75E-05 | 0 | 0.00011 | 3.42E-05 | 1.59E-05 | 0 | 0.00016 | 6.94E-05 | 5.90E-05 |

| | | | | | | | | | | | | | | | | | | |
|---|---|---|---|---|---|---|---|---|---|---|---|---|---|---|---|---|---|---|
| Nr4a2 | 0 | 2.43E-05 | 9.41E-05 | 0 | 4.24E-05 | 5.05E-05 | 3.71E-05 | 0 | 3.30E-05 | 0 | 0 | 0 | 0 | 9.43E-05 | 4.55E-05 | 0 | 0.00038 | 9.71E-05 |
| Nt5c3 | 0.00011 | 2.87E-05 | 4.82E-05 | 0 | 5.20E-06 | 5.01E-05 | 0 | 0 | 3.30E-05 | 3.41E-05 | 1.96E-06 | 4.92E-05 | 2.52E-06 | 3.75E-06 | 0 | 5.32E-05 | 2.10E-05 | 4.49E-05 |
| Morc3 | 7.47E-05 | 0.00011 | 8.26E-05 | 2.66E-05 | 2.50E-05 | 4.42E-05 | 2.10E-07 | 0 | 3.30E-05 | 3.58E-05 | 3.31E-05 | 6.70E-05 | 6.15E-05 | 4.50E-06 | 0.00015 | 8.67E-05 | 6.16E-05 | 2.91E-05 |
| Sgsm3 | 2.82E-05 | 4.31E-05 | 2.51E-05 | 0 | 4.91E-05 | 6.34E-06 | 3.64E-05 | 7.69E-06 | 3.30E-05 | 2.55E-05 | 2.36E-05 | 1.78E-06 | 0 | 1.02E-05 | 5.70E-05 | 5.12E-06 | 0 | 3.53E-05 |
| Kbtbd4 | 2.77E-05 | 4.63E-05 | 2.44E-05 | 1.83E-05 | 3.11E-05 | 5.82E-08 | 2.05E-05 | 1.33E-05 | 3.30E-05 | 1.49E-05 | 0 | 0 | 0 | 2.92E-05 | 4.49E-05 | 4.41E-05 | 0 | 9.52E-06 |
| Mgp | 0 | 0.00016 | 0 | 0 | 0 | 0 | 0 | 0.00106 | 3.30E-05 | 0 | 0 | 4.04E-05 | 0 | 0 | 0 | 0 | 0 | 0.00263 |
| Rexo1 | 2.70E-05 | 4.56E-05 | 2.08E-05 | 2.50E-05 | 1.51E-05 | 0 | 0 | 2.72E-05 | 3.30E-05 | 1.05E-05 | 1.90E-05 | 1.48E-05 | 0 | 1.95E-05 | 0.00012 | 2.86E-05 | 0.00012 | 3.64E-06 |
| Tstd2 | 0.00011 | 8.95E-05 | 7.56E-05 | 2.24E-05 | 3.61E-05 | 3.14E-05 | 4.63E-05 | 0 | 3.29E-05 | 1.95E-05 | 3.34E-05 | 4.38E-05 | 1.89E-06 | 1.79E-05 | 2.99E-05 | 5.33E-05 | 1.26E-06 | 3.93E-05 |
| Camsap1 | 1.16E-05 | 4.80E-05 | 5.60E-05 | 1.96E-05 | 1.27E-05 | 0 | 0 | 0 | 3.29E-05 | 1.17E-05 | 3.84E-05 | 3.96E-05 | 4.49E-06 | 3.50E-05 | 6.37E-05 | 2.25E-05 | 5.54E-05 | 3.21E-05 |
| Neurl4 | 3.90E-05 | 4.95E-05 | 3.22E-05 | 1.26E-05 | 2.09E-05 | 0 | 6.42E-06 | 0 | 3.29E-05 | 8.31E-06 | 2.06E-05 | 5.24E-05 | 4.10E-06 | 3.03E-05 | 8.27E-05 | 2.63E-05 | 0 | 1.46E-05 |
| Dpp4 | 0 | 2.36E-05 | 0 | 0 | 1.44E-05 | 0 | 0.00017 | 0 | 3.29E-05 | 0.00089 | 0 | 0 | 0.00019 | 0 | 0 | 0 | 0 | 0 |
| Adck5 | 1.30E-05 | 3.80E-05 | 3.48E-05 | 4.56E-05 | 6.75E-06 | 0 | 4.00E-05 | 1.25E-05 | 3.29E-05 | 1.23E-05 | 1.44E-05 | 1.38E-05 | 0 | 1.79E-05 | 7.26E-05 | 1.84E-05 | 0 | 1.50E-05 |
| Emx2 | 0 | 4.16E-05 | 0 | 6.64E-05 | 4.16E-05 | 0 | 3.30E-05 | 0 | 3.29E-05 | 0.00017 | 9.02E-05 | 0 | 0.00011 | 0 | 2.86E-05 | 3.73E-05 | 0 | 0 |
| Armcx3 | 0.00032 | 0.00012 | 8.25E-05 | 0.00027 | 0.00021 | 0.00028 | 0.00036 | 0.00025 | 3.28E-05 | 0.00012 | 0.00011 | 4.70E-05 | 0.00011 | 0.0001 | 0 | 6.93E-05 | 0.00033 | 4.88E-05 |
| Grsf1 | 0.00017 | 0.00015 | 0.00028 | 0.0002 | 0.00015 | 0.00019 | 0.0003 | 0.00011 | 3.28E-05 | 0 | 8.05E-05 | 0.00014 | 8.96E-05 | 0.00018 | 2.89E-05 | 0.00012 | 3.84E-05 | 6.87E-05 |
| Map3k20 | 0.00016 | 0.00014 | 0.00012 | 5.36E-05 | 0.00015 | 9.52E-05 | 6.55E-05 | 0 | 3.28E-05 | 7.65E-05 | 7.61E-05 | 3.70E-05 | 0.00011 | 0 | 0 | 0.00023 | 6.56E-05 | 1.14E-05 |
| Rpa3 | 0.00011 | 1.88E-05 | 5.00E-05 | 2.90E-05 | 0 | 6.41E-05 | 2.71E-06 | 2.91E-05 | 3.28E-05 | 3.21E-05 | 0 | 2.28E-05 | 3.38E-05 | 9.37E-06 | 0 | 0.0003 | 0 | 1.63E-05 |
| Cnst | 9.27E-06 | 6.47E-05 | 2.76E-05 | 1.31E-05 | 4.12E-05 | 3.78E-05 | 3.50E-05 | 0 | 3.28E-05 | 2.70E-05 | 0.00011 | 5.92E-05 | 0 | 3.47E-05 | 0.00011 | 4.53E-05 | 9.83E-05 | 3.41E-05 |
| Tmem80 | 1.58E-05 | 3.22E-05 | 5.94E-07 | 1.13E-07 | 2.59E-05 | 6.25E-06 | 0 | 1.17E-05 | 3.28E-05 | 2.94E-06 | 3.85E-06 | 1.93E-05 | 0 | 0 | 2.02E-05 | 1.14E-05 | 0 | 2.83E-05 |
| Coq4 | 4.57E-05 | 1.25E-05 | 3.36E-05 | 4.46E-05 | 1.61E-05 | 4.43E-06 | 1.06E-05 | 0 | 3.28E-05 | 1.69E-06 | 4.99E-06 | 6.45E-06 | 8.71E-06 | 0 | 2.55E-06 | 1.87E-06 | 2.72E-05 | 1.71E-05 |
| Wdr92 | 6.67E-05 | 2.99E-05 | 3.87E-05 | 9.76E-06 | 2.03E-05 | 8.10E-07 | 6.32E-06 | 8.87E-06 | 3.28E-05 | 8.00E-06 | 6.23E-06 | 0 | 2.91E-06 | 8.79E-06 | 2.72E-05 | 2.10E-05 | 0 | 1.64E-05 |
| Mettl16 | 6.62E-05 | 5.94E-05 | 0.00013 | 8.87E-05 | 0 | 0 | 0 | 0 | 3.28E-05 | 2.15E-05 | 3.66E-05 | 1.96E-05 | 6.39E-05 | 0 | 4.74E-05 | 0.0001 | 8.15E-06 | 2.86E-05 |
| Jmjd4 | 1.67E-05 | 4.48E-05 | 1.45E-05 | 1.67E-05 | 8.01E-06 | 0 | 3.57E-05 | 0 | 3.28E-05 | 1.30E-05 | 4.21E-05 | 0 | 2.35E-06 | 3.17E-05 | 0 | 2.42E-05 | 0 | 4.50E-05 |
| Aldh18a1 | 0.00018 | 1.49E-05 | 0.00026 | 0.0004 | 7.34E-05 | 0.0001 | 0.00052 | 8.95E-05 | 3.27E-05 | 0.00013 | 7.68E-05 | 4.24E-05 | 4.63E-05 | 0.00012 | 2.68E-05 | 5.74E-05 | 0 | 0.00013 |
| Mmut | 6.75E-05 | 2.79E-05 | 3.37E-06 | 0 | 2.79E-05 | 4.40E-05 | 0 | 1.08E-05 | 3.27E-05 | 5.22E-05 | 1.34E-05 | 1.52E-05 | 1.45E-05 | 0 | 2.69E-05 | 1.42E-05 | 0 | 4.06E-05 |
| Rnf141 | 4.98E-05 | 5.09E-05 | 1.54E-05 | 0 | 7.05E-06 | 3.91E-05 | 0 | 0 | 3.27E-05 | 4.14E-05 | 2.19E-05 | 4.78E-05 | 2.39E-05 | 0 | 1.55E-05 | 7.18E-05 | 0 | 5.71E-05 |
| Alkbh1 | 9.16E-05 | 3.59E-05 | 6.62E-05 | 8.89E-05 | 4.76E-05 | 3.83E-05 | 1.19E-05 | 4.50E-05 | 3.27E-05 | 4.63E-06 | 2.09E-06 | 0 | 0 | 0 | 0.00051 | 2.95E-05 | 0 | 0 |
| Coasy | 2.13E-05 | 7.16E-05 | 1.93E-05 | 0.00011 | 3.05E-05 | 2.92E-05 | 4.85E-05 | 5.04E-05 | 3.27E-05 | 3.26E-05 | 9.82E-06 | 4.43E-05 | 2.23E-05 | 5.25E-05 | 1.14E-05 | 3.35E-05 | 0 | 4.12E-05 |
| Cldn11 | 0 | 0 | 0 | 1.72E-05 | 0 | 1.86E-05 | 0 | 7.34E-07 | 3.27E-05 | 0 | 0 | 7.74E-05 | 0 | 0 | 0 | 1.94E-05 | 0.00013 | 0 |
| Rwdd2b | 3.21E-05 | 4.22E-06 | 1.19E-05 | 2.14E-05 | 2.30E-06 | 8.33E-07 | 3.99E-05 | 3.90E-07 | 3.27E-05 | 2.93E-06 | 1.48E-05 | 0 | 0 | 3.03E-05 | 0 | 1.39E-05 | 0 | 5.24E-06 |
| Aplf | 7.53E-06 | 2.04E-05 | 1.68E-05 | 0 | 1.41E-05 | 0 | 1.25E-05 | 3.65E-06 | 3.27E-05 | 2.06E-05 | 1.84E-05 | 2.63E-05 | 0 | 6.02E-06 | 2.68E-05 | 0.00013 | 0 | 1.75E-05 |
| Nrn1 | 2.60E-05 | 6.16E-06 | 0 | 0.00013 | 0 | 0 | 0 | 3.97E-05 | 3.27E-05 | 0 | 1.98E-06 | 0.00023 | 8.30E-05 | 0.00024 | 7.42E-05 | 0 | 6.81E-05 | 2.74E-05 |
| Pbxip1 | 0 | 0.00014 | 0 | 0.0001 | 0.00018 | 0.00011 | 9.78E-05 | 3.70E-05 | 3.26E-05 | 0.00018 | 0.0002 | 9.57E-05 | 0.00015 | 0.00011 | 6.14E-05 | 0 | 0 | 0.00018 |

| | | | | | | | | | | | | | | | | | | |
|---|---|---|---|---|---|---|---|---|---|---|---|---|---|---|---|---|---|---|
| Usp30 | 3.54E-05 | 7.68E-05 | 3.89E-05 | 2.85E-05 | 3.83E-05 | 2.94E-05 | 1.41E-05 | 3.86E-05 | 3.26E-05 | 2.30E-05 | 2.94E-05 | 4.47E-05 | 2.45E-05 | 2.45E-05 | 3.82E-05 | 1.91E-05 | 0 | 1.65E-05 |
| Mdk | 0 | 0.00016 | 0 | 0.00014 | 0.00048 | 0 | 0 | 0.00026 | 3.26E-05 | 0 | 0.00018 | 0 | 7.29E-05 | 0.00054 | 0.00015 | 3.60E-05 | 0 | 0.00017 |
| Dcps | 5.02E-05 | 1.63E-05 | 7.01E-06 | 0.00011 | 1.23E-05 | 0 | 2.99E-05 | 1.63E-05 | 3.26E-05 | 3.38E-05 | 0 | 0 | 4.04E-06 | 0 | 0 | 0.00015 | 3.63E-05 | 6.36E-05 |
| Drd1 | 0 | 2.21E-06 | 0 | 0 | 0 | 5.08E-05 | 0 | 0 | 3.25E-05 | 0 | 0 | 0 | 7.46E-06 | 0 | 0 | 0 | 0 | 0 |
| Tusc1 | 0 | 1.51E-05 | 0 | 5.36E-05 | 0 | 1.54E-05 | 1.10E-06 | 2.73E-05 | 3.25E-05 | 1.74E-05 | 2.13E-05 | 1.76E-06 | 1.47E-05 | 2.90E-05 | 3.35E-05 | 7.26E-06 | 1.09E-05 | 9.03E-06 |
| Znrf2 | 0 | 0.00011 | 8.82E-05 | 0.00015 | 2.07E-06 | 0 | 0 | 6.59E-05 | 3.25E-05 | 0.00017 | 2.03E-05 | 3.73E-05 | 6.25E-05 | 3.66E-05 | 4.79E-05 | 9.13E-05 | 0.00011 | 7.55E-05 |
| Nans | 0.00046 | 6.21E-06 | 0.00028 | 4.43E-05 | 6.70E-05 | 0.00017 | 0.0003 | 0.00017 | 3.24E-05 | 0 | 0.00011 | 0.00018 | 0 | 0 | 0 | 0.00028 | 0 | 0 |
| Ccnc | 0.00011 | 6.68E-05 | 0.00013 | 2.03E-05 | 1.55E-05 | 5.79E-05 | 0.0001 | 2.74E-05 | 3.24E-05 | 2.22E-05 | 4.28E-05 | 5.13E-06 | 1.34E-05 | 1.46E-06 | 1.29E-06 | 7.84E-05 | 2.69E-05 | 5.45E-05 |
| Cstad | 5.01E-06 | 2.09E-05 | 0 | 4.27E-06 | 7.53E-05 | 5.39E-05 | 0 | 1.68E-05 | 3.24E-05 | 0 | 4.48E-05 | 0 | 2.27E-05 | 0 | 9.24E-05 | 0 | 0 | 4.55E-05 |
| Galm | 1.12E-05 | 4.26E-05 | 0 | 0 | 2.18E-05 | 5.29E-05 | 0 | 0 | 3.24E-05 | 7.06E-05 | 0 | 2.75E-05 | 0 | 1.98E-05 | 0 | 4.53E-05 | 0 | 0 |
| Nudt8 | 4.75E-05 | 6.33E-05 | 4.32E-05 | 5.59E-05 | 4.97E-05 | 3.32E-05 | 1.89E-05 | 3.15E-05 | 3.24E-05 | 1.82E-05 | 1.87E-05 | 3.54E-05 | 3.97E-05 | 6.04E-05 | 0.0001 | 1.65E-06 | 0 | 1.43E-05 |
| Zswim7 | 5.45E-05 | 1.50E-05 | 1.82E-05 | 2.06E-05 | 1.66E-05 | 2.79E-05 | 0 | 2.94E-05 | 3.24E-05 | 7.52E-06 | 3.16E-05 | 3.35E-05 | 0 | 0 | 0 | 2.94E-05 | 0 | 0 |
| Ccdc22 | 1.85E-05 | 7.36E-05 | 3.78E-05 | 7.74E-05 | 2.06E-05 | 9.07E-06 | 0 | 4.83E-05 | 3.24E-05 | 2.92E-05 | 1.34E-05 | 9.40E-06 | 0 | 7.83E-05 | 4.99E-05 | 8.37E-05 | 2.47E-06 | 3.84E-05 |
| Tsr3 | 4.79E-05 | 7.52E-05 | 4.80E-05 | 0.0001 | 6.88E-06 | 0 | 7.35E-06 | 3.55E-05 | 3.24E-05 | 4.33E-05 | 3.73E-05 | 8.19E-05 | 4.20E-05 | 6.36E-05 | 0.00011 | 3.45E-05 | 0 | 9.87E-06 |
| Gpatch4 | 5.00E-05 | 9.64E-06 | 0.00012 | 7.34E-06 | 0 | 0 | 0 | 0 | 3.24E-05 | 1.82E-05 | 1.48E-05 | 0 | 1.05E-05 | 0 | 7.33E-05 | 8.37E-06 | 0 | 0 |
| Atg3 | 0.00019 | 9.05E-05 | 0.00013 | 8.51E-05 | 1.30E-05 | 8.83E-05 | 5.55E-05 | 0 | 3.23E-05 | 6.00E-05 | 5.12E-06 | 9.33E-06 | 4.54E-05 | 6.76E-05 | 6.80E-05 | 7.97E-05 | 0.00012 | 5.11E-05 |
| Lgals8 | 0.00017 | 0.00016 | 6.10E-05 | 8.89E-05 | 8.85E-05 | 6.35E-05 | 4.87E-05 | 6.09E-05 | 3.23E-05 | 0.00011 | 2.04E-05 | 7.01E-05 | 4.77E-05 | 7.31E-05 | 5.42E-05 | 1.52E-05 | 0 | 0.00015 |
| Sil1 | 4.38E-05 | 4.00E-05 | 3.23E-05 | 0.00016 | 8.51E-05 | 5.42E-05 | 4.70E-05 | 8.89E-05 | 3.23E-05 | 7.00E-05 | 2.03E-05 | 0.00012 | 0.00017 | 5.76E-05 | 0 | 4.99E-05 | 6.20E-05 | 8.63E-05 |
| Tab1 | 3.83E-05 | 5.39E-05 | 3.25E-05 | 3.91E-05 | 5.28E-05 | 3.09E-05 | 3.07E-05 | 2.97E-05 | 3.23E-05 | 2.01E-05 | 1.78E-05 | 4.65E-05 | 1.39E-05 | 2.89E-05 | 0.0001 | 3.17E-05 | 0 | 2.73E-05 |
| Akip1 | 0.00011 | 4.92E-05 | 3.67E-05 | 9.39E-05 | 3.82E-05 | 2.24E-05 | 5.94E-05 | 0 | 3.23E-05 | 3.12E-05 | 4.88E-05 | 2.84E-05 | 7.48E-05 | 0 | 0 | 0.00012 | 0 | 4.20E-05 |
| Kdm1b | 4.67E-05 | 3.34E-05 | 1.60E-05 | 9.24E-06 | 1.31E-05 | 1.45E-05 | 6.82E-06 | 0 | 3.23E-05 | 1.64E-05 | 6.21E-06 | 1.95E-06 | 1.19E-05 | 2.46E-05 | 0 | 1.74E-05 | 5.51E-06 | 3.30E-05 |
| Tmed5 | 4.38E-05 | 9.88E-05 | 0.00024 | 0.00017 | 6.33E-06 | 1.06E-05 | 0 | 2.48E-05 | 3.23E-05 | 8.24E-05 | 2.34E-05 | 7.06E-06 | 0.00017 | 2.85E-05 | 2.65E-05 | 9.41E-05 | 5.38E-05 | 6.68E-05 |
| Med19 | 3.10E-05 | 3.15E-05 | 1.89E-05 | 4.84E-05 | 1.58E-05 | 5.82E-06 | 0 | 4.11E-06 | 3.23E-05 | 2.24E-05 | 1.19E-05 | 1.99E-05 | 0 | 0 | 0 | 7.24E-05 | 0 | 1.40E-05 |
| Plpp6 | 1.75E-05 | 6.05E-05 | 7.18E-06 | 4.31E-05 | 1.17E-05 | 0 | 1.71E-05 | 1.81E-05 | 3.23E-05 | 2.12E-05 | 2.34E-05 | 1.06E-05 | 1.82E-05 | 8.93E-07 | 0 | 0 | 0 | 3.85E-05 |
| Nek8 | 2.31E-06 | 4.55E-05 | 0 | 5.79E-06 | 5.94E-06 | 0 | 2.31E-06 | 2.03E-05 | 3.23E-05 | 5.10E-06 | 1.23E-06 | 8.72E-06 | 0 | 8.63E-06 | 5.97E-05 | 1.14E-05 | 0 | 1.85E-05 |
| Bet1l | 0.00012 | 0.00013 | 9.78E-05 | 0.00029 | 0.00011 | 0.00012 | 0.00024 | 7.31E-05 | 3.22E-05 | 0.00011 | 7.43E-05 | 0.00011 | 9.27E-06 | 9.48E-05 | 0.00017 | 0 | 0 | 9.72E-05 |
| Gpbp1l1 | 0.0001 | 0.00017 | 3.81E-05 | 1.54E-05 | 8.48E-05 | 0.00012 | 4.53E-05 | 1.41E-05 | 3.22E-05 | 4.89E-05 | 4.79E-05 | 9.15E-05 | 1.68E-05 | 8.43E-05 | 0.00012 | 5.12E-05 | 6.66E-05 | 1.49E-05 |
| Ece2 | 2.46E-05 | 4.94E-06 | 1.31E-05 | 1.54E-05 | 2.83E-05 | 7.30E-05 | 5.01E-05 | 8.77E-06 | 3.22E-05 | 3.18E-06 | 0 | 3.84E-05 | 1.37E-05 | 0 | 6.28E-05 | 2.00E-05 | 0 | 0.00012 |
| Btrc | 9.09E-05 | 8.42E-05 | 1.32E-05 | 0 | 5.06E-05 | 6.40E-05 | 2.61E-05 | 3.46E-05 | 3.22E-05 | 1.62E-05 | 0 | 3.91E-05 | 2.69E-05 | 0 | 6.65E-05 | 2.73E-06 | 2.34E-05 | 7.67E-06 |
| Triap1 | 6.94E-05 | 5.54E-05 | 3.23E-05 | 3.95E-05 | 3.98E-05 | 4.99E-05 | 0 | 1.41E-05 | 3.22E-05 | 2.52E-05 | 4.29E-05 | 3.44E-05 | 6.13E-06 | 1.36E-05 | 1.33E-05 | 5.65E-05 | 0 | 2.49E-05 |
| Etfdh | 0.00015 | 6.67E-05 | 1.93E-05 | 4.69E-05 | 5.28E-05 | 3.98E-05 | 0.00011 | 0 | 3.22E-05 | 8.09E-05 | 4.00E-05 | 4.69E-05 | 5.00E-05 | 2.74E-05 | 9.93E-06 | 0.00012 | 1.39E-05 | 7.44E-05 |
| Ppip5k2 | 0.0001 | 5.99E-05 | 4.30E-05 | 3.55E-06 | 3.68E-05 | 3.61E-05 | 5.46E-06 | 0 | 3.22E-05 | 6.28E-05 | 3.61E-06 | 1.88E-05 | 1.21E-05 | 0 | 5.83E-05 | 5.79E-05 | 4.61E-05 | 1.72E-05 |
| Kansl3 | 0 | 0.0002 | 0.00016 | 4.64E-05 | 0.00011 | 3.12E-05 | 6.48E-05 | 2.34E-05 | 3.22E-05 | 6.42E-05 | 0.00011 | 8.88E-05 | 4.68E-06 | 3.22E-05 | 0.00014 | 0.00012 | 0 | 4.95E-05 |

| | | | | | | | | | | | | | | | | | | |
|---|---|---|---|---|---|---|---|---|---|---|---|---|---|---|---|---|---|---|
| Pmm1 | 0.00012 | 0.00019 | 0.00013 | 0.00012 | 0.00011 | 3.02E-05 | 3.39E-05 | 2.34E-05 | 3.22E-05 | 3.77E-05 | 5.74E-05 | 0.00019 | 6.21E-05 | 7.78E-05 | 8.87E-05 | 1.84E-05 | 5.87E-05 | 0.0002 |
| Kiss1r | 0 | 1.87E-05 | 0 | 1.21E-05 | 1.16E-05 | 1.43E-05 | 0 | 8.79E-06 | 3.22E-05 | 0 | 0 | 2.09E-05 | 1.90E-05 | 0 | 1.22E-05 | 0 | 1.77E-05 | 1.73E-05 |
| Muc1 | 1.35E-05 | 2.04E-05 | 0 | 1.19E-05 | 2.08E-05 | 5.99E-07 | 0 | 5.49E-06 | 3.22E-05 | 1.80E-05 | 1.35E-05 | 5.58E-06 | 1.58E-05 | 0 | 4.90E-05 | 2.53E-05 | 5.72E-05 | 1.23E-05 |
| Spryd4 | 3.40E-05 | 2.79E-05 | 5.85E-05 | 4.36E-05 | 1.81E-06 | 0 | 0 | 0 | 3.22E-05 | 1.27E-05 | 9.93E-06 | 0 | 9.24E-06 | 5.88E-06 | 2.25E-06 | 0 | 0 | 8.81E-06 |
| Cdc25a | 2.91E-05 | 2.70E-06 | 9.61E-05 | 3.15E-05 | 2.48E-05 | 0 | 2.32E-05 | 0 | 3.22E-05 | 0 | 1.78E-05 | 0 | 2.23E-05 | 5.57E-05 | 4.49E-05 | 0.00011 | 0 | 0 |
| Ereg | 0 | 0 | 0.00013 | 0 | 0 | 0.0004 | 0 | 0 | 3.21E-05 | 0 | 0 | 5.30E-05 | 0 | 6.25E-05 | 0 | 0 | 0.00037 | 0 |
| Klf7 | 0 | 0.00012 | 0 | 0 | 0 | 0.00019 | 0 | 7.58E-05 | 3.21E-05 | 6.38E-05 | 1.18E-05 | 0.00039 | 0 | 9.16E-05 | 0.00026 | 4.81E-05 | 0.00012 | 0.00028 |
| Washc4 | 0.00018 | 0.00014 | 1.15E-05 | 0 | 5.63E-05 | 0.00011 | 3.49E-05 | 7.29E-06 | 3.21E-05 | 5.28E-05 | 3.63E-05 | 0.00018 | 0 | 4.85E-05 | 9.12E-05 | 6.70E-05 | 3.43E-05 | 3.28E-05 |
| Cntln | 4.92E-05 | 5.55E-05 | 0 | 1.83E-05 | 4.05E-05 | 5.92E-05 | 1.42E-05 | 5.53E-05 | 3.21E-05 | 4.22E-05 | 1.53E-05 | 1.83E-05 | 0 | 0 | 0.00015 | 0.00032 | 0 | 3.13E-05 |
| Rab40c | 5.31E-05 | 0.00015 | 6.68E-05 | 0.00011 | 5.11E-05 | 5.40E-05 | 5.73E-05 | 4.55E-05 | 3.21E-05 | 1.69E-05 | 5.28E-05 | 6.43E-05 | 4.31E-05 | 5.63E-05 | 4.46E-05 | 1.01E-05 | 4.24E-05 | 2.66E-05 |
| Lcor | 7.91E-05 | 0.00013 | 8.83E-05 | 3.17E-05 | 4.60E-05 | 5.10E-05 | 3.33E-05 | 1.55E-05 | 3.21E-05 | 5.93E-05 | 7.70E-05 | 9.79E-05 | 1.68E-05 | 4.43E-05 | 0.00025 | 7.71E-05 | 0.00016 | 3.22E-05 |
| Adgrb3 | 0 | 4.68E-06 | 0 | 0 | 4.88E-06 | 4.11E-05 | 0 | 2.27E-06 | 3.21E-05 | 0 | 0 | 2.63E-06 | 0 | 0 | 0 | 0 | 0 | 4.10E-06 |
| Fktn | 4.20E-05 | 3.78E-05 | 3.55E-05 | 0 | 5.13E-05 | 3.92E-05 | 2.16E-07 | 3.21E-05 | 3.21E-05 | 3.02E-05 | 4.30E-05 | 1.82E-05 | 1.99E-06 | 2.10E-05 | 1.13E-05 | 2.55E-05 | 0 | 3.23E-06 |
| Zfp142 | 5.83E-05 | 8.85E-05 | 0.0001 | 3.09E-05 | 2.97E-05 | 3.09E-05 | 5.47E-05 | 3.62E-05 | 3.21E-05 | 0 | 0 | 0 | 8.59E-07 | 2.44E-05 | 4.78E-05 | 1.26E-05 | 0.00015 | 2.01E-05 |
| Fastkd3 | 2.67E-05 | 3.56E-05 | 5.96E-05 | 3.55E-06 | 1.33E-05 | 2.16E-05 | 0 | 0 | 3.21E-05 | 1.98E-06 | 3.65E-06 | 1.12E-05 | 1.72E-06 | 0 | 2.82E-05 | 3.84E-05 | 0 | 0 |
| Tmem184c | 0.00013 | 7.52E-05 | 3.07E-05 | 3.91E-05 | 7.62E-05 | 7.09E-05 | 2.32E-05 | 4.24E-05 | 3.20E-05 | 1.46E-05 | 0 | 1.49E-05 | 1.67E-05 | 2.22E-05 | 0 | 4.73E-05 | 0 | 4.39E-05 |
| Rrm1 | 0.00012 | 3.81E-06 | 0.00015 | 0 | 0.0001 | 4.81E-05 | 6.09E-05 | 0 | 3.20E-05 | 8.24E-05 | 0.00034 | 5.68E-07 | 2.01E-05 | 5.59E-05 | 0 | 0.0014 | 0 | 3.96E-05 |
| Lrrc8e | 1.75E-06 | 1.94E-05 | 2.53E-05 | 2.61E-05 | 1.68E-05 | 3.25E-05 | 9.41E-06 | 2.34E-05 | 3.20E-05 | 0 | 1.65E-05 | 6.01E-05 | 9.37E-06 | 2.53E-05 | 2.19E-05 | 2.09E-05 | 0 | 3.26E-05 |
| Unc119 | 1.80E-05 | 3.16E-05 | 0 | 0 | 2.97E-05 | 2.61E-05 | 1.13E-05 | 4.42E-06 | 3.20E-05 | 0 | 2.98E-05 | 3.08E-05 | 0 | 3.23E-05 | 0 | 0 | 2.51E-05 | 1.63E-05 |
| Trim46 | 5.62E-05 | 7.92E-05 | 2.74E-05 | 3.12E-05 | 1.88E-05 | 2.26E-05 | 0.0001 | 7.18E-05 | 3.20E-05 | 0 | 1.41E-06 | 2.30E-05 | 1.77E-05 | 0 | 0.00011 | 0 | 0 | 6.33E-05 |
| Cbr1 | 8.43E-06 | 4.04E-05 | 6.46E-06 | 1.92E-05 | 1.53E-05 | 2.80E-06 | 2.16E-06 | 1.40E-05 | 3.20E-05 | 1.52E-05 | 0 | 2.88E-06 | 6.30E-06 | 1.70E-05 | 8.31E-07 | 0 | 3.18E-06 | 6.83E-06 |
| DEPRECATED | 8.44E-06 | 1.24E-05 | 0 | 0 | 1.52E-05 | 0 | 3.41E-05 | 0 | 3.20E-05 | 1.23E-05 | 0 | 1.40E-05 | 2.61E-06 | 0 | 0 | 8.26E-06 | 0 | 9.03E-06 |
| Pomgnt2 | 0 | 4.68E-05 | 3.20E-08 | 3.04E-05 | 3.91E-05 | 0 | 2.95E-05 | 2.77E-05 | 3.20E-05 | 1.09E-05 | 0 | 1.79E-05 | 1.13E-06 | 1.76E-05 | 0 | 0 | 4.08E-05 | 1.36E-05 |
| Snap23 | 0.00019 | 5.17E-05 | 6.01E-05 | 2.68E-05 | 7.25E-05 | 0.00012 | 0.00013 | 0 | 3.19E-05 | 1.82E-05 | 1.11E-05 | 3.68E-05 | 0 | 7.02E-05 | 0 | 8.45E-06 | 0 | 0 |
| Serhl | 4.39E-05 | 6.24E-05 | 0 | 7.14E-05 | 7.41E-05 | 6.96E-05 | 4.03E-05 | 1.30E-05 | 3.19E-05 | 4.23E-05 | 0 | 3.90E-05 | 2.19E-05 | 4.45E-05 | 6.02E-05 | 0.00012 | 0 | 0.0001 |
| Cep104 | 8.07E-05 | 5.76E-05 | 8.98E-05 | 3.94E-05 | 6.23E-05 | 4.53E-05 | 2.81E-05 | 1.21E-05 | 3.19E-05 | 1.81E-05 | 4.53E-05 | 2.19E-05 | 0 | 4.06E-05 | 3.38E-05 | 5.15E-05 | 0 | 1.84E-05 |
| Dcun1d4 | 4.51E-05 | 4.95E-05 | 0 | 0 | 4.53E-05 | 2.70E-05 | 0 | 0 | 3.19E-05 | 9.87E-06 | 1.81E-05 | 1.75E-05 | 5.84E-05 | 4.97E-05 | 7.81E-05 | 0 | 1.26E-05 | 2.66E-05 |
| Rab30 | 4.65E-05 | 0 | 0 | 0 | 7.01E-06 | 0 | 4.26E-05 | 0 | 3.19E-05 | 0.00013 | 3.98E-05 | 1.64E-06 | 3.64E-05 | 2.51E-05 | 0 | 0 | 1.15E-06 | 0.00014 |
| Lpcat4 | 0 | 3.69E-05 | 4.11E-05 | 7.13E-05 | 2.90E-05 | 0 | 1.78E-06 | 0 | 3.19E-05 | 2.54E-05 | 9.25E-05 | 0 | 2.87E-05 | 5.74E-05 | 1.43E-06 | 3.20E-05 | 4.18E-05 | 7.85E-05 |
| Fam118a | 3.90E-05 | 3.54E-05 | 2.33E-05 | 0 | 0 | 0 | 0 | 7.87E-06 | 3.19E-05 | 2.10E-05 | 0.00015 | 0.0001 | 0 | 4.73E-05 | 8.20E-05 | 6.67E-05 | 1.53E-06 | 0 |
| Scyl2 | 6.64E-05 | 0.00013 | 4.26E-05 | 9.25E-05 | 6.11E-05 | 6.63E-05 | 9.77E-06 | 4.49E-06 | 3.18E-05 | 4.96E-05 | 8.54E-05 | 6.40E-05 | 0 | 7.65E-05 | 0 | 9.33E-05 | 8.80E-05 | 0.00011 |
| Zdhhc17 | 0.00011 | 8.24E-05 | 3.97E-05 | 0 | 5.45E-05 | 5.93E-05 | 0 | 3.56E-05 | 3.18E-05 | 1.99E-05 | 0 | 4.56E-05 | 4.37E-06 | 1.53E-05 | 0.00012 | 1.56E-05 | 5.31E-05 | 5.68E-05 |
| Gfer | 6.42E-05 | 7.63E-05 | 0.00012 | 0.00015 | 3.96E-05 | 3.04E-05 | 1.58E-05 | 0 | 3.18E-05 | 1.37E-05 | 2.85E-05 | 1.08E-05 | 5.46E-05 | 5.87E-07 | 6.03E-06 | 7.82E-05 | 3.65E-05 | 0 |

| | | | | | | | | | | | | | | | | | | |
|---|---|---|---|---|---|---|---|---|---|---|---|---|---|---|---|---|---|---|
| Atpaf2 | 5.79E-05 | 4.94E-05 | 4.51E-05 | 8.15E-05 | 3.11E-05 | 2.43E-05 | 2.65E-05 | 0 | 3.18E-05 | 0 | 2.85E-06 | 4.75E-05 | 3.46E-06 | 5.19E-05 | 3.46E-05 | 5.04E-05 | 5.55E-07 | 1.94E-05 |
| Thumpd1 | 0.00014 | 8.47E-05 | 0.00013 | 3.50E-05 | 3.68E-05 | 1.72E-05 | 1.64E-07 | 0 | 3.18E-05 | 2.58E-05 | 1.03E-05 | 2.20E-05 | 0 | 0 | 9.90E-05 | 4.17E-05 | 0 | 5.33E-05 |
| Ap1g2 | 2.43E-05 | 3.78E-05 | 8.98E-06 | 2.47E-05 | 0 | 7.05E-06 | 1.41E-05 | 0 | 3.18E-05 | 3.64E-05 | 4.87E-06 | 3.26E-05 | 2.40E-05 | 2.19E-05 | 7.67E-05 | 0 | 0 | 3.96E-05 |
| Cln8 | 0 | 5.16E-05 | 2.89E-05 | 5.98E-06 | 4.37E-05 | 0 | 6.06E-05 | 0 | 3.18E-05 | 5.87E-05 | 3.72E-05 | 3.49E-06 | 7.53E-05 | 3.97E-05 | 4.19E-06 | 8.97E-06 | 0 | 6.74E-05 |
| Fam98a | 0.00019 | 2.33E-06 | 0.00025 | 0.00028 | 5.14E-05 | 0 | 0.00024 | 1.24E-05 | 3.18E-05 | 6.84E-05 | 8.95E-05 | 0 | 5.93E-05 | 5.36E-05 | 8.08E-05 | 9.32E-05 | 0 | 7.18E-05 |
| Pcdhga5 | 5.63E-05 | 9.45E-05 | 0 | 0 | 5.91E-05 | 7.75E-05 | 5.56E-05 | 5.50E-05 | 3.17E-05 | 2.33E-05 | 3.16E-05 | 1.78E-06 | 4.75E-05 | 3.10E-05 | 0.00011 | 0 | 3.81E-05 | 2.37E-05 |
| Camkk2 | 5.22E-05 | 9.28E-05 | 0.00013 | 9.31E-05 | 1.22E-05 | 4.99E-05 | 6.73E-05 | 0 | 3.17E-05 | 6.41E-05 | 5.77E-05 | 0 | 9.85E-05 | 6.84E-05 | 3.10E-05 | 0.0001 | 0 | 0.00015 |
| Ago3 | 7.85E-05 | 9.41E-05 | 0.0001 | 0 | 4.66E-05 | 4.56E-05 | 7.52E-05 | 6.07E-05 | 3.17E-05 | 3.94E-05 | 3.41E-05 | 3.63E-05 | 9.04E-07 | 4.48E-05 | 8.51E-05 | 9.07E-06 | 8.24E-05 | 1.33E-05 |
| Polr3f | 5.26E-05 | 3.75E-05 | 1.67E-05 | 8.42E-06 | 8.51E-06 | 4.46E-05 | 6.34E-06 | 1.15E-05 | 3.17E-05 | 0 | 1.30E-05 | 6.56E-06 | 0 | 1.65E-05 | 2.33E-05 | 3.94E-05 | 0 | 0 |
| Abitram | 8.69E-05 | 1.73E-05 | 6.77E-05 | 3.31E-05 | 1.70E-05 | 1.40E-05 | 2.88E-05 | 2.59E-05 | 3.17E-05 | 0 | 0 | 3.73E-06 | 0 | 1.95E-05 | 4.42E-05 | 0 | 0 | 8.07E-06 |
| Timm8a1 | 8.78E-05 | 1.48E-05 | 0.00019 | 0 | 0 | 7.67E-06 | 0 | 0 | 3.17E-05 | 0 | 0 | 5.63E-05 | 6.10E-06 | 4.19E-06 | 0 | 0.00015 | 2.39E-06 | 7.94E-06 |
| Als2 | 4.41E-06 | 3.58E-05 | 1.53E-05 | 4.01E-05 | 4.24E-06 | 0 | 0 | 2.59E-05 | 3.17E-05 | 2.42E-05 | 1.23E-05 | 0 | 0 | 3.40E-05 | 1.44E-05 | 5.90E-05 | 5.49E-05 | 2.77E-05 |
| Brat1 | 1.47E-05 | 5.14E-05 | 0 | 6.86E-05 | 5.39E-05 | 0 | 0 | 1.97E-05 | 3.17E-05 | 3.31E-05 | 2.19E-05 | 3.28E-05 | 0 | 2.50E-05 | 3.79E-05 | 3.61E-05 | 0 | 3.84E-05 |
| Tgds | 0.00015 | 1.57E-05 | 4.42E-05 | 2.64E-06 | 2.79E-05 | 8.24E-05 | 8.45E-05 | 6.11E-05 | 3.16E-05 | 1.16E-05 | 5.68E-05 | 4.92E-05 | 4.76E-05 | 3.55E-05 | 3.19E-06 | 1.73E-05 | 0 | 5.51E-07 |
| Coa6 | 8.53E-05 | 3.83E-05 | 2.96E-05 | 6.72E-06 | 1.15E-05 | 5.34E-05 | 1.29E-05 | 1.91E-05 | 3.16E-05 | 2.03E-05 | 3.31E-08 | 2.26E-05 | 1.85E-05 | 0 | 0 | 6.67E-05 | 0 | 4.47E-06 |
| N4bp2 | 7.13E-05 | 1.36E-05 | 3.12E-05 | 0 | 0 | 5.17E-05 | 6.96E-05 | 4.74E-05 | 3.16E-05 | 0 | 3.35E-05 | 6.17E-05 | 2.40E-05 | 1.95E-05 | 5.55E-05 | 3.61E-05 | 1.14E-06 | 3.63E-06 |
| Mapk8 | 7.04E-05 | 5.86E-05 | 4.36E-05 | 2.78E-05 | 2.28E-05 | 4.30E-05 | 0 | 2.18E-05 | 3.16E-05 | 1.36E-05 | 1.46E-06 | 6.27E-05 | 0 | 0 | 0.00012 | 5.12E-05 | 0.00011 | 3.08E-05 |
| Zc3hav1 | 0.00015 | 0.00015 | 5.58E-05 | 8.76E-05 | 2.31E-05 | 3.29E-05 | 0 | 6.18E-05 | 3.16E-05 | 0.0001 | 0 | 0.00012 | 0 | 2.28E-05 | 7.30E-05 | 0.00018 | 0.00021 | 6.30E-05 |
| Exd2 | 1.67E-05 | 7.69E-05 | 2.18E-05 | 4.99E-05 | 4.46E-05 | 9.68E-06 | 1.25E-05 | 4.82E-05 | 3.16E-05 | 3.38E-05 | 2.58E-05 | 5.92E-05 | 0 | 1.40E-05 | 8.67E-05 | 6.49E-05 | 1.48E-05 | 5.21E-05 |
| Sars2 | 2.54E-05 | 2.68E-05 | 3.69E-05 | 6.22E-05 | 4.84E-06 | 3.15E-06 | 5.42E-06 | 8.84E-09 | 3.16E-05 | 4.91E-06 | 2.08E-05 | 0 | 0 | 2.05E-05 | 0.00011 | 5.50E-05 | 0 | 3.96E-06 |
| Trmt12 | 3.38E-05 | 3.27E-05 | 1.39E-05 | 2.54E-05 | 2.42E-05 | 2.94E-06 | 4.71E-06 | 0 | 3.16E-05 | 0 | 1.11E-05 | 0 | 0 | 1.32E-05 | 1.67E-05 | 6.54E-06 | 0 | 8.59E-06 |
| Slc35f5 | 0.00029 | 0.00019 | 0 | 0 | 0.00017 | 0.00024 | 0.00023 | 3.78E-05 | 3.15E-05 | 0.00013 | 0 | 0.00011 | 0.00019 | 6.14E-05 | 0 | 0 | 0.00012 | 0.00011 |
| Xrn1 | 9.55E-05 | 6.40E-05 | 0 | 0 | 4.28E-05 | 8.38E-05 | 1.42E-05 | 2.84E-05 | 3.15E-05 | 1.75E-05 | 1.22E-05 | 4.68E-05 | 3.98E-05 | 2.01E-05 | 0.00013 | 6.11E-05 | 0 | 5.23E-05 |
| Nup93 | 9.80E-05 | 4.43E-05 | 6.68E-05 | 0 | 5.13E-05 | 2.97E-05 | 7.28E-05 | 0.00012 | 3.15E-05 | 6.05E-05 | 0 | 1.97E-05 | 0.00014 | 0 | 0.00014 | 0.00023 | 2.76E-05 | 5.72E-05 |
| Actr6 | 3.20E-05 | 2.90E-05 | 2.42E-05 | 9.94E-06 | 1.32E-05 | 2.18E-05 | 0 | 8.61E-06 | 3.15E-05 | 5.12E-06 | 5.63E-06 | 1.95E-05 | 0 | 3.05E-05 | 0 | 4.71E-05 | 0 | 1.46E-05 |
| B3gntl1 | 1.57E-05 | 3.68E-05 | 0 | 2.29E-05 | 1.72E-05 | 1.00E-05 | 2.33E-05 | 5.14E-05 | 3.15E-05 | 6.44E-05 | 0 | 4.22E-06 | 0 | 0 | 0.00012 | 4.18E-05 | 7.23E-06 | 4.39E-05 |
| Accs | 5.75E-06 | 4.73E-05 | 0 | 3.46E-06 | 2.36E-05 | 4.05E-06 | 1.23E-05 | 1.72E-05 | 3.15E-05 | 1.41E-05 | 9.95E-06 | 4.41E-06 | 0 | 3.55E-05 | 1.38E-05 | 2.24E-05 | 0 | 4.84E-05 |
| Ppp1r3b | 1.71E-05 | 2.09E-05 | 0 | 1.34E-05 | 6.58E-05 | 0 | 0 | 1.63E-05 | 3.15E-05 | 0 | 2.13E-05 | 0.00048 | 1.46E-05 | 2.33E-05 | 3.05E-05 | 3.14E-06 | 0 | 0 |
| Rxylt1 | 0.00026 | 0.00014 | 0.00026 | 0.00019 | 0.0001 | 0.00015 | 0.00015 | 0.00011 | 3.14E-05 | 7.65E-05 | 8.18E-05 | 7.94E-05 | 0.00014 | 6.07E-05 | 8.55E-05 | 5.91E-05 | 3.47E-05 | 0 |
| Cog5 | 0.00019 | 8.41E-05 | 2.94E-05 | 0 | 8.12E-05 | 6.64E-05 | 8.98E-05 | 0 | 3.14E-05 | 3.40E-05 | 1.62E-05 | 2.03E-05 | 1.12E-05 | 1.28E-07 | 0.00014 | 0 | 7.18E-05 | 2.65E-05 |
| Uggt2 | 3.53E-05 | 3.03E-05 | 8.93E-05 | 7.29E-05 | 7.60E-05 | 5.22E-05 | 5.67E-05 | 0.00011 | 3.14E-05 | 2.29E-06 | 2.95E-05 | 0 | 3.86E-05 | 0.00012 | 0.00021 | 0 | 6.74E-05 | 3.31E-05 |
| Stam | 0.0001 | 0.0002 | 7.10E-05 | 0 | 2.50E-05 | 4.55E-05 | 4.76E-06 | 3.21E-06 | 3.14E-05 | 9.33E-05 | 7.87E-05 | 0.0002 | 0 | 4.70E-06 | 7.92E-05 | 0.00011 | 7.42E-05 | 2.58E-05 |
| Tspan6 | 0.00037 | 3.98E-06 | 0 | 0 | 2.14E-05 | 4.11E-05 | 0.00011 | 0.0002 | 3.14E-05 | 0 | 0.00024 | 0 | 8.69E-05 | 0.00013 | 0 | 9.34E-05 | 0 | 0.00012 |

| | | | | | | | | | | | | | | | | | | |
|---|---|---|---|---|---|---|---|---|---|---|---|---|---|---|---|---|---|---|
| Sat2 | 2.61E-05 | 1.92E-05 | 0 | 2.05E-05 | 5.10E-05 | 2.65E-05 | 0 | 6.10E-06 | 3.14E-05 | 2.30E-05 | 4.01E-07 | 0 | 1.60E-05 | 1.82E-05 | 2.76E-05 | 0 | 0 | 4.14E-05 |
| Shprh | 6.26E-05 | 9.51E-05 | 9.95E-06 | 2.33E-05 | 4.90E-05 | 2.19E-05 | 0 | 2.29E-05 | 3.14E-05 | 2.23E-05 | 0 | 5.17E-05 | 0 | 4.21E-05 | 0.00011 | 2.47E-05 | 0 | 1.30E-05 |
| Terf1 | 1.47E-05 | 4.39E-05 | 0 | 0 | 1.96E-05 | 1.65E-05 | 4.47E-05 | 0 | 3.14E-05 | 2.46E-05 | 0 | 1.00E-05 | 0 | 4.78E-06 | 0.00019 | 0.00034 | 0 | 1.31E-05 |
| Srbd1 | 2.97E-05 | 4.15E-05 | 2.72E-05 | 5.90E-05 | 2.93E-05 | 1.20E-05 | 4.50E-06 | 0 | 3.14E-05 | 1.53E-05 | 0 | 1.94E-05 | 0 | 0 | 8.29E-06 | 0.00018 | 6.23E-05 | 2.77E-05 |
| Mdc1 | 3.26E-05 | 4.39E-05 | 4.26E-05 | 4.76E-05 | 9.74E-06 | 1.38E-06 | 0 | 0 | 3.14E-05 | 1.95E-05 | 1.30E-05 | 2.38E-05 | 4.03E-05 | 0 | 6.96E-05 | 0.00028 | 4.00E-05 | 8.24E-06 |
| Ogfod3 | 3.55E-06 | 1.33E-05 | 0 | 3.29E-05 | 0 | 1.26E-06 | 2.89E-05 | 0 | 3.14E-05 | 8.51E-06 | 2.68E-06 | 1.56E-05 | 4.54E-05 | 3.64E-05 | 7.77E-06 | 3.30E-05 | 6.45E-06 | 4.34E-05 |
| Ecm1 | 0 | 7.22E-05 | 0.00017 | 0.00044 | 0.00042 | 0.00091 | 0.00092 | 0.00082 | 3.13E-05 | 0.00104 | 0.00017 | 0.00094 | 0.00091 | 0.00085 | 0.00037 | 0 | 3.66E-05 | 0.00067 |
| Amigo1 | 1.27E-06 | 5.55E-05 | 0 | 0 | 4.58E-05 | 4.83E-05 | 0 | 0 | 3.13E-05 | 5.46E-05 | 0 | 0 | 3.55E-05 | 2.19E-05 | 0 | 1.36E-06 | 0 | 3.75E-05 |
| Lin37 | 3.77E-05 | 4.09E-05 | 0 | 2.83E-05 | 1.92E-05 | 2.68E-05 | 1.99E-05 | 1.89E-05 | 3.13E-05 | 2.47E-05 | 1.92E-05 | 2.24E-05 | 0 | 2.16E-05 | 8.61E-05 | 3.39E-05 | 0 | 6.36E-05 |
| Adcy9 | 2.16E-06 | 3.96E-05 | 1.32E-05 | 8.67E-06 | 2.15E-05 | 1.39E-05 | 0 | 0 | 3.13E-05 | 4.53E-05 | 1.22E-05 | 0 | 1.80E-05 | 1.72E-05 | 4.25E-07 | 0 | 3.51E-05 | 5.53E-05 |
| Agbl5 | 6.30E-06 | 2.99E-05 | 0 | 1.42E-05 | 3.89E-05 | 3.60E-06 | 0 | 3.80E-05 | 3.13E-05 | 0 | 0 | 1.55E-05 | 0 | 2.58E-05 | 1.38E-05 | 2.84E-05 | 0 | 0 |
| Slc12a6 | 0 | 0.00014 | 0 | 1.10E-05 | 5.72E-05 | 3.78E-05 | 2.70E-05 | 6.12E-05 | 3.12E-05 | 0.00012 | 5.81E-05 | 6.70E-05 | 2.52E-05 | 1.38E-05 | 8.48E-06 | 4.45E-05 | 8.23E-05 | 4.56E-05 |
| Tsc22d2 | 0.00021 | 0.00014 | 6.58E-05 | 0 | 4.80E-05 | 2.36E-05 | 0 | 0 | 3.12E-05 | 3.71E-05 | 0.0003 | 0.00022 | 8.31E-05 | 0 | 9.30E-05 | 8.64E-05 | 0.00071 | 3.16E-05 |
| Pdss2 | 2.08E-05 | 8.22E-06 | 5.11E-06 | 0 | 1.69E-05 | 2.92E-06 | 0 | 3.09E-06 | 3.12E-05 | 1.13E-05 | 1.10E-05 | 1.36E-05 | 1.06E-05 | 0 | 1.22E-05 | 1.32E-05 | 4.25E-05 | 0 |
| Arl10 | 0 | 3.76E-05 | 3.69E-05 | 7.25E-05 | 5.20E-06 | 0 | 4.92E-06 | 8.77E-06 | 3.12E-05 | 1.43E-05 | 3.54E-05 | 1.37E-05 | 1.34E-05 | 0 | 2.33E-05 | 2.20E-05 | 5.74E-05 | 3.72E-06 |
| Ampd2 | 0.00011 | 5.24E-05 | 0.00029 | 0.00028 | 1.26E-05 | 8.19E-05 | 0.00014 | 0.0005 | 3.11E-05 | 4.13E-05 | 0 | 0 | 0 | 0.00011 | 0.00033 | 1.04E-05 | 0.00023 | 5.98E-05 |
| Marf1 | 9.61E-05 | 0.00012 | 0 | 2.17E-05 | 5.02E-05 | 7.94E-05 | 2.15E-05 | 1.94E-05 | 3.11E-05 | 3.10E-05 | 1.22E-05 | 8.86E-05 | 8.76E-06 | 4.88E-05 | 0.00021 | 6.33E-05 | 1.18E-05 | 6.21E-05 |
| Zfp719 | 5.40E-05 | 2.84E-05 | 0 | 0 | 1.46E-05 | 4.68E-05 | 0 | 5.23E-06 | 3.11E-05 | 0 | 0 | 1.67E-05 | 0 | 5.09E-06 | 2.43E-05 | 4.84E-06 | 0 | 0 |
| Otulin | 0.0001 | 9.58E-05 | 0.00016 | 0.00018 | 4.35E-05 | 4.44E-05 | 0.00015 | 0 | 3.11E-05 | 2.21E-05 | 5.19E-05 | 0 | 6.77E-05 | 6.50E-05 | 6.13E-05 | 9.38E-05 | 6.21E-05 | 6.34E-05 |
| Dusp23 | 3.36E-06 | 1.99E-05 | 0 | 0.00011 | 5.40E-06 | 2.20E-05 | 4.45E-05 | 5.62E-05 | 3.11E-05 | 5.87E-05 | 1.35E-06 | 7.50E-05 | 0 | 1.12E-05 | 1.22E-05 | 0 | 0 | 4.16E-05 |
| Spcs2 | 9.40E-05 | 1.10E-06 | 4.62E-05 | 0 | 0 | 1.95E-05 | 0 | 6.90E-06 | 3.11E-05 | 5.22E-06 | 1.05E-05 | 2.81E-05 | 0 | 3.49E-06 | 3.35E-07 | 0 | 0 | 0 |
| Elp4 | 9.47E-05 | 5.53E-05 | 2.62E-05 | 0 | 2.72E-05 | 5.91E-05 | 3.62E-05 | 0 | 3.10E-05 | 2.24E-05 | 0 | 2.04E-05 | 1.36E-05 | 0 | 2.83E-05 | 5.23E-05 | 2.42E-06 | 2.60E-05 |
| Mindy3 | 9.18E-05 | 7.55E-05 | 0.0001 | 6.30E-06 | 3.42E-05 | 5.83E-05 | 0 | 3.53E-05 | 3.10E-05 | 3.83E-05 | 9.47E-05 | 7.79E-06 | 0 | 4.56E-05 | 2.84E-05 | 4.43E-05 | 8.20E-06 | 3.38E-05 |
| Itsn2 | 8.05E-05 | 0.00013 | 7.06E-05 | 2.48E-05 | 2.77E-05 | 4.40E-05 | 0 | 2.88E-05 | 3.10E-05 | 3.27E-05 | 5.10E-05 | 9.18E-05 | 0 | 9.86E-06 | 0.00019 | 4.29E-05 | 0.00012 | 7.66E-05 |
| Bloc1s5 | 3.00E-05 | 5.39E-05 | 1.81E-05 | 9.26E-05 | 2.54E-05 | 1.96E-05 | 2.31E-05 | 1.23E-05 | 3.10E-05 | 4.42E-05 | 1.44E-05 | 2.49E-05 | 0 | 0 | 5.76E-06 | 1.77E-05 | 0 | 6.48E-05 |
| Gas2 | 1.06E-05 | 1.16E-05 | 0 | 4.43E-06 | 2.64E-07 | 3.24E-06 | 0 | 1.63E-05 | 3.10E-05 | 0 | 3.09E-06 | 2.80E-05 | 1.47E-05 | 3.94E-05 | 7.57E-06 | 2.61E-05 | 0 | 1.63E-05 |
| Ube2l6 | 0.0001 | 5.64E-05 | 0 | 9.00E-05 | 0 | 0 | 2.90E-06 | 1.23E-05 | 3.10E-05 | 0.00029 | 0 | 6.31E-05 | 0.00017 | 6.30E-06 | 0 | 4.09E-05 | 0 | 0.00041 |
| Lepr | 0 | 5.46E-05 | 0 | 0 | 0 | 0 | 0 | 0 | 3.10E-05 | 0 | 0 | 0 | 0 | 1.29E-05 | 0 | 0 | 0 | 0.0001 |
| Crocc | 0 | 2.78E-05 | 0 | 3.96E-05 | 2.18E-05 | 0 | 0 | 7.11E-06 | 3.10E-05 | 1.86E-05 | 4.28E-06 | 6.50E-05 | 1.13E-06 | 0 | 0.00011 | 6.89E-05 | 2.13E-05 | 2.84E-05 |
| Arsa | 0 | 0.00015 | 0 | 0.00021 | 0.00017 | 0.00017 | 0.00027 | 0.00017 | 3.09E-05 | 0.00027 | 4.81E-05 | 0.00023 | 0.00014 | 6.82E-05 | 0 | 0 | 5.25E-05 | 0.00029 |
| Fuom | 0 | 1.63E-05 | 0 | 4.20E-05 | 2.53E-05 | 3.06E-05 | 4.77E-05 | 0 | 3.09E-05 | 1.99E-05 | 1.52E-05 | 0 | 5.43E-05 | 4.47E-05 | 0 | 5.42E-05 | 0 | 6.06E-05 |
| Supv3l1 | 5.68E-05 | 3.03E-05 | 7.19E-05 | 4.17E-05 | 1.16E-05 | 1.39E-05 | 3.82E-06 | 0 | 3.09E-05 | 1.37E-05 | 8.27E-06 | 0 | 0 | 2.06E-05 | 1.96E-05 | 4.80E-05 | 0 | 1.35E-05 |
| Rrp9 | 2.22E-05 | 2.58E-05 | 0.00013 | 4.37E-05 | 0 | 0 | 0 | 0 | 3.09E-05 | 1.08E-05 | 3.99E-05 | 0 | 7.06E-06 | 0 | 7.48E-06 | 3.59E-05 | 0 | 0 |

| | | | | | | | | | | | | | | | | | | |
|---|---|---|---|---|---|---|---|---|---|---|---|---|---|---|---|---|---|---|
| Kbtbd2 | 0.00014 | 0.00017 | 8.72E-05 | 0 | 0.00011 | 0.00017 | 3.48E-05 | 8.16E-05 | 3.08E-05 | 3.43E-05 | 0.00011 | 0.00017 | 0 | 0.00011 | 0 | 0.0002 | 0.00011 | 7.48E-05 |
| Kat6b | 0.00011 | 0.00016 | 8.53E-07 | 0 | 0.0001 | 0.00016 | 4.19E-05 | 1.28E-05 | 3.08E-05 | 5.37E-05 | 5.58E-05 | 0.0002 | 0 | 7.32E-05 | 0.00024 | 0 | 0 | 8.09E-05 |
| Sf3b5 | 0.00024 | 7.12E-05 | 0.00027 | 0.00049 | 9.82E-05 | 6.14E-05 | 0.00021 | 8.34E-05 | 3.08E-05 | 4.82E-05 | 7.14E-05 | 4.58E-05 | 3.05E-05 | 2.27E-05 | 0 | 0.00035 | 0 | 7.12E-05 |
| Inpp5e | 3.68E-05 | 8.16E-05 | 5.14E-06 | 0 | 5.02E-06 | 3.71E-05 | 8.97E-06 | 0 | 3.08E-05 | 8.79E-06 | 4.94E-06 | 3.94E-05 | 0 | 7.04E-06 | 5.33E-05 | 1.42E-05 | 6.76E-05 | 2.86E-05 |
| Tcaf2 | 1.95E-05 | 1.43E-05 | 0 | 6.68E-05 | 9.77E-06 | 2.80E-05 | 0 | 1.95E-05 | 3.08E-05 | 5.31E-05 | 1.24E-05 | 8.30E-06 | 5.63E-05 | 1.38E-05 | 0 | 6.49E-05 | 5.65E-05 | 6.89E-05 |
| Arhgap8 | 6.44E-06 | 0 | 6.23E-08 | 9.23E-06 | 0 | 2.36E-05 | 0 | 0 | 3.08E-05 | 0 | 0 | 0 | 2.63E-06 | 2.77E-06 | 0 | 1.02E-05 | 0 | 0 |
| Abcd4 | 2.65E-05 | 5.47E-05 | 2.66E-06 | 2.86E-05 | 8.84E-06 | 1.85E-05 | 6.03E-05 | 2.88E-05 | 3.08E-05 | 3.52E-05 | 0 | 1.67E-05 | 3.79E-05 | 5.43E-06 | 4.61E-05 | 0 | 0 | 3.78E-05 |
| Kctd18 | 1.50E-05 | 4.22E-05 | 1.31E-05 | 4.60E-05 | 3.80E-05 | 1.63E-05 | 4.05E-05 | 2.00E-05 | 3.08E-05 | 1.66E-05 | 2.79E-05 | 8.07E-06 | 1.69E-05 | 2.45E-05 | 0.00016 | 4.58E-05 | 0 | 2.83E-05 |
| Fbxo46 | 0 | 5.37E-05 | 2.72E-05 | 6.90E-05 | 2.82E-05 | 0 | 2.21E-06 | 1.42E-06 | 3.08E-05 | 3.35E-05 | 4.65E-05 | 2.51E-05 | 0 | 1.63E-05 | 5.11E-05 | 1.06E-05 | 0 | 3.54E-05 |
| Eipr1 | 2.63E-05 | 3.33E-05 | 3.69E-05 | 0.00011 | 3.29E-05 | 0 | 1.91E-05 | 1.08E-05 | 3.08E-05 | 3.91E-05 | 2.02E-05 | 6.48E-06 | 1.89E-05 | 1.87E-05 | 4.27E-05 | 3.59E-05 | 1.50E-05 | 3.65E-05 |
| Zfyve16 | 0.00013 | 7.50E-05 | 2.63E-05 | 0 | 3.28E-05 | 4.51E-05 | 4.53E-05 | 0 | 3.07E-05 | 5.03E-05 | 1.51E-06 | 5.24E-05 | 4.01E-06 | 0 | 7.39E-05 | 2.25E-05 | 2.40E-06 | 9.20E-06 |
| Tubgcp6 | 0 | 6.28E-05 | 1.55E-05 | 5.94E-05 | 2.64E-05 | 4.30E-05 | 1.56E-05 | 1.33E-05 | 3.07E-05 | 2.87E-05 | 3.59E-05 | 0 | 1.36E-05 | 1.88E-05 | 0.00011 | 0.00015 | 0 | 3.97E-05 |
| Lmbr1 | 4.40E-05 | 6.94E-05 | 9.27E-06 | 0 | 4.64E-05 | 3.80E-05 | 1.81E-05 | 1.45E-05 | 3.07E-05 | 0.00014 | 5.51E-06 | 2.58E-05 | 7.33E-05 | 2.90E-05 | 0 | 0 | 0 | 9.65E-05 |
| Msrb2 | 2.25E-05 | 1.98E-05 | 0 | 0 | 2.46E-05 | 2.47E-05 | 0 | 1.56E-05 | 3.07E-05 | 2.79E-05 | 0 | 0 | 3.29E-05 | 5.20E-05 | 7.60E-06 | 5.32E-06 | 0 | 0 |
| Sap130 | 8.98E-05 | 0.00014 | 8.08E-05 | 7.47E-05 | 6.52E-05 | 2.43E-05 | 6.80E-05 | 4.82E-05 | 3.07E-05 | 3.85E-05 | 1.74E-05 | 0.00012 | 4.15E-05 | 1.80E-05 | 0.00014 | 0.0002 | 0.00013 | 4.41E-05 |
| Nup107 | 2.84E-05 | 9.25E-06 | 0.00013 | 9.69E-05 | 1.54E-05 | 1.80E-05 | 5.39E-05 | 1.72E-05 | 3.07E-05 | 3.80E-05 | 3.17E-05 | 2.38E-05 | 1.17E-05 | 2.99E-05 | 9.96E-06 | 0.00049 | 1.51E-05 | 2.95E-05 |
| Zfp428 | 0 | 2.08E-06 | 4.11E-05 | 5.20E-05 | 2.01E-05 | 1.72E-05 | 0 | 5.49E-05 | 3.07E-05 | 0 | 1.30E-05 | 1.31E-05 | 0 | 3.57E-05 | 8.69E-06 | 3.92E-05 | 2.69E-05 | 7.05E-06 |
| Txnrd3 | 1.15E-05 | 2.34E-05 | 1.68E-05 | 0 | 0 | 1.46E-05 | 0 | 0 | 3.07E-05 | 2.35E-05 | 6.24E-06 | 6.07E-05 | 0 | 3.25E-06 | 0 | 0 | 0 | 1.59E-05 |
| Mknk1 | 5.03E-05 | 6.81E-05 | 0 | 1.90E-05 | 2.03E-05 | 1.36E-05 | 6.29E-05 | 0 | 3.07E-05 | 3.73E-05 | 1.81E-05 | 1.20E-05 | 1.43E-05 | 3.56E-06 | 3.10E-05 | 0 | 6.28E-05 | 1.40E-05 |
| Mbtd1 | 9.75E-05 | 0.00012 | 2.60E-05 | 0 | 5.42E-05 | 0.00011 | 2.68E-05 | 9.11E-05 | 3.06E-05 | 0 | 6.75E-06 | 9.21E-05 | 0 | 4.58E-05 | 0.00015 | 1.38E-05 | 0.00015 | 6.63E-06 |
| Tasor | 0.00013 | 0.00012 | 0 | 0 | 9.87E-05 | 0.0001 | 2.01E-05 | 2.69E-05 | 3.06E-05 | 4.37E-05 | 6.55E-05 | 0.00011 | 3.78E-05 | 1.64E-05 | 5.04E-05 | 5.48E-05 | 2.74E-05 | 3.13E-06 |
| Wdr75 | 0.00018 | 1.95E-05 | 0.00019 | 0 | 0 | 8.02E-05 | 5.27E-06 | 0 | 3.06E-05 | 0 | 2.65E-05 | 0 | 0 | 0 | 2.56E-05 | 6.61E-05 | 0 | 0 |
| Ascc1 | 6.43E-05 | 6.63E-05 | 0 | 3.72E-05 | 1.80E-05 | 7.52E-05 | 5.77E-05 | 0 | 3.06E-05 | 2.12E-05 | 2.14E-05 | 8.15E-05 | 0 | 4.12E-05 | 0 | 3.24E-05 | 9.21E-06 | 2.08E-05 |
| Snapc1 | 0.00012 | 2.50E-05 | 4.78E-05 | 0 | 1.79E-05 | 5.28E-05 | 0 | 2.04E-05 | 3.06E-05 | 9.28E-06 | 1.03E-05 | 9.21E-05 | 9.37E-05 | 0 | 0.00011 | 7.02E-06 | 0 | 1.30E-05 |
| Bod1l | 8.26E-05 | 7.97E-05 | 3.58E-05 | 1.81E-05 | 5.67E-05 | 5.08E-05 | 5.48E-06 | 3.31E-05 | 3.06E-05 | 1.54E-05 | 0 | 6.63E-05 | 2.51E-06 | 4.93E-06 | 0.00011 | 5.84E-05 | 2.98E-05 | 3.22E-05 |
| Zdhhc1 | 0 | 0.00012 | 0 | 6.66E-05 | 0.00012 | 5.04E-05 | 6.94E-05 | 5.81E-05 | 3.06E-05 | 0.00014 | 5.64E-05 | 4.16E-05 | 1.20E-05 | 0.00013 | 5.90E-05 | 0 | 9.53E-05 | 0.00017 |
| Nthl1 | 2.47E-05 | 3.57E-05 | 0 | 2.60E-05 | 1.44E-05 | 9.89E-06 | 0 | 2.16E-05 | 3.06E-05 | 0 | 1.05E-05 | 1.24E-05 | 1.89E-05 | 3.10E-06 | 9.73E-05 | 1.01E-05 | 4.26E-06 | 9.34E-06 |
| Nde1 | 7.29E-05 | 6.02E-05 | 6.06E-05 | 4.62E-05 | 2.54E-05 | 7.83E-06 | 1.44E-05 | 2.12E-05 | 3.06E-05 | 4.32E-05 | 7.25E-05 | 0 | 2.41E-05 | 1.75E-05 | 0.00018 | 0.00043 | 0 | 8.88E-06 |
| Meak7 | 0 | 2.71E-05 | 0 | 2.88E-05 | 4.14E-05 | 6.85E-06 | 4.50E-05 | 3.97E-05 | 3.06E-05 | 2.60E-05 | 3.57E-06 | 2.61E-05 | 7.57E-06 | 3.69E-06 | 5.35E-05 | 5.27E-06 | 0 | 4.16E-05 |
| Hook2 | 1.45E-05 | 8.58E-05 | 0 | 6.40E-05 | 1.80E-05 | 0 | 2.74E-05 | 0 | 3.06E-05 | 2.16E-05 | 2.33E-06 | 0.0001 | 5.06E-05 | 2.61E-05 | 0.00013 | 2.78E-05 | 4.31E-05 | 1.15E-05 |
| Vps37d | 0 | 3.78E-05 | 0 | 2.60E-05 | 2.33E-05 | 0 | 7.79E-06 | 0 | 3.06E-05 | 3.40E-06 | 0 | 2.93E-05 | 0 | 1.49E-05 | 5.53E-05 | 0 | 0 | 2.81E-05 |
| Zfp397 | 5.74E-05 | 5.58E-05 | 7.21E-06 | 0 | 3.20E-05 | 5.02E-05 | 0 | 6.01E-06 | 3.05E-05 | 3.26E-05 | 1.17E-05 | 7.21E-05 | 9.29E-06 | 3.28E-05 | 0.00023 | 3.20E-05 | 0 | 3.72E-05 |
| Thnsl2 | 4.17E-05 | 6.07E-05 | 1.77E-06 | 1.09E-05 | 1.24E-05 | 4.39E-05 | 6.85E-05 | 2.77E-05 | 3.05E-05 | 3.38E-05 | 4.70E-06 | 3.49E-05 | 0 | 1.78E-05 | 0 | 0 | 3.03E-06 | 4.47E-05 |

| | | | | | | | | | | | | | | | | | | |
|---|---|---|---|---|---|---|---|---|---|---|---|---|---|---|---|---|---|---|
| Mfsd4b4 | 1.52E-06 | 2.80E-05 | 1.96E-06 | 1.25E-05 | 1.36E-05 | 2.61E-06 | 1.66E-05 | 2.41E-05 | 3.05E-05 | 1.04E-05 | 1.66E-06 | 4.09E-06 | 2.60E-05 | 1.33E-05 | 3.87E-05 | 4.11E-07 | 0 | 2.74E-05 |
| Pcmtd1 | 5.23E-05 | 0.00013 | 0 | 0 | 5.84E-05 | 9.91E-05 | 6.72E-05 | 1.62E-05 | 3.04E-05 | 4.00E-05 | 3.34E-05 | 1.92E-05 | 2.25E-05 | 4.24E-05 | 0 | 1.84E-05 | 0 | 0.00013 |
| Sox13 | 0 | 2.99E-05 | 3.81E-05 | 0 | 6.89E-06 | 1.51E-05 | 8.28E-06 | 6.98E-06 | 3.04E-05 | 1.00E-04 | 1.25E-05 | 0 | 0 | 0 | 1.20E-05 | 0 | 1.22E-05 | 0 |
| Dbr1 | 4.58E-05 | 4.13E-05 | 3.57E-05 | 0 | 1.84E-05 | 1.04E-05 | 0 | 1.58E-06 | 3.04E-05 | 2.35E-05 | 1.42E-05 | 8.92E-06 | 0 | 2.74E-05 | 1.01E-05 | 8.22E-05 | 0 | 5.47E-06 |
| Crebzf | 1.12E-05 | 0.00011 | 6.15E-05 | 0.00011 | 5.69E-05 | 0 | 0 | 6.66E-05 | 3.04E-05 | 8.42E-05 | 4.59E-05 | 0.00013 | 0 | 1.97E-05 | 0.00098 | 6.73E-05 | 9.22E-05 | 6.46E-05 |
| Pwwp2a | 9.58E-05 | 0.00015 | 4.00E-05 | 6.75E-05 | 6.94E-05 | 7.07E-05 | 0 | 5.13E-05 | 3.03E-05 | 1.80E-05 | 4.74E-06 | 5.68E-05 | 0 | 7.88E-05 | 0.0003 | 0.00013 | 6.58E-05 | 7.47E-06 |
| Zfp949 | 2.70E-05 | 4.22E-05 | 0 | 5.24E-07 | 2.10E-05 | 3.27E-05 | 2.68E-05 | 1.77E-05 | 3.03E-05 | 1.75E-05 | 1.32E-05 | 0 | 0 | 1.22E-05 | 4.20E-06 | 0 | 0 | 1.91E-05 |
| Kmt5c | 1.51E-05 | 7.41E-05 | 0 | 0.00013 | 7.06E-05 | 3.09E-05 | 4.36E-05 | 0.00013 | 3.03E-05 | 6.51E-05 | 1.78E-05 | 6.60E-05 | 0 | 2.50E-05 | 0.00037 | 6.85E-05 | 3.65E-05 | 6.20E-05 |
| Dkk2 | 0 | 0.00022 | 0 | 0 | 1.21E-06 | 0 | 0 | 0 | 3.03E-05 | 0.0001 | 0 | 0 | 0 | 0.00022 | 0 | 0 | 0 | 0 |
| Tut1 | 9.23E-06 | 4.10E-05 | 6.26E-05 | 6.67E-05 | 1.15E-05 | 0 | 5.00E-06 | 5.54E-06 | 3.03E-05 | 2.37E-05 | 0 | 3.83E-05 | 0 | 1.42E-05 | 0.0001 | 0 | 0 | 2.13E-05 |
| Erich1 | 1.20E-05 | 2.62E-05 | 1.65E-05 | 0 | 9.60E-06 | 5.31E-06 | 0 | 3.03E-06 | 3.02E-05 | 1.16E-05 | 1.57E-05 | 1.08E-05 | 1.65E-06 | 1.77E-05 | 3.10E-05 | 1.02E-05 | 4.92E-05 | 9.22E-06 |
| Slc35e4 | 0 | 7.00E-05 | 0.00011 | 8.07E-05 | 3.96E-05 | 0 | 0 | 0.00011 | 3.02E-05 | 3.78E-05 | 4.80E-05 | 1.65E-05 | 0 | 4.60E-05 | 4.04E-05 | 3.90E-06 | 9.20E-05 | 9.76E-05 |
| Mtg1 | 2.75E-05 | 2.55E-05 | 2.11E-05 | 4.04E-05 | 4.30E-06 | 0 | 4.52E-06 | 1.49E-05 | 3.02E-05 | 2.36E-05 | 2.99E-05 | 5.64E-06 | 0 | 2.18E-06 | 0 | 8.72E-06 | 1.66E-05 | 2.88E-05 |
| Il20ra | 1.17E-05 | 0 | 7.40E-06 | 0 | 0.00015 | 3.58E-05 | 5.21E-05 | 2.18E-05 | 3.01E-05 | 0 | 3.28E-05 | 0 | 0 | 0.00013 | 2.43E-05 | 1.65E-05 | 0 | 0 |
| Prrg1 | 7.84E-05 | 2.03E-06 | 3.66E-05 | 0 | 0 | 2.11E-05 | 3.16E-05 | 0 | 3.01E-05 | 6.20E-05 | 0 | 7.85E-05 | 5.18E-05 | 1.49E-05 | 0 | 9.22E-05 | 1.97E-05 | 3.71E-05 |
| Impa2 | 0 | 1.90E-05 | 1.47E-05 | 6.40E-06 | 1.13E-05 | 1.41E-05 | 0 | 1.06E-05 | 3.01E-05 | 0 | 4.43E-05 | 2.06E-05 | 0 | 4.09E-05 | 0 | 9.96E-05 | 0 | 1.84E-05 |
| Thap2 | 1.66E-05 | 7.57E-05 | 2.76E-05 | 0.00016 | 5.04E-05 | 0.00014 | 7.52E-05 | 5.52E-05 | 3.00E-05 | 2.80E-05 | 2.70E-05 | 5.33E-05 | 2.13E-06 | 0.00015 | 7.51E-05 | 5.52E-05 | 0 | 7.45E-05 |
| Zfp157 | 4.36E-05 | 4.54E-05 | 0 | 2.90E-06 | 2.11E-05 | 2.10E-05 | 6.54E-06 | 1.57E-06 | 3.00E-05 | 1.43E-06 | 0 | 1.38E-05 | 0 | 2.78E-05 | 6.57E-05 | 5.47E-06 | 0 | 3.54E-06 |
| Acy3 | 0 | 3.92E-05 | 0 | 2.14E-05 | 3.43E-06 | 1.11E-05 | 0 | 3.89E-05 | 3.00E-05 | 6.41E-05 | 4.25E-07 | 0 | 0 | 1.25E-05 | 0 | 0 | 0 | 3.22E-05 |
| Gstcd | 6.43E-05 | 3.96E-07 | 8.07E-05 | 0 | 0 | 0 | 2.43E-05 | 1.76E-05 | 3.00E-05 | 6.59E-06 | 4.69E-05 | 1.69E-05 | 0 | 0 | 9.11E-05 | 0.00023 | 3.14E-05 | 0 |
| Safb | 0.00012 | 0.00017 | 0.00016 | 0.00019 | 2.68E-05 | 0 | 4.41E-05 | 0 | 3.00E-05 | 4.57E-05 | 1.94E-05 | 7.12E-05 | 0 | 5.76E-05 | 0.00041 | 0.00022 | 8.21E-05 | 7.33E-05 |
| Naga | 0 | 0.00018 | 0 | 0.00032 | 0.00014 | 0.00019 | 0.00021 | 0.00019 | 2.99E-05 | 0.00015 | 0 | 0.00019 | 0.00032 | 0.00017 | 7.13E-05 | 2.90E-05 | 0.00011 | 0.00029 |
| Tcta | 3.57E-05 | 0.00011 | 0 | 0.00012 | 7.09E-05 | 4.05E-05 | 4.16E-05 | 4.58E-05 | 2.99E-05 | 7.17E-05 | 4.61E-05 | 5.59E-05 | 0 | 4.18E-06 | 0.00015 | 0 | 0 | 0.00013 |
| Mppe1 | 3.37E-05 | 3.68E-06 | 9.77E-06 | 5.10E-05 | 1.84E-05 | 3.98E-05 | 7.69E-05 | 1.91E-05 | 2.99E-05 | 6.01E-05 | 0 | 0 | 0 | 5.51E-05 | 2.29E-05 | 1.08E-05 | 0 | 5.07E-05 |
| Gpn3 | 9.83E-05 | 4.84E-05 | 5.51E-05 | 1.38E-05 | 6.71E-05 | 3.45E-05 | 4.50E-05 | 0 | 2.99E-05 | 0 | 0 | 6.45E-05 | 4.80E-05 | 1.97E-05 | 6.01E-05 | 0.00014 | 0 | 0 |
| Ap5b1 | 0 | 6.24E-05 | 0 | 9.55E-05 | 3.36E-05 | 9.93E-06 | 6.12E-06 | 2.86E-05 | 2.99E-05 | 2.69E-05 | 0 | 4.12E-05 | 0 | 1.44E-05 | 5.74E-05 | 0 | 0 | 3.75E-05 |
| Zfp668 | 2.70E-05 | 5.05E-05 | 6.16E-05 | 2.68E-05 | 2.81E-05 | 9.05E-06 | 1.78E-05 | 2.17E-05 | 2.99E-05 | 1.71E-05 | 3.71E-05 | 8.21E-07 | 0 | 9.09E-06 | 1.96E-05 | 2.97E-05 | 0 | 1.55E-05 |
| Lysmd2 | 6.72E-05 | 5.89E-05 | 2.18E-05 | 2.65E-05 | 2.66E-05 | 0 | 0 | 0 | 2.99E-05 | 0.00022 | 3.21E-05 | 0 | 0.00016 | 6.76E-05 | 0 | 4.26E-05 | 0 | 0.00018 |
| Abhd12 | 0.00011 | 0.00015 | 6.46E-05 | 0.00012 | 0.00018 | 0.00018 | 6.28E-05 | 0.00011 | 2.98E-05 | 6.64E-05 | 0.00012 | 0 | 0.00013 | 0.00012 | 5.74E-05 | 0.00018 | 0 | 7.43E-05 |
| Arv1 | 6.98E-06 | 3.94E-05 | 3.01E-05 | 1.86E-05 | 9.82E-06 | 1.16E-05 | 0 | 2.33E-06 | 2.98E-05 | 3.67E-06 | 0 | 3.69E-06 | 6.45E-07 | 5.03E-06 | 3.41E-05 | 6.25E-06 | 0 | 5.13E-06 |
| Rel | 8.12E-06 | 1.04E-05 | 3.11E-05 | 0 | 1.89E-05 | 8.04E-06 | 0 | 1.53E-05 | 2.98E-05 | 1.53E-06 | 0 | 6.20E-07 | 0 | 3.87E-05 | 2.92E-05 | 2.39E-05 | 0.00037 | 2.30E-05 |
| Nudt6 | 1.93E-05 | 2.29E-05 | 4.51E-06 | 2.49E-06 | 4.85E-06 | 0 | 9.58E-06 | 2.55E-06 | 2.98E-05 | 9.33E-06 | 0 | 0 | 0 | 9.76E-06 | 1.23E-05 | 3.81E-06 | 0 | 0 |
| Qrsl1 | 2.63E-05 | 1.47E-05 | 3.75E-05 | 6.46E-06 | 4.88E-06 | 0 | 0 | 0 | 2.98E-05 | 4.74E-06 | 1.42E-05 | 0 | 1.57E-05 | 1.48E-06 | 3.48E-05 | 4.08E-05 | 0 | 5.71E-06 |

| | | | | | | | | | | | | | | | | | | |
|---|---|---|---|---|---|---|---|---|---|---|---|---|---|---|---|---|---|---|
| Rcor2 | 3.12E-05 | 7.10E-07 | 9.75E-05 | 0 | 0 | 0 | 0 | 5.04E-06 | 2.98E-05 | 0 | 1.91E-05 | 0.00017 | 0 | 8.06E-05 | 3.54E-05 | 3.39E-05 | 0 | 0 |
| Tmem87b | 9.89E-05 | 0.00023 | 9.47E-05 | 0 | 0.0002 | 0.00029 | 0.00012 | 9.00E-05 | 2.97E-05 | 6.36E-05 | 2.21E-05 | 0.00016 | 0.00022 | 5.80E-05 | 0.00014 | 0 | 0.00011 | 8.96E-05 |
| Catip | 0 | 0 | 0 | 0 | 1.73E-05 | 2.32E-05 | 0 | 0 | 2.97E-05 | 1.52E-08 | 8.64E-06 | 0 | 1.05E-05 | 9.43E-07 | 0 | 0 | 1.67E-06 | 0 |
| Cplane1 | 7.83E-05 | 0.0001 | 4.82E-06 | 9.52E-06 | 2.71E-05 | 1.82E-05 | 2.62E-05 | 0 | 2.97E-05 | 2.75E-05 | 1.63E-05 | 4.28E-05 | 4.04E-06 | 1.48E-05 | 7.54E-05 | 7.10E-05 | 4.00E-05 | 2.31E-05 |
| Tti1 | 2.91E-05 | 3.91E-05 | 2.10E-05 | 1.05E-05 | 3.22E-05 | 1.23E-05 | 1.76E-05 | 1.74E-05 | 2.97E-05 | 2.87E-05 | 0 | 2.55E-05 | 4.78E-05 | 2.03E-05 | 3.56E-05 | 0.00011 | 0 | 1.95E-06 |
| Igip | 0.00011 | 5.41E-05 | 2.21E-05 | 0 | 7.80E-05 | 0.00011 | 4.51E-05 | 3.29E-05 | 2.96E-05 | 1.96E-05 | 2.24E-05 | 0.00011 | 0 | 0 | 3.63E-05 | 0 | 0 | 2.18E-05 |
| Ttc8 | 5.77E-05 | 4.64E-05 | 0 | 4.19E-05 | 5.00E-05 | 5.60E-05 | 1.38E-05 | 0 | 2.96E-05 | 6.45E-05 | 3.26E-06 | 1.53E-05 | 1.34E-05 | 3.51E-05 | 0 | 5.48E-05 | 4.55E-05 | 5.12E-05 |
| Asxl2 | 9.22E-05 | 0.00011 | 3.69E-05 | 0 | 7.14E-05 | 4.63E-05 | 9.80E-06 | 3.34E-07 | 2.96E-05 | 0 | 2.75E-05 | 0.00017 | 1.32E-06 | 2.00E-05 | 0.00019 | 2.76E-05 | 4.55E-05 | 2.89E-05 |
| Klf15 | 0 | 5.79E-05 | 0 | 0 | 3.83E-05 | 2.82E-05 | 0 | 8.01E-06 | 2.96E-05 | 8.69E-07 | 0 | 1.61E-05 | 0 | 0 | 8.04E-07 | 0 | 0 | 0 |
| Chac2 | 2.09E-05 | 8.47E-06 | 0 | 0 | 7.83E-06 | 1.60E-05 | 0 | 0 | 2.96E-05 | 1.89E-05 | 0 | 2.35E-05 | 0 | 3.35E-06 | 1.31E-06 | 3.23E-05 | 0 | 1.78E-05 |
| Pmf1 | 3.79E-05 | 0 | 4.97E-06 | 0.00013 | 0 | 0 | 0 | 0 | 2.96E-05 | 0 | 0 | 0 | 3.48E-05 | 4.34E-05 | 0 | 0.0007 | 4.91E-05 | 0 |
| Tfap2a | 0 | 0.00011 | 0 | 0 | 0 | 0 | 0 | 0.00011 | 2.96E-05 | 0 | 0 | 5.94E-06 | 0 | 0.00036 | 0 | 1.29E-05 | 0 | 0 |
| Tbl1xr1 | 0.00025 | 0.00015 | 2.85E-05 | 1.19E-05 | 6.20E-05 | 0.00014 | 0.00016 | 3.91E-05 | 2.95E-05 | 5.47E-05 | 1.37E-05 | 6.23E-05 | 6.70E-05 | 0 | 1.54E-06 | 0.00019 | 8.67E-05 | 0.00014 |
| Sgtb | 2.78E-05 | 5.05E-05 | 3.45E-05 | 2.72E-06 | 4.87E-06 | 2.68E-05 | 8.94E-06 | 0 | 2.95E-05 | 1.05E-05 | 2.57E-05 | 2.64E-05 | 0 | 0 | 4.31E-05 | 0 | 7.80E-05 | 0 |
| Ndor1 | 2.33E-05 | 0.0001 | 5.28E-05 | 0.0001 | 3.00E-05 | 2.46E-05 | 3.62E-05 | 0 | 2.95E-05 | 1.23E-05 | 0 | 0 | 1.05E-05 | 4.60E-05 | 0.00021 | 0 | 0 | 2.81E-05 |
| Them4 | 4.40E-05 | 5.72E-06 | 3.55E-06 | 0 | 0 | 1.96E-05 | 3.56E-06 | 0 | 2.95E-05 | 7.95E-06 | 0 | 1.32E-05 | 1.31E-05 | 0 | 1.87E-05 | 1.19E-07 | 9.08E-06 | 0 |
| Dmap1 | 5.97E-05 | 6.12E-05 | 2.34E-05 | 2.27E-05 | 2.71E-06 | 1.32E-05 | 6.64E-06 | 0 | 2.95E-05 | 2.93E-05 | 4.94E-06 | 5.85E-06 | 0 | 3.75E-06 | 9.35E-05 | 6.41E-05 | 4.11E-05 | 1.87E-05 |
| Alg8 | 9.78E-05 | 0 | 0.00015 | 0.00011 | 0 | 2.77E-06 | 0 | 7.61E-06 | 2.95E-05 | 0 | 8.39E-06 | 0 | 2.55E-05 | 0 | 5.28E-06 | 0.00033 | 1.55E-05 | 0 |
| Usp49 | 3.17E-05 | 1.95E-05 | 9.55E-06 | 0 | 2.91E-05 | 1.71E-06 | 3.98E-05 | 0 | 2.95E-05 | 3.12E-06 | 1.38E-05 | 4.03E-05 | 3.65E-05 | 2.88E-05 | 4.14E-05 | 6.11E-05 | 3.55E-05 | 7.47E-06 |
| Trip11 | 0.00019 | 0.00012 | 0.00014 | 1.47E-05 | 0.00011 | 8.11E-05 | 0.00015 | 8.94E-06 | 2.94E-05 | 3.51E-05 | 8.06E-05 | 6.54E-05 | 2.82E-05 | 0.00014 | 0.00026 | 4.25E-05 | 9.53E-05 | 0 |
| Tatdn1 | 0.00011 | 2.50E-05 | 3.37E-05 | 8.36E-06 | 7.33E-06 | 4.63E-05 | 3.09E-05 | 3.29E-05 | 2.94E-05 | 1.40E-05 | 4.32E-06 | 7.53E-05 | 0 | 1.47E-05 | 0.00014 | 6.29E-07 | 0 | 3.32E-05 |
| Cox11 | 5.99E-05 | 2.86E-05 | 2.59E-05 | 0 | 2.53E-05 | 4.41E-05 | 6.08E-06 | 1.76E-05 | 2.94E-05 | 7.62E-06 | 1.31E-05 | 0 | 0 | 2.10E-05 | 4.26E-05 | 3.88E-05 | 0 | 0 |
| Hlcs | 8.26E-06 | 1.86E-05 | 2.30E-06 | 1.58E-05 | 2.79E-05 | 3.90E-05 | 0 | 2.04E-05 | 2.94E-05 | 3.85E-05 | 1.92E-05 | 3.84E-05 | 2.65E-05 | 3.48E-05 | 4.03E-05 | 9.84E-07 | 1.34E-05 | 4.66E-05 |
| Inpp5b | 7.56E-05 | 0.00011 | 2.82E-05 | 0 | 8.53E-05 | 3.09E-05 | 1.84E-05 | 1.83E-05 | 2.94E-05 | 4.64E-05 | 7.03E-05 | 6.80E-05 | 0 | 0 | 7.30E-05 | 0.00011 | 4.41E-05 | 4.93E-05 |
| Crem | 7.95E-05 | 0.00011 | 2.14E-05 | 4.28E-08 | 5.64E-05 | 1.11E-05 | 3.07E-05 | 2.62E-05 | 2.94E-05 | 1.44E-05 | 1.66E-06 | 3.94E-05 | 1.14E-05 | 1.87E-06 | 9.30E-05 | 3.42E-05 | 0.0002 | 3.70E-05 |
| Slc12a9 | 8.93E-06 | 4.80E-05 | 0 | 7.69E-05 | 1.87E-05 | 7.47E-06 | 0 | 2.22E-05 | 2.94E-05 | 3.05E-05 | 1.97E-05 | 8.58E-06 | 1.85E-05 | 5.87E-05 | 1.54E-05 | 0 | 3.17E-06 | 3.96E-05 |
| Hoxa7 | 1.57E-05 | 5.55E-05 | 0 | 5.22E-05 | 8.47E-06 | 0 | 3.96E-05 | 0 | 2.94E-05 | 1.96E-05 | 0 | 2.45E-07 | 1.26E-06 | 0 | 0 | 1.57E-05 | 0 | 8.14E-06 |
| Exosc1 | 9.58E-05 | 4.82E-05 | 0.00013 | 5.47E-05 | 2.05E-05 | 0 | 0 | 0 | 2.94E-05 | 1.15E-05 | 2.40E-05 | 0 | 1.94E-05 | 2.20E-05 | 7.72E-05 | 6.61E-05 | 0 | 1.10E-05 |
| Fbxl3 | 9.75E-05 | 9.64E-05 | 2.21E-05 | 0 | 3.31E-05 | 7.10E-05 | 2.89E-05 | 2.93E-05 | 2.93E-05 | 2.92E-05 | 3.91E-05 | 6.79E-05 | 2.65E-05 | 2.34E-05 | 7.82E-05 | 2.78E-05 | 7.88E-05 | 5.06E-05 |
| Psmb3 | 0.00016 | 4.27E-05 | 0.00016 | 0.00018 | 2.90E-05 | 3.43E-05 | 0 | 0 | 2.93E-05 | 2.16E-05 | 3.55E-05 | 0 | 0 | 3.71E-05 | 0 | 8.03E-05 | 0 | 1.89E-05 |
| Rpap1 | 5.67E-05 | 3.78E-05 | 2.44E-05 | 2.00E-05 | 1.88E-07 | 7.61E-06 | 0 | 0 | 2.93E-05 | 1.67E-05 | 1.91E-05 | 8.80E-06 | 1.48E-05 | 0 | 0.00014 | 5.42E-05 | 4.33E-06 | 3.36E-05 |
| Clcn5 | 0 | 5.84E-05 | 0.00019 | 0.00011 | 8.65E-06 | 0.00016 | 3.56E-05 | 0.00058 | 2.92E-05 | 0.00026 | 0 | 0.00025 | 6.23E-05 | 0.00015 | 0 | 7.05E-05 | 0 | 0 |
| Rffl | 0 | 8.19E-05 | 9.64E-06 | 4.21E-05 | 1.91E-05 | 4.63E-05 | 0 | 0 | 2.92E-05 | 2.78E-05 | 0 | 0 | 0.00011 | 0.00014 | 8.41E-05 | 0 | 7.23E-05 | 0 |

| | | | | | | | | | | | | | | | | | | |
|---|---|---|---|---|---|---|---|---|---|---|---|---|---|---|---|---|---|---|
| Metrn | 4.90E-05 | 2.63E-05 | 2.20E-06 | 9.42E-05 | 2.18E-05 | 1.10E-05 | 0 | 2.78E-05 | 2.92E-05 | 1.11E-05 | 0 | 0 | 0 | 5.35E-05 | 0 | 0.00025 | 1.66E-05 | 2.57E-05 |
| Rtn4rl1 | 5.84E-06 | 1.39E-05 | 0 | 4.29E-06 | 0 | 1.05E-05 | 1.82E-06 | 0 | 2.92E-05 | 5.95E-05 | 1.81E-05 | 0 | 0 | 6.10E-05 | 0 | 8.80E-06 | 0 | 3.28E-05 |
| Rfc3 | 1.95E-05 | 0 | 4.39E-05 | 7.21E-06 | 1.85E-05 | 9.65E-06 | 2.33E-06 | 1.71E-05 | 2.92E-05 | 1.08E-05 | 9.44E-06 | 0 | 2.28E-05 | 2.56E-05 | 0 | 0.00028 | 0 | 1.11E-05 |
| Zfp691 | 1.82E-05 | 3.82E-05 | 1.69E-05 | 4.17E-05 | 6.04E-05 | 0.0001 | 0 | 0.00014 | 2.91E-05 | 1.75E-05 | 0 | 1.80E-06 | 9.55E-06 | 0 | 3.22E-05 | 0 | 0 | 0 |
| Gpatch8 | 0.0001 | 0.0001 | 2.81E-05 | 0 | 4.45E-05 | 9.72E-05 | 4.24E-05 | 6.16E-06 | 2.91E-05 | 4.63E-05 | 3.07E-06 | 8.09E-05 | 3.06E-05 | 2.82E-05 | 0.00016 | 4.37E-05 | 6.31E-05 | 3.09E-05 |
| L3mbtl3 | 0.00012 | 0.00011 | 8.01E-05 | 0 | 0.00017 | 8.57E-05 | 5.22E-05 | 0.00011 | 2.91E-05 | 0 | 5.53E-05 | 0 | 0 | 0 | 0.00012 | 0 | 0 | 6.85E-05 |
| Pan2 | 1.24E-05 | 7.49E-05 | 0 | 5.44E-05 | 2.60E-05 | 5.21E-05 | 1.58E-05 | 5.59E-05 | 2.91E-05 | 3.49E-05 | 8.41E-06 | 5.30E-05 | 0 | 2.80E-05 | 0.00019 | 0.00012 | 7.06E-05 | 3.89E-05 |
| Cc2d1b | 3.12E-05 | 7.38E-05 | 2.18E-05 | 7.45E-05 | 1.96E-05 | 2.00E-05 | 2.43E-05 | 1.56E-06 | 2.91E-05 | 4.20E-05 | 1.41E-06 | 0 | 0 | 2.92E-05 | 9.26E-05 | 5.15E-05 | 1.06E-06 | 6.04E-05 |
| Phc1 | 1.81E-05 | 8.39E-05 | 1.53E-05 | 6.19E-05 | 4.14E-05 | 1.48E-05 | 2.48E-05 | 3.63E-05 | 2.91E-05 | 1.93E-05 | 3.98E-05 | 0 | 1.29E-05 | 4.19E-05 | 4.57E-05 | 7.03E-05 | 7.49E-05 | 2.54E-05 |
| Aldh5a1 | 2.19E-05 | 1.81E-05 | 0 | 1.88E-05 | 0 | 0 | 1.58E-05 | 7.35E-06 | 2.91E-05 | 2.30E-05 | 0 | 5.84E-06 | 6.48E-06 | 2.61E-05 | 0 | 3.28E-05 | 0 | 2.99E-05 |
| Zbtb12 | 2.49E-05 | 3.24E-05 | 5.90E-06 | 2.93E-05 | 8.95E-06 | 0 | 1.46E-05 | 0 | 2.91E-05 | 0 | 2.02E-05 | 2.60E-05 | 2.55E-06 | 3.41E-06 | 3.20E-05 | 3.26E-05 | 1.92E-05 | 0 |
| Birc2 | 2.82E-05 | 3.91E-05 | 0 | 0 | 2.36E-05 | 0 | 7.74E-06 | 0 | 2.91E-05 | 9.65E-06 | 1.24E-05 | 2.55E-05 | 4.73E-05 | 1.90E-05 | 5.14E-05 | 5.09E-05 | 4.92E-05 | 3.83E-05 |
| Cdca7 | 1.28E-05 | 0 | 0.00011 | 3.74E-05 | 0 | 0 | 0 | 0 | 2.91E-05 | 0 | 0 | 0 | 3.03E-06 | 0 | 0 | 9.56E-05 | 0 | 0 |
| Pknox1 | 8.02E-05 | 6.45E-05 | 2.98E-05 | 0 | 2.47E-05 | 0.00012 | 0 | 5.43E-05 | 2.90E-05 | 0 | 0 | 0.00011 | 0 | 1.27E-05 | 6.97E-06 | 0.00013 | 5.44E-05 | 2.63E-05 |
| C2cd2 | 6.32E-06 | 2.61E-05 | 3.79E-05 | 0 | 1.60E-05 | 2.70E-05 | 2.00E-05 | 4.10E-05 | 2.90E-05 | 4.87E-05 | 3.24E-05 | 0 | 1.15E-05 | 4.54E-06 | 0 | 7.48E-05 | 0 | 4.07E-05 |
| Bend3 | 7.86E-06 | 2.86E-05 | 9.88E-05 | 2.87E-05 | 1.34E-05 | 1.86E-05 | 0 | 3.54E-05 | 2.90E-05 | 1.74E-06 | 3.19E-05 | 0 | 2.36E-05 | 3.07E-05 | 3.60E-05 | 1.85E-05 | 1.36E-05 | 0 |
| Top1mt | 3.04E-05 | 2.24E-05 | 4.79E-05 | 3.96E-05 | 2.87E-05 | 1.67E-05 | 5.90E-06 | 8.81E-06 | 2.90E-05 | 0 | 1.83E-05 | 0 | 3.18E-06 | 1.45E-06 | 2.70E-06 | 1.50E-05 | 0 | 1.54E-05 |
| Tmem229b | 8.08E-06 | 3.69E-05 | 0 | 3.36E-06 | 8.27E-05 | 1.60E-05 | 0 | 8.36E-05 | 2.90E-05 | 1.72E-05 | 0 | 2.92E-05 | 0 | 2.37E-05 | 0 | 0 | 0 | 4.73E-05 |
| Fitm2 | 2.35E-05 | 4.35E-05 | 9.42E-05 | 2.71E-05 | 3.48E-05 | 0 | 2.78E-05 | 0 | 2.90E-05 | 3.36E-05 | 0 | 0 | 3.76E-05 | 4.41E-05 | 1.73E-05 | 1.43E-05 | 0 | 0 |
| Unc93a | 0 | 2.16E-05 | 0 | 0 | 6.39E-06 | 0 | 1.48E-05 | 0 | 2.90E-05 | 0 | 0 | 0 | 3.57E-05 | 0 | 0 | 0 | 0 | 0 |
| Timm9 | 8.17E-05 | 2.83E-05 | 7.95E-05 | 0 | 2.58E-06 | 4.04E-05 | 0 | 0 | 2.89E-05 | 2.30E-06 | 4.62E-06 | 6.95E-06 | 0 | 0 | 5.19E-05 | 4.00E-05 | 0 | 0 |
| Clasp2 | 8.09E-05 | 8.73E-05 | 5.13E-05 | 0 | 2.62E-05 | 2.68E-05 | 4.21E-05 | 0 | 2.89E-05 | 4.02E-05 | 5.02E-05 | 5.29E-05 | 0 | 0 | 0.00012 | 0.00014 | 9.93E-05 | 3.09E-05 |
| Stxbp3 | 0 | 7.94E-05 | 5.55E-05 | 0.00013 | 0 | 4.18E-06 | 1.80E-05 | 1.49E-05 | 2.89E-05 | 7.14E-05 | 6.00E-05 | 3.85E-05 | 8.47E-06 | 3.09E-05 | 0.00018 | 4.23E-05 | 0 | 4.11E-05 |
| Ddx31 | 4.05E-05 | 4.83E-05 | 5.09E-05 | 8.48E-06 | 4.62E-06 | 0 | 0 | 0 | 2.89E-05 | 1.22E-05 | 9.14E-06 | 0 | 0 | 1.51E-06 | 4.24E-05 | 6.62E-06 | 0 | 1.71E-05 |
| Stx5a | 0.00047 | 0.0002 | 0.00013 | 7.48E-05 | 0.00013 | 0.00026 | 0.00047 | 8.58E-05 | 2.88E-05 | 1.40E-05 | 8.11E-05 | 0.00013 | 6.76E-05 | 0.00012 | 0.00037 | 2.06E-05 | 0.00023 | 0 |
| Fbxo7 | 3.78E-05 | 6.03E-05 | 3.48E-05 | 1.11E-05 | 3.03E-05 | 3.34E-05 | 0 | 1.33E-05 | 2.88E-05 | 2.71E-05 | 0 | 1.80E-05 | 2.85E-05 | 0 | 6.33E-05 | 3.90E-05 | 2.42E-05 | 4.81E-05 |
| Ttc38 | 1.97E-05 | 2.63E-05 | 0 | 0 | 1.43E-05 | 2.21E-05 | 1.08E-05 | 1.49E-05 | 2.88E-05 | 2.80E-05 | 0 | 0 | 0 | 2.86E-07 | 0 | 0 | 0 | 2.54E-05 |
| Prodh | 6.28E-06 | 1.59E-05 | 4.35E-07 | 1.06E-05 | 0 | 2.14E-05 | 0 | 0 | 2.88E-05 | 0 | 1.04E-06 | 2.00E-05 | 0 | 1.08E-05 | 2.22E-05 | 0 | 2.38E-05 | 9.33E-06 |
| Snapc2 | 4.42E-05 | 5.70E-05 | 4.45E-05 | 6.67E-05 | 1.79E-05 | 0 | 1.33E-05 | 2.02E-05 | 2.88E-05 | 1.97E-05 | 3.00E-05 | 0.0001 | 9.59E-06 | 1.33E-05 | 4.75E-05 | 7.54E-05 | 0 | 4.48E-05 |
| Lsm8 | 0.00015 | 0.0001 | 0.00021 | 0 | 9.86E-05 | 0.00013 | 0 | 0.00019 | 2.87E-05 | 3.05E-05 | 8.73E-05 | 4.83E-05 | 1.49E-05 | 0 | 5.36E-05 | 0.0005 | 0 | 8.37E-06 |
| Cuedc1 | 1.15E-05 | 0.00015 | 5.43E-05 | 1.93E-05 | 0.00023 | 0.00011 | 0.00027 | 0 | 2.87E-05 | 0.00016 | 0.0002 | 4.49E-05 | 6.89E-05 | 1.76E-05 | 0.00034 | 6.22E-05 | 0.00018 | 2.72E-05 |
| Wars2 | 2.85E-05 | 3.53E-05 | 2.27E-05 | 5.68E-06 | 1.56E-05 | 1.51E-05 | 2.07E-06 | 3.79E-06 | 2.87E-05 | 3.92E-06 | 3.38E-06 | 1.45E-05 | 1.43E-05 | 0 | 4.39E-05 | 1.66E-05 | 3.32E-05 | 1.63E-05 |
| Cnp | 1.45E-05 | 1.15E-05 | 0 | 1.43E-07 | 3.55E-06 | 1.16E-05 | 0 | 0 | 2.87E-05 | 6.18E-05 | 2.29E-05 | 2.47E-06 | 0 | 3.24E-06 | 0 | 6.92E-06 | 0 | 4.93E-05 |

| | | | | | | | | | | | | | | | | | | |
|---|---|---|---|---|---|---|---|---|---|---|---|---|---|---|---|---|---|---|
| Ptrhd1 | 3.21E-05 | 1.07E-05 | 1.82E-05 | 6.96E-05 | 6.22E-06 | 1.80E-06 | 1.99E-05 | 1.33E-05 | 2.87E-05 | 4.54E-06 | 1.16E-05 | 1.16E-05 | 1.53E-05 | 0 | 0 | 2.99E-05 | 9.54E-07 | 1.09E-05 |
| Greb1l | 0 | 4.37E-07 | 0 | 0 | 0 | 0 | 3.35E-05 | 0 | 2.87E-05 | 3.13E-05 | 0 | 0 | 4.16E-05 | 0 | 0 | 0 | 0 | 3.59E-06 |
| Dhx34 | 0 | 4.72E-05 | 3.69E-05 | 6.26E-05 | 7.94E-06 | 0 | 3.21E-05 | 2.06E-05 | 2.87E-05 | 3.59E-05 | 0 | 3.33E-05 | 0 | 1.68E-06 | 1.69E-05 | 1.18E-05 | 3.21E-05 | 3.01E-05 |
| Ndc1 | 4.37E-05 | 1.43E-06 | 0.00016 | 5.12E-05 | 0 | 4.95E-05 | 0 | 4.90E-05 | 2.86E-05 | 0 | 0 | 0 | 0 | 4.01E-05 | 0 | 0.00083 | 0 | 0 |
| Polr2c | 6.49E-05 | 4.26E-05 | 6.65E-05 | 0 | 7.18E-06 | 3.02E-05 | 0 | 5.76E-06 | 2.86E-05 | 8.82E-06 | 3.83E-05 | 2.39E-05 | 0 | 0 | 5.99E-05 | 6.42E-05 | 1.84E-05 | 0 |
| Asf1a | 8.96E-05 | 5.36E-05 | 4.67E-05 | 8.21E-06 | 2.17E-05 | 2.93E-05 | 0 | 0 | 2.86E-05 | 3.34E-06 | 2.16E-05 | 0 | 1.40E-05 | 0 | 6.34E-05 | 0.00013 | 5.97E-06 | 0 |
| Ptrh2 | 5.33E-05 | 2.48E-05 | 4.85E-05 | 3.15E-05 | 9.36E-06 | 1.20E-05 | 0 | 0 | 2.86E-05 | 2.19E-05 | 1.07E-05 | 1.45E-05 | 1.37E-05 | 0 | 0 | 9.76E-05 | 0 | 1.90E-05 |
| Fastkd5 | 4.31E-05 | 3.03E-05 | 2.47E-05 | 4.14E-06 | 1.97E-05 | 3.22E-06 | 0 | 0 | 2.86E-05 | 7.52E-06 | 1.01E-05 | 1.14E-05 | 0 | 0 | 5.23E-05 | 1.57E-05 | 0 | 9.88E-06 |
| Dhx29 | 6.22E-05 | 2.43E-05 | 7.75E-05 | 1.39E-05 | 2.01E-05 | 0 | 0 | 0 | 2.86E-05 | 3.00E-05 | 0 | 0 | 1.14E-05 | 1.51E-06 | 2.75E-05 | 4.82E-05 | 0 | 3.59E-05 |
| Asah2 | 5.83E-05 | 2.96E-05 | 0 | 0 | 5.10E-05 | 8.78E-05 | 0 | 7.90E-06 | 2.85E-05 | 1.83E-05 | 0 | 8.50E-06 | 7.56E-05 | 0 | 0 | 0 | 4.10E-05 | 4.45E-05 |
| Tha1 | 1.26E-05 | 8.85E-06 | 3.08E-06 | 5.02E-05 | 0 | 3.66E-05 | 4.09E-05 | 0 | 2.85E-05 | 5.24E-05 | 0 | 0 | 5.43E-06 | 3.72E-05 | 7.06E-05 | 0 | 0 | 0 |
| Akna | 1.59E-05 | 2.20E-05 | 0 | 0 | 0 | 2.59E-05 | 0 | 8.09E-06 | 2.85E-05 | 2.66E-05 | 0 | 1.45E-05 | 0 | 1.98E-05 | 9.29E-05 | 2.85E-06 | 5.97E-06 | 2.31E-05 |
| Pex11a | 0 | 2.02E-05 | 0 | 7.05E-06 | 2.62E-05 | 2.56E-05 | 0 | 0 | 2.85E-05 | 0 | 0 | 2.59E-05 | 3.20E-05 | 2.60E-05 | 3.04E-06 | 0 | 0 | 5.54E-07 |
| Mtm1 | 1.33E-05 | 5.05E-05 | 0 | 0 | 0 | 1.19E-05 | 0 | 0 | 2.85E-05 | 9.90E-06 | 0 | 5.96E-05 | 0 | 0 | 1.88E-05 | 2.09E-06 | 8.71E-07 | 0 |
| Ctu1 | 1.32E-05 | 4.18E-05 | 3.74E-05 | 6.31E-05 | 1.18E-05 | 0 | 0 | 2.26E-05 | 2.85E-05 | 1.54E-05 | 2.56E-05 | 0 | 1.21E-06 | 9.15E-06 | 4.33E-05 | 1.61E-05 | 0 | 1.25E-05 |
| Abcb10 | 4.62E-06 | 4.24E-05 | 2.26E-05 | 4.47E-05 | 2.11E-05 | 0 | 1.00E-05 | 0 | 2.85E-05 | 1.86E-05 | 9.82E-05 | 1.87E-05 | 2.52E-05 | 1.35E-05 | 3.12E-05 | 1.78E-05 | 0 | 1.53E-05 |
| Ddr1 | 0 | 8.75E-05 | 1.24E-05 | 7.40E-05 | 3.36E-05 | 0 | 0 | 0 | 2.85E-05 | 4.80E-05 | 0.00014 | 5.80E-05 | 2.59E-05 | 6.87E-05 | 3.72E-05 | 0 | 0 | 7.92E-05 |
| Inka1 | 1.36E-05 | 1.70E-05 | 5.89E-05 | 3.76E-05 | 6.50E-06 | 6.92E-05 | 0 | 0.00014 | 2.84E-05 | 0 | 0 | 0 | 0 | 0.00022 | 3.80E-05 | 3.11E-06 | 0 | 7.10E-06 |
| Trak2 | 0.00011 | 0.00016 | 0.00028 | 9.54E-05 | 0.00017 | 3.49E-05 | 0.00011 | 1.91E-05 | 2.84E-05 | 8.89E-05 | 0.00019 | 5.68E-05 | 6.07E-05 | 0.00024 | 0.00019 | 0.00021 | 4.27E-07 | 7.83E-05 |
| Dnal4 | 1.26E-05 | 7.55E-05 | 3.70E-05 | 0.00017 | 7.29E-05 | 3.04E-05 | 2.35E-05 | 6.31E-05 | 2.84E-05 | 3.97E-05 | 2.36E-05 | 4.02E-05 | 0 | 4.40E-05 | 0 | 3.26E-05 | 4.71E-05 | 4.75E-05 |
| Ak6 | 8.77E-05 | 6.88E-05 | 0.00012 | 9.03E-05 | 1.97E-05 | 2.21E-05 | 0 | 1.01E-05 | 2.84E-05 | 1.31E-05 | 1.20E-05 | 3.42E-05 | 0 | 3.80E-05 | 4.68E-05 | 0.00017 | 9.88E-05 | 2.94E-06 |
| Pcdhb20 | 6.88E-06 | 3.85E-05 | 0 | 0 | 4.35E-05 | 2.07E-05 | 0 | 1.65E-05 | 2.84E-05 | 8.81E-06 | 2.65E-05 | 7.64E-06 | 2.27E-05 | 6.84E-06 | 3.23E-05 | 0 | 0 | 2.36E-05 |
| Ints8 | 0.00011 | 7.94E-05 | 7.37E-05 | 5.43E-05 | 0 | 2.01E-05 | 0 | 0 | 2.84E-05 | 4.03E-05 | 5.09E-05 | 2.33E-05 | 0 | 1.29E-05 | 9.59E-05 | 4.82E-05 | 1.15E-05 | 3.99E-05 |
| Zfp518a | 3.05E-05 | 2.11E-05 | 0 | 0 | 8.53E-06 | 1.88E-05 | 0 | 1.02E-05 | 2.84E-05 | 1.67E-06 | 0 | 1.02E-05 | 0 | 3.16E-06 | 2.85E-05 | 1.89E-05 | 0 | 1.24E-06 |
| Gtf2e1 | 9.54E-05 | 3.41E-05 | 7.38E-05 | 2.57E-05 | 1.47E-05 | 1.56E-05 | 4.36E-05 | 0 | 2.84E-05 | 3.10E-05 | 8.98E-06 | 0 | 1.10E-05 | 0 | 2.18E-05 | 3.81E-05 | 0 | 1.61E-05 |
| Mras | 5.45E-05 | 8.63E-05 | 3.28E-05 | 0 | 8.47E-05 | 8.36E-06 | 0.00016 | 0 | 2.84E-05 | 4.07E-05 | 7.73E-05 | 1.26E-06 | 0 | 0 | 0 | 5.52E-05 | 1.89E-05 | 8.61E-05 |
| Lrrc4c | 8.36E-06 | 0 | 6.78E-06 | 0 | 2.21E-05 | 4.67E-06 | 2.21E-05 | 1.66E-05 | 2.84E-05 | 0 | 6.46E-05 | 0 | 0 | 0 | 0 | 1.80E-05 | 0 | 0 |
| Gla | 1.68E-06 | 0.00014 | 0 | 4.70E-05 | 0 | 0.00021 | 0 | 4.98E-05 | 2.83E-05 | 0 | 0 | 0.00011 | 5.92E-05 | 8.79E-05 | 0 | 0.00013 | 0.00021 | 0 |
| Slc49a4 | 0 | 0.00024 | 3.78E-05 | 0.00014 | 0.00015 | 0.00013 | 8.81E-05 | 0.00012 | 2.83E-05 | 0.00019 | 8.70E-05 | 8.01E-05 | 7.08E-05 | 0.00018 | 0 | 0 | 9.59E-05 | 0.00025 |
| Ikzf5 | 8.11E-05 | 0.00011 | 0.00012 | 3.34E-06 | 7.16E-05 | 8.37E-05 | 8.94E-05 | 4.96E-05 | 2.83E-05 | 6.14E-05 | 6.91E-05 | 2.55E-05 | 5.73E-06 | 5.72E-05 | 4.05E-05 | 0.00015 | 0.00016 | 3.62E-05 |
| Eef1akmt2 | 8.82E-05 | 9.82E-05 | 8.28E-05 | 0.00015 | 5.86E-05 | 3.70E-05 | 9.12E-05 | 5.47E-05 | 2.83E-05 | 6.87E-05 | 4.04E-05 | 4.98E-05 | 1.58E-05 | 0 | 0.00017 | 8.54E-05 | 0 | 7.17E-05 |
| Noc3l | 4.55E-05 | 3.74E-05 | 6.98E-05 | 0 | 6.07E-06 | 2.68E-05 | 0 | 0 | 2.83E-05 | 2.73E-05 | 0 | 3.30E-07 | 1.53E-05 | 0 | 3.47E-05 | 1.64E-05 | 0 | 9.01E-06 |
| Hddc3 | 4.18E-05 | 3.74E-05 | 0 | 2.10E-05 | 1.72E-05 | 2.68E-05 | 1.51E-05 | 2.49E-05 | 2.83E-05 | 0 | 1.33E-05 | 6.83E-06 | 6.74E-06 | 0 | 2.96E-05 | 1.39E-05 | 0 | 1.85E-05 |

| | | | | | | | | | | | | | | | | | | |
|---|---|---|---|---|---|---|---|---|---|---|---|---|---|---|---|---|---|---|
| Prr5l | 0 | 1.12E-05 | 8.20E-06 | 6.44E-06 | 4.46E-05 | 2.62E-05 | 4.02E-05 | 0 | 2.83E-05 | 6.07E-06 | 0 | 1.69E-05 | 0 | 6.63E-05 | 2.32E-05 | 3.06E-07 | 0 | 0 |
| Zfp707 | 9.97E-06 | 5.43E-05 | 2.57E-06 | 2.65E-05 | 6.49E-06 | 1.06E-05 | 1.02E-06 | 0 | 2.83E-05 | 6.31E-06 | 3.49E-06 | 2.64E-05 | 0 | 2.28E-05 | 6.44E-05 | 2.89E-05 | 0 | 2.83E-05 |
| Fam3c | 0.00022 | 6.19E-05 | 0.00024 | 0.00015 | 0.00019 | 0.00031 | 0.00031 | 7.63E-05 | 2.82E-05 | 0.00014 | 0.00029 | 2.63E-06 | 0.0002 | 0.00026 | 0 | 0.00023 | 0 | 0.00014 |
| Pan3 | 0.00012 | 0.00014 | 2.60E-05 | 0 | 5.28E-05 | 9.60E-05 | 3.66E-05 | 0 | 2.82E-05 | 1.65E-05 | 1.15E-05 | 0.00011 | 2.30E-05 | 2.16E-05 | 0.00025 | 1.67E-05 | 0.00014 | 0 |
| Pcdhb21 | 3.74E-06 | 2.25E-05 | 0 | 0 | 2.80E-05 | 2.67E-05 | 2.75E-06 | 3.80E-05 | 2.82E-05 | 1.80E-05 | 1.58E-05 | 0 | 2.91E-05 | 2.44E-06 | 1.94E-05 | 0 | 0 | 3.48E-05 |
| Ighmbp2 | 3.52E-05 | 2.91E-05 | 2.99E-05 | 3.01E-05 | 1.94E-06 | 1.21E-05 | 8.67E-06 | 1.74E-05 | 2.82E-05 | 1.94E-05 | 3.17E-05 | 1.71E-05 | 0 | 2.72E-05 | 5.22E-05 | 5.19E-06 | 0 | 2.27E-05 |
| Mars2 | 0 | 3.46E-05 | 3.37E-05 | 1.53E-05 | 1.46E-06 | 6.64E-06 | 0 | 0 | 2.82E-05 | 4.47E-06 | 3.11E-05 | 0 | 2.60E-06 | 2.92E-05 | 2.69E-05 | 3.28E-05 | 0 | 2.13E-05 |
| Fam133b | 1.88E-05 | 5.89E-05 | 4.62E-05 | 3.45E-05 | 3.10E-05 | 1.68E-07 | 0 | 2.12E-05 | 2.82E-05 | 1.22E-05 | 7.29E-06 | 2.91E-05 | 0 | 3.79E-06 | 0.00014 | 7.03E-05 | 0.00011 | 2.33E-05 |
| Lgals7 | 0 | 0 | 8.71E-06 | 0 | 0 | 0 | 3.30E-06 | 8.08E-05 | 2.82E-05 | 0 | 0 | 0 | 0 | 0 | 0 | 1.76E-06 | 0 | 2.14E-05 |
| Bmt2 | 8.97E-05 | 0.00013 | 2.59E-05 | 4.25E-06 | 8.41E-05 | 6.12E-05 | 3.14E-05 | 0 | 2.81E-05 | 5.08E-05 | 1.75E-05 | 0.00025 | 3.00E-06 | 2.81E-05 | 0.00022 | 2.54E-05 | 0.00018 | 6.31E-06 |
| Sox4 | 0 | 0.00036 | 8.18E-05 | 0.00047 | 0.00024 | 5.80E-05 | 0 | 0.00025 | 2.81E-05 | 0 | 0.00036 | 0.00016 | 0.00017 | 0.00034 | 0.00045 | 0 | 0.00048 | 0.00016 |
| Riok2 | 9.51E-05 | 0.00011 | 0.00017 | 9.06E-05 | 2.96E-05 | 5.19E-05 | 0 | 0 | 2.81E-05 | 1.37E-05 | 0 | 6.32E-05 | 3.60E-05 | 4.91E-06 | 8.89E-05 | 5.78E-05 | 0 | 1.73E-05 |
| Fam118b | 5.84E-05 | 5.53E-05 | 2.51E-05 | 0 | 4.86E-05 | 1.07E-05 | 2.11E-05 | 0 | 2.81E-05 | 9.38E-06 | 0 | 0 | 7.60E-06 | 0 | 1.53E-05 | 4.73E-05 | 0 | 1.83E-05 |
| Entpd6 | 4.30E-05 | 0 | 0.00011 | 0.00012 | 0 | 0 | 6.04E-05 | 1.34E-05 | 2.81E-05 | 8.12E-06 | 0 | 0 | 6.78E-05 | 0 | 7.04E-06 | 8.94E-05 | 0 | 0 |
| Lmnb1 | 0 | 0 | 0.00011 | 0.00015 | 0 | 0 | 0 | 2.42E-05 | 2.81E-05 | 0.00014 | 5.25E-05 | 0 | 0 | 0 | 7.92E-05 | 0.00145 | 0 | 0 |
| Coa7 | 3.48E-05 | 2.85E-05 | 4.01E-05 | 6.49E-05 | 0 | 0 | 0 | 0 | 2.81E-05 | 0 | 2.16E-05 | 0 | 3.70E-06 | 3.50E-05 | 2.00E-05 | 4.75E-05 | 0 | 2.31E-06 |
| Mettl6 | 6.80E-05 | 6.31E-05 | 4.04E-05 | 0 | 2.45E-05 | 4.26E-05 | 6.33E-06 | 0 | 2.80E-05 | 0 | 0 | 6.49E-05 | 4.30E-05 | 4.80E-05 | 8.40E-05 | 2.05E-05 | 0 | 0 |
| Tmem60 | 6.36E-05 | 4.84E-05 | 2.46E-05 | 0 | 2.51E-05 | 3.57E-05 | 3.36E-08 | 9.87E-08 | 2.80E-05 | 1.39E-05 | 9.34E-06 | 0.0001 | 3.72E-06 | 2.43E-06 | 0 | 1.18E-05 | 0 | 1.68E-05 |
| Dnajc2 | 0.0001 | 3.37E-05 | 0.0001 | 0 | 1.37E-05 | 3.37E-05 | 0 | 1.25E-06 | 2.80E-05 | 0 | 3.44E-06 | 2.16E-05 | 0 | 0 | 4.59E-05 | 9.68E-05 | 0 | 6.66E-06 |
| Msantd4 | 5.41E-05 | 3.00E-05 | 2.34E-05 | 0 | 3.20E-05 | 2.37E-05 | 0 | 0 | 2.80E-05 | 1.04E-05 | 6.40E-06 | 2.79E-05 | 0 | 2.22E-05 | 0 | 2.21E-05 | 0 | 3.17E-05 |
| Alkbh6 | 5.17E-05 | 7.46E-05 | 3.73E-05 | 0.00011 | 3.72E-05 | 1.92E-05 | 0 | 1.27E-07 | 2.80E-05 | 7.92E-06 | 2.82E-05 | 7.05E-05 | 0 | 0 | 9.72E-06 | 0 | 3.76E-05 | 1.34E-05 |
| Trappc2 | 5.27E-05 | 2.39E-05 | 3.69E-05 | 0 | 1.65E-05 | 1.45E-05 | 6.76E-06 | 5.76E-07 | 2.80E-05 | 3.64E-05 | 3.56E-05 | 1.56E-05 | 2.30E-05 | 0 | 0 | 2.23E-05 | 0 | 1.69E-05 |
| Zfp382 | 1.03E-05 | 2.18E-05 | 0 | 0 | 1.10E-05 | 1.42E-05 | 6.75E-06 | 8.83E-07 | 2.80E-05 | 1.33E-05 | 7.37E-06 | 2.44E-05 | 0 | 0 | 2.51E-05 | 9.10E-05 | 0 | 1.10E-05 |
| Zcchc4 | 2.67E-05 | 4.36E-05 | 1.46E-05 | 8.53E-06 | 9.37E-06 | 1.42E-05 | 0 | 4.66E-06 | 2.80E-05 | 4.19E-06 | 1.51E-05 | 2.55E-05 | 0 | 2.62E-06 | 8.76E-05 | 2.61E-05 | 0 | 0 |
| C2cd3 | 2.77E-05 | 4.64E-05 | 0 | 2.70E-05 | 2.22E-05 | 1.26E-05 | 3.25E-05 | 4.15E-05 | 2.80E-05 | 1.22E-05 | 2.53E-06 | 2.17E-05 | 0 | 1.57E-06 | 9.69E-06 | 0.0001 | 3.86E-05 | 1.61E-05 |
| Zbtb43 | 2.60E-05 | 0.00011 | 1.76E-05 | 2.20E-05 | 2.41E-05 | 0 | 0 | 0 | 2.80E-05 | 2.11E-05 | 1.68E-05 | 9.40E-05 | 0 | 5.24E-05 | 4.66E-05 | 1.37E-05 | 0 | 0 |
| Odf2 | 0.00017 | 0.00017 | 0.00011 | 2.06E-05 | 3.89E-05 | 5.56E-05 | 5.69E-05 | 6.65E-06 | 2.79E-05 | 3.62E-05 | 3.09E-05 | 0.0001 | 0 | 6.11E-05 | 0.0005 | 0.00047 | 0.00021 | 0 |
| Lactb2 | 6.29E-05 | 4.15E-05 | 9.47E-06 | 0 | 4.18E-05 | 5.14E-05 | 0 | 0 | 2.79E-05 | 3.09E-05 | 1.39E-05 | 2.94E-05 | 1.72E-05 | 0 | 3.96E-05 | 0 | 0 | 3.32E-05 |
| Slc35e2 | 5.44E-05 | 6.13E-05 | 2.55E-05 | 0 | 1.92E-05 | 2.79E-05 | 3.91E-05 | 2.09E-05 | 2.79E-05 | 9.18E-06 | 1.31E-05 | 2.65E-05 | 4.82E-06 | 1.33E-05 | 1.47E-05 | 4.23E-05 | 7.99E-05 | 4.34E-05 |
| Cwf19l1 | 2.96E-05 | 3.30E-05 | 3.99E-05 | 0 | 1.15E-05 | 1.01E-05 | 0 | 0 | 2.79E-05 | 5.13E-06 | 3.31E-06 | 0 | 0 | 2.27E-05 | 4.14E-05 | 5.84E-05 | 0 | 0 |
| Ell2 | 3.63E-06 | 2.77E-05 | 0.00014 | 7.62E-06 | 2.47E-05 | 7.95E-06 | 0 | 0 | 2.79E-05 | 0 | 8.71E-05 | 4.70E-05 | 0.00018 | 0.00021 | 9.48E-05 | 0 | 0.00027 | 0 |
| Cchcr1 | 0 | 2.29E-05 | 4.94E-05 | 7.85E-05 | 0 | 0 | 1.58E-05 | 2.25E-05 | 2.79E-05 | 3.46E-05 | 0 | 8.20E-06 | 7.76E-05 | 1.71E-05 | 0.00018 | 0.00022 | 8.74E-05 | 2.08E-05 |
| Cdan1 | 0 | 5.99E-05 | 3.98E-05 | 1.05E-05 | 2.48E-05 | 0 | 3.74E-05 | 3.35E-05 | 2.79E-05 | 2.72E-05 | 3.33E-05 | 0 | 0 | 5.71E-05 | 0.00011 | 0.00017 | 1.09E-05 | 1.99E-05 |

| | | | | | | | | | | | | | | | | | | |
|---|---|---|---|---|---|---|---|---|---|---|---|---|---|---|---|---|---|---|
| Zcchc14 | 0.00016 | 0.00039 | 0.00014 | 2.42E-05 | 0.0002 | 0.0003 | 7.21E-05 | 0.0002 | 2.78E-05 | 4.95E-05 | 9.86E-05 | 0 | 0.00022 | 0.0004 | 0.00023 | 0.00012 | 0.00024 | 0.0002 |
| Mthfs | 4.87E-05 | 1.76E-05 | 0 | 0 | 0 | 4.30E-05 | 0 | 0 | 2.78E-05 | 0 | 2.43E-06 | 1.70E-05 | 8.87E-06 | 1.60E-05 | 1.59E-05 | 5.54E-06 | 0 | 5.17E-06 |
| Fam13b | 6.18E-05 | 0.00015 | 6.62E-05 | 2.65E-06 | 7.50E-05 | 2.87E-05 | 0 | 0 | 2.78E-05 | 3.26E-05 | 5.38E-05 | 0.00016 | 0 | 2.88E-05 | 0.00014 | 4.94E-05 | 9.22E-05 | 3.50E-06 |
| Ngrn | 7.60E-05 | 5.59E-05 | 7.72E-06 | 5.77E-05 | 3.45E-05 | 2.57E-05 | 2.47E-06 | 6.31E-06 | 2.78E-05 | 8.97E-06 | 1.03E-05 | 7.91E-06 | 0 | 3.49E-06 | 8.84E-05 | 0 | 0 | 1.10E-05 |
| Dolpp1 | 2.07E-05 | 5.16E-06 | 4.30E-05 | 1.69E-05 | 1.67E-05 | 7.77E-06 | 0 | 2.32E-05 | 2.78E-05 | 3.92E-05 | 2.42E-05 | 2.04E-06 | 0 | 1.48E-06 | 0 | 1.40E-05 | 0 | 2.81E-05 |
| Ly75 | 0 | 0 | 4.01E-06 | 0 | 0 | 0.00011 | 0 | 0 | 2.77E-05 | 0 | 0.0003 | 0 | 6.79E-06 | 0.00019 | 8.89E-06 | 3.30E-05 | 0 | 0 |
| Usp33 | 0.00015 | 0.00015 | 5.79E-05 | 0 | 9.05E-05 | 8.14E-05 | 5.86E-05 | 1.95E-05 | 2.77E-05 | 3.64E-05 | 0.00011 | 6.91E-05 | 3.47E-05 | 5.41E-05 | 0.00026 | 0.00011 | 9.47E-05 | 5.58E-05 |
| Cep350 | 0.00011 | 0.00013 | 8.06E-05 | 5.25E-05 | 9.66E-05 | 7.38E-05 | 6.87E-05 | 5.79E-07 | 2.77E-05 | 4.90E-05 | 3.45E-05 | 0.0001 | 6.48E-06 | 5.32E-05 | 0.00021 | 0.00012 | 0.00013 | 4.01E-05 |
| Gstk1 | 5.30E-05 | 1.65E-05 | 1.49E-06 | 2.00E-05 | 9.34E-06 | 1.42E-05 | 1.23E-05 | 2.11E-05 | 2.77E-05 | 4.98E-05 | 0 | 2.88E-05 | 7.16E-05 | 0 | 0 | 4.76E-05 | 0 | 0 |
| Nudt7 | 5.96E-05 | 1.85E-05 | 7.00E-06 | 5.52E-05 | 3.97E-05 | 0 | 3.96E-05 | 1.83E-05 | 2.77E-05 | 4.27E-05 | 4.81E-05 | 3.75E-05 | 3.55E-06 | 0 | 2.57E-05 | 0 | 0 | 6.66E-05 |
| Asap2 | 1.63E-05 | 6.10E-05 | 5.22E-05 | 0 | 0 | 0 | 0 | 8.52E-05 | 2.77E-05 | 8.52E-05 | 5.06E-05 | 4.87E-05 | 0 | 4.83E-05 | 6.14E-05 | 6.46E-05 | 2.38E-05 | 3.66E-05 |
| Kif7 | 9.81E-06 | 4.41E-05 | 0 | 7.90E-05 | 2.98E-05 | 0 | 6.94E-06 | 1.71E-05 | 2.77E-05 | 1.24E-05 | 9.56E-06 | 0 | 2.14E-05 | 3.25E-05 | 7.98E-05 | 0 | 1.19E-05 | 5.39E-05 |
| Tmub2 | 0.00013 | 0.00016 | 0 | 0.0001 | 9.46E-05 | 0.00016 | 9.46E-05 | 5.62E-05 | 2.76E-05 | 8.36E-05 | 5.44E-05 | 0.00013 | 3.14E-05 | 8.29E-05 | 4.45E-06 | 6.34E-05 | 0 | 8.78E-05 |
| Zfp53 | 4.91E-05 | 4.53E-05 | 2.79E-06 | 1.11E-05 | 5.90E-07 | 5.72E-05 | 0 | 7.31E-05 | 2.76E-05 | 0 | 8.82E-05 | 6.51E-05 | 0 | 3.64E-06 | 1.81E-05 | 0 | 0 | 2.15E-05 |
| Pfkm | 3.85E-05 | 3.01E-05 | 1.36E-05 | 5.69E-05 | 8.28E-05 | 3.79E-05 | 7.55E-05 | 2.93E-05 | 2.76E-05 | 3.22E-05 | 2.87E-05 | 6.36E-05 | 2.98E-05 | 3.18E-05 | 4.94E-06 | 0 | 7.52E-05 | 4.93E-05 |
| Acd | 4.39E-05 | 9.32E-05 | 1.21E-05 | 9.80E-05 | 4.16E-05 | 3.47E-05 | 5.01E-05 | 0 | 2.76E-05 | 1.17E-05 | 1.59E-06 | 4.33E-05 | 2.18E-05 | 6.10E-05 | 0 | 6.54E-05 | 3.58E-05 | 5.00E-05 |
| Tent4b | 5.33E-05 | 5.80E-05 | 4.98E-05 | 0 | 3.26E-05 | 3.29E-05 | 0 | 4.15E-06 | 2.76E-05 | 0 | 3.29E-05 | 3.28E-05 | 0 | 2.98E-05 | 2.14E-05 | 2.01E-05 | 6.06E-05 | 0 |
| Bckdha | 9.04E-06 | 8.09E-05 | 0 | 8.65E-05 | 3.86E-05 | 2.46E-05 | 5.62E-05 | 2.51E-05 | 2.76E-05 | 7.76E-06 | 7.28E-06 | 4.57E-05 | 3.22E-05 | 5.46E-05 | 3.51E-06 | 0 | 0 | 5.62E-05 |
| Cyp2r1 | 2.84E-05 | 4.75E-06 | 0 | 0 | 1.51E-05 | 1.87E-05 | 8.15E-06 | 2.08E-05 | 2.76E-05 | 2.21E-06 | 0 | 0 | 1.65E-06 | 0 | 9.40E-06 | 2.51E-05 | 0 | 0 |
| Zfp27 | 3.10E-05 | 1.73E-05 | 3.96E-06 | 0 | 2.17E-05 | 1.39E-05 | 2.15E-05 | 1.60E-05 | 2.76E-05 | 2.44E-05 | 0 | 8.28E-06 | 1.17E-05 | 9.15E-06 | 2.13E-05 | 3.45E-06 | 0 | 8.86E-06 |
| Gcat | 2.91E-05 | 6.91E-06 | 2.98E-05 | 4.59E-05 | 2.48E-06 | 0 | 1.31E-05 | 0 | 2.76E-05 | 0 | 0 | 1.28E-05 | 5.79E-06 | 0 | 0 | 7.01E-05 | 0 | 0 |
| Sgf29 | 3.83E-05 | 4.13E-05 | 7.40E-05 | 4.18E-05 | 2.90E-05 | 0 | 0 | 8.89E-06 | 2.76E-05 | 1.64E-05 | 6.78E-06 | 0 | 5.08E-07 | 1.32E-05 | 7.20E-06 | 0.0001 | 0 | 1.11E-05 |
| Ppil4 | 0.00012 | 7.81E-05 | 5.05E-05 | 0 | 4.90E-05 | 7.14E-05 | 2.09E-05 | 0 | 2.75E-05 | 3.41E-05 | 0 | 7.92E-06 | 4.62E-05 | 3.89E-05 | 0.00017 | 7.93E-05 | 0 | 1.85E-06 |
| Mcm9 | 4.17E-05 | 6.79E-05 | 2.38E-05 | 0 | 2.89E-05 | 2.69E-05 | 3.42E-05 | 1.41E-05 | 2.75E-05 | 2.76E-05 | 0 | 3.95E-05 | 1.20E-05 | 0 | 7.49E-05 | 0.00013 | 0 | 9.56E-06 |
| Pde7a | 2.55E-05 | 2.52E-05 | 7.14E-06 | 0 | 2.48E-05 | 1.53E-05 | 2.97E-05 | 0 | 2.75E-05 | 2.65E-05 | 4.56E-05 | 2.44E-05 | 4.69E-05 | 1.15E-05 | 3.20E-05 | 6.12E-05 | 1.48E-05 | 5.10E-05 |
| Sec24a | 0.00025 | 9.91E-05 | 0.00021 | 0.00011 | 4.96E-05 | 0.00013 | 0.00023 | 0.00012 | 2.74E-05 | 4.87E-05 | 3.59E-05 | 6.59E-05 | 5.16E-06 | 9.46E-05 | 0.00017 | 6.96E-05 | 0.00091 | 0 |
| Cdk8 | 0.00012 | 0.00013 | 0.00014 | 2.73E-05 | 3.45E-05 | 4.79E-05 | 2.74E-05 | 2.28E-05 | 2.74E-05 | 2.97E-05 | 0.0001 | 4.91E-05 | 1.76E-05 | 4.55E-05 | 0.00014 | 5.68E-05 | 9.28E-05 | 2.64E-05 |
| Commd9 | 5.41E-05 | 3.18E-05 | 9.66E-05 | 0.00021 | 0.00011 | 4.78E-05 | 0.00011 | 1.08E-05 | 2.74E-05 | 3.64E-06 | 0 | 0 | 1.68E-05 | 8.56E-05 | 0 | 0 | 0 | 0 |
| Aftph | 7.81E-05 | 6.74E-05 | 3.86E-06 | 0 | 3.15E-05 | 4.06E-05 | 6.67E-07 | 0 | 2.74E-05 | 1.39E-05 | 1.87E-05 | 2.12E-05 | 0 | 0 | 3.27E-05 | 1.54E-05 | 1.17E-05 | 0 |
| Ogfod1 | 5.13E-05 | 4.55E-05 | 5.44E-05 | 0 | 4.18E-05 | 3.21E-05 | 0 | 1.84E-05 | 2.74E-05 | 3.14E-05 | 3.41E-05 | 2.22E-05 | 2.72E-05 | 0 | 6.26E-05 | 3.96E-05 | 9.93E-06 | 0 |
| Hddc2 | 5.91E-05 | 3.58E-05 | 1.74E-05 | 6.06E-05 | 5.17E-06 | 2.75E-05 | 0 | 1.08E-05 | 2.74E-05 | 7.01E-06 | 3.87E-05 | 3.54E-05 | 0 | 3.61E-05 | 0 | 1.66E-05 | 0 | 2.15E-05 |
| Cwc25 | 4.51E-05 | 8.29E-05 | 3.05E-05 | 7.36E-05 | 3.93E-05 | 2.69E-05 | 1.05E-05 | 0 | 2.74E-05 | 3.65E-05 | 2.47E-05 | 7.23E-05 | 2.51E-05 | 3.62E-05 | 0.00014 | 2.30E-05 | 9.48E-06 | 3.33E-05 |
| Prpf4 | 3.10E-05 | 4.41E-05 | 8.28E-05 | 4.99E-06 | 1.80E-05 | 2.65E-05 | 0 | 7.28E-06 | 2.74E-05 | 4.35E-06 | 1.47E-05 | 6.75E-06 | 8.54E-06 | 7.77E-06 | 1.72E-05 | 0.00015 | 0 | 1.19E-05 |

| | | | | | | | | | | | | | | | | | | |
|---|---|---|---|---|---|---|---|---|---|---|---|---|---|---|---|---|---|---|
| Tango6 | 4.08E-05 | 7.11E-05 | 4.44E-05 | 6.01E-05 | 1.01E-05 | 1.62E-05 | 0 | 3.36E-05 | 2.74E-05 | 2.93E-05 | 9.83E-06 | 1.32E-06 | 2.52E-05 | 0 | 0.00011 | 2.11E-05 | 0 | 2.42E-05 |
| Dffb | 6.47E-06 | 2.71E-05 | 2.59E-05 | 5.68E-05 | 2.58E-05 | 1.53E-05 | 5.25E-06 | 1.82E-05 | 2.74E-05 | 6.08E-06 | 0 | 3.87E-06 | 0 | 2.34E-05 | 5.55E-05 | 1.57E-06 | 0 | 2.83E-05 |
| Ift81 | 3.43E-05 | 5.85E-05 | 0 | 2.79E-05 | 3.92E-05 | 1.22E-05 | 9.69E-06 | 0 | 2.74E-05 | 2.58E-05 | 3.05E-05 | 5.60E-06 | 2.28E-05 | 7.50E-06 | 5.36E-05 | 2.49E-05 | 1.28E-06 | 3.68E-05 |
| Mtmr12 | 1.26E-06 | 5.79E-05 | 3.18E-05 | 4.49E-05 | 2.05E-05 | 0 | 9.89E-06 | 4.64E-05 | 2.74E-05 | 3.42E-05 | 2.55E-05 | 2.71E-05 | 3.57E-06 | 3.77E-05 | 1.24E-05 | 7.10E-05 | 7.73E-05 | 4.27E-05 |
| Nop9 | 8.99E-06 | 4.21E-05 | 7.44E-05 | 7.54E-05 | 2.03E-06 | 0 | 0 | 0 | 2.74E-05 | 6.05E-06 | 2.94E-05 | 0 | 0 | 2.19E-05 | 4.26E-05 | 8.43E-05 | 0 | 8.31E-06 |
| Cenpt | 4.58E-06 | 3.61E-05 | 3.46E-05 | 2.05E-05 | 0 | 0 | 0 | 2.05E-05 | 2.74E-05 | 3.29E-05 | 3.96E-05 | 3.52E-05 | 0 | 0 | 5.96E-05 | 0.00023 | 0 | 2.66E-05 |
| Helz2 | 7.22E-05 | 8.47E-05 | 3.01E-05 | 0.0002 | 3.14E-05 | 0 | 4.29E-05 | 0.00011 | 2.74E-05 | 0.00019 | 2.50E-05 | 4.42E-05 | 0.00014 | 3.50E-05 | 0.00019 | 3.42E-05 | 0.00022 | 5.84E-05 |
| Klhl7 | 9.47E-05 | 0.00012 | 6.05E-05 | 3.15E-05 | 9.18E-05 | 8.07E-05 | 7.14E-05 | 4.92E-05 | 2.73E-05 | 3.65E-05 | 8.42E-06 | 4.20E-05 | 7.69E-05 | 2.90E-05 | 0 | 0 | 0.00011 | 1.60E-05 |
| DEPRECATED | 0.00011 | 1.37E-05 | 4.02E-05 | 1.43E-05 | 4.21E-05 | 1.09E-05 | 5.29E-05 | 3.48E-05 | 2.73E-05 | 4.53E-06 | 2.11E-05 | 2.31E-05 | 2.18E-05 | 0 | 0 | 5.42E-05 | 0 | 1.24E-05 |
| Caskin2 | 1.61E-05 | 4.90E-05 | 1.21E-05 | 9.59E-05 | 2.69E-05 | 7.74E-06 | 1.39E-05 | 0 | 2.73E-05 | 3.26E-05 | 4.99E-05 | 0 | 5.03E-05 | 7.10E-05 | 8.73E-05 | 2.30E-05 | 0 | 4.67E-05 |
| Taf6l | 1.82E-05 | 4.49E-05 | 2.01E-05 | 2.62E-05 | 1.15E-05 | 9.79E-07 | 6.71E-06 | 3.50E-06 | 2.73E-05 | 3.59E-06 | 1.25E-05 | 6.61E-06 | 1.66E-05 | 3.70E-05 | 0 | 6.10E-05 | 0 | 8.37E-06 |
| Prkg1 | 7.76E-06 | 3.53E-05 | 0 | 0 | 0.00012 | 7.94E-05 | 1.01E-05 | 0 | 2.72E-05 | 3.05E-05 | 7.16E-05 | 8.37E-07 | 1.72E-05 | 1.92E-05 | 0 | 0 | 0 | 3.14E-06 |
| Pdcd5 | 0.00015 | 3.96E-05 | 8.98E-05 | 0.00011 | 0 | 7.34E-05 | 0 | 2.98E-05 | 2.72E-05 | 2.30E-05 | 4.64E-05 | 5.40E-05 | 0 | 3.15E-05 | 4.29E-05 | 8.51E-05 | 4.31E-05 | 2.61E-05 |
| Fam117b | 6.91E-05 | 0.00014 | 2.90E-06 | 0.00015 | 0 | 6.63E-05 | 1.95E-05 | 0.00015 | 2.72E-05 | 0 | 7.38E-05 | 0.0009 | 0 | 0.0001 | 6.61E-05 | 0 | 0.00018 | 0.00013 |
| St7l | 6.46E-05 | 5.31E-05 | 3.18E-05 | 0 | 5.21E-05 | 3.84E-05 | 1.94E-05 | 1.21E-06 | 2.72E-05 | 1.07E-05 | 2.35E-05 | 4.38E-05 | 0 | 3.12E-05 | 6.29E-05 | 2.53E-05 | 0 | 7.63E-06 |
| Cttnbp2nl | 0 | 0.00017 | 0.00014 | 5.49E-05 | 8.33E-05 | 3.48E-05 | 2.06E-06 | 9.13E-05 | 2.72E-05 | 5.17E-05 | 0.00024 | 0.00021 | 5.52E-05 | 0.00013 | 0.00017 | 5.09E-05 | 9.37E-05 | 3.93E-05 |
| Itprid2 | 2.77E-05 | 0.00011 | 4.62E-05 | 8.74E-05 | 6.41E-05 | 2.36E-05 | 0 | 2.37E-05 | 2.72E-05 | 2.88E-05 | 1.53E-05 | 8.79E-05 | 0 | 4.87E-05 | 0.00013 | 2.96E-05 | 7.17E-05 | 0.00017 |
| Lysmd4 | 3.93E-05 | 4.95E-05 | 0 | 4.10E-05 | 4.44E-05 | 1.45E-05 | 1.30E-05 | 0 | 2.72E-05 | 3.57E-05 | 2.08E-05 | 2.04E-06 | 0 | 6.72E-06 | 4.88E-05 | 1.60E-06 | 0 | 4.27E-05 |
| Tmem17 | 2.26E-05 | 1.01E-05 | 2.21E-06 | 4.20E-06 | 1.23E-05 | 4.30E-06 | 7.76E-06 | 0 | 2.72E-05 | 1.10E-05 | 1.19E-06 | 0 | 0 | 4.12E-06 | 0 | 2.02E-05 | 4.24E-06 | 3.82E-06 |
| Dipk2a | 7.18E-05 | 4.80E-05 | 0 | 0 | 1.14E-05 | 0 | 0 | 0 | 2.72E-05 | 0.00012 | 0.00014 | 0.00045 | 0 | 3.93E-05 | 0 | 5.94E-05 | 0.00019 | 0 |
| Mplkip | 0.0001 | 3.29E-05 | 1.10E-05 | 0 | 2.87E-05 | 6.74E-05 | 2.83E-06 | 2.95E-05 | 2.71E-05 | 5.56E-06 | 2.57E-05 | 4.14E-05 | 0 | 1.09E-05 | 6.46E-05 | 3.53E-05 | 4.89E-05 | 0 |
| Slc6a9 | 0 | 6.89E-05 | 3.53E-05 | 7.96E-05 | 0 | 6.30E-05 | 7.69E-05 | 2.36E-05 | 2.71E-05 | 7.30E-05 | 7.30E-06 | 0 | 9.48E-05 | 0 | 7.31E-05 | 0 | 0 | 2.02E-05 |
| Zfp526 | 2.15E-05 | 4.87E-05 | 0 | 4.09E-05 | 3.56E-05 | 4.53E-05 | 7.21E-05 | 5.37E-05 | 2.71E-05 | 5.76E-05 | 2.52E-05 | 2.84E-05 | 1.28E-05 | 6.38E-05 | 0.00011 | 2.53E-05 | 0 | 4.67E-05 |
| Slc35b3 | 6.70E-05 | 5.25E-05 | 5.13E-05 | 7.83E-06 | 2.25E-05 | 4.19E-05 | 8.24E-06 | 5.72E-05 | 2.71E-05 | 2.94E-05 | 0 | 5.64E-05 | 0 | 6.66E-06 | 7.65E-05 | 4.87E-06 | 0 | 1.61E-05 |
| Nkap | 4.06E-05 | 4.19E-05 | 3.01E-05 | 0 | 3.16E-05 | 3.75E-05 | 9.89E-06 | 1.03E-05 | 2.71E-05 | 1.19E-05 | 0 | 2.95E-05 | 0 | 2.13E-05 | 2.99E-05 | 5.02E-05 | 0 | 2.07E-05 |
| Zfp398 | 1.20E-05 | 5.84E-05 | 1.68E-05 | 1.18E-05 | 6.31E-05 | 3.72E-05 | 6.97E-05 | 0 | 2.71E-05 | 4.85E-05 | 3.57E-05 | 2.25E-06 | 2.93E-05 | 3.29E-05 | 2.99E-05 | 0 | 0 | 1.31E-05 |
| Nup205 | 6.48E-05 | 2.42E-05 | 7.05E-05 | 0 | 2.61E-05 | 3.52E-05 | 1.37E-05 | 0 | 2.71E-05 | 1.81E-05 | 1.58E-05 | 1.30E-05 | 0 | 0 | 1.87E-05 | 0.00024 | 0 | 1.05E-05 |
| Eral1 | 5.23E-05 | 3.71E-05 | 2.16E-05 | 1.24E-05 | 1.22E-05 | 2.72E-05 | 4.73E-06 | 8.02E-06 | 2.71E-05 | 2.00E-05 | 2.75E-05 | 9.68E-06 | 0 | 0 | 0 | 4.20E-05 | 0 | 1.37E-05 |
| F8a | 2.57E-06 | 2.08E-05 | 2.54E-05 | 3.83E-05 | 1.08E-05 | 0 | 2.25E-05 | 2.75E-05 | 2.71E-05 | 1.40E-05 | 0 | 0 | 2.85E-06 | 2.52E-05 | 1.37E-05 | 0 | 0 | 1.79E-05 |
| Acat1 | 0.00028 | 4.39E-05 | 0.00014 | 0.00018 | 5.12E-05 | 0.00011 | 0.00028 | 0 | 2.70E-05 | 0.0002 | 1.59E-06 | 4.92E-05 | 0.00024 | 0 | 0 | 0.00022 | 0 | 0.00013 |
| Ccdc122 | 5.54E-05 | 4.62E-05 | 0 | 0 | 3.12E-05 | 8.20E-05 | 0 | 2.91E-06 | 2.70E-05 | 3.10E-06 | 0 | 0 | 4.38E-05 | 0 | 0 | 3.29E-05 | 0 | 1.18E-06 |
| Zfp654 | 9.20E-05 | 5.49E-05 | 0 | 0 | 1.21E-05 | 8.04E-05 | 1.52E-05 | 3.31E-06 | 2.70E-05 | 1.94E-05 | 0 | 0.00019 | 1.51E-05 | 0 | 2.89E-05 | 7.96E-07 | 2.88E-05 | 0 |
| Prkar1b | 0 | 2.46E-05 | 2.97E-06 | 2.87E-05 | 8.11E-05 | 7.64E-05 | 0 | 0 | 2.70E-05 | 2.65E-05 | 5.24E-05 | 0.0001 | 2.57E-05 | 1.32E-05 | 0 | 0 | 0 | 1.92E-05 |

| | | | | | | | | | | | | | | | | | | |
|---|---|---|---|---|---|---|---|---|---|---|---|---|---|---|---|---|---|---|
| Morc2a | 0.0001 | 0.00014 | 0.00015 | 0.00013 | 7.73E-05 | 7.19E-05 | 1.07E-05 | 2.15E-05 | 2.70E-05 | 5.06E-05 | 6.75E-05 | 2.07E-05 | 3.89E-05 | 2.44E-05 | 0.00019 | 9.84E-05 | 0.00018 | 0 |
| Srr | 5.08E-05 | 5.69E-05 | 0 | 0 | 4.91E-05 | 5.54E-05 | 0 | 0 | 2.70E-05 | 3.13E-05 | 1.95E-05 | 4.73E-05 | 8.64E-06 | 0 | 1.43E-05 | 0 | 0 | 4.06E-05 |
| Hagh | 8.21E-05 | 8.90E-05 | 4.98E-06 | 3.72E-05 | 4.84E-05 | 2.70E-05 | 3.58E-05 | 2.38E-05 | 2.70E-05 | 4.29E-05 | 2.31E-05 | 4.44E-05 | 0 | 2.82E-05 | 0 | 5.34E-07 | 4.99E-05 | 2.30E-05 |
| Dhx57 | 8.96E-05 | 4.87E-05 | 1.10E-05 | 0 | 1.29E-05 | 1.50E-05 | 4.99E-05 | 0 | 2.70E-05 | 4.28E-05 | 0 | 4.70E-05 | 7.12E-07 | 2.24E-05 | 0.00015 | 1.38E-05 | 9.88E-06 | 4.21E-05 |
| Rogdi | 9.32E-06 | 5.53E-05 | 0 | 5.51E-05 | 1.20E-05 | 1.10E-05 | 0 | 0 | 2.70E-05 | 5.01E-05 | 2.54E-05 | 0 | 4.14E-05 | 3.53E-05 | 1.89E-05 | 4.17E-05 | 1.27E-05 | 2.60E-05 |
| Pdik1l | 1.95E-05 | 3.74E-05 | 1.68E-05 | 3.90E-05 | 2.62E-05 | 3.74E-06 | 1.07E-05 | 1.42E-05 | 2.70E-05 | 1.54E-05 | 2.76E-05 | 9.71E-06 | 2.85E-05 | 1.53E-05 | 1.88E-05 | 9.44E-05 | 0 | 5.81E-06 |
| Usp20 | 0 | 5.56E-05 | 0 | 6.76E-05 | 2.55E-05 | 0 | 0 | 1.96E-05 | 2.70E-05 | 1.22E-05 | 1.36E-05 | 1.11E-05 | 0 | 6.23E-06 | 4.72E-05 | 0 | 1.69E-06 | 7.95E-06 |
| Chrnb1 | 0 | 3.56E-05 | 0 | 0 | 0.00013 | 0 | 0 | 4.71E-05 | 2.70E-05 | 0 | 6.70E-05 | 2.83E-05 | 1.96E-05 | 1.73E-06 | 3.23E-05 | 0 | 2.93E-05 | 2.06E-05 |
| Yju2 | 3.12E-05 | 3.78E-05 | 2.94E-05 | 2.69E-05 | 1.37E-05 | 0 | 2.49E-05 | 0 | 2.70E-05 | 2.14E-05 | 9.87E-06 | 1.91E-05 | 1.25E-05 | 1.90E-05 | 2.08E-05 | 3.36E-05 | 0 | 0 |
| Frs2 | 7.13E-05 | 0.00016 | 7.31E-05 | 0.00018 | 0.00011 | 0.00012 | 4.40E-05 | 3.91E-05 | 2.69E-05 | 4.30E-05 | 0.00024 | 9.36E-05 | 2.96E-05 | 5.22E-05 | 0.00024 | 4.84E-05 | 0.00012 | 0.00012 |
| Emp2 | 2.77E-05 | 0.00016 | 4.30E-05 | 0 | 8.72E-05 | 9.68E-05 | 0 | 0 | 2.69E-05 | 0 | 0.00016 | 0.0003 | 0 | 0.00015 | 0 | 0.00017 | 0 | 0.00017 |
| Ezh1 | 5.50E-05 | 0.00019 | 0 | 4.46E-06 | 7.63E-05 | 8.21E-05 | 2.49E-05 | 1.51E-05 | 2.69E-05 | 5.63E-05 | 0 | 0.00011 | 0 | 1.41E-05 | 0.00011 | 2.02E-06 | 0 | 5.73E-05 |
| Smurf1 | 2.53E-05 | 0.00022 | 0.00029 | 0 | 0.00012 | 6.23E-05 | 0 | 0.0003 | 2.69E-05 | 0.00021 | 0.00031 | 0.00027 | 3.24E-06 | 0.00016 | 0.00012 | 1.65E-05 | 0.00013 | 8.28E-05 |
| Zkscan1 | 6.44E-05 | 0.0001 | 2.30E-05 | 3.94E-05 | 4.64E-05 | 6.07E-05 | 2.41E-05 | 4.72E-06 | 2.69E-05 | 2.07E-05 | 2.40E-05 | 3.31E-05 | 2.11E-05 | 0 | 7.72E-05 | 0 | 0 | 3.86E-05 |
| Scrn3 | 3.29E-05 | 2.16E-05 | 0 | 7.44E-05 | 3.96E-05 | 3.37E-05 | 2.64E-05 | 2.58E-05 | 2.69E-05 | 2.91E-05 | 8.66E-06 | 2.42E-05 | 0 | 0 | 2.07E-05 | 7.43E-05 | 0 | 4.62E-05 |
| Fbxl4 | 2.39E-05 | 5.61E-05 | 1.34E-05 | 1.43E-05 | 3.85E-05 | 3.07E-05 | 2.16E-05 | 0 | 2.69E-05 | 3.33E-05 | 1.05E-05 | 4.41E-05 | 1.97E-05 | 2.11E-05 | 2.65E-05 | 0 | 2.98E-06 | 3.95E-05 |
| Zfp790 | 4.08E-05 | 4.85E-05 | 1.33E-05 | 6.89E-06 | 2.32E-05 | 2.94E-05 | 7.38E-06 | 9.00E-06 | 2.69E-05 | 1.77E-05 | 1.87E-05 | 1.13E-05 | 0 | 1.89E-05 | 3.80E-05 | 6.21E-06 | 0 | 3.09E-05 |
| Crnkl1 | 9.94E-05 | 0.00011 | 8.19E-05 | 8.97E-05 | 3.62E-05 | 2.47E-05 | 1.06E-05 | 0 | 2.69E-05 | 2.49E-05 | 4.61E-05 | 0 | 3.57E-05 | 7.03E-05 | 5.75E-05 | 8.18E-05 | 0 | 1.82E-05 |
| Parp11 | 3.75E-05 | 2.82E-05 | 5.05E-06 | 4.47E-06 | 2.59E-05 | 1.99E-05 | 0 | 2.77E-05 | 2.69E-05 | 5.60E-05 | 1.98E-05 | 6.21E-06 | 1.54E-05 | 0 | 1.84E-05 | 0 | 0 | 5.25E-05 |
| Glis1 | 0 | 0 | 1.23E-05 | 1.44E-05 | 1.60E-05 | 1.98E-05 | 1.55E-06 | 0 | 2.69E-05 | 0 | 1.51E-05 | 3.31E-05 | 0 | 4.44E-05 | 2.81E-05 | 0 | 3.74E-05 | 4.98E-06 |
| Reep4 | 1.52E-05 | 0.00012 | 2.81E-05 | 2.26E-05 | 1.53E-05 | 1.87E-05 | 0.00012 | 6.84E-05 | 2.69E-05 | 5.55E-05 | 0.00011 | 2.44E-05 | 0 | 5.73E-05 | 7.79E-05 | 0.00067 | 0 | 4.97E-05 |
| Tbc1d25 | 1.54E-05 | 3.49E-05 | 8.12E-06 | 0 | 5.59E-08 | 1.60E-07 | 4.73E-05 | 0 | 2.69E-05 | 1.41E-05 | 6.93E-05 | 0 | 2.87E-05 | 2.46E-05 | 7.01E-05 | 0 | 0 | 4.78E-05 |
| Mospd1 | 0.00012 | 6.84E-05 | 0 | 7.41E-06 | 1.52E-05 | 0.00014 | 1.45E-05 | 4.35E-05 | 2.68E-05 | 0 | 0 | 4.83E-06 | 6.71E-05 | 0.00012 | 0 | 2.09E-06 | 0 | 7.31E-06 |
| Asb3 | 6.45E-05 | 6.18E-05 | 0 | 0 | 1.80E-05 | 6.83E-05 | 6.58E-05 | 1.18E-05 | 2.68E-05 | 4.84E-05 | 5.18E-06 | 2.37E-05 | 0 | 2.51E-05 | 7.20E-05 | 5.80E-05 | 0 | 2.82E-05 |
| Akirin1 | 0.00014 | 8.28E-05 | 0.00011 | 0 | 4.97E-05 | 4.91E-05 | 7.49E-06 | 0 | 2.68E-05 | 8.52E-06 | 3.44E-05 | 6.71E-06 | 0 | 0 | 0 | 0 | 4.60E-05 | 0 |
| Pde12 | 9.37E-05 | 2.70E-05 | 8.38E-05 | 0 | 0 | 4.85E-05 | 0 | 7.32E-06 | 2.68E-05 | 4.04E-06 | 1.13E-05 | 7.28E-06 | 0 | 0 | 1.15E-05 | 5.00E-05 | 5.25E-06 | 0 |
| Tbc1d24 | 0 | 5.36E-05 | 0 | 1.97E-05 | 5.01E-05 | 3.46E-05 | 2.87E-05 | 0 | 2.68E-05 | 2.35E-05 | 0 | 4.83E-05 | 5.39E-05 | 6.44E-05 | 6.17E-06 | 7.80E-06 | 0 | 3.63E-05 |
| Trmt6 | 4.71E-05 | 4.75E-05 | 0.00011 | 7.68E-05 | 2.21E-05 | 3.14E-05 | 0 | 1.34E-05 | 2.68E-05 | 1.52E-05 | 1.57E-05 | 1.32E-05 | 1.69E-05 | 9.24E-06 | 1.93E-05 | 8.01E-05 | 0 | 1.66E-05 |
| Phpt1 | 1.11E-05 | 1.51E-05 | 2.26E-05 | 3.71E-05 | 1.67E-05 | 1.23E-05 | 0 | 1.53E-05 | 2.68E-05 | 0 | 1.33E-05 | 1.06E-05 | 3.27E-05 | 8.80E-06 | 0 | 0 | 2.39E-05 | 2.38E-05 |
| Vps26c | 0.00016 | 0.00015 | 8.70E-06 | 7.66E-05 | 2.62E-05 | 0.0001 | 8.83E-05 | 3.29E-05 | 2.67E-05 | 7.87E-05 | 0 | 4.49E-05 | 1.58E-05 | 8.03E-05 | 2.90E-05 | 4.71E-05 | 0 | 2.83E-05 |
| Pde6d | 0.00013 | 6.64E-05 | 1.51E-05 | 2.15E-05 | 3.80E-05 | 0.0001 | 0.0001 | 3.14E-05 | 2.67E-05 | 7.75E-05 | 3.18E-05 | 7.47E-05 | 1.14E-05 | 1.57E-05 | 0 | 0.00019 | 0 | 0.0001 |
| Sncb | 0 | 0 | 0 | 0 | 0 | 4.12E-05 | 0 | 0 | 2.67E-05 | 0 | 0 | 0.00024 | 0 | 0 | 0 | 0 | 0 | 0 |
| Nepro | 4.94E-05 | 6.30E-05 | 0.00011 | 1.84E-05 | 2.35E-05 | 3.41E-05 | 0 | 1.86E-05 | 2.67E-05 | 0 | 2.24E-05 | 0 | 2.58E-05 | 3.93E-05 | 6.10E-05 | 6.60E-05 | 5.14E-05 | 0 |

| | | | | | | | | | | | | | | | | | | |
|---|---|---|---|---|---|---|---|---|---|---|---|---|---|---|---|---|---|---|
| Pus3 | 2.85E-05 | 2.20E-05 | 2.23E-05 | 0 | 1.13E-05 | 1.53E-05 | 0 | 0 | 2.67E-05 | 3.47E-06 | 0 | 5.20E-06 | 0 | 1.57E-05 | 3.58E-05 | 0 | 0 | 0 |
| Pigh | 2.24E-05 | 2.67E-05 | 0 | 0 | 1.19E-05 | 1.36E-05 | 3.28E-06 | 7.93E-06 | 2.67E-05 | 1.63E-05 | 1.84E-05 | 1.75E-05 | 1.19E-05 | 3.06E-06 | 0 | 2.52E-05 | 0 | 2.58E-05 |
| Fam151b | 1.28E-05 | 3.39E-05 | 1.14E-07 | 1.01E-05 | 4.56E-06 | 3.89E-06 | 0 | 2.01E-06 | 2.67E-05 | 1.08E-05 | 0 | 1.62E-05 | 0 | 0 | 6.02E-06 | 9.92E-06 | 0 | 5.78E-06 |
| Prorp | 2.01E-05 | 3.13E-05 | 1.78E-05 | 1.17E-05 | 3.11E-06 | 2.88E-06 | 3.04E-05 | 0 | 2.67E-05 | 9.00E-06 | 4.24E-06 | 1.43E-05 | 0 | 2.55E-06 | 6.29E-06 | 0 | 3.40E-05 | 1.04E-05 |
| Il1b | 0 | 7.97E-05 | 4.79E-05 | 0.00013 | 2.97E-05 | 0 | 0 | 0 | 2.67E-05 | 8.54E-05 | 3.21E-05 | 4.03E-05 | 7.19E-05 | 8.29E-06 | 0.00068 | 7.39E-05 | 0.00014 | 6.84E-05 |
| Armcx6 | 2.07E-05 | 4.65E-05 | 0 | 6.19E-05 | 2.06E-06 | 0 | 4.34E-05 | 5.39E-05 | 2.67E-05 | 3.45E-06 | 0 | 1.82E-05 | 1.34E-05 | 0 | 0 | 3.92E-05 | 6.75E-05 | 1.66E-05 |
| Insig2 | 0.00011 | 0.00012 | 0 | 0 | 5.66E-05 | 0.00013 | 1.51E-05 | 0 | 2.66E-05 | 7.75E-05 | 3.05E-05 | 0.00015 | 7.53E-07 | 6.79E-06 | 0 | 4.19E-06 | 0 | 2.69E-05 |
| Fam129a | 4.00E-05 | 0.0002 | 0 | 0 | 0 | 0.00012 | 0.00012 | 1.37E-05 | 2.66E-05 | 0.00025 | 0 | 0.00021 | 0 | 0 | 0 | 0 | 2.26E-06 | 8.00E-05 |
| Taf15 | 6.64E-05 | 3.07E-05 | 2.50E-05 | 0 | 0 | 2.31E-05 | 0 | 0 | 2.66E-05 | 0 | 1.17E-05 | 5.03E-05 | 0 | 1.28E-05 | 6.32E-05 | 8.41E-05 | 0 | 0 |
| Dapp1 | 5.61E-05 | 6.84E-06 | 0 | 2.20E-05 | 1.86E-05 | 1.68E-05 | 9.13E-05 | 0 | 2.66E-05 | 2.52E-05 | 0 | 5.84E-06 | 6.62E-06 | 1.40E-05 | 3.00E-06 | 1.90E-05 | 1.29E-05 | 2.22E-05 |
| Slc7a1 | 4.60E-05 | 2.66E-05 | 0.0002 | 0 | 0 | 1.35E-05 | 0 | 2.45E-06 | 2.66E-05 | 3.08E-05 | 0.00011 | 0.00014 | 0 | 5.51E-06 | 0 | 4.27E-05 | 0 | 3.55E-05 |
| Pam16 | 2.28E-05 | 2.36E-05 | 3.27E-05 | 9.37E-05 | 2.75E-05 | 0 | 0 | 1.48E-05 | 2.66E-05 | 0 | 7.19E-06 | 2.11E-05 | 6.62E-06 | 7.90E-06 | 7.47E-06 | 1.80E-05 | 4.07E-05 | 1.23E-06 |
| Ccdc126 | 6.10E-05 | 2.50E-05 | 2.15E-05 | 2.95E-05 | 4.50E-05 | 3.63E-05 | 2.60E-06 | 5.33E-05 | 2.65E-05 | 1.71E-05 | 5.46E-05 | 2.21E-05 | 0 | 1.37E-05 | 3.56E-07 | 3.19E-05 | 0 | 2.36E-05 |
| Klhl20 | 2.54E-05 | 4.74E-05 | 2.45E-05 | 2.83E-05 | 3.50E-05 | 3.21E-05 | 1.97E-05 | 4.42E-05 | 2.65E-05 | 1.93E-05 | 1.98E-05 | 2.82E-06 | 3.21E-05 | 5.83E-06 | 2.68E-08 | 5.38E-05 | 0 | 2.60E-05 |
| Rdm1 | 6.88E-06 | 3.66E-05 | 0 | 0 | 4.10E-06 | 2.15E-05 | 6.35E-06 | 2.57E-05 | 2.65E-05 | 1.17E-05 | 3.75E-06 | 2.56E-05 | 0 | 2.68E-07 | 0 | 7.37E-05 | 0 | 1.26E-05 |
| Wdpcp | 2.51E-05 | 6.57E-05 | 0 | 7.34E-06 | 2.59E-05 | 2.04E-05 | 4.98E-06 | 0 | 2.65E-05 | 3.46E-05 | 1.77E-07 | 2.00E-05 | 1.25E-05 | 3.20E-05 | 4.14E-05 | 6.78E-05 | 3.02E-05 | 1.43E-05 |
| Pex1 | 2.97E-05 | 4.35E-05 | 1.06E-05 | 2.29E-05 | 3.06E-05 | 1.37E-05 | 1.25E-06 | 9.47E-06 | 2.65E-05 | 3.60E-06 | 4.17E-06 | 0 | 2.53E-05 | 1.87E-05 | 1.43E-05 | 0 | 2.56E-05 | 9.29E-06 |
| Fra10ac1 | 5.17E-05 | 2.35E-05 | 5.51E-05 | 5.67E-05 | 1.71E-05 | 1.25E-05 | 1.41E-05 | 8.29E-06 | 2.65E-05 | 2.11E-05 | 3.76E-05 | 1.11E-05 | 1.25E-05 | 0 | 7.56E-05 | 4.90E-05 | 0 | 0 |
| Dennd4a | 0.00014 | 1.61E-05 | 4.49E-05 | 0 | 0.00022 | 0 | 0.00042 | 0 | 2.65E-05 | 0 | 0.00031 | 0 | 0.00027 | 0.00021 | 0.00011 | 1.29E-05 | 0 | 0.00026 |
| Trp53rka | 3.20E-05 | 5.32E-05 | 8.70E-05 | 6.06E-05 | 3.56E-05 | 0 | 0 | 8.52E-06 | 2.65E-05 | 9.83E-06 | 9.19E-06 | 6.78E-05 | 1.24E-06 | 1.60E-05 | 2.03E-05 | 7.67E-06 | 0 | 1.21E-05 |
| Hc | 0 | 5.87E-05 | 0 | 0 | 0.00029 | 0.00085 | 0 | 0 | 2.64E-05 | 0 | 0 | 0 | 0 | 0 | 0 | 0 | 0 | 6.92E-05 |
| Clcn7 | 4.89E-06 | 0.00016 | 4.60E-05 | 0.0001 | 0.00013 | 0.00014 | 6.52E-05 | 0.0001 | 2.64E-05 | 8.24E-05 | 3.81E-05 | 0 | 7.28E-05 | 0.00011 | 4.30E-05 | 0.00012 | 0 | 0.0001 |
| Zranb1 | 8.65E-05 | 0.00017 | 6.86E-05 | 9.97E-06 | 3.47E-05 | 8.24E-05 | 4.22E-05 | 1.22E-05 | 2.64E-05 | 1.72E-05 | 2.24E-05 | 0.0001 | 0 | 3.18E-05 | 0.0002 | 9.14E-05 | 0.00024 | 3.96E-06 |
| AI597479 | 7.26E-05 | 4.99E-05 | 3.58E-05 | 1.83E-05 | 3.64E-05 | 8.07E-05 | 3.59E-05 | 1.19E-05 | 2.64E-05 | 2.59E-05 | 0 | 7.45E-06 | 0 | 0 | 9.63E-05 | 8.99E-06 | 0 | 8.30E-06 |
| Spata6 | 7.80E-05 | 6.38E-05 | 3.86E-05 | 7.03E-07 | 3.50E-05 | 6.88E-05 | 8.74E-06 | 8.76E-05 | 2.64E-05 | 8.17E-05 | 3.57E-05 | 0.0001 | 0 | 4.32E-05 | 7.01E-05 | 4.72E-05 | 0 | 0.00011 |
| Ebag9 | 9.71E-05 | 5.33E-05 | 3.29E-05 | 0 | 2.95E-05 | 6.08E-05 | 3.84E-05 | 2.33E-05 | 2.64E-05 | 1.38E-05 | 8.85E-06 | 7.58E-06 | 0 | 0 | 0.00012 | 4.34E-05 | 3.15E-05 | 2.01E-05 |
| Trim13 | 2.67E-05 | 1.68E-05 | 0 | 0 | 0 | 5.83E-05 | 0 | 0 | 2.64E-05 | 1.70E-05 | 0 | 0 | 0 | 7.98E-06 | 0 | 0 | 0 | 1.95E-05 |
| Tk2 | 2.85E-05 | 9.06E-05 | 1.45E-05 | 2.76E-05 | 3.76E-05 | 2.13E-05 | 5.74E-06 | 9.18E-07 | 2.64E-05 | 1.71E-05 | 2.13E-05 | 3.99E-05 | 1.05E-05 | 0 | 3.59E-05 | 0 | 0 | 2.40E-05 |
| Zfp212 | 2.09E-05 | 7.47E-05 | 0 | 1.94E-05 | 1.67E-05 | 1.99E-05 | 1.97E-05 | 2.14E-05 | 2.64E-05 | 2.16E-05 | 3.87E-05 | 2.97E-05 | 0 | 3.97E-05 | 6.88E-05 | 4.70E-05 | 0 | 1.80E-05 |
| Zdhhc13 | 4.62E-05 | 1.39E-05 | 5.34E-05 | 0 | 8.58E-06 | 1.90E-05 | 0 | 2.23E-05 | 2.64E-05 | 2.30E-06 | 7.67E-06 | 2.32E-05 | 9.94E-06 | 1.35E-06 | 5.90E-06 | 4.21E-05 | 0 | 0 |
| Atpsckmt | 9.76E-06 | 2.13E-05 | 2.74E-05 | 7.05E-06 | 3.30E-05 | 7.53E-06 | 1.85E-05 | 4.11E-05 | 2.64E-05 | 3.73E-05 | 0 | 1.92E-05 | 2.37E-05 | 1.90E-05 | 8.00E-05 | 6.80E-05 | 0 | 2.04E-05 |
| Piezo1 | 2.12E-05 | 0.00029 | 0.00038 | 0.00012 | 0.00016 | 0.00026 | 5.35E-05 | 0.00025 | 2.63E-05 | 5.33E-05 | 0.0003 | 9.11E-05 | 0 | 0.00035 | 0.00024 | 0.00016 | 0.00036 | 0.00015 |
| Rngtt | 5.17E-05 | 3.58E-05 | 3.85E-05 | 0 | 1.74E-05 | 3.27E-05 | 0 | 2.35E-06 | 2.63E-05 | 1.51E-05 | 3.21E-05 | 1.32E-05 | 6.67E-06 | 0 | 1.02E-05 | 7.46E-05 | 5.38E-05 | 1.12E-05 |

| | | | | | | | | | | | | | | | | | | |
|---|---|---|---|---|---|---|---|---|---|---|---|---|---|---|---|---|---|---|
| Llph | 4.15E-05 | 8.18E-06 | 2.92E-05 | 4.16E-06 | 5.04E-06 | 1.74E-05 | 6.05E-06 | 0 | 2.63E-05 | 1.71E-06 | 5.79E-07 | 7.22E-06 | 1.79E-05 | 0 | 7.77E-06 | 5.94E-06 | 0 | 0 |
| Adal | 4.24E-05 | 2.70E-05 | 9.68E-06 | 4.37E-06 | 2.06E-05 | 1.49E-05 | 4.75E-06 | 8.92E-06 | 2.63E-05 | 1.01E-06 | 5.65E-06 | 0 | 7.38E-07 | 9.23E-06 | 0 | 2.74E-05 | 0 | 1.90E-05 |
| Rabep2 | 1.02E-05 | 6.21E-05 | 1.50E-05 | 0.00012 | 1.64E-05 | 0 | 2.60E-05 | 0.0001 | 2.63E-05 | 5.01E-05 | 9.27E-06 | 1.02E-05 | 4.87E-05 | 5.72E-05 | 0.00011 | 9.81E-06 | 2.10E-05 | 3.33E-05 |
| Yipf4 | 0.00029 | 0.00016 | 6.64E-05 | 0 | 0.00012 | 0.00029 | 0.00018 | 4.90E-05 | 2.62E-05 | 7.75E-05 | 5.87E-05 | 0.00015 | 3.88E-05 | 5.39E-05 | 0 | 8.23E-05 | 4.03E-05 | 8.17E-05 |
| Ebf1 | 0.00014 | 0.00016 | 4.27E-05 | 0 | 0 | 0.00013 | 0 | 0.00052 | 2.62E-05 | 0.00033 | 0 | 4.19E-06 | 7.45E-05 | 5.71E-05 | 0.00053 | 9.77E-05 | 0.00054 | 6.10E-05 |
| Bdp1 | 3.23E-05 | 6.54E-05 | 8.94E-05 | 0 | 2.71E-05 | 4.45E-05 | 2.66E-05 | 3.70E-05 | 2.62E-05 | 3.05E-05 | 4.24E-05 | 6.19E-06 | 1.90E-05 | 2.99E-05 | 3.91E-05 | 5.06E-05 | 9.22E-05 | 2.90E-05 |
| Slc6a6 | 3.58E-05 | 0.00026 | 8.93E-05 | 4.97E-05 | 8.01E-05 | 3.08E-05 | 0.00013 | 0 | 2.62E-05 | 0 | 0.00034 | 0 | 0.00012 | 0.00035 | 0 | 1.67E-05 | 6.27E-05 | 0.00015 |
| Jrkl | 7.79E-05 | 5.21E-05 | 6.51E-06 | 2.62E-05 | 0 | 2.53E-05 | 1.00E-05 | 3.41E-05 | 2.62E-05 | 6.25E-06 | 8.51E-07 | 2.02E-05 | 0 | 9.60E-06 | 5.85E-05 | 3.16E-05 | 0 | 1.93E-05 |
| Trim65 | 1.90E-05 | 3.26E-05 | 9.90E-06 | 1.33E-05 | 1.95E-05 | 1.88E-05 | 0 | 0 | 2.62E-05 | 0 | 4.18E-06 | 6.80E-06 | 0 | 2.71E-05 | 2.29E-05 | 3.25E-06 | 0 | 5.84E-05 |
| Dctd | 5.13E-05 | 0 | 0.00011 | 3.23E-05 | 0 | 0 | 0 | 0 | 2.62E-05 | 0 | 4.38E-06 | 0 | 3.34E-05 | 0 | 1.06E-05 | 4.95E-05 | 0 | 0 |
| Tbc1d7 | 6.18E-05 | 2.07E-05 | 2.03E-05 | 3.92E-05 | 2.66E-05 | 2.94E-05 | 1.85E-05 | 4.80E-05 | 2.61E-05 | 3.22E-05 | 2.98E-05 | 9.13E-06 | 0 | 0 | 0 | 5.39E-05 | 0 | 3.31E-05 |
| Lrp12 | 9.91E-05 | 0.00012 | 5.68E-05 | 0 | 3.61E-05 | 1.57E-05 | 0 | 7.26E-05 | 2.61E-05 | 0 | 5.58E-05 | 0 | 9.83E-05 | 4.52E-05 | 0 | 8.20E-05 | 9.84E-05 | 0.00013 |
| Cryab | 0 | 1.13E-05 | 0 | 0 | 0 | 0 | 0 | 0 | 2.61E-05 | 0.00022 | 0.00043 | 0.00039 | 0.00016 | 0.00011 | 0 | 0 | 0 | 0.00042 |
| Apln | 0 | 0 | 0 | 0 | 0 | 0 | 0 | 0 | 2.61E-05 | 0 | 0 | 0.00112 | 0.0009 | 0 | 0 | 0 | 0 | 0 |
| Hdhd3 | 1.59E-05 | 1.63E-05 | 5.17E-06 | 3.73E-05 | 1.27E-05 | 0 | 0 | 2.07E-05 | 2.61E-05 | 1.10E-05 | 1.23E-05 | 0 | 3.23E-06 | 4.33E-06 | 5.18E-05 | 0 | 0 | 1.88E-05 |
| Acap3 | 0.00012 | 0.00016 | 0.00015 | 4.65E-05 | 0.00014 | 0.00013 | 6.55E-05 | 8.12E-05 | 2.60E-05 | 0.00014 | 0.00011 | 1.99E-05 | 0 | 0.00015 | 0.00026 | 3.28E-05 | 5.36E-05 | 0.00012 |
| Spns1 | 8.47E-06 | 7.30E-05 | 1.11E-05 | 7.31E-05 | 5.67E-05 | 4.71E-05 | 3.00E-05 | 5.06E-05 | 2.60E-05 | 1.91E-05 | 2.62E-05 | 1.81E-05 | 1.18E-05 | 7.55E-05 | 0.0001 | 0 | 0 | 5.00E-05 |
| Tmem241 | 5.53E-05 | 4.04E-05 | 2.69E-06 | 1.84E-05 | 3.16E-05 | 2.04E-05 | 6.47E-05 | 4.47E-05 | 2.60E-05 | 5.59E-05 | 0 | 2.92E-05 | 2.95E-05 | 0 | 3.78E-05 | 0 | 0 | 2.83E-05 |
| Arsg | 3.42E-06 | 4.41E-05 | 0 | 9.94E-06 | 1.98E-05 | 1.07E-05 | 1.20E-06 | 1.57E-05 | 2.60E-05 | 2.65E-05 | 0 | 1.47E-05 | 2.48E-05 | 2.48E-05 | 9.39E-06 | 0 | 0 | 3.98E-05 |
| Ttc1 | 9.36E-05 | 9.07E-05 | 6.25E-05 | 6.94E-05 | 2.66E-05 | 1.02E-05 | 1.28E-05 | 0 | 2.60E-05 | 1.25E-05 | 2.43E-05 | 4.27E-05 | 0 | 3.21E-05 | 6.08E-05 | 6.12E-05 | 0 | 2.87E-05 |
| Cntrob | 0 | 1.99E-05 | 0 | 2.62E-05 | 2.62E-05 | 8.76E-06 | 0 | 8.61E-06 | 2.60E-05 | 2.10E-05 | 1.42E-06 | 1.14E-05 | 0 | 0 | 9.58E-05 | 0.00028 | 2.46E-05 | 2.14E-05 |
| Prdm15 | 1.65E-05 | 2.10E-05 | 2.65E-05 | 0 | 1.63E-07 | 2.80E-06 | 7.16E-06 | 1.67E-05 | 2.60E-05 | 5.12E-06 | 1.57E-06 | 0 | 0 | 1.77E-05 | 0.00011 | 0 | 2.17E-05 | 6.88E-06 |
| Exosc7 | 5.33E-05 | 4.05E-05 | 0.00011 | 5.52E-05 | 1.22E-05 | 0 | 0 | 0 | 2.60E-05 | 8.88E-05 | 0 | 3.60E-07 | 2.08E-05 | 4.01E-06 | 0.00013 | 0.00012 | 0 | 2.78E-05 |
| Prpf4b | 0.00028 | 0.00021 | 0.00014 | 0 | 8.93E-05 | 0.00013 | 1.57E-05 | 0 | 2.59E-05 | 2.37E-05 | 3.51E-06 | 8.92E-05 | 0 | 2.41E-05 | 0.00055 | 0.00016 | 0.00014 | 0 |
| Tceanc2 | 9.27E-05 | 5.10E-05 | 1.95E-05 | 3.40E-05 | 4.38E-05 | 5.03E-05 | 7.82E-06 | 0 | 2.59E-05 | 2.19E-05 | 3.16E-06 | 3.03E-05 | 0 | 1.39E-05 | 8.00E-05 | 4.71E-05 | 2.73E-05 | 4.06E-05 |
| Saal1 | 5.39E-05 | 1.82E-05 | 5.19E-05 | 4.10E-05 | 3.79E-05 | 4.85E-05 | 4.53E-06 | 1.06E-05 | 2.59E-05 | 3.26E-05 | 2.42E-05 | 3.38E-05 | 0.00011 | 1.29E-05 | 5.70E-07 | 0.00012 | 8.50E-06 | 0 |
| Tyw5 | 5.88E-05 | 2.19E-05 | 1.77E-05 | 0 | 8.12E-06 | 2.33E-05 | 0 | 0 | 2.59E-05 | 3.80E-06 | 9.51E-06 | 2.04E-05 | 0 | 0 | 8.87E-05 | 2.12E-05 | 0 | 1.68E-05 |
| Uevld | 3.95E-05 | 3.43E-05 | 0 | 0 | 7.48E-06 | 1.44E-05 | 0 | 0 | 2.59E-05 | 1.94E-05 | 1.05E-05 | 4.88E-05 | 0 | 4.37E-06 | 0 | 0.00011 | 0 | 2.34E-05 |
| Phlpp2 | 0 | 4.98E-05 | 3.75E-05 | 2.08E-05 | 2.90E-05 | 1.37E-05 | 0 | 0 | 2.59E-05 | 5.12E-05 | 3.04E-05 | 0 | 5.38E-05 | 2.64E-05 | 4.08E-05 | 6.75E-05 | 0.00015 | 3.72E-05 |
| Fahd1 | 4.58E-05 | 3.52E-05 | 1.40E-05 | 0.00012 | 9.70E-06 | 0 | 2.15E-05 | 3.37E-06 | 2.59E-05 | 4.43E-06 | 0 | 3.15E-05 | 1.65E-05 | 1.67E-05 | 5.48E-05 | 0 | 0 | 0 |
| Als2cl | 0 | 2.16E-05 | 0 | 0 | 0 | 0 | 0 | 3.80E-07 | 2.59E-05 | 0 | 2.04E-07 | 0 | 4.67E-05 | 0 | 4.96E-05 | 9.12E-06 | 0 | 2.26E-05 |
| Trim23 | 7.18E-05 | 4.49E-05 | 1.86E-05 | 1.30E-06 | 2.46E-05 | 5.56E-05 | 8.00E-06 | 1.07E-05 | 2.58E-05 | 2.02E-05 | 3.10E-05 | 3.53E-05 | 0 | 3.08E-05 | 2.44E-05 | 3.07E-05 | 0 | 3.12E-05 |
| Sh3bp5l | 6.68E-06 | 8.21E-05 | 2.74E-06 | 0.00011 | 5.01E-05 | 3.75E-05 | 7.18E-06 | 0 | 2.58E-05 | 6.88E-05 | 9.61E-05 | 0 | 5.66E-06 | 3.89E-05 | 6.62E-05 | 2.17E-05 | 0 | 6.01E-05 |

| | | | | | | | | | | | | | | | | | | |
|---|---|---|---|---|---|---|---|---|---|---|---|---|---|---|---|---|---|---|
| Clybl | 6.03E-05 | 3.34E-06 | 0 | 5.39E-05 | 1.45E-05 | 1.75E-05 | 5.28E-05 | 1.46E-05 | 2.58E-05 | 3.61E-05 | 2.26E-05 | 0.00033 | 0.0001 | 1.85E-05 | 0 | 0 | 0 | 3.39E-05 |
| Cby1 | 3.71E-05 | 4.67E-05 | 0 | 0.00015 | 4.48E-05 | 9.93E-06 | 2.06E-05 | 3.00E-05 | 2.58E-05 | 6.34E-05 | 4.75E-06 | 0 | 0 | 3.40E-05 | 0 | 0.00013 | 0 | 6.01E-05 |
| Zfp446 | 6.07E-06 | 1.06E-05 | 2.97E-06 | 2.51E-05 | 0 | 7.01E-06 | 0 | 9.60E-06 | 2.58E-05 | 1.94E-05 | 9.59E-06 | 1.39E-05 | 3.43E-06 | 1.25E-05 | 2.24E-05 | 6.78E-06 | 1.52E-05 | 1.23E-05 |
| Med26 | 0 | 4.89E-05 | 2.21E-05 | 5.52E-05 | 1.39E-05 | 5.42E-06 | 1.43E-05 | 0 | 2.58E-05 | 2.46E-05 | 3.35E-05 | 3.90E-05 | 9.98E-06 | 2.96E-05 | 4.16E-05 | 3.57E-05 | 6.51E-05 | 2.19E-05 |
| Pank1 | 5.53E-06 | 1.13E-05 | 3.23E-06 | 1.99E-06 | 3.01E-05 | 0 | 0 | 1.07E-05 | 2.58E-05 | 2.58E-06 | 1.06E-05 | 0 | 0 | 2.51E-05 | 0 | 2.91E-05 | 0 | 1.01E-05 |
| Mylk | 0 | 9.68E-05 | 0 | 0 | 0.00065 | 8.12E-05 | 0 | 0 | 2.57E-05 | 0 | 0.00033 | 0 | 0 | 7.14E-05 | 0 | 0 | 0 | 0.00016 |
| Hjurp | 0 | 5.12E-05 | 2.75E-05 | 0.00011 | 4.13E-05 | 7.76E-05 | 0.00025 | 0.00018 | 2.57E-05 | 0.00014 | 0 | 0.00029 | 0.00013 | 0.00015 | 0.0008 | 0.00125 | 0 | 0.00012 |
| Pcsk7 | 3.81E-05 | 6.90E-05 | 8.25E-05 | 5.98E-05 | 7.83E-05 | 6.90E-05 | 3.03E-05 | 6.86E-05 | 2.57E-05 | 3.64E-05 | 3.81E-05 | 2.48E-05 | 2.66E-06 | 6.66E-05 | 7.99E-05 | 5.91E-05 | 0 | 4.83E-05 |
| Gpr21 | 1.30E-05 | 3.42E-05 | 0 | 8.44E-06 | 3.94E-05 | 1.59E-05 | 3.47E-06 | 3.98E-06 | 2.57E-05 | 1.40E-05 | 3.22E-06 | 5.05E-06 | 0 | 1.12E-05 | 4.52E-05 | 8.17E-06 | 0 | 2.70E-05 |
| Slc25a29 | 1.96E-06 | 9.31E-06 | 2.22E-06 | 1.25E-05 | 6.60E-06 | 4.68E-06 | 0 | 2.19E-05 | 2.57E-05 | 7.97E-06 | 4.75E-06 | 2.16E-06 | 1.72E-05 | 2.03E-05 | 0 | 0 | 0 | 1.00E-05 |
| Top3a | 1.25E-05 | 2.64E-05 | 1.21E-05 | 1.89E-05 | 1.38E-05 | 3.92E-06 | 9.49E-06 | 1.78E-05 | 2.57E-05 | 2.27E-05 | 1.17E-05 | 1.34E-05 | 3.51E-06 | 0 | 2.68E-05 | 8.82E-05 | 0 | 1.39E-05 |
| Lgi3 | 3.85E-05 | 0 | 9.02E-06 | 2.24E-05 | 0 | 5.90E-07 | 5.37E-05 | 2.47E-05 | 2.57E-05 | 0.00012 | 1.93E-05 | 0 | 0 | 1.31E-05 | 6.96E-06 | 3.04E-05 | 0 | 0 |
| Tcof1 | 5.84E-05 | 4.80E-05 | 0.00029 | 0.00014 | 4.13E-05 | 0 | 0 | 0 | 2.56E-05 | 6.52E-05 | 0 | 0 | 7.02E-05 | 0 | 9.93E-05 | 0.00036 | 7.64E-05 | 3.89E-05 |
| Rnf146 | 0.00016 | 0.00017 | 5.74E-05 | 0 | 8.67E-05 | 0.00011 | 4.13E-05 | 0 | 2.55E-05 | 3.28E-05 | 4.69E-05 | 8.19E-05 | 0 | 6.98E-05 | 0.00016 | 2.45E-05 | 0 | 0 |
| Chd6 | 3.08E-05 | 0.00019 | 0 | 5.29E-05 | 0.00011 | 9.25E-05 | 7.72E-06 | 2.68E-05 | 2.55E-05 | 4.51E-05 | 6.33E-05 | 6.69E-05 | 3.17E-05 | 6.94E-05 | 0.00015 | 6.63E-05 | 8.34E-05 | 4.55E-05 |
| Phf1 | 0 | 0.00014 | 0 | 0.00013 | 2.87E-05 | 5.34E-05 | 0 | 5.78E-05 | 2.55E-05 | 3.58E-05 | 0 | 8.24E-05 | 1.56E-05 | 5.31E-05 | 9.47E-05 | 0 | 4.98E-05 | 8.16E-05 |
| Uba5 | 0.00027 | 8.56E-05 | 0.00024 | 0.00016 | 4.26E-05 | 4.79E-05 | 0.00019 | 2.91E-05 | 2.55E-05 | 5.61E-06 | 5.94E-05 | 4.89E-05 | 3.44E-06 | 2.56E-05 | 0.00012 | 0 | 0 | 3.99E-06 |
| Sinhcaf | 1.54E-05 | 1.42E-05 | 3.61E-05 | 0 | 8.54E-06 | 4.19E-05 | 0 | 2.79E-06 | 2.55E-05 | 0 | 5.73E-05 | 3.77E-05 | 0 | 2.19E-05 | 1.44E-05 | 3.21E-05 | 0 | 9.94E-06 |
| Ophn1 | 4.71E-05 | 6.33E-05 | 0 | 0 | 1.97E-05 | 1.84E-05 | 0 | 0 | 2.55E-05 | 3.15E-05 | 0 | 2.45E-07 | 0 | 7.97E-06 | 4.80E-05 | 0 | 1.64E-05 | 7.51E-05 |
| Zfp770 | 3.51E-05 | 7.22E-06 | 1.89E-05 | 0 | 6.33E-06 | 5.08E-06 | 0 | 2.47E-07 | 2.55E-05 | 5.90E-06 | 0 | 9.13E-06 | 0 | 8.52E-06 | 1.22E-05 | 1.54E-05 | 0 | 0 |
| Tyw3 | 1.55E-05 | 2.62E-05 | 1.13E-05 | 4.05E-05 | 1.87E-05 | 3.38E-06 | 0 | 0 | 2.55E-05 | 0 | 1.17E-05 | 0 | 0 | 6.26E-06 | 8.04E-05 | 1.44E-05 | 0 | 1.85E-05 |
| Dnph1 | 0 | 0 | 2.93E-05 | 1.74E-05 | 0 | 2.36E-06 | 0 | 0 | 2.55E-05 | 0 | 9.80E-06 | 0 | 8.51E-06 | 9.86E-06 | 0 | 0.00016 | 0 | 0 |
| Nmi | 4.62E-05 | 5.32E-06 | 2.66E-05 | 2.89E-05 | 0 | 0 | 7.14E-06 | 0 | 2.55E-05 | 0.0001 | 0 | 0 | 0 | 1.48E-05 | 0 | 5.43E-05 | 0 | 0.00011 |
| Smug1 | 2.16E-05 | 2.81E-05 | 1.36E-05 | 6.88E-05 | 4.54E-05 | 2.09E-05 | 3.68E-05 | 4.66E-05 | 2.54E-05 | 1.59E-05 | 8.90E-06 | 3.22E-05 | 0 | 1.13E-05 | 6.31E-05 | 0 | 0 | 3.17E-05 |
| Slc25a38 | 2.90E-05 | 4.19E-05 | 3.91E-05 | 4.35E-05 | 1.94E-05 | 1.89E-05 | 1.28E-05 | 2.40E-05 | 2.54E-05 | 1.25E-05 | 6.06E-06 | 6.15E-06 | 4.01E-06 | 0 | 1.93E-05 | 3.67E-06 | 6.13E-05 | 4.85E-06 |
| Mtg2 | 2.18E-05 | 4.36E-05 | 3.54E-05 | 4.68E-05 | 1.03E-05 | 1.69E-05 | 0 | 2.43E-05 | 2.54E-05 | 1.48E-05 | 1.24E-05 | 6.45E-06 | 1.02E-05 | 0 | 3.09E-05 | 2.01E-05 | 0 | 1.55E-05 |
| Klhl12 | 2.97E-05 | 7.75E-05 | 2.50E-05 | 8.29E-05 | 5.03E-05 | 1.36E-05 | 3.21E-06 | 3.48E-05 | 2.54E-05 | 1.71E-05 | 3.48E-05 | 1.99E-05 | 4.09E-05 | 3.67E-05 | 0.00014 | 4.57E-05 | 3.13E-05 | 2.93E-05 |
| Eda | 7.24E-06 | 2.19E-05 | 0 | 2.45E-06 | 0 | 2.45E-06 | 9.33E-06 | 5.08E-06 | 2.54E-05 | 5.04E-05 | 1.07E-05 | 1.27E-05 | 2.66E-05 | 0 | 0 | 1.44E-05 | 0 | 5.47E-05 |
| Rbmx2 | 3.64E-06 | 1.40E-05 | 2.15E-05 | 0 | 2.37E-06 | 2.37E-06 | 0 | 0 | 2.54E-05 | 1.27E-05 | 2.10E-05 | 3.90E-06 | 1.64E-05 | 1.94E-05 | 2.25E-05 | 0.00014 | 0 | 5.68E-06 |
| Jade3 | 1.48E-05 | 1.88E-05 | 0 | 0 | 4.25E-08 | 0 | 3.42E-05 | 0 | 2.54E-05 | 0 | 7.89E-06 | 2.57E-06 | 1.57E-05 | 3.96E-05 | 8.22E-06 | 3.26E-05 | 4.67E-06 | 0 |
| Cpeb4 | 0.00013 | 0.00011 | 6.04E-05 | 0 | 9.80E-05 | 8.53E-05 | 0 | 4.54E-05 | 2.53E-05 | 1.03E-05 | 4.71E-05 | 0.00013 | 0 | 4.81E-05 | 4.68E-05 | 0 | 0.00046 | 2.05E-06 |
| Comtd1 | 0 | 7.77E-05 | 4.47E-05 | 0.00012 | 0 | 5.36E-05 | 0 | 7.29E-06 | 2.53E-05 | 8.70E-06 | 2.73E-05 | 5.29E-05 | 7.26E-06 | 3.64E-05 | 4.07E-05 | 0 | 0 | 5.20E-05 |
| Fam199x | 3.27E-05 | 4.78E-05 | 2.38E-05 | 0 | 0 | 3.15E-05 | 1.53E-05 | 0 | 2.53E-05 | 1.11E-05 | 1.96E-05 | 2.18E-05 | 8.56E-06 | 3.91E-05 | 5.02E-07 | 0 | 0 | 2.10E-05 |

| | | | | | | | | | | | | | | | | | | |
|---|---|---|---|---|---|---|---|---|---|---|---|---|---|---|---|---|---|---|
| Zfp945 | 1.86E-05 | 3.60E-05 | 0 | 0 | 9.38E-06 | 2.59E-05 | 0 | 0 | 2.53E-05 | 1.86E-05 | 0 | 6.95E-07 | 0 | 1.30E-05 | 2.46E-05 | 0 | 0 | 7.61E-07 |
| Cwc27 | 6.97E-05 | 4.11E-05 | 5.30E-05 | 3.31E-05 | 0 | 2.09E-05 | 2.93E-05 | 0 | 2.53E-05 | 1.68E-05 | 1.44E-05 | 2.54E-05 | 0 | 0 | 9.50E-05 | 6.14E-05 | 3.57E-05 | 1.00E-05 |
| Gpr173 | 3.52E-06 | 1.36E-05 | 0 | 2.35E-05 | 2.87E-05 | 0 | 4.40E-06 | 1.56E-05 | 2.53E-05 | 1.41E-05 | 0 | 1.10E-05 | 1.18E-05 | 9.05E-06 | 4.85E-05 | 4.66E-07 | 0 | 1.07E-05 |
| Snx2 | 0.00029 | 0.00024 | 5.20E-05 | 6.18E-05 | 0.00013 | 0.00013 | 0 | 0.00011 | 2.52E-05 | 0.00014 | 0.00017 | 0.00021 | 1.88E-05 | 5.59E-05 | 0.00013 | 0.00012 | 0 | 6.90E-05 |
| Slc25a37 | 6.97E-06 | 0.00012 | 0.00016 | 0 | 2.24E-05 | 8.74E-05 | 0 | 8.26E-05 | 2.52E-05 | 0 | 0 | 0.0001 | 0 | 6.36E-05 | 0.00035 | 0 | 0.00019 | 0 |
| Tmem104 | 2.95E-05 | 8.27E-05 | 0.00012 | 0.0002 | 9.89E-05 | 5.78E-05 | 0.00019 | 7.69E-05 | 2.52E-05 | 5.04E-05 | 0 | 0 | 7.01E-05 | 0 | 7.26E-05 | 0 | 0 | 1.50E-05 |
| Eif2s3x | 0.00013 | 3.98E-05 | 0.00012 | 3.91E-06 | 0 | 4.31E-05 | 0 | 0 | 2.52E-05 | 0 | 1.05E-05 | 4.29E-05 | 0 | 0 | 0 | 4.78E-05 | 4.57E-05 | 2.02E-05 |
| Fuz | 3.78E-05 | 4.87E-05 | 0 | 5.06E-05 | 2.96E-05 | 3.70E-05 | 1.32E-06 | 9.50E-06 | 2.52E-05 | 6.44E-05 | 1.82E-06 | 8.67E-06 | 0 | 0 | 3.08E-05 | 0 | 0 | 8.66E-05 |
| Crybg3 | 1.60E-05 | 9.19E-05 | 8.60E-06 | 0 | 5.03E-05 | 3.18E-05 | 0 | 0 | 2.52E-05 | 1.64E-05 | 5.92E-05 | 0 | 1.35E-06 | 6.09E-05 | 4.81E-05 | 0.00018 | 3.29E-05 | 4.58E-05 |
| Brcc3 | 5.25E-05 | 2.75E-05 | 1.51E-05 | 0 | 3.95E-06 | 1.31E-05 | 0 | 0 | 2.52E-05 | 2.16E-05 | 5.16E-06 | 3.16E-05 | 4.88E-06 | 0 | 8.74E-05 | 3.59E-05 | 0 | 0 |
| Taf3 | 0 | 7.74E-05 | 6.14E-05 | 5.42E-05 | 2.99E-05 | 1.30E-05 | 0 | 2.46E-06 | 2.52E-05 | 2.25E-05 | 4.23E-05 | 3.27E-07 | 2.11E-05 | 3.40E-05 | 8.62E-06 | 8.17E-05 | 6.04E-05 | 0 |
| Zfp511 | 6.85E-05 | 7.86E-05 | 6.30E-05 | 0.00012 | 1.26E-05 | 1.03E-05 | 1.30E-05 | 0 | 2.52E-05 | 9.92E-06 | 1.58E-05 | 7.31E-05 | 1.64E-05 | 0 | 0.00014 | 4.08E-05 | 8.85E-06 | 1.45E-05 |
| Elac1 | 1.94E-05 | 4.98E-05 | 2.35E-05 | 2.47E-05 | 4.57E-05 | 8.63E-06 | 0 | 2.18E-05 | 2.52E-05 | 1.65E-05 | 6.16E-06 | 3.02E-05 | 3.21E-05 | 4.08E-05 | 3.12E-05 | 2.51E-06 | 0 | 8.06E-06 |
| Hps4 | 1.06E-05 | 3.44E-05 | 1.00E-05 | 1.82E-05 | 3.97E-05 | 5.82E-06 | 0 | 1.86E-05 | 2.52E-05 | 1.62E-05 | 2.35E-06 | 1.64E-05 | 5.83E-06 | 0 | 9.51E-05 | 6.27E-06 | 1.25E-05 | 2.62E-05 |
| Ift52 | 0.00011 | 0.00011 | 6.95E-07 | 0.00013 | 7.01E-05 | 9.51E-05 | 8.45E-05 | 3.45E-05 | 2.51E-05 | 4.11E-05 | 2.58E-05 | 9.24E-05 | 3.16E-05 | 7.91E-05 | 8.07E-05 | 4.23E-05 | 0 | 7.53E-05 |
| Sox5 | 0 | 2.80E-05 | 0 | 6.28E-06 | 0 | 2.80E-05 | 1.18E-05 | 7.73E-05 | 2.51E-05 | 3.47E-05 | 0 | 2.42E-05 | 7.89E-05 | 0 | 5.08E-05 | 0 | 6.32E-05 | 0 |
| Smim27 | 2.47E-05 | 1.78E-05 | 8.86E-06 | 1.38E-05 | 2.54E-05 | 2.60E-05 | 7.36E-05 | 3.93E-05 | 2.51E-05 | 3.49E-05 | 0 | 2.96E-05 | 1.25E-05 | 3.93E-06 | 0 | 4.54E-06 | 0 | 5.32E-05 |
| Hps6 | 8.64E-06 | 2.06E-05 | 8.51E-06 | 2.83E-05 | 1.46E-05 | 1.67E-05 | 0 | 3.74E-06 | 2.51E-05 | 2.19E-05 | 1.98E-05 | 1.06E-05 | 0 | 2.32E-05 | 4.23E-05 | 7.31E-06 | 0 | 4.21E-05 |
| Mlh3 | 1.76E-05 | 3.03E-05 | 0 | 4.22E-05 | 4.63E-05 | 3.04E-06 | 2.54E-05 | 1.56E-05 | 2.51E-05 | 3.27E-05 | 2.40E-05 | 7.66E-06 | 0 | 4.19E-06 | 5.76E-05 | 5.30E-06 | 0 | 1.69E-05 |
| Fkbpl | 2.92E-05 | 2.41E-05 | 7.98E-06 | 1.43E-06 | 1.19E-05 | 2.25E-06 | 4.64E-06 | 0 | 2.51E-05 | 0 | 0 | 4.01E-06 | 1.25E-06 | 1.45E-06 | 1.36E-05 | 8.38E-06 | 0 | 0 |
| Adcy6 | 3.46E-05 | 0.00016 | 7.97E-05 | 0.00016 | 0.00024 | 0.00026 | 4.88E-05 | 0.00038 | 2.50E-05 | 6.25E-05 | 0.00024 | 0.00018 | 8.66E-05 | 0.00022 | 0.00028 | 2.74E-05 | 5.77E-05 | 7.31E-05 |
| Med31 | 6.90E-05 | 5.59E-05 | 4.26E-05 | 1.68E-05 | 3.93E-05 | 5.75E-05 | 0 | 1.26E-05 | 2.50E-05 | 1.76E-05 | 5.28E-05 | 9.06E-05 | 0 | 4.13E-05 | 1.09E-05 | 8.62E-05 | 2.81E-05 | 3.69E-05 |
| Bola1 | 6.73E-05 | 4.26E-05 | 5.54E-05 | 0.00014 | 2.46E-05 | 4.41E-05 | 3.88E-05 | 2.06E-05 | 2.50E-05 | 3.75E-06 | 0 | 8.72E-06 | 3.97E-05 | 1.04E-06 | 1.45E-05 | 7.11E-05 | 6.89E-06 | 1.61E-05 |
| Lfng | 0 | 5.48E-06 | 0 | 3.50E-05 | 1.50E-05 | 3.47E-05 | 0 | 0 | 2.50E-05 | 0 | 0.00014 | 0 | 0 | 6.18E-05 | 2.46E-05 | 3.27E-05 | 0 | 0 |
| Baz1a | 8.99E-05 | 7.26E-05 | 0.00019 | 3.12E-05 | 4.77E-05 | 3.18E-05 | 0 | 0 | 2.50E-05 | 2.81E-05 | 6.44E-05 | 0 | 1.11E-05 | 0.00012 | 0.0001 | 0.00011 | 0.00028 | 0 |
| Tmcc2 | 0 | 3.71E-05 | 6.06E-05 | 4.08E-05 | 5.87E-05 | 1.02E-05 | 2.86E-05 | 2.48E-05 | 2.50E-05 | 1.57E-05 | 8.53E-05 | 8.46E-06 | 0 | 4.16E-05 | 6.93E-05 | 5.81E-05 | 7.22E-05 | 4.10E-05 |
| Sdr39u1 | 5.46E-05 | 1.61E-05 | 3.32E-06 | 5.25E-05 | 2.74E-05 | 0 | 2.02E-05 | 0 | 2.50E-05 | 0 | 3.60E-05 | 0 | 4.06E-05 | 3.24E-05 | 0 | 1.92E-05 | 0 | 6.31E-05 |
| Wac | 0.00025 | 0.00017 | 8.36E-05 | 0 | 6.14E-05 | 0.00016 | 0 | 0 | 2.49E-05 | 2.24E-05 | 4.24E-05 | 0.00012 | 0 | 0 | 0.0001 | 9.80E-05 | 0.00015 | 0 |
| Cyb561 | 0 | 0.0001 | 4.40E-05 | 0 | 0.00027 | 8.29E-05 | 0.00011 | 0 | 2.49E-05 | 8.86E-05 | 0 | 0 | 0.00017 | 1.45E-05 | 3.80E-05 | 0 | 0 | 0.00021 |
| B4galt7 | 0.00014 | 5.84E-05 | 8.83E-05 | 7.10E-06 | 9.00E-05 | 7.94E-05 | 9.78E-05 | 7.03E-05 | 2.49E-05 | 0 | 2.94E-06 | 2.05E-05 | 1.92E-05 | 5.50E-05 | 0 | 0 | 0 | 5.14E-05 |
| Trnt1 | 0.00013 | 3.79E-05 | 0.0001 | 0 | 2.11E-05 | 5.16E-05 | 1.95E-05 | 0 | 2.49E-05 | 6.56E-06 | 3.92E-06 | 3.56E-05 | 5.08E-05 | 0 | 0.00015 | 3.25E-05 | 0 | 1.97E-05 |
| Spef1 | 1.00E-05 | 8.49E-05 | 8.05E-05 | 8.36E-05 | 0.00015 | 4.56E-05 | 6.62E-05 | 4.88E-05 | 2.49E-05 | 1.78E-05 | 3.42E-05 | 0.00011 | 0 | 1.98E-05 | 0.00013 | 0.00013 | 0.00013 | 2.63E-05 |
| Pdk1 | 2.07E-05 | 0 | 0 | 5.09E-05 | 1.94E-05 | 3.70E-05 | 0.0002 | 7.16E-05 | 2.49E-05 | 7.73E-05 | 0 | 0.001 | 0.00016 | 0 | 0 | 0 | 5.73E-05 | 0 |

| | | | | | | | | | | | | | | | | | | |
|---|---|---|---|---|---|---|---|---|---|---|---|---|---|---|---|---|---|---|
| Nxt2 | 5.27E-05 | 2.07E-05 | 6.83E-06 | 0 | 0 | 2.63E-05 | 0 | 0 | 2.49E-05 | 1.75E-05 | 0 | 1.42E-05 | 9.47E-06 | 0 | 0 | 5.52E-05 | 0 | 9.76E-06 |
| Zfp90 | 2.81E-05 | 2.24E-05 | 0 | 0 | 1.56E-05 | 6.97E-06 | 1.28E-06 | 0 | 2.49E-05 | 3.39E-05 | 2.18E-05 | 2.18E-05 | 0 | 0 | 4.29E-05 | 1.17E-05 | 0 | 2.65E-05 |
| Srl | 8.18E-06 | 3.15E-06 | 0 | 0 | 4.54E-06 | 4.88E-06 | 0 | 9.92E-06 | 2.49E-05 | 0 | 3.58E-05 | 0 | 2.08E-06 | 1.46E-05 | 0 | 2.02E-05 | 0 | 0 |
| Emc9 | 1.02E-05 | 4.96E-05 | 6.30E-06 | 3.21E-05 | 2.50E-05 | 0 | 8.65E-06 | 2.08E-05 | 2.49E-05 | 8.28E-05 | 2.06E-06 | 2.41E-05 | 4.17E-06 | 0 | 0 | 7.45E-06 | 0 | 6.37E-05 |
| Ccdc137 | 1.62E-05 | 3.66E-05 | 8.78E-05 | 2.63E-05 | 8.52E-06 | 0 | 0 | 1.31E-05 | 2.49E-05 | 1.80E-06 | 0 | 8.40E-05 | 0 | 0 | 9.22E-05 | 3.61E-05 | 0 | 1.11E-05 |
| Atg4d | 8.58E-06 | 6.17E-05 | 1.94E-05 | 6.30E-05 | 2.79E-05 | 0 | 6.92E-06 | 3.81E-05 | 2.49E-05 | 1.57E-05 | 3.98E-05 | 3.67E-05 | 2.90E-06 | 9.97E-06 | 5.28E-05 | 4.45E-05 | 1.46E-06 | 1.02E-05 |
| Bfar | 3.77E-05 | 6.65E-05 | 5.39E-05 | 7.81E-05 | 0.00011 | 6.68E-05 | 0.00024 | 0.00012 | 2.48E-05 | 8.47E-05 | 2.88E-05 | 2.82E-05 | 8.32E-05 | 7.77E-05 | 4.41E-05 | 0 | 1.44E-05 | 0.00011 |
| Tmem248 | 0.00015 | 8.25E-05 | 0.00029 | 0.0002 | 0.0001 | 6.22E-05 | 0.00017 | 0.00014 | 2.48E-05 | 6.38E-05 | 0.00013 | 7.05E-05 | 0.00012 | 7.46E-05 | 3.23E-07 | 0.0001 | 0 | 2.44E-05 |
| Zxdb | 6.68E-05 | 6.31E-05 | 2.93E-05 | 0 | 2.78E-05 | 5.74E-05 | 0 | 2.05E-05 | 2.48E-05 | 1.81E-06 | 7.26E-06 | 7.45E-05 | 0 | 2.09E-05 | 1.99E-05 | 1.78E-05 | 3.42E-06 | 1.93E-05 |
| Tmem185a | 6.46E-05 | 2.63E-05 | 5.88E-05 | 5.31E-06 | 5.66E-05 | 3.90E-05 | 3.51E-06 | 5.04E-05 | 2.48E-05 | 3.02E-05 | 8.72E-06 | 1.62E-05 | 0 | 5.30E-05 | 0 | 1.42E-05 | 0 | 6.19E-05 |
| Tcf24 | 1.32E-05 | 1.07E-05 | 1.47E-06 | 0 | 2.06E-05 | 3.74E-05 | 0 | 3.58E-05 | 2.48E-05 | 9.80E-05 | 9.49E-08 | 0 | 1.13E-05 | 0 | 0 | 0 | 0 | 0 |
| Trp53i13 | 2.68E-05 | 2.06E-05 | 1.72E-05 | 0.00023 | 3.18E-05 | 3.65E-05 | 0.00023 | 3.75E-05 | 2.48E-05 | 2.95E-05 | 2.28E-05 | 3.16E-05 | 6.66E-05 | 4.04E-05 | 2.47E-05 | 7.46E-05 | 5.07E-05 | 7.96E-05 |
| Zfp626 | 1.31E-05 | 2.68E-05 | 0 | 3.95E-06 | 2.31E-05 | 2.72E-05 | 3.20E-06 | 1.83E-05 | 2.48E-05 | 1.82E-05 | 2.47E-05 | 1.08E-05 | 2.88E-06 | 1.07E-05 | 3.77E-05 | 1.35E-05 | 0 | 5.11E-06 |
| Tex9 | 1.68E-05 | 1.41E-05 | 0 | 0 | 4.65E-06 | 2.42E-05 | 0 | 0 | 2.48E-05 | 1.28E-05 | 0 | 0 | 0 | 3.55E-05 | 5.93E-05 | 2.41E-05 | 0 | 0 |
| Clip4 | 0 | 1.60E-05 | 0 | 0 | 0 | 1.16E-05 | 9.01E-05 | 0 | 2.48E-05 | 0.00044 | 0 | 0 | 3.62E-05 | 0 | 0 | 0 | 0 | 1.00E-04 |
| Syndig1 | 0 | 0 | 0 | 0 | 0 | 5.32E-06 | 0 | 0 | 2.48E-05 | 0 | 1.32E-05 | 0 | 4.02E-05 | 4.07E-06 | 0 | 0 | 0 | 2.45E-05 |
| Zfp825 | 1.13E-05 | 1.46E-05 | 0 | 0 | 1.05E-05 | 0 | 0 | 7.39E-06 | 2.48E-05 | 1.58E-05 | 9.90E-06 | 5.88E-06 | 0 | 6.44E-06 | 0 | 2.13E-05 | 0 | 2.35E-05 |
| Phlda3 | 2.16E-05 | 0.00021 | 0.00022 | 0 | 3.01E-05 | 0 | 3.73E-05 | 0.00029 | 2.48E-05 | 5.74E-05 | 0.00016 | 3.00E-05 | 0 | 0 | 3.51E-05 | 3.59E-05 | 0 | 0.00042 |
| Rtf1 | 7.74E-05 | 0.00011 | 9.70E-05 | 1.52E-05 | 5.25E-05 | 7.32E-05 | 4.91E-05 | 2.79E-05 | 2.47E-05 | 2.92E-05 | 5.21E-05 | 3.24E-05 | 5.55E-05 | 2.51E-05 | 0 | 8.97E-05 | 0 | 4.16E-05 |
| Bsdc1 | 5.75E-05 | 0.00013 | 6.28E-05 | 9.23E-05 | 8.35E-05 | 6.28E-05 | 8.34E-05 | 8.99E-05 | 2.47E-05 | 3.95E-05 | 5.83E-05 | 0.00012 | 4.93E-05 | 5.74E-05 | 8.89E-05 | 0 | 7.60E-05 | 9.08E-06 |
| Mis12 | 6.77E-05 | 1.79E-05 | 2.29E-05 | 0 | 2.39E-05 | 4.67E-05 | 0 | 0 | 2.47E-05 | 0 | 0 | 1.89E-05 | 0 | 2.65E-05 | 5.81E-06 | 0.00022 | 0 | 0 |
| St6galnac2 | 0 | 0 | 0 | 0 | 0 | 1.09E-05 | 0 | 3.86E-05 | 2.47E-05 | 7.97E-06 | 1.11E-05 | 0 | 5.84E-05 | 1.87E-05 | 0 | 2.29E-05 | 6.33E-06 | 2.37E-05 |
| Zbtb7b | 0 | 0.00011 | 2.49E-05 | 0.0002 | 7.46E-05 | 1.24E-06 | 3.31E-05 | 9.34E-05 | 2.47E-05 | 8.37E-05 | 8.62E-05 | 0.00019 | 8.26E-06 | 0 | 0.00015 | 0 | 3.43E-05 | 3.89E-05 |
| Parg | 0.00013 | 8.18E-05 | 4.27E-05 | 0 | 3.87E-05 | 6.64E-05 | 2.26E-05 | 0 | 2.46E-05 | 1.04E-05 | 2.01E-05 | 2.29E-05 | 3.16E-05 | 5.25E-05 | 5.17E-05 | 5.40E-05 | 1.83E-05 | 0 |
| Hs3st3b1 | 0 | 1.26E-05 | 0 | 0 | 7.59E-05 | 0 | 0 | 5.63E-05 | 2.46E-05 | 6.59E-06 | 0 | 4.30E-05 | 2.03E-05 | 0 | 1.82E-05 | 0 | 0 | 1.01E-05 |
| Naaladl2 | 0 | 2.74E-05 | 0 | 0 | 3.02E-05 | 3.85E-05 | 2.32E-05 | 0 | 2.45E-05 | 7.54E-05 | 1.29E-05 | 0 | 1.80E-05 | 5.20E-05 | 1.65E-06 | 0 | 0 | 4.95E-05 |
| Fam149b | 0.00011 | 7.28E-05 | 4.47E-05 | 4.15E-05 | 6.79E-05 | 3.19E-05 | 0.00011 | 7.53E-05 | 2.45E-05 | 3.33E-05 | 4.80E-05 | 2.02E-05 | 3.17E-05 | 2.93E-05 | 0.00012 | 7.41E-05 | 0 | 5.60E-05 |
| Dus3l | 0.0001 | 0.00013 | 0.00011 | 0.00024 | 3.18E-05 | 2.89E-05 | 6.43E-05 | 1.55E-05 | 2.45E-05 | 1.39E-05 | 6.86E-06 | 1.73E-05 | 1.02E-05 | 0 | 0.00046 | 5.46E-05 | 7.74E-05 | 1.10E-05 |
| Nsun3 | 1.84E-05 | 2.18E-05 | 9.56E-06 | 9.14E-06 | 0 | 4.90E-06 | 0 | 1.70E-05 | 2.45E-05 | 1.97E-05 | 0 | 2.43E-05 | 3.93E-06 | 2.35E-06 | 0 | 9.38E-06 | 0 | 1.38E-05 |
| Zbtb18 | 1.33E-05 | 2.76E-05 | 2.98E-05 | 4.16E-06 | 1.30E-05 | 0 | 0 | 5.27E-05 | 2.45E-05 | 5.30E-06 | 8.47E-05 | 4.76E-05 | 4.02E-05 | 0 | 6.82E-05 | 9.42E-05 | 2.74E-06 | 1.95E-05 |
| Zswim1 | 2.68E-05 | 2.62E-05 | 2.05E-05 | 7.24E-07 | 1.06E-05 | 0 | 0 | 0 | 2.45E-05 | 1.21E-05 | 1.67E-06 | 0 | 0 | 2.35E-06 | 1.75E-05 | 1.24E-05 | 0 | 1.79E-05 |
| Chrd | 0 | 0 | 0 | 0 | 5.61E-05 | 6.15E-05 | 0.00011 | 0 | 2.44E-05 | 0.00011 | 8.90E-06 | 3.63E-05 | 0.00013 | 2.73E-05 | 4.57E-05 | 0 | 0 | 0.00013 |
| Zbtb6 | 7.72E-05 | 3.27E-05 | 2.42E-05 | 0 | 1.60E-05 | 3.75E-05 | 1.07E-05 | 0 | 2.44E-05 | 3.00E-06 | 0 | 1.42E-05 | 0 | 8.22E-06 | 4.36E-05 | 7.17E-06 | 0 | 2.00E-05 |

| | | | | | | | | | | | | | | | | | | |
|---|---|---|---|---|---|---|---|---|---|---|---|---|---|---|---|---|---|---|
| Bpnt1 | 8.46E-05 | 7.04E-05 | 0.00019 | 7.86E-05 | 4.11E-05 | 3.51E-05 | 0 | 6.39E-05 | 2.44E-05 | 3.42E-05 | 5.36E-05 | 0 | 7.90E-06 | 2.40E-05 | 2.12E-05 | 6.41E-05 | 2.00E-05 | 2.14E-05 |
| Usp27x | 2.09E-05 | 2.75E-05 | 1.56E-05 | 2.95E-06 | 1.26E-05 | 1.73E-05 | 0 | 0 | 2.44E-05 | 1.07E-06 | 0 | 0 | 7.52E-06 | 1.01E-05 | 1.38E-05 | 0 | 0 | 5.44E-06 |
| Arhgef2 | 8.24E-05 | 0.00039 | 0.00034 | 3.74E-05 | 0.00016 | 0 | 0.00011 | 0.00044 | 2.44E-05 | 8.17E-05 | 9.95E-05 | 0.00037 | 0 | 6.48E-05 | 0.00051 | 0.00031 | 6.66E-05 | 0.00012 |
| H2-Q4 | 0 | 2.39E-05 | 0 | 0 | 0 | 0 | 0 | 0 | 2.44E-05 | 3.69E-05 | 0 | 2.46E-05 | 7.60E-05 | 1.03E-05 | 0 | 4.43E-05 | 5.32E-05 | 2.69E-05 |
| Zfp579 | 0 | 7.35E-05 | 0 | 4.55E-05 | 1.77E-05 | 0 | 0 | 1.35E-06 | 2.44E-05 | 3.18E-05 | 0 | 2.09E-05 | 0 | 3.80E-05 | 0.00015 | 0 | 2.94E-05 | 1.17E-05 |
| Mettl2 | 3.46E-05 | 5.52E-05 | 5.31E-05 | 2.31E-05 | 1.31E-05 | 0 | 1.18E-05 | 8.43E-06 | 2.44E-05 | 1.52E-05 | 1.00E-05 | 2.47E-05 | 1.00E-05 | 7.18E-06 | 9.69E-06 | 3.42E-05 | 1.17E-05 | 2.19E-05 |
| Zhx3 | 3.65E-05 | 7.87E-05 | 7.03E-05 | 0 | 0.00029 | 9.06E-05 | 0.00014 | 2.19E-05 | 2.43E-05 | 6.43E-05 | 0.00013 | 1.85E-05 | 2.43E-05 | 1.89E-06 | 5.91E-05 | 5.32E-05 | 0 | 4.11E-05 |
| Ciao2b | 9.98E-05 | 3.03E-05 | 6.44E-05 | 1.53E-05 | 1.40E-05 | 2.96E-05 | 4.41E-06 | 1.07E-05 | 2.43E-05 | 3.88E-06 | 4.75E-05 | 4.38E-05 | 0 | 0 | 3.66E-05 | 4.23E-05 | 3.21E-05 | 0 |
| Stk19 | 6.06E-06 | 7.22E-05 | 4.62E-05 | 5.80E-05 | 2.55E-05 | 2.24E-05 | 0 | 2.11E-05 | 2.43E-05 | 3.83E-05 | 3.56E-05 | 1.41E-05 | 1.50E-05 | 3.50E-05 | 8.37E-05 | 3.88E-05 | 6.34E-05 | 3.14E-05 |
| DEPRECATED | 3.53E-06 | 0 | 2.70E-05 | 0 | 1.67E-05 | 9.42E-06 | 2.81E-06 | 4.22E-05 | 2.43E-05 | 2.57E-05 | 1.67E-05 | 2.63E-06 | 4.07E-05 | 2.98E-05 | 6.01E-06 | 8.01E-06 | 0 | 1.96E-05 |
| Ston2 | 2.41E-05 | 6.35E-05 | 0 | 0 | 2.41E-05 | 0 | 2.11E-05 | 7.84E-05 | 2.43E-05 | 0.00017 | 0 | 1.69E-05 | 0 | 3.59E-05 | 0 | 3.97E-05 | 0 | 0 |
| Pcdhgb2 | 1.96E-05 | 0.00012 | 0 | 4.75E-05 | 0.00013 | 0.00012 | 7.49E-05 | 8.22E-05 | 2.42E-05 | 4.34E-05 | 0 | 3.54E-05 | 8.77E-05 | 4.71E-06 | 0.00014 | 0 | 0 | 9.73E-06 |
| Mmgt1 | 0.00012 | 1.52E-05 | 6.24E-05 | 0 | 1.64E-05 | 5.02E-05 | 2.67E-05 | 3.06E-05 | 2.42E-05 | 2.31E-06 | 4.17E-05 | 1.03E-05 | 1.49E-06 | 6.58E-06 | 0 | 0.00017 | 6.28E-05 | 0 |
| Taf8 | 5.32E-05 | 0.0001 | 1.30E-05 | 8.27E-05 | 5.62E-05 | 4.00E-05 | 6.35E-05 | 3.48E-05 | 2.42E-05 | 3.32E-05 | 0 | 4.22E-05 | 1.81E-05 | 3.24E-05 | 0.00014 | 7.46E-05 | 1.79E-05 | 2.22E-05 |
| Ints9 | 1.57E-05 | 3.65E-05 | 1.99E-05 | 4.15E-05 | 1.25E-05 | 2.30E-05 | 1.31E-06 | 2.29E-05 | 2.42E-05 | 3.49E-05 | 3.17E-06 | 1.97E-05 | 1.09E-05 | 1.27E-05 | 0 | 6.21E-05 | 0 | 5.28E-05 |
| Trim59 | 2.84E-05 | 1.21E-05 | 0 | 0 | 0 | 2.30E-05 | 2.65E-06 | 0 | 2.42E-05 | 0 | 3.06E-05 | 1.69E-05 | 0 | 0 | 0 | 0.00042 | 4.73E-06 | 2.54E-05 |
| Ankrd54 | 0 | 6.28E-05 | 7.25E-05 | 0.00012 | 8.80E-05 | 0 | 5.27E-05 | 5.02E-05 | 2.42E-05 | 3.72E-05 | 0 | 1.79E-05 | 1.57E-05 | 7.16E-06 | 8.16E-05 | 0.00012 | 8.82E-05 | 7.67E-05 |
| Prkca | 6.48E-05 | 4.60E-05 | 6.53E-05 | 0 | 4.09E-05 | 0.0001 | 9.89E-05 | 0 | 2.41E-05 | 0.00012 | 8.90E-06 | 0 | 8.53E-06 | 4.35E-05 | 9.78E-06 | 3.69E-05 | 1.73E-05 | 7.44E-05 |
| Odr4 | 0.00015 | 3.11E-05 | 7.14E-06 | 0 | 3.69E-05 | 9.36E-05 | 6.39E-05 | 3.05E-05 | 2.41E-05 | 2.98E-05 | 0 | 8.17E-05 | 2.84E-05 | 8.86E-07 | 4.05E-05 | 1.23E-05 | 1.07E-05 | 4.99E-05 |
| Tsg101 | 0.00016 | 1.00E-04 | 0.00012 | 0.00016 | 3.24E-05 | 5.99E-05 | 8.93E-06 | 0 | 2.41E-05 | 1.55E-05 | 0.00012 | 0.00011 | 4.40E-05 | 1.86E-05 | 0.00013 | 0.00011 | 0 | 0 |
| Klhl15 | 3.48E-05 | 2.81E-05 | 3.41E-05 | 9.28E-06 | 2.75E-05 | 5.41E-05 | 0 | 2.69E-05 | 2.41E-05 | 1.30E-05 | 0 | 5.03E-05 | 0 | 2.18E-05 | 3.25E-05 | 1.14E-05 | 8.95E-05 | 3.06E-05 |
| Rnf165 | 0 | 0 | 0 | 0 | 1.75E-06 | 4.44E-05 | 0 | 0 | 2.41E-05 | 4.33E-05 | 2.89E-05 | 0.00023 | 4.91E-06 | 3.45E-06 | 3.61E-05 | 4.84E-06 | 9.91E-05 | 1.25E-05 |
| Chchd5 | 5.38E-05 | 7.43E-05 | 0 | 4.39E-05 | 1.74E-05 | 3.68E-05 | 4.43E-05 | 2.12E-05 | 2.41E-05 | 0 | 0 | 7.93E-05 | 0 | 2.50E-05 | 1.49E-05 | 5.57E-05 | 2.45E-05 | 2.91E-05 |
| Mob3c | 3.81E-05 | 2.39E-05 | 1.44E-05 | 0 | 2.98E-05 | 9.12E-06 | 5.00E-06 | 2.78E-05 | 2.41E-05 | 1.08E-05 | 0 | 4.25E-05 | 1.01E-05 | 3.16E-06 | 9.29E-05 | 0 | 0.00011 | 1.69E-05 |
| Fcsk | 0 | 4.32E-05 | 0 | 4.40E-05 | 4.09E-05 | 8.13E-06 | 9.11E-06 | 0 | 2.41E-05 | 2.53E-05 | 1.69E-05 | 0 | 1.04E-05 | 4.37E-05 | 3.70E-05 | 0 | 0 | 3.01E-05 |
| Chrna1 | 0 | 9.79E-06 | 2.03E-06 | 3.34E-06 | 0 | 4.04E-06 | 0 | 2.86E-06 | 2.41E-05 | 0 | 0 | 0 | 0 | 2.10E-05 | 0 | 7.06E-06 | 0 | 0 |
| Acot9 | 2.30E-05 | 1.32E-05 | 2.80E-05 | 0 | 6.17E-06 | 2.59E-06 | 0 | 0 | 2.41E-05 | 1.57E-05 | 2.00E-05 | 1.55E-05 | 4.91E-06 | 1.48E-05 | 0 | 1.12E-05 | 1.70E-05 | 1.83E-05 |
| BC067074 | 0 | 1.69E-05 | 0 | 0 | 4.16E-05 | 2.06E-06 | 3.96E-05 | 0 | 2.41E-05 | 0 | 0 | 0 | 3.02E-05 | 0 | 0 | 0 | 0 | 1.55E-06 |
| Cd38 | 3.90E-06 | 2.16E-05 | 0 | 0 | 0 | 0 | 0 | 0.00013 | 2.41E-05 | 0 | 0 | 0 | 3.78E-05 | 5.06E-05 | 0 | 0 | 0 | 3.54E-05 |
| Uqcrq | 0.0002 | 2.53E-05 | 0.00021 | 0.00015 | 5.14E-05 | 4.11E-05 | 6.18E-05 | 2.16E-05 | 2.40E-05 | 0 | 0.0001 | 0 | 1.05E-05 | 4.59E-05 | 0 | 8.76E-05 | 0 | 0 |
| Ipo11 | 0.00011 | 2.85E-05 | 3.23E-05 | 0 | 1.06E-06 | 3.78E-05 | 0 | 0 | 2.40E-05 | 1.75E-07 | 0 | 2.94E-05 | 0 | 0 | 2.21E-05 | 0.00013 | 2.06E-05 | 0 |
| Rmdn3 | 2.81E-05 | 5.46E-05 | 9.08E-05 | 3.42E-05 | 6.21E-05 | 2.41E-05 | 2.91E-06 | 4.46E-05 | 2.40E-05 | 3.56E-05 | 2.36E-05 | 4.43E-05 | 1.74E-05 | 3.74E-07 | 8.29E-06 | 5.00E-05 | 1.39E-05 | 4.22E-05 |
| Ankrd26 | 1.18E-05 | 1.56E-05 | 1.35E-05 | 0 | 3.39E-06 | 3.69E-06 | 0 | 1.04E-05 | 2.40E-05 | 2.73E-06 | 1.26E-05 | 4.84E-06 | 0 | 6.77E-06 | 1.08E-05 | 3.13E-05 | 5.72E-05 | 1.56E-06 |

| | | | | | | | | | | | | | | | | | | |
|---|---|---|---|---|---|---|---|---|---|---|---|---|---|---|---|---|---|---|
| Bud13 | 2.82E-05 | 3.42E-05 | 4.17E-05 | 4.93E-05 | 7.91E-06 | 0 | 2.56E-05 | 8.56E-06 | 2.40E-05 | 1.92E-05 | 1.27E-06 | 3.32E-05 | 5.24E-06 | 2.06E-05 | 2.09E-05 | 5.88E-05 | 0 | 1.71E-05 |
| Nolc1 | 8.75E-05 | 2.63E-05 | 0.00025 | 6.65E-05 | 0 | 0 | 0 | 0 | 2.40E-05 | 5.21E-06 | 3.91E-06 | 0 | 6.91E-05 | 1.51E-05 | 3.07E-05 | 0.00012 | 0.00014 | 0 |
| Pcdhga6 | 4.29E-05 | 0.00011 | 0 | 0 | 6.70E-05 | 8.79E-05 | 1.65E-05 | 2.31E-05 | 2.39E-05 | 3.10E-05 | 3.91E-05 | 9.20E-06 | 5.10E-05 | 6.58E-05 | 8.22E-05 | 0 | 2.19E-05 | 2.19E-05 |
| Grhpr | 9.37E-05 | 8.95E-06 | 1.89E-05 | 4.34E-05 | 8.05E-05 | 5.20E-05 | 7.43E-05 | 0.00013 | 2.39E-05 | 1.01E-05 | 2.57E-05 | 0.00046 | 1.74E-05 | 3.54E-05 | 0 | 0 | 0 | 0 |
| Zfp335 | 6.51E-05 | 0.0001 | 8.55E-05 | 5.65E-05 | 2.90E-05 | 3.34E-05 | 1.47E-05 | 1.43E-05 | 2.39E-05 | 2.22E-05 | 0 | 6.17E-05 | 0 | 4.60E-05 | 0.00026 | 1.54E-05 | 3.87E-05 | 1.36E-05 |
| Ilkap | 6.10E-05 | 0.0001 | 6.12E-05 | 3.13E-06 | 2.93E-05 | 2.93E-05 | 0 | 0 | 2.39E-05 | 2.21E-05 | 4.52E-05 | 4.01E-05 | 0 | 3.62E-05 | 0.00017 | 4.40E-05 | 0 | 9.08E-06 |
| Pcdhgb5 | 0 | 7.86E-05 | 0 | 4.55E-05 | 4.68E-05 | 1.71E-05 | 5.01E-05 | 4.62E-05 | 2.39E-05 | 1.53E-05 | 3.40E-05 | 1.31E-05 | 2.27E-05 | 5.37E-05 | 0.0001 | 0 | 8.44E-05 | 2.18E-05 |
| Mettl25 | 6.71E-06 | 1.96E-05 | 6.42E-06 | 2.28E-06 | 1.70E-05 | 8.67E-06 | 9.68E-06 | 1.12E-05 | 2.39E-05 | 0 | 9.33E-06 | 1.21E-05 | 1.49E-05 | 1.31E-05 | 0 | 3.49E-06 | 0 | 1.09E-05 |
| Tmem218 | 3.44E-06 | 4.35E-05 | 0 | 6.89E-05 | 4.78E-05 | 4.65E-06 | 0 | 5.06E-05 | 2.39E-05 | 3.59E-05 | 0 | 3.09E-05 | 3.97E-05 | 2.36E-05 | 2.19E-05 | 5.87E-05 | 0 | 4.42E-05 |
| Prdm10 | 1.57E-05 | 3.23E-05 | 1.38E-05 | 9.88E-07 | 7.77E-06 | 1.70E-06 | 6.91E-06 | 2.57E-05 | 2.39E-05 | 1.80E-05 | 1.40E-05 | 1.06E-05 | 0 | 1.57E-05 | 5.03E-05 | 3.69E-05 | 4.27E-05 | 1.45E-05 |
| Nmrk1 | 1.90E-05 | 3.89E-05 | 0 | 1.09E-05 | 3.87E-05 | 2.81E-07 | 3.30E-05 | 0 | 2.39E-05 | 5.36E-06 | 3.02E-05 | 1.95E-05 | 0 | 1.52E-05 | 6.24E-05 | 6.90E-06 | 0 | 0 |
| Pkig | 0 | 0.0002 | 1.41E-05 | 0.00061 | 0.00045 | 0.00023 | 0.00019 | 0.00039 | 2.38E-05 | 0 | 0.0002 | 4.30E-05 | 0.00022 | 1.33E-05 | 0.00011 | 0.00022 | 0.00023 | 0 |
| St7 | 7.91E-05 | 5.15E-05 | 0.00011 | 0 | 9.38E-05 | 4.27E-05 | 0.00015 | 2.27E-05 | 2.38E-05 | 6.15E-05 | 5.46E-05 | 3.00E-05 | 0 | 2.99E-05 | 0 | 6.55E-05 | 4.25E-05 | 7.39E-05 |
| Pfkfb2 | 3.01E-05 | 5.97E-05 | 7.90E-06 | 2.44E-06 | 6.03E-05 | 3.31E-05 | 2.42E-05 | 2.53E-06 | 2.38E-05 | 2.22E-05 | 1.11E-05 | 1.14E-05 | 8.99E-06 | 1.64E-05 | 5.03E-05 | 0 | 4.10E-05 | 2.50E-05 |
| Xpnpep3 | 3.98E-05 | 4.13E-05 | 2.90E-05 | 1.87E-05 | 1.01E-05 | 2.39E-05 | 1.05E-05 | 8.52E-06 | 2.38E-05 | 0 | 0 | 1.64E-05 | 5.42E-06 | 0 | 3.76E-05 | 3.49E-05 | 0 | 1.39E-06 |
| Bcl2l13 | 0.00011 | 8.59E-05 | 9.22E-05 | 9.80E-05 | 5.20E-05 | 2.34E-05 | 9.47E-05 | 0 | 2.38E-05 | 1.81E-05 | 2.26E-05 | 1.92E-05 | 0 | 1.49E-05 | 2.23E-05 | 0.00012 | 0.00011 | 1.73E-05 |
| Rbm4b | 2.19E-05 | 4.30E-05 | 9.05E-06 | 3.23E-05 | 2.09E-05 | 1.19E-05 | 0 | 2.98E-05 | 2.38E-05 | 0 | 1.60E-05 | 3.13E-05 | 5.11E-07 | 3.42E-05 | 1.22E-05 | 1.93E-05 | 8.84E-05 | 1.00E-05 |
| Zkscan5 | 2.31E-05 | 3.37E-05 | 8.19E-06 | 1.03E-05 | 1.65E-05 | 3.24E-06 | 2.06E-06 | 1.54E-05 | 2.38E-05 | 1.17E-05 | 2.46E-05 | 1.32E-05 | 3.58E-06 | 7.32E-07 | 0 | 1.16E-05 | 2.36E-08 | 3.65E-06 |
| Zfp1 | 4.58E-05 | 5.90E-05 | 9.00E-06 | 0 | 1.72E-05 | 3.84E-05 | 8.57E-07 | 2.43E-05 | 2.37E-05 | 8.23E-06 | 1.62E-07 | 4.93E-05 | 5.92E-06 | 0 | 9.12E-05 | 0 | 0 | 2.14E-05 |
| Acad11 | 4.84E-05 | 8.70E-05 | 0 | 3.92E-05 | 3.51E-05 | 3.46E-05 | 5.61E-06 | 0 | 2.37E-05 | 3.81E-05 | 1.17E-05 | 0 | 5.47E-05 | 2.80E-05 | 2.91E-05 | 0 | 0 | 3.64E-05 |
| Elmod3 | 3.73E-05 | 3.26E-05 | 0 | 2.14E-05 | 3.73E-05 | 2.93E-05 | 2.10E-05 | 1.33E-05 | 2.37E-05 | 1.03E-05 | 2.88E-06 | 2.91E-05 | 1.88E-05 | 2.64E-05 | 4.00E-05 | 0 | 6.42E-05 | 1.78E-05 |
| Lrrc20 | 1.08E-05 | 4.75E-05 | 5.33E-05 | 4.88E-06 | 3.41E-05 | 2.64E-05 | 4.61E-05 | 5.39E-05 | 2.37E-05 | 4.70E-05 | 5.97E-05 | 8.29E-06 | 2.70E-05 | 3.13E-05 | 0 | 5.78E-05 | 1.37E-05 | 6.78E-06 |
| Zfx | 6.71E-05 | 3.47E-05 | 0 | 0 | 4.17E-06 | 2.45E-05 | 7.85E-06 | 0 | 2.37E-05 | 5.25E-06 | 0 | 2.98E-05 | 0 | 1.18E-05 | 4.68E-05 | 2.29E-05 | 1.48E-05 | 0 |
| L2hgdh | 8.16E-06 | 1.94E-05 | 3.24E-05 | 9.23E-06 | 1.80E-05 | 0 | 2.13E-05 | 0 | 2.37E-05 | 0 | 1.56E-05 | 0 | 4.02E-05 | 0 | 0 | 2.69E-05 | 1.60E-05 | 0 |
| Dcaf1 | 8.91E-05 | 8.43E-05 | 5.56E-05 | 0 | 1.44E-05 | 4.78E-05 | 0 | 0 | 2.36E-05 | 0 | 0 | 8.36E-05 | 0 | 0 | 6.17E-05 | 8.19E-05 | 5.25E-05 | 5.08E-06 |
| Myorg | 0 | 4.71E-05 | 0 | 0.00014 | 6.94E-05 | 1.18E-05 | 0.00012 | 0 | 2.36E-05 | 0.00012 | 7.08E-06 | 4.09E-05 | 8.49E-06 | 0 | 0 | 0 | 8.77E-05 | 4.41E-05 |
| Zbtb49 | 6.84E-06 | 3.96E-05 | 4.99E-06 | 2.62E-06 | 2.43E-05 | 1.07E-05 | 7.30E-06 | 1.43E-05 | 2.36E-05 | 1.50E-05 | 0 | 7.54E-06 | 0 | 1.56E-05 | 4.72E-05 | 5.12E-06 | 0 | 1.43E-05 |
| Npl | 0 | 1.67E-05 | 0 | 5.41E-06 | 4.74E-05 | 4.62E-06 | 2.55E-07 | 5.04E-05 | 2.36E-05 | 2.26E-06 | 0 | 0 | 1.23E-06 | 0 | 0 | 8.22E-06 | 0 | 0 |
| Lbhd1 | 1.73E-05 | 1.54E-05 | 2.27E-05 | 1.67E-05 | 1.62E-05 | 0 | 0 | 1.86E-05 | 2.36E-05 | 1.62E-05 | 0 | 0 | 1.64E-05 | 2.62E-06 | 0 | 8.70E-06 | 1.19E-05 | 0 |
| Ocel1 | 0 | 1.54E-05 | 0 | 3.26E-05 | 0 | 0 | 9.16E-06 | 0 | 2.36E-05 | 9.47E-06 | 9.53E-06 | 2.03E-05 | 2.48E-05 | 1.93E-05 | 4.50E-05 | 0 | 8.69E-07 | 1.32E-05 |
| Gpc3 | 5.63E-05 | 4.16E-05 | 0 | 0 | 0.00035 | 0.00033 | 0 | 0 | 2.35E-05 | 0.00103 | 0.0002 | 0.00066 | 0.00151 | 0.00061 | 0 | 9.10E-05 | 0.00023 | 0.00028 |
| Vamp7 | 0.00027 | 3.73E-05 | 0 | 0 | 1.10E-05 | 0.00011 | 0.0001 | 1.87E-05 | 2.35E-05 | 1.68E-07 | 0 | 2.23E-05 | 4.43E-05 | 1.97E-05 | 8.91E-07 | 0 | 6.71E-05 | 8.03E-06 |
| Rnf113a2 | 2.03E-05 | 4.32E-05 | 1.27E-05 | 9.71E-05 | 3.83E-05 | 5.21E-05 | 4.49E-05 | 4.22E-05 | 2.35E-05 | 2.38E-05 | 3.24E-05 | 7.15E-06 | 4.20E-05 | 5.85E-05 | 9.19E-05 | 3.36E-05 | 8.13E-06 | 2.83E-05 |

| | | | | | | | | | | | | | | | | | | |
|---|---|---|---|---|---|---|---|---|---|---|---|---|---|---|---|---|---|---|
| Smg9 | 4.69E-05 | 0.0001 | 9.44E-05 | 4.45E-05 | 6.69E-05 | 4.39E-05 | 2.20E-05 | 8.34E-06 | 2.35E-05 | 8.62E-06 | 1.01E-05 | 6.23E-05 | 0 | 6.08E-05 | 0.00013 | 3.47E-05 | 4.21E-05 | 2.02E-05 |
| Sprtn | 4.37E-05 | 6.29E-05 | 7.50E-05 | 1.46E-05 | 4.80E-05 | 4.26E-05 | 2.61E-05 | 1.61E-05 | 2.35E-05 | 3.02E-05 | 8.08E-05 | 0 | 9.04E-06 | 1.53E-05 | 6.20E-06 | 4.29E-05 | 0 | 2.21E-05 |
| Rnf185 | 6.01E-05 | 0.0001 | 6.75E-05 | 7.94E-05 | 7.15E-05 | 2.91E-05 | 4.04E-05 | 4.90E-05 | 2.35E-05 | 1.10E-05 | 2.43E-05 | 3.25E-05 | 7.09E-06 | 3.86E-05 | 4.51E-05 | 8.72E-07 | 2.55E-05 | 1.33E-05 |
| Cab39l | 9.41E-05 | 1.79E-05 | 2.46E-05 | 0 | 0 | 2.27E-05 | 1.28E-05 | 0 | 2.35E-05 | 5.23E-05 | 0 | 6.88E-06 | 3.90E-05 | 0 | 0 | 6.21E-05 | 0 | 5.12E-05 |
| Clpx | 6.97E-05 | 7.92E-05 | 6.58E-05 | 3.11E-05 | 7.66E-05 | 1.06E-05 | 3.21E-05 | 0 | 2.35E-05 | 2.20E-05 | 3.77E-05 | 6.55E-05 | 2.12E-05 | 5.24E-05 | 9.18E-05 | 4.93E-05 | 0 | 4.08E-05 |
| Krba1 | 0 | 3.77E-05 | 0 | 1.48E-05 | 3.48E-05 | 9.13E-06 | 0 | 1.30E-06 | 2.35E-05 | 0 | 2.55E-06 | 1.16E-05 | 0 | 2.81E-05 | 3.56E-05 | 0 | 0 | 2.98E-05 |
| Grin2d | 1.35E-05 | 2.41E-05 | 3.69E-05 | 3.62E-05 | 1.76E-05 | 7.77E-06 | 0.00011 | 5.22E-05 | 2.35E-05 | 6.11E-05 | 3.18E-05 | 3.04E-05 | 3.83E-05 | 1.50E-05 | 0.00012 | 3.54E-06 | 3.48E-05 | 8.51E-05 |
| Lgals4 | 7.60E-07 | 3.36E-05 | 0 | 0 | 2.01E-05 | 1.67E-06 | 0 | 2.02E-05 | 2.35E-05 | 3.37E-05 | 0 | 3.98E-05 | 1.61E-05 | 0 | 0.00014 | 0 | 7.40E-05 | 6.34E-05 |
| Rtn4ip1 | 1.79E-05 | 9.42E-06 | 2.17E-05 | 6.00E-07 | 0 | 1.48E-07 | 0 | 4.73E-06 | 2.35E-05 | 6.63E-06 | 6.99E-06 | 0 | 7.85E-06 | 8.10E-06 | 2.14E-05 | 1.95E-05 | 0 | 6.01E-06 |
| Esco1 | 7.80E-05 | 6.28E-05 | 2.28E-05 | 0 | 2.63E-05 | 6.63E-05 | 2.53E-06 | 3.83E-05 | 2.34E-05 | 2.10E-05 | 1.09E-05 | 3.44E-05 | 0 | 1.66E-05 | 1.26E-05 | 0 | 0 | 0 |
| Yipf6 | 0.00014 | 4.00E-05 | 7.44E-05 | 1.01E-05 | 2.20E-05 | 4.78E-05 | 1.56E-05 | 3.54E-05 | 2.34E-05 | 8.54E-06 | 4.05E-05 | 2.97E-05 | 2.43E-05 | 3.18E-05 | 9.63E-05 | 2.02E-05 | 5.38E-06 | 3.70E-05 |
| Taf12 | 0.00014 | 5.59E-05 | 0.00011 | 0.00011 | 1.92E-05 | 2.90E-05 | 9.34E-05 | 7.93E-06 | 2.34E-05 | 3.35E-05 | 5.03E-05 | 0 | 2.78E-05 | 6.89E-07 | 0 | 0.00013 | 0 | 3.48E-05 |
| Zic4 | 0 | 1.10E-05 | 0 | 0 | 0 | 9.56E-06 | 0 | 7.81E-06 | 2.34E-05 | 3.48E-06 | 0 | 1.27E-05 | 0 | 2.95E-05 | 2.97E-05 | 0 | 0 | 1.53E-06 |
| Sox7 | 0 | 0 | 1.49E-05 | 0 | 0 | 0 | 0 | 0 | 2.34E-05 | 0 | 0 | 0 | 9.93E-06 | 2.43E-05 | 0 | 5.72E-06 | 0 | 6.08E-06 |
| Fastkd1 | 1.45E-05 | 2.07E-05 | 2.11E-05 | 0 | 8.67E-07 | 0 | 0 | 0 | 2.34E-05 | 1.07E-05 | 6.07E-06 | 7.77E-06 | 0 | 4.29E-06 | 2.14E-05 | 8.92E-06 | 4.97E-07 | 2.27E-05 |
| Trak1 | 8.31E-05 | 0.00022 | 0.00011 | 4.76E-05 | 8.47E-05 | 8.83E-05 | 6.85E-05 | 0.00024 | 2.33E-05 | 0.00023 | 0.00012 | 0.00023 | 0 | 0.00012 | 0.00026 | 6.98E-05 | 0.00019 | 0.00016 |
| Fggy | 1.39E-05 | 1.09E-05 | 0 | 0 | 1.57E-05 | 1.51E-05 | 8.79E-06 | 1.86E-05 | 2.33E-05 | 0 | 1.30E-05 | 3.88E-05 | 4.95E-05 | 3.97E-06 | 4.51E-06 | 0 | 0 | 4.67E-06 |
| Lca5 | 4.20E-05 | 1.53E-05 | 0 | 0 | 4.78E-06 | 7.95E-06 | 2.87E-05 | 8.00E-06 | 2.33E-05 | 3.13E-06 | 8.10E-06 | 2.76E-05 | 0 | 0 | 6.51E-05 | 4.46E-05 | 4.96E-05 | 1.83E-05 |
| Zfp329 | 2.72E-05 | 4.03E-05 | 5.89E-06 | 1.59E-05 | 4.63E-05 | 7.31E-06 | 0 | 1.44E-05 | 2.33E-05 | 1.25E-05 | 3.92E-06 | 0 | 0 | 3.99E-05 | 4.86E-05 | 0 | 0 | 8.74E-07 |
| Tmem170b | 0 | 5.97E-05 | 3.70E-07 | 1.95E-05 | 3.16E-05 | 0 | 0 | 3.72E-05 | 2.33E-05 | 2.83E-05 | 5.02E-05 | 0.0002 | 2.75E-06 | 0 | 2.53E-05 | 0 | 7.70E-05 | 4.27E-05 |
| Elp6 | 3.29E-05 | 2.93E-05 | 4.37E-05 | 3.87E-05 | 6.62E-06 | 0 | 0 | 3.47E-06 | 2.33E-05 | 2.71E-06 | 7.54E-06 | 3.18E-05 | 2.66E-06 | 0 | 1.39E-05 | 4.73E-05 | 0 | 6.91E-06 |
| Gpt2 | 0 | 5.25E-05 | 3.25E-05 | 7.47E-05 | 2.81E-05 | 3.21E-05 | 6.09E-05 | 2.50E-05 | 2.32E-05 | 0 | 0 | 5.35E-05 | 0 | 3.45E-05 | 2.55E-05 | 0 | 0 | 1.47E-05 |
| Ube2t | 0 | 0 | 0 | 0 | 2.88E-07 | 3.06E-05 | 0 | 0 | 2.32E-05 | 0 | 2.54E-05 | 4.50E-07 | 0 | 0 | 0 | 0.00026 | 0 | 0 |
| Nxt1 | 2.85E-05 | 4.31E-05 | 4.36E-05 | 3.11E-05 | 2.02E-05 | 1.49E-05 | 2.03E-05 | 1.86E-05 | 2.32E-05 | 3.01E-05 | 1.11E-05 | 0 | 9.89E-07 | 7.41E-06 | 0 | 0.00028 | 0 | 0 |
| Recql5 | 6.44E-06 | 4.06E-05 | 1.29E-05 | 2.24E-05 | 2.90E-05 | 3.09E-06 | 0 | 1.69E-05 | 2.32E-05 | 1.81E-05 | 1.37E-05 | 9.37E-06 | 6.26E-06 | 2.32E-05 | 7.21E-05 | 1.55E-05 | 0 | 2.45E-05 |
| Chek2 | 8.60E-06 | 3.40E-06 | 0 | 0 | 9.59E-06 | 0 | 1.73E-05 | 5.93E-06 | 2.32E-05 | 2.21E-05 | 1.38E-05 | 2.52E-05 | 0 | 1.61E-05 | 1.11E-05 | 0.00013 | 0 | 2.48E-05 |
| Brf2 | 4.05E-06 | 3.52E-05 | 9.64E-06 | 6.11E-05 | 2.78E-06 | 0 | 2.69E-06 | 3.49E-06 | 2.32E-05 | 1.41E-05 | 2.45E-05 | 5.29E-06 | 0 | 0 | 3.12E-06 | 2.05E-05 | 0 | 1.45E-05 |
| Gpcpd1 | 7.99E-05 | 8.97E-05 | 3.64E-05 | 0 | 0.00011 | 0.00028 | 1.70E-05 | 0 | 2.31E-05 | 2.18E-05 | 0 | 0 | 5.63E-05 | 9.59E-05 | 5.56E-05 | 1.58E-05 | 0 | 0 |
| Cmip | 5.83E-05 | 0.00028 | 0.00023 | 9.80E-05 | 0.00023 | 0.00027 | 0.00022 | 0.00024 | 2.31E-05 | 0.00015 | 0.00014 | 0.00053 | 0 | 0.00014 | 0.0003 | 0.00014 | 0.00025 | 0.00017 |
| Mzt1 | 0.00019 | 9.18E-05 | 9.53E-05 | 0 | 4.08E-05 | 9.13E-05 | 4.21E-05 | 3.08E-06 | 2.31E-05 | 0 | 5.20E-05 | 0.00019 | 0 | 0 | 0 | 0.0003 | 0 | 1.57E-05 |
| Uckl1 | 4.94E-05 | 8.69E-05 | 2.20E-05 | 7.44E-05 | 7.79E-05 | 8.39E-05 | 3.05E-05 | 5.09E-05 | 2.31E-05 | 2.77E-05 | 4.26E-05 | 4.21E-05 | 0 | 5.62E-05 | 0.00012 | 0 | 1.56E-05 | 2.33E-05 |
| Traf6 | 1.66E-05 | 9.02E-05 | 6.33E-05 | 2.18E-05 | 6.78E-05 | 6.32E-05 | 0 | 3.81E-06 | 2.31E-05 | 2.42E-05 | 5.93E-05 | 0.00021 | 0 | 5.14E-05 | 1.68E-05 | 3.81E-05 | 0 | 2.93E-05 |
| Tgfbrap1 | 3.32E-05 | 9.77E-05 | 1.43E-05 | 2.04E-05 | 6.08E-05 | 3.35E-05 | 1.74E-05 | 1.53E-05 | 2.31E-05 | 5.72E-05 | 4.83E-05 | 0 | 3.08E-05 | 5.28E-05 | 9.11E-05 | 0 | 0 | 5.79E-05 |

| | | | | | | | | | | | | | | | | | | |
|---|---|---|---|---|---|---|---|---|---|---|---|---|---|---|---|---|---|---|
| Nsrp1 | 0.0001 | 8.20E-05 | 3.61E-05 | 2.99E-05 | 2.78E-05 | 2.83E-05 | 4.06E-05 | 0 | 2.31E-05 | 2.05E-05 | 0 | 4.90E-05 | 0 | 3.17E-05 | 0.00019 | 4.22E-05 | 0 | 1.26E-05 |
| Sp4 | 2.42E-05 | 4.41E-05 | 3.35E-06 | 0 | 2.66E-06 | 2.59E-05 | 0 | 0 | 2.31E-05 | 0 | 6.01E-06 | 1.88E-05 | 0 | 0 | 5.08E-05 | 2.49E-05 | 6.89E-05 | 0 |
| Myo1h | 0 | 0 | 0 | 0 | 7.84E-06 | 6.59E-06 | 0 | 0 | 2.31E-05 | 0 | 0 | 8.01E-05 | 0 | 1.38E-05 | 4.34E-05 | 3.22E-07 | 0 | 5.77E-06 |
| Phf3 | 0.00012 | 0.00015 | 2.30E-05 | 0 | 5.16E-05 | 0.00012 | 4.64E-05 | 0 | 2.30E-05 | 1.90E-05 | 2.08E-05 | 7.51E-05 | 0 | 4.08E-05 | 8.05E-05 | 0.00011 | 0 | 3.20E-05 |
| Arhgap5 | 0.00016 | 2.80E-05 | 7.56E-06 | 0 | 1.68E-05 | 0.00011 | 6.86E-05 | 0 | 2.30E-05 | 2.08E-05 | 7.99E-06 | 0.00013 | 0 | 0 | 0 | 2.29E-05 | 0 | 2.15E-05 |
| Rnmt | 0.00013 | 6.82E-05 | 0.00011 | 5.93E-05 | 4.00E-05 | 5.55E-05 | 8.23E-05 | 6.36E-05 | 2.30E-05 | 0 | 7.14E-06 | 6.09E-05 | 2.86E-05 | 5.20E-05 | 0.00018 | 0 | 0 | 0 |
| Mettl27 | 0 | 6.99E-05 | 0 | 1.66E-05 | 3.14E-05 | 2.79E-05 | 1.25E-05 | 0 | 2.30E-05 | 3.75E-05 | 0 | 0 | 6.39E-06 | 2.66E-05 | 6.57E-06 | 0 | 0 | 1.48E-05 |
| Paox | 0 | 1.40E-05 | 9.09E-06 | 2.79E-05 | 2.08E-05 | 1.48E-05 | 8.20E-06 | 4.30E-05 | 2.30E-05 | 2.75E-05 | 2.85E-05 | 4.01E-06 | 1.11E-05 | 0 | 2.58E-05 | 2.31E-05 | 1.27E-06 | 1.60E-05 |
| Itpripl1 | 4.26E-06 | 2.50E-05 | 2.82E-05 | 2.82E-05 | 0 | 1.22E-05 | 0 | 1.14E-05 | 2.30E-05 | 1.37E-05 | 4.38E-06 | 3.35E-05 | 0 | 3.48E-07 | 0 | 8.53E-05 | 0 | 2.66E-05 |
| Vmac | 2.66E-05 | 4.52E-05 | 0 | 4.85E-05 | 6.57E-05 | 3.08E-05 | 2.42E-05 | 1.14E-05 | 2.29E-05 | 6.14E-06 | 0 | 3.92E-05 | 1.77E-05 | 2.46E-05 | 8.29E-07 | 0 | 1.80E-05 | 5.85E-05 |
| Ankrd49 | 3.72E-05 | 2.79E-05 | 5.29E-05 | 4.54E-06 | 8.23E-06 | 2.07E-05 | 5.21E-06 | 0 | 2.29E-05 | 1.35E-05 | 3.46E-05 | 0 | 0 | 0 | 4.03E-05 | 6.31E-05 | 0 | 2.37E-05 |
| Zfp111 | 2.67E-05 | 1.16E-05 | 0 | 0 | 2.39E-05 | 1.87E-05 | 6.28E-06 | 4.16E-06 | 2.29E-05 | 1.70E-05 | 2.39E-06 | 5.28E-06 | 0 | 0 | 2.82E-05 | 4.70E-05 | 0 | 2.38E-05 |
| Slc19a2 | 2.23E-05 | 5.82E-05 | 3.10E-05 | 0 | 4.33E-06 | 1.74E-05 | 0 | 0 | 2.29E-05 | 1.93E-05 | 0 | 5.95E-05 | 4.15E-05 | 0 | 0 | 2.79E-05 | 0 | 0 |
| Alpk1 | 2.22E-05 | 1.43E-05 | 0 | 0 | 1.51E-05 | 1.29E-05 | 0 | 1.36E-05 | 2.29E-05 | 0 | 1.37E-05 | 2.28E-05 | 2.63E-05 | 2.05E-05 | 3.83E-05 | 0 | 0 | 5.53E-05 |
| Ppwd1 | 1.64E-05 | 3.50E-05 | 3.61E-05 | 5.29E-05 | 1.83E-05 | 7.67E-06 | 4.77E-06 | 1.45E-05 | 2.29E-05 | 1.40E-05 | 1.92E-08 | 1.71E-05 | 7.38E-08 | 1.87E-05 | 1.96E-05 | 0.00013 | 0 | 1.53E-05 |
| Rc3h1 | 0.00014 | 0.00023 | 5.07E-05 | 4.28E-05 | 0.00011 | 0.00013 | 4.27E-05 | 4.31E-05 | 2.28E-05 | 9.64E-05 | 7.36E-05 | 0.00014 | 2.73E-05 | 0.00012 | 0.00018 | 0.0001 | 0.00039 | 7.15E-05 |
| Foxn2 | 0.00017 | 0.0001 | 0.0002 | 0.00012 | 4.01E-05 | 8.99E-05 | 4.01E-06 | 0 | 2.28E-05 | 2.02E-05 | 1.53E-05 | 2.62E-05 | 0 | 0.00011 | 0.00012 | 6.36E-05 | 0.00024 | 5.74E-05 |
| Mbd1 | 3.66E-05 | 0.00011 | 5.61E-05 | 5.20E-05 | 7.40E-05 | 3.02E-05 | 3.65E-05 | 4.15E-05 | 2.28E-05 | 5.07E-05 | 4.26E-05 | 8.27E-05 | 8.79E-05 | 5.09E-05 | 0.00015 | 7.45E-05 | 0.00015 | 3.47E-05 |
| Ppara | 0 | 6.18E-06 | 0 | 0 | 5.25E-06 | 1.53E-05 | 0 | 0 | 2.28E-05 | 6.94E-06 | 0 | 2.88E-06 | 9.53E-06 | 1.80E-05 | 0 | 0 | 3.62E-06 | 0 |
| Rfc5 | 2.27E-05 | 3.61E-06 | 3.18E-05 | 0 | 9.94E-06 | 1.03E-05 | 0 | 1.39E-06 | 2.28E-05 | 3.30E-06 | 0 | 4.90E-06 | 0 | 1.28E-05 | 0 | 0.00039 | 0 | 1.06E-05 |
| Trmt44 | 1.66E-06 | 3.13E-05 | 2.13E-05 | 3.62E-05 | 1.30E-05 | 9.46E-06 | 0 | 4.94E-06 | 2.28E-05 | 2.32E-05 | 1.02E-05 | 2.14E-05 | 0 | 2.09E-05 | 8.11E-05 | 3.13E-05 | 0 | 1.68E-05 |
| Pard3b | 0 | 5.60E-05 | 0 | 0 | 1.67E-05 | 0 | 8.89E-06 | 3.24E-05 | 2.28E-05 | 4.96E-05 | 0 | 1.10E-05 | 0 | 9.61E-06 | 9.47E-06 | 2.97E-05 | 3.36E-06 | 0.00014 |
| Trmt10a | 3.35E-05 | 2.20E-05 | 0.00012 | 1.83E-05 | 0 | 0 | 1.77E-06 | 0 | 2.28E-05 | 2.88E-06 | 0 | 0 | 2.46E-05 | 0 | 6.99E-05 | 9.13E-05 | 2.97E-05 | 0 |
| Mcc | 5.07E-05 | 0 | 0.00015 | 0 | 6.87E-06 | 0 | 0 | 7.22E-05 | 2.28E-05 | 0.0003 | 2.78E-05 | 0 | 1.75E-05 | 0.00011 | 0.00014 | 8.31E-05 | 2.06E-05 | 0.00016 |
| Acot8 | 1.66E-05 | 4.50E-06 | 1.58E-05 | 4.73E-05 | 1.48E-05 | 0 | 8.65E-06 | 1.47E-06 | 2.28E-05 | 1.65E-05 | 1.24E-05 | 4.36E-06 | 0 | 7.38E-06 | 0 | 1.09E-05 | 2.31E-05 | 4.05E-05 |
| Mtln | 1.43E-05 | 3.19E-05 | 5.56E-05 | 6.96E-05 | 2.14E-05 | 0 | 0 | 9.63E-06 | 2.28E-05 | 3.10E-05 | 3.74E-05 | 4.21E-05 | 0 | 0 | 1.19E-05 | 0 | 0 | 7.03E-05 |
| Ldah | 9.53E-05 | 6.72E-05 | 4.47E-05 | 5.18E-06 | 6.99E-05 | 7.49E-05 | 1.79E-05 | 0 | 2.27E-05 | 2.99E-05 | 0 | 2.03E-05 | 8.27E-06 | 3.22E-05 | 4.27E-05 | 2.37E-05 | 0 | 0 |
| Xiap | 8.15E-05 | 1.71E-05 | 0 | 0 | 0 | 6.84E-05 | 0 | 0 | 2.27E-05 | 0 | 0 | 3.89E-05 | 0 | 0 | 0 | 1.46E-05 | 0 | 6.15E-06 |
| Snrnp48 | 9.66E-05 | 0.00012 | 4.90E-05 | 3.25E-05 | 3.51E-05 | 5.30E-05 | 3.80E-05 | 2.92E-06 | 2.27E-05 | 2.18E-05 | 0 | 6.91E-05 | 0 | 4.18E-05 | 0.0003 | 0.00011 | 8.39E-05 | 0 |
| Slc4a1ap | 5.72E-05 | 9.23E-05 | 5.04E-05 | 7.00E-05 | 3.59E-05 | 4.52E-05 | 3.40E-05 | 0 | 2.27E-05 | 9.28E-06 | 3.32E-05 | 2.33E-05 | 0 | 3.88E-05 | 5.95E-05 | 5.13E-05 | 0 | 3.67E-06 |
| Terf2ip | 7.26E-05 | 7.19E-05 | 1.84E-05 | 1.91E-05 | 4.46E-05 | 3.24E-05 | 2.92E-05 | 0 | 2.27E-05 | 3.09E-05 | 9.83E-07 | 3.51E-05 | 7.00E-06 | 9.20E-06 | 2.36E-05 | 1.58E-05 | 0 | 2.83E-05 |
| Ap4b1 | 4.35E-05 | 3.27E-05 | 2.84E-05 | 2.66E-05 | 2.00E-05 | 1.30E-05 | 3.23E-06 | 0 | 2.27E-05 | 6.52E-06 | 1.34E-06 | 1.40E-05 | 0 | 1.49E-05 | 2.16E-05 | 2.94E-05 | 0 | 7.68E-06 |
| Ccdc30 | 0 | 2.61E-05 | 0 | 0 | 2.75E-05 | 5.08E-06 | 1.07E-05 | 6.28E-06 | 2.27E-05 | 2.59E-06 | 0 | 0 | 0 | 0 | 1.76E-05 | 0 | 0 | 7.46E-06 |

| | | | | | | | | | | | | | | | | | | |
|---|---|---|---|---|---|---|---|---|---|---|---|---|---|---|---|---|---|---|
| Vstm2b | 0 | 7.15E-06 | 4.84E-06 | 1.77E-06 | 2.39E-07 | 1.99E-06 | 1.29E-05 | 0 | 2.27E-05 | 0 | 0 | 0 | 9.58E-06 | 3.49E-06 | 0 | 0 | 3.42E-06 | 5.75E-06 |
| Traf3ip1 | 2.09E-05 | 3.91E-05 | 4.29E-06 | 1.47E-06 | 2.43E-05 | 0 | 0 | 0 | 2.27E-05 | 1.02E-05 | 3.77E-05 | 0 | 6.97E-06 | 1.03E-05 | 0.00015 | 3.92E-05 | 4.05E-05 | 1.78E-05 |
| Coq5 | 5.81E-05 | 4.74E-05 | 5.45E-05 | 0 | 4.05E-05 | 0 | 1.68E-05 | 0 | 2.27E-05 | 8.96E-06 | 2.04E-05 | 1.31E-05 | 1.54E-05 | 0 | 4.61E-05 | 2.99E-05 | 0 | 6.00E-06 |
| Coil | 1.73E-05 | 2.92E-05 | 1.61E-05 | 2.88E-05 | 1.72E-05 | 0 | 9.37E-06 | 1.10E-05 | 2.27E-05 | 1.68E-05 | 6.11E-06 | 0 | 0 | 1.35E-05 | 7.73E-06 | 3.76E-05 | 0 | 2.23E-05 |
| Stk35 | 1.54E-05 | 3.81E-05 | 5.11E-05 | 3.12E-05 | 1.32E-05 | 0 | 0 | 0 | 2.27E-05 | 2.90E-05 | 3.03E-05 | 0 | 1.53E-05 | 4.44E-05 | 0.00011 | 8.42E-05 | 7.64E-05 | 0 |
| Immp1l | 0.00013 | 7.25E-05 | 2.90E-05 | 6.11E-05 | 6.58E-05 | 9.31E-05 | 4.18E-05 | 3.28E-05 | 2.26E-05 | 2.81E-05 | 0 | 5.62E-05 | 1.26E-05 | 1.66E-05 | 7.13E-06 | 7.13E-05 | 8.81E-06 | 3.22E-05 |
| Specc1l | 0.00012 | 0.00014 | 5.93E-05 | 1.72E-05 | 8.69E-05 | 6.77E-05 | 0.00015 | 4.05E-05 | 2.26E-05 | 7.33E-05 | 5.14E-05 | 0.00011 | 0 | 4.98E-05 | 0.00018 | 0.00013 | 0.00017 | 0.00013 |
| Med7 | 7.24E-05 | 4.71E-05 | 4.01E-05 | 0 | 2.64E-05 | 2.85E-05 | 5.02E-06 | 1.58E-05 | 2.26E-05 | 2.16E-05 | 2.47E-05 | 1.00E-05 | 3.03E-05 | 2.12E-05 | 4.45E-05 | 3.82E-05 | 0 | 1.50E-05 |
| Dgkq | 0 | 9.43E-05 | 2.50E-05 | 9.36E-05 | 4.57E-05 | 2.40E-05 | 0 | 2.86E-06 | 2.26E-05 | 4.29E-05 | 3.38E-05 | 1.13E-05 | 5.99E-06 | 5.60E-05 | 0.00017 | 0 | 0 | 4.62E-05 |
| Rad9a | 3.26E-05 | 7.80E-05 | 1.76E-05 | 5.28E-05 | 0 | 2.32E-05 | 0 | 2.01E-05 | 2.26E-05 | 4.18E-06 | 4.80E-05 | 3.08E-05 | 1.78E-05 | 7.62E-05 | 6.96E-05 | 0.00023 | 0 | 2.56E-05 |
| Nmnat1 | 3.11E-05 | 1.09E-05 | 3.26E-06 | 0 | 4.18E-06 | 2.41E-06 | 2.32E-06 | 0 | 2.26E-05 | 2.63E-06 | 2.80E-06 | 1.39E-05 | 0 | 1.48E-05 | 5.81E-05 | 4.44E-06 | 0 | 1.05E-05 |
| Syne2 | 9.66E-07 | 5.43E-05 | 0 | 0 | 0 | 0 | 5.98E-05 | 0 | 2.26E-05 | 0.00014 | 3.54E-05 | 3.94E-05 | 9.06E-06 | 7.74E-06 | 2.13E-05 | 0.00025 | 4.50E-05 | 0.0001 |
| Sema3f | 0 | 0 | 0 | 0 | 4.01E-05 | 0 | 0 | 1.67E-05 | 2.26E-05 | 0 | 4.68E-05 | 0 | 8.39E-05 | 0.00018 | 4.44E-06 | 2.66E-05 | 0 | 8.70E-05 |
| Picalm | 0.00026 | 0.00019 | 0.00019 | 0.0002 | 0.00042 | 0.00032 | 6.25E-05 | 0.00027 | 2.25E-05 | 0.00015 | 0.0009 | 0.00026 | 0.00015 | 0.00059 | 0 | 0.0001 | 0.0004 | 0.0002 |
| Cercam | 0 | 5.66E-05 | 0 | 0.00031 | 0.00028 | 0.00028 | 0.00106 | 0.00061 | 2.25E-05 | 0.00048 | 0.00028 | 0.00069 | 0.00044 | 0.00032 | 0 | 0 | 0 | 0.00045 |
| Mmachc | 4.58E-05 | 3.80E-05 | 5.59E-05 | 2.01E-05 | 2.29E-05 | 1.83E-05 | 0 | 0 | 2.25E-05 | 1.48E-05 | 1.80E-05 | 4.68E-06 | 2.12E-05 | 0 | 0 | 6.87E-05 | 0 | 0 |
| Pus10 | 0.0001 | 4.10E-05 | 2.94E-05 | 1.24E-05 | 1.75E-05 | 1.53E-05 | 2.90E-05 | 0 | 2.25E-05 | 1.89E-07 | 1.74E-06 | 1.59E-06 | 1.95E-06 | 2.98E-05 | 0.00012 | 3.90E-05 | 8.41E-05 | 1.46E-05 |
| Batf3 | 0 | 1.69E-05 | 0 | 0 | 0 | 9.65E-06 | 0 | 2.20E-05 | 2.25E-05 | 0 | 0 | 0 | 0 | 0 | 0 | 3.87E-06 | 0 | 2.20E-05 |
| Gga3 | 3.16E-05 | 5.75E-05 | 5.40E-05 | 3.16E-05 | 8.23E-06 | 8.74E-06 | 2.59E-05 | 0 | 2.25E-05 | 2.94E-05 | 5.65E-05 | 5.10E-05 | 1.40E-05 | 2.47E-05 | 9.32E-05 | 3.18E-05 | 1.77E-05 | 8.06E-06 |
| Ercc6 | 3.02E-05 | 3.56E-05 | 5.28E-05 | 1.23E-05 | 1.31E-05 | 8.53E-06 | 2.86E-05 | 5.63E-06 | 2.25E-05 | 1.29E-05 | 6.10E-05 | 9.86E-06 | 0 | 3.76E-05 | 0.00011 | 0 | 4.19E-05 | 0 |
| Thap1 | 3.38E-05 | 3.32E-05 | 5.57E-05 | 3.49E-05 | 6.93E-06 | 5.74E-06 | 4.99E-06 | 0 | 2.25E-05 | 3.15E-06 | 2.60E-05 | 1.31E-05 | 9.57E-06 | 1.30E-05 | 4.14E-05 | 3.12E-05 | 4.90E-05 | 2.26E-05 |
| Tmem181a | 4.73E-05 | 0.00011 | 4.96E-05 | 6.69E-05 | 8.56E-05 | 7.66E-05 | 7.48E-05 | 3.43E-05 | 2.24E-05 | 6.50E-05 | 0 | 0 | 6.21E-05 | 9.38E-05 | 0.00011 | 2.74E-05 | 4.77E-05 | 3.29E-05 |
| Gstz1 | 7.49E-05 | 4.71E-05 | 1.33E-06 | 3.46E-05 | 2.65E-05 | 2.27E-05 | 9.52E-05 | 0 | 2.24E-05 | 9.97E-06 | 0 | 3.40E-05 | 2.54E-05 | 7.77E-06 | 7.47E-05 | 3.99E-05 | 0 | 4.23E-05 |
| Fbxw9 | 2.15E-05 | 9.70E-05 | 8.16E-06 | 3.79E-05 | 2.99E-05 | 1.99E-05 | 0 | 1.57E-06 | 2.24E-05 | 4.31E-05 | 3.44E-05 | 9.80E-05 | 3.90E-05 | 0 | 2.00E-05 | 0 | 2.44E-05 | 7.75E-05 |
| Xpc | 3.52E-05 | 9.44E-05 | 0 | 2.48E-05 | 1.32E-05 | 1.90E-05 | 0 | 0 | 2.24E-05 | 2.23E-08 | 5.33E-07 | 2.85E-05 | 8.75E-06 | 2.76E-05 | 2.52E-05 | 0 | 1.37E-06 | 3.99E-05 |
| Zfp105 | 1.26E-05 | 2.96E-05 | 2.22E-05 | 6.23E-06 | 2.37E-05 | 1.77E-05 | 0 | 1.20E-05 | 2.24E-05 | 1.95E-06 | 1.66E-05 | 1.06E-05 | 3.22E-06 | 3.42E-06 | 4.51E-05 | 1.26E-05 | 0 | 0 |
| Sv2c | 0 | 0 | 0 | 0 | 0 | 1.28E-05 | 0 | 0 | 2.24E-05 | 1.57E-06 | 0 | 0 | 0 | 4.28E-05 | 0 | 0 | 0 | 0 |
| Hdac9 | 0 | 0 | 0 | 0 | 0 | 9.36E-06 | 0 | 5.93E-06 | 2.24E-05 | 0 | 0 | 0.00017 | 0 | 0 | 0 | 0 | 0 | 1.17E-05 |
| Golga1 | 6.19E-05 | 2.51E-05 | 1.07E-05 | 0 | 1.02E-05 | 7.74E-06 | 8.29E-06 | 0 | 2.24E-05 | 1.55E-05 | 6.26E-06 | 3.46E-05 | 0 | 0 | 0.00014 | 2.75E-05 | 0 | 2.77E-05 |
| Reg3g | 0 | 0 | 4.00E-05 | 0 | 0 | 0 | 0 | 6.31E-05 | 2.24E-05 | 0 | 0 | 0 | 0.00038 | 0 | 0 | 7.06E-05 | 0 | 0.00022 |
| Gigyf1 | 5.32E-06 | 0.00019 | 0 | 6.48E-05 | 8.32E-05 | 7.60E-05 | 3.63E-05 | 0 | 2.23E-05 | 3.35E-05 | 9.44E-06 | 0.00013 | 1.38E-05 | 9.60E-05 | 0.00036 | 4.20E-05 | 5.40E-05 | 4.83E-05 |
| Cep120 | 8.11E-05 | 0.00018 | 5.40E-05 | 0 | 6.99E-05 | 5.16E-05 | 5.71E-05 | 3.59E-05 | 2.23E-05 | 2.43E-05 | 4.85E-05 | 6.62E-05 | 2.01E-05 | 4.63E-05 | 0.00013 | 0.00014 | 4.91E-05 | 2.64E-05 |
| Rad17 | 8.55E-05 | 4.27E-05 | 5.69E-05 | 2.50E-05 | 3.95E-05 | 4.00E-05 | 1.32E-05 | 1.75E-06 | 2.23E-05 | 1.30E-05 | 3.48E-05 | 2.11E-05 | 8.80E-06 | 2.34E-05 | 5.86E-05 | 7.38E-05 | 0 | 4.79E-06 |

| | | | | | | | | | | | | | | | | | | |
|---|---|---|---|---|---|---|---|---|---|---|---|---|---|---|---|---|---|---|
| Zfp60 | 4.67E-05 | 3.44E-05 | 0 | 0 | 2.43E-05 | 1.50E-05 | 5.89E-06 | 2.76E-05 | 2.23E-05 | 2.81E-05 | 1.62E-05 | 1.22E-05 | 0 | 0 | 4.22E-05 | 1.54E-05 | 0 | 3.83E-05 |
| Zfp775 | 1.04E-05 | 4.28E-05 | 0 | 1.51E-05 | 1.13E-05 | 8.77E-06 | 1.19E-06 | 0 | 2.23E-05 | 1.66E-05 | 9.53E-06 | 3.19E-05 | 2.32E-05 | 1.27E-05 | 9.91E-06 | 1.53E-05 | 1.30E-05 | 1.62E-06 |
| Arhgap10 | 8.10E-05 | 0.00014 | 0 | 0 | 6.41E-05 | 8.78E-05 | 0 | 0 | 2.22E-05 | 0.00026 | 2.68E-05 | 0.00023 | 0.00017 | 0 | 8.72E-05 | 0.00014 | 0.00013 | 0.00012 |
| Zfp580 | 4.28E-06 | 7.00E-05 | 0 | 5.38E-05 | 8.65E-05 | 5.20E-05 | 0 | 5.56E-05 | 2.22E-05 | 1.10E-05 | 5.98E-05 | 5.98E-05 | 0 | 3.60E-05 | 0 | 1.53E-05 | 0 | 6.19E-05 |
| Krt10 | 3.59E-05 | 7.46E-06 | 0 | 4.73E-06 | 7.52E-06 | 2.84E-05 | 5.87E-06 | 1.77E-05 | 2.22E-05 | 7.70E-05 | 0 | 0 | 0 | 8.19E-06 | 0 | 1.31E-05 | 0 | 0 |
| Ska2 | 1.46E-05 | 3.90E-05 | 0 | 4.69E-05 | 3.26E-05 | 1.41E-05 | 0 | 1.88E-05 | 2.22E-05 | 2.26E-05 | 7.44E-05 | 2.19E-05 | 2.34E-06 | 3.81E-06 | 0.00011 | 0.00053 | 0 | 1.56E-05 |
| Il18rap | 4.75E-07 | 0 | 2.16E-05 | 0 | 3.60E-06 | 0 | 1.17E-06 | 1.05E-05 | 2.22E-05 | 0 | 0 | 0 | 1.67E-05 | 4.26E-06 | 1.61E-05 | 1.50E-05 | 0 | 3.20E-06 |
| Wrb | 5.00E-05 | 3.46E-05 | 1.64E-05 | 5.70E-06 | 1.03E-05 | 0 | 0 | 4.50E-05 | 2.22E-05 | 3.30E-05 | 8.74E-05 | 1.18E-05 | 0 | 0 | 1.92E-05 | 3.25E-05 | 0 | 5.33E-05 |
| Zscan26 | 0.00012 | 0.00019 | 2.63E-05 | 4.69E-05 | 0.00012 | 0.00011 | 6.95E-05 | 9.94E-05 | 2.21E-05 | 7.02E-05 | 8.61E-06 | 0.00012 | 2.68E-05 | 9.92E-05 | 0.00034 | 2.78E-05 | 0 | 8.68E-05 |
| Zfp598 | 0.00013 | 0.00022 | 0.00028 | 9.11E-05 | 8.59E-05 | 0.0001 | 0 | 0.00015 | 2.21E-05 | 7.89E-05 | 3.93E-05 | 6.74E-05 | 7.73E-06 | 2.78E-05 | 0.00023 | 8.23E-05 | 0.00018 | 0.00011 |
| Txnl4a | 0.00027 | 6.78E-05 | 0.00015 | 6.25E-05 | 2.50E-05 | 8.39E-05 | 4.75E-05 | 0 | 2.21E-05 | 9.21E-06 | 9.98E-05 | 6.35E-05 | 0 | 2.05E-05 | 1.89E-05 | 9.68E-05 | 3.30E-05 | 2.30E-05 |
| Pcdhb22 | 3.46E-05 | 3.44E-05 | 0 | 0 | 5.77E-05 | 5.67E-05 | 0 | 3.70E-05 | 2.21E-05 | 1.22E-05 | 1.51E-05 | 1.66E-05 | 2.31E-05 | 2.95E-05 | 1.35E-06 | 0 | 0 | 2.25E-05 |
| Hsdl2 | 5.58E-05 | 6.31E-05 | 0 | 2.21E-05 | 3.42E-05 | 4.14E-05 | 5.17E-05 | 7.92E-06 | 2.21E-05 | 5.46E-05 | 1.41E-05 | 5.60E-06 | 1.41E-05 | 1.51E-05 | 5.17E-07 | 3.37E-05 | 3.89E-05 | 7.45E-05 |
| Zfp715 | 8.73E-05 | 8.43E-05 | 7.44E-06 | 0 | 2.32E-05 | 4.10E-05 | 2.91E-06 | 1.35E-06 | 2.21E-05 | 0 | 0 | 3.63E-05 | 0 | 6.07E-06 | 0.0002 | 3.36E-07 | 3.91E-05 | 1.95E-06 |
| Plekhg5 | 1.13E-05 | 4.95E-05 | 2.67E-05 | 1.10E-05 | 6.37E-05 | 3.73E-05 | 0 | 2.63E-05 | 2.21E-05 | 0 | 2.00E-05 | 0 | 1.84E-05 | 2.00E-05 | 0.00019 | 1.93E-06 | 4.05E-06 | 0.00012 |
| Uxt | 0.00011 | 3.12E-05 | 9.02E-05 | 8.19E-05 | 1.89E-05 | 3.22E-05 | 2.81E-05 | 2.68E-05 | 2.21E-05 | 7.23E-06 | 4.00E-05 | 8.50E-06 | 0 | 3.61E-05 | 0 | 5.20E-05 | 0 | 2.75E-05 |
| Cars2 | 6.50E-05 | 7.13E-05 | 5.93E-05 | 4.35E-06 | 1.42E-05 | 1.38E-05 | 1.44E-05 | 2.71E-06 | 2.21E-05 | 4.55E-05 | 4.63E-06 | 1.67E-05 | 6.89E-06 | 6.94E-06 | 0.00014 | 8.38E-05 | 1.18E-05 | 4.20E-05 |
| Psmb4 | 2.17E-05 | 1.96E-05 | 5.12E-05 | 0.0001 | 1.35E-05 | 0 | 0 | 2.38E-05 | 2.21E-05 | 5.64E-06 | 2.49E-05 | 6.19E-06 | 0 | 1.90E-05 | 0 | 2.66E-05 | 0 | 7.89E-06 |
| Adgre5 | 0 | 6.50E-05 | 0.00029 | 2.71E-05 | 0 | 0.00044 | 0 | 0.00094 | 2.20E-05 | 0 | 6.86E-05 | 3.76E-05 | 0 | 0.00034 | 0.00012 | 9.89E-05 | 1.81E-05 | 0.00014 |
| Chmp1b | 0.00013 | 0.00023 | 7.78E-05 | 0.00011 | 0.00013 | 0.00016 | 0 | 2.74E-05 | 2.20E-05 | 6.81E-05 | 0.00011 | 0.0001 | 3.32E-05 | 0.00011 | 0.00012 | 8.30E-06 | 2.62E-05 | 6.80E-05 |
| Pigg | 1.58E-05 | 1.05E-05 | 1.31E-05 | 1.27E-05 | 3.30E-05 | 3.92E-05 | 0 | 3.48E-05 | 2.20E-05 | 3.58E-05 | 3.21E-05 | 2.77E-05 | 0 | 6.83E-06 | 2.35E-05 | 1.80E-05 | 0 | 3.86E-05 |
| Mrgbp | 3.24E-05 | 5.12E-05 | 2.26E-05 | 1.77E-05 | 2.88E-05 | 3.49E-05 | 5.72E-06 | 4.65E-06 | 2.20E-05 | 1.98E-05 | 2.52E-05 | 0 | 0 | 2.10E-05 | 7.68E-05 | 5.64E-05 | 0 | 0 |
| Mthfd2l | 1.39E-05 | 3.99E-06 | 8.29E-06 | 0 | 0 | 1.75E-05 | 0 | 0 | 2.20E-05 | 2.30E-06 | 8.95E-06 | 2.85E-06 | 0 | 9.26E-06 | 0 | 3.77E-06 | 5.03E-06 | 0 |
| Ndufb3 | 4.05E-05 | 2.48E-05 | 2.55E-05 | 4.52E-05 | 1.17E-05 | 1.12E-05 | 1.48E-06 | 4.31E-06 | 2.20E-05 | 1.61E-05 | 2.24E-05 | 9.24E-06 | 4.41E-06 | 1.57E-06 | 0 | 3.86E-05 | 0 | 3.28E-05 |
| Zfp629 | 5.14E-06 | 5.55E-05 | 1.51E-05 | 4.69E-05 | 1.57E-05 | 8.57E-06 | 0 | 5.83E-06 | 2.20E-05 | 7.90E-06 | 1.13E-05 | 7.50E-06 | 0 | 2.13E-05 | 0.00012 | 8.02E-06 | 0 | 1.11E-05 |
| Pusl1 | 2.21E-05 | 3.18E-05 | 7.64E-05 | 5.45E-05 | 2.15E-07 | 6.96E-06 | 4.15E-05 | 0 | 2.20E-05 | 1.47E-05 | 2.91E-05 | 3.23E-06 | 4.91E-07 | 2.60E-05 | 3.68E-05 | 0 | 0 | 4.14E-05 |
| Phykpl | 2.96E-05 | 4.70E-06 | 1.03E-05 | 1.93E-05 | 9.47E-06 | 6.58E-06 | 1.96E-05 | 1.94E-05 | 2.20E-05 | 9.90E-06 | 0 | 6.54E-05 | 0 | 4.88E-06 | 5.70E-05 | 0 | 0 | 1.46E-05 |
| Slc26a6 | 1.39E-05 | 3.70E-05 | 5.94E-06 | 2.23E-05 | 2.81E-05 | 0 | 2.92E-06 | 2.27E-05 | 2.20E-05 | 2.87E-05 | 1.26E-05 | 0 | 0 | 1.90E-05 | 2.84E-05 | 1.60E-05 | 0 | 1.80E-05 |
| Gbp5 | 1.46E-05 | 7.98E-06 | 0 | 0 | 0 | 0 | 0 | 0 | 2.20E-05 | 5.66E-05 | 0 | 0 | 6.13E-06 | 2.37E-05 | 0 | 1.06E-05 | 0 | 4.00E-05 |
| Rora | 1.97E-05 | 0.00013 | 0 | 0 | 0.00022 | 0.00033 | 0.00011 | 8.12E-05 | 2.19E-05 | 0.00012 | 0.00015 | 0.00055 | 0.00014 | 0.00029 | 0 | 0 | 0.00022 | 7.04E-05 |
| Slf2 | 0.0001 | 6.97E-05 | 2.89E-05 | 0 | 7.64E-05 | 0.00011 | 0 | 3.17E-06 | 2.19E-05 | 0 | 8.48E-06 | 5.77E-05 | 6.65E-06 | 5.69E-05 | 0.00016 | 0.00014 | 3.96E-05 | 0 |
| Zfp512 | 3.87E-05 | 2.48E-05 | 2.39E-06 | 6.47E-06 | 1.56E-05 | 2.79E-05 | 2.96E-05 | 2.57E-05 | 2.19E-05 | 3.10E-05 | 2.76E-06 | 3.52E-05 | 2.18E-05 | 3.81E-06 | 5.01E-05 | 6.74E-05 | 0 | 4.69E-05 |
| Haus1 | 6.26E-05 | 2.45E-05 | 4.70E-07 | 5.53E-05 | 1.80E-05 | 2.21E-05 | 8.74E-06 | 0 | 2.19E-05 | 2.06E-05 | 1.59E-05 | 3.81E-05 | 0 | 1.78E-05 | 8.39E-07 | 0.00031 | 0 | 1.66E-05 |

| | | | | | | | | | | | | | | | | | | |
|---|---|---|---|---|---|---|---|---|---|---|---|---|---|---|---|---|---|---|
| Spidr | 6.45E-05 | 2.55E-05 | 1.29E-05 | 0 | 2.69E-06 | 1.96E-05 | 3.40E-05 | 1.09E-05 | 2.19E-05 | 7.56E-05 | 1.61E-05 | 0.0003 | 0.00011 | 1.00E-05 | 5.87E-05 | 0.00024 | 3.18E-06 | 3.70E-05 |
| Oplah | 0 | 6.31E-05 | 0 | 1.42E-05 | 2.05E-05 | 1.79E-06 | 0 | 0 | 2.19E-05 | 6.66E-05 | 7.32E-06 | 1.69E-05 | 3.07E-05 | 3.71E-05 | 2.88E-05 | 0 | 0 | 5.20E-05 |
| Gins2 | 2.16E-05 | 0 | 8.70E-05 | 8.27E-05 | 0 | 3.04E-07 | 7.39E-06 | 0 | 2.19E-05 | 3.17E-05 | 2.40E-05 | 2.76E-06 | 4.75E-05 | 0 | 0 | 0.00051 | 0 | 0 |
| Msh3 | 3.26E-05 | 3.50E-05 | 2.37E-05 | 5.14E-05 | 7.42E-06 | 0 | 2.09E-08 | 7.42E-06 | 2.19E-05 | 0 | 0 | 1.04E-05 | 0 | 3.29E-05 | 0.00012 | 7.94E-05 | 4.60E-05 | 1.53E-05 |
| Rnf13 | 0.00023 | 0.00021 | 0 | 0 | 7.18E-05 | 0.00021 | 0 | 0 | 2.18E-05 | 0.00011 | 7.03E-07 | 0.00023 | 1.18E-05 | 0 | 6.16E-05 | 1.40E-05 | 4.43E-05 | 0.00012 |
| Ghr | 7.79E-05 | 0.00011 | 0 | 0 | 8.42E-06 | 0.00019 | 0 | 0.00011 | 2.18E-05 | 0.0002 | 0 | 0.0001 | 0.00012 | 0 | 0 | 0 | 0 | 0.00033 |
| Rragc | 0.00022 | 0.00018 | 3.36E-05 | 0 | 0.0001 | 0.00018 | 0.00011 | 0 | 2.18E-05 | 4.37E-05 | 1.42E-05 | 0.00013 | 0 | 7.19E-05 | 1.28E-05 | 6.18E-05 | 0.00019 | 9.51E-05 |
| Zfp444 | 5.42E-06 | 6.46E-05 | 0 | 6.45E-05 | 4.19E-05 | 4.64E-06 | 0 | 1.23E-05 | 2.18E-05 | 3.60E-07 | 2.15E-05 | 5.40E-05 | 0 | 4.72E-05 | 4.60E-05 | 1.97E-05 | 0 | 1.60E-05 |
| Tigd5 | 5.11E-06 | 2.56E-05 | 9.19E-06 | 5.03E-05 | 2.09E-05 | 0 | 0 | 3.17E-06 | 2.18E-05 | 2.23E-05 | 0 | 0 | 0 | 2.45E-05 | 2.87E-05 | 1.46E-05 | 0 | 2.08E-05 |
| Brms1 | 6.17E-05 | 8.60E-05 | 9.88E-05 | 0.00011 | 2.52E-05 | 0 | 9.47E-06 | 0 | 2.18E-05 | 3.28E-05 | 5.33E-05 | 2.91E-05 | 0 | 2.39E-05 | 0.00014 | 6.05E-05 | 0 | 6.93E-06 |
| Gpd2 | 1.95E-05 | 4.24E-05 | 4.70E-05 | 2.55E-07 | 2.61E-05 | 4.05E-05 | 0 | 0 | 2.17E-05 | 1.37E-05 | 3.39E-05 | 0 | 8.98E-07 | 8.71E-05 | 2.19E-05 | 3.36E-05 | 1.12E-05 | 2.73E-05 |
| Utp14a | 6.83E-05 | 5.84E-05 | 0.00012 | 2.74E-05 | 7.38E-07 | 3.85E-05 | 0 | 0 | 2.17E-05 | 1.40E-05 | 0 | 0 | 2.26E-05 | 2.37E-05 | 2.29E-05 | 5.21E-05 | 0 | 2.55E-05 |
| Mcm3ap | 1.96E-05 | 2.66E-05 | 7.85E-06 | 4.03E-06 | 4.32E-06 | 2.93E-05 | 2.24E-05 | 1.40E-05 | 2.17E-05 | 2.34E-05 | 1.32E-05 | 7.19E-06 | 0 | 1.84E-05 | 1.66E-05 | 3.15E-05 | 1.44E-05 | 1.91E-05 |
| Magohb | 5.92E-05 | 6.12E-07 | 4.88E-05 | 8.68E-06 | 4.73E-06 | 2.30E-05 | 0 | 0 | 2.17E-05 | 0 | 2.23E-05 | 7.33E-06 | 7.03E-06 | 3.95E-06 | 1.37E-05 | 4.44E-05 | 0 | 0 |
| Ythdc2 | 3.78E-05 | 1.78E-05 | 3.56E-05 | 0 | 3.71E-06 | 7.14E-06 | 0 | 0 | 2.17E-05 | 1.69E-05 | 1.65E-05 | 0 | 0 | 8.12E-06 | 3.07E-05 | 3.88E-05 | 3.42E-05 | 2.14E-07 |
| Slc12a7 | 2.82E-06 | 3.75E-05 | 0 | 0 | 2.21E-05 | 0 | 0 | 0 | 2.17E-05 | 0 | 3.86E-05 | 0 | 3.89E-05 | 2.65E-05 | 1.95E-05 | 4.28E-05 | 3.88E-06 | 2.61E-05 |
| Nexn | 3.83E-06 | 0 | 0 | 0 | 0 | 0 | 0 | 0 | 2.17E-05 | 0 | 0.0003 | 0 | 0 | 8.95E-05 | 0 | 0 | 0 | 1.75E-05 |
| Shc2 | 0 | 0.00015 | 5.20E-05 | 0.00015 | 8.41E-05 | 0.00011 | 9.59E-05 | 2.85E-05 | 2.16E-05 | 0.00023 | 5.09E-05 | 6.16E-05 | 0.00011 | 7.47E-06 | 5.39E-05 | 0 | 4.93E-05 | 2.30E-05 |
| Cops4 | 0.00029 | 9.55E-05 | 0.0001 | 7.34E-05 | 3.87E-05 | 7.79E-05 | 7.47E-05 | 0 | 2.16E-05 | 0 | 5.98E-06 | 5.45E-05 | 0 | 3.46E-06 | 4.07E-05 | 0.0001 | 4.32E-05 | 6.27E-06 |
| C1qtnf7 | 0 | 4.57E-05 | 0 | 0 | 0 | 3.29E-05 | 0 | 0 | 2.16E-05 | 0.00014 | 0 | 0 | 2.28E-05 | 0 | 0 | 0 | 0 | 0.00018 |
| Pdp2 | 2.54E-05 | 3.77E-06 | 0 | 2.16E-05 | 9.88E-06 | 1.81E-05 | 0 | 0 | 2.16E-05 | 8.13E-06 | 8.07E-08 | 0 | 0 | 2.59E-05 | 0 | 0 | 7.05E-05 | 0 |
| Cbx2 | 8.63E-06 | 2.14E-05 | 0 | 5.41E-06 | 0 | 1.12E-05 | 0 | 2.13E-05 | 2.16E-05 | 2.23E-05 | 2.54E-05 | 0 | 0 | 3.25E-05 | 4.37E-06 | 4.86E-05 | 0 | 6.61E-06 |
| Disc1 | 0 | 9.42E-06 | 0 | 4.77E-06 | 9.05E-07 | 9.69E-07 | 0 | 3.35E-05 | 2.16E-05 | 5.11E-07 | 1.36E-06 | 2.15E-05 | 0 | 1.14E-05 | 3.27E-05 | 0 | 0 | 4.83E-06 |
| Rpf2 | 4.29E-05 | 7.02E-06 | 0.00011 | 6.05E-05 | 0 | 0 | 0 | 0 | 2.16E-05 | 1.71E-05 | 0 | 0 | 3.05E-05 | 0 | 0 | 2.84E-05 | 0 | 4.62E-07 |
| Ppp1r13b | 9.01E-06 | 4.68E-05 | 4.12E-05 | 2.76E-06 | 5.72E-05 | 0 | 0 | 2.05E-05 | 2.16E-05 | 1.18E-05 | 8.30E-05 | 3.48E-05 | 0 | 0 | 4.51E-05 | 0 | 0 | 5.46E-05 |
| Fetub | 0 | 0 | 0 | 0 | 0 | 0.00014 | 0 | 0 | 2.15E-05 | 0 | 0 | 0 | 1.71E-05 | 0 | 0 | 0 | 0 | 1.68E-05 |
| Gpbp1 | 0.00019 | 9.83E-05 | 7.13E-05 | 0 | 2.90E-05 | 8.92E-05 | 4.63E-05 | 1.40E-05 | 2.15E-05 | 0 | 3.37E-05 | 8.45E-05 | 0 | 0 | 0.00012 | 0.00012 | 0.0002 | 7.46E-07 |
| Cog7 | 5.59E-05 | 8.50E-05 | 4.06E-05 | 8.93E-05 | 3.95E-05 | 5.75E-05 | 2.47E-05 | 1.99E-05 | 2.15E-05 | 4.24E-05 | 3.06E-05 | 4.58E-05 | 1.02E-05 | 1.03E-05 | 4.47E-05 | 0 | 0 | 2.69E-05 |
| Ift172 | 8.21E-05 | 6.23E-05 | 0 | 0 | 1.17E-05 | 3.79E-05 | 2.82E-05 | 1.01E-06 | 2.15E-05 | 1.91E-05 | 0 | 3.46E-05 | 1.64E-06 | 0 | 0.00018 | 4.44E-05 | 6.63E-06 | 2.73E-05 |
| Cd109 | 4.30E-05 | 0 | 0 | 0 | 0 | 2.49E-05 | 0.00011 | 0 | 2.15E-05 | 5.66E-05 | 3.57E-06 | 8.14E-05 | 0 | 0.00015 | 0 | 5.26E-05 | 0 | 6.00E-05 |
| Glmn | 4.25E-05 | 2.98E-05 | 3.17E-06 | 0 | 0 | 1.47E-05 | 0 | 1.17E-07 | 2.15E-05 | 5.91E-06 | 2.75E-06 | 2.77E-05 | 0 | 2.18E-06 | 1.46E-05 | 1.30E-05 | 0 | 9.66E-06 |
| Apbb3 | 4.38E-06 | 3.02E-05 | 0 | 1.24E-05 | 2.14E-06 | 0 | 0 | 0 | 2.15E-05 | 4.93E-06 | 2.93E-05 | 1.98E-05 | 0 | 2.24E-05 | 5.18E-05 | 0 | 0.00011 | 9.22E-06 |
| Secisbp2 | 8.15E-05 | 7.42E-05 | 0.00011 | 3.51E-05 | 5.00E-06 | 0 | 0 | 0 | 2.15E-05 | 1.85E-05 | 3.08E-05 | 3.12E-05 | 3.78E-05 | 1.37E-05 | 0.00018 | 0 | 8.39E-05 | 1.47E-05 |

| | | | | | | | | | | | | | | | | | | |
|---|---|---|---|---|---|---|---|---|---|---|---|---|---|---|---|---|---|---|
| Vps37a | 0.00013 | 0.00011 | 9.40E-05 | 0 | 5.55E-05 | 8.01E-05 | 0 | 0 | 2.14E-05 | 8.20E-06 | 6.72E-05 | 0.00014 | 9.37E-06 | 4.46E-05 | 0.00011 | 9.46E-06 | 2.56E-05 | 2.81E-05 |
| Yae1d1 | 8.36E-05 | 3.78E-05 | 6.23E-05 | 0 | 2.61E-05 | 3.67E-05 | 0 | 0 | 2.14E-05 | 2.31E-05 | 0 | 2.40E-05 | 0 | 0 | 0 | 2.47E-05 | 0 | 7.30E-06 |
| Rad1 | 5.02E-05 | 3.56E-05 | 4.06E-05 | 1.74E-05 | 5.86E-06 | 1.83E-05 | 3.28E-05 | 0 | 2.14E-05 | 1.75E-05 | 1.73E-06 | 0 | 0 | 6.15E-06 | 1.61E-05 | 6.66E-05 | 0 | 1.83E-06 |
| Hykk | 4.56E-05 | 2.19E-05 | 0 | 2.03E-05 | 1.94E-05 | 1.71E-05 | 6.42E-05 | 0 | 2.14E-05 | 5.27E-05 | 5.76E-06 | 0 | 1.11E-05 | 2.84E-06 | 1.45E-05 | 0 | 1.70E-06 | 2.88E-05 |
| Ndufaf4 | 7.46E-05 | 3.57E-05 | 0.00011 | 2.65E-05 | 1.38E-05 | 1.40E-05 | 0 | 0 | 2.14E-05 | 3.19E-06 | 3.00E-06 | 0 | 1.01E-05 | 0 | 5.49E-05 | 3.37E-05 | 0 | 1.52E-06 |
| Eri2 | 1.14E-05 | 1.79E-05 | 0 | 0 | 4.12E-05 | 7.21E-06 | 3.86E-06 | 1.62E-05 | 2.14E-05 | 1.13E-05 | 0 | 6.72E-06 | 0 | 8.83E-07 | 4.03E-05 | 0.00021 | 0 | 1.66E-05 |
| Abt1 | 2.94E-05 | 5.47E-05 | 5.59E-05 | 5.27E-05 | 1.17E-05 | 6.80E-06 | 0 | 1.70E-06 | 2.14E-05 | 3.09E-06 | 1.74E-05 | 1.11E-05 | 7.76E-07 | 2.09E-05 | 3.22E-05 | 3.94E-05 | 0 | 1.11E-05 |
| Dcaf4 | 2.71E-05 | 3.22E-05 | 1.31E-06 | 0 | 7.53E-06 | 3.49E-06 | 1.49E-05 | 1.48E-05 | 2.14E-05 | 3.59E-05 | 0 | 3.15E-05 | 0 | 0 | 2.95E-05 | 0 | 3.80E-06 | 3.39E-05 |
| Cables2 | 0 | 7.03E-05 | 5.69E-05 | 5.39E-05 | 5.20E-05 | 2.84E-06 | 5.95E-05 | 4.11E-05 | 2.14E-05 | 2.31E-05 | 0 | 8.61E-06 | 5.65E-05 | 1.87E-05 | 0.00014 | 5.02E-05 | 7.99E-05 | 4.46E-05 |
| Ttc5 | 3.12E-06 | 2.42E-05 | 0 | 2.75E-05 | 1.10E-05 | 0 | 1.99E-06 | 7.92E-06 | 2.14E-05 | 1.16E-05 | 0 | 5.59E-06 | 0 | 2.35E-07 | 3.77E-05 | 1.70E-05 | 1.18E-05 | 1.41E-05 |
| Rpap3 | 1.24E-05 | 1.72E-05 | 7.41E-06 | 1.99E-05 | 0 | 0 | 0 | 3.14E-06 | 2.14E-05 | 1.96E-06 | 0 | 6.23E-06 | 0 | 0 | 5.73E-06 | 8.72E-06 | 1.58E-05 | 7.18E-06 |
| Map3k12 | 1.92E-05 | 5.77E-05 | 4.28E-06 | 1.69E-05 | 4.64E-05 | 3.62E-05 | 5.29E-05 | 7.59E-05 | 2.13E-05 | 4.26E-05 | 3.10E-05 | 3.04E-05 | 4.87E-06 | 5.99E-05 | 4.89E-05 | 8.80E-05 | 0.00011 | 7.39E-05 |
| Bcap29 | 8.11E-05 | 1.86E-05 | 3.77E-05 | 1.15E-05 | 0 | 3.29E-05 | 0 | 1.09E-05 | 2.13E-05 | 3.07E-05 | 2.30E-05 | 1.94E-05 | 2.53E-05 | 7.13E-06 | 7.95E-06 | 1.24E-05 | 0 | 1.96E-05 |
| Nanp | 8.06E-05 | 3.70E-05 | 8.57E-06 | 0 | 2.66E-05 | 3.13E-05 | 1.88E-05 | 6.59E-07 | 2.13E-05 | 1.74E-06 | 2.86E-06 | 0 | 0 | 2.22E-05 | 2.06E-06 | 4.16E-05 | 0 | 1.22E-05 |
| Ndufb2 | 0.00011 | 0 | 7.18E-05 | 0.00018 | 3.95E-05 | 2.79E-05 | 4.81E-05 | 0 | 2.13E-05 | 2.63E-06 | 7.49E-05 | 0 | 5.85E-05 | 5.07E-05 | 0 | 6.72E-05 | 0 | 4.75E-05 |
| Cep192 | 3.32E-05 | 5.21E-05 | 4.73E-05 | 0 | 2.31E-05 | 1.98E-05 | 0 | 5.01E-05 | 2.13E-05 | 4.22E-05 | 9.66E-06 | 3.23E-05 | 0 | 3.26E-06 | 4.56E-05 | 0.00052 | 4.63E-05 | 3.85E-05 |
| Dnajc27 | 2.39E-05 | 3.64E-05 | 2.25E-05 | 0 | 1.53E-06 | 7.68E-06 | 0 | 5.04E-06 | 2.13E-05 | 3.99E-07 | 6.46E-06 | 1.41E-05 | 0 | 0 | 2.17E-05 | 2.04E-05 | 2.79E-05 | 0 |
| Cradd | 3.52E-05 | 3.04E-05 | 4.75E-06 | 6.20E-06 | 5.81E-05 | 5.77E-06 | 4.03E-05 | 7.60E-05 | 2.13E-05 | 2.99E-05 | 2.32E-05 | 0 | 7.72E-07 | 0 | 2.50E-05 | 1.62E-05 | 0 | 3.91E-05 |
| Lrrc75a | 2.58E-05 | 0.00012 | 4.27E-06 | 0.00012 | 2.17E-05 | 5.18E-06 | 3.31E-05 | 4.44E-05 | 2.13E-05 | 0.0002 | 7.42E-05 | 2.03E-05 | 8.00E-05 | 1.77E-05 | 6.25E-06 | 6.13E-05 | 3.34E-05 | 0.00025 |
| Radil | 0 | 3.31E-05 | 0 | 0 | 3.36E-05 | 0 | 3.03E-05 | 5.62E-06 | 2.13E-05 | 0.00013 | 3.17E-05 | 8.25E-05 | 0 | 3.78E-05 | 7.64E-05 | 9.10E-06 | 0 | 0.00012 |
| Exosc2 | 2.92E-05 | 1.85E-05 | 9.01E-05 | 6.20E-05 | 1.44E-05 | 0 | 0 | 8.24E-06 | 2.13E-05 | 1.34E-05 | 1.07E-06 | 0 | 0 | 0 | 5.40E-05 | 0.00014 | 0 | 1.57E-06 |
| Ccdc149 | 0 | 2.40E-05 | 0 | 3.74E-05 | 0 | 0 | 8.64E-05 | 0 | 2.13E-05 | 7.52E-05 | 2.06E-05 | 0 | 6.80E-05 | 6.23E-06 | 2.61E-06 | 0 | 0 | 2.77E-05 |
| Nedd9 | 2.65E-05 | 2.66E-05 | 4.33E-05 | 0 | 8.82E-05 | 8.77E-05 | 6.37E-05 | 0 | 2.12E-05 | 0 | 0.00037 | 0 | 4.42E-05 | 0.00013 | 0 | 0 | 6.07E-05 | 0.0002 |
| Gcdh | 3.93E-05 | 4.44E-05 | 0 | 3.68E-05 | 4.48E-05 | 4.35E-05 | 0 | 0 | 2.12E-05 | 2.74E-05 | 7.25E-06 | 2.40E-05 | 5.06E-05 | 1.40E-05 | 2.57E-05 | 5.44E-05 | 3.11E-05 | 3.57E-05 |
| Taf13 | 8.16E-05 | 2.97E-05 | 3.97E-05 | 1.43E-05 | 2.27E-05 | 3.46E-05 | 2.10E-05 | 5.80E-05 | 2.12E-05 | 0 | 3.41E-05 | 3.22E-05 | 0 | 2.02E-06 | 6.44E-07 | 3.39E-05 | 6.02E-05 | 0 |
| Zfp202 | 3.56E-06 | 1.37E-05 | 1.77E-05 | 1.55E-05 | 1.70E-05 | 9.33E-06 | 0 | 1.35E-06 | 2.12E-05 | 1.25E-05 | 0 | 0 | 4.57E-06 | 4.90E-06 | 3.05E-05 | 0 | 0 | 2.89E-06 |
| Mocos | 0 | 3.98E-05 | 0 | 0 | 3.50E-05 | 6.21E-06 | 0 | 0 | 2.12E-05 | 6.29E-05 | 0 | 6.09E-05 | 2.73E-05 | 3.72E-05 | 0 | 0 | 0 | 4.43E-05 |
| Hoxa6 | 1.13E-06 | 2.95E-05 | 0 | 2.46E-05 | 1.67E-05 | 2.14E-06 | 3.14E-06 | 5.80E-06 | 2.12E-05 | 1.09E-05 | 0 | 0 | 0 | 0 | 4.28E-05 | 1.41E-05 | 0 | 2.68E-05 |
| Tfrc | 1.78E-05 | 0 | 1.33E-05 | 0 | 0 | 0 | 0 | 0 | 2.12E-05 | 0 | 1.56E-06 | 7.42E-05 | 0 | 0 | 0 | 1.41E-05 | 6.66E-05 | 0 |
| Osbpl6 | 1.01E-05 | 2.17E-05 | 1.29E-05 | 4.28E-06 | 7.73E-06 | 0 | 1.11E-05 | 5.48E-06 | 2.12E-05 | 4.99E-05 | 2.54E-06 | 0 | 0 | 1.40E-05 | 1.31E-05 | 3.47E-05 | 5.82E-06 | 0 |
| Suox | 1.55E-05 | 3.78E-05 | 4.03E-06 | 1.66E-05 | 1.53E-05 | 0 | 5.08E-05 | 0 | 2.12E-05 | 7.46E-05 | 3.29E-05 | 1.88E-06 | 0 | 8.84E-06 | 0 | 0 | 1.41E-06 | 5.54E-05 |
| Lrrc61 | 8.04E-06 | 2.74E-05 | 0 | 2.44E-05 | 2.40E-05 | 0 | 5.36E-06 | 2.12E-05 | 2.12E-05 | 4.13E-06 | 0 | 4.74E-06 | 0 | 2.01E-05 | 0 | 0 | 0 | 1.84E-05 |
| Zw10 | 9.66E-05 | 2.76E-05 | 6.36E-05 | 0 | 1.48E-05 | 3.33E-05 | 6.55E-05 | 0 | 2.11E-05 | 3.86E-07 | 2.80E-05 | 0 | 1.09E-05 | 5.04E-05 | 0.00011 | 0.00019 | 2.59E-05 | 0 |

| | | | | | | | | | | | | | | | | | | |
|---|---|---|---|---|---|---|---|---|---|---|---|---|---|---|---|---|---|---|
| Cers4 | 0 | 3.18E-05 | 0 | 0 | 2.89E-05 | 2.56E-05 | 0 | 5.43E-05 | 2.11E-05 | 0 | 4.67E-06 | 5.46E-06 | 0 | 3.56E-05 | 5.40E-06 | 0 | 0 | 2.15E-05 |
| Zfyve9 | 3.28E-05 | 8.74E-05 | 5.30E-05 | 0 | 2.02E-05 | 2.15E-05 | 1.92E-05 | 0 | 2.11E-05 | 4.31E-05 | 2.40E-05 | 4.34E-05 | 0 | 2.74E-05 | 2.30E-05 | 2.74E-05 | 2.46E-05 | 4.62E-05 |
| Ctc1 | 0 | 4.18E-05 | 0 | 5.23E-05 | 8.29E-06 | 5.68E-06 | 1.93E-06 | 3.26E-05 | 2.11E-05 | 1.13E-05 | 0 | 2.93E-05 | 6.35E-06 | 0 | 5.13E-05 | 0.0002 | 4.49E-06 | 1.70E-05 |
| Prx | 0 | 1.78E-05 | 2.91E-07 | 0 | 0 | 0 | 0 | 0 | 2.11E-05 | 5.53E-06 | 1.54E-05 | 1.01E-05 | 1.88E-07 | 0 | 8.30E-05 | 5.42E-05 | 2.51E-05 | 3.98E-05 |
| Rap1gap2 | 0 | 0 | 7.96E-05 | 0 | 3.83E-05 | 0 | 0 | 0 | 2.11E-05 | 0 | 2.87E-05 | 1.04E-05 | 0 | 4.25E-05 | 1.09E-05 | 1.63E-05 | 1.76E-05 | 2.39E-05 |
| Hacd2 | 3.40E-05 | 2.33E-07 | 3.19E-05 | 0 | 2.58E-05 | 0 | 4.98E-05 | 0 | 2.11E-05 | 6.59E-06 | 2.43E-05 | 1.17E-05 | 0 | 0 | 0 | 2.06E-05 | 1.44E-05 | 1.79E-05 |
| Fan1 | 1.66E-05 | 2.50E-05 | 0 | 6.48E-06 | 1.67E-05 | 0 | 1.18E-05 | 0 | 2.11E-05 | 1.23E-05 | 1.01E-05 | 2.08E-06 | 7.27E-06 | 1.02E-05 | 2.46E-05 | 1.00E-05 | 0 | 2.13E-05 |
| Heatr5a | 0.00018 | 2.94E-05 | 0.00029 | 0.00022 | 3.71E-05 | 0.00015 | 0.00038 | 0.00017 | 2.10E-05 | 0.00011 | 0.00026 | 0.0002 | 0 | 7.54E-05 | 0.00025 | 0.00011 | 4.80E-06 | 6.87E-05 |
| Mdm2 | 0.00011 | 0.00026 | 0.00017 | 0 | 5.14E-05 | 5.97E-05 | 0 | 8.22E-05 | 2.10E-05 | 2.30E-05 | 4.95E-05 | 0.00029 | 0 | 0 | 0.00037 | 7.38E-05 | 0.00061 | 4.25E-05 |
| Dop1b | 3.59E-06 | 9.77E-05 | 2.62E-05 | 0 | 1.06E-05 | 4.42E-05 | 0 | 3.15E-05 | 2.10E-05 | 6.49E-05 | 3.90E-05 | 0.0001 | 4.30E-05 | 0 | 9.75E-05 | 0 | 1.56E-05 | 8.10E-05 |
| Rap2a | 0.00011 | 2.76E-05 | 0.00011 | 0 | 9.99E-05 | 3.73E-05 | 0 | 0.00037 | 2.10E-05 | 4.94E-05 | 0.00014 | 0.00014 | 0 | 0.00019 | 2.23E-05 | 6.11E-05 | 0 | 9.75E-05 |
| Clpb | 7.41E-05 | 5.74E-05 | 3.79E-05 | 2.04E-06 | 2.16E-05 | 3.70E-05 | 9.00E-06 | 0 | 2.10E-05 | 2.30E-05 | 7.60E-06 | 3.99E-05 | 8.05E-06 | 0 | 7.02E-06 | 0.00012 | 5.60E-05 | 5.31E-05 |
| Xpo4 | 8.50E-05 | 4.83E-05 | 9.60E-05 | 6.80E-06 | 4.59E-06 | 3.30E-05 | 6.57E-06 | 7.54E-06 | 2.10E-05 | 1.17E-05 | 3.85E-06 | 6.70E-06 | 9.05E-06 | 2.35E-05 | 6.20E-05 | 0.00015 | 3.71E-05 | 2.32E-05 |
| Lamtor4 | 6.01E-05 | 8.88E-05 | 0 | 0.00012 | 6.26E-05 | 1.33E-05 | 5.22E-05 | 0 | 2.10E-05 | 2.52E-05 | 3.18E-05 | 1.66E-05 | 0 | 2.53E-05 | 0 | 1.10E-05 | 0 | 7.76E-05 |
| Zfp870 | 3.16E-05 | 2.01E-05 | 0 | 0 | 3.18E-05 | 1.29E-05 | 6.88E-06 | 5.72E-06 | 2.10E-05 | 1.88E-05 | 2.96E-07 | 0 | 0 | 2.80E-06 | 4.10E-05 | 6.37E-07 | 0 | 1.59E-05 |
| Zfp160 | 1.99E-05 | 2.73E-05 | 0 | 2.07E-08 | 7.43E-06 | 6.79E-06 | 0 | 1.34E-06 | 2.10E-05 | 5.06E-06 | 1.60E-05 | 9.34E-06 | 6.37E-06 | 0 | 5.05E-05 | 8.44E-06 | 0 | 5.19E-08 |
| Itga2b | 9.21E-07 | 0 | 5.46E-06 | 0 | 0 | 5.36E-06 | 9.38E-06 | 0 | 2.10E-05 | 5.26E-05 | 7.98E-06 | 1.10E-05 | 0 | 0 | 3.50E-05 | 0 | 2.15E-05 | 0 |
| St8sia4 | 2.37E-07 | 2.64E-06 | 0 | 0 | 1.08E-06 | 2.25E-06 | 0 | 3.93E-06 | 2.10E-05 | 1.96E-06 | 0 | 0 | 1.15E-06 | 7.01E-06 | 6.62E-06 | 9.42E-07 | 0 | 7.01E-06 |
| Tvp23a | 2.15E-05 | 4.94E-06 | 1.10E-05 | 2.90E-05 | 0 | 0 | 2.01E-05 | 3.37E-06 | 2.10E-05 | 4.54E-05 | 1.19E-05 | 4.88E-07 | 2.40E-05 | 0 | 7.96E-06 | 2.95E-05 | 0 | 3.67E-05 |
| Stx11 | 2.59E-05 | 7.67E-05 | 7.04E-05 | 0 | 0 | 0 | 5.98E-05 | 4.37E-07 | 2.10E-05 | 4.89E-05 | 0 | 8.28E-05 | 5.84E-05 | 2.06E-05 | 5.13E-06 | 1.36E-05 | 8.66E-05 | 4.73E-05 |
| Map3k1 | 3.54E-05 | 5.89E-05 | 3.20E-05 | 4.69E-06 | 0 | 0.00013 | 1.01E-05 | 9.63E-05 | 2.09E-05 | 9.24E-06 | 0 | 0.00012 | 7.21E-05 | 1.66E-05 | 7.62E-05 | 4.29E-05 | 5.54E-05 | 1.58E-05 |
| Dph3 | 0.00018 | 3.56E-05 | 6.66E-05 | 4.25E-06 | 3.33E-05 | 9.78E-05 | 6.84E-05 | 5.71E-05 | 2.09E-05 | 2.49E-05 | 0.00011 | 4.59E-05 | 0 | 4.32E-05 | 9.24E-06 | 9.89E-05 | 2.80E-06 | 9.96E-06 |
| Suz12 | 0.00013 | 6.20E-05 | 6.12E-05 | 0 | 2.55E-05 | 9.58E-05 | 0 | 8.34E-06 | 2.09E-05 | 0 | 8.03E-06 | 0.00018 | 0 | 2.31E-05 | 0 | 0.00031 | 1.32E-05 | 0 |
| Sec16b | 0.00013 | 0 | 4.23E-05 | 0.00022 | 6.26E-05 | 6.91E-05 | 0.00022 | 0.0001 | 2.09E-05 | 7.02E-05 | 0 | 0 | 0 | 6.20E-05 | 0.00022 | 1.85E-05 | 0 | 0.00012 |
| Mthfsl | 2.78E-05 | 1.18E-05 | 0 | 0 | 5.67E-06 | 1.70E-05 | 0 | 0 | 2.09E-05 | 3.16E-06 | 5.62E-06 | 1.00E-05 | 1.09E-05 | 8.80E-06 | 0 | 2.13E-05 | 0 | 0 |
| E2f8 | 0 | 0 | 0 | 0 | 0 | 1.50E-05 | 0 | 3.18E-05 | 2.09E-05 | 0 | 0 | 0 | 0 | 0 | 7.32E-05 | 0.00045 | 0 | 0 |
| Rnf20 | 0.00015 | 0.00015 | 0.00012 | 9.52E-05 | 5.88E-05 | 8.28E-06 | 3.04E-05 | 0 | 2.09E-05 | 2.87E-05 | 8.48E-05 | 6.55E-05 | 0 | 5.93E-06 | 0.0001 | 7.80E-05 | 3.64E-05 | 3.34E-05 |
| Coprs | 1.67E-05 | 3.01E-05 | 0 | 4.58E-06 | 2.98E-05 | 4.68E-06 | 0 | 0 | 2.09E-05 | 2.43E-05 | 8.18E-06 | 0 | 2.69E-05 | 2.87E-05 | 0 | 1.08E-05 | 0 | 1.91E-06 |
| Gprin1 | 0 | 0 | 2.60E-06 | 0 | 6.44E-06 | 1.89E-06 | 0 | 0 | 2.09E-05 | 0 | 1.03E-06 | 0 | 9.09E-07 | 6.87E-06 | 0 | 5.54E-06 | 8.92E-06 | 0 |
| Pstpip1 | 0 | 0 | 0 | 0 | 0.00013 | 0 | 0.00019 | 0 | 2.09E-05 | 0 | 0.00027 | 7.99E-05 | 0 | 0 | 3.46E-05 | 1.13E-05 | 0 | 0 |
| Vezf1 | 0.00019 | 0.00015 | 5.91E-06 | 0 | 8.90E-05 | 0.00017 | 0.00018 | 0 | 2.08E-05 | 4.53E-06 | 9.47E-05 | 0.00011 | 4.25E-05 | 3.82E-05 | 2.50E-05 | 8.93E-05 | 0.00023 | 0 |
| Brox | 5.53E-05 | 0.00011 | 4.37E-05 | 2.26E-05 | 5.93E-05 | 9.89E-05 | 8.02E-05 | 7.12E-05 | 2.08E-05 | 4.95E-05 | 4.93E-05 | 8.26E-05 | 0 | 3.28E-05 | 4.09E-05 | 4.27E-05 | 4.65E-05 | 0.0001 |
| Wee1 | 6.03E-06 | 2.64E-05 | 5.41E-06 | 0 | 1.00E-05 | 2.17E-05 | 0 | 0 | 2.08E-05 | 1.97E-05 | 1.55E-05 | 1.55E-05 | 0 | 1.47E-06 | 6.95E-06 | 0.00032 | 4.19E-06 | 2.07E-05 |

| | | | | | | | | | | | | | | | | | | |
|---|---|---|---|---|---|---|---|---|---|---|---|---|---|---|---|---|---|---|
| Cdkl3 | 6.12E-05 | 4.57E-05 | 2.25E-05 | 2.14E-05 | 1.72E-05 | 1.02E-05 | 4.12E-06 | 2.98E-05 | 2.08E-05 | 0 | 1.85E-05 | 4.74E-06 | 7.17E-06 | 0 | 8.64E-05 | 5.04E-06 | 0 | 1.71E-05 |
| Pkd1 | 0 | 3.21E-05 | 2.82E-06 | 6.43E-05 | 4.38E-05 | 3.82E-06 | 2.40E-05 | 4.87E-05 | 2.08E-05 | 3.39E-05 | 3.90E-05 | 2.20E-05 | 0 | 3.79E-06 | 0.00015 | 0 | 1.09E-05 | 6.94E-05 |
| C1galt1c1 | 0.00018 | 7.15E-05 | 2.75E-05 | 0 | 4.88E-05 | 0.00013 | 3.80E-05 | 4.24E-05 | 2.07E-05 | 4.22E-05 | 4.96E-05 | 3.65E-05 | 0 | 4.72E-06 | 0 | 2.62E-05 | 4.42E-05 | 2.14E-05 |
| Ndfip2 | 7.25E-05 | 7.82E-05 | 7.46E-05 | 0 | 0.00011 | 9.31E-05 | 3.12E-06 | 6.96E-05 | 2.07E-05 | 3.43E-05 | 9.52E-05 | 7.28E-05 | 6.00E-07 | 7.61E-05 | 0 | 2.53E-05 | 3.26E-05 | 2.69E-05 |
| Zfp236 | 8.77E-05 | 9.04E-05 | 5.78E-05 | 0 | 4.02E-05 | 5.11E-05 | 2.49E-05 | 6.39E-05 | 2.07E-05 | 1.33E-05 | 3.19E-05 | 9.07E-05 | 0 | 3.86E-05 | 5.41E-05 | 4.42E-05 | 0.00015 | 3.67E-05 |
| Armcx5 | 2.65E-05 | 4.82E-05 | 2.54E-05 | 7.17E-06 | 1.59E-05 | 1.69E-05 | 0 | 2.09E-06 | 2.07E-05 | 6.20E-07 | 5.12E-06 | 1.54E-06 | 0 | 1.19E-05 | 4.92E-05 | 1.53E-05 | 0 | 7.71E-06 |
| Eif2ak3 | 5.91E-05 | 7.02E-05 | 7.75E-05 | 0.00011 | 3.02E-05 | 1.26E-05 | 0.0002 | 9.88E-05 | 2.07E-05 | 0.00012 | 2.57E-06 | 1.73E-05 | 3.58E-05 | 4.15E-05 | 0.00033 | 5.40E-05 | 0 | 8.93E-05 |
| Fli1 | 2.92E-05 | 1.19E-05 | 4.74E-05 | 0 | 2.98E-05 | 5.84E-07 | 7.38E-06 | 4.39E-05 | 2.07E-05 | 0 | 1.17E-05 | 0 | 0 | 1.35E-05 | 1.63E-05 | 8.21E-05 | 3.28E-05 | 6.24E-05 |
| Ints2 | 8.09E-05 | 1.18E-05 | 6.99E-05 | 0 | 0 | 0 | 2.26E-05 | 0 | 2.07E-05 | 1.29E-05 | 1.63E-05 | 1.07E-05 | 1.04E-05 | 0 | 7.75E-06 | 3.42E-05 | 0.00011 | 0 |
| Noa1 | 4.64E-05 | 2.92E-05 | 6.68E-05 | 2.15E-05 | 1.08E-05 | 0 | 0 | 3.27E-06 | 2.07E-05 | 0 | 2.02E-05 | 7.26E-06 | 0 | 1.73E-05 | 1.85E-05 | 2.72E-05 | 0 | 1.76E-05 |
| Rorb | 6.77E-06 | 0 | 0 | 0 | 0 | 0 | 6.44E-06 | 0 | 2.07E-05 | 3.54E-05 | 0 | 4.53E-08 | 0 | 0 | 0 | 5.26E-06 | 0 | 0 |
| Rtl6 | 2.10E-06 | 2.76E-05 | 0 | 9.84E-06 | 1.55E-05 | 0 | 2.17E-05 | 3.02E-06 | 2.07E-05 | 2.30E-05 | 2.24E-06 | 1.97E-05 | 9.41E-06 | 8.47E-06 | 1.02E-06 | 0 | 0 | 1.49E-05 |
| Haus2 | 9.04E-05 | 2.29E-05 | 2.01E-05 | 0 | 1.01E-05 | 6.08E-05 | 0 | 0 | 2.06E-05 | 1.22E-05 | 2.33E-05 | 3.32E-05 | 0 | 0 | 0 | 5.05E-05 | 1.57E-05 | 2.75E-06 |
| Tbc1d12 | 5.48E-05 | 7.29E-05 | 1.28E-05 | 0 | 1.52E-05 | 3.10E-05 | 0 | 2.19E-05 | 2.06E-05 | 8.47E-05 | 3.48E-05 | 3.39E-05 | 0 | 2.88E-05 | 3.86E-05 | 6.47E-06 | 0.00011 | 2.30E-05 |
| Gphn | 5.05E-05 | 1.25E-05 | 1.28E-05 | 0 | 3.33E-08 | 2.99E-05 | 0 | 1.81E-05 | 2.06E-05 | 2.26E-05 | 5.86E-06 | 7.92E-05 | 2.29E-06 | 8.44E-06 | 7.21E-05 | 4.34E-05 | 3.82E-05 | 3.24E-05 |
| Nfkbib | 7.11E-05 | 0.00012 | 0.00019 | 0.00012 | 2.71E-05 | 1.30E-05 | 0 | 0 | 2.06E-05 | 1.34E-06 | 1.69E-05 | 8.71E-05 | 0 | 5.56E-05 | 6.41E-05 | 2.46E-05 | 0.00034 | 0 |
| Neil1 | 2.45E-06 | 2.90E-05 | 0 | 2.43E-05 | 2.48E-05 | 1.20E-05 | 0 | 3.97E-06 | 2.06E-05 | 2.45E-05 | 3.53E-06 | 2.38E-06 | 4.74E-06 | 1.92E-05 | 2.86E-05 | 0 | 1.01E-05 | 3.49E-05 |
| Ccdc74a | 0 | 6.66E-06 | 0 | 0 | 0 | 6.46E-06 | 0 | 0 | 2.06E-05 | 2.95E-05 | 0 | 0 | 0 | 9.62E-06 | 0 | 1.45E-05 | 0 | 5.62E-05 |
| B9d2 | 5.50E-05 | 4.26E-05 | 5.86E-05 | 3.57E-05 | 1.82E-05 | 5.17E-06 | 1.86E-05 | 4.37E-05 | 2.06E-05 | 2.71E-05 | 1.59E-05 | 3.51E-05 | 0 | 3.15E-05 | 1.17E-06 | 4.81E-05 | 8.46E-06 | 1.34E-05 |
| Gabarap | 0.00057 | 0.00027 | 0.00024 | 0.00022 | 0.00015 | 0.00046 | 0.00077 | 6.11E-05 | 2.05E-05 | 0.0002 | 0.00019 | 0.00027 | 0.0003 | 0.00036 | 0 | 1.12E-05 | 0 | 0.00019 |
| Rasa3 | 5.46E-05 | 0.00024 | 0 | 0 | 0.00026 | 0.00016 | 0 | 0.0002 | 2.05E-05 | 0 | 0.00023 | 0 | 0.00042 | 0.00051 | 0.00017 | 0.00014 | 9.98E-05 | 0.00035 |
| Foxo1 | 5.64E-05 | 0.00013 | 1.43E-05 | 0 | 0 | 8.28E-05 | 7.53E-07 | 6.02E-05 | 2.05E-05 | 0.00014 | 3.56E-06 | 0.00011 | 2.94E-06 | 6.88E-05 | 0 | 5.79E-05 | 0.00011 | 0.00017 |
| Polh | 1.91E-05 | 7.32E-06 | 0 | 0 | 1.06E-05 | 2.54E-05 | 1.43E-05 | 0 | 2.05E-05 | 2.72E-05 | 2.29E-05 | 1.36E-05 | 2.25E-05 | 0 | 0 | 0.00031 | 0 | 3.03E-05 |
| Zc3h13 | 8.14E-05 | 0.00011 | 3.90E-05 | 6.13E-05 | 4.95E-05 | 2.48E-05 | 2.46E-05 | 1.10E-05 | 2.05E-05 | 1.08E-05 | 2.73E-05 | 5.26E-05 | 0 | 2.78E-05 | 9.51E-05 | 7.30E-05 | 0 | 1.07E-05 |
| Zfp950 | 1.13E-05 | 2.00E-05 | 0 | 6.46E-06 | 2.60E-05 | 1.70E-05 | 1.13E-05 | 0 | 2.05E-05 | 9.37E-06 | 0 | 0 | 0 | 0 | 9.25E-05 | 0 | 0 | 2.11E-05 |
| Sco1 | 5.44E-05 | 2.97E-05 | 4.22E-05 | 0 | 3.01E-06 | 7.26E-06 | 0 | 4.62E-06 | 2.05E-05 | 2.20E-05 | 1.02E-05 | 1.58E-05 | 0 | 0 | 3.82E-06 | 2.61E-05 | 0 | 1.79E-05 |
| Armh3 | 4.62E-05 | 3.74E-05 | 3.05E-05 | 1.62E-05 | 2.28E-05 | 7.22E-06 | 1.56E-05 | 0 | 2.05E-05 | 1.85E-05 | 6.34E-06 | 6.08E-06 | 0 | 0 | 1.62E-05 | 1.29E-05 | 4.22E-05 | 1.90E-05 |
| Lipt2 | 1.44E-05 | 5.30E-06 | 2.04E-05 | 2.57E-05 | 4.88E-06 | 6.50E-06 | 7.04E-06 | 7.90E-06 | 2.05E-05 | 0 | 2.46E-05 | 0 | 6.04E-06 | 1.56E-05 | 9.40E-07 | 2.21E-06 | 0 | 0 |
| Fbxl8 | 3.23E-06 | 1.26E-05 | 1.09E-07 | 6.18E-05 | 1.14E-05 | 0 | 5.13E-05 | 6.24E-05 | 2.05E-05 | 4.30E-05 | 2.60E-05 | 6.78E-05 | 1.47E-05 | 2.59E-05 | 0 | 4.12E-05 | 1.96E-05 | 5.77E-05 |
| Tmem177 | 1.81E-06 | 1.31E-05 | 0 | 0 | 1.89E-05 | 0 | 0 | 0 | 2.05E-05 | 1.03E-05 | 3.20E-06 | 2.56E-07 | 1.95E-05 | 7.26E-06 | 0 | 2.84E-05 | 0 | 9.01E-06 |
| Vps53 | 3.41E-05 | 6.29E-05 | 3.09E-05 | 0.00015 | 2.68E-05 | 0 | 3.89E-05 | 1.96E-05 | 2.05E-05 | 4.69E-05 | 3.32E-05 | 2.79E-05 | 2.32E-05 | 1.35E-05 | 0.00014 | 0 | 9.37E-05 | 7.11E-05 |
| Btbd3 | 6.24E-05 | 5.71E-05 | 1.35E-05 | 5.17E-05 | 1.41E-06 | 0.0002 | 0 | 8.98E-05 | 2.04E-05 | 2.75E-05 | 9.88E-05 | 0.0002 | 0 | 5.39E-05 | 1.37E-05 | 1.52E-05 | 0 | 2.60E-05 |
| Ubn1 | 0.00012 | 0.00015 | 6.08E-05 | 7.42E-05 | 0.00012 | 9.94E-05 | 0.00012 | 8.85E-05 | 2.04E-05 | 5.34E-05 | 1.73E-05 | 6.68E-05 | 0 | 0.00011 | 0.00031 | 0.00014 | 0.00017 | 6.86E-05 |

| | | | | | | | | | | | | | | | | | | |
|---|---|---|---|---|---|---|---|---|---|---|---|---|---|---|---|---|---|---|
| Slc5a6 | 5.53E-06 | 2.25E-05 | 3.52E-05 | 5.41E-06 | 0 | 9.86E-05 | 0 | 1.29E-06 | 2.04E-05 | 9.23E-06 | 5.10E-05 | 0 | 4.62E-06 | 8.58E-05 | 4.76E-06 | 0 | 0 | 9.08E-06 |
| Bbs2 | 3.12E-06 | 5.40E-05 | 0 | 8.46E-06 | 1.04E-05 | 2.47E-05 | 4.05E-06 | 1.54E-05 | 2.04E-05 | 2.88E-05 | 0 | 2.24E-05 | 2.83E-05 | 1.87E-05 | 4.03E-05 | 4.29E-06 | 0 | 1.43E-05 |
| Cplx2 | 0 | 2.25E-05 | 0 | 3.70E-06 | 5.41E-05 | 1.47E-05 | 0 | 3.42E-05 | 2.04E-05 | 0 | 2.55E-05 | 0 | 0 | 7.29E-05 | 7.25E-06 | 0 | 0 | 1.48E-06 |
| Zfp689 | 8.91E-06 | 1.91E-05 | 1.62E-05 | 2.44E-05 | 1.57E-05 | 1.23E-05 | 3.82E-06 | 0 | 2.04E-05 | 6.30E-06 | 5.30E-06 | 3.24E-07 | 0 | 0 | 2.97E-05 | 3.97E-05 | 0 | 7.53E-06 |
| Macf1 | 0.00017 | 0.0002 | 9.30E-05 | 0 | 0.00017 | 0.00012 | 0.0001 | 8.81E-05 | 2.03E-05 | 3.58E-05 | 0.00018 | 0.00011 | 7.08E-05 | 2.30E-05 | 0.00019 | 4.30E-05 | 0.00012 | 0.00016 |
| Khnyn | 2.20E-05 | 0.00014 | 1.88E-05 | 1.27E-05 | 7.10E-05 | 4.42E-05 | 0 | 3.19E-05 | 2.03E-05 | 2.24E-05 | 1.33E-05 | 0.00035 | 0 | 8.59E-05 | 0.00025 | 3.61E-05 | 2.01E-06 | 8.41E-05 |
| Zfp955b | 3.72E-05 | 2.31E-05 | 0 | 0 | 1.62E-05 | 2.65E-05 | 0 | 2.71E-06 | 2.03E-05 | 2.01E-05 | 0 | 2.45E-05 | 0 | 9.46E-07 | 1.43E-05 | 0 | 0 | 1.16E-05 |
| Rnf26 | 1.61E-05 | 2.15E-05 | 1.19E-05 | 6.71E-05 | 2.35E-05 | 1.77E-05 | 5.35E-05 | 4.65E-05 | 2.03E-05 | 8.34E-06 | 3.31E-05 | 0 | 1.11E-05 | 2.56E-05 | 6.55E-06 | 0.00018 | 4.12E-06 | 5.47E-06 |
| Mier1 | 0.00017 | 0.00028 | 9.49E-05 | 0.00015 | 0.00012 | 0.00015 | 0.00011 | 2.79E-05 | 2.02E-05 | 0.00011 | 2.62E-05 | 0.00016 | 0 | 0.00012 | 0.00032 | 0.0002 | 0.00015 | 6.05E-05 |
| Asns | 0.0002 | 1.32E-05 | 0.00031 | 4.90E-05 | 0 | 7.46E-05 | 4.32E-05 | 0 | 2.02E-05 | 6.64E-05 | 5.57E-05 | 0.00019 | 7.00E-05 | 3.39E-05 | 0 | 5.17E-05 | 0 | 0 |
| Poc1b | 7.56E-05 | 3.84E-05 | 1.29E-05 | 2.47E-07 | 2.50E-05 | 4.26E-05 | 7.00E-06 | 1.89E-05 | 2.02E-05 | 1.63E-05 | 1.58E-05 | 2.35E-06 | 0 | 3.38E-05 | 0 | 0.00019 | 3.51E-05 | 4.95E-06 |
| Nbn | 7.49E-05 | 4.35E-05 | 4.50E-05 | 0 | 0 | 3.40E-05 | 1.44E-05 | 0 | 2.02E-05 | 2.69E-05 | 7.12E-06 | 2.72E-05 | 1.80E-05 | 0 | 9.59E-06 | 0.00013 | 0 | 4.02E-05 |
| Wdr44 | 3.52E-05 | 2.89E-05 | 6.88E-06 | 6.02E-06 | 3.02E-06 | 3.15E-05 | 1.95E-05 | 2.01E-05 | 2.02E-05 | 4.36E-05 | 9.64E-06 | 5.33E-06 | 0 | 0 | 4.92E-05 | 2.29E-05 | 6.85E-05 | 4.53E-05 |
| Cbll1 | 3.91E-05 | 5.52E-05 | 3.68E-05 | 6.27E-06 | 2.44E-05 | 3.07E-05 | 0 | 1.74E-05 | 2.02E-05 | 2.56E-06 | 2.80E-05 | 1.99E-05 | 0 | 1.38E-05 | 5.44E-07 | 4.13E-05 | 0 | 7.05E-06 |
| Tspyl5 | 6.57E-08 | 1.26E-05 | 0 | 0 | 1.75E-05 | 9.90E-06 | 0 | 1.85E-05 | 2.02E-05 | 1.31E-05 | 1.73E-06 | 0 | 0 | 0 | 1.06E-05 | 0 | 0 | 3.81E-06 |
| Sirt7 | 1.72E-06 | 6.15E-05 | 2.07E-05 | 8.61E-05 | 9.61E-07 | 0 | 0 | 4.12E-05 | 2.02E-05 | 5.56E-05 | 2.22E-05 | 1.14E-06 | 0 | 1.57E-05 | 0.00022 | 8.45E-06 | 7.82E-05 | 4.37E-05 |
| Dusp10 | 1.56E-06 | 1.26E-05 | 2.51E-05 | 0 | 0 | 0 | 1.63E-05 | 0 | 2.02E-05 | 0.00011 | 4.80E-05 | 8.97E-05 | 4.81E-05 | 2.21E-05 | 0 | 2.67E-05 | 0.00039 | 0 |
| Jmjd7 | 6.44E-06 | 4.17E-05 | 2.49E-06 | 5.43E-05 | 0 | 0 | 0 | 5.32E-06 | 2.02E-05 | 2.52E-05 | 8.83E-06 | 1.35E-05 | 9.16E-06 | 0 | 1.14E-05 | 3.31E-05 | 3.33E-05 | 0 |
| Efs | 1.19E-05 | 9.63E-05 | 5.68E-05 | 0.00024 | 2.17E-05 | 0.00011 | 0.00014 | 0 | 2.01E-05 | 0.00018 | 6.39E-05 | 3.92E-05 | 7.54E-05 | 1.01E-05 | 0.0001 | 9.79E-05 | 5.41E-05 | 2.31E-05 |
| Msl3 | 0.00015 | 4.72E-05 | 3.49E-05 | 0 | 6.07E-05 | 9.96E-05 | 3.36E-05 | 3.03E-05 | 2.01E-05 | 3.58E-05 | 4.63E-05 | 8.59E-05 | 4.31E-05 | 7.20E-06 | 2.53E-05 | 9.71E-05 | 0 | 3.01E-05 |
| Arfrp1 | 8.26E-05 | 0.00011 | 8.24E-05 | 0 | 2.54E-05 | 4.13E-05 | 1.21E-05 | 0 | 2.01E-05 | 4.14E-05 | 1.09E-06 | 5.70E-05 | 0 | 3.01E-05 | 0.00023 | 5.02E-05 | 3.19E-05 | 4.11E-05 |
| Adck1 | 3.98E-05 | 3.89E-05 | 3.60E-05 | 8.19E-05 | 2.52E-05 | 1.90E-05 | 8.29E-06 | 1.67E-05 | 2.01E-05 | 1.60E-05 | 2.35E-05 | 0 | 1.92E-05 | 0 | 4.02E-05 | 2.50E-05 | 3.33E-05 | 4.06E-05 |
| Vrk2 | 2.60E-05 | 1.12E-05 | 1.68E-05 | 0 | 7.89E-06 | 1.70E-05 | 2.25E-05 | 0 | 2.01E-05 | 2.55E-05 | 0 | 0 | 5.05E-06 | 0 | 0 | 0.00012 | 2.95E-05 | 2.47E-05 |
| Kif16b | 2.75E-06 | 2.57E-05 | 4.92E-07 | 0 | 2.09E-05 | 1.01E-05 | 1.31E-05 | 0 | 2.01E-05 | 1.69E-05 | 9.01E-06 | 2.19E-05 | 1.44E-05 | 1.88E-05 | 3.60E-07 | 2.54E-05 | 2.41E-05 | 3.39E-05 |
| Ints4 | 4.34E-05 | 1.87E-05 | 1.82E-05 | 0 | 1.29E-05 | 8.30E-06 | 2.30E-05 | 0 | 2.01E-05 | 2.74E-05 | 4.80E-06 | 0 | 0 | 0 | 7.46E-05 | 2.73E-05 | 0 | 1.91E-05 |
| Plpp7 | 0 | 3.33E-05 | 0 | 3.75E-05 | 4.26E-05 | 7.00E-06 | 1.50E-05 | 2.57E-05 | 2.01E-05 | 1.99E-05 | 1.05E-05 | 3.36E-05 | 1.09E-05 | 0 | 8.14E-06 | 2.73E-05 | 0 | 2.48E-05 |
| Cxadr | 0 | 2.52E-05 | 3.39E-08 | 0 | 0 | 0 | 0 | 0 | 2.01E-05 | 0 | 1.86E-05 | 3.00E-05 | 8.70E-06 | 1.72E-05 | 2.13E-05 | 2.30E-07 | 1.28E-05 | 2.03E-06 |
| Wdr34 | 1.11E-05 | 4.20E-05 | 4.67E-07 | 5.70E-05 | 1.57E-05 | 0 | 0 | 4.11E-06 | 2.01E-05 | 2.02E-05 | 0 | 2.58E-05 | 8.80E-06 | 1.93E-06 | 0 | 4.59E-05 | 0 | 1.36E-05 |
| Spata5 | 4.43E-05 | 9.01E-05 | 8.25E-05 | 2.05E-05 | 2.87E-05 | 0 | 0 | 2.03E-05 | 2.01E-05 | 2.34E-05 | 1.80E-06 | 0 | 2.49E-05 | 6.39E-05 | 0.00012 | 3.84E-05 | 1.64E-05 | 1.55E-05 |
| Rictor | 0.0001 | 0.00014 | 1.79E-05 | 0 | 7.19E-05 | 0.0002 | 0 | 6.45E-05 | 2.00E-05 | 0 | 3.45E-05 | 0.00019 | 0 | 0.00011 | 0.00018 | 5.76E-05 | 0.00012 | 0 |
| Cnot10 | 0.00016 | 0.00011 | 6.23E-05 | 0 | 3.57E-05 | 7.65E-05 | 2.94E-05 | 0 | 2.00E-05 | 1.11E-05 | 0 | 3.32E-05 | 0 | 7.72E-06 | 0.0002 | 0.00018 | 2.99E-05 | 2.02E-06 |
| Micu3 | 2.96E-05 | 5.52E-05 | 3.65E-05 | 8.31E-07 | 2.71E-05 | 5.06E-05 | 8.35E-06 | 4.53E-05 | 2.00E-05 | 1.11E-05 | 4.39E-05 | 8.37E-05 | 0 | 1.19E-05 | 8.46E-05 | 0 | 6.31E-05 | 3.37E-05 |
| Zfp182 | 3.12E-05 | 3.35E-05 | 0 | 4.32E-06 | 2.06E-05 | 7.40E-06 | 5.00E-06 | 2.68E-06 | 2.00E-05 | 2.24E-05 | 6.26E-06 | 1.14E-05 | 4.32E-06 | 7.33E-06 | 3.76E-05 | 0 | 0 | 9.55E-06 |

| | | | | | | | | | | | | | | | | | | |
|---|---|---|---|---|---|---|---|---|---|---|---|---|---|---|---|---|---|---|
| Zfp830 | 4.42E-05 | 7.79E-05 | 5.18E-05 | 6.43E-05 | 3.31E-05 | 3.40E-08 | 1.50E-06 | 1.24E-05 | 2.00E-05 | 2.44E-05 | 2.89E-05 | 3.02E-05 | 0 | 6.17E-06 | 0.00011 | 4.92E-05 | 0 | 0 |
| Gimap9 | 8.03E-06 | 0 | 0 | 8.13E-06 | 0 | 0 | 0 | 1.48E-05 | 2.00E-05 | 4.09E-06 | 0 | 0 | 1.13E-06 | 0 | 0 | 8.20E-06 | 0 | 1.81E-05 |
| Ctse | 0 | 0 | 0 | 4.23E-05 | 7.67E-06 | 0 | 0 | 3.90E-05 | 2.00E-05 | 2.34E-05 | 8.10E-06 | 1.78E-05 | 9.15E-06 | 4.14E-06 | 8.31E-05 | 3.66E-06 | 2.00E-06 | 3.75E-05 |
| Tent2 | 0.00023 | 0.00013 | 0.00012 | 0 | 9.85E-05 | 0.00013 | 0.00011 | 0 | 1.99E-05 | 3.86E-05 | 7.45E-05 | 9.31E-05 | 0.00011 | 2.69E-05 | 5.53E-05 | 0.00013 | 9.74E-06 | 6.24E-05 |
| Hspb11 | 0.0001 | 4.39E-05 | 0 | 3.20E-05 | 0 | 5.92E-05 | 5.86E-05 | 2.13E-05 | 1.99E-05 | 1.08E-05 | 0 | 0.0001 | 3.29E-05 | 0 | 0 | 5.43E-05 | 9.44E-06 | 4.94E-05 |
| Mlph | 0 | 8.85E-06 | 0 | 8.55E-06 | 2.94E-05 | 4.40E-05 | 0 | 1.14E-06 | 1.99E-05 | 0 | 0 | 2.24E-06 | 0 | 3.60E-05 | 9.88E-06 | 3.14E-06 | 0 | 0 |
| Alg6 | 1.36E-05 | 7.81E-06 | 9.28E-07 | 1.82E-06 | 1.22E-05 | 9.50E-06 | 7.78E-06 | 1.50E-05 | 1.99E-05 | 1.18E-05 | 1.73E-05 | 2.22E-06 | 0 | 9.59E-06 | 0 | 6.48E-05 | 0 | 1.60E-05 |
| Itga4 | 0 | 1.26E-06 | 0 | 0 | 1.33E-06 | 8.54E-07 | 0 | 0 | 1.99E-05 | 2.66E-06 | 1.67E-05 | 1.58E-06 | 0 | 9.12E-06 | 0 | 3.95E-06 | 0 | 0 |
| Stat2 | 5.90E-05 | 6.84E-05 | 0 | 3.66E-06 | 0 | 0 | 0 | 0 | 1.99E-05 | 0.00024 | 0 | 1.01E-06 | 8.12E-05 | 2.21E-05 | 0.00014 | 7.59E-06 | 2.45E-05 | 0.00014 |
| Enpp5 | 0 | 4.50E-05 | 0 | 0 | 2.19E-05 | 0 | 0 | 0 | 1.99E-05 | 1.21E-05 | 0 | 7.58E-06 | 6.29E-05 | 2.58E-05 | 0 | 0 | 0 | 3.65E-05 |
| Wdr19 | 4.07E-06 | 1.36E-05 | 0 | 0 | 9.72E-05 | 0 | 2.33E-05 | 8.62E-05 | 1.99E-05 | 2.36E-05 | 6.96E-05 | 0.00011 | 5.08E-06 | 0 | 0 | 0 | 0 | 5.33E-05 |
| Arrdc3 | 0 | 0.00018 | 1.14E-06 | 0 | 0 | 0.0002 | 0 | 0 | 1.98E-05 | 0.00015 | 0 | 0.00018 | 0.00032 | 0.00018 | 0.00011 | 4.12E-05 | 0 | 0.0001 |
| Gtdc1 | 3.49E-05 | 3.62E-05 | 7.03E-06 | 0 | 0 | 6.78E-05 | 0 | 7.15E-06 | 1.98E-05 | 2.09E-05 | 1.55E-05 | 3.17E-05 | 1.23E-05 | 4.13E-06 | 2.14E-05 | 8.48E-05 | 2.28E-05 | 4.42E-05 |
| Zfp110 | 0.00011 | 8.79E-05 | 3.98E-05 | 3.51E-05 | 5.03E-05 | 6.32E-05 | 4.78E-05 | 4.71E-06 | 1.98E-05 | 5.89E-05 | 0 | 0 | 0 | 0 | 7.84E-05 | 2.65E-05 | 0 | 1.39E-05 |
| Polm | 0 | 3.94E-05 | 0 | 1.30E-05 | 4.38E-05 | 5.74E-05 | 3.71E-05 | 0 | 1.98E-05 | 4.59E-05 | 7.46E-05 | 9.32E-05 | 3.05E-05 | 8.90E-05 | 0.0002 | 0 | 0 | 0 |
| Rfwd3 | 5.97E-05 | 4.39E-05 | 2.31E-05 | 0 | 3.16E-05 | 5.17E-05 | 0 | 8.59E-06 | 1.98E-05 | 0 | 2.44E-05 | 2.55E-05 | 2.18E-05 | 1.07E-05 | 0 | 0.00037 | 4.06E-07 | 0 |
| Pycard | 4.72E-05 | 3.20E-06 | 0 | 0 | 0 | 3.97E-05 | 0 | 0 | 1.98E-05 | 0.00023 | 0 | 0 | 0.00014 | 0 | 0 | 4.13E-05 | 0 | 0.00017 |
| Utp3 | 9.19E-05 | 0.00011 | 9.62E-05 | 0.00014 | 1.40E-05 | 2.34E-05 | 0 | 1.98E-05 | 1.98E-05 | 3.06E-05 | 2.47E-05 | 7.36E-05 | 0 | 4.88E-05 | 0.00018 | 0.00022 | 0 | 0 |
| Atl2 | 6.67E-05 | 0.0001 | 8.52E-05 | 3.43E-06 | 3.23E-05 | 2.09E-05 | 5.52E-05 | 0 | 1.98E-05 | 6.75E-05 | 9.89E-05 | 5.48E-05 | 7.85E-05 | 1.94E-05 | 4.53E-05 | 0.00025 | 0.00033 | 0 |
| Rpap2 | 5.19E-05 | 3.35E-05 | 0 | 3.77E-05 | 1.99E-05 | 8.92E-06 | 0 | 0 | 1.98E-05 | 1.03E-05 | 0 | 3.52E-05 | 0 | 5.15E-06 | 5.03E-05 | 1.44E-05 | 0 | 2.50E-05 |
| Hoxb4 | 2.21E-06 | 3.43E-05 | 3.67E-06 | 7.14E-06 | 3.00E-05 | 6.12E-06 | 1.17E-05 | 5.03E-05 | 1.98E-05 | 2.14E-05 | 1.17E-05 | 2.46E-06 | 1.27E-05 | 0 | 2.07E-05 | 1.56E-05 | 0 | 2.19E-05 |
| Kptn | 1.14E-05 | 2.12E-05 | 0 | 4.70E-05 | 1.59E-05 | 1.13E-06 | 1.43E-05 | 2.19E-05 | 1.98E-05 | 2.33E-05 | 1.56E-05 | 1.63E-05 | 5.95E-06 | 2.49E-05 | 0 | 1.18E-05 | 7.69E-05 | 4.17E-05 |
| Slc5a3 | 0 | 1.40E-05 | 0 | 0 | 0 | 0 | 0 | 0 | 1.98E-05 | 0 | 0 | 3.17E-05 | 0.00026 | 3.34E-05 | 0 | 0 | 0.00215 | 0 |
| Bcl2 | 0 | 0.00022 | 0 | 0 | 3.43E-05 | 0 | 0 | 0 | 1.98E-05 | 5.77E-06 | 0.00011 | 0 | 0 | 0.00027 | 0.00014 | 7.29E-05 | 9.32E-05 | 5.19E-05 |
| Tgoln1 | 0.00021 | 0.00019 | 0.00021 | 0 | 0.00016 | 0.00026 | 0.00018 | 0.00014 | 1.97E-05 | 0.00011 | 7.48E-05 | 0.00027 | 0.00021 | 0.00015 | 0.00014 | 0 | 0.00038 | 4.45E-05 |
| Zc2hc1a | 0.00011 | 6.08E-05 | 0 | 0 | 4.58E-05 | 7.69E-05 | 0 | 0 | 1.97E-05 | 5.40E-05 | 0.00011 | 7.19E-05 | 0.00014 | 7.32E-05 | 4.73E-05 | 0 | 0 | 0.00015 |
| Timm21 | 0.00011 | 4.63E-05 | 1.68E-05 | 0 | 1.43E-05 | 6.59E-05 | 1.80E-05 | 7.73E-06 | 1.97E-05 | 6.40E-08 | 0 | 2.84E-05 | 0 | 1.43E-05 | 0 | 8.09E-05 | 0 | 0 |
| Rundc1 | 2.04E-05 | 7.97E-05 | 3.15E-05 | 6.28E-05 | 4.54E-05 | 2.24E-05 | 2.30E-05 | 4.65E-05 | 1.97E-05 | 4.09E-05 | 6.87E-06 | 4.90E-05 | 1.01E-06 | 5.12E-06 | 0.00014 | 1.83E-05 | 1.12E-05 | 2.18E-05 |
| Cib1 | 0.00021 | 6.99E-05 | 0.00011 | 3.58E-05 | 4.32E-05 | 1.78E-05 | 9.12E-05 | 1.70E-05 | 1.97E-05 | 4.33E-05 | 2.81E-05 | 0 | 1.54E-05 | 2.67E-05 | 0 | 0 | 0 | 9.41E-05 |
| Inip | 1.84E-05 | 1.36E-05 | 1.80E-05 | 0 | 5.68E-06 | 8.20E-06 | 1.52E-06 | 6.82E-06 | 1.97E-05 | 2.26E-06 | 4.60E-06 | 0 | 0 | 0 | 2.17E-06 | 3.62E-05 | 0 | 0 |
| Gmip | 7.93E-06 | 2.98E-05 | 2.16E-05 | 1.12E-05 | 7.03E-06 | 4.12E-07 | 7.23E-06 | 0 | 1.97E-05 | 4.41E-05 | 0 | 1.44E-06 | 1.01E-05 | 0 | 5.68E-05 | 8.61E-06 | 4.47E-05 | 2.64E-05 |
| Zpr1 | 7.24E-05 | 2.13E-05 | 0.00031 | 0.00014 | 0 | 0 | 0.00027 | 0.00013 | 1.97E-05 | 0 | 0 | 0.00019 | 3.63E-05 | 0 | 0.00012 | 0 | 0 | 3.38E-05 |
| Ybx1 | 0.00044 | 0.00016 | 0.0003 | 0.00109 | 0.00038 | 0.00071 | 0.00039 | 0.0007 | 1.96E-05 | 0.00015 | 0.00026 | 0.00019 | 0 | 0.00015 | 8.16E-05 | 0.00043 | 0.00031 | 4.34E-05 |

| | | | | | | | | | | | | | | | | | | |
|---|---|---|---|---|---|---|---|---|---|---|---|---|---|---|---|---|---|---|
| Miip | 9.82E-06 | 9.06E-05 | 0 | 0 | 3.91E-05 | 9.83E-05 | 0 | 6.58E-05 | 1.96E-05 | 4.20E-05 | 0 | 2.14E-05 | 1.28E-05 | 1.99E-05 | 7.82E-05 | 0.00027 | 0 | 1.43E-05 |
| Fbxo3 | 0.00011 | 0.00014 | 3.09E-05 | 4.79E-05 | 0.0001 | 8.56E-05 | 3.75E-05 | 0 | 1.96E-05 | 4.67E-05 | 0.0001 | 5.74E-05 | 2.79E-05 | 8.16E-05 | 9.98E-05 | 5.53E-05 | 0 | 8.37E-05 |
| Mtpap | 0.00011 | 8.81E-05 | 6.74E-05 | 0 | 2.07E-05 | 7.15E-05 | 8.66E-06 | 0 | 1.96E-05 | 3.22E-05 | 2.44E-05 | 6.25E-05 | 0 | 4.50E-05 | 0.0002 | 4.83E-05 | 0 | 1.37E-05 |
| Ptpn14 | 2.22E-06 | 4.19E-05 | 3.09E-05 | 0 | 4.49E-05 | 3.79E-05 | 3.11E-05 | 1.46E-05 | 1.96E-05 | 0 | 0.00012 | 0 | 0 | 7.66E-05 | 1.45E-05 | 7.22E-05 | 0 | 0.00011 |
| Bnip1 | 9.62E-05 | 3.65E-05 | 4.97E-05 | 4.33E-05 | 0 | 3.06E-05 | 4.44E-05 | 0 | 1.96E-05 | 1.68E-05 | 4.49E-05 | 5.39E-05 | 1.56E-05 | 1.79E-05 | 8.15E-05 | 3.36E-05 | 2.32E-05 | 2.79E-05 |
| Ttll4 | 3.51E-05 | 5.56E-05 | 8.20E-05 | 4.58E-05 | 1.70E-05 | 1.89E-05 | 1.78E-05 | 1.16E-05 | 1.96E-05 | 9.41E-06 | 5.94E-06 | 0 | 2.92E-05 | 5.49E-06 | 8.90E-05 | 6.89E-05 | 7.15E-05 | 0 |
| Lrrc8d | 6.13E-06 | 2.49E-05 | 2.89E-05 | 1.94E-06 | 0 | 1.19E-05 | 0 | 0 | 1.96E-05 | 8.79E-05 | 1.68E-05 | 0 | 5.17E-06 | 2.70E-05 | 6.70E-06 | 5.22E-06 | 0 | 6.69E-06 |
| Usp54 | 1.57E-05 | 3.81E-05 | 1.45E-05 | 0 | 1.31E-05 | 6.10E-06 | 0 | 1.32E-05 | 1.96E-05 | 2.20E-05 | 5.73E-05 | 2.23E-05 | 7.15E-06 | 2.38E-05 | 8.24E-05 | 5.63E-05 | 4.69E-05 | 2.09E-05 |
| Atat1 | 3.59E-05 | 7.54E-05 | 0 | 6.85E-05 | 2.17E-05 | 0 | 2.78E-05 | 1.89E-05 | 1.96E-05 | 3.71E-05 | 2.21E-05 | 7.19E-06 | 1.43E-05 | 0 | 0.00011 | 0 | 3.30E-05 | 2.55E-05 |
| Cyp2j6 | 5.90E-05 | 3.13E-05 | 0 | 1.75E-06 | 4.67E-05 | 5.87E-05 | 0 | 2.63E-05 | 1.95E-05 | 4.83E-05 | 4.54E-06 | 0 | 3.44E-05 | 2.00E-05 | 2.65E-05 | 0 | 0 | 6.36E-05 |
| Arl16 | 7.42E-05 | 3.99E-05 | 1.12E-05 | 5.82E-05 | 2.61E-05 | 1.37E-05 | 7.12E-05 | 1.23E-05 | 1.95E-05 | 2.27E-05 | 0 | 1.53E-05 | 0 | 4.51E-05 | 4.71E-05 | 2.05E-05 | 0 | 2.22E-05 |
| Wdr5b | 3.47E-05 | 6.49E-06 | 1.49E-05 | 6.20E-06 | 9.81E-06 | 1.29E-05 | 2.20E-05 | 2.68E-09 | 1.95E-05 | 1.73E-05 | 9.41E-06 | 0 | 6.85E-06 | 1.22E-05 | 3.22E-05 | 2.42E-05 | 1.02E-06 | 4.12E-05 |
| Isoc2a | 4.02E-05 | 8.05E-06 | 2.18E-05 | 3.85E-05 | 2.31E-05 | 8.36E-06 | 2.31E-05 | 2.04E-05 | 1.95E-05 | 2.57E-05 | 1.86E-06 | 9.89E-06 | 1.33E-05 | 0 | 0 | 8.08E-06 | 5.28E-06 | 2.09E-05 |
| Med17 | 4.66E-05 | 4.53E-05 | 4.99E-05 | 3.65E-05 | 1.90E-05 | 4.54E-06 | 2.38E-05 | 2.01E-05 | 1.95E-05 | 2.56E-05 | 9.80E-06 | 2.28E-05 | 8.54E-06 | 4.73E-05 | 5.70E-05 | 2.71E-06 | 0 | 2.15E-05 |
| Ap3m1 | 0.00011 | 1.60E-05 | 3.96E-05 | 0 | 0 | 0 | 2.74E-05 | 3.41E-06 | 1.95E-05 | 4.21E-05 | 0 | 8.52E-06 | 0 | 0 | 4.88E-05 | 0 | 0 | 6.73E-06 |
| Qser1 | 7.18E-05 | 6.76E-05 | 3.26E-05 | 0 | 2.25E-05 | 7.13E-05 | 4.79E-05 | 7.64E-06 | 1.94E-05 | 1.02E-06 | 1.63E-05 | 5.00E-05 | 0 | 2.51E-05 | 2.45E-05 | 9.54E-05 | 3.00E-06 | 2.25E-05 |
| Dusp9 | 0 | 0 | 4.99E-05 | 0 | 3.02E-05 | 4.62E-05 | 0 | 2.62E-05 | 1.94E-05 | 0 | 0 | 0.00012 | 0 | 3.19E-05 | 0 | 0 | 0 | 0 |
| Ngdn | 0.00012 | 8.76E-05 | 0.00011 | 7.22E-05 | 2.43E-05 | 2.97E-05 | 2.09E-05 | 0 | 1.94E-05 | 1.69E-05 | 2.11E-05 | 2.70E-05 | 0 | 8.00E-06 | 3.46E-05 | 0.00013 | 3.29E-05 | 0 |
| Zfp113 | 1.10E-05 | 3.48E-05 | 0 | 1.00E-05 | 2.00E-05 | 1.45E-05 | 1.42E-05 | 0 | 1.94E-05 | 1.53E-05 | 2.46E-06 | 0 | 0 | 2.99E-05 | 5.89E-05 | 0 | 0 | 1.38E-05 |
| Hdx | 1.05E-05 | 2.02E-05 | 1.72E-05 | 1.84E-05 | 8.50E-06 | 1.43E-05 | 0 | 1.39E-05 | 1.94E-05 | 8.78E-06 | 1.42E-05 | 7.16E-06 | 1.24E-05 | 4.68E-06 | 5.73E-05 | 0 | 5.35E-05 | 1.59E-05 |
| Rsad1 | 6.32E-06 | 3.26E-05 | 1.41E-05 | 7.92E-05 | 1.97E-05 | 5.63E-06 | 2.41E-05 | 1.30E-05 | 1.94E-05 | 1.17E-05 | 0 | 1.10E-05 | 3.36E-06 | 1.48E-05 | 3.87E-05 | 0 | 0 | 2.37E-05 |
| Agtr2 | 0 | 0 | 0 | 0 | 0 | 3.67E-06 | 0 | 9.13E-06 | 1.94E-05 | 7.89E-05 | 0 | 0 | 1.43E-05 | 0 | 0 | 0 | 0 | 0 |
| Zscan29 | 1.47E-05 | 6.08E-05 | 2.13E-05 | 2.19E-05 | 2.48E-05 | 5.91E-07 | 1.11E-05 | 8.59E-06 | 1.94E-05 | 3.36E-05 | 5.53E-06 | 1.00E-05 | 0 | 2.86E-05 | 3.09E-05 | 1.05E-05 | 3.36E-06 | 1.96E-05 |
| Abca5 | 5.41E-05 | 4.61E-05 | 3.94E-05 | 0 | 9.71E-06 | 0 | 5.19E-05 | 4.83E-05 | 1.94E-05 | 7.11E-05 | 5.19E-05 | 0.00011 | 5.66E-05 | 8.30E-06 | 3.44E-05 | 0 | 0 | 0 |
| Zbtb48 | 9.40E-06 | 1.64E-05 | 1.52E-05 | 1.89E-05 | 6.15E-06 | 0 | 0 | 1.66E-05 | 1.94E-05 | 1.93E-06 | 2.80E-06 | 5.24E-06 | 4.13E-06 | 2.64E-06 | 7.16E-05 | 1.46E-05 | 0 | 2.26E-05 |
| Tfap4 | 1.14E-05 | 5.17E-05 | 6.22E-05 | 0 | 0 | 2.81E-05 | 0 | 0 | 1.93E-05 | 0 | 0 | 5.70E-05 | 0 | 1.07E-05 | 0 | 7.73E-05 | 0 | 4.25E-07 |
| Tmem69 | 5.66E-05 | 1.76E-05 | 1.84E-05 | 0 | 0 | 1.81E-05 | 0 | 0 | 1.93E-05 | 1.07E-05 | 0 | 2.50E-07 | 4.88E-06 | 0 | 0 | 2.50E-05 | 0 | 0 |
| Helq | 9.75E-06 | 1.16E-05 | 1.58E-05 | 1.67E-05 | 1.68E-05 | 1.41E-05 | 1.08E-05 | 2.62E-06 | 1.93E-05 | 1.54E-05 | 3.85E-06 | 0 | 0 | 1.55E-05 | 5.32E-05 | 4.58E-05 | 2.12E-05 | 8.33E-06 |
| Wdr66 | 1.33E-06 | 2.03E-05 | 0 | 0 | 0 | 6.95E-06 | 1.80E-05 | 0 | 1.93E-05 | 3.09E-05 | 0 | 0 | 6.88E-05 | 2.96E-06 | 5.55E-05 | 0 | 7.73E-06 | 0.00013 |
| Zfyve27 | 0 | 0.00013 | 6.70E-05 | 0.00012 | 6.76E-05 | 7.77E-07 | 7.51E-06 | 4.46E-05 | 1.93E-05 | 4.91E-05 | 6.74E-05 | 0.00012 | 4.91E-05 | 8.02E-05 | 0.00024 | 0 | 4.58E-05 | 3.12E-05 |
| Cfp | 0 | 6.24E-05 | 0 | 7.00E-05 | 1.40E-05 | 0 | 7.20E-06 | 3.36E-05 | 1.93E-05 | 1.78E-05 | 9.53E-07 | 1.02E-05 | 0 | 1.49E-05 | 0.00021 | 1.41E-05 | 3.10E-05 | 6.62E-06 |
| Fnbp1l | 2.71E-05 | 4.61E-05 | 1.69E-05 | 0 | 8.66E-05 | 5.59E-05 | 0 | 0.00011 | 1.92E-05 | 0 | 9.70E-05 | 5.01E-05 | 0 | 6.96E-05 | 3.15E-05 | 8.91E-05 | 3.02E-05 | 3.76E-05 |
| Ptpn3 | 0 | 5.51E-05 | 6.60E-05 | 0 | 3.55E-05 | 4.51E-05 | 0.00011 | 1.37E-05 | 1.92E-05 | 1.67E-05 | 0 | 0 | 0 | 2.15E-05 | 0.00017 | 0 | 6.21E-05 | 8.14E-05 |

| | | | | | | | | | | | | | | | | | | |
|---|---|---|---|---|---|---|---|---|---|---|---|---|---|---|---|---|---|---|
| Cenpv | 3.08E-05 | 3.06E-05 | 9.37E-06 | 0 | 0 | 1.62E-05 | 0 | 0 | 1.92E-05 | 2.76E-05 | 1.97E-05 | 0 | 1.13E-05 | 5.50E-05 | 0 | 4.59E-05 | 0 | 2.37E-06 |
| Gmpr | 1.94E-06 | 9.27E-06 | 0 | 0 | 0 | 1.57E-05 | 0 | 1.61E-05 | 1.92E-05 | 0 | 0 | 0 | 0 | 1.23E-05 | 0 | 0 | 0 | 6.48E-08 |
| Trmt5 | 5.17E-05 | 2.14E-05 | 2.40E-05 | 7.48E-06 | 4.81E-06 | 1.52E-05 | 1.67E-06 | 0 | 1.92E-05 | 1.27E-05 | 1.36E-06 | 0 | 0 | 0 | 8.89E-06 | 2.82E-05 | 0 | 4.18E-06 |
| Tert | 1.10E-05 | 1.96E-05 | 0 | 0 | 9.91E-06 | 9.90E-06 | 1.98E-05 | 3.27E-05 | 1.92E-05 | 1.50E-05 | 0 | 1.24E-05 | 1.45E-05 | 4.43E-06 | 3.41E-05 | 0 | 0 | 1.08E-05 |
| Azin2 | 0 | 0.00015 | 0 | 0.00011 | 0.0004 | 0.00017 | 0.00036 | 0 | 1.91E-05 | 7.31E-05 | 0.00047 | 0.0001 | 0.00053 | 0.00028 | 0.00011 | 0 | 0 | 7.36E-05 |
| St6galnac4 | 9.42E-05 | 7.86E-05 | 0.00019 | 0.00011 | 7.67E-05 | 2.20E-05 | 0 | 0.00012 | 1.91E-05 | 2.30E-05 | 5.20E-05 | 0 | 7.47E-05 | 0 | 0 | 0 | 0 | 9.81E-05 |
| Eps15l1 | 2.59E-05 | 3.92E-05 | 4.20E-05 | 1.28E-05 | 2.58E-05 | 2.12E-05 | 2.17E-05 | 9.61E-06 | 1.91E-05 | 3.87E-05 | 2.00E-05 | 4.15E-05 | 4.41E-05 | 2.09E-05 | 0.00016 | 3.63E-07 | 0 | 2.08E-05 |
| Mtrf1l | 3.58E-05 | 5.02E-05 | 2.18E-05 | 1.03E-05 | 1.84E-05 | 1.99E-05 | 5.91E-06 | 1.43E-05 | 1.91E-05 | 2.58E-05 | 0 | 2.43E-05 | 1.81E-05 | 5.48E-06 | 0.00015 | 4.34E-05 | 0 | 1.55E-05 |
| Fdxacb1 | 1.76E-05 | 1.64E-05 | 1.81E-05 | 0 | 3.24E-06 | 1.40E-05 | 0 | 1.15E-06 | 1.91E-05 | 4.89E-06 | 1.43E-05 | 2.64E-06 | 0 | 1.84E-05 | 0 | 1.59E-05 | 0 | 1.54E-05 |
| Zfp983 | 1.30E-05 | 1.36E-06 | 3.04E-06 | 0 | 6.49E-06 | 1.13E-05 | 3.20E-06 | 0 | 1.91E-05 | 1.25E-05 | 0 | 1.16E-06 | 4.56E-06 | 8.25E-06 | 0 | 6.07E-07 | 0 | 5.08E-06 |
| Nprl2 | 3.22E-05 | 2.08E-05 | 3.42E-06 | 2.74E-05 | 9.61E-06 | 1.71E-06 | 0 | 0 | 1.91E-05 | 1.17E-05 | 8.38E-06 | 1.63E-05 | 1.45E-05 | 8.73E-06 | 1.87E-05 | 1.26E-05 | 0 | 1.83E-05 |
| Gng7 | 0 | 7.49E-06 | 0 | 0 | 1.45E-05 | 0 | 4.44E-06 | 2.12E-06 | 1.91E-05 | 4.81E-05 | 0 | 1.13E-05 | 3.31E-05 | 9.52E-06 | 0 | 0 | 0 | 1.01E-05 |
| Ubr2 | 0.00018 | 0.00031 | 0.00019 | 6.60E-05 | 0.00014 | 0.00014 | 0.0001 | 9.69E-05 | 1.90E-05 | 5.11E-05 | 0 | 0.00026 | 0 | 0.00011 | 0.00066 | 0.00013 | 0.00025 | 1.07E-05 |
| Ctdp1 | 4.95E-05 | 7.87E-05 | 9.88E-05 | 0 | 7.28E-05 | 2.56E-05 | 0 | 1.81E-05 | 1.90E-05 | 2.14E-07 | 0.00014 | 7.48E-05 | 0 | 3.13E-05 | 6.07E-05 | 6.46E-05 | 3.08E-05 | 0 |
| Stxbp4 | 6.66E-05 | 5.14E-05 | 0 | 0 | 5.20E-05 | 2.14E-05 | 2.15E-06 | 5.94E-06 | 1.90E-05 | 7.56E-06 | 0 | 2.70E-06 | 7.65E-06 | 0 | 6.52E-05 | 3.96E-05 | 0 | 1.76E-05 |
| Fam20c | 0 | 5.12E-05 | 7.40E-05 | 4.30E-05 | 0 | 1.14E-05 | 0 | 0.00023 | 1.90E-05 | 0.0001 | 0 | 6.44E-06 | 2.76E-05 | 7.08E-05 | 5.07E-05 | 4.20E-05 | 0.00018 | 0.00015 |
| Hpgd | 2.22E-05 | 4.13E-06 | 0 | 0 | 0 | 2.22E-06 | 0 | 0 | 1.90E-05 | 6.06E-05 | 0 | 0 | 1.19E-05 | 0.00011 | 0 | 0 | 0 | 4.34E-05 |
| Hcar2 | 0 | 1.19E-05 | 1.48E-05 | 7.82E-06 | 0 | 0 | 0 | 0 | 1.90E-05 | 1.01E-05 | 0 | 2.30E-05 | 8.58E-06 | 2.04E-05 | 9.46E-05 | 1.46E-05 | 3.76E-05 | 8.39E-06 |
| Med4 | 6.31E-05 | 7.83E-05 | 5.78E-05 | 2.78E-05 | 1.41E-05 | 5.60E-05 | 0 | 2.43E-05 | 1.89E-05 | 2.24E-05 | 3.82E-05 | 3.57E-05 | 4.55E-05 | 3.04E-05 | 8.48E-05 | 0.00023 | 0 | 1.60E-05 |
| Wdr47 | 1.76E-05 | 3.12E-05 | 1.87E-05 | 0 | 1.63E-05 | 8.08E-06 | 1.66E-05 | 0 | 1.89E-05 | 8.18E-06 | 2.71E-05 | 1.86E-05 | 4.71E-06 | 1.11E-05 | 1.79E-05 | 4.41E-05 | 7.88E-05 | 2.61E-05 |
| Smim24 | 0 | 2.35E-05 | 0 | 2.17E-08 | 2.19E-05 | 0 | 0 | 6.90E-06 | 1.89E-05 | 8.33E-06 | 0 | 2.61E-05 | 8.01E-06 | 7.59E-06 | 4.51E-06 | 0 | 1.57E-05 | 2.56E-05 |
| Cat | 0.00015 | 0.00021 | 0 | 0.00022 | 0 | 8.05E-05 | 2.30E-05 | 0.00021 | 1.88E-05 | 0.00019 | 0.0001 | 5.80E-05 | 0 | 5.47E-05 | 0 | 1.99E-05 | 0 | 0.00027 |
| Nrde2 | 1.21E-05 | 3.38E-05 | 3.87E-06 | 4.47E-05 | 7.96E-06 | 1.71E-05 | 2.50E-05 | 1.17E-05 | 1.88E-05 | 2.55E-05 | 0 | 1.66E-06 | 0 | 7.26E-06 | 8.93E-05 | 3.68E-05 | 0 | 6.57E-06 |
| Acad10 | 2.14E-06 | 1.50E-05 | 0 | 1.02E-05 | 9.74E-06 | 1.28E-06 | 0 | 0 | 1.88E-05 | 2.79E-05 | 7.06E-06 | 9.96E-06 | 9.54E-06 | 1.27E-05 | 0 | 0 | 0 | 1.92E-05 |
| Ficd | 2.18E-05 | 2.62E-05 | 4.59E-05 | 1.80E-05 | 3.86E-05 | 2.75E-08 | 2.50E-05 | 1.09E-05 | 1.88E-05 | 1.71E-05 | 2.68E-05 | 4.67E-05 | 0 | 2.43E-05 | 0 | 0 | 0 | 3.39E-05 |
| Rhobtb2 | 2.65E-05 | 4.56E-05 | 1.21E-05 | 2.64E-05 | 2.38E-05 | 0 | 3.06E-05 | 4.03E-05 | 1.88E-05 | 5.89E-05 | 0 | 0 | 1.77E-05 | 5.80E-06 | 3.02E-05 | 3.58E-05 | 1.81E-05 | 1.09E-05 |
| Sbno1 | 0.00015 | 0.00018 | 0.00015 | 5.71E-05 | 9.73E-05 | 9.07E-05 | 3.17E-05 | 3.82E-05 | 1.87E-05 | 5.84E-07 | 4.85E-05 | 0.00013 | 0 | 5.20E-05 | 0.0001 | 0.00011 | 0.00017 | 4.76E-05 |
| Baz2a | 0 | 0.00015 | 7.27E-05 | 0.00021 | 6.04E-05 | 7.98E-05 | 0 | 0.0002 | 1.87E-05 | 5.76E-05 | 6.21E-05 | 3.62E-05 | 0 | 0.0001 | 0.00032 | 0 | 0.00022 | 7.40E-06 |
| Tollip | 7.88E-05 | 0.00017 | 0.00013 | 0.00016 | 7.93E-05 | 6.05E-05 | 1.12E-05 | 0.00014 | 1.87E-05 | 9.07E-05 | 8.11E-05 | 0.0002 | 0 | 5.08E-05 | 6.57E-05 | 0.0001 | 0.00016 | 9.02E-05 |
| Arfip2 | 0.00018 | 0.00012 | 8.38E-05 | 0.00021 | 0.00012 | 6.00E-05 | 0.00038 | 3.04E-05 | 1.87E-05 | 0.0001 | 1.86E-05 | 2.66E-05 | 3.96E-05 | 0 | 0.00046 | 0 | 1.60E-05 | 0.00012 |
| Tbccd1 | 2.99E-05 | 3.02E-05 | 5.04E-06 | 2.01E-05 | 2.04E-05 | 1.32E-05 | 1.61E-05 | 1.02E-06 | 1.87E-05 | 2.18E-05 | 1.37E-05 | 6.47E-06 | 0 | 3.00E-05 | 0 | 3.31E-05 | 0 | 5.22E-06 |
| Enox2 | 2.32E-05 | 1.83E-05 | 0 | 7.82E-07 | 2.44E-06 | 1.21E-05 | 7.35E-06 | 1.39E-05 | 1.87E-05 | 3.55E-05 | 1.63E-05 | 2.69E-05 | 1.09E-06 | 4.18E-06 | 4.53E-06 | 1.34E-05 | 0 | 1.76E-05 |
| Angptl4 | 0.00015 | 0 | 0.00068 | 0.00013 | 0 | 0 | 0.00069 | 0.00035 | 1.87E-05 | 0.00043 | 0 | 0.00015 | 0.00172 | 0.0004 | 0.00017 | 0.00017 | 0 | 0.00053 |

| | | | | | | | | | | | | | | | | | | |
|---|---|---|---|---|---|---|---|---|---|---|---|---|---|---|---|---|---|---|
| Psph | 8.43E-05 | 9.31E-06 | 0.00011 | 3.58E-05 | 0 | 0 | 2.12E-06 | 0 | 1.87E-05 | 3.20E-05 | 0 | 4.46E-05 | 0 | 0 | 6.80E-05 | 4.08E-05 | 0 | 4.10E-05 |
| Rtkn | 7.72E-06 | 2.10E-05 | 6.46E-05 | 6.30E-05 | 2.50E-05 | 0 | 1.59E-05 | 0.00012 | 1.87E-05 | 1.73E-05 | 3.09E-06 | 0.00015 | 0 | 0 | 0.00012 | 1.03E-05 | 4.89E-06 | 0.00012 |
| Ubxn2b | 4.57E-05 | 2.31E-05 | 0 | 0 | 1.56E-05 | 0 | 0 | 2.32E-05 | 1.87E-05 | 1.75E-05 | 0 | 0 | 0 | 0 | 8.61E-05 | 2.60E-05 | 0 | 2.68E-05 |
| DEPRECATEI | 6.24E-05 | 7.37E-05 | 4.37E-05 | 0 | 2.45E-05 | 4.57E-05 | 0 | 0 | 1.86E-05 | 6.42E-07 | 1.11E-06 | 4.83E-05 | 0 | 2.48E-06 | 0.00013 | 1.74E-05 | 0.00017 | 2.14E-05 |
| Kif3c | 2.09E-05 | 6.74E-05 | 0 | 2.25E-05 | 3.32E-05 | 3.64E-05 | 0 | 4.86E-05 | 1.86E-05 | 8.18E-06 | 9.59E-05 | 3.49E-05 | 0 | 6.29E-05 | 6.19E-05 | 5.04E-06 | 7.29E-05 | 5.72E-05 |
| Stk4 | 4.29E-05 | 3.29E-05 | 2.75E-05 | 8.52E-06 | 2.90E-05 | 3.36E-05 | 0 | 4.18E-06 | 1.86E-05 | 1.85E-05 | 3.26E-05 | 5.41E-05 | 0 | 3.00E-05 | 0 | 4.48E-05 | 3.03E-05 | 3.62E-05 |
| Tlr3 | 2.94E-05 | 8.20E-06 | 0 | 0 | 0 | 2.67E-05 | 0 | 1.15E-05 | 1.86E-05 | 6.50E-05 | 0 | 0 | 4.01E-05 | 0 | 0 | 7.22E-07 | 0 | 4.99E-05 |
| Gemin8 | 4.94E-05 | 9.13E-06 | 1.64E-05 | 0 | 3.85E-06 | 1.44E-05 | 0 | 0 | 1.86E-05 | 0 | 0 | 8.31E-07 | 3.71E-06 | 8.80E-06 | 0 | 3.86E-05 | 1.27E-05 | 0 |
| Lrfn1 | 0 | 9.46E-06 | 1.93E-06 | 9.46E-06 | 3.87E-05 | 1.30E-05 | 0 | 0 | 1.86E-05 | 8.67E-06 | 1.61E-05 | 0 | 2.50E-05 | 2.60E-05 | 0 | 7.14E-06 | 1.54E-05 | 9.54E-06 |
| Ran | 4.39E-05 | 0 | 4.48E-05 | 0 | 0 | 8.64E-06 | 0 | 0 | 1.86E-05 | 0 | 1.81E-05 | 0 | 0 | 0 | 0 | 8.19E-05 | 0 | 0 |
| Pdss1 | 2.02E-05 | 1.32E-05 | 7.18E-05 | 0 | 0 | 4.38E-06 | 0 | 0 | 1.86E-05 | 7.98E-06 | 5.27E-06 | 9.21E-06 | 4.12E-06 | 0 | 3.68E-05 | 0.0001 | 0 | 3.10E-06 |
| Afg1l | 1.48E-05 | 2.84E-05 | 2.11E-05 | 1.48E-05 | 6.03E-06 | 3.01E-06 | 2.48E-05 | 1.21E-05 | 1.86E-05 | 3.84E-06 | 4.94E-06 | 6.02E-06 | 9.80E-06 | 0 | 2.10E-05 | 3.39E-06 | 1.65E-06 | 0 |
| Synrg | 3.27E-05 | 7.08E-05 | 1.50E-05 | 2.54E-05 | 4.13E-05 | 1.10E-06 | 1.62E-05 | 1.80E-06 | 1.86E-05 | 4.81E-05 | 7.37E-06 | 3.71E-05 | 0 | 0 | 8.52E-06 | 0 | 2.86E-05 | 4.28E-05 |
| Herc3 | 0 | 2.23E-05 | 0 | 0 | 1.70E-05 | 0 | 0 | 0 | 1.86E-05 | 0 | 3.34E-05 | 3.89E-05 | 0 | 0 | 0 | 5.06E-08 | 1.67E-05 | 6.33E-06 |
| Ahnak2 | 0 | 0.00061 | 0 | 2.24E-05 | 0.00029 | 0.00021 | 0.00011 | 0 | 1.85E-05 | 0.00037 | 5.70E-05 | 0.00033 | 0.00014 | 0.00012 | 0.00013 | 0 | 0 | 0.00047 |
| Rbm41 | 5.02E-05 | 6.66E-05 | 1.79E-05 | 0 | 1.41E-05 | 2.04E-05 | 0 | 1.37E-05 | 1.85E-05 | 1.06E-06 | 3.86E-05 | 3.84E-05 | 0 | 2.44E-05 | 0.00012 | 1.72E-05 | 5.35E-05 | 0 |
| BC024978 | 8.52E-06 | 3.36E-05 | 3.29E-05 | 4.01E-05 | 5.28E-05 | 2.04E-05 | 0 | 4.79E-05 | 1.85E-05 | 2.28E-05 | 1.77E-05 | 6.15E-06 | 3.52E-05 | 9.23E-06 | 8.43E-05 | 0 | 8.40E-05 | 2.94E-06 |
| Sdhaf1 | 3.24E-05 | 4.50E-05 | 6.82E-05 | 0.00017 | 4.10E-05 | 1.07E-05 | 8.95E-06 | 2.50E-05 | 1.85E-05 | 2.80E-05 | 2.29E-05 | 0 | 0 | 6.36E-05 | 2.26E-05 | 0 | 0 | 4.25E-05 |
| Pir | 1.59E-05 | 2.80E-05 | 0 | 0 | 6.87E-06 | 1.93E-06 | 0 | 0 | 1.85E-05 | 6.10E-05 | 0 | 6.79E-06 | 3.28E-05 | 4.24E-06 | 0 | 2.81E-05 | 4.38E-07 | 6.47E-05 |
| Fbxo24 | 0 | 6.70E-06 | 0 | 1.08E-05 | 1.01E-05 | 0 | 1.65E-06 | 0 | 1.85E-05 | 1.75E-05 | 1.34E-05 | 3.86E-05 | 3.34E-06 | 0 | 4.59E-05 | 0 | 0 | 1.37E-05 |
| Uso1 | 0.00032 | 8.71E-05 | 0.00033 | 0.00024 | 4.51E-05 | 7.54E-05 | 0.00031 | 7.41E-05 | 1.84E-05 | 4.31E-05 | 9.14E-05 | 0.00015 | 7.09E-05 | 3.63E-05 | 0.00029 | 1.10E-05 | 9.50E-05 | 8.35E-06 |
| Rgs7bp | 0 | 0 | 1.51E-05 | 0 | 0 | 2.31E-05 | 0 | 0 | 1.84E-05 | 6.86E-06 | 0 | 0 | 0 | 2.85E-05 | 0 | 0 | 0 | 0 |
| Mctp2 | 0 | 0 | 0 | 0 | 0 | 7.82E-06 | 0 | 0 | 1.84E-05 | 0 | 0 | 3.40E-06 | 5.35E-05 | 3.35E-06 | 0 | 0 | 0 | 4.46E-05 |
| Mettl17 | 1.10E-05 | 2.23E-05 | 1.62E-05 | 2.06E-05 | 1.91E-05 | 7.59E-06 | 1.59E-05 | 5.72E-06 | 1.84E-05 | 1.13E-05 | 5.39E-07 | 1.23E-06 | 0 | 5.86E-06 | 4.84E-05 | 5.11E-06 | 0 | 2.20E-05 |
| Tinf2 | 4.01E-05 | 5.69E-05 | 4.19E-05 | 0 | 1.81E-05 | 6.65E-06 | 9.16E-06 | 0 | 1.84E-05 | 1.28E-05 | 3.52E-06 | 3.83E-06 | 3.24E-05 | 7.29E-06 | 0.00014 | 0.00038 | 2.47E-05 | 1.35E-05 |
| Aar2 | 1.72E-05 | 2.03E-05 | 9.16E-06 | 2.26E-05 | 7.58E-06 | 6.58E-06 | 0 | 8.07E-06 | 1.84E-05 | 1.57E-05 | 1.57E-05 | 1.02E-06 | 0 | 1.62E-05 | 6.10E-06 | 1.64E-05 | 2.89E-06 | 5.51E-07 |
| Ube2cbp | 2.23E-05 | 7.86E-07 | 2.69E-05 | 7.30E-06 | 0 | 6.03E-06 | 0 | 0 | 1.84E-05 | 0 | 0 | 0 | 0 | 6.19E-06 | 0 | 4.01E-09 | 0 | 0 |
| Pigl | 2.79E-06 | 1.53E-05 | 8.00E-06 | 1.33E-05 | 6.74E-06 | 1.66E-06 | 0 | 1.63E-05 | 1.84E-05 | 5.71E-06 | 9.05E-06 | 0 | 7.86E-06 | 2.82E-06 | 0 | 1.35E-05 | 0 | 5.33E-06 |
| Wasf1 | 1.04E-06 | 2.36E-05 | 1.34E-05 | 3.07E-05 | 0 | 0 | 0 | 7.56E-06 | 1.84E-05 | 4.00E-05 | 2.51E-05 | 1.01E-05 | 0 | 8.25E-05 | 3.39E-06 | 2.61E-05 | 9.14E-06 | 2.51E-06 |
| Sf1 | 4.90E-05 | 1.80E-05 | 4.13E-05 | 2.40E-06 | 6.86E-06 | 0 | 0 | 0 | 1.84E-05 | 0 | 9.81E-07 | 3.48E-05 | 0 | 0 | 3.16E-05 | 2.39E-05 | 2.65E-05 | 0 |
| Gpr35 | 0 | 0 | 1.01E-05 | 0 | 2.14E-05 | 0.00016 | 3.93E-05 | 1.06E-05 | 1.83E-05 | 0 | 0 | 0.00056 | 5.34E-05 | 0 | 0 | 0 | 0 | 0 |
| Cacnb1 | 8.41E-07 | 4.07E-05 | 3.05E-05 | 0 | 4.35E-05 | 7.30E-05 | 0 | 5.84E-06 | 1.83E-05 | 1.20E-05 | 3.54E-05 | 0.00012 | 0 | 0 | 6.72E-05 | 0 | 0 | 3.21E-05 |
| Zfp68 | 0.00012 | 8.20E-05 | 2.33E-05 | 0 | 2.14E-05 | 5.36E-05 | 1.94E-05 | 0 | 1.83E-05 | 5.77E-06 | 7.21E-07 | 4.20E-05 | 5.27E-06 | 7.63E-06 | 4.52E-05 | 3.37E-05 | 0 | 2.19E-05 |

| | | | | | | | | | | | | | | | | | | |
|---|---|---|---|---|---|---|---|---|---|---|---|---|---|---|---|---|---|---|
| Mfap1b | 3.60E-05 | 4.27E-05 | 2.43E-05 | 5.20E-05 | 2.37E-05 | 1.68E-05 | 0 | 1.60E-05 | 1.83E-05 | 1.98E-05 | 1.62E-05 | 3.55E-05 | 1.22E-05 | 1.14E-05 | 1.39E-05 | 5.10E-06 | 0 | 2.42E-05 |
| Stx1a | 8.12E-07 | 7.52E-05 | 2.69E-05 | 3.92E-05 | 4.95E-05 | 5.18E-06 | 0 | 0.0001 | 1.83E-05 | 1.76E-05 | 5.75E-05 | 4.98E-06 | 0 | 0 | 0.00036 | 0 | 1.13E-07 | 1.43E-05 |
| Tgfbr3l | 0 | 2.30E-05 | 1.86E-06 | 8.36E-06 | 7.97E-06 | 0 | 3.84E-06 | 0 | 1.83E-05 | 6.49E-06 | 0 | 3.65E-05 | 2.33E-06 | 0 | 6.88E-05 | 0 | 0 | 1.13E-05 |
| Tnik | 0 | 1.09E-05 | 0 | 0 | 1.38E-05 | 0 | 0 | 5.72E-05 | 1.83E-05 | 8.11E-05 | 5.43E-05 | 1.28E-05 | 2.29E-05 | 1.41E-05 | 2.23E-06 | 0 | 8.22E-06 | 0 |
| Snapc4 | 3.59E-05 | 6.08E-05 | 4.34E-05 | 1.05E-05 | 0 | 0 | 0 | 0 | 1.83E-05 | 1.35E-05 | 2.49E-06 | 4.88E-05 | 5.30E-06 | 1.45E-05 | 0.00036 | 3.35E-05 | 4.40E-05 | 2.18E-05 |
| Ccnl2 | 0.00016 | 0.00042 | 7.85E-05 | 9.65E-05 | 0.00031 | 0.0002 | 0.00016 | 0.00013 | 1.82E-05 | 0.00013 | 0.0001 | 0.00033 | 3.72E-05 | 0.00018 | 0.00111 | 5.03E-05 | 0.00041 | 0.00012 |
| Tmem192 | 8.07E-05 | 7.70E-05 | 0 | 1.64E-05 | 2.55E-05 | 4.79E-05 | 3.27E-05 | 0 | 1.82E-05 | 5.36E-06 | 0 | 3.34E-05 | 1.34E-05 | 9.59E-05 | 0 | 1.45E-05 | 0 | 1.55E-05 |
| Eed | 0.0001 | 8.96E-05 | 7.63E-05 | 0 | 4.13E-06 | 3.80E-05 | 0 | 0 | 1.82E-05 | 2.19E-05 | 4.59E-05 | 6.62E-05 | 7.38E-06 | 0 | 5.07E-05 | 0.00013 | 0 | 0 |
| Ddx20 | 3.27E-05 | 5.49E-05 | 3.16E-05 | 0 | 1.75E-06 | 2.68E-05 | 0 | 0 | 1.82E-05 | 8.32E-06 | 6.35E-06 | 8.39E-06 | 1.98E-06 | 2.61E-06 | 5.86E-05 | 0.00015 | 0 | 0 |
| Tab3 | 1.73E-05 | 3.49E-05 | 1.83E-05 | 0 | 1.17E-05 | 2.56E-05 | 2.38E-05 | 1.44E-05 | 1.82E-05 | 7.04E-06 | 2.11E-05 | 1.51E-05 | 0 | 0 | 3.67E-05 | 4.39E-06 | 7.47E-05 | 0 |
| Ttc30b | 2.91E-05 | 3.55E-05 | 0 | 2.31E-05 | 2.16E-05 | 1.87E-05 | 3.65E-05 | 0 | 1.82E-05 | 3.72E-05 | 0 | 5.45E-06 | 9.55E-06 | 8.46E-08 | 1.71E-05 | 1.44E-06 | 1.68E-05 | 7.93E-05 |
| Il7 | 9.35E-06 | 0 | 7.87E-07 | 0 | 0 | 1.04E-05 | 0 | 6.07E-06 | 1.82E-05 | 0 | 0 | 0 | 2.62E-08 | 1.10E-05 | 0 | 0 | 0 | 5.17E-06 |
| Cdc7 | 9.26E-06 | 9.72E-06 | 0 | 0 | 3.49E-06 | 4.22E-06 | 2.98E-05 | 0 | 1.82E-05 | 2.50E-05 | 0 | 0 | 5.49E-05 | 3.00E-06 | 4.03E-05 | 0.00029 | 0 | 0 |
| Bivm | 0.00012 | 3.21E-05 | 3.22E-06 | 0 | 6.75E-06 | 5.77E-05 | 7.11E-06 | 2.43E-05 | 1.81E-05 | 0 | 1.32E-05 | 4.34E-05 | 0 | 0 | 4.05E-05 | 4.70E-07 | 0 | 0 |
| Mettl4 | 7.48E-05 | 5.81E-05 | 4.48E-05 | 7.13E-05 | 6.00E-05 | 3.74E-05 | 7.23E-05 | 1.84E-05 | 1.81E-05 | 2.04E-05 | 3.40E-05 | 3.78E-05 | 1.89E-05 | 1.47E-05 | 0 | 0.00011 | 4.60E-06 | 8.16E-06 |
| Cspp1 | 2.93E-05 | 8.35E-05 | 3.85E-06 | 2.00E-05 | 4.29E-05 | 2.72E-05 | 2.02E-05 | 0 | 1.81E-05 | 3.01E-05 | 2.16E-05 | 2.80E-06 | 2.66E-05 | 1.70E-05 | 0.00031 | 0.00011 | 5.40E-05 | 2.22E-05 |
| Rnf157 | 9.11E-06 | 4.21E-05 | 0 | 0 | 9.95E-05 | 2.49E-05 | 2.45E-05 | 2.43E-05 | 1.81E-05 | 0 | 3.67E-05 | 1.92E-05 | 0 | 5.63E-05 | 0 | 0.00013 | 0 | 0 |
| Zfp169 | 2.39E-05 | 3.81E-05 | 0 | 1.23E-05 | 3.85E-05 | 2.15E-05 | 1.26E-05 | 0 | 1.81E-05 | 7.47E-06 | 3.90E-06 | 1.38E-05 | 0 | 1.35E-05 | 6.76E-05 | 0 | 0 | 3.34E-05 |
| Ddit4l | 0 | 0 | 0 | 1.31E-05 | 3.63E-06 | 0 | 0.00013 | 0.00019 | 1.81E-05 | 0 | 0 | 0 | 0 | 0 | 0 | 1.39E-05 | 0 | 0.00012 |
| Map1lc3b | 0.00046 | 0.0006 | 0 | 0 | 0.0004 | 0.00062 | 0.00029 | 0.00016 | 1.80E-05 | 0.00033 | 5.04E-05 | 0.0004 | 0.0004 | 0.00025 | 0 | 0 | 0.0005 | 0.00024 |
| Ccnt2 | 0.00013 | 0.00016 | 5.28E-05 | 0 | 9.16E-05 | 0.00011 | 0 | 3.87E-05 | 1.80E-05 | 2.33E-05 | 3.04E-05 | 0.00011 | 0 | 1.35E-05 | 0.00051 | 1.80E-05 | 0 | 6.54E-06 |
| Kdm4b | 5.32E-05 | 0.00013 | 0 | 6.01E-05 | 9.72E-05 | 6.45E-05 | 0 | 3.61E-05 | 1.80E-05 | 2.73E-05 | 7.18E-06 | 0.00037 | 3.11E-05 | 2.62E-05 | 0.00011 | 0 | 2.33E-05 | 3.66E-05 |
| Baz2b | 6.24E-05 | 0.00013 | 2.05E-05 | 2.64E-05 | 2.75E-05 | 4.89E-05 | 4.22E-06 | 7.22E-06 | 1.80E-05 | 1.89E-05 | 4.37E-05 | 3.13E-05 | 0 | 2.99E-05 | 9.31E-05 | 2.70E-05 | 8.95E-05 | 1.18E-05 |
| Tshz1 | 4.41E-05 | 7.80E-05 | 8.83E-06 | 4.01E-05 | 4.07E-05 | 3.54E-05 | 4.34E-05 | 1.39E-05 | 1.80E-05 | 8.46E-05 | 2.03E-05 | 2.80E-05 | 5.92E-05 | 0 | 2.49E-05 | 0.00016 | 6.12E-05 | 2.80E-05 |
| Slc35d1 | 4.76E-05 | 7.44E-05 | 8.12E-05 | 0 | 3.46E-05 | 2.70E-05 | 4.44E-06 | 2.09E-05 | 1.80E-05 | 1.94E-05 | 5.07E-05 | 3.19E-08 | 2.43E-05 | 2.73E-06 | 4.36E-05 | 3.20E-05 | 1.14E-05 | 2.68E-05 |
| Paxbp1 | 5.56E-05 | 0.00017 | 0.0001 | 8.28E-06 | 7.97E-05 | 2.70E-05 | 3.79E-06 | 3.20E-06 | 1.80E-05 | 2.30E-05 | 1.46E-05 | 7.82E-05 | 0 | 2.31E-05 | 0.00058 | 0.00011 | 0.00018 | 2.63E-05 |
| Nt5m | 5.96E-05 | 7.04E-05 | 7.72E-05 | 6.39E-05 | 4.26E-05 | 1.73E-05 | 3.96E-05 | 1.92E-06 | 1.80E-05 | 2.94E-05 | 2.16E-05 | 2.55E-05 | 6.36E-06 | 1.30E-05 | 2.36E-05 | 4.67E-05 | 0 | 4.05E-05 |
| Mblac1 | 1.67E-06 | 4.30E-06 | 7.26E-06 | 4.43E-05 | 2.13E-05 | 6.55E-06 | 1.08E-06 | 1.38E-05 | 1.80E-05 | 1.04E-05 | 2.89E-05 | 0 | 1.69E-05 | 1.97E-05 | 1.79E-05 | 8.78E-06 | 0 | 1.25E-05 |
| Lpar4 | 4.15E-05 | 5.69E-06 | 0 | 0 | 9.66E-06 | 5.77E-06 | 0 | 0 | 1.80E-05 | 0.00013 | 2.82E-05 | 0 | 0 | 0 | 0 | 1.92E-05 | 0 | 0 |
| Mfsd7a | 0 | 3.05E-05 | 0 | 0 | 0 | 4.02E-06 | 1.43E-05 | 0 | 1.80E-05 | 0.00011 | 0 | 2.04E-06 | 7.67E-06 | 0 | 1.79E-05 | 0 | 0 | 4.59E-05 |
| Pkp4 | 3.04E-05 | 2.76E-05 | 5.71E-05 | 0 | 0 | 1.89E-06 | 0 | 0 | 1.80E-05 | 0 | 3.39E-05 | 6.24E-07 | 0 | 5.40E-05 | 0 | 0.00016 | 1.59E-05 | 0 |
| Proser3 | 5.18E-07 | 1.85E-05 | 0 | 8.16E-07 | 2.12E-06 | 9.77E-07 | 0 | 9.86E-07 | 1.80E-05 | 2.92E-06 | 0 | 9.63E-06 | 0 | 8.83E-06 | 3.63E-05 | 7.07E-05 | 0 | 5.51E-06 |
| Setmar | 0 | 3.86E-06 | 0 | 0 | 1.27E-05 | 0 | 1.20E-05 | 0 | 1.80E-05 | 0 | 3.27E-06 | 0 | 0 | 2.86E-05 | 8.91E-06 | 0 | 0 | 1.26E-05 |

| | | | | | | | | | | | | | | | | | | |
|---|---|---|---|---|---|---|---|---|---|---|---|---|---|---|---|---|---|---|
| Osbpl7 | 4.26E-06 | 4.25E-05 | 1.28E-05 | 2.00E-05 | 2.10E-05 | 0 | 6.96E-06 | 8.13E-06 | 1.80E-05 | 2.29E-05 | 1.17E-05 | 0 | 0 | 9.85E-06 | 3.89E-05 | 0 | 1.95E-05 | 0 |
| Phf21a | 9.10E-05 | 0.00023 | 0 | 8.04E-05 | 0.00012 | 0.00017 | 0.00013 | 4.80E-05 | 1.79E-05 | 5.19E-05 | 0 | 0.00021 | 5.65E-05 | 5.43E-05 | 0.00021 | 0 | 0.00014 | 5.85E-05 |
| Pdrg1 | 9.88E-05 | 8.45E-05 | 0.00035 | 0.00012 | 0.00023 | 0.00016 | 0.00024 | 0.00034 | 1.79E-05 | 8.15E-05 | 1.48E-05 | 6.51E-05 | 1.09E-05 | 5.23E-05 | 2.14E-05 | 0 | 3.75E-05 | 0.00016 |
| Fbxo8 | 0.00013 | 8.29E-05 | 0 | 0 | 6.07E-05 | 0.00011 | 6.45E-05 | 1.38E-05 | 1.79E-05 | 4.19E-05 | 1.27E-05 | 3.34E-05 | 2.15E-05 | 1.78E-05 | 3.54E-05 | 2.16E-05 | 0 | 3.17E-05 |
| Mre11a | 5.02E-05 | 2.97E-05 | 4.48E-05 | 4.73E-07 | 4.80E-06 | 4.16E-05 | 0 | 6.21E-06 | 1.79E-05 | 2.96E-05 | 1.87E-05 | 1.97E-05 | 4.10E-05 | 1.75E-05 | 5.08E-05 | 0.00025 | 0 | 7.74E-06 |
| Dennd1b | 3.36E-05 | 6.77E-05 | 4.78E-06 | 1.52E-05 | 6.93E-05 | 3.66E-05 | 0 | 8.43E-05 | 1.79E-05 | 4.24E-05 | 1.55E-05 | 7.42E-05 | 5.41E-06 | 6.62E-05 | 2.64E-05 | 1.90E-05 | 5.24E-05 | 5.95E-05 |
| Ncbp1 | 0.00015 | 0.00011 | 0.00021 | 3.54E-05 | 3.97E-05 | 2.75E-05 | 1.29E-05 | 5.00E-06 | 1.79E-05 | 3.27E-05 | 2.40E-05 | 0 | 3.02E-05 | 3.42E-05 | 7.49E-05 | 0.00021 | 3.20E-05 | 3.26E-05 |
| C9orf72 | 1.75E-05 | 3.46E-05 | 2.36E-05 | 0 | 7.75E-06 | 2.07E-05 | 0 | 0 | 1.79E-05 | 6.75E-06 | 2.84E-05 | 2.84E-05 | 1.28E-05 | 1.07E-05 | 1.88E-05 | 0 | 1.22E-05 | 3.46E-06 |
| Thsd4 | 0 | 2.33E-06 | 0 | 0 | 0 | 1.32E-05 | 0 | 0 | 1.79E-05 | 3.24E-05 | 0 | 8.87E-06 | 0 | 0 | 0 | 0 | 3.65E-06 | 0 |
| Primpol | 2.66E-05 | 1.22E-05 | 0 | 3.55E-06 | 1.83E-05 | 1.22E-05 | 1.35E-05 | 3.98E-05 | 1.79E-05 | 1.65E-05 | 0 | 2.51E-05 | 1.44E-05 | 0 | 4.08E-06 | 2.77E-05 | 0 | 1.68E-05 |
| Fbxo45 | 2.98E-05 | 3.51E-05 | 5.11E-05 | 0 | 7.36E-06 | 6.66E-06 | 2.03E-05 | 2.41E-05 | 1.79E-05 | 5.86E-06 | 2.45E-05 | 3.29E-05 | 1.75E-05 | 1.11E-05 | 4.68E-05 | 9.11E-05 | 7.40E-06 | 2.03E-05 |
| Sft2d3 | 2.00E-06 | 1.47E-05 | 9.20E-06 | 3.64E-05 | 1.83E-07 | 2.28E-06 | 6.02E-06 | 1.87E-05 | 1.79E-05 | 1.10E-06 | 1.14E-05 | 1.06E-05 | 0 | 2.28E-07 | 3.93E-05 | 0 | 0 | 1.67E-06 |
| Zfp7 | 1.26E-06 | 6.57E-06 | 1.20E-06 | 5.31E-07 | 0 | 1.78E-06 | 0 | 3.47E-06 | 1.79E-05 | 2.22E-06 | 8.47E-06 | 1.31E-05 | 0 | 7.17E-06 | 2.95E-05 | 8.44E-07 | 0 | 0 |
| Nfs1 | 7.90E-05 | 6.79E-05 | 5.26E-05 | 4.58E-05 | 2.47E-05 | 5.21E-05 | 1.83E-05 | 1.20E-05 | 1.78E-05 | 2.72E-05 | 7.22E-06 | 4.92E-05 | 0 | 1.83E-06 | 6.79E-05 | 3.01E-05 | 0 | 2.48E-05 |
| Bcor | 7.25E-05 | 0.00013 | 2.70E-05 | 8.40E-05 | 7.49E-05 | 4.99E-05 | 5.32E-05 | 8.15E-06 | 1.78E-05 | 1.21E-05 | 8.05E-05 | 0.00015 | 4.68E-05 | 5.08E-05 | 1.18E-05 | 8.35E-05 | 5.24E-05 | 2.32E-06 |
| Umad1 | 4.22E-05 | 4.14E-05 | 2.02E-05 | 5.12E-07 | 2.46E-05 | 3.91E-05 | 1.37E-05 | 2.33E-05 | 1.78E-05 | 4.90E-05 | 2.25E-05 | 1.29E-05 | 7.94E-05 | 0 | 6.48E-05 | 7.83E-05 | 2.93E-05 | 1.49E-05 |
| DEPRECATEI | 4.55E-05 | 2.61E-05 | 2.18E-06 | 0 | 1.80E-05 | 3.47E-05 | 1.08E-05 | 1.51E-05 | 1.78E-05 | 9.47E-06 | 1.92E-05 | 2.67E-05 | 0 | 1.72E-05 | 3.99E-05 | 1.37E-05 | 0 | 0 |
| Arhgap39 | 7.90E-06 | 5.20E-05 | 1.77E-05 | 3.98E-05 | 2.92E-05 | 1.41E-05 | 0 | 2.75E-05 | 1.78E-05 | 7.82E-07 | 4.08E-05 | 1.89E-05 | 0 | 3.39E-05 | 4.55E-05 | 0 | 1.03E-05 | 1.23E-05 |
| Gpatch1 | 1.61E-05 | 2.06E-05 | 1.06E-05 | 0 | 9.38E-06 | 7.08E-06 | 0 | 0 | 1.78E-05 | 1.02E-05 | 5.48E-07 | 1.16E-05 | 5.20E-06 | 8.80E-06 | 2.76E-05 | 3.92E-06 | 0 | 0 |
| Ctf1 | 2.93E-06 | 0 | 0 | 2.42E-05 | 2.70E-05 | 0 | 4.58E-05 | 1.89E-05 | 1.78E-05 | 0 | 7.60E-06 | 0 | 9.34E-06 | 1.90E-06 | 0 | 3.12E-06 | 0 | 2.55E-05 |
| Sar1b | 0.00036 | 0.00014 | 0.00021 | 0.00014 | 0.00012 | 0.00022 | 0.00013 | 5.14E-05 | 1.77E-05 | 5.99E-05 | 7.85E-05 | 9.70E-05 | 0.00017 | 7.18E-05 | 0 | 4.89E-05 | 0 | 0 |
| Otud5 | 0.00014 | 0.00013 | 6.84E-05 | 5.82E-06 | 3.84E-05 | 6.93E-05 | 4.77E-05 | 1.60E-05 | 1.77E-05 | 2.82E-05 | 3.71E-05 | 9.13E-05 | 0 | 1.61E-05 | 0.00036 | 5.13E-05 | 5.47E-05 | 2.68E-06 |
| Apaf1 | 0.00014 | 0.00015 | 4.78E-05 | 1.24E-05 | 5.89E-05 | 5.79E-05 | 0.00014 | 0.00012 | 1.77E-05 | 6.86E-05 | 7.31E-06 | 0.00016 | 0 | 5.75E-05 | 0.0002 | 0.00019 | 0.00022 | 4.32E-05 |
| Ezh2 | 4.69E-05 | 3.76E-05 | 6.27E-05 | 0 | 3.46E-05 | 2.78E-05 | 0 | 8.71E-06 | 1.77E-05 | 0 | 0 | 0 | 0 | 2.21E-05 | 0 | 0.00038 | 4.77E-05 | 7.66E-06 |
| Phf7 | 2.48E-05 | 1.89E-05 | 0 | 1.08E-05 | 1.60E-05 | 2.60E-05 | 7.85E-06 | 5.49E-07 | 1.77E-05 | 0 | 0 | 8.76E-06 | 0 | 0 | 0 | 2.97E-05 | 0 | 0 |
| Mcts2 | 7.12E-05 | 4.51E-05 | 5.11E-05 | 0 | 0 | 2.12E-05 | 0 | 7.51E-06 | 1.77E-05 | 0 | 5.32E-06 | 3.75E-05 | 0 | 1.93E-05 | 0.00016 | 7.08E-06 | 3.48E-05 | 1.03E-05 |
| Hoxd4 | 0 | 1.42E-05 | 0 | 0 | 1.48E-05 | 1.62E-05 | 0 | 0 | 1.77E-05 | 6.83E-06 | 6.85E-06 | 0 | 0 | 6.64E-05 | 3.50E-05 | 0 | 0 | 6.93E-05 |
| Omd | 1.17E-05 | 3.54E-08 | 0 | 0 | 0 | 0 | 0 | 1.27E-06 | 1.77E-05 | 2.43E-05 | 4.42E-07 | 0 | 0 | 4.09E-06 | 0 | 0 | 0 | 4.01E-05 |
| Herc6 | 2.13E-05 | 2.47E-05 | 0 | 0 | 1.96E-06 | 0 | 0 | 0 | 1.77E-05 | 8.54E-05 | 0 | 0 | 2.62E-05 | 1.63E-05 | 2.30E-05 | 8.04E-05 | 0 | 6.22E-05 |
| Rilp | 0 | 1.58E-05 | 1.58E-05 | 0 | 5.88E-06 | 0 | 4.85E-07 | 7.10E-06 | 1.77E-05 | 2.22E-05 | 0 | 5.25E-06 | 1.52E-05 | 2.50E-06 | 2.28E-05 | 0 | 0 | 3.72E-05 |
| Ppp4r3b | 0.00019 | 0.00011 | 3.04E-05 | 0 | 3.25E-05 | 0.00015 | 4.21E-05 | 4.47E-05 | 1.76E-05 | 1.97E-05 | 1.28E-05 | 0.00011 | 0 | 3.73E-05 | 5.94E-05 | 5.81E-05 | 0 | 5.74E-06 |
| Epc2 | 0.00013 | 0.00012 | 3.10E-05 | 0 | 2.58E-05 | 6.06E-05 | 2.57E-05 | 3.05E-05 | 1.76E-05 | 2.82E-06 | 0 | 6.72E-05 | 0 | 2.92E-05 | 1.25E-05 | 8.46E-05 | 0.00012 | 1.42E-05 |
| Tmem39b | 5.13E-05 | 5.63E-05 | 6.84E-05 | 8.37E-05 | 0 | 3.71E-05 | 0 | 2.47E-05 | 1.76E-05 | 1.33E-05 | 7.47E-06 | 2.02E-05 | 4.13E-05 | 3.66E-06 | 1.85E-05 | 6.07E-05 | 0.0001 | 4.73E-06 |

| | | | | | | | | | | | | | | | | | | |
|---|---|---|---|---|---|---|---|---|---|---|---|---|---|---|---|---|---|---|
| Trdmt1 | 3.27E-05 | 5.19E-06 | 2.07E-06 | 0 | 7.73E-06 | 1.14E-05 | 0 | 0 | 1.76E-05 | 1.32E-06 | 2.02E-05 | 3.19E-06 | 0 | 4.94E-06 | 2.50E-05 | 4.17E-05 | 0 | 1.44E-05 |
| Ajuba | 7.06E-06 | 5.69E-05 | 3.26E-05 | 7.45E-05 | 5.48E-05 | 0 | 3.78E-05 | 0 | 1.76E-05 | 0 | 0.00037 | 1.43E-05 | 3.32E-05 | 0.00012 | 8.68E-05 | 0 | 0 | 4.13E-05 |
| Notum | 0 | 7.18E-05 | 0 | 0 | 0 | 0 | 0 | 0 | 1.76E-05 | 0 | 0 | 0 | 0 | 9.60E-06 | 0 | 0 | 0 | 0 |
| Dhx32 | 4.20E-05 | 2.53E-05 | 5.22E-05 | 2.47E-05 | 8.65E-05 | 0 | 8.77E-05 | 3.57E-05 | 1.76E-05 | 6.94E-05 | 4.65E-05 | 1.27E-05 | 1.86E-05 | 0 | 2.78E-05 | 5.98E-06 | 0 | 9.02E-07 |
| Nvl | 0.00017 | 0.00012 | 0.00016 | 6.29E-05 | 6.16E-05 | 6.41E-05 | 4.81E-05 | 2.43E-05 | 1.75E-05 | 2.33E-05 | 3.10E-05 | 7.35E-05 | 6.27E-05 | 4.08E-05 | 0.00021 | 0.00015 | 0 | 4.96E-05 |
| Mill2 | 0 | 4.29E-05 | 0 | 0 | 0 | 4.53E-05 | 0 | 0 | 1.75E-05 | 0.00011 | 0 | 2.96E-05 | 0 | 0 | 6.38E-06 | 0 | 0 | 7.89E-05 |
| Hdac8 | 6.29E-05 | 7.06E-05 | 0 | 0 | 4.69E-05 | 2.88E-05 | 2.06E-05 | 1.79E-05 | 1.75E-05 | 1.07E-05 | 0 | 3.77E-05 | 2.56E-05 | 1.70E-05 | 8.62E-05 | 0 | 1.23E-05 | 9.00E-06 |
| Hus1 | 2.66E-05 | 6.83E-06 | 1.05E-05 | 0 | 9.19E-06 | 1.11E-05 | 1.51E-06 | 0 | 1.75E-05 | 0 | 0 | 0 | 0 | 2.10E-05 | 1.87E-06 | 2.60E-05 | 0 | 0 |
| Bloc1s3 | 0 | 2.40E-05 | 0 | 7.05E-05 | 3.00E-05 | 7.45E-06 | 0 | 7.46E-06 | 1.75E-05 | 1.22E-05 | 1.61E-05 | 1.87E-05 | 0 | 1.23E-05 | 4.24E-05 | 0 | 0 | 1.49E-05 |
| Atp11b | 3.56E-05 | 7.96E-05 | 3.30E-05 | 2.22E-05 | 5.16E-05 | 3.70E-06 | 0 | 2.16E-05 | 1.75E-05 | 1.69E-05 | 2.91E-05 | 0.00015 | 0 | 1.12E-05 | 6.48E-05 | 5.76E-05 | 4.98E-05 | 8.95E-05 |
| Tmem220 | 4.90E-06 | 7.65E-06 | 0 | 4.20E-06 | 1.65E-06 | 3.65E-06 | 5.76E-06 | 0 | 1.75E-05 | 2.02E-05 | 0 | 1.09E-05 | 2.82E-05 | 2.90E-06 | 0 | 0 | 7.75E-06 | 2.22E-05 |
| Pigw | 1.80E-05 | 5.20E-06 | 2.63E-05 | 4.46E-06 | 6.26E-06 | 1.92E-06 | 0 | 0 | 1.75E-05 | 2.07E-06 | 3.23E-06 | 0 | 0 | 5.32E-06 | 2.20E-05 | 0 | 0 | 0 |
| Cntfr | 6.34E-06 | 2.73E-05 | 2.16E-05 | 0 | 5.99E-05 | 0 | 0.00014 | 2.49E-05 | 1.75E-05 | 0 | 0.0001 | 0 | 0 | 0 | 0 | 2.67E-06 | 1.85E-05 | 0 |
| Mlh1 | 2.74E-05 | 9.95E-06 | 4.34E-05 | 4.63E-05 | 1.17E-06 | 0 | 2.43E-05 | 0 | 1.75E-05 | 1.90E-05 | 0 | 0 | 1.04E-06 | 0 | 8.09E-05 | 8.80E-05 | 1.84E-05 | 0 |
| Rassf3 | 1.95E-05 | 3.69E-05 | 1.57E-05 | 1.35E-05 | 0 | 0 | 0 | 1.58E-05 | 1.75E-05 | 1.15E-05 | 4.44E-05 | 2.73E-05 | 0 | 0 | 0 | 0.00012 | 0 | 0 |
| Jph1 | 0 | 6.87E-06 | 3.11E-06 | 0 | 0 | 0 | 0 | 0 | 1.75E-05 | 3.68E-06 | 4.77E-05 | 2.09E-05 | 3.77E-06 | 0 | 8.02E-07 | 5.84E-06 | 3.63E-05 | 0 |
| Zfp280c | 6.26E-05 | 4.29E-05 | 1.56E-05 | 0 | 0 | 3.17E-05 | 0 | 0 | 1.74E-05 | 0 | 0 | 2.29E-05 | 0 | 1.53E-05 | 0.00012 | 1.78E-05 | 2.46E-05 | 6.03E-06 |
| Sirt3 | 3.07E-06 | 4.64E-05 | 0 | 3.71E-05 | 3.39E-05 | 1.40E-05 | 3.66E-06 | 1.30E-05 | 1.74E-05 | 2.66E-05 | 1.12E-05 | 3.03E-05 | 0 | 1.43E-05 | 3.06E-05 | 1.52E-05 | 3.52E-05 | 5.66E-05 |
| Exoc1 | 0.00011 | 0.00014 | 0.00012 | 5.08E-05 | 8.14E-06 | 1.07E-05 | 3.01E-05 | 5.67E-05 | 1.74E-05 | 4.92E-05 | 4.63E-05 | 1.70E-05 | 2.09E-05 | 2.15E-05 | 0.00015 | 7.31E-05 | 1.79E-05 | 4.03E-05 |
| Rpusd3 | 1.46E-05 | 1.73E-05 | 2.65E-05 | 3.35E-05 | 1.55E-05 | 8.08E-06 | 0 | 0 | 1.74E-05 | 1.37E-05 | 5.71E-06 | 0 | 0 | 1.72E-05 | 1.24E-05 | 9.53E-06 | 0 | 1.48E-05 |
| Zfp213 | 2.18E-05 | 3.05E-05 | 3.76E-05 | 2.50E-05 | 5.07E-06 | 5.39E-06 | 4.89E-06 | 1.71E-05 | 1.74E-05 | 7.07E-06 | 1.24E-05 | 1.07E-05 | 4.67E-06 | 5.04E-06 | 3.13E-05 | 6.14E-05 | 0 | 1.77E-05 |
| Actr5 | 1.61E-05 | 5.64E-05 | 2.91E-05 | 2.49E-05 | 2.38E-05 | 2.27E-06 | 1.60E-05 | 1.59E-06 | 1.74E-05 | 1.45E-05 | 0 | 5.60E-06 | 0 | 1.70E-05 | 2.79E-05 | 2.03E-05 | 0 | 1.97E-05 |
| Dock6 | 9.99E-06 | 0.00011 | 0 | 0 | 6.49E-06 | 0 | 0 | 0.00022 | 1.74E-05 | 2.71E-05 | 1.56E-05 | 6.64E-05 | 0 | 0 | 0.00023 | 0 | 3.50E-05 | 0.00023 |
| Mmp28 | 0 | 2.33E-05 | 0 | 0 | 0 | 0 | 0 | 9.71E-06 | 1.74E-05 | 0 | 0 | 0 | 0 | 1.68E-05 | 1.82E-06 | 0 | 0 | 3.49E-05 |
| Polr1e | 5.40E-05 | 3.12E-05 | 7.99E-05 | 3.02E-05 | 5.37E-06 | 0 | 0 | 0 | 1.74E-05 | 1.16E-06 | 6.49E-06 | 0 | 0 | 4.98E-06 | 1.82E-05 | 2.69E-05 | 0 | 0 |
| Cacul1 | 0.0001 | 9.77E-05 | 2.86E-05 | 1.77E-05 | 7.11E-05 | 5.08E-05 | 3.09E-05 | 3.04E-05 | 1.73E-05 | 3.52E-05 | 5.08E-05 | 9.66E-05 | 1.96E-06 | 3.43E-05 | 5.96E-05 | 5.69E-05 | 0.00012 | 0 |
| D6Wsu163e | 5.64E-05 | 2.26E-05 | 2.15E-05 | 2.96E-05 | 2.22E-05 | 2.02E-05 | 2.13E-05 | 0 | 1.73E-05 | 3.98E-05 | 8.15E-06 | 0 | 5.26E-06 | 1.69E-05 | 0 | 2.64E-05 | 0 | 4.26E-05 |
| Adat2 | 3.08E-05 | 3.52E-05 | 2.62E-05 | 2.97E-05 | 0 | 3.35E-06 | 0 | 0 | 1.73E-05 | 3.30E-05 | 0 | 4.02E-05 | 0 | 0 | 1.92E-05 | 3.73E-05 | 1.21E-05 | 3.51E-05 |
| Stmn1 | 5.04E-05 | 1.13E-05 | 0 | 0.00024 | 0 | 0 | 0 | 0 | 1.73E-05 | 0 | 0 | 1.77E-05 | 0 | 2.59E-05 | 0 | 0.00191 | 0 | 3.02E-05 |
| Tnfrsf10b | 2.76E-05 | 9.14E-05 | 0.00014 | 3.09E-05 | 1.33E-05 | 0 | 2.15E-05 | 7.42E-06 | 1.73E-05 | 1.02E-06 | 3.35E-05 | 5.66E-05 | 6.42E-05 | 1.09E-05 | 4.81E-06 | 1.85E-05 | 0 | 3.03E-06 |
| Nlgn3 | 0 | 3.19E-06 | 0 | 0 | 0 | 0 | 0 | 0 | 1.73E-05 | 3.19E-06 | 0 | 2.80E-06 | 0 | 3.18E-06 | 0 | 5.91E-06 | 1.96E-05 | 3.75E-05 |
| Smc4 | 0.0001 | 2.20E-05 | 0 | 0 | 2.59E-05 | 0.00014 | 2.30E-05 | 0 | 1.72E-05 | 1.62E-05 | 1.37E-05 | 5.17E-05 | 0 | 0 | 0.00012 | 0.00093 | 7.54E-06 | 9.00E-05 |
| Slc4a7 | 3.59E-05 | 3.85E-05 | 8.41E-05 | 0 | 6.50E-06 | 6.82E-05 | 0 | 2.33E-05 | 1.72E-05 | 0 | 3.07E-05 | 5.58E-05 | 0 | 2.32E-05 | 7.54E-05 | 3.77E-05 | 0.00012 | 0 |

| | | | | | | | | | | | | | | | | | | |
|---|---|---|---|---|---|---|---|---|---|---|---|---|---|---|---|---|---|---|
| Dffa | 5.31E-05 | 3.82E-05 | 6.38E-05 | 6.30E-05 | 3.15E-05 | 5.77E-05 | 4.68E-05 | 0 | 1.72E-05 | 1.97E-05 | 6.93E-05 | 0 | 0 | 3.06E-05 | 3.71E-05 | 0 | 0 | 1.15E-05 |
| Csnk1g1 | 8.94E-05 | 6.91E-05 | 6.91E-05 | 0 | 0 | 5.49E-05 | 3.47E-05 | 3.40E-05 | 1.72E-05 | 1.40E-05 | 2.55E-05 | 5.31E-05 | 1.42E-05 | 8.23E-06 | 5.31E-05 | 1.00E-05 | 6.28E-05 | 3.27E-05 |
| Zc3h12c | 2.47E-05 | 4.87E-05 | 0.0001 | 0 | 2.21E-05 | 4.62E-05 | 0 | 3.97E-06 | 1.72E-05 | 0 | 3.63E-06 | 0 | 0 | 5.63E-05 | 0 | 3.45E-05 | 0.0001 | 2.00E-05 |
| Slc35d2 | 3.17E-05 | 2.47E-05 | 0 | 0 | 0 | 1.50E-05 | 1.37E-05 | 1.78E-05 | 1.72E-05 | 2.03E-05 | 0 | 1.27E-05 | 6.25E-06 | 0 | 0 | 8.13E-06 | 0 | 3.02E-05 |
| Colca2 | 6.11E-06 | 8.27E-06 | 0 | 0 | 4.66E-06 | 0 | 0 | 2.20E-06 | 1.72E-05 | 2.19E-05 | 0 | 0 | 5.28E-06 | 1.79E-06 | 0 | 4.84E-07 | 0 | 3.82E-06 |
| Slc25a47 | 4.42E-06 | 2.13E-05 | 1.48E-05 | 1.51E-05 | 0 | 0 | 2.05E-06 | 9.32E-06 | 1.72E-05 | 3.14E-06 | 1.82E-05 | 0 | 5.14E-06 | 1.71E-06 | 3.23E-05 | 1.17E-07 | 6.83E-05 | 0 |
| Il1rn | 0 | 1.65E-05 | 1.65E-06 | 7.19E-07 | 1.29E-05 | 0 | 0 | 4.13E-05 | 1.72E-05 | 1.94E-05 | 3.26E-05 | 1.95E-06 | 5.24E-05 | 5.43E-05 | 0.00016 | 5.36E-05 | 4.93E-05 | 4.64E-05 |
| Cfap20 | 0.00018 | 0.00013 | 7.46E-05 | 0 | 5.91E-05 | 0.00014 | 0.00015 | 3.24E-05 | 1.71E-05 | 4.14E-05 | 0 | 6.14E-05 | 1.44E-05 | 4.49E-05 | 0.00011 | 0.00017 | 9.81E-05 | 5.23E-05 |
| Dnajc4 | 0.0001 | 8.28E-05 | 1.68E-05 | 6.79E-05 | 7.65E-05 | 8.60E-05 | 6.91E-05 | 2.46E-05 | 1.71E-05 | 5.52E-05 | 3.49E-05 | 4.33E-05 | 6.36E-05 | 2.42E-05 | 7.52E-05 | 0.00011 | 0 | 1.65E-05 |
| Rabgap1l | 6.72E-05 | 0.00016 | 1.01E-05 | 0 | 0.00014 | 6.83E-05 | 0.00015 | 2.46E-05 | 1.71E-05 | 0.00022 | 0.00011 | 0.00012 | 0 | 5.23E-05 | 0.00028 | 2.54E-05 | 0.00022 | 0.00016 |
| Sema6d | 1.66E-05 | 9.20E-05 | 4.66E-05 | 0 | 0 | 5.32E-05 | 0 | 2.81E-05 | 1.71E-05 | 5.98E-05 | 0 | 0.00026 | 4.28E-05 | 8.49E-05 | 0 | 5.66E-05 | 0 | 2.59E-05 |
| Zbtb24 | 8.60E-05 | 7.92E-05 | 0 | 1.44E-06 | 2.96E-05 | 4.57E-05 | 2.84E-05 | 2.57E-05 | 1.71E-05 | 2.10E-05 | 0 | 6.92E-06 | 4.76E-06 | 0 | 0.00015 | 7.10E-06 | 0 | 2.41E-05 |
| Dut | 0.00011 | 1.51E-05 | 0.00014 | 0.00012 | 0 | 3.86E-05 | 0 | 4.97E-05 | 1.71E-05 | 1.76E-06 | 3.11E-05 | 4.87E-05 | 5.56E-06 | 1.42E-06 | 2.70E-07 | 0.00065 | 6.67E-05 | 7.10E-05 |
| Gorasp1 | 7.37E-05 | 4.14E-05 | 8.53E-05 | 0.00017 | 4.58E-05 | 3.76E-05 | 0.00013 | 2.13E-05 | 1.71E-05 | 2.69E-05 | 1.79E-05 | 1.11E-05 | 3.01E-05 | 3.10E-05 | 2.78E-05 | 0 | 0 | 3.27E-05 |
| Hps3 | 2.60E-05 | 2.56E-05 | 1.85E-05 | 0 | 2.91E-05 | 2.21E-05 | 0 | 0 | 1.71E-05 | 3.32E-05 | 2.16E-05 | 0 | 1.02E-05 | 1.46E-05 | 0 | 9.84E-06 | 0 | 3.33E-05 |
| Mtmr14 | 1.48E-05 | 5.44E-05 | 0 | 8.17E-06 | 8.69E-06 | 1.85E-05 | 0 | 1.05E-06 | 1.71E-05 | 1.99E-05 | 0 | 2.76E-05 | 0 | 9.62E-06 | 4.46E-05 | 7.34E-05 | 1.89E-05 | 0 |
| Rapgef2 | 7.60E-05 | 0.00012 | 5.75E-05 | 4.03E-05 | 5.59E-05 | 1.76E-05 | 2.45E-05 | 7.08E-06 | 1.71E-05 | 6.22E-05 | 0.00015 | 0.00012 | 0.0001 | 1.84E-05 | 0.00017 | 4.38E-05 | 0.00034 | 3.12E-05 |
| Mrto4 | 6.00E-05 | 9.65E-06 | 8.02E-05 | 0 | 0 | 9.13E-06 | 0 | 0 | 1.71E-05 | 0 | 0 | 9.14E-06 | 0 | 0 | 0 | 4.04E-05 | 0 | 0 |
| Mus81 | 1.46E-05 | 5.73E-05 | 4.85E-06 | 1.67E-05 | 8.60E-06 | 4.21E-06 | 0 | 2.38E-05 | 1.71E-05 | 1.74E-05 | 4.98E-05 | 2.50E-05 | 0 | 1.71E-05 | 8.84E-05 | 3.28E-05 | 0 | 6.21E-06 |
| Clcn6 | 2.12E-06 | 2.79E-05 | 0 | 1.01E-05 | 5.56E-06 | 0 | 0 | 1.68E-05 | 1.71E-05 | 1.75E-05 | 4.49E-06 | 2.74E-05 | 7.39E-07 | 2.08E-05 | 1.01E-05 | 6.87E-06 | 1.30E-05 | 1.44E-05 |
| Slc22a21 | 0 | 5.19E-06 | 8.73E-06 | 3.56E-06 | 9.27E-06 | 0 | 0 | 2.62E-05 | 1.71E-05 | 9.19E-06 | 2.28E-05 | 9.09E-05 | 1.70E-06 | 1.68E-05 | 2.23E-05 | 3.71E-06 | 0 | 2.66E-05 |
| Vstm5 | 0 | 0 | 0 | 0 | 0 | 0 | 4.66E-06 | 0 | 1.71E-05 | 1.22E-05 | 8.52E-06 | 8.33E-05 | 0 | 0 | 0 | 3.44E-06 | 0 | 3.70E-05 |
| Nif3l1 | 6.99E-05 | 2.39E-05 | 6.77E-05 | 1.73E-05 | 7.08E-06 | 0 | 3.39E-05 | 0 | 1.71E-05 | 1.95E-05 | 3.79E-06 | 0 | 2.26E-05 | 7.88E-06 | 0 | 4.57E-05 | 0 | 1.70E-05 |
| Ccdc59 | 0.00019 | 0.00012 | 0.00017 | 0.00016 | 4.68E-05 | 3.67E-05 | 0 | 1.16E-05 | 1.70E-05 | 0 | 0 | 0.0001 | 0 | 0 | 0.00019 | 8.63E-05 | 3.74E-05 | 1.76E-05 |
| Etv3 | 2.98E-05 | 7.52E-05 | 2.85E-05 | 3.28E-05 | 3.14E-05 | 3.26E-05 | 2.71E-05 | 4.43E-05 | 1.70E-05 | 5.20E-05 | 3.52E-05 | 5.01E-05 | 1.18E-05 | 2.26E-05 | 6.69E-05 | 2.36E-05 | 1.81E-05 | 2.79E-05 |
| Abhd13 | 2.15E-05 | 0.0001 | 9.18E-05 | 8.07E-05 | 9.15E-05 | 3.05E-05 | 3.76E-05 | 7.40E-05 | 1.70E-05 | 3.04E-05 | 0.00011 | 2.90E-05 | 4.60E-05 | 5.94E-05 | 0 | 0.00011 | 0 | 8.94E-05 |
| Eef2k | 4.27E-05 | 5.92E-05 | 7.73E-06 | 0.00012 | 0 | 1.49E-05 | 6.33E-07 | 0 | 1.70E-05 | 0.00018 | 5.90E-05 | 0.00011 | 0 | 9.86E-05 | 0 | 0 | 1.17E-05 | 0.00022 |
| Lztfl1 | 0.0001 | 4.58E-05 | 3.84E-05 | 6.21E-06 | 1.15E-05 | 2.51E-06 | 5.13E-05 | 8.50E-06 | 1.70E-05 | 5.16E-05 | 0 | 3.31E-06 | 0 | 6.32E-06 | 4.01E-05 | 0 | 0 | 1.79E-05 |
| Luc7l3 | 0.00014 | 0.00015 | 0.00012 | 0 | 1.58E-05 | 0 | 0 | 0 | 1.70E-05 | 2.62E-05 | 3.83E-05 | 5.26E-05 | 0 | 1.03E-05 | 0.00063 | 0.00017 | 8.79E-05 | 0 |
| Rabl3 | 6.84E-05 | 1.49E-05 | 2.81E-05 | 4.76E-06 | 7.91E-06 | 1.57E-05 | 4.13E-05 | 1.91E-05 | 1.69E-05 | 9.69E-06 | 0 | 2.92E-06 | 0 | 0 | 0.0001 | 2.24E-05 | 0 | 0 |
| Ephx1 | 0 | 0.00012 | 0 | 1.30E-05 | 0 | 9.51E-06 | 0 | 3.13E-05 | 1.69E-05 | 0.00026 | 4.35E-05 | 0.00017 | 0 | 0 | 3.73E-05 | 0 | 0 | 0.00018 |
| Stard9 | 0 | 8.57E-05 | 0 | 0 | 6.25E-05 | 5.68E-05 | 1.44E-06 | 0 | 1.68E-05 | 8.24E-05 | 0 | 1.59E-05 | 0 | 0 | 6.64E-05 | 6.22E-06 | 3.79E-05 | 4.13E-05 |
| Prpf18 | 0.00014 | 0.00014 | 4.57E-05 | 1.33E-05 | 6.59E-05 | 4.51E-05 | 1.28E-05 | 2.09E-05 | 1.68E-05 | 1.02E-05 | 4.07E-05 | 5.03E-05 | 1.93E-05 | 4.33E-05 | 0.00015 | 9.48E-05 | 0 | 6.04E-05 |

| | | | | | | | | | | | | | | | | | | |
|---|---|---|---|---|---|---|---|---|---|---|---|---|---|---|---|---|---|---|
| Nsmce3 | 5.40E-05 | 0.00011 | 8.93E-05 | 0.00015 | 2.51E-05 | 3.26E-05 | 0 | 3.98E-05 | 1.68E-05 | 4.71E-05 | 8.03E-05 | 0.00012 | 0 | 0.0001 | 4.21E-05 | 0.00012 | 6.98E-07 | 7.21E-05 |
| Mis18a | 0 | 2.08E-05 | 0 | 1.32E-05 | 0 | 3.04E-05 | 2.11E-06 | 2.98E-06 | 1.68E-05 | 2.44E-05 | 1.78E-05 | 5.00E-05 | 4.03E-06 | 1.86E-05 | 0 | 0.00053 | 2.48E-05 | 2.15E-05 |
| Kbtbd7 | 2.38E-05 | 2.37E-05 | 0 | 8.81E-07 | 4.64E-05 | 1.60E-05 | 4.69E-05 | 0 | 1.68E-05 | 0 | 1.61E-05 | 3.21E-05 | 0 | 8.60E-06 | 0 | 1.97E-05 | 3.79E-05 | 2.12E-05 |
| Zfp619 | 2.07E-06 | 1.09E-05 | 1.37E-06 | 4.36E-07 | 7.30E-06 | 4.95E-06 | 6.21E-06 | 1.10E-05 | 1.68E-05 | 7.58E-06 | 1.41E-05 | 1.09E-06 | 9.22E-06 | 0 | 4.31E-05 | 0 | 5.85E-05 | 0 |
| Tmem82 | 0 | 1.74E-05 | 0 | 1.28E-05 | 0 | 2.38E-06 | 0 | 7.94E-06 | 1.68E-05 | 0 | 0 | 3.03E-06 | 9.38E-06 | 0 | 0 | 0 | 1.02E-05 | 0 |
| Cln3 | 2.44E-06 | 7.79E-05 | 0 | 4.36E-05 | 3.93E-05 | 1.63E-06 | 1.60E-05 | 4.07E-05 | 1.68E-05 | 2.48E-05 | 0 | 9.78E-06 | 5.67E-06 | 1.26E-05 | 0 | 0 | 0 | 1.96E-05 |
| Nkrf | 2.36E-05 | 2.65E-05 | 3.25E-05 | 0 | 1.09E-06 | 1.16E-06 | 0 | 0 | 1.68E-05 | 2.57E-06 | 0 | 5.52E-06 | 0 | 0 | 4.62E-05 | 1.27E-05 | 3.46E-05 | 0 |
| Lsm6 | 0.00011 | 0 | 5.61E-05 | 7.44E-05 | 8.47E-07 | 0 | 0 | 8.35E-06 | 1.68E-05 | 5.46E-06 | 3.12E-05 | 0 | 1.29E-05 | 1.61E-05 | 0 | 0.00021 | 2.06E-05 | 3.83E-05 |
| Elovl5 | 0.00035 | 7.22E-05 | 8.42E-05 | 0 | 7.30E-05 | 0.00011 | 0.00027 | 9.84E-05 | 1.67E-05 | 0.00018 | 8.51E-05 | 0 | 2.85E-05 | 3.80E-05 | 0 | 0.00031 | 0 | 0.00016 |
| Tmem126a | 0.00019 | 3.75E-05 | 0.0001 | 9.40E-05 | 2.77E-05 | 5.87E-05 | 9.84E-06 | 2.10E-05 | 1.67E-05 | 2.40E-05 | 9.86E-07 | 0 | 2.89E-05 | 3.72E-05 | 4.40E-05 | 6.33E-05 | 0 | 2.21E-05 |
| Tmem219 | 5.80E-05 | 3.54E-05 | 2.42E-05 | 9.60E-05 | 8.07E-05 | 4.54E-05 | 7.62E-05 | 7.73E-05 | 1.67E-05 | 9.05E-05 | 5.06E-07 | 3.40E-05 | 2.59E-05 | 0 | 0 | 5.20E-06 | 0 | 0.00012 |
| Ankrd16 | 1.24E-05 | 8.65E-05 | 2.85E-05 | 7.06E-05 | 2.38E-05 | 3.61E-05 | 4.53E-05 | 0 | 1.67E-05 | 3.24E-05 | 5.47E-05 | 1.11E-05 | 5.95E-05 | 0.00011 | 0.00019 | 0 | 2.44E-05 | 3.11E-05 |
| Kat8 | 8.48E-05 | 6.95E-05 | 4.65E-05 | 5.68E-05 | 2.10E-06 | 2.42E-05 | 3.76E-05 | 0 | 1.67E-05 | 3.41E-06 | 9.38E-06 | 3.68E-05 | 0 | 3.71E-05 | 5.16E-05 | 2.63E-05 | 0 | 2.06E-05 |
| Pacsin3 | 0 | 4.72E-05 | 1.90E-05 | 6.89E-05 | 5.81E-05 | 2.24E-05 | 6.34E-06 | 0 | 1.67E-05 | 0 | 3.33E-05 | 6.24E-05 | 9.08E-05 | 7.57E-05 | 5.65E-05 | 0 | 0 | 5.21E-05 |
| Tmem120b | 3.31E-05 | 0 | 1.03E-05 | 1.65E-05 | 1.66E-05 | 1.13E-05 | 9.21E-05 | 0 | 1.67E-05 | 1.92E-05 | 2.84E-05 | 2.89E-05 | 3.27E-06 | 4.04E-05 | 0 | 0 | 2.09E-05 | 2.81E-05 |
| Zfp575 | 1.34E-06 | 5.28E-06 | 1.55E-06 | 1.23E-05 | 3.97E-06 | 5.72E-06 | 1.35E-06 | 1.10E-05 | 1.67E-05 | 1.08E-05 | 1.86E-05 | 6.80E-06 | 0 | 6.00E-06 | 0 | 1.24E-05 | 0 | 5.13E-06 |
| Elmo1 | 1.16E-05 | 1.10E-05 | 0 | 0 | 0 | 0 | 2.71E-05 | 0 | 1.67E-05 | 0.00013 | 0 | 3.44E-05 | 0 | 0 | 3.83E-06 | 0 | 6.01E-06 | 0 |
| Aopep | 5.81E-06 | 5.59E-05 | 0 | 0 | 8.13E-05 | 0 | 0 | 2.76E-05 | 1.67E-05 | 0 | 6.08E-05 | 0 | 0 | 6.92E-05 | 0.00012 | 0 | 0 | 4.52E-05 |
| Vbp1 | 0.00022 | 0.00013 | 0.00013 | 0.00015 | 3.60E-05 | 0.0001 | 3.56E-07 | 3.87E-05 | 1.66E-05 | 3.94E-05 | 3.82E-05 | 0.00011 | 0 | 3.37E-05 | 0.00025 | 0.00033 | 0.00011 | 4.93E-05 |
| Manea | 0.00013 | 4.42E-05 | 0 | 0 | 6.29E-05 | 9.49E-05 | 1.04E-05 | 0 | 1.66E-05 | 4.86E-05 | 2.96E-05 | 1.22E-06 | 0 | 1.17E-07 | 0 | 2.39E-05 | 1.17E-05 | 2.11E-05 |
| Uvrag | 0.00011 | 9.96E-05 | 3.60E-05 | 9.26E-07 | 2.65E-05 | 6.32E-05 | 1.45E-05 | 3.95E-05 | 1.66E-05 | 2.06E-05 | 0 | 0 | 8.84E-06 | 0.00013 | 0.00019 | 6.02E-05 | 0.00015 | 6.40E-05 |
| Pcdhga3 | 3.95E-05 | 3.89E-05 | 0 | 0 | 6.04E-05 | 6.21E-05 | 1.17E-05 | 4.85E-05 | 1.66E-05 | 1.54E-05 | 0 | 1.61E-05 | 2.17E-05 | 2.03E-05 | 0 | 0 | 0 | 5.02E-06 |
| Cpt2 | 0.0001 | 6.94E-05 | 8.27E-06 | 0.00013 | 5.21E-06 | 4.41E-05 | 9.94E-05 | 2.27E-05 | 1.66E-05 | 7.71E-05 | 4.66E-05 | 2.63E-05 | 6.25E-05 | 2.25E-05 | 0 | 9.24E-05 | 0 | 0.00013 |
| Hsbp1l1 | 0 | 0 | 1.64E-06 | 0 | 1.35E-05 | 2.44E-05 | 0 | 8.47E-06 | 1.66E-05 | 0 | 5.35E-05 | 0 | 0 | 0 | 1.98E-05 | 3.96E-06 | 0 | 0 |
| Edrf1 | 4.79E-05 | 5.21E-05 | 2.67E-05 | 2.32E-06 | 2.79E-05 | 1.35E-06 | 2.16E-05 | 0 | 1.66E-05 | 6.48E-06 | 1.92E-05 | 2.27E-05 | 7.68E-06 | 1.02E-05 | 6.28E-05 | 1.95E-05 | 0 | 3.34E-06 |
| Zdhhc15 | 0 | 3.81E-06 | 0 | 2.22E-06 | 1.47E-05 | 0 | 1.27E-05 | 8.65E-05 | 1.66E-05 | 3.06E-05 | 2.06E-05 | 0 | 9.77E-07 | 0 | 0 | 0.00016 | 0 | 3.46E-05 |
| Nasp | 6.96E-05 | 7.73E-06 | 0.00017 | 0.00016 | 0 | 0 | 0 | 0 | 1.66E-05 | 3.11E-05 | 0 | 4.20E-06 | 9.00E-05 | 1.78E-05 | 9.24E-05 | 0.00066 | 0.00029 | 0 |
| Sec62 | 0.0003 | 0.00032 | 0.00029 | 0.00033 | 0.00031 | 0.00041 | 0.00047 | 0.00032 | 1.65E-05 | 0.00022 | 0.00022 | 0.00016 | 0.00022 | 0.00024 | 0 | 0.00015 | 0.00021 | 0.00024 |
| Bcl7a | 8.08E-06 | 3.63E-05 | 1.40E-05 | 0 | 1.71E-05 | 9.27E-05 | 0 | 0 | 1.65E-05 | 0 | 4.68E-05 | 0 | 0 | 0.00016 | 0.00025 | 0.00011 | 0 | 0 |
| Fam210b | 2.80E-05 | 4.68E-05 | 0 | 0 | 8.01E-05 | 8.39E-05 | 2.29E-05 | 2.81E-05 | 1.65E-05 | 0 | 1.76E-05 | 4.01E-05 | 1.76E-05 | 4.09E-05 | 0 | 0 | 0 | 2.09E-06 |
| Samd14 | 0 | 0.00013 | 1.50E-05 | 0.00013 | 8.36E-05 | 8.23E-05 | 2.68E-05 | 4.78E-05 | 1.65E-05 | 4.16E-05 | 5.43E-05 | 0.0002 | 3.25E-05 | 4.11E-05 | 2.58E-05 | 9.03E-05 | 4.82E-05 | 0.00012 |
| B4galt4 | 1.58E-05 | 3.67E-05 | 0 | 0 | 0.00015 | 7.98E-05 | 5.27E-05 | 0 | 1.65E-05 | 6.94E-05 | 0 | 0 | 5.76E-05 | 6.01E-05 | 0 | 0 | 0 | 9.32E-05 |
| Cacfd1 | 7.48E-05 | 0.00012 | 0.00013 | 0.00032 | 0.00015 | 6.69E-05 | 0.00021 | 0.00018 | 1.65E-05 | 4.63E-05 | 2.54E-05 | 0.00018 | 3.07E-05 | 0.00019 | 0.00021 | 0 | 0 | 4.74E-05 |

| | | | | | | | | | | | | | | | | | | |
|---|---|---|---|---|---|---|---|---|---|---|---|---|---|---|---|---|---|---|
| Usp6nl | 4.85E-05 | 5.47E-05 | 2.55E-05 | 0 | 3.49E-05 | 5.29E-05 | 0 | 4.50E-05 | 1.65E-05 | 2.02E-05 | 5.59E-05 | 3.40E-05 | 3.11E-05 | 4.30E-05 | 2.25E-05 | 8.26E-06 | 9.05E-05 | 1.67E-05 |
| Dcp1b | 4.78E-06 | 4.85E-05 | 2.40E-06 | 4.85E-06 | 2.80E-05 | 1.11E-05 | 0 | 0 | 1.65E-05 | 2.75E-05 | 4.61E-06 | 2.06E-05 | 1.12E-06 | 0 | 9.59E-06 | 0 | 4.85E-05 | 3.00E-05 |
| Asb13 | 9.05E-06 | 1.18E-05 | 0 | 0 | 1.23E-05 | 2.99E-06 | 0 | 0 | 1.65E-05 | 2.76E-05 | 4.96E-06 | 0 | 1.14E-05 | 0 | 2.31E-05 | 0 | 0 | 2.86E-05 |
| Fam83f | 0 | 3.94E-07 | 0 | 2.49E-06 | 0 | 0 | 0 | 0 | 1.65E-05 | 9.70E-05 | 0 | 9.89E-06 | 4.66E-05 | 0 | 0 | 0 | 4.90E-06 | 0 |
| Chd1l | 8.16E-06 | 4.47E-05 | 1.74E-05 | 3.78E-05 | 1.01E-05 | 0 | 8.94E-06 | 1.72E-05 | 1.65E-05 | 2.62E-05 | 2.28E-05 | 0 | 0 | 7.15E-06 | 1.68E-05 | 7.50E-05 | 1.92E-05 | 2.96E-05 |
| Klhl29 | 9.03E-06 | 1.76E-06 | 1.10E-08 | 0 | 1.11E-05 | 0 | 0 | 0 | 1.65E-05 | 2.22E-05 | 7.59E-05 | 0 | 2.89E-05 | 8.07E-05 | 0 | 2.93E-05 | 0 | 4.81E-05 |
| Rexo5 | 1.76E-05 | 2.19E-05 | 0 | 0 | 1.17E-05 | 0 | 0 | 0 | 1.65E-05 | 8.11E-06 | 0 | 0 | 0 | 0 | 7.85E-05 | 0.00012 | 0 | 3.12E-06 |
| Tet2 | 9.36E-05 | 8.41E-05 | 2.69E-05 | 0 | 4.82E-05 | 7.88E-05 | 5.12E-05 | 4.52E-05 | 1.64E-05 | 4.13E-06 | 0 | 0.00017 | 0 | 1.60E-05 | 0.00019 | 0 | 0.00017 | 1.62E-05 |
| Rnf103 | 0 | 0.00018 | 2.43E-05 | 0.00014 | 0.00013 | 4.84E-05 | 3.87E-05 | 4.69E-05 | 1.64E-05 | 0.00013 | 0.00012 | 0.0002 | 3.59E-05 | 8.72E-05 | 0.00014 | 0.00011 | 0.00023 | 0.00011 |
| Xkr8 | 4.06E-06 | 3.80E-05 | 1.34E-05 | 2.84E-05 | 1.17E-05 | 1.67E-05 | 0 | 3.69E-05 | 1.64E-05 | 3.09E-05 | 0 | 1.52E-05 | 3.90E-05 | 1.72E-05 | 0 | 0 | 0 | 3.02E-05 |
| Macrod1 | 0 | 4.22E-05 | 0 | 2.85E-06 | 2.04E-05 | 1.76E-06 | 3.98E-06 | 0 | 1.64E-05 | 2.03E-05 | 8.48E-06 | 1.09E-05 | 0.00011 | 9.59E-05 | 9.80E-06 | 0 | 0 | 2.27E-05 |
| Brd3os | 2.35E-05 | 2.90E-05 | 1.48E-05 | 0 | 2.12E-05 | 0 | 0 | 2.42E-05 | 1.64E-05 | 0 | 0 | 1.58E-05 | 0 | 0 | 6.82E-06 | 0 | 1.18E-06 | 0 |
| Pex16 | 1.12E-05 | 4.33E-05 | 2.45E-05 | 6.90E-05 | 2.75E-05 | 0 | 0 | 1.25E-05 | 1.64E-05 | 2.25E-05 | 1.23E-05 | 2.76E-05 | 9.54E-06 | 1.94E-05 | 6.08E-05 | 0 | 5.67E-06 | 3.19E-05 |
| Foxp1 | 0.00022 | 0.00022 | 0.00025 | 0 | 0.00042 | 0.00012 | 0.00054 | 0.00021 | 1.63E-05 | 0.00022 | 0.00052 | 0.00053 | 0.00037 | 0 | 0.00045 | 0.0002 | 0.00036 | 0.00021 |
| Mapk8ip1 | 0 | 0.00011 | 0 | 2.58E-05 | 5.32E-05 | 8.52E-05 | 0 | 0.00013 | 1.63E-05 | 9.03E-05 | 0 | 0.00012 | 0 | 4.45E-06 | 0 | 0 | 0 | 7.13E-05 |
| Ing4 | 0.00012 | 0.00017 | 0 | 0.00015 | 0.00013 | 7.98E-05 | 0.00017 | 6.98E-05 | 1.63E-05 | 6.84E-05 | 0 | 8.62E-05 | 0 | 0.00015 | 0.00026 | 1.73E-05 | 0 | 0.00011 |
| Mbtps2 | 9.56E-05 | 8.90E-05 | 6.00E-05 | 0 | 2.18E-05 | 7.60E-05 | 1.14E-06 | 4.36E-06 | 1.63E-05 | 4.81E-06 | 3.95E-05 | 5.36E-05 | 1.24E-06 | 3.36E-05 | 0 | 4.56E-05 | 4.01E-05 | 8.72E-06 |
| Hebp2 | 8.65E-05 | 2.66E-05 | 1.09E-06 | 5.77E-05 | 5.93E-05 | 6.47E-05 | 0.00015 | 0 | 1.63E-05 | 0.0001 | 7.71E-05 | 2.42E-05 | 9.16E-05 | 0 | 0 | 0 | 0 | 7.77E-05 |
| Eri1 | 8.52E-05 | 6.32E-05 | 9.40E-05 | 0 | 8.69E-06 | 4.56E-05 | 4.69E-05 | 0 | 1.63E-05 | 1.72E-05 | 3.71E-05 | 5.51E-05 | 2.30E-05 | 5.85E-05 | 0 | 0.00038 | 0 | 1.01E-05 |
| Amn1 | 6.02E-05 | 6.55E-05 | 4.80E-06 | 0 | 1.30E-05 | 2.74E-05 | 5.04E-07 | 2.86E-05 | 1.63E-05 | 0 | 7.19E-06 | 5.59E-05 | 1.06E-05 | 1.70E-06 | 2.67E-05 | 1.72E-05 | 0 | 1.03E-05 |
| Calhm6 | 4.24E-06 | 0 | 0 | 0 | 0 | 7.81E-06 | 0 | 0 | 1.63E-05 | 1.09E-05 | 0 | 2.30E-06 | 2.50E-05 | 0 | 0 | 0 | 0 | 1.12E-05 |
| Phlpp1 | 2.10E-05 | 3.97E-05 | 3.07E-05 | 0 | 3.87E-05 | 4.96E-06 | 0 | 3.41E-06 | 1.63E-05 | 3.50E-05 | 0.00016 | 1.56E-05 | 1.46E-06 | 8.50E-05 | 5.83E-05 | 2.86E-05 | 0.00012 | 1.19E-05 |
| Pkn2 | 0.00022 | 0.0002 | 9.97E-05 | 0 | 1.86E-05 | 0.00011 | 0 | 0 | 1.62E-05 | 5.64E-05 | 5.98E-05 | 0.00016 | 0 | 0 | 8.99E-05 | 0.00021 | 0.00018 | 5.52E-05 |
| Synj1 | 6.84E-05 | 7.38E-05 | 2.14E-05 | 0 | 3.50E-05 | 5.55E-05 | 7.36E-06 | 2.04E-06 | 1.62E-05 | 5.27E-05 | 2.21E-05 | 3.89E-05 | 0 | 0 | 3.84E-05 | 3.96E-05 | 0 | 5.41E-05 |
| Rell1 | 0 | 9.57E-05 | 3.75E-05 | 0 | 3.42E-05 | 4.44E-05 | 0 | 8.23E-05 | 1.62E-05 | 4.78E-05 | 0.00028 | 0.00017 | 0 | 5.55E-05 | 0 | 0.00021 | 4.95E-05 | 6.75E-05 |
| Zfp646 | 6.38E-05 | 8.69E-05 | 4.07E-05 | 0 | 3.18E-05 | 2.52E-05 | 4.67E-06 | 1.36E-05 | 1.62E-05 | 3.08E-06 | 1.12E-05 | 2.07E-05 | 0 | 0 | 0.00011 | 1.18E-05 | 0 | 1.55E-05 |
| St6galnac3 | 0 | 5.54E-06 | 0 | 0 | 3.02E-05 | 5.12E-06 | 0 | 0 | 1.62E-05 | 0 | 0 | 0 | 0 | 1.26E-05 | 1.42E-06 | 0 | 0 | 0 |
| Faap24 | 2.32E-05 | 2.42E-05 | 3.80E-05 | 5.52E-06 | 4.69E-06 | 0 | 1.77E-06 | 7.83E-06 | 1.62E-05 | 1.60E-05 | 9.89E-06 | 7.44E-06 | 7.04E-06 | 1.40E-06 | 0 | 0.00018 | 0 | 2.87E-05 |
| Ksr1 | 2.82E-05 | 0.00011 | 0 | 0 | 4.35E-05 | 8.27E-05 | 0 | 0 | 1.61E-05 | 8.27E-05 | 2.44E-05 | 5.76E-05 | 0.00027 | 0.00011 | 7.31E-05 | 1.39E-05 | 0.0001 | 0.00012 |
| Lcorl | 3.28E-05 | 4.50E-05 | 0 | 0 | 1.77E-05 | 1.78E-05 | 1.33E-05 | 1.25E-05 | 1.61E-05 | 9.06E-06 | 0 | 2.91E-05 | 0 | 0 | 1.65E-05 | 0.0001 | 4.58E-05 | 9.92E-07 |
| Spin4 | 1.63E-05 | 0 | 1.59E-05 | 1.84E-05 | 1.28E-05 | 1.55E-05 | 1.95E-05 | 1.89E-05 | 1.61E-05 | 5.24E-05 | 0 | 8.92E-07 | 0 | 1.13E-05 | 0 | 2.70E-05 | 2.10E-05 | 2.72E-05 |
| Nudt14 | 1.00E-05 | 9.51E-06 | 0 | 1.83E-05 | 3.65E-06 | 1.12E-05 | 5.29E-06 | 3.35E-06 | 1.61E-05 | 7.48E-06 | 4.19E-06 | 0 | 1.09E-05 | 1.71E-05 | 1.09E-05 | 1.52E-06 | 0 | 9.75E-06 |
| Lipe | 2.62E-05 | 5.95E-05 | 0 | 1.76E-05 | 3.43E-05 | 7.51E-06 | 0 | 0 | 1.61E-05 | 4.37E-05 | 0 | 0 | 5.41E-05 | 2.75E-06 | 1.42E-05 | 0 | 0 | 3.56E-05 |

| | | | | | | | | | | | | | | | | | | |
|---|---|---|---|---|---|---|---|---|---|---|---|---|---|---|---|---|---|---|
| Smyd4 | 0 | 2.25E-05 | 0 | 2.04E-06 | 1.06E-05 | 8.94E-07 | 1.60E-05 | 1.37E-05 | 1.61E-05 | 1.13E-05 | 6.12E-06 | 5.19E-06 | 2.22E-06 | 1.15E-05 | 3.62E-06 | 6.56E-05 | 1.62E-05 | 1.31E-05 |
| Ggn | 0 | 9.31E-06 | 0 | 1.30E-05 | 9.11E-06 | 0 | 4.78E-06 | 1.03E-05 | 1.61E-05 | 1.81E-05 | 0 | 4.99E-05 | 0 | 0 | 0 | 6.79E-06 | 8.96E-06 | 8.70E-06 |
| Prpf39 | 0.00014 | 0.00021 | 7.66E-05 | 0 | 0.0001 | 0.00013 | 0 | 4.20E-05 | 1.60E-05 | 3.76E-05 | 0 | 0.00013 | 0 | 8.63E-05 | 0.00073 | 1.97E-05 | 9.70E-05 | 4.22E-05 |
| Etaa1 | 2.64E-05 | 2.02E-05 | 5.87E-06 | 7.97E-06 | 8.97E-06 | 1.63E-05 | 1.70E-05 | 1.94E-05 | 1.60E-05 | 4.43E-06 | 6.26E-06 | 1.15E-06 | 3.66E-06 | 1.96E-05 | 3.44E-05 | 0.00016 | 0 | 1.02E-05 |
| Nat2 | 3.15E-05 | 2.56E-05 | 0 | 4.41E-06 | 9.02E-06 | 7.15E-06 | 0 | 0 | 1.60E-05 | 2.46E-05 | 4.34E-05 | 1.18E-05 | 0 | 2.02E-05 | 0 | 6.12E-06 | 0 | 2.27E-05 |
| Setd4 | 0 | 1.97E-05 | 7.52E-06 | 4.25E-05 | 3.51E-06 | 0 | 0 | 2.21E-05 | 1.60E-05 | 6.61E-06 | 0 | 6.85E-07 | 0 | 6.90E-06 | 6.90E-05 | 0 | 0 | 1.44E-05 |
| Maml2 | 4.08E-05 | 6.34E-05 | 0 | 0 | 5.56E-05 | 6.02E-05 | 4.19E-05 | 3.16E-06 | 1.59E-05 | 0 | 2.21E-05 | 5.33E-06 | 3.19E-06 | 5.34E-06 | 2.61E-06 | 8.62E-06 | 9.61E-05 | 2.95E-05 |
| Mfsd8 | 4.85E-05 | 2.95E-05 | 1.83E-05 | 4.29E-06 | 2.05E-05 | 5.19E-05 | 2.95E-06 | 2.74E-05 | 1.59E-05 | 2.75E-05 | 0 | 3.80E-05 | 0 | 0 | 0 | 3.82E-06 | 0 | 2.08E-05 |
| Zfp81 | 4.66E-05 | 1.83E-05 | 0 | 0 | 1.19E-05 | 3.02E-05 | 6.76E-06 | 0 | 1.59E-05 | 7.68E-06 | 0 | 2.16E-05 | 0 | 0 | 0 | 2.32E-05 | 0 | 0 |
| Egfl7 | 6.81E-06 | 0 | 0 | 2.74E-05 | 2.10E-06 | 1.06E-05 | 0 | 1.49E-05 | 1.59E-05 | 0 | 4.03E-05 | 1.14E-05 | 0 | 3.51E-05 | 0 | 2.96E-05 | 0 | 0 |
| Piga | 2.59E-05 | 1.78E-05 | 2.53E-05 | 0 | 0 | 8.43E-06 | 0 | 0 | 1.59E-05 | 3.28E-05 | 0 | 0.00014 | 0 | 0 | 1.63E-05 | 7.97E-05 | 4.54E-05 | 0 |
| Wrap53 | 2.64E-05 | 3.88E-05 | 3.54E-05 | 5.70E-05 | 7.24E-06 | 4.74E-06 | 5.15E-06 | 0 | 1.59E-05 | 1.49E-06 | 0 | 0 | 7.57E-06 | 5.35E-06 | 0.00011 | 0.00015 | 4.35E-05 | 0 |
| Tom1 | 1.20E-05 | 9.19E-05 | 4.75E-05 | 0.00011 | 3.12E-05 | 0 | 5.09E-06 | 3.21E-05 | 1.59E-05 | 6.19E-05 | 3.33E-05 | 0.00015 | 1.46E-06 | 6.11E-05 | 0 | 0 | 5.44E-05 | 5.20E-05 |
| Nme6 | 2.21E-05 | 1.86E-05 | 2.73E-05 | 2.18E-05 | 1.00E-05 | 0 | 8.78E-06 | 1.47E-05 | 1.59E-05 | 0 | 0 | 2.73E-05 | 8.19E-06 | 2.24E-05 | 1.24E-05 | 3.26E-05 | 0 | 0 |
| Nudt18 | 1.08E-05 | 6.96E-05 | 5.23E-06 | 4.58E-05 | 7.50E-05 | 0 | 0.00014 | 3.41E-06 | 1.59E-05 | 8.15E-05 | 7.34E-05 | 8.89E-05 | 0 | 4.65E-06 | 0 | 0 | 0 | 0.00013 |
| Gtf3c4 | 8.35E-05 | 0.00015 | 0.00013 | 7.23E-05 | 0.00011 | 3.90E-05 | 6.79E-05 | 3.78E-05 | 1.58E-05 | 4.38E-05 | 2.54E-05 | 4.92E-05 | 0 | 0 | 0 | 8.52E-05 | 0 | 6.94E-05 |
| Zbtb5 | 3.07E-05 | 3.89E-05 | 2.23E-05 | 0 | 3.49E-05 | 3.74E-05 | 2.02E-05 | 3.93E-06 | 1.58E-05 | 2.66E-05 | 1.72E-05 | 5.78E-06 | 0 | 3.04E-05 | 2.52E-05 | 4.49E-05 | 0 | 2.91E-05 |
| Thap3 | 4.40E-05 | 6.21E-05 | 0 | 2.88E-05 | 2.34E-05 | 3.54E-05 | 2.39E-05 | 4.51E-07 | 1.58E-05 | 2.17E-05 | 0 | 6.39E-06 | 0 | 0 | 0.00013 | 1.20E-05 | 1.92E-05 | 2.09E-05 |
| Cfap97 | 5.46E-05 | 5.29E-05 | 1.49E-05 | 2.16E-05 | 3.55E-05 | 3.46E-05 | 6.16E-05 | 3.83E-06 | 1.58E-05 | 1.87E-05 | 0 | 3.83E-05 | 0 | 1.76E-05 | 1.33E-05 | 3.03E-05 | 0 | 5.18E-05 |
| Gstm4 | 3.27E-05 | 9.94E-06 | 0 | 1.85E-05 | 1.61E-05 | 3.20E-05 | 2.57E-06 | 2.73E-05 | 1.58E-05 | 7.65E-05 | 1.79E-05 | 1.78E-05 | 1.86E-05 | 0 | 0 | 0 | 2.31E-06 | 5.66E-05 |
| Zfp408 | 3.34E-05 | 6.37E-05 | 1.40E-05 | 5.72E-05 | 1.68E-05 | 1.31E-05 | 0 | 7.14E-06 | 1.58E-05 | 0 | 0 | 3.36E-05 | 0 | 2.60E-05 | 9.72E-05 | 3.02E-06 | 0 | 2.05E-05 |
| Cipc | 3.99E-05 | 3.80E-05 | 4.03E-06 | 0 | 1.82E-05 | 5.46E-06 | 0 | 0 | 1.58E-05 | 0 | 5.61E-06 | 1.64E-06 | 0 | 0 | 0 | 0 | 0 | 0 |
| Mtcp1 | 1.49E-06 | 1.73E-05 | 0 | 0 | 1.45E-05 | 2.07E-06 | 0 | 0 | 1.58E-05 | 1.08E-05 | 0 | 0 | 0 | 7.05E-07 | 7.82E-05 | 2.47E-05 | 0 | 4.00E-06 |
| Polr3a | 1.07E-05 | 4.23E-05 | 2.60E-05 | 4.86E-05 | 1.58E-05 | 0 | 3.61E-05 | 0 | 1.58E-05 | 1.36E-05 | 1.10E-05 | 1.69E-05 | 2.42E-06 | 2.47E-05 | 0.00013 | 3.19E-05 | 2.01E-05 | 1.24E-06 |
| Vps39 | 6.18E-05 | 0.0001 | 3.99E-05 | 0 | 6.73E-05 | 8.04E-05 | 3.13E-05 | 2.38E-05 | 1.57E-05 | 3.73E-05 | 3.67E-05 | 0.00011 | 3.35E-06 | 1.23E-05 | 6.83E-05 | 3.32E-05 | 3.76E-06 | 4.90E-05 |
| Dop1a | 7.39E-05 | 7.39E-05 | 3.35E-05 | 0 | 4.13E-05 | 5.73E-05 | 2.47E-05 | 2.02E-05 | 1.57E-05 | 3.20E-05 | 0 | 3.31E-05 | 4.65E-05 | 2.64E-05 | 6.84E-05 | 0 | 2.43E-05 | 6.45E-06 |
| Zup1 | 6.49E-05 | 5.31E-05 | 0 | 0 | 1.91E-05 | 3.75E-05 | 1.09E-07 | 0 | 1.57E-05 | 2.85E-05 | 0 | 4.60E-05 | 0 | 2.31E-05 | 0.00014 | 7.87E-06 | 0 | 3.05E-05 |
| Osbpl11 | 5.61E-05 | 0.00011 | 6.74E-05 | 4.62E-05 | 1.07E-05 | 2.24E-05 | 3.36E-05 | 0 | 1.57E-05 | 2.61E-05 | 5.62E-05 | 5.89E-06 | 6.59E-07 | 4.38E-05 | 9.47E-05 | 7.90E-05 | 8.84E-05 | 3.33E-05 |
| Btbd8 | 1.14E-05 | 3.56E-05 | 0 | 0 | 1.52E-06 | 1.70E-05 | 0 | 5.87E-06 | 1.57E-05 | 1.52E-05 | 1.29E-05 | 2.07E-05 | 0 | 0 | 4.34E-05 | 4.39E-05 | 0 | 4.56E-06 |
| Fastkd2 | 5.07E-05 | 1.36E-05 | 3.04E-05 | 0 | 1.57E-05 | 1.49E-05 | 0 | 0 | 1.57E-05 | 0 | 0 | 1.26E-05 | 0 | 1.98E-06 | 4.50E-05 | 1.26E-05 | 0 | 7.15E-06 |
| Tasp1 | 3.06E-05 | 3.01E-05 | 2.12E-05 | 4.54E-06 | 6.64E-06 | 1.41E-05 | 1.77E-07 | 0 | 1.57E-05 | 5.99E-06 | 1.38E-05 | 0 | 7.05E-06 | 9.23E-06 | 3.72E-05 | 1.09E-05 | 0 | 0 |
| Agbl3 | 1.73E-05 | 0 | 9.53E-06 | 0 | 8.63E-06 | 1.25E-05 | 7.96E-06 | 2.20E-05 | 1.57E-05 | 0 | 3.76E-05 | 2.60E-05 | 0 | 1.67E-05 | 9.74E-06 | 1.23E-05 | 0.00015 | 0 |
| Afmid | 2.09E-07 | 4.19E-05 | 9.32E-06 | 1.58E-05 | 9.40E-06 | 5.66E-06 | 0 | 0 | 1.57E-05 | 2.17E-05 | 0 | 1.67E-05 | 2.42E-05 | 0 | 0.0001 | 4.42E-05 | 3.06E-05 | 2.60E-05 |

| | | | | | | | | | | | | | | | | | | |
|---|---|---|---|---|---|---|---|---|---|---|---|---|---|---|---|---|---|---|
| Zdhhc24 | 1.61E-06 | 2.74E-05 | 1.06E-05 | 1.85E-05 | 1.42E-05 | 4.05E-06 | 0 | 2.16E-05 | 1.57E-05 | 2.79E-05 | 2.27E-05 | 0 | 1.16E-05 | 7.26E-06 | 3.07E-05 | 2.59E-08 | 0 | 1.79E-05 |
| Abcf3 | 4.94E-05 | 6.48E-05 | 7.04E-05 | 0.00011 | 2.23E-05 | 0 | 3.13E-05 | 2.00E-05 | 1.57E-05 | 3.15E-05 | 2.67E-05 | 2.92E-05 | 4.95E-05 | 4.00E-05 | 0.00016 | 5.80E-06 | 0 | 2.03E-05 |
| Echdc2 | 1.24E-05 | 2.99E-05 | 2.33E-05 | 3.31E-05 | 0 | 0 | 5.95E-05 | 0 | 1.57E-05 | 0 | 3.91E-05 | 0 | 0.00011 | 9.98E-05 | 8.27E-05 | 4.70E-05 | 1.28E-06 | 0.00012 |
| Mafk | 0 | 0.00014 | 2.01E-05 | 2.02E-05 | 1.10E-05 | 4.69E-05 | 0 | 3.80E-05 | 1.56E-05 | 9.65E-05 | 9.02E-05 | 6.82E-05 | 0 | 6.75E-05 | 5.62E-05 | 0 | 0.00036 | 9.09E-05 |
| Zfp563 | 1.12E-05 | 1.36E-05 | 0 | 4.24E-06 | 2.23E-05 | 1.24E-05 | 0 | 3.71E-06 | 1.56E-05 | 1.70E-05 | 0 | 0 | 2.67E-07 | 0 | 8.23E-06 | 1.20E-06 | 0 | 5.90E-06 |
| Ntn3 | 0 | 1.23E-05 | 0 | 0 | 5.26E-05 | 1.07E-05 | 1.03E-05 | 0 | 1.56E-05 | 0 | 1.60E-05 | 0 | 1.95E-05 | 8.77E-05 | 2.21E-06 | 0 | 0 | 2.77E-05 |
| Taf1b | 7.40E-05 | 8.37E-05 | 2.59E-05 | 3.98E-05 | 2.25E-05 | 7.20E-06 | 0 | 0 | 1.56E-05 | 7.21E-06 | 2.25E-05 | 1.48E-05 | 7.63E-06 | 1.24E-05 | 0.00013 | 3.86E-05 | 0 | 1.08E-05 |
| Zfp59 | 4.52E-06 | 1.17E-05 | 6.15E-06 | 6.14E-06 | 7.68E-06 | 1.99E-06 | 0 | 2.67E-06 | 1.56E-05 | 1.14E-05 | 4.77E-06 | 6.97E-06 | 0 | 0 | 1.92E-05 | 5.99E-07 | 0 | 4.72E-06 |
| Coro1b | 7.31E-06 | 1.67E-05 | 8.03E-06 | 2.30E-05 | 1.03E-05 | 0 | 0 | 8.94E-06 | 1.56E-05 | 9.88E-06 | 2.04E-05 | 1.71E-06 | 2.36E-06 | 1.20E-05 | 0 | 1.49E-07 | 0 | 8.17E-06 |
| Pcdh10 | 0 | 1.17E-05 | 0 | 0 | 1.84E-06 | 0 | 0 | 0 | 1.56E-05 | 0 | 0 | 1.19E-06 | 0 | 0 | 0 | 1.79E-07 | 0 | 3.81E-05 |
| Chuk | 7.44E-05 | 8.86E-05 | 0.00012 | 9.50E-05 | 5.57E-05 | 7.72E-05 | 0 | 0 | 1.55E-05 | 2.20E-05 | 9.28E-05 | 0 | 3.31E-05 | 6.79E-05 | 9.49E-05 | 5.33E-05 | 0.00013 | 0 |
| Mcee | 6.46E-05 | 3.63E-05 | 0 | 6.15E-06 | 3.16E-05 | 5.34E-05 | 4.69E-06 | 0 | 1.55E-05 | 3.48E-05 | 7.14E-06 | 2.61E-05 | 3.00E-05 | 1.29E-05 | 2.21E-05 | 3.11E-06 | 0 | 4.39E-05 |
| Sfmbt1 | 1.70E-05 | 0.0001 | 0.00014 | 0 | 8.08E-05 | 5.12E-05 | 2.42E-05 | 3.23E-05 | 1.55E-05 | 1.13E-05 | 0 | 2.48E-05 | 5.53E-06 | 7.45E-05 | 1.03E-05 | 0.00016 | 2.39E-05 | 0 |
| Tyw1 | 2.73E-05 | 9.22E-05 | 8.19E-05 | 7.11E-05 | 5.30E-05 | 3.36E-05 | 2.80E-05 | 5.42E-05 | 1.55E-05 | 6.37E-05 | 2.94E-05 | 3.55E-05 | 5.22E-05 | 0 | 0.00016 | 0.00013 | 3.04E-05 | 3.61E-05 |
| Ccdc115 | 4.69E-05 | 3.17E-05 | 4.36E-05 | 6.94E-05 | 4.02E-05 | 3.01E-05 | 0 | 1.17E-05 | 1.55E-05 | 2.02E-05 | 1.74E-05 | 0.00011 | 0 | 1.13E-05 | 7.86E-05 | 6.18E-06 | 0 | 1.99E-05 |
| Mto1 | 2.33E-05 | 4.79E-05 | 7.73E-05 | 7.18E-05 | 4.86E-05 | 2.77E-05 | 8.84E-06 | 0 | 1.55E-05 | 3.39E-05 | 6.05E-06 | 6.02E-08 | 2.33E-05 | 3.20E-05 | 3.52E-05 | 2.77E-05 | 0 | 3.76E-05 |
| Dusp16 | 2.46E-05 | 6.15E-05 | 6.42E-05 | 0 | 6.54E-06 | 2.36E-05 | 0 | 3.15E-05 | 1.55E-05 | 7.75E-05 | 0.0001 | 6.47E-05 | 1.53E-05 | 2.80E-05 | 1.17E-05 | 2.39E-05 | 0.00015 | 3.32E-06 |
| Wdyhv1 | 3.67E-05 | 3.31E-05 | 2.01E-05 | 3.06E-05 | 4.15E-05 | 1.54E-05 | 1.39E-05 | 0 | 1.55E-05 | 2.22E-05 | 3.21E-05 | 1.51E-05 | 1.89E-05 | 5.97E-05 | 2.43E-05 | 5.69E-05 | 0 | 1.24E-05 |
| Tspyl3 | 0 | 1.99E-05 | 0 | 3.47E-05 | 1.71E-05 | 1.00E-05 | 0 | 3.29E-05 | 1.55E-05 | 1.37E-05 | 0 | 0 | 0 | 1.28E-05 | 2.64E-05 | 0 | 0 | 3.50E-05 |
| Bag2 | 2.70E-06 | 5.09E-06 | 5.62E-06 | 5.24E-06 | 0 | 0 | 1.74E-05 | 0 | 1.55E-05 | 7.86E-06 | 8.60E-05 | 8.16E-05 | 1.16E-05 | 1.78E-05 | 0 | 0 | 0 | 1.54E-05 |
| Abhd5 | 9.11E-05 | 0.0001 | 6.27E-05 | 1.26E-05 | 2.48E-05 | 0.00012 | 2.66E-05 | 0 | 1.54E-05 | 7.04E-05 | 4.02E-06 | 9.65E-05 | 0.00019 | 5.64E-05 | 0 | 0 | 0.00046 | 1.87E-05 |
| Kat2b | 6.01E-05 | 6.26E-05 | 0 | 0 | 1.58E-05 | 4.97E-05 | 0 | 5.61E-06 | 1.54E-05 | 2.47E-05 | 0 | 0 | 1.71E-05 | 0 | 0 | 2.80E-05 | 5.70E-05 | 3.52E-05 |
| Tbp | 9.00E-05 | 6.79E-05 | 5.01E-05 | 0 | 3.92E-05 | 3.66E-05 | 0 | 0 | 1.54E-05 | 7.79E-06 | 8.91E-07 | 5.92E-05 | 5.18E-06 | 3.10E-05 | 0.00012 | 9.19E-05 | 0 | 0 |
| Dnajb3 | 0 | 7.34E-06 | 5.80E-06 | 3.81E-05 | 3.16E-05 | 3.38E-06 | 9.08E-06 | 1.31E-05 | 1.54E-05 | 2.81E-06 | 1.33E-05 | 2.05E-05 | 0 | 8.14E-06 | 9.76E-07 | 1.60E-05 | 0 | 1.82E-05 |
| Ints5 | 8.48E-06 | 2.14E-05 | 2.76E-05 | 2.55E-05 | 1.39E-06 | 0 | 2.84E-06 | 0 | 1.54E-05 | 1.12E-05 | 1.39E-05 | 1.54E-06 | 0 | 9.60E-06 | 9.15E-06 | 1.24E-05 | 0 | 1.18E-06 |
| Dph2 | 1.31E-05 | 1.51E-05 | 5.41E-05 | 2.14E-05 | 0 | 0 | 0 | 0 | 1.54E-05 | 1.25E-05 | 3.40E-06 | 0 | 3.02E-05 | 1.52E-06 | 2.82E-05 | 1.50E-05 | 1.39E-05 | 1.30E-05 |
| Gfpt1 | 0.00036 | 5.59E-05 | 0.0002 | 5.96E-05 | 8.00E-05 | 0.00014 | 0.00029 | 0.00016 | 1.53E-05 | 4.83E-05 | 4.28E-05 | 0.0002 | 1.66E-06 | 4.87E-06 | 0.00028 | 5.07E-05 | 0.00021 | 0 |
| Pdk2 | 7.16E-06 | 8.25E-05 | 0 | 4.63E-05 | 8.67E-05 | 6.04E-05 | 6.83E-05 | 3.30E-05 | 1.53E-05 | 6.47E-05 | 7.77E-06 | 1.78E-05 | 2.22E-05 | 5.31E-05 | 0 | 0 | 0 | 9.36E-05 |
| Ipo7 | 7.66E-05 | 1.91E-05 | 7.73E-05 | 3.35E-06 | 0 | 4.76E-05 | 0 | 0 | 1.53E-05 | 2.16E-06 | 3.47E-06 | 2.77E-05 | 0 | 0 | 5.32E-07 | 5.16E-05 | 5.55E-06 | 0 |
| Flrt1 | 0 | 0 | 0 | 0 | 6.89E-05 | 1.57E-05 | 0 | 0 | 1.53E-05 | 1.75E-07 | 2.39E-05 | 0 | 0 | 7.18E-05 | 1.40E-06 | 1.83E-05 | 0 | 0 |
| Naa80 | 6.23E-05 | 0.00011 | 0 | 0.00014 | 1.62E-05 | 1.51E-05 | 7.84E-05 | 4.55E-05 | 1.53E-05 | 4.83E-05 | 0 | 4.17E-05 | 1.46E-06 | 1.68E-05 | 1.93E-05 | 1.25E-05 | 3.28E-08 | 6.70E-05 |
| Letm2 | 3.39E-06 | 2.07E-05 | 0 | 0 | 6.64E-06 | 1.45E-05 | 0 | 6.92E-06 | 1.53E-05 | 6.55E-06 | 0 | 3.70E-05 | 0 | 0 | 1.66E-06 | 0 | 0 | 1.39E-05 |
| Rbm8a | 5.03E-05 | 3.65E-05 | 3.04E-05 | 2.96E-05 | 1.35E-05 | 8.77E-06 | 0 | 4.42E-06 | 1.53E-05 | 8.17E-06 | 7.25E-06 | 9.68E-06 | 0 | 1.68E-05 | 1.74E-05 | 5.58E-05 | 1.44E-05 | 1.24E-05 |

| | | | | | | | | | | | | | | | | | | |
|---|---|---|---|---|---|---|---|---|---|---|---|---|---|---|---|---|---|---|
| Zfp322a | 3.42E-05 | 2.97E-05 | 0 | 0 | 1.74E-05 | 7.97E-06 | 0 | 0 | 1.53E-05 | 6.22E-06 | 1.39E-05 | 2.36E-06 | 0 | 3.14E-05 | 5.20E-05 | 0 | 0 | 1.71E-06 |
| Zfp763 | 9.61E-06 | 1.06E-05 | 0 | 0 | 1.45E-05 | 6.00E-06 | 0 | 4.35E-06 | 1.53E-05 | 1.22E-05 | 0 | 1.19E-06 | 0 | 0 | 8.93E-06 | 1.69E-05 | 0 | 5.58E-06 |
| Zfp12 | 3.64E-06 | 3.27E-05 | 1.39E-05 | 3.67E-06 | 1.02E-05 | 2.82E-06 | 0 | 0 | 1.53E-05 | 2.67E-06 | 9.85E-06 | 1.80E-05 | 0 | 1.26E-05 | 6.24E-05 | 0 | 0 | 1.52E-05 |
| Fxn | 8.14E-06 | 1.06E-06 | 2.91E-06 | 1.85E-05 | 2.79E-06 | 0 | 6.92E-06 | 8.35E-06 | 1.53E-05 | 0 | 2.52E-06 | 0 | 0 | 9.78E-06 | 3.15E-07 | 3.03E-05 | 0 | 4.56E-06 |
| Rnf2 | 0.0002 | 0.00011 | 7.50E-05 | 0 | 4.10E-05 | 0.00018 | 6.24E-05 | 5.17E-05 | 1.52E-05 | 0 | 0 | 9.44E-05 | 0 | 0 | 6.77E-05 | 7.09E-05 | 9.77E-05 | 0 |
| Pyroxd1 | 6.26E-05 | 4.29E-05 | 6.35E-05 | 2.04E-05 | 2.99E-05 | 9.06E-05 | 6.58E-06 | 3.53E-05 | 1.52E-05 | 0 | 6.08E-06 | 1.73E-05 | 0 | 8.31E-06 | 0.00011 | 1.26E-06 | 0 | 0 |
| Dxo | 7.13E-06 | 7.84E-05 | 8.05E-05 | 7.86E-05 | 2.05E-05 | 5.76E-05 | 4.54E-05 | 7.18E-05 | 1.52E-05 | 7.50E-05 | 5.02E-05 | 1.14E-05 | 5.63E-05 | 5.84E-05 | 7.35E-05 | 0.00019 | 3.23E-05 | 5.73E-05 |
| Pex3 | 0.0001 | 3.05E-05 | 1.39E-05 | 0 | 2.20E-05 | 3.36E-05 | 6.32E-07 | 0 | 1.52E-05 | 1.38E-05 | 0 | 2.79E-05 | 0 | 1.05E-06 | 0 | 1.86E-05 | 0 | 1.45E-05 |
| Pou6f1 | 0 | 6.33E-05 | 0 | 0 | 5.56E-05 | 2.80E-05 | 4.21E-06 | 0 | 1.52E-05 | 1.49E-05 | 2.44E-05 | 6.94E-05 | 3.57E-05 | 1.83E-05 | 2.61E-05 | 0 | 0 | 4.78E-05 |
| Yars2 | 5.83E-05 | 2.87E-05 | 0.0001 | 3.24E-05 | 6.88E-06 | 2.09E-05 | 0 | 0 | 1.52E-05 | 1.65E-05 | 4.94E-05 | 4.05E-05 | 0 | 2.19E-06 | 9.45E-06 | 5.59E-05 | 1.96E-05 | 0 |
| Tspyl4 | 3.07E-05 | 1.70E-05 | 7.37E-06 | 4.86E-06 | 5.07E-07 | 1.25E-05 | 0 | 1.58E-05 | 1.52E-05 | 4.81E-05 | 0 | 0 | 0 | 0 | 9.19E-06 | 0 | 0 | 2.59E-05 |
| Zswim9 | 7.05E-06 | 1.97E-05 | 2.13E-06 | 1.11E-05 | 2.19E-05 | 5.51E-06 | 4.72E-06 | 9.56E-06 | 1.52E-05 | 4.57E-06 | 4.90E-06 | 6.13E-06 | 5.36E-06 | 1.98E-07 | 3.01E-05 | 1.12E-05 | 1.23E-05 | 6.60E-06 |
| Npy1r | 9.14E-06 | 3.25E-06 | 0 | 0 | 1.14E-05 | 0 | 0 | 0 | 1.52E-05 | 4.46E-05 | 2.73E-07 | 0 | 9.04E-06 | 0 | 0 | 0 | 0 | 2.99E-05 |
| Dusp28 | 0 | 4.43E-05 | 1.99E-05 | 4.11E-05 | 3.96E-05 | 0 | 5.42E-05 | 1.07E-05 | 1.52E-05 | 0 | 0 | 3.51E-05 | 3.36E-05 | 1.04E-06 | 4.18E-05 | 2.29E-05 | 1.08E-05 | 0 |
| Slco5a1 | 0 | 9.60E-06 | 0 | 0 | 0.0002 | 0.00038 | 0.00017 | 0 | 1.51E-05 | 0 | 0 | 0.00021 | 0.00019 | 2.75E-05 | 7.10E-05 | 0 | 4.79E-05 | 0 |
| Iqgap1 | 0.00024 | 0.00035 | 0.00048 | 0.00026 | 0.00016 | 0.00021 | 0.00065 | 0.00033 | 1.51E-05 | 0.00025 | 0.00069 | 0.00098 | 9.37E-05 | 0.00013 | 0.00052 | 0.00027 | 0.00052 | 0.00069 |
| Ccnl1 | 0.00025 | 0.00021 | 7.16E-05 | 0 | 6.32E-05 | 0.00021 | 0 | 0.00011 | 1.51E-05 | 0 | 9.15E-06 | 0.00016 | 0 | 4.99E-05 | 0.00071 | 3.32E-05 | 0.00211 | 7.65E-05 |
| Stag2 | 0.00023 | 0.00014 | 0 | 0 | 2.49E-05 | 0.00017 | 4.78E-05 | 0 | 1.51E-05 | 1.86E-05 | 0 | 0.00015 | 9.94E-06 | 0 | 6.53E-05 | 0.00017 | 0.00017 | 0 |
| Usp31 | 1.44E-05 | 9.80E-05 | 0.00017 | 1.49E-05 | 0 | 5.15E-05 | 3.36E-06 | 1.71E-05 | 1.51E-05 | 1.25E-05 | 6.42E-05 | 7.37E-05 | 0 | 5.16E-05 | 5.81E-05 | 8.35E-05 | 6.55E-05 | 0 |
| Ythdc1 | 6.56E-05 | 0.00011 | 4.79E-05 | 3.63E-05 | 4.11E-05 | 4.27E-05 | 0 | 1.01E-05 | 1.51E-05 | 1.96E-05 | 5.60E-05 | 8.84E-05 | 0 | 1.96E-05 | 0.00021 | 8.51E-05 | 0.00014 | 0 |
| Endog | 6.56E-05 | 1.42E-05 | 2.01E-05 | 0.00014 | 2.19E-06 | 2.59E-05 | 8.33E-05 | 2.41E-05 | 1.51E-05 | 2.48E-05 | 6.19E-05 | 9.61E-05 | 0 | 9.31E-05 | 0 | 0 | 0 | 0 |
| Rubcnl | 0 | 8.99E-06 | 0 | 2.13E-06 | 2.41E-06 | 2.21E-05 | 4.37E-06 | 9.47E-06 | 1.51E-05 | 7.63E-06 | 0 | 2.99E-05 | 0 | 0 | 2.10E-06 | 0 | 0 | 0 |
| Lcmt2 | 2.36E-05 | 2.89E-05 | 1.69E-05 | 0 | 1.33E-05 | 1.35E-05 | 3.46E-06 | 6.41E-06 | 1.51E-05 | 1.17E-05 | 2.08E-05 | 0 | 0 | 6.52E-06 | 2.22E-05 | 1.28E-05 | 0 | 1.53E-05 |
| Zfp784 | 9.26E-07 | 1.28E-05 | 0 | 3.84E-06 | 7.75E-06 | 2.35E-06 | 2.24E-06 | 4.59E-06 | 1.51E-05 | 5.95E-06 | 4.15E-07 | 0 | 4.59E-06 | 8.90E-06 | 2.65E-06 | 0 | 0 | 1.58E-05 |
| Eepd1 | 0 | 3.88E-05 | 0 | 0 | 2.33E-05 | 0 | 1.91E-05 | 0 | 1.51E-05 | 0 | 0 | 0 | 0 | 6.55E-05 | 2.43E-05 | 1.10E-06 | 1.76E-05 | 0 |
| Dcun1d2 | 4.07E-05 | 3.49E-05 | 2.56E-05 | 4.61E-06 | 2.60E-05 | 0 | 2.00E-05 | 0 | 1.51E-05 | 1.85E-05 | 3.74E-06 | 8.17E-06 | 0 | 0 | 2.34E-05 | 3.00E-05 | 0 | 2.80E-05 |
| Ddx59 | 5.04E-06 | 6.21E-06 | 4.50E-06 | 1.05E-05 | 5.56E-06 | 0 | 6.78E-06 | 6.19E-06 | 1.51E-05 | 5.61E-06 | 1.04E-05 | 0 | 0 | 5.25E-06 | 1.54E-05 | 3.41E-05 | 8.38E-06 | 5.80E-06 |
| Eif1ax | 0.00028 | 5.78E-05 | 0.00014 | 0 | 0 | 0.00015 | 3.17E-05 | 0 | 1.50E-05 | 0 | 6.09E-06 | 4.83E-05 | 0 | 0 | 0 | 6.02E-05 | 2.22E-05 | 4.68E-05 |
| Nbeal1 | 0.00011 | 0.00013 | 5.18E-06 | 0 | 6.11E-05 | 0.00012 | 6.89E-05 | 3.66E-05 | 1.50E-05 | 7.19E-05 | 0.00022 | 4.81E-05 | 5.02E-05 | 5.62E-05 | 0.00012 | 0 | 0.00013 | 0.00016 |
| Fam161a | 0 | 1.80E-05 | 1.14E-05 | 0 | 3.73E-06 | 9.35E-06 | 2.96E-05 | 0 | 1.50E-05 | 7.00E-06 | 0 | 2.02E-06 | 1.85E-05 | 9.34E-06 | 5.43E-05 | 0 | 1.65E-05 | 4.53E-06 |
| Dmac2l | 2.95E-05 | 2.23E-05 | 0 | 0 | 1.16E-05 | 4.90E-06 | 0 | 3.92E-06 | 1.50E-05 | 2.09E-05 | 5.09E-06 | 1.86E-05 | 3.56E-06 | 0 | 6.29E-06 | 1.73E-06 | 0 | 9.62E-06 |
| Mettl7a1 | 0 | 4.68E-05 | 0 | 0 | 1.60E-05 | 4.65E-06 | 0 | 0 | 1.50E-05 | 7.13E-05 | 0 | 0 | 1.10E-05 | 0 | 2.12E-05 | 0 | 0 | 3.20E-05 |
| Alkbh4 | 6.29E-07 | 2.07E-05 | 3.22E-06 | 3.14E-05 | 1.30E-05 | 0 | 0 | 2.43E-05 | 1.50E-05 | 1.40E-05 | 3.12E-06 | 5.22E-06 | 0 | 2.55E-05 | 3.57E-05 | 1.34E-05 | 0 | 1.11E-05 |

| | | | | | | | | | | | | | | | | | | |
|---|---|---|---|---|---|---|---|---|---|---|---|---|---|---|---|---|---|---|
| Snrnp25 | 4.43E-05 | 1.06E-05 | 0 | 6.52E-05 | 9.43E-06 | 0 | 2.37E-05 | 0 | 1.50E-05 | 1.09E-05 | 3.82E-06 | 5.37E-05 | 0 | 0 | 0 | 0.00013 | 0 | 1.88E-05 |
| Apba1 | 2.00E-05 | 4.69E-05 | 1.86E-06 | 0 | 0.0001 | 8.60E-05 | 6.44E-05 | 0 | 1.49E-05 | 8.86E-05 | 5.11E-05 | 4.89E-05 | 5.03E-05 | 6.94E-05 | 0 | 1.04E-05 | 3.79E-05 | 9.12E-05 |
| Ulk1 | 2.13E-05 | 0.00021 | 1.34E-06 | 5.10E-05 | 0.00011 | 8.01E-05 | 0.00012 | 6.11E-05 | 1.49E-05 | 5.51E-05 | 8.40E-05 | 0.0003 | 8.96E-05 | 0.00023 | 0.00013 | 7.64E-05 | 0.00019 | 0.00013 |
| Lyn | 9.73E-05 | 2.44E-05 | 3.08E-05 | 0 | 3.54E-06 | 6.17E-05 | 0 | 0.00012 | 1.49E-05 | 0 | 0.00017 | 0 | 0 | 0.0002 | 2.97E-05 | 3.50E-05 | 6.21E-05 | 0 |
| Tbc1d23 | 7.20E-05 | 6.09E-05 | 4.50E-05 | 2.76E-05 | 2.60E-05 | 2.51E-05 | 0.00016 | 1.92E-05 | 1.49E-05 | 3.67E-05 | 6.03E-05 | 5.23E-05 | 3.43E-05 | 1.33E-05 | 9.28E-05 | 2.33E-06 | 6.44E-05 | 7.28E-06 |
| Armc9 | 4.41E-05 | 3.84E-05 | 2.60E-05 | 3.35E-05 | 3.59E-05 | 2.24E-05 | 3.07E-05 | 1.58E-05 | 1.49E-05 | 1.92E-05 | 1.37E-05 | 0 | 5.29E-06 | 1.36E-05 | 4.43E-05 | 0 | 4.68E-05 | 3.69E-05 |
| Zfp937 | 1.50E-05 | 1.66E-05 | 0 | 0 | 9.40E-06 | 1.44E-05 | 2.51E-06 | 1.18E-06 | 1.49E-05 | 8.47E-06 | 0 | 0 | 1.03E-05 | 0 | 0 | 2.57E-06 | 0 | 0 |
| Ttll11 | 0 | 1.41E-06 | 4.77E-06 | 0 | 1.26E-05 | 1.34E-05 | 0 | 4.56E-06 | 1.49E-05 | 0 | 2.61E-05 | 5.24E-05 | 0 | 1.41E-05 | 9.08E-06 | 0 | 0 | 0 |
| Zfp882 | 1.34E-05 | 2.13E-05 | 1.71E-06 | 1.50E-06 | 1.22E-05 | 3.16E-06 | 0 | 6.66E-07 | 1.49E-05 | 2.82E-06 | 0 | 1.38E-06 | 0 | 1.30E-06 | 0 | 2.32E-06 | 0 | 3.15E-06 |
| Daam1 | 7.99E-05 | 3.65E-05 | 0 | 0 | 6.44E-06 | 0.00018 | 0 | 0 | 1.48E-05 | 0.00021 | 0.00026 | 0.00034 | 4.79E-05 | 0 | 0 | 0 | 8.85E-06 | 1.73E-05 |
| Etfrf1 | 9.05E-05 | 8.95E-05 | 0 | 0.00011 | 6.13E-05 | 0.00012 | 1.27E-05 | 0.0001 | 1.48E-05 | 3.10E-05 | 0 | 7.96E-05 | 9.16E-05 | 4.27E-05 | 0 | 0 | 0 | 4.59E-06 |
| Plekhg3 | 7.11E-05 | 0.00011 | 0.00011 | 0 | 6.54E-05 | 5.89E-05 | 3.50E-05 | 0.0001 | 1.48E-05 | 5.96E-05 | 0.00014 | 3.11E-05 | 0 | 2.22E-05 | 0.00015 | 0.0001 | 0.00012 | 0.00021 |
| Zkscan8 | 4.82E-05 | 4.96E-05 | 5.62E-06 | 2.44E-05 | 5.93E-05 | 4.86E-05 | 0 | 1.72E-05 | 1.48E-05 | 2.26E-05 | 1.48E-05 | 0 | 9.11E-06 | 6.27E-07 | 6.42E-05 | 0 | 0 | 2.12E-05 |
| Slc17a9 | 4.17E-06 | 0 | 2.80E-05 | 3.27E-05 | 0 | 3.48E-05 | 4.05E-05 | 2.22E-05 | 1.48E-05 | 0 | 2.46E-05 | 1.48E-05 | 0 | 3.27E-05 | 3.34E-05 | 0 | 0 | 0 |
| Uchl5 | 9.76E-05 | 4.88E-05 | 3.09E-05 | 6.44E-06 | 1.85E-05 | 2.73E-05 | 1.20E-06 | 3.77E-06 | 1.48E-05 | 0 | 1.01E-05 | 3.49E-05 | 0 | 0 | 3.42E-05 | 0.00027 | 9.98E-06 | 0 |
| Zfp764 | 1.66E-05 | 2.86E-05 | 7.41E-06 | 1.74E-07 | 2.41E-05 | 2.26E-05 | 4.35E-05 | 1.22E-05 | 1.48E-05 | 9.67E-06 | 0 | 5.44E-06 | 2.12E-06 | 1.84E-05 | 0 | 1.31E-05 | 0 | 1.60E-05 |
| Rpusd2 | 7.37E-06 | 1.59E-05 | 2.79E-05 | 5.33E-06 | 1.21E-05 | 1.38E-05 | 0 | 0 | 1.48E-05 | 6.92E-06 | 1.08E-05 | 0 | 0 | 2.26E-05 | 1.69E-05 | 5.13E-06 | 0 | 3.72E-06 |
| Kyat3 | 3.20E-05 | 7.37E-06 | 4.02E-06 | 1.48E-05 | 0 | 6.44E-06 | 0 | 0 | 1.48E-05 | 3.29E-06 | 0 | 3.40E-07 | 5.12E-06 | 0 | 0 | 1.28E-05 | 4.33E-06 | 0 |
| Rpgrip1l | 1.15E-05 | 3.00E-05 | 0 | 1.56E-05 | 2.05E-05 | 9.58E-07 | 6.50E-06 | 4.29E-06 | 1.48E-05 | 2.53E-05 | 1.29E-05 | 0 | 0 | 6.80E-06 | 8.17E-06 | 3.61E-05 | 0 | 1.41E-05 |
| Map3k11 | 1.51E-05 | 5.72E-05 | 3.24E-05 | 5.72E-05 | 2.24E-05 | 0 | 0 | 3.72E-05 | 1.48E-05 | 1.47E-05 | 1.77E-05 | 1.93E-05 | 0 | 2.61E-05 | 0.00016 | 3.07E-05 | 9.00E-05 | 5.31E-05 |
| Osbpl10 | 3.53E-06 | 0 | 2.27E-05 | 4.91E-06 | 2.82E-05 | 4.78E-05 | 5.54E-05 | 2.20E-05 | 1.47E-05 | 0 | 0 | 7.49E-05 | 0 | 0 | 2.78E-05 | 2.85E-06 | 6.77E-06 | 2.17E-05 |
| Dclre1b | 1.51E-05 | 2.56E-05 | 3.27E-05 | 2.21E-05 | 0 | 1.28E-05 | 0 | 0 | 1.47E-05 | 1.50E-05 | 1.21E-05 | 0 | 5.82E-06 | 8.59E-06 | 0 | 9.11E-05 | 0 | 1.59E-05 |
| H2-Q10 | 0 | 0 | 0 | 0 | 0 | 7.89E-06 | 0 | 7.50E-06 | 1.47E-05 | 0 | 0 | 2.93E-05 | 3.87E-05 | 4.80E-06 | 1.86E-05 | 0 | 0 | 0 |
| BC052040 | 0 | 8.73E-06 | 0 | 2.18E-05 | 3.50E-06 | 5.79E-06 | 7.66E-06 | 0 | 1.47E-05 | 7.46E-06 | 0 | 1.34E-05 | 0 | 0 | 0 | 7.45E-05 | 5.91E-06 | 2.74E-06 |
| Fchsd1 | 0 | 4.51E-05 | 3.79E-07 | 1.14E-05 | 7.57E-06 | 5.66E-06 | 0 | 8.93E-06 | 1.47E-05 | 4.43E-05 | 2.87E-05 | 2.75E-05 | 1.17E-06 | 0 | 4.21E-05 | 0 | 1.98E-05 | 4.71E-05 |
| Gba2 | 2.13E-05 | 3.00E-05 | 6.64E-06 | 5.13E-06 | 1.84E-05 | 2.95E-06 | 1.95E-07 | 3.24E-05 | 1.47E-05 | 1.82E-05 | 2.63E-05 | 6.29E-06 | 1.47E-06 | 1.71E-05 | 5.48E-05 | 3.70E-05 | 5.12E-06 | 8.27E-06 |
| Spire2 | 0 | 0 | 2.72E-06 | 1.94E-06 | 0 | 9.28E-07 | 0 | 0 | 1.47E-05 | 1.73E-06 | 1.10E-05 | 6.08E-06 | 0 | 0 | 0 | 3.81E-06 | 4.65E-06 | 1.87E-05 |
| Raver2 | 1.85E-06 | 0 | 0 | 0 | 1.82E-05 | 0 | 4.35E-05 | 4.75E-05 | 1.47E-05 | 0 | 2.98E-07 | 0 | 0 | 0 | 0 | 2.79E-05 | 0 | 4.26E-06 |
| Zfp605 | 3.20E-06 | 4.14E-06 | 0 | 0 | 8.44E-06 | 0 | 4.54E-06 | 1.46E-06 | 1.47E-05 | 1.07E-05 | 9.94E-07 | 9.49E-06 | 1.57E-06 | 0 | 0 | 4.09E-06 | 9.16E-06 | 5.09E-06 |
| Trim62 | 4.79E-06 | 5.05E-05 | 2.42E-05 | 1.56E-05 | 8.05E-06 | 0 | 2.94E-05 | 0 | 1.47E-05 | 1.46E-06 | 2.42E-05 | 0 | 1.44E-06 | 4.84E-05 | 2.50E-05 | 1.07E-05 | 1.93E-05 | 5.10E-06 |
| Lins1 | 2.96E-05 | 1.89E-05 | 3.30E-05 | 1.67E-05 | 1.06E-05 | 0 | 6.60E-07 | 0 | 1.47E-05 | 1.71E-05 | 2.91E-05 | 1.80E-05 | 7.48E-06 | 1.47E-05 | 1.68E-05 | 5.71E-07 | 1.19E-05 | 3.77E-07 |
| Fam8a1 | 0.00031 | 0.00012 | 7.44E-05 | 0 | 7.98E-05 | 0.00022 | 0.00026 | 0.00015 | 1.46E-05 | 0.00011 | 0.00013 | 0.00019 | 0.0001 | 3.68E-05 | 0 | 0 | 7.40E-06 | 0.0001 |
| Zfp623 | 4.19E-05 | 3.73E-05 | 8.95E-06 | 0 | 2.92E-05 | 2.13E-05 | 1.17E-05 | 0 | 1.46E-05 | 6.72E-06 | 6.65E-06 | 9.82E-06 | 0 | 0 | 3.38E-05 | 0 | 0 | 1.45E-05 |

| | | | | | | | | | | | | | | | | | | |
|---|---|---|---|---|---|---|---|---|---|---|---|---|---|---|---|---|---|---|
| H2-D1 | 1.18E-05 | 8.90E-05 | 6.03E-05 | 0 | 0 | 1.34E-05 | 0 | 1.70E-05 | 1.46E-05 | 0.00027 | 0 | 3.49E-05 | 0.00033 | 0.00013 | 0 | 1.36E-05 | 7.84E-05 | 0.00022 |
| Purg | 0 | 3.20E-05 | 1.21E-07 | 3.39E-05 | 1.27E-05 | 1.07E-05 | 6.74E-06 | 2.24E-05 | 1.46E-05 | 7.96E-06 | 2.50E-05 | 0 | 0 | 3.88E-05 | 0.00011 | 7.33E-06 | 4.34E-05 | 3.94E-05 |
| Mnat1 | 0.00011 | 6.27E-05 | 2.49E-05 | 0.00011 | 2.71E-05 | 8.77E-06 | 8.69E-05 | 1.35E-05 | 1.46E-05 | 2.25E-05 | 3.72E-05 | 5.20E-05 | 2.10E-05 | 6.48E-06 | 0.00017 | 4.26E-05 | 0 | 1.71E-05 |
| Mms22l | 9.47E-06 | 0 | 3.44E-05 | 0 | 7.44E-06 | 8.31E-06 | 1.34E-06 | 0 | 1.46E-05 | 4.47E-06 | 0 | 0 | 7.09E-06 | 1.99E-05 | 1.49E-05 | 0.00031 | 0 | 0 |
| Map10 | 9.42E-06 | 1.20E-05 | 3.57E-06 | 0 | 8.60E-06 | 6.25E-06 | 3.31E-06 | 5.39E-06 | 1.46E-05 | 2.15E-06 | 6.52E-06 | 1.10E-05 | 2.32E-06 | 0 | 0 | 3.18E-06 | 0 | 1.01E-05 |
| Akap17b | 1.16E-05 | 1.77E-06 | 9.97E-07 | 0 | 0 | 5.79E-06 | 1.05E-05 | 0 | 1.46E-05 | 1.35E-05 | 3.61E-06 | 0 | 0 | 1.12E-05 | 1.22E-06 | 0 | 6.35E-06 | 2.15E-06 |
| Hemk1 | 1.57E-05 | 2.86E-05 | 0 | 2.37E-05 | 7.99E-06 | 4.54E-06 | 0 | 8.20E-06 | 1.46E-05 | 6.18E-06 | 2.30E-06 | 9.95E-06 | 4.91E-06 | 0 | 6.85E-05 | 1.47E-06 | 1.50E-06 | 4.95E-06 |
| Pcdhb14 | 1.41E-05 | 1.92E-05 | 0 | 0 | 2.59E-05 | 0 | 0 | 2.24E-06 | 1.46E-05 | 1.30E-06 | 8.07E-07 | 9.76E-06 | 9.46E-06 | 9.14E-06 | 9.28E-06 | 0 | 4.99E-06 | 1.75E-05 |
| Dtd1 | 1.27E-05 | 2.62E-05 | 1.10E-05 | 3.06E-05 | 4.49E-05 | 0 | 0 | 0 | 1.46E-05 | 4.54E-05 | 7.99E-06 | 4.80E-07 | 5.07E-05 | 1.87E-05 | 0 | 0 | 1.54E-05 | 2.58E-05 |
| Tmem209 | 6.98E-05 | 5.66E-05 | 7.43E-05 | 5.15E-08 | 1.58E-05 | 4.35E-05 | 0 | 4.30E-05 | 1.45E-05 | 1.74E-05 | 0 | 3.33E-05 | 6.12E-06 | 9.72E-06 | 1.85E-05 | 0.00022 | 0 | 1.70E-05 |
| Nudt13 | 9.92E-06 | 2.65E-05 | 0 | 1.29E-05 | 1.02E-05 | 2.43E-05 | 0 | 4.84E-06 | 1.45E-05 | 3.11E-05 | 0 | 1.14E-05 | 1.68E-06 | 0 | 4.72E-05 | 2.26E-05 | 9.26E-07 | 3.89E-05 |
| Shld3 | 2.77E-05 | 1.14E-05 | 0 | 0 | 1.47E-05 | 8.88E-06 | 0 | 0 | 1.45E-05 | 0 | 0 | 0 | 0 | 8.26E-06 | 2.17E-05 | 2.61E-05 | 0 | 0 |
| Pabpc4l | 1.37E-05 | 5.91E-06 | 0 | 1.45E-05 | 2.73E-05 | 8.66E-06 | 3.58E-05 | 1.37E-05 | 1.45E-05 | 0 | 5.88E-06 | 0 | 0 | 2.51E-06 | 7.64E-06 | 0 | 0 | 7.51E-06 |
| Tmub1 | 1.15E-05 | 3.23E-05 | 0 | 2.67E-05 | 1.03E-05 | 0 | 2.87E-05 | 0 | 1.45E-05 | 1.18E-05 | 3.27E-05 | 4.01E-05 | 0 | 1.59E-05 | 0 | 0 | 0 | 1.73E-05 |
| Ift122 | 2.54E-05 | 4.60E-05 | 0 | 4.71E-05 | 2.37E-05 | 0 | 2.35E-05 | 1.65E-05 | 1.45E-05 | 3.62E-05 | 0 | 1.40E-05 | 5.25E-05 | 2.21E-05 | 4.00E-05 | 0 | 2.49E-05 | 4.60E-05 |
| Rint1 | 8.04E-05 | 4.71E-05 | 8.41E-05 | 0.0001 | 2.76E-05 | 0 | 7.17E-05 | 2.86E-05 | 1.45E-05 | 2.44E-05 | 3.73E-05 | 2.14E-05 | 0 | 2.67E-05 | 0.00016 | 4.96E-05 | 0 | 2.48E-05 |
| Bmp2k | 0.00014 | 9.91E-05 | 1.90E-05 | 0 | 7.33E-05 | 9.45E-05 | 0 | 5.67E-05 | 1.44E-05 | 4.38E-05 | 5.00E-05 | 5.65E-05 | 0 | 0.0002 | 4.21E-05 | 6.40E-05 | 0.00016 | 1.97E-05 |
| Pyroxd2 | 0 | 6.72E-05 | 0 | 3.21E-05 | 4.12E-05 | 5.97E-05 | 8.44E-05 | 5.88E-05 | 1.44E-05 | 0.00019 | 0 | 4.91E-05 | 6.35E-05 | 0 | 0.00013 | 0 | 1.36E-05 | 5.02E-05 |
| Crim1 | 0 | 0.00011 | 0 | 0 | 6.90E-06 | 5.74E-05 | 3.40E-06 | 0 | 1.44E-05 | 5.90E-05 | 0.0004 | 7.16E-05 | 9.84E-05 | 3.04E-05 | 0 | 0 | 7.02E-05 | 0 |
| Mycbp | 0.00014 | 0 | 0.00015 | 0.00011 | 5.66E-06 | 3.11E-05 | 6.20E-05 | 9.67E-05 | 1.44E-05 | 2.26E-05 | 0.0001 | 1.42E-05 | 0 | 7.63E-05 | 1.39E-05 | 0.00013 | 0 | 0 |
| Fbxo27 | 1.66E-05 | 1.80E-05 | 0 | 0 | 1.65E-05 | 1.54E-05 | 2.26E-05 | 3.04E-06 | 1.44E-05 | 1.35E-05 | 4.84E-07 | 2.09E-05 | 2.24E-05 | 1.02E-05 | 7.52E-06 | 0 | 0 | 3.28E-05 |
| Slc37a4 | 1.09E-06 | 9.71E-06 | 0 | 2.40E-05 | 7.48E-06 | 1.47E-05 | 5.07E-06 | 3.58E-05 | 1.44E-05 | 3.27E-05 | 2.12E-05 | 3.97E-05 | 0 | 9.03E-06 | 5.21E-06 | 0 | 0 | 3.00E-05 |
| Fam241a | 3.41E-06 | 2.02E-05 | 1.55E-05 | 1.77E-06 | 0 | 1.35E-05 | 0 | 3.12E-06 | 1.44E-05 | 0 | 0 | 5.79E-06 | 3.15E-05 | 5.54E-05 | 8.84E-06 | 0 | 5.16E-06 | 1.73E-05 |
| Zfp273 | 9.14E-06 | 7.97E-06 | 0 | 0 | 2.05E-06 | 7.86E-06 | 0 | 3.54E-06 | 1.44E-05 | 0 | 0 | 0 | 0 | 3.29E-06 | 1.47E-06 | 0 | 0 | 2.17E-06 |
| Syt11 | 9.59E-06 | 2.00E-05 | 0 | 0 | 1.53E-05 | 0 | 2.61E-05 | 0 | 1.44E-05 | 6.96E-05 | 1.35E-05 | 3.83E-05 | 2.03E-05 | 0 | 0 | 1.35E-05 | 0 | 0.00011 |
| Stim1 | 0 | 0.00011 | 0 | 1.85E-06 | 0.00012 | 8.45E-05 | 3.32E-06 | 3.30E-05 | 1.43E-05 | 0.00015 | 0.00014 | 5.51E-05 | 0.00011 | 5.93E-05 | 0 | 0.00018 | 4.29E-05 | 0.00019 |
| Stradb | 4.74E-05 | 5.41E-05 | 8.71E-06 | 9.61E-06 | 2.48E-05 | 4.66E-05 | 3.05E-06 | 0 | 1.43E-05 | 4.07E-06 | 8.08E-06 | 4.05E-06 | 0 | 1.78E-05 | 2.59E-05 | 1.35E-06 | 0 | 0 |
| Mfsd3 | 2.05E-06 | 1.95E-05 | 4.27E-06 | 3.54E-05 | 2.18E-05 | 3.55E-05 | 2.28E-05 | 5.00E-05 | 1.43E-05 | 1.34E-05 | 2.60E-05 | 4.48E-06 | 2.83E-05 | 2.30E-05 | 1.42E-05 | 0 | 0 | 2.96E-05 |
| Upf3b | 1.18E-05 | 6.87E-05 | 0 | 3.12E-05 | 1.60E-05 | 1.99E-05 | 0 | 0 | 1.43E-05 | 0 | 3.00E-05 | 1.16E-05 | 3.10E-05 | 2.29E-05 | 9.54E-05 | 8.66E-05 | 0 | 0 |
| Bbs12 | 1.70E-05 | 1.48E-05 | 7.75E-06 | 0 | 2.19E-06 | 8.25E-06 | 0 | 9.52E-06 | 1.43E-05 | 8.45E-06 | 0 | 0 | 0 | 2.53E-06 | 2.36E-05 | 9.45E-06 | 0 | 0 |
| Ppargc1b | 8.13E-07 | 7.31E-06 | 3.81E-05 | 2.74E-06 | 1.13E-05 | 7.42E-06 | 0 | 2.29E-05 | 1.43E-05 | 0 | 2.49E-05 | 0 | 1.74E-05 | 2.13E-05 | 2.43E-05 | 0 | 1.01E-05 | 0 |
| Setdb2 | 4.82E-05 | 9.98E-06 | 0 | 0 | 0 | 7.29E-06 | 0 | 0 | 1.43E-05 | 2.66E-05 | 0 | 0 | 7.98E-06 | 7.08E-06 | 3.89E-05 | 5.58E-05 | 0 | 2.25E-05 |
| Brd8 | 0.00017 | 0.00016 | 2.58E-05 | 5.13E-05 | 0 | 6.01E-06 | 0.00011 | 1.65E-05 | 1.43E-05 | 7.13E-05 | 0 | 4.08E-05 | 4.86E-05 | 0 | 0.0007 | 0.00047 | 4.38E-05 | 8.84E-06 |

| | | | | | | | | | | | | | | | | | | |
|---|---|---|---|---|---|---|---|---|---|---|---|---|---|---|---|---|---|---|
| C1rl | 1.27E-05 | 1.36E-05 | 0 | 1.64E-05 | 4.40E-05 | 0 | 5.45E-06 | 0 | 1.43E-05 | 7.72E-05 | 7.38E-05 | 5.48E-06 | 7.44E-06 | 1.62E-05 | 0 | 9.54E-06 | 0 | 2.45E-05 |
| Tiam1 | 5.40E-06 | 0 | 6.36E-05 | 1.60E-05 | 0 | 0 | 0.00013 | 0 | 1.43E-05 | 0.00017 | 2.74E-05 | 0 | 0.00015 | 0 | 0 | 7.51E-05 | 4.76E-05 | 0.00027 |
| Sos2 | 7.61E-05 | 0.0001 | 1.95E-05 | 0 | 2.64E-05 | 8.46E-05 | 0 | 9.67E-06 | 1.42E-05 | 1.57E-05 | 0 | 6.80E-05 | 0 | 0 | 5.58E-05 | 5.91E-06 | 0.00014 | 0 |
| Ss18l2 | 0.00013 | 1.10E-05 | 7.23E-05 | 2.04E-05 | 2.37E-05 | 6.34E-05 | 5.98E-05 | 4.09E-05 | 1.42E-05 | 0 | 2.22E-05 | 2.02E-05 | 0 | 0 | 4.20E-05 | 2.05E-05 | 8.93E-06 | 0 |
| Pcdhb19 | 6.43E-06 | 4.52E-05 | 0 | 9.66E-06 | 4.45E-05 | 3.21E-05 | 0 | 2.14E-05 | 1.42E-05 | 6.04E-07 | 1.17E-05 | 3.21E-06 | 2.85E-05 | 2.93E-05 | 3.24E-05 | 0 | 0 | 9.28E-06 |
| Slc25a26 | 3.43E-06 | 8.92E-06 | 3.05E-06 | 1.19E-05 | 1.03E-05 | 6.51E-06 | 0 | 7.15E-06 | 1.42E-05 | 7.80E-06 | 8.24E-07 | 6.96E-06 | 7.57E-06 | 1.11E-05 | 4.44E-06 | 6.63E-06 | 3.37E-06 | 2.18E-05 |
| Dnaaf5 | 2.82E-07 | 2.05E-05 | 2.62E-05 | 3.68E-05 | 1.39E-05 | 0 | 0 | 9.86E-06 | 1.42E-05 | 1.42E-05 | 1.70E-05 | 5.98E-06 | 0 | 1.85E-05 | 3.44E-05 | 1.17E-05 | 5.38E-06 | 1.05E-05 |
| Bmp6 | 0 | 1.81E-05 | 0 | 0 | 6.38E-06 | 0 | 1.42E-05 | 0 | 1.42E-05 | 0.00013 | 6.47E-06 | 5.00E-05 | 0 | 0 | 1.31E-05 | 7.36E-06 | 0 | 0 |
| Mphosph6 | 0.00012 | 0 | 0.00019 | 0.00014 | 0 | 0 | 7.13E-05 | 5.94E-05 | 1.42E-05 | 1.83E-05 | 9.11E-05 | 0.00012 | 1.92E-05 | 2.46E-05 | 9.86E-06 | 0.0001 | 0 | 0 |
| Nle1 | 3.90E-05 | 2.50E-05 | 0.00014 | 0.00012 | 0 | 0 | 0 | 2.27E-05 | 1.42E-05 | 8.53E-06 | 5.59E-06 | 2.17E-05 | 5.04E-05 | 3.66E-06 | 0.00012 | 1.09E-05 | 3.23E-05 | 0 |
| Prrc1 | 0.00028 | 7.22E-05 | 0.0004 | 0.00066 | 0.00018 | 0.00028 | 0.00068 | 0.00022 | 1.41E-05 | 0.00019 | 0.00025 | 0.00022 | 0.00024 | 0.00029 | 0.00028 | 0 | 0.00016 | 0.00014 |
| Hps5 | 4.10E-05 | 4.90E-05 | 1.40E-06 | 4.56E-05 | 3.12E-05 | 4.09E-05 | 2.81E-05 | 1.37E-05 | 1.41E-05 | 2.87E-05 | 1.75E-05 | 2.69E-05 | 2.96E-05 | 1.92E-05 | 6.26E-05 | 4.03E-05 | 4.63E-05 | 4.91E-05 |
| Tmem141 | 1.24E-05 | 9.44E-06 | 0 | 0 | 1.26E-05 | 3.95E-06 | 8.87E-06 | 0 | 1.41E-05 | 2.55E-05 | 5.27E-06 | 6.06E-06 | 8.56E-06 | 1.06E-05 | 0 | 0 | 0 | 1.61E-05 |
| Ifit2 | 3.84E-06 | 1.14E-05 | 0 | 0 | 0 | 1.77E-06 | 0 | 2.95E-06 | 1.41E-05 | 8.56E-05 | 0 | 0 | 3.17E-06 | 0 | 0 | 6.44E-05 | 0 | 3.46E-05 |
| Ap3m2 | 0 | 5.68E-05 | 0 | 6.88E-06 | 8.20E-06 | 0 | 4.74E-05 | 7.03E-05 | 1.41E-05 | 0.00015 | 6.18E-06 | 4.40E-05 | 0 | 0 | 0 | 3.14E-05 | 4.66E-06 | 4.81E-05 |
| Clk4 | 8.94E-05 | 0.0002 | 6.83E-06 | 4.19E-05 | 0.0001 | 0.00012 | 4.93E-05 | 0 | 1.40E-05 | 2.02E-05 | 1.29E-06 | 9.17E-05 | 1.38E-05 | 0.00013 | 0.00041 | 4.07E-06 | 0.00012 | 5.42E-05 |
| Tmem140 | 0 | 0.00027 | 0 | 8.05E-05 | 8.26E-05 | 5.59E-05 | 0 | 6.61E-05 | 1.40E-05 | 7.11E-05 | 0 | 0 | 0 | 0.00012 | 0 | 4.30E-06 | 0 | 0.00022 |
| Rab40b | 0 | 5.64E-06 | 0 | 7.69E-05 | 1.19E-05 | 4.84E-05 | 1.70E-05 | 1.22E-05 | 1.40E-05 | 0.00012 | 0 | 0 | 5.16E-05 | 4.06E-05 | 2.52E-06 | 0 | 0 | 0.00011 |
| Sec11c | 0.00019 | 1.53E-05 | 0.00017 | 0.00018 | 2.56E-05 | 3.19E-05 | 1.76E-05 | 9.14E-05 | 1.40E-05 | 3.63E-05 | 1.20E-05 | 3.07E-05 | 7.37E-05 | 2.70E-05 | 0.00013 | 8.23E-05 | 0 | 6.38E-05 |
| Zfp287 | 1.03E-05 | 1.61E-05 | 0 | 1.38E-05 | 0 | 1.45E-05 | 1.23E-05 | 2.73E-05 | 1.40E-05 | 1.50E-05 | 2.13E-05 | 8.43E-06 | 2.94E-06 | 3.06E-05 | 4.49E-05 | 0 | 0 | 2.14E-05 |
| Rpa2 | 1.08E-05 | 0 | 0.00011 | 5.03E-05 | 5.11E-06 | 0 | 3.69E-05 | 0 | 1.40E-05 | 1.33E-05 | 1.54E-05 | 0 | 3.08E-05 | 1.21E-05 | 0 | 0.00048 | 0 | 2.98E-05 |
| Gpr155 | 3.42E-06 | 2.63E-05 | 2.91E-06 | 0 | 3.67E-05 | 2.68E-05 | 0 | 3.89E-06 | 1.39E-05 | 2.31E-05 | 0 | 1.19E-05 | 0 | 0 | 0 | 0 | 0 | 1.23E-05 |
| Scaper | 2.77E-05 | 3.68E-05 | 0 | 0 | 1.42E-05 | 1.33E-05 | 2.49E-05 | 4.67E-06 | 1.39E-05 | 1.55E-05 | 1.35E-05 | 3.17E-05 | 2.88E-06 | 4.66E-06 | 0 | 2.65E-05 | 5.03E-05 | 1.26E-07 |
| Thumpd2 | 1.85E-05 | 4.47E-05 | 6.16E-07 | 1.27E-05 | 2.18E-05 | 7.33E-06 | 0 | 0 | 1.39E-05 | 4.48E-06 | 0 | 0 | 1.88E-05 | 0 | 2.96E-05 | 1.16E-05 | 5.95E-06 | 1.32E-05 |
| Ss18l1 | 7.38E-06 | 1.33E-05 | 1.65E-05 | 3.55E-06 | 1.07E-05 | 6.04E-06 | 4.64E-06 | 1.19E-05 | 1.39E-05 | 9.82E-06 | 8.73E-06 | 2.20E-05 | 0 | 4.80E-06 | 4.68E-05 | 0 | 2.99E-05 | 2.55E-05 |
| Il1rl1 | 0 | 0 | 0.00034 | 0 | 0 | 0 | 0 | 0 | 1.39E-05 | 0 | 0 | 0 | 0.0003 | 5.43E-05 | 0 | 6.40E-05 | 0.0003 | 0 |
| Vmo1 | 0 | 1.01E-05 | 0 | 2.09E-05 | 8.27E-06 | 0 | 5.08E-06 | 1.84E-05 | 1.39E-05 | 1.91E-06 | 3.77E-06 | 0 | 1.91E-06 | 8.58E-06 | 9.10E-06 | 0 | 0 | 7.33E-06 |
| Nxpe4 | 1.12E-05 | 1.20E-05 | 0 | 0 | 9.78E-06 | 0 | 1.32E-05 | 7.65E-06 | 1.39E-05 | 2.05E-05 | 7.01E-07 | 4.90E-06 | 3.65E-07 | 0 | 0 | 0 | 8.04E-06 | 4.56E-06 |
| Rftn2 | 5.45E-05 | 2.98E-05 | 0 | 5.90E-05 | 0.00011 | 0.00011 | 0.00014 | 6.00E-06 | 1.38E-05 | 0 | 1.89E-05 | 0 | 0 | 0.00017 | 2.48E-05 | 1.35E-05 | 0 | 7.49E-05 |
| Klf12 | 2.79E-05 | 2.57E-05 | 0 | 0 | 0 | 3.57E-05 | 3.84E-05 | 6.64E-05 | 1.38E-05 | 7.00E-05 | 8.87E-06 | 2.30E-05 | 0 | 4.10E-05 | 1.96E-05 | 0 | 3.46E-05 | 0 |
| Orc5 | 5.45E-05 | 3.31E-05 | 7.40E-06 | 0 | 8.78E-06 | 2.92E-05 | 8.41E-07 | 1.05E-05 | 1.38E-05 | 2.17E-05 | 0 | 4.35E-05 | 4.37E-05 | 5.24E-06 | 1.80E-05 | 7.05E-05 | 2.51E-05 | 7.27E-06 |
| Hint2 | 4.64E-05 | 5.22E-05 | 0 | 3.22E-05 | 2.92E-05 | 2.27E-05 | 6.75E-06 | 0 | 1.38E-05 | 2.68E-05 | 2.66E-05 | 1.28E-05 | 2.06E-05 | 1.75E-05 | 1.14E-05 | 8.73E-06 | 2.99E-05 | 0 |
| Cep76 | 4.42E-06 | 1.31E-05 | 6.38E-05 | 5.15E-05 | 9.88E-06 | 2.16E-05 | 3.78E-07 | 1.95E-05 | 1.38E-05 | 1.35E-05 | 3.20E-05 | 1.53E-05 | 0 | 1.33E-05 | 0 | 0.00018 | 0 | 2.57E-05 |

| | | | | | | | | | | | | | | | | | | |
|---|---|---|---|---|---|---|---|---|---|---|---|---|---|---|---|---|---|---|
| Klhl2 | 3.15E-05 | 6.06E-05 | 3.17E-05 | 0 | 7.38E-06 | 1.51E-05 | 0 | 2.68E-06 | 1.38E-05 | 0 | 8.14E-05 | 9.10E-05 | 0 | 5.23E-06 | 5.93E-07 | 4.15E-05 | 1.39E-05 | 1.82E-05 |
| Pctp | 2.12E-05 | 1.53E-05 | 0 | 3.03E-05 | 1.72E-05 | 2.32E-06 | 1.46E-05 | 1.74E-06 | 1.38E-05 | 1.17E-05 | 0 | 0 | 0 | 1.58E-06 | 2.22E-07 | 1.64E-05 | 0 | 0 |
| Zik1 | 6.32E-06 | 7.42E-06 | 4.56E-06 | 0 | 1.22E-05 | 0 | 0 | 0 | 1.38E-05 | 3.17E-06 | 6.44E-06 | 0 | 2.27E-06 | 2.53E-06 | 8.88E-07 | 4.10E-05 | 0 | 5.20E-06 |
| Zfp748 | 1.92E-05 | 1.69E-05 | 0 | 0 | 1.81E-05 | 1.94E-05 | 0 | 0 | 1.37E-05 | 1.32E-05 | 3.63E-06 | 6.62E-06 | 0 | 0 | 1.00E-05 | 8.17E-06 | 0 | 1.72E-05 |
| Zcchc10 | 5.16E-05 | 3.63E-05 | 4.75E-05 | 8.50E-06 | 2.44E-05 | 1.33E-05 | 0 | 0 | 1.37E-05 | 9.97E-06 | 9.23E-06 | 1.87E-05 | 9.54E-06 | 0 | 6.59E-05 | 4.57E-05 | 0 | 1.56E-05 |
| Zfp341 | 6.18E-06 | 7.64E-06 | 0 | 0 | 3.19E-06 | 1.67E-06 | 0 | 1.08E-05 | 1.37E-05 | 9.67E-06 | 8.97E-06 | 9.32E-06 | 0 | 5.70E-06 | 7.98E-06 | 4.45E-06 | 2.85E-05 | 5.97E-06 |
| Nupl2 | 2.90E-05 | 1.54E-05 | 1.82E-05 | 2.92E-05 | 1.58E-05 | 0 | 0 | 0 | 1.37E-05 | 3.78E-06 | 6.33E-06 | 4.62E-06 | 0 | 0 | 1.92E-05 | 5.52E-06 | 0 | 8.21E-07 |
| Atp23 | 3.35E-05 | 1.58E-05 | 2.90E-05 | 1.97E-05 | 4.15E-06 | 0 | 4.38E-06 | 0 | 1.37E-05 | 1.09E-05 | 0 | 0 | 1.42E-05 | 1.57E-05 | 1.85E-05 | 6.98E-06 | 0 | 4.25E-07 |
| Fam129b | 6.18E-05 | 0.00027 | 0.00084 | 0.00024 | 0.00014 | 0.00058 | 8.85E-05 | 2.87E-05 | 1.36E-05 | 1.51E-05 | 0.00059 | 0.00082 | 0.00026 | 0.00027 | 0.00032 | 0.00011 | 5.43E-05 | 0.00034 |
| Frk | 9.92E-05 | 6.34E-05 | 2.95E-05 | 0 | 8.59E-05 | 4.76E-05 | 0.00014 | 1.47E-05 | 1.36E-05 | 0 | 2.90E-05 | 0 | 3.98E-05 | 0 | 9.54E-05 | 9.09E-05 | 0 | 9.65E-06 |
| Mgat4a | 4.44E-09 | 1.05E-05 | 0 | 0 | 2.23E-05 | 1.82E-05 | 0 | 2.16E-05 | 1.36E-05 | 3.24E-06 | 5.96E-06 | 6.76E-06 | 0 | 1.11E-05 | 0 | 2.01E-05 | 2.30E-06 | 7.53E-05 |
| Cptp | 6.48E-05 | 3.36E-05 | 5.39E-05 | 7.99E-06 | 2.16E-05 | 1.80E-05 | 1.73E-05 | 2.24E-05 | 1.36E-05 | 1.75E-05 | 1.35E-05 | 1.93E-05 | 8.19E-06 | 8.63E-06 | 0 | 4.43E-05 | 0 | 1.36E-05 |
| Fndc10 | 0 | 4.02E-05 | 1.77E-05 | 3.36E-05 | 3.28E-05 | 1.38E-05 | 1.90E-05 | 2.10E-05 | 1.36E-05 | 1.43E-05 | 0 | 0 | 1.16E-06 | 0 | 1.58E-05 | 1.01E-05 | 0 | 2.79E-05 |
| Gas2l3 | 0 | 2.32E-05 | 0 | 0 | 4.92E-06 | 5.54E-06 | 0 | 0 | 1.36E-05 | 0 | 0 | 5.59E-05 | 0 | 0 | 0 | 0.00035 | 0 | 0 |
| Hhex | 0 | 7.05E-06 | 2.70E-06 | 9.60E-06 | 0 | 2.49E-06 | 0 | 7.51E-06 | 1.36E-05 | 2.90E-06 | 0 | 7.29E-06 | 0 | 0 | 2.69E-05 | 1.30E-05 | 6.27E-06 | 4.47E-06 |
| Cenpj | 1.44E-05 | 7.64E-06 | 0 | 0 | 0 | 0 | 1.30E-05 | 0 | 1.36E-05 | 1.94E-05 | 0 | 0 | 3.34E-06 | 0 | 7.77E-05 | 0.00011 | 0 | 7.99E-06 |
| Ufm1 | 0.00046 | 7.30E-05 | 0.00034 | 0.00024 | 5.57E-05 | 0.00013 | 0.0003 | 3.93E-05 | 1.35E-05 | 3.51E-05 | 0.00011 | 6.05E-05 | 3.61E-05 | 2.33E-05 | 0 | 8.28E-06 | 0 | 9.17E-06 |
| Ttc23 | 6.36E-05 | 0.00012 | 0 | 6.73E-06 | 7.45E-05 | 7.46E-05 | 0 | 1.64E-05 | 1.35E-05 | 6.24E-05 | 0 | 3.04E-05 | 1.64E-05 | 0 | 4.00E-05 | 4.66E-05 | 0 | 4.09E-05 |
| Oard1 | 8.42E-05 | 8.80E-05 | 3.14E-05 | 0.00012 | 4.48E-05 | 5.03E-05 | 5.11E-05 | 2.13E-05 | 1.35E-05 | 2.44E-05 | 0 | 2.29E-05 | 0 | 3.44E-05 | 0 | 0.00023 | 0 | 3.02E-05 |
| Rbm12b2 | 5.74E-05 | 8.59E-05 | 1.84E-05 | 0 | 4.77E-05 | 4.11E-05 | 0 | 2.55E-05 | 1.35E-05 | 3.37E-05 | 1.24E-05 | 5.48E-05 | 0 | 2.33E-05 | 0.00017 | 4.62E-05 | 0 | 2.82E-05 |
| Foxf1 | 0 | 5.42E-06 | 3.05E-05 | 0 | 1.97E-06 | 3.70E-05 | 0 | 9.50E-06 | 1.35E-05 | 0 | 8.40E-06 | 0 | 0 | 6.58E-06 | 7.65E-06 | 1.95E-06 | 2.29E-05 | 1.92E-05 |
| Lin52 | 1.33E-05 | 4.95E-05 | 0 | 1.69E-05 | 5.04E-05 | 3.63E-05 | 2.17E-05 | 2.12E-05 | 1.35E-05 | 1.14E-05 | 1.81E-05 | 2.87E-06 | 5.31E-07 | 2.52E-05 | 1.30E-05 | 7.30E-05 | 1.01E-05 | 1.55E-05 |
| Ankrd6 | 0 | 2.95E-07 | 0 | 0 | 0 | 2.78E-06 | 0 | 6.45E-06 | 1.35E-05 | 0 | 0 | 2.35E-05 | 0 | 5.53E-05 | 1.36E-06 | 0 | 0 | 2.10E-05 |
| Dnaic2 | 0 | 7.32E-06 | 5.81E-07 | 1.58E-06 | 6.58E-06 | 2.46E-06 | 1.02E-05 | 2.58E-05 | 1.35E-05 | 3.01E-05 | 0 | 7.68E-06 | 7.33E-06 | 0 | 0 | 0 | 9.56E-07 | 1.12E-05 |
| Upp1 | 0 | 1.70E-06 | 5.02E-06 | 1.60E-07 | 0 | 2.09E-06 | 0 | 5.48E-06 | 1.35E-05 | 0 | 0 | 1.08E-05 | 0 | 4.24E-07 | 0 | 5.59E-06 | 0 | 5.48E-06 |
| Il17d | 0 | 2.11E-05 | 1.04E-05 | 0 | 4.52E-07 | 0 | 0 | 0 | 1.35E-05 | 2.23E-05 | 8.56E-06 | 7.63E-06 | 2.47E-06 | 0 | 0 | 1.69E-05 | 1.46E-06 | 5.48E-05 |
| Ddx55 | 2.15E-05 | 5.81E-05 | 2.10E-05 | 3.81E-05 | 0 | 0 | 0 | 8.33E-07 | 1.35E-05 | 2.21E-05 | 7.97E-06 | 2.60E-05 | 0 | 0 | 9.63E-05 | 2.32E-05 | 0 | 9.92E-06 |
| Jpt2 | 0.00027 | 0 | 0.00025 | 7.32E-05 | 0 | 0 | 0.00017 | 0 | 1.35E-05 | 4.54E-05 | 0 | 0 | 1.18E-05 | 0 | 0 | 0.00051 | 0 | 0 |
| Slc2a8 | 0 | 7.74E-05 | 0 | 1.20E-05 | 6.87E-05 | 4.84E-05 | 3.90E-05 | 2.81E-06 | 1.34E-05 | 4.50E-05 | 0 | 7.33E-05 | 2.85E-05 | 6.50E-05 | 5.36E-05 | 1.35E-05 | 0 | 4.72E-05 |
| Slf1 | 3.17E-05 | 9.73E-06 | 1.47E-05 | 0 | 0 | 3.00E-05 | 1.76E-06 | 1.51E-06 | 1.34E-05 | 3.60E-06 | 0 | 7.02E-06 | 6.35E-07 | 2.40E-05 | 1.45E-05 | 0.00013 | 9.12E-06 | 1.92E-06 |
| Dok4 | 3.99E-05 | 4.43E-05 | 2.34E-06 | 4.32E-05 | 3.59E-05 | 2.89E-05 | 4.70E-05 | 5.87E-06 | 1.34E-05 | 1.76E-05 | 1.67E-05 | 3.49E-05 | 1.61E-05 | 1.53E-05 | 0 | 4.81E-06 | 0 | 5.18E-05 |
| Plekhm3 | 1.90E-05 | 5.22E-05 | 1.47E-05 | 0 | 3.49E-05 | 1.22E-05 | 1.54E-05 | 0 | 1.34E-05 | 3.46E-06 | 3.00E-05 | 4.34E-05 | 2.78E-05 | 5.19E-05 | 1.51E-05 | 8.23E-07 | 4.07E-05 | 2.91E-05 |
| Lratd2 | 0 | 6.20E-06 | 0 | 1.77E-06 | 6.49E-06 | 6.96E-06 | 0 | 0 | 1.34E-05 | 0 | 9.53E-05 | 1.57E-05 | 0 | 1.52E-05 | 8.36E-06 | 0 | 2.55E-06 | 0 |

| | | | | | | | | | | | | | | | | | | |
|---|---|---|---|---|---|---|---|---|---|---|---|---|---|---|---|---|---|---|
| Pgm3 | 0.00015 | 0 | 0.00025 | 0.00032 | 0 | 0 | 0.00021 | 9.57E-05 | 1.34E-05 | 0.00013 | 2.72E-05 | 1.51E-05 | 2.98E-05 | 0 | 0.00013 | 9.02E-06 | 0 | 1.38E-05 |
| Aasdh | 9.04E-06 | 2.36E-05 | 0 | 1.47E-05 | 1.70E-05 | 0 | 0 | 9.70E-06 | 1.34E-05 | 4.88E-06 | 2.77E-06 | 1.74E-05 | 6.19E-07 | 1.66E-06 | 4.73E-05 | 1.16E-07 | 0 | 1.45E-05 |
| Kdm4c | 4.55E-05 | 5.79E-05 | 2.67E-05 | 3.04E-05 | 5.63E-05 | 7.10E-05 | 3.45E-05 | 3.29E-05 | 1.33E-05 | 5.17E-05 | 6.03E-05 | 8.06E-05 | 3.69E-05 | 1.24E-05 | 1.32E-05 | 2.11E-05 | 0.00012 | 6.20E-05 |
| Maco1 | 0.00023 | 0.00012 | 0.00025 | 0.00013 | 7.43E-05 | 6.90E-05 | 0.0002 | 0 | 1.33E-05 | 2.00E-05 | 6.82E-05 | 0.00015 | 2.12E-05 | 0.0001 | 0.00019 | 4.90E-05 | 5.55E-05 | 0 |
| Swt1 | 2.91E-05 | 3.12E-05 | 1.53E-05 | 0 | 1.49E-05 | 3.57E-05 | 1.66E-05 | 0 | 1.33E-05 | 2.30E-06 | 0 | 1.18E-05 | 6.20E-06 | 6.00E-07 | 1.95E-06 | 1.60E-05 | 1.93E-05 | 0 |
| Ccdc34 | 6.67E-05 | 3.55E-05 | 0 | 0 | 6.67E-06 | 2.39E-05 | 0 | 2.59E-06 | 1.33E-05 | 3.30E-05 | 0 | 3.17E-05 | 0 | 6.86E-06 | 8.41E-06 | 0.00062 | 0 | 5.34E-05 |
| Tsc22d3 | 0 | 8.40E-05 | 0 | 4.21E-05 | 6.89E-05 | 1.99E-05 | 0 | 6.64E-05 | 1.33E-05 | 3.80E-05 | 2.92E-05 | 2.88E-05 | 0 | 4.22E-05 | 0 | 0 | 5.84E-05 | 0.00016 |
| Zfp820 | 8.61E-06 | 1.66E-05 | 0 | 3.84E-06 | 2.12E-06 | 1.87E-05 | 1.01E-05 | 1.14E-05 | 1.33E-05 | 1.88E-05 | 0 | 0 | 0 | 1.22E-05 | 9.79E-06 | 1.01E-05 | 0 | 4.36E-06 |
| Tpmt | 4.54E-05 | 2.64E-05 | 0 | 4.51E-06 | 6.59E-06 | 6.79E-06 | 0 | 2.96E-06 | 1.33E-05 | 1.56E-05 | 7.02E-06 | 0 | 1.37E-05 | 0 | 0 | 4.12E-05 | 7.54E-06 | 2.03E-05 |
| Pald1 | 0 | 4.95E-05 | 0 | 7.64E-06 | 2.28E-05 | 3.40E-06 | 0 | 3.33E-05 | 1.33E-05 | 0 | 3.46E-05 | 1.74E-07 | 0 | 0 | 5.42E-05 | 5.43E-05 | 2.04E-07 | 2.01E-05 |
| Wdr73 | 3.12E-05 | 4.72E-05 | 1.83E-05 | 4.36E-05 | 6.75E-06 | 0 | 0 | 0 | 1.33E-05 | 9.40E-06 | 0 | 0 | 0 | 1.19E-05 | 3.69E-05 | 2.60E-05 | 0 | 0 |
| Ccbe1 | 0 | 4.28E-05 | 0.00029 | 0.00023 | 0.00055 | 0 | 0.00017 | 6.27E-05 | 1.33E-05 | 0 | 0.00053 | 0.00046 | 0.00058 | 0.00097 | 0.00105 | 7.98E-05 | 1.32E-05 | 0 |
| Itpkc | 1.16E-05 | 8.58E-05 | 8.93E-05 | 7.07E-05 | 4.14E-05 | 0 | 0 | 1.31E-05 | 1.33E-05 | 5.67E-06 | 0 | 8.32E-06 | 1.50E-05 | 9.00E-05 | 3.94E-05 | 6.22E-06 | 0.00032 | 5.75E-05 |
| Faap20 | 3.89E-05 | 0.00011 | 0 | 0.00025 | 0.0001 | 0.00014 | 8.70E-05 | 9.37E-05 | 1.32E-05 | 8.46E-05 | 4.67E-05 | 0.00016 | 6.40E-05 | 6.99E-05 | 1.28E-05 | 6.82E-05 | 0 | 6.65E-05 |
| Ccdc157 | 2.27E-05 | 4.68E-05 | 1.16E-05 | 5.80E-05 | 2.54E-05 | 2.10E-05 | 8.08E-05 | 5.32E-05 | 1.32E-05 | 3.06E-05 | 0 | 9.58E-06 | 7.31E-05 | 0 | 8.24E-05 | 0 | 1.16E-05 | 6.85E-05 |
| Rmc1 | 1.08E-05 | 8.04E-05 | 6.61E-05 | 8.87E-05 | 2.43E-05 | 1.41E-05 | 4.18E-05 | 0 | 1.32E-05 | 7.80E-05 | 6.82E-06 | 3.76E-05 | 0 | 5.38E-05 | 5.11E-05 | 1.24E-05 | 4.37E-05 | 3.12E-05 |
| Dennd2c | 0 | 2.92E-05 | 5.21E-06 | 0 | 0 | 9.09E-06 | 3.90E-05 | 0 | 1.32E-05 | 2.73E-05 | 0 | 3.41E-05 | 1.53E-05 | 2.16E-05 | 6.68E-07 | 9.78E-06 | 0 | 0 |
| Arid3b | 0 | 1.04E-05 | 3.41E-06 | 1.52E-05 | 0 | 1.75E-06 | 0 | 1.40E-06 | 1.32E-05 | 0 | 0 | 1.71E-05 | 0 | 4.44E-08 | 3.56E-06 | 6.37E-06 | 4.75E-06 | 0 |
| Fam217b | 1.77E-05 | 4.87E-06 | 1.50E-06 | 1.07E-06 | 0.00016 | 1.18E-06 | 8.65E-05 | 3.49E-05 | 1.32E-05 | 0 | 6.71E-05 | 0 | 4.38E-05 | 7.40E-05 | 0 | 9.41E-07 | 0 | 3.61E-06 |
| Armc6 | 6.83E-05 | 1.97E-05 | 0.00012 | 5.76E-05 | 0 | 0 | 3.89E-05 | 0 | 1.32E-05 | 5.04E-05 | 4.25E-05 | 0 | 0 | 0 | 2.90E-06 | 2.80E-05 | 0 | 3.38E-05 |
| Cnot7 | 0.00015 | 7.19E-05 | 7.65E-05 | 0 | 4.54E-05 | 7.98E-05 | 1.69E-05 | 5.16E-06 | 1.31E-05 | 0 | 5.93E-05 | 0.00024 | 0 | 3.53E-05 | 6.16E-05 | 0.00012 | 4.78E-05 | 0 |
| Ssh3 | 0 | 6.91E-05 | 0 | 2.04E-05 | 5.92E-06 | 2.57E-05 | 4.49E-05 | 0 | 1.31E-05 | 2.24E-05 | 6.41E-06 | 3.22E-05 | 0.00014 | 0.0002 | 0 | 0 | 0 | 0.00017 |
| Baiap2 | 9.78E-06 | 1.58E-05 | 9.96E-05 | 0 | 0 | 0 | 0.00011 | 0 | 1.31E-05 | 0.00012 | 0 | 3.83E-05 | 0.00021 | 3.40E-05 | 0 | 0 | 0.00158 | 6.54E-05 |
| Cd2ap | 0.00012 | 0.00013 | 4.21E-05 | 5.08E-05 | 0.0001 | 0.00011 | 0.0001 | 0.00013 | 1.30E-05 | 4.77E-05 | 7.48E-05 | 0.00013 | 3.85E-05 | 0 | 0.00022 | 5.49E-05 | 6.66E-05 | 2.83E-05 |
| Zcchc7 | 9.14E-05 | 5.09E-05 | 3.72E-05 | 0 | 0 | 4.63E-05 | 4.72E-05 | 0 | 1.30E-05 | 3.30E-05 | 2.21E-05 | 7.19E-05 | 3.36E-06 | 5.93E-06 | 0.00016 | 0 | 0.00016 | 1.31E-05 |
| Rnd2 | 0 | 2.87E-05 | 0 | 1.62E-05 | 2.92E-05 | 1.73E-05 | 5.75E-06 | 5.23E-06 | 1.30E-05 | 4.16E-05 | 1.88E-05 | 9.63E-06 | 0 | 1.22E-05 | 0 | 0 | 2.44E-05 | 2.94E-05 |
| Pip4k2a | 2.81E-05 | 3.81E-05 | 5.21E-05 | 2.27E-06 | 4.90E-05 | 1.45E-05 | 7.35E-05 | 1.93E-05 | 1.30E-05 | 5.77E-05 | 0.00016 | 1.45E-05 | 3.11E-05 | 4.63E-05 | 0 | 5.06E-05 | 0.00014 | 0 |
| Castor1 | 2.87E-05 | 2.10E-05 | 1.65E-05 | 2.60E-05 | 4.58E-07 | 1.08E-05 | 5.10E-05 | 5.63E-06 | 1.30E-05 | 7.21E-06 | 1.35E-05 | 3.91E-05 | 2.69E-05 | 0 | 0 | 0 | 0 | 0.00013 |
| Gnb1l | 1.36E-05 | 8.70E-06 | 2.25E-05 | 1.80E-05 | 1.08E-05 | 2.09E-06 | 1.37E-05 | 1.12E-05 | 1.30E-05 | 1.09E-05 | 1.50E-06 | 1.06E-06 | 1.20E-05 | 1.97E-05 | 4.94E-06 | 4.99E-05 | 0 | 0 |
| Gdpgp1 | 7.82E-06 | 1.62E-05 | 5.41E-07 | 1.39E-05 | 1.45E-05 | 0 | 8.57E-07 | 7.22E-06 | 1.30E-05 | 1.26E-05 | 0 | 0 | 0 | 1.13E-05 | 9.06E-06 | 0 | 0 | 2.03E-06 |
| Ccdc85c | 4.46E-06 | 4.30E-05 | 2.46E-05 | 4.62E-05 | 0 | 0 | 1.72E-05 | 0 | 1.30E-05 | 6.31E-06 | 1.04E-05 | 6.11E-06 | 8.27E-06 | 2.26E-05 | 3.09E-05 | 5.52E-05 | 1.07E-05 | 7.73E-06 |
| Sec61g | 0.00042 | 0 | 0.00023 | 0.00059 | 0 | 6.17E-05 | 0.00027 | 9.07E-05 | 1.29E-05 | 0 | 7.57E-05 | 0.00017 | 1.91E-06 | 5.18E-05 | 0 | 0.00017 | 9.28E-05 | 0 |
| Ptgfr | 2.66E-05 | 2.21E-05 | 0 | 0 | 0 | 4.50E-05 | 0 | 0 | 1.29E-05 | 5.08E-05 | 0 | 0 | 0.00017 | 4.55E-05 | 0 | 0 | 0 | 0 |

| | | | | | | | | | | | | | | | | | | |
|---|---|---|---|---|---|---|---|---|---|---|---|---|---|---|---|---|---|---|
| Rhbdl3 | 4.20E-06 | 1.52E-05 | 7.52E-07 | 3.69E-05 | 1.58E-05 | 3.08E-05 | 0 | 0 | 1.29E-05 | 1.73E-08 | 2.77E-05 | 0 | 5.16E-05 | 3.21E-05 | 2.15E-06 | 2.17E-05 | 0 | 5.98E-05 |
| Zbtb40 | 2.99E-05 | 5.73E-05 | 3.63E-05 | 1.11E-05 | 3.01E-05 | 1.58E-05 | 9.03E-06 | 0 | 1.29E-05 | 1.74E-05 | 0 | 1.12E-05 | 1.42E-06 | 3.69E-05 | 0.00011 | 3.69E-05 | 0 | 1.90E-05 |
| Drp2 | 0 | 2.45E-06 | 0 | 2.19E-06 | 5.86E-07 | 6.03E-06 | 1.68E-05 | 0 | 1.29E-05 | 6.69E-06 | 1.43E-05 | 0 | 3.07E-06 | 9.31E-06 | 0 | 1.83E-05 | 0 | 3.12E-05 |
| Tsen2 | 3.76E-05 | 2.05E-05 | 4.84E-05 | 4.81E-05 | 5.41E-06 | 0 | 1.06E-05 | 6.31E-06 | 1.29E-05 | 2.96E-05 | 0 | 0 | 2.66E-05 | 0 | 1.63E-05 | 5.85E-05 | 0 | 1.89E-05 |
| Kdsr | 2.94E-05 | 0.00021 | 8.11E-05 | 9.58E-06 | 0.00011 | 0.0001 | 3.89E-05 | 0 | 1.28E-05 | 0.00013 | 3.13E-05 | 8.10E-05 | 7.15E-05 | 0.00017 | 5.64E-05 | 2.35E-05 | 0 | 7.86E-05 |
| Slc10a3 | 0.00015 | 3.70E-05 | 9.51E-05 | 0 | 4.91E-05 | 6.47E-05 | 0.00011 | 1.70E-05 | 1.28E-05 | 4.00E-05 | 0 | 0 | 7.91E-05 | 1.95E-05 | 0 | 0 | 0 | 5.69E-05 |
| Mga | 5.27E-05 | 0.00014 | 3.90E-05 | 0.00011 | 3.32E-05 | 5.10E-05 | 4.72E-05 | 5.65E-05 | 1.28E-05 | 3.15E-05 | 0 | 0.00013 | 5.91E-05 | 4.78E-05 | 0.00029 | 4.97E-06 | 0.00012 | 4.62E-05 |
| R3hcc1l | 6.11E-05 | 0.00014 | 1.27E-05 | 1.89E-05 | 1.42E-05 | 1.52E-05 | 3.98E-05 | 0 | 1.28E-05 | 4.11E-05 | 0 | 8.94E-05 | 2.27E-05 | 0 | 0.00011 | 4.57E-05 | 1.73E-05 | 4.45E-05 |
| Cutc | 5.04E-05 | 3.72E-06 | 1.56E-05 | 6.52E-06 | 0 | 7.84E-06 | 8.95E-06 | 4.41E-06 | 1.28E-05 | 2.32E-05 | 0 | 2.67E-07 | 3.78E-06 | 0 | 4.09E-06 | 1.93E-05 | 0 | 2.51E-05 |
| Fmnl1 | 0 | 2.03E-05 | 0 | 3.80E-06 | 0 | 0 | 0 | 0 | 1.28E-05 | 3.32E-05 | 1.50E-06 | 0 | 0 | 0 | 3.87E-05 | 1.11E-05 | 1.06E-05 | 2.06E-05 |
| Agbl2 | 1.25E-06 | 1.93E-06 | 0 | 0 | 0 | 0 | 0 | 0 | 1.28E-05 | 4.42E-06 | 0 | 2.08E-06 | 0 | 0 | 6.04E-06 | 3.85E-05 | 0 | 3.68E-06 |
| Cyb5rl | 6.18E-06 | 1.61E-05 | 1.00E-05 | 2.68E-05 | 1.22E-05 | 0 | 0 | 0 | 1.28E-05 | 6.22E-06 | 1.29E-05 | 6.13E-06 | 2.77E-05 | 0 | 1.34E-05 | 2.72E-06 | 1.68E-05 | 6.29E-06 |
| Thoc2 | 0.00015 | 0.00013 | 3.05E-05 | 0 | 4.25E-05 | 0.00014 | 3.72E-05 | 3.25E-05 | 1.27E-05 | 2.11E-05 | 3.78E-06 | 8.41E-05 | 0 | 2.65E-05 | 0.00019 | 3.02E-05 | 0.00018 | 1.93E-05 |
| Arid4b | 9.42E-05 | 0.00017 | 5.15E-05 | 6.72E-06 | 9.11E-05 | 0.00011 | 3.95E-05 | 9.18E-06 | 1.27E-05 | 3.86E-05 | 3.36E-05 | 9.64E-05 | 2.02E-05 | 7.89E-05 | 0.00021 | 3.48E-05 | 0.00021 | 1.58E-05 |
| Calhm5 | 0 | 8.13E-06 | 0 | 7.04E-06 | 1.86E-05 | 4.93E-05 | 0 | 0 | 1.27E-05 | 6.44E-05 | 8.29E-05 | 0 | 3.49E-05 | 0 | 0 | 0 | 0 | 7.15E-05 |
| Gpr89 | 6.28E-05 | 4.37E-05 | 1.87E-06 | 6.42E-06 | 4.95E-05 | 3.18E-05 | 5.07E-06 | 1.40E-06 | 1.27E-05 | 2.30E-05 | 9.35E-06 | 3.76E-05 | 1.29E-05 | 0 | 7.00E-06 | 3.61E-05 | 1.84E-05 | 2.31E-05 |
| Wipf2 | 1.93E-05 | 4.83E-05 | 1.94E-05 | 0 | 2.25E-05 | 2.99E-05 | 5.05E-06 | 3.87E-05 | 1.27E-05 | 2.14E-05 | 2.31E-05 | 4.79E-05 | 0 | 1.31E-05 | 1.33E-05 | 1.23E-05 | 9.81E-06 | 1.61E-05 |
| Lhx8 | 0 | 9.20E-06 | 0 | 1.63E-05 | 0 | 1.22E-05 | 5.44E-05 | 0 | 1.27E-05 | 1.26E-05 | 0 | 0 | 5.00E-05 | 0 | 8.94E-07 | 5.34E-06 | 0 | 0.00015 |
| Gatb | 2.70E-05 | 2.37E-05 | 7.33E-06 | 1.91E-05 | 4.37E-06 | 7.59E-06 | 3.83E-06 | 1.37E-06 | 1.27E-05 | 4.49E-06 | 6.75E-06 | 2.15E-06 | 0 | 0 | 2.81E-05 | 7.24E-06 | 2.40E-05 | 3.84E-06 |
| Mterf1b | 4.42E-06 | 1.89E-06 | 0 | 5.55E-07 | 5.54E-06 | 6.06E-06 | 0 | 9.95E-06 | 1.27E-05 | 5.45E-06 | 1.66E-06 | 9.38E-07 | 0 | 5.18E-06 | 7.03E-06 | 7.48E-06 | 0 | 1.00E-05 |
| Mrrf | 6.06E-05 | 7.10E-05 | 7.96E-05 | 6.70E-05 | 1.55E-05 | 2.53E-06 | 3.96E-05 | 1.01E-05 | 1.27E-05 | 4.86E-05 | 4.95E-05 | 3.71E-05 | 1.78E-05 | 0 | 0 | 4.43E-05 | 0 | 3.92E-05 |
| Ankrd9 | 0 | 7.92E-06 | 7.08E-06 | 3.18E-05 | 7.33E-06 | 0 | 0 | 0 | 1.27E-05 | 1.78E-05 | 1.58E-05 | 5.97E-06 | 1.56E-05 | 4.34E-06 | 4.82E-06 | 1.03E-05 | 4.28E-05 | 1.14E-05 |
| Spem1 | 0 | 0 | 8.34E-07 | 1.20E-05 | 0 | 0 | 0 | 0 | 1.27E-05 | 0 | 0 | 9.21E-06 | 1.75E-05 | 0 | 0 | 8.77E-06 | 9.35E-05 | 0 |
| Cttnbp2 | 0 | 2.02E-05 | 3.17E-05 | 0 | 0.00011 | 5.73E-05 | 0 | 0.00026 | 1.26E-05 | 0 | 0 | 0 | 0 | 0.00014 | 0.00011 | 2.08E-05 | 0 | 3.58E-05 |
| Pttg1 | 9.27E-05 | 0.00013 | 1.80E-05 | 5.93E-05 | 6.37E-05 | 5.33E-05 | 0 | 0 | 1.26E-05 | 0.00013 | 1.73E-05 | 3.02E-05 | 6.61E-05 | 4.30E-05 | 5.23E-05 | 9.67E-05 | 0.00012 | 0.00011 |
| Pcnx4 | 7.04E-05 | 3.39E-05 | 1.63E-05 | 0 | 4.13E-05 | 3.14E-05 | 0 | 1.27E-05 | 1.26E-05 | 5.13E-07 | 7.49E-06 | 4.15E-05 | 0 | 0 | 0 | 3.46E-06 | 3.22E-05 | 0 |
| Zfp386 | 6.32E-05 | 4.43E-05 | 0 | 0 | 2.12E-06 | 3.03E-05 | 0 | 0 | 1.26E-05 | 3.56E-05 | 0 | 2.26E-05 | 0 | 0 | 3.75E-05 | 3.80E-06 | 0 | 1.31E-05 |
| Pibf1 | 4.71E-05 | 3.69E-05 | 3.96E-06 | 0 | 1.66E-05 | 2.97E-05 | 0 | 0 | 1.26E-05 | 1.74E-05 | 6.79E-06 | 2.04E-05 | 7.51E-06 | 9.84E-06 | 6.55E-05 | 7.76E-06 | 7.09E-05 | 1.10E-05 |
| Lurap1 | 0 | 3.01E-05 | 0 | 8.69E-06 | 1.78E-05 | 2.08E-05 | 0 | 2.76E-05 | 1.26E-05 | 0 | 0 | 4.49E-06 | 0 | 2.05E-05 | 2.18E-05 | 0 | 0 | 1.14E-06 |
| Eefsec | 3.92E-05 | 3.78E-05 | 3.91E-05 | 3.88E-05 | 1.55E-05 | 1.91E-05 | 4.06E-06 | 7.34E-06 | 1.26E-05 | 1.42E-05 | 3.47E-06 | 0 | 1.13E-05 | 0 | 8.56E-06 | 0.0001 | 1.61E-05 | 0 |
| Tcea2 | 0 | 1.85E-05 | 0 | 0 | 1.21E-05 | 1.56E-05 | 0 | 1.07E-06 | 1.26E-05 | 1.25E-05 | 9.63E-06 | 0 | 5.58E-06 | 1.93E-05 | 2.48E-05 | 0 | 0 | 1.64E-05 |
| Nfxl1 | 0.00013 | 7.23E-05 | 7.63E-05 | 4.79E-05 | 4.42E-05 | 8.56E-06 | 0.00013 | 0 | 1.26E-05 | 0 | 6.36E-06 | 9.17E-05 | 3.30E-07 | 2.62E-05 | 0.0002 | 5.59E-05 | 0 | 0 |
| Ndufaf5 | 6.13E-05 | 3.67E-06 | 2.66E-05 | 5.02E-05 | 4.84E-06 | 7.96E-06 | 0 | 6.11E-06 | 1.26E-05 | 1.59E-05 | 0 | 0 | 0 | 0 | 3.49E-05 | 4.54E-05 | 3.33E-06 | 1.48E-05 |

| | | | | | | | | | | | | | | | | | | |
|---|---|---|---|---|---|---|---|---|---|---|---|---|---|---|---|---|---|---|
| Slc29a2 | 7.58E-06 | 1.19E-06 | 2.29E-05 | 2.21E-06 | 0 | 5.02E-06 | 0 | 2.54E-05 | 1.26E-05 | 9.81E-07 | 3.15E-06 | 4.45E-05 | 0 | 3.42E-05 | 8.04E-06 | 0 | 5.09E-06 | 0 |
| Ap5z1 | 9.92E-06 | 9.09E-05 | 1.16E-05 | 4.22E-05 | 2.62E-05 | 2.90E-07 | 0 | 2.91E-05 | 1.26E-05 | 1.22E-05 | 6.16E-06 | 4.31E-05 | 1.21E-05 | 2.36E-05 | 9.81E-05 | 3.63E-06 | 0 | 2.38E-05 |
| Trim45 | 3.25E-06 | 9.02E-06 | 0 | 1.98E-05 | 2.20E-06 | 0 | 0 | 0 | 1.26E-05 | 2.24E-05 | 2.82E-05 | 1.50E-05 | 0 | 0 | 1.78E-07 | 1.51E-05 | 1.42E-05 | 1.26E-05 |
| Xaf1 | 7.78E-05 | 2.65E-05 | 0 | 0 | 0 | 0 | 0 | 0 | 1.26E-05 | 0.00022 | 2.51E-06 | 3.15E-05 | 3.45E-05 | 0 | 4.30E-05 | 4.66E-05 | 0 | 0.00024 |
| Tab2 | 0.00029 | 0.00021 | 0.00014 | 0 | 0.00018 | 0.00022 | 0.00014 | 8.47E-05 | 1.25E-05 | 3.31E-05 | 0.00023 | 0.00022 | 0 | 0.00019 | 2.98E-05 | 0.00018 | 7.15E-05 | 6.16E-05 |
| Ly96 | 7.26E-05 | 5.83E-05 | 0 | 0 | 1.84E-05 | 9.83E-05 | 1.30E-05 | 2.40E-05 | 1.25E-05 | 2.51E-05 | 0 | 1.50E-05 | 5.42E-05 | 0 | 0 | 0 | 0 | 2.32E-05 |
| Surf1 | 4.53E-05 | 0.0001 | 3.98E-05 | 0.00012 | 6.17E-05 | 6.43E-05 | 6.52E-05 | 4.06E-05 | 1.25E-05 | 3.95E-05 | 2.36E-05 | 0.00012 | 0 | 4.58E-05 | 4.03E-05 | 1.00E-06 | 0.00015 | 0.00011 |
| Hoxc4 | 8.66E-06 | 2.98E-05 | 0 | 1.94E-06 | 1.04E-05 | 3.34E-05 | 6.52E-05 | 0 | 1.25E-05 | 2.37E-05 | 2.39E-05 | 1.09E-05 | 1.87E-06 | 2.85E-05 | 7.13E-05 | 1.90E-05 | 0 | 8.25E-05 |
| Hbs1l | 5.24E-05 | 8.91E-05 | 6.44E-05 | 5.68E-05 | 3.70E-05 | 1.71E-05 | 0 | 0 | 1.25E-05 | 9.65E-06 | 4.34E-05 | 8.67E-05 | 0 | 6.03E-06 | 0.00015 | 1.64E-05 | 8.21E-05 | 1.47E-05 |
| Trpt1 | 4.80E-06 | 1.91E-05 | 0 | 2.70E-06 | 4.45E-06 | 1.33E-05 | 0 | 1.15E-05 | 1.25E-05 | 8.91E-06 | 1.64E-05 | 1.46E-05 | 7.58E-06 | 1.38E-05 | 1.17E-05 | 6.39E-06 | 1.23E-06 | 1.09E-05 |
| AW209491 | 4.78E-05 | 2.01E-05 | 2.75E-05 | 0 | 1.24E-05 | 1.04E-05 | 2.59E-05 | 0 | 1.25E-05 | 9.46E-06 | 1.19E-06 | 7.71E-06 | 0 | 0 | 3.36E-05 | 0 | 0 | 4.43E-06 |
| Dmxl2 | 0 | 4.53E-05 | 3.31E-06 | 1.24E-05 | 3.54E-05 | 8.86E-06 | 0 | 5.12E-06 | 1.25E-05 | 1.51E-05 | 0 | 2.37E-05 | 0 | 2.17E-05 | 2.83E-05 | 2.01E-06 | 1.65E-05 | 1.92E-05 |
| Cenpw | 2.38E-05 | 0 | 2.81E-05 | 0 | 0 | 3.80E-06 | 0 | 8.26E-06 | 1.25E-05 | 0 | 0 | 0 | 0 | 5.98E-06 | 0 | 0.00039 | 0 | 0 |
| Scn2a | 8.15E-07 | 3.69E-07 | 0 | 0 | 4.77E-06 | 2.56E-06 | 1.16E-09 | 0 | 1.25E-05 | 2.20E-05 | 0 | 1.65E-06 | 0 | 0 | 2.84E-05 | 1.81E-06 | 5.17E-06 | 2.40E-05 |
| Cntnap1 | 0 | 2.27E-05 | 0 | 0 | 0 | 6.43E-07 | 0 | 2.55E-05 | 1.25E-05 | 2.15E-06 | 0 | 1.34E-05 | 0 | 0 | 2.54E-05 | 0 | 1.68E-06 | 3.17E-06 |
| Aqp2 | 0 | 0 | 1.64E-05 | 0 | 5.56E-05 | 0 | 1.45E-05 | 3.99E-05 | 1.25E-05 | 0 | 0 | 0 | 0 | 0 | 1.94E-05 | 0 | 0 | 8.57E-05 |
| Atp6v1d | 0.00013 | 0.0002 | 8.18E-05 | 0.00014 | 0.0001 | 0.00011 | 6.13E-05 | 0.0001 | 1.24E-05 | 3.02E-05 | 7.51E-05 | 6.26E-05 | 1.47E-05 | 0.00013 | 7.06E-05 | 7.41E-05 | 0 | 7.60E-05 |
| Mars | 0.00016 | 0.00016 | 0.00026 | 6.60E-05 | 6.43E-05 | 0.00011 | 7.57E-05 | 3.04E-05 | 1.24E-05 | 5.90E-05 | 0.00012 | 0.0001 | 0 | 5.58E-05 | 0.00023 | 0.00023 | 0 | 2.92E-05 |
| Zmym5 | 7.95E-05 | 0.00012 | 0 | 3.59E-05 | 4.04E-05 | 7.94E-05 | 0 | 2.71E-05 | 1.24E-05 | 1.87E-05 | 0 | 7.13E-05 | 0 | 5.03E-05 | 0.00018 | 3.51E-05 | 0.00021 | 1.56E-05 |
| Tmem129 | 4.72E-05 | 6.25E-05 | 6.28E-05 | 0 | 6.71E-05 | 6.25E-05 | 0.00013 | 4.03E-05 | 1.24E-05 | 4.46E-05 | 3.12E-05 | 2.46E-06 | 1.85E-05 | 0 | 0 | 0.00014 | 0 | 4.31E-05 |
| Lrrc58 | 0 | 0.00017 | 9.79E-05 | 0.0003 | 8.83E-05 | 5.08E-05 | 0 | 8.22E-05 | 1.24E-05 | 1.14E-05 | 9.26E-05 | 5.93E-05 | 3.79E-05 | 3.47E-05 | 0.00037 | 1.94E-05 | 0.00017 | 0.00013 |
| Pcdhgb4 | 1.25E-05 | 6.92E-05 | 0 | 1.52E-05 | 4.98E-05 | 4.52E-05 | 5.11E-05 | 3.17E-05 | 1.24E-05 | 2.03E-05 | 3.52E-05 | 2.51E-05 | 0 | 3.27E-05 | 0.00014 | 0 | 6.93E-05 | 1.20E-05 |
| Ninl | 0 | 2.32E-05 | 5.84E-06 | 0 | 2.65E-05 | 2.71E-05 | 0 | 1.40E-06 | 1.24E-05 | 0 | 1.60E-05 | 4.33E-06 | 0 | 5.45E-05 | 2.39E-05 | 1.52E-05 | 2.27E-05 | 3.33E-05 |
| Abraxas1 | 2.10E-05 | 2.06E-05 | 9.40E-06 | 0 | 1.24E-05 | 1.74E-05 | 0 | 1.14E-05 | 1.24E-05 | 1.72E-05 | 4.19E-06 | 1.46E-05 | 0 | 3.08E-06 | 2.57E-05 | 6.80E-05 | 0 | 5.05E-06 |
| Piwil4 | 7.67E-06 | 1.99E-07 | 0 | 0 | 0 | 1.73E-05 | 0 | 0.00011 | 1.24E-05 | 0 | 0 | 0 | 0 | 2.11E-05 | 0 | 2.82E-05 | 0 | 0 |
| Zfp35 | 3.73E-05 | 2.75E-05 | 5.34E-06 | 2.37E-05 | 1.26E-05 | 9.05E-06 | 0 | 0 | 1.24E-05 | 1.86E-05 | 3.11E-06 | 8.04E-07 | 0 | 1.47E-05 | 6.22E-06 | 3.66E-05 | 0 | 1.44E-05 |
| Ankrd24 | 5.32E-06 | 5.36E-06 | 2.72E-06 | 3.80E-06 | 0 | 3.58E-06 | 3.33E-06 | 1.24E-06 | 1.24E-05 | 2.02E-06 | 0 | 6.83E-06 | 2.03E-06 | 5.74E-06 | 4.26E-06 | 0 | 0 | 9.38E-06 |
| Gtf2e2 | 0.0002 | 3.93E-05 | 0.00015 | 2.74E-05 | 5.13E-05 | 6.12E-05 | 0.00016 | 7.93E-05 | 1.23E-05 | 0 | 4.48E-05 | 0.00035 | 0 | 0 | 1.50E-05 | 0.00016 | 0 | 0 |
| Ap5m1 | 7.40E-05 | 8.91E-05 | 1.85E-05 | 0 | 7.19E-05 | 6.08E-05 | 0 | 1.67E-06 | 1.23E-05 | 3.35E-05 | 2.23E-05 | 3.20E-05 | 2.38E-06 | 1.13E-05 | 7.32E-05 | 3.21E-05 | 0 | 3.57E-06 |
| Asnsd1 | 0.00015 | 8.92E-05 | 8.54E-05 | 0.00011 | 4.31E-05 | 5.39E-05 | 0.00013 | 5.61E-05 | 1.23E-05 | 9.68E-05 | 0 | 0.00013 | 4.21E-05 | 0 | 0 | 0.00026 | 0 | 2.25E-05 |
| Cdkal1 | 9.84E-05 | 7.85E-05 | 2.63E-05 | 0 | 3.40E-05 | 5.16E-05 | 1.98E-05 | 1.05E-05 | 1.23E-05 | 1.31E-05 | 9.74E-08 | 2.70E-05 | 0 | 0 | 0.00017 | 7.87E-05 | 2.21E-06 | 2.30E-05 |
| Pou2f1 | 4.58E-05 | 5.60E-05 | 0 | 0 | 1.35E-05 | 3.19E-05 | 1.24E-05 | 0 | 1.23E-05 | 1.38E-05 | 1.89E-05 | 3.23E-05 | 0 | 2.76E-05 | 7.91E-05 | 3.64E-05 | 7.55E-05 | 0 |
| Tbc1d32 | 1.10E-05 | 1.47E-05 | 0 | 1.87E-06 | 0 | 1.30E-05 | 6.53E-06 | 1.26E-05 | 1.23E-05 | 2.50E-05 | 0 | 3.91E-06 | 8.83E-06 | 0 | 3.72E-06 | 1.98E-05 | 0 | 1.23E-05 |

| | | | | | | | | | | | | | | | | | | |
|---|---|---|---|---|---|---|---|---|---|---|---|---|---|---|---|---|---|---|
| Sirt5 | 5.79E-06 | 3.15E-05 | 0 | 1.07E-05 | 2.31E-05 | 1.08E-05 | 1.89E-05 | 1.89E-06 | 1.23E-05 | 3.71E-05 | 1.71E-05 | 1.49E-05 | 0 | 1.02E-05 | 1.94E-05 | 2.57E-07 | 0 | 2.80E-05 |
| Zfp982 | 1.03E-05 | 2.14E-06 | 0 | 0 | 7.84E-07 | 9.74E-06 | 5.91E-07 | 0 | 1.23E-05 | 5.89E-06 | 0 | 6.30E-06 | 6.99E-06 | 0 | 6.82E-06 | 1.33E-06 | 0 | 2.82E-06 |
| Ipp | 4.88E-06 | 2.36E-05 | 9.49E-06 | 2.42E-05 | 1.88E-05 | 0 | 1.80E-05 | 5.22E-07 | 1.23E-05 | 1.07E-05 | 2.21E-05 | 1.25E-06 | 2.97E-05 | 5.25E-06 | 0 | 2.61E-05 | 0 | 9.27E-06 |
| Pheta1 | 0 | 4.93E-05 | 1.92E-05 | 5.82E-05 | 3.07E-05 | 5.10E-06 | 2.41E-05 | 1.39E-05 | 1.22E-05 | 2.07E-05 | 8.97E-06 | 2.46E-05 | 2.55E-05 | 3.42E-05 | 4.65E-05 | 0 | 0.00011 | 6.30E-06 |
| BC005561 | 2.16E-05 | 4.75E-05 | 3.43E-05 | 7.19E-06 | 9.68E-06 | 4.26E-06 | 0 | 2.44E-05 | 1.22E-05 | 1.87E-05 | 2.45E-06 | 3.72E-05 | 0 | 3.68E-05 | 0.00016 | 1.88E-05 | 9.05E-05 | 0 |
| Prokr1 | 1.70E-06 | 4.59E-06 | 0 | 1.21E-05 | 0 | 2.84E-06 | 0 | 0 | 1.22E-05 | 2.37E-05 | 0 | 0 | 1.39E-05 | 1.14E-06 | 0 | 3.62E-06 | 7.32E-07 | 0 |
| Ercc8 | 8.96E-06 | 1.21E-05 | 1.99E-05 | 0 | 1.08E-05 | 1.64E-06 | 0 | 0 | 1.22E-05 | 6.71E-06 | 7.08E-06 | 0 | 0 | 1.64E-06 | 6.35E-06 | 1.29E-05 | 0 | 1.18E-07 |
| Ccdc120 | 0 | 1.76E-05 | 3.87E-06 | 1.84E-05 | 6.51E-06 | 2.61E-07 | 1.40E-05 | 0 | 1.22E-05 | 3.09E-06 | 1.19E-05 | 1.83E-05 | 9.16E-06 | 9.24E-06 | 2.80E-05 | 2.32E-05 | 0 | 5.40E-06 |
| Pop1 | 1.09E-05 | 0 | 5.87E-05 | 3.55E-05 | 0 | 0 | 0 | 1.65E-06 | 1.22E-05 | 0 | 1.27E-05 | 0 | 0 | 2.40E-06 | 2.88E-05 | 2.32E-05 | 0 | 3.62E-06 |
| Polr1b | 1.77E-05 | 1.79E-05 | 5.63E-05 | 1.22E-05 | 0 | 0 | 0 | 0 | 1.22E-05 | 8.83E-06 | 7.28E-06 | 0 | 0 | 0 | 1.86E-05 | 1.89E-05 | 1.13E-05 | 0 |
| Fam185a | 1.78E-05 | 1.26E-05 | 4.41E-05 | 4.14E-05 | 0 | 0 | 8.08E-05 | 0 | 1.22E-05 | 1.82E-05 | 1.89E-05 | 0 | 0 | 0 | 9.34E-05 | 0 | 0 | 1.19E-06 |
| Zfp143 | 4.95E-05 | 4.61E-05 | 2.05E-05 | 0 | 4.48E-06 | 2.77E-05 | 0 | 0 | 1.21E-05 | 5.93E-06 | 3.49E-06 | 3.85E-05 | 0 | 6.76E-06 | 4.83E-05 | 4.03E-05 | 3.60E-06 | 2.09E-05 |
| Cfap69 | 7.84E-06 | 2.36E-05 | 7.81E-06 | 1.52E-05 | 8.63E-06 | 1.92E-05 | 0 | 0 | 1.21E-05 | 7.68E-06 | 0 | 3.99E-05 | 3.30E-05 | 2.71E-05 | 7.45E-05 | 1.35E-05 | 2.30E-05 | 0 |
| Zfat | 1.82E-05 | 1.22E-05 | 5.95E-06 | 0 | 0 | 9.58E-06 | 1.32E-05 | 0 | 1.21E-05 | 1.24E-05 | 7.03E-06 | 8.60E-06 | 4.11E-06 | 0 | 2.77E-05 | 3.14E-06 | 1.97E-05 | 2.36E-06 |
| Paxx | 3.31E-05 | 5.31E-05 | 0 | 4.38E-05 | 5.32E-05 | 7.24E-06 | 2.98E-05 | 0 | 1.21E-05 | 0.00015 | 2.56E-05 | 2.82E-05 | 0 | 0 | 1.19E-05 | 0 | 0 | 0.00013 |
| Slamf8 | 0 | 0 | 0 | 3.32E-06 | 0 | 0 | 0 | 0 | 1.21E-05 | 8.80E-07 | 0 | 0 | 5.26E-05 | 0 | 0 | 4.52E-06 | 0 | 3.75E-05 |
| Maob | 0 | 2.59E-05 | 0 | 0 | 5.22E-06 | 0 | 0 | 0 | 1.21E-05 | 8.24E-05 | 0 | 0 | 9.43E-05 | 0 | 0 | 0 | 0 | 2.18E-05 |
| Pole | 0 | 0 | 2.34E-05 | 0 | 0 | 0 | 0 | 0 | 1.21E-05 | 5.69E-06 | 6.11E-06 | 0 | 2.02E-05 | 1.39E-05 | 0 | 0.00052 | 0 | 0 |
| Fxyd5 | 0.00034 | 5.96E-05 | 0.00023 | 0 | 0.00025 | 0.00051 | 0.00053 | 0.00022 | 1.20E-05 | 0.00038 | 0 | 0.00045 | 0.0014 | 0.00045 | 0 | 0 | 0 | 0.00023 |
| Tmem208 | 0.00023 | 5.69E-05 | 0.00022 | 0.00025 | 0.00013 | 0.0001 | 0.00034 | 0.00011 | 1.20E-05 | 2.81E-05 | 5.68E-05 | 0 | 0.00014 | 9.81E-05 | 0.00012 | 1.30E-05 | 0 | 9.76E-05 |
| Ppm1k | 6.42E-06 | 4.74E-05 | 0 | 2.84E-05 | 6.83E-05 | 4.24E-05 | 1.03E-05 | 3.05E-05 | 1.20E-05 | 3.15E-05 | 0 | 3.07E-05 | 0 | 0 | 2.08E-05 | 3.11E-05 | 0 | 2.63E-05 |
| Vps50 | 6.66E-05 | 4.11E-05 | 3.89E-05 | 0 | 2.90E-05 | 3.02E-05 | 0 | 4.08E-07 | 1.20E-05 | 1.36E-05 | 5.57E-06 | 1.71E-05 | 0 | 2.06E-05 | 6.24E-05 | 1.80E-05 | 0 | 1.59E-05 |
| Papolg | 4.08E-05 | 3.83E-05 | 1.82E-05 | 0 | 3.51E-05 | 2.76E-05 | 1.68E-05 | 0 | 1.20E-05 | 1.15E-05 | 5.94E-07 | 5.91E-06 | 4.82E-06 | 8.28E-06 | 5.23E-05 | 1.74E-05 | 0 | 6.65E-06 |
| Fancm | 1.02E-05 | 7.66E-06 | 1.05E-05 | 0 | 2.73E-06 | 2.35E-05 | 9.75E-06 | 5.66E-06 | 1.20E-05 | 9.23E-06 | 0 | 0 | 1.30E-05 | 2.32E-05 | 8.34E-06 | 0.00019 | 2.05E-05 | 4.94E-06 |
| Slc46a1 | 4.49E-05 | 4.34E-05 | 2.25E-05 | 6.05E-05 | 8.20E-05 | 2.22E-05 | 0.00016 | 0 | 1.20E-05 | 0 | 1.91E-05 | 1.55E-05 | 4.18E-05 | 8.17E-06 | 0 | 0 | 0 | 1.28E-05 |
| Nek4 | 3.51E-05 | 2.47E-05 | 1.03E-05 | 3.99E-06 | 9.05E-06 | 2.04E-05 | 7.31E-06 | 0 | 1.20E-05 | 0 | 0 | 5.73E-06 | 2.30E-06 | 0 | 3.17E-05 | 1.94E-05 | 0 | 7.10E-06 |
| Dclre1a | 2.60E-05 | 3.41E-05 | 7.81E-06 | 5.06E-06 | 7.55E-06 | 1.05E-05 | 0 | 0 | 1.20E-05 | 2.56E-05 | 0 | 1.50E-05 | 0 | 7.86E-06 | 1.29E-05 | 6.29E-05 | 0 | 1.34E-05 |
| Nhej1 | 9.63E-06 | 1.75E-05 | 9.63E-06 | 2.03E-05 | 0 | 7.93E-06 | 6.03E-07 | 3.08E-06 | 1.20E-05 | 5.36E-06 | 5.38E-06 | 6.04E-06 | 0 | 0 | 2.08E-05 | 1.87E-05 | 3.19E-05 | 7.42E-06 |
| Mamstr | 0 | 2.40E-06 | 0 | 1.20E-05 | 0 | 4.75E-06 | 0 | 0 | 1.20E-05 | 6.73E-05 | 0 | 1.92E-05 | 0 | 0 | 0 | 7.38E-05 | 0 | 0.00011 |
| Rgl2 | 0 | 0.00011 | 6.13E-06 | 0.00016 | 6.21E-05 | 4.04E-06 | 7.97E-06 | 5.39E-06 | 1.20E-05 | 4.30E-05 | 3.19E-06 | 4.74E-05 | 0 | 7.15E-05 | 0.00019 | 0 | 1.13E-05 | 6.65E-05 |
| Slc16a6 | 0 | 1.69E-06 | 2.79E-05 | 0 | 0 | 0 | 9.62E-06 | 0 | 1.20E-05 | 1.86E-06 | 0 | 0 | 3.82E-05 | 0 | 5.38E-07 | 0 | 0 | 0 |
| Satb2 | 1.11E-05 | 1.14E-05 | 2.02E-05 | 0 | 0 | 0 | 0 | 2.71E-05 | 1.20E-05 | 0 | 1.03E-05 | 5.86E-06 | 0 | 6.13E-05 | 7.99E-06 | 6.53E-07 | 1.60E-05 | 8.97E-06 |
| Polrmt | 3.94E-06 | 1.28E-05 | 1.14E-05 | 3.40E-05 | 8.98E-06 | 0 | 0 | 5.25E-06 | 1.20E-05 | 3.65E-06 | 0 | 0 | 0 | 2.58E-05 | 2.57E-05 | 3.88E-06 | 0 | 0 |

| | | | | | | | | | | | | | | | | | | |
|---|---|---|---|---|---|---|---|---|---|---|---|---|---|---|---|---|---|---|
| Xpnpep2 | 0 | 0 | 0 | 0 | 0 | 0 | 0 | 0 | 1.20E-05 | 0.00012 | 0 | 1.04E-05 | 0.00048 | 0 | 0 | 0 | 0 | 0 |
| Mogs | 7.21E-05 | 4.76E-05 | 0.00026 | 0.00031 | 5.10E-05 | 0.00013 | 0.00012 | 0.00013 | 1.19E-05 | 3.37E-05 | 7.44E-05 | 3.85E-05 | 9.67E-05 | 2.60E-05 | 0.0003 | 0.00015 | 0 | 0 |
| Sbk1 | 1.92E-05 | 9.68E-05 | 1.61E-05 | 0 | 7.07E-05 | 5.63E-05 | 0 | 4.27E-05 | 1.19E-05 | 0 | 2.62E-05 | 3.03E-05 | 3.43E-05 | 5.87E-05 | 0 | 2.81E-05 | 1.13E-05 | 1.46E-05 |
| Mmp8 | 0 | 0 | 0 | 0 | 3.18E-05 | 2.49E-05 | 0 | 0 | 1.19E-05 | 0 | 0 | 0 | 1.94E-05 | 7.46E-05 | 0 | 7.63E-06 | 0 | 0 |
| Rtel1 | 0 | 6.87E-06 | 1.95E-05 | 2.32E-05 | 9.03E-06 | 8.19E-06 | 0 | 3.23E-06 | 1.19E-05 | 6.08E-06 | 0 | 5.14E-06 | 1.07E-05 | 9.51E-06 | 9.79E-06 | 0.00016 | 2.58E-05 | 1.61E-05 |
| Cryl1 | 9.38E-06 | 3.10E-05 | 0 | 5.20E-05 | 2.60E-05 | 6.75E-06 | 3.52E-05 | 0 | 1.19E-05 | 6.24E-05 | 1.75E-05 | 1.78E-06 | 4.03E-05 | 4.11E-05 | 0 | 9.09E-05 | 2.32E-05 | 8.35E-05 |
| Safb2 | 8.72E-05 | 0.00015 | 0.00011 | 3.15E-05 | 6.46E-05 | 0 | 2.09E-05 | 0 | 1.19E-05 | 3.90E-05 | 1.96E-05 | 0.00012 | 1.13E-05 | 0 | 0.00062 | 0.00014 | 0.00019 | 5.54E-05 |
| Kcnj2 | 8.68E-06 | 1.63E-05 | 0 | 0 | 2.58E-05 | 0 | 4.15E-06 | 0 | 1.19E-05 | 2.01E-05 | 2.19E-05 | 1.82E-05 | 0.00011 | 0 | 0 | 0 | 0 | 7.81E-06 |
| Zbtb41 | 0.00011 | 4.63E-05 | 7.05E-05 | 0 | 5.64E-05 | 7.96E-05 | 5.30E-05 | 3.66E-05 | 1.18E-05 | 1.60E-05 | 2.66E-05 | 3.68E-05 | 4.42E-05 | 0 | 3.94E-06 | 0.00015 | 3.38E-05 | 1.63E-05 |
| Rpf1 | 0.00012 | 0.00012 | 0.00018 | 0.00012 | 1.65E-05 | 1.58E-05 | 0 | 1.30E-05 | 1.18E-05 | 0 | 3.56E-05 | 9.61E-06 | 0 | 1.57E-05 | 0.00016 | 0.00014 | 0 | 0 |
| Lrrc29 | 3.20E-06 | 1.73E-05 | 0 | 1.33E-05 | 0 | 4.75E-07 | 0 | 1.38E-05 | 1.18E-05 | 1.37E-06 | 2.61E-06 | 2.59E-06 | 0 | 3.00E-06 | 3.67E-08 | 0 | 0 | 0 |
| Pard6g | 3.47E-05 | 0.00011 | 1.89E-05 | 0 | 4.23E-05 | 4.78E-05 | 4.07E-05 | 0.00012 | 1.17E-05 | 0 | 6.63E-05 | 0 | 0 | 9.58E-05 | 0 | 6.16E-05 | 3.20E-05 | 7.63E-05 |
| Lst1 | 1.26E-05 | 3.04E-05 | 0 | 1.95E-05 | 1.43E-05 | 2.41E-05 | 1.47E-05 | 1.76E-06 | 1.17E-05 | 9.54E-06 | 1.93E-05 | 8.55E-06 | 3.46E-05 | 1.33E-05 | 1.67E-05 | 0 | 0 | 6.63E-05 |
| Efna1 | 2.54E-05 | 2.89E-05 | 4.42E-06 | 1.62E-05 | 1.08E-05 | 2.34E-05 | 6.59E-05 | 0 | 1.17E-05 | 0 | 0 | 4.76E-05 | 1.38E-05 | 2.38E-05 | 9.60E-06 | 0 | 7.28E-06 | 8.09E-07 |
| Clcn2 | 0 | 2.23E-05 | 0 | 1.91E-06 | 2.76E-05 | 1.81E-05 | 3.29E-06 | 0 | 1.17E-05 | 4.01E-06 | 1.51E-05 | 2.70E-05 | 1.69E-05 | 2.62E-05 | 5.52E-05 | 0 | 5.44E-06 | 1.96E-05 |
| Ndufaf6 | 2.32E-05 | 2.35E-05 | 4.01E-05 | 2.00E-05 | 1.27E-06 | 1.36E-05 | 4.23E-07 | 0 | 1.17E-05 | 8.74E-06 | 1.25E-05 | 0 | 0 | 0 | 4.15E-05 | 3.16E-05 | 0 | 1.56E-06 |
| Mettl8 | 3.28E-05 | 2.75E-05 | 1.73E-05 | 0 | 8.96E-06 | 1.11E-05 | 8.64E-07 | 0 | 1.17E-05 | 1.01E-05 | 3.67E-06 | 3.42E-06 | 0 | 2.44E-07 | 2.86E-05 | 1.68E-05 | 1.89E-06 | 1.09E-05 |
| Zfp839 | 0 | 3.77E-05 | 3.18E-06 | 2.00E-05 | 2.45E-05 | 6.81E-06 | 0 | 1.43E-05 | 1.17E-05 | 9.29E-06 | 5.51E-06 | 2.72E-05 | 3.03E-06 | 2.44E-06 | 6.27E-05 | 0 | 0 | 1.59E-05 |
| Mterf2 | 3.82E-06 | 1.31E-05 | 0 | 0 | 5.14E-06 | 4.84E-06 | 0 | 0 | 1.17E-05 | 0 | 3.72E-07 | 1.00E-05 | 0 | 6.01E-06 | 2.81E-06 | 0 | 0 | 1.67E-05 |
| Cep290 | 2.55E-05 | 1.36E-05 | 9.61E-06 | 0 | 6.42E-06 | 2.80E-06 | 0 | 0 | 1.17E-05 | 8.64E-07 | 1.53E-06 | 2.13E-05 | 6.01E-06 | 0 | 2.10E-05 | 4.63E-05 | 1.66E-05 | 4.40E-07 |
| Scel | 3.43E-06 | 0 | 0 | 0 | 9.01E-06 | 0 | 0 | 0 | 1.17E-05 | 0 | 2.31E-05 | 0 | 3.57E-06 | 1.99E-05 | 0 | 0 | 0 | 3.53E-05 |
| Rnf144b | 0 | 1.31E-05 | 0 | 0 | 3.20E-05 | 0 | 0 | 0 | 1.17E-05 | 0 | 1.61E-06 | 3.69E-06 | 0 | 1.46E-05 | 0 | 0 | 0 | 1.07E-05 |
| Ptger1 | 9.26E-07 | 0 | 2.97E-07 | 1.24E-05 | 0 | 0 | 2.69E-06 | 1.28E-05 | 1.17E-05 | 0 | 0 | 2.54E-05 | 2.88E-05 | 0 | 0 | 4.82E-06 | 5.57E-05 | 4.91E-05 |
| Nfkbie | 2.16E-05 | 1.65E-05 | 7.30E-05 | 0 | 6.37E-06 | 2.53E-05 | 0 | 4.29E-05 | 1.16E-05 | 0 | 0 | 4.42E-06 | 4.88E-05 | 3.92E-05 | 8.02E-06 | 0 | 0.00016 | 9.13E-05 |
| Phospho2 | 5.76E-05 | 1.43E-05 | 1.49E-05 | 0 | 6.82E-06 | 2.05E-05 | 2.33E-05 | 7.92E-06 | 1.16E-05 | 3.32E-06 | 6.91E-06 | 4.77E-06 | 0 | 7.43E-06 | 7.85E-06 | 8.89E-06 | 0 | 1.08E-05 |
| Tmem199 | 4.14E-05 | 3.06E-05 | 3.88E-05 | 1.88E-05 | 1.57E-05 | 1.44E-05 | 0 | 1.97E-05 | 1.16E-05 | 0 | 0 | 0 | 1.22E-05 | 0 | 4.26E-06 | 1.25E-05 | 0 | 0 |
| Lingo1 | 0 | 2.96E-05 | 2.56E-05 | 0 | 0 | 1.36E-05 | 0 | 2.28E-05 | 1.16E-05 | 0 | 4.51E-06 | 0.00015 | 0 | 3.07E-05 | 0 | 2.24E-05 | 0 | 3.53E-05 |
| Ttc30a1 | 1.46E-05 | 4.71E-06 | 3.48E-06 | 5.85E-06 | 5.94E-06 | 1.30E-05 | 1.29E-05 | 3.63E-06 | 1.16E-05 | 0 | 3.66E-06 | 1.15E-05 | 7.62E-06 | 1.48E-05 | 3.58E-06 | 2.13E-05 | 6.45E-07 | 8.43E-06 |
| Tcp11l1 | 0 | 3.19E-05 | 2.30E-06 | 0 | 4.35E-05 | 1.19E-05 | 5.45E-06 | 0 | 1.16E-05 | 2.39E-05 | 1.55E-05 | 2.77E-05 | 3.83E-05 | 2.66E-05 | 0 | 1.59E-05 | 8.34E-06 | 2.35E-05 |
| Cyb5r1 | 8.76E-05 | 0.00013 | 0.00019 | 0 | 7.30E-06 | 8.20E-06 | 0 | 0 | 1.16E-05 | 0 | 0.00018 | 0.00039 | 0 | 4.69E-05 | 0.00018 | 0 | 0 | 0.00014 |
| Zfp568 | 1.52E-05 | 2.82E-05 | 2.93E-05 | 1.62E-05 | 1.45E-05 | 0 | 0 | 1.67E-05 | 1.16E-05 | 1.26E-05 | 9.03E-05 | 3.07E-05 | 0 | 4.85E-05 | 7.27E-05 | 1.63E-05 | 1.35E-05 | 0 |
| Zfp618 | 0 | 2.83E-05 | 0 | 0 | 4.38E-05 | 0.00016 | 0 | 0.00015 | 1.15E-05 | 0 | 8.40E-05 | 0.00014 | 0.00016 | 0.00013 | 0.00023 | 0 | 0.00014 | 0 |
| Rpp30 | 7.46E-05 | 1.71E-05 | 4.87E-05 | 0 | 0 | 3.17E-05 | 0 | 0 | 1.15E-05 | 6.98E-06 | 0 | 3.41E-05 | 7.82E-06 | 0 | 0 | 0.00015 | 0 | 4.94E-06 |

| | | | | | | | | | | | | | | | | | | |
|---|---|---|---|---|---|---|---|---|---|---|---|---|---|---|---|---|---|---|
| Det1 | 3.53E-05 | 3.63E-05 | 4.10E-05 | 3.10E-05 | 1.51E-05 | 2.62E-05 | 6.94E-06 | 0 | 1.15E-05 | 1.68E-05 | 1.72E-05 | 0 | 2.42E-07 | 5.36E-06 | 4.28E-06 | 2.81E-05 | 0 | 2.10E-05 |
| Txlng | 3.88E-05 | 4.95E-05 | 6.21E-06 | 0 | 1.02E-05 | 1.43E-05 | 2.56E-05 | 0 | 1.15E-05 | 1.58E-05 | 0 | 3.82E-05 | 2.04E-05 | 0 | 0.00014 | 0 | 1.46E-05 | 0 |
| S100a8 | 0 | 1.80E-05 | 0 | 0 | 0 | 0 | 0 | 0 | 1.15E-05 | 1.02E-05 | 0 | 6.30E-06 | 1.25E-05 | 1.04E-05 | 9.62E-05 | 3.28E-05 | 3.80E-05 | 6.19E-06 |
| Atp6v0d1 | 0.00037 | 0.00016 | 0.00015 | 6.12E-05 | 0.00016 | 0.00022 | 0.00029 | 7.38E-05 | 1.14E-05 | 7.41E-05 | 7.34E-05 | 0.00011 | 8.24E-05 | 0.00017 | 0 | 0 | 3.31E-05 | 0.00011 |
| Naalad2 | 1.84E-05 | 4.37E-05 | 0 | 0 | 1.19E-05 | 6.93E-05 | 4.62E-05 | 0.00017 | 1.14E-05 | 0.00014 | 0 | 2.75E-05 | 2.80E-05 | 0 | 0 | 0 | 0 | 0.00051 |
| Tmem67 | 6.71E-05 | 1.26E-05 | 1.54E-07 | 1.36E-05 | 6.34E-06 | 2.97E-05 | 3.48E-05 | 5.11E-06 | 1.14E-05 | 3.59E-06 | 0 | 2.81E-05 | 1.84E-05 | 9.36E-06 | 1.96E-06 | 1.64E-05 | 6.30E-05 | 0 |
| Zfp51 | 3.51E-05 | 2.14E-05 | 6.05E-06 | 0 | 1.14E-05 | 1.88E-05 | 2.21E-06 | 3.74E-07 | 1.14E-05 | 1.91E-07 | 0 | 1.92E-05 | 0 | 8.87E-06 | 2.73E-05 | 0 | 0 | 2.51E-05 |
| Kctd7 | 1.87E-05 | 2.77E-05 | 0 | 3.45E-05 | 2.81E-05 | 1.79E-05 | 1.56E-06 | 3.34E-05 | 1.14E-05 | 1.87E-05 | 0 | 0 | 0 | 1.91E-05 | 1.37E-06 | 1.34E-05 | 2.85E-05 | 1.11E-05 |
| Ercc2 | 1.70E-05 | 3.35E-05 | 4.27E-05 | 8.70E-06 | 1.06E-05 | 1.30E-05 | 3.70E-06 | 1.31E-05 | 1.14E-05 | 3.43E-05 | 2.89E-05 | 9.20E-06 | 2.65E-05 | 3.60E-06 | 6.77E-05 | 4.41E-05 | 0 | 0 |
| Zfp128 | 9.49E-06 | 2.41E-05 | 0 | 0 | 1.42E-05 | 2.75E-06 | 0 | 0 | 1.14E-05 | 1.84E-06 | 1.08E-05 | 3.62E-06 | 4.49E-06 | 6.35E-07 | 1.61E-05 | 1.48E-06 | 0 | 4.45E-06 |
| Galnt12 | 1.75E-05 | 1.40E-06 | 2.89E-05 | 1.19E-05 | 1.33E-06 | 0 | 3.74E-06 | 5.58E-05 | 1.14E-05 | 8.65E-05 | 5.32E-06 | 2.74E-05 | 5.23E-05 | 0 | 0 | 4.66E-05 | 3.91E-06 | 0 |
| Nek1 | 6.21E-05 | 5.03E-05 | 3.86E-06 | 4.76E-06 | 1.75E-05 | 0 | 9.66E-05 | 1.69E-05 | 1.14E-05 | 5.09E-05 | 4.23E-06 | 0.00016 | 0 | 0 | 7.11E-05 | 2.15E-05 | 2.40E-05 | 5.12E-05 |
| Zmat1 | 9.00E-06 | 1.22E-05 | 0 | 0 | 1.01E-05 | 0 | 1.13E-05 | 0 | 1.14E-05 | 3.92E-06 | 4.76E-06 | 4.56E-06 | 0 | 0 | 1.25E-05 | 2.39E-05 | 1.07E-05 | 2.01E-06 |
| Shroom3 | 0 | 2.78E-06 | 0 | 0 | 3.73E-05 | 0 | 6.34E-07 | 0 | 1.14E-05 | 1.37E-06 | 2.80E-05 | 1.26E-06 | 6.81E-06 | 3.83E-06 | 0 | 8.16E-06 | 0 | 5.91E-06 |
| Dnajc17 | 1.75E-05 | 2.02E-05 | 1.25E-05 | 1.89E-05 | 6.16E-06 | 0 | 0 | 0 | 1.14E-05 | 4.84E-06 | 0 | 8.72E-06 | 0 | 5.18E-06 | 1.10E-05 | 1.14E-05 | 1.84E-06 | 2.22E-05 |
| Ankhd1 | 7.28E-05 | 8.63E-05 | 3.36E-05 | 0 | 1.58E-05 | 4.52E-05 | 2.54E-05 | 2.23E-05 | 1.13E-05 | 0 | 0 | 5.95E-05 | 3.39E-05 | 0 | 6.97E-05 | 4.69E-05 | 2.30E-05 | 7.68E-06 |
| Tedc2 | 0 | 2.98E-05 | 2.17E-05 | 0 | 4.16E-05 | 4.14E-05 | 1.35E-05 | 0 | 1.13E-05 | 2.76E-06 | 1.20E-06 | 1.09E-05 | 1.79E-05 | 5.41E-05 | 7.62E-05 | 0.00021 | 1.85E-05 | 1.56E-05 |
| Prdm4 | 6.87E-05 | 8.76E-05 | 6.95E-05 | 1.08E-05 | 1.40E-05 | 3.21E-05 | 1.12E-05 | 1.19E-05 | 1.13E-05 | 4.42E-06 | 6.82E-05 | 7.43E-05 | 1.85E-05 | 3.16E-05 | 2.68E-05 | 5.50E-05 | 1.76E-05 | 2.15E-05 |
| Plekha8 | 5.76E-05 | 2.58E-05 | 7.45E-06 | 1.18E-05 | 1.11E-05 | 2.36E-05 | 1.56E-05 | 0 | 1.13E-05 | 1.54E-05 | 1.32E-05 | 1.14E-05 | 1.71E-05 | 0 | 1.88E-06 | 1.39E-05 | 2.99E-06 | 7.83E-06 |
| Ccdc66 | 4.27E-05 | 2.31E-05 | 0 | 5.24E-06 | 1.69E-05 | 1.94E-05 | 1.86E-05 | 0 | 1.13E-05 | 2.68E-05 | 1.01E-05 | 7.67E-06 | 8.66E-06 | 3.82E-06 | 3.76E-05 | 2.84E-05 | 0 | 8.50E-06 |
| Il2ra | 0 | 0 | 2.38E-05 | 0 | 4.39E-05 | 1.59E-05 | 2.01E-05 | 0 | 1.13E-05 | 0 | 0 | 0 | 0 | 5.16E-05 | 0 | 0 | 0 | 0 |
| Dpyd | 0 | 2.38E-05 | 0 | 0 | 1.06E-05 | 9.12E-06 | 0 | 0 | 1.13E-05 | 3.23E-05 | 0 | 5.63E-06 | 8.74E-07 | 0 | 2.68E-05 | 0 | 0 | 1.98E-05 |
| Atp6v1g2 | 6.30E-06 | 6.67E-06 | 0 | 1.54E-05 | 1.85E-05 | 5.87E-06 | 0 | 6.00E-06 | 1.13E-05 | 6.99E-07 | 0 | 0 | 0 | 1.14E-05 | 2.67E-05 | 0 | 0 | 0 |
| Orai2 | 0 | 4.30E-05 | 0 | 5.98E-05 | 1.32E-05 | 0 | 0 | 3.19E-05 | 1.12E-05 | 1.89E-05 | 0 | 1.52E-05 | 2.29E-05 | 3.13E-05 | 1.93E-05 | 0 | 4.48E-06 | 3.44E-05 |
| Zscan25 | 0 | 2.21E-05 | 1.32E-05 | 2.92E-05 | 3.66E-07 | 0 | 1.75E-05 | 5.32E-06 | 1.12E-05 | 6.08E-06 | 6.71E-06 | 7.62E-06 | 1.25E-05 | 0 | 4.41E-05 | 0 | 1.08E-05 | 1.47E-05 |
| Nadsyn1 | 1.11E-05 | 2.62E-05 | 2.99E-06 | 4.80E-06 | 0 | 0 | 2.67E-06 | 1.15E-05 | 1.12E-05 | 7.44E-06 | 1.45E-05 | 6.61E-06 | 6.47E-06 | 5.92E-06 | 5.65E-05 | 9.32E-06 | 0 | 1.18E-05 |
| Scai | 4.14E-05 | 6.93E-05 | 0 | 0 | 4.47E-05 | 2.64E-05 | 3.73E-05 | 0 | 1.11E-05 | 2.32E-05 | 5.56E-06 | 1.43E-05 | 0 | 1.76E-05 | 0.00012 | 2.95E-05 | 0.0001 | 1.68E-05 |
| Tmem205 | 3.55E-05 | 1.61E-05 | 0 | 4.30E-05 | 3.74E-05 | 2.16E-05 | 3.16E-05 | 1.53E-05 | 1.11E-05 | 2.93E-05 | 8.84E-06 | 1.48E-05 | 0 | 0 | 3.89E-05 | 0 | 0 | 7.59E-05 |
| Slc22a15 | 8.77E-06 | 2.03E-05 | 4.37E-06 | 0 | 3.64E-05 | 1.45E-05 | 1.51E-05 | 7.56E-06 | 1.11E-05 | 2.15E-05 | 1.23E-05 | 2.99E-06 | 5.56E-06 | 7.19E-06 | 4.52E-06 | 8.03E-06 | 3.35E-05 | 1.07E-05 |
| Inpp4b | 0 | 5.17E-06 | 1.85E-06 | 5.73E-06 | 5.02E-07 | 1.14E-06 | 0 | 8.60E-07 | 1.11E-05 | 1.53E-06 | 0 | 9.22E-07 | 1.81E-06 | 4.41E-06 | 5.76E-06 | 9.32E-07 | 6.45E-06 | 1.32E-06 |
| Orc6 | 7.49E-05 | 0 | 6.47E-05 | 0 | 0 | 3.42E-05 | 5.85E-06 | 0 | 1.10E-05 | 9.31E-06 | 7.08E-06 | 0 | 1.69E-05 | 1.24E-05 | 3.14E-05 | 0.00026 | 3.12E-05 | 0 |
| Kcp | 0 | 7.01E-06 | 0 | 0 | 6.97E-06 | 9.39E-06 | 0 | 1.74E-05 | 1.10E-05 | 2.79E-06 | 0 | 0 | 0 | 6.62E-06 | 3.04E-05 | 0 | 0 | 0 |
| N4bp3 | 0 | 1.19E-05 | 9.93E-06 | 0 | 0 | 4.01E-06 | 0 | 8.02E-06 | 1.10E-05 | 0 | 0 | 1.18E-05 | 0 | 4.28E-06 | 0 | 2.50E-05 | 2.15E-05 | 1.61E-05 |

| | | | | | | | | | | | | | | | | | | |
|---|---|---|---|---|---|---|---|---|---|---|---|---|---|---|---|---|---|---|
| Cdkn2aip | 2.38E-05 | 8.22E-05 | 4.15E-05 | 4.23E-05 | 1.37E-05 | 1.55E-06 | 5.91E-06 | 0 | 1.10E-05 | 9.73E-06 | 0 | 9.10E-05 | 0 | 9.92E-06 | 8.58E-05 | 4.17E-05 | 0.00041 | 1.96E-05 |
| Gap43 | 5.30E-06 | 0 | 3.67E-06 | 0 | 3.18E-07 | 0 | 1.12E-05 | 0 | 1.10E-05 | 4.70E-05 | 1.37E-07 | 0 | 0 | 5.87E-06 | 0 | 1.86E-05 | 0 | 0 |
| Fam174b | 1.96E-05 | 1.15E-05 | 2.08E-05 | 1.68E-05 | 0 | 0 | 0 | 5.49E-05 | 1.10E-05 | 2.30E-05 | 0 | 0 | 0 | 0.00018 | 0 | 0 | 0 | 3.95E-05 |
| Tsacc | 5.51E-06 | 6.38E-06 | 5.43E-06 | 4.11E-06 | 5.35E-06 | 0 | 0 | 0 | 1.10E-05 | 1.11E-05 | 0 | 3.34E-06 | 0 | 6.99E-06 | 5.68E-06 | 4.63E-08 | 0 | 7.82E-06 |
| Mapre2 | 0.00016 | 0.00011 | 2.68E-05 | 0 | 6.79E-05 | 0.00011 | 1.22E-05 | 0 | 1.09E-05 | 0.00011 | 0.0001 | 0.00021 | 0 | 3.91E-05 | 2.83E-05 | 3.80E-05 | 0.00011 | 3.71E-05 |
| Tmem42 | 1.80E-05 | 6.68E-07 | 0 | 0.00012 | 5.98E-05 | 4.96E-05 | 0.00015 | 2.31E-05 | 1.09E-05 | 1.65E-05 | 5.60E-06 | 7.39E-05 | 5.54E-05 | 2.91E-05 | 0 | 0 | 0 | 1.42E-05 |
| Socs7 | 8.81E-06 | 5.06E-05 | 4.61E-05 | 0 | 1.63E-05 | 2.27E-05 | 2.08E-07 | 0 | 1.09E-05 | 2.02E-06 | 2.93E-05 | 8.19E-06 | 1.02E-05 | 8.42E-06 | 2.94E-05 | 3.14E-05 | 4.78E-05 | 8.36E-06 |
| Banp | 1.74E-05 | 2.49E-05 | 1.52E-05 | 0 | 2.24E-05 | 1.62E-05 | 7.32E-06 | 7.49E-06 | 1.09E-05 | 1.06E-05 | 1.51E-05 | 2.19E-05 | 0 | 0 | 8.99E-05 | 2.18E-05 | 0 | 2.90E-06 |
| Tmem144 | 3.34E-06 | 1.23E-06 | 0 | 0 | 9.09E-06 | 7.81E-06 | 0 | 2.03E-05 | 1.09E-05 | 0 | 0 | 0 | 0 | 1.07E-05 | 0 | 4.03E-05 | 0 | 0 |
| Pex10 | 1.72E-05 | 1.35E-05 | 2.13E-06 | 1.34E-05 | 8.69E-07 | 6.35E-06 | 0 | 5.97E-06 | 1.09E-05 | 1.61E-05 | 1.72E-06 | 5.81E-06 | 9.09E-07 | 9.85E-07 | 2.53E-05 | 2.48E-05 | 0 | 6.72E-06 |
| Rapgefl1 | 2.74E-07 | 1.57E-05 | 0 | 3.04E-06 | 0 | 5.29E-06 | 1.35E-05 | 0 | 1.09E-05 | 4.79E-06 | 0 | 3.18E-06 | 9.81E-06 | 3.58E-05 | 2.08E-05 | 0 | 1.11E-05 | 0 |
| Trmt10b | 6.88E-06 | 2.65E-05 | 0 | 5.16E-06 | 1.08E-05 | 3.44E-06 | 7.39E-06 | 1.86E-06 | 1.09E-05 | 1.55E-05 | 1.87E-06 | 1.96E-05 | 0 | 0 | 1.85E-05 | 7.87E-06 | 0 | 1.74E-05 |
| Tmc4 | 0 | 2.05E-05 | 4.81E-06 | 2.63E-05 | 4.83E-06 | 0 | 6.43E-06 | 1.03E-05 | 1.09E-05 | 3.60E-07 | 0 | 2.37E-05 | 2.01E-06 | 0 | 6.07E-05 | 3.98E-06 | 4.71E-05 | 7.77E-06 |
| Id4 | 0 | 0 | 0 | 0 | 0 | 0 | 0 | 0 | 1.09E-05 | 3.62E-06 | 6.51E-05 | 1.12E-05 | 0 | 5.17E-05 | 2.62E-05 | 1.56E-06 | 0.00013 | 0.00025 |
| Ccnj | 0 | 8.11E-06 | 4.53E-06 | 9.58E-06 | 3.32E-06 | 0 | 5.45E-08 | 5.89E-06 | 1.09E-05 | 1.94E-06 | 9.19E-06 | 1.08E-06 | 0 | 0 | 2.37E-05 | 0 | 0 | 5.85E-06 |
| Chek1 | 1.62E-05 | 0 | 3.75E-05 | 0 | 0 | 0 | 1.48E-05 | 0 | 1.09E-05 | 0 | 1.04E-05 | 7.81E-06 | 0 | 0 | 1.04E-05 | 0.00018 | 8.03E-05 | 0 |
| Frmd8 | 0 | 0.00023 | 2.59E-05 | 0.00027 | 3.66E-05 | 0.00013 | 0 | 0.0003 | 1.08E-05 | 7.06E-05 | 0.00015 | 9.06E-05 | 0 | 0.00011 | 6.23E-05 | 0.00015 | 3.66E-05 | 7.56E-05 |
| Ints12 | 7.04E-05 | 6.63E-05 | 4.68E-05 | 3.18E-05 | 3.61E-06 | 3.53E-05 | 2.12E-05 | 1.12E-05 | 1.08E-05 | 2.16E-05 | 2.25E-05 | 5.60E-05 | 0 | 1.17E-05 | 4.58E-05 | 9.65E-05 | 0 | 2.53E-06 |
| Zfp933 | 2.01E-05 | 3.57E-05 | 2.50E-06 | 7.61E-07 | 3.23E-05 | 2.87E-05 | 1.34E-06 | 0 | 1.08E-05 | 1.05E-05 | 0 | 4.67E-06 | 1.13E-05 | 1.15E-05 | 2.36E-05 | 5.56E-06 | 0 | 2.74E-05 |
| Lyrm4 | 3.46E-05 | 3.03E-05 | 1.87E-05 | 0 | 3.02E-05 | 2.34E-05 | 0 | 0 | 1.08E-05 | 1.57E-05 | 3.60E-06 | 7.96E-06 | 1.24E-05 | 1.75E-05 | 0 | 3.54E-05 | 2.56E-05 | 0 |
| Rhot2 | 3.45E-05 | 5.91E-05 | 1.87E-05 | 1.66E-05 | 2.42E-05 | 2.08E-05 | 0 | 1.67E-06 | 1.08E-05 | 1.27E-05 | 0 | 1.25E-05 | 0 | 7.26E-06 | 8.29E-05 | 1.89E-05 | 3.91E-05 | 9.46E-06 |
| DEPRECATED | 6.70E-06 | 2.17E-05 | 0 | 2.03E-05 | 3.06E-05 | 1.89E-05 | 2.07E-05 | 4.57E-05 | 1.08E-05 | 1.99E-05 | 2.13E-05 | 0 | 3.50E-06 | 4.54E-05 | 8.83E-05 | 0 | 0 | 3.99E-05 |
| Neto2 | 0 | 3.91E-06 | 1.13E-05 | 0 | 1.16E-05 | 1.11E-05 | 0 | 1.21E-05 | 1.08E-05 | 0 | 0 | 1.17E-05 | 0 | 0 | 1.51E-06 | 8.71E-05 | 3.62E-05 | 0 |
| Wdhd1 | 0 | 0 | 3.35E-05 | 3.20E-06 | 0 | 2.40E-06 | 0 | 0 | 1.08E-05 | 7.30E-06 | 4.96E-06 | 0 | 5.82E-06 | 1.10E-05 | 4.98E-06 | 0.00026 | 2.64E-06 | 0 |
| Tmem81 | 4.60E-06 | 1.81E-05 | 7.61E-06 | 0 | 1.79E-05 | 1.96E-06 | 2.13E-06 | 7.36E-06 | 1.08E-05 | 3.32E-06 | 0 | 5.42E-06 | 1.35E-05 | 0 | 4.20E-05 | 1.60E-05 | 0 | 3.11E-06 |
| Rpp38 | 1.61E-05 | 1.41E-05 | 1.76E-05 | 7.90E-06 | 0 | 5.83E-07 | 0 | 1.85E-06 | 1.08E-05 | 8.46E-07 | 0 | 1.55E-05 | 0 | 0 | 1.88E-05 | 0 | 0 | 0 |
| Zfp692 | 1.24E-05 | 1.61E-05 | 1.26E-05 | 5.17E-06 | 9.91E-06 | 0 | 0 | 0 | 1.08E-05 | 1.45E-05 | 1.02E-06 | 1.62E-05 | 0 | 1.47E-06 | 0.00016 | 1.80E-05 | 2.57E-05 | 2.04E-06 |
| Eml1 | 7.65E-05 | 6.26E-05 | 9.98E-05 | 8.59E-05 | 0.00011 | 0.00024 | 0.00012 | 0 | 1.07E-05 | 8.92E-05 | 8.40E-05 | 9.06E-05 | 0 | 0.00022 | 3.78E-05 | 5.53E-05 | 0 | 0.00028 |
| Tmem267 | 4.55E-05 | 1.03E-05 | 8.24E-05 | 3.62E-05 | 1.45E-06 | 5.26E-05 | 3.30E-06 | 2.66E-05 | 1.07E-05 | 6.51E-05 | 0 | 0 | 0 | 5.38E-05 | 0 | 3.46E-05 | 6.50E-05 | 5.37E-05 |
| Spg11 | 7.73E-05 | 0.00011 | 4.15E-05 | 5.78E-05 | 4.97E-05 | 5.00E-05 | 0 | 0 | 1.07E-05 | 3.36E-05 | 1.82E-05 | 4.34E-05 | 0 | 0 | 5.47E-05 | 2.46E-05 | 7.34E-05 | 4.25E-05 |
| Mettl24 | 6.84E-06 | 0 | 4.16E-08 | 0 | 9.97E-06 | 4.31E-05 | 0 | 3.34E-05 | 1.07E-05 | 0 | 3.34E-05 | 1.18E-05 | 0 | 7.61E-05 | 0 | 1.15E-05 | 0 | 3.80E-07 |
| Mon1a | 5.07E-05 | 5.75E-05 | 1.46E-05 | 5.96E-05 | 3.72E-05 | 3.25E-05 | 3.37E-05 | 0 | 1.07E-05 | 2.46E-05 | 2.80E-05 | 0 | 4.91E-05 | 3.97E-05 | 3.46E-05 | 0 | 0 | 4.54E-05 |
| Iqcb1 | 2.30E-05 | 2.85E-05 | 0 | 0 | 6.49E-06 | 1.59E-05 | 1.68E-05 | 0 | 1.07E-05 | 1.55E-07 | 0 | 1.35E-05 | 1.12E-05 | 2.18E-05 | 2.83E-05 | 4.16E-06 | 3.16E-05 | 5.64E-06 |

| | | | | | | | | | | | | | | | | | | |
|---|---|---|---|---|---|---|---|---|---|---|---|---|---|---|---|---|---|---|
| Zc3h10 | 1.43E-05 | 5.67E-05 | 7.90E-06 | 2.65E-05 | 2.07E-05 | 1.12E-05 | 3.04E-06 | 1.34E-05 | 1.07E-05 | 1.08E-05 | 2.70E-05 | 1.57E-05 | 0 | 1.94E-05 | 3.67E-05 | 3.01E-05 | 0 | 3.50E-05 |
| Apoa2 | 2.77E-06 | 9.89E-07 | 1.35E-06 | 3.74E-06 | 1.09E-06 | 4.92E-06 | 0 | 9.05E-06 | 1.07E-05 | 2.85E-07 | 0 | 0 | 3.49E-07 | 1.35E-05 | 0 | 2.85E-06 | 0 | 1.52E-06 |
| Zfp449 | 3.92E-05 | 2.04E-05 | 0 | 6.65E-06 | 1.15E-05 | 3.20E-06 | 4.44E-05 | 0 | 1.07E-05 | 3.24E-05 | 1.36E-05 | 2.75E-06 | 0 | 0 | 1.91E-05 | 3.07E-05 | 0 | 2.43E-05 |
| Slc25a35 | 0 | 1.01E-05 | 0 | 0 | 6.62E-07 | 0 | 0 | 0 | 1.07E-05 | 2.20E-05 | 5.11E-06 | 0 | 3.09E-06 | 4.10E-06 | 0 | 0 | 0 | 9.63E-06 |
| Mtrr | 1.10E-05 | 1.14E-05 | 2.57E-05 | 5.91E-06 | 6.04E-06 | 0 | 1.77E-06 | 0 | 1.07E-05 | 5.04E-06 | 1.02E-05 | 6.02E-06 | 1.54E-06 | 0 | 2.51E-05 | 3.56E-05 | 2.21E-06 | 5.95E-06 |
| Golgb1 | 0.00024 | 0.00014 | 7.20E-05 | 8.58E-05 | 9.10E-05 | 0.00012 | 0.00028 | 4.60E-05 | 1.06E-05 | 5.84E-05 | 0 | 0.0002 | 3.73E-05 | 0 | 0.0005 | 0 | 4.02E-05 | 4.66E-05 |
| Rnf138 | 0.0001 | 8.72E-05 | 9.21E-05 | 5.21E-05 | 8.88E-05 | 5.82E-05 | 0 | 8.40E-05 | 1.06E-05 | 5.02E-05 | 3.64E-06 | 5.28E-05 | 4.40E-05 | 2.69E-05 | 7.88E-05 | 2.75E-05 | 6.82E-05 | 5.55E-05 |
| Fbxo44 | 0 | 3.75E-05 | 0 | 1.10E-05 | 3.72E-05 | 2.83E-05 | 0 | 0 | 1.06E-05 | 0 | 7.51E-06 | 3.15E-05 | 0 | 2.71E-05 | 3.97E-05 | 0 | 0 | 1.54E-06 |
| Wdfy1 | 0 | 0.00012 | 0 | 3.26E-05 | 4.73E-05 | 2.67E-05 | 0 | 0 | 1.06E-05 | 3.82E-05 | 0.00037 | 0.00014 | 0 | 0.00012 | 0.0005 | 0 | 0.00013 | 0 |
| Cep89 | 2.38E-05 | 4.76E-05 | 0 | 1.38E-05 | 9.54E-06 | 1.08E-05 | 0 | 2.41E-05 | 1.06E-05 | 2.34E-05 | 0 | 2.10E-05 | 0 | 0 | 4.93E-05 | 0.00022 | 0 | 1.41E-05 |
| Phkg2 | 2.49E-05 | 4.74E-05 | 1.14E-05 | 6.92E-05 | 1.24E-05 | 3.26E-06 | 9.61E-06 | 0 | 1.06E-05 | 2.25E-05 | 2.89E-05 | 5.78E-05 | 0 | 3.38E-06 | 4.20E-05 | 2.05E-05 | 3.00E-05 | 2.44E-05 |
| Bbs7 | 1.95E-05 | 8.79E-06 | 0 | 0 | 3.91E-06 | 0 | 8.34E-06 | 0 | 1.06E-05 | 1.73E-05 | 8.97E-07 | 0 | 1.13E-05 | 2.79E-06 | 2.77E-05 | 2.04E-05 | 0 | 6.26E-06 |
| Nfkbil1 | 0 | 3.32E-05 | 0 | 4.71E-05 | 9.14E-06 | 0 | 0 | 2.13E-05 | 1.06E-05 | 1.27E-05 | 6.82E-06 | 2.00E-05 | 0 | 0 | 5.13E-05 | 9.72E-06 | 0 | 5.38E-06 |
| Enpp4 | 1.10E-05 | 7.18E-06 | 0 | 0 | 9.56E-06 | 0 | 0 | 6.85E-06 | 1.06E-05 | 9.34E-06 | 0 | 7.55E-06 | 3.89E-05 | 1.79E-05 | 0 | 6.02E-06 | 0 | 1.33E-05 |
| Pcdhga8 | 9.36E-06 | 6.00E-05 | 0 | 0 | 4.85E-05 | 5.18E-05 | 4.33E-05 | 1.89E-05 | 1.05E-05 | 2.60E-05 | 4.25E-05 | 4.78E-06 | 3.54E-06 | 1.89E-05 | 0.0001 | 0 | 4.54E-05 | 6.98E-06 |
| Tmc7 | 0 | 0 | 2.22E-05 | 0 | 0 | 2.42E-05 | 0 | 4.96E-05 | 1.05E-05 | 4.18E-06 | 0 | 0 | 2.00E-05 | 1.66E-05 | 5.72E-06 | 0 | 9.75E-06 | 2.82E-05 |
| Eno3 | 1.15E-05 | 2.48E-05 | 0 | 2.12E-05 | 9.12E-06 | 1.23E-05 | 1.55E-05 | 0 | 1.05E-05 | 1.42E-05 | 1.14E-05 | 0 | 3.00E-06 | 0 | 1.55E-05 | 6.07E-06 | 7.22E-05 | 0 |
| Rassf4 | 0 | 0 | 0 | 1.22E-05 | 0 | 3.72E-06 | 0 | 0 | 1.05E-05 | 3.27E-06 | 4.43E-08 | 0 | 6.85E-06 | 1.63E-06 | 2.38E-05 | 8.92E-06 | 1.57E-05 | 1.33E-06 |
| Ebf3 | 1.13E-06 | 6.26E-06 | 0 | 8.23E-06 | 3.71E-06 | 0 | 3.24E-05 | 0 | 1.05E-05 | 0.00017 | 1.53E-06 | 0 | 0.00011 | 0 | 0 | 2.90E-05 | 1.13E-05 | 0 |
| Stk39 | 9.93E-05 | 1.82E-05 | 8.88E-05 | 0 | 9.83E-06 | 0 | 5.72E-05 | 6.14E-05 | 1.05E-05 | 7.98E-05 | 2.43E-05 | 7.76E-05 | 0 | 0 | 5.97E-05 | 2.98E-05 | 8.97E-05 | 0.00017 |
| Ssx2ip | 1.87E-05 | 6.05E-06 | 1.39E-05 | 0 | 2.09E-05 | 0 | 2.42E-05 | 1.63E-05 | 1.05E-05 | 3.42E-05 | 7.07E-05 | 5.10E-05 | 1.71E-05 | 0 | 0 | 7.16E-05 | 4.95E-07 | 0 |
| Zfp292 | 7.70E-05 | 0.00018 | 0 | 7.26E-05 | 9.76E-05 | 0.00011 | 0.00016 | 0 | 1.04E-05 | 6.35E-05 | 3.20E-05 | 0.00019 | 5.85E-05 | 7.37E-05 | 0.00022 | 0 | 0.00013 | 0 |
| Atg12 | 0.00012 | 9.31E-05 | 3.00E-05 | 3.21E-05 | 6.25E-05 | 5.68E-05 | 1.68E-05 | 4.07E-05 | 1.04E-05 | 1.05E-05 | 2.67E-06 | 0.00015 | 0 | 3.49E-05 | 5.16E-05 | 4.84E-05 | 9.66E-05 | 0 |
| Atg10 | 1.96E-05 | 2.90E-05 | 0 | 2.19E-05 | 6.13E-05 | 3.21E-05 | 2.60E-05 | 3.15E-05 | 1.04E-05 | 3.02E-05 | 3.44E-06 | 1.57E-05 | 5.69E-05 | 2.54E-05 | 1.10E-06 | 3.45E-05 | 0 | 3.50E-05 |
| Dctpp1 | 3.53E-05 | 0 | 5.30E-05 | 5.53E-05 | 7.20E-06 | 1.11E-05 | 1.08E-05 | 0 | 1.04E-05 | 2.07E-06 | 0 | 3.21E-06 | 6.62E-06 | 0 | 0 | 0.00027 | 2.71E-05 | 0 |
| Jrk | 1.05E-05 | 1.26E-05 | 9.68E-06 | 0 | 1.58E-06 | 0 | 0 | 0 | 1.04E-05 | 8.37E-07 | 3.19E-07 | 0 | 0 | 2.09E-06 | 4.08E-05 | 0 | 0 | 1.01E-05 |
| Foxm1 | 1.66E-05 | 4.36E-07 | 0 | 0 | 0 | 0 | 0 | 1.34E-06 | 1.04E-05 | 0 | 0 | 0 | 0 | 0 | 5.52E-05 | 0.00083 | 0 | 0 |
| Trp53rkb | 2.77E-07 | 3.00E-06 | 2.54E-06 | 8.01E-06 | 8.17E-06 | 0 | 0 | 1.86E-06 | 1.04E-05 | 2.80E-06 | 1.09E-06 | 0 | 0 | 0 | 0 | 7.26E-06 | 0 | 2.23E-06 |
| Anapc7 | 0.00016 | 0.00014 | 0.00013 | 3.78E-05 | 8.82E-05 | 8.59E-05 | 0.00011 | 0 | 1.03E-05 | 5.53E-06 | 4.44E-05 | 0.0001 | 2.82E-05 | 0.00011 | 0.0001 | 0.00013 | 0 | 5.92E-06 |
| Zfp148 | 0.00012 | 5.23E-05 | 1.93E-05 | 0 | 1.90E-05 | 6.91E-05 | 1.03E-05 | 8.00E-06 | 1.03E-05 | 1.96E-06 | 1.08E-05 | 3.59E-05 | 0 | 0 | 4.17E-05 | 4.38E-05 | 0.00011 | 2.07E-05 |
| Zfp821 | 2.60E-05 | 9.47E-05 | 4.48E-05 | 5.62E-05 | 5.18E-05 | 4.87E-05 | 5.59E-06 | 0 | 1.03E-05 | 1.39E-05 | 6.09E-05 | 8.23E-05 | 0 | 6.53E-05 | 0.00022 | 4.20E-05 | 9.32E-05 | 7.03E-05 |
| E4f1 | 3.82E-05 | 6.14E-05 | 1.04E-05 | 3.27E-05 | 6.76E-06 | 3.15E-05 | 1.67E-06 | 5.70E-06 | 1.03E-05 | 2.92E-05 | 1.17E-05 | 3.45E-05 | 0 | 0 | 7.82E-05 | 2.72E-05 | 5.28E-05 | 0 |
| Tefm | 5.80E-05 | 1.77E-05 | 2.18E-05 | 2.73E-05 | 1.21E-06 | 2.57E-05 | 0 | 1.74E-05 | 1.03E-05 | 5.44E-07 | 6.11E-06 | 0 | 0 | 1.27E-05 | 7.74E-07 | 2.59E-06 | 0 | 0 |

| | | | | | | | | | | | | | | | | | | |
|---|---|---|---|---|---|---|---|---|---|---|---|---|---|---|---|---|---|---|
| Fem1c | 1.67E-05 | 2.89E-05 | 1.03E-05 | 2.01E-05 | 3.06E-05 | 2.22E-05 | 1.89E-05 | 0 | 1.03E-05 | 8.68E-06 | 2.15E-05 | 1.36E-05 | 4.43E-06 | 2.11E-05 | 0 | 5.27E-05 | 4.34E-05 | 2.16E-05 |
| Znhit3 | 1.84E-05 | 1.74E-05 | 4.55E-05 | 5.74E-05 | 6.35E-06 | 1.28E-05 | 0 | 0 | 1.03E-05 | 2.95E-06 | 1.87E-05 | 2.31E-05 | 0 | 2.29E-05 | 1.93E-07 | 0.00012 | 0 | 0 |
| Cstf2 | 5.01E-05 | 2.83E-05 | 0.00012 | 8.07E-05 | 2.36E-05 | 0 | 0 | 1.31E-05 | 1.03E-05 | 0 | 2.09E-05 | 0.00014 | 0 | 4.53E-05 | 3.06E-05 | 0.00013 | 0 | 3.06E-05 |
| Bmyc | 7.51E-06 | 4.65E-05 | 0 | 7.00E-05 | 1.01E-05 | 0 | 0 | 0 | 1.03E-05 | 5.23E-05 | 0 | 6.57E-05 | 1.67E-05 | 9.59E-05 | 8.36E-06 | 1.57E-05 | 0 | 7.72E-05 |
| Prkch | 9.99E-06 | 7.46E-06 | 0 | 1.44E-06 | 1.68E-05 | 0 | 6.21E-05 | 0 | 1.03E-05 | 1.87E-05 | 0 | 7.40E-06 | 3.61E-05 | 6.02E-07 | 1.16E-05 | 2.82E-05 | 5.86E-06 | 0 |
| Krt18 | 0 | 3.60E-05 | 1.63E-05 | 5.13E-05 | 1.96E-05 | 0 | 0 | 2.24E-05 | 1.03E-05 | 4.46E-05 | 3.07E-05 | 4.73E-05 | 2.00E-05 | 0 | 0.00024 | 3.36E-05 | 2.26E-05 | 3.83E-05 |
| Epb41l5 | 0 | 2.30E-05 | 0 | 0 | 1.10E-05 | 0 | 0 | 0 | 1.03E-05 | 0 | 0 | 5.28E-06 | 0 | 6.08E-06 | 5.96E-06 | 0 | 9.55E-06 | 1.74E-06 |
| Aldoc | 0 | 0 | 0 | 0 | 0 | 0 | 0 | 0 | 1.03E-05 | 0 | 0 | 0.00013 | 0 | 4.36E-06 | 1.36E-05 | 0 | 0 | 1.81E-06 |
| Cyb5d2 | 5.64E-05 | 4.83E-05 | 0 | 2.92E-06 | 2.25E-05 | 3.18E-05 | 2.72E-05 | 3.79E-05 | 1.02E-05 | 3.52E-05 | 5.09E-06 | 1.51E-05 | 1.12E-05 | 2.92E-05 | 0 | 2.81E-05 | 0 | 9.14E-06 |
| Nptx2 | 0 | 0 | 0 | 0 | 0 | 2.70E-05 | 0 | 0 | 1.02E-05 | 0 | 0.00017 | 6.69E-05 | 0 | 0 | 0 | 3.08E-06 | 0 | 0 |
| Asxl3 | 0 | 5.52E-06 | 0 | 0 | 1.22E-05 | 1.31E-05 | 0 | 0 | 1.02E-05 | 0 | 1.18E-05 | 7.30E-06 | 0 | 0 | 6.24E-06 | 0 | 0 | 7.28E-06 |
| Coq6 | 3.48E-05 | 3.48E-05 | 2.46E-05 | 2.36E-05 | 1.04E-06 | 6.93E-06 | 6.02E-07 | 0 | 1.02E-05 | 6.54E-07 | 0 | 1.19E-05 | 0 | 7.26E-06 | 6.93E-06 | 4.80E-05 | 0 | 2.37E-06 |
| Cxcl3 | 0 | 0 | 1.35E-05 | 3.07E-06 | 0 | 5.23E-06 | 0 | 0 | 1.02E-05 | 0 | 4.93E-06 | 0 | 2.86E-05 | 8.53E-06 | 4.51E-06 | 3.20E-06 | 1.13E-06 | 0 |
| Cep41 | 1.94E-05 | 0 | 1.06E-05 | 0 | 1.43E-06 | 0 | 5.40E-06 | 1.09E-06 | 1.02E-05 | 0 | 4.97E-06 | 6.30E-07 | 0 | 0 | 2.91E-05 | 2.60E-05 | 0 | 0 |
| Mboat1 | 2.63E-06 | 0 | 0 | 0 | 0 | 0 | 2.38E-06 | 0 | 1.02E-05 | 2.11E-06 | 8.03E-06 | 0 | 1.32E-06 | 0 | 5.70E-06 | 8.14E-05 | 0 | 4.28E-05 |
| Zfp94 | 1.90E-06 | 1.13E-05 | 5.40E-07 | 0 | 2.99E-06 | 0 | 5.52E-06 | 2.68E-07 | 1.02E-05 | 3.15E-06 | 5.78E-06 | 8.59E-06 | 0 | 0 | 5.07E-06 | 2.59E-06 | 0 | 1.79E-06 |
| Timeless | 0 | 0 | 4.05E-05 | 0 | 0 | 0 | 0 | 0 | 1.02E-05 | 0 | 2.71E-06 | 0 | 4.81E-06 | 2.48E-06 | 0 | 0.00015 | 0 | 1.04E-05 |
| Pold3 | 3.32E-05 | 2.40E-05 | 2.42E-05 | 0 | 1.02E-05 | 3.43E-05 | 2.78E-06 | 0 | 1.01E-05 | 1.21E-05 | 1.25E-06 | 6.73E-06 | 8.53E-06 | 1.98E-05 | 7.82E-05 | 0.0002 | 2.74E-05 | 4.71E-06 |
| DEPRECATED | 0 | 2.42E-05 | 0 | 0 | 2.57E-05 | 2.24E-05 | 0 | 2.47E-05 | 1.01E-05 | 1.33E-05 | 0 | 0 | 0 | 1.35E-05 | 5.17E-05 | 0 | 0 | 2.23E-06 |
| Tbc1d31 | 1.37E-05 | 1.05E-05 | 2.86E-06 | 0 | 9.92E-06 | 1.86E-05 | 5.80E-06 | 1.48E-05 | 1.01E-05 | 1.31E-05 | 3.33E-06 | 1.21E-05 | 0 | 0 | 2.46E-05 | 0.00017 | 1.53E-05 | 9.39E-07 |
| Donson | 1.04E-05 | 2.06E-05 | 4.75E-05 | 0 | 1.83E-05 | 1.83E-05 | 1.08E-05 | 5.18E-06 | 1.01E-05 | 1.97E-05 | 2.56E-05 | 3.13E-05 | 4.54E-05 | 3.13E-05 | 6.98E-05 | 0.00031 | 0 | 3.09E-05 |
| Dhrs11 | 3.38E-05 | 4.48E-05 | 9.86E-06 | 2.16E-05 | 3.03E-05 | 5.68E-06 | 5.54E-06 | 2.55E-05 | 1.01E-05 | 1.32E-05 | 9.94E-06 | 3.60E-05 | 0 | 0 | 2.49E-05 | 0 | 1.26E-05 | 4.67E-05 |
| Zfp472 | 1.08E-05 | 1.51E-05 | 0 | 0 | 3.75E-06 | 5.64E-06 | 1.24E-05 | 8.26E-06 | 1.01E-05 | 7.05E-06 | 1.98E-05 | 0 | 0 | 8.41E-06 | 2.90E-05 | 1.10E-05 | 0 | 4.22E-06 |
| Garnl3 | 0 | 1.17E-05 | 1.30E-06 | 8.41E-06 | 1.98E-06 | 3.35E-06 | 1.87E-06 | 0 | 1.01E-05 | 6.14E-06 | 3.39E-06 | 2.17E-06 | 3.74E-05 | 0 | 2.68E-07 | 0 | 1.48E-05 | 0 |
| Lyrm9 | 6.63E-06 | 0 | 0 | 0 | 7.05E-06 | 1.21E-06 | 1.40E-05 | 0 | 1.01E-05 | 4.08E-06 | 0 | 6.42E-06 | 6.87E-06 | 0 | 0 | 7.07E-06 | 0 | 0 |
| Cgas | 8.85E-06 | 1.48E-06 | 3.69E-05 | 6.02E-06 | 0 | 7.18E-07 | 0 | 0 | 1.01E-05 | 1.50E-05 | 0 | 0 | 2.83E-05 | 8.97E-06 | 3.94E-05 | 0.00012 | 0 | 0 |
| Cdh6 | 0 | 1.64E-06 | 9.08E-07 | 0 | 0 | 0 | 0 | 0 | 1.01E-05 | 3.01E-06 | 0 | 2.82E-05 | 6.27E-06 | 2.75E-05 | 2.26E-05 | 1.25E-06 | 0 | 1.28E-06 |
| Ss18 | 0.00022 | 0.0002 | 0.00024 | 2.55E-05 | 0.00025 | 0.00012 | 0 | 0.00026 | 1.00E-05 | 4.39E-05 | 0.00013 | 7.67E-05 | 0 | 0.0001 | 0 | 0.00023 | 0.00031 | 5.27E-05 |
| Faxc | 9.72E-06 | 1.87E-05 | 4.51E-06 | 0 | 9.36E-06 | 2.71E-05 | 1.00E-05 | 0 | 1.00E-05 | 0 | 2.91E-05 | 0 | 4.79E-07 | 4.66E-05 | 0 | 2.09E-05 | 8.42E-06 | 1.69E-05 |
| Rcc1l | 5.65E-05 | 3.66E-05 | 0.00016 | 0.00011 | 4.46E-05 | 1.63E-05 | 2.77E-05 | 2.09E-05 | 1.00E-05 | 0 | 3.25E-05 | 0 | 9.24E-06 | 1.53E-05 | 4.29E-05 | 4.16E-05 | 4.66E-06 | 5.45E-05 |
| Rab42 | 1.57E-05 | 8.26E-06 | 1.39E-06 | 1.59E-05 | 0 | 2.15E-06 | 9.24E-07 | 4.29E-06 | 1.00E-05 | 4.94E-06 | 3.23E-06 | 0 | 0 | 2.14E-06 | 0 | 0 | 0 | 1.17E-06 |
| Larp1b | 3.99E-05 | 3.49E-05 | 5.77E-05 | 0 | 3.84E-06 | 0 | 1.67E-05 | 0 | 1.00E-05 | 0 | 9.50E-05 | 2.54E-05 | 0 | 2.10E-05 | 9.08E-05 | 2.23E-05 | 7.74E-05 | 1.45E-05 |
| Ifitm5 | 0 | 2.56E-05 | 1.61E-06 | 2.15E-05 | 0 | 0 | 0 | 0 | 1.00E-05 | 7.46E-06 | 0 | 0 | 0 | 1.61E-05 | 5.53E-05 | 2.03E-05 | 0 | 0 |

| | | | | | | | | | | | | | | | | | | |
|---|---|---|---|---|---|---|---|---|---|---|---|---|---|---|---|---|---|---|
| Pigf | 3.45E-05 | 1.02E-05 | 4.60E-06 | 1.09E-05 | 1.69E-05 | 2.44E-05 | 0 | 2.21E-05 | 9.99E-06 | 4.97E-06 | 6.59E-06 | 5.98E-06 | 0 | 1.04E-05 | 1.26E-05 | 5.33E-05 | 0 | 1.19E-05 |
| Zfp30 | 1.22E-05 | 2.56E-05 | 0 | 1.52E-06 | 1.55E-05 | 1.09E-05 | 2.28E-05 | 0 | 9.99E-06 | 6.56E-06 | 0 | 1.09E-06 | 3.70E-07 | 1.21E-05 | 9.02E-06 | 0 | 0 | 9.38E-06 |
| Trim34a | 1.48E-05 | 3.22E-05 | 0 | 1.13E-05 | 6.70E-06 | 0 | 0 | 2.00E-06 | 9.97E-06 | 7.57E-05 | 0 | 1.56E-05 | 5.08E-06 | 0 | 1.70E-05 | 0 | 0 | 7.00E-05 |
| Serpinb9 | 5.86E-05 | 5.90E-06 | 0 | 0 | 0.00013 | 5.58E-05 | 4.58E-05 | 6.10E-05 | 9.96E-06 | 6.60E-05 | 6.24E-05 | 0 | 2.65E-05 | 0 | 0 | 0 | 0 | 0.0003 |
| Hoxa3 | 0 | 3.22E-05 | 4.03E-05 | 3.45E-05 | 6.22E-05 | 0 | 0.00012 | 0.00019 | 9.90E-06 | 1.93E-05 | 0 | 3.01E-05 | 0 | 0 | 9.36E-05 | 3.57E-05 | 0 | 0 |
| Nr2c1 | 7.18E-05 | 7.70E-05 | 1.40E-05 | 2.47E-05 | 4.33E-05 | 4.49E-05 | 6.54E-05 | 3.25E-05 | 9.89E-06 | 2.53E-05 | 2.30E-05 | 3.87E-05 | 1.93E-05 | 4.30E-05 | 3.22E-05 | 2.75E-05 | 3.54E-05 | 3.54E-05 |
| Phtf2 | 2.28E-05 | 1.64E-05 | 2.09E-05 | 0 | 5.97E-06 | 2.59E-07 | 2.05E-06 | 6.56E-06 | 9.89E-06 | 1.11E-05 | 3.87E-05 | 0 | 0 | 2.21E-05 | 0 | 2.04E-05 | 3.62E-05 | 0 |
| Smcr8 | 8.56E-05 | 0.00018 | 9.13E-05 | 3.14E-05 | 6.80E-05 | 0.00012 | 1.32E-05 | 2.55E-05 | 9.88E-06 | 0 | 0 | 3.61E-05 | 0 | 4.60E-05 | 0.00017 | 4.53E-05 | 0.00068 | 1.62E-05 |
| Lcat | 7.42E-06 | 1.16E-05 | 3.48E-05 | 1.16E-05 | 4.01E-06 | 0 | 1.13E-05 | 3.46E-06 | 9.88E-06 | 3.85E-06 | 0 | 0 | 0 | 0 | 9.60E-05 | 5.57E-06 | 8.02E-05 | 2.23E-05 |
| C1rb | 4.90E-08 | 1.11E-05 | 0 | 0 | 1.28E-05 | 1.64E-05 | 0 | 0 | 9.87E-06 | 5.75E-05 | 0 | 5.34E-06 | 0.00014 | 7.76E-06 | 0 | 0 | 0 | 2.86E-05 |
| Pola2 | 0 | 1.99E-05 | 2.57E-05 | 2.48E-05 | 6.44E-06 | 9.93E-06 | 3.32E-05 | 0 | 9.87E-06 | 3.45E-05 | 0 | 2.21E-05 | 1.60E-05 | 2.22E-05 | 0 | 0.00033 | 0 | 4.71E-05 |
| Hoxb7 | 1.70E-05 | 9.10E-07 | 9.23E-06 | 2.36E-05 | 3.75E-05 | 4.66E-05 | 4.21E-05 | 2.53E-05 | 9.86E-06 | 6.23E-06 | 1.18E-05 | 0 | 9.59E-06 | 0 | 1.78E-05 | 2.44E-06 | 0 | 0 |
| Trmt2b | 4.92E-05 | 3.09E-05 | 0 | 0 | 4.11E-05 | 3.78E-05 | 3.28E-05 | 8.43E-06 | 9.84E-06 | 3.23E-05 | 2.52E-05 | 1.76E-05 | 0 | 2.67E-05 | 9.83E-06 | 8.39E-05 | 0 | 4.34E-05 |
| Dmac2 | 6.68E-05 | 6.44E-05 | 6.55E-06 | 3.08E-05 | 3.17E-05 | 2.70E-05 | 7.54E-05 | 6.34E-06 | 9.84E-06 | 0 | 5.19E-06 | 2.11E-05 | 7.22E-06 | 2.75E-05 | 5.19E-05 | 0 | 0 | 6.58E-06 |
| Mtmr10 | 3.79E-05 | 3.16E-05 | 1.53E-05 | 0 | 1.07E-05 | 2.45E-05 | 0 | 0 | 9.84E-06 | 1.32E-05 | 1.30E-05 | 9.24E-06 | 9.54E-06 | 1.56E-06 | 1.21E-05 | 0 | 8.03E-06 | 2.25E-05 |
| Tmem14a | 9.70E-06 | 1.37E-05 | 0 | 7.13E-06 | 1.73E-05 | 4.28E-06 | 0 | 3.63E-06 | 9.83E-06 | 2.69E-05 | 0 | 2.36E-05 | 1.23E-05 | 0 | 6.07E-06 | 4.00E-06 | 0 | 3.39E-05 |
| Wdr12 | 2.73E-05 | 1.03E-05 | 7.70E-05 | 5.97E-05 | 3.38E-06 | 0 | 0 | 0 | 9.83E-06 | 4.78E-06 | 0 | 0 | 0 | 0 | 9.69E-06 | 3.64E-05 | 0 | 2.93E-06 |
| Kcnt2 | 5.91E-06 | 5.45E-06 | 0 | 0 | 0 | 3.25E-06 | 5.61E-06 | 0 | 9.82E-06 | 3.79E-05 | 0 | 0 | 6.30E-06 | 0 | 1.96E-05 | 0 | 0 | 0.00011 |
| Nr1h3 | 0 | 7.75E-05 | 0 | 0 | 0 | 8.25E-05 | 0 | 9.13E-05 | 9.78E-06 | 2.26E-05 | 0 | 0 | 0 | 0 | 0 | 0 | 0 | 3.27E-05 |
| Plcl2 | 9.20E-06 | 6.01E-06 | 2.37E-07 | 2.03E-05 | 8.66E-06 | 0 | 1.49E-05 | 5.25E-06 | 9.78E-06 | 1.21E-05 | 0 | 0 | 7.10E-06 | 1.75E-05 | 2.47E-05 | 3.56E-05 | 1.54E-05 | 1.71E-05 |
| Shroom2 | 0 | 1.87E-05 | 0 | 0 | 1.06E-05 | 0 | 0 | 0 | 9.78E-06 | 0 | 2.90E-05 | 0 | 0 | 2.37E-06 | 0 | 1.35E-05 | 0 | 7.42E-06 |
| Hspbap1 | 2.00E-05 | 6.29E-05 | 4.82E-05 | 3.10E-05 | 1.41E-05 | 1.37E-05 | 0 | 0 | 9.76E-06 | 3.95E-05 | 2.73E-05 | 0 | 1.08E-05 | 3.03E-05 | 6.98E-05 | 2.38E-06 | 0 | 5.01E-05 |
| Ccdc159 | 4.39E-06 | 1.07E-05 | 0 | 1.60E-06 | 1.20E-06 | 3.95E-06 | 1.70E-05 | 1.79E-06 | 9.75E-06 | 8.32E-06 | 3.66E-06 | 8.62E-06 | 8.36E-06 | 0 | 4.13E-05 | 0 | 9.89E-06 | 1.40E-05 |
| Nell2 | 0 | 0 | 8.32E-06 | 0 | 0 | 0 | 1.16E-05 | 0 | 9.75E-06 | 0 | 2.40E-05 | 0 | 8.39E-06 | 1.09E-05 | 0 | 1.82E-05 | 0 | 0 |
| Senp8 | 1.29E-05 | 4.00E-06 | 0 | 0 | 0 | 0 | 1.27E-05 | 1.64E-06 | 9.73E-06 | 7.96E-06 | 5.75E-06 | 6.20E-06 | 0 | 3.03E-06 | 7.03E-06 | 1.22E-05 | 0 | 7.48E-06 |
| Zfp108 | 5.71E-06 | 8.35E-06 | 0 | 0 | 1.31E-05 | 9.92E-07 | 0 | 2.86E-07 | 9.72E-06 | 2.57E-06 | 1.99E-07 | 0 | 0 | 5.90E-07 | 0 | 0 | 0 | 6.85E-06 |
| Atf2 | 0.00021 | 0.00022 | 5.05E-05 | 0 | 8.41E-05 | 0.00018 | 0.0001 | 4.86E-05 | 9.71E-06 | 5.57E-05 | 3.63E-05 | 0.00016 | 0 | 6.85E-05 | 0.00012 | 0.00018 | 0.00028 | 3.42E-05 |
| Tprn | 2.20E-05 | 4.18E-05 | 4.80E-05 | 2.04E-05 | 2.83E-06 | 0 | 2.68E-05 | 0 | 9.71E-06 | 5.68E-06 | 0 | 0 | 2.58E-07 | 3.07E-05 | 6.21E-05 | 0 | 0 | 3.96E-05 |
| Chd9 | 0.0001 | 6.26E-05 | 0.00012 | 3.19E-05 | 0.00011 | 0.0001 | 0.00016 | 6.06E-05 | 9.69E-06 | 8.41E-05 | 5.07E-05 | 6.23E-05 | 0 | 0 | 6.38E-05 | 4.36E-05 | 0.00021 | 2.80E-05 |
| Zfp39 | 2.79E-05 | 2.19E-05 | 6.31E-06 | 0 | 2.88E-05 | 1.93E-05 | 2.03E-05 | 0 | 9.69E-06 | 1.72E-05 | 0 | 3.91E-06 | 0 | 1.93E-06 | 4.67E-05 | 2.03E-05 | 0 | 1.97E-05 |
| Dagla | 0 | 1.69E-05 | 4.20E-05 | 0 | 0 | 0 | 0 | 3.85E-05 | 9.69E-06 | 5.65E-05 | 0 | 2.18E-05 | 2.18E-06 | 2.74E-06 | 0 | 0 | 0 | 6.52E-05 |
| Klc3 | 0 | 7.72E-06 | 0 | 0 | 4.03E-05 | 1.89E-06 | 1.68E-05 | 0 | 9.68E-06 | 0 | 6.94E-05 | 6.37E-06 | 0 | 1.24E-05 | 2.04E-05 | 0 | 0 | 0 |
| Slc38a9 | 1.61E-05 | 6.54E-05 | 0 | 1.58E-05 | 2.07E-05 | 5.18E-05 | 1.76E-05 | 0 | 9.66E-06 | 3.46E-05 | 0 | 1.45E-05 | 9.56E-07 | 2.97E-05 | 6.42E-05 | 0 | 0 | 1.43E-05 |

| | | | | | | | | | | | | | | | | | | |
|---|---|---|---|---|---|---|---|---|---|---|---|---|---|---|---|---|---|---|
| Ap4s1 | 0.0001 | 3.23E-05 | 3.38E-05 | 3.02E-07 | 0 | 6.57E-05 | 6.92E-06 | 0 | 9.64E-06 | 1.77E-05 | 6.11E-06 | 0 | 6.76E-06 | 0 | 2.55E-05 | 5.40E-06 | 0 | 1.81E-05 |
| Zfp952 | 3.33E-05 | 2.53E-05 | 0 | 0 | 2.28E-05 | 2.14E-05 | 8.21E-06 | 0 | 9.64E-06 | 1.23E-05 | 0 | 0 | 0 | 0 | 6.66E-05 | 0 | 0 | 2.11E-05 |
| Shf | 0 | 5.25E-05 | 9.03E-05 | 1.27E-05 | 2.25E-05 | 2.08E-06 | 7.30E-05 | 5.88E-05 | 9.64E-06 | 2.96E-05 | 7.92E-05 | 3.20E-05 | 0 | 0 | 8.50E-05 | 1.11E-05 | 5.05E-05 | 9.57E-05 |
| Dmbt1 | 0 | 0 | 0 | 0 | 3.89E-07 | 0 | 0 | 7.69E-06 | 9.64E-06 | 8.23E-06 | 1.24E-05 | 8.88E-06 | 5.17E-06 | 0 | 3.27E-05 | 2.51E-06 | 0 | 7.09E-06 |
| Ramac | 0.00016 | 0.00011 | 9.93E-06 | 5.96E-07 | 5.72E-05 | 0.00013 | 0 | 2.46E-05 | 9.60E-06 | 2.00E-05 | 4.96E-05 | 5.13E-05 | 0 | 3.08E-05 | 1.12E-05 | 3.52E-05 | 0 | 3.46E-05 |
| Zc3h12b | 1.43E-05 | 8.12E-06 | 1.19E-07 | 0 | 2.34E-05 | 1.98E-06 | 5.95E-06 | 9.72E-06 | 9.59E-06 | 9.89E-06 | 2.88E-06 | 4.83E-06 | 1.21E-05 | 1.05E-06 | 0 | 0 | 0 | 1.42E-05 |
| Snx29 | 0 | 0.00019 | 0 | 1.56E-05 | 0.00012 | 0.00012 | 0.0001 | 9.57E-06 | 9.57E-06 | 0.00017 | 0 | 4.84E-05 | 8.87E-05 | 4.64E-05 | 0.00026 | 3.25E-06 | 0.00018 | 0.00011 |
| Tbc1d17 | 4.22E-05 | 0.00017 | 0 | 0.00014 | 5.78E-05 | 4.82E-05 | 6.66E-05 | 2.73E-05 | 9.57E-06 | 3.99E-05 | 0 | 5.81E-05 | 6.16E-06 | 2.14E-05 | 0.00024 | 0 | 0 | 6.67E-05 |
| Armc8 | 8.05E-05 | 7.29E-05 | 6.88E-05 | 0 | 4.30E-05 | 3.31E-05 | 5.91E-06 | 0 | 9.57E-06 | 2.18E-05 | 5.24E-06 | 2.47E-05 | 0 | 9.67E-06 | 3.67E-05 | 5.90E-05 | 0 | 2.28E-06 |
| Dhrs7b | 0.00011 | 4.67E-05 | 5.40E-05 | 3.63E-05 | 7.63E-06 | 3.21E-05 | 1.57E-05 | 2.02E-06 | 9.57E-06 | 4.01E-05 | 3.79E-05 | 1.50E-05 | 5.09E-06 | 2.40E-05 | 0 | 0.00017 | 6.38E-05 | 0 |
| Zfp760 | 1.53E-05 | 1.95E-05 | 0 | 0 | 1.05E-05 | 1.99E-05 | 0 | 0 | 9.56E-06 | 6.81E-06 | 0 | 0 | 0 | 0 | 1.91E-05 | 1.22E-05 | 0 | 0 |
| Lrrc27 | 3.03E-06 | 1.64E-06 | 0 | 3.77E-07 | 3.96E-06 | 5.02E-06 | 9.97E-06 | 4.38E-06 | 9.55E-06 | 4.33E-06 | 6.04E-06 | 4.04E-06 | 3.16E-06 | 0 | 0 | 0 | 0 | 4.30E-06 |
| Ctns | 1.95E-05 | 9.82E-05 | 0 | 0 | 7.38E-05 | 3.94E-05 | 4.59E-05 | 8.70E-06 | 9.51E-06 | 2.75E-05 | 0 | 3.06E-05 | 0 | 8.21E-05 | 0 | 0 | 0 | 3.53E-05 |
| Ptpn4 | 1.85E-05 | 3.07E-05 | 1.43E-05 | 0 | 1.72E-06 | 5.21E-06 | 0 | 0 | 9.49E-06 | 0 | 1.04E-05 | 1.73E-05 | 2.35E-06 | 0 | 7.56E-05 | 7.93E-06 | 8.70E-06 | 0 |
| Lrch1 | 3.06E-05 | 9.63E-05 | 0 | 0 | 0 | 0 | 2.25E-05 | 1.57E-05 | 9.48E-06 | 0.0001 | 0 | 2.74E-05 | 2.67E-05 | 1.29E-06 | 2.30E-05 | 4.24E-05 | 9.06E-05 | 6.92E-05 |
| L3mbtl2 | 2.80E-05 | 3.69E-05 | 3.07E-05 | 3.83E-06 | 5.35E-06 | 1.52E-05 | 0 | 0 | 9.47E-06 | 1.99E-05 | 1.65E-05 | 3.01E-06 | 3.68E-06 | 2.89E-06 | 1.80E-05 | 9.07E-05 | 0 | 1.05E-05 |
| Epha1 | 0 | 0 | 1.77E-05 | 1.70E-05 | 0 | 4.48E-06 | 0 | 1.35E-05 | 9.46E-06 | 9.66E-05 | 2.29E-05 | 0 | 0 | 1.20E-05 | 0 | 5.21E-06 | 0 | 0 |
| Rfxank | 4.69E-05 | 6.71E-05 | 0 | 4.92E-05 | 3.35E-05 | 1.10E-05 | 3.83E-05 | 2.38E-05 | 9.44E-06 | 1.13E-05 | 0 | 1.18E-05 | 1.83E-05 | 3.85E-05 | 7.90E-05 | 1.24E-05 | 0 | 1.80E-05 |
| Phka1 | 3.22E-05 | 6.80E-06 | 0 | 0 | 2.67E-05 | 0 | 8.69E-05 | 0 | 9.44E-06 | 5.24E-05 | 0 | 0 | 4.91E-05 | 2.90E-05 | 3.88E-05 | 2.67E-05 | 0 | 4.23E-05 |
| Dglucy | 6.47E-06 | 5.73E-05 | 0 | 1.36E-06 | 3.76E-05 | 7.78E-06 | 0 | 2.30E-05 | 9.43E-06 | 2.72E-05 | 5.35E-06 | 2.14E-07 | 2.42E-05 | 0 | 0 | 0 | 0 | 1.10E-05 |
| Fads3 | 0.00015 | 0.00011 | 0.00013 | 0.00028 | 0 | 3.86E-05 | 0.00025 | 9.47E-05 | 9.42E-06 | 0.00012 | 0.00031 | 0.0001 | 0.00021 | 0.0003 | 0 | 0.00012 | 0.00021 | 0.00019 |
| Sft2d2 | 0 | 5.19E-05 | 0 | 5.46E-06 | 2.99E-05 | 0 | 3.40E-06 | 0 | 9.41E-06 | 3.38E-05 | 5.80E-06 | 2.94E-05 | 0 | 0 | 0 | 0 | 1.82E-05 | 5.46E-05 |
| Tepsin | 0 | 4.77E-05 | 2.28E-05 | 4.51E-05 | 2.81E-05 | 1.83E-05 | 2.60E-05 | 9.79E-07 | 9.40E-06 | 5.46E-05 | 3.64E-05 | 1.24E-05 | 1.50E-05 | 4.90E-05 | 9.88E-05 | 1.95E-05 | 0 | 2.62E-05 |
| Otud1 | 4.41E-07 | 9.43E-05 | 0 | 2.21E-06 | 2.59E-05 | 2.05E-05 | 0 | 0 | 9.35E-06 | 8.40E-05 | 7.62E-05 | 6.53E-05 | 0 | 3.18E-05 | 0 | 3.86E-05 | 7.51E-05 | 6.07E-05 |
| Ubac1 | 6.07E-06 | 5.57E-06 | 4.14E-06 | 2.15E-05 | 3.48E-06 | 0 | 6.01E-06 | 0 | 9.34E-06 | 3.19E-06 | 5.20E-06 | 0 | 2.91E-08 | 6.88E-06 | 2.01E-05 | 1.42E-05 | 0 | 5.88E-06 |
| Pnpla7 | 1.74E-05 | 8.14E-05 | 0 | 0 | 2.36E-06 | 1.42E-05 | 1.90E-05 | 0 | 9.29E-06 | 2.66E-06 | 0 | 0 | 3.24E-05 | 3.99E-05 | 6.43E-05 | 0 | 1.52E-05 | 4.16E-05 |
| Oxld1 | 1.01E-05 | 1.18E-05 | 0 | 1.12E-05 | 1.74E-05 | 8.91E-06 | 2.36E-06 | 1.29E-05 | 9.29E-06 | 1.20E-05 | 8.48E-07 | 7.41E-06 | 0 | 9.64E-07 | 1.51E-05 | 2.49E-06 | 0 | 5.97E-06 |
| Map3k4 | 3.30E-05 | 3.04E-05 | 7.28E-06 | 1.73E-05 | 2.97E-05 | 1.71E-06 | 2.05E-05 | 0 | 9.27E-06 | 2.78E-05 | 1.51E-06 | 6.64E-05 | 0 | 1.48E-05 | 8.60E-05 | 3.89E-05 | 1.66E-05 | 3.68E-05 |
| Otub2 | 6.56E-06 | 1.45E-05 | 1.31E-05 | 0 | 6.25E-06 | 0 | 1.81E-06 | 0 | 9.27E-06 | 1.83E-05 | 1.49E-05 | 1.58E-05 | 0 | 1.59E-05 | 1.08E-06 | 7.74E-07 | 7.44E-06 | 6.37E-06 |
| Hexim2 | 8.67E-06 | 9.15E-06 | 3.68E-06 | 1.35E-05 | 7.22E-06 | 1.10E-05 | 0 | 2.26E-06 | 9.26E-06 | 4.12E-06 | 2.23E-06 | 3.10E-07 | 0 | 1.73E-05 | 8.69E-06 | 3.66E-06 | 0 | 0 |
| Sesn1 | 2.94E-05 | 5.98E-05 | 0 | 0 | 1.89E-05 | 2.26E-05 | 1.95E-05 | 1.12E-05 | 9.25E-06 | 5.02E-05 | 2.33E-06 | 0 | 0 | 2.36E-05 | 2.32E-05 | 3.53E-05 | 0 | 9.50E-05 |
| Mkrn1 | 5.63E-05 | 0.0001 | 3.36E-05 | 5.69E-05 | 2.93E-05 | 5.38E-05 | 4.16E-05 | 0 | 9.24E-06 | 1.23E-05 | 3.12E-05 | 4.77E-05 | 9.52E-06 | 2.09E-05 | 1.04E-05 | 1.28E-05 | 0 | 2.07E-05 |
| DEPRECATED | 2.72E-05 | 3.20E-05 | 1.37E-05 | 0 | 1.25E-05 | 2.83E-05 | 1.40E-05 | 0 | 9.22E-06 | 1.56E-05 | 0 | 1.20E-05 | 0 | 0 | 0 | 1.82E-05 | 0 | 0 |

| | | | | | | | | | | | | | | | | | | |
|---|---|---|---|---|---|---|---|---|---|---|---|---|---|---|---|---|---|---|
| Eif1ad | 0.00013 | 5.99E-05 | 0.00016 | 5.20E-05 | 4.96E-05 | 0.00011 | 0 | 4.30E-05 | 9.19E-06 | 0 | 8.09E-05 | 1.40E-05 | 1.87E-05 | 4.31E-05 | 0 | 0.0003 | 0 | 0 |
| Oscp1 | 1.29E-05 | 1.95E-05 | 0 | 1.71E-06 | 1.22E-05 | 4.07E-06 | 0 | 0 | 9.18E-06 | 1.42E-05 | 0 | 1.76E-05 | 1.30E-05 | 1.02E-05 | 5.72E-06 | 7.18E-05 | 1.26E-05 | 1.16E-05 |
| Cep152 | 0 | 6.97E-06 | 2.82E-06 | 2.96E-06 | 0 | 0 | 0 | 2.74E-06 | 9.17E-06 | 0 | 0 | 1.31E-05 | 3.20E-05 | 7.20E-06 | 5.98E-05 | 0.00012 | 5.16E-05 | 4.66E-06 |
| Tnfrsf22 | 0 | 1.67E-05 | 1.28E-05 | 1.56E-05 | 4.18E-06 | 0 | 0 | 0 | 9.16E-06 | 0 | 3.77E-06 | 1.68E-05 | 8.24E-06 | 0 | 8.38E-06 | 0 | 2.79E-05 | 0 |
| Prr3 | 1.35E-05 | 1.51E-05 | 9.90E-06 | 3.71E-05 | 6.37E-06 | 0 | 0 | 0 | 9.16E-06 | 4.01E-06 | 7.63E-06 | 5.19E-06 | 0 | 0 | 5.18E-05 | 0 | 1.34E-05 | 7.82E-06 |
| Tapt1 | 6.24E-05 | 8.96E-05 | 6.91E-05 | 2.37E-05 | 5.81E-05 | 7.17E-05 | 2.48E-05 | 8.22E-05 | 9.15E-06 | 3.20E-06 | 4.46E-05 | 2.97E-05 | 1.73E-05 | 2.55E-05 | 9.70E-05 | 7.11E-05 | 8.23E-05 | 1.98E-05 |
| Dmtf1 | 0.00013 | 9.49E-05 | 4.59E-05 | 0 | 4.29E-05 | 9.13E-05 | 1.13E-05 | 4.58E-05 | 9.13E-06 | 1.77E-05 | 0 | 8.38E-05 | 0 | 0 | 0.00029 | 1.81E-05 | 8.53E-05 | 2.52E-05 |
| Brf1 | 4.53E-05 | 0.00016 | 5.05E-05 | 5.78E-05 | 0.00012 | 3.10E-05 | 1.87E-06 | 1.77E-05 | 9.12E-06 | 5.88E-05 | 2.75E-06 | 6.98E-05 | 1.83E-05 | 5.42E-06 | 0.0004 | 4.51E-05 | 0.00014 | 0.00012 |
| Zfp189 | 2.89E-05 | 3.77E-05 | 0 | 8.46E-06 | 3.01E-05 | 2.97E-05 | 1.02E-05 | 0 | 9.12E-06 | 7.53E-06 | 0 | 1.52E-05 | 2.78E-06 | 0 | 6.13E-05 | 0 | 0 | 3.28E-05 |
| Cxxc1 | 6.97E-05 | 0.00018 | 0.00019 | 0.00021 | 0.0001 | 9.60E-06 | 6.12E-05 | 3.73E-05 | 9.12E-06 | 4.47E-05 | 6.64E-05 | 2.83E-05 | 9.68E-05 | 6.57E-05 | 0.00029 | 0.00016 | 8.40E-05 | 5.76E-05 |
| Msra | 3.33E-05 | 1.32E-05 | 2.85E-05 | 3.93E-05 | 6.48E-06 | 5.05E-06 | 7.09E-06 | 3.45E-05 | 9.12E-06 | 0 | 1.78E-05 | 3.20E-05 | 1.23E-05 | 3.13E-06 | 4.58E-05 | 3.79E-05 | 1.80E-05 | 4.22E-07 |
| Nol8 | 5.79E-05 | 2.84E-05 | 5.42E-05 | 0 | 0 | 4.27E-06 | 0 | 0 | 9.11E-06 | 7.54E-06 | 0 | 0 | 0 | 0 | 3.09E-05 | 7.61E-06 | 0 | 0 |
| Nudt15 | 1.20E-05 | 3.61E-06 | 7.98E-06 | 7.26E-06 | 0 | 0 | 0 | 0 | 9.11E-06 | 5.98E-06 | 1.59E-06 | 5.23E-06 | 3.51E-06 | 1.84E-06 | 3.76E-07 | 6.82E-06 | 0 | 4.36E-06 |
| Dclre1c | 1.85E-05 | 4.35E-05 | 4.89E-07 | 1.40E-06 | 2.97E-06 | 0 | 0 | 0 | 9.10E-06 | 1.08E-05 | 2.45E-06 | 1.19E-05 | 0 | 1.13E-05 | 2.01E-05 | 2.00E-05 | 2.91E-06 | 5.01E-06 |
| Cenpl | 2.97E-06 | 1.07E-05 | 0 | 0 | 0 | 1.68E-05 | 0 | 0 | 9.09E-06 | 2.42E-06 | 0 | 1.60E-05 | 8.61E-06 | 0 | 0 | 0.00035 | 0 | 1.04E-06 |
| Lhx6 | 0 | 0 | 0 | 3.87E-06 | 1.36E-06 | 0 | 0 | 3.80E-05 | 9.09E-06 | 0 | 4.49E-06 | 4.64E-07 | 0 | 0 | 5.54E-06 | 8.52E-06 | 0 | 1.47E-05 |
| Tgtp2 | 4.60E-06 | 4.60E-06 | 0 | 0 | 0 | 0 | 0 | 0 | 9.07E-06 | 2.15E-05 | 2.17E-06 | 0 | 1.17E-05 | 0 | 0 | 2.07E-06 | 0 | 2.87E-05 |
| Trim39 | 1.00E-05 | 7.22E-05 | 6.98E-05 | 9.09E-05 | 1.77E-05 | 4.65E-06 | 2.21E-05 | 1.02E-05 | 9.06E-06 | 7.32E-06 | 2.84E-05 | 0 | 0 | 9.78E-06 | 0.00013 | 4.57E-05 | 8.26E-05 | 1.38E-06 |
| Zfp46 | 4.87E-05 | 6.75E-05 | 2.48E-06 | 6.04E-05 | 6.16E-05 | 7.35E-05 | 5.40E-05 | 6.33E-05 | 9.05E-06 | 8.51E-05 | 5.15E-05 | 5.34E-05 | 3.41E-05 | 3.20E-05 | 0.00011 | 1.31E-05 | 0 | 0.00014 |
| Ddx52 | 0.00013 | 7.20E-05 | 3.88E-05 | 0 | 1.53E-05 | 3.82E-05 | 6.71E-06 | 8.59E-06 | 9.05E-06 | 0 | 3.86E-06 | 2.13E-05 | 0 | 7.71E-06 | 0.00015 | 0.00011 | 0 | 0 |
| Dgat1 | 2.85E-05 | 4.68E-05 | 4.29E-05 | 4.14E-05 | 1.47E-05 | 3.63E-05 | 0 | 5.52E-05 | 9.05E-06 | 1.54E-05 | 2.99E-05 | 2.58E-05 | 3.08E-05 | 2.38E-05 | 3.37E-05 | 0 | 2.33E-05 | 6.81E-06 |
| Flvcr2 | 0 | 4.45E-06 | 0 | 0 | 7.02E-06 | 1.77E-05 | 1.15E-05 | 0 | 9.05E-06 | 1.45E-05 | 0 | 4.58E-06 | 0 | 3.65E-05 | 0 | 0 | 2.63E-06 | 0 |
| Mocs3 | 2.17E-05 | 3.93E-05 | 5.60E-05 | 3.88E-05 | 3.89E-06 | 8.04E-06 | 0 | 1.30E-05 | 9.04E-06 | 3.95E-06 | 2.33E-05 | 3.69E-07 | 0 | 2.99E-05 | 2.44E-05 | 5.92E-06 | 0 | 5.64E-06 |
| Mid2 | 5.07E-05 | 2.94E-05 | 0 | 0 | 4.57E-05 | 5.38E-05 | 0 | 6.44E-05 | 9.03E-06 | 0 | 3.66E-05 | 3.33E-06 | 0 | 3.05E-05 | 0 | 0 | 0 | 0.00013 |
| Mib2 | 0 | 7.01E-05 | 2.78E-06 | 3.04E-05 | 5.32E-05 | 4.66E-05 | 1.17E-05 | 2.57E-05 | 9.03E-06 | 4.96E-05 | 0 | 3.21E-05 | 0 | 1.07E-05 | 8.82E-05 | 0 | 9.61E-06 | 3.15E-05 |
| Chic1 | 0 | 2.82E-05 | 0 | 1.98E-05 | 2.46E-05 | 4.38E-05 | 0 | 4.34E-05 | 9.03E-06 | 6.93E-06 | 0 | 2.08E-05 | 0 | 0 | 5.04E-05 | 0 | 4.22E-06 | 2.54E-05 |
| Il18bp | 7.48E-06 | 1.90E-05 | 1.55E-05 | 2.24E-06 | 0 | 0 | 3.82E-06 | 5.39E-06 | 9.03E-06 | 5.74E-05 | 0 | 0 | 8.37E-05 | 0 | 0.00013 | 6.38E-06 | 7.64E-06 | 5.25E-05 |
| Tdrd3 | 0.00018 | 6.84E-05 | 3.27E-05 | 0 | 9.51E-05 | 0.00013 | 0.00018 | 0 | 9.01E-06 | 7.86E-05 | 2.29E-05 | 0 | 2.86E-05 | 0 | 0.00012 | 4.81E-05 | 0.00013 | 9.99E-05 |
| Chml | 2.84E-05 | 3.25E-05 | 1.53E-05 | 0 | 0 | 3.18E-07 | 0 | 7.95E-06 | 9.01E-06 | 5.58E-06 | 3.09E-06 | 4.24E-05 | 0 | 1.43E-05 | 3.26E-05 | 3.70E-05 | 1.47E-05 | 3.91E-06 |
| Slitrk2 | 0 | 0 | 0 | 0 | 0 | 3.05E-05 | 0 | 2.40E-06 | 9.00E-06 | 0 | 0 | 0 | 1.16E-05 | 0 | 0 | 7.95E-06 | 0 | 0 |
| Dcaf6 | 0.00012 | 0.00012 | 0 | 0 | 9.22E-05 | 0.00011 | 0.00019 | 5.49E-07 | 8.99E-06 | 6.37E-05 | 4.92E-05 | 8.11E-05 | 6.18E-05 | 9.35E-05 | 0.00029 | 0 | 0.00012 | 5.22E-05 |
| Zkscan7 | 7.46E-06 | 9.45E-06 | 0 | 0 | 1.77E-05 | 6.10E-06 | 3.93E-07 | 0 | 8.98E-06 | 1.17E-05 | 0 | 2.50E-06 | 8.91E-07 | 0 | 2.21E-05 | 0 | 0 | 1.52E-05 |
| Zfp729b | 1.86E-05 | 1.14E-05 | 0 | 0 | 3.82E-07 | 3.14E-05 | 9.28E-07 | 0 | 8.97E-06 | 1.07E-05 | 0 | 1.56E-05 | 0 | 0 | 1.41E-05 | 2.03E-07 | 0 | 9.77E-06 |

| | | | | | | | | | | | | | | | | | | |
|---|---|---|---|---|---|---|---|---|---|---|---|---|---|---|---|---|---|---|
| Stt3a | 3.27E-05 | 0 | 3.68E-05 | 2.14E-05 | 0 | 2.79E-06 | 2.76E-05 | 6.96E-06 | 8.97E-06 | 2.02E-06 | 1.66E-05 | 2.90E-06 | 9.19E-06 | 0 | 8.55E-06 | 1.73E-05 | 6.64E-06 | 0 |
| Nfib | 0.00014 | 0 | 0 | 0 | 0 | 0 | 8.24E-05 | 2.05E-05 | 8.97E-06 | 0.00029 | 0 | 0 | 0.00032 | 0 | 3.90E-05 | 0.00016 | 0 | 0.00067 |
| Spata33 | 2.98E-06 | 4.74E-06 | 0 | 6.79E-06 | 0 | 0 | 1.71E-06 | 0 | 8.97E-06 | 1.35E-06 | 0 | 0 | 3.46E-06 | 3.15E-06 | 5.39E-06 | 0 | 7.77E-06 | 8.10E-07 |
| Phactr1 | 4.22E-06 | 5.06E-06 | 0 | 4.85E-07 | 2.22E-06 | 2.13E-05 | 0 | 0 | 8.96E-06 | 9.47E-06 | 2.30E-05 | 3.65E-05 | 9.17E-06 | 0 | 0 | 6.54E-07 | 0 | 4.55E-05 |
| Zfp248 | 1.47E-05 | 1.47E-05 | 0 | 1.71E-07 | 0 | 1.10E-05 | 0 | 2.01E-06 | 8.96E-06 | 5.37E-06 | 0 | 4.16E-06 | 0 | 7.12E-06 | 1.76E-05 | 0 | 0 | 9.32E-06 |
| Ttll1 | 2.52E-05 | 2.77E-05 | 0 | 0 | 2.60E-05 | 1.10E-05 | 5.11E-06 | 1.03E-05 | 8.96E-06 | 3.77E-05 | 3.79E-06 | 4.22E-05 | 0 | 0 | 0 | 6.12E-06 | 0 | 4.49E-05 |
| Braf | 6.64E-05 | 6.46E-05 | 2.01E-05 | 0 | 2.81E-05 | 7.46E-05 | 0 | 0 | 8.94E-06 | 7.08E-06 | 4.18E-05 | 3.60E-05 | 2.68E-06 | 1.34E-05 | 7.14E-05 | 3.69E-05 | 0.00011 | 1.85E-05 |
| Rnf113a1 | 8.49E-06 | 1.93E-06 | 1.07E-05 | 1.12E-05 | 2.00E-08 | 1.92E-05 | 0 | 0 | 8.94E-06 | 1.18E-06 | 0 | 1.79E-05 | 5.81E-08 | 9.99E-06 | 5.85E-06 | 0 | 0 | 7.00E-06 |
| Slc25a15 | 1.75E-05 | 6.90E-06 | 4.52E-05 | 5.30E-05 | 0 | 9.91E-06 | 0 | 0 | 8.94E-06 | 2.65E-05 | 4.84E-06 | 0 | 0 | 1.15E-05 | 0 | 1.54E-05 | 2.38E-05 | 4.20E-05 |
| Tst | 0 | 1.01E-05 | 5.00E-06 | 3.42E-06 | 0 | 1.57E-06 | 0 | 0 | 8.94E-06 | 1.01E-05 | 0 | 0 | 2.89E-06 | 3.03E-06 | 0 | 4.82E-06 | 0 | 1.59E-05 |
| Lrrc57 | 4.22E-05 | 4.53E-05 | 2.21E-05 | 2.01E-05 | 1.61E-05 | 3.66E-05 | 0 | 5.13E-06 | 8.93E-06 | 2.56E-05 | 2.46E-05 | 2.32E-06 | 0 | 2.67E-05 | 3.56E-05 | 5.69E-05 | 8.54E-06 | 2.23E-05 |
| C2cd5 | 4.30E-05 | 2.51E-05 | 1.07E-05 | 9.79E-06 | 0 | 2.83E-05 | 6.60E-05 | 0 | 8.92E-06 | 5.05E-05 | 2.07E-05 | 2.00E-05 | 3.05E-06 | 0 | 5.33E-05 | 0.0001 | 4.68E-05 | 3.72E-05 |
| Zbtb7c | 0 | 1.41E-07 | 1.37E-05 | 0 | 0 | 0 | 0 | 3.75E-05 | 8.92E-06 | 9.18E-06 | 3.56E-06 | 0 | 0 | 0 | 0 | 3.99E-06 | 4.27E-05 | 0.00019 |
| Kantr | 6.85E-06 | 9.37E-06 | 0 | 0 | 8.30E-06 | 1.34E-06 | 5.13E-07 | 0 | 8.91E-06 | 7.36E-06 | 0 | 4.37E-06 | 0 | 4.04E-06 | 1.80E-07 | 0 | 0 | 2.46E-06 |
| Zgpat | 5.21E-05 | 0.00017 | 5.68E-05 | 5.20E-05 | 9.19E-05 | 8.55E-05 | 3.50E-05 | 3.74E-05 | 8.86E-06 | 6.46E-05 | 6.59E-05 | 1.75E-05 | 0 | 4.91E-05 | 0.00025 | 6.45E-06 | 3.88E-05 | 5.85E-05 |
| Best1 | 4.71E-06 | 1.55E-05 | 0 | 0 | 2.71E-05 | 0 | 3.55E-05 | 0 | 8.86E-06 | 8.33E-05 | 0 | 0 | 7.65E-05 | 0 | 0 | 0 | 0 | 3.47E-05 |
| Kdm8 | 1.83E-05 | 1.02E-05 | 1.86E-05 | 1.26E-05 | 6.73E-07 | 0 | 0 | 0 | 8.86E-06 | 9.23E-06 | 0 | 0 | 4.16E-06 | 0 | 4.35E-06 | 6.42E-05 | 2.24E-06 | 1.01E-05 |
| Zfp867 | 1.50E-05 | 1.14E-05 | 0 | 0 | 1.50E-06 | 8.34E-06 | 2.88E-06 | 0 | 8.82E-06 | 1.99E-06 | 4.20E-06 | 1.41E-05 | 0 | 0 | 6.73E-08 | 3.46E-06 | 0 | 8.10E-06 |
| Irf5 | 0 | 1.13E-05 | 1.52E-05 | 7.88E-07 | 1.86E-05 | 3.97E-06 | 0 | 0 | 8.81E-06 | 5.93E-06 | 2.89E-05 | 1.74E-05 | 0 | 0 | 3.84E-05 | 0 | 4.48E-06 | 2.91E-05 |
| Sgsm1 | 1.90E-06 | 0 | 6.52E-06 | 1.10E-05 | 7.88E-06 | 0 | 1.15E-05 | 4.83E-06 | 8.81E-06 | 3.78E-06 | 0 | 2.22E-06 | 2.01E-05 | 0 | 0 | 2.05E-06 | 0 | 0 |
| B3galt4 | 8.75E-06 | 1.99E-05 | 0 | 1.25E-06 | 7.36E-06 | 4.71E-06 | 1.82E-05 | 2.42E-06 | 8.78E-06 | 1.41E-05 | 0 | 3.69E-06 | 0 | 1.80E-05 | 1.48E-05 | 0 | 0 | 1.26E-05 |
| Fgfrl1 | 2.53E-05 | 7.62E-05 | 4.82E-05 | 0 | 7.83E-05 | 5.06E-05 | 0 | 4.31E-05 | 8.76E-06 | 0 | 0.00014 | 4.30E-05 | 6.33E-05 | 0.00012 | 0 | 6.18E-05 | 4.54E-05 | 7.22E-05 |
| Adam22 | 3.91E-07 | 3.19E-06 | 0 | 0 | 2.52E-06 | 0 | 3.10E-06 | 0 | 8.74E-06 | 7.82E-05 | 0 | 7.85E-06 | 6.37E-06 | 0 | 0 | 0 | 0 | 0 |
| Pnck | 0 | 0 | 0 | 0 | 4.20E-05 | 0 | 0 | 0 | 8.73E-06 | 1.20E-06 | 2.45E-05 | 2.67E-05 | 0 | 0 | 5.85E-06 | 0 | 0 | 0 |
| Trmt10c | 7.83E-05 | 2.86E-05 | 4.90E-05 | 0 | 2.80E-06 | 3.97E-05 | 0 | 0 | 8.72E-06 | 0 | 0 | 2.16E-05 | 0 | 6.42E-06 | 3.25E-05 | 3.23E-05 | 0 | 0 |
| Hoxa1 | 4.75E-06 | 9.81E-07 | 4.72E-06 | 6.45E-06 | 2.52E-06 | 5.65E-06 | 5.74E-06 | 5.98E-06 | 8.72E-06 | 6.66E-06 | 1.06E-05 | 9.75E-06 | 0 | 1.28E-06 | 1.52E-05 | 0 | 0 | 9.83E-06 |
| Tceal1 | 2.69E-05 | 2.43E-05 | 0 | 2.10E-05 | 3.41E-05 | 3.98E-05 | 4.15E-06 | 0 | 8.71E-06 | 1.04E-05 | 5.40E-06 | 0 | 1.40E-05 | 3.17E-06 | 0 | 0 | 3.12E-06 | 1.58E-05 |
| Ppp1r3c | 0 | 0 | 0 | 0 | 0 | 1.69E-05 | 0 | 0 | 8.71E-06 | 9.71E-06 | 0 | 7.14E-05 | 2.20E-05 | 0 | 0 | 0 | 0 | 1.77E-05 |
| Entpd1 | 0 | 4.78E-05 | 0 | 0 | 6.87E-05 | 0 | 0 | 2.07E-05 | 8.71E-06 | 9.89E-05 | 0 | 0 | 0 | 5.40E-05 | 1.40E-05 | 7.59E-07 | 0 | 0 |
| Pih1d2 | 3.78E-06 | 4.37E-06 | 5.21E-06 | 0 | 0 | 0 | 5.30E-06 | 5.47E-06 | 8.69E-06 | 8.84E-06 | 2.23E-06 | 2.21E-06 | 0 | 0 | 0 | 2.57E-06 | 3.97E-06 | 5.63E-06 |
| Git2 | 0.00018 | 0.00017 | 6.36E-05 | 0 | 0.0001 | 7.13E-05 | 6.72E-05 | 8.74E-05 | 8.67E-06 | 6.67E-05 | 7.90E-05 | 0.00014 | 0 | 3.75E-05 | 0.00017 | 0.00014 | 7.07E-05 | 8.70E-05 |
| Arhgef3 | 1.40E-05 | 4.18E-05 | 0 | 0 | 0 | 0 | 0 | 0 | 8.67E-06 | 0.00011 | 0 | 2.34E-05 | 0 | 0 | 1.19E-05 | 1.43E-05 | 3.58E-05 | 3.52E-05 |
| Zfp647 | 1.42E-05 | 3.18E-05 | 1.15E-05 | 3.62E-05 | 2.58E-05 | 0 | 0 | 0 | 8.67E-06 | 1.59E-05 | 1.21E-05 | 4.40E-06 | 0 | 1.40E-05 | 1.30E-05 | 7.65E-06 | 0 | 5.64E-06 |

| | | | | | | | | | | | | | | | | | | |
|---|---|---|---|---|---|---|---|---|---|---|---|---|---|---|---|---|---|---|
| Rnaset2a | 0.00014 | 0 | 0 | 0 | 3.18E-05 | 7.83E-05 | 4.68E-05 | 4.71E-05 | 8.66E-06 | 8.17E-08 | 0 | 0 | 9.24E-05 | 7.77E-05 | 0 | 6.70E-06 | 0 | 5.23E-05 |
| Lrrc3 | 0 | 2.36E-06 | 0 | 5.19E-06 | 0 | 0 | 2.50E-06 | 0 | 8.66E-06 | 0 | 2.04E-06 | 1.18E-05 | 2.73E-05 | 1.47E-05 | 0 | 1.02E-05 | 4.60E-06 | 8.96E-06 |
| Zfp426 | 5.13E-05 | 3.06E-05 | 2.99E-06 | 1.53E-05 | 3.63E-05 | 4.38E-05 | 2.98E-05 | 1.56E-05 | 8.63E-06 | 2.79E-05 | 9.76E-06 | 1.31E-05 | 0 | 2.16E-05 | 4.10E-06 | 2.34E-05 | 0 | 3.78E-05 |
| Zfp28 | 1.57E-06 | 6.40E-06 | 0 | 0 | 0 | 6.13E-06 | 0 | 7.39E-07 | 8.63E-06 | 0 | 1.71E-06 | 1.69E-05 | 4.81E-06 | 1.02E-06 | 1.95E-05 | 1.98E-06 | 0 | 1.26E-05 |
| Zfp961 | 1.31E-05 | 2.37E-06 | 4.82E-06 | 1.07E-05 | 1.51E-05 | 5.94E-06 | 0 | 5.83E-07 | 8.61E-06 | 6.74E-06 | 6.68E-06 | 0 | 0 | 1.43E-05 | 1.09E-06 | 1.53E-05 | 0 | 1.12E-05 |
| Osr1 | 4.03E-05 | 0 | 0 | 1.53E-05 | 0 | 0 | 0 | 0 | 8.60E-06 | 0.00058 | 0 | 0 | 0.00015 | 6.32E-05 | 6.41E-06 | 2.45E-05 | 0 | 0.00034 |
| Usp35 | 6.27E-06 | 4.74E-05 | 0 | 1.54E-05 | 2.49E-06 | 0 | 1.56E-05 | 1.44E-06 | 8.60E-06 | 1.53E-05 | 0 | 3.16E-05 | 0 | 3.15E-05 | 8.46E-05 | 0 | 9.87E-06 | 2.36E-05 |
| Maml3 | 3.94E-07 | 3.87E-05 | 0 | 0 | 1.84E-05 | 7.15E-05 | 0 | 3.59E-06 | 8.57E-06 | 1.17E-05 | 0 | 6.16E-05 | 0 | 1.24E-05 | 0 | 0 | 3.30E-05 | 2.57E-05 |
| Invs | 4.64E-05 | 3.84E-05 | 0 | 0 | 1.63E-05 | 8.13E-06 | 3.61E-05 | 7.99E-06 | 8.57E-06 | 9.69E-06 | 1.04E-05 | 1.89E-05 | 8.28E-06 | 0 | 2.87E-05 | 2.87E-05 | 0 | 3.59E-05 |
| Uchl3 | 6.89E-05 | 0 | 8.35E-05 | 4.98E-05 | 0 | 0 | 8.12E-06 | 0 | 8.55E-06 | 4.73E-05 | 0 | 0 | 1.81E-05 | 0 | 0 | 7.32E-05 | 2.41E-06 | 3.87E-05 |
| Rab2b | 2.34E-05 | 2.61E-05 | 1.47E-05 | 0 | 3.26E-05 | 2.61E-05 | 4.83E-05 | 1.84E-05 | 8.54E-06 | 2.20E-05 | 2.04E-06 | 7.81E-06 | 3.47E-05 | 3.03E-06 | 0 | 0 | 0 | 2.08E-05 |
| Pars2 | 9.89E-06 | 7.81E-06 | 0 | 3.70E-06 | 9.63E-06 | 3.07E-06 | 4.57E-06 | 0 | 8.54E-06 | 0 | 8.28E-06 | 0 | 0 | 6.62E-06 | 2.32E-05 | 2.20E-05 | 0 | 6.65E-06 |
| Iqck | 0 | 5.93E-06 | 0 | 2.94E-06 | 0 | 0 | 0 | 0 | 8.52E-06 | 8.00E-06 | 1.67E-06 | 0 | 4.76E-06 | 6.16E-07 | 4.95E-06 | 1.07E-05 | 1.70E-05 | 1.29E-06 |
| Ccsap | 0 | 0 | 1.76E-05 | 0 | 2.15E-05 | 4.32E-05 | 0 | 0 | 8.50E-06 | 1.08E-05 | 6.20E-05 | 2.43E-05 | 0 | 3.51E-05 | 0.00014 | 0.00027 | 0 | 0 |
| Snap29 | 0.0001 | 7.10E-05 | 4.74E-05 | 9.71E-05 | 2.79E-05 | 3.77E-05 | 0 | 4.22E-05 | 8.50E-06 | 3.98E-05 | 8.03E-07 | 5.20E-05 | 1.17E-05 | 7.21E-06 | 0 | 2.02E-05 | 1.30E-05 | 2.90E-05 |
| Cyp4f13 | 5.41E-06 | 3.76E-05 | 0 | 2.29E-06 | 2.57E-05 | 1.18E-05 | 3.83E-06 | 7.86E-06 | 8.50E-06 | 2.48E-05 | 1.98E-05 | 3.40E-05 | 3.09E-05 | 2.22E-05 | 2.64E-05 | 0 | 0 | 9.99E-06 |
| Slc2a9 | 0 | 1.89E-05 | 0 | 7.39E-07 | 0 | 2.51E-05 | 0 | 8.64E-06 | 8.49E-06 | 1.06E-05 | 0 | 0 | 0 | 3.57E-06 | 0 | 0 | 0 | 7.10E-06 |
| Pank4 | 4.10E-05 | 4.54E-05 | 4.70E-05 | 7.49E-06 | 4.84E-05 | 6.07E-05 | 0 | 2.22E-05 | 8.47E-06 | 2.22E-05 | 3.42E-05 | 2.20E-05 | 0 | 1.06E-05 | 8.59E-05 | 3.91E-05 | 4.38E-06 | 6.42E-06 |
| Cyb561d1 | 0 | 3.68E-05 | 1.53E-05 | 1.01E-05 | 5.61E-07 | 1.14E-05 | 0 | 0 | 8.47E-06 | 1.42E-05 | 1.06E-05 | 7.23E-06 | 0 | 2.34E-05 | 2.86E-05 | 8.61E-06 | 0 | 2.21E-05 |
| Cep295nl | 0 | 1.93E-05 | 0 | 0 | 2.93E-06 | 0 | 0 | 0 | 8.47E-06 | 2.88E-05 | 1.06E-05 | 0 | 0 | 4.27E-07 | 0.00012 | 0 | 4.24E-05 | 2.32E-05 |
| Gbp3 | 5.09E-05 | 1.37E-05 | 0 | 0 | 0 | 0 | 0 | 0 | 8.46E-06 | 0.00019 | 0 | 0 | 0 | 1.19E-05 | 0 | 2.92E-05 | 0 | 0.00017 |
| Castor2 | 4.80E-05 | 8.45E-05 | 0 | 4.70E-05 | 0.00019 | 9.69E-05 | 0.0001 | 4.23E-05 | 8.44E-06 | 8.28E-05 | 0.00015 | 0 | 6.15E-05 | 2.79E-05 | 0 | 1.47E-05 | 0 | 0.00021 |
| Cog3 | 0.00018 | 0.00012 | 7.15E-05 | 0 | 5.28E-05 | 8.01E-05 | 7.83E-05 | 0 | 8.44E-06 | 7.43E-06 | 0 | 0.00017 | 0 | 7.90E-06 | 0.00011 | 9.32E-06 | 6.58E-05 | 1.23E-05 |
| Agap1 | 8.74E-06 | 0.00013 | 2.87E-05 | 0 | 0.00012 | 4.78E-05 | 0 | 0.00011 | 8.42E-06 | 0.0002 | 0.00014 | 0.0002 | 0 | 0.00013 | 0.00017 | 0.00011 | 0.0001 | 1.45E-05 |
| Miga1 | 0 | 2.40E-05 | 0 | 3.87E-07 | 7.31E-06 | 0 | 0 | 0 | 8.39E-06 | 0 | 2.23E-05 | 0 | 7.71E-06 | 2.31E-05 | 8.76E-06 | 8.67E-06 | 0 | 3.32E-06 |
| Trim14 | 4.43E-08 | 1.40E-05 | 0 | 1.36E-05 | 0 | 1.29E-05 | 0 | 0 | 8.38E-06 | 9.04E-05 | 0 | 1.39E-05 | 4.18E-05 | 4.92E-06 | 1.14E-05 | 0 | 0 | 3.40E-05 |
| Ift88 | 3.62E-05 | 3.73E-05 | 0 | 4.63E-06 | 6.66E-06 | 1.27E-05 | 1.99E-06 | 0 | 8.38E-06 | 2.17E-05 | 0 | 1.08E-05 | 1.66E-06 | 9.69E-06 | 4.77E-05 | 3.92E-05 | 8.95E-06 | 2.60E-05 |
| Zfp418 | 0 | 5.29E-06 | 0 | 0 | 2.77E-06 | 4.94E-06 | 2.15E-06 | 0 | 8.38E-06 | 8.15E-06 | 1.07E-06 | 0 | 0 | 1.58E-06 | 3.83E-06 | 6.20E-06 | 0 | 6.65E-06 |
| Plek | 0 | 4.11E-05 | 1.95E-05 | 2.50E-05 | 3.67E-06 | 0 | 0 | 2.42E-06 | 8.36E-06 | 2.51E-05 | 0 | 9.08E-05 | 7.43E-06 | 4.24E-05 | 0.00011 | 2.61E-05 | 5.94E-05 | 5.06E-05 |
| Phyhd1 | 0 | 7.79E-05 | 0 | 7.47E-06 | 3.40E-05 | 8.25E-06 | 0 | 0 | 8.35E-06 | 3.19E-05 | 0 | 6.74E-05 | 0.00011 | 0 | 5.81E-05 | 0 | 8.99E-05 | 8.71E-05 |
| Msln | 2.78E-06 | 1.62E-05 | 1.11E-06 | 1.55E-05 | 4.32E-06 | 0 | 8.54E-06 | 0 | 8.34E-06 | 9.03E-06 | 0 | 2.81E-05 | 1.25E-06 | 7.80E-06 | 0.0001 | 8.77E-06 | 8.01E-06 | 2.91E-05 |
| Gstt2 | 7.34E-06 | 3.29E-05 | 0 | 7.55E-06 | 9.54E-06 | 0 | 0 | 0 | 8.33E-06 | 1.64E-05 | 3.30E-07 | 0 | 0 | 3.51E-06 | 8.77E-06 | 2.18E-06 | 8.18E-06 | 2.31E-05 |
| Gde1 | 0.00015 | 0.00017 | 0.00012 | 0.00019 | 0.00017 | 0.00015 | 6.45E-05 | 0.00024 | 8.31E-06 | 9.19E-05 | 5.11E-05 | 0.0002 | 5.25E-05 | 1.33E-05 | 0 | 9.61E-05 | 1.31E-05 | 4.91E-05 |

| | | | | | | | | | | | | | | | | | | |
|---|---|---|---|---|---|---|---|---|---|---|---|---|---|---|---|---|---|---|
| Tslp | 5.22E-06 | 0 | 4.54E-06 | 0 | 0 | 9.55E-07 | 0 | 2.81E-06 | 8.31E-06 | 2.48E-05 | 0 | 2.44E-06 | 2.84E-06 | 0 | 0 | 6.93E-06 | 4.93E-07 | 7.19E-06 |
| Cep44 | 3.86E-05 | 4.60E-05 | 0 | 4.87E-06 | 2.26E-05 | 2.71E-05 | 0 | 1.79E-05 | 8.30E-06 | 1.03E-05 | 1.06E-05 | 1.51E-05 | 0 | 0 | 5.48E-05 | 0.00011 | 0 | 0 |
| Adh7 | 2.71E-05 | 0 | 0 | 0 | 0 | 3.12E-08 | 0 | 4.31E-07 | 8.29E-06 | 9.44E-05 | 0 | 0 | 0 | 1.86E-06 | 0 | 1.54E-05 | 0 | 4.86E-05 |
| Dhrs13 | 5.04E-06 | 2.12E-06 | 3.11E-05 | 3.30E-05 | 0 | 0 | 1.92E-05 | 5.32E-06 | 8.27E-06 | 1.68E-05 | 2.55E-05 | 4.81E-06 | 0 | 2.13E-05 | 1.50E-05 | 7.95E-05 | 0 | 5.05E-06 |
| Smim8 | 6.84E-05 | 2.65E-05 | 0 | 0 | 5.88E-06 | 4.23E-05 | 4.47E-06 | 0 | 8.25E-06 | 2.10E-05 | 0 | 5.26E-05 | 1.05E-05 | 0 | 0 | 2.11E-05 | 1.21E-05 | 2.53E-05 |
| Nmral1 | 1.67E-05 | 0 | 0 | 8.99E-06 | 0 | 0 | 2.02E-05 | 0 | 8.25E-06 | 7.31E-06 | 1.83E-06 | 0 | 8.92E-06 | 2.01E-05 | 0 | 0.00018 | 0 | 1.86E-05 |
| Ranbp17 | 8.38E-06 | 1.41E-05 | 0 | 1.39E-05 | 3.41E-05 | 2.15E-05 | 1.52E-05 | 9.14E-06 | 8.24E-06 | 6.14E-06 | 2.14E-05 | 0 | 9.14E-06 | 3.49E-05 | 2.44E-05 | 6.45E-05 | 0 | 0 |
| Kiz | 5.59E-05 | 6.73E-05 | 0 | 0 | 9.27E-06 | 8.12E-05 | 3.44E-05 | 0 | 8.22E-06 | 3.13E-05 | 0 | 3.78E-05 | 2.28E-06 | 1.72E-05 | 2.82E-05 | 6.04E-05 | 2.67E-05 | 5.09E-05 |
| Mkln1 | 0.00021 | 0.00021 | 6.30E-05 | 0 | 3.52E-05 | 0.00014 | 0 | 0 | 8.21E-06 | 4.78E-05 | 6.96E-06 | 0.00011 | 5.61E-07 | 5.86E-05 | 0.00019 | 0.00011 | 0.00014 | 5.03E-05 |
| Nup160 | 5.12E-05 | 3.39E-05 | 6.60E-05 | 0 | 1.75E-05 | 7.38E-05 | 4.28E-06 | 1.81E-05 | 8.21E-06 | 2.31E-06 | 9.85E-06 | 3.69E-05 | 2.90E-05 | 1.53E-05 | 0 | 0.0003 | 0 | 2.23E-05 |
| Them6 | 1.38E-05 | 1.94E-05 | 0 | 5.73E-05 | 1.47E-05 | 1.20E-05 | 8.01E-06 | 3.07E-05 | 8.19E-06 | 0 | 1.73E-05 | 0 | 0 | 1.16E-05 | 0 | 4.94E-05 | 0 | 3.42E-05 |
| Ikzf4 | 0 | 5.63E-06 | 0 | 1.06E-05 | 1.26E-05 | 0 | 2.02E-05 | 0 | 8.18E-06 | 4.07E-06 | 3.74E-06 | 5.57E-06 | 5.47E-06 | 2.62E-05 | 0 | 0 | 1.16E-05 | 3.08E-06 |
| Gria1 | 0 | 0 | 0 | 0 | 0.00018 | 3.95E-05 | 1.66E-06 | 0 | 8.17E-06 | 0 | 0 | 0 | 0 | 0 | 0 | 0 | 0 | 0 |
| Prickle3 | 0 | 2.94E-05 | 2.74E-05 | 6.50E-05 | 6.57E-06 | 2.67E-05 | 5.12E-05 | 8.50E-05 | 8.15E-06 | 3.32E-05 | 5.02E-05 | 1.98E-05 | 1.53E-05 | 3.53E-05 | 8.93E-05 | 2.54E-05 | 0 | 2.74E-05 |
| Prim2 | 9.53E-06 | 6.20E-06 | 2.43E-05 | 0 | 9.78E-06 | 2.14E-05 | 9.11E-06 | 1.62E-05 | 8.15E-06 | 3.50E-05 | 2.37E-05 | 2.15E-05 | 2.27E-05 | 2.24E-05 | 2.22E-05 | 0.00026 | 0 | 6.51E-06 |
| Zfp407 | 5.99E-05 | 7.96E-05 | 2.45E-05 | 4.33E-05 | 2.44E-05 | 3.83E-05 | 3.01E-05 | 2.18E-05 | 8.13E-06 | 1.10E-05 | 0 | 3.57E-05 | 0 | 3.27E-05 | 0.00016 | 3.56E-05 | 0.00014 | 8.27E-06 |
| Tfb1m | 2.12E-05 | 1.67E-05 | 1.11E-05 | 0 | 0 | 0 | 9.44E-06 | 0 | 8.13E-06 | 3.67E-06 | 0 | 1.39E-05 | 3.82E-06 | 0 | 1.05E-05 | 1.76E-05 | 0 | 1.69E-05 |
| Bmpr2 | 0.00026 | 0.00024 | 0.00031 | 0 | 0.00044 | 0.0003 | 0.00036 | 0.00047 | 8.10E-06 | 0.00011 | 0.00051 | 0.00026 | 0.00079 | 0.00077 | 3.38E-05 | 1.52E-05 | 0.00012 | 0.00025 |
| Leng9 | 1.02E-05 | 2.86E-05 | 8.53E-06 | 3.27E-05 | 1.64E-05 | 1.57E-05 | 4.89E-06 | 6.21E-06 | 8.09E-06 | 1.36E-05 | 0 | 1.94E-05 | 0 | 1.45E-05 | 8.37E-05 | 0 | 9.19E-05 | 4.51E-05 |
| Ttf1 | 4.96E-05 | 5.55E-05 | 1.96E-05 | 3.92E-05 | 2.14E-05 | 1.13E-05 | 3.75E-06 | 0 | 8.09E-06 | 1.65E-05 | 4.50E-06 | 2.77E-05 | 0 | 0 | 0.00014 | 4.43E-05 | 2.03E-06 | 3.68E-05 |
| Mcph1 | 3.53E-05 | 2.02E-05 | 3.00E-05 | 0 | 1.05E-05 | 9.67E-06 | 8.08E-06 | 7.36E-07 | 8.09E-06 | 1.23E-05 | 0 | 1.35E-05 | 7.17E-06 | 3.05E-05 | 0.00012 | 0.00022 | 4.05E-05 | 0 |
| Cdk5rap1 | 3.38E-05 | 2.28E-05 | 1.24E-05 | 3.37E-05 | 2.50E-05 | 1.76E-06 | 3.37E-05 | 0 | 8.07E-06 | 2.80E-06 | 0 | 3.83E-05 | 4.02E-05 | 1.82E-05 | 5.67E-05 | 4.99E-05 | 0 | 1.20E-05 |
| Ptpdc1 | 0 | 0.0001 | 2.77E-05 | 1.87E-05 | 1.09E-05 | 8.23E-07 | 9.12E-05 | 0 | 8.06E-06 | 6.17E-05 | 2.21E-05 | 1.62E-05 | 0 | 3.42E-05 | 9.57E-06 | 0.00011 | 6.19E-05 | 7.62E-05 |
| Cntrl | 3.74E-05 | 6.58E-05 | 0 | 0 | 5.91E-06 | 7.29E-05 | 0 | 0 | 8.05E-06 | 2.18E-05 | 0 | 0 | 0 | 0 | 0.00044 | 0.00023 | 6.35E-05 | 1.44E-05 |
| Ccdc51 | 0 | 6.94E-06 | 0 | 1.54E-05 | 6.20E-06 | 0 | 0 | 1.74E-06 | 8.04E-06 | 7.98E-06 | 0 | 8.49E-06 | 2.56E-06 | 8.01E-06 | 9.11E-06 | 1.47E-05 | 1.04E-05 | 3.76E-06 |
| Zfp964 | 4.90E-06 | 1.56E-06 | 0 | 0 | 3.36E-06 | 1.46E-05 | 0 | 0 | 8.03E-06 | 0 | 0 | 5.00E-06 | 0 | 2.27E-06 | 0 | 0 | 4.42E-06 | 3.95E-06 |
| C87436 | 2.02E-05 | 4.15E-05 | 1.10E-06 | 3.15E-05 | 1.94E-05 | 1.26E-05 | 6.40E-06 | 0 | 8.03E-06 | 2.15E-05 | 6.99E-06 | 0 | 0 | 1.35E-05 | 1.26E-05 | 3.37E-05 | 0 | 1.48E-05 |
| Rapgef6 | 0.00012 | 7.51E-05 | 3.77E-05 | 0 | 7.25E-06 | 8.27E-05 | 5.42E-06 | 1.83E-05 | 8.02E-06 | 8.89E-06 | 0 | 0.00012 | 1.55E-05 | 0 | 9.04E-05 | 0.00012 | 6.38E-05 | 1.77E-05 |
| Serp1 | 0.00033 | 0 | 0.00025 | 7.79E-05 | 1.62E-06 | 4.89E-06 | 0.00018 | 5.91E-05 | 8.01E-06 | 0 | 3.55E-05 | 6.00E-05 | 0 | 0 | 3.08E-05 | 0 | 0 | 0 |
| Lrrc8b | 0 | 1.02E-05 | 7.94E-06 | 0 | 8.57E-06 | 0 | 3.64E-06 | 6.25E-06 | 8.01E-06 | 8.25E-05 | 9.52E-06 | 0 | 0 | 1.82E-06 | 0 | 8.83E-05 | 0 | 0 |
| Tesk2 | 0 | 2.26E-05 | 0 | 3.09E-05 | 2.64E-05 | 9.01E-06 | 3.21E-05 | 0 | 8.00E-06 | 1.00E-05 | 2.06E-05 | 0 | 4.54E-05 | 3.74E-05 | 6.62E-06 | 2.97E-05 | 0 | 1.24E-05 |
| Dpep2 | 0 | 0 | 0 | 0 | 1.47E-05 | 2.30E-05 | 0 | 0 | 7.98E-06 | 0 | 1.02E-06 | 1.52E-05 | 4.80E-06 | 0 | 1.34E-05 | 0 | 0 | 4.61E-06 |
| Ropn1l | 3.34E-06 | 1.06E-05 | 0 | 1.29E-05 | 5.01E-07 | 3.25E-06 | 2.93E-06 | 5.05E-07 | 7.95E-06 | 5.90E-06 | 0 | 1.06E-05 | 7.26E-06 | 0 | 0 | 4.48E-06 | 2.86E-06 | 5.62E-06 |

| | | | | | | | | | | | | | | | | | | |
|---|---|---|---|---|---|---|---|---|---|---|---|---|---|---|---|---|---|---|
| DEPRECATEI | 1.69E-05 | 2.88E-05 | 1.34E-07 | 6.32E-05 | 9.21E-07 | 3.04E-05 | 3.18E-05 | 6.03E-05 | 7.94E-06 | 4.04E-05 | 0 | 5.08E-05 | 1.05E-06 | 8.43E-06 | 0 | 0 | 0 | 2.20E-05 |
| Rnft2 | 1.60E-06 | 0 | 1.13E-05 | 1.57E-05 | 5.67E-06 | 0 | 0 | 8.11E-06 | 7.93E-06 | 9.06E-06 | 3.08E-06 | 0 | 2.75E-07 | 0 | 2.69E-06 | 1.73E-06 | 0 | 8.46E-06 |
| Lsm3 | 0.00014 | 5.63E-06 | 0.00013 | 8.48E-05 | 0 | 9.54E-05 | 1.83E-05 | 5.44E-05 | 7.91E-06 | 1.36E-05 | 1.41E-05 | 5.93E-05 | 2.12E-05 | 1.39E-05 | 6.76E-06 | 0.00048 | 7.33E-05 | 2.31E-05 |
| Tmem86b | 1.52E-06 | 3.26E-05 | 1.63E-05 | 0 | 2.44E-06 | 0 | 0 | 0 | 7.89E-06 | 1.23E-05 | 3.27E-06 | 1.60E-06 | 1.89E-05 | 1.67E-05 | 4.20E-05 | 0 | 6.24E-05 | 0 |
| Abcg2 | 8.57E-05 | 2.86E-05 | 1.32E-05 | 0 | 2.32E-05 | 6.58E-05 | 1.04E-05 | 6.77E-05 | 7.87E-06 | 2.53E-05 | 0 | 9.74E-06 | 7.87E-06 | 2.32E-06 | 0 | 5.39E-05 | 0 | 4.45E-05 |
| Atmin | 7.71E-05 | 5.51E-05 | 6.22E-05 | 4.60E-05 | 3.06E-05 | 5.16E-05 | 8.87E-06 | 3.45E-05 | 7.87E-06 | 1.35E-05 | 1.44E-05 | 1.82E-05 | 1.34E-06 | 1.39E-05 | 1.75E-05 | 0 | 3.77E-05 | 2.44E-05 |
| Ofd1 | 2.42E-05 | 1.58E-05 | 1.29E-06 | 8.85E-06 | 3.42E-06 | 6.61E-06 | 0 | 0 | 7.87E-06 | 1.27E-05 | 1.92E-05 | 1.36E-05 | 6.49E-06 | 0 | 1.98E-05 | 2.67E-05 | 0 | 0 |
| Fam13c | 1.56E-06 | 7.62E-06 | 0 | 0 | 0 | 4.36E-06 | 0 | 0 | 7.87E-06 | 6.47E-05 | 0 | 0 | 0 | 9.47E-06 | 0 | 0 | 0 | 1.45E-05 |
| Syt12 | 0 | 0 | 2.23E-05 | 5.74E-06 | 2.54E-05 | 0 | 1.48E-05 | 1.37E-05 | 7.87E-06 | 7.66E-06 | 0 | 0 | 2.48E-05 | 0 | 5.22E-06 | 5.98E-06 | 0 | 0 |
| Bves | 7.26E-07 | 0 | 0 | 0 | 0 | 8.47E-06 | 0 | 5.53E-06 | 7.86E-06 | 1.17E-06 | 1.25E-05 | 0 | 0 | 1.35E-05 | 3.39E-06 | 5.95E-06 | 0 | 0 |
| Mest | 3.11E-05 | 7.49E-06 | 0 | 5.59E-06 | 0.00013 | 0 | 0 | 0.00043 | 7.84E-06 | 8.82E-06 | 0 | 0 | 0 | 0.0002 | 0 | 4.15E-05 | 0 | 4.46E-06 |
| Slc25a19 | 1.09E-05 | 6.09E-05 | 0 | 7.58E-05 | 2.73E-05 | 0 | 0 | 4.89E-05 | 7.83E-06 | 4.60E-05 | 1.03E-05 | 2.22E-05 | 2.91E-05 | 9.09E-07 | 2.47E-05 | 3.63E-05 | 2.75E-05 | 3.51E-05 |
| Vps54 | 0.00018 | 0.00011 | 3.56E-05 | 0 | 1.75E-05 | 7.76E-05 | 0 | 0.00011 | 7.81E-06 | 1.58E-05 | 2.78E-05 | 5.16E-05 | 0 | 2.13E-05 | 0.00012 | 4.65E-05 | 4.84E-05 | 0 |
| Cep57 | 5.96E-05 | 5.37E-05 | 0 | 0 | 1.65E-05 | 3.77E-05 | 0 | 0 | 7.80E-06 | 9.92E-07 | 0 | 1.71E-05 | 0 | 3.84E-06 | 4.58E-05 | 0.00021 | 0 | 8.16E-07 |
| Synpo2 | 0 | 3.81E-06 | 0 | 0 | 0 | 0 | 0 | 0 | 7.80E-06 | 1.14E-05 | 1.18E-05 | 1.98E-05 | 0 | 3.81E-05 | 0 | 0 | 0 | 5.34E-05 |
| Zfp101 | 0 | 2.30E-05 | 0 | 0 | 9.49E-07 | 0 | 0 | 9.35E-06 | 7.80E-06 | 7.98E-08 | 3.54E-07 | 1.06E-05 | 0 | 4.27E-06 | 9.96E-05 | 0.00016 | 0 | 8.69E-06 |
| Pigb | 5.39E-05 | 4.95E-05 | 1.70E-05 | 1.91E-06 | 3.13E-05 | 4.68E-05 | 3.17E-05 | 1.53E-06 | 7.79E-06 | 1.34E-05 | 7.27E-06 | 1.30E-06 | 0 | 0 | 1.61E-05 | 2.74E-05 | 0 | 5.47E-05 |
| Ehhadh | 1.38E-05 | 1.24E-05 | 0 | 9.96E-06 | 2.26E-05 | 2.29E-05 | 5.03E-06 | 0 | 7.79E-06 | 1.76E-05 | 7.98E-06 | 9.22E-06 | 5.16E-06 | 5.64E-06 | 4.47E-06 | 0 | 0 | 5.24E-06 |
| Zfp709 | 1.24E-05 | 1.54E-05 | 0 | 0 | 1.66E-05 | 1.66E-05 | 7.59E-07 | 8.56E-07 | 7.79E-06 | 1.53E-05 | 4.52E-06 | 0 | 0 | 7.53E-06 | 5.58E-05 | 1.89E-06 | 0 | 9.82E-06 |
| Hopx | 0 | 1.65E-06 | 0 | 0 | 0 | 6.40E-06 | 0 | 0 | 7.79E-06 | 2.30E-06 | 0 | 1.29E-05 | 0 | 2.40E-06 | 2.96E-05 | 0 | 0 | 6.04E-05 |
| Trmo | 5.11E-06 | 2.44E-06 | 1.30E-05 | 0 | 8.74E-06 | 8.94E-06 | 3.22E-06 | 3.33E-06 | 7.76E-06 | 8.11E-06 | 1.09E-05 | 1.15E-05 | 2.86E-06 | 3.23E-06 | 1.47E-05 | 6.82E-06 | 0 | 0 |
| Mrm3 | 3.85E-06 | 9.10E-06 | 3.48E-05 | 8.54E-06 | 2.88E-06 | 6.61E-06 | 0 | 0 | 7.76E-06 | 6.09E-06 | 7.01E-06 | 1.89E-07 | 1.02E-05 | 0 | 0 | 1.79E-05 | 0 | 2.27E-05 |
| Cdkn2aipnl | 3.27E-05 | 7.47E-05 | 6.98E-05 | 6.51E-05 | 1.32E-05 | 1.14E-05 | 2.52E-06 | 5.51E-05 | 7.75E-06 | 7.15E-06 | 2.37E-05 | 1.08E-05 | 0 | 2.92E-05 | 0 | 0.00031 | 1.01E-05 | 5.60E-05 |
| Rmnd5a | 0.00015 | 0.0002 | 0 | 0 | 0.00014 | 0.00019 | 5.58E-05 | 4.69E-05 | 7.74E-06 | 5.80E-05 | 3.95E-05 | 0.00011 | 1.24E-06 | 5.39E-05 | 5.63E-06 | 6.69E-05 | 0.00016 | 7.10E-05 |
| Fam49a | 6.34E-06 | 2.01E-05 | 0 | 0 | 0 | 3.82E-05 | 0 | 2.48E-06 | 7.74E-06 | 7.26E-05 | 2.56E-05 | 1.72E-05 | 5.21E-05 | 2.57E-05 | 0 | 3.24E-06 | 0 | 7.07E-05 |
| Adat1 | 1.06E-05 | 2.59E-05 | 1.74E-05 | 0 | 1.95E-06 | 3.58E-07 | 0 | 0 | 7.74E-06 | 0 | 4.49E-06 | 1.35E-05 | 4.94E-06 | 0 | 6.51E-05 | 2.62E-05 | 0 | 0 |
| Pfkfb1 | 0 | 0 | 1.07E-07 | 1.85E-06 | 0 | 2.42E-06 | 0 | 6.63E-06 | 7.73E-06 | 4.84E-06 | 5.59E-06 | 6.25E-06 | 1.12E-05 | 1.32E-05 | 1.92E-05 | 4.70E-05 | 0 | 6.83E-06 |
| Ubr1 | 6.72E-05 | 0.00011 | 1.50E-05 | 7.67E-05 | 7.46E-05 | 6.65E-05 | 6.54E-06 | 0.00012 | 7.72E-06 | 3.29E-05 | 9.36E-06 | 0 | 0 | 3.36E-05 | 6.20E-05 | 0 | 2.69E-05 | 0.00012 |
| Wdr60 | 2.01E-05 | 3.37E-05 | 7.49E-06 | 1.52E-05 | 1.71E-05 | 1.13E-05 | 0 | 0 | 7.72E-06 | 2.52E-05 | 4.78E-05 | 8.24E-06 | 0 | 2.96E-06 | 1.94E-05 | 8.26E-07 | 3.78E-05 | 2.44E-05 |
| Car13 | 0.00011 | 2.68E-05 | 0.00012 | 3.38E-05 | 0 | 0 | 0 | 3.78E-05 | 7.71E-06 | 7.75E-05 | 0 | 2.34E-05 | 0.00016 | 6.38E-05 | 0 | 0.00013 | 0.0001 | 8.89E-05 |
| Zfp607a | 1.84E-05 | 2.31E-05 | 0 | 0 | 1.36E-05 | 2.25E-05 | 0 | 8.14E-06 | 7.70E-06 | 1.45E-05 | 6.29E-06 | 5.23E-06 | 0 | 1.60E-05 | 3.61E-05 | 5.88E-06 | 0 | 2.46E-05 |
| Tmem41a | 3.11E-05 | 5.30E-05 | 0.00011 | 5.60E-05 | 3.76E-05 | 4.18E-05 | 1.53E-05 | 5.60E-05 | 7.69E-06 | 6.66E-05 | 5.08E-05 | 2.52E-05 | 6.63E-05 | 8.56E-05 | 1.39E-06 | 1.68E-06 | 0 | 7.92E-05 |
| Fen1 | 6.08E-07 | 0 | 0.00013 | 2.44E-05 | 9.88E-07 | 1.19E-05 | 5.13E-05 | 1.90E-05 | 7.68E-06 | 4.72E-06 | 2.65E-05 | 0 | 3.13E-05 | 4.52E-05 | 0 | 0.00076 | 0 | 0 |

| | | | | | | | | | | | | | | | | | | |
|---|---|---|---|---|---|---|---|---|---|---|---|---|---|---|---|---|---|---|
| Rbm12b1 | 6.93E-06 | 2.53E-05 | 1.70E-05 | 1.55E-05 | 1.38E-05 | 5.34E-06 | 0 | 1.48E-05 | 7.67E-06 | 1.50E-05 | 0 | 1.86E-05 | 0 | 5.21E-06 | 5.83E-05 | 1.45E-05 | 0 | 0 |
| Mllt11 | 5.50E-07 | 3.10E-05 | 0.00013 | 6.29E-05 | 1.86E-05 | 0 | 0 | 0.00012 | 7.66E-06 | 1.20E-05 | 1.79E-05 | 0.00012 | 0 | 9.11E-06 | 6.16E-05 | 0 | 0 | 9.76E-06 |
| D1Ertd622e | 7.50E-06 | 6.66E-06 | 1.14E-05 | 0 | 0 | 1.20E-05 | 8.81E-07 | 3.48E-06 | 7.65E-06 | 0 | 3.33E-06 | 0 | 0 | 5.11E-06 | 0 | 9.19E-06 | 1.02E-05 | 2.84E-06 |
| Fbxo2 | 0 | 9.93E-06 | 0 | 5.53E-06 | 4.73E-06 | 3.41E-06 | 0 | 0 | 7.64E-06 | 0 | 0 | 0 | 0 | 2.85E-05 | 0 | 0 | 0 | 3.66E-06 |
| Brap | 0 | 3.38E-05 | 3.34E-05 | 7.72E-05 | 4.84E-05 | 3.04E-05 | 5.76E-06 | 4.94E-05 | 7.63E-06 | 2.56E-05 | 2.02E-05 | 1.81E-05 | 1.01E-05 | 3.81E-05 | 6.88E-05 | 2.85E-06 | 4.48E-06 | 3.39E-05 |
| Zfp938 | 1.26E-05 | 1.31E-05 | 1.61E-06 | 6.79E-06 | 6.53E-06 | 3.57E-06 | 3.25E-06 | 1.78E-06 | 7.62E-06 | 8.88E-06 | 1.12E-07 | 1.14E-05 | 0 | 2.24E-06 | 4.11E-05 | 0 | 0 | 1.05E-07 |
| Reep2 | 4.32E-06 | 6.41E-06 | 1.58E-07 | 0 | 0 | 0 | 0 | 5.15E-07 | 7.60E-06 | 1.14E-05 | 0 | 2.97E-06 | 0 | 2.55E-05 | 1.09E-05 | 1.69E-05 | 2.48E-05 | 1.37E-05 |
| Tmem212 | 0 | 1.75E-05 | 0 | 0 | 4.73E-05 | 2.28E-05 | 0 | 2.73E-05 | 7.59E-06 | 0 | 0 | 0 | 0 | 6.19E-06 | 3.17E-06 | 0 | 0 | 0 |
| Stard10 | 9.74E-07 | 3.39E-05 | 0 | 2.60E-05 | 2.12E-05 | 1.14E-05 | 3.30E-06 | 0 | 7.53E-06 | 7.30E-07 | 3.12E-05 | 2.05E-05 | 1.49E-05 | 4.93E-05 | 0 | 1.13E-05 | 3.50E-05 | 2.93E-05 |
| Epsti1 | 5.22E-06 | 3.30E-06 | 0 | 6.36E-06 | 0 | 0 | 1.25E-06 | 0 | 7.53E-06 | 0 | 0 | 7.95E-07 | 6.87E-06 | 2.67E-06 | 4.90E-07 | 8.93E-06 | 7.08E-06 | 8.68E-06 |
| Zc3h8 | 1.23E-05 | 3.24E-05 | 1.30E-05 | 0 | 1.54E-05 | 1.17E-05 | 1.76E-05 | 3.31E-05 | 7.52E-06 | 0 | 2.86E-06 | 2.62E-05 | 0 | 0 | 3.54E-05 | 4.53E-06 | 9.39E-06 | 0 |
| Calr3 | 0 | 3.98E-07 | 0 | 4.71E-06 | 3.63E-06 | 5.31E-07 | 6.57E-07 | 1.63E-06 | 7.52E-06 | 2.44E-06 | 3.52E-07 | 9.33E-06 | 6.50E-06 | 2.27E-06 | 0 | 0 | 3.90E-06 | 8.39E-06 |
| Rsl1 | 1.18E-05 | 8.98E-06 | 1.92E-06 | 0 | 7.77E-06 | 1.07E-05 | 0 | 0 | 7.51E-06 | 0 | 0 | 7.68E-06 | 0 | 0 | 0 | 2.26E-06 | 0 | 5.79E-06 |
| Tekt2 | 0 | 1.00E-05 | 1.60E-06 | 3.46E-06 | 0 | 6.73E-07 | 1.79E-05 | 0 | 7.51E-06 | 9.34E-06 | 0 | 1.90E-05 | 2.63E-06 | 0 | 2.62E-05 | 0 | 4.55E-05 | 3.80E-05 |
| Dnase1l1 | 1.71E-05 | 4.82E-05 | 0 | 9.92E-06 | 9.64E-07 | 3.21E-05 | 3.46E-05 | 2.60E-05 | 7.49E-06 | 5.57E-05 | 4.10E-06 | 4.91E-05 | 1.95E-05 | 1.70E-05 | 0 | 9.48E-05 | 0 | 0.00011 |
| Zmym1 | 1.12E-05 | 3.97E-05 | 2.70E-05 | 2.21E-05 | 7.63E-06 | 2.48E-06 | 1.44E-05 | 0 | 7.49E-06 | 2.06E-05 | 0 | 0 | 1.91E-05 | 2.13E-05 | 6.77E-05 | 0.00014 | 0 | 1.32E-05 |
| H2-Eb1 | 0 | 3.85E-05 | 0 | 7.15E-05 | 0 | 0 | 0 | 2.00E-05 | 7.49E-06 | 4.97E-05 | 0 | 2.04E-05 | 8.35E-06 | 1.48E-05 | 0.00032 | 2.31E-05 | 9.90E-05 | 4.83E-05 |
| Eps8l1 | 4.36E-06 | 2.94E-05 | 0 | 0 | 8.48E-06 | 4.06E-05 | 0 | 3.27E-05 | 7.48E-06 | 2.74E-06 | 0 | 1.18E-05 | 0 | 8.87E-06 | 7.39E-07 | 0 | 0 | 2.51E-06 |
| Zfp58 | 1.60E-05 | 1.47E-05 | 0 | 0 | 3.72E-06 | 7.63E-06 | 4.95E-06 | 0 | 7.48E-06 | 5.43E-06 | 0 | 6.71E-06 | 0 | 0 | 2.66E-05 | 5.32E-07 | 0 | 6.04E-06 |
| Zfp954 | 2.93E-05 | 9.29E-06 | 7.51E-06 | 0 | 1.64E-05 | 2.25E-05 | 2.78E-05 | 0 | 7.46E-06 | 1.15E-05 | 7.87E-07 | 2.59E-05 | 1.14E-05 | 7.75E-06 | 2.16E-05 | 0 | 0 | 8.65E-06 |
| DEPRECATED | 5.53E-06 | 1.05E-05 | 0 | 0 | 0 | 1.75E-05 | 1.48E-05 | 8.06E-06 | 7.44E-06 | 1.42E-05 | 5.58E-06 | 8.87E-06 | 3.27E-05 | 1.19E-05 | 4.40E-06 | 6.57E-06 | 0 | 1.76E-05 |
| Zfp868 | 2.02E-05 | 1.05E-05 | 0 | 0 | 6.21E-07 | 1.33E-05 | 0 | 2.39E-06 | 7.44E-06 | 0 | 0 | 8.99E-06 | 0 | 4.73E-07 | 2.04E-05 | 2.51E-06 | 0 | 0 |
| Ormdl1 | 2.58E-05 | 2.02E-05 | 0 | 0 | 1.50E-05 | 1.72E-05 | 0 | 7.76E-06 | 7.43E-06 | 4.21E-06 | 0 | 3.66E-05 | 0 | 2.23E-05 | 0 | 7.53E-05 | 0 | 8.86E-06 |
| Nkiras1 | 3.81E-05 | 4.35E-05 | 5.00E-05 | 2.01E-05 | 5.16E-05 | 1.32E-05 | 6.39E-05 | 2.09E-05 | 7.43E-06 | 4.40E-05 | 1.49E-05 | 0 | 2.70E-06 | 0 | 5.94E-05 | 4.68E-06 | 0 | 5.27E-05 |
| DEPRECATED | 8.70E-06 | 8.81E-06 | 1.10E-05 | 1.06E-05 | 0 | 0 | 0 | 4.26E-06 | 7.43E-06 | 7.97E-06 | 2.38E-06 | 1.03E-05 | 0 | 0 | 3.97E-06 | 4.83E-05 | 0 | 9.16E-06 |
| Serpina3f | 0 | 1.93E-06 | 5.78E-06 | 0 | 0 | 0 | 0 | 0 | 7.41E-06 | 0 | 0 | 0 | 9.55E-05 | 9.08E-05 | 0 | 0 | 0 | 1.80E-05 |
| Fbxl16 | 1.22E-05 | 2.49E-05 | 1.82E-05 | 1.85E-05 | 0 | 1.27E-05 | 8.56E-05 | 0 | 7.38E-06 | 4.86E-06 | 1.75E-05 | 2.31E-06 | 2.44E-05 | 4.12E-05 | 0 | 0 | 0 | 0 |
| Proca1 | 0 | 6.29E-06 | 0 | 5.22E-06 | 0 | 0 | 0 | 0 | 7.38E-06 | 8.56E-06 | 0 | 4.88E-06 | 1.01E-05 | 0 | 4.51E-06 | 0 | 4.96E-05 | 6.71E-06 |
| Bcdin3d | 0 | 2.05E-05 | 9.58E-06 | 8.46E-06 | 3.15E-06 | 1.39E-06 | 0 | 0 | 7.37E-06 | 5.10E-06 | 2.66E-06 | 0 | 0 | 6.54E-06 | 1.67E-05 | 2.18E-05 | 5.01E-06 | 7.70E-06 |
| Stk10 | 0 | 3.61E-05 | 2.43E-05 | 0 | 5.02E-07 | 0 | 2.15E-05 | 1.64E-05 | 7.37E-06 | 2.63E-06 | 0 | 4.91E-05 | 0 | 0 | 1.30E-05 | 7.20E-06 | 8.29E-05 | 0 |
| Lekr1 | 5.90E-06 | 7.91E-06 | 0 | 1.67E-06 | 9.01E-06 | 3.26E-06 | 1.49E-06 | 2.23E-06 | 7.36E-06 | 9.39E-06 | 8.04E-06 | 2.22E-05 | 8.41E-06 | 1.56E-06 | 3.95E-06 | 8.32E-06 | 0 | 3.50E-07 |
| Cntf | 6.10E-06 | 5.52E-06 | 5.91E-06 | 2.18E-06 | 0 | 1.87E-06 | 6.82E-06 | 1.11E-05 | 7.36E-06 | 0 | 7.00E-06 | 8.51E-06 | 0 | 8.57E-06 | 8.83E-06 | 2.73E-05 | 3.51E-05 | 0 |
| Sugct | 0 | 9.62E-06 | 0 | 3.02E-05 | 2.89E-05 | 5.07E-06 | 5.77E-06 | 1.46E-05 | 7.34E-06 | 1.48E-05 | 2.02E-06 | 8.19E-06 | 6.13E-06 | 1.27E-05 | 3.56E-05 | 0 | 2.23E-06 | 4.93E-06 |

| | | | | | | | | | | | | | | | | | | |
|---|---|---|---|---|---|---|---|---|---|---|---|---|---|---|---|---|---|---|
| Dusp22 | 2.32E-05 | 4.17E-06 | 0 | 0 | 2.71E-05 | 5.64E-05 | 0 | 1.22E-05 | 7.33E-06 | 0.00011 | 0 | 6.89E-06 | 0 | 0 | 1.59E-05 | 0 | 0 | 9.39E-05 |
| Zfp677 | 1.89E-05 | 2.23E-05 | 0 | 0 | 1.00E-06 | 1.41E-05 | 0 | 0 | 7.33E-06 | 1.05E-05 | 0 | 0 | 0 | 2.26E-06 | 2.14E-05 | 0 | 0 | 4.99E-06 |
| Anp32a | 6.10E-05 | 0.00014 | 0.00026 | 0.00088 | 0.00021 | 0 | 0.00075 | 0.00055 | 7.32E-06 | 0.00015 | 2.76E-05 | 0.00059 | 3.45E-05 | 7.00E-05 | 0.00014 | 0.00038 | 9.00E-05 | 0.00015 |
| Efna2 | 7.74E-06 | 2.28E-05 | 2.67E-06 | 2.75E-05 | 2.89E-05 | 5.56E-05 | 0 | 3.69E-05 | 7.31E-06 | 7.11E-06 | 1.75E-05 | 2.97E-06 | 0 | 2.98E-05 | 0 | 9.87E-06 | 4.71E-05 | 0 |
| Rpgr | 1.12E-05 | 1.21E-05 | 0 | 4.36E-06 | 8.85E-06 | 1.15E-05 | 0 | 7.10E-06 | 7.31E-06 | 5.15E-06 | 0 | 1.11E-05 | 1.63E-06 | 0 | 3.62E-05 | 5.80E-06 | 4.14E-06 | 7.12E-06 |
| Impdh1 | 0.00021 | 2.24E-05 | 0.00018 | 0.0001 | 9.38E-06 | 0.00015 | 0.00028 | 0 | 7.30E-06 | 4.77E-08 | 1.12E-05 | 0.00027 | 8.33E-06 | 0.00019 | 9.29E-05 | 0 | 0 | 0 |
| Gemin6 | 2.40E-05 | 2.98E-06 | 5.58E-05 | 4.36E-06 | 1.22E-05 | 1.61E-05 | 0 | 0 | 7.30E-06 | 1.13E-05 | 6.50E-06 | 0 | 3.83E-06 | 5.64E-07 | 0 | 0.00019 | 0 | 0 |
| Peg10 | 2.54E-05 | 0 | 0 | 0 | 0 | 1.37E-06 | 0 | 0 | 7.30E-06 | 2.93E-05 | 0 | 5.13E-05 | 2.57E-05 | 2.54E-06 | 0 | 1.29E-05 | 0.00034 | 0 |
| Cep72 | 0 | 3.55E-06 | 0 | 0 | 3.49E-06 | 7.75E-06 | 0 | 9.44E-08 | 7.29E-06 | 2.24E-06 | 0 | 2.72E-06 | 0 | 0 | 1.00E-05 | 0.00011 | 0 | 0 |
| Mphosph9 | 5.51E-06 | 6.99E-06 | 0 | 0 | 0 | 8.31E-07 | 0 | 3.51E-07 | 7.29E-06 | 0 | 1.36E-07 | 0 | 0 | 1.05E-05 | 3.93E-05 | 7.24E-05 | 0 | 0 |
| Itch | 0.00025 | 0.00013 | 7.88E-05 | 0 | 6.40E-05 | 0.00012 | 9.12E-05 | 0 | 7.28E-06 | 0 | 7.05E-05 | 9.39E-05 | 0 | 3.62E-05 | 8.93E-05 | 1.76E-05 | 0.0002 | 0 |
| Tmem70 | 0.00021 | 4.90E-05 | 0.00017 | 0 | 0 | 7.77E-05 | 0 | 8.45E-06 | 7.28E-06 | 0 | 0.00036 | 2.78E-05 | 0 | 0 | 0 | 8.39E-05 | 0 | 0 |
| Klhl23 | 3.33E-06 | 0 | 5.13E-06 | 0 | 0 | 3.61E-06 | 0 | 7.34E-06 | 7.28E-06 | 0 | 1.73E-05 | 0 | 0 | 3.27E-06 | 5.54E-06 | 3.42E-05 | 0 | 7.46E-06 |
| Cenpo | 0 | 1.20E-05 | 1.36E-05 | 3.40E-05 | 1.24E-05 | 5.74E-06 | 2.22E-05 | 0 | 7.27E-06 | 3.21E-05 | 0 | 1.31E-05 | 7.29E-06 | 6.52E-06 | 3.79E-05 | 0.00019 | 3.30E-06 | 1.07E-05 |
| Zbtb10 | 3.28E-05 | 0.00013 | 6.23E-05 | 0 | 1.01E-05 | 6.58E-05 | 0 | 5.85E-05 | 7.26E-06 | 0 | 3.33E-05 | 4.42E-05 | 0 | 5.37E-05 | 8.30E-05 | 3.43E-05 | 0.00035 | 2.09E-05 |
| Stk38l | 7.37E-06 | 0.00012 | 7.26E-05 | 0 | 0 | 0 | 0 | 0 | 7.26E-06 | 4.85E-05 | 0.0001 | 3.04E-05 | 2.75E-05 | 5.37E-05 | 1.94E-05 | 1.69E-05 | 7.72E-05 | 2.32E-05 |
| Trim12a | 1.83E-05 | 2.42E-05 | 0 | 0 | 1.46E-05 | 0 | 9.51E-06 | 0 | 7.24E-06 | 3.19E-05 | 0 | 0 | 0 | 0 | 1.45E-05 | 0 | 0 | 1.61E-05 |
| Fgfr3 | 0 | 1.62E-06 | 1.47E-05 | 0 | 0 | 0 | 0 | 0 | 7.24E-06 | 0 | 7.20E-06 | 0 | 0 | 2.99E-05 | 1.00E-05 | 1.21E-05 | 2.31E-07 | 1.76E-05 |
| Mettl5 | 4.85E-05 | 3.46E-05 | 3.20E-05 | 7.77E-06 | 0 | 1.08E-05 | 0 | 2.21E-05 | 7.22E-06 | 1.39E-05 | 1.07E-05 | 6.04E-06 | 0 | 0 | 0.00011 | 5.35E-05 | 0 | 2.20E-05 |
| Kctd21 | 1.14E-05 | 1.97E-05 | 5.09E-07 | 4.20E-06 | 8.64E-06 | 5.04E-06 | 1.74E-05 | 0 | 7.22E-06 | 1.25E-05 | 4.82E-06 | 1.51E-05 | 1.43E-05 | 7.19E-06 | 0 | 0 | 0 | 5.20E-06 |
| Ppp1r3d | 1.53E-06 | 1.31E-05 | 0 | 1.96E-05 | 9.62E-05 | 8.90E-07 | 3.99E-05 | 1.74E-05 | 7.21E-06 | 0 | 6.12E-05 | 4.89E-06 | 7.83E-06 | 4.51E-05 | 1.73E-06 | 0 | 0 | 2.81E-05 |
| Rpp40 | 1.77E-05 | 1.04E-05 | 2.61E-05 | 0 | 0 | 0 | 0 | 0 | 7.20E-06 | 7.50E-06 | 0 | 0 | 1.49E-05 | 0 | 2.29E-06 | 2.07E-05 | 0 | 2.87E-06 |
| Tmem175 | 9.99E-05 | 5.23E-05 | 0 | 0 | 3.15E-05 | 6.12E-05 | 5.92E-05 | 1.24E-05 | 7.18E-06 | 3.06E-05 | 1.13E-06 | 9.41E-06 | 2.73E-05 | 1.41E-05 | 0 | 3.04E-05 | 0 | 6.77E-05 |
| Bod1 | 9.12E-05 | 7.10E-05 | 4.58E-05 | 3.37E-05 | 6.85E-05 | 7.15E-05 | 6.44E-05 | 0 | 7.15E-06 | 2.63E-05 | 3.40E-05 | 4.67E-05 | 0 | 4.59E-05 | 0 | 7.47E-05 | 8.52E-06 | 2.26E-05 |
| Pepd | 9.46E-05 | 0.00012 | 0.00016 | 0.00018 | 5.91E-05 | 0 | 0.00015 | 6.71E-06 | 7.15E-06 | 3.87E-05 | 0 | 5.87E-05 | 0 | 4.88E-05 | 0.00011 | 0 | 9.96E-06 | 2.38E-05 |
| Gspt2 | 5.02E-06 | 2.66E-05 | 3.87E-06 | 3.60E-05 | 1.00E-06 | 0 | 0 | 0 | 7.15E-06 | 1.60E-05 | 0 | 0 | 5.60E-06 | 0 | 2.86E-05 | 1.67E-07 | 0 | 2.39E-05 |
| Prcp | 0.00014 | 3.22E-06 | 4.03E-05 | 0 | 0 | 6.71E-05 | 7.48E-05 | 6.05E-05 | 7.14E-06 | 8.03E-05 | 4.72E-06 | 0 | 0 | 5.93E-05 | 0 | 4.20E-05 | 0 | 4.90E-05 |
| Rnpc3 | 6.14E-05 | 4.33E-05 | 3.07E-05 | 0 | 8.40E-06 | 1.30E-05 | 0 | 0 | 7.14E-06 | 0 | 1.29E-05 | 2.06E-05 | 0 | 2.18E-06 | 0.00014 | 1.44E-05 | 0 | 0 |
| Zc4h2 | 1.29E-05 | 1.00E-05 | 0 | 0 | 1.13E-05 | 8.77E-06 | 0 | 0 | 7.14E-06 | 2.96E-05 | 2.32E-06 | 5.31E-06 | 0 | 0 | 7.12E-07 | 2.36E-05 | 1.41E-06 | 5.00E-05 |
| Nrbp2 | 1.19E-06 | 6.96E-05 | 0 | 0 | 0 | 0 | 8.26E-05 | 0 | 7.13E-06 | 5.68E-05 | 2.44E-05 | 3.84E-05 | 9.05E-05 | 2.82E-05 | 0.00014 | 1.47E-05 | 1.80E-05 | 0.00016 |
| Poll | 2.03E-05 | 2.50E-05 | 7.75E-06 | 2.16E-05 | 0 | 1.55E-05 | 1.04E-05 | 6.63E-06 | 7.10E-06 | 1.27E-05 | 0 | 1.05E-05 | 0 | 0 | 2.57E-05 | 6.51E-05 | 7.28E-06 | 0 |
| AA986860 | 7.52E-08 | 1.60E-05 | 1.43E-06 | 5.85E-06 | 4.44E-06 | 0 | 1.33E-05 | 0 | 7.10E-06 | 4.80E-07 | 0 | 8.71E-06 | 4.13E-06 | 0 | 1.55E-05 | 5.35E-06 | 1.23E-05 | 2.28E-06 |
| Rhbdd3 | 2.56E-06 | 3.58E-05 | 1.49E-05 | 3.42E-05 | 2.05E-05 | 1.54E-05 | 1.04E-05 | 2.66E-05 | 7.03E-06 | 1.89E-05 | 0 | 2.10E-05 | 1.95E-05 | 5.03E-06 | 8.76E-05 | 2.00E-05 | 0 | 2.18E-05 |

| | | | | | | | | | | | | | | | | | | |
|---|---|---|---|---|---|---|---|---|---|---|---|---|---|---|---|---|---|---|
| Wdr76 | 2.67E-07 | 5.09E-07 | 4.58E-06 | 0 | 0 | 1.85E-06 | 0 | 0 | 7.02E-06 | 1.13E-05 | 0 | 0 | 4.58E-07 | 1.79E-06 | 3.73E-05 | 0.00012 | 0 | 9.86E-06 |
| Slc25a40 | 2.78E-05 | 1.48E-05 | 0 | 0 | 1.94E-05 | 2.14E-05 | 1.59E-05 | 8.36E-06 | 7.01E-06 | 1.85E-05 | 1.24E-05 | 1.51E-05 | 1.41E-05 | 1.07E-05 | 2.29E-05 | 7.26E-05 | 0 | 1.52E-06 |
| Rccd1 | 1.12E-06 | 3.83E-06 | 3.38E-06 | 0 | 0 | 6.55E-06 | 0 | 0 | 7.01E-06 | 5.51E-06 | 3.47E-05 | 3.70E-06 | 0 | 2.74E-06 | 2.93E-05 | 6.30E-06 | 6.75E-06 | 1.01E-05 |
| Btbd11 | 8.01E-06 | 3.37E-05 | 8.55E-06 | 0 | 2.45E-06 | 0 | 9.02E-06 | 1.06E-05 | 7.01E-06 | 1.00E-05 | 0 | 0 | 0 | 1.19E-05 | 5.83E-06 | 2.52E-05 | 0 | 4.57E-06 |
| BC024063 | 5.00E-06 | 6.48E-06 | 0 | 0 | 1.17E-05 | 8.35E-06 | 0 | 1.49E-05 | 6.97E-06 | 9.18E-06 | 0 | 0 | 0 | 0 | 3.64E-05 | 0 | 0 | 6.82E-06 |
| Rnf114 | 0.00016 | 0.00014 | 3.89E-05 | 6.96E-05 | 6.46E-06 | 6.08E-05 | 0 | 7.47E-06 | 6.96E-06 | 4.68E-05 | 2.49E-05 | 0.00015 | 2.87E-05 | 0 | 6.37E-05 | 7.54E-05 | 6.79E-05 | 0 |
| Tff1 | 0 | 2.12E-06 | 1.54E-06 | 0 | 4.57E-06 | 0 | 0 | 2.53E-06 | 6.96E-06 | 2.54E-06 | 0 | 9.96E-06 | 0 | 0 | 4.22E-05 | 0 | 0 | 1.92E-06 |
| Ythdf2 | 1.29E-05 | 2.26E-05 | 3.60E-05 | 6.47E-06 | 5.35E-06 | 0 | 0 | 5.15E-06 | 6.95E-06 | 6.74E-06 | 1.46E-05 | 6.23E-06 | 3.62E-06 | 1.59E-05 | 0 | 1.52E-05 | 0 | 7.28E-06 |
| Ino80d | 7.34E-05 | 0.00017 | 3.22E-05 | 1.41E-05 | 5.93E-05 | 0.0001 | 6.61E-05 | 4.38E-05 | 6.93E-06 | 4.40E-05 | 8.03E-05 | 8.71E-05 | 1.99E-05 | 5.10E-05 | 0.00022 | 6.68E-05 | 0.00023 | 3.85E-05 |
| Pnmal2 | 0 | 1.56E-05 | 0 | 0 | 9.50E-06 | 1.16E-05 | 0 | 2.84E-05 | 6.93E-06 | 0 | 0 | 0 | 0 | 0 | 1.75E-06 | 0 | 0 | 2.31E-05 |
| Tent5b | 0 | 1.29E-06 | 0 | 0 | 0 | 0 | 0 | 0 | 6.92E-06 | 3.26E-05 | 7.68E-05 | 0 | 0 | 8.81E-06 | 1.78E-06 | 1.19E-05 | 1.55E-06 | 4.82E-06 |
| Crls1 | 0.00019 | 3.65E-05 | 8.00E-05 | 0 | 5.71E-06 | 5.51E-05 | 8.92E-05 | 0 | 6.90E-06 | 2.19E-05 | 0 | 8.46E-05 | 0 | 0 | 0 | 7.88E-06 | 0 | 7.63E-06 |
| Tctn1 | 0 | 7.13E-05 | 0 | 8.38E-05 | 5.07E-05 | 5.00E-05 | 1.18E-05 | 4.92E-05 | 6.90E-06 | 6.03E-05 | 0 | 2.54E-05 | 3.73E-05 | 4.95E-05 | 6.64E-05 | 0 | 8.75E-05 | 0.00011 |
| Pnn | 0.00011 | 0.00016 | 0.00018 | 0.00025 | 2.39E-05 | 0 | 0 | 7.60E-06 | 6.90E-06 | 3.28E-05 | 4.23E-05 | 7.94E-06 | 0 | 4.75E-05 | 0.00067 | 0.00025 | 0 | 9.24E-06 |
| Shank1 | 0 | 5.41E-06 | 0 | 0 | 6.28E-06 | 9.23E-07 | 0 | 0 | 6.87E-06 | 4.55E-06 | 0 | 1.16E-06 | 0 | 5.42E-06 | 0 | 0 | 4.48E-06 | 1.88E-05 |
| Ing2 | 3.67E-05 | 4.32E-05 | 1.88E-05 | 2.12E-05 | 7.94E-06 | 8.36E-06 | 0 | 7.77E-06 | 6.86E-06 | 0 | 0 | 0.0001 | 0 | 9.89E-06 | 9.06E-05 | 6.61E-05 | 9.44E-05 | 0 |
| Ccdc167 | 6.08E-05 | 1.11E-05 | 1.49E-05 | 6.90E-05 | 0 | 3.37E-06 | 1.27E-05 | 1.20E-05 | 6.86E-06 | 1.18E-05 | 1.14E-05 | 2.39E-05 | 0 | 0 | 5.57E-05 | 7.32E-05 | 0 | 2.07E-05 |
| F13a1 | 0 | 0 | 0 | 0 | 0 | 0 | 6.50E-06 | 7.40E-06 | 6.86E-06 | 3.01E-06 | 5.58E-06 | 5.24E-06 | 7.25E-06 | 4.07E-06 | 9.91E-06 | 4.64E-07 | 0 | 1.30E-06 |
| Zfp951 | 2.67E-05 | 1.71E-05 | 0 | 0 | 3.61E-06 | 1.88E-05 | 0 | 0 | 6.85E-06 | 4.52E-06 | 0 | 2.65E-05 | 0 | 2.10E-08 | 1.47E-05 | 3.74E-07 | 0 | 0 |
| Rasd1 | 0 | 2.51E-05 | 0 | 0 | 0 | 1.43E-05 | 0 | 0 | 6.84E-06 | 0 | 0 | 1.23E-05 | 0 | 0 | 0 | 0 | 0.0003 | 0 |
| Mks1 | 1.19E-05 | 2.92E-05 | 1.13E-05 | 5.68E-05 | 1.78E-06 | 1.21E-05 | 1.56E-05 | 2.24E-05 | 6.83E-06 | 1.19E-05 | 1.18E-06 | 1.07E-05 | 1.51E-05 | 0 | 3.25E-05 | 8.49E-06 | 0 | 1.09E-05 |
| Zbtb3 | 9.39E-06 | 5.22E-06 | 5.36E-06 | 1.73E-06 | 6.13E-06 | 5.56E-06 | 0 | 0 | 6.83E-06 | 2.73E-06 | 1.46E-05 | 0 | 6.70E-08 | 4.14E-06 | 4.05E-06 | 8.72E-06 | 0 | 1.14E-05 |
| H2-DMb1 | 0 | 2.77E-06 | 0 | 0 | 0 | 4.50E-06 | 7.04E-06 | 1.99E-06 | 6.83E-06 | 1.16E-05 | 1.26E-05 | 6.51E-06 | 2.86E-05 | 7.44E-06 | 5.94E-06 | 2.13E-06 | 5.56E-07 | 1.16E-05 |
| Slc9a4 | 2.25E-07 | 0 | 7.74E-06 | 0 | 0 | 4.32E-06 | 6.44E-06 | 1.20E-05 | 6.83E-06 | 3.26E-06 | 0 | 2.12E-06 | 0 | 0 | 0 | 0 | 0 | 0 |
| Glyctk | 0 | 5.39E-06 | 1.51E-06 | 2.71E-06 | 3.16E-06 | 0 | 1.46E-06 | 0 | 6.82E-06 | 1.38E-06 | 2.02E-06 | 3.36E-07 | 4.28E-06 | 9.90E-06 | 1.76E-05 | 3.75E-06 | 0 | 1.21E-05 |
| Pcdhb12 | 0 | 1.64E-07 | 0 | 0 | 5.23E-06 | 0 | 9.26E-06 | 6.96E-06 | 6.80E-06 | 8.67E-06 | 5.06E-06 | 0 | 5.30E-06 | 7.52E-06 | 2.87E-07 | 0 | 0 | 7.67E-06 |
| Pex11g | 0 | 2.19E-05 | 5.23E-06 | 7.24E-05 | 1.56E-05 | 9.70E-06 | 2.80E-05 | 2.53E-05 | 6.78E-06 | 1.39E-05 | 1.39E-05 | 3.95E-05 | 1.34E-05 | 1.90E-05 | 4.29E-05 | 0 | 0 | 2.16E-05 |
| Ccdc170 | 0 | 0 | 0 | 0 | 0 | 1.30E-05 | 0 | 2.39E-06 | 6.76E-06 | 4.85E-05 | 0 | 0 | 0 | 0 | 0 | 0 | 0 | 2.16E-05 |
| Zfp383 | 3.19E-05 | 9.50E-06 | 0 | 0 | 5.72E-06 | 6.59E-06 | 3.94E-05 | 0 | 6.75E-06 | 1.79E-05 | 9.48E-07 | 8.62E-07 | 1.41E-06 | 0 | 4.62E-05 | 2.47E-06 | 0 | 6.93E-06 |
| Zfp3 | 1.32E-06 | 7.08E-06 | 0 | 4.13E-06 | 7.39E-06 | 0 | 0 | 0 | 6.75E-06 | 1.08E-05 | 0 | 2.77E-06 | 3.10E-06 | 0 | 1.91E-05 | 0 | 0 | 1.46E-05 |
| Lck | 1.60E-06 | 0 | 1.34E-06 | 0 | 7.20E-07 | 3.62E-06 | 2.49E-06 | 8.55E-06 | 6.73E-06 | 0 | 0 | 5.19E-06 | 8.90E-06 | 0 | 0 | 1.95E-06 | 0 | 0 |
| Ccdc166 | 6.47E-07 | 1.55E-06 | 1.84E-06 | 1.01E-06 | 1.77E-06 | 1.56E-07 | 0 | 4.84E-06 | 6.73E-06 | 7.13E-06 | 6.18E-07 | 0 | 4.47E-06 | 7.53E-06 | 2.17E-05 | 0 | 1.83E-07 | 0 |
| Hrct1 | 7.50E-08 | 1.12E-07 | 0 | 8.23E-06 | 0 | 0 | 0 | 1.12E-05 | 6.72E-06 | 5.59E-06 | 0 | 0 | 0 | 2.77E-05 | 0 | 4.27E-07 | 0 | 1.71E-05 |

| | | | | | | | | | | | | | | | | | | |
|---|---|---|---|---|---|---|---|---|---|---|---|---|---|---|---|---|---|---|
| Acadvl | 0.00012 | 7.57E-05 | 0.00013 | 9.26E-05 | 0.00012 | 9.31E-05 | 0.00013 | 3.25E-05 | 6.71E-06 | 0.00013 | 0.00013 | 3.42E-05 | 0.00012 | 0.00014 | 7.06E-05 | 0 | 0 | 0.00013 |
| Tnfrsf1b | 0 | 0.00012 | 5.17E-05 | 0 | 3.01E-05 | 8.57E-06 | 4.73E-05 | 3.76E-05 | 6.69E-06 | 0.00025 | 1.90E-05 | 2.73E-05 | 0.00026 | 0.00013 | 3.08E-05 | 2.19E-05 | 0 | 0.00023 |
| Ogfod2 | 2.46E-05 | 4.46E-05 | 1.55E-05 | 6.23E-05 | 3.41E-05 | 1.86E-05 | 3.15E-05 | 2.07E-05 | 6.68E-06 | 1.68E-05 | 7.10E-06 | 2.98E-05 | 0 | 3.38E-05 | 1.62E-05 | 3.96E-06 | 0 | 4.27E-05 |
| Pacc1 | 1.79E-05 | 3.66E-05 | 3.63E-06 | 3.95E-06 | 2.19E-05 | 0 | 1.41E-05 | 1.49E-05 | 6.67E-06 | 4.91E-06 | 2.91E-05 | 2.87E-05 | 0 | 1.56E-05 | 0 | 1.69E-05 | 9.34E-06 | 0 |
| Cd320 | 0.00011 | 2.94E-05 | 6.34E-05 | 0.00011 | 1.49E-05 | 3.43E-05 | 8.22E-05 | 4.11E-07 | 6.66E-06 | 0.00013 | 0 | 1.90E-05 | 0.00046 | 0 | 0 | 4.01E-05 | 4.01E-05 | 0 |
| Ago4 | 1.72E-06 | 1.29E-05 | 0 | 0 | 0 | 1.56E-05 | 0 | 0 | 6.66E-06 | 7.14E-06 | 3.81E-06 | 1.90E-05 | 0 | 3.89E-06 | 1.20E-05 | 0 | 2.58E-06 | 1.48E-05 |
| Ttc26 | 6.75E-05 | 0 | 3.01E-05 | 2.86E-05 | 6.62E-06 | 1.93E-05 | 8.92E-05 | 8.07E-06 | 6.65E-06 | 0 | 0 | 0 | 0 | 9.52E-06 | 2.16E-05 | 0 | 0 | 0 |
| Adam11 | 1.91E-06 | 1.25E-05 | 0 | 3.49E-06 | 5.52E-06 | 1.47E-06 | 0 | 6.14E-06 | 6.65E-06 | 1.04E-06 | 6.10E-07 | 1.47E-06 | 0 | 3.96E-06 | 0 | 0 | 0 | 0 |
| Abat | 5.40E-06 | 1.10E-05 | 0 | 0 | 4.98E-05 | 0 | 0 | 0 | 6.64E-06 | 6.81E-06 | 8.48E-06 | 0 | 1.71E-05 | 1.21E-05 | 0 | 0 | 1.60E-05 | 2.15E-05 |
| Hacl1 | 2.06E-05 | 3.70E-05 | 0 | 1.03E-05 | 1.47E-05 | 1.75E-05 | 0 | 2.93E-07 | 6.63E-06 | 1.21E-05 | 0 | 1.00E-05 | 7.35E-06 | 9.10E-06 | 0 | 1.06E-05 | 2.22E-05 | 2.96E-05 |
| Pcdhb9 | 0 | 1.17E-06 | 0 | 0 | 1.09E-05 | 3.92E-06 | 5.13E-06 | 1.61E-06 | 6.63E-06 | 1.25E-05 | 0 | 3.86E-06 | 2.61E-05 | 1.61E-05 | 0 | 0 | 0 | 2.28E-05 |
| Kidins220 | 0.00013 | 0.0002 | 4.29E-05 | 0 | 0.00016 | 8.82E-05 | 0.00019 | 0 | 6.61E-06 | 0.00015 | 0.00015 | 0.00012 | 1.97E-05 | 5.77E-05 | 0.00026 | 2.18E-05 | 1.41E-05 | 0.00015 |
| Epg5 | 8.41E-05 | 8.68E-05 | 6.20E-05 | 0 | 7.88E-05 | 4.72E-05 | 4.77E-05 | 0 | 6.61E-06 | 2.01E-05 | 1.91E-05 | 0.00014 | 3.24E-06 | 1.13E-05 | 7.64E-05 | 6.53E-06 | 0 | 7.03E-06 |
| Gins1 | 5.26E-06 | 0 | 3.07E-05 | 0 | 0 | 0 | 2.31E-06 | 0 | 6.61E-06 | 0 | 1.29E-06 | 0 | 2.42E-05 | 2.40E-05 | 0 | 0.00031 | 0 | 0 |
| Frem1 | 0 | 6.21E-05 | 0 | 0 | 1.50E-05 | 0 | 0 | 0.00044 | 6.60E-06 | 0 | 0 | 0 | 0 | 0 | 0 | 0 | 0 | 3.74E-05 |
| Rusc1 | 2.64E-05 | 2.34E-05 | 3.01E-05 | 9.86E-05 | 9.39E-06 | 5.70E-06 | 0.00013 | 4.39E-05 | 6.57E-06 | 4.02E-05 | 6.42E-05 | 3.02E-05 | 6.06E-06 | 1.37E-05 | 3.08E-05 | 0 | 3.56E-05 | 5.66E-05 |
| Fam205a2 | 6.29E-06 | 2.26E-06 | 0 | 0 | 0 | 3.18E-06 | 8.55E-06 | 0 | 6.57E-06 | 3.48E-05 | 0 | 6.49E-06 | 4.44E-05 | 0 | 0 | 0 | 4.45E-06 | 0 |
| Rab3il1 | 2.18E-05 | 0.00025 | 0 | 0.00031 | 0.00015 | 0 | 0.00028 | 8.17E-05 | 6.56E-06 | 0.00047 | 0 | 0 | 0 | 0.00016 | 0 | 3.11E-05 | 0 | 0.00034 |
| Mtcl1 | 0 | 2.98E-05 | 0 | 3.08E-05 | 3.01E-05 | 6.23E-05 | 0 | 0.00018 | 6.55E-06 | 0 | 0 | 9.07E-06 | 0 | 1.09E-06 | 0.0001 | 0 | 3.91E-05 | 0 |
| Zscan22 | 3.61E-05 | 2.40E-05 | 7.96E-06 | 0 | 1.42E-05 | 1.76E-05 | 3.13E-06 | 1.21E-06 | 6.54E-06 | 1.12E-06 | 7.13E-06 | 1.84E-05 | 0 | 7.32E-06 | 2.57E-05 | 1.72E-05 | 0 | 6.42E-06 |
| Tceal5 | 0 | 0 | 0 | 0 | 9.39E-06 | 0 | 1.63E-06 | 0 | 6.54E-06 | 2.33E-06 | 1.60E-05 | 0 | 6.22E-06 | 0 | 0 | 4.58E-06 | 0 | 1.13E-05 |
| Ppp2r3a | 4.77E-05 | 2.87E-05 | 1.69E-05 | 0 | 2.46E-05 | 4.84E-05 | 5.57E-05 | 0 | 6.52E-06 | 3.08E-05 | 8.91E-05 | 4.31E-05 | 2.85E-05 | 2.70E-05 | 1.24E-05 | 3.72E-05 | 2.66E-05 | 6.19E-05 |
| Slc44a3 | 2.99E-06 | 6.45E-06 | 0 | 1.79E-06 | 0 | 1.40E-05 | 0 | 0 | 6.52E-06 | 4.02E-06 | 0 | 4.74E-07 | 9.30E-06 | 1.59E-06 | 0 | 1.12E-05 | 0 | 1.25E-05 |
| Chchd6 | 2.22E-05 | 6.94E-06 | 3.76E-05 | 4.37E-05 | 2.63E-05 | 0 | 2.34E-06 | 5.14E-06 | 6.52E-06 | 6.33E-05 | 0 | 3.61E-05 | 4.50E-06 | 0 | 0 | 4.88E-05 | 1.35E-05 | 3.09E-05 |
| Crispld1 | 0 | 2.03E-05 | 0 | 0 | 0 | 5.37E-07 | 0 | 0 | 6.51E-06 | 6.20E-05 | 0 | 0 | 0 | 0 | 0 | 0 | 0 | 0 |
| Zbtb2 | 4.85E-05 | 6.78E-05 | 5.46E-05 | 0 | 0 | 1.63E-05 | 0 | 2.71E-05 | 6.49E-06 | 3.97E-06 | 2.98E-05 | 0 | 0 | 3.11E-05 | 2.75E-05 | 3.15E-05 | 0.00021 | 0 |
| Timm29 | 0 | 3.08E-06 | 7.16E-06 | 2.01E-05 | 2.45E-08 | 3.13E-07 | 8.19E-06 | 3.90E-06 | 6.49E-06 | 1.07E-05 | 0 | 7.94E-07 | 2.52E-07 | 0 | 3.63E-06 | 3.11E-06 | 1.10E-05 | 0 |
| Slc37a2 | 0 | 5.46E-06 | 1.33E-05 | 0 | 1.37E-05 | 1.61E-05 | 1.56E-05 | 2.70E-05 | 6.47E-06 | 0 | 0 | 0 | 0 | 6.45E-06 | 1.54E-05 | 3.24E-05 | 0 | 1.78E-05 |
| Tial1 | 0.0003 | 0.00017 | 0.00022 | 3.68E-05 | 0.00014 | 0.00012 | 0.00024 | 0.00011 | 6.46E-06 | 3.30E-05 | 1.36E-05 | 0.00015 | 1.93E-06 | 6.31E-05 | 0.00058 | 0.00027 | 0.00028 | 2.19E-05 |
| Ankrd34a | 0 | 0 | 3.16E-05 | 8.42E-05 | 9.04E-05 | 7.55E-05 | 0 | 0 | 6.46E-06 | 0 | 8.66E-05 | 0 | 1.76E-05 | 3.48E-05 | 4.06E-05 | 0 | 0 | 0 |
| G2e3 | 3.97E-05 | 5.83E-05 | 0 | 0 | 3.30E-05 | 2.79E-05 | 2.81E-05 | 0 | 6.45E-06 | 1.82E-05 | 0 | 0.00021 | 1.51E-05 | 0 | 2.07E-05 | 0.00048 | 0 | 0 |
| Ift57 | 8.30E-05 | 6.74E-05 | 6.21E-05 | 1.67E-05 | 6.80E-05 | 0.00011 | 2.04E-05 | 8.94E-07 | 6.43E-06 | 7.96E-05 | 0 | 7.46E-05 | 3.41E-05 | 5.21E-05 | 3.01E-05 | 3.50E-05 | 6.52E-05 | 3.63E-05 |
| Tfdp2 | 4.96E-05 | 6.42E-05 | 0 | 0 | 3.93E-05 | 7.28E-05 | 1.90E-05 | 1.28E-05 | 6.42E-06 | 0 | 0 | 2.28E-05 | 4.56E-06 | 5.99E-06 | 3.62E-05 | 0.00014 | 0 | 8.04E-06 |

| | | | | | | | | | | | | | | | | | | |
|---|---|---|---|---|---|---|---|---|---|---|---|---|---|---|---|---|---|---|
| Apoo | 3.21E-05 | 0 | 9.03E-06 | 0 | 0 | 1.54E-05 | 6.43E-07 | 0 | 6.42E-06 | 0 | 3.08E-06 | 2.16E-06 | 0 | 2.77E-06 | 0 | 1.50E-05 | 0 | 1.50E-06 |
| Glce | 0.00015 | 6.72E-05 | 9.63E-05 | 0 | 0 | 4.15E-05 | 0 | 0 | 6.40E-06 | 6.75E-05 | 0 | 0.00015 | 0 | 2.20E-05 | 0 | 3.85E-05 | 0 | 0 |
| Cep295 | 3.68E-05 | 3.74E-05 | 1.46E-05 | 0 | 2.00E-05 | 2.53E-05 | 0 | 0 | 6.40E-06 | 1.27E-05 | 5.48E-07 | 2.16E-05 | 0 | 1.47E-05 | 0.00014 | 0.00029 | 0 | 0 |
| Nubpl | 1.49E-05 | 9.80E-06 | 0 | 2.41E-06 | 5.23E-06 | 2.62E-05 | 9.66E-06 | 1.11E-05 | 6.37E-06 | 0 | 1.45E-05 | 1.17E-05 | 5.17E-06 | 1.28E-05 | 4.55E-06 | 1.12E-05 | 0 | 1.14E-05 |
| Rab33a | 3.72E-09 | 0 | 0 | 2.64E-06 | 7.88E-06 | 0 | 9.41E-06 | 0 | 6.36E-06 | 0 | 1.05E-05 | 6.41E-05 | 0 | 0 | 0 | 1.87E-07 | 0 | 0 |
| Agtr1a | 0 | 0 | 0 | 0 | 0 | 0 | 0 | 0 | 6.36E-06 | 0.00029 | 6.70E-06 | 0 | 0.00044 | 0 | 0 | 0 | 0 | 6.86E-05 |
| Spice1 | 1.40E-05 | 1.67E-05 | 0 | 0 | 1.22E-06 | 1.62E-05 | 0 | 0 | 6.35E-06 | 1.63E-05 | 1.08E-05 | 2.69E-05 | 0 | 5.57E-06 | 8.79E-06 | 3.13E-05 | 1.01E-05 | 1.77E-05 |
| Mpv17l | 1.37E-05 | 8.05E-06 | 0 | 0 | 9.24E-06 | 1.13E-05 | 1.19E-05 | 7.33E-06 | 6.35E-06 | 1.30E-06 | 1.10E-05 | 0 | 1.19E-05 | 2.16E-05 | 0 | 0 | 0 | 1.46E-05 |
| Ddx60 | 1.82E-05 | 1.17E-05 | 0 | 0 | 0 | 7.36E-06 | 0 | 0 | 6.34E-06 | 3.67E-05 | 0 | 4.49E-06 | 1.40E-05 | 0 | 0 | 0 | 0 | 4.12E-05 |
| Engase | 0 | 4.43E-06 | 5.91E-06 | 3.44E-05 | 8.06E-06 | 3.74E-06 | 0 | 6.62E-06 | 6.33E-06 | 6.05E-05 | 0 | 0 | 0 | 7.41E-06 | 0 | 0 | 0 | 1.28E-05 |
| Lmo7 | 0 | 8.67E-06 | 0 | 0 | 3.41E-05 | 0 | 0 | 0 | 6.33E-06 | 0 | 6.90E-05 | 0 | 0 | 0.00012 | 5.20E-06 | 3.86E-05 | 0 | 1.79E-05 |
| Smpd5 | 0 | 1.65E-06 | 0 | 1.90E-06 | 2.90E-06 | 4.40E-07 | 0 | 2.46E-06 | 6.30E-06 | 6.30E-07 | 7.45E-06 | 7.06E-06 | 4.39E-07 | 3.15E-05 | 5.06E-06 | 0 | 0 | 1.61E-05 |
| Slc2a10 | 0 | 1.59E-08 | 9.89E-05 | 0.00019 | 8.19E-06 | 1.99E-05 | 0.00011 | 7.17E-05 | 6.29E-06 | 4.27E-05 | 4.19E-05 | 4.55E-06 | 1.36E-05 | 6.45E-05 | 7.01E-05 | 0 | 7.74E-05 | 3.74E-05 |
| Pemt | 2.52E-05 | 9.21E-06 | 3.60E-05 | 2.77E-05 | 0 | 0 | 2.88E-06 | 0 | 6.29E-06 | 2.83E-06 | 0 | 9.68E-07 | 1.88E-05 | 2.00E-06 | 0 | 6.45E-06 | 1.88E-05 | 1.09E-05 |
| Atg16l2 | 4.95E-06 | 5.01E-05 | 1.50E-05 | 0 | 5.51E-05 | 6.38E-05 | 1.31E-05 | 3.78E-05 | 6.28E-06 | 0 | 2.05E-05 | 1.76E-05 | 0 | 4.07E-05 | 0.00013 | 6.17E-06 | 0 | 6.79E-06 |
| Ppp2r3c | 7.02E-05 | 4.05E-05 | 1.18E-05 | 3.65E-06 | 1.74E-05 | 2.54E-05 | 1.15E-05 | 0 | 6.26E-06 | 2.51E-05 | 1.17E-05 | 5.93E-05 | 0 | 0 | 0 | 2.44E-05 | 0 | 6.36E-06 |
| Bfsp1 | 2.75E-05 | 6.28E-06 | 8.69E-06 | 3.00E-05 | 4.47E-06 | 2.28E-05 | 5.31E-05 | 1.25E-05 | 6.26E-06 | 2.76E-05 | 0 | 5.12E-06 | 8.48E-06 | 0 | 0 | 9.08E-06 | 1.10E-05 | 1.30E-05 |
| Orc2 | 8.28E-05 | 3.09E-05 | 6.89E-05 | 0 | 4.39E-06 | 3.68E-05 | 0 | 0 | 6.25E-06 | 0 | 3.63E-05 | 0 | 8.38E-06 | 0 | 0 | 0.0001 | 3.49E-05 | 0 |
| Eya3 | 0.0001 | 6.23E-05 | 5.10E-05 | 0 | 3.22E-05 | 4.91E-05 | 0 | 0 | 6.22E-06 | 4.93E-05 | 7.89E-06 | 4.95E-05 | 0.0001 | 1.06E-05 | 0.00012 | 6.46E-05 | 1.25E-05 | 2.67E-05 |
| Naa16 | 3.48E-05 | 6.62E-05 | 5.30E-05 | 2.63E-05 | 1.80E-05 | 9.65E-06 | 0 | 0 | 6.22E-06 | 0 | 0 | 0.00016 | 0 | 3.80E-05 | 0.00013 | 0.0001 | 0 | 1.75E-05 |
| Cav3 | 0 | 0 | 5.44E-06 | 1.22E-07 | 0 | 1.20E-06 | 0 | 5.50E-06 | 6.22E-06 | 5.36E-07 | 2.11E-05 | 0 | 1.30E-05 | 1.12E-06 | 0 | 1.81E-06 | 0 | 2.37E-05 |
| Creg2 | 0 | 1.19E-05 | 0 | 0 | 1.55E-05 | 5.50E-06 | 0 | 2.31E-05 | 6.21E-06 | 0 | 0 | 9.23E-06 | 0 | 0 | 0 | 0 | 0 | 0 |
| Slx1b | 4.47E-06 | 5.48E-06 | 3.93E-07 | 1.01E-05 | 8.90E-07 | 0 | 0 | 0 | 6.21E-06 | 1.13E-06 | 5.03E-08 | 0 | 0 | 2.11E-06 | 7.64E-06 | 0 | 0 | 2.72E-06 |
| H2-T23 | 0 | 1.68E-05 | 0 | 0 | 9.78E-06 | 2.03E-05 | 1.89E-05 | 1.74E-05 | 6.20E-06 | 4.07E-05 | 0 | 4.14E-05 | 1.99E-05 | 0 | 0 | 0 | 0 | 4.24E-05 |
| Rbak | 3.63E-05 | 4.78E-05 | 7.67E-06 | 8.41E-06 | 3.12E-05 | 1.90E-05 | 0 | 1.03E-05 | 6.19E-06 | 5.65E-06 | 0 | 3.10E-05 | 1.20E-06 | 3.69E-05 | 0.00013 | 0 | 0 | 1.06E-05 |
| Ltb4r1 | 0 | 4.13E-06 | 3.59E-06 | 0 | 1.37E-07 | 4.32E-06 | 0 | 0 | 6.19E-06 | 0 | 6.36E-07 | 1.54E-06 | 8.78E-07 | 0 | 1.07E-05 | 2.80E-06 | 2.00E-05 | 0 |
| Grb14 | 8.57E-06 | 0 | 0 | 0 | 0 | 0 | 0 | 0 | 6.18E-06 | 0 | 0 | 0 | 4.08E-05 | 3.88E-06 | 0 | 0 | 0 | 6.87E-05 |
| Iba57 | 3.80E-06 | 3.29E-05 | 1.85E-05 | 3.73E-05 | 1.47E-05 | 0 | 1.83E-06 | 3.59E-06 | 6.17E-06 | 1.02E-05 | 1.40E-05 | 2.24E-05 | 0 | 2.09E-05 | 0 | 1.89E-06 | 0 | 1.25E-06 |
| Tigar | 1.26E-05 | 6.13E-06 | 2.71E-05 | 9.60E-06 | 0 | 0 | 7.25E-06 | 0 | 6.17E-06 | 2.30E-05 | 1.49E-05 | 0 | 3.42E-05 | 2.08E-06 | 0 | 0 | 0 | 1.68E-05 |
| Mfap1a | 2.30E-05 | 2.13E-05 | 2.47E-06 | 0 | 1.94E-05 | 1.28E-05 | 3.06E-06 | 7.66E-06 | 6.15E-06 | 7.91E-06 | 4.71E-06 | 3.62E-06 | 3.19E-06 | 0 | 0 | 1.87E-05 | 0 | 1.26E-05 |
| Prox2 | 1.86E-06 | 6.79E-06 | 0 | 0 | 2.76E-06 | 3.47E-06 | 0 | 3.17E-06 | 6.15E-06 | 2.79E-06 | 0 | 1.83E-06 | 0 | 2.71E-06 | 0 | 8.30E-06 | 0 | 3.00E-06 |
| Neil2 | 0 | 3.09E-05 | 0 | 1.32E-06 | 1.30E-06 | 9.97E-06 | 0 | 4.04E-06 | 6.14E-06 | 5.56E-06 | 0 | 8.72E-06 | 0 | 0 | 2.81E-05 | 0 | 0 | 0 |
| Tubg2 | 3.60E-06 | 4.80E-06 | 0 | 0 | 5.24E-06 | 1.53E-05 | 0 | 4.30E-06 | 6.13E-06 | 1.33E-05 | 4.80E-06 | 5.87E-06 | 1.16E-05 | 1.64E-05 | 0 | 0 | 0 | 5.45E-06 |

| | | | | | | | | | | | | | | | | | | |
|---|---|---|---|---|---|---|---|---|---|---|---|---|---|---|---|---|---|---|
| Brd2 | 0.00022 | 0.00033 | 0.00013 | 0.00019 | 9.12E-05 | 0.00016 | 4.44E-05 | 1.52E-05 | 6.11E-06 | 0 | 0.00011 | 0.00034 | 2.96E-05 | 1.23E-05 | 0.00023 | 0.00014 | 0.00175 | 0 |
| Heph | 6.02E-06 | 6.15E-06 | 0 | 4.22E-06 | 1.52E-05 | 0 | 0 | 0 | 6.09E-06 | 4.44E-05 | 0 | 0 | 1.50E-05 | 3.43E-06 | 0 | 3.41E-05 | 0 | 6.68E-05 |
| Gtpbp2 | 3.14E-06 | 0.00021 | 0 | 0.00015 | 0.00013 | 9.61E-05 | 0.00014 | 0.00012 | 6.08E-06 | 8.03E-05 | 4.93E-05 | 0.00021 | 1.16E-05 | 0.00011 | 0.00046 | 0.00024 | 0.00013 | 8.55E-05 |
| R3hdm1 | 0.00021 | 0.00011 | 9.71E-05 | 0 | 4.93E-05 | 4.22E-05 | 0.0001 | 5.65E-05 | 6.04E-06 | 2.22E-05 | 0.00011 | 0.00016 | 0 | 0 | 0.00023 | 0.00012 | 1.73E-05 | 2.47E-05 |
| St3gal5 | 1.86E-05 | 2.00E-05 | 0 | 0 | 7.21E-05 | 2.40E-06 | 1.84E-05 | 0 | 6.04E-06 | 0.00015 | 5.41E-05 | 0 | 0 | 4.98E-05 | 0 | 9.62E-05 | 0 | 0 |
| Ptger2 | 0 | 5.49E-06 | 8.69E-06 | 0 | 1.59E-05 | 2.75E-05 | 0 | 6.03E-06 | 6.03E-06 | 0 | 1.18E-06 | 0 | 0 | 2.77E-06 | 1.98E-05 | 1.35E-06 | 0 | 8.49E-06 |
| Zgrf1 | 7.97E-06 | 0 | 1.55E-05 | 0 | 0 | 1.21E-05 | 0 | 0 | 6.00E-06 | 4.51E-06 | 0 | 0 | 0 | 4.44E-06 | 1.13E-05 | 0.00018 | 0 | 0 |
| Atad5 | 4.62E-06 | 0 | 1.84E-05 | 0 | 1.73E-06 | 8.97E-06 | 0 | 0 | 6.00E-06 | 7.12E-06 | 0 | 0 | 1.42E-05 | 1.20E-05 | 1.97E-05 | 0.00022 | 0 | 0 |
| Chaf1a | 0 | 0 | 8.67E-05 | 5.59E-05 | 0 | 0 | 0 | 0 | 5.99E-06 | 1.11E-05 | 0 | 0 | 1.41E-05 | 1.32E-05 | 9.18E-05 | 0.00052 | 0 | 0 |
| Arih2 | 8.46E-05 | 0.00022 | 0.00021 | 0.0001 | 0.00011 | 5.16E-05 | 0 | 3.07E-05 | 5.98E-06 | 2.32E-05 | 7.55E-05 | 0.00029 | 1.54E-05 | 5.55E-05 | 0.00032 | 9.59E-05 | 1.74E-05 | 1.46E-05 |
| Mmgt2 | 7.03E-05 | 8.00E-05 | 4.19E-05 | 6.75E-05 | 5.85E-05 | 2.57E-05 | 8.68E-05 | 8.40E-05 | 5.94E-06 | 5.79E-06 | 2.52E-05 | 0 | 2.09E-05 | 2.45E-05 | 0 | 3.03E-05 | 0 | 5.87E-05 |
| Myrf | 1.13E-06 | 7.16E-06 | 9.29E-06 | 6.60E-06 | 2.67E-06 | 1.45E-06 | 0 | 4.09E-06 | 5.94E-06 | 1.04E-05 | 7.80E-06 | 3.60E-06 | 1.46E-05 | 4.11E-06 | 2.59E-05 | 3.04E-06 | 1.73E-05 | 4.10E-06 |
| Mgarp | 0 | 0 | 0 | 5.10E-05 | 0 | 0 | 9.24E-05 | 0 | 5.94E-06 | 0.00013 | 0 | 0.00075 | 0.00013 | 0 | 0 | 0 | 0 | 0 |
| Acacb | 3.83E-06 | 8.03E-06 | 0 | 0 | 1.02E-05 | 1.18E-05 | 0 | 3.00E-06 | 5.91E-06 | 1.64E-06 | 0 | 1.03E-05 | 0 | 0 | 1.06E-05 | 0 | 1.80E-07 | 4.34E-05 |
| Kcnn3 | 0 | 8.91E-06 | 0 | 0 | 4.36E-05 | 3.20E-06 | 0 | 1.40E-05 | 5.89E-06 | 1.40E-05 | 0 | 8.21E-06 | 0 | 0 | 2.21E-06 | 0 | 0 | 0 |
| Zfp943 | 1.21E-06 | 2.77E-05 | 1.02E-05 | 3.99E-05 | 2.08E-05 | 2.50E-05 | 4.47E-05 | 3.34E-05 | 5.83E-06 | 1.98E-05 | 5.53E-06 | 5.37E-06 | 0 | 2.83E-05 | 1.09E-05 | 2.88E-05 | 0 | 2.29E-05 |
| Kpna3 | 0.00021 | 8.25E-05 | 8.58E-05 | 0 | 6.22E-05 | 0 | 1.65E-05 | 6.69E-05 | 5.83E-06 | 5.34E-07 | 2.45E-05 | 5.70E-05 | 0 | 2.00E-05 | 8.62E-06 | 0.00021 | 4.73E-05 | 0 |
| Tmem59l | 0 | 0 | 3.32E-05 | 0 | 4.29E-05 | 0 | 1.32E-05 | 5.91E-05 | 5.82E-06 | 0 | 0 | 0 | 0 | 0 | 9.86E-06 | 0 | 0 | 2.04E-05 |
| Scly | 3.31E-05 | 0.00012 | 0 | 0.0001 | 5.42E-05 | 4.03E-05 | 0 | 8.89E-05 | 5.81E-06 | 2.84E-05 | 3.49E-05 | 1.01E-05 | 4.09E-05 | 5.91E-05 | 1.24E-06 | 7.63E-06 | 0 | 1.29E-05 |
| Clba1 | 1.66E-05 | 4.48E-05 | 3.74E-05 | 1.28E-05 | 3.19E-05 | 2.02E-05 | 1.51E-05 | 0 | 5.81E-06 | 7.79E-06 | 1.20E-05 | 0 | 0 | 4.74E-05 | 0 | 0 | 0 | 1.60E-05 |
| Scamp1 | 5.02E-05 | 0.00011 | 4.19E-05 | 7.29E-06 | 8.26E-05 | 0.00011 | 2.97E-05 | 1.36E-05 | 5.79E-06 | 3.92E-05 | 1.25E-05 | 7.09E-05 | 4.76E-05 | 4.34E-05 | 6.65E-06 | 0 | 0 | 8.89E-05 |
| Tmem161b | 5.58E-05 | 5.18E-05 | 1.89E-05 | 0 | 2.87E-05 | 5.95E-05 | 2.47E-05 | 2.47E-05 | 5.77E-06 | 0 | 1.40E-05 | 1.38E-05 | 8.32E-06 | 4.10E-06 | 7.41E-05 | 3.04E-05 | 7.19E-06 | 1.65E-05 |
| Pcdhgb1 | 1.82E-05 | 2.23E-05 | 0 | 0 | 2.81E-05 | 4.19E-05 | 1.85E-06 | 2.15E-05 | 5.77E-06 | 7.84E-06 | 0 | 2.66E-05 | 1.59E-05 | 2.30E-05 | 3.29E-05 | 0 | 0 | 0 |
| Fcgr3 | 0 | 9.26E-06 | 1.50E-05 | 2.31E-05 | 3.18E-05 | 0 | 1.16E-05 | 7.19E-06 | 5.74E-06 | 8.12E-06 | 1.20E-05 | 3.72E-06 | 3.13E-05 | 0 | 0.0001 | 1.39E-05 | 0 | 9.71E-06 |
| Cdc14b | 2.77E-05 | 3.73E-05 | 5.91E-06 | 0 | 0 | 4.16E-05 | 3.87E-06 | 0 | 5.73E-06 | 3.41E-05 | 1.47E-06 | 3.15E-05 | 0 | 4.02E-05 | 1.47E-05 | 4.15E-06 | 2.56E-05 | 4.80E-05 |
| Slc25a42 | 0 | 1.82E-05 | 0 | 0 | 1.05E-05 | 4.55E-06 | 0 | 0 | 5.73E-06 | 3.44E-05 | 5.32E-06 | 7.43E-07 | 1.12E-05 | 9.87E-06 | 1.50E-05 | 2.49E-06 | 0 | 1.88E-05 |
| Acss1 | 0 | 1.42E-05 | 0 | 0 | 3.61E-06 | 6.17E-06 | 0 | 0 | 5.72E-06 | 0 | 0 | 6.81E-06 | 0 | 5.93E-06 | 1.35E-05 | 0 | 0 | 0 |
| Rab19 | 1.76E-06 | 0 | 0 | 6.39E-06 | 0 | 1.16E-07 | 0 | 1.64E-06 | 5.71E-06 | 1.86E-05 | 0 | 0 | 0 | 0 | 0 | 0 | 0 | 1.92E-05 |
| Rpp25 | 0 | 0 | 0 | 1.15E-05 | 1.55E-05 | 0 | 1.38E-05 | 4.34E-06 | 5.71E-06 | 0 | 4.97E-06 | 0 | 0 | 0 | 0 | 4.50E-06 | 0 | 0 |
| Rtl5 | 0 | 0 | 9.55E-06 | 1.75E-05 | 0 | 0 | 0 | 5.16E-06 | 5.70E-06 | 1.33E-05 | 0 | 2.77E-05 | 3.66E-06 | 0 | 0 | 0 | 0 | 2.14E-06 |
| Gng5 | 0.00022 | 1.81E-05 | 2.99E-05 | 0 | 1.30E-06 | 0.00013 | 4.28E-05 | 3.57E-05 | 5.68E-06 | 2.02E-05 | 0 | 0.00012 | 0 | 0 | 0 | 1.84E-05 | 5.01E-05 | 3.73E-06 |
| Parp2 | 6.34E-05 | 4.33E-05 | 4.94E-05 | 2.07E-05 | 1.73E-05 | 6.57E-05 | 1.68E-05 | 2.50E-05 | 5.68E-06 | 3.94E-05 | 2.93E-05 | 3.33E-05 | 0 | 8.16E-06 | 0.00016 | 0.00023 | 0 | 3.69E-05 |
| Zfp219 | 0 | 5.06E-05 | 9.03E-06 | 1.87E-05 | 1.71E-05 | 1.91E-05 | 0 | 0 | 5.67E-06 | 2.55E-05 | 1.13E-05 | 1.35E-05 | 1.13E-05 | 0 | 5.40E-05 | 2.97E-06 | 7.61E-06 | 2.17E-05 |

| | | | | | | | | | | | | | | | | | | |
|---|---|---|---|---|---|---|---|---|---|---|---|---|---|---|---|---|---|---|
| Chst10 | 4.18E-06 | 0 | 0 | 0 | 0 | 6.34E-07 | 1.69E-05 | 0 | 5.65E-06 | 8.13E-06 | 0 | 0 | 0 | 2.67E-05 | 5.27E-08 | 3.22E-06 | 0 | 1.06E-05 |
| Lsm5 | 5.71E-05 | 1.52E-06 | 2.92E-05 | 2.59E-05 | 0 | 1.43E-05 | 0 | 8.12E-06 | 5.64E-06 | 0 | 0 | 6.03E-06 | 2.05E-06 | 0 | 0 | 0.00013 | 0 | 5.90E-08 |
| DEPRECATED | 1.99E-06 | 0 | 0 | 4.93E-06 | 3.94E-06 | 0 | 0 | 1.00E-05 | 5.64E-06 | 0 | 1.64E-05 | 6.46E-06 | 0 | 2.03E-05 | 0 | 1.08E-05 | 7.96E-06 | 2.56E-06 |
| Zfp267 | 5.70E-05 | 3.80E-05 | 2.06E-05 | 2.89E-05 | 2.46E-05 | 3.65E-05 | 3.38E-05 | 2.46E-05 | 5.63E-06 | 2.11E-05 | 3.22E-06 | 1.12E-05 | 0 | 8.66E-06 | 2.09E-05 | 5.45E-06 | 0 | 4.11E-05 |
| Mpeg1 | 1.35E-06 | 8.86E-06 | 0 | 0 | 1.60E-06 | 9.48E-06 | 0 | 0 | 5.63E-06 | 6.82E-05 | 5.49E-06 | 7.68E-07 | 6.40E-05 | 7.51E-06 | 3.61E-05 | 3.23E-06 | 0 | 6.29E-05 |
| Wwox | 1.91E-05 | 5.36E-05 | 1.55E-05 | 2.88E-05 | 1.65E-06 | 0 | 5.46E-05 | 3.33E-05 | 5.63E-06 | 3.66E-05 | 1.77E-05 | 2.45E-05 | 0 | 0 | 0.00024 | 7.50E-05 | 0.00011 | 5.42E-05 |
| Hspb6 | 0 | 5.60E-06 | 0 | 8.61E-06 | 0 | 0 | 0 | 0 | 5.63E-06 | 0 | 0 | 0 | 0 | 1.36E-05 | 1.49E-05 | 2.70E-05 | 1.99E-05 | 7.20E-06 |
| Alkbh2 | 1.37E-05 | 1.43E-05 | 4.82E-05 | 5.99E-05 | 1.16E-05 | 0 | 0 | 3.96E-05 | 5.63E-06 | 0 | 8.46E-06 | 6.19E-05 | 8.94E-06 | 3.84E-06 | 5.86E-05 | 0 | 0 | 1.70E-05 |
| Mysm1 | 0.00013 | 0.00014 | 1.59E-05 | 0 | 2.47E-05 | 0.0001 | 4.96E-05 | 1.57E-05 | 5.62E-06 | 5.12E-06 | 0 | 8.48E-05 | 3.68E-06 | 4.19E-05 | 0.00015 | 2.80E-05 | 0 | 0 |
| Pced1b | 0 | 2.28E-05 | 0 | 0 | 5.13E-06 | 0 | 0 | 5.15E-06 | 5.60E-06 | 4.12E-05 | 5.74E-06 | 1.34E-05 | 1.29E-05 | 0 | 0 | 0 | 0 | 2.23E-05 |
| Ppih | 1.98E-05 | 0 | 1.65E-05 | 0 | 0 | 9.72E-06 | 0 | 0 | 5.59E-06 | 3.51E-06 | 0 | 0 | 0 | 3.00E-06 | 2.55E-05 | 9.08E-05 | 2.29E-08 | 0 |
| Rsph9 | 0 | 2.21E-05 | 0 | 8.47E-06 | 1.63E-05 | 0 | 0 | 1.34E-07 | 5.59E-06 | 3.26E-05 | 0 | 2.00E-05 | 6.39E-06 | 1.48E-05 | 0 | 0 | 3.53E-05 | 2.72E-05 |
| Trhr2 | 0 | 5.77E-06 | 0 | 0 | 7.37E-06 | 3.55E-06 | 0 | 0 | 5.58E-06 | 0 | 0 | 0 | 0 | 1.56E-06 | 0 | 1.43E-06 | 2.96E-05 | 4.11E-06 |
| Zfp40 | 1.32E-05 | 1.61E-05 | 0 | 0 | 1.57E-06 | 2.42E-05 | 0 | 2.63E-06 | 5.56E-06 | 4.03E-06 | 7.16E-06 | 1.42E-05 | 0 | 0 | 1.87E-05 | 1.74E-06 | 0 | 5.34E-06 |
| Pqlc2 | 2.03E-05 | 4.82E-05 | 9.73E-06 | 2.78E-05 | 3.52E-05 | 1.13E-05 | 0 | 1.34E-05 | 5.56E-06 | 2.94E-06 | 2.69E-07 | 0 | 0 | 1.31E-05 | 2.48E-05 | 0 | 0 | 1.12E-05 |
| Nat8f1 | 1.07E-06 | 1.00E-05 | 1.06E-06 | 0 | 2.98E-06 | 3.78E-06 | 6.09E-06 | 5.08E-06 | 5.54E-06 | 1.02E-05 | 0 | 9.27E-07 | 3.47E-06 | 0 | 1.17E-05 | 1.11E-07 | 0 | 1.15E-05 |
| Anapc10 | 2.41E-05 | 6.77E-07 | 0 | 0 | 0 | 1.41E-05 | 3.67E-06 | 7.19E-06 | 5.53E-06 | 3.01E-06 | 0 | 9.54E-06 | 0 | 0 | 0 | 4.19E-06 | 0 | 4.91E-07 |
| Rnf170 | 2.41E-05 | 1.81E-05 | 0 | 2.06E-05 | 1.53E-05 | 1.27E-05 | 1.54E-05 | 1.47E-05 | 5.53E-06 | 3.01E-06 | 0 | 8.48E-07 | 0 | 0 | 2.74E-05 | 4.54E-06 | 0 | 1.74E-05 |
| Arhgap26 | 6.35E-07 | 1.27E-05 | 0 | 0 | 0 | 9.32E-06 | 0 | 1.83E-05 | 5.53E-06 | 1.91E-05 | 1.91E-05 | 1.75E-05 | 0 | 1.06E-08 | 1.12E-05 | 0 | 9.97E-06 | 1.74E-05 |
| P4htm | 0 | 1.75E-05 | 0 | 1.45E-05 | 0 | 0 | 0 | 0 | 5.53E-06 | 2.91E-05 | 0 | 1.91E-05 | 4.11E-07 | 5.45E-06 | 2.63E-05 | 2.13E-06 | 3.67E-06 | 3.96E-05 |
| Pigx | 0.00015 | 6.03E-05 | 6.51E-05 | 0 | 1.61E-05 | 8.38E-05 | 6.51E-05 | 1.17E-05 | 5.52E-06 | 2.20E-05 | 1.18E-05 | 4.97E-05 | 4.98E-05 | 4.42E-05 | 0 | 2.62E-05 | 0 | 2.67E-05 |
| Bcr | 1.32E-05 | 0.0001 | 0.00017 | 4.08E-05 | 0.00013 | 6.12E-05 | 5.12E-05 | 0.00014 | 5.52E-06 | 7.39E-05 | 9.66E-05 | 5.01E-05 | 3.30E-05 | 2.52E-05 | 3.19E-05 | 1.76E-05 | 0.00018 | 0.00017 |
| Zfp141 | 2.24E-05 | 2.35E-05 | 1.17E-06 | 0 | 0 | 1.83E-05 | 0 | 1.11E-05 | 5.52E-06 | 3.21E-06 | 0 | 3.42E-05 | 3.03E-06 | 0 | 9.38E-06 | 6.74E-08 | 0 | 2.98E-06 |
| Pole3 | 0.00014 | 2.66E-05 | 0.00011 | 0.00011 | 3.72E-05 | 3.46E-05 | 9.97E-05 | 5.45E-07 | 5.51E-06 | 0 | 7.33E-06 | 7.19E-06 | 3.15E-05 | 4.39E-06 | 2.30E-05 | 0.0003 | 1.29E-05 | 0 |
| Zfp454 | 4.92E-06 | 1.40E-07 | 0 | 2.06E-06 | 5.76E-06 | 0 | 0 | 0 | 5.51E-06 | 3.50E-06 | 0 | 0 | 1.86E-06 | 2.72E-06 | 2.02E-05 | 2.05E-06 | 0 | 0 |
| Atad2b | 5.56E-05 | 6.12E-05 | 3.43E-05 | 0 | 2.82E-05 | 3.94E-05 | 2.23E-05 | 2.37E-05 | 5.50E-06 | 1.39E-05 | 1.95E-05 | 3.37E-05 | 1.52E-05 | 7.32E-06 | 0.00015 | 3.33E-05 | 5.94E-05 | 8.42E-06 |
| Cep95 | 2.10E-05 | 4.29E-05 | 0 | 1.04E-05 | 3.94E-06 | 3.83E-06 | 0 | 1.03E-05 | 5.50E-06 | 0 | 0 | 5.90E-05 | 0 | 1.28E-05 | 9.17E-05 | 3.22E-05 | 6.45E-07 | 5.72E-06 |
| Hyls1 | 0 | 2.41E-05 | 0 | 0 | 5.92E-06 | 0 | 2.89E-06 | 6.05E-06 | 5.50E-06 | 1.69E-05 | 0 | 1.62E-05 | 0 | 1.50E-05 | 2.38E-05 | 0.0002 | 4.93E-05 | 1.50E-07 |
| Diablo | 7.36E-05 | 3.24E-05 | 2.27E-05 | 0 | 1.88E-05 | 3.05E-05 | 0 | 0 | 5.49E-06 | 1.45E-05 | 1.39E-05 | 4.56E-05 | 3.20E-07 | 1.58E-06 | 6.15E-05 | 1.31E-05 | 0 | 0 |
| Zfand4 | 1.80E-06 | 1.37E-06 | 0 | 0 | 0 | 3.11E-06 | 0 | 1.38E-06 | 5.49E-06 | 7.06E-06 | 0 | 0 | 0 | 3.64E-06 | 0 | 5.46E-05 | 0 | 0 |
| Gsta4 | 5.70E-07 | 1.89E-05 | 0 | 5.09E-06 | 8.09E-06 | 0 | 0 | 5.37E-07 | 5.49E-06 | 8.57E-06 | 7.48E-06 | 7.23E-06 | 0 | 0 | 6.41E-05 | 1.03E-05 | 1.05E-05 | 2.75E-05 |
| Fahd2a | 0 | 7.17E-06 | 0 | 7.21E-06 | 2.40E-06 | 0 | 0 | 0 | 5.48E-06 | 1.16E-05 | 0 | 0 | 1.16E-05 | 2.40E-06 | 4.70E-06 | 0 | 0 | 6.91E-06 |
| Rnf167 | 7.76E-05 | 0.00011 | 0 | 0 | 4.91E-05 | 7.67E-05 | 2.88E-05 | 1.69E-05 | 5.41E-06 | 6.76E-05 | 0 | 3.70E-05 | 9.46E-06 | 5.18E-05 | 7.81E-05 | 2.21E-05 | 0 | 4.36E-05 |

| | | | | | | | | | | | | | | | | | | |
|---|---|---|---|---|---|---|---|---|---|---|---|---|---|---|---|---|---|---|
| Mutyh | 2.79E-06 | 4.60E-06 | 0 | 9.47E-06 | 0 | 3.31E-06 | 0 | 0 | 5.41E-06 | 6.64E-06 | 9.29E-06 | 4.17E-07 | 0 | 0 | 2.72E-05 | 4.75E-05 | 0 | 1.10E-05 |
| Egflam | 0 | 0.00012 | 0 | 0 | 0 | 8.51E-05 | 0 | 0 | 5.38E-06 | 0 | 0 | 0 | 0 | 0.00041 | 0 | 0 | 0 | 0.00013 |
| Kif3a | 4.90E-05 | 0.00012 | 6.07E-06 | 4.88E-05 | 1.31E-05 | 0 | 2.24E-05 | 0 | 5.38E-06 | 2.42E-05 | 2.39E-05 | 4.91E-05 | 0 | 3.18E-05 | 8.52E-05 | 2.11E-05 | 5.32E-05 | 6.52E-05 |
| Tubgcp5 | 7.13E-05 | 4.67E-05 | 3.00E-05 | 0 | 7.00E-06 | 5.52E-05 | 5.65E-06 | 0 | 5.37E-06 | 1.58E-05 | 0 | 3.66E-06 | 0 | 0 | 8.98E-05 | 2.75E-05 | 0 | 1.17E-05 |
| Acer2 | 1.93E-06 | 3.75E-06 | 0 | 6.10E-06 | 0 | 1.71E-06 | 0 | 0 | 5.37E-06 | 3.13E-06 | 0 | 1.34E-05 | 7.43E-06 | 1.62E-07 | 0 | 3.27E-06 | 2.51E-05 | 0 |
| Zfp358 | 0 | 3.71E-05 | 0 | 7.18E-05 | 2.62E-05 | 2.18E-05 | 9.56E-06 | 3.43E-05 | 5.36E-06 | 2.20E-05 | 1.49E-05 | 3.86E-05 | 4.30E-06 | 2.23E-05 | 2.04E-06 | 7.01E-07 | 0 | 3.57E-05 |
| Abca2 | 0 | 6.23E-05 | 1.33E-05 | 9.64E-07 | 3.21E-05 | 0 | 0 | 2.89E-06 | 5.36E-06 | 0.00012 | 0 | 1.23E-05 | 0 | 0 | 0 | 6.51E-06 | 1.47E-06 | 0.00011 |
| Rmdn1 | 2.39E-05 | 3.72E-05 | 0 | 0 | 2.89E-05 | 2.31E-05 | 0 | 2.44E-05 | 5.35E-06 | 2.27E-05 | 0 | 5.69E-06 | 0 | 0 | 4.26E-05 | 0 | 0 | 1.79E-05 |
| Zfp367 | 0 | 0 | 5.65E-05 | 0 | 2.82E-06 | 0 | 0 | 0 | 5.35E-06 | 1.70E-05 | 1.51E-05 | 8.90E-06 | 3.29E-05 | 1.81E-05 | 0 | 0.00037 | 4.36E-05 | 0 |
| Lrrc40 | 4.19E-05 | 8.75E-06 | 1.51E-05 | 0 | 1.04E-05 | 2.87E-05 | 2.44E-06 | 0 | 5.34E-06 | 2.56E-06 | 0 | 2.27E-05 | 8.98E-06 | 0 | 8.54E-06 | 0.00013 | 0 | 6.03E-06 |
| Zfp991 | 6.78E-06 | 1.35E-05 | 1.17E-06 | 3.38E-06 | 3.57E-06 | 1.06E-05 | 1.41E-05 | 4.85E-06 | 5.34E-06 | 1.80E-06 | 0 | 2.80E-06 | 1.25E-05 | 0 | 4.07E-05 | 2.01E-06 | 3.70E-06 | 7.15E-06 |
| Mkks | 2.95E-05 | 7.92E-06 | 1.04E-05 | 0 | 1.11E-06 | 9.82E-06 | 1.13E-05 | 1.71E-05 | 5.34E-06 | 4.28E-06 | 0 | 1.22E-05 | 0 | 5.33E-06 | 3.00E-06 | 2.21E-05 | 0 | 1.45E-05 |
| DEPRECATED | 5.42E-05 | 3.51E-05 | 2.25E-05 | 4.68E-06 | 2.85E-05 | 4.11E-05 | 3.83E-05 | 1.55E-05 | 5.33E-06 | 2.17E-05 | 1.35E-05 | 1.14E-05 | 0 | 1.18E-05 | 1.44E-05 | 4.78E-05 | 0 | 2.03E-05 |
| Ralgps1 | 0 | 3.10E-05 | 0 | 0 | 0 | 9.02E-06 | 0 | 7.18E-06 | 5.32E-06 | 2.79E-05 | 0 | 6.00E-05 | 0 | 0 | 1.78E-05 | 5.99E-07 | 0 | 0 |
| Uros | 3.85E-05 | 9.35E-06 | 9.39E-06 | 2.11E-05 | 1.87E-05 | 8.75E-06 | 0 | 1.99E-05 | 5.32E-06 | 1.07E-05 | 5.97E-06 | 0 | 0 | 5.62E-07 | 3.81E-05 | 1.74E-05 | 2.77E-05 | 0 |
| Slc27a6 | 2.68E-06 | 0 | 0 | 7.06E-06 | 0 | 9.96E-06 | 0 | 8.38E-06 | 5.31E-06 | 3.68E-06 | 1.24E-05 | 0 | 1.01E-05 | 0 | 0 | 2.69E-06 | 0 | 1.32E-05 |
| Arc | 0 | 0 | 0 | 0 | 0 | 0 | 0 | 0 | 5.30E-06 | 0 | 0 | 4.45E-06 | 5.29E-05 | 0 | 0 | 0 | 0.0006 | 0 |
| Sms | 1.34E-06 | 4.71E-05 | 3.83E-05 | 8.62E-06 | 0 | 5.87E-05 | 0 | 0 | 5.29E-06 | 1.93E-05 | 2.95E-05 | 0.00013 | 0 | 6.19E-05 | 3.78E-06 | 8.16E-05 | 0 | 4.27E-06 |
| Dnah8 | 2.57E-06 | 8.93E-07 | 0 | 0 | 2.84E-06 | 0 | 9.05E-07 | 0 | 5.29E-06 | 1.26E-05 | 0 | 3.59E-06 | 0 | 0 | 0 | 0 | 0 | 1.53E-05 |
| Sobp | 3.63E-05 | 6.98E-05 | 4.86E-05 | 0 | 8.33E-06 | 2.37E-05 | 2.09E-05 | 0 | 5.28E-06 | 5.29E-05 | 4.86E-05 | 0 | 9.26E-05 | 4.83E-05 | 5.88E-05 | 0 | 7.43E-05 | 4.95E-05 |
| Sdsl | 0 | 2.79E-07 | 3.48E-06 | 1.40E-05 | 0 | 0 | 1.37E-06 | 3.10E-06 | 5.28E-06 | 2.57E-05 | 0 | 0 | 3.26E-05 | 0 | 3.11E-06 | 0 | 0 | 1.07E-05 |
| Gsap | 0 | 3.22E-05 | 2.86E-05 | 0 | 3.23E-05 | 7.81E-05 | 0 | 0 | 5.25E-06 | 0 | 0 | 3.42E-05 | 3.17E-05 | 0.00013 | 4.01E-05 | 0 | 0 | 1.21E-05 |
| Tnnt1 | 1.26E-05 | 0 | 4.05E-06 | 1.09E-05 | 0 | 1.60E-05 | 1.18E-05 | 2.71E-05 | 5.25E-06 | 0 | 2.72E-06 | 1.28E-05 | 0 | 0 | 0 | 4.91E-06 | 0 | 8.07E-06 |
| Isca1 | 2.69E-06 | 3.12E-06 | 5.88E-07 | 3.64E-06 | 3.40E-06 | 1.80E-09 | 7.63E-06 | 0 | 5.25E-06 | 0 | 0 | 1.25E-05 | 4.86E-06 | 2.52E-06 | 6.17E-06 | 9.92E-06 | 7.19E-05 | 4.70E-06 |
| Lrrc1 | 2.80E-05 | 3.08E-05 | 0 | 0 | 0 | 0 | 4.29E-06 | 0 | 5.23E-06 | 1.84E-05 | 5.89E-06 | 9.25E-06 | 0 | 1.48E-07 | 0 | 1.92E-05 | 0 | 8.02E-06 |
| Pick1 | 8.70E-07 | 2.50E-05 | 0 | 6.03E-06 | 9.35E-06 | 0 | 2.79E-05 | 0 | 5.22E-06 | 1.37E-05 | 3.50E-06 | 1.57E-05 | 0 | 0 | 4.60E-05 | 0 | 0 | 4.78E-05 |
| Neu1 | 0 | 0.0001 | 0 | 6.99E-05 | 8.45E-05 | 2.42E-05 | 4.42E-05 | 1.08E-05 | 5.18E-06 | 4.25E-05 | 2.61E-05 | 3.37E-05 | 1.77E-05 | 8.34E-05 | 2.27E-05 | 0 | 1.76E-05 | 6.50E-05 |
| Zfp235 | 3.83E-06 | 5.41E-06 | 0 | 0 | 6.24E-06 | 0 | 1.69E-06 | 2.65E-06 | 5.18E-06 | 6.22E-06 | 2.28E-06 | 0 | 2.25E-06 | 0 | 4.55E-06 | 0 | 0 | 1.46E-06 |
| C1ql1 | 3.69E-07 | 1.09E-07 | 3.28E-06 | 0 | 7.02E-06 | 0 | 1.82E-06 | 6.25E-06 | 5.17E-06 | 0 | 0 | 0 | 5.14E-06 | 2.22E-05 | 0 | 2.03E-05 | 0 | 0 |
| Itpr2 | 6.49E-05 | 7.68E-05 | 0 | 0 | 2.62E-05 | 0.00013 | 1.97E-06 | 0.0002 | 5.16E-06 | 3.97E-05 | 3.45E-05 | 2.52E-05 | 0 | 7.73E-05 | 8.31E-05 | 0 | 5.68E-05 | 3.94E-05 |
| Slc16a13 | 0 | 9.74E-06 | 0 | 0 | 4.00E-06 | 0 | 0 | 0 | 5.15E-06 | 0 | 7.84E-06 | 4.03E-06 | 0 | 4.08E-06 | 7.34E-06 | 0 | 2.21E-06 | 7.04E-06 |
| AI987944 | 9.79E-06 | 9.07E-06 | 0 | 0 | 4.62E-06 | 6.43E-06 | 0 | 0 | 5.13E-06 | 6.68E-06 | 3.94E-06 | 7.40E-06 | 0 | 0 | 0 | 0 | 2.76E-06 | 8.67E-06 |
| Rxfp3 | 0 | 0 | 0 | 0 | 0.00015 | 1.26E-05 | 2.78E-05 | 0 | 5.12E-06 | 0 | 2.65E-05 | 0 | 0 | 4.28E-06 | 0 | 0 | 0 | 0 |

| | | | | | | | | | | | | | | | | | | |
|---|---|---|---|---|---|---|---|---|---|---|---|---|---|---|---|---|---|---|
| Dus4l | 1.39E-05 | 5.58E-06 | 1.84E-05 | 0 | 0 | 2.82E-06 | 0 | 0 | 5.12E-06 | 8.49E-06 | 0 | 1.17E-06 | 5.38E-06 | 1.34E-06 | 1.87E-06 | 8.86E-06 | 0 | 0 |
| Kifc2 | 0 | 2.00E-05 | 0 | 0 | 1.34E-06 | 0 | 0 | 0 | 5.10E-06 | 5.55E-06 | 0 | 0 | 0 | 9.20E-06 | 4.41E-05 | 0 | 0 | 1.21E-05 |
| Dnaaf4 | 6.25E-06 | 3.95E-06 | 9.52E-07 | 0 | 6.36E-06 | 3.36E-06 | 1.47E-06 | 9.78E-07 | 5.09E-06 | 4.19E-06 | 2.49E-06 | 4.58E-06 | 4.98E-06 | 0 | 0 | 1.11E-06 | 7.16E-06 | 0 |
| Il10ra | 0 | 2.79E-06 | 3.24E-06 | 3.42E-06 | 3.89E-08 | 0 | 6.50E-07 | 0 | 5.08E-06 | 4.06E-06 | 1.36E-06 | 2.91E-06 | 8.92E-06 | 4.96E-06 | 2.29E-05 | 2.29E-06 | 0 | 1.68E-06 |
| Flywch2 | 0 | 2.98E-05 | 0 | 5.10E-06 | 4.01E-05 | 1.42E-05 | 1.42E-05 | 0 | 5.05E-06 | 1.12E-05 | 3.76E-06 | 1.43E-05 | 1.66E-05 | 3.77E-05 | 4.59E-06 | 0 | 0 | 0 |
| Paqr3 | 1.03E-05 | 1.79E-05 | 2.51E-05 | 1.12E-05 | 7.13E-06 | 2.20E-05 | 1.00E-05 | 1.92E-05 | 5.04E-06 | 4.77E-06 | 1.17E-05 | 0 | 1.35E-05 | 2.53E-05 | 2.93E-05 | 3.17E-06 | 0 | 5.24E-06 |
| Mnd1 | 3.68E-06 | 0 | 1.75E-06 | 7.72E-06 | 0 | 0 | 0 | 0 | 5.04E-06 | 4.95E-06 | 0 | 1.37E-06 | 0 | 4.85E-06 | 0 | 4.70E-05 | 1.25E-05 | 5.62E-06 |
| U2af1l4 | 3.96E-06 | 5.68E-06 | 7.85E-06 | 2.48E-06 | 1.81E-06 | 0 | 3.46E-07 | 3.81E-06 | 5.03E-06 | 3.78E-06 | 0 | 0 | 0 | 0 | 3.63E-05 | 1.83E-06 | 0 | 0 |
| DEPRECATED | 0 | 7.38E-05 | 7.33E-05 | 1.24E-05 | 5.27E-05 | 8.17E-06 | 0 | 3.64E-05 | 5.02E-06 | 7.14E-06 | 0.0001 | 3.74E-05 | 0 | 2.54E-05 | 5.83E-05 | 6.55E-05 | 7.41E-05 | 0 |
| Cda | 0 | 0 | 0 | 0 | 0 | 2.09E-05 | 0 | 0 | 5.01E-06 | 0 | 0 | 0.00017 | 1.49E-05 | 0 | 0 | 0 | 0 | 4.15E-05 |
| Vwc2 | 0 | 0 | 0 | 0 | 0 | 9.48E-06 | 0 | 0 | 5.01E-06 | 1.99E-05 | 0 | 0 | 2.81E-05 | 0 | 0 | 0 | 0 | 1.99E-05 |
| Pde3b | 2.46E-05 | 1.03E-05 | 3.59E-05 | 9.57E-06 | 0 | 8.68E-05 | 0 | 6.48E-05 | 5.00E-06 | 0 | 0 | 1.64E-05 | 0 | 7.21E-05 | 6.43E-05 | 0.00011 | 0 | 4.10E-05 |
| Xpa | 0.00011 | 0.00012 | 6.08E-06 | 4.70E-05 | 4.78E-05 | 0.00012 | 2.47E-05 | 4.81E-06 | 4.99E-06 | 2.94E-05 | 1.06E-05 | 8.60E-05 | 0 | 6.36E-05 | 7.02E-05 | 0 | 0 | 4.87E-05 |
| Zfp948 | 7.58E-05 | 0 | 3.82E-05 | 0 | 0 | 3.85E-05 | 0 | 2.64E-06 | 4.99E-06 | 2.40E-06 | 0 | 6.15E-05 | 0 | 0 | 8.12E-05 | 0.0001 | 6.76E-05 | 8.77E-06 |
| Kcne1l | 0 | 0 | 0 | 2.23E-06 | 5.17E-06 | 0 | 1.22E-05 | 8.49E-06 | 4.98E-06 | 2.30E-05 | 0 | 8.37E-06 | 1.46E-05 | 4.25E-07 | 0 | 7.01E-06 | 2.26E-06 | 0 |
| Adcyap1r1 | 0 | 0 | 0 | 0 | 1.74E-05 | 0 | 0 | 0.00036 | 4.97E-06 | 2.28E-05 | 0 | 2.31E-05 | 0.0001 | 0 | 0 | 0 | 0 | 0.00039 |
| Cyp2u1 | 4.24E-06 | 1.49E-05 | 1.42E-05 | 0 | 6.15E-06 | 0 | 2.00E-05 | 9.40E-06 | 4.93E-06 | 2.49E-05 | 0 | 0 | 1.37E-05 | 0 | 0 | 0 | 0 | 3.55E-05 |
| Creb3l2 | 0.00011 | 0.00028 | 0.00039 | 0 | 0.00053 | 0.00037 | 0.00036 | 0.00086 | 4.92E-06 | 0.00024 | 0.00034 | 0.00043 | 0.00018 | 0.00023 | 0.00014 | 0.00016 | 0.00025 | 0.00021 |
| Tmem170 | 6.37E-06 | 1.88E-05 | 1.66E-05 | 6.75E-06 | 0 | 5.03E-06 | 1.93E-05 | 2.97E-07 | 4.92E-06 | 5.40E-06 | 1.30E-05 | 1.10E-05 | 1.74E-05 | 2.22E-05 | 2.54E-05 | 0 | 1.45E-05 | 1.32E-05 |
| Acbd3 | 0.00032 | 0.00018 | 0.00029 | 0.00022 | 8.83E-05 | 0.00019 | 0.00034 | 0.00015 | 4.89E-06 | 7.19E-05 | 0.0001 | 0.00024 | 2.72E-05 | 3.92E-05 | 0.0004 | 0.00012 | 0.00028 | 6.45E-05 |
| Nhlrc1 | 0 | 2.12E-06 | 7.50E-07 | 3.67E-06 | 7.28E-07 | 5.51E-07 | 7.71E-06 | 3.61E-06 | 4.88E-06 | 1.05E-05 | 0 | 5.45E-06 | 4.97E-07 | 1.79E-06 | 2.95E-06 | 0 | 7.11E-07 | 2.06E-05 |
| Ftl1-ps1 | 2.05E-07 | 1.29E-05 | 4.04E-06 | 4.55E-06 | 1.46E-06 | 9.75E-07 | 0 | 0 | 4.87E-06 | 2.35E-06 | 4.04E-06 | 1.17E-05 | 0 | 2.74E-06 | 0 | 0 | 3.34E-05 | 5.84E-06 |
| Trim11 | 0 | 4.82E-05 | 1.58E-05 | 2.05E-05 | 1.18E-05 | 0 | 0 | 0 | 4.87E-06 | 5.35E-05 | 5.29E-07 | 1.19E-08 | 9.46E-06 | 0 | 0.00015 | 0 | 7.26E-06 | 1.73E-05 |
| Ggact | 3.29E-05 | 1.52E-05 | 2.07E-06 | 1.91E-05 | 2.23E-05 | 1.43E-05 | 4.04E-05 | 0 | 4.85E-06 | 0 | 1.41E-05 | 0 | 3.94E-05 | 3.46E-05 | 0 | 7.61E-06 | 0 | 2.98E-05 |
| Fam228b | 4.61E-06 | 7.62E-06 | 3.09E-06 | 1.17E-06 | 0 | 0 | 7.71E-06 | 0 | 4.84E-06 | 1.37E-05 | 0 | 0 | 7.08E-07 | 4.98E-06 | 3.50E-05 | 0 | 8.39E-06 | 1.69E-05 |
| AU041133 | 1.08E-05 | 1.01E-05 | 0 | 0 | 3.25E-06 | 4.02E-06 | 0 | 8.55E-07 | 4.81E-06 | 1.16E-05 | 0 | 6.31E-06 | 1.56E-06 | 0 | 4.38E-06 | 1.56E-06 | 0 | 1.66E-08 |
| Oxnad1 | 2.50E-05 | 1.43E-05 | 1.81E-05 | 3.92E-06 | 9.54E-08 | 4.34E-06 | 1.55E-05 | 3.85E-05 | 4.80E-06 | 1.04E-05 | 5.15E-06 | 1.78E-05 | 0 | 0 | 1.49E-05 | 2.50E-05 | 0 | 3.22E-07 |
| Zfp934 | 1.18E-05 | 0 | 0 | 0 | 1.83E-06 | 4.85E-06 | 3.90E-06 | 0 | 4.79E-06 | 5.28E-06 | 5.92E-06 | 5.21E-06 | 3.15E-06 | 0 | 5.72E-06 | 4.64E-05 | 0 | 7.65E-06 |
| Nlrc3 | 0 | 1.07E-05 | 0 | 0 | 0 | 0 | 1.24E-05 | 4.67E-05 | 4.78E-06 | 0 | 0 | 3.23E-06 | 1.47E-05 | 0 | 0 | 0 | 0 | 0 |
| Ptcd1 | 4.85E-05 | 6.78E-05 | 3.19E-05 | 3.37E-05 | 2.34E-05 | 4.79E-05 | 9.57E-06 | 0 | 4.75E-06 | 2.46E-05 | 0 | 9.33E-06 | 1.92E-06 | 0 | 5.14E-05 | 1.93E-05 | 0 | 1.67E-05 |
| Mpzl3 | 3.30E-06 | 5.15E-06 | 0 | 0 | 1.59E-05 | 6.90E-06 | 0 | 2.01E-05 | 4.75E-06 | 2.63E-06 | 0 | 0 | 1.94E-06 | 0 | 0 | 1.94E-06 | 7.45E-06 | 1.35E-06 |
| Mllt10 | 6.60E-05 | 9.49E-05 | 3.41E-05 | 2.36E-06 | 5.53E-05 | 7.19E-05 | 2.05E-05 | 1.92E-05 | 4.74E-06 | 3.83E-05 | 4.77E-05 | 0.00013 | 2.83E-05 | 2.82E-05 | 0.0001 | 9.22E-05 | 0.00019 | 4.36E-05 |
| Kif1a | 0 | 7.21E-06 | 0 | 0 | 0 | 1.11E-05 | 0 | 0 | 4.71E-06 | 0 | 0 | 4.25E-06 | 0 | 9.76E-05 | 0 | 0 | 0 | 1.12E-05 |

| | | | | | | | | | | | | | | | | | | |
|---|---|---|---|---|---|---|---|---|---|---|---|---|---|---|---|---|---|---|
| Churc1 | 6.98E-05 | 3.40E-05 | 2.36E-05 | 6.16E-05 | 2.62E-05 | 4.72E-05 | 6.28E-05 | 3.47E-05 | 4.69E-06 | 2.51E-05 | 2.96E-05 | 1.53E-05 | 5.06E-06 | 3.18E-05 | 0 | 2.52E-05 | 0 | 2.98E-05 |
| Fntb | 6.18E-05 | 1.60E-05 | 6.07E-05 | 9.09E-05 | 8.03E-06 | 2.39E-05 | 7.67E-05 | 4.03E-05 | 4.69E-06 | 1.00E-05 | 2.36E-05 | 0.00012 | 0 | 1.17E-05 | 2.43E-05 | 0.00013 | 0 | 5.06E-05 |
| Clspn | 0 | 0 | 1.73E-05 | 0 | 6.62E-06 | 3.02E-06 | 0 | 0 | 4.68E-06 | 0 | 0 | 0 | 1.26E-05 | 0 | 1.58E-05 | 0.00051 | 0 | 0 |
| St8sia2 | 0 | 4.29E-06 | 0 | 0 | 0 | 0 | 0 | 0.00019 | 4.67E-06 | 0 | 4.77E-05 | 3.51E-05 | 0 | 0.0001 | 0 | 2.92E-05 | 0 | 3.17E-05 |
| Hsdl1 | 0 | 6.41E-05 | 0 | 8.15E-06 | 7.90E-07 | 0 | 1.06E-05 | 0 | 4.66E-06 | 2.81E-05 | 1.23E-05 | 2.70E-05 | 1.22E-05 | 1.43E-05 | 1.58E-05 | 5.85E-07 | 0 | 1.33E-05 |
| Suv39h2 | 5.64E-06 | 0 | 1.85E-05 | 0 | 0 | 0 | 0 | 0 | 4.64E-06 | 0 | 0 | 0 | 0 | 0 | 0 | 8.95E-05 | 1.11E-05 | 2.18E-06 |
| Ccdc171 | 3.69E-06 | 2.32E-05 | 0 | 0 | 0 | 8.51E-06 | 0 | 2.05E-05 | 4.63E-06 | 2.88E-05 | 0 | 9.12E-06 | 3.22E-06 | 0 | 4.36E-05 | 0 | 0 | 3.18E-05 |
| Pcbp3 | 1.50E-05 | 2.81E-05 | 8.21E-06 | 0 | 0 | 0 | 0 | 3.96E-05 | 4.63E-06 | 6.80E-05 | 0 | 1.46E-05 | 8.94E-05 | 0 | 5.71E-05 | 7.27E-05 | 3.53E-05 | 5.13E-05 |
| Zfp109 | 1.38E-05 | 7.48E-06 | 0 | 7.73E-07 | 2.26E-06 | 9.28E-06 | 1.13E-07 | 0 | 4.62E-06 | 1.19E-06 | 1.28E-06 | 3.55E-06 | 0 | 5.66E-07 | 7.43E-06 | 0 | 0 | 0 |
| Ccl27b | 1.90E-06 | 0 | 0 | 0 | 6.86E-07 | 5.58E-06 | 4.74E-06 | 0 | 4.62E-06 | 2.59E-06 | 0 | 9.98E-06 | 1.29E-05 | 0 | 0 | 0 | 0 | 0 |
| Arl4c | 0 | 0.00012 | 0.00022 | 0 | 0.00023 | 0 | 0.00042 | 5.09E-05 | 4.61E-06 | 0 | 4.93E-05 | 0.00015 | 0 | 7.50E-05 | 0.00011 | 0.00017 | 0 | 0 |
| Mamld1 | 2.94E-06 | 2.30E-05 | 7.45E-06 | 0 | 7.78E-07 | 2.64E-05 | 0 | 2.29E-05 | 4.59E-06 | 0 | 2.32E-05 | 1.75E-05 | 0 | 1.88E-05 | 0 | 7.25E-06 | 0 | 2.26E-05 |
| Mrm1 | 7.60E-06 | 2.46E-05 | 3.05E-06 | 2.44E-05 | 1.43E-07 | 9.44E-06 | 0 | 1.81E-05 | 4.58E-06 | 9.77E-06 | 1.55E-05 | 3.73E-05 | 0 | 4.61E-06 | 9.61E-06 | 0 | 1.27E-05 | 1.11E-05 |
| Osgepl1 | 2.97E-05 | 3.85E-06 | 1.19E-07 | 0 | 0 | 8.39E-06 | 0 | 1.88E-05 | 4.58E-06 | 5.82E-06 | 0 | 4.39E-05 | 0 | 0 | 0 | 2.18E-05 | 0 | 6.88E-06 |
| Lmln | 5.62E-07 | 0 | 3.44E-06 | 6.78E-06 | 0 | 0 | 8.19E-06 | 0 | 4.58E-06 | 3.49E-06 | 8.56E-06 | 3.01E-05 | 2.53E-06 | 8.22E-06 | 0 | 6.27E-06 | 9.43E-06 | 4.22E-06 |
| Chid1 | 0.00022 | 2.03E-05 | 5.39E-05 | 9.74E-05 | 2.78E-05 | 9.38E-05 | 0.00014 | 8.79E-05 | 4.57E-06 | 1.24E-05 | 2.24E-05 | 6.34E-05 | 0.00023 | 0 | 0 | 5.50E-05 | 0 | 3.43E-05 |
| Zfp697 | 0 | 4.11E-05 | 1.84E-05 | 1.71E-05 | 2.67E-05 | 1.34E-05 | 4.80E-06 | 7.22E-06 | 4.55E-06 | 0 | 3.31E-05 | 1.43E-05 | 5.56E-06 | 1.76E-05 | 0.00012 | 0 | 4.59E-05 | 2.55E-05 |
| Il18 | 0 | 0 | 0 | 0 | 0 | 0 | 0 | 6.55E-06 | 4.55E-06 | 0.00023 | 0 | 0 | 0 | 0 | 0 | 0 | 0 | 0 |
| Bbs4 | 1.41E-05 | 2.76E-05 | 0 | 0 | 1.62E-05 | 3.03E-05 | 1.55E-06 | 3.24E-06 | 4.54E-06 | 9.16E-06 | 0 | 1.69E-05 | 0 | 1.97E-06 | 3.20E-06 | 1.23E-05 | 0 | 7.58E-06 |
| Mdm1 | 7.28E-06 | 3.24E-05 | 0 | 0 | 3.86E-06 | 0 | 0 | 1.24E-05 | 4.54E-06 | 9.55E-06 | 0 | 5.89E-06 | 0 | 5.28E-06 | 4.30E-05 | 0.00016 | 6.54E-06 | 0 |
| Osgin2 | 3.72E-05 | 2.90E-05 | 5.01E-05 | 0 | 2.96E-05 | 4.33E-05 | 5.00E-06 | 0 | 4.53E-06 | 0 | 8.81E-05 | 0 | 0 | 0 | 1.19E-05 | 2.51E-05 | 1.75E-05 | 2.40E-06 |
| Kmt2e | 0.00022 | 0.00026 | 2.22E-07 | 0 | 0.0001 | 0.00025 | 0 | 0.00011 | 4.50E-06 | 0 | 0 | 0.00022 | 0 | 9.29E-05 | 0.00026 | 5.85E-05 | 0.00026 | 0.00012 |
| Pcdhb18 | 9.62E-06 | 2.38E-05 | 0 | 0 | 2.05E-05 | 1.63E-05 | 0 | 1.18E-05 | 4.50E-06 | 6.47E-06 | 5.60E-06 | 7.90E-06 | 4.71E-06 | 0 | 1.96E-05 | 0 | 0 | 2.10E-05 |
| Zfp874b | 4.97E-06 | 1.85E-05 | 0 | 0 | 3.85E-06 | 1.51E-05 | 0 | 1.35E-08 | 4.50E-06 | 3.64E-06 | 2.20E-06 | 2.00E-06 | 0 | 2.72E-07 | 5.18E-06 | 0 | 0 | 7.70E-06 |
| Ets2 | 0 | 0.00023 | 0.00014 | 2.81E-05 | 6.99E-05 | 0.00012 | 0 | 0 | 4.48E-06 | 2.27E-06 | 0.00019 | 0.00024 | 0 | 0.00061 | 0.0001 | 3.25E-05 | 0.00027 | 7.33E-05 |
| Ccl12 | 0 | 0 | 0 | 0 | 1.44E-05 | 0 | 0 | 3.70E-05 | 4.47E-06 | 9.94E-07 | 0 | 0 | 2.22E-06 | 3.94E-06 | 0 | 1.11E-05 | 0 | 2.77E-06 |
| Lsr | 0 | 1.34E-05 | 0 | 8.10E-06 | 0 | 0 | 0 | 0 | 4.46E-06 | 6.00E-06 | 0 | 1.57E-05 | 4.16E-05 | 0 | 2.35E-05 | 0 | 0 | 1.78E-05 |
| Ccdc186 | 0.00013 | 0.00011 | 3.99E-05 | 0.00017 | 3.11E-05 | 5.44E-05 | 0.00013 | 2.90E-05 | 4.45E-06 | 2.14E-05 | 4.64E-05 | 0.00013 | 0 | 4.73E-05 | 0.00026 | 7.90E-05 | 0.00017 | 7.04E-05 |
| Tmem35b | 0 | 3.92E-05 | 0 | 1.67E-05 | 1.52E-05 | 5.67E-06 | 2.44E-07 | 4.56E-06 | 4.43E-06 | 1.85E-05 | 0 | 5.41E-05 | 3.75E-05 | 0 | 0 | 2.28E-06 | 0 | 3.67E-05 |
| Mrnip | 6.18E-07 | 1.10E-05 | 0 | 0 | 4.47E-06 | 1.73E-06 | 0 | 0 | 4.43E-06 | 5.33E-06 | 0 | 4.57E-06 | 0 | 5.49E-07 | 1.01E-05 | 5.83E-05 | 0 | 0 |
| Simc1 | 1.86E-05 | 2.53E-05 | 4.72E-05 | 5.41E-06 | 0 | 0 | 0 | 0 | 4.43E-06 | 2.28E-06 | 0 | 2.93E-05 | 6.62E-06 | 9.66E-06 | 3.95E-06 | 2.16E-05 | 0.00011 | 0 |
| Zfp442 | 1.11E-05 | 1.74E-06 | 8.68E-07 | 0 | 4.11E-07 | 4.73E-06 | 0 | 5.55E-07 | 4.42E-06 | 2.19E-06 | 0 | 3.50E-06 | 0 | 0 | 0 | 0 | 0 | 0 |
| Rita1 | 1.50E-05 | 3.07E-05 | 1.55E-05 | 2.36E-05 | 9.32E-06 | 0 | 0 | 0 | 4.42E-06 | 7.73E-06 | 4.81E-07 | 0 | 0 | 0 | 2.53E-05 | 0 | 2.84E-05 | 1.94E-06 |

| | | | | | | | | | | | | | | | | | | |
|---|---|---|---|---|---|---|---|---|---|---|---|---|---|---|---|---|---|---|
| Tpcn1 | 8.73E-05 | 0.00011 | 7.29E-05 | 0.00011 | 7.12E-07 | 1.76E-05 | 0.00014 | 2.70E-05 | 4.41E-06 | 0.00013 | 0 | 4.34E-07 | 0 | 2.37E-05 | 9.05E-05 | 2.95E-05 | 1.69E-05 | 7.18E-05 |
| Nup43 | 1.41E-05 | 2.53E-06 | 3.89E-05 | 0 | 0 | 5.98E-06 | 0 | 0 | 4.41E-06 | 9.26E-06 | 9.05E-06 | 0 | 0 | 1.42E-06 | 0 | 0.00011 | 0 | 2.37E-06 |
| Abcb9 | 3.38E-06 | 1.29E-06 | 0 | 1.09E-05 | 2.78E-06 | 2.03E-06 | 1.61E-05 | 1.37E-05 | 4.40E-06 | 2.87E-05 | 2.14E-06 | 0 | 4.01E-06 | 2.00E-05 | 9.69E-06 | 0 | 0 | 1.59E-05 |
| Txnl4b | 3.49E-05 | 2.30E-05 | 3.45E-06 | 7.04E-06 | 5.31E-06 | 3.73E-05 | 1.13E-05 | 2.56E-05 | 4.38E-06 | 4.67E-06 | 0 | 1.85E-05 | 8.73E-07 | 1.93E-05 | 2.83E-06 | 1.87E-05 | 0 | 0 |
| Arhgef9 | 8.20E-06 | 1.16E-05 | 0 | 0 | 3.80E-06 | 3.47E-06 | 0 | 0 | 4.38E-06 | 1.48E-05 | 0 | 0 | 0 | 4.47E-08 | 2.01E-05 | 0 | 0 | 9.89E-06 |
| Nos1ap | 0 | 7.63E-06 | 0 | 0 | 0 | 0 | 0 | 0 | 4.38E-06 | 0 | 7.33E-06 | 9.58E-06 | 8.23E-06 | 9.74E-07 | 1.62E-05 | 3.52E-06 | 1.04E-07 | 1.26E-05 |
| Cd302 | 0.00034 | 4.81E-05 | 0 | 0 | 0.00017 | 0.00095 | 0 | 0.00014 | 4.37E-06 | 0.00013 | 0 | 0 | 0.00054 | 0.0004 | 0 | 0 | 0 | 8.76E-05 |
| Zmym6 | 2.37E-05 | 3.35E-05 | 0 | 0 | 2.73E-05 | 3.41E-05 | 1.03E-05 | 7.51E-06 | 4.37E-06 | 1.24E-05 | 0 | 3.23E-05 | 0 | 6.81E-06 | 3.53E-05 | 0 | 0 | 2.22E-05 |
| Map3k3 | 8.75E-05 | 0.00021 | 0.00013 | 0 | 0.0001 | 0.00015 | 0.00011 | 5.29E-05 | 4.36E-06 | 9.95E-05 | 6.39E-05 | 9.67E-05 | 5.89E-05 | 6.44E-05 | 0.00018 | 0.00028 | 0 | 0.00015 |
| Dock11 | 5.49E-05 | 3.00E-05 | 2.07E-05 | 0 | 0 | 5.30E-05 | 0 | 3.21E-05 | 4.36E-06 | 7.95E-05 | 0 | 4.32E-05 | 0 | 0 | 0 | 5.63E-05 | 1.98E-05 | 0 |
| Hus1b | 3.26E-08 | 9.68E-06 | 4.66E-07 | 0 | 1.73E-06 | 0 | 0 | 0 | 4.36E-06 | 6.40E-06 | 1.08E-06 | 1.21E-06 | 0 | 1.20E-06 | 2.37E-05 | 6.05E-09 | 0 | 2.82E-07 |
| Gpr52 | 1.19E-06 | 5.47E-06 | 0 | 0 | 1.62E-06 | 9.10E-07 | 3.70E-06 | 3.83E-06 | 4.34E-06 | 1.13E-06 | 7.80E-06 | 0 | 1.41E-06 | 4.64E-07 | 0 | 2.45E-06 | 4.68E-06 | 3.74E-06 |
| Zfp65 | 2.70E-05 | 1.87E-05 | 0 | 0 | 1.21E-05 | 2.36E-05 | 4.81E-06 | 0 | 4.33E-06 | 1.06E-05 | 0 | 3.27E-06 | 0 | 3.68E-06 | 1.93E-05 | 0 | 0 | 2.71E-05 |
| Tigd2 | 2.97E-05 | 6.53E-05 | 0 | 7.06E-06 | 1.27E-05 | 4.11E-05 | 0 | 1.75E-05 | 4.30E-06 | 1.70E-05 | 0 | 9.58E-05 | 0 | 0 | 3.74E-05 | 8.89E-06 | 0 | 2.26E-05 |
| Ormdl3 | 1.41E-05 | 9.08E-05 | 0.00012 | 0.00026 | 0.00014 | 7.92E-05 | 0.00021 | 0.00011 | 4.29E-06 | 1.48E-05 | 7.79E-05 | 3.13E-05 | 0.00016 | 8.50E-05 | 0.00019 | 6.82E-06 | 1.17E-05 | 4.43E-05 |
| Mtrf1 | 1.95E-05 | 1.08E-05 | 5.39E-06 | 9.34E-06 | 2.51E-06 | 1.00E-05 | 0 | 1.07E-06 | 4.29E-06 | 6.05E-06 | 6.58E-06 | 3.26E-06 | 0 | 0 | 7.77E-08 | 1.10E-05 | 0 | 1.49E-05 |
| Zkscan4 | 6.62E-07 | 6.50E-06 | 0 | 3.03E-06 | 9.76E-06 | 9.59E-06 | 0 | 9.36E-07 | 4.29E-06 | 4.18E-06 | 7.62E-09 | 2.89E-06 | 4.84E-06 | 0 | 7.53E-06 | 0 | 0 | 4.58E-06 |
| Ccdc174 | 7.82E-05 | 8.92E-05 | 4.53E-05 | 2.84E-05 | 5.04E-05 | 3.88E-05 | 4.00E-05 | 3.42E-05 | 4.28E-06 | 9.37E-05 | 1.81E-05 | 5.70E-05 | 9.75E-06 | 0 | 0.00015 | 5.96E-05 | 6.88E-05 | 0 |
| Znrd1as | 1.50E-05 | 1.74E-05 | 7.06E-07 | 0 | 8.16E-08 | 4.40E-06 | 1.05E-05 | 6.54E-07 | 4.25E-06 | 7.93E-06 | 0 | 1.46E-05 | 0 | 6.23E-08 | 4.53E-06 | 1.90E-05 | 0 | 2.59E-06 |
| Pakap | 0 | 7.60E-05 | 2.45E-05 | 0 | 0 | 0.0003 | 0 | 0.00026 | 4.24E-06 | 0 | 0.00068 | 0 | 0 | 0.00028 | 0.00018 | 2.89E-05 | 3.48E-05 | 0 |
| Stard8 | 2.14E-05 | 4.79E-05 | 0 | 2.89E-06 | 2.01E-06 | 0 | 0 | 2.96E-05 | 4.24E-06 | 8.01E-06 | 0 | 9.72E-06 | 0 | 0 | 0 | 0.00017 | 1.64E-06 | 8.71E-05 |
| Zfp37 | 3.35E-05 | 1.71E-05 | 0 | 0 | 1.47E-05 | 1.84E-05 | 7.18E-06 | 0 | 4.23E-06 | 1.08E-05 | 0 | 1.18E-05 | 0 | 1.08E-05 | 6.11E-09 | 1.00E-05 | 0 | 0 |
| Flt4 | 0 | 6.56E-06 | 1.01E-05 | 0 | 2.54E-06 | 0 | 0 | 0 | 4.23E-06 | 0 | 2.08E-05 | 0 | 1.09E-05 | 0 | 0 | 1.40E-05 | 0 | 0 |
| Acot1 | 1.10E-05 | 2.15E-05 | 0 | 1.61E-05 | 2.95E-06 | 0 | 0 | 0 | 4.20E-06 | 2.33E-06 | 0 | 9.10E-06 | 5.91E-06 | 2.18E-05 | 0 | 9.85E-06 | 0 | 0 |
| Tagap1 | 1.50E-05 | 1.82E-05 | 7.06E-06 | 4.73E-06 | 8.57E-06 | 1.63E-05 | 0 | 0 | 4.19E-06 | 8.99E-06 | 0 | 2.47E-07 | 0 | 7.78E-06 | 3.89E-05 | 2.55E-05 | 0 | 0 |
| Ankrd23 | 3.12E-06 | 2.92E-05 | 2.29E-06 | 6.45E-05 | 1.66E-05 | 1.61E-05 | 1.53E-05 | 2.29E-05 | 4.19E-06 | 8.05E-06 | 0 | 9.85E-06 | 6.25E-07 | 4.83E-06 | 8.24E-05 | 8.17E-07 | 1.19E-05 | 7.86E-06 |
| Ccdc69 | 1.56E-06 | 0 | 1.75E-06 | 9.48E-07 | 3.42E-07 | 1.37E-06 | 5.31E-06 | 1.36E-05 | 4.17E-06 | 4.38E-06 | 0 | 0 | 6.36E-06 | 1.69E-06 | 0 | 1.45E-05 | 0 | 0 |
| Ptprf | 6.19E-06 | 4.83E-05 | 3.50E-05 | 0 | 0 | 4.07E-06 | 0 | 9.84E-06 | 4.16E-06 | 1.66E-06 | 4.27E-05 | 6.32E-05 | 0 | 0.0002 | 6.11E-05 | 1.98E-05 | 0 | 0.00035 |
| Furin | 0.00018 | 0.00038 | 0.00058 | 0.00049 | 0.00052 | 0.00057 | 0.00037 | 0.00032 | 4.15E-06 | 9.97E-05 | 0.00066 | 0.00061 | 0.00022 | 0.00057 | 0.00055 | 9.81E-05 | 8.30E-05 | 0.00027 |
| Izumo4 | 8.05E-07 | 2.37E-05 | 0 | 0 | 3.96E-05 | 2.20E-05 | 2.12E-05 | 0 | 4.15E-06 | 1.72E-05 | 1.37E-05 | 1.07E-05 | 1.48E-05 | 9.91E-06 | 9.21E-05 | 0 | 0 | 4.60E-05 |
| Smad6 | 0 | 6.06E-06 | 0 | 1.25E-05 | 0 | 0 | 7.28E-07 | 0 | 4.13E-06 | 2.21E-06 | 0 | 0 | 8.98E-06 | 6.88E-06 | 4.42E-05 | 1.03E-05 | 0 | 1.55E-05 |
| Cacna1d | 0 | 0 | 0 | 0 | 0 | 0 | 0 | 0 | 4.13E-06 | 0 | 0 | 0 | 0.00013 | 2.31E-05 | 1.94E-05 | 3.65E-06 | 0 | 0 |
| Camta1 | 4.29E-05 | 3.62E-05 | 2.59E-06 | 1.21E-05 | 4.57E-05 | 5.92E-05 | 0 | 6.84E-05 | 4.12E-06 | 0 | 2.93E-05 | 4.71E-05 | 9.34E-06 | 3.43E-05 | 6.46E-05 | 3.17E-05 | 2.92E-05 | 1.48E-05 |

| | | | | | | | | | | | | | | | | | | |
|---|---|---|---|---|---|---|---|---|---|---|---|---|---|---|---|---|---|---|
| Tef | 4.37E-05 | 0.00012 | 0 | 6.79E-05 | 1.12E-06 | 0 | 0 | 0 | 4.11E-06 | 0 | 4.47E-05 | 4.05E-05 | 0 | 0.0001 | 0.00018 | 0 | 0.0001 | 5.18E-05 |
| Fmc1 | 7.76E-05 | 1.63E-05 | 4.65E-05 | 5.50E-05 | 0 | 3.29E-05 | 1.72E-05 | 1.22E-06 | 4.10E-06 | 2.84E-05 | 0 | 1.22E-05 | 0 | 0 | 0 | 5.21E-05 | 3.06E-05 | 2.80E-05 |
| Kif18a | 7.92E-07 | 5.80E-06 | 0 | 0 | 0 | 0 | 0 | 0 | 4.09E-06 | 0 | 0 | 0 | 0 | 0 | 1.70E-05 | 0.00025 | 0 | 4.67E-06 |
| Mgme1 | 1.48E-05 | 4.29E-05 | 3.10E-05 | 0 | 8.29E-06 | 1.86E-05 | 7.91E-06 | 2.66E-05 | 4.08E-06 | 2.75E-05 | 5.70E-06 | 6.40E-05 | 0 | 0 | 1.77E-05 | 6.04E-05 | 1.64E-05 | 1.91E-05 |
| Plxnc1 | 0 | 0 | 0 | 0 | 8.57E-07 | 0 | 1.57E-05 | 7.13E-06 | 4.08E-06 | 0 | 5.93E-06 | 0 | 0 | 0 | 0 | 1.42E-06 | 0 | 1.87E-05 |
| Fsip1 | 3.22E-07 | 0 | 0 | 0 | 4.66E-07 | 0 | 0 | 0 | 4.07E-06 | 0 | 7.64E-06 | 1.22E-05 | 1.22E-05 | 7.18E-06 | 0 | 5.55E-06 | 1.36E-05 | 0 |
| Rbm48 | 2.26E-05 | 2.95E-05 | 0 | 8.37E-06 | 6.92E-06 | 2.84E-05 | 0 | 1.38E-06 | 4.03E-06 | 2.24E-06 | 0 | 1.06E-05 | 0 | 0 | 7.75E-05 | 4.57E-05 | 0 | 2.87E-07 |
| Anapc15 | 2.14E-05 | 0 | 7.79E-05 | 5.46E-05 | 0 | 0 | 0 | 1.80E-05 | 4.03E-06 | 1.26E-05 | 5.39E-05 | 3.79E-06 | 2.83E-06 | 1.08E-05 | 0 | 0.00016 | 0 | 1.67E-05 |
| Hace1 | 5.33E-05 | 2.56E-05 | 0 | 0 | 2.26E-05 | 5.72E-05 | 0 | 1.84E-05 | 4.02E-06 | 0 | 1.32E-06 | 4.42E-05 | 0 | 0 | 0 | 3.44E-05 | 4.35E-05 | 1.08E-05 |
| Wdr78 | 6.24E-06 | 5.01E-06 | 0 | 0 | 1.87E-06 | 8.82E-07 | 1.28E-06 | 1.22E-05 | 4.02E-06 | 6.65E-06 | 0 | 6.61E-06 | 0 | 0 | 2.83E-05 | 1.13E-05 | 1.48E-05 | 0 |
| Itgb7 | 0 | 9.52E-06 | 3.47E-06 | 3.39E-06 | 0 | 0 | 0 | 0 | 4.02E-06 | 2.19E-05 | 0 | 6.53E-06 | 0 | 0 | 5.21E-06 | 0 | 7.26E-06 | 1.19E-05 |
| Icam5 | 0 | 2.42E-06 | 0 | 1.34E-05 | 0 | 0 | 8.58E-06 | 0 | 4.02E-06 | 0 | 0 | 0 | 2.88E-05 | 0 | 2.27E-05 | 0 | 5.09E-05 | 0 |
| Mettl22 | 9.95E-06 | 8.60E-06 | 5.68E-06 | 6.51E-06 | 4.37E-06 | 7.89E-06 | 7.52E-06 | 4.75E-06 | 4.00E-06 | 5.73E-06 | 0 | 1.28E-05 | 0 | 1.33E-06 | 2.47E-05 | 0 | 0 | 1.28E-05 |
| Ggct | 2.81E-06 | 0 | 7.29E-05 | 0 | 0 | 0 | 2.49E-05 | 5.41E-05 | 4.00E-06 | 0.0001 | 0 | 0 | 7.64E-05 | 0 | 8.93E-07 | 2.75E-05 | 0 | 9.37E-05 |
| Rab26 | 2.44E-06 | 6.27E-06 | 2.53E-06 | 2.37E-06 | 0 | 0 | 0 | 3.35E-07 | 4.00E-06 | 7.18E-07 | 6.01E-06 | 2.22E-06 | 0 | 3.80E-06 | 1.02E-05 | 2.77E-06 | 0 | 7.10E-06 |
| Tmem191c | 0 | 8.40E-06 | 0 | 4.98E-06 | 3.53E-06 | 0 | 0 | 0 | 3.99E-06 | 0 | 0 | 1.70E-05 | 0 | 8.79E-06 | 5.43E-05 | 0 | 9.65E-06 | 5.23E-06 |
| Trappc6a | 0 | 8.79E-05 | 0 | 0.00017 | 7.80E-05 | 5.82E-05 | 0 | 1.73E-05 | 3.97E-06 | 5.93E-05 | 3.23E-05 | 0.00039 | 0.00013 | 4.92E-05 | 0 | 1.28E-05 | 0 | 3.14E-05 |
| Eme2 | 0 | 1.97E-05 | 1.52E-06 | 2.32E-05 | 6.54E-06 | 0 | 0 | 0 | 3.95E-06 | 2.40E-06 | 2.29E-06 | 6.43E-06 | 1.17E-05 | 0 | 5.62E-05 | 0 | 0 | 3.30E-06 |
| Nebl | 0 | 6.11E-07 | 0 | 0 | 1.28E-05 | 0 | 5.85E-06 | 3.85E-06 | 3.94E-06 | 0 | 1.42E-05 | 0 | 0 | 0 | 0 | 0 | 0 | 6.28E-06 |
| Zfp661 | 6.40E-06 | 8.82E-06 | 0 | 7.12E-06 | 1.46E-05 | 1.08E-06 | 1.09E-05 | 4.47E-06 | 3.93E-06 | 6.12E-06 | 3.24E-07 | 0 | 0 | 2.90E-07 | 1.89E-05 | 0 | 0 | 3.96E-06 |
| 1-Mar | 2.51E-07 | 9.00E-07 | 0 | 1.29E-06 | 3.87E-06 | 0 | 0 | 3.51E-06 | 3.91E-06 | 0 | 1.82E-05 | 1.26E-07 | 2.56E-05 | 7.95E-07 | 0 | 1.94E-06 | 5.18E-06 | 1.32E-06 |
| Gtf2h1 | 0.00014 | 0.0001 | 1.80E-05 | 0 | 1.72E-05 | 8.18E-05 | 0 | 0 | 3.90E-06 | 0 | 0 | 0.00011 | 0 | 3.00E-05 | 0.00012 | 2.83E-05 | 9.07E-05 | 0 |
| Zfp82 | 9.14E-06 | 1.12E-05 | 1.31E-06 | 1.88E-05 | 0 | 1.90E-06 | 1.58E-06 | 0 | 3.88E-06 | 6.60E-06 | 3.04E-06 | 9.43E-06 | 0 | 4.83E-06 | 1.65E-06 | 0 | 0 | 1.21E-05 |
| Ulk4 | 6.08E-06 | 7.92E-06 | 0 | 4.24E-06 | 4.12E-06 | 1.19E-06 | 0 | 6.44E-06 | 3.88E-06 | 5.59E-06 | 0 | 1.10E-05 | 0 | 0 | 0 | 3.90E-06 | 8.19E-06 | 1.45E-05 |
| Rasl10b | 0 | 1.58E-06 | 0 | 9.70E-06 | 3.00E-05 | 0 | 2.20E-06 | 0 | 3.88E-06 | 7.03E-06 | 0 | 0 | 7.57E-06 | 0 | 6.94E-06 | 0 | 1.40E-06 | 3.48E-07 |
| Sh3d21 | 0 | 1.46E-05 | 0 | 9.54E-06 | 2.83E-06 | 8.45E-06 | 1.92E-06 | 0 | 3.86E-06 | 8.35E-06 | 3.36E-06 | 1.13E-05 | 0 | 2.86E-05 | 0 | 0 | 6.16E-05 | 3.22E-05 |
| Phf19 | 0 | 0 | 0 | 0 | 1.47E-06 | 0 | 0 | 0 | 3.86E-06 | 5.73E-06 | 0 | 0 | 0 | 0 | 0 | 7.19E-05 | 0 | 0 |
| Vsig10l | 0 | 3.33E-05 | 0 | 7.29E-07 | 0 | 1.78E-05 | 0 | 0 | 3.84E-06 | 3.02E-06 | 0 | 0 | 2.15E-05 | 5.26E-05 | 0 | 0 | 0 | 4.67E-06 |
| Nutf2 | 0 | 6.76E-06 | 4.72E-06 | 1.93E-06 | 1.86E-06 | 0 | 0 | 0 | 3.83E-06 | 2.51E-07 | 7.77E-06 | 0 | 0 | 4.77E-06 | 3.32E-05 | 6.12E-06 | 0 | 0 |
| Mapkapk5 | 3.48E-05 | 4.98E-05 | 7.81E-05 | 4.48E-05 | 2.34E-05 | 0 | 4.52E-06 | 3.00E-06 | 3.83E-06 | 1.23E-05 | 3.84E-05 | 2.08E-05 | 0 | 2.01E-06 | 6.57E-05 | 3.95E-05 | 6.14E-05 | 0 |
| Ccdc14 | 0 | 6.20E-06 | 0 | 0 | 6.99E-07 | 3.62E-06 | 0 | 0 | 3.81E-06 | 1.38E-06 | 0 | 0 | 0 | 1.50E-06 | 3.18E-05 | 4.45E-05 | 0 | 2.15E-06 |
| Gpr19 | 0 | 1.60E-05 | 0 | 7.03E-06 | 0 | 1.32E-05 | 1.00E-05 | 1.64E-05 | 3.79E-06 | 1.59E-05 | 0 | 1.27E-05 | 0 | 4.13E-06 | 9.41E-06 | 4.36E-05 | 3.02E-05 | 2.15E-05 |
| Cracr2a | 0 | 0 | 0 | 0 | 2.00E-06 | 8.13E-06 | 2.54E-06 | 8.58E-06 | 3.79E-06 | 1.15E-05 | 0 | 0 | 9.06E-06 | 1.32E-05 | 8.61E-06 | 0 | 0 | 4.47E-05 |

| | | | | | | | | | | | | | | | | | | |
|---|---|---|---|---|---|---|---|---|---|---|---|---|---|---|---|---|---|---|
| Gpr18 | 0 | 4.61E-06 | 0 | 0 | 0 | 3.81E-06 | 5.00E-06 | 3.41E-05 | 3.79E-06 | 0 | 0 | 0 | 8.28E-06 | 0 | 3.54E-08 | 2.48E-06 | 0 | 0 |
| Rap1gap | 0 | 1.48E-05 | 0 | 0 | 6.75E-07 | 0 | 0 | 0 | 3.79E-06 | 0 | 3.04E-05 | 2.59E-05 | 8.49E-06 | 1.20E-05 | 7.18E-06 | 0 | 0 | 4.89E-06 |
| Abcc4 | 1.93E-05 | 9.51E-06 | 4.98E-05 | 1.73E-05 | 1.46E-05 | 6.55E-07 | 8.05E-05 | 1.27E-05 | 3.78E-06 | 1.78E-05 | 5.94E-06 | 0 | 3.76E-05 | 2.41E-06 | 8.55E-06 | 0 | 0 | 0 |
| Irak2 | 0 | 1.99E-05 | 0 | 2.19E-05 | 2.97E-05 | 0 | 0 | 0 | 3.78E-06 | 1.35E-05 | 1.98E-05 | 2.52E-05 | 1.64E-06 | 2.25E-05 | 3.85E-06 | 0 | 8.64E-06 | 2.14E-05 |
| Gin1 | 2.06E-05 | 2.10E-05 | 7.33E-08 | 2.34E-05 | 2.03E-06 | 3.79E-06 | 2.52E-06 | 1.75E-05 | 3.74E-06 | 1.27E-05 | 1.77E-06 | 1.03E-06 | 0 | 0 | 1.03E-05 | 2.06E-05 | 0 | 1.43E-05 |
| Dmrt2 | 0 | 2.24E-05 | 0 | 0 | 0 | 0 | 7.44E-05 | 0 | 3.74E-06 | 0.00013 | 0 | 0 | 8.09E-05 | 8.20E-06 | 0 | 0 | 0 | 0 |
| Mtus1 | 0 | 8.17E-05 | 0 | 0 | 0.00022 | 0.00013 | 5.31E-05 | 0 | 3.70E-06 | 0 | 0.0001 | 4.63E-05 | 0 | 0.0001 | 8.47E-06 | 0 | 3.87E-05 | 6.64E-05 |
| Rapgef3 | 0 | 1.43E-05 | 0 | 0 | 0 | 0 | 0 | 0 | 3.70E-06 | 9.68E-06 | 0 | 5.62E-07 | 7.65E-06 | 3.94E-05 | 0 | 0 | 3.33E-06 | 3.39E-05 |
| Rhbdd1 | 4.44E-05 | 0.0001 | 6.25E-05 | 3.69E-05 | 7.43E-05 | 3.25E-05 | 3.77E-05 | 6.83E-05 | 3.69E-06 | 5.56E-05 | 6.80E-05 | 0.00012 | 5.45E-05 | 2.29E-05 | 5.75E-05 | 5.98E-05 | 0 | 4.48E-05 |
| Ramp3 | 0 | 0 | 0 | 0 | 0 | 4.66E-05 | 0 | 0 | 3.68E-06 | 0 | 0 | 9.53E-06 | 0 | 0 | 9.52E-08 | 8.91E-06 | 3.26E-05 | 8.28E-06 |
| Ptch2 | 0 | 3.10E-05 | 0 | 6.67E-05 | 5.19E-05 | 3.05E-05 | 0 | 5.98E-05 | 3.67E-06 | 0.00015 | 0 | 0 | 0 | 0 | 7.84E-05 | 0 | 0 | 0.0004 |
| Lmtk2 | 3.88E-05 | 4.18E-05 | 3.05E-05 | 0 | 8.05E-06 | 1.34E-05 | 0 | 0 | 3.67E-06 | 0 | 2.08E-06 | 3.01E-05 | 0 | 0 | 0 | 2.02E-05 | 3.73E-05 | 0 |
| Wdcp | 0 | 2.81E-06 | 7.29E-06 | 9.63E-06 | 8.23E-06 | 0 | 0 | 5.41E-06 | 3.67E-06 | 6.97E-06 | 6.26E-06 | 8.85E-06 | 7.60E-07 | 2.05E-06 | 0 | 1.09E-05 | 2.12E-06 | 1.11E-05 |
| Clec4n | 0 | 7.52E-06 | 1.13E-05 | 8.82E-06 | 1.77E-06 | 0 | 0 | 0 | 3.67E-06 | 6.44E-06 | 6.66E-07 | 7.66E-06 | 1.33E-05 | 1.05E-05 | 4.46E-05 | 1.19E-05 | 1.53E-06 | 9.06E-06 |
| Smim14 | 0.00019 | 0.00026 | 4.53E-05 | 0.00025 | 0.00023 | 0.0002 | 0.00032 | 6.94E-05 | 3.64E-06 | 0.00019 | 0.00013 | 0.00028 | 0 | 4.95E-05 | 1.86E-05 | 0 | 0 | 0.00022 |
| Cyth1 | 1.37E-05 | 0.00018 | 8.76E-05 | 6.92E-05 | 0.00015 | 2.32E-05 | 2.93E-05 | 7.39E-05 | 3.64E-06 | 2.65E-05 | 3.97E-05 | 1.88E-05 | 0 | 0.00011 | 0.00014 | 0 | 9.01E-05 | 0 |
| Fbxl22 | 2.54E-07 | 1.19E-05 | 9.46E-06 | 0 | 0 | 0 | 1.60E-05 | 3.68E-06 | 3.64E-06 | 1.02E-07 | 3.03E-06 | 1.07E-05 | 1.07E-05 | 2.00E-05 | 6.94E-06 | 0 | 6.37E-05 | 7.55E-07 |
| Trmu | 2.41E-05 | 3.26E-05 | 3.27E-05 | 2.61E-05 | 5.04E-06 | 0 | 0 | 4.43E-07 | 3.64E-06 | 0 | 4.01E-06 | 9.00E-06 | 0 | 0 | 5.22E-05 | 7.19E-06 | 1.63E-05 | 4.79E-06 |
| Dgke | 1.43E-05 | 2.25E-05 | 2.17E-05 | 0 | 1.91E-05 | 2.16E-05 | 0 | 0 | 3.63E-06 | 7.14E-07 | 1.68E-05 | 1.44E-05 | 0 | 8.01E-06 | 2.00E-05 | 0 | 0 | 3.31E-05 |
| Spa17 | 5.83E-06 | 4.88E-06 | 5.26E-06 | 1.13E-05 | 0 | 0 | 0 | 0 | 3.63E-06 | 6.04E-06 | 0 | 6.66E-06 | 0 | 0 | 0 | 6.66E-06 | 0 | 7.03E-06 |
| Kctd1 | 7.10E-06 | 4.17E-05 | 0 | 6.61E-07 | 0 | 0 | 0 | 2.78E-05 | 3.62E-06 | 0 | 0 | 0 | 0 | 0.00016 | 0 | 0 | 0 | 4.81E-05 |
| Adamts8 | 0 | 2.21E-06 | 1.19E-05 | 0 | 0 | 0 | 0 | 3.32E-06 | 3.61E-06 | 1.42E-05 | 0 | 0 | 1.14E-06 | 0 | 0 | 3.19E-06 | 5.22E-06 | 1.67E-05 |
| Zfp688 | 3.50E-05 | 6.88E-05 | 5.37E-06 | 5.01E-05 | 3.73E-05 | 5.35E-05 | 2.14E-05 | 7.40E-06 | 3.59E-06 | 6.85E-06 | 2.63E-07 | 1.13E-05 | 1.47E-05 | 2.48E-05 | 7.22E-05 | 0 | 0 | 0 |
| Fam122b | 7.09E-06 | 9.83E-06 | 0 | 1.59E-06 | 1.11E-05 | 9.87E-06 | 0 | 5.79E-06 | 3.57E-06 | 1.82E-05 | 0 | 9.22E-06 | 0 | 0 | 3.40E-06 | 8.49E-05 | 1.72E-05 | 2.80E-07 |
| Cep19 | 4.52E-05 | 3.44E-05 | 0 | 7.65E-06 | 3.22E-05 | 7.71E-06 | 2.24E-05 | 8.00E-06 | 3.57E-06 | 2.69E-05 | 7.28E-07 | 4.39E-06 | 0 | 0 | 2.09E-06 | 8.87E-06 | 1.11E-05 | 2.07E-05 |
| Cdc14a | 1.01E-05 | 2.25E-05 | 0 | 6.76E-06 | 0 | 8.17E-06 | 0 | 3.48E-07 | 3.56E-06 | 8.25E-06 | 0 | 0 | 0 | 0 | 5.94E-06 | 3.15E-06 | 1.64E-05 | 7.41E-06 |
| Loxl4 | 0 | 0 | 0.00011 | 3.48E-05 | 0 | 0 | 0.00013 | 0 | 3.56E-06 | 0.00035 | 0 | 9.20E-06 | 0.00027 | 0.00018 | 0 | 0 | 0 | 0.00012 |
| Nek3 | 1.94E-06 | 1.33E-05 | 0 | 0 | 0 | 4.86E-06 | 5.27E-06 | 9.06E-06 | 3.55E-06 | 8.45E-06 | 0 | 1.10E-06 | 1.34E-05 | 0 | 1.54E-05 | 2.03E-05 | 3.44E-05 | 7.24E-06 |
| Zfp959 | 4.90E-06 | 1.49E-07 | 6.12E-06 | 0 | 2.22E-06 | 5.88E-06 | 5.06E-06 | 2.72E-06 | 3.54E-06 | 3.64E-06 | 0 | 7.35E-06 | 0 | 9.50E-06 | 0 | 5.80E-06 | 0 | 0 |
| Map9 | 2.88E-06 | 4.19E-06 | 7.32E-06 | 3.41E-06 | 0 | 0 | 2.01E-06 | 7.26E-06 | 3.54E-06 | 9.67E-06 | 0 | 0 | 6.43E-06 | 0 | 0 | 0 | 2.25E-06 | 5.75E-06 |
| Sass6 | 1.11E-05 | 0 | 0 | 0 | 1.33E-05 | 0 | 7.29E-06 | 7.60E-06 | 3.52E-06 | 4.02E-06 | 1.05E-06 | 0 | 5.59E-06 | 1.14E-05 | 2.57E-05 | 0.00029 | 0 | 3.34E-06 |
| Phf8 | 8.77E-05 | 0.00011 | 8.86E-05 | 0 | 5.80E-05 | 4.99E-05 | 7.58E-05 | 3.47E-05 | 3.51E-06 | 3.51E-05 | 1.61E-05 | 4.88E-05 | 8.60E-05 | 5.51E-05 | 0.00015 | 6.38E-06 | 5.67E-05 | 0 |
| Lmnb2 | 0 | 0 | 2.50E-05 | 4.82E-05 | 0 | 0 | 0 | 2.13E-05 | 3.50E-06 | 2.26E-05 | 0 | 0 | 0 | 0 | 4.94E-05 | 0.00032 | 2.45E-05 | 0 |

| | | | | | | | | | | | | | | | | | | |
|---|---|---|---|---|---|---|---|---|---|---|---|---|---|---|---|---|---|---|
| Ankzf1 | 2.37E-05 | 2.18E-05 | 7.04E-06 | 1.55E-05 | 2.67E-05 | 4.09E-05 | 1.80E-05 | 1.75E-05 | 3.49E-06 | 2.34E-06 | 1.16E-05 | 3.95E-05 | 0 | 1.03E-05 | 1.46E-05 | 0 | 0 | 1.28E-05 |
| Clmn | 0 | 6.22E-06 | 0 | 0 | 3.14E-05 | 0 | 3.60E-05 | 0 | 3.49E-06 | 7.97E-06 | 1.21E-05 | 3.65E-05 | 0 | 0 | 6.23E-06 | 2.03E-05 | 4.06E-06 | 2.19E-06 |
| Bmp2 | 0 | 1.32E-05 | 0 | 0 | 0 | 0 | 0 | 0 | 3.49E-06 | 7.20E-06 | 0 | 1.95E-05 | 1.25E-05 | 0 | 0 | 0 | 0.0001 | 0 |
| Ak4 | 3.35E-05 | 0 | 5.90E-05 | 0 | 0 | 0 | 4.52E-05 | 0 | 3.46E-06 | 1.86E-05 | 0 | 0.00056 | 2.67E-05 | 0 | 0 | 0 | 0 | 0 |
| Irf8 | 0 | 1.58E-05 | 0 | 0 | 0 | 0 | 0 | 0 | 3.45E-06 | 1.89E-06 | 0 | 3.52E-07 | 0 | 0 | 2.10E-05 | 3.41E-06 | 1.38E-05 | 1.76E-05 |
| Zfp438 | 4.36E-06 | 2.71E-05 | 7.96E-06 | 0 | 1.32E-05 | 1.10E-05 | 5.11E-06 | 6.80E-06 | 3.44E-06 | 8.67E-06 | 0 | 1.84E-05 | 0 | 0 | 8.46E-06 | 5.60E-06 | 3.44E-05 | 7.82E-06 |
| Akap6 | 0 | 0 | 1.77E-07 | 0 | 7.97E-06 | 7.34E-06 | 0 | 0 | 3.44E-06 | 3.42E-07 | 2.52E-05 | 0 | 0 | 1.12E-05 | 1.93E-05 | 2.03E-06 | 0 | 0 |
| Mid1 | 0 | 6.74E-07 | 0 | 9.30E-06 | 6.50E-06 | 0 | 0 | 0 | 3.42E-06 | 6.70E-06 | 0 | 0 | 4.28E-06 | 0 | 1.34E-05 | 3.29E-06 | 5.90E-06 | 2.70E-06 |
| Zfp597 | 5.27E-05 | 2.54E-05 | 1.78E-05 | 0 | 9.30E-06 | 2.55E-05 | 1.33E-05 | 1.27E-05 | 3.40E-06 | 8.76E-06 | 8.73E-06 | 2.09E-05 | 3.82E-06 | 0 | 3.49E-05 | 2.94E-05 | 0 | 2.37E-06 |
| Cdkn1b | 0.00019 | 0.00035 | 0 | 0.00014 | 0.00017 | 0.00025 | 0.00015 | 0 | 3.39E-06 | 0.00016 | 0.00011 | 0.00063 | 2.59E-05 | 0 | 0.00025 | 0.00029 | 0 | 0.00022 |
| Mtmr2 | 8.01E-06 | 0 | 7.53E-06 | 0 | 7.64E-06 | 2.65E-06 | 4.96E-06 | 1.59E-06 | 3.38E-06 | 1.38E-06 | 2.78E-06 | 3.59E-06 | 0 | 0 | 0 | 8.00E-06 | 0 | 0 |
| Mns1 | 0 | 0 | 0 | 3.61E-06 | 0 | 2.69E-07 | 0 | 3.22E-07 | 3.38E-06 | 0 | 0 | 0 | 0 | 0 | 0 | 0.00019 | 0 | 9.19E-07 |
| Ubap2 | 0.00011 | 0.00011 | 0.00018 | 0.00021 | 4.32E-05 | 7.27E-05 | 0.00016 | 9.98E-05 | 3.37E-06 | 7.07E-05 | 0.00014 | 9.42E-05 | 8.92E-05 | 6.95E-05 | 0.00021 | 0.00037 | 0.00016 | 7.69E-05 |
| Wnk3 | 9.02E-06 | 0 | 5.36E-06 | 0 | 0 | 1.61E-06 | 6.22E-06 | 0 | 3.37E-06 | 0 | 0 | 9.36E-06 | 0 | 5.81E-06 | 0 | 3.96E-06 | 4.15E-06 | 0 |
| Siah1b | 7.89E-06 | 4.09E-06 | 2.32E-05 | 1.49E-05 | 1.39E-06 | 8.25E-06 | 0 | 3.81E-06 | 3.36E-06 | 7.66E-06 | 4.28E-06 | 0 | 1.47E-05 | 1.32E-05 | 5.34E-05 | 0.00013 | 0 | 1.30E-05 |
| Trim36 | 0 | 0 | 1.28E-05 | 0 | 9.23E-06 | 2.82E-05 | 0 | 1.49E-05 | 3.35E-06 | 0 | 5.20E-06 | 0 | 0 | 0 | 0 | 7.66E-07 | 1.63E-05 | 0 |
| Eppk1 | 0 | 1.58E-05 | 5.26E-08 | 6.03E-06 | 0 | 0 | 0 | 0 | 3.35E-06 | 3.26E-05 | 1.15E-05 | 1.39E-06 | 0 | 9.77E-06 | 6.21E-06 | 0 | 0 | 3.89E-05 |
| Zbtb42 | 1.07E-06 | 3.25E-05 | 3.64E-06 | 3.81E-06 | 2.63E-06 | 8.19E-06 | 0 | 3.70E-06 | 3.33E-06 | 4.70E-06 | 2.89E-05 | 1.95E-05 | 0 | 0 | 1.18E-05 | 0 | 1.83E-05 | 1.49E-05 |
| Cdkn3 | 1.91E-05 | 0 | 0 | 0 | 0 | 0 | 0 | 0 | 3.32E-06 | 4.63E-06 | 0 | 0 | 0 | 0 | 0 | 0.00029 | 0 | 0 |
| Cycs | 1.90E-05 | 2.11E-06 | 1.42E-05 | 0 | 0 | 3.83E-06 | 0 | 0 | 3.29E-06 | 0 | 5.60E-06 | 0 | 3.68E-06 | 0 | 0 | 1.98E-05 | 1.48E-05 | 0 |
| Atg13 | 0.00012 | 0.00011 | 1.80E-05 | 7.17E-05 | 6.81E-05 | 9.17E-05 | 0.00014 | 2.24E-05 | 3.28E-06 | 2.08E-05 | 4.35E-05 | 8.04E-05 | 1.68E-05 | 6.28E-05 | 6.65E-05 | 0 | 1.32E-05 | 2.04E-05 |
| Cenps | 0 | 0 | 2.27E-05 | 0 | 0 | 0 | 0 | 0 | 3.28E-06 | 0 | 3.86E-06 | 0 | 7.31E-06 | 3.63E-06 | 2.33E-06 | 0.0002 | 0 | 5.07E-06 |
| Frat2 | 0 | 3.71E-05 | 0 | 0 | 2.25E-05 | 6.19E-06 | 0 | 0 | 3.26E-06 | 0 | 1.89E-06 | 5.69E-05 | 0 | 3.80E-05 | 9.38E-05 | 3.33E-05 | 0.00012 | 0 |
| Cep85l | 4.48E-05 | 8.13E-05 | 1.95E-05 | 0 | 1.47E-05 | 1.14E-05 | 2.95E-05 | 1.43E-05 | 3.23E-06 | 5.71E-05 | 6.78E-06 | 3.33E-05 | 2.55E-05 | 1.11E-05 | 5.71E-05 | 2.03E-05 | 0.00014 | 0 |
| Prr22 | 1.40E-07 | 0 | 3.57E-06 | 4.74E-06 | 0 | 0 | 0 | 6.29E-06 | 3.23E-06 | 2.31E-06 | 0 | 0 | 1.00E-05 | 5.32E-06 | 5.38E-06 | 1.49E-06 | 0 | 5.24E-06 |
| Map3k13 | 0 | 2.44E-06 | 0 | 2.58E-06 | 6.17E-06 | 0 | 6.40E-06 | 3.89E-06 | 3.21E-06 | 2.35E-06 | 0 | 6.94E-06 | 2.54E-07 | 3.06E-06 | 4.20E-06 | 0 | 3.00E-06 | 1.47E-06 |
| BC017158 | 5.33E-05 | 1.48E-05 | 3.34E-05 | 7.39E-05 | 4.77E-05 | 9.02E-05 | 7.97E-05 | 7.64E-05 | 3.20E-06 | 2.69E-05 | 2.29E-05 | 1.31E-05 | 3.46E-05 | 6.80E-05 | 3.35E-05 | 8.27E-07 | 0 | 2.94E-05 |
| Pitpnm1 | 5.45E-06 | 9.52E-05 | 4.07E-05 | 7.06E-05 | 2.24E-05 | 5.00E-05 | 0 | 0 | 3.20E-06 | 5.92E-06 | 1.94E-05 | 0 | 0 | 0.00012 | 0.00011 | 1.25E-05 | 8.30E-05 | 4.90E-05 |
| Fzd6 | 4.24E-06 | 6.09E-06 | 0 | 0 | 0 | 0 | 7.81E-05 | 0 | 3.20E-06 | 3.03E-05 | 0 | 3.10E-06 | 0 | 0 | 0 | 1.77E-06 | 0 | 1.99E-05 |
| Zfp316 | 0 | 2.98E-05 | 0 | 2.99E-05 | 9.92E-06 | 0 | 0 | 0 | 3.20E-06 | 8.03E-06 | 1.18E-05 | 3.21E-05 | 0 | 1.35E-05 | 1.94E-06 | 0 | 0 | 3.25E-05 |
| Fam177a | 0 | 1.16E-05 | 1.14E-05 | 6.49E-06 | 0 | 0 | 0 | 3.62E-06 | 3.18E-06 | 1.18E-05 | 1.26E-05 | 2.00E-06 | 0 | 1.04E-05 | 2.72E-05 | 1.07E-05 | 0 | 1.61E-05 |
| Adam1a | 0 | 6.70E-06 | 1.13E-05 | 0 | 0 | 0 | 0 | 7.07E-06 | 3.18E-06 | 5.59E-06 | 6.24E-06 | 5.54E-07 | 0 | 5.25E-06 | 1.78E-05 | 6.73E-06 | 5.32E-05 | 0 |
| Ticam1 | 4.87E-05 | 2.78E-05 | 1.56E-05 | 2.93E-05 | 1.77E-05 | 2.24E-05 | 9.57E-06 | 1.52E-05 | 3.17E-06 | 2.45E-05 | 9.96E-06 | 5.56E-05 | 0 | 2.07E-07 | 3.61E-05 | 0 | 0 | 8.12E-06 |

| | | | | | | | | | | | | | | | | | | |
|---|---|---|---|---|---|---|---|---|---|---|---|---|---|---|---|---|---|---|
| Tes3-ps | 1.76E-05 | 2.33E-06 | 5.03E-06 | 0 | 6.97E-06 | 3.86E-06 | 2.23E-06 | 2.44E-06 | 3.16E-06 | 1.40E-06 | 0 | 1.06E-05 | 0 | 0 | 0 | 0 | 1.53E-05 | 4.93E-06 |
| Zfp93 | 5.24E-06 | 1.32E-05 | 0 | 3.57E-06 | 2.81E-06 | 3.74E-06 | 5.86E-06 | 0 | 3.16E-06 | 1.45E-06 | 3.37E-06 | 3.69E-06 | 3.00E-06 | 1.82E-06 | 0 | 0 | 0 | 6.73E-06 |
| Ttc32 | 1.72E-05 | 8.41E-06 | 0 | 1.03E-05 | 1.75E-06 | 1.06E-05 | 7.73E-07 | 0 | 3.14E-06 | 6.52E-06 | 1.71E-06 | 6.45E-06 | 0 | 1.60E-05 | 0 | 3.19E-05 | 0 | 8.49E-08 |
| Rmi2 | 0 | 0 | 6.37E-06 | 6.64E-06 | 0 | 0 | 0 | 0 | 3.12E-06 | 0 | 0 | 0 | 3.47E-07 | 0 | 4.15E-06 | 7.05E-05 | 0 | 0 |
| Ppp1r14c | 2.55E-06 | 4.47E-06 | 0 | 0 | 4.07E-06 | 0 | 4.30E-07 | 0 | 3.11E-06 | 1.99E-06 | 1.63E-06 | 2.61E-06 | 0 | 0 | 1.79E-05 | 0 | 0 | 9.38E-06 |
| Pot1a | 3.51E-05 | 2.73E-05 | 0 | 1.93E-06 | 2.86E-06 | 4.42E-05 | 1.82E-06 | 2.31E-05 | 3.10E-06 | 8.99E-06 | 1.76E-06 | 2.78E-05 | 9.44E-06 | 0 | 1.35E-05 | 5.42E-05 | 0 | 4.49E-06 |
| Zfp369 | 1.83E-05 | 8.85E-05 | 6.21E-06 | 3.37E-05 | 3.03E-05 | 2.49E-05 | 1.95E-05 | 0 | 3.10E-06 | 2.15E-05 | 1.53E-05 | 1.40E-05 | 1.87E-05 | 3.41E-05 | 3.93E-05 | 0 | 4.48E-06 | 8.96E-06 |
| Rgs17 | 5.51E-06 | 7.47E-06 | 1.27E-06 | 0 | 0 | 0 | 1.12E-05 | 0 | 3.10E-06 | 1.65E-05 | 0 | 2.15E-05 | 3.76E-07 | 0 | 0 | 0 | 2.66E-05 | 0 |
| Mcpt2 | 2.59E-06 | 1.67E-05 | 0 | 0 | 4.43E-06 | 0 | 2.27E-06 | 0 | 3.09E-06 | 2.56E-05 | 0 | 2.53E-05 | 0 | 0 | 0.00023 | 9.49E-06 | 4.60E-05 | 2.78E-05 |
| Taf4 | 3.15E-05 | 0.00012 | 5.20E-05 | 3.62E-05 | 4.25E-05 | 2.36E-05 | 0 | 3.82E-05 | 3.08E-06 | 2.30E-05 | 2.66E-05 | 4.92E-05 | 0 | 2.77E-05 | 0.00019 | 0.00015 | 0.00012 | 1.41E-05 |
| Ccdc142 | 1.27E-05 | 2.04E-05 | 1.36E-06 | 0 | 1.30E-05 | 0 | 5.20E-06 | 5.92E-06 | 3.08E-06 | 0 | 0 | 0 | 0 | 2.13E-06 | 7.83E-05 | 7.52E-06 | 0 | 0 |
| Ccdc17 | 0 | 7.96E-06 | 1.43E-07 | 0 | 5.21E-06 | 1.34E-06 | 6.29E-06 | 2.15E-06 | 3.07E-06 | 2.39E-06 | 0 | 7.95E-06 | 4.16E-06 | 3.60E-06 | 1.33E-05 | 0 | 0 | 2.81E-06 |
| Rab6b | 8.52E-06 | 0.00011 | 1.05E-05 | 0 | 6.12E-05 | 4.71E-05 | 0 | 0.00015 | 3.06E-06 | 9.09E-05 | 3.90E-05 | 0.0001 | 0 | 0 | 0 | 0 | 0 | 1.71E-05 |
| Zfp955a | 2.04E-05 | 9.21E-06 | 0 | 0 | 1.03E-05 | 1.49E-05 | 0 | 0 | 3.05E-06 | 1.11E-05 | 0 | 1.21E-05 | 0 | 0 | 1.00E-05 | 0 | 0 | 1.28E-06 |
| Frs3 | 1.20E-05 | 1.72E-05 | 1.54E-06 | 0 | 1.82E-05 | 1.73E-05 | 7.85E-06 | 1.59E-06 | 3.03E-06 | 8.30E-07 | 0 | 2.38E-05 | 1.96E-05 | 0 | 2.51E-05 | 4.95E-06 | 0 | 0 |
| Trim5 | 7.53E-06 | 8.43E-06 | 0 | 0 | 0 | 3.41E-06 | 0 | 0 | 3.03E-06 | 1.26E-05 | 0 | 0 | 0 | 0 | 0 | 0 | 0 | 8.92E-06 |
| Adck2 | 3.72E-05 | 4.55E-05 | 4.12E-06 | 3.70E-05 | 4.92E-05 | 3.34E-05 | 2.92E-05 | 2.27E-05 | 3.02E-06 | 5.90E-05 | 1.39E-05 | 1.78E-05 | 4.30E-05 | 0 | 6.64E-05 | 1.89E-06 | 0 | 4.44E-05 |
| Shb | 0 | 5.84E-05 | 3.26E-05 | 0.00011 | 7.22E-05 | 0 | 1.89E-05 | 5.12E-05 | 3.01E-06 | 0.00017 | 6.42E-05 | 0.00012 | 0.00013 | 0 | 0.00019 | 1.26E-05 | 0.00021 | 0.0001 |
| Ccdc191 | 6.32E-06 | 2.39E-05 | 0 | 8.29E-06 | 0 | 0 | 1.54E-05 | 9.00E-08 | 3.01E-06 | 9.89E-06 | 0 | 2.49E-05 | 0 | 0 | 0.00011 | 1.56E-05 | 4.12E-05 | 8.17E-06 |
| Xylb | 6.98E-06 | 8.82E-06 | 2.17E-06 | 4.50E-06 | 1.71E-06 | 4.14E-06 | 6.49E-06 | 0 | 3.00E-06 | 3.38E-06 | 0 | 0 | 3.08E-06 | 1.37E-05 | 0 | 2.13E-05 | 1.33E-05 | 0 |
| Gpx2 | 0 | 1.97E-06 | 1.89E-06 | 0 | 8.29E-06 | 1.76E-06 | 0 | 3.20E-07 | 3.00E-06 | 1.98E-06 | 1.48E-05 | 2.59E-06 | 0 | 0 | 3.34E-07 | 3.28E-06 | 8.05E-07 | 4.25E-06 |
| Itgb3bp | 2.36E-05 | 1.48E-06 | 0 | 0 | 0 | 1.69E-07 | 0 | 0 | 3.00E-06 | 1.92E-05 | 0 | 4.30E-06 | 0 | 0 | 0 | 6.32E-05 | 0 | 6.15E-06 |
| Osgin1 | 0 | 1.40E-05 | 6.47E-06 | 1.34E-05 | 0 | 0 | 5.10E-06 | 2.11E-06 | 3.00E-06 | 1.11E-05 | 0 | 2.77E-06 | 0 | 0 | 0 | 0 | 5.58E-05 | 1.53E-05 |
| Sirt4 | 2.89E-05 | 4.16E-05 | 0 | 2.87E-05 | 1.19E-05 | 1.82E-05 | 0 | 1.18E-05 | 2.99E-06 | 8.23E-06 | 0 | 8.79E-06 | 6.95E-06 | 2.50E-05 | 6.21E-05 | 0 | 0 | 3.72E-05 |
| Gpn2 | 3.66E-05 | 5.47E-05 | 0.00015 | 3.61E-05 | 1.76E-05 | 0 | 2.95E-05 | 2.39E-06 | 2.99E-06 | 1.69E-05 | 4.46E-05 | 0 | 8.72E-06 | 1.31E-05 | 2.37E-05 | 8.17E-05 | 6.50E-05 | 3.99E-05 |
| Zfp942 | 3.21E-05 | 1.31E-05 | 0 | 0 | 1.13E-05 | 2.00E-05 | 7.43E-07 | 5.03E-06 | 2.97E-06 | 4.75E-06 | 0 | 1.43E-05 | 0 | 0 | 1.40E-05 | 6.48E-07 | 0 | 3.22E-06 |
| Alyref | 0 | 3.20E-05 | 1.71E-05 | 1.59E-05 | 0 | 1.78E-06 | 0 | 3.45E-05 | 2.96E-06 | 1.33E-05 | 0 | 0.00012 | 0 | 0 | 6.16E-05 | 0.00012 | 0.00064 | 0 |
| Lyrm1 | 8.28E-06 | 1.04E-05 | 2.44E-06 | 0 | 1.17E-05 | 8.38E-06 | 1.37E-05 | 0 | 2.95E-06 | 6.52E-06 | 2.20E-06 | 2.92E-06 | 5.68E-06 | 0 | 0 | 9.28E-06 | 0 | 7.03E-06 |
| Gen1 | 0 | 0 | 0 | 0 | 1.87E-06 | 7.52E-07 | 0 | 0 | 2.95E-06 | 0 | 0 | 0 | 0 | 0 | 9.64E-06 | 0.00024 | 0 | 0 |
| Ralgapa2 | 1.04E-05 | 3.86E-05 | 0 | 0 | 3.92E-05 | 5.01E-05 | 0 | 0 | 2.94E-06 | 1.04E-05 | 0 | 0.0001 | 0 | 0 | 5.63E-05 | 9.52E-06 | 4.14E-05 | 1.06E-05 |
| Tnfsf18 | 0 | 0 | 0.00013 | 0 | 0 | 0 | 0.00056 | 0 | 2.94E-06 | 0 | 0 | 0 | 0.00014 | 0 | 0 | 0 | 0 | 0.00014 |
| Zfp846 | 2.90E-05 | 2.09E-05 | 5.09E-06 | 0 | 5.22E-06 | 2.86E-05 | 7.08E-06 | 0 | 2.93E-06 | 4.06E-06 | 0 | 3.70E-06 | 8.16E-08 | 0 | 6.80E-06 | 2.03E-05 | 0 | 1.52E-05 |
| Myo5b | 0 | 0 | 0 | 0 | 3.32E-05 | 3.94E-06 | 0 | 0 | 2.93E-06 | 6.42E-06 | 0 | 1.12E-05 | 0 | 0 | 1.02E-05 | 1.65E-07 | 1.34E-05 | 8.87E-06 |

| | | | | | | | | | | | | | | | | | | |
|---|---|---|---|---|---|---|---|---|---|---|---|---|---|---|---|---|---|---|
| Fgf7 | 6.26E-05 | 0 | 0.00013 | 0 | 0 | 1.91E-05 | 0 | 0 | 2.92E-06 | 0.00032 | 0 | 0 | 0.00187 | 0.00024 | 0.00014 | 0 | 0 | 0.00051 |
| Dclk2 | 0 | 8.81E-06 | 0 | 0 | 6.80E-06 | 0 | 1.75E-06 | 0 | 2.92E-06 | 0 | 0 | 1.58E-05 | 1.35E-08 | 7.25E-06 | 2.70E-05 | 1.98E-05 | 0 | 1.27E-05 |
| Prss36 | 1.30E-06 | 2.91E-06 | 3.29E-06 | 1.07E-05 | 1.03E-06 | 0 | 1.01E-05 | 5.35E-06 | 2.92E-06 | 1.03E-05 | 0 | 5.59E-06 | 9.61E-08 | 0 | 2.87E-06 | 0 | 0 | 3.00E-05 |
| Ttc33 | 7.35E-05 | 0.00013 | 0 | 3.11E-05 | 9.95E-05 | 0.0001 | 4.18E-05 | 3.06E-05 | 2.91E-06 | 2.44E-05 | 1.35E-05 | 8.06E-05 | 2.37E-06 | 1.20E-05 | 0.00016 | 3.48E-05 | 0 | 9.73E-05 |
| Ager | 0 | 9.29E-06 | 0 | 5.83E-06 | 2.96E-06 | 9.49E-06 | 0 | 2.45E-05 | 2.91E-06 | 0 | 0 | 0 | 0 | 0 | 3.29E-05 | 8.87E-06 | 1.10E-05 | 0 |
| Sh2d4a | 1.88E-06 | 7.29E-06 | 0 | 0 | 0 | 7.63E-06 | 0 | 0 | 2.91E-06 | 8.24E-06 | 0 | 0 | 0 | 0 | 0 | 3.60E-06 | 0 | 2.31E-05 |
| Metap1d | 3.33E-05 | 8.77E-05 | 0 | 6.45E-07 | 4.29E-05 | 7.93E-05 | 0 | 2.89E-05 | 2.90E-06 | 2.63E-05 | 0 | 3.01E-05 | 3.27E-05 | 1.28E-05 | 0.00027 | 2.58E-05 | 4.28E-05 | 2.42E-05 |
| Tonsl | 0 | 0 | 1.62E-05 | 2.40E-06 | 4.69E-07 | 0 | 0 | 0 | 2.88E-06 | 3.19E-06 | 0 | 0 | 6.80E-07 | 0 | 1.30E-05 | 0.00015 | 1.35E-05 | 9.96E-06 |
| Zfp788 | 2.31E-05 | 2.33E-05 | 0 | 0 | 1.62E-07 | 1.80E-05 | 0 | 3.81E-06 | 2.87E-06 | 0 | 0 | 1.15E-05 | 0 | 0 | 4.44E-05 | 0 | 0 | 6.12E-07 |
| Cd59b | 1.50E-06 | 0 | 0 | 0 | 1.46E-05 | 1.01E-06 | 7.09E-07 | 0 | 2.87E-06 | 2.55E-07 | 1.20E-05 | 0 | 2.64E-06 | 0 | 0 | 0 | 9.32E-07 | 4.56E-06 |
| Arrdc2 | 0 | 1.75E-05 | 0 | 3.18E-05 | 0 | 2.59E-05 | 0 | 2.39E-05 | 2.85E-06 | 4.12E-06 | 6.28E-06 | 3.74E-05 | 0 | 2.38E-05 | 3.44E-05 | 0 | 0.00015 | 2.61E-05 |
| Syne1 | 1.75E-05 | 4.89E-05 | 0 | 0 | 7.71E-05 | 0.00011 | 0 | 3.37E-05 | 2.84E-06 | 7.12E-06 | 4.34E-05 | 7.48E-05 | 0 | 0.00011 | 6.93E-05 | 6.36E-06 | 9.33E-06 | 7.72E-05 |
| Dnajc12 | 5.62E-06 | 4.94E-06 | 0 | 3.83E-06 | 1.20E-06 | 2.40E-05 | 0 | 7.48E-06 | 2.81E-06 | 1.42E-06 | 0 | 0 | 0 | 1.71E-05 | 0 | 4.47E-07 | 0 | 0 |
| Napb | 3.68E-06 | 1.00E-05 | 0 | 0 | 1.23E-05 | 0 | 4.68E-06 | 0 | 2.81E-06 | 6.31E-06 | 0 | 7.24E-06 | 0 | 0 | 2.99E-05 | 0 | 1.29E-06 | 3.52E-06 |
| Socs2 | 0 | 6.49E-06 | 0.00015 | 0.00013 | 0.00018 | 2.67E-06 | 0 | 0.00022 | 2.80E-06 | 2.09E-05 | 0.00014 | 0 | 0 | 0 | 0.00014 | 0 | 0 | 0.00011 |
| Btc | 4.41E-07 | 1.31E-05 | 3.96E-07 | 3.05E-06 | 0 | 7.13E-08 | 0 | 2.13E-06 | 2.77E-06 | 0 | 3.77E-07 | 1.79E-05 | 0 | 0 | 3.03E-05 | 8.33E-06 | 3.10E-07 | 1.33E-05 |
| Dennd6b | 7.90E-06 | 1.97E-05 | 2.61E-06 | 0 | 8.36E-06 | 0 | 9.86E-06 | 0 | 2.77E-06 | 0 | 0 | 1.24E-06 | 0 | 0 | 7.39E-05 | 6.86E-06 | 1.11E-05 | 0 |
| Zfp808 | 0 | 5.68E-06 | 0 | 1.01E-05 | 2.99E-06 | 0 | 1.22E-07 | 0 | 2.76E-06 | 7.92E-06 | 0 | 0 | 9.09E-06 | 1.23E-06 | 3.40E-05 | 3.82E-05 | 0 | 0 |
| Spag4 | 0 | 2.39E-06 | 5.91E-06 | 0 | 0 | 8.99E-06 | 0 | 1.89E-06 | 2.75E-06 | 0 | 0 | 0 | 5.93E-06 | 4.64E-06 | 4.17E-06 | 7.92E-06 | 1.41E-05 | 7.12E-06 |
| Zfp963 | 2.61E-06 | 2.64E-05 | 0 | 3.26E-05 | 6.62E-06 | 7.39E-06 | 2.69E-06 | 2.19E-05 | 2.75E-06 | 7.17E-06 | 5.13E-06 | 0 | 0 | 1.59E-05 | 9.71E-06 | 0 | 0 | 1.39E-05 |
| Bdh1 | 7.57E-06 | 1.96E-06 | 3.74E-08 | 0 | 0 | 0 | 5.53E-08 | 0 | 2.75E-06 | 0 | 2.59E-05 | 0 | 7.66E-06 | 1.87E-05 | 0 | 0 | 0 | 4.52E-06 |
| Samsn1 | 0 | 3.70E-06 | 0 | 2.12E-06 | 3.95E-07 | 0 | 0 | 1.11E-06 | 2.75E-06 | 5.68E-06 | 3.71E-06 | 5.66E-06 | 8.26E-07 | 0 | 9.30E-06 | 1.07E-05 | 6.92E-06 | 8.78E-06 |
| Nxph3 | 0 | 7.56E-06 | 9.29E-06 | 0 | 0 | 0 | 0 | 0 | 2.74E-06 | 0 | 0 | 1.28E-05 | 6.65E-06 | 0 | 4.43E-07 | 6.36E-06 | 0 | 0 |
| Cfap298 | 4.71E-05 | 5.64E-05 | 5.91E-05 | 7.69E-05 | 0 | 0 | 0 | 0 | 2.71E-06 | 5.24E-05 | 1.45E-05 | 0 | 0 | 0 | 0 | 7.14E-05 | 3.83E-05 | 4.86E-05 |
| Sephs2 | 0.00011 | 0.0001 | 0.00011 | 8.42E-05 | 7.59E-05 | 8.19E-05 | 2.50E-06 | 0 | 2.68E-06 | 1.76E-05 | 2.29E-05 | 0 | 0.00011 | 9.37E-05 | 0 | 0.00012 | 0.00011 | 1.86E-06 |
| Snx16 | 3.19E-05 | 3.81E-05 | 8.64E-06 | 0 | 6.88E-06 | 2.65E-05 | 0 | 0 | 2.68E-06 | 6.89E-06 | 2.24E-05 | 2.34E-05 | 0 | 3.16E-06 | 5.79E-06 | 7.70E-06 | 0 | 0 |
| Gnpda1 | 1.66E-05 | 5.71E-05 | 0 | 1.58E-05 | 3.85E-05 | 9.33E-06 | 0 | 0 | 2.68E-06 | 0 | 0 | 1.69E-05 | 0 | 5.52E-05 | 3.49E-05 | 4.61E-06 | 0 | 4.01E-05 |
| Tulp1 | 0 | 5.66E-06 | 0 | 4.69E-07 | 2.03E-05 | 0 | 3.93E-06 | 0 | 2.68E-06 | 0 | 1.81E-05 | 0 | 1.53E-05 | 2.22E-05 | 0 | 0 | 2.80E-06 | 3.26E-05 |
| Tuba4a | 0 | 0 | 0 | 0 | 0 | 0 | 0 | 0 | 2.68E-06 | 0 | 0 | 4.25E-05 | 8.00E-06 | 1.13E-05 | 0 | 0 | 0.00013 | 0 |
| Cpeb3 | 4.43E-06 | 1.55E-05 | 0 | 4.47E-05 | 0 | 9.01E-06 | 4.42E-05 | 1.41E-05 | 2.67E-06 | 2.55E-05 | 0 | 0 | 9.23E-06 | 4.38E-06 | 3.30E-05 | 0 | 1.05E-05 | 4.17E-05 |
| Nexmif | 2.41E-06 | 2.61E-06 | 2.89E-06 | 9.76E-07 | 9.93E-07 | 0 | 5.09E-06 | 0 | 2.67E-06 | 2.12E-05 | 0 | 0 | 4.03E-05 | 0 | 0 | 1.02E-05 | 5.89E-06 | 0 |
| H2-Aa | 0 | 4.12E-05 | 6.80E-06 | 5.71E-05 | 9.05E-06 | 0 | 0 | 4.84E-06 | 2.66E-06 | 3.37E-05 | 1.68E-05 | 1.61E-05 | 3.32E-05 | 1.63E-06 | 0.00017 | 2.95E-05 | 3.64E-05 | 2.93E-05 |
| Dsn1 | 1.76E-05 | 1.01E-05 | 0 | 0 | 1.26E-05 | 2.13E-05 | 2.45E-05 | 5.45E-08 | 2.65E-06 | 0 | 0 | 9.28E-06 | 1.30E-05 | 1.66E-06 | 2.30E-05 | 0.00021 | 2.68E-05 | 1.47E-05 |

| | | | | | | | | | | | | | | | | | | |
|---|---|---|---|---|---|---|---|---|---|---|---|---|---|---|---|---|---|---|
| Pcdhb7 | 3.16E-06 | 1.96E-06 | 0 | 0 | 6.88E-06 | 9.48E-06 | 2.61E-06 | 3.06E-06 | 2.65E-06 | 3.00E-06 | 7.05E-06 | 0 | 2.98E-05 | 4.69E-06 | 0 | 3.39E-07 | 0 | 1.85E-06 |
| Nceh1 | 0 | 6.97E-05 | 0 | 0 | 0 | 4.75E-07 | 0 | 0 | 2.64E-06 | 8.29E-07 | 0 | 0 | 0 | 5.60E-05 | 0 | 1.25E-05 | 0 | 1.45E-05 |
| Dnaaf3 | 0 | 1.57E-05 | 0 | 1.04E-05 | 5.17E-06 | 0 | 0 | 9.14E-06 | 2.63E-06 | 9.55E-07 | 0 | 1.75E-05 | 8.30E-06 | 0 | 4.39E-05 | 0 | 0 | 9.34E-06 |
| Trim30d | 1.45E-05 | 1.60E-05 | 0 | 0 | 0 | 0 | 0 | 0 | 2.62E-06 | 0.00013 | 0 | 1.82E-06 | 1.87E-05 | 0 | 4.42E-05 | 0 | 0 | 2.38E-05 |
| Emid1 | 0 | 1.22E-05 | 0 | 0 | 3.79E-06 | 7.86E-08 | 0 | 0 | 2.61E-06 | 0 | 0 | 0 | 0 | 3.88E-05 | 1.32E-05 | 9.79E-06 | 0 | 2.12E-05 |
| Fxyd2 | 4.38E-06 | 1.02E-06 | 0 | 0 | 0 | 0 | 0 | 8.06E-08 | 2.60E-06 | 1.10E-05 | 0 | 3.88E-06 | 1.46E-05 | 0 | 0 | 0 | 0 | 9.99E-06 |
| Chd2 | 9.68E-05 | 0.00012 | 2.22E-05 | 3.34E-05 | 4.58E-05 | 5.44E-05 | 8.52E-05 | 1.95E-06 | 2.59E-06 | 6.88E-05 | 0 | 0.00011 | 0 | 0 | 0.0004 | 7.40E-05 | 0.00036 | 1.90E-05 |
| Mad2l1 | 2.84E-05 | 0 | 1.87E-05 | 0 | 0 | 8.32E-06 | 0 | 1.47E-05 | 2.56E-06 | 0 | 2.94E-07 | 0 | 0 | 7.09E-06 | 0 | 0.00077 | 0 | 0 |
| Nog | 0 | 0 | 8.78E-06 | 0 | 0 | 7.98E-06 | 0 | 2.10E-06 | 2.56E-06 | 0 | 5.40E-06 | 0 | 0 | 1.53E-05 | 1.42E-05 | 9.61E-06 | 0 | 8.39E-05 |
| Hic2 | 1.30E-05 | 2.00E-05 | 2.71E-05 | 0 | 2.74E-05 | 2.03E-05 | 0 | 1.01E-05 | 2.55E-06 | 6.60E-06 | 6.00E-06 | 1.97E-05 | 0 | 2.33E-05 | 2.66E-05 | 2.22E-06 | 7.46E-05 | 0 |
| Peli3 | 0 | 1.86E-05 | 1.08E-06 | 0 | 1.81E-06 | 5.52E-06 | 0 | 0 | 2.55E-06 | 3.02E-06 | 1.54E-06 | 8.58E-06 | 1.07E-05 | 1.75E-05 | 8.91E-06 | 0 | 0 | 1.51E-06 |
| Ptprj | 0 | 0 | 0.00031 | 0 | 0 | 0.00016 | 4.52E-05 | 0.00014 | 2.52E-06 | 0.00016 | 0 | 7.47E-05 | 3.95E-05 | 0 | 2.69E-05 | 2.36E-05 | 0 | 0 |
| 1-Sep | 1.45E-06 | 1.18E-05 | 0 | 1.40E-05 | 2.07E-05 | 2.13E-05 | 6.49E-07 | 1.37E-05 | 2.50E-06 | 9.96E-06 | 2.79E-07 | 5.82E-07 | 1.98E-05 | 7.67E-06 | 2.53E-06 | 1.25E-05 | 1.57E-06 | 2.13E-05 |
| Pak1 | 4.87E-05 | 0 | 7.94E-05 | 0 | 4.45E-05 | 0 | 2.62E-05 | 1.75E-05 | 2.49E-06 | 0 | 8.18E-05 | 3.94E-05 | 0.00013 | 2.14E-05 | 4.94E-05 | 7.19E-05 | 2.85E-05 | 0 |
| Kbtbd6 | 2.33E-06 | 0 | 1.97E-06 | 0 | 2.54E-06 | 0 | 4.11E-06 | 3.09E-08 | 2.49E-06 | 0 | 6.92E-07 | 0 | 1.86E-07 | 6.41E-07 | 2.30E-06 | 4.40E-05 | 1.68E-06 | 0 |
| Abcb7 | 7.59E-05 | 4.82E-05 | 4.42E-05 | 1.62E-05 | 4.47E-06 | 1.19E-05 | 0 | 6.26E-06 | 2.47E-06 | 9.73E-06 | 0 | 3.59E-06 | 3.53E-06 | 0 | 3.95E-05 | 8.35E-05 | 6.38E-05 | 5.89E-06 |
| Zfp276 | 1.76E-05 | 5.27E-05 | 1.93E-05 | 4.22E-05 | 1.23E-05 | 9.43E-06 | 2.02E-05 | 9.16E-06 | 2.47E-06 | 2.51E-05 | 1.91E-05 | 1.99E-05 | 0 | 3.72E-05 | 0.0002 | 0 | 4.50E-05 | 2.99E-05 |
| Zfp595 | 4.72E-06 | 1.61E-05 | 0 | 2.69E-06 | 0 | 5.23E-06 | 4.56E-06 | 6.07E-06 | 2.47E-06 | 6.15E-06 | 1.35E-05 | 0 | 0 | 1.27E-06 | 1.72E-05 | 1.18E-05 | 0 | 4.13E-06 |
| Sorl1 | 0 | 0 | 0 | 3.25E-06 | 8.29E-07 | 0 | 0 | 0 | 2.46E-06 | 2.85E-06 | 6.96E-06 | 4.27E-07 | 6.50E-06 | 0 | 3.90E-05 | 7.09E-06 | 9.36E-06 | 1.12E-05 |
| Pcdhga10 | 4.35E-05 | 2.83E-05 | 1.19E-05 | 1.24E-05 | 6.58E-05 | 7.21E-05 | 4.73E-05 | 4.44E-05 | 2.45E-06 | 0 | 1.92E-05 | 0 | 2.96E-05 | 2.89E-05 | 0.00015 | 0 | 0 | 3.90E-06 |
| Pex26 | 0 | 3.39E-05 | 4.71E-06 | 5.04E-05 | 2.61E-05 | 6.05E-05 | 7.53E-06 | 0 | 2.45E-06 | 2.01E-05 | 2.53E-05 | 3.25E-05 | 0 | 3.17E-05 | 9.99E-05 | 0 | 0 | 3.41E-05 |
| Dnajc28 | 2.52E-06 | 1.47E-05 | 0 | 0 | 6.48E-06 | 6.88E-07 | 3.93E-06 | 0 | 2.45E-06 | 6.46E-06 | 0 | 0 | 1.20E-06 | 5.84E-06 | 0 | 0 | 0 | 2.39E-06 |
| Matn3 | 7.86E-06 | 0 | 0 | 0 | 2.39E-06 | 0 | 2.84E-06 | 0 | 2.45E-06 | 0 | 5.53E-05 | 0 | 0 | 5.62E-06 | 0 | 6.59E-05 | 0 | 1.95E-07 |
| Aunip | 0 | 0 | 2.47E-06 | 0 | 0 | 0 | 0 | 0 | 2.45E-06 | 0 | 0 | 0 | 4.47E-06 | 3.60E-06 | 0 | 8.59E-05 | 0 | 0 |
| Samd13 | 3.77E-08 | 0 | 1.19E-05 | 5.00E-06 | 1.59E-06 | 0 | 1.01E-06 | 1.11E-05 | 2.44E-06 | 0 | 6.87E-07 | 2.62E-06 | 9.97E-06 | 0 | 6.75E-06 | 8.53E-07 | 1.81E-05 | 0 |
| Wnk4 | 0 | 3.25E-06 | 0 | 0 | 0 | 0 | 6.38E-06 | 6.28E-06 | 2.44E-06 | 8.76E-06 | 0 | 4.32E-05 | 8.65E-06 | 0 | 4.46E-05 | 0 | 2.12E-05 | 0 |
| Shld1 | 4.11E-06 | 6.87E-06 | 0 | 7.22E-06 | 9.07E-06 | 7.02E-06 | 3.14E-06 | 2.18E-06 | 2.43E-06 | 1.40E-05 | 4.20E-06 | 3.74E-06 | 5.93E-06 | 0 | 0 | 0 | 0 | 1.97E-05 |
| Angptl8 | 0 | 7.38E-06 | 1.14E-06 | 0 | 0 | 1.55E-05 | 7.89E-06 | 0 | 2.42E-06 | 2.03E-05 | 0 | 2.45E-05 | 2.17E-05 | 0 | 0 | 0 | 0 | 0 |
| Zfp61 | 9.58E-06 | 3.15E-05 | 0 | 2.36E-05 | 1.14E-05 | 8.55E-06 | 5.85E-06 | 1.22E-05 | 2.41E-06 | 1.30E-05 | 1.06E-05 | 8.94E-06 | 0 | 1.21E-05 | 4.17E-05 | 6.08E-06 | 8.22E-06 | 4.96E-06 |
| Wdr62 | 0 | 2.65E-07 | 7.01E-07 | 1.15E-05 | 0 | 0 | 2.45E-06 | 4.77E-06 | 2.41E-06 | 7.12E-06 | 0 | 0 | 0 | 0 | 3.23E-06 | 0.00014 | 5.05E-06 | 3.26E-06 |
| Alg13 | 8.76E-06 | 1.28E-05 | 3.61E-06 | 5.03E-06 | 5.53E-06 | 4.86E-06 | 1.23E-06 | 1.53E-06 | 2.40E-06 | 3.82E-07 | 4.43E-06 | 2.44E-05 | 4.93E-06 | 8.89E-06 | 1.88E-05 | 7.68E-05 | 3.34E-06 | 0 |
| Rdh12 | 0 | 2.00E-06 | 0 | 0 | 1.48E-06 | 4.37E-06 | 0 | 0 | 2.39E-06 | 0 | 0 | 2.13E-06 | 1.01E-05 | 6.29E-06 | 1.44E-05 | 2.71E-05 | 0 | 4.84E-06 |
| Pla2g4b | 1.97E-06 | 1.23E-05 | 6.74E-08 | 1.82E-05 | 1.22E-06 | 0 | 2.41E-06 | 0 | 2.38E-06 | 7.80E-06 | 0 | 1.89E-06 | 1.38E-05 | 0 | 2.52E-05 | 4.57E-06 | 6.26E-05 | 3.78E-07 |

| | | | | | | | | | | | | | | | | | | |
|---|---|---|---|---|---|---|---|---|---|---|---|---|---|---|---|---|---|---|
| Mlycd | 1.43E-05 | 6.09E-05 | 5.44E-05 | 0.00012 | 7.11E-05 | 4.80E-05 | 1.70E-07 | 1.73E-05 | 2.37E-06 | 6.19E-05 | 4.79E-05 | 0 | 3.09E-05 | 3.87E-05 | 4.43E-05 | 1.40E-05 | 1.10E-05 | 1.39E-05 |
| Zswim3 | 2.30E-05 | 1.89E-05 | 3.40E-06 | 9.81E-06 | 1.02E-05 | 7.36E-06 | 3.12E-05 | 0 | 2.35E-06 | 9.37E-06 | 0 | 5.14E-05 | 0 | 2.08E-05 | 6.79E-06 | 2.19E-05 | 7.83E-05 | 5.37E-06 |
| Nxf1 | 0.00026 | 0.00025 | 0.00018 | 0 | 8.27E-05 | 0.00013 | 0 | 1.11E-05 | 2.33E-06 | 0 | 1.54E-05 | 0.0002 | 0 | 5.10E-05 | 0.00033 | 0.0002 | 0.00032 | 9.71E-06 |
| Ranbp6 | 2.78E-05 | 4.32E-05 | 8.23E-06 | 9.73E-07 | 2.71E-05 | 3.10E-05 | 0 | 2.59E-06 | 2.33E-06 | 1.90E-05 | 9.88E-06 | 5.74E-06 | 0 | 9.92E-07 | 1.51E-05 | 1.62E-05 | 0 | 1.78E-05 |
| Morc4 | 5.91E-05 | 1.35E-05 | 3.33E-05 | 3.90E-06 | 2.00E-05 | 2.64E-05 | 0.00011 | 4.19E-07 | 2.33E-06 | 1.84E-05 | 1.77E-05 | 2.09E-05 | 0 | 1.46E-05 | 0 | 7.85E-05 | 3.65E-05 | 2.41E-05 |
| Mss51 | 4.96E-06 | 1.84E-05 | 3.33E-06 | 2.38E-06 | 1.46E-06 | 3.91E-06 | 3.43E-06 | 0 | 2.33E-06 | 4.48E-06 | 5.13E-06 | 1.97E-05 | 7.12E-06 | 3.86E-06 | 4.24E-05 | 2.15E-05 | 6.29E-05 | 0 |
| Abhd15 | 2.66E-06 | 2.85E-06 | 8.61E-06 | 1.36E-05 | 0 | 0 | 4.98E-06 | 0 | 2.33E-06 | 7.41E-06 | 0 | 0 | 1.27E-05 | 6.11E-06 | 0 | 1.73E-05 | 5.50E-06 | 8.71E-06 |
| Ttc41 | 2.82E-06 | 7.03E-06 | 0 | 1.05E-06 | 0 | 0 | 2.62E-06 | 1.01E-06 | 2.33E-06 | 1.93E-06 | 0 | 1.56E-05 | 0 | 0 | 9.07E-06 | 3.47E-06 | 2.23E-06 | 0 |
| Ccdc130 | 2.95E-05 | 4.67E-05 | 1.53E-05 | 0 | 8.64E-06 | 2.27E-06 | 9.99E-06 | 1.33E-05 | 2.31E-06 | 8.81E-06 | 5.62E-06 | 1.65E-05 | 0 | 1.22E-05 | 9.46E-05 | 2.29E-05 | 0 | 5.88E-06 |
| Otud3 | 2.21E-05 | 4.49E-05 | 5.22E-05 | 0 | 2.27E-05 | 3.70E-05 | 2.74E-07 | 0 | 2.30E-06 | 5.83E-06 | 9.75E-06 | 2.68E-05 | 2.80E-06 | 2.38E-05 | 2.88E-05 | 0 | 2.72E-05 | 1.72E-06 |
| Trmt13 | 4.07E-05 | 4.57E-05 | 4.25E-05 | 3.89E-05 | 1.12E-05 | 2.39E-05 | 2.41E-05 | 6.09E-06 | 2.30E-06 | 1.39E-05 | 5.74E-06 | 2.00E-05 | 1.03E-05 | 9.04E-06 | 0.00021 | 0 | 1.20E-05 | 1.54E-05 |
| Wdr90 | 0 | 1.49E-05 | 0 | 8.28E-06 | 1.49E-05 | 1.05E-05 | 0 | 0 | 2.30E-06 | 1.14E-05 | 0 | 1.59E-05 | 0 | 2.25E-06 | 6.92E-05 | 0.00022 | 0 | 5.69E-06 |
| Armc2 | 6.98E-06 | 4.29E-06 | 0 | 1.44E-06 | 7.68E-06 | 4.13E-06 | 5.67E-06 | 0 | 2.28E-06 | 0 | 4.92E-06 | 1.50E-05 | 4.28E-06 | 6.54E-07 | 1.19E-05 | 1.64E-05 | 0 | 9.66E-06 |
| Dtwd1 | 4.09E-05 | 3.43E-07 | 1.69E-05 | 2.93E-05 | 0 | 0 | 0 | 0 | 2.28E-06 | 5.83E-05 | 0 | 2.81E-06 | 9.17E-05 | 7.47E-07 | 4.28E-05 | 2.56E-05 | 7.33E-06 | 1.44E-05 |
| Grik2 | 0 | 1.59E-06 | 0 | 2.12E-05 | 7.54E-07 | 0.00024 | 0 | 1.07E-05 | 2.26E-06 | 0 | 0 | 0 | 0 | 0 | 1.76E-05 | 0 | 0 | 0 |
| Dnajc19 | 5.91E-05 | 6.32E-06 | 2.26E-05 | 4.66E-06 | 3.46E-06 | 2.58E-05 | 1.15E-05 | 7.55E-06 | 2.26E-06 | 9.91E-06 | 7.96E-06 | 0 | 0 | 7.11E-07 | 8.69E-06 | 3.09E-05 | 2.32E-05 | 2.92E-05 |
| Hspe1-rs1 | 1.60E-05 | 0 | 2.37E-06 | 3.32E-06 | 0 | 1.56E-06 | 6.52E-06 | 0 | 2.26E-06 | 0 | 0 | 3.62E-06 | 9.68E-06 | 8.28E-07 | 0 | 3.90E-06 | 4.80E-05 | 0 |
| Epb41 | 1.06E-05 | 2.01E-05 | 7.65E-06 | 0 | 1.33E-05 | 6.13E-05 | 8.43E-07 | 1.84E-06 | 2.25E-06 | 0 | 2.05E-05 | 7.86E-06 | 0 | 5.54E-05 | 1.52E-05 | 7.73E-05 | 2.68E-05 | 2.53E-05 |
| Ccdc84 | 1.43E-05 | 6.15E-05 | 4.46E-06 | 4.31E-05 | 2.44E-05 | 4.01E-05 | 1.23E-05 | 0 | 2.25E-06 | 1.25E-05 | 1.02E-05 | 3.59E-05 | 0 | 3.52E-05 | 0.00012 | 1.69E-05 | 0 | 9.08E-06 |
| Dnhd1 | 1.10E-05 | 7.33E-06 | 0 | 0 | 1.20E-05 | 1.08E-05 | 2.33E-05 | 0 | 2.24E-06 | 0 | 0 | 1.02E-05 | 1.98E-05 | 1.23E-06 | 2.91E-05 | 0 | 0 | 1.22E-05 |
| Ing1 | 0.00013 | 0.00016 | 0.00012 | 6.35E-05 | 8.53E-05 | 0.00013 | 8.99E-05 | 0.00019 | 2.23E-06 | 0 | 7.73E-05 | 0.00021 | 7.73E-05 | 0.0001 | 0.00023 | 0.00036 | 0.00025 | 1.91E-05 |
| Hyal1 | 0 | 2.78E-05 | 0 | 0 | 3.79E-06 | 0 | 3.29E-06 | 0 | 2.22E-06 | 1.56E-05 | 0 | 2.64E-05 | 1.05E-05 | 1.02E-05 | 2.16E-05 | 0 | 0 | 4.93E-05 |
| Avl9 | 2.36E-05 | 8.43E-05 | 1.68E-05 | 1.84E-05 | 2.77E-05 | 3.09E-05 | 4.30E-06 | 1.66E-05 | 2.21E-06 | 2.97E-06 | 1.92E-05 | 7.13E-05 | 1.44E-05 | 1.10E-05 | 3.24E-05 | 2.18E-06 | 9.42E-06 | 8.66E-06 |
| Acod1 | 0 | 7.68E-06 | 4.84E-06 | 7.09E-06 | 0 | 0 | 0 | 2.13E-06 | 2.21E-06 | 6.85E-06 | 0 | 0 | 1.28E-05 | 4.88E-06 | 4.27E-05 | 1.64E-05 | 2.37E-05 | 1.14E-05 |
| Fsd1l | 1.09E-05 | 1.29E-05 | 5.67E-06 | 0 | 0 | 0 | 0 | 0 | 2.21E-06 | 1.38E-05 | 3.33E-07 | 4.35E-05 | 0 | 1.16E-05 | 1.24E-05 | 2.22E-05 | 1.07E-05 | 2.24E-05 |
| Zbtb17 | 3.70E-05 | 6.57E-05 | 6.10E-05 | 5.99E-05 | 1.71E-05 | 1.10E-05 | 0 | 1.21E-05 | 2.17E-06 | 1.21E-06 | 4.64E-05 | 2.04E-05 | 0 | 1.88E-06 | 0.00011 | 2.15E-05 | 5.25E-05 | 0 |
| Pradc1 | 1.47E-05 | 4.35E-06 | 2.05E-06 | 7.91E-06 | 1.59E-05 | 2.40E-05 | 1.92E-05 | 1.15E-05 | 2.16E-06 | 4.06E-05 | 1.13E-05 | 2.57E-05 | 0 | 4.79E-06 | 0 | 0.00016 | 1.29E-06 | 2.31E-05 |
| Zfp974 | 1.17E-06 | 7.71E-06 | 0 | 0 | 8.38E-07 | 3.20E-07 | 4.59E-06 | 4.91E-06 | 2.14E-06 | 4.42E-06 | 3.05E-06 | 3.42E-06 | 0 | 4.83E-07 | 0 | 2.86E-06 | 0 | 3.38E-06 |
| Cbx8 | 0 | 6.71E-06 | 0 | 2.88E-05 | 2.21E-06 | 0 | 4.71E-06 | 1.34E-05 | 2.14E-06 | 6.28E-06 | 0 | 3.93E-06 | 1.23E-05 | 8.77E-06 | 2.43E-05 | 0 | 1.51E-05 | 1.05E-05 |
| Ippk | 1.25E-05 | 4.46E-05 | 8.78E-05 | 5.45E-05 | 8.59E-06 | 0 | 1.36E-05 | 6.63E-06 | 2.13E-06 | 6.53E-06 | 3.81E-05 | 1.04E-05 | 1.15E-06 | 1.83E-05 | 1.08E-05 | 1.70E-05 | 8.50E-06 | 1.23E-05 |
| Mical2 | 0 | 0.00014 | 0.00071 | 4.07E-05 | 0.00058 | 0.00015 | 0.00042 | 0.0004 | 2.12E-06 | 0 | 0.00115 | 0.00048 | 0 | 0.00016 | 0.00056 | 0 | 0.00029 | 0.00041 |
| Fmn1 | 3.52E-05 | 4.91E-05 | 1.36E-05 | 0 | 0.00025 | 0.00012 | 0.00019 | 0 | 2.12E-06 | 1.79E-05 | 8.60E-05 | 0 | 7.00E-05 | 0.00012 | 0.00017 | 0 | 0.00014 | 0 |
| Atxn7l2 | 1.76E-05 | 3.89E-05 | 0 | 4.10E-05 | 0 | 0 | 0 | 7.79E-06 | 2.12E-06 | 0 | 1.12E-07 | 0.00015 | 0 | 1.95E-05 | 2.77E-05 | 2.08E-05 | 0 | 1.25E-05 |

| | | | | | | | | | | | | | | | | | | |
|---|---|---|---|---|---|---|---|---|---|---|---|---|---|---|---|---|---|---|
| Slc9a5 | 0 | 0 | 7.91E-06 | 1.22E-05 | 1.58E-05 | 1.62E-05 | 0 | 0 | 2.10E-06 | 4.71E-06 | 3.05E-05 | 1.70E-05 | 0 | 4.81E-05 | 5.05E-05 | 3.95E-05 | 0 | 2.06E-05 |
| Mbd4 | 1.29E-05 | 1.91E-05 | 0 | 0 | 5.28E-06 | 2.49E-06 | 8.23E-07 | 3.20E-06 | 2.07E-06 | 4.26E-06 | 0 | 0 | 6.16E-06 | 0 | 4.39E-05 | 4.07E-05 | 0 | 8.90E-06 |
| Mansc1 | 0 | 1.90E-05 | 0 | 0 | 0 | 5.90E-07 | 0 | 0 | 2.07E-06 | 6.93E-05 | 0 | 0 | 0 | 1.68E-05 | 0 | 0 | 0 | 5.90E-05 |
| Taf5 | 2.31E-05 | 3.25E-05 | 1.69E-05 | 0 | 1.03E-05 | 0 | 5.10E-06 | 0 | 2.07E-06 | 0 | 5.00E-06 | 3.03E-05 | 8.99E-06 | 1.12E-05 | 3.81E-05 | 0.00013 | 0 | 2.66E-06 |
| Hspb2 | 0 | 4.81E-06 | 0 | 1.01E-06 | 1.37E-05 | 0 | 0 | 0 | 2.06E-06 | 2.14E-06 | 7.87E-06 | 2.05E-05 | 0 | 1.60E-05 | 2.48E-06 | 0 | 0 | 4.05E-05 |
| Tsga10 | 1.41E-05 | 1.66E-05 | 0 | 0 | 0 | 1.36E-05 | 0 | 0 | 2.05E-06 | 1.08E-05 | 0 | 2.16E-06 | 0 | 2.22E-06 | 9.36E-06 | 3.11E-05 | 0 | 2.76E-06 |
| Zcwpw2 | 0 | 5.05E-06 | 0 | 0 | 1.20E-05 | 1.12E-05 | 0 | 1.09E-05 | 2.04E-06 | 6.79E-06 | 0 | 3.81E-06 | 0 | 0 | 2.56E-05 | 0 | 2.14E-05 | 9.66E-06 |
| Tob1 | 7.75E-06 | 7.91E-05 | 0 | 2.48E-05 | 0 | 0 | 0.00014 | 0 | 2.04E-06 | 9.91E-05 | 0 | 0.0002 | 9.30E-05 | 3.12E-05 | 0 | 0 | 0.00222 | 0 |
| Snx5 | 0.00018 | 3.87E-05 | 7.83E-05 | 0 | 0 | 7.08E-05 | 2.23E-05 | 0 | 2.03E-06 | 0 | 0 | 5.98E-05 | 1.15E-05 | 0 | 1.27E-05 | 5.71E-05 | 5.54E-05 | 1.96E-05 |
| Smarca1 | 7.71E-05 | 1.42E-05 | 0 | 8.19E-06 | 5.71E-06 | 1.06E-05 | 3.98E-05 | 6.54E-05 | 2.02E-06 | 2.11E-05 | 0 | 0 | 4.49E-05 | 0 | 1.18E-05 | 0 | 3.91E-05 | 5.24E-05 |
| Fignl1 | 0 | 0 | 2.36E-05 | 0 | 0 | 0 | 0 | 0 | 2.01E-06 | 2.83E-06 | 3.82E-06 | 0 | 5.88E-06 | 0 | 0 | 0.0003 | 0 | 0 |
| Inpp5d | 0 | 3.72E-06 | 1.22E-06 | 1.54E-08 | 0 | 0 | 0 | 0 | 1.99E-06 | 4.82E-06 | 0 | 3.07E-06 | 5.78E-07 | 0 | 2.81E-05 | 1.24E-05 | 1.75E-05 | 1.10E-05 |
| Rinl | 0 | 1.14E-06 | 1.35E-06 | 9.62E-06 | 2.87E-06 | 0 | 1.08E-05 | 2.54E-06 | 1.99E-06 | 3.26E-06 | 0 | 7.54E-06 | 0 | 0 | 1.24E-05 | 0 | 6.94E-06 | 1.11E-05 |
| Spaca1 | 1.57E-05 | 0 | 0 | 0 | 1.21E-05 | 1.84E-06 | 3.75E-06 | 1.48E-06 | 1.97E-06 | 2.18E-06 | 7.13E-06 | 3.19E-07 | 1.29E-05 | 0 | 3.99E-06 | 7.45E-06 | 0 | 0 |
| Ubxn2a | 4.28E-05 | 4.92E-05 | 6.78E-05 | 1.39E-05 | 8.28E-05 | 8.34E-05 | 4.56E-05 | 2.82E-05 | 1.96E-06 | 2.61E-05 | 7.28E-05 | 1.05E-05 | 5.71E-05 | 6.42E-05 | 0 | 1.88E-05 | 3.45E-05 | 0.0001 |
| Fhit | 0 | 2.32E-06 | 0 | 0 | 3.29E-06 | 7.31E-07 | 0 | 3.82E-06 | 1.96E-06 | 3.23E-05 | 0 | 0 | 3.33E-07 | 0 | 0 | 6.00E-07 | 0 | 0 |
| Mkrn3 | 1.90E-06 | 8.62E-06 | 0 | 6.30E-06 | 4.75E-06 | 0 | 0 | 5.31E-06 | 1.96E-06 | 0 | 5.45E-06 | 4.70E-06 | 0 | 0 | 4.99E-06 | 6.18E-06 | 0 | 0 |
| Neurl3 | 0 | 9.21E-06 | 0 | 2.40E-05 | 0 | 0 | 0 | 2.82E-06 | 1.94E-06 | 6.44E-06 | 0 | 0 | 1.94E-06 | 0 | 2.35E-05 | 1.43E-05 | 3.30E-05 | 4.45E-06 |
| Gdf9 | 0 | 4.18E-07 | 0 | 0 | 0 | 0 | 9.03E-07 | 0 | 1.94E-06 | 1.66E-06 | 0 | 2.67E-05 | 1.29E-05 | 0 | 3.09E-06 | 0 | 5.29E-05 | 0 |
| Inafm2 | 0 | 5.29E-05 | 8.17E-05 | 0.00013 | 6.89E-05 | 6.70E-05 | 0 | 0.00011 | 1.93E-06 | 1.03E-05 | 4.90E-05 | 4.71E-05 | 0 | 2.15E-05 | 0.00013 | 0 | 3.77E-05 | 0 |
| Slc52a3 | 0 | 0 | 0 | 1.28E-05 | 0 | 0 | 4.44E-05 | 0 | 1.93E-06 | 0 | 2.97E-05 | 0 | 2.46E-05 | 0 | 0 | 6.18E-06 | 0 | 0.0001 |
| Rhbdl1 | 0 | 1.64E-05 | 0 | 3.10E-05 | 7.52E-06 | 5.96E-06 | 0 | 1.60E-05 | 1.92E-06 | 1.25E-05 | 0 | 4.34E-06 | 0 | 0 | 8.10E-05 | 0 | 2.35E-05 | 1.04E-05 |
| Ogt | 0.00017 | 0.0004 | 0 | 0 | 0.00015 | 0.00017 | 0 | 5.83E-05 | 1.91E-06 | 5.88E-05 | 0 | 0.00048 | 0 | 8.10E-05 | 0.00067 | 0 | 0 | 0.00015 |
| Slc26a11 | 1.09E-07 | 9.31E-05 | 1.14E-06 | 3.03E-05 | 9.16E-05 | 7.69E-05 | 1.52E-05 | 5.58E-05 | 1.89E-06 | 1.77E-05 | 0 | 3.87E-05 | 1.24E-05 | 6.36E-05 | 8.27E-05 | 0 | 0 | 1.74E-05 |
| Smox | 2.26E-05 | 0.00013 | 9.56E-05 | 0 | 1.71E-05 | 0 | 0 | 0.00021 | 1.89E-06 | 1.73E-06 | 0 | 0.00038 | 0 | 0 | 8.57E-05 | 0 | 0 | 5.46E-05 |
| Ccnb1 | 0 | 0 | 0 | 0 | 0 | 1.26E-06 | 0 | 0 | 1.88E-06 | 0 | 0 | 0 | 0 | 0 | 0 | 8.59E-05 | 0 | 0 |
| Lilr4b | 0 | 1.06E-05 | 0 | 2.26E-05 | 2.33E-06 | 0 | 1.34E-07 | 0 | 1.87E-06 | 1.66E-06 | 1.47E-06 | 1.68E-06 | 1.01E-05 | 0 | 2.31E-05 | 1.22E-05 | 6.67E-06 | 1.94E-06 |
| Pola1 | 0 | 0 | 2.83E-05 | 0 | 0 | 1.39E-05 | 0 | 3.30E-06 | 1.86E-06 | 2.79E-05 | 0 | 0 | 3.98E-05 | 1.03E-05 | 6.70E-05 | 0.00045 | 1.10E-05 | 2.49E-05 |
| Neurl1a | 0 | 5.74E-07 | 1.85E-07 | 3.89E-06 | 4.53E-06 | 6.21E-06 | 0 | 1.22E-05 | 1.86E-06 | 6.67E-06 | 1.77E-05 | 0 | 0 | 6.00E-06 | 4.66E-06 | 0 | 0 | 5.61E-06 |
| Pik3cg | 0 | 3.72E-06 | 0 | 3.08E-07 | 0 | 1.61E-06 | 0 | 6.37E-06 | 1.86E-06 | 6.88E-07 | 3.83E-06 | 0 | 0 | 2.29E-05 | 3.33E-06 | 5.27E-06 | 3.33E-06 | 0 |
| Ranbp9 | 9.96E-05 | 0.00017 | 4.41E-05 | 5.04E-06 | 6.15E-05 | 0.00011 | 2.20E-05 | 5.98E-05 | 1.84E-06 | 5.56E-05 | 7.06E-05 | 5.04E-05 | 0.00012 | 8.90E-06 | 0.00017 | 0.00011 | 0.00015 | 5.36E-05 |
| Coq3 | 5.40E-05 | 1.39E-05 | 6.83E-05 | 7.99E-05 | 0 | 0 | 3.50E-06 | 4.71E-06 | 1.84E-06 | 0 | 2.31E-05 | 0 | 2.81E-06 | 1.38E-05 | 2.10E-05 | 3.42E-05 | 5.40E-05 | 1.40E-05 |
| Zfp2 | 3.75E-06 | 1.58E-05 | 0 | 0 | 2.47E-06 | 0 | 0 | 0 | 1.84E-06 | 2.16E-06 | 0 | 0 | 0 | 7.63E-06 | 0 | 0 | 0 | 5.69E-06 |

| | | | | | | | | | | | | | | | | | | |
|---|---|---|---|---|---|---|---|---|---|---|---|---|---|---|---|---|---|---|
| Cytip | 0 | 4.32E-06 | 4.25E-06 | 1.62E-05 | 0 | 0 | 0 | 0 | 1.84E-06 | 9.35E-06 | 2.23E-06 | 0 | 1.64E-06 | 0 | 1.49E-05 | 7.30E-06 | 2.18E-05 | 3.22E-06 |
| Acer3 | 6.90E-05 | 7.19E-06 | 4.29E-05 | 0 | 0 | 5.07E-05 | 4.39E-05 | 0 | 1.83E-06 | 1.91E-05 | 1.61E-05 | 9.78E-06 | 7.35E-06 | 9.83E-06 | 1.43E-05 | 3.06E-05 | 1.54E-05 | 0 |
| Cbarp | 0 | 2.94E-05 | 3.61E-06 | 0 | 0 | 2.15E-05 | 0 | 2.68E-05 | 1.83E-06 | 0 | 5.46E-06 | 3.33E-05 | 0 | 5.01E-06 | 0 | 0 | 2.61E-05 | 4.21E-06 |
| Cmbl | 0 | 2.53E-06 | 0 | 0 | 1.45E-05 | 5.82E-07 | 0 | 1.74E-05 | 1.83E-06 | 1.06E-05 | 0 | 3.41E-06 | 0 | 0 | 0 | 0 | 0 | 3.34E-05 |
| Exog | 2.62E-05 | 4.72E-06 | 3.44E-05 | 2.01E-05 | 8.52E-06 | 0 | 1.09E-05 | 0 | 1.83E-06 | 9.52E-06 | 1.40E-05 | 1.76E-05 | 1.67E-05 | 4.20E-06 | 7.00E-05 | 7.31E-05 | 0 | 1.03E-05 |
| Zfp729a | 1.03E-05 | 1.31E-05 | 0 | 0 | 2.05E-06 | 7.30E-06 | 0 | 1.17E-06 | 1.82E-06 | 1.86E-06 | 0 | 2.56E-06 | 0 | 0 | 2.88E-06 | 2.94E-06 | 0 | 4.67E-06 |
| Carns1 | 0 | 1.48E-05 | 0 | 1.67E-05 | 0 | 0 | 0 | 1.87E-05 | 1.81E-06 | 3.95E-06 | 7.18E-06 | 3.41E-06 | 0 | 4.64E-06 | 1.07E-06 | 0 | 3.07E-06 | 1.01E-05 |
| Myo19 | 3.43E-05 | 1.61E-05 | 0.00011 | 0 | 5.92E-06 | 0 | 2.00E-05 | 0 | 1.80E-06 | 1.17E-05 | 2.03E-05 | 0 | 1.64E-05 | 1.24E-05 | 9.84E-05 | 7.36E-05 | 0 | 2.94E-05 |
| Tigd3 | 0 | 7.47E-06 | 1.61E-06 | 1.28E-07 | 0 | 0 | 6.92E-07 | 0 | 1.79E-06 | 7.12E-06 | 0 | 1.88E-05 | 3.46E-06 | 0 | 1.51E-05 | 1.22E-05 | 5.74E-05 | 1.68E-06 |
| Noxo1 | 0 | 4.65E-06 | 0 | 0 | 2.83E-06 | 0 | 0 | 2.24E-06 | 1.77E-06 | 2.04E-05 | 0 | 7.67E-06 | 0 | 0 | 1.91E-06 | 2.41E-06 | 0 | 5.13E-06 |
| Nkain4 | 3.80E-06 | 4.83E-06 | 0 | 0 | 0 | 2.13E-06 | 6.85E-06 | 9.17E-06 | 1.76E-06 | 1.28E-05 | 0 | 0 | 1.15E-06 | 0 | 6.73E-06 | 3.17E-06 | 0 | 5.32E-06 |
| Xrcc3 | 1.14E-06 | 2.28E-06 | 3.00E-06 | 0 | 0 | 0 | 0 | 0 | 1.76E-06 | 3.03E-06 | 0 | 0 | 6.17E-06 | 0 | 1.22E-05 | 4.21E-05 | 0 | 1.05E-06 |
| Lsm11 | 3.09E-06 | 2.51E-05 | 5.65E-05 | 1.64E-05 | 0 | 1.01E-05 | 8.95E-06 | 0 | 1.74E-06 | 1.39E-05 | 4.98E-05 | 4.99E-05 | 6.56E-06 | 2.62E-05 | 0 | 0.00011 | 0 | 0 |
| Tma16 | 3.51E-05 | 4.29E-05 | 7.39E-05 | 6.57E-07 | 3.98E-06 | 1.96E-05 | 0 | 0 | 1.73E-06 | 0 | 0 | 1.62E-05 | 2.18E-05 | 0 | 2.12E-05 | 0 | 5.92E-06 | 0 |
| Ephx3 | 0 | 1.98E-06 | 2.20E-06 | 0 | 0 | 4.35E-07 | 5.05E-07 | 2.93E-07 | 1.73E-06 | 2.49E-07 | 0 | 6.11E-06 | 3.23E-06 | 5.88E-07 | 0 | 3.17E-06 | 6.02E-05 | 0 |
| Glcci1 | 3.46E-05 | 1.35E-05 | 0 | 0 | 0 | 4.09E-05 | 3.38E-05 | 5.99E-06 | 1.72E-06 | 3.99E-06 | 6.30E-06 | 2.18E-05 | 3.42E-05 | 1.61E-05 | 3.57E-05 | 0 | 2.78E-05 | 0 |
| Klhdc8b | 0 | 4.85E-05 | 0 | 2.92E-05 | 1.09E-05 | 3.09E-05 | 0 | 4.19E-05 | 1.72E-06 | 4.22E-05 | 1.99E-06 | 2.06E-06 | 0 | 1.48E-05 | 5.22E-05 | 0 | 0 | 3.83E-05 |
| Srfbp1 | 5.47E-05 | 7.42E-05 | 5.41E-05 | 0 | 6.11E-06 | 1.51E-05 | 0 | 0 | 1.72E-06 | 2.05E-05 | 1.28E-05 | 2.45E-05 | 1.82E-05 | 1.05E-05 | 0.00014 | 3.05E-06 | 0 | 0 |
| Carf | 1.47E-05 | 1.04E-05 | 0 | 0 | 8.02E-06 | 1.18E-05 | 1.61E-05 | 2.99E-07 | 1.72E-06 | 2.83E-06 | 0 | 2.37E-06 | 2.78E-06 | 0 | 1.39E-05 | 0 | 7.18E-06 | 0 |
| Tsc1 | 5.13E-05 | 0.00013 | 2.93E-05 | 2.91E-05 | 6.54E-05 | 5.51E-05 | 2.85E-05 | 7.75E-06 | 1.71E-06 | 0 | 0 | 4.68E-05 | 2.81E-05 | 1.52E-05 | 0.00014 | 3.93E-05 | 0.00016 | 0 |
| E2f2 | 0 | 0 | 0 | 0 | 4.62E-07 | 1.19E-05 | 1.47E-05 | 4.68E-06 | 1.70E-06 | 0 | 0 | 0 | 5.68E-06 | 1.86E-06 | 3.31E-07 | 8.09E-05 | 0 | 0 |
| Atg14 | 2.17E-05 | 4.77E-05 | 0 | 0 | 1.31E-06 | 1.27E-05 | 0 | 3.52E-06 | 1.68E-06 | 0 | 0 | 2.30E-05 | 0 | 3.08E-06 | 6.26E-05 | 5.19E-06 | 2.84E-05 | 5.16E-06 |
| Tmem151a | 0 | 9.11E-06 | 0 | 5.21E-06 | 0 | 1.45E-06 | 1.02E-06 | 4.71E-06 | 1.67E-06 | 0 | 1.30E-06 | 7.69E-06 | 9.53E-06 | 0 | 1.66E-05 | 0 | 0 | 0 |
| Slc22a17 | 0 | 4.65E-05 | 0 | 0.00024 | 7.13E-05 | 5.00E-05 | 0.00015 | 6.45E-05 | 1.65E-06 | 2.04E-05 | 0 | 0 | 1.22E-05 | 0.00011 | 0.00028 | 0 | 1.48E-05 | 6.11E-05 |
| Mettl15 | 1.10E-05 | 2.09E-05 | 0 | 0 | 5.54E-06 | 9.05E-06 | 0 | 0 | 1.64E-06 | 3.53E-06 | 0 | 1.38E-05 | 0 | 0 | 6.98E-06 | 9.01E-05 | 1.65E-05 | 2.95E-05 |
| Ybey | 8.75E-06 | 1.15E-05 | 0 | 1.52E-05 | 7.46E-07 | 6.41E-06 | 0 | 1.39E-06 | 1.64E-06 | 0 | 4.75E-06 | 2.64E-06 | 0 | 1.54E-06 | 1.01E-05 | 1.11E-05 | 0 | 0 |
| Atl1 | 3.19E-06 | 1.35E-05 | 0 | 0 | 5.25E-05 | 2.57E-06 | 2.76E-05 | 4.77E-06 | 1.64E-06 | 3.53E-06 | 1.34E-05 | 2.13E-05 | 0 | 2.31E-05 | 4.74E-05 | 0 | 0 | 0 |
| P2ry10b | 9.82E-06 | 0 | 0 | 0 | 1.41E-05 | 0 | 6.93E-05 | 0 | 1.64E-06 | 4.69E-05 | 0 | 0 | 0 | 0 | 0 | 4.98E-06 | 0 | 0 |
| Zfp869 | 2.08E-05 | 1.51E-05 | 1.29E-06 | 0 | 0 | 1.99E-05 | 0 | 2.49E-07 | 1.63E-06 | 5.63E-07 | 0 | 3.21E-05 | 0 | 3.09E-06 | 0 | 1.29E-05 | 0 | 0 |
| Ttll7 | 1.47E-06 | 6.09E-06 | 9.75E-06 | 1.76E-06 | 4.52E-06 | 0 | 7.20E-06 | 0 | 1.63E-06 | 8.72E-07 | 0 | 8.56E-06 | 1.21E-06 | 4.16E-06 | 0 | 8.70E-07 | 0 | 0 |
| Ndrg4 | 3.54E-06 | 0 | 0.00014 | 3.53E-05 | 0 | 5.10E-05 | 0.00031 | 0 | 1.62E-06 | 0.00034 | 0.00017 | 7.53E-05 | 0 | 0.00012 | 0 | 0 | 0 | 0.00023 |
| Saxo2 | 0 | 9.73E-06 | 0 | 4.38E-06 | 5.86E-06 | 3.43E-06 | 0 | 5.82E-06 | 1.61E-06 | 1.07E-05 | 0 | 7.07E-06 | 0 | 0 | 3.52E-07 | 0 | 0 | 1.05E-07 |
| Cetn4 | 1.72E-05 | 5.65E-07 | 0 | 1.56E-05 | 1.63E-05 | 0 | 0 | 2.23E-05 | 1.61E-06 | 6.19E-05 | 0 | 1.32E-05 | 3.30E-05 | 7.63E-06 | 0 | 2.83E-05 | 0 | 0 |

| | | | | | | | | | | | | | | | | | | |
|---|---|---|---|---|---|---|---|---|---|---|---|---|---|---|---|---|---|---|
| Dock10 | 7.15E-07 | 3.15E-05 | 0 | 0 | 0 | 0 | 0 | 4.22E-06 | 1.61E-06 | 2.16E-05 | 0 | 2.25E-06 | 0 | 0 | 7.62E-06 | 1.62E-05 | 6.28E-06 | 4.10E-05 |
| Nrbf2 | 0.00015 | 0.00016 | 0.00013 | 0.00014 | 9.80E-05 | 4.79E-05 | 4.08E-05 | 5.05E-05 | 1.58E-06 | 9.64E-06 | 7.28E-05 | 2.98E-05 | 7.73E-05 | 6.53E-05 | 8.89E-05 | 0 | 0 | 4.09E-05 |
| Stx1b | 0 | 1.04E-05 | 0 | 0 | 3.00E-06 | 0 | 0 | 1.67E-05 | 1.58E-06 | 3.28E-05 | 0 | 5.57E-06 | 0 | 0 | 3.58E-05 | 0 | 0 | 5.26E-06 |
| Taf1a | 2.78E-05 | 4.84E-05 | 5.86E-05 | 3.64E-05 | 2.99E-06 | 0 | 0 | 0 | 1.58E-06 | 6.23E-07 | 1.37E-07 | 1.66E-06 | 0 | 5.72E-06 | 0.00016 | 1.90E-05 | 0 | 1.48E-05 |
| Bbs10 | 7.49E-06 | 1.12E-05 | 1.08E-06 | 3.12E-06 | 0 | 5.47E-06 | 1.39E-05 | 2.93E-06 | 1.57E-06 | 5.14E-06 | 0 | 1.64E-05 | 2.03E-06 | 0 | 0 | 1.05E-05 | 2.02E-05 | 8.63E-06 |
| Rad51c | 3.96E-06 | 0 | 0 | 0 | 0 | 4.08E-06 | 0 | 0 | 1.56E-06 | 0 | 1.89E-06 | 0 | 0 | 0 | 1.02E-05 | 6.73E-05 | 0 | 0 |
| Dsp | 0 | 8.99E-06 | 4.08E-06 | 0 | 1.69E-06 | 0 | 0 | 7.08E-06 | 1.56E-06 | 6.05E-07 | 1.30E-05 | 1.26E-05 | 0 | 0 | 7.71E-07 | 0 | 0 | 7.42E-07 |
| Elf3 | 0 | 2.00E-06 | 0 | 1.44E-05 | 1.85E-06 | 0 | 1.56E-06 | 2.74E-06 | 1.55E-06 | 1.47E-06 | 0 | 2.23E-06 | 0 | 0 | 2.99E-05 | 0 | 1.13E-05 | 1.57E-06 |
| Zfp780b | 1.17E-05 | 6.59E-06 | 0 | 0 | 2.42E-06 | 9.85E-06 | 8.70E-06 | 0 | 1.54E-06 | 4.71E-06 | 0 | 4.39E-06 | 0 | 0 | 1.48E-05 | 0 | 0 | 0 |
| Hspa12a | 1.64E-05 | 2.37E-05 | 0 | 0 | 1.67E-05 | 0 | 0 | 6.95E-05 | 1.53E-06 | 9.37E-05 | 2.04E-05 | 1.30E-06 | 0.0001 | 0 | 1.94E-06 | 3.85E-05 | 0 | 0.00017 |
| Pbp2 | 4.24E-06 | 2.34E-06 | 1.55E-06 | 0 | 2.24E-05 | 0 | 0 | 5.23E-05 | 1.52E-06 | 0 | 0 | 2.50E-05 | 0 | 0 | 1.61E-05 | 9.09E-06 | 0 | 2.95E-05 |
| Pomc | 3.98E-06 | 1.80E-05 | 0 | 8.50E-06 | 0 | 3.26E-06 | 0 | 2.14E-05 | 1.51E-06 | 0 | 4.05E-06 | 6.12E-06 | 6.40E-06 | 0 | 1.40E-05 | 2.69E-07 | 1.06E-05 | 1.19E-06 |
| Stil | 5.87E-06 | 0 | 0 | 0 | 0 | 0 | 0 | 0 | 1.51E-06 | 0 | 7.28E-06 | 0 | 0 | 9.94E-06 | 0 | 0.00031 | 0 | 0 |
| Mfsd9 | 9.47E-07 | 2.02E-05 | 0 | 2.24E-05 | 2.87E-05 | 0 | 1.78E-05 | 2.18E-06 | 1.50E-06 | 2.74E-05 | 0 | 6.39E-05 | 0 | 0 | 0 | 0 | 0 | 2.42E-05 |
| Camkmt | 1.12E-05 | 1.15E-05 | 1.58E-06 | 4.58E-06 | 7.42E-06 | 2.17E-06 | 9.93E-06 | 0 | 1.47E-06 | 1.21E-05 | 3.17E-06 | 8.21E-07 | 0 | 0 | 0 | 1.90E-05 | 2.54E-06 | 1.02E-05 |
| Wdr63 | 1.89E-06 | 1.08E-06 | 0 | 1.81E-06 | 1.41E-05 | 8.18E-06 | 0 | 1.62E-05 | 1.46E-06 | 0 | 0 | 6.05E-07 | 0 | 5.12E-06 | 8.80E-07 | 9.27E-07 | 0 | 0 |
| Papolb | 0 | 2.89E-06 | 0 | 0 | 0 | 0 | 0 | 1.32E-06 | 1.46E-06 | 1.20E-05 | 2.99E-06 | 9.54E-06 | 0 | 1.05E-05 | 1.61E-05 | 3.33E-06 | 0 | 3.50E-06 |
| Pcdhb16 | 7.17E-06 | 4.77E-05 | 0 | 0 | 6.01E-05 | 7.11E-05 | 0 | 2.75E-05 | 1.45E-06 | 1.79E-06 | 4.19E-05 | 0 | 3.69E-05 | 2.59E-05 | 0 | 0 | 3.20E-06 | 4.82E-05 |
| Gne | 9.20E-05 | 2.02E-05 | 4.39E-05 | 2.01E-05 | 4.33E-06 | 8.33E-06 | 8.09E-05 | 0 | 1.45E-06 | 3.42E-05 | 2.52E-06 | 0 | 0 | 0 | 1.04E-05 | 0 | 0 | 4.35E-05 |
| Agpat5 | 0.00011 | 2.07E-05 | 9.44E-05 | 2.83E-06 | 1.63E-05 | 2.78E-05 | 1.43E-06 | 2.87E-06 | 1.44E-06 | 8.38E-06 | 3.80E-05 | 0.00021 | 2.29E-05 | 0 | 0 | 0.0001 | 0 | 3.87E-05 |
| Pidd1 | 0 | 1.02E-05 | 1.08E-05 | 8.99E-06 | 0 | 1.64E-06 | 0 | 6.82E-06 | 1.44E-06 | 2.30E-06 | 0 | 0 | 1.70E-05 | 0 | 5.49E-06 | 6.29E-05 | 2.71E-06 | 0 |
| Tex29 | 0 | 2.18E-06 | 2.74E-06 | 0 | 0 | 0 | 5.11E-06 | 0 | 1.44E-06 | 3.01E-06 | 0 | 2.53E-06 | 2.10E-05 | 0 | 0 | 1.63E-06 | 6.88E-05 | 0 |
| Rwdd3 | 3.28E-06 | 7.99E-06 | 0 | 1.53E-06 | 6.71E-06 | 5.41E-06 | 1.60E-05 | 1.46E-05 | 1.43E-06 | 9.02E-07 | 0 | 3.75E-06 | 0 | 2.72E-06 | 2.05E-05 | 0 | 0 | 1.94E-05 |
| Tmem246 | 0 | 1.27E-05 | 0 | 0 | 2.32E-06 | 2.27E-06 | 0 | 0 | 1.43E-06 | 3.38E-07 | 5.52E-07 | 2.31E-06 | 0 | 3.47E-05 | 0 | 0 | 0 | 2.19E-05 |
| Harbi1 | 3.92E-05 | 2.89E-05 | 1.20E-05 | 2.79E-05 | 2.11E-06 | 5.25E-06 | 1.29E-05 | 1.46E-05 | 1.42E-06 | 7.27E-06 | 1.29E-05 | 1.29E-05 | 0 | 1.63E-05 | 3.44E-05 | 0 | 0 | 1.17E-05 |
| Efcab2 | 5.71E-06 | 6.76E-06 | 4.07E-06 | 3.19E-06 | 1.87E-06 | 2.67E-06 | 0 | 0 | 1.42E-06 | 7.00E-06 | 7.25E-06 | 3.43E-07 | 4.92E-07 | 0 | 0 | 2.21E-06 | 4.09E-06 | 0 |
| Anxa10 | 0 | 8.32E-06 | 0 | 1.73E-05 | 6.80E-06 | 0 | 1.29E-06 | 5.50E-06 | 1.40E-06 | 3.42E-06 | 0 | 2.37E-05 | 4.12E-06 | 0 | 2.06E-05 | 0 | 0 | 9.34E-06 |
| Ptpn6 | 0 | 5.42E-06 | 1.07E-06 | 1.06E-05 | 0 | 0 | 0 | 2.96E-06 | 1.40E-06 | 5.93E-06 | 1.37E-07 | 1.12E-05 | 1.56E-06 | 1.88E-06 | 0 | 5.01E-06 | 2.19E-05 | 1.40E-05 |
| Cldn34c1 | 1.14E-05 | 2.21E-05 | 0 | 0 | 2.49E-05 | 2.40E-05 | 0 | 9.75E-07 | 1.39E-06 | 0 | 0 | 1.76E-05 | 0 | 3.20E-06 | 0 | 1.38E-05 | 0 | 1.23E-06 |
| Tssk6 | 0 | 1.86E-06 | 5.95E-06 | 0 | 3.83E-06 | 4.04E-06 | 3.70E-06 | 0 | 1.39E-06 | 2.24E-06 | 4.34E-06 | 4.29E-06 | 0 | 0 | 0 | 3.42E-06 | 0 | 4.21E-06 |
| Scp2 | 0.0002 | 8.47E-05 | 1.75E-05 | 8.45E-05 | 1.34E-05 | 0.00012 | 0.00011 | 8.38E-06 | 1.38E-06 | 8.51E-06 | 2.54E-05 | 7.45E-05 | 0.00016 | 2.35E-05 | 0 | 3.68E-05 | 0 | 0.00018 |
| Nipal2 | 7.12E-06 | 1.24E-05 | 0 | 0 | 0 | 3.06E-06 | 0 | 5.38E-06 | 1.38E-06 | 4.70E-05 | 0 | 0 | 2.17E-05 | 1.42E-05 | 0 | 0 | 0 | 3.50E-05 |
| Ntn4 | 4.07E-05 | 0 | 8.03E-05 | 0 | 7.70E-05 | 3.24E-05 | 0 | 0.00029 | 1.35E-06 | 0.00016 | 0.00076 | 0 | 0 | 0 | 0 | 0 | 0 | 7.29E-05 |

| | | | | | | | | | | | | | | | | | | |
|---|---|---|---|---|---|---|---|---|---|---|---|---|---|---|---|---|---|---|
| Podxl2 | 0 | 6.89E-05 | 0 | 0 | 0.00012 | 1.15E-05 | 1.50E-05 | 4.56E-05 | 1.35E-06 | 0.0001 | 4.79E-05 | 1.80E-05 | 0.00015 | 0.0001 | 0 | 7.20E-07 | 1.52E-05 | 8.43E-05 |
| Tnfsf4 | 0 | 0 | 1.57E-05 | 0 | 0 | 0 | 4.23E-05 | 7.05E-06 | 1.35E-06 | 0 | 0 | 0 | 8.25E-06 | 0 | 0 | 1.19E-05 | 0 | 1.26E-05 |
| Rtl10 | 0 | 1.96E-06 | 4.52E-06 | 1.08E-05 | 4.53E-06 | 0 | 2.31E-06 | 0 | 1.34E-06 | 9.78E-07 | 3.59E-07 | 0 | 3.86E-06 | 0 | 4.67E-07 | 8.50E-06 | 0 | 9.58E-06 |
| Dars2 | 8.23E-06 | 8.76E-06 | 8.57E-06 | 2.06E-05 | 1.41E-05 | 2.87E-05 | 3.19E-05 | 1.24E-05 | 1.32E-06 | 2.92E-05 | 1.27E-05 | 1.18E-05 | 2.48E-05 | 3.70E-06 | 1.55E-05 | 0.00018 | 0 | 2.85E-05 |
| Slc12a8 | 0 | 0 | 7.10E-06 | 0 | 4.81E-06 | 0 | 3.51E-05 | 1.13E-06 | 1.29E-06 | 5.21E-05 | 0 | 8.93E-07 | 0 | 0 | 0 | 1.48E-05 | 0 | 0 |
| Mgmt | 1.66E-05 | 3.01E-05 | 0 | 0 | 3.00E-06 | 1.25E-05 | 8.43E-06 | 0 | 1.27E-06 | 3.67E-05 | 0 | 1.19E-05 | 6.22E-05 | 0 | 3.45E-06 | 6.96E-06 | 0 | 7.18E-06 |
| Cdc6 | 2.95E-07 | 0 | 5.41E-05 | 0 | 0 | 1.10E-06 | 0 | 0 | 1.26E-06 | 0 | 0 | 0 | 1.56E-05 | 2.88E-06 | 0 | 0.00016 | 0 | 0 |
| Dennd5b | 3.84E-05 | 3.97E-05 | 2.59E-05 | 0 | 5.53E-05 | 2.47E-05 | 2.56E-05 | 5.05E-05 | 1.25E-06 | 1.22E-05 | 4.54E-05 | 6.21E-06 | 0 | 3.03E-05 | 1.58E-05 | 5.14E-05 | 4.13E-05 | 5.55E-06 |
| Palb2 | 1.04E-06 | 3.77E-06 | 4.68E-06 | 0 | 0 | 7.53E-07 | 1.88E-06 | 0 | 1.25E-06 | 1.28E-06 | 0 | 3.98E-06 | 4.51E-06 | 1.41E-06 | 7.83E-06 | 7.74E-05 | 0 | 3.18E-06 |
| Cmtr2 | 1.89E-05 | 5.95E-06 | 1.08E-05 | 9.13E-06 | 5.98E-06 | 0 | 0 | 3.81E-06 | 1.25E-06 | 1.28E-05 | 0 | 6.77E-06 | 6.42E-06 | 9.84E-06 | 0 | 3.55E-07 | 5.04E-08 | 5.88E-06 |
| Kin | 4.21E-05 | 3.83E-05 | 2.46E-05 | 7.18E-06 | 1.20E-05 | 0 | 0 | 0 | 1.24E-06 | 2.36E-05 | 0 | 2.62E-05 | 0 | 0 | 0.00024 | 3.89E-05 | 6.99E-06 | 5.67E-06 |
| Fsbp | 0 | 0 | 0 | 0 | 3.64E-06 | 4.55E-06 | 2.65E-06 | 0 | 1.22E-06 | 3.34E-06 | 6.06E-06 | 0 | 8.24E-06 | 0 | 1.83E-06 | 7.98E-05 | 3.44E-06 | 4.32E-06 |
| Gatm | 0 | 8.14E-06 | 6.89E-07 | 1.16E-05 | 0 | 0 | 0 | 0 | 1.21E-06 | 2.28E-06 | 7.78E-06 | 1.01E-05 | 0 | 0 | 1.23E-05 | 1.48E-05 | 2.71E-06 | 8.63E-06 |
| Dnaja4 | 3.27E-05 | 7.18E-05 | 6.27E-05 | 0 | 7.11E-05 | 4.15E-05 | 4.91E-05 | 5.71E-06 | 1.20E-06 | 1.17E-05 | 5.43E-05 | 2.82E-05 | 0 | 4.13E-05 | 1.84E-06 | 1.03E-06 | 0.00016 | 3.10E-05 |
| Zfp781 | 1.09E-05 | 8.46E-06 | 0 | 0 | 2.75E-06 | 7.70E-06 | 0 | 7.80E-06 | 1.19E-06 | 5.16E-06 | 0 | 3.46E-06 | 0 | 0 | 0 | 0 | 0 | 6.40E-06 |
| Cd3eap | 2.61E-05 | 2.25E-05 | 8.30E-05 | 4.70E-05 | 1.73E-05 | 0 | 0 | 4.22E-06 | 1.19E-06 | 2.55E-08 | 3.27E-05 | 3.94E-05 | 2.81E-05 | 8.71E-06 | 5.65E-05 | 0 | 0 | 9.40E-06 |
| Map4k1 | 0 | 1.17E-05 | 0 | 0 | 0 | 0 | 0 | 2.17E-05 | 1.19E-06 | 0 | 0 | 0 | 0 | 0 | 3.01E-06 | 1.23E-05 | 0 | 0 |
| Optn | 0 | 1.99E-05 | 0 | 0 | 0 | 0 | 0 | 0 | 1.18E-06 | 4.81E-05 | 0 | 1.87E-05 | 2.28E-05 | 3.03E-05 | 0 | 3.84E-05 | 0 | 8.10E-05 |
| Fam120c | 1.72E-05 | 2.15E-05 | 0 | 0 | 1.64E-05 | 8.17E-06 | 1.63E-05 | 1.10E-05 | 1.17E-06 | 8.20E-06 | 0 | 6.18E-06 | 2.71E-05 | 0 | 1.21E-05 | 1.86E-05 | 3.29E-05 | 1.81E-05 |
| Osm | 0 | 5.54E-06 | 5.46E-07 | 0 | 3.62E-07 | 3.52E-08 | 0 | 2.20E-06 | 1.17E-06 | 6.52E-06 | 2.12E-06 | 4.80E-06 | 4.06E-06 | 6.65E-06 | 4.15E-05 | 9.74E-06 | 1.48E-05 | 3.58E-06 |
| Nrcam | 2.13E-06 | 0 | 0 | 0 | 0 | 0 | 0 | 0 | 1.15E-06 | 0.00023 | 0 | 6.76E-07 | 0.0001 | 0 | 0 | 0 | 0 | 0 |
| Acyp1 | 1.95E-05 | 3.68E-05 | 6.22E-06 | 8.55E-07 | 2.68E-05 | 5.00E-05 | 0 | 2.07E-06 | 1.14E-06 | 4.89E-05 | 1.86E-05 | 3.83E-05 | 0 | 1.42E-05 | 0 | 2.20E-05 | 0 | 3.91E-06 |
| Mad2l1bp | 3.89E-05 | 5.65E-05 | 2.43E-05 | 7.49E-05 | 2.27E-05 | 1.59E-05 | 3.07E-06 | 1.36E-05 | 1.14E-06 | 2.47E-05 | 2.72E-05 | 6.22E-05 | 0 | 1.20E-05 | 2.13E-05 | 0.00012 | 1.17E-05 | 5.45E-05 |
| Mblac2 | 2.42E-06 | 2.32E-05 | 0 | 0 | 5.39E-06 | 2.70E-06 | 0 | 8.79E-06 | 1.14E-06 | 9.92E-06 | 4.49E-06 | 1.34E-05 | 0 | 0 | 2.46E-05 | 0 | 0 | 4.51E-06 |
| Ccdc18 | 0 | 2.42E-06 | 0 | 0 | 0 | 6.45E-08 | 0 | 0 | 1.14E-06 | 1.93E-06 | 0 | 0 | 0 | 0 | 1.73E-05 | 7.03E-05 | 0 | 0 |
| Enkd1 | 0 | 1.01E-05 | 3.07E-05 | 5.69E-05 | 6.35E-06 | 0 | 0 | 0 | 1.13E-06 | 1.97E-05 | 4.29E-06 | 2.39E-05 | 1.35E-05 | 1.00E-05 | 7.06E-05 | 0.00018 | 0 | 1.31E-05 |
| Nt5dc3 | 2.02E-05 | 4.78E-05 | 9.16E-05 | 0 | 1.28E-05 | 0 | 2.57E-05 | 0 | 1.13E-06 | 2.46E-05 | 3.44E-05 | 4.77E-05 | 4.07E-05 | 5.89E-06 | 4.46E-05 | 1.27E-05 | 0.0001 | 2.19E-05 |
| Parpbp | 0 | 7.71E-06 | 0 | 0 | 0 | 0 | 0 | 0 | 1.13E-06 | 1.63E-07 | 0 | 0 | 0 | 0 | 0 | 0.00048 | 0 | 0 |
| Spi1 | 0 | 5.31E-06 | 4.82E-06 | 1.37E-05 | 0 | 0 | 0 | 0 | 1.12E-06 | 8.20E-06 | 0 | 1.49E-07 | 0 | 1.23E-05 | 2.28E-05 | 8.66E-06 | 1.75E-05 | 1.46E-05 |
| Phyh | 7.43E-05 | 5.92E-05 | 0 | 1.68E-07 | 0 | 3.55E-05 | 8.53E-05 | 0 | 1.11E-06 | 8.27E-05 | 5.51E-06 | 4.78E-05 | 0.0002 | 1.61E-05 | 0 | 0 | 0 | 7.21E-05 |
| Akr1c13 | 3.01E-06 | 4.84E-06 | 0 | 1.60E-06 | 0 | 0 | 0 | 0 | 1.11E-06 | 3.17E-06 | 4.23E-06 | 1.44E-06 | 0 | 2.53E-06 | 1.38E-05 | 0 | 5.23E-06 | 1.42E-05 |
| Nphp4 | 2.01E-06 | 1.17E-05 | 2.37E-06 | 0 | 2.26E-05 | 8.34E-06 | 2.88E-08 | 1.32E-05 | 1.10E-06 | 2.72E-06 | 8.54E-06 | 0 | 0 | 3.20E-06 | 2.50E-05 | 4.25E-05 | 6.55E-06 | 3.45E-07 |
| Dyrk1b | 2.96E-05 | 0.00013 | 0 | 1.25E-05 | 5.27E-05 | 7.41E-05 | 5.27E-06 | 1.64E-05 | 1.09E-06 | 5.05E-05 | 0 | 6.85E-05 | 0 | 5.18E-05 | 6.77E-05 | 0 | 0 | 4.34E-05 |

| | | | | | | | | | | | | | | | | | | |
|---|---|---|---|---|---|---|---|---|---|---|---|---|---|---|---|---|---|---|
| Lrp11 | 1.86E-06 | 1.78E-05 | 3.28E-06 | 0 | 3.73E-05 | 2.14E-05 | 6.38E-06 | 0 | 1.08E-06 | 1.60E-05 | 3.11E-05 | 1.74E-05 | 1.66E-05 | 3.38E-05 | 7.79E-06 | 1.12E-05 | 0 | 2.00E-06 |
| Cap2 | 4.24E-06 | 0 | 0 | 0 | 0 | 2.05E-05 | 0 | 3.01E-05 | 1.06E-06 | 0 | 2.86E-05 | 0 | 0 | 3.07E-05 | 4.46E-07 | 0 | 0 | 0 |
| Tm4sf19 | 0 | 0 | 8.67E-06 | 0 | 3.28E-06 | 7.12E-06 | 0 | 6.65E-06 | 1.05E-06 | 0 | 2.78E-06 | 0 | 0 | 4.96E-07 | 0 | 3.19E-06 | 8.23E-06 | 0 |
| Xkr4 | 0 | 0 | 0 | 0 | 1.12E-05 | 6.68E-06 | 8.78E-06 | 0 | 1.05E-06 | 2.78E-06 | 4.45E-06 | 0 | 1.61E-05 | 0 | 6.01E-06 | 0 | 0 | 0 |
| Uchl4 | 5.28E-06 | 2.07E-06 | 7.98E-06 | 4.44E-06 | 0 | 0 | 0 | 0 | 1.01E-06 | 6.63E-06 | 1.32E-07 | 0 | 0 | 0 | 6.68E-06 | 7.61E-06 | 0 | 0 |
| Sdhaf3 | 1.44E-05 | 2.00E-05 | 1.62E-05 | 5.45E-06 | 6.73E-06 | 1.06E-05 | 3.46E-06 | 1.19E-05 | 1.00E-06 | 1.33E-05 | 9.47E-06 | 0 | 0 | 0 | 0 | 5.91E-06 | 3.13E-05 | 1.29E-06 |
| Retreg1 | 0 | 5.18E-06 | 0 | 0 | 1.69E-06 | 0 | 0 | 0 | 9.98E-07 | 0 | 0 | 6.96E-06 | 0 | 1.84E-05 | 0 | 3.77E-06 | 5.85E-06 | 3.46E-05 |
| Trim30a | 1.76E-05 | 9.33E-06 | 0 | 0 | 0 | 3.60E-06 | 0 | 4.74E-06 | 9.88E-07 | 0.00021 | 0 | 6.32E-06 | 8.40E-05 | 0 | 2.61E-05 | 1.20E-05 | 1.74E-07 | 6.09E-05 |
| Trim24 | 0.00013 | 0.00011 | 2.94E-06 | 0 | 1.67E-06 | 5.28E-05 | 0 | 0 | 9.87E-07 | 3.24E-05 | 8.53E-05 | 3.53E-05 | 0 | 7.84E-06 | 0.00019 | 5.53E-05 | 8.32E-05 | 0 |
| Haus8 | 6.00E-06 | 1.99E-05 | 2.50E-05 | 1.75E-05 | 0 | 0 | 0 | 7.42E-06 | 9.74E-07 | 1.30E-05 | 0 | 4.51E-05 | 0 | 0 | 1.18E-05 | 0.00033 | 9.08E-05 | 2.32E-05 |
| Nipal1 | 0 | 2.08E-06 | 1.27E-05 | 0 | 0 | 2.63E-05 | 0 | 0 | 9.67E-07 | 0 | 4.85E-05 | 5.63E-05 | 0 | 5.79E-06 | 1.53E-05 | 0 | 0 | 7.68E-06 |
| Lonrf3 | 0 | 3.80E-05 | 0 | 0 | 0 | 0 | 0 | 0 | 9.61E-07 | 5.91E-06 | 0 | 0.00014 | 0 | 0 | 2.44E-05 | 0.00012 | 0.00019 | 0 |
| Rasa2 | 5.60E-05 | 8.15E-05 | 8.72E-05 | 0 | 8.06E-05 | 2.52E-05 | 0 | 1.64E-05 | 9.56E-07 | 0 | 4.01E-05 | 7.78E-05 | 0 | 6.26E-05 | 9.52E-05 | 6.62E-06 | 1.54E-05 | 1.09E-05 |
| Plxnb1 | 0 | 3.18E-05 | 0 | 2.16E-07 | 3.19E-05 | 0 | 0 | 3.20E-05 | 9.30E-07 | 0 | 5.34E-05 | 0 | 2.24E-06 | 4.65E-05 | 3.08E-05 | 4.69E-05 | 0 | 0 |
| Adap1 | 0 | 2.99E-07 | 7.14E-06 | 5.74E-06 | 0 | 0 | 0 | 0 | 9.29E-07 | 3.36E-05 | 0 | 2.82E-06 | 6.75E-06 | 2.81E-06 | 9.02E-06 | 2.59E-06 | 0 | 3.70E-05 |
| Ganc | 2.60E-05 | 1.67E-05 | 0 | 0 | 8.65E-06 | 3.02E-05 | 7.86E-06 | 6.90E-06 | 9.16E-07 | 2.41E-05 | 0 | 1.04E-05 | 0 | 0 | 2.12E-05 | 2.40E-06 | 1.43E-05 | 1.63E-05 |
| Cystm1 | 1.69E-06 | 3.53E-05 | 3.88E-06 | 9.94E-05 | 0 | 0 | 0 | 1.00E-05 | 9.16E-07 | 1.70E-05 | 3.98E-06 | 8.08E-05 | 0 | 0 | 0.00015 | 2.25E-05 | 7.55E-05 | 2.88E-05 |
| Fam214a | 0 | 0.00015 | 2.90E-05 | 0 | 0.00017 | 0 | 4.21E-05 | 0.00023 | 9.13E-07 | 4.55E-05 | 0 | 8.78E-05 | 0 | 0.00019 | 0.00011 | 0 | 6.95E-05 | 2.16E-05 |
| Ssc4d | 9.70E-06 | 3.68E-06 | 0 | 1.07E-05 | 0 | 7.83E-06 | 0 | 9.43E-06 | 9.05E-07 | 1.11E-06 | 0 | 1.31E-05 | 8.55E-06 | 0 | 0 | 2.21E-05 | 1.46E-06 | 9.17E-06 |
| Srgap1 | 0 | 5.66E-05 | 7.91E-05 | 0 | 0 | 0 | 0 | 0 | 8.98E-07 | 0.00022 | 0 | 1.86E-05 | 6.81E-05 | 0 | 6.99E-05 | 1.48E-05 | 4.40E-05 | 1.12E-05 |
| C1qa | 0 | 1.78E-05 | 7.80E-06 | 4.16E-05 | 7.32E-06 | 0 | 3.23E-06 | 7.07E-06 | 8.95E-07 | 7.25E-06 | 6.20E-06 | 3.34E-05 | 1.35E-05 | 8.22E-06 | 0.00012 | 1.20E-06 | 0 | 1.78E-05 |
| Usp15 | 0.00016 | 0.00013 | 0.00013 | 5.66E-05 | 3.71E-05 | 5.10E-05 | 2.96E-05 | 2.13E-05 | 8.94E-07 | 4.96E-05 | 4.40E-05 | 8.73E-05 | 0 | 0 | 8.11E-05 | 8.34E-05 | 0.00012 | 5.29E-05 |
| Cdh1 | 3.99E-06 | 1.33E-05 | 0 | 2.45E-05 | 8.77E-06 | 0 | 0 | 5.22E-06 | 8.78E-07 | 9.92E-06 | 1.04E-06 | 1.19E-05 | 2.43E-06 | 0 | 3.31E-05 | 8.71E-06 | 7.38E-06 | 2.14E-05 |
| Creb3l4 | 5.28E-06 | 7.84E-06 | 0 | 7.82E-07 | 3.91E-06 | 2.30E-06 | 9.66E-06 | 0 | 8.76E-07 | 4.52E-06 | 3.25E-06 | 5.17E-06 | 1.64E-05 | 1.11E-06 | 1.02E-05 | 0 | 0 | 0 |
| Ermard | 5.35E-06 | 2.56E-06 | 0 | 0 | 4.39E-06 | 6.49E-06 | 1.11E-06 | 0 | 8.74E-07 | 2.08E-07 | 0 | 9.22E-06 | 0 | 0 | 0 | 3.55E-06 | 0 | 1.01E-05 |
| Zfand1 | 3.35E-05 | 2.56E-05 | 0 | 1.03E-05 | 2.24E-05 | 3.64E-05 | 3.38E-05 | 2.60E-06 | 8.72E-07 | 9.72E-06 | 4.91E-06 | 2.47E-05 | 9.51E-06 | 0 | 7.34E-05 | 0 | 0 | 9.49E-06 |
| Tmem11 | 3.12E-05 | 0.00013 | 0.00025 | 0.00028 | 3.41E-05 | 5.96E-05 | 3.37E-05 | 6.94E-05 | 8.71E-07 | 5.06E-05 | 3.59E-05 | 0.00012 | 4.87E-07 | 4.96E-05 | 0.00013 | 8.93E-05 | 0.00013 | 4.09E-05 |
| Btbd10 | 1.75E-05 | 5.03E-05 | 4.43E-05 | 1.49E-06 | 1.71E-05 | 7.63E-06 | 3.68E-06 | 4.67E-06 | 8.64E-07 | 3.85E-05 | 2.36E-05 | 2.77E-06 | 3.66E-09 | 0 | 2.76E-06 | 6.27E-05 | 5.69E-05 | 2.99E-05 |
| Cdkl1 | 0 | 0 | 0 | 1.24E-06 | 5.60E-06 | 1.15E-05 | 9.11E-06 | 0 | 8.49E-07 | 0 | 2.26E-05 | 1.01E-06 | 1.27E-05 | 5.55E-06 | 4.85E-06 | 0 | 1.22E-05 | 0 |
| Cdk15 | 0 | 2.04E-05 | 0 | 0 | 0 | 1.07E-05 | 1.45E-06 | 0 | 8.30E-07 | 0 | 0 | 2.70E-05 | 0 | 2.19E-05 | 0 | 2.15E-06 | 0 | 3.68E-06 |
| Adam4 | 1.65E-05 | 1.55E-05 | 5.60E-06 | 0 | 4.16E-06 | 0 | 7.98E-06 | 2.76E-06 | 8.29E-07 | 6.56E-06 | 0 | 2.63E-06 | 0 | 1.55E-06 | 3.03E-05 | 0 | 0 | 5.50E-06 |
| Adgrg6 | 0 | 6.31E-06 | 9.13E-06 | 0 | 0 | 0 | 3.80E-07 | 5.40E-06 | 8.27E-07 | 8.66E-05 | 2.59E-06 | 6.21E-06 | 0 | 0 | 1.15E-05 | 3.60E-06 | 0 | 0 |
| Hectd3 | 7.26E-05 | 0.00011 | 1.87E-05 | 0.00013 | 5.04E-05 | 5.33E-05 | 0.0001 | 4.49E-05 | 8.13E-07 | 5.51E-05 | 0 | 1.05E-05 | 0 | 7.19E-05 | 0.00017 | 5.21E-06 | 3.25E-05 | 0.00018 |

| | | | | | | | | | | | | | | | | | | |
|---|---|---|---|---|---|---|---|---|---|---|---|---|---|---|---|---|---|---|
| Pld4 | 0 | 1.57E-06 | 8.00E-07 | 1.79E-05 | 0 | 0 | 0 | 0 | 8.06E-07 | 4.14E-06 | 0 | 0 | 6.44E-06 | 0 | 5.48E-05 | 1.51E-06 | 3.01E-05 | 3.41E-06 |
| Ezr | 0 | 6.96E-05 | 8.33E-06 | 6.14E-05 | 0 | 1.20E-05 | 0 | 2.06E-05 | 8.01E-07 | 9.31E-06 | 0 | 3.50E-05 | 0 | 0 | 0.0001 | 3.61E-05 | 2.67E-05 | 5.01E-05 |
| Zswim4 | 0 | 0.00013 | 7.73E-05 | 9.76E-05 | 7.94E-05 | 7.35E-05 | 0 | 8.76E-05 | 7.98E-07 | 0 | 0.00011 | 0.00016 | 0 | 6.87E-05 | 0.00028 | 0 | 0.00056 | 1.38E-05 |
| Mab21l1 | 0 | 0 | 0 | 1.55E-06 | 0 | 0 | 2.25E-05 | 7.14E-06 | 7.79E-07 | 6.01E-05 | 0 | 0 | 0 | 0 | 0 | 1.27E-06 | 0 | 0 |
| Lilrb4a | 0 | 2.21E-05 | 8.84E-06 | 4.87E-05 | 0 | 0 | 0 | 0 | 7.73E-07 | 7.44E-06 | 0 | 1.93E-05 | 1.87E-06 | 0 | 7.11E-05 | 2.02E-05 | 1.26E-06 | 1.30E-05 |
| Wfikkn1 | 0 | 1.13E-05 | 2.46E-07 | 0 | 0 | 8.09E-07 | 0 | 0 | 7.72E-07 | 4.30E-06 | 0 | 5.32E-06 | 3.95E-06 | 0 | 4.18E-05 | 5.43E-06 | 5.30E-05 | 8.91E-07 |
| Pcdhga2 | 0 | 1.10E-05 | 0 | 0 | 5.27E-06 | 7.69E-06 | 0 | 1.76E-06 | 7.39E-07 | 2.52E-06 | 0 | 0 | 3.29E-06 | 1.26E-05 | 2.11E-05 | 0 | 0 | 1.29E-06 |
| Usp16 | 0.00022 | 6.40E-05 | 0.0002 | 5.07E-05 | 5.41E-05 | 6.95E-05 | 0.00011 | 2.80E-05 | 7.33E-07 | 8.00E-06 | 1.97E-05 | 1.69E-05 | 5.60E-05 | 7.98E-05 | 0.00018 | 1.10E-05 | 0.00021 | 0 |
| Pms1 | 1.60E-05 | 1.28E-05 | 0 | 0 | 0 | 3.92E-06 | 0 | 5.41E-06 | 7.21E-07 | 3.57E-06 | 0 | 1.77E-05 | 0 | 0 | 1.81E-05 | 5.94E-05 | 0 | 7.58E-07 |
| Liph | 0 | 4.57E-07 | 0 | 0 | 0 | 6.38E-05 | 0 | 3.13E-06 | 6.97E-07 | 0 | 0 | 2.57E-05 | 0 | 1.55E-05 | 0 | 1.34E-05 | 0 | 0 |
| Kcnn1 | 1.70E-06 | 5.50E-06 | 0 | 4.50E-06 | 1.41E-05 | 6.52E-06 | 6.93E-06 | 0 | 6.91E-07 | 9.70E-06 | 6.48E-06 | 2.46E-06 | 1.09E-05 | 6.10E-06 | 2.01E-05 | 0 | 0 | 1.02E-05 |
| Dmtn | 0 | 1.07E-06 | 3.16E-06 | 6.12E-06 | 7.74E-06 | 0 | 4.69E-06 | 1.62E-06 | 6.78E-07 | 4.83E-06 | 2.61E-06 | 6.04E-06 | 0 | 3.38E-06 | 3.82E-06 | 0 | 6.07E-06 | 1.86E-05 |
| Shld2 | 1.40E-05 | 1.88E-05 | 0 | 2.07E-06 | 0 | 1.37E-05 | 0 | 0 | 6.71E-07 | 7.55E-06 | 4.00E-06 | 0 | 4.37E-07 | 5.51E-06 | 1.56E-05 | 1.31E-05 | 0 | 9.10E-06 |
| Cage1 | 0 | 1.26E-05 | 6.58E-06 | 3.15E-06 | 4.33E-06 | 1.26E-05 | 0 | 0 | 6.63E-07 | 6.05E-06 | 2.05E-06 | 1.40E-06 | 0 | 0 | 2.51E-07 | 0 | 1.46E-05 | 3.38E-06 |
| Pdcd2l | 3.40E-05 | 8.07E-05 | 6.66E-05 | 2.22E-05 | 8.72E-06 | 0 | 0 | 0 | 6.54E-07 | 1.97E-05 | 2.18E-05 | 6.02E-05 | 0 | 5.21E-06 | 0.00019 | 3.56E-05 | 7.52E-05 | 1.56E-05 |
| Upk1a | 6.14E-07 | 0 | 2.44E-06 | 3.68E-06 | 1.45E-06 | 3.71E-06 | 1.89E-06 | 8.58E-06 | 6.47E-07 | 0 | 1.04E-06 | 4.07E-06 | 2.89E-06 | 1.64E-06 | 1.42E-05 | 0 | 3.10E-06 | 4.11E-06 |
| Gal3st4 | 0 | 4.43E-06 | 0 | 0 | 4.02E-05 | 0 | 3.50E-05 | 0 | 6.40E-07 | 6.68E-06 | 4.31E-06 | 0 | 0 | 0 | 0 | 0 | 0 | 2.11E-05 |
| Cobll1 | 0 | 3.58E-05 | 0 | 0 | 0 | 0 | 0 | 3.38E-05 | 6.40E-07 | 0 | 0 | 3.00E-05 | 0 | 2.72E-05 | 6.36E-06 | 1.76E-05 | 0 | 3.58E-05 |
| Hoxb5 | 5.48E-07 | 0 | 0 | 0 | 5.34E-06 | 8.66E-08 | 4.52E-06 | 0 | 6.36E-07 | 1.54E-06 | 8.84E-06 | 0 | 0 | 0 | 3.49E-05 | 1.42E-06 | 0 | 0 |
| Hoxa2 | 7.86E-06 | 1.85E-05 | 3.00E-06 | 3.35E-06 | 4.83E-06 | 0 | 3.40E-06 | 6.86E-06 | 6.23E-07 | 3.56E-06 | 0 | 6.99E-06 | 0 | 0 | 8.74E-06 | 7.34E-06 | 0 | 2.51E-06 |
| Plag1 | 1.90E-06 | 1.09E-05 | 2.88E-06 | 0 | 0 | 4.61E-06 | 4.33E-07 | 3.38E-06 | 6.19E-07 | 2.27E-06 | 0 | 5.56E-06 | 0 | 3.52E-06 | 5.14E-06 | 3.26E-06 | 1.07E-05 | 5.96E-07 |
| Klhl8 | 7.89E-06 | 1.69E-05 | 0 | 7.23E-06 | 1.06E-05 | 5.73E-06 | 1.99E-06 | 2.08E-05 | 5.97E-07 | 1.10E-05 | 4.92E-06 | 2.54E-06 | 0 | 4.02E-06 | 5.18E-06 | 0 | 0 | 2.15E-05 |
| Cmc2 | 8.67E-06 | 7.15E-06 | 1.45E-05 | 0.00021 | 3.68E-05 | 2.86E-05 | 3.55E-05 | 3.12E-05 | 5.81E-07 | 3.25E-05 | 1.42E-08 | 0 | 1.30E-05 | 0 | 0 | 0.00049 | 0 | 4.61E-05 |
| Zscan2 | 6.32E-06 | 2.26E-05 | 0 | 0 | 1.89E-05 | 9.91E-06 | 0 | 8.84E-06 | 5.78E-07 | 0 | 4.36E-06 | 1.85E-05 | 0 | 9.75E-06 | 1.59E-05 | 6.43E-06 | 0 | 1.46E-05 |
| Efr3b | 2.28E-06 | 2.39E-06 | 9.34E-06 | 5.50E-06 | 5.14E-06 | 0 | 5.01E-06 | 1.63E-06 | 5.30E-07 | 1.41E-07 | 3.11E-06 | 1.40E-05 | 2.70E-05 | 0 | 2.75E-06 | 1.56E-05 | 0 | 1.06E-05 |
| C1ql3 | 0 | 1.87E-06 | 0 | 0 | 0 | 0 | 1.63E-05 | 0 | 5.19E-07 | 0 | 1.00E-05 | 1.89E-05 | 1.27E-05 | 0 | 0 | 5.84E-07 | 2.28E-06 | 0 |
| Prob1 | 0 | 8.02E-06 | 0 | 6.17E-06 | 1.10E-06 | 5.69E-07 | 1.45E-06 | 1.88E-06 | 5.16E-07 | 0 | 1.37E-06 | 0 | 1.70E-06 | 6.00E-06 | 0 | 6.79E-06 | 1.03E-05 | 0 |
| Gp1bb | 0 | 8.38E-07 | 2.34E-06 | 3.33E-06 | 2.82E-06 | 0 | 4.80E-06 | 2.67E-06 | 5.13E-07 | 6.57E-07 | 9.16E-06 | 6.20E-06 | 2.14E-06 | 4.18E-07 | 0 | 1.55E-06 | 9.60E-07 | 6.42E-06 |
| Zfp976 | 3.02E-06 | 6.80E-06 | 1.97E-06 | 0 | 6.32E-06 | 8.41E-07 | 1.10E-06 | 0 | 4.98E-07 | 7.30E-06 | 0 | 0 | 1.34E-06 | 1.78E-06 | 4.03E-06 | 1.16E-06 | 0 | 9.53E-06 |
| Scml2 | 0 | 0 | 0 | 0 | 0 | 1.73E-06 | 1.08E-05 | 0 | 4.94E-07 | 2.52E-05 | 0 | 2.16E-06 | 8.50E-06 | 0 | 0 | 7.36E-05 | 1.84E-06 | 0 |
| Map7d3 | 1.40E-05 | 2.80E-06 | 0 | 3.45E-07 | 6.31E-06 | 4.30E-06 | 0 | 0 | 4.90E-07 | 4.48E-06 | 6.20E-06 | 1.12E-07 | 5.27E-06 | 5.98E-06 | 0 | 1.69E-05 | 0 | 6.77E-06 |
| Pou5f2 | 0 | 2.09E-05 | 0 | 4.70E-06 | 9.53E-06 | 9.97E-06 | 0 | 0 | 4.89E-07 | 3.53E-05 | 1.65E-05 | 4.67E-06 | 2.81E-06 | 0 | 0.00014 | 1.15E-05 | 6.76E-05 | 2.79E-05 |
| Prkdc | 1.53E-05 | 1.48E-05 | 4.24E-06 | 1.19E-05 | 1.66E-06 | 2.46E-05 | 7.22E-06 | 0 | 4.79E-07 | 1.62E-05 | 3.33E-06 | 0 | 1.74E-07 | 7.67E-07 | 1.99E-05 | 1.48E-05 | 2.00E-06 | 2.30E-05 |

| | | | | | | | | | | | | | | | | | | |
|---|---|---|---|---|---|---|---|---|---|---|---|---|---|---|---|---|---|---|
| Bbs1 | 2.41E-06 | 2.18E-05 | 0 | 1.29E-05 | 1.90E-05 | 1.45E-05 | 0 | 1.10E-05 | 4.67E-07 | 3.10E-05 | 0 | 1.73E-06 | 3.53E-06 | 2.00E-05 | 1.03E-05 | 2.00E-05 | 0 | 1.28E-05 |
| Atp13a3 | 0.00028 | 0.00016 | 0.00033 | 0 | 0.00016 | 0.00026 | 0 | 2.32E-05 | 4.51E-07 | 4.48E-05 | 0.0006 | 0.00017 | 2.06E-06 | 0.00027 | 0 | 0.00018 | 0.0003 | 0 |
| Zfp940 | 3.09E-06 | 2.72E-06 | 0 | 6.79E-06 | 1.12E-05 | 1.76E-06 | 1.46E-06 | 0 | 4.31E-07 | 1.01E-06 | 4.47E-06 | 0 | 0 | 9.76E-06 | 7.89E-06 | 0 | 0 | 0 |
| Homez | 1.49E-05 | 1.91E-05 | 5.04E-06 | 0 | 1.62E-05 | 1.86E-05 | 1.63E-05 | 6.40E-06 | 4.07E-07 | 4.53E-06 | 6.35E-07 | 5.63E-06 | 5.90E-06 | 5.95E-06 | 4.55E-07 | 0 | 0 | 5.55E-06 |
| Aph1b | 1.18E-05 | 9.57E-06 | 3.31E-06 | 2.14E-06 | 6.41E-06 | 6.51E-06 | 1.79E-05 | 4.85E-06 | 3.98E-07 | 2.40E-06 | 1.14E-05 | 6.21E-06 | 8.94E-06 | 4.03E-06 | 2.07E-05 | 0 | 0 | 4.08E-06 |
| S100a14 | 6.30E-08 | 0 | 2.32E-06 | 1.27E-05 | 0 | 0 | 0 | 1.24E-05 | 3.95E-07 | 1.55E-05 | 0 | 5.84E-06 | 8.10E-07 | 0 | 8.67E-06 | 7.88E-06 | 1.16E-05 | 1.36E-05 |
| Pirb | 0 | 2.95E-06 | 0 | 1.14E-05 | 1.85E-07 | 0 | 0 | 0 | 3.78E-07 | 7.34E-06 | 0 | 1.89E-06 | 0 | 9.00E-06 | 3.48E-05 | 4.76E-06 | 1.16E-05 | 0 |
| Tmigd3 | 0 | 1.94E-06 | 0 | 0 | 8.99E-05 | 4.68E-05 | 3.26E-05 | 0 | 3.77E-07 | 0 | 0 | 0 | 4.83E-05 | 2.26E-05 | 0 | 8.10E-06 | 0 | 0 |
| Ccdc189 | 6.54E-06 | 6.80E-06 | 0 | 1.07E-05 | 1.10E-06 | 2.72E-06 | 0 | 8.25E-06 | 3.57E-07 | 2.06E-05 | 0 | 1.91E-06 | 0 | 9.33E-07 | 2.00E-05 | 0 | 2.52E-05 | 6.80E-06 |
| Zranb3 | 2.77E-06 | 0 | 0 | 0 | 2.34E-07 | 7.73E-07 | 1.42E-05 | 5.51E-06 | 3.54E-07 | 1.83E-06 | 1.58E-05 | 7.31E-06 | 0 | 0 | 2.43E-05 | 0.00011 | 0 | 8.53E-06 |
| Lysmd1 | 1.32E-06 | 1.07E-05 | 2.67E-06 | 1.88E-05 | 1.46E-06 | 8.47E-06 | 1.26E-05 | 9.11E-06 | 3.48E-07 | 3.97E-06 | 1.75E-06 | 4.90E-06 | 7.20E-06 | 1.59E-07 | 1.78E-06 | 0 | 0 | 1.22E-06 |
| Zfp54 | 9.04E-06 | 6.13E-06 | 0 | 0 | 3.66E-08 | 9.04E-06 | 0 | 0 | 3.44E-07 | 3.92E-06 | 3.50E-07 | 1.45E-06 | 0 | 0 | 2.93E-05 | 1.02E-05 | 0 | 3.07E-06 |
| Hecw2 | 1.36E-06 | 1.61E-05 | 1.24E-05 | 0 | 0 | 0 | 1.58E-05 | 0 | 3.34E-07 | 5.23E-05 | 2.74E-06 | 0 | 1.30E-05 | 0 | 1.26E-05 | 6.71E-06 | 3.42E-06 | 2.28E-05 |
| Cul9 | 0 | 1.61E-05 | 0 | 1.20E-05 | 8.19E-06 | 4.73E-06 | 0 | 0 | 3.29E-07 | 3.48E-05 | 0 | 8.25E-06 | 4.39E-06 | 1.01E-05 | 1.67E-05 | 0 | 0 | 6.74E-06 |
| Met | 0 | 4.71E-06 | 0 | 0 | 0 | 0 | 0 | 0 | 3.25E-07 | 2.92E-05 | 6.28E-06 | 1.13E-06 | 1.22E-05 | 0 | 0 | 4.33E-06 | 7.86E-07 | 1.28E-05 |
| Stk38 | 0.00021 | 0.00032 | 2.79E-05 | 0.00019 | 0.00026 | 0.00013 | 0.00057 | 0.00023 | 3.18E-07 | 0.00035 | 5.02E-05 | 0.00025 | 0.00046 | 6.36E-05 | 0.00085 | 0.00018 | 9.95E-05 | 0 |
| Triqk | 4.61E-05 | 7.39E-06 | 0 | 0 | 1.40E-05 | 1.93E-05 | 3.15E-05 | 8.21E-06 | 3.18E-07 | 4.65E-06 | 2.86E-05 | 2.75E-05 | 0 | 1.95E-05 | 0 | 0 | 2.42E-06 | 8.65E-06 |
| Mgst2 | 0 | 1.04E-05 | 0 | 5.40E-06 | 5.46E-06 | 0 | 0 | 0 | 3.17E-07 | 7.53E-07 | 0 | 1.61E-06 | 9.88E-06 | 0 | 8.13E-06 | 2.98E-06 | 2.18E-06 | 6.73E-07 |
| Tnpo2 | 0.00021 | 0.00024 | 0.00028 | 0.00012 | 0.00013 | 0.00012 | 0.00018 | 0.00012 | 3.15E-07 | 6.92E-05 | 9.59E-05 | 0.00014 | 0 | 5.10E-05 | 0.00021 | 0.00019 | 0.00011 | 7.69E-05 |
| Hat1 | 5.43E-05 | 2.79E-05 | 0.00013 | 0 | 3.33E-06 | 5.82E-05 | 0 | 3.66E-05 | 3.12E-07 | 2.19E-05 | 0 | 4.19E-05 | 5.77E-05 | 8.72E-06 | 0 | 0.00057 | 2.11E-05 | 1.27E-06 |
| Bcl2a1b | 0 | 5.05E-06 | 0 | 1.22E-05 | 5.31E-06 | 0 | 0 | 0 | 3.08E-07 | 9.88E-06 | 4.50E-06 | 4.46E-06 | 1.04E-05 | 0 | 5.56E-05 | 1.12E-05 | 0 | 2.16E-05 |
| Zcwpw1 | 0 | 2.49E-06 | 0 | 0 | 0 | 5.29E-06 | 0 | 4.66E-06 | 2.99E-07 | 3.44E-07 | 0 | 1.62E-05 | 0 | 0 | 6.81E-06 | 3.16E-05 | 1.91E-05 | 0 |
| Zfp873 | 6.53E-06 | 1.41E-05 | 0 | 0 | 2.46E-06 | 0 | 2.66E-06 | 0 | 2.96E-07 | 5.14E-06 | 2.03E-06 | 0 | 1.39E-06 | 7.85E-06 | 1.54E-05 | 7.73E-07 | 0 | 1.43E-06 |
| Kmt2c | 9.67E-05 | 9.53E-05 | 2.60E-05 | 0 | 2.31E-05 | 8.05E-05 | 3.27E-05 | 3.00E-07 | 2.51E-07 | 3.88E-05 | 8.03E-06 | 6.84E-05 | 2.30E-05 | 0 | 2.58E-05 | 1.96E-05 | 0.00015 | 1.84E-05 |
| Pnldc1 | 0 | 1.30E-06 | 0 | 5.01E-06 | 3.14E-06 | 4.45E-06 | 4.69E-06 | 1.73E-06 | 2.44E-07 | 2.37E-06 | 0 | 4.69E-06 | 7.57E-06 | 0 | 0 | 4.02E-06 | 0 | 7.13E-06 |
| Cd300lf | 0 | 6.03E-06 | 0 | 2.33E-05 | 0 | 0 | 0 | 0 | 2.43E-07 | 2.27E-06 | 0 | 3.91E-06 | 2.51E-06 | 2.53E-06 | 1.87E-05 | 1.09E-05 | 1.79E-05 | 4.44E-06 |
| Abhd3 | 0 | 1.23E-05 | 0 | 0 | 0 | 5.73E-06 | 3.54E-06 | 0 | 2.25E-07 | 1.81E-05 | 2.39E-06 | 0 | 6.49E-06 | 4.25E-06 | 0 | 0 | 0 | 1.11E-05 |
| Ubox5 | 0 | 6.20E-06 | 2.05E-06 | 1.99E-06 | 0 | 2.24E-06 | 7.56E-06 | 0 | 2.25E-07 | 0 | 2.63E-06 | 4.43E-06 | 0 | 7.26E-06 | 3.35E-06 | 4.79E-07 | 2.67E-06 | 1.67E-07 |
| Ubald2 | 7.05E-05 | 0.00039 | 2.64E-05 | 0.0003 | 0.00019 | 0.00021 | 0 | 0.00038 | 2.09E-07 | 0.00012 | 5.15E-05 | 0.00045 | 0.00025 | 0.0002 | 3.79E-05 | 0.00068 | 0.00015 | 0.00013 |
| Rgs1 | 0 | 5.37E-06 | 0 | 1.84E-06 | 5.82E-07 | 0 | 0 | 4.67E-06 | 1.94E-07 | 5.67E-06 | 0 | 1.36E-05 | 0 | 0 | 4.19E-05 | 1.03E-05 | 1.27E-05 | 1.21E-05 |
| Poc1a | 8.61E-06 | 8.09E-06 | 0 | 1.12E-05 | 0 | 7.83E-06 | 0 | 0 | 1.63E-07 | 5.82E-06 | 0 | 9.46E-06 | 7.01E-06 | 9.76E-06 | 3.74E-06 | 0.00024 | 0 | 8.37E-06 |
| Dna2 | 0 | 0 | 4.08E-06 | 0 | 0 | 0 | 0 | 0 | 1.37E-07 | 0 | 0 | 0 | 0 | 0 | 1.57E-05 | 9.04E-05 | 0 | 2.84E-07 |
| Srp54b | 0.00027 | 6.82E-05 | 0.00022 | 0 | 7.40E-06 | 0.00011 | 0.00021 | 5.49E-05 | 1.35E-07 | 3.81E-05 | 7.27E-05 | 0.0001 | 1.41E-05 | 0 | 6.27E-05 | 7.72E-05 | 8.53E-05 | 7.44E-05 |

| | | | | | | | | | | | | | | | | | | |
|---|---|---|---|---|---|---|---|---|---|---|---|---|---|---|---|---|---|---|
| Zfp956 | 4.69E-06 | 1.79E-05 | 0 | 4.75E-06 | 6.12E-06 | 1.14E-06 | 1.46E-05 | 0 | 1.34E-07 | 7.08E-06 | 0 | 1.32E-06 | 2.42E-05 | 0 | 4.86E-05 | 1.04E-05 | 2.28E-05 | 0 |
| Depdc7 | 1.99E-07 | 5.83E-06 | 0 | 7.77E-08 | 1.29E-06 | 5.03E-06 | 0 | 0 | 1.31E-07 | 6.29E-08 | 0 | 6.23E-06 | 5.58E-06 | 7.29E-06 | 3.22E-06 | 1.27E-05 | 1.39E-05 | 2.91E-06 |
| Zc3h3 | 1.81E-05 | 6.90E-05 | 3.76E-05 | 4.07E-05 | 2.30E-05 | 3.88E-05 | 2.28E-05 | 0 | 1.12E-07 | 2.51E-05 | 0 | 5.85E-05 | 0 | 4.49E-06 | 8.85E-05 | 4.71E-07 | 0.00011 | 1.52E-05 |
| Fjx1 | 0 | 1.08E-05 | 0.00014 | 0 | 0.00011 | 0 | 0 | 0.00049 | 1.10E-07 | 0 | 0 | 6.25E-05 | 0 | 0 | 0 | 0 | 0.0002 | 0 |
| Zfp560 | 6.75E-05 | 5.57E-05 | 0 | 0 | 5.23E-05 | 4.92E-05 | 4.11E-05 | 2.62E-05 | 9.71E-08 | 1.75E-05 | 2.71E-05 | 3.19E-05 | 3.00E-05 | 0 | 0 | 8.83E-06 | 0 | 1.24E-05 |
| Coq8a | 0 | 2.15E-05 | 0 | 6.47E-06 | 5.40E-06 | 3.23E-06 | 0 | 0 | 7.57E-08 | 2.34E-05 | 0 | 0 | 1.46E-06 | 5.70E-06 | 1.08E-05 | 0 | 0 | 2.08E-05 |
| Mettl14 | 3.09E-05 | 3.91E-05 | 0 | 0 | 1.94E-05 | 7.73E-06 | 0 | 0 | 7.30E-08 | 3.40E-05 | 1.42E-05 | 3.66E-05 | 0 | 0 | 0 | 7.79E-05 | 0 | 2.28E-05 |
| Nsun7 | 0 | 3.21E-06 | 3.73E-06 | 2.75E-06 | 0 | 0 | 0 | 0 | 7.17E-08 | 7.05E-06 | 0 | 0 | 1.16E-05 | 0 | 1.75E-05 | 0 | 4.12E-06 | 1.95E-05 |
| Gpt | 1.98E-06 | 1.22E-05 | 1.11E-05 | 1.30E-05 | 4.46E-06 | 0 | 3.15E-06 | 1.48E-06 | 4.24E-08 | 8.04E-06 | 0 | 1.00E-06 | 6.43E-06 | 0 | 4.89E-05 | 1.54E-05 | 1.74E-05 | 1.11E-05 |
| Actn3 | 0 | 1.87E-06 | 1.97E-06 | 3.30E-06 | 4.03E-06 | 0 | 0 | 0 | 4.10E-08 | 5.67E-06 | 9.94E-06 | 0 | 1.32E-05 | 6.23E-06 | 0 | 2.04E-07 | 0 | 0 |
| Sp110 | 3.32E-06 | 5.78E-07 | 0 | 0 | 0 | 0 | 0 | 0 | 4.02E-08 | 4.76E-05 | 0 | 0 | 3.37E-06 | 0 | 0 | 5.13E-06 | 0 | 8.06E-06 |
| Cox18 | 8.24E-05 | 2.45E-06 | 7.37E-05 | 5.54E-05 | 0 | 2.94E-06 | 2.52E-05 | 0 | 3.58E-08 | 1.02E-06 | 1.57E-05 | 0 | 1.78E-05 | 3.95E-07 | 2.20E-06 | 3.78E-05 | 8.48E-05 | 0 |
| Piwil2 | 0 | 2.59E-07 | 8.92E-06 | 5.77E-06 | 3.95E-06 | 1.48E-06 | 1.27E-05 | 0 | 2.34E-08 | 0 | 0 | 1.68E-05 | 9.58E-06 | 2.46E-05 | 3.49E-06 | 0 | 4.05E-05 | 2.14E-06 |
| Kif24 | 0 | 0 | 0 | 0 | 4.00E-07 | 1.01E-06 | 0 | 0 | 2.10E-08 | 3.12E-06 | 0 | 2.13E-06 | 0 | 6.14E-06 | 0 | 7.94E-05 | 0 | 0 |
| Ccdc138 | 6.44E-06 | 0 | 1.72E-06 | 0 | 0 | 4.65E-06 | 0 | 0 | 7.37E-09 | 1.24E-05 | 0 | 1.40E-05 | 7.96E-07 | 4.20E-06 | 1.18E-05 | 2.25E-05 | 4.13E-05 | 0 |
| Fn1 | 4.11E-05 | 0.00038 | 0.00017 | 0.00088 | 0.00165 | 0.00222 | 0.00172 | 0.00261 | 0 | 0.0021 | 0.00175 | 0.00129 | 0.00137 | 0.00056 | 0.00226 | 0.00028 | 0.00074 | 0.00141 |
| Col4a1 | 0 | 0 | 0 | 0.00044 | 0 | 0.00169 | 0 | 0.00236 | 0 | 0.00111 | 4.29E-05 | 0 | 0.00177 | 0.00223 | 0.00151 | 0.00017 | 0 | 0.00134 |
| Col7a1 | 0 | 0 | 0 | 0 | 0.00214 | 0.0015 | 0 | 0 | 0 | 0 | 0.00083 | 0.00039 | 0 | 0.00191 | 0.00113 | 0 | 0 | 0 |
| Wnt5a | 0 | 0 | 0.00044 | 0 | 0.00113 | 0.00146 | 0 | 0.00025 | 0 | 0 | 0.00139 | 0.00039 | 0 | 0.00111 | 0.00042 | 7.40E-05 | 0 | 0 |
| Col5a3 | 0 | 0 | 0.001 | 0.00058 | 0.00054 | 0.00136 | 0.00108 | 0.00238 | 0 | 0.00063 | 0.00041 | 0.00074 | 0 | 0.00102 | 0.00219 | 0.00025 | 0 | 0 |
| Loxl2 | 0.00048 | 0 | 0.00055 | 0.00047 | 0.00074 | 0.00132 | 0.00153 | 0.00079 | 0 | 0.00098 | 0.00153 | 0.00118 | 0 | 0.00107 | 6.12E-05 | 0.00034 | 0.00014 | 0.00013 |
| Col4a2 | 0 | 0 | 0 | 0.0004 | 0 | 0.0013 | 0 | 0.00174 | 0 | 0.00081 | 7.03E-05 | 0 | 0.00131 | 0.00194 | 0.00174 | 0 | 0 | 0.00122 |
| Tbx3 | 0 | 0 | 5.40E-05 | 0 | 0.00069 | 0.00115 | 0 | 5.55E-05 | 0 | 0 | 0.00038 | 0.00019 | 0 | 0.00011 | 0.00026 | 0 | 0 | 0 |
| Htra1 | 0.00013 | 0.00023 | 0 | 0 | 0.00117 | 0.00103 | 0.00043 | 0 | 0 | 0.00066 | 0.00034 | 0.00065 | 3.95E-05 | 0.00045 | 0 | 0 | 0.00023 | 1.55E-05 |
| Cd53 | 0 | 0 | 0 | 0 | 0.00078 | 0.00097 | 0 | 0.00058 | 0 | 0 | 0.00016 | 5.57E-05 | 0 | 9.23E-06 | 0 | 0.00012 | 0 | 0 |
| Pmp22 | 0 | 0 | 0 | 0.0001 | 0 | 0.00093 | 0 | 0 | 0 | 0.00088 | 0.0006 | 0.00014 | 0.00092 | 0.00072 | 0 | 0.00049 | 0 | 0.00061 |
| Pxdn | 0.00011 | 0.00015 | 0.00075 | 0 | 0.00067 | 0.00087 | 0.00129 | 0.00129 | 0 | 0.00066 | 0.00032 | 0.00064 | 9.21E-05 | 0.00084 | 0.00039 | 0.00022 | 0.00025 | 0 |
| Creb3l1 | 0.00021 | 0 | 0.00093 | 0.00103 | 0.00057 | 0.00083 | 0.00142 | 0.00047 | 0 | 0.00045 | 0.00064 | 0.00041 | 0.00073 | 0.00063 | 0.00068 | 0.00036 | 0.00021 | 0.00018 |
| Fkbp9 | 0.00037 | 0 | 0.00029 | 0.00081 | 0.00031 | 0.00082 | 0.0014 | 0.00048 | 0 | 0.00065 | 0.00054 | 7.82E-05 | 0.001 | 0.00105 | 0.00041 | 0.00029 | 0.00028 | 0.00073 |
| Tnfrsf21 | 0 | 0.00015 | 0 | 0 | 0.00048 | 0.00076 | 0 | 0 | 0 | 0 | 0 | 0 | 0 | 0.00096 | 0 | 0 | 0 | 0 |
| Ctsh | 0.00058 | 0 | 0.00026 | 0.0003 | 0 | 0.00075 | 0.00158 | 0.00059 | 0 | 0.00053 | 0 | 0.00026 | 0.00141 | 0.00086 | 0 | 0.00018 | 0 | 0 |
| Antxr1 | 0.00011 | 0.00057 | 0.00041 | 0 | 0.00128 | 0.00074 | 0.00143 | 0.00089 | 0 | 0.00061 | 0.00026 | 0.00012 | 0.00073 | 0.00124 | 0.0008 | 8.01E-05 | 0.00034 | 0.00048 |
| Tmed3 | 0.00036 | 0 | 0.0004 | 0.00057 | 0.00078 | 0.00074 | 0.00129 | 0.00076 | 0 | 0.00045 | 0.00054 | 0.00042 | 0.00113 | 0.00069 | 0 | 0.00019 | 0 | 0.00045 |

| | | | | | | | | | | | | | | | | | | |
|---|---|---|---|---|---|---|---|---|---|---|---|---|---|---|---|---|---|---|
| DEPRECATEI | 0 | 0.00014 | 0 | 0 | 0.00085 | 0.00069 | 0.00029 | 0 | 0 | 0.00047 | 0.00025 | 0.00044 | 0 | 0.00023 | 0 | 0 | 0.00014 | 3.12E-05 |
| Pcsk5 | 1.67E-05 | 0.00016 | 0.00039 | 0.0003 | 0.00094 | 0.00068 | 0.0016 | 0.00073 | 0 | 0.00013 | 0.00018 | 7.65E-06 | 0.00197 | 0.00054 | 0.00104 | 4.26E-06 | 0.00028 | 0 |
| Rabac1 | 0.00026 | 0.00038 | 0.00028 | 0.00094 | 0.00049 | 0.00067 | 0.00087 | 0.00048 | 0 | 0.00035 | 0.00053 | 0.00061 | 0.0005 | 0.00055 | 0 | 0 | 0.00022 | 0.00043 |
| Stk26 | 0 | 0 | 0 | 0 | 0.00044 | 0.00067 | 0 | 0 | 0 | 0 | 0.00033 | 0 | 0 | 8.98E-06 | 0 | 0 | 0 | 0 |
| Lox | 0.00039 | 0 | 0.00053 | 0.00091 | 0.00082 | 0.00065 | 0.00225 | 0.00021 | 0 | 0.00091 | 0.00161 | 0.0015 | 0.00251 | 0.00041 | 0.00034 | 0.00038 | 0 | 0.00057 |
| Tcn2 | 0 | 0.0004 | 0 | 0.00017 | 0.00051 | 0.00065 | 9.04E-05 | 0.00041 | 0 | 0.00045 | 0.00011 | 0.00028 | 0.00068 | 0.00052 | 0 | 0 | 0 | 0.00051 |
| Lpl | 0 | 0.0003 | 0 | 0 | 5.56E-05 | 0.00063 | 0 | 0.00065 | 0 | 0.0003 | 0 | 0 | 0.00086 | 0.00025 | 0 | 0 | 0 | 0.00026 |
| Tcim | 0 | 0 | 5.70E-05 | 0 | 0.00025 | 0.00062 | 0 | 8.18E-05 | 0 | 0 | 0 | 0.00017 | 0 | 0.00037 | 0.00013 | 0 | 0.00243 | 0 |
| Cygb | 0 | 0.00025 | 0 | 0.00056 | 0.00013 | 0.0006 | 0 | 0.00204 | 0 | 0.00088 | 0 | 0 | 0.00061 | 0 | 6.43E-05 | 0 | 0 | 0.00183 |
| Ace | 0 | 0.00032 | 3.35E-05 | 0 | 0.00081 | 0.00058 | 0 | 0.00015 | 0 | 0.0006 | 0 | 0 | 0.00183 | 0 | 0 | 0 | 0 | 0.00086 |
| Mmd | 3.75E-05 | 0.0003 | 0.00021 | 0 | 0.00028 | 0.00058 | 1.11E-05 | 0.00054 | 0 | 0 | 0.00013 | 0.00021 | 0 | 0 | 0 | 0.00015 | 0 | 0 |
| A4galt | 0 | 0.00014 | 0 | 0.00013 | 0.00042 | 0.00057 | 0.00022 | 6.06E-05 | 0 | 2.24E-05 | 0 | 0.00053 | 0.00044 | 0.00036 | 0 | 0 | 0 | 0.00025 |
| P4ha1 | 0.00028 | 0 | 0.00036 | 0.00069 | 0.00017 | 0.00057 | 0.00089 | 0.00064 | 0 | 0.00028 | 0.00023 | 0.00171 | 0.00042 | 2.35E-05 | 0.00026 | 0.00015 | 0.00066 | 0 |
| Ext1 | 0.00028 | 0.00015 | 0.00034 | 0 | 0.00028 | 0.00056 | 0.0003 | 0.00064 | 0 | 0.0007 | 0.00055 | 0.00014 | 0.00046 | 0.00068 | 0.00046 | 0.00013 | 0.00042 | 0.00033 |
| Ptprn | 0 | 0 | 0.00027 | 0 | 0.00018 | 0.00056 | 0 | 0.00096 | 0 | 0 | 0.00032 | 0.00032 | 0 | 0.00025 | 0.00022 | 0 | 0 | 0 |
| Fermt2 | 0.0004 | 0.00021 | 0.00034 | 0 | 0.0003 | 0.00054 | 0 | 0.00044 | 0 | 8.18E-05 | 0.00072 | 0.00088 | 0 | 0.00041 | 3.82E-05 | 0.00011 | 0.00037 | 0 |
| Ddit4 | 0 | 1.23E-05 | 0.00015 | 6.35E-05 | 0.0002 | 0.00054 | 0.00013 | 0.00037 | 0 | 0 | 0 | 0.00147 | 0.0002 | 0.00016 | 0.00025 | 0 | 0.00171 | 0 |
| Sulf1 | 0.00011 | 0 | 0.00034 | 0 | 0.00107 | 0.00049 | 0.00161 | 0.00104 | 0 | 0.00087 | 0.00055 | 0.00021 | 0.0009 | 0.00016 | 6.07E-05 | 5.71E-05 | 0 | 0 |
| Creg1 | 7.01E-05 | 0.00017 | 0.00013 | 9.96E-05 | 0.00047 | 0.00047 | 0.00042 | 0.0002 | 0 | 0.00022 | 0 | 0.00012 | 0.00031 | 6.18E-05 | 0 | 4.66E-05 | 0 | 0.00018 |
| Kdelr3 | 0.00056 | 0 | 0.00044 | 0.00087 | 0.00022 | 0.00046 | 0.00111 | 0.00047 | 0 | 0.00015 | 0.00019 | 0.00035 | 8.93E-05 | 0.00045 | 0.0002 | 0.0001 | 0 | 0.00011 |
| Nrp1 | 0.00022 | 2.12E-05 | 0.00025 | 0 | 0.00035 | 0.00046 | 3.04E-05 | 0.0002 | 0 | 0.00025 | 0 | 0 | 0.00019 | 0.00017 | 0 | 0.00013 | 9.39E-05 | 0.00052 |
| Vkorc1 | 0.00033 | 0 | 0.00014 | 0.00087 | 0.00029 | 0.00045 | 0.00121 | 0.00057 | 0 | 0.00026 | 7.71E-05 | 0.00015 | 0.00077 | 0.00026 | 1.92E-05 | 0.00018 | 6.82E-05 | 0.00032 |
| Igf2r | 2.97E-05 | 0.00023 | 0.00024 | 0.00012 | 0.00035 | 0.00045 | 0.00043 | 0.00057 | 0 | 0.0004 | 1.93E-05 | 9.41E-05 | 0.00015 | 0.00013 | 8.25E-05 | 0.00011 | 0.00017 | 0.00012 |
| Ets1 | 5.51E-05 | 0.00012 | 0.00021 | 1.09E-05 | 7.49E-05 | 0.00045 | 0 | 0.00105 | 0 | 0 | 0.00024 | 0.00036 | 0 | 0.00043 | 7.92E-05 | 0.00013 | 0.00026 | 2.27E-05 |
| Emilin1 | 6.60E-05 | 0 | 0.00013 | 0.00048 | 0.00047 | 0.00044 | 0.00052 | 0.0007 | 0 | 0.00013 | 0.00074 | 0.00066 | 0.0007 | 0.00049 | 0.00057 | 0.0001 | 0 | 0.00053 |
| Sod3 | 0.00011 | 0 | 0.00083 | 0.00086 | 0.00051 | 0.00044 | 0.0015 | 0.00073 | 0 | 0.00106 | 0.00053 | 0 | 0.00234 | 0.00103 | 0.0008 | 0.00033 | 0 | 0.00096 |
| Sh3glb1 | 0.00063 | 0.00025 | 0.00032 | 5.82E-05 | 0.00015 | 0.00044 | 0.00043 | 0.00032 | 0 | 0.00023 | 0.00025 | 0.00047 | 0.00014 | 0.00014 | 9.64E-05 | 0.00013 | 0.00011 | 0.00017 |
| Camk2n1 | 0 | 0.00043 | 8.71E-05 | 0.00023 | 0.00053 | 0.00044 | 0 | 0.00043 | 0 | 3.80E-05 | 0.00019 | 0.00048 | 0 | 0.00018 | 0.00015 | 0 | 0.00011 | 4.15E-05 |
| Ece1 | 0.00011 | 0.00023 | 0.00029 | 0 | 0.00013 | 0.00044 | 8.61E-05 | 0.0002 | 0 | 0.0005 | 0.00043 | 0.00019 | 0.0001 | 0.00039 | 4.72E-05 | 0 | 0.00015 | 0.00027 |
| Htra3 | 4.73E-05 | 0.00018 | 0 | 0 | 0.00025 | 0.00044 | 0.00075 | 0.00063 | 0 | 0.00174 | 0.00013 | 0.00021 | 3.43E-05 | 0.00047 | 4.62E-05 | 0 | 0 | 0.00035 |
| Cadm4 | 0 | 0 | 0 | 0 | 0 | 0.00043 | 0 | 0.00058 | 0 | 0 | 0 | 2.61E-05 | 0 | 0 | 0 | 0 | 0 | 0 |
| Sned1 | 0 | 0.00014 | 0 | 0 | 0.00037 | 0.00043 | 0 | 0.00031 | 0 | 0.00125 | 6.09E-05 | 7.72E-05 | 0.00088 | 0.00023 | 0.00069 | 0 | 0 | 0.00085 |
| Aldh1a2 | 0 | 0 | 0 | 0 | 0.00024 | 0.00043 | 0 | 0 | 0 | 0 | 0.0001 | 0 | 0 | 0 | 0 | 0 | 0 | 0 |

| | | | | | | | | | | | | | | | | | | |
|---|---|---|---|---|---|---|---|---|---|---|---|---|---|---|---|---|---|---|
| Hist1h1e | 1.58E-05 | 0.00017 | 0 | 0 | 0.0001 | 0.00043 | 3.32E-05 | 1.95E-05 | 0 | 0.0001 | 0 | 3.22E-05 | 0.00016 | 0.00013 | 0.00015 | 0.00188 | 0 | 2.35E-05 |
| Fkbp7 | 0.00064 | 0 | 5.81E-05 | 7.58E-05 | 0.00019 | 0.00043 | 0.00087 | 0.00028 | 0 | 8.78E-05 | 9.99E-05 | 0 | 0.00031 | 0.00015 | 0 | 7.47E-05 | 0 | 6.18E-05 |
| Tmem119 | 0 | 4.98E-05 | 0.00018 | 0.00045 | 0.0011 | 0.00042 | 0.00069 | 0.00042 | 0 | 1.77E-05 | 0.00042 | 0.00091 | 0.00048 | 0.00051 | 0.00037 | 7.84E-05 | 0 | 0.00053 |
| Itgb3 | 0 | 2.61E-05 | 0.00019 | 2.12E-06 | 0.00038 | 0.00042 | 0 | 0.00088 | 0 | 0.00026 | 0.00039 | 0.00039 | 0 | 0.00029 | 0.0001 | 0 | 0 | 0 |
| Ucn2 | 0 | 0 | 0 | 0 | 0.00065 | 0.00042 | 0 | 0 | 0 | 0 | 0.00022 | 7.86E-05 | 0 | 0.00082 | 0.00038 | 0 | 0 | 0 |
| Hbp1 | 0.00021 | 0.0005 | 0 | 3.05E-05 | 0.00035 | 0.00042 | 0.00019 | 4.94E-05 | 0 | 0.00011 | 4.31E-05 | 0.00031 | 0.00013 | 0.00037 | 0.00023 | 4.19E-05 | 0 | 0.00016 |
| Hspa1b | 0.00011 | 9.24E-05 | 0 | 0 | 8.06E-05 | 0.00041 | 0.00019 | 7.76E-05 | 0 | 0.00012 | 4.58E-05 | 0 | 0 | 0.00014 | 0 | 0 | 0.00508 | 0.00047 |
| Cbfa2t3 | 0 | 6.87E-05 | 0 | 0 | 0 | 0.0004 | 0 | 0.00066 | 0 | 0 | 0 | 0 | 0 | 7.84E-05 | 1.82E-05 | 0 | 0 | 0 |
| Tcp11l2 | 3.55E-05 | 0.00039 | 0 | 0 | 0.00026 | 0.0004 | 0 | 0.00012 | 0 | 0.0001 | 0 | 0.00034 | 0 | 8.13E-05 | 0 | 0 | 0.00026 | 6.81E-05 |
| Gabarapl1 | 1.08E-06 | 0.00052 | 0.00011 | 0.00039 | 0.00036 | 0.00039 | 0.00018 | 8.01E-05 | 0 | 0.00011 | 3.89E-05 | 0.00031 | 0.00028 | 0.00011 | 0 | 0 | 0.00021 | 0.00024 |
| Zfp532 | 7.59E-05 | 0.00022 | 0.00014 | 0 | 0.00025 | 0.00039 | 0 | 0.00039 | 0 | 6.23E-06 | 0.00027 | 0.00037 | 0 | 0.00033 | 0.00028 | 4.78E-05 | 0.00025 | 9.53E-06 |
| Ccni | 0.00029 | 0.00036 | 0.00011 | 0.00022 | 0.0003 | 0.00038 | 3.94E-05 | 0.00031 | 0 | 0.0002 | 0.00032 | 0.00066 | 0.00024 | 0.00012 | 0 | 0.00016 | 1.84E-05 | 0.0003 |
| Jun | 6.78E-05 | 0.00013 | 0.00011 | 0.00025 | 0.00023 | 0.00038 | 0.00061 | 0.00051 | 0 | 0.00053 | 0.00016 | 0.0005 | 0.00065 | 0.00046 | 0.00023 | 0.00022 | 0.01039 | 0.00047 |
| Crispld2 | 0 | 0 | 0.00066 | 0 | 0.00037 | 0.00035 | 0.00096 | 0 | 0 | 0 | 0.00132 | 0 | 0.00025 | 0.00246 | 0.00035 | 6.47E-05 | 0 | 0.0002 |
| Rnf144a | 0 | 0 | 0 | 7.83E-05 | 3.17E-05 | 0.00035 | 0.00071 | 0 | 0 | 8.25E-05 | 0 | 0.0004 | 0.0005 | 1.66E-05 | 3.88E-05 | 1.28E-05 | 0.00013 | 3.40E-05 |
| Hspa1a | 0.00014 | 5.71E-05 | 0 | 0 | 8.49E-05 | 0.00035 | 7.79E-05 | 4.46E-05 | 0 | 5.15E-05 | 0 | 0 | 0 | 9.88E-05 | 0 | 0 | 0.00511 | 0.00025 |
| Nrep | 0 | 0 | 6.38E-06 | 0.00048 | 0.00109 | 0.00034 | 8.46E-05 | 0.00245 | 0 | 0 | 0.00023 | 0 | 0 | 0.00103 | 0.00013 | 0.0001 | 0 | 0 |
| Nsd3 | 0.00038 | 0.00042 | 0.00011 | 0 | 0.00026 | 0.00034 | 0.0001 | 6.91E-05 | 0 | 0.00012 | 4.14E-05 | 0.00036 | 8.27E-05 | 6.40E-05 | 0.00078 | 0.00028 | 0.00059 | 4.16E-05 |
| Tmem106b | 0.00032 | 0.00023 | 0 | 0 | 0.00024 | 0.00034 | 0.00013 | 3.40E-05 | 0 | 0.00015 | 8.87E-05 | 4.56E-05 | 0.00017 | 0.00014 | 0 | 0 | 0 | 0.00019 |
| Prdm1 | 0 | 1.44E-05 | 0 | 0 | 0.00048 | 0.00034 | 0 | 0 | 0 | 0 | 4.67E-05 | 0.00027 | 0 | 0.00094 | 0.00029 | 0 | 0 | 0 |
| Tgfb2 | 0 | 0.00015 | 9.99E-05 | 0 | 0.00021 | 0.00033 | 0.00029 | 0.00053 | 0 | 3.30E-05 | 0.00061 | 0 | 0 | 0.00026 | 0 | 0 | 0.00012 | 0 |
| Rab31 | 9.82E-05 | 0.00039 | 0 | 0.00011 | 0.00087 | 0.00033 | 0.00023 | 0.00039 | 0 | 0 | 0.00038 | 0.00026 | 0.00031 | 0.00076 | 0 | 8.67E-05 | 0.00014 | 0.00058 |
| Pla2g15 | 0 | 0.00012 | 5.31E-06 | 0.00018 | 0.00017 | 0.00033 | 0.00037 | 0 | 0 | 0.00013 | 0.00045 | 0.00014 | 0.00038 | 0.00016 | 0 | 0.00012 | 0 | 9.96E-05 |
| Pld1 | 3.00E-05 | 8.11E-05 | 0 | 0 | 0.00013 | 0.00033 | 0 | 0.00019 | 0 | 3.59E-05 | 0 | 3.92E-05 | 3.98E-05 | 1.78E-05 | 0 | 0 | 6.86E-05 | 0 |
| Fndc3a | 9.37E-05 | 0.00028 | 2.47E-05 | 7.63E-05 | 9.85E-05 | 0.00032 | 1.54E-05 | 5.34E-05 | 0 | 0.00017 | 2.77E-05 | 0.00057 | 0.0003 | 0 | 4.61E-05 | 6.40E-05 | 0.00023 | 0 |
| Anpep | 0.00013 | 0.00034 | 0 | 0 | 0.00031 | 0.00032 | 0.00101 | 2.13E-05 | 0 | 0.00067 | 0 | 0.00013 | 0.00041 | 0 | 0 | 0 | 0 | 0.00042 |
| Timp3 | 0 | 0 | 0.00074 | 0 | 0 | 0.00032 | 0.00061 | 0.0006 | 0 | 0.0005 | 0.00189 | 0.00064 | 0 | 0 | 0 | 4.08E-05 | 0 | 0.00118 |
| Ypel3 | 0 | 0.00067 | 0 | 9.52E-05 | 0.00041 | 0.00032 | 0.00027 | 0 | 0 | 0.0002 | 0 | 0.00025 | 0.00022 | 0.00028 | 0.00059 | 0 | 3.50E-05 | 0.00023 |
| Nr1d1 | 6.46E-05 | 0.00023 | 0 | 9.43E-05 | 0.0001 | 0.00032 | 0.00012 | 0 | 0 | 3.02E-05 | 0.0002 | 3.38E-05 | 0.00011 | 0.00016 | 6.94E-05 | 2.45E-05 | 0.00019 | 0.00011 |
| Serf2 | 0.00014 | 4.99E-05 | 0.0001 | 0.00101 | 0.00051 | 0.0003 | 0.00106 | 0.00044 | 0 | 0.00037 | 0.00048 | 0.0002 | 0.00049 | 0.00059 | 0.00037 | 9.13E-05 | 0.0003 | 0.00046 |
| Tmem38b | 0 | 0.00015 | 0 | 0 | 4.72E-05 | 0.00029 | 0 | 0 | 0 | 7.78E-05 | 0 | 0.00013 | 0.00042 | 0.00013 | 0 | 0 | 0 | 8.44E-06 |
| Prnp | 0 | 0.00021 | 0.00011 | 8.01E-05 | 0.00049 | 0.00029 | 0.00038 | 0 | 0 | 0.00082 | 0.00029 | 0.00018 | 0.00085 | 0.00039 | 0 | 0 | 4.07E-05 | 0.00068 |
| 4-Sep | 0 | 0.00013 | 0 | 0 | 0.00022 | 0.00029 | 0 | 0.00045 | 0 | 0 | 9.34E-05 | 0 | 0 | 0.00015 | 0 | 0 | 0 | 7.12E-05 |

| | | | | | | | | | | | | | | | | | | |
|---|---|---|---|---|---|---|---|---|---|---|---|---|---|---|---|---|---|---|
| Frrs1 | 1.88E-05 | 0.0002 | 0.00014 | 0 | 0.00021 | 0.00029 | 8.77E-05 | 5.36E-05 | 0 | 4.90E-06 | 3.52E-06 | 9.24E-05 | 8.64E-05 | 0 | 9.80E-06 | 2.54E-05 | 0 | 0.0001 |
| Mmp11 | 0 | 0.00092 | 0 | 0 | 0.00077 | 0.00029 | 0 | 0 | 0 | 0 | 0 | 0 | 0 | 0.00127 | 0.00014 | 0 | 0 | 0 |
| Ptprk | 0 | 9.74E-05 | 0 | 0 | 7.29E-05 | 0.00028 | 0 | 9.42E-06 | 0 | 3.15E-06 | 5.46E-05 | 0.00012 | 0 | 5.47E-05 | 0 | 3.56E-05 | 9.74E-05 | 0 |
| B3gnt3 | 0.0001 | 1.04E-05 | 0 | 2.32E-05 | 4.29E-05 | 0.00028 | 1.42E-05 | 0.00023 | 0 | 1.92E-05 | 0.00015 | 0.00035 | 5.07E-05 | 2.08E-05 | 0 | 0 | 3.31E-05 | 0.0001 |
| Clk1 | 0.0003 | 0.00045 | 0 | 0 | 0.00014 | 0.00028 | 0.00011 | 0 | 0 | 3.39E-05 | 0 | 0.00028 | 0 | 0.00015 | 0.00071 | 1.49E-05 | 0.00095 | 7.15E-05 |
| Klhl24 | 4.38E-05 | 0.0004 | 0 | 0 | 0.0002 | 0.00028 | 6.23E-05 | 5.43E-05 | 0 | 1.34E-05 | 0 | 0.00017 | 0 | 0.00022 | 8.74E-05 | 0 | 4.55E-05 | 5.26E-05 |
| Serpinb6a | 0.00057 | 0.00038 | 0.00042 | 0.00058 | 0 | 0.00028 | 0.00029 | 6.63E-05 | 0 | 0.00082 | 0.00019 | 0.00054 | 0.00043 | 6.17E-05 | 0 | 0.00022 | 0.0001 | 0.0005 |
| Phldb1 | 0 | 0.00022 | 0.00027 | 7.52E-05 | 8.49E-05 | 0.00028 | 7.31E-05 | 0.00036 | 0 | 0.00036 | 0.00031 | 0.00011 | 0 | 0.00016 | 0.00056 | 9.08E-05 | 0.00028 | 0.00036 |
| Plac8 | 0.00048 | 2.02E-05 | 0 | 0.00015 | 2.30E-05 | 0.00028 | 0.00013 | 0 | 0 | 0.001 | 0.00057 | 0.00021 | 0.00141 | 0 | 1.65E-05 | 6.12E-05 | 0 | 0.00041 |
| Adgrl3 | 0 | 0 | 0 | 0 | 0.00019 | 0.00027 | 0 | 0.00011 | 0 | 0 | 2.50E-05 | 0.00011 | 0 | 2.83E-06 | 0 | 0 | 0 | 0 |
| Lmo4 | 9.30E-05 | 0.0002 | 0.0002 | 5.60E-05 | 0.00025 | 0.00027 | 0 | 0 | 0 | 0.00013 | 0.0002 | 5.60E-05 | 0.00042 | 0.00049 | 8.51E-06 | 0.00016 | 0.00016 | 0.00031 |
| Prxl2c | 0.00023 | 0.00014 | 0 | 0 | 0.00016 | 0.00027 | 0.00023 | 4.68E-05 | 0 | 0.00012 | 0 | 0.00011 | 0.00013 | 0.00013 | 0 | 1.42E-05 | 0 | 2.83E-05 |
| Dusp1 | 8.35E-05 | 2.89E-05 | 0 | 0 | 0 | 0.00027 | 0.00027 | 0 | 0 | 0.00056 | 4.76E-06 | 0.00022 | 0.00032 | 0.00029 | 0 | 0 | 0.00818 | 0.00014 |
| Nipbl | 0.00028 | 0.00018 | 6.38E-05 | 0 | 7.57E-05 | 0.00027 | 4.88E-05 | 0 | 0 | 1.03E-05 | 1.20E-05 | 6.51E-05 | 0 | 2.21E-05 | 5.44E-05 | 7.64E-05 | 0.00028 | 0 |
| Pfkfb4 | 8.03E-06 | 0 | 0.00011 | 0 | 8.84E-05 | 0.00027 | 5.51E-05 | 0.00011 | 0 | 8.51E-05 | 8.83E-05 | 0.00013 | 0 | 0 | 0.00034 | 0 | 0 | 0 |
| Adamts7 | 0 | 0.00019 | 0 | 0 | 0.00028 | 0.00027 | 0 | 0.00021 | 0 | 0 | 3.23E-05 | 0 | 0.00028 | 0.00058 | 3.15E-05 | 0 | 0 | 3.86E-05 |
| Galnt10 | 0.00011 | 2.30E-05 | 1.47E-05 | 2.03E-05 | 0.00032 | 0.00026 | 0.00024 | 0.00011 | 0 | 8.87E-05 | 0.00045 | 8.14E-05 | 0.00049 | 0.00013 | 4.92E-05 | 4.13E-05 | 0 | 0.00029 |
| Dusp4 | 0 | 9.03E-05 | 0 | 0 | 0 | 0.00026 | 0 | 0.00029 | 0 | 0 | 0 | 0.00081 | 0 | 0.00019 | 0.00015 | 0 | 0 | 0 |
| Gucy1a1 | 0 | 0 | 0 | 0 | 0 | 0.00026 | 0 | 0.00023 | 0 | 4.39E-07 | 0 | 0 | 0 | 0 | 0 | 0 | 0 | 0.00014 |
| Aoc3 | 0 | 0 | 0 | 0.00039 | 0 | 0.00026 | 0.00033 | 0.00047 | 0 | 0.00012 | 0.00065 | 0.00057 | 0.00047 | 0 | 6.07E-05 | 0 | 0 | 0.00023 |
| Fam110c | 0 | 0 | 0 | 3.19E-05 | 1.14E-05 | 0.00026 | 0 | 0 | 0 | 0 | 0 | 0 | 0.00021 | 0.00014 | 0.00015 | 0 | 0 | 0 |
| Spock1 | 0 | 1.57E-05 | 0 | 0 | 0 | 0.00025 | 0 | 3.69E-06 | 0 | 0 | 0 | 0 | 0 | 0 | 0 | 0 | 0 | 0 |
| Cd82 | 0 | 0.0001 | 6.38E-05 | 0 | 0.0001 | 0.00025 | 0 | 1.32E-05 | 0 | 0 | 0 | 0.00042 | 0 | 0.00018 | 1.78E-05 | 0 | 0 | 0.00018 |
| Dusp6 | 0 | 8.66E-05 | 3.65E-05 | 8.28E-05 | 0.0003 | 0.00025 | 0 | 0.00096 | 0 | 5.24E-05 | 0.00038 | 0 | 0 | 0.0003 | 0.00042 | 3.19E-06 | 0.00014 | 0 |
| Bpgm | 0 | 0 | 4.19E-05 | 0 | 0.00115 | 0.00025 | 0 | 0 | 0 | 0 | 0.00076 | 0.0004 | 0.00023 | 0.00124 | 6.51E-05 | 3.36E-05 | 0.00027 | 0.00034 |
| Pla2g7 | 0 | 3.90E-05 | 0 | 0 | 0.00091 | 0.00025 | 0 | 0 | 0 | 0 | 0.00018 | 0 | 0 | 0.00036 | 0 | 0 | 0 | 0 |
| Col24a1 | 4.22E-06 | 0 | 0 | 0 | 0.00015 | 0.00025 | 0.00021 | 0 | 0 | 4.96E-07 | 2.97E-06 | 6.09E-05 | 0.00023 | 0.00026 | 0.0001 | 0 | 4.19E-05 | 0 |
| Tmem150a | 0 | 0.00012 | 1.12E-05 | 0.00011 | 0.00017 | 0.00024 | 0.00018 | 6.23E-05 | 0 | 7.03E-05 | 0.00018 | 0.00011 | 0.00035 | 0.00017 | 0 | 0 | 0 | 0.0003 |
| Moxd1 | 0 | 0 | 0 | 0 | 0.00163 | 0.00024 | 0 | 0.00038 | 0 | 0 | 0.00036 | 0 | 0 | 0 | 0 | 0 | 0 | 0 |
| Tmem108 | 0 | 3.96E-05 | 0 | 0 | 9.76E-05 | 0.00024 | 0 | 7.58E-06 | 0 | 0 | 0 | 4.05E-05 | 0 | 0 | 0 | 0 | 0 | 0 |
| Mxra7 | 0 | 0.00011 | 7.24E-05 | 0.00055 | 0.00031 | 0.00024 | 0 | 0.00044 | 0 | 0.00028 | 0.00067 | 0 | 0.00012 | 7.76E-05 | 0.0005 | 3.82E-05 | 0 | 0.00077 |
| Cux1 | 0.00015 | 8.33E-05 | 2.03E-05 | 0 | 0.00027 | 0.00024 | 0 | 0.00015 | 0 | 0 | 0.00022 | 0.00032 | 0 | 0.00032 | 0.00031 | 8.83E-05 | 5.11E-05 | 2.29E-05 |
| Cpne3 | 0.00022 | 0.00023 | 5.76E-05 | 0 | 0.00043 | 0.00023 | 0 | 0 | 0 | 3.12E-05 | 7.22E-05 | 0.0001 | 0.00012 | 4.00E-05 | 0 | 6.45E-05 | 3.61E-05 | 4.48E-05 |

| | | | | | | | | | | | | | | | | | | |
|---|---|---|---|---|---|---|---|---|---|---|---|---|---|---|---|---|---|---|
| F3 | 6.39E-05 | 0 | 5.17E-06 | 0 | 0 | 0.00023 | 0 | 0 | 0 | 2.09E-05 | 0.00034 | 6.15E-05 | 0.00058 | 3.36E-05 | 0 | 0 | 0.00035 | 0.00068 |
| Nr4a3 | 0 | 0 | 5.87E-05 | 6.82E-06 | 5.49E-06 | 0.00023 | 0 | 2.50E-05 | 0 | 0 | 0 | 0 | 0 | 3.01E-05 | 2.32E-05 | 0 | 0.00092 | 3.77E-05 |
| Trappc6b | 0.00022 | 0.00014 | 0.00017 | 8.63E-05 | 9.08E-05 | 0.00022 | 0.00011 | 0.00014 | 0 | 4.45E-05 | 0.00017 | 0.00017 | 2.84E-05 | 8.16E-05 | 0.00024 | 8.95E-05 | 0 | 0.00011 |
| Ankrd44 | 0 | 3.13E-05 | 0 | 0 | 2.45E-05 | 0.00022 | 0 | 0 | 0 | 7.08E-06 | 0 | 0.00057 | 0 | 2.25E-05 | 5.16E-06 | 2.91E-05 | 0 | 5.03E-05 |
| Tbc1d15 | 0.00021 | 0.00026 | 8.46E-05 | 0.0001 | 7.82E-05 | 0.00022 | 0.00019 | 2.27E-06 | 0 | 0 | 0 | 0.00017 | 3.20E-05 | 0.00014 | 0.00018 | 0.00017 | 0.00018 | 0 |
| Abca1 | 0 | 0.00028 | 0 | 0.00012 | 0.00015 | 0.00022 | 0 | 0 | 0 | 0.00032 | 0 | 6.48E-05 | 0.00039 | 0.0002 | 0.00016 | 0 | 0.00016 | 0.00011 |
| Ccng1 | 0.00025 | 0.00033 | 0.00024 | 7.26E-05 | 0.00012 | 0.00021 | 0.00029 | 8.14E-05 | 0 | 0.0002 | 7.25E-05 | 6.13E-05 | 0.00019 | 0 | 0 | 1.51E-05 | 0 | 8.64E-05 |
| Unc5b | 0 | 0 | 9.35E-05 | 0 | 0.00032 | 0.00021 | 0 | 0.0011 | 0 | 0 | 0.00028 | 0.00042 | 0 | 0.00087 | 0.00016 | 0 | 0 | 4.38E-05 |
| Ndufa4l2 | 0 | 0 | 0 | 0 | 0 | 0.00021 | 0.00035 | 0.00026 | 0 | 0.00013 | 0 | 0.00199 | 0.00037 | 0 | 0 | 0 | 0.00014 | 0 |
| Stom | 4.25E-05 | 0.00019 | 8.33E-05 | 0 | 0 | 0.00021 | 0 | 0.00012 | 0 | 0.00012 | 0.00019 | 0.00024 | 6.63E-05 | 0 | 0 | 9.55E-05 | 0 | 0.00013 |
| Ass1 | 0 | 0 | 0.0009 | 0 | 1.27E-05 | 0.00021 | 0 | 0.00064 | 0 | 0 | 0 | 0.00024 | 0.00014 | 0.00108 | 0 | 0 | 0 | 0.00016 |
| Ccpg1 | 0 | 0.00035 | 0 | 5.32E-06 | 0.00031 | 0.00021 | 0.00027 | 0.00025 | 0 | 0.00027 | 0.00015 | 0.00016 | 0.00025 | 0.00017 | 0 | 0 | 0 | 0.00037 |
| Zbtb34 | 2.73E-05 | 9.83E-05 | 3.38E-05 | 1.19E-05 | 0.00013 | 0.00021 | 2.31E-06 | 0.00024 | 0 | 1.39E-05 | 1.69E-05 | 3.79E-05 | 4.76E-05 | 0.00015 | 8.71E-05 | 0 | 7.19E-05 | 0 |
| Akt3 | 0.00014 | 0.00018 | 4.17E-05 | 0 | 0.00013 | 0.00021 | 3.83E-05 | 5.00E-05 | 0 | 1.12E-05 | 0.00018 | 0.00022 | 0 | 7.08E-05 | 0.00012 | 7.44E-05 | 0.00012 | 0.00012 |
| Ralgds | 7.88E-05 | 0.00022 | 3.67E-05 | 4.30E-05 | 0 | 0.0002 | 4.64E-05 | 0 | 0 | 5.89E-05 | 2.36E-06 | 4.26E-05 | 0.00024 | 0.00012 | 5.48E-05 | 0 | 0.00013 | 8.70E-06 |
| Gng11 | 0.00015 | 9.82E-05 | 9.72E-05 | 0.00019 | 0.00073 | 0.0002 | 0 | 0.00193 | 0 | 0 | 0.00023 | 0.00043 | 0 | 0.00075 | 0 | 3.15E-05 | 0 | 0.00011 |
| Myo1d | 0 | 0.00013 | 8.42E-05 | 0 | 0.00041 | 0.0002 | 4.13E-05 | 9.85E-05 | 0 | 0 | 0.00039 | 5.84E-05 | 0 | 0.00025 | 2.08E-05 | 0 | 7.04E-05 | 0.00044 |
| Nupr1 | 0 | 0.00033 | 0.00035 | 1.31E-05 | 0.00023 | 0.0002 | 0.0005 | 0.00012 | 0 | 0.00035 | 0 | 0.00069 | 0 | 0.00033 | 0.0003 | 4.83E-06 | 0.00065 | 0.00061 |
| Eif1 | 0.00039 | 0.00026 | 0.00024 | 0 | 4.77E-05 | 0.0002 | 4.37E-05 | 0 | 0 | 3.36E-06 | 8.67E-05 | 0.00038 | 0 | 1.39E-06 | 0 | 0.00017 | 0.00079 | 0 |
| Uaca | 0.00011 | 8.54E-05 | 0 | 0.0001 | 0.00035 | 0.0002 | 5.88E-07 | 0 | 0 | 0.00012 | 0.00019 | 0.00029 | 0.00032 | 0.00024 | 0.00021 | 0.00013 | 0 | 6.21E-05 |
| Saa3 | 0 | 0 | 0.00035 | 0 | 0 | 0.0002 | 0 | 0 | 0 | 5.74E-05 | 0 | 0 | 0.00484 | 0.00053 | 0 | 5.43E-05 | 0 | 0 |
| Fzd7 | 0 | 0.00018 | 0 | 0.0002 | 0.00015 | 0.0002 | 0.0002 | 0 | 0 | 0.00017 | 0.00032 | 0.00043 | 0.00017 | 0.00013 | 4.51E-05 | 1.87E-05 | 0.00017 | 9.68E-05 |
| Rnf181 | 0.00014 | 0.0001 | 2.63E-05 | 1.12E-05 | 0.0001 | 0.0002 | 1.75E-05 | 7.17E-05 | 0 | 5.99E-05 | 0.0001 | 0.0003 | 4.94E-05 | 2.76E-05 | 5.33E-05 | 0.00017 | 0 | 0.00015 |
| Cnot6l | 0.00021 | 0.00021 | 3.96E-05 | 0 | 0.00011 | 0.0002 | 5.86E-05 | 6.89E-05 | 0 | 5.85E-05 | 2.59E-05 | 0.00015 | 0 | 0 | 3.72E-05 | 0.0002 | 7.08E-05 | 3.18E-05 |
| Nudt9 | 1.14E-05 | 0.00015 | 0.00016 | 0.00019 | 0.00019 | 0.0002 | 6.00E-05 | 0.00031 | 0 | 8.85E-05 | 0.00013 | 7.61E-05 | 9.29E-05 | 0.00015 | 0 | 0.00016 | 1.85E-05 | 0.00018 |
| Vsir | 0 | 0.00021 | 0 | 0 | 0.00022 | 0.00019 | 0 | 0 | 0 | 0.00015 | 0.00014 | 0.00021 | 0 | 6.08E-05 | 9.48E-05 | 0 | 0 | 0.00033 |
| Emb | 0.00023 | 0 | 0.00018 | 0.00032 | 0 | 0.00019 | 4.17E-05 | 0.00034 | 0 | 0.0003 | 0 | 0 | 0.00204 | 0.00058 | 0 | 0.00015 | 0 | 0.00079 |
| Csgalnact2 | 0.00013 | 0.00012 | 0.00016 | 0 | 9.54E-05 | 0.00019 | 0.00025 | 7.69E-05 | 0 | 0.00021 | 7.04E-05 | 6.35E-05 | 1.28E-05 | 2.84E-05 | 0 | 0.00013 | 1.19E-05 | 0.00012 |
| Tnfrsf9 | 0 | 0 | 0 | 0 | 0 | 0.00019 | 0 | 0 | 0 | 0 | 0 | 0.00053 | 0 | 0 | 0 | 0 | 0 | 0 |
| Ubr3 | 0.00019 | 0.00026 | 0.00018 | 7.64E-05 | 0.00015 | 0.00019 | 0.00011 | 5.40E-05 | 0 | 4.64E-05 | 9.07E-05 | 0.00015 | 2.86E-05 | 7.75E-05 | 0.00014 | 5.42E-05 | 0.0002 | 9.90E-05 |
| Rin2 | 7.49E-05 | 0.00033 | 0 | 0 | 0.00011 | 0.00019 | 0.00013 | 0 | 0 | 7.57E-05 | 0 | 0 | 0.00028 | 0.0003 | 0 | 0 | 0 | 9.15E-05 |
| Supt4a | 0.0002 | 0.00025 | 1.27E-05 | 0.0003 | 0.00018 | 0.00019 | 0.0002 | 5.70E-05 | 0 | 2.94E-06 | 0.00012 | 0.00018 | 1.37E-05 | 0.00018 | 6.36E-07 | 0.00012 | 4.50E-05 | 2.52E-05 |
| Xylt1 | 0 | 0.0002 | 0 | 0 | 2.08E-05 | 0.00019 | 8.75E-05 | 0 | 0 | 9.87E-05 | 0 | 0.00012 | 6.90E-06 | 0.00024 | 6.11E-05 | 7.06E-05 | 0.00024 | 0.00015 |

| | | | | | | | | | | | | | | | | | | |
|---|---|---|---|---|---|---|---|---|---|---|---|---|---|---|---|---|---|---|
| Ccng2 | 0 | 0.00026 | 0 | 0 | 0.00015 | 0.00019 | 0 | 0.00012 | 0 | 0.00012 | 1.50E-05 | 0.00077 | 9.43E-05 | 0.00016 | 0.00019 | 0.00037 | 0.00017 | 8.31E-05 |
| Stard13 | 0 | 2.44E-05 | 7.69E-05 | 4.55E-05 | 0 | 0.00019 | 0 | 0.00062 | 0 | 1.99E-05 | 0.00017 | 0.0001 | 0 | 0 | 4.49E-05 | 4.27E-05 | 0.0001 | 0.0001 |
| Heyl | 0 | 0 | 0.00014 | 0 | 0 | 0.00019 | 0 | 0.00143 | 0 | 0 | 0 | 0 | 0 | 0.00031 | 0 | 8.75E-05 | 0 | 9.13E-05 |
| Bet1 | 0.00027 | 3.39E-05 | 0.00011 | 0.0004 | 0.00019 | 0.00019 | 0.00031 | 0.00026 | 0 | 8.07E-05 | 0.0001 | 1.80E-05 | 8.73E-05 | 0.00013 | 7.60E-05 | 0.00011 | 0 | 2.16E-05 |
| Ctsf | 0 | 0.00023 | 0 | 0 | 0.00038 | 0.00018 | 6.88E-05 | 4.34E-05 | 0 | 0.0001 | 4.50E-05 | 0.0002 | 0.00018 | 0.00017 | 0 | 0 | 0 | 0.00013 |
| Calcrl | 0 | 9.49E-05 | 0 | 0 | 0 | 0.00018 | 0 | 0.00022 | 0 | 0.00017 | 0 | 0.00021 | 0 | 0 | 0 | 0 | 0.00011 | 8.39E-06 |
| Igf2bp2 | 1.26E-05 | 0.00017 | 7.04E-05 | 0 | 0 | 0.00018 | 0 | 0 | 0 | 0 | 0 | 0.00051 | 0.00012 | 0.00017 | 0 | 4.62E-05 | 0 | 0 |
| Npc1 | 7.16E-05 | 0.00023 | 6.56E-05 | 0.00017 | 0.00017 | 0.00018 | 0.00026 | 8.11E-05 | 0 | 7.34E-05 | 8.76E-05 | 0.00015 | 2.99E-06 | 0.00024 | 0.00015 | 9.16E-05 | 3.56E-05 | 5.14E-05 |
| Rbp1 | 0 | 0 | 0.00028 | 0.00028 | 0.00092 | 0.00018 | 0.00025 | 0.0012 | 0 | 0 | 0.001 | 0.00026 | 0.00045 | 0 | 0 | 3.16E-05 | 0 | 0.00156 |
| Mamdc2 | 0 | 3.93E-05 | 0 | 0 | 0.00034 | 0.00018 | 0 | 0 | 0 | 0 | 0.00016 | 8.45E-05 | 0 | 0.00059 | 0 | 0 | 0 | 0 |
| Commd3 | 0.0003 | 0.00018 | 3.50E-05 | 0.00013 | 0.00011 | 0.00018 | 5.29E-05 | 6.81E-05 | 0 | 2.27E-05 | 5.86E-05 | 6.16E-05 | 9.65E-05 | 4.43E-05 | 0 | 0.00013 | 0 | 0 |
| Hmga2 | 0 | 5.05E-05 | 5.14E-05 | 0 | 0 | 0.00018 | 0 | 0 | 0 | 0 | 0 | 0.00018 | 0 | 0.00017 | 0.00022 | 2.04E-06 | 0 | 0 |
| Adm | 0 | 0 | 0 | 0 | 0.00035 | 0.00018 | 0 | 0 | 0 | 0.00074 | 3.69E-05 | 0.00146 | 0.00078 | 0 | 0 | 0 | 0 | 0 |
| Msx1 | 0 | 0 | 9.02E-05 | 0.00015 | 4.14E-05 | 0.00017 | 0 | 0.00081 | 0 | 0 | 2.33E-05 | 0 | 0 | 0 | 0.0001 | 0 | 2.17E-05 | 5.73E-05 |
| Efhd2 | 3.59E-06 | 9.02E-05 | 0.00029 | 0.00011 | 0.00022 | 0.00017 | 1.14E-05 | 0.00033 | 0 | 0.00015 | 0.00033 | 0.00018 | 2.47E-05 | 0.00019 | 4.64E-05 | 8.84E-05 | 0 | 0 |
| Sik1 | 0 | 4.26E-05 | 0 | 0 | 0 | 0.00017 | 0 | 0.00012 | 0 | 0 | 7.17E-05 | 0.00072 | 0 | 0 | 0.00017 | 0 | 0.00111 | 0.00014 |
| Ctsc | 0.00015 | 2.54E-05 | 0 | 0 | 0.00095 | 0.00017 | 0 | 0.00224 | 0 | 0 | 0.00027 | 0 | 0 | 0.00044 | 0.00036 | 4.28E-05 | 0 | 0 |
| Rbpms2 | 0 | 4.93E-06 | 9.84E-05 | 2.63E-05 | 0.00013 | 0.00017 | 0 | 0.00026 | 0 | 0 | 0.00026 | 3.92E-05 | 0 | 0.00032 | 1.95E-05 | 5.87E-05 | 0 | 0 |
| Mgat5 | 0 | 6.64E-05 | 5.89E-05 | 0 | 0.00014 | 0.00017 | 7.64E-05 | 0.00025 | 0 | 3.96E-05 | 0 | 0 | 0 | 0.00016 | 1.77E-05 | 4.45E-05 | 3.27E-05 | 4.68E-05 |
| Rhbdf2 | 0 | 6.92E-05 | 5.57E-05 | 0 | 2.54E-05 | 0.00017 | 0 | 0.00048 | 0 | 0 | 0 | 0.00016 | 0 | 0.00018 | 2.14E-05 | 0 | 0 | 0 |
| Gadd45b | 1.63E-05 | 4.77E-05 | 0.00027 | 0.00016 | 8.49E-05 | 0.00017 | 0.0002 | 0 | 0 | 0 | 0.00053 | 0.00041 | 0.00045 | 0.00021 | 0 | 0 | 0.00347 | 0.00015 |
| Kdm7a | 7.85E-05 | 0.00027 | 4.25E-06 | 0 | 3.51E-05 | 0.00017 | 4.02E-06 | 4.83E-05 | 0 | 2.28E-06 | 0 | 8.54E-05 | 0 | 0.00015 | 0.00014 | 0 | 0.00025 | 0 |
| Wdr45 | 4.88E-05 | 0.00021 | 0 | 5.06E-05 | 0.00011 | 0.00017 | 0.00015 | 5.06E-05 | 0 | 4.65E-05 | 2.01E-05 | 0.0001 | 5.51E-05 | 0.00016 | 3.62E-05 | 0 | 0 | 9.48E-05 |
| Bmi1 | 0.00015 | 8.97E-05 | 0.00011 | 1.87E-06 | 9.49E-05 | 0.00017 | 2.56E-05 | 5.51E-05 | 0 | 0 | 8.56E-05 | 0 | 4.03E-05 | 9.71E-05 | 6.63E-05 | 0.00012 | 5.60E-05 | 0 |
| Rab15 | 1.02E-05 | 0 | 2.31E-05 | 4.61E-05 | 0 | 0.00017 | 2.29E-06 | 0.00019 | 0 | 4.17E-06 | 0.00013 | 0.00017 | 0 | 4.26E-05 | 0 | 8.18E-05 | 0 | 2.53E-05 |
| Arfgef1 | 0.00023 | 0.00012 | 0.00011 | 0 | 3.32E-05 | 0.00016 | 8.06E-05 | 0 | 0 | 6.06E-06 | 8.25E-05 | 9.74E-05 | 7.95E-06 | 4.60E-06 | 0.00018 | 1.83E-05 | 0.00012 | 1.31E-05 |
| Sqstm1 | 0.00015 | 0.00044 | 0.00013 | 0.0005 | 0.00013 | 0.00016 | 8.16E-05 | 0.00012 | 0 | 3.31E-05 | 4.67E-05 | 0.00051 | 0 | 0.00026 | 0.00015 | 0 | 0.00246 | 0.00021 |
| Gm2a | 0 | 0.00015 | 0 | 0.0002 | 0.00025 | 0.00016 | 0.00027 | 0.00021 | 0 | 0.00023 | 0.00013 | 0.00046 | 0.00049 | 0 | 4.15E-05 | 0 | 0 | 0.00027 |
| Cd80 | 0 | 4.14E-05 | 0.0001 | 0 | 1.87E-05 | 0.00016 | 0 | 4.57E-05 | 0 | 2.53E-05 | 3.87E-05 | 0 | 0 | 0 | 5.40E-05 | 0 | 0 | 0 |
| Dtx3 | 0 | 0.00037 | 0 | 0.0002 | 0.00029 | 0.00016 | 0.00014 | 0.0003 | 0 | 0.00016 | 6.63E-05 | 0.00018 | 6.56E-05 | 9.17E-05 | 0.00087 | 0 | 0.00016 | 0.00028 |
| Pamr1 | 0 | 0.00016 | 0 | 0 | 0 | 0.00016 | 0 | 0 | 0 | 0.00044 | 0 | 1.19E-05 | 0 | 0 | 0 | 0 | 0 | 0 |
| Aif1l | 0 | 0 | 0 | 0 | 0.00027 | 0.00016 | 0 | 0 | 0 | 0.00022 | 0.0002 | 5.95E-05 | 2.42E-05 | 0 | 0 | 5.41E-07 | 0 | 0 |
| Sun2 | 0 | 0.00028 | 0 | 3.59E-05 | 9.65E-05 | 0.00016 | 0 | 0 | 0 | 0 | 0.00011 | 0.00029 | 0 | 0.00024 | 0.00017 | 0.00039 | 2.13E-05 | 0.00013 |

| | | | | | | | | | | | | | | | | | | |
|---|---|---|---|---|---|---|---|---|---|---|---|---|---|---|---|---|---|---|
| Usp8 | 0.0002 | 0.00021 | 9.64E-05 | 0.0002 | 0.00012 | 0.00016 | 0.00015 | 5.33E-05 | 0 | 0.00011 | 0 | 0.00018 | 0.00015 | 6.67E-05 | 0.00037 | 0.00011 | 1.49E-06 | 6.26E-05 |
| Plau | 0 | 5.62E-05 | 0 | 9.30E-05 | 4.68E-05 | 0.00016 | 0 | 0.00192 | 0 | 0.00074 | 0.00067 | 0.00031 | 0.00046 | 0.0008 | 0 | 0.00017 | 0 | 0 |
| Plpp5 | 0.0002 | 2.81E-05 | 0.00025 | 0.00052 | 0.00024 | 0.00016 | 0.0007 | 0.00055 | 0 | 0.00016 | 0.00016 | 0.00051 | 0.00022 | 0.00011 | 0.00011 | 0 | 0.00015 | 0.00016 |
| Itga5 | 0 | 1.67E-05 | 0.00107 | 2.20E-05 | 0.00018 | 0.00016 | 0.0001 | 0.00086 | 0 | 0.00028 | 0.00145 | 0.00057 | 0.00019 | 0.00042 | 0.00042 | 9.45E-05 | 0.00036 | 0.0001 |
| Sh3pxd2a | 0.00012 | 0.00034 | 0.00041 | 0.00037 | 0.00035 | 0.00016 | 0.00079 | 0.0001 | 0 | 6.82E-06 | 0.00018 | 0.00015 | 7.50E-05 | 0.00028 | 0.00036 | 0.00025 | 1.39E-05 | 0.00034 |
| Golga5 | 0.00028 | 9.87E-05 | 0.0001 | 0 | 5.93E-05 | 0.00016 | 0.00022 | 0 | 0 | 7.16E-06 | 3.69E-05 | 1.54E-05 | 3.70E-05 | 1.11E-05 | 0.00025 | 2.70E-05 | 0.00013 | 0 |
| Hexb | 0.00022 | 0.00019 | 2.69E-07 | 0 | 0.00022 | 0.00015 | 0.00018 | 0.00017 | 0 | 0.00016 | 5.54E-05 | 0.00044 | 0.00032 | 0.0002 | 0 | 9.04E-05 | 0 | 0.00036 |
| Zfp36 | 5.84E-06 | 1.50E-05 | 0 | 0.00033 | 0.00016 | 0.00015 | 0 | 0.00011 | 0 | 0.0002 | 0.00018 | 8.93E-05 | 0.00043 | 0.00024 | 0.0003 | 3.09E-06 | 0.00776 | 0.00013 |
| Ehd3 | 0 | 0 | 6.90E-05 | 1.62E-05 | 0.00014 | 0.00015 | 0 | 8.16E-07 | 0 | 0 | 5.41E-05 | 0 | 0 | 0.00018 | 9.62E-06 | 0 | 0 | 1.37E-06 |
| Qprt | 0 | 5.88E-05 | 0 | 9.62E-05 | 0.00011 | 0.00015 | 0 | 0.0002 | 0 | 6.56E-06 | 0 | 2.06E-05 | 4.37E-07 | 0 | 0 | 0 | 0 | 0 |
| Kdm5a | 0.00016 | 0.00021 | 2.54E-05 | 1.58E-05 | 9.05E-05 | 0.00015 | 0.00017 | 8.18E-05 | 0 | 0 | 0 | 0.00027 | 2.27E-05 | 8.02E-05 | 0.00027 | 0 | 0.00017 | 5.08E-05 |
| Zc3h12a | 0 | 3.66E-05 | 0.00014 | 0 | 6.65E-05 | 0.00015 | 0 | 0.00013 | 0 | 3.45E-05 | 0 | 0.00023 | 0.00034 | 0.00037 | 0.00024 | 0 | 0.00021 | 0 |
| Herpud2 | 0.00018 | 0.00013 | 1.66E-05 | 0 | 8.91E-05 | 0.00015 | 9.65E-05 | 5.86E-05 | 0 | 5.51E-05 | 6.09E-05 | 0.00015 | 0 | 0 | 0 | 2.37E-05 | 0.00012 | 8.35E-05 |
| Pcdh1 | 0 | 1.99E-05 | 0 | 0 | 0 | 0.00015 | 0 | 0.00015 | 0 | 8.38E-05 | 0 | 0 | 0 | 9.09E-05 | 0 | 0 | 0 | 0 |
| Csnk1g3 | 0.00024 | 0.00017 | 0.00014 | 0 | 9.27E-05 | 0.00015 | 7.33E-05 | 4.25E-05 | 0 | 2.19E-05 | 6.27E-05 | 0.00015 | 1.32E-05 | 4.77E-05 | 0.00016 | 0.00011 | 2.02E-05 | 0 |
| Zfp469 | 0 | 1.34E-05 | 0.00039 | 2.30E-05 | 0.0003 | 0.00015 | 0.00033 | 7.42E-05 | 0 | 4.68E-05 | 0.00038 | 0.00037 | 7.96E-06 | 0.0005 | 0.00045 | 0 | 0 | 1.67E-05 |
| Slc29a1 | 1.37E-05 | 1.27E-05 | 0 | 0 | 0 | 0.00015 | 0 | 0.00034 | 0 | 0.00023 | 0 | 0 | 0 | 7.66E-06 | 0 | 0.00018 | 0 | 0.00022 |
| Fbxl20 | 9.14E-05 | 0.00022 | 0 | 0 | 0.00011 | 0.00015 | 9.33E-05 | 5.02E-06 | 0 | 2.53E-05 | 2.66E-05 | 0.00014 | 0 | 6.21E-05 | 4.25E-06 | 0 | 0 | 3.68E-05 |
| Cnppd1 | 0 | 0.00035 | 5.48E-05 | 0.00026 | 9.87E-05 | 0.00015 | 4.64E-05 | 4.68E-05 | 0 | 0.00014 | 5.35E-05 | 0.00028 | 2.06E-05 | 0.00021 | 0.00013 | 6.18E-06 | 0.00025 | 0.00013 |
| Agrn | 0 | 0.00018 | 0 | 3.99E-05 | 0.00021 | 0.00015 | 0 | 0.00036 | 0 | 0.00014 | 0.00023 | 0.00017 | 8.49E-05 | 0.00011 | 0.00023 | 0 | 0.00011 | 9.26E-05 |
| Pold4 | 7.66E-05 | 0.0001 | 6.40E-05 | 0.00022 | 9.06E-05 | 0.00015 | 0.00013 | 0.00034 | 0 | 3.92E-05 | 1.57E-05 | 0.0002 | 1.20E-05 | 0.00013 | 1.93E-05 | 0 | 3.69E-05 | 0.0001 |
| Ssh1 | 5.56E-05 | 0.00017 | 0.00021 | 0 | 0.00022 | 0.00015 | 0.0001 | 0.00015 | 0 | 0.00012 | 0.00027 | 0.00017 | 0 | 0.00014 | 0.00015 | 0 | 3.46E-05 | 0.00011 |
| Cdkn1c | 0 | 2.59E-05 | 0 | 5.08E-06 | 4.26E-05 | 0.00015 | 0 | 0 | 0 | 0.00015 | 1.89E-05 | 6.80E-05 | 0 | 6.79E-06 | 7.15E-05 | 0 | 0 | 0 |
| Olfr810 | 0 | 0 | 0 | 0 | 5.50E-05 | 0.00014 | 0 | 0.00016 | 0 | 0 | 0 | 0 | 0 | 0 | 2.29E-05 | 0 | 0 | 0 |
| Boc | 0 | 0.00012 | 4.71E-05 | 1.96E-06 | 0.00015 | 0.00014 | 2.64E-05 | 0 | 0 | 5.77E-05 | 6.72E-05 | 9.03E-05 | 5.28E-06 | 0.00018 | 4.34E-05 | 9.73E-06 | 0.00014 | 0.00022 |
| Ugp2 | 0.00037 | 0 | 8.16E-05 | 0 | 0 | 0.00014 | 0.0001 | 0 | 0 | 0.00036 | 0 | 0.00036 | 0 | 0 | 0 | 9.15E-05 | 0 | 7.44E-06 |
| Hsd17b11 | 8.03E-05 | 0.00032 | 0 | 0 | 0.00015 | 0.00014 | 0.0002 | 0.00013 | 0 | 0.00018 | 9.93E-05 | 4.80E-05 | 0.00011 | 0 | 0 | 2.94E-05 | 0.00015 | 0.00023 |
| Crebrf | 1.05E-05 | 0.00024 | 0 | 0 | 5.78E-05 | 0.00014 | 0 | 0 | 0 | 3.82E-05 | 0 | 0.00011 | 0 | 2.98E-05 | 4.15E-05 | 0 | 0.00013 | 3.80E-05 |
| Pqlc1 | 0 | 3.68E-05 | 0.00026 | 0.00018 | 0.00024 | 0.00014 | 0.0001 | 0.00036 | 0 | 0 | 0.00035 | 0.00014 | 5.97E-05 | 2.61E-05 | 0.00037 | 0 | 0 | 0 |
| Trappc8 | 0.00019 | 0.00013 | 8.71E-05 | 0 | 8.94E-05 | 0.00014 | 3.39E-05 | 0 | 0 | 2.61E-05 | 1.15E-05 | 0.0001 | 0 | 3.33E-05 | 3.93E-05 | 0.0001 | 1.30E-05 | 3.76E-05 |
| Dcaf10 | 0.0001 | 0.00013 | 0.00022 | 6.58E-05 | 0.00012 | 0.00014 | 0.00018 | 0.00012 | 0 | 7.27E-05 | 0.00014 | 8.92E-05 | 4.46E-05 | 7.03E-05 | 0 | 6.08E-05 | 2.55E-05 | 7.04E-05 |
| Nr1d2 | 0.00011 | 0.00018 | 0 | 0 | 5.89E-05 | 0.00014 | 2.00E-05 | 0 | 0 | 0 | 5.71E-05 | 4.81E-05 | 0 | 0.00014 | 8.82E-05 | 4.08E-05 | 0.00053 | 0 |
| Gem | 0 | 0.00011 | 0.00012 | 0.00011 | 0 | 0.00014 | 0 | 0 | 0 | 0 | 0 | 0 | 0 | 0.00015 | 0 | 0 | 0.00292 | 0.00016 |

| | | | | | | | | | | | | | | | | | | |
|---|---|---|---|---|---|---|---|---|---|---|---|---|---|---|---|---|---|---|
| Loxl3 | 0.0002 | 0 | 0.00016 | 0.0008 | 0.00053 | 0.00013 | 0.00057 | 0.00112 | 0 | 4.61E-05 | 0.00073 | 2.57E-05 | 0.00064 | 0.00028 | 0.00095 | 0.00029 | 0.00025 | 0.0002 |
| Hivep1 | 5.30E-05 | 0.00013 | 0.00012 | 0 | 8.65E-05 | 0.00013 | 4.98E-05 | 0 | 0 | 0.00014 | 0.00018 | 5.39E-05 | 0 | 2.18E-05 | 1.04E-05 | 0 | 0.00023 | 2.16E-05 |
| Rnf145 | 2.67E-05 | 0.00013 | 0.00012 | 0.00022 | 0.00014 | 0.00013 | 0.00021 | 0.00016 | 0 | 0.00023 | 0.00015 | 0.00023 | 0 | 8.79E-05 | 4.99E-06 | 0.00014 | 0.00018 | 0.00015 |
| Hilpda | 9.25E-05 | 6.74E-05 | 8.55E-05 | 2.54E-05 | 0 | 0.00013 | 6.67E-06 | 0 | 0 | 0 | 0 | 0.00107 | 0 | 3.98E-05 | 8.50E-05 | 2.76E-05 | 0.0002 | 2.32E-05 |
| Smc5 | 0.00015 | 0.00016 | 2.21E-05 | 0 | 6.39E-05 | 0.00013 | 7.21E-05 | 0 | 0 | 2.05E-06 | 0.00027 | 7.28E-05 | 0 | 0.00022 | 0.00055 | 0.00029 | 0.00017 | 0 |
| Itga10 | 0 | 0 | 0 | 0 | 0.00011 | 0.00013 | 0 | 2.43E-05 | 0 | 3.90E-06 | 4.69E-05 | 0 | 0 | 0 | 3.80E-05 | 0 | 5.30E-06 | 0 |
| Fnip1 | 0.00014 | 0.00021 | 0 | 0 | 7.24E-05 | 0.00013 | 8.89E-05 | 0 | 0 | 0 | 0 | 0.00017 | 1.67E-05 | 7.03E-05 | 0.00013 | 5.96E-05 | 0.00046 | 0 |
| Pdia5 | 0.00029 | 0 | 0.00022 | 0.00034 | 0 | 0.00013 | 0.00098 | 0.00038 | 0 | 2.78E-05 | 0.00013 | 0.0002 | 0.00047 | 0.0002 | 2.61E-05 | 0.00015 | 0 | 0.00024 |
| Tns1 | 8.53E-06 | 0.00012 | 2.09E-05 | 0 | 7.21E-06 | 0.00013 | 0 | 0.00022 | 0 | 0.00025 | 0.00048 | 6.23E-06 | 0 | 0 | 2.72E-05 | 3.41E-05 | 0 | 0.00015 |
| Tmem189 | 0.0001 | 6.46E-05 | 0.00015 | 6.15E-05 | 7.42E-05 | 0.00013 | 0.00018 | 8.10E-05 | 0 | 3.13E-05 | 0.00013 | 0.0002 | 2.61E-05 | 0.00012 | 0.00018 | 0.0002 | 2.49E-05 | 7.40E-05 |
| Pck2 | 7.58E-05 | 0.00018 | 2.86E-05 | 0.00042 | 1.22E-05 | 0.00013 | 0.00053 | 0.00038 | 0 | 7.11E-05 | 6.61E-05 | 0.0003 | 1.82E-05 | 0.00028 | 0.00043 | 0.00034 | 0 | 0.00013 |
| Htatip2 | 0.0001 | 9.73E-05 | 2.89E-05 | 8.50E-05 | 6.41E-05 | 0.00013 | 3.86E-05 | 0 | 0 | 9.27E-05 | 4.67E-05 | 4.70E-05 | 9.29E-05 | 0.0001 | 0 | 2.48E-05 | 0 | 0.00011 |
| Fcrlb | 0 | 0 | 5.48E-06 | 4.18E-06 | 8.07E-05 | 0.00013 | 0 | 0 | 0 | 0 | 0 | 0 | 1.18E-05 | 0.00015 | 6.07E-05 | 1.37E-05 | 0 | 0 |
| Nes | 0 | 0 | 0 | 0 | 1.91E-05 | 0.00013 | 0 | 0.0007 | 0 | 0 | 0.00019 | 9.60E-05 | 0 | 3.39E-05 | 0 | 2.33E-05 | 0 | 0 |
| Fnip2 | 0 | 0.00021 | 6.74E-05 | 0 | 7.13E-05 | 0.00013 | 1.24E-05 | 0 | 0 | 2.78E-06 | 0 | 0.00012 | 0.00017 | 0.00013 | 0.00012 | 1.22E-05 | 7.61E-05 | 0 |
| Ufsp2 | 0.00033 | 0 | 0.00018 | 0.00023 | 8.87E-05 | 0.00013 | 0.00038 | 6.83E-05 | 0 | 3.37E-05 | 0 | 8.19E-05 | 5.38E-05 | 7.95E-05 | 3.54E-05 | 8.57E-05 | 0 | 0 |
| Cln5 | 3.97E-05 | 0.00019 | 0.00013 | 0.00033 | 0.0002 | 0.00013 | 0.00019 | 0.00019 | 0 | 0.00017 | 4.09E-05 | 0.00014 | 1.23E-05 | 0.00021 | 0 | 6.17E-05 | 0 | 0.00019 |
| Akap13 | 0.0002 | 0.00032 | 0.00028 | 0 | 0.00026 | 0.00013 | 0.00023 | 0.00019 | 0 | 0.00018 | 0.00014 | 0.00023 | 0.00017 | 0.00031 | 0.00064 | 5.39E-05 | 0.0004 | 0.00029 |
| Tnfrsf23 | 0.00021 | 2.90E-05 | 0.00045 | 0.00016 | 0.00023 | 0.00012 | 0.00077 | 0.00013 | 0 | 4.59E-05 | 0.00015 | 0.00058 | 0.00023 | 0.00037 | 3.01E-05 | 2.04E-05 | 0 | 0 |
| Gpx3 | 0.00052 | 6.66E-05 | 0.00012 | 0.00121 | 0.00085 | 0.00012 | 0.00227 | 0.00121 | 0 | 0.00144 | 0.00107 | 0 | 0.00349 | 0.00176 | 0 | 0.00033 | 0 | 0.00059 |
| Dnajc25 | 0.00038 | 0 | 0.00028 | 0.00012 | 4.09E-05 | 0.00012 | 0.00048 | 3.69E-05 | 0 | 6.53E-05 | 7.07E-05 | 1.24E-05 | 0.00016 | 6.62E-05 | 9.34E-05 | 5.97E-05 | 0 | 5.94E-05 |
| Parvb | 0 | 2.68E-05 | 0 | 0 | 5.27E-05 | 0.00012 | 0 | 9.25E-05 | 0 | 0 | 0 | 0.00017 | 0 | 0 | 0 | 0 | 0 | 0 |
| Tex264 | 8.32E-06 | 0.00022 | 0.00014 | 0.00038 | 0.00015 | 0.00012 | 0.00034 | 0.00017 | 0 | 0.00028 | 0.0002 | 0.00038 | 0.00014 | 0.00023 | 7.89E-05 | 4.98E-05 | 4.33E-05 | 0.00021 |
| Gucy1b1 | 0 | 0 | 0 | 0 | 0 | 0.00012 | 0 | 0.0002 | 0 | 1.06E-05 | 0 | 0 | 0 | 0 | 0 | 7.69E-06 | 0 | 6.48E-05 |
| Dpp7 | 2.45E-05 | 0.00014 | 0 | 0 | 8.52E-05 | 0.00012 | 1.34E-05 | 4.61E-06 | 0 | 3.62E-05 | 0 | 5.04E-05 | 0.00013 | 7.50E-05 | 2.09E-06 | 0 | 0 | 8.72E-05 |
| Steap4 | 0 | 0 | 0 | 0 | 0 | 0.00012 | 0 | 0 | 0 | 0.00071 | 0 | 0 | 0.00078 | 0 | 0 | 0 | 0 | 0.0004 |
| Naglu | 0 | 0.00014 | 0 | 9.81E-06 | 8.72E-05 | 0.00012 | 0.00014 | 0 | 0 | 0.0002 | 0 | 0.00021 | 0.00057 | 0 | 0 | 1.28E-05 | 0 | 0.00018 |
| Ttc39b | 0.0001 | 0.00011 | 0.00016 | 0 | 0 | 0.00012 | 0 | 0.00015 | 0 | 0 | 7.26E-05 | 0 | 0.00016 | 8.74E-05 | 0.00021 | 4.79E-05 | 0 | 4.72E-05 |
| Dhx40 | 0.0002 | 0.00013 | 4.15E-06 | 1.20E-05 | 0 | 0.00012 | 0.00013 | 8.35E-06 | 0 | 5.60E-06 | 0 | 0.0002 | 4.19E-05 | 8.18E-05 | 4.02E-05 | 5.09E-05 | 0.00017 | 2.47E-05 |
| Vgll4 | 5.49E-05 | 0.00034 | 2.28E-05 | 0.0001 | 0.00026 | 0.00012 | 0.0003 | 0.0002 | 0 | 0 | 0.00014 | 0.0001 | 5.99E-05 | 0.00052 | 0.00015 | 0.00014 | 0.00027 | 0.00016 |
| Clstn2 | 0 | 0 | 0 | 0 | 0.00043 | 0.00012 | 0 | 0 | 0 | 0 | 0.00036 | 0 | 0 | 0.0003 | 0 | 0 | 0 | 0 |
| Cdk19 | 8.81E-05 | 0.00018 | 0 | 0 | 4.59E-05 | 0.00012 | 6.63E-05 | 5.99E-05 | 0 | 5.53E-05 | 0 | 6.69E-05 | 0 | 1.77E-05 | 1.50E-05 | 0.00025 | 9.72E-05 | 2.41E-05 |
| Mt3 | 0 | 0 | 0 | 0 | 0 | 0.00011 | 0 | 0 | 0 | 0 | 0 | 2.06E-05 | 0 | 0 | 0 | 0 | 0 | 0 |

| | | | | | | | | | | | | | | | | | | |
|---|---|---|---|---|---|---|---|---|---|---|---|---|---|---|---|---|---|---|
| Etl4 | 0 | 8.69E-05 | 0 | 0 | 0.00013 | 0.00011 | 0 | 5.42E-05 | 0 | 0 | 2.37E-05 | 0 | 0 | 8.88E-06 | 0 | 0 | 0 | 0.0001 |
| Hivep2 | 0.00014 | 0.00016 | 0.00019 | 0 | 3.20E-05 | 0.00011 | 0 | 6.54E-05 | 0 | 0.00013 | 6.39E-05 | 0.00021 | 0 | 2.74E-05 | 6.28E-05 | 4.18E-05 | 0.00026 | 4.04E-05 |
| Ostm1 | 0.00015 | 0.0002 | 6.54E-05 | 0 | 0.00013 | 0.00011 | 0.00015 | 2.03E-05 | 0 | 3.83E-05 | 1.22E-05 | 0 | 4.33E-05 | 2.46E-05 | 0 | 0 | 0 | 0.00011 |
| Gid4 | 0.00011 | 9.28E-05 | 5.31E-05 | 2.23E-06 | 7.42E-05 | 0.00011 | 7.93E-05 | 7.58E-06 | 0 | 9.12E-06 | 6.24E-07 | 0.00014 | 0 | 1.46E-05 | 0 | 5.16E-05 | 5.88E-06 | 2.18E-05 |
| Zbtb46 | 7.74E-06 | 5.51E-05 | 2.60E-05 | 0 | 3.16E-05 | 0.00011 | 0 | 0.0002 | 0 | 0 | 1.86E-05 | 0 | 0 | 0 | 2.84E-05 | 0 | 1.38E-05 | 5.13E-06 |
| Mxi1 | 2.93E-05 | 0.00014 | 0 | 2.50E-05 | 0.00011 | 0.00011 | 7.84E-05 | 3.50E-05 | 0 | 3.07E-05 | 2.31E-05 | 0.00081 | 0 | 8.14E-05 | 0.00014 | 2.20E-05 | 0 | 0.0001 |
| Got1 | 4.84E-06 | 8.68E-05 | 0.00026 | 0.00024 | 9.25E-05 | 0.00011 | 7.68E-05 | 0.00021 | 0 | 2.22E-05 | 0.00012 | 0.00021 | 0 | 0 | 6.01E-05 | 7.21E-05 | 8.52E-05 | 1.23E-05 |
| Npas2 | 3.38E-05 | 4.77E-05 | 0 | 0 | 0.00023 | 0.00011 | 0.00019 | 4.73E-05 | 0 | 0.00011 | 5.89E-05 | 3.44E-05 | 0.00018 | 0 | 7.08E-05 | 3.66E-05 | 0 | 0.00014 |
| Zbed4 | 2.33E-05 | 7.43E-06 | 2.35E-05 | 2.53E-06 | 5.88E-05 | 0.00011 | 1.59E-05 | 0 | 0 | 0 | 2.74E-05 | 0 | 0 | 0 | 6.47E-05 | 8.17E-06 | 1.34E-05 | 0 |
| Itga7 | 0 | 0 | 0 | 0 | 5.33E-06 | 0.00011 | 0 | 3.06E-05 | 0 | 0 | 6.13E-06 | 0.00035 | 0 | 1.46E-05 | 4.05E-05 | 6.58E-06 | 0 | 2.22E-05 |
| Tmtc1 | 0 | 3.87E-05 | 0 | 0 | 4.87E-05 | 0.00011 | 0 | 0 | 0 | 1.10E-05 | 6.31E-05 | 0 | 0 | 0.00012 | 0 | 0 | 0 | 0.00029 |
| Slc17a5 | 4.67E-05 | 9.04E-05 | 0 | 0 | 8.50E-05 | 0.00011 | 0.0001 | 0 | 0 | 1.30E-05 | 1.59E-05 | 0 | 7.05E-05 | 0.00017 | 0 | 0 | 0 | 2.20E-05 |
| Ier3ip1 | 0.00021 | 4.06E-05 | 0.00036 | 0.00019 | 0.00011 | 0.00011 | 0.00048 | 0.00037 | 0 | 7.04E-05 | 0.00028 | 0.0002 | 9.58E-05 | 8.84E-05 | 1.45E-05 | 8.62E-05 | 3.75E-05 | 2.25E-05 |
| Gas7 | 0 | 0.00016 | 3.77E-05 | 0 | 3.72E-05 | 0.00011 | 0.00036 | 0 | 0 | 0.00118 | 0 | 0.00012 | 0 | 0 | 0 | 7.42E-05 | 0 | 0.00061 |
| Rb1cc1 | 0.00015 | 0.0002 | 1.83E-05 | 0 | 6.78E-05 | 0.00011 | 6.37E-05 | 0 | 0 | 5.86E-05 | 8.17E-05 | 0.00015 | 0 | 8.60E-05 | 0.00011 | 3.78E-05 | 0 | 6.17E-05 |
| Usp3 | 0.0001 | 0.00012 | 3.36E-05 | 2.75E-05 | 5.37E-05 | 0.00011 | 5.11E-05 | 3.51E-05 | 0 | 1.85E-05 | 0 | 4.06E-05 | 7.21E-06 | 5.89E-05 | 0.00013 | 0 | 5.92E-05 | 3.14E-06 |
| Tgfb1 | 0 | 0.00013 | 0.00068 | 0.00044 | 0.00022 | 0.00011 | 0 | 0.0017 | 0 | 0 | 0.00026 | 0.00055 | 0 | 0.0007 | 0.00031 | 9.31E-05 | 0.00023 | 2.92E-06 |
| Cyp4v3 | 3.73E-05 | 8.21E-05 | 0 | 1.82E-05 | 5.69E-05 | 0.00011 | 0.00012 | 0 | 0 | 2.27E-05 | 0 | 4.16E-05 | 9.52E-05 | 0 | 0 | 0 | 0 | 5.69E-05 |
| Klf10 | 3.17E-05 | 0.00016 | 0.00023 | 1.26E-05 | 0 | 0.00011 | 0 | 0 | 0 | 5.14E-05 | 0 | 2.64E-05 | 0 | 3.92E-05 | 0 | 4.85E-05 | 0.00045 | 0.00025 |
| Atp1b2 | 0 | 4.20E-06 | 0 | 0 | 9.53E-05 | 0.0001 | 0 | 5.17E-05 | 0 | 1.23E-05 | 0.0001 | 0 | 6.81E-05 | 0 | 4.33E-06 | 0 | 0 | 1.23E-05 |
| Dlg4 | 0.00018 | 0.0001 | 0.00015 | 0 | 0.00012 | 0.0001 | 1.51E-05 | 0.00011 | 0 | 4.73E-05 | 4.80E-05 | 0.00017 | 0 | 0 | 0.00027 | 0 | 0.00025 | 5.08E-05 |
| Clock | 0.00012 | 8.69E-05 | 0.00014 | 0 | 6.11E-05 | 0.0001 | 7.85E-05 | 0 | 0 | 6.09E-05 | 2.97E-05 | 2.37E-05 | 2.92E-05 | 1.60E-05 | 6.13E-06 | 9.06E-05 | 7.95E-05 | 3.38E-05 |
| Serf1 | 0.00011 | 1.66E-05 | 0 | 1.18E-06 | 3.09E-05 | 0.0001 | 0.00021 | 6.81E-05 | 0 | 6.29E-06 | 1.48E-05 | 3.94E-05 | 1.81E-05 | 4.78E-06 | 0 | 0 | 1.53E-05 | 8.22E-05 |
| Ebf4 | 0 | 4.80E-05 | 0 | 0 | 8.38E-05 | 0.0001 | 0 | 0 | 0 | 0 | 2.15E-05 | 6.64E-05 | 0 | 4.85E-05 | 7.63E-05 | 0 | 0 | 0 |
| Col4a3bp | 4.10E-05 | 0.00016 | 0 | 6.53E-05 | 0.00021 | 0.0001 | 1.17E-05 | 6.37E-05 | 0 | 4.45E-05 | 0.00014 | 0.00011 | 0 | 0.00017 | 0.00028 | 5.79E-05 | 0.00014 | 9.95E-05 |
| Dbndd2 | 0 | 3.10E-05 | 3.80E-06 | 1.46E-05 | 0 | 0.0001 | 0 | 0.00047 | 0 | 9.87E-05 | 0 | 4.75E-05 | 0 | 0.00021 | 0 | 0 | 0 | 0.00037 |
| Cacna2d3 | 0 | 0 | 0 | 0 | 8.51E-05 | 0.0001 | 0 | 0 | 0 | 0 | 0 | 2.59E-06 | 0 | 2.43E-05 | 2.63E-06 | 1.81E-06 | 0 | 0 |
| Mxd1 | 0 | 8.40E-05 | 0 | 4.35E-05 | 3.75E-05 | 0.0001 | 0 | 3.35E-05 | 0 | 9.67E-06 | 0 | 9.05E-05 | 0 | 0.00015 | 0.00011 | 1.04E-05 | 0.00041 | 0 |
| Ankrd35 | 0 | 3.86E-06 | 0 | 5.13E-05 | 0.00015 | 0.0001 | 3.82E-07 | 5.16E-05 | 0 | 5.67E-06 | 0 | 0 | 0 | 1.14E-05 | 3.61E-05 | 0 | 4.25E-06 | 1.40E-05 |
| Cir1 | 5.84E-05 | 0.00012 | 0 | 1.08E-05 | 4.92E-05 | 0.0001 | 3.12E-05 | 0 | 0 | 1.32E-05 | 2.89E-06 | 6.16E-05 | 8.23E-06 | 2.12E-05 | 6.44E-05 | 1.36E-05 | 0 | 2.05E-05 |
| Lpin1 | 9.21E-06 | 0.00015 | 0 | 0 | 2.02E-05 | 0.0001 | 0 | 2.67E-05 | 0 | 0 | 0 | 8.89E-05 | 0 | 0 | 0 | 0 | 0 | 3.61E-05 |
| Olfml2b | 0 | 0 | 0.00012 | 0.0001 | 0.00019 | 0.0001 | 0.00027 | 0.00027 | 0 | 0.00054 | 0.00054 | 0 | 0 | 0.00051 | 3.92E-07 | 0.00013 | 0 | 0.00068 |
| Tmem200b | 0 | 2.48E-05 | 6.89E-05 | 7.84E-05 | 0.00013 | 0.0001 | 0 | 0 | 0 | 0 | 4.92E-05 | 0 | 0 | 1.49E-05 | 9.31E-05 | 2.36E-05 | 0 | 2.33E-05 |

| | | | | | | | | | | | | | | | | | | |
|---|---|---|---|---|---|---|---|---|---|---|---|---|---|---|---|---|---|---|
| Samd8 | 0.00012 | 0.00011 | 8.07E-05 | 0 | 2.35E-05 | 0.0001 | 0 | 5.44E-05 | 0 | 2.83E-06 | 4.13E-05 | 9.81E-05 | 0 | 1.16E-05 | 0 | 4.11E-05 | 0.00011 | 1.30E-06 |
| Edil3 | 7.68E-06 | 0 | 0 | 0 | 0.00025 | 9.95E-05 | 2.16E-06 | 0.00013 | 0 | 0 | 2.40E-05 | 6.42E-05 | 0 | 2.48E-05 | 0 | 0 | 0 | 0 |
| Vps13d | 6.91E-05 | 0.00018 | 9.86E-05 | 0 | 9.87E-05 | 9.86E-05 | 5.02E-05 | 0.0001 | 0 | 0.00012 | 0.00011 | 0.00015 | 1.78E-05 | 1.43E-05 | 0.00027 | 2.18E-05 | 6.40E-05 | 5.83E-05 |
| Fgd6 | 0 | 3.63E-05 | 0 | 0 | 7.85E-05 | 9.82E-05 | 0 | 0.00021 | 0 | 0 | 5.00E-05 | 8.05E-05 | 0 | 0.00013 | 3.31E-05 | 0 | 0 | 2.71E-05 |
| Cyb561a3 | 1.98E-05 | 0.00011 | 0 | 0.00011 | 0.00016 | 9.77E-05 | 8.07E-06 | 0.00015 | 0 | 7.22E-05 | 3.66E-05 | 5.89E-05 | 0 | 0 | 5.15E-05 | 3.57E-05 | 0 | 9.24E-05 |
| Dgka | 5.76E-05 | 0.00012 | 3.05E-05 | 5.37E-05 | 9.94E-05 | 9.74E-05 | 4.49E-05 | 1.82E-05 | 0 | 1.67E-05 | 4.47E-05 | 7.43E-06 | 0 | 1.29E-05 | 0.0001 | 0 | 4.52E-05 | 0.00018 |
| Zfp9 | 0.00027 | 3.18E-05 | 0.00015 | 6.61E-05 | 1.99E-05 | 9.71E-05 | 0.00032 | 4.31E-05 | 0 | 7.02E-06 | 0 | 0.00019 | 2.59E-05 | 7.09E-05 | 0.00026 | 1.85E-05 | 9.20E-05 | 1.51E-05 |
| Il13ra2 | 3.62E-06 | 0 | 2.13E-05 | 0 | 0 | 9.67E-05 | 0 | 0 | 0 | 0 | 0 | 0 | 0.00038 | 3.36E-05 | 0 | 1.13E-05 | 0 | 0 |
| Mfhas1 | 1.08E-05 | 7.06E-05 | 0.00029 | 0.00013 | 0.00018 | 9.63E-05 | 7.66E-05 | 0.00013 | 0 | 0.00026 | 0.0004 | 0.00012 | 4.90E-05 | 0.00027 | 0.00019 | 6.37E-05 | 5.18E-05 | 6.74E-06 |
| Mast4 | 0 | 0.00022 | 0.00016 | 0 | 0.00057 | 9.63E-05 | 8.62E-05 | 0.00049 | 0 | 0.0001 | 0.00021 | 0.00019 | 0 | 7.24E-05 | 0.00026 | 8.50E-05 | 0.00014 | 0 |
| Rnf169 | 6.47E-05 | 0.00022 | 4.65E-05 | 4.23E-05 | 7.43E-05 | 9.56E-05 | 3.38E-05 | 4.81E-05 | 0 | 1.98E-05 | 0 | 2.16E-05 | 4.08E-05 | 0 | 0.00022 | 3.54E-05 | 1.26E-05 | 2.32E-05 |
| Plcg1 | 1.46E-05 | 0.00021 | 4.21E-05 | 7.47E-05 | 0.00043 | 9.50E-05 | 0.00011 | 0.00019 | 0 | 6.87E-05 | 0.00024 | 0.00017 | 4.65E-05 | 0.00016 | 0.00032 | 0 | 2.05E-07 | 0.00014 |
| Dact1 | 0 | 0.00015 | 0 | 0.00022 | 0.00079 | 9.49E-05 | 2.14E-05 | 0 | 0 | 0.00041 | 0.00034 | 0.00019 | 0 | 0.00027 | 0.00023 | 0 | 0.00031 | 0.00031 |
| Bicral | 6.03E-05 | 0.00012 | 0 | 0 | 4.65E-05 | 9.42E-05 | 2.33E-05 | 2.31E-05 | 0 | 3.57E-05 | 2.69E-05 | 6.95E-05 | 0 | 1.84E-05 | 1.70E-05 | 1.19E-05 | 0.00014 | 7.27E-05 |
| Des | 0 | 0 | 0 | 0 | 0 | 9.41E-05 | 0 | 4.09E-05 | 0 | 1.02E-05 | 0.00031 | 0 | 8.98E-06 | 0 | 0 | 0 | 0 | 0 |
| Slco4a1 | 0 | 0.00013 | 0 | 0 | 0.00015 | 9.41E-05 | 0 | 0.0002 | 0 | 0 | 0 | 0.00015 | 0 | 0 | 6.28E-06 | 0 | 1.29E-05 | 0 |
| Mkx | 0 | 1.78E-05 | 2.35E-05 | 0 | 9.42E-05 | 9.33E-05 | 0.0002 | 1.59E-05 | 0 | 0 | 0 | 0 | 0.00012 | 0.00023 | 0 | 4.00E-06 | 0 | 0 |
| Bach1 | 0.00011 | 0.00025 | 0.00041 | 4.51E-05 | 0.00013 | 9.33E-05 | 0.00036 | 0.00047 | 0 | 0.00011 | 0 | 0.00019 | 0.00024 | 0.00018 | 0.00024 | 0 | 0.00042 | 6.52E-05 |
| Nbas | 0.00021 | 5.97E-06 | 0.00022 | 0.00017 | 3.65E-05 | 9.31E-05 | 0.00042 | 7.14E-05 | 0 | 0.0001 | 1.24E-05 | 0 | 0.00013 | 8.88E-06 | 0.00031 | 0 | 9.11E-06 | 1.73E-05 |
| Gprc5b | 0 | 3.43E-05 | 0 | 0 | 0.00011 | 9.28E-05 | 0 | 8.65E-06 | 0 | 0.00052 | 4.40E-05 | 0 | 0 | 0.0003 | 3.25E-06 | 0.00017 | 0 | 0.00027 |
| Mif | 0.0002 | 4.94E-05 | 0.00023 | 0.00044 | 0.00014 | 9.18E-05 | 0.00024 | 6.76E-05 | 0 | 3.14E-06 | 2.16E-05 | 0.00145 | 0.00043 | 0.00021 | 0 | 0.00015 | 0.00033 | 0 |
| Trib2 | 2.57E-05 | 9.60E-05 | 4.33E-06 | 0 | 4.36E-05 | 9.15E-05 | 0 | 9.42E-05 | 0 | 0 | 4.70E-05 | 8.52E-05 | 0 | 5.85E-05 | 0 | 0 | 4.09E-06 | 0.00025 |
| Yeats4 | 0.00016 | 2.76E-05 | 4.00E-05 | 0 | 2.35E-05 | 9.14E-05 | 1.66E-05 | 0 | 0 | 0 | 1.84E-05 | 4.36E-05 | 0 | 7.89E-06 | 6.02E-05 | 0.00012 | 0 | 0 |
| Spock3 | 0 | 0 | 0 | 0 | 0.00043 | 9.13E-05 | 0 | 9.73E-05 | 0 | 0 | 0.00018 | 6.30E-06 | 0 | 0 | 0.0001 | 0 | 0 | 0 |
| Cog6 | 0.00016 | 2.18E-05 | 8.16E-05 | 0.0001 | 4.40E-05 | 9.06E-05 | 0.00031 | 8.09E-05 | 0 | 1.38E-05 | 7.23E-05 | 2.35E-05 | 7.19E-05 | 1.92E-05 | 5.43E-05 | 0 | 1.15E-05 | 0 |
| Fpgt | 8.22E-05 | 8.11E-05 | 1.86E-05 | 1.50E-07 | 9.83E-05 | 8.96E-05 | 0 | 4.13E-05 | 0 | 0 | 3.37E-05 | 0.00012 | 3.70E-06 | 0 | 3.87E-06 | 1.93E-05 | 0 | 6.30E-06 |
| Cpt1a | 3.83E-05 | 0.00014 | 0 | 0.00013 | 9.09E-05 | 8.93E-05 | 0.00017 | 0 | 0 | 0.00026 | 4.52E-05 | 0 | 0.00025 | 0.0002 | 0 | 6.33E-05 | 0 | 0.0003 |
| Mtss2 | 0 | 0.00022 | 0 | 0 | 0.00016 | 8.92E-05 | 0.00015 | 0 | 0 | 0 | 0.0002 | 0 | 0 | 0.00046 | 0.00013 | 8.14E-06 | 1.88E-05 | 0 |
| Il2rg | 0 | 2.19E-06 | 0 | 0 | 0 | 8.88E-05 | 0 | 0.0005 | 0 | 3.15E-06 | 8.97E-05 | 8.18E-06 | 0 | 0.00017 | 0 | 0 | 0 | 0 |
| Col25a1 | 0 | 0 | 0 | 0 | 0 | 8.85E-05 | 0 | 0 | 0 | 0 | 0 | 2.19E-05 | 0 | 2.55E-05 | 9.83E-06 | 0 | 0 | 0 |
| DEPRECATEI | 5.18E-05 | 6.94E-05 | 0 | 0 | 1.98E-05 | 8.77E-05 | 0 | 0 | 0 | 0 | 0 | 0.00045 | 0 | 2.15E-05 | 6.66E-05 | 3.53E-06 | 0 | 3.00E-05 |
| Appl2 | 5.16E-05 | 0.00021 | 0 | 0 | 7.20E-05 | 8.76E-05 | 2.62E-06 | 3.91E-06 | 0 | 7.03E-05 | 0 | 0.00017 | 0 | 0 | 0.00013 | 0 | 8.42E-05 | 9.34E-05 |
| Elmod2 | 9.40E-05 | 3.46E-05 | 2.68E-05 | 0 | 2.37E-05 | 8.73E-05 | 8.55E-05 | 4.56E-05 | 0 | 2.08E-05 | 0 | 3.38E-05 | 0.00012 | 6.28E-06 | 1.61E-05 | 4.97E-05 | 1.93E-05 | 1.75E-05 |

| | | | | | | | | | | | | | | | | | | |
|---|---|---|---|---|---|---|---|---|---|---|---|---|---|---|---|---|---|---|
| Tomm7 | 0.00017 | 0.00014 | 0.00018 | 0.00031 | 6.84E-05 | 8.66E-05 | 0.00023 | 0.00012 | 0 | 5.08E-05 | 1.54E-05 | 0.0001 | 0.00011 | 1.31E-05 | 0 | 0.00012 | 0.00014 | 9.48E-05 |
| Prorsd1 | 0.00018 | 1.33E-06 | 2.11E-05 | 9.17E-05 | 4.52E-06 | 8.57E-05 | 0.00013 | 3.97E-06 | 0 | 6.42E-06 | 0 | 0 | 0 | 0 | 3.71E-05 | 0 | 0 | 0 |
| Tuba8 | 0 | 0 | 3.46E-07 | 0 | 0 | 8.51E-05 | 0 | 0 | 0 | 0 | 2.39E-06 | 0 | 0 | 0 | 0 | 5.79E-06 | 0 | 0 |
| Tnrc6a | 0.0002 | 0.00017 | 5.31E-05 | 0 | 5.12E-05 | 8.46E-05 | 8.78E-05 | 0 | 0 | 2.81E-05 | 6.92E-05 | 1.72E-05 | 6.92E-05 | 3.62E-06 | 0.00023 | 0.00013 | 0.00031 | 6.19E-05 |
| Clip1 | 0.00011 | 0.00012 | 0.0002 | 0 | 0 | 8.38E-05 | 6.43E-05 | 0 | 0 | 0.00029 | 9.66E-05 | 0 | 0.00012 | 0.00034 | 0.00027 | 4.38E-05 | 0.00022 | 6.48E-05 |
| Ppt2 | 2.74E-05 | 6.59E-05 | 1.43E-05 | 0.00014 | 6.80E-05 | 8.36E-05 | 3.19E-05 | 5.87E-05 | 0 | 2.74E-05 | 8.19E-05 | 5.53E-05 | 0 | 9.77E-05 | 0 | 5.62E-05 | 0 | 1.53E-05 |
| Gramd3 | 0.00013 | 1.53E-05 | 0.00013 | 3.98E-05 | 7.80E-05 | 8.35E-05 | 0.00022 | 0.00012 | 0 | 0.00012 | 0 | 0 | 6.84E-05 | 0.00015 | 4.41E-05 | 2.68E-05 | 8.53E-05 | 0 |
| Ufl1 | 0.0002 | 8.10E-05 | 9.37E-05 | 1.08E-05 | 9.71E-05 | 8.32E-05 | 0.00011 | 6.51E-05 | 0 | 0 | 1.85E-05 | 0.00012 | 0 | 0 | 8.78E-05 | 1.81E-05 | 0 | 1.36E-05 |
| Tars | 0.0004 | 9.83E-05 | 0.00047 | 0.00041 | 7.26E-05 | 8.28E-05 | 0.00028 | 1.82E-05 | 0 | 1.17E-05 | 0.00012 | 9.59E-05 | 0 | 5.13E-05 | 0.00027 | 0.00019 | 0 | 5.90E-05 |
| Crbn | 0.00011 | 0.00011 | 0 | 0 | 5.36E-05 | 8.27E-05 | 2.18E-05 | 0 | 0 | 2.06E-05 | 1.30E-05 | 8.69E-05 | 0 | 5.09E-05 | 7.05E-05 | 1.81E-05 | 0 | 5.25E-05 |
| Jph2 | 0 | 1.61E-05 | 0 | 0 | 0.00011 | 8.26E-05 | 0 | 0 | 0 | 0 | 0.00057 | 9.98E-05 | 0 | 0 | 0.00012 | 0 | 0 | 0 |
| Wdr48 | 0.00014 | 0.00015 | 6.39E-05 | 1.33E-05 | 5.44E-05 | 8.25E-05 | 0.00012 | 1.11E-05 | 0 | 1.17E-05 | 0 | 7.97E-05 | 0 | 4.53E-06 | 0.00017 | 0.00012 | 0 | 0 |
| Rab32 | 3.10E-05 | 6.36E-05 | 4.92E-05 | 0 | 0 | 8.23E-05 | 0 | 0 | 0 | 0.00043 | 2.64E-05 | 3.95E-05 | 0 | 4.93E-05 | 0 | 4.08E-05 | 0 | 0.0001 |
| Trpm4 | 0 | 5.73E-05 | 3.80E-05 | 6.84E-05 | 3.38E-06 | 8.13E-05 | 0.00015 | 8.10E-05 | 0 | 4.19E-05 | 4.05E-05 | 7.07E-05 | 3.53E-05 | 7.78E-05 | 0.00016 | 4.11E-05 | 0 | 5.54E-05 |
| Rtn2 | 3.96E-06 | 5.67E-05 | 0 | 0 | 0.00016 | 8.09E-05 | 2.25E-05 | 0 | 0 | 6.07E-05 | 5.64E-05 | 1.38E-05 | 8.36E-05 | 7.94E-05 | 0 | 0 | 0 | 4.05E-05 |
| Insyn1 | 0 | 0.00011 | 0.0002 | 5.15E-05 | 0.0002 | 8.02E-05 | 0.0003 | 0 | 0 | 0.00022 | 0.00017 | 3.62E-05 | 0 | 2.62E-05 | 3.81E-05 | 0 | 2.80E-05 | 0.00012 |
| Rbm33 | 0.00017 | 0.00016 | 3.39E-05 | 0 | 5.47E-05 | 8.00E-05 | 6.23E-05 | 1.03E-05 | 0 | 3.77E-05 | 0 | 0.00012 | 0 | 2.90E-05 | 0.00073 | 8.18E-05 | 0.00022 | 0 |
| Mertk | 0 | 1.12E-05 | 0 | 3.71E-05 | 5.39E-05 | 7.97E-05 | 9.29E-05 | 1.65E-05 | 0 | 0 | 0 | 2.50E-05 | 0 | 1.32E-05 | 2.40E-05 | 0 | 0 | 0 |
| Flt1 | 0 | 0 | 0 | 0 | 0 | 7.94E-05 | 0 | 4.50E-05 | 0 | 0 | 8.08E-05 | 0.00034 | 0 | 0.00036 | 3.31E-06 | 7.27E-06 | 6.88E-05 | 1.62E-05 |
| Smad7 | 7.92E-05 | 8.94E-05 | 0.00011 | 0 | 0.00022 | 7.92E-05 | 0 | 0 | 0 | 1.15E-05 | 5.72E-05 | 0.00061 | 5.09E-05 | 0.00026 | 0.00014 | 2.77E-05 | 0.00118 | 8.30E-05 |
| Jade1 | 0 | 7.02E-05 | 8.14E-05 | 0.00013 | 0.00015 | 7.88E-05 | 9.68E-05 | 0 | 0 | 3.15E-05 | 0.00023 | 2.41E-05 | 0 | 0 | 6.25E-05 | 0.00015 | 0 | 0.00022 |
| Ap4m1 | 6.26E-05 | 7.61E-05 | 2.93E-05 | 0.00011 | 3.64E-05 | 7.88E-05 | 8.11E-05 | 3.25E-05 | 0 | 3.74E-05 | 0 | 4.05E-05 | 4.06E-05 | 3.67E-05 | 6.72E-05 | 2.09E-05 | 0.00014 | 4.01E-05 |
| Smim4 | 9.75E-05 | 7.91E-05 | 3.23E-05 | 0.00019 | 6.30E-05 | 7.86E-05 | 3.05E-05 | 1.03E-05 | 0 | 3.39E-05 | 2.77E-05 | 4.58E-05 | 1.99E-05 | 0 | 2.64E-05 | 6.28E-05 | 0 | 8.18E-05 |
| Pde4a | 0 | 7.54E-05 | 1.73E-05 | 0 | 3.41E-05 | 7.80E-05 | 0 | 0.00012 | 0 | 2.13E-05 | 0.00011 | 0.00011 | 0 | 8.78E-05 | 2.32E-05 | 3.51E-05 | 2.67E-06 | 4.55E-05 |
| Fbxo30 | 6.67E-05 | 7.52E-05 | 5.98E-05 | 0 | 1.66E-05 | 7.77E-05 | 0 | 5.55E-05 | 0 | 3.27E-05 | 0.00016 | 0.00015 | 0 | 5.27E-05 | 0 | 3.88E-05 | 0.00094 | 0 |
| Praf2 | 0.00035 | 0 | 0.00022 | 0.00032 | 3.69E-05 | 7.74E-05 | 0.00068 | 0.00012 | 0 | 7.92E-05 | 0.00016 | 6.88E-05 | 7.96E-05 | 7.48E-05 | 0 | 0 | 0 | 0.00011 |
| Cr1l | 6.31E-05 | 2.41E-05 | 9.82E-06 | 8.78E-05 | 0 | 7.71E-05 | 1.61E-05 | 6.99E-05 | 0 | 5.96E-05 | 1.04E-05 | 4.33E-05 | 7.76E-05 | 3.99E-05 | 0 | 4.46E-05 | 0 | 7.36E-05 |
| Zdhhc14 | 0 | 2.32E-05 | 0 | 0 | 5.99E-05 | 7.66E-05 | 7.00E-05 | 0 | 0 | 0 | 3.45E-05 | 0.00019 | 6.51E-05 | 0.00013 | 0.00012 | 0 | 0 | 7.93E-05 |
| Blzf1 | 0.00013 | 3.74E-05 | 8.39E-05 | 5.18E-05 | 2.78E-05 | 7.66E-05 | 0.00012 | 3.16E-05 | 0 | 2.68E-05 | 5.85E-05 | 3.78E-05 | 0 | 5.30E-05 | 7.11E-05 | 0 | 5.97E-05 | 2.63E-05 |
| Tmem98 | 5.65E-05 | 0.00017 | 0 | 0.0001 | 0.0003 | 7.61E-05 | 0.00022 | 0 | 0 | 0 | 0.00026 | 0 | 0.00027 | 0 | 0 | 0.00019 | 0 | 0.00019 |
| Cryz | 4.99E-05 | 6.66E-05 | 0 | 7.55E-05 | 5.54E-05 | 7.56E-05 | 2.47E-05 | 6.79E-06 | 0 | 1.47E-05 | 0 | 9.95E-05 | 8.36E-05 | 0 | 0 | 2.65E-05 | 0 | 5.07E-05 |
| Tbpl1 | 0.00018 | 5.99E-05 | 4.90E-05 | 0 | 2.56E-05 | 7.49E-05 | 4.18E-05 | 3.15E-05 | 0 | 7.53E-06 | 0 | 0.00011 | 0 | 0 | 7.38E-05 | 5.91E-05 | 0.00011 | 0 |
| Trim33 | 0.00015 | 9.73E-05 | 5.63E-05 | 0 | 6.15E-05 | 7.49E-05 | 1.95E-05 | 0 | 0 | 0 | 2.95E-05 | 0.00012 | 0 | 0 | 0.00012 | 5.35E-05 | 0 | 1.12E-05 |

| | | | | | | | | | | | | | | | | | | |
|---|---|---|---|---|---|---|---|---|---|---|---|---|---|---|---|---|---|---|
| Idua | 0 | 0.00011 | 0 | 0 | 6.49E-05 | 7.47E-05 | 2.71E-05 | 0 | 0 | 6.35E-05 | 3.12E-05 | 2.13E-05 | 0.00011 | 5.78E-05 | 4.30E-05 | 0 | 0 | 6.31E-05 |
| Chst13 | 0 | 0 | 0 | 2.01E-05 | 7.19E-05 | 7.44E-05 | 0 | 1.50E-06 | 0 | 0 | 9.15E-05 | 0 | 0 | 1.69E-05 | 3.64E-05 | 0 | 0 | 0 |
| Ap1s2 | 7.69E-05 | 0.0001 | 0 | 0 | 5.32E-05 | 7.39E-05 | 0 | 7.70E-06 | 0 | 0 | 0.00013 | 6.93E-05 | 0 | 0.00024 | 2.33E-05 | 7.48E-05 | 1.26E-05 | 0.00014 |
| Hist1h2bc | 0.00038 | 0.00023 | 3.84E-05 | 0.00013 | 0 | 7.39E-05 | 9.13E-05 | 3.43E-05 | 0 | 0.00019 | 6.98E-05 | 0.00053 | 0.00024 | 0 | 5.90E-05 | 0 | 0.00018 | 0.00037 |
| Tshz2 | 0 | 4.96E-05 | 0 | 3.03E-05 | 0 | 7.37E-05 | 8.74E-05 | 0.00019 | 0 | 0.00032 | 0 | 0 | 0.00053 | 0 | 5.36E-05 | 0 | 0 | 0.00027 |
| Fosb | 0 | 0 | 0 | 2.58E-05 | 3.64E-05 | 7.36E-05 | 0.00039 | 6.68E-05 | 0 | 4.17E-05 | 0 | 0.00012 | 0.00033 | 0 | 6.28E-05 | 5.78E-05 | 0.0111 | 0.00021 |
| Slc25a16 | 7.45E-05 | 2.86E-05 | 6.59E-05 | 9.78E-05 | 3.64E-05 | 7.30E-05 | 8.85E-05 | 0 | 0 | 3.93E-05 | 5.25E-05 | 0 | 5.59E-05 | 9.58E-05 | 0 | 3.91E-05 | 0 | 5.80E-05 |
| Gle1 | 0.00011 | 7.46E-05 | 0.00012 | 0.00012 | 5.17E-05 | 7.30E-05 | 0.00013 | 9.46E-06 | 0 | 5.95E-06 | 4.02E-05 | 1.02E-05 | 6.13E-05 | 4.18E-05 | 9.27E-05 | 0.00029 | 4.04E-06 | 1.06E-05 |
| Slc43a1 | 0.00015 | 3.70E-05 | 3.75E-05 | 8.40E-05 | 1.68E-05 | 7.28E-05 | 0.0002 | 6.25E-05 | 0 | 1.09E-05 | 2.33E-05 | 0 | 0.00015 | 0.00015 | 8.01E-05 | 2.83E-05 | 0 | 0 |
| Grem2 | 5.00E-05 | 0.00018 | 0.00069 | 0 | 0.00057 | 7.23E-05 | 0.00015 | 0.00049 | 0 | 0 | 0.00105 | 0.00025 | 0.00109 | 0.00056 | 0.00017 | 0 | 0 | 6.02E-05 |
| Thrb | 0 | 7.91E-05 | 0 | 0 | 1.87E-05 | 7.17E-05 | 0 | 2.84E-06 | 0 | 1.46E-05 | 0 | 2.00E-06 | 0 | 8.47E-06 | 1.12E-05 | 0 | 6.24E-05 | 0 |
| Slc27a1 | 0 | 0.00019 | 0 | 7.12E-05 | 0.00016 | 7.16E-05 | 0 | 0 | 0 | 0 | 0 | 0.00013 | 0 | 0.00023 | 0.00019 | 0 | 0 | 3.07E-05 |
| Glipr1 | 0 | 2.31E-05 | 0 | 0 | 6.47E-05 | 7.11E-05 | 0 | 6.61E-06 | 0 | 0 | 0.00014 | 4.88E-06 | 0 | 0 | 0 | 3.61E-06 | 4.87E-07 | 0 |
| Tlr4 | 2.16E-05 | 0.00012 | 0 | 0 | 3.78E-06 | 7.06E-05 | 0 | 1.00E-05 | 0 | 0.00017 | 2.06E-05 | 0 | 3.48E-05 | 7.82E-06 | 0 | 0 | 0 | 8.06E-05 |
| Sdk2 | 0 | 3.13E-05 | 0 | 0 | 5.80E-05 | 7.03E-05 | 0 | 3.80E-05 | 0 | 0 | 0 | 0 | 0 | 9.29E-05 | 0 | 0 | 5.69E-06 | 0 |
| Iglon5 | 0 | 0.00011 | 0 | 0 | 0.0002 | 7.01E-05 | 0 | 0 | 0 | 0.00012 | 6.37E-05 | 1.69E-05 | 5.61E-05 | 0.00014 | 0 | 0 | 0 | 0 |
| Ttc39c | 5.25E-05 | 3.58E-05 | 0.00011 | 0 | 0 | 7.00E-05 | 1.60E-05 | 2.62E-05 | 0 | 0.0001 | 5.03E-05 | 0 | 0 | 0 | 9.79E-05 | 2.36E-05 | 7.38E-05 | 7.83E-05 |
| Enox1 | 0 | 5.82E-05 | 4.47E-05 | 0 | 0.00011 | 6.91E-05 | 4.65E-05 | 3.49E-05 | 0 | 8.49E-06 | 1.76E-05 | 5.11E-05 | 0 | 9.40E-05 | 5.28E-05 | 1.69E-05 | 4.59E-05 | 4.09E-05 |
| Galnt7 | 7.20E-05 | 5.08E-05 | 3.73E-05 | 0 | 4.75E-05 | 6.89E-05 | 3.90E-05 | 1.11E-05 | 0 | 0 | 2.96E-05 | 5.15E-06 | 0 | 3.05E-05 | 2.76E-05 | 2.52E-05 | 5.64E-05 | 0 |
| Fkbp11 | 0.0004 | 0 | 0.00049 | 0.00056 | 0 | 6.77E-05 | 0.00083 | 9.50E-05 | 0 | 0 | 8.98E-05 | 6.37E-05 | 3.87E-05 | 0.00015 | 6.38E-05 | 4.01E-05 | 0 | 0 |
| Zfp120 | 6.15E-05 | 3.25E-05 | 0 | 0 | 3.38E-05 | 6.74E-05 | 3.39E-06 | 6.05E-06 | 0 | 7.61E-07 | 0 | 3.38E-05 | 6.84E-06 | 5.69E-06 | 0 | 1.31E-05 | 0 | 0 |
| Brinp1 | 0 | 1.64E-05 | 0 | 0 | 5.29E-05 | 6.74E-05 | 0 | 6.44E-05 | 0 | 0 | 6.00E-05 | 4.26E-05 | 0 | 0 | 0 | 0 | 0 | 0 |
| Cep170b | 0 | 0.00015 | 8.41E-05 | 2.12E-05 | 6.53E-05 | 6.72E-05 | 2.76E-05 | 0 | 0 | 0 | 0.00015 | 0.00018 | 9.31E-05 | 0.00017 | 0.00016 | 0 | 3.24E-05 | 9.56E-05 |
| Eno2 | 0 | 0.00012 | 0 | 1.44E-05 | 4.89E-05 | 6.66E-05 | 0 | 0 | 0 | 0 | 0 | 4.26E-05 | 7.72E-05 | 1.09E-05 | 0.00016 | 0 | 9.92E-05 | 7.67E-05 |
| Twnk | 3.41E-05 | 3.36E-05 | 3.46E-05 | 0 | 5.18E-05 | 6.66E-05 | 3.66E-05 | 0 | 0 | 0 | 2.32E-05 | 2.39E-05 | 0 | 0 | 4.52E-05 | 1.49E-05 | 1.56E-05 | 0 |
| Notch3 | 0 | 0 | 0 | 0 | 0 | 6.65E-05 | 0 | 0.00035 | 0 | 0 | 0 | 0.00016 | 0 | 0 | 1.02E-05 | 0 | 0 | 8.03E-05 |
| Atg4c | 6.10E-05 | 7.71E-05 | 3.13E-05 | 0 | 6.21E-05 | 6.63E-05 | 5.04E-05 | 2.02E-05 | 0 | 3.20E-05 | 8.78E-06 | 9.24E-05 | 0 | 0 | 6.93E-05 | 6.77E-06 | 0 | 4.81E-05 |
| Gga2 | 0.00023 | 0.00013 | 9.84E-05 | 0.00019 | 5.72E-05 | 6.61E-05 | 0.00024 | 0 | 0 | 9.58E-05 | 0 | 1.07E-05 | 0.00011 | 3.31E-05 | 4.97E-05 | 8.94E-06 | 9.38E-05 | 0 |
| Cdh2 | 2.13E-05 | 1.04E-05 | 3.44E-05 | 8.56E-06 | 0 | 6.59E-05 | 0.00012 | 3.04E-05 | 0 | 2.29E-05 | 0 | 0.00015 | 0 | 0 | 0 | 0.00019 | 6.05E-05 | 0.00013 |
| Tent5a | 0.00017 | 0 | 0.00035 | 0 | 0 | 6.56E-05 | 0.0002 | 3.35E-05 | 0 | 6.86E-05 | 7.15E-05 | 0.00046 | 0.0007 | 0.00024 | 5.78E-05 | 0 | 5.17E-05 | 0.00011 |
| Slc2a13 | 8.36E-05 | 1.91E-05 | 7.09E-05 | 3.10E-06 | 3.62E-06 | 6.53E-05 | 0.00018 | 4.50E-06 | 0 | 1.73E-05 | 2.57E-06 | 0 | 3.34E-05 | 6.55E-05 | 8.86E-05 | 0 | 9.84E-05 | 6.75E-05 |
| Klhl28 | 5.27E-05 | 7.17E-05 | 3.59E-05 | 3.02E-05 | 4.32E-05 | 6.52E-05 | 3.98E-05 | 2.25E-05 | 0 | 2.53E-05 | 1.52E-05 | 8.95E-05 | 5.30E-06 | 1.15E-05 | 2.10E-05 | 3.43E-05 | 0 | 1.43E-05 |
| Hoxb2 | 2.60E-05 | 1.03E-05 | 0.00017 | 0.00022 | 0.00011 | 6.51E-05 | 8.02E-05 | 0.00024 | 0 | 8.55E-06 | 0.00018 | 0 | 0 | 0 | 0 | 0.00011 | 3.67E-05 | 0 |

| | | | | | | | | | | | | | | | | | | |
|---|---|---|---|---|---|---|---|---|---|---|---|---|---|---|---|---|---|---|
| C2cd2l | 2.62E-05 | 7.17E-05 | 0.00014 | 1.10E-05 | 7.19E-05 | 6.49E-05 | 4.91E-05 | 8.16E-05 | 0 | 3.38E-05 | 9.30E-05 | 4.15E-05 | 6.74E-06 | 5.04E-05 | 8.82E-05 | 0.00013 | 0.00011 | 4.15E-05 |
| Jam2 | 0 | 1.66E-05 | 0 | 0 | 0 | 6.48E-05 | 0 | 0 | 0 | 0.00019 | 0 | 5.20E-05 | 0.0002 | 4.37E-05 | 0 | 0 | 0 | 0.00027 |
| Slpi | 0 | 0 | 7.32E-05 | 0 | 0 | 6.47E-05 | 3.49E-05 | 0 | 0 | 0 | 6.78E-06 | 0 | 0.00039 | 0.00019 | 0 | 3.90E-05 | 1.74E-05 | 0.00011 |
| Zfp800 | 8.95E-05 | 6.37E-05 | 6.15E-05 | 9.24E-06 | 3.32E-05 | 6.45E-05 | 2.12E-05 | 7.83E-06 | 0 | 1.77E-05 | 4.20E-06 | 3.87E-05 | 0 | 2.59E-05 | 6.77E-05 | 1.49E-05 | 3.29E-05 | 1.81E-05 |
| Tmem251 | 7.04E-05 | 8.24E-05 | 3.55E-05 | 0 | 3.50E-05 | 6.44E-05 | 1.89E-05 | 0 | 0 | 1.84E-06 | 0 | 2.10E-05 | 0 | 3.92E-05 | 0 | 3.29E-05 | 0.00018 | 7.44E-06 |
| Guk1 | 0.00018 | 8.19E-05 | 0.00018 | 0.00021 | 2.62E-05 | 6.42E-05 | 0.00026 | 0 | 0 | 0.0001 | 0.00013 | 2.20E-05 | 4.15E-05 | 5.01E-05 | 0 | 0 | 1.54E-05 | 0.00011 |
| Ccl11 | 1.58E-05 | 0 | 0.00023 | 0 | 0 | 6.38E-05 | 0 | 0.00077 | 0 | 0.00085 | 0 | 0 | 0.00085 | 0 | 0 | 0 | 0 | 0.00028 |
| Plekhb2 | 0 | 7.05E-05 | 4.81E-05 | 0.0001 | 0.00013 | 6.36E-05 | 4.40E-05 | 5.13E-05 | 0 | 9.89E-05 | 0.00027 | 0.0003 | 0 | 0 | 1.99E-05 | 4.90E-05 | 0 | 0 |
| Fam193b | 3.58E-05 | 0.00025 | 1.74E-06 | 5.80E-05 | 0.00013 | 6.34E-05 | 7.83E-05 | 4.89E-05 | 0 | 6.23E-05 | 2.87E-05 | 9.37E-05 | 0 | 6.57E-05 | 0.00051 | 6.13E-05 | 0.00017 | 7.02E-05 |
| Med27 | 8.59E-05 | 4.47E-05 | 0.00012 | 6.18E-05 | 2.55E-05 | 6.27E-05 | 0 | 0 | 0 | 1.75E-05 | 5.12E-05 | 0 | 2.65E-05 | 4.82E-05 | 8.85E-05 | 4.91E-05 | 2.19E-05 | 5.34E-05 |
| Prps2 | 4.63E-05 | 4.67E-05 | 3.48E-05 | 0 | 6.89E-05 | 6.24E-05 | 0 | 0 | 0 | 3.76E-05 | 0 | 0 | 3.89E-05 | 1.38E-06 | 0 | 9.06E-05 | 0 | 4.98E-05 |
| Zfp346 | 2.55E-05 | 8.52E-05 | 3.32E-05 | 1.67E-05 | 4.89E-05 | 6.21E-05 | 7.32E-06 | 0 | 0 | 2.57E-05 | 8.61E-06 | 1.26E-05 | 2.07E-05 | 4.63E-05 | 6.57E-06 | 5.70E-05 | 0 | 0 |
| Rflnb | 0 | 3.33E-05 | 0.00013 | 0.0001 | 0.00033 | 6.20E-05 | 0.00022 | 0.00019 | 0 | 0 | 0.00024 | 0 | 0 | 0.00021 | 2.23E-05 | 5.20E-05 | 0 | 3.52E-05 |
| Igsf3 | 0 | 0.00012 | 0.00011 | 0 | 0.0002 | 6.18E-05 | 8.92E-05 | 0.00058 | 0 | 0 | 7.56E-05 | 0.00012 | 0 | 0 | 4.70E-05 | 0 | 8.84E-06 | 0.00017 |
| Rin3 | 0 | 2.09E-05 | 0 | 0 | 7.62E-06 | 6.16E-05 | 0 | 8.48E-05 | 0 | 9.89E-05 | 0.00011 | 0 | 0 | 8.03E-05 | 0.00013 | 7.66E-05 | 0 | 0 |
| Garem1 | 9.67E-06 | 4.40E-05 | 2.15E-05 | 0 | 7.27E-05 | 6.16E-05 | 5.39E-05 | 0 | 0 | 6.64E-06 | 4.66E-05 | 0 | 0 | 3.28E-05 | 3.60E-06 | 9.82E-06 | 5.96E-05 | 4.35E-05 |
| Fhod1 | 0 | 6.89E-05 | 2.19E-05 | 7.11E-05 | 4.00E-05 | 6.09E-05 | 0 | 2.53E-05 | 0 | 2.12E-06 | 0.00011 | 1.20E-06 | 0 | 0.00016 | 0.0001 | 4.88E-05 | 0 | 8.04E-05 |
| Tmem204 | 0 | 0.00018 | 0 | 0 | 0.00028 | 6.07E-05 | 0 | 0.00042 | 0 | 0 | 7.90E-05 | 2.17E-05 | 0 | 0.00067 | 0 | 0 | 0 | 0.00012 |
| Gucy1a2 | 3.99E-06 | 0 | 0 | 0 | 0 | 6.05E-05 | 0 | 8.63E-05 | 0 | 0 | 0 | 0 | 0 | 0 | 0 | 8.45E-06 | 0 | 9.54E-05 |
| Il11 | 0 | 0 | 0.0002 | 0 | 0 | 6.04E-05 | 0 | 3.10E-05 | 0 | 0 | 6.71E-05 | 0.00017 | 0 | 0 | 8.41E-06 | 0 | 0.00013 | 0 |
| Acvr2b | 9.19E-07 | 3.42E-05 | 0 | 0 | 2.07E-05 | 6.02E-05 | 0 | 2.15E-05 | 0 | 0 | 0 | 3.85E-05 | 0 | 3.54E-05 | 6.00E-05 | 0 | 4.01E-06 | 0 |
| Upf2 | 0.00011 | 8.53E-05 | 1.37E-05 | 0 | 3.17E-05 | 5.97E-05 | 6.38E-05 | 0 | 0 | 8.18E-06 | 2.66E-05 | 8.22E-05 | 4.19E-05 | 2.42E-05 | 0.00021 | 3.20E-05 | 7.53E-05 | 0 |
| Abcg1 | 0 | 2.75E-05 | 1.88E-05 | 0 | 0.00014 | 5.95E-05 | 0 | 3.65E-05 | 0 | 1.27E-05 | 3.11E-05 | 4.82E-05 | 0 | 0.00015 | 2.30E-05 | 0 | 6.11E-06 | 2.87E-05 |
| Irak1bp1 | 5.02E-05 | 6.70E-05 | 0 | 3.96E-05 | 0.00011 | 5.95E-05 | 0 | 0 | 0 | 1.10E-05 | 5.07E-05 | 0 | 1.99E-05 | 0.00012 | 0 | 1.28E-05 | 0 | 3.11E-05 |
| Ak5 | 0 | 0 | 0 | 0 | 0.00013 | 5.94E-05 | 0 | 0 | 0 | 0 | 0.00028 | 0 | 2.90E-05 | 0.00012 | 1.12E-05 | 0 | 0 | 0.00013 |
| Fos | 0 | 0 | 0 | 8.26E-05 | 3.02E-05 | 5.90E-05 | 0.00056 | 0.00017 | 0 | 0.00022 | 0 | 9.22E-05 | 0.00033 | 0 | 0 | 9.75E-05 | 0.01253 | 0.00019 |
| Pithd1 | 8.34E-05 | 6.12E-05 | 6.01E-05 | 7.16E-05 | 4.99E-05 | 5.88E-05 | 7.18E-05 | 2.37E-05 | 0 | 0 | 2.76E-05 | 4.11E-05 | 0 | 6.45E-05 | 0 | 0.00018 | 1.13E-05 | 0 |
| Ccdc163 | 1.14E-05 | 5.15E-05 | 0 | 1.21E-05 | 4.22E-05 | 5.87E-05 | 6.50E-06 | 3.54E-05 | 0 | 1.92E-05 | 0 | 9.79E-05 | 6.08E-05 | 2.40E-05 | 6.82E-05 | 9.27E-05 | 5.57E-05 | 0 |
| Ackr3 | 0 | 0 | 0 | 0 | 0.00012 | 5.82E-05 | 0.00018 | 0 | 0 | 0.00096 | 0 | 0.0004 | 0.00035 | 0 | 0 | 1.95E-05 | 0.00013 | 0 |
| Pcdhga7 | 2.08E-05 | 7.21E-05 | 0 | 0 | 8.78E-05 | 5.79E-05 | 4.26E-05 | 3.64E-05 | 0 | 3.32E-05 | 2.53E-05 | 2.24E-05 | 1.41E-05 | 1.94E-05 | 0.00016 | 0 | 6.71E-05 | 2.85E-05 |
| Acot13 | 0.00011 | 8.51E-05 | 0 | 7.31E-05 | 4.27E-05 | 5.79E-05 | 8.95E-05 | 1.18E-05 | 0 | 7.35E-06 | 0 | 7.06E-05 | 2.81E-05 | 5.29E-05 | 0 | 7.56E-06 | 0 | 4.50E-05 |
| Pycr1 | 0.00019 | 0 | 1.38E-05 | 0.0003 | 6.40E-05 | 5.78E-05 | 0.0009 | 0.00013 | 0 | 3.28E-05 | 0.0001 | 0.0001 | 0 | 3.17E-06 | 9.41E-05 | 0 | 0 | 0.00016 |
| Gpr146 | 0 | 4.02E-05 | 0 | 0 | 1.82E-05 | 5.78E-05 | 0 | 3.86E-05 | 0 | 1.43E-05 | 0 | 0.00013 | 1.92E-05 | 0 | 1.02E-05 | 0 | 0 | 4.39E-05 |

| | | | | | | | | | | | | | | | | | | |
|---|---|---|---|---|---|---|---|---|---|---|---|---|---|---|---|---|---|---|
| Kalrn | 0 | 6.33E-05 | 7.62E-05 | 0 | 0.00043 | 5.76E-05 | 0 | 2.80E-05 | 0 | 0 | 0.00046 | 8.91E-05 | 0 | 0.00034 | 0.00038 | 0 | 0.00018 | 6.25E-05 |
| Angpt2 | 0 | 1.03E-05 | 0 | 0 | 0.0001 | 5.69E-05 | 0 | 0 | 0 | 0 | 0 | 0 | 0 | 3.36E-05 | 0 | 0 | 0 | 0 |
| Fcho2 | 8.80E-05 | 0.00011 | 0 | 0 | 2.07E-05 | 5.69E-05 | 0 | 0 | 0 | 0 | 2.10E-08 | 8.02E-05 | 0 | 5.15E-06 | 0.00022 | 2.82E-05 | 4.89E-05 | 0 |
| Dysf | 0 | 0 | 0 | 0.00012 | 5.62E-05 | 5.67E-05 | 6.04E-05 | 3.60E-06 | 0 | 0 | 0.00036 | 6.93E-06 | 6.57E-05 | 8.55E-05 | 1.65E-05 | 0 | 4.12E-06 | 0 |
| Slc38a7 | 0 | 0.00012 | 1.80E-06 | 3.87E-05 | 5.96E-05 | 5.64E-05 | 1.60E-05 | 0 | 0 | 2.21E-05 | 3.03E-05 | 2.65E-05 | 4.53E-06 | 6.43E-05 | 3.47E-05 | 0 | 0 | 1.84E-05 |
| Hmgxb4 | 5.49E-05 | 4.62E-05 | 4.74E-05 | 0 | 5.82E-05 | 5.63E-05 | 8.96E-05 | 5.80E-05 | 0 | 1.77E-05 | 4.50E-05 | 3.67E-05 | 1.34E-05 | 1.33E-05 | 1.75E-05 | 0.00017 | 5.82E-05 | 5.66E-05 |
| Uap1 | 0.00024 | 4.48E-05 | 0.00051 | 0 | 7.33E-05 | 5.61E-05 | 0.0003 | 0 | 0 | 0.0009 | 7.32E-05 | 0.00012 | 0.00012 | 3.49E-05 | 0.00014 | 8.50E-05 | 0.00055 | 0.00011 |
| Spn | 0 | 0 | 0 | 0 | 0.0001 | 5.57E-05 | 0 | 0.00017 | 0 | 0 | 0.0001 | 3.80E-05 | 0 | 0 | 2.81E-05 | 0 | 0 | 0 |
| Rnf135 | 2.48E-05 | 6.72E-05 | 0 | 5.14E-05 | 8.03E-05 | 5.57E-05 | 0 | 0.00012 | 0 | 4.10E-05 | 0 | 4.35E-06 | 0 | 8.84E-05 | 5.21E-06 | 0 | 0 | 1.87E-05 |
| Sipa1l2 | 0 | 7.90E-05 | 0.0001 | 0 | 0 | 5.56E-05 | 0 | 0.00012 | 0 | 0 | 2.33E-05 | 0.00023 | 0.00011 | 5.71E-05 | 4.60E-05 | 5.83E-05 | 0 | 3.46E-05 |
| Hcfc2 | 0.00015 | 5.66E-05 | 7.24E-05 | 0 | 6.25E-05 | 5.54E-05 | 6.48E-05 | 0 | 0 | 5.52E-05 | 2.48E-05 | 0.00012 | 0 | 2.79E-05 | 0.0001 | 4.74E-05 | 0 | 2.89E-06 |
| Ptpre | 0 | 2.82E-05 | 4.41E-06 | 0 | 3.71E-05 | 5.50E-05 | 0 | 6.68E-05 | 0 | 0 | 0 | 8.30E-05 | 0 | 4.36E-05 | 0.0001 | 0 | 0 | 0 |
| Inpp5k | 2.32E-05 | 6.43E-05 | 0 | 1.19E-06 | 3.55E-05 | 5.47E-05 | 3.15E-05 | 0 | 0 | 3.68E-05 | 2.77E-05 | 4.43E-05 | 1.75E-05 | 1.54E-05 | 0 | 0 | 1.23E-05 | 4.87E-05 |
| Cnnm3 | 5.38E-06 | 0.00011 | 1.88E-07 | 7.87E-06 | 6.40E-05 | 5.47E-05 | 3.15E-05 | 3.53E-05 | 0 | 6.63E-05 | 2.43E-05 | 6.91E-05 | 4.65E-05 | 5.21E-05 | 6.32E-05 | 5.43E-05 | 0 | 6.12E-05 |
| Mylip | 0 | 0.00018 | 0 | 5.63E-05 | 5.02E-05 | 5.47E-05 | 0 | 0 | 0 | 0.00024 | 2.44E-05 | 0 | 8.14E-05 | 0.00014 | 0 | 7.79E-06 | 0.0005 | 0 |
| Stk11ip | 2.49E-05 | 0.00011 | 1.30E-05 | 3.79E-05 | 4.18E-05 | 5.31E-05 | 4.20E-05 | 0 | 0 | 2.46E-05 | 1.17E-05 | 8.38E-06 | 1.86E-05 | 3.61E-05 | 0.00011 | 1.94E-05 | 1.16E-05 | 6.02E-05 |
| Il10 | 0 | 1.32E-06 | 0 | 0 | 0 | 5.29E-05 | 0 | 5.82E-06 | 0 | 0 | 0 | 7.33E-05 | 0 | 0 | 0 | 0 | 0 | 0 |
| Nrm | 0 | 8.36E-06 | 0 | 0.00015 | 9.33E-07 | 5.24E-05 | 5.09E-05 | 0 | 0 | 4.02E-05 | 4.53E-05 | 3.65E-05 | 0 | 5.42E-05 | 0 | 0.00048 | 0 | 2.40E-05 |
| Zfp992 | 4.23E-05 | 3.52E-05 | 0 | 0 | 1.29E-05 | 5.22E-05 | 1.98E-05 | 0 | 0 | 1.47E-05 | 0 | 3.48E-05 | 1.94E-05 | 0 | 0 | 0 | 0 | 2.30E-05 |
| Pla2g12a | 0.0001 | 2.36E-05 | 3.57E-05 | 0 | 1.72E-05 | 5.22E-05 | 0.00013 | 0 | 0 | 5.15E-06 | 0 | 0.00025 | 2.54E-06 | 1.42E-05 | 0 | 4.44E-06 | 0 | 3.02E-05 |
| Dnajb9 | 0.00021 | 0.00011 | 0 | 0 | 0 | 5.20E-05 | 0 | 0 | 0 | 6.36E-05 | 0 | 0.00042 | 0.00016 | 0 | 0 | 0 | 0.00125 | 1.10E-05 |
| Adamtsl4 | 0 | 0 | 3.06E-05 | 0 | 0 | 5.19E-05 | 0 | 0 | 0 | 0.00039 | 0 | 1.20E-05 | 0 | 3.74E-05 | 6.97E-05 | 0 | 0 | 0.00015 |
| Trib1 | 0 | 0 | 0.00018 | 0 | 3.63E-06 | 5.15E-05 | 0 | 3.81E-05 | 0 | 6.68E-05 | 3.24E-05 | 7.49E-05 | 0.0001 | 0.00017 | 0 | 0 | 0.00134 | 0 |
| Papss2 | 0.0002 | 0 | 0.00032 | 0.00013 | 5.34E-05 | 5.09E-05 | 0.00045 | 8.05E-05 | 0 | 2.83E-05 | 1.84E-05 | 0.00012 | 6.21E-05 | 2.48E-05 | 9.81E-06 | 4.15E-05 | 0 | 0.00013 |
| Mtf2 | 6.09E-05 | 6.06E-05 | 2.42E-05 | 0 | 2.56E-05 | 5.08E-05 | 0 | 3.06E-05 | 0 | 0 | 0 | 4.59E-05 | 2.03E-05 | 5.42E-05 | 8.94E-05 | 0.00012 | 0 | 0 |
| Slc8a1 | 7.52E-05 | 0 | 0 | 0 | 2.18E-05 | 5.07E-05 | 0.00025 | 0 | 0 | 0.00023 | 0.00051 | 0 | 0.00028 | 2.01E-05 | 9.72E-05 | 0.00015 | 0 | 0.00023 |
| Tfpi2 | 0.00021 | 0 | 0.00047 | 0 | 0.00056 | 5.06E-05 | 0.00021 | 0.00153 | 0 | 1.43E-05 | 0.00031 | 0 | 3.03E-05 | 0.00046 | 0 | 7.00E-06 | 0 | 8.99E-06 |
| Smim26 | 8.71E-05 | 4.59E-05 | 4.89E-05 | 0 | 3.80E-05 | 5.04E-05 | 0 | 9.39E-06 | 0 | 1.28E-05 | 1.26E-05 | 1.62E-05 | 0 | 1.67E-06 | 4.60E-06 | 2.79E-05 | 1.58E-05 | 0 |
| Jmy | 1.81E-05 | 0.00022 | 4.58E-05 | 0.00018 | 0.00017 | 4.94E-05 | 0.00024 | 6.64E-05 | 0 | 3.87E-05 | 0.00015 | 0.00032 | 6.29E-05 | 0.00018 | 0.0003 | 5.45E-05 | 5.68E-05 | 9.24E-05 |
| Lin54 | 1.40E-05 | 3.59E-05 | 2.19E-05 | 0 | 2.55E-05 | 4.92E-05 | 2.01E-05 | 0 | 0 | 1.31E-05 | 7.63E-06 | 4.21E-05 | 6.79E-07 | 6.09E-06 | 1.90E-05 | 0.00038 | 5.31E-05 | 0 |
| Rad18 | 0 | 0 | 6.00E-05 | 0 | 0 | 4.90E-05 | 3.58E-05 | 2.49E-05 | 0 | 1.49E-05 | 2.94E-05 | 5.94E-05 | 1.70E-05 | 8.26E-05 | 0 | 0.00063 | 0 | 0 |
| Zfp609 | 5.66E-05 | 0.00018 | 0.00012 | 2.81E-05 | 0.00014 | 4.90E-05 | 0 | 8.26E-05 | 0 | 0 | 2.55E-05 | 0.00013 | 2.44E-06 | 0.00011 | 0.0001 | 5.69E-05 | 0.00019 | 0 |
| Clk3 | 0.0001 | 0.00014 | 0.00012 | 6.73E-05 | 1.59E-05 | 4.89E-05 | 6.42E-05 | 0 | 0 | 3.36E-05 | 3.50E-05 | 0.00017 | 5.39E-06 | 2.56E-05 | 0.00029 | 0.00014 | 8.89E-05 | 5.00E-05 |

| | | | | | | | | | | | | | | | | | | |
|---|---|---|---|---|---|---|---|---|---|---|---|---|---|---|---|---|---|---|
| Rfesd | 5.94E-05 | 1.82E-05 | 0 | 0 | 1.45E-05 | 4.88E-05 | 0 | 2.29E-05 | 0 | 0 | 0 | 4.20E-05 | 0 | 0 | 0 | 8.25E-07 | 0 | 7.62E-06 |
| Plxdc1 | 0 | 0.00011 | 0 | 0 | 0 | 4.87E-05 | 0 | 7.17E-06 | 0 | 0 | 0 | 7.52E-05 | 0 | 3.70E-05 | 0 | 0 | 0 | 0 |
| Slc22a5 | 3.87E-05 | 7.25E-05 | 0 | 0 | 1.42E-05 | 4.85E-05 | 0 | 0 | 0 | 4.69E-05 | 3.19E-05 | 5.86E-05 | 3.35E-05 | 6.28E-06 | 2.06E-05 | 0 | 4.46E-05 | 3.90E-05 |
| Ppp3cc | 3.46E-05 | 4.45E-05 | 3.09E-05 | 0 | 2.33E-05 | 4.84E-05 | 0 | 9.07E-06 | 0 | 0 | 6.61E-06 | 5.27E-05 | 1.04E-05 | 1.93E-05 | 0 | 1.90E-05 | 8.70E-05 | 0 |
| Parp6 | 7.48E-05 | 8.41E-05 | 1.28E-05 | 0 | 4.51E-05 | 4.82E-05 | 1.01E-05 | 2.07E-05 | 0 | 4.36E-05 | 2.61E-05 | 0.00011 | 5.86E-06 | 1.68E-06 | 0.00015 | 0 | 0 | 5.15E-05 |
| Casp8 | 0.0001 | 2.61E-05 | 7.92E-05 | 0 | 4.47E-05 | 4.81E-05 | 4.97E-05 | 0.00021 | 0 | 4.59E-06 | 0 | 0 | 6.71E-05 | 0.00018 | 9.84E-05 | 9.67E-05 | 0 | 5.64E-05 |
| Slc31a2 | 4.76E-06 | 0.00011 | 0 | 0 | 5.14E-05 | 4.78E-05 | 0 | 1.15E-05 | 0 | 0.0002 | 3.75E-05 | 3.67E-05 | 0 | 7.32E-05 | 6.01E-06 | 0 | 0 | 6.27E-05 |
| Dzip1l | 1.16E-05 | 5.08E-05 | 5.59E-05 | 4.82E-05 | 0.00017 | 4.74E-05 | 0 | 5.44E-05 | 0 | 0 | 0.00024 | 0 | 4.60E-05 | 0.00016 | 0.00054 | 0.00015 | 0.00011 | 5.67E-05 |
| Wnt5b | 0 | 0 | 8.17E-06 | 0 | 2.61E-05 | 4.72E-05 | 4.07E-05 | 0 | 0 | 1.64E-05 | 9.72E-05 | 0 | 0 | 0.00018 | 1.29E-06 | 7.35E-05 | 0 | 0 |
| Angel2 | 8.71E-05 | 6.23E-05 | 6.50E-06 | 2.11E-05 | 2.68E-05 | 4.70E-05 | 6.47E-05 | 1.36E-05 | 0 | 9.19E-06 | 1.77E-05 | 4.18E-05 | 0 | 2.75E-05 | 4.03E-05 | 7.24E-05 | 0 | 2.03E-05 |
| Myo1e | 0 | 0.00016 | 3.64E-05 | 0 | 0.00029 | 4.68E-05 | 5.64E-05 | 2.11E-05 | 0 | 0 | 0.00035 | 0.00011 | 0 | 6.20E-05 | 2.47E-05 | 6.90E-05 | 1.68E-05 | 0.00021 |
| Atm | 6.79E-05 | 5.79E-05 | 0 | 0 | 0 | 4.68E-05 | 0 | 0 | 0 | 7.21E-06 | 0 | 2.09E-05 | 3.29E-06 | 0 | 4.64E-05 | 0.00012 | 8.12E-06 | 1.08E-05 |
| Scfd2 | 0.00011 | 3.06E-05 | 5.74E-05 | 3.31E-05 | 5.26E-05 | 4.67E-05 | 9.92E-05 | 0 | 0 | 4.53E-05 | 0 | 0 | 7.59E-05 | 2.66E-05 | 0.00018 | 0 | 3.01E-05 | 6.20E-05 |
| Wdr81 | 6.67E-05 | 0.00018 | 5.44E-06 | 3.68E-05 | 4.63E-05 | 4.64E-05 | 5.82E-05 | 4.18E-05 | 0 | 3.54E-08 | 0 | 0.00017 | 0 | 9.68E-05 | 6.49E-05 | 0 | 0 | 4.08E-05 |
| Borcs5 | 5.57E-05 | 4.13E-05 | 2.54E-05 | 1.61E-05 | 4.71E-05 | 4.61E-05 | 3.02E-05 | 0 | 0 | 7.29E-05 | 1.61E-05 | 0 | 0 | 6.84E-07 | 1.29E-05 | 2.59E-05 | 0 | 4.47E-05 |
| Unc13a | 0 | 0 | 0 | 0 | 7.10E-06 | 4.59E-05 | 0 | 2.50E-05 | 0 | 0 | 2.29E-06 | 0.00021 | 0 | 0 | 1.59E-05 | 0 | 0 | 0 |
| Rcsd1 | 0 | 2.15E-05 | 0 | 0 | 4.49E-06 | 4.59E-05 | 0 | 3.67E-05 | 0 | 0 | 1.93E-06 | 0 | 0 | 2.00E-05 | 0 | 0 | 8.36E-06 | 0 |
| Plxdc2 | 0.00012 | 0.0003 | 0 | 0 | 0.00011 | 4.58E-05 | 0 | 8.67E-05 | 0 | 0.00024 | 0 | 0 | 0.00028 | 0.00014 | 0 | 0 | 9.29E-06 | 0.00047 |
| Polb | 0.00014 | 5.25E-05 | 0.0001 | 0 | 4.22E-05 | 4.58E-05 | 0.00026 | 0 | 0 | 0.00024 | 0.00024 | 6.00E-05 | 0.00011 | 3.51E-05 | 2.53E-05 | 4.02E-05 | 0 | 0 |
| Ncapd3 | 3.22E-05 | 3.62E-05 | 0 | 0 | 1.75E-05 | 4.58E-05 | 0 | 6.71E-07 | 0 | 2.10E-06 | 0 | 2.56E-05 | 6.81E-06 | 1.88E-05 | 6.70E-05 | 0.00042 | 2.00E-06 | 0 |
| Stx3 | 1.30E-05 | 4.83E-05 | 2.62E-06 | 0 | 4.81E-05 | 4.56E-05 | 0 | 1.92E-06 | 0 | 0 | 3.40E-05 | 2.18E-05 | 0 | 4.80E-05 | 5.85E-05 | 0 | 0.00011 | 0 |
| Heca | 0 | 0.0003 | 1.40E-05 | 2.76E-05 | 6.64E-05 | 4.49E-05 | 0.00011 | 2.37E-05 | 0 | 0.00024 | 0.00011 | 0.0002 | 2.23E-05 | 0.00011 | 0.00026 | 0.00013 | 0.00028 | 0.00012 |
| Prkg2 | 1.66E-05 | 0 | 6.11E-05 | 0 | 0 | 4.48E-05 | 0 | 7.04E-05 | 0 | 1.05E-05 | 6.85E-06 | 0 | 0 | 5.61E-05 | 0 | 5.97E-05 | 0 | 0 |
| Zfp282 | 2.21E-05 | 7.29E-05 | 3.47E-05 | 4.08E-05 | 4.55E-05 | 4.40E-05 | 4.53E-05 | 1.92E-05 | 0 | 4.41E-05 | 3.41E-05 | 4.33E-07 | 1.46E-05 | 4.03E-05 | 6.86E-05 | 3.81E-05 | 8.03E-06 | 4.61E-05 |
| Polk | 3.94E-05 | 5.47E-05 | 0 | 2.25E-05 | 0.00012 | 4.39E-05 | 0 | 6.67E-05 | 0 | 1.25E-05 | 6.47E-05 | 6.55E-05 | 0 | 6.95E-05 | 9.45E-05 | 3.24E-05 | 0 | 4.01E-05 |
| Lrmda | 1.72E-05 | 1.94E-05 | 0 | 0 | 9.75E-06 | 4.39E-05 | 0 | 1.33E-05 | 0 | 8.88E-06 | 0 | 7.71E-05 | 6.29E-05 | 5.05E-05 | 4.07E-05 | 0 | 1.16E-05 | 0 |
| Trp53inp1 | 5.42E-06 | 0.00035 | 0 | 0 | 6.06E-05 | 4.36E-05 | 0 | 0 | 0 | 3.19E-05 | 0 | 2.83E-05 | 0 | 0 | 3.29E-05 | 0 | 2.14E-05 | 4.59E-05 |
| Stk17b | 0.00012 | 0 | 5.08E-05 | 0 | 4.09E-05 | 4.34E-05 | 2.46E-05 | 0.00021 | 0 | 0 | 6.69E-05 | 5.67E-05 | 0 | 7.66E-05 | 1.41E-06 | 9.02E-05 | 8.61E-05 | 5.01E-05 |
| Herpud1 | 0.00031 | 0.00021 | 0.00022 | 9.80E-05 | 4.21E-05 | 4.34E-05 | 0.0003 | 0 | 0 | 8.69E-05 | 7.33E-05 | 0.00038 | 0 | 0.00014 | 0.0002 | 0 | 0.00018 | 0.00017 |
| Nhlrc3 | 3.25E-05 | 3.23E-05 | 1.83E-05 | 0 | 1.13E-05 | 4.34E-05 | 2.19E-05 | 5.29E-06 | 0 | 4.47E-06 | 0 | 4.92E-05 | 0 | 7.40E-06 | 0 | 4.61E-06 | 0 | 6.34E-06 |
| Pbx3 | 2.33E-05 | 4.46E-06 | 1.56E-05 | 0 | 2.32E-05 | 4.33E-05 | 2.76E-05 | 3.53E-05 | 0 | 4.61E-06 | 2.05E-05 | 0 | 0 | 2.14E-05 | 7.87E-05 | 0.00016 | 4.40E-05 | 0 |
| Zc3h6 | 0 | 8.46E-05 | 0 | 2.68E-05 | 6.44E-05 | 4.33E-05 | 2.18E-05 | 1.49E-05 | 0 | 3.59E-05 | 1.48E-05 | 5.21E-05 | 2.52E-05 | 2.33E-05 | 0.00013 | 4.08E-05 | 5.04E-06 | 5.97E-05 |
| Tceal3 | 0 | 0 | 7.57E-07 | 9.80E-06 | 2.01E-05 | 4.33E-05 | 0 | 0 | 0 | 0 | 0 | 0 | 0 | 7.20E-05 | 3.60E-06 | 4.17E-06 | 0 | 0 |

| | | | | | | | | | | | | | | | | | | |
|---|---|---|---|---|---|---|---|---|---|---|---|---|---|---|---|---|---|---|
| Arid4a | 3.81E-05 | 6.72E-05 | 1.23E-05 | 0 | 8.54E-06 | 4.32E-05 | 0 | 3.72E-06 | 0 | 1.58E-05 | 3.15E-05 | 2.40E-05 | 0 | 0 | 5.28E-05 | 4.66E-05 | 0.00011 | 0 |
| Nfatc2ip | 1.53E-05 | 2.13E-05 | 2.36E-05 | 0 | 3.82E-05 | 4.31E-05 | 0 | 1.12E-05 | 0 | 3.94E-06 | 0 | 0 | 1.33E-05 | 3.39E-05 | 0 | 0.00019 | 1.65E-05 | 0 |
| Hax1 | 0.00017 | 0.00011 | 0.00012 | 5.52E-05 | 4.35E-05 | 4.29E-05 | 4.42E-05 | 0 | 0 | 4.12E-05 | 0.00012 | 0.0002 | 4.29E-05 | 0 | 4.98E-05 | 5.90E-05 | 4.85E-05 | 0 |
| Gmeb2 | 7.40E-05 | 9.71E-05 | 5.58E-05 | 0 | 1.53E-05 | 4.27E-05 | 0 | 1.16E-05 | 0 | 0 | 3.48E-05 | 7.72E-05 | 0 | 3.68E-05 | 0.00015 | 7.88E-05 | 9.14E-05 | 0 |
| Zfand2a | 4.88E-05 | 5.54E-05 | 5.53E-05 | 0 | 3.30E-05 | 4.27E-05 | 0 | 1.52E-05 | 0 | 6.56E-06 | 1.59E-05 | 6.09E-05 | 0 | 1.16E-05 | 2.27E-05 | 9.70E-05 | 7.65E-05 | 3.03E-05 |
| Flcn | 5.22E-05 | 0.00028 | 0 | 0 | 5.05E-05 | 4.25E-05 | 2.58E-05 | 1.63E-05 | 0 | 4.11E-05 | 0 | 6.67E-05 | 0 | 0.00011 | 0.00021 | 1.02E-05 | 0.00012 | 8.62E-05 |
| Ercc5 | 5.53E-05 | 6.52E-05 | 1.95E-05 | 1.39E-05 | 1.82E-05 | 4.25E-05 | 2.34E-05 | 0 | 0 | 3.50E-05 | 3.54E-06 | 1.78E-05 | 1.47E-05 | 0 | 4.67E-05 | 3.23E-05 | 0 | 6.94E-06 |
| Slc25a45 | 1.92E-05 | 5.33E-05 | 0 | 3.11E-05 | 2.44E-05 | 4.21E-05 | 2.02E-05 | 4.58E-05 | 0 | 5.33E-05 | 1.01E-05 | 7.00E-06 | 3.67E-05 | 1.04E-05 | 0 | 0 | 0 | 5.58E-05 |
| Gata6 | 0 | 0 | 3.20E-05 | 0 | 0 | 4.19E-05 | 6.39E-05 | 0 | 0 | 0.00011 | 0 | 0.00011 | 0 | 0 | 0 | 9.79E-06 | 8.21E-06 | 0.0002 |
| Ccdc93 | 7.44E-05 | 0.0001 | 3.73E-05 | 3.26E-05 | 7.39E-05 | 4.17E-05 | 7.74E-05 | 0 | 0 | 4.32E-05 | 1.08E-05 | 1.90E-05 | 3.78E-05 | 4.94E-05 | 6.92E-05 | 0 | 3.12E-05 | 5.20E-05 |
| Tfb2m | 0.00011 | 6.27E-05 | 4.66E-05 | 0 | 3.26E-05 | 4.16E-05 | 2.35E-05 | 0 | 0 | 0 | 8.43E-06 | 3.75E-05 | 0 | 0 | 3.91E-05 | 0 | 1.10E-05 | 6.16E-06 |
| Mpp6 | 8.46E-05 | 0 | 0.00016 | 4.31E-07 | 0.0001 | 4.13E-05 | 0.0003 | 0 | 0 | 0.00028 | 9.92E-05 | 0.00014 | 2.20E-05 | 2.26E-05 | 1.79E-05 | 0.00017 | 0 | 3.36E-05 |
| Med14 | 0.00016 | 4.64E-05 | 0.00011 | 0 | 5.94E-06 | 4.13E-05 | 0 | 5.92E-05 | 0 | 0 | 5.37E-05 | 7.96E-06 | 0 | 0 | 8.74E-05 | 0.00015 | 0.00013 | 6.12E-05 |
| Ints6l | 4.01E-05 | 3.81E-05 | 6.59E-06 | 0 | 1.41E-05 | 4.11E-05 | 0 | 1.20E-05 | 0 | 0 | 2.21E-05 | 2.84E-05 | 0 | 0 | 7.54E-05 | 8.53E-06 | 7.49E-05 | 5.64E-06 |
| Kdm3a | 7.52E-05 | 0.00023 | 0 | 0 | 7.43E-05 | 4.09E-05 | 2.63E-05 | 1.06E-05 | 0 | 6.61E-05 | 0 | 0.00047 | 2.54E-05 | 0 | 0.00035 | 7.32E-05 | 0.00024 | 0 |
| Rb1 | 3.18E-05 | 7.56E-05 | 0 | 0 | 3.37E-05 | 4.08E-05 | 0 | 0 | 0 | 8.84E-06 | 9.57E-06 | 5.59E-05 | 0 | 3.52E-05 | 2.90E-05 | 5.24E-05 | 2.79E-05 | 5.61E-05 |
| Unkl | 1.22E-05 | 5.95E-05 | 0 | 5.26E-05 | 3.49E-05 | 4.05E-05 | 0 | 9.50E-08 | 0 | 2.61E-05 | 4.72E-06 | 5.17E-05 | 2.91E-05 | 3.11E-05 | 8.47E-05 | 1.13E-05 | 5.63E-05 | 1.49E-05 |
| Klhl21 | 2.75E-05 | 0.0002 | 0.00013 | 0.00011 | 7.12E-05 | 4.04E-05 | 0 | 0 | 0 | 0 | 1.53E-05 | 0.00041 | 4.39E-05 | 0.00011 | 0.00011 | 0 | 0.00115 | 0 |
| Tmc6 | 5.77E-06 | 7.74E-05 | 4.18E-05 | 6.81E-05 | 8.07E-06 | 4.03E-05 | 0.00019 | 0.00015 | 0 | 6.08E-05 | 0 | 4.15E-05 | 8.19E-06 | 1.59E-05 | 0.00012 | 0 | 0 | 0.00013 |
| Slc23a2 | 3.64E-05 | 0.0001 | 7.45E-05 | 0 | 3.09E-05 | 3.99E-05 | 0 | 2.93E-05 | 0 | 6.41E-05 | 0 | 1.77E-05 | 8.55E-05 | 0.0001 | 5.97E-05 | 7.19E-06 | 4.75E-05 | 0.0001 |
| Gramd1b | 0 | 9.31E-05 | 1.65E-05 | 1.43E-06 | 0 | 3.96E-05 | 0 | 0 | 0 | 1.72E-05 | 0 | 0.00019 | 0 | 0 | 0 | 2.31E-05 | 4.45E-05 | 0 |
| Aktip | 8.84E-05 | 0.00013 | 1.43E-05 | 0 | 5.65E-05 | 3.95E-05 | 6.25E-05 | 1.05E-05 | 0 | 6.81E-05 | 9.02E-05 | 0.00011 | 1.86E-05 | 0 | 0 | 2.53E-05 | 3.70E-05 | 0.00011 |
| Sesn2 | 4.01E-06 | 9.14E-05 | 0.0001 | 3.02E-05 | 3.36E-05 | 3.95E-05 | 0 | 0 | 0 | 0 | 0 | 0.00016 | 3.89E-05 | 2.74E-05 | 3.28E-05 | 0 | 8.48E-05 | 0 |
| Dll1 | 0 | 2.23E-05 | 0 | 2.12E-06 | 8.36E-05 | 3.94E-05 | 0 | 0 | 0 | 0 | 0 | 0 | 0 | 0.00027 | 0 | 0 | 0 | 0 |
| Acbd5 | 0.00014 | 0.00015 | 2.90E-05 | 4.07E-05 | 4.01E-05 | 3.91E-05 | 5.40E-05 | 0 | 0 | 4.00E-05 | 4.88E-05 | 7.61E-05 | 0 | 6.91E-06 | 0.00012 | 1.97E-05 | 4.24E-05 | 9.03E-05 |
| Il17rd | 0 | 0.00012 | 0 | 0 | 6.00E-05 | 3.91E-05 | 0 | 1.79E-05 | 0 | 2.24E-05 | 3.07E-05 | 1.52E-05 | 0 | 4.52E-05 | 0 | 2.18E-05 | 1.13E-05 | 6.04E-05 |
| Cdc42bpa | 9.69E-05 | 0.00012 | 0.00025 | 0 | 0.00049 | 3.88E-05 | 0.00017 | 0.00023 | 0 | 6.20E-05 | 0.00075 | 0.00055 | 0 | 0.00032 | 0.00014 | 0.00011 | 0.00013 | 4.20E-05 |
| Ero1l | 0.00018 | 0.0001 | 0.00012 | 3.41E-05 | 0 | 3.87E-05 | 1.72E-05 | 0.0001 | 0 | 2.50E-05 | 2.78E-05 | 0.00213 | 6.37E-05 | 0 | 0.00017 | 4.30E-05 | 0.00021 | 5.48E-05 |
| Bmf | 0 | 0.00025 | 0 | 6.40E-05 | 0.00021 | 3.87E-05 | 0 | 0 | 0 | 8.20E-06 | 1.22E-06 | 0 | 0 | 0 | 5.21E-05 | 0 | 0 | 4.45E-05 |
| Zfp180 | 4.57E-05 | 8.62E-05 | 8.75E-05 | 3.53E-05 | 2.91E-05 | 3.87E-05 | 7.74E-06 | 0 | 0 | 7.22E-06 | 1.09E-05 | 0 | 6.95E-06 | 1.58E-05 | 0.0001 | 7.96E-05 | 5.85E-05 | 1.56E-05 |
| Pgm2l1 | 7.55E-06 | 6.94E-05 | 0 | 0 | 4.42E-05 | 3.86E-05 | 0 | 0 | 0 | 4.01E-06 | 0.00023 | 5.79E-05 | 0 | 1.82E-05 | 0 | 6.01E-05 | 1.32E-05 | 1.59E-06 |
| Chd7 | 0 | 1.87E-05 | 0 | 0 | 7.59E-06 | 3.85E-05 | 0 | 1.95E-05 | 0 | 0 | 0 | 7.94E-05 | 0 | 0.00013 | 3.51E-05 | 0 | 6.10E-05 | 0 |
| Oser1 | 0.00011 | 0.0002 | 4.52E-05 | 4.21E-05 | 3.00E-05 | 3.84E-05 | 6.73E-06 | 1.33E-05 | 0 | 0 | 2.72E-05 | 0.00026 | 0 | 5.97E-05 | 9.86E-05 | 0.00019 | 0.00041 | 6.31E-05 |

| | | | | | | | | | | | | | | | | | | |
|---|---|---|---|---|---|---|---|---|---|---|---|---|---|---|---|---|---|---|
| Pcdh15 | 0 | 3.52E-06 | 0 | 0 | 9.12E-05 | 3.84E-05 | 0 | 0 | 0 | 0 | 1.46E-05 | 8.53E-06 | 0 | 0 | 0 | 0 | 0 | 0 |
| Egr3 | 0 | 0 | 0.00032 | 0 | 0 | 3.84E-05 | 0 | 0.00064 | 0 | 0 | 0 | 0.00014 | 0 | 0 | 0 | 0 | 0.00213 | 0 |
| Cep97 | 1.74E-05 | 4.81E-05 | 0 | 0 | 2.75E-05 | 3.83E-05 | 0 | 1.06E-05 | 0 | 5.85E-06 | 1.05E-06 | 0 | 0 | 2.43E-05 | 0 | 3.12E-05 | 0 | 4.94E-06 |
| Tor3a | 6.10E-07 | 6.12E-05 | 0 | 9.32E-05 | 1.45E-05 | 3.80E-05 | 0 | 1.32E-05 | 0 | 0.00015 | 0 | 0 | 8.97E-05 | 2.29E-05 | 2.70E-05 | 6.19E-06 | 3.10E-05 | 0.00011 |
| Rufy2 | 6.52E-05 | 8.16E-05 | 8.62E-06 | 0 | 3.89E-05 | 3.79E-05 | 2.64E-05 | 0 | 0 | 1.97E-05 | 9.87E-06 | 2.80E-05 | 0 | 2.75E-05 | 0.00014 | 3.55E-06 | 0 | 0 |
| Pacs2 | 6.25E-06 | 7.82E-05 | 1.98E-05 | 2.34E-06 | 5.69E-05 | 3.78E-05 | 1.14E-05 | 3.83E-05 | 0 | 8.01E-05 | 2.00E-05 | 4.61E-05 | 0 | 1.64E-05 | 0 | 1.22E-05 | 5.65E-06 | 9.08E-05 |
| Cgrrf1 | 0.0001 | 7.30E-05 | 2.80E-05 | 1.19E-05 | 6.49E-05 | 3.75E-05 | 2.65E-05 | 8.05E-06 | 0 | 5.97E-05 | 3.73E-05 | 3.80E-05 | 1.54E-05 | 4.97E-05 | 2.53E-05 | 4.17E-05 | 8.46E-06 | 3.59E-05 |
| Elf4 | 0.00012 | 7.94E-05 | 0.00016 | 0 | 5.76E-05 | 3.71E-05 | 0.00024 | 0.0001 | 0 | 0.00011 | 6.54E-05 | 0.00011 | 0 | 0 | 0.00018 | 0.00012 | 0 | 2.94E-05 |
| P4ha2 | 0.00024 | 0 | 0.0003 | 0.00053 | 4.89E-05 | 3.68E-05 | 0.00119 | 0.00013 | 0 | 0.00017 | 0.00038 | 0.00169 | 0.00034 | 9.93E-05 | 0.00035 | 5.80E-05 | 5.60E-05 | 0 |
| Parp16 | 3.10E-05 | 2.49E-05 | 4.68E-05 | 2.96E-05 | 5.26E-05 | 3.67E-05 | 6.46E-05 | 2.71E-05 | 0 | 1.82E-05 | 4.96E-06 | 8.09E-05 | 0 | 3.45E-05 | 1.71E-05 | 0 | 1.93E-05 | 3.81E-05 |
| Hhipl1 | 0 | 0.0001 | 7.04E-05 | 0.00016 | 6.94E-05 | 3.67E-05 | 0.0005 | 0 | 0 | 7.68E-05 | 0.00054 | 0 | 0.00022 | 0.00012 | 0.00018 | 2.46E-05 | 2.90E-05 | 0.00019 |
| Rasgrf1 | 0 | 4.29E-05 | 0 | 0 | 0 | 3.64E-05 | 0 | 0.00012 | 0 | 0 | 0 | 9.60E-05 | 0 | 0 | 1.98E-05 | 0 | 0 | 0 |
| Asphd2 | 0 | 1.34E-05 | 0 | 7.21E-06 | 4.68E-05 | 3.62E-05 | 0 | 0 | 0 | 1.61E-05 | 0.0001 | 0 | 3.90E-06 | 0 | 1.63E-05 | 0 | 0 | 0 |
| Gpr135 | 3.41E-05 | 1.55E-05 | 1.48E-05 | 5.07E-05 | 3.55E-05 | 3.61E-05 | 0.0001 | 5.55E-06 | 0 | 8.54E-06 | 2.43E-05 | 2.89E-05 | 4.19E-05 | 1.55E-05 | 3.46E-05 | 0 | 1.08E-05 | 3.75E-05 |
| Otud7a | 0 | 1.18E-07 | 0 | 0 | 9.76E-06 | 3.60E-05 | 0 | 3.14E-05 | 0 | 0 | 1.85E-05 | 3.78E-05 | 0 | 0 | 1.21E-05 | 0 | 0 | 0 |
| Rmi1 | 1.77E-05 | 2.70E-05 | 2.51E-05 | 2.61E-05 | 2.96E-05 | 3.55E-05 | 2.75E-05 | 2.38E-05 | 0 | 2.27E-05 | 9.66E-06 | 3.33E-05 | 0 | 1.40E-05 | 1.34E-05 | 8.24E-05 | 0 | 3.82E-05 |
| Gab2 | 0 | 0.00016 | 4.57E-06 | 5.18E-05 | 7.36E-05 | 3.55E-05 | 0 | 0.00017 | 0 | 8.21E-05 | 8.94E-05 | 4.14E-05 | 0 | 4.24E-06 | 8.04E-05 | 1.07E-05 | 0.00021 | 0.00024 |
| Pcnt | 0 | 7.70E-05 | 1.39E-05 | 6.62E-05 | 3.48E-05 | 3.54E-05 | 2.85E-05 | 1.77E-05 | 0 | 4.37E-05 | 2.43E-05 | 4.03E-05 | 0 | 2.53E-05 | 0.00014 | 0.00038 | 7.03E-05 | 2.75E-05 |
| Cyp2d22 | 0 | 0.00013 | 0 | 0.00019 | 0.0002 | 3.51E-05 | 8.95E-05 | 0.00018 | 0 | 4.69E-05 | 0 | 4.87E-05 | 2.67E-05 | 0 | 0.00011 | 0 | 0 | 9.65E-05 |
| Mtmr1 | 3.22E-05 | 4.14E-05 | 0 | 0 | 4.78E-05 | 3.49E-05 | 4.40E-05 | 0 | 0 | 6.49E-05 | 0 | 3.87E-05 | 3.89E-05 | 0 | 0 | 0 | 5.38E-06 | 3.98E-05 |
| Tcf19 | 0 | 0 | 3.18E-05 | 3.91E-05 | 2.77E-05 | 3.49E-05 | 5.95E-06 | 5.29E-05 | 0 | 3.68E-05 | 0 | 0 | 0.00013 | 4.63E-05 | 0 | 0.00098 | 0 | 5.96E-05 |
| Sars | 4.26E-05 | 0.00019 | 0.0002 | 0.00033 | 1.89E-05 | 3.48E-05 | 3.30E-05 | 0.00013 | 0 | 4.99E-05 | 3.19E-05 | 0.00033 | 0 | 7.60E-05 | 0.00013 | 1.40E-06 | 0 | 7.43E-05 |
| Nfatc1 | 1.36E-05 | 1.31E-05 | 0.00024 | 0 | 1.82E-05 | 3.47E-05 | 0.00016 | 4.77E-06 | 0 | 0 | 0.00012 | 0.0001 | 0 | 0.0001 | 5.94E-05 | 3.75E-05 | 0.00029 | 0.00013 |
| Tacc2 | 1.80E-05 | 0.00012 | 0.0001 | 1.24E-05 | 8.48E-05 | 3.47E-05 | 0 | 8.29E-06 | 0 | 9.54E-05 | 0.00013 | 6.08E-05 | 0 | 6.69E-05 | 9.16E-05 | 3.92E-05 | 5.75E-06 | 4.00E-05 |
| Cnnm2 | 6.64E-06 | 0.0001 | 0 | 9.77E-06 | 9.65E-05 | 3.46E-05 | 1.89E-05 | 0 | 0 | 0 | 8.58E-05 | 4.45E-05 | 1.58E-05 | 1.58E-05 | 6.30E-05 | 1.30E-07 | 5.01E-05 | 2.96E-06 |
| Pcbd2 | 9.97E-05 | 4.05E-05 | 0 | 6.55E-05 | 4.52E-06 | 3.44E-05 | 2.92E-05 | 4.69E-05 | 0 | 2.85E-05 | 2.47E-07 | 2.38E-05 | 1.66E-05 | 4.03E-05 | 0 | 0.00012 | 0 | 6.45E-05 |
| Evi2a | 4.30E-05 | 0 | 0 | 2.17E-05 | 1.16E-05 | 3.43E-05 | 5.18E-05 | 5.71E-05 | 0 | 1.60E-05 | 0 | 3.30E-05 | 0 | 3.47E-06 | 0 | 0 | 0 | 0 |
| Slc9b2 | 0 | 0 | 0 | 0 | 0 | 3.43E-05 | 5.15E-05 | 0 | 0 | 0 | 0 | 0.00016 | 6.42E-05 | 0 | 0 | 0 | 0 | 1.04E-05 |
| Trim68 | 1.49E-05 | 4.29E-05 | 1.32E-05 | 2.36E-05 | 2.01E-05 | 3.42E-05 | 5.09E-05 | 3.76E-06 | 0 | 2.01E-05 | 4.79E-06 | 2.43E-05 | 1.78E-05 | 0 | 7.27E-05 | 3.03E-05 | 0 | 3.50E-05 |
| Daam2 | 0 | 2.33E-05 | 0 | 0 | 4.03E-05 | 3.41E-05 | 0 | 0 | 0 | 0 | 4.02E-05 | 0 | 0 | 4.50E-05 | 1.72E-05 | 0 | 0 | 0 |
| Eid2b | 2.92E-06 | 3.52E-05 | 0 | 4.57E-05 | 2.99E-05 | 3.39E-05 | 0 | 1.42E-05 | 0 | 1.33E-05 | 1.48E-05 | 1.20E-05 | 2.48E-05 | 1.69E-05 | 2.12E-05 | 5.42E-06 | 0 | 4.71E-06 |
| Synj2 | 4.55E-06 | 0.00011 | 8.60E-05 | 0 | 0.00017 | 3.38E-05 | 0.00013 | 0 | 0 | 1.74E-05 | 0.00021 | 0 | 6.37E-05 | 7.01E-05 | 0.00014 | 0 | 5.31E-05 | 0.00019 |
| Slx4ip | 2.99E-05 | 1.96E-05 | 1.06E-05 | 1.66E-05 | 1.25E-05 | 3.35E-05 | 1.50E-05 | 4.78E-05 | 0 | 9.88E-06 | 9.87E-06 | 1.88E-05 | 0 | 0 | 0 | 4.30E-05 | 4.65E-05 | 1.44E-05 |

| | | | | | | | | | | | | | | | | | | |
|---|---|---|---|---|---|---|---|---|---|---|---|---|---|---|---|---|---|---|
| Krt80 | 0 | 0 | 0 | 2.36E-05 | 0 | 3.33E-05 | 0 | 0 | 0 | 5.33E-05 | 6.10E-05 | 7.28E-05 | 0.00013 | 0 | 0 | 4.21E-05 | 0 | 0 |
| Aagab | 7.05E-05 | 0.00013 | 6.26E-05 | 8.14E-05 | 7.13E-05 | 3.33E-05 | 7.64E-05 | 6.79E-05 | 0 | 5.57E-05 | 0.00012 | 1.37E-05 | 3.95E-05 | 0 | 0.00013 | 8.14E-05 | 6.14E-06 | 4.84E-05 |
| Rufy3 | 4.18E-05 | 8.35E-05 | 8.91E-05 | 3.21E-05 | 4.11E-06 | 3.30E-05 | 2.10E-05 | 1.18E-05 | 0 | 2.18E-05 | 1.38E-05 | 2.71E-05 | 2.83E-05 | 0 | 0.00012 | 6.64E-05 | 2.07E-05 | 1.41E-05 |
| Nudt5 | 0.00018 | 6.84E-07 | 0.00015 | 0.00015 | 2.26E-05 | 3.28E-05 | 9.61E-05 | 2.93E-05 | 0 | 1.20E-05 | 0 | 3.65E-05 | 4.39E-05 | 1.28E-05 | 0 | 0.00026 | 3.83E-05 | 0 |
| Tctn3 | 0 | 5.40E-05 | 3.40E-05 | 8.59E-05 | 4.90E-05 | 3.28E-05 | 8.80E-05 | 4.36E-05 | 0 | 7.55E-05 | 0 | 0 | 3.41E-05 | 5.34E-05 | 7.70E-05 | 8.41E-05 | 0 | 1.52E-05 |
| Cklf | 0 | 2.45E-05 | 1.06E-05 | 3.10E-05 | 1.15E-05 | 3.28E-05 | 5.11E-06 | 2.65E-05 | 0 | 1.58E-05 | 3.19E-05 | 2.36E-05 | 9.89E-06 | 4.20E-06 | 1.24E-05 | 9.47E-05 | 1.42E-05 | 1.61E-05 |
| Lrrc28 | 3.57E-05 | 4.85E-05 | 0 | 6.57E-06 | 2.94E-05 | 3.26E-05 | 5.70E-05 | 1.55E-05 | 0 | 1.61E-05 | 9.18E-06 | 0 | 4.23E-06 | 3.73E-05 | 2.20E-06 | 1.11E-05 | 2.93E-05 | 2.15E-05 |
| Fbxo32 | 0 | 0.00012 | 0 | 0 | 0.0001 | 3.25E-05 | 0 | 0 | 0 | 0 | 0.00017 | 0.0002 | 0 | 0.00018 | 0 | 0 | 0 | 0 |
| Taf7 | 4.55E-05 | 4.26E-05 | 2.44E-06 | 0 | 5.68E-06 | 3.25E-05 | 0 | 2.10E-05 | 0 | 0 | 0 | 3.26E-05 | 0 | 0 | 2.11E-05 | 1.15E-05 | 0.00015 | 1.14E-05 |
| Zfp617 | 4.67E-05 | 2.70E-05 | 8.34E-06 | 0 | 3.62E-06 | 3.22E-05 | 0 | 9.33E-06 | 0 | 5.36E-06 | 1.59E-06 | 1.58E-05 | 0 | 1.79E-05 | 3.05E-05 | 1.30E-05 | 0 | 2.59E-06 |
| Zbtb25 | 7.70E-05 | 4.62E-05 | 8.85E-06 | 0 | 7.21E-05 | 3.22E-05 | 3.97E-05 | 5.85E-06 | 0 | 0 | 4.08E-05 | 4.11E-05 | 0 | 4.03E-06 | 0.00011 | 6.08E-05 | 0 | 3.47E-05 |
| Ccdc117 | 6.53E-05 | 0.00011 | 1.84E-05 | 3.76E-05 | 6.67E-05 | 3.20E-05 | 6.16E-05 | 1.83E-05 | 0 | 4.82E-07 | 5.63E-06 | 3.99E-05 | 0 | 3.53E-05 | 7.13E-05 | 6.21E-05 | 8.48E-05 | 8.95E-06 |
| Nup35 | 6.46E-05 | 4.30E-05 | 3.81E-05 | 0 | 1.82E-05 | 3.20E-05 | 0 | 0 | 0 | 2.48E-05 | 1.23E-05 | 2.99E-06 | 4.10E-06 | 0 | 1.90E-05 | 0.00021 | 0 | 2.15E-06 |
| Natd1 | 1.55E-06 | 7.69E-05 | 0 | 4.02E-05 | 6.86E-05 | 3.19E-05 | 2.29E-05 | 4.45E-05 | 0 | 2.91E-05 | 0 | 6.12E-05 | 1.02E-05 | 4.84E-05 | 0 | 3.50E-05 | 0 | 4.54E-05 |
| Nudcd1 | 0.00016 | 0 | 0.00016 | 3.37E-06 | 0 | 3.17E-05 | 8.49E-05 | 0 | 0 | 2.34E-06 | 0 | 0 | 0 | 0 | 0 | 0.00014 | 0.00012 | 0 |
| Mapk12 | 9.43E-06 | 2.81E-05 | 2.86E-05 | 1.62E-05 | 1.23E-05 | 3.17E-05 | 0.00017 | 0 | 0 | 9.94E-05 | 1.69E-05 | 0 | 5.70E-05 | 0 | 2.53E-05 | 2.07E-05 | 2.23E-06 | 0 |
| Rcan3 | 5.79E-06 | 5.27E-05 | 1.55E-05 | 5.93E-05 | 9.10E-05 | 3.17E-05 | 0.00012 | 2.05E-05 | 0 | 7.35E-05 | 5.94E-05 | 0.0001 | 0 | 8.47E-05 | 3.48E-05 | 7.51E-05 | 0 | 7.88E-05 |
| Itsn1 | 0.00024 | 0.00014 | 0 | 0 | 7.22E-05 | 3.17E-05 | 3.67E-05 | 0 | 0 | 0.00012 | 0.00013 | 5.92E-05 | 0.00039 | 1.18E-05 | 0.00032 | 9.70E-05 | 0.0002 | 0.00037 |
| Vsig10 | 0 | 9.05E-05 | 0 | 0 | 6.06E-05 | 3.16E-05 | 0 | 0 | 0 | 6.55E-05 | 6.02E-05 | 3.98E-05 | 3.75E-05 | 2.64E-05 | 3.93E-05 | 9.70E-05 | 0 | 2.07E-05 |
| Suco | 0.00011 | 8.56E-05 | 6.74E-05 | 0 | 2.82E-05 | 3.15E-05 | 3.85E-05 | 3.50E-07 | 0 | 9.44E-07 | 1.76E-05 | 0.00018 | 0 | 0 | 0.00015 | 0.00012 | 0.00011 | 0 |
| Cacng7 | 0 | 2.86E-05 | 7.06E-06 | 0 | 0.00023 | 3.14E-05 | 0.00015 | 0 | 0 | 1.76E-05 | 8.53E-05 | 3.47E-05 | 0.0001 | 0 | 0 | 0 | 5.41E-05 | 0 |
| Bace2 | 2.79E-05 | 0 | 4.51E-06 | 0 | 9.13E-06 | 3.08E-05 | 0.00011 | 0 | 0 | 5.70E-05 | 0 | 2.85E-06 | 0 | 8.58E-05 | 0 | 2.40E-05 | 0 | 0.00025 |
| Abtb1 | 0 | 0.00012 | 0 | 8.39E-05 | 5.64E-05 | 3.06E-05 | 0 | 0 | 0 | 4.68E-05 | 0 | 6.77E-05 | 0 | 7.17E-05 | 0.00015 | 0 | 0 | 4.43E-05 |
| Hectd2 | 2.89E-05 | 1.51E-05 | 4.96E-05 | 0 | 1.07E-05 | 3.05E-05 | 0 | 1.01E-05 | 0 | 0 | 1.62E-06 | 8.30E-05 | 0 | 0 | 3.62E-06 | 6.39E-06 | 0 | 3.10E-05 |
| Chchd1 | 0.0002 | 2.70E-05 | 0.0001 | 0.00011 | 1.22E-05 | 3.02E-05 | 3.60E-05 | 0 | 0 | 1.24E-05 | 3.46E-05 | 6.34E-06 | 2.36E-05 | 8.29E-06 | 0.00013 | 0.00011 | 0 | 0 |
| Fam126b | 2.36E-05 | 3.34E-05 | 2.61E-06 | 0 | 1.22E-05 | 3.01E-05 | 0 | 0 | 0 | 1.01E-05 | 4.08E-05 | 4.59E-05 | 0 | 1.03E-05 | 6.90E-05 | 2.16E-05 | 3.40E-05 | 1.80E-06 |
| Klc4 | 3.37E-05 | 9.35E-05 | 0 | 8.05E-05 | 6.71E-05 | 3.00E-05 | 6.86E-05 | 4.62E-05 | 0 | 1.28E-05 | 1.17E-05 | 3.35E-05 | 0 | 4.48E-05 | 0.00012 | 0 | 4.21E-05 | 2.18E-05 |
| Entpd2 | 0 | 6.07E-05 | 0 | 0 | 3.07E-08 | 2.97E-05 | 0 | 0.00018 | 0 | 0.00039 | 0 | 0 | 7.26E-05 | 0 | 0 | 0 | 0 | 0.00062 |
| Arhgap28 | 1.60E-05 | 1.37E-05 | 0 | 0 | 9.09E-06 | 2.96E-05 | 6.62E-05 | 0 | 0 | 3.35E-05 | 1.42E-05 | 4.18E-05 | 4.63E-05 | 2.38E-05 | 1.58E-07 | 1.44E-05 | 1.10E-05 | 3.63E-05 |
| Tifa | 1.82E-05 | 6.68E-06 | 0 | 0 | 2.95E-06 | 2.96E-05 | 0 | 5.49E-07 | 0 | 0 | 0 | 9.70E-06 | 3.75E-05 | 3.67E-05 | 0 | 0 | 4.13E-05 | 2.99E-05 |
| Vrk1 | 2.56E-05 | 2.16E-05 | 4.66E-05 | 0 | 1.39E-05 | 2.95E-05 | 4.15E-06 | 9.80E-08 | 0 | 1.70E-05 | 1.30E-05 | 1.46E-05 | 2.75E-05 | 3.17E-05 | 3.92E-05 | 0.00047 | 7.02E-06 | 3.67E-05 |
| Sec14l2 | 0 | 9.28E-06 | 2.81E-06 | 2.57E-06 | 0 | 2.95E-05 | 0 | 0 | 0 | 2.42E-06 | 1.90E-06 | 7.23E-06 | 0 | 4.52E-05 | 1.70E-05 | 0 | 0 | 2.61E-06 |
| Filip1 | 0 | 2.82E-06 | 0 | 0 | 0 | 2.89E-05 | 0 | 3.39E-05 | 0 | 6.32E-06 | 0 | 5.99E-05 | 0 | 0 | 0 | 0 | 0 | 1.60E-05 |

| | | | | | | | | | | | | | | | | | | |
|---|---|---|---|---|---|---|---|---|---|---|---|---|---|---|---|---|---|---|
| Golga4 | 0.00013 | 0.00017 | 0.00013 | 3.07E-05 | 0.00011 | 2.86E-05 | 0.00019 | 0 | 0 | 2.84E-05 | 4.73E-06 | 0.00017 | 0 | 3.58E-05 | 0.00043 | 0 | 0.0002 | 6.15E-05 |
| Mkrn2 | 3.74E-05 | 5.34E-05 | 3.08E-05 | 5.53E-05 | 3.12E-05 | 2.85E-05 | 1.36E-05 | 2.20E-05 | 0 | 2.39E-05 | 2.24E-05 | 4.54E-05 | 0 | 2.05E-05 | 0 | 4.40E-05 | 5.85E-06 | 2.83E-05 |
| Pcdhgb7 | 5.47E-06 | 1.22E-05 | 5.58E-06 | 6.35E-06 | 7.17E-06 | 2.85E-05 | 5.76E-06 | 1.90E-05 | 0 | 4.37E-06 | 1.98E-06 | 0 | 9.65E-06 | 1.56E-05 | 3.06E-06 | 0 | 7.25E-06 | 6.26E-06 |
| Ypel2 | 0 | 0.00011 | 0 | 0 | 8.03E-05 | 2.85E-05 | 0 | 0 | 0 | 3.82E-06 | 0 | 3.51E-05 | 0 | 2.81E-05 | 0 | 0 | 0 | 4.48E-06 |
| Arfgef2 | 2.39E-05 | 8.83E-05 | 1.21E-05 | 0.00011 | 3.88E-05 | 2.83E-05 | 4.31E-05 | 0 | 0 | 7.07E-05 | 4.89E-05 | 0 | 0.0001 | 4.82E-05 | 9.30E-05 | 0 | 0 | 1.14E-05 |
| Isoc2b | 2.37E-05 | 2.92E-05 | 0 | 4.32E-05 | 4.38E-05 | 2.81E-05 | 8.72E-05 | 5.16E-05 | 0 | 3.81E-05 | 0 | 7.03E-05 | 0 | 0 | 0 | 0 | 0 | 5.61E-05 |
| Nucb2 | 0.00028 | 0 | 8.01E-05 | 0.00023 | 0 | 2.79E-05 | 0.00041 | 0.00018 | 0 | 0.00029 | 0 | 0.00023 | 0.00026 | 0 | 0 | 2.96E-05 | 0 | 0.00028 |
| Pnpla2 | 0 | 0.00016 | 0 | 8.11E-05 | 6.07E-05 | 2.78E-05 | 0 | 1.49E-05 | 0 | 3.33E-05 | 5.49E-05 | 4.18E-05 | 0 | 1.28E-05 | 1.25E-05 | 0 | 6.85E-05 | 6.59E-05 |
| Trnp1 | 0 | 3.57E-05 | 0 | 2.17E-05 | 4.56E-05 | 2.78E-05 | 0 | 0 | 0 | 3.07E-05 | 2.35E-06 | 0 | 2.54E-05 | 1.94E-05 | 1.38E-05 | 0 | 0 | 1.83E-06 |
| Ankrd37 | 0 | 4.27E-05 | 0 | 0 | 1.96E-05 | 2.76E-05 | 0 | 0 | 0 | 0 | 0 | 0.00059 | 0 | 0 | 3.39E-05 | 0 | 0 | 0 |
| Tchh | 0 | 1.70E-07 | 0 | 2.88E-06 | 3.46E-06 | 2.76E-05 | 0 | 1.38E-05 | 0 | 0 | 0 | 6.27E-05 | 0 | 9.84E-06 | 3.81E-05 | 0 | 0 | 0 |
| Cldn34c2 | 1.36E-05 | 1.54E-05 | 0 | 0 | 1.62E-05 | 2.75E-05 | 0 | 3.95E-06 | 0 | 0 | 0 | 9.03E-06 | 0 | 0 | 0 | 3.94E-07 | 0 | 0 |
| Msantd2 | 6.75E-05 | 7.23E-05 | 3.53E-05 | 0 | 4.70E-05 | 2.74E-05 | 0 | 2.75E-05 | 0 | 2.32E-05 | 0 | 4.39E-05 | 0 | 0 | 0.00017 | 9.86E-05 | 0.00015 | 1.93E-05 |
| Alms1 | 2.71E-05 | 4.50E-05 | 0 | 0 | 6.18E-06 | 2.72E-05 | 2.28E-05 | 0 | 0 | 8.60E-06 | 6.74E-06 | 1.40E-05 | 5.43E-05 | 0 | 3.89E-05 | 8.07E-05 | 9.57E-06 | 0 |
| Dhfr | 2.76E-05 | 0 | 2.22E-05 | 0 | 0 | 2.69E-05 | 3.94E-06 | 0 | 0 | 0 | 0 | 1.10E-05 | 2.52E-05 | 8.07E-06 | 0 | 0.00058 | 4.46E-05 | 0 |
| Tmem107 | 2.12E-05 | 1.39E-05 | 1.07E-05 | 5.42E-05 | 0 | 2.69E-05 | 4.98E-05 | 4.88E-05 | 0 | 4.08E-05 | 7.88E-06 | 2.81E-05 | 0 | 0 | 0 | 0.00012 | 0 | 4.62E-05 |
| Hsf4 | 0 | 8.96E-06 | 2.90E-07 | 1.48E-05 | 9.90E-06 | 2.69E-05 | 6.32E-06 | 0 | 0 | 4.90E-07 | 1.83E-05 | 5.96E-05 | 3.04E-05 | 0 | 0 | 2.86E-06 | 3.05E-05 | 1.39E-05 |
| Batf | 2.43E-06 | 1.53E-06 | 0 | 3.16E-06 | 1.89E-05 | 2.67E-05 | 0 | 1.10E-05 | 0 | 0 | 0 | 5.80E-06 | 8.83E-06 | 3.63E-05 | 0 | 0 | 0 | 1.33E-07 |
| Kcnk6 | 3.02E-05 | 0 | 4.21E-05 | 7.98E-05 | 0 | 2.67E-05 | 0.00012 | 0 | 0 | 5.14E-05 | 0 | 1.96E-05 | 0 | 5.48E-05 | 2.67E-05 | 0 | 0 | 3.48E-05 |
| Sclt1 | 2.41E-05 | 2.13E-05 | 0 | 0 | 7.92E-06 | 2.67E-05 | 0 | 1.67E-05 | 0 | 8.81E-06 | 1.76E-05 | 7.26E-06 | 0 | 0 | 4.85E-05 | 0.00011 | 2.78E-05 | 5.32E-06 |
| Dusp18 | 4.84E-05 | 0 | 3.13E-05 | 6.31E-06 | 0 | 2.65E-05 | 2.36E-05 | 3.16E-05 | 0 | 1.83E-05 | 0 | 0 | 0 | 0 | 0 | 0 | 0.0002 | 3.62E-05 |
| Fas | 1.82E-05 | 3.15E-05 | 8.52E-05 | 1.38E-05 | 0 | 2.63E-05 | 2.58E-05 | 0 | 0 | 8.64E-05 | 4.87E-06 | 0 | 0.00014 | 0 | 1.41E-05 | 0 | 2.21E-05 | 0.00017 |
| Stxbp2 | 7.13E-06 | 1.44E-05 | 3.48E-07 | 7.40E-06 | 8.39E-06 | 2.63E-05 | 0 | 0 | 0 | 0 | 1.99E-05 | 5.95E-06 | 0 | 4.10E-05 | 4.64E-05 | 1.45E-06 | 9.49E-06 | 0 |
| Uimc1 | 5.89E-05 | 5.05E-05 | 1.96E-05 | 0 | 4.17E-05 | 2.62E-05 | 8.81E-06 | 0 | 0 | 1.05E-05 | 2.29E-06 | 1.79E-05 | 3.14E-05 | 3.26E-05 | 0.00012 | 7.44E-05 | 6.48E-05 | 2.11E-05 |
| Atad2 | 2.20E-05 | 0 | 4.56E-05 | 0 | 0 | 2.61E-05 | 1.23E-05 | 0 | 0 | 1.94E-06 | 4.67E-06 | 0 | 1.58E-05 | 1.93E-05 | 1.26E-05 | 0.00063 | 2.75E-05 | 2.62E-06 |
| Casp8ap2 | 7.17E-05 | 2.79E-05 | 3.94E-05 | 0 | 1.67E-05 | 2.60E-05 | 4.94E-06 | 0 | 0 | 9.90E-06 | 1.36E-05 | 0 | 1.77E-06 | 1.86E-05 | 4.37E-05 | 0.0001 | 0 | 4.22E-06 |
| Pygl | 0 | 0 | 0 | 0.00012 | 0 | 2.60E-05 | 0.00023 | 7.23E-05 | 0 | 0.00032 | 0 | 0.00063 | 0.00017 | 0 | 0 | 0 | 0 | 1.22E-05 |
| Tmem231 | 1.85E-05 | 2.33E-05 | 0 | 0 | 3.12E-05 | 2.58E-05 | 7.10E-06 | 0 | 0 | 1.90E-05 | 7.60E-06 | 1.41E-05 | 1.39E-05 | 1.07E-05 | 0 | 1.64E-05 | 0 | 1.07E-05 |
| Hoxb6 | 0 | 1.15E-05 | 3.30E-06 | 3.20E-05 | 2.16E-05 | 2.56E-05 | 3.27E-05 | 3.88E-05 | 0 | 2.14E-05 | 2.31E-05 | 0 | 9.63E-06 | 0 | 6.48E-05 | 2.98E-05 | 0 | 0 |
| Cdr2 | 7.08E-05 | 2.07E-05 | 0.00028 | 0.00012 | 1.59E-05 | 2.55E-05 | 0.00026 | 7.47E-05 | 0 | 4.32E-05 | 0.00018 | 0 | 1.04E-05 | 6.18E-05 | 4.45E-05 | 0.00016 | 0.00027 | 4.13E-05 |
| Fbxo17 | 3.81E-05 | 0 | 8.40E-05 | 0.00023 | 1.38E-05 | 2.53E-05 | 0.00022 | 0 | 0 | 7.78E-05 | 8.52E-05 | 0 | 9.07E-05 | 0.00015 | 2.61E-05 | 2.74E-05 | 0 | 2.47E-05 |
| Krt7 | 0 | 9.86E-06 | 0 | 6.29E-05 | 3.52E-05 | 2.51E-05 | 0 | 0 | 0 | 2.61E-05 | 6.07E-05 | 6.29E-05 | 0 | 0 | 0.00014 | 6.37E-06 | 4.95E-05 | 2.01E-05 |
| Aaas | 6.14E-05 | 3.67E-05 | 0.0001 | 5.56E-05 | 0 | 2.50E-05 | 2.59E-06 | 9.95E-06 | 0 | 1.92E-05 | 6.40E-06 | 0 | 2.85E-05 | 1.26E-05 | 6.97E-05 | 0.00058 | 1.62E-07 | 5.70E-07 |

| | | | | | | | | | | | | | | | | | | |
|---|---|---|---|---|---|---|---|---|---|---|---|---|---|---|---|---|---|---|
| Rgl3 | 0 | 2.29E-05 | 0 | 8.82E-06 | 1.55E-05 | 2.50E-05 | 2.57E-05 | 0 | 0 | 4.78E-06 | 0 | 2.04E-05 | 2.06E-05 | 0 | 5.50E-05 | 0 | 0 | 6.02E-06 |
| Gtf2ird2 | 2.48E-05 | 3.07E-05 | 3.03E-06 | 2.49E-05 | 8.05E-06 | 2.49E-05 | 3.26E-05 | 1.20E-05 | 0 | 2.57E-06 | 0 | 3.01E-06 | 6.37E-06 | 9.70E-06 | 5.85E-05 | 1.49E-05 | 0 | 7.87E-06 |
| Tceanc | 2.35E-05 | 1.44E-05 | 0 | 0 | 1.90E-05 | 2.47E-05 | 2.49E-06 | 0 | 0 | 6.69E-06 | 5.14E-06 | 1.08E-05 | 1.93E-05 | 0 | 0 | 8.70E-06 | 0 | 8.60E-06 |
| Adam19 | 0 | 9.34E-05 | 0.00019 | 9.55E-05 | 0.00054 | 2.46E-05 | 0.00047 | 0.00048 | 0 | 0.00029 | 0.00092 | 0.0005 | 0.00062 | 0.00028 | 0.00031 | 0 | 0.00016 | 0.00011 |
| Gpr68 | 0 | 8.41E-06 | 1.85E-05 | 0 | 0 | 2.46E-05 | 0 | 2.72E-05 | 0 | 0 | 0 | 2.15E-05 | 0 | 1.35E-05 | 0 | 0 | 0 | 1.60E-05 |
| Sowahc | 1.47E-05 | 6.12E-05 | 0 | 1.41E-05 | 6.86E-06 | 2.46E-05 | 0 | 2.89E-06 | 0 | 3.11E-05 | 0 | 4.06E-05 | 4.15E-05 | 1.26E-05 | 0 | 4.40E-05 | 0.00016 | 3.50E-05 |
| Srgap3 | 0 | 2.04E-05 | 1.26E-05 | 0 | 1.81E-05 | 2.42E-05 | 0 | 0 | 0 | 0 | 0.00011 | 0.0003 | 0 | 3.48E-05 | 1.24E-05 | 0 | 0 | 0 |
| Cmc4 | 1.82E-05 | 2.86E-05 | 3.19E-06 | 6.26E-06 | 2.35E-05 | 2.39E-05 | 2.80E-06 | 9.88E-06 | 0 | 1.41E-05 | 2.94E-06 | 4.89E-06 | 1.09E-06 | 6.10E-06 | 4.61E-05 | 4.20E-05 | 4.42E-06 | 2.51E-05 |
| Rfxap | 9.67E-05 | 8.33E-05 | 6.44E-05 | 2.87E-05 | 4.72E-05 | 2.38E-05 | 9.02E-05 | 2.68E-05 | 0 | 3.04E-05 | 1.81E-05 | 0.00013 | 0 | 1.98E-05 | 3.83E-05 | 4.82E-05 | 0 | 0 |
| Cdk5rap2 | 0 | 4.63E-05 | 0 | 1.62E-06 | 3.36E-05 | 2.36E-05 | 2.44E-05 | 1.14E-05 | 0 | 7.29E-05 | 6.86E-06 | 1.33E-05 | 2.09E-06 | 3.04E-05 | 4.03E-05 | 0.00034 | 5.48E-05 | 5.88E-05 |
| Arntl | 1.15E-05 | 5.56E-05 | 6.99E-05 | 0 | 7.13E-05 | 2.36E-05 | 5.97E-05 | 2.34E-05 | 0 | 9.58E-05 | 8.32E-05 | 8.79E-05 | 3.00E-07 | 5.64E-06 | 7.84E-05 | 3.30E-05 | 0 | 0.00011 |
| Sema4g | 0 | 1.27E-05 | 0 | 0 | 5.24E-05 | 2.34E-05 | 0 | 0 | 0 | 0 | 2.01E-05 | 8.14E-05 | 0 | 0 | 1.46E-05 | 0 | 0 | 0 |
| Nmnat2 | 0 | 0 | 3.41E-05 | 3.15E-06 | 0 | 2.33E-05 | 0 | 4.95E-05 | 0 | 3.54E-05 | 0 | 1.29E-05 | 0 | 0 | 0 | 0 | 0 | 9.50E-06 |
| Rrm2b | 5.05E-05 | 2.76E-05 | 2.03E-06 | 0 | 0 | 2.33E-05 | 0 | 1.34E-06 | 0 | 1.49E-05 | 0 | 5.83E-05 | 0 | 0 | 0 | 3.70E-06 | 6.89E-05 | 6.44E-06 |
| Atp2b4 | 0 | 0 | 4.95E-05 | 0 | 0 | 2.32E-05 | 2.36E-07 | 0 | 0 | 0.0002 | 0.00013 | 0 | 0 | 0.00015 | 2.25E-05 | 2.31E-05 | 0 | 0.00022 |
| Klhdc1 | 9.08E-06 | 8.88E-06 | 0 | 0 | 0 | 2.32E-05 | 8.53E-06 | 0 | 0 | 7.76E-06 | 0 | 2.38E-05 | 1.09E-05 | 0 | 1.12E-05 | 1.25E-06 | 0 | 0 |
| Cd1d1 | 2.01E-05 | 2.68E-05 | 1.24E-05 | 0 | 0 | 2.32E-05 | 2.46E-06 | 0 | 0 | 0 | 0 | 0 | 1.48E-05 | 3.51E-05 | 0 | 1.14E-05 | 0 | 1.82E-07 |
| Madd | 2.49E-05 | 2.84E-05 | 0 | 2.35E-05 | 0 | 2.31E-05 | 1.08E-05 | 9.96E-06 | 0 | 1.21E-05 | 0 | 0 | 0 | 9.16E-05 | 3.07E-05 | 0 | 0 | 2.32E-05 |
| Haus3 | 5.88E-05 | 3.86E-05 | 1.69E-05 | 0 | 5.63E-06 | 2.31E-05 | 1.08E-05 | 6.63E-05 | 0 | 0 | 0 | 1.51E-05 | 0 | 0 | 0.00017 | 0.00018 | 6.38E-05 | 0 |
| Odf2l | 2.80E-05 | 4.07E-05 | 1.22E-05 | 1.59E-05 | 1.03E-05 | 2.29E-05 | 2.06E-05 | 4.76E-05 | 0 | 1.01E-05 | 2.28E-05 | 2.39E-05 | 5.00E-06 | 1.79E-05 | 0.00016 | 0.00019 | 0 | 0 |
| Atoh8 | 0 | 5.23E-06 | 0 | 2.55E-06 | 6.17E-06 | 2.28E-05 | 1.83E-05 | 0 | 0 | 7.03E-06 | 8.69E-05 | 1.22E-05 | 4.85E-05 | 0 | 2.52E-05 | 4.42E-05 | 0 | 0.00012 |
| Usp53 | 0 | 3.54E-05 | 0 | 0 | 5.23E-06 | 2.28E-05 | 0 | 0 | 0 | 1.99E-05 | 6.41E-05 | 0.00026 | 0 | 6.40E-05 | 0 | 0 | 7.02E-05 | 7.41E-05 |
| Gmds | 0.0001 | 8.12E-06 | 2.33E-05 | 0.00014 | 0 | 2.27E-05 | 8.27E-05 | 0 | 0 | 0.00015 | 3.14E-05 | 3.56E-06 | 0.00014 | 0 | 2.38E-05 | 0.00011 | 9.38E-05 | 5.24E-05 |
| Paip2b | 7.31E-05 | 5.37E-05 | 2.66E-05 | 4.41E-05 | 1.75E-05 | 2.26E-05 | 6.81E-07 | 0 | 0 | 1.49E-05 | 2.20E-06 | 0.0001 | 8.68E-06 | 1.27E-05 | 0 | 0.00015 | 4.18E-05 | 1.87E-05 |
| Ptpru | 9.37E-06 | 1.83E-06 | 5.60E-06 | 0 | 2.55E-05 | 2.26E-05 | 3.59E-05 | 1.14E-05 | 0 | 0 | 5.95E-06 | 7.62E-06 | 0 | 6.12E-05 | 5.54E-05 | 3.23E-05 | 0 | 0 |
| Nsun6 | 2.83E-05 | 2.18E-05 | 0 | 0 | 0 | 2.26E-05 | 1.42E-05 | 0 | 0 | 1.30E-05 | 3.88E-06 | 2.86E-05 | 7.46E-06 | 1.45E-05 | 3.48E-05 | 3.96E-05 | 1.18E-05 | 3.50E-06 |
| Adamts3 | 2.40E-05 | 0 | 2.17E-05 | 0 | 7.60E-05 | 2.26E-05 | 4.65E-05 | 0 | 0 | 2.88E-05 | 1.51E-06 | 0 | 0 | 0 | 2.61E-05 | 3.67E-05 | 0 | 0.00011 |
| Myo7a | 0 | 0 | 0.00016 | 3.59E-05 | 0.00034 | 2.25E-05 | 0.00065 | 0 | 0 | 0 | 0.00041 | 0 | 0.00046 | 0.00048 | 0.00025 | 1.65E-05 | 0 | 0.00023 |
| Muc13 | 0 | 0 | 0 | 0 | 6.93E-06 | 2.24E-05 | 4.39E-06 | 4.69E-05 | 0 | 2.10E-05 | 7.85E-05 | 4.76E-05 | 5.28E-05 | 0 | 0 | 1.07E-05 | 0.00013 | 0 |
| Tafa1 | 0 | 0 | 0 | 0 | 4.78E-05 | 2.23E-05 | 0 | 3.11E-06 | 0 | 0 | 0 | 2.71E-06 | 0 | 0 | 0 | 0 | 3.24E-06 | 3.13E-06 |
| F2rl3 | 0 | 0 | 0 | 0 | 2.43E-05 | 2.21E-05 | 0 | 0.0003 | 0 | 0 | 0 | 0 | 0 | 0 | 0 | 0 | 0 | 0 |
| Zfp524 | 4.32E-05 | 6.38E-05 | 0 | 6.90E-05 | 2.04E-05 | 2.20E-05 | 2.24E-05 | 2.10E-05 | 0 | 7.33E-06 | 0 | 5.76E-05 | 0 | 7.72E-06 | 0 | 1.92E-05 | 0 | 1.11E-05 |
| Zfp758 | 3.73E-05 | 2.54E-05 | 0 | 0 | 0 | 2.20E-05 | 4.06E-06 | 0 | 0 | 1.66E-07 | 0 | 5.08E-06 | 4.57E-07 | 7.30E-06 | 1.54E-05 | 0 | 0 | 0 |

| | | | | | | | | | | | | | | | | | | |
|---|---|---|---|---|---|---|---|---|---|---|---|---|---|---|---|---|---|---|
| Zfp984 | 2.48E-05 | 1.24E-05 | 0 | 0 | 0 | 2.19E-05 | 0 | 2.18E-06 | 0 | 1.84E-07 | 0 | 1.16E-05 | 0 | 0 | 0 | 6.81E-07 | 0 | 2.66E-06 |
| Eda2r | 0 | 8.96E-05 | 2.32E-05 | 0 | 0 | 2.18E-05 | 0 | 1.11E-05 | 0 | 2.17E-05 | 0 | 0 | 5.11E-06 | 0 | 0 | 0 | 6.06E-06 | 2.37E-05 |
| Haus6 | 1.32E-05 | 1.41E-05 | 6.27E-05 | 2.04E-06 | 0 | 2.18E-05 | 9.33E-06 | 1.78E-05 | 0 | 7.72E-06 | 3.22E-05 | 1.57E-05 | 1.50E-05 | 1.06E-05 | 0 | 0.00037 | 0 | 0 |
| Atr | 4.79E-05 | 3.19E-05 | 2.10E-05 | 0 | 2.87E-06 | 2.16E-05 | 0 | 0 | 0 | 4.56E-06 | 0 | 3.94E-05 | 1.22E-05 | 0 | 3.70E-05 | 7.38E-05 | 2.01E-05 | 4.70E-06 |
| Ogdhl | 0 | 2.21E-05 | 0 | 0 | 7.85E-05 | 2.15E-05 | 0 | 0 | 0 | 0 | 1.53E-05 | 0 | 0 | 4.45E-05 | 0 | 0 | 0 | 0 |
| Nim1k | 8.46E-06 | 1.12E-05 | 0 | 2.33E-06 | 0 | 2.13E-05 | 2.06E-05 | 0 | 0 | 1.25E-05 | 0 | 4.43E-06 | 1.61E-05 | 2.03E-05 | 6.27E-06 | 0 | 0 | 7.10E-06 |
| Fbxl7 | 2.44E-07 | 4.27E-05 | 9.30E-06 | 2.62E-06 | 1.20E-05 | 2.09E-05 | 2.35E-06 | 1.68E-05 | 0 | 3.66E-05 | 0 | 0 | 0 | 6.41E-05 | 3.39E-05 | 7.68E-06 | 0 | 2.95E-05 |
| Emsy | 0.00013 | 7.38E-05 | 5.26E-05 | 0 | 3.25E-05 | 2.09E-05 | 5.09E-05 | 1.71E-06 | 0 | 0 | 0 | 7.38E-05 | 0 | 2.51E-05 | 0.00022 | 0.00016 | 0 | 0 |
| Ube2ql1 | 0 | 5.93E-05 | 0 | 1.94E-05 | 0.00019 | 2.09E-05 | 0 | 6.59E-05 | 0 | 0 | 9.60E-05 | 6.61E-05 | 0 | 0 | 7.26E-05 | 4.34E-06 | 5.29E-05 | 0 |
| Hltf | 5.64E-05 | 4.57E-05 | 6.61E-06 | 1.00E-05 | 2.32E-05 | 2.09E-05 | 8.18E-05 | 1.73E-05 | 0 | 1.85E-05 | 0 | 1.26E-06 | 3.89E-05 | 0 | 0.00015 | 5.29E-05 | 0 | 3.15E-05 |
| Tmem132e | 3.75E-06 | 0 | 1.95E-05 | 0 | 8.35E-05 | 2.08E-05 | 8.21E-06 | 1.70E-05 | 0 | 0 | 6.72E-05 | 0 | 0 | 1.24E-05 | 5.83E-06 | 6.74E-06 | 0 | 0 |
| Mturn | 0 | 9.57E-05 | 0 | 0 | 3.66E-05 | 2.08E-05 | 0 | 0 | 0 | 0.00014 | 2.01E-05 | 6.67E-05 | 4.50E-05 | 3.44E-05 | 2.35E-05 | 0 | 0 | 0 |
| Tlcd1 | 3.83E-05 | 3.74E-05 | 0 | 2.01E-05 | 1.07E-06 | 2.07E-05 | 5.30E-05 | 0 | 0 | 3.90E-06 | 0 | 5.72E-06 | 1.65E-05 | 3.78E-06 | 2.92E-05 | 0 | 8.63E-06 | 2.38E-05 |
| Fam214b | 4.13E-07 | 0.00014 | 5.83E-05 | 8.85E-05 | 7.29E-05 | 2.07E-05 | 1.20E-05 | 1.80E-05 | 0 | 4.83E-05 | 8.90E-05 | 0.00011 | 0 | 6.20E-05 | 0.00011 | 0 | 6.86E-05 | 8.18E-05 |
| Nat9 | 5.86E-05 | 3.75E-05 | 1.08E-05 | 5.53E-05 | 1.95E-05 | 2.06E-05 | 4.61E-05 | 6.86E-05 | 0 | 3.02E-05 | 1.56E-05 | 3.03E-05 | 0 | 0 | 3.51E-05 | 0 | 0 | 4.73E-05 |
| Fam234b | 2.96E-05 | 1.37E-05 | 1.54E-05 | 2.86E-05 | 0.0001 | 2.06E-05 | 6.39E-06 | 5.65E-05 | 0 | 3.39E-05 | 2.41E-05 | 0 | 1.12E-05 | 1.37E-05 | 0 | 0 | 3.94E-05 | 1.56E-05 |
| Mrgpre | 0 | 2.95E-05 | 0 | 0 | 2.84E-06 | 2.05E-05 | 0 | 2.72E-05 | 0 | 2.00E-05 | 0 | 1.84E-05 | 0 | 1.08E-05 | 0 | 0 | 0 | 8.43E-06 |
| Stard6 | 8.51E-06 | 4.54E-06 | 0 | 0 | 4.81E-06 | 2.04E-05 | 0 | 8.79E-06 | 0 | 8.00E-06 | 0 | 3.92E-06 | 0 | 1.00E-05 | 0 | 1.24E-05 | 0 | 0 |
| Alyref2 | 8.44E-06 | 7.64E-06 | 8.93E-07 | 4.50E-05 | 1.96E-05 | 2.02E-05 | 2.84E-05 | 1.59E-05 | 0 | 9.69E-06 | 0 | 1.48E-07 | 0 | 4.49E-06 | 1.30E-05 | 1.07E-05 | 0 | 9.35E-06 |
| Cd244a | 0 | 0 | 3.51E-05 | 0 | 0 | 2.02E-05 | 0 | 9.86E-05 | 0 | 0 | 0 | 0 | 0 | 6.40E-05 | 1.56E-05 | 2.54E-05 | 0 | 0 |
| Gstm1 | 0 | 2.77E-05 | 0 | 0 | 0 | 2.01E-05 | 0 | 1.34E-05 | 0 | 0.0003 | 0 | 3.76E-05 | 0 | 1.32E-05 | 0 | 0 | 0 | 5.18E-05 |
| Smc2 | 2.60E-05 | 0 | 2.28E-05 | 0 | 0 | 1.99E-05 | 3.32E-06 | 2.17E-07 | 0 | 9.84E-06 | 0 | 0 | 0 | 0 | 0 | 0.00102 | 0 | 0 |
| Bnip3 | 0 | 2.57E-05 | 0 | 0 | 3.50E-05 | 1.99E-05 | 0 | 0 | 0 | 2.44E-05 | 1.24E-07 | 0.00072 | 5.06E-05 | 3.35E-06 | 0 | 0 | 0 | 0 |
| P2ry1 | 0 | 1.92E-05 | 0 | 0 | 1.57E-05 | 1.95E-05 | 0 | 4.67E-05 | 0 | 0 | 0 | 0 | 0 | 0 | 0 | 7.16E-06 | 4.75E-06 | 0 |
| Glp1r | 0 | 6.91E-06 | 1.03E-05 | 2.03E-05 | 0.00033 | 1.95E-05 | 0 | 0 | 0 | 0 | 0 | 0.00012 | 0 | 0 | 5.26E-05 | 0 | 0 | 1.25E-05 |
| Negr1 | 0 | 9.64E-06 | 0 | 0 | 3.10E-05 | 1.95E-05 | 0 | 0 | 0 | 0 | 2.78E-06 | 0 | 0 | 0 | 0 | 0 | 0 | 0 |
| Slc16a12 | 1.25E-05 | 2.59E-05 | 0 | 0 | 0 | 1.94E-05 | 0 | 3.92E-05 | 0 | 9.51E-05 | 0 | 0 | 6.30E-05 | 0 | 0 | 0 | 0 | 0 |
| Tstd3 | 9.06E-05 | 0 | 7.01E-06 | 1.69E-05 | 0 | 1.94E-05 | 2.30E-05 | 3.71E-05 | 0 | 3.61E-05 | 0 | 5.64E-06 | 1.19E-05 | 0 | 0 | 8.10E-06 | 0 | 0 |
| Tnfsf13b | 0 | 0 | 0 | 0 | 0 | 1.93E-05 | 0 | 0 | 0 | 8.74E-05 | 0 | 4.21E-05 | 8.25E-05 | 0 | 0 | 0 | 0 | 0.00044 |
| Snx30 | 6.14E-06 | 0.00011 | 6.49E-06 | 5.52E-05 | 8.24E-05 | 1.92E-05 | 0 | 0 | 0 | 1.81E-05 | 0.0003 | 6.03E-05 | 9.81E-05 | 2.88E-05 | 0 | 0 | 0.00012 | 1.85E-05 |
| Adcy1 | 0 | 0 | 0 | 0 | 1.57E-05 | 1.90E-05 | 0 | 5.46E-06 | 0 | 0 | 0.00014 | 2.63E-05 | 0 | 0 | 6.15E-05 | 0 | 0 | 0 |
| Cnih2 | 0 | 0 | 0 | 1.09E-06 | 1.54E-05 | 1.89E-05 | 0 | 1.54E-05 | 0 | 0 | 1.48E-05 | 9.93E-06 | 0 | 0 | 7.30E-06 | 1.87E-05 | 0 | 0 |
| Gpank1 | 1.58E-05 | 5.95E-05 | 2.75E-05 | 6.14E-05 | 1.76E-05 | 1.88E-05 | 0 | 4.57E-06 | 0 | 1.54E-05 | 4.42E-06 | 1.55E-05 | 0 | 2.94E-05 | 0 | 5.25E-05 | 0 | 2.41E-05 |

| | | | | | | | | | | | | | | | | | | |
|---|---|---|---|---|---|---|---|---|---|---|---|---|---|---|---|---|---|---|
| Nktr | 0.00025 | 0.00022 | 0.00013 | 0 | 6.19E-05 | 1.88E-05 | 0.0002 | 6.86E-06 | 0 | 2.29E-05 | 6.51E-06 | 0.0002 | 0 | 2.42E-05 | 0.00114 | 7.07E-05 | 0.00059 | 1.36E-05 |
| Ccpg1os | 3.37E-05 | 1.81E-05 | 0 | 1.70E-05 | 1.36E-06 | 1.86E-05 | 2.66E-05 | 1.46E-05 | 0 | 4.06E-06 | 0 | 1.52E-05 | 9.40E-06 | 2.87E-06 | 1.55E-06 | 2.39E-07 | 0 | 1.86E-05 |
| Kcnab1 | 0 | 0 | 0 | 0 | 1.75E-05 | 1.85E-05 | 0 | 0 | 0 | 0 | 4.10E-06 | 0 | 0 | 2.10E-05 | 0 | 0 | 0 | 2.07E-05 |
| Gng4 | 3.74E-06 | 6.25E-06 | 1.96E-05 | 0 | 0 | 1.85E-05 | 2.96E-05 | 0 | 0 | 1.47E-05 | 2.70E-06 | 5.65E-06 | 0 | 0 | 3.58E-07 | 0 | 2.35E-05 | 2.97E-06 |
| Crebl2 | 0 | 0.0001 | 0 | 3.06E-05 | 5.18E-05 | 1.85E-05 | 5.62E-06 | 0 | 0 | 8.16E-05 | 1.31E-05 | 7.84E-05 | 1.19E-05 | 6.95E-06 | 1.15E-05 | 0 | 4.30E-06 | 5.00E-05 |
| Gfod1 | 0 | 2.55E-05 | 2.70E-05 | 0 | 4.18E-05 | 1.84E-05 | 0 | 0 | 0 | 0 | 0.00014 | 5.20E-05 | 0 | 0 | 1.79E-05 | 0 | 0 | 0 |
| Sord | 0 | 5.47E-05 | 0 | 5.41E-06 | 3.28E-05 | 1.82E-05 | 0 | 0 | 0 | 5.10E-05 | 0 | 1.21E-05 | 2.01E-05 | 1.06E-05 | 0 | 3.52E-05 | 0 | 1.25E-05 |
| Tmem62 | 1.53E-05 | 3.10E-05 | 6.42E-06 | 0 | 0 | 1.82E-05 | 0 | 1.42E-05 | 0 | 7.53E-06 | 3.56E-06 | 5.65E-06 | 0 | 0 | 1.41E-06 | 0 | 0 | 2.09E-05 |
| Pex12 | 4.81E-05 | 3.68E-05 | 1.83E-05 | 4.78E-05 | 1.39E-06 | 1.82E-05 | 2.77E-05 | 0 | 0 | 4.81E-07 | 2.70E-06 | 1.73E-05 | 1.23E-05 | 0 | 0 | 2.41E-06 | 1.53E-05 | 7.07E-06 |
| Doc2g | 0 | 2.60E-05 | 7.79E-06 | 1.81E-05 | 2.67E-05 | 1.80E-05 | 0 | 1.36E-06 | 0 | 2.81E-06 | 3.93E-06 | 1.47E-05 | 1.54E-05 | 1.64E-05 | 5.39E-05 | 0 | 6.96E-07 | 0 |
| DEPRECATED | 2.40E-05 | 4.92E-07 | 0 | 6.08E-06 | 1.79E-05 | 1.80E-05 | 1.98E-05 | 1.30E-05 | 0 | 1.03E-05 | 5.90E-06 | 1.55E-05 | 3.90E-06 | 2.29E-05 | 0 | 2.79E-06 | 0 | 0 |
| Plekhg1 | 0 | 5.45E-05 | 0 | 0 | 2.29E-05 | 1.78E-05 | 1.70E-05 | 3.99E-05 | 0 | 0.00014 | 5.94E-05 | 4.10E-05 | 0 | 1.09E-05 | 0 | 0 | 4.49E-05 | 6.89E-05 |
| Cx3cl1 | 1.22E-06 | 0 | 0 | 0 | 0 | 1.78E-05 | 0 | 0.00024 | 0 | 0 | 0.00011 | 0 | 0 | 7.22E-05 | 0 | 5.10E-05 | 0 | 0.00023 |
| Gnao1 | 9.99E-07 | 1.97E-05 | 0 | 0 | 1.10E-06 | 1.77E-05 | 0 | 1.33E-05 | 0 | 0 | 2.80E-05 | 8.46E-06 | 0 | 2.78E-05 | 1.28E-05 | 0 | 1.51E-05 | 3.99E-05 |
| Pias4 | 2.92E-05 | 8.98E-05 | 3.98E-05 | 5.58E-05 | 1.80E-05 | 1.75E-05 | 2.28E-05 | 2.89E-06 | 0 | 6.01E-06 | 1.66E-05 | 6.18E-05 | 4.28E-06 | 1.56E-05 | 0.00011 | 5.58E-05 | 7.77E-06 | 1.81E-05 |
| Zfp946 | 3.54E-05 | 4.93E-08 | 0 | 7.67E-06 | 4.59E-06 | 1.75E-05 | 1.91E-05 | 0 | 0 | 1.33E-05 | 0 | 0 | 0 | 0 | 6.81E-06 | 4.84E-05 | 0 | 1.34E-05 |
| Map3k8 | 0 | 8.44E-05 | 0 | 0 | 2.11E-05 | 1.75E-05 | 0 | 0 | 0 | 0.00012 | 5.42E-05 | 1.62E-05 | 0.0002 | 0.00011 | 8.17E-05 | 0 | 8.44E-05 | 4.38E-06 |
| Nemp2 | 3.86E-06 | 6.78E-06 | 0 | 0 | 3.30E-06 | 1.75E-05 | 0 | 3.69E-05 | 0 | 1.36E-06 | 0 | 0 | 0 | 3.72E-06 | 6.30E-06 | 3.30E-05 | 0 | 8.93E-06 |
| Thpo | 0 | 5.46E-06 | 0 | 0 | 1.61E-05 | 1.74E-05 | 7.00E-06 | 0 | 0 | 1.11E-05 | 1.41E-06 | 1.42E-05 | 3.47E-06 | 2.71E-07 | 9.96E-06 | 0 | 0 | 2.35E-05 |
| Pcdhb15 | 1.04E-06 | 1.43E-05 | 2.72E-07 | 0 | 1.42E-05 | 1.74E-05 | 0 | 1.06E-05 | 0 | 4.04E-06 | 9.50E-06 | 5.65E-06 | 4.34E-06 | 0 | 7.89E-07 | 0 | 1.55E-05 | 4.71E-06 |
| Ccrl2 | 0 | 1.47E-05 | 1.53E-05 | 2.91E-05 | 4.65E-05 | 1.73E-05 | 0 | 5.47E-05 | 0 | 1.46E-05 | 0 | 2.64E-05 | 0 | 0 | 1.79E-05 | 2.38E-05 | 0.00011 | 2.00E-05 |
| Cenpq | 1.16E-05 | 2.66E-06 | 0 | 0 | 7.96E-06 | 1.73E-05 | 0 | 0 | 0 | 1.11E-06 | 0 | 1.58E-05 | 0 | 0 | 0 | 0.00029 | 0 | 0 |
| Pcdhga1 | 0 | 1.67E-05 | 0 | 1.15E-06 | 1.27E-05 | 1.73E-05 | 8.03E-07 | 1.96E-05 | 0 | 4.44E-06 | 0 | 1.25E-05 | 2.29E-05 | 3.74E-05 | 4.00E-05 | 0 | 0 | 3.21E-05 |
| Foxs1 | 0 | 0 | 2.43E-05 | 0 | 0 | 1.71E-05 | 0 | 0.00037 | 0 | 0 | 0.00018 | 2.86E-05 | 0 | 3.50E-06 | 1.28E-05 | 0 | 0 | 2.43E-05 |
| Hist4h4 | 0 | 1.84E-05 | 0 | 0 | 0 | 1.71E-05 | 0 | 0 | 0 | 2.43E-06 | 0 | 1.00E-05 | 0 | 0 | 7.50E-07 | 1.87E-05 | 9.12E-05 | 0 |
| Cxcl16 | 1.77E-05 | 0 | 0.00011 | 0 | 0.00017 | 1.70E-05 | 0 | 7.68E-05 | 0 | 0 | 0.00017 | 5.19E-06 | 0.00038 | 0.00019 | 0 | 0 | 2.05E-05 | 0.00033 |
| Pop5 | 4.89E-05 | 1.87E-05 | 3.93E-05 | 6.57E-05 | 4.58E-05 | 1.69E-05 | 0 | 1.52E-05 | 0 | 7.77E-06 | 5.31E-06 | 0 | 0 | 1.62E-05 | 0 | 4.24E-05 | 0 | 1.80E-05 |
| Eef1akmt4 | 9.83E-05 | 0 | 0.00018 | 0.0002 | 1.50E-05 | 1.68E-05 | 0.0001 | 0.00011 | 0 | 0 | 4.56E-05 | 0 | 1.56E-05 | 6.33E-05 | 0 | 0 | 2.12E-05 | 0 |
| Gpatch3 | 1.63E-05 | 4.67E-05 | 2.01E-05 | 3.07E-05 | 0 | 1.67E-05 | 0 | 0 | 0 | 0 | 2.84E-06 | 0 | 0 | 1.85E-05 | 2.63E-05 | 1.05E-05 | 2.96E-05 | 0 |
| Adap2 | 0 | 4.98E-05 | 0 | 0 | 4.42E-05 | 1.67E-05 | 0 | 4.94E-05 | 0 | 0 | 1.91E-06 | 0 | 0 | 0.0001 | 1.17E-05 | 0 | 0 | 0 |
| Ank1 | 0 | 0 | 3.11E-05 | 0 | 0 | 1.66E-05 | 0 | 4.12E-05 | 0 | 3.37E-06 | 0 | 0 | 0 | 0 | 0 | 2.66E-05 | 1.87E-05 | 0 |
| Zscan12 | 2.47E-05 | 3.88E-05 | 1.25E-05 | 1.34E-05 | 1.43E-05 | 1.65E-05 | 1.67E-05 | 5.39E-06 | 0 | 5.69E-06 | 2.77E-06 | 1.35E-05 | 0 | 0 | 4.26E-05 | 1.35E-05 | 0 | 8.75E-06 |
| Dot1l | 7.46E-05 | 9.79E-05 | 0.00018 | 0 | 6.18E-05 | 1.65E-05 | 0 | 0 | 0 | 7.15E-06 | 0.00016 | 7.39E-05 | 0 | 5.18E-05 | 0.00018 | 0.00011 | 0.00026 | 0 |

| | | | | | | | | | | | | | | | | | | |
|---|---|---|---|---|---|---|---|---|---|---|---|---|---|---|---|---|---|---|
| Fmo1 | 0 | 3.57E-05 | 0 | 0 | 9.14E-06 | 1.63E-05 | 0 | 0 | 0 | 0.00011 | 0 | 0 | 0 | 7.76E-05 | 0 | 0 | 0 | 0.00031 |
| Uhrf2 | 0.00014 | 0.00024 | 0 | 0.00012 | 0 | 1.63E-05 | 7.46E-05 | 0 | 0 | 8.99E-05 | 0 | 0.00025 | 0.00027 | 0 | 0.00032 | 0.00044 | 0.00027 | 0.00018 |
| Slc25a27 | 0 | 9.47E-05 | 0 | 0 | 3.04E-05 | 1.63E-05 | 0 | 0 | 0 | 8.60E-06 | 0 | 0 | 0 | 7.31E-06 | 0.00013 | 0 | 0 | 1.48E-05 |
| Atg4a | 1.39E-05 | 2.11E-05 | 1.23E-05 | 0 | 8.35E-06 | 1.63E-05 | 2.22E-05 | 0 | 0 | 1.18E-05 | 4.41E-06 | 4.40E-06 | 0 | 1.58E-05 | 0 | 7.38E-06 | 0 | 2.56E-05 |
| Jsrp1 | 0 | 0 | 0 | 9.71E-07 | 1.70E-06 | 1.59E-05 | 0 | 3.28E-05 | 0 | 0 | 1.85E-05 | 1.89E-05 | 0 | 0 | 0 | 0 | 0 | 0 |
| Fuca2 | 1.69E-05 | 2.76E-05 | 0 | 0 | 1.22E-05 | 1.59E-05 | 9.96E-05 | 0 | 0 | 4.41E-05 | 5.78E-06 | 1.75E-06 | 2.19E-05 | 3.63E-05 | 0 | 2.19E-05 | 0 | 8.08E-05 |
| Zfp712 | 4.67E-06 | 5.20E-06 | 0 | 0 | 1.02E-05 | 1.58E-05 | 0 | 0 | 0 | 9.50E-06 | 3.62E-08 | 2.65E-06 | 0 | 0 | 1.35E-07 | 0 | 1.21E-05 | 1.05E-05 |
| Tcaim | 8.01E-06 | 1.36E-05 | 3.76E-06 | 2.19E-05 | 1.44E-05 | 1.58E-05 | 0 | 1.07E-06 | 0 | 6.30E-06 | 0 | 0 | 2.06E-05 | 8.24E-06 | 1.00E-05 | 7.85E-06 | 4.22E-06 | 1.33E-05 |
| Aff3 | 0 | 2.53E-06 | 0 | 0 | 8.11E-05 | 1.57E-05 | 0 | 0 | 0 | 5.78E-05 | 5.99E-05 | 3.35E-05 | 0 | 4.03E-05 | 2.89E-06 | 1.65E-05 | 0 | 0.0001 |
| Mtbp | 2.98E-05 | 1.36E-05 | 2.07E-05 | 0 | 1.06E-05 | 1.57E-05 | 0 | 0 | 0 | 8.43E-06 | 3.56E-06 | 3.21E-05 | 2.26E-05 | 1.41E-05 | 2.08E-05 | 0.00022 | 0 | 0 |
| H2-Ke6 | 4.99E-05 | 9.72E-05 | 1.60E-05 | 4.62E-05 | 3.39E-05 | 1.56E-05 | 8.49E-05 | 0 | 0 | 4.88E-05 | 0 | 3.06E-06 | 2.40E-06 | 0 | 8.74E-05 | 2.67E-06 | 8.51E-05 | 4.28E-05 |
| Cgref1 | 2.47E-05 | 0 | 8.52E-05 | 7.67E-05 | 0 | 1.56E-05 | 0.00052 | 0 | 0 | 4.05E-05 | 0 | 0 | 6.40E-05 | 4.34E-05 | 0 | 0 | 0 | 5.94E-05 |
| Pcdhb5 | 1.75E-05 | 6.72E-06 | 0 | 0 | 1.12E-05 | 1.55E-05 | 1.02E-05 | 1.02E-05 | 0 | 2.63E-07 | 0 | 0 | 5.82E-06 | 2.11E-05 | 0 | 0 | 2.49E-06 | 2.94E-06 |
| Ankrd42 | 2.86E-05 | 2.85E-05 | 7.22E-06 | 7.49E-06 | 5.65E-06 | 1.55E-05 | 1.34E-05 | 0 | 0 | 5.71E-07 | 1.61E-05 | 2.59E-05 | 1.74E-05 | 7.11E-06 | 0 | 1.18E-05 | 8.30E-05 | 2.82E-06 |
| Rnaseh2b | 0.00011 | 1.03E-05 | 3.96E-05 | 0 | 2.53E-05 | 1.54E-05 | 8.63E-05 | 9.68E-06 | 0 | 1.23E-05 | 3.22E-05 | 0 | 0 | 0 | 0 | 0.00027 | 0 | 8.82E-06 |
| Rgs4 | 2.59E-06 | 0 | 0 | 0 | 0 | 1.54E-05 | 1.92E-05 | 0 | 0 | 7.42E-06 | 7.44E-05 | 9.54E-06 | 0 | 0 | 0 | 0 | 0 | 5.23E-05 |
| Htr2b | 0 | 2.01E-05 | 0 | 0 | 0.0004 | 1.53E-05 | 0 | 4.47E-07 | 0 | 0 | 0 | 5.53E-05 | 0 | 0 | 0 | 0 | 0 | 0 |
| Cbx7 | 0 | 5.58E-05 | 0 | 0 | 0 | 1.53E-05 | 0 | 0 | 0 | 2.74E-05 | 0 | 2.77E-05 | 0 | 1.81E-05 | 2.69E-05 | 0 | 1.32E-05 | 3.87E-06 |
| Pcgf1 | 4.19E-05 | 4.97E-05 | 4.91E-05 | 1.86E-05 | 2.75E-05 | 1.53E-05 | 1.55E-05 | 2.10E-05 | 0 | 2.86E-06 | 1.62E-05 | 2.33E-05 | 2.27E-05 | 2.59E-05 | 8.75E-06 | 0 | 0 | 0 |
| Aplp1 | 0 | 3.37E-05 | 0 | 0 | 0.00011 | 1.51E-05 | 0 | 7.27E-05 | 0 | 0 | 0.00011 | 0.00028 | 0 | 0.00014 | 2.64E-05 | 0 | 0 | 0 |
| Rbbp8 | 6.64E-05 | 1.21E-05 | 8.89E-05 | 0 | 2.39E-06 | 1.49E-05 | 2.08E-05 | 1.09E-05 | 0 | 1.04E-05 | 0 | 4.98E-05 | 0 | 2.22E-05 | 7.07E-05 | 0.00023 | 5.79E-05 | 6.50E-06 |
| Apold1 | 0 | 7.87E-06 | 0 | 0 | 0 | 1.48E-05 | 0 | 2.67E-05 | 0 | 0 | 0 | 1.14E-05 | 0 | 4.75E-07 | 2.04E-05 | 5.62E-06 | 3.19E-05 | 0 |
| Gabrb3 | 0 | 0 | 0 | 0 | 0.00011 | 1.47E-05 | 5.80E-05 | 0 | 0 | 0 | 8.70E-05 | 0 | 0 | 0 | 0 | 1.35E-05 | 0 | 0 |
| Rundc3a | 0 | 2.36E-05 | 0 | 0 | 7.70E-06 | 1.47E-05 | 0 | 5.20E-06 | 0 | 5.92E-07 | 3.12E-06 | 3.32E-06 | 0 | 0 | 6.20E-06 | 0 | 1.64E-05 | 0 |
| Akap1 | 5.02E-05 | 8.03E-05 | 7.65E-05 | 0 | 0 | 1.47E-05 | 0 | 0 | 0 | 1.97E-05 | 5.66E-05 | 1.26E-05 | 1.17E-05 | 0 | 0 | 1.97E-05 | 5.37E-05 | 2.26E-05 |
| Xrcc2 | 7.35E-06 | 1.42E-05 | 3.96E-06 | 0 | 0 | 1.47E-05 | 0 | 0 | 0 | 4.21E-06 | 5.12E-06 | 0 | 0 | 0 | 3.65E-05 | 0.00017 | 0 | 2.18E-06 |
| Prr7 | 3.05E-06 | 0 | 7.24E-05 | 9.06E-05 | 0 | 1.47E-05 | 0.00016 | 0 | 0 | 0 | 5.48E-05 | 0 | 0 | 0 | 0 | 6.53E-06 | 5.01E-05 | 2.51E-05 |
| Nemp1 | 3.49E-06 | 1.46E-05 | 0 | 0 | 5.41E-06 | 1.46E-05 | 0 | 0 | 0 | 2.45E-06 | 0 | 3.25E-05 | 5.71E-06 | 0 | 0 | 0.00015 | 0 | 0 |
| Fbxo41 | 0 | 1.21E-05 | 0 | 9.67E-06 | 0 | 1.46E-05 | 0 | 1.65E-06 | 0 | 0 | 7.55E-06 | 1.81E-05 | 0 | 1.23E-05 | 1.28E-05 | 0 | 0 | 1.93E-06 |
| Tspyl2 | 9.77E-05 | 0.00012 | 4.93E-05 | 2.02E-05 | 1.75E-05 | 1.45E-05 | 4.80E-05 | 0 | 0 | 1.41E-05 | 1.01E-05 | 0.00011 | 0 | 3.49E-06 | 0.00053 | 0 | 7.49E-05 | 3.55E-05 |
| Ripk2 | 4.39E-05 | 3.85E-05 | 5.47E-05 | 0 | 4.84E-05 | 1.45E-05 | 7.90E-06 | 0 | 0 | 0 | 9.16E-05 | 4.81E-05 | 5.97E-05 | 0 | 0 | 9.40E-06 | 0.00013 | 2.53E-05 |
| Bid | 8.32E-06 | 4.09E-05 | 0 | 0 | 0 | 1.44E-05 | 0 | 0 | 0 | 2.99E-06 | 2.25E-06 | 0 | 2.97E-05 | 6.72E-05 | 0 | 2.03E-06 | 0 | 2.26E-05 |
| Asrgl1 | 3.03E-05 | 1.49E-05 | 5.63E-07 | 1.83E-05 | 1.39E-05 | 1.43E-05 | 5.37E-08 | 9.37E-06 | 0 | 1.37E-05 | 1.46E-06 | 0 | 1.04E-05 | 1.76E-05 | 0 | 4.14E-05 | 2.72E-05 | 2.93E-05 |

| | | | | | | | | | | | | | | | | | | |
|---|---|---|---|---|---|---|---|---|---|---|---|---|---|---|---|---|---|---|
| Zscan20 | 4.73E-06 | 1.98E-05 | 1.53E-06 | 2.54E-05 | 2.80E-07 | 1.43E-05 | 1.68E-05 | 1.02E-05 | 0 | 8.75E-06 | 0 | 1.14E-05 | 0 | 1.05E-05 | 4.90E-05 | 6.44E-06 | 0 | 4.40E-06 |
| Brca2 | 5.36E-06 | 1.27E-08 | 0 | 0 | 4.29E-06 | 1.43E-05 | 0 | 3.61E-06 | 0 | 6.97E-06 | 0 | 0 | 2.55E-06 | 3.25E-06 | 0 | 0.00013 | 0 | 0 |
| Zfp52 | 5.24E-05 | 4.94E-05 | 8.79E-06 | 0 | 9.16E-06 | 1.42E-05 | 3.17E-05 | 0 | 0 | 0 | 4.48E-05 | 7.10E-05 | 0 | 2.50E-05 | 6.90E-05 | 1.69E-05 | 0 | 6.39E-06 |
| Mettl26 | 0 | 3.61E-05 | 0 | 3.81E-05 | 3.33E-05 | 1.42E-05 | 0 | 9.38E-06 | 0 | 3.85E-06 | 1.86E-06 | 2.20E-05 | 0 | 1.84E-05 | 2.80E-05 | 2.25E-06 | 1.59E-05 | 2.94E-05 |
| Ddb2 | 6.51E-06 | 4.14E-05 | 0 | 2.06E-06 | 1.53E-05 | 1.41E-05 | 1.29E-06 | 0 | 0 | 8.29E-06 | 0 | 1.44E-05 | 0 | 1.18E-05 | 3.53E-05 | 1.61E-05 | 1.66E-06 | 4.24E-06 |
| Slc25a14 | 2.69E-05 | 3.29E-05 | 2.04E-05 | 3.09E-05 | 1.31E-05 | 1.41E-05 | 3.10E-05 | 2.83E-05 | 0 | 1.07E-05 | 0 | 1.63E-06 | 6.91E-07 | 1.41E-05 | 0 | 4.91E-05 | 0 | 7.36E-06 |
| Il12rb1 | 0 | 1.24E-05 | 0 | 6.11E-06 | 0 | 1.41E-05 | 0 | 2.26E-05 | 0 | 0 | 0 | 1.81E-07 | 1.21E-06 | 8.17E-06 | 0 | 0 | 5.31E-05 | 0 |
| Kyat1 | 9.39E-06 | 2.46E-05 | 1.59E-05 | 1.13E-05 | 5.69E-06 | 1.39E-05 | 0 | 1.67E-05 | 0 | 1.96E-05 | 1.16E-05 | 1.11E-05 | 0 | 8.51E-06 | 3.26E-05 | 0 | 7.21E-06 | 5.17E-06 |
| Abhd14a | 5.96E-05 | 6.51E-05 | 2.87E-05 | 0.00011 | 4.87E-05 | 1.39E-05 | 9.94E-05 | 3.14E-05 | 0 | 5.88E-05 | 2.63E-05 | 1.82E-05 | 8.46E-05 | 9.38E-05 | 1.18E-05 | 0 | 0 | 6.67E-05 |
| Rhebl1 | 4.75E-06 | 3.29E-05 | 0 | 0 | 2.63E-06 | 1.38E-05 | 1.41E-05 | 0 | 0 | 0 | 1.33E-05 | 1.55E-05 | 0 | 1.45E-05 | 2.16E-05 | 4.52E-05 | 9.43E-06 | 4.02E-06 |
| Sap25 | 0 | 5.52E-05 | 0 | 2.92E-05 | 1.37E-05 | 1.37E-05 | 0 | 1.93E-05 | 0 | 1.23E-05 | 0 | 0 | 0 | 2.00E-05 | 0.00011 | 5.05E-05 | 0 | 3.07E-05 |
| Cd46 | 5.00E-06 | 2.49E-05 | 0 | 2.21E-07 | 1.59E-06 | 1.37E-05 | 0 | 1.43E-05 | 0 | 8.61E-06 | 0 | 1.31E-05 | 0 | 0 | 2.73E-05 | 0 | 0 | 1.14E-05 |
| Hist1h1d | 0 | 0 | 0 | 0 | 6.24E-06 | 1.37E-05 | 0 | 0 | 0 | 0 | 0 | 0 | 0 | 0 | 0 | 0.00053 | 0 | 0 |
| Slc43a2 | 1.25E-06 | 2.27E-05 | 0 | 1.47E-06 | 2.59E-06 | 1.36E-05 | 0 | 8.34E-06 | 0 | 5.36E-06 | 0 | 0 | 0 | 3.85E-05 | 4.94E-05 | 5.47E-06 | 3.29E-05 | 2.89E-06 |
| Psrc1 | 0 | 8.44E-05 | 1.12E-05 | 0 | 0 | 1.36E-05 | 0 | 3.38E-06 | 0 | 0 | 0 | 0 | 2.55E-05 | 0 | 2.50E-05 | 0.00013 | 0 | 1.35E-05 |
| Itpkb | 0 | 8.90E-05 | 1.88E-05 | 0.00012 | 8.56E-05 | 1.34E-05 | 4.38E-06 | 7.45E-05 | 0 | 3.62E-05 | 1.60E-05 | 2.54E-05 | 0 | 0.00012 | 0.00013 | 1.01E-05 | 0.00012 | 0.00012 |
| Enpep | 0 | 0 | 0 | 0 | 0 | 1.34E-05 | 0 | 0.0002 | 0 | 0.00035 | 0 | 4.81E-06 | 0.00017 | 0 | 0 | 0 | 0 | 0 |
| Zfp958 | 9.15E-06 | 2.85E-05 | 0 | 0 | 8.70E-06 | 1.34E-05 | 0 | 2.05E-06 | 0 | 8.73E-06 | 0 | 1.75E-05 | 1.70E-06 | 0 | 0 | 2.25E-05 | 0 | 1.19E-05 |
| Coa4 | 2.08E-05 | 1.80E-05 | 4.16E-06 | 2.82E-05 | 1.38E-05 | 1.34E-05 | 0 | 6.58E-06 | 0 | 1.31E-05 | 6.26E-06 | 2.22E-05 | 0 | 1.74E-05 | 0 | 1.25E-05 | 0 | 7.69E-06 |
| Cep112 | 6.17E-06 | 1.86E-05 | 0 | 0 | 5.19E-06 | 1.33E-05 | 0 | 0 | 0 | 1.45E-05 | 0 | 1.36E-05 | 0 | 0 | 0 | 4.50E-05 | 0 | 0 |
| Zfp994 | 1.54E-05 | 6.30E-06 | 0 | 0 | 3.08E-06 | 1.33E-05 | 1.87E-06 | 0 | 0 | 5.85E-06 | 0 | 6.69E-06 | 0 | 0 | 0 | 9.99E-07 | 0 | 0 |
| Bbs9 | 0 | 5.00E-05 | 0 | 9.91E-06 | 2.98E-05 | 1.33E-05 | 2.01E-07 | 2.01E-06 | 0 | 2.53E-05 | 0 | 7.92E-06 | 7.00E-06 | 1.10E-05 | 1.06E-06 | 2.23E-05 | 2.39E-05 | 6.41E-05 |
| Vamp5 | 0 | 0 | 0 | 2.96E-06 | 0 | 1.33E-05 | 0 | 0 | 0 | 4.34E-06 | 0.00011 | 2.14E-05 | 2.07E-06 | 9.66E-05 | 5.83E-06 | 0 | 0 | 9.09E-05 |
| Zfp804a | 0 | 0 | 3.35E-06 | 0 | 2.60E-05 | 1.32E-05 | 0 | 0 | 0 | 0 | 1.43E-05 | 0 | 0 | 0 | 0 | 0 | 0 | 0 |
| Pitpnm2 | 8.66E-06 | 2.73E-05 | 1.40E-05 | 0 | 0 | 1.32E-05 | 0 | 0 | 0 | 1.74E-05 | 3.54E-05 | 0 | 0 | 1.75E-06 | 1.38E-05 | 6.67E-05 | 0 | 4.85E-05 |
| Lyrm7 | 1.57E-05 | 3.98E-06 | 5.81E-06 | 0 | 3.96E-06 | 1.31E-05 | 0 | 3.08E-06 | 0 | 2.11E-06 | 0 | 7.91E-06 | 5.13E-06 | 0 | 0 | 1.61E-05 | 0 | 7.76E-06 |
| Ess2 | 1.57E-05 | 2.36E-05 | 1.26E-05 | 1.68E-05 | 1.31E-05 | 1.30E-05 | 0 | 1.21E-06 | 0 | 1.77E-05 | 6.01E-06 | 6.66E-06 | 8.50E-06 | 1.75E-05 | 9.53E-05 | 4.45E-05 | 0 | 2.11E-05 |
| Rad9b | 6.55E-07 | 5.52E-06 | 3.49E-07 | 1.22E-06 | 6.11E-06 | 1.29E-05 | 1.07E-06 | 4.75E-06 | 0 | 2.26E-06 | 0 | 7.43E-06 | 0 | 9.73E-07 | 7.30E-06 | 6.72E-06 | 0 | 0 |
| Tyms | 3.67E-05 | 0 | 5.81E-05 | 0 | 0 | 1.29E-05 | 0 | 0 | 0 | 0 | 0 | 0 | 5.76E-05 | 0 | 0 | 0.00076 | 0 | 1.64E-05 |
| Olfml2a | 0 | 0.00015 | 0 | 0 | 0 | 1.29E-05 | 0 | 0.00011 | 0 | 0 | 0 | 0 | 0 | 0 | 0 | 0 | 0 | 8.59E-05 |
| Cfap100 | 0 | 0 | 0 | 0 | 2.66E-05 | 1.28E-05 | 0 | 0 | 0 | 0 | 2.59E-05 | 0 | 0 | 1.30E-05 | 4.44E-05 | 0 | 0 | 0 |
| Zfp944 | 3.28E-06 | 8.54E-06 | 4.23E-06 | 7.39E-06 | 1.17E-05 | 1.28E-05 | 1.44E-05 | 1.72E-06 | 0 | 1.16E-05 | 5.56E-06 | 2.30E-06 | 8.36E-06 | 9.34E-06 | 1.55E-05 | 1.75E-06 | 0 | 1.04E-05 |
| Lig4 | 3.56E-05 | 1.30E-05 | 1.55E-05 | 0 | 1.49E-05 | 1.26E-05 | 1.17E-05 | 1.56E-06 | 0 | 6.47E-06 | 0 | 0 | 0 | 1.29E-06 | 6.08E-06 | 1.24E-05 | 0 | 0 |

| | | | | | | | | | | | | | | | | | | |
|---|---|---|---|---|---|---|---|---|---|---|---|---|---|---|---|---|---|---|
| Efhc1 | 2.64E-05 | 2.14E-05 | 0 | 7.84E-07 | 3.95E-06 | 1.26E-05 | 3.18E-05 | 2.53E-05 | 0 | 7.23E-06 | 0 | 2.67E-05 | 7.55E-06 | 0 | 1.42E-05 | 0 | 0 | 9.69E-06 |
| Spata1 | 1.64E-05 | 1.07E-05 | 0 | 0 | 9.60E-06 | 1.25E-05 | 0 | 1.04E-05 | 0 | 0 | 0 | 1.74E-05 | 7.72E-06 | 2.04E-06 | 7.92E-06 | 0 | 0 | 1.25E-05 |
| Camk2a | 0 | 0 | 3.52E-06 | 0 | 2.20E-05 | 1.23E-05 | 5.71E-05 | 0 | 0 | 0 | 8.88E-06 | 0 | 7.34E-05 | 1.05E-05 | 3.40E-05 | 0 | 0 | 1.16E-05 |
| Sfxn4 | 9.41E-06 | 6.56E-06 | 0 | 0 | 2.07E-06 | 1.22E-05 | 0 | 0 | 0 | 1.25E-05 | 0 | 1.08E-05 | 0 | 0 | 2.32E-05 | 1.82E-06 | 0 | 9.04E-06 |
| Rasal2 | 5.80E-05 | 4.09E-05 | 1.37E-05 | 0 | 2.41E-05 | 1.21E-05 | 5.94E-05 | 2.59E-05 | 0 | 1.07E-05 | 4.10E-07 | 2.95E-05 | 4.56E-06 | 0 | 4.53E-05 | 2.50E-05 | 0.0001 | 6.32E-05 |
| Blm | 7.31E-06 | 2.51E-06 | 3.01E-06 | 0 | 0 | 1.21E-05 | 0 | 0 | 0 | 6.55E-07 | 2.07E-06 | 1.51E-05 | 0 | 0 | 3.28E-05 | 0.00029 | 0 | 0 |
| Thsd1 | 1.90E-05 | 7.18E-06 | 8.26E-06 | 0 | 0 | 1.21E-05 | 0 | 5.20E-05 | 0 | 3.34E-05 | 1.75E-07 | 1.46E-06 | 3.31E-05 | 5.12E-06 | 7.99E-07 | 3.22E-06 | 5.92E-06 | 6.70E-05 |
| Hcn4 | 0 | 0 | 0 | 0 | 0 | 1.20E-05 | 0 | 0 | 0 | 0.00015 | 0 | 0.00011 | 0 | 0 | 0 | 0 | 0 | 0 |
| Ypel4 | 0 | 8.27E-05 | 0 | 0 | 1.29E-05 | 1.19E-05 | 3.29E-05 | 0 | 0 | 3.13E-05 | 0 | 3.94E-06 | 0 | 2.30E-05 | 3.96E-05 | 0 | 0 | 1.84E-06 |
| Mastl | 0 | 0 | 0 | 0 | 0 | 1.17E-05 | 0 | 0 | 0 | 1.43E-07 | 0 | 3.98E-06 | 0 | 0 | 0 | 0.0004 | 0 | 0 |
| Uqcrh | 1.53E-05 | 9.90E-06 | 1.55E-05 | 1.84E-05 | 5.37E-06 | 1.16E-05 | 2.14E-05 | 4.93E-06 | 0 | 3.14E-06 | 0 | 6.49E-06 | 6.46E-06 | 2.22E-05 | 0 | 5.74E-06 | 1.87E-05 | 5.76E-06 |
| Mdga1 | 2.00E-05 | 0 | 0.00015 | 5.02E-05 | 0 | 1.16E-05 | 0.00018 | 0.00017 | 0 | 0 | 2.39E-05 | 3.37E-05 | 3.51E-05 | 0.00016 | 0.00015 | 7.15E-05 | 5.97E-05 | 1.70E-05 |
| Fscn3 | 0 | 1.84E-06 | 0 | 9.55E-07 | 1.90E-05 | 1.16E-05 | 0 | 0 | 0 | 0 | 8.34E-06 | 0 | 0 | 0 | 1.07E-05 | 4.01E-06 | 2.23E-05 | 0 |
| Pecr | 6.10E-05 | 1.80E-05 | 6.23E-05 | 2.71E-05 | 0 | 1.14E-05 | 3.37E-05 | 0 | 0 | 1.57E-05 | 0 | 6.60E-06 | 2.19E-05 | 0 | 0 | 0 | 0 | 4.88E-06 |
| Fam83a | 0 | 0 | 3.22E-05 | 0 | 1.90E-05 | 1.14E-05 | 0 | 4.70E-06 | 0 | 0 | 4.57E-09 | 0 | 0 | 8.28E-06 | 0 | 4.91E-07 | 0 | 1.46E-06 |
| Pclaf | 0 | 0 | 3.36E-05 | 0 | 1.28E-05 | 1.14E-05 | 0 | 3.01E-06 | 0 | 0 | 0 | 0 | 1.91E-05 | 3.84E-05 | 0 | 0.00119 | 0 | 1.78E-05 |
| Kazald1 | 9.72E-06 | 2.98E-06 | 0 | 1.77E-05 | 6.44E-06 | 1.13E-05 | 4.47E-05 | 0 | 0 | 2.90E-05 | 0 | 4.11E-06 | 0 | 2.82E-05 | 0 | 0 | 6.60E-06 | 3.85E-05 |
| Slfn2 | 8.90E-06 | 2.39E-05 | 9.67E-06 | 0 | 0 | 1.12E-05 | 0 | 0 | 0 | 0.00014 | 0.00012 | 0 | 0.00011 | 1.12E-05 | 4.95E-06 | 2.84E-05 | 0 | 2.68E-05 |
| Serp2 | 6.71E-06 | 1.08E-05 | 8.60E-06 | 6.03E-06 | 3.03E-05 | 1.11E-05 | 2.96E-05 | 3.01E-05 | 0 | 0 | 3.17E-06 | 2.48E-06 | 0 | 3.00E-05 | 7.51E-06 | 1.22E-05 | 0 | 0 |
| Abcc3 | 0 | 1.99E-05 | 0 | 5.17E-05 | 6.63E-05 | 1.11E-05 | 3.91E-05 | 2.15E-05 | 0 | 3.77E-05 | 5.05E-06 | 0 | 9.29E-05 | 0 | 2.03E-05 | 0 | 1.00E-05 | 0 |
| Cib2 | 0 | 8.32E-06 | 0 | 0 | 1.65E-05 | 1.10E-05 | 0 | 0 | 0 | 0 | 1.38E-05 | 0 | 0 | 2.56E-05 | 0 | 0 | 0 | 9.45E-06 |
| Slc15a3 | 0 | 1.13E-05 | 3.19E-05 | 4.51E-05 | 9.69E-06 | 1.10E-05 | 0 | 5.54E-05 | 0 | 6.36E-06 | 6.39E-06 | 1.38E-05 | 8.55E-05 | 3.78E-05 | 0.0001 | 1.27E-05 | 0 | 9.15E-05 |
| Frat1 | 0 | 1.97E-05 | 0 | 0 | 1.31E-05 | 1.09E-05 | 4.77E-07 | 0 | 0 | 0 | 0 | 6.73E-06 | 0 | 1.96E-05 | 0 | 0 | 1.83E-05 | 0 |
| Nppc | 0 | 0 | 0 | 0 | 0 | 1.09E-05 | 0 | 5.92E-05 | 0 | 0 | 0 | 0 | 0 | 0 | 0 | 0 | 4.88E-05 | 3.14E-05 |
| Ppp1r3f | 0 | 8.52E-06 | 6.07E-07 | 1.52E-05 | 2.87E-05 | 1.08E-05 | 0 | 1.57E-05 | 0 | 0 | 2.57E-05 | 9.41E-06 | 5.60E-06 | 1.20E-05 | 5.78E-06 | 1.57E-05 | 3.41E-05 | 8.94E-06 |
| Eif3j1 | 2.11E-05 | 8.36E-06 | 1.74E-06 | 0 | 0 | 1.08E-05 | 0 | 2.39E-06 | 0 | 3.37E-06 | 0 | 7.97E-06 | 0 | 0 | 7.23E-06 | 8.57E-06 | 5.40E-05 | 0 |
| Inava | 0 | 0 | 0.00011 | 5.69E-05 | 0.00023 | 1.06E-05 | 0 | 0.00017 | 0 | 0 | 0.00013 | 0 | 0 | 0.00014 | 0.00014 | 1.65E-06 | 0 | 0 |
| Dck | 2.04E-05 | 0 | 3.70E-06 | 0 | 0 | 1.05E-05 | 0 | 0 | 0 | 8.78E-06 | 4.80E-06 | 4.92E-05 | 7.30E-06 | 1.09E-05 | 0 | 0.00031 | 4.75E-06 | 5.44E-06 |
| Aff2 | 3.94E-05 | 0 | 0 | 0 | 3.65E-05 | 1.05E-05 | 0 | 0 | 0 | 3.13E-05 | 2.97E-05 | 0 | 6.59E-05 | 1.56E-05 | 4.66E-05 | 0 | 0 | 7.49E-05 |
| Ccdc112 | 0 | 1.89E-05 | 1.77E-05 | 6.28E-06 | 1.72E-05 | 1.03E-05 | 4.65E-05 | 1.77E-05 | 0 | 1.05E-05 | 4.95E-05 | 7.32E-06 | 4.30E-06 | 0 | 3.48E-05 | 2.40E-05 | 1.81E-05 | 3.28E-05 |
| Fgf1 | 0 | 1.74E-06 | 0 | 0 | 0 | 1.03E-05 | 0 | 5.31E-05 | 0 | 1.59E-05 | 0 | 5.51E-05 | 0 | 0 | 0 | 0 | 0 | 0.00018 |
| Lin9 | 1.32E-05 | 1.25E-05 | 9.77E-06 | 0 | 4.97E-06 | 1.02E-05 | 0 | 3.00E-06 | 0 | 0 | 4.81E-06 | 4.39E-06 | 0 | 1.90E-05 | 0 | 0.00023 | 0 | 0 |
| Zfp119a | 2.36E-05 | 3.60E-05 | 0 | 0 | 8.87E-06 | 1.02E-05 | 0 | 5.18E-06 | 0 | 0 | 0 | 1.16E-05 | 0 | 0 | 1.16E-05 | 0 | 0 | 4.89E-07 |

| | | | | | | | | | | | | | | | | | | |
|---|---|---|---|---|---|---|---|---|---|---|---|---|---|---|---|---|---|---|
| Fbxo36 | 1.16E-05 | 2.11E-06 | 0 | 1.97E-05 | 0 | 1.01E-05 | 2.36E-05 | 0 | 0 | 7.11E-06 | 0 | 5.21E-06 | 1.93E-05 | 6.99E-06 | 0 | 0 | 1.15E-06 | 2.17E-05 |
| Poli | 9.23E-06 | 1.67E-05 | 0 | 2.32E-06 | 3.14E-06 | 1.01E-05 | 1.66E-05 | 1.78E-05 | 0 | 1.53E-05 | 0 | 3.93E-05 | 0 | 0 | 3.58E-05 | 5.30E-06 | 3.64E-06 | 1.50E-05 |
| Areg | 0 | 0 | 2.33E-06 | 0 | 0 | 1.00E-05 | 0 | 2.75E-05 | 0 | 3.53E-06 | 0 | 2.39E-05 | 0 | 0 | 3.39E-06 | 0 | 7.62E-05 | 2.44E-06 |
| Zfp112 | 4.94E-06 | 7.55E-06 | 2.02E-06 | 0 | 1.14E-05 | 1.00E-05 | 0 | 6.59E-06 | 0 | 4.98E-06 | 1.29E-06 | 8.20E-06 | 0 | 0 | 2.70E-07 | 0 | 0 | 6.88E-06 |
| Ncald | 5.42E-07 | 7.06E-06 | 0 | 0 | 7.82E-06 | 9.97E-06 | 0 | 5.27E-05 | 0 | 2.69E-05 | 0 | 0 | 0 | 2.24E-05 | 0 | 0 | 0 | 7.41E-05 |
| Mtfp1 | 6.26E-07 | 8.32E-06 | 5.66E-06 | 1.04E-05 | 9.14E-06 | 9.80E-06 | 0 | 0 | 0 | 0 | 3.78E-07 | 0 | 0 | 1.13E-05 | 7.31E-06 | 1.61E-05 | 0 | 5.82E-06 |
| Whamm | 5.58E-05 | 7.78E-05 | 1.32E-05 | 5.09E-06 | 6.63E-06 | 9.62E-06 | 0 | 0 | 0 | 3.21E-05 | 1.40E-06 | 0.00011 | 0 | 0 | 0.00012 | 0 | 0 | 1.22E-05 |
| Immp2l | 1.17E-05 | 2.18E-05 | 1.14E-05 | 0 | 0 | 9.48E-06 | 0 | 0 | 0 | 2.17E-05 | 0 | 2.37E-05 | 9.41E-06 | 6.78E-06 | 4.03E-05 | 1.54E-05 | 1.17E-05 | 0 |
| Ebf2 | 0 | 0 | 8.64E-06 | 0 | 0 | 9.40E-06 | 1.53E-05 | 7.21E-05 | 0 | 0.00025 | 0 | 3.00E-05 | 0 | 0 | 0 | 1.06E-05 | 0 | 0.00035 |
| Rsph3b | 2.07E-05 | 3.19E-05 | 7.35E-06 | 7.64E-05 | 2.34E-06 | 9.25E-06 | 9.79E-05 | 6.01E-07 | 0 | 6.96E-06 | 1.31E-06 | 1.95E-05 | 2.24E-05 | 4.16E-07 | 0 | 3.40E-05 | 0 | 9.88E-06 |
| Prss53 | 0 | 0 | 0 | 1.65E-05 | 2.57E-06 | 9.12E-06 | 4.95E-05 | 3.64E-06 | 0 | 1.00E-05 | 0 | 9.67E-06 | 9.80E-07 | 0 | 6.22E-05 | 0 | 0 | 7.64E-06 |
| Card6 | 5.61E-06 | 1.68E-05 | 0 | 0 | 2.62E-05 | 9.10E-06 | 0 | 3.36E-05 | 0 | 0 | 0 | 0 | 0 | 8.92E-06 | 0 | 5.87E-06 | 0 | 6.47E-06 |
| Zfp772 | 1.10E-05 | 1.42E-05 | 1.35E-06 | 0 | 1.22E-05 | 9.03E-06 | 1.78E-06 | 0 | 0 | 4.44E-06 | 2.75E-06 | 2.34E-06 | 7.31E-06 | 0 | 3.78E-05 | 0 | 0 | 4.05E-06 |
| Deptor | 1.70E-05 | 4.12E-06 | 0 | 0 | 0 | 8.96E-06 | 0 | 0 | 0 | 3.13E-05 | 0 | 0 | 2.99E-05 | 0 | 0 | 0 | 0 | 0.00016 |
| Zfp583 | 9.77E-06 | 1.41E-05 | 1.23E-06 | 0 | 2.89E-06 | 8.94E-06 | 7.63E-06 | 0 | 0 | 7.59E-07 | 3.61E-06 | 1.87E-06 | 0 | 0 | 0 | 0 | 0 | 8.17E-06 |
| Musk | 1.05E-06 | 1.10E-06 | 0 | 1.17E-06 | 1.76E-05 | 8.94E-06 | 0 | 0 | 0 | 4.76E-06 | 1.08E-05 | 0 | 3.20E-05 | 0 | 3.20E-07 | 0 | 0 | 3.98E-05 |
| Pot1b | 8.59E-06 | 1.79E-05 | 0 | 0 | 1.15E-05 | 8.93E-06 | 8.00E-06 | 0 | 0 | 1.30E-05 | 0 | 7.81E-06 | 0 | 4.91E-06 | 2.31E-05 | 3.21E-05 | 0 | 1.19E-05 |
| Hist1h4i | 0 | 0 | 0 | 0 | 0 | 8.89E-06 | 0 | 0 | 0 | 0 | 0 | 1.59E-05 | 9.96E-06 | 0 | 0 | 0.00027 | 0.00011 | 1.43E-05 |
| Efcc1 | 0 | 0 | 1.75E-06 | 0 | 1.63E-06 | 8.70E-06 | 8.89E-06 | 0 | 0 | 3.17E-05 | 0 | 6.74E-06 | 0 | 5.55E-06 | 7.47E-06 | 0 | 0 | 4.42E-05 |
| Nudt16 | 2.61E-06 | 4.13E-05 | 0 | 4.60E-05 | 2.47E-05 | 8.69E-06 | 4.83E-06 | 5.71E-06 | 0 | 3.17E-05 | 1.99E-05 | 5.15E-05 | 0 | 2.06E-05 | 5.32E-05 | 4.14E-06 | 0 | 1.14E-05 |
| Rnf227 | 0 | 1.25E-05 | 0 | 1.09E-05 | 3.66E-07 | 8.67E-06 | 0 | 2.18E-06 | 0 | 0 | 0 | 0 | 1.86E-06 | 1.64E-05 | 2.95E-05 | 4.77E-05 | 1.03E-05 | 1.08E-06 |
| Srrm4 | 0 | 4.63E-06 | 0 | 1.06E-05 | 2.13E-06 | 8.65E-06 | 2.90E-05 | 0 | 0 | 1.55E-05 | 2.73E-06 | 7.06E-06 | 0 | 0 | 5.25E-05 | 0 | 5.36E-05 | 1.16E-06 |
| Zkscan14 | 0 | 3.68E-05 | 0 | 3.38E-05 | 1.79E-05 | 8.53E-06 | 0 | 8.99E-07 | 0 | 1.45E-05 | 7.45E-06 | 1.19E-05 | 0 | 1.50E-05 | 3.68E-05 | 0 | 0 | 2.83E-05 |
| Abhd18 | 1.27E-05 | 2.05E-05 | 0 | 0 | 2.77E-05 | 8.52E-06 | 2.65E-06 | 5.85E-07 | 0 | 5.10E-06 | 0 | 0.00012 | 0 | 1.20E-05 | 2.40E-05 | 0 | 0 | 8.19E-06 |
| Malt1 | 3.89E-05 | 3.97E-05 | 0 | 0 | 7.24E-06 | 8.52E-06 | 3.46E-06 | 1.61E-06 | 0 | 0 | 8.30E-06 | 4.15E-05 | 0 | 0 | 4.49E-05 | 6.39E-06 | 2.38E-05 | 6.15E-07 |
| Rttn | 5.18E-06 | 6.83E-06 | 0 | 0 | 1.51E-05 | 8.52E-06 | 7.03E-06 | 0 | 0 | 4.99E-06 | 0 | 9.01E-06 | 0 | 6.48E-06 | 3.51E-05 | 0.00013 | 0 | 0 |
| Ccl25 | 5.98E-06 | 6.87E-07 | 1.51E-05 | 0 | 7.98E-06 | 8.43E-06 | 0 | 1.66E-05 | 0 | 7.83E-06 | 0 | 1.25E-05 | 2.45E-05 | 0 | 1.48E-05 | 4.75E-07 | 1.86E-06 | 0 |
| S100b | 0 | 0 | 0 | 0 | 3.14E-06 | 8.40E-06 | 0 | 2.69E-07 | 0 | 1.32E-06 | 0 | 4.30E-06 | 0 | 0 | 0 | 0 | 0 | 0.00012 |
| Csmd1 | 0 | 0 | 0 | 0 | 0 | 8.35E-06 | 0 | 0 | 0 | 2.11E-05 | 0 | 0 | 0 | 0 | 0 | 0 | 0 | 0.00014 |
| Ccp110 | 2.62E-05 | 5.01E-05 | 0 | 0 | 1.85E-05 | 8.31E-06 | 0 | 3.68E-06 | 0 | 0 | 0 | 1.99E-05 | 0 | 1.28E-05 | 6.92E-05 | 0.00015 | 2.27E-05 | 0 |
| Taf4b | 2.63E-05 | 1.33E-05 | 3.91E-05 | 0 | 0 | 8.29E-06 | 0 | 0 | 0 | 0 | 0 | 0 | 1.20E-05 | 2.95E-05 | 0 | 1.67E-05 | 0.00019 | 0 |
| Zfp11 | 1.95E-05 | 2.22E-05 | 1.00E-06 | 1.47E-05 | 1.30E-05 | 8.26E-06 | 1.78E-06 | 4.42E-06 | 0 | 9.19E-06 | 4.19E-06 | 1.94E-06 | 6.92E-06 | 8.58E-06 | 8.80E-06 | 8.87E-06 | 0 | 5.88E-06 |
| Nat8f4 | 1.34E-07 | 6.00E-06 | 1.32E-06 | 6.40E-07 | 6.90E-06 | 8.26E-06 | 1.37E-06 | 9.87E-08 | 0 | 5.18E-06 | 0 | 0 | 5.59E-07 | 5.72E-06 | 0 | 2.54E-06 | 0 | 2.64E-06 |

| | | | | | | | | | | | | | | | | | | |
|---|---|---|---|---|---|---|---|---|---|---|---|---|---|---|---|---|---|---|
| Serac1 | 1.43E-06 | 4.54E-05 | 0 | 0 | 9.88E-06 | 8.23E-06 | 0 | 1.00E-05 | 0 | 4.47E-06 | 0 | 1.66E-05 | 0 | 1.38E-05 | 8.17E-06 | 1.13E-06 | 0 | 4.65E-06 |
| Cep135 | 1.85E-05 | 2.70E-05 | 4.56E-06 | 0 | 1.57E-05 | 8.19E-06 | 0 | 1.61E-05 | 0 | 0 | 0 | 1.69E-05 | 0 | 0 | 0.0001 | 0.0001 | 1.74E-05 | 0 |
| Zfp667 | 8.53E-06 | 9.16E-06 | 1.61E-06 | 5.74E-07 | 3.20E-06 | 8.06E-06 | 2.38E-06 | 2.07E-06 | 0 | 0 | 4.15E-06 | 4.60E-06 | 0 | 0 | 1.18E-05 | 0 | 0 | 1.28E-06 |
| Zfp174 | 1.15E-05 | 1.63E-05 | 0 | 0 | 1.12E-05 | 8.05E-06 | 8.51E-07 | 0 | 0 | 2.33E-06 | 1.28E-06 | 4.89E-06 | 0 | 6.61E-07 | 1.81E-05 | 0 | 0 | 3.31E-06 |
| Gramd1c | 0 | 6.01E-06 | 0 | 0 | 2.53E-06 | 8.03E-06 | 0 | 1.13E-05 | 0 | 9.78E-08 | 0 | 7.79E-06 | 0 | 0 | 1.25E-05 | 1.80E-05 | 0 | 8.55E-06 |
| Fbxo5 | 0 | 8.71E-06 | 0 | 0 | 4.79E-06 | 8.02E-06 | 0 | 0 | 0 | 0 | 0 | 0 | 0 | 0 | 0 | 0.0004 | 0 | 0 |
| Pard6a | 1.16E-05 | 3.76E-05 | 5.07E-06 | 4.56E-05 | 1.97E-05 | 7.90E-06 | 2.06E-05 | 0 | 0 | 5.28E-06 | 4.71E-06 | 1.72E-05 | 0 | 7.84E-06 | 0 | 4.57E-05 | 2.53E-05 | 7.75E-06 |
| Bcl6b | 0 | 9.48E-06 | 1.37E-05 | 0 | 1.44E-05 | 7.90E-06 | 0 | 4.34E-05 | 0 | 0 | 1.47E-05 | 9.29E-06 | 0 | 9.77E-06 | 1.86E-06 | 1.85E-05 | 8.88E-06 | 1.37E-06 |
| Phldb3 | 0 | 1.91E-05 | 0 | 0 | 3.32E-05 | 7.74E-06 | 0 | 0 | 0 | 0 | 2.09E-05 | 8.04E-06 | 0 | 4.73E-05 | 9.15E-06 | 0 | 0 | 4.47E-06 |
| Ccdc77 | 9.96E-06 | 9.00E-06 | 0 | 0 | 9.74E-07 | 7.67E-06 | 3.87E-06 | 6.01E-06 | 0 | 5.08E-06 | 0 | 6.16E-06 | 0 | 0 | 6.93E-05 | 8.12E-05 | 0 | 3.83E-06 |
| C7 | 0 | 0 | 0 | 0 | 2.59E-06 | 7.66E-06 | 0 | 0 | 0 | 8.55E-06 | 0 | 0 | 0 | 0 | 0 | 0 | 0 | 0.00012 |
| Pcdhgc4 | 0 | 3.17E-05 | 0 | 4.75E-05 | 1.79E-05 | 7.57E-06 | 0 | 1.25E-05 | 0 | 2.47E-05 | 0 | 0 | 8.62E-06 | 2.53E-06 | 0.00011 | 1.93E-05 | 3.65E-05 | 2.59E-05 |
| Klhl3 | 0 | 0 | 0 | 0 | 2.74E-06 | 7.55E-06 | 0 | 0 | 0 | 0 | 0 | 0.0002 | 0 | 0 | 2.52E-05 | 0 | 0 | 0 |
| Ccl19 | 1.49E-06 | 0 | 9.39E-06 | 0 | 0 | 7.49E-06 | 0 | 0 | 0 | 9.43E-05 | 0 | 0 | 0.00026 | 0 | 0 | 0 | 0 | 0.0007 |
| Inha | 0 | 2.35E-05 | 0 | 4.27E-05 | 5.33E-05 | 7.46E-06 | 4.66E-05 | 0 | 0 | 1.16E-05 | 2.36E-06 | 0.00013 | 4.59E-06 | 4.86E-05 | 0.00011 | 0 | 0 | 8.49E-06 |
| Wdr53 | 7.78E-06 | 0 | 5.23E-06 | 5.68E-06 | 3.02E-06 | 7.44E-06 | 9.51E-06 | 0 | 0 | 3.13E-06 | 3.02E-06 | 0 | 9.24E-07 | 5.74E-06 | 0 | 1.18E-05 | 0 | 4.97E-06 |
| Procr | 4.14E-06 | 1.69E-05 | 9.09E-05 | 3.80E-05 | 4.10E-05 | 7.37E-06 | 0 | 3.37E-05 | 0 | 0.00022 | 1.11E-05 | 3.73E-05 | 9.45E-05 | 2.52E-05 | 0 | 0 | 0 | 0 |
| Serpinb6b | 0 | 0 | 0 | 0 | 2.55E-05 | 7.29E-06 | 0 | 0 | 0 | 1.60E-05 | 1.10E-05 | 9.77E-07 | 0 | 0 | 0 | 0 | 0 | 4.41E-05 |
| Rfc4 | 2.59E-05 | 8.02E-06 | 3.46E-05 | 2.83E-06 | 0 | 7.24E-06 | 5.68E-06 | 0 | 0 | 9.93E-06 | 0 | 7.75E-06 | 4.49E-06 | 2.23E-06 | 0 | 0.00031 | 0 | 3.70E-06 |
| Tkfc | 3.39E-06 | 1.30E-05 | 4.79E-06 | 6.13E-05 | 2.11E-05 | 7.18E-06 | 1.83E-05 | 3.72E-05 | 0 | 1.05E-05 | 7.11E-06 | 9.14E-06 | 6.98E-06 | 2.93E-06 | 0 | 2.68E-05 | 0 | 1.94E-05 |
| Gins3 | 1.27E-06 | 0 | 1.20E-05 | 0 | 7.24E-06 | 7.14E-06 | 1.32E-05 | 1.90E-06 | 0 | 8.26E-06 | 3.29E-06 | 0 | 1.53E-05 | 1.33E-05 | 1.27E-07 | 0.00019 | 0 | 1.22E-05 |
| Rec114 | 0 | 0 | 0 | 2.75E-06 | 1.46E-05 | 7.10E-06 | 0 | 0 | 0 | 1.16E-05 | 3.62E-05 | 1.63E-05 | 0 | 0 | 0 | 0 | 0 | 0 |
| Snx24 | 0 | 9.87E-06 | 1.42E-05 | 3.08E-06 | 7.45E-05 | 7.07E-06 | 0 | 3.66E-05 | 0 | 9.84E-06 | 0.00014 | 6.00E-05 | 3.95E-06 | 0 | 0 | 2.19E-05 | 0 | 0 |
| Anxa9 | 8.97E-07 | 7.47E-06 | 0 | 1.07E-06 | 9.88E-06 | 7.06E-06 | 8.47E-06 | 6.80E-06 | 0 | 8.64E-06 | 0 | 9.40E-06 | 1.85E-05 | 1.16E-06 | 2.58E-05 | 0 | 0 | 3.92E-06 |
| Haus4 | 1.34E-05 | 9.20E-05 | 0 | 1.09E-05 | 4.32E-05 | 7.01E-06 | 7.91E-05 | 3.11E-06 | 0 | 0 | 0 | 0 | 0 | 8.42E-05 | 0 | 0.00065 | 2.74E-05 | 2.84E-05 |
| Gdpd5 | 0 | 7.68E-05 | 0 | 0 | 0.00017 | 7.00E-06 | 0 | 0 | 0 | 6.82E-05 | 0.00029 | 2.56E-05 | 0 | 0.00021 | 0 | 0 | 0 | 6.57E-05 |
| Prrg2 | 3.38E-05 | 2.77E-05 | 0 | 6.54E-05 | 8.70E-06 | 6.99E-06 | 1.63E-06 | 1.73E-05 | 0 | 2.42E-05 | 2.59E-05 | 4.76E-05 | 3.76E-05 | 4.60E-06 | 1.41E-05 | 0 | 2.19E-05 | 2.61E-05 |
| Dnmt3b | 5.44E-06 | 1.12E-05 | 1.43E-05 | 4.36E-06 | 6.94E-06 | 6.97E-06 | 8.57E-06 | 3.74E-08 | 0 | 4.55E-07 | 0 | 4.98E-07 | 0 | 0 | 2.21E-05 | 1.25E-05 | 1.25E-05 | 0 |
| Tcfl5 | 9.39E-06 | 0 | 2.91E-05 | 5.12E-06 | 3.87E-06 | 6.94E-06 | 7.20E-05 | 1.16E-06 | 0 | 1.33E-05 | 0 | 2.24E-05 | 0 | 0 | 0 | 1.60E-05 | 0 | 0 |
| Aqp1 | 0 | 3.69E-05 | 7.94E-05 | 0 | 0 | 6.92E-06 | 6.24E-05 | 0 | 0 | 4.78E-05 | 0 | 2.09E-05 | 0 | 5.69E-05 | 4.55E-05 | 9.30E-07 | 1.48E-05 | 0.00041 |
| Polr2l | 5.18E-08 | 1.28E-05 | 0 | 3.47E-07 | 6.04E-06 | 6.91E-06 | 0 | 0 | 0 | 0 | 4.55E-06 | 6.69E-08 | 1.08E-05 | 1.62E-06 | 0 | 4.76E-06 | 4.96E-05 | 8.90E-06 |
| Prag1 | 0 | 0 | 9.27E-05 | 0 | 4.15E-05 | 6.89E-06 | 3.60E-06 | 3.88E-05 | 0 | 0 | 2.67E-05 | 2.11E-05 | 0 | 0 | 3.91E-05 | 0 | 0 | 2.23E-05 |
| Lyplal1 | 3.07E-05 | 3.28E-05 | 0 | 2.55E-06 | 0 | 6.85E-06 | 4.65E-05 | 5.12E-05 | 0 | 1.92E-05 | 3.49E-06 | 2.24E-05 | 0 | 0 | 0 | 0 | 0 | 9.72E-06 |

| | | | | | | | | | | | | | | | | | | |
|---|---|---|---|---|---|---|---|---|---|---|---|---|---|---|---|---|---|---|
| Asic1 | 7.46E-08 | 0 | 1.47E-06 | 7.83E-07 | 1.84E-05 | 6.84E-06 | 0 | 0 | 0 | 0 | 1.36E-05 | 0 | 4.26E-07 | 1.65E-05 | 9.19E-07 | 0 | 0 | 5.65E-06 |
| Zfp551 | 6.38E-06 | 1.08E-05 | 0 | 0 | 1.15E-05 | 6.82E-06 | 0 | 0 | 0 | 6.40E-06 | 0 | 5.70E-06 | 0 | 0 | 1.13E-05 | 0 | 0 | 7.48E-06 |
| Shank2 | 0 | 3.98E-06 | 0 | 0 | 2.46E-06 | 6.79E-06 | 0 | 0 | 0 | 0 | 3.06E-05 | 2.89E-05 | 0 | 0 | 8.75E-06 | 0 | 0 | 0 |
| Hoxb9 | 0 | 5.38E-06 | 2.68E-06 | 0 | 1.42E-05 | 6.77E-06 | 0 | 0 | 0 | 0 | 3.59E-05 | 1.98E-05 | 0 | 0 | 4.83E-06 | 0 | 0 | 0 |
| BC055324 | 1.90E-06 | 1.31E-06 | 0 | 0 | 0 | 6.70E-06 | 0 | 0 | 0 | 2.70E-06 | 0 | 1.74E-05 | 1.62E-06 | 0 | 0 | 0.00016 | 0 | 0 |
| Rnf128 | 6.62E-06 | 3.18E-06 | 4.31E-06 | 0 | 0 | 6.70E-06 | 0 | 8.62E-08 | 0 | 0 | 3.05E-06 | 3.50E-05 | 0 | 3.84E-05 | 7.42E-07 | 0 | 0 | 0 |
| Agap2 | 0 | 4.74E-05 | 0 | 0 | 3.16E-05 | 6.69E-06 | 0 | 0.00017 | 0 | 0 | 1.14E-05 | 0 | 0 | 0 | 4.15E-06 | 0 | 0 | 0 |
| Hoxb8 | 1.34E-06 | 2.07E-06 | 3.45E-06 | 1.12E-05 | 2.13E-05 | 6.69E-06 | 0 | 1.80E-05 | 0 | 0 | 2.81E-05 | 0 | 0 | 2.31E-06 | 8.62E-06 | 0 | 0 | 0 |
| Nat14 | 3.04E-06 | 1.67E-05 | 4.75E-06 | 6.39E-05 | 2.55E-05 | 6.68E-06 | 4.35E-06 | 3.72E-05 | 0 | 1.07E-05 | 6.86E-06 | 6.84E-06 | 0 | 1.39E-05 | 3.61E-05 | 1.45E-06 | 1.80E-05 | 1.68E-05 |
| Mycn | 0 | 0 | 1.07E-05 | 9.52E-07 | 2.27E-06 | 6.64E-06 | 0 | 1.36E-06 | 0 | 0 | 6.36E-06 | 1.29E-05 | 0 | 1.38E-05 | 3.32E-06 | 1.73E-05 | 5.13E-06 | 0 |
| Styx | 2.37E-06 | 6.76E-06 | 0 | 0 | 0 | 6.63E-06 | 0 | 1.22E-06 | 0 | 0 | 0 | 4.47E-06 | 0 | 0 | 0 | 2.92E-06 | 1.68E-05 | 0 |
| Acsl1 | 1.38E-05 | 5.58E-05 | 1.11E-05 | 1.11E-05 | 0 | 6.63E-06 | 0 | 2.85E-05 | 0 | 4.18E-06 | 8.37E-06 | 8.25E-06 | 3.86E-05 | 0 | 0 | 1.34E-05 | 2.76E-05 | 0 |
| Glt8d2 | 9.38E-06 | 0 | 0 | 0 | 4.81E-06 | 6.63E-06 | 0 | 8.54E-06 | 0 | 6.84E-05 | 2.88E-06 | 0 | 0 | 1.93E-05 | 0 | 0 | 0 | 5.69E-05 |
| Pcdhb11 | 0 | 4.82E-06 | 0 | 1.82E-06 | 7.15E-06 | 6.63E-06 | 8.47E-06 | 5.97E-06 | 0 | 5.86E-06 | 4.17E-07 | 0 | 3.51E-06 | 5.94E-06 | 0 | 0 | 0 | 3.61E-06 |
| Rbks | 5.36E-06 | 4.40E-06 | 0 | 7.92E-06 | 4.46E-06 | 6.55E-06 | 0 | 1.02E-05 | 0 | 7.89E-06 | 0 | 0 | 1.93E-05 | 0 | 5.23E-06 | 7.98E-06 | 0 | 0 |
| Dipk1a | 2.29E-05 | 1.95E-05 | 0 | 0 | 0 | 6.54E-06 | 0 | 0 | 0 | 6.89E-05 | 0 | 1.76E-05 | 9.45E-05 | 0 | 0 | 5.85E-05 | 0 | 2.78E-05 |
| Fv1 | 0 | 1.66E-05 | 9.79E-06 | 0 | 4.95E-06 | 6.54E-06 | 0 | 3.48E-06 | 0 | 6.37E-06 | 2.64E-07 | 0 | 0 | 5.40E-06 | 4.63E-06 | 1.99E-05 | 0 | 5.64E-06 |
| Agmo | 5.86E-06 | 7.84E-06 | 0 | 0 | 0 | 6.51E-06 | 0 | 3.08E-05 | 0 | 1.71E-05 | 0 | 0 | 0 | 0 | 0 | 0 | 0 | 0.0002 |
| Carmil3 | 0 | 1.92E-05 | 0 | 0 | 0 | 6.46E-06 | 0 | 0 | 0 | 0 | 0 | 5.58E-06 | 0 | 2.85E-07 | 2.95E-05 | 0 | 0 | 0 |
| Zfp41 | 0 | 1.76E-05 | 0 | 4.28E-06 | 1.44E-05 | 6.39E-06 | 1.09E-05 | 0 | 0 | 7.64E-06 | 5.17E-06 | 0 | 0 | 1.28E-06 | 3.04E-05 | 6.76E-05 | 0 | 1.58E-05 |
| Zfp429 | 6.27E-06 | 1.06E-05 | 1.68E-07 | 0 | 7.72E-08 | 6.32E-06 | 7.92E-06 | 2.84E-06 | 0 | 2.27E-06 | 0 | 3.51E-06 | 1.72E-06 | 4.10E-06 | 0 | 5.33E-06 | 0 | 2.32E-06 |
| Aoc2 | 4.74E-06 | 5.41E-05 | 6.31E-05 | 1.72E-05 | 1.31E-05 | 6.27E-06 | 0 | 8.70E-06 | 0 | 7.31E-06 | 0 | 1.43E-05 | 6.55E-06 | 0 | 6.50E-05 | 1.42E-05 | 0.00011 | 0 |
| Tac1 | 0 | 2.78E-06 | 7.45E-08 | 6.19E-06 | 0 | 6.24E-06 | 1.16E-05 | 8.96E-07 | 0 | 0 | 0 | 0 | 1.16E-05 | 0 | 2.06E-06 | 0 | 8.57E-05 | 1.73E-06 |
| Pask | 0 | 0 | 0 | 0 | 5.89E-06 | 6.24E-06 | 0 | 3.06E-06 | 0 | 4.87E-06 | 7.03E-06 | 3.29E-06 | 1.45E-05 | 9.74E-07 | 9.09E-06 | 0.00021 | 0 | 7.30E-07 |
| Scn2b | 0 | 1.39E-05 | 0 | 0 | 1.57E-05 | 6.21E-06 | 0 | 2.62E-05 | 0 | 0 | 0 | 0 | 0 | 0 | 0 | 0 | 0 | 0 |
| Zfp420 | 1.08E-05 | 1.49E-05 | 1.95E-06 | 2.06E-06 | 8.83E-06 | 6.19E-06 | 0 | 0 | 0 | 1.27E-05 | 7.30E-06 | 1.47E-05 | 1.24E-06 | 4.51E-06 | 6.53E-06 | 2.54E-07 | 0 | 1.04E-06 |
| Serpinb8 | 0 | 9.98E-06 | 0 | 0 | 0 | 6.14E-06 | 0 | 1.10E-05 | 0 | 2.04E-05 | 0 | 9.81E-06 | 0 | 0 | 0 | 3.82E-05 | 0 | 3.30E-05 |
| Nr3c2 | 0 | 9.51E-06 | 0 | 0 | 0 | 6.12E-06 | 0 | 1.16E-05 | 0 | 6.19E-06 | 0 | 0 | 0 | 0 | 1.43E-05 | 0 | 0 | 1.81E-05 |
| Tet1 | 8.19E-06 | 4.66E-07 | 0 | 0 | 9.73E-06 | 6.09E-06 | 0 | 1.08E-05 | 0 | 7.64E-07 | 0 | 1.74E-05 | 0 | 7.86E-06 | 2.80E-06 | 1.48E-06 | 0 | 6.07E-06 |
| Samd11 | 0 | 9.02E-07 | 0 | 0 | 4.55E-06 | 5.98E-06 | 0 | 8.91E-06 | 0 | 2.59E-06 | 3.54E-06 | 1.92E-05 | 0 | 0 | 0 | 0 | 1.58E-05 | 0 |
| Naip2 | 0 | 0 | 2.00E-06 | 7.63E-06 | 0 | 5.98E-06 | 4.47E-06 | 4.48E-06 | 0 | 6.39E-06 | 0 | 1.14E-06 | 1.28E-05 | 0 | 4.70E-07 | 3.27E-07 | 1.82E-06 | 6.78E-06 |
| Cdk18 | 5.30E-07 | 3.11E-06 | 8.32E-06 | 1.55E-05 | 0 | 5.94E-06 | 0 | 2.91E-06 | 0 | 2.21E-06 | 1.24E-05 | 2.87E-06 | 0 | 6.52E-06 | 1.39E-05 | 0 | 0 | 8.38E-07 |
| Zfp78 | 7.49E-06 | 0 | 0 | 0 | 2.24E-06 | 5.94E-06 | 0 | 5.42E-06 | 0 | 1.56E-05 | 4.52E-06 | 9.70E-06 | 0 | 2.55E-07 | 9.66E-06 | 0 | 0 | 3.60E-06 |

| | | | | | | | | | | | | | | | | | | |
|---|---|---|---|---|---|---|---|---|---|---|---|---|---|---|---|---|---|---|
| Cit | 0 | 0 | 0 | 0 | 6.80E-05 | 5.89E-06 | 0 | 0 | 0 | 0 | 3.52E-05 | 2.88E-05 | 0 | 0 | 0.00019 | 0.00053 | 1.79E-05 | 0 |
| Cdnf | 0 | 9.81E-06 | 0 | 2.30E-06 | 5.55E-06 | 5.89E-06 | 5.76E-06 | 0 | 0 | 4.02E-06 | 3.60E-06 | 4.70E-06 | 0 | 0 | 0 | 5.78E-06 | 0 | 8.63E-06 |
| Nr6a1 | 0 | 2.87E-05 | 9.33E-06 | 1.30E-05 | 0 | 5.88E-06 | 7.81E-06 | 1.33E-05 | 0 | 5.48E-06 | 5.89E-06 | 1.57E-06 | 0 | 2.50E-06 | 1.64E-05 | 0 | 7.40E-05 | 3.13E-06 |
| Bard1 | 0 | 0 | 5.33E-06 | 0 | 0 | 5.86E-06 | 0 | 0 | 0 | 0 | 0 | 0 | 2.51E-06 | 3.64E-06 | 0 | 0.00021 | 4.62E-06 | 3.74E-07 |
| Arhgap19 | 0 | 6.32E-06 | 0 | 0 | 5.43E-07 | 5.85E-06 | 0 | 0 | 0 | 0 | 0 | 3.83E-06 | 0 | 0 | 0 | 0.0003 | 0 | 0 |
| Tmem37 | 0 | 8.36E-07 | 0 | 0 | 4.26E-06 | 5.76E-06 | 0 | 1.06E-07 | 0 | 0 | 0 | 3.65E-05 | 0 | 4.05E-06 | 1.83E-05 | 7.38E-07 | 0 | 3.94E-06 |
| Adssl1 | 0 | 3.09E-05 | 0 | 0 | 0 | 5.71E-06 | 0 | 0 | 0 | 0 | 1.15E-06 | 3.56E-05 | 0 | 3.97E-05 | 1.69E-05 | 0 | 5.84E-05 | 0 |
| Oacyl | 0 | 0 | 9.76E-06 | 0 | 0 | 5.70E-06 | 0 | 2.46E-05 | 0 | 0 | 0 | 0 | 0 | 1.31E-05 | 1.64E-06 | 6.47E-06 | 0 | 0 |
| Fmn2 | 0 | 0 | 0 | 0 | 1.21E-05 | 5.62E-06 | 0 | 0 | 0 | 1.67E-06 | 2.94E-05 | 0 | 0 | 1.41E-05 | 1.51E-05 | 2.09E-05 | 1.65E-05 | 0 |
| Plk4 | 3.41E-05 | 0 | 1.32E-06 | 0 | 6.67E-06 | 5.62E-06 | 3.21E-05 | 0 | 0 | 0 | 5.09E-06 | 0 | 7.98E-07 | 0 | 4.15E-05 | 0.00072 | 2.40E-05 | 1.49E-06 |
| St3gal4 | 0.00012 | 0.00013 | 0.00015 | 1.51E-05 | 0.00017 | 5.53E-06 | 0.00034 | 9.05E-05 | 0 | 0.0002 | 0 | 0 | 0.00034 | 0.00018 | 0 | 0.00016 | 5.19E-05 | 0.00021 |
| Haspin | 0 | 5.75E-07 | 0 | 0 | 9.02E-06 | 5.25E-06 | 0 | 0 | 0 | 2.36E-06 | 0 | 1.04E-06 | 0 | 0 | 2.91E-06 | 0.0002 | 0 | 0 |
| Drd4 | 0 | 0 | 7.96E-06 | 1.71E-05 | 5.32E-06 | 5.21E-06 | 1.98E-05 | 0 | 0 | 0 | 1.25E-05 | 0 | 2.55E-06 | 1.57E-06 | 0 | 0 | 0 | 0 |
| Usp13 | 1.85E-06 | 7.77E-06 | 4.38E-05 | 0 | 5.23E-05 | 5.20E-06 | 4.35E-05 | 0 | 0 | 6.39E-06 | 7.62E-05 | 0.00036 | 2.89E-06 | 2.67E-05 | 0.00018 | 5.50E-05 | 2.17E-05 | 0 |
| Zfp458 | 7.03E-06 | 6.46E-06 | 0 | 0 | 2.39E-06 | 5.20E-06 | 1.91E-06 | 0 | 0 | 0 | 1.30E-06 | 8.81E-07 | 5.32E-08 | 0 | 0 | 6.38E-07 | 0 | 3.77E-06 |
| H2-DMa | 0 | 2.87E-06 | 0 | 2.88E-05 | 8.18E-06 | 5.16E-06 | 2.59E-06 | 1.07E-05 | 0 | 3.82E-06 | 3.60E-06 | 1.60E-05 | 2.53E-05 | 0 | 3.28E-05 | 0 | 3.59E-05 | 1.10E-05 |
| Fam129c | 0 | 3.52E-06 | 1.37E-06 | 0 | 6.18E-06 | 5.08E-06 | 0 | 3.06E-06 | 0 | 2.12E-06 | 2.80E-06 | 5.67E-06 | 4.65E-06 | 9.75E-06 | 1.69E-05 | 0 | 1.05E-05 | 2.82E-06 |
| Fcor | 1.90E-07 | 2.01E-06 | 0 | 5.11E-07 | 1.99E-08 | 5.05E-06 | 0 | 0 | 0 | 2.72E-06 | 0 | 9.36E-06 | 3.76E-05 | 3.66E-06 | 0 | 0 | 0 | 0 |
| Fancg | 0 | 2.11E-05 | 9.41E-07 | 6.97E-06 | 8.48E-06 | 5.01E-06 | 3.17E-06 | 0 | 0 | 1.20E-05 | 6.79E-06 | 7.15E-06 | 8.03E-06 | 9.33E-06 | 4.62E-05 | 8.67E-05 | 3.36E-06 | 1.91E-05 |
| Trpc1 | 1.59E-05 | 4.30E-05 | 6.37E-05 | 0 | 2.13E-06 | 5.00E-06 | 5.46E-06 | 3.38E-05 | 0 | 2.06E-05 | 4.79E-05 | 0.00013 | 0 | 3.88E-05 | 4.19E-05 | 0 | 3.57E-05 | 1.92E-05 |
| Pkmyt1 | 0 | 0 | 4.47E-06 | 5.16E-06 | 0 | 4.98E-06 | 0 | 0 | 0 | 3.70E-06 | 1.43E-05 | 0 | 1.76E-05 | 3.58E-06 | 1.89E-05 | 0.00044 | 4.32E-06 | 4.02E-07 |
| P3h2 | 3.83E-06 | 4.94E-06 | 0 | 6.85E-06 | 1.77E-06 | 4.93E-06 | 0 | 8.20E-06 | 0 | 1.41E-06 | 4.38E-07 | 3.42E-07 | 2.18E-05 | 7.96E-06 | 0 | 4.11E-06 | 0 | 2.26E-06 |
| Mylpf | 5.08E-06 | 1.42E-06 | 0 | 0 | 1.35E-06 | 4.89E-06 | 8.66E-06 | 1.81E-06 | 0 | 0 | 0 | 2.63E-06 | 0 | 1.34E-06 | 0 | 1.41E-05 | 0 | 0 |
| Lrtm1 | 0 | 0 | 0 | 0 | 6.33E-05 | 4.85E-06 | 7.36E-06 | 0 | 0 | 0 | 2.77E-05 | 0 | 0 | 0 | 0 | 0 | 0 | 0 |
| Catsper2 | 5.63E-06 | 2.96E-06 | 6.78E-06 | 1.35E-05 | 4.98E-06 | 4.84E-06 | 4.44E-06 | 7.81E-06 | 0 | 2.53E-06 | 0 | 8.47E-06 | 8.17E-06 | 0 | 1.71E-05 | 5.71E-07 | 8.62E-06 | 1.23E-05 |
| Tmem151b | 0 | 1.71E-05 | 0 | 0 | 3.68E-05 | 4.82E-06 | 0 | 0 | 0 | 0 | 7.91E-06 | 0 | 0 | 2.74E-05 | 1.36E-05 | 0 | 9.74E-06 | 0 |
| Ms4a6d | 3.12E-06 | 1.81E-06 | 5.00E-06 | 4.43E-07 | 1.17E-05 | 4.81E-06 | 0 | 6.10E-05 | 0 | 7.33E-06 | 0 | 0 | 9.52E-06 | 1.38E-05 | 0 | 1.76E-05 | 0 | 0 |
| Fam227a | 1.88E-06 | 7.11E-06 | 0 | 7.74E-07 | 5.47E-06 | 4.80E-06 | 0 | 4.69E-06 | 0 | 0 | 0 | 1.07E-06 | 0 | 2.80E-06 | 9.67E-06 | 0 | 0 | 5.75E-07 |
| Pik3r3 | 2.21E-05 | 1.78E-05 | 0 | 7.64E-06 | 0 | 4.75E-06 | 4.73E-07 | 2.96E-05 | 0 | 0 | 1.48E-05 | 0 | 1.06E-05 | 6.13E-06 | 0 | 1.90E-05 | 2.89E-05 | 0.00011 |
| Atf4 | 1.44E-06 | 0.00012 | 0.0001 | 0.00021 | 4.27E-05 | 4.74E-06 | 0 | 8.81E-06 | 0 | 3.01E-05 | 0.00011 | 0.00027 | 0 | 4.62E-05 | 0.00022 | 2.31E-05 | 0.00099 | 0 |
| Kcnmb4 | 0 | 1.34E-05 | 0 | 0 | 3.63E-05 | 4.61E-06 | 0 | 7.80E-06 | 0 | 0 | 4.06E-05 | 5.50E-06 | 0 | 0 | 2.59E-08 | 0 | 3.25E-05 | 0 |
| Oit3 | 0 | 4.39E-06 | 0 | 0 | 3.13E-05 | 4.55E-06 | 0 | 0 | 0 | 0 | 0 | 1.10E-05 | 0 | 1.46E-06 | 0 | 0 | 2.62E-06 | 0 |
| Mfsd6 | 0 | 7.08E-06 | 0 | 0 | 0 | 4.44E-06 | 1.12E-05 | 2.34E-05 | 0 | 6.22E-05 | 0 | 0 | 0 | 8.47E-06 | 0 | 9.64E-06 | 0 | 8.71E-05 |

| | | | | | | | | | | | | | | | | | | |
|---|---|---|---|---|---|---|---|---|---|---|---|---|---|---|---|---|---|---|
| Tmem198 | 1.27E-06 | 3.16E-06 | 5.10E-06 | 1.81E-05 | 4.74E-06 | 4.41E-06 | 1.44E-07 | 1.36E-05 | 0 | 0 | 0 | 2.61E-06 | 6.40E-06 | 2.83E-06 | 6.13E-06 | 0 | 1.05E-05 | 0 |
| Cep57l1 | 7.39E-06 | 3.15E-06 | 0 | 0 | 0 | 4.41E-06 | 0 | 0 | 0 | 6.59E-06 | 0 | 1.53E-05 | 0 | 0 | 3.50E-05 | 8.70E-05 | 0 | 7.61E-06 |
| Tlr5 | 0 | 1.39E-06 | 0 | 0 | 0 | 4.41E-06 | 4.57E-05 | 0 | 0 | 8.46E-05 | 0 | 2.92E-05 | 2.12E-05 | 2.49E-06 | 0 | 0 | 0 | 0 |
| Ccdc28a | 0 | 3.44E-05 | 0 | 7.26E-06 | 1.49E-05 | 4.41E-06 | 3.21E-05 | 1.46E-05 | 0 | 2.37E-05 | 0 | 3.25E-05 | 1.81E-05 | 0 | 2.23E-05 | 0 | 0 | 2.67E-05 |
| Tns4 | 0 | 4.16E-06 | 0 | 6.90E-06 | 9.09E-06 | 4.39E-06 | 0 | 5.81E-06 | 0 | 1.31E-05 | 5.11E-06 | 9.33E-06 | 1.36E-05 | 6.57E-06 | 5.63E-05 | 5.54E-06 | 1.71E-05 | 3.97E-05 |
| Kntc1 | 0 | 0 | 0 | 0 | 0 | 4.33E-06 | 0 | 0 | 0 | 0 | 0 | 0 | 0 | 0 | 0 | 0.00029 | 0 | 0 |
| Fam229b | 1.15E-05 | 6.20E-06 | 0 | 1.12E-05 | 5.87E-06 | 4.24E-06 | 1.60E-05 | 4.66E-06 | 0 | 1.27E-05 | 1.73E-06 | 1.10E-06 | 0 | 0 | 0 | 1.03E-05 | 0 | 1.13E-05 |
| Sdccag8 | 7.58E-05 | 5.64E-05 | 1.39E-06 | 0 | 0 | 4.23E-06 | 5.00E-05 | 5.02E-05 | 0 | 0.0001 | 0 | 0.00012 | 1.36E-05 | 0 | 0.00015 | 3.27E-05 | 0 | 2.33E-05 |
| Morn1 | 0 | 2.32E-05 | 0 | 9.94E-06 | 1.08E-05 | 4.22E-06 | 0 | 0 | 0 | 1.05E-05 | 0 | 5.74E-06 | 1.96E-06 | 0 | 2.46E-05 | 1.73E-06 | 3.90E-06 | 4.33E-06 |
| Lipt1 | 1.48E-05 | 6.65E-06 | 0 | 4.05E-06 | 1.25E-06 | 4.17E-06 | 0 | 0 | 0 | 1.12E-05 | 1.07E-05 | 4.20E-07 | 0 | 2.10E-06 | 0 | 3.27E-06 | 0 | 2.66E-06 |
| Zbed5 | 2.44E-05 | 0 | 1.33E-05 | 6.39E-06 | 9.73E-06 | 4.17E-06 | 4.21E-05 | 2.47E-05 | 0 | 0 | 0 | 6.66E-06 | 0 | 2.39E-05 | 3.61E-05 | 0 | 3.61E-05 | 6.12E-06 |
| Kcnab3 | 4.93E-07 | 1.29E-05 | 0 | 8.31E-06 | 6.00E-06 | 4.15E-06 | 1.06E-05 | 2.60E-06 | 0 | 0 | 0 | 0 | 0 | 1.28E-05 | 1.31E-06 | 1.71E-05 | 0 | 0 |
| Pde2a | 0 | 0 | 0 | 0 | 0 | 4.12E-06 | 0 | 7.82E-05 | 0 | 1.70E-05 | 4.81E-05 | 4.98E-05 | 6.30E-06 | 0.00011 | 0 | 0 | 0 | 5.59E-05 |
| Mmaa | 6.44E-06 | 3.70E-06 | 0 | 0 | 5.80E-06 | 4.11E-06 | 0 | 0 | 0 | 1.29E-05 | 5.18E-06 | 0 | 6.39E-06 | 3.74E-06 | 2.30E-05 | 1.06E-05 | 0 | 6.02E-06 |
| Ovgp1 | 1.26E-07 | 1.03E-05 | 2.22E-06 | 3.15E-06 | 1.64E-06 | 4.06E-06 | 0 | 1.37E-05 | 0 | 6.35E-06 | 0 | 6.71E-06 | 1.44E-05 | 5.69E-06 | 1.94E-05 | 1.33E-06 | 5.01E-05 | 0 |
| Dnaaf2 | 2.35E-05 | 5.00E-06 | 1.68E-05 | 0 | 8.87E-06 | 4.01E-06 | 0 | 0 | 0 | 6.83E-06 | 3.17E-06 | 2.02E-05 | 1.62E-06 | 0 | 1.10E-05 | 0.00012 | 0 | 0 |
| Xkrx | 0 | 0 | 0 | 0 | 3.39E-05 | 4.00E-06 | 0 | 0 | 0 | 1.02E-06 | 6.13E-05 | 0 | 0 | 0 | 0 | 0 | 0 | 0 |
| Zfp827 | 0 | 3.32E-05 | 1.64E-05 | 2.40E-05 | 9.56E-05 | 3.92E-06 | 9.83E-06 | 0 | 0 | 5.16E-05 | 0.00025 | 6.60E-05 | 0 | 5.71E-05 | 0.00013 | 0 | 2.57E-06 | 0 |
| Siglecg | 0 | 0 | 0 | 0 | 0 | 3.90E-06 | 0 | 0 | 0 | 0 | 4.68E-05 | 0 | 0 | 7.01E-06 | 2.05E-06 | 5.12E-07 | 0 | 0 |
| Pde4dip | 9.10E-05 | 6.42E-05 | 0.00011 | 0.00015 | 7.16E-05 | 3.89E-06 | 0.00025 | 7.17E-05 | 0 | 3.19E-05 | 5.38E-05 | 0.00024 | 0 | 7.50E-05 | 0.00032 | 3.82E-05 | 0 | 0.00013 |
| Ubxn11 | 0 | 1.43E-05 | 0 | 9.34E-06 | 8.34E-07 | 3.83E-06 | 8.37E-06 | 0 | 0 | 3.07E-06 | 2.86E-06 | 2.35E-06 | 1.47E-05 | 0 | 1.88E-05 | 0 | 1.14E-05 | 0 |
| Zwilch | 6.85E-07 | 0 | 0 | 0 | 0 | 3.78E-06 | 0 | 0 | 0 | 2.94E-06 | 0 | 5.20E-06 | 2.58E-06 | 0 | 2.82E-05 | 0.00042 | 0 | 0 |
| Shpk | 0 | 2.31E-05 | 0 | 7.70E-06 | 1.09E-05 | 3.77E-06 | 5.71E-06 | 2.42E-06 | 0 | 3.23E-05 | 8.39E-06 | 4.31E-06 | 1.01E-05 | 0 | 2.65E-06 | 0 | 0 | 4.32E-06 |
| Cenpn | 6.86E-06 | 1.27E-06 | 0 | 0 | 0 | 3.69E-06 | 0 | 0 | 0 | 1.65E-06 | 0 | 3.77E-07 | 0 | 6.26E-07 | 0 | 0.00046 | 0 | 0 |
| Mmp16 | 0 | 3.62E-05 | 0 | 0 | 7.79E-05 | 3.68E-06 | 0 | 0 | 0 | 0 | 0 | 0 | 0 | 1.34E-05 | 0 | 0 | 0 | 1.71E-05 |
| Cmas | 0.00017 | 2.20E-05 | 0.00013 | 0.00022 | 3.78E-05 | 3.68E-06 | 0.00044 | 0.00017 | 0 | 4.34E-05 | 5.92E-05 | 0.00015 | 5.01E-05 | 0 | 0.0001 | 0 | 0 | 3.80E-05 |
| Vdr | 1.59E-05 | 1.65E-05 | 8.91E-06 | 0 | 0 | 3.66E-06 | 0 | 1.53E-06 | 0 | 0 | 1.48E-05 | 0 | 5.10E-05 | 0.00013 | 2.22E-05 | 0 | 0 | 0.00011 |
| Phgdh | 2.27E-05 | 9.01E-06 | 4.51E-05 | 2.33E-05 | 1.57E-06 | 3.60E-06 | 0 | 0 | 0 | 5.02E-06 | 1.48E-05 | 1.80E-05 | 0 | 7.75E-06 | 0 | 7.77E-05 | 0 | 2.57E-05 |
| Zfp874a | 1.23E-05 | 1.28E-05 | 0 | 0 | 9.46E-06 | 3.60E-06 | 0 | 0 | 0 | 5.66E-06 | 2.06E-06 | 0 | 5.74E-06 | 3.98E-07 | 0 | 0 | 0 | 1.58E-07 |
| AW554918 | 3.02E-05 | 4.00E-05 | 0 | 0 | 0 | 3.57E-06 | 0 | 2.63E-06 | 0 | 2.13E-05 | 1.24E-05 | 2.31E-05 | 0 | 1.02E-05 | 0.00022 | 0.00011 | 0 | 5.04E-06 |
| Mipol1 | 1.08E-05 | 1.22E-05 | 5.82E-06 | 1.66E-06 | 1.04E-05 | 3.56E-06 | 5.24E-06 | 0 | 0 | 8.73E-06 | 2.77E-06 | 8.71E-06 | 9.43E-06 | 0 | 2.71E-05 | 6.32E-06 | 2.16E-05 | 5.58E-06 |
| Dbf4 | 1.53E-05 | 2.07E-05 | 3.48E-06 | 0 | 1.71E-05 | 3.51E-06 | 2.30E-05 | 0 | 0 | 0 | 0 | 0 | 0 | 2.38E-05 | 3.70E-05 | 0.00059 | 1.26E-05 | 0 |
| Nudt12 | 9.57E-08 | 1.10E-05 | 3.71E-06 | 1.71E-06 | 1.40E-06 | 3.47E-06 | 0 | 9.42E-06 | 0 | 4.97E-06 | 0 | 0 | 7.92E-06 | 3.31E-07 | 0 | 5.85E-07 | 0 | 1.06E-05 |

| | | | | | | | | | | | | | | | | | | |
|---|---|---|---|---|---|---|---|---|---|---|---|---|---|---|---|---|---|---|
| Tarm1 | 0 | 0 | 0 | 4.01E-06 | 7.69E-06 | 3.44E-06 | 0 | 0 | 0 | 9.16E-07 | 3.99E-05 | 1.04E-06 | 5.44E-06 | 2.34E-08 | 2.62E-06 | 6.58E-06 | 0 | 0 |
| Cenpu | 0 | 0 | 0 | 0 | 0 | 3.39E-06 | 0 | 0 | 0 | 0 | 0 | 0 | 0 | 2.48E-06 | 1.23E-05 | 0.00016 | 0 | 0 |
| Gk5 | 3.89E-05 | 1.61E-06 | 2.70E-05 | 0 | 0 | 3.35E-06 | 2.47E-05 | 5.35E-06 | 0 | 1.12E-05 | 0 | 5.80E-05 | 3.16E-05 | 1.27E-05 | 4.26E-05 | 0 | 0.00019 | 0 |
| H2-T24 | 0 | 4.49E-06 | 0 | 0 | 2.36E-06 | 3.26E-06 | 0 | 1.24E-06 | 0 | 0.00011 | 0 | 4.85E-07 | 1.33E-05 | 1.48E-06 | 0 | 0 | 0 | 1.84E-05 |
| Ccdc15 | 8.85E-06 | 1.92E-05 | 2.42E-06 | 0 | 6.53E-06 | 3.20E-06 | 1.34E-05 | 2.03E-06 | 0 | 3.10E-06 | 0 | 3.75E-06 | 0 | 0 | 1.89E-05 | 9.92E-05 | 0 | 4.56E-06 |
| Csf2rb | 0 | 1.09E-05 | 0 | 0 | 0 | 3.17E-06 | 0 | 0 | 0 | 4.58E-07 | 0 | 9.30E-07 | 1.62E-05 | 3.62E-06 | 5.26E-05 | 1.09E-05 | 1.31E-05 | 6.50E-05 |
| Zfp747 | 9.09E-06 | 1.28E-05 | 0 | 3.20E-06 | 1.06E-05 | 3.17E-06 | 0 | 3.18E-06 | 0 | 6.69E-06 | 5.77E-06 | 1.78E-05 | 0 | 4.94E-06 | 3.10E-05 | 8.22E-06 | 0 | 8.23E-06 |
| Shroom1 | 1.13E-06 | 3.10E-06 | 6.21E-06 | 2.31E-05 | 1.52E-05 | 3.15E-06 | 2.13E-05 | 2.96E-06 | 0 | 0 | 2.13E-05 | 1.17E-05 | 3.27E-06 | 4.74E-06 | 3.68E-05 | 0 | 5.29E-06 | 5.83E-06 |
| Incenp | 0 | 1.74E-05 | 0 | 0 | 1.29E-05 | 3.10E-06 | 0 | 0 | 0 | 0 | 0 | 0 | 9.29E-06 | 3.43E-06 | 8.16E-05 | 0.00083 | 0 | 0 |
| Hcn2 | 4.02E-06 | 7.67E-06 | 8.34E-06 | 0 | 1.20E-06 | 3.10E-06 | 1.49E-05 | 2.17E-06 | 0 | 2.46E-05 | 0 | 1.92E-05 | 1.73E-05 | 0 | 0 | 5.83E-05 | 0 | 0 |
| Acat3 | 0 | 0 | 0 | 0 | 5.17E-05 | 3.09E-06 | 0 | 0 | 0 | 0 | 0 | 0 | 0.00024 | 8.93E-05 | 0 | 0 | 0 | 0 |
| Ptgs2 | 0 | 0 | 0.0002 | 0 | 4.16E-05 | 2.99E-06 | 0 | 0.00017 | 0 | 5.60E-06 | 0.00013 | 0.00073 | 0 | 0 | 3.81E-05 | 0 | 0.00163 | 1.66E-05 |
| C1s2 | 0 | 5.69E-06 | 0 | 0 | 0 | 2.94E-06 | 0 | 9.40E-07 | 0 | 1.93E-05 | 0 | 0 | 3.14E-05 | 0 | 0 | 0 | 0 | 3.32E-05 |
| Tinagl1 | 0 | 1.54E-05 | 0 | 1.63E-05 | 9.56E-06 | 2.89E-06 | 0 | 0 | 0 | 7.18E-06 | 6.52E-05 | 2.32E-05 | 0 | 0 | 4.85E-05 | 0 | 0 | 4.94E-06 |
| Mcm7 | 0 | 2.32E-05 | 0.00012 | 0.00014 | 3.02E-06 | 2.87E-06 | 2.53E-06 | 1.94E-05 | 0 | 1.36E-05 | 1.98E-06 | 0 | 3.00E-05 | 2.66E-05 | 1.74E-05 | 0.00071 | 3.32E-05 | 0 |
| Nmb | 2.76E-06 | 1.42E-06 | 2.40E-06 | 0 | 0 | 2.86E-06 | 0 | 0 | 0 | 6.60E-06 | 0 | 1.60E-06 | 0 | 2.36E-06 | 1.47E-05 | 2.01E-05 | 6.69E-06 | 8.47E-06 |
| Gpr137c | 0 | 6.62E-06 | 0 | 6.01E-06 | 4.58E-06 | 2.70E-06 | 0 | 7.31E-06 | 0 | 0 | 1.11E-05 | 0 | 0 | 1.61E-05 | 1.07E-05 | 8.53E-06 | 5.82E-05 | 0 |
| Inca1 | 0 | 1.19E-05 | 0 | 0 | 6.73E-06 | 2.69E-06 | 0 | 0 | 0 | 2.23E-06 | 4.95E-06 | 6.58E-06 | 1.26E-05 | 0 | 1.97E-06 | 0 | 0 | 0 |
| Cd37 | 0 | 1.80E-06 | 2.71E-06 | 7.46E-06 | 3.74E-06 | 2.65E-06 | 0 | 1.58E-06 | 0 | 2.83E-06 | 0 | 1.94E-06 | 5.32E-07 | 0 | 1.20E-05 | 1.64E-06 | 6.13E-06 | 6.36E-06 |
| Cdkl5 | 0 | 3.93E-06 | 0 | 0 | 1.25E-05 | 2.64E-06 | 9.09E-06 | 0 | 0 | 3.68E-06 | 2.08E-05 | 1.66E-05 | 7.08E-06 | 0 | 0 | 1.30E-05 | 0 | 0 |
| Ercc6l | 8.10E-06 | 0 | 2.84E-06 | 0 | 0 | 2.64E-06 | 0 | 0 | 0 | 0 | 0 | 0 | 0 | 0 | 0 | 0.00029 | 0 | 0 |
| Slc23a3 | 0 | 2.10E-05 | 0 | 0 | 0 | 2.64E-06 | 0 | 0 | 0 | 0 | 0 | 3.58E-05 | 0 | 0 | 0 | 0 | 0.00015 | 0 |
| Il2rb | 0 | 2.85E-06 | 2.64E-06 | 1.33E-07 | 1.30E-06 | 2.63E-06 | 3.91E-06 | 7.21E-06 | 0 | 9.02E-07 | 2.66E-06 | 1.10E-06 | 2.62E-06 | 3.76E-06 | 6.90E-06 | 1.41E-06 | 1.08E-06 | 4.45E-06 |
| Lnx2 | 4.58E-05 | 5.80E-05 | 4.16E-05 | 0.00012 | 7.94E-05 | 2.62E-06 | 8.99E-05 | 4.86E-05 | 0 | 2.55E-05 | 5.44E-05 | 3.69E-05 | 3.30E-05 | 1.21E-05 | 2.83E-05 | 1.96E-05 | 0 | 1.25E-05 |
| Cdc45 | 0 | 0 | 5.99E-05 | 0 | 1.24E-06 | 2.58E-06 | 1.33E-05 | 0 | 0 | 0 | 1.14E-05 | 8.04E-06 | 5.20E-06 | 1.32E-05 | 0 | 0.00048 | 6.87E-07 | 0 |
| Tdrkh | 0 | 0 | 0 | 0 | 0 | 2.56E-06 | 0 | 8.27E-06 | 0 | 5.50E-06 | 1.83E-05 | 9.53E-06 | 0 | 0 | 0 | 2.13E-05 | 0 | 0 |
| Adam5 | 6.96E-06 | 4.95E-07 | 0 | 0 | 0 | 2.55E-06 | 0 | 0 | 0 | 1.04E-05 | 0 | 0 | 4.69E-06 | 0 | 0 | 0 | 0 | 1.79E-05 |
| Efcab7 | 1.10E-05 | 8.20E-06 | 0 | 5.93E-07 | 0 | 2.54E-06 | 4.76E-06 | 0 | 0 | 2.59E-06 | 0 | 3.85E-06 | 6.86E-07 | 0 | 8.95E-06 | 1.98E-05 | 3.25E-06 | 6.88E-06 |
| Ikzf1 | 0 | 1.09E-06 | 4.65E-06 | 0 | 0 | 2.54E-06 | 1.23E-08 | 0 | 0 | 7.17E-06 | 4.68E-06 | 2.77E-06 | 4.69E-06 | 1.95E-06 | 1.15E-05 | 4.11E-06 | 3.47E-07 | 6.81E-06 |
| Zfp14 | 9.89E-06 | 1.97E-05 | 1.79E-06 | 0 | 9.24E-06 | 2.54E-06 | 6.33E-07 | 1.70E-06 | 0 | 6.42E-06 | 1.74E-06 | 2.04E-07 | 0 | 0 | 1.05E-05 | 4.44E-06 | 0 | 1.15E-05 |
| Ncapg2 | 0 | 0 | 2.31E-05 | 0 | 2.75E-06 | 2.52E-06 | 0 | 0 | 0 | 0 | 0 | 1.28E-05 | 1.14E-05 | 1.46E-06 | 1.03E-05 | 0.00073 | 6.57E-05 | 4.85E-07 |
| Ctnnal1 | 0 | 9.30E-06 | 0 | 0 | 0 | 2.51E-06 | 0 | 0 | 0 | 1.27E-06 | 0 | 1.70E-05 | 0 | 0 | 6.92E-06 | 0 | 1.28E-05 | 3.18E-05 |
| Dixdc1 | 0 | 7.69E-06 | 0 | 0 | 0 | 2.46E-06 | 0 | 0 | 0 | 1.76E-06 | 4.01E-06 | 6.06E-05 | 0 | 2.16E-05 | 2.42E-05 | 0 | 0 | 1.55E-05 |

| | | | | | | | | | | | | | | | | | | |
|---|---|---|---|---|---|---|---|---|---|---|---|---|---|---|---|---|---|---|
| Dhtkd1 | 4.76E-06 | 2.38E-05 | 0 | 0 | 2.13E-06 | 2.41E-06 | 0 | 0 | 0 | 1.48E-07 | 0 | 0 | 0 | 1.48E-06 | 3.33E-05 | 0 | 0 | 0 |
| Igflr1 | 0 | 4.14E-05 | 0 | 3.69E-05 | 1.94E-06 | 2.34E-06 | 0 | 1.83E-05 | 0 | 9.52E-09 | 0 | 2.01E-05 | 0 | 0 | 7.69E-05 | 2.38E-05 | 2.44E-05 | 8.65E-06 |
| Trbc2 | 0 | 9.51E-06 | 0 | 0 | 2.59E-05 | 2.33E-06 | 0 | 0 | 0 | 0 | 1.70E-05 | 0 | 0 | 0 | 1.13E-06 | 0 | 0 | 4.64E-07 |
| Fam171b | 2.56E-05 | 4.66E-06 | 8.46E-06 | 7.70E-06 | 0 | 2.28E-06 | 4.11E-05 | 1.89E-06 | 0 | 5.06E-05 | 2.29E-05 | 0 | 0 | 4.34E-05 | 0 | 1.91E-05 | 0 | 1.79E-05 |
| Naif1 | 1.34E-05 | 1.81E-05 | 0 | 1.47E-05 | 6.78E-06 | 2.25E-06 | 9.90E-06 | 3.11E-06 | 0 | 3.26E-06 | 8.33E-06 | 1.42E-05 | 0 | 8.35E-06 | 0 | 4.08E-05 | 0 | 5.42E-06 |
| Neurl2 | 0 | 3.48E-05 | 0 | 2.27E-05 | 1.36E-05 | 2.24E-06 | 0 | 0 | 0 | 0 | 0 | 3.82E-06 | 0 | 2.32E-06 | 0 | 0 | 1.82E-05 | 2.30E-05 |
| Nars2 | 3.42E-05 | 1.26E-05 | 2.64E-05 | 2.63E-05 | 1.39E-05 | 2.22E-06 | 0 | 6.24E-06 | 0 | 9.69E-06 | 3.91E-06 | 2.62E-05 | 0 | 0 | 5.48E-05 | 0 | 0 | 1.04E-05 |
| Ppp4r4 | 1.61E-06 | 1.65E-05 | 0 | 0 | 3.86E-06 | 2.11E-06 | 0 | 0 | 0 | 6.37E-06 | 0 | 6.42E-06 | 0 | 3.74E-07 | 6.32E-06 | 7.62E-06 | 6.30E-07 | 0 |
| Dmd | 0 | 1.72E-05 | 0 | 0 | 9.75E-05 | 2.09E-06 | 1.74E-05 | 0 | 0 | 2.75E-05 | 0.00031 | 0 | 1.10E-05 | 0 | 0 | 2.90E-05 | 0 | 9.50E-05 |
| Sfn | 0 | 1.78E-05 | 3.29E-07 | 9.05E-06 | 0 | 2.08E-06 | 1.46E-05 | 6.11E-06 | 0 | 1.52E-05 | 3.48E-05 | 0 | 0 | 0 | 4.17E-05 | 3.41E-06 | 2.51E-05 | 1.36E-05 |
| Platr25 | 1.58E-05 | 2.51E-06 | 0 | 0 | 4.54E-07 | 2.05E-06 | 3.87E-06 | 0 | 0 | 1.06E-05 | 0 | 4.06E-06 | 0 | 2.90E-06 | 2.02E-05 | 3.40E-05 | 0 | 3.71E-06 |
| Tubd1 | 1.49E-05 | 6.96E-06 | 0 | 9.39E-06 | 0 | 2.04E-06 | 2.98E-06 | 5.70E-06 | 0 | 1.97E-05 | 8.43E-06 | 0 | 8.92E-06 | 0 | 0 | 6.19E-05 | 0 | 2.18E-05 |
| Zfp558 | 5.05E-06 | 6.14E-06 | 0 | 0 | 3.77E-06 | 2.01E-06 | 4.44E-06 | 0 | 0 | 1.96E-06 | 0 | 5.29E-06 | 4.10E-06 | 0 | 0 | 4.76E-06 | 6.46E-07 | 9.51E-07 |
| AI429214 | 1.75E-05 | 1.57E-05 | 0 | 1.75E-06 | 0 | 2.00E-06 | 0 | 3.02E-06 | 0 | 2.05E-05 | 0 | 3.65E-06 | 0 | 0 | 1.63E-05 | 0 | 0 | 5.66E-06 |
| Slfn8 | 4.59E-05 | 0 | 0 | 0 | 0 | 2.00E-06 | 1.50E-05 | 0 | 0 | 0.00014 | 0 | 8.77E-06 | 9.04E-05 | 0 | 0 | 6.42E-05 | 0 | 6.88E-05 |
| Zfp354c | 5.04E-05 | 1.69E-05 | 0 | 7.63E-05 | 5.50E-05 | 1.99E-06 | 0.00024 | 4.55E-05 | 0 | 1.43E-05 | 3.22E-05 | 0 | 5.71E-05 | 3.04E-05 | 1.64E-05 | 0 | 0 | 0.00012 |
| Aif1 | 0 | 3.58E-07 | 0 | 1.93E-05 | 1.81E-07 | 1.98E-06 | 0 | 6.88E-06 | 0 | 2.55E-06 | 6.32E-06 | 4.87E-06 | 3.07E-06 | 3.30E-06 | 1.66E-05 | 1.59E-06 | 2.08E-05 | 4.10E-06 |
| Mfap3l | 3.13E-06 | 0 | 0 | 0 | 0 | 1.98E-06 | 0 | 0 | 0 | 3.44E-05 | 0 | 0 | 0 | 5.71E-06 | 0 | 4.01E-06 | 0 | 7.09E-05 |
| Prim1 | 7.16E-06 | 0 | 1.75E-05 | 0 | 0 | 1.97E-06 | 1.21E-07 | 0 | 0 | 2.07E-06 | 0 | 0 | 4.71E-06 | 2.11E-06 | 0 | 0.00034 | 0 | 0 |
| Tlr6 | 9.08E-06 | 0 | 0 | 0 | 0 | 1.92E-06 | 1.67E-05 | 4.05E-06 | 0 | 6.12E-06 | 1.72E-06 | 8.61E-06 | 0 | 0 | 4.87E-06 | 0 | 0 | 1.49E-05 |
| Kif21b | 0 | 2.87E-05 | 3.09E-05 | 0 | 0 | 1.90E-06 | 0 | 0.00013 | 0 | 0 | 2.70E-05 | 0.00038 | 0 | 5.83E-05 | 0.00022 | 0 | 0 | 0 |
| Amt | 0 | 1.87E-05 | 0 | 3.99E-06 | 7.07E-06 | 1.85E-06 | 0 | 3.61E-06 | 0 | 0 | 2.98E-06 | 1.69E-05 | 0 | 0 | 1.91E-05 | 1.54E-05 | 2.78E-06 | 5.20E-06 |
| Tpk1 | 1.03E-05 | 2.20E-05 | 0 | 0 | 1.17E-06 | 1.83E-06 | 5.23E-07 | 0 | 0 | 7.72E-07 | 1.97E-05 | 1.57E-05 | 3.12E-06 | 8.28E-06 | 2.40E-05 | 7.42E-06 | 0 | 7.54E-06 |
| Zmynd15 | 0 | 3.39E-06 | 6.72E-06 | 0 | 2.69E-05 | 1.73E-06 | 0 | 2.76E-06 | 0 | 8.19E-07 | 3.93E-06 | 0 | 3.99E-05 | 7.88E-06 | 4.02E-05 | 0 | 1.58E-05 | 5.25E-05 |
| Rdh5 | 0 | 0 | 1.24E-05 | 3.72E-06 | 3.34E-07 | 1.70E-06 | 4.72E-06 | 2.39E-05 | 0 | 9.93E-06 | 7.96E-07 | 8.05E-07 | 0 | 9.87E-07 | 1.98E-05 | 0 | 0 | 1.30E-05 |
| Xkr5 | 0 | 0 | 0 | 0 | 0 | 1.69E-06 | 0 | 0 | 0 | 0 | 0 | 0 | 0 | 0 | 2.00E-06 | 8.73E-05 | 0 | 0 |
| Ablim3 | 0 | 2.80E-06 | 1.14E-06 | 0 | 0 | 1.68E-06 | 5.22E-06 | 2.58E-06 | 0 | 2.10E-05 | 0 | 5.78E-06 | 0 | 0 | 0 | 0 | 3.51E-06 | 2.61E-05 |
| Zfp607b | 1.70E-06 | 9.03E-07 | 1.76E-06 | 4.04E-06 | 3.14E-06 | 1.65E-06 | 0 | 1.88E-05 | 0 | 3.51E-06 | 0 | 2.51E-05 | 0 | 0 | 0 | 1.02E-05 | 0 | 3.54E-06 |
| Tpd52l1 | 2.06E-06 | 9.96E-06 | 0 | 1.52E-05 | 0 | 1.58E-06 | 0 | 2.26E-07 | 0 | 8.28E-08 | 3.62E-05 | 1.00E-05 | 0 | 7.10E-06 | 4.16E-06 | 3.75E-06 | 0 | 0 |
| Mcm8 | 0 | 0 | 2.24E-05 | 9.88E-08 | 0 | 1.57E-06 | 0 | 0 | 0 | 1.98E-06 | 1.25E-06 | 1.84E-07 | 0 | 0 | 0 | 0.00016 | 7.78E-07 | 0 |
| Hist1h2af | 0 | 0 | 0 | 0 | 0 | 1.56E-06 | 0 | 0 | 0 | 0 | 0 | 0 | 0 | 0 | 0 | 0.00018 | 0 | 0 |
| Oip5 | 0 | 0 | 0 | 0 | 0 | 1.56E-06 | 0 | 0 | 0 | 2.60E-06 | 0 | 0 | 0 | 0 | 0 | 0.00015 | 0 | 0 |
| Sgo2a | 0 | 0 | 0 | 0 | 0 | 1.55E-06 | 0 | 0 | 0 | 0 | 0 | 1.41E-06 | 0 | 0 | 0 | 0.00045 | 0 | 0 |

| | | | | | | | | | | | | | | | | | | |
|---|---|---|---|---|---|---|---|---|---|---|---|---|---|---|---|---|---|---|
| Efna3 | 0 | 2.72E-06 | 1.34E-05 | 0 | 1.67E-05 | 1.55E-06 | 3.43E-06 | 0 | 0 | 0 | 5.52E-06 | 0 | 0 | 9.50E-06 | 0 | 1.12E-06 | 0 | 4.96E-06 |
| Kif9 | 8.16E-07 | 4.13E-06 | 0 | 5.65E-06 | 3.03E-06 | 1.54E-06 | 2.71E-06 | 0 | 0 | 9.62E-06 | 0 | 2.42E-06 | 6.15E-06 | 0 | 0 | 0 | 4.06E-06 | 8.44E-06 |
| Spx | 0 | 0 | 7.38E-06 | 9.77E-06 | 0 | 1.48E-06 | 1.62E-06 | 1.30E-05 | 0 | 5.54E-06 | 4.43E-06 | 1.28E-05 | 0 | 0 | 2.65E-05 | 3.05E-06 | 2.54E-05 | 0 |
| Brip1 | 0 | 0 | 0 | 0 | 0 | 1.46E-06 | 3.90E-06 | 0 | 0 | 0 | 0 | 0 | 1.55E-05 | 0 | 3.52E-05 | 0.00034 | 1.75E-06 | 0 |
| Chchd10 | 0 | 0.00038 | 0 | 0 | 0.00023 | 1.45E-06 | 0 | 0 | 0 | 0 | 0 | 0 | 0 | 0.00028 | 0 | 0 | 0 | 0 |
| Tspan12 | 0 | 1.09E-06 | 0 | 2.52E-05 | 0 | 1.41E-06 | 0 | 3.80E-05 | 0 | 1.47E-05 | 0 | 8.63E-05 | 4.30E-05 | 7.25E-05 | 0 | 4.65E-06 | 0 | 5.53E-05 |
| Sdcbp2 | 0 | 1.82E-05 | 6.22E-06 | 7.44E-06 | 1.01E-06 | 1.36E-06 | 0 | 0 | 0 | 4.34E-06 | 4.21E-06 | 8.14E-06 | 0 | 0 | 1.97E-05 | 4.15E-06 | 6.27E-05 | 2.54E-06 |
| Mpzl2 | 0 | 1.32E-05 | 0 | 0 | 2.23E-05 | 1.36E-06 | 0 | 4.99E-05 | 0 | 0 | 0 | 0 | 0 | 0 | 0 | 0 | 0 | 0 |
| Cyth4 | 0 | 5.05E-06 | 0 | 5.57E-06 | 5.24E-07 | 1.35E-06 | 0 | 3.26E-06 | 0 | 1.08E-05 | 1.49E-06 | 0 | 0 | 6.62E-07 | 3.14E-05 | 3.61E-06 | 3.61E-05 | 1.50E-06 |
| Hist2h4 | 0 | 8.46E-06 | 0 | 0 | 0 | 1.35E-06 | 0 | 0 | 0 | 4.46E-06 | 0 | 2.10E-05 | 0 | 0 | 1.56E-05 | 7.46E-06 | 1.48E-05 | 8.24E-06 |
| Nipal4 | 0 | 0 | 0 | 0 | 2.50E-05 | 1.29E-06 | 0 | 0 | 0 | 0 | 6.35E-05 | 1.64E-05 | 0 | 1.30E-05 | 0 | 0 | 0 | 0 |
| Myzap | 0 | 3.23E-06 | 6.15E-06 | 8.49E-06 | 0 | 1.21E-06 | 5.48E-07 | 0 | 0 | 1.44E-05 | 1.19E-06 | 8.05E-06 | 6.30E-06 | 1.58E-06 | 0 | 7.34E-06 | 0 | 3.99E-07 |
| Frg2f1 | 0 | 2.01E-05 | 1.94E-06 | 6.49E-06 | 8.40E-06 | 1.20E-06 | 8.22E-06 | 0 | 0 | 0 | 0 | 6.04E-06 | 1.13E-05 | 0 | 1.98E-05 | 2.95E-06 | 0 | 1.14E-05 |
| Rnf180 | 5.85E-08 | 5.61E-06 | 3.18E-06 | 4.79E-06 | 1.81E-07 | 1.19E-06 | 8.01E-06 | 4.76E-05 | 0 | 1.27E-05 | 0 | 0 | 4.39E-06 | 0 | 2.21E-05 | 0 | 3.35E-06 | 1.45E-05 |
| Fmo5 | 1.48E-06 | 3.37E-05 | 0 | 0 | 4.85E-06 | 1.18E-06 | 0 | 0 | 0 | 5.57E-05 | 0 | 1.62E-05 | 0 | 4.34E-08 | 0 | 0 | 0 | 4.47E-05 |
| Cenpi | 2.00E-06 | 0 | 0 | 0 | 0 | 1.14E-06 | 0 | 0 | 0 | 0 | 0 | 0 | 0 | 0 | 1.85E-05 | 0.00036 | 0 | 0 |
| Csdc2 | 0 | 8.16E-06 | 0 | 0 | 1.04E-05 | 1.12E-06 | 0 | 8.01E-06 | 0 | 0 | 1.08E-05 | 0 | 0 | 8.11E-06 | 5.99E-06 | 0 | 0 | 3.23E-05 |
| Spata2l | 0 | 9.38E-06 | 0 | 2.89E-05 | 1.79E-05 | 1.09E-06 | 5.58E-06 | 1.38E-05 | 0 | 9.05E-06 | 4.42E-06 | 5.66E-06 | 0 | 3.29E-06 | 1.43E-05 | 0 | 4.02E-06 | 8.64E-06 |
| Cenpp | 3.46E-06 | 0 | 0 | 0 | 0 | 1.07E-06 | 0 | 0 | 0 | 0 | 0 | 0 | 0 | 5.70E-06 | 0 | 0.00018 | 0 | 0 |
| Chaf1b | 0 | 0 | 2.41E-05 | 0 | 0 | 1.04E-06 | 0 | 0 | 0 | 4.47E-06 | 0 | 0 | 2.15E-05 | 1.22E-05 | 0 | 0.00024 | 0 | 0 |
| Ephx2 | 7.20E-06 | 7.85E-06 | 0 | 0 | 0 | 1.01E-06 | 1.71E-07 | 0 | 0 | 9.82E-06 | 0 | 8.70E-06 | 2.90E-05 | 1.05E-05 | 6.03E-06 | 0 | 4.60E-06 | 5.05E-05 |
| Zfp738 | 3.33E-06 | 1.23E-05 | 0 | 0 | 4.32E-06 | 9.83E-07 | 0 | 0 | 0 | 2.44E-07 | 2.53E-06 | 5.23E-06 | 0 | 1.34E-06 | 0 | 0 | 0 | 5.54E-06 |
| Fancd2 | 0 | 3.78E-07 | 0 | 0 | 4.89E-06 | 9.80E-07 | 2.63E-06 | 3.49E-06 | 0 | 5.35E-06 | 0 | 2.12E-06 | 8.04E-06 | 0 | 0 | 0.00023 | 1.79E-06 | 0 |
| Brca1 | 0 | 0 | 0 | 0 | 0 | 9.61E-07 | 0 | 0 | 0 | 0 | 0 | 0 | 5.79E-06 | 0 | 0 | 0.00024 | 0 | 0 |
| D3Ertd751e | 2.83E-06 | 2.87E-06 | 0 | 0 | 4.19E-06 | 9.51E-07 | 0 | 6.58E-06 | 0 | 9.31E-06 | 0 | 0 | 3.09E-06 | 4.68E-06 | 1.20E-05 | 1.54E-05 | 0 | 4.71E-06 |
| Fam72a | 4.23E-06 | 0 | 1.03E-05 | 0 | 0 | 9.49E-07 | 0 | 8.42E-07 | 0 | 6.40E-06 | 0 | 7.09E-06 | 0 | 0 | 1.46E-06 | 5.90E-05 | 4.52E-05 | 0 |
| Mreg | 0 | 4.56E-06 | 3.42E-06 | 1.86E-06 | 0 | 9.34E-07 | 0 | 0 | 0 | 0 | 0 | 1.35E-05 | 7.87E-06 | 3.84E-06 | 0 | 1.04E-05 | 1.65E-05 | 0 |
| Armh4 | 9.62E-06 | 0 | 0 | 1.34E-06 | 0 | 9.29E-07 | 0.00027 | 5.00E-06 | 0 | 7.11E-05 | 0 | 0.00027 | 0.00016 | 0 | 0 | 1.56E-05 | 0 | 0.00019 |
| Psmc3ip | 4.07E-06 | 0 | 1.49E-05 | 0 | 0 | 9.15E-07 | 0 | 0 | 0 | 0 | 0 | 0 | 8.22E-07 | 1.20E-05 | 7.46E-06 | 0.00018 | 0 | 9.78E-06 |
| Slfn9 | 5.13E-06 | 0 | 2.55E-06 | 0 | 0 | 8.76E-07 | 0 | 0 | 0 | 1.56E-05 | 0 | 0 | 2.32E-05 | 0 | 0 | 0.00024 | 0 | 0 |
| Arhgef6 | 2.09E-06 | 1.14E-05 | 0 | 0 | 6.19E-07 | 8.54E-07 | 0 | 0 | 0 | 0 | 8.00E-06 | 0 | 0 | 3.42E-05 | 0 | 0 | 0 | 0.0001 |
| Mybl1 | 0 | 6.81E-06 | 0 | 0 | 0 | 8.31E-07 | 0 | 0 | 0 | 9.73E-06 | 0 | 3.10E-06 | 0 | 0 | 0 | 8.95E-05 | 1.65E-05 | 0 |
| Wfdc17 | 0 | 0 | 0 | 0 | 0 | 8.26E-07 | 0 | 1.85E-07 | 0 | 5.18E-06 | 1.28E-06 | 3.04E-06 | 2.40E-05 | 0 | 1.81E-05 | 2.74E-06 | 1.69E-05 | 0 |

| | | | | | | | | | | | | | | | | | | |
|---|---|---|---|---|---|---|---|---|---|---|---|---|---|---|---|---|---|---|
| Phf11c | 0 | 4.85E-06 | 2.48E-06 | 7.10E-06 | 0 | 8.09E-07 | 6.91E-06 | 2.62E-06 | 0 | 1.89E-05 | 2.00E-06 | 0 | 6.07E-06 | 0 | 6.01E-06 | 1.70E-05 | 2.79E-06 | 7.64E-06 |
| Bnc2 | 0.00019 | 0.00017 | 0.00023 | 0 | 0.00026 | 8.04E-07 | 0.00053 | 0.00014 | 0 | 9.68E-05 | 0.00034 | 8.62E-05 | 0.00039 | 0 | 0.0003 | 1.80E-05 | 0.00023 | 8.79E-05 |
| Cenpk | 3.77E-06 | 0 | 1.80E-07 | 0 | 1.63E-06 | 7.03E-07 | 0 | 0 | 0 | 7.34E-06 | 1.73E-06 | 0 | 5.77E-07 | 0 | 1.10E-05 | 0.00017 | 0 | 0 |
| Fam160a1 | 3.66E-06 | 5.32E-06 | 0 | 0 | 2.38E-05 | 5.78E-07 | 0 | 5.36E-05 | 0 | 0 | 4.03E-05 | 6.62E-06 | 5.61E-06 | 0.0001 | 2.44E-05 | 0 | 5.06E-05 | 3.10E-05 |
| Rcan2 | 0 | 1.27E-05 | 0 | 0 | 5.09E-05 | 5.37E-07 | 0 | 0 | 0 | 0.00021 | 0 | 2.18E-05 | 9.21E-05 | 2.85E-05 | 0 | 0 | 0 | 0 |
| Zfr2 | 0 | 1.41E-05 | 0 | 4.52E-06 | 2.22E-05 | 5.30E-07 | 0 | 0 | 0 | 2.51E-05 | 0 | 2.92E-05 | 7.92E-06 | 0 | 3.13E-05 | 0 | 0 | 1.89E-05 |
| Ttll3 | 0 | 1.24E-05 | 4.54E-07 | 0 | 3.62E-06 | 5.01E-07 | 0 | 1.08E-06 | 0 | 0 | 8.41E-06 | 2.07E-05 | 0 | 0 | 4.34E-05 | 0 | 3.96E-05 | 8.79E-06 |
| Rnf125 | 5.85E-05 | 6.26E-05 | 0.0001 | 0 | 0 | 4.82E-07 | 5.16E-05 | 0.00021 | 0 | 0 | 0 | 0 | 0.00011 | 9.41E-05 | 0 | 3.74E-05 | 3.63E-05 | 2.11E-05 |
| Bmp8a | 0 | 9.53E-07 | 0 | 0 | 0 | 4.73E-07 | 0 | 2.22E-05 | 0 | 0.00015 | 0 | 5.01E-06 | 0 | 1.65E-06 | 0 | 0 | 0 | 0 |
| Car15 | 0 | 1.52E-05 | 1.87E-06 | 1.06E-05 | 1.28E-05 | 4.46E-07 | 0 | 0 | 0 | 1.09E-05 | 0 | 1.46E-05 | 3.68E-07 | 2.36E-06 | 1.31E-05 | 7.80E-06 | 2.31E-05 | 2.31E-06 |
| Nanos1 | 0 | 0 | 6.39E-06 | 5.17E-07 | 0 | 4.39E-07 | 3.08E-06 | 0 | 0 | 0 | 1.53E-06 | 2.34E-06 | 2.74E-06 | 4.01E-06 | 0 | 3.48E-05 | 7.18E-06 | 0 |
| Galnt5 | 1.88E-05 | 0 | 1.92E-06 | 1.01E-05 | 0 | 4.17E-07 | 6.24E-05 | 9.05E-06 | 0 | 0 | 0 | 0 | 0 | 1.40E-05 | 4.76E-07 | 1.98E-06 | 0 | 0 |
| Abcg4 | 0 | 5.62E-06 | 0 | 0 | 6.03E-07 | 4.01E-07 | 1.10E-06 | 0 | 0 | 0 | 2.50E-06 | 3.64E-06 | 0 | 2.13E-05 | 2.14E-06 | 0 | 5.76E-06 | 2.38E-05 |
| Mapk1ip1 | 1.62E-05 | 9.18E-05 | 0 | 4.71E-05 | 7.75E-06 | 3.99E-07 | 1.93E-05 | 0 | 0 | 2.72E-05 | 1.42E-05 | 4.97E-05 | 0 | 1.31E-05 | 0.00011 | 0 | 1.63E-05 | 3.73E-05 |
| Nol4l | 0 | 1.31E-05 | 3.17E-06 | 0 | 0 | 3.98E-07 | 0 | 7.31E-06 | 0 | 0 | 6.87E-06 | 0 | 0 | 1.16E-05 | 1.15E-05 | 0 | 0 | 7.81E-06 |
| Map4k2 | 0 | 2.90E-05 | 0 | 3.65E-05 | 2.58E-05 | 3.51E-07 | 0 | 0 | 0 | 6.44E-07 | 4.25E-06 | 1.21E-05 | 0 | 3.87E-05 | 4.63E-05 | 3.89E-05 | 4.70E-05 | 1.14E-05 |
| Pcdhb13 | 3.42E-06 | 5.85E-06 | 4.75E-07 | 0 | 7.03E-06 | 3.23E-07 | 0 | 2.28E-06 | 0 | 1.68E-07 | 2.99E-06 | 1.96E-06 | 9.38E-06 | 3.04E-06 | 1.14E-06 | 0 | 0 | 0 |
| Enkur | 0 | 6.93E-07 | 0 | 0 | 0 | 3.22E-07 | 1.28E-05 | 9.65E-07 | 0 | 7.33E-06 | 0 | 3.75E-06 | 0 | 0 | 0 | 2.84E-06 | 1.90E-05 | 8.72E-06 |
| Sec22c | 1.38E-06 | 1.48E-05 | 8.29E-06 | 1.79E-05 | 1.27E-05 | 2.96E-07 | 0 | 2.52E-05 | 0 | 2.41E-05 | 0 | 6.20E-06 | 0 | 0 | 0 | 2.27E-05 | 0 | 2.72E-05 |
| Cox6b2 | 2.91E-07 | 4.23E-05 | 2.41E-05 | 0 | 1.31E-05 | 2.86E-07 | 0 | 1.93E-05 | 0 | 0 | 3.33E-05 | 2.24E-05 | 0 | 7.67E-06 | 0 | 5.00E-07 | 7.22E-05 | 3.94E-05 |
| Tesmin | 0 | 1.17E-07 | 0 | 0 | 3.77E-07 | 2.57E-07 | 8.38E-06 | 0 | 0 | 1.87E-06 | 0 | 0 | 1.39E-05 | 3.15E-06 | 2.30E-06 | 6.19E-06 | 1.94E-05 | 4.87E-06 |
| Fgd4 | 9.07E-06 | 7.23E-05 | 0 | 0 | 1.69E-05 | 2.40E-07 | 1.95E-05 | 1.52E-05 | 0 | 3.25E-05 | 0 | 2.37E-05 | 0 | 0 | 3.06E-05 | 6.30E-06 | 4.33E-06 | 4.24E-05 |
| Fam221a | 2.84E-06 | 0 | 0 | 6.00E-06 | 2.86E-06 | 2.29E-07 | 0 | 2.44E-06 | 0 | 0 | 0 | 9.60E-06 | 4.79E-06 | 1.04E-05 | 0 | 2.65E-05 | 4.98E-06 | 4.17E-06 |
| Mlf1 | 1.67E-06 | 0 | 0 | 0 | 0 | 2.18E-07 | 0 | 0 | 0 | 9.50E-07 | 0 | 8.21E-07 | 6.88E-06 | 0 | 0 | 0.00011 | 6.22E-05 | 1.45E-05 |
| Car2 | 0 | 1.05E-05 | 1.72E-06 | 0 | 0 | 1.47E-07 | 0 | 0 | 0 | 3.76E-06 | 5.14E-05 | 1.16E-05 | 0 | 0 | 2.16E-05 | 5.71E-06 | 2.20E-05 | 0 |
| Ovca2 | 3.66E-06 | 1.90E-05 | 1.25E-05 | 2.75E-05 | 1.31E-05 | 1.43E-07 | 0 | 1.74E-05 | 0 | 4.70E-07 | 4.02E-06 | 0 | 1.44E-05 | 4.30E-06 | 0 | 0 | 1.14E-06 | 0 |
| Mfrp | 0 | 0 | 0 | 0 | 0 | 1.17E-07 | 0 | 1.85E-06 | 0 | 0 | 0 | 1.88E-05 | 0 | 0 | 0 | 0 | 7.46E-05 | 3.47E-06 |
| Cenph | 0 | 0 | 1.46E-05 | 0 | 0 | 1.17E-07 | 0 | 0 | 0 | 0 | 0 | 0 | 4.10E-06 | 0 | 0 | 0.00044 | 0 | 0 |
| Rad51b | 0 | 0 | 0 | 0 | 0 | 7.73E-08 | 0 | 0 | 0 | 1.24E-06 | 0 | 6.04E-06 | 0 | 0 | 4.63E-05 | 0.00017 | 0 | 0 |
| Fign | 4.67E-08 | 0 | 0 | 0 | 0 | 7.25E-08 | 0 | 0 | 0 | 5.29E-06 | 0 | 0 | 2.46E-05 | 0 | 1.07E-05 | 7.16E-06 | 0 | 4.65E-05 |
| Cacna1a | 0 | 2.05E-05 | 0 | 1.72E-05 | 0 | 7.09E-08 | 7.71E-06 | 1.14E-05 | 0 | 1.63E-05 | 0 | 5.89E-07 | 0 | 0 | 6.58E-06 | 0 | 2.55E-05 | 1.94E-05 |
| G0s2 | 0 | 3.26E-05 | 0 | 8.32E-06 | 2.73E-05 | 4.24E-08 | 0 | 2.07E-05 | 0 | 1.06E-05 | 0 | 2.43E-05 | 1.52E-05 | 0 | 5.67E-05 | 1.94E-05 | 0 | 5.03E-05 |
| Tnni3 | 0 | 7.71E-06 | 0 | 0 | 5.10E-06 | 1.02E-08 | 0 | 7.94E-07 | 0 | 4.37E-06 | 0 | 1.28E-05 | 0 | 0 | 4.85E-05 | 0 | 3.55E-05 | 6.70E-06 |

| | | | | | | | | | | | | | | | | | | |
|---|---|---|---|---|---|---|---|---|---|---|---|---|---|---|---|---|---|---|
| Mmp13 | 0 | 0 | 0 | 0 | 0.00367 | 0 | 0 | 0.00105 | 0 | 0 | 0 | 0 | 0 | 0.00283 | 0 | 0 | 0 | 0 |
| Mmp9 | 0 | 0 | 0 | 0 | 0 | 0 | 0 | 0.00155 | 0 | 0 | 0 | 0 | 0 | 0.00206 | 0 | 0 | 0 | 0 |
| Tnn | 0 | 0 | 0 | 0 | 0 | 0 | 0.00032 | 0 | 0 | 0 | 0.00112 | 0 | 0 | 0.00193 | 0 | 0 | 0 | 0 |
| Lrrc15 | 0 | 0 | 0.00021 | 0 | 0.00177 | 0 | 0.00167 | 0.00015 | 0 | 0 | 0.00253 | 9.28E-05 | 0 | 0.00146 | 0 | 6.34E-05 | 1.66E-05 | 0 |
| Hp | 0 | 0 | 1.20E-05 | 0 | 0 | 0 | 0 | 0 | 0 | 0 | 0 | 0 | 0.00197 | 0.00123 | 0 | 0.00012 | 0 | 0.00023 |
| Tgfbi | 9.30E-06 | 0 | 6.79E-05 | 4.23E-05 | 0 | 0 | 0.00106 | 0 | 0 | 0 | 0.00039 | 0 | 0 | 0.00122 | 0 | 0.00022 | 0 | 0.00024 |
| Srpx2 | 0.0003 | 0 | 3.06E-06 | 0 | 0.00077 | 0 | 0.00071 | 0.0005 | 0 | 0 | 0.00053 | 0 | 0.00068 | 0.00116 | 0 | 0.00018 | 0 | 0.0004 |
| Smoc2 | 0 | 7.71E-05 | 0 | 0 | 0 | 0 | 0 | 0.00116 | 0 | 0.00016 | 0 | 0 | 0.00065 | 0.00104 | 0 | 0 | 0 | 0.0009 |
| Tagln | 0 | 0 | 3.32E-05 | 0.00018 | 0.00015 | 0 | 0 | 7.00E-05 | 0 | 0 | 0.00477 | 0 | 0 | 0.00091 | 0.00015 | 0.00022 | 0 | 0 |
| Tln2 | 0.00013 | 0 | 0.00043 | 1.47E-05 | 2.71E-05 | 0 | 0.00028 | 0.00058 | 0 | 3.32E-05 | 0.00055 | 7.08E-06 | 0 | 0.00068 | 0.00023 | 0.00018 | 2.32E-05 | 0.00025 |
| Ntn1 | 0 | 7.02E-05 | 0 | 0 | 0 | 0 | 0 | 0 | 0 | 0.00019 | 0 | 0 | 0.00019 | 0.00064 | 0 | 0 | 0 | 0.00068 |
| Acta2 | 0 | 0 | 0.00022 | 0.0004 | 0 | 0 | 7.94E-06 | 0.00068 | 0 | 0 | 0.00447 | 0 | 0 | 0.00059 | 0 | 0.00046 | 0 | 0 |
| Mmp10 | 0 | 0 | 0 | 0 | 0.00079 | 0 | 0 | 0.00023 | 0 | 0 | 9.66E-06 | 0.00033 | 0 | 0.00057 | 0 | 0 | 0 | 0 |
| Ltbp2 | 0 | 0 | 0.00044 | 0 | 0 | 0 | 1.98E-05 | 0 | 0 | 0 | 0.00077 | 0 | 0.00128 | 0.00047 | 0 | 7.21E-05 | 0 | 0.00054 |
| Gask1b | 0.0002 | 3.98E-05 | 0.0002 | 7.98E-05 | 0.00107 | 0 | 0.00015 | 0.00049 | 0 | 0 | 0.0007 | 0 | 2.10E-06 | 0.0004 | 0 | 0.00016 | 0 | 0.0004 |
| Cspg4 | 0 | 6.76E-05 | 0 | 0 | 7.23E-05 | 0 | 0 | 0.00092 | 0 | 0 | 0.00042 | 0.00018 | 0 | 0.00039 | 0 | 0 | 0 | 0 |
| Peg3 | 9.45E-05 | 0 | 0 | 5.48E-05 | 6.57E-05 | 0 | 0.00014 | 0 | 0 | 5.12E-06 | 9.58E-05 | 0 | 0 | 0.00038 | 0.00011 | 0 | 7.78E-05 | 0.0003 |
| Gpnmb | 0 | 0.00025 | 0 | 0 | 0 | 0 | 0 | 0 | 0 | 2.00E-05 | 0 | 0 | 0.00027 | 0.00036 | 0 | 0 | 0 | 0.00018 |
| Slc20a1 | 0 | 3.91E-05 | 0.00029 | 0 | 0 | 0 | 0 | 0.00134 | 0 | 0 | 0 | 6.19E-05 | 0 | 0.00036 | 0.00011 | 1.18E-05 | 0.00103 | 3.45E-05 |
| Adamts16 | 0 | 0 | 0 | 0 | 0.0004 | 0 | 0 | 7.44E-05 | 0 | 3.70E-05 | 0.00062 | 0 | 0 | 0.00033 | 0.00029 | 0 | 0 | 0 |
| Tes | 8.16E-05 | 3.85E-05 | 0.00031 | 0 | 2.07E-05 | 0 | 0 | 0.00017 | 0 | 0 | 0.00055 | 0.00044 | 0 | 0.00031 | 0 | 6.62E-05 | 0 | 0 |
| Sdc3 | 0 | 2.60E-05 | 0 | 0.00013 | 0 | 0 | 0 | 0.00033 | 0 | 8.30E-05 | 0.00014 | 0.00058 | 2.06E-05 | 0.0003 | 0 | 6.67E-05 | 6.41E-05 | 0.00019 |
| Itga11 | 0 | 2.16E-05 | 0 | 0 | 0 | 0 | 0.0001 | 0 | 0 | 0.00045 | 0.00034 | 3.24E-05 | 0.00062 | 0.00029 | 0 | 0 | 7.34E-06 | 0.00033 |
| Spsb1 | 0 | 0.00011 | 0.00017 | 0 | 0.00024 | 0 | 0.0003 | 0.00026 | 0 | 0.0004 | 0.00019 | 0 | 0.00078 | 0.00028 | 0 | 0 | 0 | 0.00013 |
| Hspb1 | 0 | 0.00019 | 0 | 0 | 0 | 0 | 0 | 0 | 0 | 0.00017 | 0.00036 | 9.35E-07 | 1.14E-05 | 0.00027 | 0 | 0 | 0.00073 | 0.00017 |
| Cdc42ep3 | 0 | 2.76E-05 | 0 | 0 | 0 | 0 | 0 | 0 | 0 | 0 | 0.00063 | 0.00016 | 0 | 0.00026 | 0 | 0 | 0 | 4.79E-05 |
| Lrrc55 | 0 | 0 | 0.00012 | 0 | 3.81E-05 | 0 | 0 | 0.00041 | 0 | 0 | 8.98E-05 | 0 | 0 | 0.00026 | 0 | 7.76E-05 | 0 | 0 |
| Pdgfrl | 4.10E-05 | 0 | 1.63E-05 | 0 | 7.07E-05 | 0 | 0.00037 | 5.68E-05 | 0 | 0 | 0.00013 | 0 | 0 | 0.00025 | 0 | 7.34E-05 | 0 | 0.0008 |
| Apoe | 0 | 0.00035 | 1.71E-05 | 7.33E-05 | 0 | 0 | 0 | 0 | 0 | 0.00014 | 0 | 3.84E-05 | 0.00052 | 0.00025 | 0.00073 | 8.98E-06 | 0.00012 | 0.00089 |
| Runx3 | 0 | 2.93E-05 | 0 | 0 | 0 | 0 | 0 | 0 | 0 | 0 | 0 | 1.97E-05 | 0 | 0.00025 | 0 | 0 | 0 | 2.24E-05 |
| Podn | 0 | 0 | 0 | 0 | 0 | 0 | 0.0003 | 0 | 0 | 0.00024 | 0.00025 | 0 | 0.00162 | 0.00024 | 0 | 0 | 0 | 0.0014 |
| Pcolce2 | 9.95E-06 | 0 | 0.00016 | 0 | 0 | 0 | 0 | 0.00011 | 0 | 0.00074 | 0 | 0 | 6.80E-05 | 0.00024 | 0 | 0.00013 | 0 | 0.00054 |
| Pappa | 0 | 4.09E-05 | 0 | 0 | 0 | 0 | 8.48E-05 | 0 | 0 | 0 | 0.00036 | 0 | 3.33E-05 | 0.00023 | 0 | 0 | 0 | 3.55E-05 |

| | | | | | | | | | | | | | | | | | | |
|---|---|---|---|---|---|---|---|---|---|---|---|---|---|---|---|---|---|---|
| Prokr2 | 0 | 0 | 0.00038 | 0 | 0.00015 | 0 | 6.04E-05 | 0.00041 | 0 | 0 | 1.15E-05 | 0.00013 | 9.07E-05 | 0.00023 | 0 | 1.74E-05 | 0 | 0 |
| Arsi | 0 | 0 | 7.40E-05 | 0.00042 | 0.00015 | 0 | 0.00079 | 0 | 0 | 0.00093 | 9.76E-05 | 0 | 0.00066 | 0.00023 | 0 | 0.00025 | 0 | 0.0004 |
| Jag1 | 0 | 0 | 0.00033 | 0 | 0 | 0 | 1.58E-05 | 0.00093 | 0 | 0.00013 | 0.00014 | 0 | 0 | 0.00023 | 0.00019 | 0 | 0.00024 | 0.00028 |
| Syt13 | 0 | 3.81E-06 | 0 | 0 | 0 | 0 | 0 | 0 | 0 | 0 | 0 | 0 | 0 | 0.00022 | 0 | 0 | 0 | 9.40E-05 |
| Ctla2a | 0 | 0 | 0 | 0 | 0 | 0 | 0 | 0 | 0 | 8.21E-05 | 2.94E-05 | 0 | 0.00027 | 0.00022 | 0 | 1.48E-05 | 0 | 0 |
| Lamc3 | 0 | 7.98E-05 | 0 | 0 | 0 | 0 | 0 | 0 | 0 | 0 | 0 | 3.79E-05 | 0 | 0.00022 | 0 | 0 | 0 | 0 |
| Bcl11b | 0 | 0 | 0 | 0 | 0 | 0 | 0 | 0 | 0 | 0 | 2.35E-05 | 0 | 0 | 0.00021 | 0 | 5.53E-06 | 0 | 3.26E-05 |
| Tmem254c | 0 | 3.91E-05 | 0 | 0.00016 | 0 | 0 | 6.33E-05 | 0 | 0 | 0.00034 | 6.85E-05 | 0 | 2.68E-05 | 0.00021 | 0 | 4.33E-06 | 0 | 0.00028 |
| Itprip | 0 | 7.36E-05 | 7.93E-05 | 6.73E-05 | 4.47E-05 | 0 | 0 | 9.69E-05 | 0 | 0 | 3.39E-05 | 0.0002 | 0 | 0.00021 | 0.00021 | 7.26E-05 | 8.40E-05 | 0 |
| Cpe | 0 | 0.00026 | 0 | 0 | 0.0001 | 0 | 0 | 0 | 0 | 0 | 0 | 0.00015 | 0 | 0.00021 | 0 | 0 | 7.68E-05 | 0.00015 |
| Vax2 | 0 | 1.31E-05 | 7.84E-06 | 3.05E-05 | 0 | 0 | 0 | 1.76E-05 | 0 | 0 | 4.42E-05 | 3.56E-06 | 0 | 0.0002 | 3.36E-05 | 0 | 0 | 2.23E-05 |
| Bmp4 | 0 | 1.34E-05 | 0 | 0 | 0 | 0 | 0 | 0 | 0 | 4.07E-05 | 0 | 0 | 0 | 0.0002 | 0 | 0 | 0 | 2.09E-05 |
| Cd40 | 0 | 4.38E-05 | 0 | 0 | 0.00023 | 0 | 0 | 0.00021 | 0 | 0 | 0 | 5.66E-05 | 0 | 0.0002 | 0 | 0 | 0 | 0 |
| Dpep1 | 0 | 0 | 0 | 0 | 0.0004 | 0 | 0.00038 | 0 | 0 | 0.00077 | 0 | 1.35E-05 | 0.00181 | 0.0002 | 0 | 0 | 0 | 0.00019 |
| Gfra1 | 0 | 0 | 0 | 8.04E-05 | 0 | 0 | 0 | 0.0004 | 0 | 0 | 0 | 0 | 0.00116 | 0.0002 | 0 | 0.00011 | 0 | 0.00041 |
| Slc6a17 | 0 | 0 | 0 | 0 | 0 | 0 | 1.07E-05 | 0 | 0 | 5.09E-06 | 1.78E-05 | 0 | 0 | 0.0002 | 7.21E-06 | 1.05E-05 | 0 | 0 |
| Arhgef19 | 0 | 0 | 0 | 0.00013 | 0.0001 | 0 | 0.00017 | 0 | 0 | 0 | 0.00016 | 2.35E-05 | 5.25E-05 | 0.00019 | 0.00014 | 0 | 0 | 0 |
| Pianp | 0 | 3.14E-05 | 0 | 0 | 0 | 0 | 0 | 0 | 0 | 3.01E-06 | 0 | 0 | 0 | 0.00019 | 0 | 0 | 0 | 5.82E-05 |
| Klhl33 | 0 | 2.08E-05 | 0 | 0 | 9.50E-05 | 0 | 0 | 0 | 0 | 1.53E-05 | 3.98E-06 | 0 | 0 | 0.00019 | 2.58E-05 | 0 | 0 | 6.45E-05 |
| Fabp4 | 1.70E-06 | 0 | 0 | 1.72E-06 | 0 | 0 | 0 | 0 | 0 | 2.70E-05 | 1.86E-05 | 0 | 0.00029 | 0.00019 | 0 | 2.93E-06 | 3.64E-05 | 0 |
| Sema3g | 0 | 5.10E-06 | 0 | 0 | 0 | 0 | 0 | 0 | 0 | 2.01E-07 | 0 | 0 | 0 | 0.00018 | 0 | 0 | 0 | 1.54E-05 |
| Itga8 | 0 | 1.71E-05 | 0 | 0 | 0 | 0 | 0 | 0 | 0 | 0 | 9.62E-05 | 0 | 0 | 0.00018 | 0 | 0 | 0 | 2.89E-06 |
| Cyp1b1 | 0.00014 | 0 | 0 | 5.38E-05 | 0 | 0 | 0 | 3.67E-05 | 0 | 0 | 0 | 0 | 0.00154 | 0.00018 | 0 | 3.91E-06 | 0 | 0.00161 |
| Ier5 | 0 | 0.00012 | 0 | 0.00035 | 0.0002 | 0 | 0 | 0 | 0 | 1.68E-05 | 0.00026 | 0.00017 | 0.00013 | 0.00018 | 0.00079 | 2.19E-05 | 0.00312 | 0 |
| Ednrb | 0 | 2.42E-05 | 0 | 0 | 0 | 0 | 0 | 0 | 0 | 0 | 0 | 1.00E-05 | 0 | 0.00018 | 0 | 0 | 0 | 6.46E-05 |
| Nox4 | 0 | 0 | 0 | 0 | 0.00038 | 0 | 0 | 0 | 0 | 0.00035 | 0.00027 | 0 | 2.55E-05 | 0.00018 | 0 | 0 | 0 | 0.00034 |
| Cfb | 0 | 0 | 0.00023 | 0 | 0 | 0 | 0.00038 | 0.00023 | 0 | 0.00114 | 0 | 0 | 0.00242 | 0.00017 | 0 | 5.41E-05 | 0 | 0.00111 |
| Serinc2 | 0 | 0 | 0.00013 | 6.52E-05 | 0.00039 | 0 | 1.30E-05 | 0.00039 | 0 | 0 | 0.00033 | 0 | 0 | 0.00017 | 0 | 0 | 0 | 0 |
| Smtn | 2.00E-05 | 0 | 0.00021 | 0.00014 | 0 | 0 | 0.00029 | 0.00029 | 0 | 0.00015 | 0.00055 | 0.00045 | 0 | 0.00017 | 0.00057 | 0.00044 | 0.00016 | 0.0002 |
| Nav2 | 0 | 2.99E-05 | 0 | 0 | 0 | 0 | 0 | 0 | 0 | 0 | 0 | 0 | 0 | 0.00017 | 0 | 0 | 0 | 1.67E-05 |
| Rgs3 | 0 | 0.00022 | 0.00016 | 0.00011 | 0.00022 | 0 | 0.00017 | 0.00044 | 0 | 0.00016 | 0.00027 | 0 | 0.00044 | 0.00016 | 9.86E-05 | 0 | 0.00032 | 0.00015 |
| Tmem254a | 3.54E-06 | 2.84E-05 | 0 | 7.76E-05 | 0 | 0 | 7.76E-05 | 4.21E-06 | 0 | 0.00027 | 5.49E-05 | 0 | 1.65E-05 | 0.00016 | 0 | 3.95E-06 | 0 | 0.0002 |
| Stac2 | 0 | 1.65E-05 | 2.72E-05 | 0 | 0 | 0 | 3.05E-05 | 0 | 0 | 1.52E-05 | 0 | 0.00012 | 0 | 0.00016 | 0 | 0 | 0 | 0.00012 |

| | | | | | | | | | | | | | | | | | | |
|---|---|---|---|---|---|---|---|---|---|---|---|---|---|---|---|---|---|---|
| S1pr3 | 8.46E-05 | 7.11E-05 | 2.16E-05 | 7.17E-05 | 9.30E-05 | 0 | 0 | 0.00037 | 0 | 8.62E-05 | 0.00037 | 0.00057 | 0.00012 | 0.00016 | 0 | 4.53E-05 | 0.00018 | 0.00017 |
| Alx4 | 0 | 4.40E-05 | 0 | 0 | 0 | 0 | 0 | 0 | 0 | 0 | 0 | 0 | 0 | 0.00016 | 0 | 0 | 0 | 3.81E-05 |
| Parm1 | 0 | 0 | 9.56E-06 | 0 | 0 | 0 | 0 | 0 | 0 | 0.00013 | 5.52E-05 | 0 | 0.00046 | 0.00015 | 0 | 0 | 0 | 0.00017 |
| Col11a1 | 0 | 0 | 0 | 0 | 0 | 0 | 6.59E-05 | 0 | 0 | 0 | 3.94E-05 | 0 | 0 | 0.00015 | 0 | 0 | 0 | 0.00019 |
| Mn1 | 0 | 8.31E-07 | 0 | 0 | 6.54E-05 | 0 | 1.10E-06 | 0 | 0 | 7.00E-05 | 0.00017 | 0 | 0.00013 | 0.00015 | 0 | 0 | 0 | 0.00021 |
| Creb3l3 | 3.81E-05 | 0 | 4.38E-05 | 0.00013 | 0 | 0 | 0.00045 | 0 | 0 | 0 | 0 | 0 | 0.00015 | 0.00015 | 0 | 0 | 0 | 0 |
| Mgll | 0 | 0 | 0 | 0 | 0 | 0 | 0 | 0 | 0 | 0.00045 | 0.00028 | 0.00012 | 0 | 0.00015 | 0 | 0 | 0 | 2.52E-05 |
| Lrrn2 | 0 | 0 | 5.02E-05 | 0 | 0 | 0 | 0 | 5.34E-05 | 0 | 0 | 6.63E-05 | 0 | 3.87E-05 | 0.00015 | 0 | 0 | 0 | 0.00022 |
| Sox11 | 0 | 0 | 1.36E-05 | 0 | 0 | 0 | 0 | 0 | 0 | 6.54E-06 | 0 | 5.54E-06 | 4.52E-05 | 0.00015 | 0 | 7.41E-05 | 0.00015 | 0 |
| Gzmd | 0 | 0 | 0 | 0 | 9.83E-06 | 0 | 0 | 2.83E-05 | 0 | 0 | 4.55E-05 | 0 | 0 | 0.00014 | 0 | 1.69E-05 | 0 | 0 |
| Smpdl3a | 1.78E-06 | 5.35E-05 | 0 | 0 | 8.59E-05 | 0 | 3.67E-05 | 0 | 0 | 0.00015 | 0 | 0 | 0.00021 | 0.00014 | 0 | 0 | 0 | 0.00012 |
| Chst2 | 0 | 2.29E-06 | 5.39E-05 | 0.00015 | 0.0001 | 0 | 0 | 0.0004 | 0 | 0 | 0.00025 | 0 | 0 | 0.00014 | 3.18E-05 | 0.00011 | 0.00013 | 5.02E-05 |
| Bmp3 | 0 | 5.57E-05 | 0 | 0 | 0 | 0 | 0 | 0 | 0 | 0.00034 | 0 | 0 | 0 | 0.00013 | 0 | 0 | 0 | 0 |
| Endod1 | 0 | 3.17E-05 | 2.42E-05 | 0 | 8.18E-05 | 0 | 0 | 0.00027 | 0 | 3.21E-06 | 9.89E-05 | 0 | 0 | 0.00013 | 0 | 8.68E-05 | 0 | 0.00017 |
| Bok | 0 | 0 | 0 | 5.73E-05 | 0 | 0 | 0 | 0 | 0 | 1.38E-05 | 0.00026 | 0 | 0 | 0.00013 | 0 | 0.00028 | 0 | 0.00027 |
| Mfap4 | 0 | 0 | 0 | 0.00012 | 0.00089 | 0 | 0.00076 | 0 | 0 | 0 | 0.00041 | 0 | 0.00077 | 0.00013 | 0 | 0 | 0 | 0.00082 |
| Kdm6b | 0 | 0.00013 | 0.00034 | 0.00042 | 3.75E-05 | 0 | 0 | 0.00014 | 0 | 4.65E-05 | 0 | 0.00027 | 0 | 0.00013 | 0.00036 | 0 | 0.00179 | 0 |
| Tnfaip3 | 1.10E-05 | 0 | 0.00016 | 0 | 0 | 0 | 0 | 0 | 0 | 0 | 0 | 0 | 0.00013 | 0.00013 | 0 | 2.57E-05 | 0.0013 | 0 |
| Ncam1 | 7.34E-05 | 0 | 0.00022 | 2.71E-05 | 0.0007 | 0 | 0.00057 | 3.37E-05 | 0 | 0 | 0.00126 | 0.00035 | 0 | 0.00013 | 0.00038 | 0.00034 | 0.0001 | 0.00017 |
| Fgf2 | 1.25E-05 | 9.97E-05 | 0.00012 | 0 | 0.00015 | 0 | 0 | 9.52E-05 | 0 | 0.00039 | 0.0001 | 6.07E-05 | 0.00031 | 0.00012 | 0 | 0 | 0 | 5.65E-05 |
| Prelp | 0 | 0 | 0 | 0 | 0 | 0 | 0.0004 | 0 | 0 | 0.00133 | 0 | 0 | 0.00198 | 0.00012 | 0 | 0 | 0 | 0.00115 |
| Pdlim1 | 0 | 0 | 0.00016 | 2.63E-05 | 0 | 0 | 0 | 0.00023 | 0 | 0.0005 | 0.00034 | 0.00053 | 0 | 0.00012 | 0 | 0.00014 | 0 | 0.0004 |
| Rasl11b | 0 | 9.22E-05 | 0 | 0 | 0.00022 | 0 | 0.00013 | 0 | 0 | 0 | 0.00031 | 0 | 0 | 0.00012 | 0 | 0 | 0.00021 | 0.00041 |
| Tmem95 | 0 | 0 | 0.00021 | 0 | 0 | 0 | 0 | 0.00021 | 0 | 0 | 9.70E-07 | 0.00041 | 0 | 0.00011 | 0.00025 | 0 | 0.00083 | 0 |
| Mical3 | 0 | 0.00013 | 2.11E-05 | 3.31E-05 | 4.81E-05 | 0 | 2.22E-05 | 1.27E-05 | 0 | 9.22E-06 | 0.0001 | 4.57E-05 | 2.63E-05 | 0.00011 | 0.00014 | 0.00011 | 2.49E-05 | 0.00018 |
| Plac9a | 0 | 0 | 0 | 4.03E-05 | 0 | 0 | 0 | 0 | 0 | 0.00048 | 0 | 0 | 0 | 0.00011 | 0 | 0 | 0 | 0.00027 |
| Kcnab2 | 0 | 8.36E-06 | 0 | 0 | 0 | 0 | 0 | 0 | 0 | 2.63E-06 | 0 | 4.08E-05 | 0 | 0.00011 | 0 | 3.01E-05 | 0 | 2.06E-05 |
| Ifi30 | 0.00011 | 5.19E-05 | 5.55E-05 | 0.00013 | 2.87E-06 | 0 | 0.00013 | 0 | 0 | 0 | 2.81E-05 | 0.00016 | 0 | 0.0001 | 3.88E-05 | 0.00021 | 7.88E-05 | 0.00018 |
| Itgbl1 | 0 | 3.01E-06 | 0 | 0 | 7.00E-05 | 0 | 0.00027 | 0 | 0 | 0.00014 | 5.22E-07 | 0 | 0 | 0.0001 | 0 | 0 | 0 | 0.00081 |
| Gpr84 | 0 | 0 | 3.87E-05 | 0 | 0 | 0 | 0 | 1.47E-05 | 0 | 0 | 0 | 6.05E-06 | 1.68E-05 | 9.98E-05 | 0 | 0 | 1.11E-06 | 0 |
| Hhip | 0 | 0 | 0 | 0 | 3.34E-05 | 0 | 0 | 0 | 0 | 0 | 0 | 0 | 0 | 9.87E-05 | 0 | 0 | 0 | 0.00011 |
| Ndst3 | 0 | 0 | 7.66E-06 | 0 | 0 | 0 | 0 | 0 | 0 | 1.66E-06 | 0 | 0 | 0 | 9.81E-05 | 0 | 2.16E-05 | 0 | 0 |
| Sema6c | 0 | 5.59E-05 | 0 | 6.93E-05 | 5.86E-05 | 0 | 0.00025 | 0 | 0 | 7.70E-05 | 0 | 3.34E-05 | 0.00014 | 9.76E-05 | 0.00011 | 0 | 1.46E-05 | 5.70E-05 |

| | | | | | | | | | | | | | | | | | | |
|---|---|---|---|---|---|---|---|---|---|---|---|---|---|---|---|---|---|---|
| Pde1b | 0 | 1.77E-05 | 5.68E-05 | 0 | 0 | 0 | 0 | 4.08E-05 | 0 | 0 | 4.91E-05 | 3.59E-05 | 2.58E-06 | 9.59E-05 | 0 | 0 | 1.03E-05 | 1.43E-05 |
| Ptx3 | 0 | 0 | 0 | 8.13E-05 | 0 | 0 | 0 | 0 | 0 | 0.00083 | 8.27E-05 | 2.80E-05 | 0.00229 | 9.56E-05 | 0 | 0.00011 | 7.55E-05 | 0 |
| Fabp5 | 4.86E-05 | 0 | 0 | 0 | 3.64E-05 | 0 | 0 | 3.05E-05 | 0 | 5.69E-06 | 0.00057 | 0.00053 | 0 | 9.09E-05 | 0 | 8.92E-05 | 0 | 0 |
| Slc2a1 | 0 | 2.91E-05 | 0.00025 | 0.00028 | 0.00014 | 0 | 0 | 0.00029 | 0 | 6.60E-05 | 0.00018 | 0.00211 | 7.50E-05 | 9.07E-05 | 0.00019 | 0 | 0.00069 | 0 |
| Fmo2 | 0 | 2.68E-05 | 0 | 0 | 0 | 0 | 0 | 0 | 0 | 2.33E-05 | 0 | 7.02E-05 | 0 | 9.05E-05 | 0 | 0 | 0 | 0.00011 |
| Cemip | 0 | 3.43E-05 | 0 | 0 | 1.49E-05 | 0 | 0 | 0 | 0 | 2.36E-05 | 6.95E-05 | 0 | 0.00089 | 8.87E-05 | 0 | 0 | 0 | 0 |
| Ramp1 | 0 | 3.12E-05 | 0 | 0 | 2.50E-05 | 0 | 0 | 0 | 0 | 0 | 0.00023 | 0 | 0 | 8.59E-05 | 0 | 0 | 0 | 0 |
| Wnt11 | 0 | 0 | 0.00024 | 0 | 0.00017 | 0 | 0 | 0.00036 | 0 | 9.93E-06 | 0.00021 | 0.00017 | 0 | 8.49E-05 | 0 | 0 | 1.72E-05 | 2.69E-05 |
| Klhdc8a | 0 | 0 | 3.91E-05 | 0 | 0 | 0 | 0 | 2.14E-06 | 0 | 1.00E-05 | 8.45E-05 | 0 | 0 | 8.46E-05 | 0 | 6.31E-06 | 0 | 1.66E-05 |
| Hoxd9 | 0 | 1.24E-05 | 0 | 0 | 0 | 0 | 3.68E-06 | 0 | 0 | 1.33E-05 | 7.88E-06 | 0 | 0 | 8.37E-05 | 0 | 0 | 0 | 8.20E-05 |
| Thbs3 | 3.33E-05 | 0 | 6.56E-05 | 0 | 7.84E-05 | 0 | 0.00073 | 0.00018 | 0 | 0.00088 | 0.00012 | 0 | 0.00034 | 8.32E-05 | 0.00042 | 3.79E-05 | 0 | 0.0009 |
| Schip1 | 0 | 6.63E-05 | 0.00022 | 0 | 0.00016 | 0 | 0.00027 | 0.00018 | 0 | 0 | 0.00034 | 8.43E-05 | 0 | 8.17E-05 | 0 | 0.00012 | 6.02E-05 | 7.62E-05 |
| Fzd4 | 0 | 0 | 0 | 8.75E-05 | 0 | 0 | 0 | 2.67E-05 | 0 | 0.00044 | 0 | 0 | 0.00012 | 8.12E-05 | 2.04E-05 | 7.79E-05 | 0 | 0.00055 |
| Ccdc6 | 0.00012 | 0.00025 | 0.00027 | 0.00036 | 0.0002 | 0 | 0.00019 | 0.00034 | 0 | 0.00024 | 0.0006 | 0.00034 | 0 | 7.87E-05 | 0.00032 | 0.00021 | 7.49E-05 | 0.00018 |
| Piezo2 | 0 | 6.44E-05 | 0 | 0 | 0.00041 | 0 | 0 | 0.00041 | 0 | 0 | 0.00031 | 0.00018 | 0 | 7.79E-05 | 6.18E-05 | 0 | 0 | 3.89E-05 |
| Shmt2 | 4.60E-05 | 9.30E-05 | 0.00049 | 0.00039 | 0 | 0 | 0.00016 | 0.00014 | 0 | 5.81E-05 | 0 | 0.00046 | 0 | 7.64E-05 | 0.00015 | 0.00065 | 0.00025 | 8.85E-06 |
| Rnf150 | 6.87E-06 | 4.28E-05 | 0 | 5.09E-05 | 8.75E-05 | 0 | 0.00017 | 0 | 0 | 9.34E-06 | 0.00012 | 0 | 0.00043 | 7.53E-05 | 3.86E-05 | 2.67E-05 | 0.0002 | 0.00034 |
| Lrig3 | 4.83E-05 | 5.70E-05 | 0 | 0 | 0 | 0 | 0.00019 | 0 | 0 | 0 | 7.82E-05 | 0 | 1.81E-05 | 7.46E-05 | 0 | 4.21E-05 | 1.14E-05 | 0.00021 |
| Aplnr | 0 | 0 | 1.58E-06 | 0 | 3.48E-05 | 0 | 0 | 6.92E-05 | 0 | 2.26E-06 | 0 | 0 | 1.20E-05 | 7.39E-05 | 2.58E-05 | 2.27E-05 | 0 | 0 |
| Car4 | 0 | 0 | 2.96E-05 | 0 | 0 | 0 | 0 | 0 | 0 | 0 | 0 | 0 | 0.00024 | 7.34E-05 | 0 | 3.27E-05 | 1.13E-05 | 0 |
| Il1a | 0 | 0 | 1.70E-05 | 0 | 0 | 0 | 0 | 0 | 0 | 9.42E-06 | 1.40E-05 | 2.44E-06 | 1.57E-05 | 7.19E-05 | 6.58E-05 | 7.24E-06 | 5.40E-05 | 0 |
| Tspan7 | 0 | 2.73E-05 | 0 | 1.50E-05 | 9.78E-06 | 0 | 0 | 0.00011 | 0 | 2.15E-05 | 0 | 5.94E-07 | 2.55E-05 | 7.17E-05 | 0 | 1.94E-05 | 0 | 3.85E-05 |
| Igfbp2 | 0 | 0 | 0 | 0 | 0 | 0 | 0 | 0 | 0 | 0 | 0.00012 | 0 | 0 | 7.13E-05 | 0 | 0 | 0 | 3.43E-05 |
| Zfp521 | 0 | 5.20E-05 | 0 | 0 | 0 | 0 | 0.00035 | 0 | 0 | 0.00021 | 0 | 0 | 0.00011 | 7.08E-05 | 0.00016 | 2.02E-05 | 0 | 0.00045 |
| Slc7a2 | 0 | 0 | 4.11E-06 | 1.31E-05 | 0 | 0 | 0 | 4.28E-05 | 0 | 0 | 0.00011 | 4.87E-05 | 0.00011 | 7.08E-05 | 2.35E-05 | 0 | 0 | 0.00047 |
| Rasl12 | 0 | 0 | 0.00017 | 4.67E-05 | 0.00013 | 0 | 0.0001 | 0.00017 | 0 | 0 | 0.00019 | 9.70E-05 | 0.00014 | 7.03E-05 | 1.63E-05 | 0 | 0 | 4.26E-05 |
| F5 | 0 | 2.16E-05 | 0 | 0 | 0 | 0 | 0 | 0 | 0 | 0 | 0 | 0 | 0 | 6.96E-05 | 0 | 0 | 0 | 3.84E-05 |
| Prex1 | 0 | 7.03E-05 | 4.40E-06 | 3.92E-05 | 2.87E-05 | 0 | 0.00018 | 0 | 0 | 0.0001 | 0 | 0 | 8.48E-05 | 6.96E-05 | 0 | 2.13E-05 | 6.22E-06 | 0 |
| Ctxn1 | 0 | 7.02E-05 | 0 | 5.48E-05 | 0 | 0 | 0 | 0.00017 | 0 | 0 | 0.00012 | 0.00013 | 0 | 6.91E-05 | 6.72E-05 | 0 | 0 | 5.24E-05 |
| Dock9 | 0 | 2.11E-05 | 0 | 0 | 0 | 0 | 0 | 0 | 0 | 1.06E-05 | 5.09E-06 | 1.11E-05 | 0 | 6.59E-05 | 0 | 0 | 0 | 0.00027 |
| Mcam | 0 | 5.14E-05 | 0 | 0 | 0 | 0 | 0 | 0.00049 | 0 | 0 | 0.00012 | 0 | 0 | 6.47E-05 | 0 | 0 | 0 | 0 |
| Ucp2 | 0 | 1.53E-05 | 1.19E-05 | 7.31E-05 | 0 | 0 | 0 | 5.68E-05 | 0 | 2.63E-05 | 4.71E-05 | 9.92E-07 | 1.52E-05 | 6.33E-05 | 0.00012 | 6.67E-05 | 1.02E-05 | 0.00019 |
| Susd2 | 0 | 0 | 0 | 5.09E-06 | 0 | 0 | 0 | 5.81E-05 | 0 | 7.28E-05 | 0 | 1.17E-06 | 2.34E-05 | 6.32E-05 | 0 | 0 | 0 | 0.00017 |

| | | | | | | | | | | | | | | | | | | |
|---|---|---|---|---|---|---|---|---|---|---|---|---|---|---|---|---|---|---|
| Cxcl2 | 0 | 0 | 9.56E-05 | 0 | 9.50E-06 | 0 | 0 | 0 | 0 | 3.22E-05 | 1.33E-05 | 0 | 0.00016 | 6.31E-05 | 0.00036 | 7.85E-05 | 0.00071 | 0 |
| Ppm1l | 0 | 4.75E-05 | 4.51E-06 | 0 | 0 | 0 | 0 | 0 | 0 | 0 | 0 | 8.49E-06 | 0 | 6.24E-05 | 2.12E-07 | 0 | 5.59E-06 | 0 |
| Pawr | 0 | 3.86E-05 | 0 | 0 | 0.00011 | 0 | 0 | 0 | 0 | 0 | 0.0002 | 0 | 0 | 6.15E-05 | 0 | 0 | 0 | 2.60E-05 |
| Eif2ak4 | 6.05E-05 | 9.56E-05 | 0.0001 | 0.0001 | 8.94E-05 | 0 | 0 | 1.53E-05 | 0 | 0 | 4.20E-05 | 8.20E-05 | 3.76E-05 | 6.13E-05 | 0.0002 | 3.65E-05 | 5.36E-05 | 2.64E-05 |
| Kcnmb1 | 0 | 0 | 0 | 0 | 2.58E-05 | 0 | 0 | 9.47E-05 | 0 | 0 | 0.00016 | 7.23E-06 | 0 | 6.11E-05 | 0 | 0 | 0 | 0 |
| Rab11fip4 | 0 | 4.11E-06 | 0 | 0 | 3.00E-05 | 0 | 0 | 0 | 0 | 0 | 3.04E-05 | 1.09E-05 | 0 | 6.11E-05 | 0 | 0 | 0 | 0 |
| Garem2 | 0 | 0 | 2.87E-05 | 3.94E-06 | 0 | 0 | 2.02E-05 | 2.58E-05 | 0 | 0 | 2.07E-05 | 0 | 1.03E-05 | 6.10E-05 | 2.00E-05 | 1.60E-05 | 0 | 0 |
| Lysmd3 | 0.00016 | 2.44E-05 | 0 | 8.03E-05 | 0 | 0 | 0.00026 | 0 | 0 | 6.78E-05 | 9.70E-06 | 1.31E-05 | 7.76E-05 | 6.05E-05 | 7.52E-05 | 2.82E-05 | 0.00042 | 0 |
| Ooep | 0 | 0 | 0 | 0 | 0 | 0 | 7.82E-06 | 0 | 0 | 8.70E-07 | 0 | 0 | 0 | 5.98E-05 | 0 | 0 | 0 | 1.13E-05 |
| Foxo3 | 0 | 0.00019 | 0 | 0.00018 | 9.51E-05 | 0 | 0 | 0 | 0 | 2.15E-05 | 0 | 0.00025 | 0 | 5.92E-05 | 0.00057 | 1.69E-05 | 0.00063 | 7.39E-05 |
| Megf10 | 0 | 9.74E-06 | 0 | 0 | 1.50E-05 | 0 | 0 | 0 | 0 | 0 | 0 | 0 | 0 | 5.90E-05 | 0 | 0 | 0 | 6.10E-05 |
| Cpox | 8.18E-05 | 1.66E-05 | 0.00019 | 0.00014 | 0 | 0 | 0.00015 | 0 | 0 | 2.45E-06 | 8.93E-06 | 8.16E-05 | 2.72E-05 | 5.76E-05 | 0 | 0.00012 | 3.49E-05 | 1.07E-05 |
| Nfatc2 | 0 | 0 | 3.51E-05 | 0 | 0 | 0 | 0 | 0 | 0 | 1.84E-05 | 1.54E-06 | 2.87E-06 | 0 | 5.75E-05 | 3.67E-06 | 4.69E-06 | 3.08E-05 | 3.23E-05 |
| Micall2 | 0 | 1.07E-05 | 1.57E-05 | 0 | 8.69E-07 | 0 | 0 | 3.02E-05 | 0 | 0 | 4.10E-05 | 2.97E-05 | 1.76E-05 | 5.75E-05 | 3.87E-05 | 0 | 0 | 3.80E-05 |
| Gadd45a | 0 | 0.00037 | 0 | 0 | 3.15E-05 | 0 | 0 | 0.00015 | 0 | 4.16E-05 | 0 | 0.00026 | 7.52E-06 | 5.58E-05 | 7.66E-05 | 0 | 0.00067 | 0 |
| Dusp5 | 0 | 0 | 0.00013 | 0 | 0 | 0 | 0 | 0.00025 | 0 | 0 | 0 | 0.00045 | 0 | 5.56E-05 | 0.0002 | 0 | 0.00108 | 0 |
| Cpeb1 | 0 | 9.07E-05 | 3.14E-05 | 0 | 0 | 0 | 1.83E-05 | 0 | 0 | 3.67E-06 | 0 | 8.70E-05 | 6.90E-05 | 5.54E-05 | 4.42E-05 | 0 | 0 | 0.0001 |
| Lrrc8c | 3.94E-05 | 2.71E-05 | 0.00016 | 0 | 0 | 0 | 6.20E-05 | 1.23E-05 | 0 | 0.00013 | 0.00012 | 5.39E-05 | 0 | 5.53E-05 | 0 | 7.39E-05 | 0 | 0 |
| Sh2b2 | 0 | 8.43E-05 | 4.63E-05 | 6.31E-05 | 0 | 0 | 0 | 6.66E-05 | 0 | 0 | 0 | 0.00017 | 1.02E-05 | 5.47E-05 | 0.00017 | 0 | 0.00012 | 0.00011 |
| Trim47 | 0 | 1.86E-05 | 0.00013 | 3.97E-05 | 0 | 0 | 0 | 0 | 0 | 0.00011 | 0 | 0 | 1.41E-05 | 5.46E-05 | 6.03E-05 | 9.41E-05 | 1.62E-05 | 0.00034 |
| Plk3 | 0 | 2.09E-05 | 0.00016 | 0 | 9.80E-05 | 0 | 0 | 0.00025 | 0 | 0 | 3.98E-05 | 2.92E-05 | 0 | 5.44E-05 | 7.60E-05 | 0 | 0.00104 | 0 |
| Flrt3 | 1.71E-05 | 0 | 2.70E-05 | 5.65E-06 | 0 | 0 | 4.71E-06 | 8.16E-07 | 0 | 0 | 0 | 7.05E-05 | 2.75E-05 | 5.43E-05 | 1.81E-05 | 2.10E-05 | 1.31E-06 | 0 |
| Cxcr6 | 0 | 0 | 0 | 0 | 5.07E-06 | 0 | 3.67E-06 | 0 | 0 | 0 | 2.67E-05 | 0 | 0 | 5.19E-05 | 0 | 0 | 0 | 0 |
| Sertad4 | 3.72E-06 | 1.51E-05 | 0 | 9.90E-05 | 4.50E-06 | 0 | 0.00014 | 0 | 0 | 0 | 0.00016 | 0 | 0 | 5.12E-05 | 0 | 8.58E-05 | 0 | 0.00063 |
| Zim1 | 0 | 0 | 0 | 0 | 1.63E-05 | 0 | 0 | 0 | 0 | 2.01E-06 | 0 | 0 | 0 | 5.09E-05 | 0 | 1.49E-06 | 0 | 0.00013 |
| Rasa4 | 0 | 6.04E-06 | 0 | 0 | 0 | 0 | 6.54E-06 | 0 | 0 | 2.37E-05 | 0 | 4.60E-05 | 3.78E-05 | 5.07E-05 | 0 | 0 | 0 | 7.24E-05 |
| Afap1l2 | 0 | 2.25E-05 | 0 | 0 | 2.52E-08 | 0 | 0 | 1.60E-05 | 0 | 9.08E-05 | 0 | 0 | 0 | 5.06E-05 | 0 | 0 | 0 | 3.24E-05 |
| Ctla2b | 0 | 0 | 3.88E-07 | 0 | 9.00E-07 | 0 | 1.04E-05 | 1.20E-06 | 0 | 2.68E-06 | 0 | 0 | 4.22E-05 | 5.05E-05 | 0 | 0 | 0 | 0 |
| Sorcs2 | 0 | 0 | 0.00011 | 4.97E-05 | 0.00048 | 0 | 0.0006 | 0 | 0 | 0 | 0.00085 | 1.83E-05 | 0.00011 | 5.00E-05 | 0.00028 | 8.65E-05 | 1.63E-05 | 0.00065 |
| Gzmc | 0 | 0 | 0 | 0 | 3.12E-05 | 0 | 0 | 6.82E-06 | 0 | 0 | 0.00012 | 0 | 0 | 4.96E-05 | 0 | 1.00E-05 | 0 | 0 |
| Cadm3 | 4.46E-05 | 0 | 2.23E-05 | 0 | 0 | 0 | 0 | 0 | 0 | 0.00109 | 0 | 5.01E-05 | 5.64E-05 | 4.96E-05 | 0 | 0 | 0 | 6.51E-05 |
| H2-Q6 | 0 | 0 | 3.27E-05 | 0 | 0 | 0 | 0 | 0 | 0 | 7.41E-05 | 0 | 0.00019 | 0.0002 | 4.93E-05 | 3.54E-06 | 4.63E-05 | 0 | 6.83E-05 |
| Pmaip1 | 0 | 0.00011 | 0 | 0 | 0.00022 | 0 | 0 | 0.00022 | 0 | 0 | 8.58E-05 | 7.27E-05 | 0 | 4.88E-05 | 1.49E-06 | 6.02E-05 | 0.00018 | 0 |

| | | | | | | | | | | | | | | | | | | |
|---|---|---|---|---|---|---|---|---|---|---|---|---|---|---|---|---|---|---|
| Nuak2 | 0 | 0 | 5.29E-05 | 8.12E-05 | 0 | 0 | 0 | 5.20E-05 | 0 | 7.11E-05 | 0.00016 | 0 | 0.00012 | 4.74E-05 | 3.65E-05 | 8.27E-06 | 6.10E-05 | 4.69E-05 |
| Fkbp1b | 8.49E-06 | 3.51E-06 | 0 | 4.24E-05 | 2.00E-05 | 0 | 4.43E-05 | 4.10E-06 | 0 | 0 | 2.90E-05 | 4.26E-05 | 0 | 4.74E-05 | 0 | 7.02E-07 | 0 | 7.25E-05 |
| Magi1 | 5.98E-06 | 5.39E-05 | 0 | 8.50E-06 | 1.74E-05 | 0 | 0 | 0 | 0 | 6.80E-06 | 0 | 0 | 2.31E-05 | 4.63E-05 | 8.37E-05 | 2.80E-05 | 5.69E-05 | 5.98E-05 |
| Acot7 | 7.52E-05 | 1.06E-05 | 0.00022 | 5.06E-05 | 4.14E-05 | 0 | 0.00012 | 0.0003 | 0 | 0 | 0.00022 | 0.00017 | 0 | 4.45E-05 | 0 | 0.00016 | 2.54E-05 | 0.00028 |
| Megf11 | 0 | 0 | 0 | 0 | 1.94E-05 | 0 | 0 | 9.22E-05 | 0 | 0 | 4.90E-05 | 0 | 0 | 4.44E-05 | 0 | 0 | 0 | 4.88E-05 |
| Tmem44 | 0 | 5.00E-06 | 0 | 1.66E-06 | 9.84E-07 | 0 | 0 | 0 | 0 | 0 | 4.77E-05 | 0 | 0 | 4.43E-05 | 1.82E-05 | 0 | 0 | 0 |
| Egln3 | 0 | 0 | 0 | 0 | 0 | 0 | 0 | 0 | 0 | 0 | 0 | 0.00139 | 0 | 4.38E-05 | 0 | 0 | 0 | 0 |
| Cars | 7.10E-05 | 0.00012 | 0.00018 | 0.00012 | 0 | 0 | 0.00013 | 1.06E-05 | 0 | 5.98E-05 | 6.13E-05 | 0.00022 | 0 | 4.36E-05 | 0.00042 | 0.00019 | 0 | 7.49E-05 |
| Clcf1 | 0 | 6.21E-05 | 0.00021 | 4.50E-05 | 0 | 0 | 0 | 1.57E-06 | 0 | 0 | 0 | 0 | 0 | 4.26E-05 | 0.00014 | 0 | 0.00017 | 2.33E-05 |
| Aldh1l2 | 4.83E-05 | 0 | 0.00022 | 0.00023 | 0 | 0 | 0.00088 | 1.14E-05 | 0 | 0 | 0 | 0.0004 | 0 | 4.25E-05 | 2.72E-05 | 9.25E-05 | 0 | 0 |
| Slc35g2 | 1.72E-05 | 0 | 3.89E-05 | 0 | 0 | 0 | 3.64E-05 | 0 | 0 | 1.46E-05 | 2.33E-05 | 0 | 2.85E-05 | 4.12E-05 | 0 | 1.49E-06 | 0 | 0 |
| Pfkfb3 | 3.97E-05 | 3.40E-05 | 4.70E-05 | 0 | 7.83E-05 | 0 | 1.65E-05 | 0 | 0 | 3.48E-05 | 1.30E-05 | 0.00071 | 2.50E-05 | 4.09E-05 | 7.66E-05 | 7.49E-06 | 0.00013 | 0 |
| Zfp423 | 0 | 0 | 2.89E-05 | 7.42E-06 | 3.84E-05 | 0 | 0.0002 | 0 | 0 | 0.00014 | 0 | 0 | 0 | 4.07E-05 | 5.41E-05 | 0 | 0 | 0.00017 |
| Tmem8b | 0 | 2.44E-05 | 0 | 6.35E-06 | 1.20E-05 | 0 | 0 | 0 | 0 | 3.41E-05 | 1.42E-05 | 9.47E-06 | 2.72E-05 | 4.07E-05 | 2.82E-05 | 0 | 0 | 9.09E-06 |
| Itpk1 | 3.95E-06 | 4.36E-05 | 0 | 1.93E-05 | 0 | 0 | 0 | 1.50E-05 | 0 | 0 | 0 | 0.00022 | 2.88E-05 | 4.03E-05 | 0.00015 | 1.34E-05 | 0 | 3.94E-05 |
| Prex2 | 0 | 2.58E-06 | 0 | 0 | 0 | 0 | 0 | 0 | 0 | 0.0001 | 0 | 0 | 0 | 4.03E-05 | 0 | 6.50E-06 | 0 | 6.58E-05 |
| Fmod | 0 | 0 | 0 | 0 | 0 | 0 | 0.00013 | 0 | 0 | 0 | 0.00016 | 0 | 0 | 3.99E-05 | 0 | 4.85E-05 | 0 | 0.00063 |
| Nabp1 | 3.31E-05 | 0 | 2.51E-05 | 0 | 2.25E-05 | 0 | 0 | 0.0001 | 0 | 0 | 2.85E-05 | 0.00012 | 3.24E-05 | 3.93E-05 | 0 | 4.43E-05 | 0.00034 | 0 |
| Gstt3 | 7.42E-07 | 1.62E-05 | 0 | 3.65E-05 | 9.87E-07 | 0 | 8.12E-05 | 0 | 0 | 0.00015 | 0 | 0 | 7.06E-05 | 3.82E-05 | 0 | 4.33E-06 | 0 | 0.00016 |
| Cdh3 | 0 | 0 | 0 | 0 | 0 | 0 | 0 | 0 | 0 | 0 | 0.00016 | 0 | 0 | 3.71E-05 | 0 | 6.54E-06 | 0 | 0 |
| Rcan1 | 0 | 0 | 1.08E-05 | 0 | 0.00019 | 0 | 0.00069 | 0 | 0 | 0.00021 | 0.00015 | 0.00026 | 2.00E-05 | 3.68E-05 | 0 | 0 | 0 | 0 |
| Ebpl | 8.24E-05 | 6.45E-05 | 0 | 8.24E-05 | 0.00019 | 0 | 8.10E-05 | 7.47E-05 | 0 | 5.12E-05 | 0.00015 | 3.31E-05 | 4.24E-05 | 3.61E-05 | 0 | 0 | 0 | 8.07E-05 |
| Scara5 | 0 | 0 | 0 | 0 | 0 | 0 | 0 | 0 | 0 | 0.00091 | 0 | 0 | 0 | 3.58E-05 | 0 | 0 | 0 | 0.00017 |
| Caln1 | 0 | 0 | 0 | 0 | 4.92E-05 | 0 | 0 | 6.27E-05 | 0 | 0 | 0 | 0 | 0 | 3.53E-05 | 7.90E-06 | 0 | 0 | 0 |
| Plekhg4 | 4.16E-07 | 0 | 0 | 4.41E-06 | 2.09E-05 | 0 | 4.51E-05 | 0 | 0 | 0 | 9.12E-06 | 0 | 0 | 3.39E-05 | 2.08E-05 | 0 | 0 | 1.92E-05 |
| Trp53i11 | 0 | 7.64E-05 | 0 | 0 | 2.02E-05 | 0 | 0 | 3.28E-05 | 0 | 0 | 1.74E-05 | 6.59E-05 | 0 | 3.37E-05 | 0 | 1.95E-05 | 0 | 7.54E-06 |
| Pdk4 | 3.41E-06 | 8.01E-06 | 0 | 0 | 0 | 0 | 0 | 0 | 0 | 1.40E-05 | 0 | 0 | 9.63E-06 | 3.32E-05 | 0 | 1.27E-05 | 1.29E-05 | 5.84E-06 |
| Nlrc5 | 0 | 2.40E-05 | 0 | 0 | 0 | 0 | 0 | 0 | 0 | 7.62E-05 | 0 | 0 | 6.36E-06 | 3.28E-05 | 1.85E-05 | 0 | 4.23E-06 | 5.56E-05 |
| Relt | 0 | 0 | 4.82E-05 | 1.90E-05 | 0 | 0 | 0 | 0 | 0 | 0 | 5.13E-05 | 0 | 1.59E-05 | 3.23E-05 | 1.01E-05 | 6.60E-05 | 0.00015 | 0 |
| Bcl2l11 | 0 | 0.00012 | 0 | 3.16E-05 | 8.35E-05 | 0 | 0 | 7.69E-05 | 0 | 0 | 2.70E-05 | 8.27E-05 | 0 | 3.22E-05 | 0.00017 | 3.82E-05 | 0.00089 | 0 |
| Fam189a2 | 0 | 3.37E-05 | 0 | 0 | 0 | 0 | 0 | 9.03E-06 | 0 | 0 | 3.06E-05 | 3.44E-07 | 0 | 3.21E-05 | 0 | 0 | 0 | 3.12E-05 |
| Snx32 | 0 | 3.66E-05 | 0 | 0 | 3.12E-05 | 0 | 0 | 0 | 0 | 2.71E-05 | 0 | 0 | 0 | 3.17E-05 | 0.00013 | 0 | 0 | 3.65E-05 |
| Mmp17 | 0 | 4.28E-06 | 0 | 0 | 1.08E-05 | 0 | 0 | 2.26E-05 | 0 | 0 | 0.00014 | 3.72E-05 | 0 | 3.16E-05 | 0 | 1.80E-05 | 0 | 1.12E-06 |

| | | | | | | | | | | | | | | | | | | |
|---|---|---|---|---|---|---|---|---|---|---|---|---|---|---|---|---|---|---|
| Zfpm1 | 0 | 8.46E-05 | 0.00016 | 0 | 7.45E-05 | 0 | 5.99E-05 | 8.35E-05 | 0 | 8.42E-06 | 7.51E-05 | 0.00018 | 2.21E-05 | 3.11E-05 | 4.42E-05 | 0 | 0 | 2.69E-05 |
| Tom1l1 | 1.14E-05 | 2.36E-05 | 0 | 2.22E-05 | 2.33E-05 | 0 | 0 | 0 | 0 | 0 | 2.25E-05 | 1.38E-05 | 4.61E-06 | 3.09E-05 | 0 | 9.66E-07 | 1.25E-05 | 5.39E-05 |
| Tbxa2r | 0 | 0 | 0 | 1.55E-06 | 0 | 0 | 0 | 0 | 0 | 1.56E-06 | 3.39E-05 | 0 | 0 | 3.08E-05 | 0 | 0 | 0 | 0.0001 |
| Sgk3 | 3.78E-05 | 1.49E-06 | 9.18E-06 | 8.12E-06 | 2.10E-05 | 0 | 4.89E-05 | 5.08E-05 | 0 | 3.58E-05 | 1.70E-05 | 0 | 0 | 3.06E-05 | 2.52E-06 | 4.47E-05 | 2.14E-06 | 5.08E-05 |
| Tpbgl | 0 | 0 | 0 | 2.20E-05 | 0 | 0 | 0 | 0 | 0 | 0.00011 | 0 | 0 | 0 | 3.06E-05 | 0 | 1.47E-06 | 0 | 4.23E-05 |
| Epha2 | 0 | 2.12E-05 | 0 | 0 | 1.91E-05 | 0 | 0 | 1.08E-05 | 0 | 5.54E-06 | 2.46E-05 | 9.71E-06 | 2.33E-06 | 3.04E-05 | 6.24E-05 | 4.63E-07 | 0.00012 | 1.17E-05 |
| Cnnm4 | 0 | 3.24E-05 | 5.58E-05 | 4.52E-05 | 0.00019 | 0 | 0.00011 | 0 | 0 | 0 | 0.00017 | 0.00012 | 0 | 3.00E-05 | 6.89E-05 | 7.38E-05 | 0.00017 | 3.92E-05 |
| Cyp4f16 | 3.60E-06 | 4.29E-05 | 4.92E-05 | 3.62E-05 | 2.52E-05 | 0 | 2.78E-05 | 0.00013 | 0 | 2.03E-05 | 2.68E-05 | 3.33E-05 | 3.96E-06 | 3.00E-05 | 2.96E-05 | 2.30E-05 | 0 | 1.55E-05 |
| Adam23 | 6.83E-05 | 0 | 0 | 7.13E-05 | 0 | 0 | 0 | 0 | 0 | 0.00041 | 3.62E-05 | 0 | 0.00069 | 2.98E-05 | 0 | 0.00016 | 0 | 0 |
| Rtn4r | 0 | 5.51E-06 | 0 | 0 | 0 | 0 | 0 | 0 | 0 | 0 | 0 | 0 | 4.47E-06 | 2.91E-05 | 0 | 8.41E-06 | 0 | 3.18E-05 |
| Shroom4 | 0 | 2.90E-06 | 0 | 0 | 2.25E-05 | 0 | 0 | 1.91E-05 | 0 | 2.44E-06 | 9.57E-05 | 3.37E-05 | 0 | 2.90E-05 | 4.08E-05 | 5.81E-06 | 1.68E-05 | 2.37E-05 |
| Foxd2 | 0 | 0 | 0 | 0 | 0 | 0 | 0 | 6.89E-06 | 0 | 0 | 0 | 7.16E-06 | 0 | 2.82E-05 | 7.28E-06 | 0 | 0 | 5.90E-05 |
| Eif5a2 | 3.13E-05 | 2.07E-05 | 3.85E-05 | 0 | 0 | 0 | 3.36E-05 | 9.62E-07 | 0 | 0 | 1.85E-05 | 7.42E-06 | 1.01E-05 | 2.80E-05 | 0 | 1.75E-05 | 1.88E-05 | 3.06E-05 |
| Slc35d3 | 0 | 4.97E-06 | 1.46E-05 | 0 | 2.78E-07 | 0 | 0 | 0 | 0 | 0 | 3.14E-07 | 1.89E-05 | 0 | 2.77E-05 | 5.50E-06 | 0 | 9.08E-06 | 0 |
| Dok5 | 5.32E-06 | 1.25E-05 | 2.59E-05 | 0 | 0 | 0 | 1.19E-06 | 0 | 0 | 0 | 8.81E-06 | 4.05E-05 | 0 | 2.75E-05 | 1.65E-05 | 0 | 0 | 3.01E-06 |
| Gpr157 | 0 | 2.61E-05 | 3.46E-06 | 2.29E-05 | 0 | 0 | 8.42E-06 | 0 | 0 | 0 | 1.51E-05 | 4.47E-05 | 0 | 2.70E-05 | 7.65E-06 | 6.12E-08 | 0 | 2.76E-06 |
| Gstt1 | 3.12E-06 | 1.13E-06 | 0 | 0 | 0 | 0 | 0 | 0 | 0 | 0.00031 | 0 | 0 | 0.0002 | 2.59E-05 | 0 | 0 | 0 | 0.00033 |
| Ppp1r13l | 0 | 1.16E-05 | 9.64E-05 | 6.16E-05 | 5.57E-05 | 0 | 4.34E-05 | 8.84E-06 | 0 | 2.53E-06 | 0.00014 | 0.00023 | 0 | 2.56E-05 | 5.75E-05 | 4.63E-06 | 7.43E-05 | 0.00011 |
| Ulk3 | 5.82E-06 | 2.96E-05 | 1.26E-05 | 3.03E-06 | 0 | 0 | 2.11E-05 | 0 | 0 | 2.65E-05 | 1.81E-05 | 0.00011 | 8.88E-07 | 2.52E-05 | 0.00014 | 0 | 0 | 3.44E-05 |
| Hr | 0 | 0 | 0 | 0 | 0 | 0 | 0 | 0 | 0 | 3.12E-05 | 0 | 5.87E-08 | 0 | 2.47E-05 | 0 | 0 | 0 | 0.00013 |
| Cish | 0 | 4.25E-05 | 5.10E-05 | 3.27E-05 | 0 | 0 | 0 | 0 | 0 | 3.70E-06 | 1.12E-06 | 5.44E-06 | 4.53E-05 | 2.44E-05 | 2.85E-05 | 0 | 0.00013 | 2.55E-06 |
| Vcan | 0.00024 | 6.55E-06 | 0.00041 | 0.0005 | 0.00044 | 0 | 0.00022 | 0.00085 | 0 | 0.00064 | 0.00011 | 4.43E-05 | 0.00027 | 2.41E-05 | 0.00019 | 0.00025 | 0.0005 | 0.00066 |
| Tmod2 | 8.97E-06 | 8.60E-06 | 1.27E-06 | 0 | 0 | 0 | 0 | 0 | 0 | 0.0001 | 0 | 0 | 2.16E-05 | 2.39E-05 | 0 | 3.79E-05 | 0 | 0.00019 |
| Anxa8 | 0 | 2.02E-05 | 0 | 0 | 0 | 0 | 5.68E-05 | 0 | 0 | 0.0001 | 0 | 0 | 0.0002 | 2.38E-05 | 0 | 0 | 0 | 0.00036 |
| Arid5a | 0 | 5.06E-05 | 3.34E-05 | 4.88E-05 | 0 | 0 | 0 | 4.06E-05 | 0 | 0 | 0 | 2.84E-05 | 3.71E-05 | 2.35E-05 | 9.78E-05 | 2.23E-05 | 0.0002 | 1.71E-05 |
| Elfn1 | 1.02E-05 | 0 | 0.00011 | 2.40E-05 | 1.39E-06 | 0 | 0 | 0.00011 | 0 | 2.62E-05 | 8.85E-05 | 0.00011 | 0 | 2.35E-05 | 0 | 2.12E-05 | 0 | 2.44E-05 |
| Lyz2 | 0 | 3.13E-05 | 5.46E-06 | 0.00018 | 8.98E-06 | 0 | 0 | 0 | 0 | 3.37E-05 | 0 | 3.33E-05 | 8.27E-05 | 2.33E-05 | 0.00037 | 1.64E-05 | 3.54E-05 | 5.35E-05 |
| Wdr54 | 0 | 6.07E-06 | 0 | 4.88E-06 | 2.76E-06 | 0 | 0 | 2.01E-06 | 0 | 2.10E-06 | 1.41E-05 | 0.00011 | 0 | 2.32E-05 | 1.18E-05 | 1.80E-05 | 0 | 1.32E-05 |
| Me2 | 9.01E-05 | 2.26E-05 | 7.04E-05 | 0 | 6.06E-05 | 0 | 0 | 0 | 0 | 5.80E-05 | 7.69E-05 | 0.0002 | 0 | 2.31E-05 | 0 | 0.00018 | 0 | 3.67E-05 |
| Sox18 | 0 | 4.02E-07 | 1.81E-05 | 0 | 0 | 0 | 0 | 0 | 0 | 0 | 0 | 0 | 0 | 2.29E-05 | 0 | 8.64E-06 | 0 | 5.19E-05 |
| Arhgap33 | 0 | 2.16E-05 | 0 | 1.31E-05 | 0 | 0 | 3.45E-05 | 0 | 0 | 1.73E-05 | 0 | 1.14E-05 | 0 | 2.28E-05 | 5.80E-05 | 8.26E-05 | 6.36E-06 | 6.24E-06 |
| Cd274 | 0 | 2.91E-05 | 0 | 5.09E-07 | 0 | 0 | 0 | 3.40E-06 | 0 | 2.21E-05 | 0 | 1.56E-05 | 2.07E-05 | 2.28E-05 | 2.25E-05 | 1.01E-05 | 1.68E-05 | 1.32E-05 |
| 3-Mar | 0 | 0 | 1.67E-05 | 0.00011 | 0 | 0 | 0 | 0 | 0 | 0.00016 | 4.60E-05 | 3.24E-05 | 0.00028 | 2.27E-05 | 3.49E-05 | 0 | 0.00015 | 0.00017 |

| | | | | | | | | | | | | | | | | | | |
|---|---|---|---|---|---|---|---|---|---|---|---|---|---|---|---|---|---|---|
| Trpv4 | 1.81E-06 | 0 | 1.56E-05 | 5.30E-06 | 0 | 0 | 5.88E-05 | 0 | 0 | 3.14E-05 | 4.87E-06 | 0 | 9.11E-06 | 2.26E-05 | 2.31E-05 | 2.04E-06 | 1.16E-06 | 1.15E-05 |
| Sh3bp2 | 9.27E-06 | 3.87E-05 | 2.52E-05 | 3.04E-05 | 0 | 0 | 1.88E-06 | 1.66E-05 | 0 | 3.02E-05 | 1.46E-05 | 4.07E-06 | 4.63E-05 | 2.20E-05 | 4.09E-05 | 0 | 3.28E-05 | 1.24E-05 |
| Cd55 | 1.44E-05 | 0 | 0 | 0 | 0 | 0 | 0 | 0 | 0 | 0.00064 | 0 | 0 | 0 | 2.17E-05 | 0 | 0 | 0 | 0 |
| Arhgef26 | 0 | 1.30E-05 | 0 | 0 | 1.88E-05 | 0 | 0 | 1.66E-05 | 0 | 0 | 1.20E-05 | 0 | 0 | 2.14E-05 | 0 | 9.71E-06 | 5.33E-06 | 5.77E-06 |
| Hip1r | 0 | 1.74E-05 | 2.23E-05 | 6.84E-06 | 0 | 0 | 7.06E-06 | 1.48E-07 | 0 | 9.55E-06 | 1.32E-05 | 1.48E-05 | 0 | 2.13E-05 | 2.84E-05 | 2.53E-05 | 0 | 1.74E-05 |
| Vash1 | 0 | 0 | 1.26E-05 | 0 | 8.46E-05 | 0 | 2.74E-06 | 5.70E-05 | 0 | 5.01E-05 | 5.39E-05 | 0 | 0 | 2.12E-05 | 1.59E-05 | 1.67E-05 | 0 | 0 |
| Adarb1 | 0 | 1.99E-05 | 3.35E-05 | 0 | 1.60E-06 | 0 | 0.00017 | 0 | 0 | 0 | 8.29E-05 | 2.20E-05 | 0.00028 | 2.11E-05 | 0 | 4.25E-06 | 1.67E-05 | 1.59E-05 |
| Per2 | 0 | 7.61E-06 | 0 | 1.09E-05 | 1.05E-05 | 0 | 2.57E-05 | 0 | 0 | 4.75E-05 | 1.40E-05 | 3.19E-05 | 0 | 2.07E-05 | 5.50E-06 | 0 | 7.91E-05 | 1.17E-05 |
| Cks2 | 3.98E-05 | 0 | 3.12E-05 | 5.15E-05 | 0 | 0 | 0 | 2.99E-06 | 0 | 0 | 7.82E-06 | 8.36E-05 | 0 | 2.03E-05 | 0 | 0.00091 | 0.00021 | 0 |
| Nyap1 | 0 | 1.49E-05 | 0 | 6.56E-06 | 1.37E-05 | 0 | 0 | 0 | 0 | 0 | 5.68E-06 | 1.43E-05 | 0 | 1.99E-05 | 1.19E-05 | 0 | 7.83E-06 | 1.35E-05 |
| Wnt7b | 0 | 0 | 0 | 8.59E-06 | 1.86E-05 | 0 | 0 | 2.55E-05 | 0 | 8.98E-07 | 3.93E-06 | 6.43E-06 | 0 | 1.99E-05 | 4.39E-06 | 0 | 0 | 0 |
| Ackr1 | 0 | 0 | 0 | 0 | 0 | 0 | 0 | 0 | 0 | 0.00058 | 0 | 0 | 0 | 1.96E-05 | 0 | 0 | 0 | 0 |
| Strbp | 7.04E-06 | 2.51E-05 | 0 | 5.27E-06 | 6.73E-06 | 0 | 1.33E-05 | 0 | 0 | 9.83E-06 | 5.28E-06 | 0 | 1.40E-05 | 1.96E-05 | 0 | 3.30E-05 | 0 | 1.07E-05 |
| Btbd19 | 0 | 2.62E-05 | 2.70E-05 | 5.01E-05 | 1.93E-06 | 0 | 5.82E-06 | 6.01E-05 | 0 | 4.05E-05 | 4.45E-06 | 2.65E-05 | 1.55E-05 | 1.91E-05 | 5.27E-05 | 0 | 0.00017 | 2.99E-08 |
| Abcc10 | 0 | 7.22E-05 | 0 | 9.54E-05 | 2.55E-05 | 0 | 8.50E-06 | 2.35E-07 | 0 | 3.96E-05 | 4.12E-06 | 2.13E-05 | 2.78E-05 | 1.90E-05 | 0.00018 | 0 | 3.91E-05 | 8.00E-06 |
| Hspa2 | 0 | 2.98E-05 | 0 | 0 | 0 | 0 | 5.16E-05 | 1.30E-05 | 0 | 3.22E-05 | 0.00014 | 0.00016 | 7.73E-06 | 1.88E-05 | 0 | 0 | 0.00011 | 0.0001 |
| Patj | 0 | 1.68E-05 | 0 | 8.99E-06 | 1.38E-05 | 0 | 0 | 0 | 0 | 1.41E-05 | 4.75E-06 | 3.24E-06 | 2.07E-05 | 1.88E-05 | 3.23E-05 | 1.02E-06 | 2.07E-05 | 3.54E-05 |
| D6Ertd527e | 4.72E-06 | 0 | 2.52E-06 | 0 | 0 | 0 | 2.48E-05 | 1.66E-06 | 0 | 0 | 0 | 1.16E-05 | 1.05E-05 | 1.84E-05 | 0 | 9.16E-07 | 8.31E-05 | 0 |
| Dio2 | 0 | 2.66E-05 | 0 | 0 | 0 | 0 | 0 | 0 | 0 | 8.09E-05 | 0 | 0 | 2.38E-05 | 1.84E-05 | 0 | 0 | 0 | 0.00013 |
| Map2k6 | 5.59E-08 | 8.21E-06 | 1.41E-06 | 4.90E-06 | 0 | 0 | 0 | 4.84E-05 | 0 | 2.27E-05 | 0 | 0.00011 | 0 | 1.83E-05 | 2.56E-05 | 1.93E-05 | 0 | 2.36E-05 |
| Ptafr | 0 | 0 | 5.03E-06 | 5.16E-06 | 0 | 0 | 0 | 8.61E-06 | 0 | 4.77E-06 | 1.13E-07 | 5.91E-06 | 5.97E-06 | 1.81E-05 | 4.47E-05 | 4.82E-06 | 2.36E-05 | 4.34E-06 |
| Fam89a | 0 | 0 | 1.29E-05 | 5.43E-06 | 0 | 0 | 0 | 5.62E-06 | 0 | 0 | 1.07E-06 | 5.04E-06 | 0 | 1.81E-05 | 0 | 2.71E-06 | 0 | 0 |
| Abhd6 | 0 | 1.51E-05 | 3.43E-07 | 5.43E-06 | 1.33E-05 | 0 | 2.72E-05 | 0 | 0 | 2.28E-05 | 1.78E-05 | 1.45E-05 | 0 | 1.81E-05 | 3.39E-06 | 2.60E-07 | 0 | 1.72E-05 |
| Kcnk12 | 0 | 7.41E-07 | 1.02E-05 | 6.79E-07 | 7.76E-06 | 0 | 0 | 0 | 0 | 0 | 3.99E-05 | 2.12E-06 | 0 | 1.76E-05 | 0 | 0 | 0 | 0 |
| Tbkbp1 | 0 | 0.00011 | 0 | 1.67E-05 | 2.59E-05 | 0 | 0 | 3.26E-05 | 0 | 7.92E-05 | 0 | 0.00011 | 4.27E-05 | 1.76E-05 | 0.00013 | 0 | 4.81E-05 | 2.77E-05 |
| Taf1c | 0 | 2.38E-05 | 1.31E-05 | 2.88E-05 | 1.09E-06 | 0 | 0 | 0 | 0 | 6.44E-06 | 5.87E-06 | 1.41E-06 | 2.09E-06 | 1.73E-05 | 5.78E-05 | 4.93E-06 | 0 | 1.14E-06 |
| Cacna1h | 0 | 5.35E-06 | 0 | 0 | 8.28E-06 | 0 | 0 | 0 | 0 | 0 | 7.82E-06 | 6.73E-05 | 0 | 1.71E-05 | 1.66E-05 | 0 | 0 | 0 |
| Smyd3 | 0 | 5.58E-05 | 0 | 5.74E-05 | 7.10E-06 | 0 | 4.53E-06 | 1.09E-05 | 0 | 2.87E-05 | 0 | 5.21E-05 | 9.15E-06 | 1.69E-05 | 0.00012 | 5.79E-05 | 3.50E-05 | 3.48E-05 |
| Slc38a4 | 2.87E-05 | 0 | 3.30E-05 | 3.67E-05 | 0 | 0 | 0.00014 | 0 | 0 | 0 | 0 | 0 | 0 | 1.66E-05 | 0 | 8.95E-05 | 0 | 0 |
| Adam8 | 0 | 6.94E-06 | 1.34E-05 | 1.06E-05 | 0 | 0 | 0 | 0 | 0 | 1.21E-05 | 0 | 2.13E-05 | 0 | 1.64E-05 | 4.48E-05 | 1.30E-05 | 2.02E-05 | 2.97E-06 |
| Sdf2l1 | 0.00023 | 3.32E-05 | 0.00015 | 0.00031 | 3.34E-05 | 0 | 0 | 2.10E-05 | 0 | 0.00024 | 7.50E-05 | 0.00018 | 9.26E-05 | 1.64E-05 | 1.42E-06 | 7.62E-05 | 0 | 3.69E-05 |
| Lrp8 | 3.18E-09 | 0 | 1.78E-05 | 9.16E-06 | 0 | 0 | 1.82E-05 | 0 | 0 | 0 | 4.59E-06 | 0 | 1.36E-05 | 1.62E-05 | 0 | 4.55E-05 | 1.27E-05 | 0 |
| Clec4d | 0 | 0 | 9.35E-07 | 9.30E-07 | 0 | 0 | 0 | 2.31E-06 | 0 | 8.28E-06 | 2.38E-06 | 4.36E-06 | 9.72E-06 | 1.61E-05 | 4.73E-05 | 1.47E-05 | 3.15E-05 | 3.60E-06 |

| | | | | | | | | | | | | | | | | | | |
|---|---|---|---|---|---|---|---|---|---|---|---|---|---|---|---|---|---|---|
| Gjb3 | 0 | 0 | 5.29E-05 | 0 | 0 | 0 | 1.41E-05 | 0 | 0 | 1.46E-06 | 2.94E-05 | 1.22E-05 | 0 | 1.57E-05 | 1.71E-05 | 0 | 0 | 2.13E-05 |
| Cldn18 | 0 | 2.74E-05 | 0 | 4.47E-05 | 6.45E-06 | 0 | 0 | 1.47E-05 | 0 | 2.02E-05 | 4.25E-06 | 4.15E-05 | 0 | 1.56E-05 | 9.38E-05 | 2.30E-05 | 4.59E-05 | 2.58E-05 |
| Per3 | 3.00E-06 | 3.49E-05 | 0 | 1.02E-05 | 5.77E-06 | 0 | 0 | 1.30E-06 | 0 | 0 | 2.06E-05 | 1.48E-05 | 0 | 1.55E-05 | 4.24E-05 | 0 | 0 | 2.90E-05 |
| Fabp3 | 3.78E-06 | 4.58E-06 | 0 | 0 | 6.67E-06 | 0 | 1.02E-05 | 0 | 0 | 0 | 1.45E-06 | 0 | 0 | 1.54E-05 | 0 | 1.45E-06 | 7.78E-06 | 1.34E-06 |
| Lcp1 | 0 | 2.00E-05 | 8.93E-06 | 5.57E-05 | 0 | 0 | 0 | 0 | 0 | 1.90E-05 | 5.61E-07 | 1.99E-05 | 1.35E-05 | 1.53E-05 | 9.38E-05 | 2.82E-05 | 1.80E-05 | 1.78E-05 |
| Rbm38 | 0 | 2.18E-05 | 0 | 0 | 0 | 0 | 0 | 0 | 0 | 0 | 0 | 3.80E-05 | 0 | 1.52E-05 | 1.42E-05 | 1.23E-05 | 0.00036 | 0 |
| Nfkbid | 0 | 2.72E-06 | 0 | 2.56E-05 | 0 | 0 | 0 | 0 | 0 | 3.68E-06 | 5.15E-06 | 1.17E-06 | 0 | 1.52E-05 | 4.73E-05 | 1.29E-05 | 6.12E-05 | 1.38E-05 |
| Lef1 | 0 | 5.91E-05 | 0 | 0 | 8.80E-07 | 0 | 0 | 3.20E-05 | 0 | 0 | 0 | 6.96E-06 | 0 | 1.49E-05 | 0 | 0 | 4.47E-06 | 0 |
| Fam131a | 0 | 2.47E-05 | 0 | 0 | 1.75E-05 | 0 | 0 | 0 | 0 | 5.13E-06 | 1.31E-05 | 1.72E-06 | 0 | 1.46E-05 | 2.80E-05 | 0 | 1.69E-05 | 1.38E-05 |
| Cables1 | 0 | 1.53E-05 | 0 | 0 | 0 | 0 | 0 | 7.12E-06 | 0 | 4.89E-05 | 5.42E-07 | 1.42E-05 | 0 | 1.46E-05 | 0 | 3.66E-06 | 0 | 7.84E-05 |
| Arrb2 | 2.76E-05 | 6.43E-05 | 1.73E-05 | 3.70E-05 | 2.92E-06 | 0 | 0 | 5.22E-05 | 0 | 1.33E-05 | 0 | 0 | 1.05E-05 | 1.46E-05 | 0.00011 | 6.90E-05 | 4.68E-05 | 3.65E-05 |
| Slc40a1 | 0 | 2.43E-05 | 0 | 0 | 0 | 0 | 7.23E-06 | 0 | 0 | 0 | 2.63E-06 | 0 | 0 | 1.43E-05 | 0 | 0 | 0 | 1.92E-05 |
| Lmo2 | 0 | 0 | 0 | 0 | 0 | 0 | 0 | 0 | 0 | 0.00012 | 0 | 3.60E-06 | 0 | 1.43E-05 | 1.41E-05 | 0 | 0 | 8.16E-05 |
| Nrtn | 0 | 9.36E-06 | 0 | 4.71E-06 | 8.13E-06 | 0 | 0 | 0 | 0 | 0 | 5.50E-06 | 0 | 1.25E-06 | 1.43E-05 | 1.70E-05 | 1.70E-08 | 1.21E-05 | 7.34E-06 |
| Ccl9 | 4.51E-06 | 0 | 9.85E-06 | 2.97E-06 | 0 | 0 | 0 | 0 | 0 | 8.32E-06 | 2.20E-05 | 0 | 3.38E-05 | 1.43E-05 | 1.86E-05 | 1.70E-05 | 2.14E-06 | 6.89E-05 |
| Baiap2l1 | 0 | 5.87E-06 | 1.74E-05 | 9.93E-06 | 3.77E-06 | 0 | 0 | 6.12E-06 | 0 | 3.49E-06 | 6.87E-05 | 0.00013 | 0 | 1.41E-05 | 4.21E-05 | 0 | 0 | 6.20E-06 |
| Adgrf5 | 0 | 1.05E-05 | 0 | 0 | 1.79E-05 | 0 | 0 | 1.06E-05 | 0 | 3.06E-05 | 5.15E-05 | 1.55E-07 | 0 | 1.41E-05 | 0 | 5.93E-06 | 0 | 0 |
| Cd14 | 0 | 1.94E-05 | 0 | 0.0001 | 1.15E-05 | 0 | 0 | 0 | 0 | 1.54E-05 | 0 | 1.01E-05 | 7.67E-05 | 1.39E-05 | 0.00017 | 4.02E-05 | 2.89E-05 | 7.09E-05 |
| Mmp12 | 0 | 1.92E-05 | 7.45E-06 | 3.38E-05 | 1.38E-05 | 0 | 0 | 2.07E-05 | 0 | 1.72E-05 | 1.20E-05 | 5.02E-05 | 1.27E-06 | 1.38E-05 | 6.21E-05 | 0 | 9.51E-06 | 2.30E-06 |
| Cd200 | 0 | 1.02E-05 | 0 | 0 | 0 | 0 | 0 | 9.89E-05 | 0 | 1.54E-05 | 0 | 6.97E-05 | 0 | 1.37E-05 | 0 | 0 | 0 | 0.00022 |
| Sh3bgrl2 | 0 | 1.37E-05 | 1.39E-05 | 5.70E-07 | 0 | 0 | 1.70E-05 | 1.83E-05 | 0 | 0 | 1.08E-05 | 1.04E-05 | 0 | 1.35E-05 | 1.11E-05 | 2.25E-05 | 0 | 2.10E-05 |
| Ptprc | 0 | 5.85E-06 | 3.40E-06 | 3.34E-06 | 0 | 0 | 0 | 0 | 0 | 7.76E-06 | 0 | 5.34E-06 | 1.24E-06 | 1.35E-05 | 4.81E-05 | 1.40E-05 | 9.71E-06 | 1.05E-05 |
| Syce2 | 0 | 0 | 0 | 2.60E-05 | 0 | 0 | 1.04E-05 | 0 | 0 | 0 | 2.00E-07 | 0.00021 | 3.26E-06 | 1.34E-05 | 0 | 0.00029 | 0 | 0 |
| B3gnt8 | 0 | 0 | 1.08E-06 | 1.47E-05 | 1.36E-05 | 0 | 4.67E-06 | 7.74E-05 | 0 | 0 | 1.64E-05 | 5.37E-06 | 0 | 1.33E-05 | 1.12E-06 | 2.76E-05 | 0 | 8.75E-05 |
| Intu | 1.89E-06 | 4.23E-05 | 3.44E-06 | 3.13E-05 | 0 | 0 | 5.91E-06 | 0 | 0 | 1.57E-05 | 1.27E-05 | 0 | 6.39E-06 | 1.33E-05 | 8.14E-05 | 3.59E-05 | 3.87E-05 | 7.53E-06 |
| Chac1 | 0 | 2.67E-06 | 1.76E-05 | 0 | 0 | 0 | 8.47E-06 | 0 | 0 | 3.36E-06 | 7.33E-06 | 2.04E-05 | 0 | 1.33E-05 | 0 | 0 | 0 | 6.15E-06 |
| Slc27a3 | 0 | 9.67E-06 | 0 | 8.23E-06 | 0 | 0 | 1.50E-05 | 0 | 0 | 1.91E-05 | 0 | 8.14E-06 | 0 | 1.33E-05 | 0 | 0 | 0 | 2.16E-06 |
| Ung | 7.99E-06 | 0 | 9.65E-05 | 3.05E-05 | 0 | 0 | 3.89E-05 | 0 | 0 | 0 | 3.37E-06 | 0 | 8.89E-06 | 1.33E-05 | 0 | 8.06E-05 | 0 | 0 |
| Rac2 | 0 | 5.83E-06 | 0 | 2.66E-05 | 0 | 0 | 0 | 1.83E-06 | 0 | 1.03E-05 | 0 | 1.12E-05 | 0 | 1.32E-05 | 5.55E-05 | 1.90E-05 | 1.10E-05 | 3.34E-05 |
| Dgat2 | 1.07E-06 | 0 | 5.98E-06 | 5.73E-06 | 0 | 0 | 6.85E-06 | 0 | 0 | 2.00E-06 | 0 | 6.85E-05 | 3.08E-05 | 1.32E-05 | 0 | 0 | 0.00012 | 0 |
| Cxcr4 | 0 | 3.03E-05 | 0 | 1.03E-06 | 0 | 0 | 0 | 0 | 0 | 2.59E-06 | 0 | 6.12E-06 | 0 | 1.32E-05 | 8.14E-05 | 3.01E-05 | 1.21E-05 | 4.18E-05 |
| Rab27a | 0 | 1.84E-05 | 0 | 0 | 0 | 0 | 0 | 0 | 0 | 0 | 0 | 5.64E-06 | 0 | 1.31E-05 | 0 | 1.99E-05 | 0 | 1.22E-05 |
| Npas4 | 0 | 0 | 0 | 0 | 0 | 0 | 0 | 0 | 0 | 0 | 4.40E-05 | 0 | 0 | 1.31E-05 | 1.34E-05 | 0 | 0 | 0 |

| | | | | | | | | | | | | | | | | | | |
|---|---|---|---|---|---|---|---|---|---|---|---|---|---|---|---|---|---|---|
| Pnpo | 1.35E-05 | 5.02E-05 | 1.60E-05 | 5.93E-05 | 2.23E-06 | 0 | 3.07E-05 | 0 | 0 | 1.33E-05 | 4.37E-05 | 2.49E-06 | 0 | 1.29E-05 | 0 | 0 | 0 | 1.58E-05 |
| Mpp2 | 0 | 0 | 0 | 9.88E-06 | 1.43E-05 | 0 | 0 | 0.0003 | 0 | 0 | 5.34E-05 | 0.00038 | 0 | 1.27E-05 | 2.92E-05 | 0 | 0 | 4.34E-06 |
| Rfx3 | 1.61E-05 | 2.97E-05 | 1.31E-05 | 0 | 7.61E-06 | 0 | 4.18E-06 | 2.98E-05 | 0 | 0 | 0 | 2.98E-05 | 1.34E-06 | 1.25E-05 | 0.00015 | 3.02E-05 | 8.47E-05 | 4.15E-05 |
| Gpc2 | 0 | 2.37E-05 | 0 | 1.60E-05 | 1.23E-05 | 0 | 0 | 0 | 0 | 0 | 1.90E-05 | 0 | 2.23E-06 | 1.25E-05 | 3.95E-05 | 7.51E-06 | 0 | 0 |
| Arap3 | 0 | 1.63E-05 | 0 | 0 | 0 | 0 | 0 | 6.87E-05 | 0 | 2.98E-06 | 5.69E-06 | 0 | 0 | 1.25E-05 | 2.25E-05 | 0 | 0 | 9.77E-06 |
| Zdhhc2 | 4.83E-06 | 0 | 0 | 3.21E-06 | 0 | 0 | 4.52E-07 | 0 | 0 | 2.00E-06 | 1.10E-06 | 0 | 0 | 1.23E-05 | 4.89E-06 | 1.76E-05 | 4.76E-06 | 1.34E-05 |
| Sap30 | 5.21E-07 | 1.88E-05 | 0 | 0 | 3.71E-05 | 0 | 0 | 0 | 0 | 0 | 6.51E-05 | 0.00028 | 0 | 1.22E-05 | 1.55E-07 | 0.00012 | 0 | 0 |
| Hils1 | 0 | 0 | 0 | 9.67E-06 | 4.60E-06 | 0 | 3.18E-06 | 0 | 0 | 0 | 3.64E-06 | 4.95E-05 | 7.39E-06 | 1.22E-05 | 3.27E-05 | 1.08E-05 | 0 | 0 |
| Tmem136 | 0 | 2.62E-05 | 0 | 0 | 3.06E-05 | 0 | 0 | 2.31E-05 | 0 | 1.52E-06 | 5.08E-06 | 1.54E-05 | 0 | 1.22E-05 | 0 | 0 | 0 | 1.50E-05 |
| Mustn1 | 0 | 7.14E-06 | 0 | 0 | 0 | 0 | 0 | 0 | 0 | 0.00017 | 0 | 0 | 0 | 1.20E-05 | 0 | 0 | 0 | 0.00012 |
| Tnnt2 | 0 | 4.94E-06 | 5.04E-06 | 2.42E-05 | 0 | 0 | 0 | 0 | 0 | 6.25E-06 | 1.42E-05 | 6.66E-06 | 0 | 1.18E-05 | 4.39E-05 | 8.04E-06 | 2.51E-05 | 1.89E-06 |
| Bcam | 0 | 1.83E-05 | 1.09E-06 | 5.89E-07 | 0 | 0 | 0 | 0 | 0 | 3.12E-06 | 0 | 9.06E-06 | 0 | 1.18E-05 | 1.70E-05 | 8.52E-07 | 0 | 3.63E-05 |
| Acan | 1.67E-06 | 4.61E-06 | 0 | 1.46E-05 | 0 | 0 | 5.12E-06 | 0 | 0 | 7.45E-06 | 0 | 3.27E-06 | 0 | 1.17E-05 | 2.35E-05 | 0 | 0 | 1.59E-05 |
| Car8 | 0 | 0 | 0 | 0 | 4.94E-06 | 0 | 0 | 0 | 0 | 0.00021 | 0 | 0 | 0 | 1.16E-05 | 0 | 0 | 0 | 0 |
| Tbx6 | 0 | 3.99E-05 | 0 | 0 | 9.73E-06 | 0 | 0 | 0 | 0 | 0 | 0 | 3.89E-05 | 0 | 1.16E-05 | 4.62E-05 | 0 | 1.90E-05 | 1.72E-07 |
| Irf7 | 2.79E-05 | 3.44E-06 | 0 | 3.78E-05 | 0 | 0 | 0 | 0 | 0 | 0.00057 | 6.89E-07 | 2.69E-05 | 0.00015 | 1.15E-05 | 1.17E-05 | 0 | 3.43E-05 | 0.00018 |
| Pkn3 | 0 | 0 | 5.06E-06 | 1.35E-05 | 4.08E-06 | 0 | 0 | 0 | 0 | 0 | 0 | 4.81E-06 | 0 | 1.15E-05 | 2.12E-05 | 9.19E-05 | 5.81E-05 | 6.91E-06 |
| Snrpn | 0 | 8.18E-06 | 0 | 0 | 0 | 0 | 2.49E-06 | 0 | 0 | 3.10E-05 | 0 | 5.57E-06 | 0 | 1.14E-05 | 0 | 0 | 0 | 5.46E-05 |
| Ppm1j | 0 | 0 | 5.91E-06 | 0 | 6.42E-06 | 0 | 1.55E-06 | 3.66E-05 | 0 | 7.44E-07 | 2.47E-05 | 8.11E-06 | 7.44E-06 | 1.13E-05 | 2.40E-05 | 0 | 0 | 1.53E-06 |
| Arhgap44 | 0 | 2.34E-05 | 1.71E-05 | 4.76E-06 | 0 | 0 | 3.19E-05 | 0 | 0 | 3.67E-05 | 1.78E-05 | 0 | 1.03E-05 | 1.12E-05 | 1.98E-05 | 0 | 0 | 4.91E-05 |
| Tyrobp | 0 | 1.47E-05 | 0 | 4.83E-05 | 0 | 0 | 0 | 0 | 0 | 2.14E-05 | 0 | 1.03E-05 | 0 | 1.12E-05 | 0.00014 | 3.29E-05 | 6.07E-05 | 3.14E-05 |
| Pdzd2 | 0 | 4.82E-06 | 0 | 0 | 0 | 0 | 0 | 0 | 0 | 0.00016 | 5.50E-06 | 0 | 0 | 1.12E-05 | 0 | 0 | 0 | 6.61E-05 |
| Angel1 | 2.05E-07 | 2.51E-05 | 1.25E-05 | 1.22E-05 | 2.12E-05 | 0 | 4.47E-06 | 0 | 0 | 1.45E-05 | 4.26E-06 | 0 | 0 | 1.11E-05 | 1.78E-05 | 1.02E-05 | 1.76E-05 | 0 |
| Prkab2 | 0 | 1.40E-05 | 0 | 0 | 0 | 0 | 0 | 1.98E-05 | 0 | 6.59E-05 | 0.00011 | 0.00023 | 0 | 1.11E-05 | 0.00012 | 0 | 0 | 3.30E-05 |
| Tdp2 | 8.33E-06 | 5.68E-05 | 1.32E-05 | 0.00012 | 1.13E-05 | 0 | 1.15E-07 | 2.70E-05 | 0 | 2.32E-05 | 0 | 0.00016 | 5.19E-05 | 1.10E-05 | 0.00025 | 0.00024 | 0.00017 | 0 |
| Hist2h3c1 | 3.01E-06 | 2.82E-05 | 0 | 0 | 0 | 0 | 2.03E-06 | 4.86E-06 | 0 | 2.99E-05 | 1.02E-05 | 9.87E-05 | 3.51E-05 | 1.10E-05 | 0 | 0 | 0.00016 | 2.17E-05 |
| Ldlrad4 | 1.10E-05 | 7.02E-05 | 0.00012 | 0 | 8.57E-05 | 0 | 0.00024 | 0.00023 | 0 | 0.0002 | 0.00026 | 3.48E-05 | 9.81E-06 | 1.09E-05 | 8.74E-05 | 0 | 0.00012 | 0.00035 |
| Kcng2 | 0 | 0 | 7.79E-07 | 1.92E-06 | 6.96E-06 | 0 | 2.51E-05 | 0 | 0 | 0 | 5.45E-06 | 2.83E-05 | 0 | 1.08E-05 | 1.56E-05 | 0 | 0 | 0 |
| Snx20 | 0 | 5.40E-07 | 1.66E-06 | 6.26E-06 | 3.19E-07 | 0 | 0 | 0 | 0 | 8.76E-06 | 0 | 0 | 0 | 1.07E-05 | 4.50E-05 | 1.21E-05 | 1.04E-05 | 4.05E-06 |
| Pak3 | 6.83E-06 | 0 | 3.81E-06 | 1.73E-06 | 0 | 0 | 0 | 0 | 0 | 2.61E-06 | 4.47E-06 | 0 | 4.00E-05 | 1.07E-05 | 0 | 2.63E-06 | 7.93E-06 | 1.29E-05 |
| Csf3r | 0 | 4.59E-06 | 0 | 1.50E-05 | 6.36E-07 | 0 | 0 | 0 | 0 | 2.14E-06 | 0 | 0 | 8.84E-06 | 1.06E-05 | 2.12E-05 | 3.13E-05 | 1.12E-05 | 1.12E-05 |
| Smn1 | 3.08E-05 | 2.02E-05 | 2.93E-05 | 1.26E-05 | 5.38E-06 | 0 | 0 | 3.34E-06 | 0 | 0 | 4.68E-06 | 5.48E-06 | 1.82E-06 | 1.05E-05 | 2.92E-05 | 2.18E-05 | 2.07E-06 | 0 |
| Arnt2 | 0 | 8.66E-06 | 0 | 0 | 7.31E-07 | 0 | 0 | 0 | 0 | 7.48E-06 | 0 | 0 | 3.48E-05 | 1.04E-05 | 1.12E-07 | 7.06E-06 | 0 | 0 |

| | | | | | | | | | | | | | | | | | | |
|---|---|---|---|---|---|---|---|---|---|---|---|---|---|---|---|---|---|---|
| Mapk8ip3 | 7.92E-05 | 0.00018 | 5.67E-05 | 0 | 1.18E-05 | 0 | 0 | 7.98E-05 | 0 | 0 | 0 | 0.00031 | 0 | 1.03E-05 | 0.0006 | 0 | 0.00018 | 4.17E-05 |
| Gemin2 | 1.42E-05 | 2.47E-05 | 2.76E-05 | 4.77E-05 | 6.28E-06 | 0 | 1.82E-06 | 1.66E-05 | 0 | 1.51E-05 | 7.46E-06 | 6.67E-06 | 0 | 1.03E-05 | 1.97E-05 | 0.0001 | 0 | 2.04E-05 |
| Grk4 | 0 | 1.87E-05 | 0 | 0 | 1.84E-05 | 0 | 1.67E-05 | 0 | 0 | 2.21E-05 | 0 | 1.57E-06 | 0 | 1.02E-05 | 5.94E-05 | 0 | 0 | 1.99E-05 |
| Hells | 2.50E-05 | 0 | 0.00014 | 0 | 0 | 0 | 0 | 0 | 0 | 0 | 8.90E-08 | 0 | 4.94E-05 | 1.01E-05 | 0 | 0.00047 | 0 | 0 |
| Gmnn | 0 | 0 | 3.52E-05 | 0 | 0 | 0 | 0 | 0 | 0 | 0 | 2.58E-05 | 0 | 2.96E-05 | 1.01E-05 | 0 | 0.00069 | 0 | 0 |
| Rab38 | 7.14E-06 | 0 | 3.30E-06 | 9.29E-06 | 1.47E-06 | 0 | 0 | 4.50E-05 | 0 | 3.15E-05 | 0 | 0 | 0 | 1.00E-05 | 0 | 2.50E-05 | 0 | 2.47E-05 |
| Dtwd2 | 2.57E-05 | 2.64E-05 | 1.54E-05 | 0 | 1.30E-05 | 0 | 2.32E-05 | 2.53E-05 | 0 | 1.16E-05 | 0 | 2.57E-06 | 6.56E-06 | 1.00E-05 | 1.25E-05 | 4.79E-06 | 1.28E-05 | 7.35E-06 |
| Hvcn1 | 0 | 2.79E-05 | 0 | 1.37E-06 | 0 | 0 | 0 | 0 | 0 | 0 | 0 | 7.76E-06 | 3.30E-05 | 9.81E-06 | 0 | 2.89E-06 | 0 | 4.39E-05 |
| Ppp1r14a | 0 | 9.50E-06 | 0 | 0 | 0 | 0 | 0 | 4.15E-05 | 0 | 0 | 3.91E-06 | 0 | 0 | 9.78E-06 | 0 | 0 | 0 | 2.17E-05 |
| Srgn | 0 | 5.05E-05 | 1.88E-05 | 0.00012 | 3.86E-06 | 0 | 0 | 0 | 0 | 0.00029 | 0 | 3.97E-05 | 0.00024 | 9.76E-06 | 0.00031 | 4.97E-05 | 1.21E-05 | 0.0004 |
| Il20rb | 3.28E-06 | 6.64E-06 | 1.68E-05 | 2.40E-05 | 0 | 0 | 9.82E-06 | 1.42E-06 | 0 | 1.33E-05 | 7.76E-06 | 3.08E-06 | 4.54E-05 | 9.65E-06 | 2.27E-05 | 1.10E-05 | 9.09E-05 | 9.38E-06 |
| Skil | 0.00012 | 0.00015 | 0.00023 | 0 | 0.00035 | 0 | 0.00013 | 0.00027 | 0 | 0 | 0.00029 | 0.00012 | 0 | 9.60E-06 | 0.00047 | 0 | 0.00132 | 0.00018 |
| Hck | 0 | 3.97E-07 | 7.34E-06 | 0 | 0 | 0 | 0 | 0 | 0 | 2.99E-06 | 6.68E-07 | 1.57E-06 | 0 | 9.52E-06 | 1.78E-05 | 4.77E-06 | 2.06E-05 | 5.56E-05 |
| Mybl2 | 0 | 0 | 3.68E-05 | 0 | 0 | 0 | 0 | 7.75E-05 | 0 | 0 | 0 | 0 | 0 | 9.41E-06 | 0 | 0.00065 | 0 | 0 |
| Tarsl2 | 5.40E-08 | 1.56E-06 | 0 | 0 | 0 | 0 | 1.36E-05 | 0 | 0 | 1.07E-05 | 0 | 2.18E-06 | 0 | 9.18E-06 | 0 | 1.07E-05 | 0 | 6.23E-06 |
| Hspa4l | 1.15E-05 | 4.50E-05 | 0 | 0 | 1.67E-05 | 0 | 1.15E-05 | 0 | 0 | 4.89E-05 | 5.09E-05 | 5.89E-05 | 3.86E-05 | 9.13E-06 | 1.65E-05 | 5.64E-05 | 3.71E-05 | 7.26E-05 |
| N4bp2l1 | 0 | 3.87E-05 | 0 | 0 | 0 | 0 | 0 | 0 | 0 | 7.61E-06 | 0 | 3.34E-06 | 1.30E-05 | 8.77E-06 | 0 | 0 | 8.17E-06 | 0 |
| Bin2 | 0 | 0 | 0 | 1.55E-05 | 0 | 0 | 0 | 3.92E-06 | 0 | 9.14E-06 | 0 | 1.20E-06 | 0 | 8.77E-06 | 2.47E-05 | 0 | 7.40E-06 | 9.56E-06 |
| Adgrb2 | 0 | 1.31E-05 | 0 | 1.67E-06 | 3.62E-06 | 0 | 1.15E-06 | 0 | 0 | 0 | 0 | 0 | 0 | 8.72E-06 | 1.48E-05 | 2.17E-05 | 7.64E-06 | 0 |
| Scrn1 | 0 | 2.04E-05 | 0 | 4.79E-06 | 0 | 0 | 2.70E-05 | 0 | 0 | 0.00012 | 0 | 0 | 9.35E-06 | 8.63E-06 | 0 | 0 | 0 | 3.31E-05 |
| Chadl | 0 | 9.18E-06 | 0 | 1.11E-05 | 4.93E-06 | 0 | 1.00E-05 | 0 | 0 | 1.20E-05 | 5.68E-07 | 0 | 1.40E-05 | 8.63E-06 | 7.33E-06 | 0 | 1.20E-05 | 1.33E-05 |
| Ust | 9.83E-05 | 8.44E-05 | 0.00015 | 0 | 0.00016 | 0 | 0.00016 | 3.21E-05 | 0 | 0.00016 | 0.00014 | 3.82E-05 | 2.75E-05 | 8.48E-06 | 0 | 0 | 0.00013 | 0 |
| Lnx1 | 0 | 0 | 0 | 1.29E-05 | 3.14E-06 | 0 | 4.61E-06 | 0 | 0 | 8.49E-06 | 0 | 0 | 3.02E-05 | 8.40E-06 | 5.78E-06 | 8.75E-06 | 1.09E-05 | 0 |
| Gdpd1 | 0 | 2.45E-05 | 0 | 0 | 1.70E-06 | 0 | 0 | 0 | 0 | 0 | 0 | 6.54E-06 | 5.55E-06 | 8.39E-06 | 3.57E-06 | 0 | 0 | 0 |
| Spock2 | 0 | 0 | 0 | 1.53E-05 | 8.04E-07 | 0 | 1.60E-05 | 0 | 0 | 8.23E-06 | 1.03E-06 | 8.96E-06 | 0 | 8.20E-06 | 6.57E-06 | 0 | 3.24E-06 | 7.58E-05 |
| Lonrf1 | 0 | 2.79E-05 | 4.91E-06 | 0 | 6.10E-06 | 0 | 0 | 0 | 0 | 0 | 0 | 0.0002 | 0 | 8.20E-06 | 3.43E-05 | 0 | 0 | 3.46E-05 |
| Bambi | 3.56E-06 | 2.78E-08 | 2.49E-05 | 0 | 9.22E-06 | 0 | 0.00013 | 0 | 0 | 0 | 5.73E-05 | 3.43E-05 | 0.00012 | 8.17E-06 | 0 | 4.19E-05 | 0.00019 | 0 |
| Coro1a | 0 | 1.18E-05 | 7.44E-06 | 1.74E-05 | 3.09E-07 | 0 | 0 | 0 | 0 | 2.68E-06 | 1.64E-06 | 0 | 8.57E-07 | 8.14E-06 | 5.75E-05 | 2.00E-05 | 3.20E-05 | 1.98E-05 |
| Stk36 | 1.59E-05 | 2.19E-05 | 0 | 1.23E-06 | 1.84E-05 | 0 | 1.02E-05 | 2.64E-06 | 0 | 1.08E-05 | 1.51E-06 | 1.35E-05 | 8.77E-06 | 8.10E-06 | 3.14E-05 | 0 | 0 | 4.60E-06 |
| Adrb2 | 0 | 0 | 9.81E-07 | 0 | 0 | 0 | 1.99E-06 | 0 | 0 | 5.03E-06 | 4.53E-07 | 5.94E-06 | 5.55E-06 | 8.07E-06 | 2.71E-05 | 2.02E-05 | 8.69E-05 | 2.43E-06 |
| Nlrp3 | 0 | 6.92E-06 | 3.47E-06 | 1.24E-05 | 2.89E-06 | 0 | 3.33E-06 | 4.86E-07 | 0 | 4.52E-06 | 2.11E-06 | 6.41E-06 | 0 | 8.05E-06 | 2.88E-05 | 1.93E-05 | 2.54E-05 | 1.23E-05 |
| S100a9 | 0 | 1.77E-05 | 0 | 0 | 0 | 0 | 0 | 0 | 0 | 1.67E-05 | 8.37E-06 | 1.74E-05 | 2.00E-05 | 7.94E-06 | 0.00012 | 3.07E-05 | 4.21E-05 | 1.37E-05 |
| Zfp566 | 1.31E-05 | 2.73E-05 | 2.10E-05 | 0 | 0 | 0 | 0 | 0 | 0 | 7.39E-06 | 0 | 8.94E-06 | 0 | 7.93E-06 | 6.12E-05 | 0 | 0 | 1.28E-05 |

| | | | | | | | | | | | | | | | | | | |
|---|---|---|---|---|---|---|---|---|---|---|---|---|---|---|---|---|---|---|
| Clec4e | 0 | 0 | 6.35E-06 | 7.07E-06 | 0 | 0 | 0 | 0 | 0 | 4.65E-06 | 2.78E-06 | 4.59E-06 | 3.16E-07 | 7.76E-06 | 2.03E-05 | 5.14E-06 | 1.52E-05 | 1.24E-05 |
| Rfx2 | 0 | 1.59E-05 | 9.39E-06 | 0 | 0 | 0 | 4.71E-06 | 0 | 0 | 8.38E-05 | 0 | 1.41E-06 | 0 | 7.75E-06 | 1.98E-05 | 3.53E-05 | 3.63E-05 | 3.19E-05 |
| Abcb1b | 0 | 9.25E-07 | 3.51E-05 | 0 | 0 | 0 | 0 | 0 | 0 | 1.44E-05 | 0 | 0 | 3.88E-05 | 7.72E-06 | 0 | 2.00E-05 | 1.86E-05 | 1.31E-05 |
| Acsbg1 | 0 | 0 | 3.65E-06 | 0 | 0 | 0 | 0 | 0.00012 | 0 | 0 | 1.01E-05 | 6.60E-05 | 0 | 7.67E-06 | 0 | 1.24E-05 | 0 | 1.33E-05 |
| C5ar1 | 0 | 9.01E-07 | 1.35E-06 | 3.64E-05 | 0 | 0 | 0 | 0 | 0 | 3.98E-06 | 0 | 3.41E-06 | 5.64E-06 | 7.66E-06 | 4.31E-05 | 1.23E-05 | 3.01E-05 | 3.20E-06 |
| Myl4 | 5.28E-06 | 7.12E-06 | 1.15E-05 | 1.46E-05 | 1.21E-05 | 0 | 2.63E-05 | 1.92E-07 | 0 | 9.82E-06 | 5.07E-06 | 0 | 1.18E-05 | 7.61E-06 | 0 | 3.33E-05 | 3.36E-06 | 1.78E-05 |
| Lpar2 | 0 | 1.31E-05 | 0 | 1.10E-05 | 1.13E-06 | 0 | 1.66E-06 | 0 | 0 | 7.21E-07 | 2.73E-06 | 0 | 2.25E-05 | 7.52E-06 | 4.07E-06 | 2.89E-06 | 2.75E-05 | 7.03E-06 |
| Cd300a | 0 | 3.59E-06 | 0 | 3.23E-06 | 0 | 0 | 0 | 3.25E-06 | 0 | 4.72E-06 | 2.65E-06 | 0 | 0 | 7.44E-06 | 4.40E-05 | 1.08E-05 | 3.08E-06 | 3.04E-06 |
| Galnt18 | 0 | 0 | 6.54E-06 | 0 | 0 | 0 | 0 | 0 | 0 | 7.72E-05 | 0 | 0 | 0 | 7.38E-06 | 1.17E-06 | 0 | 0 | 8.29E-05 |
| Whrn | 0 | 2.17E-05 | 2.42E-05 | 6.24E-06 | 5.21E-05 | 0 | 2.07E-05 | 6.27E-05 | 0 | 0 | 1.79E-05 | 8.66E-06 | 0 | 7.29E-06 | 5.13E-05 | 0 | 6.49E-05 | 8.03E-06 |
| Ncf2 | 0 | 1.28E-05 | 2.56E-06 | 0 | 0 | 0 | 5.63E-06 | 0 | 0 | 3.82E-06 | 7.43E-06 | 4.83E-06 | 1.18E-05 | 7.21E-06 | 5.17E-05 | 1.29E-05 | 0 | 7.04E-07 |
| Ndrg2 | 0 | 6.30E-05 | 0 | 2.78E-06 | 2.98E-05 | 0 | 0 | 0 | 0 | 1.56E-05 | 7.02E-05 | 0.00025 | 0.00024 | 7.16E-06 | 0 | 0 | 0 | 0.00021 |
| Arhgap27 | 0 | 1.02E-05 | 0 | 8.99E-06 | 0 | 0 | 0 | 0 | 0 | 4.84E-06 | 0 | 1.85E-05 | 1.98E-06 | 7.12E-06 | 4.69E-06 | 3.45E-06 | 3.95E-05 | 4.21E-06 |
| Eef1aknmt | 4.30E-05 | 5.24E-05 | 7.80E-05 | 7.14E-05 | 1.14E-05 | 0 | 0 | 0 | 0 | 1.63E-05 | 0 | 0 | 1.58E-05 | 7.07E-06 | 3.95E-05 | 1.30E-05 | 0 | 4.56E-05 |
| Chchd4 | 3.27E-05 | 3.00E-05 | 0.00011 | 2.22E-05 | 1.46E-06 | 0 | 0 | 0 | 0 | 0 | 1.67E-07 | 0 | 3.58E-07 | 7.02E-06 | 0 | 4.29E-05 | 0 | 0 |
| Slc11a1 | 0 | 1.01E-05 | 8.63E-06 | 1.74E-05 | 0 | 0 | 0 | 0 | 0 | 5.44E-06 | 0 | 5.44E-06 | 0 | 6.92E-06 | 3.72E-05 | 6.23E-06 | 6.76E-06 | 1.35E-05 |
| Hlx | 0 | 2.10E-05 | 1.12E-06 | 2.44E-06 | 0 | 0 | 0 | 0 | 0 | 0 | 3.80E-06 | 2.29E-06 | 0 | 6.87E-06 | 2.28E-05 | 1.90E-05 | 9.84E-06 | 7.28E-06 |
| Arhgef28 | 0 | 2.68E-05 | 0 | 0 | 8.07E-05 | 0 | 3.93E-05 | 0 | 0 | 2.65E-05 | 8.36E-05 | 7.65E-05 | 5.41E-05 | 6.81E-06 | 9.90E-06 | 0 | 0 | 0 |
| C1qtnf3 | 0 | 0 | 0 | 0 | 0.00087 | 0 | 0.00107 | 0 | 0 | 0 | 0.00015 | 0 | 0 | 6.80E-06 | 0 | 0 | 0 | 0.00076 |
| Tnip2 | 1.88E-05 | 0.00017 | 5.17E-05 | 9.02E-05 | 1.44E-05 | 0 | 2.38E-05 | 3.58E-05 | 0 | 0.00012 | 0 | 7.92E-05 | 7.76E-05 | 6.52E-06 | 0.00043 | 0 | 0.00013 | 6.06E-05 |
| Tanc2 | 0 | 1.43E-05 | 1.72E-05 | 8.48E-08 | 6.68E-06 | 0 | 6.15E-05 | 0 | 0 | 5.91E-05 | 6.04E-05 | 3.36E-05 | 7.41E-06 | 6.43E-06 | 0 | 7.94E-06 | 1.34E-05 | 1.76E-06 |
| Slc24a3 | 0 | 0 | 0 | 0 | 5.74E-05 | 0 | 2.49E-05 | 0 | 0 | 8.39E-06 | 6.55E-05 | 0 | 1.58E-05 | 6.33E-06 | 6.98E-06 | 0 | 0 | 8.00E-06 |
| Ncapd2 | 0 | 1.57E-05 | 0 | 1.39E-05 | 0 | 0 | 2.47E-05 | 0 | 0 | 3.30E-05 | 0 | 2.63E-06 | 1.04E-06 | 6.32E-06 | 8.63E-05 | 0.00144 | 2.86E-05 | 2.77E-05 |
| Tnf | 0 | 8.16E-06 | 0 | 5.14E-05 | 0 | 0 | 0 | 0 | 0 | 9.95E-06 | 0 | 4.35E-06 | 1.54E-05 | 6.30E-06 | 0.0001 | 1.39E-05 | 7.54E-05 | 2.05E-05 |
| Apol9b | 0 | 0 | 1.51E-05 | 0 | 3.69E-05 | 0 | 2.43E-05 | 2.99E-05 | 0 | 1.51E-05 | 5.35E-06 | 7.28E-06 | 5.81E-06 | 6.27E-06 | 0 | 3.24E-05 | 0 | 1.12E-05 |
| Trpv2 | 0 | 2.37E-06 | 6.97E-09 | 7.59E-06 | 0 | 0 | 1.91E-06 | 1.06E-05 | 0 | 6.32E-06 | 7.27E-05 | 7.91E-06 | 0 | 6.25E-06 | 7.44E-06 | 0 | 0 | 3.09E-05 |
| Epop | 0 | 0 | 3.21E-05 | 8.35E-06 | 0 | 0 | 0 | 0 | 0 | 3.17E-06 | 0 | 1.92E-05 | 7.83E-06 | 6.23E-06 | 4.99E-06 | 2.24E-05 | 3.63E-05 | 0 |
| Entpd4 | 6.94E-06 | 0 | 0 | 1.30E-05 | 5.86E-06 | 0 | 1.05E-05 | 4.16E-06 | 0 | 4.10E-06 | 3.38E-06 | 1.10E-05 | 6.79E-06 | 6.20E-06 | 1.01E-05 | 0 | 4.19E-06 | 0 |
| Abca3 | 0 | 3.93E-05 | 0 | 0 | 1.64E-05 | 0 | 0 | 0 | 0 | 6.40E-05 | 0 | 3.45E-05 | 1.63E-05 | 6.16E-06 | 3.22E-06 | 0 | 0 | 7.85E-05 |
| Mtfmt | 0 | 8.14E-07 | 0 | 4.05E-05 | 7.73E-06 | 0 | 0 | 1.42E-05 | 0 | 1.20E-05 | 0 | 0 | 5.29E-06 | 6.05E-06 | 1.81E-05 | 2.45E-06 | 0 | 0 |
| Tram1l1 | 8.02E-06 | 0 | 4.97E-06 | 3.85E-06 | 0 | 0 | 1.95E-05 | 2.53E-06 | 0 | 6.11E-06 | 4.37E-06 | 0 | 0 | 6.00E-06 | 1.88E-06 | 8.72E-06 | 0 | 1.32E-06 |
| Foxc2 | 0 | 2.67E-06 | 0 | 0 | 0 | 0 | 0 | 3.75E-05 | 0 | 0 | 0 | 2.19E-05 | 0 | 5.94E-06 | 0 | 0 | 0 | 7.28E-05 |
| Homer2 | 3.11E-07 | 0 | 2.24E-05 | 0 | 0 | 0 | 8.57E-06 | 0 | 0 | 1.02E-05 | 0 | 0 | 2.35E-06 | 5.94E-06 | 0 | 3.40E-05 | 0 | 3.88E-05 |

| | | | | | | | | | | | | | | | | | | |
|---|---|---|---|---|---|---|---|---|---|---|---|---|---|---|---|---|---|---|
| Gata3 | 0 | 0 | 1.82E-05 | 4.07E-06 | 2.50E-06 | 0 | 0 | 1.71E-05 | 0 | 0 | 1.96E-05 | 1.45E-06 | 6.81E-06 | 5.92E-06 | 1.16E-05 | 2.78E-06 | 3.39E-06 | 0 |
| Ydjc | 1.78E-05 | 0 | 6.61E-05 | 9.37E-05 | 0 | 0 | 6.72E-05 | 0 | 0 | 0 | 1.87E-05 | 4.74E-06 | 0 | 5.89E-06 | 3.38E-05 | 0 | 0 | 2.35E-05 |
| Mroh3 | 0 | 0 | 6.25E-06 | 0 | 3.64E-06 | 0 | 0 | 4.45E-06 | 0 | 0 | 1.01E-05 | 0 | 0 | 5.89E-06 | 1.52E-05 | 0 | 2.12E-05 | 0 |
| Igkc | 0 | 5.69E-06 | 6.51E-07 | 7.23E-06 | 0 | 0 | 0 | 0 | 0 | 1.63E-06 | 2.93E-06 | 1.53E-06 | 3.30E-06 | 5.76E-06 | 2.01E-05 | 2.25E-06 | 0 | 1.10E-05 |
| Nrros | 0 | 7.56E-06 | 0 | 1.36E-05 | 0 | 0 | 0 | 4.25E-06 | 0 | 4.89E-06 | 0 | 6.76E-07 | 0 | 5.75E-06 | 5.02E-06 | 1.25E-05 | 1.10E-05 | 0 |
| Arg2 | 0 | 0 | 0 | 1.52E-05 | 0 | 0 | 5.03E-07 | 0 | 0 | 2.74E-06 | 0 | 9.57E-07 | 4.03E-06 | 5.74E-06 | 1.87E-05 | 7.71E-06 | 7.40E-06 | 0 |
| Mogat2 | 9.05E-07 | 0 | 0 | 5.03E-06 | 0 | 0 | 1.20E-05 | 0 | 0 | 0 | 7.43E-06 | 0 | 1.88E-05 | 5.69E-06 | 0 | 1.35E-06 | 0 | 0 |
| Agt | 0 | 1.51E-06 | 0 | 0 | 0 | 0 | 0 | 0 | 0 | 4.48E-06 | 0 | 0 | 3.25E-05 | 5.66E-06 | 0 | 0 | 0 | 8.36E-05 |
| Clec2l | 0 | 8.23E-07 | 0 | 4.68E-06 | 0 | 0 | 6.58E-06 | 0 | 0 | 2.37E-06 | 0 | 5.85E-06 | 1.11E-05 | 5.65E-06 | 0 | 1.88E-05 | 2.35E-05 | 0 |
| Esam | 0 | 7.43E-06 | 0 | 1.33E-05 | 0 | 0 | 0 | 4.23E-06 | 0 | 0 | 7.70E-06 | 9.57E-06 | 0 | 5.55E-06 | 7.37E-06 | 7.32E-06 | 3.65E-05 | 0 |
| Net1 | 0 | 4.92E-05 | 5.14E-05 | 4.01E-05 | 0 | 0 | 1.41E-05 | 0 | 0 | 1.71E-05 | 9.09E-05 | 0.0001 | 0 | 5.51E-06 | 0.00017 | 0.00023 | 0 | 8.31E-06 |
| Cpt1b | 5.39E-07 | 7.08E-06 | 0 | 0 | 0 | 0 | 7.05E-07 | 4.20E-06 | 0 | 5.28E-06 | 0 | 1.18E-05 | 1.25E-05 | 5.48E-06 | 6.41E-05 | 1.07E-05 | 2.44E-05 | 0 |
| Cpn2 | 0 | 2.53E-07 | 0 | 8.61E-06 | 1.90E-05 | 0 | 0 | 0 | 0 | 0 | 4.81E-05 | 1.37E-05 | 2.19E-06 | 5.43E-06 | 0 | 0 | 2.48E-05 | 0 |
| Rab11fip1 | 0 | 9.79E-06 | 0 | 1.72E-05 | 3.83E-07 | 0 | 0 | 0 | 0 | 3.88E-06 | 2.04E-05 | 7.13E-06 | 0 | 5.41E-06 | 2.72E-05 | 6.20E-06 | 1.86E-05 | 7.34E-08 |
| Klhl30 | 0 | 1.46E-06 | 0 | 0 | 5.94E-05 | 0 | 0 | 0 | 0 | 0 | 0.00012 | 0 | 0 | 5.24E-06 | 0 | 0 | 0 | 0 |
| Slc14a1 | 0 | 0 | 0 | 9.65E-06 | 3.76E-06 | 0 | 0 | 8.19E-08 | 0 | 6.68E-06 | 0 | 0 | 1.76E-06 | 5.23E-06 | 3.98E-05 | 2.97E-06 | 2.44E-05 | 3.80E-06 |
| Cabp1 | 0 | 5.12E-06 | 0 | 0 | 0 | 0 | 7.20E-07 | 1.97E-05 | 0 | 2.23E-06 | 1.36E-06 | 4.23E-06 | 0 | 5.21E-06 | 5.26E-05 | 0 | 1.74E-06 | 2.44E-05 |
| Adcy5 | 0 | 1.18E-05 | 0 | 0 | 6.98E-06 | 0 | 0.00015 | 0 | 0 | 6.83E-05 | 2.33E-05 | 0 | 0.00011 | 5.20E-06 | 0 | 0 | 0 | 0.00011 |
| Itgb2 | 0 | 2.10E-05 | 0 | 4.20E-05 | 5.72E-06 | 0 | 0 | 4.36E-06 | 0 | 9.84E-06 | 0 | 1.30E-05 | 8.51E-06 | 5.18E-06 | 8.48E-05 | 1.06E-05 | 6.41E-06 | 1.87E-05 |
| Fhl4 | 5.89E-06 | 0 | 2.56E-06 | 0 | 0 | 0 | 6.18E-06 | 0 | 0 | 0 | 0 | 1.64E-05 | 5.42E-06 | 5.18E-06 | 0 | 3.82E-06 | 7.04E-05 | 7.21E-07 |
| Sort1 | 2.35E-07 | 2.07E-05 | 0 | 0 | 0 | 0 | 0 | 0 | 0 | 4.55E-06 | 9.90E-06 | 0 | 0 | 5.16E-06 | 8.30E-06 | 6.63E-06 | 0 | 1.09E-05 |
| Bend5 | 0 | 3.86E-06 | 0 | 0 | 0 | 0 | 0 | 0 | 0 | 3.74E-06 | 0 | 5.26E-05 | 0 | 5.08E-06 | 2.08E-06 | 2.82E-06 | 0 | 4.43E-06 |
| Cyp46a1 | 0 | 8.12E-06 | 0 | 0 | 1.27E-05 | 0 | 6.95E-06 | 0 | 0 | 0 | 0 | 1.66E-06 | 1.38E-05 | 4.96E-06 | 7.37E-05 | 0 | 1.81E-05 | 6.21E-06 |
| Ccl3 | 0 | 1.96E-05 | 4.01E-06 | 3.42E-05 | 6.35E-06 | 0 | 0 | 0 | 0 | 1.77E-05 | 3.80E-06 | 2.28E-05 | 2.87E-05 | 4.93E-06 | 0.00026 | 2.38E-05 | 1.69E-05 | 2.59E-05 |
| Sema3e | 0 | 0 | 0 | 2.85E-06 | 0 | 0 | 0 | 7.27E-06 | 0 | 7.34E-05 | 9.62E-06 | 3.09E-05 | 0 | 4.90E-06 | 2.22E-05 | 0 | 0 | 0 |
| Gpr132 | 0 | 6.79E-06 | 3.22E-06 | 7.12E-06 | 0 | 0 | 0 | 0 | 0 | 1.75E-06 | 0 | 1.67E-06 | 6.42E-07 | 4.90E-06 | 3.69E-05 | 7.39E-06 | 7.67E-06 | 2.20E-06 |
| Cd52 | 0 | 6.22E-06 | 1.42E-07 | 4.43E-06 | 0 | 0 | 0 | 0 | 0 | 1.20E-05 | 2.69E-06 | 1.12E-05 | 1.05E-05 | 4.76E-06 | 0.00016 | 2.84E-05 | 5.10E-05 | 2.01E-05 |
| Igsf6 | 0 | 1.72E-06 | 0 | 6.09E-06 | 0 | 0 | 0 | 2.37E-06 | 0 | 0 | 0 | 0 | 8.61E-06 | 4.54E-06 | 4.61E-05 | 4.27E-06 | 2.46E-06 | 3.15E-06 |
| Ccdc116 | 3.45E-06 | 0 | 1.26E-05 | 2.81E-06 | 0 | 0 | 2.52E-05 | 0 | 0 | 0 | 0 | 3.21E-06 | 0 | 4.44E-06 | 1.12E-05 | 1.07E-06 | 1.71E-05 | 3.44E-06 |
| Pik3cb | 0 | 1.94E-05 | 6.85E-06 | 0 | 0 | 0 | 0 | 0 | 0 | 0 | 7.43E-06 | 6.15E-06 | 0 | 4.44E-06 | 1.09E-05 | 1.18E-05 | 0 | 8.19E-06 |
| Tspan8 | 0 | 1.28E-05 | 0 | 1.13E-05 | 1.29E-05 | 0 | 0 | 0 | 0 | 1.59E-05 | 3.10E-06 | 2.81E-05 | 0 | 4.22E-06 | 5.47E-05 | 1.18E-05 | 5.86E-06 | 3.08E-05 |
| Coro6 | 0 | 9.93E-07 | 0 | 0 | 0 | 0 | 0 | 1.46E-06 | 0 | 7.97E-07 | 1.39E-05 | 9.58E-06 | 7.17E-06 | 4.21E-06 | 8.08E-06 | 0 | 2.77E-05 | 2.12E-06 |
| Eya2 | 0 | 1.00E-05 | 0 | 6.90E-06 | 0 | 0 | 0 | 0 | 0 | 6.57E-07 | 0 | 0 | 0 | 4.20E-06 | 1.50E-05 | 0 | 0 | 0.0001 |

| | | | | | | | | | | | | | | | | | | |
|---|---|---|---|---|---|---|---|---|---|---|---|---|---|---|---|---|---|---|
| Wfdc2 | 0 | 7.80E-06 | 0 | 3.29E-05 | 1.14E-05 | 0 | 0 | 5.19E-06 | 0 | 2.36E-05 | 8.04E-06 | 4.64E-05 | 9.04E-06 | 4.17E-06 | 0.00013 | 1.69E-05 | 1.13E-05 | 3.35E-05 |
| Klhl35 | 4.50E-06 | 0 | 0 | 2.65E-06 | 0 | 0 | 3.86E-06 | 1.40E-05 | 0 | 0 | 6.90E-06 | 9.73E-06 | 0 | 4.13E-06 | 3.88E-07 | 0 | 0 | 4.92E-06 |
| Mboat2 | 0 | 0 | 0 | 0 | 1.32E-05 | 0 | 0 | 2.23E-06 | 0 | 0 | 1.64E-05 | 0.0001 | 0 | 4.10E-06 | 0 | 0 | 0 | 0 |
| Arvcf | 0 | 1.91E-05 | 0 | 2.47E-06 | 0 | 0 | 0 | 0 | 0 | 7.96E-05 | 1.32E-05 | 0 | 1.23E-05 | 4.06E-06 | 2.39E-06 | 0 | 0 | 8.06E-05 |
| Tbc1d9 | 0 | 7.51E-07 | 2.87E-06 | 9.00E-06 | 1.42E-06 | 0 | 2.32E-06 | 0 | 0 | 3.11E-06 | 2.15E-06 | 9.60E-07 | 1.12E-05 | 4.04E-06 | 0 | 0 | 2.41E-05 | 7.76E-06 |
| Dbp | 0 | 7.71E-05 | 0 | 0 | 1.90E-05 | 0 | 0 | 3.09E-05 | 0 | 0 | 4.19E-05 | 0.00011 | 0 | 3.99E-06 | 1.50E-05 | 0 | 0 | 2.22E-05 |
| Ms4a4a | 0 | 2.19E-06 | 2.34E-06 | 1.51E-05 | 1.54E-06 | 0 | 0 | 0 | 0 | 4.66E-06 | 0 | 0 | 9.30E-06 | 3.91E-06 | 4.02E-05 | 0 | 8.74E-06 | 3.06E-07 |
| Cry1 | 3.34E-05 | 3.13E-05 | 7.41E-05 | 2.98E-06 | 2.37E-05 | 0 | 0 | 1.41E-05 | 0 | 4.74E-05 | 3.77E-05 | 1.08E-05 | 0 | 3.89E-06 | 4.53E-05 | 1.13E-05 | 6.37E-05 | 6.91E-06 |
| Perp | 0 | 4.14E-06 | 8.50E-07 | 5.95E-06 | 0 | 0 | 0 | 0 | 0 | 5.32E-06 | 3.76E-06 | 7.09E-06 | 0 | 3.85E-06 | 2.06E-05 | 2.81E-06 | 0 | 5.87E-06 |
| Ticrr | 0 | 0 | 0 | 0 | 0 | 0 | 0 | 2.73E-06 | 0 | 0 | 0 | 0 | 0 | 3.84E-06 | 0 | 0.00018 | 0 | 0 |
| Spns2 | 0 | 0 | 0 | 3.99E-06 | 8.09E-07 | 0 | 0 | 0 | 0 | 2.86E-06 | 9.14E-06 | 3.63E-05 | 8.94E-07 | 3.76E-06 | 0 | 2.56E-08 | 0 | 2.94E-05 |
| Sh3yl1 | 1.67E-05 | 1.13E-05 | 4.01E-06 | 1.70E-05 | 1.66E-05 | 0 | 4.71E-05 | 1.65E-05 | 0 | 1.63E-05 | 2.10E-05 | 0.00011 | 1.80E-05 | 3.74E-06 | 4.17E-05 | 3.31E-06 | 0 | 3.06E-05 |
| Ccr1 | 0 | 1.28E-05 | 4.40E-06 | 4.93E-05 | 0 | 0 | 0 | 0 | 0 | 3.68E-06 | 0 | 1.43E-05 | 4.84E-06 | 3.68E-06 | 4.26E-05 | 1.27E-05 | 0 | 0 |
| Itga3 | 0 | 1.96E-05 | 0 | 2.47E-05 | 2.25E-05 | 0 | 0 | 0 | 0 | 1.57E-05 | 2.78E-06 | 1.56E-05 | 0 | 3.63E-06 | 7.39E-05 | 1.63E-05 | 2.74E-05 | 2.36E-05 |
| Tro | 1.48E-05 | 0 | 2.43E-05 | 0 | 6.07E-06 | 0 | 8.96E-05 | 1.70E-05 | 0 | 0 | 1.12E-05 | 1.60E-05 | 0 | 3.62E-06 | 0 | 2.33E-06 | 0 | 1.10E-05 |
| Hist1h4a | 5.19E-07 | 0 | 0 | 0 | 0 | 0 | 0 | 0 | 0 | 1.72E-06 | 1.07E-06 | 4.28E-05 | 0 | 3.61E-06 | 8.85E-06 | 6.82E-06 | 7.11E-05 | 0 |
| Ly86 | 0 | 6.50E-06 | 1.13E-06 | 1.10E-05 | 1.18E-06 | 0 | 0 | 0 | 0 | 6.45E-06 | 1.51E-06 | 3.29E-06 | 1.74E-06 | 3.58E-06 | 1.90E-05 | 1.67E-06 | 7.84E-06 | 0 |
| Csf1r | 0 | 4.29E-06 | 0 | 3.36E-05 | 0 | 0 | 0 | 0 | 0 | 2.37E-06 | 0 | 2.39E-05 | 0 | 3.54E-06 | 5.64E-05 | 3.24E-06 | 9.06E-05 | 3.81E-06 |
| Frmd4b | 0 | 7.84E-06 | 0 | 0 | 0 | 0 | 0 | 0 | 0 | 4.48E-05 | 0 | 0 | 0 | 3.50E-06 | 3.21E-06 | 9.25E-06 | 0 | 2.52E-05 |
| Cldn15 | 0 | 6.64E-06 | 0 | 0 | 0 | 0 | 0 | 0 | 0 | 6.05E-06 | 0 | 0 | 7.40E-05 | 3.49E-06 | 4.53E-06 | 0 | 0 | 3.91E-05 |
| Arap2 | 0 | 2.26E-05 | 0 | 0 | 7.17E-06 | 0 | 0 | 1.05E-05 | 0 | 0 | 1.01E-05 | 1.22E-05 | 0 | 3.49E-06 | 0 | 0 | 0 | 3.48E-05 |
| Card10 | 0 | 2.73E-05 | 0 | 0 | 0 | 0 | 0 | 0 | 0 | 0.00012 | 0 | 0 | 0 | 3.49E-06 | 4.89E-05 | 0 | 0 | 3.22E-05 |
| Crybg1 | 0 | 4.38E-06 | 0 | 6.77E-06 | 8.05E-07 | 0 | 0 | 5.26E-07 | 0 | 4.50E-06 | 1.36E-06 | 3.91E-06 | 0 | 3.40E-06 | 1.12E-05 | 5.72E-06 | 4.78E-06 | 7.14E-06 |
| Pik3c2b | 0 | 0 | 0 | 0 | 0 | 0 | 0 | 0 | 0 | 2.27E-05 | 0 | 0 | 0 | 3.33E-06 | 3.63E-05 | 0 | 0 | 5.45E-05 |
| Fam71e1 | 0 | 7.01E-06 | 0 | 3.00E-06 | 7.00E-06 | 0 | 0 | 9.10E-06 | 0 | 1.70E-06 | 1.24E-06 | 1.04E-05 | 0 | 3.24E-06 | 9.79E-06 | 0 | 0 | 3.58E-06 |
| Rasgef1b | 0 | 0 | 1.47E-06 | 0 | 0 | 0 | 0 | 2.44E-05 | 0 | 9.28E-06 | 0 | 0 | 0 | 3.14E-06 | 1.60E-05 | 1.07E-05 | 0.00012 | 4.01E-05 |
| Asap3 | 3.93E-05 | 5.97E-05 | 0 | 1.30E-05 | 1.25E-05 | 0 | 0.0001 | 0 | 0 | 0.00019 | 0 | 0 | 0.00031 | 3.08E-06 | 0.00014 | 0 | 0 | 0.0003 |
| Ripor2 | 0 | 6.82E-06 | 0 | 0 | 0 | 0 | 0 | 0 | 0 | 3.76E-06 | 0 | 2.42E-05 | 0 | 3.07E-06 | 4.03E-06 | 3.01E-06 | 0 | 5.71E-05 |
| Ccr5 | 0 | 7.83E-06 | 0 | 1.40E-05 | 0 | 0 | 0 | 3.52E-06 | 0 | 2.74E-06 | 0 | 4.05E-06 | 9.41E-08 | 3.03E-06 | 2.78E-05 | 4.19E-06 | 4.41E-06 | 3.04E-06 |
| DEPRECATED | 4.89E-06 | 2.35E-05 | 2.93E-05 | 1.24E-05 | 2.17E-05 | 0 | 4.44E-06 | 1.08E-06 | 0 | 1.25E-05 | 3.81E-05 | 4.78E-05 | 2.71E-05 | 2.88E-06 | 1.35E-05 | 7.55E-07 | 0.0002 | 3.00E-05 |
| Elmo3 | 0 | 6.32E-06 | 0 | 5.02E-06 | 0 | 0 | 9.18E-06 | 2.22E-07 | 0 | 8.19E-06 | 0 | 1.05E-06 | 7.09E-06 | 2.83E-06 | 2.66E-05 | 0 | 0 | 5.62E-06 |
| Acrbp | 1.44E-06 | 2.54E-05 | 0 | 1.84E-06 | 0 | 0 | 0 | 0 | 0 | 0 | 6.82E-07 | 1.91E-05 | 1.46E-05 | 2.81E-06 | 7.47E-05 | 5.53E-06 | 1.47E-05 | 8.82E-06 |
| Palmd | 1.81E-06 | 6.54E-06 | 1.29E-06 | 0 | 0 | 0 | 3.19E-06 | 1.20E-06 | 0 | 3.02E-05 | 6.36E-06 | 1.36E-06 | 0 | 2.78E-06 | 5.39E-06 | 2.08E-06 | 6.20E-06 | 0 |

| | | | | | | | | | | | | | | | | | | |
|---|---|---|---|---|---|---|---|---|---|---|---|---|---|---|---|---|---|---|
| Lad1 | 0 | 3.17E-06 | 0 | 1.24E-05 | 1.22E-05 | 0 | 0 | 3.91E-06 | 0 | 3.40E-06 | 0 | 1.33E-05 | 6.53E-06 | 2.71E-06 | 1.95E-05 | 3.05E-06 | 2.08E-05 | 1.19E-05 |
| Rtp4 | 6.47E-05 | 2.42E-05 | 0 | 0 | 0 | 0 | 0 | 0 | 0 | 0.00025 | 0 | 0 | 6.82E-05 | 2.70E-06 | 0 | 1.59E-05 | 0 | 0.00021 |
| Zfp622 | 6.23E-05 | 0.00012 | 9.35E-05 | 7.93E-05 | 1.13E-05 | 0 | 4.81E-05 | 0 | 0 | 1.44E-05 | 2.54E-05 | 0.00011 | 0 | 2.69E-06 | 0.00014 | 3.90E-05 | 0.00018 | 6.06E-06 |
| Pms2 | 1.47E-05 | 1.33E-05 | 4.12E-06 | 0 | 7.87E-07 | 0 | 8.38E-06 | 0 | 0 | 6.63E-06 | 0 | 2.13E-05 | 0 | 2.63E-06 | 4.74E-05 | 3.83E-05 | 4.64E-06 | 0 |
| Fcer1g | 0 | 1.30E-05 | 0 | 4.99E-05 | 2.64E-06 | 0 | 0 | 0 | 0 | 2.45E-05 | 1.21E-05 | 7.43E-06 | 1.38E-05 | 2.61E-06 | 0.00021 | 2.71E-05 | 6.16E-05 | 2.34E-05 |
| Npnt | 5.17E-07 | 7.37E-06 | 3.35E-06 | 9.10E-06 | 1.15E-05 | 0 | 0 | 4.99E-06 | 0 | 1.20E-05 | 2.05E-05 | 2.85E-06 | 1.72E-05 | 2.61E-06 | 4.00E-05 | 1.95E-06 | 3.81E-05 | 1.39E-05 |
| Cavin2 | 3.92E-06 | 0 | 0 | 0 | 0 | 0 | 0 | 0 | 0 | 0.00014 | 4.93E-05 | 0 | 0 | 2.48E-06 | 0 | 3.37E-05 | 0 | 8.57E-05 |
| Camkk1 | 9.72E-06 | 0 | 3.55E-05 | 0 | 1.88E-05 | 0 | 8.13E-05 | 3.09E-06 | 0 | 6.27E-05 | 5.17E-06 | 0 | 0 | 2.48E-06 | 3.50E-05 | 1.88E-05 | 1.32E-06 | 4.64E-05 |
| Gprc5a | 0 | 0 | 5.12E-06 | 2.11E-05 | 0 | 0 | 0 | 0 | 0 | 1.13E-05 | 0 | 7.59E-06 | 1.15E-05 | 2.48E-06 | 4.53E-05 | 1.69E-05 | 2.41E-05 | 2.82E-06 |
| Lmo1 | 2.20E-06 | 1.63E-07 | 3.42E-06 | 6.00E-06 | 0 | 0 | 1.05E-05 | 0 | 0 | 8.75E-06 | 0 | 0 | 1.10E-05 | 2.44E-06 | 0 | 8.74E-06 | 4.55E-06 | 3.60E-05 |
| Sfmbt2 | 0 | 9.51E-06 | 0 | 0 | 1.02E-05 | 0 | 0 | 5.63E-07 | 0 | 0 | 1.13E-05 | 2.59E-05 | 7.78E-06 | 2.42E-06 | 5.03E-06 | 0 | 0 | 0 |
| Kank3 | 4.52E-07 | 1.52E-05 | 2.55E-07 | 0 | 0 | 0 | 1.08E-05 | 0 | 0 | 1.62E-05 | 2.48E-06 | 1.22E-05 | 1.68E-06 | 2.32E-06 | 1.21E-05 | 3.62E-06 | 0 | 1.19E-05 |
| Thsd7a | 0 | 0 | 0 | 0 | 0 | 0 | 0 | 0 | 0 | 6.33E-05 | 0 | 7.57E-06 | 9.78E-05 | 2.28E-06 | 0 | 0 | 0 | 3.66E-05 |
| Gjb4 | 0 | 5.10E-07 | 1.49E-05 | 3.57E-06 | 0 | 0 | 5.81E-06 | 0 | 0 | 7.30E-07 | 5.61E-06 | 1.24E-05 | 0 | 2.27E-06 | 2.20E-05 | 5.59E-06 | 9.67E-06 | 9.84E-06 |
| Atf7ip2 | 0 | 0 | 0 | 0 | 0 | 0 | 0 | 0 | 0 | 0 | 6.82E-06 | 0 | 6.45E-06 | 2.24E-06 | 0 | 4.28E-05 | 1.05E-05 | 0 |
| Hdc | 0 | 0 | 2.50E-06 | 1.73E-05 | 0 | 0 | 0 | 0 | 0 | 2.75E-06 | 0 | 9.25E-06 | 0 | 2.19E-06 | 1.91E-05 | 1.42E-05 | 1.33E-05 | 1.81E-05 |
| Ctsw | 0 | 0 | 2.18E-06 | 0 | 0 | 0 | 6.53E-06 | 3.85E-05 | 0 | 0 | 0 | 7.46E-06 | 5.68E-06 | 2.16E-06 | 0 | 4.37E-06 | 0 | 5.78E-06 |
| Slc4a11 | 0 | 0 | 4.49E-07 | 1.51E-06 | 0 | 0 | 0 | 0 | 0 | 1.23E-06 | 0 | 1.44E-05 | 2.34E-05 | 2.14E-06 | 6.83E-07 | 5.54E-07 | 4.25E-05 | 6.38E-06 |
| Pcyox1l | 1.39E-05 | 8.17E-07 | 5.97E-05 | 2.58E-05 | 0 | 0 | 0 | 0 | 0 | 1.53E-05 | 2.56E-05 | 2.48E-06 | 1.61E-06 | 2.13E-06 | 2.86E-05 | 0.0001 | 0 | 1.91E-05 |
| Ltn1 | 0 | 3.14E-05 | 1.44E-05 | 0.00011 | 6.81E-06 | 0 | 0 | 1.56E-05 | 0 | 0 | 0 | 4.86E-06 | 0 | 2.10E-06 | 1.42E-05 | 2.41E-05 | 0 | 6.59E-06 |
| Lims2 | 0 | 0 | 0 | 0 | 0 | 0 | 0 | 0 | 0 | 0.00026 | 9.87E-05 | 2.88E-05 | 0 | 2.06E-06 | 4.04E-06 | 0 | 0 | 0 |
| Mpp7 | 3.25E-06 | 3.61E-05 | 4.98E-06 | 0 | 2.90E-05 | 0 | 3.76E-06 | 0 | 0 | 3.23E-05 | 2.04E-05 | 0 | 2.31E-05 | 2.02E-06 | 3.04E-05 | 4.65E-05 | 2.92E-05 | 0 |
| Ano1 | 0 | 2.05E-05 | 0 | 1.73E-05 | 5.42E-06 | 0 | 0 | 0 | 0 | 7.35E-06 | 0 | 1.76E-05 | 0 | 2.01E-06 | 9.80E-05 | 8.81E-06 | 6.95E-06 | 3.03E-05 |
| Zfp286 | 5.56E-06 | 9.59E-06 | 0 | 8.22E-06 | 1.15E-05 | 0 | 2.35E-07 | 0 | 0 | 2.04E-06 | 7.30E-06 | 0 | 7.09E-06 | 1.97E-06 | 9.87E-06 | 0 | 0 | 1.02E-05 |
| Lama5 | 0 | 8.12E-06 | 0 | 1.99E-05 | 0 | 0 | 0 | 0 | 0 | 3.65E-06 | 3.61E-06 | 4.65E-06 | 2.09E-06 | 1.95E-06 | 1.08E-05 | 3.72E-07 | 8.15E-06 | 2.84E-06 |
| Rrad | 0 | 0 | 0 | 3.48E-06 | 0 | 0 | 3.64E-05 | 0 | 0 | 2.20E-05 | 0 | 0 | 0.00017 | 1.83E-06 | 0 | 0 | 0.00029 | 4.93E-05 |
| Laptm5 | 0 | 1.50E-05 | 0 | 0.00012 | 0 | 0 | 0 | 0 | 0 | 1.17E-05 | 0 | 3.79E-05 | 7.31E-06 | 1.82E-06 | 0.00016 | 3.49E-05 | 3.96E-05 | 2.28E-05 |
| Prss22 | 0 | 1.20E-06 | 0 | 8.66E-06 | 2.59E-06 | 0 | 5.28E-06 | 0 | 0 | 8.80E-07 | 0 | 7.12E-06 | 1.19E-05 | 1.72E-06 | 1.82E-05 | 1.41E-06 | 1.04E-05 | 0 |
| Selplg | 0 | 2.45E-06 | 1.43E-06 | 7.05E-06 | 0 | 0 | 0 | 0 | 0 | 1.77E-06 | 9.73E-08 | 2.33E-06 | 9.99E-06 | 1.72E-06 | 4.95E-06 | 6.93E-06 | 2.60E-05 | 3.87E-06 |
| Tchp | 2.53E-05 | 3.56E-05 | 1.82E-05 | 0 | 0 | 0 | 2.65E-05 | 0 | 0 | 1.72E-05 | 0 | 0 | 2.42E-05 | 1.68E-06 | 0.00027 | 3.16E-05 | 0 | 1.41E-05 |
| Opn3 | 8.32E-06 | 7.01E-06 | 1.03E-05 | 0 | 2.52E-06 | 0 | 0 | 1.09E-05 | 0 | 2.06E-05 | 4.07E-06 | 1.39E-05 | 0 | 1.67E-06 | 0 | 2.07E-05 | 0 | 0 |
| Exo1 | 0 | 0 | 2.40E-05 | 0 | 0 | 0 | 0 | 0 | 0 | 0 | 0 | 0 | 0 | 1.60E-06 | 0 | 0.00029 | 0 | 0 |
| Tusc2 | 1.85E-05 | 7.64E-05 | 7.84E-06 | 5.06E-05 | 1.37E-05 | 0 | 7.74E-06 | 0 | 0 | 3.54E-05 | 0 | 0 | 1.28E-05 | 1.57E-06 | 3.76E-05 | 2.95E-05 | 0.00015 | 4.76E-05 |

| | | | | | | | | | | | | | | | | | | |
|---|---|---|---|---|---|---|---|---|---|---|---|---|---|---|---|---|---|---|
| Spsb4 | 0 | 0 | 1.74E-05 | 0 | 0 | 0 | 2.04E-05 | 1.32E-06 | 0 | 2.90E-05 | 0 | 0 | 0 | 1.57E-06 | 0 | 2.03E-05 | 0 | 0 |
| Wdr25 | 7.66E-06 | 7.77E-06 | 4.50E-06 | 5.32E-06 | 2.03E-06 | 0 | 0 | 7.69E-06 | 0 | 2.83E-06 | 2.38E-06 | 3.45E-06 | 0 | 1.55E-06 | 8.61E-06 | 2.92E-06 | 9.62E-06 | 3.98E-06 |
| Tnnc1 | 1.29E-06 | 9.28E-06 | 0 | 0 | 0 | 0 | 0 | 0 | 0 | 8.40E-06 | 1.11E-05 | 6.26E-05 | 0 | 1.54E-06 | 1.18E-05 | 0 | 0 | 0 |
| Recql4 | 0 | 0 | 5.88E-06 | 0 | 0 | 0 | 0 | 0 | 0 | 2.83E-07 | 0 | 0 | 0 | 1.48E-06 | 1.10E-05 | 7.58E-05 | 1.23E-05 | 0 |
| Clec14a | 0 | 9.19E-06 | 0 | 0 | 0 | 0 | 0 | 0 | 0 | 1.68E-05 | 0 | 0 | 0 | 1.41E-06 | 0 | 0 | 0 | 6.07E-05 |
| C1qtnf4 | 0 | 1.25E-05 | 0 | 0 | 6.44E-06 | 0 | 0 | 2.09E-06 | 0 | 0 | 6.65E-06 | 1.27E-05 | 0 | 1.25E-06 | 6.66E-06 | 0 | 0 | 0 |
| Tuft1 | 1.58E-07 | 2.55E-05 | 2.01E-06 | 0 | 8.15E-06 | 0 | 0 | 4.21E-07 | 0 | 0 | 1.32E-05 | 1.54E-05 | 0 | 1.12E-06 | 5.47E-05 | 4.04E-06 | 1.58E-06 | 3.36E-06 |
| H2-Q7 | 1.24E-05 | 0 | 3.11E-05 | 0 | 0 | 0 | 0 | 0 | 0 | 4.16E-05 | 0 | 0.00025 | 0.00019 | 1.07E-06 | 0 | 8.84E-05 | 0 | 1.55E-05 |
| Xndc1 | 0 | 1.85E-05 | 1.46E-05 | 4.08E-05 | 2.91E-06 | 0 | 2.28E-05 | 0 | 0 | 2.31E-05 | 0 | 0 | 9.48E-06 | 9.69E-07 | 5.37E-05 | 7.10E-05 | 1.21E-06 | 8.79E-06 |
| Tmem154 | 0 | 1.22E-05 | 0 | 0 | 0 | 0 | 2.70E-05 | 0 | 0 | 7.67E-06 | 3.68E-06 | 0 | 0 | 9.14E-07 | 0 | 0 | 0 | 1.12E-05 |
| Cd24a | 0 | 5.20E-05 | 1.32E-05 | 1.73E-05 | 1.03E-05 | 0 | 0 | 0 | 0 | 1.70E-05 | 2.26E-06 | 1.81E-05 | 1.62E-05 | 9.05E-07 | 8.99E-05 | 2.22E-05 | 5.60E-05 | 3.24E-05 |
| Lrrc75b | 0 | 0 | 1.27E-05 | 1.13E-08 | 0 | 0 | 2.91E-05 | 0 | 0 | 0 | 0 | 0 | 2.17E-06 | 8.94E-07 | 0 | 0 | 0 | 9.11E-06 |
| Itgax | 0 | 3.28E-06 | 2.91E-06 | 3.91E-06 | 0 | 0 | 0 | 0 | 0 | 3.64E-06 | 3.82E-06 | 3.08E-06 | 0 | 8.84E-07 | 3.50E-05 | 9.94E-06 | 1.61E-05 | 5.48E-06 |
| Dock8 | 0 | 1.90E-07 | 3.01E-08 | 0 | 9.23E-06 | 0 | 0 | 0 | 0 | 3.29E-06 | 6.71E-06 | 1.86E-05 | 0 | 8.77E-07 | 3.94E-05 | 4.09E-06 | 0 | 9.93E-06 |
| Clcn3 | 1.96E-05 | 2.30E-05 | 1.73E-05 | 0 | 8.92E-07 | 0 | 3.78E-05 | 1.55E-05 | 0 | 3.57E-05 | 5.77E-07 | 0.0001 | 0 | 8.37E-07 | 4.73E-05 | 2.19E-06 | 0 | 3.89E-05 |
| Doc2b | 0 | 6.68E-06 | 0 | 0 | 0 | 0 | 3.51E-06 | 2.95E-06 | 0 | 4.41E-06 | 0 | 6.32E-06 | 6.07E-06 | 7.91E-07 | 3.85E-06 | 0 | 0 | 2.94E-05 |
| Mettl18 | 6.05E-06 | 1.15E-05 | 0 | 3.50E-06 | 2.13E-06 | 0 | 0 | 4.85E-06 | 0 | 4.10E-08 | 0 | 8.02E-07 | 3.35E-06 | 5.60E-07 | 0 | 1.30E-05 | 9.24E-06 | 2.89E-06 |
| Ccdc57 | 0 | 1.19E-05 | 2.00E-06 | 0 | 1.79E-06 | 0 | 0 | 0 | 0 | 1.13E-05 | 0 | 3.51E-05 | 0 | 5.55E-07 | 7.13E-05 | 1.89E-05 | 2.57E-07 | 1.09E-05 |
| Pik3ap1 | 0 | 8.42E-06 | 0 | 3.14E-05 | 2.44E-06 | 0 | 0 | 0 | 0 | 3.16E-06 | 0 | 3.80E-06 | 0 | 5.40E-07 | 4.13E-05 | 2.07E-05 | 1.73E-05 | 5.75E-06 |
| Krtcap3 | 0 | 2.56E-06 | 2.05E-07 | 4.67E-06 | 4.51E-06 | 0 | 0 | 5.65E-06 | 0 | 3.04E-06 | 0 | 3.53E-06 | 6.39E-08 | 5.36E-07 | 2.28E-05 | 8.93E-06 | 0 | 3.45E-06 |
| Arhgap11a | 0 | 8.75E-06 | 0 | 0 | 9.14E-07 | 0 | 4.81E-06 | 3.03E-06 | 0 | 0 | 0 | 6.52E-06 | 0 | 5.18E-07 | 0 | 0.00056 | 0 | 0 |
| Cep70 | 7.69E-07 | 7.31E-06 | 0 | 0 | 5.17E-07 | 0 | 0 | 0 | 0 | 2.19E-06 | 0 | 0 | 2.68E-06 | 5.13E-07 | 7.02E-06 | 3.41E-05 | 0 | 1.21E-05 |
| Nbeal2 | 0 | 1.26E-05 | 0 | 1.70E-05 | 0 | 0 | 7.24E-06 | 0 | 0 | 1.36E-05 | 0 | 0 | 0 | 4.56E-07 | 1.84E-05 | 0 | 1.40E-05 | 1.75E-06 |
| Krt23 | 0 | 0 | 0 | 0 | 5.59E-06 | 0 | 0 | 2.84E-06 | 0 | 0.00011 | 0 | 0 | 0 | 4.38E-07 | 3.46E-06 | 0 | 0 | 0 |
| Ufsp1 | 5.85E-06 | 6.90E-06 | 4.72E-06 | 6.12E-06 | 2.56E-06 | 0 | 0 | 0 | 0 | 2.73E-07 | 4.15E-06 | 0 | 3.13E-06 | 4.21E-07 | 1.87E-06 | 2.41E-06 | 0 | 4.31E-06 |
| Cys1 | 0 | 0 | 0 | 0 | 6.57E-06 | 0 | 0 | 0 | 0 | 9.10E-06 | 6.51E-07 | 0 | 0 | 3.81E-07 | 1.69E-05 | 0 | 0 | 8.23E-05 |
| Igf2 | 0 | 4.16E-05 | 0 | 4.20E-06 | 0 | 0 | 0 | 0 | 0 | 0 | 0 | 0 | 0 | 1.80E-07 | 1.40E-05 | 7.13E-06 | 3.60E-05 | 6.04E-06 |
| Prl | 0 | 1.34E-06 | 0 | 1.27E-05 | 0 | 0 | 0 | 0 | 0 | 1.84E-06 | 6.55E-06 | 4.41E-06 | 0 | 1.44E-07 | 3.52E-05 | 4.42E-06 | 1.21E-05 | 0 |
| F11r | 0 | 1.30E-05 | 9.31E-07 | 4.38E-05 | 0 | 0 | 0 | 7.03E-07 | 0 | 2.10E-05 | 7.93E-06 | 1.23E-05 | 5.82E-06 | 1.31E-07 | 7.22E-05 | 4.93E-06 | 3.62E-05 | 9.00E-05 |
| Gipr | 0 | 0 | 0 | 0 | 0 | 0 | 0 | 0 | 0 | 0 | 0 | 0.00017 | 0 | 1.27E-07 | 2.24E-05 | 0 | 0 | 0 |
| Chn2 | 5.04E-09 | 5.02E-06 | 1.32E-06 | 0 | 9.23E-07 | 0 | 2.07E-05 | 0 | 0 | 1.49E-05 | 1.13E-06 | 0 | 0 | 1.25E-07 | 0 | 5.56E-06 | 0 | 0 |
| Cd68 | 0 | 1.43E-05 | 0 | 0 | 0 | 0 | 0 | 0 | 0 | 1.75E-05 | 0 | 0.00012 | 7.68E-06 | 6.78E-08 | 1.47E-05 | 1.54E-05 | 0.00043 | 0 |
| Zcchc12 | 0 | 0 | 6.63E-06 | 0 | 0 | 0 | 1.56E-05 | 3.51E-06 | 0 | 0 | 0 | 4.57E-06 | 7.95E-06 | 6.38E-08 | 3.78E-06 | 4.16E-06 | 1.99E-05 | 0 |

| | | | | | | | | | | | | | | | | | | |
|---|---|---|---|---|---|---|---|---|---|---|---|---|---|---|---|---|---|---|
| Ms4a7 | 0 | 2.10E-07 | 3.71E-07 | 5.83E-06 | 1.56E-06 | 0 | 0 | 0 | 0 | 6.76E-06 | 5.35E-06 | 2.93E-06 | 2.82E-06 | 4.45E-08 | 2.69E-05 | 1.03E-06 | 1.19E-05 | 0 |
| Foxg1 | 0 | 1.85E-06 | 0 | 3.15E-05 | 0 | 0 | 1.98E-06 | 2.02E-06 | 0 | 1.12E-06 | 2.31E-06 | 3.04E-06 | 0 | 4.24E-08 | 3.08E-05 | 1.04E-06 | 1.28E-05 | 1.05E-06 |
| Cdiptos | 3.18E-06 | 7.40E-07 | 9.06E-07 | 0 | 1.20E-06 | 0 | 7.70E-07 | 2.54E-06 | 0 | 9.88E-06 | 0 | 5.55E-06 | 5.29E-06 | 4.13E-08 | 1.23E-06 | 0 | 4.39E-06 | 1.32E-05 |
| Msc | 0 | 0 | 0 | 0 | 0 | 0 | 0 | 3.21E-05 | 0 | 8.42E-06 | 0 | 0 | 0 | 0 | 0 | 0 | 0 | 6.16E-05 |
| Bend6 | 1.59E-05 | 0 | 7.41E-06 | 0 | 0 | 0 | 2.63E-05 | 0 | 0 | 2.42E-05 | 0 | 1.98E-05 | 2.68E-06 | 0 | 0 | 3.57E-06 | 0 | 1.74E-05 |
| Zap70 | 0 | 2.34E-06 | 0 | 0 | 1.89E-05 | 0 | 0 | 4.15E-06 | 0 | 0 | 9.33E-06 | 1.61E-05 | 0 | 0 | 0 | 0 | 0 | 4.28E-06 |
| Tbc1d8 | 0 | 6.11E-06 | 0 | 0 | 0 | 0 | 0 | 9.38E-05 | 0 | 7.93E-06 | 0 | 2.28E-05 | 0 | 0 | 0 | 1.53E-06 | 0 | 3.35E-05 |
| Tex30 | 2.73E-05 | 1.23E-05 | 3.92E-05 | 0 | 0 | 0 | 0 | 0 | 0 | 2.89E-06 | 0 | 3.39E-05 | 0 | 0 | 0 | 0.00015 | 9.84E-06 | 0 |
| Tmeff2 | 0 | 0 | 0 | 0 | 0 | 0 | 0 | 0 | 0 | 0.00079 | 0 | 0 | 0.00061 | 0 | 0 | 9.91E-05 | 0 | 0.00084 |
| Aox3 | 0 | 0 | 0 | 0 | 0 | 0 | 0 | 0 | 0 | 0.00024 | 0 | 0 | 7.34E-05 | 0 | 0 | 0 | 0 | 0 |
| Fzd5 | 0 | 1.60E-05 | 0 | 7.90E-05 | 0 | 0 | 1.18E-05 | 1.45E-05 | 0 | 0 | 9.61E-05 | 0.00017 | 0 | 0 | 0 | 3.86E-05 | 0.00047 | 0.00014 |
| Igfbp5 | 0 | 0 | 0.00033 | 0 | 0 | 0 | 0 | 0.00044 | 0 | 0.00065 | 0 | 0 | 0 | 0 | 4.31E-05 | 0 | 0 | 0.00059 |
| Gpbar1 | 0 | 0 | 4.73E-05 | 0 | 0 | 0 | 3.73E-06 | 2.63E-05 | 0 | 0 | 0 | 0 | 0 | 0 | 0 | 3.58E-06 | 0 | 0 |
| Kcne4 | 0 | 2.25E-05 | 0 | 0 | 0 | 0 | 1.85E-05 | 8.49E-05 | 0 | 0 | 0 | 5.21E-06 | 0.00027 | 0 | 0 | 0 | 0.0002 | 0.00027 |
| Col4a4 | 0 | 0 | 0 | 0 | 3.43E-06 | 0 | 0 | 1.02E-06 | 0 | 3.69E-05 | 0 | 0 | 0 | 0 | 2.87E-06 | 2.90E-06 | 0 | 3.58E-05 |
| Slc19a3 | 0 | 0 | 0 | 0 | 0 | 0 | 3.23E-05 | 0 | 0 | 2.30E-06 | 0 | 1.37E-05 | 2.37E-05 | 0 | 0 | 1.03E-06 | 0 | 0 |
| Sphkap | 5.23E-06 | 0 | 0 | 0 | 0 | 0 | 0 | 0 | 0 | 1.77E-05 | 0 | 0 | 2.01E-05 | 0 | 0 | 7.96E-06 | 0 | 1.47E-05 |
| Efhd1 | 0 | 0 | 0 | 0 | 0 | 0 | 0 | 0 | 0 | 0.00051 | 0 | 0 | 0 | 0 | 0 | 0 | 0 | 0 |
| Ngef | 0 | 0 | 2.20E-05 | 0 | 0 | 0 | 2.22E-05 | 0 | 0 | 2.96E-06 | 0 | 0 | 0 | 0 | 0 | 0 | 0 | 0.00014 |
| Gbx2 | 0 | 0 | 1.66E-05 | 3.68E-05 | 0 | 0 | 0 | 1.49E-05 | 0 | 0 | 5.96E-05 | 0 | 1.82E-05 | 0 | 5.95E-05 | 2.83E-05 | 0 | 0.00013 |
| Erfe | 0 | 0 | 0 | 0 | 0 | 0 | 6.08E-05 | 0 | 0 | 2.38E-05 | 3.60E-05 | 1.27E-06 | 0 | 0 | 0 | 0.00011 | 0 | 0 |
| Agxt | 3.46E-06 | 0 | 0 | 0 | 0 | 0 | 0 | 0 | 0 | 2.88E-05 | 0 | 0 | 2.58E-05 | 0 | 0 | 2.44E-06 | 0 | 0 |
| Crocc2 | 0 | 0 | 0 | 0 | 4.69E-06 | 0 | 2.67E-07 | 0 | 0 | 3.57E-05 | 0 | 0 | 1.56E-05 | 0 | 4.49E-05 | 1.00E-05 | 1.07E-06 | 3.13E-06 |
| Serpinb2 | 0 | 0 | 6.16E-05 | 0 | 0 | 0 | 0 | 0 | 0 | 0.00016 | 0 | 0.00022 | 0.00038 | 0 | 0 | 1.75E-05 | 0 | 0 |
| Gpr39 | 0 | 0 | 3.03E-05 | 0 | 9.91E-05 | 0 | 0 | 3.47E-05 | 0 | 2.95E-05 | 0.00013 | 2.49E-05 | 2.31E-05 | 0 | 0 | 0 | 0 | 0.00016 |
| Slc45a3 | 0 | 4.67E-06 | 3.07E-06 | 1.16E-06 | 0 | 0 | 0 | 2.22E-06 | 0 | 0 | 3.31E-06 | 1.89E-05 | 0 | 0 | 9.77E-06 | 1.02E-05 | 0 | 2.23E-05 |
| Lgr6 | 0 | 0 | 2.17E-05 | 0 | 0 | 0 | 2.93E-05 | 0 | 0 | 0 | 3.26E-05 | 3.77E-05 | 1.36E-06 | 0 | 0 | 8.96E-06 | 0 | 0 |
| Lmod1 | 0 | 0 | 0 | 0 | 0 | 0 | 0 | 0 | 0 | 0 | 0.00014 | 0 | 0 | 0 | 0 | 0 | 0 | 0 |
| Kif14 | 0 | 0 | 0 | 0 | 0 | 0 | 0 | 0 | 0 | 0 | 0 | 0 | 0 | 0 | 2.58E-05 | 0.00049 | 0 | 0 |
| Aspm | 0 | 1.98E-06 | 0 | 0 | 0 | 0 | 0 | 0 | 0 | 4.72E-06 | 0 | 0 | 0 | 0 | 2.89E-05 | 0.00055 | 0 | 0 |
| Prg4 | 0 | 0 | 0 | 0 | 0 | 0 | 6.20E-05 | 0 | 0 | 0.00143 | 0 | 0 | 0.00177 | 0 | 0 | 0 | 0 | 3.92E-06 |
| Lamc2 | 9.88E-09 | 5.02E-06 | 0 | 6.94E-06 | 3.23E-06 | 0 | 0 | 4.50E-06 | 0 | 2.13E-05 | 9.35E-06 | 1.81E-05 | 2.74E-05 | 0 | 4.40E-05 | 7.35E-06 | 3.51E-05 | 0 |
| Ralgps2 | 6.21E-06 | 1.51E-05 | 1.05E-06 | 0 | 2.28E-05 | 0 | 0 | 2.03E-05 | 0 | 0 | 2.14E-05 | 5.77E-06 | 0 | 0 | 0 | 1.19E-05 | 0 | 6.54E-05 |

| | | | | | | | | | | | | | | | | | | |
|---|---|---|---|---|---|---|---|---|---|---|---|---|---|---|---|---|---|---|
| Angptl1 | 0 | 0 | 0 | 0 | 8.37E-06 | 0 | 3.28E-05 | 0 | 0 | 0 | 1.00E-05 | 0 | 0 | 0 | 0 | 4.25E-06 | 0 | 0.00015 |
| Ccdc181 | 3.22E-06 | 5.67E-06 | 0 | 0 | 0 | 0 | 1.37E-05 | 0 | 0 | 2.26E-05 | 0 | 1.13E-05 | 0 | 0 | 0 | 1.03E-05 | 0 | 6.70E-07 |
| Atp1b1 | 0 | 1.17E-05 | 0 | 8.49E-06 | 0 | 0 | 0 | 0 | 0 | 6.89E-05 | 9.83E-06 | 5.07E-06 | 0 | 0 | 3.06E-05 | 1.35E-06 | 2.25E-06 | 7.94E-06 |
| Lmx1a | 0 | 0 | 8.67E-06 | 0 | 0 | 0 | 1.20E-05 | 7.86E-06 | 0 | 4.90E-05 | 0 | 9.04E-05 | 7.84E-06 | 0 | 0 | 1.61E-05 | 0 | 1.26E-05 |
| Nuf2 | 7.49E-06 | 0 | 0 | 0 | 0 | 0 | 0 | 0 | 0 | 0 | 0 | 0 | 0 | 0 | 0 | 0.0007 | 0 | 0 |
| Cfap45 | 0 | 9.58E-07 | 0 | 0 | 0 | 0 | 0 | 7.91E-06 | 0 | 5.60E-06 | 0 | 8.00E-06 | 1.57E-05 | 0 | 0 | 6.20E-06 | 2.77E-05 | 7.34E-06 |
| Spta1 | 0 | 0 | 0 | 0 | 0 | 0 | 0 | 0 | 0 | 9.06E-05 | 0 | 0 | 1.62E-05 | 0 | 1.14E-05 | 0 | 0 | 0 |
| Cenpf | 0 | 2.72E-07 | 0 | 0 | 0 | 0 | 0 | 0 | 0 | 7.22E-06 | 0 | 0 | 0 | 0 | 4.35E-05 | 0.00063 | 0 | 0 |
| Nsl1 | 0 | 0 | 0 | 0 | 0 | 0 | 0 | 0 | 0 | 0 | 0 | 0 | 0 | 0 | 4.73E-05 | 0.00033 | 0 | 0 |
| Fam71a | 0 | 0 | 0 | 0 | 0 | 0 | 0 | 0 | 0 | 1.24E-05 | 0 | 4.79E-05 | 2.28E-05 | 0 | 0 | 0 | 0.00012 | 0 |
| Atf3 | 0 | 0 | 0 | 0 | 0 | 0 | 0 | 0 | 0 | 0.00014 | 0 | 8.85E-05 | 0.00025 | 0 | 0 | 0 | 0.00271 | 0 |
| Dtl | 1.19E-05 | 0 | 3.22E-05 | 0 | 0 | 0 | 0 | 0 | 0 | 0 | 0 | 0 | 1.05E-05 | 0 | 0 | 0.00017 | 0 | 0 |
| Nek2 | 0 | 0 | 0 | 0 | 0 | 0 | 0 | 0 | 0 | 0 | 0 | 0 | 0 | 0 | 8.42E-05 | 0.00065 | 0 | 0 |
| Camk1g | 0 | 1.15E-05 | 0 | 0 | 2.86E-05 | 0 | 0 | 2.37E-05 | 0 | 0 | 0 | 0 | 0 | 0 | 0 | 0 | 0 | 2.47E-06 |
| Cd34 | 0 | 0 | 5.85E-05 | 0 | 0 | 0 | 0.00047 | 0 | 0 | 0.00191 | 0 | 0 | 0.00012 | 0 | 0 | 4.94E-05 | 0 | 0.00085 |
| Mcm10 | 0 | 0 | 2.56E-05 | 0 | 0 | 0 | 0 | 0 | 0 | 0 | 0 | 0 | 0 | 0 | 0 | 0.00026 | 0 | 0 |
| Camk1d | 0 | 4.69E-06 | 4.13E-06 | 0 | 1.13E-06 | 0 | 4.67E-06 | 0 | 0 | 3.58E-05 | 1.36E-06 | 4.99E-06 | 0 | 0 | 1.81E-06 | 2.68E-06 | 0 | 2.42E-05 |
| Itih5 | 0 | 0 | 0 | 0 | 0.00029 | 0 | 0.00026 | 0.00029 | 0 | 0.00109 | 0 | 0 | 0 | 0 | 0 | 0 | 0 | 0 |
| Mrc1 | 0 | 6.82E-06 | 0 | 2.75E-05 | 2.08E-06 | 0 | 3.60E-07 | 0 | 0 | 1.17E-05 | 0 | 5.79E-06 | 6.57E-06 | 0 | 7.38E-05 | 7.08E-06 | 7.22E-06 | 7.54E-06 |
| Thnsl1 | 1.35E-05 | 0 | 0 | 0 | 7.88E-06 | 0 | 2.83E-05 | 0 | 0 | 1.06E-05 | 8.89E-08 | 1.14E-05 | 0 | 0 | 0 | 0 | 0 | 3.04E-05 |
| Nrarp | 0 | 0 | 3.94E-05 | 0 | 0 | 0 | 0 | 8.87E-05 | 0 | 0 | 0 | 0 | 0 | 0 | 0 | 2.13E-05 | 3.33E-05 | 0 |
| Stpg3 | 0 | 0 | 0 | 0 | 0 | 0 | 0 | 0 | 0 | 2.85E-06 | 0 | 2.51E-05 | 5.52E-05 | 0 | 0 | 0 | 0.00017 | 0 |
| Sapcd2 | 0 | 0 | 0 | 0 | 0 | 0 | 0 | 0 | 0 | 0 | 0 | 0 | 0 | 0 | 3.27E-05 | 0.00022 | 0 | 0 |
| Lcn2 | 0 | 0 | 0 | 0 | 0 | 0 | 0 | 0 | 0 | 0.00011 | 0 | 0 | 0.00045 | 0 | 0 | 0 | 0 | 0 |
| Traf1 | 0 | 4.96E-06 | 9.62E-06 | 3.92E-06 | 0 | 0 | 0 | 0 | 0 | 4.46E-06 | 0 | 3.07E-06 | 1.34E-05 | 0 | 2.00E-05 | 5.24E-07 | 2.43E-05 | 2.69E-05 |
| Acvr2a | 9.46E-05 | 2.88E-05 | 0 | 5.78E-05 | 0 | 0 | 2.38E-05 | 0 | 0 | 0.0001 | 7.72E-05 | 2.99E-05 | 0.00016 | 0 | 0 | 3.16E-05 | 0.00011 | 0.00034 |
| Kif5c | 0 | 5.48E-06 | 3.77E-05 | 0 | 1.69E-06 | 0 | 8.44E-06 | 2.29E-06 | 0 | 0 | 6.32E-06 | 2.90E-05 | 0 | 0 | 4.41E-06 | 0 | 0 | 0 |
| Tnfaip6 | 0 | 0 | 0.00054 | 5.71E-06 | 0 | 0 | 7.84E-05 | 0 | 0 | 0.00036 | 0 | 0 | 0.00032 | 0 | 0 | 0.00014 | 0.00092 | 7.02E-06 |
| Galnt13 | 1.59E-06 | 4.09E-07 | 0 | 7.78E-07 | 0 | 0 | 0 | 0 | 0 | 4.55E-06 | 0 | 0 | 4.39E-05 | 0 | 0 | 4.42E-06 | 9.69E-07 | 0 |
| Gca | 0 | 0 | 0 | 8.72E-06 | 0 | 0 | 0 | 0 | 0 | 3.43E-05 | 0 | 3.49E-06 | 1.37E-05 | 0 | 0 | 8.63E-06 | 1.03E-05 | 0 |
| Scn7a | 3.88E-05 | 0 | 0 | 0 | 0 | 0 | 0 | 0 | 0 | 0.00071 | 0 | 0 | 0.00065 | 0 | 0 | 0 | 0 | 0.00045 |
| B3galt1 | 3.59E-06 | 1.01E-05 | 0 | 0 | 0 | 0 | 0 | 1.57E-05 | 0 | 6.18E-06 | 0 | 1.26E-06 | 0 | 0 | 0 | 0 | 0 | 2.18E-05 |
| Nostrin | 0 | 0 | 0 | 0 | 0 | 0 | 0 | 2.41E-05 | 0 | 3.80E-05 | 0 | 4.04E-06 | 0 | 0 | 0 | 3.27E-06 | 0 | 5.27E-05 |

| | | | | | | | | | | | | | | | | | | |
|---|---|---|---|---|---|---|---|---|---|---|---|---|---|---|---|---|---|---|
| Spc25 | 0 | 0 | 0 | 0 | 0 | 0 | 0 | 0 | 0 | 0 | 0 | 0 | 0 | 0 | 0 | 0.00076 | 0 | 0 |
| Dhrs9 | 0 | 0 | 0 | 0 | 5.58E-06 | 0 | 0 | 1.09E-05 | 0 | 8.77E-05 | 0 | 1.27E-05 | 0 | 0 | 0 | 0 | 0 | 0 |
| Itga6 | 0 | 5.48E-05 | 0 | 0 | 0 | 0 | 0 | 9.91E-05 | 0 | 0 | 0 | 0.00029 | 0 | 0 | 5.77E-05 | 1.86E-06 | 0 | 0.00012 |
| Ccdc141 | 0 | 6.49E-06 | 2.08E-05 | 4.13E-06 | 0 | 0 | 9.16E-06 | 2.46E-06 | 0 | 3.11E-05 | 0 | 1.12E-05 | 1.27E-05 | 0 | 2.23E-05 | 1.95E-05 | 8.20E-05 | 4.62E-05 |
| Zfp385b | 1.22E-05 | 0 | 0 | 0 | 0 | 0 | 3.86E-05 | 0 | 0 | 4.01E-06 | 0 | 7.22E-05 | 0.00016 | 0 | 0 | 2.91E-06 | 2.66E-08 | 4.45E-05 |
| Selenoh | 7.79E-05 | 0 | 4.96E-05 | 0.00036 | 0 | 0 | 0 | 0 | 0 | 0 | 0 | 0 | 0 | 0 | 0 | 0.0008 | 1.99E-05 | 0 |
| Tspan18 | 0 | 0 | 0.00015 | 0 | 0 | 0 | 0.0004 | 0 | 0 | 1.25E-05 | 0 | 9.37E-05 | 0 | 0 | 0 | 0 | 0 | 0.00033 |
| Abtb2 | 0 | 0 | 7.11E-05 | 4.78E-06 | 0 | 0 | 3.40E-05 | 0.00011 | 0 | 0 | 9.87E-06 | 2.71E-05 | 0 | 0 | 1.04E-05 | 2.53E-06 | 0 | 5.88E-05 |
| Cd59a | 0 | 1.77E-05 | 0 | 0 | 0.00021 | 0 | 0 | 0 | 0 | 0 | 0.00028 | 0 | 0 | 0 | 0 | 0 | 0 | 1.52E-05 |
| Wt1 | 0 | 0 | 0 | 0 | 1.93E-06 | 0 | 0 | 0 | 0 | 0 | 3.91E-05 | 6.81E-05 | 0 | 0 | 1.73E-05 | 0 | 0 | 0 |
| Ano3 | 0 | 2.05E-05 | 0 | 0 | 0 | 0 | 1.10E-06 | 0 | 0 | 6.57E-05 | 0 | 0 | 3.50E-06 | 0 | 0 | 0 | 0 | 0 |
| Bub1b | 0 | 4.47E-06 | 0 | 0 | 0 | 0 | 0 | 0 | 0 | 2.76E-06 | 0 | 0 | 0 | 0 | 0 | 0.00072 | 0 | 0 |
| Knstrn | 4.19E-05 | 0 | 0 | 3.59E-05 | 0 | 0 | 0 | 0 | 0 | 1.59E-06 | 0 | 0 | 0 | 0 | 4.12E-05 | 0.00089 | 0 | 0 |
| Knl1 | 0 | 0 | 0 | 0 | 0 | 0 | 0 | 0 | 0 | 0 | 0 | 0 | 0 | 0 | 3.25E-05 | 0.00046 | 0 | 0 |
| Rad51 | 0 | 0 | 1.32E-05 | 0 | 0 | 0 | 0 | 0 | 0 | 0 | 0 | 0 | 1.56E-05 | 0 | 0 | 0.00049 | 0 | 0 |
| Dll4 | 0 | 8.36E-06 | 1.48E-06 | 1.57E-05 | 5.48E-06 | 0 | 0 | 0 | 0 | 3.02E-06 | 2.40E-06 | 0 | 9.74E-07 | 0 | 3.57E-05 | 5.84E-06 | 1.63E-05 | 0 |
| Nusap1 | 0 | 0 | 0 | 0 | 0 | 0 | 0 | 0 | 0 | 2.07E-06 | 0 | 0 | 0 | 0 | 0 | 0.0007 | 0 | 0 |
| Ppip5k1 | 0 | 3.84E-05 | 0 | 9.56E-06 | 1.12E-05 | 0 | 5.56E-06 | 0 | 0 | 4.52E-05 | 1.38E-05 | 9.80E-06 | 2.11E-06 | 0 | 2.46E-05 | 2.40E-05 | 1.40E-05 | 1.42E-05 |
| Trim69 | 0 | 0 | 0 | 0 | 0 | 0 | 0 | 0 | 0 | 8.28E-07 | 0 | 3.62E-05 | 4.56E-05 | 0 | 0 | 0 | 0.00035 | 0 |
| Duoxa1 | 0 | 0 | 0 | 0 | 0 | 0 | 3.78E-05 | 0 | 0 | 0.00015 | 0 | 0 | 0 | 0 | 0 | 0 | 0 | 0 |
| Duox1 | 0 | 0 | 0 | 1.43E-06 | 0 | 0 | 1.95E-05 | 0 | 0 | 0.00017 | 0 | 0 | 0 | 0 | 0 | 7.85E-07 | 0 | 0 |
| AA467197 | 0 | 0 | 3.12E-05 | 0 | 0 | 0 | 3.42E-06 | 3.70E-06 | 0 | 4.36E-05 | 1.00E-05 | 0.00016 | 0 | 0 | 3.79E-05 | 0 | 0 | 4.47E-06 |
| Ncaph | 0 | 0 | 0 | 0 | 0 | 0 | 0 | 0 | 0 | 0 | 0 | 0 | 0 | 0 | 5.92E-05 | 0.00076 | 0 | 0 |
| Dusp2 | 0 | 3.22E-06 | 0 | 1.70E-05 | 0 | 0 | 0 | 0 | 0 | 0 | 0 | 0 | 0 | 0 | 2.21E-05 | 1.67E-06 | 4.08E-05 | 0 |
| Bub1 | 0 | 0 | 0 | 0 | 0 | 0 | 0 | 0 | 0 | 0 | 0 | 1.59E-06 | 0 | 0 | 0 | 0.00043 | 0 | 0 |
| Ckap2l | 0 | 0 | 0 | 0 | 0 | 0 | 0 | 0 | 0 | 7.38E-06 | 0 | 0 | 0 | 0 | 0 | 0.00088 | 0 | 0 |
| Oxt | 0 | 0 | 0 | 0 | 0 | 0 | 0 | 0 | 0 | 0 | 0 | 7.88E-06 | 2.45E-05 | 0 | 0 | 0 | 5.80E-05 | 0 |
| Lzts3 | 0 | 1.53E-05 | 1.03E-05 | 3.89E-05 | 1.09E-05 | 0 | 0 | 0 | 0 | 1.35E-05 | 1.41E-05 | 4.75E-06 | 0 | 0 | 2.81E-05 | 2.03E-05 | 2.32E-05 | 0 |
| Cdc25b | 0 | 5.22E-05 | 0 | 0 | 0 | 0 | 0 | 0 | 0 | 2.03E-05 | 0 | 0 | 0 | 0 | 0.00016 | 0.00065 | 0 | 5.40E-05 |
| Plcb4 | 1.85E-05 | 0 | 3.24E-06 | 0 | 3.57E-06 | 0 | 2.98E-05 | 0 | 0 | 8.84E-05 | 1.13E-05 | 0 | 0 | 0 | 3.20E-05 | 1.31E-05 | 1.07E-05 | 7.70E-05 |
| Macrod2 | 1.85E-05 | 1.07E-05 | 0 | 0 | 3.25E-05 | 0 | 3.36E-05 | 0 | 0 | 1.96E-05 | 1.39E-05 | 5.23E-06 | 0 | 0 | 3.43E-05 | 3.78E-05 | 0 | 2.49E-05 |
| Cst7 | 0 | 0 | 0 | 0 | 2.99E-05 | 0 | 0 | 3.24E-05 | 0 | 0 | 4.95E-06 | 3.93E-06 | 0 | 0 | 0 | 0 | 0 | 0 |
| Trib3 | 0 | 3.85E-06 | 2.44E-07 | 0 | 0 | 0 | 0 | 0 | 0 | 2.61E-05 | 0 | 0.00023 | 0 | 0 | 2.07E-05 | 0 | 0 | 1.45E-05 |

| | | | | | | | | | | | | | | | | | | |
|---|---|---|---|---|---|---|---|---|---|---|---|---|---|---|---|---|---|---|
| Cox4i2 | 0 | 0 | 0 | 0 | 0 | 0 | 0 | 0 | 0 | 0 | 0 | 0.00014 | 1.02E-05 | 0 | 0 | 0 | 0 | 1.11E-06 |
| Tpx2 | 0 | 0 | 5.89E-07 | 0 | 0 | 0 | 0 | 1.19E-05 | 0 | 6.04E-06 | 0 | 5.46E-06 | 0 | 0 | 2.30E-05 | 0.00151 | 0 | 0 |
| Mylk2 | 0 | 0 | 2.09E-05 | 0 | 0 | 0 | 0 | 3.45E-05 | 0 | 0 | 0 | 5.86E-06 | 0 | 0 | 0 | 1.04E-05 | 0 | 0 |
| Snta1 | 0 | 3.18E-05 | 1.78E-05 | 0.00024 | 4.22E-05 | 0 | 3.48E-05 | 3.41E-05 | 0 | 1.02E-05 | 0.00013 | 0.00019 | 0.00019 | 0 | 5.61E-05 | 6.48E-05 | 0 | 0.00011 |
| Actl10 | 0 | 2.24E-06 | 0 | 0 | 0 | 0 | 3.45E-06 | 0 | 0 | 3.98E-07 | 0 | 0 | 1.90E-05 | 0 | 0 | 0 | 6.00E-05 | 0 |
| Ahcy | 2.26E-05 | 6.65E-06 | 0 | 0 | 0 | 0 | 3.71E-05 | 0 | 0 | 0 | 2.77E-05 | 0 | 4.49E-05 | 0 | 0 | 4.51E-05 | 3.32E-05 | 0 |
| Cnbd2 | 1.72E-08 | 8.90E-06 | 0 | 0 | 2.30E-06 | 0 | 7.27E-08 | 1.24E-05 | 0 | 6.61E-06 | 0 | 5.97E-06 | 2.84E-06 | 0 | 1.39E-05 | 0 | 1.25E-05 | 8.88E-06 |
| Snhg11 | 0 | 1.46E-05 | 0 | 0 | 0 | 0 | 0 | 0 | 0 | 0.0001 | 0 | 7.88E-06 | 0.0001 | 0 | 7.30E-05 | 0 | 0 | 0 |
| Fam83d | 0 | 2.74E-06 | 0 | 0 | 6.10E-08 | 0 | 0 | 0 | 0 | 0 | 0 | 0 | 0 | 0 | 0 | 0.00032 | 0 | 1.06E-05 |
| L3mbtl1 | 0 | 0 | 2.01E-06 | 0 | 0 | 0 | 8.51E-07 | 0 | 0 | 0 | 0 | 1.86E-05 | 9.10E-06 | 0 | 0 | 0 | 5.78E-05 | 0 |
| Ccn5 | 0 | 0 | 0 | 0 | 0 | 0 | 0.00156 | 0 | 0 | 0.00066 | 0 | 0 | 0.00296 | 0 | 0 | 0 | 0 | 0.0012 |
| Matn4 | 0 | 0 | 0 | 0 | 0 | 0 | 0 | 0 | 0 | 3.55E-06 | 0 | 1.99E-05 | 6.82E-05 | 0 | 0 | 0 | 0.00016 | 0 |
| Ube2c | 0 | 0 | 0 | 0 | 0 | 0 | 0 | 0 | 0 | 0 | 0 | 2.10E-05 | 0 | 0 | 1.38E-05 | 0.00137 | 0 | 0 |
| Pltp | 0 | 0.0003 | 0 | 0 | 7.49E-05 | 0 | 0 | 8.19E-05 | 0 | 0.00011 | 0 | 0 | 2.04E-05 | 0 | 1.19E-05 | 0 | 7.00E-05 | 0.00097 |
| Sulf2 | 0 | 7.44E-05 | 0 | 0 | 0 | 0 | 0.00091 | 0 | 0 | 0.00057 | 0 | 0 | 0 | 0 | 0 | 2.96E-05 | 0 | 0.00035 |
| Ptgis | 0 | 0 | 0.00019 | 8.90E-05 | 0 | 0 | 0.00082 | 0 | 0 | 0.00049 | 0 | 0.0003 | 0.00074 | 0 | 0 | 1.76E-05 | 0 | 0 |
| Aurka | 0 | 0 | 0 | 0 | 0 | 0 | 0 | 0 | 0 | 0 | 0 | 0 | 0 | 0 | 3.22E-05 | 0.00068 | 0 | 0 |
| Col9a3 | 0 | 2.28E-06 | 3.90E-07 | 6.14E-06 | 2.82E-06 | 0 | 4.44E-06 | 3.29E-06 | 0 | 3.27E-06 | 0 | 2.47E-06 | 0 | 0 | 8.45E-06 | 5.64E-06 | 3.07E-05 | 7.19E-07 |
| Col20a1 | 0 | 0 | 8.23E-06 | 5.14E-06 | 0 | 0 | 1.63E-05 | 0 | 0 | 0 | 0 | 3.52E-05 | 2.89E-05 | 0 | 2.28E-06 | 0 | 0.00011 | 0 |
| Pkia | 8.45E-06 | 0 | 0 | 0 | 0 | 0 | 0 | 0 | 0 | 6.19E-05 | 0 | 0 | 0 | 0 | 0 | 8.02E-06 | 0 | 0.00012 |
| Hey1 | 0 | 0 | 0.00012 | 0 | 0 | 0 | 0 | 0.00026 | 0 | 0 | 0 | 0 | 0 | 0 | 0 | 3.47E-05 | 0 | 0 |
| Tpd52 | 0 | 1.42E-05 | 1.61E-05 | 2.03E-05 | 0 | 0 | 4.30E-07 | 0 | 0 | 0 | 0 | 3.16E-05 | 0 | 0 | 3.26E-05 | 9.42E-06 | 2.85E-06 | 9.64E-06 |
| Ect2 | 0 | 5.19E-06 | 0 | 0 | 0 | 0 | 0 | 0 | 0 | 4.72E-06 | 0 | 3.51E-06 | 0 | 0 | 0 | 0.00085 | 0 | 0 |
| Mecom | 1.67E-06 | 7.39E-06 | 6.13E-06 | 2.15E-05 | 0 | 0 | 1.18E-05 | 2.89E-06 | 0 | 6.39E-06 | 1.03E-05 | 1.41E-05 | 5.18E-06 | 0 | 8.54E-06 | 8.83E-06 | 4.04E-06 | 1.81E-06 |
| Ccna2 | 0 | 0 | 0 | 0 | 0 | 0 | 0 | 0 | 0 | 4.70E-06 | 0 | 2.05E-06 | 0 | 0 | 0 | 0.00179 | 0 | 0 |
| Trpc3 | 5.09E-08 | 1.98E-05 | 0 | 0 | 0 | 0 | 0 | 5.75E-06 | 0 | 0 | 0 | 0 | 7.25E-05 | 0 | 0 | 3.27E-06 | 2.48E-05 | 0 |
| Slc7a11 | 0 | 3.85E-06 | 5.36E-06 | 4.59E-05 | 0 | 0 | 0 | 0 | 0 | 3.08E-06 | 0 | 1.83E-05 | 2.09E-05 | 0 | 4.69E-05 | 1.09E-06 | 1.43E-05 | 6.39E-06 |
| Noct | 2.93E-05 | 0 | 0.00019 | 0 | 0 | 0 | 0 | 0 | 0 | 4.44E-05 | 0.00011 | 0.00014 | 0.00015 | 0 | 0 | 3.32E-05 | 0.00033 | 0 |
| Pdgfc | 0 | 0 | 8.18E-05 | 0 | 0 | 0 | 0.00016 | 0 | 0 | 0.00011 | 0.00018 | 1.54E-06 | 0 | 0 | 0 | 3.58E-05 | 0 | 0 |
| Lrat | 0 | 0 | 0 | 0 | 0.00013 | 0 | 0 | 0 | 0 | 0 | 3.47E-06 | 0 | 4.88E-06 | 0 | 0 | 0 | 0 | 0 |
| Dchs2 | 0 | 5.05E-05 | 0 | 0 | 0.00017 | 0 | 0 | 0 | 0 | 0 | 0 | 5.14E-05 | 0 | 0 | 0 | 0 | 0 | 0 |
| Crabp2 | 0 | 0 | 9.05E-05 | 1.84E-05 | 0 | 0 | 0.00015 | 1.43E-05 | 0 | 0 | 0.00012 | 0 | 0 | 0 | 1.00E-06 | 0 | 0 | 6.97E-05 |
| Iqgap3 | 0 | 0 | 0 | 0 | 0 | 0 | 0 | 0 | 0 | 0 | 0 | 7.02E-06 | 0 | 0 | 4.83E-05 | 0.00065 | 0 | 0 |

| | | | | | | | | | | | | | | | | | | |
|---|---|---|---|---|---|---|---|---|---|---|---|---|---|---|---|---|---|---|
| Sema4a | 0 | 0 | 0 | 0 | 0 | 0 | 0 | 0 | 0 | 0 | 0 | 4.10E-05 | 0.00016 | 0 | 0 | 0 | 0.00053 | 0 |
| Cks1b | 8.10E-05 | 0 | 7.96E-05 | 5.04E-05 | 0 | 0 | 0 | 0 | 0 | 2.72E-05 | 0 | 0 | 2.47E-05 | 0 | 0 | 0.00064 | 0 | 0 |
| Npr1 | 0 | 1.23E-05 | 0 | 0 | 0 | 0 | 3.91E-05 | 0 | 0 | 0.00049 | 0 | 0 | 5.79E-05 | 0 | 1.46E-05 | 0 | 0 | 0 |
| Sprr1a | 0 | 1.68E-05 | 0 | 3.16E-05 | 1.04E-05 | 0 | 3.42E-06 | 1.20E-05 | 0 | 1.55E-05 | 0 | 4.28E-05 | 0 | 0 | 0.00012 | 3.13E-05 | 3.38E-05 | 0 |
| Selenbp1 | 0 | 0 | 0 | 0 | 0 | 0 | 0 | 0 | 0 | 6.09E-05 | 0 | 0.0003 | 1.11E-05 | 0 | 0 | 0 | 0 | 3.92E-05 |
| Ctss | 0 | 3.40E-05 | 0 | 0.0002 | 1.44E-05 | 0 | 0 | 0 | 0 | 3.66E-05 | 1.14E-06 | 2.76E-05 | 4.35E-05 | 0 | 0.00035 | 2.78E-05 | 7.23E-05 | 4.25E-05 |
| Ciart | 0 | 9.11E-06 | 0 | 0 | 0 | 0 | 0 | 0 | 0 | 0 | 0 | 8.27E-05 | 0 | 0 | 1.97E-05 | 0 | 0 | 0 |
| Adam30 | 0 | 0 | 3.11E-06 | 0 | 0 | 0 | 3.14E-06 | 0 | 0 | 0 | 0 | 1.08E-05 | 3.93E-05 | 0 | 0 | 0 | 5.75E-05 | 0 |
| Tent5c | 0 | 0 | 0 | 0 | 0 | 0 | 0 | 0 | 0 | 2.15E-06 | 0 | 2.55E-05 | 1.02E-05 | 0 | 0 | 0 | 0.00016 | 2.62E-05 |
| Cd101 | 0 | 0 | 3.79E-06 | 0 | 0 | 0 | 4.92E-06 | 0 | 0 | 0 | 0 | 0 | 1.96E-05 | 0 | 0 | 0 | 3.34E-05 | 0 |
| Ngf | 5.02E-06 | 0 | 1.88E-05 | 0 | 0 | 0 | 6.23E-08 | 0 | 0 | 0.00015 | 0 | 0 | 0 | 0 | 0 | 1.40E-05 | 9.63E-06 | 5.81E-05 |
| Tspan2 | 0 | 2.90E-06 | 0 | 0 | 0 | 0 | 0 | 0 | 0 | 0 | 4.38E-05 | 0 | 0 | 0 | 9.65E-06 | 5.47E-06 | 0 | 1.43E-05 |
| Ampd1 | 0 | 0 | 4.39E-06 | 0 | 0 | 0 | 0 | 0 | 0 | 0 | 0 | 1.55E-05 | 1.70E-05 | 0 | 0 | 4.51E-06 | 0.00014 | 0 |
| Amy1 | 0 | 4.04E-06 | 0 | 0 | 0 | 0 | 0 | 0 | 0 | 6.21E-05 | 0 | 0 | 3.01E-05 | 0 | 0 | 0 | 0 | 3.79E-05 |
| Vcam1 | 4.77E-05 | 0 | 0 | 0 | 0 | 0 | 0.00016 | 0 | 0 | 0.00014 | 0 | 0 | 0.0008 | 0 | 0 | 0.0001 | 6.20E-05 | 0.00056 |
| Gpr88 | 0 | 0 | 0 | 3.50E-06 | 0 | 0 | 1.51E-06 | 0 | 0 | 2.32E-07 | 0 | 0 | 2.45E-05 | 0 | 0 | 0 | 0 | 9.87E-05 |
| Bcar3 | 0 | 1.57E-05 | 7.44E-07 | 1.01E-06 | 2.94E-05 | 0 | 1.20E-05 | 0 | 0 | 0 | 6.05E-05 | 0 | 1.85E-05 | 0 | 1.42E-05 | 0 | 0 | 1.13E-05 |
| Prss12 | 0 | 1.52E-05 | 0 | 0 | 2.90E-05 | 0 | 3.14E-05 | 1.01E-05 | 0 | 5.79E-05 | 3.86E-05 | 4.00E-05 | 0 | 0 | 0 | 0 | 0 | 6.02E-05 |
| Cfi | 0 | 0 | 0 | 0 | 0 | 0 | 1.25E-05 | 1.58E-06 | 0 | 0 | 0 | 0 | 1.19E-05 | 0 | 0 | 0 | 0 | 0.00029 |
| Mcub | 0.00013 | 0 | 6.22E-05 | 0 | 0 | 0 | 0.00022 | 0 | 0 | 0.00017 | 0 | 0.00016 | 1.43E-05 | 0 | 0 | 4.62E-05 | 0 | 2.43E-05 |
| Cenpe | 0 | 0 | 0 | 0 | 0 | 0 | 0 | 0 | 0 | 0 | 0 | 0 | 0 | 0 | 7.93E-05 | 0.00066 | 0 | 0 |
| Bdh2 | 0 | 1.61E-05 | 0 | 0 | 0 | 0 | 0 | 0 | 0 | 2.02E-05 | 0 | 2.63E-06 | 6.32E-06 | 0 | 0 | 0 | 0 | 4.36E-06 |
| Adh1 | 2.20E-06 | 0 | 0 | 0 | 0 | 0 | 0 | 0 | 0 | 0.00023 | 0 | 0 | 3.43E-06 | 0 | 0 | 0 | 0 | 0.00011 |
| Uox | 0 | 0 | 1.62E-05 | 0 | 0 | 0 | 3.29E-05 | 6.81E-06 | 0 | 1.23E-07 | 0 | 0 | 0 | 0 | 0 | 1.52E-05 | 0 | 0 |
| Ifi44 | 6.71E-06 | 0 | 0 | 0 | 0 | 0 | 0 | 0 | 0 | 5.02E-05 | 0 | 0 | 1.64E-05 | 0 | 0 | 0 | 0 | 2.79E-05 |
| Depdc1a | 0 | 0 | 0 | 0 | 0 | 0 | 0 | 0 | 0 | 0 | 0 | 0 | 0 | 0 | 0 | 0.00059 | 0 | 0 |
| Ccne2 | 5.92E-06 | 0 | 4.30E-05 | 0 | 0 | 0 | 0 | 0 | 0 | 0 | 0 | 6.53E-06 | 3.39E-06 | 0 | 0 | 0.00026 | 0 | 0 |
| Rad54b | 0 | 0 | 0 | 0 | 0 | 0 | 0 | 0 | 0 | 1.11E-07 | 0 | 0 | 0 | 0 | 4.24E-05 | 0.00016 | 0 | 0 |
| Cdh17 | 0 | 3.34E-06 | 0 | 1.41E-05 | 0 | 0 | 0 | 0 | 0 | 8.47E-06 | 0 | 2.62E-05 | 0 | 0 | 4.92E-05 | 3.07E-06 | 0 | 9.43E-06 |
| Bach2 | 0 | 9.59E-05 | 0 | 0 | 3.54E-05 | 0 | 7.11E-05 | 0 | 0 | 0 | 7.63E-05 | 5.39E-05 | 7.56E-07 | 0 | 3.45E-07 | 0 | 0.00013 | 0 |
| Ddx58 | 0.0001 | 4.37E-05 | 3.32E-05 | 3.19E-05 | 4.92E-05 | 0 | 6.60E-05 | 5.27E-05 | 0 | 0.00014 | 0 | 0 | 0.0001 | 0 | 6.26E-05 | 1.96E-06 | 2.15E-05 | 0.00019 |
| Fam219a | 7.01E-06 | 0.00013 | 7.57E-05 | 4.34E-05 | 0 | 0 | 2.12E-05 | 1.11E-06 | 0 | 4.74E-05 | 2.12E-05 | 0.00015 | 4.71E-05 | 0 | 8.79E-05 | 9.69E-06 | 0 | 4.39E-05 |
| Enho | 0 | 2.36E-05 | 0 | 0 | 0.00014 | 0 | 0 | 0.00021 | 0 | 0 | 0 | 1.15E-05 | 0 | 0 | 0 | 0 | 0 | 0 |

| | | | | | | | | | | | | | | | | | | |
|---|---|---|---|---|---|---|---|---|---|---|---|---|---|---|---|---|---|---|
| Ccl21b | 0 | 1.59E-06 | 0 | 0 | 0 | 0 | 1.03E-05 | 0 | 0 | 1.92E-07 | 0 | 1.02E-05 | 1.28E-05 | 0 | 0 | 0 | 4.42E-05 | 0 |
| Ccl21a | 0 | 1.67E-06 | 0 | 0 | 0 | 0 | 2.12E-06 | 0 | 0 | 0 | 0 | 1.82E-05 | 2.52E-05 | 0 | 0 | 0 | 2.84E-05 | 0 |
| Phf24 | 0 | 0 | 0 | 0 | 0 | 0 | 0 | 0 | 0 | 0.00011 | 0 | 1.16E-05 | 5.71E-05 | 0 | 1.85E-05 | 0 | 7.69E-06 | 0 |
| Arhgef39 | 0 | 3.06E-07 | 0 | 0 | 0 | 0 | 0 | 0 | 0 | 0 | 0 | 1.47E-05 | 0 | 0 | 2.48E-05 | 0.00023 | 3.32E-06 | 0 |
| Car9 | 0 | 0 | 0 | 0 | 0 | 0 | 8.65E-06 | 0 | 0 | 0 | 0 | 0.00014 | 0 | 0 | 1.99E-06 | 0 | 0 | 0 |
| Ccin | 0 | 0 | 1.85E-06 | 0 | 0 | 0 | 5.14E-06 | 0 | 0 | 2.00E-05 | 0 | 2.00E-05 | 2.53E-05 | 0 | 0 | 5.10E-06 | 7.78E-05 | 0 |
| Melk | 0 | 0 | 0 | 5.11E-06 | 0 | 0 | 0 | 0 | 0 | 0 | 0 | 0 | 0 | 0 | 0 | 0.00054 | 0 | 0 |
| Tmod1 | 0 | 8.81E-06 | 0 | 0 | 0 | 0 | 0 | 0 | 0 | 9.57E-05 | 0 | 8.60E-06 | 5.38E-05 | 0 | 0 | 0 | 0 | 0 |
| Coro2a | 0 | 3.67E-06 | 6.44E-08 | 7.49E-06 | 1.13E-06 | 0 | 0 | 0 | 0 | 3.45E-06 | 3.05E-06 | 2.05E-05 | 1.01E-05 | 0 | 2.11E-05 | 3.65E-05 | 1.19E-05 | 2.60E-06 |
| Tbc1d2 | 0 | 1.24E-05 | 0 | 2.25E-07 | 3.88E-06 | 0 | 2.01E-05 | 0 | 0 | 0 | 2.44E-05 | 1.15E-07 | 8.17E-06 | 0 | 0 | 0 | 0 | 5.14E-06 |
| Tmeff1 | 0 | 3.66E-05 | 0.00014 | 0 | 0.00029 | 0 | 6.79E-05 | 0 | 0 | 0 | 0.00042 | 0.00015 | 0.00013 | 0 | 0 | 0 | 0.00014 | 0 |
| Cavin4 | 0 | 0 | 2.82E-06 | 2.92E-06 | 9.12E-06 | 0 | 0 | 0 | 0 | 0 | 2.67E-05 | 4.14E-05 | 5.68E-06 | 0 | 9.42E-06 | 0 | 1.14E-05 | 0 |
| Klf4 | 0 | 0 | 0 | 0 | 0 | 0 | 0 | 0 | 0 | 0.00089 | 0 | 0 | 0.00022 | 0 | 0 | 0 | 0.00389 | 0.00028 |
| Txndc8 | 0 | 0 | 0 | 0 | 0 | 0 | 0 | 0 | 0 | 0 | 0 | 2.50E-06 | 5.95E-05 | 0 | 0 | 0 | 3.73E-05 | 0 |
| Olfr267 | 0 | 0 | 3.40E-06 | 0 | 0 | 0 | 0 | 0 | 0 | 2.11E-06 | 0 | 1.19E-05 | 1.94E-05 | 0 | 0 | 1.77E-06 | 5.56E-05 | 0 |
| Megf9 | 8.52E-06 | 3.26E-05 | 0 | 0 | 6.22E-06 | 0 | 1.74E-05 | 0 | 0 | 0.00025 | 4.33E-05 | 0 | 9.00E-05 | 0 | 0 | 6.50E-05 | 3.92E-05 | 0 |
| Tle1 | 9.29E-06 | 5.73E-05 | 4.46E-06 | 8.96E-06 | 3.47E-05 | 0 | 5.53E-05 | 3.09E-05 | 0 | 4.08E-05 | 1.76E-05 | 5.71E-06 | 7.60E-05 | 0 | 2.20E-05 | 3.61E-06 | 5.72E-05 | 0 |
| Lurap1l | 1.83E-06 | 0 | 0 | 0 | 0 | 0 | 0 | 0 | 0 | 5.82E-05 | 2.70E-05 | 0 | 0 | 0 | 0 | 1.13E-06 | 0 | 7.15E-06 |
| Tek | 0 | 2.40E-05 | 0 | 0 | 0 | 0 | 0 | 0 | 0 | 0.00037 | 0 | 0 | 0 | 0 | 0 | 0 | 0 | 0 |
| Cyp2j9 | 2.34E-06 | 2.21E-05 | 0 | 0 | 0 | 0 | 0 | 0 | 0 | 1.27E-05 | 0 | 0 | 7.02E-06 | 0 | 0 | 3.79E-06 | 0 | 2.36E-05 |
| Fyb2 | 0 | 0 | 0 | 6.45E-06 | 0 | 0 | 5.07E-05 | 0 | 0 | 4.49E-05 | 8.34E-06 | 1.70E-06 | 1.66E-05 | 0 | 0 | 1.67E-05 | 0 | 0 |
| Orc1 | 0 | 0 | 2.11E-05 | 0 | 0 | 0 | 0 | 0 | 0 | 0 | 0 | 0 | 1.25E-05 | 0 | 0 | 8.46E-05 | 7.97E-06 | 0 |
| Dmrta2 | 0 | 0 | 0 | 1.18E-05 | 0 | 0 | 4.37E-06 | 2.09E-06 | 0 | 0 | 0 | 3.26E-06 | 0 | 0 | 1.92E-05 | 3.17E-07 | 1.80E-05 | 3.78E-06 |
| Trabd2b | 2.11E-06 | 2.20E-05 | 2.72E-05 | 0 | 0 | 0 | 0 | 0 | 0 | 9.10E-05 | 0 | 2.31E-05 | 0 | 0 | 0 | 4.64E-05 | 6.00E-06 | 0.00051 |
| Rad54l | 0 | 0 | 0 | 0 | 2.10E-06 | 0 | 0 | 0 | 0 | 0 | 0 | 0 | 1.20E-06 | 0 | 0 | 0.00036 | 0 | 0 |
| Kif2c | 0 | 0 | 0 | 0 | 0 | 0 | 0 | 0 | 0 | 0 | 0 | 0 | 0 | 0 | 4.96E-05 | 0.00085 | 0 | 0 |
| Cdc20 | 0 | 0 | 7.50E-05 | 1.84E-05 | 0 | 0 | 0 | 7.64E-06 | 0 | 4.32E-06 | 0 | 0 | 0 | 0 | 0 | 0.00109 | 0 | 0 |
| Kcnq4 | 0 | 1.09E-05 | 0 | 4.81E-06 | 1.60E-06 | 0 | 0 | 2.80E-06 | 0 | 4.58E-05 | 1.04E-09 | 2.96E-06 | 1.63E-05 | 0 | 0 | 0 | 0 | 2.77E-05 |
| Mfsd2a | 0 | 0 | 8.71E-06 | 0 | 0 | 0 | 0 | 0 | 0 | 0 | 1.38E-05 | 0 | 2.36E-05 | 0 | 0 | 0 | 1.22E-05 | 0 |
| Cdca8 | 1.17E-06 | 0 | 0 | 0 | 0 | 0 | 0 | 4.54E-07 | 0 | 2.84E-07 | 0 | 3.07E-06 | 0 | 0 | 1.27E-05 | 0.00097 | 0 | 0 |
| Col8a2 | 0 | 0 | 0 | 0 | 0 | 0 | 3.07E-05 | 0 | 0 | 0 | 0.00012 | 0 | 0 | 0 | 0 | 0 | 0 | 0.00034 |
| Gjb5 | 0 | 0 | 0.00012 | 0 | 0 | 0 | 0.00012 | 0 | 0 | 0 | 0.00012 | 1.17E-05 | 0 | 0 | 0 | 8.05E-06 | 0 | 0.00011 |
| A3galt2 | 0 | 3.95E-06 | 0 | 1.70E-06 | 0 | 0 | 0 | 6.88E-06 | 0 | 0 | 0 | 3.20E-06 | 0 | 0 | 4.93E-06 | 1.99E-06 | 6.47E-05 | 0 |

| | | | | | | | | | | | | | | | | | | |
|---|---|---|---|---|---|---|---|---|---|---|---|---|---|---|---|---|---|---|
| Tmem54 | 0 | 0 | 1.71E-06 | 4.84E-06 | 0 | 0 | 0 | 0 | 0 | 0 | 7.05E-06 | 3.56E-06 | 2.61E-05 | 0 | 0 | 7.42E-06 | 7.22E-05 | 0 |
| C77080 | 0 | 6.52E-05 | 0 | 3.01E-05 | 3.96E-05 | 0 | 4.36E-05 | 0 | 0 | 0 | 1.22E-05 | 0.00017 | 0 | 0 | 4.93E-05 | 1.61E-05 | 0 | 0.00012 |
| Rcc1 | 4.17E-05 | 0 | 0.00038 | 0.00024 | 0 | 0 | 4.72E-05 | 0 | 0 | 5.05E-06 | 0 | 0 | 5.68E-05 | 0 | 0.00012 | 0.00064 | 0.00036 | 0 |
| Smpdl3b | 1.51E-06 | 0 | 1.14E-05 | 5.89E-06 | 0 | 0 | 1.86E-05 | 0 | 0 | 0 | 0 | 0 | 1.90E-05 | 0 | 0 | 2.70E-05 | 0 | 5.03E-06 |
| Fgr | 0 | 4.49E-06 | 1.98E-06 | 1.09E-05 | 0 | 0 | 0 | 0 | 0 | 4.06E-06 | 0 | 0 | 0 | 0 | 4.52E-05 | 1.60E-05 | 3.35E-07 | 9.21E-06 |
| Cd164l2 | 0 | 0 | 0 | 8.14E-07 | 0 | 0 | 0 | 4.57E-06 | 0 | 3.32E-06 | 1.31E-06 | 1.03E-05 | 1.17E-05 | 0 | 1.35E-06 | 0 | 3.75E-05 | 0 |
| Gale | 3.32E-05 | 0 | 6.41E-05 | 6.69E-05 | 0 | 0 | 6.01E-05 | 0 | 0 | 3.54E-06 | 3.28E-05 | 0 | 0 | 0 | 1.11E-06 | 1.46E-06 | 0 | 3.30E-06 |
| Tcea3 | 0 | 0 | 0 | 0 | 0 | 0 | 7.48E-05 | 0 | 0 | 9.35E-05 | 0 | 0 | 0.00012 | 0 | 0 | 0 | 0 | 0.00023 |
| C1qb | 0 | 5.30E-06 | 0 | 0.0001 | 4.79E-07 | 0 | 0 | 0 | 0 | 1.60E-05 | 0 | 2.37E-05 | 1.94E-05 | 0 | 0.0003 | 1.04E-06 | 6.80E-05 | 1.98E-05 |
| C1qc | 0 | 3.12E-06 | 0 | 4.71E-05 | 0 | 0 | 0 | 3.55E-06 | 0 | 9.28E-06 | 0 | 5.08E-06 | 0 | 0 | 7.56E-05 | 2.84E-06 | 3.33E-05 | 5.26E-06 |
| Alpl | 0 | 0 | 0 | 0 | 0 | 0 | 0 | 0 | 0 | 6.72E-06 | 0 | 0 | 0 | 0 | 0 | 0 | 0 | 0.00027 |
| Sh2d5 | 0 | 4.09E-07 | 4.66E-06 | 5.66E-06 | 0 | 0 | 0 | 0 | 0 | 6.51E-07 | 0 | 1.82E-07 | 8.89E-06 | 0 | 3.08E-06 | 4.13E-06 | 4.90E-05 | 0 |
| Pla2g2e | 0 | 2.54E-05 | 1.01E-05 | 0 | 0 | 0 | 0 | 0 | 0 | 1.31E-05 | 0 | 0 | 6.04E-05 | 0 | 0 | 0 | 0 | 5.71E-06 |
| Hspb7 | 0 | 0 | 0 | 1.60E-05 | 0 | 0 | 2.73E-05 | 0 | 0 | 0 | 1.27E-05 | 0 | 9.64E-05 | 0 | 0 | 0 | 2.58E-06 | 3.24E-06 |
| Pramef12 | 0 | 0 | 0 | 0 | 2.94E-05 | 0 | 0 | 0 | 0 | 5.10E-07 | 7.56E-05 | 0 | 0 | 0 | 0 | 0 | 0 | 5.96E-06 |
| Srm | 0.0004 | 0 | 0.00077 | 0.00032 | 0 | 0 | 0.00031 | 0 | 0 | 2.72E-05 | 0 | 0 | 0.00012 | 0 | 3.84E-06 | 2.76E-05 | 0 | 0 |
| Pik3cd | 0 | 0 | 0 | 7.19E-05 | 0 | 0 | 4.28E-06 | 8.65E-05 | 0 | 8.02E-06 | 0 | 9.36E-06 | 4.33E-05 | 0 | 6.27E-05 | 6.37E-06 | 4.21E-05 | 5.21E-05 |
| Slc25a33 | 3.01E-05 | 1.90E-05 | 3.25E-05 | 4.80E-06 | 1.56E-06 | 0 | 0 | 5.29E-06 | 0 | 4.27E-06 | 9.05E-06 | 2.40E-06 | 0 | 0 | 0 | 0 | 0.00013 | 0 |
| Cfap74 | 2.68E-05 | 0 | 3.64E-05 | 1.46E-05 | 0 | 0 | 3.32E-05 | 1.16E-05 | 0 | 2.00E-06 | 0 | 2.07E-06 | 0 | 0 | 1.96E-05 | 0 | 0.00015 | 0 |
| Vwa1 | 0 | 7.92E-06 | 0 | 0 | 0 | 0 | 4.94E-05 | 0 | 0 | 1.65E-05 | 3.24E-05 | 1.05E-05 | 4.15E-05 | 0 | 0 | 1.34E-05 | 2.45E-05 | 1.48E-05 |
| Isg15 | 0.0001 | 0 | 7.57E-06 | 7.60E-05 | 0 | 0 | 0 | 0 | 0 | 0.00048 | 0 | 6.00E-05 | 0.00016 | 0 | 0 | 7.56E-05 | 0.00018 | 0.00031 |
| Abcb1a | 0 | 1.84E-06 | 6.39E-06 | 3.63E-06 | 0 | 0 | 0 | 2.03E-05 | 0 | 6.22E-06 | 0 | 1.48E-06 | 8.95E-06 | 0 | 0 | 3.02E-06 | 5.49E-06 | 5.32E-06 |
| Sema3c | 0 | 0 | 0 | 0 | 0 | 0 | 8.26E-05 | 0 | 0 | 0.00118 | 5.23E-05 | 0 | 8.56E-05 | 0 | 0 | 3.02E-05 | 2.68E-05 | 0 |
| Napepld | 8.72E-06 | 1.44E-05 | 4.88E-06 | 1.18E-05 | 0 | 0 | 7.25E-06 | 0 | 0 | 1.87E-05 | 1.47E-06 | 5.28E-07 | 2.40E-05 | 0 | 0 | 8.95E-06 | 0 | 0 |
| Atg9b | 0 | 0 | 0 | 8.27E-06 | 0 | 0 | 3.25E-06 | 0 | 0 | 0 | 0 | 3.09E-05 | 0 | 0 | 2.08E-05 | 0 | 0 | 5.91E-06 |
| Actr3b | 0 | 0 | 0 | 5.44E-07 | 0 | 0 | 8.19E-06 | 0 | 0 | 1.30E-05 | 7.96E-06 | 4.25E-06 | 0 | 0 | 0 | 0 | 8.38E-06 | 4.08E-05 |
| Dpp6 | 0 | 0 | 1.10E-06 | 0 | 3.69E-05 | 0 | 3.07E-05 | 6.60E-06 | 0 | 0 | 0 | 2.53E-05 | 0 | 0 | 0 | 0 | 0 | 0 |
| Il6 | 0 | 0 | 7.50E-05 | 0 | 0 | 0 | 0 | 0 | 0 | 5.58E-05 | 0 | 0 | 0.00039 | 0 | 0 | 0 | 0.00035 | 0 |
| Cenpa | 1.77E-05 | 0 | 0 | 6.65E-05 | 0 | 0 | 0 | 1.14E-05 | 0 | 0 | 0 | 0 | 0 | 0 | 7.18E-05 | 0.00077 | 0 | 0 |
| Spon2 | 0 | 0 | 0 | 2.25E-05 | 0 | 0 | 0.00039 | 0 | 0 | 0.00117 | 0 | 4.11E-05 | 0 | 0 | 0 | 0 | 0 | 0.00046 |
| Tacc3 | 0 | 0 | 4.66E-05 | 0.00014 | 0 | 0 | 0 | 1.75E-05 | 0 | 0 | 0 | 1.54E-08 | 0 | 0 | 4.46E-05 | 0.00141 | 0 | 0 |
| Zfyve28 | 0 | 0 | 2.81E-05 | 5.77E-06 | 0 | 0 | 4.91E-05 | 2.48E-05 | 0 | 5.21E-06 | 0 | 0 | 0 | 0 | 2.27E-05 | 0 | 0 | 6.57E-05 |
| Msantd1 | 0 | 0 | 1.21E-05 | 6.51E-06 | 0 | 0 | 5.05E-06 | 0 | 0 | 1.83E-05 | 0 | 9.46E-06 | 1.17E-05 | 0 | 6.83E-05 | 3.47E-07 | 0.00013 | 0 |

| | | | | | | | | | | | | | | | | | | |
|---|---|---|---|---|---|---|---|---|---|---|---|---|---|---|---|---|---|---|
| Hgfac | 0 | 4.69E-06 | 0 | 1.46E-05 | 0 | 0 | 0 | 0 | 0 | 1.45E-06 | 2.67E-08 | 5.67E-06 | 1.26E-05 | 0 | 7.49E-06 | 5.77E-06 | 2.85E-05 | 5.81E-06 |
| Cpeb2 | 0 | 4.32E-05 | 4.89E-05 | 3.12E-05 | 1.41E-05 | 0 | 0 | 7.10E-06 | 0 | 1.89E-05 | 7.79E-05 | 8.52E-05 | 4.55E-05 | 0 | 6.43E-05 | 0 | 0.00019 | 5.01E-05 |
| Ldb2 | 3.60E-05 | 3.54E-05 | 0 | 0 | 7.93E-06 | 0 | 0 | 3.54E-05 | 0 | 6.36E-05 | 1.85E-05 | 0 | 3.47E-05 | 0 | 0 | 3.71E-05 | 0 | 5.44E-05 |
| Ncapg | 0 | 0 | 0 | 0 | 0 | 0 | 0 | 0 | 0 | 0 | 0 | 0 | 0 | 0 | 9.20E-05 | 0.00063 | 0 | 0 |
| Sel1l3 | 0 | 0 | 0 | 0 | 1.75E-06 | 0 | 0 | 0 | 0 | 0.00039 | 0 | 9.09E-07 | 0.00011 | 0 | 0 | 0 | 0 | 0 |
| Tbc1d1 | 2.29E-05 | 3.47E-05 | 0.00014 | 0 | 0.0001 | 0 | 0 | 0.00012 | 0 | 2.89E-05 | 0.0004 | 0.00016 | 0 | 0 | 0 | 0.00014 | 9.62E-05 | 0.00014 |
| Rbm47 | 0 | 1.23E-05 | 1.59E-05 | 1.61E-05 | 0 | 0 | 0 | 0 | 0 | 3.36E-06 | 3.12E-05 | 0 | 7.72E-06 | 0 | 2.52E-05 | 5.60E-06 | 2.17E-05 | 0 |
| Uchl1 | 0 | 0 | 2.66E-05 | 0 | 0 | 0 | 1.88E-06 | 5.73E-06 | 0 | 3.75E-05 | 1.46E-05 | 2.42E-05 | 0 | 0 | 0 | 1.93E-06 | 0 | 3.22E-05 |
| Atp8a1 | 0 | 2.13E-06 | 0 | 0 | 1.44E-05 | 0 | 0 | 9.51E-07 | 0 | 7.91E-06 | 0 | 0 | 3.62E-06 | 0 | 5.34E-05 | 9.49E-06 | 2.18E-05 | 1.07E-05 |
| Tec | 2.48E-06 | 1.76E-05 | 0 | 0 | 0 | 0 | 0 | 6.22E-05 | 0 | 0 | 1.85E-05 | 5.21E-05 | 0 | 0 | 1.88E-05 | 1.01E-05 | 0 | 6.27E-05 |
| Kdr | 1.90E-06 | 4.30E-06 | 0 | 0 | 0 | 0 | 2.51E-05 | 0 | 0 | 4.51E-05 | 2.39E-06 | 0 | 0 | 0 | 0 | 2.93E-06 | 0 | 0 |
| Slc4a4 | 0 | 9.07E-06 | 0 | 4.71E-06 | 0 | 0 | 0 | 0 | 0 | 0.00029 | 0 | 0 | 5.38E-05 | 0 | 2.12E-05 | 9.82E-06 | 4.71E-05 | 0 |
| Pf4 | 0 | 4.83E-06 | 9.31E-06 | 3.07E-05 | 3.09E-06 | 0 | 0 | 5.28E-06 | 0 | 9.07E-06 | 0 | 1.03E-05 | 4.43E-09 | 0 | 4.88E-05 | 3.79E-06 | 2.78E-05 | 0 |
| Cxcl1 | 0 | 0 | 0.00019 | 0 | 0 | 0 | 0 | 0 | 0 | 0 | 0 | 0 | 0.0006 | 0 | 0 | 9.94E-05 | 0.00201 | 0 |
| Cdkl2 | 2.81E-06 | 3.94E-06 | 2.88E-06 | 0 | 0 | 0 | 5.68E-06 | 1.63E-06 | 0 | 9.79E-06 | 0 | 8.38E-06 | 5.33E-06 | 0 | 0 | 0 | 3.54E-06 | 2.65E-05 |
| Cxcl10 | 0 | 0 | 0 | 0 | 0 | 0 | 0 | 0 | 0 | 5.03E-05 | 0 | 0 | 0.00028 | 0 | 0 | 0.00011 | 0.0004 | 0 |
| Stbd1 | 0 | 0 | 0 | 1.41E-05 | 0 | 0 | 0 | 5.06E-06 | 0 | 1.56E-05 | 2.38E-05 | 8.77E-05 | 0 | 0 | 2.01E-05 | 2.16E-06 | 0 | 1.26E-05 |
| Cxcl13 | 0 | 0 | 0 | 0 | 0 | 0 | 0 | 0 | 0 | 0.00026 | 0 | 0 | 0 | 0 | 0 | 0 | 0 | 0 |
| Anxa3 | 8.79E-05 | 0 | 4.74E-05 | 0.00012 | 0 | 0 | 0 | 0 | 0 | 0.0015 | 0.00043 | 9.41E-05 | 0.00035 | 0 | 0.00015 | 5.71E-05 | 4.62E-05 | 0 |
| Prdm8 | 0 | 3.76E-07 | 0 | 0 | 2.95E-06 | 0 | 0 | 0 | 0 | 6.19E-05 | 0 | 0 | 0 | 0 | 0 | 4.29E-06 | 0 | 0 |
| Sparcl1 | 0 | 9.98E-05 | 0 | 0 | 0 | 0 | 0 | 0.00138 | 0 | 0 | 0 | 0 | 0 | 0 | 0 | 0 | 0 | 0.00055 |
| Ephx4 | 3.66E-06 | 0 | 0 | 0 | 0 | 0 | 6.20E-06 | 0 | 0 | 3.46E-05 | 0 | 4.84E-07 | 0 | 0 | 0 | 6.88E-07 | 0 | 0 |
| Grk3 | 0 | 2.54E-05 | 0 | 0 | 1.99E-05 | 0 | 0 | 2.33E-05 | 0 | 0 | 0 | 0 | 0 | 0 | 0 | 0 | 0 | 6.51E-07 |
| Cmklr1 | 0 | 0 | 3.08E-05 | 0 | 0 | 0 | 0 | 0.00083 | 0 | 0 | 9.72E-06 | 9.13E-05 | 0 | 0 | 0 | 0 | 0 | 0.00022 |
| Oasl2 | 3.24E-05 | 6.54E-05 | 0 | 6.78E-05 | 8.49E-06 | 0 | 0 | 0.00018 | 0 | 0.00047 | 0 | 5.85E-06 | 0.00015 | 0 | 7.36E-05 | 5.55E-05 | 0 | 0.00024 |
| Oasl1 | 1.30E-05 | 0 | 1.10E-05 | 9.00E-06 | 0 | 0 | 0 | 0 | 0 | 4.65E-05 | 0 | 4.33E-06 | 2.74E-06 | 0 | 0 | 5.59E-06 | 5.71E-06 | 1.61E-05 |
| Slc8b1 | 0 | 7.20E-05 | 0 | 3.15E-05 | 0 | 0 | 0 | 0 | 0 | 9.92E-05 | 0 | 5.12E-06 | 4.83E-05 | 0 | 3.11E-05 | 0 | 3.73E-05 | 7.94E-05 |
| Oas2 | 8.43E-06 | 0 | 0 | 0 | 0 | 0 | 0 | 0 | 0 | 0.00024 | 0 | 0 | 7.41E-05 | 0 | 7.29E-06 | 0 | 0 | 0.00013 |
| Oas3 | 0 | 0 | 0 | 0 | 0 | 0 | 0 | 0 | 0 | 7.52E-05 | 0 | 0 | 3.94E-05 | 0 | 0 | 5.48E-06 | 0 | 0 |
| Oas1g | 1.08E-06 | 0 | 1.46E-07 | 8.42E-07 | 0 | 0 | 0 | 0 | 0 | 3.04E-05 | 0 | 0 | 1.11E-05 | 0 | 2.56E-06 | 2.48E-06 | 0 | 1.26E-05 |
| Oas1a | 1.54E-05 | 0 | 1.52E-05 | 1.65E-05 | 0 | 0 | 5.50E-07 | 0 | 0 | 0.0002 | 0 | 0 | 5.00E-05 | 0 | 0 | 1.05E-05 | 0 | 7.02E-05 |
| P2rx7 | 0 | 2.77E-05 | 1.13E-06 | 0 | 6.35E-06 | 0 | 1.16E-05 | 0 | 0 | 1.90E-05 | 2.58E-05 | 0 | 8.01E-06 | 0 | 2.92E-05 | 2.04E-05 | 0 | 7.09E-05 |
| Rhof | 3.37E-06 | 1.04E-05 | 1.84E-06 | 9.95E-06 | 0 | 0 | 1.48E-06 | 0 | 0 | 0 | 1.07E-05 | 3.66E-06 | 3.63E-06 | 0 | 1.73E-05 | 0 | 5.27E-05 | 7.93E-06 |

| | | | | | | | | | | | | | | | | | | |
|---|---|---|---|---|---|---|---|---|---|---|---|---|---|---|---|---|---|---|
| Glt1d1 | 0 | 0 | 0 | 0 | 0 | 0 | 0 | 0 | 0 | 0 | 0 | 0.00013 | 0 | 0 | 0 | 0 | 0 | 0 |
| Adgrd1 | 0 | 0 | 0.00016 | 0 | 0 | 0 | 0.00058 | 0 | 0 | 0.00132 | 5.70E-05 | 9.82E-05 | 0.00086 | 0 | 0 | 5.01E-06 | 0 | 0.00064 |
| Ncf1 | 0 | 1.29E-05 | 0 | 2.06E-05 | 0 | 0 | 0 | 0 | 0 | 6.92E-06 | 0 | 4.97E-06 | 8.38E-06 | 0 | 3.31E-05 | 1.19E-05 | 3.98E-05 | 3.05E-06 |
| Lat2 | 0 | 5.04E-07 | 0 | 5.71E-06 | 0 | 0 | 0 | 0 | 0 | 2.95E-06 | 0 | 8.54E-06 | 0 | 0 | 4.36E-05 | 6.08E-06 | 0.00011 | 0 |
| Vgf | 0 | 0 | 0 | 1.62E-06 | 0 | 0 | 0 | 0 | 0 | 0 | 0 | 1.48E-05 | 0.00011 | 0 | 0 | 3.40E-06 | 0 | 0 |
| Serpine1 | 0 | 0 | 0.00078 | 0 | 0 | 0 | 0 | 0 | 0 | 0 | 0.00155 | 0.00194 | 0.00234 | 0 | 0 | 0 | 0.00326 | 0 |
| Tfr2 | 0 | 0 | 0 | 6.29E-06 | 0.00016 | 0 | 5.95E-05 | 7.22E-06 | 0 | 0 | 0.00012 | 1.60E-05 | 1.68E-05 | 0 | 2.21E-05 | 0 | 0 | 1.28E-05 |
| Pilra | 0 | 6.63E-06 | 0 | 0 | 0 | 0 | 0 | 0.00012 | 0 | 5.83E-06 | 0 | 3.32E-06 | 7.27E-06 | 0 | 0 | 0 | 0 | 3.03E-05 |
| Pilrb2 | 0 | 2.22E-07 | 0 | 4.97E-07 | 0 | 0 | 0 | 3.04E-05 | 0 | 3.85E-06 | 6.71E-07 | 3.41E-07 | 1.21E-05 | 0 | 1.50E-06 | 4.60E-07 | 1.17E-06 | 1.22E-05 |
| Pdgfa | 0 | 7.77E-06 | 3.13E-05 | 0 | 4.52E-05 | 0 | 0 | 0.00028 | 0 | 0 | 0.00046 | 0.00013 | 0 | 0 | 0 | 0 | 0 | 9.73E-05 |
| Amz1 | 0 | 0 | 0 | 0 | 0 | 0 | 0 | 0 | 0 | 0 | 0 | 8.30E-05 | 0 | 0 | 0 | 0 | 0 | 4.71E-06 |
| Rasl11a | 0 | 0 | 2.77E-05 | 1.67E-06 | 0.00018 | 0 | 0 | 0 | 0 | 0 | 0.00042 | 0.00029 | 0.00191 | 0 | 0 | 0 | 0 | 0 |
| Tex26 | 0 | 0 | 0 | 5.77E-06 | 3.56E-06 | 0 | 2.80E-05 | 0 | 0 | 0 | 1.48E-05 | 2.38E-05 | 3.47E-05 | 0 | 0 | 2.32E-06 | 3.64E-06 | 0 |
| Fry | 0 | 2.77E-06 | 0 | 0 | 4.10E-05 | 0 | 0 | 7.08E-07 | 0 | 1.67E-06 | 2.89E-05 | 0 | 0 | 0 | 8.33E-06 | 0 | 0 | 3.83E-05 |
| Samd9l | 8.45E-05 | 0 | 0 | 0 | 0 | 0 | 1.48E-05 | 0 | 0 | 0.00011 | 0 | 8.46E-05 | 1.21E-05 | 0 | 0 | 1.57E-05 | 0 | 0.0001 |
| Ppp1r9a | 0 | 4.13E-06 | 0 | 0 | 0 | 0 | 0 | 0 | 0 | 4.17E-05 | 0 | 0 | 3.30E-06 | 0 | 1.66E-05 | 0 | 0 | 0 |
| Col28a1 | 0 | 0 | 0 | 0 | 0 | 0 | 0 | 0 | 0 | 0.00025 | 0 | 0 | 0.00057 | 0 | 0 | 0 | 0 | 0.00015 |
| Wnt16 | 0 | 4.10E-05 | 0 | 0 | 0 | 0 | 4.35E-05 | 0 | 0 | 0.00043 | 0 | 0 | 0.00014 | 0 | 0 | 0 | 0 | 0 |
| Tmem229a | 0 | 0 | 0 | 0 | 0 | 0 | 0 | 0 | 0 | 0.00011 | 0 | 0 | 0 | 0 | 0 | 0 | 0 | 0 |
| Fam71f2 | 0 | 0 | 4.95E-06 | 1.08E-05 | 0 | 0 | 0 | 0 | 0 | 0 | 0 | 5.60E-05 | 0 | 0 | 6.65E-06 | 3.01E-06 | 7.28E-05 | 0 |
| Ccdc136 | 0 | 1.81E-05 | 0 | 0 | 2.96E-06 | 0 | 1.68E-05 | 0 | 0 | 3.94E-06 | 0 | 2.63E-05 | 0 | 0 | 3.12E-05 | 0 | 1.59E-05 | 1.32E-05 |
| Plxna4 | 0 | 0 | 3.54E-06 | 2.44E-05 | 0 | 0 | 0.00019 | 0 | 0 | 6.83E-05 | 8.66E-06 | 1.39E-05 | 3.03E-05 | 0 | 0 | 4.36E-05 | 0 | 0.00025 |
| Ptn | 1.76E-05 | 2.23E-05 | 0 | 0 | 0 | 0 | 1.64E-05 | 0 | 0 | 0 | 0 | 0 | 0 | 0 | 0 | 7.54E-06 | 0 | 8.68E-05 |
| Ephb6 | 0 | 6.09E-05 | 0 | 0 | 3.67E-05 | 0 | 0 | 1.97E-05 | 0 | 0 | 5.00E-05 | 2.03E-05 | 0 | 0 | 5.18E-05 | 0 | 0 | 1.87E-05 |
| Tas2r143 | 0 | 0 | 0 | 0 | 0 | 0 | 0 | 0 | 0 | 1.84E-06 | 0 | 2.94E-06 | 1.74E-05 | 0 | 0 | 0 | 5.77E-05 | 0 |
| Tas2r135 | 0 | 0 | 0 | 0 | 0 | 0 | 0 | 0 | 0 | 1.96E-06 | 0 | 3.05E-06 | 9.21E-06 | 0 | 0 | 0 | 7.08E-05 | 0 |
| Tas2r126 | 0 | 0 | 1.92E-06 | 0 | 0 | 0 | 0 | 0 | 0 | 9.81E-06 | 0 | 0 | 1.94E-05 | 0 | 0 | 0 | 8.79E-05 | 0 |
| Hoxa4 | 0 | 8.58E-06 | 4.18E-05 | 3.10E-05 | 1.50E-05 | 0 | 4.53E-05 | 6.63E-05 | 0 | 4.02E-06 | 0 | 1.11E-05 | 0 | 0 | 7.54E-06 | 1.43E-05 | 0 | 0 |
| Creb5 | 0 | 0 | 1.16E-05 | 0 | 0 | 0 | 0 | 0 | 0 | 0.00046 | 6.57E-05 | 4.47E-06 | 0 | 0 | 2.29E-05 | 1.35E-05 | 0 | 0 |
| Tril | 0 | 3.65E-05 | 0 | 9.45E-05 | 0 | 0 | 0 | 1.70E-05 | 0 | 8.64E-05 | 0 | 0 | 0 | 0 | 0 | 8.41E-05 | 0 | 0.00021 |
| Sh2d6 | 0 | 6.20E-08 | 5.68E-06 | 0 | 0 | 0 | 0 | 0 | 0 | 4.03E-05 | 0 | 0 | 0 | 0 | 3.16E-06 | 0 | 7.19E-06 | 3.85E-07 |
| Gcfc2 | 1.56E-05 | 3.79E-05 | 6.28E-07 | 0 | 1.33E-05 | 0 | 1.72E-05 | 0 | 0 | 0 | 0 | 2.54E-05 | 1.05E-05 | 0 | 0.00012 | 5.82E-05 | 1.19E-06 | 0 |
| Hk2 | 7.38E-05 | 0 | 0.00046 | 8.09E-05 | 9.55E-05 | 0 | 6.15E-05 | 0.00014 | 0 | 6.10E-05 | 0.00015 | 0.00094 | 0.00014 | 0 | 0.00028 | 3.01E-05 | 0.00076 | 0 |

| | | | | | | | | | | | | | | | | | | |
|---|---|---|---|---|---|---|---|---|---|---|---|---|---|---|---|---|---|---|
| M1ap | 0 | 0 | 0 | 7.94E-06 | 3.19E-05 | 0 | 0 | 1.46E-05 | 0 | 1.75E-06 | 4.02E-05 | 0 | 0 | 0 | 0 | 2.04E-05 | 0 | 0 |
| Mthfd2 | 0.00012 | 0 | 0.00037 | 0 | 0 | 0 | 0.0001 | 0 | 0 | 0 | 8.29E-05 | 0.00015 | 0 | 0 | 0 | 0.00032 | 0 | 0 |
| Actg2 | 0 | 0 | 0 | 0 | 0 | 0 | 0 | 0 | 0 | 0 | 0.00121 | 0 | 0 | 0 | 0 | 0 | 0 | 0 |
| Add2 | 0 | 5.27E-06 | 0 | 0 | 0 | 0 | 0 | 0.00014 | 0 | 0 | 0 | 4.51E-05 | 0 | 0 | 0 | 0 | 0 | 1.51E-05 |
| Fgd5 | 0 | 0 | 0 | 0 | 0.00013 | 0 | 4.15E-05 | 9.01E-05 | 0 | 1.26E-05 | 1.02E-05 | 1.08E-05 | 0 | 0 | 0 | 2.36E-06 | 0 | 0.00031 |
| Lmcd1 | 0 | 0 | 0 | 0 | 0 | 0 | 0 | 0 | 0 | 1.32E-05 | 0 | 7.86E-05 | 0.0005 | 0 | 0 | 0 | 0 | 0 |
| Ret | 0 | 0 | 3.25E-05 | 0 | 0 | 0 | 0 | 1.50E-05 | 0 | 5.26E-05 | 0 | 4.37E-05 | 0 | 0 | 0 | 0 | 0 | 0 |
| Usp18 | 2.34E-05 | 0 | 0 | 0 | 0 | 0 | 0 | 0 | 0 | 0.00021 | 0 | 1.06E-05 | 5.95E-05 | 0 | 0 | 1.14E-05 | 0 | 5.03E-05 |
| C3ar1 | 0 | 1.13E-05 | 2.92E-06 | 1.34E-05 | 2.30E-06 | 0 | 0 | 0 | 0 | 5.27E-06 | 0 | 0 | 3.22E-06 | 0 | 3.44E-05 | 2.36E-06 | 0 | 2.09E-06 |
| Cdca3 | 4.80E-06 | 0 | 2.59E-05 | 0.00017 | 0 | 0 | 0 | 0 | 0 | 0 | 0 | 0 | 0 | 0 | 7.29E-05 | 0.00157 | 0 | 0 |
| Gpr162 | 0 | 0 | 0 | 6.36E-06 | 0 | 0 | 0 | 8.74E-07 | 0 | 0 | 0 | 2.84E-06 | 2.57E-05 | 0 | 0 | 0 | 5.94E-05 | 0 |
| Lag3 | 0 | 0 | 0 | 0 | 0 | 0 | 0 | 0 | 0 | 0 | 0 | 4.50E-05 | 1.54E-05 | 0 | 2.86E-05 | 6.39E-06 | 0.00089 | 0 |
| Rad51ap1 | 0 | 0 | 0 | 0 | 1.03E-06 | 0 | 0 | 0 | 0 | 0 | 0 | 0 | 0 | 0 | 0 | 0.00042 | 0 | 0 |
| Olr1 | 0 | 2.10E-06 | 3.44E-05 | 0 | 1.48E-05 | 0 | 0 | 3.31E-05 | 0 | 0 | 0 | 0 | 1.99E-05 | 0 | 2.10E-05 | 2.68E-06 | 4.20E-05 | 6.01E-05 |
| Plbd1 | 0 | 1.35E-06 | 0 | 0 | 0 | 0 | 0 | 0 | 0 | 5.24E-06 | 1.63E-06 | 6.27E-06 | 4.16E-06 | 0 | 6.67E-05 | 1.05E-05 | 6.39E-06 | 2.64E-06 |
| Kcnj8 | 0 | 2.96E-05 | 0 | 0 | 0 | 0 | 0.00015 | 0 | 0 | 7.65E-05 | 0 | 2.35E-05 | 7.73E-05 | 0 | 0 | 0 | 0 | 6.16E-05 |
| Bhlhe41 | 0 | 0 | 0.00021 | 0 | 0 | 0 | 0 | 0.00082 | 0 | 0 | 1.90E-05 | 0.00013 | 0 | 0 | 0 | 0 | 0 | 0.00027 |
| Prkcg | 0 | 0 | 1.60E-05 | 0 | 0 | 0 | 0 | 0 | 0 | 0 | 1.35E-05 | 1.54E-05 | 0 | 0 | 0 | 7.29E-06 | 0.00022 | 6.37E-06 |
| Syt5 | 0 | 0 | 2.41E-05 | 0 | 0 | 0 | 0 | 2.20E-05 | 0 | 0 | 0 | 0.00015 | 0 | 0 | 0 | 0 | 0 | 6.38E-06 |
| Tmem238 | 0 | 0 | 2.32E-05 | 2.98E-05 | 0 | 0 | 1.84E-05 | 0 | 0 | 2.86E-05 | 0 | 3.66E-05 | 3.11E-05 | 0 | 0 | 1.04E-05 | 7.08E-05 | 2.56E-05 |
| Ssc5d | 0 | 4.78E-06 | 0 | 0.00012 | 2.75E-05 | 0 | 0.00056 | 0 | 0 | 0.00074 | 0 | 0 | 0.00042 | 0 | 0.00025 | 0 | 5.39E-05 | 0.00028 |
| Zscan18 | 3.14E-07 | 8.56E-06 | 0 | 0 | 1.76E-06 | 0 | 7.58E-07 | 4.45E-06 | 0 | 1.28E-07 | 1.47E-06 | 4.08E-06 | 0 | 0 | 1.29E-05 | 0 | 1.75E-05 | 6.46E-06 |
| Rnf225 | 0 | 0 | 1.03E-05 | 3.14E-06 | 0 | 0 | 0 | 9.25E-07 | 0 | 0 | 0 | 0 | 8.33E-06 | 0 | 1.03E-05 | 1.02E-06 | 7.60E-05 | 0 |
| Lig1 | 0 | 0 | 3.20E-05 | 0.00011 | 7.54E-05 | 0 | 1.88E-06 | 0.00017 | 0 | 6.88E-06 | 0 | 0 | 6.37E-05 | 0 | 0 | 0.00068 | 4.41E-05 | 4.18E-05 |
| Slc8a2 | 0 | 8.06E-06 | 0 | 0 | 0 | 0 | 0 | 0 | 0 | 0 | 0 | 5.87E-06 | 2.08E-07 | 0 | 8.70E-07 | 0 | 7.29E-05 | 2.42E-06 |
| Bbc3 | 0 | 9.39E-05 | 3.35E-06 | 2.02E-05 | 0 | 0 | 0 | 0 | 0 | 3.94E-05 | 0 | 3.30E-05 | 0 | 0 | 0.00011 | 0 | 5.83E-05 | 2.85E-05 |
| Dmpk | 0 | 6.86E-05 | 0 | 1.07E-05 | 2.28E-05 | 0 | 0 | 0 | 0 | 0.00027 | 9.35E-05 | 0 | 0.00018 | 0 | 0.00018 | 0 | 1.38E-05 | 0.00012 |
| Ppm1n | 0 | 0 | 0 | 0 | 0 | 0 | 0 | 0 | 0 | 3.79E-07 | 0 | 8.56E-05 | 4.45E-05 | 0 | 0 | 1.38E-05 | 9.89E-05 | 0 |
| Pvr | 5.70E-05 | 0.00014 | 0.0002 | 0 | 1.29E-05 | 0 | 4.84E-05 | 0 | 0 | 8.34E-05 | 0.00017 | 0.0006 | 0.00021 | 0 | 0 | 0 | 0.00013 | 0 |
| Kcnn4 | 0 | 1.16E-05 | 0 | 1.11E-05 | 4.27E-07 | 0 | 0 | 0 | 0 | 3.97E-06 | 0 | 5.32E-06 | 0 | 0 | 3.80E-05 | 7.00E-06 | 1.40E-05 | 9.82E-06 |
| Cd79a | 0 | 0 | 1.18E-05 | 0 | 0 | 0 | 0 | 0 | 0 | 0 | 0 | 0 | 5.70E-05 | 0 | 0 | 3.87E-06 | 0.00017 | 0 |
| Grik5 | 0 | 6.38E-05 | 0 | 6.00E-06 | 4.25E-05 | 0 | 2.13E-05 | 0 | 0 | 3.68E-05 | 0 | 2.28E-05 | 4.00E-05 | 0 | 7.95E-05 | 0 | 3.85E-06 | 1.13E-05 |
| Dedd2 | 0 | 3.70E-06 | 0 | 1.28E-05 | 0 | 0 | 0 | 0 | 0 | 4.77E-06 | 0 | 0 | 8.99E-05 | 0 | 5.01E-05 | 0 | 0.00043 | 0 |

| | | | | | | | | | | | | | | | | | | |
|---|---|---|---|---|---|---|---|---|---|---|---|---|---|---|---|---|---|---|
| Cyp2s1 | 0 | 6.05E-06 | 0 | 0 | 0 | 0 | 0 | 0.00011 | 0 | 0 | 0 | 2.48E-05 | 0 | 0 | 1.37E-05 | 0 | 0 | 0 |
| Ltbp4 | 0 | 0 | 0 | 0 | 0 | 0 | 0 | 0.00028 | 0 | 0.00027 | 0 | 5.95E-05 | 0 | 0 | 0.00015 | 0 | 0 | 0.00147 |
| Spint2 | 1.47E-06 | 3.54E-06 | 0 | 9.34E-06 | 3.38E-06 | 0 | 0 | 1.33E-06 | 0 | 1.02E-05 | 1.52E-05 | 1.38E-05 | 6.85E-06 | 0 | 4.21E-05 | 3.81E-06 | 4.30E-05 | 2.09E-05 |
| Dpf1 | 0 | 0 | 1.55E-06 | 6.15E-07 | 0 | 0 | 0 | 0 | 0 | 0 | 0 | 3.89E-05 | 0 | 0 | 7.80E-06 | 5.98E-05 | 0 | 5.22E-06 |
| Cox7a1 | 0 | 0 | 0 | 1.26E-05 | 0 | 0 | 1.53E-06 | 1.11E-05 | 0 | 7.30E-06 | 0 | 6.47E-05 | 3.77E-07 | 0 | 0 | 0 | 6.86E-06 | 0 |
| Haus5 | 0 | 5.92E-06 | 0 | 2.02E-06 | 0 | 0 | 0 | 0 | 0 | 5.02E-06 | 0 | 1.50E-06 | 0 | 0 | 5.64E-05 | 0.0001 | 6.86E-06 | 5.76E-07 |
| Sbsn | 0 | 0 | 0 | 0 | 0 | 0 | 0.00041 | 0 | 0 | 0.00091 | 0 | 0.00021 | 0.00025 | 0 | 0 | 0 | 0 | 0 |
| Dmkn | 0 | 0 | 0 | 0 | 0 | 0 | 1.43E-06 | 0 | 0 | 0.00039 | 0 | 7.13E-06 | 0 | 0 | 0 | 0 | 0 | 0 |
| Fgf21 | 0 | 0 | 1.23E-05 | 0 | 0 | 0 | 2.73E-05 | 1.96E-05 | 0 | 0 | 2.15E-05 | 5.20E-05 | 0 | 0 | 0 | 0 | 0 | 0 |
| Sec1 | 0 | 1.81E-06 | 0 | 0 | 6.23E-06 | 0 | 0 | 3.13E-05 | 0 | 0 | 1.94E-06 | 1.56E-05 | 0 | 0 | 0 | 0 | 0 | 6.53E-07 |
| Car11 | 0 | 5.91E-05 | 0 | 0 | 3.75E-06 | 0 | 0 | 0.00023 | 0 | 0 | 0 | 0.00017 | 0 | 0 | 0 | 0 | 0 | 4.60E-06 |
| Ccdc114 | 3.20E-06 | 1.23E-05 | 0 | 3.13E-06 | 1.16E-05 | 0 | 1.14E-05 | 1.07E-06 | 0 | 9.94E-06 | 0 | 2.17E-05 | 2.03E-06 | 0 | 6.46E-05 | 7.89E-06 | 4.36E-05 | 1.99E-05 |
| Saa1 | 0 | 0 | 0 | 0 | 0 | 0 | 0 | 0 | 0 | 8.68E-05 | 0 | 0 | 0.00016 | 0 | 0 | 0 | 0 | 0 |
| Pcsk6 | 0 | 0 | 0 | 0 | 0 | 0 | 0 | 0 | 0 | 0.00047 | 1.22E-05 | 0 | 0 | 0 | 2.26E-05 | 0 | 0 | 0 |
| Aldh1a3 | 0 | 6.48E-06 | 0 | 0 | 3.65E-06 | 0 | 0 | 6.90E-05 | 0 | 0.00027 | 2.55E-05 | 0 | 0 | 0 | 3.13E-06 | 0 | 0 | 1.62E-05 |
| Synm | 0 | 0 | 0 | 0 | 0 | 0 | 5.72E-05 | 0 | 0 | 2.50E-05 | 0 | 0 | 0 | 0 | 0 | 0 | 0 | 0.00011 |
| Fanci | 0 | 0 | 0 | 0 | 0 | 0 | 1.58E-05 | 9.41E-06 | 0 | 4.49E-06 | 0 | 1.81E-05 | 0 | 0 | 0 | 0.00027 | 3.62E-05 | 0 |
| Prc1 | 0 | 0 | 0 | 0 | 0 | 0 | 0 | 0 | 0 | 0 | 0 | 0 | 0 | 0 | 9.01E-05 | 0.00123 | 0 | 0 |
| Sh3gl3 | 0 | 0 | 0 | 1.03E-07 | 0 | 0 | 0 | 5.95E-06 | 0 | 3.02E-05 | 7.14E-06 | 2.46E-05 | 0 | 0 | 0 | 0 | 0 | 0 |
| Adamtsl3 | 0 | 0 | 4.01E-06 | 0 | 0 | 0 | 0 | 0 | 0 | 7.38E-06 | 0 | 0 | 9.68E-05 | 0 | 0 | 1.94E-05 | 0 | 0.00029 |
| Sytl2 | 0 | 0 | 0 | 0 | 0 | 0 | 0 | 8.70E-05 | 0 | 0.00021 | 0.0003 | 0 | 0 | 0 | 0 | 0 | 0 | 0 |
| Ddias | 7.06E-06 | 1.32E-06 | 0 | 0 | 0 | 0 | 0 | 0 | 0 | 0 | 0 | 3.97E-06 | 0 | 0 | 0 | 0.00012 | 2.99E-05 | 0 |
| Kctd14 | 4.28E-06 | 0 | 8.91E-06 | 2.42E-06 | 0 | 0 | 2.51E-05 | 0 | 0 | 0 | 0 | 0 | 7.33E-06 | 0 | 0 | 0 | 0 | 1.47E-06 |
| Slco2b1 | 0 | 2.57E-05 | 0 | 0 | 0 | 0 | 0 | 0 | 0 | 3.88E-05 | 0 | 0 | 0 | 0 | 0 | 0 | 0 | 0.00011 |
| Neu3 | 1.99E-06 | 4.78E-06 | 4.40E-07 | 8.19E-06 | 1.78E-05 | 0 | 0 | 5.35E-07 | 0 | 7.26E-06 | 1.76E-06 | 5.72E-06 | 2.28E-05 | 0 | 1.30E-05 | 0 | 0 | 3.17E-05 |
| P2ry6 | 0 | 1.88E-07 | 0 | 1.76E-06 | 0 | 0 | 0 | 0 | 0 | 0 | 0 | 4.37E-05 | 6.59E-05 | 0 | 1.48E-05 | 0 | 2.01E-05 | 9.51E-06 |
| Lrrc51 | 7.04E-06 | 7.81E-06 | 0 | 1.85E-06 | 1.11E-07 | 0 | 0 | 3.26E-07 | 0 | 6.79E-06 | 0 | 1.56E-05 | 9.55E-06 | 0 | 3.62E-06 | 1.05E-06 | 0 | 0 |
| Hpx | 0 | 7.82E-06 | 0 | 0 | 0 | 0 | 0 | 0 | 0 | 0 | 3.19E-06 | 1.54E-05 | 6.13E-06 | 0 | 2.24E-05 | 0 | 4.66E-05 | 6.18E-06 |
| Dchs1 | 0 | 0.00018 | 0 | 0.00019 | 0.00045 | 0 | 0.0003 | 0 | 0 | 0.00014 | 9.46E-05 | 9.32E-05 | 0.00053 | 0 | 2.89E-05 | 0 | 0.00011 | 0 |
| Gvin1 | 3.20E-05 | 0 | 0 | 0 | 0 | 0 | 0 | 4.15E-06 | 0 | 0.00018 | 0 | 0 | 6.66E-05 | 0 | 1.49E-05 | 0 | 0 | 0.00011 |
| Scube2 | 1.54E-06 | 0 | 0 | 0 | 0 | 0 | 3.16E-05 | 0 | 0 | 0 | 2.23E-06 | 0 | 7.65E-06 | 0 | 0 | 3.34E-06 | 3.64E-06 | 2.73E-05 |
| Swap70 | 5.43E-05 | 0.00016 | 0 | 0.00014 | 6.84E-05 | 0 | 7.72E-06 | 4.73E-06 | 0 | 0.00012 | 0.00019 | 8.04E-05 | 0.00028 | 0 | 0.00026 | 5.11E-05 | 4.98E-05 | 4.34E-05 |
| Mrvi1 | 0 | 0 | 0 | 0 | 0 | 0 | 0 | 0 | 0 | 5.66E-07 | 9.30E-05 | 0 | 0 | 0 | 0 | 0 | 0 | 5.11E-06 |

| | | | | | | | | | | | | | | | | | | |
|---|---|---|---|---|---|---|---|---|---|---|---|---|---|---|---|---|---|---|
| Insc | 0 | 6.77E-06 | 0 | 0 | 2.97E-05 | 0 | 0 | 0 | 0 | 0 | 5.93E-05 | 0 | 0 | 0 | 0 | 0 | 0 | 8.65E-06 |
| Syt17 | 0 | 2.22E-05 | 0 | 0 | 3.71E-05 | 0 | 7.89E-05 | 0 | 0 | 8.24E-07 | 0 | 4.00E-05 | 9.32E-05 | 0 | 0 | 0 | 0 | 0 |
| Plk1 | 0 | 0 | 0 | 0 | 0 | 0 | 0 | 1.08E-05 | 0 | 0 | 0 | 0 | 0 | 0 | 4.77E-06 | 0.00116 | 0 | 0 |
| Il21r | 0 | 0 | 0 | 8.78E-06 | 0 | 0 | 0 | 0 | 0 | 0 | 0 | 9.58E-06 | 5.72E-05 | 0 | 0 | 4.31E-08 | 4.88E-05 | 0 |
| Apobr | 0 | 4.42E-06 | 2.76E-07 | 2.38E-05 | 1.09E-05 | 0 | 0 | 1.62E-05 | 0 | 9.49E-06 | 0 | 2.78E-05 | 0 | 0 | 9.07E-06 | 0 | 6.16E-05 | 9.16E-06 |
| Hirip3 | 2.60E-06 | 2.55E-05 | 5.87E-05 | 1.36E-05 | 0 | 0 | 0 | 0 | 0 | 2.72E-05 | 0 | 0 | 4.03E-06 | 0 | 0.0002 | 0.00076 | 5.08E-05 | 0 |
| Kif22 | 0 | 0 | 0 | 0 | 0 | 0 | 0 | 0 | 0 | 0 | 0 | 0 | 0 | 0 | 6.19E-05 | 0.00064 | 0 | 0 |
| Itgam | 0 | 5.97E-06 | 4.42E-06 | 1.74E-05 | 0 | 0 | 0 | 0 | 0 | 1.74E-06 | 1.04E-06 | 1.45E-05 | 1.23E-05 | 0 | 4.38E-05 | 1.48E-05 | 1.58E-05 | 1.18E-05 |
| Mki67 | 0 | 0 | 0 | 0 | 0 | 0 | 0 | 0 | 0 | 1.99E-07 | 0 | 0 | 0 | 0 | 6.01E-05 | 0.00193 | 0 | 0 |
| Sprn | 0 | 5.57E-06 | 0 | 2.13E-05 | 1.68E-05 | 0 | 2.53E-05 | 0 | 0 | 7.91E-06 | 3.67E-06 | 3.30E-06 | 2.69E-05 | 0 | 1.56E-05 | 0 | 3.68E-06 | 2.12E-05 |
| Lrrc56 | 0 | 2.48E-05 | 0 | 2.17E-05 | 1.25E-05 | 0 | 2.49E-06 | 5.17E-06 | 0 | 5.64E-06 | 0 | 1.91E-06 | 7.23E-06 | 0 | 2.35E-05 | 2.63E-05 | 4.90E-06 | 6.61E-06 |
| Eps8l2 | 0 | 4.14E-06 | 0 | 0 | 1.08E-05 | 0 | 1.35E-06 | 0 | 0 | 1.80E-06 | 7.85E-06 | 1.21E-05 | 0 | 0 | 0 | 0 | 0 | 9.94E-06 |
| Cend1 | 1.42E-06 | 0 | 3.68E-06 | 1.71E-06 | 1.08E-06 | 0 | 2.12E-05 | 0 | 0 | 0 | 6.37E-06 | 0 | 2.55E-06 | 0 | 0 | 0 | 7.07E-06 | 2.02E-05 |
| Muc2 | 0 | 0 | 0 | 0 | 0 | 0 | 0 | 0 | 0 | 6.52E-07 | 0 | 0.0003 | 5.44E-05 | 0 | 1.30E-05 | 0 | 3.41E-05 | 0 |
| Dusp8 | 0 | 0 | 4.52E-05 | 0 | 0 | 0 | 0 | 0 | 0 | 4.40E-06 | 7.72E-05 | 1.44E-05 | 0 | 0 | 0 | 0 | 0.00031 | 0 |
| Ifitm10 | 0 | 4.42E-06 | 0 | 6.48E-06 | 0 | 0 | 0 | 0 | 0 | 0 | 0 | 1.34E-05 | 0 | 0 | 9.03E-06 | 0 | 0.0001 | 4.42E-05 |
| Lsp1 | 0.00011 | 2.58E-05 | 0.00045 | 0 | 0.0007 | 0 | 0.00133 | 0.00069 | 0 | 0.00084 | 0.00061 | 0 | 0.00035 | 0 | 0.00023 | 0.00024 | 0 | 0.0003 |
| Tnnt3 | 0 | 1.50E-06 | 0 | 0 | 5.33E-06 | 0 | 2.55E-05 | 0 | 0 | 2.02E-05 | 0 | 0 | 2.52E-05 | 0 | 6.30E-05 | 0 | 8.03E-05 | 1.15E-05 |
| Camsap3 | 0 | 0 | 0 | 8.33E-06 | 0 | 0 | 0 | 0 | 0 | 7.85E-05 | 0 | 0 | 0 | 0 | 0 | 0 | 0 | 6.47E-05 |
| Prr36 | 5.82E-07 | 1.93E-06 | 2.44E-06 | 2.67E-06 | 0 | 0 | 1.75E-06 | 6.06E-06 | 0 | 0 | 2.75E-06 | 1.61E-05 | 1.76E-06 | 0 | 7.85E-06 | 3.90E-06 | 4.29E-06 | 3.39E-06 |
| Shcbp1 | 1.43E-06 | 0 | 0 | 0 | 0 | 0 | 0 | 0 | 0 | 0 | 0 | 0 | 0 | 0 | 0 | 0.00054 | 0 | 0 |
| Mcf2l | 0 | 0 | 5.59E-05 | 0 | 0 | 0 | 1.28E-05 | 1.76E-05 | 0 | 8.40E-05 | 0 | 1.56E-05 | 0 | 0 | 0 | 6.73E-06 | 0 | 0 |
| F10 | 0 | 2.43E-06 | 0 | 1.04E-05 | 0 | 0 | 0 | 0 | 0 | 1.51E-06 | 0 | 1.37E-06 | 5.82E-06 | 0 | 3.83E-05 | 8.19E-06 | 5.81E-06 | 5.79E-06 |
| Grtp1 | 0 | 2.22E-05 | 0 | 0 | 0 | 0 | 0 | 3.83E-05 | 0 | 0 | 0 | 8.16E-06 | 0 | 0 | 0 | 0 | 0 | 3.31E-06 |
| Gas6 | 0 | 0.00011 | 0 | 0 | 1.16E-05 | 0 | 0 | 0 | 0 | 0.00035 | 2.87E-05 | 0 | 0.00086 | 0 | 0 | 0 | 0 | 0.00206 |
| Kbtbd11 | 0 | 0 | 0 | 1.69E-05 | 0 | 0 | 4.92E-06 | 0 | 0 | 0 | 0 | 0.00011 | 1.36E-05 | 0 | 0 | 0 | 0 | 1.55E-05 |
| Ckap2 | 6.87E-06 | 2.31E-05 | 0 | 0 | 0 | 0 | 0 | 0 | 0 | 2.69E-05 | 0 | 7.40E-06 | 0 | 0 | 4.21E-05 | 0.0008 | 0 | 1.89E-05 |
| Star | 0 | 3.08E-06 | 1.25E-05 | 0 | 0 | 0 | 3.27E-05 | 0 | 0 | 0 | 0 | 1.82E-05 | 1.59E-05 | 0 | 1.06E-05 | 0 | 1.19E-05 | 0 |
| Dlc1 | 5.22E-05 | 7.62E-05 | 3.44E-05 | 3.34E-05 | 0.00021 | 0 | 4.21E-05 | 0.00017 | 0 | 0.00046 | 0.00043 | 0.00046 | 9.11E-05 | 0 | 6.41E-05 | 7.71E-05 | 0.00032 | 0.00027 |
| Msr1 | 0 | 3.50E-06 | 0 | 1.23E-06 | 0 | 0 | 0 | 3.87E-05 | 0 | 8.84E-06 | 0 | 5.52E-06 | 7.04E-06 | 0 | 2.69E-05 | 6.52E-06 | 0 | 0 |
| Neil3 | 0 | 0 | 0 | 0 | 0 | 0 | 0 | 0 | 0 | 0 | 0 | 0 | 0 | 0 | 0 | 0.00043 | 0 | 0 |
| Hapln4 | 0 | 0 | 4.14E-05 | 3.62E-05 | 2.17E-07 | 0 | 0.00016 | 0 | 0 | 0 | 0 | 0 | 4.55E-06 | 0 | 1.85E-05 | 0 | 0 | 0 |
| Lrrc25 | 0 | 5.04E-06 | 0 | 1.01E-05 | 0 | 0 | 0 | 0 | 0 | 1.90E-06 | 1.36E-06 | 6.41E-06 | 1.87E-06 | 0 | 1.67E-05 | 3.52E-06 | 8.63E-06 | 7.93E-06 |

| | | | | | | | | | | | | | | | | | | |
|---|---|---|---|---|---|---|---|---|---|---|---|---|---|---|---|---|---|---|
| Gdf15 | 0 | 8.35E-06 | 0 | 2.28E-05 | 0 | 0 | 0 | 0 | 0 | 0 | 0 | 1.87E-06 | 1.45E-05 | 0 | 2.61E-05 | 1.64E-06 | 4.10E-05 | 0 |
| Iqcn | 0 | 0 | 0 | 0 | 0 | 0 | 0 | 0 | 0 | 0 | 0 | 0.00025 | 0.00055 | 0 | 0 | 0 | 0.00177 | 0 |
| Pde4c | 0 | 0 | 0 | 0 | 0 | 0 | 0 | 0 | 0 | 0 | 0 | 0.00011 | 0.00026 | 0 | 0 | 0 | 0.00051 | 0 |
| Ankle1 | 0 | 0 | 0 | 0 | 0 | 0 | 0 | 0 | 0 | 0 | 0 | 0 | 0 | 0 | 0.00017 | 0.00028 | 0 | 0 |
| Plvap | 0 | 3.40E-05 | 0 | 0 | 0 | 0 | 0 | 8.46E-05 | 0 | 5.21E-06 | 0 | 0 | 0 | 0 | 0 | 1.80E-05 | 0 | 5.59E-05 |
| Bst2 | 0.0002 | 4.70E-05 | 4.33E-05 | 0.00018 | 0 | 0 | 0 | 0 | 0 | 0.0009 | 0 | 9.38E-05 | 0.00058 | 0 | 0 | 7.41E-05 | 0.00023 | 0.00068 |
| Klf2 | 0 | 0 | 0 | 4.55E-05 | 0 | 0 | 0 | 0.00011 | 0 | 0.00049 | 3.55E-05 | 2.22E-05 | 0 | 0 | 7.02E-05 | 0 | 0.00192 | 0.00037 |
| Mcm5 | 0 | 0 | 0.00016 | 0 | 0 | 0 | 0 | 0 | 0 | 1.01E-05 | 0 | 0 | 1.62E-05 | 0 | 0 | 0.00056 | 0 | 0 |
| Asf1b | 0 | 0 | 2.27E-05 | 0 | 0 | 0 | 0 | 7.79E-07 | 0 | 1.19E-06 | 0 | 0 | 0 | 0 | 0 | 0.00064 | 0 | 0 |
| Dand5 | 0 | 0 | 0 | 1.20E-05 | 0 | 0 | 0 | 0 | 0 | 5.04E-07 | 0 | 6.21E-05 | 6.46E-05 | 0 | 0 | 0 | 0.00033 | 0 |
| Adcy7 | 0 | 6.11E-05 | 0.00014 | 0 | 0.00032 | 0 | 0.0002 | 0.00038 | 0 | 0.00016 | 0.00023 | 0 | 0 | 0 | 0.0001 | 5.12E-05 | 0 | 0.0005 |
| Lpcat2 | 0 | 2.16E-05 | 0 | 1.40E-06 | 0 | 0 | 7.86E-06 | 0 | 0 | 6.48E-06 | 0 | 0 | 2.71E-05 | 0 | 1.32E-05 | 0 | 1.81E-05 | 5.89E-06 |
| Ccl17 | 0 | 1.56E-06 | 0 | 2.68E-05 | 6.49E-08 | 0 | 0 | 1.72E-07 | 0 | 3.36E-06 | 0 | 6.79E-06 | 1.01E-05 | 0 | 7.91E-06 | 4.99E-06 | 0 | 9.53E-06 |
| Ces2g | 0 | 0 | 0 | 0 | 0 | 0 | 7.96E-05 | 0 | 0 | 0.00057 | 0 | 0 | 0.00026 | 0 | 0 | 3.34E-05 | 0 | 0.00037 |
| Tppp3 | 0 | 0 | 0 | 0 | 0 | 0 | 0 | 0.00015 | 0 | 0.00105 | 5.53E-05 | 0 | 0 | 0 | 0 | 2.55E-05 | 0 | 0.00067 |
| Carmil2 | 0 | 2.59E-06 | 3.90E-06 | 9.47E-07 | 0 | 0 | 0 | 0 | 0 | 0 | 0 | 2.23E-06 | 4.20E-06 | 0 | 9.45E-06 | 9.87E-06 | 2.99E-05 | 0 |
| Esrp2 | 0 | 0 | 0 | 0 | 7.83E-07 | 0 | 0 | 0 | 0 | 0 | 4.02E-05 | 0 | 0 | 0 | 2.52E-05 | 0 | 0 | 0 |
| Smpd3 | 0 | 0 | 0 | 0 | 6.96E-06 | 0 | 0 | 0 | 0 | 0.00016 | 0 | 0 | 0 | 0 | 0 | 0 | 0 | 0 |
| Nqo1 | 5.60E-06 | 3.48E-06 | 0 | 0 | 0 | 0 | 0 | 0 | 0 | 7.02E-06 | 0 | 0 | 0 | 0 | 0 | 2.78E-05 | 0 | 1.71E-05 |
| Zfhx3 | 0 | 1.93E-05 | 0 | 4.11E-05 | 0 | 0 | 0.00025 | 3.01E-05 | 0 | 0.00015 | 2.66E-05 | 2.60E-05 | 0.00026 | 0 | 2.94E-05 | 2.25E-05 | 0.00016 | 2.99E-05 |
| Sdr42e1 | 0 | 6.98E-06 | 0 | 5.64E-06 | 1.38E-06 | 0 | 0 | 0 | 0 | 2.51E-05 | 0 | 3.05E-06 | 1.11E-05 | 0 | 0 | 0 | 0 | 1.18E-06 |
| Cotl1 | 0 | 0.0001 | 2.81E-05 | 0.00028 | 9.75E-05 | 0 | 0 | 0.00081 | 0 | 0.0005 | 0.00076 | 1.44E-05 | 0.00017 | 0 | 0 | 0.00019 | 0 | 5.46E-05 |
| Sult5a1 | 0 | 5.94E-06 | 0 | 1.14E-06 | 1.04E-05 | 0 | 0 | 2.04E-05 | 0 | 8.75E-05 | 0 | 0 | 8.06E-05 | 0 | 1.81E-05 | 0 | 0 | 9.05E-05 |
| Fanca | 0 | 0 | 0 | 0 | 0 | 0 | 0 | 0 | 0 | 0 | 0 | 0 | 0 | 0 | 3.19E-05 | 0.00043 | 0 | 0 |
| Tubb3 | 6.35E-05 | 0 | 0.00011 | 0 | 0 | 0 | 0 | 0.0001 | 0 | 1.33E-05 | 0.00015 | 0 | 0 | 0 | 0 | 0.00022 | 0 | 2.80E-05 |
| Cfap300 | 2.29E-05 | 9.33E-06 | 7.90E-06 | 1.83E-05 | 8.12E-06 | 0 | 2.20E-05 | 0 | 0 | 1.82E-05 | 0 | 1.70E-05 | 2.19E-05 | 0 | 2.12E-05 | 0 | 1.47E-05 | 0 |
| Panx1 | 7.00E-05 | 3.60E-05 | 0.0001 | 0 | 3.60E-05 | 0 | 0.00018 | 8.42E-05 | 0 | 0.00013 | 2.72E-08 | 0 | 3.07E-06 | 0 | 0 | 4.66E-05 | 0 | 0.00016 |
| Olfm2 | 0 | 1.01E-05 | 0 | 0 | 0 | 0 | 0 | 4.30E-05 | 0 | 0 | 4.51E-06 | 3.30E-05 | 0 | 0 | 1.13E-06 | 0 | 6.44E-07 | 0 |
| Rdh8 | 0 | 0 | 2.24E-05 | 0 | 0 | 0 | 6.34E-06 | 2.00E-06 | 0 | 0 | 0 | 3.81E-06 | 0 | 0 | 0 | 4.78E-06 | 0 | 0 |
| Angptl6 | 0 | 1.05E-06 | 0 | 0 | 0 | 0 | 0 | 0 | 0 | 4.62E-06 | 0 | 5.67E-05 | 0 | 0 | 0 | 6.51E-06 | 0 | 1.19E-06 |
| Icam4 | 0 | 0 | 0 | 0 | 0 | 0 | 0 | 0 | 0 | 0 | 0 | 6.01E-06 | 6.18E-05 | 0 | 0 | 0 | 0.00011 | 0 |
| Zglp1 | 0 | 0 | 6.89E-06 | 9.31E-07 | 0 | 0 | 0 | 0 | 0 | 0 | 0 | 9.91E-06 | 4.41E-06 | 0 | 5.44E-06 | 2.07E-06 | 6.11E-05 | 0 |
| Spc24 | 0 | 0 | 0 | 8.05E-05 | 0 | 0 | 0 | 0 | 0 | 0 | 0 | 0 | 0 | 0 | 8.42E-06 | 0.00082 | 0 | 0 |

| | | | | | | | | | | | | | | | | | | |
|---|---|---|---|---|---|---|---|---|---|---|---|---|---|---|---|---|---|---|
| Cnn1 | 0 | 0 | 0 | 0 | 0 | 0 | 0 | 0 | 0 | 0 | 0.00032 | 0 | 0 | 0 | 0 | 0 | 0 | 0 |
| Acp5 | 0 | 0 | 0 | 8.26E-05 | 5.92E-06 | 0 | 0.00011 | 2.97E-05 | 0 | 2.46E-05 | 0 | 7.54E-06 | 1.33E-05 | 0 | 7.52E-05 | 1.12E-05 | 0 | 0.0002 |
| Bmper | 8.04E-06 | 0 | 0 | 0 | 0 | 0 | 0.00016 | 0 | 0 | 0.00036 | 0 | 0 | 0.00043 | 0 | 0 | 4.83E-05 | 0 | 0.00051 |
| Igsf9b | 0 | 1.05E-05 | 4.95E-06 | 0 | 0 | 0 | 0 | 0 | 0 | 2.94E-05 | 0 | 4.86E-07 | 3.12E-06 | 0 | 0 | 0 | 0 | 0 |
| Opcml | 7.09E-06 | 0 | 7.27E-06 | 0 | 0 | 0 | 0 | 0 | 0 | 0.00013 | 0 | 0 | 0 | 0 | 0 | 3.11E-06 | 0 | 0 |
| Barx2 | 0 | 0 | 0 | 0 | 0 | 0 | 6.73E-05 | 0 | 0 | 6.84E-05 | 0 | 2.21E-05 | 0 | 0 | 0 | 2.61E-06 | 0 | 0 |
| Cadm1 | 0 | 8.56E-06 | 0 | 1.74E-05 | 0 | 0 | 0 | 8.18E-06 | 0 | 1.20E-05 | 1.28E-06 | 7.73E-06 | 4.97E-06 | 0 | 2.58E-05 | 2.38E-06 | 1.45E-05 | 0 |
| Zbtb16 | 0 | 6.37E-06 | 0 | 0 | 0 | 0 | 0 | 0 | 0 | 0.0002 | 0 | 0 | 0 | 0 | 1.66E-05 | 0 | 0 | 0 |
| Arhgap20 | 0 | 0 | 0 | 0 | 0 | 0 | 0 | 0 | 0 | 0.0002 | 0 | 4.99E-06 | 0 | 0 | 0 | 0 | 0 | 3.37E-05 |
| Islr2 | 0 | 2.10E-07 | 0 | 4.74E-05 | 0.00018 | 0 | 2.66E-05 | 7.98E-05 | 0 | 0.00035 | 3.60E-05 | 0 | 0 | 0 | 2.93E-05 | 0 | 0 | 0 |
| Kif23 | 0 | 6.22E-06 | 0 | 0 | 0 | 0 | 0 | 0 | 0 | 0 | 0 | 6.67E-06 | 0 | 0 | 0 | 0.00081 | 0 | 0 |
| Coro2b | 0 | 2.45E-07 | 1.17E-05 | 4.66E-05 | 0 | 0 | 6.10E-05 | 1.23E-05 | 0 | 0.00014 | 0 | 2.19E-05 | 2.10E-05 | 0 | 6.93E-06 | 2.17E-05 | 1.56E-05 | 6.97E-05 |
| Calml4 | 6.65E-07 | 0 | 0 | 0 | 0 | 0 | 2.39E-06 | 0 | 0 | 8.23E-05 | 0 | 1.82E-05 | 0 | 0 | 0 | 0 | 0 | 2.08E-05 |
| Cilp | 0 | 0 | 3.42E-05 | 0 | 9.77E-05 | 0 | 0.00055 | 3.25E-05 | 0 | 0 | 0.00016 | 0 | 0 | 0 | 2.22E-05 | 7.15E-05 | 0 | 0.00189 |
| Pif1 | 0 | 0 | 0 | 0 | 0 | 0 | 0 | 0 | 0 | 0 | 0 | 0 | 0 | 0 | 4.11E-05 | 0.00023 | 0 | 0 |
| Car12 | 0 | 0 | 0.0002 | 4.65E-05 | 0 | 0 | 0.00044 | 0 | 0 | 0 | 3.31E-05 | 0 | 0 | 0 | 8.02E-06 | 1.70E-05 | 0 | 0 |
| Ccnb2 | 9.35E-05 | 0 | 3.04E-06 | 0.00012 | 0 | 0 | 0 | 0 | 0 | 0 | 0 | 0 | 0 | 0 | 0 | 0.00115 | 0 | 0 |
| Ttk | 0 | 0 | 0 | 0 | 0 | 0 | 0 | 0 | 0 | 0 | 0 | 0 | 0 | 0 | 8.04E-06 | 0.00046 | 0 | 0 |
| Prss35 | 0 | 0 | 0 | 0 | 0.00031 | 0 | 4.14E-05 | 0 | 0 | 1.92E-05 | 0.00025 | 0 | 0 | 0 | 0 | 0 | 0 | 0 |
| Snap91 | 0 | 0 | 0 | 0 | 3.02E-05 | 0 | 6.79E-06 | 0 | 0 | 0 | 0 | 1.02E-05 | 0 | 0 | 0 | 0 | 0 | 0 |
| Plscr2 | 1.34E-05 | 0 | 0 | 0 | 0 | 0 | 0 | 0 | 0 | 0.00019 | 0 | 0 | 0.00017 | 0 | 0 | 2.93E-05 | 0 | 9.19E-05 |
| Plod2 | 0.00024 | 0 | 6.01E-05 | 0 | 0 | 0 | 0.00073 | 0.0002 | 0 | 0.00039 | 0 | 0.00083 | 0.00036 | 0 | 0 | 0.00012 | 0 | 0.00026 |
| Acy1 | 1.93E-05 | 1.47E-05 | 2.48E-05 | 4.00E-05 | 5.14E-06 | 0 | 1.55E-05 | 9.63E-10 | 0 | 2.63E-05 | 0 | 1.25E-08 | 4.74E-06 | 0 | 1.69E-05 | 3.52E-05 | 6.57E-06 | 1.40E-05 |
| Gpr62 | 0 | 3.49E-06 | 0 | 7.40E-07 | 0 | 0 | 0 | 0 | 0 | 1.15E-06 | 0 | 3.95E-06 | 6.55E-06 | 0 | 0 | 2.28E-06 | 6.11E-05 | 1.46E-06 |
| Sema3b | 0 | 2.44E-05 | 0 | 6.53E-05 | 4.03E-05 | 0 | 0.00029 | 0 | 0 | 0.00012 | 0 | 5.57E-05 | 0.00021 | 0 | 0 | 0 | 0 | 0 |
| Traip | 7.65E-07 | 0 | 4.38E-06 | 0 | 0 | 0 | 8.59E-07 | 0 | 0 | 5.99E-06 | 0 | 0 | 1.02E-06 | 0 | 5.16E-05 | 0.0002 | 0 | 1.77E-06 |
| Gmppb | 0.00027 | 0 | 0.00031 | 0.00021 | 0 | 0 | 0.00026 | 0.0001 | 0 | 0 | 0.00011 | 0 | 0 | 0 | 0.0001 | 0 | 0 | 2.10E-05 |
| Amigo3 | 1.31E-05 | 0 | 0.00012 | 8.37E-05 | 0 | 0 | 9.63E-05 | 6.49E-05 | 0 | 0 | 2.74E-05 | 0 | 0 | 0 | 3.30E-05 | 0 | 8.99E-05 | 2.23E-05 |
| Cspg5 | 0 | 0 | 1.87E-06 | 6.79E-06 | 0 | 0 | 0 | 0 | 0 | 1.33E-06 | 0 | 0 | 1.65E-05 | 0 | 0 | 1.55E-06 | 5.01E-05 | 0 |
| Prss46 | 0 | 0 | 8.63E-06 | 0 | 0 | 0 | 6.81E-06 | 0 | 0 | 0 | 5.04E-05 | 3.23E-06 | 1.60E-05 | 0 | 0 | 0 | 0 | 0 |
| Klhl40 | 0 | 0 | 0 | 0 | 0 | 0 | 0 | 0 | 0 | 0 | 0 | 7.39E-06 | 2.22E-05 | 0 | 0 | 0 | 0.00011 | 0 |
| Kif15 | 0 | 0 | 0 | 0 | 0 | 0 | 0 | 0 | 0 | 0 | 0 | 0 | 0 | 0 | 4.92E-05 | 0.00058 | 0 | 0 |
| Clec3b | 0 | 0 | 0 | 0 | 0 | 0 | 0 | 0 | 0 | 0.00168 | 0 | 0 | 0 | 0 | 0 | 0 | 0 | 0.00032 |

| | | | | | | | | | | | | | | | | | | |
|---|---|---|---|---|---|---|---|---|---|---|---|---|---|---|---|---|---|---|
| Epm2a | 0 | 7.48E-06 | 0 | 0 | 2.39E-05 | 0 | 0 | 1.49E-05 | 0 | 2.75E-06 | 0 | 0.00017 | 0 | 0 | 0 | 0 | 0 | 3.14E-05 |
| Plagl1 | 0 | 0 | 0 | 0.00011 | 0 | 0 | 0.00049 | 0 | 0 | 0.00041 | 0 | 0 | 0.00011 | 0 | 0 | 6.65E-05 | 0.0003 | 0 |
| Cited2 | 0 | 0 | 0 | 0 | 0 | 0 | 0 | 0 | 0 | 0.00018 | 0.00014 | 0.00016 | 0.00016 | 0 | 0 | 0 | 0.00071 | 0 |
| Nhsl1 | 0 | 5.97E-05 | 4.82E-05 | 0 | 9.49E-05 | 0 | 3.02E-06 | 0.00032 | 0 | 0.0003 | 1.50E-05 | 4.12E-05 | 0 | 0 | 0 | 5.19E-05 | 7.70E-05 | 0 |
| Map7 | 0 | 1.55E-05 | 0 | 2.13E-06 | 0 | 0 | 1.85E-06 | 0 | 0 | 1.18E-05 | 0 | 1.31E-05 | 1.66E-05 | 0 | 1.32E-05 | 0 | 0 | 1.20E-05 |
| Mtfr2 | 0 | 0 | 4.97E-07 | 0 | 0 | 0 | 0 | 4.23E-07 | 0 | 4.13E-07 | 0 | 1.84E-05 | 0 | 0 | 8.73E-06 | 0.00024 | 9.82E-05 | 0 |
| Ccn2 | 0 | 0 | 0 | 0 | 0 | 0 | 0 | 0 | 0 | 0.00022 | 0.00263 | 0 | 0.00035 | 0 | 0 | 0 | 8.81E-05 | 0.00106 |
| Enpp1 | 0 | 0 | 9.17E-05 | 0 | 0 | 0 | 0.00026 | 0.00058 | 0 | 0 | 0.00014 | 0.00012 | 6.46E-05 | 0 | 0 | 1.01E-05 | 0 | 0.00013 |
| Enpp3 | 4.11E-05 | 5.86E-06 | 0 | 4.03E-05 | 1.27E-05 | 0 | 7.88E-05 | 4.96E-05 | 0 | 2.83E-05 | 0 | 0 | 3.65E-05 | 0 | 0 | 6.40E-05 | 0 | 2.38E-06 |
| Arg1 | 0 | 0 | 3.36E-06 | 0.0002 | 0.00011 | 0 | 0.00012 | 0.00039 | 0 | 0 | 0 | 0.00024 | 0 | 0 | 0.00017 | 0 | 0 | 0 |
| Tube1 | 0 | 0 | 0 | 0 | 0 | 0 | 0 | 0 | 0 | 5.18E-06 | 0 | 2.57E-05 | 3.78E-05 | 0 | 2.65E-05 | 0.00012 | 9.79E-05 | 0 |
| Slc16a10 | 0 | 4.35E-05 | 1.52E-05 | 0 | 2.63E-05 | 0 | 9.10E-06 | 0 | 0 | 4.98E-05 | 8.04E-06 | 1.42E-05 | 5.22E-05 | 0 | 1.00E-05 | 4.58E-05 | 0.00026 | 0 |
| Sim1 | 0 | 0 | 0 | 3.30E-06 | 2.39E-06 | 0 | 2.18E-05 | 0 | 0 | 3.90E-06 | 9.62E-06 | 4.52E-07 | 1.81E-05 | 0 | 0 | 4.95E-06 | 6.21E-06 | 0 |
| Ppa1 | 0.00027 | 0 | 0.0006 | 6.96E-05 | 0 | 0 | 0.00047 | 0 | 0 | 0.00035 | 8.60E-05 | 0 | 0 | 0 | 0 | 0.00013 | 0.00033 | 0 |
| Tspan15 | 0 | 5.56E-06 | 0 | 1.20E-05 | 0 | 0 | 0 | 0 | 0 | 2.05E-06 | 3.17E-06 | 4.12E-06 | 0 | 0 | 1.04E-05 | 9.22E-06 | 0 | 7.04E-06 |
| Zfp365 | 2.27E-06 | 1.54E-07 | 1.06E-05 | 1.88E-06 | 0 | 0 | 0 | 0 | 0 | 3.17E-05 | 0 | 5.27E-06 | 0 | 0 | 0 | 7.44E-07 | 1.48E-06 | 2.17E-05 |
| Slc16a9 | 0 | 0 | 0 | 0 | 0 | 0 | 4.61E-05 | 0 | 0 | 1.99E-05 | 0 | 0 | 0 | 0 | 0 | 0 | 0 | 6.45E-05 |
| Adora2a | 0 | 0 | 0 | 0 | 0 | 0 | 0 | 0 | 0 | 7.64E-07 | 0 | 2.39E-05 | 3.37E-05 | 0 | 0 | 4.56E-07 | 3.30E-05 | 7.31E-05 |
| Ggt5 | 0 | 4.94E-05 | 0 | 0.00012 | 0 | 0 | 0 | 0 | 0 | 0.00055 | 0 | 0 | 0.00066 | 0 | 0 | 0 | 0 | 0 |
| Gzmm | 0 | 0 | 0 | 0 | 0 | 0 | 0 | 0 | 0 | 0 | 0 | 1.78E-05 | 1.61E-05 | 0 | 0 | 1.15E-06 | 0.00013 | 0 |
| Misp | 0 | 8.81E-06 | 0 | 5.21E-06 | 0 | 0 | 7.69E-06 | 0 | 0 | 6.71E-06 | 0 | 3.48E-06 | 1.60E-05 | 0 | 1.29E-05 | 1.33E-06 | 3.65E-05 | 5.36E-06 |
| Elane | 0 | 0 | 0 | 2.81E-06 | 0 | 0 | 0 | 0 | 0 | 8.46E-07 | 0 | 2.28E-05 | 2.23E-05 | 0 | 0 | 0 | 6.63E-05 | 0 |
| Arhgap45 | 1.02E-05 | 1.84E-06 | 5.54E-06 | 2.43E-05 | 0 | 0 | 3.56E-05 | 8.40E-06 | 0 | 1.83E-05 | 0 | 1.39E-05 | 0 | 0 | 4.27E-05 | 4.26E-06 | 8.51E-06 | 6.64E-05 |
| Klf16 | 0 | 5.62E-05 | 0.00011 | 0.00022 | 0 | 0 | 0 | 5.32E-05 | 0 | 2.69E-05 | 4.26E-05 | 4.62E-05 | 0 | 0 | 0.00013 | 5.34E-06 | 0.00037 | 5.30E-05 |
| Eid3 | 0 | 0 | 1.31E-05 | 5.11E-06 | 0 | 0 | 0 | 0 | 0 | 0 | 0 | 0 | 0 | 0 | 0.00011 | 3.13E-05 | 0.00018 | 4.87E-06 |
| E2f7 | 0 | 0 | 0 | 0 | 0 | 0 | 0 | 0 | 0 | 9.72E-07 | 0 | 0 | 0 | 0 | 8.91E-05 | 0.00039 | 0 | 0 |
| Csrp2 | 0 | 0 | 0.0002 | 4.08E-05 | 0 | 0 | 0.00157 | 0.00035 | 0 | 0.00051 | 0.00064 | 0.0002 | 0.00012 | 0 | 0 | 0.00024 | 0 | 0.00069 |
| Ptprb | 0 | 0 | 0 | 0 | 1.07E-05 | 0 | 0 | 1.90E-05 | 0 | 0 | 1.52E-05 | 1.79E-06 | 0 | 0 | 0 | 0 | 0 | 0.00012 |
| Tafa2 | 0 | 0 | 0 | 0 | 0 | 0 | 0 | 0 | 0 | 0 | 0 | 0.00023 | 0 | 0 | 0 | 0 | 0 | 0 |
| Avil | 0 | 1.98E-05 | 0 | 2.25E-05 | 0 | 0 | 6.92E-06 | 0 | 0 | 2.59E-05 | 0 | 3.52E-05 | 3.58E-06 | 0 | 6.21E-05 | 7.48E-06 | 0.00011 | 2.64E-05 |
| Eef1akmt3 | 3.89E-05 | 0 | 5.67E-05 | 9.03E-05 | 0 | 0 | 0.00014 | 1.24E-05 | 0 | 0 | 0 | 2.00E-05 | 3.81E-05 | 0 | 6.42E-05 | 0 | 0 | 1.83E-06 |
| Cyp27b1 | 0 | 1.63E-06 | 2.78E-06 | 3.41E-06 | 0 | 0 | 0 | 0 | 0 | 5.42E-07 | 1.39E-06 | 1.22E-05 | 1.16E-05 | 0 | 0 | 4.73E-07 | 2.42E-05 | 6.74E-07 |
| 9-Mar | 0 | 4.83E-05 | 0 | 4.24E-06 | 3.16E-05 | 0 | 0 | 0 | 0 | 4.54E-06 | 0 | 7.97E-05 | 0 | 0 | 0 | 0 | 0 | 8.53E-06 |

| | | | | | | | | | | | | | | | | | | |
|---|---|---|---|---|---|---|---|---|---|---|---|---|---|---|---|---|---|---|
| Ddit3 | 0 | 0.00014 | 4.66E-05 | 5.37E-05 | 0 | 0 | 0 | 0 | 0 | 3.82E-05 | 0 | 0.0005 | 0 | 0 | 0.00023 | 1.81E-05 | 0.00069 | 0 |
| Stac3 | 2.18E-06 | 3.06E-06 | 2.18E-06 | 5.09E-06 | 0 | 0 | 0 | 3.07E-06 | 0 | 0 | 0 | 8.50E-08 | 8.75E-07 | 0 | 2.89E-05 | 1.25E-07 | 3.67E-05 | 0 |
| Nxph4 | 0 | 0 | 0 | 4.34E-06 | 0 | 0 | 1.03E-05 | 0 | 0 | 0 | 0 | 0.00031 | 5.33E-05 | 0 | 0 | 0 | 7.01E-05 | 0 |
| Gls2 | 0 | 0 | 1.95E-05 | 0 | 0 | 0 | 2.86E-05 | 0 | 0 | 0 | 2.12E-06 | 4.47E-05 | 0 | 0 | 9.81E-06 | 1.33E-05 | 0 | 7.19E-06 |
| Myl6b | 0 | 1.39E-05 | 0 | 1.69E-06 | 1.05E-05 | 0 | 2.52E-05 | 5.99E-06 | 0 | 3.28E-06 | 0 | 0 | 2.17E-05 | 0 | 1.35E-05 | 1.35E-06 | 0 | 1.29E-05 |
| Ormdl2 | 1.47E-05 | 2.61E-05 | 5.77E-06 | 0 | 0 | 0 | 0 | 0 | 0 | 4.45E-05 | 0 | 2.52E-05 | 2.58E-06 | 0 | 0 | 1.03E-05 | 0 | 2.53E-05 |
| Pik3ip1 | 0 | 0.00014 | 0 | 3.74E-06 | 1.16E-05 | 0 | 0 | 0 | 0 | 8.03E-05 | 3.02E-05 | 4.50E-05 | 2.37E-06 | 0 | 5.11E-05 | 0 | 0 | 0.00017 |
| Osbp2 | 0 | 6.53E-06 | 0 | 0 | 6.98E-05 | 0 | 0 | 4.31E-05 | 0 | 3.98E-06 | 0 | 1.15E-05 | 0 | 0 | 0 | 0 | 3.31E-05 | 0 |
| Lif | 0 | 0 | 0.00025 | 0 | 0 | 0 | 9.59E-05 | 0 | 0 | 0 | 0 | 8.38E-05 | 0.0003 | 0 | 0 | 0 | 0.00016 | 4.81E-05 |
| Myo1g | 0 | 9.18E-07 | 0 | 1.47E-05 | 0 | 0 | 0 | 0 | 0 | 1.55E-06 | 3.66E-07 | 3.35E-06 | 4.77E-06 | 0 | 1.44E-05 | 4.89E-06 | 2.26E-05 | 5.40E-06 |
| Nacad | 0 | 0 | 0 | 0 | 0 | 0 | 0 | 0 | 0 | 6.37E-06 | 3.37E-05 | 6.48E-05 | 0 | 0 | 0 | 0 | 0 | 1.61E-05 |
| Fbxo48 | 0 | 0 | 0 | 0 | 0 | 0 | 0 | 0 | 0 | 2.26E-06 | 0 | 1.53E-05 | 0.0001 | 0 | 0 | 0.00013 | 0.00013 | 0 |
| Slc1a4 | 1.21E-05 | 4.17E-06 | 6.67E-05 | 3.68E-05 | 3.72E-05 | 0 | 0.00032 | 1.61E-05 | 0 | 6.94E-05 | 0.00013 | 0.00017 | 0 | 0 | 0 | 0 | 0 | 7.17E-05 |
| Ccdc85a | 0 | 0 | 0 | 4.19E-06 | 0 | 0 | 0 | 0 | 0 | 8.04E-05 | 0 | 0 | 4.11E-05 | 0 | 0 | 0 | 0 | 0 |
| Eml6 | 9.11E-06 | 1.66E-06 | 8.85E-07 | 0 | 0 | 0 | 4.03E-06 | 6.08E-06 | 0 | 2.56E-05 | 0 | 1.20E-05 | 0 | 0 | 2.71E-06 | 0 | 0 | 1.15E-05 |
| Fgf18 | 0 | 0 | 4.08E-05 | 0 | 0 | 0 | 7.88E-05 | 0 | 0 | 0.00012 | 0 | 0.00015 | 4.89E-06 | 0 | 0 | 0 | 0.00016 | 0.00019 |
| Spdl1 | 0 | 0 | 0 | 0 | 0 | 0 | 0 | 0 | 0 | 0 | 0 | 0 | 0 | 0 | 2.33E-05 | 0.00044 | 0 | 0 |
| Hmmr | 0 | 0 | 0 | 0 | 0 | 0 | 0 | 1.49E-06 | 0 | 2.44E-06 | 0 | 0 | 0 | 0 | 2.46E-05 | 0.00103 | 0 | 0 |
| Trim7 | 0 | 1.41E-05 | 3.04E-06 | 0 | 0 | 0 | 3.93E-06 | 0 | 0 | 1.12E-06 | 9.57E-06 | 9.46E-06 | 1.66E-05 | 0 | 8.48E-06 | 0 | 0 | 0 |
| Ltc4s | 0 | 2.66E-05 | 0 | 9.30E-06 | 0 | 0 | 0 | 0 | 0 | 1.74E-05 | 0 | 5.92E-06 | 1.24E-05 | 0 | 0 | 0 | 5.02E-06 | 2.99E-05 |
| Acsl6 | 0 | 1.11E-05 | 0 | 0 | 0 | 0 | 0 | 0 | 0 | 3.64E-06 | 2.56E-05 | 1.24E-05 | 6.97E-06 | 0 | 1.27E-05 | 0 | 0 | 0 |
| Gjc2 | 0 | 9.30E-07 | 0 | 1.91E-06 | 4.32E-06 | 0 | 0 | 5.87E-06 | 0 | 8.87E-06 | 0 | 2.14E-05 | 0 | 0 | 0 | 0 | 3.65E-06 | 3.53E-05 |
| Wnt9a | 0 | 3.88E-05 | 3.67E-05 | 0 | 0.00011 | 0 | 0.00058 | 6.91E-05 | 0 | 0 | 2.63E-05 | 0 | 0 | 0 | 1.18E-05 | 0 | 0 | 0.00034 |
| Shmt1 | 0 | 0 | 1.84E-05 | 0 | 0 | 0 | 0 | 0 | 0 | 0 | 0 | 0 | 8.09E-06 | 0 | 0 | 7.38E-05 | 0 | 0 |
| Rcvrn | 0 | 0 | 0 | 0 | 0 | 0 | 2.88E-06 | 0 | 0 | 2.67E-05 | 0 | 2.52E-06 | 0 | 0 | 0 | 0 | 8.55E-05 | 1.19E-05 |
| Pik3r5 | 0 | 3.04E-06 | 0 | 0 | 0 | 0 | 0 | 0 | 0 | 4.11E-05 | 0 | 1.04E-05 | 0 | 0 | 1.39E-05 | 1.60E-06 | 0 | 8.50E-05 |
| Aurkb | 0 | 0 | 0 | 0 | 0 | 0 | 0 | 0 | 0 | 0 | 0 | 0 | 0 | 0 | 4.24E-05 | 0.00062 | 0 | 0 |
| Tmem88 | 0 | 0 | 0.0001 | 0 | 0 | 0 | 0 | 0 | 0 | 0 | 0 | 0 | 6.18E-05 | 0 | 0 | 0 | 0.0011 | 0 |
| Tnk1 | 0 | 0 | 0 | 1.32E-05 | 0 | 0 | 1.05E-05 | 0 | 0 | 2.33E-05 | 0 | 8.99E-06 | 6.51E-05 | 0 | 0 | 0 | 1.58E-05 | 4.78E-05 |
| Mgl2 | 0 | 1.54E-06 | 0 | 9.92E-06 | 2.78E-06 | 0 | 2.24E-06 | 3.62E-06 | 0 | 2.78E-06 | 0 | 4.13E-07 | 2.67E-06 | 0 | 1.29E-05 | 4.90E-06 | 1.32E-05 | 0 |
| Pimreg | 0 | 0 | 0 | 0 | 0 | 0 | 0 | 0 | 0 | 0 | 0 | 0 | 0 | 0 | 9.20E-05 | 0.00086 | 0 | 0 |
| Smtnl2 | 0 | 0 | 0 | 0 | 0 | 0 | 0 | 0 | 0 | 1.69E-06 | 0 | 0.00029 | 0 | 0 | 0 | 0 | 0 | 1.39E-05 |
| Fam57a | 7.24E-05 | 0 | 7.95E-05 | 1.77E-06 | 0 | 0 | 3.19E-05 | 0 | 0 | 3.34E-05 | 3.96E-05 | 0.00012 | 0 | 0 | 0 | 0.00012 | 0 | 4.26E-05 |

| | | | | | | | | | | | | | | | | | | |
|---|---|---|---|---|---|---|---|---|---|---|---|---|---|---|---|---|---|---|
| Spag5 | 0 | 0 | 0 | 0 | 0 | 0 | 0 | 0 | 0 | 0 | 0 | 0 | 0 | 0 | 0.00022 | 0.00059 | 0 | 0 |
| Vtn | 0 | 0 | 0 | 0 | 0 | 0 | 0 | 0 | 0 | 1.87E-05 | 0 | 0 | 4.60E-05 | 0 | 0 | 0 | 0 | 0.00011 |
| Nos2 | 0 | 0 | 0 | 0 | 0 | 0 | 5.60E-05 | 0 | 0 | 1.48E-05 | 0 | 0.00035 | 0.00011 | 0 | 0 | 0 | 0 | 0 |
| Cdk5r1 | 0 | 3.41E-06 | 2.23E-06 | 4.94E-06 | 9.41E-08 | 0 | 0 | 9.95E-07 | 0 | 0 | 2.08E-05 | 2.76E-05 | 0 | 0 | 3.83E-05 | 4.85E-06 | 5.19E-05 | 0 |
| Ccl2 | 5.20E-06 | 0 | 0.0002 | 0 | 0 | 0 | 0 | 0 | 0 | 3.19E-05 | 0 | 0 | 0.00068 | 0 | 0 | 0.00013 | 0.00066 | 4.69E-05 |
| Ccl7 | 0.00012 | 0 | 0.0002 | 0 | 0 | 0 | 5.26E-05 | 0 | 0 | 0.00025 | 5.84E-06 | 0 | 0.00122 | 0 | 0 | 8.79E-05 | 0.00104 | 0 |
| Ccl8 | 1.50E-06 | 2.35E-06 | 1.96E-05 | 0 | 5.04E-05 | 0 | 0.00027 | 0.00016 | 0 | 0.00024 | 0 | 0 | 0.00081 | 0 | 0 | 3.24E-05 | 0 | 0 |
| Unc45b | 0 | 2.13E-05 | 0 | 0 | 5.38E-07 | 0 | 2.01E-05 | 0 | 0 | 0.00013 | 0 | 0 | 8.16E-05 | 0 | 4.10E-05 | 0 | 0 | 0 |
| Ccl5 | 0 | 5.67E-06 | 1.02E-05 | 0 | 0 | 0 | 0 | 0 | 0 | 7.92E-06 | 0 | 0 | 1.72E-05 | 0 | 1.58E-05 | 1.18E-05 | 1.47E-05 | 7.29E-06 |
| Ccl6 | 0 | 4.51E-06 | 0 | 2.85E-05 | 2.34E-07 | 0 | 0 | 2.37E-06 | 0 | 1.22E-05 | 0 | 9.99E-06 | 4.76E-06 | 0 | 5.41E-05 | 9.69E-06 | 4.42E-06 | 7.27E-06 |
| Ccl4 | 0 | 6.91E-06 | 1.90E-06 | 2.70E-05 | 3.52E-06 | 0 | 0 | 3.81E-06 | 0 | 1.05E-05 | 0 | 4.54E-06 | 7.30E-06 | 0 | 5.79E-05 | 1.62E-05 | 3.98E-05 | 6.26E-06 |
| Dusp14 | 9.63E-06 | 1.81E-08 | 1.04E-05 | 0 | 2.19E-05 | 0 | 3.92E-06 | 0 | 0 | 4.63E-06 | 2.61E-07 | 0 | 0 | 0 | 0 | 0 | 1.55E-06 | 5.48E-07 |
| Prr11 | 0 | 0 | 0 | 0 | 0 | 0 | 0 | 0 | 0 | 0 | 0 | 6.66E-06 | 0 | 0 | 0 | 0.00103 | 0 | 0 |
| Eme1 | 0 | 0 | 0 | 0 | 0 | 0 | 0 | 0 | 0 | 6.38E-07 | 0 | 0 | 0 | 0 | 6.75E-05 | 0.00013 | 0 | 0 |
| Ngfr | 3.42E-06 | 0 | 0 | 4.02E-05 | 0 | 0 | 0 | 4.46E-05 | 0 | 0.00021 | 0 | 0 | 3.24E-05 | 0 | 0 | 2.29E-05 | 0 | 0 |
| Sp6 | 0 | 0 | 0 | 0 | 2.24E-05 | 0 | 0 | 7.06E-05 | 0 | 0 | 1.64E-05 | 5.31E-05 | 0 | 0 | 0 | 0 | 0 | 0 |
| Top2a | 0 | 0 | 0 | 0 | 0 | 0 | 0 | 0 | 0 | 0 | 0 | 0 | 0 | 0 | 0.00011 | 0.00213 | 0 | 0 |
| Krt20 | 0 | 0 | 0 | 1.23E-05 | 2.31E-06 | 0 | 6.21E-06 | 5.00E-06 | 0 | 3.34E-05 | 6.07E-06 | 2.01E-06 | 0 | 0 | 3.43E-05 | 5.42E-06 | 0 | 2.86E-05 |
| Krt19 | 0 | 0 | 0 | 0.00011 | 0 | 0 | 0 | 0 | 0 | 5.47E-05 | 0 | 0.00037 | 2.21E-06 | 0 | 0.00014 | 1.40E-05 | 0 | 3.04E-05 |
| Jup | 0 | 0.00021 | 0 | 0 | 4.18E-05 | 0 | 0 | 0.00052 | 0 | 0 | 9.47E-05 | 1.14E-05 | 0 | 0 | 0 | 0 | 0 | 0.0001 |
| Dhx58 | 1.46E-05 | 8.28E-07 | 0 | 5.57E-06 | 0 | 0 | 0 | 0 | 0 | 0.00016 | 0 | 2.20E-06 | 3.44E-05 | 0 | 7.99E-06 | 2.33E-05 | 0 | 3.93E-05 |
| Arl4d | 0 | 0 | 0 | 0 | 0 | 0 | 0 | 0 | 0 | 0.0001 | 0 | 7.00E-06 | 0 | 0 | 0 | 0 | 0.00023 | 0 |
| Meox1 | 0 | 0 | 3.90E-05 | 0 | 0 | 0 | 0 | 4.25E-05 | 0 | 0 | 0 | 0 | 0 | 0 | 0 | 8.65E-06 | 0 | 8.88E-05 |
| BC030867 | 0 | 0 | 6.98E-08 | 0 | 0 | 0 | 0 | 0 | 0 | 0 | 0 | 7.38E-06 | 0 | 0 | 1.22E-05 | 0.0005 | 0 | 0 |
| Kif18b | 0 | 0 | 0 | 0 | 0 | 0 | 0 | 0 | 0 | 0 | 0 | 0 | 0 | 0 | 1.42E-05 | 0.00066 | 0 | 0 |
| Icam2 | 0 | 0 | 1.04E-05 | 0 | 0 | 0 | 0 | 0 | 0 | 0 | 0 | 1.23E-05 | 2.38E-05 | 0 | 0 | 0 | 6.51E-05 | 0 |
| Pecam1 | 0 | 0 | 0 | 1.07E-05 | 0 | 0 | 0 | 0 | 0 | 3.90E-06 | 0 | 3.12E-05 | 6.89E-05 | 0 | 0 | 3.33E-06 | 0.00027 | 0 |
| Polg2 | 0 | 0 | 0 | 4.29E-07 | 0 | 0 | 0 | 0 | 0 | 2.00E-06 | 0 | 0.00028 | 0 | 0 | 0 | 0 | 0.00231 | 0 |
| Abca8b | 0 | 8.36E-06 | 0 | 0 | 0 | 0 | 0 | 2.41E-05 | 0 | 4.13E-05 | 0 | 0 | 0 | 0 | 0 | 0 | 0 | 5.56E-05 |
| Abca8a | 0 | 6.79E-05 | 0 | 0 | 0 | 0 | 0 | 0.00026 | 0 | 0.00012 | 0 | 0 | 0 | 0 | 0 | 0 | 0 | 0.00022 |
| Abca6 | 0 | 2.01E-05 | 0 | 0 | 0 | 0 | 0 | 1.78E-05 | 0 | 0.00017 | 0 | 0 | 0 | 0 | 0 | 0 | 0 | 3.13E-05 |
| Sox9 | 0 | 0 | 1.31E-05 | 0.00013 | 0 | 0 | 1.50E-05 | 2.09E-05 | 0 | 0 | 0 | 0.0002 | 4.07E-05 | 0 | 0 | 5.27E-05 | 0.00038 | 0.00033 |
| Gprc5c | 0 | 1.90E-05 | 0 | 0 | 0 | 0 | 0 | 5.07E-05 | 0 | 0 | 0 | 1.79E-06 | 0 | 0 | 0 | 0 | 0 | 7.65E-06 |

| | | | | | | | | | | | | | | | | | | |
|---|---|---|---|---|---|---|---|---|---|---|---|---|---|---|---|---|---|---|
| Fdxr | 1.92E-05 | 2.84E-05 | 0 | 0.00021 | 3.79E-05 | 0 | 0.00025 | 0.00014 | 0 | 0 | 0 | 0 | 0 | 0 | 0.00011 | 2.56E-05 | 9.81E-06 | 1.28E-05 |
| Fads6 | 0 | 1.69E-05 | 0 | 1.47E-05 | 0 | 0 | 2.82E-05 | 0 | 0 | 7.27E-06 | 0 | 0 | 3.14E-05 | 0 | 0 | 0 | 0 | 1.72E-06 |
| Armc7 | 3.78E-06 | 2.07E-05 | 1.57E-05 | 2.25E-05 | 1.17E-05 | 0 | 2.71E-07 | 8.54E-06 | 0 | 4.03E-06 | 3.08E-06 | 1.04E-05 | 6.93E-06 | 0 | 1.14E-05 | 6.03E-06 | 0 | 1.71E-05 |
| Itgb4 | 0 | 1.70E-05 | 0 | 3.09E-05 | 0 | 0 | 0 | 0 | 0 | 1.22E-05 | 4.17E-06 | 1.29E-05 | 0 | 0 | 6.62E-05 | 7.13E-06 | 4.48E-06 | 2.79E-05 |
| Galr2 | 0 | 2.66E-05 | 0 | 0 | 0 | 0 | 0 | 1.06E-05 | 0 | 0 | 0 | 4.03E-05 | 0 | 0 | 0 | 0 | 0 | 3.59E-05 |
| Tmc8 | 0 | 4.60E-06 | 1.02E-06 | 9.62E-07 | 0 | 0 | 2.13E-06 | 4.21E-06 | 0 | 2.10E-06 | 0 | 1.38E-05 | 3.71E-06 | 0 | 2.20E-05 | 0 | 1.70E-05 | 3.18E-06 |
| Tk1 | 0 | 0 | 0 | 3.56E-05 | 0 | 0 | 0 | 0 | 0 | 0 | 0 | 0 | 0 | 0 | 0 | 0.00125 | 0 | 0 |
| Birc5 | 2.52E-06 | 0 | 1.10E-05 | 2.60E-05 | 0 | 0 | 7.22E-09 | 0 | 0 | 0 | 0 | 0 | 0 | 0 | 2.48E-05 | 0.00143 | 0 | 0 |
| Ppp1r27 | 0 | 0 | 0 | 0 | 0 | 0 | 0 | 0 | 0 | 0 | 0 | 3.04E-05 | 1.45E-05 | 0 | 2.33E-06 | 0 | 4.67E-05 | 0 |
| Apob | 0 | 1.21E-05 | 0 | 4.98E-05 | 2.05E-06 | 0 | 0 | 1.74E-05 | 0 | 1.03E-05 | 5.88E-06 | 8.77E-06 | 1.59E-05 | 0 | 4.48E-05 | 1.46E-06 | 2.30E-06 | 7.03E-06 |
| Greb1 | 4.52E-06 | 1.95E-05 | 0 | 7.71E-06 | 0 | 0 | 0 | 0 | 0 | 6.82E-05 | 0 | 0 | 5.14E-05 | 0 | 2.08E-05 | 2.72E-07 | 0 | 3.92E-05 |
| Rrm2 | 0 | 0 | 0.00018 | 0 | 0 | 0 | 0 | 0 | 0 | 0 | 1.90E-05 | 0 | 0 | 0 | 4.99E-05 | 0.00172 | 0 | 0 |
| Rsad2 | 5.18E-06 | 0 | 0 | 0 | 0 | 0 | 0 | 0 | 0 | 7.63E-05 | 0 | 0 | 2.92E-05 | 0 | 0 | 5.40E-06 | 0 | 5.18E-06 |
| Cmpk2 | 0 | 0 | 0 | 0 | 0 | 0 | 0 | 0 | 0 | 9.90E-05 | 0 | 0 | 0 | 0 | 0 | 2.80E-06 | 0 | 2.85E-05 |
| Tspan13 | 0 | 3.21E-06 | 0 | 5.77E-06 | 0 | 0 | 3.44E-05 | 0 | 0 | 6.44E-06 | 1.86E-06 | 0 | 0 | 0 | 1.16E-05 | 7.31E-07 | 0 | 3.09E-05 |
| Meox2 | 0 | 0 | 0 | 0 | 0 | 0 | 0 | 0 | 0 | 4.77E-05 | 0 | 0 | 0 | 0 | 0 | 2.49E-06 | 0 | 0.00066 |
| Scin | 0 | 2.05E-06 | 2.42E-05 | 0 | 0 | 0 | 5.21E-07 | 3.58E-06 | 0 | 1.58E-05 | 0 | 0 | 1.72E-05 | 0 | 0 | 6.72E-06 | 2.27E-05 | 3.92E-05 |
| Lsmem1 | 0 | 0 | 0 | 0 | 0 | 0 | 0 | 0 | 0 | 0 | 0 | 0.00011 | 6.04E-05 | 0 | 0 | 0 | 0.00063 | 0 |
| Stxbp6 | 0 | 0 | 0 | 0 | 0 | 0 | 0 | 0 | 0 | 0.00022 | 0 | 0 | 0 | 0 | 0 | 6.03E-06 | 0 | 2.23E-05 |
| Mis18bp1 | 0 | 0 | 0 | 0 | 0 | 0 | 0 | 0 | 0 | 0 | 0 | 0 | 0 | 0 | 5.13E-05 | 0.00028 | 0 | 0 |
| Lrr1 | 0 | 0 | 5.05E-06 | 0 | 0 | 0 | 0 | 0 | 0 | 2.40E-07 | 0 | 0 | 0 | 0 | 0 | 0.00013 | 0 | 0 |
| Rtn1 | 0 | 2.72E-06 | 0 | 4.11E-06 | 0 | 0 | 0 | 0 | 0 | 3.46E-05 | 0 | 0 | 2.83E-05 | 0 | 2.89E-06 | 2.60E-06 | 0 | 0 |
| Six1 | 0 | 6.28E-06 | 0 | 4.41E-05 | 4.07E-06 | 0 | 0.00011 | 1.56E-05 | 0 | 0.00017 | 0 | 1.89E-05 | 0.00014 | 0 | 0 | 4.94E-05 | 0 | 0 |
| Six4 | 2.27E-06 | 0 | 3.66E-05 | 0 | 1.48E-05 | 0 | 0.00012 | 8.12E-06 | 0 | 5.10E-06 | 0 | 0 | 6.44E-05 | 0 | 0 | 2.06E-05 | 1.77E-05 | 0 |
| Slc38a6 | 0 | 6.16E-05 | 0 | 0.00012 | 3.63E-05 | 0 | 9.43E-05 | 0.00011 | 0 | 4.36E-05 | 0 | 4.02E-06 | 3.11E-05 | 0 | 8.81E-05 | 0 | 8.86E-05 | 2.37E-05 |
| Tmem30b | 0 | 1.93E-06 | 0 | 2.65E-06 | 0 | 0 | 8.24E-05 | 0 | 0 | 0.00011 | 0 | 0 | 0.00012 | 0 | 0 | 0 | 0 | 0 |
| Galnt16 | 0 | 0 | 4.74E-05 | 0 | 0 | 0 | 0.00043 | 0 | 0 | 0.00121 | 2.50E-05 | 0 | 0.00053 | 0 | 0 | 8.78E-05 | 0 | 0 |
| Ttc9 | 0 | 0 | 1.05E-05 | 0 | 0 | 0 | 0 | 3.12E-05 | 0 | 0 | 1.16E-06 | 3.31E-05 | 0 | 0 | 0 | 5.46E-08 | 0 | 0 |
| Papln | 0 | 0 | 0 | 0 | 0 | 0 | 0 | 1.71E-06 | 0 | 0.0001 | 0 | 6.41E-06 | 0 | 0 | 2.30E-06 | 0 | 0 | 0 |
| Acot2 | 1.41E-05 | 4.58E-05 | 6.19E-06 | 5.86E-05 | 0 | 0 | 3.89E-06 | 1.78E-05 | 0 | 2.98E-05 | 0 | 0.00012 | 0 | 0 | 0 | 0 | 0 | 5.81E-05 |
| Acot4 | 1.72E-06 | 0 | 1.79E-05 | 0 | 0 | 0 | 2.16E-05 | 0 | 0 | 1.77E-05 | 0 | 2.25E-05 | 2.91E-05 | 0 | 0 | 0 | 0 | 3.56E-06 |
| Pnma1 | 0 | 0 | 0 | 1.50E-06 | 0 | 0 | 9.16E-06 | 0 | 0 | 2.51E-06 | 0 | 1.11E-05 | 1.17E-05 | 0 | 2.72E-06 | 1.89E-06 | 2.97E-05 | 0 |
| Pgf | 0 | 0 | 0 | 0 | 0 | 0 | 0 | 0 | 0 | 0 | 0.00023 | 0.00104 | 0 | 0 | 0 | 0 | 0 | 0 |

| | | | | | | | | | | | | | | | | | | |
|---|---|---|---|---|---|---|---|---|---|---|---|---|---|---|---|---|---|---|
| Cep128 | 0 | 5.38E-06 | 0 | 0 | 0 | 0 | 0 | 0 | 0 | 1.17E-05 | 0 | 2.52E-07 | 0 | 0 | 0.00013 | 0.00035 | 0 | 1.20E-06 |
| Efcab11 | 0 | 0 | 0 | 0 | 0 | 0 | 0 | 0 | 0 | 4.62E-06 | 0 | 0 | 0 | 0 | 4.15E-05 | 0.00026 | 0 | 0 |
| Ifi27l2a | 4.45E-05 | 0 | 0 | 0 | 0 | 0 | 0 | 0 | 0 | 0.00073 | 0 | 0 | 0 | 0 | 0 | 7.44E-06 | 0 | 0.0002 |
| Serpina3n | 0 | 0 | 0.00045 | 0 | 0 | 0 | 0.00052 | 0 | 0 | 0.00163 | 0 | 0 | 0.00426 | 0 | 0 | 0.00025 | 0 | 0.00119 |
| Mok | 5.52E-06 | 1.10E-05 | 0 | 9.30E-06 | 2.00E-06 | 0 | 0 | 0 | 0 | 1.58E-06 | 0 | 9.65E-06 | 0 | 0 | 1.82E-05 | 7.41E-06 | 0 | 6.49E-06 |
| Ckb | 0 | 0 | 0 | 1.29E-05 | 3.22E-06 | 0 | 0 | 0 | 0 | 9.98E-05 | 0.00013 | 3.65E-05 | 0 | 0 | 0 | 0 | 0 | 9.54E-05 |
| Crip2 | 0 | 6.63E-06 | 0.00011 | 1.07E-05 | 0 | 0 | 0 | 0.00012 | 0 | 0.00017 | 0 | 0.00011 | 0 | 0 | 0 | 1.62E-05 | 0 | 0.00032 |
| Tedc1 | 0 | 0 | 1.53E-05 | 4.17E-05 | 0 | 0 | 1.79E-05 | 0 | 0 | 0 | 0 | 0 | 2.87E-06 | 0 | 3.72E-05 | 0.00056 | 0.00013 | 8.39E-06 |
| Cdca7l | 0 | 0 | 1.55E-05 | 0 | 0 | 0 | 9.61E-07 | 0 | 0 | 0 | 0 | 0 | 4.33E-06 | 0 | 0 | 0.00013 | 0 | 5.17E-06 |
| Akr1c14 | 1.86E-06 | 3.68E-06 | 0 | 0 | 0 | 0 | 0 | 0 | 0 | 0.00018 | 0 | 0 | 0 | 0 | 0 | 0 | 0 | 0 |
| Akr1c18 | 0 | 0 | 0 | 0 | 0.00019 | 0 | 0 | 4.46E-05 | 0 | 0.00028 | 0 | 0 | 0 | 0 | 0 | 0 | 0 | 0 |
| Pfkp | 6.16E-05 | 8.87E-05 | 0.00017 | 3.72E-05 | 0 | 0 | 9.56E-05 | 0.00011 | 0 | 0.00035 | 0.00018 | 0.00092 | 0 | 0 | 0.00011 | 9.27E-05 | 2.74E-05 | 0.00014 |
| Ero1lb | 1.46E-05 | 3.87E-05 | 3.77E-06 | 0 | 1.18E-06 | 0 | 0 | 0 | 0 | 0 | 4.34E-06 | 5.18E-05 | 2.73E-05 | 0 | 1.34E-05 | 5.02E-05 | 0 | 9.91E-06 |
| Sfrp4 | 0 | 0 | 0 | 0 | 0 | 0 | 1.81E-05 | 2.59E-05 | 0 | 0.00116 | 0 | 0 | 0 | 0 | 0 | 0 | 0 | 0.0009 |
| Hist1h2bn | 0 | 0 | 0 | 0 | 0 | 0 | 0 | 0 | 0 | 0 | 0 | 0 | 1.84E-07 | 0 | 3.50E-05 | 0.0013 | 6.78E-05 | 0 |
| Hist1h1b | 0 | 0 | 0 | 0 | 0 | 0 | 0 | 0 | 0 | 0 | 0 | 0 | 0 | 0 | 0 | 0.00179 | 0 | 0 |
| Hist1h2ap | 0 | 0 | 0 | 0 | 0 | 0 | 0 | 0 | 0 | 0 | 0 | 2.41E-06 | 5.10E-07 | 0 | 0 | 0.00213 | 3.35E-05 | 0 |
| Hist1h2ae | 0 | 0 | 0 | 0 | 0 | 0 | 0 | 0 | 0 | 0 | 0 | 0 | 0 | 0 | 0 | 0.00129 | 0 | 0 |
| Hist1h3d | 0 | 0 | 0 | 0 | 0 | 0 | 3.01E-06 | 0 | 0 | 0 | 0 | 9.81E-06 | 1.33E-06 | 0 | 0 | 6.43E-05 | 5.09E-05 | 0 |
| Hist1h2be | 7.19E-06 | 7.56E-06 | 1.31E-06 | 0 | 0 | 0 | 0 | 0 | 0 | 1.12E-06 | 0 | 5.15E-06 | 0 | 0 | 0 | 2.67E-05 | 4.79E-05 | 1.34E-06 |
| Hist1h2ac | 0 | 1.66E-06 | 0 | 0 | 0 | 0 | 0 | 0 | 0 | 0 | 0 | 2.88E-05 | 2.82E-06 | 0 | 0 | 2.10E-05 | 4.55E-05 | 0 |
| Hist1h1t | 0 | 0 | 0 | 0 | 0 | 0 | 0 | 0 | 0 | 0 | 0 | 1.13E-05 | 0 | 0 | 0 | 0.00012 | 3.06E-05 | 0 |
| Hist1h3c | 0 | 0 | 0 | 0 | 0 | 0 | 0 | 0 | 0 | 0 | 0 | 0 | 0 | 0 | 0 | 0.00026 | 0 | 0 |
| Hist1h3a | 0 | 0 | 0 | 0 | 0 | 0 | 0 | 0 | 0 | 0 | 0 | 0 | 4.04E-06 | 0 | 0 | 0.0005 | 3.87E-05 | 0 |
| Hist1h1a | 0 | 0 | 0 | 0 | 0 | 0 | 0 | 0 | 0 | 0 | 0 | 0 | 2.41E-06 | 0 | 0 | 0.00098 | 6.06E-06 | 0 |
| Cmah | 0 | 0 | 0 | 2.52E-05 | 0 | 0 | 7.72E-05 | 0 | 0 | 0.00084 | 0.00018 | 0 | 4.80E-05 | 0 | 0 | 2.62E-05 | 0 | 0 |
| Irf4 | 0 | 0 | 0 | 0 | 0 | 0 | 0 | 0 | 0 | 0.00011 | 0 | 0 | 0 | 0 | 0 | 0 | 0 | 0 |
| Foxf2 | 0 | 3.20E-06 | 1.34E-05 | 7.45E-06 | 0 | 0 | 0 | 0 | 0 | 5.32E-06 | 1.13E-06 | 1.59E-05 | 1.57E-05 | 0 | 3.48E-06 | 0 | 9.73E-06 | 0 |
| Foxc1 | 2.80E-07 | 0 | 0 | 5.92E-06 | 0 | 0 | 0 | 1.08E-05 | 0 | 0 | 1.05E-05 | 0 | 1.31E-05 | 0 | 3.93E-05 | 4.14E-05 | 7.80E-05 | 0.00012 |
| Serpinb1a | 4.69E-05 | 0 | 2.53E-05 | 0 | 0 | 0 | 8.67E-05 | 0 | 0 | 0.00033 | 0 | 0 | 0.00033 | 0 | 0 | 0.00014 | 0 | 0.00029 |
| Tubb2a | 0.00015 | 0 | 0.00033 | 0.00046 | 0 | 0 | 0.00049 | 5.26E-05 | 0 | 0.00023 | 0.00033 | 0 | 0.0002 | 0 | 0 | 0.00048 | 0.0003 | 0.00056 |
| Tubb2b | 0 | 0 | 0.00037 | 0.00021 | 0 | 0 | 0.00017 | 0.0002 | 0 | 0.00036 | 0.00034 | 0 | 0 | 0 | 0 | 0.00023 | 0 | 0.00046 |
| Edn1 | 0 | 0 | 0 | 0 | 0 | 0 | 0 | 0 | 0 | 8.28E-05 | 7.90E-05 | 0 | 0 | 0 | 0 | 0 | 0 | 0 |

| | | | | | | | | | | | | | | | | | | |
|---|---|---|---|---|---|---|---|---|---|---|---|---|---|---|---|---|---|---|
| Fgd3 | 0 | 6.56E-06 | 0 | 0 | 0 | 0 | 0 | 0 | 0 | 2.28E-06 | 0.00015 | 0 | 0 | 0 | 6.65E-06 | 0 | 0 | 0 |
| Ecm2 | 5.34E-05 | 1.07E-05 | 0 | 0 | 8.09E-06 | 0 | 2.95E-05 | 5.38E-05 | 0 | 4.76E-05 | 2.14E-05 | 0 | 0 | 0 | 0 | 0 | 0 | 0.00017 |
| Aspn | 0.00016 | 0 | 0 | 0 | 0 | 0 | 0.00065 | 0.00062 | 0 | 0.00031 | 0.00028 | 0 | 0 | 0 | 0 | 0 | 0 | 0.00031 |
| Ogn | 0.0001 | 0 | 0 | 1.72E-05 | 0 | 0 | 3.19E-05 | 0 | 0 | 0.00102 | 0 | 0 | 0.00017 | 0 | 0 | 0 | 0 | 6.19E-05 |
| Nxnl2 | 1.58E-06 | 0 | 0 | 0 | 0 | 0 | 0 | 9.72E-07 | 0 | 2.93E-05 | 0 | 1.37E-06 | 0 | 0 | 1.42E-05 | 0 | 3.13E-05 | 0 |
| Sema4d | 0 | 7.12E-06 | 0 | 2.23E-05 | 0 | 0 | 0 | 0 | 0 | 2.97E-05 | 0 | 5.58E-06 | 2.16E-05 | 0 | 5.37E-05 | 1.15E-05 | 1.67E-05 | 7.04E-06 |
| Gadd45g | 0 | 5.75E-05 | 0 | 0.00042 | 0 | 0 | 0.00112 | 0 | 0 | 0.00037 | 0.00137 | 0.00038 | 0.00155 | 0 | 0.00038 | 8.35E-05 | 0.00107 | 0 |
| Syk | 0 | 0 | 0 | 3.60E-05 | 0 | 0 | 0 | 0 | 0 | 0.0004 | 0 | 0 | 0 | 0 | 4.54E-05 | 5.94E-05 | 0 | 0 |
| Mxd3 | 0 | 0 | 0 | 0 | 0 | 0 | 0 | 0 | 0 | 0 | 0 | 0 | 0 | 0 | 4.14E-05 | 0.00087 | 0 | 0 |
| Dapk1 | 0 | 8.43E-05 | 0 | 0 | 0 | 0 | 0 | 0 | 0 | 0.00041 | 0 | 0 | 0.00026 | 0 | 2.63E-05 | 0 | 0 | 0.00032 |
| Srd5a1 | 2.29E-06 | 1.04E-05 | 0 | 9.82E-06 | 0 | 0 | 3.13E-06 | 1.81E-05 | 0 | 0 | 0 | 1.71E-05 | 2.62E-06 | 0 | 0 | 3.63E-05 | 9.08E-06 | 0 |
| Trip13 | 1.27E-05 | 0 | 1.79E-05 | 0 | 0 | 0 | 0 | 0 | 0 | 0 | 0 | 0 | 0 | 0 | 0 | 0.0004 | 0 | 0 |
| Polr3g | 0 | 0 | 2.93E-05 | 4.14E-06 | 0 | 0 | 4.34E-06 | 0 | 0 | 0 | 2.68E-07 | 7.54E-05 | 2.17E-06 | 0 | 0 | 2.28E-05 | 0 | 9.56E-07 |
| Thbs4 | 0 | 0 | 0.00016 | 0.0001 | 0 | 0 | 0.00046 | 0 | 0 | 0 | 0 | 0 | 0.00028 | 0 | 0 | 9.63E-05 | 0 | 0.00035 |
| Cmya5 | 0 | 8.12E-06 | 0 | 0 | 0 | 0 | 4.66E-05 | 0 | 0 | 7.17E-06 | 0 | 2.16E-05 | 0.00018 | 0 | 0 | 0 | 0 | 2.12E-05 |
| Iqgap2 | 0 | 0 | 7.75E-05 | 0 | 0 | 0 | 5.06E-05 | 0 | 0 | 4.29E-05 | 0 | 0 | 0 | 0 | 0 | 7.82E-05 | 0 | 8.29E-05 |
| Map1b | 0.00018 | 7.73E-05 | 0.00012 | 0 | 0 | 0 | 6.25E-05 | 0.00013 | 0 | 4.63E-05 | 0.00033 | 0.00016 | 0 | 0 | 0 | 0 | 0.00021 | 0.00016 |
| Adamts6 | 1.11E-06 | 3.87E-06 | 2.35E-05 | 0 | 0.0002 | 0 | 7.65E-05 | 7.37E-06 | 0 | 3.09E-05 | 0.00066 | 0.00012 | 0 | 0 | 0.00013 | 0 | 0 | 2.91E-05 |
| Depdc1b | 0 | 0 | 0 | 0 | 0 | 0 | 0 | 0 | 0 | 0 | 0 | 0 | 0 | 0 | 3.22E-05 | 0.00015 | 0 | 0 |
| Plk2 | 2.58E-05 | 1.53E-05 | 0 | 0 | 0 | 0 | 0 | 0 | 0 | 0 | 7.35E-05 | 0.00017 | 0 | 0 | 0 | 0 | 0.00074 | 7.38E-05 |
| Ankrd55 | 1.41E-06 | 0 | 0 | 1.48E-06 | 0 | 0 | 0 | 0 | 0 | 2.15E-05 | 0 | 0 | 2.53E-05 | 0 | 0 | 0 | 0 | 5.45E-05 |
| Ccno | 0 | 0 | 0 | 1.28E-06 | 0 | 0 | 0 | 0 | 0 | 0 | 0 | 7.10E-06 | 2.37E-05 | 0 | 0 | 2.35E-06 | 7.44E-05 | 0 |
| Esm1 | 0 | 0 | 0 | 0 | 0 | 0 | 0 | 0 | 0 | 0 | 0 | 2.91E-05 | 0 | 0 | 0 | 0 | 0 | 0.00032 |
| Itga2 | 0 | 0 | 1.73E-05 | 0 | 0 | 0 | 8.51E-06 | 0 | 0 | 4.65E-05 | 0 | 7.93E-05 | 5.89E-05 | 0 | 0 | 7.06E-06 | 0 | 0 |
| Kcnk5 | 0 | 0 | 2.09E-06 | 0 | 0 | 0 | 0 | 0 | 0 | 9.56E-05 | 0 | 1.58E-05 | 1.99E-06 | 0 | 0 | 5.22E-05 | 0.00015 | 0 |
| Kcnma1 | 0 | 0 | 0 | 0 | 4.84E-05 | 0 | 0.00025 | 0 | 0 | 0 | 0.00014 | 0 | 9.37E-05 | 0 | 0 | 0 | 0 | 0.0003 |
| Stab1 | 0 | 0 | 2.11E-05 | 2.05E-05 | 2.47E-05 | 0 | 8.96E-06 | 0 | 0 | 0 | 0 | 2.55E-05 | 1.03E-05 | 0 | 7.12E-05 | 0 | 1.70E-05 | 0 |
| Arhgap22 | 0 | 1.51E-05 | 0 | 0 | 0 | 0 | 0 | 0 | 0 | 2.01E-05 | 0 | 7.44E-05 | 0 | 0 | 0 | 1.17E-06 | 0 | 0 |
| Gdf10 | 0 | 0 | 0 | 0 | 0 | 0 | 0 | 0 | 0 | 3.58E-07 | 0 | 0 | 0 | 0 | 0 | 0 | 0 | 0.00093 |
| Dlgap5 | 0 | 0 | 0 | 0 | 0 | 0 | 0 | 0 | 0 | 3.04E-06 | 0 | 0 | 0 | 0 | 4.11E-05 | 0.00075 | 0 | 0 |
| Pnp2 | 0 | 0 | 0 | 0 | 0 | 0 | 0 | 0 | 0 | 4.96E-05 | 0 | 3.18E-05 | 9.13E-05 | 0 | 0 | 0 | 0.00016 | 0 |
| Trac | 0 | 0 | 0 | 0 | 0 | 0 | 0 | 1.80E-05 | 0 | 0 | 3.50E-05 | 0 | 0 | 0 | 0 | 1.62E-05 | 0 | 4.49E-05 |
| Slc7a7 | 8.18E-06 | 5.49E-05 | 1.65E-05 | 0 | 3.92E-05 | 0 | 3.78E-05 | 0.00015 | 0 | 5.28E-05 | 4.67E-05 | 3.74E-05 | 0 | 0 | 0 | 0 | 1.24E-05 | 8.66E-05 |

| | | | | | | | | | | | | | | | | | | |
|---|---|---|---|---|---|---|---|---|---|---|---|---|---|---|---|---|---|---|
| Cdh24 | 0 | 5.50E-06 | 0 | 1.81E-05 | 9.08E-06 | 0 | 0 | 2.78E-05 | 0 | 0 | 1.29E-05 | 3.11E-08 | 0 | 0 | 4.03E-05 | 9.43E-05 | 3.89E-05 | 0 |
| Ppp1r3e | 0 | 3.85E-06 | 0 | 1.94E-05 | 0 | 0 | 3.16E-06 | 1.06E-05 | 0 | 0 | 7.84E-06 | 5.24E-05 | 2.10E-06 | 0 | 6.62E-07 | 0 | 0 | 7.46E-06 |
| Tssk4 | 4.09E-06 | 7.62E-06 | 2.04E-06 | 1.10E-05 | 1.51E-06 | 0 | 0 | 0 | 0 | 3.84E-06 | 0 | 0 | 1.41E-05 | 0 | 4.65E-05 | 1.63E-05 | 5.11E-06 | 0 |
| Mcpt1 | 1.87E-06 | 0 | 0 | 6.96E-05 | 0 | 0 | 1.78E-05 | 0.00012 | 0 | 1.77E-05 | 0 | 3.11E-05 | 0 | 0 | 9.10E-05 | 9.71E-06 | 1.19E-05 | 0 |
| Ska3 | 0 | 0 | 0 | 0 | 0 | 0 | 0 | 0 | 0 | 0 | 0 | 1.53E-06 | 0 | 0 | 1.45E-05 | 0.00032 | 0 | 0 |
| Phf11b | 1.84E-05 | 0 | 0 | 0 | 0 | 0 | 0 | 0 | 0 | 0.00013 | 0 | 0 | 2.75E-05 | 0 | 0 | 8.65E-06 | 0 | 2.37E-05 |
| Phf11d | 6.80E-05 | 0 | 0 | 0 | 0 | 0 | 0 | 0 | 0 | 0.00026 | 0 | 5.53E-06 | 0.00016 | 0 | 2.89E-06 | 2.32E-05 | 0 | 0.00013 |
| Fam124a | 0 | 2.65E-05 | 0 | 0 | 0 | 0 | 0 | 0 | 0 | 6.43E-05 | 0 | 0 | 5.84E-05 | 0 | 0 | 0 | 0 | 3.90E-05 |
| Defb48 | 0 | 0 | 0 | 0 | 0 | 0 | 0 | 0.00011 | 0 | 0 | 0 | 0 | 6.93E-06 | 0 | 0 | 1.60E-05 | 0 | 1.27E-06 |
| Fam167a | 0 | 2.05E-05 | 3.29E-05 | 0 | 4.22E-05 | 0 | 0 | 0.00012 | 0 | 3.84E-05 | 0 | 0 | 0 | 0 | 0 | 1.93E-05 | 0 | 2.26E-05 |
| Pbk | 0 | 0 | 0 | 0 | 0 | 0 | 0 | 0 | 0 | 0 | 0 | 0 | 0 | 0 | 0 | 0.0009 | 0 | 0 |
| Esco2 | 0 | 0 | 0 | 0 | 0 | 0 | 0 | 0 | 0 | 0 | 0 | 0 | 0 | 0 | 0 | 0.00039 | 0 | 0 |
| Clu | 0 | 0.00032 | 0 | 0.00089 | 0.00018 | 0 | 0 | 0.00014 | 0 | 0.00042 | 4.37E-05 | 0.00031 | 0.00087 | 0 | 0.00189 | 0.00014 | 0.00057 | 0.00029 |
| Ptk2b | 0 | 1.42E-05 | 5.15E-05 | 1.04E-06 | 0 | 0 | 7.83E-05 | 0.0001 | 0 | 0.0001 | 0.00019 | 0.00011 | 0 | 0 | 3.45E-05 | 0 | 0 | 0.00016 |
| Stmn4 | 0 | 0 | 0 | 0 | 0 | 0 | 0 | 2.07E-05 | 0 | 3.46E-06 | 0.00033 | 6.96E-05 | 0 | 0 | 0 | 0 | 0 | 0 |
| Cdca2 | 0 | 0 | 0 | 0 | 0 | 0 | 0 | 0 | 0 | 0 | 0 | 0 | 0 | 0 | 1.74E-05 | 0.00048 | 0 | 0 |
| Dok2 | 0 | 6.13E-06 | 0 | 1.28E-05 | 1.87E-06 | 0 | 0 | 3.51E-06 | 0 | 3.37E-06 | 5.58E-06 | 8.35E-07 | 0 | 0 | 5.19E-06 | 2.97E-06 | 8.84E-07 | 2.22E-06 |
| Gfra2 | 0 | 0 | 2.61E-05 | 0 | 0 | 0 | 0.00011 | 1.28E-05 | 0 | 0.00033 | 0 | 5.07E-05 | 0 | 0 | 0 | 8.01E-05 | 0 | 0.00026 |
| Htr2a | 6.21E-05 | 0 | 0.00013 | 0 | 0 | 0 | 3.45E-05 | 0 | 0 | 9.85E-05 | 0.0003 | 0.00019 | 0 | 0 | 0 | 0 | 2.57E-05 | 0 |
| Diaph3 | 2.10E-05 | 0 | 1.75E-06 | 0 | 0 | 0 | 0 | 0 | 0 | 2.09E-05 | 0 | 0 | 0 | 0 | 0 | 0.00059 | 0 | 1.99E-06 |
| Bora | 5.94E-06 | 8.07E-06 | 6.05E-06 | 0 | 5.12E-06 | 0 | 0 | 1.15E-05 | 0 | 0 | 1.07E-05 | 1.02E-06 | 0 | 0 | 0.00012 | 0.00046 | 0 | 0 |
| Kctd12 | 0 | 0 | 0 | 0.00023 | 0 | 0 | 1.74E-05 | 4.39E-05 | 0 | 8.45E-05 | 0.0002 | 0 | 2.04E-05 | 0 | 9.23E-05 | 0.00011 | 0.00062 | 0.00044 |
| Cldn10 | 0 | 0 | 0 | 0 | 0 | 0 | 0.00049 | 0 | 0 | 9.79E-05 | 0 | 0 | 0.00014 | 0 | 0 | 0 | 0 | 0 |
| Slc1a3 | 0 | 0 | 0 | 0 | 0 | 0 | 0.0001 | 0 | 0 | 0 | 0 | 0 | 0.00036 | 0 | 0 | 0 | 0 | 0.00037 |
| Npr3 | 0 | 3.06E-06 | 0 | 0 | 1.24E-05 | 0 | 0 | 0 | 0 | 2.60E-06 | 4.89E-05 | 6.59E-06 | 0 | 0 | 0 | 0 | 0 | 2.20E-05 |
| Cdh10 | 4.96E-07 | 0 | 5.22E-06 | 1.33E-06 | 1.58E-05 | 0 | 7.19E-06 | 2.79E-06 | 0 | 0 | 1.17E-05 | 0 | 0 | 0 | 0 | 3.38E-06 | 6.28E-07 | 0 |
| Zfpm2 | 5.20E-06 | 0 | 0 | 0 | 0 | 0 | 7.03E-05 | 0 | 0 | 0.0001 | 0 | 0 | 4.38E-05 | 0 | 0 | 9.59E-06 | 2.61E-05 | 0.00011 |
| Ccn3 | 0 | 0 | 0 | 0 | 0 | 0 | 0.00036 | 0 | 0 | 0.00085 | 8.73E-06 | 0 | 1.30E-05 | 0 | 0 | 1.46E-05 | 0 | 0.00072 |
| Dscc1 | 0 | 0 | 1.56E-05 | 0 | 0 | 0 | 0 | 0 | 0 | 5.49E-06 | 0 | 0 | 8.16E-06 | 0 | 0 | 0.00019 | 0 | 5.01E-06 |
| Col14a1 | 2.66E-05 | 0 | 0 | 6.14E-06 | 0 | 0 | 0.00081 | 4.74E-05 | 0 | 0.00147 | 5.70E-05 | 0 | 0.00126 | 0 | 0 | 0.0001 | 1.73E-05 | 0.00091 |
| Mtss1 | 0 | 5.41E-05 | 0 | 0 | 0 | 0 | 0 | 0 | 0 | 0 | 0 | 0.00012 | 0 | 0 | 2.11E-05 | 0 | 0 | 5.33E-05 |
| Kcnq3 | 0 | 1.07E-06 | 0 | 0 | 1.99E-05 | 0 | 0 | 9.51E-05 | 0 | 0 | 0 | 2.57E-05 | 0 | 0 | 0 | 0 | 0 | 0 |
| Dennd3 | 2.20E-06 | 2.84E-05 | 0 | 4.57E-05 | 3.39E-05 | 0 | 7.63E-05 | 0 | 0 | 0 | 4.34E-05 | 0 | 2.21E-05 | 0 | 3.90E-05 | 0 | 4.87E-05 | 4.35E-05 |

| | | | | | | | | | | | | | | | | | | |
|---|---|---|---|---|---|---|---|---|---|---|---|---|---|---|---|---|---|---|
| Ly6c1 | 3.80E-05 | 0 | 0 | 0 | 0 | 0 | 0 | 0 | 0 | 0.00147 | 0 | 0 | 0.00017 | 0 | 0 | 0 | 0 | 0.00033 |
| Mb | 0 | 0 | 3.27E-06 | 0 | 0 | 0 | 7.07E-08 | 0 | 0 | 4.46E-07 | 0 | 5.44E-06 | 2.28E-05 | 0 | 0 | 0 | 6.49E-05 | 0 |
| Apol6 | 1.19E-05 | 0 | 0 | 5.80E-06 | 0 | 0 | 0 | 0 | 0 | 0.00012 | 0 | 0 | 3.69E-05 | 0 | 1.13E-06 | 1.41E-05 | 0 | 9.55E-06 |
| Apol7a | 0 | 0 | 0 | 0 | 9.11E-06 | 0 | 1.15E-05 | 0 | 0 | 1.16E-05 | 4.12E-06 | 1.82E-06 | 0 | 0 | 1.18E-05 | 0 | 0 | 4.61E-06 |
| Apol9a | 0 | 3.45E-06 | 7.27E-06 | 0 | 7.29E-05 | 0 | 1.89E-05 | 0 | 0 | 2.55E-05 | 7.20E-06 | 2.36E-05 | 1.16E-05 | 0 | 0 | 2.80E-05 | 0 | 0 |
| Apol8 | 0 | 0 | 5.23E-05 | 0 | 0 | 0 | 2.78E-05 | 0 | 0 | 0 | 0.00013 | 0.00013 | 3.56E-05 | 0 | 3.44E-05 | 0 | 0.00086 | 0 |
| Ncf4 | 0 | 0 | 5.63E-06 | 8.45E-06 | 0 | 0 | 0 | 2.53E-07 | 0 | 2.96E-06 | 3.62E-06 | 3.43E-06 | 0 | 0 | 2.29E-05 | 9.26E-07 | 3.61E-06 | 9.25E-07 |
| Csf2rb2 | 0 | 5.64E-06 | 0 | 5.18E-06 | 0 | 0 | 0 | 3.57E-06 | 0 | 2.19E-05 | 0 | 0 | 2.96E-06 | 0 | 6.77E-06 | 0 | 5.99E-06 | 8.07E-05 |
| Lgals2 | 3.70E-06 | 8.53E-06 | 2.10E-06 | 8.93E-06 | 1.16E-06 | 0 | 0 | 0 | 0 | 2.09E-06 | 1.67E-06 | 7.95E-06 | 2.01E-06 | 0 | 1.40E-05 | 9.01E-07 | 7.96E-06 | 4.90E-06 |
| Pdxp | 0 | 0 | 2.58E-05 | 1.03E-06 | 1.14E-05 | 0 | 0 | 1.53E-05 | 0 | 8.29E-07 | 1.18E-05 | 0.00025 | 0 | 0 | 1.94E-05 | 2.61E-05 | 0 | 3.73E-06 |
| Triobp | 5.37E-05 | 6.02E-05 | 0 | 0.00015 | 7.33E-06 | 0 | 6.53E-05 | 5.56E-05 | 0 | 0.00035 | 4.63E-05 | 0.00105 | 0 | 0 | 0.00012 | 0.00026 | 3.95E-05 | 9.21E-05 |
| Cenpm | 0 | 0 | 7.26E-06 | 9.63E-05 | 0 | 0 | 0 | 0 | 0 | 0 | 0 | 0 | 0 | 0 | 0 | 0.00063 | 0 | 0 |
| Scube1 | 0 | 3.91E-05 | 0 | 0 | 4.03E-05 | 0 | 0 | 9.60E-05 | 0 | 0 | 8.37E-06 | 3.71E-05 | 0 | 0 | 0 | 0 | 0 | 6.59E-06 |
| Gtse1 | 0 | 1.90E-05 | 0 | 0 | 0 | 0 | 0 | 0 | 0 | 5.63E-06 | 0 | 0 | 2.30E-06 | 0 | 6.21E-05 | 0.00085 | 0 | 0 |
| Tafa5 | 0 | 6.77E-06 | 7.81E-06 | 0 | 3.45E-05 | 0 | 0 | 8.95E-06 | 0 | 0 | 2.09E-05 | 2.65E-05 | 0 | 0 | 0 | 0 | 0 | 0 |
| Adm2 | 0 | 0 | 0 | 0 | 0 | 0 | 0 | 0 | 0 | 0 | 0 | 0.00033 | 0 | 0 | 0 | 0 | 0 | 0 |
| Pdzrn4 | 0 | 0 | 0 | 0 | 1.84E-05 | 0 | 2.45E-05 | 0 | 0 | 8.91E-06 | 0 | 0 | 3.03E-05 | 0 | 0 | 0 | 3.17E-06 | 0 |
| Dbx2 | 0 | 0 | 1.08E-05 | 0 | 0 | 0 | 3.35E-05 | 0 | 0 | 9.79E-06 | 0 | 0 | 9.43E-06 | 0 | 0 | 0 | 0 | 4.85E-06 |
| Slc38a1 | 0 | 1.84E-06 | 2.08E-06 | 1.03E-05 | 3.97E-06 | 0 | 0 | 2.72E-06 | 0 | 0 | 2.36E-06 | 4.96E-06 | 0 | 0 | 5.07E-06 | 1.37E-05 | 7.10E-06 | 1.67E-06 |
| Amigo2 | 6.89E-07 | 7.74E-06 | 0 | 0 | 0 | 0 | 0 | 0 | 0 | 2.88E-05 | 0 | 0 | 1.99E-05 | 0 | 0 | 0 | 3.23E-06 | 7.38E-06 |
| Rnd1 | 0 | 0 | 5.98E-05 | 0 | 0 | 0 | 0 | 0 | 0 | 0 | 1.50E-05 | 3.91E-05 | 0.00033 | 0 | 0 | 2.94E-06 | 9.84E-05 | 0 |
| Wnt10b | 0 | 0 | 4.14E-07 | 0 | 0 | 0 | 0 | 0 | 0 | 6.15E-05 | 1.09E-05 | 0 | 0 | 0 | 0 | 0 | 0 | 0 |
| Troap | 0 | 0 | 0 | 0 | 0 | 0 | 0 | 0 | 0 | 0 | 0 | 0 | 0 | 0 | 6.78E-05 | 0.00039 | 0 | 0 |
| Racgap1 | 0 | 0 | 0 | 7.39E-05 | 0 | 0 | 0 | 0 | 0 | 0 | 0 | 0 | 0 | 0 | 1.17E-05 | 0.00136 | 0 | 5.47E-06 |
| Gpd1 | 0 | 8.30E-06 | 4.35E-06 | 8.12E-06 | 3.47E-06 | 0 | 7.06E-07 | 0 | 0 | 0 | 0 | 5.15E-07 | 3.51E-06 | 0 | 1.11E-05 | 4.83E-06 | 1.13E-05 | 0 |
| Cela1 | 0 | 2.61E-06 | 0 | 1.41E-06 | 9.10E-06 | 0 | 4.82E-05 | 0 | 0 | 4.68E-05 | 0 | 1.60E-05 | 1.14E-05 | 0 | 0 | 0 | 0 | 0 |
| Galnt6 | 0 | 0 | 0 | 0 | 0 | 0 | 0.00019 | 0 | 0 | 0.00012 | 0 | 6.92E-05 | 0 | 0 | 0 | 0 | 0 | 0 |
| Krt8 | 0 | 5.14E-05 | 0 | 0.00023 | 6.82E-06 | 0 | 0 | 1.32E-05 | 0 | 5.55E-05 | 6.03E-06 | 6.45E-05 | 5.86E-06 | 0 | 0.00055 | 4.44E-05 | 9.74E-05 | 7.13E-05 |
| Soat2 | 0 | 4.14E-06 | 3.03E-06 | 0 | 0 | 0 | 0 | 0 | 0 | 1.72E-06 | 0 | 4.69E-05 | 0 | 0 | 2.31E-05 | 3.73E-06 | 0 | 0 |
| Espl1 | 0 | 0 | 0 | 0 | 0 | 0 | 0 | 0 | 0 | 0 | 0 | 0 | 0 | 0 | 4.21E-05 | 0.00044 | 0 | 0 |
| Npff | 0 | 2.42E-05 | 0 | 1.06E-05 | 0 | 0 | 0 | 0 | 0 | 1.06E-05 | 0 | 3.09E-05 | 0 | 0 | 5.85E-05 | 0 | 8.64E-05 | 4.25E-06 |
| Nckap1l | 0 | 2.57E-06 | 2.00E-06 | 2.33E-05 | 0 | 0 | 0 | 1.53E-05 | 0 | 3.79E-06 | 0 | 2.26E-05 | 0 | 0 | 4.61E-05 | 9.33E-06 | 1.25E-05 | 8.12E-06 |
| Ppl | 0 | 1.17E-05 | 0 | 0 | 0 | 0 | 0 | 0 | 0 | 7.29E-05 | 0 | 0 | 1.61E-05 | 0 | 3.66E-07 | 0 | 0 | 8.33E-06 |

| | | | | | | | | | | | | | | | | | | |
|---|---|---|---|---|---|---|---|---|---|---|---|---|---|---|---|---|---|---|
| Ypel1 | 0 | 2.21E-05 | 0 | 0 | 0 | 0 | 0 | 0 | 0 | 8.07E-06 | 0 | 3.11E-05 | 3.31E-05 | 0 | 0 | 0 | 9.16E-05 | 0 |
| St6gal1 | 9.29E-06 | 7.22E-05 | 0 | 0 | 0 | 0 | 0 | 4.59E-05 | 0 | 7.13E-05 | 0 | 1.08E-05 | 0 | 0 | 0 | 6.80E-06 | 0 | 5.91E-05 |
| Hcls1 | 0 | 3.69E-06 | 2.71E-06 | 1.64E-05 | 2.77E-06 | 0 | 0 | 0 | 0 | 5.39E-06 | 4.05E-06 | 7.36E-06 | 0 | 0 | 2.65E-05 | 1.71E-05 | 2.46E-05 | 3.86E-06 |
| Polq | 0 | 0 | 0 | 0 | 0 | 0 | 0 | 0 | 0 | 0 | 0 | 0 | 0 | 0 | 0 | 0.00021 | 0 | 0 |
| Tagln3 | 0 | 0 | 0 | 4.44E-06 | 0 | 0 | 4.19E-06 | 0 | 0 | 0 | 1.69E-05 | 2.96E-06 | 4.38E-05 | 0 | 0 | 0 | 0.00014 | 0 |
| Plcxd2 | 3.75E-06 | 0 | 6.42E-06 | 0 | 0 | 0 | 0 | 0 | 0 | 6.84E-06 | 9.68E-06 | 9.52E-06 | 8.54E-06 | 0 | 0 | 0 | 3.66E-06 | 0 |
| Cip2a | 3.74E-05 | 0 | 1.68E-05 | 0 | 0 | 0 | 0 | 0 | 0 | 0 | 0 | 0 | 0 | 0 | 8.16E-05 | 0.0009 | 0 | 0 |
| Alcam | 5.07E-07 | 2.82E-05 | 2.76E-05 | 0 | 1.57E-06 | 0 | 0 | 1.72E-05 | 0 | 0 | 2.48E-05 | 2.97E-05 | 0 | 0 | 2.02E-05 | 2.24E-05 | 6.70E-06 | 9.88E-06 |
| Col8a1 | 5.55E-05 | 0 | 0 | 0 | 0.00076 | 0 | 0.00058 | 0.00064 | 0 | 0 | 0.00209 | 0 | 0.00059 | 0 | 0 | 0.00029 | 0 | 0.00062 |
| Nrip1 | 0.00016 | 9.34E-05 | 3.34E-05 | 0 | 0.00022 | 0 | 0.0002 | 0.00037 | 0 | 8.04E-05 | 8.32E-05 | 0.00027 | 2.92E-05 | 0 | 2.76E-05 | 3.65E-06 | 0.00011 | 5.43E-05 |
| Kcnj15 | 0 | 0 | 0 | 0 | 2.38E-05 | 0 | 0.00019 | 0 | 0 | 0 | 6.73E-05 | 0 | 0.00014 | 0 | 0 | 5.26E-06 | 0 | 0.00013 |
| Igsf5 | 1.67E-06 | 8.19E-06 | 0 | 0 | 4.20E-05 | 0 | 0 | 9.10E-08 | 0 | 0 | 0 | 0 | 0 | 0 | 1.17E-05 | 9.51E-06 | 0 | 0 |
| Dact2 | 0 | 1.61E-05 | 3.68E-07 | 0 | 1.15E-05 | 0 | 0 | 2.56E-05 | 0 | 5.28E-05 | 0 | 1.51E-05 | 0 | 0 | 0 | 0 | 0 | 0 |
| Has1 | 0 | 0 | 0.00023 | 4.33E-05 | 0 | 0 | 0 | 0 | 0 | 0.00027 | 0 | 0 | 2.67E-05 | 0 | 0 | 0 | 0.001 | 0.00012 |
| Mmp25 | 0 | 0 | 6.78E-06 | 5.93E-06 | 0 | 0 | 5.50E-06 | 1.70E-05 | 0 | 0 | 0 | 1.83E-05 | 2.58E-05 | 0 | 0 | 1.15E-05 | 0.00013 | 3.59E-06 |
| Ccnf | 0 | 0 | 0 | 0 | 0 | 0 | 0 | 0 | 0 | 0 | 0 | 2.08E-06 | 0 | 0 | 6.01E-05 | 0.00037 | 0 | 0 |
| Dnase1l2 | 2.23E-07 | 6.35E-06 | 5.06E-07 | 5.18E-07 | 2.99E-06 | 0 | 0 | 0 | 0 | 1.21E-06 | 1.68E-06 | 6.56E-06 | 0 | 0 | 3.64E-05 | 1.42E-06 | 8.31E-06 | 5.97E-06 |
| Tpsb2 | 0 | 2.60E-06 | 0 | 1.15E-05 | 0 | 0 | 0 | 0 | 0 | 9.55E-06 | 7.56E-07 | 3.75E-06 | 0 | 0 | 4.00E-05 | 4.56E-06 | 2.67E-05 | 1.24E-06 |
| Chtf18 | 0 | 0 | 0 | 1.29E-05 | 0 | 0 | 0 | 0 | 0 | 0 | 0 | 0 | 0 | 0 | 2.32E-05 | 0.00037 | 0 | 5.74E-06 |
| Mcrip2 | 1.81E-05 | 7.86E-06 | 6.97E-05 | 8.03E-05 | 0 | 0 | 0 | 2.57E-05 | 0 | 2.89E-05 | 0 | 2.30E-05 | 2.87E-05 | 0 | 0 | 0 | 9.40E-06 | 6.02E-08 |
| Arhgdig | 0 | 0 | 4.28E-05 | 0 | 0 | 0 | 3.52E-05 | 0 | 0 | 9.62E-06 | 4.38E-05 | 5.21E-05 | 4.27E-06 | 0 | 0 | 7.88E-06 | 0 | 2.49E-05 |
| Rgs11 | 0 | 1.75E-05 | 0 | 0 | 2.75E-05 | 0 | 0 | 0 | 0 | 0 | 0 | 0.00025 | 0 | 0 | 6.41E-05 | 0 | 0 | 0 |
| Neurl1b | 0 | 2.66E-06 | 0 | 0 | 0 | 0 | 0 | 0.00015 | 0 | 0 | 0 | 0 | 0 | 0 | 9.48E-06 | 0.00027 | 0 | 0.00011 |
| Cdkn1a | 0 | 0.00041 | 0.00013 | 0 | 0 | 0 | 0 | 0 | 0 | 0.00014 | 0 | 0.00114 | 7.15E-05 | 0 | 0.00015 | 0 | 0.00085 | 0 |
| Pi16 | 0 | 0 | 0 | 0 | 0 | 0 | 0.00033 | 0 | 0 | 0.00238 | 0 | 0 | 0.00147 | 0 | 0 | 0 | 0 | 0.00023 |
| Rsph1 | 7.62E-06 | 0 | 5.62E-06 | 4.00E-06 | 2.90E-06 | 0 | 2.42E-05 | 6.54E-06 | 0 | 5.19E-06 | 0 | 4.68E-06 | 0 | 0 | 0 | 3.60E-06 | 0 | 1.43E-05 |
| Slc37a1 | 0 | 5.88E-06 | 0 | 7.61E-06 | 0 | 0 | 0 | 2.18E-06 | 0 | 3.42E-07 | 0 | 3.04E-05 | 0 | 0 | 6.26E-06 | 7.34E-07 | 0 | 5.89E-07 |
| Pde9a | 0 | 0 | 0 | 0 | 0 | 0 | 0 | 9.97E-05 | 0 | 0 | 0 | 0 | 0 | 0 | 0 | 2.68E-05 | 0 | 9.53E-06 |
| Zfp811 | 0 | 1.01E-05 | 0 | 0 | 2.24E-06 | 0 | 4.02E-07 | 0 | 0 | 5.37E-06 | 0 | 1.03E-05 | 0 | 0 | 1.70E-05 | 1.20E-05 | 0 | 9.83E-06 |
| Kifc1 | 0 | 0 | 0 | 0 | 0 | 0 | 0 | 0 | 0 | 6.31E-06 | 0 | 0 | 0 | 0 | 0.00012 | 0.00095 | 0 | 0 |
| Col11a2 | 0 | 1.62E-06 | 0 | 0 | 1.04E-06 | 0 | 1.03E-05 | 7.60E-06 | 0 | 0 | 0 | 1.34E-05 | 0 | 0 | 6.53E-05 | 0 | 5.34E-05 | 1.95E-06 |
| H2-Ab1 | 0 | 0 | 0 | 0.00019 | 0 | 0 | 0 | 2.54E-05 | 0 | 2.68E-05 | 0 | 0.0007 | 0 | 0 | 0.00031 | 2.13E-05 | 0 | 5.90E-05 |
| Tnxb | 0 | 0 | 0 | 0 | 0 | 0 | 0 | 0 | 0 | 0.00185 | 0 | 0 | 0.00084 | 0 | 0 | 0 | 0 | 0.00028 |

| | | | | | | | | | | | | | | | | | | |
|---|---|---|---|---|---|---|---|---|---|---|---|---|---|---|---|---|---|---|
| C4b | 0 | 0 | 9.01E-05 | 0 | 0 | 0 | 0.00078 | 0 | 0 | 0.00187 | 0 | 0 | 0.00294 | 0 | 0.0001 | 0 | 0 | 0.001 |
| C4a | 0 | 2.96E-05 | 0 | 0 | 0 | 0 | 5.78E-05 | 0 | 0 | 0.00091 | 0 | 0 | 0.00145 | 0 | 0 | 0 | 0 | 0.00036 |
| Lrrc73 | 9.16E-06 | 1.11E-05 | 5.84E-06 | 4.90E-05 | 1.28E-05 | 0 | 3.61E-06 | 1.83E-05 | 0 | 2.45E-06 | 9.35E-06 | 3.70E-06 | 1.20E-06 | 0 | 1.70E-05 | 0 | 2.53E-05 | 1.41E-05 |
| Efhb | 1.00E-06 | 1.82E-06 | 0 | 0 | 2.84E-06 | 0 | 0 | 4.65E-06 | 0 | 9.18E-06 | 0 | 8.69E-06 | 0 | 0 | 1.09E-05 | 3.57E-06 | 1.36E-05 | 9.03E-06 |
| Sgo1 | 0 | 0 | 0 | 0 | 2.20E-06 | 0 | 0 | 0 | 0 | 0 | 0 | 0 | 0 | 0 | 2.63E-05 | 0.00033 | 0 | 0 |
| Lrg1 | 0 | 1.30E-05 | 0 | 0 | 0 | 0 | 0 | 0 | 0 | 3.14E-05 | 0 | 0 | 9.02E-05 | 0 | 3.97E-06 | 0 | 0 | 0.00015 |
| Uhrf1 | 0 | 0 | 0.00016 | 3.59E-05 | 0 | 0 | 0 | 0 | 0 | 7.30E-06 | 0 | 0 | 3.84E-05 | 0 | 0 | 0.00091 | 0 | 0 |
| Tubb4a | 0 | 1.43E-05 | 0 | 0 | 0 | 0 | 0 | 0 | 0 | 0.00024 | 0 | 9.72E-05 | 0 | 0 | 0 | 0 | 0 | 0 |
| C3 | 0 | 0 | 0.00017 | 2.23E-05 | 0 | 0 | 0.00022 | 0.00011 | 0 | 0.00208 | 0 | 0 | 0.00364 | 0 | 0 | 0 | 0 | 0.001 |
| Vav1 | 0 | 8.66E-07 | 0 | 7.37E-06 | 1.48E-06 | 0 | 0 | 0 | 0 | 8.27E-07 | 0 | 1.70E-06 | 1.51E-06 | 0 | 1.58E-05 | 8.92E-06 | 1.72E-05 | 1.02E-05 |
| Ddx11 | 0 | 7.05E-06 | 0 | 3.63E-06 | 0 | 0 | 0 | 0 | 0 | 7.22E-06 | 0 | 0 | 9.85E-06 | 0 | 7.94E-05 | 0.0002 | 0 | 0 |
| Lama1 | 0 | 0 | 0 | 0 | 0.0001 | 0 | 0 | 2.64E-05 | 0 | 0 | 0.00013 | 0 | 0 | 0 | 0 | 0 | 0 | 0 |
| Dlgap1 | 0 | 0 | 0 | 4.99E-06 | 1.05E-05 | 0 | 0 | 0 | 0 | 4.83E-05 | 0 | 2.48E-05 | 0.00015 | 0 | 0 | 0 | 6.15E-06 | 6.57E-06 |
| Ndc80 | 0 | 0 | 0 | 0 | 0 | 0 | 0 | 0 | 0 | 0 | 0 | 0 | 0 | 0 | 5.33E-05 | 0.00068 | 0 | 0 |
| Vit | 0 | 0 | 0 | 0 | 0 | 0 | 0 | 0 | 0 | 0.00011 | 0 | 0 | 0 | 0 | 0 | 0 | 0 | 0.00019 |
| Morn2 | 1.15E-05 | 6.42E-06 | 0 | 1.59E-05 | 0 | 0 | 2.02E-05 | 3.61E-05 | 0 | 1.59E-05 | 0 | 9.08E-06 | 7.17E-07 | 0 | 0 | 1.18E-05 | 0 | 1.94E-05 |
| Stpg4 | 0 | 0 | 0 | 0 | 0 | 0 | 0 | 0 | 0 | 0 | 0 | 2.98E-05 | 1.02E-07 | 0 | 0 | 2.46E-06 | 0.0001 | 0 |
| Ankrd29 | 2.22E-06 | 0 | 0 | 0 | 0 | 0 | 2.13E-05 | 0 | 0 | 3.82E-05 | 0 | 0 | 0 | 0 | 0 | 1.21E-05 | 0 | 1.63E-05 |
| Cabyr | 0 | 2.92E-06 | 2.91E-06 | 0 | 0 | 0 | 0 | 0 | 0 | 7.68E-07 | 0 | 0 | 1.28E-06 | 0 | 6.33E-06 | 2.43E-06 | 6.24E-05 | 4.92E-06 |
| Camk4 | 0 | 0 | 0 | 0 | 2.19E-06 | 0 | 8.76E-05 | 0 | 0 | 0 | 6.38E-05 | 0 | 0 | 0 | 0 | 0 | 0 | 5.11E-05 |
| Kif20a | 3.20E-05 | 0 | 3.80E-06 | 3.13E-05 | 0 | 0 | 0 | 0 | 0 | 0 | 0 | 0 | 0 | 0 | 6.42E-06 | 0.00083 | 0 | 0 |
| Cdc25c | 5.15E-06 | 0 | 0 | 0 | 0 | 0 | 0 | 0 | 0 | 0 | 0 | 0 | 0 | 0 | 8.81E-06 | 0.00037 | 0 | 0 |
| Hbegf | 0 | 0 | 0.00019 | 0 | 0 | 0 | 0 | 0 | 0 | 0 | 0.00012 | 0.00048 | 0 | 0 | 0 | 0 | 0.00059 | 0 |
| Prelid2 | 1.96E-05 | 0 | 0 | 0 | 0 | 0 | 4.08E-05 | 0 | 0 | 4.24E-06 | 0 | 0.00037 | 1.73E-05 | 0 | 0 | 4.91E-06 | 0 | 0 |
| Arl14epl | 0 | 6.86E-06 | 0 | 3.47E-06 | 0 | 0 | 0 | 2.74E-06 | 0 | 2.50E-06 | 0 | 2.05E-06 | 1.99E-05 | 0 | 0 | 0 | 1.69E-05 | 0 |
| Prdm6 | 0 | 0 | 9.66E-07 | 0 | 3.54E-06 | 0 | 0 | 0 | 0 | 2.85E-05 | 0 | 0 | 0 | 0 | 0 | 0 | 0 | 8.66E-05 |
| Adamts19 | 7.63E-07 | 0 | 5.39E-07 | 0 | 0 | 0 | 0 | 0 | 0 | 9.12E-05 | 0 | 0 | 0 | 0 | 1.18E-06 | 2.58E-06 | 0 | 0 |
| Minar2 | 0 | 0 | 0 | 0 | 0 | 0 | 2.09E-05 | 0 | 0 | 3.67E-05 | 0 | 0 | 4.07E-06 | 0 | 0 | 1.28E-07 | 0 | 0 |
| Myoz3 | 0 | 0 | 3.97E-06 | 0 | 0 | 0 | 1.10E-06 | 0 | 0 | 0 | 0 | 1.22E-05 | 2.56E-05 | 0 | 9.29E-06 | 0 | 8.28E-05 | 0 |
| Cd74 | 0 | 5.18E-05 | 0 | 0.00076 | 0 | 0 | 0 | 0 | 0 | 0.0001 | 0 | 4.02E-05 | 3.45E-05 | 0 | 0.00101 | 2.90E-05 | 0.00023 | 0.00015 |
| Onecut2 | 0 | 1.18E-05 | 0 | 2.07E-05 | 9.64E-06 | 0 | 4.99E-06 | 0 | 0 | 1.45E-05 | 1.14E-06 | 1.08E-05 | 4.69E-06 | 0 | 6.77E-05 | 4.83E-06 | 1.73E-05 | 1.23E-05 |
| Mc5r | 0 | 3.37E-06 | 1.90E-06 | 0 | 0 | 0 | 0 | 0 | 0 | 2.58E-06 | 0 | 0 | 1.28E-05 | 0 | 0 | 0 | 6.96E-05 | 0 |
| Ccdc68 | 0 | 2.89E-06 | 0 | 0 | 0 | 0 | 6.82E-06 | 0 | 0 | 0.00013 | 1.58E-05 | 4.01E-06 | 0 | 0 | 0 | 8.37E-07 | 0 | 0 |

| | | | | | | | | | | | | | | | | | | |
|---|---|---|---|---|---|---|---|---|---|---|---|---|---|---|---|---|---|---|
| Rab27b | 0 | 0 | 0 | 0 | 0 | 0 | 3.52E-05 | 0 | 0 | 4.82E-05 | 1.25E-06 | 5.35E-06 | 0 | 0 | 2.24E-05 | 1.43E-05 | 0 | 0 |
| Ska1 | 0 | 0 | 0 | 0 | 0 | 0 | 0 | 0 | 0 | 0 | 0 | 0 | 0 | 0 | 2.63E-06 | 0.00041 | 0 | 0 |
| Pstpip2 | 7.68E-06 | 0 | 9.87E-07 | 1.31E-05 | 0 | 0 | 2.40E-05 | 0 | 0 | 2.91E-06 | 6.39E-06 | 4.78E-07 | 5.47E-06 | 0 | 0 | 4.93E-06 | 0 | 0 |
| Mbp | 0 | 2.02E-05 | 0 | 5.76E-06 | 0 | 0 | 0 | 0 | 0 | 4.53E-05 | 0 | 0 | 5.31E-05 | 0 | 4.90E-07 | 0 | 0 | 3.31E-05 |
| Ankrd13d | 0 | 7.15E-06 | 0 | 0 | 0 | 0 | 0 | 3.56E-06 | 0 | 1.90E-06 | 0 | 1.50E-05 | 1.54E-05 | 0 | 1.79E-05 | 0 | 4.65E-05 | 0 |
| Cdca5 | 0 | 0 | 0 | 0 | 0 | 0 | 0 | 0 | 0 | 0 | 0 | 0 | 0 | 0 | 1.78E-05 | 0.00051 | 0 | 0 |
| Pygm | 0 | 5.42E-06 | 5.36E-06 | 0 | 0 | 0 | 0 | 0 | 0 | 0 | 0 | 1.98E-05 | 8.42E-06 | 0 | 0 | 3.02E-06 | 0.00017 | 9.23E-06 |
| Ccdc88b | 0 | 3.77E-06 | 8.40E-06 | 6.85E-06 | 6.90E-06 | 0 | 9.35E-06 | 7.77E-06 | 0 | 1.34E-05 | 1.10E-05 | 1.13E-05 | 1.93E-05 | 0 | 2.01E-05 | 5.80E-06 | 0 | 9.63E-06 |
| Fermt3 | 0 | 0 | 1.78E-06 | 7.19E-06 | 1.60E-06 | 0 | 0 | 0 | 0 | 4.76E-06 | 0 | 5.83E-06 | 0 | 0 | 3.22E-05 | 4.03E-06 | 1.88E-05 | 1.30E-05 |
| Lrrn4cl | 0 | 0.00022 | 0 | 7.10E-05 | 0 | 0 | 0.00014 | 0 | 0 | 0.00125 | 0 | 0 | 0.00014 | 0 | 0 | 0 | 9.66E-05 | 6.87E-05 |
| Vwce | 0 | 0 | 4.03E-07 | 9.26E-06 | 0 | 0 | 0 | 0 | 0 | 9.79E-07 | 0 | 1.05E-05 | 1.08E-05 | 0 | 0 | 2.74E-06 | 6.14E-05 | 0 |
| Psat1 | 5.38E-05 | 0 | 0.00014 | 6.23E-05 | 0 | 0 | 2.78E-05 | 0 | 0 | 4.55E-05 | 0 | 5.73E-05 | 4.69E-06 | 0 | 0 | 0.00029 | 0 | 0 |
| Gcnt1 | 8.77E-06 | 0 | 1.38E-05 | 0 | 0 | 0 | 9.69E-07 | 4.29E-06 | 0 | 0 | 0 | 3.28E-06 | 0 | 0 | 2.40E-05 | 9.19E-06 | 0 | 0 |
| Trpm6 | 0 | 1.69E-06 | 0 | 0 | 0 | 0 | 5.61E-05 | 0 | 0 | 0 | 0 | 1.42E-07 | 0 | 0 | 0 | 9.72E-07 | 0 | 5.64E-06 |
| Aldh1a1 | 0 | 0 | 0 | 0 | 0 | 0 | 0 | 0 | 0 | 0.00044 | 0 | 0 | 0.00018 | 0 | 0 | 0 | 0 | 0 |
| Tjp2 | 1.11E-05 | 4.66E-05 | 0.00011 | 0 | 9.88E-05 | 0 | 0 | 4.06E-05 | 0 | 0 | 7.46E-05 | 8.34E-05 | 0 | 0 | 8.22E-05 | 6.02E-05 | 0 | 4.83E-05 |
| Vldlr | 0 | 0 | 0 | 0 | 0 | 0 | 0 | 0 | 0 | 3.03E-05 | 0 | 0.00053 | 0 | 0 | 0 | 0 | 0 | 1.41E-05 |
| Gldc | 9.26E-06 | 0 | 0 | 4.34E-06 | 0 | 0 | 0 | 0 | 0 | 6.36E-05 | 0 | 6.98E-07 | 7.67E-05 | 0 | 0 | 2.58E-06 | 0 | 4.64E-06 |
| Ifit3 | 1.37E-05 | 0 | 0 | 0 | 0 | 0 | 0 | 0 | 0 | 0.00017 | 0 | 0 | 1.69E-05 | 0 | 0 | 0 | 0 | 0.00013 |
| Ifit1 | 2.30E-05 | 0 | 0 | 0 | 0 | 0 | 0 | 0 | 0 | 0.00017 | 0 | 0 | 3.17E-05 | 0 | 0 | 0 | 0 | 5.39E-05 |
| Kif20b | 0 | 0 | 0 | 0 | 0 | 0 | 0 | 0 | 0 | 0 | 0 | 0 | 0 | 0 | 9.92E-05 | 0.00041 | 0 | 0 |
| Ankrd1 | 0 | 0 | 0 | 5.97E-06 | 0 | 0 | 0 | 0 | 0 | 1.29E-05 | 0.00015 | 7.50E-06 | 2.33E-05 | 0 | 9.75E-05 | 1.70E-05 | 0 | 1.23E-06 |
| Kif11 | 0 | 0 | 0 | 0 | 0 | 0 | 0 | 0 | 0 | 0 | 0 | 0 | 0 | 0 | 4.21E-05 | 0.00087 | 0 | 0 |
| Exoc6 | 6.26E-06 | 1.22E-05 | 1.48E-06 | 0 | 0 | 0 | 2.10E-06 | 1.05E-05 | 0 | 3.52E-06 | 1.01E-05 | 2.74E-05 | 0 | 0 | 0 | 2.58E-06 | 0 | 8.25E-06 |
| Cep55 | 0 | 0 | 0 | 0 | 0 | 0 | 0 | 0 | 0 | 0 | 0 | 0 | 0 | 0 | 6.65E-06 | 0.00051 | 0 | 0 |
| Rbp4 | 0 | 0 | 0 | 5.09E-05 | 0 | 0 | 9.37E-05 | 0 | 0 | 1.44E-05 | 0 | 0.00021 | 0.00026 | 0 | 0 | 2.98E-05 | 8.07E-05 | 0 |
| Slc35g1 | 7.28E-07 | 2.63E-07 | 6.13E-06 | 0 | 0 | 0 | 0 | 0 | 0 | 2.64E-05 | 0 | 2.61E-06 | 7.66E-06 | 0 | 0 | 2.92E-05 | 0 | 0 |
| Plce1 | 0 | 1.27E-05 | 0 | 0 | 0 | 0 | 1.28E-05 | 1.06E-05 | 0 | 0.00022 | 1.97E-05 | 3.19E-05 | 6.00E-05 | 0 | 7.13E-06 | 3.96E-05 | 4.58E-05 | 7.80E-05 |
| Sorbs1 | 0 | 5.93E-05 | 0 | 0 | 0 | 0 | 0 | 0 | 0 | 0 | 0.00028 | 0 | 0 | 0 | 1.87E-05 | 1.94E-06 | 0 | 3.44E-05 |
| Slit1 | 0 | 0 | 0 | 0 | 7.62E-05 | 0 | 0 | 0.00011 | 0 | 0 | 4.51E-05 | 0.00042 | 0 | 0 | 5.19E-05 | 0 | 0 | 0 |
| Hoga1 | 0 | 9.46E-06 | 0 | 7.78E-06 | 0 | 0 | 1.12E-05 | 0 | 0 | 3.82E-05 | 0 | 4.39E-05 | 1.79E-05 | 0 | 9.43E-06 | 0 | 4.37E-06 | 2.20E-05 |
| Gsto2 | 0 | 0 | 0 | 0 | 0 | 0 | 0 | 0 | 0 | 0 | 0 | 6.74E-05 | 6.81E-05 | 0 | 0 | 0 | 0.0002 | 0 |
| Csf2ra | 0 | 6.82E-06 | 4.99E-06 | 8.27E-05 | 0 | 0 | 5.35E-05 | 1.70E-05 | 0 | 8.38E-05 | 0 | 2.14E-06 | 5.04E-05 | 0 | 0.00018 | 2.72E-05 | 4.24E-05 | 5.03E-05 |

| | | | | | | | | | | | | | | | | | | |
|---|---|---|---|---|---|---|---|---|---|---|---|---|---|---|---|---|---|---|
| Was | 0 | 3.34E-06 | 0 | 9.19E-06 | 1.05E-06 | 0 | 0 | 1.45E-05 | 0 | 1.86E-07 | 2.66E-06 | 0 | 0 | 0 | 1.64E-05 | 6.62E-06 | 1.34E-05 | 0 |
| Cybb | 0 | 4.17E-06 | 0 | 1.49E-05 | 1.25E-06 | 0 | 0 | 0 | 0 | 1.77E-06 | 3.51E-06 | 1.29E-06 | 4.08E-06 | 0 | 2.57E-05 | 0 | 6.56E-06 | 1.14E-06 |
| Klhl13 | 7.78E-05 | 2.30E-05 | 0 | 0 | 0 | 0 | 0 | 0 | 0 | 0.00017 | 0 | 0 | 2.78E-05 | 0 | 0 | 2.15E-05 | 0 | 9.80E-05 |
| Fhl1 | 0 | 0 | 0 | 0 | 0 | 0 | 5.47E-06 | 0 | 0 | 0.00086 | 0.0006 | 0 | 0 | 0 | 0 | 0 | 0 | 0.0008 |
| Prrg3 | 0 | 3.78E-05 | 0 | 0 | 0 | 0 | 0 | 0 | 0 | 0.00026 | 0 | 1.36E-05 | 0.00019 | 0 | 0 | 0 | 0 | 0.00011 |
| Gabre | 0 | 0 | 0 | 0 | 1.37E-05 | 0 | 0 | 0 | 0 | 0.00011 | 0 | 6.83E-05 | 0.00013 | 0 | 6.76E-05 | 0 | 7.97E-06 | 1.11E-05 |
| Gabra3 | 2.32E-05 | 0 | 0 | 0 | 0 | 0 | 3.12E-05 | 0 | 0 | 0.0003 | 0 | 0 | 2.60E-05 | 0 | 0 | 0 | 0 | 0.00023 |
| Kif4 | 0 | 0 | 0 | 0 | 0 | 0 | 0 | 0 | 0 | 0 | 0 | 0 | 0 | 0 | 9.33E-05 | 0.00066 | 0 | 0 |
| Gdpd2 | 0 | 1.76E-06 | 0 | 0 | 0 | 0 | 0 | 0 | 0 | 0 | 0 | 0 | 0.0001 | 0 | 0 | 0 | 0 | 4.66E-05 |
| Rtl3 | 0 | 0 | 0 | 0 | 1.35E-06 | 0 | 0 | 0 | 0 | 0.00016 | 1.39E-06 | 0 | 0 | 0 | 0 | 1.82E-07 | 0 | 0 |
| Pcdh11x | 0 | 0 | 0 | 0 | 0 | 0 | 0 | 0 | 0 | 4.54E-05 | 0 | 0 | 0 | 0 | 4.75E-06 | 0 | 0 | 4.42E-05 |
| Tmem35a | 2.06E-07 | 5.18E-06 | 0 | 0 | 3.07E-06 | 0 | 0 | 0 | 0 | 4.75E-07 | 2.43E-05 | 0 | 2.11E-06 | 0 | 0 | 1.34E-06 | 0 | 2.08E-05 |
| Nrk | 0 | 0 | 0 | 0 | 0 | 0 | 8.05E-07 | 0 | 0 | 0.00011 | 2.21E-08 | 0 | 0 | 0 | 0 | 1.85E-05 | 0 | 0 |
| Pwwp3b | 7.24E-06 | 0 | 9.63E-07 | 5.86E-07 | 0 | 0 | 1.01E-05 | 0 | 0 | 1.49E-05 | 0 | 1.12E-05 | 0 | 0 | 0 | 1.79E-05 | 0.00013 | 0 |
| Cldn2 | 0 | 1.08E-05 | 0 | 1.78E-05 | 1.32E-06 | 0 | 0 | 0 | 0 | 3.83E-06 | 2.88E-06 | 5.24E-06 | 0 | 0 | 5.96E-05 | 5.71E-06 | 2.25E-05 | 3.35E-05 |
| Prps1 | 4.45E-05 | 7.56E-06 | 7.09E-05 | 6.96E-05 | 0 | 0 | 0 | 0 | 0 | 1.11E-05 | 0 | 0 | 0 | 0 | 0 | 9.15E-05 | 3.53E-05 | 3.00E-06 |
| Col4a5 | 0 | 0 | 0 | 0 | 0 | 0 | 4.19E-05 | 0 | 0 | 8.89E-05 | 1.25E-05 | 0 | 1.10E-05 | 0 | 0 | 1.88E-06 | 0 | 2.74E-05 |
| Chrdl1 | 0 | 0 | 0 | 0 | 0 | 0 | 4.78E-05 | 0 | 0 | 0.00072 | 0 | 0 | 0.0004 | 0 | 0 | 0 | 0 | 0.00063 |
| Capn6 | 1.22E-05 | 0 | 0 | 0 | 3.62E-05 | 0 | 2.77E-05 | 1.63E-05 | 0 | 0 | 3.04E-05 | 0 | 0 | 0 | 0 | 0 | 0 | 1.51E-06 |
| Bclaf3 | 5.47E-05 | 5.22E-05 | 2.89E-05 | 0 | 0 | 0 | 0 | 4.07E-05 | 0 | 7.84E-06 | 5.37E-06 | 4.31E-05 | 0 | 0 | 0.00013 | 9.04E-05 | 3.16E-05 | 3.25E-05 |
| Map3k15 | 0 | 2.73E-06 | 0 | 0 | 3.75E-06 | 0 | 2.56E-06 | 4.77E-06 | 0 | 2.03E-06 | 2.43E-06 | 1.81E-05 | 0 | 0 | 0 | 0 | 1.57E-05 | 0 |
| Adgrg2 | 4.37E-07 | 0 | 6.98E-06 | 0 | 0 | 0 | 2.69E-05 | 0 | 0 | 9.14E-05 | 1.57E-06 | 0 | 0 | 0 | 0 | 4.15E-06 | 0 | 0 |
| Reps2 | 0 | 1.56E-05 | 0 | 0 | 0 | 0 | 0 | 2.97E-06 | 0 | 4.87E-05 | 0 | 0 | 3.88E-05 | 0 | 0 | 3.20E-06 | 0 | 5.20E-05 |
| Car5b | 9.08E-06 | 2.03E-05 | 0 | 0 | 0 | 0 | 1.65E-05 | 0 | 0 | 4.36E-05 | 0 | 0 | 0 | 0 | 0 | 0 | 0 | 9.75E-05 |
| Fancb | 5.33E-06 | 0 | 7.90E-06 | 0 | 0 | 0 | 0 | 0 | 0 | 8.24E-06 | 0 | 0 | 8.77E-07 | 0 | 0 | 0.00016 | 0 | 0 |
| DEPRECATED | 0 | 0 | 0 | 9.57E-07 | 0 | 0 | 0 | 0 | 0 | 7.50E-05 | 0 | 0 | 0 | 0 | 5.14E-06 | 0 | 0 | 0 |
| DEPRECATED | 0 | 0 | 0.00067 | 0 | 0 | 0 | 9.64E-05 | 0 | 0 | 0.00042 | 0 | 0 | 0.00043 | 0 | 0 | 0.00018 | 0.00111 | 3.37E-05 |
