## Supplementary Table 2 for "Discovery of a Linked Constellation of Gene Expression Revealed by Local Editing of Fibroblasts in Tumors"

Differentially expressed genes (DEGs) for each cell subtype identified withn major cell types. Major cell types grouped by sheet.

| | p_val | avg_log2FC | pct.1 | pct.2 | p_val_adj | cluster | gene | clustering_1 |
|---|---|---|---|---|---|---|---|---|
| Moxd1 | 0 | 2.65877543 | 0.735 | 0.214 | 0 | 0 | Moxd1 | myCAF_1 |
| Mmp13 | 0 | 2.67031861 | 0.79 | 0.29 | 0 | 0 | Mmp13 | myCAF_1 |
| Podnl1 | 0 | 1.895699 | 0.782 | 0.283 | 0 | 0 | Podnl1 | myCAF_1 |
| Col7a1 | 0 | 2.0325552 | 0.94 | 0.46 | 0 | 0 | Col7a1 | myCAF_1 |
| Adamts16 | 0 | 2.19147118 | 0.676 | 0.201 | 0 | 0 | Adamts16 | myCAF_1 |
| Stra6 | 0 | 1.47010792 | 0.943 | 0.473 | 0 | 0 | Stra6 | myCAF_1 |
| Ucn2 | 0 | 1.94446323 | 0.707 | 0.243 | 0 | 0 | Ucn2 | myCAF_1 |
| Fhod3 | 0 | 1.95190659 | 0.638 | 0.191 | 0 | 0 | Fhod3 | myCAF_1 |
| Stk26 | 0 | 1.70455988 | 0.649 | 0.215 | 0 | 0 | Stk26 | myCAF_1 |
| Pla2g7 | 0 | 2.75294714 | 0.654 | 0.228 | 0 | 0 | Pla2g7 | myCAF_1 |
| Tbx3 | 0 | 1.300483 | 0.801 | 0.377 | 0 | 0 | Tbx3 | myCAF_1 |
| Brinp3 | 0 | 1.49913693 | 0.798 | 0.38 | 0 | 0 | Brinp3 | myCAF_1 |
| Grem1 | 0 | 1.76932403 | 0.672 | 0.266 | 0 | 0 | Grem1 | myCAF_1 |
| Aqp5 | 0 | 2.27396689 | 0.572 | 0.166 | 0 | 0 | Aqp5 | myCAF_1 |
| Mylk | 0 | 1.92351545 | 0.621 | 0.221 | 0 | 0 | Mylk | myCAF_1 |
| Serinc2 | 0 | 1.54215115 | 0.67 | 0.271 | 0 | 0 | Serinc2 | myCAF_1 |
| Clstn2 | 0 | 2.7757586 | 0.54 | 0.142 | 0 | 0 | Clstn2 | myCAF_1 |
| Myl9 | 0 | 1.91944255 | 0.79 | 0.398 | 0 | 0 | Myl9 | myCAF_1 |
| Prdm1 | 0 | 1.16785296 | 0.683 | 0.3 | 0 | 0 | Prdm1 | myCAF_1 |
| Col23a1 | 0 | 1.10734644 | 0.917 | 0.535 | 0 | 0 | Col23a1 | myCAF_1 |
| Ctsc | 0 | 1.04837381 | 0.925 | 0.544 | 0 | 0 | Ctsc | myCAF_1 |
| Wnt2b | 0 | 1.80455149 | 0.759 | 0.379 | 0 | 0 | Wnt2b | myCAF_1 |
| Tagln | 0 | 2.00849743 | 0.821 | 0.443 | 0 | 0 | Tagln | myCAF_1 |
| Piezo2 | 0 | 1.10222972 | 0.632 | 0.261 | 0 | 0 | Piezo2 | myCAF_1 |
| Hs3st1 | 0 | 1.36049756 | 0.871 | 0.509 | 0 | 0 | Hs3st1 | myCAF_1 |
| Adra2a | 0 | 1.04237554 | 0.677 | 0.323 | 0 | 0 | Adra2a | myCAF_1 |
| Rflna | 0 | 1.21435277 | 0.69 | 0.339 | 0 | 0 | Rflna | myCAF_1 |
| Tenm3 | 0 | 1.32230942 | 0.749 | 0.4 | 0 | 0 | Tenm3 | myCAF_1 |
| Nrep | 0 | 1.00855075 | 0.943 | 0.594 | 0 | 0 | Nrep | myCAF_1 |
| Cpxm2 | 0 | 1.38948049 | 0.918 | 0.58 | 0 | 0 | Cpxm2 | myCAF_1 |
| Inhba | 0 | 0.82618664 | 0.952 | 0.619 | 0 | 0 | Inhba | myCAF_1 |
| Mdk | 0 | 1.05743174 | 0.742 | 0.416 | 0 | 0 | Mdk | myCAF_1 |
| Kalrn | 0 | 1.18046614 | 0.731 | 0.406 | 0 | 0 | Kalrn | myCAF_1 |
| Rem1 | 0 | 1.05114144 | 0.679 | 0.355 | 0 | 0 | Rem1 | myCAF_1 |
| Nuak1 | 0 | 1.59777294 | 0.853 | 0.533 | 0 | 0 | Nuak1 | myCAF_1 |
| Gask1b | 0 | 1.36463201 | 0.931 | 0.612 | 0 | 0 | Gask1b | myCAF_1 |
| Nkd2 | 0 | 1.12878112 | 0.809 | 0.491 | 0 | 0 | Nkd2 | myCAF_1 |
| Mamdc2 | 0 | 1.66243594 | 0.515 | 0.207 | 0 | 0 | Mamdc2 | myCAF_1 |
| Cd53 | 0 | 1.35585256 | 0.629 | 0.321 | 0 | 0 | Cd53 | myCAF_1 |
| Lrrc15 | 0 | 1.92113244 | 0.973 | 0.671 | 0 | 0 | Lrrc15 | myCAF_1 |
| Tnfrsf21 | 0 | 1.32153443 | 0.565 | 0.267 | 0 | 0 | Tnfrsf21 | myCAF_1 |
| Mmp10 | 0 | 1.69915558 | 0.444 | 0.149 | 0 | 0 | Mmp10 | myCAF_1 |
| Tnfrsf26 | 0 | 0.77396207 | 0.836 | 0.543 | 0 | 0 | Tnfrsf26 | myCAF_1 |
| Htr2b | 0 | 2.4133547 | 0.364 | 0.073 | 0 | 0 | Htr2b | myCAF_1 |
| Wnt5a | 0 | 1.58597616 | 0.962 | 0.671 | 0 | 0 | Wnt5a | myCAF_1 |
| Bpgm | 0 | 1.32857155 | 0.873 | 0.585 | 0 | 0 | Bpgm | myCAF_1 |
| Syde2 | 0 | 1.33777672 | 0.594 | 0.308 | 0 | 0 | Syde2 | myCAF_1 |
| Tmem86a | 0 | 1.00069814 | 0.902 | 0.617 | 0 | 0 | Tmem86a | myCAF_1 |
| Tanc1 | 0 | 1.09391342 | 0.812 | 0.541 | 0 | 0 | Tanc1 | myCAF_1 |
| Spock3 | 0 | 2.00993083 | 0.369 | 0.101 | 0 | 0 | Spock3 | myCAF_1 |
| Acta2 | 0 | 1.6519612 | 0.803 | 0.535 | 0 | 0 | Acta2 | myCAF_1 |
| Igfbp3 | 0 | 1.51182392 | 0.918 | 0.653 | 0 | 0 | Igfbp3 | myCAF_1 |
| Ncam1 | 0 | 0.97836973 | 0.906 | 0.646 | 0 | 0 | Ncam1 | myCAF_1 |
| Crabp1 | 0 | 0.62206517 | 0.941 | 0.683 | 0 | 0 | Crabp1 | myCAF_1 |
| Sbspon | 0 | 1.87170984 | 0.366 | 0.11 | 0 | 0 | Sbspon | myCAF_1 |
| Sema7a | 0 | 0.81406559 | 0.977 | 0.722 | 0 | 0 | Sema7a | myCAF_1 |
| C1qtnf3 | 0 | 1.13858333 | 0.506 | 0.251 | 0 | 0 | C1qtnf3 | myCAF_1 |
| Hs6st2 | 0 | 1.02720728 | 0.731 | 0.477 | 0 | 0 | Hs6st2 | myCAF_1 |
| Ccbe1 | 0 | 0.58389627 | 0.908 | 0.659 | 0 | 0 | Ccbe1 | myCAF_1 |
| Fbln7 | 0 | 1.05276844 | 0.559 | 0.312 | 0 | 0 | Fbln7 | myCAF_1 |
| Kif26b | 0 | 0.97353409 | 0.959 | 0.72 | 0 | 0 | Kif26b | myCAF_1 |
| Fbln5 | 0 | 1.29821662 | 0.818 | 0.583 | 0 | 0 | Fbln5 | myCAF_1 |
| Tnn | 0 | 1.70022604 | 0.433 | 0.199 | 0 | 0 | Tnn | myCAF_1 |
| Scarf2 | 0 | 0.9469122 | 0.86 | 0.626 | 0 | 0 | Scarf2 | myCAF_1 |
| Ptk7 | 0 | 1.13626 | 0.986 | 0.753 | 0 | 0 | Ptk7 | myCAF_1 |
| Pdlim2 | 0 | 0.87127548 | 0.805 | 0.573 | 0 | 0 | Pdlim2 | myCAF_1 |
| Col27a1 | 0 | 0.89316483 | 0.958 | 0.732 | 0 | 0 | Col27a1 | myCAF_1 |
| Spats2l | 0 | 0.77935528 | 0.888 | 0.663 | 0 | 0 | Spats2l | myCAF_1 |
| Tfpi2 | 0 | 0.7716135 | 0.887 | 0.663 | 0 | 0 | Tfpi2 | myCAF_1 |
| Fcgr2b | 0 | 1.01857214 | 0.528 | 0.305 | 0 | 0 | Fcgr2b | myCAF_1 |
| Ston1 | 0 | 0.9465465 | 0.869 | 0.647 | 0 | 0 | Ston1 | myCAF_1 |
| Ermp1 | 0 | 1.02968062 | 0.879 | 0.658 | 0 | 0 | Ermp1 | myCAF_1 |
| Mmp11 | 0 | 1.11276124 | 0.75 | 0.53 | 0 | 0 | Mmp11 | myCAF_1 |
| Serpine2 | 0 | 1.10610569 | 0.974 | 0.755 | 0 | 0 | Serpine2 | myCAF_1 |
| Crispld2 | 0 | 1.0288674 | 0.905 | 0.691 | 0 | 0 | Crispld2 | myCAF_1 |
| Col8a1 | 0 | 0.90685795 | 0.717 | 0.505 | 0 | 0 | Col8a1 | myCAF_1 |
| Megf6 | 0 | 1.57298147 | 0.347 | 0.142 | 0 | 0 | Megf6 | myCAF_1 |
| Ccn4 | 0 | 0.94007554 | 0.919 | 0.716 | 0 | 0 | Ccn4 | myCAF_1 |
| Prss35 | 0 | 2.56940435 | 0.272 | 0.07 | 0 | 0 | Prss35 | myCAF_1 |
| Actg2 | 0 | 2.72019888 | 0.279 | 0.078 | 0 | 0 | Actg2 | myCAF_1 |
| Dusp6 | 0 | 0.81485143 | 0.725 | 0.525 | 0 | 0 | Dusp6 | myCAF_1 |
| Dact1 | 0 | 0.76874519 | 0.828 | 0.628 | 0 | 0 | Dact1 | myCAF_1 |
| Gng11 | 0 | 0.56993641 | 0.942 | 0.744 | 0 | 0 | Gng11 | myCAF_1 |
| Cd59a | 0 | 2.13087527 | 0.295 | 0.1 | 0 | 0 | Cd59a | myCAF_1 |
| Lbh | 0 | 1.34352315 | 0.945 | 0.752 | 0 | 0 | Lbh | myCAF_1 |
| Tpm2 | 0 | 1.02423406 | 0.898 | 0.707 | 0 | 0 | Tpm2 | myCAF_1 |
| Hs6st1 | 0 | 0.86435403 | 0.803 | 0.615 | 0 | 0 | Hs6st1 | myCAF_1 |
| Srpx2 | 0 | 0.81551119 | 0.909 | 0.723 | 0 | 0 | Srpx2 | myCAF_1 |
| Fbn2 | 0 | 0.91031904 | 0.805 | 0.621 | 0 | 0 | Fbn2 | myCAF_1 |
| Dkk3 | 0 | 1.29780233 | 0.964 | 0.782 | 0 | 0 | Dkk3 | myCAF_1 |
| Maf | 0 | 1.11994719 | 0.954 | 0.778 | 0 | 0 | Maf | myCAF_1 |
| Lgalsl | 0 | 0.9606957 | 0.826 | 0.652 | 0 | 0 | Lgalsl | myCAF_1 |
| Chchd10 | 0 | 1.08985077 | 0.316 | 0.144 | 0 | 0 | Chchd10 | myCAF_1 |
| Uck2 | 0 | 1.29946959 | 0.991 | 0.832 | 0 | 0 | Uck2 | myCAF_1 |
| Spon1 | 0 | 1.23065218 | 0.995 | 0.841 | 0 | 0 | Spon1 | myCAF_1 |
| Ecrg4 | 0 | 0.92520841 | 0.498 | 0.345 | 0 | 0 | Ecrg4 | myCAF_1 |
| Cnn2 | 0 | 0.9617309 | 0.942 | 0.789 | 0 | 0 | Cnn2 | myCAF_1 |
| Mmp9 | 0 | 0.51705285 | 0.427 | 0.277 | 0 | 0 | Mmp9 | myCAF_1 |
| Dhrs3 | 0 | 0.66324612 | 0.832 | 0.682 | 0 | 0 | Dhrs3 | myCAF_1 |
| Fzd1 | 0 | 0.84629135 | 0.97 | 0.826 | 0 | 0 | Fzd1 | myCAF_1 |
| Pla1a | 0 | 0.74250696 | 0.982 | 0.838 | 0 | 0 | Pla1a | myCAF_1 |
| Fndc1 | 0 | 0.7353566 | 0.909 | 0.765 | 0 | 0 | Fndc1 | myCAF_1 |
| Fam20a | 0 | 0.79219213 | 0.898 | 0.755 | 0 | 0 | Fam20a | myCAF_1 |
| Rbp1 | 0 | 0.57593369 | 0.923 | 0.781 | 0 | 0 | Rbp1 | myCAF_1 |
| Csrp1 | 0 | 1.22832952 | 0.97 | 0.829 | 0 | 0 | Csrp1 | myCAF_1 |
| Cfh | 0 | 0.94900388 | 0.956 | 0.817 | 0 | 0 | Cfh | myCAF_1 |
| Tnc | 0 | 0.53948303 | 1 | 0.864 | 0 | 0 | Tnc | myCAF_1 |
| Mical2 | 0 | 0.59519601 | 0.939 | 0.804 | 0 | 0 | Mical2 | myCAF_1 |
| Twist2 | 0 | 0.6054222 | 0.904 | 0.772 | 0 | 0 | Twist2 | myCAF_1 |
| Islr | 0 | 0.66133914 | 0.903 | 0.777 | 0 | 0 | Islr | myCAF_1 |
| Rnf149 | 0 | 0.64515558 | 0.982 | 0.859 | 0 | 0 | Rnf149 | myCAF_1 |
| Adam12 | 0 | 0.68640676 | 0.96 | 0.838 | 0 | 0 | Adam12 | myCAF_1 |
| Dcakd | 0 | 0.85797263 | 0.942 | 0.822 | 0 | 0 | Dcakd | myCAF_1 |
| Thbs1 | 0 | 0.73061383 | 0.911 | 0.796 | 0 | 0 | Thbs1 | myCAF_1 |
| DEPRECATED-l | 0 | 0.65746761 | 0.79 | 0.677 | 0 | 0 | DEPRECATED-l | myCAF_1 |
| Adamts12 | 0 | 1.25755215 | 0.99 | 0.881 | 0 | 0 | Adamts12 | myCAF_1 |
| Sdc1 | 0 | 0.35095993 | 0.99 | 0.885 | 0 | 0 | Sdc1 | myCAF_1 |
| Rab31 | 0 | 0.85501861 | 0.929 | 0.825 | 0 | 0 | Rab31 | myCAF_1 |
| Akr1b8 | 0 | 0.68021197 | 0.953 | 0.856 | 0 | 0 | Akr1b8 | myCAF_1 |
| Fzd2 | 0 | 0.72072675 | 0.919 | 0.825 | 0 | 0 | Fzd2 | myCAF_1 |
| Cdh11 | 0 | 0.76765888 | 0.981 | 0.888 | 0 | 0 | Cdh11 | myCAF_1 |
| Adamts4 | 0 | 0.54092373 | 0.972 | 0.88 | 0 | 0 | Adamts4 | myCAF_1 |
| Spp1 | 0 | 0.87696167 | 0.993 | 0.905 | 0 | 0 | Spp1 | myCAF_1 |
| Palld | 0 | 1.14278572 | 0.978 | 0.891 | 0 | 0 | Palld | myCAF_1 |
| Kremen1 | 0 | 0.48497498 | 0.965 | 0.878 | 0 | 0 | Kremen1 | myCAF_1 |
| Postn | 0 | 0.94552493 | 0.998 | 0.921 | 0 | 0 | Postn | myCAF_1 |
| Htra1 | 0 | 0.66355227 | 0.881 | 0.805 | 0 | 0 | Htra1 | myCAF_1 |
| Actn1 | 0 | 0.48380047 | 0.96 | 0.884 | 0 | 0 | Actn1 | myCAF_1 |
| Tcf4 | 0 | 0.50908137 | 0.975 | 0.9 | 0 | 0 | Tcf4 | myCAF_1 |
| Ccn2 | 0 | 0.96745604 | 0.506 | 0.433 | 0 | 0 | Ccn2 | myCAF_1 |
| Col12a1 | 0 | 1.22996327 | 0.998 | 0.928 | 0 | 0 | Col12a1 | myCAF_1 |
| Pdgfrb | 0 | 0.47304522 | 0.992 | 0.925 | 0 | 0 | Pdgfrb | myCAF_1 |
| Ccn1 | 0 | 0.42054193 | 0.946 | 0.882 | 0 | 0 | Ccn1 | myCAF_1 |
| Tpm1 | 0 | 1.34115951 | 0.993 | 0.93 | 0 | 0 | Tpm1 | myCAF_1 |
| Esyt2 | 0 | 0.61300611 | 0.915 | 0.857 | 0 | 0 | Esyt2 | myCAF_1 |
| Mfge8 | 0 | 0.76021166 | 0.984 | 0.932 | 0 | 0 | Mfge8 | myCAF_1 |
| C1qtnf6 | 0 | 0.47111809 | 0.981 | 0.931 | 0 | 0 | C1qtnf6 | myCAF_1 |
| Flna | 0 | 0.620492 | 0.982 | 0.935 | 0 | 0 | Flna | myCAF_1 |
| Egfp | 0 | 0.43825585 | 0.966 | 0.919 | 0 | 0 | Egfp | myCAF_1 |
| Pdgfra | 0 | 0.57085664 | 0.974 | 0.932 | 0 | 0 | Pdgfra | myCAF_1 |
| Prkar1a | 0 | 0.6062479 | 0.973 | 0.931 | 0 | 0 | Prkar1a | myCAF_1 |
| Mxra8 | 0 | 0.56710396 | 0.992 | 0.951 | 0 | 0 | Mxra8 | myCAF_1 |
| Pmepa1 | 0 | 1.06854266 | 0.995 | 0.955 | 0 | 0 | Pmepa1 | myCAF_1 |
| Itgb5 | 0 | 0.71026208 | 0.989 | 0.954 | 0 | 0 | Itgb5 | myCAF_1 |
| Loxl2 | 0 | 0.37542048 | 0.987 | 0.954 | 0 | 0 | Loxl2 | myCAF_1 |
| Angptl2 | 0 | 1.08173008 | 0.995 | 0.966 | 0 | 0 | Angptl2 | myCAF_1 |
| Antxr1 | 0 | 0.61186927 | 0.99 | 0.961 | 0 | 0 | Antxr1 | myCAF_1 |
| Mrc2 | 0 | 0.52444828 | 0.992 | 0.963 | 0 | 0 | Mrc2 | myCAF_1 |
| Mtch1 | 0 | 0.61352881 | 0.992 | 0.964 | 0 | 0 | Mtch1 | myCAF_1 |
| Ctsk | 0 | 0.39319209 | 0.991 | 0.964 | 0 | 0 | Ctsk | myCAF_1 |
| Thbs2 | 0 | 0.44197548 | 0.994 | 0.968 | 0 | 0 | Thbs2 | myCAF_1 |
| Ppic | 0 | 0.78320799 | 0.994 | 0.968 | 0 | 0 | Ppic | myCAF_1 |
| Itm2c | 0 | 0.37583937 | 0.988 | 0.965 | 0 | 0 | Itm2c | myCAF_1 |
| Col5a2 | 0 | 0.65475372 | 0.997 | 0.976 | 0 | 0 | Col5a2 | myCAF_1 |
| Itgb1 | 0 | 0.46830652 | 0.991 | 0.97 | 0 | 0 | Itgb1 | myCAF_1 |
| Fgfr1 | 0 | 0.69247163 | 0.992 | 0.972 | 0 | 0 | Fgfr1 | myCAF_1 |
| Eln | 0 | 0.4860434 | 0.558 | 0.54 | 0 | 0 | Eln | myCAF_1 |
| Adamts2 | 0 | 0.62035535 | 0.996 | 0.978 | 0 | 0 | Adamts2 | myCAF_1 |
| Cald1 | 0 | 0.42410165 | 0.995 | 0.979 | 0 | 0 | Cald1 | myCAF_1 |
| Mmp14 | 0 | 0.30525399 | 0.997 | 0.981 | 0 | 0 | Mmp14 | myCAF_1 |
| Col6a3 | 0 | 0.6642283 | 0.999 | 0.984 | 0 | 0 | Col6a3 | myCAF_1 |
| Col6a1 | 0 | 0.6082337 | 1 | 0.986 | 0 | 0 | Col6a1 | myCAF_1 |
| App | 0 | 0.306621 | 0.996 | 0.982 | 0 | 0 | App | myCAF_1 |
| Tmsb4x | 0 | 0.47924914 | 0.996 | 0.982 | 0 | 0 | Tmsb4x | myCAF_1 |
| Col6a2 | 0 | 0.6228196 | 0.999 | 0.987 | 0 | 0 | Col6a2 | myCAF_1 |
| Bgn | 0 | 0.3794213 | 0.999 | 0.987 | 0 | 0 | Bgn | myCAF_1 |
| Igfbp7 | 0 | 0.47734602 | 1 | 0.99 | 0 | 0 | Igfbp7 | myCAF_1 |
| Fn1 | 0 | 0.26352407 | 0.999 | 0.99 | 0 | 0 | Fn1 | myCAF_1 |
| Col5a1 | 0 | 0.50133789 | 0.997 | 0.988 | 0 | 0 | Col5a1 | myCAF_1 |
| Col1a2 | 0 | 0.35170474 | 1 | 0.996 | 0 | 0 | Col1a2 | myCAF_1 |
| Col1a1 | 0 | 0.43897356 | 1 | 0.997 | 0 | 0 | Col1a1 | myCAF_1 |
| Plaat3 | 8.68E-293 | 0.95903621 | 0.799 | 0.541 | 1.68E-288 | 0 | Plaat3 | myCAF_1 |
| Eid1 | 2.42E-293 | 0.38809624 | 0.98 | 0.947 | 4.68E-289 | 0 | Eid1 | myCAF_1 |
| Apcdd1 | 3.72E-293 | 0.77154303 | 0.659 | 0.395 | 7.19E-289 | 0 | Apcdd1 | myCAF_1 |
| Glp1r | 1.23E-291 | 1.58812371 | 0.277 | 0.129 | 2.39E-287 | 0 | Glp1r | myCAF_1 |
| Nkd1 | 2.88E-290 | 1.0819472 | 0.525 | 0.298 | 5.58E-286 | 0 | Nkd1 | myCAF_1 |
| Tmem119 | 1.12E-288 | 0.45549951 | 0.962 | 0.855 | 2.17E-284 | 0 | Tmem119 | myCAF_1 |
| Tgfb2 | 2.13E-288 | 0.77689616 | 0.767 | 0.545 | 4.11E-284 | 0 | Tgfb2 | myCAF_1 |
| Myh9 | 2.02E-284 | 0.45605384 | 0.966 | 0.913 | 3.90E-280 | 0 | Myh9 | myCAF_1 |
| Tgfb3 | 8.96E-284 | 0.71416782 | 0.857 | 0.634 | 1.73E-279 | 0 | Tgfb3 | myCAF_1 |
| Fnbp1 | 2.44E-283 | 0.60556438 | 0.902 | 0.767 | 4.72E-279 | 0 | Fnbp1 | myCAF_1 |
| Tmem204 | 3.56E-283 | 0.87159361 | 0.544 | 0.262 | 6.88E-279 | 0 | Tmem204 | myCAF_1 |
| Fbln2 | 6.12E-278 | 0.28952724 | 0.996 | 0.979 | 1.18E-273 | 0 | Fbln2 | myCAF_1 |
| Cacna1c | 1.25E-275 | 0.70871799 | 0.883 | 0.662 | 2.42E-271 | 0 | Cacna1c | myCAF_1 |
| Mfap2 | 1.51E-271 | 0.41913766 | 0.979 | 0.914 | 2.93E-267 | 0 | Mfap2 | myCAF_1 |
| Ociad2 | 1.67E-270 | 0.829942 | 0.683 | 0.404 | 3.24E-266 | 0 | Ociad2 | myCAF_1 |
| Gna14 | 4.63E-269 | 1.25763582 | 0.515 | 0.233 | 8.96E-265 | 0 | Gna14 | myCAF_1 |
| Lpp | 1.24E-266 | 0.44363956 | 0.957 | 0.914 | 2.41E-262 | 0 | Lpp | myCAF_1 |
| Picalm | 2.36E-264 | 0.59089903 | 0.931 | 0.864 | 4.56E-260 | 0 | Picalm | myCAF_1 |
| Myl12a | 3.18E-263 | 0.3958413 | 0.975 | 0.942 | 6.16E-259 | 0 | Myl12a | myCAF_1 |
| Grem2 | 1.42E-261 | 0.5531018 | 0.914 | 0.776 | 2.74E-257 | 0 | Grem2 | myCAF_1 |
| Nudt4 | 1.62E-258 | 0.5294683 | 0.952 | 0.855 | 3.13E-254 | 0 | Nudt4 | myCAF_1 |
| Ntn4 | 6.89E-258 | 1.06717251 | 0.547 | 0.371 | 1.33E-253 | 0 | Ntn4 | myCAF_1 |
| Gpr176 | 2.13E-254 | 0.7320163 | 0.834 | 0.635 | 4.12E-250 | 0 | Gpr176 | myCAF_1 |
| Adamts6 | 2.38E-252 | 1.15937418 | 0.54 | 0.288 | 4.59E-248 | 0 | Adamts6 | myCAF_1 |
| Eva1b | 3.29E-248 | 0.55459435 | 0.931 | 0.804 | 6.36E-244 | 0 | Eva1b | myCAF_1 |
| Cemip2 | 4.85E-246 | 0.64354016 | 0.791 | 0.584 | 9.37E-242 | 0 | Cemip2 | myCAF_1 |
| Trp53inp2 | 8.16E-246 | 0.77720106 | 0.824 | 0.647 | 1.58E-241 | 0 | Trp53inp2 | myCAF_1 |
| Arxes2 | 4.39E-244 | 1.28245083 | 0.589 | 0.304 | 8.49E-240 | 0 | Arxes2 | myCAF_1 |
| Jph2 | 2.24E-242 | 1.32368429 | 0.429 | 0.211 | 4.34E-238 | 0 | Jph2 | myCAF_1 |
| Cd63 | 1.40E-236 | 0.29428039 | 0.996 | 0.985 | 2.71E-232 | 0 | Cd63 | myCAF_1 |
| Vmp1 | 3.29E-236 | 0.39669455 | 0.982 | 0.933 | 6.36E-232 | 0 | Vmp1 | myCAF_1 |
| Htr1b | 4.01E-234 | 1.08479411 | 0.584 | 0.298 | 7.76E-230 | 0 | Htr1b | myCAF_1 |
| Myo1d | 8.16E-233 | 0.90603635 | 0.76 | 0.495 | 1.58E-228 | 0 | Myo1d | myCAF_1 |
| Cdc42bpa | 1.76E-228 | 0.59330465 | 0.863 | 0.737 | 3.40E-224 | 0 | Cdc42bpa | myCAF_1 |
| Sorcs2 | 1.96E-226 | 0.70950314 | 0.792 | 0.574 | 3.78E-222 | 0 | Sorcs2 | myCAF_1 |
| Sulf1 | 5.98E-223 | 0.39010116 | 0.961 | 0.905 | 1.16E-218 | 0 | Sulf1 | myCAF_1 |
| Ybx3 | 9.04E-221 | 0.3325676 | 0.974 | 0.92 | 1.75E-216 | 0 | Ybx3 | myCAF_1 |
| Unc5b | 3.53E-213 | 0.59156587 | 0.83 | 0.526 | 6.82E-209 | 0 | Unc5b | myCAF_1 |
| Parva | 3.27E-209 | 0.47303954 | 0.961 | 0.898 | 6.32E-205 | 0 | Parva | myCAF_1 |
| Cdc42ep3 | 5.77E-208 | 1.0894803 | 0.355 | 0.183 | 1.12E-203 | 0 | Cdc42ep3 | myCAF_1 |
| Ptgfrn | 1.49E-206 | 0.42760728 | 0.98 | 0.945 | 2.87E-202 | 0 | Ptgfrn | myCAF_1 |
| Plcg1 | 8.98E-206 | 0.80878994 | 0.801 | 0.61 | 1.74E-201 | 0 | Plcg1 | myCAF_1 |
| Myh10 | 1.93E-205 | 0.63191456 | 0.806 | 0.691 | 3.73E-201 | 0 | Myh10 | myCAF_1 |
| Prickle1 | 5.69E-204 | 0.57191412 | 0.877 | 0.736 | 1.10E-199 | 0 | Prickle1 | myCAF_1 |
| Gpc1 | 3.16E-203 | 0.28115182 | 0.99 | 0.95 | 6.11E-199 | 0 | Gpc1 | myCAF_1 |
| Gdpd5 | 6.30E-203 | 1.25171618 | 0.425 | 0.211 | 1.22E-198 | 0 | Gdpd5 | myCAF_1 |
| Edil3 | 1.85E-202 | 1.34083158 | 0.362 | 0.139 | 3.58E-198 | 0 | Edil3 | myCAF_1 |
| Mast4 | 2.00E-202 | 0.65423757 | 0.823 | 0.627 | 3.88E-198 | 0 | Mast4 | myCAF_1 |
| Loxl3 | 2.07E-197 | 0.45940256 | 0.971 | 0.894 | 3.99E-193 | 0 | Loxl3 | myCAF_1 |
| Atp1a3 | 7.02E-197 | 1.31770081 | 0.389 | 0.17 | 1.36E-192 | 0 | Atp1a3 | myCAF_1 |
| Msrb3 | 3.26E-196 | 0.65975222 | 0.837 | 0.711 | 6.31E-192 | 0 | Msrb3 | myCAF_1 |
| Itgb3 | 6.38E-196 | 0.57103263 | 0.815 | 0.621 | 1.23E-191 | 0 | Itgb3 | myCAF_1 |
| Tspan3 | 2.43E-195 | 0.39356181 | 0.977 | 0.943 | 4.70E-191 | 0 | Tspan3 | myCAF_1 |
| Ak1 | 4.11E-195 | 0.45724159 | 0.929 | 0.842 | 7.96E-191 | 0 | Ak1 | myCAF_1 |
| Dstn | 7.49E-193 | 0.44456906 | 0.919 | 0.884 | 1.45E-188 | 0 | Dstn | myCAF_1 |
| Rflnb | 1.72E-190 | 0.94230807 | 0.671 | 0.409 | 3.33E-186 | 0 | Rflnb | myCAF_1 |
| Inava | 2.23E-190 | 1.32324309 | 0.548 | 0.264 | 4.31E-186 | 0 | Inava | myCAF_1 |
| Furin | 3.06E-190 | 0.39780683 | 0.977 | 0.932 | 5.91E-186 | 0 | Furin | myCAF_1 |
| Rai14 | 5.31E-190 | 0.42256964 | 0.922 | 0.805 | 1.03E-185 | 0 | Rai14 | myCAF_1 |
| Qsox1 | 9.26E-189 | 0.43202413 | 0.964 | 0.921 | 1.79E-184 | 0 | Qsox1 | myCAF_1 |
| Serinc3 | 4.31E-187 | 0.36509206 | 0.976 | 0.946 | 8.32E-183 | 0 | Serinc3 | myCAF_1 |
| Rgma | 6.04E-187 | 0.59445487 | 0.79 | 0.664 | 1.17E-182 | 0 | Rgma | myCAF_1 |
| Camk2n1 | 7.28E-187 | 0.45526042 | 0.862 | 0.721 | 1.41E-182 | 0 | Camk2n1 | myCAF_1 |
| Pdzrn3 | 1.14E-182 | 0.53502026 | 0.938 | 0.83 | 2.21E-178 | 0 | Pdzrn3 | myCAF_1 |
| Odc1 | 3.14E-180 | 0.49002736 | 0.945 | 0.858 | 6.07E-176 | 0 | Odc1 | myCAF_1 |
| Ptprn | 5.00E-180 | 0.60454073 | 0.798 | 0.472 | 9.66E-176 | 0 | Ptprn | myCAF_1 |
| Rnf122 | 2.37E-178 | 0.88323583 | 0.659 | 0.49 | 4.58E-174 | 0 | Rnf122 | myCAF_1 |
| Abhd2 | 1.61E-175 | 0.35549009 | 0.946 | 0.879 | 3.12E-171 | 0 | Abhd2 | myCAF_1 |
| Bace1 | 4.44E-173 | 0.55071751 | 0.915 | 0.799 | 8.58E-169 | 0 | Bace1 | myCAF_1 |
| Tpm4 | 2.17E-172 | 0.31939994 | 0.982 | 0.954 | 4.20E-168 | 0 | Tpm4 | myCAF_1 |
| Clec2d | 2.49E-172 | 0.34137647 | 0.93 | 0.815 | 4.82E-168 | 0 | Clec2d | myCAF_1 |
| Pqlc1 | 4.10E-170 | 0.71277868 | 0.745 | 0.575 | 7.94E-166 | 0 | Pqlc1 | myCAF_1 |
| Gxylt2 | 6.54E-170 | 0.65895724 | 0.818 | 0.632 | 1.26E-165 | 0 | Gxylt2 | myCAF_1 |
| Entpd7 | 2.82E-168 | 0.67709929 | 0.834 | 0.674 | 5.45E-164 | 0 | Entpd7 | myCAF_1 |
| Cd40 | 5.05E-166 | 1.07786021 | 0.315 | 0.124 | 9.77E-162 | 0 | Cd40 | myCAF_1 |
| Mfap4 | 2.12E-164 | 0.61597355 | 0.808 | 0.533 | 4.10E-160 | 0 | Mfap4 | myCAF_1 |
| Adamts7 | 3.78E-163 | 0.79462577 | 0.691 | 0.468 | 7.30E-159 | 0 | Adamts7 | myCAF_1 |
| Pthlh | 3.06E-160 | 1.23019551 | 0.282 | 0.136 | 5.92E-156 | 0 | Pthlh | myCAF_1 |
| Myl6 | 4.97E-160 | 0.4715978 | 0.913 | 0.833 | 9.61E-156 | 0 | Myl6 | myCAF_1 |
| Ttyh3 | 8.56E-159 | 0.49993199 | 0.902 | 0.808 | 1.65E-154 | 0 | Ttyh3 | myCAF_1 |
| Sphk1 | 1.69E-158 | 0.63589192 | 0.841 | 0.65 | 3.26E-154 | 0 | Sphk1 | myCAF_1 |
| Foxn3 | 1.95E-157 | 0.59288278 | 0.807 | 0.676 | 3.77E-153 | 0 | Foxn3 | myCAF_1 |
| Crlf1 | 3.50E-157 | 0.67301884 | 0.559 | 0.331 | 6.76E-153 | 0 | Crlf1 | myCAF_1 |
| Azin2 | 3.75E-157 | 0.62371796 | 0.8 | 0.646 | 7.24E-153 | 0 | Azin2 | myCAF_1 |
| Map1lc3a | 2.27E-156 | 0.43248785 | 0.934 | 0.863 | 4.39E-152 | 0 | Map1lc3a | myCAF_1 |
| Inpp5a | 5.87E-156 | 0.58306524 | 0.814 | 0.714 | 1.14E-151 | 0 | Inpp5a | myCAF_1 |
| Pitpnc1 | 1.85E-154 | 0.74592778 | 0.719 | 0.496 | 3.58E-150 | 0 | Pitpnc1 | myCAF_1 |
| Cacnb3 | 3.99E-153 | 0.57426937 | 0.871 | 0.749 | 7.71E-149 | 0 | Cacnb3 | myCAF_1 |
| Rbpms2 | 6.74E-153 | 0.84293448 | 0.577 | 0.329 | 1.30E-148 | 0 | Rbpms2 | myCAF_1 |
| Tfr2 | 9.45E-153 | 1.53708028 | 0.289 | 0.106 | 1.83E-148 | 0 | Tfr2 | myCAF_1 |
| Gas1 | 8.12E-152 | 0.37565779 | 0.934 | 0.823 | 1.57E-147 | 0 | Gas1 | myCAF_1 |
| Slc9a3r2 | 1.25E-150 | 0.94469825 | 0.517 | 0.291 | 2.43E-146 | 0 | Slc9a3r2 | myCAF_1 |
| Irf2bp2 | 2.81E-150 | 0.30591479 | 0.972 | 0.932 | 5.43E-146 | 0 | Irf2bp2 | myCAF_1 |
| Nxn | 4.08E-150 | 0.37986469 | 0.973 | 0.899 | 7.89E-146 | 0 | Nxn | myCAF_1 |
| Actn4 | 7.98E-150 | 0.31506121 | 0.968 | 0.942 | 1.54E-145 | 0 | Actn4 | myCAF_1 |
| Syn1 | 2.22E-148 | 1.32888546 | 0.32 | 0.14 | 4.30E-144 | 0 | Syn1 | myCAF_1 |
| Adam19 | 6.41E-148 | 0.42045006 | 0.898 | 0.825 | 1.24E-143 | 0 | Adam19 | myCAF_1 |
| Ehd3 | 2.12E-146 | 1.49607851 | 0.363 | 0.172 | 4.10E-142 | 0 | Ehd3 | myCAF_1 |
| Mtss2 | 1.91E-144 | 0.73087226 | 0.59 | 0.386 | 3.68E-140 | 0 | Mtss2 | myCAF_1 |
| Ak5 | 6.26E-144 | 1.42168349 | 0.386 | 0.174 | 1.21E-139 | 0 | Ak5 | myCAF_1 |
| Myo1e | 1.07E-140 | 0.74625939 | 0.594 | 0.397 | 2.06E-136 | 0 | Myo1e | myCAF_1 |
| Slc6a8 | 5.66E-139 | 0.36378608 | 0.904 | 0.835 | 1.09E-134 | 0 | Slc6a8 | myCAF_1 |
| Laptm4b | 1.65E-138 | 0.69053155 | 0.767 | 0.605 | 3.18E-134 | 0 | Laptm4b | myCAF_1 |
| Limk2 | 3.00E-138 | 0.79068287 | 0.666 | 0.484 | 5.81E-134 | 0 | Limk2 | myCAF_1 |
| Il20ra | 7.48E-138 | 1.47214419 | 0.338 | 0.132 | 1.45E-133 | 0 | Il20ra | myCAF_1 |
| Pdgfa | 1.35E-134 | 0.8490733 | 0.353 | 0.218 | 2.61E-130 | 0 | Pdgfa | myCAF_1 |
| Filip1l | 1.75E-132 | 0.91275531 | 0.464 | 0.329 | 3.38E-128 | 0 | Filip1l | myCAF_1 |
| Mxra7 | 1.85E-132 | 0.42526931 | 0.882 | 0.795 | 3.58E-128 | 0 | Mxra7 | myCAF_1 |
| Aif1l | 4.85E-131 | 1.06505686 | 0.401 | 0.252 | 9.38E-127 | 0 | Aif1l | myCAF_1 |
| Gpm6b | 1.69E-129 | 0.51222035 | 0.895 | 0.766 | 3.27E-125 | 0 | Gpm6b | myCAF_1 |
| Csnk1g2 | 7.66E-129 | 0.39870997 | 0.931 | 0.874 | 1.48E-124 | 0 | Csnk1g2 | myCAF_1 |
| Kazn | 2.04E-128 | 0.67756186 | 0.562 | 0.359 | 3.94E-124 | 0 | Kazn | myCAF_1 |
| Vcl | 3.64E-128 | 0.31053882 | 0.889 | 0.85 | 7.04E-124 | 0 | Vcl | myCAF_1 |
| Gng2 | 7.87E-127 | 0.85501367 | 0.473 | 0.259 | 1.52E-122 | 0 | Gng2 | myCAF_1 |
| Avpr1a | 1.16E-126 | 0.61533422 | 0.779 | 0.586 | 2.25E-122 | 0 | Avpr1a | myCAF_1 |
| Tln1 | 2.05E-126 | 0.34404995 | 0.959 | 0.915 | 3.97E-122 | 0 | Tln1 | myCAF_1 |
| Robo1 | 3.28E-126 | 0.55440533 | 0.671 | 0.402 | 6.33E-122 | 0 | Robo1 | myCAF_1 |
| Ptprm | 5.86E-126 | 0.65575686 | 0.769 | 0.58 | 1.13E-121 | 0 | Ptprm | myCAF_1 |
| Pakap | 1.46E-125 | 0.62203594 | 0.632 | 0.412 | 2.82E-121 | 0 | Pakap | myCAF_1 |
| Stx6 | 2.62E-125 | 0.57262371 | 0.802 | 0.667 | 5.07E-121 | 0 | Stx6 | myCAF_1 |
| Ilk | 1.38E-123 | 0.36662866 | 0.948 | 0.905 | 2.68E-119 | 0 | Ilk | myCAF_1 |
| Shc4 | 4.30E-123 | 0.65609849 | 0.622 | 0.431 | 8.31E-119 | 0 | Shc4 | myCAF_1 |
| Zfp469 | 4.53E-123 | 0.55448885 | 0.822 | 0.661 | 8.76E-119 | 0 | Zfp469 | myCAF_1 |
| Fat4 | 2.07E-122 | 1.18366892 | 0.366 | 0.185 | 4.01E-118 | 0 | Fat4 | myCAF_1 |
| Man2a2 | 4.76E-122 | 0.47257248 | 0.92 | 0.812 | 9.20E-118 | 0 | Man2a2 | myCAF_1 |
| Cacna1g | 9.83E-122 | 0.68597041 | 0.743 | 0.505 | 1.90E-117 | 0 | Cacna1g | myCAF_1 |
| Bmf | 6.95E-120 | 0.83964274 | 0.41 | 0.224 | 1.34E-115 | 0 | Bmf | myCAF_1 |
| Mtus1 | 9.14E-120 | 0.90750943 | 0.442 | 0.247 | 1.77E-115 | 0 | Mtus1 | myCAF_1 |
| Tmeff1 | 1.90E-119 | 0.84535871 | 0.557 | 0.351 | 3.68E-115 | 0 | Tmeff1 | myCAF_1 |
| Tbc1d2b | 2.40E-119 | 0.39310109 | 0.928 | 0.847 | 4.64E-115 | 0 | Tbc1d2b | myCAF_1 |
| Ddah1 | 3.10E-119 | 0.40249828 | 0.937 | 0.824 | 6.00E-115 | 0 | Ddah1 | myCAF_1 |
| Fhl2 | 1.16E-116 | 0.71953188 | 0.598 | 0.39 | 2.24E-112 | 0 | Fhl2 | myCAF_1 |
| Socs2 | 2.91E-116 | 0.86813477 | 0.523 | 0.314 | 5.62E-112 | 0 | Socs2 | myCAF_1 |
| Ube2ql1 | 1.29E-114 | 0.9893748 | 0.36 | 0.172 | 2.49E-110 | 0 | Ube2ql1 | myCAF_1 |
| Masp1 | 2.26E-113 | 0.54209671 | 0.406 | 0.335 | 4.37E-109 | 0 | Masp1 | myCAF_1 |
| Lhfp | 4.04E-112 | 0.25913471 | 0.992 | 0.966 | 7.81E-108 | 0 | Lhfp | myCAF_1 |
| Matn2 | 2.74E-111 | 0.59316729 | 0.776 | 0.625 | 5.29E-107 | 0 | Matn2 | myCAF_1 |
| Arg1 | 4.28E-111 | 0.58468983 | 0.326 | 0.217 | 8.28E-107 | 0 | Arg1 | myCAF_1 |
| Sox4 | 5.11E-110 | 0.33574194 | 0.832 | 0.714 | 9.88E-106 | 0 | Sox4 | myCAF_1 |
| 4-Sep | 6.81E-110 | 0.66137279 | 0.544 | 0.305 | 1.32E-105 | 0 | 4-Sep | myCAF_1 |
| Arxes1 | 9.07E-110 | 1.52710367 | 0.28 | 0.11 | 1.75E-105 | 0 | Arxes1 | myCAF_1 |
| Nynrin | 9.75E-110 | 0.57814651 | 0.775 | 0.634 | 1.89E-105 | 0 | Nynrin | myCAF_1 |
| B4galnt1 | 1.10E-107 | 0.6013504 | 0.697 | 0.471 | 2.14E-103 | 0 | B4galnt1 | myCAF_1 |
| B3gnt9 | 2.97E-107 | 0.42656018 | 0.924 | 0.866 | 5.75E-103 | 0 | B3gnt9 | myCAF_1 |
| Pfn2 | 1.25E-105 | 0.86776536 | 0.444 | 0.299 | 2.42E-101 | 0 | Pfn2 | myCAF_1 |
| Pmaip1 | 1.35E-104 | 0.65252943 | 0.4 | 0.221 | 2.61E-100 | 0 | Pmaip1 | myCAF_1 |
| Fndc3b | 1.58E-104 | 0.27282545 | 0.986 | 0.955 | 3.05E-100 | 0 | Fndc3b | myCAF_1 |
| Hc | 2.33E-104 | 0.72736386 | 0.438 | 0.218 | 4.50E-100 | 0 | Hc | myCAF_1 |
| DEPRECATED-l | 5.61E-104 | 0.52532486 | 0.716 | 0.581 | 1.08E-99 | 0 | DEPRECATED-l | myCAF_1 |
| Oaz2 | 7.62E-104 | 0.42515924 | 0.893 | 0.819 | 1.47E-99 | 0 | Oaz2 | myCAF_1 |
| Elk3 | 8.17E-104 | 0.52310121 | 0.876 | 0.771 | 1.58E-99 | 0 | Elk3 | myCAF_1 |
| Cacng7 | 2.32E-102 | 0.90485557 | 0.327 | 0.217 | 4.48E-98 | 0 | Cacng7 | myCAF_1 |
| Adcy3 | 2.91E-102 | 0.435328 | 0.811 | 0.683 | 5.63E-98 | 0 | Adcy3 | myCAF_1 |
| Pkd2 | 7.73E-102 | 0.36455878 | 0.944 | 0.898 | 1.50E-97 | 0 | Pkd2 | myCAF_1 |
| Bex3 | 2.30E-101 | 0.65544559 | 0.698 | 0.512 | 4.44E-97 | 0 | Bex3 | myCAF_1 |
| Zhx3 | 2.38E-100 | 0.78704823 | 0.578 | 0.432 | 4.60E-96 | 0 | Zhx3 | myCAF_1 |
| Pex19 | 2.60E-100 | 0.57925884 | 0.746 | 0.621 | 5.02E-96 | 0 | Pex19 | myCAF_1 |
| Cux1 | 4.86E-100 | 0.49559541 | 0.756 | 0.585 | 9.39E-96 | 0 | Cux1 | myCAF_1 |
| Tmcc3 | 5.29E-99 | 0.68962444 | 0.592 | 0.419 | 1.02E-94 | 0 | Tmcc3 | myCAF_1 |
| Eva1a | 6.36E-99 | 1.14191465 | 0.355 | 0.171 | 1.23E-94 | 0 | Eva1a | myCAF_1 |
| Colec12 | 1.63E-98 | 0.28764233 | 0.957 | 0.893 | 3.16E-94 | 0 | Colec12 | myCAF_1 |
| Mgat3 | 4.95E-98 | 0.67351476 | 0.573 | 0.367 | 9.58E-94 | 0 | Mgat3 | myCAF_1 |
| Nox4 | 1.37E-97 | 0.5163034 | 0.456 | 0.342 | 2.65E-93 | 0 | Nox4 | myCAF_1 |
| Golm1 | 4.34E-97 | 0.51573576 | 0.856 | 0.718 | 8.38E-93 | 0 | Golm1 | myCAF_1 |
| Nmnat3 | 1.06E-96 | 1.11637608 | 0.408 | 0.234 | 2.05E-92 | 0 | Nmnat3 | myCAF_1 |
| Dcbld1 | 2.30E-94 | 0.76028652 | 0.549 | 0.313 | 4.44E-90 | 0 | Dcbld1 | myCAF_1 |
| Pkig | 6.30E-94 | 0.46604651 | 0.852 | 0.735 | 1.22E-89 | 0 | Pkig | myCAF_1 |
| Dynlrb1 | 2.50E-93 | 0.3313165 | 0.961 | 0.928 | 4.83E-89 | 0 | Dynlrb1 | myCAF_1 |
| Fance | 5.67E-93 | 0.61085303 | 0.69 | 0.537 | 1.10E-88 | 0 | Fance | myCAF_1 |
| C1qtnf2 | 5.41E-92 | 0.50263715 | 0.776 | 0.648 | 1.05E-87 | 0 | C1qtnf2 | myCAF_1 |
| Adgrl1 | 7.49E-92 | 0.6182263 | 0.738 | 0.541 | 1.45E-87 | 0 | Adgrl1 | myCAF_1 |
| Ninj1 | 7.68E-92 | 0.4351247 | 0.728 | 0.579 | 1.48E-87 | 0 | Ninj1 | myCAF_1 |
| Shisa4 | 3.17E-91 | 0.79496067 | 0.497 | 0.351 | 6.12E-87 | 0 | Shisa4 | myCAF_1 |
| Arhgap31 | 7.93E-91 | 0.61940967 | 0.763 | 0.6 | 1.53E-86 | 0 | Arhgap31 | myCAF_1 |
| Frzb | 1.87E-90 | 0.86645959 | 0.362 | 0.179 | 3.62E-86 | 0 | Frzb | myCAF_1 |
| Fbxo32 | 7.15E-90 | 0.66345883 | 0.367 | 0.214 | 1.38E-85 | 0 | Fbxo32 | myCAF_1 |
| Nfatc4 | 9.29E-90 | 0.46164113 | 0.895 | 0.793 | 1.80E-85 | 0 | Nfatc4 | myCAF_1 |
| Fmn1 | 1.01E-89 | 0.70336065 | 0.546 | 0.395 | 1.95E-85 | 0 | Fmn1 | myCAF_1 |
| Irf2bpl | 2.77E-89 | 0.49402374 | 0.852 | 0.745 | 5.35E-85 | 0 | Irf2bpl | myCAF_1 |
| Fibin | 1.19E-87 | 0.29437893 | 0.828 | 0.741 | 2.30E-83 | 0 | Fibin | myCAF_1 |
| Adgrl3 | 1.50E-87 | 0.88132352 | 0.377 | 0.178 | 2.90E-83 | 0 | Adgrl3 | myCAF_1 |
| Sfxn3 | 3.13E-87 | 0.36582746 | 0.878 | 0.799 | 6.06E-83 | 0 | Sfxn3 | myCAF_1 |
| Rftn1 | 9.05E-87 | 0.46072533 | 0.871 | 0.756 | 1.75E-82 | 0 | Rftn1 | myCAF_1 |
| Arhgef19 | 8.28E-86 | 0.93692164 | 0.464 | 0.269 | 1.60E-81 | 0 | Arhgef19 | myCAF_1 |
| Dock4 | 3.91E-85 | 0.79086057 | 0.478 | 0.333 | 7.56E-81 | 0 | Dock4 | myCAF_1 |
| Ajuba | 5.87E-85 | 0.83317528 | 0.456 | 0.301 | 1.13E-80 | 0 | Ajuba | myCAF_1 |
| Ankrd35 | 6.44E-85 | 1.19785096 | 0.285 | 0.135 | 1.24E-80 | 0 | Ankrd35 | myCAF_1 |
| Lats2 | 1.43E-84 | 0.43227472 | 0.86 | 0.772 | 2.77E-80 | 0 | Lats2 | myCAF_1 |
| Otulinl | 1.66E-84 | 1.07704176 | 0.297 | 0.151 | 3.20E-80 | 0 | Otulinl | myCAF_1 |
| Rasl11b | 2.74E-84 | 0.61805597 | 0.474 | 0.314 | 5.30E-80 | 0 | Rasl11b | myCAF_1 |
| Prxl2a | 3.50E-84 | 1.11155618 | 0.312 | 0.165 | 6.78E-80 | 0 | Prxl2a | myCAF_1 |
| Slc27a1 | 5.88E-84 | 0.6874843 | 0.484 | 0.312 | 1.14E-79 | 0 | Slc27a1 | myCAF_1 |
| Efnb1 | 6.21E-84 | 0.5122343 | 0.705 | 0.534 | 1.20E-79 | 0 | Efnb1 | myCAF_1 |
| Fam217b | 8.38E-84 | 1.17252945 | 0.345 | 0.181 | 1.62E-79 | 0 | Fam217b | myCAF_1 |
| Dmd | 3.19E-83 | 1.06761976 | 0.294 | 0.176 | 6.16E-79 | 0 | Dmd | myCAF_1 |
| Snai2 | 5.07E-83 | 0.59553245 | 0.631 | 0.468 | 9.80E-79 | 0 | Snai2 | myCAF_1 |
| Igdcc4 | 4.03E-82 | 0.5789081 | 0.652 | 0.507 | 7.79E-78 | 0 | Igdcc4 | myCAF_1 |
| Cpne3 | 7.78E-82 | 0.63802163 | 0.731 | 0.619 | 1.51E-77 | 0 | Cpne3 | myCAF_1 |
| Dysf | 1.09E-81 | 0.94897147 | 0.396 | 0.249 | 2.10E-77 | 0 | Dysf | myCAF_1 |
| Unc5a | 2.19E-81 | 0.89401059 | 0.438 | 0.25 | 4.24E-77 | 0 | Unc5a | myCAF_1 |
| Etl4 | 5.88E-81 | 0.92761947 | 0.305 | 0.149 | 1.14E-76 | 0 | Etl4 | myCAF_1 |
| DEPRECATED-l | 1.01E-80 | 0.61577256 | 0.354 | 0.27 | 1.95E-76 | 0 | DEPRECATED-l | myCAF_1 |
| Tns3 | 1.08E-80 | 0.32722151 | 0.841 | 0.596 | 2.10E-76 | 0 | Tns3 | myCAF_1 |
| Ctsf | 1.74E-80 | 0.56506489 | 0.629 | 0.511 | 3.36E-76 | 0 | Ctsf | myCAF_1 |
| Gcnt4 | 1.04E-79 | 0.55717312 | 0.551 | 0.298 | 2.02E-75 | 0 | Gcnt4 | myCAF_1 |
| Ankrd34a | 4.81E-79 | 1.16310669 | 0.303 | 0.154 | 9.30E-75 | 0 | Ankrd34a | myCAF_1 |
| Pde11a | 6.86E-79 | 0.73982625 | 0.293 | 0.202 | 1.33E-74 | 0 | Pde11a | myCAF_1 |
| Dok1 | 1.23E-78 | 0.47859696 | 0.805 | 0.659 | 2.38E-74 | 0 | Dok1 | myCAF_1 |
| Wipi1 | 2.41E-78 | 0.3520329 | 0.946 | 0.901 | 4.67E-74 | 0 | Wipi1 | myCAF_1 |
| Zfp532 | 8.79E-78 | 0.43056693 | 0.823 | 0.694 | 1.70E-73 | 0 | Zfp532 | myCAF_1 |
| Cttnbp2 | 3.57E-76 | 0.94360283 | 0.36 | 0.197 | 6.90E-72 | 0 | Cttnbp2 | myCAF_1 |
| Atxn1 | 5.37E-76 | 0.50782403 | 0.697 | 0.565 | 1.04E-71 | 0 | Atxn1 | myCAF_1 |
| Psd | 8.51E-76 | 0.99831983 | 0.287 | 0.152 | 1.64E-71 | 0 | Psd | myCAF_1 |
| Plekhf1 | 1.84E-75 | 0.65473698 | 0.632 | 0.477 | 3.56E-71 | 0 | Plekhf1 | myCAF_1 |
| Pls3 | 8.02E-75 | 0.39366708 | 0.882 | 0.807 | 1.55E-70 | 0 | Pls3 | myCAF_1 |
| Etv4 | 3.34E-74 | 0.55661309 | 0.635 | 0.435 | 6.46E-70 | 0 | Etv4 | myCAF_1 |
| Myo7a | 4.59E-73 | 0.57801108 | 0.776 | 0.586 | 8.88E-69 | 0 | Myo7a | myCAF_1 |
| Ccnd2 | 5.76E-73 | 0.29985238 | 0.78 | 0.666 | 1.11E-68 | 0 | Ccnd2 | myCAF_1 |
| Pde4b | 7.60E-73 | 0.54181043 | 0.579 | 0.357 | 1.47E-68 | 0 | Pde4b | myCAF_1 |
| Akap7 | 1.27E-72 | 0.80590907 | 0.426 | 0.286 | 2.46E-68 | 0 | Akap7 | myCAF_1 |
| Nck2 | 2.17E-72 | 0.64007969 | 0.555 | 0.382 | 4.20E-68 | 0 | Nck2 | myCAF_1 |
| Itga5 | 3.34E-72 | 0.28284552 | 0.937 | 0.876 | 6.46E-68 | 0 | Itga5 | myCAF_1 |
| Wls | 7.28E-72 | 0.36722387 | 0.936 | 0.863 | 1.41E-67 | 0 | Wls | myCAF_1 |
| Dzip1l | 7.49E-72 | 0.54050062 | 0.608 | 0.434 | 1.45E-67 | 0 | Dzip1l | myCAF_1 |
| Tspan4 | 8.80E-72 | 0.34753421 | 0.97 | 0.915 | 1.70E-67 | 0 | Tspan4 | myCAF_1 |
| Abcg1 | 1.30E-71 | 0.90877001 | 0.309 | 0.175 | 2.52E-67 | 0 | Abcg1 | myCAF_1 |
| Iglon5 | 2.72E-71 | 0.75750101 | 0.355 | 0.228 | 5.27E-67 | 0 | Iglon5 | myCAF_1 |
| Zcchc24 | 3.16E-70 | 0.27245026 | 0.965 | 0.924 | 6.12E-66 | 0 | Zcchc24 | myCAF_1 |
| Aopep | 1.10E-69 | 0.85742572 | 0.284 | 0.143 | 2.13E-65 | 0 | Aopep | myCAF_1 |
| Ccdc102a | 1.54E-69 | 0.45414792 | 0.808 | 0.69 | 2.98E-65 | 0 | Ccdc102a | myCAF_1 |
| Sgpl1 | 2.73E-69 | 0.41143179 | 0.809 | 0.72 | 5.28E-65 | 0 | Sgpl1 | myCAF_1 |
| Kctd10 | 1.25E-68 | 0.29850754 | 0.913 | 0.859 | 2.43E-64 | 0 | Kctd10 | myCAF_1 |
| Yap1 | 2.09E-68 | 0.36186627 | 0.893 | 0.825 | 4.04E-64 | 0 | Yap1 | myCAF_1 |
| Cspg4 | 7.48E-68 | 0.28192078 | 0.526 | 0.308 | 1.45E-63 | 0 | Cspg4 | myCAF_1 |
| Marveld1 | 8.66E-68 | 0.30772203 | 0.933 | 0.876 | 1.68E-63 | 0 | Marveld1 | myCAF_1 |
| Chrnb1 | 2.91E-67 | 0.97134108 | 0.273 | 0.145 | 5.62E-63 | 0 | Chrnb1 | myCAF_1 |
| Tnk2 | 3.91E-67 | 0.52570705 | 0.558 | 0.433 | 7.56E-63 | 0 | Tnk2 | myCAF_1 |
| Gnb4 | 4.19E-67 | 0.54232778 | 0.617 | 0.488 | 8.10E-63 | 0 | Gnb4 | myCAF_1 |
| Snx9 | 4.45E-67 | 0.33600696 | 0.953 | 0.904 | 8.60E-63 | 0 | Snx9 | myCAF_1 |
| Rcn2 | 1.35E-66 | 0.35363298 | 0.928 | 0.867 | 2.62E-62 | 0 | Rcn2 | myCAF_1 |
| H1fx | 3.06E-66 | 0.6095658 | 0.567 | 0.367 | 5.92E-62 | 0 | H1fx | myCAF_1 |
| Tmem268 | 3.09E-66 | 0.99005997 | 0.367 | 0.224 | 5.98E-62 | 0 | Tmem268 | myCAF_1 |
| Rhod | 5.02E-66 | 0.77304949 | 0.479 | 0.326 | 9.70E-62 | 0 | Rhod | myCAF_1 |
| Porcn | 6.13E-66 | 0.77465982 | 0.477 | 0.317 | 1.18E-61 | 0 | Porcn | myCAF_1 |
| Kifc3 | 7.28E-66 | 0.70316636 | 0.565 | 0.384 | 1.41E-61 | 0 | Kifc3 | myCAF_1 |
| Lrrc55 | 2.14E-64 | 0.82826987 | 0.335 | 0.181 | 4.15E-60 | 0 | Lrrc55 | myCAF_1 |
| Fut8 | 2.92E-64 | 0.48483044 | 0.706 | 0.571 | 5.65E-60 | 0 | Fut8 | myCAF_1 |
| Leng8 | 3.34E-64 | 0.37213419 | 0.853 | 0.749 | 6.46E-60 | 0 | Leng8 | myCAF_1 |
| Hmcn1 | 4.24E-64 | 0.78378655 | 0.349 | 0.206 | 8.20E-60 | 0 | Hmcn1 | myCAF_1 |
| Wdfy1 | 4.67E-64 | 0.46472247 | 0.534 | 0.393 | 9.02E-60 | 0 | Wdfy1 | myCAF_1 |
| Rtn2 | 5.19E-63 | 0.76492737 | 0.346 | 0.216 | 1.00E-58 | 0 | Rtn2 | myCAF_1 |
| Zfp827 | 1.05E-62 | 0.86751151 | 0.385 | 0.229 | 2.03E-58 | 0 | Zfp827 | myCAF_1 |
| Wnt11 | 2.59E-62 | 0.493154 | 0.475 | 0.285 | 5.01E-58 | 0 | Wnt11 | myCAF_1 |
| Calcoco1 | 1.00E-61 | 0.31427042 | 0.696 | 0.576 | 1.94E-57 | 0 | Calcoco1 | myCAF_1 |
| Bmpr2 | 1.10E-61 | 0.33767669 | 0.929 | 0.881 | 2.12E-57 | 0 | Bmpr2 | myCAF_1 |
| Sc5d | 2.37E-61 | 0.5104595 | 0.577 | 0.423 | 4.58E-57 | 0 | Sc5d | myCAF_1 |
| Mcrip1 | 2.60E-60 | 0.31469278 | 0.937 | 0.885 | 5.03E-56 | 0 | Mcrip1 | myCAF_1 |
| Chst12 | 4.66E-60 | 0.33668968 | 0.921 | 0.859 | 9.01E-56 | 0 | Chst12 | myCAF_1 |
| Pde4d | 5.66E-60 | 0.58255177 | 0.626 | 0.467 | 1.10E-55 | 0 | Pde4d | myCAF_1 |
| Ednra | 1.33E-59 | 0.29593775 | 0.776 | 0.513 | 2.57E-55 | 0 | Ednra | myCAF_1 |
| Ankrd12 | 1.71E-59 | 0.6736494 | 0.495 | 0.357 | 3.32E-55 | 0 | Ankrd12 | myCAF_1 |
| Tmem200b | 2.46E-59 | 0.83604552 | 0.351 | 0.216 | 4.76E-55 | 0 | Tmem200b | myCAF_1 |
| Lzts2 | 6.32E-59 | 0.31080076 | 0.852 | 0.786 | 1.22E-54 | 0 | Lzts2 | myCAF_1 |
| Lpin3 | 8.33E-59 | 0.66423463 | 0.526 | 0.412 | 1.61E-54 | 0 | Lpin3 | myCAF_1 |
| Msi2 | 1.42E-58 | 0.39308457 | 0.691 | 0.557 | 2.75E-54 | 0 | Msi2 | myCAF_1 |
| Adcy6 | 2.29E-58 | 0.38761369 | 0.805 | 0.68 | 4.43E-54 | 0 | Adcy6 | myCAF_1 |
| Ttc28 | 2.81E-57 | 0.47340663 | 0.662 | 0.555 | 5.43E-53 | 0 | Ttc28 | myCAF_1 |
| Emilin1 | 3.40E-57 | 0.25650662 | 0.958 | 0.908 | 6.57E-53 | 0 | Emilin1 | myCAF_1 |
| Vgll4 | 4.12E-57 | 0.36244354 | 0.81 | 0.7 | 7.97E-53 | 0 | Vgll4 | myCAF_1 |
| Arhgap42 | 6.03E-57 | 0.34622559 | 0.85 | 0.746 | 1.17E-52 | 0 | Arhgap42 | myCAF_1 |
| Scarb1 | 8.98E-57 | 0.49748588 | 0.613 | 0.442 | 1.74E-52 | 0 | Scarb1 | myCAF_1 |
| Kctd5 | 2.54E-56 | 0.52748706 | 0.714 | 0.599 | 4.91E-52 | 0 | Kctd5 | myCAF_1 |
| Draxin | 8.15E-56 | 0.55688009 | 0.552 | 0.372 | 1.58E-51 | 0 | Draxin | myCAF_1 |
| Ccdc9b | 1.21E-55 | 0.6215926 | 0.459 | 0.316 | 2.33E-51 | 0 | Ccdc9b | myCAF_1 |
| Arhgef17 | 1.50E-55 | 0.40072822 | 0.771 | 0.645 | 2.91E-51 | 0 | Arhgef17 | myCAF_1 |
| Prkg1 | 1.74E-55 | 0.96004084 | 0.286 | 0.167 | 3.37E-51 | 0 | Prkg1 | myCAF_1 |
| Ppfibp1 | 2.28E-55 | 0.31280561 | 0.905 | 0.841 | 4.41E-51 | 0 | Ppfibp1 | myCAF_1 |
| Phldb2 | 4.94E-55 | 0.35190742 | 0.835 | 0.764 | 9.54E-51 | 0 | Phldb2 | myCAF_1 |
| Col4a3bp | 9.10E-55 | 0.44423111 | 0.632 | 0.506 | 1.76E-50 | 0 | Col4a3bp | myCAF_1 |
| M6pr | 2.80E-54 | 0.36412428 | 0.849 | 0.789 | 5.42E-50 | 0 | M6pr | myCAF_1 |
| Bcar1 | 4.05E-54 | 0.30300256 | 0.783 | 0.683 | 7.83E-50 | 0 | Bcar1 | myCAF_1 |
| Actr1b | 5.09E-54 | 0.31838384 | 0.937 | 0.878 | 9.84E-50 | 0 | Actr1b | myCAF_1 |
| Cyp51 | 9.24E-54 | 0.32647564 | 0.718 | 0.581 | 1.79E-49 | 0 | Cyp51 | myCAF_1 |
| Pcsk9 | 1.40E-53 | 0.64454768 | 0.447 | 0.315 | 2.71E-49 | 0 | Pcsk9 | myCAF_1 |
| Ier5l | 1.41E-53 | 0.37512629 | 0.812 | 0.698 | 2.73E-49 | 0 | Ier5l | myCAF_1 |
| Stox2 | 8.14E-53 | 0.62112263 | 0.451 | 0.307 | 1.57E-48 | 0 | Stox2 | myCAF_1 |
| Uqcrc1 | 9.10E-53 | 0.27934231 | 0.959 | 0.918 | 1.76E-48 | 0 | Uqcrc1 | myCAF_1 |
| Plekha5 | 4.18E-52 | 0.5441793 | 0.627 | 0.492 | 8.08E-48 | 0 | Plekha5 | myCAF_1 |
| Snx8 | 5.01E-52 | 0.59714627 | 0.552 | 0.412 | 9.69E-48 | 0 | Snx8 | myCAF_1 |
| Abracl | 6.31E-52 | 0.40008456 | 0.777 | 0.693 | 1.22E-47 | 0 | Abracl | myCAF_1 |

| | | | | | | | | |
|---|---|---|---|---|---|---|---|---|
| Aplp1 | 6.59E-52 | 0.52534071 | 0.325 | 0.199 | 1.27E-47 | 0 | Aplp1 | myCAF_1 |
| Me1 | 7.34E-52 | 0.60124284 | 0.451 | 0.354 | 1.42E-47 | 0 | Me1 | myCAF_1 |
| L3mbtl3 | 1.29E-51 | 0.50924813 | 0.542 | 0.415 | 2.49E-47 | 0 | L3mbtl3 | myCAF_1 |
| Fut4 | 6.16E-51 | 0.54414126 | 0.508 | 0.349 | 1.19E-46 | 0 | Fut4 | myCAF_1 |
| Tpp1 | 7.04E-51 | 0.35353165 | 0.825 | 0.754 | 1.36E-46 | 0 | Tpp1 | myCAF_1 |
| Ifitm1 | 9.12E-51 | 0.26776516 | 0.467 | 0.337 | 1.76E-46 | 0 | Ifitm1 | myCAF_1 |
| Paqr7 | 1.36E-50 | 0.63533079 | 0.432 | 0.305 | 2.63E-46 | 0 | Paqr7 | myCAF_1 |
| Pde5a | 5.51E-50 | 0.45921454 | 0.592 | 0.407 | 1.07E-45 | 0 | Pde5a | myCAF_1 |
| Kansl2 | 1.10E-49 | 0.3533147 | 0.873 | 0.78 | 2.13E-45 | 0 | Kansl2 | myCAF_1 |
| Ebpl | 3.22E-49 | 0.59900526 | 0.491 | 0.388 | 6.22E-45 | 0 | Ebpl | myCAF_1 |
| Klhl24 | 6.33E-49 | 0.38664493 | 0.648 | 0.515 | 1.22E-44 | 0 | Klhl24 | myCAF_1 |
| Mid1ip1 | 2.46E-48 | 0.4069139 | 0.703 | 0.571 | 4.76E-44 | 0 | Mid1ip1 | myCAF_1 |
| Irak1bp1 | 4.63E-48 | 0.79589289 | 0.356 | 0.245 | 8.96E-44 | 0 | Irak1bp1 | myCAF_1 |
| Tet3 | 8.95E-48 | 0.37162407 | 0.613 | 0.492 | 1.73E-43 | 0 | Tet3 | myCAF_1 |
| a | 9.27E-48 | 0.54421117 | 0.425 | 0.267 | 1.79E-43 | 0 | a | myCAF_1 |
| Dip2a | 1.03E-47 | 0.36657838 | 0.73 | 0.6 | 1.98E-43 | 0 | Dip2a | myCAF_1 |
| Cask | 1.39E-47 | 0.4295501 | 0.77 | 0.673 | 2.68E-43 | 0 | Cask | myCAF_1 |
| Pink1 | 1.44E-47 | 0.27435155 | 0.79 | 0.689 | 2.78E-43 | 0 | Pink1 | myCAF_1 |
| Cnnm2 | 1.72E-47 | 0.52429436 | 0.306 | 0.218 | 3.33E-43 | 0 | Cnnm2 | myCAF_1 |
| Rida | 6.62E-47 | 0.70696665 | 0.323 | 0.21 | 1.28E-42 | 0 | Rida | myCAF_1 |
| Synj2 | 7.64E-47 | 0.58673343 | 0.542 | 0.407 | 1.48E-42 | 0 | Synj2 | myCAF_1 |
| Lactb | 9.86E-47 | 0.79941245 | 0.368 | 0.243 | 1.91E-42 | 0 | Lactb | myCAF_1 |
| Arhgef5 | 1.79E-46 | 0.78464209 | 0.307 | 0.183 | 3.46E-42 | 0 | Arhgef5 | myCAF_1 |
| Nectin2 | 2.19E-45 | 0.33957068 | 0.831 | 0.72 | 4.23E-41 | 0 | Nectin2 | myCAF_1 |
| Nlk | 2.85E-45 | 0.61687044 | 0.467 | 0.34 | 5.51E-41 | 0 | Nlk | myCAF_1 |
| Tmem9 | 3.30E-45 | 0.36070717 | 0.806 | 0.718 | 6.39E-41 | 0 | Tmem9 | myCAF_1 |
| Zbtb34 | 3.31E-45 | 0.47167612 | 0.538 | 0.406 | 6.40E-41 | 0 | Zbtb34 | myCAF_1 |
| Efhd2 | 3.76E-45 | 0.40204653 | 0.747 | 0.639 | 7.27E-41 | 0 | Efhd2 | myCAF_1 |
| Svbp | 1.09E-44 | 0.52320119 | 0.611 | 0.487 | 2.10E-40 | 0 | Svbp | myCAF_1 |
| Apbb2 | 1.61E-44 | 0.33162164 | 0.802 | 0.688 | 3.12E-40 | 0 | Apbb2 | myCAF_1 |
| Agrn | 2.10E-44 | 0.38782465 | 0.667 | 0.575 | 4.07E-40 | 0 | Agrn | myCAF_1 |
| Src | 2.56E-44 | 0.3247985 | 0.812 | 0.734 | 4.96E-40 | 0 | Src | myCAF_1 |
| Tead3 | 2.96E-44 | 0.37121675 | 0.743 | 0.639 | 5.73E-40 | 0 | Tead3 | myCAF_1 |
| Atp13a3 | 4.20E-44 | 0.36894728 | 0.804 | 0.742 | 8.12E-40 | 0 | Atp13a3 | myCAF_1 |
| Pstpip1 | 5.24E-44 | 0.69553962 | 0.335 | 0.209 | 1.01E-39 | 0 | Pstpip1 | myCAF_1 |
| Arid3a | 8.31E-44 | 0.45572951 | 0.506 | 0.368 | 1.61E-39 | 0 | Arid3a | myCAF_1 |
| Qprt | 8.31E-44 | 0.62987399 | 0.283 | 0.194 | 1.61E-39 | 0 | Qprt | myCAF_1 |
| Desi1 | 1.06E-43 | 0.44545444 | 0.735 | 0.636 | 2.05E-39 | 0 | Desi1 | myCAF_1 |
| Nedd9 | 1.09E-43 | 0.68672066 | 0.474 | 0.337 | 2.10E-39 | 0 | Nedd9 | myCAF_1 |
| Gna12 | 2.03E-43 | 0.30459642 | 0.907 | 0.818 | 3.93E-39 | 0 | Gna12 | myCAF_1 |
| Smim1 | 7.36E-43 | 0.68510911 | 0.342 | 0.231 | 1.42E-38 | 0 | Smim1 | myCAF_1 |
| Shc1 | 3.27E-42 | 0.27973096 | 0.918 | 0.868 | 6.32E-38 | 0 | Shc1 | myCAF_1 |
| Mdfic | 1.03E-41 | 0.29655475 | 0.824 | 0.748 | 1.99E-37 | 0 | Mdfic | myCAF_1 |
| Gcnt2 | 1.40E-41 | 0.62415914 | 0.499 | 0.365 | 2.70E-37 | 0 | Gcnt2 | myCAF_1 |
| Spg20 | 6.10E-41 | 0.43848223 | 0.736 | 0.63 | 1.18E-36 | 0 | Spg20 | myCAF_1 |
| Ror2 | 6.27E-41 | 0.33793829 | 0.856 | 0.754 | 1.21E-36 | 0 | Ror2 | myCAF_1 |
| Zfp810 | 1.31E-40 | 0.68148461 | 0.378 | 0.254 | 2.53E-36 | 0 | Zfp810 | myCAF_1 |
| Pear1 | 1.85E-40 | 0.31170464 | 0.761 | 0.565 | 3.58E-36 | 0 | Pear1 | myCAF_1 |
| Cyth1 | 2.02E-40 | 0.51226778 | 0.524 | 0.404 | 3.91E-36 | 0 | Cyth1 | myCAF_1 |
| Spry4 | 6.99E-40 | 0.39817213 | 0.517 | 0.397 | 1.35E-35 | 0 | Spry4 | myCAF_1 |
| Sike1 | 9.14E-40 | 0.38602994 | 0.755 | 0.67 | 1.77E-35 | 0 | Sike1 | myCAF_1 |
| Hbp1 | 1.16E-39 | 0.26149919 | 0.847 | 0.78 | 2.25E-35 | 0 | Hbp1 | myCAF_1 |
| Slc25a51 | 2.11E-39 | 0.27852057 | 0.877 | 0.829 | 4.08E-35 | 0 | Slc25a51 | myCAF_1 |
| Cyb561 | 2.23E-39 | 0.54365889 | 0.463 | 0.347 | 4.32E-35 | 0 | Cyb561 | myCAF_1 |
| Fzd3 | 2.37E-39 | 0.70669967 | 0.307 | 0.187 | 4.58E-35 | 0 | Fzd3 | myCAF_1 |
| Osbpl5 | 2.64E-39 | 0.30470395 | 0.761 | 0.682 | 5.10E-35 | 0 | Osbpl5 | myCAF_1 |
| Tceal9 | 2.09E-38 | 0.27785626 | 0.967 | 0.927 | 4.04E-34 | 0 | Tceal9 | myCAF_1 |
| Tmem132a | 2.20E-38 | 0.46336787 | 0.514 | 0.411 | 4.25E-34 | 0 | Tmem132a | myCAF_1 |
| Ly75 | 2.54E-38 | 0.75677563 | 0.282 | 0.178 | 4.90E-34 | 0 | Ly75 | myCAF_1 |
| Fgd6 | 2.96E-38 | 0.63004454 | 0.339 | 0.218 | 5.72E-34 | 0 | Fgd6 | myCAF_1 |
| Mmd | 6.78E-38 | 0.29669792 | 0.738 | 0.614 | 1.31E-33 | 0 | Mmd | myCAF_1 |
| Wwtr1 | 3.90E-37 | 0.26047291 | 0.916 | 0.877 | 7.55E-33 | 0 | Wwtr1 | myCAF_1 |
| Cyba | 4.13E-37 | 0.32585576 | 0.849 | 0.782 | 7.98E-33 | 0 | Cyba | myCAF_1 |
| Gria3 | 1.36E-36 | 0.57709646 | 0.538 | 0.399 | 2.64E-32 | 0 | Gria3 | myCAF_1 |
| Prkd1 | 2.25E-36 | 0.56931813 | 0.377 | 0.247 | 4.35E-32 | 0 | Prkd1 | myCAF_1 |
| Samd4 | 3.26E-36 | 0.32163756 | 0.733 | 0.66 | 6.30E-32 | 0 | Samd4 | myCAF_1 |
| Lman2l | 3.81E-36 | 0.58255389 | 0.403 | 0.3 | 7.36E-32 | 0 | Lman2l | myCAF_1 |
| Mgat5 | 3.85E-36 | 0.50258448 | 0.459 | 0.368 | 7.44E-32 | 0 | Mgat5 | myCAF_1 |
| Ptar1 | 4.04E-36 | 0.29265199 | 0.849 | 0.793 | 7.81E-32 | 0 | Ptar1 | myCAF_1 |
| Etv1 | 4.74E-36 | 0.73800729 | 0.253 | 0.161 | 9.17E-32 | 0 | Etv1 | myCAF_1 |
| Ptprd | 8.57E-36 | 0.69967094 | 0.293 | 0.191 | 1.66E-31 | 0 | Ptprd | myCAF_1 |
| Tcp11l2 | 1.91E-35 | 0.2507395 | 0.647 | 0.539 | 3.70E-31 | 0 | Tcp11l2 | myCAF_1 |
| Gsto1 | 3.46E-35 | 0.34158372 | 0.916 | 0.816 | 6.68E-31 | 0 | Gsto1 | myCAF_1 |
| Pofut2 | 6.91E-35 | 0.25944854 | 0.963 | 0.924 | 1.34E-30 | 0 | Pofut2 | myCAF_1 |
| Igsf3 | 6.94E-35 | 0.34848768 | 0.588 | 0.462 | 1.34E-30 | 0 | Igsf3 | myCAF_1 |
| Abhd8 | 2.47E-34 | 0.49266763 | 0.666 | 0.543 | 4.78E-30 | 0 | Abhd8 | myCAF_1 |
| Lipo3 | 3.94E-34 | 0.58146353 | 0.403 | 0.273 | 7.61E-30 | 0 | Lipo3 | myCAF_1 |
| Castor2 | 4.56E-34 | 0.48728902 | 0.522 | 0.416 | 8.82E-30 | 0 | Castor2 | myCAF_1 |
| Cped1 | 6.13E-34 | 0.25097453 | 0.87 | 0.789 | 1.19E-29 | 0 | Cped1 | myCAF_1 |
| Fam214a | 1.32E-33 | 0.40363064 | 0.47 | 0.361 | 2.56E-29 | 0 | Fam214a | myCAF_1 |
| Slc25a1 | 1.86E-33 | 0.26711546 | 0.801 | 0.722 | 3.60E-29 | 0 | Slc25a1 | myCAF_1 |
| Fmnl3 | 2.88E-33 | 0.30791438 | 0.837 | 0.698 | 5.56E-29 | 0 | Fmnl3 | myCAF_1 |
| Pgm2l1 | 3.38E-33 | 0.51892778 | 0.279 | 0.202 | 6.53E-29 | 0 | Pgm2l1 | myCAF_1 |
| Slc44a2 | 5.18E-33 | 0.30991178 | 0.862 | 0.781 | 1.00E-28 | 0 | Slc44a2 | myCAF_1 |
| Pcdhgc5 | 2.09E-32 | 0.53527792 | 0.446 | 0.338 | 4.04E-28 | 0 | Pcdhgc5 | myCAF_1 |
| Med10 | 2.10E-32 | 0.31214545 | 0.828 | 0.739 | 4.05E-28 | 0 | Med10 | myCAF_1 |
| Rab20 | 3.73E-32 | 0.40514764 | 0.629 | 0.488 | 7.22E-28 | 0 | Rab20 | myCAF_1 |
| Arl2 | 6.46E-32 | 0.39897612 | 0.627 | 0.525 | 1.25E-27 | 0 | Arl2 | myCAF_1 |
| Zfp809 | 8.93E-32 | 0.52346155 | 0.488 | 0.384 | 1.73E-27 | 0 | Zfp809 | myCAF_1 |
| Nmt1 | 1.06E-31 | 0.32905072 | 0.896 | 0.81 | 2.05E-27 | 0 | Nmt1 | myCAF_1 |
| Nrg1 | 1.08E-31 | 0.45911124 | 0.466 | 0.321 | 2.09E-27 | 0 | Nrg1 | myCAF_1 |
| Ssh1 | 1.13E-31 | 0.29430023 | 0.726 | 0.633 | 2.19E-27 | 0 | Ssh1 | myCAF_1 |
| Arid5b | 1.15E-31 | 0.2750083 | 0.777 | 0.674 | 2.22E-27 | 0 | Arid5b | myCAF_1 |
| Vash2 | 1.16E-31 | 0.63277434 | 0.443 | 0.327 | 2.25E-27 | 0 | Vash2 | myCAF_1 |
| Adamts9 | 1.58E-31 | 0.51515784 | 0.344 | 0.258 | 3.05E-27 | 0 | Adamts9 | myCAF_1 |
| Myo10 | 1.61E-31 | 0.33533499 | 0.656 | 0.507 | 3.10E-27 | 0 | Myo10 | myCAF_1 |
| Spats2 | 1.63E-31 | 0.27167892 | 0.764 | 0.687 | 3.16E-27 | 0 | Spats2 | myCAF_1 |
| Il34 | 3.12E-31 | 0.32613107 | 0.335 | 0.231 | 6.03E-27 | 0 | Il34 | myCAF_1 |
| Pmpcb | 5.31E-31 | 0.39385731 | 0.723 | 0.641 | 1.03E-26 | 0 | Pmpcb | myCAF_1 |
| Pik3r1 | 5.44E-31 | 0.28569376 | 0.784 | 0.681 | 1.05E-26 | 0 | Pik3r1 | myCAF_1 |
| Fdps | 7.11E-31 | 0.25132946 | 0.626 | 0.54 | 1.37E-26 | 0 | Fdps | myCAF_1 |
| Snx30 | 8.03E-31 | 0.40168478 | 0.413 | 0.34 | 1.55E-26 | 0 | Snx30 | myCAF_1 |
| Rgs10 | 8.39E-31 | 0.45559691 | 0.422 | 0.34 | 1.62E-26 | 0 | Rgs10 | myCAF_1 |
| Mef2a | 1.13E-30 | 0.35719545 | 0.689 | 0.602 | 2.18E-26 | 0 | Mef2a | myCAF_1 |
| Cnnm4 | 3.28E-30 | 0.51062605 | 0.463 | 0.347 | 6.34E-26 | 0 | Cnnm4 | myCAF_1 |
| Zfp281 | 5.89E-30 | 0.38276892 | 0.69 | 0.57 | 1.14E-25 | 0 | Zfp281 | myCAF_1 |
| Arhgef7 | 7.71E-30 | 0.25228558 | 0.886 | 0.802 | 1.49E-25 | 0 | Arhgef7 | myCAF_1 |
| Dynlt3 | 9.27E-30 | 0.31303759 | 0.738 | 0.662 | 1.79E-25 | 0 | Dynlt3 | myCAF_1 |
| Lyn | 1.04E-29 | 0.53958828 | 0.441 | 0.316 | 2.00E-25 | 0 | Lyn | myCAF_1 |
| Rftn2 | 1.37E-29 | 0.50532836 | 0.432 | 0.33 | 2.65E-25 | 0 | Rftn2 | myCAF_1 |
| Kif1c | 1.56E-29 | 0.31034566 | 0.788 | 0.715 | 3.01E-25 | 0 | Kif1c | myCAF_1 |
| Nr2f6 | 1.62E-29 | 0.37295403 | 0.661 | 0.556 | 3.14E-25 | 0 | Nr2f6 | myCAF_1 |
| Hoxc6 | 2.10E-29 | 0.31465913 | 0.788 | 0.706 | 4.05E-25 | 0 | Hoxc6 | myCAF_1 |
| Tmem98 | 2.16E-29 | 0.43714218 | 0.645 | 0.55 | 4.18E-25 | 0 | Tmem98 | myCAF_1 |
| Serinc5 | 2.28E-29 | 0.39416186 | 0.739 | 0.625 | 4.41E-25 | 0 | Serinc5 | myCAF_1 |
| En1 | 2.31E-29 | 0.33145781 | 0.656 | 0.565 | 4.47E-25 | 0 | En1 | myCAF_1 |
| Polk | 2.52E-29 | 0.49736062 | 0.381 | 0.271 | 4.88E-25 | 0 | Polk | myCAF_1 |
| Plekho1 | 2.56E-29 | 0.35381865 | 0.577 | 0.479 | 4.96E-25 | 0 | Plekho1 | myCAF_1 |
| Enah | 2.87E-29 | 0.2964762 | 0.874 | 0.791 | 5.55E-25 | 0 | Enah | myCAF_1 |
| Specc1 | 3.02E-29 | 0.47567512 | 0.372 | 0.29 | 5.84E-25 | 0 | Specc1 | myCAF_1 |
| Foxs1 | 3.78E-29 | 0.49309556 | 0.269 | 0.167 | 7.32E-25 | 0 | Foxs1 | myCAF_1 |
| Myo1b | 4.01E-29 | 0.29942603 | 0.832 | 0.692 | 7.76E-25 | 0 | Myo1b | myCAF_1 |
| Tes | 7.93E-29 | 0.2906908 | 0.646 | 0.524 | 1.53E-24 | 0 | Tes | myCAF_1 |
| Ap2s1 | 1.04E-28 | 0.33642883 | 0.842 | 0.767 | 2.01E-24 | 0 | Ap2s1 | myCAF_1 |
| Epas1 | 1.47E-28 | 0.25819569 | 0.775 | 0.605 | 2.85E-24 | 0 | Epas1 | myCAF_1 |
| St3gal3 | 1.75E-28 | 0.53209101 | 0.458 | 0.354 | 3.39E-24 | 0 | St3gal3 | myCAF_1 |
| Tead1 | 1.77E-28 | 0.25952424 | 0.894 | 0.807 | 3.43E-24 | 0 | Tead1 | myCAF_1 |
| Casc4 | 1.94E-28 | 0.38125415 | 0.67 | 0.555 | 3.75E-24 | 0 | Casc4 | myCAF_1 |
| Cmtm4 | 2.03E-28 | 0.46011201 | 0.539 | 0.414 | 3.93E-24 | 0 | Cmtm4 | myCAF_1 |
| Hoxd8 | 4.62E-28 | 0.40639693 | 0.478 | 0.377 | 8.93E-24 | 0 | Hoxd8 | myCAF_1 |
| Peg3 | 4.62E-28 | 0.45263597 | 0.392 | 0.306 | 8.94E-24 | 0 | Peg3 | myCAF_1 |
| Galnt10 | 6.46E-28 | 0.38507706 | 0.787 | 0.689 | 1.25E-23 | 0 | Galnt10 | myCAF_1 |
| Nrp2 | 7.53E-28 | 0.35455361 | 0.499 | 0.392 | 1.46E-23 | 0 | Nrp2 | myCAF_1 |
| Cyp2d22 | 1.41E-27 | 0.41763927 | 0.473 | 0.385 | 2.73E-23 | 0 | Cyp2d22 | myCAF_1 |
| Hmg20b | 1.82E-27 | 0.26911434 | 0.792 | 0.699 | 3.51E-23 | 0 | Hmg20b | myCAF_1 |
| Zdhhc16 | 1.92E-27 | 0.4302268 | 0.696 | 0.565 | 3.71E-23 | 0 | Zdhhc16 | myCAF_1 |
| Wtip | 3.79E-27 | 0.30655287 | 0.79 | 0.707 | 7.33E-23 | 0 | Wtip | myCAF_1 |
| Ddhd2 | 4.47E-27 | 0.4558771 | 0.542 | 0.444 | 8.64E-23 | 0 | Ddhd2 | myCAF_1 |
| Smim10l1 | 1.20E-26 | 0.30969358 | 0.783 | 0.705 | 2.31E-22 | 0 | Smim10l1 | myCAF_1 |
| Parp8 | 1.26E-26 | 0.32659006 | 0.647 | 0.568 | 2.43E-22 | 0 | Parp8 | myCAF_1 |
| Rab7 | 2.00E-26 | 0.27295082 | 0.85 | 0.802 | 3.87E-22 | 0 | Rab7 | myCAF_1 |
| Ice1 | 2.20E-26 | 0.3687434 | 0.634 | 0.563 | 4.25E-22 | 0 | Ice1 | myCAF_1 |
| Ttll5 | 2.69E-26 | 0.63559533 | 0.372 | 0.275 | 5.20E-22 | 0 | Ttll5 | myCAF_1 |
| Cdip1 | 2.82E-26 | 0.34121162 | 0.595 | 0.495 | 5.46E-22 | 0 | Cdip1 | myCAF_1 |
| Abhd4 | 3.94E-26 | 0.25901182 | 0.802 | 0.732 | 7.62E-22 | 0 | Abhd4 | myCAF_1 |
| Clcn4 | 4.36E-26 | 0.30979905 | 0.781 | 0.712 | 8.43E-22 | 0 | Clcn4 | myCAF_1 |
| Pld1 | 5.96E-26 | 0.50059684 | 0.444 | 0.351 | 1.15E-21 | 0 | Pld1 | myCAF_1 |
| Smo | 1.38E-25 | 0.26602587 | 0.891 | 0.827 | 2.67E-21 | 0 | Smo | myCAF_1 |
| B4galt4 | 2.17E-25 | 0.60507009 | 0.306 | 0.239 | 4.20E-21 | 0 | B4galt4 | myCAF_1 |
| As3mt | 2.38E-25 | 0.54593624 | 0.424 | 0.319 | 4.60E-21 | 0 | As3mt | myCAF_1 |
| Hoxc5 | 2.50E-25 | 0.66029925 | 0.281 | 0.196 | 4.83E-21 | 0 | Hoxc5 | myCAF_1 |
| Chst2 | 3.00E-25 | 0.35592469 | 0.497 | 0.358 | 5.79E-21 | 0 | Chst2 | myCAF_1 |
| Epb41l1 | 4.72E-25 | 0.27342571 | 0.616 | 0.536 | 9.13E-21 | 0 | Epb41l1 | myCAF_1 |
| Mcoln2 | 6.36E-25 | 0.34028304 | 0.597 | 0.517 | 1.23E-20 | 0 | Mcoln2 | myCAF_1 |
| Armcx4 | 7.99E-25 | 0.35844233 | 0.684 | 0.587 | 1.54E-20 | 0 | Armcx4 | myCAF_1 |
| Rad12 | 1.42E-24 | 0.49298486 | 0.5 | 0.345 | 2.75E-20 | 0 | Rad12 | myCAF_1 |
| F2rl1 | 1.51E-24 | 0.36063836 | 0.707 | 0.553 | 2.93E-20 | 0 | F2rl1 | myCAF_1 |
| Kctd15 | 3.15E-24 | 0.39569297 | 0.524 | 0.405 | 6.08E-20 | 0 | Kctd15 | myCAF_1 |
| Rnf135 | 5.13E-24 | 0.47308751 | 0.32 | 0.242 | 9.91E-20 | 0 | Rnf135 | myCAF_1 |
| Wwc2 | 6.64E-24 | 0.31973929 | 0.79 | 0.706 | 1.28E-19 | 0 | Wwc2 | myCAF_1 |
| Cdc42ep1 | 6.84E-24 | 0.31652519 | 0.632 | 0.542 | 1.32E-19 | 0 | Cdc42ep1 | myCAF_1 |
| Rhbdf2 | 7.45E-24 | 0.30738241 | 0.414 | 0.294 | 1.44E-19 | 0 | Rhbdf2 | myCAF_1 |
| Ldlrap1 | 9.73E-24 | 0.35598313 | 0.635 | 0.54 | 1.88E-19 | 0 | Ldlrap1 | myCAF_1 |
| Ube2f | 1.16E-23 | 0.31161876 | 0.771 | 0.692 | 2.24E-19 | 0 | Ube2f | myCAF_1 |
| Dgcr6 | 1.37E-23 | 0.33727272 | 0.646 | 0.561 | 2.64E-19 | 0 | Dgcr6 | myCAF_1 |
| Commd9 | 1.43E-23 | 0.50745366 | 0.472 | 0.385 | 2.77E-19 | 0 | Commd9 | myCAF_1 |
| Tm7sf3 | 2.03E-23 | 0.27730038 | 0.856 | 0.789 | 3.92E-19 | 0 | Tm7sf3 | myCAF_1 |
| Adcy7 | 2.19E-23 | 0.27905281 | 0.664 | 0.539 | 4.23E-19 | 0 | Adcy7 | myCAF_1 |
| Etv6 | 2.35E-23 | 0.32751877 | 0.74 | 0.648 | 4.55E-19 | 0 | Etv6 | myCAF_1 |
| Slc6a6 | 3.30E-23 | 0.31991586 | 0.652 | 0.531 | 6.38E-19 | 0 | Slc6a6 | myCAF_1 |
| Uaca | 3.41E-23 | 0.28398004 | 0.719 | 0.648 | 6.59E-19 | 0 | Uaca | myCAF_1 |
| Lims1 | 5.88E-23 | 0.26335023 | 0.882 | 0.838 | 1.14E-18 | 0 | Lims1 | myCAF_1 |
| Pkn1 | 6.81E-23 | 0.28762812 | 0.622 | 0.529 | 1.32E-18 | 0 | Pkn1 | myCAF_1 |
| Nhsl2 | 7.93E-23 | 0.42221955 | 0.406 | 0.318 | 1.53E-18 | 0 | Nhsl2 | myCAF_1 |
| Lpcat1 | 9.12E-23 | 0.41960211 | 0.577 | 0.481 | 1.76E-18 | 0 | Lpcat1 | myCAF_1 |
| Cept1 | 1.15E-22 | 0.34397127 | 0.791 | 0.714 | 2.23E-18 | 0 | Cept1 | myCAF_1 |
| Slc26a11 | 2.43E-22 | 0.5553251 | 0.322 | 0.236 | 4.69E-18 | 0 | Slc26a11 | myCAF_1 |
| Peli2 | 3.76E-22 | 0.32784268 | 0.568 | 0.458 | 7.26E-18 | 0 | Peli2 | myCAF_1 |
| Fam210b | 5.03E-22 | 0.58356943 | 0.28 | 0.198 | 9.72E-18 | 0 | Fam210b | myCAF_1 |
| Patz1 | 7.16E-22 | 0.36305699 | 0.523 | 0.43 | 1.38E-17 | 0 | Patz1 | myCAF_1 |
| Tesk1 | 8.98E-22 | 0.31581629 | 0.686 | 0.6 | 1.74E-17 | 0 | Tesk1 | myCAF_1 |
| Gpr107 | 9.85E-22 | 0.27897853 | 0.759 | 0.7 | 1.90E-17 | 0 | Gpr107 | myCAF_1 |
| Syne1 | 1.81E-21 | 0.45027929 | 0.328 | 0.234 | 3.50E-17 | 0 | Syne1 | myCAF_1 |
| Smim3 | 1.83E-21 | 0.35955706 | 0.755 | 0.622 | 3.54E-17 | 0 | Smim3 | myCAF_1 |
| Klhl22 | 2.28E-21 | 0.37358932 | 0.636 | 0.557 | 4.40E-17 | 0 | Klhl22 | myCAF_1 |
| Rap2a | 3.43E-21 | 0.31688351 | 0.579 | 0.487 | 6.63E-17 | 0 | Rap2a | myCAF_1 |
| Mfhas1 | 3.48E-21 | 0.32892042 | 0.75 | 0.653 | 6.73E-17 | 0 | Mfhas1 | myCAF_1 |
| Tbc1d16 | 6.53E-21 | 0.45400175 | 0.351 | 0.264 | 1.26E-16 | 0 | Tbc1d16 | myCAF_1 |
| Pgpep1 | 6.86E-21 | 0.33639647 | 0.484 | 0.418 | 1.33E-16 | 0 | Pgpep1 | myCAF_1 |
| Pfkfb4 | 1.03E-20 | 0.38220234 | 0.451 | 0.379 | 1.99E-16 | 0 | Pfkfb4 | myCAF_1 |
| Wbp1 | 1.34E-20 | 0.26109797 | 0.535 | 0.448 | 2.59E-16 | 0 | Wbp1 | myCAF_1 |
| Scap | 1.64E-20 | 0.26901131 | 0.721 | 0.645 | 3.17E-16 | 0 | Scap | myCAF_1 |
| Lclat1 | 1.91E-20 | 0.41679679 | 0.662 | 0.565 | 3.69E-16 | 0 | Lclat1 | myCAF_1 |
| Prr14 | 2.28E-20 | 0.32218824 | 0.63 | 0.55 | 4.41E-16 | 0 | Prr14 | myCAF_1 |
| Inka1 | 2.46E-20 | 0.53896589 | 0.285 | 0.206 | 4.75E-16 | 0 | Inka1 | myCAF_1 |
| Pcdhga7 | 3.09E-20 | 0.32634351 | 0.275 | 0.22 | 5.98E-16 | 0 | Pcdhga7 | myCAF_1 |
| Meis1 | 5.29E-20 | 0.32691961 | 0.664 | 0.552 | 1.02E-15 | 0 | Meis1 | myCAF_1 |
| Msantd3 | 5.90E-20 | 0.48814626 | 0.425 | 0.338 | 1.14E-15 | 0 | Msantd3 | myCAF_1 |
| Col24a1 | 7.27E-20 | 0.49330278 | 0.486 | 0.372 | 1.40E-15 | 0 | Col24a1 | myCAF_1 |
| Dnm3 | 7.52E-20 | 0.46705389 | 0.47 | 0.376 | 1.45E-15 | 0 | Dnm3 | myCAF_1 |
| Sh3rf3 | 7.53E-20 | 0.44601179 | 0.331 | 0.252 | 1.46E-15 | 0 | Sh3rf3 | myCAF_1 |
| Lrfn4 | 9.50E-20 | 0.40818436 | 0.525 | 0.423 | 1.84E-15 | 0 | Lrfn4 | myCAF_1 |
| Fah | 1.13E-19 | 0.45626546 | 0.569 | 0.484 | 2.18E-15 | 0 | Fah | myCAF_1 |
| Enox1 | 1.22E-19 | 0.4364204 | 0.321 | 0.238 | 2.36E-15 | 0 | Enox1 | myCAF_1 |
| Limd2 | 1.29E-19 | 0.3510643 | 0.649 | 0.538 | 2.49E-15 | 0 | Limd2 | myCAF_1 |
| Gdf11 | 1.38E-19 | 0.45278543 | 0.377 | 0.29 | 2.66E-15 | 0 | Gdf11 | myCAF_1 |
| Slco5a1 | 1.43E-19 | 0.43903108 | 0.509 | 0.374 | 2.76E-15 | 0 | Slco5a1 | myCAF_1 |
| Bmpr1b | 1.44E-19 | 0.62160686 | 0.305 | 0.204 | 2.79E-15 | 0 | Bmpr1b | myCAF_1 |
| Wdr19 | 2.80E-19 | 0.48107867 | 0.261 | 0.194 | 5.41E-15 | 0 | Wdr19 | myCAF_1 |
| Fhod1 | 5.01E-19 | 0.47373486 | 0.348 | 0.261 | 9.69E-15 | 0 | Fhod1 | myCAF_1 |
| Stx1a | 7.24E-19 | 0.30881722 | 0.337 | 0.256 | 1.40E-14 | 0 | Stx1a | myCAF_1 |
| Amacr | 7.45E-19 | 0.59576615 | 0.283 | 0.211 | 1.44E-14 | 0 | Amacr | myCAF_1 |
| Crim1 | 7.76E-19 | 0.35313781 | 0.385 | 0.325 | 1.50E-14 | 0 | Crim1 | myCAF_1 |
| Blcap | 7.78E-19 | 0.25027872 | 0.69 | 0.616 | 1.50E-14 | 0 | Blcap | myCAF_1 |
| BC034090 | 1.05E-18 | 0.36257743 | 0.531 | 0.443 | 2.03E-14 | 0 | BC034090 | myCAF_1 |
| Plekhn1 | 1.47E-18 | 0.28326753 | 0.479 | 0.381 | 2.84E-14 | 0 | Plekhn1 | myCAF_1 |
| Gcc1 | 1.72E-18 | 0.27680409 | 0.538 | 0.441 | 3.32E-14 | 0 | Gcc1 | myCAF_1 |
| Dcaf8 | 1.82E-18 | 0.26088866 | 0.749 | 0.695 | 3.52E-14 | 0 | Dcaf8 | myCAF_1 |
| Sema3a | 2.42E-18 | 0.33566713 | 0.378 | 0.22 | 4.68E-14 | 0 | Sema3a | myCAF_1 |
| Cyhr1 | 3.03E-18 | 0.31715286 | 0.579 | 0.498 | 5.87E-14 | 0 | Cyhr1 | myCAF_1 |
| Amn | 5.81E-18 | 0.42981049 | 0.337 | 0.21 | 1.12E-13 | 0 | Amn | myCAF_1 |
| Zfp651 | 5.92E-18 | 0.28549585 | 0.66 | 0.578 | 1.14E-13 | 0 | Zfp651 | myCAF_1 |
| Lrch2 | 6.79E-18 | 0.49291491 | 0.306 | 0.225 | 1.31E-13 | 0 | Lrch2 | myCAF_1 |
| Taf9b | 6.97E-18 | 0.58565814 | 0.312 | 0.237 | 1.35E-13 | 0 | Taf9b | myCAF_1 |
| Slc17a5 | 7.47E-18 | 0.46854404 | 0.384 | 0.311 | 1.44E-13 | 0 | Slc17a5 | myCAF_1 |
| Atg16l2 | 2.17E-17 | 0.48291833 | 0.252 | 0.175 | 4.20E-13 | 0 | Atg16l2 | myCAF_1 |
| Rufy1 | 2.56E-17 | 0.36824645 | 0.606 | 0.492 | 4.95E-13 | 0 | Rufy1 | myCAF_1 |
| Ephb2 | 2.58E-17 | 0.30084497 | 0.707 | 0.576 | 4.99E-13 | 0 | Ephb2 | myCAF_1 |
| Pxdc1 | 2.80E-17 | 0.32380764 | 0.624 | 0.515 | 5.42E-13 | 0 | Pxdc1 | myCAF_1 |
| Dennd4a | 3.25E-17 | 0.41289313 | 0.666 | 0.53 | 6.29E-13 | 0 | Dennd4a | myCAF_1 |
| Ccdc91 | 4.57E-17 | 0.44859061 | 0.494 | 0.408 | 8.84E-13 | 0 | Ccdc91 | myCAF_1 |
| Atp13a2 | 5.16E-17 | 0.3960931 | 0.33 | 0.248 | 9.97E-13 | 0 | Atp13a2 | myCAF_1 |
| Dhcr7 | 5.86E-17 | 0.43901687 | 0.346 | 0.269 | 1.13E-12 | 0 | Dhcr7 | myCAF_1 |
| Tjap1 | 7.25E-17 | 0.27687015 | 0.682 | 0.617 | 1.40E-12 | 0 | Tjap1 | myCAF_1 |
| S1pr2 | 7.62E-17 | 0.26036834 | 0.81 | 0.74 | 1.47E-12 | 0 | S1pr2 | myCAF_1 |
| Gli3 | 8.08E-17 | 0.35666937 | 0.671 | 0.557 | 1.56E-12 | 0 | Gli3 | myCAF_1 |
| Ttpal | 8.30E-17 | 0.37224527 | 0.557 | 0.467 | 1.60E-12 | 0 | Ttpal | myCAF_1 |
| Zfp866 | 8.65E-17 | 0.33638825 | 0.438 | 0.355 | 1.67E-12 | 0 | Zfp866 | myCAF_1 |
| Dlg5 | 9.83E-17 | 0.2658736 | 0.735 | 0.653 | 1.90E-12 | 0 | Dlg5 | myCAF_1 |
| Snx7 | 1.17E-16 | 0.37251566 | 0.702 | 0.608 | 2.26E-12 | 0 | Snx7 | myCAF_1 |
| Srpk2 | 1.19E-16 | 0.27984728 | 0.602 | 0.531 | 2.31E-12 | 0 | Srpk2 | myCAF_1 |
| Zfp523 | 1.89E-16 | 0.36398829 | 0.497 | 0.423 | 3.64E-12 | 0 | Zfp523 | myCAF_1 |
| Tmem47 | 2.01E-16 | 0.38134632 | 0.409 | 0.323 | 3.88E-12 | 0 | Tmem47 | myCAF_1 |
| Zfc3h1 | 2.28E-16 | 0.30438825 | 0.573 | 0.479 | 4.41E-12 | 0 | Zfc3h1 | myCAF_1 |
| Garem1 | 4.10E-16 | 0.60813987 | 0.252 | 0.188 | 7.93E-12 | 0 | Garem1 | myCAF_1 |
| Rin1 | 4.98E-16 | 0.26856535 | 0.507 | 0.384 | 9.63E-12 | 0 | Rin1 | myCAF_1 |
| Epn2 | 5.07E-16 | 0.26550537 | 0.533 | 0.45 | 9.80E-12 | 0 | Epn2 | myCAF_1 |
| Pcdhga10 | 7.62E-16 | 0.53799636 | 0.278 | 0.214 | 1.47E-11 | 0 | Pcdhga10 | myCAF_1 |
| Klc2 | 7.67E-16 | 0.31297448 | 0.439 | 0.358 | 1.48E-11 | 0 | Klc2 | myCAF_1 |
| Tln2 | 1.00E-15 | 0.27842327 | 0.79 | 0.68 | 1.93E-11 | 0 | Tln2 | myCAF_1 |
| Prr5 | 1.32E-15 | 0.34526968 | 0.686 | 0.579 | 2.56E-11 | 0 | Prr5 | myCAF_1 |
| Tmem123 | 1.42E-15 | 0.28398251 | 0.704 | 0.641 | 2.75E-11 | 0 | Tmem123 | myCAF_1 |
| Fam234b | 1.82E-15 | 0.53725278 | 0.265 | 0.197 | 3.51E-11 | 0 | Fam234b | myCAF_1 |
| Epm2aip1 | 3.44E-15 | 0.27796354 | 0.594 | 0.513 | 6.65E-11 | 0 | Epm2aip1 | myCAF_1 |
| Ric1 | 4.04E-15 | 0.27736138 | 0.684 | 0.611 | 7.82E-11 | 0 | Ric1 | myCAF_1 |
| Mn1 | 5.27E-15 | 0.55331189 | 0.291 | 0.207 | 1.02E-10 | 0 | Mn1 | myCAF_1 |
| Zfp57 | 6.02E-15 | 0.45098642 | 0.267 | 0.186 | 1.16E-10 | 0 | Zfp57 | myCAF_1 |
| Cdyl2 | 7.49E-15 | 0.30789186 | 0.483 | 0.388 | 1.45E-10 | 0 | Cdyl2 | myCAF_1 |
| Ctns | 8.10E-15 | 0.40788017 | 0.281 | 0.215 | 1.57E-10 | 0 | Ctns | myCAF_1 |
| Cdc42bpg | 9.85E-15 | 0.38162461 | 0.268 | 0.202 | 1.91E-10 | 0 | Cdc42bpg | myCAF_1 |
| Slc38a7 | 1.12E-14 | 0.40076053 | 0.286 | 0.218 | 2.16E-10 | 0 | Slc38a7 | myCAF_1 |
| Speg | 1.37E-14 | 0.40388104 | 0.299 | 0.226 | 2.64E-10 | 0 | Speg | myCAF_1 |
| Creg1 | 1.39E-14 | 0.25842669 | 0.801 | 0.756 | 2.68E-10 | 0 | Creg1 | myCAF_1 |
| Mxi1 | 1.66E-14 | 0.30072577 | 0.459 | 0.346 | 3.20E-10 | 0 | Mxi1 | myCAF_1 |
| Spata13 | 2.10E-14 | 0.54703003 | 0.347 | 0.247 | 4.06E-10 | 0 | Spata13 | myCAF_1 |
| Hsd17b7 | 2.43E-14 | 0.41856908 | 0.284 | 0.221 | 4.70E-10 | 0 | Hsd17b7 | myCAF_1 |
| Rnf41 | 2.51E-14 | 0.27792051 | 0.54 | 0.476 | 4.85E-10 | 0 | Rnf41 | myCAF_1 |
| Unc45a | 2.88E-14 | 0.26656069 | 0.535 | 0.456 | 5.57E-10 | 0 | Unc45a | myCAF_1 |
| Fnbp1l | 3.10E-14 | 0.34483676 | 0.369 | 0.269 | 6.00E-10 | 0 | Fnbp1l | myCAF_1 |
| Angpt4 | 4.32E-14 | 0.27720501 | 0.527 | 0.425 | 8.35E-10 | 0 | Angpt4 | myCAF_1 |
| Pigyl | 4.56E-14 | 0.43773525 | 0.337 | 0.272 | 8.82E-10 | 0 | Pigyl | myCAF_1 |
| Pbxip1 | 4.97E-14 | 0.27136536 | 0.602 | 0.541 | 9.61E-10 | 0 | Pbxip1 | myCAF_1 |
| Msi1 | 5.36E-14 | 0.45812779 | 0.333 | 0.242 | 1.04E-09 | 0 | Msi1 | myCAF_1 |
| Chic2 | 5.95E-14 | 0.28130846 | 0.693 | 0.621 | 1.15E-09 | 0 | Chic2 | myCAF_1 |
| Mvd | 7.11E-14 | 0.3954285 | 0.275 | 0.207 | 1.38E-09 | 0 | Mvd | myCAF_1 |
| Prokr2 | 8.40E-14 | 0.37782358 | 0.52 | 0.362 | 1.62E-09 | 0 | Prokr2 | myCAF_1 |
| Kank1 | 1.89E-13 | 0.28981413 | 0.441 | 0.355 | 3.66E-09 | 0 | Kank1 | myCAF_1 |
| Efna4 | 2.10E-13 | 0.39287382 | 0.283 | 0.217 | 4.06E-09 | 0 | Efna4 | myCAF_1 |
| Rnf19b | 2.23E-13 | 0.25128991 | 0.941 | 0.861 | 4.32E-09 | 0 | Rnf19b | myCAF_1 |
| Fkrp | 2.24E-13 | 0.25394341 | 0.625 | 0.552 | 4.33E-09 | 0 | Fkrp | myCAF_1 |
| Pskh1 | 2.67E-13 | 0.29592685 | 0.458 | 0.383 | 5.17E-09 | 0 | Pskh1 | myCAF_1 |
| Rab4a | 2.93E-13 | 0.32257442 | 0.304 | 0.238 | 5.66E-09 | 0 | Rab4a | myCAF_1 |
| Scamp5 | 2.99E-13 | 0.34286403 | 0.342 | 0.281 | 5.77E-09 | 0 | Scamp5 | myCAF_1 |
| Dlg3 | 4.35E-13 | 0.31210598 | 0.418 | 0.34 | 8.41E-09 | 0 | Dlg3 | myCAF_1 |
| Sall2 | 4.52E-13 | 0.41685521 | 0.341 | 0.271 | 8.73E-09 | 0 | Sall2 | myCAF_1 |
| Spns1 | 4.59E-13 | 0.34876407 | 0.346 | 0.274 | 8.87E-09 | 0 | Spns1 | myCAF_1 |
| Zfhx4 | 4.87E-13 | 0.3189628 | 0.573 | 0.502 | 9.41E-09 | 0 | Zfhx4 | myCAF_1 |
| Dnajb5 | 5.26E-13 | 0.38862005 | 0.355 | 0.292 | 1.02E-08 | 0 | Dnajb5 | myCAF_1 |
| Inafm2 | 5.60E-13 | 0.30081634 | 0.387 | 0.312 | 1.08E-08 | 0 | Inafm2 | myCAF_1 |
| Zfp580 | 6.45E-13 | 0.43442769 | 0.307 | 0.243 | 1.25E-08 | 0 | Zfp580 | myCAF_1 |
| Frrs1 | 6.96E-13 | 0.32109861 | 0.56 | 0.48 | 1.35E-08 | 0 | Frrs1 | myCAF_1 |
| Def6 | 7.46E-13 | 0.34876497 | 0.344 | 0.277 | 1.44E-08 | 0 | Def6 | myCAF_1 |
| Adcy4 | 8.20E-13 | 0.29215736 | 0.335 | 0.261 | 1.59E-08 | 0 | Adcy4 | myCAF_1 |
| Kif3c | 8.23E-13 | 0.34480918 | 0.302 | 0.249 | 1.59E-08 | 0 | Kif3c | myCAF_1 |
| Ttc14 | 8.28E-13 | 0.27269776 | 0.552 | 0.482 | 1.60E-08 | 0 | Ttc14 | myCAF_1 |
| Uckl1 | 8.62E-13 | 0.33937639 | 0.429 | 0.354 | 1.67E-08 | 0 | Uckl1 | myCAF_1 |
| Snf8 | 8.83E-13 | 0.25619521 | 0.808 | 0.731 | 1.71E-08 | 0 | Snf8 | myCAF_1 |
| Atp5md | 1.45E-12 | 0.2865511 | 0.703 | 0.636 | 2.81E-08 | 0 | Atp5md | myCAF_1 |
| Ric8b | 2.11E-12 | 0.29895374 | 0.548 | 0.471 | 4.08E-08 | 0 | Ric8b | myCAF_1 |
| Hoxb3 | 2.31E-12 | 0.32120047 | 0.292 | 0.226 | 4.46E-08 | 0 | Hoxb3 | myCAF_1 |
| Ccdc107 | 2.59E-12 | 0.34223952 | 0.511 | 0.442 | 5.00E-08 | 0 | Ccdc107 | myCAF_1 |
| Mtfr1 | 2.68E-12 | 0.29611271 | 0.562 | 0.489 | 5.19E-08 | 0 | Mtfr1 | myCAF_1 |
| Hes6 | 3.19E-12 | 0.27788309 | 0.377 | 0.309 | 6.17E-08 | 0 | Hes6 | myCAF_1 |
| Cstf2t | 3.56E-12 | 0.29078012 | 0.591 | 0.511 | 6.88E-08 | 0 | Cstf2t | myCAF_1 |
| Kansl1l | 4.04E-12 | 0.38885215 | 0.344 | 0.274 | 7.81E-08 | 0 | Kansl1l | myCAF_1 |
| Spaca6 | 4.85E-12 | 0.25425117 | 0.459 | 0.378 | 9.37E-08 | 0 | Spaca6 | myCAF_1 |
| Bicra | 5.01E-12 | 0.30713367 | 0.427 | 0.37 | 9.68E-08 | 0 | Bicra | myCAF_1 |
| Slc46a1 | 5.88E-12 | 0.49770175 | 0.341 | 0.277 | 1.14E-07 | 0 | Slc46a1 | myCAF_1 |
| Helb | 6.32E-12 | 0.37540173 | 0.429 | 0.352 | 1.22E-07 | 0 | Helb | myCAF_1 |
| Ptpn14 | 6.59E-12 | 0.40985223 | 0.292 | 0.223 | 1.27E-07 | 0 | Ptpn14 | myCAF_1 |
| Pcdhb17 | 6.74E-12 | 0.41579194 | 0.277 | 0.215 | 1.30E-07 | 0 | Pcdhb17 | myCAF_1 |
| Fryl | 7.04E-12 | 0.29695546 | 0.567 | 0.487 | 1.36E-07 | 0 | Fryl | myCAF_1 |
| Ccs | 8.24E-12 | 0.27254549 | 0.433 | 0.37 | 1.59E-07 | 0 | Ccs | myCAF_1 |
| Klc4 | 1.01E-11 | 0.25639211 | 0.368 | 0.302 | 1.95E-07 | 0 | Klc4 | myCAF_1 |
| Aldh9a1 | 1.02E-11 | 0.26977158 | 0.677 | 0.637 | 1.97E-07 | 0 | Aldh9a1 | myCAF_1 |
| Endod1 | 1.13E-11 | 0.36489279 | 0.364 | 0.269 | 2.19E-07 | 0 | Endod1 | myCAF_1 |
| Utp25 | 1.23E-11 | 0.51725261 | 0.317 | 0.26 | 2.37E-07 | 0 | Utp25 | myCAF_1 |
| Mkx | 1.52E-11 | 0.53516177 | 0.328 | 0.232 | 2.94E-07 | 0 | Mkx | myCAF_1 |
| Vangl2 | 1.55E-11 | 0.37856135 | 0.427 | 0.356 | 3.00E-07 | 0 | Vangl2 | myCAF_1 |
| Rad54l2 | 1.59E-11 | 0.29152672 | 0.553 | 0.473 | 3.08E-07 | 0 | Rad54l2 | myCAF_1 |
| Uggt2 | 1.69E-11 | 0.42325064 | 0.454 | 0.383 | 3.26E-07 | 0 | Uggt2 | myCAF_1 |
| Schip1 | 2.29E-11 | 0.32050789 | 0.606 | 0.483 | 4.43E-07 | 0 | Schip1 | myCAF_1 |
| Ndfip2 | 2.47E-11 | 0.27583929 | 0.465 | 0.405 | 4.78E-07 | 0 | Ndfip2 | myCAF_1 |
| Gpr137 | 2.62E-11 | 0.27028559 | 0.464 | 0.408 | 5.06E-07 | 0 | Gpr137 | myCAF_1 |
| Hoxb2 | 2.62E-11 | 0.37214031 | 0.543 | 0.445 | 5.06E-07 | 0 | Hoxb2 | myCAF_1 |
| Jade1 | 2.70E-11 | 0.26052009 | 0.499 | 0.412 | 5.22E-07 | 0 | Jade1 | myCAF_1 |
| Cyb561a3 | 2.84E-11 | 0.35032792 | 0.46 | 0.389 | 5.49E-07 | 0 | Cyb561a3 | myCAF_1 |
| Slc22a17 | 3.60E-11 | 0.30759172 | 0.453 | 0.381 | 6.96E-07 | 0 | Slc22a17 | myCAF_1 |
| Efcab14 | 5.31E-11 | 0.32771308 | 0.614 | 0.553 | 1.03E-06 | 0 | Efcab14 | myCAF_1 |
| Txndc16 | 6.00E-11 | 0.27979767 | 0.431 | 0.364 | 1.16E-06 | 0 | Txndc16 | myCAF_1 |
| Runx1t1 | 6.88E-11 | 0.3623992 | 0.337 | 0.278 | 1.33E-06 | 0 | Runx1t1 | myCAF_1 |
| Il2rg | 7.41E-11 | 0.312218 | 0.281 | 0.186 | 1.43E-06 | 0 | Il2rg | myCAF_1 |
| Alkbh1 | 1.16E-10 | 0.26630758 | 0.445 | 0.368 | 2.25E-06 | 0 | Alkbh1 | myCAF_1 |
| Hoxc10 | 1.24E-10 | 0.25931364 | 0.607 | 0.523 | 2.39E-06 | 0 | Hoxc10 | myCAF_1 |
| Dok6 | 1.27E-10 | 0.33425025 | 0.256 | 0.201 | 2.45E-06 | 0 | Dok6 | myCAF_1 |
| Xxylt1 | 1.52E-10 | 0.28931172 | 0.667 | 0.582 | 2.93E-06 | 0 | Xxylt1 | myCAF_1 |
| Fgfrl1 | 1.62E-10 | 0.37559863 | 0.403 | 0.327 | 3.14E-06 | 0 | Fgfrl1 | myCAF_1 |
| Rnpep | 1.64E-10 | 0.27698619 | 0.488 | 0.427 | 3.16E-06 | 0 | Rnpep | myCAF_1 |
| Pcdhgb6 | 2.12E-10 | 0.35279923 | 0.258 | 0.218 | 4.10E-06 | 0 | Pcdhgb6 | myCAF_1 |
| Wdr35 | 2.13E-10 | 0.38463695 | 0.351 | 0.29 | 4.12E-06 | 0 | Wdr35 | myCAF_1 |
| Ormdl3 | 2.80E-10 | 0.25534393 | 0.623 | 0.574 | 5.42E-06 | 0 | Ormdl3 | myCAF_1 |
| Pbx2 | 2.81E-10 | 0.25259284 | 0.638 | 0.573 | 5.44E-06 | 0 | Pbx2 | myCAF_1 |
| Ppt2 | 3.14E-10 | 0.32929434 | 0.416 | 0.347 | 6.06E-06 | 0 | Ppt2 | myCAF_1 |
| D16Ertd472e | 4.36E-10 | 0.33488024 | 0.39 | 0.322 | 8.43E-06 | 0 | D16Ertd472e | myCAF_1 |
| Asb8 | 5.39E-10 | 0.33502457 | 0.573 | 0.498 | 1.04E-05 | 0 | Asb8 | myCAF_1 |
| Tbx18 | 5.51E-10 | 0.33634067 | 0.382 | 0.339 | 1.07E-05 | 0 | Tbx18 | myCAF_1 |
| Dnal1 | 6.56E-10 | 0.36264089 | 0.3 | 0.244 | 1.27E-05 | 0 | Dnal1 | myCAF_1 |
| Rnf157 | 6.81E-10 | 0.46593088 | 0.252 | 0.187 | 1.32E-05 | 0 | Rnf157 | myCAF_1 |
| Ak2 | 7.79E-10 | 0.26048251 | 0.933 | 0.861 | 1.51E-05 | 0 | Ak2 | myCAF_1 |
| Hdhd2 | 8.79E-10 | 0.33154053 | 0.441 | 0.384 | 1.70E-05 | 0 | Hdhd2 | myCAF_1 |
| Zic1 | 1.04E-09 | 0.26239315 | 0.321 | 0.255 | 2.01E-05 | 0 | Zic1 | myCAF_1 |
| Fpgt | 1.10E-09 | 0.34451117 | 0.363 | 0.303 | 2.12E-05 | 0 | Fpgt | myCAF_1 |
| Hhipl1 | 1.91E-09 | 0.2503407 | 0.637 | 0.576 | 3.69E-05 | 0 | Hhipl1 | myCAF_1 |
| Sbk1 | 1.99E-09 | 0.39022631 | 0.281 | 0.231 | 3.86E-05 | 0 | Sbk1 | myCAF_1 |
| Pip5k1a | 2.22E-09 | 0.25746294 | 0.657 | 0.581 | 4.30E-05 | 0 | Pip5k1a | myCAF_1 |
| Arntl2 | 2.70E-09 | 0.34624662 | 0.287 | 0.227 | 5.21E-05 | 0 | Arntl2 | myCAF_1 |
| Zfp691 | 2.83E-09 | 0.38517017 | 0.281 | 0.229 | 5.47E-05 | 0 | Zfp691 | myCAF_1 |
| Polm | 2.84E-09 | 0.32767534 | 0.306 | 0.248 | 5.49E-05 | 0 | Polm | myCAF_1 |
| Agpat2 | 4.72E-09 | 0.30405184 | 0.564 | 0.484 | 9.13E-05 | 0 | Agpat2 | myCAF_1 |
| Hebp1 | 5.75E-09 | 0.26740805 | 0.398 | 0.346 | 0.0001111 | 0 | Hebp1 | myCAF_1 |
| Akr1b10 | 5.99E-09 | 0.29641313 | 0.391 | 0.338 | 0.00011581 | 0 | Akr1b10 | myCAF_1 |
| Tjp2 | 6.12E-09 | 0.25474723 | 0.324 | 0.261 | 0.00011834 | 0 | Tjp2 | myCAF_1 |
| Ptprk | 6.46E-09 | 0.30882858 | 0.343 | 0.277 | 0.00012481 | 0 | Ptprk | myCAF_1 |
| Ddr1 | 6.53E-09 | 0.27242961 | 0.317 | 0.273 | 0.00012618 | 0 | Ddr1 | myCAF_1 |
| Rnf123 | 9.06E-09 | 0.29333605 | 0.361 | 0.303 | 0.00017508 | 0 | Rnf123 | myCAF_1 |
| Vmac | 1.07E-08 | 0.25953073 | 0.252 | 0.206 | 0.0002069 | 0 | Vmac | myCAF_1 |
| Alg2 | 1.07E-08 | 0.27370207 | 0.433 | 0.372 | 0.00020741 | 0 | Alg2 | myCAF_1 |
| Fndc4 | 1.13E-08 | 0.32241513 | 0.491 | 0.417 | 0.00021797 | 0 | Fndc4 | myCAF_1 |
| Mmab | 1.43E-08 | 0.36667817 | 0.287 | 0.232 | 0.00027737 | 0 | Mmab | myCAF_1 |
| Abtb1 | 1.66E-08 | 0.28349072 | 0.293 | 0.246 | 0.0003208 | 0 | Abtb1 | myCAF_1 |
| Shank3 | 1.75E-08 | 0.33494259 | 0.354 | 0.294 | 0.00033919 | 0 | Shank3 | myCAF_1 |
| Tnfsf13 | 1.91E-08 | 0.3085363 | 0.293 | 0.247 | 0.00036875 | 0 | Tnfsf13 | myCAF_1 |
| Rasa2 | 1.96E-08 | 0.27217348 | 0.357 | 0.306 | 0.00037936 | 0 | Rasa2 | myCAF_1 |
| Itpr1 | 2.36E-08 | 0.28878873 | 0.271 | 0.224 | 0.00045675 | 0 | Itpr1 | myCAF_1 |
| Phlpp1 | 2.96E-08 | 0.35333468 | 0.309 | 0.251 | 0.00057158 | 0 | Phlpp1 | myCAF_1 |
| Tmem70 | 3.54E-08 | 0.25061875 | 0.514 | 0.486 | 0.00068518 | 0 | Tmem70 | myCAF_1 |
| Smim13 | 3.81E-08 | 0.2826997 | 0.419 | 0.365 | 0.00073745 | 0 | Smim13 | myCAF_1 |
| Tmcc2 | 3.83E-08 | 0.32031882 | 0.316 | 0.26 | 0.00073977 | 0 | Tmcc2 | myCAF_1 |
| Dbt | 3.84E-08 | 0.29375261 | 0.323 | 0.294 | 0.00074322 | 0 | Dbt | myCAF_1 |
| Cxcl16 | 3.92E-08 | 0.39669673 | 0.47 | 0.391 | 0.0007582 | 0 | Cxcl16 | myCAF_1 |
| Mex3a | 5.21E-08 | 0.28422195 | 0.298 | 0.234 | 0.00100761 | 0 | Mex3a | myCAF_1 |
| Gpcpd1 | 5.90E-08 | 0.2930252 | 0.447 | 0.375 | 0.00114077 | 0 | Gpcpd1 | myCAF_1 |
| Dffa | 8.61E-08 | 0.32227801 | 0.333 | 0.286 | 0.00166491 | 0 | Dffa | myCAF_1 |
| Ptger4 | 8.63E-08 | 0.37615564 | 0.366 | 0.274 | 0.00166865 | 0 | Ptger4 | myCAF_1 |
| Zswim6 | 1.21E-07 | 0.25626441 | 0.575 | 0.5 | 0.00233746 | 0 | Zswim6 | myCAF_1 |
| Dpy19l3 | 1.57E-07 | 0.34914089 | 0.296 | 0.247 | 0.00303531 | 0 | Dpy19l3 | myCAF_1 |
| Copg2 | 2.62E-07 | 0.26076974 | 0.301 | 0.253 | 0.00506799 | 0 | Copg2 | myCAF_1 |
| Lrfn3 | 3.21E-07 | 0.42396144 | 0.253 | 0.212 | 0.00621548 | 0 | Lrfn3 | myCAF_1 |
| Mad2l2 | 3.29E-07 | 0.38467529 | 0.365 | 0.295 | 0.00636905 | 0 | Mad2l2 | myCAF_1 |
| Slc52a2 | 4.19E-07 | 0.36988973 | 0.288 | 0.24 | 0.00809894 | 0 | Slc52a2 | myCAF_1 |
| Plekhg5 | 4.23E-07 | 0.29399074 | 0.279 | 0.222 | 0.00817338 | 0 | Plekhg5 | myCAF_1 |
| B4galt6 | 7.12E-07 | 0.25195974 | 0.481 | 0.409 | 0.01376204 | 0 | B4galt6 | myCAF_1 |
| Bcl7a | 8.26E-07 | 0.26440215 | 0.281 | 0.218 | 0.01597932 | 0 | Bcl7a | myCAF_1 |
| Pheta2 | 1.27E-06 | 0.33940454 | 0.465 | 0.405 | 0.02458895 | 0 | Pheta2 | myCAF_1 |
| Pdk2 | 1.47E-06 | 0.31951409 | 0.318 | 0.274 | 0.02842451 | 0 | Pdk2 | myCAF_1 |
| Map11 | 1.84E-06 | 0.27441148 | 0.366 | 0.314 | 0.03554091 | 0 | Map11 | myCAF_1 |
| Gk | 1.88E-06 | 0.34477022 | 0.384 | 0.319 | 0.0364403 | 0 | Gk | myCAF_1 |
| Zkscan8 | 1.90E-06 | 0.29370263 | 0.251 | 0.211 | 0.03682897 | 0 | Zkscan8 | myCAF_1 |
| Gdnf | 2.34E-06 | 0.38609364 | 0.273 | 0.193 | 0.04523754 | 0 | Gdnf | myCAF_1 |
| Irx5 | 2.38E-06 | 0.32991085 | 0.41 | 0.337 | 0.04600073 | 0 | Irx5 | myCAF_1 |
| Nlrx1 | 4.50E-06 | 0.2739001 | 0.325 | 0.28 | 0.0869621 | 0 | Nlrx1 | myCAF_1 |
| Zdhhc12 | 4.62E-06 | 0.29619947 | 0.282 | 0.239 | 0.08925172 | 0 | Zdhhc12 | myCAF_1 |
| Zfp507 | 4.64E-06 | 0.27977517 | 0.428 | 0.363 | 0.08974973 | 0 | Zfp507 | myCAF_1 |
| Ethe1 | 4.88E-06 | 0.26677952 | 0.477 | 0.419 | 0.09428047 | 0 | Ethe1 | myCAF_1 |
| Sphk2 | 5.40E-06 | 0.26462943 | 0.375 | 0.328 | 0.10433065 | 0 | Sphk2 | myCAF_1 |
| Crppa | 5.55E-06 | 0.2878906 | 0.258 | 0.219 | 0.10725005 | 0 | Crppa | myCAF_1 |
| B3gnt2 | 5.71E-06 | 0.26931439 | 0.448 | 0.382 | 0.11041534 | 0 | B3gnt2 | myCAF_1 |
| B4galt7 | 7.29E-06 | 0.28400176 | 0.466 | 0.423 | 0.14099404 | 0 | B4galt7 | myCAF_1 |
| Tor4a | 1.09E-05 | 0.25004817 | 0.397 | 0.3 | 0.21022867 | 0 | Tor4a | myCAF_1 |
| Smco4 | 1.17E-05 | 0.31418431 | 0.597 | 0.509 | 0.22711081 | 0 | Smco4 | myCAF_1 |
| Bmi1 | 1.87E-05 | 0.26181164 | 0.554 | 0.485 | 0.36197272 | 0 | Bmi1 | myCAF_1 |
| Ctdp1 | 2.09E-05 | 0.32915164 | 0.396 | 0.338 | 0.40330215 | 0 | Ctdp1 | myCAF_1 |
| Acsf2 | 2.76E-05 | 0.28359643 | 0.378 | 0.323 | 0.5327098 | 0 | Acsf2 | myCAF_1 |
| D11Wsu47e | 3.51E-05 | 0.29801144 | 0.317 | 0.275 | 0.67767253 | 0 | D11Wsu47e | myCAF_1 |
| Eif2ak4 | 3.63E-05 | 0.25461788 | 0.446 | 0.41 | 0.70178112 | 0 | Eif2ak4 | myCAF_1 |
| Chsy3 | 3.76E-05 | 0.26762959 | 0.425 | 0.368 | 0.72782148 | 0 | Chsy3 | myCAF_1 |
| Dgkq | 3.83E-05 | 0.25465951 | 0.312 | 0.267 | 0.73988022 | 0 | Dgkq | myCAF_1 |
| Eogt | 6.05E-05 | 0.28300094 | 0.273 | 0.235 | 1 | 0 | Eogt | myCAF_1 |
| Fbxl6 | 6.91E-05 | 0.25043621 | 0.479 | 0.433 | 1 | 0 | Fbxl6 | myCAF_1 |
| Tyro3 | 7.85E-05 | 0.38954261 | 0.271 | 0.224 | 1 | 0 | Tyro3 | myCAF_1 |
| Brsk1 | 7.98E-05 | 0.2606475 | 0.295 | 0.247 | 1 | 0 | Brsk1 | myCAF_1 |
| Sgsh | 9.77E-05 | 0.25703269 | 0.364 | 0.315 | 1 | 0 | Sgsh | myCAF_1 |
| Elk1 | 0.00011497 | 0.26824052 | 0.44 | 0.395 | 1 | 0 | Elk1 | myCAF_1 |
| Csgalnact1 | 0.00012409 | 0.25787705 | 0.829 | 0.736 | 1 | 0 | Csgalnact1 | myCAF_1 |
| Srek1ip1 | 0.00013775 | 0.27143763 | 0.324 | 0.291 | 1 | 0 | Srek1ip1 | myCAF_1 |
| Pdzd4 | 0.00026957 | 0.29218414 | 0.361 | 0.321 | 1 | 0 | Pdzd4 | myCAF_1 |
| Tgfbi | 0.00027275 | 0.32803268 | 0.573 | 0.457 | 1 | 0 | Tgfbi | myCAF_1 |
| Mlycd | 0.00028242 | 0.29335227 | 0.343 | 0.308 | 1 | 0 | Mlycd | myCAF_1 |
| Tmtc4 | 0.00028638 | 0.35911192 | 0.433 | 0.346 | 1 | 0 | Tmtc4 | myCAF_1 |
| Usp46 | 0.00029799 | 0.26386321 | 0.383 | 0.344 | 1 | 0 | Usp46 | myCAF_1 |
| Lgr5 | 0.00031677 | 0.33924462 | 0.413 | 0.283 | 1 | 0 | Lgr5 | myCAF_1 |
| DEPRECATED-l | 0.00045395 | 0.27932109 | 0.298 | 0.263 | 1 | 0 | DEPRECATED-l | myCAF_1 |
| Pgap3 | 0.00045966 | 0.32879022 | 0.257 | 0.226 | 1 | 0 | Pgap3 | myCAF_1 |
| Galnt7 | 0.00047332 | 0.32707451 | 0.303 | 0.265 | 1 | 0 | Galnt7 | myCAF_1 |
| Siah1a | 0.00047722 | 0.28712306 | 0.266 | 0.233 | 1 | 0 | Siah1a | myCAF_1 |
| Lnx2 | 0.00052815 | 0.30368541 | 0.385 | 0.348 | 1 | 0 | Lnx2 | myCAF_1 |
| Zbtb46 | 0.0006583 | 0.25862621 | 0.254 | 0.206 | 1 | 0 | Zbtb46 | myCAF_1 |
| Ccdc32 | 0.00074354 | 0.28240715 | 0.278 | 0.25 | 1 | 0 | Ccdc32 | myCAF_1 |
| Maml2 | 0.00098865 | 0.27801304 | 0.256 | 0.219 | 1 | 0 | Maml2 | myCAF_1 |
| Camk2n2 | 0.00114799 | 0.33433329 | 0.303 | 0.257 | 1 | 0 | Camk2n2 | myCAF_1 |
| Pptc7 | 0.00123474 | 0.26633523 | 0.309 | 0.279 | 1 | 0 | Pptc7 | myCAF_1 |
| Zfp462 | 0.00127024 | 0.28428598 | 0.386 | 0.338 | 1 | 0 | Zfp462 | myCAF_1 |
| Ccdc126 | 0.00159972 | 0.27398099 | 0.28 | 0.258 | 1 | 0 | Ccdc126 | myCAF_1 |
| Sharpin | 0.00212606 | 0.25581117 | 0.385 | 0.347 | 1 | 0 | Sharpin | myCAF_1 |
| Tmem68 | 0.00559668 | 0.25410541 | 0.35 | 0.317 | 1 | 0 | Tmem68 | myCAF_1 |
| Peg12 | 0.00660308 | 0.32454169 | 0.26 | 0.219 | 1 | 0 | Peg12 | myCAF_1 |
| Acox3 | 0.00836881 | 0.25358188 | 0.372 | 0.335 | 1 | 0 | Acox3 | myCAF_1 |
| Ccn5 | 0 | 2.26236665 | 0.916 | 0.356 | 0 | 1 | Ccn5 | infl_CAF |
| Prelp | 0 | 1.22017114 | 0.889 | 0.348 | 0 | 1 | Prelp | infl_CAF |
| C4b | 0 | 1.36465731 | 0.95 | 0.462 | 0 | 1 | C4b | infl_CAF |
| C4a | 0 | 1.51480807 | 0.788 | 0.302 | 0 | 1 | C4a | infl_CAF |
| C3 | 0 | 1.10773875 | 0.939 | 0.455 | 0 | 1 | C3 | infl_CAF |
| Serpina3n | 0 | 1.63522874 | 0.966 | 0.512 | 0 | 1 | Serpina3n | infl_CAF |
| Prg4 | 0 | 0.99992437 | 0.679 | 0.242 | 0 | 1 | Prg4 | infl_CAF |
| Ptgis | 0 | 1.42610975 | 0.763 | 0.362 | 0 | 1 | Ptgis | infl_CAF |
| Chl1 | 0 | 1.44298621 | 0.725 | 0.326 | 0 | 1 | Chl1 | infl_CAF |
| Col14a1 | 0 | 0.85823717 | 0.882 | 0.488 | 0 | 1 | Col14a1 | infl_CAF |
| Lgi2 | 0 | 1.88718765 | 0.925 | 0.533 | 0 | 1 | Lgi2 | infl_CAF |
| Fgf7 | 0 | 1.76109578 | 0.815 | 0.425 | 0 | 1 | Fgf7 | infl_CAF |
| Vcam1 | 0 | 1.62089649 | 0.668 | 0.279 | 0 | 1 | Vcam1 | infl_CAF |
| Atf3 | 0 | 1.72702222 | 0.627 | 0.249 | 0 | 1 | Atf3 | infl_CAF |
| Saa3 | 0 | 2.43473528 | 0.711 | 0.34 | 0 | 1 | Saa3 | infl_CAF |

| | | | | | | | | |
|---|---|---|---|---|---|---|---|---|
| Ccl7 | 0 | 1.59200349 | 0.7 | 0.332 | 0 | 1 | Ccl7 | infl_CAF |
| Podn | 0 | 1.73462817 | 0.862 | 0.5 | 0 | 1 | Podn | infl_CAF |
| Dpt | 0 | 0.43192614 | 0.953 | 0.593 | 0 | 1 | Dpt | infl_CAF |
| Cfb | 0 | 1.32283806 | 0.933 | 0.584 | 0 | 1 | Cfb | infl_CAF |
| Il33 | 0 | 0.67397605 | 0.684 | 0.34 | 0 | 1 | Il33 | infl_CAF |
| Klf4 | 0 | 1.51284552 | 0.737 | 0.399 | 0 | 1 | Klf4 | infl_CAF |
| Cyp1b1 | 0 | 1.9753792 | 0.677 | 0.341 | 0 | 1 | Cyp1b1 | infl_CAF |
| Slfn5 | 0 | 0.72756534 | 0.82 | 0.491 | 0 | 1 | Slfn5 | infl_CAF |
| Csf1 | 0 | 1.13593268 | 0.849 | 0.521 | 0 | 1 | Csf1 | infl_CAF |
| C2 | 0 | 1.2886436 | 0.754 | 0.427 | 0 | 1 | C2 | infl_CAF |
| Iqcn | 0 | 2.15649791 | 0.464 | 0.141 | 0 | 1 | Iqcn | infl_CAF |
| Dpep1 | 0 | 1.36772021 | 0.875 | 0.552 | 0 | 1 | Dpep1 | infl_CAF |
| Gas6 | 0 | 1.09701343 | 0.621 | 0.3 | 0 | 1 | Gas6 | infl_CAF |
| Mmp3 | 0 | 1.29051932 | 0.874 | 0.554 | 0 | 1 | Mmp3 | infl_CAF |
| Ptx3 | 0 | 1.83839665 | 0.65 | 0.338 | 0 | 1 | Ptx3 | infl_CAF |
| Gfra1 | 0 | 1.85287449 | 0.603 | 0.297 | 0 | 1 | Gfra1 | infl_CAF |
| Plagl1 | 0 | 1.42529115 | 0.526 | 0.223 | 0 | 1 | Plagl1 | infl_CAF |
| Icam1 | 0 | 1.41408894 | 0.744 | 0.443 | 0 | 1 | Icam1 | infl_CAF |
| Slc1a3 | 0 | 2.14974155 | 0.415 | 0.114 | 0 | 1 | Slc1a3 | infl_CAF |
| Ltbp2 | 0 | 1.69047191 | 0.674 | 0.374 | 0 | 1 | Ltbp2 | infl_CAF |
| Ccl8 | 0 | 1.42447395 | 0.617 | 0.318 | 0 | 1 | Ccl8 | infl_CAF |
| Fosb | 0 | 1.21492636 | 0.908 | 0.61 | 0 | 1 | Fosb | infl_CAF |
| Ccl2 | 0 | 1.8762365 | 0.468 | 0.17 | 0 | 1 | Ccl2 | infl_CAF |
| Cxcl1 | 0 | 2.1707711 | 0.479 | 0.185 | 0 | 1 | Cxcl1 | infl_CAF |
| Rasl11a | 0 | 2.00432519 | 0.709 | 0.419 | 0 | 1 | Rasl11a | infl_CAF |
| Apod | 0 | 0.65769145 | 0.864 | 0.578 | 0 | 1 | Apod | infl_CAF |
| Itga11 | 0 | 1.21772728 | 0.583 | 0.297 | 0 | 1 | Itga11 | infl_CAF |
| Cyp7b1 | 0 | 1.33345352 | 0.805 | 0.528 | 0 | 1 | Cyp7b1 | infl_CAF |
| Lrrc32 | 0 | 1.71776643 | 0.928 | 0.655 | 0 | 1 | Lrrc32 | infl_CAF |
| Slc5a3 | 0 | 1.52944616 | 0.574 | 0.301 | 0 | 1 | Slc5a3 | infl_CAF |
| Cemip | 0 | 2.46368802 | 0.428 | 0.159 | 0 | 1 | Cemip | infl_CAF |
| Tmem88 | 0 | 2.05425419 | 0.393 | 0.13 | 0 | 1 | Tmem88 | infl_CAF |
| Btg2 | 0 | 0.83159197 | 0.845 | 0.587 | 0 | 1 | Btg2 | infl_CAF |
| Abi3bp | 0 | 1.79549539 | 0.938 | 0.681 | 0 | 1 | Abi3bp | infl_CAF |
| Baiap2 | 0 | 1.349643 | 0.655 | 0.399 | 0 | 1 | Baiap2 | infl_CAF |
| DEPRECATED-( | 0 | 1.32820242 | 0.675 | 0.427 | 0 | 1 | DEPRECATED-( | infl_CAF |
| Tnfaip6 | 0 | 1.32337356 | 0.627 | 0.38 | 0 | 1 | Tnfaip6 | infl_CAF |
| Hspa1a | 0 | 0.34924166 | 0.75 | 0.505 | 0 | 1 | Hspa1a | infl_CAF |
| Lif | 0 | 2.34013215 | 0.391 | 0.147 | 0 | 1 | Lif | infl_CAF |
| Cxcl5 | 0 | 0.7893042 | 0.548 | 0.308 | 0 | 1 | Cxcl5 | infl_CAF |
| Xpnpep2 | 0 | 2.1157025 | 0.349 | 0.11 | 0 | 1 | Xpnpep2 | infl_CAF |
| Angptl4 | 0 | 1.15763561 | 0.962 | 0.724 | 0 | 1 | Angptl4 | infl_CAF |
| Apln | 0 | 1.54586707 | 0.5 | 0.265 | 0 | 1 | Apln | infl_CAF |
| Tob1 | 0 | 1.084608 | 0.677 | 0.446 | 0 | 1 | Tob1 | infl_CAF |
| Hp | 0 | 2.07101791 | 0.467 | 0.237 | 0 | 1 | Hp | infl_CAF |
| Serpine1 | 0 | 1.64701424 | 0.905 | 0.675 | 0 | 1 | Serpine1 | infl_CAF |
| Fos | 0 | 1.2204085 | 0.932 | 0.705 | 0 | 1 | Fos | infl_CAF |
| Hspa1b | 0 | 0.28924758 | 0.783 | 0.562 | 0 | 1 | Hspa1b | infl_CAF |
| Scara3 | 0 | 0.85895313 | 0.839 | 0.622 | 0 | 1 | Scara3 | infl_CAF |
| Nr4a1 | 0 | 1.07485023 | 0.81 | 0.596 | 0 | 1 | Nr4a1 | infl_CAF |
| Irx1 | 0 | 0.80616447 | 0.772 | 0.565 | 0 | 1 | Irx1 | infl_CAF |
| Slc16a1 | 0 | 0.74901656 | 0.887 | 0.68 | 0 | 1 | Slc16a1 | infl_CAF |
| Emb | 0 | 1.09674244 | 0.896 | 0.69 | 0 | 1 | Emb | infl_CAF |
| Cxcl12 | 0 | 1.91337913 | 0.969 | 0.765 | 0 | 1 | Cxcl12 | infl_CAF |
| Il6 | 0 | 2.37290077 | 0.318 | 0.116 | 0 | 1 | Il6 | infl_CAF |
| Mfap41 | 0 | 0.88537246 | 0.777 | 0.578 | 0 | 1 | Mfap4 | infl_CAF |
| Tnfsf18 | 0 | 1.80173092 | 0.305 | 0.109 | 0 | 1 | Tnfsf18 | infl_CAF |
| Has1 | 0 | 1.22961828 | 0.416 | 0.222 | 0 | 1 | Has1 | infl_CAF |
| Nfkbiz | 0 | 1.13608785 | 0.827 | 0.633 | 0 | 1 | Nfkbiz | infl_CAF |
| Thbs4 | 0 | 2.3362525 | 0.318 | 0.127 | 0 | 1 | Thbs4 | infl_CAF |
| Gadd45g | 0 | 1.26750372 | 0.925 | 0.736 | 0 | 1 | Gadd45g | infl_CAF |
| Dusp1 | 0 | 1.09158822 | 0.889 | 0.701 | 0 | 1 | Dusp1 | infl_CAF |
| Il15ra | 0 | 0.79622119 | 0.824 | 0.639 | 0 | 1 | Il15ra | infl_CAF |
| Pnp | 0 | 0.80021816 | 0.872 | 0.692 | 0 | 1 | Pnp | infl_CAF |
| Rnd1 | 0 | 2.17390684 | 0.282 | 0.102 | 0 | 1 | Rnd1 | infl_CAF |
| Tnfaip3 | 0 | 1.32513347 | 0.44 | 0.263 | 0 | 1 | Tnfaip3 | infl_CAF |
| Myc | 0 | 1.01648686 | 0.705 | 0.53 | 0 | 1 | Myc | infl_CAF |
| Nfkbia | 0 | 1.34448389 | 0.916 | 0.746 | 0 | 1 | Nfkbia | infl_CAF |
| Il1rl1 | 0 | 1.60291182 | 0.344 | 0.174 | 0 | 1 | Il1rl1 | infl_CAF |
| Rrad | 0 | 2.21114583 | 0.254 | 0.085 | 0 | 1 | Rrad | infl_CAF |
| Gem | 0 | 0.87875153 | 0.664 | 0.496 | 0 | 1 | Gem | infl_CAF |
| Gadd45b | 0 | 0.86579836 | 0.905 | 0.74 | 0 | 1 | Gadd45b | infl_CAF |
| Dnajb1 | 0 | 1.33114947 | 0.802 | 0.638 | 0 | 1 | Dnajb1 | infl_CAF |
| Prdx6 | 0 | 0.88724062 | 0.832 | 0.669 | 0 | 1 | Prdx6 | infl_CAF |
| Ugcg | 0 | 0.91720113 | 0.923 | 0.761 | 0 | 1 | Ugcg | infl_CAF |
| Cilp | 0 | 0.87507371 | 0.398 | 0.236 | 0 | 1 | Cilp | infl_CAF |
| Sgk1 | 0 | 0.67110632 | 0.817 | 0.659 | 0 | 1 | Sgk1 | infl_CAF |
| Dio3 | 0 | 0.7823612 | 0.503 | 0.349 | 0 | 1 | Dio3 | infl_CAF |
| Cxcl14 | 0 | 1.62216602 | 0.882 | 0.729 | 0 | 1 | Cxcl14 | infl_CAF |
| Rarres2 | 0 | 1.88735687 | 0.97 | 0.824 | 0 | 1 | Rarres2 | infl_CAF |
| Mfap5 | 0 | 0.67847202 | 0.967 | 0.824 | 0 | 1 | Mfap5 | infl_CAF |
| Sdc4 | 0 | 0.75314747 | 0.881 | 0.741 | 0 | 1 | Sdc4 | infl_CAF |
| Gpc3 | 0 | 0.65578924 | 0.878 | 0.748 | 0 | 1 | Gpc3 | infl_CAF |
| Cthrc1 | 0 | 0.84284162 | 0.921 | 0.793 | 0 | 1 | Cthrc1 | infl_CAF |
| Sfrp2 | 0 | 1.01011054 | 0.944 | 0.817 | 0 | 1 | Sfrp2 | infl_CAF |
| Svep1 | 0 | 1.15192918 | 0.958 | 0.831 | 0 | 1 | Svep1 | infl_CAF |
| Ier3 | 0 | 1.09617333 | 0.902 | 0.777 | 0 | 1 | Ier3 | infl_CAF |
| Slc16a2 | 0 | 0.79411724 | 0.829 | 0.705 | 0 | 1 | Slc16a2 | infl_CAF |
| Sfrp1 | 0 | 1.11213261 | 0.812 | 0.689 | 0 | 1 | Sfrp1 | infl_CAF |
| Dclk1 | 0 | 0.73188616 | 0.908 | 0.785 | 0 | 1 | Dclk1 | infl_CAF |
| Zfp36 | 0 | 1.12623873 | 0.922 | 0.803 | 0 | 1 | Zfp36 | infl_CAF |
| Mt2 | 0 | 1.61492424 | 0.98 | 0.861 | 0 | 1 | Mt2 | infl_CAF |
| Ifrd1 | 0 | 0.85529258 | 0.844 | 0.727 | 0 | 1 | Ifrd1 | infl_CAF |
| Ace | 0 | 0.95716421 | 0.835 | 0.72 | 0 | 1 | Ace | infl_CAF |
| H2-K1 | 0 | 0.73402506 | 0.911 | 0.802 | 0 | 1 | H2-K1 | infl_CAF |
| Tgfbr3 | 0 | 0.78808466 | 0.945 | 0.837 | 0 | 1 | Tgfbr3 | infl_CAF |
| Ier2 | 0 | 0.6833777 | 0.885 | 0.78 | 0 | 1 | Ier2 | infl_CAF |
| Ier5 | 0 | 0.69213518 | 0.84 | 0.738 | 0 | 1 | Ier5 | infl_CAF |
| Ndufa4l2 | 0 | 0.50862119 | 0.51 | 0.413 | 0 | 1 | Ndufa4l2 | infl_CAF |
| Gfpt2 | 0 | 0.55163427 | 0.921 | 0.828 | 0 | 1 | Gfpt2 | infl_CAF |
| Egr1 | 0 | 0.94427997 | 0.966 | 0.877 | 0 | 1 | Egr1 | infl_CAF |
| Socs3 | 0 | 0.57007546 | 0.866 | 0.778 | 0 | 1 | Socs3 | infl_CAF |
| C1ra | 0 | 0.82205736 | 0.955 | 0.868 | 0 | 1 | C1ra | infl_CAF |
| Cxcl2 | 0 | 1.18031685 | 0.36 | 0.275 | 0 | 1 | Cxcl2 | infl_CAF |
| Sod2 | 0 | 0.56707125 | 0.851 | 0.769 | 0 | 1 | Sod2 | infl_CAF |
| Lgmn | 0 | 0.58422809 | 0.932 | 0.852 | 0 | 1 | Lgmn | infl_CAF |
| Ccn11 | 0 | 0.48885859 | 0.967 | 0.887 | 0 | 1 | Ccn1 | infl_CAF |
| Thbs11 | 0 | 0.39900704 | 0.895 | 0.816 | 0 | 1 | Thbs1 | infl_CAF |
| Mt1 | 0 | 1.52225069 | 0.978 | 0.902 | 0 | 1 | Mt1 | infl_CAF |
| Jun | 0 | 1.03343872 | 0.974 | 0.899 | 0 | 1 | Jun | infl_CAF |
| Egfr | 0 | 0.58870088 | 0.939 | 0.864 | 0 | 1 | Egfr | infl_CAF |
| Rhob | 0 | 0.38289887 | 0.924 | 0.85 | 0 | 1 | Rhob | infl_CAF |
| Gns | 0 | 0.68774219 | 0.95 | 0.876 | 0 | 1 | Gns | infl_CAF |
| Il6st | 0 | 0.53980905 | 0.957 | 0.883 | 0 | 1 | Il6st | infl_CAF |
| Gpx7 | 0 | 0.58090093 | 0.936 | 0.864 | 0 | 1 | Gpx7 | infl_CAF |
| Phlda1 | 0 | 0.56817773 | 0.669 | 0.597 | 0 | 1 | Phlda1 | infl_CAF |
| Enc1 | 0 | 0.69588694 | 0.863 | 0.793 | 0 | 1 | Enc1 | infl_CAF |
| Sod3 | 0 | 0.87782983 | 0.989 | 0.92 | 0 | 1 | Sod3 | infl_CAF |
| Fam162a | 0 | 0.60025494 | 0.839 | 0.77 | 0 | 1 | Fam162a | infl_CAF |
| Igfbp4 | 0 | 0.44894231 | 0.819 | 0.885 | 0 | 1 | Igfbp4 | infl_CAF |
| Clu | 0 | 0.49265339 | 0.941 | 0.875 | 0 | 1 | Clu | infl_CAF |
| Itm2a | 0 | 0.65455115 | 0.842 | 0.78 | 0 | 1 | Itm2a | infl_CAF |
| Glis3 | 0 | 0.6677163 | 0.936 | 0.876 | 0 | 1 | Glis3 | infl_CAF |
| Fxyd5 | 0 | 0.68638232 | 0.916 | 0.857 | 0 | 1 | Fxyd5 | infl_CAF |
| C1s1 | 0 | 0.51982977 | 0.975 | 0.916 | 0 | 1 | C1s1 | infl_CAF |
| Junb | 0 | 0.74157913 | 0.978 | 0.923 | 0 | 1 | Junb | infl_CAF |
| Lgals3bp | 0 | 0.4517036 | 0.958 | 0.905 | 0 | 1 | Lgals3bp | infl_CAF |
| Twist1 | 0 | 0.47413558 | 0.95 | 0.897 | 0 | 1 | Twist1 | infl_CAF |
| Il4ra | 0 | 1.09861019 | 0.975 | 0.924 | 0 | 1 | Il4ra | infl_CAF |
| Nfix | 0 | 0.50198617 | 0.969 | 0.918 | 0 | 1 | Nfix | infl_CAF |
| Man2a1 | 0 | 0.67929626 | 0.971 | 0.92 | 0 | 1 | Man2a1 | infl_CAF |
| Gpx3 | 0 | 1.09464795 | 0.992 | 0.943 | 0 | 1 | Gpx3 | infl_CAF |
| Pdpn | 0 | 0.68003108 | 0.966 | 0.919 | 0 | 1 | Pdpn | infl_CAF |
| Ctsh | 0 | 0.73641278 | 0.953 | 0.907 | 0 | 1 | Ctsh | infl_CAF |
| Fbln1 | 0 | 0.52933237 | 0.957 | 0.916 | 0 | 1 | Fbln1 | infl_CAF |
| Ubc | 0 | 0.52146902 | 0.968 | 0.929 | 0 | 1 | Ubc | infl_CAF |
| Pcsk5 | 0 | 0.86778446 | 0.963 | 0.927 | 0 | 1 | Pcsk5 | infl_CAF |
| Dcn | 0 | 0.30102934 | 0.986 | 0.952 | 0 | 1 | Dcn | infl_CAF |
| Rnase4 | 0 | 0.7593207 | 0.975 | 0.941 | 0 | 1 | Rnase4 | infl_CAF |
| Cpxm1 | 0 | 0.94403139 | 0.985 | 0.952 | 0 | 1 | Cpxm1 | infl_CAF |
| Vasn | 0 | 0.49449564 | 0.966 | 0.933 | 0 | 1 | Vasn | infl_CAF |
| Jund | 0 | 0.72327124 | 0.988 | 0.958 | 0 | 1 | Jund | infl_CAF |
| Slc39a14 | 0 | 0.70666946 | 0.977 | 0.949 | 0 | 1 | Slc39a14 | infl_CAF |
| Lox | 0 | 0.89467793 | 0.988 | 0.96 | 0 | 1 | Lox | infl_CAF |
| Nbl1 | 0 | 0.78828088 | 0.948 | 0.92 | 0 | 1 | Nbl1 | infl_CAF |
| Npc2 | 0 | 0.5001464 | 0.984 | 0.959 | 0 | 1 | Npc2 | infl_CAF |
| Timp1 | 0 | 0.94914191 | 0.985 | 0.961 | 0 | 1 | Timp1 | infl_CAF |
| Serping1 | 0 | 0.58204502 | 0.984 | 0.961 | 0 | 1 | Serping1 | infl_CAF |
| Mxra81 | 0 | 0.4333674 | 0.978 | 0.96 | 0 | 1 | Mxra8 | infl_CAF |
| Fkbp9 | 0 | 0.38023987 | 0.98 | 0.962 | 0 | 1 | Fkbp9 | infl_CAF |
| Mmp2 | 0 | 0.89078369 | 0.989 | 0.972 | 0 | 1 | Mmp2 | infl_CAF |
| Fth1 | 0 | 0.61260504 | 0.997 | 0.982 | 0 | 1 | Fth1 | infl_CAF |
| Efemp2 | 0 | 0.36918302 | 0.973 | 0.963 | 0 | 1 | Efemp2 | infl_CAF |
| Pcolce | 0 | 0.45138877 | 0.991 | 0.982 | 0 | 1 | Pcolce | infl_CAF |
| Rcn3 | 0 | 0.38247628 | 0.988 | 0.98 | 0 | 1 | Rcn3 | infl_CAF |
| Aebp1 | 0 | 0.41096057 | 0.99 | 0.982 | 0 | 1 | Aebp1 | infl_CAF |
| Col3a1 | 0 | 0.65720234 | 1 | 0.996 | 0 | 1 | Col3a1 | infl_CAF |
| Thbs21 | 0 | 0.30373069 | 0.979 | 0.975 | 0 | 1 | Thbs2 | infl_CAF |
| Col1a21 | 0 | 0.30817858 | 0.999 | 0.996 | 0 | 1 | Col1a2 | infl_CAF |
| Maged1 | 1.07E-293 | 0.25140092 | 0.973 | 0.971 | 2.06E-289 | 1 | Maged1 | infl_CAF |
| Chpf | 1.21E-293 | 0.54448814 | 0.948 | 0.88 | 2.35E-289 | 1 | Chpf | infl_CAF |
| Tent5a | 1.77E-293 | 0.75133311 | 0.753 | 0.602 | 3.42E-289 | 1 | Tent5a | infl_CAF |
| Il11ra1 | 3.62E-293 | 0.95287639 | 0.769 | 0.533 | 7.00E-289 | 1 | Il11ra1 | infl_CAF |
| Lgr51 | 3.03E-293 | 1.22433821 | 0.453 | 0.293 | 5.86E-289 | 1 | Lgr5 | infl_CAF |
| P4ha2 | 4.63E-293 | 0.37846808 | 0.935 | 0.869 | 8.95E-289 | 1 | P4ha2 | infl_CAF |
| Rnf144a | 5.68E-293 | 0.86858905 | 0.739 | 0.533 | 1.10E-288 | 1 | Rnf144a | infl_CAF |
| Pam | 8.95E-293 | 0.68354889 | 0.865 | 0.695 | 1.73E-288 | 1 | Pam | infl_CAF |
| Adam23 | 1.23E-292 | 1.14594021 | 0.62 | 0.31 | 2.38E-288 | 1 | Adam23 | infl_CAF |
| Vkorc1 | 4.14E-292 | 0.49094032 | 0.961 | 0.915 | 8.00E-288 | 1 | Vkorc1 | infl_CAF |
| Mgst1 | 2.02E-288 | 0.55099801 | 0.926 | 0.8 | 3.91E-284 | 1 | Mgst1 | infl_CAF |
| Has2 | 3.80E-288 | 0.70260078 | 0.616 | 0.507 | 7.34E-284 | 1 | Has2 | infl_CAF |
| Sema4a | 1.16E-286 | 2.02635638 | 0.262 | 0.101 | 2.23E-282 | 1 | Sema4a | infl_CAF |
| Adgrd1 | 2.80E-285 | 0.65465454 | 0.877 | 0.415 | 5.41E-281 | 1 | Adgrd1 | infl_CAF |
| Cstb | 3.56E-283 | 0.4943865 | 0.937 | 0.885 | 6.89E-279 | 1 | Cstb | infl_CAF |
| Fxyd6 | 2.36E-282 | 0.68075076 | 0.75 | 0.62 | 4.56E-278 | 1 | Fxyd6 | infl_CAF |
| Ifi27 | 4.51E-282 | 0.65320842 | 0.893 | 0.79 | 8.71E-278 | 1 | Ifi27 | infl_CAF |
| Plod3 | 1.23E-281 | 0.43507611 | 0.971 | 0.934 | 2.38E-277 | 1 | Plod3 | infl_CAF |
| Rnf19b1 | 3.25E-280 | 0.33340095 | 0.921 | 0.877 | 6.28E-276 | 1 | Rnf19b | infl_CAF |
| Pkdcc | 3.79E-278 | 0.85046487 | 0.872 | 0.671 | 7.33E-274 | 1 | Pkdcc | infl_CAF |
| Plac8 | 9.67E-278 | 0.41531826 | 0.877 | 0.715 | 1.87E-273 | 1 | Plac8 | infl_CAF |
| Mknk2 | 2.34E-276 | 0.7955562 | 0.755 | 0.629 | 4.53E-272 | 1 | Mknk2 | infl_CAF |
| Adgra2 | 9.38E-275 | 0.61577323 | 0.959 | 0.879 | 1.81E-270 | 1 | Adgra2 | infl_CAF |
| Mat2a | 7.76E-268 | 0.54286859 | 0.868 | 0.806 | 1.50E-263 | 1 | Mat2a | infl_CAF |
| Tmem176b | 4.88E-264 | 0.55805184 | 0.85 | 0.696 | 9.43E-260 | 1 | Tmem176b | infl_CAF |
| Ch25h | 1.89E-260 | 1.48529204 | 0.329 | 0.175 | 3.65E-256 | 1 | Ch25h | infl_CAF |
| Flrt2 | 3.82E-260 | 0.60320133 | 0.833 | 0.614 | 7.38E-256 | 1 | Flrt2 | infl_CAF |
| Cited2 | 1.31E-258 | 1.12111908 | 0.408 | 0.239 | 2.52E-254 | 1 | Cited2 | infl_CAF |
| Polg2 | 3.76E-258 | 0.95156598 | 0.571 | 0.272 | 7.27E-254 | 1 | Polg2 | infl_CAF |
| Tsku | 1.04E-252 | 0.53720855 | 0.926 | 0.847 | 2.01E-248 | 1 | Tsku | infl_CAF |
| Arsi | 1.55E-252 | 0.67095641 | 0.857 | 0.609 | 3.00E-248 | 1 | Arsi | infl_CAF |
| Ccn3 | 4.58E-252 | 0.652952 | 0.494 | 0.211 | 8.85E-248 | 1 | Ccn3 | infl_CAF |
| Tm4sf1 | 5.61E-252 | 0.63587821 | 0.846 | 0.709 | 1.08E-247 | 1 | Tm4sf1 | infl_CAF |
| Sertad1 | 4.91E-251 | 0.62781176 | 0.775 | 0.62 | 9.50E-247 | 1 | Sertad1 | infl_CAF |
| Kng2 | 9.89E-251 | 1.30809516 | 0.425 | 0.195 | 1.91E-246 | 1 | Kng2 | infl_CAF |
| Slpi | 6.88E-249 | 1.26071574 | 0.324 | 0.165 | 1.33E-244 | 1 | Slpi | infl_CAF |
| Lsmem1 | 7.11E-249 | 2.07786591 | 0.262 | 0.073 | 1.38E-244 | 1 | Lsmem1 | infl_CAF |
| Rab3a | 7.56E-248 | 1.53746158 | 0.285 | 0.197 | 1.46E-243 | 1 | Rab3a | infl_CAF |
| Pdia5 | 3.72E-247 | 0.57803771 | 0.89 | 0.792 | 7.19E-243 | 1 | Pdia5 | infl_CAF |
| Nsg1 | 1.62E-246 | 0.89584106 | 0.814 | 0.554 | 3.12E-242 | 1 | Nsg1 | infl_CAF |
| Egr3 | 9.23E-243 | 0.46746052 | 0.504 | 0.436 | 1.78E-238 | 1 | Egr3 | infl_CAF |
| Il1r1 | 1.60E-239 | 0.38483401 | 0.95 | 0.897 | 3.08E-235 | 1 | Il1r1 | infl_CAF |
| Rbp4 | 7.74E-239 | 1.52038169 | 0.359 | 0.153 | 1.50E-234 | 1 | Rbp4 | infl_CAF |
| Fxyd1 | 9.90E-239 | 0.78523555 | 0.821 | 0.596 | 1.91E-234 | 1 | Fxyd1 | infl_CAF |
| Nop2 | 8.72E-238 | 1.00798495 | 0.639 | 0.463 | 1.69E-233 | 1 | Nop2 | infl_CAF |
| Kcne4 | 2.10E-236 | 1.517778 | 0.324 | 0.115 | 4.06E-232 | 1 | Kcne4 | infl_CAF |
| Bmper | 1.02E-233 | 1.18273473 | 0.514 | 0.218 | 1.97E-229 | 1 | Bmper | infl_CAF |
| Agtr1a | 1.45E-233 | 1.39697676 | 0.425 | 0.154 | 2.80E-229 | 1 | Agtr1a | infl_CAF |
| Pnrc1 | 2.22E-231 | 0.35042515 | 0.911 | 0.868 | 4.29E-227 | 1 | Pnrc1 | infl_CAF |
| Ifi205 | 8.54E-231 | 0.44322212 | 0.854 | 0.59 | 1.65E-226 | 1 | Ifi205 | infl_CAF |
| Col28a1 | 2.06E-230 | 1.46276607 | 0.302 | 0.124 | 3.98E-226 | 1 | Col28a1 | infl_CAF |
| Ube2v2 | 8.35E-230 | 0.60665791 | 0.54 | 0.544 | 1.61E-225 | 1 | Ube2v2 | infl_CAF |
| F3 | 5.09E-229 | 0.83141478 | 0.564 | 0.415 | 9.83E-225 | 1 | F3 | infl_CAF |
| Medag | 1.65E-228 | 0.48777569 | 0.902 | 0.799 | 3.19E-224 | 1 | Medag | infl_CAF |
| Prdx5 | 7.65E-228 | 0.30887272 | 0.984 | 0.96 | 1.48E-223 | 1 | Prdx5 | infl_CAF |
| Ggt5 | 8.53E-223 | 1.11783385 | 0.536 | 0.231 | 1.65E-218 | 1 | Ggt5 | infl_CAF |
| Cldn10 | 3.80E-221 | 1.54252493 | 0.432 | 0.175 | 7.35E-217 | 1 | Cldn10 | infl_CAF |
| Kdm6b | 2.63E-220 | 0.55658123 | 0.763 | 0.696 | 5.09E-216 | 1 | Kdm6b | infl_CAF |
| Jak2 | 8.67E-220 | 0.46946101 | 0.877 | 0.799 | 1.68E-215 | 1 | Jak2 | infl_CAF |
| B4galt5 | 2.55E-216 | 0.47079577 | 0.882 | 0.824 | 4.92E-212 | 1 | B4galt5 | infl_CAF |
| Fst | 5.18E-216 | 0.7682847 | 0.745 | 0.51 | 1.00E-211 | 1 | Fst | infl_CAF |
| Klf2 | 1.36E-214 | 0.60702898 | 0.588 | 0.379 | 2.63E-210 | 1 | Klf2 | infl_CAF |
| Trib1 | 1.41E-214 | 0.72670163 | 0.531 | 0.398 | 2.73E-210 | 1 | Trib1 | infl_CAF |
| Plod2 | 4.60E-214 | 0.59485847 | 0.797 | 0.621 | 8.89E-210 | 1 | Plod2 | infl_CAF |
| Slc38a2 | 4.95E-214 | 0.3474693 | 0.939 | 0.912 | 9.58E-210 | 1 | Slc38a2 | infl_CAF |
| Cp | 1.98E-212 | 0.81908185 | 0.533 | 0.379 | 3.83E-208 | 1 | Cp | infl_CAF |
| Col16a1 | 1.24E-210 | 0.28392219 | 0.977 | 0.96 | 2.39E-206 | 1 | Col16a1 | infl_CAF |
| Ssc5d | 2.34E-210 | 0.86936061 | 0.777 | 0.458 | 4.53E-206 | 1 | Ssc5d | infl_CAF |
| Pknox2 | 2.51E-210 | 1.03758557 | 0.645 | 0.4 | 4.86E-206 | 1 | Pknox2 | infl_CAF |
| Zfp36l2 | 2.60E-204 | 0.42147071 | 0.908 | 0.823 | 5.02E-200 | 1 | Zfp36l2 | infl_CAF |
| Ptgs2 | 3.98E-201 | 0.27064647 | 0.503 | 0.422 | 7.70E-197 | 1 | Ptgs2 | infl_CAF |
| Il3ra | 1.08E-197 | 0.55404957 | 0.918 | 0.822 | 2.08E-193 | 1 | Il3ra | infl_CAF |
| Fmnl2 | 1.09E-196 | 1.01476367 | 0.66 | 0.379 | 2.10E-192 | 1 | Fmnl2 | infl_CAF |
| Ppp1r15a | 2.94E-196 | 0.45377804 | 0.795 | 0.609 | 5.68E-192 | 1 | Ppp1r15a | infl_CAF |
| Ssr2 | 2.37E-195 | 0.30568021 | 0.968 | 0.948 | 4.58E-191 | 1 | Ssr2 | infl_CAF |
| Bag3 | 1.14E-193 | 0.47967234 | 0.891 | 0.806 | 2.21E-189 | 1 | Bag3 | infl_CAF |
| Ank3 | 2.00E-192 | 1.08643148 | 0.536 | 0.319 | 3.87E-188 | 1 | Ank3 | infl_CAF |
| Fgl2 | 5.82E-192 | 0.45669526 | 0.745 | 0.561 | 1.13E-187 | 1 | Fgl2 | infl_CAF |
| Creb3l3 | 1.92E-190 | 1.09372174 | 0.491 | 0.284 | 3.71E-186 | 1 | Creb3l3 | infl_CAF |
| Spsb1 | 1.82E-189 | 0.62389274 | 0.76 | 0.547 | 3.52E-185 | 1 | Spsb1 | infl_CAF |
| Rnf150 | 4.69E-185 | 0.99958547 | 0.641 | 0.391 | 9.07E-181 | 1 | Rnf150 | infl_CAF |
| Ppp1r10 | 4.95E-184 | 0.6585247 | 0.653 | 0.536 | 9.58E-180 | 1 | Ppp1r10 | infl_CAF |
| Rdh10 | 1.01E-183 | 0.51554572 | 0.763 | 0.688 | 1.95E-179 | 1 | Rdh10 | infl_CAF |
| Vegfd | 3.25E-183 | 0.57441442 | 0.664 | 0.446 | 6.27E-179 | 1 | Vegfd | infl_CAF |
| Naglu | 2.11E-182 | 0.92750026 | 0.66 | 0.453 | 4.07E-178 | 1 | Naglu | infl_CAF |
| Sema3b | 3.37E-180 | 1.31686084 | 0.491 | 0.254 | 6.51E-176 | 1 | Sema3b | infl_CAF |
| Klf6 | 1.22E-176 | 0.33544601 | 0.905 | 0.856 | 2.36E-172 | 1 | Klf6 | infl_CAF |
| Scn7a | 9.35E-176 | 0.8127443 | 0.542 | 0.247 | 1.81E-171 | 1 | Scn7a | infl_CAF |
| Adarb1 | 2.86E-172 | 1.36036051 | 0.46 | 0.212 | 5.53E-168 | 1 | Adarb1 | infl_CAF |
| Csrp2 | 4.46E-172 | 0.5967752 | 0.863 | 0.642 | 8.62E-168 | 1 | Csrp2 | infl_CAF |
| Adamts1 | 4.68E-172 | 0.43348266 | 0.745 | 0.62 | 9.04E-168 | 1 | Adamts1 | infl_CAF |
| Birc3 | 4.88E-171 | 0.67947071 | 0.716 | 0.577 | 9.44E-167 | 1 | Birc3 | infl_CAF |
| Adamtsl1 | 9.92E-171 | 0.5441424 | 0.878 | 0.635 | 1.92E-166 | 1 | Adamtsl1 | infl_CAF |
| Adamts15 | 4.79E-170 | 0.68889469 | 0.58 | 0.401 | 9.25E-166 | 1 | Adamts15 | infl_CAF |
| Zfp36l1 | 1.25E-168 | 0.31967677 | 0.882 | 0.859 | 2.42E-164 | 1 | Zfp36l1 | infl_CAF |
| H6pd | 2.25E-166 | 0.34991358 | 0.938 | 0.901 | 4.36E-162 | 1 | H6pd | infl_CAF |
| Agfg1 | 2.57E-166 | 0.49222546 | 0.889 | 0.809 | 4.97E-162 | 1 | Agfg1 | infl_CAF |
| Snai1 | 6.29E-162 | 0.35727887 | 0.872 | 0.791 | 1.22E-157 | 1 | Snai1 | infl_CAF |
| Comt | 4.00E-161 | 0.59619457 | 0.893 | 0.711 | 7.74E-157 | 1 | Comt | infl_CAF |
| Car12 | 8.92E-160 | 1.08921988 | 0.297 | 0.209 | 1.72E-155 | 1 | Car12 | infl_CAF |
| Csgalnact11 | 8.12E-159 | 0.47380628 | 0.842 | 0.746 | 1.57E-154 | 1 | Csgalnact1 | infl_CAF |
| Zfhx3 | 8.63E-159 | 1.17227468 | 0.547 | 0.291 | 1.67E-154 | 1 | Zfhx3 | infl_CAF |
| Abcc9 | 2.52E-158 | 0.9083279 | 0.393 | 0.194 | 4.88E-154 | 1 | Abcc9 | infl_CAF |
| Adamts5 | 1.06E-156 | 0.51671138 | 0.696 | 0.502 | 2.06E-152 | 1 | Adamts5 | infl_CAF |
| Dchs1 | 6.27E-156 | 0.70948642 | 0.744 | 0.528 | 1.21E-151 | 1 | Dchs1 | infl_CAF |
| Tmem176a | 8.62E-156 | 0.41557752 | 0.799 | 0.656 | 1.67E-151 | 1 | Tmem176a | infl_CAF |
| Cebpd | 1.64E-155 | 0.39446631 | 0.878 | 0.786 | 3.16E-151 | 1 | Cebpd | infl_CAF |
| Pim1 | 3.83E-155 | 0.40467876 | 0.862 | 0.772 | 7.40E-151 | 1 | Pim1 | infl_CAF |
| Dram1 | 5.22E-155 | 0.44680113 | 0.926 | 0.829 | 1.01E-150 | 1 | Dram1 | infl_CAF |
| Aldh1l2 | 6.57E-152 | 0.6730938 | 0.693 | 0.51 | 1.27E-147 | 1 | Aldh1l2 | infl_CAF |
| Lag3 | 1.05E-150 | 1.11075663 | 0.301 | 0.138 | 2.02E-146 | 1 | Lag3 | infl_CAF |
| Runx1 | 1.94E-150 | 0.2753356 | 0.953 | 0.889 | 3.74E-146 | 1 | Runx1 | infl_CAF |
| Chst11 | 1.20E-148 | 0.37053785 | 0.861 | 0.799 | 2.32E-144 | 1 | Chst11 | infl_CAF |
| Arhgap29 | 1.27E-148 | 0.61668063 | 0.751 | 0.56 | 2.46E-144 | 1 | Arhgap29 | infl_CAF |
| Irf1 | 2.62E-148 | 0.43320509 | 0.722 | 0.647 | 5.07E-144 | 1 | Irf1 | infl_CAF |
| Cdkn1a | 8.49E-148 | 0.39260088 | 0.494 | 0.411 | 1.64E-143 | 1 | Cdkn1a | infl_CAF |
| Ddah11 | 1.34E-145 | 0.33247625 | 0.919 | 0.844 | 2.59E-141 | 1 | Ddah1 | infl_CAF |
| Clec11a | 2.46E-144 | 0.53344111 | 0.63 | 0.549 | 4.76E-140 | 1 | Clec11a | infl_CAF |
| Cgref1 | 1.17E-143 | 0.99711309 | 0.495 | 0.298 | 2.27E-139 | 1 | Cgref1 | infl_CAF |
| Galnt2 | 1.47E-141 | 0.35675646 | 0.944 | 0.912 | 2.84E-137 | 1 | Galnt2 | infl_CAF |
| Cd320 | 2.79E-139 | 0.85950004 | 0.623 | 0.415 | 5.39E-135 | 1 | Cd320 | infl_CAF |
| Tgm2 | 1.60E-138 | 0.54586543 | 0.62 | 0.448 | 3.10E-134 | 1 | Tgm2 | infl_CAF |
| Parm1 | 7.59E-138 | 1.0376092 | 0.356 | 0.194 | 1.47E-133 | 1 | Parm1 | infl_CAF |
| Mcfd2 | 3.16E-137 | 0.29709926 | 0.95 | 0.923 | 6.10E-133 | 1 | Mcfd2 | infl_CAF |
| Bcat1 | 6.01E-137 | 0.41209027 | 0.895 | 0.801 | 1.16E-132 | 1 | Bcat1 | infl_CAF |
| Sox9 | 6.75E-137 | 0.85148356 | 0.365 | 0.211 | 1.30E-132 | 1 | Sox9 | infl_CAF |
| Dedd2 | 2.57E-133 | 1.87570927 | 0.253 | 0.094 | 4.97E-129 | 1 | Dedd2 | infl_CAF |
| Apol8 | 6.22E-133 | 0.8579257 | 0.429 | 0.205 | 1.20E-128 | 1 | Apol8 | infl_CAF |
| Fkbp11 | 1.27E-131 | 0.4232339 | 0.847 | 0.761 | 2.46E-127 | 1 | Fkbp11 | infl_CAF |
| Homer1 | 5.80E-131 | 0.50995522 | 0.585 | 0.489 | 1.12E-126 | 1 | Homer1 | infl_CAF |
| Galnt17 | 1.70E-130 | 0.72415493 | 0.696 | 0.488 | 3.29E-126 | 1 | Galnt17 | infl_CAF |
| Myo7a1 | 2.47E-130 | 0.49749359 | 0.765 | 0.615 | 4.77E-126 | 1 | Myo7a | infl_CAF |
| Rac3 | 9.42E-130 | 0.9908622 | 0.49 | 0.296 | 1.82E-125 | 1 | Rac3 | infl_CAF |
| mt-Nd6 | 1.00E-129 | 0.33861167 | 0.825 | 0.668 | 1.94E-125 | 1 | mt-Nd6 | infl_CAF |
| Prdx4 | 6.66E-129 | 0.40229771 | 0.859 | 0.786 | 1.29E-124 | 1 | Prdx4 | infl_CAF |
| Anpep | 2.10E-126 | 0.42231501 | 0.868 | 0.753 | 4.06E-122 | 1 | Anpep | infl_CAF |
| Lsp1 | 2.27E-125 | 0.27129083 | 0.904 | 0.847 | 4.38E-121 | 1 | Lsp1 | infl_CAF |
| Rpn1 | 1.49E-124 | 0.26813285 | 0.956 | 0.936 | 2.87E-120 | 1 | Rpn1 | infl_CAF |
| Slc9a3r1 | 2.71E-124 | 0.76576293 | 0.644 | 0.432 | 5.23E-120 | 1 | Slc9a3r1 | infl_CAF |
| Nectin21 | 7.73E-124 | 0.44975699 | 0.826 | 0.736 | 1.49E-119 | 1 | Nectin2 | infl_CAF |
| Srm | 1.84E-122 | 0.50299443 | 0.786 | 0.681 | 3.56E-118 | 1 | Srm | infl_CAF |
| Lifr | 9.97E-122 | 0.57666042 | 0.709 | 0.601 | 1.93E-117 | 1 | Lifr | infl_CAF |
| Tcf7l2 | 1.88E-120 | 0.83109408 | 0.52 | 0.344 | 3.63E-116 | 1 | Tcf7l2 | infl_CAF |
| Rab8b | 6.73E-119 | 0.59257426 | 0.765 | 0.609 | 1.30E-114 | 1 | Rab8b | infl_CAF |
| Hmox1 | 8.84E-119 | 0.2995636 | 0.759 | 0.695 | 1.71E-114 | 1 | Hmox1 | infl_CAF |
| Nod1 | 7.16E-118 | 0.61953139 | 0.741 | 0.517 | 1.39E-113 | 1 | Nod1 | infl_CAF |
| Nid2 | 9.63E-118 | 0.55495254 | 0.489 | 0.443 | 1.86E-113 | 1 | Nid2 | infl_CAF |
| Il17ra | 2.57E-117 | 0.51219275 | 0.787 | 0.642 | 4.97E-113 | 1 | Il17ra | infl_CAF |
| Socs1 | 3.98E-117 | 0.5817246 | 0.659 | 0.507 | 7.69E-113 | 1 | Socs1 | infl_CAF |
| Shisa5 | 2.07E-116 | 0.31777709 | 0.956 | 0.91 | 4.00E-112 | 1 | Shisa5 | infl_CAF |
| Klhl21 | 2.78E-114 | 0.39719255 | 0.649 | 0.571 | 5.37E-110 | 1 | Klhl21 | infl_CAF |
| Slc38a10 | 9.11E-114 | 0.31985063 | 0.952 | 0.915 | 1.76E-109 | 1 | Slc38a10 | infl_CAF |
| Fam102a | 1.81E-113 | 0.74021771 | 0.668 | 0.507 | 3.50E-109 | 1 | Fam102a | infl_CAF |
| Prnp | 2.87E-113 | 0.37043628 | 0.899 | 0.83 | 5.55E-109 | 1 | Prnp | infl_CAF |
| Amotl2 | 4.43E-113 | 0.3782424 | 0.839 | 0.761 | 8.57E-109 | 1 | Amotl2 | infl_CAF |
| Dnajb9 | 3.38E-112 | 0.50695711 | 0.698 | 0.546 | 6.53E-108 | 1 | Dnajb9 | infl_CAF |
| Atp8b1 | 5.88E-112 | 0.64015055 | 0.767 | 0.586 | 1.14E-107 | 1 | Atp8b1 | infl_CAF |
| Mif | 7.54E-112 | 0.29579252 | 0.84 | 0.786 | 1.46E-107 | 1 | Mif | infl_CAF |
| Cmtm3 | 8.78E-112 | 0.25712128 | 0.94 | 0.914 | 1.70E-107 | 1 | Cmtm3 | infl_CAF |
| Cbx4 | 2.55E-111 | 0.69334566 | 0.512 | 0.391 | 4.93E-107 | 1 | Cbx4 | infl_CAF |
| Irs2 | 1.57E-110 | 0.40817843 | 0.712 | 0.597 | 3.03E-106 | 1 | Irs2 | infl_CAF |
| Mmp23 | 7.77E-109 | 0.35527889 | 0.899 | 0.834 | 1.50E-104 | 1 | Mmp23 | infl_CAF |
| Cadm2 | 8.90E-109 | 1.11450161 | 0.329 | 0.163 | 1.72E-104 | 1 | Cadm2 | infl_CAF |
| Lrrc17 | 1.30E-108 | 0.29963044 | 0.741 | 0.72 | 2.51E-104 | 1 | Lrrc17 | infl_CAF |
| Ampd3 | 2.56E-108 | 0.53830948 | 0.636 | 0.461 | 4.95E-104 | 1 | Ampd3 | infl_CAF |
| Loxl4 | 1.97E-107 | 0.86050656 | 0.517 | 0.288 | 3.80E-103 | 1 | Loxl4 | infl_CAF |
| Gpc4 | 2.50E-107 | 0.5571395 | 0.708 | 0.588 | 4.83E-103 | 1 | Gpc4 | infl_CAF |
| Ifnar2 | 8.31E-107 | 0.34273435 | 0.895 | 0.834 | 1.61E-102 | 1 | Ifnar2 | infl_CAF |
| Creld1 | 4.93E-106 | 0.6459954 | 0.735 | 0.592 | 9.54E-102 | 1 | Creld1 | infl_CAF |
| Gpnmb | 1.32E-105 | 0.87843247 | 0.322 | 0.201 | 2.54E-101 | 1 | Gpnmb | infl_CAF |
| Srxn1 | 4.47E-105 | 0.67170034 | 0.519 | 0.447 | 8.64E-101 | 1 | Srxn1 | infl_CAF |
| Tcea3 | 1.21E-104 | 1.68004858 | 0.302 | 0.101 | 2.33E-100 | 1 | Tcea3 | infl_CAF |
| Il13ra1 | 9.81E-102 | 0.29635038 | 0.878 | 0.825 | 1.90E-97 | 1 | Il13ra1 | infl_CAF |
| Susd6 | 3.70E-100 | 0.30898273 | 0.928 | 0.869 | 7.15E-96 | 1 | Susd6 | infl_CAF |
| St3gal1 | 3.83E-100 | 0.49673469 | 0.733 | 0.549 | 7.40E-96 | 1 | St3gal1 | infl_CAF |
| Ahr | 7.01E-100 | 0.74996677 | 0.522 | 0.3 | 1.35E-95 | 1 | Ahr | infl_CAF |
| Tspan18 | 4.71E-99 | 0.95995012 | 0.348 | 0.181 | 9.10E-95 | 1 | Tspan18 | infl_CAF |
| Anxa8 | 8.15E-99 | 0.94258665 | 0.274 | 0.119 | 1.58E-94 | 1 | Anxa8 | infl_CAF |
| Noct | 6.03E-98 | 0.84853956 | 0.428 | 0.288 | 1.17E-93 | 1 | Noct | infl_CAF |
| Kcnma1 | 9.28E-98 | 1.05197351 | 0.292 | 0.162 | 1.79E-93 | 1 | Kcnma1 | infl_CAF |
| Sulf2 | 1.26E-97 | 0.62276046 | 0.627 | 0.368 | 2.43E-93 | 1 | Sulf2 | infl_CAF |
| 3-Mar | 3.03E-97 | 1.01899368 | 0.463 | 0.266 | 5.85E-93 | 1 | 3-Mar | infl_CAF |
| Ints6 | 1.19E-96 | 0.60367968 | 0.646 | 0.501 | 2.31E-92 | 1 | Ints6 | infl_CAF |
| Mycbp2 | 1.07E-95 | 0.37317465 | 0.841 | 0.757 | 2.08E-91 | 1 | Mycbp2 | infl_CAF |
| Tgfbi1 | 1.18E-95 | 0.31903809 | 0.545 | 0.48 | 2.29E-91 | 1 | Tgfbi | infl_CAF |
| Dnm31 | 1.83E-95 | 0.64498757 | 0.522 | 0.378 | 3.55E-91 | 1 | Dnm3 | infl_CAF |
| Ccl11 | 2.34E-95 | 0.34950249 | 0.589 | 0.398 | 4.52E-91 | 1 | Ccl11 | infl_CAF |
| Trf | 3.15E-95 | 0.64516519 | 0.426 | 0.345 | 6.09E-91 | 1 | Trf | infl_CAF |
| Rpn2 | 3.17E-95 | 0.28648348 | 0.939 | 0.903 | 6.14E-91 | 1 | Rpn2 | infl_CAF |
| Bnc2 | 3.54E-95 | 0.43184563 | 0.847 | 0.739 | 6.85E-91 | 1 | Bnc2 | infl_CAF |
| Lpgat1 | 8.34E-95 | 0.44162105 | 0.871 | 0.797 | 1.61E-90 | 1 | Lpgat1 | infl_CAF |
| Zfp385b | 1.95E-94 | 1.54589682 | 0.255 | 0.099 | 3.77E-90 | 1 | Zfp385b | infl_CAF |
| Lgals9 | 5.03E-94 | 0.3943626 | 0.8 | 0.656 | 9.72E-90 | 1 | Lgals9 | infl_CAF |
| Cdr2l | 7.36E-91 | 0.27873797 | 0.897 | 0.84 | 1.42E-86 | 1 | Cdr2l | infl_CAF |
| Tiam1 | 8.99E-91 | 0.96114149 | 0.475 | 0.26 | 1.74E-86 | 1 | Tiam1 | infl_CAF |
| Zbtb38 | 9.71E-91 | 0.44030623 | 0.783 | 0.678 | 1.88E-86 | 1 | Zbtb38 | infl_CAF |
| Dennd4a1 | 2.26E-90 | 0.44213663 | 0.668 | 0.549 | 4.37E-86 | 1 | Dennd4a | infl_CAF |
| Kctd12 | 7.20E-90 | 0.60270922 | 0.461 | 0.349 | 1.39E-85 | 1 | Kctd12 | infl_CAF |
| Lpcat3 | 1.26E-89 | 0.51636078 | 0.761 | 0.636 | 2.44E-85 | 1 | Lpcat3 | infl_CAF |
| Tapbp | 5.83E-89 | 0.34562044 | 0.895 | 0.799 | 1.13E-84 | 1 | Tapbp | infl_CAF |
| Bambi | 1.94E-87 | 0.99165561 | 0.306 | 0.174 | 3.74E-83 | 1 | Bambi | infl_CAF |
| Nfia | 2.96E-87 | 0.51093407 | 0.744 | 0.564 | 5.73E-83 | 1 | Nfia | infl_CAF |
| Plpp1 | 7.47E-87 | 0.40723879 | 0.702 | 0.611 | 1.44E-82 | 1 | Plpp1 | infl_CAF |
| Mvp | 1.12E-86 | 0.32431599 | 0.884 | 0.811 | 2.16E-82 | 1 | Mvp | infl_CAF |
| Tshz2 | 1.05E-85 | 0.70087729 | 0.606 | 0.394 | 2.02E-81 | 1 | Tshz2 | infl_CAF |
| H2-D1 | 1.09E-85 | 0.65461584 | 0.51 | 0.346 | 2.10E-81 | 1 | H2-D1 | infl_CAF |
| Tmeff2 | 1.05E-84 | 0.59135586 | 0.621 | 0.286 | 2.03E-80 | 1 | Tmeff2 | infl_CAF |
| Fmr1 | 1.53E-83 | 0.43381752 | 0.775 | 0.706 | 2.96E-79 | 1 | Fmr1 | infl_CAF |
| Bst2 | 2.71E-83 | 0.38094241 | 0.759 | 0.518 | 5.24E-79 | 1 | Bst2 | infl_CAF |
| Slc10a6 | 5.35E-83 | 1.02286224 | 0.395 | 0.205 | 1.03E-78 | 1 | Slc10a6 | infl_CAF |
| Nfkb2 | 8.68E-83 | 0.40067474 | 0.838 | 0.757 | 1.68E-78 | 1 | Nfkb2 | infl_CAF |
| Zbp1 | 7.71E-82 | 0.4812169 | 0.658 | 0.475 | 1.49E-77 | 1 | Zbp1 | infl_CAF |
| Flnc | 4.59E-79 | 0.31073678 | 0.838 | 0.79 | 8.88E-75 | 1 | Flnc | infl_CAF |
| Glul | 7.04E-79 | 0.35340037 | 0.687 | 0.671 | 1.36E-74 | 1 | Glul | infl_CAF |
| Pcdhgc3 | 9.67E-79 | 0.32989748 | 0.895 | 0.83 | 1.87E-74 | 1 | Pcdhgc3 | infl_CAF |
| Mark1 | 1.11E-78 | 0.34366732 | 0.804 | 0.754 | 2.14E-74 | 1 | Mark1 | infl_CAF |
| Itgbl1 | 1.23E-78 | 0.6355113 | 0.35 | 0.219 | 2.37E-74 | 1 | Itgbl1 | infl_CAF |
| Dst | 1.40E-78 | 0.37869761 | 0.854 | 0.766 | 2.71E-74 | 1 | Dst | infl_CAF |
| Ppp3ca | 2.67E-78 | 0.38811935 | 0.806 | 0.703 | 5.16E-74 | 1 | Ppp3ca | infl_CAF |
| Asap3 | 4.22E-78 | 0.74679211 | 0.534 | 0.33 | 8.17E-74 | 1 | Asap3 | infl_CAF |
| Plaur | 7.64E-78 | 0.29398562 | 0.854 | 0.758 | 1.48E-73 | 1 | Plaur | infl_CAF |
| Smad3 | 1.29E-77 | 0.47606038 | 0.745 | 0.612 | 2.49E-73 | 1 | Smad3 | infl_CAF |
| Rgs3 | 1.54E-77 | 0.35902459 | 0.753 | 0.674 | 2.98E-73 | 1 | Rgs3 | infl_CAF |
| Pim3 | 1.65E-77 | 0.39134517 | 0.534 | 0.394 | 3.19E-73 | 1 | Pim3 | infl_CAF |
| Bach1 | 2.34E-77 | 0.29968858 | 0.829 | 0.786 | 4.53E-73 | 1 | Bach1 | infl_CAF |
| Bdkrb1 | 3.17E-77 | 0.78387182 | 0.402 | 0.297 | 6.13E-73 | 1 | Bdkrb1 | infl_CAF |
| Nr4a3 | 4.18E-77 | 0.30857857 | 0.276 | 0.225 | 8.09E-73 | 1 | Nr4a3 | infl_CAF |
| Cdon | 2.63E-76 | 0.51127033 | 0.756 | 0.66 | 5.08E-72 | 1 | Cdon | infl_CAF |
| Npdc1 | 9.67E-76 | 0.40162018 | 0.902 | 0.798 | 1.87E-71 | 1 | Npdc1 | infl_CAF |
| Nucb2 | 2.47E-75 | 0.42259994 | 0.8 | 0.663 | 4.78E-71 | 1 | Nucb2 | infl_CAF |
| Ern1 | 3.15E-74 | 0.47075587 | 0.647 | 0.505 | 6.08E-70 | 1 | Ern1 | infl_CAF |
| Tubb2a | 5.17E-74 | 0.46037294 | 0.826 | 0.695 | 1.00E-69 | 1 | Tubb2a | infl_CAF |
| Adcy5 | 2.73E-73 | 1.14710618 | 0.303 | 0.137 | 5.28E-69 | 1 | Adcy5 | infl_CAF |
| Sema6c | 4.90E-73 | 0.81753948 | 0.475 | 0.316 | 9.47E-69 | 1 | Sema6c | infl_CAF |
| Man1a | 5.47E-73 | 0.28964875 | 0.923 | 0.842 | 1.06E-68 | 1 | Man1a | infl_CAF |
| Erf | 8.95E-73 | 0.35169886 | 0.782 | 0.717 | 1.73E-68 | 1 | Erf | infl_CAF |
| Steap4 | 1.87E-72 | 0.45027426 | 0.504 | 0.273 | 3.61E-68 | 1 | Steap4 | infl_CAF |
| Mapk6 | 3.10E-72 | 0.25810767 | 0.833 | 0.791 | 5.99E-68 | 1 | Mapk6 | infl_CAF |
| Arl5b | 3.24E-72 | 0.45885646 | 0.555 | 0.457 | 6.26E-68 | 1 | Arl5b | infl_CAF |
| Hsd11b1 | 6.23E-72 | 0.36655394 | 0.578 | 0.44 | 1.20E-67 | 1 | Hsd11b1 | infl_CAF |
| Zfp521 | 7.12E-72 | 0.83302715 | 0.572 | 0.337 | 1.38E-67 | 1 | Zfp521 | infl_CAF |
| Ipo4 | 7.99E-72 | 0.36139674 | 0.874 | 0.8 | 1.54E-67 | 1 | Ipo4 | infl_CAF |
| Nenf | 9.23E-72 | 0.25346345 | 0.941 | 0.909 | 1.79E-67 | 1 | Nenf | infl_CAF |
| Mdfi | 4.63E-71 | 0.57201217 | 0.608 | 0.469 | 8.96E-67 | 1 | Mdfi | infl_CAF |
| Rab29 | 5.77E-71 | 0.75298208 | 0.501 | 0.337 | 1.12E-66 | 1 | Rab29 | infl_CAF |
| Angptl41 | 1.76E-69 | 0.50177157 | 0.57 | 0.43 | 3.40E-65 | 1 | Angptl4 | infl_CAF |
| Lysmd3 | 1.03E-68 | 0.57046627 | 0.596 | 0.432 | 1.99E-64 | 1 | Lysmd3 | infl_CAF |
| Chka | 1.55E-68 | 0.55738846 | 0.425 | 0.309 | 2.99E-64 | 1 | Chka | infl_CAF |
| Slc7a2 | 5.62E-68 | 0.73769961 | 0.267 | 0.179 | 1.09E-63 | 1 | Slc7a2 | infl_CAF |
| Hbegf | 6.62E-68 | 0.41475285 | 0.337 | 0.238 | 1.28E-63 | 1 | Hbegf | infl_CAF |
| Glis2 | 1.26E-67 | 0.37053789 | 0.835 | 0.748 | 2.44E-63 | 1 | Glis2 | infl_CAF |
| Pdgfrl | 1.76E-67 | 0.45165942 | 0.415 | 0.364 | 3.39E-63 | 1 | Pdgfrl | infl_CAF |
| Csrnp1 | 2.45E-67 | 0.33473098 | 0.538 | 0.508 | 4.73E-63 | 1 | Csrnp1 | infl_CAF |
| Tspo | 1.78E-66 | 0.28391645 | 0.913 | 0.856 | 3.43E-62 | 1 | Tspo | infl_CAF |
| Ang | 8.89E-66 | 0.65354308 | 0.56 | 0.4 | 1.72E-61 | 1 | Ang | infl_CAF |
| Slc9a9 | 1.41E-65 | 1.12996408 | 0.297 | 0.16 | 2.73E-61 | 1 | Slc9a9 | infl_CAF |
| Nr4a2 | 2.37E-65 | 0.53822461 | 0.286 | 0.224 | 4.58E-61 | 1 | Nr4a2 | infl_CAF |
| Pycr1 | 2.38E-65 | 0.40204014 | 0.734 | 0.631 | 4.59E-61 | 1 | Pycr1 | infl_CAF |
| Wnt9a | 2.55E-65 | 0.55500112 | 0.382 | 0.289 | 4.92E-61 | 1 | Wnt9a | infl_CAF |
| Ndrg2 | 2.71E-65 | 0.69267175 | 0.364 | 0.238 | 5.23E-61 | 1 | Ndrg2 | infl_CAF |
| H2-Q6 | 4.70E-65 | 0.65434535 | 0.267 | 0.149 | 9.09E-61 | 1 | H2-Q6 | infl_CAF |
| Abhd5 | 8.17E-65 | 0.51794614 | 0.57 | 0.459 | 1.58E-60 | 1 | Abhd5 | infl_CAF |
| Azin21 | 1.42E-64 | 0.31002567 | 0.758 | 0.677 | 2.74E-60 | 1 | Azin2 | infl_CAF |
| Serpinb1a | 1.88E-64 | 0.77535751 | 0.392 | 0.226 | 3.64E-60 | 1 | Serpinb1a | infl_CAF |
| Neo1 | 1.92E-64 | 0.33295417 | 0.694 | 0.642 | 3.72E-60 | 1 | Neo1 | infl_CAF |
| Oat | 3.26E-63 | 0.26118688 | 0.933 | 0.891 | 6.29E-59 | 1 | Oat | infl_CAF |
| Itsn1 | 6.19E-63 | 0.51933678 | 0.718 | 0.64 | 1.20E-58 | 1 | Itsn1 | infl_CAF |
| Dab2 | 9.64E-63 | 0.41150962 | 0.65 | 0.551 | 1.86E-58 | 1 | Dab2 | infl_CAF |
| Ces2g | 1.00E-62 | 0.64185042 | 0.461 | 0.202 | 1.93E-58 | 1 | Ces2g | infl_CAF |
| Setbp1 | 8.99E-62 | 0.41717072 | 0.675 | 0.561 | 1.74E-57 | 1 | Setbp1 | infl_CAF |
| Obsl1 | 3.71E-61 | 0.42036629 | 0.756 | 0.657 | 7.18E-57 | 1 | Obsl1 | infl_CAF |
| Tll1 | 7.69E-61 | 0.87505734 | 0.306 | 0.172 | 1.49E-56 | 1 | Tll1 | infl_CAF |
| Bin1 | 6.35E-60 | 0.29749912 | 0.812 | 0.739 | 1.23E-55 | 1 | Bin1 | infl_CAF |
| Sigmar1 | 5.93E-59 | 0.34505664 | 0.892 | 0.801 | 1.15E-54 | 1 | Sigmar1 | infl_CAF |
| Pabpc4 | 7.72E-59 | 0.29363818 | 0.857 | 0.798 | 1.49E-54 | 1 | Pabpc4 | infl_CAF |
| Gch1 | 1.04E-58 | 0.64052245 | 0.354 | 0.23 | 2.01E-54 | 1 | Gch1 | infl_CAF |
| Pgm1 | 3.81E-58 | 0.29998247 | 0.729 | 0.648 | 7.37E-54 | 1 | Pgm1 | infl_CAF |
| Fam110a | 5.40E-58 | 0.7015457 | 0.513 | 0.373 | 1.04E-53 | 1 | Fam110a | infl_CAF |
| Ell2 | 7.63E-58 | 0.55944522 | 0.458 | 0.338 | 1.48E-53 | 1 | Ell2 | infl_CAF |
| Hspb8 | 1.30E-57 | 0.36970489 | 0.742 | 0.638 | 2.52E-53 | 1 | Hspb8 | infl_CAF |
| Il16 | 1.57E-57 | 0.87318109 | 0.327 | 0.167 | 3.03E-53 | 1 | Il16 | infl_CAF |
| Armh4 | 2.22E-56 | 0.78858557 | 0.454 | 0.275 | 4.29E-52 | 1 | Armh4 | infl_CAF |
| Rcan1 | 2.52E-56 | 0.40470661 | 0.425 | 0.365 | 4.87E-52 | 1 | Rcan1 | infl_CAF |
| Kif21a | 3.46E-56 | 0.91320639 | 0.31 | 0.15 | 6.70E-52 | 1 | Kif21a | infl_CAF |
| Pvr | 6.34E-56 | 0.33905897 | 0.633 | 0.506 | 1.23E-51 | 1 | Pvr | infl_CAF |
| Plk2 | 4.82E-55 | 0.28097059 | 0.29 | 0.2 | 9.31E-51 | 1 | Plk2 | infl_CAF |
| Cpne2 | 7.47E-55 | 0.82985037 | 0.415 | 0.263 | 1.44E-50 | 1 | Cpne2 | infl_CAF |
| Dapk1 | 1.22E-54 | 0.71924557 | 0.428 | 0.234 | 2.37E-50 | 1 | Dapk1 | infl_CAF |
| Dusp10 | 2.96E-54 | 0.52880663 | 0.315 | 0.198 | 5.73E-50 | 1 | Dusp10 | infl_CAF |
| Dnajc25 | 8.71E-54 | 0.33504188 | 0.77 | 0.696 | 1.68E-49 | 1 | Dnajc25 | infl_CAF |
| Pth1r | 9.62E-54 | 0.2541255 | 0.832 | 0.778 | 1.86E-49 | 1 | Pth1r | infl_CAF |
| Ccl19 | 6.83E-53 | 0.64709988 | 0.285 | 0.158 | 1.32E-48 | 1 | Ccl19 | infl_CAF |
| Praf2 | 9.56E-53 | 0.29737802 | 0.808 | 0.73 | 1.85E-48 | 1 | Praf2 | infl_CAF |
| Zpr1 | 1.99E-52 | 0.48214876 | 0.612 | 0.496 | 3.84E-48 | 1 | Zpr1 | infl_CAF |
| Rcl1 | 3.66E-52 | 0.59579381 | 0.467 | 0.351 | 7.09E-48 | 1 | Rcl1 | infl_CAF |
| Med13l | 7.71E-52 | 0.25002092 | 0.908 | 0.856 | 1.49E-47 | 1 | Med13l | infl_CAF |
| Nfib | 8.04E-52 | 0.64337629 | 0.612 | 0.398 | 1.55E-47 | 1 | Nfib | infl_CAF |
| Cpne8 | 1.85E-51 | 0.49276921 | 0.6 | 0.407 | 3.57E-47 | 1 | Cpne8 | infl_CAF |
| Gcsh | 1.15E-50 | 0.42148086 | 0.724 | 0.577 | 2.23E-46 | 1 | Gcsh | infl_CAF |
| Park7 | 1.38E-50 | 0.27087726 | 0.834 | 0.773 | 2.67E-46 | 1 | Park7 | infl_CAF |
| H2-Q7 | 6.84E-50 | 0.64186376 | 0.265 | 0.156 | 1.32E-45 | 1 | H2-Q7 | infl_CAF |
| Map3k14 | 8.00E-50 | 0.50253343 | 0.503 | 0.364 | 1.55E-45 | 1 | Map3k14 | infl_CAF |
| Steap1 | 1.53E-49 | 0.60223699 | 0.562 | 0.41 | 2.97E-45 | 1 | Steap1 | infl_CAF |

| | | | | | | | | |
|---|---|---|---|---|---|---|---|---|
| St3gal4 | 1.90E-48 | 0.39766802 | 0.714 | 0.606 | 3.67E-44 | 1 | St3gal4 | infl_CAF |
| Xdh | 2.17E-48 | 0.25317288 | 0.806 | 0.708 | 4.19E-44 | 1 | Xdh | infl_CAF |
| Pygl | 3.82E-48 | 0.38247151 | 0.567 | 0.404 | 7.39E-44 | 1 | Pygl | infl_CAF |
| Fzd5 | 1.15E-47 | 0.56732353 | 0.371 | 0.243 | 2.23E-43 | 1 | Fzd5 | infl_CAF |
| Porcn1 | 2.81E-47 | 0.3171462 | 0.398 | 0.356 | 5.42E-43 | 1 | Porcn | infl_CAF |
| Gm2a | 8.41E-47 | 0.28415985 | 0.754 | 0.658 | 1.63E-42 | 1 | Gm2a | infl_CAF |
| Nkain1 | 8.73E-47 | 0.53355613 | 0.53 | 0.412 | 1.69E-42 | 1 | Nkain1 | infl_CAF |
| Smpdl3a | 1.08E-46 | 0.59620041 | 0.347 | 0.226 | 2.09E-42 | 1 | Smpdl3a | infl_CAF |
| Nbas | 2.52E-46 | 0.3834112 | 0.733 | 0.633 | 4.87E-42 | 1 | Nbas | infl_CAF |
| Ctps | 4.33E-46 | 0.39318402 | 0.635 | 0.531 | 8.37E-42 | 1 | Ctps | infl_CAF |
| Six1 | 1.52E-45 | 0.89120023 | 0.316 | 0.17 | 2.94E-41 | 1 | Six1 | infl_CAF |
| Ppp1r14b | 1.85E-45 | 0.29126875 | 0.815 | 0.736 | 3.57E-41 | 1 | Ppp1r14b | infl_CAF |
| Slc30a1 | 2.21E-45 | 0.58625034 | 0.562 | 0.423 | 4.26E-41 | 1 | Slc30a1 | infl_CAF |
| Atxn7l1 | 8.59E-45 | 0.37913565 | 0.644 | 0.542 | 1.66E-40 | 1 | Atxn7l1 | infl_CAF |
| Fas | 1.09E-44 | 0.77049593 | 0.339 | 0.223 | 2.10E-40 | 1 | Fas | infl_CAF |
| Nfasc | 2.00E-44 | 0.4695334 | 0.518 | 0.362 | 3.87E-40 | 1 | Nfasc | infl_CAF |
| Gstt1 | 2.46E-44 | 0.7721446 | 0.327 | 0.172 | 4.76E-40 | 1 | Gstt1 | infl_CAF |
| Tmem243 | 7.00E-44 | 0.72908701 | 0.309 | 0.213 | 1.35E-39 | 1 | Tmem243 | infl_CAF |
| Sbsn | 4.08E-43 | 0.35728244 | 0.596 | 0.325 | 7.88E-39 | 1 | Sbsn | infl_CAF |
| Sh3d19 | 8.40E-43 | 0.31750654 | 0.862 | 0.77 | 1.62E-38 | 1 | Sh3d19 | infl_CAF |
| Slc1a5 | 1.24E-42 | 0.31937984 | 0.772 | 0.633 | 2.41E-38 | 1 | Slc1a5 | infl_CAF |
| Hhipl11 | 1.31E-42 | 0.38398709 | 0.682 | 0.575 | 2.53E-38 | 1 | Hhipl1 | infl_CAF |
| Sdk1 | 1.53E-42 | 0.41646613 | 0.622 | 0.509 | 2.95E-38 | 1 | Sdk1 | infl_CAF |
| Chd3 | 1.73E-42 | 0.33271723 | 0.803 | 0.717 | 3.34E-38 | 1 | Chd3 | infl_CAF |
| Ppa1 | 3.73E-42 | 0.32770942 | 0.757 | 0.64 | 7.20E-38 | 1 | Ppa1 | infl_CAF |
| Plpbp | 4.63E-42 | 0.3293427 | 0.641 | 0.537 | 8.96E-38 | 1 | Plpbp | infl_CAF |
| Ccnyl1 | 4.98E-42 | 0.41817007 | 0.534 | 0.447 | 9.63E-38 | 1 | Ccnyl1 | infl_CAF |
| Tmem150a | 1.20E-41 | 0.29242484 | 0.658 | 0.578 | 2.32E-37 | 1 | Tmem150a | infl_CAF |
| Phyhd1 | 1.71E-40 | 0.56768804 | 0.269 | 0.178 | 3.31E-36 | 1 | Phyhd1 | infl_CAF |
| Celf2 | 3.39E-40 | 0.49493528 | 0.648 | 0.474 | 6.55E-36 | 1 | Celf2 | infl_CAF |
| St5 | 1.17E-39 | 0.37692875 | 0.679 | 0.564 | 2.26E-35 | 1 | St5 | infl_CAF |
| Sirpa | 5.48E-39 | 0.31882538 | 0.812 | 0.704 | 1.06E-34 | 1 | Sirpa | infl_CAF |
| Mylip | 7.70E-39 | 0.28914897 | 0.469 | 0.366 | 1.49E-34 | 1 | Mylip | infl_CAF |
| Pxylp1 | 9.16E-39 | 0.27745532 | 0.72 | 0.635 | 1.77E-34 | 1 | Pxylp1 | infl_CAF |
| Zdhhc9 | 1.41E-38 | 0.31132844 | 0.744 | 0.658 | 2.72E-34 | 1 | Zdhhc9 | infl_CAF |
| Cmas | 2.04E-38 | 0.29411525 | 0.69 | 0.607 | 3.94E-34 | 1 | Cmas | infl_CAF |
| Acvr2a | 3.36E-38 | 0.53225076 | 0.467 | 0.351 | 6.49E-34 | 1 | Acvr2a | infl_CAF |
| Phyh | 3.62E-38 | 0.55818126 | 0.406 | 0.291 | 6.99E-34 | 1 | Phyh | infl_CAF |
| Ndn | 6.91E-38 | 0.28323144 | 0.802 | 0.636 | 1.34E-33 | 1 | Ndn | infl_CAF |
| Gga2 | 8.69E-38 | 0.29074723 | 0.675 | 0.589 | 1.68E-33 | 1 | Gga2 | infl_CAF |
| Eps8 | 9.25E-38 | 0.48306214 | 0.5 | 0.343 | 1.79E-33 | 1 | Eps8 | infl_CAF |
| Pdlim5 | 9.86E-38 | 0.27120007 | 0.735 | 0.65 | 1.91E-33 | 1 | Pdlim5 | infl_CAF |
| Fgf18 | 9.98E-38 | 0.70224826 | 0.286 | 0.143 | 1.93E-33 | 1 | Fgf18 | infl_CAF |
| Galk2 | 1.76E-37 | 0.50250849 | 0.605 | 0.504 | 3.41E-33 | 1 | Galk2 | infl_CAF |
| Papss2 | 2.19E-37 | 0.34929259 | 0.705 | 0.61 | 4.23E-33 | 1 | Papss2 | infl_CAF |
| Ssh3 | 2.91E-37 | 0.53511916 | 0.354 | 0.23 | 5.63E-33 | 1 | Ssh3 | infl_CAF |
| Clcf1 | 1.76E-36 | 0.37677197 | 0.318 | 0.228 | 3.41E-32 | 1 | Clcf1 | infl_CAF |
| Dmpk | 2.22E-36 | 0.79801584 | 0.363 | 0.244 | 4.30E-32 | 1 | Dmpk | infl_CAF |
| Prrx2 | 4.74E-36 | 0.2822827 | 0.78 | 0.673 | 9.17E-32 | 1 | Prrx2 | infl_CAF |
| Layn | 6.46E-36 | 0.33758827 | 0.609 | 0.514 | 1.25E-31 | 1 | Layn | infl_CAF |
| Ddit3 | 1.71E-35 | 0.27879534 | 0.509 | 0.378 | 3.31E-31 | 1 | Ddit3 | infl_CAF |
| Galt | 4.63E-35 | 0.6608917 | 0.452 | 0.318 | 8.95E-31 | 1 | Galt | infl_CAF |
| Man2b2 | 7.14E-35 | 0.25900979 | 0.829 | 0.771 | 1.38E-30 | 1 | Man2b2 | infl_CAF |
| Stat5a | 1.96E-34 | 0.37978867 | 0.52 | 0.429 | 3.79E-30 | 1 | Stat5a | infl_CAF |
| Babam2 | 2.77E-34 | 0.35853293 | 0.593 | 0.518 | 5.36E-30 | 1 | Babam2 | infl_CAF |
| Thbs3 | 3.20E-34 | 0.3305939 | 0.841 | 0.651 | 6.19E-30 | 1 | Thbs3 | infl_CAF |
| Spry2 | 3.40E-34 | 0.31889437 | 0.668 | 0.568 | 6.57E-30 | 1 | Spry2 | infl_CAF |
| Pak1 | 6.36E-34 | 0.53668681 | 0.36 | 0.261 | 1.23E-29 | 1 | Pak1 | infl_CAF |
| Iigp1 | 1.96E-33 | 0.47338693 | 0.271 | 0.152 | 3.78E-29 | 1 | Iigp1 | infl_CAF |
| Guk1 | 2.40E-33 | 0.29241595 | 0.671 | 0.591 | 4.63E-29 | 1 | Guk1 | infl_CAF |
| Ccdc149 | 2.55E-33 | 0.79740222 | 0.257 | 0.142 | 4.93E-29 | 1 | Ccdc149 | infl_CAF |
| Notch2 | 8.37E-33 | 0.34068186 | 0.665 | 0.567 | 1.62E-28 | 1 | Notch2 | infl_CAF |
| Eef1akmt3 | 9.21E-33 | 0.63548772 | 0.341 | 0.22 | 1.78E-28 | 1 | Eef1akmt3 | infl_CAF |
| Eya4 | 1.03E-32 | 0.82069452 | 0.278 | 0.169 | 2.00E-28 | 1 | Eya4 | infl_CAF |
| Ttc12 | 2.52E-32 | 0.71582774 | 0.25 | 0.154 | 4.87E-28 | 1 | Ttc12 | infl_CAF |
| Efnb2 | 8.09E-32 | 0.25724909 | 0.602 | 0.491 | 1.56E-27 | 1 | Efnb2 | infl_CAF |
| Gda | 1.04E-31 | 0.35676075 | 0.703 | 0.49 | 2.01E-27 | 1 | Gda | infl_CAF |
| Scn1b | 3.28E-31 | 0.25405387 | 0.809 | 0.701 | 6.34E-27 | 1 | Scn1b | infl_CAF |
| Alas1 | 3.73E-31 | 0.47479615 | 0.509 | 0.43 | 7.21E-27 | 1 | Alas1 | infl_CAF |
| Pigs | 8.37E-31 | 0.28215227 | 0.772 | 0.682 | 1.62E-26 | 1 | Pigs | infl_CAF |
| Cpeb1 | 9.03E-31 | 0.62559608 | 0.266 | 0.164 | 1.75E-26 | 1 | Cpeb1 | infl_CAF |
| Cpt1a | 1.73E-30 | 0.34897159 | 0.632 | 0.52 | 3.34E-26 | 1 | Cpt1a | infl_CAF |
| Acvr1b | 2.52E-30 | 0.32339103 | 0.578 | 0.482 | 4.86E-26 | 1 | Acvr1b | infl_CAF |
| Gstt3 | 3.05E-30 | 0.65008309 | 0.285 | 0.173 | 5.90E-26 | 1 | Gstt3 | infl_CAF |
| Prrg3 | 5.07E-30 | 0.6449734 | 0.274 | 0.146 | 9.81E-26 | 1 | Prrg3 | infl_CAF |
| Car13 | 5.44E-30 | 0.39332456 | 0.487 | 0.372 | 1.05E-25 | 1 | Car13 | infl_CAF |
| Slc16a10 | 6.18E-30 | 0.56644705 | 0.253 | 0.161 | 1.19E-25 | 1 | Slc16a10 | infl_CAF |
| Tnfrsf1b | 6.71E-30 | 0.43357794 | 0.52 | 0.385 | 1.30E-25 | 1 | Tnfrsf1b | infl_CAF |
| Arl15 | 1.24E-29 | 0.36026054 | 0.62 | 0.517 | 2.40E-25 | 1 | Arl15 | infl_CAF |
| Erap1 | 2.02E-29 | 0.49190858 | 0.434 | 0.336 | 3.91E-25 | 1 | Erap1 | infl_CAF |
| Ebp | 2.21E-29 | 0.31446032 | 0.599 | 0.556 | 4.27E-25 | 1 | Ebp | infl_CAF |
| Galnt16 | 2.79E-29 | 0.39262991 | 0.711 | 0.306 | 5.40E-25 | 1 | Galnt16 | infl_CAF |
| Pde1a | 1.17E-28 | 0.63776871 | 0.3 | 0.205 | 2.26E-24 | 1 | Pde1a | infl_CAF |
| Nrbp2 | 3.30E-28 | 0.47419967 | 0.331 | 0.222 | 6.39E-24 | 1 | Nrbp2 | infl_CAF |
| Ralgds | 4.58E-28 | 0.28576387 | 0.558 | 0.472 | 8.86E-24 | 1 | Ralgds | infl_CAF |
| Mgarp | 5.14E-28 | 0.39953256 | 0.378 | 0.264 | 9.94E-24 | 1 | Mgarp | infl_CAF |
| Ripor3 | 1.17E-27 | 0.32960392 | 0.539 | 0.453 | 2.27E-23 | 1 | Ripor3 | infl_CAF |
| Tssc4 | 1.53E-27 | 0.36992564 | 0.638 | 0.526 | 2.97E-23 | 1 | Tssc4 | infl_CAF |
| Grasp | 1.72E-27 | 0.34303642 | 0.531 | 0.469 | 3.33E-23 | 1 | Grasp | infl_CAF |
| Rab23 | 1.79E-27 | 0.36559117 | 0.6 | 0.484 | 3.45E-23 | 1 | Rab23 | infl_CAF |
| Plk3 | 1.93E-27 | 0.27780443 | 0.478 | 0.402 | 3.73E-23 | 1 | Plk3 | infl_CAF |
| Rnf213 | 2.13E-27 | 0.25039123 | 0.558 | 0.44 | 4.11E-23 | 1 | Rnf213 | infl_CAF |
| Echdc2 | 2.71E-27 | 0.66756776 | 0.338 | 0.231 | 5.24E-23 | 1 | Echdc2 | infl_CAF |
| Htra2 | 2.71E-27 | 0.32541294 | 0.735 | 0.638 | 5.24E-23 | 1 | Htra2 | infl_CAF |
| Mkx1 | 3.19E-27 | 0.36644545 | 0.316 | 0.248 | 6.16E-23 | 1 | Mkx | infl_CAF |
| Acvrl1 | 3.39E-27 | 0.26880137 | 0.776 | 0.69 | 6.55E-23 | 1 | Acvrl1 | infl_CAF |
| Spryd7 | 4.83E-27 | 0.5455808 | 0.436 | 0.325 | 9.34E-23 | 1 | Spryd7 | infl_CAF |
| Chid1 | 9.98E-27 | 0.29050552 | 0.605 | 0.527 | 1.93E-22 | 1 | Chid1 | infl_CAF |
| Aldh18a1 | 1.47E-26 | 0.26215415 | 0.789 | 0.719 | 2.85E-22 | 1 | Aldh18a1 | infl_CAF |
| Man2b1 | 1.54E-26 | 0.26234936 | 0.751 | 0.661 | 2.98E-22 | 1 | Man2b1 | infl_CAF |
| Mapk12 | 1.74E-26 | 0.71768946 | 0.319 | 0.209 | 3.36E-22 | 1 | Mapk12 | infl_CAF |
| Nelfb | 5.73E-26 | 0.25488697 | 0.691 | 0.646 | 1.11E-21 | 1 | Nelfb | infl_CAF |
| Srgap1 | 6.96E-26 | 0.47909884 | 0.317 | 0.203 | 1.35E-21 | 1 | Srgap1 | infl_CAF |
| Zfp354c | 1.52E-25 | 0.38564113 | 0.416 | 0.313 | 2.95E-21 | 1 | Zfp354c | infl_CAF |
| Ibtk | 1.55E-25 | 0.25162787 | 0.658 | 0.592 | 3.00E-21 | 1 | Ibtk | infl_CAF |
| Swap70 | 2.53E-25 | 0.34071898 | 0.595 | 0.503 | 4.89E-21 | 1 | Swap70 | infl_CAF |
| Cog6 | 3.68E-25 | 0.27485895 | 0.599 | 0.517 | 7.11E-21 | 1 | Cog6 | infl_CAF |
| Cers6 | 5.01E-25 | 0.30385808 | 0.602 | 0.483 | 9.68E-21 | 1 | Cers6 | infl_CAF |
| Fbxo17 | 2.13E-24 | 0.37563418 | 0.513 | 0.402 | 4.12E-20 | 1 | Fbxo17 | infl_CAF |
| Ksr1 | 2.39E-24 | 0.39668486 | 0.456 | 0.361 | 4.62E-20 | 1 | Ksr1 | infl_CAF |
| Pter | 2.43E-24 | 0.6785526 | 0.267 | 0.176 | 4.69E-20 | 1 | Pter | infl_CAF |
| Tmem104 | 1.59E-23 | 0.35124911 | 0.572 | 0.465 | 3.07E-19 | 1 | Tmem104 | infl_CAF |
| Prr7 | 4.52E-23 | 0.64413887 | 0.286 | 0.202 | 8.74E-19 | 1 | Prr7 | infl_CAF |
| Qpctl | 5.54E-23 | 0.42865934 | 0.527 | 0.431 | 1.07E-18 | 1 | Qpctl | infl_CAF |
| Prex1 | 5.65E-23 | 0.60041428 | 0.351 | 0.234 | 1.09E-18 | 1 | Prex1 | infl_CAF |
| B4galt2 | 8.40E-23 | 0.30176923 | 0.675 | 0.584 | 1.62E-18 | 1 | B4galt2 | infl_CAF |
| Gk5 | 1.28E-22 | 0.61304233 | 0.256 | 0.171 | 2.47E-18 | 1 | Gk5 | infl_CAF |
| Psen2 | 1.64E-22 | 0.28835046 | 0.485 | 0.398 | 3.16E-18 | 1 | Psen2 | infl_CAF |
| Rasl121 | 1.82E-22 | 0.25328172 | 0.469 | 0.374 | 3.53E-18 | 1 | Rasl12 | infl_CAF |
| Ggct | 2.25E-22 | 0.5660862 | 0.261 | 0.16 | 4.35E-18 | 1 | Ggct | infl_CAF |
| Vegfb | 2.35E-22 | 0.40129218 | 0.531 | 0.449 | 4.55E-18 | 1 | Vegfb | infl_CAF |
| Pltp | 3.38E-22 | 0.2876248 | 0.421 | 0.293 | 6.53E-18 | 1 | Pltp | infl_CAF |
| Prkag2 | 4.58E-22 | 0.51549957 | 0.383 | 0.295 | 8.85E-18 | 1 | Prkag2 | infl_CAF |
| Mcub | 2.67E-21 | 0.36730181 | 0.454 | 0.338 | 5.16E-17 | 1 | Mcub | infl_CAF |
| Tmed5 | 3.33E-21 | 0.42255827 | 0.573 | 0.465 | 6.44E-17 | 1 | Tmed5 | infl_CAF |
| Zbtb11 | 3.41E-21 | 0.30172665 | 0.499 | 0.415 | 6.60E-17 | 1 | Zbtb11 | infl_CAF |
| Stx11 | 8.86E-21 | 0.44973251 | 0.345 | 0.25 | 1.71E-16 | 1 | Stx11 | infl_CAF |
| Rab3d | 1.14E-20 | 0.33543998 | 0.458 | 0.345 | 2.21E-16 | 1 | Rab3d | infl_CAF |
| Slc6a9 | 1.53E-20 | 0.35994328 | 0.333 | 0.257 | 2.97E-16 | 1 | Slc6a9 | infl_CAF |
| Sertad4 | 1.83E-20 | 0.37085149 | 0.334 | 0.258 | 3.55E-16 | 1 | Sertad4 | infl_CAF |
| Kcnd1 | 2.71E-20 | 0.43788008 | 0.339 | 0.241 | 5.24E-16 | 1 | Kcnd1 | infl_CAF |
| Pgm3 | 4.00E-20 | 0.26610562 | 0.667 | 0.559 | 7.73E-16 | 1 | Pgm3 | infl_CAF |
| Cpeb2 | 4.34E-20 | 0.41273611 | 0.324 | 0.226 | 8.38E-16 | 1 | Cpeb2 | infl_CAF |
| Shb | 4.35E-20 | 0.26677014 | 0.494 | 0.399 | 8.42E-16 | 1 | Shb | infl_CAF |
| Trp53i13 | 4.44E-20 | 0.36899767 | 0.508 | 0.418 | 8.57E-16 | 1 | Trp53i13 | infl_CAF |
| Gnai1 | 5.11E-20 | 0.39539881 | 0.343 | 0.267 | 9.88E-16 | 1 | Gnai1 | infl_CAF |
| Ptpn2 | 5.90E-20 | 0.33740634 | 0.534 | 0.44 | 1.14E-15 | 1 | Ptpn2 | infl_CAF |
| Ltv1 | 6.17E-20 | 0.43844798 | 0.506 | 0.425 | 1.19E-15 | 1 | Ltv1 | infl_CAF |
| Hebp2 | 6.85E-20 | 0.36457749 | 0.457 | 0.365 | 1.32E-15 | 1 | Hebp2 | infl_CAF |
| Capn5 | 1.20E-19 | 0.34411791 | 0.383 | 0.278 | 2.31E-15 | 1 | Capn5 | infl_CAF |
| Nol9 | 1.79E-19 | 0.29220306 | 0.455 | 0.386 | 3.46E-15 | 1 | Nol9 | infl_CAF |
| Zmat3 | 1.83E-19 | 0.35690472 | 0.521 | 0.455 | 3.55E-15 | 1 | Zmat3 | infl_CAF |
| Iffo1 | 2.48E-19 | 0.27612242 | 0.391 | 0.325 | 4.79E-15 | 1 | Iffo1 | infl_CAF |
| Trmt61a | 3.17E-19 | 0.41486527 | 0.38 | 0.303 | 6.13E-15 | 1 | Trmt61a | infl_CAF |
| Snta1 | 4.08E-19 | 0.36487013 | 0.471 | 0.387 | 7.89E-15 | 1 | Snta1 | infl_CAF |
| Lrrk2 | 6.34E-19 | 0.34532842 | 0.46 | 0.379 | 1.22E-14 | 1 | Lrrk2 | infl_CAF |
| Ccdc157 | 7.58E-19 | 0.43026491 | 0.337 | 0.255 | 1.47E-14 | 1 | Ccdc157 | infl_CAF |
| Tmem106a | 8.13E-19 | 0.4311054 | 0.413 | 0.317 | 1.57E-14 | 1 | Tmem106a | infl_CAF |
| Tmem38b | 8.81E-19 | 0.29720871 | 0.503 | 0.41 | 1.70E-14 | 1 | Tmem38b | infl_CAF |
| Gmds | 2.49E-18 | 0.42928927 | 0.496 | 0.392 | 4.82E-14 | 1 | Gmds | infl_CAF |
| Agpat3 | 2.97E-18 | 0.34953565 | 0.612 | 0.479 | 5.74E-14 | 1 | Agpat3 | infl_CAF |
| Cxcl161 | 3.70E-18 | 0.25821686 | 0.491 | 0.398 | 7.15E-14 | 1 | Cxcl16 | infl_CAF |
| Lrig3 | 5.22E-18 | 0.39295783 | 0.343 | 0.26 | 1.01E-13 | 1 | Lrig3 | infl_CAF |
| Entpd6 | 7.41E-18 | 0.54763414 | 0.359 | 0.265 | 1.43E-13 | 1 | Entpd6 | infl_CAF |
| Bcl2l1 | 7.82E-18 | 0.29757812 | 0.365 | 0.315 | 1.51E-13 | 1 | Bcl2l1 | infl_CAF |
| Ptgr1 | 7.94E-18 | 0.26511853 | 0.533 | 0.47 | 1.53E-13 | 1 | Ptgr1 | infl_CAF |
| Map3k8 | 8.64E-18 | 0.38789043 | 0.336 | 0.253 | 1.67E-13 | 1 | Map3k8 | infl_CAF |
| Slc46a11 | 1.02E-17 | 0.30275565 | 0.361 | 0.282 | 1.97E-13 | 1 | Slc46a1 | infl_CAF |
| Tob2 | 4.07E-17 | 0.36772803 | 0.446 | 0.378 | 7.86E-13 | 1 | Tob2 | infl_CAF |
| Hsph1 | 6.26E-17 | 0.36671967 | 0.497 | 0.4 | 1.21E-12 | 1 | Hsph1 | infl_CAF |
| Nt5dc3 | 1.83E-16 | 0.29816285 | 0.317 | 0.233 | 3.54E-12 | 1 | Nt5dc3 | infl_CAF |
| Traf3 | 3.34E-16 | 0.27538397 | 0.469 | 0.387 | 6.46E-12 | 1 | Traf3 | infl_CAF |
| Nfkbie | 3.66E-16 | 0.52916996 | 0.259 | 0.193 | 7.08E-12 | 1 | Nfkbie | infl_CAF |
| Npm3 | 4.42E-16 | 0.2942819 | 0.545 | 0.446 | 8.55E-12 | 1 | Npm3 | infl_CAF |
| Casp4 | 5.35E-16 | 0.27109321 | 0.527 | 0.434 | 1.03E-11 | 1 | Casp4 | infl_CAF |
| Alyref | 5.95E-16 | 0.30718578 | 0.282 | 0.249 | 1.15E-11 | 1 | Alyref | infl_CAF |
| Tmc6 | 8.60E-16 | 0.27097439 | 0.425 | 0.351 | 1.66E-11 | 1 | Tmc6 | infl_CAF |
| Itpkc | 2.51E-15 | 0.3428684 | 0.353 | 0.311 | 4.85E-11 | 1 | Itpkc | infl_CAF |
| Slc1a4 | 3.38E-15 | 0.33967955 | 0.424 | 0.336 | 6.53E-11 | 1 | Slc1a4 | infl_CAF |
| Olfml1 | 3.91E-15 | 0.46232764 | 0.364 | 0.237 | 7.56E-11 | 1 | Olfml1 | infl_CAF |
| Mn11 | 6.69E-15 | 0.37206324 | 0.28 | 0.221 | 1.29E-10 | 1 | Mn1 | infl_CAF |
| Kcnk6 | 7.74E-15 | 0.4248778 | 0.285 | 0.216 | 1.50E-10 | 1 | Kcnk6 | infl_CAF |
| Procr | 9.30E-15 | 0.36596297 | 0.337 | 0.245 | 1.80E-10 | 1 | Procr | infl_CAF |
| Serpina3i | 1.08E-14 | 0.27255128 | 0.382 | 0.302 | 2.09E-10 | 1 | Serpina3i | infl_CAF |
| Fbxl18 | 1.13E-14 | 0.47072934 | 0.253 | 0.199 | 2.18E-10 | 1 | Fbxl18 | infl_CAF |
| Prickle2 | 1.27E-14 | 0.29138804 | 0.419 | 0.356 | 2.46E-10 | 1 | Prickle2 | infl_CAF |
| Plpp2 | 1.66E-14 | 0.32797086 | 0.349 | 0.274 | 3.21E-10 | 1 | Plpp2 | infl_CAF |
| Nol6 | 2.03E-14 | 0.31659639 | 0.581 | 0.492 | 3.93E-10 | 1 | Nol6 | infl_CAF |
| Fpgs | 2.47E-14 | 0.28524375 | 0.53 | 0.441 | 4.78E-10 | 1 | Fpgs | infl_CAF |
| Mccc1 | 5.26E-14 | 0.30100453 | 0.301 | 0.258 | 1.02E-09 | 1 | Mccc1 | infl_CAF |
| Nuak2 | 6.00E-14 | 0.38825772 | 0.291 | 0.229 | 1.16E-09 | 1 | Nuak2 | infl_CAF |
| Ddx58 | 9.28E-14 | 0.33784716 | 0.43 | 0.352 | 1.79E-09 | 1 | Ddx58 | infl_CAF |
| Grk5 | 2.61E-13 | 0.28883419 | 0.536 | 0.436 | 5.04E-09 | 1 | Grk5 | infl_CAF |
| Lysmd2 | 4.98E-13 | 0.2619983 | 0.418 | 0.334 | 9.64E-09 | 1 | Lysmd2 | infl_CAF |
| Nedd4l | 5.30E-13 | 0.36096843 | 0.344 | 0.274 | 1.02E-08 | 1 | Nedd4l | infl_CAF |
| Amigo3 | 5.73E-13 | 0.29480128 | 0.283 | 0.232 | 1.11E-08 | 1 | Amigo3 | infl_CAF |
| Jam2 | 9.54E-13 | 0.47567865 | 0.268 | 0.191 | 1.84E-08 | 1 | Jam2 | infl_CAF |
| Entpd5 | 1.22E-12 | 0.26895009 | 0.483 | 0.413 | 2.35E-08 | 1 | Entpd5 | infl_CAF |
| Rhbdd2 | 1.33E-12 | 0.38974381 | 0.355 | 0.328 | 2.57E-08 | 1 | Rhbdd2 | infl_CAF |
| Myorg | 3.09E-12 | 0.34012664 | 0.354 | 0.281 | 5.98E-08 | 1 | Myorg | infl_CAF |
| Gpr27 | 3.24E-12 | 0.37171658 | 0.359 | 0.274 | 6.26E-08 | 1 | Gpr27 | infl_CAF |
| Nckap5l | 4.26E-12 | 0.2621138 | 0.333 | 0.267 | 8.23E-08 | 1 | Nckap5l | infl_CAF |
| Chrdl1 | 8.96E-12 | 0.29419673 | 0.491 | 0.226 | 1.73E-07 | 1 | Chrdl1 | infl_CAF |
| Ddt | 9.71E-12 | 0.25429202 | 0.437 | 0.37 | 1.88E-07 | 1 | Ddt | infl_CAF |
| Sqor | 9.94E-12 | 0.31820985 | 0.408 | 0.291 | 1.92E-07 | 1 | Sqor | infl_CAF |
| Aldh1l1 | 1.21E-11 | 0.36323591 | 0.381 | 0.318 | 2.34E-07 | 1 | Aldh1l1 | infl_CAF |
| Csf2ra | 1.33E-11 | 0.29336681 | 0.254 | 0.188 | 2.57E-07 | 1 | Csf2ra | infl_CAF |
| Fkbp5 | 1.82E-11 | 0.29859232 | 0.581 | 0.507 | 3.53E-07 | 1 | Fkbp5 | infl_CAF |
| Abhd11 | 2.09E-11 | 0.26891671 | 0.505 | 0.443 | 4.03E-07 | 1 | Abhd11 | infl_CAF |
| Gpr135 | 2.15E-11 | 0.31695069 | 0.285 | 0.232 | 4.16E-07 | 1 | Gpr135 | infl_CAF |
| Slc25a25 | 2.29E-11 | 0.25431431 | 0.392 | 0.331 | 4.43E-07 | 1 | Slc25a25 | infl_CAF |
| Tnfrsf10b | 3.46E-11 | 0.36025754 | 0.336 | 0.279 | 6.69E-07 | 1 | Tnfrsf10b | infl_CAF |
| Wnt16 | 4.99E-11 | 0.43803749 | 0.253 | 0.115 | 9.64E-07 | 1 | Wnt16 | infl_CAF |
| Cryzl2 | 5.02E-11 | 0.30711167 | 0.315 | 0.252 | 9.71E-07 | 1 | Cryzl2 | infl_CAF |
| Abhd17c | 5.26E-11 | 0.25852762 | 0.522 | 0.452 | 1.02E-06 | 1 | Abhd17c | infl_CAF |
| Chrd | 5.65E-11 | 0.27728286 | 0.308 | 0.239 | 1.09E-06 | 1 | Chrd | infl_CAF |
| Dzip1 | 7.77E-11 | 0.25169153 | 0.425 | 0.335 | 1.50E-06 | 1 | Dzip1 | infl_CAF |
| Smarca1 | 1.16E-10 | 0.30355861 | 0.256 | 0.199 | 2.25E-06 | 1 | Smarca1 | infl_CAF |
| Rpf2 | 1.33E-10 | 0.41029468 | 0.255 | 0.198 | 2.57E-06 | 1 | Rpf2 | infl_CAF |
| Fzd8 | 1.62E-10 | 0.31229046 | 0.331 | 0.276 | 3.14E-06 | 1 | Fzd8 | infl_CAF |
| Repin1 | 2.14E-10 | 0.28152528 | 0.385 | 0.323 | 4.14E-06 | 1 | Repin1 | infl_CAF |
| Sfxn2 | 6.98E-10 | 0.29757211 | 0.388 | 0.307 | 1.35E-05 | 1 | Sfxn2 | infl_CAF |
| Cnpy3 | 1.05E-09 | 0.30431988 | 0.651 | 0.57 | 2.04E-05 | 1 | Cnpy3 | infl_CAF |
| Tmem106c | 1.30E-09 | 0.37817059 | 0.347 | 0.277 | 2.51E-05 | 1 | Tmem106c | infl_CAF |
| Emx2 | 1.41E-09 | 0.32733549 | 0.316 | 0.248 | 2.72E-05 | 1 | Emx2 | infl_CAF |
| Hfe | 1.42E-09 | 0.30888467 | 0.266 | 0.201 | 2.74E-05 | 1 | Hfe | infl_CAF |
| Rusc1 | 2.21E-09 | 0.26655709 | 0.354 | 0.286 | 4.27E-05 | 1 | Rusc1 | infl_CAF |
| Uprt | 3.02E-09 | 0.27793148 | 0.534 | 0.451 | 5.83E-05 | 1 | Uprt | infl_CAF |
| Fdxr | 4.39E-09 | 0.2528613 | 0.374 | 0.33 | 8.48E-05 | 1 | Fdxr | infl_CAF |
| Rab30 | 5.22E-09 | 0.2970722 | 0.257 | 0.196 | 0.00010094 | 1 | Rab30 | infl_CAF |
| Pde8a | 1.29E-08 | 0.26158605 | 0.451 | 0.312 | 0.0002489 | 1 | Pde8a | infl_CAF |
| Nle1 | 1.58E-08 | 0.29859109 | 0.352 | 0.289 | 0.00030603 | 1 | Nle1 | infl_CAF |
| Slc4a3 | 1.86E-08 | 0.44279875 | 0.288 | 0.23 | 0.00036004 | 1 | Slc4a3 | infl_CAF |
| Cyb561d2 | 2.04E-08 | 0.3059117 | 0.287 | 0.232 | 0.00039455 | 1 | Cyb561d2 | infl_CAF |
| Sgms2 | 2.20E-08 | 0.26952445 | 0.433 | 0.36 | 0.00042545 | 1 | Sgms2 | infl_CAF |
| Mthfr | 2.84E-08 | 0.26076523 | 0.436 | 0.373 | 0.00054882 | 1 | Mthfr | infl_CAF |
| Nolc1 | 2.90E-08 | 0.25459747 | 0.461 | 0.376 | 0.00056135 | 1 | Nolc1 | infl_CAF |
| Fes | 5.31E-08 | 0.29021505 | 0.443 | 0.369 | 0.00102706 | 1 | Fes | infl_CAF |
| Lhpp | 7.11E-08 | 0.53468132 | 0.265 | 0.206 | 0.00137512 | 1 | Lhpp | infl_CAF |
| Ftsj3 | 7.55E-08 | 0.35582104 | 0.351 | 0.28 | 0.00145947 | 1 | Ftsj3 | infl_CAF |
| Bahcc1 | 9.44E-08 | 0.38291545 | 0.371 | 0.312 | 0.00182608 | 1 | Bahcc1 | infl_CAF |
| Tbkbp1 | 1.01E-07 | 0.25569361 | 0.265 | 0.219 | 0.00195878 | 1 | Tbkbp1 | infl_CAF |
| Slc35c2 | 1.03E-07 | 0.32689635 | 0.518 | 0.449 | 0.00199825 | 1 | Slc35c2 | infl_CAF |
| Mapkbp1 | 2.59E-07 | 0.29855562 | 0.305 | 0.252 | 0.00501316 | 1 | Mapkbp1 | infl_CAF |
| Abcd4 | 6.42E-07 | 0.3332165 | 0.255 | 0.209 | 0.01240533 | 1 | Abcd4 | infl_CAF |
| Tle2 | 6.57E-07 | 0.35385466 | 0.295 | 0.241 | 0.01271164 | 1 | Tle2 | infl_CAF |
| Enpep | 3.73E-06 | 0.31744539 | 0.277 | 0.186 | 0.07208409 | 1 | Enpep | infl_CAF |
| Tapbpl | 4.77E-06 | 0.28564154 | 0.334 | 0.282 | 0.09218252 | 1 | Tapbpl | infl_CAF |
| Gyg | 5.94E-06 | 0.28697227 | 0.423 | 0.371 | 0.11484954 | 1 | Gyg | infl_CAF |
| Cln8 | 1.47E-05 | 0.28232809 | 0.264 | 0.218 | 0.28380517 | 1 | Cln8 | infl_CAF |
| Txnrd2 | 1.76E-05 | 0.30001589 | 0.312 | 0.264 | 0.33982784 | 1 | Txnrd2 | infl_CAF |
| Erlin1 | 0.0001053 | 0.25625457 | 0.338 | 0.278 | 1 | 1 | Erlin1 | infl_CAF |
| Tcaf2 | 0.00051605 | 0.2689586 | 0.253 | 0.213 | 1 | 1 | Tcaf2 | infl_CAF |
| Trim47 | 0.00093258 | 0.29506751 | 0.294 | 0.238 | 1 | 1 | Trim47 | infl_CAF |
| Tor3a | 0.0022227 | 0.26095818 | 0.273 | 0.239 | 1 | 1 | Tor3a | infl_CAF |
| Ccna2 | 0 | 6.22786906 | 0.866 | 0.049 | 0 | 2 | Ccna2 | myCAF_prolif |
| Mki67 | 0 | 5.69299042 | 0.863 | 0.059 | 0 | 2 | Mki67 | myCAF_prolif |
| Tpx2 | 0 | 5.61034487 | 0.836 | 0.051 | 0 | 2 | Tpx2 | myCAF_prolif |
| Birc5 | 0 | 5.19550953 | 0.84 | 0.064 | 0 | 2 | Birc5 | myCAF_prolif |
| Anln | 0 | 4.85984263 | 0.878 | 0.112 | 0 | 2 | Anln | myCAF_prolif |
| Rrm2 | 0 | 5.44324575 | 0.874 | 0.112 | 0 | 2 | Rrm2 | myCAF_prolif |
| Racgap1 | 0 | 4.88784106 | 0.841 | 0.082 | 0 | 2 | Racgap1 | myCAF_prolif |
| Top2a | 0 | 6.33846062 | 0.837 | 0.079 | 0 | 2 | Top2a | myCAF_prolif |
| Tk1 | 0 | 5.61226819 | 0.799 | 0.046 | 0 | 2 | Tk1 | myCAF_prolif |
| Tacc3 | 0 | 4.17114375 | 0.882 | 0.133 | 0 | 2 | Tacc3 | myCAF_prolif |
| Cdca3 | 0 | 4.27774087 | 0.839 | 0.098 | 0 | 2 | Cdca3 | myCAF_prolif |
| Ncapd2 | 0 | 4.36599689 | 0.868 | 0.133 | 0 | 2 | Ncapd2 | myCAF_prolif |
| Kifc1 | 0 | 5.84848941 | 0.759 | 0.032 | 0 | 2 | Kifc1 | myCAF_prolif |
| Cdca8 | 0 | 5.65313156 | 0.754 | 0.035 | 0 | 2 | Cdca8 | myCAF_prolif |
| Spc24 | 0 | 4.72450399 | 0.759 | 0.049 | 0 | 2 | Spc24 | myCAF_prolif |
| Prr11 | 0 | 6.19885169 | 0.731 | 0.027 | 0 | 2 | Prr11 | myCAF_prolif |
| Pclaf | 0 | 4.41794557 | 0.806 | 0.104 | 0 | 2 | Pclaf | myCAF_prolif |
| Plk1 | 0 | 6.08878775 | 0.734 | 0.033 | 0 | 2 | Plk1 | myCAF_prolif |
| Cip2a | 0 | 4.47221994 | 0.773 | 0.078 | 0 | 2 | Cip2a | myCAF_prolif |
| Ckap2l | 0 | 5.76177586 | 0.715 | 0.026 | 0 | 2 | Ckap2l | myCAF_prolif |
| Hmmr | 0 | 5.8835839 | 0.711 | 0.023 | 0 | 2 | Hmmr | myCAF_prolif |
| Pimreg | 0 | 6.42067284 | 0.703 | 0.016 | 0 | 2 | Pimreg | myCAF_prolif |
| Cdc20 | 0 | 5.23517454 | 0.746 | 0.06 | 0 | 2 | Cdc20 | myCAF_prolif |
| Stmn1 | 0 | 3.81049549 | 0.906 | 0.223 | 0 | 2 | Stmn1 | myCAF_prolif |
| Mad2l1 | 0 | 4.0065449 | 0.772 | 0.089 | 0 | 2 | Mad2l1 | myCAF_prolif |
| Smc2 | 0 | 4.30450218 | 0.789 | 0.108 | 0 | 2 | Smc2 | myCAF_prolif |
| Ube2c | 0 | 5.98676714 | 0.726 | 0.046 | 0 | 2 | Ube2c | myCAF_prolif |
| Dlgap5 | 0 | 6.47553025 | 0.696 | 0.016 | 0 | 2 | Dlgap5 | myCAF_prolif |
| Ect2 | 0 | 5.6666818 | 0.709 | 0.031 | 0 | 2 | Ect2 | myCAF_prolif |
| Gtse1 | 0 | 4.70176893 | 0.727 | 0.05 | 0 | 2 | Gtse1 | myCAF_prolif |
| Uhrf1 | 0 | 4.37597014 | 0.782 | 0.107 | 0 | 2 | Uhrf1 | myCAF_prolif |
| Prc1 | 0 | 6.73586121 | 0.701 | 0.029 | 0 | 2 | Prc1 | myCAF_prolif |
| Asf1b | 0 | 5.27080814 | 0.693 | 0.028 | 0 | 2 | Asf1b | myCAF_prolif |
| Kif2c | 0 | 6.90644533 | 0.674 | 0.011 | 0 | 2 | Kif2c | myCAF_prolif |
| Mxd3 | 0 | 5.31611486 | 0.694 | 0.035 | 0 | 2 | Mxd3 | myCAF_prolif |
| Pbk | 0 | 7.8140422 | 0.665 | 0.007 | 0 | 2 | Pbk | myCAF_prolif |
| Knstrn | 0 | 4.23506873 | 0.714 | 0.066 | 0 | 2 | Knstrn | myCAF_prolif |
| Ncapg2 | 0 | 4.37176095 | 0.717 | 0.07 | 0 | 2 | Ncapg2 | myCAF_prolif |
| Kif4 | 0 | 5.86617379 | 0.669 | 0.023 | 0 | 2 | Kif4 | myCAF_prolif |
| Cdk1 | 0 | 4.57989909 | 0.836 | 0.19 | 0 | 2 | Cdk1 | myCAF_prolif |
| Kif20a | 0 | 4.43574281 | 0.703 | 0.058 | 0 | 2 | Kif20a | myCAF_prolif |
| Ncaph | 0 | 5.72499811 | 0.672 | 0.028 | 0 | 2 | Ncaph | myCAF_prolif |
| Foxm1 | 0 | 4.38217985 | 0.718 | 0.074 | 0 | 2 | Foxm1 | myCAF_prolif |
| Lmnb1 | 0 | 3.52301875 | 0.913 | 0.271 | 0 | 2 | Lmnb1 | myCAF_prolif |
| Ndc80 | 0 | 7.09265035 | 0.649 | 0.01 | 0 | 2 | Ndc80 | myCAF_prolif |
| Kif22 | 0 | 5.7619165 | 0.658 | 0.021 | 0 | 2 | Kif22 | myCAF_prolif |
| Cenpm | 0 | 4.167365 | 0.694 | 0.058 | 0 | 2 | Cenpm | myCAF_prolif |
| Bub1b | 0 | 5.39828255 | 0.66 | 0.025 | 0 | 2 | Bub1b | myCAF_prolif |
| Ncapg | 0 | 6.42599454 | 0.647 | 0.013 | 0 | 2 | Ncapg | myCAF_prolif |
| Ckap2 | 0 | 4.33833347 | 0.711 | 0.078 | 0 | 2 | Ckap2 | myCAF_prolif |
| Ccnb2 | 0 | 3.53485688 | 0.735 | 0.104 | 0 | 2 | Ccnb2 | myCAF_prolif |
| Spag5 | 0 | 5.70140325 | 0.639 | 0.016 | 0 | 2 | Spag5 | myCAF_prolif |
| Tedc1 | 0 | 3.81435085 | 0.701 | 0.078 | 0 | 2 | Tedc1 | myCAF_prolif |
| Plk4 | 0 | 3.7067891 | 0.737 | 0.115 | 0 | 2 | Plk4 | myCAF_prolif |
| Kif11 | 0 | 6.61240699 | 0.634 | 0.012 | 0 | 2 | Kif11 | myCAF_prolif |
| Kif15 | 0 | 6.3324654 | 0.629 | 0.013 | 0 | 2 | Kif15 | myCAF_prolif |
| Nek2 | 0 | 6.51562449 | 0.628 | 0.013 | 0 | 2 | Nek2 | myCAF_prolif |
| Spc25 | 0 | 6.31240365 | 0.632 | 0.02 | 0 | 2 | Spc25 | myCAF_prolif |
| Hirip3 | 0 | 3.18761168 | 0.771 | 0.163 | 0 | 2 | Hirip3 | myCAF_prolif |
| Cenpe | 0 | 5.29304558 | 0.624 | 0.017 | 0 | 2 | Cenpe | myCAF_prolif |
| Tyms | 0 | 3.50645435 | 0.746 | 0.139 | 0 | 2 | Tyms | myCAF_prolif |
| Tmpo | 0 | 2.94663085 | 0.831 | 0.224 | 0 | 2 | Tmpo | myCAF_prolif |
| Gmnn | 0 | 4.46864779 | 0.67 | 0.064 | 0 | 2 | Gmnn | myCAF_prolif |
| Nuf2 | 0 | 5.29483062 | 0.63 | 0.024 | 0 | 2 | Nuf2 | myCAF_prolif |
| Mybl2 | 0 | 4.869417 | 0.642 | 0.036 | 0 | 2 | Mybl2 | myCAF_prolif |
| Kif23 | 0 | 5.15763715 | 0.636 | 0.031 | 0 | 2 | Kif23 | myCAF_prolif |
| BC030867 | 0 | 6.14544754 | 0.615 | 0.012 | 0 | 2 | BC030867 | myCAF_prolif |
| Cks2 | 0 | 3.14356073 | 0.796 | 0.196 | 0 | 2 | Cks2 | myCAF_prolif |
| Melk | 0 | 5.8733898 | 0.612 | 0.015 | 0 | 2 | Melk | myCAF_prolif |
| Shcbp1 | 0 | 5.68365606 | 0.606 | 0.014 | 0 | 2 | Shcbp1 | myCAF_prolif |
| Incenp | 0 | 4.15230033 | 0.688 | 0.096 | 0 | 2 | Incenp | myCAF_prolif |
| Cenpa | 0 | 4.01631545 | 0.649 | 0.065 | 0 | 2 | Cenpa | myCAF_prolif |
| Dnajc9 | 0 | 2.83861198 | 0.837 | 0.255 | 0 | 2 | Dnajc9 | myCAF_prolif |
| Pmf1 | 0 | 2.67292771 | 0.765 | 0.184 | 0 | 2 | Pmf1 | myCAF_prolif |
| Clspn | 0 | 4.99906576 | 0.615 | 0.034 | 0 | 2 | Clspn | myCAF_prolif |
| Kif18b | 0 | 7.53548038 | 0.587 | 0.006 | 0 | 2 | Kif18b | myCAF_prolif |
| Diaph3 | 0 | 4.2695176 | 0.631 | 0.052 | 0 | 2 | Diaph3 | myCAF_prolif |
| Ndc1 | 0 | 2.72247914 | 0.827 | 0.25 | 0 | 2 | Ndc1 | myCAF_prolif |
| Pole | 0 | 4.40053817 | 0.622 | 0.047 | 0 | 2 | Pole | myCAF_prolif |
| Fen1 | 0 | 3.22454478 | 0.759 | 0.184 | 0 | 2 | Fen1 | myCAF_prolif |
| Cep55 | 0 | 6.34773344 | 0.585 | 0.01 | 0 | 2 | Cep55 | myCAF_prolif |
| Ttk | 0 | 5.89840806 | 0.577 | 0.01 | 0 | 2 | Ttk | myCAF_prolif |
| Selenoh | 0 | 2.41442995 | 0.84 | 0.275 | 0 | 2 | Selenoh | myCAF_prolif |
| Mcm5 | 0 | 3.86044778 | 0.649 | 0.085 | 0 | 2 | Mcm5 | myCAF_prolif |
| Cenpn | 0 | 4.62953462 | 0.592 | 0.029 | 0 | 2 | Cenpn | myCAF_prolif |
| Dbf4 | 0 | 3.44419059 | 0.672 | 0.11 | 0 | 2 | Dbf4 | myCAF_prolif |
| Arhgap11a | 0 | 4.59195739 | 0.594 | 0.036 | 0 | 2 | Arhgap11a | myCAF_prolif |
| Rad51 | 0 | 5.73184509 | 0.577 | 0.019 | 0 | 2 | Rad51 | myCAF_prolif |
| Hist1h2ap | 0 | 6.52414424 | 0.646 | 0.089 | 0 | 2 | Hist1h2ap | myCAF_prolif |
| Cks1b | 0 | 2.68734954 | 0.763 | 0.207 | 0 | 2 | Cks1b | myCAF_prolif |
| Rad18 | 0 | 2.84165681 | 0.735 | 0.179 | 0 | 2 | Rad18 | myCAF_prolif |
| Depdc1a | 0 | 6.72477848 | 0.566 | 0.01 | 0 | 2 | Depdc1a | myCAF_prolif |
| Aurka | 0 | 5.98044842 | 0.573 | 0.018 | 0 | 2 | Aurka | myCAF_prolif |
| Cdca5 | 0 | 6.33817542 | 0.567 | 0.012 | 0 | 2 | Cdca5 | myCAF_prolif |
| Cdc45 | 0 | 3.884565 | 0.622 | 0.069 | 0 | 2 | Cdc45 | myCAF_prolif |
| Aurkb | 0 | 6.42232549 | 0.566 | 0.014 | 0 | 2 | Aurkb | myCAF_prolif |
| Chaf1a | 0 | 3.16530251 | 0.676 | 0.129 | 0 | 2 | Chaf1a | myCAF_prolif |
| Cenph | 0 | 5.41863892 | 0.567 | 0.021 | 0 | 2 | Cenph | myCAF_prolif |
| Atad2 | 0 | 3.37282059 | 0.665 | 0.123 | 0 | 2 | Atad2 | myCAF_prolif |
| Fanca | 0 | 5.08345878 | 0.562 | 0.021 | 0 | 2 | Fanca | myCAF_prolif |
| Iqgap3 | 0 | 5.22563898 | 0.563 | 0.029 | 0 | 2 | Iqgap3 | myCAF_prolif |
| Trip13 | 0 | 4.56899144 | 0.565 | 0.031 | 0 | 2 | Trip13 | myCAF_prolif |
| Dhfr | 0 | 3.54129344 | 0.635 | 0.104 | 0 | 2 | Dhfr | myCAF_prolif |
| Hist1h1b | 0 | 6.8272357 | 0.584 | 0.053 | 0 | 2 | Hist1h1b | myCAF_prolif |
| Cit | 0 | 3.07256973 | 0.652 | 0.124 | 0 | 2 | Cit | myCAF_prolif |
| Spdl1 | 0 | 5.6320905 | 0.545 | 0.017 | 0 | 2 | Spdl1 | myCAF_prolif |
| Pkmyt1 | 0 | 4.15285355 | 0.569 | 0.044 | 0 | 2 | Pkmyt1 | myCAF_prolif |
| Zwilch | 0 | 4.49050473 | 0.556 | 0.033 | 0 | 2 | Zwilch | myCAF_prolif |
| Espl1 | 0 | 6.11988294 | 0.531 | 0.01 | 0 | 2 | Espl1 | myCAF_prolif |
| Cenpf | 0 | 6.43960159 | 0.534 | 0.014 | 0 | 2 | Cenpf | myCAF_prolif |
| Knl1 | 0 | 6.40036504 | 0.528 | 0.008 | 0 | 2 | Knl1 | myCAF_prolif |
| Aspm | 0 | 6.1268375 | 0.53 | 0.012 | 0 | 2 | Aspm | myCAF_prolif |
| Bub1 | 0 | 5.82093395 | 0.53 | 0.012 | 0 | 2 | Bub1 | myCAF_prolif |
| Cenpw | 0 | 3.35233278 | 0.584 | 0.066 | 0 | 2 | Cenpw | myCAF_prolif |
| Rad51ap1 | 0 | 6.70919928 | 0.525 | 0.008 | 0 | 2 | Rad51ap1 | myCAF_prolif |
| Mis18a | 0 | 2.8143949 | 0.645 | 0.129 | 0 | 2 | Mis18a | myCAF_prolif |
| Parpbp | 0 | 5.2838219 | 0.535 | 0.022 | 0 | 2 | Parpbp | myCAF_prolif |
| Rangap1 | 0 | 2.49058865 | 0.873 | 0.36 | 0 | 2 | Rangap1 | myCAF_prolif |
| Gins2 | 0 | 2.61183066 | 0.699 | 0.186 | 0 | 2 | Gins2 | myCAF_prolif |
| Aaas | 0 | 2.21008452 | 0.767 | 0.256 | 0 | 2 | Aaas | myCAF_prolif |
| Ccdc34 | 0 | 2.54431512 | 0.703 | 0.195 | 0 | 2 | Ccdc34 | myCAF_prolif |
| Mcm7 | 0 | 2.89879419 | 0.719 | 0.212 | 0 | 2 | Mcm7 | myCAF_prolif |
| Haus4 | 0 | 2.29636357 | 0.722 | 0.216 | 0 | 2 | Haus4 | myCAF_prolif |
| Vrk1 | 0 | 2.38177559 | 0.68 | 0.178 | 0 | 2 | Vrk1 | myCAF_prolif |
| Kif20b | 0 | 5.45723606 | 0.513 | 0.011 | 0 | 2 | Kif20b | myCAF_prolif |
| Nusap1 | 0 | 6.36593592 | 0.516 | 0.015 | 0 | 2 | Nusap1 | myCAF_prolif |
| Cdca2 | 0 | 5.51948744 | 0.515 | 0.014 | 0 | 2 | Cdca2 | myCAF_prolif |
| Tcf19 | 0 | 3.36292165 | 0.711 | 0.21 | 0 | 2 | Tcf19 | myCAF_prolif |
| E2f8 | 0 | 3.94149241 | 0.546 | 0.051 | 0 | 2 | E2f8 | myCAF_prolif |
| Mcm3 | 0 | 3.37795214 | 0.642 | 0.148 | 0 | 2 | Mcm3 | myCAF_prolif |
| Topbp1 | 0 | 2.39152194 | 0.727 | 0.236 | 0 | 2 | Topbp1 | myCAF_prolif |
| Pola1 | 0 | 3.11497911 | 0.587 | 0.096 | 0 | 2 | Pola1 | myCAF_prolif |
| Rfc5 | 0 | 2.84194033 | 0.592 | 0.102 | 0 | 2 | Rfc5 | myCAF_prolif |
| E2f7 | 0 | 6.04967547 | 0.503 | 0.013 | 0 | 2 | E2f7 | myCAF_prolif |
| Chtf18 | 0 | 4.8695409 | 0.506 | 0.02 | 0 | 2 | Chtf18 | myCAF_prolif |
| Cdc25c | 0 | 5.208555 | 0.502 | 0.016 | 0 | 2 | Cdc25c | myCAF_prolif |
| Hist1h2bn | 0 | 5.53024534 | 0.561 | 0.076 | 0 | 2 | Hist1h2bn | myCAF_prolif |
| Sgo2a | 0 | 5.44130626 | 0.501 | 0.016 | 0 | 2 | Sgo2a | myCAF_prolif |
| Mastl | 0 | 5.24046564 | 0.501 | 0.018 | 0 | 2 | Mastl | myCAF_prolif |
| Rpa2 | 0 | 2.78384668 | 0.645 | 0.162 | 0 | 2 | Rpa2 | myCAF_prolif |
| Bora | 0 | 3.42306429 | 0.57 | 0.087 | 0 | 2 | Bora | myCAF_prolif |
| Smc4 | 0 | 2.40920684 | 0.808 | 0.329 | 0 | 2 | Smc4 | myCAF_prolif |
| Cenpi | 0 | 5.11166108 | 0.498 | 0.019 | 0 | 2 | Cenpi | myCAF_prolif |
| Haus6 | 0 | 2.49044059 | 0.608 | 0.131 | 0 | 2 | Haus6 | myCAF_prolif |
| Hells | 0 | 3.48218622 | 0.587 | 0.111 | 0 | 2 | Hells | myCAF_prolif |
| Pold1 | 0 | 2.4530193 | 0.656 | 0.184 | 0 | 2 | Pold1 | myCAF_prolif |
| Rad54l | 0 | 5.58463583 | 0.478 | 0.012 | 0 | 2 | Rad54l | myCAF_prolif |
| Ska2 | 0 | 2.34898701 | 0.682 | 0.217 | 0 | 2 | Ska2 | myCAF_prolif |
| Cep192 | 0 | 2.04123339 | 0.713 | 0.248 | 0 | 2 | Cep192 | myCAF_prolif |
| Lig1 | 0 | 2.69126888 | 0.678 | 0.217 | 0 | 2 | Lig1 | myCAF_prolif |
| Exosc8 | 0 | 2.00037653 | 0.799 | 0.338 | 0 | 2 | Exosc8 | myCAF_prolif |
| Rbl1 | 0 | 3.09392801 | 0.563 | 0.102 | 0 | 2 | Rbl1 | myCAF_prolif |
| Prim1 | 0 | 4.23150228 | 0.496 | 0.035 | 0 | 2 | Prim1 | myCAF_prolif |
| Hist1h2ae | 0 | 6.11042697 | 0.504 | 0.046 | 0 | 2 | Hist1h2ae | myCAF_prolif |
| Cdc25b | 0 | 3.48662154 | 0.577 | 0.12 | 0 | 2 | Cdc25b | myCAF_prolif |
| Nsl1 | 0 | 5.90507833 | 0.466 | 0.01 | 0 | 2 | Nsl1 | myCAF_prolif |
| Nup85 | 0 | 2.0912964 | 0.775 | 0.321 | 0 | 2 | Nup85 | myCAF_prolif |
| Usp1 | 0 | 2.0260772 | 0.763 | 0.31 | 0 | 2 | Usp1 | myCAF_prolif |
| Nup107 | 0 | 1.89512462 | 0.754 | 0.301 | 0 | 2 | Nup107 | myCAF_prolif |
| Mms22l | 0 | 3.28677937 | 0.523 | 0.072 | 0 | 2 | Mms22l | myCAF_prolif |
| Kpna2 | 0 | 3.4115173 | 0.633 | 0.184 | 0 | 2 | Kpna2 | myCAF_prolif |
| Suv39h1 | 0 | 1.76335415 | 0.734 | 0.285 | 0 | 2 | Suv39h1 | myCAF_prolif |
| Rrm1 | 0 | 2.68247278 | 0.902 | 0.454 | 0 | 2 | Rrm1 | myCAF_prolif |
| Ulbp1 | 0 | 2.80482909 | 0.589 | 0.141 | 0 | 2 | Ulbp1 | myCAF_prolif |
| Gins1 | 0 | 3.44984762 | 0.5 | 0.053 | 0 | 2 | Gins1 | myCAF_prolif |
| Blm | 0 | 3.57141768 | 0.491 | 0.044 | 0 | 2 | Blm | myCAF_prolif |
| Troap | 0 | 6.43261831 | 0.453 | 0.008 | 0 | 2 | Troap | myCAF_prolif |
| Hat1 | 0 | 2.23450912 | 0.701 | 0.256 | 0 | 2 | Hat1 | myCAF_prolif |
| Cep128 | 0 | 3.87688485 | 0.49 | 0.046 | 0 | 2 | Cep128 | myCAF_prolif |
| Cdkn2c | 0 | 2.77965246 | 0.704 | 0.26 | 0 | 2 | Cdkn2c | myCAF_prolif |
| Brip1 | 0 | 4.58062798 | 0.473 | 0.029 | 0 | 2 | Brip1 | myCAF_prolif |
| Lmnb2 | 0 | 2.76042749 | 0.536 | 0.093 | 0 | 2 | Lmnb2 | myCAF_prolif |
| Ska3 | 0 | 5.48081737 | 0.454 | 0.013 | 0 | 2 | Ska3 | myCAF_prolif |
| Slc25a10 | 0 | 1.67071265 | 0.74 | 0.3 | 0 | 2 | Slc25a10 | myCAF_prolif |
| Nasp | 0 | 2.0846897 | 0.811 | 0.374 | 0 | 2 | Nasp | myCAF_prolif |
| H2afx | 0 | 2.79154074 | 0.871 | 0.435 | 0 | 2 | H2afx | myCAF_prolif |
| Exo1 | 0 | 5.92742917 | 0.444 | 0.009 | 0 | 2 | Exo1 | myCAF_prolif |
| Fbxo5 | 0 | 4.37957061 | 0.461 | 0.026 | 0 | 2 | Fbxo5 | myCAF_prolif |
| Hmgb3 | 0 | 1.81330465 | 0.742 | 0.307 | 0 | 2 | Hmgb3 | myCAF_prolif |
| Mcm2 | 0 | 2.69017482 | 0.661 | 0.227 | 0 | 2 | Mcm2 | myCAF_prolif |
| Hist1h1a | 0 | 6.22536086 | 0.459 | 0.025 | 0 | 2 | Hist1h1a | myCAF_prolif |
| Rfc4 | 0 | 2.78932721 | 0.521 | 0.088 | 0 | 2 | Rfc4 | myCAF_prolif |
| Cmc2 | 0 | 1.88110765 | 0.683 | 0.252 | 0 | 2 | Cmc2 | myCAF_prolif |
| Nrm | 0 | 1.94956552 | 0.652 | 0.221 | 0 | 2 | Nrm | myCAF_prolif |
| Sgo1 | 0 | 5.44679468 | 0.444 | 0.013 | 0 | 2 | Sgo1 | myCAF_prolif |
| Rpa1 | 0 | 2.29105881 | 0.724 | 0.295 | 0 | 2 | Rpa1 | myCAF_prolif |
| Ercc6l | 0 | 4.51068934 | 0.453 | 0.024 | 0 | 2 | Ercc6l | myCAF_prolif |
| Stil | 0 | 4.68204164 | 0.451 | 0.024 | 0 | 2 | Stil | myCAF_prolif |
| Rcc1 | 0 | 1.85287743 | 0.856 | 0.43 | 0 | 2 | Rcc1 | myCAF_prolif |
| Kntc1 | 0 | 5.54980671 | 0.435 | 0.01 | 0 | 2 | Kntc1 | myCAF_prolif |
| Ezh2 | 0 | 2.01390366 | 0.627 | 0.202 | 0 | 2 | Ezh2 | myCAF_prolif |
| Zfp367 | 0 | 3.42063237 | 0.504 | 0.079 | 0 | 2 | Zfp367 | myCAF_prolif |
| Mcm6 | 0 | 2.74025774 | 0.696 | 0.272 | 0 | 2 | Mcm6 | myCAF_prolif |
| Fam111a | 0 | 1.99328321 | 0.628 | 0.205 | 0 | 2 | Fam111a | myCAF_prolif |
| Tinf2 | 0 | 1.92200127 | 0.644 | 0.222 | 0 | 2 | Tinf2 | myCAF_prolif |
| Sass6 | 0 | 3.18023831 | 0.479 | 0.059 | 0 | 2 | Sass6 | myCAF_prolif |
| Lin54 | 0 | 2.05013671 | 0.591 | 0.171 | 0 | 2 | Lin54 | myCAF_prolif |
| Mcm4 | 0 | 2.26373425 | 0.699 | 0.28 | 0 | 2 | Mcm4 | myCAF_prolif |
| Trim59 | 0 | 2.72761791 | 0.534 | 0.116 | 0 | 2 | Trim59 | myCAF_prolif |
| Mcm10 | 0 | 5.60105314 | 0.427 | 0.01 | 0 | 2 | Mcm10 | myCAF_prolif |
| Dut | 0 | 1.99303149 | 0.784 | 0.368 | 0 | 2 | Dut | myCAF_prolif |
| Haus8 | 0 | 2.32257832 | 0.534 | 0.12 | 0 | 2 | Haus8 | myCAF_prolif |
| Fignl1 | 0 | 4.33287502 | 0.441 | 0.027 | 0 | 2 | Fignl1 | myCAF_prolif |
| Ncapd3 | 0 | 2.18207975 | 0.575 | 0.162 | 0 | 2 | Ncapd3 | myCAF_prolif |
| Pold2 | 0 | 1.79832492 | 0.623 | 0.211 | 0 | 2 | Pold2 | myCAF_prolif |
| Cenpq | 0 | 3.35371806 | 0.458 | 0.048 | 0 | 2 | Cenpq | myCAF_prolif |
| Fanci | 0 | 3.60456832 | 0.44 | 0.031 | 0 | 2 | Fanci | myCAF_prolif |
| Nup37 | 0 | 1.74859297 | 0.686 | 0.28 | 0 | 2 | Nup37 | myCAF_prolif |
| Alg8 | 0 | 1.5861026 | 0.697 | 0.291 | 0 | 2 | Alg8 | myCAF_prolif |
| Sac3d1 | 0 | 1.43693341 | 0.713 | 0.314 | 0 | 2 | Sac3d1 | myCAF_prolif |
| Hist1h3a | 0 | 4.85697043 | 0.424 | 0.027 | 0 | 2 | Hist1h3a | myCAF_prolif |
| Ppil1 | 0 | 1.54037152 | 0.736 | 0.341 | 0 | 2 | Ppil1 | myCAF_prolif |
| Ccnf | 0 | 4.8995808 | 0.412 | 0.018 | 0 | 2 | Ccnf | myCAF_prolif |
| Wdhd1 | 0 | 3.45733461 | 0.439 | 0.046 | 0 | 2 | Wdhd1 | myCAF_prolif |
| Tipin | 0 | 1.86514729 | 0.756 | 0.364 | 0 | 2 | Tipin | myCAF_prolif |
| Cse1l | 0 | 1.72590142 | 0.79 | 0.399 | 0 | 2 | Cse1l | myCAF_prolif |
| Lsm3 | 0 | 1.49832526 | 0.778 | 0.393 | 0 | 2 | Lsm3 | myCAF_prolif |

| | | | | | | | | |
|---|---|---|---|---|---|---|---|---|
| Ccne1 | 0 | 2.70853995 | 0.523 | 0.139 | 0 | 2 | Ccne1 | myCAF_prolif |
| Pcnt | 0 | 1.52850287 | 0.661 | 0.277 | 0 | 2 | Pcnt | myCAF_prolif |
| Tubg1 | 0 | 1.69731096 | 0.74 | 0.356 | 0 | 2 | Tubg1 | myCAF_prolif |
| Reep4 | 0 | 2.06449772 | 0.747 | 0.364 | 0 | 2 | Reep4 | myCAF_prolif |
| Cenpl | 0 | 3.43102496 | 0.437 | 0.058 | 0 | 2 | Cenpl | myCAF_prolif |
| Rfc3 | 0 | 2.31682172 | 0.509 | 0.13 | 0 | 2 | Rfc3 | myCAF_prolif |
| Skp2 | 0 | 2.31401168 | 0.508 | 0.13 | 0 | 2 | Skp2 | myCAF_prolif |
| Nup155 | 0 | 1.53361443 | 0.693 | 0.315 | 0 | 2 | Nup155 | myCAF_prolif |
| Pola2 | 0 | 2.4299727 | 0.509 | 0.132 | 0 | 2 | Pola2 | myCAF_prolif |
| Lin9 | 0 | 2.719469 | 0.435 | 0.059 | 0 | 2 | Lin9 | myCAF_prolif |
| Wee1 | 0 | 2.46054816 | 0.483 | 0.108 | 0 | 2 | Wee1 | myCAF_prolif |
| Donson | 0 | 1.86497113 | 0.549 | 0.176 | 0 | 2 | Donson | myCAF_prolif |
| Dctpp1 | 0 | 1.98390292 | 0.527 | 0.154 | 0 | 2 | Dctpp1 | myCAF_prolif |
| Hist1h1d | 0 | 5.37905255 | 0.394 | 0.021 | 0 | 2 | Hist1h1d | myCAF_prolif |
| Prim2 | 0 | 2.05559976 | 0.505 | 0.133 | 0 | 2 | Prim2 | myCAF_prolif |
| Rfwd3 | 0 | 1.65671179 | 0.599 | 0.227 | 0 | 2 | Rfwd3 | myCAF_prolif |
| Psat1 | 0 | 1.6722774 | 0.579 | 0.207 | 0 | 2 | Psat1 | myCAF_prolif |
| Ankle1 | 0 | 5.72273902 | 0.382 | 0.013 | 0 | 2 | Ankle1 | myCAF_prolif |
| Mis18bp1 | 0 | 5.72636395 | 0.378 | 0.011 | 0 | 2 | Mis18bp1 | myCAF_prolif |
| Cdc7 | 0 | 2.52750786 | 0.466 | 0.1 | 0 | 2 | Cdc7 | myCAF_prolif |
| Mad1l1 | 0 | 1.64479816 | 0.613 | 0.247 | 0 | 2 | Mad1l1 | myCAF_prolif |
| Cdkn2aipnl | 0 | 1.41197247 | 0.625 | 0.259 | 0 | 2 | Cdkn2aipnl | myCAF_prolif |
| Mtfr2 | 0 | 3.28787743 | 0.403 | 0.038 | 0 | 2 | Mtfr2 | myCAF_prolif |
| Poc1a | 0 | 2.87618627 | 0.422 | 0.057 | 0 | 2 | Poc1a | myCAF_prolif |
| Polh | 0 | 2.03836188 | 0.499 | 0.134 | 0 | 2 | Polh | myCAF_prolif |
| Cdkn3 | 0 | 4.03847683 | 0.391 | 0.027 | 0 | 2 | Cdkn3 | myCAF_prolif |
| Nde1 | 0 | 1.6354986 | 0.71 | 0.346 | 0 | 2 | Nde1 | myCAF_prolif |
| Lsm2 | 0 | 1.72721169 | 0.851 | 0.487 | 0 | 2 | Lsm2 | myCAF_prolif |
| Arhgap19 | 0 | 4.41761421 | 0.385 | 0.021 | 0 | 2 | Arhgap19 | myCAF_prolif |
| Nup160 | 0 | 1.38995707 | 0.601 | 0.239 | 0 | 2 | Nup160 | myCAF_prolif |
| Jpt2 | 0 | 1.54901849 | 0.799 | 0.44 | 0 | 2 | Jpt2 | myCAF_prolif |
| Ccsap | 0 | 1.91924024 | 0.492 | 0.133 | 0 | 2 | Ccsap | myCAF_prolif |
| Ube2t | 0 | 3.31093734 | 0.399 | 0.041 | 0 | 2 | Ube2t | myCAF_prolif |
| Fam83d | 0 | 4.48489882 | 0.38 | 0.022 | 0 | 2 | Fam83d | myCAF_prolif |
| Dck | 0 | 2.77487943 | 0.441 | 0.083 | 0 | 2 | Dck | myCAF_prolif |
| Cdk5rap2 | 0 | 1.74496918 | 0.544 | 0.187 | 0 | 2 | Cdk5rap2 | myCAF_prolif |
| Eri1 | 0 | 1.42623199 | 0.68 | 0.324 | 0 | 2 | Eri1 | myCAF_prolif |
| Efcab11 | 0 | 4.81369449 | 0.375 | 0.019 | 0 | 2 | Efcab11 | myCAF_prolif |
| Slfn9 | 0 | 4.41086412 | 0.372 | 0.019 | 0 | 2 | Slfn9 | myCAF_prolif |
| Haus1 | 0 | 1.48016343 | 0.565 | 0.212 | 0 | 2 | Haus1 | myCAF_prolif |
| Lbr | 0 | 1.45255436 | 0.824 | 0.472 | 0 | 2 | Lbr | myCAF_prolif |
| Gins4 | 0 | 1.3488523 | 0.693 | 0.341 | 0 | 2 | Gins4 | myCAF_prolif |
| Atad5 | 0 | 3.35268399 | 0.397 | 0.046 | 0 | 2 | Atad5 | myCAF_prolif |
| Chaf1b | 0 | 4.02502336 | 0.381 | 0.03 | 0 | 2 | Chaf1b | myCAF_prolif |
| Mdc1 | 0 | 1.54103343 | 0.573 | 0.222 | 0 | 2 | Mdc1 | myCAF_prolif |
| Orc6 | 0 | 1.83264021 | 0.547 | 0.197 | 0 | 2 | Orc6 | myCAF_prolif |
| Gstcd | 0 | 1.49227449 | 0.57 | 0.221 | 0 | 2 | Gstcd | myCAF_prolif |
| Terf1 | 0 | 2.07510338 | 0.538 | 0.191 | 0 | 2 | Terf1 | myCAF_prolif |
| Rpa3 | 0 | 1.21122732 | 0.649 | 0.302 | 0 | 2 | Rpa3 | myCAF_prolif |
| Syce2 | 0 | 2.1180116 | 0.451 | 0.104 | 0 | 2 | Syce2 | myCAF_prolif |
| Cep295 | 0 | 1.83034758 | 0.527 | 0.18 | 0 | 2 | Cep295 | myCAF_prolif |
| Dnmt1 | 0 | 1.96480833 | 0.808 | 0.461 | 0 | 2 | Dnmt1 | myCAF_prolif |
| Cenps | 0 | 4.04377547 | 0.374 | 0.028 | 0 | 2 | Cenps | myCAF_prolif |
| Fancd2 | 0 | 3.92971869 | 0.368 | 0.022 | 0 | 2 | Fancd2 | myCAF_prolif |
| Mre11a | 0 | 1.40314828 | 0.56 | 0.216 | 0 | 2 | Mre11a | myCAF_prolif |
| Umps | 0 | 1.28132781 | 0.691 | 0.347 | 0 | 2 | Umps | myCAF_prolif |
| Xrcc1 | 0 | 1.19480287 | 0.683 | 0.34 | 0 | 2 | Xrcc1 | myCAF_prolif |
| Rnaseh2b | 0 | 1.31714166 | 0.585 | 0.242 | 0 | 2 | Rnaseh2b | myCAF_prolif |
| Rbmx | 0 | 1.32933374 | 0.67 | 0.328 | 0 | 2 | Rbmx | myCAF_prolif |
| Uchl5 | 0 | 1.31371349 | 0.587 | 0.245 | 0 | 2 | Uchl5 | myCAF_prolif |
| Mtbp | 0 | 1.8059842 | 0.461 | 0.12 | 0 | 2 | Mtbp | myCAF_prolif |
| Rfc2 | 0 | 1.6550615 | 0.801 | 0.461 | 0 | 2 | Rfc2 | myCAF_prolif |
| Gas2l3 | 0 | 3.34603303 | 0.392 | 0.052 | 0 | 2 | Gas2l3 | myCAF_prolif |
| Cdk2 | 0 | 1.70835452 | 0.717 | 0.377 | 0 | 2 | Cdk2 | myCAF_prolif |
| Dsn1 | 0 | 2.20390032 | 0.434 | 0.095 | 0 | 2 | Dsn1 | myCAF_prolif |
| Nup133 | 0 | 1.23002289 | 0.587 | 0.248 | 0 | 2 | Nup133 | myCAF_prolif |
| Larp7 | 0 | 1.16169016 | 0.718 | 0.38 | 0 | 2 | Larp7 | myCAF_prolif |
| Rbbp8 | 0 | 1.47600605 | 0.567 | 0.229 | 0 | 2 | Rbbp8 | myCAF_prolif |
| Hpf1 | 0 | 1.34980111 | 0.664 | 0.327 | 0 | 2 | Hpf1 | myCAF_prolif |
| Kif18a | 0 | 4.21805227 | 0.359 | 0.022 | 0 | 2 | Kif18a | myCAF_prolif |
| Mthfd2 | 0 | 1.35083 | 0.709 | 0.372 | 0 | 2 | Mthfd2 | myCAF_prolif |
| Anp32b | 0 | 1.15857176 | 0.727 | 0.391 | 0 | 2 | Anp32b | myCAF_prolif |
| Haus7 | 0 | 1.35578859 | 0.66 | 0.325 | 0 | 2 | Haus7 | myCAF_prolif |
| Gins3 | 0 | 2.94526612 | 0.382 | 0.048 | 0 | 2 | Gins3 | myCAF_prolif |
| Tcof1 | 0 | 1.32575156 | 0.751 | 0.418 | 0 | 2 | Tcof1 | myCAF_prolif |
| Nxt1 | 0 | 1.48401378 | 0.532 | 0.2 | 0 | 2 | Nxt1 | myCAF_prolif |
| Ifrd2 | 0 | 1.32436772 | 0.689 | 0.357 | 0 | 2 | Ifrd2 | myCAF_prolif |
| Tfdp2 | 0 | 1.70985062 | 0.501 | 0.17 | 0 | 2 | Tfdp2 | myCAF_prolif |
| Pole2 | 0 | 1.8575134 | 0.477 | 0.146 | 0 | 2 | Pole2 | myCAF_prolif |
| Ckap5 | 0 | 1.71094728 | 0.818 | 0.488 | 0 | 2 | Ckap5 | myCAF_prolif |
| Traip | 0 | 4.14534179 | 0.354 | 0.025 | 0 | 2 | Traip | myCAF_prolif |
| Nedd1 | 0 | 1.10727844 | 0.672 | 0.344 | 0 | 2 | Nedd1 | myCAF_prolif |
| Lnpk | 0 | 1.13504548 | 0.744 | 0.416 | 0 | 2 | Lnpk | myCAF_prolif |
| Smchd1 | 0 | 1.15409028 | 0.747 | 0.419 | 0 | 2 | Smchd1 | myCAF_prolif |
| Lsm6 | 0 | 1.25405938 | 0.593 | 0.265 | 0 | 2 | Lsm6 | myCAF_prolif |
| Cdt1 | 0 | 2.06097835 | 0.52 | 0.193 | 0 | 2 | Cdt1 | myCAF_prolif |
| Bok | 0 | 1.46111866 | 0.515 | 0.19 | 0 | 2 | Bok | myCAF_prolif |
| Lyar | 0 | 1.2146713 | 0.711 | 0.386 | 0 | 2 | Lyar | myCAF_prolif |
| Ccne2 | 0 | 4.33344378 | 0.352 | 0.03 | 0 | 2 | Ccne2 | myCAF_prolif |
| Pole3 | 0 | 1.14886113 | 0.718 | 0.396 | 0 | 2 | Pole3 | myCAF_prolif |
| Mcph1 | 0 | 1.58910604 | 0.489 | 0.167 | 0 | 2 | Mcph1 | myCAF_prolif |
| Bard1 | 0 | 4.48405279 | 0.335 | 0.014 | 0 | 2 | Bard1 | myCAF_prolif |
| Cdkn2d | 0 | 1.88359268 | 0.659 | 0.338 | 0 | 2 | Cdkn2d | myCAF_prolif |
| Nsd2 | 0 | 1.63428162 | 0.861 | 0.54 | 0 | 2 | Nsd2 | myCAF_prolif |
| Gemin6 | 0 | 2.02185872 | 0.428 | 0.108 | 0 | 2 | Gemin6 | myCAF_prolif |
| G2e3 | 0 | 1.6893257 | 0.574 | 0.255 | 0 | 2 | G2e3 | myCAF_prolif |
| Nup205 | 0 | 1.34220193 | 0.547 | 0.23 | 0 | 2 | Nup205 | myCAF_prolif |
| Slc43a3 | 0 | 0.7415141 | 0.813 | 0.496 | 0 | 2 | Slc43a3 | myCAF_prolif |
| Cbx5 | 0 | 1.62015548 | 0.81 | 0.494 | 0 | 2 | Cbx5 | myCAF_prolif |
| Metrn | 0 | 1.26059049 | 0.534 | 0.218 | 0 | 2 | Metrn | myCAF_prolif |
| Pask | 0 | 3.68744247 | 0.347 | 0.031 | 0 | 2 | Pask | myCAF_prolif |
| Chek1 | 0 | 2.14986628 | 0.403 | 0.089 | 0 | 2 | Chek1 | myCAF_prolif |
| Nptxr | 0 | 1.27428736 | 0.515 | 0.201 | 0 | 2 | Nptxr | myCAF_prolif |
| Psmc3ip | 0 | 3.80203913 | 0.339 | 0.027 | 0 | 2 | Psmc3ip | myCAF_prolif |
| Aasdl | 0 | 1.02961691 | 0.739 | 0.427 | 0 | 2 | Aasdl | myCAF_prolif |
| Fancm | 0 | 2.03129877 | 0.398 | 0.087 | 0 | 2 | Fancm | myCAF_prolif |
| Dcaf15 | 0 | 1.21073885 | 0.676 | 0.367 | 0 | 2 | Dcaf15 | myCAF_prolif |
| Cntln | 0 | 1.47167863 | 0.6 | 0.291 | 0 | 2 | Cntln | myCAF_prolif |
| Ncaph2 | 0 | 1.06448394 | 0.772 | 0.463 | 0 | 2 | Ncaph2 | myCAF_prolif |
| Gpsm2 | 0 | 1.29912508 | 0.663 | 0.355 | 0 | 2 | Gpsm2 | myCAF_prolif |
| Ruvbl2 | 0 | 1.1179728 | 0.718 | 0.41 | 0 | 2 | Ruvbl2 | myCAF_prolif |
| Cnot9 | 0 | 1.17453301 | 0.701 | 0.395 | 0 | 2 | Cnot9 | myCAF_prolif |
| Cntrob | 0 | 1.96723307 | 0.441 | 0.136 | 0 | 2 | Cntrob | myCAF_prolif |
| Timm50 | 0 | 1.11942385 | 0.778 | 0.474 | 0 | 2 | Timm50 | myCAF_prolif |
| Rnaseh2c | 0 | 1.0767554 | 0.81 | 0.507 | 0 | 2 | Rnaseh2c | myCAF_prolif |
| Snrnp40 | 0 | 0.99661365 | 0.75 | 0.447 | 0 | 2 | Snrnp40 | myCAF_prolif |
| Nsmce4a | 0 | 1.21967261 | 0.781 | 0.479 | 0 | 2 | Nsmce4a | myCAF_prolif |
| Nucks1 | 0 | 1.62822937 | 0.843 | 0.541 | 0 | 2 | Nucks1 | myCAF_prolif |
| Hspa14 | 0 | 1.08813585 | 0.793 | 0.492 | 0 | 2 | Hspa14 | myCAF_prolif |
| H2afz | 0 | 1.95304155 | 0.889 | 0.588 | 0 | 2 | H2afz | myCAF_prolif |
| Rad21 | 0 | 1.80444458 | 0.879 | 0.578 | 0 | 2 | Rad21 | myCAF_prolif |
| Myef2 | 0 | 1.31663443 | 0.808 | 0.508 | 0 | 2 | Myef2 | myCAF_prolif |
| Zgrf1 | 0 | 3.32866388 | 0.337 | 0.037 | 0 | 2 | Zgrf1 | myCAF_prolif |
| Snrpa1 | 0 | 1.07428779 | 0.778 | 0.478 | 0 | 2 | Snrpa1 | myCAF_prolif |
| Cenpc1 | 0 | 1.20399507 | 0.622 | 0.323 | 0 | 2 | Cenpc1 | myCAF_prolif |
| Wdr90 | 0 | 2.31325858 | 0.373 | 0.074 | 0 | 2 | Wdr90 | myCAF_prolif |
| Paxip1 | 0 | 1.25377394 | 0.66 | 0.362 | 0 | 2 | Paxip1 | myCAF_prolif |
| Pfas | 0 | 1.31786238 | 0.527 | 0.23 | 0 | 2 | Pfas | myCAF_prolif |
| Pelp1 | 0 | 0.85668856 | 0.715 | 0.419 | 0 | 2 | Pelp1 | myCAF_prolif |
| Mis12 | 0 | 1.4001938 | 0.484 | 0.188 | 0 | 2 | Mis12 | myCAF_prolif |
| Ppp2r5d | 0 | 1.05149126 | 0.763 | 0.467 | 0 | 2 | Ppp2r5d | myCAF_prolif |
| Arpin | 0 | 1.07396415 | 0.726 | 0.431 | 0 | 2 | Arpin | myCAF_prolif |
| Ddx39 | 0 | 1.07147208 | 0.814 | 0.519 | 0 | 2 | Ddx39 | myCAF_prolif |
| Dek | 0 | 1.7188733 | 0.891 | 0.597 | 0 | 2 | Dek | myCAF_prolif |
| Eif1ad | 0 | 1.00197792 | 0.708 | 0.415 | 0 | 2 | Eif1ad | myCAF_prolif |
| Dkc1 | 0 | 1.2156904 | 0.567 | 0.276 | 0 | 2 | Dkc1 | myCAF_prolif |
| Timm8a1 | 0 | 1.10389231 | 0.539 | 0.249 | 0 | 2 | Timm8a1 | myCAF_prolif |
| Dtl | 0 | 4.21025383 | 0.315 | 0.025 | 0 | 2 | Dtl | myCAF_prolif |
| Anp32e | 0 | 1.67344182 | 0.92 | 0.631 | 0 | 2 | Anp32e | myCAF_prolif |
| Suz12 | 0 | 0.95132379 | 0.649 | 0.36 | 0 | 2 | Suz12 | myCAF_prolif |
| Pop4 | 0 | 0.99169522 | 0.648 | 0.36 | 0 | 2 | Pop4 | myCAF_prolif |
| Timeless | 0 | 3.06690638 | 0.32 | 0.033 | 0 | 2 | Timeless | myCAF_prolif |
| G3bp1 | 0 | 0.93548289 | 0.805 | 0.518 | 0 | 2 | G3bp1 | myCAF_prolif |
| Neurl1b | 0 | 2.76773716 | 0.353 | 0.066 | 0 | 2 | Neurl1b | myCAF_prolif |
| Ssrp1 | 0 | 1.26971654 | 0.804 | 0.518 | 0 | 2 | Ssrp1 | myCAF_prolif |
| Nup35 | 0 | 1.32042828 | 0.481 | 0.195 | 0 | 2 | Nup35 | myCAF_prolif |
| Arhgef39 | 0 | 4.46911635 | 0.304 | 0.018 | 0 | 2 | Arhgef39 | myCAF_prolif |
| Hjurp | 0 | 2.12449114 | 0.916 | 0.631 | 0 | 2 | Hjurp | myCAF_prolif |
| Rhno1 | 0 | 1.03377834 | 0.704 | 0.422 | 0 | 2 | Rhno1 | myCAF_prolif |
| Cdc6 | 0 | 3.86864698 | 0.311 | 0.029 | 0 | 2 | Cdc6 | myCAF_prolif |
| Nop58 | 0 | 1.066877 | 0.784 | 0.504 | 0 | 2 | Nop58 | myCAF_prolif |
| Nudt5 | 0 | 0.93785016 | 0.724 | 0.444 | 0 | 2 | Nudt5 | myCAF_prolif |
| Parp1 | 0 | 1.17958015 | 0.653 | 0.374 | 0 | 2 | Parp1 | myCAF_prolif |
| Prpf31 | 0 | 0.89914102 | 0.698 | 0.419 | 0 | 2 | Prpf31 | myCAF_prolif |
| Dtymk | 0 | 1.47083026 | 0.862 | 0.584 | 0 | 2 | Dtymk | myCAF_prolif |
| Snrpd1 | 0 | 1.46854644 | 0.861 | 0.583 | 0 | 2 | Snrpd1 | myCAF_prolif |
| Lsm8 | 0 | 1.25758116 | 0.807 | 0.53 | 0 | 2 | Lsm8 | myCAF_prolif |
| Khdrbs3 | 0 | 0.9127494 | 0.653 | 0.379 | 0 | 2 | Khdrbs3 | myCAF_prolif |
| Tnfaip61 | 0 | 0.66491253 | 0.649 | 0.378 | 0 | 2 | Tnfaip6 | myCAF_prolif |
| Mthfd1 | 0 | 1.1174632 | 0.516 | 0.245 | 0 | 2 | Mthfd1 | myCAF_prolif |
| DEPRECATED-l | 0 | 0.70159345 | 0.696 | 0.425 | 0 | 2 | DEPRECATED-l | myCAF_prolif |
| Ilf2 | 0 | 1.06688368 | 0.818 | 0.547 | 0 | 2 | Ilf2 | myCAF_prolif |
| Tmx2 | 0 | 0.76612709 | 0.687 | 0.417 | 0 | 2 | Tmx2 | myCAF_prolif |
| Banf1 | 0 | 0.85906452 | 0.701 | 0.431 | 0 | 2 | Banf1 | myCAF_prolif |
| Gle1 | 0 | 0.878496 | 0.734 | 0.465 | 0 | 2 | Gle1 | myCAF_prolif |
| Hist1h4i | 0 | 3.18194951 | 0.319 | 0.051 | 0 | 2 | Hist1h4i | myCAF_prolif |
| Psip1 | 0 | 1.5258768 | 0.904 | 0.639 | 0 | 2 | Psip1 | myCAF_prolif |
| Grk6 | 0 | 0.93221484 | 0.768 | 0.503 | 0 | 2 | Grk6 | myCAF_prolif |
| Pbdc1 | 0 | 0.8722498 | 0.768 | 0.503 | 0 | 2 | Pbdc1 | myCAF_prolif |
| Med30 | 0 | 0.92746173 | 0.733 | 0.468 | 0 | 2 | Med30 | myCAF_prolif |
| Phf5a | 0 | 0.85456783 | 0.787 | 0.523 | 0 | 2 | Phf5a | myCAF_prolif |
| Thop1 | 0 | 0.77311728 | 0.68 | 0.418 | 0 | 2 | Thop1 | myCAF_prolif |
| Slc7a5 | 0 | 0.67175929 | 0.753 | 0.491 | 0 | 2 | Slc7a5 | myCAF_prolif |
| Dpy30 | 0 | 0.86918342 | 0.669 | 0.408 | 0 | 2 | Dpy30 | myCAF_prolif |
| E2f1 | 0 | 1.47680469 | 0.508 | 0.247 | 0 | 2 | E2f1 | myCAF_prolif |
| Tubb2b | 0 | 0.84450196 | 0.712 | 0.451 | 0 | 2 | Tubb2b | myCAF_prolif |
| Ppp1r8 | 0 | 0.81180639 | 0.68 | 0.42 | 0 | 2 | Ppp1r8 | myCAF_prolif |
| Pa2g4 | 0 | 1.02690508 | 0.728 | 0.469 | 0 | 2 | Pa2g4 | myCAF_prolif |
| Ndufaf2 | 0 | 0.85622694 | 0.751 | 0.492 | 0 | 2 | Ndufaf2 | myCAF_prolif |
| Actl6a | 0 | 0.84110143 | 0.696 | 0.437 | 0 | 2 | Actl6a | myCAF_prolif |
| Arl6ip1 | 0 | 1.71190262 | 0.562 | 0.305 | 0 | 2 | Arl6ip1 | myCAF_prolif |
| Bub3 | 0 | 1.25338469 | 0.734 | 0.477 | 0 | 2 | Bub3 | myCAF_prolif |
| Ubr7 | 0 | 0.82851797 | 0.666 | 0.41 | 0 | 2 | Ubr7 | myCAF_prolif |
| Pcna | 0 | 2.10786835 | 0.83 | 0.575 | 0 | 2 | Pcna | myCAF_prolif |
| U2af1 | 0 | 0.76803194 | 0.756 | 0.501 | 0 | 2 | U2af1 | myCAF_prolif |
| Ttc9c | 0 | 0.87671257 | 0.716 | 0.462 | 0 | 2 | Ttc9c | myCAF_prolif |
| Gjc1 | 0 | 0.73981847 | 0.691 | 0.438 | 0 | 2 | Gjc1 | myCAF_prolif |
| Phip | 0 | 0.85709413 | 0.736 | 0.484 | 0 | 2 | Phip | myCAF_prolif |
| Tmem97 | 0 | 0.98708476 | 0.823 | 0.571 | 0 | 2 | Tmem97 | myCAF_prolif |
| Brd8 | 0 | 1.24809398 | 0.804 | 0.552 | 0 | 2 | Brd8 | myCAF_prolif |
| Sephs1 | 0 | 0.95374905 | 0.796 | 0.545 | 0 | 2 | Sephs1 | myCAF_prolif |
| Fst1 | 0 | 0.84879511 | 0.761 | 0.51 | 0 | 2 | Fst | myCAF_prolif |
| Smc3 | 0 | 0.90618908 | 0.741 | 0.49 | 0 | 2 | Smc3 | myCAF_prolif |
| Cacybp | 0 | 0.8197464 | 0.731 | 0.481 | 0 | 2 | Cacybp | myCAF_prolif |
| Cdca4 | 0 | 1.11276189 | 0.775 | 0.525 | 0 | 2 | Cdca4 | myCAF_prolif |
| Ctcf | 0 | 0.88514328 | 0.73 | 0.481 | 0 | 2 | Ctcf | myCAF_prolif |
| Nop56 | 0 | 0.93408188 | 0.785 | 0.537 | 0 | 2 | Nop56 | myCAF_prolif |
| Tubb3 | 0 | 1.40334668 | 0.413 | 0.166 | 0 | 2 | Tubb3 | myCAF_prolif |
| Rbm15 | 0 | 0.98334504 | 0.706 | 0.459 | 0 | 2 | Rbm15 | myCAF_prolif |
| Has21 | 0 | 0.95316227 | 0.73 | 0.485 | 0 | 2 | Has2 | myCAF_prolif |
| Fkbp51 | 0 | 0.79377481 | 0.724 | 0.479 | 0 | 2 | Fkbp5 | myCAF_prolif |
| Sf3a3 | 0 | 0.84493746 | 0.736 | 0.492 | 0 | 2 | Sf3a3 | myCAF_prolif |
| Cep89 | 0 | 1.89828072 | 0.383 | 0.139 | 0 | 2 | Cep89 | myCAF_prolif |
| Ripk3 | 0 | 0.87348288 | 0.842 | 0.598 | 0 | 2 | Ripk3 | myCAF_prolif |
| Wbp11 | 0 | 0.80824737 | 0.735 | 0.492 | 0 | 2 | Wbp11 | myCAF_prolif |
| Nsg11 | 0 | 0.5561042 | 0.801 | 0.559 | 0 | 2 | Nsg1 | myCAF_prolif |
| Supt16 | 0 | 0.94868032 | 0.849 | 0.607 | 0 | 2 | Supt16 | myCAF_prolif |
| Dio31 | 0 | 0.87851739 | 0.578 | 0.336 | 0 | 2 | Dio3 | myCAF_prolif |
| Mzt1 | 0 | 0.76804432 | 0.708 | 0.466 | 0 | 2 | Mzt1 | myCAF_prolif |
| Pnn | 0 | 0.88971328 | 0.797 | 0.557 | 0 | 2 | Pnn | myCAF_prolif |
| Nubp1 | 0 | 0.80111676 | 0.784 | 0.544 | 0 | 2 | Nubp1 | myCAF_prolif |
| Ncbp1 | 0 | 0.70168722 | 0.699 | 0.459 | 0 | 2 | Ncbp1 | myCAF_prolif |
| Prelid3b | 0 | 0.78483396 | 0.656 | 0.417 | 0 | 2 | Prelid3b | myCAF_prolif |
| Shmt2 | 0 | 1.20192027 | 0.901 | 0.663 | 0 | 2 | Shmt2 | myCAF_prolif |
| Nt5dc2 | 0 | 1.20664083 | 0.897 | 0.659 | 0 | 2 | Nt5dc2 | myCAF_prolif |
| Fzr1 | 0 | 1.30627175 | 0.725 | 0.488 | 0 | 2 | Fzr1 | myCAF_prolif |
| Nin | 0 | 0.84397742 | 0.731 | 0.494 | 0 | 2 | Nin | myCAF_prolif |
| Alad | 0 | 0.78943242 | 0.728 | 0.492 | 0 | 2 | Alad | myCAF_prolif |
| H1fx1 | 0 | 0.92016602 | 0.622 | 0.386 | 0 | 2 | H1fx | myCAF_prolif |
| Cinp | 0 | 0.80968351 | 0.768 | 0.532 | 0 | 2 | Cinp | myCAF_prolif |
| Hist1h1e | 0 | 3.2671643 | 0.755 | 0.52 | 0 | 2 | Hist1h1e | myCAF_prolif |
| Gar1 | 0 | 0.74014253 | 0.776 | 0.542 | 0 | 2 | Gar1 | myCAF_prolif |
| Psmd14 | 0 | 0.76454176 | 0.776 | 0.543 | 0 | 2 | Psmd14 | myCAF_prolif |
| Cdca7 | 0 | 2.01878658 | 0.327 | 0.095 | 0 | 2 | Cdca7 | myCAF_prolif |
| Snrpe | 0 | 0.79499072 | 0.783 | 0.551 | 0 | 2 | Snrpe | myCAF_prolif |
| Tsc41 | 0 | 0.68524429 | 0.739 | 0.507 | 0 | 2 | Tsc4 | myCAF_prolif |
| Kmt5a | 0 | 1.03342124 | 0.842 | 0.611 | 0 | 2 | Kmt5a | myCAF_prolif |
| Srrt | 0 | 0.93116907 | 0.828 | 0.598 | 0 | 2 | Srrt | myCAF_prolif |
| Xpo1 | 0 | 1.10489423 | 0.825 | 0.596 | 0 | 2 | Xpo1 | myCAF_prolif |
| Gart | 0 | 0.76573198 | 0.698 | 0.469 | 0 | 2 | Gart | myCAF_prolif |
| Odf2 | 0 | 1.236072 | 0.806 | 0.578 | 0 | 2 | Odf2 | myCAF_prolif |
| Kctd9 | 0 | 0.71516922 | 0.68 | 0.452 | 0 | 2 | Kctd9 | myCAF_prolif |
| Cdr2 | 0 | 0.66633936 | 0.745 | 0.517 | 0 | 2 | Cdr2 | myCAF_prolif |
| Rfc1 | 0 | 0.97478142 | 0.62 | 0.394 | 0 | 2 | Rfc1 | myCAF_prolif |
| Cad | 0 | 0.67043852 | 0.813 | 0.588 | 0 | 2 | Cad | myCAF_prolif |
| Nup62 | 0 | 0.74371942 | 0.756 | 0.531 | 0 | 2 | Nup62 | myCAF_prolif |
| Nme1 | 0 | 0.88172605 | 0.846 | 0.621 | 0 | 2 | Nme1 | myCAF_prolif |
| Baz1b | 0 | 0.90611994 | 0.808 | 0.585 | 0 | 2 | Baz1b | myCAF_prolif |
| Ube2s | 0 | 1.77589061 | 0.917 | 0.694 | 0 | 2 | Ube2s | myCAF_prolif |
| Slbp | 0 | 1.28801594 | 0.775 | 0.554 | 0 | 2 | Slbp | myCAF_prolif |
| Tent4a | 0 | 0.76409098 | 0.722 | 0.501 | 0 | 2 | Tent4a | myCAF_prolif |
| Cox7a2 | 0 | 0.75986022 | 0.796 | 0.577 | 0 | 2 | Cox7a2 | myCAF_prolif |
| Cdk2ap1 | 0 | 0.70992626 | 0.815 | 0.598 | 0 | 2 | Cdk2ap1 | myCAF_prolif |
| Kpnb1 | 0 | 0.9639128 | 0.847 | 0.631 | 0 | 2 | Kpnb1 | myCAF_prolif |
| Snrpg | 0 | 0.70735545 | 0.787 | 0.572 | 0 | 2 | Snrpg | myCAF_prolif |
| Lsm4 | 0 | 0.80881806 | 0.857 | 0.642 | 0 | 2 | Lsm4 | myCAF_prolif |
| Sf3a2 | 0 | 0.63093684 | 0.764 | 0.549 | 0 | 2 | Sf3a2 | myCAF_prolif |
| Acta21 | 0 | 0.40539899 | 0.792 | 0.578 | 0 | 2 | Acta2 | myCAF_prolif |
| Smc1a | 0 | 0.91472745 | 0.841 | 0.627 | 0 | 2 | Smc1a | myCAF_prolif |
| Ilf3 | 0 | 0.63252019 | 0.772 | 0.559 | 0 | 2 | Ilf3 | myCAF_prolif |
| Srprb | 0 | 0.75300144 | 0.821 | 0.608 | 0 | 2 | Srprb | myCAF_prolif |
| Sf3b4 | 0 | 0.60905638 | 0.762 | 0.55 | 0 | 2 | Sf3b4 | myCAF_prolif |
| Gxylt1 | 0 | 0.57700675 | 0.772 | 0.56 | 0 | 2 | Gxylt1 | myCAF_prolif |
| Pfdn6 | 0 | 0.60094894 | 0.792 | 0.581 | 0 | 2 | Pfdn6 | myCAF_prolif |
| Ada | 0 | 0.74137201 | 0.643 | 0.432 | 0 | 2 | Ada | myCAF_prolif |
| Gtf2a2 | 0 | 0.60849122 | 0.727 | 0.517 | 0 | 2 | Gtf2a2 | myCAF_prolif |
| Uxs1 | 0 | 0.61328662 | 0.791 | 0.582 | 0 | 2 | Uxs1 | myCAF_prolif |
| Sf3a1 | 0 | 0.70252623 | 0.825 | 0.616 | 0 | 2 | Sf3a1 | myCAF_prolif |
| Cdc27 | 0 | 0.66966402 | 0.759 | 0.55 | 0 | 2 | Cdc27 | myCAF_prolif |
| Nup153 | 0 | 0.72933462 | 0.699 | 0.491 | 0 | 2 | Nup153 | myCAF_prolif |
| Cnih4 | 0 | 0.59670398 | 0.74 | 0.534 | 0 | 2 | Cnih4 | myCAF_prolif |
| Stard7 | 0 | 0.63357576 | 0.744 | 0.538 | 0 | 2 | Stard7 | myCAF_prolif |
| Nudcd2 | 0 | 0.71547724 | 0.813 | 0.607 | 0 | 2 | Nudcd2 | myCAF_prolif |
| Psmb8 | 0 | 0.54704362 | 0.699 | 0.493 | 0 | 2 | Psmb8 | myCAF_prolif |
| Nap1l1 | 0 | 0.82435086 | 0.801 | 0.596 | 0 | 2 | Nap1l1 | myCAF_prolif |
| Gfra11 | 0 | 0.38426665 | 0.521 | 0.316 | 0 | 2 | Gfra1 | myCAF_prolif |
| Vars | 0 | 1.03581335 | 0.893 | 0.689 | 0 | 2 | Vars | myCAF_prolif |
| Vbp1 | 0 | 0.74462762 | 0.799 | 0.596 | 0 | 2 | Vbp1 | myCAF_prolif |
| Nhp2 | 0 | 0.93415257 | 0.846 | 0.643 | 0 | 2 | Nhp2 | myCAF_prolif |
| Ndufa5 | 0 | 0.70726757 | 0.781 | 0.579 | 0 | 2 | Ndufa5 | myCAF_prolif |
| Prmt1 | 0 | 0.64399077 | 0.78 | 0.578 | 0 | 2 | Prmt1 | myCAF_prolif |
| Impdh2 | 0 | 0.87452042 | 0.859 | 0.657 | 0 | 2 | Impdh2 | myCAF_prolif |
| Uhrf2 | 0 | 0.7351467 | 0.815 | 0.613 | 0 | 2 | Uhrf2 | myCAF_prolif |
| Smc6 | 0 | 0.70896015 | 0.78 | 0.581 | 0 | 2 | Smc6 | myCAF_prolif |
| Trim28 | 0 | 0.64871687 | 0.806 | 0.609 | 0 | 2 | Trim28 | myCAF_prolif |
| Lmf2 | 0 | 0.64690889 | 0.828 | 0.631 | 0 | 2 | Lmf2 | myCAF_prolif |
| Sae1 | 0 | 1.28060219 | 0.903 | 0.709 | 0 | 2 | Sae1 | myCAF_prolif |
| Gcsh1 | 0 | 0.58754515 | 0.765 | 0.571 | 0 | 2 | Gcsh | myCAF_prolif |
| Foxk2 | 0 | 0.59070333 | 0.78 | 0.587 | 0 | 2 | Foxk2 | myCAF_prolif |
| Srpk1 | 0 | 0.74319128 | 0.83 | 0.637 | 0 | 2 | Srpk1 | myCAF_prolif |
| Emd | 0 | 0.57964265 | 0.773 | 0.581 | 0 | 2 | Emd | myCAF_prolif |
| Hspd1 | 0 | 0.89969773 | 0.817 | 0.625 | 0 | 2 | Hspd1 | myCAF_prolif |
| Acp1 | 0 | 0.66125193 | 0.753 | 0.561 | 0 | 2 | Acp1 | myCAF_prolif |
| H2afv | 0 | 1.25154585 | 0.897 | 0.706 | 0 | 2 | H2afv | myCAF_prolif |
| Tlnrd1 | 0 | 0.70520707 | 0.843 | 0.653 | 0 | 2 | Tlnrd1 | myCAF_prolif |
| Hdac2 | 0 | 0.55656674 | 0.746 | 0.556 | 0 | 2 | Hdac2 | myCAF_prolif |
| Il1rl11 | 0 | 1.01923079 | 0.361 | 0.172 | 0 | 2 | Il1rl1 | myCAF_prolif |
| Ltbp21 | 0 | 0.52350031 | 0.584 | 0.395 | 0 | 2 | Ltbp2 | myCAF_prolif |
| Snrpc | 0 | 0.69175684 | 0.862 | 0.674 | 0 | 2 | Snrpc | myCAF_prolif |
| Taf1d | 0 | 0.59334869 | 0.777 | 0.59 | 0 | 2 | Taf1d | myCAF_prolif |
| Set | 0 | 0.65939986 | 0.815 | 0.628 | 0 | 2 | Set | myCAF_prolif |
| Htra21 | 0 | 0.53939915 | 0.809 | 0.624 | 0 | 2 | Htra2 | myCAF_prolif |
| Eif2s1 | 0 | 0.71039455 | 0.791 | 0.606 | 0 | 2 | Eif2s1 | myCAF_prolif |
| Pxylp11 | 0 | 0.55952194 | 0.803 | 0.62 | 0 | 2 | Pxylp1 | myCAF_prolif |
| Cdc123 | 0 | 0.69983418 | 0.834 | 0.651 | 0 | 2 | Cdc123 | myCAF_prolif |
| Ppa11 | 0 | 0.58261574 | 0.813 | 0.63 | 0 | 2 | Ppa1 | myCAF_prolif |
| Ssna1 | 0 | 0.74792655 | 0.871 | 0.689 | 0 | 2 | Ssna1 | myCAF_prolif |
| Tgfbi2 | 0 | 0.51236012 | 0.643 | 0.461 | 0 | 2 | Tgfbi | myCAF_prolif |
| Srsf3 | 0 | 0.3899346 | 0.73 | 0.548 | 0 | 2 | Srsf3 | myCAF_prolif |
| Jpt1 | 0 | 1.35394868 | 0.9 | 0.719 | 0 | 2 | Jpt1 | myCAF_prolif |
| Idh2 | 0 | 1.04886899 | 0.899 | 0.719 | 0 | 2 | Idh2 | myCAF_prolif |
| Trp53 | 0 | 0.69131319 | 0.832 | 0.652 | 0 | 2 | Trp53 | myCAF_prolif |
| Tubb4b | 0 | 1.44788559 | 0.919 | 0.74 | 0 | 2 | Tubb4b | myCAF_prolif |
| Srsf10 | 0 | 0.69993951 | 0.834 | 0.655 | 0 | 2 | Srsf10 | myCAF_prolif |
| Hnrnpd | 0 | 0.84386618 | 0.887 | 0.709 | 0 | 2 | Hnrnpd | myCAF_prolif |
| Smarca5 | 0 | 0.59763615 | 0.68 | 0.502 | 0 | 2 | Smarca5 | myCAF_prolif |
| Trim27 | 0 | 0.65656376 | 0.796 | 0.618 | 0 | 2 | Trim27 | myCAF_prolif |
| Srsf4 | 0 | 0.68353151 | 0.827 | 0.65 | 0 | 2 | Srsf4 | myCAF_prolif |
| Eftud2 | 0 | 0.85986387 | 0.873 | 0.697 | 0 | 2 | Eftud2 | myCAF_prolif |
| Hars | 0 | 0.59040428 | 0.803 | 0.627 | 0 | 2 | Hars | myCAF_prolif |
| Hprt | 0 | 0.56322776 | 0.775 | 0.599 | 0 | 2 | Hprt | myCAF_prolif |
| Ung | 0 | 2.33641834 | 0.266 | 0.09 | 0 | 2 | Ung | myCAF_prolif |
| Tm4sf11 | 0 | 0.86706981 | 0.879 | 0.704 | 0 | 2 | Tm4sf1 | myCAF_prolif |
| Ccl21 | 0 | 0.85056559 | 0.368 | 0.193 | 0 | 2 | Ccl2 | myCAF_prolif |
| Csrp21 | 0 | 0.34583921 | 0.827 | 0.652 | 0 | 2 | Csrp2 | myCAF_prolif |
| Snrpa | 0 | 0.4713552 | 0.794 | 0.62 | 0 | 2 | Snrpa | myCAF_prolif |
| Smtn | 0 | 0.7877851 | 0.869 | 0.695 | 0 | 2 | Smtn | myCAF_prolif |
| Ubap2 | 0 | 0.76872117 | 0.824 | 0.65 | 0 | 2 | Ubap2 | myCAF_prolif |
| Elovl5 | 0 | 0.58110381 | 0.792 | 0.619 | 0 | 2 | Elovl5 | myCAF_prolif |
| Tmem109 | 0 | 0.68013776 | 0.844 | 0.671 | 0 | 2 | Tmem109 | myCAF_prolif |
| Tuba1b | 0 | 1.89187572 | 0.94 | 0.768 | 0 | 2 | Tuba1b | myCAF_prolif |
| Pole4 | 0 | 0.62223839 | 0.787 | 0.616 | 0 | 2 | Pole4 | myCAF_prolif |
| Ciao2a | 0 | 0.58915184 | 0.869 | 0.698 | 0 | 2 | Ciao2a | myCAF_prolif |
| Cdc23 | 0 | 0.53899629 | 0.78 | 0.609 | 0 | 2 | Cdc23 | myCAF_prolif |
| Slc20a1 | 0 | 0.41931234 | 0.701 | 0.53 | 0 | 2 | Slc20a1 | myCAF_prolif |
| Mtap | 0 | 0.64652493 | 0.812 | 0.642 | 0 | 2 | Mtap | myCAF_prolif |
| Eef1d | 0 | 0.56968427 | 0.83 | 0.66 | 0 | 2 | Eef1d | myCAF_prolif |
| Arl6ip6 | 0 | 0.6675285 | 0.665 | 0.496 | 0 | 2 | Arl6ip6 | myCAF_prolif |
| Cbx3 | 0 | 0.7174919 | 0.791 | 0.622 | 0 | 2 | Cbx3 | myCAF_prolif |
| Lrrc551 | 0 | 0.96569548 | 0.367 | 0.198 | 0 | 2 | Lrrc55 | myCAF_prolif |
| Csf11 | 0 | 0.2738691 | 0.719 | 0.55 | 0 | 2 | Csf1 | myCAF_prolif |
| Sf3b5 | 0 | 0.73429611 | 0.872 | 0.704 | 0 | 2 | Sf3b5 | myCAF_prolif |
| Ebna1bp2 | 0 | 0.60007384 | 0.818 | 0.65 | 0 | 2 | Ebna1bp2 | myCAF_prolif |
| Ppm1g | 0 | 0.85483416 | 0.846 | 0.678 | 0 | 2 | Ppm1g | myCAF_prolif |
| Tomm40 | 0 | 0.54079865 | 0.819 | 0.651 | 0 | 2 | Tomm40 | myCAF_prolif |
| Ing1 | 0 | 0.64990868 | 0.808 | 0.641 | 0 | 2 | Ing1 | myCAF_prolif |
| Dok11 | 0 | 0.59118391 | 0.841 | 0.674 | 0 | 2 | Dok1 | myCAF_prolif |
| F2rl11 | 0 | 0.78756938 | 0.736 | 0.571 | 0 | 2 | F2rl1 | myCAF_prolif |
| Snrpb | 0 | 0.5893568 | 0.826 | 0.661 | 0 | 2 | Snrpb | myCAF_prolif |
| Abce1 | 0 | 0.56083488 | 0.717 | 0.552 | 0 | 2 | Abce1 | myCAF_prolif |
| Pi4k2b | 0 | 0.46154484 | 0.789 | 0.625 | 0 | 2 | Pi4k2b | myCAF_prolif |
| Immt | 0 | 0.49006772 | 0.773 | 0.609 | 0 | 2 | Immt | myCAF_prolif |
| Cox5a | 0 | 0.63827392 | 0.863 | 0.699 | 0 | 2 | Cox5a | myCAF_prolif |
| Srsf2 | 0 | 0.54474785 | 0.87 | 0.706 | 0 | 2 | Srsf2 | myCAF_prolif |
| Creld2 | 0 | 0.4030232 | 0.809 | 0.645 | 0 | 2 | Creld2 | myCAF_prolif |
| Psmc2 | 0 | 0.56188909 | 0.842 | 0.678 | 0 | 2 | Psmc2 | myCAF_prolif |
| Notch1 | 0 | 0.49475286 | 0.659 | 0.497 | 0 | 2 | Notch1 | myCAF_prolif |
| Idh3a | 0 | 0.61181969 | 0.804 | 0.642 | 0 | 2 | Idh3a | myCAF_prolif |
| Hnrnpab | 0 | 0.82808871 | 0.889 | 0.727 | 0 | 2 | Hnrnpab | myCAF_prolif |
| Psmd12 | 0 | 0.54952185 | 0.811 | 0.649 | 0 | 2 | Psmd12 | myCAF_prolif |
| Fmr11 | 0 | 0.56282994 | 0.853 | 0.691 | 0 | 2 | Fmr1 | myCAF_prolif |
| Hcfc1 | 0 | 0.5273779 | 0.819 | 0.657 | 0 | 2 | Hcfc1 | myCAF_prolif |
| Ipo9 | 0 | 0.84105997 | 0.861 | 0.7 | 0 | 2 | Ipo9 | myCAF_prolif |
| Dcbld2 | 0 | 0.47138534 | 0.848 | 0.687 | 0 | 2 | Dcbld2 | myCAF_prolif |
| Tuba1c | 0 | 1.43921939 | 0.913 | 0.753 | 0 | 2 | Tuba1c | myCAF_prolif |
| Chchd3 | 0 | 0.59291444 | 0.82 | 0.66 | 0 | 2 | Chchd3 | myCAF_prolif |
| Mmp31 | 0 | 0.28045211 | 0.742 | 0.583 | 0 | 2 | Mmp3 | myCAF_prolif |
| Tubb2a1 | 0 | 0.92202098 | 0.851 | 0.692 | 0 | 2 | Tubb2a | myCAF_prolif |
| Ddx39b | 0 | 0.74742138 | 0.85 | 0.691 | 0 | 2 | Ddx39b | myCAF_prolif |
| Atp5g3 | 0 | 0.46705675 | 0.781 | 0.623 | 0 | 2 | Atp5g3 | myCAF_prolif |
| Ank | 0 | 0.35482038 | 0.707 | 0.55 | 0 | 2 | Ank | myCAF_prolif |
| Tra2b | 0 | 0.48411931 | 0.813 | 0.656 | 0 | 2 | Tra2b | myCAF_prolif |
| Rnf126 | 0 | 0.54846557 | 0.853 | 0.697 | 0 | 2 | Rnf126 | myCAF_prolif |
| Cdk4 | 0 | 0.57443869 | 0.83 | 0.674 | 0 | 2 | Cdk4 | myCAF_prolif |
| Prpf19 | 0 | 0.5699201 | 0.839 | 0.683 | 0 | 2 | Prpf19 | myCAF_prolif |
| Slc25a5 | 0 | 0.61562578 | 0.756 | 0.601 | 0 | 2 | Slc25a5 | myCAF_prolif |
| Cnot6 | 0 | 0.56008189 | 0.783 | 0.629 | 0 | 2 | Cnot6 | myCAF_prolif |
| Sumo2 | 0 | 0.5160171 | 0.72 | 0.566 | 0 | 2 | Sumo2 | myCAF_prolif |
| Cnot1 | 0 | 0.59634569 | 0.798 | 0.645 | 0 | 2 | Cnot1 | myCAF_prolif |
| Cpsf6 | 0 | 0.66825152 | 0.857 | 0.704 | 0 | 2 | Cpsf6 | myCAF_prolif |
| Hint1 | 0 | 0.5856894 | 0.782 | 0.629 | 0 | 2 | Hint1 | myCAF_prolif |
| Tubb6 | 0 | 1.6170842 | 0.944 | 0.791 | 0 | 2 | Tubb6 | myCAF_prolif |
| Txnrd1 | 0 | 0.86185546 | 0.888 | 0.736 | 0 | 2 | Txnrd1 | myCAF_prolif |
| Evl | 0 | 0.63097915 | 0.843 | 0.692 | 0 | 2 | Evl | myCAF_prolif |
| Tln21 | 0 | 0.53132494 | 0.838 | 0.687 | 0 | 2 | Tln2 | myCAF_prolif |
| Zwint | 0 | 0.62515659 | 0.807 | 0.657 | 0 | 2 | Zwint | myCAF_prolif |
| Mdh2 | 0 | 0.43932384 | 0.832 | 0.682 | 0 | 2 | Mdh2 | myCAF_prolif |
| Nudt21 | 0 | 0.63226204 | 0.824 | 0.674 | 0 | 2 | Nudt21 | myCAF_prolif |
| Akr1b3 | 0 | 0.4485577 | 0.85 | 0.701 | 0 | 2 | Akr1b3 | myCAF_prolif |
| Sgk11 | 0 | 0.3708672 | 0.811 | 0.662 | 0 | 2 | Sgk1 | myCAF_prolif |
| Cxcl11 | 0 | 0.84705371 | 0.361 | 0.212 | 0 | 2 | Cxcl1 | myCAF_prolif |
| Ppp4c | 0 | 0.43379278 | 0.811 | 0.663 | 0 | 2 | Ppp4c | myCAF_prolif |
| Myh101 | 0 | 0.31353309 | 0.848 | 0.7 | 0 | 2 | Myh10 | myCAF_prolif |
| Magoh | 0 | 0.57533829 | 0.801 | 0.654 | 0 | 2 | Magoh | myCAF_prolif |
| Tfdp1 | 0 | 1.3098424 | 0.909 | 0.762 | 0 | 2 | Tfdp1 | myCAF_prolif |
| Anapc1 | 0 | 0.48037424 | 0.845 | 0.699 | 0 | 2 | Anapc1 | myCAF_prolif |
| Fubp1 | 0 | 0.56721692 | 0.859 | 0.713 | 0 | 2 | Fubp1 | myCAF_prolif |
| Gatad2a | 0 | 0.53619314 | 0.841 | 0.695 | 0 | 2 | Gatad2a | myCAF_prolif |
| Tmx1 | 0 | 0.80554527 | 0.893 | 0.747 | 0 | 2 | Tmx1 | myCAF_prolif |
| Psmc4 | 0 | 0.52158767 | 0.816 | 0.67 | 0 | 2 | Psmc4 | myCAF_prolif |
| Cct6a | 0 | 0.56737018 | 0.836 | 0.692 | 0 | 2 | Cct6a | myCAF_prolif |
| Smarca4 | 0 | 0.70023696 | 0.887 | 0.743 | 0 | 2 | Smarca4 | myCAF_prolif |
| Polr2f | 0 | 0.4783377 | 0.832 | 0.688 | 0 | 2 | Polr2f | myCAF_prolif |
| Srm1 | 0 | 0.69781587 | 0.819 | 0.675 | 0 | 2 | Srm | myCAF_prolif |
| Ptges3 | 0 | 0.659501 | 0.829 | 0.685 | 0 | 2 | Ptges3 | myCAF_prolif |
| Psmb7 | 0 | 0.49888413 | 0.816 | 0.673 | 0 | 2 | Psmb7 | myCAF_prolif |
| Cct3 | 0 | 0.52947433 | 0.828 | 0.685 | 0 | 2 | Cct3 | myCAF_prolif |
| Atic | 0 | 0.48593939 | 0.831 | 0.689 | 0 | 2 | Atic | myCAF_prolif |
| Pitrm1 | 0 | 0.61686783 | 0.816 | 0.674 | 0 | 2 | Pitrm1 | myCAF_prolif |
| Jag1 | 0 | 0.49100804 | 0.647 | 0.506 | 0 | 2 | Jag1 | myCAF_prolif |
| Nans | 0 | 0.61883358 | 0.841 | 0.7 | 0 | 2 | Nans | myCAF_prolif |
| Gspt1 | 0 | 0.63568878 | 0.869 | 0.728 | 0 | 2 | Gspt1 | myCAF_prolif |
| Txlna | 0 | 0.57351423 | 0.872 | 0.732 | 0 | 2 | Txlna | myCAF_prolif |
| Stip1 | 0 | 0.73622078 | 0.836 | 0.696 | 0 | 2 | Stip1 | myCAF_prolif |
| Cops8 | 0 | 0.52988821 | 0.842 | 0.703 | 0 | 2 | Cops8 | myCAF_prolif |
| Uba2 | 0 | 0.65930862 | 0.843 | 0.704 | 0 | 2 | Uba2 | myCAF_prolif |
| Mcmbp | 0 | 0.50764191 | 0.798 | 0.66 | 0 | 2 | Mcmbp | myCAF_prolif |
| Acat1 | 0 | 0.40204911 | 0.788 | 0.65 | 0 | 2 | Acat1 | myCAF_prolif |
| Srsf7 | 0 | 0.89488799 | 0.913 | 0.776 | 0 | 2 | Srsf7 | myCAF_prolif |
| Ube2k | 0 | 0.45804898 | 0.805 | 0.669 | 0 | 2 | Ube2k | myCAF_prolif |
| Psma1 | 0 | 0.46542741 | 0.767 | 0.631 | 0 | 2 | Psma1 | myCAF_prolif |
| Arglu1 | 0 | 1.21290913 | 0.914 | 0.778 | 0 | 2 | Arglu1 | myCAF_prolif |
| Lrrc171 | 0 | 0.35966908 | 0.835 | 0.701 | 0 | 2 | Lrrc17 | myCAF_prolif |
| Gtf3c2 | 0 | 0.51538037 | 0.83 | 0.696 | 0 | 2 | Gtf3c2 | myCAF_prolif |
| Ccl71 | 0 | 0.40928848 | 0.508 | 0.374 | 0 | 2 | Ccl7 | myCAF_prolif |
| Srsf1 | 0 | 1.02005398 | 0.934 | 0.8 | 0 | 2 | Srsf1 | myCAF_prolif |
| Psmc1 | 0 | 0.46276512 | 0.792 | 0.658 | 0 | 2 | Psmc1 | myCAF_prolif |
| Fkbp2 | 0 | 0.71454659 | 0.909 | 0.775 | 0 | 2 | Fkbp2 | myCAF_prolif |
| Nop10 | 0 | 0.44026018 | 0.813 | 0.679 | 0 | 2 | Nop10 | myCAF_prolif |
| Angptl41 | 0 | 0.36380343 | 0.877 | 0.744 | 0 | 2 | Angptl4 | myCAF_prolif |
| Psmd13 | 0 | 0.45735224 | 0.849 | 0.717 | 0 | 2 | Psmd13 | myCAF_prolif |
| Commd1 | 0 | 0.54539431 | 0.863 | 0.731 | 0 | 2 | Commd1 | myCAF_prolif |
| Fibin1 | 0 | 0.38973058 | 0.876 | 0.745 | 0 | 2 | Fibin | myCAF_prolif |
| Magt1 | 0 | 0.456022 | 0.866 | 0.735 | 0 | 2 | Magt1 | myCAF_prolif |
| Smarcc1 | 0 | 0.56304521 | 0.833 | 0.703 | 0 | 2 | Smarcc1 | myCAF_prolif |
| Ranbp1 | 0 | 1.24621029 | 0.92 | 0.79 | 0 | 2 | Ranbp1 | myCAF_prolif |
| Cct7 | 0 | 0.5227274 | 0.842 | 0.713 | 0 | 2 | Cct7 | myCAF_prolif |
| Spcs3 | 0 | 0.50331638 | 0.841 | 0.713 | 0 | 2 | Spcs3 | myCAF_prolif |
| Pck2 | 0 | 0.34422566 | 0.876 | 0.748 | 0 | 2 | Pck2 | myCAF_prolif |
| Cyc1 | 0 | 0.53318175 | 0.861 | 0.733 | 0 | 2 | Cyc1 | myCAF_prolif |
| Naa50 | 0 | 0.90861565 | 0.904 | 0.776 | 0 | 2 | Naa50 | myCAF_prolif |
| Ipo5 | 0 | 0.78088245 | 0.87 | 0.743 | 0 | 2 | Ipo5 | myCAF_prolif |
| Ddx21 | 0 | 0.68295021 | 0.876 | 0.75 | 0 | 2 | Ddx21 | myCAF_prolif |
| Get4 | 0 | 0.4218233 | 0.806 | 0.681 | 0 | 2 | Get4 | myCAF_prolif |
| Atp1b3 | 0 | 0.42142012 | 0.846 | 0.721 | 0 | 2 | Atp1b3 | myCAF_prolif |
| Sec61g | 0 | 0.46968161 | 0.863 | 0.739 | 0 | 2 | Sec61g | myCAF_prolif |
| mt-Nd1 | 0 | 1.39741672 | 0.789 | 0.665 | 0 | 2 | mt-Nd1 | myCAF_prolif |
| Bcat11 | 0 | 0.68009236 | 0.92 | 0.797 | 0 | 2 | Bcat1 | myCAF_prolif |
| Hp1 | 0 | 0.43715313 | 0.38 | 0.257 | 0 | 2 | Hp | myCAF_prolif |
| Thoc7 | 0 | 0.42119173 | 0.854 | 0.731 | 0 | 2 | Thoc7 | myCAF_prolif |
| Rdh101 | 0 | 0.25199697 | 0.803 | 0.681 | 0 | 2 | Rdh10 | myCAF_prolif |
| Heyl | 0 | 0.75946568 | 0.451 | 0.329 | 0 | 2 | Heyl | myCAF_prolif |
| Dhx15 | 0 | 0.74129516 | 0.894 | 0.772 | 0 | 2 | Dhx15 | myCAF_prolif |
| Ddx23 | 0 | 0.5286639 | 0.88 | 0.758 | 0 | 2 | Ddx23 | myCAF_prolif |
| Tmem43 | 0 | 0.29356092 | 0.854 | 0.733 | 0 | 2 | Tmem43 | myCAF_prolif |
| Thbs41 | 0 | 0.80439223 | 0.261 | 0.141 | 0 | 2 | Thbs4 | myCAF_prolif |
| Eif4e | 0 | 0.55172049 | 0.863 | 0.743 | 0 | 2 | Eif4e | myCAF_prolif |
| Eif3b | 0 | 0.44720335 | 0.856 | 0.736 | 0 | 2 | Eif3b | myCAF_prolif |
| Akirin2 | 0 | 0.57683093 | 0.808 | 0.689 | 0 | 2 | Akirin2 | myCAF_prolif |
| Tsn | 0 | 0.54668309 | 0.88 | 0.761 | 0 | 2 | Tsn | myCAF_prolif |
| Top1 | 0 | 0.47565645 | 0.793 | 0.674 | 0 | 2 | Top1 | myCAF_prolif |
| Emg1 | 0 | 0.54400033 | 0.865 | 0.746 | 0 | 2 | Emg1 | myCAF_prolif |
| Rdx | 0 | 0.51334666 | 0.832 | 0.713 | 0 | 2 | Rdx | myCAF_prolif |
| Txnl1 | 0 | 0.56202267 | 0.866 | 0.747 | 0 | 2 | Txnl1 | myCAF_prolif |
| Ecrg41 | 0 | 0.43179194 | 0.488 | 0.37 | 0 | 2 | Ecrg4 | myCAF_prolif |
| Mast2 | 0 | 0.45637437 | 0.864 | 0.746 | 0 | 2 | Mast2 | myCAF_prolif |
| Atpif1 | 0 | 0.43295762 | 0.851 | 0.733 | 0 | 2 | Atpif1 | myCAF_prolif |
| Slit2 | 0 | 0.26039952 | 0.742 | 0.624 | 0 | 2 | Slit2 | myCAF_prolif |
| Drg1 | 0 | 0.43394166 | 0.861 | 0.743 | 0 | 2 | Drg1 | myCAF_prolif |
| Slc35a4 | 0 | 0.43209465 | 0.849 | 0.731 | 0 | 2 | Slc35a4 | myCAF_prolif |
| Snrpd3 | 0 | 0.65042736 | 0.87 | 0.753 | 0 | 2 | Snrpd3 | myCAF_prolif |
| Cthrc11 | 0 | 0.44831715 | 0.913 | 0.796 | 0 | 2 | Cthrc1 | myCAF_prolif |

| | | | | | | | | |
|---|---|---|---|---|---|---|---|---|
| Rnps1 | 0 | 0.69342519 | 0.907 | 0.79 | 0 | 2 | Rnps1 | myCAF_prolif |
| Cnbp | 0 | 0.50088881 | 0.858 | 0.742 | 0 | 2 | Cnbp | myCAF_prolif |
| Tcp1 | 0 | 0.74606878 | 0.907 | 0.791 | 0 | 2 | Tcp1 | myCAF_prolif |
| Ppp1r14b1 | 0 | 0.40034891 | 0.847 | 0.731 | 0 | 2 | Ppp1r14b | myCAF_prolif |
| Prdx41 | 0 | 0.58208717 | 0.896 | 0.78 | 0 | 2 | Prdx4 | myCAF_prolif |
| Ass1 | 0 | 0.54779234 | 0.642 | 0.527 | 0 | 2 | Ass1 | myCAF_prolif |
| Nudc | 0 | 0.99452682 | 0.918 | 0.803 | 0 | 2 | Nudc | myCAF_prolif |
| Edem1 | 0 | 0.45167245 | 0.873 | 0.758 | 0 | 2 | Edem1 | myCAF_prolif |
| Kars | 0 | 0.53355827 | 0.872 | 0.757 | 0 | 2 | Kars | myCAF_prolif |
| Cct2 | 0 | 0.40472268 | 0.87 | 0.755 | 0 | 2 | Cct2 | myCAF_prolif |
| Cisd2 | 0 | 0.44491137 | 0.83 | 0.718 | 0 | 2 | Cisd2 | myCAF_prolif |
| Tpm21 | 0 | 0.3711026 | 0.856 | 0.744 | 0 | 2 | Tpm2 | myCAF_prolif |
| Ubtf | 0 | 0.43623614 | 0.846 | 0.734 | 0 | 2 | Ubtf | myCAF_prolif |
| Eif3d | 0 | 0.51439053 | 0.866 | 0.754 | 0 | 2 | Eif3d | myCAF_prolif |
| Tial1 | 0 | 0.49342647 | 0.861 | 0.75 | 0 | 2 | Tial1 | myCAF_prolif |
| Snrpf | 0 | 0.80247657 | 0.905 | 0.795 | 0 | 2 | Snrpf | myCAF_prolif |
| Hnrnph1 | 0 | 0.47319319 | 0.827 | 0.718 | 0 | 2 | Hnrnph1 | myCAF_prolif |
| Basp1 | 0 | 0.33435922 | 0.855 | 0.746 | 0 | 2 | Basp1 | myCAF_prolif |
| Srf | 0 | 0.42017575 | 0.844 | 0.735 | 0 | 2 | Srf | myCAF_prolif |
| Tardbp | 0 | 0.62723245 | 0.87 | 0.762 | 0 | 2 | Tardbp | myCAF_prolif |
| Plaur1 | 0 | 0.64224089 | 0.865 | 0.757 | 0 | 2 | Plaur | myCAF_prolif |
| Slc35e1 | 0 | 0.51652126 | 0.872 | 0.764 | 0 | 2 | Slc35e1 | myCAF_prolif |
| Ddx1 | 0 | 0.5028456 | 0.855 | 0.747 | 0 | 2 | Ddx1 | myCAF_prolif |
| Psmd2 | 0 | 0.46998504 | 0.866 | 0.759 | 0 | 2 | Psmd2 | myCAF_prolif |
| Ncl | 0 | 0.6309237 | 0.871 | 0.765 | 0 | 2 | Ncl | myCAF_prolif |
| Hdgf | 0 | 0.65398226 | 0.904 | 0.798 | 0 | 2 | Hdgf | myCAF_prolif |
| Tsen34 | 0 | 0.63984278 | 0.888 | 0.782 | 0 | 2 | Tsen34 | myCAF_prolif |
| Scaf11 | 0 | 0.35763924 | 0.842 | 0.736 | 0 | 2 | Scaf11 | myCAF_prolif |
| Hif1a | 0 | 0.40879561 | 0.854 | 0.749 | 0 | 2 | Hif1a | myCAF_prolif |
| Dlst | 0 | 0.47828104 | 0.875 | 0.771 | 0 | 2 | Dlst | myCAF_prolif |
| Tmco1 | 0 | 0.36813548 | 0.851 | 0.748 | 0 | 2 | Tmco1 | myCAF_prolif |
| Psma6 | 0 | 0.35908152 | 0.868 | 0.766 | 0 | 2 | Psma6 | myCAF_prolif |
| Tars | 0 | 0.51259946 | 0.873 | 0.771 | 0 | 2 | Tars | myCAF_prolif |
| Ndufb8 | 0 | 0.39738837 | 0.86 | 0.758 | 0 | 2 | Ndufb8 | myCAF_prolif |
| Caprin1 | 0 | 0.41036231 | 0.817 | 0.716 | 0 | 2 | Caprin1 | myCAF_prolif |
| Wdr82 | 0 | 0.3623977 | 0.861 | 0.76 | 0 | 2 | Wdr82 | myCAF_prolif |
| Purb | 0 | 0.48330254 | 0.888 | 0.787 | 0 | 2 | Purb | myCAF_prolif |
| Psme3 | 0 | 0.53159533 | 0.879 | 0.779 | 0 | 2 | Psme3 | myCAF_prolif |
| Fkbp111 | 0 | 0.47697027 | 0.859 | 0.759 | 0 | 2 | Fkbp11 | myCAF_prolif |
| Psma5 | 0 | 0.36319145 | 0.839 | 0.741 | 0 | 2 | Psma5 | myCAF_prolif |
| Ndufb6 | 0 | 0.32873517 | 0.857 | 0.759 | 0 | 2 | Ndufb6 | myCAF_prolif |
| Ivns1abp | 0 | 0.31123698 | 0.788 | 0.691 | 0 | 2 | Ivns1abp | myCAF_prolif |
| Kif5b | 0 | 0.3348632 | 0.853 | 0.756 | 0 | 2 | Kif5b | myCAF_prolif |
| H2afy | 0 | 0.78568269 | 0.926 | 0.83 | 0 | 2 | H2afy | myCAF_prolif |
| Atp5a1 | 0 | 0.35912031 | 0.851 | 0.755 | 0 | 2 | Atp5a1 | myCAF_prolif |
| Mmp91 | 0 | 0.42233819 | 0.4 | 0.305 | 0 | 2 | Mmp9 | myCAF_prolif |
| Atf1 | 0 | 0.36161715 | 0.859 | 0.765 | 0 | 2 | Atf1 | myCAF_prolif |
| Anapc11 | 0 | 0.83967929 | 0.906 | 0.813 | 0 | 2 | Anapc11 | myCAF_prolif |
| Aars | 0 | 0.3627896 | 0.853 | 0.76 | 0 | 2 | Aars | myCAF_prolif |
| Psmc5 | 0 | 0.44000684 | 0.892 | 0.799 | 0 | 2 | Psmc5 | myCAF_prolif |
| Ubald2 | 0 | 0.66823787 | 0.881 | 0.788 | 0 | 2 | Ubald2 | myCAF_prolif |
| mt-Nd4l | 0 | 1.52530722 | 0.877 | 0.784 | 0 | 2 | mt-Nd4l | myCAF_prolif |
| Marcksl1 | 0 | 0.46478497 | 0.879 | 0.787 | 0 | 2 | Marcksl1 | myCAF_prolif |
| Tra2a | 0 | 0.26025394 | 0.853 | 0.761 | 0 | 2 | Tra2a | myCAF_prolif |
| Tbcb | 0 | 0.27142071 | 0.87 | 0.778 | 0 | 2 | Tbcb | myCAF_prolif |
| Psmd1 | 0 | 0.37489219 | 0.859 | 0.768 | 0 | 2 | Psmd1 | myCAF_prolif |
| Sdhb | 0 | 0.30094252 | 0.868 | 0.777 | 0 | 2 | Sdhb | myCAF_prolif |
| Etfa | 0 | 0.39223112 | 0.874 | 0.783 | 0 | 2 | Etfa | myCAF_prolif |
| Arhgef2 | 0 | 0.36317802 | 0.865 | 0.775 | 0 | 2 | Arhgef2 | myCAF_prolif |
| Rbbp4 | 0 | 0.66480306 | 0.875 | 0.785 | 0 | 2 | Rbbp4 | myCAF_prolif |
| Sdhd | 0 | 0.3681893 | 0.873 | 0.783 | 0 | 2 | Sdhd | myCAF_prolif |
| Pdia51 | 0 | 0.37508564 | 0.884 | 0.794 | 0 | 2 | Pdia5 | myCAF_prolif |
| Psmb2 | 0 | 0.45541771 | 0.885 | 0.796 | 0 | 2 | Psmb2 | myCAF_prolif |
| Atp5o | 0 | 0.48996177 | 0.875 | 0.786 | 0 | 2 | Atp5o | myCAF_prolif |
| mt-Co1 | 0 | 1.15157012 | 0.894 | 0.805 | 0 | 2 | mt-Co1 | myCAF_prolif |
| mt-Cytb | 0 | 1.52946443 | 0.894 | 0.806 | 0 | 2 | mt-Cytb | myCAF_prolif |
| Gps1 | 0 | 0.38593583 | 0.844 | 0.757 | 0 | 2 | Gps1 | myCAF_prolif |
| Sec61b | 0 | 0.29577472 | 0.844 | 0.758 | 0 | 2 | Sec61b | myCAF_prolif |
| Bclaf1 | 0 | 0.41409081 | 0.861 | 0.775 | 0 | 2 | Bclaf1 | myCAF_prolif |
| Eif4a1 | 0 | 0.47831545 | 0.882 | 0.796 | 0 | 2 | Eif4a1 | myCAF_prolif |
| Cdv3 | 0 | 0.30998283 | 0.823 | 0.739 | 0 | 2 | Cdv3 | myCAF_prolif |
| Metap2 | 0 | 0.3593385 | 0.852 | 0.768 | 0 | 2 | Metap2 | myCAF_prolif |
| Psmc6 | 0 | 0.44274755 | 0.863 | 0.779 | 0 | 2 | Psmc6 | myCAF_prolif |
| Snrnp70 | 0 | 0.71302539 | 0.928 | 0.845 | 0 | 2 | Snrnp70 | myCAF_prolif |
| Clic1 | 0 | 0.4472666 | 0.902 | 0.819 | 0 | 2 | Clic1 | myCAF_prolif |
| Elavl1 | 0 | 0.51137998 | 0.894 | 0.811 | 0 | 2 | Elavl1 | myCAF_prolif |
| Ndufv1 | 0 | 0.35721522 | 0.861 | 0.778 | 0 | 2 | Ndufv1 | myCAF_prolif |
| Zfp91 | 0 | 0.30890592 | 0.834 | 0.751 | 0 | 2 | Zfp91 | myCAF_prolif |
| Rbm3 | 0 | 0.68453872 | 0.917 | 0.835 | 0 | 2 | Rbm3 | myCAF_prolif |
| Mdh1 | 0 | 0.36512025 | 0.859 | 0.777 | 0 | 2 | Mdh1 | myCAF_prolif |
| U2af2 | 0 | 0.56248786 | 0.919 | 0.838 | 0 | 2 | U2af2 | myCAF_prolif |
| Gatad1 | 0 | 0.3626866 | 0.859 | 0.778 | 0 | 2 | Gatad1 | myCAF_prolif |
| Cfdp1 | 0 | 0.39330912 | 0.873 | 0.792 | 0 | 2 | Cfdp1 | myCAF_prolif |
| Serbp1 | 0 | 0.61858494 | 0.897 | 0.817 | 0 | 2 | Serbp1 | myCAF_prolif |
| Eif3l | 0 | 0.50345232 | 0.912 | 0.832 | 0 | 2 | Eif3l | myCAF_prolif |
| Uggt1 | 0 | 0.31323131 | 0.863 | 0.783 | 0 | 2 | Uggt1 | myCAF_prolif |
| Prdx1 | 0 | 0.35553477 | 0.884 | 0.804 | 0 | 2 | Prdx1 | myCAF_prolif |
| Ywhah | 0 | 0.63252339 | 0.892 | 0.813 | 0 | 2 | Ywhah | myCAF_prolif |
| Adam121 | 0 | 0.25038239 | 0.939 | 0.86 | 0 | 2 | Adam12 | myCAF_prolif |
| Rbm17 | 0 | 0.52882985 | 0.903 | 0.826 | 0 | 2 | Rbm17 | myCAF_prolif |
| Sf3b3 | 0 | 0.47894716 | 0.89 | 0.813 | 0 | 2 | Sf3b3 | myCAF_prolif |
| Polr2a | 0 | 0.2735098 | 0.847 | 0.77 | 0 | 2 | Polr2a | myCAF_prolif |
| Ptma | 0 | 0.88209603 | 0.933 | 0.857 | 0 | 2 | Ptma | myCAF_prolif |
| Uqcrfs1 | 0 | 0.37199237 | 0.838 | 0.762 | 0 | 2 | Uqcrfs1 | myCAF_prolif |
| Ahsa1 | 0 | 0.68945651 | 0.897 | 0.822 | 0 | 2 | Ahsa1 | myCAF_prolif |
| Tm9sf1 | 0 | 0.43434977 | 0.888 | 0.813 | 0 | 2 | Tm9sf1 | myCAF_prolif |
| Cand1 | 0 | 0.33090077 | 0.85 | 0.775 | 0 | 2 | Cand1 | myCAF_prolif |
| Tfpi21 | 0 | 0.3313557 | 0.79 | 0.716 | 0 | 2 | Tfpi2 | myCAF_prolif |
| Maz | 0 | 0.48117516 | 0.896 | 0.822 | 0 | 2 | Maz | myCAF_prolif |
| Pabpc41 | 0 | 0.26703419 | 0.87 | 0.796 | 0 | 2 | Pabpc4 | myCAF_prolif |
| Hnrnpf | 0 | 0.55562262 | 0.877 | 0.803 | 0 | 2 | Hnrnpf | myCAF_prolif |
| Dhx9 | 0 | 0.53631797 | 0.876 | 0.803 | 0 | 2 | Dhx9 | myCAF_prolif |
| Mgat2 | 0 | 0.26784941 | 0.851 | 0.778 | 0 | 2 | Mgat2 | myCAF_prolif |
| mt-Atp8 | 0 | 1.41251106 | 0.903 | 0.831 | 0 | 2 | mt-Atp8 | myCAF_prolif |
| Ssb | 0 | 0.64972602 | 0.917 | 0.846 | 0 | 2 | Ssb | myCAF_prolif |
| Dynll2 | 0 | 0.36097698 | 0.895 | 0.824 | 0 | 2 | Dynll2 | myCAF_prolif |
| Chd4 | 0 | 0.27334141 | 0.876 | 0.805 | 0 | 2 | Chd4 | myCAF_prolif |
| Cwc15 | 0 | 0.32897588 | 0.878 | 0.807 | 0 | 2 | Cwc15 | myCAF_prolif |
| Eif4g1 | 0 | 0.37571894 | 0.865 | 0.794 | 0 | 2 | Eif4g1 | myCAF_prolif |
| Sfpq | 0 | 0.45971787 | 0.916 | 0.846 | 0 | 2 | Sfpq | myCAF_prolif |
| Hnrnpa1 | 0 | 0.82648766 | 0.931 | 0.861 | 0 | 2 | Hnrnpa1 | myCAF_prolif |
| Snu13 | 0 | 0.47046439 | 0.885 | 0.817 | 0 | 2 | Snu13 | myCAF_prolif |
| Dynll1 | 0 | 0.34838804 | 0.867 | 0.799 | 0 | 2 | Dynll1 | myCAF_prolif |
| Pdap1 | 0 | 0.50122496 | 0.925 | 0.858 | 0 | 2 | Pdap1 | myCAF_prolif |
| Ppia | 0 | 0.54941875 | 0.902 | 0.835 | 0 | 2 | Ppia | myCAF_prolif |
| Eif3g | 0 | 0.3075763 | 0.864 | 0.797 | 0 | 2 | Eif3g | myCAF_prolif |
| Tmem14c | 0 | 0.28329337 | 0.856 | 0.789 | 0 | 2 | Tmem14c | myCAF_prolif |
| Lsp11 | 0 | 0.29559511 | 0.912 | 0.846 | 0 | 2 | Lsp1 | myCAF_prolif |
| Ncln | 0 | 0.25818607 | 0.887 | 0.821 | 0 | 2 | Ncln | myCAF_prolif |
| Rbbp7 | 0 | 0.79227298 | 0.924 | 0.858 | 0 | 2 | Rbbp7 | myCAF_prolif |
| Gna11 | 0 | 0.3057287 | 0.874 | 0.808 | 0 | 2 | Gna11 | myCAF_prolif |
| Got2 | 0 | 0.3903014 | 0.896 | 0.831 | 0 | 2 | Got2 | myCAF_prolif |
| Arhgef40 | 0 | 0.35545091 | 0.916 | 0.851 | 0 | 2 | Arhgef40 | myCAF_prolif |
| Ccnd3 | 0 | 0.33350865 | 0.864 | 0.799 | 0 | 2 | Ccnd3 | myCAF_prolif |
| Cox5b | 0 | 0.35629175 | 0.909 | 0.845 | 0 | 2 | Cox5b | myCAF_prolif |
| Nfic | 0 | 0.45694854 | 0.897 | 0.833 | 0 | 2 | Nfic | myCAF_prolif |
| Ube2m | 0 | 0.35185525 | 0.897 | 0.834 | 0 | 2 | Ube2m | myCAF_prolif |
| Khdrbs1 | 0 | 0.31481368 | 0.892 | 0.83 | 0 | 2 | Khdrbs1 | myCAF_prolif |
| Gpx71 | 0 | 0.60128615 | 0.928 | 0.867 | 0 | 2 | Gpx7 | myCAF_prolif |
| Mfap51 | 0 | 0.27073715 | 0.9 | 0.839 | 0 | 2 | Mfap5 | myCAF_prolif |
| mt-Atp6 | 0 | 1.20854494 | 0.905 | 0.844 | 0 | 2 | mt-Atp6 | myCAF_prolif |
| Ndufc2 | 0 | 0.37470467 | 0.899 | 0.839 | 0 | 2 | Ndufc2 | myCAF_prolif |
| Khsrp | 0 | 0.40489258 | 0.912 | 0.852 | 0 | 2 | Khsrp | myCAF_prolif |
| Rnf4 | 0 | 0.50978469 | 0.915 | 0.856 | 0 | 2 | Rnf4 | myCAF_prolif |
| Anp32a | 0 | 0.44164689 | 0.922 | 0.863 | 0 | 2 | Anp32a | myCAF_prolif |
| Pdlim7 | 0 | 0.32227507 | 0.913 | 0.855 | 0 | 2 | Pdlim7 | myCAF_prolif |
| Tkt | 0 | 0.26961274 | 0.87 | 0.812 | 0 | 2 | Tkt | myCAF_prolif |
| Rad23b | 0 | 0.35816175 | 0.877 | 0.82 | 0 | 2 | Rad23b | myCAF_prolif |
| Golt1b | 0 | 0.52509219 | 0.91 | 0.853 | 0 | 2 | Golt1b | myCAF_prolif |
| Mta1 | 0 | 0.26985749 | 0.878 | 0.821 | 0 | 2 | Mta1 | myCAF_prolif |
| Ubqln1 | 0 | 0.2693644 | 0.876 | 0.819 | 0 | 2 | Ubqln1 | myCAF_prolif |
| Mapre1 | 0 | 0.51492509 | 0.901 | 0.845 | 0 | 2 | Mapre1 | myCAF_prolif |
| Psma4 | 0 | 0.44212819 | 0.899 | 0.844 | 0 | 2 | Psma4 | myCAF_prolif |
| Cnih1 | 0 | 0.29894471 | 0.879 | 0.824 | 0 | 2 | Cnih1 | myCAF_prolif |
| Cd44 | 0 | 0.64622024 | 0.923 | 0.869 | 0 | 2 | Cd44 | myCAF_prolif |
| Tgfb1 | 0 | 0.41021716 | 0.887 | 0.833 | 0 | 2 | Tgfb1 | myCAF_prolif |
| Tnc1 | 0 | 0.45212103 | 0.949 | 0.895 | 0 | 2 | Tnc | myCAF_prolif |
| Cox8a | 0 | 0.29427329 | 0.908 | 0.855 | 0 | 2 | Cox8a | myCAF_prolif |
| Actr1a | 0 | 0.32892091 | 0.896 | 0.843 | 0 | 2 | Actr1a | myCAF_prolif |
| Rlim | 0 | 0.44080242 | 0.908 | 0.855 | 0 | 2 | Rlim | myCAF_prolif |
| Hnrnpu | 0 | 0.47844402 | 0.904 | 0.852 | 0 | 2 | Hnrnpu | myCAF_prolif |
| Cenpb | 0 | 0.35576002 | 0.923 | 0.871 | 0 | 2 | Cenpb | myCAF_prolif |
| 11-Sep | 0 | 0.29904364 | 0.901 | 0.849 | 0 | 2 | 11-Sep | myCAF_prolif |
| Dnaja2 | 0 | 0.255378 | 0.877 | 0.825 | 0 | 2 | Dnaja2 | myCAF_prolif |
| Lims11 | 0 | 0.26945827 | 0.894 | 0.842 | 0 | 2 | Lims1 | myCAF_prolif |
| Hsp90aa1 | 0 | 0.26572333 | 0.893 | 0.841 | 0 | 2 | Hsp90aa1 | myCAF_prolif |
| Sf3b2 | 0 | 0.38049268 | 0.905 | 0.853 | 0 | 2 | Sf3b2 | myCAF_prolif |
| Ak21 | 0 | 0.51199221 | 0.925 | 0.874 | 0 | 2 | Ak2 | myCAF_prolif |
| Hnrnpl | 0 | 0.61761166 | 0.918 | 0.867 | 0 | 2 | Hnrnpl | myCAF_prolif |
| Cox6b1 | 0 | 0.25137691 | 0.892 | 0.841 | 0 | 2 | Cox6b1 | myCAF_prolif |
| Hnrnpm | 0 | 0.54006289 | 0.931 | 0.88 | 0 | 2 | Hnrnpm | myCAF_prolif |
| Bzw1 | 0 | 0.33241848 | 0.896 | 0.846 | 0 | 2 | Bzw1 | myCAF_prolif |
| Cul3 | 0 | 0.32869843 | 0.889 | 0.839 | 0 | 2 | Cul3 | myCAF_prolif |
| Psmd7 | 0 | 0.39375365 | 0.903 | 0.854 | 0 | 2 | Psmd7 | myCAF_prolif |
| Cggbp1 | 0 | 0.4221328 | 0.898 | 0.849 | 0 | 2 | Cggbp1 | myCAF_prolif |
| Nono | 0 | 0.63164792 | 0.928 | 0.879 | 0 | 2 | Nono | myCAF_prolif |
| Dnajc10 | 0 | 0.49514668 | 0.911 | 0.863 | 0 | 2 | Dnajc10 | myCAF_prolif |
| Psmc3 | 0 | 0.47678759 | 0.923 | 0.875 | 0 | 2 | Psmc3 | myCAF_prolif |
| Txn1 | 0 | 0.51475673 | 0.899 | 0.851 | 0 | 2 | Txn1 | myCAF_prolif |
| Tbrg1 | 0 | 0.25991218 | 0.903 | 0.857 | 0 | 2 | Tbrg1 | myCAF_prolif |
| Ube2z | 0 | 0.28797567 | 0.899 | 0.854 | 0 | 2 | Ube2z | myCAF_prolif |
| Pabpn1 | 0 | 0.46486457 | 0.937 | 0.892 | 0 | 2 | Pabpn1 | myCAF_prolif |
| Rtn3 | 0 | 0.31159873 | 0.887 | 0.842 | 0 | 2 | Rtn3 | myCAF_prolif |
| Tmem258 | 0 | 0.34446106 | 0.904 | 0.859 | 0 | 2 | Tmem258 | myCAF_prolif |
| Hp1bp3 | 0 | 0.56377756 | 0.922 | 0.878 | 0 | 2 | Hp1bp3 | myCAF_prolif |
| Eif5a | 0 | 0.59756227 | 0.927 | 0.883 | 0 | 2 | Eif5a | myCAF_prolif |
| Arf6 | 0 | 0.37614498 | 0.909 | 0.865 | 0 | 2 | Arf6 | myCAF_prolif |
| mt-Co2 | 0 | 1.30719021 | 0.942 | 0.898 | 0 | 2 | mt-Co2 | myCAF_prolif |
| Pafah1b2 | 0 | 0.28811706 | 0.896 | 0.853 | 0 | 2 | Pafah1b2 | myCAF_prolif |
| Timm13 | 0 | 0.32316455 | 0.913 | 0.87 | 0 | 2 | Timm13 | myCAF_prolif |
| Ogdh | 0 | 0.31621356 | 0.91 | 0.867 | 0 | 2 | Ogdh | myCAF_prolif |
| Capn2 | 0 | 0.25314808 | 0.9 | 0.858 | 0 | 2 | Capn2 | myCAF_prolif |
| Nme2 | 0 | 0.32494568 | 0.868 | 0.826 | 0 | 2 | Nme2 | myCAF_prolif |
| Syncrip | 0 | 0.62885284 | 0.942 | 0.9 | 0 | 2 | Syncrip | myCAF_prolif |
| Ptbp3 | 0 | 0.31088501 | 0.906 | 0.865 | 0 | 2 | Ptbp3 | myCAF_prolif |
| Prrc2a | 0 | 0.39888709 | 0.914 | 0.873 | 0 | 2 | Prrc2a | myCAF_prolif |
| Calm2 | 0 | 0.31085604 | 0.872 | 0.831 | 0 | 2 | Calm2 | myCAF_prolif |
| Dazap1 | 0 | 0.45940386 | 0.939 | 0.898 | 0 | 2 | Dazap1 | myCAF_prolif |
| Ywhae | 0 | 0.44133095 | 0.93 | 0.89 | 0 | 2 | Ywhae | myCAF_prolif |
| Loxl31 | 0 | 0.3706873 | 0.95 | 0.91 | 0 | 2 | Loxl3 | myCAF_prolif |
| Dnajb11 | 0 | 0.3292176 | 0.903 | 0.865 | 0 | 2 | Dnajb11 | myCAF_prolif |
| Actn11 | 0 | 0.32841787 | 0.938 | 0.9 | 0 | 2 | Actn1 | myCAF_prolif |
| Chtop | 0 | 0.26634215 | 0.901 | 0.865 | 0 | 2 | Chtop | myCAF_prolif |
| Mettl9 | 0 | 0.27099107 | 0.895 | 0.859 | 0 | 2 | Mettl9 | myCAF_prolif |
| Erp29 | 0 | 0.26306073 | 0.916 | 0.882 | 0 | 2 | Erp29 | myCAF_prolif |
| H1f0 | 0 | 0.55225393 | 0.914 | 0.88 | 0 | 2 | H1f0 | myCAF_prolif |
| Atp5b | 0 | 0.43916661 | 0.911 | 0.878 | 0 | 2 | Atp5b | myCAF_prolif |
| Itga51 | 0 | 0.28861013 | 0.921 | 0.888 | 0 | 2 | Itga5 | myCAF_prolif |
| P4ha3 | 0 | 0.25038114 | 0.943 | 0.91 | 0 | 2 | P4ha3 | myCAF_prolif |
| Sdha | 0 | 0.34797733 | 0.914 | 0.882 | 0 | 2 | Sdha | myCAF_prolif |
| Ncor2 | 0 | 0.4136405 | 0.919 | 0.888 | 0 | 2 | Ncor2 | myCAF_prolif |
| Sod31 | 0 | 0.33845892 | 0.958 | 0.927 | 0 | 2 | Sod3 | myCAF_prolif |
| Fkbp1a | 0 | 0.26111986 | 0.921 | 0.892 | 0 | 2 | Fkbp1a | myCAF_prolif |
| Stt3b | 0 | 0.35134991 | 0.904 | 0.875 | 0 | 2 | Stt3b | myCAF_prolif |
| Dnajc3 | 0 | 0.31539639 | 0.923 | 0.895 | 0 | 2 | Dnajc3 | myCAF_prolif |
| Qk | 0 | 0.33469172 | 0.918 | 0.89 | 0 | 2 | Qk | myCAF_prolif |
| Ganab | 0 | 0.26043184 | 0.92 | 0.892 | 0 | 2 | Ganab | myCAF_prolif |
| Lrrc59 | 0 | 0.75501921 | 0.949 | 0.921 | 0 | 2 | Lrrc59 | myCAF_prolif |
| Rer1 | 0 | 0.29351743 | 0.889 | 0.862 | 0 | 2 | Rer1 | myCAF_prolif |
| Puf60 | 0 | 0.35884702 | 0.928 | 0.901 | 0 | 2 | Puf60 | myCAF_prolif |
| Cfl1 | 0 | 0.3026307 | 0.953 | 0.98 | 0 | 2 | Cfl1 | myCAF_prolif |
| Crip1 | 0 | 0.37397849 | 0.944 | 0.917 | 0 | 2 | Crip1 | myCAF_prolif |
| Sec13 | 0 | 0.3576683 | 0.928 | 0.902 | 0 | 2 | Sec13 | myCAF_prolif |
| Fus | 0 | 0.34941143 | 0.916 | 0.89 | 0 | 2 | Fus | myCAF_prolif |
| Map4k4 | 0 | 0.27492379 | 0.918 | 0.893 | 0 | 2 | Map4k4 | myCAF_prolif |
| Srp9 | 0 | 0.27137321 | 0.907 | 0.882 | 0 | 2 | Srp9 | myCAF_prolif |
| Hnrnpa2b1 | 0 | 0.69905399 | 0.913 | 0.888 | 0 | 2 | Hnrnpa2b1 | myCAF_prolif |
| Pcbp1 | 0 | 0.31685063 | 0.901 | 0.876 | 0 | 2 | Pcbp1 | myCAF_prolif |
| Rnf187 | 0 | 0.25227357 | 0.926 | 0.901 | 0 | 2 | Rnf187 | myCAF_prolif |
| Cbfb | 0 | 0.56460717 | 0.947 | 0.923 | 0 | 2 | Cbfb | myCAF_prolif |
| Kdelr2 | 0 | 0.34146268 | 0.943 | 0.966 | 0 | 2 | Kdelr2 | myCAF_prolif |
| Stub1 | 0 | 0.54187311 | 0.937 | 0.914 | 0 | 2 | Stub1 | myCAF_prolif |
| Sec61a1 | 0 | 0.29821903 | 0.899 | 0.877 | 0 | 2 | Sec61a1 | myCAF_prolif |
| Calm1 | 0 | 0.29993682 | 0.922 | 0.9 | 0 | 2 | Calm1 | myCAF_prolif |
| Srsf9 | 0 | 0.32165608 | 0.925 | 0.904 | 0 | 2 | Srsf9 | myCAF_prolif |
| Hnrnpk | 0 | 0.29841202 | 0.906 | 0.885 | 0 | 2 | Hnrnpk | myCAF_prolif |
| Nedd8 | 0 | 0.3013038 | 0.926 | 0.905 | 0 | 2 | Nedd8 | myCAF_prolif |
| Mydgf | 0 | 0.30155866 | 0.914 | 0.894 | 0 | 2 | Mydgf | myCAF_prolif |
| Srsf6 | 0 | 0.32717924 | 0.946 | 0.926 | 0 | 2 | Srsf6 | myCAF_prolif |
| Wdr1 | 0 | 0.35370591 | 0.904 | 0.885 | 0 | 2 | Wdr1 | myCAF_prolif |
| Npm1 | 0 | 0.64047915 | 0.941 | 0.922 | 0 | 2 | Npm1 | myCAF_prolif |
| Cops6 | 0 | 0.26771726 | 0.893 | 0.875 | 0 | 2 | Cops6 | myCAF_prolif |
| 7-Sep | 0 | 0.26585796 | 0.915 | 0.897 | 0 | 2 | 7-Sep | myCAF_prolif |
| Ssr4 | 0 | 0.33771464 | 0.906 | 0.888 | 0 | 2 | Ssr4 | myCAF_prolif |
| Gorasp2 | 0 | 0.26922258 | 0.907 | 0.89 | 0 | 2 | Gorasp2 | myCAF_prolif |
| Raly | 0 | 0.29095581 | 0.929 | 0.912 | 0 | 2 | Raly | myCAF_prolif |
| Vcp | 0 | 0.32089641 | 0.946 | 0.963 | 0 | 2 | Vcp | myCAF_prolif |
| Map4 | 0 | 0.28756742 | 0.915 | 0.898 | 0 | 2 | Map4 | myCAF_prolif |
| Mbd3 | 0 | 0.2916758 | 0.928 | 0.911 | 0 | 2 | Mbd3 | myCAF_prolif |
| Hnrnpul2 | 0 | 0.25057769 | 0.91 | 0.893 | 0 | 2 | Hnrnpul2 | myCAF_prolif |
| Plp2 | 0 | 0.40938673 | 0.935 | 0.918 | 0 | 2 | Plp2 | myCAF_prolif |
| Rpn21 | 0 | 0.37648121 | 0.923 | 0.907 | 0 | 2 | Rpn2 | myCAF_prolif |
| Gnai3 | 0 | 0.30153265 | 0.919 | 0.903 | 0 | 2 | Gnai3 | myCAF_prolif |
| Copg1 | 0 | 0.27076534 | 0.927 | 0.911 | 0 | 2 | Copg1 | myCAF_prolif |
| Rcn31 | 0 | 0.26630499 | 0.968 | 0.984 | 0 | 2 | Rcn3 | myCAF_prolif |
| Snx3 | 0 | 0.25667152 | 0.918 | 0.902 | 0 | 2 | Snx3 | myCAF_prolif |
| Reep5 | 0 | 0.51969304 | 0.937 | 0.921 | 0 | 2 | Reep5 | myCAF_prolif |
| Atn1 | 0 | 0.30819178 | 0.918 | 0.903 | 0 | 2 | Atn1 | myCAF_prolif |
| Tpm41 | 0 | 0.27342762 | 0.949 | 0.964 | 0 | 2 | Tpm4 | myCAF_prolif |
| Hspa8 | 0 | 0.40388324 | 0.949 | 0.964 | 0 | 2 | Hspa8 | myCAF_prolif |
| Ddost | 0 | 0.3145024 | 0.937 | 0.952 | 0 | 2 | Ddost | myCAF_prolif |
| Lmna | 0 | 0.42401861 | 0.968 | 0.982 | 0 | 2 | Lmna | myCAF_prolif |
| Snd1 | 0 | 0.33487462 | 0.949 | 0.935 | 0 | 2 | Snd1 | myCAF_prolif |
| Hnrnpul1 | 0 | 0.61485735 | 0.949 | 0.935 | 0 | 2 | Hnrnpul1 | myCAF_prolif |
| Tuba1a | 0 | 0.71311453 | 0.951 | 0.937 | 0 | 2 | Tuba1a | myCAF_prolif |
| Mlf2 | 0 | 0.40195258 | 0.942 | 0.931 | 0 | 2 | Mlf2 | myCAF_prolif |
| Ptbp1 | 0 | 0.44920041 | 0.942 | 0.932 | 0 | 2 | Ptbp1 | myCAF_prolif |
| Ssr21 | 0 | 0.42526201 | 0.944 | 0.953 | 0 | 2 | Ssr2 | myCAF_prolif |
| Uqcrc11 | 0 | 0.30187171 | 0.937 | 0.928 | 0 | 2 | Uqcrc1 | myCAF_prolif |
| Actn41 | 0 | 0.38820926 | 0.956 | 0.948 | 0 | 2 | Actn4 | myCAF_prolif |
| Atp1a1 | 0 | 0.28107381 | 0.936 | 0.929 | 0 | 2 | Atp1a1 | myCAF_prolif |
| Ckap4 | 0 | 0.86762332 | 0.97 | 0.977 | 0 | 2 | Ckap4 | myCAF_prolif |
| Timp11 | 0 | 0.90471518 | 0.959 | 0.966 | 0 | 2 | Timp1 | myCAF_prolif |
| Tceal8 | 0 | 0.26921032 | 0.945 | 0.938 | 0 | 2 | Tceal8 | myCAF_prolif |
| Tceal91 | 0 | 0.31632863 | 0.944 | 0.937 | 0 | 2 | Tceal9 | myCAF_prolif |
| Mlec | 0 | 0.34610282 | 0.939 | 0.945 | 0 | 2 | Mlec | myCAF_prolif |
| Trir | 0 | 0.27844522 | 0.922 | 0.916 | 0 | 2 | Trir | myCAF_prolif |
| Actg1 | 0 | 0.44803176 | 0.956 | 0.962 | 0 | 2 | Actg1 | myCAF_prolif |
| Ssr3 | 0 | 0.33765628 | 0.941 | 0.935 | 0 | 2 | Ssr3 | myCAF_prolif |
| Ugdh | 0 | 0.25972053 | 0.941 | 0.935 | 0 | 2 | Ugdh | myCAF_prolif |
| Plod31 | 0 | 0.25842936 | 0.936 | 0.942 | 0 | 2 | Plod3 | myCAF_prolif |
| Manf | 0 | 0.46346614 | 0.935 | 0.941 | 0 | 2 | Manf | myCAF_prolif |
| Creb3l1 | 0 | 0.44445346 | 0.958 | 0.963 | 0 | 2 | Creb3l1 | myCAF_prolif |
| Ctbp1 | 0 | 0.25001757 | 0.919 | 0.914 | 0 | 2 | Ctbp1 | myCAF_prolif |
| Arf1 | 0 | 0.30876273 | 0.939 | 0.944 | 0 | 2 | Arf1 | myCAF_prolif |
| Ssr1 | 0 | 0.63545977 | 0.947 | 0.952 | 0 | 2 | Ssr1 | myCAF_prolif |
| Ybx1 | 0 | 0.29449959 | 0.946 | 0.942 | 0 | 2 | Ybx1 | myCAF_prolif |
| Pfn1 | 0 | 0.35914945 | 0.956 | 0.96 | 0 | 2 | Pfn1 | myCAF_prolif |
| Arhgdia | 0 | 0.30609251 | 0.947 | 0.943 | 0 | 2 | Arhgdia | myCAF_prolif |
| Msn | 0 | 0.32076041 | 0.963 | 0.967 | 0 | 2 | Msn | myCAF_prolif |
| Tagln2 | 0 | 0.73658787 | 0.965 | 0.968 | 0 | 2 | Tagln2 | myCAF_prolif |
| Ostc | 0 | 0.28092629 | 0.935 | 0.933 | 0 | 2 | Ostc | myCAF_prolif |
| Ptms | 0 | 0.5311399 | 0.989 | 0.991 | 0 | 2 | Ptms | myCAF_prolif |
| Eif4g2 | 0 | 0.27237252 | 0.931 | 0.933 | 0 | 2 | Eif4g2 | myCAF_prolif |
| Col5a3 | 0 | 0.25359841 | 0.953 | 0.951 | 0 | 2 | Col5a3 | myCAF_prolif |
| Oaf | 0 | 0.39470225 | 0.936 | 0.934 | 0 | 2 | Oaf | myCAF_prolif |
| Tubb5 | 0 | 1.29521122 | 0.949 | 0.947 | 0 | 2 | Tubb5 | myCAF_prolif |
| Lman1 | 0 | 0.34660955 | 0.947 | 0.949 | 0 | 2 | Lman1 | myCAF_prolif |
| Rpn11 | 0 | 0.41111644 | 0.939 | 0.94 | 0 | 2 | Rpn1 | myCAF_prolif |
| Akt1 | 0 | 0.28190786 | 0.953 | 0.952 | 0 | 2 | Akt1 | myCAF_prolif |
| Ndufb7 | 5.84E-293 | 0.34778754 | 0.858 | 0.73 | 1.13E-294 | 2 | Ndufb7 | myCAF_prolif |
| Nelfe | 1.13E-293 | 0.88795495 | 0.709 | 0.446 | 2.18E-293 | 2 | Nelfe | myCAF_prolif |
| Mapk1 | 2.56E-295 | 0.25237602 | 0.866 | 0.805 | 4.95E-293 | 2 | Mapk1 | myCAF_prolif |
| Ranbp3 | 1.02E-293 | 0.65114545 | 0.794 | 0.622 | 1.97E-293 | 2 | Ranbp3 | myCAF_prolif |
| Stx12 | 3.62E-293 | 0.28982796 | 0.838 | 0.774 | 7.00E-293 | 2 | Stx12 | myCAF_prolif |
| Ppp5c | 6.93E-293 | 0.68090903 | 0.737 | 0.521 | 1.34E-293 | 2 | Ppp5c | myCAF_prolif |
| Tmem183a | 1.12E-293 | 0.44648004 | 0.78 | 0.594 | 2.17E-293 | 2 | Tmem183a | myCAF_prolif |
| Pspc1 | 1.82E-293 | 0.75061886 | 0.718 | 0.461 | 3.51E-295 | 2 | Pspc1 | myCAF_prolif |
| Tubgcp3 | 2.29E-293 | 0.89284455 | 0.635 | 0.386 | 4.42E-293 | 2 | Tubgcp3 | myCAF_prolif |
| Ppp1r7 | 3.97E-293 | 0.64272712 | 0.729 | 0.505 | 7.67E-293 | 2 | Ppp1r7 | myCAF_prolif |
| Ccdc88a | 4.18E-293 | 0.60863312 | 0.723 | 0.498 | 8.08E-293 | 2 | Ccdc88a | myCAF_prolif |
| Haspin | 5.03E-293 | 3.99002495 | 0.328 | 0.02 | 9.73E-293 | 2 | Haspin | myCAF_prolif |
| Tedc2 | 8.52E-294 | 1.66376483 | 0.435 | 0.149 | 1.65E-293 | 2 | Tedc2 | myCAF_prolif |
| Dnph1 | 1.45E-293 | 2.40498935 | 0.33 | 0.056 | 2.80E-293 | 2 | Dnph1 | myCAF_prolif |
| Rbm14 | 7.70E-293 | 0.44771372 | 0.854 | 0.709 | 1.49E-293 | 2 | Rbm14 | myCAF_prolif |
| Ddx11 | 1.52E-293 | 3.04232488 | 0.346 | 0.039 | 2.94E-294 | 2 | Ddx11 | myCAF_prolif |
| Bud31 | 9.04E-293 | 0.33226507 | 0.831 | 0.733 | 1.75E-293 | 2 | Bud31 | myCAF_prolif |
| Nolc11 | 2.87E-293 | 0.94995852 | 0.604 | 0.348 | 5.54E-293 | 2 | Nolc1 | myCAF_prolif |
| Actr3 | 4.89E-293 | 0.36468307 | 0.778 | 0.669 | 9.45E-293 | 2 | Actr3 | myCAF_prolif |
| Dcun1d5 | 4.03E-294 | 0.3200001 | 0.832 | 0.737 | 7.79E-293 | 2 | Dcun1d5 | myCAF_prolif |
| Steap11 | 7.69E-293 | 0.7030147 | 0.645 | 0.395 | 1.49E-293 | 2 | Steap1 | myCAF_prolif |
| Igsf8 | 1.33E-293 | 0.25587175 | 0.851 | 0.733 | 2.58E-293 | 2 | Igsf8 | myCAF_prolif |
| Pacsin2 | 1.40E-293 | 0.56102901 | 0.775 | 0.554 | 2.70E-293 | 2 | Pacsin2 | myCAF_prolif |
| Atp5f1 | 9.64E-293 | 0.28928455 | 0.825 | 0.733 | 1.86E-293 | 2 | Atp5f1 | myCAF_prolif |
| Nup50 | 1.38E-293 | 0.69776615 | 0.71 | 0.493 | 2.68E-293 | 2 | Nup50 | myCAF_prolif |
| Pkdcc1 | 3.71E-293 | 0.29871556 | 0.826 | 0.682 | 7.17E-293 | 2 | Pkdcc | myCAF_prolif |
| Nup93 | 1.63E-293 | 0.92801322 | 0.677 | 0.414 | 3.15E-293 | 2 | Nup93 | myCAF_prolif |
| Slc35b2 | 2.18E-293 | 0.28483014 | 0.875 | 0.789 | 4.21E-293 | 2 | Slc35b2 | myCAF_prolif |
| Prep | 2.33E-293 | 0.37072267 | 0.795 | 0.668 | 4.50E-293 | 2 | Prep | myCAF_prolif |
| Cenpo | 3.69E-293 | 1.85025984 | 0.403 | 0.103 | 7.14E-293 | 2 | Cenpo | myCAF_prolif |
| Dscc1 | 4.37E-293 | 5.10947393 | 0.325 | 0.012 | 8.46E-293 | 2 | Dscc1 | myCAF_prolif |
| Usp14 | 5.22E-293 | 0.61885691 | 0.682 | 0.463 | 1.01E-292 | 2 | Usp14 | myCAF_prolif |
| Ppp1cb | 5.58E-293 | 0.35187853 | 0.885 | 0.8 | 1.08E-292 | 2 | Ppp1cb | myCAF_prolif |
| Gen1 | 3.60E-293 | 5.02438262 | 0.357 | 0.013 | 6.96E-292 | 2 | Gen1 | myCAF_prolif |
| Acin1 | 5.42E-293 | 0.46305748 | 0.851 | 0.744 | 1.05E-291 | 2 | Acin1 | myCAF_prolif |
| Nid21 | 6.51E-293 | 0.59583996 | 0.642 | 0.413 | 1.26E-291 | 2 | Nid2 | myCAF_prolif |
| Cxcl21 | 7.01E-293 | 0.8898694 | 0.385 | 0.271 | 1.36E-291 | 2 | Cxcl2 | myCAF_prolif |
| Naa20 | 1.24E-293 | 0.32106148 | 0.8 | 0.676 | 2.40E-291 | 2 | Naa20 | myCAF_prolif |
| Nmral1 | 1.50E-293 | 2.28460805 | 0.348 | 0.068 | 2.90E-291 | 2 | Nmral1 | myCAF_prolif |
| Tnpo1 | 4.98E-293 | 0.4011214 | 0.833 | 0.717 | 9.62E-291 | 2 | Tnpo1 | myCAF_prolif |
| Zfp207 | 8.59E-293 | 0.2909002 | 0.882 | 0.816 | 1.66E-290 | 2 | Zfp207 | myCAF_prolif |
| Hnrnph2 | 1.22E-294 | 0.34590707 | 0.833 | 0.711 | 2.36E-290 | 2 | Hnrnph2 | myCAF_prolif |
| Blmh | 1.01E-293 | 0.36290043 | 0.819 | 0.692 | 1.95E-289 | 2 | Blmh | myCAF_prolif |
| Pdia5a | 2.78E-293 | 0.60410934 | 0.814 | 0.635 | 5.37E-289 | 2 | Pdia5a | myCAF_prolif |
| DEPRECATED-I | 4.39E-293 | 1.30938729 | 0.494 | 0.183 | 8.49E-289 | 2 | DEPRECATED-I | myCAF_prolif |
| Fbl | 8.88E-293 | 0.61703165 | 0.726 | 0.509 | 1.72E-288 | 2 | Fbl | myCAF_prolif |
| Rabggtb | 1.02E-292 | 0.42769068 | 0.806 | 0.654 | 1.97E-288 | 2 | Rabggtb | myCAF_prolif |
| Mapk1ip1l | 4.42E-292 | 0.27489071 | 0.874 | 0.798 | 8.55E-288 | 2 | Mapk1ip1l | myCAF_prolif |
| Hnrnpr | 6.55E-292 | 0.45978672 | 0.751 | 0.59 | 1.27E-287 | 2 | Hnrnpr | myCAF_prolif |
| Polr2e | 2.29E-291 | 0.28729396 | 0.832 | 0.748 | 4.42E-287 | 2 | Polr2e | myCAF_prolif |
| Ahctf1 | 7.82E-291 | 0.65481886 | 0.706 | 0.487 | 1.51E-286 | 2 | Ahctf1 | myCAF_prolif |
| Cenpt | 2.84E-290 | 1.66186958 | 0.476 | 0.165 | 5.49E-286 | 2 | Cenpt | myCAF_prolif |
| Eri2 | 9.74E-290 | 1.91858266 | 0.399 | 0.11 | 1.88E-285 | 2 | Eri2 | myCAF_prolif |
| Exosc10 | 2.92E-289 | 0.59596371 | 0.738 | 0.518 | 5.64E-285 | 2 | Exosc10 | myCAF_prolif |
| Cchcr1 | 5.39E-289 | 1.22932179 | 0.54 | 0.235 | 1.04E-284 | 2 | Cchcr1 | myCAF_prolif |
| Kpna3 | 6.06E-289 | 0.70937978 | 0.644 | 0.43 | 1.17E-284 | 2 | Kpna3 | myCAF_prolif |
| Gsr | 6.85E-289 | 0.25844459 | 0.816 | 0.701 | 1.32E-284 | 2 | Gsr | myCAF_prolif |
| Sumo3 | 7.62E-289 | 0.29304556 | 0.862 | 0.774 | 1.47E-284 | 2 | Sumo3 | myCAF_prolif |
| Xrcc2 | 2.76E-288 | 2.63973381 | 0.335 | 0.052 | 5.34E-284 | 2 | Xrcc2 | myCAF_prolif |
| Zdhhc5 | 5.97E-288 | 0.25245437 | 0.855 | 0.809 | 1.15E-283 | 2 | Zdhhc5 | myCAF_prolif |
| Timm17a | 9.45E-288 | 0.38504001 | 0.798 | 0.676 | 1.83E-283 | 2 | Timm17a | myCAF_prolif |
| Pdlim4 | 1.76E-287 | 0.33231026 | 0.709 | 0.585 | 3.41E-283 | 2 | Pdlim4 | myCAF_prolif |
| Ndufv2 | 2.64E-287 | 0.34198426 | 0.815 | 0.715 | 5.10E-283 | 2 | Ndufv2 | myCAF_prolif |
| Faap24 | 4.47E-287 | 1.72581117 | 0.414 | 0.124 | 8.65E-283 | 2 | Faap24 | myCAF_prolif |
| Nup188 | 7.13E-287 | 0.64501835 | 0.701 | 0.467 | 1.38E-282 | 2 | Nup188 | myCAF_prolif |
| Phb2 | 3.98E-286 | 0.48599739 | 0.845 | 0.702 | 7.70E-282 | 2 | Phb2 | myCAF_prolif |
| Mcm8 | 6.34E-286 | 4.04887642 | 0.327 | 0.02 | 1.23E-281 | 2 | Mcm8 | myCAF_prolif |
| Pold3 | 6.90E-286 | 1.47434556 | 0.428 | 0.159 | 1.33E-281 | 2 | Pold3 | myCAF_prolif |
| Tonsl | 1.93E-285 | 3.50480895 | 0.307 | 0.025 | 3.74E-281 | 2 | Tonsl | myCAF_prolif |
| Fam3c | 5.42E-285 | 0.28363403 | 0.847 | 0.757 | 1.05E-280 | 2 | Fam3c | myCAF_prolif |
| Fancb | 5.87E-285 | 4.08487727 | 0.317 | 0.019 | 1.13E-280 | 2 | Fancb | myCAF_prolif |
| Mogs | 7.55E-285 | 0.44518659 | 0.773 | 0.603 | 1.46E-280 | 2 | Mogs | myCAF_prolif |
| Asnsd1 | 2.16E-284 | 0.64777325 | 0.727 | 0.503 | 4.17E-280 | 2 | Asnsd1 | myCAF_prolif |
| Tex30 | 3.45E-284 | 1.85313946 | 0.377 | 0.084 | 6.67E-280 | 2 | Tex30 | myCAF_prolif |
| Slc30a5 | 7.10E-284 | 0.33305503 | 0.82 | 0.7 | 1.37E-279 | 2 | Slc30a5 | myCAF_prolif |
| Smc5 | 9.34E-284 | 0.72102996 | 0.76 | 0.595 | 1.81E-279 | 2 | Smc5 | myCAF_prolif |
| Aimp2 | 2.31E-283 | 0.71859033 | 0.68 | 0.445 | 4.47E-279 | 2 | Aimp2 | myCAF_prolif |
| Zmynd19 | 6.82E-283 | 0.75450622 | 0.568 | 0.382 | 1.32E-278 | 2 | Zmynd19 | myCAF_prolif |
| Il1b | 7.12E-283 | 1.01616298 | 0.453 | 0.372 | 1.38E-278 | 2 | Il1b | myCAF_prolif |
| Soga1 | 3.87E-282 | 0.60122066 | 0.76 | 0.602 | 7.47E-278 | 2 | Soga1 | myCAF_prolif |
| Lrwd1 | 8.02E-282 | 1.0031892 | 0.602 | 0.302 | 1.55E-277 | 2 | Lrwd1 | myCAF_prolif |
| Hnrnpa0 | 1.10E-281 | 0.35453785 | 0.88 | 0.773 | 2.13E-277 | 2 | Hnrnpa0 | myCAF_prolif |
| Rassf1 | 2.70E-281 | 0.27988591 | 0.841 | 0.743 | 5.22E-277 | 2 | Rassf1 | myCAF_prolif |
| Mars | 3.42E-281 | 0.52829759 | 0.775 | 0.602 | 6.61E-277 | 2 | Mars | myCAF_prolif |
| Ier3ip1 | 3.50E-281 | 0.28311531 | 0.854 | 0.76 | 6.76E-277 | 2 | Ier3ip1 | myCAF_prolif |
| Brca1 | 7.32E-281 | 5.32427945 | 0.37 | 0.009 | 1.41E-276 | 2 | Brca1 | myCAF_prolif |
| Smg5 | 1.53E-280 | 0.63070778 | 0.765 | 0.546 | 2.96E-276 | 2 | Smg5 | myCAF_prolif |
| Nab1 | 4.90E-280 | 0.35915262 | 0.801 | 0.683 | 9.47E-276 | 2 | Nab1 | myCAF_prolif |
| Bckdk | 8.16E-280 | 0.36726131 | 0.744 | 0.606 | 1.58E-275 | 2 | Bckdk | myCAF_prolif |
| Ppp3cb | 8.74E-280 | 0.28844432 | 0.848 | 0.761 | 1.69E-275 | 2 | Ppp3cb | myCAF_prolif |
| Hyls1 | 1.33E-279 | 2.02788097 | 0.308 | 0.081 | 2.58E-275 | 2 | Hyls1 | myCAF_prolif |
| Ccdc124 | 4.97E-279 | 0.37835737 | 0.879 | 0.753 | 9.62E-275 | 2 | Ccdc124 | myCAF_prolif |
| Isy1 | 1.80E-278 | 0.49383051 | 0.769 | 0.553 | 3.48E-274 | 2 | Isy1 | myCAF_prolif |
| Fkbp4 | 2.71E-278 | 0.61646998 | 0.703 | 0.473 | 5.24E-274 | 2 | Fkbp4 | myCAF_prolif |
| Stk11 | 2.76E-278 | 0.4049328 | 0.854 | 0.746 | 5.33E-274 | 2 | Stk11 | myCAF_prolif |
| Pkp4 | 1.12E-277 | 1.56525997 | 0.441 | 0.138 | 2.17E-273 | 2 | Pkp4 | myCAF_prolif |
| Cnep1r1 | 3.04E-277 | 0.33735357 | 0.791 | 0.669 | 5.87E-273 | 2 | Cnep1r1 | myCAF_prolif |
| Tmem165 | 4.78E-277 | 0.27371298 | 0.847 | 0.774 | 9.25E-273 | 2 | Tmem165 | myCAF_prolif |
| Dnajc251 | 6.49E-277 | 0.32850299 | 0.808 | 0.69 | 1.25E-272 | 2 | Dnajc25 | myCAF_prolif |
| Eif4h | 1.81E-276 | 0.44679395 | 0.812 | 0.678 | 3.50E-272 | 2 | Eif4h | myCAF_prolif |
| Prkaca | 2.38E-276 | 0.26941181 | 0.846 | 0.769 | 4.60E-272 | 2 | Prkaca | myCAF_prolif |
| Recql | 4.02E-276 | 1.06121841 | 0.634 | 0.37 | 7.78E-272 | 2 | Recql | myCAF_prolif |
| Etf1 | 6.89E-276 | 0.41007124 | 0.746 | 0.584 | 1.33E-271 | 2 | Etf1 | myCAF_prolif |
| Gmps | 9.69E-276 | 0.54445135 | 0.716 | 0.498 | 1.87E-271 | 2 | Gmps | myCAF_prolif |
| Bola2 | 1.27E-275 | 0.27595878 | 0.834 | 0.745 | 2.46E-271 | 2 | Bola2 | myCAF_prolif |
| Hdac1 | 1.44E-275 | 0.39055111 | 0.816 | 0.681 | 2.79E-271 | 2 | Hdac1 | myCAF_prolif |
| Anapc4 | 1.86E-275 | 0.34535752 | 0.822 | 0.703 | 3.60E-271 | 2 | Anapc4 | myCAF_prolif |
| Tmtc41 | 1.90E-275 | 0.81277328 | 0.573 | 0.331 | 3.68E-271 | 2 | Tmtc4 | myCAF_prolif |
| Atp5k | 6.42E-275 | 0.34886451 | 0.815 | 0.686 | 1.24E-270 | 2 | Atp5k | myCAF_prolif |
| Rac1 | 1.06E-274 | 0.31703695 | 0.841 | 0.745 | 2.05E-270 | 2 | Rac1 | myCAF_prolif |
| Eloc | 1.35E-274 | 0.46791534 | 0.725 | 0.553 | 2.60E-270 | 2 | Eloc | myCAF_prolif |
| Ska1 | 1.43E-274 | 6.8397768 | 0.503 | 0.006 | 2.76E-270 | 2 | Ska1 | myCAF_prolif |
| Clcc1 | 1.62E-274 | 0.75654558 | 0.668 | 0.435 | 3.13E-270 | 2 | Clcc1 | myCAF_prolif |
| Thrap3 | 2.35E-274 | 0.54636438 | 0.786 | 0.607 | 4.55E-270 | 2 | Thrap3 | myCAF_prolif |
| Ss18 | 2.55E-274 | 0.25548477 | 0.809 | 0.703 | 4.92E-270 | 2 | Ss18 | myCAF_prolif |
| Bcl7c | 5.02E-274 | 0.40815122 | 0.843 | 0.702 | 9.71E-270 | 2 | Bcl7c | myCAF_prolif |
| Hmgn1 | 5.69E-274 | 0.4693802 | 0.788 | 0.6 | 1.10E-269 | 2 | Hmgn1 | myCAF_prolif |
| Casp7 | 1.08E-273 | 1.08990628 | 0.497 | 0.244 | 2.09E-269 | 2 | Casp7 | myCAF_prolif |
| Abracl1 | 1.52E-273 | 0.34227531 | 0.815 | 0.698 | 2.94E-269 | 2 | Abracl | myCAF_prolif |
| Fh1 | 1.54E-273 | 0.66463027 | 0.691 | 0.476 | 2.97E-269 | 2 | Fh1 | myCAF_prolif |
| Prkar2a | 1.94E-273 | 0.8039153 | 0.64 | 0.399 | 3.75E-269 | 2 | Prkar2a | myCAF_prolif |
| Tmem209 | 3.19E-273 | 1.14047713 | 0.544 | 0.261 | 6.16E-269 | 2 | Tmem209 | myCAF_prolif |
| Srrm1 | 3.86E-273 | 0.42340888 | 0.807 | 0.661 | 7.47E-269 | 2 | Srrm1 | myCAF_prolif |
| Mfsd14b | 5.15E-273 | 0.31482686 | 0.77 | 0.656 | 9.95E-269 | 2 | Mfsd14b | myCAF_prolif |
| Fam98b | 5.99E-273 | 0.52861391 | 0.759 | 0.562 | 1.16E-268 | 2 | Fam98b | myCAF_prolif |
| Wdr77 | 3.52E-272 | 0.47685763 | 0.716 | 0.547 | 6.81E-268 | 2 | Wdr77 | myCAF_prolif |
| Gemin7 | 5.28E-272 | 0.33065908 | 0.827 | 0.713 | 1.02E-267 | 2 | Gemin7 | myCAF_prolif |
| Luc7l2 | 6.07E-272 | 0.27381866 | 0.916 | 0.896 | 1.17E-267 | 2 | Luc7l2 | myCAF_prolif |
| Pea15a | 7.50E-272 | 0.33688587 | 0.844 | 0.713 | 1.45E-267 | 2 | Pea15a | myCAF_prolif |
| Nudcd1 | 1.09E-271 | 0.82906231 | 0.594 | 0.334 | 2.10E-267 | 2 | Nudcd1 | myCAF_prolif |
| Gars | 2.44E-271 | 0.48254796 | 0.708 | 0.526 | 4.71E-267 | 2 | Gars | myCAF_prolif |
| Atp10a | 2.53E-271 | 0.71568876 | 0.615 | 0.365 | 4.88E-267 | 2 | Atp10a | myCAF_prolif |
| Mlip | 4.96E-271 | 1.2298207 | 0.466 | 0.252 | 9.59E-267 | 2 | Mlip | myCAF_prolif |
| Atl2 | 7.72E-271 | 0.6920969 | 0.675 | 0.436 | 1.49E-266 | 2 | Atl2 | myCAF_prolif |
| Pih1d1 | 9.34E-271 | 0.74600881 | 0.674 | 0.42 | 1.81E-266 | 2 | Pih1d1 | myCAF_prolif |
| Ino80e | 1.19E-270 | 0.58810054 | 0.751 | 0.524 | 2.30E-266 | 2 | Ino80e | myCAF_prolif |
| Med9 | 1.85E-270 | 0.68384469 | 0.699 | 0.458 | 3.58E-266 | 2 | Med9 | myCAF_prolif |
| Usp5 | 3.66E-270 | 0.37697478 | 0.839 | 0.723 | 7.08E-266 | 2 | Usp5 | myCAF_prolif |
| Kras | 4.12E-270 | 0.43430933 | 0.761 | 0.572 | 7.96E-266 | 2 | Kras | myCAF_prolif |
| Ubl4a | 6.22E-270 | 0.44536042 | 0.795 | 0.611 | 1.20E-265 | 2 | Ubl4a | myCAF_prolif |
| Fam98a | 1.46E-268 | 0.54515301 | 0.744 | 0.575 | 2.83E-264 | 2 | Fam98a | myCAF_prolif |
| Med4 | 3.21E-268 | 0.8861809 | 0.611 | 0.32 | 6.21E-264 | 2 | Med4 | myCAF_prolif |
| Lsm7 | 9.66E-268 | 0.38690389 | 0.812 | 0.67 | 1.87E-263 | 2 | Lsm7 | myCAF_prolif |
| Txndc12 | 1.30E-267 | 0.44479249 | 0.766 | 0.614 | 2.52E-263 | 2 | Txndc12 | myCAF_prolif |
| Dynlt1f | 2.06E-267 | 0.56458084 | 0.703 | 0.493 | 3.98E-263 | 2 | Dynlt1f | myCAF_prolif |
| Tpr | 1.04E-266 | 0.36142554 | 0.787 | 0.676 | 2.01E-262 | 2 | Tpr | myCAF_prolif |
| BC055324 | 3.03E-266 | 3.54402046 | 0.304 | 0.025 | 5.87E-262 | 2 | BC055324 | myCAF_prolif |
| Txn2 | 4.11E-266 | 0.27016595 | 0.797 | 0.717 | 7.95E-262 | 2 | Txn2 | myCAF_prolif |
| Hnrnpdl | 5.41E-266 | 0.60731786 | 0.728 | 0.515 | 1.05E-261 | 2 | Hnrnpdl | myCAF_prolif |
| Hdac3 | 1.23E-265 | 0.32292677 | 0.804 | 0.692 | 2.39E-261 | 2 | Hdac3 | myCAF_prolif |
| Wdr5 | 1.62E-265 | 0.6092343 | 0.698 | 0.479 | 3.14E-261 | 2 | Wdr5 | myCAF_prolif |
| Nacc1 | 1.97E-265 | 0.49698257 | 0.752 | 0.547 | 3.82E-261 | 2 | Nacc1 | myCAF_prolif |
| Nmt2 | 4.12E-265 | 0.38882318 | 0.77 | 0.627 | 7.97E-261 | 2 | Nmt2 | myCAF_prolif |
| Psmd3 | 5.27E-265 | 0.28997566 | 0.804 | 0.698 | 1.02E-260 | 2 | Psmd3 | myCAF_prolif |
| Kif14 | 6.63E-265 | 7.02058338 | 0.534 | 0.004 | 1.28E-260 | 2 | Kif14 | myCAF_prolif |
| Enkd1 | 6.85E-265 | 1.99610735 | 0.409 | 0.117 | 1.32E-260 | 2 | Enkd1 | myCAF_prolif |
| Acot7 | 1.10E-264 | 0.54769753 | 0.687 | 0.488 | 2.12E-260 | 2 | Acot7 | myCAF_prolif |
| Cenpk | 1.12E-264 | 4.10069953 | 0.309 | 0.022 | 2.16E-260 | 2 | Cenpk | myCAF_prolif |
| Rab231 | 1.60E-264 | 0.55480406 | 0.667 | 0.472 | 3.08E-260 | 2 | Rab23 | myCAF_prolif |
| Cct4 | 2.46E-264 | 0.32434729 | 0.76 | 0.655 | 4.76E-260 | 2 | Cct4 | myCAF_prolif |
| Myc1 | 3.37E-264 | 0.33845183 | 0.672 | 0.538 | 6.52E-260 | 2 | Myc | myCAF_prolif |
| Pom121 | 3.65E-264 | 0.56417732 | 0.711 | 0.493 | 7.06E-260 | 2 | Pom121 | myCAF_prolif |
| Tcerg1 | 6.78E-264 | 0.7873416 | 0.708 | 0.477 | 1.31E-259 | 2 | Tcerg1 | myCAF_prolif |
| Cenpx | 7.19E-264 | 0.62178484 | 0.687 | 0.454 | 1.39E-259 | 2 | Cenpx | myCAF_prolif |
| Casp3 | 1.22E-263 | 0.50108488 | 0.733 | 0.529 | 2.36E-259 | 2 | Casp3 | myCAF_prolif |
| Ankrd52 | 1.56E-263 | 0.4357543 | 0.819 | 0.675 | 3.02E-259 | 2 | Ankrd52 | myCAF_prolif |
| Rad9a | 2.16E-263 | 0.97548214 | 0.551 | 0.254 | 4.18E-259 | 2 | Rad9a | myCAF_prolif |
| Rnf168 | 3.68E-263 | 0.88803619 | 0.557 | 0.319 | 7.12E-259 | 2 | Rnf168 | myCAF_prolif |
| Mns1 | 5.54E-263 | 4.65234119 | 0.315 | 0.014 | 1.07E-258 | 2 | Mns1 | myCAF_prolif |
| Adgre5 | 7.14E-263 | 0.30897124 | 0.707 | 0.596 | 1.38E-258 | 2 | Adgre5 | myCAF_prolif |
| Mesd | 1.73E-262 | 0.25637375 | 0.839 | 0.742 | 3.34E-258 | 2 | Mesd | myCAF_prolif |
| Stat1 | 1.99E-262 | 0.42324244 | 0.727 | 0.523 | 3.85E-258 | 2 | Stat1 | myCAF_prolif |

| | | | | | | | | |
|---|---|---|---|---|---|---|---|---|
| Cox17 | 4.34E-262 | 0.41452087 | 0.777 | 0.634 | 8.38E-258 | 2 | Cox17 | myCAF_prolif |
| Lclat11 | 4.63E-262 | 0.41560259 | 0.759 | 0.561 | 8.96E-258 | 2 | Lclat1 | myCAF_prolif |
| Npm31 | 8.29E-262 | 0.77104607 | 0.661 | 0.424 | 1.60E-257 | 2 | Npm3 | myCAF_prolif |
| Parp2 | 2.11E-261 | 0.9981689 | 0.577 | 0.301 | 4.09E-257 | 2 | Parp2 | myCAF_prolif |
| Pgp | 2.67E-261 | 0.47903591 | 0.749 | 0.538 | 5.17E-257 | 2 | Pgp | myCAF_prolif |
| Msh6 | 5.19E-261 | 1.3043454 | 0.396 | 0.184 | 1.00E-256 | 2 | Msh6 | myCAF_prolif |
| Dpy19l1 | 6.30E-261 | 0.34324993 | 0.782 | 0.655 | 1.22E-256 | 2 | Dpy19l1 | myCAF_prolif |
| Cep250 | 1.26E-260 | 1.45914365 | 0.501 | 0.199 | 2.43E-256 | 2 | Cep250 | myCAF_prolif |
| Stard8 | 1.44E-260 | 1.67706816 | 0.328 | 0.104 | 2.78E-256 | 2 | Stard8 | myCAF_prolif |
| Psmb6 | 1.77E-260 | 0.39116184 | 0.79 | 0.638 | 3.43E-256 | 2 | Psmb6 | myCAF_prolif |
| Erf1 | 3.23E-260 | 0.25125509 | 0.832 | 0.708 | 6.25E-256 | 2 | Erf | myCAF_prolif |
| Dcp2 | 3.27E-260 | 0.64742921 | 0.672 | 0.44 | 6.33E-256 | 2 | Dcp2 | myCAF_prolif |
| Denr | 4.77E-260 | 0.36730166 | 0.818 | 0.699 | 9.23E-256 | 2 | Denr | myCAF_prolif |
| Zw10 | 1.32E-259 | 0.92536013 | 0.555 | 0.299 | 2.54E-255 | 2 | Zw10 | myCAF_prolif |
| Fgd1 | 4.47E-259 | 0.71628965 | 0.657 | 0.408 | 8.65E-255 | 2 | Fgd1 | myCAF_prolif |
| Ndufa4 | 1.55E-258 | 0.41573565 | 0.739 | 0.572 | 3.00E-254 | 2 | Ndufa4 | myCAF_prolif |
| Cgas | 2.61E-258 | 2.32449181 | 0.322 | 0.07 | 5.05E-254 | 2 | Cgas | myCAF_prolif |
| Nomo1 | 1.17E-257 | 0.40995194 | 0.742 | 0.589 | 2.26E-253 | 2 | Nomo1 | myCAF_prolif |
| Pml | 1.60E-257 | 0.538753 | 0.725 | 0.521 | 3.10E-253 | 2 | Pml | myCAF_prolif |
| Myg1 | 2.59E-257 | 0.8018628 | 0.604 | 0.33 | 5.00E-253 | 2 | Myg1 | myCAF_prolif |
| Cep76 | 2.95E-257 | 1.43108706 | 0.459 | 0.158 | 5.71E-253 | 2 | Cep76 | myCAF_prolif |
| Gpaa1 | 6.63E-257 | 0.54305653 | 0.727 | 0.481 | 1.28E-252 | 2 | Gpaa1 | myCAF_prolif |
| Mybbp1a | 3.49E-256 | 0.32488143 | 0.76 | 0.644 | 6.75E-252 | 2 | Mybbp1a | myCAF_prolif |
| Slc35b4 | 6.64E-256 | 0.32160264 | 0.779 | 0.656 | 1.28E-251 | 2 | Slc35b4 | myCAF_prolif |
| Polr3k | 6.94E-256 | 0.5210529 | 0.729 | 0.516 | 1.34E-251 | 2 | Polr3k | myCAF_prolif |
| Cpox | 7.99E-256 | 0.62195168 | 0.609 | 0.383 | 1.55E-251 | 2 | Cpox | myCAF_prolif |
| Nfatc2ip | 1.61E-255 | 1.4978884 | 0.415 | 0.13 | 3.11E-251 | 2 | Nfatc2ip | myCAF_prolif |
| Ndufc1 | 1.96E-255 | 0.37933216 | 0.815 | 0.69 | 3.78E-251 | 2 | Ndufc1 | myCAF_prolif |
| Hspe1 | 5.71E-255 | 0.41981423 | 0.742 | 0.591 | 1.10E-250 | 2 | Hspe1 | myCAF_prolif |
| Eif6 | 2.21E-254 | 0.4290977 | 0.801 | 0.643 | 4.27E-250 | 2 | Eif6 | myCAF_prolif |
| Tcirg1 | 2.43E-254 | 0.4755289 | 0.777 | 0.595 | 4.70E-250 | 2 | Tcirg1 | myCAF_prolif |
| Anapc15 | 4.43E-254 | 1.44410305 | 0.47 | 0.162 | 8.57E-250 | 2 | Anapc15 | myCAF_prolif |
| Hmgn5 | 7.55E-254 | 1.23053336 | 0.506 | 0.221 | 1.46E-249 | 2 | Hmgn5 | myCAF_prolif |
| Dnajc8 | 4.65E-253 | 0.47537291 | 0.756 | 0.605 | 9.00E-249 | 2 | Dnajc8 | myCAF_prolif |
| Sf3b6 | 5.59E-253 | 0.27704185 | 0.801 | 0.705 | 1.08E-248 | 2 | Sf3b6 | myCAF_prolif |
| Cct8 | 6.62E-253 | 0.3369962 | 0.759 | 0.639 | 1.28E-248 | 2 | Cct8 | myCAF_prolif |
| Map3k20 | 9.83E-253 | 0.43402706 | 0.718 | 0.55 | 1.90E-248 | 2 | Map3k20 | myCAF_prolif |
| Psmd6 | 1.47E-252 | 0.27648265 | 0.777 | 0.68 | 2.84E-248 | 2 | Psmd6 | myCAF_prolif |
| Pcmt1 | 1.64E-252 | 0.36204711 | 0.758 | 0.614 | 3.18E-248 | 2 | Pcmt1 | myCAF_prolif |
| Timm23 | 1.96E-252 | 0.4437873 | 0.747 | 0.574 | 3.78E-248 | 2 | Timm23 | myCAF_prolif |
| Cenpp | 2.19E-252 | 4.45666046 | 0.287 | 0.012 | 4.23E-248 | 2 | Cenpp | myCAF_prolif |
| Map3k3 | 7.96E-252 | 0.41844197 | 0.747 | 0.593 | 1.54E-247 | 2 | Map3k3 | myCAF_prolif |
| Adam10 | 1.02E-251 | 0.26661135 | 0.797 | 0.676 | 1.97E-247 | 2 | Adam10 | myCAF_prolif |
| Abcf2 | 2.17E-251 | 0.60685473 | 0.674 | 0.463 | 4.20E-247 | 2 | Abcf2 | myCAF_prolif |
| Tnfaip31 | 4.69E-251 | 0.53510529 | 0.373 | 0.279 | 9.07E-247 | 2 | Tnfaip3 | myCAF_prolif |
| Lars | 8.92E-251 | 0.35886562 | 0.745 | 0.575 | 1.73E-246 | 2 | Lars | myCAF_prolif |
| Noc2l | 1.65E-250 | 0.32125394 | 0.741 | 0.621 | 3.20E-246 | 2 | Noc2l | myCAF_prolif |
| Gnl3 | 6.88E-250 | 0.36017187 | 0.768 | 0.649 | 1.33E-245 | 2 | Gnl3 | myCAF_prolif |
| Vps36 | 4.73E-249 | 0.57053965 | 0.716 | 0.504 | 9.15E-245 | 2 | Vps36 | myCAF_prolif |
| Wrnip1 | 6.81E-249 | 0.4683584 | 0.729 | 0.53 | 1.32E-244 | 2 | Wrnip1 | myCAF_prolif |
| Upf1 | 1.52E-248 | 0.34294805 | 0.837 | 0.698 | 2.95E-244 | 2 | Upf1 | myCAF_prolif |
| Rad51b | 2.17E-248 | 4.58266795 | 0.294 | 0.015 | 4.19E-244 | 2 | Rad51b | myCAF_prolif |
| Strap | 3.81E-248 | 0.43809042 | 0.739 | 0.582 | 7.38E-244 | 2 | Strap | myCAF_prolif |
| Usp47 | 4.19E-248 | 0.27247938 | 0.825 | 0.737 | 8.11E-244 | 2 | Usp47 | myCAF_prolif |
| Tdp1 | 5.78E-248 | 1.01247638 | 0.518 | 0.24 | 1.12E-243 | 2 | Tdp1 | myCAF_prolif |
| Plaa | 7.89E-248 | 0.3749349 | 0.771 | 0.617 | 1.53E-243 | 2 | Plaa | myCAF_prolif |
| Net1 | 9.48E-248 | 1.06634692 | 0.499 | 0.191 | 1.83E-243 | 2 | Net1 | myCAF_prolif |
| Rrn3 | 9.79E-248 | 0.63080537 | 0.727 | 0.526 | 1.89E-243 | 2 | Rrn3 | myCAF_prolif |
| Nsun2 | 4.03E-247 | 0.36027769 | 0.747 | 0.62 | 7.79E-243 | 2 | Nsun2 | myCAF_prolif |
| Rtel1 | 1.78E-246 | 1.85355751 | 0.353 | 0.082 | 3.44E-242 | 2 | Rtel1 | myCAF_prolif |
| Fads2 | 2.19E-246 | 0.42466173 | 0.709 | 0.525 | 4.24E-242 | 2 | Fads2 | myCAF_prolif |
| Utp6 | 3.59E-246 | 0.58037021 | 0.691 | 0.479 | 6.94E-242 | 2 | Utp6 | myCAF_prolif |
| Nap1l4 | 5.83E-246 | 0.49360453 | 0.781 | 0.629 | 1.13E-241 | 2 | Nap1l4 | myCAF_prolif |
| Cdc16 | 2.18E-245 | 0.28374969 | 0.789 | 0.684 | 4.21E-241 | 2 | Cdc16 | myCAF_prolif |
| Pak1ip1 | 2.85E-245 | 0.50344102 | 0.705 | 0.523 | 5.50E-241 | 2 | Pak1ip1 | myCAF_prolif |
| Sun2 | 4.98E-245 | 0.51494212 | 0.627 | 0.487 | 9.63E-241 | 2 | Sun2 | myCAF_prolif |
| Tsr2 | 6.43E-245 | 0.69326852 | 0.657 | 0.399 | 1.24E-240 | 2 | Tsr2 | myCAF_prolif |
| Coq7 | 7.08E-245 | 0.87525573 | 0.558 | 0.298 | 1.37E-240 | 2 | Coq7 | myCAF_prolif |
| Ndufab1 | 1.30E-244 | 0.52932045 | 0.69 | 0.485 | 2.51E-240 | 2 | Ndufab1 | myCAF_prolif |
| Runx2 | 1.34E-244 | 0.4597242 | 0.788 | 0.647 | 2.58E-240 | 2 | Runx2 | myCAF_prolif |
| Ywhaq | 1.85E-244 | 0.39493037 | 0.74 | 0.605 | 3.57E-240 | 2 | Ywhaq | myCAF_prolif |
| Syne2 | 1.97E-244 | 1.18217548 | 0.479 | 0.209 | 3.80E-240 | 2 | Syne2 | myCAF_prolif |
| Trak2 | 2.15E-244 | 0.51582382 | 0.794 | 0.646 | 4.15E-240 | 2 | Trak2 | myCAF_prolif |
| Mtrex | 4.33E-244 | 0.74550239 | 0.627 | 0.386 | 8.37E-240 | 2 | Mtrex | myCAF_prolif |
| Grk2 | 9.19E-244 | 0.56039096 | 0.734 | 0.555 | 1.78E-239 | 2 | Grk2 | myCAF_prolif |
| Cs | 1.24E-243 | 0.53007991 | 0.751 | 0.563 | 2.40E-239 | 2 | Cs | myCAF_prolif |
| Anapc5 | 1.41E-243 | 0.587808 | 0.696 | 0.522 | 2.72E-239 | 2 | Anapc5 | myCAF_prolif |
| Tbc1d31 | 1.51E-243 | 2.0186791 | 0.354 | 0.089 | 2.92E-239 | 2 | Tbc1d31 | myCAF_prolif |
| Uqcc2 | 2.35E-243 | 0.30960457 | 0.81 | 0.68 | 4.55E-239 | 2 | Uqcc2 | myCAF_prolif |
| Cep57 | 4.06E-243 | 1.29538687 | 0.442 | 0.181 | 7.85E-239 | 2 | Cep57 | myCAF_prolif |
| Stoml2 | 7.24E-243 | 0.45855969 | 0.756 | 0.578 | 1.40E-238 | 2 | Stoml2 | myCAF_prolif |
| Prpf38a | 3.23E-242 | 0.669079 | 0.651 | 0.417 | 6.25E-238 | 2 | Prpf38a | myCAF_prolif |
| Ifi30 | 4.14E-242 | 0.62337788 | 0.641 | 0.418 | 8.00E-238 | 2 | Ifi30 | myCAF_prolif |
| Gsk3a | 6.71E-242 | 0.27899168 | 0.835 | 0.72 | 1.30E-237 | 2 | Gsk3a | myCAF_prolif |
| Erh | 1.10E-241 | 0.94810304 | 0.582 | 0.316 | 2.13E-237 | 2 | Erh | myCAF_prolif |
| Eif3c | 4.38E-241 | 0.38011261 | 0.815 | 0.711 | 8.46E-237 | 2 | Eif3c | myCAF_prolif |
| Wdr62 | 5.08E-241 | 3.39586412 | 0.295 | 0.023 | 9.82E-237 | 2 | Wdr62 | myCAF_prolif |
| Zc3h4 | 5.89E-241 | 0.50294556 | 0.785 | 0.599 | 1.14E-236 | 2 | Zc3h4 | myCAF_prolif |
| Tmed1 | 2.10E-240 | 0.27904846 | 0.81 | 0.68 | 4.05E-236 | 2 | Tmed1 | myCAF_prolif |
| Utp11 | 4.08E-240 | 0.44501594 | 0.786 | 0.631 | 7.90E-236 | 2 | Utp11 | myCAF_prolif |
| Fam110a1 | 1.40E-239 | 0.74678427 | 0.585 | 0.36 | 2.71E-235 | 2 | Fam110a | myCAF_prolif |
| Nudt3 | 1.90E-239 | 0.29684897 | 0.888 | 0.807 | 3.67E-235 | 2 | Nudt3 | myCAF_prolif |
| Rabl6 | 2.07E-239 | 0.35189622 | 0.827 | 0.725 | 4.00E-235 | 2 | Rabl6 | myCAF_prolif |
| Stard4 | 2.17E-239 | 0.56824199 | 0.642 | 0.403 | 4.19E-235 | 2 | Stard4 | myCAF_prolif |
| Psmg2 | 2.35E-239 | 0.81231819 | 0.599 | 0.336 | 4.55E-235 | 2 | Psmg2 | myCAF_prolif |
| Spg7 | 3.21E-239 | 0.35522871 | 0.789 | 0.647 | 6.20E-235 | 2 | Spg7 | myCAF_prolif |
| Utp3 | 4.90E-239 | 0.73587466 | 0.683 | 0.427 | 9.47E-235 | 2 | Utp3 | myCAF_prolif |
| Rcbtb2 | 6.25E-239 | 0.52205248 | 0.723 | 0.541 | 1.21E-234 | 2 | Rcbtb2 | myCAF_prolif |
| Smu1 | 1.08E-238 | 0.41325766 | 0.766 | 0.62 | 2.09E-234 | 2 | Smu1 | myCAF_prolif |
| Pnrc2 | 3.43E-238 | 0.41955089 | 0.78 | 0.673 | 6.63E-234 | 2 | Pnrc2 | myCAF_prolif |
| Alg10b | 2.16E-237 | 0.31954381 | 0.794 | 0.653 | 4.17E-233 | 2 | Alg10b | myCAF_prolif |
| Safb | 6.94E-237 | 0.64683933 | 0.752 | 0.57 | 1.34E-232 | 2 | Safb | myCAF_prolif |
| Hypk | 1.18E-236 | 0.39549697 | 0.799 | 0.63 | 2.28E-232 | 2 | Hypk | myCAF_prolif |
| Gfm1 | 1.53E-236 | 0.62278969 | 0.657 | 0.427 | 2.96E-232 | 2 | Gfm1 | myCAF_prolif |
| Mmgt1 | 4.29E-236 | 0.87930783 | 0.555 | 0.296 | 8.30E-232 | 2 | Mmgt1 | myCAF_prolif |
| Oard1 | 6.12E-236 | 0.83523008 | 0.613 | 0.346 | 1.18E-231 | 2 | Oard1 | myCAF_prolif |
| Cyb5b | 1.28E-235 | 0.34198545 | 0.805 | 0.649 | 2.48E-231 | 2 | Cyb5b | myCAF_prolif |
| Suds3 | 1.44E-235 | 0.32224117 | 0.849 | 0.755 | 2.79E-231 | 2 | Suds3 | myCAF_prolif |
| Myo5a | 2.07E-235 | 0.31459441 | 0.786 | 0.654 | 4.00E-231 | 2 | Myo5a | myCAF_prolif |
| Rbmx2 | 2.37E-235 | 1.80968949 | 0.353 | 0.085 | 4.58E-231 | 2 | Rbmx2 | myCAF_prolif |
| Ranbp2 | 5.21E-235 | 0.28717423 | 0.815 | 0.73 | 1.01E-230 | 2 | Ranbp2 | myCAF_prolif |
| Zcchc8 | 7.42E-235 | 0.86171288 | 0.628 | 0.361 | 1.44E-230 | 2 | Zcchc8 | myCAF_prolif |
| Tiprl | 1.27E-234 | 0.29551005 | 0.818 | 0.699 | 2.45E-230 | 2 | Tiprl | myCAF_prolif |
| Zbed3 | 6.31E-234 | 0.72413182 | 0.638 | 0.447 | 1.22E-229 | 2 | Zbed3 | myCAF_prolif |
| Kpna4 | 4.85E-233 | 0.43092353 | 0.758 | 0.597 | 9.38E-229 | 2 | Kpna4 | myCAF_prolif |
| Serpinb1a1 | 9.67E-233 | 0.66256668 | 0.366 | 0.233 | 1.87E-228 | 2 | Serpinb1a | myCAF_prolif |
| Hspa4 | 2.63E-232 | 0.41845425 | 0.753 | 0.602 | 5.08E-228 | 2 | Hspa4 | myCAF_prolif |
| Nemp1 | 1.09E-231 | 2.14620575 | 0.297 | 0.05 | 2.11E-227 | 2 | Nemp1 | myCAF_prolif |
| Rad50 | 1.21E-231 | 0.85833524 | 0.578 | 0.322 | 2.35E-227 | 2 | Rad50 | myCAF_prolif |
| Obsl11 | 2.09E-231 | 0.37091817 | 0.792 | 0.651 | 4.04E-227 | 2 | Obsl1 | myCAF_prolif |
| Cpsf4 | 2.17E-231 | 0.7908376 | 0.577 | 0.332 | 4.19E-227 | 2 | Cpsf4 | myCAF_prolif |
| Cbx1 | 2.25E-231 | 1.00406989 | 0.643 | 0.385 | 4.34E-227 | 2 | Cbx1 | myCAF_prolif |
| Nfyb | 2.94E-231 | 0.66728329 | 0.653 | 0.436 | 5.68E-227 | 2 | Nfyb | myCAF_prolif |
| Pds5b | 8.80E-231 | 0.67120261 | 0.632 | 0.425 | 1.70E-226 | 2 | Pds5b | myCAF_prolif |
| Brix1 | 8.83E-231 | 0.57438076 | 0.696 | 0.506 | 1.71E-226 | 2 | Brix1 | myCAF_prolif |
| Cisd1 | 2.63E-230 | 0.70621693 | 0.626 | 0.403 | 5.09E-226 | 2 | Cisd1 | myCAF_prolif |
| Tdp2 | 5.38E-230 | 0.93334756 | 0.592 | 0.302 | 1.04E-225 | 2 | Tdp2 | myCAF_prolif |
| Matr3 | 7.21E-230 | 0.49732387 | 0.797 | 0.648 | 1.39E-225 | 2 | Matr3 | myCAF_prolif |
| Zdhhc161 | 1.27E-229 | 0.48976353 | 0.757 | 0.573 | 2.46E-225 | 2 | Zdhhc16 | myCAF_prolif |
| Naa40 | 3.45E-229 | 0.890162 | 0.518 | 0.275 | 6.67E-225 | 2 | Naa40 | myCAF_prolif |
| Nutf2-ps1 | 8.63E-229 | 0.68004402 | 0.67 | 0.436 | 1.67E-224 | 2 | Nutf2-ps1 | myCAF_prolif |
| Fut10 | 1.41E-228 | 0.75961203 | 0.597 | 0.364 | 2.72E-224 | 2 | Fut10 | myCAF_prolif |
| Tnfsf11 | 3.09E-228 | 0.3920439 | 0.477 | 0.279 | 5.98E-224 | 2 | Tnfsf11 | myCAF_prolif |
| Haus3 | 3.16E-228 | 1.24793801 | 0.497 | 0.216 | 6.12E-224 | 2 | Haus3 | myCAF_prolif |
| Trmt112 | 6.40E-228 | 0.28175445 | 0.847 | 0.761 | 1.24E-223 | 2 | Trmt112 | myCAF_prolif |
| Papss21 | 6.48E-228 | 0.33650159 | 0.747 | 0.602 | 1.25E-223 | 2 | Papss2 | myCAF_prolif |
| Cox6a1 | 6.69E-228 | 0.40519773 | 0.804 | 0.655 | 1.29E-223 | 2 | Cox6a1 | myCAF_prolif |
| Bzw2 | 1.99E-227 | 0.55545639 | 0.686 | 0.468 | 3.85E-223 | 2 | Bzw2 | myCAF_prolif |
| Mpdu1 | 2.82E-227 | 0.26922828 | 0.846 | 0.74 | 5.45E-223 | 2 | Mpdu1 | myCAF_prolif |
| Zc3h18 | 4.05E-227 | 0.62196848 | 0.701 | 0.488 | 7.82E-223 | 2 | Zc3h18 | myCAF_prolif |
| Raet1d | 5.95E-227 | 2.14416226 | 0.259 | 0.049 | 1.15E-222 | 2 | Raet1d | myCAF_prolif |
| Etaa1 | 2.08E-226 | 1.46070548 | 0.389 | 0.126 | 4.03E-222 | 2 | Etaa1 | myCAF_prolif |
| Ddias | 3.36E-226 | 3.20361613 | 0.259 | 0.024 | 6.49E-222 | 2 | Ddias | myCAF_prolif |
| Rpp30 | 4.65E-226 | 1.23637293 | 0.449 | 0.182 | 8.98E-222 | 2 | Rpp30 | myCAF_prolif |
| Fkbp3 | 1.04E-225 | 0.53667803 | 0.647 | 0.437 | 2.02E-221 | 2 | Fkbp3 | myCAF_prolif |
| Fbxo48 | 1.45E-225 | 1.75605165 | 0.297 | 0.049 | 2.81E-221 | 2 | Fbxo48 | myCAF_prolif |
| Ctc1 | 2.61E-225 | 1.44190194 | 0.423 | 0.145 | 5.05E-221 | 2 | Ctc1 | myCAF_prolif |
| Lsm12 | 4.19E-225 | 0.27499801 | 0.804 | 0.69 | 8.11E-221 | 2 | Lsm12 | myCAF_prolif |
| Med14 | 6.47E-225 | 0.70474032 | 0.587 | 0.372 | 1.25E-220 | 2 | Med14 | myCAF_prolif |
| Zdhhc15 | 1.12E-224 | 1.64131233 | 0.355 | 0.1 | 2.17E-220 | 2 | Zdhhc15 | myCAF_prolif |
| Pbrm1 | 1.13E-224 | 0.37589056 | 0.768 | 0.628 | 2.18E-220 | 2 | Pbrm1 | myCAF_prolif |
| Dars2 | 3.83E-224 | 1.52649004 | 0.404 | 0.135 | 7.40E-220 | 2 | Dars2 | myCAF_prolif |
| Senp1 | 4.76E-224 | 0.9232044 | 0.57 | 0.315 | 9.20E-220 | 2 | Senp1 | myCAF_prolif |
| Map2k3 | 6.11E-224 | 0.36088735 | 0.811 | 0.688 | 1.18E-219 | 2 | Map2k3 | myCAF_prolif |
| Gadd45gip1 | 6.24E-224 | 0.46271085 | 0.773 | 0.605 | 1.21E-219 | 2 | Gadd45gip1 | myCAF_prolif |
| mt-Co3 | 8.60E-224 | 1.11142669 | 0.882 | 0.81 | 1.66E-219 | 2 | mt-Co3 | myCAF_prolif |
| Rars | 1.66E-223 | 0.51924858 | 0.713 | 0.533 | 3.20E-219 | 2 | Rars | myCAF_prolif |
| Pphln1 | 3.93E-223 | 0.4770102 | 0.713 | 0.529 | 7.59E-219 | 2 | Pphln1 | myCAF_prolif |
| Ddx46 | 3.42E-222 | 0.55303736 | 0.76 | 0.594 | 6.61E-218 | 2 | Ddx46 | myCAF_prolif |
| Rab35 | 5.29E-222 | 0.36526071 | 0.822 | 0.701 | 1.02E-217 | 2 | Rab35 | myCAF_prolif |
| Tbrg4 | 1.75E-221 | 0.60073116 | 0.668 | 0.464 | 3.38E-217 | 2 | Tbrg4 | myCAF_prolif |
| Vkorc1l1 | 3.50E-221 | 0.36226036 | 0.72 | 0.578 | 6.78E-217 | 2 | Vkorc1l1 | myCAF_prolif |
| Exosc2 | 3.95E-221 | 1.36517178 | 0.465 | 0.182 | 7.64E-217 | 2 | Exosc2 | myCAF_prolif |
| Arpp19 | 8.03E-221 | 0.59580766 | 0.675 | 0.471 | 1.55E-216 | 2 | Arpp19 | myCAF_prolif |
| Pradc1 | 1.33E-220 | 1.49132152 | 0.373 | 0.115 | 2.57E-216 | 2 | Pradc1 | myCAF_prolif |
| Mthfd10 | 2.50E-220 | 0.3061551 | 0.778 | 0.612 | 4.83E-216 | 2 | Mthfd10 | myCAF_prolif |
| Abhd17c1 | 3.19E-220 | 0.55109463 | 0.653 | 0.427 | 6.17E-216 | 2 | Abhd17c | myCAF_prolif |
| DEPRECATED-I | 3.94E-220 | 0.37323502 | 0.803 | 0.698 | 7.62E-216 | 2 | DEPRECATED-I | myCAF_prolif |
| Mpzl1 | 4.05E-220 | 0.42355534 | 0.679 | 0.524 | 7.83E-216 | 2 | Mpzl1 | myCAF_prolif |
| Htatsf5a | 6.58E-220 | 0.29927755 | 0.839 | 0.724 | 1.27E-215 | 2 | Htatsf5a | myCAF_prolif |
| Amd1 | 9.30E-220 | 0.84821279 | 0.549 | 0.296 | 1.80E-215 | 2 | Amd1 | myCAF_prolif |
| Dld | 9.96E-220 | 0.33623402 | 0.749 | 0.63 | 1.93E-215 | 2 | Dld | myCAF_prolif |
| Tead2 | 3.12E-219 | 0.51107929 | 0.622 | 0.425 | 6.03E-215 | 2 | Tead2 | myCAF_prolif |
| Adsl | 3.33E-219 | 0.64906948 | 0.633 | 0.402 | 6.43E-215 | 2 | Adsl | myCAF_prolif |
| Uqcr10 | 3.57E-219 | 0.48878498 | 0.746 | 0.561 | 6.91E-215 | 2 | Uqcr10 | myCAF_prolif |
| Zfp101 | 3.91E-219 | 2.13554729 | 0.314 | 0.067 | 7.57E-215 | 2 | Zfp101 | myCAF_prolif |
| Ccdc25 | 5.69E-219 | 0.6028504 | 0.668 | 0.46 | 1.10E-214 | 2 | Ccdc25 | myCAF_prolif |
| Hdgfl2 | 6.09E-219 | 0.51619076 | 0.765 | 0.579 | 1.18E-214 | 2 | Hdgfl2 | myCAF_prolif |
| Eif4a3 | 7.08E-219 | 0.54071566 | 0.668 | 0.465 | 1.37E-214 | 2 | Eif4a3 | myCAF_prolif |
| E2f6 | 8.62E-219 | 0.84108116 | 0.517 | 0.286 | 1.67E-214 | 2 | E2f6 | myCAF_prolif |
| Hras | 9.72E-219 | 0.43482274 | 0.719 | 0.552 | 1.88E-214 | 2 | Hras | myCAF_prolif |
| Hspa13 | 4.01E-218 | 0.29317841 | 0.747 | 0.594 | 7.75E-214 | 2 | Hspa13 | myCAF_prolif |
| Itpa | 4.51E-218 | 0.62724828 | 0.683 | 0.46 | 8.71E-214 | 2 | Itpa | myCAF_prolif |
| Tmem248 | 4.67E-218 | 0.31921646 | 0.777 | 0.656 | 9.04E-214 | 2 | Tmem248 | myCAF_prolif |
| Mycbp | 1.16E-217 | 0.60393288 | 0.654 | 0.425 | 2.24E-213 | 2 | Mycbp | myCAF_prolif |
| Ccar1 | 1.75E-217 | 0.54346291 | 0.773 | 0.63 | 3.39E-213 | 2 | Ccar1 | myCAF_prolif |
| Ppp1r11 | 2.18E-217 | 0.40727207 | 0.75 | 0.592 | 4.21E-213 | 2 | Ppp1r11 | myCAF_prolif |
| E2f4 | 2.26E-217 | 0.26076354 | 0.838 | 0.746 | 4.37E-213 | 2 | E2f4 | myCAF_prolif |
| Nploc4 | 4.41E-217 | 0.27709812 | 0.797 | 0.68 | 8.53E-213 | 2 | Nploc4 | myCAF_prolif |
| Msh2 | 6.52E-217 | 0.93087139 | 0.431 | 0.24 | 1.26E-212 | 2 | Msh2 | myCAF_prolif |
| Rock2 | 8.82E-217 | 0.25003362 | 0.843 | 0.77 | 1.71E-212 | 2 | Rock2 | myCAF_prolif |
| Ebp1 | 9.51E-217 | 0.41456768 | 0.73 | 0.53 | 1.84E-212 | 2 | Ebp | myCAF_prolif |
| Chst21 | 1.41E-216 | 0.51948703 | 0.579 | 0.363 | 2.74E-212 | 2 | Chst2 | myCAF_prolif |
| 10-Sep | 1.86E-216 | 0.65464076 | 0.61 | 0.399 | 3.59E-212 | 2 | 10-Sep | myCAF_prolif |
| Blvra | 3.16E-216 | 0.80709328 | 0.566 | 0.304 | 6.11E-212 | 2 | Blvra | myCAF_prolif |
| Cntrl | 3.64E-216 | 1.71932975 | 0.508 | 0.256 | 7.04E-212 | 2 | Cntrl | myCAF_prolif |
| Cetn3 | 8.55E-216 | 0.35909703 | 0.778 | 0.62 | 1.65E-211 | 2 | Cetn3 | myCAF_prolif |
| Prokr21 | 8.73E-216 | 0.46430697 | 0.548 | 0.381 | 1.69E-211 | 2 | Prokr2 | myCAF_prolif |
| Cpsf2 | 1.75E-215 | 0.61054991 | 0.68 | 0.457 | 3.38E-211 | 2 | Cpsf2 | myCAF_prolif |
| Cd2bp2 | 1.93E-215 | 0.4425749 | 0.809 | 0.674 | 3.73E-211 | 2 | Cd2bp2 | myCAF_prolif |
| Chd1 | 3.05E-215 | 0.55761822 | 0.742 | 0.561 | 5.90E-211 | 2 | Chd1 | myCAF_prolif |
| Aldh1l21 | 3.35E-215 | 0.28591654 | 0.678 | 0.515 | 6.47E-211 | 2 | Aldh1l2 | myCAF_prolif |
| Bcas2 | 5.96E-215 | 0.39909599 | 0.766 | 0.62 | 1.15E-210 | 2 | Bcas2 | myCAF_prolif |
| Rnf26 | 6.40E-215 | 1.16151851 | 0.474 | 0.205 | 1.24E-210 | 2 | Rnf26 | myCAF_prolif |
| Poc1b | 8.67E-215 | 1.02021917 | 0.483 | 0.223 | 1.68E-210 | 2 | Poc1b | myCAF_prolif |
| Tnpo3 | 2.43E-214 | 0.40293168 | 0.771 | 0.609 | 4.69E-210 | 2 | Tnpo3 | myCAF_prolif |
| Cyth2 | 2.76E-214 | 0.34801784 | 0.83 | 0.71 | 5.33E-210 | 2 | Cyth2 | myCAF_prolif |
| Pitpnb | 3.83E-214 | 0.30138262 | 0.799 | 0.663 | 7.41E-210 | 2 | Pitpnb | myCAF_prolif |
| Hnrnpll | 1.13E-213 | 0.26015544 | 0.77 | 0.646 | 2.19E-209 | 2 | Hnrnpll | myCAF_prolif |
| Bak1 | 1.23E-213 | 0.26606626 | 0.842 | 0.745 | 2.37E-209 | 2 | Bak1 | myCAF_prolif |
| Ppat | 1.97E-213 | 0.80244904 | 0.488 | 0.268 | 3.80E-209 | 2 | Ppat | myCAF_prolif |
| Rnaseh2a | 2.46E-213 | 0.68908969 | 0.623 | 0.38 | 4.75E-209 | 2 | Rnaseh2a | myCAF_prolif |
| Setd1a | 3.66E-213 | 0.36285372 | 0.776 | 0.622 | 7.08E-209 | 2 | Setd1a | myCAF_prolif |
| Slc20a2 | 4.93E-213 | 0.44582431 | 0.666 | 0.479 | 9.52E-209 | 2 | Slc20a2 | myCAF_prolif |
| Pithd1 | 7.12E-213 | 0.74068342 | 0.592 | 0.346 | 1.38E-208 | 2 | Pithd1 | myCAF_prolif |
| Nup98 | 1.12E-212 | 0.35203338 | 0.703 | 0.571 | 2.17E-208 | 2 | Nup98 | myCAF_prolif |
| Ctdspl2 | 2.64E-212 | 0.59947145 | 0.643 | 0.444 | 5.10E-208 | 2 | Ctdspl2 | myCAF_prolif |
| Adss | 2.65E-212 | 0.34112805 | 0.771 | 0.626 | 5.12E-208 | 2 | Adss | myCAF_prolif |
| Nup43 | 4.77E-212 | 2.04250375 | 0.299 | 0.063 | 9.23E-208 | 2 | Nup43 | myCAF_prolif |
| Rad54b | 1.13E-210 | 5.23252776 | 0.265 | 0.01 | 2.18E-206 | 2 | Rad54b | myCAF_prolif |
| Pknox21 | 1.99E-210 | 0.48580504 | 0.596 | 0.413 | 3.86E-206 | 2 | Pknox2 | myCAF_prolif |
| Tmem189 | 3.65E-210 | 0.49442253 | 0.764 | 0.584 | 7.07E-206 | 2 | Tmem189 | myCAF_prolif |
| Ube3c | 2.52E-209 | 0.38076284 | 0.76 | 0.62 | 4.87E-205 | 2 | Ube3c | myCAF_prolif |
| Scrib | 2.53E-209 | 0.71320107 | 0.561 | 0.313 | 4.88E-205 | 2 | Scrib | myCAF_prolif |
| Prkag1 | 3.21E-209 | 0.36062519 | 0.735 | 0.58 | 6.20E-205 | 2 | Prkag1 | myCAF_prolif |
| Pomp | 5.18E-209 | 0.25586792 | 0.832 | 0.726 | 1.00E-204 | 2 | Pomp | myCAF_prolif |
| Nsmce1 | 9.66E-209 | 0.85836702 | 0.516 | 0.256 | 1.87E-204 | 2 | Nsmce1 | myCAF_prolif |
| Mllt1 | 4.31E-208 | 0.47093382 | 0.728 | 0.529 | 8.33E-204 | 2 | Mllt1 | myCAF_prolif |
| Zc3h15 | 1.07E-207 | 0.33841782 | 0.778 | 0.627 | 2.06E-203 | 2 | Zc3h15 | myCAF_prolif |
| Coq2 | 1.88E-207 | 0.48903761 | 0.707 | 0.526 | 3.63E-203 | 2 | Coq2 | myCAF_prolif |
| Mmadhc | 9.42E-207 | 0.31067719 | 0.766 | 0.632 | 1.82E-202 | 2 | Mmadhc | myCAF_prolif |
| Larp4 | 9.57E-207 | 0.32486413 | 0.753 | 0.636 | 1.85E-202 | 2 | Larp4 | myCAF_prolif |
| Tnpo2 | 1.62E-206 | 0.27034409 | 0.836 | 0.726 | 3.13E-202 | 2 | Tnpo2 | myCAF_prolif |
| Siva1 | 3.45E-206 | 0.79403627 | 0.527 | 0.303 | 6.67E-202 | 2 | Siva1 | myCAF_prolif |
| Nop21 | 6.33E-206 | 0.39385396 | 0.63 | 0.467 | 1.22E-201 | 2 | Nop2 | myCAF_prolif |
| Rdh11 | 5.93E-205 | 0.79564357 | 0.557 | 0.306 | 1.15E-200 | 2 | Rdh11 | myCAF_prolif |
| Dxo | 7.45E-205 | 0.66042611 | 0.602 | 0.363 | 1.44E-200 | 2 | Dxo | myCAF_prolif |
| Wdr46 | 1.61E-204 | 0.56095034 | 0.625 | 0.425 | 3.11E-200 | 2 | Wdr46 | myCAF_prolif |
| Spidr | 6.51E-204 | 0.58987219 | 0.577 | 0.337 | 1.26E-199 | 2 | Spidr | myCAF_prolif |
| Cnot3 | 7.31E-204 | 0.43312834 | 0.805 | 0.651 | 1.41E-199 | 2 | Cnot3 | myCAF_prolif |
| Rbm25 | 7.91E-204 | 0.30264139 | 0.811 | 0.714 | 1.53E-199 | 2 | Rbm25 | myCAF_prolif |
| Plekha3 | 1.19E-203 | 0.33099357 | 0.757 | 0.604 | 2.29E-199 | 2 | Plekha3 | myCAF_prolif |
| Eif2b1 | 1.83E-203 | 0.45632494 | 0.68 | 0.477 | 3.53E-199 | 2 | Eif2b1 | myCAF_prolif |
| Adam231 | 2.21E-203 | 0.43868084 | 0.546 | 0.329 | 4.27E-199 | 2 | Adam23 | myCAF_prolif |
| Atp5g1 | 5.97E-203 | 0.44254955 | 0.759 | 0.585 | 1.15E-198 | 2 | Atp5g1 | myCAF_prolif |
| Mdm1 | 7.18E-203 | 2.10805525 | 0.31 | 0.075 | 1.39E-198 | 2 | Mdm1 | myCAF_prolif |
| Trim37 | 3.54E-202 | 0.95319985 | 0.451 | 0.191 | 6.84E-198 | 2 | Trim37 | myCAF_prolif |
| Prpf4 | 3.82E-202 | 1.05498493 | 0.463 | 0.195 | 7.39E-198 | 2 | Prpf4 | myCAF_prolif |
| Cars | 1.87E-201 | 0.50435468 | 0.722 | 0.528 | 3.62E-197 | 2 | Cars | myCAF_prolif |
| Nfya | 2.47E-201 | 0.39077841 | 0.666 | 0.496 | 4.77E-197 | 2 | Nfya | myCAF_prolif |
| Brms1l | 3.25E-201 | 0.80777099 | 0.692 | 0.469 | 6.29E-197 | 2 | Brms1l | myCAF_prolif |
| Zcrb1 | 6.21E-201 | 0.46868533 | 0.725 | 0.533 | 1.20E-196 | 2 | Zcrb1 | myCAF_prolif |
| Ndufs8 | 7.12E-201 | 0.2626939 | 0.78 | 0.655 | 1.38E-196 | 2 | Ndufs8 | myCAF_prolif |
| Yaf2 | 8.11E-201 | 0.45044812 | 0.729 | 0.546 | 1.57E-196 | 2 | Yaf2 | myCAF_prolif |
| Noc4l | 8.57E-201 | 0.89259326 | 0.511 | 0.251 | 1.66E-196 | 2 | Noc4l | myCAF_prolif |
| Opa3 | 1.48E-200 | 0.5565599 | 0.624 | 0.4 | 2.87E-196 | 2 | Opa3 | myCAF_prolif |
| Slf1 | 2.94E-200 | 1.45342311 | 0.347 | 0.105 | 5.68E-196 | 2 | Slf1 | myCAF_prolif |
| Fundc2 | 3.05E-200 | 0.2515933 | 0.816 | 0.715 | 5.91E-196 | 2 | Fundc2 | myCAF_prolif |
| Cd74 | 7.49E-200 | 0.74448469 | 0.618 | 0.514 | 1.45E-195 | 2 | Cd74 | myCAF_prolif |
| Arpc5l | 8.24E-200 | 0.42077452 | 0.724 | 0.555 | 1.59E-195 | 2 | Arpc5l | myCAF_prolif |
| Clasp1 | 1.45E-199 | 0.36043547 | 0.765 | 0.626 | 2.81E-195 | 2 | Clasp1 | myCAF_prolif |
| Acsl5 | 2.86E-199 | 0.40970372 | 0.748 | 0.6 | 5.53E-195 | 2 | Acsl5 | myCAF_prolif |
| Zmym1 | 3.18E-199 | 1.43609714 | 0.377 | 0.136 | 6.15E-195 | 2 | Zmym1 | myCAF_prolif |
| Sec11c | 3.61E-199 | 0.49130932 | 0.687 | 0.499 | 6.98E-195 | 2 | Sec11c | myCAF_prolif |
| Tspan6 | 4.29E-199 | 0.29162384 | 0.692 | 0.552 | 8.29E-195 | 2 | Tspan6 | myCAF_prolif |
| Hsd17b12 | 6.20E-199 | 0.40904356 | 0.702 | 0.553 | 1.20E-194 | 2 | Hsd17b12 | myCAF_prolif |
| Dnaaf2 | 1.49E-198 | 1.8565328 | 0.301 | 0.074 | 2.88E-194 | 2 | Dnaaf2 | myCAF_prolif |
| Gtf2e2 | 1.86E-198 | 0.33155392 | 0.698 | 0.536 | 3.60E-194 | 2 | Gtf2e2 | myCAF_prolif |
| Idh3g | 2.76E-198 | 0.2974457 | 0.804 | 0.691 | 5.34E-194 | 2 | Idh3g | myCAF_prolif |
| Oxct1 | 2.82E-198 | 0.38409619 | 0.677 | 0.525 | 5.46E-194 | 2 | Oxct1 | myCAF_prolif |
| B3galnt1 | 4.47E-198 | 0.49158337 | 0.672 | 0.448 | 8.65E-194 | 2 | B3galnt1 | myCAF_prolif |
| Naa15 | 5.66E-198 | 0.40668612 | 0.706 | 0.548 | 1.09E-193 | 2 | Naa15 | myCAF_prolif |
| Hoxb21 | 1.04E-197 | 0.49354126 | 0.635 | 0.441 | 2.00E-193 | 2 | Hoxb2 | myCAF_prolif |
| Trappc1 | 1.11E-197 | 0.43479018 | 0.705 | 0.538 | 2.14E-193 | 2 | Trappc1 | myCAF_prolif |
| Pkn2 | 1.41E-197 | 0.32053294 | 0.708 | 0.564 | 2.72E-193 | 2 | Pkn2 | myCAF_prolif |
| Lsm5 | 2.98E-197 | 1.47485293 | 0.372 | 0.127 | 5.77E-193 | 2 | Lsm5 | myCAF_prolif |
| Thoc3 | 5.44E-197 | 0.62057729 | 0.597 | 0.349 | 1.05E-192 | 2 | Thoc3 | myCAF_prolif |
| Ccm2 | 1.26E-196 | 0.26917826 | 0.804 | 0.658 | 2.43E-192 | 2 | Ccm2 | myCAF_prolif |
| Grpel1 | 1.29E-196 | 0.32336644 | 0.782 | 0.672 | 2.49E-192 | 2 | Grpel1 | myCAF_prolif |
| Lsm14b | 1.33E-196 | 0.30659644 | 0.851 | 0.765 | 2.56E-192 | 2 | Lsm14b | myCAF_prolif |
| Cenpu | 1.92E-196 | 4.97552877 | 0.275 | 0.009 | 3.70E-192 | 2 | Cenpu | myCAF_prolif |
| Ddx19a | 3.12E-196 | 0.52640877 | 0.734 | 0.533 | 6.02E-192 | 2 | Ddx19a | myCAF_prolif |
| Rad23a | 5.49E-196 | 0.26568536 | 0.804 | 0.677 | 1.06E-191 | 2 | Rad23a | myCAF_prolif |
| Ppp6r3 | 9.43E-196 | 0.40232751 | 0.773 | 0.646 | 1.82E-191 | 2 | Ppp6r3 | myCAF_prolif |
| Phf13 | 1.74E-195 | 0.4222565 | 0.704 | 0.523 | 3.36E-191 | 2 | Phf13 | myCAF_prolif |
| Cdan1 | 4.19E-195 | 1.04700351 | 0.464 | 0.212 | 8.11E-191 | 2 | Cdan1 | myCAF_prolif |
| Ubqln4 | 4.58E-195 | 0.4997243 | 0.747 | 0.578 | 8.86E-191 | 2 | Ubqln4 | myCAF_prolif |
| Xpo4 | 9.99E-195 | 0.76289449 | 0.518 | 0.285 | 1.93E-190 | 2 | Xpo4 | myCAF_prolif |
| Elp3 | 1.93E-194 | 0.26982727 | 0.784 | 0.666 | 3.73E-190 | 2 | Elp3 | myCAF_prolif |
| Odf2l | 1.48E-193 | 1.33407213 | 0.456 | 0.208 | 2.86E-189 | 2 | Odf2l | myCAF_prolif |
| Ammecr1 | 2.19E-193 | 0.33198915 | 0.713 | 0.559 | 4.23E-189 | 2 | Ammecr1 | myCAF_prolif |
| Tube1 | 2.22E-193 | 2.4532255 | 0.277 | 0.048 | 4.29E-189 | 2 | Tube1 | myCAF_prolif |
| Polq | 3.59E-193 | 5.75472334 | 0.351 | 0.005 | 6.95E-189 | 2 | Polq | myCAF_prolif |
| Aimp1 | 4.22E-193 | 0.40683671 | 0.751 | 0.589 | 8.16E-189 | 2 | Aimp1 | myCAF_prolif |
| Med18 | 9.13E-193 | 1.38451752 | 0.386 | 0.145 | 1.77E-188 | 2 | Med18 | myCAF_prolif |
| Taf1 | 1.32E-192 | 0.55867672 | 0.679 | 0.484 | 2.56E-188 | 2 | Taf1 | myCAF_prolif |
| Pbx3 | 3.43E-192 | 1.40675901 | 0.415 | 0.158 | 6.64E-188 | 2 | Pbx3 | myCAF_prolif |
| Ppm1m | 1.25E-191 | 0.37137806 | 0.745 | 0.586 | 2.41E-187 | 2 | Ppm1m | myCAF_prolif |
| Car121 | 1.39E-191 | 0.69074732 | 0.365 | 0.196 | 2.68E-187 | 2 | Car12 | myCAF_prolif |
| Cdc25a | 1.71E-191 | 1.18461442 | 0.449 | 0.194 | 3.31E-187 | 2 | Cdc25a | myCAF_prolif |
| Srbd1 | 2.07E-191 | 1.05591584 | 0.466 | 0.206 | 4.01E-187 | 2 | Srbd1 | myCAF_prolif |
| Cnpy4 | 3.21E-191 | 0.34688943 | 0.75 | 0.577 | 6.20E-187 | 2 | Cnpy4 | myCAF_prolif |
| Polr2b | 3.35E-191 | 0.36124779 | 0.658 | 0.486 | 6.47E-187 | 2 | Polr2b | myCAF_prolif |
| Ticrr | 4.60E-191 | 5.6234914 | 0.324 | 0.006 | 8.90E-187 | 2 | Ticrr | myCAF_prolif |
| Gtf3c5 | 4.76E-191 | 0.68021412 | 0.584 | 0.361 | 9.21E-187 | 2 | Gtf3c5 | myCAF_prolif |
| Snrpd2 | 1.53E-190 | 0.5425359 | 0.658 | 0.451 | 2.95E-186 | 2 | Snrpd2 | myCAF_prolif |
| Tmtc3 | 1.70E-190 | 0.37917971 | 0.718 | 0.535 | 3.29E-186 | 2 | Tmtc3 | myCAF_prolif |
| Pak4 | 2.00E-190 | 1.0589173 | 0.437 | 0.203 | 3.87E-186 | 2 | Pak4 | myCAF_prolif |
| Yy1 | 3.27E-190 | 0.39173673 | 0.894 | 0.838 | 6.33E-186 | 2 | Yy1 | myCAF_prolif |
| Tomm70a | 3.72E-190 | 0.38708465 | 0.623 | 0.47 | 7.19E-186 | 2 | Tomm70a | myCAF_prolif |
| Prpf40a | 4.78E-190 | 0.29366017 | 0.748 | 0.6 | 9.23E-186 | 2 | Prpf40a | myCAF_prolif |
| Slah1b | 7.43E-190 | 1.80269587 | 0.316 | 0.082 | 1.44E-185 | 2 | Slah1b | myCAF_prolif |
| Cluh | 1.54E-189 | 0.39694673 | 0.658 | 0.492 | 2.97E-185 | 2 | Cluh | myCAF_prolif |
| Yif1b | 1.59E-189 | 0.28549092 | 0.777 | 0.64 | 3.07E-185 | 2 | Yif1b | myCAF_prolif |
| Pin1 | 2.57E-189 | 0.71867462 | 0.575 | 0.362 | 4.97E-185 | 2 | Pin1 | myCAF_prolif |
| Ehbp1l1 | 2.79E-189 | 0.29330147 | 0.788 | 0.662 | 5.40E-185 | 2 | Ehbp1l1 | myCAF_prolif |
| Grwd1 | 3.83E-189 | 0.79945135 | 0.479 | 0.259 | 7.41E-185 | 2 | Grwd1 | myCAF_prolif |
| Rtca | 5.65E-189 | 0.36526165 | 0.722 | 0.53 | 1.09E-184 | 2 | Rtca | myCAF_prolif |
| Coro1c | 6.26E-189 | 0.37192819 | 0.701 | 0.566 | 1.21E-184 | 2 | Coro1c | myCAF_prolif |
| Ak6 | 1.08E-188 | 0.70657673 | 0.602 | 0.367 | 2.09E-184 | 2 | Ak6 | myCAF_prolif |
| Sap130 | 1.26E-188 | 0.42839454 | 0.664 | 0.498 | 2.43E-184 | 2 | Sap130 | myCAF_prolif |
| Sema6b | 1.36E-188 | 0.91715318 | 0.458 | 0.28 | 2.62E-184 | 2 | Sema6b | myCAF_prolif |
| Ash2l | 2.21E-188 | 0.37577026 | 0.736 | 0.568 | 4.28E-184 | 2 | Ash2l | myCAF_prolif |
| Tufm | 2.97E-188 | 0.32233607 | 0.718 | 0.562 | 5.73E-184 | 2 | Tufm | myCAF_prolif |
| Ccdc82 | 6.76E-188 | 1.40872316 | 0.559 | 0.349 | 1.31E-183 | 2 | Ccdc82 | myCAF_prolif |
| Cdh2 | 7.16E-188 | 0.76377134 | 0.447 | 0.236 | 1.38E-183 | 2 | Cdh2 | myCAF_prolif |
| Rprd1b | 7.59E-188 | 0.74200786 | 0.597 | 0.381 | 1.47E-183 | 2 | Rprd1b | myCAF_prolif |
| Spout1 | 1.44E-187 | 0.46523383 | 0.656 | 0.454 | 2.79E-183 | 2 | Spout1 | myCAF_prolif |
| Mphosph6 | 1.62E-187 | 0.55395236 | 0.649 | 0.421 | 3.13E-183 | 2 | Mphosph6 | myCAF_prolif |
| Cyp39a1 | 1.36E-186 | 1.09146757 | 0.377 | 0.209 | 2.63E-182 | 2 | Cyp39a1 | myCAF_prolif |
| Dis3 | 1.53E-186 | 0.91151739 | 0.474 | 0.224 | 2.96E-182 | 2 | Dis3 | myCAF_prolif |
| Ddx20 | 2.00E-186 | 1.202729 | 0.399 | 0.16 | 3.86E-182 | 2 | Ddx20 | myCAF_prolif |
| Psmd9 | 3.81E-186 | 0.37608145 | 0.72 | 0.544 | 7.37E-182 | 2 | Psmd9 | myCAF_prolif |
| Dpagt1 | 4.01E-186 | 0.3606343 | 0.722 | 0.552 | 7.76E-182 | 2 | Dpagt1 | myCAF_prolif |
| Rif1 | 2.77E-185 | 1.02941187 | 0.442 | 0.214 | 5.35E-181 | 2 | Rif1 | myCAF_prolif |
| Hmbs | 1.80E-184 | 0.45568046 | 0.659 | 0.458 | 3.48E-180 | 2 | Hmbs | myCAF_prolif |
| Sfmbt1 | 3.84E-184 | 0.71658751 | 0.503 | 0.305 | 7.43E-180 | 2 | Sfmbt1 | myCAF_prolif |
| Eef1e1 | 6.04E-184 | 0.8531063 | 0.489 | 0.291 | 1.17E-179 | 2 | Eef1e1 | myCAF_prolif |
| Esco2 | 5.27E-183 | 7.46817154 | 0.424 | 0.003 | 1.02E-178 | 2 | Esco2 | myCAF_prolif |
| Uqcrb | 5.69E-183 | 0.3575136 | 0.701 | 0.551 | 1.10E-178 | 2 | Uqcrb | myCAF_prolif |
| H2afy2 | 7.42E-183 | 0.68753575 | 0.559 | 0.349 | 1.44E-178 | 2 | H2afy2 | myCAF_prolif |
| Ints7 | 1.04E-182 | 0.54537865 | 0.532 | 0.363 | 2.02E-178 | 2 | Ints7 | myCAF_prolif |
| Pank2 | 1.51E-182 | 0.52226777 | 0.72 | 0.548 | 2.93E-178 | 2 | Pank2 | myCAF_prolif |
| Cops5 | 2.01E-182 | 0.32946224 | 0.79 | 0.663 | 3.89E-178 | 2 | Cops5 | myCAF_prolif |
| Ctps1 | 2.19E-182 | 0.27385606 | 0.671 | 0.525 | 4.23E-178 | 2 | Ctps | myCAF_prolif |
| Elof1 | 4.11E-182 | 0.4934248 | 0.646 | 0.439 | 7.95E-178 | 2 | Elof1 | myCAF_prolif |
| Has11 | 6.57E-182 | 0.39359432 | 0.362 | 0.235 | 1.27E-177 | 2 | Has1 | myCAF_prolif |
| Syne3 | 1.30E-181 | 0.62425883 | 0.555 | 0.349 | 2.52E-177 | 2 | Syne3 | myCAF_prolif |
| mt-Nd5 | 1.88E-181 | 1.80413412 | 0.767 | 0.633 | 3.63E-177 | 2 | mt-Nd5 | myCAF_prolif |
| Uqcrq | 3.49E-181 | 0.48570132 | 0.689 | 0.5 | 6.75E-177 | 2 | Uqcrq | myCAF_prolif |
| Baz1a | 4.37E-181 | 0.5053552 | 0.628 | 0.441 | 8.44E-177 | 2 | Baz1a | myCAF_prolif |
| Adgra3 | 8.66E-181 | 0.34307997 | 0.653 | 0.477 | 1.67E-176 | 2 | Adgra3 | myCAF_prolif |
| Cops2 | 2.73E-180 | 0.34336419 | 0.761 | 0.634 | 5.28E-176 | 2 | Cops2 | myCAF_prolif |
| Pno1 | 5.15E-180 | 0.54439003 | 0.587 | 0.382 | 9.95E-176 | 2 | Pno1 | myCAF_prolif |
| Dcaf13 | 5.61E-180 | 0.39607271 | 0.673 | 0.495 | 1.08E-175 | 2 | Dcaf13 | myCAF_prolif |
| Taf12 | 2.44E-179 | 0.48059147 | 0.618 | 0.418 | 4.72E-175 | 2 | Taf12 | myCAF_prolif |
| Arl4c | 4.22E-179 | 0.40663072 | 0.629 | 0.457 | 8.17E-175 | 2 | Arl4c | myCAF_prolif |
| Serp1 | 5.28E-179 | 0.33810884 | 0.682 | 0.535 | 1.02E-174 | 2 | Serp1 | myCAF_prolif |
| Pdss1 | 7.29E-179 | 1.40346115 | 0.328 | 0.113 | 1.41E-174 | 2 | Pdss1 | myCAF_prolif |
| Csnk2b | 8.42E-179 | 0.28274017 | 0.759 | 0.615 | 1.63E-174 | 2 | Csnk2b | myCAF_prolif |
| Dhrs4 | 1.52E-178 | 0.32155043 | 0.765 | 0.612 | 2.93E-174 | 2 | Dhrs4 | myCAF_prolif |
| Mpdz | 1.53E-178 | 0.55431122 | 0.646 | 0.429 | 2.96E-174 | 2 | Mpdz | myCAF_prolif |
| Ergic2 | 1.78E-178 | 0.46537766 | 0.686 | 0.494 | 3.43E-174 | 2 | Ergic2 | myCAF_prolif |
| Luc7l3 | 4.31E-178 | 0.76674866 | 0.699 | 0.485 | 8.33E-174 | 2 | Luc7l3 | myCAF_prolif |
| Schip11 | 4.92E-178 | 0.41417931 | 0.656 | 0.492 | 9.52E-174 | 2 | Schip1 | myCAF_prolif |
| Pxk | 9.61E-178 | 0.30672336 | 0.803 | 0.686 | 1.86E-173 | 2 | Pxk | myCAF_prolif |
| Nsmce2 | 1.79E-177 | 0.47495661 | 0.687 | 0.499 | 3.47E-173 | 2 | Nsmce2 | myCAF_prolif |
| Lrrc40 | 2.67E-177 | 1.45360674 | 0.358 | 0.122 | 5.16E-173 | 2 | Lrrc40 | myCAF_prolif |
| Rttn | 3.82E-177 | 2.23588126 | 0.278 | 0.054 | 7.40E-173 | 2 | Rttn | myCAF_prolif |
| Tbl3 | 5.90E-177 | 0.60063556 | 0.576 | 0.352 | 1.14E-172 | 2 | Tbl3 | myCAF_prolif |
| Rpf1 | 3.61E-176 | 0.46491282 | 0.647 | 0.425 | 6.98E-172 | 2 | Rpf1 | myCAF_prolif |
| Cherp | 8.75E-176 | 0.44800768 | 0.78 | 0.613 | 1.69E-171 | 2 | Cherp | myCAF_prolif |
| C1qbp | 9.63E-176 | 0.59018039 | 0.553 | 0.382 | 1.86E-171 | 2 | C1qbp | myCAF_prolif |
| Cdc34 | 5.26E-175 | 0.32915093 | 0.764 | 0.597 | 1.02E-170 | 2 | Cdc34 | myCAF_prolif |
| Brd3 | 5.96E-175 | 0.46589254 | 0.671 | 0.471 | 1.15E-170 | 2 | Brd3 | myCAF_prolif |
| Poglut2 | 6.41E-175 | 0.31303827 | 0.753 | 0.609 | 1.24E-170 | 2 | Poglut2 | myCAF_prolif |
| Wdr75 | 1.34E-174 | 0.54917759 | 0.541 | 0.362 | 2.59E-170 | 2 | Wdr75 | myCAF_prolif |
| Slc29a1 | 1.66E-174 | 0.53590364 | 0.502 | 0.268 | 3.21E-170 | 2 | Slc29a1 | myCAF_prolif |
| Ddx27 | 2.12E-174 | 0.54157532 | 0.625 | 0.428 | 4.10E-170 | 2 | Ddx27 | myCAF_prolif |
| Arf2 | 3.31E-174 | 0.35322816 | 0.678 | 0.522 | 6.40E-170 | 2 | Arf2 | myCAF_prolif |
| Nop16 | 6.23E-174 | 0.5202014 | 0.608 | 0.411 | 1.20E-169 | 2 | Nop16 | myCAF_prolif |
| Arhgef1 | 1.15E-173 | 0.26047863 | 0.87 | 0.805 | 2.23E-169 | 2 | Arhgef1 | myCAF_prolif |
| Cdc5l | 1.68E-173 | 0.49003247 | 0.622 | 0.412 | 3.25E-169 | 2 | Cdc5l | myCAF_prolif |
| Cdk11b | 2.17E-173 | 0.49125018 | 0.76 | 0.589 | 4.19E-169 | 2 | Cdk11b | myCAF_prolif |
| Chchd1 | 4.16E-173 | 0.74604459 | 0.616 | 0.415 | 8.05E-169 | 2 | Chchd1 | myCAF_prolif |
| Ibtk1 | 5.96E-173 | 0.32577787 | 0.732 | 0.579 | 1.15E-168 | 2 | Ibtk | myCAF_prolif |
| Dhrs7b | 7.02E-173 | 0.77669965 | 0.549 | 0.303 | 1.36E-168 | 2 | Dhrs7b | myCAF_prolif |
| Ppp4r3a | 9.19E-173 | 0.37774069 | 0.67 | 0.481 | 1.78E-168 | 2 | Ppp4r3a | myCAF_prolif |
| Ube2n | 1.06E-172 | 0.57941135 | 0.646 | 0.441 | 2.04E-168 | 2 | Ube2n | myCAF_prolif |
| Paics | 1.45E-172 | 0.40060077 | 0.63 | 0.464 | 2.81E-168 | 2 | Paics | myCAF_prolif |
| Wrap53 | 3.59E-172 | 1.18126674 | 0.424 | 0.179 | 6.95E-168 | 2 | Wrap53 | myCAF_prolif |
| Larp4b | 4.47E-172 | 0.30230546 | 0.763 | 0.615 | 8.65E-168 | 2 | Larp4b | myCAF_prolif |
| Elp5 | 4.92E-172 | 0.30429126 | 0.758 | 0.616 | 9.51E-168 | 2 | Elp5 | myCAF_prolif |
| Snrpb2 | 8.56E-172 | 0.38019737 | 0.68 | 0.508 | 1.66E-167 | 2 | Snrpb2 | myCAF_prolif |
| Cep83 | 1.96E-171 | 0.95275921 | 0.497 | 0.252 | 3.79E-167 | 2 | Cep83 | myCAF_prolif |
| Hist1h3c | 2.90E-171 | 7.03400737 | 0.294 | 0.003 | 5.62E-167 | 2 | Hist1h3c | myCAF_prolif |
| Car131 | 3.57E-171 | 0.67487041 | 0.587 | 0.354 | 6.89E-167 | 2 | Car13 | myCAF_prolif |
| Otud6b | 4.57E-171 | 0.54126326 | 0.568 | 0.349 | 8.84E-167 | 2 | Otud6b | myCAF_prolif |
| Efnb21 | 6.06E-171 | 0.28778708 | 0.633 | 0.486 | 1.17E-166 | 2 | Efnb2 | myCAF_prolif |
| Ddx56 | 6.83E-171 | 0.40592745 | 0.657 | 0.475 | 1.32E-166 | 2 | Ddx56 | myCAF_prolif |
| Gtf3a | 8.75E-171 | 0.4689439 | 0.678 | 0.494 | 1.69E-166 | 2 | Gtf3a | myCAF_prolif |
| Brd9 | 1.14E-170 | 0.87852126 | 0.538 | 0.319 | 2.21E-166 | 2 | Brd9 | myCAF_prolif |
| Xrn2 | 1.73E-170 | 0.32968455 | 0.735 | 0.615 | 3.35E-166 | 2 | Xrn2 | myCAF_prolif |
| Erg28 | 1.89E-170 | 0.29248957 | 0.707 | 0.548 | 3.66E-166 | 2 | Erg28 | myCAF_prolif |
| Smyd5 | 2.22E-170 | 0.43972555 | 0.609 | 0.453 | 4.29E-166 | 2 | Smyd5 | myCAF_prolif |
| Zdhhc18 | 6.87E-170 | 0.37958049 | 0.756 | 0.617 | 1.33E-165 | 2 | Zdhhc18 | myCAF_prolif |
| Sin3a | 6.75E-169 | 0.41414126 | 0.684 | 0.519 | 1.31E-164 | 2 | Sin3a | myCAF_prolif |
| Rbmxl1 | 9.15E-169 | 0.64270651 | 0.6 | 0.368 | 1.77E-164 | 2 | Rbmxl1 | myCAF_prolif |
| Wdr36 | 1.92E-168 | 0.42527199 | 0.631 | 0.453 | 3.72E-164 | 2 | Wdr36 | myCAF_prolif |
| Smpd4 | 1.95E-168 | 0.60616356 | 0.561 | 0.324 | 3.78E-164 | 2 | Smpd4 | myCAF_prolif |
| Nelfcd | 2.88E-168 | 0.66189081 | 0.538 | 0.295 | 5.56E-164 | 2 | Nelfcd | myCAF_prolif |
| Thap12 | 3.04E-168 | 0.35613145 | 0.713 | 0.544 | 5.87E-164 | 2 | Thap12 | myCAF_prolif |
| Commd10 | 5.22E-168 | 0.61730579 | 0.685 | 0.49 | 1.01E-163 | 2 | Commd10 | myCAF_prolif |
| Ube2e3 | 6.58E-168 | 0.31090995 | 0.713 | 0.551 | 1.27E-163 | 2 | Ube2e3 | myCAF_prolif |
| Tmem701 | 1.65E-167 | 0.49332127 | 0.635 | 0.466 | 3.18E-163 | 2 | Tmem70 | myCAF_prolif |
| Dyrk1a | 1.65E-167 | 0.34063453 | 0.676 | 0.519 | 3.19E-163 | 2 | Dyrk1a | myCAF_prolif |
| Gabpa | 1.99E-167 | 0.44691985 | 0.699 | 0.515 | 3.85E-163 | 2 | Gabpa | myCAF_prolif |
| Emsy | 2.57E-167 | 0.72532222 | 0.55 | 0.356 | 4.97E-163 | 2 | Emsy | myCAF_prolif |
| Slirp | 2.59E-167 | 0.33672931 | 0.707 | 0.557 | 5.01E-163 | 2 | Slirp | myCAF_prolif |
| Mpp6 | 3.23E-167 | 0.36327349 | 0.658 | 0.464 | 6.24E-163 | 2 | Mpp6 | myCAF_prolif |
| Ptgr11 | 4.87E-167 | 0.46151536 | 0.639 | 0.449 | 9.42E-163 | 2 | Ptgr1 | myCAF_prolif |
| Fam57a | 6.31E-167 | 0.92588204 | 0.444 | 0.223 | 1.22E-162 | 2 | Fam57a | myCAF_prolif |
| Exosc7 | 1.09E-166 | 0.87739441 | 0.491 | 0.278 | 2.11E-162 | 2 | Exosc7 | myCAF_prolif |
| Plrg1 | 1.68E-166 | 0.33847438 | 0.694 | 0.519 | 3.25E-162 | 2 | Plrg1 | myCAF_prolif |
| Trappc13 | 2.49E-166 | 0.62013437 | 0.609 | 0.415 | 4.82E-162 | 2 | Trappc13 | myCAF_prolif |
| Cdk19 | 4.04E-166 | 0.53365699 | 0.584 | 0.422 | 7.82E-162 | 2 | Cdk19 | myCAF_prolif |
| Farsb | 5.54E-166 | 0.47062707 | 0.619 | 0.433 | 1.07E-161 | 2 | Farsb | myCAF_prolif |
| Ipo11 | 1.29E-165 | 0.7263023 | 0.459 | 0.234 | 2.50E-161 | 2 | Ipo11 | myCAF_prolif |
| Ufd1 | 5.93E-165 | 0.3783444 | 0.751 | 0.609 | 1.15E-160 | 2 | Ufd1 | myCAF_prolif |
| Cox6c | 6.90E-165 | 0.30238214 | 0.786 | 0.642 | 1.33E-160 | 2 | Cox6c | myCAF_prolif |
| Ampd2 | 9.93E-165 | 0.28776444 | 0.788 | 0.656 | 1.92E-160 | 2 | Ampd2 | myCAF_prolif |
| Gtpbp1 | 1.05E-164 | 0.41795643 | 0.632 | 0.457 | 2.03E-160 | 2 | Gtpbp1 | myCAF_prolif |
| Emc1 | 1.14E-164 | 0.3673519 | 0.732 | 0.582 | 2.20E-160 | 2 | Emc1 | myCAF_prolif |
| Cep152 | 1.44E-164 | 1.71147979 | 0.277 | 0.067 | 2.79E-160 | 2 | Cep152 | myCAF_prolif |
| Usp39 | 6.25E-164 | 0.54001487 | 0.604 | 0.395 | 1.21E-159 | 2 | Usp39 | myCAF_prolif |
| Pmpca | 7.62E-164 | 0.47332231 | 0.673 | 0.497 | 1.47E-159 | 2 | Pmpca | myCAF_prolif |
| Pcyox1l | 9.92E-164 | 1.59597125 | 0.328 | 0.102 | 1.92E-159 | 2 | Pcyox1l | myCAF_prolif |
| Orc2 | 1.78E-163 | 0.819635 | 0.422 | 0.213 | 3.44E-159 | 2 | Orc2 | myCAF_prolif |
| Bhlhe41 | 7.85E-163 | 0.39998932 | 0.402 | 0.283 | 1.52E-158 | 2 | Bhlhe41 | myCAF_prolif |
| Zbtb41 | 9.50E-163 | 0.56567994 | 0.578 | 0.363 | 1.84E-158 | 2 | Zbtb41 | myCAF_prolif |
| Timm22 | 1.12E-162 | 0.33490453 | 0.703 | 0.533 | 2.17E-158 | 2 | Timm22 | myCAF_prolif |
| Tomm7 | 1.13E-162 | 0.26285158 | 0.797 | 0.678 | 2.18E-158 | 2 | Tomm7 | myCAF_prolif |
| Tmed51 | 1.37E-162 | 0.4403132 | 0.636 | 0.453 | 2.65E-158 | 2 | Tmed5 | myCAF_prolif |
| Cxxc1 | 1.60E-162 | 0.36395318 | 0.741 | 0.593 | 3.10E-158 | 2 | Cxxc1 | myCAF_prolif |
| Srp54b | 2.12E-162 | 0.27780421 | 0.719 | 0.594 | 4.11E-158 | 2 | Srp54b | myCAF_prolif |
| Cnot11 | 2.53E-162 | 0.42176494 | 0.649 | 0.449 | 4.89E-158 | 2 | Cnot11 | myCAF_prolif |
| Dcps | 6.37E-162 | 0.83694177 | 0.476 | 0.226 | 1.23E-157 | 2 | Dcps | myCAF_prolif |
| Nfyc | 1.04E-161 | 0.51435541 | 0.637 | 0.44 | 2.02E-157 | 2 | Nfyc | myCAF_prolif |
| Nvl | 1.73E-161 | 0.32026857 | 0.713 | 0.54 | 3.34E-157 | 2 | Nvl | myCAF_prolif |
| Sart3 | 3.02E-161 | 0.66711321 | 0.522 | 0.282 | 5.84E-157 | 2 | Sart3 | myCAF_prolif |
| Fam216a | 3.13E-161 | 0.79623189 | 0.513 | 0.299 | 6.05E-157 | 2 | Fam216a | myCAF_prolif |
| Klhdc4 | 4.30E-161 | 0.6947749 | 0.588 | 0.361 | 8.31E-157 | 2 | Klhdc4 | myCAF_prolif |
| Lancl2 | 6.33E-161 | 0.7449188 | 0.478 | 0.245 | 1.22E-156 | 2 | Lancl2 | myCAF_prolif |
| Actr10 | 6.75E-161 | 0.32469358 | 0.768 | 0.622 | 1.31E-156 | 2 | Actr10 | myCAF_prolif |
| Entpd61 | 8.27E-161 | 0.76839643 | 0.476 | 0.243 | 1.60E-156 | 2 | Entpd6 | myCAF_prolif |
| Bola3 | 1.24E-160 | 0.49893757 | 0.623 | 0.417 | 2.41E-156 | 2 | Bola3 | myCAF_prolif |
| Slc4a2 | 1.29E-160 | 0.31663719 | 0.816 | 0.692 | 2.50E-156 | 2 | Slc4a2 | myCAF_prolif |
| Stk16 | 1.43E-160 | 0.38361421 | 0.773 | 0.626 | 2.77E-156 | 2 | Stk16 | myCAF_prolif |
| Taf5 | 2.00E-160 | 1.28049619 | 0.33 | 0.118 | 3.87E-156 | 2 | Taf5 | myCAF_prolif |
| Slc35b1 | 2.90E-160 | 0.2783795 | 0.742 | 0.582 | 5.62E-156 | 2 | Slc35b1 | myCAF_prolif |
| Znhit3 | 2.92E-160 | 1.23195614 | 0.38 | 0.142 | 5.65E-156 | 2 | Znhit3 | myCAF_prolif |
| Cnot10 | 4.21E-160 | 0.53489527 | 0.616 | 0.421 | 8.14E-156 | 2 | Cnot10 | myCAF_prolif |

| | | | | | | | | |
|---|---|---|---|---|---|---|---|---|
| Cers61 | 5.23E-160 | 0.55338965 | 0.655 | 0.474 | 1.01E-155 | 2 | Cers6 | myCAF_prolif |
| Hmgxb4 | 6.02E-160 | 0.63317108 | 0.547 | 0.321 | 1.16E-155 | 2 | Hmgxb4 | myCAF_prolif |
| Ube2l3 | 1.35E-159 | 0.37235419 | 0.721 | 0.554 | 2.61E-155 | 2 | Ube2l3 | myCAF_prolif |
| Pdzd8 | 1.76E-159 | 0.29458551 | 0.67 | 0.512 | 3.39E-155 | 2 | Pdzd8 | myCAF_prolif |
| Aifm1 | 1.77E-159 | 0.60766531 | 0.552 | 0.336 | 3.42E-155 | 2 | Aifm1 | myCAF_prolif |
| Ppie | 2.03E-159 | 0.56033927 | 0.631 | 0.406 | 3.92E-155 | 2 | Ppie | myCAF_prolif |
| Rnf121 | 2.48E-159 | 0.48650748 | 0.682 | 0.508 | 4.80E-155 | 2 | Rnf121 | myCAF_prolif |
| Iws1 | 5.08E-159 | 0.36585366 | 0.635 | 0.467 | 9.82E-155 | 2 | Iws1 | myCAF_prolif |
| Lpcat11 | 6.83E-159 | 0.44784531 | 0.673 | 0.476 | 1.32E-154 | 2 | Lpcat1 | myCAF_prolif |
| Psmg1 | 1.17E-158 | 0.47928912 | 0.64 | 0.447 | 2.26E-154 | 2 | Psmg1 | myCAF_prolif |
| Selenot | 1.62E-158 | 0.3176146 | 0.717 | 0.616 | 3.14E-154 | 2 | Selenot | myCAF_prolif |
| Tsen15 | 1.88E-158 | 0.75419126 | 0.489 | 0.248 | 3.64E-154 | 2 | Tsen15 | myCAF_prolif |
| Cfap20 | 1.90E-158 | 0.29706491 | 0.696 | 0.538 | 3.67E-154 | 2 | Cfap20 | myCAF_prolif |
| Ppwd1 | 4.10E-158 | 1.08277375 | 0.404 | 0.169 | 7.93E-154 | 2 | Ppwd1 | myCAF_prolif |
| Tmem881 | 9.12E-158 | 0.58528347 | 0.257 | 0.16 | 1.76E-153 | 2 | Tmem88 | myCAF_prolif |
| Vegfc | 1.21E-157 | 0.28237331 | 0.644 | 0.517 | 2.35E-153 | 2 | Vegfc | myCAF_prolif |
| Slc8a1 | 5.04E-157 | 0.32194439 | 0.663 | 0.495 | 9.75E-153 | 2 | Slc8a1 | myCAF_prolif |
| Poc5 | 5.18E-157 | 0.99611079 | 0.4 | 0.19 | 1.00E-152 | 2 | Poc5 | myCAF_prolif |
| Ints14 | 6.21E-157 | 0.36946316 | 0.663 | 0.488 | 1.20E-152 | 2 | Ints14 | myCAF_prolif |
| Tsr1 | 1.82E-156 | 0.40267929 | 0.731 | 0.616 | 3.52E-152 | 2 | Tsr1 | myCAF_prolif |
| Ncbp2 | 3.64E-156 | 0.30310699 | 0.782 | 0.635 | 7.04E-152 | 2 | Ncbp2 | myCAF_prolif |
| Smg7 | 4.08E-156 | 0.28057207 | 0.687 | 0.538 | 7.88E-152 | 2 | Smg7 | myCAF_prolif |
| Ccp110 | 8.94E-156 | 1.29378663 | 0.33 | 0.131 | 1.73E-151 | 2 | Ccp110 | myCAF_prolif |
| Yars | 1.17E-155 | 0.34834088 | 0.734 | 0.595 | 2.26E-151 | 2 | Yars | myCAF_prolif |
| Pde6d | 1.19E-155 | 0.38682236 | 0.609 | 0.431 | 2.31E-151 | 2 | Pde6d | myCAF_prolif |
| Derl2 | 1.24E-155 | 0.25519803 | 0.753 | 0.627 | 2.40E-151 | 2 | Derl2 | myCAF_prolif |
| Aplf | 1.75E-155 | 1.17800625 | 0.344 | 0.116 | 3.38E-151 | 2 | Aplf | myCAF_prolif |
| Cstf2 | 5.51E-155 | 0.6585526 | 0.52 | 0.296 | 1.07E-150 | 2 | Cstf2 | myCAF_prolif |
| Olp5 | 7.82E-155 | 5.44641901 | 0.268 | 0.005 | 1.51E-150 | 2 | Olp5 | myCAF_prolif |
| Cct5 | 8.30E-155 | 0.43912776 | 0.641 | 0.463 | 1.61E-150 | 2 | Cct5 | myCAF_prolif |
| Asns | 1.33E-154 | 0.3193472 | 0.648 | 0.479 | 2.57E-150 | 2 | Asns | myCAF_prolif |
| Nelfa | 1.83E-154 | 0.46785559 | 0.684 | 0.52 | 3.54E-150 | 2 | Nelfa | myCAF_prolif |
| Phf10 | 2.59E-154 | 0.25494266 | 0.752 | 0.624 | 5.01E-150 | 2 | Phf10 | myCAF_prolif |
| Lif1 | 3.03E-154 | 0.48802607 | 0.267 | 0.175 | 5.86E-150 | 2 | Lif | myCAF_prolif |
| Taf6 | 3.49E-154 | 0.56433809 | 0.587 | 0.378 | 6.74E-150 | 2 | Taf6 | myCAF_prolif |
| Tubgcp2 | 4.25E-154 | 0.75323315 | 0.479 | 0.264 | 8.22E-150 | 2 | Tubgcp2 | myCAF_prolif |
| Vdac3 | 1.04E-153 | 0.46340847 | 0.696 | 0.52 | 2.00E-149 | 2 | Vdac3 | myCAF_prolif |
| Wdr18 | 1.11E-153 | 0.49741217 | 0.637 | 0.451 | 2.15E-149 | 2 | Wdr18 | myCAF_prolif |
| Plagl11 | 2.19E-153 | 0.37093941 | 0.411 | 0.249 | 4.24E-149 | 2 | Plagl1 | myCAF_prolif |
| Psma2 | 2.43E-153 | 0.29543678 | 0.723 | 0.577 | 4.70E-149 | 2 | Psma2 | myCAF_prolif |
| St51 | 4.19E-153 | 0.5032336 | 0.717 | 0.558 | 8.10E-149 | 2 | St5 | myCAF_prolif |
| Prps1l3 | 4.36E-153 | 0.45915332 | 0.635 | 0.424 | 8.43E-149 | 2 | Prps1l3 | myCAF_prolif |
| Zdhhc6 | 7.89E-153 | 0.41355175 | 0.663 | 0.478 | 1.52E-148 | 2 | Zdhhc6 | myCAF_prolif |
| Cstf3 | 1.01E-152 | 0.59698854 | 0.539 | 0.323 | 1.96E-148 | 2 | Cstf3 | myCAF_prolif |
| Abhd10 | 1.20E-152 | 1.0314136 | 0.406 | 0.183 | 2.31E-148 | 2 | Abhd10 | myCAF_prolif |
| Alg5 | 1.27E-152 | 0.34240879 | 0.712 | 0.542 | 2.45E-148 | 2 | Alg5 | myCAF_prolif |
| Glrx5 | 1.48E-152 | 0.28890712 | 0.754 | 0.614 | 2.86E-148 | 2 | Glrx5 | myCAF_prolif |
| Memo1 | 4.47E-152 | 0.27717571 | 0.711 | 0.571 | 8.64E-148 | 2 | Memo1 | myCAF_prolif |
| D10Wsu102e | 1.29E-151 | 0.46666518 | 0.559 | 0.355 | 2.50E-147 | 2 | D10Wsu102e | myCAF_prolif |
| Ik | 1.74E-151 | 0.25362799 | 0.849 | 0.762 | 3.37E-147 | 2 | Ik | myCAF_prolif |
| Vrk2 | 2.01E-151 | 1.37775542 | 0.346 | 0.121 | 3.89E-147 | 2 | Vrk2 | myCAF_prolif |
| Gpn3 | 2.42E-151 | 0.56094935 | 0.539 | 0.331 | 4.67E-147 | 2 | Gpn3 | myCAF_prolif |
| Prmt5 | 3.16E-151 | 0.4354012 | 0.638 | 0.44 | 6.11E-147 | 2 | Prmt5 | myCAF_prolif |
| Meaf6 | 3.33E-151 | 0.71246306 | 0.601 | 0.432 | 6.43E-147 | 2 | Meaf6 | myCAF_prolif |
| Nbn | 7.27E-151 | 0.70617136 | 0.44 | 0.228 | 1.41E-146 | 2 | Nbn | myCAF_prolif |
| Prps1 | 1.38E-150 | 1.08859212 | 0.382 | 0.162 | 2.68E-146 | 2 | Prps1 | myCAF_prolif |
| Polr1c | 8.68E-150 | 0.34106607 | 0.708 | 0.542 | 1.68E-145 | 2 | Polr1c | myCAF_prolif |
| Psma3 | 2.38E-149 | 0.32659521 | 0.704 | 0.608 | 4.61E-145 | 2 | Psma3 | myCAF_prolif |
| Sap18b | 6.67E-149 | 0.33107728 | 0.717 | 0.551 | 1.29E-144 | 2 | Sap18b | myCAF_prolif |
| Casp8ap2 | 7.32E-149 | 1.13517782 | 0.396 | 0.189 | 1.42E-144 | 2 | Casp8ap2 | myCAF_prolif |
| Tnfaip1 | 7.52E-149 | 0.35149268 | 0.724 | 0.555 | 1.45E-144 | 2 | Tnfaip1 | myCAF_prolif |
| Csnk2a2 | 8.42E-149 | 0.27775819 | 0.748 | 0.628 | 1.63E-144 | 2 | Csnk2a2 | myCAF_prolif |
| Chek2 | 9.10E-149 | 1.55373785 | 0.292 | 0.09 | 1.76E-144 | 2 | Chek2 | myCAF_prolif |
| Stk17b | 9.56E-149 | 0.40188285 | 0.523 | 0.344 | 1.85E-144 | 2 | Stk17b | myCAF_prolif |
| Dgcr8 | 1.29E-148 | 0.73265894 | 0.534 | 0.327 | 2.49E-144 | 2 | Dgcr8 | myCAF_prolif |
| Tubgcp4 | 1.65E-148 | 0.61893772 | 0.629 | 0.413 | 3.20E-144 | 2 | Tubgcp4 | myCAF_prolif |
| Leo1 | 1.90E-148 | 0.59043309 | 0.636 | 0.423 | 3.67E-144 | 2 | Leo1 | myCAF_prolif |
| Nup54 | 7.59E-148 | 0.53845818 | 0.515 | 0.311 | 1.47E-143 | 2 | Nup54 | myCAF_prolif |
| Me2 | 9.20E-148 | 0.5712934 | 0.545 | 0.319 | 1.78E-143 | 2 | Me2 | myCAF_prolif |
| Rcbtb1 | 1.49E-147 | 0.28169957 | 0.683 | 0.508 | 2.87E-143 | 2 | Rcbtb1 | myCAF_prolif |
| Psma7 | 1.54E-147 | 0.26511147 | 0.673 | 0.545 | 2.98E-143 | 2 | Psma7 | myCAF_prolif |
| Ints11 | 3.53E-147 | 0.48784611 | 0.588 | 0.413 | 6.82E-143 | 2 | Ints11 | myCAF_prolif |
| Xpo7 | 1.10E-146 | 0.34988002 | 0.689 | 0.511 | 2.12E-142 | 2 | Xpo7 | myCAF_prolif |
| Ppig | 1.35E-146 | 0.31103351 | 0.671 | 0.494 | 2.61E-142 | 2 | Ppig | myCAF_prolif |
| Tcf3 | 3.30E-146 | 0.35686931 | 0.696 | 0.522 | 6.39E-142 | 2 | Tcf3 | myCAF_prolif |
| Sart1 | 2.51E-145 | 0.25827323 | 0.698 | 0.563 | 4.85E-141 | 2 | Sart1 | myCAF_prolif |
| Spag7 | 3.06E-145 | 0.29761305 | 0.709 | 0.568 | 5.91E-141 | 2 | Spag7 | myCAF_prolif |
| Usp10 | 3.13E-145 | 0.58530439 | 0.572 | 0.362 | 6.05E-141 | 2 | Usp10 | myCAF_prolif |
| Ylpm1 | 3.61E-145 | 0.46857748 | 0.689 | 0.507 | 6.98E-141 | 2 | Ylpm1 | myCAF_prolif |
| Pcdh19 | 5.24E-145 | 0.32459639 | 0.57 | 0.423 | 1.01E-140 | 2 | Pcdh19 | myCAF_prolif |
| Xpo5 | 6.53E-145 | 0.38242381 | 0.623 | 0.462 | 1.26E-140 | 2 | Xpo5 | myCAF_prolif |
| Exosc3 | 7.32E-145 | 0.38010313 | 0.62 | 0.434 | 1.41E-140 | 2 | Exosc3 | myCAF_prolif |
| Cfl2 | 8.86E-145 | 0.29592777 | 0.703 | 0.563 | 1.71E-140 | 2 | Cfl2 | myCAF_prolif |
| Ruvbl1 | 1.03E-144 | 0.53980649 | 0.584 | 0.396 | 2.00E-140 | 2 | Ruvbl1 | myCAF_prolif |
| Cmklr1 | 1.44E-144 | 0.31552803 | 0.361 | 0.243 | 2.79E-140 | 2 | Cmklr1 | myCAF_prolif |
| Mettl16 | 1.56E-144 | 0.65220392 | 0.535 | 0.314 | 3.02E-140 | 2 | Mettl16 | myCAF_prolif |
| Xrcc6 | 2.97E-144 | 0.73449843 | 0.466 | 0.262 | 5.73E-140 | 2 | Xrcc6 | myCAF_prolif |
| Dele1 | 4.60E-144 | 0.50807802 | 0.617 | 0.406 | 8.90E-140 | 2 | Dele1 | myCAF_prolif |
| Gmds1 | 5.76E-144 | 0.49908988 | 0.572 | 0.378 | 1.11E-139 | 2 | Gmds | myCAF_prolif |
| Gabpb1 | 7.51E-144 | 0.58909955 | 0.528 | 0.319 | 1.45E-139 | 2 | Gabpb1 | myCAF_prolif |
| Kpna6 | 8.43E-144 | 0.2668625 | 0.723 | 0.564 | 1.63E-139 | 2 | Kpna6 | myCAF_prolif |
| Cops3 | 1.11E-143 | 0.52970205 | 0.604 | 0.413 | 2.15E-139 | 2 | Cops3 | myCAF_prolif |
| Dnm1l | 2.02E-143 | 0.27471133 | 0.727 | 0.592 | 3.91E-139 | 2 | Dnm1l | myCAF_prolif |
| Mdga1 | 2.20E-143 | 0.68442428 | 0.596 | 0.362 | 4.26E-139 | 2 | Mdga1 | myCAF_prolif |
| Farsa | 2.22E-143 | 0.3405111 | 0.67 | 0.504 | 4.29E-139 | 2 | Farsa | myCAF_prolif |
| Ift80 | 3.29E-143 | 0.79635117 | 0.473 | 0.267 | 6.37E-139 | 2 | Ift80 | myCAF_prolif |
| Dhx38 | 4.78E-143 | 0.42236913 | 0.727 | 0.588 | 9.24E-139 | 2 | Dhx38 | myCAF_prolif |
| Ngdn | 9.12E-143 | 0.50284666 | 0.573 | 0.375 | 1.76E-138 | 2 | Ngdn | myCAF_prolif |
| Hikeshi | 1.13E-142 | 0.37931846 | 0.697 | 0.534 | 2.18E-138 | 2 | Hikeshi | myCAF_prolif |
| Pif1 | 1.32E-142 | 7.38433897 | 0.272 | 0.003 | 2.55E-138 | 2 | Pif1 | myCAF_prolif |
| Casp8 | 2.34E-142 | 0.49956388 | 0.595 | 0.398 | 4.52E-138 | 2 | Casp8 | myCAF_prolif |
| Rbm28 | 3.12E-142 | 0.44179633 | 0.639 | 0.448 | 6.03E-138 | 2 | Rbm28 | myCAF_prolif |
| Rassf3 | 4.09E-142 | 1.14366396 | 0.297 | 0.119 | 7.92E-138 | 2 | Rassf3 | myCAF_prolif |
| Bud23 | 4.49E-142 | 0.35914048 | 0.648 | 0.444 | 8.68E-138 | 2 | Bud23 | myCAF_prolif |
| Arid1a | 4.90E-142 | 0.27870992 | 0.706 | 0.553 | 9.48E-138 | 2 | Arid1a | myCAF_prolif |
| Naa38 | 5.92E-142 | 0.26084324 | 0.747 | 0.619 | 1.15E-137 | 2 | Naa38 | myCAF_prolif |
| Rbm22 | 6.68E-142 | 0.36845263 | 0.704 | 0.553 | 1.29E-137 | 2 | Rbm22 | myCAF_prolif |
| Lrrc8c | 1.01E-141 | 0.55614085 | 0.435 | 0.271 | 1.96E-137 | 2 | Lrrc8c | myCAF_prolif |
| Trim471 | 2.62E-141 | 0.56199743 | 0.418 | 0.214 | 5.06E-137 | 2 | Trim47 | myCAF_prolif |
| Irf2bp1 | 4.34E-141 | 0.32286499 | 0.636 | 0.449 | 8.40E-137 | 2 | Irf2bp1 | myCAF_prolif |
| Fopnl | 4.54E-141 | 0.60350265 | 0.558 | 0.335 | 8.77E-137 | 2 | Fopnl | myCAF_prolif |
| Ran | 1.81E-140 | 1.31116087 | 0.313 | 0.107 | 3.51E-136 | 2 | Ran | myCAF_prolif |
| Snai21 | 1.94E-140 | 0.29078406 | 0.622 | 0.494 | 3.75E-136 | 2 | Snai2 | myCAF_prolif |
| Rexo5 | 1.99E-140 | 1.8806214 | 0.277 | 0.085 | 3.85E-136 | 2 | Rexo5 | myCAF_prolif |
| Phf6 | 3.43E-140 | 0.71860041 | 0.44 | 0.242 | 6.63E-136 | 2 | Phf6 | myCAF_prolif |
| Fip1l1 | 3.47E-140 | 0.28739381 | 0.687 | 0.544 | 6.70E-136 | 2 | Fip1l1 | myCAF_prolif |
| Elf4 | 3.50E-140 | 0.38736904 | 0.671 | 0.525 | 6.77E-136 | 2 | Elf4 | myCAF_prolif |
| Ciapin1 | 4.01E-140 | 0.62442059 | 0.556 | 0.357 | 7.75E-136 | 2 | Ciapin1 | myCAF_prolif |
| Med8 | 4.09E-140 | 0.35031901 | 0.675 | 0.523 | 7.91E-136 | 2 | Med8 | myCAF_prolif |
| Eed | 4.88E-140 | 0.41667489 | 0.52 | 0.319 | 9.43E-136 | 2 | Eed | myCAF_prolif |
| Ptcd3 | 5.20E-140 | 0.41715956 | 0.553 | 0.363 | 1.01E-135 | 2 | Ptcd3 | myCAF_prolif |
| Hsph11 | 1.29E-139 | 0.36700406 | 0.566 | 0.388 | 2.50E-135 | 2 | Hsph1 | myCAF_prolif |
| Asf1a | 1.34E-139 | 0.70813193 | 0.465 | 0.252 | 2.59E-135 | 2 | Asf1a | myCAF_prolif |
| C2cd2l | 2.41E-139 | 0.36298372 | 0.576 | 0.398 | 4.66E-135 | 2 | C2cd2l | myCAF_prolif |
| Nfib1 | 4.44E-139 | 0.34694767 | 0.606 | 0.401 | 8.59E-135 | 2 | Nfib | myCAF_prolif |
| Tamm41 | 4.62E-139 | 1.09635668 | 0.385 | 0.169 | 8.93E-135 | 2 | Tamm41 | myCAF_prolif |
| Eef1akmt1 | 5.12E-139 | 0.54118159 | 0.595 | 0.383 | 9.89E-135 | 2 | Eef1akmt1 | myCAF_prolif |
| Ppih | 8.42E-139 | 2.02001414 | 0.256 | 0.059 | 1.63E-134 | 2 | Ppih | myCAF_prolif |
| Zc3hav1 | 1.15E-138 | 0.28464393 | 0.682 | 0.503 | 2.23E-134 | 2 | Zc3hav1 | myCAF_prolif |
| Enoph1 | 1.38E-138 | 0.58599934 | 0.588 | 0.373 | 2.68E-134 | 2 | Enoph1 | myCAF_prolif |
| Eif4enif1 | 1.96E-138 | 0.44640986 | 0.603 | 0.394 | 3.79E-134 | 2 | Eif4enif1 | myCAF_prolif |
| Slpi1 | 2.20E-138 | 0.33668552 | 0.288 | 0.173 | 4.26E-134 | 2 | Slpi | myCAF_prolif |
| Cstf1 | 2.54E-138 | 0.53496274 | 0.504 | 0.309 | 4.90E-134 | 2 | Cstf1 | myCAF_prolif |
| Psmb3 | 3.63E-138 | 0.419553 | 0.638 | 0.433 | 7.01E-134 | 2 | Psmb3 | myCAF_prolif |
| Fxr2 | 4.60E-138 | 0.39953684 | 0.713 | 0.566 | 8.89E-134 | 2 | Fxr2 | myCAF_prolif |
| Rell1 | 4.88E-138 | 0.3674436 | 0.566 | 0.375 | 9.43E-134 | 2 | Rell1 | myCAF_prolif |
| Adrm1 | 1.58E-137 | 0.53319149 | 0.627 | 0.427 | 3.05E-133 | 2 | Adrm1 | myCAF_prolif |
| Usp37 | 7.40E-137 | 0.65278834 | 0.439 | 0.24 | 1.43E-132 | 2 | Usp37 | myCAF_prolif |
| Ndufs1 | 7.78E-137 | 0.36957707 | 0.68 | 0.511 | 1.50E-132 | 2 | Ndufs1 | myCAF_prolif |
| Fmnl21 | 1.18E-136 | 0.39103478 | 0.568 | 0.401 | 2.29E-132 | 2 | Fmnl2 | myCAF_prolif |
| Prpf4b | 1.35E-136 | 0.32608807 | 0.763 | 0.658 | 2.62E-132 | 2 | Prpf4b | myCAF_prolif |
| Stard3nl | 1.97E-136 | 0.25059265 | 0.732 | 0.608 | 3.81E-132 | 2 | Stard3nl | myCAF_prolif |
| Rbx1 | 2.37E-136 | 0.33144816 | 0.691 | 0.526 | 4.57E-132 | 2 | Rbx1 | myCAF_prolif |
| Slc35a3 | 2.47E-136 | 0.49834613 | 0.558 | 0.374 | 4.78E-132 | 2 | Slc35a3 | myCAF_prolif |
| Naa10 | 2.86E-136 | 0.68149917 | 0.485 | 0.269 | 5.52E-132 | 2 | Naa10 | myCAF_prolif |
| Dnttip2 | 3.99E-136 | 0.32949802 | 0.633 | 0.472 | 7.71E-132 | 2 | Dnttip2 | myCAF_prolif |
| Akap8 | 4.57E-136 | 0.37454569 | 0.689 | 0.541 | 8.84E-132 | 2 | Akap8 | myCAF_prolif |
| Gtf2f1 | 1.81E-135 | 0.50736401 | 0.656 | 0.484 | 3.50E-131 | 2 | Gtf2f1 | myCAF_prolif |
| Arhgap21 | 3.10E-135 | 0.27894869 | 0.685 | 0.551 | 5.99E-131 | 2 | Arhgap21 | myCAF_prolif |
| Arl6ip4 | 3.37E-135 | 0.40119379 | 0.692 | 0.534 | 6.52E-131 | 2 | Arl6ip4 | myCAF_prolif |
| Ppp1r12c | 4.90E-135 | 0.26077201 | 0.799 | 0.687 | 9.48E-131 | 2 | Ppp1r12c | myCAF_prolif |
| Supt20 | 5.02E-135 | 0.29322632 | 0.762 | 0.649 | 9.71E-131 | 2 | Supt20 | myCAF_prolif |
| Reps1 | 1.20E-134 | 0.29650653 | 0.654 | 0.469 | 2.33E-130 | 2 | Reps1 | myCAF_prolif |
| Ccnyl11 | 1.38E-134 | 0.38500814 | 0.601 | 0.435 | 2.66E-130 | 2 | Ccnyl1 | myCAF_prolif |
| Pwp1 | 1.47E-134 | 0.43788894 | 0.515 | 0.331 | 2.84E-130 | 2 | Pwp1 | myCAF_prolif |
| Crybg3 | 2.64E-134 | 0.81838578 | 0.451 | 0.23 | 5.11E-130 | 2 | Crybg3 | myCAF_prolif |
| Ints3 | 2.96E-134 | 0.31861859 | 0.687 | 0.524 | 5.72E-130 | 2 | Ints3 | myCAF_prolif |
| Pgm31 | 2.07E-133 | 0.33598486 | 0.712 | 0.552 | 4.00E-129 | 2 | Pgm3 | myCAF_prolif |
| Clasp2 | 7.09E-133 | 0.63712515 | 0.539 | 0.355 | 1.37E-128 | 2 | Clasp2 | myCAF_prolif |
| Tmco4 | 7.24E-133 | 0.43113205 | 0.513 | 0.309 | 1.40E-128 | 2 | Tmco4 | myCAF_prolif |
| Fntb | 7.70E-133 | 0.59348285 | 0.523 | 0.319 | 1.49E-128 | 2 | Fntb | myCAF_prolif |
| Mical1 | 1.45E-132 | 0.33101392 | 0.659 | 0.531 | 2.80E-128 | 2 | Mical1 | myCAF_prolif |
| Trmt10a | 1.47E-132 | 1.10499429 | 0.392 | 0.186 | 2.84E-128 | 2 | Trmt10a | myCAF_prolif |
| Snrnp25 | 1.48E-132 | 1.10541703 | 0.377 | 0.162 | 2.85E-128 | 2 | Snrnp25 | myCAF_prolif |
| Srsf11 | 1.66E-132 | 0.29335621 | 0.808 | 0.719 | 3.21E-128 | 2 | Srsf11 | myCAF_prolif |
| Ppp2r1b | 1.99E-132 | 0.50899577 | 0.596 | 0.405 | 3.85E-128 | 2 | Ppp2r1b | myCAF_prolif |
| Bcl2l12 | 2.30E-132 | 0.78673057 | 0.501 | 0.263 | 4.45E-128 | 2 | Bcl2l12 | myCAF_prolif |
| Tbl2 | 2.46E-132 | 0.34845406 | 0.7 | 0.527 | 4.76E-128 | 2 | Tbl2 | myCAF_prolif |
| Sapcd2 | 3.47E-132 | 5.75408611 | 0.294 | 0.002 | 6.70E-128 | 2 | Sapcd2 | myCAF_prolif |
| Commd2 | 5.27E-132 | 0.4380096 | 0.615 | 0.427 | 1.02E-127 | 2 | Commd2 | myCAF_prolif |
| Aven | 6.87E-132 | 0.51012792 | 0.532 | 0.326 | 1.33E-127 | 2 | Aven | myCAF_prolif |
| Snx11 | 9.58E-132 | 0.67847158 | 0.571 | 0.389 | 1.85E-127 | 2 | Snx11 | myCAF_prolif |
| Pgam5 | 1.31E-131 | 0.26288599 | 0.66 | 0.497 | 2.52E-127 | 2 | Pgam5 | myCAF_prolif |
| Txnl4a | 1.90E-131 | 0.30405526 | 0.67 | 0.522 | 3.68E-127 | 2 | Txnl4a | myCAF_prolif |
| Srek1 | 5.88E-131 | 0.38361495 | 0.795 | 0.678 | 1.14E-126 | 2 | Srek1 | myCAF_prolif |
| Trap1 | 1.14E-130 | 0.33498841 | 0.639 | 0.475 | 2.21E-126 | 2 | Trap1 | myCAF_prolif |
| Otulin | 1.33E-130 | 0.27259354 | 0.679 | 0.509 | 2.58E-126 | 2 | Otulin | myCAF_prolif |
| U2surp | 1.71E-130 | 0.38021441 | 0.675 | 0.498 | 3.31E-126 | 2 | U2surp | myCAF_prolif |
| Tmem18 | 2.96E-130 | 0.41267902 | 0.534 | 0.34 | 5.73E-126 | 2 | Tmem18 | myCAF_prolif |
| Dnajc2 | 6.89E-130 | 0.66382764 | 0.463 | 0.266 | 1.33E-125 | 2 | Dnajc2 | myCAF_prolif |
| Snx27 | 7.28E-130 | 0.36725705 | 0.597 | 0.419 | 1.41E-125 | 2 | Snx27 | myCAF_prolif |
| Nqo2 | 1.13E-129 | 0.27094498 | 0.613 | 0.45 | 2.19E-125 | 2 | Nqo2 | myCAF_prolif |
| Paip2b | 1.73E-129 | 0.74807434 | 0.461 | 0.258 | 3.34E-125 | 2 | Paip2b | myCAF_prolif |
| Zfp948 | 2.92E-129 | 1.11400708 | 0.392 | 0.188 | 5.65E-125 | 2 | Zfp948 | myCAF_prolif |
| Slc25a28 | 3.76E-129 | 0.55552643 | 0.603 | 0.402 | 7.27E-125 | 2 | Slc25a28 | myCAF_prolif |
| Psmd5 | 6.50E-129 | 0.31848488 | 0.608 | 0.442 | 1.26E-124 | 2 | Psmd5 | myCAF_prolif |
| Dnajc7 | 9.11E-129 | 0.25241577 | 0.73 | 0.604 | 1.76E-124 | 2 | Dnajc7 | myCAF_prolif |
| Rexo4 | 1.84E-128 | 0.6438478 | 0.587 | 0.392 | 3.56E-124 | 2 | Rexo4 | myCAF_prolif |
| Apmap | 4.91E-128 | 0.28164517 | 0.635 | 0.447 | 9.49E-124 | 2 | Apmap | myCAF_prolif |
| Ing5 | 1.74E-127 | 0.66081691 | 0.497 | 0.289 | 3.36E-123 | 2 | Ing5 | myCAF_prolif |
| Sgf29 | 1.88E-127 | 0.70429036 | 0.413 | 0.202 | 3.63E-123 | 2 | Sgf29 | myCAF_prolif |
| Tubgcp6 | 2.39E-127 | 0.78737615 | 0.457 | 0.227 | 4.62E-123 | 2 | Tubgcp6 | myCAF_prolif |
| Scoc | 2.95E-127 | 0.6568882 | 0.485 | 0.276 | 5.71E-123 | 2 | Scoc | myCAF_prolif |
| Ints10 | 3.29E-127 | 0.62920455 | 0.504 | 0.283 | 6.36E-123 | 2 | Ints10 | myCAF_prolif |
| Emc8 | 8.14E-127 | 0.74259989 | 0.518 | 0.301 | 1.57E-122 | 2 | Emc8 | myCAF_prolif |
| Ipo8 | 9.30E-127 | 0.35278396 | 0.634 | 0.49 | 1.80E-122 | 2 | Ipo8 | myCAF_prolif |
| Ttll12 | 9.77E-127 | 0.53624317 | 0.566 | 0.361 | 1.89E-122 | 2 | Ttll12 | myCAF_prolif |
| Ppif | 9.82E-127 | 0.60479095 | 0.427 | 0.219 | 1.90E-122 | 2 | Ppif | myCAF_prolif |
| Pfdn4 | 1.00E-126 | 0.45285723 | 0.551 | 0.356 | 1.94E-122 | 2 | Pfdn4 | myCAF_prolif |
| Tfam | 1.56E-126 | 0.56324204 | 0.601 | 0.406 | 3.02E-122 | 2 | Tfam | myCAF_prolif |
| Cenpj | 1.71E-126 | 2.09316273 | 0.251 | 0.073 | 3.31E-122 | 2 | Cenpj | myCAF_prolif |
| Mpv17l2 | 3.08E-126 | 0.47782987 | 0.605 | 0.413 | 5.95E-122 | 2 | Mpv17l2 | myCAF_prolif |
| Gemin2 | 3.94E-126 | 0.96361021 | 0.322 | 0.12 | 7.62E-122 | 2 | Gemin2 | myCAF_prolif |
| Prkg2 | 4.55E-126 | 1.02700888 | 0.265 | 0.114 | 8.80E-122 | 2 | Prkg2 | myCAF_prolif |
| Vcam11 | 8.19E-126 | 0.42172601 | 0.513 | 0.314 | 1.58E-121 | 2 | Vcam1 | myCAF_prolif |
| Zfp706 | 9.11E-126 | 0.38556174 | 0.599 | 0.415 | 1.76E-121 | 2 | Zfp706 | myCAF_prolif |
| Taf2 | 9.77E-126 | 0.30700988 | 0.64 | 0.471 | 1.89E-121 | 2 | Taf2 | myCAF_prolif |
| Bdkrb11 | 9.89E-126 | 0.40112063 | 0.439 | 0.291 | 1.91E-121 | 2 | Bdkrb1 | myCAF_prolif |
| Noct1 | 1.39E-125 | 0.34332178 | 0.435 | 0.288 | 2.69E-121 | 2 | Noct | myCAF_prolif |
| Rnf6 | 2.41E-125 | 0.29406856 | 0.74 | 0.619 | 4.65E-121 | 2 | Rnf6 | myCAF_prolif |
| Gpn2 | 3.14E-125 | 0.60107878 | 0.482 | 0.273 | 6.07E-121 | 2 | Gpn2 | myCAF_prolif |
| Traf2 | 4.56E-125 | 0.25949991 | 0.602 | 0.436 | 8.83E-121 | 2 | Traf2 | myCAF_prolif |
| Impa2 | 4.59E-125 | 0.97084891 | 0.278 | 0.103 | 8.88E-121 | 2 | Impa2 | myCAF_prolif |
| Med21 | 6.92E-125 | 0.32689046 | 0.675 | 0.548 | 1.34E-120 | 2 | Med21 | myCAF_prolif |
| Cops4 | 8.39E-125 | 0.27942805 | 0.67 | 0.53 | 1.62E-120 | 2 | Cops4 | myCAF_prolif |
| Pias3 | 9.73E-125 | 0.42460301 | 0.689 | 0.545 | 1.88E-120 | 2 | Pias3 | myCAF_prolif |
| Ap1ar | 1.07E-124 | 0.30089448 | 0.67 | 0.512 | 2.07E-120 | 2 | Ap1ar | myCAF_prolif |
| Stat5a1 | 1.15E-124 | 0.43540891 | 0.575 | 0.419 | 2.22E-120 | 2 | Stat5a | myCAF_prolif |
| Ube2j2 | 1.15E-124 | 0.26505705 | 0.675 | 0.523 | 2.22E-120 | 2 | Ube2j2 | myCAF_prolif |
| Dctd | 1.76E-124 | 0.86210582 | 0.347 | 0.162 | 3.40E-120 | 2 | Dctd | myCAF_prolif |
| Phgdh | 1.79E-124 | 1.0520148 | 0.26 | 0.102 | 3.45E-120 | 2 | Phgdh | myCAF_prolif |
| Gltp | 2.39E-124 | 0.29435012 | 0.667 | 0.534 | 4.62E-120 | 2 | Gltp | myCAF_prolif |
| Ppp2r2d | 4.03E-124 | 0.36359061 | 0.543 | 0.377 | 7.79E-120 | 2 | Ppp2r2d | myCAF_prolif |
| Nol7 | 5.07E-124 | 0.41718121 | 0.573 | 0.402 | 9.80E-120 | 2 | Nol7 | myCAF_prolif |
| Taf10 | 7.85E-124 | 0.42647278 | 0.607 | 0.412 | 1.52E-119 | 2 | Taf10 | myCAF_prolif |
| Clu1 | 1.55E-123 | 0.29049584 | 0.924 | 0.879 | 3.00E-119 | 2 | Clu | myCAF_prolif |
| Nfatc1 | 1.86E-123 | 0.35015931 | 0.541 | 0.382 | 3.59E-119 | 2 | Nfatc1 | myCAF_prolif |
| Uchl3 | 1.91E-123 | 0.88103264 | 0.392 | 0.203 | 3.69E-119 | 2 | Uchl3 | myCAF_prolif |
| Sp2 | 2.74E-123 | 0.55958373 | 0.481 | 0.279 | 5.29E-119 | 2 | Sp2 | myCAF_prolif |
| Tbc1d1 | 3.85E-123 | 0.2972981 | 0.609 | 0.429 | 7.45E-119 | 2 | Tbc1d1 | myCAF_prolif |
| Sephs2 | 4.99E-123 | 0.34121948 | 0.597 | 0.426 | 9.65E-119 | 2 | Sephs2 | myCAF_prolif |
| Atm | 6.85E-123 | 0.8695574 | 0.364 | 0.178 | 1.32E-118 | 2 | Atm | myCAF_prolif |
| Sclt1 | 6.87E-123 | 1.28042804 | 0.313 | 0.113 | 1.33E-118 | 2 | Sclt1 | myCAF_prolif |
| Ttc13 | 6.94E-123 | 0.4279898 | 0.568 | 0.38 | 1.34E-118 | 2 | Ttc13 | myCAF_prolif |
| Clpp | 1.05E-122 | 0.37159699 | 0.626 | 0.468 | 2.02E-118 | 2 | Clpp | myCAF_prolif |
| Ado | 1.91E-122 | 0.30555136 | 0.627 | 0.446 | 3.69E-118 | 2 | Ado | myCAF_prolif |
| Fadd | 2.19E-122 | 0.6352947 | 0.539 | 0.349 | 4.23E-118 | 2 | Fadd | myCAF_prolif |
| Prmt7 | 2.94E-122 | 0.52943349 | 0.497 | 0.307 | 5.69E-118 | 2 | Prmt7 | myCAF_prolif |
| Dot1l | 3.13E-122 | 0.32751096 | 0.616 | 0.436 | 6.06E-118 | 2 | Dot1l | myCAF_prolif |
| Lzic | 3.78E-122 | 0.38006688 | 0.592 | 0.421 | 7.30E-118 | 2 | Lzic | myCAF_prolif |
| Lum11 | 3.88E-122 | 0.9802762 | 0.353 | 0.14 | 7.50E-118 | 2 | Lum11 | myCAF_prolif |
| Armt1 | 5.47E-122 | 0.67238456 | 0.457 | 0.261 | 1.06E-117 | 2 | Armt1 | myCAF_prolif |
| Yeats4 | 5.71E-122 | 0.62034342 | 0.491 | 0.322 | 1.10E-117 | 2 | Yeats4 | myCAF_prolif |
| Bpnt1 | 6.58E-122 | 0.29409173 | 0.566 | 0.398 | 1.27E-117 | 2 | Bpnt1 | myCAF_prolif |
| Kat7 | 7.51E-122 | 0.31623174 | 0.644 | 0.483 | 1.45E-117 | 2 | Kat7 | myCAF_prolif |
| Drosha | 9.93E-122 | 0.31476789 | 0.612 | 0.446 | 1.92E-117 | 2 | Drosha | myCAF_prolif |
| Akip1 | 1.17E-121 | 0.46914041 | 0.544 | 0.359 | 2.26E-117 | 2 | Akip1 | myCAF_prolif |
| Gtf2f2 | 1.42E-121 | 0.3976154 | 0.56 | 0.391 | 2.74E-117 | 2 | Gtf2f2 | myCAF_prolif |
| Exosc9 | 1.44E-121 | 0.66697885 | 0.561 | 0.351 | 2.79E-117 | 2 | Exosc9 | myCAF_prolif |
| Prpf38b | 1.86E-121 | 0.46349462 | 0.725 | 0.582 | 3.59E-117 | 2 | Prpf38b | myCAF_prolif |
| Tmem129 | 2.05E-121 | 0.59122278 | 0.509 | 0.329 | 3.97E-117 | 2 | Tmem129 | myCAF_prolif |
| Ptprj | 2.21E-121 | 0.37719841 | 0.508 | 0.357 | 4.28E-117 | 2 | Ptprj | myCAF_prolif |
| Gtf3c6 | 2.45E-121 | 0.43117372 | 0.663 | 0.499 | 4.74E-117 | 2 | Gtf3c6 | myCAF_prolif |
| Wdr74 | 2.61E-121 | 0.43418365 | 0.574 | 0.393 | 5.04E-117 | 2 | Wdr74 | myCAF_prolif |
| Abcf1 | 3.02E-121 | 0.31452456 | 0.676 | 0.526 | 5.84E-117 | 2 | Abcf1 | myCAF_prolif |
| Gpbp1 | 6.43E-121 | 0.37664424 | 0.642 | 0.477 | 1.24E-116 | 2 | Gpbp1 | myCAF_prolif |
| Auts2 | 1.10E-120 | 0.52291906 | 0.653 | 0.532 | 2.13E-116 | 2 | Auts2 | myCAF_prolif |
| Fam92a | 2.10E-120 | 0.54088457 | 0.585 | 0.394 | 4.06E-116 | 2 | Fam92a | myCAF_prolif |
| Camkk2 | 2.75E-120 | 0.31256169 | 0.592 | 0.419 | 5.31E-116 | 2 | Camkk2 | myCAF_prolif |
| Pycr2 | 2.97E-120 | 0.36097839 | 0.623 | 0.448 | 5.73E-116 | 2 | Pycr2 | myCAF_prolif |
| Slc39a10 | 4.12E-120 | 0.47750507 | 0.457 | 0.293 | 7.97E-116 | 2 | Slc39a10 | myCAF_prolif |
| Exosc5 | 4.86E-120 | 0.3839861 | 0.593 | 0.413 | 9.40E-116 | 2 | Exosc5 | myCAF_prolif |
| Osbpl1a | 5.17E-120 | 0.37418082 | 0.53 | 0.36 | 1.00E-115 | 2 | Osbpl1a | myCAF_prolif |
| Upf3a | 1.26E-119 | 0.32058605 | 0.678 | 0.533 | 2.44E-115 | 2 | Upf3a | myCAF_prolif |
| Ptrh2 | 2.83E-119 | 0.72295865 | 0.398 | 0.194 | 5.48E-115 | 2 | Ptrh2 | myCAF_prolif |
| Cand2 | 4.55E-119 | 1.00548748 | 0.32 | 0.132 | 8.79E-115 | 2 | Cand2 | myCAF_prolif |
| Bahcc11 | 7.20E-119 | 0.43288471 | 0.477 | 0.292 | 1.39E-114 | 2 | Bahcc1 | myCAF_prolif |
| Nipsnap3b | 9.69E-119 | 0.34596423 | 0.552 | 0.382 | 1.87E-114 | 2 | Nipsnap3b | myCAF_prolif |
| Bysl | 9.79E-119 | 0.5958155 | 0.531 | 0.325 | 1.89E-114 | 2 | Bysl | myCAF_prolif |
| Dclre1b | 1.61E-118 | 1.01184492 | 0.304 | 0.112 | 3.11E-114 | 2 | Dclre1b | myCAF_prolif |
| Phtf1 | 2.22E-118 | 0.29135825 | 0.662 | 0.505 | 4.29E-114 | 2 | Phtf1 | myCAF_prolif |
| Slain2 | 2.45E-118 | 0.3434694 | 0.648 | 0.492 | 4.74E-114 | 2 | Slain2 | myCAF_prolif |
| Clns1a | 3.49E-118 | 0.37135741 | 0.639 | 0.461 | 6.75E-114 | 2 | Clns1a | myCAF_prolif |
| Zranb2 | 6.53E-118 | 0.46647907 | 0.628 | 0.455 | 1.26E-113 | 2 | Zranb2 | myCAF_prolif |
| Ift27 | 7.48E-118 | 0.27100805 | 0.687 | 0.491 | 1.45E-113 | 2 | Ift27 | myCAF_prolif |
| Toe1 | 1.30E-117 | 0.75154538 | 0.454 | 0.255 | 2.52E-113 | 2 | Toe1 | myCAF_prolif |
| Nav3 | 1.46E-117 | 0.28936565 | 0.651 | 0.537 | 2.82E-113 | 2 | Nav3 | myCAF_prolif |
| Atad3a | 1.49E-117 | 0.40095691 | 0.544 | 0.346 | 2.87E-113 | 2 | Atad3a | myCAF_prolif |
| Cln6 | 1.92E-117 | 0.37581725 | 0.513 | 0.339 | 3.71E-113 | 2 | Cln6 | myCAF_prolif |
| Adk | 3.98E-117 | 0.26536355 | 0.625 | 0.475 | 7.70E-113 | 2 | Adk | myCAF_prolif |
| Nup88 | 5.58E-117 | 0.40541058 | 0.565 | 0.374 | 1.08E-112 | 2 | Nup88 | myCAF_prolif |
| Mlh1 | 6.08E-117 | 1.118144 | 0.356 | 0.148 | 1.17E-112 | 2 | Mlh1 | myCAF_prolif |
| Clpb | 7.73E-117 | 0.51274093 | 0.463 | 0.258 | 1.49E-112 | 2 | Clpb | myCAF_prolif |
| Npat | 1.08E-116 | 0.88350137 | 0.409 | 0.217 | 2.08E-112 | 2 | Npat | myCAF_prolif |
| Saal1 | 1.66E-116 | 0.51148935 | 0.462 | 0.277 | 3.20E-112 | 2 | Saal1 | myCAF_prolif |
| Zfp131 | 3.14E-116 | 0.48111775 | 0.697 | 0.536 | 6.08E-112 | 2 | Zfp131 | myCAF_prolif |
| Rpia | 3.32E-116 | 0.46694738 | 0.459 | 0.294 | 6.42E-112 | 2 | Rpia | myCAF_prolif |
| Apex1 | 3.80E-116 | 0.25200366 | 0.635 | 0.496 | 7.35E-112 | 2 | Apex1 | myCAF_prolif |
| Rgs12 | 2.25E-115 | 0.31833269 | 0.618 | 0.459 | 4.35E-111 | 2 | Rgs12 | myCAF_prolif |
| Tmem138 | 5.60E-115 | 0.60791507 | 0.465 | 0.283 | 1.08E-110 | 2 | Tmem138 | myCAF_prolif |
| Polr3h | 5.65E-115 | 0.44791445 | 0.551 | 0.376 | 1.09E-110 | 2 | Polr3h | myCAF_prolif |
| Tmem107 | 6.07E-115 | 0.81913383 | 0.386 | 0.18 | 1.17E-110 | 2 | Tmem107 | myCAF_prolif |
| Armc1 | 6.35E-115 | 0.28713752 | 0.646 | 0.489 | 1.23E-110 | 2 | Armc1 | myCAF_prolif |
| Dhrs13 | 6.57E-115 | 1.27688004 | 0.259 | 0.093 | 1.27E-110 | 2 | Dhrs13 | myCAF_prolif |
| Dhdds | 7.60E-115 | 0.40052419 | 0.651 | 0.486 | 1.47E-110 | 2 | Dhdds | myCAF_prolif |
| Spata131 | 7.85E-115 | 0.53035823 | 0.409 | 0.25 | 1.52E-110 | 2 | Spata13 | myCAF_prolif |
| Thyn1 | 7.99E-115 | 0.68605164 | 0.449 | 0.242 | 1.54E-110 | 2 | Thyn1 | myCAF_prolif |
| Gtpbp4 | 8.03E-115 | 0.26609861 | 0.666 | 0.531 | 1.55E-110 | 2 | Gtpbp4 | myCAF_prolif |
| Crlf3 | 1.67E-114 | 0.44601703 | 0.509 | 0.304 | 3.24E-110 | 2 | Crlf3 | myCAF_prolif |
| Timm10 | 4.28E-114 | 0.36895407 | 0.552 | 0.388 | 8.28E-110 | 2 | Timm10 | myCAF_prolif |
| Tdg | 9.75E-114 | 0.51482005 | 0.541 | 0.327 | 1.88E-109 | 2 | Tdg | myCAF_prolif |
| Eefsec | 1.10E-113 | 0.85526116 | 0.382 | 0.177 | 2.13E-109 | 2 | Eefsec | myCAF_prolif |
| S1pr1 | 2.48E-113 | 0.6808637 | 0.28 | 0.137 | 4.80E-109 | 2 | S1pr1 | myCAF_prolif |
| Mtf2 | 3.56E-113 | 0.79921984 | 0.435 | 0.251 | 6.88E-109 | 2 | Mtf2 | myCAF_prolif |
| Exosc1 | 8.57E-113 | 0.57637158 | 0.489 | 0.314 | 1.66E-108 | 2 | Exosc1 | myCAF_prolif |
| Slf2 | 1.11E-112 | 0.65142269 | 0.507 | 0.358 | 2.15E-108 | 2 | Slf2 | myCAF_prolif |
| Bcl2l13 | 1.27E-112 | 0.41063902 | 0.607 | 0.42 | 2.46E-108 | 2 | Bcl2l13 | myCAF_prolif |
| Tsg101 | 2.17E-112 | 0.32793877 | 0.689 | 0.525 | 4.19E-108 | 2 | Tsg101 | myCAF_prolif |
| Bbip1 | 2.60E-112 | 0.46577387 | 0.525 | 0.349 | 5.02E-108 | 2 | Bbip1 | myCAF_prolif |
| Erc1 | 3.62E-112 | 0.36020924 | 0.738 | 0.619 | 7.01E-108 | 2 | Erc1 | myCAF_prolif |
| Rock1 | 5.37E-112 | 0.25599196 | 0.714 | 0.619 | 1.04E-107 | 2 | Rock1 | myCAF_prolif |
| Senp2 | 5.63E-112 | 0.29330757 | 0.575 | 0.389 | 1.09E-107 | 2 | Senp2 | myCAF_prolif |
| Tyw1 | 7.70E-112 | 0.46265029 | 0.543 | 0.369 | 1.49E-107 | 2 | Tyw1 | myCAF_prolif |
| Hccs | 1.13E-111 | 0.47254895 | 0.535 | 0.335 | 2.18E-107 | 2 | Hccs | myCAF_prolif |
| Myo19 | 1.17E-111 | 0.92733368 | 0.373 | 0.185 | 2.26E-107 | 2 | Myo19 | myCAF_prolif |
| Gabpb2 | 1.71E-111 | 0.5285408 | 0.528 | 0.34 | 3.31E-107 | 2 | Gabpb2 | myCAF_prolif |
| Trmt6 | 1.80E-111 | 0.64585667 | 0.476 | 0.266 | 3.48E-107 | 2 | Trmt6 | myCAF_prolif |
| Alg14 | 2.60E-111 | 0.2508985 | 0.621 | 0.472 | 5.03E-107 | 2 | Alg14 | myCAF_prolif |
| Eny2 | 2.90E-111 | 0.5038041 | 0.544 | 0.339 | 5.60E-107 | 2 | Eny2 | myCAF_prolif |
| Gpn1 | 3.37E-111 | 0.54422028 | 0.499 | 0.304 | 6.52E-107 | 2 | Gpn1 | myCAF_prolif |
| Tarbp2 | 1.26E-110 | 0.58363577 | 0.477 | 0.277 | 2.44E-106 | 2 | Tarbp2 | myCAF_prolif |
| Pfdn2 | 1.86E-110 | 0.30656561 | 0.687 | 0.544 | 3.60E-106 | 2 | Pfdn2 | myCAF_prolif |
| Dnlz | 5.43E-110 | 0.27459882 | 0.663 | 0.524 | 1.05E-105 | 2 | Dnlz | myCAF_prolif |
| Tbc1d10a | 9.25E-110 | 0.31348715 | 0.737 | 0.588 | 1.79E-105 | 2 | Tbc1d10a | myCAF_prolif |
| Pde8b | 1.04E-109 | 0.73178864 | 0.38 | 0.183 | 2.01E-105 | 2 | Pde8b | myCAF_prolif |
| Safb2 | 1.53E-109 | 0.53734507 | 0.677 | 0.519 | 2.95E-105 | 2 | Safb2 | myCAF_prolif |
| Clk3 | 1.84E-109 | 0.27980442 | 0.667 | 0.507 | 3.57E-105 | 2 | Clk3 | myCAF_prolif |
| Tspan181 | 3.25E-109 | 0.39232565 | 0.297 | 0.193 | 6.28E-105 | 2 | Tspan18 | myCAF_prolif |
| Otud4 | 3.88E-109 | 0.27108899 | 0.577 | 0.41 | 7.49E-105 | 2 | Otud4 | myCAF_prolif |
| Apip | 6.59E-109 | 0.42037282 | 0.504 | 0.321 | 1.27E-104 | 2 | Apip | myCAF_prolif |
| Ftsj1 | 7.08E-109 | 0.27624401 | 0.591 | 0.425 | 1.37E-104 | 2 | Ftsj1 | myCAF_prolif |
| Sltm | 1.15E-108 | 0.55023314 | 0.672 | 0.527 | 2.22E-104 | 2 | Sltm | myCAF_prolif |
| Ch25h1 | 1.26E-108 | 0.389154 | 0.304 | 0.181 | 2.43E-104 | 2 | Ch25h | myCAF_prolif |
| Ccnc | 1.92E-108 | 0.4088029 | 0.539 | 0.392 | 3.71E-104 | 2 | Ccnc | myCAF_prolif |
| Gtf2h4 | 2.28E-108 | 0.43250154 | 0.529 | 0.336 | 4.40E-104 | 2 | Gtf2h4 | myCAF_prolif |
| Bop1 | 3.09E-108 | 0.49376446 | 0.549 | 0.348 | 5.98E-104 | 2 | Bop1 | myCAF_prolif |
| Eif2b4 | 5.26E-108 | 0.46425734 | 0.567 | 0.381 | 1.02E-103 | 2 | Eif2b4 | myCAF_prolif |
| Setdb1 | 5.97E-108 | 0.36347306 | 0.57 | 0.402 | 1.15E-103 | 2 | Setdb1 | myCAF_prolif |
| Neil3 | 1.10E-107 | 8.97308796 | 0.466 | 0.001 | 2.13E-103 | 2 | Neil3 | myCAF_prolif |
| Zrsr2 | 2.62E-107 | 0.48650939 | 0.639 | 0.453 | 5.06E-103 | 2 | Zrsr2 | myCAF_prolif |
| Plscr1 | 2.91E-107 | 0.49033713 | 0.452 | 0.276 | 5.62E-103 | 2 | Plscr1 | myCAF_prolif |
| Mtr | 3.01E-107 | 0.65310417 | 0.492 | 0.294 | 5.83E-103 | 2 | Mtr | myCAF_prolif |
| Fam171a1 | 5.62E-107 | 0.25181232 | 0.67 | 0.509 | 1.09E-102 | 2 | Fam171a1 | myCAF_prolif |
| Taf4 | 5.99E-107 | 0.48961955 | 0.531 | 0.342 | 1.16E-102 | 2 | Taf4 | myCAF_prolif |
| Fam76b | 6.56E-107 | 0.65837331 | 0.62 | 0.447 | 1.27E-102 | 2 | Fam76b | myCAF_prolif |
| Tshz1 | 1.04E-106 | 0.52133165 | 0.484 | 0.308 | 2.00E-102 | 2 | Tshz1 | myCAF_prolif |
| Lrr1 | 1.20E-106 | 6.57527969 | 0.254 | 0.002 | 2.33E-102 | 2 | Lrr1 | myCAF_prolif |
| Exog | 3.44E-106 | 1.08110528 | 0.32 | 0.128 | 6.65E-102 | 2 | Exog | myCAF_prolif |
| Rnf125 | 4.10E-106 | 0.31382725 | 0.456 | 0.293 | 7.93E-102 | 2 | Rnf125 | myCAF_prolif |
| Wars | 5.53E-106 | 0.27884574 | 0.643 | 0.465 | 1.07E-101 | 2 | Wars | myCAF_prolif |
| Tbc1d8b | 5.76E-106 | 0.34047763 | 0.601 | 0.43 | 1.11E-101 | 2 | Tbc1d8b | myCAF_prolif |
| Hdgfl3 | 6.55E-106 | 0.35290916 | 0.488 | 0.322 | 1.27E-101 | 2 | Hdgfl3 | myCAF_prolif |
| Relt | 6.81E-106 | 1.22578048 | 0.284 | 0.105 | 1.32E-101 | 2 | Relt | myCAF_prolif |
| C2cd5 | 1.13E-105 | 0.69762712 | 0.38 | 0.207 | 2.19E-101 | 2 | C2cd5 | myCAF_prolif |
| Fli1 | 1.29E-105 | 0.9004249 | 0.326 | 0.159 | 2.50E-101 | 2 | Fli1 | myCAF_prolif |
| Rnf5 | 2.71E-105 | 0.45879845 | 0.564 | 0.376 | 5.24E-101 | 2 | Rnf5 | myCAF_prolif |
| Zfp330 | 6.54E-105 | 0.33870137 | 0.664 | 0.501 | 1.26E-100 | 2 | Zfp330 | myCAF_prolif |
| Nt5c3b | 1.54E-104 | 0.26586946 | 0.553 | 0.407 | 2.99E-100 | 2 | Nt5c3b | myCAF_prolif |
| Brd7 | 1.81E-104 | 0.25484926 | 0.644 | 0.484 | 3.49E-100 | 2 | Brd7 | myCAF_prolif |
| Kctd121 | 3.12E-104 | 0.28585661 | 0.509 | 0.341 | 6.04E-100 | 2 | Kctd12 | myCAF_prolif |
| Uba6 | 3.66E-104 | 0.30354158 | 0.525 | 0.366 | 7.08E-100 | 2 | Uba6 | myCAF_prolif |
| Emc6 | 4.51E-104 | 0.26020041 | 0.611 | 0.451 | 8.72E-100 | 2 | Emc6 | myCAF_prolif |
| Nosip | 1.64E-103 | 0.34353558 | 0.667 | 0.517 | 3.17E-99 | 2 | Nosip | myCAF_prolif |
| Nsf | 3.10E-103 | 0.31492863 | 0.645 | 0.482 | 6.00E-99 | 2 | Nsf | myCAF_prolif |
| Cep135 | 3.19E-103 | 1.62582494 | 0.285 | 0.108 | 6.16E-99 | 2 | Cep135 | myCAF_prolif |
| Mcm9 | 4.84E-103 | 0.58940225 | 0.406 | 0.229 | 9.37E-99 | 2 | Mcm9 | myCAF_prolif |
| Tpgs2 | 6.21E-103 | 0.57550189 | 0.423 | 0.237 | 1.20E-98 | 2 | Tpgs2 | myCAF_prolif |
| Gmeb1 | 7.63E-103 | 0.37232109 | 0.49 | 0.311 | 1.48E-98 | 2 | Gmeb1 | myCAF_prolif |
| Tfdp2 | 8.32E-103 | 0.7432538 | 0.383 | 0.217 | 1.61E-98 | 2 | Tfdp2 | myCAF_prolif |
| Zcchc17 | 1.70E-102 | 0.38200732 | 0.536 | 0.365 | 3.28E-98 | 2 | Zcchc17 | myCAF_prolif |
| Pcbd2 | 1.75E-102 | 0.5385041 | 0.47 | 0.283 | 3.38E-98 | 2 | Pcbd2 | myCAF_prolif |
| Snx5 | 2.18E-102 | 0.29465041 | 0.483 | 0.347 | 4.21E-98 | 2 | Snx5 | myCAF_prolif |
| Vrk3 | 2.51E-102 | 0.33503585 | 0.681 | 0.543 | 4.85E-98 | 2 | Vrk3 | myCAF_prolif |
| Atp11c | 2.81E-102 | 0.51502636 | 0.463 | 0.275 | 5.43E-98 | 2 | Atp11c | myCAF_prolif |
| Morc4 | 2.96E-102 | 0.5474028 | 0.384 | 0.217 | 5.72E-98 | 2 | Morc4 | myCAF_prolif |
| Zfp414 | 8.73E-102 | 0.25106035 | 0.655 | 0.511 | 1.69E-97 | 2 | Zfp414 | myCAF_prolif |
| H2-Q71 | 8.75E-102 | 0.42826423 | 0.298 | 0.15 | 1.69E-97 | 2 | H2-Q7 | myCAF_prolif |
| Kif2a | 8.75E-102 | 0.4579036 | 0.568 | 0.402 | 1.69E-97 | 2 | Kif2a | myCAF_prolif |
| Zpr11 | 2.48E-101 | 0.25334629 | 0.632 | 0.493 | 4.79E-97 | 2 | Zpr1 | myCAF_prolif |
| Abcb7 | 3.76E-101 | 0.5942438 | 0.403 | 0.212 | 7.28E-97 | 2 | Abcb7 | myCAF_prolif |
| Gps2 | 4.49E-101 | 0.54026176 | 0.532 | 0.351 | 8.68E-97 | 2 | Gps2 | myCAF_prolif |
| Tmem201 | 5.72E-101 | 0.45546649 | 0.477 | 0.292 | 1.11E-96 | 2 | Tmem201 | myCAF_prolif |
| Ddx49 | 7.08E-101 | 0.30347624 | 0.644 | 0.5 | 1.37E-96 | 2 | Ddx49 | myCAF_prolif |
| Prrg1 | 2.09E-100 | 0.57418706 | 0.413 | 0.236 | 4.04E-96 | 2 | Prrg1 | myCAF_prolif |
| Pwwp3a | 2.31E-100 | 0.68247385 | 0.538 | 0.389 | 4.46E-96 | 2 | Pwwp3a | myCAF_prolif |
| Clock | 2.52E-100 | 0.35500327 | 0.581 | 0.43 | 4.87E-96 | 2 | Clock | myCAF_prolif |
| Bmi11 | 2.94E-100 | 0.40139396 | 0.633 | 0.48 | 5.69E-96 | 2 | Bmi1 | myCAF_prolif |
| Prmt3 | 4.25E-100 | 0.51655464 | 0.462 | 0.287 | 8.22E-96 | 2 | Prmt3 | myCAF_prolif |
| Carnmt1 | 4.45E-100 | 0.44177208 | 0.468 | 0.3 | 8.61E-96 | 2 | Carnmt1 | myCAF_prolif |
| Sacs | 4.85E-100 | 0.44715319 | 0.441 | 0.272 | 9.38E-96 | 2 | Sacs | myCAF_prolif |
| Ctu2 | 6.93E-100 | 0.4937689 | 0.486 | 0.308 | 1.34E-95 | 2 | Ctu2 | myCAF_prolif |
| Usp31 | 7.99E-100 | 0.43969505 | 0.485 | 0.325 | 1.54E-95 | 2 | Usp31 | myCAF_prolif |
| Agpat5 | 1.32E-99 | 0.4355269 | 0.499 | 0.321 | 2.55E-95 | 2 | Agpat5 | myCAF_prolif |
| Nsfl1c | 1.48E-99 | 0.41311856 | 0.706 | 0.562 | 2.86E-95 | 2 | Nsfl1c | myCAF_prolif |
| Rrs1 | 3.11E-99 | 0.35397277 | 0.525 | 0.366 | 6.01E-95 | 2 | Rrs1 | myCAF_prolif |
| Tyro31 | 4.20E-99 | 0.58341401 | 0.384 | 0.209 | 8.11E-95 | 2 | Tyro3 | myCAF_prolif |
| Pqbp1 | 4.56E-99 | 0.27558888 | 0.716 | 0.583 | 8.81E-95 | 2 | Pqbp1 | myCAF_prolif |
| Rcl11 | 6.73E-99 | 0.27394525 | 0.517 | 0.343 | 1.30E-94 | 2 | Rcl1 | myCAF_prolif |
| Ecd | 9.06E-99 | 0.46449828 | 0.561 | 0.373 | 1.75E-94 | 2 | Ecd | myCAF_prolif |
| Sfswap | 1.36E-98 | 0.32935754 | 0.665 | 0.538 | 2.63E-94 | 2 | Sfswap | myCAF_prolif |
| Map3k5 | 1.36E-98 | 0.78462657 | 0.38 | 0.193 | 2.63E-94 | 2 | Map3k5 | myCAF_prolif |
| Snrnp35 | 1.47E-98 | 0.68569411 | 0.451 | 0.288 | 2.84E-94 | 2 | Snrnp35 | myCAF_prolif |
| Znrd2 | 1.63E-98 | 0.41319631 | 0.508 | 0.34 | 3.15E-94 | 2 | Znrd2 | myCAF_prolif |
| B4galt61 | 1.81E-98 | 0.28491474 | 0.565 | 0.403 | 3.51E-94 | 2 | B4galt6 | myCAF_prolif |
| Lmbrd2 | 4.14E-98 | 0.34758778 | 0.502 | 0.328 | 8.00E-94 | 2 | Lmbrd2 | myCAF_prolif |
| Kat14 | 5.10E-98 | 0.28420635 | 0.622 | 0.456 | 9.87E-94 | 2 | Kat14 | myCAF_prolif |
| Gtpbp2 | 6.08E-98 | 0.25616326 | 0.703 | 0.612 | 1.18E-93 | 2 | Gtpbp2 | myCAF_prolif |
| Ebf1 | 1.21E-97 | 0.26729137 | 0.763 | 0.655 | 2.33E-93 | 2 | Ebf1 | myCAF_prolif |
| Rab13 | 1.77E-97 | 0.33939552 | 0.642 | 0.503 | 3.43E-93 | 2 | Rab13 | myCAF_prolif |
| Urb2 | 2.03E-97 | 0.47961953 | 0.445 | 0.278 | 3.92E-93 | 2 | Urb2 | myCAF_prolif |
| Ankrd54 | 2.04E-97 | 0.35334281 | 0.511 | 0.332 | 3.95E-93 | 2 | Ankrd54 | myCAF_prolif |
| Fancf | 3.36E-97 | 0.66870917 | 0.375 | 0.196 | 6.49E-93 | 2 | Fancf | myCAF_prolif |
| Mak16 | 4.13E-97 | 0.44040892 | 0.519 | 0.359 | 7.99E-93 | 2 | Mak16 | myCAF_prolif |
| Jade11 | 4.48E-97 | 0.30448561 | 0.57 | 0.411 | 8.67E-93 | 2 | Jade1 | myCAF_prolif |
| Tmem126a | 5.04E-97 | 0.37938638 | 0.581 | 0.422 | 9.75E-93 | 2 | Tmem126a | myCAF_prolif |
| Ints13 | 7.70E-97 | 0.28133452 | 0.538 | 0.38 | 1.49E-92 | 2 | Ints13 | myCAF_prolif |
| L3mbtl2 | 9.26E-97 | 0.78702308 | 0.299 | 0.137 | 1.79E-92 | 2 | L3mbtl2 | myCAF_prolif |
| Mcc | 1.15E-96 | 0.47772473 | 0.527 | 0.36 | 2.23E-92 | 2 | Mcc | myCAF_prolif |
| Mpp1 | 1.73E-96 | 0.57059667 | 0.518 | 0.367 | 3.34E-92 | 2 | Mpp1 | myCAF_prolif |
| Nudt19 | 1.93E-96 | 0.35896302 | 0.489 | 0.328 | 3.74E-92 | 2 | Nudt19 | myCAF_prolif |
| Ssbp1 | 2.21E-96 | 0.33814386 | 0.587 | 0.411 | 4.27E-92 | 2 | Ssbp1 | myCAF_prolif |
| Tmem245 | 3.95E-96 | 0.53638818 | 0.561 | 0.378 | 7.64E-92 | 2 | Tmem245 | myCAF_prolif |
| Pus1 | 5.78E-96 | 0.2805524 | 0.566 | 0.416 | 1.12E-91 | 2 | Pus1 | myCAF_prolif |
| AW554918 | 6.57E-96 | 1.47906377 | 0.332 | 0.155 | 1.27E-91 | 2 | AW554918 | myCAF_prolif |
| Fam45a | 8.87E-96 | 0.36716362 | 0.614 | 0.436 | 1.71E-91 | 2 | Fam45a | myCAF_prolif |
| mt-Nd3 | 1.32E-95 | 1.39299019 | 0.576 | 0.459 | 2.56E-91 | 2 | mt-Nd3 | myCAF_prolif |
| Pdcd5 | 1.34E-95 | 0.32048385 | 0.591 | 0.412 | 2.59E-91 | 2 | Pdcd5 | myCAF_prolif |
| Psmg4 | 2.16E-95 | 0.25471182 | 0.546 | 0.411 | 4.17E-91 | 2 | Psmg4 | myCAF_prolif |
| Tex10 | 2.92E-95 | 0.27590819 | 0.553 | 0.402 | 5.65E-91 | 2 | Tex10 | myCAF_prolif |
| Ewsr1 | 3.55E-95 | 0.36773284 | 0.544 | 0.347 | 6.87E-91 | 2 | Ewsr1 | myCAF_prolif |
| Tfap4 | 5.37E-95 | 0.81640892 | 0.275 | 0.127 | 1.04E-90 | 2 | Tfap4 | myCAF_prolif |
| Rbm10 | 7.44E-95 | 0.62126551 | 0.415 | 0.226 | 1.44E-90 | 2 | Rbm10 | myCAF_prolif |
| Ddx52 | 8.23E-95 | 0.67436402 | 0.487 | 0.308 | 1.59E-90 | 2 | Ddx52 | myCAF_prolif |
| Slc25a251 | 1.16E-94 | 0.3485316 | 0.47 | 0.316 | 2.24E-90 | 2 | Slc25a25 | myCAF_prolif |
| Pfdn1 | 1.28E-94 | 0.4764816 | 0.642 | 0.482 | 2.48E-90 | 2 | Pfdn1 | myCAF_prolif |
| Mphosph10 | 1.43E-94 | 0.65876816 | 0.473 | 0.288 | 2.76E-90 | 2 | Mphosph10 | myCAF_prolif |
| Stk35 | 2.58E-94 | 0.73825989 | 0.393 | 0.204 | 4.99E-90 | 2 | Stk35 | myCAF_prolif |
| Ankrd13c | 7.87E-94 | 0.30488629 | 0.654 | 0.517 | 1.52E-89 | 2 | Ankrd13c | myCAF_prolif |
| Tnrc6a | 8.39E-94 | 0.28334995 | 0.687 | 0.564 | 1.62E-89 | 2 | Tnrc6a | myCAF_prolif |
| Faf1 | 8.58E-94 | 0.34134098 | 0.676 | 0.558 | 1.66E-89 | 2 | Faf1 | myCAF_prolif |
| Peg121 | 8.59E-94 | 0.63853588 | 0.375 | 0.203 | 1.66E-89 | 2 | Peg12 | myCAF_prolif |
| Stim2 | 1.49E-93 | 0.34860527 | 0.546 | 0.388 | 2.89E-89 | 2 | Stim2 | myCAF_prolif |
| Fnbp4 | 1.90E-93 | 0.4012654 | 0.615 | 0.439 | 3.67E-89 | 2 | Fnbp4 | myCAF_prolif |
| Dzip1l1 | 4.26E-93 | 0.66764804 | 0.628 | 0.456 | 8.23E-89 | 2 | Dzip1l | myCAF_prolif |
| Nrf1 | 5.80E-93 | 0.48516197 | 0.443 | 0.27 | 1.12E-88 | 2 | Nrf1 | myCAF_prolif |
| Pak11 | 6.79E-93 | 0.66674806 | 0.43 | 0.248 | 1.31E-88 | 2 | Pak1 | myCAF_prolif |
| Eef1akmt4 | 1.24E-92 | 0.30589477 | 0.556 | 0.413 | 2.39E-88 | 2 | Eef1akmt4 | myCAF_prolif |
| Polg | 1.87E-92 | 0.41170745 | 0.584 | 0.411 | 3.61E-88 | 2 | Polg | myCAF_prolif |
| Topors | 2.04E-92 | 0.30821224 | 0.6 | 0.444 | 3.94E-88 | 2 | Topors | myCAF_prolif |
| Polr1a | 2.16E-92 | 0.34784461 | 0.482 | 0.336 | 4.17E-88 | 2 | Polr1a | myCAF_prolif |
| Ddx18 | 2.88E-92 | 0.33978413 | 0.558 | 0.389 | 5.56E-88 | 2 | Ddx18 | myCAF_prolif |
| Ppan | 3.39E-92 | 0.26903287 | 0.566 | 0.423 | 6.56E-88 | 2 | Ppan | myCAF_prolif |
| Mtmr4 | 4.85E-92 | 0.28605616 | 0.569 | 0.408 | 9.38E-88 | 2 | Mtmr4 | myCAF_prolif |
| Heatr3 | 8.20E-92 | 0.31055365 | 0.497 | 0.341 | 1.59E-87 | 2 | Heatr3 | myCAF_prolif |
| Nop9 | 1.42E-91 | 0.71335334 | 0.374 | 0.186 | 2.75E-87 | 2 | Nop9 | myCAF_prolif |
| Diaph1 | 1.57E-91 | 0.31139717 | 0.603 | 0.424 | 3.04E-87 | 2 | Diaph1 | myCAF_prolif |
| Cep44 | 1.69E-91 | 0.89719937 | 0.328 | 0.162 | 3.27E-87 | 2 | Cep44 | myCAF_prolif |
| Chchd4 | 2.18E-91 | 0.75511932 | 0.304 | 0.144 | 4.21E-87 | 2 | Chchd4 | myCAF_prolif |
| Pdgfc | 2.43E-91 | 0.56311736 | 0.295 | 0.175 | 4.71E-87 | 2 | Pdgfc | myCAF_prolif |
| Sfxn21 | 2.76E-91 | 0.53946217 | 0.472 | 0.291 | 5.33E-87 | 2 | Sfxn2 | myCAF_prolif |
| Terf2 | 2.78E-91 | 0.47125022 | 0.532 | 0.375 | 5.38E-87 | 2 | Terf2 | myCAF_prolif |
| Csrnp2 | 2.88E-91 | 0.52666863 | 0.453 | 0.293 | 5.57E-87 | 2 | Csrnp2 | myCAF_prolif |

| | | | | | | | | |
|---|---|---|---|---|---|---|---|---|
| Cby1 | 3.27E-91 | 0.54028524 | 0.455 | 0.28 | 6.33E-87 | 2 | Cby1 | myCAF_prolif |
| Kat2a | 4.51E-91 | 0.31405243 | 0.556 | 0.395 | 8.73E-87 | 2 | Kat2a | myCAF_prolif |
| Nol10 | 4.84E-91 | 0.54312718 | 0.444 | 0.278 | 9.36E-87 | 2 | Nol10 | myCAF_prolif |
| Gfer | 9.53E-91 | 0.3650967 | 0.537 | 0.363 | 1.84E-86 | 2 | Gfer | myCAF_prolif |
| Ptges2 | 1.44E-90 | 0.39521093 | 0.541 | 0.362 | 2.79E-86 | 2 | Ptges2 | myCAF_prolif |
| Bclaf3 | 1.68E-90 | 0.74466477 | 0.387 | 0.206 | 3.25E-86 | 2 | Bclaf3 | myCAF_prolif |
| Fbxo45 | 2.03E-90 | 0.6857796 | 0.371 | 0.194 | 3.92E-86 | 2 | Fbxo45 | myCAF_prolif |
| Aqr | 2.41E-90 | 0.25440847 | 0.574 | 0.408 | 4.65E-86 | 2 | Aqr | myCAF_prolif |
| Cmtr1 | 3.35E-90 | 0.2795859 | 0.654 | 0.501 | 6.48E-86 | 2 | Cmtr1 | myCAF_prolif |
| Utp18 | 4.17E-90 | 0.33249164 | 0.486 | 0.321 | 8.06E-86 | 2 | Utp18 | myCAF_prolif |
| Zfp518b | 4.86E-90 | 0.40345411 | 0.442 | 0.297 | 9.40E-86 | 2 | Zfp518b | myCAF_prolif |
| Bmpr1b1 | 5.16E-90 | 0.54252849 | 0.349 | 0.211 | 9.97E-86 | 2 | Bmpr1b | myCAF_prolif |
| Ftsj31 | 5.45E-90 | 0.4917849 | 0.432 | 0.265 | 1.05E-85 | 2 | Ftsj3 | myCAF_prolif |
| Rnf1571 | 5.65E-90 | 0.6237018 | 0.329 | 0.181 | 1.09E-85 | 2 | Rnf157 | myCAF_prolif |
| Camk2n21 | 7.16E-90 | 0.55652387 | 0.416 | 0.241 | 1.38E-85 | 2 | Camk2n2 | myCAF_prolif |
| Mettl4 | 7.58E-90 | 0.33303964 | 0.494 | 0.33 | 1.46E-85 | 2 | Mettl4 | myCAF_prolif |
| Dis3l | 8.03E-90 | 0.36024266 | 0.463 | 0.304 | 1.55E-85 | 2 | Dis3l | myCAF_prolif |
| Dnajc15 | 1.07E-89 | 0.2713824 | 0.566 | 0.413 | 2.07E-85 | 2 | Dnajc15 | myCAF_prolif |
| Polr2d | 1.64E-89 | 0.33524738 | 0.553 | 0.389 | 3.16E-85 | 2 | Polr2d | myCAF_prolif |
| Bcl7a1 | 2.12E-89 | 1.07688423 | 0.368 | 0.211 | 4.10E-85 | 2 | Bcl7a | myCAF_prolif |
| Rapgef6 | 2.24E-89 | 0.43467295 | 0.51 | 0.355 | 4.33E-85 | 2 | Rapgef6 | myCAF_prolif |
| Pmm2 | 4.34E-89 | 0.28427516 | 0.627 | 0.49 | 8.39E-85 | 2 | Pmm2 | myCAF_prolif |
| Utp20 | 5.84E-89 | 0.44475362 | 0.506 | 0.337 | 1.13E-84 | 2 | Utp20 | myCAF_prolif |
| Rnf40 | 9.70E-89 | 0.2821105 | 0.68 | 0.548 | 1.88E-84 | 2 | Rnf40 | myCAF_prolif |
| Bag4 | 1.10E-88 | 0.38780152 | 0.488 | 0.319 | 2.13E-84 | 2 | Bag4 | myCAF_prolif |
| Prpf3 | 1.11E-88 | 0.50378463 | 0.51 | 0.323 | 2.14E-84 | 2 | Prpf3 | myCAF_prolif |
| Dbr1 | 1.30E-88 | 0.44200446 | 0.328 | 0.178 | 2.52E-84 | 2 | Dbr1 | myCAF_prolif |
| Lrfn41 | 1.67E-88 | 0.35184737 | 0.604 | 0.423 | 3.23E-84 | 2 | Lrfn4 | myCAF_prolif |
| Uevld | 2.37E-88 | 0.72834316 | 0.291 | 0.144 | 4.57E-84 | 2 | Uevld | myCAF_prolif |
| Katnb1 | 2.66E-88 | 0.46543187 | 0.441 | 0.262 | 5.14E-84 | 2 | Katnb1 | myCAF_prolif |
| Nol11 | 5.99E-88 | 0.39961505 | 0.487 | 0.33 | 1.16E-83 | 2 | Nol11 | myCAF_prolif |
| Ndufb2 | 8.69E-88 | 0.31088595 | 0.553 | 0.39 | 1.68E-83 | 2 | Ndufb2 | myCAF_prolif |
| Pigu | 3.35E-87 | 0.42998798 | 0.58 | 0.407 | 6.47E-83 | 2 | Pigu | myCAF_prolif |
| Ttl1 | 3.39E-87 | 0.65399595 | 0.367 | 0.2 | 6.54E-83 | 2 | Ttl1 | myCAF_prolif |
| Aldh16a1 | 3.55E-87 | 0.76143651 | 0.396 | 0.218 | 6.86E-83 | 2 | Aldh16a1 | myCAF_prolif |
| Vangl1 | 3.96E-87 | 0.26703231 | 0.553 | 0.449 | 7.66E-83 | 2 | Vangl1 | myCAF_prolif |
| Ppm1d | 5.26E-87 | 0.36864422 | 0.462 | 0.28 | 1.02E-82 | 2 | Ppm1d | myCAF_prolif |
| Alyref1 | 6.12E-87 | 0.25054176 | 0.425 | 0.221 | 1.18E-82 | 2 | Alyref | myCAF_prolif |
| Suco | 1.16E-86 | 0.44003796 | 0.549 | 0.391 | 2.25E-82 | 2 | Suco | myCAF_prolif |
| Cnot7 | 1.42E-86 | 0.26041463 | 0.589 | 0.442 | 2.74E-82 | 2 | Cnot7 | myCAF_prolif |
| Ptger41 | 1.59E-86 | 0.27106974 | 0.411 | 0.279 | 3.08E-82 | 2 | Ptger4 | myCAF_prolif |
| Erlin11 | 1.65E-86 | 0.45838296 | 0.425 | 0.261 | 3.18E-82 | 2 | Erlin1 | myCAF_prolif |
| Ercc1 | 1.82E-86 | 0.2507691 | 0.658 | 0.512 | 3.53E-82 | 2 | Ercc1 | myCAF_prolif |
| Hnrnph3 | 2.76E-86 | 0.47094744 | 0.596 | 0.435 | 5.33E-82 | 2 | Hnrnph3 | myCAF_prolif |
| Esf1 | 3.12E-86 | 0.48202559 | 0.477 | 0.297 | 6.03E-82 | 2 | Esf1 | myCAF_prolif |
| Abhd13 | 3.76E-86 | 0.25378342 | 0.566 | 0.408 | 7.28E-82 | 2 | Abhd13 | myCAF_prolif |
| mt-Nd2 | 8.49E-86 | 1.46035282 | 0.582 | 0.462 | 1.64E-81 | 2 | mt-Nd2 | myCAF_prolif |
| Slc7a1 | 9.99E-86 | 0.33087164 | 0.45 | 0.302 | 1.93E-81 | 2 | Slc7a1 | myCAF_prolif |
| Htatsf1 | 1.38E-85 | 0.32871111 | 0.598 | 0.441 | 2.67E-81 | 2 | Htatsf1 | myCAF_prolif |
| Nf2 | 1.40E-85 | 0.2958377 | 0.782 | 0.671 | 2.71E-81 | 2 | Nf2 | myCAF_prolif |
| Mrto4 | 1.71E-85 | 0.76199788 | 0.303 | 0.147 | 3.31E-81 | 2 | Mrto4 | myCAF_prolif |
| Bhlhb9 | 1.88E-85 | 0.41373011 | 0.444 | 0.287 | 3.64E-81 | 2 | Bhlhb9 | myCAF_prolif |
| Itpripl1 | 2.05E-85 | 1.02641797 | 0.261 | 0.111 | 3.97E-81 | 2 | Itpripl1 | myCAF_prolif |
| Usp2 | 2.25E-85 | 0.61793645 | 0.309 | 0.174 | 4.35E-81 | 2 | Usp2 | myCAF_prolif |
| Endod11 | 5.86E-85 | 0.43787018 | 0.451 | 0.266 | 1.13E-80 | 2 | Endod1 | myCAF_prolif |
| C2cd3 | 1.00E-84 | 0.86104997 | 0.346 | 0.188 | 1.93E-80 | 2 | C2cd3 | myCAF_prolif |
| Pdik1l | 1.08E-84 | 0.702033 | 0.356 | 0.174 | 2.10E-80 | 2 | Pdik1l | myCAF_prolif |
| Med6 | 1.29E-84 | 0.31026289 | 0.514 | 0.341 | 2.50E-80 | 2 | Med6 | myCAF_prolif |
| Pdk3 | 1.47E-84 | 0.2818661 | 0.588 | 0.436 | 2.84E-80 | 2 | Pdk3 | myCAF_prolif |
| Dph5 | 1.66E-84 | 0.63218651 | 0.361 | 0.19 | 3.21E-80 | 2 | Dph5 | myCAF_prolif |
| Traf31 | 2.41E-84 | 0.33667452 | 0.54 | 0.373 | 4.65E-80 | 2 | Traf3 | myCAF_prolif |
| Dzip3 | 2.99E-84 | 0.56489525 | 0.391 | 0.225 | 5.78E-80 | 2 | Dzip3 | myCAF_prolif |
| Ttll4 | 3.02E-84 | 0.46562576 | 0.431 | 0.252 | 5.85E-80 | 2 | Ttll4 | myCAF_prolif |
| Rpp25l | 6.22E-84 | 0.27905408 | 0.509 | 0.355 | 1.20E-79 | 2 | Rpp25l | myCAF_prolif |
| Nsun5 | 1.49E-83 | 0.66866973 | 0.421 | 0.243 | 2.87E-79 | 2 | Nsun5 | myCAF_prolif |
| Zbtb18 | 1.67E-83 | 0.71759109 | 0.364 | 0.194 | 3.22E-79 | 2 | Zbtb18 | myCAF_prolif |
| Atg5 | 3.36E-83 | 0.25066356 | 0.591 | 0.445 | 6.49E-79 | 2 | Atg5 | myCAF_prolif |
| Rwdd4a | 6.69E-83 | 0.35203506 | 0.565 | 0.398 | 1.29E-78 | 2 | Rwdd4a | myCAF_prolif |
| Cpsf1 | 1.73E-82 | 0.309277 | 0.551 | 0.4 | 3.35E-78 | 2 | Cpsf1 | myCAF_prolif |
| Timm17b | 3.22E-82 | 0.2727514 | 0.604 | 0.444 | 6.23E-78 | 2 | Timm17b | myCAF_prolif |
| Rbm27 | 3.44E-82 | 0.3469139 | 0.562 | 0.403 | 6.66E-78 | 2 | Rbm27 | myCAF_prolif |
| Vps37b | 5.05E-82 | 0.35425691 | 0.459 | 0.304 | 9.76E-78 | 2 | Vps37b | myCAF_prolif |
| Gcat | 5.25E-82 | 1.04065286 | 0.275 | 0.118 | 1.01E-77 | 2 | Gcat | myCAF_prolif |
| Ubiad1 | 5.26E-82 | 0.46621259 | 0.469 | 0.298 | 1.02E-77 | 2 | Ubiad1 | myCAF_prolif |
| Rcc1l | 5.77E-82 | 0.4245417 | 0.499 | 0.329 | 1.12E-77 | 2 | Rcc1l | myCAF_prolif |
| Evc | 7.20E-82 | 0.34801205 | 0.511 | 0.353 | 1.39E-77 | 2 | Evc | myCAF_prolif |
| Ccdc86 | 1.29E-81 | 0.65506746 | 0.484 | 0.326 | 2.49E-77 | 2 | Ccdc86 | myCAF_prolif |
| Fam204a | 1.93E-81 | 0.4161562 | 0.542 | 0.37 | 3.73E-77 | 2 | Fam204a | myCAF_prolif |
| Pym1 | 2.02E-81 | 0.4741287 | 0.448 | 0.28 | 3.90E-77 | 2 | Pym1 | myCAF_prolif |
| Fgfr1op | 2.42E-81 | 0.57230182 | 0.428 | 0.26 | 4.69E-77 | 2 | Fgfr1op | myCAF_prolif |
| Eva1a1 | 3.46E-81 | 0.48605667 | 0.327 | 0.205 | 6.69E-77 | 2 | Eva1a | myCAF_prolif |
| Apex2 | 5.86E-81 | 0.71439014 | 0.478 | 0.29 | 1.13E-76 | 2 | Apex2 | myCAF_prolif |
| Lcorl | 8.70E-81 | 0.58405223 | 0.306 | 0.154 | 1.68E-76 | 2 | Lcorl | myCAF_prolif |
| Mlst8 | 1.32E-80 | 0.48238299 | 0.414 | 0.245 | 2.55E-76 | 2 | Mlst8 | myCAF_prolif |
| Tars2 | 2.74E-80 | 0.77896222 | 0.337 | 0.175 | 5.30E-76 | 2 | Tars2 | myCAF_prolif |
| Cep85 | 3.54E-80 | 0.61599515 | 0.455 | 0.29 | 6.84E-76 | 2 | Cep85 | myCAF_prolif |
| Slc10a7 | 3.77E-80 | 0.29048904 | 0.576 | 0.444 | 7.29E-76 | 2 | Slc10a7 | myCAF_prolif |
| Dimt1 | 4.27E-80 | 0.93676849 | 0.402 | 0.216 | 8.26E-76 | 2 | Dimt1 | myCAF_prolif |
| Cox10 | 1.66E-79 | 0.64711978 | 0.425 | 0.247 | 3.20E-75 | 2 | Cox10 | myCAF_prolif |
| Nufip1 | 1.90E-79 | 0.41816782 | 0.501 | 0.322 | 3.67E-75 | 2 | Nufip1 | myCAF_prolif |
| Mad2l1bp | 2.34E-79 | 0.50347903 | 0.41 | 0.25 | 4.53E-75 | 2 | Mad2l1bp | myCAF_prolif |
| Nudt1 | 2.35E-79 | 1.08737783 | 0.267 | 0.114 | 4.54E-75 | 2 | Nudt1 | myCAF_prolif |
| Nol12 | 3.44E-79 | 0.49141181 | 0.588 | 0.415 | 6.65E-75 | 2 | Nol12 | myCAF_prolif |
| Ipo7 | 7.39E-79 | 0.43574321 | 0.365 | 0.204 | 1.43E-74 | 2 | Ipo7 | myCAF_prolif |
| Enpp3 | 1.04E-78 | 0.81061852 | 0.294 | 0.153 | 2.02E-74 | 2 | Enpp3 | myCAF_prolif |
| Ncoa5 | 1.06E-78 | 0.37242508 | 0.594 | 0.452 | 2.04E-74 | 2 | Ncoa5 | myCAF_prolif |
| Nae1 | 1.17E-78 | 0.49662528 | 0.521 | 0.361 | 2.27E-74 | 2 | Nae1 | myCAF_prolif |
| Clasrp | 1.37E-78 | 0.60603747 | 0.62 | 0.466 | 2.65E-74 | 2 | Clasrp | myCAF_prolif |
| Shkbp1 | 1.53E-78 | 0.49309208 | 0.481 | 0.314 | 2.96E-74 | 2 | Shkbp1 | myCAF_prolif |
| Las1l | 2.12E-78 | 0.30910681 | 0.545 | 0.386 | 4.10E-74 | 2 | Las1l | myCAF_prolif |
| Scaf4 | 3.64E-78 | 0.26507091 | 0.586 | 0.445 | 7.04E-74 | 2 | Scaf4 | myCAF_prolif |
| Cklf | 4.29E-78 | 0.81115083 | 0.287 | 0.131 | 8.29E-74 | 2 | Cklf | myCAF_prolif |
| Tia1 | 4.52E-78 | 0.40377775 | 0.573 | 0.416 | 8.74E-74 | 2 | Tia1 | myCAF_prolif |
| Tmem186 | 4.99E-78 | 0.48338263 | 0.453 | 0.267 | 9.66E-74 | 2 | Tmem186 | myCAF_prolif |
| Zfp280b | 5.20E-78 | 0.46235359 | 0.451 | 0.274 | 1.00E-73 | 2 | Zfp280b | myCAF_prolif |
| Nln | 5.71E-78 | 0.32325147 | 0.503 | 0.359 | 1.10E-73 | 2 | Nln | myCAF_prolif |
| Kti12 | 6.79E-78 | 0.37017082 | 0.509 | 0.341 | 1.31E-73 | 2 | Kti12 | myCAF_prolif |
| Dhodh | 7.43E-78 | 0.44644097 | 0.391 | 0.227 | 1.44E-73 | 2 | Dhodh | myCAF_prolif |
| Pdhx | 8.57E-78 | 0.47068133 | 0.443 | 0.273 | 1.66E-73 | 2 | Pdhx | myCAF_prolif |
| Tysnd1 | 9.89E-78 | 0.27751253 | 0.569 | 0.399 | 1.91E-73 | 2 | Tysnd1 | myCAF_prolif |
| Depdc5 | 1.54E-77 | 0.45852081 | 0.456 | 0.294 | 2.98E-73 | 2 | Depdc5 | myCAF_prolif |
| Mex3a1 | 1.62E-77 | 0.51511816 | 0.38 | 0.227 | 3.14E-73 | 2 | Mex3a | myCAF_prolif |
| Dnm2 | 2.34E-77 | 0.2663707 | 0.787 | 0.692 | 4.52E-73 | 2 | Dnm2 | myCAF_prolif |
| Pycrl | 2.42E-77 | 0.52490051 | 0.431 | 0.255 | 4.68E-73 | 2 | Pycrl | myCAF_prolif |
| Top3a | 3.62E-77 | 0.80705342 | 0.289 | 0.131 | 6.99E-73 | 2 | Top3a | myCAF_prolif |
| Knop1 | 4.20E-77 | 0.66663309 | 0.456 | 0.29 | 8.12E-73 | 2 | Knop1 | myCAF_prolif |
| Alms1 | 4.66E-77 | 0.64880428 | 0.301 | 0.144 | 9.01E-73 | 2 | Alms1 | myCAF_prolif |
| Smarcd1 | 4.78E-77 | 0.33610757 | 0.529 | 0.384 | 9.25E-73 | 2 | Smarcd1 | myCAF_prolif |
| Fam49b | 5.31E-77 | 0.2746398 | 0.465 | 0.333 | 1.03E-72 | 2 | Fam49b | myCAF_prolif |
| Gtpbp10 | 7.96E-77 | 0.50444962 | 0.394 | 0.227 | 1.54E-72 | 2 | Gtpbp10 | myCAF_prolif |
| Rbpms21 | 1.04E-76 | 0.27819081 | 0.504 | 0.381 | 2.02E-72 | 2 | Rbpms2 | myCAF_prolif |
| Rpp21 | 1.81E-76 | 0.41779712 | 0.562 | 0.406 | 3.50E-72 | 2 | Rpp21 | myCAF_prolif |
| Cars2 | 2.08E-76 | 0.51017096 | 0.443 | 0.272 | 4.02E-72 | 2 | Cars2 | myCAF_prolif |
| Eif2s3x | 3.24E-76 | 0.36308319 | 0.454 | 0.314 | 6.26E-72 | 2 | Eif2s3x | myCAF_prolif |
| Ncbp3 | 3.32E-76 | 0.27366966 | 0.589 | 0.454 | 6.43E-72 | 2 | Ncbp3 | myCAF_prolif |
| Ttc4 | 5.93E-76 | 0.50582647 | 0.517 | 0.366 | 1.15E-71 | 2 | Ttc4 | myCAF_prolif |
| Fam107b | 6.97E-76 | 0.4742267 | 0.445 | 0.284 | 1.35E-71 | 2 | Fam107b | myCAF_prolif |
| Nepro | 8.06E-76 | 0.38648515 | 0.446 | 0.272 | 1.56E-71 | 2 | Nepro | myCAF_prolif |
| Tsfm | 1.06E-75 | 0.36502481 | 0.556 | 0.407 | 2.04E-71 | 2 | Tsfm | myCAF_prolif |
| Ints12 | 1.09E-75 | 0.56096033 | 0.43 | 0.274 | 2.11E-71 | 2 | Ints12 | myCAF_prolif |
| Msh3 | 1.80E-75 | 0.66857785 | 0.343 | 0.18 | 3.48E-71 | 2 | Msh3 | myCAF_prolif |
| Ino80c | 2.12E-75 | 0.27919085 | 0.521 | 0.385 | 4.11E-71 | 2 | Ino80c | myCAF_prolif |
| Tiam11 | 2.66E-75 | 0.27118976 | 0.45 | 0.267 | 5.14E-71 | 2 | Tiam1 | myCAF_prolif |
| Wdr43 | 3.13E-75 | 0.47870885 | 0.517 | 0.37 | 6.06E-71 | 2 | Wdr43 | myCAF_prolif |
| Cfap298 | 4.88E-75 | 0.47645314 | 0.377 | 0.219 | 9.44E-71 | 2 | Cfap298 | myCAF_prolif |
| Cep78 | 5.27E-75 | 0.90324049 | 0.325 | 0.169 | 1.02E-70 | 2 | Cep78 | myCAF_prolif |
| Atr | 5.49E-75 | 0.50253982 | 0.308 | 0.15 | 1.06E-70 | 2 | Atr | myCAF_prolif |
| Mtx1 | 7.63E-75 | 0.41821667 | 0.461 | 0.288 | 1.47E-70 | 2 | Mtx1 | myCAF_prolif |
| Ndufaf4 | 7.96E-75 | 0.42678075 | 0.392 | 0.233 | 1.54E-70 | 2 | Ndufaf4 | myCAF_prolif |
| Rad1 | 8.16E-75 | 0.53289834 | 0.336 | 0.181 | 1.58E-70 | 2 | Rad1 | myCAF_prolif |
| Eps15 | 9.57E-75 | 0.27786831 | 0.684 | 0.572 | 1.85E-70 | 2 | Eps15 | myCAF_prolif |
| Fpgs1 | 9.98E-75 | 0.25107551 | 0.569 | 0.435 | 1.93E-70 | 2 | Fpgs | myCAF_prolif |
| Chmp6 | 1.25E-74 | 0.28726969 | 0.602 | 0.459 | 2.41E-70 | 2 | Chmp6 | myCAF_prolif |
| Cox18 | 4.55E-74 | 0.52319399 | 0.398 | 0.223 | 8.79E-70 | 2 | Cox18 | myCAF_prolif |
| Timm9 | 4.58E-74 | 0.70835779 | 0.347 | 0.216 | 8.85E-70 | 2 | Timm9 | myCAF_prolif |
| Psph | 4.66E-74 | 0.58075612 | 0.396 | 0.243 | 9.01E-70 | 2 | Psph | myCAF_prolif |
| Lsg1 | 4.80E-74 | 0.2597427 | 0.515 | 0.345 | 9.29E-70 | 2 | Lsg1 | myCAF_prolif |
| Nktr | 5.42E-74 | 0.29606593 | 0.809 | 0.736 | 1.05E-69 | 2 | Nktr | myCAF_prolif |
| Fbxo171 | 5.43E-74 | 0.31116435 | 0.54 | 0.398 | 1.05E-69 | 2 | Fbxo17 | myCAF_prolif |
| Ucp2 | 6.18E-74 | 0.71713619 | 0.282 | 0.173 | 1.19E-69 | 2 | Ucp2 | myCAF_prolif |
| Mipep | 7.74E-74 | 0.54012094 | 0.403 | 0.238 | 1.50E-69 | 2 | Mipep | myCAF_prolif |
| Bms1 | 7.80E-74 | 0.37466193 | 0.46 | 0.296 | 1.51E-69 | 2 | Bms1 | myCAF_prolif |
| Polr2i | 8.45E-74 | 0.47665636 | 0.498 | 0.34 | 1.63E-69 | 2 | Polr2i | myCAF_prolif |
| Pde12 | 1.08E-73 | 0.47755167 | 0.391 | 0.229 | 2.09E-69 | 2 | Pde12 | myCAF_prolif |
| Hira | 2.75E-73 | 0.26836179 | 0.543 | 0.378 | 5.31E-69 | 2 | Hira | myCAF_prolif |
| Cab39l | 4.01E-73 | 0.43228668 | 0.375 | 0.222 | 7.75E-69 | 2 | Cab39l | myCAF_prolif |
| Bola1 | 4.83E-73 | 0.31133817 | 0.454 | 0.31 | 9.33E-69 | 2 | Bola1 | myCAF_prolif |
| Nrg11 | 8.32E-73 | 0.36805986 | 0.466 | 0.343 | 1.61E-68 | 2 | Nrg1 | myCAF_prolif |
| Nup214 | 1.74E-72 | 0.25433002 | 0.533 | 0.373 | 3.37E-68 | 2 | Nup214 | myCAF_prolif |
| Rnaseh1 | 1.76E-72 | 0.25092301 | 0.48 | 0.328 | 3.40E-68 | 2 | Rnaseh1 | myCAF_prolif |
| Yars2 | 3.18E-72 | 0.45394209 | 0.4 | 0.237 | 6.14E-68 | 2 | Yars2 | myCAF_prolif |
| Gmcl1 | 3.38E-72 | 0.26728354 | 0.518 | 0.383 | 6.54E-68 | 2 | Gmcl1 | myCAF_prolif |
| Parl | 4.46E-72 | 0.34585711 | 0.505 | 0.343 | 8.62E-68 | 2 | Parl | myCAF_prolif |
| Foxred1 | 4.55E-72 | 0.44706455 | 0.509 | 0.348 | 8.79E-68 | 2 | Foxred1 | myCAF_prolif |
| Lrrcc1 | 4.67E-72 | 0.92124972 | 0.378 | 0.218 | 9.03E-68 | 2 | Lrrcc1 | myCAF_prolif |
| Pomt1 | 5.26E-72 | 0.78405345 | 0.35 | 0.19 | 1.02E-67 | 2 | Pomt1 | myCAF_prolif |
| Med24 | 5.46E-72 | 0.26606551 | 0.552 | 0.395 | 1.06E-67 | 2 | Med24 | myCAF_prolif |
| Uxt | 5.67E-72 | 0.36576967 | 0.492 | 0.335 | 1.10E-67 | 2 | Uxt | myCAF_prolif |
| Utp14a | 5.83E-72 | 0.44562129 | 0.422 | 0.263 | 1.13E-67 | 2 | Utp14a | myCAF_prolif |
| Gatc | 6.12E-72 | 0.59711731 | 0.385 | 0.21 | 1.18E-67 | 2 | Gatc | myCAF_prolif |
| Tctn3 | 6.24E-72 | 0.36718761 | 0.433 | 0.276 | 1.21E-67 | 2 | Tctn3 | myCAF_prolif |
| Hspbp1 | 1.07E-71 | 0.32106178 | 0.516 | 0.362 | 2.08E-67 | 2 | Hspbp1 | myCAF_prolif |
| Eif2b3 | 1.44E-71 | 0.38582817 | 0.425 | 0.273 | 2.79E-67 | 2 | Eif2b3 | myCAF_prolif |
| B3galt6 | 2.29E-71 | 0.35108739 | 0.459 | 0.299 | 4.43E-67 | 2 | B3galt6 | myCAF_prolif |
| Spata24 | 7.50E-71 | 0.88224743 | 0.263 | 0.118 | 1.45E-66 | 2 | Spata24 | myCAF_prolif |
| Armc6 | 2.32E-70 | 0.43765478 | 0.424 | 0.256 | 4.49E-66 | 2 | Armc6 | myCAF_prolif |
| Zfand2a | 2.74E-70 | 0.45015948 | 0.408 | 0.246 | 5.30E-66 | 2 | Zfand2a | myCAF_prolif |
| Snx25 | 2.97E-70 | 0.28410478 | 0.535 | 0.405 | 5.74E-66 | 2 | Snx25 | myCAF_prolif |
| Krr1 | 7.53E-70 | 0.28573487 | 0.459 | 0.298 | 1.46E-65 | 2 | Krr1 | myCAF_prolif |
| Fam149a | 8.95E-70 | 0.27736939 | 0.606 | 0.467 | 1.73E-65 | 2 | Fam149a | myCAF_prolif |
| Paqr4 | 1.15E-69 | 0.58175804 | 0.295 | 0.167 | 2.22E-65 | 2 | Paqr4 | myCAF_prolif |
| Ccdc167 | 1.55E-69 | 0.52763868 | 0.356 | 0.19 | 3.00E-65 | 2 | Ccdc167 | myCAF_prolif |
| Cactin | 2.46E-69 | 0.61004081 | 0.415 | 0.242 | 4.75E-65 | 2 | Cactin | myCAF_prolif |
| Aatf | 2.53E-69 | 0.41825578 | 0.458 | 0.309 | 4.89E-65 | 2 | Aatf | myCAF_prolif |
| Slx4 | 2.76E-69 | 0.53223406 | 0.405 | 0.242 | 5.34E-65 | 2 | Slx4 | myCAF_prolif |
| Tsen2 | 3.13E-69 | 0.66015676 | 0.312 | 0.17 | 6.05E-65 | 2 | Tsen2 | myCAF_prolif |
| Rrp9 | 3.86E-69 | 0.60410218 | 0.332 | 0.185 | 7.47E-65 | 2 | Rrp9 | myCAF_prolif |
| Snrnp48 | 3.94E-69 | 0.42692534 | 0.575 | 0.412 | 7.61E-65 | 2 | Snrnp48 | myCAF_prolif |
| Prps2 | 4.79E-69 | 0.31387002 | 0.368 | 0.217 | 9.26E-65 | 2 | Prps2 | myCAF_prolif |
| Cops7b | 4.95E-69 | 0.38700994 | 0.454 | 0.3 | 9.57E-65 | 2 | Cops7b | myCAF_prolif |
| Frk | 6.50E-69 | 0.33314801 | 0.494 | 0.354 | 1.26E-64 | 2 | Frk | myCAF_prolif |
| Wdr61 | 7.28E-69 | 0.27699893 | 0.621 | 0.477 | 1.41E-64 | 2 | Wdr61 | myCAF_prolif |
| Mpi | 7.30E-69 | 0.3148476 | 0.456 | 0.307 | 1.41E-64 | 2 | Mpi | myCAF_prolif |
| Slc25a32 | 8.10E-69 | 0.26110451 | 0.432 | 0.297 | 1.57E-64 | 2 | Slc25a32 | myCAF_prolif |
| Zfp180 | 1.05E-68 | 0.46042743 | 0.439 | 0.281 | 2.04E-64 | 2 | Zfp180 | myCAF_prolif |
| Mlkl | 1.17E-68 | 0.28537516 | 0.491 | 0.35 | 2.26E-64 | 2 | Mlkl | myCAF_prolif |
| Cep164 | 1.40E-68 | 0.47870405 | 0.454 | 0.284 | 2.70E-64 | 2 | Cep164 | myCAF_prolif |
| Pan2 | 5.65E-68 | 0.44636033 | 0.445 | 0.29 | 1.09E-63 | 2 | Pan2 | myCAF_prolif |
| Rangrf | 7.19E-68 | 0.35437692 | 0.441 | 0.298 | 1.39E-63 | 2 | Rangrf | myCAF_prolif |
| Gemin5 | 1.20E-67 | 0.26089382 | 0.442 | 0.299 | 2.32E-63 | 2 | Gemin5 | myCAF_prolif |
| Kri1 | 1.40E-67 | 0.48202428 | 0.541 | 0.403 | 2.70E-63 | 2 | Kri1 | myCAF_prolif |
| Dcaf1 | 2.44E-67 | 0.28059457 | 0.437 | 0.299 | 4.71E-63 | 2 | Dcaf1 | myCAF_prolif |
| Sox12 | 2.55E-67 | 0.45453376 | 0.372 | 0.259 | 4.92E-63 | 2 | Sox12 | myCAF_prolif |
| Wdr12 | 2.64E-67 | 0.7457217 | 0.27 | 0.128 | 5.11E-63 | 2 | Wdr12 | myCAF_prolif |
| Tada2a | 3.58E-67 | 0.41936467 | 0.446 | 0.284 | 6.93E-63 | 2 | Tada2a | myCAF_prolif |
| Hsf1 | 4.01E-67 | 0.28959921 | 0.563 | 0.438 | 7.75E-63 | 2 | Hsf1 | myCAF_prolif |
| Taf15 | 4.40E-67 | 0.68539514 | 0.341 | 0.193 | 8.50E-63 | 2 | Taf15 | myCAF_prolif |
| Polr2c | 4.72E-67 | 0.49733628 | 0.403 | 0.234 | 9.12E-63 | 2 | Polr2c | myCAF_prolif |
| Rpf21 | 5.45E-67 | 0.57444442 | 0.344 | 0.181 | 1.05E-62 | 2 | Rpf2 | myCAF_prolif |
| Sertad41 | 6.30E-67 | 0.3165504 | 0.38 | 0.25 | 1.22E-62 | 2 | Sertad4 | myCAF_prolif |
| Dus1l | 6.73E-67 | 0.42476467 | 0.493 | 0.33 | 1.30E-62 | 2 | Dus1l | myCAF_prolif |
| Ppid | 7.80E-67 | 0.40019535 | 0.402 | 0.257 | 1.51E-62 | 2 | Ppid | myCAF_prolif |
| Rmi1 | 8.55E-67 | 0.5035079 | 0.332 | 0.18 | 1.65E-62 | 2 | Rmi1 | myCAF_prolif |
| Nip7 | 1.06E-66 | 0.30535956 | 0.497 | 0.358 | 2.06E-62 | 2 | Nip7 | myCAF_prolif |
| Mphosph8 | 1.44E-66 | 0.34812395 | 0.434 | 0.287 | 2.79E-62 | 2 | Mphosph8 | myCAF_prolif |
| Med11 | 2.36E-66 | 0.27241923 | 0.505 | 0.349 | 4.57E-62 | 2 | Med11 | myCAF_prolif |
| Msantd2 | 2.46E-66 | 0.35645665 | 0.442 | 0.305 | 4.76E-62 | 2 | Msantd2 | myCAF_prolif |
| Rint1 | 5.88E-66 | 0.36596295 | 0.518 | 0.354 | 1.14E-61 | 2 | Rint1 | myCAF_prolif |
| Ptpdc1 | 6.25E-66 | 0.3288368 | 0.378 | 0.244 | 1.21E-61 | 2 | Ptpdc1 | myCAF_prolif |
| Samd1 | 7.32E-66 | 0.44425976 | 0.382 | 0.236 | 1.41E-61 | 2 | Samd1 | myCAF_prolif |
| Piga | 9.05E-66 | 0.46058782 | 0.309 | 0.148 | 1.75E-61 | 2 | Piga | myCAF_prolif |
| Lyn1 | 9.14E-66 | 0.33185776 | 0.447 | 0.324 | 1.77E-61 | 2 | Lyn | myCAF_prolif |
| Zfp213 | 1.35E-65 | 0.67155326 | 0.287 | 0.147 | 2.62E-61 | 2 | Zfp213 | myCAF_prolif |
| Cc2d2a | 2.18E-65 | 0.59929951 | 0.384 | 0.239 | 4.22E-61 | 2 | Cc2d2a | myCAF_prolif |
| Rabepk | 2.37E-65 | 0.3376616 | 0.403 | 0.259 | 4.59E-61 | 2 | Rabepk | myCAF_prolif |
| Paxbp1 | 2.47E-65 | 0.3933581 | 0.613 | 0.477 | 4.78E-61 | 2 | Paxbp1 | myCAF_prolif |
| E2f5 | 2.50E-65 | 0.55597336 | 0.361 | 0.221 | 4.83E-61 | 2 | E2f5 | myCAF_prolif |
| Med22 | 3.78E-65 | 0.25595228 | 0.624 | 0.48 | 7.31E-61 | 2 | Med22 | myCAF_prolif |
| Tut4 | 4.08E-65 | 0.35349662 | 0.528 | 0.365 | 7.88E-61 | 2 | Tut4 | myCAF_prolif |
| Phf20l1 | 4.21E-65 | 0.31268655 | 0.659 | 0.522 | 8.14E-61 | 2 | Phf20l1 | myCAF_prolif |
| Ccdc22 | 4.84E-65 | 0.3688931 | 0.42 | 0.258 | 9.35E-61 | 2 | Ccdc22 | myCAF_prolif |
| Magohb | 2.01E-64 | 0.64134749 | 0.311 | 0.162 | 3.88E-60 | 2 | Magohb | myCAF_prolif |
| Exoc1 | 2.76E-64 | 0.30980611 | 0.591 | 0.438 | 5.33E-60 | 2 | Exoc1 | myCAF_prolif |
| Ccdc134 | 2.85E-64 | 0.3358622 | 0.42 | 0.283 | 5.51E-60 | 2 | Ccdc134 | myCAF_prolif |
| Crnkl1 | 3.55E-64 | 0.2551938 | 0.541 | 0.395 | 6.87E-60 | 2 | Crnkl1 | myCAF_prolif |
| Gkap1 | 3.66E-64 | 0.26097471 | 0.506 | 0.378 | 7.08E-60 | 2 | Gkap1 | myCAF_prolif |
| Psmg3 | 4.33E-64 | 0.30060348 | 0.484 | 0.321 | 8.37E-60 | 2 | Psmg3 | myCAF_prolif |
| Dnajc11 | 5.45E-64 | 0.26355282 | 0.485 | 0.332 | 1.05E-59 | 2 | Dnajc11 | myCAF_prolif |
| Qser1 | 5.69E-64 | 0.34835933 | 0.45 | 0.289 | 1.10E-59 | 2 | Qser1 | myCAF_prolif |
| Chd1l | 6.58E-64 | 0.52856445 | 0.285 | 0.146 | 1.27E-59 | 2 | Chd1l | myCAF_prolif |
| Ptpn21 | 8.27E-64 | 0.27349448 | 0.568 | 0.418 | 1.60E-59 | 2 | Ptpn21 | myCAF_prolif |
| Agfg2 | 1.47E-63 | 0.37854553 | 0.344 | 0.204 | 2.83E-59 | 2 | Agfg2 | myCAF_prolif |
| Ccdc61 | 1.64E-63 | 0.74289055 | 0.339 | 0.191 | 3.17E-59 | 2 | Ccdc61 | myCAF_prolif |
| Rad17 | 1.89E-63 | 0.40087772 | 0.445 | 0.294 | 3.65E-59 | 2 | Rad17 | myCAF_prolif |
| Ogg1 | 2.17E-63 | 0.8158209 | 0.322 | 0.167 | 4.20E-59 | 2 | Ogg1 | myCAF_prolif |
| Trappc10 | 2.44E-63 | 0.27326517 | 0.53 | 0.385 | 4.71E-59 | 2 | Trappc10 | myCAF_prolif |
| Trnt1 | 2.65E-63 | 0.41160518 | 0.478 | 0.349 | 5.13E-59 | 2 | Trnt1 | myCAF_prolif |
| Bambi1 | 3.36E-63 | 0.42414471 | 0.309 | 0.175 | 6.51E-59 | 2 | Bambi | myCAF_prolif |
| Tmem158 | 4.30E-63 | 0.3305033 | 0.381 | 0.223 | 8.31E-59 | 2 | Tmem158 | myCAF_prolif |
| Timm21 | 4.36E-63 | 0.49336756 | 0.412 | 0.257 | 8.43E-59 | 2 | Timm21 | myCAF_prolif |
| Ercc6l2 | 4.48E-63 | 0.29268379 | 0.41 | 0.253 | 8.66E-59 | 2 | Ercc6l2 | myCAF_prolif |
| Nif3l1 | 4.72E-63 | 0.45269379 | 0.366 | 0.21 | 9.12E-59 | 2 | Nif3l1 | myCAF_prolif |
| Mecr | 6.23E-63 | 0.34646566 | 0.424 | 0.276 | 1.20E-58 | 2 | Mecr | myCAF_prolif |
| Clcf11 | 1.14E-62 | 0.62284551 | 0.338 | 0.225 | 2.19E-58 | 2 | Clcf1 | myCAF_prolif |
| Timmdc1 | 1.27E-62 | 0.30091423 | 0.542 | 0.403 | 2.45E-58 | 2 | Timmdc1 | myCAF_prolif |
| Arrb2 | 1.36E-62 | 0.67980561 | 0.343 | 0.191 | 2.63E-58 | 2 | Arrb2 | myCAF_prolif |
| Nom1 | 2.34E-62 | 0.26721437 | 0.525 | 0.367 | 4.52E-58 | 2 | Nom1 | myCAF_prolif |
| Wnt5b | 2.93E-62 | 0.61420347 | 0.25 | 0.153 | 5.67E-58 | 2 | Wnt5b | myCAF_prolif |
| Tmem237 | 2.97E-62 | 0.46498919 | 0.355 | 0.206 | 5.73E-58 | 2 | Tmem237 | myCAF_prolif |
| Rpusd1 | 4.24E-62 | 0.60350011 | 0.318 | 0.178 | 8.20E-58 | 2 | Rpusd1 | myCAF_prolif |
| St3gal5 | 6.02E-62 | 0.31501757 | 0.292 | 0.168 | 1.16E-57 | 2 | St3gal5 | myCAF_prolif |
| Ehmt1 | 9.67E-62 | 0.28183016 | 0.706 | 0.588 | 1.87E-57 | 2 | Ehmt1 | myCAF_prolif |
| Elfn1 | 1.22E-61 | 0.32746743 | 0.318 | 0.205 | 2.36E-57 | 2 | Elfn1 | myCAF_prolif |
| Tbp | 1.50E-61 | 0.31973123 | 0.444 | 0.3 | 2.91E-57 | 2 | Tbp | myCAF_prolif |
| Kdm2b | 1.74E-61 | 0.36081299 | 0.42 | 0.259 | 3.37E-57 | 2 | Kdm2b | myCAF_prolif |
| Rbm19 | 4.75E-61 | 0.36805585 | 0.453 | 0.326 | 9.18E-57 | 2 | Rbm19 | myCAF_prolif |
| Cav2 | 4.99E-61 | 0.29140765 | 0.385 | 0.263 | 9.65E-57 | 2 | Cav2 | myCAF_prolif |
| DEPRECATED-l | 6.04E-61 | 0.29997112 | 0.438 | 0.285 | 1.17E-56 | 2 | DEPRECATED-l | myCAF_prolif |
| Pus7 | 6.72E-61 | 0.32414831 | 0.378 | 0.243 | 1.30E-56 | 2 | Pus7 | myCAF_prolif |
| Cx3cl1 | 6.86E-61 | 0.39291816 | 0.265 | 0.167 | 1.33E-56 | 2 | Cx3cl1 | myCAF_prolif |
| Ankrd49 | 8.92E-61 | 0.77010852 | 0.32 | 0.171 | 1.73E-56 | 2 | Ankrd49 | myCAF_prolif |
| mt-Nd4 | 1.10E-60 | 1.04789988 | 0.659 | 0.577 | 2.12E-56 | 2 | mt-Nd4 | myCAF_prolif |
| Eef1akmt2 | 1.63E-60 | 0.27312669 | 0.628 | 0.486 | 3.16E-56 | 2 | Eef1akmt2 | myCAF_prolif |
| Vti1a | 1.68E-60 | 0.4116203 | 0.629 | 0.514 | 3.25E-56 | 2 | Vti1a | myCAF_prolif |
| Cwf19l1 | 1.71E-60 | 0.56940099 | 0.281 | 0.137 | 3.31E-56 | 2 | Cwf19l1 | myCAF_prolif |
| Prkrip1 | 3.90E-60 | 0.61352783 | 0.392 | 0.236 | 7.55E-56 | 2 | Prkrip1 | myCAF_prolif |
| Trappc12 | 4.43E-60 | 0.34444256 | 0.546 | 0.42 | 8.57E-56 | 2 | Trappc12 | myCAF_prolif |
| Vac14 | 5.24E-60 | 0.29738259 | 0.476 | 0.332 | 1.01E-55 | 2 | Vac14 | myCAF_prolif |
| Rnf219 | 6.41E-60 | 0.56256705 | 0.287 | 0.149 | 1.24E-55 | 2 | Rnf219 | myCAF_prolif |
| Rngtt | 7.01E-60 | 0.5448743 | 0.363 | 0.207 | 1.35E-55 | 2 | Rngtt | myCAF_prolif |
| Thoc1 | 1.01E-59 | 0.31374636 | 0.489 | 0.344 | 1.95E-55 | 2 | Thoc1 | myCAF_prolif |
| C2cd2 | 1.29E-59 | 0.554888 | 0.32 | 0.171 | 2.49E-55 | 2 | C2cd2 | myCAF_prolif |
| Atpsckmt | 1.75E-59 | 0.49174245 | 0.327 | 0.178 | 3.38E-55 | 2 | Atpsckmt | myCAF_prolif |
| Med19 | 1.88E-59 | 0.48456637 | 0.321 | 0.168 | 3.63E-55 | 2 | Med19 | myCAF_prolif |
| Pnpt1 | 2.60E-59 | 0.27525601 | 0.423 | 0.279 | 5.02E-55 | 2 | Pnpt1 | myCAF_prolif |
| Dmac1 | 3.07E-59 | 0.27832103 | 0.52 | 0.376 | 5.94E-55 | 2 | Dmac1 | myCAF_prolif |
| Coq3 | 3.12E-59 | 0.38226741 | 0.347 | 0.196 | 6.03E-55 | 2 | Coq3 | myCAF_prolif |
| Taf3 | 3.14E-59 | 0.39274161 | 0.373 | 0.225 | 6.06E-55 | 2 | Taf3 | myCAF_prolif |
| Cdk12 | 3.59E-59 | 0.26985969 | 0.622 | 0.518 | 6.95E-55 | 2 | Cdk12 | myCAF_prolif |
| Miga2 | 3.78E-59 | 0.35498236 | 0.442 | 0.282 | 7.30E-55 | 2 | Miga2 | myCAF_prolif |
| Ciz1 | 7.29E-59 | 0.31336598 | 0.663 | 0.536 | 1.41E-54 | 2 | Ciz1 | myCAF_prolif |
| Pwwp2a | 8.42E-59 | 0.28342609 | 0.597 | 0.464 | 1.63E-54 | 2 | Pwwp2a | myCAF_prolif |
| Cryl1 | 9.47E-59 | 0.44569469 | 0.338 | 0.195 | 1.83E-54 | 2 | Cryl1 | myCAF_prolif |
| Ints2 | 9.62E-59 | 0.4777159 | 0.365 | 0.221 | 1.86E-54 | 2 | Ints2 | myCAF_prolif |
| Eif2ak3 | 1.28E-58 | 0.25374228 | 0.618 | 0.484 | 2.48E-54 | 2 | Eif2ak3 | myCAF_prolif |
| Fam149b | 2.37E-58 | 0.25475613 | 0.556 | 0.432 | 4.58E-54 | 2 | Fam149b | myCAF_prolif |
| Naa16 | 2.45E-58 | 0.45699653 | 0.416 | 0.26 | 4.73E-54 | 2 | Naa16 | myCAF_prolif |
| Cdyl | 3.96E-58 | 0.2607281 | 0.63 | 0.512 | 7.66E-54 | 2 | Cdyl | myCAF_prolif |
| Cwc27 | 7.44E-58 | 0.59095866 | 0.387 | 0.254 | 1.44E-53 | 2 | Cwc27 | myCAF_prolif |
| Nudt2 | 7.71E-58 | 0.31264727 | 0.411 | 0.282 | 1.49E-53 | 2 | Nudt2 | myCAF_prolif |
| Pcbp3 | 1.32E-57 | 0.6503688 | 0.297 | 0.157 | 2.56E-53 | 2 | Pcbp3 | myCAF_prolif |
| B3glct | 1.37E-57 | 0.30681452 | 0.497 | 0.355 | 2.64E-53 | 2 | B3glct | myCAF_prolif |
| Sugp2 | 1.56E-57 | 0.29220549 | 0.539 | 0.412 | 3.01E-53 | 2 | Sugp2 | myCAF_prolif |
| Suclg1 | 2.15E-57 | 0.31045543 | 0.588 | 0.469 | 4.16E-53 | 2 | Suclg1 | myCAF_prolif |
| Fnbp1l1 | 2.16E-57 | 0.35886531 | 0.397 | 0.279 | 4.17E-53 | 2 | Fnbp1l | myCAF_prolif |
| Mmachc | 2.20E-57 | 0.5499911 | 0.333 | 0.196 | 4.26E-53 | 2 | Mmachc | myCAF_prolif |
| Rufy3 | 2.47E-57 | 0.43651422 | 0.419 | 0.28 | 4.77E-53 | 2 | Rufy3 | myCAF_prolif |
| Osbpl3 | 2.98E-57 | 0.35125051 | 0.487 | 0.358 | 5.76E-53 | 2 | Osbpl3 | myCAF_prolif |
| Dhx29 | 3.39E-57 | 0.51833368 | 0.351 | 0.208 | 6.56E-53 | 2 | Dhx29 | myCAF_prolif |
| Phka2 | 6.30E-57 | 0.59601663 | 0.33 | 0.192 | 1.22E-52 | 2 | Phka2 | myCAF_prolif |
| Nmi | 7.21E-57 | 0.32212484 | 0.298 | 0.169 | 1.39E-52 | 2 | Nmi | myCAF_prolif |
| Ciao2b | 8.92E-57 | 0.4922173 | 0.416 | 0.274 | 1.72E-52 | 2 | Ciao2b | myCAF_prolif |
| Atrip | 1.03E-56 | 0.74450482 | 0.318 | 0.173 | 1.99E-52 | 2 | Atrip | myCAF_prolif |
| Nabp1 | 1.22E-56 | 0.30199319 | 0.321 | 0.214 | 2.36E-52 | 2 | Nabp1 | myCAF_prolif |
| Wdr351 | 1.48E-56 | 0.41412656 | 0.425 | 0.285 | 2.86E-52 | 2 | Wdr35 | myCAF_prolif |
| Wdr70 | 4.33E-56 | 0.42559597 | 0.471 | 0.312 | 8.38E-52 | 2 | Wdr70 | myCAF_prolif |
| Slfn8 | 5.70E-56 | 0.40049962 | 0.275 | 0.152 | 1.10E-51 | 2 | Slfn8 | myCAF_prolif |
| Tmem39b | 8.27E-56 | 0.27915615 | 0.427 | 0.284 | 1.60E-51 | 2 | Tmem39b | myCAF_prolif |
| Heatr6 | 1.81E-55 | 0.28599723 | 0.462 | 0.327 | 3.50E-51 | 2 | Heatr6 | myCAF_prolif |
| Ssx2ip | 2.29E-55 | 0.73193481 | 0.278 | 0.144 | 4.44E-51 | 2 | Ssx2ip | myCAF_prolif |
| Zbtb14 | 2.81E-55 | 0.25307073 | 0.397 | 0.279 | 5.44E-51 | 2 | Zbtb14 | myCAF_prolif |
| Ndufaf5 | 4.89E-55 | 0.75522445 | 0.301 | 0.16 | 9.46E-51 | 2 | Ndufaf5 | myCAF_prolif |
| Twistnb | 6.44E-55 | 0.33960548 | 0.402 | 0.263 | 1.24E-50 | 2 | Twistnb | myCAF_prolif |
| Angpt1 | 1.02E-54 | 0.4427674 | 0.308 | 0.189 | 1.97E-50 | 2 | Angpt1 | myCAF_prolif |
| Srgn | 1.20E-54 | 0.27441817 | 0.408 | 0.325 | 2.33E-50 | 2 | Srgn | myCAF_prolif |
| Lrrc20 | 1.84E-54 | 0.26729167 | 0.373 | 0.236 | 3.55E-50 | 2 | Lrrc20 | myCAF_prolif |
| Psmb9 | 2.27E-54 | 0.32196651 | 0.297 | 0.195 | 4.40E-50 | 2 | Psmb9 | myCAF_prolif |
| Asap2 | 2.40E-54 | 0.39974206 | 0.383 | 0.248 | 4.64E-50 | 2 | Asap2 | myCAF_prolif |
| Brpf1 | 2.54E-54 | 0.44323 | 0.472 | 0.324 | 4.90E-50 | 2 | Brpf1 | myCAF_prolif |
| Serhl | 5.88E-54 | 0.39960951 | 0.463 | 0.333 | 1.14E-49 | 2 | Serhl | myCAF_prolif |
| Cep162 | 9.58E-54 | 0.55520202 | 0.288 | 0.16 | 1.85E-49 | 2 | Cep162 | myCAF_prolif |
| Zdhhc21 | 1.33E-53 | 0.52925243 | 0.338 | 0.198 | 2.57E-49 | 2 | Zdhhc21 | myCAF_prolif |
| Coa7 | 1.42E-53 | 0.55307179 | 0.304 | 0.166 | 2.75E-49 | 2 | Coa7 | myCAF_prolif |
| Naa35 | 1.56E-53 | 0.32542037 | 0.455 | 0.33 | 3.01E-49 | 2 | Naa35 | myCAF_prolif |
| Ncoa7 | 1.63E-53 | 0.33534783 | 0.412 | 0.273 | 3.14E-49 | 2 | Ncoa7 | myCAF_prolif |
| Abca9 | 2.73E-53 | 0.4532336 | 0.344 | 0.205 | 5.27E-49 | 2 | Abca9 | myCAF_prolif |
| Gpatch4 | 5.59E-53 | 0.71271758 | 0.305 | 0.177 | 1.08E-48 | 2 | Gpatch4 | myCAF_prolif |
| Ppil4 | 6.99E-53 | 0.44512187 | 0.503 | 0.376 | 1.35E-48 | 2 | Ppil4 | myCAF_prolif |
| Zdhhc13 | 8.32E-53 | 0.46200495 | 0.305 | 0.163 | 1.61E-48 | 2 | Zdhhc13 | myCAF_prolif |
| Cdkal1 | 9.52E-53 | 0.46223377 | 0.436 | 0.311 | 1.84E-48 | 2 | Cdkal1 | myCAF_prolif |
| Upf3b | 1.09E-52 | 0.73835771 | 0.309 | 0.171 | 2.12E-48 | 2 | Upf3b | myCAF_prolif |
| Taf6l | 1.15E-52 | 0.44221528 | 0.275 | 0.145 | 2.22E-48 | 2 | Taf6l | myCAF_prolif |
| Btf3l4 | 3.06E-52 | 0.27989391 | 0.431 | 0.291 | 5.91E-48 | 2 | Btf3l4 | myCAF_prolif |
| Nle11 | 3.62E-52 | 0.41135967 | 0.411 | 0.278 | 7.00E-48 | 2 | Nle1 | myCAF_prolif |
| Fastkd3 | 4.61E-52 | 0.55022484 | 0.267 | 0.149 | 8.91E-48 | 2 | Fastkd3 | myCAF_prolif |
| Cptp | 4.73E-52 | 0.25911775 | 0.356 | 0.218 | 9.15E-48 | 2 | Cptp | myCAF_prolif |
| Pop5 | 7.52E-52 | 0.31391975 | 0.322 | 0.192 | 1.45E-47 | 2 | Pop5 | myCAF_prolif |
| Hltf | 2.04E-51 | 0.50463253 | 0.384 | 0.248 | 3.94E-47 | 2 | Hltf | myCAF_prolif |
| Rin3 | 2.68E-51 | 0.46615093 | 0.312 | 0.188 | 5.17E-47 | 2 | Rin3 | myCAF_prolif |
| Bloc1s4 | 2.78E-51 | 0.34117047 | 0.427 | 0.274 | 5.37E-47 | 2 | Bloc1s4 | myCAF_prolif |
| Usp49 | 3.34E-51 | 0.50076621 | 0.29 | 0.177 | 6.46E-47 | 2 | Usp49 | myCAF_prolif |
| Fbf1 | 3.43E-51 | 0.48532518 | 0.516 | 0.382 | 6.63E-47 | 2 | Fbf1 | myCAF_prolif |
| Fmc1 | 3.49E-51 | 0.57286738 | 0.365 | 0.219 | 6.74E-47 | 2 | Fmc1 | myCAF_prolif |
| Eogt1 | 4.06E-51 | 0.4368777 | 0.353 | 0.225 | 7.85E-47 | 2 | Eogt | myCAF_prolif |
| Gclc | 4.23E-51 | 0.3543648 | 0.401 | 0.269 | 8.17E-47 | 2 | Gclc | myCAF_prolif |
| 6-Sep | 7.50E-51 | 0.58669276 | 0.279 | 0.148 | 1.45E-46 | 2 | 6-Sep | myCAF_prolif |
| Gtf2e1 | 8.35E-51 | 0.36288411 | 0.416 | 0.271 | 1.61E-46 | 2 | Gtf2e1 | myCAF_prolif |
| Zfp428 | 1.34E-50 | 0.53441114 | 0.259 | 0.141 | 2.59E-46 | 2 | Zfp428 | myCAF_prolif |
| Eif4ebp3 | 2.63E-50 | 0.26990901 | 0.399 | 0.269 | 5.09E-46 | 2 | Eif4ebp3 | myCAF_prolif |
| Bud13 | 3.11E-50 | 0.37867098 | 0.323 | 0.187 | 6.02E-46 | 2 | Bud13 | myCAF_prolif |
| Dnase1l1 | 3.22E-50 | 0.49617758 | 0.319 | 0.196 | 6.22E-46 | 2 | Dnase1l1 | myCAF_prolif |
| Bend3 | 3.60E-50 | 0.33640571 | 0.306 | 0.189 | 6.97E-46 | 2 | Bend3 | myCAF_prolif |
| Fam133b | 4.35E-50 | 0.50656705 | 0.382 | 0.24 | 8.41E-46 | 2 | Fam133b | myCAF_prolif |
| Gas8 | 4.96E-50 | 0.37219629 | 0.414 | 0.281 | 9.59E-46 | 2 | Gas8 | myCAF_prolif |
| Ss18l2 | 5.19E-50 | 0.26385706 | 0.463 | 0.324 | 1.00E-45 | 2 | Ss18l2 | myCAF_prolif |
| Galnt12 | 7.60E-50 | 0.54877208 | 0.253 | 0.143 | 1.47E-45 | 2 | Galnt12 | myCAF_prolif |
| Ate1 | 7.70E-50 | 0.51224562 | 0.452 | 0.332 | 1.49E-45 | 2 | Ate1 | myCAF_prolif |
| Snapc2 | 8.47E-50 | 0.25535391 | 0.427 | 0.28 | 1.64E-45 | 2 | Snapc2 | myCAF_prolif |
| Ing2 | 1.24E-49 | 0.47165423 | 0.329 | 0.204 | 2.40E-45 | 2 | Ing2 | myCAF_prolif |
| Wwox | 1.27E-49 | 0.64439155 | 0.403 | 0.254 | 2.46E-45 | 2 | Wwox | myCAF_prolif |
| Gtdc1 | 1.49E-49 | 0.30708102 | 0.286 | 0.174 | 2.87E-45 | 2 | Gtdc1 | myCAF_prolif |
| Trmt2b | 1.51E-49 | 0.3206987 | 0.33 | 0.208 | 2.92E-45 | 2 | Trmt2b | myCAF_prolif |
| Mettl5 | 1.53E-49 | 0.46942357 | 0.315 | 0.174 | 2.95E-45 | 2 | Mettl5 | myCAF_prolif |
| Map3k12 | 1.74E-49 | 0.33567819 | 0.428 | 0.294 | 3.37E-45 | 2 | Map3k12 | myCAF_prolif |
| Sars2 | 1.84E-49 | 0.73006837 | 0.305 | 0.174 | 3.56E-45 | 2 | Sars2 | myCAF_prolif |
| Eya3 | 1.94E-49 | 0.28546237 | 0.462 | 0.35 | 3.75E-45 | 2 | Eya3 | myCAF_prolif |
| Mgme1 | 2.90E-49 | 0.38502243 | 0.28 | 0.155 | 5.60E-45 | 2 | Mgme1 | myCAF_prolif |
| Supv3l1 | 3.65E-49 | 0.40126607 | 0.355 | 0.219 | 7.05E-45 | 2 | Supv3l1 | myCAF_prolif |
| Uimc1 | 4.46E-49 | 0.48926433 | 0.376 | 0.251 | 8.63E-45 | 2 | Uimc1 | myCAF_prolif |
| Umad1 | 5.84E-49 | 0.49920311 | 0.375 | 0.248 | 1.13E-44 | 2 | Umad1 | myCAF_prolif |
| Slc39a8 | 9.14E-49 | 0.43927428 | 0.274 | 0.155 | 1.77E-44 | 2 | Slc39a8 | myCAF_prolif |
| Vcpkmt | 9.99E-49 | 0.26839447 | 0.411 | 0.267 | 1.93E-44 | 2 | Vcpkmt | myCAF_prolif |
| Polr2k | 1.08E-48 | 0.38663952 | 0.363 | 0.234 | 2.08E-44 | 2 | Polr2k | myCAF_prolif |
| Orc5 | 1.16E-48 | 0.42546164 | 0.31 | 0.187 | 2.23E-44 | 2 | Orc5 | myCAF_prolif |
| Phlpp2 | 1.84E-48 | 0.26011922 | 0.361 | 0.225 | 3.55E-44 | 2 | Phlpp2 | myCAF_prolif |
| Polr1e | 2.70E-48 | 0.50040304 | 0.311 | 0.177 | 5.21E-44 | 2 | Polr1e | myCAF_prolif |
| Nadk2 | 3.11E-48 | 0.27875984 | 0.377 | 0.247 | 6.01E-44 | 2 | Nadk2 | myCAF_prolif |
| Lrig31 | 3.19E-48 | 0.35101048 | 0.368 | 0.255 | 6.17E-44 | 2 | Lrig3 | myCAF_prolif |
| Bcs1l | 4.11E-48 | 0.26671122 | 0.384 | 0.248 | 7.95E-44 | 2 | Bcs1l | myCAF_prolif |
| Maip1 | 5.61E-48 | 0.38338399 | 0.375 | 0.25 | 1.08E-43 | 2 | Maip1 | myCAF_prolif |
| B9d2 | 5.98E-48 | 0.28221189 | 0.372 | 0.244 | 1.16E-43 | 2 | B9d2 | myCAF_prolif |
| Pdcd7 | 9.89E-48 | 0.43684808 | 0.444 | 0.294 | 1.91E-43 | 2 | Pdcd7 | myCAF_prolif |
| Zc3h12c | 1.17E-47 | 0.25426381 | 0.34 | 0.218 | 2.27E-43 | 2 | Zc3h12c | myCAF_prolif |
| Il1r2 | 1.46E-47 | 0.27942061 | 0.348 | 0.26 | 2.83E-43 | 2 | Il1r2 | myCAF_prolif |
| Cspp1 | 2.23E-47 | 0.61649164 | 0.428 | 0.289 | 4.31E-43 | 2 | Cspp1 | myCAF_prolif |
| Rnf8 | 7.42E-47 | 0.36057485 | 0.356 | 0.217 | 1.43E-42 | 2 | Rnf8 | myCAF_prolif |
| Caap1 | 2.52E-46 | 0.38694767 | 0.401 | 0.267 | 4.87E-42 | 2 | Caap1 | myCAF_prolif |
| Dnmt3l | 3.89E-46 | 0.45952438 | 0.284 | 0.169 | 7.51E-42 | 2 | Dnmt3l | myCAF_prolif |
| Mapkapk5 | 4.65E-46 | 0.45147421 | 0.361 | 0.234 | 8.99E-42 | 2 | Mapkapk5 | myCAF_prolif |
| Hoxa3 | 5.66E-46 | 0.27085568 | 0.359 | 0.246 | 1.09E-41 | 2 | Hoxa3 | myCAF_prolif |
| Sgk3 | 8.78E-46 | 0.55661473 | 0.297 | 0.164 | 1.70E-41 | 2 | Sgk3 | myCAF_prolif |
| Slc4a7 | 1.06E-45 | 0.34128196 | 0.363 | 0.246 | 2.05E-41 | 2 | Slc4a7 | myCAF_prolif |
| Cdkn2a | 2.45E-45 | 0.53500961 | 0.256 | 0.16 | 4.73E-41 | 2 | Cdkn2a | myCAF_prolif |
| Kctd13 | 3.68E-45 | 0.26308042 | 0.38 | 0.257 | 7.11E-41 | 2 | Kctd13 | myCAF_prolif |
| Ggct1 | 3.99E-45 | 0.58938794 | 0.276 | 0.158 | 7.71E-41 | 2 | Ggct | myCAF_prolif |
| Srgap2 | 4.08E-45 | 0.26458767 | 0.563 | 0.47 | 7.88E-41 | 2 | Srgap2 | myCAF_prolif |
| Btbd10 | 4.61E-45 | 0.49003044 | 0.299 | 0.165 | 8.90E-41 | 2 | Btbd10 | myCAF_prolif |
| Ythdc1 | 4.67E-45 | 0.25357749 | 0.506 | 0.388 | 9.02E-41 | 2 | Ythdc1 | myCAF_prolif |
| Tjp21 | 4.98E-45 | 0.36996355 | 0.375 | 0.26 | 9.62E-41 | 2 | Tjp2 | myCAF_prolif |
| Usp45 | 5.45E-45 | 0.29002164 | 0.539 | 0.418 | 1.05E-40 | 2 | Usp45 | myCAF_prolif |
| Ift140 | 7.54E-45 | 0.37010028 | 0.334 | 0.208 | 1.46E-40 | 2 | Ift140 | myCAF_prolif |
| Coa6 | 7.97E-45 | 0.34911944 | 0.37 | 0.241 | 1.54E-40 | 2 | Coa6 | myCAF_prolif |
| Ccnd1 | 1.02E-44 | 0.39038864 | 0.274 | 0.189 | 1.98E-40 | 2 | Ccnd1 | myCAF_prolif |
| Mtmr12 | 1.04E-44 | 0.27476964 | 0.335 | 0.22 | 2.01E-40 | 2 | Mtmr12 | myCAF_prolif |
| Noa1 | 1.14E-44 | 0.35376296 | 0.296 | 0.175 | 2.20E-40 | 2 | Noa1 | myCAF_prolif |
| Fra10ac1 | 2.51E-44 | 0.4890606 | 0.362 | 0.233 | 4.85E-40 | 2 | Fra10ac1 | myCAF_prolif |
| Fdxr1 | 4.49E-44 | 0.30029462 | 0.446 | 0.316 | 8.67E-40 | 2 | Fdxr | myCAF_prolif |
| Armc8 | 5.33E-44 | 0.27635333 | 0.399 | 0.272 | 1.03E-39 | 2 | Armc8 | myCAF_prolif |
| Rsrc1 | 7.08E-44 | 0.40986255 | 0.434 | 0.306 | 1.37E-39 | 2 | Rsrc1 | myCAF_prolif |
| Tril | 7.46E-44 | 0.38601517 | 0.251 | 0.142 | 1.44E-39 | 2 | Tril | myCAF_prolif |
| Dhx37 | 8.38E-44 | 0.31457527 | 0.381 | 0.257 | 1.62E-39 | 2 | Dhx37 | myCAF_prolif |
| Rpap1 | 9.07E-44 | 0.61960225 | 0.337 | 0.211 | 1.75E-39 | 2 | Rpap1 | myCAF_prolif |
| Gtf2h2 | 1.21E-43 | 0.2826807 | 0.376 | 0.253 | 2.34E-39 | 2 | Gtf2h2 | myCAF_prolif |
| Nxt2 | 1.37E-43 | 0.41427703 | 0.256 | 0.137 | 2.65E-39 | 2 | Nxt2 | myCAF_prolif |
| Taco1 | 1.52E-43 | 0.64999818 | 0.318 | 0.19 | 2.94E-39 | 2 | Taco1 | myCAF_prolif |
| Fbxo34 | 2.13E-43 | 0.33224185 | 0.369 | 0.251 | 4.12E-39 | 2 | Fbxo34 | myCAF_prolif |
| Ercc2 | 2.49E-43 | 0.5392796 | 0.29 | 0.165 | 4.81E-39 | 2 | Ercc2 | myCAF_prolif |
| Rbm8a | 3.89E-43 | 0.58623465 | 0.299 | 0.178 | 7.53E-39 | 2 | Rbm8a | myCAF_prolif |
| Mapk11 | 4.37E-43 | 0.462782 | 0.279 | 0.155 | 8.46E-39 | 2 | Mapk11 | myCAF_prolif |
| Wdr37 | 4.60E-43 | 0.36215758 | 0.478 | 0.364 | 8.90E-39 | 2 | Wdr37 | myCAF_prolif |
| Zcchc10 | 6.94E-43 | 0.39485595 | 0.314 | 0.193 | 1.34E-38 | 2 | Zcchc10 | myCAF_prolif |
| Uvrag | 1.06E-42 | 0.34650051 | 0.513 | 0.407 | 2.05E-38 | 2 | Uvrag | myCAF_prolif |
| Scrn3 | 1.11E-42 | 0.3003566 | 0.335 | 0.213 | 2.15E-38 | 2 | Scrn3 | myCAF_prolif |
| Pde7a | 1.38E-42 | 0.28937317 | 0.316 | 0.191 | 2.67E-38 | 2 | Pde7a | myCAF_prolif |
| Dennd5b | 1.74E-42 | 0.43997701 | 0.328 | 0.218 | 3.35E-38 | 2 | Dennd5b | myCAF_prolif |
| Elp6 | 2.24E-42 | 0.37423537 | 0.27 | 0.159 | 4.32E-38 | 2 | Elp6 | myCAF_prolif |
| Zc3hc1 | 2.82E-42 | 0.3112699 | 0.361 | 0.239 | 5.46E-38 | 2 | Zc3hc1 | myCAF_prolif |
| Wrn | 3.20E-42 | 0.27691459 | 0.397 | 0.266 | 6.19E-38 | 2 | Wrn | myCAF_prolif |
| Zbtb45 | 3.37E-42 | 0.44394584 | 0.369 | 0.243 | 6.51E-38 | 2 | Zbtb45 | myCAF_prolif |
| Ints9 | 5.06E-42 | 0.31598014 | 0.297 | 0.177 | 9.78E-38 | 2 | Ints9 | myCAF_prolif |
| Tbc1d19 | 6.03E-42 | 0.27226497 | 0.496 | 0.382 | 1.17E-37 | 2 | Tbc1d19 | myCAF_prolif |
| Sms | 6.32E-42 | 0.27730783 | 0.287 | 0.173 | 1.22E-37 | 2 | Sms | myCAF_prolif |
| Mov10 | 8.67E-42 | 0.25632912 | 0.412 | 0.287 | 1.68E-37 | 2 | Mov10 | myCAF_prolif |
| Rbm33 | 9.16E-42 | 0.50726589 | 0.673 | 0.572 | 1.77E-37 | 2 | Rbm33 | myCAF_prolif |
| Acad9 | 1.28E-41 | 0.41879622 | 0.427 | 0.304 | 2.48E-37 | 2 | Acad9 | myCAF_prolif |
| N4bp2 | 1.31E-41 | 0.28495191 | 0.389 | 0.267 | 2.52E-37 | 2 | N4bp2 | myCAF_prolif |
| Bhlhe22 | 2.87E-41 | 0.42417961 | 0.273 | 0.164 | 5.54E-37 | 2 | Bhlhe22 | myCAF_prolif |
| Als2 | 3.98E-41 | 0.48056298 | 0.265 | 0.148 | 7.70E-37 | 2 | Als2 | myCAF_prolif |
| Hoxc9 | 7.11E-41 | 0.28487849 | 0.427 | 0.322 | 1.37E-36 | 2 | Hoxc9 | myCAF_prolif |
| Gpatch11 | 9.12E-41 | 0.32715498 | 0.467 | 0.34 | 1.76E-36 | 2 | Gpatch11 | myCAF_prolif |
| Chst1 | 1.02E-40 | 0.28487837 | 0.264 | 0.169 | 1.97E-36 | 2 | Chst1 | myCAF_prolif |
| Acat2 | 1.48E-40 | 0.29126936 | 0.306 | 0.204 | 2.86E-36 | 2 | Acat2 | myCAF_prolif |
| Tmcc21 | 1.58E-40 | 0.38409983 | 0.384 | 0.255 | 3.06E-36 | 2 | Tmcc2 | myCAF_prolif |
| Gmeb2 | 3.49E-40 | 0.38885286 | 0.445 | 0.324 | 6.74E-36 | 2 | Gmeb2 | myCAF_prolif |
| Alkbh11 | 4.41E-40 | 0.52796951 | 0.488 | 0.371 | 8.52E-36 | 2 | Alkbh1 | myCAF_prolif |
| BC003965 | 4.71E-40 | 0.41233013 | 0.444 | 0.328 | 9.11E-36 | 2 | BC003965 | myCAF_prolif |
| Tmem681 | 7.13E-40 | 0.26041284 | 0.425 | 0.307 | 1.38E-35 | 2 | Tmem68 | myCAF_prolif |
| Chchd6 | 7.46E-40 | 0.36196718 | 0.277 | 0.162 | 1.44E-35 | 2 | Chchd6 | myCAF_prolif |
| Lpin2 | 1.01E-39 | 0.29303772 | 0.431 | 0.313 | 1.95E-35 | 2 | Lpin2 | myCAF_prolif |
| Ddx51 | 1.09E-39 | 0.29250589 | 0.308 | 0.196 | 2.10E-35 | 2 | Ddx51 | myCAF_prolif |
| Ttc1 | 1.14E-39 | 0.29033988 | 0.47 | 0.342 | 2.21E-35 | 2 | Ttc1 | myCAF_prolif |
| Ptpn141 | 1.41E-39 | 0.40423238 | 0.336 | 0.225 | 2.73E-35 | 2 | Ptpn14 | myCAF_prolif |

| | | | | | | | | |
|---|---|---|---|---|---|---|---|---|
| Prr71 | 1.45E-39 | 0.376549 | 0.31 | 0.199 | 2.80E-35 | 2 | Prr7 | myCAF_prolif |
| Prkce | 1.90E-39 | 0.30365918 | 0.384 | 0.265 | 3.67E-35 | 2 | Prkce | myCAF_prolif |
| Nop14 | 2.80E-39 | 0.36953131 | 0.335 | 0.212 | 5.42E-35 | 2 | Nop14 | myCAF_prolif |
| Tbc1d7 | 3.50E-39 | 0.30856975 | 0.361 | 0.234 | 6.77E-35 | 2 | Tbc1d7 | myCAF_prolif |
| Lrrc57 | 4.48E-39 | 0.26410784 | 0.314 | 0.194 | 8.67E-35 | 2 | Lrrc57 | myCAF_prolif |
| Ccnq | 8.23E-39 | 0.25764368 | 0.33 | 0.217 | 1.59E-34 | 2 | Ccnq | myCAF_prolif |
| Casp2 | 1.21E-38 | 0.26966172 | 0.338 | 0.224 | 2.33E-34 | 2 | Casp2 | myCAF_prolif |
| Dnajc19 | 1.93E-38 | 0.54368499 | 0.261 | 0.149 | 3.74E-34 | 2 | Dnajc19 | myCAF_prolif |
| Dmap1 | 4.41E-38 | 0.30079733 | 0.343 | 0.235 | 8.53E-34 | 2 | Dmap1 | myCAF_prolif |
| Acer3 | 4.90E-38 | 0.4498541 | 0.319 | 0.201 | 9.47E-34 | 2 | Acer3 | myCAF_prolif |
| Six11 | 5.02E-38 | 0.44456828 | 0.268 | 0.181 | 9.71E-34 | 2 | Six1 | myCAF_prolif |
| Rbm34 | 1.31E-37 | 0.43210837 | 0.364 | 0.24 | 2.53E-33 | 2 | Rbm34 | myCAF_prolif |
| Zfp512 | 1.41E-37 | 0.26007177 | 0.306 | 0.199 | 2.72E-33 | 2 | Zfp512 | myCAF_prolif |
| Tmem143 | 1.97E-37 | 0.46150032 | 0.293 | 0.191 | 3.81E-33 | 2 | Tmem143 | myCAF_prolif |
| Pus10 | 2.33E-37 | 0.31023885 | 0.398 | 0.274 | 4.51E-33 | 2 | Pus10 | myCAF_prolif |
| Tcaf21 | 4.44E-37 | 0.32777558 | 0.315 | 0.201 | 8.58E-33 | 2 | Tcaf2 | myCAF_prolif |
| Haus2 | 4.47E-37 | 0.33586845 | 0.341 | 0.221 | 8.64E-33 | 2 | Haus2 | myCAF_prolif |
| Ttc27 | 5.39E-37 | 0.37601258 | 0.332 | 0.219 | 1.04E-32 | 2 | Ttc27 | myCAF_prolif |
| Gon4l | 6.36E-37 | 0.26340671 | 0.441 | 0.319 | 1.23E-32 | 2 | Gon4l | myCAF_prolif |
| Fuom | 7.11E-37 | 0.29016818 | 0.276 | 0.169 | 1.37E-32 | 2 | Fuom | myCAF_prolif |
| Cdk5rap1 | 8.85E-37 | 0.40506582 | 0.282 | 0.175 | 1.71E-32 | 2 | Cdk5rap1 | myCAF_prolif |
| Abcg2 | 1.37E-36 | 0.25314861 | 0.364 | 0.245 | 2.66E-32 | 2 | Abcg2 | myCAF_prolif |
| Ptcd2 | 4.61E-36 | 0.35737232 | 0.302 | 0.194 | 8.91E-32 | 2 | Ptcd2 | myCAF_prolif |
| Faap100 | 7.64E-36 | 0.26677759 | 0.32 | 0.203 | 1.48E-31 | 2 | Faap100 | myCAF_prolif |
| Actr6 | 7.79E-36 | 0.4805332 | 0.261 | 0.151 | 1.51E-31 | 2 | Actr6 | myCAF_prolif |
| Lin52 | 1.22E-35 | 0.52818201 | 0.279 | 0.172 | 2.36E-31 | 2 | Lin52 | myCAF_prolif |
| Pinx1 | 3.07E-35 | 0.43062824 | 0.273 | 0.159 | 5.93E-31 | 2 | Pinx1 | myCAF_prolif |
| Arfrp1 | 4.99E-35 | 0.27369438 | 0.473 | 0.366 | 9.65E-31 | 2 | Arfrp1 | myCAF_prolif |
| Wdyhv1 | 7.60E-35 | 0.4004621 | 0.32 | 0.211 | 1.47E-30 | 2 | Wdyhv1 | myCAF_prolif |
| Supt3 | 8.85E-35 | 0.28113477 | 0.348 | 0.246 | 1.71E-30 | 2 | Supt3 | myCAF_prolif |
| Klhl25 | 9.07E-35 | 0.39115653 | 0.306 | 0.2 | 1.75E-30 | 2 | Klhl25 | myCAF_prolif |
| Amigo31 | 1.09E-34 | 0.32335046 | 0.332 | 0.223 | 2.10E-30 | 2 | Amigo3 | myCAF_prolif |
| Thoc5 | 1.52E-34 | 0.32984215 | 0.413 | 0.312 | 2.93E-30 | 2 | Thoc5 | myCAF_prolif |
| Pnkp | 2.09E-34 | 0.31929911 | 0.411 | 0.302 | 4.05E-30 | 2 | Pnkp | myCAF_prolif |
| Afdn | 5.62E-34 | 0.30559768 | 0.437 | 0.355 | 1.09E-29 | 2 | Afdn | myCAF_prolif |
| Nemf | 9.40E-34 | 0.29433689 | 0.545 | 0.44 | 1.82E-29 | 2 | Nemf | myCAF_prolif |
| Pop7 | 1.38E-33 | 0.37209587 | 0.291 | 0.181 | 2.67E-29 | 2 | Pop7 | myCAF_prolif |
| Pnisr | 1.74E-33 | 0.27798426 | 0.676 | 0.602 | 3.36E-29 | 2 | Pnisr | myCAF_prolif |
| Lca5 | 1.86E-33 | 0.50058635 | 0.254 | 0.149 | 3.59E-29 | 2 | Lca5 | myCAF_prolif |
| Ctss | 9.15E-33 | 0.54140493 | 0.315 | 0.24 | 1.77E-28 | 2 | Ctss | myCAF_prolif |
| Krt8 | 9.28E-33 | 0.74891806 | 0.382 | 0.31 | 1.80E-28 | 2 | Krt8 | myCAF_prolif |
| Ccdc137 | 9.74E-33 | 0.42546764 | 0.303 | 0.192 | 1.88E-28 | 2 | Ccdc137 | myCAF_prolif |
| Slc25a14 | 1.92E-32 | 0.69649158 | 0.254 | 0.152 | 3.71E-28 | 2 | Slc25a14 | myCAF_prolif |
| mt-Nd61 | 1.93E-32 | 1.29241317 | 0.72 | 0.691 | 3.72E-28 | 2 | mt-Nd6 | myCAF_prolif |
| Crebzf | 2.26E-32 | 0.56717642 | 0.606 | 0.508 | 4.38E-28 | 2 | Crebzf | myCAF_prolif |
| Utp23 | 4.06E-32 | 0.37097765 | 0.298 | 0.189 | 7.85E-28 | 2 | Utp23 | myCAF_prolif |
| Mxra | 4.30E-32 | 0.5493644 | 0.263 | 0.164 | 8.31E-28 | 2 | Mxra | myCAF_prolif |
| Lrrc45 | 4.35E-32 | 0.28222751 | 0.439 | 0.339 | 8.41E-28 | 2 | Lrrc45 | myCAF_prolif |
| Gale | 4.56E-32 | 0.55721747 | 0.256 | 0.146 | 8.82E-28 | 2 | Gale | myCAF_prolif |
| Trmt10c | 5.88E-32 | 0.43767752 | 0.289 | 0.191 | 1.14E-27 | 2 | Trmt10c | myCAF_prolif |
| Zc3hav1l | 6.33E-32 | 0.28689093 | 0.392 | 0.282 | 1.22E-27 | 2 | Zc3hav1l | myCAF_prolif |
| Brcc3 | 6.47E-32 | 0.25241949 | 0.267 | 0.168 | 1.25E-27 | 2 | Brcc3 | myCAF_prolif |
| Gpr161 | 6.87E-32 | 0.28424467 | 0.389 | 0.279 | 1.33E-27 | 2 | Gpr161 | myCAF_prolif |
| Sfxn5 | 9.45E-32 | 0.37836361 | 0.342 | 0.237 | 1.83E-27 | 2 | Sfxn5 | myCAF_prolif |
| Setd6 | 1.26E-31 | 0.58098493 | 0.284 | 0.18 | 2.43E-27 | 2 | Setd6 | myCAF_prolif |
| Mcat | 3.69E-31 | 0.28392675 | 0.36 | 0.243 | 7.13E-27 | 2 | Mcat | myCAF_prolif |
| Ing3 | 4.48E-31 | 0.25789225 | 0.297 | 0.197 | 8.66E-27 | 2 | Ing3 | myCAF_prolif |
| Prpsap2 | 6.66E-31 | 0.32468167 | 0.293 | 0.187 | 1.29E-26 | 2 | Prpsap2 | myCAF_prolif |
| Usp54 | 8.81E-31 | 0.50889112 | 0.276 | 0.177 | 1.70E-26 | 2 | Usp54 | myCAF_prolif |
| Trub1 | 9.39E-31 | 0.50334568 | 0.343 | 0.247 | 1.81E-26 | 2 | Trub1 | myCAF_prolif |
| Dusp19 | 9.78E-31 | 0.28730877 | 0.278 | 0.181 | 1.89E-26 | 2 | Dusp19 | myCAF_prolif |
| Crocc | 1.65E-30 | 0.60260283 | 0.25 | 0.15 | 3.19E-26 | 2 | Crocc | myCAF_prolif |
| Ptrhd1 | 5.30E-30 | 0.30544965 | 0.254 | 0.156 | 1.02E-25 | 2 | Ptrhd1 | myCAF_prolif |
| Zfp668 | 1.17E-29 | 0.27579345 | 0.306 | 0.209 | 2.27E-25 | 2 | Zfp668 | myCAF_prolif |
| Plagl2 | 1.32E-29 | 0.27268571 | 0.294 | 0.196 | 2.55E-25 | 2 | Plagl2 | myCAF_prolif |
| Gstk1 | 2.31E-29 | 0.2866307 | 0.278 | 0.178 | 4.47E-25 | 2 | Gstk1 | myCAF_prolif |
| Slx4ip | 6.08E-29 | 0.27474593 | 0.258 | 0.153 | 1.18E-24 | 2 | Slx4ip | myCAF_prolif |
| Amer1 | 6.73E-29 | 0.38602351 | 0.284 | 0.193 | 1.30E-24 | 2 | Amer1 | myCAF_prolif |
| Kmt5c | 9.09E-29 | 0.30428905 | 0.495 | 0.406 | 1.76E-24 | 2 | Kmt5c | myCAF_prolif |
| Rbm43 | 2.45E-28 | 0.27396514 | 0.29 | 0.203 | 4.73E-24 | 2 | Rbm43 | myCAF_prolif |
| Thap1 | 3.53E-28 | 0.42636736 | 0.301 | 0.19 | 6.83E-24 | 2 | Thap1 | myCAF_prolif |
| Lgi3 | 4.01E-28 | 0.33218003 | 0.258 | 0.153 | 7.76E-24 | 2 | Lgi3 | myCAF_prolif |
| Gcc2 | 4.21E-28 | 0.26260063 | 0.439 | 0.339 | 8.14E-24 | 2 | Gcc2 | myCAF_prolif |
| Kin | 5.25E-28 | 0.60282609 | 0.294 | 0.19 | 1.02E-23 | 2 | Kin | myCAF_prolif |
| Pdcd2l | 6.35E-28 | 0.29847565 | 0.36 | 0.258 | 1.23E-23 | 2 | Pdcd2l | myCAF_prolif |
| Iffo11 | 2.83E-27 | 0.56619704 | 0.418 | 0.32 | 5.47E-23 | 2 | Iffo1 | myCAF_prolif |
| Itpr11 | 5.89E-27 | 0.31782146 | 0.329 | 0.219 | 1.14E-22 | 2 | Itpr1 | myCAF_prolif |
| Nme7 | 1.97E-26 | 0.26259485 | 0.342 | 0.246 | 3.80E-22 | 2 | Nme7 | myCAF_prolif |
| Wrb | 2.03E-26 | 0.30375228 | 0.296 | 0.205 | 3.92E-22 | 2 | Wrb | myCAF_prolif |
| Smyd3 | 5.30E-26 | 0.48244774 | 0.263 | 0.172 | 1.03E-21 | 2 | Smyd3 | myCAF_prolif |
| Nkap | 5.76E-26 | 0.3438097 | 0.297 | 0.201 | 1.11E-21 | 2 | Nkap | myCAF_prolif |
| Mtrf1l | 6.11E-26 | 0.43726407 | 0.306 | 0.202 | 1.18E-21 | 2 | Mtrf1l | myCAF_prolif |
| Rhobtb2 | 6.82E-26 | 0.27942784 | 0.287 | 0.192 | 1.32E-21 | 2 | Rhobtb2 | myCAF_prolif |
| Wdpcp | 7.70E-26 | 0.32433604 | 0.266 | 0.189 | 1.49E-21 | 2 | Wdpcp | myCAF_prolif |
| Dido1 | 8.16E-26 | 0.27379177 | 0.46 | 0.366 | 1.58E-21 | 2 | Dido1 | myCAF_prolif |
| Csf2ra1 | 2.35E-25 | 0.5716582 | 0.285 | 0.183 | 4.54E-21 | 2 | Csf2ra | myCAF_prolif |
| Luc7l | 2.97E-25 | 0.3767199 | 0.459 | 0.361 | 5.74E-21 | 2 | Luc7l | myCAF_prolif |
| Cbll1 | 3.54E-25 | 0.29311335 | 0.284 | 0.196 | 6.85E-21 | 2 | Cbll1 | myCAF_prolif |
| Zfp830 | 4.55E-25 | 0.38119449 | 0.368 | 0.273 | 8.81E-21 | 2 | Zfp830 | myCAF_prolif |
| Ippk | 5.77E-25 | 0.30598202 | 0.277 | 0.19 | 1.11E-20 | 2 | Ippk | myCAF_prolif |
| Rabl2 | 2.67E-24 | 0.29111317 | 0.353 | 0.261 | 5.17E-20 | 2 | Rabl2 | myCAF_prolif |
| Osgin2 | 3.19E-24 | 0.3064584 | 0.271 | 0.189 | 6.17E-20 | 2 | Osgin2 | myCAF_prolif |
| Stxbp3 | 3.80E-24 | 0.27622361 | 0.418 | 0.313 | 7.34E-20 | 2 | Stxbp3 | myCAF_prolif |
| Chd2 | 3.88E-24 | 0.29633577 | 0.537 | 0.476 | 7.51E-20 | 2 | Chd2 | myCAF_prolif |
| Xpnpep3 | 4.33E-24 | 0.33994882 | 0.253 | 0.172 | 8.38E-20 | 2 | Xpnpep3 | myCAF_prolif |
| Armcx6 | 1.04E-23 | 0.264932 | 0.268 | 0.179 | 2.00E-19 | 2 | Armcx6 | myCAF_prolif |
| Ahi1 | 1.23E-23 | 0.27372395 | 0.301 | 0.218 | 2.38E-19 | 2 | Ahi1 | myCAF_prolif |
| Slc2a13 | 1.59E-23 | 0.26688967 | 0.425 | 0.338 | 3.08E-19 | 2 | Slc2a13 | myCAF_prolif |
| Mast3 | 1.69E-23 | 0.28939192 | 0.368 | 0.279 | 3.27E-19 | 2 | Mast3 | myCAF_prolif |
| Begain | 2.41E-23 | 0.27811267 | 0.327 | 0.243 | 4.67E-19 | 2 | Begain | myCAF_prolif |
| Gpd2 | 8.40E-23 | 0.34268133 | 0.269 | 0.188 | 1.62E-18 | 2 | Gpd2 | myCAF_prolif |
| Ttf1 | 1.19E-22 | 0.33285702 | 0.319 | 0.225 | 2.30E-18 | 2 | Ttf1 | myCAF_prolif |
| Cry1 | 1.41E-22 | 0.26668251 | 0.275 | 0.191 | 2.72E-18 | 2 | Cry1 | myCAF_prolif |
| Slc35e3 | 2.33E-22 | 0.28413023 | 0.37 | 0.28 | 4.50E-18 | 2 | Slc35e3 | myCAF_prolif |
| Fn3krp | 2.64E-22 | 0.32057442 | 0.29 | 0.203 | 5.11E-18 | 2 | Fn3krp | myCAF_prolif |
| Taf1a | 2.76E-22 | 0.60246404 | 0.263 | 0.172 | 5.34E-18 | 2 | Taf1a | myCAF_prolif |
| Tyw5 | 3.33E-22 | 0.28121396 | 0.254 | 0.166 | 6.43E-18 | 2 | Tyw5 | myCAF_prolif |
| Nsrp1 | 5.10E-22 | 0.27480038 | 0.442 | 0.343 | 9.85E-18 | 2 | Nsrp1 | myCAF_prolif |
| Tecpr2 | 7.18E-22 | 0.40763615 | 0.293 | 0.195 | 1.39E-17 | 2 | Tecpr2 | myCAF_prolif |
| Lyz2 | 1.10E-21 | 0.57733193 | 0.313 | 0.246 | 2.13E-17 | 2 | Lyz2 | myCAF_prolif |
| Asb3 | 1.80E-21 | 0.30534358 | 0.377 | 0.278 | 3.49E-17 | 2 | Asb3 | myCAF_prolif |
| Hdac10 | 1.86E-21 | 0.27938387 | 0.282 | 0.201 | 3.59E-17 | 2 | Hdac10 | myCAF_prolif |
| Taf1b | 2.79E-21 | 0.28032973 | 0.352 | 0.264 | 5.40E-17 | 2 | Taf1b | myCAF_prolif |
| Ppp1r35 | 7.09E-21 | 0.3344255 | 0.381 | 0.295 | 1.37E-16 | 2 | Ppp1r35 | myCAF_prolif |
| Krit1 | 1.77E-20 | 0.27265405 | 0.433 | 0.35 | 3.43E-16 | 2 | Krit1 | myCAF_prolif |
| Tubgcp5 | 2.76E-20 | 0.27561844 | 0.294 | 0.21 | 5.34E-16 | 2 | Tubgcp5 | myCAF_prolif |
| Inpp1 | 9.29E-20 | 0.25908942 | 0.342 | 0.266 | 1.80E-15 | 2 | Inpp1 | myCAF_prolif |
| Atad2b | 1.14E-19 | 0.25544836 | 0.363 | 0.274 | 2.21E-15 | 2 | Atad2b | myCAF_prolif |
| Kctd18 | 3.20E-19 | 0.41242072 | 0.325 | 0.235 | 6.20E-15 | 2 | Kctd18 | myCAF_prolif |
| Tchp | 7.00E-19 | 0.92556104 | 0.255 | 0.166 | 1.35E-14 | 2 | Tchp | myCAF_prolif |
| Tgs1 | 2.44E-18 | 0.39492257 | 0.285 | 0.209 | 4.72E-14 | 2 | Tgs1 | myCAF_prolif |
| Polr3a | 2.50E-18 | 0.31932586 | 0.272 | 0.186 | 4.83E-14 | 2 | Polr3a | myCAF_prolif |
| Adck5 | 2.85E-17 | 0.25580139 | 0.251 | 0.182 | 5.51E-13 | 2 | Adck5 | myCAF_prolif |
| Pex7 | 3.53E-17 | 0.44214228 | 0.299 | 0.211 | 6.83E-13 | 2 | Pex7 | myCAF_prolif |
| Rnpc3 | 6.44E-17 | 0.34969552 | 0.261 | 0.185 | 1.25E-12 | 2 | Rnpc3 | myCAF_prolif |
| Vps9d1 | 2.82E-16 | 0.2938294 | 0.333 | 0.257 | 5.46E-12 | 2 | Vps9d1 | myCAF_prolif |
| Snapc4 | 1.15E-15 | 0.29555836 | 0.34 | 0.249 | 2.23E-11 | 2 | Snapc4 | myCAF_prolif |
| B3gntl1 | 1.23E-15 | 0.36150295 | 0.27 | 0.192 | 2.37E-11 | 2 | B3gntl1 | myCAF_prolif |
| Ift172 | 2.59E-15 | 0.50899902 | 0.346 | 0.264 | 5.01E-11 | 2 | Ift172 | myCAF_prolif |
| Ppp1r21 | 1.32E-14 | 0.49705809 | 0.284 | 0.214 | 2.55E-10 | 2 | Ppp1r21 | myCAF_prolif |
| Jrkl | 2.79E-14 | 0.25204982 | 0.298 | 0.23 | 5.40E-10 | 2 | Jrkl | myCAF_prolif |
| Golga1 | 1.05E-13 | 0.28666988 | 0.265 | 0.194 | 2.04E-09 | 2 | Golga1 | myCAF_prolif |
| Clp1 | 1.79E-13 | 0.3035483 | 0.256 | 0.198 | 3.46E-09 | 2 | Clp1 | myCAF_prolif |
| Nt5dc31 | 7.19E-13 | 0.3369871 | 0.311 | 0.235 | 1.39E-08 | 2 | Nt5dc3 | myCAF_prolif |
| Tspyl2 | 5.35E-05 | 0.28599093 | 0.457 | 0.423 | 1 | 2 | Tspyl2 | myCAF_prolif |
| Ccnt2 | 6.84E-05 | 0.26563431 | 0.544 | 0.509 | 1 | 2 | Ccnt2 | myCAF_prolif |
| Hc1 | 0 | 2.47784012 | 0.736 | 0.225 | 0 | 3 | Hc | Col18a1 |
| Pde7b | 0 | 2.6918714 | 0.689 | 0.192 | 0 | 3 | Pde7b | Col18a1 |
| Gcnt41 | 0 | 2.20719676 | 0.813 | 0.316 | 0 | 3 | Gcnt4 | Col18a1 |
| Lratd1 | 0 | 3.75737364 | 0.578 | 0.103 | 0 | 3 | Lratd1 | Col18a1 |
| Ltbp1 | 0 | 1.87930346 | 0.788 | 0.33 | 0 | 3 | Ltbp1 | Col18a1 |
| Prr16 | 0 | 1.92296143 | 0.827 | 0.372 | 0 | 3 | Prr16 | Col18a1 |
| Bmp7 | 0 | 3.0052373 | 0.52 | 0.086 | 0 | 3 | Bmp7 | Col18a1 |
| Ociad21 | 0 | 1.73300313 | 0.861 | 0.438 | 0 | 3 | Ociad2 | Col18a1 |
| Sema3a1 | 0 | 2.51246956 | 0.631 | 0.22 | 0 | 3 | Sema3a | Col18a1 |
| Cbfa2t3 | 0 | 1.50622355 | 0.621 | 0.244 | 0 | 3 | Cbfa2t3 | Col18a1 |
| Rspo4 | 0 | 4.24470186 | 0.396 | 0.023 | 0 | 3 | Rspo4 | Col18a1 |
| Fcgr2b1 | 0 | 2.32301069 | 0.699 | 0.329 | 0 | 3 | Fcgr2b | Col18a1 |
| Rarres1 | 0 | 2.66072285 | 0.508 | 0.139 | 0 | 3 | Rarres1 | Col18a1 |
| Ptgir | 0 | 1.95244989 | 0.655 | 0.293 | 0 | 3 | Ptgir | Col18a1 |
| Dusp4 | 0 | 1.52445749 | 0.644 | 0.282 | 0 | 3 | Dusp4 | Col18a1 |
| a1 | 0 | 1.70460697 | 0.63 | 0.273 | 0 | 3 | a | Col18a1 |
| Amn2 | 0 | 2.02259405 | 0.561 | 0.207 | 0 | 3 | Amn | Col18a1 |
| Col18a1 | 0 | 2.94200495 | 0.963 | 0.609 | 0 | 3 | Col18a1 | Col18a1 |
| Chst15 | 0 | 1.60520631 | 0.793 | 0.441 | 0 | 3 | Chst15 | Col18a1 |
| Ereg | 0 | 2.73049257 | 0.502 | 0.15 | 0 | 3 | Ereg | Col18a1 |
| Arhgdib | 0 | 1.83148095 | 0.928 | 0.577 | 0 | 3 | Arhgdib | Col18a1 |
| Tbx31 | 0 | 1.29155886 | 0.809 | 0.462 | 0 | 3 | Tbx3 | Col18a1 |
| Cyp26b1 | 0 | 1.59321972 | 0.812 | 0.467 | 0 | 3 | Cyp26b1 | Col18a1 |
| Inhbb | 0 | 1.89559349 | 0.953 | 0.611 | 0 | 3 | Inhbb | Col18a1 |
| Draxin1 | 0 | 1.55225259 | 0.722 | 0.387 | 0 | 3 | Draxin | Col18a1 |
| Pde5a1 | 0 | 1.58327149 | 0.757 | 0.424 | 0 | 3 | Pde5a | Col18a1 |
| Brinp31 | 0 | 1.35740206 | 0.791 | 0.465 | 0 | 3 | Brinp3 | Col18a1 |
| Pgap1 | 0 | 2.36558988 | 0.541 | 0.216 | 0 | 3 | Pgap1 | Col18a1 |
| Fosl1 | 0 | 2.07517739 | 0.664 | 0.344 | 0 | 3 | Fosl1 | Col18a1 |
| Il1rap | 0 | 1.62801703 | 0.797 | 0.48 | 0 | 3 | Il1rap | Col18a1 |
| Tmem26 | 0 | 2.28077785 | 0.452 | 0.137 | 0 | 3 | Tmem26 | Col18a1 |
| Cadm4 | 0 | 2.24521206 | 0.435 | 0.127 | 0 | 3 | Cadm4 | Col18a1 |
| Sema5a | 0 | 1.77783152 | 0.76 | 0.459 | 0 | 3 | Sema5a | Col18a1 |
| Mitf | 0 | 1.78146108 | 0.765 | 0.467 | 0 | 3 | Mitf | Col18a1 |
| Tiparp | 0 | 1.89122243 | 0.772 | 0.475 | 0 | 3 | Tiparp | Col18a1 |
| Rgs2 | 0 | 2.11347234 | 0.476 | 0.179 | 0 | 3 | Rgs2 | Col18a1 |
| Ndufa4l21 | 0 | 1.46229931 | 0.694 | 0.397 | 0 | 3 | Ndufa4l2 | Col18a1 |
| Epas11 | 0 | 1.48396651 | 0.918 | 0.622 | 0 | 3 | Epas1 | Col18a1 |
| Slc2a3 | 0 | 1.726644 | 0.592 | 0.298 | 0 | 3 | Slc2a3 | Col18a1 |
| Vstm4 | 0 | 1.28905931 | 0.76 | 0.467 | 0 | 3 | Vstm4 | Col18a1 |
| Ifitm11 | 0 | 1.34166664 | 0.634 | 0.342 | 0 | 3 | Ifitm1 | Col18a1 |
| Masp11 | 0 | 1.42454835 | 0.615 | 0.323 | 0 | 3 | Masp1 | Col18a1 |
| Itga1 | 0 | 1.78655977 | 0.885 | 0.598 | 0 | 3 | Itga1 | Col18a1 |
| Pde4b1 | 0 | 1.63769449 | 0.674 | 0.391 | 0 | 3 | Pde4b | Col18a1 |
| Cxcl51 | 0 | 2.17747328 | 0.601 | 0.319 | 0 | 3 | Cxcl5 | Col18a1 |
| DEPRECATED-I | 0 | 1.33400931 | 0.539 | 0.264 | 0 | 3 | DEPRECATED-I | Col18a1 |
| Cnksr3 | 0 | 1.72178088 | 0.551 | 0.277 | 0 | 3 | Cnksr3 | Col18a1 |
| Lgals3 | 0 | 1.31680716 | 0.761 | 0.487 | 0 | 3 | Lgals3 | Col18a1 |
| Pakap.1 | 0 | 1.33523811 | 0.908 | 0.636 | 0 | 3 | Pakap.1 | Col18a1 |
| Inhba1 | 0 | 0.84769812 | 0.956 | 0.686 | 0 | 3 | Inhba | Col18a1 |
| Tcim | 0 | 1.17035526 | 0.777 | 0.51 | 0 | 3 | Tcim | Col18a1 |
| Spock1 | 0 | 3.9487572 | 0.295 | 0.028 | 0 | 3 | Spock1 | Col18a1 |
| Serpina3g | 0 | 1.29866227 | 0.822 | 0.557 | 0 | 3 | Serpina3g | Col18a1 |
| Rgs16 | 0 | 1.07318596 | 0.726 | 0.466 | 0 | 3 | Rgs16 | Col18a1 |
| Crabp11 | 0 | 2.06891249 | 0.987 | 0.73 | 0 | 3 | Crabp1 | Col18a1 |
| Hgf | 0 | 1.74551942 | 0.577 | 0.322 | 0 | 3 | Hgf | Col18a1 |
| Cpxm21 | 0 | 0.72322889 | 0.903 | 0.65 | 0 | 3 | Cpxm2 | Col18a1 |
| Eif2s2 | 0 | 1.62993064 | 0.883 | 0.633 | 0 | 3 | Eif2s2 | Col18a1 |
| Cav1 | 0 | 1.65373864 | 0.855 | 0.605 | 0 | 3 | Cav1 | Col18a1 |
| Tnfrsf211 | 0 | 1.36529254 | 0.572 | 0.327 | 0 | 3 | Tnfrsf21 | Col18a1 |
| Slco2a1 | 0 | 1.45258228 | 0.604 | 0.366 | 0 | 3 | Slco2a1 | Col18a1 |
| Xdh1 | 0 | 1.01279527 | 0.936 | 0.699 | 0 | 3 | Xdh | Col18a1 |
| Tbx2 | 0 | 2.0408215 | 0.406 | 0.169 | 0 | 3 | Tbx2 | Col18a1 |
| Tns31 | 0 | 1.04382807 | 0.877 | 0.641 | 0 | 3 | Tns3 | Col18a1 |
| Tnfsf9 | 0 | 1.94722515 | 0.479 | 0.244 | 0 | 3 | Tnfsf9 | Col18a1 |
| Olfm1 | 0 | 1.21275183 | 0.785 | 0.553 | 0 | 3 | Olfm1 | Col18a1 |
| Cd531 | 0 | 0.89694283 | 0.612 | 0.386 | 0 | 3 | Cd53 | Col18a1 |
| Ankrd44 | 0 | 2.07832465 | 0.443 | 0.219 | 0 | 3 | Ankrd44 | Col18a1 |
| Ptger3 | 0 | 1.88369043 | 0.358 | 0.135 | 0 | 3 | Ptger3 | Col18a1 |
| Ldlr | 0 | 1.13996129 | 0.812 | 0.59 | 0 | 3 | Ldlr | Col18a1 |
| Klf5 | 0 | 2.06809057 | 0.46 | 0.24 | 0 | 3 | Klf5 | Col18a1 |
| Mme | 0 | 2.75689446 | 0.301 | 0.083 | 0 | 3 | Mme | Col18a1 |
| S100a4 | 0 | 1.35132904 | 0.73 | 0.512 | 0 | 3 | S100a4 | Col18a1 |
| Gucy1a1 | 0 | 2.16427366 | 0.334 | 0.117 | 0 | 3 | Gucy1a1 | Col18a1 |
| Calcrl | 0 | 2.22091215 | 0.434 | 0.22 | 0 | 3 | Calcrl | Col18a1 |
| Hsd11b11 | 0 | 1.29808236 | 0.653 | 0.44 | 0 | 3 | Hsd11b1 | Col18a1 |
| Glul1 | 0 | 1.21415703 | 0.863 | 0.65 | 0 | 3 | Glul | Col18a1 |
| Peak1 | 0 | 1.34408671 | 0.881 | 0.672 | 0 | 3 | Peak1 | Col18a1 |
| Btg1 | 0 | 1.25175346 | 0.902 | 0.696 | 0 | 3 | Btg1 | Col18a1 |
| Id2 | 0 | 1.18824953 | 0.811 | 0.607 | 0 | 3 | Id2 | Col18a1 |
| Rgcc | 0 | 1.38556367 | 0.818 | 0.616 | 0 | 3 | Rgcc | Col18a1 |
| Spats2l1 | 0 | 1.17054567 | 0.907 | 0.706 | 0 | 3 | Spats2l | Col18a1 |
| Cd302 | 0 | 1.32773621 | 0.893 | 0.705 | 0 | 3 | Cd302 | Col18a1 |
| Ntrk2 | 0 | 1.37226307 | 0.461 | 0.275 | 0 | 3 | Ntrk2 | Col18a1 |
| Stc1 | 0 | 2.96785117 | 0.289 | 0.103 | 0 | 3 | Stc1 | Col18a1 |
| Ddit4 | 0 | 1.1793957 | 0.843 | 0.657 | 0 | 3 | Ddit4 | Col18a1 |
| Sik1 | 0 | 1.41644665 | 0.603 | 0.428 | 0 | 3 | Sik1 | Col18a1 |
| Serpine21 | 0 | 1.04310303 | 0.974 | 0.799 | 0 | 3 | Serpine2 | Col18a1 |
| Sema7a1 | 0 | 0.53580702 | 0.947 | 0.778 | 0 | 3 | Sema7a | Col18a1 |
| Itga7 | 0 | 2.28108604 | 0.305 | 0.139 | 0 | 3 | Itga7 | Col18a1 |
| Col15a1 | 0 | 0.84979543 | 0.944 | 0.78 | 0 | 3 | Col15a1 | Col18a1 |
| Igf1 | 0 | 1.16910985 | 0.77 | 0.609 | 0 | 3 | Igf1 | Col18a1 |
| Pgk1 | 0 | 1.16868003 | 0.936 | 0.775 | 0 | 3 | Pgk1 | Col18a1 |
| mt-Nd11 | 0 | 0.47726599 | 0.824 | 0.668 | 0 | 3 | mt-Nd1 | Col18a1 |
| Vegfa | 0 | 0.99476935 | 0.835 | 0.68 | 0 | 3 | Vegfa | Col18a1 |
| Ndrg1 | 0 | 1.42605736 | 0.797 | 0.651 | 0 | 3 | Ndrg1 | Col18a1 |
| Hmgcs1 | 0 | 1.25166881 | 0.852 | 0.715 | 0 | 3 | Hmgcs1 | Col18a1 |
| Twist21 | 0 | 1.41090482 | 0.931 | 0.795 | 0 | 3 | Twist2 | Col18a1 |
| Cfh1 | 0 | 0.94918418 | 0.978 | 0.842 | 0 | 3 | Cfh | Col18a1 |
| Hilpda | 0 | 1.27277962 | 0.619 | 0.483 | 0 | 3 | Hilpda | Col18a1 |
| Cpz | 0 | 1.23645062 | 0.746 | 0.615 | 0 | 3 | Cpz | Col18a1 |
| Sod21 | 0 | 0.94308628 | 0.9 | 0.769 | 0 | 3 | Sod2 | Col18a1 |
| Lhfpl2 | 0 | 0.99070407 | 0.946 | 0.817 | 0 | 3 | Lhfpl2 | Col18a1 |
| Ero1l | 0 | 1.18102636 | 0.797 | 0.676 | 0 | 3 | Ero1l | Col18a1 |
| mt-Co31 | 0 | 0.37745632 | 0.922 | 0.809 | 0 | 3 | mt-Co3 | Col18a1 |
| Spon11 | 0 | 0.50358834 | 0.985 | 0.874 | 0 | 3 | Spon1 | Col18a1 |
| Mt21 | 0 | 0.36160277 | 0.979 | 0.87 | 0 | 3 | Mt2 | Col18a1 |
| Lum | 0 | 1.5556444 | 0.944 | 0.836 | 0 | 3 | Lum | Col18a1 |
| Osmr | 0 | 1.17931529 | 0.972 | 0.865 | 0 | 3 | Osmr | Col18a1 |
| Pla1a1 | 0 | 0.90696109 | 0.975 | 0.868 | 0 | 3 | Pla1a | Col18a1 |
| Rnd3 | 0 | 1.02499783 | 0.923 | 0.817 | 0 | 3 | Rnd3 | Col18a1 |
| Cyth3 | 0 | 0.82641566 | 0.955 | 0.85 | 0 | 3 | Cyth3 | Col18a1 |
| Nfe2l2 | 0 | 0.96446704 | 0.963 | 0.865 | 0 | 3 | Nfe2l2 | Col18a1 |
| Insig1 | 0 | 0.92091769 | 0.811 | 0.715 | 0 | 3 | Insig1 | Col18a1 |
| Sat1 | 0 | 0.97885456 | 0.92 | 0.826 | 0 | 3 | Sat1 | Col18a1 |
| Tbc1d2b1 | 0 | 0.94740333 | 0.953 | 0.86 | 0 | 3 | Tbc1d2b | Col18a1 |
| Gpi1 | 0 | 0.9364499 | 0.937 | 0.845 | 0 | 3 | Gpi1 | Col18a1 |
| Colec121 | 0 | 1.01677871 | 0.991 | 0.901 | 0 | 3 | Colec12 | Col18a1 |
| mt-Atp61 | 0 | 0.35874938 | 0.933 | 0.844 | 0 | 3 | mt-Atp6 | Col18a1 |
| Sdc11 | 0 | 1.35468013 | 0.994 | 0.906 | 0 | 3 | Sdc1 | Col18a1 |
| Pten | 0 | 1.05200144 | 0.965 | 0.877 | 0 | 3 | Pten | Col18a1 |
| Kremen11 | 0 | 0.79756498 | 0.98 | 0.893 | 0 | 3 | Kremen1 | Col18a1 |
| Gas11 | 0 | 0.66703477 | 0.932 | 0.846 | 0 | 3 | Gas1 | Col18a1 |
| Cd9 | 0 | 0.82095824 | 0.975 | 0.893 | 0 | 3 | Cd9 | Col18a1 |
| Cdh111 | 0 | 0.66191654 | 0.985 | 0.906 | 0 | 3 | Cdh11 | Col18a1 |
| Plod1 | 0 | 0.96892492 | 0.963 | 0.885 | 0 | 3 | Plod1 | Col18a1 |
| Tnfrsf11b | 0 | 1.41772501 | 0.436 | 0.361 | 0 | 3 | Tnfrsf11b | Col18a1 |
| Errfi1 | 0 | 0.77369074 | 0.852 | 0.777 | 0 | 3 | Errfi1 | Col18a1 |
| Rassf8 | 0 | 1.13576257 | 0.977 | 0.903 | 0 | 3 | Rassf8 | Col18a1 |
| Tcf41 | 0 | 0.95625461 | 0.984 | 0.914 | 0 | 3 | Tcf4 | Col18a1 |
| Tnfaip2 | 0 | 0.93332789 | 0.969 | 0.901 | 0 | 3 | Tnfaip2 | Col18a1 |
| 9-Sep | 0 | 0.89104557 | 0.983 | 0.92 | 0 | 3 | 9-Sep | Col18a1 |
| Ybx31 | 0 | 0.87197569 | 0.99 | 0.929 | 0 | 3 | Ybx3 | Col18a1 |
| Egfp1 | 0 | 0.69846193 | 0.982 | 0.926 | 0 | 3 | Egfp | Col18a1 |
| Pdgfrb1 | 0 | 0.67854809 | 0.992 | 0.938 | 0 | 3 | Pdgfrb | Col18a1 |
| Col12a11 | 0 | 0.58446216 | 0.996 | 0.943 | 0 | 3 | Col12a1 | Col18a1 |
| Spp11 | 0 | 0.61325025 | 0.976 | 0.925 | 0 | 3 | Spp1 | Col18a1 |
| Lamp2 | 0 | 0.94870045 | 0.991 | 0.942 | 0 | 3 | Lamp2 | Col18a1 |
| Pkm | 0 | 0.68165497 | 0.995 | 0.95 | 0 | 3 | Pkm | Col18a1 |
| Dcn1 | 0 | 1.53948774 | 0.998 | 0.953 | 0 | 3 | Dcn | Col18a1 |
| Cpxm11 | 0 | 0.73865134 | 0.991 | 0.953 | 0 | 3 | Cpxm1 | Col18a1 |
| Gpc11 | 0 | 0.55390917 | 0.994 | 0.958 | 0 | 3 | Gpc1 | Col18a1 |
| Ctsl | 0 | 0.97424698 | 0.993 | 0.961 | 0 | 3 | Ctsl | Col18a1 |
| Pros1 | 0 | 0.86294955 | 0.987 | 0.956 | 0 | 3 | Pros1 | Col18a1 |
| Hspa1b1 | 0 | 0.69150314 | 0.586 | 0.603 | 0 | 3 | Hspa1b | Col18a1 |
| S100a6 | 0 | 1.01259833 | 0.998 | 0.982 | 0 | 3 | S100a6 | Col18a1 |
| Adamts21 | 0 | 0.62398064 | 0.998 | 0.982 | 0 | 3 | Adamts2 | Col18a1 |
| Egr11 | 0 | 0.50274951 | 0.907 | 0.891 | 0 | 3 | Egr1 | Col18a1 |
| Ctsb | 0 | 0.54486358 | 0.997 | 0.984 | 0 | 3 | Ctsb | Col18a1 |
| Col6a31 | 0 | 0.82639536 | 0.998 | 0.987 | 0 | 3 | Col6a3 | Col18a1 |
| Thy1 | 0 | 0.80432178 | 0.992 | 0.981 | 0 | 3 | Thy1 | Col18a1 |
| Col6a11 | 0 | 1.01509663 | 1 | 0.989 | 0 | 3 | Col6a1 | Col18a1 |
| Tpt1 | 0 | 0.85863807 | 0.992 | 0.981 | 0 | 3 | Tpt1 | Col18a1 |
| Col6a21 | 0 | 0.88839687 | 0.997 | 0.989 | 0 | 3 | Col6a2 | Col18a1 |
| Eef1a1 | 0 | 0.56014252 | 0.997 | 0.99 | 0 | 3 | Eef1a1 | Col18a1 |
| Dusp12 | 0 | 0.81953315 | 0.734 | 0.734 | 0 | 3 | Dusp1 | Col18a1 |
| Nr2f2 | 6.78E-293 | 1.57332914 | 0.671 | 0.37 | 1.31E-288 | 3 | Nr2f2 | Col18a1 |
| Spred1 | 4.11E-293 | 0.72853362 | 0.97 | 0.926 | 7.95E-289 | 3 | Spred1 | Col18a1 |
| Tnfrsf19 | 1.00E-293 | 1.17399856 | 0.53 | 0.292 | 1.94E-289 | 3 | Tnfrsf19 | Col18a1 |
| Icosl | 5.72E-293 | 2.49464302 | 0.365 | 0.106 | 1.11E-288 | 3 | Icosl | Col18a1 |
| Hspa1a1 | 1.18E-293 | 0.75466514 | 0.545 | 0.548 | 2.28E-289 | 3 | Hspa1a | Col18a1 |
| Sqle | 3.84E-293 | 1.41568902 | 0.616 | 0.363 | 7.41E-289 | 3 | Sqle | Col18a1 |
| Stk261 | 1.47E-292 | 1.37321594 | 0.608 | 0.308 | 2.83E-288 | 3 | Stk26 | Col18a1 |
| Tspan9 | 1.63E-287 | 1.19398475 | 0.831 | 0.62 | 3.15E-283 | 3 | Tspan9 | Col18a1 |
| Il1r11 | 2.42E-282 | 0.71614623 | 0.969 | 0.898 | 4.67E-278 | 3 | Il1r1 | Col18a1 |
| Ahr1 | 4.23E-280 | 1.57856997 | 0.553 | 0.312 | 8.17E-276 | 3 | Ahr | Col18a1 |
| Lama4 | 8.43E-271 | 0.71900227 | 0.97 | 0.919 | 1.63E-266 | 3 | Lama4 | Col18a1 |
| Fau | 1.68E-269 | 0.59342847 | 0.991 | 0.972 | 3.26E-265 | 3 | Fau | Col18a1 |
| Zmiz1 | 5.36E-268 | 0.88479672 | 0.936 | 0.835 | 1.04E-263 | 3 | Zmiz1 | Col18a1 |
| Prrx1 | 1.13E-266 | 0.64574221 | 0.992 | 0.961 | 2.19E-262 | 3 | Prrx1 | Col18a1 |
| Fetub | 5.78E-266 | 3.66538123 | 0.25 | 0.034 | 1.12E-261 | 3 | Fetub | Col18a1 |
| Scpep1 | 4.88E-264 | 0.70290146 | 0.982 | 0.941 | 9.43E-260 | 3 | Scpep1 | Col18a1 |
| Grik2 | 1.52E-261 | 2.71282649 | 0.315 | 0.074 | 2.94E-257 | 3 | Grik2 | Col18a1 |
| Hmga2 | 1.91E-257 | 1.64968921 | 0.382 | 0.178 | 3.69E-253 | 3 | Hmga2 | Col18a1 |
| Rasa1 | 6.88E-257 | 1.17467404 | 0.848 | 0.631 | 1.33E-252 | 3 | Rasa1 | Col18a1 |
| Gpm6b1 | 9.15E-255 | 0.81262984 | 0.883 | 0.794 | 1.77E-250 | 3 | Gpm6b | Col18a1 |
| Nt5e | 3.68E-253 | 1.57055963 | 0.415 | 0.252 | 7.12E-249 | 3 | Nt5e | Col18a1 |
| Tmem108 | 1.95E-252 | 2.08965374 | 0.352 | 0.112 | 3.78E-248 | 3 | Tmem108 | Col18a1 |
| Tshz3 | 3.15E-250 | 1.2003532 | 0.893 | 0.636 | 6.10E-246 | 3 | Tshz3 | Col18a1 |
| Smim41 | 3.70E-249 | 2.02210733 | 0.318 | 0.088 | 7.15E-245 | 3 | Smim41 | Col18a1 |
| Tpbg | 5.73E-248 | 0.74799077 | 0.962 | 0.903 | 1.11E-243 | 3 | Tpbg | Col18a1 |
| Asah1 | 9.77E-247 | 1.05987294 | 0.883 | 0.7 | 1.89E-242 | 3 | Asah1 | Col18a1 |
| Hmox11 | 1.33E-245 | 1.0003113 | 0.796 | 0.695 | 2.57E-241 | 3 | Hmox1 | Col18a1 |
| Ldha | 8.57E-244 | 0.60609837 | 0.991 | 0.95 | 1.66E-239 | 3 | Ldha | Col18a1 |
| Ednra1 | 1.19E-240 | 0.85994649 | 0.783 | 0.565 | 2.30E-236 | 3 | Ednra | Col18a1 |
| Tnfrsf261 | 9.94E-237 | 1.00139024 | 0.864 | 0.599 | 1.92E-232 | 3 | Tnfrsf26 | Col18a1 |
| Glrx | 2.10E-233 | 1.26567808 | 0.725 | 0.504 | 4.06E-229 | 3 | Glrx | Col18a1 |
| Psap | 1.75E-232 | 0.63167678 | 0.984 | 0.945 | 3.38E-228 | 3 | Psap | Col18a1 |
| Plekha6 | 1.47E-231 | 1.81421099 | 0.386 | 0.124 | 2.85E-227 | 3 | Plekha6 | Col18a1 |
| Ugcg1 | 4.22E-231 | 0.78539749 | 0.842 | 0.783 | 8.16E-227 | 3 | Ugcg | Col18a1 |
| Cd47 | 3.89E-229 | 0.70331625 | 0.968 | 0.911 | 7.51E-225 | 3 | Cd47 | Col18a1 |
| Slc16a11 | 5.22E-227 | 0.87106577 | 0.82 | 0.703 | 1.01E-222 | 3 | Slc16a1 | Col18a1 |
| Ifi207 | 4.62E-225 | 1.06477061 | 0.822 | 0.541 | 8.94E-221 | 3 | Ifi207 | Col18a1 |
| Btg21 | 4.00E-224 | 0.88890141 | 0.674 | 0.627 | 7.74E-220 | 3 | Btg2 | Col18a1 |
| Ptpn1 | 8.43E-222 | 1.00361002 | 0.855 | 0.668 | 1.63E-217 | 3 | Ptpn1 | Col18a1 |
| Il15 | 3.77E-221 | 2.43556718 | 0.289 | 0.083 | 7.29E-217 | 3 | Il15 | Col18a1 |
| Sipa1l1 | 4.66E-221 | 1.42923242 | 0.735 | 0.423 | 9.02E-217 | 3 | Sipa1l1 | Col18a1 |
| Fam78b | 1.46E-219 | 1.26595856 | 0.721 | 0.479 | 2.82E-215 | 3 | Fam78b | Col18a1 |
| Sc5d1 | 1.01E-218 | 1.30250232 | 0.696 | 0.439 | 1.95E-214 | 3 | Sc5d | Col18a1 |
| Arrb1 | 1.09E-216 | 1.37486614 | 0.549 | 0.311 | 2.11E-212 | 3 | Arrb1 | Col18a1 |
| Lhfp1 | 4.02E-215 | 0.47991886 | 0.993 | 0.971 | 7.78E-211 | 3 | Lhfp | Col18a1 |
| Il161 | 9.72E-214 | 1.81668259 | 0.39 | 0.171 | 1.88E-209 | 3 | Il16 | Col18a1 |
| Sdc2 | 2.94E-210 | 0.79561356 | 0.938 | 0.845 | 5.69E-206 | 3 | Sdc2 | Col18a1 |
| Nr4a11 | 3.49E-208 | 0.72811958 | 0.689 | 0.626 | 6.74E-204 | 3 | Nr4a1 | Col18a1 |
| Flt1 | 5.22E-208 | 1.35571969 | 0.264 | 0.155 | 1.01E-203 | 3 | Flt1 | Col18a1 |
| Sav1 | 9.68E-208 | 0.97219119 | 0.914 | 0.757 | 1.87E-203 | 3 | Sav1 | Col18a1 |
| Klf7 | 4.29E-206 | 1.26997893 | 0.63 | 0.41 | 8.29E-202 | 3 | Klf7 | Col18a1 |
| Egln3 | 1.10E-203 | 1.01946562 | 0.312 | 0.178 | 2.13E-199 | 3 | Egln3 | Col18a1 |
| P2ry2 | 9.69E-203 | 1.50788033 | 0.528 | 0.27 | 1.87E-198 | 3 | P2ry2 | Col18a1 |
| Il331 | 5.73E-201 | 0.72211912 | 0.413 | 0.399 | 1.11E-196 | 3 | Il33 | Col18a1 |
| Cebpb | 5.79E-201 | 0.40248903 | 0.986 | 0.96 | 1.12E-196 | 3 | Cebpb | Col18a1 |
| Ptk71 | 6.19E-197 | 0.55757325 | 0.98 | 0.801 | 1.20E-192 | 3 | Ptk7 | Col18a1 |
| Apcdd11 | 8.05E-196 | 0.9829401 | 0.674 | 0.446 | 1.56E-191 | 3 | Apcdd1 | Col18a1 |
| Tacc1 | 4.77E-195 | 0.78776285 | 0.945 | 0.833 | 9.23E-191 | 3 | Tacc1 | Col18a1 |
| Zfp36l21 | 1.04E-191 | 0.70849841 | 0.921 | 0.827 | 2.00E-187 | 3 | Zfp36l2 | Col18a1 |
| Junb1 | 1.04E-187 | 0.50682427 | 0.947 | 0.931 | 2.00E-183 | 3 | Junb | Col18a1 |
| Gk1 | 2.39E-186 | 1.52873272 | 0.577 | 0.308 | 4.61E-182 | 3 | Gk | Col18a1 |
| Aldoa | 5.29E-186 | 0.61583991 | 0.966 | 0.915 | 1.02E-181 | 3 | Aldoa | Col18a1 |
| Pear11 | 9.08E-186 | 0.92137775 | 0.812 | 0.598 | 1.76E-181 | 3 | Pear1 | Col18a1 |
| Tfpi | 1.66E-185 | 0.84554186 | 0.892 | 0.797 | 3.20E-181 | 3 | Tfpi | Col18a1 |
| Mgst3 | 8.38E-185 | 1.15907854 | 0.601 | 0.376 | 1.62E-180 | 3 | Mgst3 | Col18a1 |
| Gpr35 | 7.11E-184 | 1.45207864 | 0.491 | 0.207 | 1.37E-179 | 3 | Gpr35 | Col18a1 |
| Tcp11l21 | 8.09E-183 | 0.95954245 | 0.78 | 0.545 | 1.57E-178 | 3 | Tcp11l2 | Col18a1 |
| Cblb | 1.12E-182 | 0.7728627 | 0.896 | 0.778 | 2.17E-178 | 3 | Cblb | Col18a1 |
| Nptn | 1.31E-181 | 0.70024552 | 0.974 | 0.906 | 2.53E-177 | 3 | Nptn | Col18a1 |
| Ccnd21 | 5.37E-180 | 0.82000304 | 0.82 | 0.684 | 1.04E-175 | 3 | Ccnd2 | Col18a1 |
| Slc2a1 | 1.50E-179 | 0.62393417 | 0.787 | 0.697 | 2.90E-175 | 3 | Slc2a1 | Col18a1 |
| Nlgn2 | 2.86E-177 | 0.69369717 | 0.977 | 0.908 | 5.53E-173 | 3 | Nlgn2 | Col18a1 |
| Egr2 | 3.39E-177 | 0.93337826 | 0.696 | 0.466 | 6.56E-173 | 3 | Egr2 | Col18a1 |
| Rasgrp2 | 4.07E-177 | 2.42944079 | 0.293 | 0.077 | 7.88E-173 | 3 | Rasgrp2 | Col18a1 |
| Nr3c1 | 3.13E-176 | 0.96496318 | 0.856 | 0.676 | 6.05E-172 | 3 | Nr3c1 | Col18a1 |
| Litaf | 4.32E-176 | 0.64074104 | 0.977 | 0.924 | 8.36E-172 | 3 | Litaf | Col18a1 |
| Tmem45a | 1.77E-174 | 0.82987885 | 0.872 | 0.737 | 3.41E-170 | 3 | Tmem45a | Col18a1 |
| Tnfsf111 | 2.15E-174 | 1.23523024 | 0.378 | 0.304 | 4.15E-170 | 3 | Tnfsf11 | Col18a1 |
| Pi15 | 2.73E-174 | 1.41214451 | 0.38 | 0.229 | 5.27E-170 | 3 | Pi15 | Col18a1 |
| Ipmk | 1.03E-170 | 1.32318431 | 0.642 | 0.411 | 1.99E-166 | 3 | Ipmk | Col18a1 |
| A4galt | 1.75E-168 | 0.84567556 | 0.812 | 0.661 | 3.39E-164 | 3 | A4galt | Col18a1 |
| Fndc3a | 1.75E-167 | 0.97456235 | 0.791 | 0.674 | 3.38E-163 | 3 | Fndc3a | Col18a1 |
| Msmo1 | 2.20E-167 | 1.273813 | 0.48 | 0.314 | 4.25E-163 | 3 | Msmo1 | Col18a1 |
| Sept111 | 3.66E-167 | 0.73550125 | 0.962 | 0.845 | 7.07E-163 | 3 | 11-Sep | Col18a1 |
| Adgrl31 | 9.43E-165 | 1.4989437 | 0.443 | 0.21 | 1.82E-160 | 3 | Adgrl3 | Col18a1 |
| Megf61 | 6.03E-163 | 1.13345414 | 0.314 | 0.188 | 1.17E-158 | 3 | Megf6 | Col18a1 |
| Adcy31 | 4.12E-162 | 0.90305643 | 0.878 | 0.7 | 7.97E-158 | 3 | Adcy3 | Col18a1 |
| Nr2f1 | 7.03E-162 | 1.45385274 | 0.466 | 0.246 | 1.36E-157 | 3 | Nr2f1 | Col18a1 |
| Fdps1 | 1.25E-161 | 0.95477085 | 0.678 | 0.551 | 2.41E-157 | 3 | Fdps | Col18a1 |
| Cpq | 1.56E-160 | 1.2099651 | 0.707 | 0.453 | 3.01E-156 | 3 | Cpq | Col18a1 |
| Htra11 | 1.60E-160 | 0.5849826 | 0.928 | 0.815 | 3.10E-156 | 3 | Htra1 | Col18a1 |
| Nbea | 6.06E-158 | 1.39271186 | 0.599 | 0.344 | 1.17E-153 | 3 | Nbea | Col18a1 |
| Pink11 | 5.72E-156 | 0.80302493 | 0.882 | 0.698 | 1.11E-151 | 3 | Pink1 | Col18a1 |
| Kitl | 9.65E-156 | 1.6421508 | 0.422 | 0.193 | 1.87E-151 | 3 | Kitl | Col18a1 |
| Cept11 | 3.27E-155 | 0.85724279 | 0.859 | 0.721 | 6.32E-151 | 3 | Cept1 | Col18a1 |
| Ecrg42 | 6.88E-155 | 0.85834841 | 0.43 | 0.384 | 1.33E-150 | 3 | Ecrg4 | Col18a1 |
| Ak3 | 4.60E-153 | 0.78231516 | 0.886 | 0.771 | 8.89E-149 | 3 | Ak3 | Col18a1 |
| Plcb1 | 3.83E-152 | 1.59711106 | 0.41 | 0.211 | 7.41E-148 | 3 | Plcb1 | Col18a1 |
| Fosl2 | 6.77E-152 | 0.69357209 | 0.936 | 0.86 | 1.31E-147 | 3 | Fosl2 | Col18a1 |
| Zeb2 | 1.46E-151 | 0.58566526 | 0.976 | 0.92 | 2.82E-147 | 3 | Zeb2 | Col18a1 |
| Tspan41 | 3.23E-151 | 0.50477624 | 0.975 | 0.925 | 6.24E-147 | 3 | Tspan4 | Col18a1 |
| Rbms1 | 5.00E-150 | 0.66753028 | 0.972 | 0.904 | 9.66E-146 | 3 | Rbms1 | Col18a1 |
| Jarid2 | 6.80E-150 | 1.05718724 | 0.623 | 0.425 | 1.31E-145 | 3 | Jarid2 | Col18a1 |
| Ddx5 | 4.75E-149 | 0.84543711 | 0.909 | 0.81 | 9.18E-145 | 3 | Ddx5 | Col18a1 |
| Jak21 | 6.72E-149 | 0.77888495 | 0.917 | 0.799 | 1.30E-144 | 3 | Jak2 | Col18a1 |
| Sh2b3 | 7.34E-149 | 0.98070045 | 0.77 | 0.541 | 1.42E-144 | 3 | Sh2b3 | Col18a1 |
| Lin7a | 1.80E-147 | 1.78157165 | 0.365 | 0.134 | 3.47E-143 | 3 | Lin7a | Col18a1 |
| Rem11 | 2.88E-146 | 0.87159761 | 0.657 | 0.423 | 5.58E-142 | 3 | Rem1 | Col18a1 |
| Ptprk1 | 7.32E-144 | 1.48958647 | 0.524 | 0.267 | 1.41E-139 | 3 | Ptprk | Col18a1 |
| Ror1 | 6.34E-143 | 1.26058129 | 0.609 | 0.381 | 1.23E-138 | 3 | Ror1 | Col18a1 |
| Mmp19 | 1.92E-142 | 0.56892761 | 0.965 | 0.929 | 3.71E-138 | 3 | Mmp19 | Col18a1 |
| Prkar1a1 | 5.28E-142 | 0.5220527 | 0.977 | 0.939 | 1.02E-137 | 3 | Prkar1a | Col18a1 |
| Frzb1 | 1.01E-140 | 1.28579896 | 0.412 | 0.21 | 1.96E-136 | 3 | Frzb | Col18a1 |
| Ccnd11 | 3.64E-140 | 1.23240391 | 0.309 | 0.19 | 7.03E-136 | 3 | Ccnd1 | Col18a1 |
| Cd82 | 4.19E-140 | 1.12964285 | 0.598 | 0.379 | 8.09E-136 | 3 | Cd82 | Col18a1 |
| Ptprg | 7.88E-139 | 1.14534998 | 0.701 | 0.457 | 1.52E-134 | 3 | Ptprg | Col18a1 |
| Pfkl | 1.72E-138 | 0.70275865 | 0.829 | 0.683 | 3.33E-134 | 3 | Pfkl | Col18a1 |
| Pik3r11 | 2.05E-138 | 0.73990697 | 0.831 | 0.696 | 3.97E-134 | 3 | Pik3r1 | Col18a1 |
| Sdc41 | 3.23E-138 | 0.67087366 | 0.886 | 0.75 | 6.24E-134 | 3 | Sdc4 | Col18a1 |
| Klf9 | 8.38E-138 | 0.55843305 | 0.915 | 0.847 | 1.62E-133 | 3 | Klf9 | Col18a1 |
| Sp3 | 8.95E-138 | 0.91752898 | 0.883 | 0.733 | 1.73E-133 | 3 | Sp3 | Col18a1 |
| Ggta1 | 1.82E-137 | 1.12564641 | 0.697 | 0.475 | 3.51E-133 | 3 | Ggta1 | Col18a1 |
| Tmem30a | 5.43E-135 | 0.87230194 | 0.877 | 0.731 | 1.05E-130 | 3 | Tmem30a | Col18a1 |
| Igf2bp2 | 4.14E-134 | 1.16065812 | 0.572 | 0.381 | 8.00E-130 | 3 | Igf2bp2 | Col18a1 |
| Slco5a11 | 1.11E-133 | 1.19561445 | 0.614 | 0.388 | 2.14E-129 | 3 | Slco5a1 | Col18a1 |
| Slc44a1 | 5.61E-133 | 0.81202935 | 0.91 | 0.796 | 1.08E-128 | 3 | Slc44a1 | Col18a1 |
| Fam110c | 2.17E-132 | 1.35543881 | 0.356 | 0.171 | 4.19E-128 | 3 | Fam110c | Col18a1 |
| Gli2 | 2.21E-132 | 1.23702037 | 0.62 | 0.375 | 4.28E-128 | 3 | Gli2 | Col18a1 |
| Pcdh1 | 1.06E-130 | 1.6843898 | 0.311 | 0.124 | 2.05E-126 | 3 | Pcdh1 | Col18a1 |
| Cd81 | 1.01E-129 | 0.46461706 | 0.984 | 0.96 | 1.96E-125 | 3 | Cd81 | Col18a1 |
| Pgr | 2.99E-128 | 1.46669411 | 0.278 | 0.096 | 5.77E-124 | 3 | Pgr | Col18a1 |
| Ecscr | 1.78E-127 | 1.08709475 | 0.693 | 0.489 | 3.43E-123 | 3 | Ecscr | Col18a1 |
| Cdo1 | 2.40E-127 | 1.42974566 | 0.291 | 0.172 | 4.63E-123 | 3 | Cdo1 | Col18a1 |
| Nr4a31 | 3.97E-127 | 1.40990799 | 0.311 | 0.224 | 7.68E-123 | 3 | Nr4a3 | Col18a1 |
| Bhlhe40 | 5.12E-127 | 0.57825623 | 0.924 | 0.845 | 9.91E-123 | 3 | Bhlhe40 | Col18a1 |
| Tpi1 | 2.53E-126 | 0.6284522 | 0.927 | 0.829 | 4.90E-122 | 3 | Tpi1 | Col18a1 |
| Fgf10 | 1.89E-124 | 1.86756265 | 0.318 | 0.108 | 3.66E-120 | 3 | Fgf10 | Col18a1 |
| BC028528 | 6.82E-124 | 1.17807591 | 0.596 | 0.369 | 1.32E-119 | 3 | BC028528 | Col18a1 |
| Gbe1 | 1.45E-123 | 1.31118967 | 0.422 | 0.253 | 2.80E-119 | 3 | Gbe1 | Col18a1 |
| Cyp511 | 4.00E-123 | 0.83111194 | 0.725 | 0.608 | 7.73E-119 | 3 | Cyp51 | Col18a1 |
| Tgfbr31 | 1.11E-122 | 0.52650094 | 0.964 | 0.842 | 2.14E-118 | 3 | Tgfbr3 | Col18a1 |
| Srebf2 | 1.69E-122 | 0.64861046 | 0.901 | 0.79 | 3.26E-118 | 3 | Srebf2 | Col18a1 |
| Tpst1 | 1.72E-122 | 0.8112457 | 0.886 | 0.778 | 3.32E-118 | 3 | Tpst1 | Col18a1 |
| Ptprn1 | 4.59E-122 | 0.71804367 | 0.786 | 0.54 | 8.88E-118 | 3 | Ptprn | Col18a1 |
| P4ha1 | 5.71E-121 | 0.55813961 | 0.981 | 0.931 | 1.10E-116 | 3 | P4ha1 | Col18a1 |
| Gsr1 | 5.82E-121 | 0.81637768 | 0.852 | 0.703 | 1.12E-116 | 3 | Gsr | Col18a1 |
| Pitpnc11 | 1.93E-120 | 0.95680194 | 0.707 | 0.543 | 3.73E-116 | 3 | Pitpnc1 | Col18a1 |
| Lxn | 2.12E-120 | 1.12754555 | 0.687 | 0.468 | 4.10E-116 | 3 | Lxn | Col18a1 |
| Scd2 | 6.48E-120 | 0.66205459 | 0.867 | 0.76 | 1.25E-115 | 3 | Scd2 | Col18a1 |
| Tgfbr1 | 1.26E-119 | 0.85988399 | 0.884 | 0.732 | 2.44E-115 | 3 | Tgfbr1 | Col18a1 |
| Dram2 | 1.30E-118 | 1.12727136 | 0.691 | 0.465 | 2.51E-114 | 3 | Dram2 | Col18a1 |
| Adamtsl5 | 1.46E-118 | 0.94850982 | 0.702 | 0.473 | 2.83E-114 | 3 | Adamtsl5 | Col18a1 |
| Lrig1 | 4.68E-118 | 0.97561099 | 0.787 | 0.587 | 9.05E-114 | 3 | Lrig1 | Col18a1 |
| Ankrd37 | 1.78E-116 | 1.24581504 | 0.309 | 0.152 | 3.44E-112 | 3 | Ankrd37 | Col18a1 |
| Mmd1 | 1.89E-116 | 0.78466937 | 0.792 | 0.633 | 3.66E-112 | 3 | Mmd | Col18a1 |
| Ptgs21 | 2.95E-116 | 0.86149554 | 0.483 | 0.43 | 5.71E-112 | 3 | Ptgs2 | Col18a1 |
| Ccdc85b | 4.53E-116 | 0.84662266 | 0.664 | 0.481 | 8.76E-112 | 3 | Ccdc85b | Col18a1 |
| Cadm21 | 6.76E-116 | 1.55258665 | 0.366 | 0.17 | 1.31E-111 | 3 | Cadm2 | Col18a1 |
| Inf2 | 5.32E-115 | 0.67259126 | 0.879 | 0.721 | 1.03E-110 | 3 | Inf2 | Col18a1 |
| Man2a21 | 7.19E-115 | 0.61973466 | 0.923 | 0.834 | 1.39E-110 | 3 | Man2a2 | Col18a1 |
| Kazn1 | 1.03E-113 | 0.85942066 | 0.62 | 0.393 | 2.00E-109 | 3 | Kazn | Col18a1 |
| Fryl1 | 2.02E-113 | 1.03492156 | 0.696 | 0.487 | 3.91E-109 | 3 | Fryl | Col18a1 |
| Polg21 | 2.07E-113 | 0.91799197 | 0.355 | 0.32 | 4.01E-109 | 3 | Polg2 | Col18a1 |
| Wnt5a1 | 7.33E-113 | 0.44811033 | 0.944 | 0.732 | 1.42E-108 | 3 | Wnt5a | Col18a1 |
| Arsj | 1.10E-111 | 2.02346595 | 0.255 | 0.098 | 2.13E-107 | 3 | Arsj | Col18a1 |
| Adm | 9.46E-111 | 0.56873854 | 0.538 | 0.447 | 1.83E-106 | 3 | Adm | Col18a1 |
| Idi1 | 3.63E-110 | 1.67247499 | 0.314 | 0.145 | 7.02E-106 | 3 | Idi1 | Col18a1 |
| Tmem59 | 6.87E-109 | 0.58123806 | 0.962 | 0.908 | 1.33E-104 | 3 | Tmem59 | Col18a1 |
| Hivep3 | 1.10E-107 | 0.82797948 | 0.686 | 0.525 | 2.13E-103 | 3 | Hivep3 | Col18a1 |
| Atxn7l11 | 3.39E-107 | 0.828448 | 0.692 | 0.543 | 6.56E-103 | 3 | Atxn7l1 | Col18a1 |

| | | | | | | | | |
|---|---|---|---|---|---|---|---|---|
| P2rx4 | 1.93E-106 | 0.82679957 | 0.783 | 0.615 | 3.73E-102 | 3 | P2rx4 | Col18a1 |
| B3gnt3 | 2.30E-106 | 1.0044282 | 0.618 | 0.438 | 4.45E-102 | 3 | B3gnt3 | Col18a1 |
| Stc2 | 3.46E-106 | 1.17530992 | 0.347 | 0.179 | 6.69E-102 | 3 | Stc2 | Col18a1 |
| Gab1 | 1.73E-105 | 1.09927093 | 0.547 | 0.385 | 3.35E-101 | 3 | Gab1 | Col18a1 |
| Hdac5 | 8.84E-105 | 0.72914302 | 0.826 | 0.664 | 1.71E-100 | 3 | Hdac5 | Col18a1 |
| Pld11 | 2.00E-104 | 1.13071026 | 0.572 | 0.354 | 3.87E-100 | 3 | Pld1 | Col18a1 |
| Adra2a1 | 1.59E-103 | 0.59998391 | 0.553 | 0.411 | 3.08E-99 | 3 | Adra2a | Col18a1 |
| Art3 | 3.18E-103 | 1.42454508 | 0.328 | 0.166 | 6.15E-99 | 3 | Art3 | Col18a1 |
| Limd1 | 1.53E-102 | 0.69060447 | 0.856 | 0.787 | 2.96E-98 | 3 | Limd1 | Col18a1 |
| Fam117b | 5.37E-102 | 0.76376397 | 0.703 | 0.566 | 1.04E-97 | 3 | Fam117b | Col18a1 |
| Eps81 | 9.16E-102 | 1.17233062 | 0.56 | 0.346 | 1.77E-97 | 3 | Eps8 | Col18a1 |
| Scarb11 | 1.86E-101 | 1.03059841 | 0.707 | 0.465 | 3.59E-97 | 3 | Scarb1 | Col18a1 |
| Arrdc3 | 3.72E-101 | 0.77742555 | 0.516 | 0.358 | 7.20E-97 | 3 | Arrdc3 | Col18a1 |
| Serinc1 | 4.91E-101 | 0.65895116 | 0.945 | 0.859 | 9.49E-97 | 3 | Serinc1 | Col18a1 |
| Hpse | 5.26E-101 | 1.30587361 | 0.541 | 0.312 | 1.02E-96 | 3 | Hpse | Col18a1 |
| Bnip3 | 1.15E-100 | 1.11515977 | 0.399 | 0.219 | 2.22E-96 | 3 | Bnip3 | Col18a1 |
| Anxa7 | 1.62E-98 | 0.72143199 | 0.911 | 0.796 | 3.14E-94 | 3 | Anxa7 | Col18a1 |
| Dio32 | 2.97E-98 | 0.83701722 | 0.433 | 0.369 | 5.73E-94 | 3 | Dio3 | Col18a1 |
| Eif4a2 | 7.54E-97 | 0.59927628 | 0.937 | 0.871 | 1.46E-92 | 3 | Eif4a2 | Col18a1 |
| Lamc1 | 8.02E-97 | 0.41688958 | 0.993 | 0.966 | 1.55E-92 | 3 | Lamc1 | Col18a1 |
| Prkx | 9.95E-97 | 1.25650085 | 0.481 | 0.276 | 1.92E-92 | 3 | Prkx | Col18a1 |
| Pnrc11 | 2.60E-96 | 0.5233633 | 0.933 | 0.868 | 5.02E-92 | 3 | Pnrc1 | Col18a1 |
| Btbd3 | 5.68E-95 | 1.13319956 | 0.562 | 0.362 | 1.10E-90 | 3 | Btbd3 | Col18a1 |
| Eif3f | 6.21E-95 | 0.67138658 | 0.947 | 0.864 | 1.20E-90 | 3 | Eif3f | Col18a1 |
| Tspan17 | 8.13E-95 | 0.84559718 | 0.629 | 0.464 | 1.57E-90 | 3 | Tspan17 | Col18a1 |
| Slit21 | 9.83E-95 | 0.71063126 | 0.812 | 0.622 | 1.90E-90 | 3 | Slit2 | Col18a1 |
| Tmem51 | 1.78E-94 | 1.68695793 | 0.278 | 0.098 | 3.44E-90 | 3 | Tmem51 | Col18a1 |
| Nr1h3 | 2.18E-94 | 1.58212971 | 0.267 | 0.109 | 4.21E-90 | 3 | Nr1h3 | Col18a1 |
| Xbp1 | 7.56E-94 | 0.49873376 | 0.987 | 0.945 | 1.46E-89 | 3 | Xbp1 | Col18a1 |
| Arhgap421 | 8.99E-94 | 0.57944379 | 0.876 | 0.764 | 1.74E-89 | 3 | Arhgap42 | Col18a1 |
| Fstl3 | 1.19E-92 | 0.57651416 | 0.817 | 0.679 | 2.30E-88 | 3 | Fstl3 | Col18a1 |
| Sep-41 | 2.91E-91 | 0.83605936 | 0.566 | 0.351 | 5.63E-87 | 3 | 4-Sep | Col18a1 |
| Ctsk1 | 4.21E-91 | 0.29251463 | 0.993 | 0.969 | 8.14E-87 | 3 | Ctsk | Col18a1 |
| Il13ra11 | 5.66E-91 | 0.69835909 | 0.932 | 0.822 | 1.10E-86 | 3 | Il13ra1 | Col18a1 |
| Soat1 | 9.35E-91 | 0.60915374 | 0.929 | 0.834 | 1.81E-86 | 3 | Soat1 | Col18a1 |
| Slc16a3 | 1.51E-90 | 0.57109951 | 0.781 | 0.636 | 2.92E-86 | 3 | Slc16a3 | Col18a1 |
| Nid1 | 1.57E-90 | 0.289067 | 0.997 | 0.979 | 3.03E-86 | 3 | Nid1 | Col18a1 |
| Ube2v21 | 2.01E-90 | 1.00885594 | 0.641 | 0.531 | 3.88E-86 | 3 | Ube2v2 | Col18a1 |
| Sesn3 | 2.46E-90 | 0.8365383 | 0.684 | 0.509 | 4.75E-86 | 3 | Sesn3 | Col18a1 |
| Acss3 | 2.18E-89 | 1.51542277 | 0.345 | 0.16 | 4.22E-85 | 3 | Acss3 | Col18a1 |
| Egln1 | 5.87E-89 | 0.50781318 | 0.913 | 0.861 | 1.13E-84 | 3 | Egln1 | Col18a1 |
| Stk24 | 2.39E-88 | 0.74891831 | 0.873 | 0.783 | 4.63E-84 | 3 | Stk24 | Col18a1 |
| Fam129b | 3.12E-88 | 0.57108391 | 0.945 | 0.873 | 6.04E-84 | 3 | Fam129b | Col18a1 |
| Nfil3 | 7.54E-88 | 0.78429243 | 0.766 | 0.635 | 1.46E-83 | 3 | Nfil3 | Col18a1 |
| Steap2 | 1.76E-87 | 0.78266704 | 0.796 | 0.658 | 3.40E-83 | 3 | Steap2 | Col18a1 |
| Lipo31 | 2.71E-87 | 1.22632624 | 0.492 | 0.288 | 5.24E-83 | 3 | Lipo3 | Col18a1 |
| Pcgf5 | 4.54E-87 | 1.14457695 | 0.488 | 0.308 | 8.77E-83 | 3 | Pcgf5 | Col18a1 |
| Laptm4a | 5.16E-87 | 0.53773026 | 0.979 | 0.917 | 9.98E-83 | 3 | Laptm4a | Col18a1 |
| Gpc6 | 6.87E-87 | 0.93796542 | 0.655 | 0.519 | 1.33E-82 | 3 | Gpc6 | Col18a1 |
| Vdac2 | 8.06E-87 | 0.91736387 | 0.793 | 0.606 | 1.56E-82 | 3 | Vdac2 | Col18a1 |
| Gla | 1.66E-86 | 1.170264 | 0.513 | 0.335 | 3.20E-82 | 3 | Gla | Col18a1 |
| Rnf1491 | 6.01E-86 | 0.38407269 | 0.971 | 0.886 | 1.16E-81 | 3 | Rnf149 | Col18a1 |
| Adgrl2 | 1.11E-85 | 1.07851118 | 0.657 | 0.422 | 2.14E-81 | 3 | Adgrl2 | Col18a1 |
| Map1lc3b | 1.90E-85 | 0.53263632 | 0.962 | 0.892 | 3.67E-81 | 3 | Map1lc3b | Col18a1 |
| Map4k41 | 1.98E-84 | 0.53974264 | 0.957 | 0.889 | 3.83E-80 | 3 | Map4k4 | Col18a1 |
| Cyp39a11 | 2.79E-83 | 1.24206494 | 0.426 | 0.213 | 5.40E-79 | 3 | Cyp39a1 | Col18a1 |
| Parvb | 1.06E-82 | 1.37635818 | 0.326 | 0.133 | 2.06E-78 | 3 | Parvb | Col18a1 |
| Gna121 | 1.05E-81 | 0.59910516 | 0.93 | 0.833 | 2.03E-77 | 3 | Gna12 | Col18a1 |
| Lbh1 | 1.06E-81 | 0.39995138 | 0.895 | 0.797 | 2.04E-77 | 3 | Lbh | Col18a1 |
| Dnajb91 | 1.80E-81 | 0.84348953 | 0.616 | 0.567 | 3.49E-77 | 3 | Dnajb9 | Col18a1 |
| Use1 | 2.65E-81 | 0.66504385 | 0.909 | 0.794 | 5.11E-77 | 3 | Use1 | Col18a1 |
| Svil | 4.67E-81 | 0.58644383 | 0.904 | 0.795 | 9.02E-77 | 3 | Svil | Col18a1 |
| Tspan5 | 6.45E-81 | 0.99689797 | 0.633 | 0.459 | 1.25E-76 | 3 | Tspan5 | Col18a1 |
| Tmem38b1 | 1.89E-80 | 0.94321329 | 0.579 | 0.407 | 3.65E-76 | 3 | Tmem38b | Col18a1 |
| Rbms2 | 3.37E-80 | 0.64579539 | 0.925 | 0.843 | 6.51E-76 | 3 | Rbms2 | Col18a1 |
| Ets1 | 4.90E-80 | 0.60296783 | 0.826 | 0.691 | 9.47E-76 | 3 | Ets1 | Col18a1 |
| Rnf24 | 6.73E-80 | 1.09259575 | 0.617 | 0.394 | 1.30E-75 | 3 | Rnf24 | Col18a1 |
| Shc11 | 1.81E-79 | 0.54519045 | 0.949 | 0.874 | 3.50E-75 | 3 | Shc1 | Col18a1 |
| Pdk1 | 7.27E-79 | 0.82027856 | 0.58 | 0.415 | 1.41E-74 | 3 | Pdk1 | Col18a1 |
| B4galnt11 | 9.09E-79 | 0.8085018 | 0.745 | 0.511 | 1.76E-74 | 3 | B4galnt1 | Col18a1 |
| Myo1b1 | 3.29E-78 | 0.7155011 | 0.885 | 0.713 | 6.35E-74 | 3 | Myo1b | Col18a1 |
| Gpcpd11 | 9.02E-78 | 1.08440188 | 0.529 | 0.38 | 1.74E-73 | 3 | Gpcpd1 | Col18a1 |
| Apbb21 | 2.49E-77 | 0.6983581 | 0.856 | 0.704 | 4.82E-73 | 3 | Apbb2 | Col18a1 |
| Sema6a | 2.70E-77 | 1.66206939 | 0.299 | 0.118 | 5.21E-73 | 3 | Sema6a | Col18a1 |
| Hs3st11 | 6.82E-77 | 0.50251764 | 0.771 | 0.595 | 1.32E-72 | 3 | Hs3st1 | Col18a1 |
| Jak1 | 7.06E-77 | 0.56023289 | 0.945 | 0.893 | 1.37E-72 | 3 | Jak1 | Col18a1 |
| DEPRECATED-l | 1.61E-76 | 1.13859536 | 0.472 | 0.281 | 3.11E-72 | 3 | DEPRECATED-l | Col18a1 |
| Pla2r1 | 1.63E-76 | 1.02338674 | 0.586 | 0.373 | 3.14E-72 | 3 | Pla2r1 | Col18a1 |
| Gng8 | 1.93E-76 | 1.25742232 | 0.324 | 0.149 | 3.74E-72 | 3 | Gng8 | Col18a1 |
| Spry41 | 2.60E-76 | 0.82931671 | 0.599 | 0.411 | 5.03E-72 | 3 | Spry4 | Col18a1 |
| Bnip3l | 7.04E-76 | 0.98220597 | 0.598 | 0.389 | 1.36E-71 | 3 | Bnip3l | Col18a1 |
| Cyb5a | 7.49E-76 | 0.73017738 | 0.841 | 0.661 | 1.45E-71 | 3 | Cyb5a | Col18a1 |
| Cdkn1c | 8.19E-76 | 1.05976763 | 0.265 | 0.15 | 1.58E-71 | 3 | Cdkn1c | Col18a1 |
| Trps1 | 1.39E-75 | 0.78889097 | 0.44 | 0.231 | 2.69E-71 | 3 | Trps1 | Col18a1 |
| Slc6a81 | 1.92E-75 | 0.47630201 | 0.899 | 0.85 | 3.71E-71 | 3 | Slc6a8 | Col18a1 |
| Meis11 | 2.17E-75 | 0.66557496 | 0.728 | 0.567 | 4.20E-71 | 3 | Meis1 | Col18a1 |
| Slco4a1 | 4.93E-75 | 1.00825357 | 0.319 | 0.171 | 9.53E-71 | 3 | Slco4a1 | Col18a1 |
| mt-Nd41 | 2.50E-74 | 0.31168242 | 0.685 | 0.579 | 4.83E-70 | 3 | mt-Nd4 | Col18a1 |
| Phlda11 | 3.40E-74 | 0.68312495 | 0.742 | 0.593 | 6.57E-70 | 3 | Phlda1 | Col18a1 |
| Cebpa | 5.15E-74 | 0.50035482 | 0.752 | 0.616 | 9.95E-70 | 3 | Cebpa | Col18a1 |
| Map1lc3a1 | 1.63E-73 | 0.51001518 | 0.958 | 0.875 | 3.15E-69 | 3 | Map1lc3a | Col18a1 |
| Mcoln21 | 1.88E-73 | 0.71917847 | 0.705 | 0.52 | 3.64E-69 | 3 | Mcoln2 | Col18a1 |
| Tpp11 | 2.80E-73 | 0.60075253 | 0.867 | 0.763 | 5.41E-69 | 3 | Tpp1 | Col18a1 |
| Frmd6 | 1.07E-72 | 0.77808708 | 0.765 | 0.606 | 2.07E-68 | 3 | Frmd6 | Col18a1 |
| Zic11 | 2.50E-72 | 1.0110019 | 0.407 | 0.258 | 4.83E-68 | 3 | Zic1 | Col18a1 |
| S100a1 | 9.56E-72 | 0.63383071 | 0.819 | 0.688 | 1.85E-67 | 3 | S100a1 | Col18a1 |
| Cacna1g1 | 1.01E-71 | 0.66707061 | 0.707 | 0.558 | 1.95E-67 | 3 | Cacna1g | Col18a1 |
| Stat3 | 1.12E-71 | 0.54589615 | 0.979 | 0.903 | 2.16E-67 | 3 | Stat3 | Col18a1 |
| Tspan11 | 3.50E-71 | 0.55393094 | 0.874 | 0.719 | 6.77E-67 | 3 | Tspan11 | Col18a1 |
| Cdkn1a1 | 3.77E-71 | 0.44258852 | 0.529 | 0.413 | 7.28E-67 | 3 | Cdkn1a | Col18a1 |
| Cask1 | 5.58E-71 | 0.72249218 | 0.824 | 0.686 | 1.08E-66 | 3 | Cask | Col18a1 |
| Wdr59 | 1.50E-70 | 1.29213334 | 0.371 | 0.203 | 2.90E-66 | 3 | Wdr59 | Col18a1 |
| Chka1 | 3.71E-70 | 1.14082297 | 0.417 | 0.318 | 7.16E-66 | 3 | Chka | Col18a1 |
| Mxi1 | 4.59E-70 | 0.72556326 | 0.643 | 0.494 | 8.87E-66 | 3 | Mxi1 | Col18a1 |
| Fdft1 | 6.23E-70 | 0.75548071 | 0.617 | 0.49 | 1.21E-65 | 3 | Fdft1 | Col18a1 |
| DEPRECATED-l | 1.47E-69 | 0.52275115 | 0.832 | 0.695 | 2.85E-65 | 3 | DEPRECATED-l | Col18a1 |
| Lmbrd1 | 1.74E-69 | 0.92521937 | 0.671 | 0.483 | 3.36E-65 | 3 | Lmbrd1 | Col18a1 |
| Ier21 | 1.91E-69 | 0.50953535 | 0.829 | 0.794 | 3.69E-65 | 3 | Ier2 | Col18a1 |
| Dlg2 | 5.09E-69 | 1.43830335 | 0.338 | 0.173 | 9.84E-65 | 3 | Dlg2 | Col18a1 |
| Iqsec1 | 5.23E-69 | 0.69688222 | 0.795 | 0.679 | 1.01E-64 | 3 | Iqsec1 | Col18a1 |
| Prxl2a1 | 5.95E-69 | 1.16948141 | 0.343 | 0.191 | 1.15E-64 | 3 | Prxl2a | Col18a1 |
| Adgre51 | 3.81E-68 | 0.69670421 | 0.815 | 0.589 | 7.36E-64 | 3 | Adgre5 | Col18a1 |
| Txnip | 7.39E-68 | 0.40138349 | 0.868 | 0.844 | 1.43E-63 | 3 | Txnip | Col18a1 |
| Egflam | 1.24E-67 | 1.08474314 | 0.286 | 0.127 | 2.40E-63 | 3 | Egflam | Col18a1 |
| Nfasc1 | 2.13E-67 | 0.94557447 | 0.556 | 0.368 | 4.11E-63 | 3 | Nfasc | Col18a1 |
| Aoc3 | 7.92E-67 | 0.54154343 | 0.647 | 0.647 | 1.53E-62 | 3 | Aoc3 | Col18a1 |
| Nsa2 | 1.78E-66 | 1.02061883 | 0.72 | 0.492 | 3.45E-62 | 3 | Nsa2 | Col18a1 |
| Nsdhl | 2.50E-66 | 1.08209551 | 0.406 | 0.29 | 4.82E-62 | 3 | Nsdhl | Col18a1 |
| Axin2 | 4.21E-66 | 0.78071092 | 0.664 | 0.478 | 8.13E-62 | 3 | Axin2 | Col18a1 |
| Tbl1x | 1.05E-65 | 0.81206624 | 0.829 | 0.662 | 2.03E-61 | 3 | Tbl1x | Col18a1 |
| Sntb1 | 2.19E-65 | 1.6318108 | 0.262 | 0.11 | 4.24E-61 | 3 | Sntb1 | Col18a1 |
| Ifitm2 | 3.95E-65 | 0.35359423 | 0.99 | 0.973 | 7.63E-61 | 3 | Ifitm2 | Col18a1 |
| Gng12 | 1.16E-64 | 0.53601429 | 0.949 | 0.866 | 2.25E-60 | 3 | Gng12 | Col18a1 |
| Dusp5 | 1.21E-64 | 0.74016147 | 0.456 | 0.388 | 2.33E-60 | 3 | Dusp5 | Col18a1 |
| Dnaja1 | 1.66E-64 | 0.68340193 | 0.811 | 0.761 | 3.20E-60 | 3 | Dnaja1 | Col18a1 |
| Arhgef71 | 2.71E-64 | 0.58960646 | 0.917 | 0.815 | 5.24E-60 | 3 | Arhgef7 | Col18a1 |
| Ryk | 5.54E-64 | 0.54625016 | 0.96 | 0.893 | 1.07E-59 | 3 | Ryk | Col18a1 |
| Lss | 8.52E-64 | 1.20262862 | 0.302 | 0.175 | 1.65E-59 | 3 | Lss | Col18a1 |
| Fez1 | 1.14E-63 | 0.84741974 | 0.555 | 0.383 | 2.21E-59 | 3 | Fez1 | Col18a1 |
| Htr1b1 | 3.50E-62 | 0.88409424 | 0.563 | 0.359 | 6.77E-58 | 3 | Htr1b | Col18a1 |
| Id3 | 4.19E-62 | 0.48814453 | 0.69 | 0.658 | 8.10E-58 | 3 | Id3 | Col18a1 |
| Hes1 | 4.42E-62 | 0.49318153 | 0.714 | 0.7 | 8.55E-58 | 3 | Hes1 | Col18a1 |
| Clk1 | 8.89E-62 | 0.54615395 | 0.893 | 0.805 | 1.72E-57 | 3 | Clk1 | Col18a1 |
| Kdm5b | 2.09E-61 | 0.63421067 | 0.818 | 0.687 | 4.05E-57 | 3 | Kdm5b | Col18a1 |
| Bsg | 5.17E-61 | 0.52834728 | 0.954 | 0.884 | 9.99E-57 | 3 | Bsg | Col18a1 |
| Lpl | 7.43E-61 | 0.42209445 | 0.689 | 0.574 | 1.44E-56 | 3 | Lpl | Col18a1 |
| Dtnb | 1.57E-60 | 0.89314547 | 0.548 | 0.365 | 3.04E-56 | 3 | Dtnb | Col18a1 |
| Adamts14 | 1.60E-60 | 0.74342635 | 0.6 | 0.494 | 3.09E-56 | 3 | Adamts14 | Col18a1 |
| Ripor31 | 7.93E-60 | 0.73390012 | 0.629 | 0.448 | 1.53E-55 | 3 | Ripor3 | Col18a1 |
| Isyna1 | 1.25E-59 | 0.77458356 | 0.54 | 0.466 | 2.42E-55 | 3 | Isyna1 | Col18a1 |
| Ctnnb1 | 1.68E-59 | 0.43772118 | 0.982 | 0.95 | 3.25E-55 | 3 | Ctnnb1 | Col18a1 |
| Angpt42 | 6.39E-59 | 0.69388245 | 0.563 | 0.441 | 1.24E-54 | 3 | Angpt4 | Col18a1 |
| Pde11a1 | 7.52E-59 | 1.02530529 | 0.316 | 0.218 | 1.45E-54 | 3 | Pde11a | Col18a1 |
| Pfkfb41 | 7.71E-59 | 0.86423872 | 0.538 | 0.382 | 1.49E-54 | 3 | Pfkfb4 | Col18a1 |
| Rap2c | 8.57E-59 | 0.85601785 | 0.669 | 0.518 | 1.66E-54 | 3 | Rap2c | Col18a1 |
| Dap | 1.09E-58 | 0.55345393 | 0.962 | 0.885 | 2.11E-54 | 3 | Dap | Col18a1 |
| Tle3 | 1.42E-58 | 0.55356511 | 0.878 | 0.792 | 2.74E-54 | 3 | Tle3 | Col18a1 |
| Ankrd28 | 1.56E-58 | 0.78272495 | 0.658 | 0.486 | 3.01E-54 | 3 | Ankrd28 | Col18a1 |
| Pcbp2 | 2.00E-58 | 0.68778625 | 0.914 | 0.777 | 3.88E-54 | 3 | Pcbp2 | Col18a1 |
| Slk | 2.39E-58 | 0.71691929 | 0.841 | 0.664 | 4.62E-54 | 3 | Slk | Col18a1 |
| Rin11 | 2.52E-58 | 0.74389957 | 0.571 | 0.401 | 4.88E-54 | 3 | Rin1 | Col18a1 |
| Cav21 | 4.00E-58 | 1.11198383 | 0.439 | 0.263 | 7.73E-54 | 3 | Cav2 | Col18a1 |
| Gda1 | 4.17E-58 | 0.54136676 | 0.751 | 0.499 | 8.05E-54 | 3 | Gda | Col18a1 |
| Rgma1 | 4.46E-58 | 0.54198633 | 0.808 | 0.687 | 8.63E-54 | 3 | Rgma | Col18a1 |
| Eif3h | 1.46E-57 | 0.81498076 | 0.817 | 0.649 | 2.83E-53 | 3 | Eif3h | Col18a1 |
| Abhd21 | 3.11E-57 | 0.40543384 | 0.951 | 0.892 | 6.01E-53 | 3 | Abhd2 | Col18a1 |
| Dgkz | 7.03E-57 | 0.74079036 | 0.638 | 0.569 | 1.36E-52 | 3 | Dgkz | Col18a1 |
| Rora | 8.72E-57 | 0.65990077 | 0.755 | 0.625 | 1.69E-52 | 3 | Rora | Col18a1 |
| Zfp395 | 9.93E-57 | 0.79597808 | 0.488 | 0.336 | 1.92E-52 | 3 | Zfp395 | Col18a1 |
| Raly1 | 1.02E-56 | 0.51112944 | 0.969 | 0.908 | 1.96E-52 | 3 | Raly | Col18a1 |
| Capn1 | 1.27E-56 | 0.68483627 | 0.814 | 0.654 | 2.45E-52 | 3 | Capn1 | Col18a1 |
| Glg1 | 4.03E-56 | 0.51390958 | 0.962 | 0.91 | 7.79E-52 | 3 | Glg1 | Col18a1 |
| Eva1b1 | 4.72E-56 | 0.43411057 | 0.912 | 0.832 | 9.12E-52 | 3 | Eva1b | Col18a1 |
| Sdcbp | 8.83E-56 | 0.48955496 | 0.971 | 0.906 | 1.71E-51 | 3 | Sdcbp | Col18a1 |
| Azin1 | 9.15E-56 | 0.96638308 | 0.659 | 0.498 | 1.77E-51 | 3 | Azin1 | Col18a1 |
| Lpin1 | 1.06E-55 | 1.17298941 | 0.343 | 0.179 | 2.06E-51 | 3 | Lpin1 | Col18a1 |
| Stom | 1.47E-55 | 0.93386637 | 0.614 | 0.46 | 2.84E-51 | 3 | Stom | Col18a1 |
| Dpysl2 | 2.20E-55 | 0.61562954 | 0.884 | 0.761 | 4.26E-51 | 3 | Dpysl2 | Col18a1 |
| Tcf20 | 6.50E-55 | 0.75461824 | 0.743 | 0.573 | 1.26E-50 | 3 | Tcf20 | Col18a1 |
| Arhgap35 | 6.51E-55 | 0.77350225 | 0.809 | 0.639 | 1.26E-50 | 3 | Arhgap35 | Col18a1 |
| Daam1 | 1.14E-54 | 0.86743833 | 0.515 | 0.411 | 2.21E-50 | 3 | Daam1 | Col18a1 |
| Spin1 | 1.27E-54 | 0.65606243 | 0.863 | 0.746 | 2.46E-50 | 3 | Spin1 | Col18a1 |
| Prickle11 | 2.40E-54 | 0.61068371 | 0.893 | 0.763 | 4.64E-50 | 3 | Prickle1 | Col18a1 |
| Ctsa | 1.20E-53 | 0.42951574 | 0.959 | 0.907 | 2.33E-49 | 3 | Ctsa | Col18a1 |
| Abcc91 | 1.52E-53 | 0.92183846 | 0.325 | 0.217 | 2.95E-49 | 3 | Abcc9 | Col18a1 |
| Epb41l11 | 3.43E-53 | 0.73261658 | 0.73 | 0.538 | 6.63E-49 | 3 | Epb41l1 | Col18a1 |
| Mapk3 | 6.24E-53 | 0.70341125 | 0.828 | 0.692 | 1.21E-48 | 3 | Mapk3 | Col18a1 |
| Rbms3 | 2.48E-52 | 0.44403457 | 0.938 | 0.864 | 4.79E-48 | 3 | Rbms3 | Col18a1 |
| Chm | 3.57E-52 | 0.84577968 | 0.589 | 0.411 | 6.91E-48 | 3 | Chm | Col18a1 |
| Fermt2 | 3.78E-52 | 0.49666243 | 0.951 | 0.863 | 7.30E-48 | 3 | Fermt2 | Col18a1 |
| Notch3 | 5.12E-52 | 0.95883217 | 0.315 | 0.179 | 9.89E-48 | 3 | Notch3 | Col18a1 |
| Cdkn2a1 | 7.52E-52 | 0.89083094 | 0.267 | 0.164 | 1.45E-47 | 3 | Cdkn2a | Col18a1 |
| Slit3 | 7.89E-52 | 0.32079332 | 0.99 | 0.965 | 1.53E-47 | 3 | Slit3 | Col18a1 |
| Ston11 | 9.36E-52 | 0.55173537 | 0.857 | 0.693 | 1.81E-47 | 3 | Ston1 | Col18a1 |
| Emilin2 | 9.99E-52 | 0.46474329 | 0.943 | 0.745 | 1.93E-47 | 3 | Emilin2 | Col18a1 |
| Hexim1 | 1.16E-51 | 0.66670341 | 0.754 | 0.675 | 2.24E-47 | 3 | Hexim1 | Col18a1 |
| Clip2 | 1.41E-51 | 0.5389437 | 0.836 | 0.703 | 2.72E-47 | 3 | Clip2 | Col18a1 |
| Naca | 1.80E-51 | 0.6037354 | 0.937 | 0.825 | 3.47E-47 | 3 | Naca | Col18a1 |
| Ubxn6 | 1.04E-50 | 0.78002538 | 0.706 | 0.535 | 2.01E-46 | 3 | Ubxn6 | Col18a1 |
| Hbp11 | 2.50E-50 | 0.49269727 | 0.889 | 0.788 | 4.83E-46 | 3 | Hbp1 | Col18a1 |
| Acp2 | 3.71E-50 | 0.94304895 | 0.556 | 0.41 | 7.17E-46 | 3 | Acp2 | Col18a1 |
| Plec | 3.90E-50 | 0.31225569 | 0.991 | 0.968 | 7.54E-46 | 3 | Plec | Col18a1 |
| Adcy2 | 6.01E-50 | 0.82450965 | 0.531 | 0.394 | 1.16E-45 | 3 | Adcy2 | Col18a1 |
| Rnf13 | 6.79E-50 | 0.71528638 | 0.708 | 0.552 | 1.31E-45 | 3 | Rnf13 | Col18a1 |
| Spg201 | 7.54E-50 | 0.6616158 | 0.782 | 0.646 | 1.46E-45 | 3 | Spg20 | Col18a1 |
| Col5a21 | 7.69E-50 | 0.31624902 | 0.996 | 0.98 | 1.49E-45 | 3 | Col5a2 | Col18a1 |
| Atp7a | 1.37E-49 | 0.81812202 | 0.618 | 0.484 | 2.65E-45 | 3 | Atp7a | Col18a1 |
| Tcf12 | 1.76E-49 | 0.59802361 | 0.916 | 0.801 | 3.41E-45 | 3 | Tcf12 | Col18a1 |
| Kmt2e | 1.87E-49 | 0.62177434 | 0.781 | 0.627 | 3.62E-45 | 3 | Kmt2e | Col18a1 |
| Casp6 | 2.12E-49 | 0.66575232 | 0.788 | 0.667 | 4.10E-45 | 3 | Casp6 | Col18a1 |
| Ptpn5 | 3.14E-49 | 1.0979227 | 0.253 | 0.105 | 6.08E-45 | 3 | Ptpn5 | Col18a1 |
| Nfat5 | 3.82E-49 | 0.54054327 | 0.872 | 0.766 | 7.39E-45 | 3 | Nfat5 | Col18a1 |
| Pbx1 | 5.29E-49 | 0.50398326 | 0.879 | 0.791 | 1.02E-44 | 3 | Pbx1 | Col18a1 |
| Mark11 | 5.65E-49 | 0.5579411 | 0.866 | 0.749 | 1.09E-44 | 3 | Mark1 | Col18a1 |
| Zfp36l11 | 6.39E-49 | 0.44487196 | 0.907 | 0.857 | 1.23E-44 | 3 | Zfp36l1 | Col18a1 |
| Ank2 | 8.85E-49 | 0.79842112 | 0.607 | 0.462 | 1.71E-44 | 3 | Ank2 | Col18a1 |
| Eif3e | 1.08E-48 | 0.87623697 | 0.64 | 0.452 | 2.10E-44 | 3 | Eif3e | Col18a1 |
| Sbspon1 | 1.31E-48 | 0.99309755 | 0.315 | 0.169 | 2.53E-44 | 3 | Sbspon | Col18a1 |
| Fam162a1 | 1.66E-48 | 0.48500591 | 0.868 | 0.771 | 3.21E-44 | 3 | Fam162a | Col18a1 |
| Pcca | 2.24E-48 | 0.69813727 | 0.644 | 0.515 | 4.32E-44 | 3 | Pcca | Col18a1 |
| Klhl241 | 4.04E-48 | 0.61973527 | 0.685 | 0.537 | 7.81E-44 | 3 | Klhl24 | Col18a1 |
| Pcsk91 | 4.57E-48 | 0.83244861 | 0.455 | 0.34 | 8.83E-44 | 3 | Pcsk9 | Col18a1 |
| Epb41l3 | 7.81E-48 | 0.63544912 | 0.856 | 0.716 | 1.51E-43 | 3 | Epb41l3 | Col18a1 |
| Slc39a6 | 9.62E-48 | 0.76639445 | 0.702 | 0.617 | 1.86E-43 | 3 | Slc39a6 | Col18a1 |
| Ube2q2 | 1.07E-47 | 0.84041499 | 0.654 | 0.483 | 2.08E-43 | 3 | Ube2q2 | Col18a1 |
| Kcnip3 | 1.60E-47 | 1.5008923 | 0.251 | 0.117 | 3.10E-43 | 3 | Kcnip3 | Col18a1 |
| Mgarp1 | 1.85E-47 | 0.77150215 | 0.387 | 0.271 | 3.58E-43 | 3 | Mgarp | Col18a1 |
| Dok61 | 2.41E-47 | 1.14454452 | 0.361 | 0.199 | 4.67E-43 | 3 | Dok6 | Col18a1 |
| Itfg1 | 3.55E-47 | 0.64013748 | 0.824 | 0.712 | 6.87E-43 | 3 | Itfg1 | Col18a1 |
| Akap71 | 4.85E-47 | 0.78948505 | 0.457 | 0.311 | 9.38E-43 | 3 | Akap7 | Col18a1 |
| Fam129a | 6.04E-47 | 0.8490713 | 0.46 | 0.356 | 1.17E-42 | 3 | Fam129a | Col18a1 |
| Cdc42 | 9.38E-47 | 0.50078895 | 0.95 | 0.891 | 1.81E-42 | 3 | Cdc42 | Col18a1 |
| Lamb1 | 9.59E-47 | 0.32706724 | 0.975 | 0.946 | 1.85E-42 | 3 | Lamb1 | Col18a1 |
| 1-Jun | 1.04E-46 | 0.28266388 | 0.889 | 0.915 | 2.01E-42 | 3 | Jun | Col18a1 |
| Ptpn9 | 1.59E-46 | 0.60888218 | 0.849 | 0.73 | 3.07E-42 | 3 | Ptpn9 | Col18a1 |
| Slc50a1 | 2.70E-46 | 0.60448282 | 0.856 | 0.756 | 5.23E-42 | 3 | Slc50a1 | Col18a1 |
| Oxr1 | 5.90E-46 | 0.75338825 | 0.604 | 0.454 | 1.14E-41 | 3 | Oxr1 | Col18a1 |
| Ogt | 1.12E-45 | 0.52368775 | 0.808 | 0.688 | 2.16E-41 | 3 | Ogt | Col18a1 |
| Hmgcr | 1.99E-45 | 0.62298239 | 0.684 | 0.626 | 3.84E-41 | 3 | Hmgcr | Col18a1 |
| Arrdc4 | 6.19E-45 | 0.63032944 | 0.614 | 0.509 | 1.20E-40 | 3 | Arrdc4 | Col18a1 |
| Fam43a | 1.11E-44 | 0.58305887 | 0.632 | 0.497 | 2.15E-40 | 3 | Fam43a | Col18a1 |
| Arl5c | 1.31E-44 | 1.16148219 | 0.264 | 0.149 | 2.53E-40 | 3 | Arl5c | Col18a1 |
| Pttg1ip | 1.66E-44 | 0.43509214 | 0.957 | 0.915 | 3.20E-40 | 3 | Pttg1ip | Col18a1 |
| Dcbld11 | 1.92E-44 | 0.78014543 | 0.578 | 0.357 | 3.71E-40 | 3 | Dcbld1 | Col18a1 |
| Rere | 2.17E-44 | 0.51377806 | 0.855 | 0.724 | 4.20E-40 | 3 | Rere | Col18a1 |
| Vwa5a | 2.31E-44 | 0.56042835 | 0.833 | 0.723 | 4.47E-40 | 3 | Vwa5a | Col18a1 |
| Ppp1r15a1 | 2.85E-44 | 0.56900947 | 0.696 | 0.635 | 5.51E-40 | 3 | Ppp1r15a | Col18a1 |
| Asph | 3.53E-44 | 0.41113678 | 0.961 | 0.921 | 6.83E-40 | 3 | Asph | Col18a1 |
| Aldh6a1 | 8.10E-44 | 0.96505896 | 0.499 | 0.326 | 1.57E-39 | 3 | Aldh6a1 | Col18a1 |
| Wipf1 | 3.29E-43 | 0.47714651 | 0.912 | 0.848 | 6.36E-39 | 3 | Wipf1 | Col18a1 |
| Pcdh18 | 6.21E-43 | 0.87437009 | 0.503 | 0.358 | 1.20E-38 | 3 | Pcdh18 | Col18a1 |
| Unc5a1 | 6.40E-43 | 0.8660799 | 0.436 | 0.288 | 1.24E-38 | 3 | Unc5a | Col18a1 |
| Rictor | 6.46E-43 | 0.75835169 | 0.665 | 0.482 | 1.25E-38 | 3 | Rictor | Col18a1 |
| Ifrd11 | 8.80E-43 | 0.57242324 | 0.811 | 0.74 | 1.70E-38 | 3 | Ifrd1 | Col18a1 |
| Cp1 | 9.08E-43 | 0.68740882 | 0.503 | 0.394 | 1.75E-38 | 3 | Cp | Col18a1 |
| Atp6v1c1 | 1.88E-42 | 0.72235575 | 0.664 | 0.517 | 3.64E-38 | 3 | Atp6v1c1 | Col18a1 |
| Alkbh5 | 1.90E-42 | 0.48911356 | 0.937 | 0.868 | 3.66E-38 | 3 | Alkbh5 | Col18a1 |
| Zfp654 | 2.58E-42 | 1.00758825 | 0.429 | 0.282 | 4.99E-38 | 3 | Zfp654 | Col18a1 |
| Mapkapk2 | 2.82E-42 | 0.41890421 | 0.962 | 0.901 | 5.46E-38 | 3 | Mapkapk2 | Col18a1 |
| Thra | 3.52E-42 | 0.6758113 | 0.759 | 0.598 | 6.80E-38 | 3 | Thra | Col18a1 |
| Ebf4 | 4.41E-42 | 0.92733343 | 0.252 | 0.135 | 8.54E-38 | 3 | Ebf4 | Col18a1 |
| Efemp21 | 1.05E-41 | 0.40224407 | 0.993 | 0.961 | 2.04E-37 | 3 | Efemp2 | Col18a1 |
| Rab14 | 1.17E-41 | 0.52722767 | 0.937 | 0.844 | 2.26E-37 | 3 | Rab14 | Col18a1 |
| Eif4e3 | 1.21E-41 | 0.66031393 | 0.68 | 0.543 | 2.34E-37 | 3 | Eif4e3 | Col18a1 |
| Ctnnd1 | 1.61E-41 | 0.42127116 | 0.946 | 0.885 | 3.11E-37 | 3 | Ctnnd1 | Col18a1 |
| Cdh13 | 3.22E-41 | 0.81312088 | 0.387 | 0.259 | 6.22E-37 | 3 | Cdh13 | Col18a1 |
| Rarg | 4.51E-41 | 0.51535327 | 0.88 | 0.776 | 8.72E-37 | 3 | Rarg | Col18a1 |
| Ccnl1 | 5.42E-41 | 0.55357506 | 0.867 | 0.763 | 1.05E-36 | 3 | Ccnl1 | Col18a1 |
| Pabpc1 | 1.39E-40 | 0.54748 | 0.962 | 0.867 | 2.69E-36 | 3 | Pabpc1 | Col18a1 |
| Cox7a2l | 1.65E-40 | 0.56299822 | 0.893 | 0.782 | 3.19E-36 | 3 | Cox7a2l | Col18a1 |
| Mid1ip11 | 2.24E-40 | 0.50071692 | 0.692 | 0.599 | 4.33E-36 | 3 | Mid1ip1 | Col18a1 |
| Aldh1l11 | 3.34E-40 | 0.6619667 | 0.447 | 0.314 | 6.47E-36 | 3 | Aldh1l1 | Col18a1 |
| Aff4 | 3.45E-40 | 0.52839556 | 0.91 | 0.825 | 6.67E-36 | 3 | Aff4 | Col18a1 |
| Dhcr24 | 3.65E-40 | 0.85346267 | 0.458 | 0.318 | 7.06E-36 | 3 | Dhcr24 | Col18a1 |
| Gbp2 | 4.10E-40 | 0.92607185 | 0.256 | 0.271 | 7.92E-36 | 3 | Gbp2 | Col18a1 |
| Il6ra | 4.12E-40 | 0.92367135 | 0.283 | 0.161 | 7.97E-36 | 3 | Il6ra | Col18a1 |
| Apln1 | 4.29E-40 | 0.50426025 | 0.486 | 0.284 | 8.29E-36 | 3 | Apln | Col18a1 |
| Igf1r | 7.87E-40 | 0.51880199 | 0.86 | 0.746 | 1.52E-35 | 3 | Igf1r | Col18a1 |
| Pqlc3 | 8.23E-40 | 1.04066601 | 0.392 | 0.249 | 1.59E-35 | 3 | Pqlc3 | Col18a1 |
| Crebrf | 1.75E-39 | 0.79791079 | 0.462 | 0.322 | 3.37E-35 | 3 | Crebrf | Col18a1 |
| Kdm7a | 5.80E-39 | 0.63983385 | 0.605 | 0.451 | 1.12E-34 | 3 | Kdm7a | Col18a1 |
| Map3k1 | 6.77E-39 | 0.91778611 | 0.464 | 0.316 | 1.31E-34 | 3 | Map3k1 | Col18a1 |
| Alg1 | 7.22E-39 | 0.81997035 | 0.456 | 0.318 | 1.40E-34 | 3 | Alg1 | Col18a1 |
| Eaf1 | 7.23E-39 | 0.62793186 | 0.796 | 0.668 | 1.40E-34 | 3 | Eaf1 | Col18a1 |
| Ilrun | 7.24E-39 | 0.55854518 | 0.889 | 0.78 | 1.40E-34 | 3 | Ilrun | Col18a1 |
| Rab201 | 1.52E-38 | 0.56189936 | 0.662 | 0.512 | 2.94E-34 | 3 | Rab20 | Col18a1 |
| Robo11 | 1.90E-38 | 0.50454481 | 0.636 | 0.461 | 3.67E-34 | 3 | Robo1 | Col18a1 |
| Fbxl5 | 2.16E-38 | 0.75088337 | 0.666 | 0.504 | 4.18E-34 | 3 | Fbxl5 | Col18a1 |
| Gsk3b | 2.27E-38 | 0.5561038 | 0.89 | 0.793 | 4.39E-34 | 3 | Gsk3b | Col18a1 |
| Fam126a | 3.18E-38 | 0.74688217 | 0.67 | 0.534 | 6.16E-34 | 3 | Fam126a | Col18a1 |
| Pde1a1 | 3.35E-38 | 1.17783482 | 0.355 | 0.205 | 6.47E-34 | 3 | Pde1a | Col18a1 |
| Ifngr2 | 4.02E-38 | 0.54396107 | 0.863 | 0.743 | 7.78E-34 | 3 | Ifngr2 | Col18a1 |
| Gramd4 | 5.08E-38 | 0.56823644 | 0.626 | 0.5 | 9.83E-34 | 3 | Gramd4 | Col18a1 |
| Acadsb | 1.50E-37 | 0.5456837 | 0.812 | 0.684 | 2.91E-33 | 3 | Acadsb | Col18a1 |
| Arid5b1 | 1.73E-37 | 0.55833179 | 0.797 | 0.692 | 3.34E-33 | 3 | Arid5b | Col18a1 |
| Ccng2 | 2.13E-37 | 0.53620781 | 0.72 | 0.607 | 4.11E-33 | 3 | Ccng2 | Col18a1 |
| Jak3 | 2.15E-37 | 0.63619992 | 0.491 | 0.355 | 4.16E-33 | 3 | Jak3 | Col18a1 |
| Arhgef12 | 2.77E-37 | 0.51602857 | 0.898 | 0.799 | 5.35E-33 | 3 | Arhgef12 | Col18a1 |
| Selenot1 | 3.55E-37 | 0.68781083 | 0.774 | 0.615 | 6.86E-33 | 3 | Selenot | Col18a1 |
| Cltc | 3.63E-37 | 0.55387093 | 0.896 | 0.786 | 7.02E-33 | 3 | Cltc | Col18a1 |
| Zswim61 | 3.94E-37 | 0.67253057 | 0.65 | 0.506 | 7.61E-33 | 3 | Zswim6 | Col18a1 |
| Xiap | 6.88E-37 | 1.28589409 | 0.322 | 0.167 | 1.33E-32 | 3 | Xiap | Col18a1 |
| Lamb2 | 8.00E-37 | 0.45239103 | 0.879 | 0.799 | 1.55E-32 | 3 | Lamb2 | Col18a1 |
| Gys1 | 1.12E-36 | 0.49475652 | 0.678 | 0.579 | 2.16E-32 | 3 | Gys1 | Col18a1 |
| Camk2n11 | 1.23E-36 | 0.34666963 | 0.882 | 0.747 | 2.38E-32 | 3 | Camk2n1 | Col18a1 |
| Casp1 | 1.27E-36 | 0.94156896 | 0.345 | 0.221 | 2.46E-32 | 3 | Casp1 | Col18a1 |
| Vps13c | 1.67E-36 | 0.72354763 | 0.645 | 0.499 | 3.23E-32 | 3 | Vps13c | Col18a1 |
| Sh3bgrl3 | 2.27E-36 | 0.41506377 | 0.946 | 0.896 | 4.40E-32 | 3 | Sh3bgrl3 | Col18a1 |
| Ago1 | 3.05E-36 | 0.59298758 | 0.629 | 0.494 | 5.91E-32 | 3 | Ago1 | Col18a1 |
| Stra61 | 3.68E-36 | 0.33373708 | 0.895 | 0.575 | 7.11E-32 | 3 | Stra6 | Col18a1 |
| Tnfaip8 | 3.85E-36 | 1.09441141 | 0.345 | 0.217 | 7.44E-32 | 3 | Tnfaip8 | Col18a1 |
| Klhdc2 | 4.36E-36 | 0.54945348 | 0.877 | 0.77 | 8.44E-32 | 3 | Klhdc2 | Col18a1 |
| Armcx1 | 1.63E-35 | 0.69103173 | 0.621 | 0.478 | 3.16E-31 | 3 | Armcx1 | Col18a1 |
| Prdm11 | 2.23E-35 | 0.46299251 | 0.557 | 0.394 | 4.31E-31 | 3 | Prdm1 | Col18a1 |
| Ism1 | 2.54E-35 | 1.10820968 | 0.291 | 0.154 | 4.91E-31 | 3 | Ism1 | Col18a1 |
| Fam117a | 3.40E-35 | 0.97441463 | 0.323 | 0.185 | 6.56E-31 | 3 | Fam117a | Col18a1 |
| Agpat21 | 3.66E-35 | 0.75087373 | 0.651 | 0.489 | 7.07E-31 | 3 | Agpat2 | Col18a1 |
| Cdk20 | 4.34E-35 | 0.77663712 | 0.446 | 0.313 | 8.39E-31 | 3 | Cdk20 | Col18a1 |
| Ecm1 | 9.94E-35 | 0.25184723 | 0.969 | 0.953 | 1.92E-30 | 3 | Ecm1 | Col18a1 |
| Rab12 | 1.06E-34 | 0.51834382 | 0.903 | 0.821 | 2.05E-30 | 3 | Rab12 | Col18a1 |
| Pamr1 | 2.08E-34 | 0.83764032 | 0.332 | 0.161 | 4.03E-30 | 3 | Pamr1 | Col18a1 |
| Tnks2 | 2.61E-34 | 0.60649595 | 0.853 | 0.737 | 5.05E-30 | 3 | Tnks2 | Col18a1 |
| Cacnb1 | 2.77E-34 | 0.91096024 | 0.305 | 0.178 | 5.35E-30 | 3 | Cacnb1 | Col18a1 |
| Nop53 | 4.51E-34 | 0.46236889 | 0.813 | 0.719 | 8.71E-30 | 3 | Nop53 | Col18a1 |
| Epha4 | 4.75E-34 | 1.02058759 | 0.265 | 0.18 | 9.18E-30 | 3 | Epha4 | Col18a1 |
| Ptch1 | 8.62E-34 | 1.06744206 | 0.281 | 0.216 | 1.67E-29 | 3 | Ptch1 | Col18a1 |
| Tmem50a | 1.35E-33 | 0.38170292 | 0.961 | 0.925 | 2.61E-29 | 3 | Tmem50a | Col18a1 |
| Tor4a1 | 1.78E-33 | 0.74151103 | 0.453 | 0.313 | 3.44E-29 | 3 | Tor4a | Col18a1 |
| Plppr3 | 1.87E-33 | 0.58658153 | 0.505 | 0.356 | 3.62E-29 | 3 | Plppr3 | Col18a1 |
| Fgf11 | 1.96E-33 | 0.80937553 | 0.306 | 0.181 | 3.80E-29 | 3 | Fgf11 | Col18a1 |
| Pfdn5 | 2.93E-33 | 0.50628498 | 0.87 | 0.783 | 5.66E-29 | 3 | Pfdn5 | Col18a1 |
| Qpct | 3.28E-33 | 0.60950362 | 0.503 | 0.345 | 6.34E-29 | 3 | Qpct | Col18a1 |
| Ikbke | 8.17E-33 | 0.62722712 | 0.512 | 0.377 | 1.58E-28 | 3 | Ikbke | Col18a1 |
| Serpina3i1 | 8.24E-33 | 0.73613955 | 0.432 | 0.301 | 1.59E-28 | 3 | Serpina3i | Col18a1 |
| Emp1 | 8.42E-33 | 0.34154582 | 0.994 | 0.966 | 1.63E-28 | 3 | Emp1 | Col18a1 |
| 7-Mar | 1.11E-32 | 0.59957337 | 0.783 | 0.651 | 2.15E-28 | 3 | 7-Mar | Col18a1 |
| Fzd11 | 1.96E-32 | 0.32298114 | 0.948 | 0.858 | 3.79E-28 | 3 | Fzd1 | Col18a1 |
| Slc25a36 | 2.22E-32 | 0.59579464 | 0.682 | 0.531 | 4.30E-28 | 3 | Slc25a36 | Col18a1 |
| Serinc31 | 3.56E-32 | 0.31719376 | 0.982 | 0.952 | 6.88E-28 | 3 | Serinc3 | Col18a1 |
| Zfp5321 | 4.22E-32 | 0.45279687 | 0.835 | 0.718 | 8.16E-28 | 3 | Zfp532 | Col18a1 |
| Znrf1 | 4.89E-32 | 0.53094314 | 0.78 | 0.681 | 9.45E-28 | 3 | Znrf1 | Col18a1 |
| Camk2d | 8.94E-32 | 0.49189639 | 0.777 | 0.685 | 1.73E-27 | 3 | Camk2d | Col18a1 |
| Pld3 | 1.02E-31 | 0.39411189 | 0.903 | 0.857 | 1.98E-27 | 3 | Pld3 | Col18a1 |
| Hsp90ab1 | 1.45E-31 | 0.46631306 | 0.974 | 0.914 | 2.81E-27 | 3 | Hsp90ab1 | Col18a1 |
| Ppard | 1.50E-31 | 0.56842807 | 0.736 | 0.599 | 2.90E-27 | 3 | Ppard | Col18a1 |
| Gnpda2 | 1.65E-31 | 0.73083674 | 0.492 | 0.382 | 3.18E-27 | 3 | Gnpda2 | Col18a1 |
| Frrs11 | 1.79E-31 | 0.6703961 | 0.623 | 0.488 | 3.46E-27 | 3 | Frrs1 | Col18a1 |
| Uap1l1 | 1.79E-31 | 0.63072328 | 0.623 | 0.456 | 3.47E-27 | 3 | Uap1l1 | Col18a1 |
| Tmem65 | 2.62E-31 | 0.59201242 | 0.827 | 0.724 | 5.07E-27 | 3 | Tmem65 | Col18a1 |
| Gcnt21 | 3.17E-31 | 0.66961112 | 0.54 | 0.387 | 6.13E-27 | 3 | Gcnt2 | Col18a1 |
| Grina | 3.21E-31 | 0.40513984 | 0.731 | 0.638 | 6.21E-27 | 3 | Grina | Col18a1 |
| Syngap1 | 3.34E-31 | 0.53181167 | 0.635 | 0.515 | 6.46E-27 | 3 | Syngap1 | Col18a1 |
| Atraid | 3.38E-31 | 0.508387 | 0.852 | 0.752 | 6.53E-27 | 3 | Atraid | Col18a1 |
| Pcyt2 | 5.03E-31 | 0.69974371 | 0.42 | 0.296 | 9.72E-27 | 3 | Pcyt2 | Col18a1 |
| Chsy31 | 5.22E-31 | 0.88373849 | 0.52 | 0.367 | 1.01E-26 | 3 | Chsy3 | Col18a1 |
| Stxbp1 | 8.84E-31 | 0.65053292 | 0.635 | 0.508 | 1.71E-26 | 3 | Stxbp1 | Col18a1 |
| Ap3s1 | 1.21E-30 | 1.07078468 | 0.297 | 0.17 | 2.34E-26 | 3 | Ap3s1 | Col18a1 |
| Pdlim21 | 1.22E-30 | 0.42022964 | 0.762 | 0.625 | 2.36E-26 | 3 | Pdlim2 | Col18a1 |
| Peli21 | 1.32E-30 | 0.62345169 | 0.625 | 0.473 | 2.55E-26 | 3 | Peli2 | Col18a1 |
| Sertad11 | 2.78E-30 | 0.50548431 | 0.698 | 0.64 | 5.38E-26 | 3 | Sertad1 | Col18a1 |
| Arhgef171 | 2.94E-30 | 0.4927325 | 0.801 | 0.667 | 5.69E-26 | 3 | Arhgef17 | Col18a1 |
| Fem1b | 3.15E-30 | 0.65937313 | 0.724 | 0.641 | 6.09E-26 | 3 | Fem1b | Col18a1 |
| Xylt1 | 4.29E-30 | 0.63626171 | 0.516 | 0.401 | 8.30E-26 | 3 | Xylt1 | Col18a1 |
| Tcn2 | 5.32E-30 | 0.31381749 | 0.931 | 0.863 | 1.03E-25 | 3 | Tcn2 | Col18a1 |
| Pgpep11 | 6.04E-30 | 0.58448154 | 0.539 | 0.424 | 1.17E-25 | 3 | Pgpep1 | Col18a1 |
| Taok1 | 8.92E-30 | 0.60801682 | 0.825 | 0.689 | 1.73E-25 | 3 | Taok1 | Col18a1 |
| Mettl91 | 9.94E-30 | 0.52303068 | 0.942 | 0.855 | 1.92E-25 | 3 | Mettl9 | Col18a1 |
| Prpf8 | 1.70E-29 | 0.55803172 | 0.888 | 0.77 | 3.30E-25 | 3 | Prpf8 | Col18a1 |
| Tnfsf131 | 1.79E-29 | 0.83796563 | 0.392 | 0.244 | 3.47E-25 | 3 | Tnfsf13 | Col18a1 |
| Gnai11 | 2.11E-29 | 0.91107637 | 0.398 | 0.266 | 4.08E-25 | 3 | Gnai1 | Col18a1 |
| Arntl21 | 2.44E-29 | 0.92674429 | 0.357 | 0.23 | 4.72E-25 | 3 | Arntl2 | Col18a1 |
| Rabgap1 | 2.70E-29 | 0.7017064 | 0.624 | 0.484 | 5.22E-25 | 3 | Rabgap1 | Col18a1 |
| Pmp22 | 2.91E-29 | 0.41751354 | 0.867 | 0.713 | 5.62E-25 | 3 | Pmp22 | Col18a1 |
| Insig2 | 3.59E-29 | 0.76966654 | 0.521 | 0.381 | 6.94E-25 | 3 | Insig2 | Col18a1 |
| Kng21 | 4.04E-29 | 0.70893265 | 0.291 | 0.229 | 7.81E-25 | 3 | Kng2 | Col18a1 |
| Ak4 | 4.09E-29 | 0.89797828 | 0.357 | 0.232 | 7.92E-25 | 3 | Ak4 | Col18a1 |
| Sema6d | 5.53E-29 | 0.83802454 | 0.392 | 0.273 | 1.07E-24 | 3 | Sema6d | Col18a1 |
| Stambpl1 | 6.07E-29 | 0.50965583 | 0.714 | 0.617 | 1.17E-24 | 3 | Stambpl1 | Col18a1 |
| Farp1 | 6.48E-29 | 0.48304061 | 0.877 | 0.778 | 1.25E-24 | 3 | Farp1 | Col18a1 |
| Fnbp11 | 9.23E-29 | 0.31510465 | 0.89 | 0.796 | 1.78E-24 | 3 | Fnbp1 | Col18a1 |
| Dusp11 | 9.95E-29 | 0.53115434 | 0.804 | 0.689 | 1.92E-24 | 3 | Dusp11 | Col18a1 |
| Cacna1c1 | 1.39E-28 | 0.38247901 | 0.877 | 0.708 | 2.68E-24 | 3 | Cacna1c | Col18a1 |
| Ddit31 | 1.44E-28 | 0.50898692 | 0.433 | 0.397 | 2.78E-24 | 3 | Ddit3 | Col18a1 |
| Foxj3 | 1.48E-28 | 0.67582071 | 0.546 | 0.401 | 2.86E-24 | 3 | Foxj3 | Col18a1 |
| Irf2 | 1.64E-28 | 0.72625072 | 0.537 | 0.393 | 3.17E-24 | 3 | Irf2 | Col18a1 |
| Mgat31 | 1.74E-28 | 0.56254608 | 0.571 | 0.409 | 3.37E-24 | 3 | Mgat3 | Col18a1 |
| Pmvk | 2.01E-28 | 0.83724756 | 0.373 | 0.273 | 3.89E-24 | 3 | Pmvk | Col18a1 |
| Runx21 | 2.34E-28 | 0.39114651 | 0.823 | 0.651 | 4.52E-24 | 3 | Runx2 | Col18a1 |
| Itga9 | 3.10E-28 | 0.50944813 | 0.615 | 0.507 | 5.99E-24 | 3 | Itga9 | Col18a1 |
| B4galt51 | 3.79E-28 | 0.47212681 | 0.903 | 0.825 | 7.32E-24 | 3 | B4galt5 | Col18a1 |
| Tmem87b | 4.13E-28 | 0.54197261 | 0.756 | 0.657 | 7.98E-24 | 3 | Tmem87b | Col18a1 |
| Ppp1r3b | 4.52E-28 | 0.67206506 | 0.345 | 0.246 | 8.75E-24 | 3 | Ppp1r3b | Col18a1 |
| Cds2 | 6.13E-28 | 0.57333814 | 0.686 | 0.56 | 1.19E-23 | 3 | Cds2 | Col18a1 |
| Paip2 | 8.34E-28 | 0.43054633 | 0.956 | 0.881 | 1.61E-23 | 3 | Paip2 | Col18a1 |
| Ginm1 | 9.10E-28 | 0.47093219 | 0.865 | 0.781 | 1.76E-23 | 3 | Ginm1 | Col18a1 |
| Per1 | 1.24E-27 | 0.56685333 | 0.648 | 0.535 | 2.39E-23 | 3 | Per1 | Col18a1 |
| Ubash3b | 1.24E-27 | 0.65305409 | 0.416 | 0.269 | 2.40E-23 | 3 | Ubash3b | Col18a1 |
| Mbnl2 | 1.56E-27 | 0.36256941 | 0.959 | 0.928 | 3.01E-23 | 3 | Mbnl2 | Col18a1 |
| Timp2 | 1.78E-27 | 0.2502835 | 0.982 | 0.955 | 3.44E-23 | 3 | Timp2 | Col18a1 |
| Slc30a9 | 2.05E-27 | 0.64178145 | 0.669 | 0.538 | 3.97E-23 | 3 | Slc30a9 | Col18a1 |
| Gem1 | 2.72E-27 | 0.49107379 | 0.576 | 0.519 | 5.27E-23 | 3 | Gem | Col18a1 |
| Nol3 | 2.77E-27 | 0.82289246 | 0.379 | 0.261 | 5.35E-23 | 3 | Nol3 | Col18a1 |
| Stk40 | 2.82E-27 | 0.49872686 | 0.755 | 0.644 | 5.45E-23 | 3 | Stk40 | Col18a1 |
| Rnf165 | 3.08E-27 | 0.97330419 | 0.264 | 0.153 | 5.96E-23 | 3 | Rnf165 | Col18a1 |
| Notch11 | 3.24E-27 | 0.49988486 | 0.709 | 0.501 | 6.26E-23 | 3 | Notch1 | Col18a1 |
| Tead11 | 3.62E-27 | 0.4495254 | 0.894 | 0.825 | 7.00E-23 | 3 | Tead1 | Col18a1 |
| Smad2 | 3.95E-27 | 0.65743301 | 0.737 | 0.607 | 7.63E-23 | 3 | Smad2 | Col18a1 |
| Hsd17b71 | 4.24E-27 | 0.89615174 | 0.313 | 0.23 | 8.19E-23 | 3 | Hsd17b7 | Col18a1 |
| Cnrip1 | 4.63E-27 | 0.5275145 | 0.733 | 0.599 | 8.95E-23 | 3 | Cnrip1 | Col18a1 |
| Btg3 | 4.89E-27 | 0.56682797 | 0.808 | 0.723 | 9.45E-23 | 3 | Btg3 | Col18a1 |
| Aida | 5.63E-27 | 0.6546712 | 0.794 | 0.632 | 1.09E-22 | 3 | Aida | Col18a1 |
| Erlec1 | 7.91E-27 | 0.55520334 | 0.788 | 0.692 | 1.53E-22 | 3 | Erlec1 | Col18a1 |
| Diaph2 | 9.16E-27 | 0.62719712 | 0.621 | 0.5 | 1.77E-22 | 3 | Diaph2 | Col18a1 |
| Pign | 9.84E-27 | 0.65139915 | 0.707 | 0.558 | 1.90E-22 | 3 | Pign | Col18a1 |
| Gna141 | 9.88E-27 | 0.6384969 | 0.427 | 0.301 | 1.91E-22 | 3 | Gna14 | Col18a1 |
| Slc25a37 | 1.21E-26 | 0.60903691 | 0.512 | 0.39 | 2.35E-22 | 3 | Slc25a37 | Col18a1 |
| Map3k6 | 1.32E-26 | 0.69436073 | 0.483 | 0.33 | 2.56E-22 | 3 | Map3k6 | Col18a1 |
| Vill | 1.74E-26 | 0.85588417 | 0.254 | 0.17 | 3.37E-22 | 3 | Vill | Col18a1 |
| Qprt1 | 1.74E-26 | 0.70369339 | 0.312 | 0.208 | 3.37E-22 | 3 | Qprt | Col18a1 |
| Mef2c | 3.82E-26 | 0.83574502 | 0.352 | 0.273 | 7.39E-22 | 3 | Mef2c | Col18a1 |
| Paip1 | 4.24E-26 | 0.55634797 | 0.851 | 0.735 | 8.19E-22 | 3 | Paip1 | Col18a1 |
| Zfp740 | 5.89E-26 | 0.58055907 | 0.721 | 0.588 | 1.14E-21 | 3 | Zfp740 | Col18a1 |
| Rab24 | 6.13E-26 | 0.45028729 | 0.784 | 0.674 | 1.19E-21 | 3 | Rab24 | Col18a1 |
| Mthfd1l | 7.68E-26 | 0.75352261 | 0.419 | 0.321 | 1.48E-21 | 3 | Mthfd1l | Col18a1 |
| Sgms1 | 8.29E-26 | 0.75600188 | 0.579 | 0.451 | 1.60E-21 | 3 | Sgms1 | Col18a1 |
| DEPRECATED-l | 8.29E-26 | 0.79954693 | 0.377 | 0.247 | 1.60E-21 | 3 | DEPRECATED-l | Col18a1 |
| Eid11 | 8.99E-26 | 0.25408334 | 0.989 | 0.952 | 1.74E-21 | 3 | Eid1 | Col18a1 |
| Zfp8101 | 9.89E-26 | 0.74584718 | 0.397 | 0.277 | 1.91E-21 | 3 | Zfp810 | Col18a1 |
| Insr | 1.32E-25 | 0.56469333 | 0.663 | 0.554 | 2.54E-21 | 3 | Insr | Col18a1 |
| Unc93b1 | 1.33E-25 | 0.57697436 | 0.712 | 0.599 | 2.57E-21 | 3 | Unc93b1 | Col18a1 |
| Tmem86a1 | 2.44E-25 | 0.27906713 | 0.781 | 0.69 | 4.72E-21 | 3 | Tmem86a | Col18a1 |
| Atp2b1 | 2.80E-25 | 0.60256454 | 0.821 | 0.692 | 5.42E-21 | 3 | Atp2b1 | Col18a1 |
| Sfxn1 | 3.01E-25 | 0.46913326 | 0.826 | 0.727 | 5.82E-21 | 3 | Sfxn1 | Col18a1 |
| Map2k4 | 3.56E-25 | 0.71970234 | 0.529 | 0.39 | 6.87E-21 | 3 | Map2k4 | Col18a1 |
| Appl2 | 3.77E-25 | 0.52403746 | 0.487 | 0.388 | 7.29E-21 | 3 | Appl2 | Col18a1 |
| Larp6 | 3.85E-25 | 0.63797281 | 0.442 | 0.338 | 7.44E-21 | 3 | Larp6 | Col18a1 |
| Pdcd10 | 4.37E-25 | 0.60378366 | 0.653 | 0.521 | 8.44E-21 | 3 | Pdcd10 | Col18a1 |
| Slc25a52 | 4.92E-25 | 0.68119489 | 0.745 | 0.611 | 9.51E-21 | 3 | Slc25a5 | Col18a1 |
| Atp6v1b2 | 5.01E-25 | 0.35860423 | 0.826 | 0.75 | 9.69E-21 | 3 | Atp6v1b2 | Col18a1 |
| Cdadc1 | 6.08E-25 | 0.68170545 | 0.53 | 0.401 | 1.18E-20 | 3 | Cdadc1 | Col18a1 |
| Strn3 | 7.19E-25 | 0.5370142 | 0.808 | 0.69 | 1.39E-20 | 3 | Strn3 | Col18a1 |
| Sbf2 | 7.76E-25 | 0.60831267 | 0.645 | 0.533 | 1.50E-20 | 3 | Sbf2 | Col18a1 |
| Slc25a11 | 1.12E-24 | 0.44765568 | 0.838 | 0.734 | 2.16E-20 | 3 | Slc25a1 | Col18a1 |
| Sbno2 | 1.68E-24 | 0.30558987 | 0.927 | 0.89 | 3.24E-20 | 3 | Sbno2 | Col18a1 |
| Arhgap6 | 1.93E-24 | 0.94571741 | 0.308 | 0.206 | 3.73E-20 | 3 | Arhgap6 | Col18a1 |
| Gdnf1 | 1.94E-24 | 0.75394835 | 0.312 | 0.204 | 3.75E-20 | 3 | Gdnf | Col18a1 |
| Vma21 | 1.97E-24 | 0.91488917 | 0.462 | 0.318 | 3.81E-20 | 3 | Vma21 | Col18a1 |
| Bbx | 2.16E-24 | 0.59977084 | 0.607 | 0.487 | 4.18E-20 | 3 | Bbx | Col18a1 |
| Twist11 | 3.49E-24 | 0.33627874 | 0.928 | 0.903 | 6.75E-20 | 3 | Twist1 | Col18a1 |
| Spg21 | 4.23E-24 | 0.5610386 | 0.748 | 0.652 | 8.19E-20 | 3 | Spg21 | Col18a1 |
| Hmcn11 | 4.62E-24 | 0.78800753 | 0.356 | 0.234 | 8.94E-20 | 3 | Hmcn1 | Col18a1 |
| Usp391 | 5.05E-24 | 0.7357363 | 0.532 | 0.416 | 9.76E-20 | 3 | Usp39 | Col18a1 |
| Irx3 | 8.45E-24 | 0.5889218 | 0.527 | 0.4 | 1.63E-19 | 3 | Irx3 | Col18a1 |
| Kdm4b | 8.80E-24 | 0.60459298 | 0.538 | 0.425 | 1.70E-19 | 3 | Kdm4b | Col18a1 |
| Srgap3 | 1.07E-23 | 0.74398356 | 0.254 | 0.168 | 2.08E-19 | 3 | Srgap3 | Col18a1 |
| Slc9a3r21 | 1.11E-23 | 0.54041324 | 0.476 | 0.342 | 2.15E-19 | 3 | Slc9a3r2 | Col18a1 |
| Rab6a | 1.18E-23 | 0.52736411 | 0.906 | 0.805 | 2.28E-19 | 3 | Rab6a | Col18a1 |
| Mindy4 | 1.25E-23 | 0.59827483 | 0.432 | 0.339 | 2.41E-19 | 3 | Mindy4 | Col18a1 |
| Nipbl | 1.40E-23 | 0.63027292 | 0.741 | 0.597 | 2.71E-19 | 3 | Nipbl | Col18a1 |
| Wnk1 | 1.51E-23 | 0.48921405 | 0.924 | 0.835 | 2.92E-19 | 3 | Wnk1 | Col18a1 |
| Pnpla8 | 2.46E-23 | 0.61502752 | 0.692 | 0.56 | 4.76E-19 | 3 | Pnpla8 | Col18a1 |
| Ms4a4d | 2.48E-23 | 0.4887677 | 0.76 | 0.663 | 4.79E-19 | 3 | Ms4a4d | Col18a1 |
| Rhbdf21 | 2.84E-23 | 0.50867377 | 0.464 | 0.312 | 5.50E-19 | 3 | Rhbdf2 | Col18a1 |
| Ubc1 | 2.89E-23 | 0.32412621 | 0.969 | 0.932 | 5.59E-19 | 3 | Ubc | Col18a1 |
| Plpp11 | 2.93E-23 | 0.61107694 | 0.683 | 0.62 | 5.67E-19 | 3 | Plpp1 | Col18a1 |
| Rlf | 4.41E-23 | 0.6624726 | 0.596 | 0.478 | 8.53E-19 | 3 | Rlf | Col18a1 |
| Zfp618 | 4.45E-23 | 0.49639189 | 0.479 | 0.371 | 8.61E-19 | 3 | Zfp618 | Col18a1 |
| Calu | 4.86E-23 | 0.53567373 | 0.884 | 0.787 | 9.40E-19 | 3 | Calu | Col18a1 |
| Rara | 6.01E-23 | 0.52270568 | 0.717 | 0.618 | 1.16E-18 | 3 | Rara | Col18a1 |
| Lnpep | 7.32E-23 | 0.41737055 | 0.911 | 0.838 | 1.41E-18 | 3 | Lnpep | Col18a1 |
| Hnrnpll1 | 7.94E-23 | 0.62162245 | 0.78 | 0.653 | 1.53E-18 | 3 | Hnrnpll | Col18a1 |
| Zer1 | 8.42E-23 | 0.67301216 | 0.553 | 0.436 | 1.63E-18 | 3 | Zer1 | Col18a1 |

| | | | | | | | | |
|---|---|---|---|---|---|---|---|---|
| Atp6v1h | 1.01E-22 | 0.55852481 | 0.727 | 0.62 | 1.95E-18 | 3 | Atp6v1h | Col18a1 |
| Tmbim6 | 1.01E-22 | 0.38686995 | 0.965 | 0.909 | 1.95E-18 | 3 | Tmbim6 | Col18a1 |
| Herc4 | 1.04E-22 | 0.67610141 | 0.553 | 0.428 | 2.02E-18 | 3 | Herc4 | Col18a1 |
| Gpr271 | 1.32E-22 | 0.64414322 | 0.376 | 0.278 | 2.56E-18 | 3 | Gpr27 | Col18a1 |
| Rack1 | 1.83E-22 | 0.32877672 | 0.992 | 0.973 | 3.54E-18 | 3 | Rack1 | Col18a1 |
| Klhl211 | 2.07E-22 | 0.51681726 | 0.627 | 0.579 | 4.01E-18 | 3 | Klhl21 | Col18a1 |
| Rsbn1 | 3.21E-22 | 0.68658081 | 0.525 | 0.404 | 6.21E-18 | 3 | Rsbn1 | Col18a1 |
| Adprh | 3.46E-22 | 0.31361305 | 0.97 | 0.939 | 6.69E-18 | 3 | Adprh | Col18a1 |
| Zdhhc20 | 3.55E-22 | 0.56050005 | 0.768 | 0.643 | 6.86E-18 | 3 | Zdhhc20 | Col18a1 |
| Atf5 | 3.62E-22 | 0.25478368 | 0.854 | 0.809 | 7.00E-18 | 3 | Atf5 | Col18a1 |
| Cped11 | 4.63E-22 | 0.34301041 | 0.864 | 0.806 | 8.95E-18 | 3 | Cped1 | Col18a1 |
| Scarb2 | 5.09E-22 | 0.43587717 | 0.893 | 0.805 | 9.84E-18 | 3 | Scarb2 | Col18a1 |
| Rbpj | 5.98E-22 | 0.50460389 | 0.809 | 0.679 | 1.16E-17 | 3 | Rbpj | Col18a1 |
| Rnf130 | 6.06E-22 | 0.5937016 | 0.716 | 0.568 | 1.17E-17 | 3 | Rnf130 | Col18a1 |
| Rraga | 8.81E-22 | 0.57551282 | 0.734 | 0.623 | 1.70E-17 | 3 | Rraga | Col18a1 |
| Zfp644 | 9.59E-22 | 0.733517 | 0.504 | 0.365 | 1.85E-17 | 3 | Zfp644 | Col18a1 |
| Igsf10 | 1.26E-21 | 0.46157214 | 0.825 | 0.754 | 2.43E-17 | 3 | Igsf10 | Col18a1 |
| Narf | 2.11E-21 | 0.59825193 | 0.549 | 0.439 | 4.08E-17 | 3 | Narf | Col18a1 |
| Sbds | 2.11E-21 | 0.56659441 | 0.758 | 0.651 | 4.08E-17 | 3 | Sbds | Col18a1 |
| Usp53 | 2.12E-21 | 0.8371505 | 0.264 | 0.198 | 4.11E-17 | 3 | Usp53 | Col18a1 |
| Rnf115 | 2.18E-21 | 0.70233133 | 0.583 | 0.448 | 4.21E-17 | 3 | Rnf115 | Col18a1 |
| Tkt1 | 2.61E-21 | 0.49141235 | 0.893 | 0.813 | 5.04E-17 | 3 | Tkt | Col18a1 |
| Gss | 2.63E-21 | 0.61870975 | 0.541 | 0.413 | 5.09E-17 | 3 | Gss | Col18a1 |
| Wapl | 2.92E-21 | 0.61919018 | 0.688 | 0.542 | 5.64E-17 | 3 | Wapl | Col18a1 |
| Lgals1 | 2.95E-21 | 0.45256605 | 0.942 | 0.863 | 5.70E-17 | 3 | Lgals1 | Col18a1 |
| Pip5k1a1 | 3.41E-21 | 0.53105956 | 0.712 | 0.589 | 6.59E-17 | 3 | Pip5k1a | Col18a1 |
| Fchsd2 | 4.34E-21 | 0.54514494 | 0.65 | 0.538 | 8.39E-17 | 3 | Fchsd2 | Col18a1 |
| Dipk1b | 5.41E-21 | 0.78801307 | 0.261 | 0.161 | 1.05E-16 | 3 | Dipk1b | Col18a1 |
| Mlkl1 | 5.44E-21 | 0.76941491 | 0.483 | 0.359 | 1.05E-16 | 3 | Mlkl | Col18a1 |
| Ube2b | 5.70E-21 | 0.39646257 | 0.935 | 0.874 | 1.10E-16 | 3 | Ube2b | Col18a1 |
| Rnf2 | 6.09E-21 | 0.59825171 | 0.627 | 0.499 | 1.18E-16 | 3 | Rnf2 | Col18a1 |
| Mospd1 | 7.99E-21 | 0.67067616 | 0.446 | 0.342 | 1.55E-16 | 3 | Mospd1 | Col18a1 |
| Prr141 | 8.99E-21 | 0.57850098 | 0.675 | 0.56 | 1.74E-16 | 3 | Prr14 | Col18a1 |
| Srsf31 | 1.02E-20 | 0.6608621 | 0.67 | 0.567 | 1.97E-16 | 3 | Srsf3 | Col18a1 |
| Kat6b | 1.25E-20 | 0.5090367 | 0.606 | 0.498 | 2.42E-16 | 3 | Kat6b | Col18a1 |
| Lyst | 1.29E-20 | 0.62160703 | 0.497 | 0.394 | 2.49E-16 | 3 | Lyst | Col18a1 |
| Pigp | 1.31E-20 | 0.63351974 | 0.613 | 0.496 | 2.53E-16 | 3 | Pigp | Col18a1 |
| Reep3 | 1.37E-20 | 0.46378271 | 0.856 | 0.767 | 2.65E-16 | 3 | Reep3 | Col18a1 |
| Brd2 | 1.70E-20 | 0.42869532 | 0.856 | 0.804 | 3.28E-16 | 3 | Brd2 | Col18a1 |
| Add3 | 2.15E-20 | 0.50819869 | 0.709 | 0.543 | 4.16E-16 | 3 | Add3 | Col18a1 |
| Atf7 | 2.23E-20 | 0.5080279 | 0.82 | 0.73 | 4.31E-16 | 3 | Atf7 | Col18a1 |
| Ppt1 | 2.24E-20 | 0.47020974 | 0.745 | 0.668 | 4.33E-16 | 3 | Ppt1 | Col18a1 |
| Lrp3 | 2.29E-20 | 0.52094333 | 0.523 | 0.434 | 4.44E-16 | 3 | Lrp3 | Col18a1 |
| Prrc2c | 2.31E-20 | 0.46837312 | 0.932 | 0.838 | 4.47E-16 | 3 | Prrc2c | Col18a1 |
| Rab15 | 2.45E-20 | 0.71303231 | 0.4 | 0.297 | 4.73E-16 | 3 | Rab15 | Col18a1 |
| Btbd1 | 2.49E-20 | 0.5494591 | 0.697 | 0.578 | 4.82E-16 | 3 | Btbd1 | Col18a1 |
| Sos2 | 3.08E-20 | 0.64275007 | 0.405 | 0.297 | 5.95E-16 | 3 | Sos2 | Col18a1 |
| Ggt7 | 3.24E-20 | 0.87770934 | 0.255 | 0.153 | 6.26E-16 | 3 | Ggt7 | Col18a1 |
| Mxd1 | 3.33E-20 | 0.5784102 | 0.369 | 0.298 | 6.43E-16 | 3 | Mxd1 | Col18a1 |
| Fbxl20 | 3.52E-20 | 0.59513903 | 0.569 | 0.44 | 6.80E-16 | 3 | Fbxl20 | Col18a1 |
| Elmo2 | 3.62E-20 | 0.58381047 | 0.621 | 0.516 | 7.01E-16 | 3 | Elmo2 | Col18a1 |
| Fads1 | 4.30E-20 | 0.54025447 | 0.7 | 0.616 | 8.31E-16 | 3 | Fads1 | Col18a1 |
| Dennd4c | 6.25E-20 | 0.62505676 | 0.588 | 0.45 | 1.21E-15 | 3 | Dennd4c | Col18a1 |
| Ctnna1 | 6.43E-20 | 0.41156204 | 0.953 | 0.886 | 1.24E-15 | 3 | Ctnna1 | Col18a1 |
| H2-Ab1 | 7.23E-20 | 0.26899422 | 0.385 | 0.318 | 1.40E-15 | 3 | H2-Ab1 | Col18a1 |
| Sqstm1 | 7.58E-20 | 0.35113724 | 0.905 | 0.874 | 1.47E-15 | 3 | Sqstm1 | Col18a1 |
| Frmd4a | 8.14E-20 | 0.47628149 | 0.675 | 0.585 | 1.57E-15 | 3 | Frmd4a | Col18a1 |
| Gabarapl1 | 8.52E-20 | 0.32698575 | 0.862 | 0.809 | 1.65E-15 | 3 | Gabarapl1 | Col18a1 |
| Rchy1 | 8.59E-20 | 0.60029628 | 0.659 | 0.531 | 1.66E-15 | 3 | Rchy1 | Col18a1 |
| Nnmt | 8.84E-20 | 0.48852058 | 0.748 | 0.642 | 1.71E-15 | 3 | Nnmt | Col18a1 |
| Pdcd4 | 9.39E-20 | 0.4054931 | 0.72 | 0.649 | 1.82E-15 | 3 | Pdcd4 | Col18a1 |
| Ino80 | 1.20E-19 | 0.60969978 | 0.551 | 0.436 | 2.33E-15 | 3 | Ino80 | Col18a1 |
| Dennd1a | 1.66E-19 | 0.6118982 | 0.594 | 0.479 | 3.22E-15 | 3 | Dennd1a | Col18a1 |
| Hadh | 1.83E-19 | 0.51074193 | 0.629 | 0.523 | 3.54E-15 | 3 | Hadh | Col18a1 |
| Eif4b | 1.98E-19 | 0.33731434 | 0.909 | 0.855 | 3.84E-15 | 3 | Eif4b | Col18a1 |
| Igip | 2.02E-19 | 0.63448675 | 0.462 | 0.357 | 3.90E-15 | 3 | Igip | Col18a1 |
| Hfe1 | 2.04E-19 | 0.72040364 | 0.293 | 0.202 | 3.95E-15 | 3 | Hfe | Col18a1 |
| Plekhn11 | 2.42E-19 | 0.5105332 | 0.503 | 0.398 | 4.69E-15 | 3 | Plekhn1 | Col18a1 |
| Slc39a101 | 2.53E-19 | 0.76826787 | 0.416 | 0.308 | 4.88E-15 | 3 | Slc39a10 | Col18a1 |
| Mrgprf | 2.57E-19 | 0.50399633 | 0.62 | 0.521 | 4.97E-15 | 3 | Mrgprf | Col18a1 |
| Vps33a | 2.59E-19 | 0.74430237 | 0.361 | 0.257 | 5.00E-15 | 3 | Vps33a | Col18a1 |
| Zbtb461 | 2.70E-19 | 0.76621514 | 0.293 | 0.211 | 5.22E-15 | 3 | Zbtb46 | Col18a1 |
| Cdk6 | 2.86E-19 | 0.89711871 | 0.386 | 0.27 | 5.53E-15 | 3 | Cdk6 | Col18a1 |
| Tusc3 | 3.62E-19 | 0.42991474 | 0.909 | 0.823 | 6.99E-15 | 3 | Tusc3 | Col18a1 |
| Rhobtb3 | 4.46E-19 | 0.61404053 | 0.7 | 0.579 | 8.63E-15 | 3 | Rhobtb3 | Col18a1 |
| Rab5b | 4.47E-19 | 0.56607533 | 0.661 | 0.562 | 8.65E-15 | 3 | Rab5b | Col18a1 |
| Mapk8ip1 | 4.56E-19 | 0.55870572 | 0.355 | 0.274 | 8.82E-15 | 3 | Mapk8ip1 | Col18a1 |
| DEPRECATED-I | 5.62E-19 | 0.87864061 | 0.353 | 0.238 | 1.09E-14 | 3 | DEPRECATED-I | Col18a1 |
| Zfp2811 | 5.85E-19 | 0.430735 | 0.688 | 0.595 | 1.13E-14 | 3 | Zfp281 | Col18a1 |
| Mb21d2 | 5.88E-19 | 0.97788608 | 0.284 | 0.193 | 1.14E-14 | 3 | Mb21d2 | Col18a1 |
| Il341 | 5.98E-19 | 0.58916452 | 0.315 | 0.255 | 1.16E-14 | 3 | Il34 | Col18a1 |
| Syne11 | 6.27E-19 | 0.68989404 | 0.361 | 0.248 | 1.21E-14 | 3 | Syne1 | Col18a1 |
| Mycbp21 | 6.51E-19 | 0.44525518 | 0.858 | 0.761 | 1.26E-14 | 3 | Mycbp2 | Col18a1 |
| Cdkn1b | 7.29E-19 | 0.41009709 | 0.836 | 0.763 | 1.41E-14 | 3 | Cdkn1b | Col18a1 |
| Relch | 7.34E-19 | 0.5959175 | 0.491 | 0.383 | 1.42E-14 | 3 | Relch | Col18a1 |
| Plekha2 | 7.46E-19 | 0.50803507 | 0.622 | 0.513 | 1.44E-14 | 3 | Plekha2 | Col18a1 |
| Pth1r1 | 7.79E-19 | 0.35909492 | 0.85 | 0.78 | 1.51E-14 | 3 | Pth1r | Col18a1 |
| Lrig2 | 8.46E-19 | 0.68164185 | 0.491 | 0.371 | 1.64E-14 | 3 | Lrig2 | Col18a1 |
| Yipf4 | 1.06E-18 | 0.53824527 | 0.799 | 0.672 | 2.06E-14 | 3 | Yipf4 | Col18a1 |
| Plekho2 | 1.11E-18 | 0.34020494 | 0.895 | 0.848 | 2.14E-14 | 3 | Plekho2 | Col18a1 |
| Cdyl21 | 1.20E-18 | 0.55563695 | 0.534 | 0.401 | 2.32E-14 | 3 | Cdyl2 | Col18a1 |
| Nsd3 | 1.34E-18 | 0.34593481 | 0.927 | 0.865 | 2.58E-14 | 3 | Nsd3 | Col18a1 |
| Dnajb12 | 1.50E-18 | 0.45644679 | 0.678 | 0.665 | 2.90E-14 | 3 | Dnajb1 | Col18a1 |
| Ktn1 | 1.56E-18 | 0.47783993 | 0.674 | 0.569 | 3.01E-14 | 3 | Ktn1 | Col18a1 |
| AI413582 | 1.87E-18 | 0.68970819 | 0.363 | 0.246 | 3.61E-14 | 3 | AI413582 | Col18a1 |
| Prkar1b | 2.19E-18 | 0.71298584 | 0.282 | 0.185 | 4.24E-14 | 3 | Prkar1b | Col18a1 |
| Acsf21 | 2.29E-18 | 0.63890666 | 0.427 | 0.328 | 4.43E-14 | 3 | Acsf2 | Col18a1 |
| Arhgap32 | 2.61E-18 | 0.77416492 | 0.408 | 0.314 | 5.05E-14 | 3 | Arhgap32 | Col18a1 |
| Ubtd2 | 2.72E-18 | 0.45623596 | 0.765 | 0.677 | 5.25E-14 | 3 | Ubtd2 | Col18a1 |
| Sdf4 | 2.86E-18 | 0.32136399 | 0.958 | 0.92 | 5.53E-14 | 3 | Sdf4 | Col18a1 |
| Gsap | 3.10E-18 | 0.86982278 | 0.25 | 0.151 | 5.99E-14 | 3 | Gsap | Col18a1 |
| Psd1 | 3.47E-18 | 0.60384415 | 0.268 | 0.182 | 6.71E-14 | 3 | Psd | Col18a1 |
| Irf2bp21 | 3.79E-18 | 0.26714931 | 0.975 | 0.94 | 7.32E-14 | 3 | Irf2bp2 | Col18a1 |
| Gng5 | 4.20E-18 | 0.70352471 | 0.55 | 0.406 | 8.13E-14 | 3 | Gng5 | Col18a1 |
| Casp12 | 4.48E-18 | 0.53708026 | 0.542 | 0.432 | 8.66E-14 | 3 | Casp12 | Col18a1 |
| Phf21a | 5.42E-18 | 0.43723554 | 0.683 | 0.579 | 1.05E-13 | 3 | Phf21a | Col18a1 |
| Tanc11 | 5.55E-18 | 0.40021923 | 0.748 | 0.604 | 1.07E-13 | 3 | Tanc1 | Col18a1 |
| Dcaf11 | 5.72E-18 | 0.62258409 | 0.522 | 0.406 | 1.11E-13 | 3 | Dcaf11 | Col18a1 |
| Zmym2 | 5.77E-18 | 0.57549365 | 0.589 | 0.468 | 1.12E-13 | 3 | Zmym2 | Col18a1 |
| Fhl3 | 6.41E-18 | 0.46865577 | 0.796 | 0.695 | 1.24E-13 | 3 | Fhl3 | Col18a1 |
| Dram11 | 6.66E-18 | 0.33946489 | 0.906 | 0.839 | 1.29E-13 | 3 | Dram1 | Col18a1 |
| Wdr13 | 7.98E-18 | 0.48935934 | 0.61 | 0.508 | 1.54E-13 | 3 | Wdr13 | Col18a1 |
| Stat6 | 8.33E-18 | 0.4186084 | 0.762 | 0.674 | 1.61E-13 | 3 | Stat6 | Col18a1 |
| Shc41 | 8.97E-18 | 0.44702822 | 0.564 | 0.477 | 1.73E-13 | 3 | Shc4 | Col18a1 |
| Stag2 | 9.17E-18 | 0.60348306 | 0.654 | 0.52 | 1.77E-13 | 3 | Stag2 | Col18a1 |
| Rbm18 | 1.04E-17 | 0.65442937 | 0.551 | 0.427 | 2.02E-13 | 3 | Rbm18 | Col18a1 |
| Tcf7 | 1.07E-17 | 0.80867471 | 0.265 | 0.166 | 2.07E-13 | 3 | Tcf7 | Col18a1 |
| Fbxo33 | 1.11E-17 | 0.70666429 | 0.432 | 0.32 | 2.15E-13 | 3 | Fbxo33 | Col18a1 |
| Arhgef11 | 1.18E-17 | 0.63341363 | 0.568 | 0.44 | 2.27E-13 | 3 | Arhgef11 | Col18a1 |
| Vsir | 1.23E-17 | 0.39935997 | 0.517 | 0.451 | 2.38E-13 | 3 | Vsir | Col18a1 |
| Etnk1 | 1.24E-17 | 0.45743089 | 0.819 | 0.733 | 2.40E-13 | 3 | Etnk1 | Col18a1 |
| Myo101 | 1.36E-17 | 0.46227166 | 0.667 | 0.536 | 2.63E-13 | 3 | Myo10 | Col18a1 |
| Srsf5 | 1.37E-17 | 0.2518565 | 0.953 | 0.914 | 2.64E-13 | 3 | Srsf5 | Col18a1 |
| Wipi11 | 1.40E-17 | 0.37221009 | 0.963 | 0.908 | 2.70E-13 | 3 | Wipi1 | Col18a1 |
| Zhx1 | 1.40E-17 | 0.6010729 | 0.557 | 0.433 | 2.71E-13 | 3 | Zhx1 | Col18a1 |
| Atp6v1a | 1.70E-17 | 0.40678947 | 0.772 | 0.714 | 3.30E-13 | 3 | Atp6v1a | Col18a1 |
| Selenok | 1.78E-17 | 0.38101203 | 0.892 | 0.825 | 3.44E-13 | 3 | Selenok | Col18a1 |
| Mocs1 | 1.80E-17 | 0.42828462 | 0.492 | 0.386 | 3.48E-13 | 3 | Mocs1 | Col18a1 |
| Pcmtd2 | 2.08E-17 | 0.43105233 | 0.46 | 0.356 | 4.03E-13 | 3 | Pcmtd2 | Col18a1 |
| Rfx7 | 2.38E-17 | 0.6456372 | 0.401 | 0.293 | 4.61E-13 | 3 | Rfx7 | Col18a1 |
| Hsd17b10 | 2.40E-17 | 0.46427138 | 0.788 | 0.692 | 4.64E-13 | 3 | Hsd17b10 | Col18a1 |
| Efr3a | 3.06E-17 | 0.56737592 | 0.687 | 0.577 | 5.91E-13 | 3 | Efr3a | Col18a1 |
| Nampt | 3.33E-17 | 0.63086724 | 0.492 | 0.386 | 6.43E-13 | 3 | Nampt | Col18a1 |
| Nup981 | 3.34E-17 | 0.6472714 | 0.716 | 0.577 | 6.46E-13 | 3 | Nup98 | Col18a1 |
| Trib2 | 3.40E-17 | 0.68110434 | 0.337 | 0.261 | 6.57E-13 | 3 | Trib2 | Col18a1 |
| Snx251 | 3.94E-17 | 0.60464644 | 0.523 | 0.415 | 7.62E-13 | 3 | Snx25 | Col18a1 |
| Arsk | 3.94E-17 | 0.92041332 | 0.329 | 0.221 | 7.62E-13 | 3 | Arsk | Col18a1 |
| Srebf1 | 4.04E-17 | 0.41785424 | 0.743 | 0.649 | 7.80E-13 | 3 | Srebf1 | Col18a1 |
| Smim11 | 4.36E-17 | 0.59149207 | 0.354 | 0.252 | 8.43E-13 | 3 | Smim1 | Col18a1 |
| Stard13 | 4.38E-17 | 0.56968344 | 0.514 | 0.431 | 8.46E-13 | 3 | Stard13 | Col18a1 |
| Ly96 | 4.50E-17 | 0.74607374 | 0.341 | 0.236 | 8.70E-13 | 3 | Ly96 | Col18a1 |
| Api5 | 4.56E-17 | 0.49459541 | 0.815 | 0.697 | 8.81E-13 | 3 | Api5 | Col18a1 |
| Card19 | 4.59E-17 | 0.47441887 | 0.549 | 0.474 | 8.87E-13 | 3 | Card19 | Col18a1 |
| Rnf217 | 4.79E-17 | 0.57275463 | 0.522 | 0.426 | 9.27E-13 | 3 | Rnf217 | Col18a1 |
| Oga | 4.99E-17 | 0.46735052 | 0.765 | 0.659 | 9.64E-13 | 3 | Oga | Col18a1 |
| Eml2 | 5.71E-17 | 0.57614438 | 0.4 | 0.298 | 1.10E-12 | 3 | Eml2 | Col18a1 |
| Mbtd1 | 5.83E-17 | 0.57922232 | 0.542 | 0.423 | 1.13E-12 | 3 | Mbtd1 | Col18a1 |
| Sms1 | 6.65E-17 | 0.92925242 | 0.265 | 0.182 | 1.28E-12 | 3 | Sms | Col18a1 |
| Arl6ip5 | 6.90E-17 | 0.43147672 | 0.842 | 0.763 | 1.33E-12 | 3 | Arl6ip5 | Col18a1 |
| Erp44 | 7.14E-17 | 0.53297664 | 0.832 | 0.706 | 1.38E-12 | 3 | Erp44 | Col18a1 |
| Lama2 | 7.63E-17 | 0.33018743 | 0.751 | 0.66 | 1.48E-12 | 3 | Lama2 | Col18a1 |
| Mff | 8.55E-17 | 0.49160084 | 0.762 | 0.656 | 1.65E-12 | 3 | Mff | Col18a1 |
| Eef1b2 | 9.48E-17 | 0.36809236 | 0.985 | 0.954 | 1.83E-12 | 3 | Eef1b2 | Col18a1 |
| Speg1 | 9.76E-17 | 0.66664835 | 0.338 | 0.236 | 1.89E-12 | 3 | Speg | Col18a1 |
| Rbbp6 | 1.02E-16 | 0.45216188 | 0.718 | 0.635 | 1.97E-12 | 3 | Rbbp6 | Col18a1 |
| Hnrnpr1 | 1.06E-16 | 0.58325254 | 0.735 | 0.602 | 2.04E-12 | 3 | Hnrnpr | Col18a1 |
| Syne31 | 1.08E-16 | 0.58319546 | 0.503 | 0.368 | 2.09E-12 | 3 | Syne3 | Col18a1 |
| Mapk13 | 1.09E-16 | 0.44395855 | 0.899 | 0.804 | 2.11E-12 | 3 | Mapk1 | Col18a1 |
| Ldb1 | 1.18E-16 | 0.36340729 | 0.87 | 0.797 | 2.27E-12 | 3 | Ldb1 | Col18a1 |
| Layn1 | 1.19E-16 | 0.45038038 | 0.603 | 0.522 | 2.30E-12 | 3 | Layn | Col18a1 |
| Psma31 | 1.33E-16 | 0.53776605 | 0.733 | 0.61 | 2.57E-12 | 3 | Psma3 | Col18a1 |
| Capns1 | 1.41E-16 | 0.43047734 | 0.925 | 0.838 | 2.72E-12 | 3 | Capns1 | Col18a1 |
| Mgat4b | 1.42E-16 | 0.5779144 | 0.624 | 0.521 | 2.75E-12 | 3 | Mgat4b | Col18a1 |
| Isg20 | 1.64E-16 | 0.44023894 | 0.543 | 0.471 | 3.18E-12 | 3 | Isg20 | Col18a1 |
| Dhcr71 | 1.66E-16 | 0.61666314 | 0.35 | 0.284 | 3.21E-12 | 3 | Dhcr7 | Col18a1 |
| Smyd2 | 1.68E-16 | 0.48125655 | 0.777 | 0.653 | 3.25E-12 | 3 | Smyd2 | Col18a1 |
| Ankrd121 | 1.71E-16 | 0.5687914 | 0.484 | 0.386 | 3.31E-12 | 3 | Ankrd12 | Col18a1 |
| Iffo2 | 1.79E-16 | 0.44960389 | 0.634 | 0.502 | 3.46E-12 | 3 | Iffo2 | Col18a1 |
| Decr2 | 2.01E-16 | 0.64121992 | 0.447 | 0.333 | 3.89E-12 | 3 | Decr2 | Col18a1 |
| Mndal | 2.05E-16 | 0.77279954 | 0.357 | 0.262 | 3.97E-12 | 3 | Mndal | Col18a1 |
| Timm8b | 2.20E-16 | 0.50614214 | 0.687 | 0.586 | 4.25E-12 | 3 | Timm8b | Col18a1 |
| Snx14 | 2.26E-16 | 0.66787567 | 0.463 | 0.369 | 4.36E-12 | 3 | Snx14 | Col18a1 |
| Hexa | 2.39E-16 | 0.33890181 | 0.936 | 0.882 | 4.62E-12 | 3 | Hexa | Col18a1 |
| Ifnar1 | 2.42E-16 | 0.50926144 | 0.811 | 0.686 | 4.68E-12 | 3 | Ifnar1 | Col18a1 |
| Lrif1 | 2.49E-16 | 0.89928204 | 0.268 | 0.181 | 4.82E-12 | 3 | Lrif1 | Col18a1 |
| Tbx15 | 2.64E-16 | 0.55087461 | 0.774 | 0.668 | 5.10E-12 | 3 | Tbx15 | Col18a1 |
| Irs21 | 3.38E-16 | 0.41663919 | 0.656 | 0.612 | 6.54E-12 | 3 | Irs2 | Col18a1 |
| Gtf2h1 | 3.40E-16 | 0.57909599 | 0.477 | 0.36 | 6.58E-12 | 3 | Gtf2h1 | Col18a1 |
| Map4k3 | 3.47E-16 | 0.45459667 | 0.669 | 0.559 | 6.71E-12 | 3 | Map4k3 | Col18a1 |
| Socs5 | 3.72E-16 | 0.45545883 | 0.744 | 0.668 | 7.20E-12 | 3 | Socs5 | Col18a1 |
| B2m | 4.13E-16 | 0.44179957 | 0.918 | 0.839 | 7.99E-12 | 3 | B2m | Col18a1 |
| Tcea1 | 4.39E-16 | 0.52530715 | 0.774 | 0.659 | 8.49E-12 | 3 | Tcea1 | Col18a1 |
| Rarb | 5.29E-16 | 0.65007242 | 0.312 | 0.229 | 1.02E-11 | 3 | Rarb | Col18a1 |
| Calm3 | 6.46E-16 | 0.2795334 | 0.982 | 0.956 | 1.25E-11 | 3 | Calm3 | Col18a1 |
| Rnf144a1 | 6.69E-16 | 0.49935875 | 0.623 | 0.562 | 1.29E-11 | 3 | Rnf144a | Col18a1 |
| Pfkfb3 | 6.74E-16 | 0.50122813 | 0.404 | 0.348 | 1.30E-11 | 3 | Pfkfb3 | Col18a1 |
| Crtc3 | 6.85E-16 | 0.47227125 | 0.623 | 0.527 | 1.32E-11 | 3 | Crtc3 | Col18a1 |
| AW549877 | 8.26E-16 | 0.40320785 | 0.609 | 0.522 | 1.60E-11 | 3 | AW549877 | Col18a1 |
| Tpr1 | 9.02E-16 | 0.52984282 | 0.807 | 0.68 | 1.74E-11 | 3 | Tpr | Col18a1 |
| Prrc2b | 9.06E-16 | 0.37152476 | 0.89 | 0.839 | 1.75E-11 | 3 | Prrc2b | Col18a1 |
| Ralgds1 | 9.53E-16 | 0.56940775 | 0.58 | 0.475 | 1.84E-11 | 3 | Ralgds | Col18a1 |
| Tnrc6b | 9.60E-16 | 0.50941819 | 0.671 | 0.567 | 1.86E-11 | 3 | Tnrc6b | Col18a1 |
| Ptges | 9.68E-16 | 0.4026274 | 0.782 | 0.733 | 1.87E-11 | 3 | Ptges | Col18a1 |
| Ndrg3 | 9.99E-16 | 0.58898782 | 0.531 | 0.431 | 1.93E-11 | 3 | Ndrg3 | Col18a1 |
| Ppp4r3b | 1.01E-15 | 0.60135196 | 0.585 | 0.465 | 1.96E-11 | 3 | Ppp4r3b | Col18a1 |
| Arhgap5 | 1.19E-15 | 0.74251999 | 0.474 | 0.35 | 2.30E-11 | 3 | Arhgap5 | Col18a1 |
| Tjp1 | 1.31E-15 | 0.46963973 | 0.705 | 0.609 | 2.54E-11 | 3 | Tjp1 | Col18a1 |
| Rab34 | 1.47E-15 | 0.31469648 | 0.871 | 0.82 | 2.83E-11 | 3 | Rab34 | Col18a1 |
| Smim131 | 1.52E-15 | 0.63212973 | 0.473 | 0.369 | 2.94E-11 | 3 | Smim13 | Col18a1 |
| Ggnbp2 | 1.58E-15 | 0.48389909 | 0.754 | 0.64 | 3.06E-11 | 3 | Ggnbp2 | Col18a1 |
| Agpat31 | 1.60E-15 | 0.39929252 | 0.648 | 0.484 | 3.10E-11 | 3 | Agpat3 | Col18a1 |
| Ccdc9b1 | 1.76E-15 | 0.40601308 | 0.434 | 0.348 | 3.40E-11 | 3 | Ccdc9b | Col18a1 |
| Mbtps1 | 1.80E-15 | 0.43663833 | 0.864 | 0.772 | 3.49E-11 | 3 | Mbtps1 | Col18a1 |
| Npepps | 2.04E-15 | 0.45505584 | 0.816 | 0.738 | 3.94E-11 | 3 | Npepps | Col18a1 |
| Pik3ca | 2.08E-15 | 0.46698387 | 0.732 | 0.611 | 4.02E-11 | 3 | Pik3ca | Col18a1 |
| Snx91 | 2.09E-15 | 0.30221742 | 0.96 | 0.913 | 4.04E-11 | 3 | Snx9 | Col18a1 |
| Pan3 | 2.12E-15 | 0.48915704 | 0.559 | 0.457 | 4.10E-11 | 3 | Pan3 | Col18a1 |
| Zfp6911 | 2.25E-15 | 0.63107929 | 0.33 | 0.233 | 4.36E-11 | 3 | Zfp691 | Col18a1 |
| Kdm3a | 2.71E-15 | 0.35828444 | 0.605 | 0.523 | 5.23E-11 | 3 | Kdm3a | Col18a1 |
| Pknox1 | 2.85E-15 | 0.78260844 | 0.426 | 0.32 | 5.51E-11 | 3 | Pknox1 | Col18a1 |
| Axl1 | 3.30E-15 | 0.50455868 | 0.668 | 0.563 | 6.38E-11 | 3 | Axl1 | Col18a1 |
| Anks1 | 3.38E-15 | 0.50642802 | 0.452 | 0.368 | 6.54E-11 | 3 | Anks1 | Col18a1 |
| Hgsnat | 3.39E-15 | 0.49970379 | 0.813 | 0.727 | 6.55E-11 | 3 | Hgsnat | Col18a1 |
| Calcoco11 | 3.61E-15 | 0.32422237 | 0.696 | 0.6 | 6.98E-11 | 3 | Calcoco1 | Col18a1 |
| Suclg2 | 3.65E-15 | 0.74839708 | 0.405 | 0.319 | 7.05E-11 | 3 | Suclg2 | Col18a1 |
| Ghr | 4.21E-15 | 0.48058888 | 0.564 | 0.415 | 8.15E-11 | 3 | Ghr | Col18a1 |
| Zfp592 | 4.73E-15 | 0.52517905 | 0.63 | 0.528 | 9.15E-11 | 3 | Zfp592 | Col18a1 |
| Ndufa6 | 5.46E-15 | 0.46519 | 0.879 | 0.793 | 1.06E-10 | 3 | Ndufa6 | Col18a1 |
| Cux11 | 5.74E-15 | 0.32182548 | 0.703 | 0.626 | 1.11E-10 | 3 | Cux1 | Col18a1 |
| Adam33 | 6.83E-15 | 0.48495543 | 0.302 | 0.198 | 1.32E-10 | 3 | Adam33 | Col18a1 |
| Lacc1 | 7.27E-15 | 0.70890688 | 0.307 | 0.224 | 1.41E-10 | 3 | Lacc1 | Col18a1 |
| Smad7 | 7.83E-15 | 0.46156909 | 0.674 | 0.623 | 1.51E-10 | 3 | Smad7 | Col18a1 |
| Zfand5 | 7.96E-15 | 0.40733801 | 0.914 | 0.831 | 1.54E-10 | 3 | Zfand5 | Col18a1 |
| Gabbr1 | 8.37E-15 | 0.3011855 | 0.559 | 0.464 | 1.62E-10 | 3 | Gabbr1 | Col18a1 |
| Hist3h2a | 8.42E-15 | 0.65700058 | 0.28 | 0.208 | 1.63E-10 | 3 | Hist3h2a | Col18a1 |
| Arxes21 | 8.52E-15 | 0.53787769 | 0.498 | 0.373 | 1.65E-10 | 3 | Arxes2 | Col18a1 |
| Nlk1 | 1.05E-14 | 0.57020758 | 0.468 | 0.365 | 2.03E-10 | 3 | Nlk | Col18a1 |
| Kdm6a | 1.22E-14 | 0.52744348 | 0.509 | 0.418 | 2.36E-10 | 3 | Kdm6a | Col18a1 |
| Itpr2 | 1.32E-14 | 0.52148766 | 0.43 | 0.345 | 2.55E-10 | 3 | Itpr2 | Col18a1 |
| Ccdc122 | 1.69E-14 | 0.92579135 | 0.28 | 0.194 | 3.27E-10 | 3 | Ccdc122 | Col18a1 |
| Chmp4b | 1.78E-14 | 0.43368423 | 0.911 | 0.828 | 3.45E-10 | 3 | Chmp4b | Col18a1 |
| Pum2 | 1.82E-14 | 0.44017426 | 0.916 | 0.831 | 3.53E-10 | 3 | Pum2 | Col18a1 |
| Sf3b1 | 2.13E-14 | 0.35780007 | 0.952 | 0.893 | 4.13E-10 | 3 | Sf3b1 | Col18a1 |
| Auh | 2.19E-14 | 0.36513985 | 0.67 | 0.596 | 4.23E-10 | 3 | Auh | Col18a1 |
| Casp41 | 2.28E-14 | 0.54048921 | 0.526 | 0.44 | 4.41E-10 | 3 | Casp4 | Col18a1 |
| Zfp799 | 2.46E-14 | 0.74375119 | 0.287 | 0.198 | 4.76E-10 | 3 | Zfp799 | Col18a1 |
| Mroh1 | 2.69E-14 | 0.38349115 | 0.478 | 0.386 | 5.21E-10 | 3 | Mroh1 | Col18a1 |
| Ptpra | 2.92E-14 | 0.38729685 | 0.926 | 0.853 | 5.65E-10 | 3 | Ptpra | Col18a1 |
| Khnyn | 3.25E-14 | 0.39348635 | 0.475 | 0.411 | 6.28E-10 | 3 | Khnyn | Col18a1 |
| Tomm70a1 | 3.95E-14 | 0.67387936 | 0.629 | 0.479 | 7.64E-10 | 3 | Tomm70a | Col18a1 |
| Pon2 | 5.13E-14 | 0.46836648 | 0.648 | 0.553 | 9.92E-10 | 3 | Pon2 | Col18a1 |
| Trip12 | 5.32E-14 | 0.396138 | 0.949 | 0.877 | 1.03E-09 | 3 | Trip12 | Col18a1 |
| Gng21 | 5.64E-14 | 0.45521625 | 0.448 | 0.305 | 1.09E-09 | 3 | Gng2 | Col18a1 |
| Ppp1r12b | 6.59E-14 | 0.65214421 | 0.405 | 0.292 | 1.27E-09 | 3 | Ppp1r12b | Col18a1 |
| Cachd1 | 6.77E-14 | 0.52366903 | 0.55 | 0.458 | 1.31E-09 | 3 | Cachd1 | Col18a1 |
| Ube2h | 7.27E-14 | 0.38336119 | 0.784 | 0.711 | 1.41E-09 | 3 | Ube2h | Col18a1 |
| Ddx6 | 7.50E-14 | 0.42026855 | 0.919 | 0.833 | 1.45E-09 | 3 | Ddx6 | Col18a1 |
| Capza1 | 7.74E-14 | 0.56391563 | 0.675 | 0.553 | 1.50E-09 | 3 | Capza1 | Col18a1 |
| Fnip2 | 7.77E-14 | 0.5254386 | 0.463 | 0.401 | 1.50E-09 | 3 | Fnip2 | Col18a1 |
| Zfp608 | 8.29E-14 | 0.55054615 | 0.614 | 0.492 | 1.60E-09 | 3 | Zfp608 | Col18a1 |
| Ezh1 | 8.41E-14 | 0.51292316 | 0.469 | 0.368 | 1.63E-09 | 3 | Ezh1 | Col18a1 |
| Atp6v1f | 8.82E-14 | 0.37479576 | 0.78 | 0.699 | 1.71E-09 | 3 | Atp6v1f | Col18a1 |
| Eif1 | 9.68E-14 | 0.50075828 | 0.841 | 0.745 | 1.87E-09 | 3 | Eif1 | Col18a1 |
| Grhpr | 9.87E-14 | 0.51204355 | 0.514 | 0.424 | 1.91E-09 | 3 | Grhpr | Col18a1 |
| Fibp | 1.03E-13 | 0.41069487 | 0.75 | 0.674 | 1.99E-09 | 3 | Fibp | Col18a1 |
| Eif3k | 1.27E-13 | 0.36004052 | 0.961 | 0.909 | 2.46E-09 | 3 | Eif3k | Col18a1 |
| Alg11 | 1.30E-13 | 0.70953473 | 0.448 | 0.337 | 2.52E-09 | 3 | Alg11 | Col18a1 |
| Dag1 | 1.33E-13 | 0.39869299 | 0.93 | 0.848 | 2.58E-09 | 3 | Dag1 | Col18a1 |
| Sde2 | 1.37E-13 | 0.52318413 | 0.572 | 0.48 | 2.66E-09 | 3 | Sde2 | Col18a1 |
| Mvd1 | 1.44E-13 | 0.62744183 | 0.266 | 0.221 | 2.78E-09 | 3 | Mvd | Col18a1 |
| Glt8d1 | 1.45E-13 | 0.48554729 | 0.778 | 0.667 | 2.80E-09 | 3 | Glt8d1 | Col18a1 |
| Srgap21 | 1.47E-13 | 0.45856921 | 0.585 | 0.473 | 2.85E-09 | 3 | Srgap2 | Col18a1 |
| Nrp21 | 1.49E-13 | 0.50789497 | 0.506 | 0.412 | 2.88E-09 | 3 | Nrp2 | Col18a1 |
| Efcab141 | 1.80E-13 | 0.48552805 | 0.661 | 0.56 | 3.48E-09 | 3 | Efcab14 | Col18a1 |
| Tut7 | 1.81E-13 | 0.37455463 | 0.831 | 0.752 | 3.50E-09 | 3 | Tut7 | Col18a1 |
| Ctso | 1.84E-13 | 0.46851418 | 0.574 | 0.475 | 3.55E-09 | 3 | Ctso | Col18a1 |
| Carmmt11 | 2.03E-13 | 0.6806257 | 0.417 | 0.316 | 3.92E-09 | 3 | Carmmt1 | Col18a1 |
| Sacm1l | 2.19E-13 | 0.52667235 | 0.714 | 0.603 | 4.24E-09 | 3 | Sacm1l | Col18a1 |
| Cerk | 2.23E-13 | 0.60335304 | 0.396 | 0.308 | 4.30E-09 | 3 | Cerk | Col18a1 |
| Hspa9 | 2.38E-13 | 0.45111823 | 0.736 | 0.634 | 4.60E-09 | 3 | Hspa9 | Col18a1 |
| Atrx | 2.50E-13 | 0.41028533 | 0.783 | 0.681 | 4.82E-09 | 3 | Atrx | Col18a1 |
| Zc3h12a | 2.67E-13 | 0.38217601 | 0.518 | 0.463 | 5.16E-09 | 3 | Zc3h12a | Col18a1 |
| Rab43 | 2.74E-13 | 0.40063114 | 0.786 | 0.712 | 5.29E-09 | 3 | Rab43 | Col18a1 |
| Sp1 | 2.92E-13 | 0.5470878 | 0.65 | 0.511 | 5.65E-09 | 3 | Sp1 | Col18a1 |
| Abi2 | 2.99E-13 | 0.55600494 | 0.53 | 0.423 | 5.78E-09 | 3 | Abi2 | Col18a1 |
| Pebp1 | 3.03E-13 | 0.36879578 | 0.887 | 0.817 | 5.86E-09 | 3 | Pebp1 | Col18a1 |
| Capza2 | 3.26E-13 | 0.53406238 | 0.748 | 0.616 | 6.30E-09 | 3 | Capza2 | Col18a1 |
| Tmem39a | 3.36E-13 | 0.47472464 | 0.805 | 0.711 | 6.50E-09 | 3 | Tmem39a | Col18a1 |
| Map2k1 | 3.54E-13 | 0.35382513 | 0.82 | 0.765 | 6.84E-09 | 3 | Map2k1 | Col18a1 |
| Mad2l21 | 3.67E-13 | 0.632039 | 0.411 | 0.303 | 7.10E-09 | 3 | Mad2l2 | Col18a1 |
| Hebp11 | 3.73E-13 | 0.33795719 | 0.425 | 0.353 | 7.21E-09 | 3 | Hebp1 | Col18a1 |
| Aacs | 3.74E-13 | 0.4684736 | 0.582 | 0.511 | 7.22E-09 | 3 | Aacs | Col18a1 |
| Kdm5a | 4.23E-13 | 0.41461159 | 0.708 | 0.618 | 8.19E-09 | 3 | Kdm5a | Col18a1 |
| Akt2 | 4.35E-13 | 0.40790909 | 0.82 | 0.751 | 8.41E-09 | 3 | Akt2 | Col18a1 |
| Xpa | 4.54E-13 | 0.53532785 | 0.477 | 0.393 | 8.78E-09 | 3 | Xpa | Col18a1 |
| Sdk11 | 5.33E-13 | 0.4823439 | 0.637 | 0.516 | 1.03E-08 | 3 | Sdk1 | Col18a1 |
| Klhl5 | 5.44E-13 | 0.4845647 | 0.768 | 0.663 | 1.05E-08 | 3 | Klhl5 | Col18a1 |
| Asah2 | 5.86E-13 | 0.82117436 | 0.321 | 0.232 | 1.13E-08 | 3 | Asah2 | Col18a1 |
| Paqr71 | 6.26E-13 | 0.48886533 | 0.423 | 0.332 | 1.21E-08 | 3 | Paqr7 | Col18a1 |
| Rragc | 6.74E-13 | 0.49647668 | 0.691 | 0.583 | 1.30E-08 | 3 | Rragc | Col18a1 |
| Vps35 | 6.84E-13 | 0.56857612 | 0.718 | 0.6 | 1.32E-08 | 3 | Vps35 | Col18a1 |
| Etfrf1 | 7.16E-13 | 0.45467461 | 0.483 | 0.409 | 1.38E-08 | 3 | Etfrf1 | Col18a1 |
| Cmip | 7.95E-13 | 0.3353189 | 0.853 | 0.801 | 1.54E-08 | 3 | Cmip | Col18a1 |
| Pcnx | 8.08E-13 | 0.48501784 | 0.728 | 0.618 | 1.56E-08 | 3 | Pcnx | Col18a1 |
| Nbr1 | 8.37E-13 | 0.47201233 | 0.625 | 0.523 | 1.62E-08 | 3 | Nbr1 | Col18a1 |
| Tsc2 | 8.38E-13 | 0.47947944 | 0.725 | 0.636 | 1.62E-08 | 3 | Tsc2 | Col18a1 |
| Zfp871 | 8.43E-13 | 0.52078736 | 0.643 | 0.537 | 1.63E-08 | 3 | Zfp871 | Col18a1 |
| Ube2g1 | 8.63E-13 | 0.43934563 | 0.738 | 0.636 | 1.67E-08 | 3 | Ube2g1 | Col18a1 |
| Tbx181 | 9.18E-13 | 0.59021517 | 0.435 | 0.341 | 1.77E-08 | 3 | Tbx18 | Col18a1 |
| H3f3b | 9.33E-13 | 0.43625262 | 0.96 | 0.886 | 1.80E-08 | 3 | H3f3b | Col18a1 |
| Rab5a | 1.09E-12 | 0.40352463 | 0.929 | 0.856 | 2.11E-08 | 3 | Rab5a | Col18a1 |
| Ccny | 1.11E-12 | 0.42023651 | 0.862 | 0.77 | 2.15E-08 | 3 | Ccny | Col18a1 |
| Zfp8091 | 1.28E-12 | 0.51488561 | 0.465 | 0.408 | 2.48E-08 | 3 | Zfp809 | Col18a1 |
| Fnip1 | 1.29E-12 | 0.47682358 | 0.642 | 0.536 | 2.50E-08 | 3 | Fnip1 | Col18a1 |
| Fam210b1 | 1.42E-12 | 0.64300118 | 0.308 | 0.212 | 2.75E-08 | 3 | Fam210b | Col18a1 |
| Smarca2 | 1.46E-12 | 0.49005428 | 0.694 | 0.594 | 2.82E-08 | 3 | Smarca2 | Col18a1 |
| Slc41a1 | 1.76E-12 | 0.43465776 | 0.72 | 0.657 | 3.40E-08 | 3 | Slc41a1 | Col18a1 |
| Sardh | 1.78E-12 | 0.44830211 | 0.458 | 0.362 | 3.44E-08 | 3 | Sardh | Col18a1 |
| Dennd6a | 2.10E-12 | 0.55010341 | 0.495 | 0.399 | 4.07E-08 | 3 | Dennd6a | Col18a1 |
| Ttpal1 | 2.19E-12 | 0.47208625 | 0.56 | 0.485 | 4.24E-08 | 3 | Ttpal | Col18a1 |
| Fndc3b1 | 2.38E-12 | 0.25607716 | 0.982 | 0.962 | 4.60E-08 | 3 | Fndc3b | Col18a1 |
| Atp5j | 2.49E-12 | 0.53979515 | 0.748 | 0.625 | 4.82E-08 | 3 | Atp5j | Col18a1 |
| Tet2 | 2.52E-12 | 0.57855754 | 0.502 | 0.401 | 4.87E-08 | 3 | Tet2 | Col18a1 |
| Yes1 | 2.76E-12 | 0.59998866 | 0.511 | 0.417 | 5.33E-08 | 3 | Yes1 | Col18a1 |
| Taf9b1 | 2.80E-12 | 0.75639722 | 0.32 | 0.251 | 5.42E-08 | 3 | Taf9b | Col18a1 |
| Ago2 | 2.90E-12 | 0.39697504 | 0.848 | 0.77 | 5.62E-08 | 3 | Ago2 | Col18a1 |
| Bmt2 | 2.93E-12 | 0.40108192 | 0.551 | 0.464 | 5.67E-08 | 3 | Bmt2 | Col18a1 |
| Csde1 | 3.06E-12 | 0.41782165 | 0.932 | 0.833 | 5.92E-08 | 3 | Csde1 | Col18a1 |
| Ddx50 | 3.28E-12 | 0.43079275 | 0.828 | 0.725 | 6.33E-08 | 3 | Ddx50 | Col18a1 |
| Vps41 | 3.34E-12 | 0.48170389 | 0.539 | 0.457 | 6.45E-08 | 3 | Vps41 | Col18a1 |
| Cirbp | 3.46E-12 | 0.31919652 | 0.658 | 0.573 | 6.70E-08 | 3 | Cirbp | Col18a1 |
| Zfp53 | 4.02E-12 | 0.74580007 | 0.303 | 0.22 | 7.78E-08 | 3 | Zfp53 | Col18a1 |
| Renbp | 4.03E-12 | 0.4286587 | 0.599 | 0.501 | 7.80E-08 | 3 | Renbp | Col18a1 |
| Asap1 | 4.06E-12 | 0.49798222 | 0.658 | 0.573 | 7.84E-08 | 3 | Asap1 | Col18a1 |
| Ccdc50 | 4.20E-12 | 0.40328701 | 0.705 | 0.594 | 8.12E-08 | 3 | Ccdc50 | Col18a1 |
| B4gat1 | 4.43E-12 | 0.30562841 | 0.717 | 0.646 | 8.57E-08 | 3 | B4gat1 | Col18a1 |
| Lrch3 | 4.45E-12 | 0.46639914 | 0.681 | 0.586 | 8.59E-08 | 3 | Lrch3 | Col18a1 |
| Meis2 | 4.48E-12 | 0.35564819 | 0.54 | 0.465 | 8.67E-08 | 3 | Meis2 | Col18a1 |
| Setd5 | 4.50E-12 | 0.36746053 | 0.885 | 0.816 | 8.69E-08 | 3 | Setd5 | Col18a1 |
| Tlr2 | 4.54E-12 | 0.59013957 | 0.259 | 0.178 | 8.78E-08 | 3 | Tlr2 | Col18a1 |
| Col24a11 | 4.78E-12 | 0.530964 | 0.483 | 0.395 | 9.25E-08 | 3 | Col24a1 | Col18a1 |
| Wdr26 | 5.32E-12 | 0.30811404 | 0.957 | 0.903 | 1.03E-07 | 3 | Wdr26 | Col18a1 |
| Fam110b | 5.60E-12 | 0.59027762 | 0.348 | 0.255 | 1.08E-07 | 3 | Fam110b | Col18a1 |
| Mllt6 | 5.75E-12 | 0.30493831 | 0.822 | 0.753 | 1.11E-07 | 3 | Mllt6 | Col18a1 |
| Fignl2 | 5.86E-12 | 0.40941709 | 0.416 | 0.325 | 1.13E-07 | 3 | Fignl2 | Col18a1 |
| Arl21 | 6.69E-12 | 0.33757977 | 0.619 | 0.547 | 1.29E-07 | 3 | Arl2 | Col18a1 |
| Nbdy | 6.83E-12 | 0.53774262 | 0.454 | 0.354 | 1.32E-07 | 3 | Nbdy | Col18a1 |
| Rnft1 | 6.86E-12 | 0.5192824 | 0.658 | 0.545 | 1.33E-07 | 3 | Rnft1 | Col18a1 |
| Thoc2 | 7.72E-12 | 0.51929597 | 0.592 | 0.477 | 1.49E-07 | 3 | Thoc2 | Col18a1 |
| Egr31 | 7.75E-12 | 0.50594981 | 0.533 | 0.437 | 1.50E-07 | 3 | Egr3 | Col18a1 |
| Pyroxd1 | 8.51E-12 | 0.71315172 | 0.35 | 0.27 | 1.65E-07 | 3 | Pyroxd1 | Col18a1 |
| Cfap36 | 8.54E-12 | 0.64030236 | 0.378 | 0.284 | 1.65E-07 | 3 | Cfap36 | Col18a1 |
| Gngt2 | 9.71E-12 | 0.44963914 | 0.501 | 0.399 | 1.88E-07 | 3 | Gngt2 | Col18a1 |
| Ppp2r5a | 9.91E-12 | 0.46300231 | 0.706 | 0.606 | 1.92E-07 | 3 | Ppp2r5a | Col18a1 |
| Arhgef25 | 1.01E-11 | 0.2841706 | 0.92 | 0.866 | 1.95E-07 | 3 | Arhgef25 | Col18a1 |
| Eif3m | 1.02E-11 | 0.69548374 | 0.43 | 0.332 | 1.98E-07 | 3 | Eif3m | Col18a1 |
| Pgd | 1.05E-11 | 0.36059021 | 0.812 | 0.737 | 2.02E-07 | 3 | Pgd | Col18a1 |
| Nr1d1 | 1.06E-11 | 0.36603035 | 0.594 | 0.478 | 2.04E-07 | 3 | Nr1d1 | Col18a1 |
| Slc25a321 | 1.09E-11 | 0.73171324 | 0.401 | 0.309 | 2.10E-07 | 3 | Slc25a32 | Col18a1 |
| Inpp5e | 1.13E-11 | 0.66142023 | 0.28 | 0.207 | 2.19E-07 | 3 | Inpp5e | Col18a1 |
| Sirt1 | 1.14E-11 | 0.55657265 | 0.349 | 0.294 | 2.21E-07 | 3 | Sirt1 | Col18a1 |
| Crebbp | 1.18E-11 | 0.44559758 | 0.796 | 0.694 | 2.28E-07 | 3 | Crebbp | Col18a1 |
| Cop1 | 1.23E-11 | 0.42054922 | 0.748 | 0.657 | 2.38E-07 | 3 | Cop1 | Col18a1 |
| Slc4a71 | 1.28E-11 | 0.75442522 | 0.344 | 0.255 | 2.47E-07 | 3 | Slc4a7 | Col18a1 |
| Epm2aip11 | 1.30E-11 | 0.39679119 | 0.616 | 0.527 | 2.50E-07 | 3 | Epm2aip1 | Col18a1 |
| Ccdc821 | 1.31E-11 | 0.3195631 | 0.488 | 0.371 | 2.53E-07 | 3 | Ccdc82 | Col18a1 |
| Sema4b | 1.41E-11 | 0.40477542 | 0.615 | 0.522 | 2.72E-07 | 3 | Sema4b | Col18a1 |
| Vamp3 | 1.42E-11 | 0.51918886 | 0.783 | 0.664 | 2.75E-07 | 3 | Vamp3 | Col18a1 |
| Epn21 | 1.47E-11 | 0.48273044 | 0.562 | 0.463 | 2.84E-07 | 3 | Epn2 | Col18a1 |
| Sdf2 | 1.72E-11 | 0.33443935 | 0.896 | 0.827 | 3.32E-07 | 3 | Sdf2 | Col18a1 |
| Azi2 | 1.77E-11 | 0.40005433 | 0.621 | 0.535 | 3.43E-07 | 3 | Azi2 | Col18a1 |
| Fkbp8 | 1.85E-11 | 0.31955203 | 0.957 | 0.908 | 3.58E-07 | 3 | Fkbp8 | Col18a1 |
| Prxl2b | 2.08E-11 | 0.71119295 | 0.273 | 0.195 | 4.02E-07 | 3 | Prxl2b | Col18a1 |
| Arih1 | 2.28E-11 | 0.42839989 | 0.862 | 0.765 | 4.40E-07 | 3 | Arih1 | Col18a1 |
| Rc3h2 | 2.43E-11 | 0.47675749 | 0.674 | 0.571 | 4.69E-07 | 3 | Rc3h2 | Col18a1 |
| Sgcd | 2.46E-11 | 0.58541495 | 0.39 | 0.312 | 4.76E-07 | 3 | Sgcd | Col18a1 |
| Ash1l | 2.50E-11 | 0.38922263 | 0.792 | 0.702 | 4.84E-07 | 3 | Ash1l | Col18a1 |
| Tmx3 | 2.85E-11 | 0.43833901 | 0.752 | 0.658 | 5.50E-07 | 3 | Tmx3 | Col18a1 |
| Atxn10 | 3.23E-11 | 0.39613414 | 0.893 | 0.808 | 6.24E-07 | 3 | Atxn10 | Col18a1 |
| Pdp1 | 3.33E-11 | 0.7890307 | 0.313 | 0.227 | 6.44E-07 | 3 | Pdp1 | Col18a1 |
| Fbxl17 | 3.41E-11 | 0.50500673 | 0.658 | 0.537 | 6.59E-07 | 3 | Fbxl17 | Col18a1 |
| Aldh3a2 | 3.71E-11 | 0.51392003 | 0.52 | 0.418 | 7.18E-07 | 3 | Aldh3a2 | Col18a1 |
| Tmem63a | 4.79E-11 | 0.33339268 | 0.722 | 0.639 | 9.26E-07 | 3 | Tmem63a | Col18a1 |
| Ahnak2 | 4.85E-11 | 0.26155289 | 0.757 | 0.694 | 9.37E-07 | 3 | Ahnak2 | Col18a1 |
| Rwdd1 | 5.48E-11 | 0.54957785 | 0.581 | 0.479 | 1.06E-06 | 3 | Rwdd1 | Col18a1 |
| Zfp280d | 5.77E-11 | 0.45587051 | 0.552 | 0.466 | 1.11E-06 | 3 | Zfp280d | Col18a1 |
| Srr | 6.60E-11 | 0.62924592 | 0.295 | 0.212 | 1.28E-06 | 3 | Srr | Col18a1 |
| Tmem41b | 7.10E-11 | 0.5981523 | 0.613 | 0.511 | 1.37E-06 | 3 | Tmem41b | Col18a1 |
| Zfp639 | 7.50E-11 | 0.60455994 | 0.489 | 0.404 | 1.45E-06 | 3 | Zfp639 | Col18a1 |
| Nicn1 | 7.54E-11 | 0.69489241 | 0.351 | 0.265 | 1.46E-06 | 3 | Nicn1 | Col18a1 |
| Dlg51 | 7.67E-11 | 0.26968749 | 0.727 | 0.671 | 1.48E-06 | 3 | Dlg5 | Col18a1 |
| Ep300 | 8.45E-11 | 0.39438376 | 0.707 | 0.613 | 1.63E-06 | 3 | Ep300 | Col18a1 |
| Gas | 8.91E-11 | 0.25299943 | 0.967 | 0.919 | 1.72E-06 | 3 | Gas | Col18a1 |
| Washc4 | 9.33E-11 | 0.50536706 | 0.598 | 0.5 | 1.80E-06 | 3 | Washc4 | Col18a1 |
| Fdx1 | 1.02E-10 | 0.62351957 | 0.451 | 0.357 | 1.97E-06 | 3 | Fdx1 | Col18a1 |
| Morf4l2 | 1.05E-10 | 0.4514203 | 0.947 | 0.853 | 2.03E-06 | 3 | Morf4l2 | Col18a1 |
| AI597479 | 1.07E-10 | 0.56186311 | 0.356 | 0.282 | 2.07E-06 | 3 | AI597479 | Col18a1 |
| Prmt2 | 1.09E-10 | 0.48978901 | 0.334 | 0.262 | 2.11E-06 | 3 | Prmt2 | Col18a1 |
| C1galt1c1 | 1.12E-10 | 0.5592241 | 0.512 | 0.427 | 2.16E-06 | 3 | C1galt1c1 | Col18a1 |
| Snapin | 1.40E-10 | 0.4524378 | 0.59 | 0.498 | 2.71E-06 | 3 | Snapin | Col18a1 |
| Irx2 | 1.47E-10 | 0.3255293 | 0.629 | 0.554 | 2.83E-06 | 3 | Irx2 | Col18a1 |
| Cog3 | 1.53E-10 | 0.4615099 | 0.54 | 0.473 | 2.96E-06 | 3 | Cog3 | Col18a1 |
| Ptpmt1 | 1.55E-10 | 0.53138042 | 0.576 | 0.476 | 3.00E-06 | 3 | Ptpmt1 | Col18a1 |
| Bivm | 1.60E-10 | 0.72636379 | 0.332 | 0.251 | 3.10E-06 | 3 | Bivm | Col18a1 |
| Sgpl11 | 1.64E-10 | 0.39072811 | 0.834 | 0.735 | 3.17E-06 | 3 | Sgpl1 | Col18a1 |
| Zbtb341 | 1.80E-10 | 0.41442073 | 0.538 | 0.433 | 3.47E-06 | 3 | Zbtb34 | Col18a1 |
| Agl | 1.85E-10 | 0.55753297 | 0.363 | 0.295 | 3.59E-06 | 3 | Agl | Col18a1 |
| Mob4 | 1.88E-10 | 0.50847335 | 0.592 | 0.502 | 3.64E-06 | 3 | Mob4 | Col18a1 |
| Spen | 1.94E-10 | 0.42319308 | 0.686 | 0.593 | 3.76E-06 | 3 | Spen | Col18a1 |
| Stk3 | 1.96E-10 | 0.30308454 | 0.782 | 0.716 | 3.79E-06 | 3 | Stk3 | Col18a1 |
| Erg281 | 1.97E-10 | 0.4824517 | 0.641 | 0.566 | 3.80E-06 | 3 | Erg28 | Col18a1 |
| Nf1 | 2.00E-10 | 0.42405356 | 0.747 | 0.667 | 3.86E-06 | 3 | Nf1 | Col18a1 |
| H2afy21 | 2.03E-10 | 0.55761993 | 0.458 | 0.375 | 3.92E-06 | 3 | H2afy2 | Col18a1 |
| Impact | 2.16E-10 | 0.32770653 | 0.659 | 0.593 | 4.17E-06 | 3 | Impact | Col18a1 |
| Slc7a8 | 2.24E-10 | 0.4650911 | 0.257 | 0.185 | 4.34E-06 | 3 | Slc7a8 | Col18a1 |
| Clk4 | 2.41E-10 | 0.36757946 | 0.588 | 0.514 | 4.65E-06 | 3 | Clk4 | Col18a1 |
| Selenof | 2.58E-10 | 0.39360708 | 0.829 | 0.739 | 4.98E-06 | 3 | Selenof | Col18a1 |
| Maf1 | 2.69E-10 | 0.46522606 | 0.542 | 0.458 | 5.20E-06 | 3 | Maf1 | Col18a1 |
| Traf6 | 2.89E-10 | 0.53300832 | 0.396 | 0.333 | 5.58E-06 | 3 | Traf6 | Col18a1 |
| Pim31 | 3.02E-10 | 0.47429429 | 0.468 | 0.413 | 5.85E-06 | 3 | Pim3 | Col18a1 |
| Dmxl1 | 3.04E-10 | 0.51956623 | 0.509 | 0.442 | 5.87E-06 | 3 | Dmxl1 | Col18a1 |
| Hnrnpk1 | 3.24E-10 | 0.38809112 | 0.947 | 0.881 | 6.26E-06 | 3 | Hnrnpk | Col18a1 |
| Plxnd1 | 3.36E-10 | 0.36192967 | 0.629 | 0.545 | 6.50E-06 | 3 | Plxnd1 | Col18a1 |
| Slc48a1 | 3.50E-10 | 0.32811948 | 0.529 | 0.447 | 6.78E-06 | 3 | Slc48a1 | Col18a1 |
| Bicral | 3.58E-10 | 0.58315586 | 0.409 | 0.33 | 6.93E-06 | 3 | Bicral | Col18a1 |
| Zmynd11 | 3.71E-10 | 0.36125461 | 0.908 | 0.828 | 7.18E-06 | 3 | Zmynd11 | Col18a1 |
| Mvk | 3.82E-10 | 0.40941985 | 0.497 | 0.437 | 7.39E-06 | 3 | Mvk | Col18a1 |
| Bnip2 | 4.18E-10 | 0.3828554 | 0.861 | 0.775 | 8.07E-06 | 3 | Bnip2 | Col18a1 |
| Abhd41 | 4.18E-10 | 0.28344242 | 0.817 | 0.744 | 8.08E-06 | 3 | Abhd4 | Col18a1 |
| Pakap1 | 4.55E-10 | 0.28100598 | 0.599 | 0.461 | 8.80E-06 | 3 | Pakap | Col18a1 |
| Etl41 | 4.73E-10 | 0.63491643 | 0.26 | 0.187 | 9.14E-06 | 3 | Etl4 | Col18a1 |
| Sgsh1 | 4.76E-10 | 0.46868835 | 0.399 | 0.32 | 9.20E-06 | 3 | Sgsh | Col18a1 |
| Slc25a511 | 4.81E-10 | 0.31429934 | 0.9 | 0.836 | 9.30E-06 | 3 | Slc25a51 | Col18a1 |
| Eno2 | 5.36E-10 | 0.39767089 | 0.317 | 0.239 | 1.04E-05 | 3 | Eno2 | Col18a1 |
| Slmap | 5.38E-10 | 0.40226607 | 0.66 | 0.577 | 1.04E-05 | 3 | Slmap | Col18a1 |
| Pde3b1 | 5.77E-10 | 0.64393816 | 0.288 | 0.207 | 1.12E-05 | 3 | Pde3b | Col18a1 |
| Zbtb44 | 6.08E-10 | 0.51411014 | 0.594 | 0.499 | 1.17E-05 | 3 | Zbtb44 | Col18a1 |
| Slc36a4 | 6.63E-10 | 0.4783959 | 0.633 | 0.529 | 1.28E-05 | 3 | Slc36a4 | Col18a1 |
| Mysm1 | 6.69E-10 | 0.4951877 | 0.492 | 0.402 | 1.29E-05 | 3 | Mysm1 | Col18a1 |
| Homer11 | 6.74E-10 | 0.54049588 | 0.533 | 0.502 | 1.30E-05 | 3 | Homer1 | Col18a1 |
| Prdm2 | 6.97E-10 | 0.53609929 | 0.644 | 0.564 | 1.35E-05 | 3 | Prdm2 | Col18a1 |
| Atf7ip | 7.06E-10 | 0.40504009 | 0.767 | 0.707 | 1.36E-05 | 3 | Atf7ip | Col18a1 |
| Rap1b | 7.28E-10 | 0.34356985 | 0.959 | 0.893 | 1.41E-05 | 3 | Rap1b | Col18a1 |
| Zfp638 | 7.51E-10 | 0.46639453 | 0.523 | 0.433 | 1.45E-05 | 3 | Zfp638 | Col18a1 |
| Mosmo | 7.52E-10 | 0.32727732 | 0.615 | 0.536 | 1.45E-05 | 3 | Mosmo | Col18a1 |
| Ece2 | 7.70E-10 | 0.68490631 | 0.261 | 0.203 | 1.49E-05 | 3 | Ece2 | Col18a1 |
| Sub1 | 8.17E-10 | 0.53354544 | 0.664 | 0.541 | 1.58E-05 | 3 | Sub1 | Col18a1 |
| Stxbp5 | 8.53E-10 | 0.38075391 | 0.556 | 0.495 | 1.65E-05 | 3 | Stxbp5 | Col18a1 |
| Zxdc | 8.99E-10 | 0.52916882 | 0.406 | 0.316 | 1.74E-05 | 3 | Zxdc | Col18a1 |
| Acyp2 | 9.00E-10 | 0.63309792 | 0.25 | 0.186 | 1.74E-05 | 3 | Acyp2 | Col18a1 |
| S100pbp | 9.10E-10 | 0.53465112 | 0.429 | 0.332 | 1.76E-05 | 3 | S100pbp | Col18a1 |
| Wac | 9.34E-10 | 0.49395466 | 0.684 | 0.576 | 1.81E-05 | 3 | Wac | Col18a1 |
| Mpg | 9.64E-10 | 0.37678367 | 0.601 | 0.511 | 1.86E-05 | 3 | Mpg | Col18a1 |
| Dhx40 | 9.75E-10 | 0.5442824 | 0.599 | 0.509 | 1.89E-05 | 3 | Dhx40 | Col18a1 |
| Ecpas | 9.84E-10 | 0.42814332 | 0.717 | 0.619 | 1.90E-05 | 3 | Ecpas | Col18a1 |
| Ascc3 | 1.05E-09 | 0.43376364 | 0.765 | 0.66 | 2.03E-05 | 3 | Ascc3 | Col18a1 |
| Stox21 | 1.12E-09 | 0.60911716 | 0.435 | 0.338 | 2.16E-05 | 3 | Stox2 | Col18a1 |
| Cir1 | 1.12E-09 | 0.60059434 | 0.369 | 0.292 | 2.17E-05 | 3 | Cir1 | Col18a1 |
| Zxdb | 1.12E-09 | 0.6361518 | 0.344 | 0.253 | 2.17E-05 | 3 | Zxdb | Col18a1 |
| Atxn3 | 1.20E-09 | 0.53377642 | 0.402 | 0.308 | 2.32E-05 | 3 | Atxn3 | Col18a1 |
| Dpy19l4 | 1.21E-09 | 0.64364294 | 0.408 | 0.315 | 2.33E-05 | 3 | Dpy19l4 | Col18a1 |
| Tob11 | 1.22E-09 | 0.5033462 | 0.452 | 0.49 | 2.36E-05 | 3 | Tob1 | Col18a1 |
| Ppp6r31 | 1.24E-09 | 0.46417012 | 0.76 | 0.655 | 2.40E-05 | 3 | Ppp6r3 | Col18a1 |
| Pds5b1 | 1.25E-09 | 0.52879547 | 0.541 | 0.449 | 2.41E-05 | 3 | Pds5b | Col18a1 |
| Dmtf1 | 1.27E-09 | 0.39448528 | 0.5 | 0.418 | 2.45E-05 | 3 | Dmtf1 | Col18a1 |
| Esco1 | 1.30E-09 | 0.60008048 | 0.336 | 0.257 | 2.51E-05 | 3 | Esco1 | Col18a1 |
| Ivd | 1.30E-09 | 0.49880885 | 0.467 | 0.386 | 2.52E-05 | 3 | Ivd | Col18a1 |
| Apeh | 1.32E-09 | 0.32251753 | 0.621 | 0.537 | 2.55E-05 | 3 | Apeh | Col18a1 |
| Rtraf | 1.35E-09 | 0.48065402 | 0.726 | 0.622 | 2.60E-05 | 3 | Rtraf | Col18a1 |
| Kiz | 1.40E-09 | 0.56795508 | 0.309 | 0.24 | 2.71E-05 | 3 | Kiz | Col18a1 |
| Mamdc21 | 1.50E-09 | 0.36458661 | 0.383 | 0.286 | 2.90E-05 | 3 | Mamdc2 | Col18a1 |
| Tmem168 | 1.53E-09 | 0.57830439 | 0.5 | 0.403 | 2.95E-05 | 3 | Tmem168 | Col18a1 |

| | | | | | | | | |
|---|---|---|---|---|---|---|---|---|
| Tead21 | 1.54E-09 | 0.49212544 | 0.554 | 0.445 | 2.98E-05 | 3 | Tead2 | Col18a1 |
| Fpgt1 | 1.57E-09 | 0.56621904 | 0.392 | 0.312 | 3.03E-05 | 3 | Fpgt | Col18a1 |
| AU040320 | 1.64E-09 | 0.42244912 | 0.559 | 0.482 | 3.16E-05 | 3 | AU040320 | Col18a1 |
| Pde10a | 1.76E-09 | 0.56481306 | 0.413 | 0.33 | 3.40E-05 | 3 | Pde10a | Col18a1 |
| Mecp2 | 1.86E-09 | 0.4491719 | 0.587 | 0.488 | 3.59E-05 | 3 | Mecp2 | Col18a1 |
| Vcpip1 | 2.00E-09 | 0.54892735 | 0.477 | 0.382 | 3.87E-05 | 3 | Vcpip1 | Col18a1 |
| Ociad1 | 2.01E-09 | 0.36668735 | 0.818 | 0.746 | 3.89E-05 | 3 | Ociad1 | Col18a1 |
| Dyrk1b | 2.03E-09 | 0.50717837 | 0.343 | 0.267 | 3.92E-05 | 3 | Dyrk1b | Col18a1 |
| Zc3h14 | 2.15E-09 | 0.40040695 | 0.799 | 0.703 | 4.15E-05 | 3 | Zc3h14 | Col18a1 |
| Nkd11 | 2.15E-09 | 0.3725187 | 0.452 | 0.353 | 4.15E-05 | 3 | Nkd1 | Col18a1 |
| Mpst | 2.22E-09 | 0.46266734 | 0.452 | 0.367 | 4.29E-05 | 3 | Mpst | Col18a1 |
| Rbbp9 | 2.74E-09 | 0.57533769 | 0.403 | 0.327 | 5.30E-05 | 3 | Rbbp9 | Col18a1 |
| Mapre2 | 2.85E-09 | 0.45054018 | 0.564 | 0.489 | 5.51E-05 | 3 | Mapre2 | Col18a1 |
| Pde4d1 | 2.87E-09 | 0.51798708 | 0.612 | 0.501 | 5.55E-05 | 3 | Pde4d | Col18a1 |
| Vdac1 | 2.90E-09 | 0.42457556 | 0.785 | 0.673 | 5.61E-05 | 3 | Vdac1 | Col18a1 |
| Pcdhb171 | 2.97E-09 | 0.61846944 | 0.269 | 0.228 | 5.74E-05 | 3 | Pcdhb17 | Col18a1 |
| Eci2 | 3.03E-09 | 0.47952735 | 0.565 | 0.493 | 5.86E-05 | 3 | Eci2 | Col18a1 |
| Pced1a | 3.04E-09 | 0.42624499 | 0.508 | 0.449 | 5.88E-05 | 3 | Pced1a | Col18a1 |
| Fam135a | 3.07E-09 | 0.5842898 | 0.411 | 0.321 | 5.93E-05 | 3 | Fam135a | Col18a1 |
| Eif4ebp1 | 3.21E-09 | 0.33705849 | 0.929 | 0.856 | 6.21E-05 | 3 | Eif4ebp1 | Col18a1 |
| Rab3ip | 3.29E-09 | 0.4837817 | 0.341 | 0.305 | 6.36E-05 | 3 | Rab3ip | Col18a1 |
| Fam220a.1 | 3.43E-09 | 0.49287689 | 0.591 | 0.499 | 6.64E-05 | 3 | Fam220a.1 | Col18a1 |
| Begain1 | 3.49E-09 | 0.39922994 | 0.323 | 0.249 | 6.75E-05 | 3 | Begain | Col18a1 |
| Cul2 | 3.49E-09 | 0.67052887 | 0.468 | 0.375 | 6.75E-05 | 3 | Cul2 | Col18a1 |
| Ly751 | 3.59E-09 | 0.60372265 | 0.289 | 0.198 | 6.94E-05 | 3 | Ly75 | Col18a1 |
| Usp28 | 3.77E-09 | 0.60443756 | 0.253 | 0.181 | 7.29E-05 | 3 | Usp28 | Col18a1 |
| Naf1 | 4.03E-09 | 0.64978607 | 0.379 | 0.293 | 7.80E-05 | 3 | Naf1 | Col18a1 |
| Fah1 | 4.15E-09 | 0.42923876 | 0.589 | 0.499 | 8.03E-05 | 3 | Fah | Col18a1 |
| Ube2q1 | 4.19E-09 | 0.37816761 | 0.822 | 0.73 | 8.11E-05 | 3 | Ube2q1 | Col18a1 |
| Med13 | 4.32E-09 | 0.35873506 | 0.855 | 0.768 | 8.35E-05 | 3 | Med13 | Col18a1 |
| Bri3bp | 4.46E-09 | 0.37084283 | 0.727 | 0.635 | 8.62E-05 | 3 | Bri3bp | Col18a1 |
| Mfsd14b1 | 4.60E-09 | 0.47920948 | 0.775 | 0.663 | 8.90E-05 | 3 | Mfsd14b | Col18a1 |
| Rnf181 | 4.68E-09 | 0.41794289 | 0.636 | 0.558 | 9.06E-05 | 3 | Rnf181 | Col18a1 |
| Zfp512b | 4.91E-09 | 0.44953189 | 0.622 | 0.519 | 9.50E-05 | 3 | Zfp512b | Col18a1 |
| Ro60 | 6.04E-09 | 0.56856599 | 0.475 | 0.403 | 0.00011678 | 3 | Ro60 | Col18a1 |
| Trp53inp1 | 6.11E-09 | 0.33142013 | 0.341 | 0.265 | 0.00011807 | 3 | Trp53inp1 | Col18a1 |
| Mef2d | 6.31E-09 | 0.32578915 | 0.848 | 0.777 | 0.00012195 | 3 | Mef2d | Col18a1 |
| Mplkip | 6.38E-09 | 0.53388507 | 0.362 | 0.283 | 0.00012341 | 3 | Mplkip | Col18a1 |
| Bri3 | 6.43E-09 | 0.26906991 | 0.902 | 0.855 | 0.00012423 | 3 | Bri3 | Col18a1 |
| Fkbp31 | 7.06E-09 | 0.49368573 | 0.561 | 0.46 | 0.00013648 | 3 | Fkbp3 | Col18a1 |
| Mbip | 7.38E-09 | 0.63444003 | 0.289 | 0.215 | 0.00014268 | 3 | Mbip | Col18a1 |
| Tmem164 | 7.44E-09 | 0.36424874 | 0.735 | 0.649 | 0.0001438 | 3 | Tmem164 | Col18a1 |
| Rab9 | 7.69E-09 | 0.38371048 | 0.659 | 0.574 | 0.00014877 | 3 | Rab9 | Col18a1 |
| Pcdhgb2 | 7.74E-09 | 0.39279078 | 0.324 | 0.283 | 0.0001497 | 3 | Pcdhgb2 | Col18a1 |
| Vamp2 | 8.37E-09 | 0.30972394 | 0.605 | 0.518 | 0.00016192 | 3 | Vamp2 | Col18a1 |
| Ube2d3 | 8.38E-09 | 0.45292847 | 0.839 | 0.734 | 0.00016206 | 3 | Ube2d3 | Col18a1 |
| Rmnd5a | 8.70E-09 | 0.46233219 | 0.666 | 0.56 | 0.00016825 | 3 | Rmnd5a | Col18a1 |
| Eif1ax | 8.92E-09 | 0.48244596 | 0.594 | 0.488 | 0.00017237 | 3 | Eif1ax | Col18a1 |
| Pisd | 9.35E-09 | 0.398413 | 0.696 | 0.618 | 0.00018073 | 3 | Pisd | Col18a1 |
| Wdfy2 | 9.49E-09 | 0.39326322 | 0.559 | 0.491 | 0.00018357 | 3 | Wdfy2 | Col18a1 |
| Mif1 | 9.57E-09 | 0.31362169 | 0.842 | 0.79 | 0.00018512 | 3 | Mif | Col18a1 |
| Lamtor2 | 9.61E-09 | 0.35035834 | 0.836 | 0.77 | 0.00018571 | 3 | Lamtor2 | Col18a1 |
| Manba | 1.06E-08 | 0.46576518 | 0.446 | 0.376 | 0.00020456 | 3 | Manba | Col18a1 |
| Acsl3 | 1.10E-08 | 0.67080883 | 0.406 | 0.325 | 0.00021245 | 3 | Acsl3 | Col18a1 |
| Wipi2 | 1.22E-08 | 0.38258763 | 0.678 | 0.594 | 0.0002368 | 3 | Wipi2 | Col18a1 |
| Pank3 | 1.24E-08 | 0.46686628 | 0.691 | 0.622 | 0.00024008 | 3 | Pank3 | Col18a1 |
| Ifi203 | 1.24E-08 | 0.45970208 | 0.579 | 0.442 | 0.00024015 | 3 | Ifi203 | Col18a1 |
| Gpalpp1 | 1.27E-08 | 0.50223685 | 0.562 | 0.473 | 0.00024511 | 3 | Gpalpp1 | Col18a1 |
| Fancel | 1.27E-08 | 0.27895949 | 0.638 | 0.574 | 0.00024534 | 3 | Fance | Col18a1 |
| Pgrmc1 | 1.31E-08 | 0.33749295 | 0.89 | 0.825 | 0.00025272 | 3 | Pgrmc1 | Col18a1 |
| Arhgap24 | 1.31E-08 | 0.46872713 | 0.517 | 0.435 | 0.00025298 | 3 | Arhgap24 | Col18a1 |
| Myo9b | 1.34E-08 | 0.5045486 | 0.658 | 0.558 | 0.00025941 | 3 | Myo9b | Col18a1 |
| Tnfrsf1a | 1.36E-08 | 0.26307431 | 0.978 | 0.958 | 0.00026346 | 3 | Tnfrsf1a | Col18a1 |
| Tm2d1 | 1.38E-08 | 0.50340164 | 0.432 | 0.353 | 0.00026735 | 3 | Tm2d1 | Col18a1 |
| Ddx28 | 1.45E-08 | 0.67222198 | 0.266 | 0.199 | 0.000281 | 3 | Ddx28 | Col18a1 |
| Qrich1 | 1.47E-08 | 0.35033155 | 0.846 | 0.764 | 0.00028473 | 3 | Qrich1 | Col18a1 |
| Kif21b | 1.50E-08 | 0.26618858 | 0.283 | 0.228 | 0.00029079 | 3 | Kif21b | Col18a1 |
| Fam114a2 | 1.54E-08 | 0.40721025 | 0.717 | 0.631 | 0.00029806 | 3 | Fam114a2 | Col18a1 |
| Rgmb | 1.58E-08 | 0.34718925 | 0.728 | 0.697 | 0.00030592 | 3 | Rgmb | Col18a1 |
| Galc | 1.59E-08 | 0.55328203 | 0.271 | 0.214 | 0.00030694 | 3 | Galc | Col18a1 |
| Galnt11 | 1.63E-08 | 0.41367496 | 0.477 | 0.393 | 0.00031462 | 3 | Galnt11 | Col18a1 |
| Gtpbp6 | 1.64E-08 | 0.45701931 | 0.428 | 0.345 | 0.00031665 | 3 | Gtpbp6 | Col18a1 |
| Fyttd1 | 1.84E-08 | 0.39801619 | 0.775 | 0.678 | 0.00035662 | 3 | Fyttd1 | Col18a1 |
| Mbd5 | 1.94E-08 | 0.50654897 | 0.42 | 0.344 | 0.00037449 | 3 | Mbd5 | Col18a1 |
| Rcor1 | 2.01E-08 | 0.41465132 | 0.738 | 0.629 | 0.00038828 | 3 | Rcor1 | Col18a1 |
| Xxylt11 | 2.11E-08 | 0.40078021 | 0.676 | 0.598 | 0.00040703 | 3 | Xxylt1 | Col18a1 |
| St6galnac6 | 2.12E-08 | 0.3517034 | 0.712 | 0.658 | 0.00040986 | 3 | St6galnac6 | Col18a1 |
| Mkln1 | 2.21E-08 | 0.39183568 | 0.629 | 0.568 | 0.00042768 | 3 | Mkln1 | Col18a1 |
| Minpp1 | 2.24E-08 | 0.58196009 | 0.62 | 0.505 | 0.00043252 | 3 | Minpp1 | Col18a1 |
| Cebpz | 2.26E-08 | 0.59534358 | 0.384 | 0.31 | 0.00043666 | 3 | Cebpz | Col18a1 |
| Casp9 | 2.31E-08 | 0.64834273 | 0.282 | 0.224 | 0.0004459 | 3 | Casp9 | Col18a1 |
| Shoc2 | 2.33E-08 | 0.49465286 | 0.623 | 0.536 | 0.00045057 | 3 | Shoc2 | Col18a1 |
| Ppm1a | 2.35E-08 | 0.28451378 | 0.926 | 0.874 | 0.00045348 | 3 | Ppm1a | Col18a1 |
| Snai11 | 2.37E-08 | 0.29024271 | 0.85 | 0.8 | 0.00045744 | 3 | Snai1 | Col18a1 |
| Tgoln1 | 2.37E-08 | 0.35395501 | 0.836 | 0.759 | 0.00045768 | 3 | Tgoln1 | Col18a1 |
| Haus21 | 2.39E-08 | 0.66160026 | 0.302 | 0.233 | 0.00046234 | 3 | Haus2 | Col18a1 |
| Exoc6b | 2.49E-08 | 0.34965735 | 0.629 | 0.533 | 0.00048116 | 3 | Exoc6b | Col18a1 |
| Elf1 | 2.59E-08 | 0.40798016 | 0.644 | 0.569 | 0.00050154 | 3 | Elf1 | Col18a1 |
| Dpp7 | 2.70E-08 | 0.44599985 | 0.413 | 0.325 | 0.00052258 | 3 | Dpp7 | Col18a1 |
| Pcm1 | 2.93E-08 | 0.42120104 | 0.654 | 0.581 | 0.00056594 | 3 | Pcm1 | Col18a1 |
| Ctps2 | 3.06E-08 | 0.38555469 | 0.713 | 0.621 | 0.00059118 | 3 | Ctps2 | Col18a1 |
| Uba3 | 3.10E-08 | 0.37041427 | 0.718 | 0.653 | 0.00059844 | 3 | Uba3 | Col18a1 |
| Gatd1 | 3.17E-08 | 0.36079925 | 0.728 | 0.666 | 0.00061273 | 3 | Gatd1 | Col18a1 |
| Ccnt21 | 3.20E-08 | 0.25405554 | 0.573 | 0.508 | 0.00061847 | 3 | Ccnt2 | Col18a1 |
| H2afj | 3.40E-08 | 0.32107016 | 0.335 | 0.28 | 0.00065696 | 3 | H2afj | Col18a1 |
| Ell | 3.52E-08 | 0.32712884 | 0.554 | 0.478 | 0.00068064 | 3 | Ell | Col18a1 |
| Maoa | 3.66E-08 | 0.58473835 | 0.372 | 0.296 | 0.00070724 | 3 | Maoa | Col18a1 |
| Fcho2 | 3.81E-08 | 0.45345178 | 0.39 | 0.31 | 0.00073712 | 3 | Fcho2 | Col18a1 |
| Ncoa1 | 3.83E-08 | 0.57445824 | 0.309 | 0.239 | 0.00074111 | 3 | Ncoa1 | Col18a1 |
| Dhx36 | 4.00E-08 | 0.42256428 | 0.682 | 0.573 | 0.00077339 | 3 | Dhx36 | Col18a1 |
| Sun1 | 4.12E-08 | 0.36773348 | 0.611 | 0.533 | 0.0007967 | 3 | Sun1 | Col18a1 |
| Gls | 4.30E-08 | 0.41015154 | 0.831 | 0.755 | 0.00083178 | 3 | Gls | Col18a1 |
| Fos1 | 5.08E-08 | 0.3000629 | 0.689 | 0.751 | 0.00098225 | 3 | Fos | Col18a1 |
| Zfp1 | 5.08E-08 | 0.47018477 | 0.273 | 0.201 | 0.00098273 | 3 | Zfp1 | Col18a1 |
| Aplp2 | 5.53E-08 | 0.26235208 | 0.861 | 0.819 | 0.00106888 | 3 | Aplp2 | Col18a1 |
| Ell24 | 5.80E-08 | 0.43419387 | 0.605 | 0.526 | 0.00112042 | 3 | Ell24 | Col18a1 |
| Tprgl | 5.99E-08 | 0.33288804 | 0.665 | 0.595 | 0.00115819 | 3 | Tprgl | Col18a1 |
| Fgfr1op2 | 6.01E-08 | 0.32069567 | 0.871 | 0.804 | 0.00116271 | 3 | Fgfr1op2 | Col18a1 |
| Fzd31 | 6.14E-08 | 0.76688683 | 0.273 | 0.216 | 0.0011869 | 3 | Fzd3 | Col18a1 |
| Nab2 | 6.29E-08 | 0.33713632 | 0.786 | 0.71 | 0.00121631 | 3 | Nab2 | Col18a1 |
| Arhgap17 | 6.75E-08 | 0.37757689 | 0.542 | 0.46 | 0.00130417 | 3 | Arhgap17 | Col18a1 |
| Senp5 | 7.05E-08 | 0.46257438 | 0.693 | 0.58 | 0.00136225 | 3 | Senp5 | Col18a1 |
| Ppp3r1 | 7.13E-08 | 0.39658843 | 0.842 | 0.747 | 0.00137941 | 3 | Ppp3r1 | Col18a1 |
| Ncstn | 7.45E-08 | 0.32720647 | 0.919 | 0.862 | 0.00143989 | 3 | Ncstn | Col18a1 |
| Ccdc90b | 7.93E-08 | 0.47790797 | 0.537 | 0.465 | 0.00153317 | 3 | Ccdc90b | Col18a1 |
| Kxd1 | 8.04E-08 | 0.47317708 | 0.316 | 0.258 | 0.00155483 | 3 | Kxd1 | Col18a1 |
| Ddx19b | 8.07E-08 | 0.68830648 | 0.293 | 0.232 | 0.00156048 | 3 | Ddx19b | Col18a1 |
| Rest | 8.20E-08 | 0.45770834 | 0.614 | 0.513 | 0.00158567 | 3 | Rest | Col18a1 |
| Sccpdh | 8.36E-08 | 0.4472115 | 0.584 | 0.487 | 0.00161643 | 3 | Sccpdh | Col18a1 |
| Eid2 | 8.50E-08 | 0.34897226 | 0.504 | 0.428 | 0.00164316 | 3 | Eid2 | Col18a1 |
| Tgm1 | 8.55E-08 | 0.51402135 | 0.329 | 0.259 | 0.00165244 | 3 | Tgm1 | Col18a1 |
| Slc35a1 | 8.71E-08 | 0.45398351 | 0.586 | 0.514 | 0.001684 | 3 | Slc35a1 | Col18a1 |
| Pum1 | 8.85E-08 | 0.39175078 | 0.789 | 0.701 | 0.0017108 | 3 | Pum1 | Col18a1 |
| Ttll51 | 9.00E-08 | 0.51210821 | 0.362 | 0.296 | 0.00174085 | 3 | Ttll5 | Col18a1 |
| Surf2 | 9.34E-08 | 0.47656188 | 0.31 | 0.242 | 0.00180573 | 3 | Surf2 | Col18a1 |
| C1qtnf12 | 9.79E-08 | 0.37243404 | 0.635 | 0.548 | 0.00189247 | 3 | C1qtnf12 | Col18a1 |
| Plekhf11 | 1.00E-07 | 0.38403987 | 0.585 | 0.515 | 0.00194219 | 3 | Plekhf1 | Col18a1 |
| Poglut1 | 1.11E-07 | 0.48665447 | 0.467 | 0.406 | 0.00214279 | 3 | Poglut1 | Col18a1 |
| Lactb2 | 1.18E-07 | 0.64550567 | 0.28 | 0.22 | 0.00227387 | 3 | Lactb2 | Col18a1 |
| Wdr91 | 1.21E-07 | 0.43886668 | 0.436 | 0.361 | 0.00234739 | 3 | Wdr91 | Col18a1 |
| Tmx4 | 1.22E-07 | 0.45640592 | 0.553 | 0.491 | 0.00236183 | 3 | Tmx4 | Col18a1 |
| Spata132 | 1.25E-07 | 0.53734316 | 0.337 | 0.269 | 0.00242248 | 3 | Spata13 | Col18a1 |
| Cdk13 | 1.28E-07 | 0.40943691 | 0.72 | 0.646 | 0.00246766 | 3 | Cdk13 | Col18a1 |
| Ankmy2 | 1.31E-07 | 0.50615906 | 0.397 | 0.329 | 0.0025263 | 3 | Ankmy2 | Col18a1 |
| Gria31 | 1.32E-07 | 0.41765792 | 0.5 | 0.432 | 0.0025479 | 3 | Gria3 | Col18a1 |
| Ubxn7 | 1.36E-07 | 0.26526058 | 0.691 | 0.637 | 0.00262634 | 3 | Ubxn7 | Col18a1 |
| Gamt | 1.41E-07 | 0.45251564 | 0.36 | 0.288 | 0.0027308 | 3 | Gamt | Col18a1 |
| Abhd51 | 1.42E-07 | 0.4960425 | 0.527 | 0.473 | 0.00274462 | 3 | Abhd5 | Col18a1 |
| Klhl15 | 1.46E-07 | 0.64633487 | 0.299 | 0.22 | 0.00282131 | 3 | Klhl15 | Col18a1 |
| Nckap1 | 1.52E-07 | 0.42051336 | 0.804 | 0.697 | 0.00294451 | 3 | Nckap1 | Col18a1 |
| Gskip | 1.54E-07 | 0.31986731 | 0.674 | 0.603 | 0.00297116 | 3 | Gskip | Col18a1 |
| Lamtor5 | 1.58E-07 | 0.39493743 | 0.516 | 0.441 | 0.00304967 | 3 | Lamtor5 | Col18a1 |
| Btrc | 1.59E-07 | 0.54579239 | 0.383 | 0.325 | 0.00307012 | 3 | Btrc | Col18a1 |
| Psme1 | 1.65E-07 | 0.34183872 | 0.816 | 0.739 | 0.00319049 | 3 | Psme1 | Col18a1 |
| Enah1 | 1.70E-07 | 0.31974082 | 0.883 | 0.807 | 0.00329385 | 3 | Enah | Col18a1 |
| Sipa1l2 | 1.71E-07 | 0.48667524 | 0.38 | 0.318 | 0.00330962 | 3 | Sipa1l2 | Col18a1 |
| Zfp277 | 1.82E-07 | 0.64709826 | 0.304 | 0.226 | 0.0035186 | 3 | Zfp277 | Col18a1 |
| Usf3 | 1.88E-07 | 0.40930789 | 0.475 | 0.408 | 0.00364414 | 3 | Usf3 | Col18a1 |
| Mfsd14a | 2.12E-07 | 0.46048006 | 0.508 | 0.427 | 0.00410595 | 3 | Mfsd14a | Col18a1 |
| Mindy1 | 2.22E-07 | 0.36271941 | 0.576 | 0.522 | 0.00429968 | 3 | Mindy1 | Col18a1 |
| Gjc11 | 2.24E-07 | 0.50668114 | 0.592 | 0.466 | 0.00433387 | 3 | Gjc1 | Col18a1 |
| Top2b | 2.31E-07 | 0.39292916 | 0.798 | 0.713 | 0.0044603 | 3 | Top2b | Col18a1 |
| Dbndd2 | 2.33E-07 | 0.31391372 | 0.421 | 0.342 | 0.0045003 | 3 | Dbndd2 | Col18a1 |
| Zmym5 | 2.36E-07 | 0.4209654 | 0.466 | 0.387 | 0.00457095 | 3 | Zmym5 | Col18a1 |
| Haghl | 2.37E-07 | 0.39192328 | 0.336 | 0.276 | 0.00457795 | 3 | Haghl | Col18a1 |
| Mcoln1 | 2.42E-07 | 0.4158781 | 0.524 | 0.453 | 0.00467169 | 3 | Mcoln1 | Col18a1 |
| Polr2a1 | 2.44E-07 | 0.41562141 | 0.856 | 0.774 | 0.00470893 | 3 | Polr2a | Col18a1 |
| Tlk2 | 2.46E-07 | 0.36247125 | 0.611 | 0.523 | 0.00476182 | 3 | Tlk2 | Col18a1 |
| Zfc3h11 | 2.48E-07 | 0.43230473 | 0.58 | 0.498 | 0.00479101 | 3 | Zfc3h1 | Col18a1 |
| Mst1r | 2.53E-07 | 0.37216199 | 0.295 | 0.222 | 0.0048959 | 3 | Mst1r | Col18a1 |
| Ypel5 | 2.54E-07 | 0.34786168 | 0.795 | 0.688 | 0.00491565 | 3 | Ypel5 | Col18a1 |
| Prkra | 2.65E-07 | 0.57297442 | 0.321 | 0.27 | 0.00511756 | 3 | Prkra | Col18a1 |
| Rp9 | 2.71E-07 | 0.38943624 | 0.669 | 0.584 | 0.00523605 | 3 | Rp9 | Col18a1 |
| Clybl | 2.75E-07 | 0.40252135 | 0.385 | 0.323 | 0.00531819 | 3 | Clybl | Col18a1 |
| Bcl10 | 2.89E-07 | 0.43477715 | 0.73 | 0.621 | 0.0055809 | 3 | Bcl10 | Col18a1 |
| Dynlt31 | 2.92E-07 | 0.38301271 | 0.773 | 0.673 | 0.00565499 | 3 | Dynlt3 | Col18a1 |
| Borcs8 | 2.94E-07 | 0.37270728 | 0.672 | 0.612 | 0.005686 | 3 | Borcs8 | Col18a1 |
| Cdc40 | 3.24E-07 | 0.49910132 | 0.428 | 0.355 | 0.00626549 | 3 | Cdc40 | Col18a1 |
| Tlk1 | 3.29E-07 | 0.4731813 | 0.46 | 0.387 | 0.0063546 | 3 | Tlk1 | Col18a1 |
| Cdip11 | 3.38E-07 | 0.42483117 | 0.605 | 0.514 | 0.00653727 | 3 | Cdip1 | Col18a1 |
| Tmem159 | 3.42E-07 | 0.34931666 | 0.617 | 0.526 | 0.00661393 | 3 | Tmem159 | Col18a1 |
| Fgl21 | 3.45E-07 | 0.28032914 | 0.671 | 0.583 | 0.0066659 | 3 | Fgl2 | Col18a1 |
| Parp3 | 3.47E-07 | 0.25328371 | 0.752 | 0.649 | 0.00670645 | 3 | Parp3 | Col18a1 |
| Metap1d | 3.74E-07 | 0.3810422 | 0.344 | 0.286 | 0.00722459 | 3 | Metap1d | Col18a1 |
| Nr1d2 | 3.74E-07 | 0.4623466 | 0.579 | 0.447 | 0.00723528 | 3 | Nr1d2 | Col18a1 |
| Tmem60 | 3.77E-07 | 0.58299556 | 0.3 | 0.238 | 0.00728128 | 3 | Tmem60 | Col18a1 |
| Mtor | 4.21E-07 | 0.28668575 | 0.529 | 0.447 | 0.00813653 | 3 | Mtor | Col18a1 |
| Nap1l11 | 4.21E-07 | 0.5161817 | 0.742 | 0.616 | 0.00813964 | 3 | Nap1l1 | Col18a1 |
| Snx13 | 4.30E-07 | 0.38812241 | 0.512 | 0.435 | 0.00832076 | 3 | Snx13 | Col18a1 |
| Zfp397 | 4.31E-07 | 0.42912852 | 0.353 | 0.286 | 0.00832959 | 3 | Zfp397 | Col18a1 |
| Vps11 | 4.44E-07 | 0.37448099 | 0.546 | 0.465 | 0.00858244 | 3 | Vps11 | Col18a1 |
| Cpsf3 | 4.45E-07 | 0.42072021 | 0.553 | 0.481 | 0.0085961 | 3 | Cpsf3 | Col18a1 |
| Ets2 | 4.78E-07 | 0.27198369 | 0.58 | 0.567 | 0.0092387 | 3 | Ets2 | Col18a1 |
| Phf3 | 4.90E-07 | 0.51348687 | 0.537 | 0.441 | 0.00947048 | 3 | Phf3 | Col18a1 |
| Matr31 | 5.04E-07 | 0.36754884 | 0.755 | 0.662 | 0.00974341 | 3 | Matr3 | Col18a1 |
| Hspb11 | 5.74E-07 | 0.53579689 | 0.367 | 0.31 | 0.01110446 | 3 | Hspb11 | Col18a1 |
| Rusc2 | 5.76E-07 | 0.31973982 | 0.525 | 0.489 | 0.01113018 | 3 | Rusc2 | Col18a1 |
| Xpr1 | 5.80E-07 | 0.57758099 | 0.405 | 0.328 | 0.01120992 | 3 | Xpr1 | Col18a1 |
| Arfgef1 | 6.19E-07 | 0.45791563 | 0.677 | 0.56 | 0.01196576 | 3 | Arfgef1 | Col18a1 |
| Jkamp | 6.22E-07 | 0.39939559 | 0.659 | 0.574 | 0.01203449 | 3 | Jkamp | Col18a1 |
| Herpud2 | 6.44E-07 | 0.4264767 | 0.615 | 0.538 | 0.01245484 | 3 | Herpud2 | Col18a1 |
| Tomm20 | 6.50E-07 | 0.46630965 | 0.667 | 0.556 | 0.0125625 | 3 | Tomm20 | Col18a1 |
| Ccnt1 | 6.53E-07 | 0.38923527 | 0.509 | 0.432 | 0.01262715 | 3 | Ccnt1 | Col18a1 |
| Foxo1 | 6.62E-07 | 0.37278313 | 0.44 | 0.385 | 0.01279835 | 3 | Foxo1 | Col18a1 |
| Zfp715 | 6.91E-07 | 0.37812248 | 0.332 | 0.275 | 0.01335527 | 3 | Zfp715 | Col18a1 |
| Fbf11 | 7.05E-07 | 0.3976669 | 0.475 | 0.395 | 0.01362496 | 3 | Fbf1 | Col18a1 |
| Ascc1 | 7.13E-07 | 0.55534554 | 0.367 | 0.294 | 0.01377721 | 3 | Ascc1 | Col18a1 |
| Akap8l | 7.19E-07 | 0.39089593 | 0.443 | 0.372 | 0.01390061 | 3 | Akap8l | Col18a1 |
| Fam120a | 7.47E-07 | 0.4284936 | 0.886 | 0.776 | 0.01444119 | 3 | Fam120a | Col18a1 |
| Il1rl2 | 7.72E-07 | 0.34335403 | 0.738 | 0.648 | 0.01493231 | 3 | Il1rl2 | Col18a1 |
| Esr1 | 7.75E-07 | 0.34603242 | 0.465 | 0.391 | 0.01499058 | 3 | Esr1 | Col18a1 |
| Gtdc11 | 7.82E-07 | 0.73815017 | 0.25 | 0.185 | 0.01512268 | 3 | Gtdc1 | Col18a1 |
| Atp6v0a1 | 7.97E-07 | 0.37085224 | 0.356 | 0.272 | 0.01541216 | 3 | Atp6v0a1 | Col18a1 |
| Pacs1 | 8.17E-07 | 0.4970916 | 0.572 | 0.482 | 0.01580015 | 3 | Pacs1 | Col18a1 |
| Tbc1d15 | 8.24E-07 | 0.401552 | 0.759 | 0.685 | 0.01592684 | 3 | Tbc1d15 | Col18a1 |
| Dock7 | 8.28E-07 | 0.3162348 | 0.821 | 0.752 | 0.01599947 | 3 | Dock7 | Col18a1 |
| Ngly1 | 8.30E-07 | 0.48278037 | 0.518 | 0.456 | 0.01604786 | 3 | Ngly1 | Col18a1 |
| Agtrap | 8.49E-07 | 0.31690942 | 0.388 | 0.339 | 0.01642361 | 3 | Agtrap | Col18a1 |
| Scamp1 | 8.62E-07 | 0.4688182 | 0.447 | 0.361 | 0.01666598 | 3 | Scamp1 | Col18a1 |
| Yme1l1 | 8.64E-07 | 0.33033419 | 0.804 | 0.719 | 0.0166995 | 3 | Yme1l1 | Col18a1 |
| Vps37a | 9.21E-07 | 0.39647829 | 0.522 | 0.446 | 0.01779711 | 3 | Vps37a | Col18a1 |
| Pcdh7 | 9.57E-07 | 0.42545927 | 0.662 | 0.593 | 0.01849663 | 3 | Pcdh7 | Col18a1 |
| Atp8b2 | 9.63E-07 | 0.31128179 | 0.876 | 0.804 | 0.01861416 | 3 | Atp8b2 | Col18a1 |
| Ctdspl | 1.01E-06 | 0.4912266 | 0.348 | 0.274 | 0.01951365 | 3 | Ctdspl | Col18a1 |
| Vti1b | 1.03E-06 | 0.38913179 | 0.744 | 0.666 | 0.01995512 | 3 | Vti1b | Col18a1 |
| Tmem106b | 1.04E-06 | 0.35606071 | 0.796 | 0.706 | 0.0201725 | 3 | Tmem106b | Col18a1 |
| Nrg12 | 1.05E-06 | 0.51432938 | 0.399 | 0.359 | 0.02039307 | 3 | Nrg1 | Col18a1 |
| Atrnl1 | 1.13E-06 | 0.33881987 | 0.851 | 0.75 | 0.02182587 | 3 | Atrnl1 | Col18a1 |
| Ewsr11 | 1.17E-06 | 0.512064 | 0.462 | 0.369 | 0.02265543 | 3 | Ewsr1 | Col18a1 |
| Pcdhga6 | 1.17E-06 | 0.43418595 | 0.338 | 0.281 | 0.02268814 | 3 | Pcdhga6 | Col18a1 |
| Rxra | 1.17E-06 | 0.35228185 | 0.724 | 0.651 | 0.02270424 | 3 | Rxra | Col18a1 |
| Rsbn1l | 1.21E-06 | 0.45533284 | 0.395 | 0.318 | 0.02330346 | 3 | Rsbn1l | Col18a1 |
| Patz11 | 1.32E-06 | 0.33939047 | 0.535 | 0.447 | 0.02548967 | 3 | Patz1 | Col18a1 |
| Prkd3 | 1.40E-06 | 0.41247096 | 0.686 | 0.598 | 0.0270254 | 3 | Prkd3 | Col18a1 |
| Atp6v0b | 1.42E-06 | 0.29711079 | 0.765 | 0.715 | 0.02736223 | 3 | Atp6v0b | Col18a1 |
| Crat | 1.42E-06 | 0.41046314 | 0.434 | 0.377 | 0.02748677 | 3 | Crat | Col18a1 |
| Gigyf1 | 1.47E-06 | 0.37058415 | 0.493 | 0.432 | 0.02849581 | 3 | Gigyf1 | Col18a1 |
| Smcr8 | 1.52E-06 | 0.34446148 | 0.605 | 0.522 | 0.0294734 | 3 | Smcr8 | Col18a1 |
| Ramac | 1.53E-06 | 0.47046274 | 0.511 | 0.416 | 0.0296082 | 3 | Ramac | Col18a1 |
| Sugt1 | 1.58E-06 | 0.40701083 | 0.739 | 0.626 | 0.03050768 | 3 | Sugt1 | Col18a1 |
| Xrn1 | 1.65E-06 | 0.47268267 | 0.414 | 0.346 | 0.03180953 | 3 | Xrn1 | Col18a1 |
| Snw1 | 1.69E-06 | 0.43659032 | 0.633 | 0.548 | 0.0326668 | 3 | Snw1 | Col18a1 |
| Abcc1 | 1.72E-06 | 0.39975374 | 0.643 | 0.562 | 0.03329388 | 3 | Abcc1 | Col18a1 |
| Cpeb4 | 1.97E-06 | 0.46482698 | 0.545 | 0.493 | 0.0380654 | 3 | Cpeb4 | Col18a1 |
| Atad1 | 1.97E-06 | 0.46271662 | 0.542 | 0.459 | 0.03809543 | 3 | Atad1 | Col18a1 |
| Crppa1 | 2.00E-06 | 0.57509889 | 0.278 | 0.224 | 0.03860282 | 3 | Crppa | Col18a1 |
| Twistnb1 | 2.01E-06 | 0.558965 | 0.349 | 0.279 | 0.03883594 | 3 | Twistnb | Col18a1 |
| Rsf1 | 2.03E-06 | 0.38898059 | 0.532 | 0.452 | 0.03915201 | 3 | Rsf1 | Col18a1 |
| Bmpr1a | 2.04E-06 | 0.36434314 | 0.855 | 0.769 | 0.0394499 | 3 | Bmpr1a | Col18a1 |
| Sesn2 | 2.08E-06 | 0.40987981 | 0.312 | 0.264 | 0.04029975 | 3 | Sesn2 | Col18a1 |
| Wdr45 | 2.09E-06 | 0.3376846 | 0.583 | 0.512 | 0.04036674 | 3 | Wdr45 | Col18a1 |
| Polr2j | 2.19E-06 | 0.50961783 | 0.52 | 0.439 | 0.0422857 | 3 | Polr2j | Col18a1 |
| Tspan14 | 2.22E-06 | 0.49866097 | 0.488 | 0.393 | 0.04291836 | 3 | Tspan14 | Col18a1 |
| Jmjd6 | 2.24E-06 | 0.39927718 | 0.652 | 0.59 | 0.04323117 | 3 | Jmjd6 | Col18a1 |
| Prdm5 | 2.29E-06 | 0.47384936 | 0.425 | 0.377 | 0.04431372 | 3 | Prdm5 | Col18a1 |
| Phf1 | 2.30E-06 | 0.36273772 | 0.41 | 0.335 | 0.04446945 | 3 | Phf1 | Col18a1 |
| Fam98c | 2.38E-06 | 0.35183856 | 0.458 | 0.398 | 0.04603639 | 3 | Fam98c | Col18a1 |
| Acaca | 2.51E-06 | 0.35885535 | 0.65 | 0.598 | 0.04848411 | 3 | Acaca | Col18a1 |
| Tatdn1 | 2.54E-06 | 0.48666473 | 0.381 | 0.317 | 0.04905844 | 3 | Tatdn1 | Col18a1 |
| Gid4 | 2.59E-06 | 0.38518222 | 0.47 | 0.402 | 0.05006095 | 3 | Gid4 | Col18a1 |
| Flot2 | 2.61E-06 | 0.32739523 | 0.796 | 0.719 | 0.05052324 | 3 | Flot2 | Col18a1 |
| Tank | 2.87E-06 | 0.37645243 | 0.496 | 0.435 | 0.05541403 | 3 | Tank | Col18a1 |
| Arap1 | 2.91E-06 | 0.25264524 | 0.868 | 0.809 | 0.05619617 | 3 | Arap1 | Col18a1 |
| Samd8 | 2.92E-06 | 0.50679803 | 0.484 | 0.403 | 0.05645566 | 3 | Samd8 | Col18a1 |
| Echs1 | 2.94E-06 | 0.36413849 | 0.805 | 0.714 | 0.05688501 | 3 | Echs1 | Col18a1 |
| Ywhaq1 | 2.95E-06 | 0.48725694 | 0.739 | 0.614 | 0.05700818 | 3 | Ywhaq | Col18a1 |
| Nsd1 | 2.96E-06 | 0.28918505 | 0.914 | 0.856 | 0.0572342 | 3 | Nsd1 | Col18a1 |
| Rp2 | 3.18E-06 | 0.51416265 | 0.435 | 0.373 | 0.06139251 | 3 | Rp2 | Col18a1 |
| Pgm11 | 3.22E-06 | 0.25421341 | 0.689 | 0.659 | 0.06227007 | 3 | Pgm1 | Col18a1 |
| Tut41 | 3.34E-06 | 0.3706079 | 0.448 | 0.384 | 0.06459086 | 3 | Tut4 | Col18a1 |
| Rnf38 | 3.38E-06 | 0.43502393 | 0.565 | 0.482 | 0.06526672 | 3 | Rnf38 | Col18a1 |
| Wdsub1 | 3.39E-06 | 0.57350057 | 0.312 | 0.246 | 0.0655334 | 3 | Wdsub1 | Col18a1 |
| Wbp2 | 3.42E-06 | 0.37695375 | 0.523 | 0.453 | 0.06620624 | 3 | Wbp2 | Col18a1 |
| Ncoa6 | 3.44E-06 | 0.44885744 | 0.622 | 0.55 | 0.06647107 | 3 | Ncoa6 | Col18a1 |
| Zfp280c | 3.47E-06 | 0.61442676 | 0.252 | 0.194 | 0.06706299 | 3 | Zfp280c | Col18a1 |
| Dnajc71 | 3.55E-06 | 0.39655413 | 0.691 | 0.617 | 0.0686738 | 3 | Dnajc7 | Col18a1 |
| Rars2 | 3.58E-06 | 0.51841672 | 0.287 | 0.225 | 0.06914746 | 3 | Rars2 | Col18a1 |
| Cstb1 | 3.63E-06 | 0.33274834 | 0.942 | 0.888 | 0.07014311 | 3 | Cstb | Col18a1 |
| Ulk2 | 3.72E-06 | 0.33915007 | 0.69 | 0.624 | 0.07198255 | 3 | Ulk2 | Col18a1 |
| Gnl2 | 3.78E-06 | 0.4066323 | 0.472 | 0.389 | 0.07310552 | 3 | Gnl2 | Col18a1 |
| Rab7b | 3.80E-06 | 0.55352658 | 0.366 | 0.306 | 0.07341911 | 3 | Rab7b | Col18a1 |
| Limk1 | 3.82E-06 | 0.3386437 | 0.715 | 0.661 | 0.07378208 | 3 | Limk1 | Col18a1 |
| Parp4 | 3.82E-06 | 0.42075555 | 0.427 | 0.37 | 0.07391548 | 3 | Parp4 | Col18a1 |
| Cdc42ep4 | 4.01E-06 | 0.34787605 | 0.767 | 0.695 | 0.0775533 | 3 | Cdc42ep4 | Col18a1 |
| Borcs6 | 4.02E-06 | 0.33601835 | 0.457 | 0.417 | 0.07775871 | 3 | Borcs6 | Col18a1 |
| Polr3gl | 4.02E-06 | 0.35095181 | 0.46 | 0.395 | 0.07779799 | 3 | Polr3gl | Col18a1 |
| Tpp2 | 4.18E-06 | 0.36130815 | 0.703 | 0.607 | 0.08081919 | 3 | Tpp2 | Col18a1 |
| Cwf19l2 | 4.21E-06 | 0.42189916 | 0.391 | 0.32 | 0.08132434 | 3 | Cwf19l2 | Col18a1 |
| Marf1 | 4.26E-06 | 0.43866854 | 0.492 | 0.411 | 0.08239055 | 3 | Marf1 | Col18a1 |
| Mynn | 4.46E-06 | 0.46859963 | 0.37 | 0.299 | 0.08620958 | 3 | Mynn | Col18a1 |
| Extl2 | 4.54E-06 | 0.48862342 | 0.435 | 0.358 | 0.08780296 | 3 | Extl2 | Col18a1 |
| Comtd1 | 4.60E-06 | 0.42392695 | 0.29 | 0.246 | 0.08901964 | 3 | Comtd1 | Col18a1 |
| Vav2 | 4.66E-06 | 0.4085098 | 0.386 | 0.315 | 0.09001828 | 3 | Vav2 | Col18a1 |
| Gch11 | 4.69E-06 | 0.48847658 | 0.289 | 0.247 | 0.09075544 | 3 | Gch1 | Col18a1 |
| Tor1aip2 | 4.72E-06 | 0.32744146 | 0.946 | 0.89 | 0.09117254 | 3 | Tor1aip2 | Col18a1 |
| Fes1 | 4.72E-06 | 0.31637947 | 0.447 | 0.374 | 0.09122649 | 3 | Fes | Col18a1 |
| Ugdh1 | 4.87E-06 | 0.31311735 | 0.977 | 0.931 | 0.09414793 | 3 | Ugdh | Col18a1 |
| Usp3 | 5.06E-06 | 0.34161426 | 0.466 | 0.407 | 0.09786031 | 3 | Usp3 | Col18a1 |
| Lima1 | 5.15E-06 | 0.31143627 | 0.909 | 0.834 | 0.09954978 | 3 | Lima1 | Col18a1 |
| Slc26a111 | 5.27E-06 | 0.27732132 | 0.324 | 0.254 | 0.10195274 | 3 | Slc26a11 | Col18a1 |
| Traf3ip2 | 5.52E-06 | 0.39585803 | 0.542 | 0.479 | 0.10664215 | 3 | Traf3ip2 | Col18a1 |
| Ube4a | 5.52E-06 | 0.39370527 | 0.682 | 0.594 | 0.10668012 | 3 | Ube4a | Col18a1 |
| Sri | 5.56E-06 | 0.31399157 | 0.913 | 0.839 | 0.10753584 | 3 | Sri | Col18a1 |
| Ppp2cb | 5.62E-06 | 0.28462875 | 0.912 | 0.849 | 0.10867591 | 3 | Ppp2cb | Col18a1 |
| Gorab | 5.66E-06 | 0.51828901 | 0.391 | 0.323 | 0.10934131 | 3 | Gorab | Col18a1 |
| Fut41 | 5.77E-06 | 0.30053811 | 0.462 | 0.387 | 0.11151661 | 3 | Fut4 | Col18a1 |
| Lancl1 | 5.81E-06 | 0.45770592 | 0.425 | 0.357 | 0.11227226 | 3 | Lancl1 | Col18a1 |
| Maz1 | 5.84E-06 | 0.28829451 | 0.907 | 0.825 | 0.11295022 | 3 | Maz | Col18a1 |
| Nek7 | 5.96E-06 | 0.42599834 | 0.82 | 0.723 | 0.11523599 | 3 | Nek7 | Col18a1 |
| St13 | 6.01E-06 | 0.4029663 | 0.693 | 0.607 | 0.11618242 | 3 | St13 | Col18a1 |
| Aprt | 6.04E-06 | 0.39708091 | 0.537 | 0.469 | 0.11674183 | 3 | Aprt | Col18a1 |
| Setx | 6.08E-06 | 0.50220745 | 0.417 | 0.361 | 0.11761502 | 3 | Setx | Col18a1 |
| Tmem126b | 6.23E-06 | 0.43854558 | 0.332 | 0.28 | 0.12042831 | 3 | Tmem126b | Col18a1 |
| Ptpa | 6.35E-06 | 0.32291064 | 0.812 | 0.733 | 0.12286687 | 3 | Ptpa | Col18a1 |
| Pura | 6.44E-06 | 0.26537478 | 0.867 | 0.806 | 0.1244854 | 3 | Pura | Col18a1 |
| Micu3 | 6.60E-06 | 0.4669728 | 0.318 | 0.262 | 0.12752101 | 3 | Micu3 | Col18a1 |
| Slc12a4 | 6.82E-06 | 0.35581533 | 0.817 | 0.73 | 0.13193661 | 3 | Slc12a4 | Col18a1 |
| Spred2 | 6.83E-06 | 0.38841681 | 0.661 | 0.571 | 0.13210311 | 3 | Spred2 | Col18a1 |
| Fgd61 | 7.03E-06 | 0.33418903 | 0.315 | 0.246 | 0.13596655 | 3 | Fgd6 | Col18a1 |
| Pcmtd1 | 7.09E-06 | 0.45373808 | 0.414 | 0.348 | 0.13709495 | 3 | Pcmtd1 | Col18a1 |
| Ube2d2a | 7.19E-06 | 0.34109869 | 0.901 | 0.843 | 0.13907526 | 3 | Ube2d2a | Col18a1 |
| Pdpk1 | 7.21E-06 | 0.38125655 | 0.689 | 0.612 | 0.13946177 | 3 | Pdpk1 | Col18a1 |
| Osbpl51 | 7.30E-06 | 0.25827658 | 0.768 | 0.697 | 0.1411529 | 3 | Osbpl5 | Col18a1 |
| Tgif1 | 7.30E-06 | 0.3584408 | 0.66 | 0.602 | 0.14118587 | 3 | Tgif1 | Col18a1 |
| Dhps | 7.38E-06 | 0.40404781 | 0.453 | 0.404 | 0.14275278 | 3 | Dhps | Col18a1 |
| Ip6k2 | 7.52E-06 | 0.35585562 | 0.514 | 0.442 | 0.14540093 | 3 | Ip6k2 | Col18a1 |
| Mmab1 | 7.64E-06 | 0.4677448 | 0.295 | 0.242 | 0.14765718 | 3 | Mmab | Col18a1 |
| Casd1 | 7.81E-06 | 0.35041358 | 0.536 | 0.472 | 0.15098813 | 3 | Casd1 | Col18a1 |
| Nt5c3b1 | 8.07E-06 | 0.49361781 | 0.513 | 0.421 | 0.15601598 | 3 | Nt5c3b | Col18a1 |
| Plpp21 | 8.20E-06 | 0.53796515 | 0.338 | 0.281 | 0.15844967 | 3 | Plpp2 | Col18a1 |
| Mib1 | 8.52E-06 | 0.3266573 | 0.897 | 0.818 | 0.16473971 | 3 | Mib1 | Col18a1 |
| Sgpp1 | 8.62E-06 | 0.2666832 | 0.848 | 0.811 | 0.16660368 | 3 | Sgpp1 | Col18a1 |
| Sppl3 | 8.65E-06 | 0.31846061 | 0.825 | 0.757 | 0.16720394 | 3 | Sppl3 | Col18a1 |
| Mettl23 | 8.74E-06 | 0.4604177 | 0.363 | 0.305 | 0.16906379 | 3 | Mettl23 | Col18a1 |
| Tap2 | 8.87E-06 | 0.25399749 | 0.64 | 0.599 | 0.17142638 | 3 | Tap2 | Col18a1 |
| Ch25h2 | 8.97E-06 | 0.36600489 | 0.251 | 0.196 | 0.17333186 | 3 | Ch25h | Col18a1 |
| Cdk17 | 9.32E-06 | 0.44195876 | 0.524 | 0.46 | 0.18019583 | 3 | Cdk17 | Col18a1 |
| Pard3 | 9.35E-06 | 0.32570712 | 0.64 | 0.57 | 0.18067794 | 3 | Pard3 | Col18a1 |
| Gpatch8 | 9.67E-06 | 0.43014949 | 0.486 | 0.418 | 0.18705332 | 3 | Gpatch8 | Col18a1 |
| Arpc1a | 9.82E-06 | 0.38997348 | 0.737 | 0.657 | 0.18981149 | 3 | Arpc1a | Col18a1 |
| Smox | 9.99E-06 | 0.25565342 | 0.435 | 0.365 | 0.19308405 | 3 | Smox | Col18a1 |
| Zfp263 | 1.00E-05 | 0.4565028 | 0.295 | 0.235 | 0.19326057 | 3 | Zfp263 | Col18a1 |
| Ctbs | 1.00E-05 | 0.52863782 | 0.304 | 0.25 | 0.19390121 | 3 | Ctbs | Col18a1 |
| Thnsl2 | 1.04E-05 | 0.54617214 | 0.289 | 0.227 | 0.20076665 | 3 | Thnsl2 | Col18a1 |
| Mdm2 | 1.07E-05 | 0.33161453 | 0.687 | 0.623 | 0.20623352 | 3 | Mdm2 | Col18a1 |
| Gpbp1l1 | 1.07E-05 | 0.31963819 | 0.569 | 0.499 | 0.20682994 | 3 | Gpbp1l1 | Col18a1 |
| DEPRECATED-l | 1.10E-05 | 0.43738665 | 0.356 | 0.293 | 0.21230046 | 3 | DEPRECATED-l | Col18a1 |
| Dmwd | 1.11E-05 | 0.36793131 | 0.664 | 0.581 | 0.21391985 | 3 | Dmwd | Col18a1 |
| Zfp503 | 1.12E-05 | 0.28546329 | 0.542 | 0.496 | 0.21749913 | 3 | Zfp503 | Col18a1 |
| Mgrn1 | 1.16E-05 | 0.26164103 | 0.866 | 0.828 | 0.22387969 | 3 | Mgrn1 | Col18a1 |
| Amn1 | 1.18E-05 | 0.46191968 | 0.26 | 0.205 | 0.22882899 | 3 | Amn1 | Col18a1 |
| Mtdh | 1.27E-05 | 0.29666059 | 0.833 | 0.726 | 0.2449287 | 3 | Mtdh | Col18a1 |
| Arid3a1 | 1.28E-05 | 0.37226273 | 0.474 | 0.4 | 0.24748976 | 3 | Arid3a | Col18a1 |
| Pxdc11 | 1.30E-05 | 0.32165178 | 0.621 | 0.537 | 0.25178978 | 3 | Pxdc1 | Col18a1 |
| Memo11 | 1.31E-05 | 0.42899341 | 0.689 | 0.582 | 0.25306213 | 3 | Memo1 | Col18a1 |
| Tmem131 | 1.31E-05 | 0.32145305 | 0.765 | 0.68 | 0.25336367 | 3 | Tmem131 | Col18a1 |
| Arid1b | 1.32E-05 | 0.35177195 | 0.795 | 0.693 | 0.25442339 | 3 | Arid1b | Col18a1 |
| Crbn | 1.34E-05 | 0.43986111 | 0.429 | 0.361 | 0.25857461 | 3 | Crbn | Col18a1 |
| Ggps1 | 1.35E-05 | 0.38950372 | 0.581 | 0.51 | 0.26068388 | 3 | Ggps1 | Col18a1 |
| St3gal31 | 1.37E-05 | 0.42413272 | 0.447 | 0.376 | 0.2641641 | 3 | St3gal3 | Col18a1 |
| Zbtb10 | 1.50E-05 | 0.38591184 | 0.407 | 0.351 | 0.290023 | 3 | Zbtb10 | Col18a1 |
| Tbce | 1.50E-05 | 0.45440156 | 0.41 | 0.353 | 0.29073482 | 3 | Tbce | Col18a1 |
| Vps13b | 1.51E-05 | 0.6144748 | 0.339 | 0.276 | 0.2911028 | 3 | Vps13b | Col18a1 |
| Smndc1 | 1.53E-05 | 0.42234691 | 0.544 | 0.471 | 0.29595085 | 3 | Smndc1 | Col18a1 |
| Mdm4 | 1.54E-05 | 0.39276375 | 0.576 | 0.517 | 0.29780733 | 3 | Mdm4 | Col18a1 |
| Ift74 | 1.57E-05 | 0.48990755 | 0.402 | 0.327 | 0.30330135 | 3 | Ift74 | Col18a1 |
| Axl2 | 1.58E-05 | 0.44048944 | 0.462 | 0.4 | 0.30535619 | 3 | Axl2 | Col18a1 |
| Isoc1 | 1.59E-05 | 0.59928193 | 0.356 | 0.292 | 0.30675366 | 3 | Isoc1 | Col18a1 |
| Riok3 | 1.60E-05 | 0.33539445 | 0.843 | 0.761 | 0.30985056 | 3 | Riok3 | Col18a1 |
| Cnot71 | 1.61E-05 | 0.45154879 | 0.547 | 0.456 | 0.31104959 | 3 | Cnot7 | Col18a1 |
| Cyp4v3 | 1.63E-05 | 0.46587493 | 0.361 | 0.308 | 0.31611073 | 3 | Cyp4v3 | Col18a1 |
| Akr1b101 | 1.65E-05 | 0.43200674 | 0.406 | 0.346 | 0.31906988 | 3 | Akr1b10 | Col18a1 |
| Fbxo8 | 1.67E-05 | 0.39818098 | 0.434 | 0.387 | 0.3220905 | 3 | Fbxo8 | Col18a1 |
| Manea | 1.70E-05 | 0.5664415 | 0.374 | 0.306 | 0.32834536 | 3 | Manea | Col18a1 |
| Ntan1 | 1.71E-05 | 0.45621493 | 0.52 | 0.452 | 0.33002684 | 3 | Ntan1 | Col18a1 |
| Smap1 | 1.72E-05 | 0.39125188 | 0.731 | 0.639 | 0.33269334 | 3 | Smap1 | Col18a1 |
| Sept101 | 1.79E-05 | 0.50203017 | 0.523 | 0.422 | 0.34607197 | 3 | 10-Sep | Col18a1 |
| Senp6 | 1.80E-05 | 0.32045921 | 0.722 | 0.67 | 0.34734216 | 3 | Senp6 | Col18a1 |
| Raph1 | 1.83E-05 | 0.41320803 | 0.358 | 0.307 | 0.3541876 | 3 | Raph1 | Col18a1 |
| Smco41 | 1.84E-05 | 0.42151013 | 0.603 | 0.526 | 0.35540717 | 3 | Smco4 | Col18a1 |
| Mnt | 1.88E-05 | 0.42566007 | 0.432 | 0.353 | 0.36404625 | 3 | Mnt | Col18a1 |
| Rnf146 | 1.91E-05 | 0.33303612 | 0.584 | 0.506 | 0.36857172 | 3 | Rnf146 | Col18a1 |
| Ralbp1 | 1.91E-05 | 0.28930077 | 0.788 | 0.725 | 0.36857995 | 3 | Ralbp1 | Col18a1 |
| Rad52 | 1.91E-05 | 0.52555309 | 0.3 | 0.235 | 0.36911344 | 3 | Rad52 | Col18a1 |
| Srpk21 | 1.92E-05 | 0.30069728 | 0.621 | 0.543 | 0.37149545 | 3 | Srpk2 | Col18a1 |
| Cbx6 | 1.93E-05 | 0.33362142 | 0.583 | 0.527 | 0.37252979 | 3 | Cbx6 | Col18a1 |
| Mindy2 | 1.95E-05 | 0.45540856 | 0.508 | 0.436 | 0.37672878 | 3 | Mindy2 | Col18a1 |
| Nit2 | 1.98E-05 | 0.47036379 | 0.354 | 0.304 | 0.38209893 | 3 | Nit2 | Col18a1 |
| Cdc42ep2 | 2.02E-05 | 0.36903172 | 0.462 | 0.366 | 0.39075016 | 3 | Cdc42ep2 | Col18a1 |
| Pip4k2c | 2.25E-05 | 0.26892954 | 0.594 | 0.541 | 0.43547552 | 3 | Pip4k2c | Col18a1 |
| Ap4s1 | 2.27E-05 | 0.70547741 | 0.293 | 0.232 | 0.43868202 | 3 | Ap4s1 | Col18a1 |
| Copg21 | 2.34E-05 | 0.45081858 | 0.321 | 0.261 | 0.45238755 | 3 | Copg2 | Col18a1 |
| Irx51 | 2.35E-05 | 0.45879603 | 0.397 | 0.354 | 0.45446502 | 3 | Irx5 | Col18a1 |
| Rprd2 | 2.36E-05 | 0.32265955 | 0.659 | 0.589 | 0.45554802 | 3 | Rprd2 | Col18a1 |
| Sike11 | 2.36E-05 | 0.28665819 | 0.758 | 0.687 | 0.45690889 | 3 | Sike1 | Col18a1 |
| Fam49b1 | 2.36E-05 | 0.49269462 | 0.428 | 0.346 | 0.45711238 | 3 | Fam49b | Col18a1 |
| Zdhhc14 | 2.42E-05 | 0.30898863 | 0.306 | 0.27 | 0.46722855 | 3 | Zdhhc14 | Col18a1 |
| Gatad11 | 2.43E-05 | 0.37685743 | 0.878 | 0.781 | 0.46957905 | 3 | Gatad1 | Col18a1 |
| Kcmf1 | 2.50E-05 | 0.33237645 | 0.826 | 0.755 | 0.48335199 | 3 | Kcmf1 | Col18a1 |
| Ldah | 2.53E-05 | 0.48724082 | 0.373 | 0.327 | 0.48958199 | 3 | Ldah | Col18a1 |
| Pcdhga11 | 2.57E-05 | 0.37873501 | 0.3 | 0.251 | 0.49707826 | 3 | Pcdhga11 | Col18a1 |
| Psmb1 | 2.63E-05 | 0.36842569 | 0.794 | 0.715 | 0.50907651 | 3 | Psmb1 | Col18a1 |
| Adamts91 | 2.64E-05 | 0.35216522 | 0.313 | 0.279 | 0.51065637 | 3 | Adamts9 | Col18a1 |
| Bzw11 | 2.66E-05 | 0.40639041 | 0.937 | 0.844 | 0.5145402 | 3 | Bzw1 | Col18a1 |
| Rbm7 | 2.68E-05 | 0.39453283 | 0.701 | 0.639 | 0.51780868 | 3 | Rbm7 | Col18a1 |
| Ahcyl2 | 2.88E-05 | 0.52351506 | 0.263 | 0.213 | 0.55676867 | 3 | Ahcyl2 | Col18a1 |
| Zfp317 | 2.91E-05 | 0.4445016 | 0.323 | 0.269 | 0.56200195 | 3 | Zfp317 | Col18a1 |
| Tmem87a | 2.93E-05 | 0.43886292 | 0.634 | 0.543 | 0.56713864 | 3 | Tmem87a | Col18a1 |
| Zup1 | 2.96E-05 | 0.41378053 | 0.282 | 0.225 | 0.5731841 | 3 | Zup1 | Col18a1 |
| Tmbim1 | 3.01E-05 | 0.34657392 | 0.77 | 0.688 | 0.58196401 | 3 | Tmbim1 | Col18a1 |
| Odr4 | 3.11E-05 | 0.4846342 | 0.453 | 0.378 | 0.60066606 | 3 | Odr4 | Col18a1 |
| Ppcdc | 3.18E-05 | 0.50380452 | 0.26 | 0.212 | 0.61565132 | 3 | Ppcdc | Col18a1 |
| Rpia1 | 3.22E-05 | 0.41169849 | 0.372 | 0.315 | 0.62232844 | 3 | Rpia | Col18a1 |
| Cbwd1 | 3.29E-05 | 0.40603964 | 0.278 | 0.221 | 0.63634718 | 3 | Cbwd1 | Col18a1 |
| Eif4ebp2 | 3.41E-05 | 0.32593179 | 0.865 | 0.792 | 0.65914248 | 3 | Eif4ebp2 | Col18a1 |
| Ppp2r5c | 3.42E-05 | 0.35099998 | 0.879 | 0.789 | 0.66203537 | 3 | Ppp2r5c | Col18a1 |
| Igbp1 | 3.50E-05 | 0.36813162 | 0.688 | 0.614 | 0.6765162 | 3 | Igbp1 | Col18a1 |
| Tbck | 3.52E-05 | 0.49078679 | 0.361 | 0.304 | 0.68119287 | 3 | Tbck | Col18a1 |
| Zbtb37 | 3.62E-05 | 0.3784391 | 0.279 | 0.228 | 0.70066378 | 3 | Zbtb37 | Col18a1 |
| Sec63 | 3.65E-05 | 0.32442316 | 0.848 | 0.769 | 0.70612858 | 3 | Sec63 | Col18a1 |
| Pcdhga5 | 3.66E-05 | 0.36677516 | 0.311 | 0.286 | 0.70677425 | 3 | Pcdhga5 | Col18a1 |
| Slc22a5 | 3.74E-05 | 0.39527685 | 0.257 | 0.218 | 0.72346803 | 3 | Slc22a5 | Col18a1 |
| Brwd3 | 3.76E-05 | 0.32826988 | 0.391 | 0.342 | 0.72692485 | 3 | Brwd3 | Col18a1 |
| Fam210a | 3.79E-05 | 0.39783946 | 0.476 | 0.418 | 0.7326823 | 3 | Fam210a | Col18a1 |
| Zfp655 | 3.88E-05 | 0.33135781 | 0.598 | 0.533 | 0.75104992 | 3 | Zfp655 | Col18a1 |
| Snx15 | 4.14E-05 | 0.29625675 | 0.354 | 0.309 | 0.80022294 | 3 | Snx15 | Col18a1 |
| Cmtm6 | 4.14E-05 | 0.27616597 | 0.747 | 0.69 | 0.80115262 | 3 | Cmtm6 | Col18a1 |
| Lamtor3 | 4.20E-05 | 0.33090092 | 0.866 | 0.786 | 0.81149167 | 3 | Lamtor3 | Col18a1 |
| Wdtc1 | 4.37E-05 | 0.4067699 | 0.57 | 0.501 | 0.84493276 | 3 | Wdtc1 | Col18a1 |
| Pla2g12a | 4.48E-05 | 0.43480401 | 0.384 | 0.328 | 0.86628886 | 3 | Pla2g12a | Col18a1 |
| Cnot6l | 4.52E-05 | 0.42550618 | 0.686 | 0.593 | 0.87458347 | 3 | Cnot6l | Col18a1 |
| Stau1 | 4.68E-05 | 0.30612908 | 0.823 | 0.741 | 0.905622 | 3 | Stau1 | Col18a1 |
| Ube2e31 | 4.71E-05 | 0.34453253 | 0.638 | 0.571 | 0.91074202 | 3 | Ube2e3 | Col18a1 |
| Cep63 | 4.92E-05 | 0.40460157 | 0.369 | 0.3 | 0.95049796 | 3 | Cep63 | Col18a1 |
| Pik3c2a | 4.96E-05 | 0.38785699 | 0.483 | 0.424 | 0.95984363 | 3 | Pik3c2a | Col18a1 |
| Ube3a | 5.13E-05 | 0.30426194 | 0.789 | 0.716 | 0.99240848 | 3 | Ube3a | Col18a1 |
| Gufl1 | 5.19E-05 | 0.45555046 | 0.308 | 0.257 | 1 | 3 | Gufl1 | Col18a1 |
| Chchd5 | 5.29E-05 | 0.38948917 | 0.324 | 0.274 | 1 | 3 | Chchd5 | Col18a1 |
| Jade2 | 5.30E-05 | 0.42825737 | 0.278 | 0.223 | 1 | 3 | Jade2 | Col18a1 |
| Rfx5 | 5.31E-05 | 0.55086307 | 0.265 | 0.214 | 1 | 3 | Rfx5 | Col18a1 |
| Ttc23 | 5.40E-05 | 0.38390134 | 0.369 | 0.313 | 1 | 3 | Ttc23 | Col18a1 |
| Slc25a46 | 5.41E-05 | 0.4774202 | 0.416 | 0.349 | 1 | 3 | Slc25a46 | Col18a1 |
| Abhd17b | 5.48E-05 | 0.31213869 | 0.688 | 0.607 | 1 | 3 | Abhd17b | Col18a1 |
| Akap11 | 5.52E-05 | 0.44251216 | 0.655 | 0.579 | 1 | 3 | Akap11 | Col18a1 |
| Btf3l41 | 5.84E-05 | 0.46293927 | 0.369 | 0.307 | 1 | 3 | Btf3l4 | Col18a1 |
| Cdk10 | 5.87E-05 | 0.37807283 | 0.32 | 0.259 | 1 | 3 | Cdk10 | Col18a1 |
| Mlxip | 5.94E-05 | 0.30051856 | 0.787 | 0.721 | 1 | 3 | Mlxip | Col18a1 |
| Tecr | 5.94E-05 | 0.26957191 | 0.901 | 0.837 | 1 | 3 | Tecr | Col18a1 |
| Dph6 | 5.96E-05 | 0.41156741 | 0.378 | 0.316 | 1 | 3 | Dph6 | Col18a1 |
| Pip4p1 | 5.98E-05 | 0.3454067 | 0.696 | 0.633 | 1 | 3 | Pip4p1 | Col18a1 |
| Isca2 | 5.99E-05 | 0.26728699 | 0.495 | 0.443 | 1 | 3 | Isca2 | Col18a1 |
| Slc16a21 | 6.15E-05 | 0.36468016 | 0.832 | 0.714 | 1 | 3 | Slc16a2 | Col18a1 |
| Dbi | 6.15E-05 | 0.40041611 | 0.674 | 0.605 | 1 | 3 | Dbi | Col18a1 |
| Wdr81 | 6.25E-05 | 0.32443432 | 0.441 | 0.389 | 1 | 3 | Wdr81 | Col18a1 |
| Echdc1 | 6.41E-05 | 0.53991681 | 0.273 | 0.234 | 1 | 3 | Echdc1 | Col18a1 |
| Sdhaf2 | 6.41E-05 | 0.43028903 | 0.523 | 0.461 | 1 | 3 | Sdhaf2 | Col18a1 |
| Itfg2 | 6.57E-05 | 0.41989764 | 0.405 | 0.35 | 1 | 3 | Itfg2 | Col18a1 |
| Dock41 | 6.60E-05 | 0.30792376 | 0.445 | 0.367 | 1 | 3 | Dock4 | Col18a1 |
| Vps39 | 6.62E-05 | 0.34909154 | 0.429 | 0.363 | 1 | 3 | Vps39 | Col18a1 |
| Zbtb21 | 6.72E-05 | 0.34137379 | 0.47 | 0.395 | 1 | 3 | Zbtb21 | Col18a1 |
| Zkscan6 | 6.78E-05 | 0.36276964 | 0.322 | 0.261 | 1 | 3 | Zkscan6 | Col18a1 |
| Braf | 7.00E-05 | 0.48307616 | 0.356 | 0.287 | 1 | 3 | Braf | Col18a1 |
| Trrap | 7.04E-05 | 0.37052863 | 0.765 | 0.698 | 1 | 3 | Trrap | Col18a1 |
| Vapa | 7.12E-05 | 0.41786757 | 0.619 | 0.547 | 1 | 3 | Vapa | Col18a1 |
| Vps28 | 7.26E-05 | 0.3276611 | 0.759 | 0.708 | 1 | 3 | Vps28 | Col18a1 |
| Naxe | 7.72E-05 | 0.38067038 | 0.705 | 0.627 | 1 | 3 | Naxe | Col18a1 |
| Thap2 | 7.81E-05 | 0.35441212 | 0.476 | 0.42 | 1 | 3 | Thap2 | Col18a1 |
| Gnb5 | 7.95E-05 | 0.28076873 | 0.654 | 0.602 | 1 | 3 | Gnb5 | Col18a1 |
| Akap10 | 8.08E-05 | 0.37649239 | 0.43 | 0.379 | 1 | 3 | Akap10 | Col18a1 |
| Nes | 8.22E-05 | 0.33847839 | 0.375 | 0.258 | 1 | 3 | Nes | Col18a1 |
| Pcdhgb4 | 8.27E-05 | 0.30657644 | 0.267 | 0.222 | 1 | 3 | Pcdhgb4 | Col18a1 |
| Pdcd6 | 8.42E-05 | 0.30626848 | 0.842 | 0.765 | 1 | 3 | Pdcd6 | Col18a1 |

| | | | | | | | | |
|---|---|---|---|---|---|---|---|---|
| Nipa1 | 8.48E-05 | 0.41197795 | 0.374 | 0.326 | 1 | 3 | Nipa1 | Col18a1 |
| Poldip3 | 8.67E-05 | 0.33183043 | 0.855 | 0.78 | 1 | 3 | Poldip3 | Col18a1 |
| Tubgcp51 | 8.76E-05 | 0.43694574 | 0.26 | 0.219 | 1 | 3 | Tubgcp5 | Col18a1 |
| Ttc17 | 9.04E-05 | 0.36686913 | 0.526 | 0.455 | 1 | 3 | Ttc17 | Col18a1 |
| Cryzl1 | 9.08E-05 | 0.30966322 | 0.585 | 0.524 | 1 | 3 | Cryzl1 | Col18a1 |
| Atf2 | 9.18E-05 | 0.36196793 | 0.728 | 0.64 | 1 | 3 | Atf2 | Col18a1 |
| Zfr | 9.29E-05 | 0.34081783 | 0.819 | 0.732 | 1 | 3 | Zfr | Col18a1 |
| Zranb1 | 9.32E-05 | 0.36744257 | 0.574 | 0.483 | 1 | 3 | Zranb1 | Col18a1 |
| Cbfa2t2 | 9.36E-05 | 0.41891303 | 0.538 | 0.459 | 1 | 3 | Cbfa2t2 | Col18a1 |
| Gstm5 | 9.46E-05 | 0.50768738 | 0.364 | 0.303 | 1 | 3 | Gstm5 | Col18a1 |
| Proser1 | 9.90E-05 | 0.59079162 | 0.302 | 0.255 | 1 | 3 | Proser1 | Col18a1 |
| Slc35f5 | 9.99E-05 | 0.34229203 | 0.753 | 0.682 | 1 | 3 | Slc35f5 | Col18a1 |
| Zfp26 | 0.00010115 | 0.46178892 | 0.38 | 0.314 | 1 | 3 | Zfp26 | Col18a1 |
| Phf20 | 0.00010145 | 0.46100827 | 0.448 | 0.394 | 1 | 3 | Phf20 | Col18a1 |
| Zdhhc211 | 0.00010379 | 0.54634907 | 0.259 | 0.216 | 1 | 3 | Zdhhc21 | Col18a1 |
| Slc35f6 | 0.00010633 | 0.38340139 | 0.528 | 0.477 | 1 | 3 | Slc35f6 | Col18a1 |
| Tgif2 | 0.00010826 | 0.47276607 | 0.262 | 0.224 | 1 | 3 | Tgif2 | Col18a1 |
| Kbtbd2 | 0.00011091 | 0.38749863 | 0.663 | 0.594 | 1 | 3 | Kbtbd2 | Col18a1 |
| Retsat | 0.00011135 | 0.47541341 | 0.303 | 0.255 | 1 | 3 | Retsat | Col18a1 |
| Ppp2r5e | 0.00011136 | 0.40131248 | 0.674 | 0.58 | 1 | 3 | Ppp2r5e | Col18a1 |
| Setd3 | 0.00011369 | 0.33037226 | 0.767 | 0.698 | 1 | 3 | Setd3 | Col18a1 |
| Mapkbp11 | 0.00011586 | 0.46466565 | 0.319 | 0.254 | 1 | 3 | Mapkbp1 | Col18a1 |
| Rapgef1 | 0.00011847 | 0.31880809 | 0.808 | 0.739 | 1 | 3 | Rapgef1 | Col18a1 |
| Mtmr3 | 0.00011941 | 0.39200716 | 0.549 | 0.485 | 1 | 3 | Mtmr3 | Col18a1 |
| Zfp266 | 0.00012114 | 0.34266373 | 0.65 | 0.59 | 1 | 3 | Zfp266 | Col18a1 |
| Ankrd13b | 0.00012243 | 0.42762049 | 0.3 | 0.249 | 1 | 3 | Ankrd13b | Col18a1 |
| Fhdc1 | 0.00012372 | 0.3966182 | 0.401 | 0.35 | 1 | 3 | Fhdc1 | Col18a1 |
| Prxl2c | 0.00012628 | 0.33219823 | 0.706 | 0.632 | 1 | 3 | Prxl2c | Col18a1 |
| Tmem100 | 0.00012702 | 0.37115188 | 0.512 | 0.447 | 1 | 3 | Tmem100 | Col18a1 |
| Cxxc5 | 0.00012723 | 0.30793582 | 0.865 | 0.795 | 1 | 3 | Cxxc5 | Col18a1 |
| Mtmr6 | 0.0001278 | 0.32432544 | 0.78 | 0.7 | 1 | 3 | Mtmr6 | Col18a1 |
| Arl1 | 0.00012812 | 0.3799337 | 0.914 | 0.832 | 1 | 3 | Arl1 | Col18a1 |
| Prpf40b | 0.00012905 | 0.3454232 | 0.307 | 0.259 | 1 | 3 | Prpf40b | Col18a1 |
| Snx6 | 0.00012993 | 0.36147383 | 0.648 | 0.585 | 1 | 3 | Snx6 | Col18a1 |
| Mbtps2 | 0.00013284 | 0.49553958 | 0.404 | 0.335 | 1 | 3 | Mbtps2 | Col18a1 |
| Fut11 | 0.0001329 | 0.36391062 | 0.644 | 0.564 | 1 | 3 | Fut11 | Col18a1 |
| Cbx31 | 0.00013355 | 0.44287696 | 0.757 | 0.637 | 1 | 3 | Cbx3 | Col18a1 |
| Dlg1 | 0.00013829 | 0.33188359 | 0.778 | 0.704 | 1 | 3 | Dlg1 | Col18a1 |
| Rbm221 | 0.00013971 | 0.3657598 | 0.641 | 0.57 | 1 | 3 | Rbm22 | Col18a1 |
| Dop1b | 0.00014337 | 0.30210926 | 0.318 | 0.271 | 1 | 3 | Dop1b | Col18a1 |
| Pafah1b3 | 0.00015121 | 0.3651892 | 0.61 | 0.547 | 1 | 3 | Pafah1b3 | Col18a1 |
| Arl13b | 0.00015211 | 0.40431867 | 0.568 | 0.516 | 1 | 3 | Arl13b | Col18a1 |
| Myo6 | 0.0001523 | 0.46332346 | 0.284 | 0.233 | 1 | 3 | Myo6 | Col18a1 |
| Prkab1 | 0.00015348 | 0.50574907 | 0.393 | 0.328 | 1 | 3 | Prkab1 | Col18a1 |
| Ptp4a2 | 0.00016534 | 0.32718591 | 0.754 | 0.679 | 1 | 3 | Ptp4a2 | Col18a1 |
| Prdx3 | 0.0001655 | 0.34929804 | 0.71 | 0.649 | 1 | 3 | Prdx3 | Col18a1 |
| Sun21 | 0.00016848 | 0.28572025 | 0.599 | 0.499 | 1 | 3 | Sun2 | Col18a1 |
| N4bp2l2 | 0.00016952 | 0.33343978 | 0.628 | 0.552 | 1 | 3 | N4bp2l2 | Col18a1 |
| Btbd7 | 0.00017173 | 0.36938064 | 0.687 | 0.625 | 1 | 3 | Btbd7 | Col18a1 |
| Amdhd2 | 0.00017458 | 0.26501352 | 0.457 | 0.396 | 1 | 3 | Amdhd2 | Col18a1 |
| Sorbs3 | 0.00017504 | 0.33526122 | 0.535 | 0.478 | 1 | 3 | Sorbs3 | Col18a1 |
| Camta1 | 0.00018914 | 0.44619437 | 0.304 | 0.248 | 1 | 3 | Camta1 | Col18a1 |
| Med1 | 0.00019138 | 0.43727089 | 0.662 | 0.572 | 1 | 3 | Med1 | Col18a1 |
| Acap2 | 0.00019499 | 0.28164157 | 0.736 | 0.675 | 1 | 3 | Acap2 | Col18a1 |
| Pdhb | 0.00019687 | 0.33979639 | 0.637 | 0.545 | 1 | 3 | Pdhb | Col18a1 |
| Ranbp21 | 0.00019918 | 0.34348924 | 0.823 | 0.734 | 1 | 3 | Ranbp2 | Col18a1 |
| Cd151 | 0.00020362 | 0.32198013 | 0.693 | 0.636 | 1 | 3 | Cd151 | Col18a1 |
| Men1 | 0.00020925 | 0.27888857 | 0.56 | 0.505 | 1 | 3 | Men1 | Col18a1 |
| Cd99l2 | 0.00021173 | 0.34880902 | 0.543 | 0.482 | 1 | 3 | Cd99l2 | Col18a1 |
| Gltp1 | 0.00021265 | 0.31759907 | 0.627 | 0.547 | 1 | 3 | Gltp | Col18a1 |
| Nsmaf | 0.00021288 | 0.28913493 | 0.492 | 0.419 | 1 | 3 | Nsmaf | Col18a1 |
| Dazap2 | 0.00021992 | 0.36061735 | 0.702 | 0.636 | 1 | 3 | Dazap2 | Col18a1 |
| Meis3 | 0.0002217 | 0.29493987 | 0.465 | 0.411 | 1 | 3 | Meis3 | Col18a1 |
| Cyfip1 | 0.00022505 | 0.33933954 | 0.808 | 0.727 | 1 | 3 | Cyfip1 | Col18a1 |
| Slc31a1 | 0.00023011 | 0.44692568 | 0.665 | 0.576 | 1 | 3 | Slc31a1 | Col18a1 |
| Eif1a | 0.00023015 | 0.28170783 | 0.832 | 0.794 | 1 | 3 | Eif1a | Col18a1 |
| Zfyve1 | 0.00023283 | 0.34455677 | 0.611 | 0.552 | 1 | 3 | Zfyve1 | Col18a1 |
| Ado1 | 0.00023374 | 0.34651224 | 0.535 | 0.469 | 1 | 3 | Ado | Col18a1 |
| Trim25 | 0.00023949 | 0.29723912 | 0.6 | 0.548 | 1 | 3 | Trim25 | Col18a1 |
| Rnf214 | 0.00024067 | 0.28375323 | 0.58 | 0.506 | 1 | 3 | Rnf214 | Col18a1 |
| Fam91a1 | 0.00024147 | 0.31798258 | 0.845 | 0.758 | 1 | 3 | Fam91a1 | Col18a1 |
| Mtx2 | 0.00025223 | 0.35319881 | 0.74 | 0.659 | 1 | 3 | Mtx2 | Col18a1 |
| Eapp | 0.00025256 | 0.26303646 | 0.688 | 0.626 | 1 | 3 | Eapp | Col18a1 |
| Hmgcl | 0.00025682 | 0.26879019 | 0.661 | 0.623 | 1 | 3 | Hmgcl | Col18a1 |
| Gatd3a | 0.00026344 | 0.25151628 | 0.571 | 0.524 | 1 | 3 | Gatd3a | Col18a1 |
| Dcaf14 | 0.00027298 | 0.51821593 | 0.382 | 0.314 | 1 | 3 | Dcaf1 | Col18a1 |
| Ncoa2 | 0.00027504 | 0.45102381 | 0.464 | 0.405 | 1 | 3 | Ncoa2 | Col18a1 |
| Tmem260 | 0.00027539 | 0.42960571 | 0.436 | 0.367 | 1 | 3 | Tmem260 | Col18a1 |
| Pdxk | 0.00027713 | 0.43728748 | 0.475 | 0.427 | 1 | 3 | Pdxk | Col18a1 |
| Aptx | 0.00028335 | 0.42620324 | 0.32 | 0.27 | 1 | 3 | Aptx | Col18a1 |
| Rapgef61 | 0.00028398 | 0.49006783 | 0.447 | 0.373 | 1 | 3 | Rapgef6 | Col18a1 |
| Vangl11 | 0.00028965 | 0.42289567 | 0.533 | 0.458 | 1 | 3 | Vangl1 | Col18a1 |
| Kmt2c | 0.00029326 | 0.41195494 | 0.42 | 0.353 | 1 | 3 | Kmt2c | Col18a1 |
| Rnf139 | 0.00029658 | 0.40705813 | 0.532 | 0.475 | 1 | 3 | Rnf139 | Col18a1 |
| Siah1a1 | 0.00030633 | 0.436413 | 0.28 | 0.238 | 1 | 3 | Siah1a | Col18a1 |
| Ube3b | 0.00031715 | 0.36868388 | 0.499 | 0.45 | 1 | 3 | Ube3b | Col18a1 |
| Atp11c1 | 0.00032019 | 0.4218368 | 0.366 | 0.298 | 1 | 3 | Atp11c | Col18a1 |
| Eps151 | 0.00032043 | 0.31565125 | 0.668 | 0.581 | 1 | 3 | Eps15 | Col18a1 |
| Atg4c | 0.00032844 | 0.39772734 | 0.359 | 0.315 | 1 | 3 | Atg4c | Col18a1 |
| Cnot2 | 0.00032848 | 0.32261855 | 0.736 | 0.667 | 1 | 3 | Cnot2 | Col18a1 |
| Pts | 0.00033413 | 0.38104259 | 0.27 | 0.22 | 1 | 3 | Pts | Col18a1 |
| Klhl11 | 0.00034069 | 0.44040224 | 0.542 | 0.464 | 1 | 3 | Klhl11 | Col18a1 |
| Kmt5b | 0.00034203 | 0.26360958 | 0.642 | 0.592 | 1 | 3 | Kmt5b | Col18a1 |
| Trmt11 | 0.00034584 | 0.42712867 | 0.298 | 0.253 | 1 | 3 | Trmt11 | Col18a1 |
| Slc25a24 | 0.00035012 | 0.32875587 | 0.829 | 0.747 | 1 | 3 | Slc25a24 | Col18a1 |
| Tmem234 | 0.00035395 | 0.28904786 | 0.891 | 0.818 | 1 | 3 | Tmem234 | Col18a1 |
| N4bp21 | 0.00035462 | 0.46339613 | 0.328 | 0.282 | 1 | 3 | N4bp2 | Col18a1 |
| Pex13 | 0.00035682 | 0.41679994 | 0.44 | 0.394 | 1 | 3 | Pex13 | Col18a1 |
| Wasl | 0.00035784 | 0.3280366 | 0.725 | 0.652 | 1 | 3 | Wasl | Col18a1 |
| Tprkb | 0.00035934 | 0.33611038 | 0.454 | 0.391 | 1 | 3 | Tprkb | Col18a1 |
| Strip1 | 0.00036481 | 0.33067076 | 0.417 | 0.362 | 1 | 3 | Strip1 | Col18a1 |
| Tmem179b | 0.00036792 | 0.39115123 | 0.493 | 0.445 | 1 | 3 | Tmem179b | Col18a1 |
| Mrtfb | 0.00037684 | 0.30839575 | 0.551 | 0.495 | 1 | 3 | Mrtfb | Col18a1 |
| Kifc31 | 0.00038439 | 0.31994739 | 0.499 | 0.429 | 1 | 3 | Kifc3 | Col18a1 |
| Plxna1 | 0.0003984 | 0.28073754 | 0.862 | 0.794 | 1 | 3 | Plxna1 | Col18a1 |
| Arid4b | 0.00041391 | 0.32322749 | 0.581 | 0.508 | 1 | 3 | Arid4b | Col18a1 |
| Ube2v1 | 0.00042572 | 0.30598662 | 0.933 | 0.866 | 1 | 3 | Ube2v1 | Col18a1 |
| Supt31 | 0.00042772 | 0.41374766 | 0.311 | 0.257 | 1 | 3 | Supt3 | Col18a1 |
| Elp2 | 0.00042983 | 0.45996569 | 0.456 | 0.391 | 1 | 3 | Elp2 | Col18a1 |
| Bahcc12 | 0.00043872 | 0.4814859 | 0.346 | 0.32 | 1 | 3 | Bahcc1 | Col18a1 |
| Adgrl11 | 0.000441 | 0.36205437 | 0.685 | 0.588 | 1 | 3 | Adgrl1 | Col18a1 |
| Ccni | 0.00046133 | 0.2581652 | 0.921 | 0.872 | 1 | 3 | Ccni | Col18a1 |
| Ocrl | 0.00046523 | 0.37314073 | 0.556 | 0.503 | 1 | 3 | Ocrl | Col18a1 |
| Wdr33 | 0.00047426 | 0.29078471 | 0.722 | 0.644 | 1 | 3 | Wdr33 | Col18a1 |
| Wrn1 | 0.00047821 | 0.39629992 | 0.337 | 0.282 | 1 | 3 | Wrn | Col18a1 |
| Kat71 | 0.00049256 | 0.44859996 | 0.593 | 0.499 | 1 | 3 | Kat7 | Col18a1 |
| Gzf1 | 0.00049388 | 0.35141489 | 0.433 | 0.369 | 1 | 3 | Gzf1 | Col18a1 |
| Tada1 | 0.00049906 | 0.35249109 | 0.498 | 0.437 | 1 | 3 | Tada1 | Col18a1 |
| Nadk21 | 0.00050059 | 0.37847323 | 0.318 | 0.263 | 1 | 3 | Nadk2 | Col18a1 |
| Ppp1r101 | 0.00050282 | 0.32303965 | 0.54 | 0.558 | 1 | 3 | Ppp1r10 | Col18a1 |
| Slc30a6 | 0.00050856 | 0.50664966 | 0.337 | 0.29 | 1 | 3 | Slc30a6 | Col18a1 |
| Rreb1 | 0.0005114 | 0.35783132 | 0.553 | 0.483 | 1 | 3 | Rreb1 | Col18a1 |
| Acox1 | 0.00051487 | 0.27120171 | 0.563 | 0.553 | 1 | 3 | Acox1 | Col18a1 |
| Vamp7 | 0.00053229 | 0.45262251 | 0.507 | 0.439 | 1 | 3 | Vamp7 | Col18a1 |
| Pias1 | 0.00056211 | 0.39116689 | 0.42 | 0.365 | 1 | 3 | Pias1 | Col18a1 |
| Snrk | 0.00058135 | 0.31180129 | 0.401 | 0.36 | 1 | 3 | Snrk | Col18a1 |
| Chrac1 | 0.00059619 | 0.37907838 | 0.529 | 0.467 | 1 | 3 | Chrac1 | Col18a1 |
| Fam136a | 0.00059886 | 0.47160985 | 0.451 | 0.382 | 1 | 3 | Fam136a | Col18a1 |
| Wtap | 0.00060201 | 0.3580941 | 0.642 | 0.566 | 1 | 3 | Wtap | Col18a1 |
| Pnkd | 0.00060466 | 0.49718508 | 0.387 | 0.338 | 1 | 3 | Pnkd | Col18a1 |
| Utp251 | 0.00064737 | 0.37687782 | 0.312 | 0.272 | 1 | 3 | Utp25 | Col18a1 |
| Chmp2b | 0.00065091 | 0.33696199 | 0.66 | 0.587 | 1 | 3 | Chmp2b | Col18a1 |
| Slf21 | 0.00065419 | 0.41939571 | 0.447 | 0.374 | 1 | 3 | Slf2 | Col18a1 |
| Ccdc58 | 0.00067203 | 0.36351462 | 0.439 | 0.39 | 1 | 3 | Ccdc58 | Col18a1 |
| Dhrs7 | 0.00067737 | 0.27235442 | 0.723 | 0.636 | 1 | 3 | Dhrs7 | Col18a1 |
| Ttc141 | 0.00069317 | 0.27521506 | 0.553 | 0.496 | 1 | 3 | Ttc14 | Col18a1 |
| As3mt1 | 0.0007016 | 0.32934404 | 0.392 | 0.345 | 1 | 3 | As3mt | Col18a1 |
| Sep-71 | 0.00070615 | 0.3139898 | 0.95 | 0.893 | 1 | 3 | 7-Sep | Col18a1 |
| Bdkrb12 | 0.0007094 | 0.42954878 | 0.401 | 0.305 | 1 | 3 | Bdkrb1 | Col18a1 |
| Ccnh | 0.00072851 | 0.4495866 | 0.383 | 0.344 | 1 | 3 | Ccnh | Col18a1 |
| Helz | 0.00073027 | 0.37774335 | 0.572 | 0.512 | 1 | 3 | Helz | Col18a1 |
| Mccc11 | 0.0007401 | 0.43209093 | 0.303 | 0.26 | 1 | 3 | Mccc1 | Col18a1 |
| Washc5 | 0.00075038 | 0.32990794 | 0.482 | 0.427 | 1 | 3 | Washc5 | Col18a1 |
| Mcee | 0.00076724 | 0.43982379 | 0.266 | 0.221 | 1 | 3 | Mcee | Col18a1 |
| Arsb | 0.00080071 | 0.27538962 | 0.66 | 0.631 | 1 | 3 | Arsb | Col18a1 |
| Tmed7 | 0.00080872 | 0.25067279 | 0.932 | 0.894 | 1 | 3 | Tmed7 | Col18a1 |
| Ccser2 | 0.00081184 | 0.31109187 | 0.573 | 0.512 | 1 | 3 | Ccser2 | Col18a1 |
| Ints61 | 0.00083722 | 0.39509013 | 0.554 | 0.523 | 1 | 3 | Ints6 | Col18a1 |
| DEPRECATED-I | 0.00084838 | 0.37995096 | 0.559 | 0.478 | 1 | 3 | DEPRECATED-I | Col18a1 |
| Tle21 | 0.00084901 | 0.27048065 | 0.292 | 0.245 | 1 | 3 | Tle2 | Col18a1 |
| Rab10 | 0.00088135 | 0.29799362 | 0.769 | 0.69 | 1 | 3 | Rab10 | Col18a1 |
| Thap4 | 0.00090105 | 0.43953037 | 0.484 | 0.428 | 1 | 3 | Thap4 | Col18a1 |
| Celf1 | 0.00090416 | 0.36186077 | 0.587 | 0.511 | 1 | 3 | Celf1 | Col18a1 |
| Fbxl3 | 0.00092971 | 0.40371649 | 0.443 | 0.392 | 1 | 3 | Fbxl3 | Col18a1 |
| Cyp20a1 | 0.00093041 | 0.39483883 | 0.668 | 0.598 | 1 | 3 | Cyp20a1 | Col18a1 |
| Coa61 | 0.00094632 | 0.43419061 | 0.31 | 0.256 | 1 | 3 | Coa6 | Col18a1 |
| Zfp68 | 0.00096208 | 0.4616725 | 0.367 | 0.312 | 1 | 3 | Zfp68 | Col18a1 |
| Strn4 | 0.00096472 | 0.28180162 | 0.837 | 0.773 | 1 | 3 | Strn4 | Col18a1 |
| Acvr1 | 0.00097572 | 0.30047301 | 0.862 | 0.795 | 1 | 3 | Acvr1 | Col18a1 |
| Spopl | 0.00099951 | 0.30328057 | 0.501 | 0.446 | 1 | 3 | Spopl | Col18a1 |
| Fubp3 | 0.00100122 | 0.3127056 | 0.63 | 0.565 | 1 | 3 | Fubp3 | Col18a1 |
| Slc35a5 | 0.00100243 | 0.35759874 | 0.446 | 0.385 | 1 | 3 | Slc35a5 | Col18a1 |
| Gclc1 | 0.0010138 | 0.45239556 | 0.33 | 0.286 | 1 | 3 | Gclc | Col18a1 |
| Uspl1 | 0.00104495 | 0.37189089 | 0.371 | 0.336 | 1 | 3 | Uspl1 | Col18a1 |
| Bex31 | 0.00105379 | 0.33517745 | 0.64 | 0.557 | 1 | 3 | Bex3 | Col18a1 |
| Ttc21b | 0.00109088 | 0.35770354 | 0.265 | 0.231 | 1 | 3 | Ttc21b | Col18a1 |
| Akt3 | 0.00111117 | 0.30715844 | 0.663 | 0.591 | 1 | 3 | Akt3 | Col18a1 |
| Psmf1 | 0.00111152 | 0.27966646 | 0.49 | 0.445 | 1 | 3 | Psmf1 | Col18a1 |
| Sin3a1 | 0.00112465 | 0.32394488 | 0.622 | 0.536 | 1 | 3 | Sin3a | Col18a1 |
| Agtpbp1 | 0.00116159 | 0.34111175 | 0.535 | 0.465 | 1 | 3 | Agtpbp1 | Col18a1 |
| Zkscan1 | 0.00117204 | 0.35671839 | 0.377 | 0.32 | 1 | 3 | Zkscan1 | Col18a1 |
| Fbxo6 | 0.00117728 | 0.2502255 | 0.572 | 0.523 | 1 | 3 | Fbxo6 | Col18a1 |
| Brox | 0.0011849 | 0.33318283 | 0.485 | 0.428 | 1 | 3 | Brox | Col18a1 |
| Bcl2l11 | 0.00118523 | 0.28348579 | 0.366 | 0.319 | 1 | 3 | Bcl2l1 | Col18a1 |
| Lemd3 | 0.00118574 | 0.31159842 | 0.553 | 0.515 | 1 | 3 | Lemd3 | Col18a1 |
| Tmub2 | 0.00119288 | 0.29566561 | 0.602 | 0.565 | 1 | 3 | Tmub2 | Col18a1 |
| Dusp7 | 0.00120351 | 0.30495342 | 0.819 | 0.753 | 1 | 3 | Dusp7 | Col18a1 |
| Whamm | 0.0012492 | 0.29576539 | 0.263 | 0.23 | 1 | 3 | Whamm | Col18a1 |
| Snapc1 | 0.0012766 | 0.35292458 | 0.381 | 0.338 | 1 | 3 | Snapc1 | Col18a1 |
| Ncs1 | 0.00128277 | 0.39128399 | 0.626 | 0.546 | 1 | 3 | Ncs1 | Col18a1 |
| Dld1 | 0.0013175 | 0.32243602 | 0.719 | 0.641 | 1 | 3 | Dld | Col18a1 |
| Hnrnpc | 0.00133342 | 0.4704976 | 0.648 | 0.528 | 1 | 3 | Hnrnpc | Col18a1 |
| Alyref2 | 0.00134467 | 0.32301214 | 0.257 | 0.255 | 1 | 3 | Alyref | Col18a1 |
| Arid1a1 | 0.00139044 | 0.33136164 | 0.644 | 0.57 | 1 | 3 | Arid1a | Col18a1 |
| Ppm1d1 | 0.0014128 | 0.45490287 | 0.345 | 0.306 | 1 | 3 | Ppm1d | Col18a1 |
| Dnmt3a | 0.00143946 | 0.35807563 | 0.603 | 0.542 | 1 | 3 | Dnmt3a | Col18a1 |
| Pcdhga101 | 0.00145555 | 0.32024742 | 0.262 | 0.229 | 1 | 3 | Pcdhga10 | Col18a1 |
| Patl1 | 0.00145888 | 0.26014809 | 0.579 | 0.526 | 1 | 3 | Patl1 | Col18a1 |
| Hdac11 | 0.00146055 | 0.31161171 | 0.768 | 0.696 | 1 | 3 | Hdac1 | Col18a1 |
| Kifap3 | 0.00146737 | 0.27588426 | 0.625 | 0.588 | 1 | 3 | Kifap3 | Col18a1 |
| Sik3 | 0.00149204 | 0.27220135 | 0.711 | 0.632 | 1 | 3 | Sik3 | Col18a1 |
| Phax | 0.00149874 | 0.36503931 | 0.611 | 0.568 | 1 | 3 | Phax | Col18a1 |
| Gse1 | 0.00151415 | 0.40918844 | 0.36 | 0.302 | 1 | 3 | Gse1 | Col18a1 |
| Dcaf17 | 0.00151912 | 0.34541862 | 0.259 | 0.209 | 1 | 3 | Dcaf17 | Col18a1 |
| N4bp1 | 0.00154044 | 0.30099111 | 0.674 | 0.612 | 1 | 3 | N4bp1 | Col18a1 |
| Yeats2 | 0.00159894 | 0.49822158 | 0.368 | 0.315 | 1 | 3 | Yeats2 | Col18a1 |
| Ccdc163 | 0.00163286 | 0.3220371 | 0.269 | 0.241 | 1 | 3 | Ccdc163 | Col18a1 |
| Ccdc321 | 0.00167428 | 0.44624014 | 0.296 | 0.253 | 1 | 3 | Ccdc32 | Col18a1 |
| Srek1ip11 | 0.00170332 | 0.40114759 | 0.344 | 0.295 | 1 | 3 | Srek1ip1 | Col18a1 |
| Gdi2 | 0.00170846 | 0.38647257 | 0.889 | 0.799 | 1 | 3 | Gdi2 | Col18a1 |
| Eif2a | 0.00170875 | 0.4113456 | 0.676 | 0.582 | 1 | 3 | Eif2a | Col18a1 |
| Lsm14b1 | 0.00170897 | 0.29022362 | 0.837 | 0.772 | 1 | 3 | Lsm14b | Col18a1 |
| Zcchc7 | 0.00171664 | 0.45311901 | 0.375 | 0.336 | 1 | 3 | Zcchc7 | Col18a1 |
| Yaf21 | 0.0017309 | 0.34541286 | 0.635 | 0.569 | 1 | 3 | Yaf2 | Col18a1 |
| Uhrf1bp1l | 0.00173761 | 0.36287159 | 0.536 | 0.474 | 1 | 3 | Uhrf1bp1l | Col18a1 |
| Elp4 | 0.00173849 | 0.52128758 | 0.332 | 0.292 | 1 | 3 | Elp4 | Col18a1 |
| Hacd3 | 0.00177028 | 0.28518935 | 0.614 | 0.565 | 1 | 3 | Hacd3 | Col18a1 |
| Timm211 | 0.00178053 | 0.45705788 | 0.338 | 0.276 | 1 | 3 | Timm21 | Col18a1 |
| Rock11 | 0.0017908 | 0.26274196 | 0.693 | 0.627 | 1 | 3 | Rock1 | Col18a1 |
| Ddx42 | 0.00180458 | 0.2504295 | 0.638 | 0.583 | 1 | 3 | Ddx42 | Col18a1 |
| Ubxn8 | 0.00181729 | 0.35229755 | 0.642 | 0.56 | 1 | 3 | Ubxn8 | Col18a1 |
| Gphn | 0.00182744 | 0.46781307 | 0.25 | 0.207 | 1 | 3 | Gphn | Col18a1 |
| Tbca | 0.00183216 | 0.31358471 | 0.903 | 0.83 | 1 | 3 | Tbca | Col18a1 |
| Nxf1 | 0.00184672 | 0.33859062 | 0.754 | 0.684 | 1 | 3 | Nxf1 | Col18a1 |
| Galns | 0.00186245 | 0.34618424 | 0.448 | 0.398 | 1 | 3 | Galns | Col18a1 |
| Usp38 | 0.00186391 | 0.41969382 | 0.452 | 0.381 | 1 | 3 | Usp38 | Col18a1 |
| Evi5 | 0.001898 | 0.32999924 | 0.468 | 0.426 | 1 | 3 | Evi5 | Col18a1 |
| Plekhh3 | 0.00195418 | 0.32644748 | 0.303 | 0.261 | 1 | 3 | Plekhh3 | Col18a1 |
| Sestd1 | 0.00197251 | 0.33397973 | 0.548 | 0.499 | 1 | 3 | Sestd1 | Col18a1 |
| Epc1 | 0.00197802 | 0.27483851 | 0.683 | 0.653 | 1 | 3 | Epc1 | Col18a1 |
| Smg71 | 0.00197992 | 0.36998269 | 0.635 | 0.553 | 1 | 3 | Smg7 | Col18a1 |
| Tasor | 0.00201304 | 0.35503097 | 0.508 | 0.438 | 1 | 3 | Tasor | Col18a1 |
| Spire1 | 0.00208199 | 0.4659443 | 0.432 | 0.378 | 1 | 3 | Spire1 | Col18a1 |
| Zcchc2 | 0.00209871 | 0.48218287 | 0.314 | 0.271 | 1 | 3 | Zcchc2 | Col18a1 |
| Tmem128 | 0.00210981 | 0.38249889 | 0.615 | 0.537 | 1 | 3 | Tmem128 | Col18a1 |
| Ptcd21 | 0.00211197 | 0.56280172 | 0.25 | 0.207 | 1 | 3 | Ptcd2 | Col18a1 |
| Ube2o | 0.00212114 | 0.29856506 | 0.549 | 0.503 | 1 | 3 | Ube2o | Col18a1 |
| Rcor3 | 0.00212478 | 0.31241093 | 0.468 | 0.416 | 1 | 3 | Rcor3 | Col18a1 |
| Ephb21 | 0.00217187 | 0.31997578 | 0.658 | 0.609 | 1 | 3 | Ephb2 | Col18a1 |
| Med61 | 0.00219787 | 0.50482012 | 0.43 | 0.362 | 1 | 3 | Med6 | Col18a1 |
| Ric11 | 0.00221131 | 0.31130931 | 0.693 | 0.625 | 1 | 3 | Ric1 | Col18a1 |
| Csnk1g1 | 0.00224654 | 0.40934389 | 0.394 | 0.336 | 1 | 3 | Csnk1g1 | Col18a1 |
| Ube2d1 | 0.0022583 | 0.31583265 | 0.578 | 0.523 | 1 | 3 | Ube2d1 | Col18a1 |
| Wdr11 | 0.00226597 | 0.33055005 | 0.433 | 0.374 | 1 | 3 | Wdr11 | Col18a1 |
| Ctdspl21 | 0.00226917 | 0.44358363 | 0.561 | 0.466 | 1 | 3 | Ctdspl2 | Col18a1 |
| Erbin | 0.00230336 | 0.29378889 | 0.826 | 0.761 | 1 | 3 | Erbin | Col18a1 |
| Fxr1 | 0.00232012 | 0.31387889 | 0.811 | 0.75 | 1 | 3 | Fxr1 | Col18a1 |
| Mapk14 | 0.00235548 | 0.25502132 | 0.79 | 0.722 | 1 | 3 | Mapk14 | Col18a1 |
| Rab22a | 0.00236098 | 0.25612027 | 0.788 | 0.739 | 1 | 3 | Rab22a | Col18a1 |
| Cul4a | 0.00239354 | 0.3239212 | 0.819 | 0.754 | 1 | 3 | Cul4a | Col18a1 |
| Uhmk1 | 0.00254466 | 0.35257919 | 0.533 | 0.471 | 1 | 3 | Uhmk1 | Col18a1 |
| Usp34 | 0.00256734 | 0.35599026 | 0.66 | 0.589 | 1 | 3 | Usp34 | Col18a1 |
| Polr2m | 0.00260427 | 0.39677699 | 0.596 | 0.525 | 1 | 3 | Polr2m | Col18a1 |
| Dync1i2 | 0.0026071 | 0.32652929 | 0.822 | 0.743 | 1 | 3 | Dync1i2 | Col18a1 |
| Rbm41 | 0.00270142 | 0.32278648 | 0.272 | 0.239 | 1 | 3 | Rbm41 | Col18a1 |
| Tmem173 | 0.00275635 | 0.31523815 | 0.806 | 0.761 | 1 | 3 | Tmem173 | Col18a1 |
| Aftph | 0.00276585 | 0.54211943 | 0.278 | 0.236 | 1 | 3 | Aftph | Col18a1 |
| Srsf21 | 0.00278124 | 0.32636058 | 0.782 | 0.727 | 1 | 3 | Srsf2 | Col18a1 |
| Tmem251 | 0.00279149 | 0.41525748 | 0.324 | 0.292 | 1 | 3 | Tmem251 | Col18a1 |
| Metap21 | 0.0028505 | 0.31933077 | 0.854 | 0.773 | 1 | 3 | Metap2 | Col18a1 |
| Iqcc | 0.00286302 | 0.40630476 | 0.265 | 0.225 | 1 | 3 | Iqcc | Col18a1 |
| Zkscan17 | 0.00286732 | 0.39278226 | 0.467 | 0.416 | 1 | 3 | Zkscan17 | Col18a1 |
| Emc3 | 0.00288057 | 0.40389233 | 0.763 | 0.666 | 1 | 3 | Emc3 | Col18a1 |
| Fbxw4 | 0.00292023 | 0.3433547 | 0.259 | 0.22 | 1 | 3 | Fbxw4 | Col18a1 |
| Ten1 | 0.00296643 | 0.38305825 | 0.381 | 0.329 | 1 | 3 | Ten1 | Col18a1 |
| Arpc3 | 0.00298449 | 0.33335191 | 0.801 | 0.722 | 1 | 3 | Arpc3 | Col18a1 |
| Appl1 | 0.00304866 | 0.32645621 | 0.748 | 0.662 | 1 | 3 | Appl1 | Col18a1 |
| Itgav | 0.00305159 | 0.27511715 | 0.937 | 0.863 | 1 | 3 | Itgav | Col18a1 |
| Numa1 | 0.00306373 | 0.28538066 | 0.799 | 0.714 | 1 | 3 | Numa1 | Col18a1 |
| Uqcrc2 | 0.00308339 | 0.36406369 | 0.79 | 0.701 | 1 | 3 | Uqcrc2 | Col18a1 |
| Atl3 | 0.00309537 | 0.27963485 | 0.948 | 0.879 | 1 | 3 | Atl3 | Col18a1 |
| Tmem161b | 0.00325687 | 0.35507636 | 0.273 | 0.235 | 1 | 3 | Tmem161b | Col18a1 |
| Zbtb33 | 0.00327335 | 0.38273487 | 0.423 | 0.371 | 1 | 3 | Zbtb33 | Col18a1 |
| Cryz | 0.00328157 | 0.35776074 | 0.337 | 0.301 | 1 | 3 | Cryz | Col18a1 |
| Vps4b | 0.0033208 | 0.34153855 | 0.536 | 0.481 | 1 | 3 | Vps4b | Col18a1 |
| Zbtb6 | 0.00333864 | 0.44766666 | 0.252 | 0.21 | 1 | 3 | Zbtb6 | Col18a1 |
| Rtl8b | 0.00339201 | 0.38233253 | 0.327 | 0.288 | 1 | 3 | Rtl8b | Col18a1 |
| Rev1 | 0.00346856 | 0.34435335 | 0.311 | 0.267 | 1 | 3 | Rev1 | Col18a1 |
| Vkorc1l11 | 0.00352707 | 0.41042351 | 0.666 | 0.594 | 1 | 3 | Vkorc1l1 | Col18a1 |
| Zbtb1 | 0.00353022 | 0.29114716 | 0.663 | 0.609 | 1 | 3 | Zbtb1 | Col18a1 |
| Epc2 | 0.00353068 | 0.41138232 | 0.451 | 0.397 | 1 | 3 | Epc2 | Col18a1 |
| Acox31 | 0.00354415 | 0.28095488 | 0.386 | 0.341 | 1 | 3 | Acox3 | Col18a1 |
| Lipa | 0.00365043 | 0.36988486 | 0.763 | 0.683 | 1 | 3 | Lipa | Col18a1 |
| Zswim4 | 0.00368835 | 0.28490988 | 0.553 | 0.492 | 1 | 3 | Zswim4 | Col18a1 |
| Baz2b | 0.0036982 | 0.32241583 | 0.378 | 0.327 | 1 | 3 | Baz2b | Col18a1 |
| Dcaf12 | 0.00375281 | 0.29032124 | 0.682 | 0.636 | 1 | 3 | Dcaf12 | Col18a1 |
| Hoxd81 | 0.00376821 | 0.30043236 | 0.458 | 0.4 | 1 | 3 | Hoxd8 | Col18a1 |
| Pnrc21 | 0.00382159 | 0.27034222 | 0.763 | 0.682 | 1 | 3 | Pnrc2 | Col18a1 |
| Brwd1 | 0.00386357 | 0.28624698 | 0.766 | 0.696 | 1 | 3 | Brwd1 | Col18a1 |
| Polr2k1 | 0.00390379 | 0.42020571 | 0.283 | 0.252 | 1 | 3 | Polr2k | Col18a1 |
| Cops21 | 0.00396641 | 0.32355052 | 0.715 | 0.648 | 1 | 3 | Cops2 | Col18a1 |
| Smarca51 | 0.00404302 | 0.4420065 | 0.614 | 0.521 | 1 | 3 | Smarca5 | Col18a1 |
| Shox2 | 0.00410234 | 0.2507713 | 0.432 | 0.411 | 1 | 3 | Shox2 | Col18a1 |
| Ngrn | 0.00421962 | 0.25853705 | 0.267 | 0.249 | 1 | 3 | Ngrn | Col18a1 |
| Rnf114 | 0.00426644 | 0.35490262 | 0.509 | 0.464 | 1 | 3 | Rnf114 | Col18a1 |
| Ebag9 | 0.0043653 | 0.28093442 | 0.356 | 0.314 | 1 | 3 | Ebag9 | Col18a1 |
| Adpgk | 0.00437079 | 0.33529369 | 0.8 | 0.717 | 1 | 3 | Adpgk | Col18a1 |
| Frg1 | 0.00439002 | 0.2728633 | 0.498 | 0.461 | 1 | 3 | Frg1 | Col18a1 |
| Usp13 | 0.00441366 | 0.25425543 | 0.311 | 0.286 | 1 | 3 | Usp13 | Col18a1 |
| Eef1g | 0.00441823 | 0.35160298 | 0.939 | 0.863 | 1 | 3 | Eef1g | Col18a1 |
| Dera | 0.00445656 | 0.45911407 | 0.381 | 0.348 | 1 | 3 | Dera | Col18a1 |
| L3hypdh | 0.00452059 | 0.32037418 | 0.496 | 0.464 | 1 | 3 | L3hypdh | Col18a1 |
| Vopp1 | 0.0045934 | 0.32256503 | 0.583 | 0.538 | 1 | 3 | Vopp1 | Col18a1 |
| C1d | 0.00460766 | 0.35994843 | 0.664 | 0.588 | 1 | 3 | C1d | Col18a1 |
| Nt5c3 | 0.00461839 | 0.49541998 | 0.333 | 0.283 | 1 | 3 | Nt5c3 | Col18a1 |
| Diablo | 0.00467508 | 0.43566283 | 0.255 | 0.213 | 1 | 3 | Diablo | Col18a1 |
| Tbpl1 | 0.00472801 | 0.3681668 | 0.462 | 0.414 | 1 | 3 | Tbpl1 | Col18a1 |
| Mms19 | 0.00481129 | 0.38513346 | 0.38 | 0.328 | 1 | 3 | Mms19 | Col18a1 |
| Tent4b | 0.00482601 | 0.35162801 | 0.289 | 0.252 | 1 | 3 | Tent4b | Col18a1 |
| Zbtb24 | 0.00489971 | 0.30834839 | 0.312 | 0.282 | 1 | 3 | Zbtb24 | Col18a1 |
| Ubap2l | 0.00494972 | 0.30694545 | 0.909 | 0.83 | 1 | 3 | Ubap2l | Col18a1 |
| Rnf167 | 0.00495542 | 0.25770936 | 0.376 | 0.328 | 1 | 3 | Rnf167 | Col18a1 |
| Atg12 | 0.0050218 | 0.3665823 | 0.464 | 0.411 | 1 | 3 | Atg12 | Col18a1 |
| Trappc8 | 0.00506014 | 0.38448389 | 0.574 | 0.511 | 1 | 3 | Trappc8 | Col18a1 |
| Cebpzos | 0.00508822 | 0.26794103 | 0.415 | 0.373 | 1 | 3 | Cebpzos | Col18a1 |
| Oser1 | 0.00514589 | 0.27220632 | 0.579 | 0.537 | 1 | 3 | Oser1 | Col18a1 |
| Ccz1 | 0.00516302 | 0.33214303 | 0.526 | 0.456 | 1 | 3 | Ccz1 | Col18a1 |
| Nadk | 0.00516852 | 0.29370668 | 0.806 | 0.753 | 1 | 3 | Nadk | Col18a1 |
| Fbxo31 | 0.00517145 | 0.28790788 | 0.403 | 0.37 | 1 | 3 | Fbxo31 | Col18a1 |
| Trap11 | 0.00522403 | 0.37586988 | 0.564 | 0.495 | 1 | 3 | Trap1 | Col18a1 |
| Mmut | 0.00524452 | 0.33182813 | 0.261 | 0.223 | 1 | 3 | Mmut | Col18a1 |
| Acads | 0.00525301 | 0.28718838 | 0.614 | 0.557 | 1 | 3 | Acads | Col18a1 |
| Mob3a | 0.00530263 | 0.3447668 | 0.519 | 0.483 | 1 | 3 | Mob3a | Col18a1 |
| Pelo | 0.00534257 | 0.38208179 | 0.475 | 0.47 | 1 | 3 | Pelo | Col18a1 |
| Ing31 | 0.00539842 | 0.48211441 | 0.256 | 0.208 | 1 | 3 | Ing3 | Col18a1 |
| Dock11 | 0.00546057 | 0.4788929 | 0.257 | 0.207 | 1 | 3 | Dock11 | Col18a1 |
| Ttbk2 | 0.00555916 | 0.42954828 | 0.317 | 0.267 | 1 | 3 | Ttbk2 | Col18a1 |
| Vps18 | 0.00557173 | 0.33109667 | 0.345 | 0.309 | 1 | 3 | Vps18 | Col18a1 |
| Psma71 | 0.00561848 | 0.48570519 | 0.657 | 0.555 | 1 | 3 | Psma7 | Col18a1 |
| Ythdf3 | 0.0056344 | 0.26255437 | 0.882 | 0.833 | 1 | 3 | Ythdf3 | Col18a1 |
| Agps | 0.00565659 | 0.40562714 | 0.533 | 0.456 | 1 | 3 | Agps | Col18a1 |
| M6pr1 | 0.00576753 | 0.28486379 | 0.863 | 0.799 | 1 | 3 | M6pr | Col18a1 |
| Polr3c | 0.00584568 | 0.39283122 | 0.397 | 0.336 | 1 | 3 | Polr3c | Col18a1 |
| Stag1 | 0.00588956 | 0.39933695 | 0.462 | 0.41 | 1 | 3 | Stag1 | Col18a1 |
| Esf11 | 0.00595867 | 0.40655107 | 0.373 | 0.321 | 1 | 3 | Esf1 | Col18a1 |
| Pogk | 0.00597258 | 0.3623233 | 0.33 | 0.291 | 1 | 3 | Pogk | Col18a1 |
| Hcfc1r1 | 0.00600527 | 0.30283804 | 0.84 | 0.775 | 1 | 3 | Hcfc1r1 | Col18a1 |
| Trim3 | 0.00605309 | 0.26090555 | 0.604 | 0.554 | 1 | 3 | Trim3 | Col18a1 |
| Mbd6 | 0.00612059 | 0.25522023 | 0.308 | 0.268 | 1 | 3 | Mbd6 | Col18a1 |
| Ppp1r15b | 0.00618796 | 0.29420631 | 0.714 | 0.658 | 1 | 3 | Ppp1r15b | Col18a1 |
| Ptdss2 | 0.00623634 | 0.32269892 | 0.455 | 0.404 | 1 | 3 | Ptdss2 | Col18a1 |
| Zzz3 | 0.00625259 | 0.30876793 | 0.736 | 0.677 | 1 | 3 | Zzz3 | Col18a1 |
| Elovl1 | 0.00627867 | 0.31833146 | 0.82 | 0.737 | 1 | 3 | Elovl1 | Col18a1 |
| Slc29a11 | 0.0062857 | 0.39385133 | 0.346 | 0.302 | 1 | 3 | Slc29a1 | Col18a1 |
| Cbr3 | 0.00628644 | 0.43946169 | 0.257 | 0.215 | 1 | 3 | Cbr3 | Col18a1 |
| Cox7b | 0.00639362 | 0.27492021 | 0.806 | 0.745 | 1 | 3 | Cox7b | Col18a1 |
| Aasdhppt | 0.00666252 | 0.29923219 | 0.434 | 0.402 | 1 | 3 | Aasdhppt | Col18a1 |
| Klhl28 | 0.00667596 | 0.3394639 | 0.339 | 0.309 | 1 | 3 | Klhl28 | Col18a1 |
| Inka11 | 0.00669539 | 0.28292786 | 0.273 | 0.223 | 1 | 3 | Inka1 | Col18a1 |
| Katna1 | 0.0066954 | 0.29054339 | 0.305 | 0.266 | 1 | 3 | Katna1 | Col18a1 |
| Khsrp1 | 0.0067357 | 0.25862241 | 0.907 | 0.857 | 1 | 3 | Khsrp | Col18a1 |
| Katnbl1 | 0.00675967 | 0.37414799 | 0.44 | 0.39 | 1 | 3 | Katnbl1 | Col18a1 |
| Fat1 | 0.00681605 | 0.29056653 | 0.918 | 0.864 | 1 | 3 | Fat1 | Col18a1 |
| Ptprj1 | 0.00696339 | 0.45153142 | 0.442 | 0.375 | 1 | 3 | Ptprj | Col18a1 |
| Tfcp2 | 0.00701277 | 0.36414602 | 0.326 | 0.287 | 1 | 3 | Tfcp2 | Col18a1 |
| Zc3h3 | 0.0070308 | 0.31563297 | 0.271 | 0.244 | 1 | 3 | Zc3h3 | Col18a1 |
| Sec24b | 0.00703722 | 0.2787749 | 0.732 | 0.666 | 1 | 3 | Sec24b | Col18a1 |
| Ehd4 | 0.00712364 | 0.39420564 | 0.351 | 0.34 | 1 | 3 | Ehd4 | Col18a1 |
| Pde4a | 0.00714031 | 0.26160342 | 0.334 | 0.29 | 1 | 3 | Pde4a | Col18a1 |
| Ccndbp1 | 0.00714925 | 0.26772471 | 0.546 | 0.494 | 1 | 3 | Ccndbp1 | Col18a1 |
| Ankrd13c1 | 0.00715427 | 0.33266001 | 0.588 | 0.533 | 1 | 3 | Ankrd13c | Col18a1 |
| Rhot1 | 0.00717629 | 0.36310686 | 0.483 | 0.433 | 1 | 3 | Rhot1 | Col18a1 |
| Sco2 | 0.00718725 | 0.3116122 | 0.362 | 0.315 | 1 | 3 | Sco2 | Col18a1 |
| Rfk | 0.00731855 | 0.3893778 | 0.623 | 0.58 | 1 | 3 | Rfk | Col18a1 |
| Ubr4 | 0.00735482 | 0.25081321 | 0.825 | 0.771 | 1 | 3 | Ubr4 | Col18a1 |
| Cetn2 | 0.00747532 | 0.34289798 | 0.378 | 0.332 | 1 | 3 | Cetn2 | Col18a1 |
| Pramef8 | 0.00747678 | 0.33196046 | 0.304 | 0.286 | 1 | 3 | Pramef8 | Col18a1 |
| Rcc2 | 0.00753304 | 0.26638852 | 0.721 | 0.673 | 1 | 3 | Rcc2 | Col18a1 |
| Prkci | 0.00753382 | 0.2641427 | 0.55 | 0.49 | 1 | 3 | Prkci | Col18a1 |
| Nfs1 | 0.00755518 | 0.38427874 | 0.369 | 0.32 | 1 | 3 | Nfs1 | Col18a1 |
| Jam3 | 0.00759656 | 0.28098338 | 0.671 | 0.612 | 1 | 3 | Jam3 | Col18a1 |
| Impdh1 | 0.00777037 | 0.32368112 | 0.674 | 0.594 | 1 | 3 | Impdh1 | Col18a1 |
| Atp6v0c | 0.00784329 | 0.28747423 | 0.917 | 0.86 | 1 | 3 | Atp6v0c | Col18a1 |
| Trpm7 | 0.00785609 | 0.29587867 | 0.637 | 0.575 | 1 | 3 | Trpm7 | Col18a1 |
| Rab131 | 0.00788219 | 0.28790699 | 0.571 | 0.52 | 1 | 3 | Rab13 | Col18a1 |
| Ube2e1 | 0.00789822 | 0.37490753 | 0.642 | 0.552 | 1 | 3 | Ube2e1 | Col18a1 |
| Gpbp11 | 0.00796982 | 0.2922756 | 0.558 | 0.498 | 1 | 3 | Gpbp1 | Col18a1 |
| Lonp2 | 0.0082196 | 0.3605991 | 0.495 | 0.436 | 1 | 3 | Lonp2 | Col18a1 |
| Ythdf1 | 0.00824013 | 0.38079635 | 0.702 | 0.629 | 1 | 3 | Ythdf1 | Col18a1 |
| Ing21 | 0.00832503 | 0.32020036 | 0.259 | 0.22 | 1 | 3 | Ing2 | Col18a1 |
| Trim23 | 0.00835589 | 0.52505535 | 0.312 | 0.262 | 1 | 3 | Trim23 | Col18a1 |
| Hmgb1 | 0.00837465 | 0.25235418 | 0.571 | 0.546 | 1 | 3 | Hmgb1 | Col18a1 |
| Immp1l | 0.0083899 | 0.34082851 | 0.447 | 0.409 | 1 | 3 | Immp1l | Col18a1 |
| Dbt1 | 0.00841405 | 0.33365445 | 0.346 | 0.297 | 1 | 3 | Dbt | Col18a1 |
| Qser11 | 0.0085867 | 0.44081819 | 0.355 | 0.311 | 1 | 3 | Qser1 | Col18a1 |
| Fbxl4 | 0.00865016 | 0.37138729 | 0.25 | 0.216 | 1 | 3 | Fbxl4 | Col18a1 |
| Mterf3 | 0.00882235 | 0.3605396 | 0.42 | 0.375 | 1 | 3 | Mterf3 | Col18a1 |
| S1pr21 | 0.00895089 | 0.28075108 | 0.814 | 0.754 | 1 | 3 | S1pr2 | Col18a1 |
| Ivns1abp1 | 0.00897465 | 0.36422415 | 0.784 | 0.697 | 1 | 3 | Ivns1abp | Col18a1 |
| Ptk2 | 0.00899417 | 0.29692805 | 0.719 | 0.655 | 1 | 3 | Ptk2 | Col18a1 |
| Asb31 | 0.00909872 | 0.34685464 | 0.331 | 0.29 | 1 | 3 | Asb3 | Col18a1 |
| Lrrfip2 | 0.00911411 | 0.31359399 | 0.566 | 0.512 | 1 | 3 | Lrrfip2 | Col18a1 |
| Txndc161 | 0.00916641 | 0.25567019 | 0.417 | 0.38 | 1 | 3 | Txndc16 | Col18a1 |
| Rfng | 0.0091823 | 0.32830209 | 0.518 | 0.463 | 1 | 3 | Rfng | Col18a1 |
| Psmb10 | 0.00927166 | 0.28034718 | 0.435 | 0.419 | 1 | 3 | Psmb10 | Col18a1 |
| Sertad2 | 0.00928291 | 0.3876852 | 0.506 | 0.45 | 1 | 3 | Sertad2 | Col18a1 |
| Zfp9481 | 0.00929841 | 0.50123944 | 0.257 | 0.218 | 1 | 3 | Zfp948 | Col18a1 |
| Mlx | 0.00936766 | 0.36482222 | 0.41 | 0.369 | 1 | 3 | Mlx | Col18a1 |
| Cmc1 | 0.00943705 | 0.36039621 | 0.605 | 0.552 | 1 | 3 | Cmc1 | Col18a1 |
| Taf151 | 0.00947944 | 0.3696585 | 0.262 | 0.212 | 1 | 3 | Taf15 | Col18a1 |
| Suz121 | 0.00978732 | 0.46288423 | 0.486 | 0.399 | 1 | 3 | Suz12 | Col18a1 |
| Ipo71 | 0.00980016 | 0.65329289 | 0.267 | 0.226 | 1 | 3 | Ipo7 | Col18a1 |
| Tbcc | 0.00994606 | 0.27519685 | 0.467 | 0.429 | 1 | 3 | Tbcc | Col18a1 |
| Adi1 | 0.00999746 | 0.2853128 | 0.445 | 0.385 | 1 | 3 | Adi1 | Col18a1 |
| Piezo21 | 0 | 2.14821519 | 0.723 | 0.326 | 0 | 4 | Piezo2 | myCAF_2 |
| Cspg41 | 0 | 2.27655431 | 0.711 | 0.33 | 0 | 4 | Cspg4 | myCAF_2 |
| Dusp41 | 0 | 2.51907447 | 0.656 | 0.282 | 0 | 4 | Dusp4 | myCAF_2 |
| Egln31 | 0 | 3.32245943 | 0.498 | 0.155 | 0 | 4 | Egln3 | myCAF_2 |
| Cmklr11 | 0 | 2.30225845 | 0.558 | 0.226 | 0 | 4 | Cmklr1 | myCAF_2 |
| Slc2a31 | 0 | 2.58770168 | 0.627 | 0.295 | 0 | 4 | Slc2a3 | myCAF_2 |
| Ndufa4l22 | 0 | 1.51324788 | 0.713 | 0.395 | 0 | 4 | Ndufa4l2 | myCAF_2 |
| Nes1 | 0 | 2.0074848 | 0.552 | 0.236 | 0 | 4 | Nes | myCAF_2 |
| Sparcl1 | 0 | 3.35900925 | 0.532 | 0.222 | 0 | 4 | Sparcl1 | myCAF_2 |
| Slit1 | 0 | 3.64747524 | 0.394 | 0.091 | 0 | 4 | Slit1 | myCAF_2 |
| Pgf | 0 | 4.20451497 | 0.428 | 0.131 | 0 | 4 | Pgf | myCAF_2 |
| Unc5b1 | 0 | 1.49891996 | 0.864 | 0.584 | 0 | 4 | Unc5b | myCAF_2 |
| Jup | 0 | 2.67965366 | 0.506 | 0.232 | 0 | 4 | Jup | myCAF_2 |
| Ankrd371 | 0 | 2.60677589 | 0.399 | 0.141 | 0 | 4 | Ankrd37 | myCAF_2 |
| Mmp92 | 0 | 2.11260488 | 0.548 | 0.292 | 0 | 4 | Mmp9 | myCAF_2 |
| Mpp2 | 0 | 2.35264921 | 0.472 | 0.216 | 0 | 4 | Mpp2 | myCAF_2 |
| Ctsc1 | 0 | 1.33018653 | 0.881 | 0.628 | 0 | 4 | Ctsc | myCAF_2 |
| Car11 | 0 | 3.06012181 | 0.328 | 0.088 | 0 | 4 | Car11 | myCAF_2 |
| Mcam | 0 | 2.98637847 | 0.352 | 0.119 | 0 | 4 | Mcam | myCAF_2 |
| Egr21 | 0 | 1.58164816 | 0.691 | 0.467 | 0 | 4 | Egr2 | myCAF_2 |
| Sik11 | 0 | 1.73911022 | 0.64 | 0.424 | 0 | 4 | Sik1 | myCAF_2 |
| Slc20a11 | 0 | 2.12373485 | 0.75 | 0.534 | 0 | 4 | Slc20a1 | myCAF_2 |
| Slco4a11 | 0 | 2.31359437 | 0.378 | 0.165 | 0 | 4 | Slco4a1 | myCAF_2 |
| Cadm41 | 0 | 2.01645375 | 0.348 | 0.138 | 0 | 4 | Cadm4 | myCAF_2 |
| Nrep1 | 0 | 1.22630893 | 0.878 | 0.673 | 0 | 4 | Nrep | myCAF_2 |
| Vldlr | 0 | 2.56109005 | 0.3 | 0.109 | 0 | 4 | Vldlr | myCAF_2 |
| Fabp5 | 0 | 2.5894798 | 0.435 | 0.255 | 0 | 4 | Fabp5 | myCAF_2 |
| Rgs161 | 0 | 1.58127351 | 0.653 | 0.475 | 0 | 4 | Rgs16 | myCAF_2 |
| S1pr3 | 0 | 2.01282159 | 0.652 | 0.477 | 0 | 4 | S1pr3 | myCAF_2 |
| Cdkn1a2 | 0 | 1.34203061 | 0.576 | 0.407 | 0 | 4 | Cdkn1a | myCAF_2 |
| Mmp101 | 0 | 1.74312632 | 0.38 | 0.217 | 0 | 4 | Mmp10 | myCAF_2 |
| Slc16a31 | 0 | 1.62105934 | 0.79 | 0.636 | 0 | 4 | Slc16a3 | myCAF_2 |
| Slc2a11 | 0 | 2.49999458 | 0.844 | 0.691 | 0 | 4 | Slc2a1 | myCAF_2 |
| Inhba2 | 0 | 1.06601425 | 0.851 | 0.699 | 0 | 4 | Inhba | myCAF_2 |
| Hbegf1 | 0 | 1.99386803 | 0.39 | 0.239 | 0 | 4 | Hbegf | myCAF_2 |
| Gng111 | 0 | 1.67764597 | 0.925 | 0.786 | 0 | 4 | Gng11 | myCAF_2 |
| Tgfb11 | 0 | 1.52156186 | 0.965 | 0.827 | 0 | 4 | Tgfb1 | myCAF_2 |
| Ptgs22 | 0 | 1.59843905 | 0.558 | 0.421 | 0 | 4 | Ptgs2 | myCAF_2 |
| Fam117b1 | 0 | 1.65641117 | 0.7 | 0.567 | 0 | 4 | Fam117b | myCAF_2 |
| Ank1 | 0 | 1.56477224 | 0.684 | 0.562 | 0 | 4 | Ank | myCAF_2 |
| Vegfa1 | 0 | 2.02441769 | 0.806 | 0.684 | 0 | 4 | Vegfa | myCAF_2 |
| Arg11 | 0 | 1.8330943 | 0.348 | 0.237 | 0 | 4 | Arg1 | myCAF_2 |
| Ddit41 | 0 | 1.39211303 | 0.776 | 0.666 | 0 | 4 | Ddit4 | myCAF_2 |
| Ndrg11 | 0 | 2.17475754 | 0.763 | 0.655 | 0 | 4 | Ndrg1 | myCAF_2 |
| Spry1 | 0 | 1.57446562 | 0.676 | 0.573 | 0 | 4 | Spry1 | myCAF_2 |
| Hilpda1 | 0 | 1.99183964 | 0.582 | 0.488 | 0 | 4 | Hilpda | myCAF_2 |
| Ero1l1 | 0 | 2.69273851 | 0.774 | 0.68 | 0 | 4 | Ero1l | myCAF_2 |
| Tnc2 | 0 | 0.28655422 | 0.986 | 0.894 | 0 | 4 | Tnc | myCAF_2 |
| Serpine22 | 0 | 1.04829024 | 0.894 | 0.81 | 0 | 4 | Serpine2 | myCAF_2 |
| Col15a11 | 0 | 1.33994389 | 0.87 | 0.789 | 0 | 4 | Col15a1 | myCAF_2 |
| Mgp | 0 | 1.9825215 | 0.387 | 0.331 | 0 | 4 | Mgp | myCAF_2 |
| Igfbp31 | 0 | 1.11119313 | 0.778 | 0.725 | 0 | 4 | Igfbp3 | myCAF_2 |
| Serpine11 | 0 | 1.14559369 | 0.751 | 0.711 | 0 | 4 | Serpine1 | myCAF_2 |
| Egln11 | 0 | 1.52229705 | 0.895 | 0.864 | 0 | 4 | Egln1 | myCAF_2 |
| Plau | 0 | 1.39603351 | 0.79 | 0.765 | 0 | 4 | Plau | myCAF_2 |
| P4ha11 | 0 | 1.05150395 | 0.953 | 0.935 | 0 | 4 | P4ha1 | myCAF_2 |
| P4ha21 | 0 | 1.32998701 | 0.868 | 0.882 | 0 | 4 | P4ha2 | myCAF_2 |
| Fn11 | 0 | 0.69315036 | 0.998 | 0.992 | 0 | 4 | Fn1 | myCAF_2 |
| Col6a32 | 0 | 0.26364199 | 0.99 | 0.988 | 0 | 4 | Col6a3 | myCAF_2 |
| Hk1 | 2.02E-293 | 1.43943031 | 0.831 | 0.772 | 3.90E-293 | 4 | Hk1 | myCAF_2 |
| Gpi11 | 4.44E-293 | 1.2695605 | 0.886 | 0.852 | 8.58E-293 | 4 | Gpi1 | myCAF_2 |
| Cpe | 9.85E-276 | 1.75144473 | 0.351 | 0.223 | 1.90E-271 | 4 | Cpe | myCAF_2 |
| H2-Ab11 | 3.09E-274 | 1.78998955 | 0.491 | 0.305 | 5.98E-270 | 4 | H2-Ab1 | myCAF_2 |
| Amotl1 | 3.82E-266 | 1.33683007 | 0.886 | 0.791 | 7.39E-262 | 4 | Amotl1 | myCAF_2 |
| Tmem2041 | 7.32E-259 | 1.6487786 | 0.546 | 0.319 | 1.41E-254 | 4 | Tmem204 | myCAF_2 |
| Stra62 | 3.51E-258 | 0.98949364 | 0.843 | 0.582 | 6.79E-254 | 4 | Stra6 | myCAF_2 |
| Bhlhe401 | 4.55E-255 | 1.12406752 | 0.895 | 0.849 | 8.79E-251 | 4 | Bhlhe40 | myCAF_2 |
| Heyl1 | 1.60E-254 | 1.32577307 | 0.509 | 0.329 | 3.10E-250 | 4 | Heyl | myCAF_2 |

| | | | | | | | | |
|---|---|---|---|---|---|---|---|---|
| Frem1 | 3.17E-254 | 2.46285562 | 0.327 | 0.137 | 6.13E-250 | 4 | Frem1 | myCAF_2 |
| Kctd11 | 3.29E-250 | 1.23863087 | 0.88 | 0.838 | 6.36E-246 | 4 | Kctd11 | myCAF_2 |
| Wnt111 | 3.70E-249 | 1.8073911 | 0.572 | 0.312 | 7.16E-245 | 4 | Wnt11 | myCAF_2 |
| Krt19 | 4.10E-247 | 2.10253388 | 0.3 | 0.181 | 7.92E-243 | 4 | Krt19 | myCAF_2 |
| Fermt21 | 5.81E-247 | 1.07362358 | 0.901 | 0.869 | 1.12E-242 | 4 | Fermt2 | myCAF_2 |
| Itga6 | 1.59E-246 | 2.26943226 | 0.311 | 0.122 | 3.07E-242 | 4 | Itga6 | myCAF_2 |
| Ednra2 | 2.08E-241 | 1.29069578 | 0.664 | 0.58 | 4.02E-237 | 4 | Ednra | myCAF_2 |
| Tmem1191 | 3.23E-239 | 1.03647236 | 0.894 | 0.885 | 6.25E-235 | 4 | Tmem119 | myCAF_2 |
| Rhob1 | 4.48E-236 | 1.26605966 | 0.904 | 0.858 | 8.66E-232 | 4 | Rhob | myCAF_2 |
| Bhlhe411 | 3.25E-230 | 1.80226328 | 0.564 | 0.271 | 6.28E-226 | 4 | Bhlhe41 | myCAF_2 |
| Egr12 | 7.48E-226 | 0.84748662 | 0.928 | 0.888 | 1.45E-221 | 4 | Egr1 | myCAF_2 |
| Sdc3 | 6.63E-225 | 1.68138031 | 0.64 | 0.405 | 1.28E-220 | 4 | Sdc3 | myCAF_2 |
| 2-Jun | 2.75E-221 | 0.91868071 | 0.918 | 0.911 | 5.32E-217 | 4 | Jun | myCAF_2 |
| Itgb31 | 2.82E-218 | 1.20490862 | 0.833 | 0.659 | 5.46E-214 | 4 | Itgb3 | myCAF_2 |
| Fjx1 | 3.29E-216 | 1.95768842 | 0.485 | 0.283 | 6.36E-212 | 4 | Fjx1 | myCAF_2 |
| Plod11 | 2.21E-214 | 0.9769639 | 0.896 | 0.894 | 4.28E-210 | 4 | Plod1 | myCAF_2 |
| Pfkl1 | 1.29E-208 | 1.21999186 | 0.783 | 0.689 | 2.49E-204 | 4 | Pfkl | myCAF_2 |
| Pfkp | 8.56E-207 | 1.44174522 | 0.657 | 0.622 | 1.66E-202 | 4 | Pfkp | myCAF_2 |
| Stc21 | 3.31E-200 | 1.81973919 | 0.303 | 0.185 | 6.40E-196 | 4 | Stc2 | myCAF_2 |
| Bnip31 | 1.43E-199 | 1.90708211 | 0.42 | 0.217 | 2.77E-195 | 4 | Bnip3 | myCAF_2 |
| Fstl31 | 2.52E-199 | 1.27602175 | 0.742 | 0.688 | 4.87E-195 | 4 | Fstl3 | myCAF_2 |
| Pdk11 | 4.42E-195 | 1.4932096 | 0.583 | 0.415 | 8.55E-191 | 4 | Pdk1 | myCAF_2 |
| Col6a22 | 3.71E-192 | 0.31225947 | 0.991 | 0.99 | 7.18E-188 | 4 | Col6a2 | myCAF_2 |
| Triobp | 5.33E-191 | 1.47356546 | 0.638 | 0.571 | 1.03E-186 | 4 | Triobp | myCAF_2 |
| Gys11 | 4.69E-190 | 1.4482893 | 0.656 | 0.582 | 9.06E-186 | 4 | Gys1 | myCAF_2 |
| Cbfa2t31 | 2.08E-189 | 1.80339155 | 0.517 | 0.258 | 4.01E-185 | 4 | Cbfa2t3 | myCAF_2 |
| Dusp51 | 2.08E-189 | 1.68333963 | 0.559 | 0.376 | 4.03E-185 | 4 | Dusp5 | myCAF_2 |
| Cd401 | 5.30E-187 | 1.99688997 | 0.386 | 0.154 | 1.03E-182 | 4 | Cd40 | myCAF_2 |
| Flnb | 2.16E-181 | 1.05620554 | 0.854 | 0.83 | 4.18E-177 | 4 | Flnb | myCAF_2 |
| Hk2 | 2.20E-181 | 1.32755419 | 0.756 | 0.658 | 4.26E-177 | 4 | Hk2 | myCAF_2 |
| Smox1 | 3.73E-179 | 1.79301866 | 0.536 | 0.353 | 7.21E-175 | 4 | Smox | myCAF_2 |
| Ets11 | 3.01E-178 | 1.24638841 | 0.85 | 0.688 | 5.82E-174 | 4 | Ets1 | myCAF_2 |
| Tmem95 | 6.17E-175 | 1.73857728 | 0.548 | 0.379 | 1.19E-170 | 4 | Tmem95 | myCAF_2 |
| Egr32 | 8.19E-174 | 1.29639294 | 0.626 | 0.426 | 1.58E-169 | 4 | Egr3 | myCAF_2 |
| Kif21b1 | 1.14E-173 | 1.96978995 | 0.446 | 0.208 | 2.21E-169 | 4 | Kif21b | myCAF_2 |
| Pfkfb31 | 9.47E-169 | 1.70987612 | 0.481 | 0.339 | 1.83E-164 | 4 | Pfkfb3 | myCAF_2 |
| Ptprn2 | 3.84E-168 | 1.14059899 | 0.754 | 0.544 | 7.43E-164 | 4 | Ptprn | myCAF_2 |
| Tpi11 | 8.96E-167 | 0.95853214 | 0.84 | 0.84 | 1.73E-162 | 4 | Tpi1 | myCAF_2 |
| Camk2n12 | 1.61E-166 | 1.1089525 | 0.865 | 0.749 | 3.11E-162 | 4 | Camk2n1 | myCAF_2 |
| Pdgfa1 | 3.27E-166 | 1.85093751 | 0.43 | 0.236 | 6.32E-162 | 4 | Pdgfa | myCAF_2 |
| Mxi11 | 2.67E-165 | 1.51601131 | 0.622 | 0.497 | 5.16E-161 | 4 | Mxi1 | myCAF_2 |
| Rhbdf22 | 4.50E-161 | 1.70499123 | 0.526 | 0.304 | 8.70E-157 | 4 | Rhbdf2 | myCAF_2 |
| Col5a31 | 2.42E-158 | 0.85314764 | 0.989 | 0.947 | 4.67E-154 | 4 | Col5a3 | myCAF_2 |
| Aplp11 | 6.83E-157 | 1.93423738 | 0.436 | 0.211 | 1.32E-152 | 4 | Aplp1 | myCAF_2 |
| Col4a1 | 5.89E-151 | 0.29416345 | 0.836 | 0.902 | 1.14E-146 | 4 | Col4a1 | myCAF_2 |
| Tagln21 | 2.08E-147 | 0.27743185 | 0.972 | 0.967 | 4.03E-143 | 4 | Tagln2 | myCAF_2 |
| Mif2 | 2.17E-147 | 1.06933187 | 0.831 | 0.791 | 4.20E-143 | 4 | Mif | myCAF_2 |
| Hmox12 | 4.14E-147 | 1.13273558 | 0.749 | 0.701 | 8.00E-143 | 4 | Hmox1 | myCAF_2 |
| Pgk11 | 6.09E-147 | 0.96422425 | 0.788 | 0.794 | 1.18E-142 | 4 | Pgk1 | myCAF_2 |
| Kremen12 | 1.66E-146 | 0.85584306 | 0.947 | 0.898 | 3.20E-142 | 4 | Kremen1 | myCAF_2 |
| Mgarp2 | 2.86E-146 | 1.57839861 | 0.414 | 0.268 | 5.54E-142 | 4 | Mgarp | myCAF_2 |
| Sep-42 | 1.46E-145 | 1.47984869 | 0.5 | 0.359 | 2.82E-141 | 4 | 4-Sep | myCAF_2 |
| Gbe11 | 1.59E-145 | 1.79518876 | 0.387 | 0.258 | 3.08E-141 | 4 | Gbe1 | myCAF_2 |
| Fos2 | 2.31E-143 | 0.86564095 | 0.799 | 0.738 | 4.47E-139 | 4 | Fos | myCAF_2 |
| Ddit32 | 4.18E-143 | 1.56197664 | 0.476 | 0.392 | 8.08E-139 | 4 | Ddit3 | myCAF_2 |
| Ppp1r3b1 | 2.74E-139 | 1.77808279 | 0.403 | 0.24 | 5.30E-135 | 4 | Ppp1r3b | myCAF_2 |
| Jag11 | 3.02E-139 | 1.2357014 | 0.63 | 0.517 | 5.84E-135 | 4 | Jag1 | myCAF_2 |
| Mmp131 | 5.03E-138 | 0.41971645 | 0.601 | 0.416 | 9.72E-134 | 4 | Mmp13 | myCAF_2 |
| Fosb1 | 1.05E-137 | 0.98076202 | 0.743 | 0.652 | 2.02E-133 | 4 | Fosb | myCAF_2 |
| Pkm1 | 8.08E-137 | 0.86827493 | 0.972 | 0.953 | 1.56E-132 | 4 | Pkm | myCAF_2 |
| Junb2 | 6.00E-134 | 0.89161438 | 0.956 | 0.93 | 1.16E-129 | 4 | Junb | myCAF_2 |
| Tbx21 | 1.50E-133 | 1.83455 | 0.351 | 0.177 | 2.89E-129 | 4 | Tbx2 | myCAF_2 |
| Maff | 4.72E-131 | 1.5344028 | 0.635 | 0.502 | 9.13E-127 | 4 | Maff | myCAF_2 |
| Zfp3951 | 1.85E-130 | 1.54086027 | 0.484 | 0.337 | 3.57E-126 | 4 | Zfp395 | myCAF_2 |
| Itga52 | 7.49E-126 | 0.89295524 | 0.915 | 0.891 | 1.45E-121 | 4 | Itga5 | myCAF_2 |
| Hes11 | 8.13E-124 | 1.06233608 | 0.714 | 0.7 | 1.57E-119 | 4 | Hes1 | myCAF_2 |
| Jund1 | 1.04E-123 | 0.90755182 | 0.974 | 0.962 | 2.01E-119 | 4 | Jund | myCAF_2 |
| Cygb | 1.98E-122 | 0.94211111 | 0.788 | 0.839 | 3.83E-118 | 4 | Cygb | myCAF_2 |
| Fam162a2 | 4.15E-122 | 0.98938955 | 0.791 | 0.781 | 8.02E-118 | 4 | Fam162a | myCAF_2 |
| Tcim1 | 1.52E-116 | 1.29729073 | 0.611 | 0.531 | 2.93E-112 | 4 | Tcim | myCAF_2 |
| Mmd2 | 1.56E-116 | 1.16651262 | 0.725 | 0.641 | 3.02E-112 | 4 | Mmd | myCAF_2 |
| Tpm11 | 3.15E-116 | 0.25979643 | 0.949 | 0.948 | 6.09E-112 | 4 | Tpm1 | myCAF_2 |
| Higd1a | 4.30E-116 | 1.07908039 | 0.787 | 0.788 | 8.32E-112 | 4 | Higd1a | myCAF_2 |
| Cemip21 | 4.75E-115 | 1.21134543 | 0.746 | 0.632 | 9.19E-111 | 4 | Cemip2 | myCAF_2 |
| Bsg1 | 9.48E-113 | 0.89863055 | 0.882 | 0.894 | 1.83E-108 | 4 | Bsg | myCAF_2 |
| Mmp141 | 2.29E-112 | 0.31274692 | 0.986 | 0.985 | 4.43E-108 | 4 | Mmp14 | myCAF_2 |
| Adra2a2 | 1.04E-109 | 1.17623424 | 0.592 | 0.406 | 2.02E-105 | 4 | Adra2a | myCAF_2 |
| Ttyh2 | 1.19E-109 | 1.03104541 | 0.859 | 0.823 | 2.30E-105 | 4 | Ttyh2 | myCAF_2 |
| Tpt11 | 4.85E-109 | 0.3235407 | 0.991 | 0.981 | 9.38E-105 | 4 | Tpt1 | myCAF_2 |
| Rack11 | 1.96E-108 | 0.25739445 | 0.98 | 0.974 | 3.79E-104 | 4 | Rack1 | myCAF_2 |
| Dusp61 | 2.23E-106 | 1.16481703 | 0.696 | 0.569 | 4.31E-102 | 4 | Dusp6 | myCAF_2 |
| Enpp1 | 1.20E-105 | 1.50750223 | 0.528 | 0.356 | 2.31E-101 | 4 | Enpp1 | myCAF_2 |
| Gadd45a | 7.36E-104 | 1.33453563 | 0.55 | 0.401 | 1.42E-99 | 4 | Gadd45a | myCAF_2 |
| Ninj11 | 1.47E-102 | 1.1732269 | 0.721 | 0.61 | 2.84E-98 | 4 | Ninj1 | myCAF_2 |
| Cdc42bpa1 | 1.99E-102 | 0.96428928 | 0.799 | 0.771 | 3.85E-98 | 4 | Cdc42bpa | myCAF_2 |
| Cryab | 1.09E-98 | 1.27632702 | 0.249 | 0.353 | 2.11E-94 | 4 | Cryab | myCAF_2 |
| Col6a12 | 9.28E-98 | 0.32220614 | 0.991 | 0.99 | 1.79E-93 | 4 | Col6a1 | myCAF_2 |
| Ier31 | 7.34E-97 | 0.81114928 | 0.825 | 0.796 | 1.42E-92 | 4 | Ier3 | myCAF_2 |
| Pdgfrb2 | 3.08E-96 | 0.75629969 | 0.961 | 0.942 | 5.96E-92 | 4 | Pdgfrb | myCAF_2 |
| Gpr351 | 1.44E-94 | 1.41130057 | 0.375 | 0.222 | 2.79E-90 | 4 | Gpr35 | myCAF_2 |
| Notch31 | 4.82E-93 | 1.76741039 | 0.344 | 0.176 | 9.32E-89 | 4 | Notch3 | myCAF_2 |
| Jarid21 | 7.52E-93 | 1.38410169 | 0.603 | 0.428 | 1.45E-88 | 4 | Jarid2 | myCAF_2 |
| Srgap31 | 9.05E-93 | 1.93336806 | 0.307 | 0.162 | 1.75E-88 | 4 | Srgap3 | myCAF_2 |
| Robo12 | 3.80E-90 | 1.17232481 | 0.634 | 0.462 | 7.36E-86 | 4 | Robo1 | myCAF_2 |
| Il2rg1 | 2.81E-89 | 1.57437393 | 0.368 | 0.195 | 5.43E-85 | 4 | Il2rg | myCAF_2 |
| Ankrd441 | 1.00E-88 | 1.44435927 | 0.348 | 0.231 | 1.93E-84 | 4 | Ankrd44 | myCAF_2 |
| Tnfrsf191 | 1.19E-88 | 1.05338907 | 0.378 | 0.311 | 2.31E-84 | 4 | Tnfrsf19 | myCAF_2 |
| Fbxo321 | 1.46E-88 | 1.45797868 | 0.364 | 0.246 | 2.83E-84 | 4 | Fbxo32 | myCAF_2 |
| Loxl21 | 4.55E-88 | 0.27428933 | 0.961 | 0.964 | 8.79E-84 | 4 | Loxl2 | myCAF_2 |
| Sh2b31 | 1.75E-87 | 1.26708706 | 0.706 | 0.549 | 3.38E-83 | 4 | Sh2b3 | myCAF_2 |
| Afap1 | 1.16E-86 | 1.04357439 | 0.823 | 0.782 | 2.24E-82 | 4 | Afap1 | myCAF_2 |
| Fbln71 | 7.40E-86 | 1.19839442 | 0.568 | 0.361 | 1.43E-81 | 4 | Fbln7 | myCAF_2 |
| Ak41 | 9.73E-86 | 1.58257292 | 0.368 | 0.231 | 1.88E-81 | 4 | Ak4 | myCAF_2 |
| Jak31 | 1.10E-84 | 1.48814948 | 0.549 | 0.348 | 2.12E-80 | 4 | Jak3 | myCAF_2 |
| Aldoa1 | 3.38E-84 | 0.82901464 | 0.922 | 0.921 | 6.54E-80 | 4 | Aldoa | myCAF_2 |
| Dcbld21 | 2.28E-83 | 1.09077096 | 0.754 | 0.709 | 4.41E-79 | 4 | Dcbld2 | myCAF_2 |
| Itga12 | 4.33E-83 | 0.93685608 | 0.688 | 0.623 | 8.38E-79 | 4 | Itga1 | myCAF_2 |
| Parvb1 | 4.42E-83 | 1.82020112 | 0.309 | 0.136 | 8.54E-79 | 4 | Parvb | myCAF_2 |
| App1 | 2.35E-82 | 0.27125077 | 0.988 | 0.986 | 4.55E-78 | 4 | App | myCAF_2 |
| Peak11 | 2.57E-82 | 1.05196906 | 0.767 | 0.686 | 4.97E-78 | 4 | Peak1 | myCAF_2 |
| Ier22 | 1.51E-81 | 0.95276456 | 0.844 | 0.792 | 2.92E-77 | 4 | Ier2 | myCAF_2 |
| Mxx11 | 3.46E-80 | 1.37063651 | 0.521 | 0.362 | 6.69E-76 | 4 | Mxx1 | myCAF_2 |
| Ybx32 | 7.49E-80 | 0.75157191 | 0.94 | 0.935 | 1.45E-75 | 4 | Ybx3 | myCAF_2 |
| Spock31 | 8.86E-80 | 1.24996019 | 0.295 | 0.165 | 1.71E-75 | 4 | Spock3 | myCAF_2 |
| Nectin1 | 1.36E-79 | 1.09018545 | 0.737 | 0.682 | 2.62E-75 | 4 | Nectin1 | myCAF_2 |
| Ccng21 | 2.27E-79 | 1.04541988 | 0.698 | 0.61 | 4.39E-75 | 4 | Ccng2 | myCAF_2 |
| Igsf31 | 3.87E-79 | 1.25981107 | 0.647 | 0.481 | 7.47E-75 | 4 | Igsf3 | myCAF_2 |
| Smad71 | 4.93E-78 | 1.15064213 | 0.68 | 0.622 | 9.52E-74 | 4 | Smad7 | myCAF_2 |
| Enho | 6.35E-77 | 1.90680282 | 0.277 | 0.128 | 1.23E-72 | 4 | Enho | myCAF_2 |
| Map2k11 | 1.19E-76 | 0.96055642 | 0.771 | 0.771 | 2.31E-72 | 4 | Map2k1 | myCAF_2 |
| Ermp11 | 2.05E-71 | 0.85215994 | 0.737 | 0.721 | 3.97E-67 | 4 | Ermp1 | myCAF_2 |
| Sema7a2 | 2.64E-71 | 0.74225138 | 0.934 | 0.78 | 5.10E-67 | 4 | Sema7a | myCAF_2 |
| Prelid2 | 3.76E-71 | 1.82147234 | 0.294 | 0.142 | 7.27E-67 | 4 | Prelid2 | myCAF_2 |
| Mast41 | 1.79E-70 | 1.08496445 | 0.797 | 0.67 | 3.46E-66 | 4 | Mast4 | myCAF_2 |
| Uck21 | 3.94E-70 | 0.31503878 | 0.963 | 0.868 | 7.62E-66 | 4 | Uck2 | myCAF_2 |
| Lama21 | 4.08E-70 | 0.81119041 | 0.618 | 0.676 | 7.89E-66 | 4 | Lama2 | myCAF_2 |
| Tfpi22 | 4.25E-70 | 0.98131245 | 0.81 | 0.718 | 8.21E-66 | 4 | Tfpi2 | myCAF_2 |
| Pvr1 | 1.17E-69 | 1.17408712 | 0.574 | 0.523 | 2.26E-65 | 4 | Pvr | myCAF_2 |
| Rbp11 | 2.17E-69 | 0.75752261 | 0.895 | 0.813 | 4.20E-65 | 4 | Rbp1 | myCAF_2 |
| Plekha61 | 2.57E-69 | 1.49686944 | 0.27 | 0.139 | 4.96E-65 | 4 | Plekha6 | myCAF_2 |
| Stard131 | 2.19E-68 | 1.34823156 | 0.583 | 0.423 | 4.24E-64 | 4 | Stard13 | myCAF_2 |
| Pltp1 | 9.26E-68 | 1.07107426 | 0.35 | 0.311 | 1.79E-63 | 4 | Pltp | myCAF_2 |
| Nrip1 | 1.49E-67 | 1.12171246 | 0.656 | 0.583 | 2.89E-63 | 4 | Nrip1 | myCAF_2 |
| Cyb5r1 | 4.05E-67 | 1.19887054 | 0.458 | 0.485 | 7.82E-63 | 4 | Cyb5r1 | myCAF_2 |
| Ubash3b1 | 1.19E-66 | 1.50028898 | 0.44 | 0.266 | 2.30E-62 | 4 | Ubash3b | myCAF_2 |
| Vim | 1.20E-66 | 0.52341391 | 0.999 | 0.998 | 2.33E-62 | 4 | Vim | myCAF_2 |
| Apln2 | 1.93E-65 | 1.19686225 | 0.458 | 0.288 | 3.74E-61 | 4 | Apln | myCAF_2 |
| Tspan91 | 2.16E-65 | 0.97535824 | 0.693 | 0.638 | 4.18E-61 | 4 | Tspan9 | myCAF_2 |
| Abhd22 | 2.30E-65 | 0.80174714 | 0.91 | 0.897 | 4.44E-61 | 4 | Abhd2 | myCAF_2 |
| Polg22 | 3.13E-65 | 1.24686937 | 0.402 | 0.315 | 6.04E-61 | 4 | Polg2 | myCAF_2 |
| Gramd41 | 3.47E-65 | 1.17430798 | 0.63 | 0.5 | 6.70E-61 | 4 | Gramd4 | myCAF_2 |
| Foxs11 | 6.51E-65 | 1.67907746 | 0.368 | 0.176 | 1.26E-60 | 4 | Foxs1 | myCAF_2 |
| DEPRECATED-I | 1.39E-63 | 1.45946329 | 0.383 | 0.292 | 2.69E-59 | 4 | DEPRECATED-I | myCAF_2 |
| P4hb | 2.47E-63 | 0.33426444 | 0.988 | 0.98 | 4.77E-59 | 4 | P4hb | myCAF_2 |
| Dad1 | 5.06E-63 | 0.26811868 | 0.939 | 0.951 | 9.79E-59 | 4 | Dad1 | myCAF_2 |
| Slc7a51 | 5.75E-63 | 1.25120255 | 0.613 | 0.525 | 1.11E-58 | 4 | Slc7a5 | myCAF_2 |
| Il11 | 1.33E-62 | 1.07478674 | 0.267 | 0.159 | 2.58E-58 | 4 | Il11 | myCAF_2 |
| Pgam1 | 1.51E-61 | 0.90902239 | 0.887 | 0.856 | 2.91E-57 | 4 | Pgam1 | myCAF_2 |
| Grhpr1 | 3.90E-61 | 1.22265427 | 0.534 | 0.421 | 7.53E-57 | 4 | Grhpr | myCAF_2 |
| Bcar11 | 9.49E-61 | 1.01316267 | 0.745 | 0.708 | 1.83E-56 | 4 | Bcar1 | myCAF_2 |
| Pmaip11 | 1.28E-59 | 1.41346699 | 0.389 | 0.259 | 2.47E-55 | 4 | Pmaip1 | myCAF_2 |
| Pgm12 | 1.40E-59 | 0.95587364 | 0.679 | 0.66 | 2.71E-55 | 4 | Pgm1 | myCAF_2 |
| Creb3l2 | 8.54E-59 | 0.86893759 | 0.918 | 0.895 | 1.65E-54 | 4 | Creb3l2 | myCAF_2 |
| Plekha21 | 3.85E-56 | 1.10750493 | 0.601 | 0.516 | 7.44E-52 | 4 | Plekha2 | myCAF_2 |
| Mthfd1l1 | 5.87E-56 | 1.39942827 | 0.42 | 0.321 | 1.13E-51 | 4 | Mthfd1l | myCAF_2 |
| Plp21 | 1.21E-55 | 0.28609391 | 0.916 | 0.921 | 2.34E-51 | 4 | Plp2 | myCAF_2 |
| Ccn21 | 2.86E-55 | 0.68541391 | 0.458 | 0.454 | 5.53E-51 | 4 | Ccn2 | myCAF_2 |
| Hsp90b1 | 3.00E-55 | 0.27301272 | 0.92 | 0.952 | 5.79E-51 | 4 | Hsp90b1 | myCAF_2 |
| Fap | 4.84E-55 | 1.11018249 | 0.625 | 0.563 | 9.36E-51 | 4 | Fap | myCAF_2 |
| Rflna1 | 1.38E-54 | 0.97215566 | 0.521 | 0.432 | 2.67E-50 | 4 | Rflna | myCAF_2 |
| Iqgap1 | 1.54E-54 | 0.8380462 | 0.947 | 0.938 | 2.97E-50 | 4 | Iqgap1 | myCAF_2 |
| Gng22 | 4.28E-54 | 1.30871067 | 0.469 | 0.303 | 8.27E-50 | 4 | Gng2 | myCAF_2 |
| H1f01 | 5.62E-54 | 0.31226926 | 0.864 | 0.889 | 1.09E-49 | 4 | H1f0 | myCAF_2 |
| Arrb11 | 1.67E-53 | 1.17708345 | 0.404 | 0.329 | 3.22E-49 | 4 | Arrb1 | myCAF_2 |
| Vdac11 | 1.68E-53 | 1.00543109 | 0.679 | 0.686 | 3.26E-49 | 4 | Vdac1 | myCAF_2 |
| Tcp11l22 | 3.33E-52 | 0.94744993 | 0.679 | 0.558 | 6.45E-48 | 4 | Tcp11l2 | myCAF_2 |
| Cd821 | 8.84E-52 | 1.04970606 | 0.475 | 0.395 | 1.71E-47 | 4 | Cd82 | myCAF_2 |
| Etv41 | 1.02E-51 | 1.12758162 | 0.613 | 0.478 | 1.98E-47 | 4 | Etv4 | myCAF_2 |
| Ctxn1 | 4.03E-51 | 1.34359249 | 0.398 | 0.241 | 7.79E-47 | 4 | Ctxn1 | myCAF_2 |
| Loxl32 | 1.78E-50 | 0.35502803 | 0.886 | 0.92 | 3.44E-46 | 4 | Loxl3 | myCAF_2 |
| Pla2g4a | 2.66E-50 | 1.13321073 | 0.632 | 0.553 | 5.14E-46 | 4 | Pla2g4a | myCAF_2 |
| Csrp11 | 2.69E-50 | 0.25392524 | 0.879 | 0.869 | 5.20E-46 | 4 | Csrp1 | myCAF_2 |
| Sap30 | 3.24E-50 | 1.62858828 | 0.254 | 0.163 | 6.27E-46 | 4 | Sap30 | myCAF_2 |
| Ldha1 | 4.07E-50 | 0.7069495 | 0.955 | 0.954 | 7.86E-46 | 4 | Ldha | myCAF_2 |
| Tes1 | 1.07E-49 | 1.06367341 | 0.631 | 0.551 | 2.07E-45 | 4 | Tes | myCAF_2 |
| Fosl11 | 1.92E-48 | 1.02542921 | 0.415 | 0.376 | 3.72E-44 | 4 | Fosl1 | myCAF_2 |
| Tnfrsf12a | 2.35E-48 | 0.31518969 | 0.761 | 0.824 | 4.55E-44 | 4 | Tnfrsf12a | myCAF_2 |
| Zfp5322 | 1.31E-47 | 0.94533398 | 0.8 | 0.723 | 2.54E-43 | 4 | Zfp532 | myCAF_2 |
| Cdc42ep31 | 4.21E-47 | 1.16374048 | 0.314 | 0.223 | 8.14E-43 | 4 | Cdc42ep3 | myCAF_2 |
| Prkab2 | 5.79E-47 | 1.6349935 | 0.303 | 0.186 | 1.12E-42 | 4 | Prkab2 | myCAF_2 |
| Flna1 | 1.14E-46 | 0.30252671 | 0.939 | 0.949 | 2.20E-42 | 4 | Flna | myCAF_2 |
| Itgb11 | 1.23E-46 | 0.31783711 | 0.981 | 0.975 | 2.38E-42 | 4 | Itgb1 | myCAF_2 |
| Ybx11 | 1.76E-46 | 0.32036678 | 0.931 | 0.944 | 3.41E-42 | 4 | Ybx1 | myCAF_2 |
| Ppp1r15a2 | 2.18E-46 | 1.03553488 | 0.697 | 0.635 | 4.22E-42 | 4 | Ppp1r15a | myCAF_2 |
| Klf51 | 9.15E-45 | 1.26013451 | 0.355 | 0.254 | 1.77E-40 | 4 | Klf5 | myCAF_2 |
| Notch12 | 1.08E-44 | 1.06125917 | 0.672 | 0.506 | 2.09E-40 | 4 | Notch1 | myCAF_2 |
| Il342 | 5.25E-44 | 1.17176788 | 0.356 | 0.25 | 1.02E-39 | 4 | Il34 | myCAF_2 |
| Sqstm11 | 6.63E-44 | 0.88296881 | 0.898 | 0.875 | 1.28E-39 | 4 | Sqstm1 | myCAF_2 |
| Myo1c | 3.24E-43 | 0.27442285 | 0.935 | 0.945 | 6.27E-39 | 4 | Myo1c | myCAF_2 |
| Itga71 | 1.10E-42 | 1.47143676 | 0.254 | 0.146 | 2.13E-38 | 4 | Itga7 | myCAF_2 |
| Usp131 | 2.63E-42 | 1.29768178 | 0.366 | 0.28 | 5.08E-38 | 4 | Usp13 | myCAF_2 |
| Dipk2a | 3.78E-42 | 1.12461341 | 0.425 | 0.382 | 7.32E-38 | 4 | Dipk2a | myCAF_2 |
| Ube2o1 | 3.98E-42 | 1.1872462 | 0.583 | 0.499 | 7.70E-38 | 4 | Ube2o | myCAF_2 |
| Dock6 | 7.88E-42 | 1.12264154 | 0.402 | 0.303 | 1.52E-37 | 4 | Dock6 | myCAF_2 |
| Igfbp71 | 3.10E-41 | 0.38288486 | 0.992 | 0.993 | 5.99E-37 | 4 | Igfbp7 | myCAF_2 |
| Fgd62 | 5.69E-41 | 1.48741341 | 0.388 | 0.237 | 1.10E-36 | 4 | Fgd6 | myCAF_2 |
| Dhrs31 | 8.32E-41 | 0.81570751 | 0.748 | 0.723 | 1.61E-36 | 4 | Dhrs3 | myCAF_2 |
| Tmem170b | 8.38E-41 | 1.40022624 | 0.327 | 0.211 | 1.62E-36 | 4 | Tmem170b | myCAF_2 |
| Homer12 | 4.34E-40 | 1.12549595 | 0.57 | 0.498 | 8.39E-36 | 4 | Homer1 | myCAF_2 |
| Fgf111 | 8.61E-40 | 1.40390217 | 0.296 | 0.183 | 1.66E-35 | 4 | Fgf11 | myCAF_2 |
| Skil | 1.37E-39 | 0.90163365 | 0.801 | 0.744 | 2.66E-35 | 4 | Skil | myCAF_2 |
| Itpripl2 | 1.56E-39 | 0.29560385 | 0.892 | 0.916 | 3.01E-35 | 4 | Itpripl2 | myCAF_2 |
| Myc2 | 3.10E-39 | 0.26465865 | 0.532 | 0.564 | 6.00E-35 | 4 | Myc | myCAF_2 |
| Nfkbiz1 | 9.89E-39 | 0.28001861 | 0.666 | 0.667 | 1.91E-34 | 4 | Nfkbiz | myCAF_2 |
| Zcchc241 | 6.40E-38 | 0.28667955 | 0.888 | 0.942 | 1.24E-33 | 4 | Zcchc24 | myCAF_2 |
| Fam214a1 | 1.03E-37 | 1.13134819 | 0.479 | 0.383 | 2.00E-33 | 4 | Fam214a | myCAF_2 |
| Slco2a11 | 1.38E-37 | 0.88209915 | 0.476 | 0.383 | 2.66E-33 | 4 | Slco2a1 | myCAF_2 |
| Adprh1 | 1.43E-37 | 0.25077487 | 0.921 | 0.945 | 2.76E-33 | 4 | Adprh | myCAF_2 |
| Pdlim1 | 8.12E-35 | 1.02510664 | 0.616 | 0.528 | 1.57E-30 | 4 | Pdlim1 | myCAF_2 |
| Dlc1 | 1.98E-34 | 0.88561493 | 0.716 | 0.654 | 3.82E-30 | 4 | Dlc1 | myCAF_2 |
| Thy11 | 2.49E-34 | 0.37996307 | 0.974 | 0.983 | 4.82E-30 | 4 | Thy1 | myCAF_2 |
| Ankrd50 | 2.73E-34 | 1.06318307 | 0.649 | 0.546 | 5.28E-30 | 4 | Ankrd50 | myCAF_2 |
| Arhgef21 | 7.00E-34 | 0.82278878 | 0.824 | 0.786 | 1.35E-29 | 4 | Arhgef2 | myCAF_2 |
| Pgp1 | 1.14E-33 | 0.98623845 | 0.563 | 0.574 | 2.21E-29 | 4 | Pgp | myCAF_2 |
| Htr2b1 | 1.31E-33 | 1.30899111 | 0.274 | 0.143 | 2.53E-29 | 4 | Htr2b | myCAF_2 |
| Soat11 | 2.12E-33 | 0.74539754 | 0.88 | 0.84 | 4.10E-29 | 4 | Soat1 | myCAF_2 |
| Tmem45a1 | 2.63E-33 | 0.75451966 | 0.725 | 0.756 | 5.08E-29 | 4 | Tmem45a | myCAF_2 |
| Itpk1 | 2.81E-33 | 1.29352608 | 0.27 | 0.176 | 5.43E-29 | 4 | Itpk1 | myCAF_2 |
| Smoc2 | 4.65E-33 | 0.7886725 | 0.471 | 0.481 | 9.00E-29 | 4 | Smoc2 | myCAF_2 |
| Bcl2l111 | 5.61E-33 | 1.38386471 | 0.434 | 0.33 | 1.08E-28 | 4 | Bcl2l11 | myCAF_2 |
| Fus1 | 6.33E-33 | 0.27212478 | 0.872 | 0.897 | 1.22E-28 | 4 | Fus | myCAF_2 |
| Zmiz11 | 1.30E-32 | 0.7516533 | 0.869 | 0.844 | 2.51E-28 | 4 | Zmiz1 | myCAF_2 |
| Nupr1 | 1.38E-32 | 0.73290787 | 0.795 | 0.814 | 2.67E-28 | 4 | Nupr1 | myCAF_2 |
| Sat11 | 2.28E-32 | 0.86869467 | 0.845 | 0.835 | 4.42E-28 | 4 | Sat1 | myCAF_2 |
| Nr2f11 | 2.46E-32 | 1.32556256 | 0.336 | 0.263 | 4.77E-28 | 4 | Nr2f1 | myCAF_2 |
| Adcyap1r1 | 2.83E-32 | 1.21321943 | 0.258 | 0.172 | 5.47E-28 | 4 | Adcyap1r1 | myCAF_2 |
| Tiparp1 | 2.87E-32 | 0.8775124 | 0.574 | 0.5 | 5.54E-28 | 4 | Tiparp | myCAF_2 |
| Ogt1 | 4.30E-32 | 0.91533457 | 0.732 | 0.698 | 8.32E-28 | 4 | Ogt | myCAF_2 |
| Plk31 | 1.28E-31 | 1.08887517 | 0.513 | 0.403 | 2.48E-27 | 4 | Plk3 | myCAF_2 |
| Khsrp2 | 1.88E-31 | 0.28169897 | 0.826 | 0.867 | 3.64E-27 | 4 | Khsrp | myCAF_2 |
| Son | 4.93E-31 | 0.31795399 | 0.919 | 0.926 | 9.53E-27 | 4 | Son | myCAF_2 |
| Mat2a1 | 1.05E-30 | 0.36482456 | 0.763 | 0.824 | 2.04E-26 | 4 | Mat2a | myCAF_2 |
| Gnb1 | 1.55E-30 | 0.33164393 | 0.92 | 0.945 | 3.00E-26 | 4 | Gnb1 | myCAF_2 |
| Usp531 | 3.38E-30 | 1.30880225 | 0.26 | 0.199 | 6.54E-26 | 4 | Usp53 | myCAF_2 |
| Sfpq1 | 7.54E-30 | 0.29519875 | 0.846 | 0.859 | 1.46E-25 | 4 | Sfpq | myCAF_2 |
| Mical21 | 8.77E-30 | 0.77742663 | 0.898 | 0.836 | 1.70E-25 | 4 | Mical2 | myCAF_2 |
| Capg | 9.24E-30 | 0.72414792 | 0.917 | 0.895 | 1.79E-25 | 4 | Capg | myCAF_2 |
| Lpp1 | 1.19E-29 | 0.27129495 | 0.908 | 0.929 | 2.30E-25 | 4 | Lpp | myCAF_2 |
| Pygl1 | 1.86E-29 | 0.79010921 | 0.453 | 0.431 | 3.60E-25 | 4 | Pygl | myCAF_2 |
| Adgre52 | 7.74E-29 | 0.88438923 | 0.719 | 0.601 | 1.50E-24 | 4 | Adgre5 | myCAF_2 |
| Adipor1 | 8.01E-29 | 0.28954652 | 0.904 | 0.936 | 1.55E-24 | 4 | Adipor1 | myCAF_2 |
| Vmp11 | 3.31E-28 | 0.36177829 | 0.939 | 0.948 | 6.39E-24 | 4 | Vmp1 | myCAF_2 |
| Myl12a1 | 3.82E-28 | 0.35849129 | 0.942 | 0.953 | 7.39E-24 | 4 | Myl12a | myCAF_2 |
| Mapk8ip3 | 4.84E-28 | 1.00902807 | 0.593 | 0.489 | 9.35E-24 | 4 | Mapk8ip3 | myCAF_2 |
| Agrn1 | 5.36E-28 | 0.94270315 | 0.66 | 0.594 | 1.04E-23 | 4 | Agrn | myCAF_2 |
| Kdm3a1 | 7.62E-28 | 1.00290075 | 0.583 | 0.526 | 1.47E-23 | 4 | Kdm3a | myCAF_2 |
| Tcf42 | 7.98E-28 | 0.68924986 | 0.936 | 0.92 | 1.54E-23 | 4 | Tcf4 | myCAF_2 |
| Rin12 | 1.21E-27 | 1.05957131 | 0.522 | 0.407 | 2.33E-23 | 4 | Rin1 | myCAF_2 |
| Kdm4b1 | 1.57E-27 | 1.01595225 | 0.521 | 0.427 | 3.03E-23 | 4 | Kdm4b | myCAF_2 |
| Lonp1 | 2.17E-27 | 0.90974188 | 0.665 | 0.672 | 4.19E-23 | 4 | Lonp1 | myCAF_2 |
| Rora1 | 2.88E-27 | 0.78220586 | 0.663 | 0.637 | 5.56E-23 | 4 | Rora | myCAF_2 |
| Fzd12 | 3.45E-27 | 0.27155156 | 0.868 | 0.869 | 6.66E-23 | 4 | Fzd1 | myCAF_2 |
| BC005537 | 3.86E-27 | 0.25352507 | 0.874 | 0.922 | 7.45E-23 | 4 | BC005537 | myCAF_2 |
| Jmjd61 | 4.03E-27 | 0.96347672 | 0.642 | 0.592 | 7.79E-23 | 4 | Jmjd6 | myCAF_2 |
| Sertad12 | 4.17E-27 | 0.89328268 | 0.675 | 0.643 | 8.06E-23 | 4 | Sertad1 | myCAF_2 |
| Tjp11 | 6.01E-27 | 0.91495159 | 0.667 | 0.614 | 1.16E-22 | 4 | Tjp1 | myCAF_2 |
| Smg6 | 6.41E-27 | 0.76200851 | 0.737 | 0.726 | 1.24E-22 | 4 | Smg6 | myCAF_2 |
| Ptbp31 | 1.05E-26 | 0.25881332 | 0.839 | 0.876 | 2.02E-22 | 4 | Ptbp3 | myCAF_2 |
| Khnyn1 | 1.35E-26 | 1.09614769 | 0.484 | 0.41 | 2.61E-22 | 4 | Khnyn | myCAF_2 |
| Mfge81 | 1.53E-26 | 0.62524477 | 0.959 | 0.945 | 2.96E-22 | 4 | Mfge8 | myCAF_2 |
| Myof | 1.69E-26 | 0.2741877 | 0.889 | 0.926 | 3.27E-22 | 4 | Myof | myCAF_2 |
| Igf2bp21 | 4.19E-26 | 1.03769735 | 0.446 | 0.397 | 8.10E-22 | 4 | Igf2bp2 | myCAF_2 |
| Adm1 | 4.56E-26 | 0.66462194 | 0.504 | 0.451 | 8.81E-22 | 4 | Adm | myCAF_2 |
| Sh3rf1 | 1.12E-25 | 1.15630167 | 0.541 | 0.436 | 2.17E-21 | 4 | Sh3rf1 | myCAF_2 |
| Nr4a12 | 1.21E-25 | 0.64914955 | 0.65 | 0.631 | 2.33E-21 | 4 | Nr4a1 | myCAF_2 |
| Mapk8ip11 | 2.87E-25 | 1.13724198 | 0.369 | 0.272 | 5.56E-21 | 4 | Mapk8ip1 | myCAF_2 |
| Cacna1c2 | 3.06E-25 | 0.7508735 | 0.792 | 0.718 | 5.92E-21 | 4 | Cacna1c | myCAF_2 |
| Adam191 | 3.44E-25 | 0.65012138 | 0.826 | 0.849 | 6.65E-21 | 4 | Adam19 | myCAF_2 |
| Wwtr11 | 4.69E-25 | 0.25405017 | 0.857 | 0.892 | 9.08E-21 | 4 | Wwtr1 | myCAF_2 |
| Vhl | 5.04E-25 | 0.9206076 | 0.548 | 0.512 | 9.75E-21 | 4 | Vhl | myCAF_2 |
| Nptn1 | 6.17E-25 | 0.26966686 | 0.89 | 0.916 | 1.19E-20 | 4 | Nptn | myCAF_2 |
| Ptrh1 | 1.11E-24 | 1.0152835 | 0.554 | 0.526 | 2.14E-20 | 4 | Ptrh1 | myCAF_2 |
| Zfand3 | 1.29E-24 | 0.2513886 | 0.893 | 0.917 | 2.50E-20 | 4 | Zfand3 | myCAF_2 |
| Spry42 | 1.42E-24 | 0.96611583 | 0.508 | 0.423 | 2.75E-20 | 4 | Spry4 | myCAF_2 |
| Ppme1 | 1.60E-24 | 0.86491811 | 0.682 | 0.667 | 3.10E-20 | 4 | Ppme1 | myCAF_2 |
| Apbb22 | 2.54E-24 | 0.79949143 | 0.774 | 0.715 | 4.91E-20 | 4 | Apbb2 | myCAF_2 |
| Tpd52l2 | 2.63E-24 | 0.27084169 | 0.826 | 0.862 | 5.09E-20 | 4 | Tpd52l2 | myCAF_2 |
| Dkk31 | 2.83E-24 | 0.25123023 | 0.829 | 0.836 | 5.46E-20 | 4 | Dkk3 | myCAF_2 |
| Trim35 | 3.05E-24 | 0.28280675 | 0.845 | 0.894 | 5.90E-20 | 4 | Trim35 | myCAF_2 |
| Ptch11 | 4.37E-24 | 1.00077973 | 0.254 | 0.22 | 8.45E-20 | 4 | Ptch1 | myCAF_2 |
| Srsf51 | 6.82E-24 | 0.26013946 | 0.887 | 0.922 | 1.32E-19 | 4 | Srsf5 | myCAF_2 |
| Ier5l1 | 7.19E-24 | 0.26288934 | 0.694 | 0.736 | 1.39E-19 | 4 | Ier5l | myCAF_2 |
| Mllt11 | 7.50E-24 | 1.20761834 | 0.338 | 0.252 | 1.45E-19 | 4 | Mllt11 | myCAF_2 |
| Vasp | 1.56E-23 | 0.25971792 | 0.845 | 0.877 | 3.01E-19 | 4 | Vasp | myCAF_2 |
| Dbndd21 | 2.45E-23 | 1.02295696 | 0.429 | 0.341 | 4.73E-19 | 4 | Dbndd2 | myCAF_2 |
| Sptan1 | 2.67E-23 | 0.25508012 | 0.866 | 0.902 | 5.17E-19 | 4 | Sptan1 | myCAF_2 |
| Btg11 | 2.77E-23 | 0.86980068 | 0.76 | 0.714 | 5.35E-19 | 4 | Btg1 | myCAF_2 |
| Dennd2a | 5.69E-23 | 0.90225417 | 0.651 | 0.625 | 1.10E-18 | 4 | Dennd2a | myCAF_2 |
| Ccdc85b1 | 7.02E-23 | 0.9849078 | 0.569 | 0.493 | 1.36E-18 | 4 | Ccdc85b | myCAF_2 |
| Sep-91 | 8.15E-23 | 0.67420114 | 0.95 | 0.925 | 1.58E-18 | 4 | 9-Sep | myCAF_2 |
| Rab6b | 1.14E-22 | 1.06094829 | 0.369 | 0.263 | 2.19E-18 | 4 | Rab6b | myCAF_2 |
| Rai141 | 1.90E-22 | 0.70049063 | 0.866 | 0.836 | 3.66E-18 | 4 | Rai14 | myCAF_2 |
| Crlf11 | 1.99E-22 | 0.69005994 | 0.452 | 0.391 | 3.84E-18 | 4 | Crlf1 | myCAF_2 |
| Rab151 | 2.09E-22 | 1.09129034 | 0.349 | 0.303 | 4.04E-18 | 4 | Rab15 | myCAF_2 |
| Atf51 | 2.69E-22 | 0.79981564 | 0.826 | 0.813 | 5.20E-18 | 4 | Atf5 | myCAF_2 |
| Mafb | 3.41E-22 | 0.83558353 | 0.708 | 0.658 | 6.59E-18 | 4 | Mafb | myCAF_2 |
| Mef2d1 | 4.69E-22 | 0.25326678 | 0.721 | 0.793 | 9.06E-18 | 4 | Mef2d | myCAF_2 |
| Apbb1ip | 6.11E-22 | 0.25386405 | 0.738 | 0.804 | 1.18E-17 | 4 | Apbb1ip | myCAF_2 |
| Hsp90ab11 | 8.26E-22 | 0.36537654 | 0.913 | 0.921 | 1.60E-17 | 4 | Hsp90ab1 | myCAF_2 |
| Chst111 | 8.69E-22 | 0.66292336 | 0.817 | 0.809 | 1.68E-17 | 4 | Chst11 | myCAF_2 |
| Tmod3 | 1.16E-21 | 0.27079144 | 0.86 | 0.892 | 2.24E-17 | 4 | Tmod3 | myCAF_2 |
| Chst22 | 1.24E-21 | 1.23736405 | 0.496 | 0.387 | 2.39E-17 | 4 | Chst2 | myCAF_2 |
| Asph1 | 1.69E-21 | 0.30883094 | 0.889 | 0.93 | 3.26E-17 | 4 | Asph | myCAF_2 |
| Ncor21 | 1.87E-21 | 0.28962256 | 0.888 | 0.894 | 3.62E-17 | 4 | Ncor2 | myCAF_2 |
| Cdc42se1 | 1.91E-21 | 0.27217394 | 0.786 | 0.82 | 3.69E-17 | 4 | Cdc42se1 | myCAF_2 |
| Spp12 | 2.38E-21 | 0.43865918 | 0.981 | 0.925 | 4.61E-17 | 4 | Spp1 | myCAF_2 |
| Marcksl11 | 2.57E-21 | 0.30616348 | 0.778 | 0.805 | 4.96E-17 | 4 | Marcksl1 | myCAF_2 |
| Akt1s1 | 3.42E-21 | 0.29205526 | 0.745 | 0.8 | 6.61E-17 | 4 | Akt1s1 | myCAF_2 |
| Gadd45b1 | 4.19E-21 | 0.74786774 | 0.756 | 0.77 | 8.10E-17 | 4 | Gadd45b | myCAF_2 |
| Qprt2 | 4.23E-21 | 1.1870669 | 0.281 | 0.212 | 8.18E-17 | 4 | Qprt | myCAF_2 |
| Selenok1 | 5.59E-21 | 0.26042429 | 0.789 | 0.838 | 1.08E-16 | 4 | Selenok | myCAF_2 |
| Pum21 | 5.94E-21 | 0.29445979 | 0.812 | 0.844 | 1.15E-16 | 4 | Pum2 | myCAF_2 |
| Itgav1 | 6.79E-21 | 0.71211682 | 0.895 | 0.868 | 1.31E-16 | 4 | Itgav | myCAF_2 |
| Fau1 | 1.06E-20 | 0.69904788 | 0.981 | 0.974 | 2.05E-16 | 4 | Fau | myCAF_2 |
| Mapkapk21 | 1.52E-20 | 0.30532028 | 0.884 | 0.911 | 2.93E-16 | 4 | Mapkapk2 | myCAF_2 |
| Ptk72 | 1.66E-20 | 0.27711476 | 0.905 | 0.81 | 3.20E-16 | 4 | Ptk7 | myCAF_2 |
| Adamts61 | 2.21E-20 | 0.90255497 | 0.419 | 0.354 | 4.27E-16 | 4 | Adamts6 | myCAF_2 |
| Zc3h151 | 2.36E-20 | 0.25300157 | 0.583 | 0.661 | 4.56E-16 | 4 | Zc3h15 | myCAF_2 |
| Arhgef401 | 3.54E-20 | 0.25972483 | 0.817 | 0.867 | 6.84E-16 | 4 | Arhgef40 | myCAF_2 |
| Gnai2 | 3.54E-20 | 0.36971638 | 0.9 | 0.916 | 6.85E-16 | 4 | Gnai2 | myCAF_2 |
| Sipa1l21 | 3.80E-20 | 0.9961755 | 0.387 | 0.317 | 7.34E-16 | 4 | Sipa1l2 | myCAF_2 |
| Lamb11 | 4.37E-20 | 0.36817543 | 0.957 | 0.949 | 8.45E-16 | 4 | Lamb1 | myCAF_2 |
| Pxdn | 7.38E-20 | 0.58817549 | 0.956 | 0.95 | 1.43E-15 | 4 | Pxdn | myCAF_2 |
| Mex3d | 7.38E-20 | 0.90622253 | 0.676 | 0.661 | 1.43E-15 | 4 | Mex3d | myCAF_2 |
| Oaf1 | 8.38E-20 | 0.37957029 | 0.936 | 0.934 | 1.62E-15 | 4 | Oaf | myCAF_2 |
| Cdr2l1 | 1.10E-19 | 0.30486521 | 0.81 | 0.855 | 2.13E-15 | 4 | Cdr2l | myCAF_2 |
| Col7a11 | 1.16E-19 | 0.41159999 | 0.754 | 0.581 | 2.25E-15 | 4 | Col7a1 | myCAF_2 |
| Dip2a1 | 1.25E-19 | 0.86370786 | 0.671 | 0.634 | 2.42E-15 | 4 | Dip2a | myCAF_2 |
| Tra2a1 | 1.37E-19 | 0.29820092 | 0.731 | 0.782 | 2.64E-15 | 4 | Tra2a | myCAF_2 |
| Dlgap4 | 2.21E-19 | 0.26306039 | 0.871 | 0.907 | 4.26E-15 | 4 | Dlgap4 | myCAF_2 |
| Slc9a1 | 3.14E-19 | 0.26964006 | 0.736 | 0.8 | 6.07E-15 | 4 | Slc9a1 | myCAF_2 |
| Akr1b81 | 3.21E-19 | 0.26820038 | 0.841 | 0.89 | 6.21E-15 | 4 | Akr1b8 | myCAF_2 |
| Cald11 | 4.25E-19 | 0.40490427 | 0.985 | 0.983 | 8.22E-15 | 4 | Cald1 | myCAF_2 |
| B3gnt31 | 7.71E-19 | 0.83499002 | 0.523 | 0.45 | 1.49E-14 | 4 | B3gnt3 | myCAF_2 |
| Sun11 | 9.56E-19 | 0.95093092 | 0.57 | 0.538 | 1.85E-14 | 4 | Sun1 | myCAF_2 |
| Edil31 | 1.54E-18 | 1.07706676 | 0.293 | 0.193 | 2.97E-14 | 4 | Edil3 | myCAF_2 |
| Mtcl1 | 1.72E-18 | 1.29523425 | 0.262 | 0.165 | 3.32E-14 | 4 | Mtcl1 | myCAF_2 |
| Atp1a31 | 1.95E-18 | 1.10527692 | 0.318 | 0.224 | 3.77E-14 | 4 | Atp1a3 | myCAF_2 |
| Timp3 | 2.53E-18 | 0.42978811 | 0.68 | 0.672 | 4.90E-14 | 4 | Timp3 | myCAF_2 |
| P2ry21 | 2.63E-18 | 0.98393028 | 0.359 | 0.291 | 5.08E-14 | 4 | P2ry2 | myCAF_2 |
| Card191 | 3.35E-18 | 0.89587886 | 0.537 | 0.475 | 6.48E-14 | 4 | Card19 | myCAF_2 |
| Ptprs | 5.14E-18 | 0.32890942 | 0.894 | 0.904 | 9.93E-14 | 4 | Ptprs | myCAF_2 |
| Itprip | 5.16E-18 | 1.12498334 | 0.442 | 0.35 | 9.97E-14 | 4 | Itprip | myCAF_2 |
| Ccnl1 | 5.26E-18 | 0.69671312 | 0.866 | 0.879 | 1.02E-13 | 4 | Ccnl1 | myCAF_2 |
| Pcgf51 | 5.45E-18 | 0.99731727 | 0.331 | 0.328 | 1.05E-13 | 4 | Pcgf5 | myCAF_2 |
| Rab5a1 | 5.75E-18 | 0.26618341 | 0.823 | 0.869 | 1.11E-13 | 4 | Rab5a | myCAF_2 |
| Dtx4 | 6.08E-18 | 0.92076655 | 0.577 | 0.523 | 1.18E-13 | 4 | Dtx4 | myCAF_2 |
| Serinc21 | 6.72E-18 | 0.80310687 | 0.484 | 0.376 | 1.30E-13 | 4 | Serinc2 | myCAF_2 |
| Dbn1 | 7.80E-18 | 0.30074436 | 0.855 | 0.867 | 1.51E-13 | 4 | Dbn1 | myCAF_2 |
| Mboat7 | 9.52E-18 | 0.25040077 | 0.762 | 0.822 | 1.84E-13 | 4 | Mboat7 | myCAF_2 |
| Ppp2cb1 | 1.14E-17 | 0.31690855 | 0.827 | 0.86 | 2.21E-13 | 4 | Ppp2cb | myCAF_2 |
| Ptpn12 | 1.17E-17 | 0.28197106 | 0.768 | 0.797 | 2.26E-13 | 4 | Ptpn12 | myCAF_2 |
| Fem1b1 | 1.19E-17 | 0.84718138 | 0.658 | 0.65 | 2.29E-13 | 4 | Fem1b | myCAF_2 |
| Adgrl32 | 1.71E-17 | 0.92941932 | 0.342 | 0.223 | 3.30E-13 | 4 | Adgrl3 | myCAF_2 |
| Ier3ip11 | 2.71E-17 | 0.27437699 | 0.734 | 0.781 | 5.23E-13 | 4 | Ier3ip1 | myCAF_2 |
| Plk21 | 3.17E-17 | 1.07670941 | 0.263 | 0.21 | 6.13E-13 | 4 | Plk2 | myCAF_2 |
| Mbnl21 | 4.21E-17 | 0.40729777 | 0.928 | 0.932 | 8.13E-13 | 4 | Mbnl2 | myCAF_2 |
| Srcap | 4.35E-17 | 0.25616389 | 0.751 | 0.797 | 8.42E-13 | 4 | Srcap | myCAF_2 |
| Plekhb2 | 5.54E-17 | 0.99168969 | 0.486 | 0.393 | 1.07E-12 | 4 | Plekhb2 | myCAF_2 |
| Pde4a1 | 7.52E-17 | 0.92098291 | 0.361 | 0.286 | 1.45E-12 | 4 | Pde4a | myCAF_2 |
| Plekho21 | 7.77E-17 | 0.27745293 | 0.81 | 0.859 | 1.50E-12 | 4 | Plekho2 | myCAF_2 |
| Tet31 | 8.75E-17 | 0.83639101 | 0.617 | 0.516 | 1.69E-12 | 4 | Tet3 | myCAF_2 |
| Rnf19a | 9.03E-17 | 0.82592843 | 0.624 | 0.601 | 1.74E-12 | 4 | Rnf19a | myCAF_2 |
| Rab3ip1 | 9.54E-17 | 0.96075876 | 0.36 | 0.303 | 1.85E-12 | 4 | Rab3ip | myCAF_2 |
| Wdr191 | 1.21E-16 | 1.05125783 | 0.281 | 0.205 | 2.34E-12 | 4 | Wdr19 | myCAF_2 |
| Smurf1 | 1.66E-16 | 0.80742007 | 0.73 | 0.683 | 3.22E-12 | 4 | Smurf1 | myCAF_2 |
| Sin3b | 1.67E-16 | 0.30283075 | 0.837 | 0.88 | 3.22E-12 | 4 | Sin3b | myCAF_2 |
| Tbc1d151 | 2.55E-16 | 0.25717354 | 0.634 | 0.7 | 4.92E-12 | 4 | Tbc1d15 | myCAF_2 |
| Csnk1d | 2.98E-16 | 0.31644592 | 0.848 | 0.877 | 5.76E-12 | 4 | Csnk1d | myCAF_2 |
| Lamb21 | 2.98E-16 | 0.29173412 | 0.793 | 0.81 | 5.77E-12 | 4 | Lamb2 | myCAF_2 |
| Acbd3 | 3.55E-16 | 0.26205884 | 0.769 | 0.833 | 6.86E-12 | 4 | Acbd3 | myCAF_2 |
| Trappc6a | 3.57E-16 | 1.00975794 | 0.477 | 0.404 | 6.90E-12 | 4 | Trappc6a | myCAF_2 |
| Pfdn51 | 3.99E-16 | 0.29259378 | 0.741 | 0.799 | 7.71E-12 | 4 | Pfdn5 | myCAF_2 |
| Ccdc59 | 4.24E-16 | 0.25587154 | 0.477 | 0.563 | 8.19E-12 | 4 | Ccdc59 | myCAF_2 |
| Cdyl22 | 4.48E-16 | 0.92555397 | 0.502 | 0.405 | 8.67E-12 | 4 | Cdyl2 | myCAF_2 |
| Ube2b1 | 4.88E-16 | 0.29457795 | 0.86 | 0.884 | 9.43E-12 | 4 | Ube2b | myCAF_2 |
| Ddx17 | 5.88E-16 | 0.35679834 | 0.954 | 0.953 | 1.14E-11 | 4 | Ddx17 | myCAF_2 |
| Desi11 | 9.19E-16 | 0.26313295 | 0.639 | 0.668 | 1.78E-11 | 4 | Desi1 | myCAF_2 |
| Inppl1 | 1.01E-15 | 0.25007784 | 0.745 | 0.801 | 1.95E-11 | 4 | Inppl1 | myCAF_2 |
| Sh2b2 | 1.12E-15 | 1.08415599 | 0.369 | 0.285 | 2.17E-11 | 4 | Sh2b2 | myCAF_2 |
| Ube2ql11 | 1.16E-15 | 1.20628639 | 0.3 | 0.218 | 2.24E-11 | 4 | Ube2ql1 | myCAF_2 |
| Rnf2171 | 1.25E-15 | 0.9615359 | 0.462 | 0.433 | 2.43E-11 | 4 | Rnf217 | myCAF_2 |
| Med101 | 1.26E-15 | 0.28850211 | 0.72 | 0.771 | 2.43E-11 | 4 | Med10 | myCAF_2 |
| Slc44a21 | 1.54E-15 | 0.26703148 | 0.782 | 0.807 | 2.98E-11 | 4 | Slc44a2 | myCAF_2 |
| Klf71 | 1.60E-15 | 0.83034667 | 0.523 | 0.424 | 3.10E-11 | 4 | Klf7 | myCAF_2 |
| Aamp | 1.67E-15 | 0.31192813 | 0.886 | 0.903 | 3.22E-11 | 4 | Aamp | myCAF_2 |
| Eaf11 | 1.86E-15 | 0.82135954 | 0.701 | 0.68 | 3.59E-11 | 4 | Eaf1 | myCAF_2 |
| Msi21 | 2.24E-15 | 0.86497763 | 0.671 | 0.587 | 4.34E-11 | 4 | Msi2 | myCAF_2 |
| Smim31 | 2.49E-15 | 0.25712719 | 0.646 | 0.663 | 4.81E-11 | 4 | Smim3 | myCAF_2 |
| Ctbp2 | 2.77E-15 | 0.25712413 | 0.715 | 0.76 | 5.36E-11 | 4 | Ctbp2 | myCAF_2 |
| Itgb51 | 2.96E-15 | 0.37577471 | 0.943 | 0.967 | 5.72E-11 | 4 | Itgb5 | myCAF_2 |
| Ist1 | 3.09E-15 | 0.27370622 | 0.77 | 0.819 | 5.98E-11 | 4 | Ist1 | myCAF_2 |
| Cab39 | 3.42E-15 | 0.27787528 | 0.759 | 0.81 | 6.61E-11 | 4 | Cab39 | myCAF_2 |
| Tmem50a1 | 3.43E-15 | 0.26478474 | 0.901 | 0.932 | 6.63E-11 | 4 | Tmem50a | myCAF_2 |
| Igf2r | 5.23E-15 | 0.25613716 | 0.76 | 0.827 | 1.01E-10 | 4 | Igf2r | myCAF_2 |
| Lpin11 | 7.98E-15 | 1.04275513 | 0.269 | 0.188 | 1.54E-10 | 4 | Lpin1 | myCAF_2 |
| Bnip3l1 | 9.52E-15 | 0.88873384 | 0.453 | 0.408 | 1.84E-10 | 4 | Bnip3l | myCAF_2 |
| Nudcd3 | 9.82E-15 | 0.25187829 | 0.634 | 0.699 | 1.90E-10 | 4 | Nudcd3 | myCAF_2 |
| Ubald1 | 1.06E-14 | 0.84216829 | 0.576 | 0.539 | 2.05E-10 | 4 | Ubald1 | myCAF_2 |
| Pafah1b31 | 1.08E-14 | 0.8636149 | 0.568 | 0.553 | 2.09E-10 | 4 | Pafah1b3 | myCAF_2 |
| Kctd101 | 1.18E-14 | 0.37311985 | 0.852 | 0.877 | 2.29E-10 | 4 | Kctd10 | myCAF_2 |
| Fastk | 1.28E-14 | 0.25383231 | 0.709 | 0.769 | 2.48E-10 | 4 | Fastk | myCAF_2 |
| Sdf21 | 1.32E-14 | 0.31938419 | 0.801 | 0.839 | 2.55E-10 | 4 | Sdf2 | myCAF_2 |
| Lnpep1 | 1.33E-14 | 0.27603185 | 0.821 | 0.849 | 2.57E-10 | 4 | Lnpep | myCAF_2 |
| Sars | 1.33E-14 | 0.83733941 | 0.649 | 0.617 | 2.58E-10 | 4 | Sars | myCAF_2 |
| Sel1l | 1.47E-14 | 0.2907605 | 0.815 | 0.881 | 2.84E-10 | 4 | Sel1l | myCAF_2 |
| Ctnna11 | 1.91E-14 | 0.29587267 | 0.868 | 0.896 | 3.70E-10 | 4 | Ctnna1 | myCAF_2 |
| Cmip1 | 2.37E-14 | 0.77490243 | 0.827 | 0.804 | 4.57E-10 | 4 | Cmip | myCAF_2 |
| Nck21 | 2.45E-14 | 0.88892552 | 0.511 | 0.423 | 4.74E-10 | 4 | Nck2 | myCAF_2 |
| Mcl1 | 2.57E-14 | 0.45397229 | 0.937 | 0.946 | 4.96E-10 | 4 | Mcl1 | myCAF_2 |
| Tbkbp11 | 3.22E-14 | 1.01039544 | 0.303 | 0.218 | 6.22E-10 | 4 | Tbkbp1 | myCAF_2 |
| Nrn1 | 3.65E-14 | 0.94010443 | 0.263 | 0.223 | 7.05E-10 | 4 | Nrn1 | myCAF_2 |
| Wbp1l | 4.13E-14 | 0.29970439 | 0.851 | 0.893 | 7.99E-10 | 4 | Wbp1l | myCAF_2 |
| Atxn2l | 4.70E-14 | 0.34377613 | 0.856 | 0.878 | 9.08E-10 | 4 | Atxn2l | myCAF_2 |
| Grina1 | 4.76E-14 | 0.71068351 | 0.694 | 0.643 | 9.21E-10 | 4 | Grina | myCAF_2 |
| Pcyox1 | 5.06E-14 | 0.3228449 | 0.723 | 0.798 | 9.78E-10 | 4 | Pcyox1 | myCAF_2 |
| R3hdm4 | 5.79E-14 | 0.29296887 | 0.783 | 0.844 | 1.12E-09 | 4 | R3hdm4 | myCAF_2 |
| Cyth31 | 6.34E-14 | 0.3140187 | 0.824 | 0.867 | 1.23E-09 | 4 | Cyth3 | myCAF_2 |
| Pmepa11 | 7.03E-14 | 0.59902341 | 0.986 | 0.964 | 1.36E-09 | 4 | Pmepa1 | myCAF_2 |
| Clic11 | 7.04E-14 | 0.31649973 | 0.802 | 0.837 | 1.36E-09 | 4 | Clic1 | myCAF_2 |
| Rab101 | 7.57E-14 | 0.28382337 | 0.677 | 0.701 | 1.46E-09 | 4 | Rab10 | myCAF_2 |
| Lrch21 | 8.44E-14 | 0.96230856 | 0.302 | 0.242 | 1.63E-09 | 4 | Lrch2 | myCAF_2 |
| Ascc31 | 8.75E-14 | 0.30832189 | 0.608 | 0.679 | 1.69E-09 | 4 | Ascc3 | myCAF_2 |
| Fhl21 | 1.10E-13 | 0.89353146 | 0.514 | 0.443 | 2.13E-09 | 4 | Fhl2 | myCAF_2 |
| Clcn5 | 1.14E-13 | 0.83291016 | 0.607 | 0.583 | 2.21E-09 | 4 | Clcn5 | myCAF_2 |
| Prdm12 | 1.18E-13 | 0.71008886 | 0.494 | 0.402 | 2.27E-09 | 4 | Prdm1 | myCAF_2 |
| Irs22 | 1.21E-13 | 0.9005408 | 0.637 | 0.615 | 2.34E-09 | 4 | Irs2 | myCAF_2 |
| Luc7l21 | 1.23E-13 | 0.36744445 | 0.888 | 0.9 | 2.38E-09 | 4 | Luc7l2 | myCAF_2 |
| Arfip1 | 1.35E-13 | 0.28649334 | 0.732 | 0.796 | 2.62E-09 | 4 | Arfip1 | myCAF_2 |
| Ubxn1 | 1.55E-13 | 0.25114666 | 0.764 | 0.809 | 2.99E-09 | 4 | Ubxn1 | myCAF_2 |
| Rsrp1 | 1.63E-13 | 0.62022333 | 0.909 | 0.896 | 3.16E-09 | 4 | Rsrp1 | myCAF_2 |
| Dock1 | 1.78E-13 | 0.25806095 | 0.776 | 0.819 | 3.44E-09 | 4 | Dock1 | myCAF_2 |
| Rere1 | 1.81E-13 | 0.29967302 | 0.703 | 0.743 | 3.49E-09 | 4 | Rere | myCAF_2 |
| Apol81 | 1.89E-13 | 1.05121798 | 0.287 | 0.239 | 3.65E-09 | 4 | Apol8 | myCAF_2 |
| Rusc21 | 2.12E-13 | 0.79868357 | 0.519 | 0.49 | 4.11E-09 | 4 | Rusc2 | myCAF_2 |
| Dock5 | 2.51E-13 | 0.93656202 | 0.361 | 0.3 | 4.84E-09 | 4 | Dock5 | myCAF_2 |
| Podnl11 | 3.09E-13 | 0.78012611 | 0.575 | 0.411 | 5.97E-09 | 4 | Podnl1 | myCAF_2 |
| Scd21 | 3.46E-13 | 0.64371774 | 0.802 | 0.768 | 6.68E-09 | 4 | Scd2 | myCAF_2 |
| Rbm39 | 4.04E-13 | 0.32416752 | 0.732 | 0.776 | 7.82E-09 | 4 | Rbm39 | myCAF_2 |
| Ifi301 | 8.33E-13 | 0.28316933 | 0.371 | 0.465 | 1.61E-08 | 4 | Ifi30 | myCAF_2 |
| Pex5 | 8.50E-13 | 0.27305511 | 0.58 | 0.663 | 1.64E-08 | 4 | Pex5 | myCAF_2 |
| Fblim1 | 9.70E-13 | 0.37533826 | 0.76 | 0.728 | 1.87E-08 | 4 | Fblim1 | myCAF_2 |
| Mlxip1 | 1.09E-12 | 0.26617321 | 0.688 | 0.734 | 2.11E-08 | 4 | Mlxip | myCAF_2 |
| Rnd31 | 1.11E-12 | 0.28672375 | 0.785 | 0.834 | 2.14E-08 | 4 | Rnd3 | myCAF_2 |
| Dcbld12 | 1.14E-12 | 0.78860549 | 0.447 | 0.374 | 2.20E-08 | 4 | Dcbld1 | myCAF_2 |
| Atp6v1a1 | 1.40E-12 | 0.27271389 | 0.653 | 0.728 | 2.71E-08 | 4 | Atp6v1a | myCAF_2 |
| Ap3b1 | 1.42E-12 | 0.28499336 | 0.693 | 0.732 | 2.74E-08 | 4 | Ap3b1 | myCAF_2 |
| Atp11a | 1.53E-12 | 0.705227 | 0.85 | 0.828 | 2.96E-08 | 4 | Atp11a | myCAF_2 |
| Trpc1 | 1.95E-12 | 0.96964299 | 0.278 | 0.216 | 3.77E-08 | 4 | Trpc1 | myCAF_2 |
| Endod12 | 2.24E-12 | 0.99937434 | 0.345 | 0.291 | 4.34E-08 | 4 | Endod1 | myCAF_2 |
| Src1 | 2.51E-12 | 0.30529689 | 0.709 | 0.762 | 4.84E-08 | 4 | Src | myCAF_2 |
| Ssbp4 | 2.72E-12 | 0.31050071 | 0.618 | 0.706 | 5.26E-08 | 4 | Ssbp4 | myCAF_2 |
| Slc3a2 | 2.90E-12 | 0.67825124 | 0.827 | 0.849 | 5.61E-08 | 4 | Slc3a2 | myCAF_2 |
| Zswim41 | 3.24E-12 | 1.03489576 | 0.57 | 0.49 | 6.26E-08 | 4 | Zswim4 | myCAF_2 |
| Lrrc552 | 3.28E-12 | 0.92387959 | 0.294 | 0.217 | 6.33E-08 | 4 | Lrrc55 | myCAF_2 |
| Acsl4 | 3.57E-12 | 0.32705077 | 0.728 | 0.762 | 6.90E-08 | 4 | Acsl4 | myCAF_2 |
| Calcrl1 | 3.64E-12 | 0.81412276 | 0.293 | 0.238 | 7.04E-08 | 4 | Calcrl | myCAF_2 |
| H3f3b1 | 3.75E-12 | 0.42452009 | 0.891 | 0.895 | 7.24E-08 | 4 | H3f3b | myCAF_2 |
| Prkcsh | 4.48E-12 | 0.35701594 | 0.871 | 0.897 | 8.67E-08 | 4 | Prkcsh | myCAF_2 |
| Klf91 | 4.59E-12 | 0.29213074 | 0.84 | 0.856 | 8.87E-08 | 4 | Klf9 | myCAF_2 |
| Syf2 | 4.68E-12 | 0.32532186 | 0.745 | 0.797 | 9.05E-08 | 4 | Syf2 | myCAF_2 |
| Usf2 | 4.99E-12 | 0.34446312 | 0.855 | 0.872 | 9.66E-08 | 4 | Usf2 | myCAF_2 |

| | | | | | | | | |
|---|---|---|---|---|---|---|---|---|
| Ing11 | 5.04E-12 | 0.28478395 | 0.64 | 0.673 | 9.74E-08 | 4 | Ing1 | myCAF_2 |
| Cebpa1 | 7.72E-12 | 0.2856681 | 0.561 | 0.64 | 1.49E-07 | 4 | Cebpa | myCAF_2 |
| Ppp1r13l | 8.21E-12 | 1.117472 | 0.391 | 0.319 | 1.59E-07 | 4 | Ppp1r13l | myCAF_2 |
| Tgfb1i1 | 8.84E-12 | 0.36211548 | 0.844 | 0.867 | 1.71E-07 | 4 | Tgfb1i1 | myCAF_2 |
| Arl3 | 1.05E-11 | 0.29559131 | 0.735 | 0.796 | 2.03E-07 | 4 | Arl3 | myCAF_2 |
| Nsd31 | 1.17E-11 | 0.35219368 | 0.856 | 0.874 | 2.26E-07 | 4 | Nsd3 | myCAF_2 |
| Tnfrsf23 | 1.18E-11 | 0.40567251 | 0.729 | 0.816 | 2.28E-07 | 4 | Tnfrsf23 | myCAF_2 |
| Bri31 | 1.21E-11 | 0.31000213 | 0.818 | 0.865 | 2.34E-07 | 4 | Bri3 | myCAF_2 |
| Atf4 | 1.34E-11 | 0.95793381 | 0.623 | 0.565 | 2.58E-07 | 4 | Atf4 | myCAF_2 |
| Arf61 | 1.42E-11 | 0.35050975 | 0.853 | 0.875 | 2.74E-07 | 4 | Arf6 | myCAF_2 |
| Wsb1 | 1.42E-11 | 0.71829587 | 0.827 | 0.849 | 2.74E-07 | 4 | Wsb1 | myCAF_2 |
| Elk4 | 1.45E-11 | 0.26362624 | 0.74 | 0.79 | 2.81E-07 | 4 | Elk4 | myCAF_2 |
| Sec24c | 1.59E-11 | 0.25943975 | 0.697 | 0.758 | 3.08E-07 | 4 | Sec24c | myCAF_2 |
| Islr1 | 1.83E-11 | 0.34300002 | 0.732 | 0.824 | 3.54E-07 | 4 | Islr | myCAF_2 |
| Slc29a12 | 1.91E-11 | 0.84624133 | 0.326 | 0.304 | 3.70E-07 | 4 | Slc29a1 | myCAF_2 |
| Klf61 | 2.09E-11 | 0.53253923 | 0.826 | 0.87 | 4.04E-07 | 4 | Klf6 | myCAF_2 |
| Zbtb462 | 2.43E-11 | 1.04926054 | 0.288 | 0.212 | 4.69E-07 | 4 | Zbtb46 | myCAF_2 |
| Gnl1 | 2.50E-11 | 0.31771012 | 0.746 | 0.787 | 4.84E-07 | 4 | Gnl1 | myCAF_2 |
| Srsf22 | 2.55E-11 | 0.44721935 | 0.709 | 0.736 | 4.93E-07 | 4 | Srsf2 | myCAF_2 |
| Oaz21 | 2.95E-11 | 0.29827089 | 0.791 | 0.847 | 5.71E-07 | 4 | Oaz2 | myCAF_2 |
| Slc4a21 | 3.25E-11 | 0.32514977 | 0.644 | 0.721 | 6.28E-07 | 4 | Slc4a2 | myCAF_2 |
| Golga7 | 3.34E-11 | 0.31536382 | 0.826 | 0.855 | 6.45E-07 | 4 | Golga7 | myCAF_2 |
| Rtkn | 3.62E-11 | 1.06268022 | 0.348 | 0.278 | 7.00E-07 | 4 | Rtkn | myCAF_2 |
| Pum11 | 3.75E-11 | 0.26134803 | 0.642 | 0.719 | 7.24E-07 | 4 | Pum1 | myCAF_2 |
| Mgat32 | 3.80E-11 | 0.81416702 | 0.478 | 0.421 | 7.34E-07 | 4 | Mgat3 | myCAF_2 |
| Zfp598 | 4.66E-11 | 0.31331743 | 0.622 | 0.666 | 9.01E-07 | 4 | Zfp598 | myCAF_2 |
| Fxr11 | 4.77E-11 | 0.31411519 | 0.713 | 0.762 | 9.22E-07 | 4 | Fxr1 | myCAF_2 |
| Nectin3 | 5.03E-11 | 0.25204804 | 0.58 | 0.661 | 9.72E-07 | 4 | Nectin3 | myCAF_2 |
| Klhl212 | 5.09E-11 | 0.9750291 | 0.604 | 0.582 | 9.84E-07 | 4 | Klhl21 | myCAF_2 |
| Hspa5 | 5.22E-11 | 0.42432032 | 0.888 | 0.929 | 1.01E-06 | 4 | Hspa5 | myCAF_2 |
| Shc12 | 5.52E-11 | 0.27748624 | 0.849 | 0.887 | 1.07E-06 | 4 | Shc1 | myCAF_2 |
| Vcan | 5.66E-11 | 0.45905179 | 0.847 | 0.843 | 1.09E-06 | 4 | Vcan | myCAF_2 |
| Thoc6 | 6.51E-11 | 1.03562002 | 0.476 | 0.449 | 1.26E-06 | 4 | Thoc6 | myCAF_2 |
| Fndc11 | 6.53E-11 | 0.28705836 | 0.709 | 0.819 | 1.26E-06 | 4 | Fndc1 | myCAF_2 |
| Spred3 | 6.93E-11 | 0.95535464 | 0.365 | 0.285 | 1.34E-06 | 4 | Spred3 | myCAF_2 |
| Arl8b | 8.34E-11 | 0.29407423 | 0.605 | 0.681 | 1.61E-06 | 4 | Arl8b | myCAF_2 |
| Rnf20 | 9.88E-11 | 0.25008794 | 0.441 | 0.523 | 1.91E-06 | 4 | Rnf20 | myCAF_2 |
| Atp2c1 | 1.02E-10 | 0.28340192 | 0.721 | 0.763 | 1.98E-06 | 4 | Atp2c1 | myCAF_2 |
| Ghitm | 1.03E-10 | 0.29223713 | 0.556 | 0.65 | 1.99E-06 | 4 | Ghitm | myCAF_2 |
| Cblb1 | 1.11E-10 | 0.70473755 | 0.812 | 0.788 | 2.14E-06 | 4 | Cblb | myCAF_2 |
| Kdm5b1 | 1.21E-10 | 0.68447054 | 0.734 | 0.698 | 2.34E-06 | 4 | Kdm5b | myCAF_2 |
| Hsp90aa11 | 1.23E-10 | 0.46750491 | 0.836 | 0.851 | 2.37E-06 | 4 | Hsp90aa1 | myCAF_2 |
| Arih11 | 1.30E-10 | 0.30610796 | 0.72 | 0.783 | 2.52E-06 | 4 | Arih1 | myCAF_2 |
| Myl61 | 1.36E-10 | 0.40499608 | 0.837 | 0.859 | 2.64E-06 | 4 | Myl6 | myCAF_2 |
| Sdc12 | 1.47E-10 | 0.53665608 | 0.975 | 0.909 | 2.83E-06 | 4 | Sdc1 | myCAF_2 |
| Sema6d1 | 1.47E-10 | 0.94204355 | 0.294 | 0.285 | 2.85E-06 | 4 | Sema6d | myCAF_2 |
| Slc7a6os | 1.48E-10 | 0.26862534 | 0.384 | 0.456 | 2.85E-06 | 4 | Slc7a6os | myCAF_2 |
| Prpsap1 | 1.48E-10 | 0.28647118 | 0.58 | 0.648 | 2.86E-06 | 4 | Prpsap1 | myCAF_2 |
| Agap1 | 1.60E-10 | 0.85948006 | 0.513 | 0.472 | 3.09E-06 | 4 | Agap1 | myCAF_2 |
| Ythdf31 | 1.64E-10 | 0.34917263 | 0.804 | 0.842 | 3.16E-06 | 4 | Ythdf3 | myCAF_2 |
| Mgrn11 | 1.84E-10 | 0.30698994 | 0.804 | 0.836 | 3.56E-06 | 4 | Mgrn1 | myCAF_2 |
| Rhoc | 1.89E-10 | 0.38550052 | 0.906 | 0.933 | 3.65E-06 | 4 | Rhoc | myCAF_2 |
| Fut42 | 2.14E-10 | 0.90354005 | 0.432 | 0.391 | 4.13E-06 | 4 | Fut4 | myCAF_2 |
| Akap13 | 2.20E-10 | 0.2698252 | 0.816 | 0.854 | 4.25E-06 | 4 | Akap13 | myCAF_2 |
| Zfp771 | 2.29E-10 | 0.26094008 | 0.407 | 0.492 | 4.42E-06 | 4 | Zfp771 | myCAF_2 |
| Wdr591 | 2.31E-10 | 1.11073267 | 0.27 | 0.216 | 4.46E-06 | 4 | Wdr59 | myCAF_2 |
| Usp12 | 2.45E-10 | 0.29429075 | 0.458 | 0.531 | 4.74E-06 | 4 | Usp12 | myCAF_2 |
| Smyd21 | 2.76E-10 | 0.2955824 | 0.621 | 0.673 | 5.33E-06 | 4 | Smyd2 | myCAF_2 |
| Sox91 | 2.91E-10 | 1.20053228 | 0.302 | 0.231 | 5.64E-06 | 4 | Sox9 | myCAF_2 |
| Tln22 | 3.38E-10 | 0.31290571 | 0.705 | 0.713 | 6.53E-06 | 4 | Tln2 | myCAF_2 |
| Abi1 | 3.46E-10 | 0.39166822 | 0.82 | 0.833 | 6.69E-06 | 4 | Abi1 | myCAF_2 |
| Rprd21 | 3.48E-10 | 0.28233182 | 0.528 | 0.605 | 6.73E-06 | 4 | Rprd2 | myCAF_2 |
| Frmd4a1 | 3.83E-10 | 0.83152433 | 0.663 | 0.587 | 7.41E-06 | 4 | Frmd4a | myCAF_2 |
| Colec122 | 3.94E-10 | 0.32440087 | 0.911 | 0.911 | 7.61E-06 | 4 | Colec12 | myCAF_2 |
| Nfya1 | 4.17E-10 | 0.26155079 | 0.482 | 0.529 | 8.05E-06 | 4 | Nfya | myCAF_2 |
| Dusp3 | 4.75E-10 | 0.62793228 | 0.77 | 0.755 | 9.18E-06 | 4 | Dusp3 | myCAF_2 |
| Mast21 | 5.08E-10 | 0.42559706 | 0.736 | 0.769 | 9.81E-06 | 4 | Mast2 | myCAF_2 |
| Ptk21 | 5.56E-10 | 0.30555077 | 0.616 | 0.668 | 1.07E-05 | 4 | Ptk2 | myCAF_2 |
| Hs2st1 | 6.07E-10 | 0.34759712 | 0.68 | 0.739 | 1.17E-05 | 4 | Hs2st1 | myCAF_2 |
| Lin7c | 6.08E-10 | 0.28210062 | 0.597 | 0.647 | 1.18E-05 | 4 | Lin7c | myCAF_2 |
| Atp6v1g1 | 6.39E-10 | 0.38083703 | 0.847 | 0.86 | 1.24E-05 | 4 | Atp6v1g1 | myCAF_2 |
| Rhbdf1 | 6.47E-10 | 0.35137586 | 0.712 | 0.776 | 1.25E-05 | 4 | Rhbdf1 | myCAF_2 |
| Scand1 | 6.94E-10 | 0.33574313 | 0.667 | 0.717 | 1.34E-05 | 4 | Scand1 | myCAF_2 |
| D10Wsu102e | 6.96E-10 | 0.26375071 | 0.335 | 0.395 | 1.34E-05 | 4 | D10Wsu102e | myCAF_2 |
| Sdf2l1 | 7.06E-10 | 0.30596831 | 0.455 | 0.583 | 1.36E-05 | 4 | Sdf2l1 | myCAF_2 |
| Larp4b1 | 8.05E-10 | 0.28747433 | 0.582 | 0.647 | 1.56E-05 | 4 | Larp4b | myCAF_2 |
| Yaf22 | 8.54E-10 | 0.32898956 | 0.529 | 0.582 | 1.65E-05 | 4 | Yaf2 | myCAF_2 |
| Max | 8.55E-10 | 0.293444 | 0.586 | 0.652 | 1.65E-05 | 4 | Max | myCAF_2 |
| Tbc1d11 | 8.64E-10 | 0.83746614 | 0.495 | 0.455 | 1.67E-05 | 4 | Tbc1d1 | myCAF_2 |
| Fam50a | 8.76E-10 | 0.26141543 | 0.501 | 0.572 | 1.69E-05 | 4 | Fam50a | myCAF_2 |
| Trappc5 | 9.81E-10 | 0.29517217 | 0.545 | 0.613 | 1.90E-05 | 4 | Trappc5 | myCAF_2 |
| Prkar1a2 | 9.87E-10 | 0.39764119 | 0.953 | 0.942 | 1.91E-05 | 4 | Prkar1a | myCAF_2 |
| Tmeff11 | 1.02E-09 | 0.8228929 | 0.463 | 0.405 | 1.97E-05 | 4 | Tmeff1 | myCAF_2 |
| Tbc1d9b | 1.27E-09 | 0.28148521 | 0.732 | 0.8 | 2.46E-05 | 4 | Tbc1d9b | myCAF_2 |
| Stx4a | 1.30E-09 | 0.27491241 | 0.707 | 0.742 | 2.52E-05 | 4 | Stx4a | myCAF_2 |
| Trib21 | 1.52E-09 | 0.89434695 | 0.315 | 0.263 | 2.94E-05 | 4 | Trib2 | myCAF_2 |
| Ints62 | 1.52E-09 | 0.29212736 | 0.468 | 0.534 | 2.94E-05 | 4 | Ints6 | myCAF_2 |
| Fbxo11 | 1.57E-09 | 0.26492854 | 0.611 | 0.674 | 3.03E-05 | 4 | Fbxo11 | myCAF_2 |
| Prokr22 | 1.59E-09 | 0.83222849 | 0.523 | 0.394 | 3.07E-05 | 4 | Prokr2 | myCAF_2 |
| Btbd11 | 1.63E-09 | 0.25135439 | 0.541 | 0.597 | 3.15E-05 | 4 | Btbd1 | myCAF_2 |
| Cdk131 | 1.67E-09 | 0.28309762 | 0.595 | 0.662 | 3.24E-05 | 4 | Cdk13 | myCAF_2 |
| Nsmf | 1.68E-09 | 0.28311674 | 0.631 | 0.707 | 3.25E-05 | 4 | Nsmf | myCAF_2 |
| Tmem1891 | 1.70E-09 | 0.27027936 | 0.568 | 0.619 | 3.28E-05 | 4 | Tmem189 | myCAF_2 |
| Arl6ip61 | 1.96E-09 | 0.33428092 | 0.506 | 0.527 | 3.80E-05 | 4 | Arl6ip6 | myCAF_2 |
| Setd51 | 2.05E-09 | 0.30926864 | 0.786 | 0.828 | 3.97E-05 | 4 | Setd5 | myCAF_2 |
| Klf21 | 2.09E-09 | 1.01242152 | 0.422 | 0.414 | 4.04E-05 | 4 | Klf2 | myCAF_2 |
| Rasl122 | 2.17E-09 | 0.93720919 | 0.41 | 0.388 | 4.20E-05 | 4 | Rasl12 | myCAF_2 |
| Rlf1 | 2.40E-09 | 0.84537273 | 0.524 | 0.487 | 4.65E-05 | 4 | Rlf | myCAF_2 |
| Traf4 | 2.45E-09 | 0.8381289 | 0.475 | 0.422 | 4.74E-05 | 4 | Traf4 | myCAF_2 |
| Ccdc186 | 2.51E-09 | 0.27612463 | 0.509 | 0.573 | 4.85E-05 | 4 | Ccdc186 | myCAF_2 |
| Sap1301 | 2.67E-09 | 0.29300708 | 0.476 | 0.532 | 5.16E-05 | 4 | Sap130 | myCAF_2 |
| Foxo3 | 2.69E-09 | 0.86626266 | 0.571 | 0.526 | 5.20E-05 | 4 | Foxo3 | myCAF_2 |
| Stk401 | 2.78E-09 | 0.75783484 | 0.696 | 0.652 | 5.37E-05 | 4 | Stk40 | myCAF_2 |
| Tns32 | 2.96E-09 | 0.61373165 | 0.767 | 0.655 | 5.72E-05 | 4 | Tns3 | myCAF_2 |
| Trp53bp1 | 3.16E-09 | 0.27377906 | 0.352 | 0.414 | 6.11E-05 | 4 | Trp53bp1 | myCAF_2 |
| Bicd2 | 3.49E-09 | 0.74565341 | 0.699 | 0.677 | 6.74E-05 | 4 | Bicd2 | myCAF_2 |
| Irf3 | 4.39E-09 | 0.31020078 | 0.423 | 0.497 | 8.50E-05 | 4 | Irf3 | myCAF_2 |
| Dtx3 | 4.42E-09 | 0.71742318 | 0.817 | 0.781 | 8.54E-05 | 4 | Dtx3 | myCAF_2 |
| G2e31 | 4.44E-09 | 0.42130962 | 0.294 | 0.31 | 8.58E-05 | 4 | G2e3 | myCAF_2 |
| Picalm1 | 4.87E-09 | 0.37418252 | 0.875 | 0.884 | 9.41E-05 | 4 | Picalm | myCAF_2 |
| Pgrmc2 | 4.91E-09 | 0.3867505 | 0.794 | 0.834 | 9.49E-05 | 4 | Pgrmc2 | myCAF_2 |
| Cyp512 | 4.99E-09 | 0.35406279 | 0.534 | 0.632 | 9.64E-05 | 4 | Cyp51 | myCAF_2 |
| Atp6v1c11 | 5.29E-09 | 0.25032986 | 0.474 | 0.541 | 0.00010222 | 4 | Atp6v1c1 | myCAF_2 |
| Nxf11 | 5.95E-09 | 0.37176086 | 0.661 | 0.696 | 0.00011505 | 4 | Nxf1 | myCAF_2 |
| Nhsl1 | 6.04E-09 | 0.78222128 | 0.418 | 0.352 | 0.00011672 | 4 | Nhsl1 | myCAF_2 |
| Rab11b | 7.04E-09 | 0.34007835 | 0.762 | 0.802 | 0.00013603 | 4 | Rab11b | myCAF_2 |
| Fam110b1 | 7.34E-09 | 0.97900108 | 0.301 | 0.261 | 0.00014189 | 4 | Fam110b | myCAF_2 |
| Mknk21 | 7.46E-09 | 0.42520931 | 0.612 | 0.656 | 0.00014426 | 4 | Mknk2 | myCAF_2 |
| Afg3l2 | 7.57E-09 | 0.28914477 | 0.607 | 0.653 | 0.00014635 | 4 | Afg3l2 | myCAF_2 |
| Smg72 | 8.13E-09 | 0.26687248 | 0.518 | 0.568 | 0.00015724 | 4 | Smg7 | myCAF_2 |
| Cry2 | 8.16E-09 | 0.76986345 | 0.36 | 0.333 | 0.00015773 | 4 | Cry2 | myCAF_2 |
| Ror21 | 8.28E-09 | 0.69323744 | 0.818 | 0.78 | 0.00016008 | 4 | Ror2 | myCAF_2 |
| N4bp2l21 | 9.50E-09 | 0.29686685 | 0.5 | 0.568 | 0.0001837 | 4 | N4bp2l2 | myCAF_2 |
| Clasp11 | 9.62E-09 | 0.27837838 | 0.593 | 0.656 | 0.00018595 | 4 | Clasp1 | myCAF_2 |
| Rap2a1 | 9.74E-09 | 0.84531383 | 0.567 | 0.507 | 0.00018837 | 4 | Rap2a | myCAF_2 |
| Rab11fip5 | 1.05E-08 | 0.25936585 | 0.645 | 0.713 | 0.00020358 | 4 | Rab11fip5 | myCAF_2 |
| Mtdh1 | 1.10E-08 | 0.37453705 | 0.71 | 0.741 | 0.00021243 | 4 | Mtdh | myCAF_2 |
| Hic1 | 1.12E-08 | 0.72080688 | 0.849 | 0.882 | 0.00021618 | 4 | Hic1 | myCAF_2 |
| Nabp11 | 1.13E-08 | 0.9425425 | 0.27 | 0.227 | 0.00021855 | 4 | Nabp1 | myCAF_2 |
| Tpm22 | 1.22E-08 | 0.48101387 | 0.816 | 0.756 | 0.00023499 | 4 | Tpm2 | myCAF_2 |
| Phrf1 | 1.23E-08 | 0.31437514 | 0.497 | 0.566 | 0.00023727 | 4 | Phrf1 | myCAF_2 |
| Ebf11 | 1.23E-08 | 0.61450679 | 0.648 | 0.676 | 0.00023816 | 4 | Ebf1 | myCAF_2 |
| Dgka | 1.24E-08 | 0.25242649 | 0.327 | 0.399 | 0.00024047 | 4 | Dgka | myCAF_2 |
| Crlf2 | 1.33E-08 | 0.81981896 | 0.54 | 0.464 | 0.00025737 | 4 | Crlf2 | myCAF_2 |
| Unc5a2 | 1.37E-08 | 0.80356456 | 0.354 | 0.299 | 0.00026464 | 4 | Unc5a | myCAF_2 |
| Orc4 | 1.48E-08 | 0.25428622 | 0.475 | 0.537 | 0.0002859 | 4 | Orc4 | myCAF_2 |
| Rarb1 | 1.60E-08 | 1.00445202 | 0.291 | 0.232 | 0.00030977 | 4 | Rarb | myCAF_2 |
| Runx22 | 1.71E-08 | 0.69942432 | 0.717 | 0.665 | 0.00033124 | 4 | Runx2 | myCAF_2 |
| Rabggtb1 | 1.81E-08 | 0.3334292 | 0.64 | 0.684 | 0.0003508 | 4 | Rabggtb | myCAF_2 |
| Kmt2d | 1.83E-08 | 0.33221878 | 0.687 | 0.741 | 0.0003536 | 4 | Kmt2d | myCAF_2 |
| Ctif | 1.87E-08 | 0.78624857 | 0.637 | 0.594 | 0.00036194 | 4 | Ctif | myCAF_2 |
| Rbm4 | 2.07E-08 | 0.29893272 | 0.596 | 0.649 | 0.0004001 | 4 | Rbm4 | myCAF_2 |
| Wwc21 | 2.08E-08 | 0.30033995 | 0.677 | 0.737 | 0.00040309 | 4 | Wwc2 | myCAF_2 |
| Fxr21 | 2.16E-08 | 0.26835909 | 0.547 | 0.595 | 0.00041856 | 4 | Fxr2 | myCAF_2 |
| Eloa | 2.28E-08 | 0.32573534 | 0.768 | 0.801 | 0.00044045 | 4 | Eloa | myCAF_2 |
| Tnks | 2.37E-08 | 0.33356645 | 0.658 | 0.715 | 0.0004583 | 4 | Tnks | myCAF_2 |
| Klc1 | 2.38E-08 | 0.25985189 | 0.626 | 0.681 | 0.00046086 | 4 | Klc1 | myCAF_2 |
| Gli31 | 2.40E-08 | 0.2521562 | 0.514 | 0.6 | 0.00046453 | 4 | Gli3 | myCAF_2 |
| Tgoln11 | 2.52E-08 | 0.37272029 | 0.723 | 0.773 | 0.00048778 | 4 | Tgoln1 | myCAF_2 |
| Mbd1 | 2.56E-08 | 0.29843334 | 0.407 | 0.437 | 0.00049575 | 4 | Mbd1 | myCAF_2 |
| Clybl1 | 2.59E-08 | 0.93289034 | 0.358 | 0.326 | 0.00050162 | 4 | Clybl | myCAF_2 |
| Srp68 | 3.24E-08 | 0.34644779 | 0.752 | 0.789 | 0.00062629 | 4 | Srp68 | myCAF_2 |
| Gcnt22 | 3.54E-08 | 0.26277303 | 0.384 | 0.407 | 0.00068398 | 4 | Gcnt2 | myCAF_2 |
| Aprt1 | 3.67E-08 | 0.28081123 | 0.423 | 0.484 | 0.00071045 | 4 | Aprt | myCAF_2 |
| Lum14b2 | 3.79E-08 | 0.26789228 | 0.733 | 0.785 | 0.00073189 | 4 | Lum14b | myCAF_2 |
| Qdpr | 3.81E-08 | 0.28756847 | 0.543 | 0.599 | 0.00073658 | 4 | Qdpr | myCAF_2 |
| Fhod31 | 3.82E-08 | 0.56890823 | 0.371 | 0.315 | 0.00073823 | 4 | Fhod3 | myCAF_2 |
| Fez2 | 3.84E-08 | 0.28871099 | 0.57 | 0.613 | 0.00074182 | 4 | Fez2 | myCAF_2 |
| Khdc4 | 4.05E-08 | 0.30551968 | 0.634 | 0.667 | 0.00078241 | 4 | Khdc4 | myCAF_2 |
| Brwd11 | 4.09E-08 | 0.38641877 | 0.674 | 0.708 | 0.00079048 | 4 | Brwd1 | myCAF_2 |
| Alad1 | 4.17E-08 | 0.35239304 | 0.482 | 0.537 | 0.0008064 | 4 | Alad | myCAF_2 |
| Med141 | 4.45E-08 | 0.27015976 | 0.346 | 0.415 | 0.00086045 | 4 | Med14 | myCAF_2 |
| Wasl1 | 4.47E-08 | 0.27639802 | 0.605 | 0.667 | 0.00086499 | 4 | Wasl | myCAF_2 |
| Yrdc | 4.79E-08 | 0.28792255 | 0.46 | 0.505 | 0.0009254 | 4 | Yrdc | myCAF_2 |
| Snrk1 | 4.93E-08 | 0.74054133 | 0.384 | 0.363 | 0.00095386 | 4 | Snrk | myCAF_2 |
| Gopc | 5.13E-08 | 0.83213015 | 0.573 | 0.569 | 0.00099199 | 4 | Gopc | myCAF_2 |
| Wdfy3 | 5.18E-08 | 0.28145352 | 0.757 | 0.813 | 0.00100164 | 4 | Wdfy3 | myCAF_2 |
| Rbpms22 | 5.51E-08 | 0.89619755 | 0.487 | 0.391 | 0.00106564 | 4 | Rbpms2 | myCAF_2 |
| Neo11 | 5.57E-08 | 0.35758714 | 0.602 | 0.657 | 0.00107612 | 4 | Neo1 | myCAF_2 |
| Supt5 | 5.92E-08 | 0.35905742 | 0.444 | 0.497 | 0.00114521 | 4 | Supt5 | myCAF_2 |
| Tns2 | 5.98E-08 | 0.34592218 | 0.65 | 0.737 | 0.00115666 | 4 | Tns2 | myCAF_2 |
| Rbm26 | 6.17E-08 | 0.29054741 | 0.583 | 0.639 | 0.00119376 | 4 | Rbm26 | myCAF_2 |
| Phlda12 | 7.44E-08 | 0.58826606 | 0.608 | 0.61 | 0.00143755 | 4 | Phlda1 | myCAF_2 |
| Col8a11 | 7.64E-08 | 0.63468489 | 0.625 | 0.56 | 0.00147804 | 4 | Col8a1 | myCAF_2 |
| Hcfc1r11 | 7.78E-08 | 0.34752244 | 0.769 | 0.784 | 0.00150397 | 4 | Hcfc1r1 | myCAF_2 |
| Necap2 | 7.90E-08 | 0.3242365 | 0.619 | 0.671 | 0.00152823 | 4 | Necap2 | myCAF_2 |
| Eif4ebp11 | 8.84E-08 | 0.64432579 | 0.826 | 0.869 | 0.00170894 | 4 | Eif4ebp1 | myCAF_2 |
| Ncam11 | 1.01E-07 | 0.35229613 | 0.737 | 0.72 | 0.00195761 | 4 | Ncam1 | myCAF_2 |
| Acaa2 | 1.05E-07 | 0.33114778 | 0.601 | 0.678 | 0.00203265 | 4 | Acaa2 | myCAF_2 |
| Eva1b2 | 1.15E-07 | 0.34459788 | 0.805 | 0.846 | 0.00222495 | 4 | Eva1b | myCAF_2 |
| Adpgk1 | 1.15E-07 | 0.33018794 | 0.681 | 0.732 | 0.0022311 | 4 | Adpgk | myCAF_2 |
| Grem21 | 1.36E-07 | 0.44907979 | 0.824 | 0.815 | 0.00263848 | 4 | Grem2 | myCAF_2 |
| Smad4 | 1.39E-07 | 0.35855302 | 0.848 | 0.899 | 0.00269018 | 4 | Smad4 | myCAF_2 |
| Pigm | 1.45E-07 | 0.35379106 | 0.397 | 0.441 | 0.00279665 | 4 | Pigm | myCAF_2 |
| Tmem134 | 1.69E-07 | 0.37669352 | 0.703 | 0.735 | 0.00326164 | 4 | Tmem134 | myCAF_2 |
| Shoc21 | 1.70E-07 | 0.29397455 | 0.502 | 0.551 | 0.00327938 | 4 | Shoc2 | myCAF_2 |
| Rilpl2 | 1.76E-07 | 0.36213486 | 0.348 | 0.42 | 0.00339606 | 4 | Rilpl2 | myCAF_2 |
| Phf2 | 1.85E-07 | 0.25733476 | 0.595 | 0.639 | 0.00357272 | 4 | Phf2 | myCAF_2 |
| Aff41 | 2.06E-07 | 0.62794224 | 0.818 | 0.836 | 0.00397414 | 4 | Aff4 | myCAF_2 |
| Hbp12 | 2.09E-07 | 0.28765856 | 0.761 | 0.804 | 0.00404834 | 4 | Hbp1 | myCAF_2 |
| Wdfy11 | 2.12E-07 | 0.75827792 | 0.468 | 0.43 | 0.00409854 | 4 | Wdfy1 | myCAF_2 |
| Zbtb7a | 2.18E-07 | 0.34248997 | 0.687 | 0.74 | 0.00421657 | 4 | Zbtb7a | myCAF_2 |
| Phf131 | 2.27E-07 | 0.3463971 | 0.51 | 0.558 | 0.00438145 | 4 | Phf13 | myCAF_2 |
| Trim44 | 2.31E-07 | 0.32321863 | 0.752 | 0.791 | 0.00447544 | 4 | Trim44 | myCAF_2 |
| Stk25 | 2.33E-07 | 0.37135498 | 0.827 | 0.852 | 0.0045048 | 4 | Stk25 | myCAF_2 |
| Cnot6l1 | 2.34E-07 | 0.28137679 | 0.548 | 0.61 | 0.00451813 | 4 | Cnot6l | myCAF_2 |
| Vdac21 | 2.44E-07 | 0.29413086 | 0.587 | 0.632 | 0.0047249 | 4 | Vdac2 | myCAF_2 |
| Pcgf3 | 2.77E-07 | 0.33046899 | 0.562 | 0.6 | 0.0053493 | 4 | Pcgf3 | myCAF_2 |
| Alyref3 | 3.07E-07 | 0.96187446 | 0.278 | 0.252 | 0.00593305 | 4 | Alyref | myCAF_2 |
| Trappc4 | 3.17E-07 | 0.33239263 | 0.715 | 0.756 | 0.00613683 | 4 | Trappc4 | myCAF_2 |
| Htt | 3.19E-07 | 0.2549893 | 0.418 | 0.484 | 0.00615964 | 4 | Htt | myCAF_2 |
| Atp6v1d | 3.24E-07 | 0.31982232 | 0.536 | 0.616 | 0.00627374 | 4 | Atp6v1d | myCAF_2 |
| Dnajb4 | 3.25E-07 | 0.25829768 | 0.385 | 0.432 | 0.00628554 | 4 | Dnajb4 | myCAF_2 |
| U2surp1 | 3.30E-07 | 0.36004873 | 0.484 | 0.533 | 0.0063856 | 4 | U2surp | myCAF_2 |
| Gak | 3.42E-07 | 0.28724833 | 0.503 | 0.567 | 0.00661104 | 4 | Gak | myCAF_2 |
| Ankrd541 | 3.79E-07 | 0.25511978 | 0.313 | 0.367 | 0.00733355 | 4 | Ankrd54 | myCAF_2 |
| Pkig1 | 3.86E-07 | 0.43959697 | 0.726 | 0.775 | 0.00747121 | 4 | Pkig | myCAF_2 |
| Id21 | 3.89E-07 | 0.59505272 | 0.646 | 0.627 | 0.00751683 | 4 | Id2 | myCAF_2 |
| Ube2e2 | 3.90E-07 | 0.28376888 | 0.542 | 0.59 | 0.00753672 | 4 | Ube2e2 | myCAF_2 |
| Gna13 | 3.94E-07 | 0.76212045 | 0.837 | 0.809 | 0.00761777 | 4 | Gna13 | myCAF_2 |
| Nfkb1 | 3.98E-07 | 0.35231161 | 0.653 | 0.698 | 0.00769426 | 4 | Nfkb1 | myCAF_2 |
| Fbrs | 3.99E-07 | 0.31597695 | 0.683 | 0.732 | 0.00771702 | 4 | Fbrs | myCAF_2 |
| Exoc6b1 | 4.00E-07 | 0.2514185 | 0.492 | 0.551 | 0.00773064 | 4 | Exoc6b | myCAF_2 |
| Mysm11 | 4.01E-07 | 0.25638005 | 0.356 | 0.419 | 0.00775948 | 4 | Mysm1 | myCAF_2 |
| Fnbp12 | 4.11E-07 | 0.31034622 | 0.796 | 0.808 | 0.00794636 | 4 | Fnbp1 | myCAF_2 |
| Ubn1 | 4.38E-07 | 0.32307582 | 0.564 | 0.613 | 0.00846525 | 4 | Ubn1 | myCAF_2 |
| Cbx41 | 4.77E-07 | 0.2633671 | 0.361 | 0.419 | 0.00921532 | 4 | Cbx4 | myCAF_2 |
| Galk1 | 4.94E-07 | 0.35863744 | 0.759 | 0.779 | 0.00954332 | 4 | Galk1 | myCAF_2 |
| Rab311 | 5.33E-07 | 0.35304118 | 0.86 | 0.855 | 0.01029883 | 4 | Rab31 | myCAF_2 |
| Gdpd51 | 5.41E-07 | 0.79168475 | 0.325 | 0.267 | 0.01045497 | 4 | Gdpd5 | myCAF_2 |
| Ptk2b | 5.46E-07 | 0.68696812 | 0.287 | 0.276 | 0.01055836 | 4 | Ptk2b | myCAF_2 |
| Soga11 | 5.47E-07 | 0.80188245 | 0.674 | 0.623 | 0.01057891 | 4 | Soga1 | myCAF_2 |
| Hoxb22 | 5.76E-07 | 0.34523354 | 0.427 | 0.479 | 0.01114579 | 4 | Hoxb2 | myCAF_2 |
| Kdm6b1 | 5.95E-07 | 0.744417 | 0.768 | 0.701 | 0.01150853 | 4 | Kdm6b | myCAF_2 |
| Rala | 5.98E-07 | 0.36332662 | 0.556 | 0.612 | 0.01156655 | 4 | Rala | myCAF_2 |
| Stard41 | 6.01E-07 | 0.29971053 | 0.391 | 0.449 | 0.01161679 | 4 | Stard4 | myCAF_2 |
| Sipa1l3 | 6.03E-07 | 0.80730024 | 0.629 | 0.584 | 0.01165985 | 4 | Sipa1l3 | myCAF_2 |
| Ddx421 | 6.13E-07 | 0.294858 | 0.553 | 0.594 | 0.01184277 | 4 | Ddx42 | myCAF_2 |
| Ufl1 | 6.53E-07 | 0.34003917 | 0.453 | 0.522 | 0.0126193 | 4 | Ufl1 | myCAF_2 |
| Gtf3c4 | 6.81E-07 | 0.25550761 | 0.423 | 0.489 | 0.0131602 | 4 | Gtf3c4 | myCAF_2 |
| Zdhhc201 | 7.06E-07 | 0.28663633 | 0.625 | 0.66 | 0.01365639 | 4 | Zdhhc20 | myCAF_2 |
| Frs2 | 7.16E-07 | 0.25717074 | 0.56 | 0.619 | 0.01384454 | 4 | Frs2 | myCAF_2 |
| Smap2 | 7.40E-07 | 0.34679336 | 0.592 | 0.625 | 0.01430743 | 4 | Smap2 | myCAF_2 |
| Nbr11 | 7.55E-07 | 0.26046821 | 0.476 | 0.542 | 0.01460121 | 4 | Nbr1 | myCAF_2 |
| Pi4k2a | 7.62E-07 | 0.29795013 | 0.509 | 0.582 | 0.01473591 | 4 | Pi4k2a | myCAF_2 |
| Sart31 | 7.63E-07 | 0.3075171 | 0.279 | 0.327 | 0.01475119 | 4 | Sart3 | myCAF_2 |
| Atxn7l3b | 7.67E-07 | 0.4134226 | 0.814 | 0.832 | 0.0148217 | 4 | Atxn7l3b | myCAF_2 |
| Wbp12 | 7.76E-07 | 0.26887154 | 0.43 | 0.479 | 0.01500714 | 4 | Wbp1 | myCAF_2 |
| Ip6k1 | 8.52E-07 | 0.36232844 | 0.823 | 0.858 | 0.01646613 | 4 | Ip6k1 | myCAF_2 |
| Katnbl11 | 8.54E-07 | 0.38035043 | 0.345 | 0.401 | 0.01650972 | 4 | Katnbl1 | myCAF_2 |
| Mdm21 | 8.73E-07 | 0.78913536 | 0.656 | 0.627 | 0.01688476 | 4 | Mdm2 | myCAF_2 |
| Pde3a | 8.99E-07 | 0.7353492 | 0.356 | 0.317 | 0.01737906 | 4 | Pde3a | myCAF_2 |
| Smad31 | 9.08E-07 | 0.33136922 | 0.577 | 0.642 | 0.01755791 | 4 | Smad3 | myCAF_2 |
| Smim14 | 9.32E-07 | 0.29947972 | 0.72 | 0.761 | 0.0180135 | 4 | Smim14 | myCAF_2 |
| C2cd2l1 | 9.73E-07 | 0.31379998 | 0.384 | 0.433 | 0.01881456 | 4 | C2cd2l | myCAF_2 |
| Yars1 | 9.89E-07 | 0.32632119 | 0.555 | 0.626 | 0.01911484 | 4 | Yars | myCAF_2 |
| Setdb11 | 1.06E-06 | 0.27138737 | 0.394 | 0.435 | 0.0204265 | 4 | Setdb1 | myCAF_2 |
| Srf1 | 1.11E-06 | 0.41429325 | 0.732 | 0.756 | 0.02153328 | 4 | Srf | myCAF_2 |
| Crk | 1.13E-06 | 0.45738497 | 0.788 | 0.798 | 0.02190073 | 4 | Crk | myCAF_2 |
| C1qtnf121 | 1.28E-06 | 0.31572077 | 0.515 | 0.563 | 0.02478755 | 4 | C1qtnf12 | myCAF_2 |
| Dmxl11 | 1.28E-06 | 0.26248441 | 0.404 | 0.455 | 0.02479662 | 4 | Dmxl1 | myCAF_2 |
| Zbtb342 | 1.29E-06 | 0.71397438 | 0.465 | 0.442 | 0.02493376 | 4 | Zbtb34 | myCAF_2 |
| Ptbp2 | 1.31E-06 | 0.25372169 | 0.613 | 0.659 | 0.02539843 | 4 | Ptbp2 | myCAF_2 |
| Adnp2 | 1.46E-06 | 0.31685838 | 0.437 | 0.496 | 0.02830819 | 4 | Adnp2 | myCAF_2 |
| Aspn | 1.50E-06 | 0.53964652 | 0.46 | 0.429 | 0.02904167 | 4 | Aspn | myCAF_2 |
| Pwwp2a1 | 1.55E-06 | 0.25968178 | 0.423 | 0.494 | 0.02999334 | 4 | Pwwp2a | myCAF_2 |
| Plod21 | 1.59E-06 | 0.69530372 | 0.702 | 0.646 | 0.0307717 | 4 | Plod2 | myCAF_2 |
| Cflar | 1.62E-06 | 0.27119676 | 0.57 | 0.621 | 0.03141209 | 4 | Cflar | myCAF_2 |
| Rnf21 | 1.70E-06 | 0.26747978 | 0.462 | 0.52 | 0.03284736 | 4 | Rnf2 | myCAF_2 |
| Ppp1r12c1 | 1.73E-06 | 0.37122156 | 0.651 | 0.712 | 0.03348246 | 4 | Ppp1r12c | myCAF_2 |
| Syde21 | 1.76E-06 | 0.55132361 | 0.434 | 0.386 | 0.03398529 | 4 | Syde2 | myCAF_2 |
| Ltbp4 | 1.84E-06 | 0.54936862 | 0.354 | 0.304 | 0.03550337 | 4 | Ltbp4 | myCAF_2 |
| Uhrf1bp1l1 | 1.85E-06 | 0.27185289 | 0.42 | 0.488 | 0.03571568 | 4 | Uhrf1bp1l | myCAF_2 |
| Ammecr1l | 1.88E-06 | 0.26199132 | 0.484 | 0.53 | 0.03639135 | 4 | Ammecr1l | myCAF_2 |
| Rap1gds1 | 1.88E-06 | 0.33046177 | 0.683 | 0.734 | 0.03639311 | 4 | Rap1gds1 | myCAF_2 |
| Rc3h1 | 1.91E-06 | 0.32945907 | 0.601 | 0.66 | 0.03698168 | 4 | Rc3h1 | myCAF_2 |
| Vgll41 | 1.93E-06 | 0.29847479 | 0.686 | 0.737 | 0.03731682 | 4 | Vgll4 | myCAF_2 |
| Cnrip11 | 1.93E-06 | 0.26008432 | 0.544 | 0.623 | 0.03740068 | 4 | Cnrip1 | myCAF_2 |
| Tead22 | 1.95E-06 | 0.29683103 | 0.417 | 0.463 | 0.03763402 | 4 | Tead2 | myCAF_2 |
| Med221 | 1.95E-06 | 0.30326844 | 0.446 | 0.511 | 0.03773753 | 4 | Med22 | myCAF_2 |
| B3gat3 | 2.19E-06 | 0.29633062 | 0.383 | 0.449 | 0.04238398 | 4 | B3gat3 | myCAF_2 |
| Elfn11 | 2.35E-06 | 0.8715129 | 0.251 | 0.22 | 0.04544555 | 4 | Elfn1 | myCAF_2 |
| Rsrc2 | 2.49E-06 | 0.42603048 | 0.655 | 0.701 | 0.04809464 | 4 | Rsrc2 | myCAF_2 |
| Zfp410 | 2.49E-06 | 0.32576869 | 0.44 | 0.496 | 0.04822736 | 4 | Zfp410 | myCAF_2 |
| Hs6st21 | 2.54E-06 | 0.66474652 | 0.54 | 0.553 | 0.04901871 | 4 | Hs6st2 | myCAF_2 |
| Myef21 | 2.54E-06 | 0.34063644 | 0.566 | 0.557 | 0.04905884 | 4 | Myef2 | myCAF_2 |
| Rictor1 | 2.57E-06 | 0.64351132 | 0.536 | 0.499 | 0.04966207 | 4 | Rictor | myCAF_2 |
| Zrsr21 | 2.73E-06 | 0.37373041 | 0.437 | 0.49 | 0.05283824 | 4 | Zrsr2 | myCAF_2 |
| Gnaq | 2.73E-06 | 0.35808423 | 0.655 | 0.704 | 0.05284346 | 4 | Gnaq | myCAF_2 |
| Ubn2 | 2.85E-06 | 0.25828168 | 0.422 | 0.49 | 0.05503176 | 4 | Ubn2 | myCAF_2 |
| Arid2 | 2.85E-06 | 0.67872729 | 0.645 | 0.63 | 0.05512644 | 4 | Arid2 | myCAF_2 |
| Galns1 | 2.88E-06 | 0.33521512 | 0.345 | 0.411 | 0.05577076 | 4 | Galns | myCAF_2 |
| Socs11 | 2.93E-06 | 0.48686468 | 0.496 | 0.539 | 0.05666666 | 4 | Socs1 | myCAF_2 |
| Cux12 | 3.08E-06 | 0.67906588 | 0.703 | 0.626 | 0.05955704 | 4 | Cux1 | myCAF_2 |
| Rfx71 | 3.17E-06 | 0.26061607 | 0.25 | 0.312 | 0.06131976 | 4 | Rfx7 | myCAF_2 |
| Gmeb11 | 3.24E-06 | 0.32939152 | 0.295 | 0.347 | 0.06259816 | 4 | Gmeb1 | myCAF_2 |
| Aebp2 | 3.33E-06 | 0.4654424 | 0.696 | 0.728 | 0.06444362 | 4 | Aebp2 | myCAF_2 |
| Kansl1 | 3.40E-06 | 0.26167112 | 0.67 | 0.724 | 0.06579503 | 4 | Kansl1 | myCAF_2 |
| Hoxa31 | 3.42E-06 | 0.84551342 | 0.306 | 0.26 | 0.06602573 | 4 | Hoxa3 | myCAF_2 |
| Zbtb7b | 3.63E-06 | 0.85272809 | 0.473 | 0.436 | 0.07016663 | 4 | Zbtb7b | myCAF_2 |
| Grcc10 | 3.71E-06 | 0.38319307 | 0.84 | 0.887 | 0.07167905 | 4 | Grcc10 | myCAF_2 |
| Cebpg | 3.86E-06 | 0.29301713 | 0.555 | 0.646 | 0.07454951 | 4 | Cebpg | myCAF_2 |
| Tpgs1 | 4.13E-06 | 0.36118666 | 0.547 | 0.618 | 0.07975326 | 4 | Tpgs1 | myCAF_2 |
| Tom1 | 4.14E-06 | 0.73487753 | 0.362 | 0.328 | 0.08009321 | 4 | Tom1 | myCAF_2 |
| Acot71 | 4.28E-06 | 0.83514073 | 0.58 | 0.514 | 0.08271029 | 4 | Acot7 | myCAF_2 |
| Pank31 | 4.30E-06 | 0.26740668 | 0.569 | 0.637 | 0.0830398 | 4 | Pank3 | myCAF_2 |
| Ubtd1 | 4.50E-06 | 0.35050355 | 0.61 | 0.667 | 0.08694284 | 4 | Ubtd1 | myCAF_2 |
| Timm44 | 4.58E-06 | 0.29390378 | 0.36 | 0.419 | 0.08859934 | 4 | Timm44 | myCAF_2 |
| Rnf131 | 4.59E-06 | 0.30595023 | 0.527 | 0.575 | 0.08868739 | 4 | Rnf13 | myCAF_2 |
| Mdfic1 | 4.62E-06 | 0.35576997 | 0.736 | 0.775 | 0.08929264 | 4 | Mdfic | myCAF_2 |
| Fam219a | 4.64E-06 | 0.61593244 | 0.327 | 0.29 | 0.08975942 | 4 | Fam219a | myCAF_2 |
| Fkrp1 | 4.71E-06 | 0.38247903 | 0.532 | 0.578 | 0.09103283 | 4 | Fkrp | myCAF_2 |
| Cfh2 | 4.73E-06 | 0.30317816 | 0.848 | 0.859 | 0.09140877 | 4 | Cfh | myCAF_2 |
| Pcsk7 | 4.78E-06 | 0.27342169 | 0.337 | 0.401 | 0.09244025 | 4 | Pcsk7 | myCAF_2 |
| Ccpg1 | 5.03E-06 | 0.31715899 | 0.688 | 0.745 | 0.09730324 | 4 | Ccpg1 | myCAF_2 |
| Tgfb21 | 5.12E-06 | 0.65456858 | 0.644 | 0.606 | 0.09905009 | 4 | Tgfb2 | myCAF_2 |
| Gabbr11 | 5.44E-06 | 0.72796917 | 0.537 | 0.467 | 0.10523964 | 4 | Gabbr1 | myCAF_2 |
| Ctsf1 | 5.45E-06 | 0.55645248 | 0.568 | 0.543 | 0.10544108 | 4 | Ctsf | myCAF_2 |
| Akr7a5 | 5.49E-06 | 0.26153163 | 0.44 | 0.495 | 0.10608942 | 4 | Akr7a5 | myCAF_2 |
| Tut71 | 5.56E-06 | 0.35379788 | 0.74 | 0.763 | 0.10759093 | 4 | Tut7 | myCAF_2 |
| Dhx401 | 5.95E-06 | 0.26202308 | 0.457 | 0.526 | 0.11509043 | 4 | Dhx40 | myCAF_2 |
| Pik3r2 | 6.31E-06 | 0.3276623 | 0.658 | 0.704 | 0.12204173 | 4 | Pik3r2 | myCAF_2 |
| Tbcc1 | 6.35E-06 | 0.25839714 | 0.377 | 0.44 | 0.12281093 | 4 | Tbcc | myCAF_2 |
| Vps181 | 6.45E-06 | 0.28233579 | 0.248 | 0.321 | 0.12473863 | 4 | Vps18 | myCAF_2 |
| Tbc1d10b | 6.47E-06 | 0.43722832 | 0.623 | 0.648 | 0.12503435 | 4 | Tbc1d10b | myCAF_2 |
| Pstpip11 | 6.54E-06 | 0.96122908 | 0.284 | 0.242 | 0.12636746 | 4 | Pstpip1 | myCAF_2 |
| Cyp2d221 | 6.97E-06 | 0.73837247 | 0.434 | 0.408 | 0.13471773 | 4 | Cyp2d22 | myCAF_2 |
| Purb1 | 7.65E-06 | 0.53310705 | 0.811 | 0.803 | 0.14789851 | 4 | Purb | myCAF_2 |
| Cxxc51 | 7.97E-06 | 0.59513802 | 0.794 | 0.803 | 0.15403402 | 4 | Cxxc5 | myCAF_2 |
| BC0340901 | 8.09E-06 | 0.31752117 | 0.411 | 0.476 | 0.15635305 | 4 | BC034090 | myCAF_2 |
| Oxr11 | 8.31E-06 | 0.34693922 | 0.416 | 0.478 | 0.16072181 | 4 | Oxr1 | myCAF_2 |
| Eif4a21 | 8.50E-06 | 0.3665562 | 0.839 | 0.883 | 0.16437929 | 4 | Eif4a2 | myCAF_2 |
| Kcmf11 | 8.52E-06 | 0.38930627 | 0.723 | 0.767 | 0.16477557 | 4 | Kcmf1 | myCAF_2 |
| Raph11 | 8.74E-06 | 0.38621307 | 0.263 | 0.319 | 0.16905075 | 4 | Raph1 | myCAF_2 |
| Emp11 | 8.95E-06 | 0.54109046 | 0.961 | 0.97 | 0.17298865 | 4 | Emp1 | myCAF_2 |
| Pcm11 | 9.32E-06 | 0.27651593 | 0.537 | 0.595 | 0.18026066 | 4 | Pcm1 | myCAF_2 |
| Setd31 | 9.34E-06 | 0.36509205 | 0.659 | 0.712 | 0.18061847 | 4 | Setd3 | myCAF_2 |
| Fzd51 | 9.94E-06 | 0.82769551 | 0.306 | 0.26 | 0.19219614 | 4 | Fzd5 | myCAF_2 |
| Mapkapk51 | 1.02E-05 | 0.27215916 | 0.214 | 0.261 | 0.19728353 | 4 | Mapkapk5 | myCAF_2 |
| Wdr811 | 1.11E-05 | 0.67329142 | 0.408 | 0.393 | 0.21516466 | 4 | Wdr81 | myCAF_2 |
| Stard3 | 1.14E-05 | 0.70404499 | 0.504 | 0.475 | 0.22087605 | 4 | Stard3 | myCAF_2 |
| Rnf138 | 1.14E-05 | 0.38172952 | 0.406 | 0.465 | 0.22112741 | 4 | Rnf138 | myCAF_2 |
| Jcad | 1.19E-05 | 0.26370501 | 0.387 | 0.464 | 0.22991281 | 4 | Jcad | myCAF_2 |
| Mar-71 | 1.21E-05 | 0.29615854 | 0.627 | 0.671 | 0.23388265 | 4 | 7-Mar | myCAF_2 |
| Sf3b11 | 1.23E-05 | 0.41968315 | 0.88 | 0.902 | 0.23875665 | 4 | Sf3b1 | myCAF_2 |
| Lpin21 | 1.24E-05 | 0.27976113 | 0.28 | 0.34 | 0.23983158 | 4 | Lpin2 | myCAF_2 |
| Pbx21 | 1.26E-05 | 0.29221064 | 0.542 | 0.598 | 0.24334454 | 4 | Pbx2 | myCAF_2 |
| Cdk121 | 1.33E-05 | 0.26312446 | 0.496 | 0.54 | 0.25628168 | 4 | Cdk12 | myCAF_2 |
| Washc41 | 1.38E-05 | 0.26561614 | 0.461 | 0.517 | 0.26650235 | 4 | Washc4 | myCAF_2 |
| Dstyk | 1.39E-05 | 0.28776273 | 0.482 | 0.536 | 0.26840879 | 4 | Dstyk | myCAF_2 |
| Atp6v1f1 | 1.43E-05 | 0.26303342 | 0.663 | 0.714 | 0.27688845 | 4 | Atp6v1f | myCAF_2 |
| Gss1 | 1.50E-05 | 0.3426829 | 0.382 | 0.433 | 0.289211 | 4 | Gss | myCAF_2 |
| Zmym21 | 1.51E-05 | 0.27357246 | 0.43 | 0.488 | 0.29141216 | 4 | Zmym2 | myCAF_2 |
| Ldlrad4 | 1.51E-05 | 0.34372512 | 0.528 | 0.554 | 0.29145275 | 4 | Ldlrad4 | myCAF_2 |
| Edc3 | 1.52E-05 | 0.28369497 | 0.453 | 0.51 | 0.2941053 | 4 | Edc3 | myCAF_2 |
| Suz122 | 1.53E-05 | 0.33196888 | 0.372 | 0.413 | 0.2958684 | 4 | Suz12 | myCAF_2 |
| Fam78b1 | 1.54E-05 | 0.66594834 | 0.528 | 0.503 | 0.29710172 | 4 | Fam78b | myCAF_2 |
| Zbtb111 | 1.56E-05 | 0.31630207 | 0.362 | 0.438 | 0.30097685 | 4 | Zbtb11 | myCAF_2 |
| Slc25a512 | 1.69E-05 | 0.32105109 | 0.805 | 0.848 | 0.32640282 | 4 | Slc25a51 | myCAF_2 |
| Emp2 | 1.73E-05 | 0.82497323 | 0.471 | 0.468 | 0.33415109 | 4 | Emp2 | myCAF_2 |
| Gatd3a1 | 1.78E-05 | 0.34670126 | 0.478 | 0.535 | 0.34326364 | 4 | Gatd3a | myCAF_2 |
| Yeats21 | 1.78E-05 | 0.28001803 | 0.275 | 0.326 | 0.34330159 | 4 | Yeats2 | myCAF_2 |
| Spred21 | 1.85E-05 | 0.65080966 | 0.6 | 0.579 | 0.35746472 | 4 | Spred2 | myCAF_2 |
| Sox121 | 1.87E-05 | 0.28593791 | 0.246 | 0.282 | 0.36101052 | 4 | Sox12 | myCAF_2 |
| Map2k41 | 1.92E-05 | 0.27465404 | 0.352 | 0.413 | 0.37078997 | 4 | Map2k4 | myCAF_2 |
| Orai1 | 1.93E-05 | 0.26145793 | 0.219 | 0.272 | 0.37221539 | 4 | Orai1 | myCAF_2 |
| Ttc171 | 1.93E-05 | 0.27777855 | 0.416 | 0.468 | 0.37319225 | 4 | Ttc17 | myCAF_2 |
| Pip5k1a2 | 1.95E-05 | 0.35969438 | 0.584 | 0.605 | 0.37795574 | 4 | Pip5k1a | myCAF_2 |
| Tgfbr11 | 2.02E-05 | 0.28294119 | 0.687 | 0.757 | 0.39052001 | 4 | Tgfbr1 | myCAF_2 |
| Fntb1 | 2.03E-05 | 0.39732608 | 0.314 | 0.358 | 0.39279872 | 4 | Fntb | myCAF_2 |
| Hexim11 | 2.06E-05 | 0.41649709 | 0.619 | 0.691 | 0.39895375 | 4 | Hexim1 | myCAF_2 |
| Ptprm1 | 2.07E-05 | 0.67586486 | 0.663 | 0.631 | 0.40006865 | 4 | Ptprm | myCAF_2 |
| Haus31 | 2.09E-05 | 0.29351591 | 0.229 | 0.266 | 0.40415809 | 4 | Haus3 | myCAF_2 |
| Vopp11 | 2.16E-05 | 0.31335836 | 0.499 | 0.548 | 0.41732967 | 4 | Vopp1 | myCAF_2 |
| Golga4 | 2.16E-05 | 0.30100952 | 0.563 | 0.618 | 0.41772265 | 4 | Golga4 | myCAF_2 |
| Ccdc9 | 2.26E-05 | 0.39711591 | 0.584 | 0.626 | 0.43681306 | 4 | Ccdc9 | myCAF_2 |
| Btbd2 | 2.29E-05 | 0.29445445 | 0.477 | 0.539 | 0.44280558 | 4 | Btbd2 | myCAF_2 |
| Blcap1 | 2.30E-05 | 0.68749012 | 0.641 | 0.637 | 0.44444425 | 4 | Blcap | myCAF_2 |
| Cic | 2.31E-05 | 0.30141361 | 0.633 | 0.689 | 0.4463788 | 4 | Cic | myCAF_2 |
| Phldb1 | 2.36E-05 | 0.40124633 | 0.752 | 0.776 | 0.45610156 | 4 | Phldb1 | myCAF_2 |
| Ifrd12 | 2.46E-05 | 0.54273127 | 0.762 | 0.746 | 0.47564153 | 4 | Ifrd1 | myCAF_2 |
| Ier51 | 2.52E-05 | 0.75894477 | 0.778 | 0.753 | 0.48689872 | 4 | Ier5 | myCAF_2 |
| Sil1 | 2.56E-05 | 0.27416912 | 0.423 | 0.484 | 0.4942349 | 4 | Sil1 | myCAF_2 |
| Ddah2 | 2.65E-05 | 0.31289316 | 0.648 | 0.679 | 0.51294284 | 4 | Ddah2 | myCAF_2 |
| Tnfsf91 | 2.81E-05 | 0.68343462 | 0.329 | 0.263 | 0.54419462 | 4 | Tnfsf9 | myCAF_2 |
| Sgcd1 | 2.94E-05 | 0.91604868 | 0.337 | 0.319 | 0.56923302 | 4 | Sgcd | myCAF_2 |
| Sesn21 | 2.98E-05 | 0.74407822 | 0.286 | 0.268 | 0.57651346 | 4 | Sesn2 | myCAF_2 |
| Utrn | 2.99E-05 | 0.70628467 | 0.59 | 0.578 | 0.57735464 | 4 | Utrn | myCAF_2 |
| Nudt41 | 3.10E-05 | 0.69017742 | 0.9 | 0.882 | 0.59881376 | 4 | Nudt4 | myCAF_2 |
| Fbxo30 | 3.16E-05 | 0.76637065 | 0.495 | 0.468 | 0.61110561 | 4 | Fbxo30 | myCAF_2 |
| Cyth21 | 3.17E-05 | 0.32292425 | 0.707 | 0.733 | 0.61270616 | 4 | Cyth2 | myCAF_2 |
| Pskh11 | 3.28E-05 | 0.26668541 | 0.355 | 0.411 | 0.63385152 | 4 | Pskh1 | myCAF_2 |
| Ypel3 | 3.29E-05 | 0.34640389 | 0.779 | 0.796 | 0.63616866 | 4 | Ypel3 | myCAF_2 |
| Adcy61 | 3.31E-05 | 0.66913515 | 0.741 | 0.713 | 0.64051531 | 4 | Adcy6 | myCAF_2 |
| 5-Sep | 3.58E-05 | 0.38874094 | 0.656 | 0.693 | 0.69220794 | 4 | 5-Sep | myCAF_2 |
| Ctdspl22 | 3.58E-05 | 0.37105708 | 0.461 | 0.479 | 0.69292898 | 4 | Ctdspl2 | myCAF_2 |
| Trpm4 | 3.66E-05 | 0.31386047 | 0.324 | 0.384 | 0.7074503 | 4 | Trpm4 | myCAF_2 |
| Sympk | 3.72E-05 | 0.33315669 | 0.658 | 0.7 | 0.71846817 | 4 | Sympk | myCAF_2 |
| Pdpk11 | 3.74E-05 | 0.29735975 | 0.572 | 0.627 | 0.72227133 | 4 | Pdpk1 | myCAF_2 |
| Mark4 | 3.76E-05 | 0.26338428 | 0.575 | 0.635 | 0.72699565 | 4 | Mark4 | myCAF_2 |
| Fam210a1 | 3.95E-05 | 0.37296937 | 0.368 | 0.431 | 0.76340056 | 4 | Fam210a | myCAF_2 |
| Slc7a7 | 4.04E-05 | 0.8355323 | 0.273 | 0.221 | 0.78088623 | 4 | Slc7a7 | myCAF_2 |
| Mosmo1 | 4.10E-05 | 0.2742934 | 0.49 | 0.552 | 0.79320805 | 4 | Mosmo | myCAF_2 |
| Osbpl52 | 4.17E-05 | 0.33675157 | 0.663 | 0.71 | 0.80611011 | 4 | Osbpl5 | myCAF_2 |
| Mef2a1 | 4.34E-05 | 0.60188019 | 0.608 | 0.629 | 0.83931082 | 4 | Mef2a | myCAF_2 |
| Timmdc11 | 4.35E-05 | 0.28726898 | 0.395 | 0.43 | 0.84120932 | 4 | Timmdc1 | myCAF_2 |
| Clip21 | 4.42E-05 | 0.66814822 | 0.727 | 0.717 | 0.85405721 | 4 | Clip2 | myCAF_2 |
| Bri3bp1 | 4.58E-05 | 0.39462793 | 0.631 | 0.647 | 0.88577645 | 4 | Bri3bp | myCAF_2 |
| Hdac4 | 4.63E-05 | 0.32880406 | 0.552 | 0.596 | 0.89459396 | 4 | Hdac4 | myCAF_2 |
| Ntan11 | 4.69E-05 | 0.30350826 | 0.402 | 0.466 | 0.90580288 | 4 | Ntan1 | myCAF_2 |
| Mau2 | 4.70E-05 | 0.3481251 | 0.579 | 0.619 | 0.90782882 | 4 | Mau2 | myCAF_2 |
| Plekhg3 | 4.72E-05 | 0.2673449 | 0.455 | 0.506 | 0.91174247 | 4 | Plekhg3 | myCAF_2 |
| Atp11c2 | 4.76E-05 | 0.30885994 | 0.259 | 0.312 | 0.92056177 | 4 | Atp11c | myCAF_2 |
| Zfp319 | 4.77E-05 | 0.28136285 | 0.279 | 0.327 | 0.92316665 | 4 | Zfp319 | myCAF_2 |
| Abl2 | 4.84E-05 | 0.35341156 | 0.504 | 0.57 | 0.93560971 | 4 | Abl2 | myCAF_2 |
| Gramd1a | 4.95E-05 | 0.75368041 | 0.612 | 0.601 | 0.95644941 | 4 | Gramd1a | myCAF_2 |
| Fyttd11 | 5.19E-05 | 0.37818972 | 0.645 | 0.694 | 1 | 4 | Fyttd1 | myCAF_2 |
| Jph21 | 5.32E-05 | 0.76985351 | 0.311 | 0.27 | 1 | 4 | Jph2 | myCAF_2 |
| Tob12 | 5.42E-05 | 0.40870785 | 0.435 | 0.492 | 1 | 4 | Tob1 | myCAF_2 |
| Rab21 | 5.49E-05 | 0.47132544 | 0.893 | 0.899 | 1 | 4 | Rab21 | myCAF_2 |
| Sbno1 | 5.51E-05 | 0.34313143 | 0.554 | 0.6 | 1 | 4 | Sbno1 | myCAF_2 |
| Arid4b1 | 5.58E-05 | 0.25595207 | 0.461 | 0.523 | 1 | 4 | Arid4b | myCAF_2 |
| Mark2 | 5.61E-05 | 0.27441336 | 0.43 | 0.486 | 1 | 4 | Mark2 | myCAF_2 |
| Zfp217 | 5.76E-05 | 0.29267916 | 0.584 | 0.638 | 1 | 4 | Zfp217 | myCAF_2 |
| Dcp1a | 5.89E-05 | 0.3398827 | 0.333 | 0.392 | 1 | 4 | Dcp1a | myCAF_2 |
| Tmem135 | 6.08E-05 | 0.31577339 | 0.343 | 0.4 | 1 | 4 | Tmem135 | myCAF_2 |
| Rem12 | 6.15E-05 | 0.39993155 | 0.459 | 0.448 | 1 | 4 | Rem1 | myCAF_2 |
| Dyrk2 | 6.23E-05 | 0.35041697 | 0.549 | 0.618 | 1 | 4 | Dyrk2 | myCAF_2 |
| Otud41 | 6.27E-05 | 0.28048452 | 0.405 | 0.442 | 1 | 4 | Otud4 | myCAF_2 |
| Ube2g11 | 6.31E-05 | 0.37403433 | 0.586 | 0.655 | 1 | 4 | Ube2g1 | myCAF_2 |
| Pip4p11 | 6.48E-05 | 0.31805515 | 0.593 | 0.646 | 1 | 4 | Pip4p1 | myCAF_2 |
| Ado2 | 6.52E-05 | 0.37117483 | 0.448 | 0.479 | 1 | 4 | Ado | myCAF_2 |
| Acox11 | 6.99E-05 | 0.28263994 | 0.512 | 0.559 | 1 | 4 | Acox1 | myCAF_2 |
| Rasal2 | 7.06E-05 | 0.27893967 | 0.211 | 0.257 | 1 | 4 | Rasal2 | myCAF_2 |
| Zfpm1 | 7.22E-05 | 0.82978346 | 0.391 | 0.361 | 1 | 4 | Zfpm1 | myCAF_2 |
| Zranb11 | 7.26E-05 | 0.28467259 | 0.452 | 0.499 | 1 | 4 | Zranb1 | myCAF_2 |
| Wdr451 | 7.54E-05 | 0.25667621 | 0.473 | 0.525 | 1 | 4 | Wdr45 | myCAF_2 |
| BC029722 | 7.59E-05 | 0.2560806 | 0.494 | 0.544 | 1 | 4 | BC029722 | myCAF_2 |
| Ndrg21 | 7.67E-05 | 0.58879874 | 0.292 | 0.257 | 1 | 4 | Ndrg2 | myCAF_2 |
| Rptor | 7.84E-05 | 0.29739168 | 0.437 | 0.481 | 1 | 4 | Rptor | myCAF_2 |
| Naalad2 | 7.89E-05 | 0.27188853 | 0.234 | 0.265 | 1 | 4 | Naalad2 | myCAF_2 |
| Per11 | 7.89E-05 | 0.67005358 | 0.569 | 0.545 | 1 | 4 | Per1 | myCAF_2 |
| Got1 | 7.93E-05 | 0.69371792 | 0.601 | 0.565 | 1 | 4 | Got1 | myCAF_2 |
| Zfp6551 | 7.97E-05 | 0.38096155 | 0.503 | 0.545 | 1 | 4 | Zfp655 | myCAF_2 |
| Slc27a11 | 8.14E-05 | 0.84714965 | 0.413 | 0.356 | 1 | 4 | Slc27a1 | myCAF_2 |
| Yy11 | 8.61E-05 | 0.43104225 | 0.848 | 0.847 | 1 | 4 | Yy1 | myCAF_2 |
| Nfe2l1 | 8.69E-05 | 0.40329449 | 0.858 | 0.896 | 1 | 4 | Nfe2l1 | myCAF_2 |
| Numb | 8.79E-05 | 0.28174996 | 0.443 | 0.501 | 1 | 4 | Numb | myCAF_2 |
| Cirbp1 | 8.89E-05 | 0.36196408 | 0.547 | 0.587 | 1 | 4 | Cirbp | myCAF_2 |
| Cmtm61 | 8.89E-05 | 0.37434816 | 0.665 | 0.7 | 1 | 4 | Cmtm6 | myCAF_2 |
| Tspyl1 | 9.14E-05 | 0.46105303 | 0.584 | 0.64 | 1 | 4 | Tspyl1 | myCAF_2 |
| Phc3 | 9.15E-05 | 0.26889944 | 0.487 | 0.552 | 1 | 4 | Phc3 | myCAF_2 |
| Cited21 | 9.19E-05 | 0.40792022 | 0.256 | 0.27 | 1 | 4 | Cited2 | myCAF_2 |
| Nars | 9.29E-05 | 0.36832211 | 0.618 | 0.672 | 1 | 4 | Nars | myCAF_2 |
| Ubxn71 | 9.36E-05 | 0.36057431 | 0.618 | 0.646 | 1 | 4 | Ubxn7 | myCAF_2 |
| Rnf1141 | 9.45E-05 | 0.33862477 | 0.411 | 0.476 | 1 | 4 | Rnf114 | myCAF_2 |
| Atxn31 | 9.59E-05 | 0.38915045 | 0.277 | 0.324 | 1 | 4 | Atxn3 | myCAF_2 |
| Agl1 | 9.68E-05 | 0.85458626 | 0.339 | 0.298 | 1 | 4 | Agl | myCAF_2 |
| Idnk | 9.78E-05 | 0.25674213 | 0.322 | 0.392 | 1 | 4 | Idnk | myCAF_2 |
| Mpg1 | 9.82E-05 | 0.40921559 | 0.476 | 0.527 | 1 | 4 | Mpg | myCAF_2 |
| Slc27a4 | 9.88E-05 | 0.29918774 | 0.544 | 0.607 | 1 | 4 | Slc27a4 | myCAF_2 |
| Pold4 | 0.00010317 | 0.6736379 | 0.62 | 0.604 | 1 | 4 | Pold4 | myCAF_2 |
| Atg2a | 0.00010669 | 0.3159998 | 0.426 | 0.484 | 1 | 4 | Atg2a | myCAF_2 |
| Hes61 | 0.00010817 | 0.73611405 | 0.332 | 0.329 | 1 | 4 | Hes6 | myCAF_2 |
| Gdf111 | 0.00010836 | 0.30604392 | 0.287 | 0.319 | 1 | 4 | Gdf11 | myCAF_2 |
| Gab2 | 0.00010843 | 0.73779203 | 0.437 | 0.408 | 1 | 4 | Gab2 | myCAF_2 |
| Myo1e1 | 0.00010855 | 0.68708768 | 0.501 | 0.448 | 1 | 4 | Myo1e | myCAF_2 |
| Abcg11 | 0.00011073 | 0.8197044 | 0.264 | 0.208 | 1 | 4 | Abcg1 | myCAF_2 |
| B4galnt12 | 0.00011437 | 0.67920872 | 0.596 | 0.53 | 1 | 4 | B4galnt1 | myCAF_2 |
| Gamt1 | 0.0001157 | 0.86132796 | 0.323 | 0.293 | 1 | 4 | Gamt | myCAF_2 |
| Zfp511 | 0.00011893 | 0.27569252 | 0.309 | 0.359 | 1 | 4 | Zfp511 | myCAF_2 |
| Kbtbd21 | 0.00011921 | 0.36424711 | 0.555 | 0.608 | 1 | 4 | Kbtbd2 | myCAF_2 |

| | | | | | | | | |
|---|---|---|---|---|---|---|---|---|
| Cttnbp2nl | 0.00012078 | 0.65663228 | 0.572 | 0.547 | 1 | 4 | Cttnbp2nl | myCAF_2 |
| Mecp21 | 0.00012183 | 0.38359595 | 0.455 | 0.505 | 1 | 4 | Mecp2 | myCAF_2 |
| Ldlrad3 | 0.00012268 | 0.76351284 | 0.382 | 0.364 | 1 | 4 | Ldlrad3 | myCAF_2 |
| Trmt1 | 0.00012343 | 0.27714299 | 0.305 | 0.35 | 1 | 4 | Trmt1 | myCAF_2 |
| Tex264 | 0.00012585 | 0.35744632 | 0.766 | 0.798 | 1 | 4 | Tex264 | myCAF_2 |
| Togaram1 | 0.00012719 | 0.27147344 | 0.395 | 0.452 | 1 | 4 | Togaram1 | myCAF_2 |
| Pgrmc11 | 0.0001272 | 0.37055255 | 0.782 | 0.839 | 1 | 4 | Pgrmc1 | myCAF_2 |
| Znrf11 | 0.0001274 | 0.55818669 | 0.693 | 0.692 | 1 | 4 | Znrf1 | myCAF_2 |
| Mta11 | 0.00012939 | 0.41105861 | 0.822 | 0.831 | 1 | 4 | Mta1 | myCAF_2 |
| Lrch4 | 0.00012965 | 0.33330593 | 0.46 | 0.528 | 1 | 4 | Lrch4 | myCAF_2 |
| Cd1511 | 0.00013292 | 0.33084267 | 0.594 | 0.648 | 1 | 4 | Cd151 | myCAF_2 |
| Cog4 | 0.00013332 | 0.32951259 | 0.333 | 0.381 | 1 | 4 | Cog4 | myCAF_2 |
| Tmem11 | 0.00013661 | 0.43847809 | 0.516 | 0.576 | 1 | 4 | Tmem11 | myCAF_2 |
| E2f11 | 0.00013778 | 0.48724615 | 0.289 | 0.29 | 1 | 4 | E2f1 | myCAF_2 |
| Hoxb31 | 0.00013908 | 0.93592432 | 0.284 | 0.241 | 1 | 4 | Hoxb3 | myCAF_2 |
| Spred11 | 0.00014201 | 0.62151019 | 0.941 | 0.929 | 1 | 4 | Spred1 | myCAF_2 |
| Shmt21 | 0.00014212 | 0.47749332 | 0.691 | 0.704 | 1 | 4 | Shmt2 | myCAF_2 |
| Fmnl31 | 0.00014233 | 0.45724959 | 0.764 | 0.735 | 1 | 4 | Fmnl3 | myCAF_2 |
| Sgta | 0.00014625 | 0.29759875 | 0.332 | 0.388 | 1 | 4 | Sgta | myCAF_2 |
| Irx31 | 0.00014738 | 0.80032419 | 0.467 | 0.408 | 1 | 4 | Irx3 | myCAF_2 |
| Abhd42 | 0.00014802 | 0.32498945 | 0.69 | 0.76 | 1 | 4 | Abhd4 | myCAF_2 |
| Rxrb | 0.00014897 | 0.39782493 | 0.587 | 0.623 | 1 | 4 | Rxrb | myCAF_2 |
| Ids | 0.0001492 | 0.30590544 | 0.47 | 0.505 | 1 | 4 | Ids | myCAF_2 |
| Foxk1 | 0.00015 | 0.74726124 | 0.554 | 0.532 | 1 | 4 | Foxk1 | myCAF_2 |
| Iars | 0.00015627 | 0.36079007 | 0.54 | 0.597 | 1 | 4 | Iars | myCAF_2 |
| Crtc31 | 0.00015788 | 0.33178209 | 0.482 | 0.544 | 1 | 4 | Crtc3 | myCAF_2 |
| Cables2 | 0.0001585 | 0.26507423 | 0.27 | 0.313 | 1 | 4 | Cables2 | myCAF_2 |
| Flywch1 | 0.00015851 | 0.33285902 | 0.491 | 0.547 | 1 | 4 | Flywch1 | myCAF_2 |
| Piezo1 | 0.00016024 | 0.517988 | 0.777 | 0.771 | 1 | 4 | Piezo1 | myCAF_2 |
| Socs21 | 0.00016398 | 0.31689601 | 0.341 | 0.379 | 1 | 4 | Socs2 | myCAF_2 |
| Stk241 | 0.00016819 | 0.33948306 | 0.743 | 0.799 | 1 | 4 | Stk24 | myCAF_2 |
| Zfand2b | 0.00017318 | 0.29429163 | 0.342 | 0.387 | 1 | 4 | Zfand2b | myCAF_2 |
| R3hdm1 | 0.00017541 | 0.37624021 | 0.521 | 0.576 | 1 | 4 | R3hdm1 | myCAF_2 |
| Gm2a1 | 0.00017782 | 0.57877125 | 0.646 | 0.679 | 1 | 4 | Gm2a | myCAF_2 |
| Gli21 | 0.00017865 | 0.60716643 | 0.41 | 0.401 | 1 | 4 | Gli2 | myCAF_2 |
| Zfp292 | 0.00018572 | 0.3197389 | 0.472 | 0.547 | 1 | 4 | Zfp292 | myCAF_2 |
| Mrtfa | 0.00018758 | 0.40547756 | 0.53 | 0.563 | 1 | 4 | Mrtfa | myCAF_2 |
| B4galt11 | 0.00018927 | 0.33712628 | 0.599 | 0.661 | 1 | 4 | B4galt1 | myCAF_2 |
| Ercc11 | 0.00019267 | 0.40595443 | 0.486 | 0.543 | 1 | 4 | Ercc1 | myCAF_2 |
| Hmgcr1 | 0.00019624 | 0.34554216 | 0.602 | 0.637 | 1 | 4 | Hmgcr | myCAF_2 |
| Usp311 | 0.00019752 | 0.30707471 | 0.305 | 0.358 | 1 | 4 | Usp31 | myCAF_2 |
| Ttc142 | 0.00020241 | 0.42922866 | 0.454 | 0.508 | 1 | 4 | Ttc14 | myCAF_2 |
| Mink1 | 0.00020756 | 0.75278146 | 0.403 | 0.357 | 1 | 4 | Mink1 | myCAF_2 |
| Cotl1 | 0.00020896 | 0.6527902 | 0.669 | 0.662 | 1 | 4 | Cotl1 | myCAF_2 |
| Plppr31 | 0.00022066 | 0.50818214 | 0.403 | 0.369 | 1 | 4 | Plppr3 | myCAF_2 |
| Atp7a1 | 0.00022424 | 0.25229301 | 0.437 | 0.506 | 1 | 4 | Atp7a | myCAF_2 |
| Tmem471 | 0.00022822 | 0.86266628 | 0.396 | 0.342 | 1 | 4 | Tmem47 | myCAF_2 |
| Kdm4c | 0.00022891 | 0.33530357 | 0.325 | 0.369 | 1 | 4 | Kdm4c | myCAF_2 |
| Kalrn1 | 0.00022972 | 0.51447919 | 0.523 | 0.498 | 1 | 4 | Kalrn | myCAF_2 |
| Gpr1371 | 0.00023107 | 0.27663323 | 0.378 | 0.43 | 1 | 4 | Gpr137 | myCAF_2 |
| Pakap2 | 0.00023502 | 0.51317458 | 0.539 | 0.469 | 1 | 4 | Pakap | myCAF_2 |
| Lrp5 | 0.00024142 | 0.65184546 | 0.691 | 0.7 | 1 | 4 | Lrp5 | myCAF_2 |
| Ndufa61 | 0.00024181 | 0.35499223 | 0.776 | 0.806 | 1 | 4 | Ndufa6 | myCAF_2 |
| Pex191 | 0.00024212 | 0.338152 | 0.615 | 0.662 | 1 | 4 | Pex19 | myCAF_2 |
| Pura1 | 0.00024728 | 0.43622471 | 0.797 | 0.814 | 1 | 4 | Pura | myCAF_2 |
| Plxnd11 | 0.00025384 | 0.36242416 | 0.531 | 0.557 | 1 | 4 | Plxnd1 | myCAF_2 |
| Thoc21 | 0.00025423 | 0.33904412 | 0.465 | 0.493 | 1 | 4 | Thoc2 | myCAF_2 |
| Bsdc1 | 0.00025525 | 0.25569686 | 0.434 | 0.507 | 1 | 4 | Bsdc1 | myCAF_2 |
| Hs3st12 | 0.00026054 | 0.34096569 | 0.662 | 0.609 | 1 | 4 | Hs3st1 | myCAF_2 |
| Rnf169 | 0.00026549 | 0.26789692 | 0.4 | 0.447 | 1 | 4 | Rnf169 | myCAF_2 |
| Zfp664 | 0.00026848 | 0.32190793 | 0.401 | 0.438 | 1 | 4 | Zfp664 | myCAF_2 |
| Ahdc1 | 0.00027024 | 0.70052033 | 0.695 | 0.668 | 1 | 4 | Ahdc1 | myCAF_2 |
| Cog31 | 0.00027229 | 0.41085198 | 0.427 | 0.487 | 1 | 4 | Cog3 | myCAF_2 |
| Arl4c1 | 0.00027535 | 0.43072394 | 0.479 | 0.487 | 1 | 4 | Arl4c | myCAF_2 |
| Pmm1 | 0.0002813 | 0.37980614 | 0.546 | 0.593 | 1 | 4 | Pmm1 | myCAF_2 |
| Vps111 | 0.00029161 | 0.25255905 | 0.428 | 0.48 | 1 | 4 | Vps11 | myCAF_2 |
| Fiz1 | 0.00029508 | 0.26183998 | 0.291 | 0.337 | 1 | 4 | Fiz1 | myCAF_2 |
| Insig11 | 0.0002981 | 0.41652179 | 0.696 | 0.73 | 1 | 4 | Insig1 | myCAF_2 |
| Def61 | 0.00030429 | 0.25037107 | 0.262 | 0.301 | 1 | 4 | Def6 | myCAF_2 |
| Bloc1s41 | 0.00030683 | 0.31642112 | 0.258 | 0.304 | 1 | 4 | Bloc1s4 | myCAF_2 |
| Dot1l1 | 0.0003082 | 0.35261534 | 0.423 | 0.471 | 1 | 4 | Dot1l | myCAF_2 |
| Zdhhc1 | 0.00031252 | 0.27019342 | 0.376 | 0.431 | 1 | 4 | Zdhhc1 | myCAF_2 |
| S100pbp1 | 0.00031542 | 0.26492266 | 0.3 | 0.348 | 1 | 4 | S100pbp | myCAF_2 |
| Fbxw11 | 0.00031888 | 0.30709649 | 0.679 | 0.729 | 1 | 4 | Fbxw11 | myCAF_2 |
| Imp3 | 0.00031953 | 0.35015552 | 0.565 | 0.604 | 1 | 4 | Imp3 | myCAF_2 |
| Med31 | 0.00032857 | 0.38585262 | 0.294 | 0.335 | 1 | 4 | Med31 | myCAF_2 |
| Rnf2141 | 0.00032977 | 0.31190946 | 0.464 | 0.521 | 1 | 4 | Rnf214 | myCAF_2 |
| Clk41 | 0.00033373 | 0.32642394 | 0.49 | 0.526 | 1 | 4 | Clk4 | myCAF_2 |
| Eea1 | 0.00033542 | 0.42199303 | 0.519 | 0.568 | 1 | 4 | Eea1 | myCAF_2 |
| Cdkn1b1 | 0.00033731 | 0.5800825 | 0.739 | 0.775 | 1 | 4 | Cdkn1b | myCAF_2 |
| Evl1 | 0.00036719 | 0.35852039 | 0.704 | 0.719 | 1 | 4 | Evl | myCAF_2 |
| Ampd21 | 0.0003678 | 0.48590901 | 0.629 | 0.684 | 1 | 4 | Ampd2 | myCAF_2 |
| Kdm3b | 0.00036814 | 0.38158441 | 0.591 | 0.638 | 1 | 4 | Kdm3b | myCAF_2 |
| Cdk171 | 0.000369 | 0.43020311 | 0.439 | 0.471 | 1 | 4 | Cdk17 | myCAF_2 |
| Slc35e2 | 0.00038233 | 0.26068813 | 0.227 | 0.268 | 1 | 4 | Slc35e2 | myCAF_2 |
| Clk31 | 0.00038515 | 0.37701926 | 0.477 | 0.54 | 1 | 4 | Clk3 | myCAF_2 |
| Ttc33 | 0.00038774 | 0.259591 | 0.362 | 0.412 | 1 | 4 | Ttc33 | myCAF_2 |
| Nampt1 | 0.00041066 | 0.69988939 | 0.407 | 0.396 | 1 | 4 | Nampt | myCAF_2 |
| Mical3 | 0.00041097 | 0.33951488 | 0.313 | 0.359 | 1 | 4 | Mical3 | myCAF_2 |
| Trim39 | 0.00042478 | 0.32405898 | 0.237 | 0.278 | 1 | 4 | Trim39 | myCAF_2 |
| Vps54 | 0.00042535 | 0.26062908 | 0.384 | 0.447 | 1 | 4 | Vps54 | myCAF_2 |
| Zcchc21 | 0.00043129 | 0.25807592 | 0.232 | 0.282 | 1 | 4 | Zcchc2 | myCAF_2 |
| Zfand2a1 | 0.00043938 | 0.43108898 | 0.233 | 0.278 | 1 | 4 | Zfand2a | myCAF_2 |
| Btg22 | 0.00044161 | 0.79781391 | 0.653 | 0.63 | 1 | 4 | Btg2 | myCAF_2 |
| Clk11 | 0.0004577 | 0.32397655 | 0.776 | 0.819 | 1 | 4 | Clk1 | myCAF_2 |
| Tasor1 | 0.00045839 | 0.32504491 | 0.397 | 0.452 | 1 | 4 | Tasor | myCAF_2 |
| Klhl7 | 0.00047108 | 0.27336934 | 0.411 | 0.453 | 1 | 4 | Klhl7 | myCAF_2 |
| Pcmtd21 | 0.0004794 | 0.29220924 | 0.325 | 0.373 | 1 | 4 | Pcmtd2 | myCAF_2 |
| Patz12 | 0.00048269 | 0.32264606 | 0.413 | 0.462 | 1 | 4 | Patz1 | myCAF_2 |
| Phc1 | 0.00049325 | 0.2745448 | 0.253 | 0.297 | 1 | 4 | Phc1 | myCAF_2 |
| Hgs | 0.00049541 | 0.36372117 | 0.648 | 0.693 | 1 | 4 | Hgs | myCAF_2 |
| Ywhaz | 0.00049733 | 0.37028535 | 0.426 | 0.476 | 1 | 4 | Ywhaz | myCAF_2 |
| Plcg11 | 0.00050145 | 0.56789713 | 0.673 | 0.665 | 1 | 4 | Plcg1 | myCAF_2 |
| Rnft11 | 0.00051712 | 0.44058587 | 0.497 | 0.565 | 1 | 4 | Rnft1 | myCAF_2 |
| DEPRECATED-l | 0.0005358 | 0.28075259 | 0.343 | 0.395 | 1 | 4 | DEPRECATED-l | myCAF_2 |
| Zfp6181 | 0.00055296 | 0.78251431 | 0.425 | 0.378 | 1 | 4 | Zfp618 | myCAF_2 |
| Cwc25 | 0.00055865 | 0.25924631 | 0.291 | 0.343 | 1 | 4 | Cwc25 | myCAF_2 |
| Carmil1 | 0.00056116 | 0.2638832 | 0.264 | 0.312 | 1 | 4 | Carmil1 | myCAF_2 |
| Prkx1 | 0.00056831 | 0.65945561 | 0.325 | 0.296 | 1 | 4 | Prkx | myCAF_2 |
| Map2k7 | 0.00057156 | 0.29915725 | 0.552 | 0.608 | 1 | 4 | Map2k7 | myCAF_2 |
| Klhl12 | 0.00057428 | 0.36394779 | 0.274 | 0.328 | 1 | 4 | Klhl12 | myCAF_2 |
| Zfp8711 | 0.00057718 | 0.31576461 | 0.503 | 0.554 | 1 | 4 | Zfp871 | myCAF_2 |
| Taf13 | 0.00059691 | 0.2598654 | 0.248 | 0.291 | 1 | 4 | Taf13 | myCAF_2 |
| Wdr911 | 0.00059781 | 0.63253023 | 0.378 | 0.368 | 1 | 4 | Wdr91 | myCAF_2 |
| Micu1 | 0.00062405 | 0.26776405 | 0.503 | 0.552 | 1 | 4 | Micu1 | myCAF_2 |
| Sos1 | 0.00062531 | 0.37134503 | 0.343 | 0.379 | 1 | 4 | Sos1 | myCAF_2 |
| Uba7 | 0.00065346 | 0.37280249 | 0.357 | 0.397 | 1 | 4 | Uba7 | myCAF_2 |
| Map3k11 | 0.00067563 | 0.44343472 | 0.208 | 0.255 | 1 | 4 | Map3k11 | myCAF_2 |
| Git2 | 0.0006811 | 0.44792387 | 0.58 | 0.598 | 1 | 4 | Git2 | myCAF_2 |
| Cars1 | 0.00068117 | 0.37941078 | 0.503 | 0.567 | 1 | 4 | Cars | myCAF_2 |
| Vapa1 | 0.00068377 | 0.28591188 | 0.517 | 0.56 | 1 | 4 | Vapa | myCAF_2 |
| Tex101 | 0.00068427 | 0.37378339 | 0.397 | 0.431 | 1 | 4 | Tex10 | myCAF_2 |
| Ptpn211 | 0.00069206 | 0.26300164 | 0.399 | 0.449 | 1 | 4 | Ptpn21 | myCAF_2 |
| Mapre21 | 0.00072596 | 0.36437933 | 0.439 | 0.505 | 1 | 4 | Mapre2 | myCAF_2 |
| Pcdhgb21 | 0.00074161 | 0.26472711 | 0.269 | 0.29 | 1 | 4 | Pcdhgb2 | myCAF_2 |
| Rsbn11 | 0.0007444 | 0.89067698 | 0.431 | 0.416 | 1 | 4 | Rsbn1 | myCAF_2 |
| Kmt2c1 | 0.00075363 | 0.260349 | 0.32 | 0.365 | 1 | 4 | Kmt2c | myCAF_2 |
| Clip3 | 0.00075871 | 0.32139841 | 0.71 | 0.747 | 1 | 4 | Clip3 | myCAF_2 |
| Usp22 | 0.00077461 | 0.38291917 | 0.68 | 0.751 | 1 | 4 | Usp22 | myCAF_2 |
| Rmdn3 | 0.00077535 | 0.41190824 | 0.28 | 0.308 | 1 | 4 | Rmdn3 | myCAF_2 |
| Lpcat12 | 0.0007962 | 0.32480599 | 0.477 | 0.513 | 1 | 4 | Lpcat1 | myCAF_2 |
| Fzd81 | 0.0008016 | 0.32713063 | 0.233 | 0.292 | 1 | 4 | Fzd8 | myCAF_2 |
| Ccbe11 | 0.00080177 | 0.52285735 | 0.756 | 0.729 | 1 | 4 | Ccbe1 | myCAF_2 |
| Nudt9 | 0.00081039 | 0.385176 | 0.61 | 0.674 | 1 | 4 | Nudt9 | myCAF_2 |
| Ogfod1 | 0.00082087 | 0.26832237 | 0.222 | 0.27 | 1 | 4 | Ogfod1 | myCAF_2 |
| Cdc42ep5 | 0.00084057 | 0.29158455 | 0.36 | 0.4 | 1 | 4 | Cdc42ep5 | myCAF_2 |
| Itpkc1 | 0.00084403 | 0.34820533 | 0.279 | 0.323 | 1 | 4 | Itpkc | myCAF_2 |
| Tox4 | 0.00086345 | 0.28732509 | 0.547 | 0.588 | 1 | 4 | Tox4 | myCAF_2 |
| Slc25a461 | 0.00086424 | 0.27636835 | 0.316 | 0.361 | 1 | 4 | Slc25a46 | myCAF_2 |
| Tmem2451 | 0.00088021 | 0.30689425 | 0.36 | 0.414 | 1 | 4 | Tmem245 | myCAF_2 |
| Macf1 | 0.00089638 | 0.3958548 | 0.611 | 0.662 | 1 | 4 | Macf1 | myCAF_2 |
| Itpr21 | 0.00090453 | 0.6575151 | 0.38 | 0.352 | 1 | 4 | Itpr2 | myCAF_2 |
| Ccdc117 | 0.00092709 | 0.50837194 | 0.304 | 0.348 | 1 | 4 | Ccdc117 | myCAF_2 |
| Prdm4 | 0.00093105 | 0.32349221 | 0.294 | 0.343 | 1 | 4 | Prdm4 | myCAF_2 |
| Tbpl11 | 0.00095797 | 0.28727035 | 0.37 | 0.425 | 1 | 4 | Tbpl1 | myCAF_2 |
| Kmt2a | 0.00096497 | 0.38763421 | 0.53 | 0.564 | 1 | 4 | Kmt2a | myCAF_2 |
| Gan | 0.00097446 | 0.37284881 | 0.431 | 0.463 | 1 | 4 | Gan | myCAF_2 |
| Saysd1 | 0.0009936 | 0.28236378 | 0.213 | 0.266 | 1 | 4 | Saysd1 | myCAF_2 |
| Traf61 | 0.0010039 | 0.63450988 | 0.358 | 0.338 | 1 | 4 | Traf6 | myCAF_2 |
| Brd1 | 0.00101651 | 0.38887792 | 0.654 | 0.671 | 1 | 4 | Brd1 | myCAF_2 |
| Tnip2 | 0.00102129 | 0.29638809 | 0.41 | 0.45 | 1 | 4 | Tnip2 | myCAF_2 |
| Efnb11 | 0.00102486 | 0.37162655 | 0.568 | 0.586 | 1 | 4 | Efnb1 | myCAF_2 |
| Tbc1d12 | 0.00103865 | 0.25366098 | 0.235 | 0.277 | 1 | 4 | Tbc1d12 | myCAF_2 |
| Pcnx1 | 0.00104288 | 0.34171301 | 0.593 | 0.635 | 1 | 4 | Pcnx | myCAF_2 |
| Ppp1r12a | 0.0010643 | 0.47496839 | 0.692 | 0.728 | 1 | 4 | Ppp1r12a | myCAF_2 |
| Nudt18 | 0.00107385 | 0.30137939 | 0.276 | 0.318 | 1 | 4 | Nudt18 | myCAF_2 |
| Tmem132a1 | 0.00107738 | 0.67850195 | 0.46 | 0.439 | 1 | 4 | Tmem132a | myCAF_2 |
| Pmvk1 | 0.00111811 | 0.35080103 | 0.251 | 0.288 | 1 | 4 | Pmvk | myCAF_2 |
| Hist1h2bc | 0.00112079 | 0.49477628 | 0.592 | 0.666 | 1 | 4 | Hist1h2bc | myCAF_2 |
| Arntl | 0.00114195 | 0.41984389 | 0.283 | 0.333 | 1 | 4 | Arntl | myCAF_2 |
| Map1b | 0.00116253 | 0.68263589 | 0.513 | 0.512 | 1 | 4 | Map1b | myCAF_2 |
| Fut81 | 0.0011693 | 0.59941732 | 0.613 | 0.61 | 1 | 4 | Fut8 | myCAF_2 |
| Rnf185 | 0.00118152 | 0.26216638 | 0.349 | 0.386 | 1 | 4 | Rnf185 | myCAF_2 |
| Plekhh31 | 0.00121797 | 0.63695391 | 0.3 | 0.261 | 1 | 4 | Plekhh3 | myCAF_2 |
| Etv5 | 0.00121892 | 0.62510373 | 0.626 | 0.61 | 1 | 4 | Etv5 | myCAF_2 |
| Unc119b | 0.00123573 | 0.35216601 | 0.506 | 0.558 | 1 | 4 | Unc119b | myCAF_2 |
| Ino80d | 0.00123753 | 0.32348907 | 0.471 | 0.518 | 1 | 4 | Ino80d | myCAF_2 |
| Eif2ak41 | 0.00124627 | 0.38183989 | 0.384 | 0.425 | 1 | 4 | Eif2ak4 | myCAF_2 |
| Klhl26 | 0.00125647 | 0.41721646 | 0.386 | 0.447 | 1 | 4 | Klhl26 | myCAF_2 |
| Rab5b1 | 0.00126882 | 0.27737822 | 0.529 | 0.578 | 1 | 4 | Rab5b | myCAF_2 |
| Zfand51 | 0.00127512 | 0.45952163 | 0.815 | 0.843 | 1 | 4 | Zfand5 | myCAF_2 |
| Rubcn | 0.00128471 | 0.33100202 | 0.366 | 0.422 | 1 | 4 | Rubcn | myCAF_2 |
| Stx7 | 0.00130579 | 0.62923692 | 0.709 | 0.703 | 1 | 4 | Stx7 | myCAF_2 |
| Ankrd11 | 0.00130893 | 0.41828727 | 0.717 | 0.756 | 1 | 4 | Ankrd11 | myCAF_2 |
| Prkacb | 0.0013678 | 0.69201758 | 0.805 | 0.804 | 1 | 4 | Prkacb | myCAF_2 |
| Sec24b1 | 0.00140343 | 0.41746071 | 0.64 | 0.677 | 1 | 4 | Sec24b | myCAF_2 |
| Slc25a371 | 0.00140349 | 0.66426224 | 0.427 | 0.401 | 1 | 4 | Slc25a37 | myCAF_2 |
| Tapt1 | 0.00144946 | 0.25116775 | 0.364 | 0.399 | 1 | 4 | Tapt1 | myCAF_2 |
| Tmem230 | 0.00154133 | 0.36130621 | 0.52 | 0.567 | 1 | 4 | Tmem230 | myCAF_2 |
| Slc12a6 | 0.00155303 | 0.33954663 | 0.29 | 0.348 | 1 | 4 | Slc12a6 | myCAF_2 |
| Lmbr1l | 0.00156692 | 0.25250108 | 0.386 | 0.436 | 1 | 4 | Lmbr1l | myCAF_2 |
| Ago21 | 0.001574 | 0.64594161 | 0.778 | 0.779 | 1 | 4 | Ago2 | myCAF_2 |
| Gla1 | 0.00157509 | 0.37083354 | 0.33 | 0.358 | 1 | 4 | Gla | myCAF_2 |
| Fam133b1 | 0.00158525 | 0.26031043 | 0.236 | 0.267 | 1 | 4 | Fam133b | myCAF_2 |
| Por | 0.00159124 | 0.69756484 | 0.663 | 0.636 | 1 | 4 | Por | myCAF_2 |
| Ppt21 | 0.00160312 | 0.33845035 | 0.329 | 0.372 | 1 | 4 | Ppt2 | myCAF_2 |
| Klhl281 | 0.00161501 | 0.26116584 | 0.276 | 0.317 | 1 | 4 | Klhl28 | myCAF_2 |
| Stx61 | 0.00161584 | 0.44367731 | 0.681 | 0.71 | 1 | 4 | Stx6 | myCAF_2 |
| Pan21 | 0.00161982 | 0.30286753 | 0.287 | 0.319 | 1 | 4 | Pan2 | myCAF_2 |
| Herpud1 | 0.00164335 | 0.62620695 | 0.661 | 0.74 | 1 | 4 | Herpud1 | myCAF_2 |
| Nudt8 | 0.00166307 | 0.32911651 | 0.279 | 0.327 | 1 | 4 | Nudt8 | myCAF_2 |
| Lemd31 | 0.00166782 | 0.30573826 | 0.475 | 0.525 | 1 | 4 | Lemd3 | myCAF_2 |
| Fam129b1 | 0.00168321 | 0.61418337 | 0.849 | 0.885 | 1 | 4 | Fam129b | myCAF_2 |
| Epha3 | 0.0017056 | 0.43571363 | 0.471 | 0.478 | 1 | 4 | Epha3 | myCAF_2 |
| Vps37a1 | 0.00171102 | 0.34725073 | 0.404 | 0.461 | 1 | 4 | Vps37a | myCAF_2 |
| Vps37c | 0.00174469 | 0.30712775 | 0.314 | 0.374 | 1 | 4 | Vps37c | myCAF_2 |
| Pank4 | 0.00175275 | 0.293463 | 0.23 | 0.261 | 1 | 4 | Pank4 | myCAF_2 |
| Cdh21 | 0.00176891 | 0.29025246 | 0.241 | 0.275 | 1 | 4 | Cdh2 | myCAF_2 |
| Trim31 | 0.00180436 | 0.33047559 | 0.521 | 0.564 | 1 | 4 | Trim3 | myCAF_2 |
| Srrm2 | 0.0018277 | 0.50811665 | 0.853 | 0.871 | 1 | 4 | Srrm2 | myCAF_2 |
| Cpeb21 | 0.00188354 | 0.7428132 | 0.272 | 0.24 | 1 | 4 | Cpeb2 | myCAF_2 |
| Sardh1 | 0.00195632 | 0.2757615 | 0.331 | 0.378 | 1 | 4 | Sardh | myCAF_2 |
| Foxp1 | 0.00198332 | 0.47326656 | 0.853 | 0.881 | 1 | 4 | Foxp1 | myCAF_2 |
| Ghdc | 0.0019911 | 0.34949721 | 0.26 | 0.299 | 1 | 4 | Ghdc | myCAF_2 |
| Ctnnd11 | 0.00204206 | 0.42379671 | 0.872 | 0.895 | 1 | 4 | Ctnnd1 | myCAF_2 |
| Appl21 | 0.00205689 | 0.65135259 | 0.405 | 0.398 | 1 | 4 | Appl2 | myCAF_2 |
| Gas2l1 | 0.00207593 | 0.47041149 | 0.649 | 0.666 | 1 | 4 | Gas2l1 | myCAF_2 |
| Ube2h1 | 0.00207841 | 0.37324268 | 0.68 | 0.724 | 1 | 4 | Ube2h | myCAF_2 |
| Relb | 0.00209467 | 0.40822228 | 0.468 | 0.513 | 1 | 4 | Relb | myCAF_2 |
| Frrs12 | 0.00211787 | 0.33949467 | 0.455 | 0.509 | 1 | 4 | Frrs1 | myCAF_2 |
| Kdm2a | 0.00214044 | 0.60827059 | 0.849 | 0.868 | 1 | 4 | Kdm2a | myCAF_2 |
| Nf11 | 0.00219011 | 0.38774005 | 0.654 | 0.679 | 1 | 4 | Nf1 | myCAF_2 |
| Klhl42 | 0.00222305 | 0.26183653 | 0.318 | 0.363 | 1 | 4 | Klhl42 | myCAF_2 |
| Dpysl21 | 0.00222921 | 0.52979445 | 0.769 | 0.776 | 1 | 4 | Dpysl2 | myCAF_2 |
| Ppp4r1 | 0.00223928 | 0.33736858 | 0.376 | 0.42 | 1 | 4 | Ppp4r1 | myCAF_2 |
| Zfp5801 | 0.00233085 | 0.66346692 | 0.295 | 0.258 | 1 | 4 | Zfp580 | myCAF_2 |
| Fbxl61 | 0.00235486 | 0.26328771 | 0.403 | 0.451 | 1 | 4 | Fbxl6 | myCAF_2 |
| Zfp280d1 | 0.00240444 | 0.3477083 | 0.45 | 0.479 | 1 | 4 | Zfp280d | myCAF_2 |
| Pfn21 | 0.00240808 | 0.52886775 | 0.302 | 0.346 | 1 | 4 | Pfn2 | myCAF_2 |
| Tmem115 | 0.0024548 | 0.40558955 | 0.635 | 0.689 | 1 | 4 | Tmem115 | myCAF_2 |
| Etfrf11 | 0.00262955 | 0.26588266 | 0.374 | 0.423 | 1 | 4 | Etfrf1 | myCAF_2 |
| Ubxn61 | 0.00266679 | 0.30185056 | 0.516 | 0.559 | 1 | 4 | Ubxn6 | myCAF_2 |
| Oga1 | 0.00267348 | 0.43018449 | 0.628 | 0.676 | 1 | 4 | Oga | myCAF_2 |
| Dohh | 0.00272056 | 0.42899983 | 0.408 | 0.449 | 1 | 4 | Dohh | myCAF_2 |
| Bpgm1 | 0.00273291 | 0.507803 | 0.656 | 0.671 | 1 | 4 | Bpgm | myCAF_2 |
| Pip4k2c1 | 0.00277133 | 0.41904638 | 0.519 | 0.55 | 1 | 4 | Pip4k2c | myCAF_2 |
| Atp6v0b1 | 0.00282417 | 0.44782773 | 0.681 | 0.726 | 1 | 4 | Atp6v0b | myCAF_2 |
| Akap9 | 0.00282936 | 0.33774278 | 0.505 | 0.547 | 1 | 4 | Akap9 | myCAF_2 |
| Polr2d1 | 0.0029009 | 0.40036091 | 0.386 | 0.42 | 1 | 4 | Polr2d | myCAF_2 |
| Spopl1 | 0.0029035 | 0.37175861 | 0.42 | 0.457 | 1 | 4 | Spopl | myCAF_2 |
| Arhgdib1 | 0.00291285 | 0.36436737 | 0.653 | 0.611 | 1 | 4 | Arhgdib | myCAF_2 |
| Zfp7151 | 0.00295931 | 0.31837754 | 0.252 | 0.285 | 1 | 4 | Zfp715 | myCAF_2 |
| Rbm12b2 | 0.00296097 | 0.35701183 | 0.272 | 0.309 | 1 | 4 | Rbm12b2 | myCAF_2 |
| Sirt7 | 0.00301192 | 0.38060405 | 0.22 | 0.255 | 1 | 4 | Sirt7 | myCAF_2 |
| Vstm41 | 0.00301526 | 0.30673275 | 0.445 | 0.507 | 1 | 4 | Vstm4 | myCAF_2 |
| Trip10 | 0.00302901 | 0.44345568 | 0.68 | 0.709 | 1 | 4 | Trip10 | myCAF_2 |
| Zfp609 | 0.00305505 | 0.6404319 | 0.516 | 0.496 | 1 | 4 | Zfp609 | myCAF_2 |
| Mthfd14a1 | 0.00305918 | 0.31862433 | 0.412 | 0.439 | 1 | 4 | Mthfd14a | myCAF_2 |
| Tcf201 | 0.00306289 | 0.42896826 | 0.56 | 0.596 | 1 | 4 | Tcf20 | myCAF_2 |
| Stoml1 | 0.00306731 | 0.26670265 | 0.381 | 0.418 | 1 | 4 | Stoml1 | myCAF_2 |
| Plekhn12 | 0.00309404 | 0.75727616 | 0.431 | 0.407 | 1 | 4 | Plekhn1 | myCAF_2 |
| Ntmt1 | 0.00314146 | 0.42603228 | 0.441 | 0.484 | 1 | 4 | Ntmt1 | myCAF_2 |
| Serinc32 | 0.00323214 | 0.46131669 | 0.957 | 0.955 | 1 | 4 | Serinc3 | myCAF_2 |
| Pramef81 | 0.00326309 | 0.25602785 | 0.238 | 0.295 | 1 | 4 | Pramef8 | myCAF_2 |
| St7l | 0.00334895 | 0.28111109 | 0.246 | 0.276 | 1 | 4 | St7l | myCAF_2 |
| Abcc11 | 0.0033628 | 0.61604922 | 0.582 | 0.57 | 1 | 4 | Abcc1 | myCAF_2 |
| Whamm1 | 0.00338878 | 0.72918849 | 0.254 | 0.231 | 1 | 4 | Whamm | myCAF_2 |
| Sdc21 | 0.00341934 | 0.40199393 | 0.844 | 0.857 | 1 | 4 | Sdc2 | myCAF_2 |
| Pxdc12 | 0.00348603 | 0.60766522 | 0.552 | 0.546 | 1 | 4 | Pxdc1 | myCAF_2 |
| Cd811 | 0.00355637 | 0.50968236 | 0.957 | 0.964 | 1 | 4 | Cd81 | myCAF_2 |
| Fbxl171 | 0.00358466 | 0.28248448 | 0.509 | 0.556 | 1 | 4 | Fbxl17 | myCAF_2 |
| Pdrg1 | 0.003697 | 0.43472984 | 0.639 | 0.689 | 1 | 4 | Pdrg1 | myCAF_2 |
| Slc9a3r22 | 0.00372795 | 0.73313999 | 0.377 | 0.355 | 1 | 4 | Slc9a3r2 | myCAF_2 |
| Myh91 | 0.00388256 | 0.47796442 | 0.918 | 0.93 | 1 | 4 | Myh9 | myCAF_2 |
| Tcf25 | 0.00390593 | 0.5971456 | 0.948 | 0.948 | 1 | 4 | Tcf25 | myCAF_2 |
| Hax1 | 0.00393566 | 0.37973934 | 0.486 | 0.53 | 1 | 4 | Hax1 | myCAF_2 |
| Cpt1c | 0.00393865 | 0.33412756 | 0.467 | 0.503 | 1 | 4 | Cpt1c | myCAF_2 |
| Narf1 | 0.00394384 | 0.60723366 | 0.474 | 0.448 | 1 | 4 | Narf | myCAF_2 |
| Nap1l41 | 0.00397031 | 0.39431678 | 0.623 | 0.658 | 1 | 4 | Nap1l4 | myCAF_2 |
| Msantd21 | 0.00397719 | 0.36541942 | 0.309 | 0.33 | 1 | 4 | Msantd2 | myCAF_2 |
| Lrrc8a | 0.00406636 | 0.61708395 | 0.738 | 0.715 | 1 | 4 | Lrrc8a | myCAF_2 |
| Kmt2e1 | 0.00411601 | 0.60177808 | 0.64 | 0.645 | 1 | 4 | Kmt2e | myCAF_2 |
| Kank11 | 0.0041725 | 0.26682184 | 0.332 | 0.386 | 1 | 4 | Kank1 | myCAF_2 |
| Fhod11 | 0.00421032 | 0.27208619 | 0.253 | 0.291 | 1 | 4 | Fhod1 | myCAF_2 |
| Vangl21 | 0.00425865 | 0.28404057 | 0.329 | 0.383 | 1 | 4 | Vangl2 | myCAF_2 |
| Rapgef2 | 0.00427537 | 0.4175432 | 0.457 | 0.517 | 1 | 4 | Rapgef2 | myCAF_2 |
| Hs1bp3 | 0.00429608 | 0.30717975 | 0.348 | 0.394 | 1 | 4 | Hs1bp3 | myCAF_2 |
| Zfp8271 | 0.0046014 | 0.6846789 | 0.29 | 0.273 | 1 | 4 | Zfp827 | myCAF_2 |
| Ric8b1 | 0.00462402 | 0.35073856 | 0.462 | 0.497 | 1 | 4 | Ric8b | myCAF_2 |
| Tsc22d3 | 0.00470624 | 0.81253849 | 0.271 | 0.241 | 1 | 4 | Tsc22d3 | myCAF_2 |
| Usp32 | 0.00489785 | 0.28065983 | 0.429 | 0.479 | 1 | 4 | Usp32 | myCAF_2 |
| Slc52a21 | 0.00493195 | 0.40295367 | 0.232 | 0.256 | 1 | 4 | Slc52a2 | myCAF_2 |
| Unk | 0.00501693 | 0.37865697 | 0.333 | 0.368 | 1 | 4 | Unk | myCAF_2 |
| Meis31 | 0.0050489 | 0.27327822 | 0.373 | 0.423 | 1 | 4 | Meis3 | myCAF_2 |
| Nav31 | 0.00506563 | 0.43965289 | 0.542 | 0.558 | 1 | 4 | Nav3 | myCAF_2 |
| St6galnac4 | 0.00512322 | 0.32921863 | 0.42 | 0.47 | 1 | 4 | St6galnac4 | myCAF_2 |
| Ldlrap11 | 0.00518107 | 0.39791558 | 0.534 | 0.572 | 1 | 4 | Ldlrap1 | myCAF_2 |
| Eif4b1 | 0.00535828 | 0.49262774 | 0.854 | 0.862 | 1 | 4 | Eif4b | myCAF_2 |
| Apaf1 | 0.00536467 | 0.38817583 | 0.533 | 0.589 | 1 | 4 | Apaf1 | myCAF_2 |
| Spire11 | 0.00543768 | 0.27787373 | 0.339 | 0.389 | 1 | 4 | Spire1 | myCAF_2 |
| Rcan3 | 0.00544251 | 0.35013359 | 0.311 | 0.353 | 1 | 4 | Rcan3 | myCAF_2 |
| Foxj31 | 0.00550846 | 0.50103521 | 0.384 | 0.421 | 1 | 4 | Foxj3 | myCAF_2 |
| Mast31 | 0.00554741 | 0.66021997 | 0.315 | 0.291 | 1 | 4 | Mast3 | myCAF_2 |
| Sirt11 | 0.00558541 | 0.32726071 | 0.269 | 0.304 | 1 | 4 | Sirt1 | myCAF_2 |
| Pias4 | 0.00560434 | 0.38215368 | 0.241 | 0.283 | 1 | 4 | Pias4 | myCAF_2 |
| Rnf1492 | 0.00584041 | 0.61253449 | 0.969 | 0.886 | 1 | 4 | Rnf149 | myCAF_2 |
| Zbtb211 | 0.00587085 | 0.29102203 | 0.373 | 0.407 | 1 | 4 | Zbtb21 | myCAF_2 |
| Jmjd1c | 0.00592418 | 0.44144992 | 0.555 | 0.574 | 1 | 4 | Jmjd1c | myCAF_2 |
| Nktr1 | 0.00595898 | 0.39057576 | 0.724 | 0.751 | 1 | 4 | Nktr | myCAF_2 |
| Myadm | 0.00595935 | 0.63026796 | 0.942 | 0.925 | 1 | 4 | Myadm | myCAF_2 |
| 6-Mar | 0.00609077 | 0.44124677 | 0.706 | 0.716 | 1 | 4 | 6-Mar | myCAF_2 |
| Pla2g12a1 | 0.00637457 | 0.72258689 | 0.314 | 0.337 | 1 | 4 | Pla2g12a | myCAF_2 |
| Rsbn1l1 | 0.0067329 | 0.32083611 | 0.279 | 0.333 | 1 | 4 | Rsbn1l | myCAF_2 |
| Ccdc581 | 0.00676438 | 0.33427765 | 0.358 | 0.4 | 1 | 4 | Ccdc58 | myCAF_2 |
| Wiz | 0.00696201 | 0.36880618 | 0.382 | 0.411 | 1 | 4 | Wiz | myCAF_2 |
| Limd11 | 0.0070046 | 0.57503221 | 0.779 | 0.797 | 1 | 4 | Limd1 | myCAF_2 |
| Fhdc11 | 0.00703804 | 0.40829933 | 0.345 | 0.357 | 1 | 4 | Fhdc1 | myCAF_2 |
| Fam98c1 | 0.00706695 | 0.71873985 | 0.428 | 0.402 | 1 | 4 | Fam98c | myCAF_2 |
| Zbtb17 | 0.00711376 | 0.38133481 | 0.228 | 0.266 | 1 | 4 | Zbtb17 | myCAF_2 |
| Brf1 | 0.00716498 | 0.2569209 | 0.447 | 0.487 | 1 | 4 | Brf1 | myCAF_2 |
| Dmwd1 | 0.00717301 | 0.7021143 | 0.598 | 0.589 | 1 | 4 | Dmwd | myCAF_2 |
| Ankhd1 | 0.00718141 | 0.31106636 | 0.271 | 0.31 | 1 | 4 | Ankhd1 | myCAF_2 |
| Ss181 | 0.00730608 | 0.42002528 | 0.686 | 0.724 | 1 | 4 | Ss18 | myCAF_2 |
| Fam122a | 0.00736811 | 0.49887243 | 0.356 | 0.389 | 1 | 4 | Fam122a | myCAF_2 |
| Tatdn11 | 0.00755082 | 0.38755528 | 0.282 | 0.329 | 1 | 4 | Tatdn1 | myCAF_2 |
| Leng81 | 0.00775378 | 0.48453022 | 0.768 | 0.78 | 1 | 4 | Leng8 | myCAF_2 |
| Renbp1 | 0.00796697 | 0.39348198 | 0.491 | 0.514 | 1 | 4 | Renbp | myCAF_2 |
| Mettl231 | 0.0079833 | 0.37139362 | 0.295 | 0.313 | 1 | 4 | Mettl23 | myCAF_2 |
| Elf11 | 0.00804185 | 0.37591918 | 0.711 | 0.762 | 1 | 4 | Elf1 | myCAF_2 |
| Pcgf2 | 0.00810626 | 0.30784377 | 0.487 | 0.526 | 1 | 4 | Pcgf2 | myCAF_2 |
| Ets21 | 0.0081218 | 0.3119136 | 0.532 | 0.573 | 1 | 4 | Ets2 | myCAF_2 |
| Hdac51 | 0.00818652 | 0.58680635 | 0.68 | 0.683 | 1 | 4 | Hdac5 | myCAF_2 |
| Cacul1 | 0.00833818 | 0.40834902 | 0.392 | 0.431 | 1 | 4 | Cacul1 | myCAF_2 |
| Sept112 | 0.00854672 | 0.53478141 | 0.859 | 0.858 | 1 | 4 | 11-Sep | myCAF_2 |
| Fam43a1 | 0.00868154 | 0.38306644 | 0.509 | 0.512 | 1 | 4 | Fam43a | myCAF_2 |
| Rnf1251 | 0.00875964 | 0.32939789 | 0.263 | 0.327 | 1 | 4 | Rnf125 | myCAF_2 |
| Spats21 | 0.0089171 | 0.5648017 | 0.723 | 0.708 | 1 | 4 | Spats2 | myCAF_2 |
| Nsmce3 | 0.00906253 | 0.36225516 | 0.413 | 0.468 | 1 | 4 | Nsmce3 | myCAF_2 |
| Irs1 | 0.00908486 | 0.53009471 | 0.678 | 0.699 | 1 | 4 | Irs1 | myCAF_2 |
| Brd21 | 0.00917505 | 0.6937557 | 0.804 | 0.81 | 1 | 4 | Brd2 | myCAF_2 |
| Mepce | 0.00920213 | 0.41620478 | 0.412 | 0.431 | 1 | 4 | Mepce | myCAF_2 |
| Vsir1 | 0.00924214 | 0.49291512 | 0.445 | 0.46 | 1 | 4 | Vsir | myCAF_2 |
| Limk21 | 0.00944487 | 0.41459464 | 0.52 | 0.539 | 1 | 4 | Limk2 | myCAF_2 |
| Pik3r12 | 0.00960089 | 0.41124537 | 0.643 | 0.719 | 1 | 4 | Pik3r1 | myCAF_2 |
| Ccdc9b2 | 0.0096292 | 0.26110104 | 0.326 | 0.362 | 1 | 4 | Ccdc9b | myCAF_2 |
| Slc25a12 | 0.00966324 | 0.34918528 | 0.713 | 0.749 | 1 | 4 | Slc25a1 | myCAF_2 |
| Evi5l | 0.00987861 | 0.37370212 | 0.416 | 0.438 | 1 | 4 | Evi5l | myCAF_2 |
| Pex6 | 0.00996569 | 0.41238546 | 0.446 | 0.489 | 1 | 4 | Pex6 | myCAF_2 |
| Clec3b | 0 | 5.37458777 | 0.915 | 0.084 | 0 | 5 | Clec3b | homeostatic |
| Ly6c1 | 0 | 4.45543519 | 0.962 | 0.194 | 0 | 5 | Ly6c1 | homeostatic |
| Tnxb | 0 | 4.33134424 | 0.984 | 0.238 | 0 | 5 | Tnxb | homeostatic |
| Sema3c | 0 | 4.62293168 | 0.923 | 0.189 | 0 | 5 | Sema3c | homeostatic |
| Sfrp4 | 0 | 3.45389248 | 0.872 | 0.149 | 0 | 5 | Sfrp4 | homeostatic |
| Dpp4 | 0 | 3.93272818 | 0.924 | 0.207 | 0 | 5 | Dpp4 | homeostatic |
| Cmah | 0 | 3.99795213 | 0.898 | 0.194 | 0 | 5 | Cmah | homeostatic |
| Spon2 | 0 | 3.58194798 | 0.886 | 0.193 | 0 | 5 | Spon2 | homeostatic |
| Prg41 | 0 | 3.50006687 | 0.944 | 0.262 | 0 | 5 | Prg4 | homeostatic |
| Ogn | 0 | 4.03950024 | 0.893 | 0.213 | 0 | 5 | Ogn | homeostatic |
| Ifi27l2a | 0 | 3.86341664 | 0.793 | 0.12 | 0 | 5 | Ifi27l2a | homeostatic |
| Cd34 | 0 | 4.19343212 | 0.989 | 0.323 | 0 | 5 | Cd34 | homeostatic |
| Pi16 | 0 | 4.68116418 | 0.977 | 0.313 | 0 | 5 | Pi16 | homeostatic |
| Cadm3 | 0 | 4.33984341 | 0.891 | 0.228 | 0 | 5 | Cadm3 | homeostatic |
| Galnt161 | 0 | 3.59860577 | 0.975 | 0.323 | 0 | 5 | Galnt16 | homeostatic |
| Scara5 | 0 | 4.93364188 | 0.703 | 0.052 | 0 | 5 | Scara5 | homeostatic |
| Ackr1 | 0 | 4.88700615 | 0.718 | 0.073 | 0 | 5 | Ackr1 | homeostatic |
| Npr1 | 0 | 4.05819086 | 0.76 | 0.116 | 0 | 5 | Npr1 | homeostatic |
| Cd55 | 0 | 4.85767354 | 0.746 | 0.11 | 0 | 5 | Cd55 | homeostatic |
| Itih5 | 0 | 3.36148247 | 0.916 | 0.281 | 0 | 5 | Itih5 | homeostatic |
| Ces2g1 | 0 | 3.07960619 | 0.83 | 0.195 | 0 | 5 | Ces2g | homeostatic |
| Efemp1 | 0 | 3.97416358 | 0.919 | 0.287 | 0 | 5 | Efemp1 | homeostatic |
| Lrrn4cl | 0 | 3.92660413 | 0.956 | 0.334 | 0 | 5 | Lrrn4cl | homeostatic |
| Ccn31 | 0 | 2.98726241 | 0.827 | 0.21 | 0 | 5 | Ccn3 | homeostatic |
| Tppp3 | 0 | 3.61275908 | 0.793 | 0.188 | 0 | 5 | Tppp3 | homeostatic |
| Pcsk6 | 0 | 4.82115803 | 0.671 | 0.067 | 0 | 5 | Pcsk6 | homeostatic |
| Clip4 | 0 | 3.3562075 | 0.75 | 0.146 | 0 | 5 | Clip4 | homeostatic |
| Tmeff21 | 0 | 2.64103248 | 0.898 | 0.295 | 0 | 5 | Tmeff2 | homeostatic |
| C4a1 | 0 | 2.58188124 | 0.94 | 0.337 | 0 | 5 | C4a | homeostatic |
| Scn7a1 | 0 | 3.02244836 | 0.844 | 0.249 | 0 | 5 | Scn7a | homeostatic |
| Fhl1 | 0 | 2.68466213 | 0.867 | 0.274 | 0 | 5 | Fhl1 | homeostatic |
| Chrdl11 | 0 | 2.96162286 | 0.816 | 0.224 | 0 | 5 | Chrdl1 | homeostatic |
| Sel1l3 | 0 | 4.28998198 | 0.661 | 0.071 | 0 | 5 | Sel1l3 | homeostatic |
| Steap41 | 0 | 2.86259586 | 0.853 | 0.265 | 0 | 5 | Steap4 | homeostatic |
| Creb5 | 0 | 3.96334472 | 0.695 | 0.108 | 0 | 5 | Creb5 | homeostatic |
| Wnt161 | 0 | 3.59939204 | 0.676 | 0.091 | 0 | 5 | Wnt16 | homeostatic |
| Prelp1 | 0 | 2.70588409 | 0.972 | 0.395 | 0 | 5 | Prelp | homeostatic |
| Sbsn1 | 0 | 3.6498191 | 0.9 | 0.324 | 0 | 5 | Sbsn | homeostatic |
| Ackr3 | 0 | 3.04481214 | 0.928 | 0.363 | 0 | 5 | Ackr3 | homeostatic |
| Adgrd11 | 0 | 3.01265285 | 0.989 | 0.451 | 0 | 5 | Adgrd1 | homeostatic |
| Osr1 | 0 | 3.02698031 | 0.77 | 0.232 | 0 | 5 | Osr1 | homeostatic |
| Aldh1a1 | 0 | 3.97195578 | 0.621 | 0.089 | 0 | 5 | Aldh1a1 | homeostatic |
| Syk | 0 | 3.64062962 | 0.622 | 0.093 | 0 | 5 | Syk | homeostatic |
| Mgll | 0 | 2.77333009 | 0.703 | 0.178 | 0 | 5 | Mgll | homeostatic |
| Ablim1 | 0 | 2.88554968 | 0.703 | 0.183 | 0 | 5 | Ablim1 | homeostatic |
| Plac9a | 0 | 3.57889958 | 0.652 | 0.136 | 0 | 5 | Plac9a | homeostatic |
| Ly6a | 0 | 3.61194762 | 0.992 | 0.476 | 0 | 5 | Ly6a | homeostatic |
| Pde8a1 | 0 | 2.61641781 | 0.808 | 0.294 | 0 | 5 | Pde8a | homeostatic |
| Ltbp11 | 0 | 2.07454911 | 0.849 | 0.34 | 0 | 5 | Ltbp1 | homeostatic |
| Ggt51 | 0 | 2.86408701 | 0.75 | 0.243 | 0 | 5 | Ggt5 | homeostatic |
| Irf7 | 0 | 2.99173408 | 0.69 | 0.186 | 0 | 5 | Irf7 | homeostatic |
| Dapk11 | 0 | 2.48011521 | 0.727 | 0.227 | 0 | 5 | Dapk1 | homeostatic |
| Olfml11 | 0 | 2.46264068 | 0.717 | 0.218 | 0 | 5 | Olfml1 | homeostatic |
| Il332 | 0 | 1.38309224 | 0.857 | 0.359 | 0 | 5 | Il33 | homeostatic |
| Tek | 0 | 4.23118392 | 0.565 | 0.068 | 0 | 5 | Tek | homeostatic |
| Slc4a4 | 0 | 3.59470223 | 0.59 | 0.094 | 0 | 5 | Slc4a4 | homeostatic |
| Gabra3 | 0 | 3.20525706 | 0.614 | 0.121 | 0 | 5 | Gabra3 | homeostatic |
| Efhd1 | 0 | 5.968837 | 0.511 | 0.019 | 0 | 5 | Efhd1 | homeostatic |
| C31 | 0 | 3.16140989 | 0.99 | 0.499 | 0 | 5 | C3 | homeostatic |
| C4b1 | 0 | 2.81336786 | 0.996 | 0.507 | 0 | 5 | C4b | homeostatic |
| Bmper1 | 0 | 2.11077886 | 0.713 | 0.23 | 0 | 5 | Bmper | homeostatic |
| Klf41 | 0 | 1.63335401 | 0.897 | 0.418 | 0 | 5 | Klf4 | homeostatic |
| Vegfd1 | 0 | 3.11016759 | 0.923 | 0.445 | 0 | 5 | Vegfd | homeostatic |
| Dmkn | 0 | 6.02855571 | 0.496 | 0.018 | 0 | 5 | Dmkn | homeostatic |
| Pamr11 | 0 | 3.21470995 | 0.617 | 0.14 | 0 | 5 | Pamr1 | homeostatic |
| Adamts151 | 0 | 2.32530919 | 0.864 | 0.393 | 0 | 5 | Adamts15 | homeostatic |
| Plagl12 | 0 | 2.215953 | 0.704 | 0.237 | 0 | 5 | Plagl1 | homeostatic |
| Gas7 | 0 | 3.43692733 | 0.937 | 0.471 | 0 | 5 | Gas7 | homeostatic |
| Ptx31 | 0 | 2.25753748 | 0.82 | 0.354 | 0 | 5 | Ptx3 | homeostatic |
| Col14a11 | 0 | 2.96855595 | 0.984 | 0.519 | 0 | 5 | Col14a1 | homeostatic |
| Nova1 | 0 | 2.83972106 | 0.775 | 0.312 | 0 | 5 | Nova1 | homeostatic |
| Ccl111 | 0 | 1.82659608 | 0.854 | 0.394 | 0 | 5 | Ccl11 | homeostatic |
| Ssc5d1 | 0 | 2.5130448 | 0.935 | 0.476 | 0 | 5 | Sssc5d | homeostatic |
| AW551984 | 0 | 2.46684913 | 0.836 | 0.378 | 0 | 5 | AW551984 | homeostatic |
| Anxa3 | 0 | 3.86834027 | 0.97 | 0.513 | 0 | 5 | Anxa3 | homeostatic |
| Slfn51 | 0 | 2.88337578 | 0.966 | 0.511 | 0 | 5 | Slfn5 | homeostatic |
| Prrg31 | 0 | 2.91855783 | 0.584 | 0.131 | 0 | 5 | Prrg3 | homeostatic |
| Gfra2 | 0 | 2.72777189 | 0.575 | 0.128 | 0 | 5 | Gfra2 | homeostatic |
| Ramp2 | 0 | 3.24755923 | 0.615 | 0.169 | 0 | 5 | Ramp2 | homeostatic |
| Lims2 | 0 | 3.57351732 | 0.506 | 0.061 | 0 | 5 | Lims2 | homeostatic |
| Serpinb1a2 | 0 | 2.07481616 | 0.661 | 0.218 | 0 | 5 | Serpinb1a | homeostatic |
| Gda2 | 0 | 2.40274911 | 0.934 | 0.491 | 0 | 5 | Gda | homeostatic |
| Sulf21 | 0 | 2.30573665 | 0.82 | 0.377 | 0 | 5 | Sulf2 | homeostatic |
| Gstt11 | 0 | 2.49598874 | 0.603 | 0.162 | 0 | 5 | Gstt1 | homeostatic |
| Adamts51 | 0 | 2.55246062 | 0.934 | 0.5 | 0 | 5 | Adamts5 | homeostatic |
| Ptgis1 | 0 | 1.8293336 | 0.83 | 0.396 | 0 | 5 | Ptgis | homeostatic |
| Sqor1 | 0 | 2.5018007 | 0.706 | 0.276 | 0 | 5 | Sqor | homeostatic |
| Serpina3n1 | 0 | 2.40099665 | 0.981 | 0.556 | 0 | 5 | Serpina3n | homeostatic |
| Bmp3 | 0 | 3.78254669 | 0.462 | 0.038 | 0 | 5 | Bmp3 | homeostatic |
| Gstm1 | 0 | 3.34393499 | 0.533 | 0.11 | 0 | 5 | Gstm1 | homeostatic |
| Rab32 | 0 | 2.52729807 | 0.678 | 0.257 | 0 | 5 | Rab32 | homeostatic |
| Loxl41 | 0 | 2.08317481 | 0.713 | 0.293 | 0 | 5 | Loxl4 | homeostatic |
| Pdgfd | 0 | 2.23999489 | 0.65 | 0.232 | 0 | 5 | Pdgfd | homeostatic |
| Irak3 | 0 | 2.23031677 | 0.792 | 0.375 | 0 | 5 | Irak3 | homeostatic |
| Agtr1a1 | 0 | 2.54668413 | 0.582 | 0.168 | 0 | 5 | Agtr1a | homeostatic |
| Adamtsl4 | 0 | 3.35055907 | 0.57 | 0.157 | 0 | 5 | Adamtsl4 | homeostatic |
| Ccn51 | 0 | 1.15661361 | 0.832 | 0.42 | 0 | 5 | Ccn5 | homeostatic |
| Bst21 | 0 | 2.30710722 | 0.938 | 0.526 | 0 | 5 | Bst2 | homeostatic |
| Irgm2 | 0 | 2.25106953 | 0.674 | 0.264 | 0 | 5 | Irgm2 | homeostatic |
| Gprc5b | 0 | 2.24536097 | 0.745 | 0.335 | 0 | 5 | Gprc5b | homeostatic |
| Cyp4b1 | 0 | 2.34810584 | 0.586 | 0.177 | 0 | 5 | Cyp4b1 | homeostatic |
| Enpep1 | 0 | 2.80405516 | 0.572 | 0.169 | 0 | 5 | Enpep | homeostatic |
| Oasl2 | 0 | 2.20851537 | 0.723 | 0.325 | 0 | 5 | Oasl2 | homeostatic |
| Megf9 | 0 | 2.63297513 | 0.576 | 0.179 | 0 | 5 | Megf9 | homeostatic |
| Lgi4 | 0 | 2.30203504 | 0.612 | 0.216 | 0 | 5 | Lgi4 | homeostatic |
| Pcolce2 | 0 | 2.93063602 | 0.712 | 0.319 | 0 | 5 | Pcolce2 | homeostatic |
| Stxbp6 | 0 | 4.38483402 | 0.418 | 0.028 | 0 | 5 | Stxbp6 | homeostatic |
| Lgals31 | 0 | 0.95390194 | 0.873 | 0.486 | 0 | 5 | Lgals3 | homeostatic |
| Il18 | 0 | 4.48945538 | 0.426 | 0.041 | 0 | 5 | Il18 | homeostatic |
| Slc10a61 | 0 | 2.35675513 | 0.589 | 0.207 | 0 | 5 | Slc10a6 | homeostatic |
| Chl11 | 0 | 1.5543261 | 0.745 | 0.365 | 0 | 5 | Chl1 | homeostatic |
| Oas2 | 0 | 3.15086006 | 0.463 | 0.084 | 0 | 5 | Oas2 | homeostatic |
| Phf11d | 0 | 2.44067806 | 0.563 | 0.185 | 0 | 5 | Phf11d | homeostatic |
| Ifi2051 | 0 | 3.06903496 | 0.98 | 0.606 | 0 | 5 | Ifi205 | homeostatic |
| Sp100 | 0 | 2.41197209 | 0.576 | 0.202 | 0 | 5 | Sp100 | homeostatic |
| Adam232 | 0 | 2.22093998 | 0.707 | 0.334 | 0 | 5 | Adam23 | homeostatic |
| Dpt1 | 0 | 3.35606565 | 0.997 | 0.625 | 0 | 5 | Dpt | homeostatic |
| Zbp11 | 0 | 2.11613678 | 0.848 | 0.476 | 0 | 5 | Zbp1 | homeostatic |
| Iigp11 | 0 | 1.96220583 | 0.514 | 0.142 | 0 | 5 | Iigp1 | homeostatic |
| Fzd4 | 0 | 2.34656329 | 0.607 | 0.237 | 0 | 5 | Fzd4 | homeostatic |
| Adora2b | 0 | 2.1408563 | 0.626 | 0.258 | 0 | 5 | Adora2b | homeostatic |
| Qpct1 | 0 | 2.27151678 | 0.699 | 0.332 | 0 | 5 | Qpct | homeostatic |
| Ifi2031 | 0 | 2.09379748 | 0.793 | 0.427 | 0 | 5 | Ifi203 | homeostatic |
| Cfb1 | 0 | 1.98063123 | 0.981 | 0.615 | 0 | 5 | Cfb | homeostatic |
| Nrcam | 0 | 3.60382856 | 0.415 | 0.051 | 0 | 5 | Nrcam | homeostatic |
| Csf12 | 0 | 1.82684557 | 0.911 | 0.548 | 0 | 5 | Csf1 | homeostatic |
| Ptgs1 | 0 | 2.63067203 | 0.595 | 0.233 | 0 | 5 | Ptgs1 | homeostatic |
| Wnt2 | 0 | 2.72206149 | 0.524 | 0.162 | 0 | 5 | Wnt2 | homeostatic |
| Prss23 | 0 | 2.86339651 | 0.978 | 0.617 | 0 | 5 | Prss23 | homeostatic |
| Epdr1 | 0 | 1.76090468 | 0.754 | 0.393 | 0 | 5 | Epdr1 | homeostatic |
| Aox3 | 0 | 4.61825275 | 0.388 | 0.031 | 0 | 5 | Aox3 | homeostatic |
| Duox1 | 0 | 4.33649821 | 0.385 | 0.028 | 0 | 5 | Duox1 | homeostatic |
| Cdc42ep21 | 0 | 1.8321528 | 0.703 | 0.347 | 0 | 5 | Cdc42ep2 | homeostatic |
| Isg15 | 0 | 2.16195109 | 0.614 | 0.26 | 0 | 5 | Isg15 | homeostatic |

| | | | | | | | | |
|---|---|---|---|---|---|---|---|---|
| Add31 | 0 | 2.08081176 | 0.885 | 0.533 | 0 | 5 | Add3 | homeostatic |
| Ifi204 | 0 | 2.12656758 | 0.862 | 0.513 | 0 | 5 | Ifi204 | homeostatic |
| Arhgap20 | 0 | 3.95721037 | 0.388 | 0.04 | 0 | 5 | Arhgap20 | homeostatic |
| Nsg12 | 0 | 1.60535957 | 0.91 | 0.571 | 0 | 5 | Nsg1 | homeostatic |
| Arsi1 | 0 | 2.10715184 | 0.963 | 0.625 | 0 | 5 | Arsi | homeostatic |
| Rcan2 | 0 | 2.85549471 | 0.447 | 0.11 | 0 | 5 | Rcan2 | homeostatic |
| Tubb4a | 0 | 3.39498061 | 0.397 | 0.06 | 0 | 5 | Tubb4a | homeostatic |
| Islr2 | 0 | 2.72862338 | 0.548 | 0.212 | 0 | 5 | Islr2 | homeostatic |
| Osr2 | 0 | 1.6775954 | 0.783 | 0.447 | 0 | 5 | Osr2 | homeostatic |
| Psd3 | 0 | 1.9874064 | 0.853 | 0.518 | 0 | 5 | Psd3 | homeostatic |
| Scara31 | 0 | 2.43561385 | 0.967 | 0.632 | 0 | 5 | Scara3 | homeostatic |
| Apod1 | 0 | 1.79516218 | 0.935 | 0.601 | 0 | 5 | Apod | homeostatic |
| Ampd31 | 0 | 1.62122293 | 0.798 | 0.464 | 0 | 5 | Ampd3 | homeostatic |
| Slc43a31 | 0 | 1.97575035 | 0.855 | 0.521 | 0 | 5 | Slc43a3 | homeostatic |
| Thbd | 0 | 2.59526167 | 0.708 | 0.376 | 0 | 5 | Thbd | homeostatic |
| Adm2 | 0 | 1.94475323 | 0.761 | 0.43 | 0 | 5 | Adm | homeostatic |
| Igfbp6 | 0 | 2.63312841 | 0.981 | 0.65 | 0 | 5 | Igfbp6 | homeostatic |
| Entpd2 | 0 | 1.99278266 | 0.608 | 0.278 | 0 | 5 | Entpd2 | homeostatic |
| Rtp4 | 0 | 2.23019743 | 0.49 | 0.163 | 0 | 5 | Rtp4 | homeostatic |
| Dpep11 | 0 | 1.88372016 | 0.904 | 0.582 | 0 | 5 | Dpep1 | homeostatic |
| Rab3il1 | 0 | 1.54090508 | 0.848 | 0.533 | 0 | 5 | Rab3il1 | homeostatic |
| Igfbp5 | 0 | 2.06485923 | 0.494 | 0.181 | 0 | 5 | Igfbp5 | homeostatic |
| Rtl3 | 0 | 4.60657439 | 0.334 | 0.021 | 0 | 5 | Rtl3 | homeostatic |
| Tmem1001 | 0 | 1.68317776 | 0.74 | 0.428 | 0 | 5 | Tmem100 | homeostatic |
| Lgi21 | 0 | 1.12487407 | 0.887 | 0.576 | 0 | 5 | Lgi2 | homeostatic |
| Flrt21 | 0 | 2.10626292 | 0.937 | 0.627 | 0 | 5 | Flrt2 | homeostatic |
| Abca6 | 0 | 3.8345693 | 0.355 | 0.046 | 0 | 5 | Abca6 | homeostatic |
| Adamtsl11 | 0 | 1.88086072 | 0.961 | 0.652 | 0 | 5 | Adamtsl1 | homeostatic |
| Klf22 | 0 | 1.52662599 | 0.698 | 0.39 | 0 | 5 | Klf2 | homeostatic |
| Usp18 | 0 | 2.79934557 | 0.383 | 0.079 | 0 | 5 | Usp18 | homeostatic |
| Galnt171 | 0 | 1.48481652 | 0.803 | 0.5 | 0 | 5 | Galnt17 | homeostatic |
| Car8 | 0 | 5.08331024 | 0.327 | 0.025 | 0 | 5 | Car8 | homeostatic |
| St3gal11 | 0 | 1.70984828 | 0.858 | 0.556 | 0 | 5 | St3gal1 | homeostatic |
| Adh1 | 0 | 3.93769371 | 0.343 | 0.042 | 0 | 5 | Adh1 | homeostatic |
| Zbtb16 | 0 | 4.38266693 | 0.326 | 0.029 | 0 | 5 | Zbtb16 | homeostatic |
| Galnt6 | 0 | 3.30600218 | 0.364 | 0.067 | 0 | 5 | Galnt6 | homeostatic |
| Thbs31 | 0 | 2.08586928 | 0.956 | 0.66 | 0 | 5 | Thbs3 | homeostatic |
| Nfia1 | 0 | 1.58314849 | 0.864 | 0.571 | 0 | 5 | Nfia | homeostatic |
| Akr1c18 | 0 | 2.66594596 | 0.368 | 0.077 | 0 | 5 | Akr1c18 | homeostatic |
| Lbp | 0 | 3.24875943 | 0.953 | 0.668 | 0 | 5 | Lbp | homeostatic |
| Rras2 | 0 | 2.27867014 | 0.85 | 0.568 | 0 | 5 | Rras2 | homeostatic |
| Col28a11 | 0 | 2.3549904 | 0.412 | 0.132 | 0 | 5 | Col28a1 | homeostatic |
| DEPRECATED-I | 0 | 1.17576729 | 0.725 | 0.447 | 0 | 5 | DEPRECATED-I | homeostatic |
| Rnf2131 | 0 | 1.77627525 | 0.714 | 0.438 | 0 | 5 | Rnf213 | homeostatic |
| Fyn | 0 | 1.80404006 | 0.819 | 0.543 | 0 | 5 | Fyn | homeostatic |
| Aldh1a3 | 0 | 3.31744426 | 0.343 | 0.069 | 0 | 5 | Aldh1a3 | homeostatic |
| Pdlim11 | 0 | 1.36235258 | 0.788 | 0.515 | 0 | 5 | Pdlim1 | homeostatic |
| Arhgap291 | 0 | 1.54049177 | 0.843 | 0.571 | 0 | 5 | Arhgap29 | homeostatic |
| Fxyd11 | 0 | 1.36152315 | 0.882 | 0.613 | 0 | 5 | Fxyd1 | homeostatic |
| Ifit1 | 0 | 2.77050061 | 0.313 | 0.044 | 0 | 5 | Ifit1 | homeostatic |
| Ndn1 | 0 | 1.43712221 | 0.909 | 0.643 | 0 | 5 | Ndn | homeostatic |
| Ptpn51 | 0 | 2.61885563 | 0.36 | 0.1 | 0 | 5 | Ptpn5 | homeostatic |
| Adamts11 | 0 | 1.49650729 | 0.879 | 0.62 | 0 | 5 | Adamts1 | homeostatic |
| Fam102b | 0 | 1.70375094 | 0.89 | 0.632 | 0 | 5 | Fam102b | homeostatic |
| Ifi211 | 0 | 1.86758193 | 0.882 | 0.626 | 0 | 5 | Ifi211 | homeostatic |
| Cxcl13 | 0 | 5.10839319 | 0.265 | 0.01 | 0 | 5 | Cxcl13 | homeostatic |
| Vgll3 | 0 | 2.05548251 | 0.954 | 0.701 | 0 | 5 | Vgll3 | homeostatic |
| Akr1c14 | 0 | 4.51207759 | 0.265 | 0.015 | 0 | 5 | Akr1c14 | homeostatic |
| Plac81 | 0 | 2.0330479 | 0.968 | 0.723 | 0 | 5 | Plac8 | homeostatic |
| Hcn4 | 0 | 3.42892156 | 0.284 | 0.039 | 0 | 5 | Hcn4 | homeostatic |
| Zfp385a | 0 | 1.99534058 | 0.849 | 0.606 | 0 | 5 | Zfp385a | homeostatic |
| Steap3 | 0 | 1.85971037 | 0.737 | 0.496 | 0 | 5 | Steap3 | homeostatic |
| Slco3a1 | 0 | 1.55633224 | 0.82 | 0.579 | 0 | 5 | Slco3a1 | homeostatic |
| Samhd1 | 0 | 1.50670994 | 0.777 | 0.537 | 0 | 5 | Samhd1 | homeostatic |
| Spry21 | 0 | 1.49397238 | 0.801 | 0.566 | 0 | 5 | Spry2 | homeostatic |
| Heg1 | 0 | 3.00016987 | 0.995 | 0.764 | 0 | 5 | Heg1 | homeostatic |
| Prr13 | 0 | 1.38745869 | 0.871 | 0.641 | 0 | 5 | Prr13 | homeostatic |
| Pam1 | 0 | 1.98844394 | 0.933 | 0.706 | 0 | 5 | Pam | homeostatic |
| Fgl22 | 0 | 1.39242354 | 0.797 | 0.575 | 0 | 5 | Fgl2 | homeostatic |
| Fgf2 | 0 | 1.74389838 | 0.668 | 0.447 | 0 | 5 | Fgf2 | homeostatic |
| Pgr1 | 0 | 2.69897802 | 0.319 | 0.098 | 0 | 5 | Pgr | homeostatic |
| Tgm21 | 0 | 1.72963753 | 0.679 | 0.46 | 0 | 5 | Tgm2 | homeostatic |
| Spsb11 | 0 | 1.05402616 | 0.78 | 0.566 | 0 | 5 | Spsb1 | homeostatic |
| Lgals91 | 0 | 1.50922758 | 0.878 | 0.664 | 0 | 5 | Lgals9 | homeostatic |
| Cyb5a1 | 0 | 1.36872152 | 0.877 | 0.663 | 0 | 5 | Cyb5a | homeostatic |
| Xdh2 | 0 | 1.42909411 | 0.921 | 0.708 | 0 | 5 | Xdh | homeostatic |
| Sned1 | 0 | 2.35012383 | 0.961 | 0.748 | 0 | 5 | Sned1 | homeostatic |
| Pmp221 | 0 | 1.22353519 | 0.924 | 0.713 | 0 | 5 | Pmp22 | homeostatic |
| Emilin21 | 0 | 2.26854895 | 0.961 | 0.75 | 0 | 5 | Emilin2 | homeostatic |
| Hdac7 | 0 | 1.51357826 | 0.86 | 0.653 | 0 | 5 | Hdac7 | homeostatic |
| Lcn2 | 0 | 2.4986042 | 0.266 | 0.065 | 0 | 5 | Lcn2 | homeostatic |
| Pnp1 | 0 | 1.33859741 | 0.907 | 0.707 | 0 | 5 | Pnp | homeostatic |
| Uap1 | 0 | 2.41042737 | 0.934 | 0.736 | 0 | 5 | Uap1 | homeostatic |
| Comt1 | 0 | 1.21347539 | 0.924 | 0.726 | 0 | 5 | Comt | homeostatic |
| S100a41 | 0 | 0.92382143 | 0.717 | 0.52 | 0 | 5 | S100a4 | homeostatic |
| Ms4a4d1 | 0 | 1.41613413 | 0.855 | 0.658 | 0 | 5 | Ms4a4d | homeostatic |
| Gpc31 | 0 | 1.8317235 | 0.951 | 0.755 | 0 | 5 | Gpc3 | homeostatic |
| Rhou | 0 | 1.46607353 | 0.879 | 0.685 | 0 | 5 | Rhou | homeostatic |
| Ugp2 | 0 | 1.5813671 | 0.768 | 0.575 | 0 | 5 | Ugp2 | homeostatic |
| Sirpa1 | 0 | 1.25544352 | 0.895 | 0.708 | 0 | 5 | Sirpa | homeostatic |
| Plod22 | 0 | 1.27953438 | 0.82 | 0.637 | 0 | 5 | Plod2 | homeostatic |
| Lrrc172 | 0 | 1.26354203 | 0.888 | 0.708 | 0 | 5 | Lrrc17 | homeostatic |
| Csrp22 | 0 | 0.96718922 | 0.844 | 0.666 | 0 | 5 | Csrp2 | homeostatic |
| Tmem431 | 0 | 1.38687635 | 0.915 | 0.739 | 0 | 5 | Tmem43 | homeostatic |
| Anpep1 | 0 | 1.6154845 | 0.929 | 0.759 | 0 | 5 | Anpep | homeostatic |
| Carhsp1 | 0 | 1.57683852 | 0.921 | 0.755 | 0 | 5 | Carhsp1 | homeostatic |
| Htra3 | 0 | 3.39895681 | 0.987 | 0.822 | 0 | 5 | Htra3 | homeostatic |
| Ppa12 | 0 | 1.46794819 | 0.807 | 0.647 | 0 | 5 | Ppa1 | homeostatic |
| Cebpd1 | 0 | 1.84730178 | 0.949 | 0.789 | 0 | 5 | Cebpd | homeostatic |
| Selenop | 0 | 1.65902184 | 0.881 | 0.723 | 0 | 5 | Selenop | homeostatic |
| Rarres21 | 0 | 1.2590991 | 0.984 | 0.837 | 0 | 5 | Rarres2 | homeostatic |
| Mfap52 | 0 | 1.28980449 | 0.984 | 0.837 | 0 | 5 | Mfap5 | homeostatic |
| Apbb1ip1 | 0 | 1.21087557 | 0.931 | 0.785 | 0 | 5 | Apbb1ip | homeostatic |
| Basp11 | 0 | 1.24737625 | 0.897 | 0.752 | 0 | 5 | Basp1 | homeostatic |
| Dpysl22 | 0 | 1.02202507 | 0.905 | 0.763 | 0 | 5 | Dpysl2 | homeostatic |
| Lamb22 | 0 | 1.18956483 | 0.935 | 0.797 | 0 | 5 | Lamb2 | homeostatic |
| Dpysl3 | 0 | 1.02836966 | 0.878 | 0.744 | 0 | 5 | Dpysl3 | homeostatic |
| Mgst11 | 0 | 0.98540279 | 0.945 | 0.811 | 0 | 5 | Mgst1 | homeostatic |
| Cavin3 | 0 | 1.20462203 | 0.918 | 0.785 | 0 | 5 | Cavin3 | homeostatic |
| Man1a1 | 0 | 1.68151697 | 0.975 | 0.845 | 0 | 5 | Man1a | homeostatic |
| Tgfbr32 | 0 | 1.19938192 | 0.972 | 0.846 | 0 | 5 | Tgfbr3 | homeostatic |
| S100a10 | 0 | 1.97003606 | 0.978 | 0.853 | 0 | 5 | S100a10 | homeostatic |
| Prnp1 | 0 | 1.46493497 | 0.956 | 0.832 | 0 | 5 | Prnp | homeostatic |
| Ddr2 | 0 | 1.45529174 | 0.968 | 0.85 | 0 | 5 | Ddr2 | homeostatic |
| Islr3 | 0 | 1.06476642 | 0.921 | 0.804 | 0 | 5 | Islr | homeostatic |
| S100a13 | 0 | 1.36043155 | 0.954 | 0.838 | 0 | 5 | S100a13 | homeostatic |
| Cxcl121 | 0 | 0.72853386 | 0.907 | 0.791 | 0 | 5 | Cxcl12 | homeostatic |
| C1ra1 | 0 | 1.68200591 | 0.986 | 0.874 | 0 | 5 | C1ra | homeostatic |
| Lsp12 | 0 | 0.97639229 | 0.959 | 0.847 | 0 | 5 | Lsp1 | homeostatic |
| Gfpt21 | 0 | 1.8002017 | 0.947 | 0.835 | 0 | 5 | Gfpt2 | homeostatic |
| Emp3 | 0 | 1.04811595 | 0.949 | 0.842 | 0 | 5 | Emp3 | homeostatic |
| Capn21 | 0 | 1.07578296 | 0.962 | 0.856 | 0 | 5 | Capn2 | homeostatic |
| Txnip1 | 0 | 0.98697954 | 0.939 | 0.838 | 0 | 5 | Txnip | homeostatic |
| Cavin1 | 0 | 1.82735606 | 0.991 | 0.894 | 0 | 5 | Cavin1 | homeostatic |
| Medag1 | 0 | 0.88088434 | 0.905 | 0.809 | 0 | 5 | Medag | homeostatic |
| Svep11 | 0 | 1.09239733 | 0.937 | 0.846 | 0 | 5 | Svep1 | homeostatic |
| Plpp3 | 0 | 1.8488765 | 0.93 | 0.839 | 0 | 5 | Plpp3 | homeostatic |
| Mt22 | 0 | 1.26962321 | 0.961 | 0.875 | 0 | 5 | Mt2 | homeostatic |
| Egfr1 | 0 | 1.16425352 | 0.951 | 0.871 | 0 | 5 | Egfr | homeostatic |
| Lgals3bp1 | 0 | 1.26778484 | 0.986 | 0.908 | 0 | 5 | Lgals3bp | homeostatic |
| Anxa1 | 0 | 1.22672697 | 0.98 | 0.902 | 0 | 5 | Anxa1 | homeostatic |
| B2m1 | 0 | 1.01887859 | 0.915 | 0.842 | 0 | 5 | B2m | homeostatic |
| Saa31 | 0 | 0.90974466 | 0.472 | 0.399 | 0 | 5 | Saa3 | homeostatic |
| Clec2d1 | 0 | 1.11854727 | 0.914 | 0.842 | 0 | 5 | Clec2d | homeostatic |
| Sfrp21 | 0 | 0.80661898 | 0.904 | 0.833 | 0 | 5 | Sfrp2 | homeostatic |
| S100a16 | 0 | 1.48491009 | 0.989 | 0.918 | 0 | 5 | S100a16 | homeostatic |
| C1s11 | 0 | 1.60264276 | 0.992 | 0.921 | 0 | 5 | C1s1 | homeostatic |
| Pdpn1 | 0 | 1.19919909 | 0.991 | 0.921 | 0 | 5 | Pdpn | homeostatic |
| Sod32 | 0 | 0.67621652 | 0.994 | 0.926 | 0 | 5 | Sod3 | homeostatic |
| Cd248 | 0 | 1.6948806 | 0.987 | 0.923 | 0 | 5 | Cd248 | homeostatic |
| Serpinb6a | 0 | 1.56305865 | 0.972 | 0.91 | 0 | 5 | Serpinb6a | homeostatic |
| Nfix1 | 0 | 0.99476503 | 0.982 | 0.922 | 0 | 5 | Nfix | homeostatic |
| Gja1 | 0 | 0.9159669 | 0.952 | 0.897 | 0 | 5 | Gja1 | homeostatic |
| Nbl11 | 0 | 0.87527925 | 0.975 | 0.921 | 0 | 5 | Nbl1 | homeostatic |
| Gpx31 | 0 | 1.27998899 | 1 | 0.948 | 0 | 5 | Gpx3 | homeostatic |
| Clmp | 0 | 0.79563357 | 0.97 | 0.92 | 0 | 5 | Clmp | homeostatic |
| Il6st1 | 0 | 0.79297385 | 0.937 | 0.892 | 0 | 5 | Il6st | homeostatic |
| Mt11 | 0 | 0.96843302 | 0.957 | 0.912 | 0 | 5 | Mt1 | homeostatic |
| Rnase41 | 0 | 1.33900839 | 0.987 | 0.944 | 0 | 5 | Rnase4 | homeostatic |
| Crip11 | 0 | 1.35867432 | 0.961 | 0.918 | 0 | 5 | Crip1 | homeostatic |
| Ly6e | 0 | 1.24153301 | 0.987 | 0.945 | 0 | 5 | Ly6e | homeostatic |
| Ifitm3 | 0 | 1.01412967 | 0.964 | 0.922 | 0 | 5 | Ifitm3 | homeostatic |
| Fbln11 | 0 | 0.86150086 | 0.962 | 0.92 | 0 | 5 | Fbln1 | homeostatic |
| Dcn2 | 0 | 1.02499425 | 0.996 | 0.955 | 0 | 5 | Dcn | homeostatic |
| Ahnak | 0 | 1.40087566 | 0.992 | 0.956 | 0 | 5 | Ahnak | homeostatic |
| Ecm11 | 0 | 0.93866823 | 0.987 | 0.952 | 0 | 5 | Ecm1 | homeostatic |
| Ugdh2 | 0 | 0.74087513 | 0.968 | 0.933 | 0 | 5 | Ugdh | homeostatic |
| Tgfbr2 | 0 | 1.68320782 | 0.995 | 0.961 | 0 | 5 | Tgfbr2 | homeostatic |
| Gsn | 0 | 2.67777115 | 0.999 | 0.966 | 0 | 5 | Gsn | homeostatic |
| Serping11 | 0 | 1.28608872 | 0.995 | 0.962 | 0 | 5 | Serping1 | homeostatic |
| Axl | 0 | 1.45229301 | 0.995 | 0.963 | 0 | 5 | Axl | homeostatic |
| Sulf11 | 0 | 0.93244952 | 0.951 | 0.919 | 0 | 5 | Sulf1 | homeostatic |
| Clic4 | 0 | 1.01007904 | 0.949 | 0.917 | 0 | 5 | Clic4 | homeostatic |
| Anxa2 | 0 | 1.31957069 | 0.997 | 0.968 | 0 | 5 | Anxa2 | homeostatic |
| Timp21 | 0 | 0.98219908 | 0.985 | 0.956 | 0 | 5 | Timp2 | homeostatic |
| Mmp21 | 0 | 0.58336008 | 1 | 0.972 | 0 | 5 | Mmp2 | homeostatic |
| Cd164 | 0 | 0.97253122 | 0.981 | 0.955 | 0 | 5 | Cd164 | homeostatic |
| Lox1 | 0 | 0.5979174 | 0.989 | 0.963 | 0 | 5 | Lox | homeostatic |
| Lamc11 | 0 | 0.89058149 | 0.992 | 0.967 | 0 | 5 | Lamc1 | homeostatic |
| Loxl1 | 0 | 0.58209875 | 0.995 | 0.973 | 0 | 5 | Loxl1 | homeostatic |
| Cthrc12 | 0 | 0.72205865 | 0.836 | 0.814 | 0 | 5 | Cthrc1 | homeostatic |
| Nid11 | 0 | 1.16243033 | 1 | 0.979 | 0 | 5 | Nid1 | homeostatic |
| Marcks | 0 | 0.94746274 | 0.986 | 0.969 | 0 | 5 | Marcks | homeostatic |
| Fbn1 | 0 | 1.26882445 | 0.995 | 0.979 | 0 | 5 | Fbn1 | homeostatic |
| Fth11 | 0 | 1.35300309 | 0.999 | 0.983 | 0 | 5 | Fth1 | homeostatic |
| Mmp191 | 0 | 0.62028965 | 0.947 | 0.932 | 0 | 5 | Mmp19 | homeostatic |
| Anxa5 | 0 | 0.50884465 | 0.985 | 0.974 | 0 | 5 | Anxa5 | homeostatic |
| Fstl1 | 0 | 1.17967481 | 0.999 | 0.99 | 0 | 5 | Fstl1 | homeostatic |
| Col3a11 | 0 | 0.63967658 | 1 | 0.996 | 0 | 5 | Col3a1 | homeostatic |
| Fn12 | 0 | 0.58852732 | 0.996 | 0.992 | 0 | 5 | Fn1 | homeostatic |
| Vim1 | 0 | 0.3498721 | 0.999 | 0.998 | 0 | 5 | Vim | homeostatic |
| Ccdc80 | 9.63E-298 | 0.64642902 | 0.989 | 0.954 | 1.86E-293 | 5 | Ccdc80 | homeostatic |
| Tmem254c | 2.66E-297 | 1.97057479 | 0.671 | 0.329 | 5.14E-293 | 5 | Tmem254c | homeostatic |
| Sptbn1 | 3.41E-297 | 0.78168641 | 0.984 | 0.955 | 6.60E-293 | 5 | Sptbn1 | homeostatic |
| Vcan1 | 4.65E-297 | 0.80928346 | 0.934 | 0.835 | 8.99E-293 | 5 | Vcan | homeostatic |
| Srgn1 | 2.01E-293 | 1.37869502 | 0.577 | 0.317 | 3.88E-289 | 5 | Srgn | homeostatic |
| Oas1a | 2.14E-293 | 2.88323818 | 0.485 | 0.105 | 4.14E-289 | 5 | Oas1a | homeostatic |
| Ifi2071 | 1.91E-292 | 1.21634743 | 0.759 | 0.555 | 3.69E-288 | 5 | Ifi207 | homeostatic |
| Plscr2 | 2.34E-291 | 2.81181776 | 0.377 | 0.096 | 4.53E-287 | 5 | Plscr2 | homeostatic |
| Gbp21 | 1.55E-290 | 1.75612057 | 0.556 | 0.244 | 3.00E-286 | 5 | Gbp2 | homeostatic |
| Mustn1 | 6.20E-290 | 3.50019962 | 0.286 | 0.052 | 1.20E-285 | 5 | Mustn1 | homeostatic |
| Il1r21 | 9.69E-290 | 2.18951937 | 0.539 | 0.251 | 1.87E-285 | 5 | Il1r2 | homeostatic |
| Sigmar11 | 2.01E-289 | 1.07614845 | 0.929 | 0.807 | 3.88E-285 | 5 | Sigmar1 | homeostatic |
| Gstm2 | 5.47E-287 | 1.05450229 | 0.949 | 0.847 | 1.06E-282 | 5 | Gstm2 | homeostatic |
| Ndrg4 | 2.31E-286 | 1.58523594 | 0.721 | 0.418 | 4.46E-282 | 5 | Ndrg4 | homeostatic |
| Rassf2 | 2.32E-286 | 2.05565347 | 0.58 | 0.208 | 4.48E-282 | 5 | Rassf2 | homeostatic |
| Grk51 | 8.05E-284 | 1.48190324 | 0.73 | 0.429 | 1.56E-279 | 5 | Grk5 | homeostatic |
| Cd471 | 2.54E-281 | 0.72341271 | 0.959 | 0.914 | 4.92E-277 | 5 | Cd47 | homeostatic |
| Duoxa1 | 1.33E-279 | 3.89042091 | 0.34 | 0.035 | 2.56E-275 | 5 | Duoxa1 | homeostatic |
| Gvin1 | 6.93E-279 | 2.49752393 | 0.439 | 0.098 | 1.34E-274 | 5 | Gvin1 | homeostatic |
| Npc21 | 1.85E-278 | 0.71858865 | 0.99 | 0.961 | 3.57E-274 | 5 | Npc2 | homeostatic |
| Antxr2 | 3.00E-274 | 1.11592927 | 0.864 | 0.766 | 5.80E-270 | 5 | Antxr2 | homeostatic |
| Trim30a | 5.87E-271 | 2.56224496 | 0.472 | 0.124 | 1.14E-266 | 5 | Trim30a | homeostatic |
| Pycard | 1.51E-270 | 2.10679022 | 0.486 | 0.182 | 2.92E-266 | 5 | Pycard | homeostatic |
| Ngf | 5.03E-270 | 2.95941654 | 0.34 | 0.053 | 9.72E-266 | 5 | Ngf | homeostatic |
| Il17ra1 | 2.30E-267 | 1.23234997 | 0.869 | 0.649 | 4.45E-263 | 5 | Il17ra | homeostatic |
| Tnfaip62 | 1.10E-266 | 1.20662529 | 0.701 | 0.398 | 2.13E-262 | 5 | Tnfaip6 | homeostatic |
| Bmp8a | 4.87E-266 | 3.7907551 | 0.272 | 0.035 | 9.41E-262 | 5 | Bmp8a | homeostatic |
| Celf21 | 4.98E-265 | 1.6012754 | 0.815 | 0.477 | 9.63E-261 | 5 | Celf2 | homeostatic |
| Glipr2 | 9.98E-265 | 0.93190033 | 0.92 | 0.812 | 1.93E-260 | 5 | Glipr2 | homeostatic |
| Stat2 | 2.32E-262 | 1.88644759 | 0.602 | 0.255 | 4.49E-258 | 5 | Stat2 | homeostatic |
| Tshz21 | 3.66E-262 | 1.54877338 | 0.775 | 0.4 | 7.08E-258 | 5 | Tshz2 | homeostatic |
| Ccdc68 | 1.27E-259 | 3.60737819 | 0.299 | 0.032 | 2.45E-255 | 5 | Ccdc68 | homeostatic |
| Barx2 | 1.12E-258 | 3.36313514 | 0.251 | 0.029 | 2.17E-254 | 5 | Barx2 | homeostatic |
| Ube2l6 | 7.03E-258 | 1.60825869 | 0.716 | 0.375 | 1.36E-253 | 5 | Ube2l6 | homeostatic |
| Pdzd2 | 2.87E-256 | 3.53539723 | 0.32 | 0.045 | 5.55E-252 | 5 | Pdzd2 | homeostatic |
| Dmpk1 | 5.52E-256 | 1.89964274 | 0.607 | 0.234 | 1.07E-251 | 5 | Dmpk | homeostatic |
| Itga111 | 8.09E-252 | 1.64035864 | 0.678 | 0.317 | 1.56E-247 | 5 | Itga11 | homeostatic |
| Prdx51 | 4.57E-251 | 0.7135325 | 0.989 | 0.962 | 8.83E-247 | 5 | Prdx5 | homeostatic |
| Dhx58 | 1.81E-248 | 2.80118563 | 0.345 | 0.076 | 3.50E-244 | 5 | Dhx58 | homeostatic |
| Ebf2 | 3.44E-248 | 2.22073719 | 0.519 | 0.156 | 6.64E-244 | 5 | Ebf2 | homeostatic |
| Ccl72 | 4.43E-247 | 0.88921424 | 0.613 | 0.377 | 8.56E-243 | 5 | Ccl7 | homeostatic |
| Lama41 | 4.87E-246 | 0.66267577 | 0.966 | 0.921 | 9.41E-242 | 5 | Lama4 | homeostatic |
| Rcan11 | 6.21E-246 | 1.07546646 | 0.58 | 0.357 | 1.20E-241 | 5 | Rcan1 | homeostatic |
| Aldh2 | 8.73E-242 | 0.98846176 | 0.963 | 0.879 | 1.69E-237 | 5 | Aldh2 | homeostatic |
| Tmem254a | 9.01E-242 | 2.04061676 | 0.609 | 0.265 | 1.74E-237 | 5 | Tmem254a | homeostatic |
| Ctps3 | 3.04E-239 | 1.38007939 | 0.746 | 0.531 | 5.87E-235 | 5 | Ctps | homeostatic |
| Ace1 | 2.05E-238 | 0.90379251 | 0.862 | 0.729 | 3.97E-234 | 5 | Ace | homeostatic |
| Ephx1 | 3.33E-238 | 1.96494329 | 0.504 | 0.242 | 6.43E-234 | 5 | Ephx1 | homeostatic |
| Daglb | 8.28E-236 | 2.20489694 | 0.566 | 0.204 | 1.60E-231 | 5 | Daglb | homeostatic |
| Poglut3 | 1.59E-235 | 1.3062924 | 0.774 | 0.584 | 3.08E-231 | 5 | Poglut3 | homeostatic |
| Irgm1 | 3.62E-231 | 1.53629834 | 0.591 | 0.34 | 6.99E-227 | 5 | Irgm1 | homeostatic |
| Cpne81 | 3.95E-229 | 1.58155225 | 0.749 | 0.413 | 7.64E-225 | 5 | Cpne8 | homeostatic |
| Pxn | 8.04E-227 | 1.59459097 | 0.732 | 0.401 | 1.55E-222 | 5 | Pxn | homeostatic |
| Naaa | 1.12E-226 | 1.54795914 | 0.694 | 0.386 | 2.16E-222 | 5 | Naaa | homeostatic |
| Hspa12b | 1.37E-226 | 2.22364295 | 0.459 | 0.179 | 2.66E-222 | 5 | Hspa12b | homeostatic |
| Reck | 2.00E-225 | 1.54065044 | 0.711 | 0.447 | 3.86E-221 | 5 | Reck | homeostatic |
| C1galt1 | 2.12E-225 | 1.50170289 | 0.704 | 0.468 | 4.10E-221 | 5 | C1galt1 | homeostatic |
| Xaf1 | 8.05E-224 | 1.84145471 | 0.544 | 0.221 | 1.56E-219 | 5 | Xaf1 | homeostatic |
| Klhl13 | 8.75E-222 | 2.30402826 | 0.481 | 0.166 | 1.69E-217 | 5 | Klhl13 | homeostatic |
| Pygl2 | 3.39E-220 | 1.43017116 | 0.761 | 0.403 | 6.56E-216 | 5 | Pygl | homeostatic |
| Kif21a1 | 5.73E-220 | 2.20202679 | 0.48 | 0.151 | 1.11E-215 | 5 | Kif21a | homeostatic |
| Gbp3 | 4.53E-218 | 2.02111101 | 0.406 | 0.124 | 8.75E-214 | 5 | Gbp3 | homeostatic |
| Cavin2 | 7.84E-218 | 2.6812777 | 0.331 | 0.078 | 1.52E-213 | 5 | Cavin2 | homeostatic |
| Lrp1 | 2.32E-215 | 0.41418124 | 0.996 | 0.989 | 4.48E-211 | 5 | Lrp1 | homeostatic |
| Gng121 | 7.27E-215 | 0.84446909 | 0.892 | 0.874 | 1.41E-210 | 5 | Gng12 | homeostatic |
| Opcml | 2.08E-212 | 3.94847251 | 0.272 | 0.038 | 4.02E-208 | 5 | Opcml | homeostatic |
| Phf11b | 1.94E-211 | 2.91892363 | 0.32 | 0.06 | 3.76E-207 | 5 | Phf11b | homeostatic |
| Metrnl | 6.69E-211 | 1.25801781 | 0.84 | 0.684 | 1.29E-206 | 5 | Metrnl | homeostatic |
| Tmem30b | 1.21E-208 | 2.60377389 | 0.382 | 0.076 | 2.35E-204 | 5 | Tmem30b | homeostatic |
| Tlr5 | 7.44E-208 | 2.92946872 | 0.305 | 0.048 | 1.44E-203 | 5 | Tlr5 | homeostatic |
| Ndufa4l23 | 4.58E-207 | 0.43346529 | 0.435 | 0.43 | 8.86E-203 | 5 | Ndufa4l2 | homeostatic |
| Cygb1 | 1.23E-205 | 0.67041389 | 0.943 | 0.824 | 2.38E-201 | 5 | Cygb | homeostatic |
| Anxa81 | 2.29E-203 | 1.5820107 | 0.327 | 0.13 | 4.42E-199 | 5 | Anxa8 | homeostatic |
| Dmrt2 | 2.65E-203 | 2.49492404 | 0.379 | 0.088 | 5.12E-199 | 5 | Dmrt2 | homeostatic |
| Elmo1 | 5.89E-203 | 2.62820191 | 0.355 | 0.074 | 1.14E-198 | 5 | Elmo1 | homeostatic |
| Apol6 | 1.92E-201 | 3.16373035 | 0.264 | 0.05 | 3.72E-197 | 5 | Apol6 | homeostatic |
| Mndal1 | 4.99E-201 | 1.71322188 | 0.539 | 0.248 | 9.64E-197 | 5 | Mndal | homeostatic |
| Ifih1 | 2.52E-200 | 1.62169284 | 0.591 | 0.309 | 4.88E-196 | 5 | Ifih1 | homeostatic |
| Ergic1 | 9.42E-199 | 0.82020789 | 0.939 | 0.878 | 1.82E-194 | 5 | Ergic1 | homeostatic |
| Crip2 | 1.70E-198 | 1.57430683 | 0.414 | 0.191 | 3.29E-194 | 5 | Crip2 | homeostatic |
| Dtnbp1 | 1.56E-197 | 0.92955008 | 0.937 | 0.831 | 3.01E-193 | 5 | Dtnbp1 | homeostatic |
| Plce1 | 1.49E-196 | 2.44056609 | 0.464 | 0.148 | 2.89E-192 | 5 | Plce1 | homeostatic |
| Zbtb20 | 2.68E-196 | 1.37156175 | 0.755 | 0.476 | 5.18E-192 | 5 | Zbtb20 | homeostatic |
| Idh1 | 1.70E-195 | 1.10360701 | 0.852 | 0.696 | 3.28E-191 | 5 | Idh1 | homeostatic |
| Fam129a1 | 2.24E-195 | 1.31788051 | 0.673 | 0.34 | 4.33E-191 | 5 | Fam129a | homeostatic |
| Igsf101 | 2.82E-195 | 0.99410477 | 0.888 | 0.751 | 5.45E-191 | 5 | Igsf10 | homeostatic |
| Bmp6 | 2.00E-192 | 2.58071685 | 0.279 | 0.068 | 3.86E-188 | 5 | Bmp6 | homeostatic |
| Srgap11 | 3.54E-192 | 2.02779282 | 0.53 | 0.195 | 6.85E-188 | 5 | Srgap1 | homeostatic |
| Srpx | 1.30E-191 | 0.9797286 | 0.891 | 0.76 | 2.51E-187 | 5 | Srpx | homeostatic |
| Trim30d | 3.76E-191 | 2.69923718 | 0.332 | 0.078 | 7.26E-187 | 5 | Trim30d | homeostatic |
| Sema3d | 5.96E-191 | 2.33474826 | 0.386 | 0.08 | 1.15E-186 | 5 | Sema3d | homeostatic |
| Tmem106a1 | 1.19E-190 | 1.79115987 | 0.641 | 0.306 | 2.30E-186 | 5 | Tmem106a | homeostatic |
| Ebf3 | 2.38E-189 | 2.51916973 | 0.426 | 0.116 | 4.59E-185 | 5 | Ebf3 | homeostatic |
| Ext1 | 1.52E-187 | 0.66009495 | 0.959 | 0.925 | 2.94E-183 | 5 | Ext1 | homeostatic |
| Sec14l1 | 1.85E-186 | 1.16557772 | 0.821 | 0.635 | 3.58E-182 | 5 | Sec14l1 | homeostatic |
| Mpp61 | 3.37E-186 | 1.3205543 | 0.681 | 0.48 | 6.51E-182 | 5 | Mpp6 | homeostatic |
| Rab232 | 5.79E-186 | 1.37594263 | 0.737 | 0.483 | 1.12E-181 | 5 | Rab23 | homeostatic |
| Epb41l2 | 5.98E-186 | 0.87396535 | 0.914 | 0.822 | 1.16E-181 | 5 | Epb41l2 | homeostatic |
| Irx11 | 5.43E-185 | 1.00734908 | 0.73 | 0.59 | 1.05E-180 | 5 | Irx1 | homeostatic |
| Grb10 | 1.16E-184 | 0.90432199 | 0.836 | 0.738 | 2.24E-180 | 5 | Grb10 | homeostatic |
| H2-K11 | 1.57E-183 | 0.90768424 | 0.918 | 0.812 | 3.03E-179 | 5 | H2-K1 | homeostatic |
| Pkdcc2 | 4.08E-180 | 0.93951557 | 0.83 | 0.695 | 7.90E-176 | 5 | Pkdcc | homeostatic |
| Nfib2 | 1.90E-179 | 1.31971873 | 0.749 | 0.407 | 3.67E-175 | 5 | Nfib | homeostatic |
| Man2a11 | 1.93E-178 | 0.64146845 | 0.958 | 0.926 | 3.73E-174 | 5 | Man2a1 | homeostatic |
| Esr11 | 4.51E-178 | 1.42654396 | 0.646 | 0.377 | 8.71E-174 | 5 | Esr1 | homeostatic |
| Cpq1 | 1.38E-177 | 1.30931049 | 0.707 | 0.461 | 2.67E-173 | 5 | Cpq | homeostatic |
| Cldn101 | 2.19E-176 | 1.69594966 | 0.463 | 0.198 | 4.23E-172 | 5 | Cldn10 | homeostatic |
| Fgf71 | 1.80E-175 | 0.90734112 | 0.82 | 0.464 | 3.48E-171 | 5 | Fgf7 | homeostatic |
| S100a11 | 2.67E-175 | 0.39177605 | 0.992 | 0.985 | 5.16E-171 | 5 | S100a11 | homeostatic |
| Kcnj8 | 1.26E-174 | 2.0603244 | 0.26 | 0.076 | 2.43E-170 | 5 | Kcnj8 | homeostatic |
| Mcub1 | 1.00E-173 | 1.4605476 | 0.591 | 0.337 | 1.94E-169 | 5 | Mcub | homeostatic |
| Ctsh1 | 1.84E-173 | 0.61547827 | 0.961 | 0.911 | 3.56E-169 | 5 | Ctsh | homeostatic |
| Rhoj | 3.05E-173 | 0.61299227 | 0.968 | 0.924 | 5.90E-169 | 5 | Rhoj | homeostatic |
| Ap2a2 | 7.07E-173 | 0.85196495 | 0.929 | 0.851 | 1.37E-168 | 5 | Ap2a2 | homeostatic |
| Dag11 | 8.25E-172 | 0.80009863 | 0.921 | 0.851 | 1.60E-167 | 5 | Dag1 | homeostatic |
| Ccl81 | 1.19E-171 | 1.26461621 | 0.664 | 0.344 | 2.31E-167 | 5 | Ccl8 | homeostatic |
| Jak11 | 4.20E-170 | 0.73035931 | 0.953 | 0.894 | 8.11E-166 | 5 | Jak1 | homeostatic |
| Adgrg2 | 2.46E-169 | 3.57712303 | 0.263 | 0.032 | 4.76E-165 | 5 | Adgrg2 | homeostatic |
| Six12 | 9.56E-169 | 1.94709145 | 0.478 | 0.17 | 1.85E-164 | 5 | Six1 | homeostatic |
| Slc1a51 | 1.38E-168 | 1.23384892 | 0.863 | 0.639 | 2.66E-164 | 5 | Slc1a5 | homeostatic |
| Gnpnat1 | 2.29E-168 | 1.21118867 | 0.775 | 0.624 | 4.43E-164 | 5 | Gnpnat1 | homeostatic |
| H2-D11 | 2.37E-165 | 1.44275999 | 0.624 | 0.353 | 4.59E-161 | 5 | H2-D1 | homeostatic |
| Card10 | 2.57E-165 | 2.8143364 | 0.259 | 0.051 | 4.98E-161 | 5 | Card10 | homeostatic |
| Scrn1 | 1.12E-164 | 2.91201638 | 0.325 | 0.066 | 2.16E-160 | 5 | Scrn1 | homeostatic |
| Phf24 | 2.59E-164 | 3.17006015 | 0.288 | 0.048 | 5.00E-160 | 5 | Phf24 | homeostatic |
| Irf9 | 2.93E-164 | 1.50851333 | 0.659 | 0.367 | 5.67E-160 | 5 | Irf9 | homeostatic |
| Unc45b | 8.26E-164 | 2.60679259 | 0.341 | 0.083 | 1.60E-159 | 5 | Unc45b | homeostatic |
| Ston2 | 8.56E-164 | 1.84485133 | 0.42 | 0.188 | 1.66E-159 | 5 | Ston2 | homeostatic |
| Adam15 | 7.46E-163 | 1.06861781 | 0.858 | 0.711 | 1.44E-158 | 5 | Adam15 | homeostatic |
| Atp2b4 | 8.79E-163 | 1.73865995 | 0.444 | 0.19 | 1.70E-158 | 5 | Atp2b4 | homeostatic |
| Polb | 2.74E-162 | 1.12553666 | 0.708 | 0.518 | 5.29E-158 | 5 | Polb | homeostatic |
| Il10rb | 5.49E-162 | 1.1491834 | 0.766 | 0.545 | 1.06E-157 | 5 | Il10rb | homeostatic |
| Has12 | 6.28E-162 | 1.30039258 | 0.533 | 0.231 | 1.21E-157 | 5 | Has1 | homeostatic |
| Tor1aip1 | 7.29E-162 | 0.93086898 | 0.883 | 0.78 | 1.41E-157 | 5 | Tor1aip1 | homeostatic |
| Jam21 | 2.86E-161 | 1.74320603 | 0.438 | 0.184 | 5.53E-157 | 5 | Jam2 | homeostatic |
| Hspb81 | 5.49E-159 | 0.87784548 | 0.786 | 0.645 | 1.06E-154 | 5 | Hspb8 | homeostatic |
| Rab3d1 | 1.84E-157 | 1.37176362 | 0.599 | 0.344 | 3.57E-153 | 5 | Rab3d | homeostatic |
| Nhsl11 | 5.68E-156 | 1.62773232 | 0.551 | 0.342 | 1.10E-151 | 5 | Nhsl1 | homeostatic |
| Fxyd51 | 5.67E-155 | 0.59200155 | 0.881 | 0.866 | 1.10E-150 | 5 | Fxyd5 | homeostatic |
| Shisa51 | 2.62E-154 | 0.71552802 | 0.954 | 0.915 | 5.06E-150 | 5 | Shisa5 | homeostatic |
| Mfsd7a | 8.32E-154 | 2.50922341 | 0.338 | 0.08 | 1.61E-149 | 5 | Mfsd7a | homeostatic |
| Aspn1 | 1.66E-152 | 0.86077477 | 0.673 | 0.411 | 3.21E-148 | 5 | Aspn | homeostatic |
| Sash1 | 1.28E-151 | 1.00349554 | 0.871 | 0.76 | 2.47E-147 | 5 | Sash1 | homeostatic |
| Igtp | 5.77E-151 | 1.84895336 | 0.355 | 0.14 | 1.12E-146 | 5 | Igtp | homeostatic |
| Dst1 | 4.51E-149 | 0.84539734 | 0.897 | 0.771 | 8.72E-145 | 5 | Dst | homeostatic |
| Gns1 | 2.61E-148 | 0.73128135 | 0.943 | 0.884 | 5.06E-144 | 5 | Gns | homeostatic |
| Col4a5 | 4.48E-148 | 2.93470376 | 0.284 | 0.053 | 8.65E-144 | 5 | Col4a5 | homeostatic |
| Mcc1 | 7.24E-148 | 1.48095436 | 0.619 | 0.366 | 1.40E-143 | 5 | Mcc | homeostatic |
| Ntrk21 | 5.89E-146 | 1.35149294 | 0.571 | 0.271 | 1.14E-141 | 5 | Ntrk2 | homeostatic |
| Eya41 | 5.62E-145 | 2.06777372 | 0.468 | 0.163 | 1.09E-140 | 5 | Eya4 | homeostatic |
| Procr1 | 6.38E-143 | 1.99934228 | 0.489 | 0.241 | 1.23E-138 | 5 | Procr | homeostatic |
| Cyp7b11 | 1.19E-142 | 0.92643271 | 0.604 | 0.574 | 2.29E-138 | 5 | Cyp7b1 | homeostatic |
| Fam102a1 | 3.67E-142 | 1.21745558 | 0.723 | 0.518 | 7.09E-138 | 5 | Fam102a | homeostatic |
| Dlc11 | 4.25E-141 | 0.97452849 | 0.832 | 0.646 | 8.21E-137 | 5 | Dlc1 | homeostatic |
| Dkk2 | 2.89E-140 | 0.96928456 | 0.273 | 0.124 | 5.58E-136 | 5 | Dkk2 | homeostatic |
| C1qtnf7 | 7.43E-140 | 2.22901202 | 0.363 | 0.103 | 1.44E-135 | 5 | C1qtnf7 | homeostatic |
| Rassf5 | 1.22E-139 | 1.13000616 | 0.572 | 0.374 | 2.35E-135 | 5 | Rassf5 | homeostatic |
| Gabre | 3.60E-139 | 2.00353866 | 0.327 | 0.102 | 6.96E-135 | 5 | Gabre | homeostatic |
| Klf3 | 1.91E-138 | 0.91649381 | 0.797 | 0.708 | 3.69E-134 | 5 | Klf3 | homeostatic |
| Ddah21 | 4.76E-138 | 1.07475021 | 0.836 | 0.661 | 9.20E-134 | 5 | Ddah2 | homeostatic |
| Ahnak21 | 2.25E-137 | 0.8667003 | 0.819 | 0.691 | 4.36E-133 | 5 | Ahnak2 | homeostatic |
| Lysmd21 | 4.55E-137 | 1.5890749 | 0.6 | 0.326 | 8.80E-133 | 5 | Lysmd2 | homeostatic |
| Asap31 | 1.64E-135 | 1.31778113 | 0.64 | 0.341 | 3.18E-131 | 5 | Asap3 | homeostatic |
| Nucb21 | 6.86E-135 | 0.9986602 | 0.849 | 0.673 | 1.33E-130 | 5 | Nucb2 | homeostatic |
| Mrgprf1 | 1.04E-134 | 1.24792025 | 0.77 | 0.51 | 2.02E-130 | 5 | Mrgprf | homeostatic |
| Dzip11 | 1.46E-134 | 1.6130018 | 0.608 | 0.327 | 2.82E-130 | 5 | Dzip1 | homeostatic |
| Abca1 | 5.20E-133 | 1.06947029 | 0.76 | 0.564 | 1.00E-128 | 5 | Abca1 | homeostatic |
| Dsel | 2.61E-132 | 0.91312819 | 0.772 | 0.664 | 5.04E-128 | 5 | Dsel | homeostatic |
| Ankrd46 | 2.11E-131 | 1.2101662 | 0.651 | 0.469 | 4.09E-127 | 5 | Ankrd46 | homeostatic |
| Vwa5a1 | 1.68E-130 | 0.82557477 | 0.841 | 0.725 | 3.24E-126 | 5 | Vwa5a | homeostatic |
| Eif2ak2 | 6.91E-130 | 1.2677591 | 0.643 | 0.406 | 1.34E-125 | 5 | Eif2ak2 | homeostatic |
| Twsg1 | 1.19E-128 | 0.7674259 | 0.929 | 0.864 | 2.31E-124 | 5 | Twsg1 | homeostatic |
| Lgi31 | 1.38E-128 | 1.96378601 | 0.345 | 0.154 | 2.67E-124 | 5 | Lgi3 | homeostatic |
| Slc16a12 | 2.01E-128 | 2.17099395 | 0.277 | 0.084 | 3.90E-124 | 5 | Slc16a12 | homeostatic |
| S100a61 | 2.44E-128 | 0.33307806 | 0.997 | 0.983 | 4.71E-124 | 5 | S100a6 | homeostatic |
| Sytl2 | 4.62E-128 | 2.00824104 | 0.416 | 0.157 | 8.93E-124 | 5 | Sytl2 | homeostatic |
| Tlr4 | 1.79E-126 | 1.71811778 | 0.506 | 0.222 | 3.46E-122 | 5 | Tlr4 | homeostatic |
| Vat1 | 2.89E-126 | 0.59528903 | 0.972 | 0.946 | 5.58E-122 | 5 | Vat1 | homeostatic |
| Slfn81 | 1.27E-125 | 1.94251185 | 0.41 | 0.151 | 2.45E-121 | 5 | Slfn8 | homeostatic |
| Angptl42 | 1.30E-125 | 0.60322544 | 0.912 | 0.752 | 2.51E-121 | 5 | Angptl4 | homeostatic |
| Lpar4 | 4.14E-125 | 2.25973115 | 0.329 | 0.116 | 8.01E-121 | 5 | Lpar4 | homeostatic |
| Abcd2 | 6.15E-125 | 2.06455556 | 0.396 | 0.137 | 1.19E-120 | 5 | Abcd2 | homeostatic |
| Trf1 | 7.93E-124 | 1.15675429 | 0.457 | 0.35 | 1.53E-119 | 5 | Trf | homeostatic |
| Cpne21 | 2.46E-123 | 1.46496861 | 0.541 | 0.267 | 4.75E-119 | 5 | Cpne2 | homeostatic |
| Tle5 | 3.54E-123 | 0.88561454 | 0.876 | 0.768 | 6.84E-119 | 5 | Tle5 | homeostatic |
| Palm | 5.36E-123 | 0.87387086 | 0.888 | 0.788 | 1.04E-118 | 5 | Palm | homeostatic |
| Hacd4 | 1.58E-122 | 1.47739638 | 0.567 | 0.331 | 3.06E-118 | 5 | Hacd4 | homeostatic |
| Blvrb | 3.30E-122 | 1.21607766 | 0.676 | 0.456 | 6.39E-118 | 5 | Blvrb | homeostatic |
| Tnfrsf1b1 | 1.02E-121 | 1.36333853 | 0.665 | 0.385 | 1.98E-117 | 5 | Tnfrsf1b | homeostatic |
| Arhgap10 | 3.00E-120 | 1.11771877 | 0.698 | 0.507 | 5.80E-116 | 5 | Arhgap10 | homeostatic |
| Cltb | 5.58E-118 | 0.87398215 | 0.905 | 0.794 | 1.08E-113 | 5 | Cltb | homeostatic |
| Fbxw17 | 2.12E-117 | 1.49952972 | 0.518 | 0.27 | 4.09E-113 | 5 | Fbxw17 | homeostatic |
| Nod11 | 2.20E-117 | 1.09824708 | 0.817 | 0.533 | 4.24E-113 | 5 | Nod1 | homeostatic |
| Parp14 | 2.26E-117 | 1.51172829 | 0.503 | 0.276 | 4.37E-113 | 5 | Parp14 | homeostatic |
| Slc31a2 | 9.28E-117 | 1.70903495 | 0.504 | 0.242 | 1.79E-112 | 5 | Slc31a2 | homeostatic |
| Gstt31 | 2.39E-115 | 1.78967271 | 0.449 | 0.169 | 4.61E-111 | 5 | Gstt3 | homeostatic |
| Atp10a1 | 3.30E-114 | 1.31532363 | 0.626 | 0.387 | 6.39E-110 | 5 | Atp10a | homeostatic |
| Mylip1 | 4.88E-113 | 1.29037823 | 0.59 | 0.365 | 9.44E-109 | 5 | Mylip | homeostatic |
| Tap1 | 1.28E-112 | 1.37816944 | 0.605 | 0.363 | 2.47E-108 | 5 | Tap1 | homeostatic |
| Hsd3b7 | 1.34E-112 | 1.03854938 | 0.701 | 0.49 | 2.60E-108 | 5 | Hsd3b7 | homeostatic |
| Samd9l | 5.56E-110 | 1.59383235 | 0.407 | 0.187 | 1.08E-105 | 5 | Samd9l | homeostatic |
| Zfp423 | 3.26E-109 | 1.605401 | 0.39 | 0.194 | 6.29E-105 | 5 | Zfp423 | homeostatic |
| Snhg11 | 1.71E-108 | 2.033206 | 0.286 | 0.072 | 3.30E-104 | 5 | Snhg11 | homeostatic |
| Efna5 | 1.02E-107 | 1.49639488 | 0.572 | 0.31 | 1.97E-103 | 5 | Efna5 | homeostatic |
| Plekha4 | 2.46E-107 | 1.51602916 | 0.486 | 0.227 | 4.75E-103 | 5 | Plekha4 | homeostatic |
| Stx2 | 1.21E-106 | 0.95413975 | 0.756 | 0.588 | 2.35E-102 | 5 | Stx2 | homeostatic |
| Parp9 | 1.83E-106 | 1.10795186 | 0.579 | 0.406 | 3.54E-102 | 5 | Parp9 | homeostatic |
| Nnmt1 | 1.19E-105 | 0.92338441 | 0.806 | 0.64 | 2.29E-101 | 5 | Nnmt | homeostatic |
| Il1rl21 | 5.18E-105 | 0.95709716 | 0.794 | 0.646 | 1.00E-100 | 5 | Il1rl2 | homeostatic |
| 8-Sep | 9.90E-105 | 0.69583466 | 0.954 | 0.892 | 1.91E-100 | 5 | 8-Sep | homeostatic |
| Ar | 2.41E-104 | 1.406162 | 0.594 | 0.371 | 4.67E-100 | 5 | Ar | homeostatic |
| Atp8b11 | 5.85E-104 | 0.89567881 | 0.774 | 0.604 | 1.13E-99 | 5 | Atp8b1 | homeostatic |
| Emx21 | 7.41E-104 | 1.64565568 | 0.513 | 0.237 | 1.43E-99 | 5 | Emx2 | homeostatic |
| Glud1 | 4.45E-103 | 0.6802026 | 0.916 | 0.857 | 8.60E-99 | 5 | Glud1 | homeostatic |
| 31-Mar | 6.83E-103 | 1.39500082 | 0.542 | 0.278 | 1.32E-98 | 5 | 3-Mar | homeostatic |
| Ildr2 | 8.02E-103 | 2.02345343 | 0.329 | 0.112 | 1.55E-98 | 5 | Ildr2 | homeostatic |
| Xpnpep21 | 1.25E-102 | 1.69512714 | 0.367 | 0.133 | 2.41E-98 | 5 | Xpnpep2 | homeostatic |
| Selenow | 3.55E-102 | 0.50848589 | 0.991 | 0.965 | 6.87E-98 | 5 | Selenow | homeostatic |
| Nqo21 | 4.34E-102 | 1.2656924 | 0.676 | 0.459 | 8.38E-98 | 5 | Nqo2 | homeostatic |
| Zfp5211 | 8.92E-100 | 1.1839045 | 0.648 | 0.353 | 1.72E-95 | 5 | Zfp521 | homeostatic |
| C1qtnf1 | 1.29E-99 | 0.83030495 | 0.825 | 0.74 | 2.49E-95 | 5 | C1qtnf1 | homeostatic |
| Myl12b | 9.72E-98 | 0.9633581 | 0.779 | 0.594 | 1.88E-93 | 5 | Myl12b | homeostatic |
| Fgf181 | 1.84E-97 | 1.37433431 | 0.353 | 0.151 | 3.55E-93 | 5 | Fgf18 | homeostatic |
| Pim11 | 2.52E-97 | 0.6024562 | 0.871 | 0.78 | 4.87E-93 | 5 | Pim1 | homeostatic |
| Ifi35 | 6.69E-97 | 1.45008779 | 0.51 | 0.3 | 1.29E-92 | 5 | Ifi35 | homeostatic |
| Sdk12 | 1.01E-96 | 1.08382282 | 0.75 | 0.509 | 1.94E-92 | 5 | Sdk1 | homeostatic |
| Ap3m2 | 1.82E-96 | 1.70026573 | 0.43 | 0.184 | 3.52E-92 | 5 | Ap3m2 | homeostatic |
| Pyroxd2 | 1.12E-95 | 1.37963342 | 0.536 | 0.287 | 2.16E-91 | 5 | Pyroxd2 | homeostatic |
| Tcea31 | 2.12E-95 | 1.66971604 | 0.335 | 0.119 | 4.10E-91 | 5 | Tcea3 | homeostatic |
| Plekhg1 | 6.97E-95 | 1.57718926 | 0.393 | 0.182 | 1.35E-90 | 5 | Plekhg1 | homeostatic |
| Adgra21 | 1.74E-94 | 0.61576467 | 0.937 | 0.889 | 3.37E-90 | 5 | Adgra2 | homeostatic |
| Etfbkmt | 1.87E-94 | 1.51927376 | 0.437 | 0.233 | 3.61E-90 | 5 | Etfbkmt | homeostatic |
| Socs31 | 3.50E-94 | 0.51379662 | 0.825 | 0.791 | 6.78E-90 | 5 | Socs3 | homeostatic |
| Agfg11 | 3.77E-94 | 0.62939148 | 0.877 | 0.818 | 7.28E-90 | 5 | Agfg1 | homeostatic |
| Zfpm2 | 3.80E-94 | 2.06912037 | 0.325 | 0.105 | 7.35E-90 | 5 | Zfpm2 | homeostatic |
| Sdf2l11 | 6.56E-94 | 1.03166483 | 0.756 | 0.552 | 1.27E-89 | 5 | Sdf2l1 | homeostatic |
| Cpt1a1 | 7.57E-94 | 0.99849363 | 0.753 | 0.52 | 1.46E-89 | 5 | Cpt1a | homeostatic |
| Nox41 | 9.77E-94 | 1.24188896 | 0.619 | 0.353 | 1.89E-89 | 5 | Nox4 | homeostatic |
| Gas61 | 2.60E-93 | 0.71095633 | 0.607 | 0.334 | 5.02E-89 | 5 | Gas6 | homeostatic |
| Tiam12 | 3.84E-93 | 1.45301267 | 0.536 | 0.276 | 7.42E-89 | 5 | Tiam1 | homeostatic |
| Lpgat11 | 9.28E-93 | 0.7515653 | 0.881 | 0.804 | 1.79E-88 | 5 | Lpgat1 | homeostatic |
| Plxdc2 | 5.56E-92 | 0.99567885 | 0.697 | 0.499 | 1.08E-87 | 5 | Plxdc2 | homeostatic |
| Rab301 | 6.56E-92 | 1.62750175 | 0.41 | 0.188 | 1.27E-87 | 5 | Rab30 | homeostatic |
| Scn1b1 | 7.09E-92 | 0.78854584 | 0.877 | 0.706 | 1.37E-87 | 5 | Scn1b | homeostatic |
| Agpat4 | 8.65E-92 | 1.02645767 | 0.683 | 0.513 | 1.67E-87 | 5 | Agpat4 | homeostatic |
| Parp31 | 7.51E-91 | 0.89721871 | 0.812 | 0.647 | 1.45E-86 | 5 | Parp3 | homeostatic |
| Lrrc321 | 8.39E-91 | 0.29852272 | 0.754 | 0.698 | 1.62E-86 | 5 | Lrrc32 | homeostatic |
| Arhgap211 | 2.61E-90 | 0.93700935 | 0.721 | 0.56 | 5.05E-86 | 5 | Arhgap21 | homeostatic |
| Osbpl9 | 6.67E-90 | 0.83643986 | 0.779 | 0.658 | 1.29E-85 | 5 | Osbpl9 | homeostatic |
| Rab8b1 | 1.57E-89 | 0.81304747 | 0.736 | 0.627 | 3.04E-85 | 5 | Rab8b | homeostatic |
| Capn51 | 2.16E-89 | 1.47598329 | 0.546 | 0.274 | 4.18E-85 | 5 | Capn5 | homeostatic |
| Paxx | 1.14E-88 | 1.59421696 | 0.454 | 0.24 | 2.21E-84 | 5 | Paxx | homeostatic |
| Arhgef3 | 1.46E-88 | 1.88018937 | 0.311 | 0.109 | 2.82E-84 | 5 | Arhgef3 | homeostatic |
| Rab7b1 | 1.85E-88 | 1.25675838 | 0.522 | 0.294 | 3.57E-84 | 5 | Rab7b | homeostatic |
| Pcolce1 | 3.96E-88 | 0.38206884 | 0.994 | 0.983 | 7.66E-84 | 5 | Pcolce | homeostatic |
| Dhrs71 | 7.14E-88 | 0.81677658 | 0.779 | 0.634 | 1.38E-83 | 5 | Dhrs7 | homeostatic |
| Ece1 | 2.01E-87 | 0.67774249 | 0.897 | 0.8 | 3.89E-83 | 5 | Ece1 | homeostatic |
| Il11ra11 | 5.29E-87 | 0.94798999 | 0.789 | 0.555 | 1.02E-82 | 5 | Il11ra1 | homeostatic |
| Ssbp2 | 1.26E-86 | 1.14082413 | 0.596 | 0.378 | 2.44E-82 | 5 | Ssbp2 | homeostatic |
| Pcdhgc31 | 1.82E-86 | 0.67700185 | 0.915 | 0.835 | 3.51E-82 | 5 | Pcdhgc3 | homeostatic |
| Coro2b | 2.04E-86 | 2.15358149 | 0.374 | 0.144 | 3.94E-82 | 5 | Coro2b | homeostatic |
| Arhgef10l | 2.28E-86 | 1.30997858 | 0.527 | 0.319 | 4.41E-82 | 5 | Arhgef10l | homeostatic |
| Sh3d191 | 3.00E-86 | 0.79571039 | 0.878 | 0.778 | 5.79E-82 | 5 | Sh3d19 | homeostatic |
| Tubb2b1 | 4.63E-86 | 0.904197 | 0.684 | 0.478 | 8.94E-82 | 5 | Tubb2b | homeostatic |
| St6gal1 | 9.73E-86 | 1.49608765 | 0.269 | 0.101 | 1.88E-81 | 5 | St6gal1 | homeostatic |
| Triobp1 | 2.34E-85 | 0.82852597 | 0.779 | 0.56 | 4.53E-81 | 5 | Triobp | homeostatic |
| Sult5a1 | 3.68E-85 | 2.16235159 | 0.253 | 0.079 | 7.12E-81 | 5 | Sult5a1 | homeostatic |
| Lama22 | 3.99E-85 | 0.68936417 | 0.822 | 0.656 | 7.70E-81 | 5 | Lama2 | homeostatic |
| Gngt21 | 5.48E-85 | 1.15680135 | 0.593 | 0.394 | 1.06E-80 | 5 | Gngt2 | homeostatic |
| Ifi47 | 9.35E-85 | 1.42763821 | 0.456 | 0.264 | 1.81E-80 | 5 | Ifi47 | homeostatic |
| Pfkp1 | 4.24E-84 | 0.77643537 | 0.798 | 0.61 | 8.21E-80 | 5 | Pfkp | homeostatic |
| Clip1 | 5.88E-84 | 0.85381487 | 0.773 | 0.593 | 1.14E-79 | 5 | Clip1 | homeostatic |
| Fxyd61 | 7.37E-84 | 0.73712033 | 0.631 | 0.644 | 1.43E-79 | 5 | Fxyd6 | homeostatic |
| Snx1 | 3.43E-83 | 0.7743987 | 0.817 | 0.724 | 6.62E-79 | 5 | Snx1 | homeostatic |
| Clu2 | 6.76E-83 | 0.38466952 | 0.924 | 0.883 | 1.31E-78 | 5 | Clu | homeostatic |
| Dnm1 | 1.06E-82 | 0.61217341 | 0.919 | 0.842 | 2.05E-78 | 5 | Dnm1 | homeostatic |
| Arl151 | 2.31E-82 | 1.02854704 | 0.69 | 0.521 | 4.47E-78 | 5 | Arl15 | homeostatic |
| Stat11 | 2.81E-82 | 0.97034423 | 0.73 | 0.541 | 5.44E-78 | 5 | Stat1 | homeostatic |
| Higd1a1 | 3.88E-82 | 0.34716286 | 0.778 | 0.789 | 7.50E-78 | 5 | Higd1a | homeostatic |
| Acadsb1 | 7.66E-82 | 0.80951736 | 0.799 | 0.689 | 1.48E-77 | 5 | Acadsb | homeostatic |
| Etfa1 | 1.27E-81 | 0.81171501 | 0.883 | 0.791 | 2.46E-77 | 5 | Etfa | homeostatic |

| | | | | | | | | |
|---|---|---|---|---|---|---|---|---|
| Nmt21 | 1.84E-81 | 0.91493011 | 0.78 | 0.639 | 3.56E-77 | 5 | Nmt2 | homeostatic |
| Zfhx31 | 1.08E-80 | 1.21641829 | 0.57 | 0.314 | 2.09E-76 | 5 | Zfhx3 | homeostatic |
| Adar | 1.72E-79 | 1.04548794 | 0.65 | 0.452 | 3.33E-75 | 5 | Adar | homeostatic |
| Best1 | 3.00E-79 | 2.04503345 | 0.284 | 0.104 | 5.80E-75 | 5 | Best1 | homeostatic |
| Emb1 | 3.89E-79 | 0.57455305 | 0.765 | 0.723 | 7.53E-75 | 5 | Emb | homeostatic |
| Fmnl22 | 7.35E-79 | 1.01237419 | 0.681 | 0.406 | 1.42E-74 | 5 | Fmnl2 | homeostatic |
| Tmem45a2 | 1.04E-78 | 0.50538257 | 0.84 | 0.745 | 2.01E-74 | 5 | Tmem45a | homeostatic |
| Dusp22 | 1.21E-78 | 1.71736061 | 0.307 | 0.129 | 2.33E-74 | 5 | Dusp22 | homeostatic |
| Mturn | 1.78E-78 | 1.6390877 | 0.392 | 0.177 | 3.45E-74 | 5 | Mturn | homeostatic |
| Il6ra1 | 2.47E-78 | 1.76665981 | 0.395 | 0.155 | 4.78E-74 | 5 | Il6ra | homeostatic |
| Cdkn1c1 | 4.24E-78 | 1.62177258 | 0.301 | 0.15 | 8.21E-74 | 5 | Cdkn1c | homeostatic |
| Tnfrsf1a1 | 5.77E-78 | 0.45686227 | 0.975 | 0.959 | 1.11E-73 | 5 | Tnfrsf1a | homeostatic |
| Hist1h2bc1 | 7.45E-78 | 0.54449232 | 0.764 | 0.648 | 1.44E-73 | 5 | Hist1h2bc | homeostatic |
| Abca2 | 9.37E-78 | 1.71449159 | 0.349 | 0.135 | 1.81E-73 | 5 | Abca2 | homeostatic |
| Adam331 | 1.69E-77 | 1.54451018 | 0.397 | 0.192 | 3.26E-73 | 5 | Adam33 | homeostatic |
| Mill2 | 2.74E-77 | 1.95032455 | 0.298 | 0.099 | 5.30E-73 | 5 | Mill2 | homeostatic |
| Sgce | 2.99E-77 | 0.89668539 | 0.841 | 0.707 | 5.79E-73 | 5 | Sgce | homeostatic |
| Cry21 | 1.36E-76 | 1.2628032 | 0.52 | 0.319 | 2.63E-72 | 5 | Cry2 | homeostatic |
| Mgarp3 | 2.87E-76 | 1.01249686 | 0.497 | 0.264 | 5.54E-72 | 5 | Mgarp | homeostatic |
| Tcf24 | 3.29E-76 | 1.92620922 | 0.286 | 0.11 | 6.36E-72 | 5 | Tcf24 | homeostatic |
| Creld11 | 5.51E-76 | 0.79341387 | 0.77 | 0.603 | 1.06E-71 | 5 | Creld1 | homeostatic |
| Fmo1 | 1.47E-75 | 1.53803353 | 0.312 | 0.114 | 2.84E-71 | 5 | Fmo1 | homeostatic |
| Ehbp1 | 1.43E-74 | 1.26262869 | 0.496 | 0.281 | 2.76E-70 | 5 | Ehbp1 | homeostatic |
| Lrrc75a | 9.38E-74 | 1.14434388 | 0.598 | 0.391 | 1.81E-69 | 5 | Lrrc75a | homeostatic |
| Pla2r11 | 1.96E-73 | 1.10549911 | 0.591 | 0.379 | 3.79E-69 | 5 | Pla2r1 | homeostatic |
| Shc2 | 5.92E-73 | 1.11197114 | 0.673 | 0.465 | 1.15E-68 | 5 | Shc2 | homeostatic |
| Jcad1 | 1.73E-72 | 0.92858049 | 0.642 | 0.439 | 3.35E-68 | 5 | Jcad | homeostatic |
| Rai2 | 2.61E-72 | 1.92785508 | 0.289 | 0.103 | 5.05E-68 | 5 | Rai2 | homeostatic |
| Iah1 | 3.21E-72 | 0.77992431 | 0.841 | 0.725 | 6.20E-68 | 5 | Iah1 | homeostatic |
| Sntb2 | 4.08E-72 | 0.95315045 | 0.773 | 0.626 | 7.89E-68 | 5 | Sntb2 | homeostatic |
| Dtx3l | 4.45E-72 | 1.03751499 | 0.548 | 0.386 | 8.60E-68 | 5 | Dtx3l | homeostatic |
| Sppl2a | 7.52E-72 | 0.77564115 | 0.769 | 0.664 | 1.45E-67 | 5 | Sppl2a | homeostatic |
| St3gal51 | 1.15E-71 | 1.67564949 | 0.341 | 0.175 | 2.22E-67 | 5 | St3gal5 | homeostatic |
| Cast | 1.37E-71 | 0.5354184 | 0.906 | 0.863 | 2.66E-67 | 5 | Cast | homeostatic |
| Diaph11 | 2.74E-71 | 1.1012335 | 0.627 | 0.438 | 5.30E-67 | 5 | Diaph1 | homeostatic |
| Maob | 3.61E-71 | 2.17731022 | 0.263 | 0.083 | 6.98E-67 | 5 | Maob | homeostatic |
| Eef2k | 4.06E-71 | 1.24639665 | 0.544 | 0.33 | 7.85E-67 | 5 | Eef2k | homeostatic |
| Aida1 | 3.87E-70 | 0.78340269 | 0.766 | 0.64 | 7.49E-66 | 5 | Aida | homeostatic |
| Prex11 | 8.25E-70 | 1.22494552 | 0.431 | 0.238 | 1.59E-65 | 5 | Prex1 | homeostatic |
| Pdgfc1 | 1.89E-69 | 1.40442463 | 0.326 | 0.183 | 3.64E-65 | 5 | Pdgfc | homeostatic |
| Olfml2b | 6.26E-69 | 0.79686766 | 0.844 | 0.682 | 1.21E-64 | 5 | Olfml2b | homeostatic |
| Parm11 | 6.77E-69 | 1.20671882 | 0.419 | 0.205 | 1.31E-64 | 5 | Parm1 | homeostatic |
| Ang1 | 1.78E-68 | 1.13030212 | 0.61 | 0.411 | 3.44E-64 | 5 | Ang | homeostatic |
| Tmod2 | 2.95E-68 | 1.67198203 | 0.287 | 0.108 | 5.71E-64 | 5 | Tmod2 | homeostatic |
| Amotl21 | 4.48E-68 | 0.65145679 | 0.839 | 0.769 | 8.65E-64 | 5 | Amotl2 | homeostatic |
| Slfn2 | 7.94E-68 | 1.56766441 | 0.331 | 0.155 | 1.54E-63 | 5 | Slfn2 | homeostatic |
| Ddx581 | 3.35E-67 | 0.9980505 | 0.523 | 0.352 | 6.48E-63 | 5 | Ddx58 | homeostatic |
| Trim251 | 5.13E-67 | 0.8099711 | 0.657 | 0.545 | 9.93E-63 | 5 | Trim25 | homeostatic |
| Stk38 | 2.62E-66 | 0.51332144 | 0.92 | 0.865 | 5.06E-62 | 5 | Stk38 | homeostatic |
| Rbpj1 | 8.17E-66 | 0.70409146 | 0.787 | 0.685 | 1.58E-61 | 5 | Rbpj | homeostatic |
| Smad1 | 3.16E-65 | 0.93389445 | 0.67 | 0.525 | 6.11E-61 | 5 | Smad1 | homeostatic |
| Adcy51 | 3.80E-65 | 1.50065668 | 0.332 | 0.151 | 7.34E-61 | 5 | Adcy5 | homeostatic |
| Tspan51 | 5.34E-65 | 0.91686619 | 0.647 | 0.463 | 1.03E-60 | 5 | Tspan5 | homeostatic |
| Sp140 | 5.69E-65 | 1.7789326 | 0.33 | 0.142 | 1.10E-60 | 5 | Sp140 | homeostatic |
| Ugcg2 | 7.99E-65 | 0.47110892 | 0.876 | 0.782 | 1.54E-60 | 5 | Ugcg | homeostatic |
| Radil | 2.19E-63 | 1.39720637 | 0.393 | 0.196 | 4.24E-59 | 5 | Radil | homeostatic |
| Cd3201 | 1.12E-62 | 1.06562783 | 0.615 | 0.437 | 2.17E-58 | 5 | Cd320 | homeostatic |
| Slc29a13 | 1.93E-62 | 1.19060283 | 0.478 | 0.291 | 3.74E-58 | 5 | Slc29a1 | homeostatic |
| Gan1 | 2.01E-62 | 1.03439375 | 0.538 | 0.453 | 3.89E-58 | 5 | Gan | homeostatic |
| Trip101 | 2.48E-62 | 0.72016296 | 0.817 | 0.695 | 4.80E-58 | 5 | Trip10 | homeostatic |
| Rbm431 | 2.74E-62 | 1.33910275 | 0.382 | 0.203 | 5.29E-58 | 5 | Rbm43 | homeostatic |
| Tle6 | 2.97E-62 | 1.64369303 | 0.277 | 0.106 | 5.75E-58 | 5 | Tle6 | homeostatic |
| Dcbld22 | 7.39E-62 | 0.66344722 | 0.836 | 0.702 | 1.43E-57 | 5 | Dcbld2 | homeostatic |
| Map3k81 | 1.40E-61 | 0.97436944 | 0.44 | 0.252 | 2.71E-57 | 5 | Map3k8 | homeostatic |
| Dram12 | 1.91E-61 | 0.44953424 | 0.893 | 0.842 | 3.70E-57 | 5 | Dram1 | homeostatic |
| Slc8b1 | 2.39E-61 | 1.69988942 | 0.321 | 0.124 | 4.62E-57 | 5 | Slc8b1 | homeostatic |
| Hpse1 | 2.62E-61 | 1.13077186 | 0.552 | 0.319 | 5.06E-57 | 5 | Hpse | homeostatic |
| Plcb4 | 5.21E-61 | 1.7951681 | 0.264 | 0.101 | 1.01E-56 | 5 | Plcb4 | homeostatic |
| Smurf2 | 1.84E-60 | 0.86351055 | 0.646 | 0.555 | 3.56E-56 | 5 | Smurf2 | homeostatic |
| Mmp231 | 2.37E-60 | 0.59422772 | 0.923 | 0.838 | 4.57E-56 | 5 | Mmp23 | homeostatic |
| Hspg2 | 2.81E-60 | 0.37624514 | 0.985 | 0.976 | 5.44E-56 | 5 | Hspg2 | homeostatic |
| Flt3l | 1.35E-59 | 1.79415398 | 0.261 | 0.094 | 2.60E-55 | 5 | Flt3l | homeostatic |
| Fam171a11 | 2.08E-59 | 0.96526093 | 0.665 | 0.524 | 4.02E-55 | 5 | Fam171a1 | homeostatic |
| Rbpms | 2.16E-59 | 0.63398924 | 0.831 | 0.756 | 4.18E-55 | 5 | Rbpms | homeostatic |
| Map1lc3b1 | 3.53E-59 | 0.45092123 | 0.943 | 0.895 | 6.83E-55 | 5 | Map1lc3b | homeostatic |
| Lum1 | 4.09E-59 | 0.38285145 | 0.966 | 0.838 | 7.92E-55 | 5 | Lum | homeostatic |
| Lrrc8d | 5.95E-59 | 1.67203403 | 0.268 | 0.105 | 1.15E-54 | 5 | Lrrc8d | homeostatic |
| Phactr2 | 7.25E-59 | 1.05121492 | 0.56 | 0.362 | 1.40E-54 | 5 | Phactr2 | homeostatic |
| Trim34a | 8.97E-59 | 1.63541198 | 0.268 | 0.1 | 1.73E-54 | 5 | Trim34a | homeostatic |
| Npepl1 | 1.05E-58 | 0.76963671 | 0.753 | 0.624 | 2.02E-54 | 5 | Npepl1 | homeostatic |
| Vcam12 | 3.79E-58 | 0.86697817 | 0.59 | 0.325 | 7.33E-54 | 5 | Vcam1 | homeostatic |
| Rab40b | 6.26E-58 | 1.4724766 | 0.372 | 0.182 | 1.21E-53 | 5 | Rab40b | homeostatic |
| Gulp1 | 1.42E-57 | 0.90595037 | 0.558 | 0.412 | 2.74E-53 | 5 | Gulp1 | homeostatic |
| Sox13 | 2.00E-57 | 1.67285695 | 0.299 | 0.116 | 3.86E-53 | 5 | Sox13 | homeostatic |
| Smpdl3a1 | 4.54E-57 | 1.38733766 | 0.449 | 0.229 | 8.78E-53 | 5 | Smpdl3a | homeostatic |
| Panx1 | 2.60E-56 | 0.93507157 | 0.513 | 0.356 | 5.02E-52 | 5 | Panx1 | homeostatic |
| Gmds2 | 3.88E-56 | 1.12814908 | 0.59 | 0.394 | 7.50E-52 | 5 | Gmds | homeostatic |
| Atf31 | 6.55E-56 | 0.61384446 | 0.398 | 0.308 | 1.27E-51 | 5 | Atf3 | homeostatic |
| Phyh1 | 8.84E-56 | 1.06492353 | 0.466 | 0.298 | 1.71E-51 | 5 | Phyh | homeostatic |
| Agpat32 | 1.09E-55 | 0.93042471 | 0.745 | 0.481 | 2.11E-51 | 5 | Agpat3 | homeostatic |
| Acat11 | 1.82E-55 | 0.7642121 | 0.782 | 0.663 | 3.53E-51 | 5 | Acat1 | homeostatic |
| Lmbr1 | 3.66E-55 | 1.33054888 | 0.487 | 0.28 | 7.08E-51 | 5 | Lmbr1 | homeostatic |
| Rsu1 | 3.92E-55 | 0.61998725 | 0.918 | 0.837 | 7.57E-51 | 5 | Rsu1 | homeostatic |
| Tor3a1 | 8.99E-55 | 1.31228858 | 0.431 | 0.228 | 1.74E-50 | 5 | Tor3a | homeostatic |
| Gadd45g1 | 1.09E-54 | 0.29990969 | 0.846 | 0.762 | 2.12E-50 | 5 | Gadd45g | homeostatic |
| Agtrap1 | 2.28E-54 | 1.26150254 | 0.516 | 0.329 | 4.41E-50 | 5 | Agtrap | homeostatic |
| Flnc1 | 2.62E-54 | 0.54130971 | 0.846 | 0.794 | 5.06E-50 | 5 | Flnc | homeostatic |
| Ghr1 | 2.66E-54 | 1.02112219 | 0.544 | 0.421 | 5.14E-50 | 5 | Ghr | homeostatic |
| Ech1 | 3.00E-54 | 0.67717799 | 0.802 | 0.692 | 5.80E-50 | 5 | Ech1 | homeostatic |
| Helz2 | 4.66E-54 | 0.88662391 | 0.626 | 0.505 | 9.00E-50 | 5 | Helz2 | homeostatic |
| Ankrd40 | 4.76E-54 | 0.55911749 | 0.857 | 0.813 | 9.19E-50 | 5 | Ankrd40 | homeostatic |
| Aif1l1 | 6.46E-54 | 1.00452248 | 0.5 | 0.277 | 1.25E-49 | 5 | Aif1l | homeostatic |
| Tcf7l21 | 9.47E-54 | 0.90890354 | 0.555 | 0.358 | 1.83E-49 | 5 | Tcf7l2 | homeostatic |
| Lpar1 | 1.08E-53 | 0.64149044 | 0.911 | 0.82 | 2.10E-49 | 5 | Lpar1 | homeostatic |
| Acly | 1.47E-53 | 0.62521935 | 0.887 | 0.819 | 2.85E-49 | 5 | Acly | homeostatic |
| Gata6 | 2.11E-53 | 1.26152008 | 0.364 | 0.167 | 4.07E-49 | 5 | Gata6 | homeostatic |
| Eps82 | 2.42E-53 | 0.81360118 | 0.57 | 0.352 | 4.69E-49 | 5 | Eps8 | homeostatic |
| Fam171a2 | 1.25E-52 | 1.16172073 | 0.53 | 0.336 | 2.42E-48 | 5 | Fam171a2 | homeostatic |
| Rabgap1l | 1.51E-52 | 0.64583186 | 0.664 | 0.559 | 2.92E-48 | 5 | Rabgap1l | homeostatic |
| Parp12 | 2.21E-52 | 0.90038216 | 0.59 | 0.43 | 4.28E-48 | 5 | Parp12 | homeostatic |
| Lrrc8c1 | 3.11E-52 | 1.05396467 | 0.443 | 0.285 | 6.01E-48 | 5 | Lrrc8c | homeostatic |
| Daam11 | 4.95E-52 | 0.8507374 | 0.586 | 0.408 | 9.57E-48 | 5 | Daam1 | homeostatic |
| Ifnar21 | 1.86E-51 | 0.59406667 | 0.885 | 0.841 | 3.60E-47 | 5 | Ifnar2 | homeostatic |
| Rgl1 | 4.75E-51 | 0.73936739 | 0.793 | 0.669 | 9.18E-47 | 5 | Rgl1 | homeostatic |
| B4galt1 | 5.01E-51 | 0.38517316 | 0.977 | 0.95 | 9.68E-47 | 5 | B4galt1 | homeostatic |
| Ifitm21 | 5.57E-51 | 0.3434361 | 0.994 | 0.973 | 1.08E-46 | 5 | Ifitm2 | homeostatic |
| Lgmn1 | 8.67E-51 | 0.48758131 | 0.92 | 0.861 | 1.68E-46 | 5 | Lgmn | homeostatic |
| Nmi1 | 1.01E-50 | 1.37296412 | 0.349 | 0.176 | 1.96E-46 | 5 | Nmi | homeostatic |
| Disp1 | 2.06E-50 | 1.5882408 | 0.321 | 0.152 | 3.97E-46 | 5 | Disp1 | homeostatic |
| Flot21 | 2.08E-50 | 0.64517159 | 0.82 | 0.719 | 4.01E-46 | 5 | Flot2 | homeostatic |
| Plin3 | 2.88E-50 | 0.6777354 | 0.831 | 0.708 | 5.56E-46 | 5 | Plin3 | homeostatic |
| Grn | 4.67E-50 | 0.41722902 | 0.984 | 0.962 | 9.03E-46 | 5 | Grn | homeostatic |
| Fez11 | 1.28E-49 | 1.07694638 | 0.572 | 0.387 | 2.47E-45 | 5 | Fez1 | homeostatic |
| Emc2 | 1.40E-49 | 0.83768599 | 0.701 | 0.568 | 2.70E-45 | 5 | Emc2 | homeostatic |
| Pdcd41 | 1.96E-49 | 0.58123252 | 0.784 | 0.646 | 3.79E-45 | 5 | Pdcd4 | homeostatic |
| Slc25a121 | 2.45E-49 | 1.01487079 | 0.584 | 0.427 | 4.74E-45 | 5 | Slc25a12 | homeostatic |
| H2-M3 | 2.83E-49 | 1.17430804 | 0.506 | 0.371 | 5.47E-45 | 5 | H2-M3 | homeostatic |
| Rap1b1 | 4.08E-49 | 0.43447671 | 0.921 | 0.898 | 7.89E-45 | 5 | Rap1b | homeostatic |
| Edem2 | 6.47E-49 | 0.93793732 | 0.694 | 0.534 | 1.25E-44 | 5 | Edem2 | homeostatic |
| Cd1512 | 1.07E-48 | 0.68080616 | 0.77 | 0.631 | 2.07E-44 | 5 | Cd151 | homeostatic |
| Cdon1 | 5.32E-48 | 0.67967904 | 0.735 | 0.671 | 1.03E-43 | 5 | Cdon | homeostatic |
| Snx18 | 5.86E-48 | 0.63137106 | 0.813 | 0.74 | 1.13E-43 | 5 | Snx18 | homeostatic |
| Fndc12 | 1.30E-47 | 0.51601289 | 0.923 | 0.797 | 2.51E-43 | 5 | Fndc1 | homeostatic |
| Mindy21 | 2.32E-47 | 0.83268383 | 0.579 | 0.432 | 4.49E-43 | 5 | Mindy2 | homeostatic |
| Ggct2 | 3.09E-47 | 1.45945768 | 0.341 | 0.163 | 5.97E-43 | 5 | Ggct | homeostatic |
| Ypel51 | 3.21E-47 | 0.53104635 | 0.766 | 0.694 | 6.22E-43 | 5 | Ypel5 | homeostatic |
| Fcgrt | 5.61E-47 | 0.6137339 | 0.93 | 0.89 | 1.08E-42 | 5 | Fcgrt | homeostatic |
| Eln1 | 1.81E-46 | 0.62762933 | 0.789 | 0.523 | 3.49E-42 | 5 | Eln | homeostatic |
| Mapk121 | 2.10E-46 | 1.17513389 | 0.382 | 0.214 | 4.06E-42 | 5 | Mapk12 | homeostatic |
| Abca7 | 5.32E-46 | 1.21159427 | 0.326 | 0.157 | 1.03E-41 | 5 | Abca7 | homeostatic |
| Adam17 | 8.51E-46 | 0.62550229 | 0.827 | 0.751 | 1.65E-41 | 5 | Adam17 | homeostatic |
| Chst13 | 1.42E-45 | 1.29533859 | 0.308 | 0.173 | 2.75E-41 | 5 | Chst1 | homeostatic |
| Tspo1 | 1.51E-45 | 0.52750403 | 0.918 | 0.862 | 2.93E-41 | 5 | Tspo | homeostatic |
| Slc8a11 | 2.48E-45 | 0.83007202 | 0.69 | 0.508 | 4.79E-41 | 5 | Slc8a1 | homeostatic |
| Gpam | 4.28E-45 | 0.95016074 | 0.478 | 0.356 | 8.27E-41 | 5 | Gpam | homeostatic |
| Notch21 | 6.38E-45 | 0.78085987 | 0.669 | 0.577 | 1.23E-40 | 5 | Notch2 | homeostatic |
| Elovl51 | 9.97E-45 | 0.63120068 | 0.723 | 0.641 | 1.93E-40 | 5 | Elovl5 | homeostatic |
| Efemp22 | 1.90E-44 | 0.26338982 | 0.972 | 0.964 | 3.68E-40 | 5 | Efemp2 | homeostatic |
| Fam162a3 | 2.89E-44 | 0.27890298 | 0.778 | 0.782 | 5.59E-40 | 5 | Fam162a | homeostatic |
| Rap2b | 3.34E-44 | 0.77237593 | 0.655 | 0.568 | 6.46E-40 | 5 | Rap2b | homeostatic |
| Nenf1 | 3.40E-44 | 0.5103646 | 0.948 | 0.912 | 6.58E-40 | 5 | Nenf | homeostatic |
| Sh3bp5 | 3.63E-44 | 0.90538817 | 0.572 | 0.425 | 7.01E-40 | 5 | Sh3bp5 | homeostatic |
| Sema3b1 | 8.42E-44 | 1.07682425 | 0.462 | 0.28 | 1.63E-39 | 5 | Sema3b | homeostatic |
| Dipk1a | 1.18E-43 | 1.41954337 | 0.272 | 0.127 | 2.28E-39 | 5 | Dipk1a | homeostatic |
| Herc6 | 2.75E-43 | 1.63785392 | 0.255 | 0.123 | 5.32E-39 | 5 | Herc6 | homeostatic |
| Rev3l | 3.96E-43 | 0.86522734 | 0.485 | 0.339 | 7.65E-39 | 5 | Rev3l | homeostatic |
| Fer | 6.50E-43 | 0.70684648 | 0.708 | 0.59 | 1.26E-38 | 5 | Fer | homeostatic |
| Tpcn1 | 7.92E-43 | 0.8190531 | 0.574 | 0.41 | 1.53E-38 | 5 | Tpcn1 | homeostatic |
| Gpc61 | 1.24E-42 | 0.70648218 | 0.581 | 0.53 | 2.39E-38 | 5 | Gpc6 | homeostatic |
| Fas1 | 1.34E-42 | 0.95690357 | 0.411 | 0.229 | 2.59E-38 | 5 | Fas | homeostatic |
| Tdrd7 | 3.41E-42 | 0.92667626 | 0.522 | 0.371 | 6.59E-38 | 5 | Tdrd7 | homeostatic |
| Adam9 | 4.11E-42 | 0.41058021 | 0.956 | 0.939 | 7.95E-38 | 5 | Adam9 | homeostatic |
| Sgpp11 | 6.65E-42 | 0.6758399 | 0.869 | 0.81 | 1.29E-37 | 5 | Sgpp1 | homeostatic |
| St3gal2 | 7.04E-42 | 0.66824714 | 0.796 | 0.685 | 1.36E-37 | 5 | St3gal2 | homeostatic |
| Iqsec2 | 7.88E-42 | 0.92634804 | 0.552 | 0.395 | 1.52E-37 | 5 | Iqsec2 | homeostatic |
| Capn11 | 1.10E-41 | 0.57289605 | 0.76 | 0.664 | 2.14E-37 | 5 | Capn1 | homeostatic |
| Plxna4 | 1.42E-41 | 1.05085244 | 0.272 | 0.15 | 2.74E-37 | 5 | Plxna4 | homeostatic |
| Lrch1 | 2.18E-41 | 1.25633519 | 0.415 | 0.225 | 4.22E-37 | 5 | Lrch1 | homeostatic |
| Cep170 | 2.21E-41 | 0.53955182 | 0.84 | 0.767 | 4.28E-37 | 5 | Cep170 | homeostatic |
| 8-Mar | 3.26E-41 | 0.8111465 | 0.659 | 0.508 | 6.30E-37 | 5 | 8-Mar | homeostatic |
| Zyx | 3.90E-41 | 0.49625503 | 0.953 | 0.908 | 7.53E-37 | 5 | Zyx | homeostatic |
| Ptch2 | 4.76E-41 | 0.95232008 | 0.395 | 0.222 | 9.21E-37 | 5 | Ptch2 | homeostatic |
| Dnpep | 7.10E-41 | 0.62669032 | 0.791 | 0.672 | 1.37E-36 | 5 | Dnpep | homeostatic |
| Galnt1 | 9.00E-41 | 0.59123983 | 0.896 | 0.831 | 1.74E-36 | 5 | Galnt1 | homeostatic |
| Syne21 | 9.64E-41 | 1.08231337 | 0.439 | 0.237 | 1.86E-36 | 5 | Syne2 | homeostatic |
| Ppp1r2 | 2.12E-40 | 0.59621274 | 0.736 | 0.647 | 4.10E-36 | 5 | Ppp1r2 | homeostatic |
| Glis31 | 3.52E-40 | 0.26963382 | 0.85 | 0.89 | 6.80E-36 | 5 | Glis3 | homeostatic |
| Limk11 | 5.21E-40 | 0.65124289 | 0.766 | 0.658 | 1.01E-35 | 5 | Limk1 | homeostatic |
| Hspa12a | 2.88E-39 | 1.12403285 | 0.352 | 0.178 | 5.58E-35 | 5 | Hspa12a | homeostatic |
| Pcdh191 | 4.49E-39 | 0.98663668 | 0.636 | 0.43 | 8.69E-35 | 5 | Pcdh19 | homeostatic |
| Naglu1 | 9.16E-39 | 0.80229639 | 0.664 | 0.474 | 1.77E-34 | 5 | Naglu | homeostatic |
| Ctso1 | 1.24E-38 | 0.88552818 | 0.624 | 0.474 | 2.40E-34 | 5 | Ctso | homeostatic |
| Cercam | 1.29E-38 | 0.46938343 | 0.926 | 0.873 | 2.48E-34 | 5 | Cercam | homeostatic |
| Sgsm2 | 1.88E-38 | 1.21065868 | 0.423 | 0.259 | 3.64E-34 | 5 | Sgsm2 | homeostatic |
| Zdhhc8 | 2.09E-38 | 0.59333563 | 0.777 | 0.706 | 4.04E-34 | 5 | Zdhhc8 | homeostatic |
| Elf11 | 3.76E-38 | 0.70862019 | 0.706 | 0.566 | 7.27E-34 | 5 | Elf1 | homeostatic |
| Osbpl1a1 | 3.88E-38 | 0.91448908 | 0.558 | 0.373 | 7.50E-34 | 5 | Osbpl1a | homeostatic |
| Lhpp1 | 5.30E-38 | 1.19192184 | 0.35 | 0.204 | 1.02E-33 | 5 | Lhpp | homeostatic |
| Il3ra1 | 5.81E-38 | 0.6335236 | 0.912 | 0.832 | 1.12E-33 | 5 | Il3ra | homeostatic |
| Ddhd1 | 7.16E-38 | 0.86500039 | 0.453 | 0.333 | 1.38E-33 | 5 | Ddhd1 | homeostatic |
| Nfil31 | 7.61E-38 | 0.4502298 | 0.739 | 0.642 | 1.47E-33 | 5 | Nfil3 | homeostatic |
| Gpx1 | 8.67E-38 | 0.33655491 | 0.947 | 0.913 | 1.68E-33 | 5 | Gpx1 | homeostatic |
| Arhgef10 | 9.55E-38 | 1.08096791 | 0.497 | 0.337 | 1.85E-33 | 5 | Arhgef10 | homeostatic |
| Plgrkt | 1.34E-37 | 0.70438833 | 0.561 | 0.44 | 2.60E-33 | 5 | Plgrkt | homeostatic |
| Ccng1 | 1.46E-37 | 0.61004713 | 0.784 | 0.72 | 2.82E-33 | 5 | Ccng1 | homeostatic |
| Ehd2 | 1.90E-37 | 0.43122541 | 0.971 | 0.934 | 3.67E-33 | 5 | Ehd2 | homeostatic |
| Tlr3 | 2.67E-37 | 1.42621073 | 0.254 | 0.11 | 5.16E-33 | 5 | Tlr3 | homeostatic |
| Ehd1 | 5.19E-37 | 0.70062732 | 0.778 | 0.663 | 1.00E-32 | 5 | Ehd1 | homeostatic |
| Fbxo4 | 8.28E-37 | 0.92310236 | 0.533 | 0.37 | 1.60E-32 | 5 | Fbxo4 | homeostatic |
| Chrd1 | 8.34E-37 | 0.93241848 | 0.41 | 0.236 | 1.61E-32 | 5 | Chrd | homeostatic |
| Samd4b | 9.15E-37 | 0.52441793 | 0.827 | 0.782 | 1.77E-32 | 5 | Samd4b | homeostatic |
| Shb1 | 1.94E-36 | 0.89722454 | 0.537 | 0.405 | 3.76E-32 | 5 | Shb | homeostatic |
| Ltbp3 | 2.64E-36 | 0.40980435 | 0.877 | 0.84 | 5.10E-32 | 5 | Ltbp3 | homeostatic |
| Stx16 | 3.64E-36 | 0.61307241 | 0.692 | 0.604 | 7.05E-32 | 5 | Stx16 | homeostatic |
| Naaladl2 | 7.69E-36 | 1.31581325 | 0.278 | 0.134 | 1.49E-31 | 5 | Naaladl2 | homeostatic |
| Cat | 8.60E-36 | 0.69689749 | 0.731 | 0.586 | 1.66E-31 | 5 | Cat | homeostatic |
| 2-Mar | 1.09E-35 | 0.87954292 | 0.489 | 0.344 | 2.10E-31 | 5 | 2-Mar | homeostatic |
| Cited22 | 1.36E-35 | 0.92307072 | 0.451 | 0.252 | 2.62E-31 | 5 | Cited2 | homeostatic |
| Selenon | 2.02E-35 | 0.48814708 | 0.929 | 0.888 | 3.91E-31 | 5 | Selenon | homeostatic |
| Plscr4 | 2.20E-35 | 1.04446336 | 0.461 | 0.288 | 4.25E-31 | 5 | Plscr4 | homeostatic |
| Daxx | 2.72E-35 | 0.88085883 | 0.609 | 0.5 | 5.27E-31 | 5 | Daxx | homeostatic |
| Abca91 | 4.31E-35 | 1.29230992 | 0.391 | 0.214 | 8.33E-31 | 5 | Abca9 | homeostatic |
| Rnf19a1 | 4.64E-35 | 0.56356955 | 0.692 | 0.596 | 8.97E-31 | 5 | Rnf19a | homeostatic |
| Mmp27 | 5.52E-35 | 1.56843129 | 0.273 | 0.137 | 1.07E-30 | 5 | Mmp27 | homeostatic |
| Cotl11 | 7.74E-35 | 0.6661019 | 0.832 | 0.647 | 1.50E-30 | 5 | Cotl1 | homeostatic |
| Arsa | 8.78E-35 | 0.67280044 | 0.784 | 0.655 | 1.70E-30 | 5 | Arsa | homeostatic |
| Dnajc101 | 1.35E-34 | 0.52308266 | 0.916 | 0.867 | 2.62E-30 | 5 | Dnajc10 | homeostatic |
| Kctd91 | 1.41E-34 | 0.80441141 | 0.645 | 0.476 | 2.72E-30 | 5 | Kctd9 | homeostatic |
| Abl1 | 1.53E-34 | 0.57364532 | 0.835 | 0.772 | 2.96E-30 | 5 | Abl1 | homeostatic |
| Mst1r1 | 2.79E-34 | 1.09038404 | 0.365 | 0.218 | 5.39E-30 | 5 | Mst1r | homeostatic |
| Chp1 | 3.49E-34 | 0.45585849 | 0.877 | 0.853 | 6.76E-30 | 5 | Chp1 | homeostatic |
| Ncoa71 | 3.88E-34 | 0.99124869 | 0.439 | 0.284 | 7.51E-30 | 5 | Ncoa7 | homeostatic |
| St3gal41 | 4.39E-34 | 0.62915333 | 0.74 | 0.615 | 8.50E-30 | 5 | St3gal4 | homeostatic |
| Arrdc31 | 6.32E-34 | 0.51685155 | 0.522 | 0.363 | 1.22E-29 | 5 | Arrdc3 | homeostatic |
| Tbkbp12 | 9.64E-34 | 0.93019074 | 0.359 | 0.215 | 1.86E-29 | 5 | Tbkbp1 | homeostatic |
| B3galnt11 | 9.94E-34 | 0.88261657 | 0.618 | 0.473 | 1.92E-29 | 5 | B3galnt1 | homeostatic |
| Slc25a281 | 1.15E-33 | 0.85893429 | 0.584 | 0.422 | 2.23E-29 | 5 | Slc25a28 | homeostatic |
| Ccdc1491 | 1.26E-33 | 1.36736609 | 0.293 | 0.15 | 2.44E-29 | 5 | Ccdc149 | homeostatic |
| Ebf12 | 2.51E-33 | 0.54607694 | 0.791 | 0.663 | 4.86E-29 | 5 | Ebf1 | homeostatic |
| Agap11 | 3.25E-33 | 0.67417887 | 0.57 | 0.468 | 6.29E-29 | 5 | Agap1 | homeostatic |
| Suox | 4.15E-33 | 1.20965015 | 0.287 | 0.153 | 8.02E-29 | 5 | Suox | homeostatic |
| Capns11 | 4.32E-33 | 0.45781842 | 0.872 | 0.846 | 8.35E-29 | 5 | Capns1 | homeostatic |
| Cers62 | 4.33E-33 | 0.6006789 | 0.593 | 0.496 | 8.37E-29 | 5 | Cers6 | homeostatic |
| Anxa11 | 5.36E-33 | 0.67276305 | 0.758 | 0.641 | 1.04E-28 | 5 | Anxa11 | homeostatic |
| Dbn11 | 7.46E-33 | 0.43056115 | 0.876 | 0.865 | 1.44E-28 | 5 | Dbn1 | homeostatic |
| Tbrg11 | 7.74E-33 | 0.4271993 | 0.888 | 0.862 | 1.50E-28 | 5 | Tbrg1 | homeostatic |
| Dusp101 | 1.42E-32 | 0.9009361 | 0.36 | 0.206 | 2.75E-28 | 5 | Dusp10 | homeostatic |
| Taok3 | 2.36E-32 | 0.87557645 | 0.553 | 0.405 | 4.57E-28 | 5 | Taok3 | homeostatic |
| Inpp11 | 2.87E-32 | 1.08104497 | 0.411 | 0.267 | 5.56E-28 | 5 | Inpp1 | homeostatic |
| Add1 | 3.37E-32 | 0.4444633 | 0.929 | 0.892 | 6.52E-28 | 5 | Add1 | homeostatic |
| Larp61 | 3.55E-32 | 0.79648762 | 0.485 | 0.337 | 6.87E-28 | 5 | Larp6 | homeostatic |
| G6pdx | 5.67E-32 | 0.90098412 | 0.575 | 0.438 | 1.10E-27 | 5 | G6pdx | homeostatic |
| Lynx1 | 1.28E-31 | 0.77453495 | 0.624 | 0.479 | 2.47E-27 | 5 | Lynx1 | homeostatic |
| Aldh3b1 | 1.39E-31 | 1.16403744 | 0.364 | 0.22 | 2.68E-27 | 5 | Aldh3b1 | homeostatic |
| Tns1 | 1.76E-31 | 0.72994605 | 0.6 | 0.467 | 3.40E-27 | 5 | Tns1 | homeostatic |
| Atp6v0a11 | 1.96E-31 | 0.84765152 | 0.411 | 0.27 | 3.79E-27 | 5 | Atp6v0a1 | homeostatic |
| Ltbr | 2.09E-31 | 0.47284372 | 0.864 | 0.785 | 4.04E-27 | 5 | Ltbr | homeostatic |
| Mtmr11 | 2.87E-31 | 0.77509049 | 0.539 | 0.413 | 5.54E-27 | 5 | Mtmr11 | homeostatic |
| Mocs11 | 4.09E-31 | 0.91911643 | 0.534 | 0.385 | 7.91E-27 | 5 | Mocs1 | homeostatic |
| Igsf81 | 4.11E-31 | 0.60714859 | 0.819 | 0.747 | 7.95E-27 | 5 | Igsf8 | homeostatic |
| Slc25a23 | 4.13E-31 | 1.03613121 | 0.294 | 0.173 | 7.99E-27 | 5 | Slc25a23 | homeostatic |
| Lrrc49 | 4.52E-31 | 0.98138072 | 0.404 | 0.251 | 8.74E-27 | 5 | Lrrc49 | homeostatic |
| Tmbim11 | 4.79E-31 | 0.59859383 | 0.769 | 0.691 | 9.26E-27 | 5 | Tmbim1 | homeostatic |
| Flii | 5.14E-31 | 0.57117018 | 0.737 | 0.646 | 9.94E-27 | 5 | Flii | homeostatic |
| Ppp1r14b2 | 5.22E-31 | 0.60630319 | 0.824 | 0.743 | 1.01E-26 | 5 | Ppp1r14b | homeostatic |
| Trafd1 | 6.60E-31 | 0.77547772 | 0.654 | 0.541 | 1.28E-26 | 5 | Trafd1 | homeostatic |
| Snx33 | 1.02E-30 | 0.76093262 | 0.654 | 0.521 | 1.97E-26 | 5 | Snx33 | homeostatic |
| Gypc | 1.22E-30 | 0.77851762 | 0.605 | 0.481 | 2.37E-26 | 5 | Gypc | homeostatic |
| Gm2a2 | 1.49E-30 | 0.38195513 | 0.761 | 0.667 | 2.89E-26 | 5 | Gm2a | homeostatic |
| Trim16 | 2.25E-30 | 1.09148434 | 0.296 | 0.18 | 4.36E-26 | 5 | Trim16 | homeostatic |
| Ralbp11 | 2.30E-30 | 0.53696713 | 0.794 | 0.726 | 4.45E-26 | 5 | Ralbp1 | homeostatic |
| Gstm4 | 3.12E-30 | 1.25885095 | 0.299 | 0.167 | 6.03E-26 | 5 | Gstm4 | homeostatic |
| Trim21 | 3.51E-30 | 0.97115001 | 0.312 | 0.174 | 6.79E-26 | 5 | Trim21 | homeostatic |
| Heca | 3.73E-30 | 0.67544043 | 0.73 | 0.586 | 7.21E-26 | 5 | Heca | homeostatic |
| Podn1 | 6.40E-30 | 0.53235074 | 0.754 | 0.547 | 1.24E-25 | 5 | Podn | homeostatic |
| Sptan11 | 7.79E-30 | 0.3686279 | 0.911 | 0.897 | 1.51E-25 | 5 | Sptan1 | homeostatic |
| Syt11 | 1.39E-29 | 1.20543005 | 0.265 | 0.127 | 2.69E-25 | 5 | Syt11 | homeostatic |
| Dock111 | 1.81E-29 | 1.09764767 | 0.34 | 0.201 | 3.50E-25 | 5 | Dock11 | homeostatic |
| Ddt1 | 2.07E-29 | 0.84070258 | 0.516 | 0.369 | 3.99E-25 | 5 | Ddt | homeostatic |
| Npdc11 | 6.00E-29 | 0.46322825 | 0.891 | 0.81 | 1.16E-24 | 5 | Npdc1 | homeostatic |
| Znfx1 | 6.40E-29 | 0.92502007 | 0.387 | 0.267 | 1.24E-24 | 5 | Znfx1 | homeostatic |
| Tmem1581 | 1.06E-28 | 1.17063036 | 0.326 | 0.243 | 2.04E-24 | 5 | Tmem158 | homeostatic |
| Prkca | 1.10E-28 | 0.84189509 | 0.471 | 0.335 | 2.13E-24 | 5 | Prkca | homeostatic |
| Mbd2 | 1.30E-28 | 0.34464931 | 0.951 | 0.937 | 2.52E-24 | 5 | Mbd2 | homeostatic |
| Nckap11 | 2.20E-28 | 0.559371 | 0.773 | 0.703 | 4.25E-24 | 5 | Nckap1 | homeostatic |
| Park71 | 3.43E-28 | 0.48654627 | 0.839 | 0.779 | 6.64E-24 | 5 | Park7 | homeostatic |
| Itgbl11 | 3.65E-28 | 0.76459919 | 0.402 | 0.227 | 7.06E-24 | 5 | Itgbl1 | homeostatic |
| Ift271 | 5.81E-28 | 0.74362217 | 0.637 | 0.513 | 1.12E-23 | 5 | Ift27 | homeostatic |
| Insyn1 | 6.27E-28 | 0.69185174 | 0.654 | 0.541 | 1.21E-23 | 5 | Insyn1 | homeostatic |
| Crebl2 | 7.15E-28 | 1.07943105 | 0.338 | 0.201 | 1.38E-23 | 5 | Crebl2 | homeostatic |
| Ino80c1 | 7.99E-28 | 0.8342337 | 0.549 | 0.395 | 1.54E-23 | 5 | Ino80c | homeostatic |
| Psmb91 | 1.29E-27 | 1.0378614 | 0.336 | 0.201 | 2.49E-23 | 5 | Psmb9 | homeostatic |
| Calhm2 | 2.06E-27 | 0.6378945 | 0.636 | 0.525 | 3.98E-23 | 5 | Calhm2 | homeostatic |
| Entpd51 | 2.13E-27 | 0.84597299 | 0.561 | 0.413 | 4.12E-23 | 5 | Entpd5 | homeostatic |
| Mia3 | 2.97E-27 | 0.41470968 | 0.935 | 0.906 | 5.75E-23 | 5 | Mia3 | homeostatic |
| Csf2ra2 | 3.95E-27 | 1.14633142 | 0.329 | 0.188 | 7.64E-23 | 5 | Csf2ra | homeostatic |
| Myorg1 | 6.29E-27 | 1.02757882 | 0.411 | 0.283 | 1.22E-22 | 5 | Myorg | homeostatic |
| Fap1 | 7.50E-27 | 0.57026804 | 0.697 | 0.558 | 1.45E-22 | 5 | Fap | homeostatic |
| Snapc5 | 8.96E-27 | 0.90379038 | 0.478 | 0.342 | 1.73E-22 | 5 | Snapc5 | homeostatic |
| Sdccag8 | 1.12E-26 | 0.82960665 | 0.429 | 0.292 | 2.16E-22 | 5 | Sdccag8 | homeostatic |
| Sri1 | 1.92E-26 | 0.39786467 | 0.878 | 0.844 | 3.72E-22 | 5 | Sri | homeostatic |
| Tm4sf12 | 2.57E-26 | 0.38113822 | 0.824 | 0.725 | 4.96E-22 | 5 | Tm4sf1 | homeostatic |
| Csgalnact2 | 2.71E-26 | 0.61058831 | 0.702 | 0.619 | 5.24E-22 | 5 | Csgalnact2 | homeostatic |
| Stx4a1 | 3.35E-26 | 0.51012016 | 0.81 | 0.731 | 6.47E-22 | 5 | Stx4a | homeostatic |
| Akap12 | 4.14E-26 | 0.75263564 | 0.497 | 0.411 | 8.00E-22 | 5 | Akap12 | homeostatic |
| C21 | 4.39E-26 | 0.60109989 | 0.679 | 0.467 | 8.48E-22 | 5 | C2 | homeostatic |
| Erlin12 | 5.82E-26 | 0.95278311 | 0.418 | 0.277 | 1.13E-21 | 5 | Erlin1 | homeostatic |
| Dtwd1 | 6.62E-26 | 1.18403762 | 0.294 | 0.15 | 1.28E-21 | 5 | Dtwd1 | homeostatic |
| Exoc4 | 8.24E-26 | 0.53830225 | 0.688 | 0.624 | 1.59E-21 | 5 | Exoc4 | homeostatic |
| Armh41 | 1.01E-25 | 0.57921612 | 0.405 | 0.297 | 1.96E-21 | 5 | Armh4 | homeostatic |
| Lima11 | 1.09E-25 | 0.37010935 | 0.876 | 0.84 | 2.10E-21 | 5 | Lima1 | homeostatic |
| Arhgap18 | 1.49E-25 | 1.27524189 | 0.279 | 0.157 | 2.88E-21 | 5 | Arhgap18 | homeostatic |
| Emc9 | 1.97E-25 | 1.1673306 | 0.306 | 0.167 | 3.82E-21 | 5 | Emc9 | homeostatic |
| Rnf145 | 2.43E-25 | 0.47843248 | 0.734 | 0.686 | 4.69E-21 | 5 | Rnf145 | homeostatic |
| Znrf2 | 2.63E-25 | 1.00714753 | 0.528 | 0.403 | 5.09E-21 | 5 | Znrf2 | homeostatic |
| Hspb1 | 2.87E-25 | 0.28272455 | 0.485 | 0.385 | 5.54E-21 | 5 | Hspb1 | homeostatic |
| Evi5l1 | 3.39E-25 | 0.67613105 | 0.536 | 0.426 | 6.55E-21 | 5 | Evi5l | homeostatic |
| Ids1 | 3.42E-25 | 0.64793213 | 0.589 | 0.493 | 6.61E-21 | 5 | Ids | homeostatic |
| Adk1 | 1.04E-24 | 0.7614268 | 0.627 | 0.488 | 2.01E-20 | 5 | Adk | homeostatic |
| Efs | 1.25E-24 | 0.79530667 | 0.647 | 0.483 | 2.41E-20 | 5 | Efs | homeostatic |
| Zeb1 | 1.33E-24 | 0.51712375 | 0.673 | 0.578 | 2.58E-20 | 5 | Zeb1 | homeostatic |
| Rhobtb31 | 1.47E-24 | 0.56252961 | 0.685 | 0.584 | 2.84E-20 | 5 | Rhobtb3 | homeostatic |
| Galnt121 | 1.62E-24 | 1.23860309 | 0.268 | 0.152 | 3.14E-20 | 5 | Galnt12 | homeostatic |
| Borcs5 | 1.74E-24 | 0.83776143 | 0.36 | 0.233 | 3.37E-20 | 5 | Borcs5 | homeostatic |
| Nav1 | 2.03E-24 | 0.42937478 | 0.91 | 0.852 | 3.93E-20 | 5 | Nav1 | homeostatic |
| Abca8a | 3.19E-24 | 1.09597521 | 0.255 | 0.12 | 6.17E-20 | 5 | Abca8a | homeostatic |
| Trabd2b | 3.59E-24 | 0.87342229 | 0.291 | 0.138 | 6.94E-20 | 5 | Trabd2b | homeostatic |
| Hsd17b11 | 3.66E-24 | 0.51011922 | 0.689 | 0.609 | 7.07E-20 | 5 | Hsd17b11 | homeostatic |
| Lztfl1 | 7.31E-24 | 0.82675461 | 0.374 | 0.262 | 1.41E-19 | 5 | Lztfl1 | homeostatic |
| Casp42 | 8.25E-24 | 0.52396 | 0.543 | 0.442 | 1.59E-19 | 5 | Casp4 | homeostatic |
| Calcrl2 | 9.89E-24 | 0.60646738 | 0.407 | 0.23 | 1.91E-19 | 5 | Calcrl | homeostatic |
| Bckdhb | 1.00E-23 | 0.70188853 | 0.378 | 0.252 | 1.94E-19 | 5 | Bckdhb | homeostatic |
| Lysmd31 | 1.41E-23 | 0.52118928 | 0.582 | 0.45 | 2.73E-19 | 5 | Lysmd3 | homeostatic |
| Tex2641 | 1.58E-23 | 0.44140173 | 0.845 | 0.79 | 3.06E-19 | 5 | Tex264 | homeostatic |
| Sdc42 | 3.07E-23 | 0.35078812 | 0.835 | 0.759 | 5.94E-19 | 5 | Sdc4 | homeostatic |
| Mtmr1 | 4.97E-23 | 1.04408733 | 0.315 | 0.191 | 9.62E-19 | 5 | Mtmr1 | homeostatic |
| Magi3 | 6.54E-23 | 1.00046235 | 0.363 | 0.236 | 1.26E-18 | 5 | Magi3 | homeostatic |
| Ldlrad41 | 7.58E-23 | 0.5129054 | 0.631 | 0.544 | 1.46E-18 | 5 | Ldlrad4 | homeostatic |
| Sgms21 | 1.01E-22 | 0.73703125 | 0.51 | 0.361 | 1.96E-18 | 5 | Sgms2 | homeostatic |
| Rap1a | 1.45E-22 | 0.47784643 | 0.854 | 0.802 | 2.80E-18 | 5 | Rap1a | homeostatic |
| Slco2a12 | 2.61E-22 | 0.54852328 | 0.585 | 0.376 | 5.06E-18 | 5 | Slco2a1 | homeostatic |
| Chd31 | 2.88E-22 | 0.43705076 | 0.788 | 0.727 | 5.56E-18 | 5 | Chd3 | homeostatic |
| Camk2d1 | 3.35E-22 | 0.49705121 | 0.755 | 0.69 | 6.47E-18 | 5 | Camk2d | homeostatic |
| Irak4 | 4.02E-22 | 0.7188194 | 0.53 | 0.416 | 7.78E-18 | 5 | Irak4 | homeostatic |
| Zdhhc91 | 4.83E-22 | 0.53786832 | 0.749 | 0.666 | 9.33E-18 | 5 | Zdhhc9 | homeostatic |
| Zfp36l12 | 1.11E-21 | 0.37368532 | 0.912 | 0.858 | 2.14E-17 | 5 | Zfp36l1 | homeostatic |
| Smarca21 | 1.13E-21 | 0.53485548 | 0.694 | 0.597 | 2.18E-17 | 5 | Smarca2 | homeostatic |
| Edem11 | 1.33E-21 | 0.45486811 | 0.84 | 0.772 | 2.57E-17 | 5 | Edem1 | homeostatic |
| Pid1 | 1.64E-21 | 0.69564713 | 0.35 | 0.197 | 3.17E-17 | 5 | Pid1 | homeostatic |
| Pgd1 | 1.67E-21 | 0.49495601 | 0.783 | 0.742 | 3.24E-17 | 5 | Pgd | homeostatic |
| Mxd4 | 1.72E-21 | 0.30409405 | 0.923 | 0.889 | 3.32E-17 | 5 | Mxd4 | homeostatic |
| Usp24 | 2.50E-21 | 0.57294273 | 0.617 | 0.509 | 4.83E-17 | 5 | Usp24 | homeostatic |
| Oplah | 2.67E-21 | 1.06287088 | 0.259 | 0.14 | 5.17E-17 | 5 | Oplah | homeostatic |
| Foxo11 | 2.81E-21 | 0.74610126 | 0.514 | 0.38 | 5.42E-17 | 5 | Foxo1 | homeostatic |
| Plpbp1 | 3.87E-21 | 0.49434956 | 0.61 | 0.551 | 7.48E-17 | 5 | Plpbp | homeostatic |
| Lpl1 | 4.90E-21 | 0.50209473 | 0.551 | 0.59 | 9.47E-17 | 5 | Lpl | homeostatic |
| Pttg1 | 4.97E-21 | 0.62735029 | 0.558 | 0.431 | 9.60E-17 | 5 | Pttg1 | homeostatic |
| Slc4a31 | 5.23E-21 | 0.70006004 | 0.353 | 0.23 | 1.01E-16 | 5 | Slc4a3 | homeostatic |
| Ctbs1 | 5.65E-21 | 0.96036718 | 0.371 | 0.246 | 1.09E-16 | 5 | Ctbs | homeostatic |
| Tril1 | 6.89E-21 | 1.10143311 | 0.279 | 0.149 | 1.33E-16 | 5 | Tril | homeostatic |
| Arl6ip51 | 7.31E-21 | 0.41158118 | 0.806 | 0.768 | 1.41E-16 | 5 | Arl6ip5 | homeostatic |
| Smarcd2 | 1.07E-20 | 0.59133147 | 0.622 | 0.548 | 2.08E-16 | 5 | Smarcd2 | homeostatic |
| Tbc1d121 | 1.15E-20 | 0.85842847 | 0.367 | 0.264 | 2.22E-16 | 5 | Tbc1d12 | homeostatic |
| Ap3m1 | 1.21E-20 | 0.74075249 | 0.34 | 0.236 | 2.35E-16 | 5 | Ap3m1 | homeostatic |
| Bad | 1.38E-20 | 0.61464614 | 0.726 | 0.612 | 2.67E-16 | 5 | Bad | homeostatic |
| Iglon51 | 1.96E-20 | 0.75174292 | 0.382 | 0.254 | 3.78E-16 | 5 | Iglon5 | homeostatic |
| Hebp21 | 2.01E-20 | 0.69085393 | 0.468 | 0.374 | 3.89E-16 | 5 | Hebp2 | homeostatic |
| Itgb1bp1 | 2.33E-20 | 0.64716597 | 0.586 | 0.49 | 4.50E-16 | 5 | Itgb1bp1 | homeostatic |
| Stat5b | 2.34E-20 | 0.68483139 | 0.621 | 0.478 | 4.53E-16 | 5 | Stat5b | homeostatic |
| Naprt | 3.24E-20 | 0.98739854 | 0.401 | 0.266 | 6.26E-16 | 5 | Naprt | homeostatic |
| Pik3ip1 | 3.68E-20 | 1.20625243 | 0.296 | 0.186 | 7.12E-16 | 5 | Pik3ip1 | homeostatic |
| Tram2 | 4.05E-20 | 0.47479165 | 0.849 | 0.795 | 7.83E-16 | 5 | Tram2 | homeostatic |
| Hivep2 | 4.36E-20 | 0.39281335 | 0.615 | 0.547 | 8.42E-16 | 5 | Hivep2 | homeostatic |
| Glul2 | 7.20E-20 | 0.44143936 | 0.742 | 0.668 | 1.39E-15 | 5 | Glul | homeostatic |
| Gpsm21 | 9.00E-20 | 0.70457489 | 0.558 | 0.392 | 1.74E-15 | 5 | Gpsm2 | homeostatic |
| Ccpg11 | 1.03E-19 | 0.45505485 | 0.807 | 0.732 | 1.99E-15 | 5 | Ccpg1 | homeostatic |
| Kif13b | 1.05E-19 | 0.77750881 | 0.463 | 0.335 | 2.02E-15 | 5 | Kif13b | homeostatic |
| Pts1 | 1.42E-19 | 1.0053746 | 0.324 | 0.217 | 2.74E-15 | 5 | Pts | homeostatic |
| Lrba | 1.71E-19 | 0.93756387 | 0.274 | 0.176 | 3.30E-15 | 5 | Lrba | homeostatic |
| Tmem106c1 | 2.75E-19 | 0.89856218 | 0.421 | 0.277 | 5.32E-15 | 5 | Tmem106c | homeostatic |
| Ube2e21 | 4.60E-19 | 0.4635218 | 0.646 | 0.58 | 8.88E-15 | 5 | Ube2e2 | homeostatic |
| Klf12 | 4.83E-19 | 0.85286469 | 0.265 | 0.179 | 9.33E-15 | 5 | Klf12 | homeostatic |
| Acap21 | 6.79E-19 | 0.50458545 | 0.749 | 0.676 | 1.31E-14 | 5 | Acap2 | homeostatic |
| Cav11 | 7.49E-19 | 0.28045554 | 0.751 | 0.623 | 1.45E-14 | 5 | Cav1 | homeostatic |
| Fkbp91 | 1.15E-18 | 0.29005355 | 0.981 | 0.964 | 2.23E-14 | 5 | Fkbp9 | homeostatic |
| Dtx2 | 1.21E-18 | 0.867618 | 0.421 | 0.316 | 2.34E-14 | 5 | Dtx2 | homeostatic |
| Jmjd8 | 1.38E-18 | 0.65652259 | 0.499 | 0.395 | 2.68E-14 | 5 | Jmjd8 | homeostatic |
| Anxa71 | 1.67E-18 | 0.39570158 | 0.855 | 0.805 | 3.23E-14 | 5 | Anxa7 | homeostatic |
| Atl31 | 1.70E-18 | 0.3969865 | 0.905 | 0.886 | 3.29E-14 | 5 | Atl3 | homeostatic |
| Gbp7 | 2.20E-18 | 0.74217557 | 0.293 | 0.215 | 4.26E-14 | 5 | Gbp7 | homeostatic |
| Fam114a1 | 2.54E-18 | 0.29456572 | 0.93 | 0.915 | 4.91E-14 | 5 | Fam114a1 | homeostatic |
| Ttc39c | 4.59E-18 | 0.7050067 | 0.423 | 0.302 | 8.87E-14 | 5 | Ttc39c | homeostatic |
| Acvr2a1 | 5.46E-18 | 0.73907569 | 0.522 | 0.358 | 1.06E-13 | 5 | Acvr2a | homeostatic |
| Psme11 | 5.73E-18 | 0.46606372 | 0.807 | 0.742 | 1.11E-13 | 5 | Psme1 | homeostatic |
| Pgm32 | 6.93E-18 | 0.48769626 | 0.66 | 0.571 | 1.34E-13 | 5 | Pgm3 | homeostatic |
| Nampt2 | 7.27E-18 | 0.58616168 | 0.511 | 0.387 | 1.41E-13 | 5 | Nampt | homeostatic |
| Reep31 | 8.35E-18 | 0.37623804 | 0.825 | 0.773 | 1.61E-13 | 5 | Reep3 | homeostatic |
| Dnajc31 | 8.72E-18 | 0.38605354 | 0.929 | 0.897 | 1.69E-13 | 5 | Dnajc3 | homeostatic |
| Fig4 | 9.41E-18 | 0.96523425 | 0.393 | 0.276 | 1.82E-13 | 5 | Fig4 | homeostatic |
| Mpdz1 | 2.37E-17 | 0.7581967 | 0.571 | 0.456 | 4.58E-13 | 5 | Mpdz | homeostatic |
| Rab291 | 3.62E-17 | 0.70176982 | 0.477 | 0.355 | 7.00E-13 | 5 | Rab29 | homeostatic |
| Hykk | 4.47E-17 | 0.99759669 | 0.289 | 0.185 | 8.65E-13 | 5 | Hykk | homeostatic |
| Cyb5r3 | 5.16E-17 | 0.27538883 | 0.937 | 0.945 | 9.98E-13 | 5 | Cyb5r3 | homeostatic |
| Rgmb1 | 5.24E-17 | 0.48874896 | 0.787 | 0.693 | 1.01E-12 | 5 | Rgmb | homeostatic |
| Itsn11 | 5.82E-17 | 0.36645379 | 0.712 | 0.648 | 1.12E-12 | 5 | Itsn1 | homeostatic |
| Stat61 | 6.63E-17 | 0.43664367 | 0.747 | 0.678 | 1.28E-12 | 5 | Stat6 | homeostatic |
| Pcbp31 | 6.70E-17 | 0.71410097 | 0.264 | 0.173 | 1.30E-12 | 5 | Pcbp3 | homeostatic |
| Nadk1 | 8.43E-17 | 0.36175513 | 0.813 | 0.754 | 1.63E-12 | 5 | Nadk | homeostatic |
| Bace2 | 8.51E-17 | 0.79605075 | 0.269 | 0.194 | 1.64E-12 | 5 | Bace2 | homeostatic |
| Cgref11 | 8.89E-17 | 0.53392742 | 0.401 | 0.326 | 1.72E-12 | 5 | Cgref1 | homeostatic |
| Tbc1d14 | 9.04E-17 | 0.50525794 | 0.61 | 0.519 | 1.75E-12 | 5 | Tbc1d14 | homeostatic |
| Tnfsf13b | 9.45E-17 | 0.96695634 | 0.279 | 0.136 | 1.83E-12 | 5 | Tnfsf13b | homeostatic |
| Trim2 | 1.33E-16 | 0.79035811 | 0.437 | 0.326 | 2.58E-12 | 5 | Trim2 | homeostatic |
| Slc9a91 | 1.45E-16 | 0.91854141 | 0.275 | 0.176 | 2.81E-12 | 5 | Slc9a9 | homeostatic |
| Tcn21 | 1.73E-16 | 0.36080488 | 0.909 | 0.867 | 3.35E-12 | 5 | Tcn2 | homeostatic |
| Hsf2 | 2.17E-16 | 0.99295325 | 0.35 | 0.238 | 4.20E-12 | 5 | Hsf2 | homeostatic |
| Pml1 | 2.51E-16 | 0.54986168 | 0.633 | 0.548 | 4.85E-12 | 5 | Pml | homeostatic |
| Slc44a11 | 2.57E-16 | 0.39672118 | 0.872 | 0.803 | 4.97E-12 | 5 | Slc44a1 | homeostatic |
| Ctdsp2 | 2.58E-16 | 0.31551475 | 0.986 | 0.975 | 4.99E-12 | 5 | Ctdsp2 | homeostatic |
| Snx29 | 3.10E-16 | 0.68547436 | 0.574 | 0.462 | 6.00E-12 | 5 | Snx29 | homeostatic |
| Bscl2 | 3.87E-16 | 0.5157771 | 0.741 | 0.672 | 7.48E-12 | 5 | Bscl2 | homeostatic |
| Cep85l | 4.50E-16 | 0.77062398 | 0.35 | 0.259 | 8.71E-12 | 5 | Cep85l | homeostatic |
| Tm2d2 | 7.99E-16 | 0.41947431 | 0.849 | 0.797 | 1.54E-11 | 5 | Tm2d2 | homeostatic |
| Lpcat31 | 1.24E-15 | 0.50787462 | 0.749 | 0.65 | 2.40E-11 | 5 | Lpcat3 | homeostatic |
| Mindy11 | 1.26E-15 | 0.52776075 | 0.612 | 0.521 | 2.43E-11 | 5 | Mindy1 | homeostatic |
| Swap701 | 1.29E-15 | 0.45882769 | 0.591 | 0.512 | 2.49E-11 | 5 | Swap70 | homeostatic |
| Rtn31 | 1.33E-15 | 0.41304326 | 0.883 | 0.846 | 2.57E-11 | 5 | Rtn3 | homeostatic |
| Abhd111 | 1.38E-15 | 0.68868539 | 0.547 | 0.445 | 2.67E-11 | 5 | Abhd11 | homeostatic |
| Ptpn13 | 1.40E-15 | 0.57627267 | 0.574 | 0.439 | 2.71E-11 | 5 | Ptpn13 | homeostatic |
| Snx10 | 1.42E-15 | 0.86762092 | 0.289 | 0.192 | 2.74E-11 | 5 | Snx10 | homeostatic |
| Ccdc174 | 1.48E-15 | 0.56301283 | 0.481 | 0.381 | 2.86E-11 | 5 | Ccdc174 | homeostatic |
| Fdx11 | 1.54E-15 | 0.56119516 | 0.464 | 0.359 | 2.98E-11 | 5 | Fdx1 | homeostatic |
| Kcnd11 | 1.65E-15 | 0.82665285 | 0.336 | 0.251 | 3.19E-11 | 5 | Kcnd1 | homeostatic |
| Htr2a | 2.00E-15 | 0.64073758 | 0.392 | 0.289 | 3.86E-11 | 5 | Htr2a | homeostatic |
| Slc12a61 | 2.63E-15 | 0.66431156 | 0.438 | 0.333 | 5.08E-11 | 5 | Slc12a6 | homeostatic |
| Cnih11 | 4.02E-15 | 0.40579467 | 0.871 | 0.829 | 7.77E-11 | 5 | Cnih1 | homeostatic |
| Mpst1 | 5.64E-15 | 0.62118277 | 0.48 | 0.368 | 1.09E-10 | 5 | Mpst | homeostatic |
| Naalad21 | 6.01E-15 | 0.73571617 | 0.345 | 0.254 | 1.16E-10 | 5 | Naalad2 | homeostatic |
| Dpy19l11 | 6.11E-15 | 0.4867749 | 0.758 | 0.668 | 1.18E-10 | 5 | Dpy19l1 | homeostatic |
| Slc49a4 | 6.92E-15 | 0.43919607 | 0.675 | 0.612 | 1.34E-10 | 5 | Slc49a4 | homeostatic |
| Golph3l | 7.68E-15 | 0.57547687 | 0.459 | 0.361 | 1.48E-10 | 5 | Golph3l | homeostatic |
| Fuz | 1.04E-14 | 0.72955062 | 0.307 | 0.209 | 2.01E-10 | 5 | Fuz | homeostatic |
| Fam189b | 1.05E-14 | 0.57413928 | 0.544 | 0.452 | 2.03E-10 | 5 | Fam189b | homeostatic |
| Otud1 | 1.08E-14 | 0.73416386 | 0.289 | 0.214 | 2.10E-10 | 5 | Otud1 | homeostatic |
| Rock21 | 1.10E-14 | 0.4038914 | 0.81 | 0.78 | 2.13E-10 | 5 | Rock2 | homeostatic |
| Lrp6 | 1.15E-14 | 0.3402191 | 0.874 | 0.844 | 2.23E-10 | 5 | Lrp6 | homeostatic |
| Ap1m1 | 1.26E-14 | 0.45502601 | 0.708 | 0.627 | 2.43E-10 | 5 | Ap1m1 | homeostatic |
| Abhd14b | 1.31E-14 | 0.88986934 | 0.338 | 0.241 | 2.54E-10 | 5 | Abhd14b | homeostatic |
| Il13ra12 | 1.54E-14 | 0.27201838 | 0.859 | 0.832 | 2.99E-10 | 5 | Il13ra1 | homeostatic |
| Hebp12 | 1.56E-14 | 0.46981897 | 0.48 | 0.35 | 3.02E-10 | 5 | Hebp1 | homeostatic |
| P2rx41 | 1.58E-14 | 0.49791629 | 0.726 | 0.626 | 3.05E-10 | 5 | P2rx4 | homeostatic |
| Nfasc2 | 1.65E-14 | 0.52023501 | 0.494 | 0.38 | 3.19E-10 | 5 | Nfasc | homeostatic |
| Smim20 | 1.89E-14 | 0.56516001 | 0.712 | 0.622 | 3.66E-10 | 5 | Smim20 | homeostatic |
| Jmjd1c1 | 2.67E-14 | 0.34425892 | 0.647 | 0.565 | 5.17E-10 | 5 | Jmjd1c | homeostatic |
| Xrcc5 | 2.86E-14 | 0.93579394 | 0.272 | 0.18 | 5.54E-10 | 5 | Xrcc5 | homeostatic |
| Guk11 | 3.03E-14 | 0.42857061 | 0.654 | 0.601 | 5.87E-10 | 5 | Guk1 | homeostatic |
| Mif4gd | 3.55E-14 | 0.54773072 | 0.504 | 0.411 | 6.87E-10 | 5 | Mif4gd | homeostatic |
| Rab28 | 3.66E-14 | 0.46050946 | 0.74 | 0.69 | 7.07E-10 | 5 | Rab28 | homeostatic |
| Galt1 | 3.75E-14 | 0.71136005 | 0.447 | 0.332 | 7.26E-10 | 5 | Galt | homeostatic |
| Setd7 | 3.92E-14 | 0.53045078 | 0.793 | 0.721 | 7.58E-10 | 5 | Setd7 | homeostatic |
| Acyp21 | 3.92E-14 | 0.92420524 | 0.275 | 0.186 | 7.59E-10 | 5 | Acyp2 | homeostatic |
| Slc25a30 | 5.17E-14 | 0.49008188 | 0.542 | 0.47 | 1.00E-09 | 5 | Slc25a30 | homeostatic |
| Sertad3 | 5.36E-14 | 0.74806551 | 0.404 | 0.275 | 1.04E-09 | 5 | Sertad3 | homeostatic |
| Ltbp41 | 7.20E-14 | 0.62142718 | 0.494 | 0.293 | 1.39E-09 | 5 | Ltbp4 | homeostatic |
| Man2b11 | 9.10E-14 | 0.51857956 | 0.77 | 0.668 | 1.76E-09 | 5 | Man2b1 | homeostatic |
| Pigs1 | 9.17E-14 | 0.415498 | 0.742 | 0.694 | 1.77E-09 | 5 | Pigs | homeostatic |
| Tnip21 | 9.71E-14 | 0.46623245 | 0.532 | 0.438 | 1.88E-09 | 5 | Tnip2 | homeostatic |
| Wdfy21 | 9.97E-14 | 0.55691111 | 0.581 | 0.491 | 1.93E-09 | 5 | Wdfy2 | homeostatic |
| Ust | 1.09E-13 | 0.68255587 | 0.595 | 0.493 | 2.10E-09 | 5 | Ust | homeostatic |
| St52 | 1.11E-13 | 0.42526883 | 0.635 | 0.58 | 2.16E-09 | 5 | St5 | homeostatic |
| Itm2a1 | 1.16E-13 | 0.4214185 | 0.819 | 0.788 | 2.25E-09 | 5 | Itm2a | homeostatic |
| Cab39l1 | 1.25E-13 | 0.76387775 | 0.345 | 0.239 | 2.42E-09 | 5 | Cab39l | homeostatic |

| | | | | | | | | |
|---|---|---|---|---|---|---|---|---|
| Ccdc251 | 1.30E-13 | 0.49198362 | 0.577 | 0.487 | 2.52E-09 | 5 | Ccdc25 | homeostatic |
| Sfi1 | 1.35E-13 | 0.57483771 | 0.402 | 0.327 | 2.61E-09 | 5 | Sfi1 | homeostatic |
| Fez21 | 1.66E-13 | 0.4672551 | 0.666 | 0.603 | 3.21E-09 | 5 | Fez2 | homeostatic |
| Pygb | 2.03E-13 | 0.55273682 | 0.734 | 0.656 | 3.93E-09 | 5 | Pygb | homeostatic |
| Psmb101 | 2.21E-13 | 0.5572124 | 0.495 | 0.414 | 4.28E-09 | 5 | Psmb10 | homeostatic |
| Lrp31 | 2.23E-13 | 0.64610638 | 0.548 | 0.435 | 4.32E-09 | 5 | Lrp3 | homeostatic |
| Parp11 | 2.29E-13 | 0.75719769 | 0.27 | 0.18 | 4.43E-09 | 5 | Parp11 | homeostatic |
| Rpain | 2.42E-13 | 0.74948757 | 0.358 | 0.256 | 4.68E-09 | 5 | Rpain | homeostatic |
| Chchd61 | 2.90E-13 | 0.95877349 | 0.274 | 0.172 | 5.62E-09 | 5 | Chchd6 | homeostatic |
| Snx21 | 2.96E-13 | 0.55278498 | 0.613 | 0.517 | 5.71E-09 | 5 | Snx21 | homeostatic |
| Arl4a | 3.00E-13 | 0.71215408 | 0.303 | 0.19 | 5.81E-09 | 5 | Arl4a | homeostatic |
| Cc2d2a1 | 3.39E-13 | 0.81667305 | 0.368 | 0.254 | 6.56E-09 | 5 | Cc2d2a | homeostatic |
| Rnf132 | 4.54E-13 | 0.36478064 | 0.636 | 0.564 | 8.79E-09 | 5 | Rnf13 | homeostatic |
| Man2b21 | 5.93E-13 | 0.45921954 | 0.826 | 0.777 | 1.15E-08 | 5 | Man2b2 | homeostatic |
| Fam234a | 7.72E-13 | 0.41206407 | 0.685 | 0.61 | 1.49E-08 | 5 | Fam234a | homeostatic |
| Txn11 | 7.97E-13 | 0.34428195 | 0.89 | 0.856 | 1.54E-08 | 5 | Txn1 | homeostatic |
| Gstk11 | 8.02E-13 | 0.80114794 | 0.279 | 0.187 | 1.55E-08 | 5 | Gstk1 | homeostatic |
| Npas2 | 8.37E-13 | 0.28216854 | 0.51 | 0.45 | 1.62E-08 | 5 | Npas2 | homeostatic |
| Uba71 | 9.56E-13 | 0.56135214 | 0.458 | 0.386 | 1.85E-08 | 5 | Uba7 | homeostatic |
| Vps13a | 9.81E-13 | 0.7770156 | 0.269 | 0.179 | 1.90E-08 | 5 | Vps13a | homeostatic |
| Prcp | 1.04E-12 | 0.72833776 | 0.429 | 0.338 | 2.01E-08 | 5 | Prcp | homeostatic |
| Traf3ip21 | 1.29E-12 | 0.46339048 | 0.544 | 0.481 | 2.49E-08 | 5 | Traf3ip2 | homeostatic |
| Phka1 | 1.33E-12 | 0.7942254 | 0.272 | 0.179 | 2.57E-08 | 5 | Phka1 | homeostatic |
| Ggh | 1.62E-12 | 0.45783932 | 0.6 | 0.521 | 3.14E-08 | 5 | Ggh | homeostatic |
| Arfip2 | 1.70E-12 | 0.30568881 | 0.701 | 0.663 | 3.29E-08 | 5 | Arfip2 | homeostatic |
| Prxl2c1 | 1.76E-12 | 0.3989914 | 0.678 | 0.637 | 3.40E-08 | 5 | Prxl2c | homeostatic |
| Mcu | 1.82E-12 | 0.59148422 | 0.538 | 0.456 | 3.53E-08 | 5 | Mcu | homeostatic |
| Pgm13 | 1.86E-12 | 0.30927907 | 0.735 | 0.656 | 3.60E-08 | 5 | Pgm1 | homeostatic |
| Cacna2d1 | 1.88E-12 | 0.35010682 | 0.853 | 0.821 | 3.64E-08 | 5 | Cacna2d1 | homeostatic |
| Pdlim51 | 1.93E-12 | 0.33997548 | 0.74 | 0.658 | 3.73E-08 | 5 | Pdlim5 | homeostatic |
| Tstd3 | 1.96E-12 | 0.79214303 | 0.261 | 0.185 | 3.78E-08 | 5 | Tstd3 | homeostatic |
| Etfdh | 2.03E-12 | 0.53406984 | 0.557 | 0.452 | 3.92E-08 | 5 | Etfdh | homeostatic |
| Mageh1 | 2.50E-12 | 0.5510636 | 0.552 | 0.456 | 4.83E-08 | 5 | Mageh1 | homeostatic |
| Tmem63a1 | 2.61E-12 | 0.5155087 | 0.72 | 0.642 | 5.05E-08 | 5 | Tmem63a | homeostatic |
| Snx61 | 2.63E-12 | 0.48129592 | 0.662 | 0.586 | 5.09E-08 | 5 | Snx6 | homeostatic |
| Tmem106b1 | 2.75E-12 | 0.33755121 | 0.764 | 0.712 | 5.32E-08 | 5 | Tmem106b | homeostatic |
| Insig21 | 3.15E-12 | 0.52295324 | 0.481 | 0.389 | 6.10E-08 | 5 | Insig2 | homeostatic |
| Erlin2 | 5.04E-12 | 0.42907853 | 0.668 | 0.598 | 9.74E-08 | 5 | Erlin2 | homeostatic |
| Mospd2 | 5.25E-12 | 0.58019017 | 0.431 | 0.344 | 1.01E-07 | 5 | Mospd2 | homeostatic |
| Pofut1 | 5.86E-12 | 0.48173573 | 0.753 | 0.676 | 1.13E-07 | 5 | Pofut1 | homeostatic |
| Fech | 6.50E-12 | 0.40777988 | 0.759 | 0.706 | 1.26E-07 | 5 | Fech | homeostatic |
| Tcf7l1 | 7.17E-12 | 0.65965828 | 0.495 | 0.399 | 1.39E-07 | 5 | Tcf7l1 | homeostatic |
| Phldb11 | 8.95E-12 | 0.34437133 | 0.784 | 0.773 | 1.73E-07 | 5 | Phldb1 | homeostatic |
| Rab6b1 | 1.07E-11 | 0.64104967 | 0.349 | 0.268 | 2.08E-07 | 5 | Rab6b | homeostatic |
| Clstn1 | 1.14E-11 | 0.41606781 | 0.812 | 0.755 | 2.20E-07 | 5 | Clstn1 | homeostatic |
| Anks11 | 1.23E-11 | 0.65366667 | 0.464 | 0.369 | 2.38E-07 | 5 | Anks1 | homeostatic |
| Rexo2 | 1.29E-11 | 0.34224319 | 0.912 | 0.865 | 2.49E-07 | 5 | Rexo2 | homeostatic |
| Psen21 | 1.30E-11 | 0.59908916 | 0.509 | 0.405 | 2.52E-07 | 5 | Psen2 | homeostatic |
| Ppp2r5a1 | 1.47E-11 | 0.39149474 | 0.665 | 0.613 | 2.85E-07 | 5 | Ppp2r5a | homeostatic |
| Trim472 | 1.52E-11 | 0.57286182 | 0.352 | 0.238 | 2.93E-07 | 5 | Trim47 | homeostatic |
| Bphl | 1.56E-11 | 0.6837403 | 0.382 | 0.281 | 3.02E-07 | 5 | Bphl | homeostatic |
| Rhoa | 1.71E-11 | 0.31485575 | 0.937 | 0.928 | 3.30E-07 | 5 | Rhoa | homeostatic |
| Kidins220 | 2.43E-11 | 0.32382358 | 0.674 | 0.615 | 4.71E-07 | 5 | Kidins220 | homeostatic |
| Etfb | 2.88E-11 | 0.47083868 | 0.86 | 0.788 | 5.56E-07 | 5 | Etfb | homeostatic |
| Tep1 | 3.09E-11 | 0.51432789 | 0.467 | 0.398 | 5.98E-07 | 5 | Tep1 | homeostatic |
| Ncoa4 | 3.51E-11 | 0.37752974 | 0.699 | 0.672 | 6.78E-07 | 5 | Ncoa4 | homeostatic |
| Gga21 | 4.04E-11 | 0.46221719 | 0.688 | 0.596 | 7.80E-07 | 5 | Gga2 | homeostatic |
| Lipa1 | 4.64E-11 | 0.30106573 | 0.72 | 0.69 | 8.98E-07 | 5 | Lipa | homeostatic |
| Tbc1d191 | 4.84E-11 | 0.55555574 | 0.482 | 0.393 | 9.36E-07 | 5 | Tbc1d19 | homeostatic |
| Slc6a91 | 4.89E-11 | 0.84898784 | 0.335 | 0.264 | 9.45E-07 | 5 | Slc6a9 | homeostatic |
| Vps26c | 5.14E-11 | 0.44771953 | 0.546 | 0.489 | 9.94E-07 | 5 | Vps26c | homeostatic |
| Sobp | 6.36E-11 | 0.44210031 | 0.344 | 0.275 | 1.23E-06 | 5 | Sobp | homeostatic |
| Arfgef2 | 6.45E-11 | 0.58330392 | 0.412 | 0.324 | 1.25E-06 | 5 | Arfgef2 | homeostatic |
| Zfp110 | 8.97E-11 | 0.52190806 | 0.435 | 0.363 | 1.73E-06 | 5 | Zfp110 | homeostatic |
| Trak1 | 9.57E-11 | 0.34301527 | 0.73 | 0.688 | 1.85E-06 | 5 | Trak1 | homeostatic |
| Zbtb441 | 1.15E-10 | 0.42390685 | 0.562 | 0.505 | 2.22E-06 | 5 | Zbtb44 | homeostatic |
| Vapb | 1.25E-10 | 0.42202041 | 0.822 | 0.771 | 2.41E-06 | 5 | Vapb | homeostatic |
| Igf2r1 | 1.29E-10 | 0.26761199 | 0.859 | 0.816 | 2.50E-06 | 5 | Igf2r | homeostatic |
| Ap2b1 | 1.37E-10 | 0.39883729 | 0.832 | 0.804 | 2.64E-06 | 5 | Ap2b1 | homeostatic |
| Ptprj2 | 1.38E-10 | 0.5718109 | 0.492 | 0.372 | 2.67E-06 | 5 | Ptprj | homeostatic |
| Kcnk61 | 1.42E-10 | 0.57543846 | 0.302 | 0.222 | 2.75E-06 | 5 | Kcnk6 | homeostatic |
| Stk31 | 1.61E-10 | 0.35589145 | 0.72 | 0.724 | 3.11E-06 | 5 | Stk3 | homeostatic |
| Stxbp51 | 1.80E-10 | 0.56376203 | 0.585 | 0.494 | 3.47E-06 | 5 | Stxbp5 | homeostatic |
| Adam101 | 2.28E-10 | 0.36488079 | 0.736 | 0.693 | 4.41E-06 | 5 | Adam10 | homeostatic |
| Rnf1671 | 2.40E-10 | 0.45307138 | 0.412 | 0.326 | 4.64E-06 | 5 | Rnf167 | homeostatic |
| Repin11 | 2.68E-10 | 0.42858971 | 0.424 | 0.326 | 5.19E-06 | 5 | Repin1 | homeostatic |
| Exosc71 | 2.78E-10 | 0.57138407 | 0.398 | 0.306 | 5.38E-06 | 5 | Exosc7 | homeostatic |
| Rogdi | 2.88E-10 | 0.76279967 | 0.255 | 0.183 | 5.57E-06 | 5 | Rogdi | homeostatic |
| Ank31 | 3.14E-10 | 0.38342828 | 0.43 | 0.35 | 6.07E-06 | 5 | Ank3 | homeostatic |
| Dnajb41 | 3.14E-10 | 0.43890993 | 0.511 | 0.42 | 6.08E-06 | 5 | Dnajb4 | homeostatic |
| Tnfsf12 | 4.02E-10 | 0.39201497 | 0.806 | 0.757 | 7.77E-06 | 5 | Tnfsf12 | homeostatic |
| Slc35f51 | 4.31E-10 | 0.41047523 | 0.739 | 0.685 | 8.33E-06 | 5 | Slc35f5 | homeostatic |
| Rab11fip51 | 5.09E-10 | 0.46194579 | 0.746 | 0.702 | 9.83E-06 | 5 | Rab11fip5 | homeostatic |
| Wwp1 | 5.18E-10 | 0.4384091 | 0.571 | 0.501 | 1.00E-05 | 5 | Wwp1 | homeostatic |
| Podxl2 | 5.45E-10 | 0.75210403 | 0.388 | 0.282 | 1.05E-05 | 5 | Podxl2 | homeostatic |
| Ncoa21 | 5.92E-10 | 0.4957662 | 0.505 | 0.403 | 1.14E-05 | 5 | Ncoa2 | homeostatic |
| Sema6c1 | 6.04E-10 | 0.53909112 | 0.429 | 0.336 | 1.17E-05 | 5 | Sema6c | homeostatic |
| Ssh2 | 6.40E-10 | 0.64642531 | 0.425 | 0.32 | 1.24E-05 | 5 | Ssh2 | homeostatic |
| Map3k141 | 6.80E-10 | 0.41390035 | 0.461 | 0.382 | 1.31E-05 | 5 | Map3k14 | homeostatic |
| Prrg11 | 7.31E-10 | 0.65351872 | 0.345 | 0.259 | 1.41E-05 | 5 | Prrg1 | homeostatic |
| Stard9 | 7.40E-10 | 0.77937637 | 0.273 | 0.188 | 1.43E-05 | 5 | Stard9 | homeostatic |
| Kctd17 | 7.59E-10 | 0.3064443 | 0.85 | 0.795 | 1.47E-05 | 5 | Kctd17 | homeostatic |
| Dnmbp | 8.30E-10 | 0.63963067 | 0.438 | 0.352 | 1.60E-05 | 5 | Dnmbp | homeostatic |
| Zc3hav11 | 8.35E-10 | 0.44333551 | 0.586 | 0.528 | 1.61E-05 | 5 | Zc3hav1 | homeostatic |
| Acaa21 | 9.04E-10 | 0.40639204 | 0.742 | 0.663 | 1.75E-05 | 5 | Acaa2 | homeostatic |
| Dhdh | 9.08E-10 | 0.81265378 | 0.259 | 0.184 | 1.76E-05 | 5 | Dhdh | homeostatic |
| Rin31 | 9.46E-10 | 0.63428257 | 0.254 | 0.205 | 1.83E-05 | 5 | Rin3 | homeostatic |
| Crot | 9.78E-10 | 0.56479329 | 0.398 | 0.318 | 1.89E-05 | 5 | Crot | homeostatic |
| Fn3krp1 | 1.21E-09 | 0.89227754 | 0.303 | 0.209 | 2.34E-05 | 5 | Fn3krp | homeostatic |
| Pbxip11 | 1.21E-09 | 0.488426 | 0.633 | 0.552 | 2.34E-05 | 5 | Pbxip1 | homeostatic |
| Anxa4 | 1.22E-09 | 0.36964534 | 0.76 | 0.704 | 2.35E-05 | 5 | Anxa4 | homeostatic |
| Hsd17b4 | 1.44E-09 | 0.40605926 | 0.692 | 0.613 | 2.79E-05 | 5 | Hsd17b4 | homeostatic |
| Rnh1 | 1.45E-09 | 0.35448006 | 0.864 | 0.814 | 2.80E-05 | 5 | Rnh1 | homeostatic |
| Narf2 | 1.56E-09 | 0.38398003 | 0.532 | 0.444 | 3.02E-05 | 5 | Narf | homeostatic |
| Mapre22 | 1.62E-09 | 0.37279108 | 0.558 | 0.492 | 3.13E-05 | 5 | Mapre2 | homeostatic |
| Cryl11 | 1.78E-09 | 0.68157134 | 0.298 | 0.211 | 3.44E-05 | 5 | Cryl1 | homeostatic |
| Hivep1 | 1.99E-09 | 0.45805833 | 0.562 | 0.47 | 3.85E-05 | 5 | Hivep1 | homeostatic |
| Slc1a41 | 2.10E-09 | 0.43394804 | 0.4 | 0.347 | 4.06E-05 | 5 | Slc1a4 | homeostatic |
| Kif2a1 | 2.32E-09 | 0.61006729 | 0.497 | 0.424 | 4.49E-05 | 5 | Kif2a | homeostatic |
| Zfp6081 | 2.44E-09 | 0.34042939 | 0.547 | 0.502 | 4.72E-05 | 5 | Zfp608 | homeostatic |
| Acss31 | 2.44E-09 | 1.06969515 | 0.275 | 0.172 | 4.72E-05 | 5 | Acss3 | homeostatic |
| Uchl31 | 2.47E-09 | 0.81653631 | 0.322 | 0.227 | 4.77E-05 | 5 | Uchl3 | homeostatic |
| 2-Mar | 2.70E-09 | 0.37870722 | 0.807 | 0.746 | 5.22E-05 | 5 | 2-Mar | homeostatic |
| Slae | 2.85E-09 | 0.5453364 | 0.345 | 0.256 | 5.50E-05 | 5 | Slae | homeostatic |
| Wbp1l1 | 2.93E-09 | 0.27914655 | 0.906 | 0.887 | 5.67E-05 | 5 | Wbp1l | homeostatic |
| Acsl51 | 3.21E-09 | 0.41404696 | 0.685 | 0.619 | 6.21E-05 | 5 | Acsl5 | homeostatic |
| Casp121 | 3.23E-09 | 0.50796751 | 0.49 | 0.44 | 6.25E-05 | 5 | Casp12 | homeostatic |
| Zfp2171 | 3.39E-09 | 0.29637484 | 0.684 | 0.628 | 6.55E-05 | 5 | Zfp217 | homeostatic |
| Fes2 | 3.62E-09 | 0.40534318 | 0.454 | 0.376 | 6.99E-05 | 5 | Fes | homeostatic |
| Cgnl1 | 4.03E-09 | 0.68815054 | 0.272 | 0.19 | 7.79E-05 | 5 | Cgnl1 | homeostatic |
| Arntl1 | 4.14E-09 | 0.49374095 | 0.402 | 0.321 | 8.01E-05 | 5 | Arntl | homeostatic |
| Hsdl2 | 5.04E-09 | 0.4887903 | 0.353 | 0.27 | 9.75E-05 | 5 | Hsdl2 | homeostatic |
| Akr1e1 | 6.69E-09 | 0.68558723 | 0.296 | 0.224 | 0.00012927 | 5 | Akr1e1 | homeostatic |
| Dse | 6.69E-09 | 0.35101089 | 0.859 | 0.824 | 0.00012943 | 5 | Dse | homeostatic |
| Rsrc11 | 6.73E-09 | 0.36901402 | 0.382 | 0.322 | 0.00013018 | 5 | Rsrc1 | homeostatic |
| Trp53inp11 | 7.08E-09 | 0.34297494 | 0.297 | 0.272 | 0.00013691 | 5 | Trp53inp1 | homeostatic |
| Gpd1l | 7.65E-09 | 0.77188741 | 0.272 | 0.208 | 0.00014785 | 5 | Gpd1l | homeostatic |
| Oxct11 | 8.09E-09 | 0.51613319 | 0.626 | 0.543 | 0.00015639 | 5 | Oxct1 | homeostatic |
| Cmtr11 | 8.48E-09 | 0.47708798 | 0.599 | 0.519 | 0.00016398 | 5 | Cmtr1 | homeostatic |
| Git1 | 8.63E-09 | 0.34466718 | 0.717 | 0.666 | 0.00016688 | 5 | Git1 | homeostatic |
| Prickle21 | 8.88E-09 | 0.42608778 | 0.426 | 0.361 | 0.00017175 | 5 | Prickle2 | homeostatic |
| Slc25a45 | 9.14E-09 | 0.68941115 | 0.286 | 0.195 | 0.0001767 | 5 | Slc25a45 | homeostatic |
| Cry11 | 9.98E-09 | 0.64824793 | 0.259 | 0.2 | 0.00019293 | 5 | Cry1 | homeostatic |
| Ndst2 | 1.03E-08 | 0.59634439 | 0.327 | 0.266 | 0.00019818 | 5 | Ndst2 | homeostatic |
| Tnks1bp1 | 1.10E-08 | 0.29940418 | 0.874 | 0.856 | 0.00021231 | 5 | Tnks1bp1 | homeostatic |
| Zdhhc11 | 1.22E-08 | 0.54981301 | 0.506 | 0.417 | 0.00023674 | 5 | Zdhhc1 | homeostatic |
| Ccl191 | 1.41E-08 | 0.75777451 | 0.298 | 0.17 | 0.00027325 | 5 | Ccl19 | homeostatic |
| Tshz11 | 1.52E-08 | 0.55978322 | 0.421 | 0.33 | 0.00029317 | 5 | Tshz1 | homeostatic |
| Tlcd2 | 1.55E-08 | 0.82266439 | 0.259 | 0.173 | 0.00029962 | 5 | Tlcd2 | homeostatic |
| Aldh6a11 | 1.72E-08 | 0.50838466 | 0.423 | 0.339 | 0.00033348 | 5 | Aldh6a1 | homeostatic |
| Gne | 1.74E-08 | 0.54331982 | 0.325 | 0.244 | 0.00033669 | 5 | Gne | homeostatic |
| Cpt2 | 1.78E-08 | 0.55023549 | 0.505 | 0.421 | 0.00034429 | 5 | Cpt2 | homeostatic |
| Nek1 | 1.79E-08 | 0.52498287 | 0.376 | 0.292 | 0.00034563 | 5 | Nek1 | homeostatic |
| Plekho22 | 1.85E-08 | 0.29469521 | 0.891 | 0.85 | 0.00035761 | 5 | Plekho2 | homeostatic |
| Pigt | 1.85E-08 | 0.38533031 | 0.834 | 0.789 | 0.00035782 | 5 | Pigt | homeostatic |
| H2-Ke6 | 2.13E-08 | 0.50657641 | 0.354 | 0.284 | 0.00041221 | 5 | H2-Ke6 | homeostatic |
| Hmgn3 | 2.13E-08 | 0.43588526 | 0.552 | 0.489 | 0.00041251 | 5 | Hmgn3 | homeostatic |
| Plekha22 | 2.41E-08 | 0.41943688 | 0.586 | 0.52 | 0.0004665 | 5 | Plekha2 | homeostatic |
| Ralgds2 | 2.84E-08 | 0.25292718 | 0.544 | 0.482 | 0.00054925 | 5 | Ralgds | homeostatic |
| Kif13a | 3.16E-08 | 0.35751218 | 0.504 | 0.457 | 0.00061045 | 5 | Kif13a | homeostatic |
| Eci21 | 3.41E-08 | 0.41394426 | 0.556 | 0.496 | 0.00065986 | 5 | Eci2 | homeostatic |
| Mknk1 | 3.78E-08 | 0.48288433 | 0.312 | 0.246 | 0.0007317 | 5 | Mknk1 | homeostatic |
| Ifngr1 | 4.06E-08 | 0.4554643 | 0.862 | 0.826 | 0.00078407 | 5 | Ifngr1 | homeostatic |
| Kdsr | 5.05E-08 | 0.41869698 | 0.533 | 0.483 | 0.00097651 | 5 | Kdsr | homeostatic |
| Ap4e1 | 5.07E-08 | 0.51441756 | 0.373 | 0.281 | 0.00097977 | 5 | Ap4e1 | homeostatic |
| Gyg1 | 5.26E-08 | 0.52161732 | 0.451 | 0.374 | 0.00101667 | 5 | Gyg | homeostatic |
| Zdhhc3 | 5.27E-08 | 0.43111736 | 0.834 | 0.774 | 0.00101855 | 5 | Zdhhc3 | homeostatic |
| Tpcn2 | 5.87E-08 | 0.52470064 | 0.423 | 0.348 | 0.0011342 | 5 | Tpcn2 | homeostatic |
| Fam172a | 6.85E-08 | 0.47574907 | 0.458 | 0.399 | 0.00132516 | 5 | Fam172a | homeostatic |
| Zadh2 | 7.15E-08 | 0.45198544 | 0.615 | 0.532 | 0.00138221 | 5 | Zadh2 | homeostatic |
| Mmut1 | 7.33E-08 | 0.62066406 | 0.289 | 0.222 | 0.0014173 | 5 | Mmut | homeostatic |
| Mppe1 | 7.65E-08 | 0.64677092 | 0.292 | 0.227 | 0.00147966 | 5 | Mppe1 | homeostatic |
| Sfxn11 | 7.76E-08 | 0.42552222 | 0.788 | 0.733 | 0.00150071 | 5 | Sfxn1 | homeostatic |
| Rmc1 | 7.80E-08 | 0.58804689 | 0.392 | 0.293 | 0.00150843 | 5 | Rmc1 | homeostatic |
| Ntn1 | 7.93E-08 | 0.38187443 | 0.456 | 0.29 | 0.00153226 | 5 | Ntn1 | homeostatic |
| Cyld | 8.09E-08 | 0.50056022 | 0.428 | 0.355 | 0.00156413 | 5 | Cyld | homeostatic |
| Pdxk1 | 8.12E-08 | 0.49130788 | 0.503 | 0.426 | 0.00156913 | 5 | Pdxk | homeostatic |
| Zmat5 | 8.50E-08 | 0.5285823 | 0.348 | 0.271 | 0.00164335 | 5 | Zmat5 | homeostatic |
| Smim141 | 8.73E-08 | 0.34197075 | 0.801 | 0.752 | 0.00168838 | 5 | Smim14 | homeostatic |
| Tmem241 | 8.78E-08 | 0.52905868 | 0.308 | 0.252 | 0.00169773 | 5 | Tmem241 | homeostatic |
| Pnpt11 | 9.39E-08 | 0.49436247 | 0.364 | 0.298 | 0.00181581 | 5 | Pnpt1 | homeostatic |
| Eif2ak31 | 1.01E-07 | 0.42721676 | 0.58 | 0.5 | 0.00194547 | 5 | Eif2ak3 | homeostatic |
| Dusp23 | 1.01E-07 | 0.78087478 | 0.275 | 0.221 | 0.00196033 | 5 | Dusp23 | homeostatic |
| Ext2 | 1.04E-07 | 0.2686198 | 0.855 | 0.842 | 0.00200198 | 5 | Ext2 | homeostatic |
| Synrg | 1.04E-07 | 0.69664953 | 0.282 | 0.209 | 0.00202 | 5 | Synrg | homeostatic |
| Nudt181 | 1.10E-07 | 0.48862126 | 0.401 | 0.306 | 0.002132 | 5 | Nudt18 | homeostatic |
| Adamts141 | 1.24E-07 | 0.31081346 | 0.595 | 0.498 | 0.00240608 | 5 | Adamts14 | homeostatic |
| Arsb1 | 1.31E-07 | 0.43891199 | 0.678 | 0.63 | 0.00253905 | 5 | Arsb | homeostatic |
| Bcl2l14 | 1.35E-07 | 0.60609096 | 0.386 | 0.318 | 0.00261969 | 5 | Bcl2l1 | homeostatic |
| Ttc37 | 1.36E-07 | 0.28983476 | 0.662 | 0.629 | 0.00262241 | 5 | Ttc37 | homeostatic |
| Zfyve16 | 1.42E-07 | 0.43935455 | 0.4 | 0.35 | 0.00275133 | 5 | Zfyve16 | homeostatic |
| Ints31 | 1.43E-07 | 0.44793975 | 0.603 | 0.547 | 0.00275586 | 5 | Ints3 | homeostatic |
| Rab11fip2 | 1.69E-07 | 0.36114215 | 0.494 | 0.437 | 0.00326193 | 5 | Rab11fip2 | homeostatic |
| Pxk1 | 1.71E-07 | 0.27543334 | 0.735 | 0.703 | 0.00329853 | 5 | Pxk | homeostatic |
| Abhd16a | 1.71E-07 | 0.4365806 | 0.635 | 0.574 | 0.00331399 | 5 | Abhd16a | homeostatic |
| Pacs2 | 1.75E-07 | 0.6976824 | 0.313 | 0.234 | 0.00337764 | 5 | Pacs2 | homeostatic |
| Nck1 | 1.79E-07 | 0.45802003 | 0.539 | 0.484 | 0.0034681 | 5 | Nck1 | homeostatic |
| Mgat21 | 1.82E-07 | 0.31808237 | 0.813 | 0.788 | 0.00352529 | 5 | Mgat2 | homeostatic |
| Mras | 1.93E-07 | 0.3039224 | 0.35 | 0.345 | 0.00373047 | 5 | Mras | homeostatic |
| Stx111 | 2.02E-07 | 0.37951807 | 0.341 | 0.26 | 0.00391065 | 5 | Stx11 | homeostatic |
| Sptssa | 2.08E-07 | 0.38429633 | 0.586 | 0.561 | 0.00401502 | 5 | Sptssa | homeostatic |
| Setd61 | 2.08E-07 | 0.55459756 | 0.259 | 0.192 | 0.00402423 | 5 | Setd6 | homeostatic |
| Gstm51 | 2.30E-07 | 0.49281872 | 0.372 | 0.304 | 0.00445136 | 5 | Gstm5 | homeostatic |
| Gaa1 | 2.44E-07 | 0.29760086 | 0.956 | 0.921 | 0.00472264 | 5 | Gaa | homeostatic |
| Apbb1 | 2.47E-07 | 0.52249213 | 0.339 | 0.284 | 0.00478497 | 5 | Apbb1 | homeostatic |
| Stam | 2.63E-07 | 0.40314503 | 0.563 | 0.488 | 0.00508738 | 5 | Stam | homeostatic |
| Aplp21 | 2.96E-07 | 0.33932071 | 0.858 | 0.821 | 0.00572576 | 5 | Aplp2 | homeostatic |
| Grina2 | 2.97E-07 | 0.42430986 | 0.723 | 0.641 | 0.00573654 | 5 | Grina | homeostatic |
| Psen1 | 3.77E-07 | 0.36476512 | 0.718 | 0.677 | 0.00728293 | 5 | Psen1 | homeostatic |
| Coro7 | 3.79E-07 | 0.49799804 | 0.393 | 0.316 | 0.00732235 | 5 | Coro7 | homeostatic |
| Evc1 | 3.79E-07 | 0.53831587 | 0.44 | 0.374 | 0.0073286 | 5 | Evc | homeostatic |
| Ksr11 | 5.29E-07 | 0.43087964 | 0.423 | 0.374 | 0.0102343 | 5 | Ksr1 | homeostatic |
| Dyrk1b1 | 5.52E-07 | 0.33894579 | 0.336 | 0.27 | 0.0106668 | 5 | Dyrk1b | homeostatic |
| Cul7 | 5.65E-07 | 0.31986376 | 0.702 | 0.652 | 0.01093089 | 5 | Cul7 | homeostatic |
| Vdac31 | 6.09E-07 | 0.44527377 | 0.608 | 0.544 | 0.01177659 | 5 | Vdac3 | homeostatic |
| Sesn1 | 6.22E-07 | 0.62950808 | 0.25 | 0.19 | 0.01202574 | 5 | Sesn1 | homeostatic |
| Mien1 | 6.69E-07 | 0.43091245 | 0.579 | 0.509 | 0.01293719 | 5 | Mien1 | homeostatic |
| Bin11 | 7.98E-07 | 0.3428398 | 0.774 | 0.75 | 0.01542413 | 5 | Bin1 | homeostatic |
| Letmd1 | 8.08E-07 | 0.50503858 | 0.433 | 0.365 | 0.01562908 | 5 | Letmd1 | homeostatic |
| Znhit6 | 8.21E-07 | 0.61914451 | 0.345 | 0.296 | 0.01587597 | 5 | Znhit6 | homeostatic |
| Eya31 | 8.93E-07 | 0.34103139 | 0.431 | 0.363 | 0.0172607 | 5 | Eya3 | homeostatic |
| Tm2d3 | 9.09E-07 | 0.39459003 | 0.513 | 0.451 | 0.017569 | 5 | Tm2d3 | homeostatic |
| Micu11 | 9.59E-07 | 0.37569976 | 0.589 | 0.543 | 0.01853778 | 5 | Micu1 | homeostatic |
| Abcc5 | 1.10E-06 | 0.30680185 | 0.642 | 0.604 | 0.02128362 | 5 | Abcc5 | homeostatic |
| Ppp2r3c | 1.16E-06 | 0.5083427 | 0.259 | 0.205 | 0.02238335 | 5 | Ppp2r3c | homeostatic |
| Nfic1 | 1.20E-06 | 0.26861583 | 0.867 | 0.842 | 0.02314782 | 5 | Nfic | homeostatic |
| Ikbkb | 1.25E-06 | 0.40193996 | 0.514 | 0.45 | 0.02414174 | 5 | Ikbkb | homeostatic |
| Tmem50b | 1.36E-06 | 0.46293095 | 0.499 | 0.433 | 0.02622908 | 5 | Tmem50b | homeostatic |
| Cdk14 | 1.37E-06 | 0.30139663 | 0.657 | 0.604 | 0.02648092 | 5 | Cdk14 | homeostatic |
| Cbr31 | 1.44E-06 | 0.66969587 | 0.279 | 0.214 | 0.02791189 | 5 | Cbr3 | homeostatic |
| Nrbp21 | 1.47E-06 | 0.48434878 | 0.302 | 0.235 | 0.02835155 | 5 | Nrbp2 | homeostatic |
| Ccnyl12 | 1.47E-06 | 0.36855754 | 0.538 | 0.456 | 0.02844216 | 5 | Ccnyl1 | homeostatic |
| Lrch31 | 1.52E-06 | 0.37908166 | 0.636 | 0.593 | 0.02933337 | 5 | Lrch3 | homeostatic |
| Parp10 | 1.53E-06 | 0.72413385 | 0.255 | 0.201 | 0.02955721 | 5 | Parp10 | homeostatic |
| Pdk21 | 2.06E-06 | 0.40721172 | 0.349 | 0.281 | 0.03973846 | 5 | Pdk2 | homeostatic |
| Dact11 | 2.06E-06 | 0.43642106 | 0.711 | 0.684 | 0.0398767 | 5 | Dact1 | homeostatic |
| Glce | 2.07E-06 | 0.56904791 | 0.43 | 0.369 | 0.04010809 | 5 | Glce | homeostatic |
| Lgals8 | 2.28E-06 | 0.46012075 | 0.643 | 0.561 | 0.04405552 | 5 | Lgals8 | homeostatic |
| Eef2kmt | 2.45E-06 | 0.40488008 | 0.452 | 0.388 | 0.04734598 | 5 | Eef2kmt | homeostatic |
| Dusp16 | 2.72E-06 | 0.45900797 | 0.359 | 0.291 | 0.05258952 | 5 | Dusp16 | homeostatic |
| Adamtsl51 | 2.89E-06 | 0.42100526 | 0.522 | 0.496 | 0.05589559 | 5 | Adamtsl5 | homeostatic |
| Arhgap12 | 2.92E-06 | 0.37540292 | 0.425 | 0.359 | 0.05651827 | 5 | Arhgap12 | homeostatic |
| Cd109 | 3.24E-06 | 0.5470555 | 0.272 | 0.215 | 0.06264383 | 5 | Cd109 | homeostatic |
| Nectin31 | 3.31E-06 | 0.36185006 | 0.692 | 0.648 | 0.06399188 | 5 | Nectin3 | homeostatic |
| Ripk1 | 3.40E-06 | 0.3140664 | 0.704 | 0.655 | 0.06576786 | 5 | Ripk1 | homeostatic |
| Leprot | 3.53E-06 | 0.36037703 | 0.895 | 0.858 | 0.06823786 | 5 | Leprot | homeostatic |
| Nhlrc2 | 3.58E-06 | 0.49215898 | 0.541 | 0.483 | 0.06915441 | 5 | Nhlrc2 | homeostatic |
| Sgsh2 | 3.59E-06 | 0.39943632 | 0.39 | 0.324 | 0.06933424 | 5 | Sgsh | homeostatic |
| Pccb | 3.64E-06 | 0.46523856 | 0.435 | 0.367 | 0.07035405 | 5 | Pccb | homeostatic |
| Fam111a1 | 3.79E-06 | 0.77034549 | 0.383 | 0.265 | 0.07333811 | 5 | Fam111a | homeostatic |
| Stk39 | 4.12E-06 | 0.40004024 | 0.451 | 0.367 | 0.07958932 | 5 | Stk39 | homeostatic |
| Cfap2981 | 4.67E-06 | 0.56910813 | 0.302 | 0.24 | 0.09034131 | 5 | Cfap298 | homeostatic |
| Sprytd71 | 4.76E-06 | 0.40037461 | 0.406 | 0.339 | 0.09203744 | 5 | Spryd7 | homeostatic |
| Gramd3 | 4.81E-06 | 0.38707021 | 0.598 | 0.523 | 0.09295575 | 5 | Gramd3 | homeostatic |
| Gtpbp8 | 4.94E-06 | 0.40715008 | 0.261 | 0.211 | 0.09541825 | 5 | Gtpbp8 | homeostatic |
| Fut111 | 5.69E-06 | 0.40116999 | 0.643 | 0.566 | 0.1100705 | 5 | Fut11 | homeostatic |
| Oxsm | 5.77E-06 | 0.56942778 | 0.282 | 0.226 | 0.11152676 | 5 | Oxsm | homeostatic |
| Kif1b | 5.94E-06 | 0.33587357 | 0.584 | 0.546 | 0.11483381 | 5 | Kif1b | homeostatic |
| Fst2 | 6.19E-06 | 0.52437696 | 0.73 | 0.535 | 0.11974727 | 5 | Fst | homeostatic |
| B3gntl11 | 6.21E-06 | 0.63549813 | 0.259 | 0.2 | 0.12015959 | 5 | B3gntl1 | homeostatic |
| Tango2 | 6.59E-06 | 0.46140333 | 0.382 | 0.324 | 0.12747009 | 5 | Tango2 | homeostatic |
| Atf52 | 6.96E-06 | 0.30667758 | 0.846 | 0.811 | 0.13450991 | 5 | Atf5 | homeostatic |
| Rras | 6.97E-06 | 0.26012056 | 0.881 | 0.85 | 0.13467265 | 5 | Rras | homeostatic |
| Nbas1 | 7.16E-06 | 0.3201613 | 0.674 | 0.649 | 0.13836663 | 5 | Nbas | homeostatic |
| H2-T22 | 8.12E-06 | 0.47093944 | 0.405 | 0.34 | 0.15701814 | 5 | H2-T22 | homeostatic |
| Fbxo341 | 8.17E-06 | 0.42011716 | 0.334 | 0.265 | 0.15794818 | 5 | Fbxo34 | homeostatic |
| Dpagt11 | 8.17E-06 | 0.34666423 | 0.626 | 0.576 | 0.15797635 | 5 | Dpagt1 | homeostatic |
| Aktip | 8.31E-06 | 0.37179281 | 0.447 | 0.392 | 0.16063925 | 5 | Aktip | homeostatic |
| Flot1 | 8.54E-06 | 0.26699392 | 0.807 | 0.795 | 0.16510852 | 5 | Flot1 | homeostatic |
| Fam8a1 | 8.63E-06 | 0.27434763 | 0.703 | 0.697 | 0.16684643 | 5 | Fam8a1 | homeostatic |
| Smim111 | 8.65E-06 | 0.428136 | 0.519 | 0.474 | 0.16730604 | 5 | Smim11 | homeostatic |
| Mapk8ip12 | 8.74E-06 | 0.33279649 | 0.33 | 0.278 | 0.16893281 | 5 | Mapk8ip1 | homeostatic |
| Dhx57 | 9.01E-06 | 0.43789766 | 0.353 | 0.289 | 0.17415835 | 5 | Dhx57 | homeostatic |
| Scamp51 | 1.00E-05 | 0.49146128 | 0.373 | 0.292 | 0.19411051 | 5 | Scamp5 | homeostatic |
| Tns21 | 1.01E-05 | 0.39513913 | 0.76 | 0.724 | 0.1943851 | 5 | Tns2 | homeostatic |
| Oat1 | 1.04E-05 | 0.35199942 | 0.943 | 0.894 | 0.20064641 | 5 | Oat | homeostatic |
| Alas11 | 1.08E-05 | 0.33775067 | 0.518 | 0.437 | 0.2088206 | 5 | Alas1 | homeostatic |
| Vsir2 | 1.10E-05 | 0.26432449 | 0.52 | 0.453 | 0.21181591 | 5 | Vsir | homeostatic |
| BC0297221 | 1.12E-05 | 0.51031665 | 0.61 | 0.532 | 0.21576618 | 5 | BC029722 | homeostatic |
| Taldo1 | 1.14E-05 | 0.3942724 | 0.709 | 0.645 | 0.22073563 | 5 | Taldo1 | homeostatic |
| Cnpy31 | 1.20E-05 | 0.5202829 | 0.641 | 0.58 | 0.23178387 | 5 | Cnpy3 | homeostatic |
| Mitd1 | 1.20E-05 | 0.62052421 | 0.293 | 0.223 | 0.23242643 | 5 | Mitd1 | homeostatic |
| Gsdmd | 1.23E-05 | 0.29149591 | 0.638 | 0.616 | 0.23758036 | 5 | Gsdmd | homeostatic |
| Extl21 | 1.30E-05 | 0.53567198 | 0.419 | 0.362 | 0.25122918 | 5 | Extl2 | homeostatic |
| Ptpdc11 | 1.31E-05 | 0.50589528 | 0.334 | 0.26 | 0.25267903 | 5 | Ptpdc1 | homeostatic |
| Dipk2a1 | 1.37E-05 | 0.5343109 | 0.462 | 0.38 | 0.26432404 | 5 | Dipk2a | homeostatic |
| Hfe2 | 1.37E-05 | 0.61138864 | 0.273 | 0.207 | 0.26581102 | 5 | Hfe | homeostatic |
| Ttc8 | 1.70E-05 | 0.6021709 | 0.339 | 0.283 | 0.32799467 | 5 | Ttc8 | homeostatic |
| Ano6 | 1.70E-05 | 0.2898631 | 0.829 | 0.79 | 0.32847629 | 5 | Ano6 | homeostatic |
| Syngr2 | 1.70E-05 | 0.41405526 | 0.571 | 0.512 | 0.32954809 | 5 | Syngr2 | homeostatic |
| Zfp398 | 1.76E-05 | 0.41961008 | 0.292 | 0.239 | 0.34040933 | 5 | Zfp398 | homeostatic |
| Cryzl11 | 1.77E-05 | 0.28204454 | 0.576 | 0.527 | 0.34173655 | 5 | Cryzl1 | homeostatic |
| Ttc231 | 1.86E-05 | 0.4237075 | 0.379 | 0.314 | 0.3587433 | 5 | Ttc23 | homeostatic |
| Pex2 | 1.98E-05 | 0.49764558 | 0.52 | 0.458 | 0.3836197 | 5 | Pex2 | homeostatic |
| Ly961 | 2.01E-05 | 0.34385709 | 0.293 | 0.244 | 0.38851961 | 5 | Ly96 | homeostatic |
| Taok2 | 2.18E-05 | 0.25620792 | 0.741 | 0.676 | 0.42244201 | 5 | Taok2 | homeostatic |
| Abca5 | 2.27E-05 | 0.5592376 | 0.359 | 0.296 | 0.43893654 | 5 | Abca5 | homeostatic |
| Fam168a | 2.52E-05 | 0.35392591 | 0.737 | 0.69 | 0.48709964 | 5 | Fam168a | homeostatic |
| Mfsd12 | 2.57E-05 | 0.39410559 | 0.306 | 0.248 | 0.49755441 | 5 | Mfsd12 | homeostatic |
| Cdk2ap2 | 2.58E-05 | 0.30242481 | 0.537 | 0.472 | 0.49787483 | 5 | Cdk2ap2 | homeostatic |
| Tdrd3 | 2.79E-05 | 0.27669221 | 0.558 | 0.515 | 0.53945843 | 5 | Tdrd3 | homeostatic |
| Fbxo172 | 3.08E-05 | 0.42109682 | 0.495 | 0.415 | 0.59498676 | 5 | Fbxo17 | homeostatic |
| Cfl21 | 3.34E-05 | 0.35142221 | 0.633 | 0.582 | 0.64601778 | 5 | Cfl2 | homeostatic |
| Capn7 | 3.41E-05 | 0.37494385 | 0.53 | 0.48 | 0.65921172 | 5 | Capn7 | homeostatic |
| Arnt | 3.80E-05 | 0.301145 | 0.855 | 0.82 | 0.73502598 | 5 | Arnt | homeostatic |
| Ibtk2 | 3.80E-05 | 0.36544233 | 0.648 | 0.6 | 0.73547675 | 5 | Ibtk | homeostatic |
| Qpctl1 | 4.16E-05 | 0.40267461 | 0.52 | 0.441 | 0.80426854 | 5 | Qpctl | homeostatic |
| Mat2b | 4.65E-05 | 0.37342113 | 0.618 | 0.581 | 0.89990093 | 5 | Mat2b | homeostatic |
| Pip5k2 | 4.74E-05 | 0.48752047 | 0.377 | 0.325 | 0.91576503 | 5 | Pip5k2 | homeostatic |
| Phkb | 4.86E-05 | 0.34812648 | 0.341 | 0.286 | 0.94000711 | 5 | Phkb | homeostatic |
| Car132 | 5.03E-05 | 0.41767927 | 0.471 | 0.385 | 0.97217776 | 5 | Car13 | homeostatic |
| Nt5c | 5.07E-05 | 0.30342392 | 0.662 | 0.605 | 0.98029277 | 5 | Nt5c | homeostatic |
| Idua | 5.17E-05 | 0.5150034 | 0.329 | 0.271 | 1 | 5 | Idua | homeostatic |
| Saysd11 | 5.35E-05 | 0.48529247 | 0.312 | 0.255 | 1 | 5 | Saysd1 | homeostatic |
| Dnase1l11 | 5.60E-05 | 0.51787031 | 0.265 | 0.212 | 1 | 5 | Dnase1l1 | homeostatic |
| Mapk9 | 5.62E-05 | 0.39705155 | 0.632 | 0.59 | 1 | 5 | Mapk9 | homeostatic |
| Faim | 5.86E-05 | 0.39529707 | 0.468 | 0.401 | 1 | 5 | Faim | homeostatic |
| Aamdc | 5.92E-05 | 0.41063201 | 0.409 | 0.348 | 1 | 5 | Aamdc | homeostatic |
| Spag9 | 6.01E-05 | 0.27426568 | 0.797 | 0.779 | 1 | 5 | Spag9 | homeostatic |
| Arhgap61 | 6.60E-05 | 0.46059138 | 0.277 | 0.212 | 1 | 5 | Arhgap6 | homeostatic |
| Plcb3 | 6.70E-05 | 0.29772793 | 0.712 | 0.656 | 1 | 5 | Plcb3 | homeostatic |
| Kras1 | 6.77E-05 | 0.29846606 | 0.632 | 0.601 | 1 | 5 | Kras | homeostatic |
| Glod4 | 6.78E-05 | 0.35002601 | 0.66 | 0.604 | 1 | 5 | Glod4 | homeostatic |
| Fbh1 | 7.09E-05 | 0.38357208 | 0.684 | 0.634 | 1 | 5 | Fbh1 | homeostatic |
| Rhobtb21 | 7.22E-05 | 0.68223552 | 0.263 | 0.203 | 1 | 5 | Rhobtb2 | homeostatic |
| Rnf1142 | 7.37E-05 | 0.28231778 | 0.486 | 0.467 | 1 | 5 | Rnf114 | homeostatic |
| Lrp10 | 7.56E-05 | 0.27844998 | 0.918 | 0.889 | 1 | 5 | Lrp10 | homeostatic |
| Tusc2 | 7.80E-05 | 0.41561652 | 0.26 | 0.204 | 1 | 5 | Tusc2 | homeostatic |
| LTO1 | 7.86E-05 | 0.3770018 | 0.345 | 0.303 | 1 | 5 | LTO1 | homeostatic |
| Ankfy1 | 7.87E-05 | 0.3379763 | 0.546 | 0.515 | 1 | 5 | Ankfy1 | homeostatic |
| Rp21 | 8.18E-05 | 0.36558621 | 0.428 | 0.375 | 1 | 5 | Rp2 | homeostatic |
| Cant1 | 8.27E-05 | 0.35929312 | 0.766 | 0.74 | 1 | 5 | Cant1 | homeostatic |
| Prkag21 | 8.72E-05 | 0.50488445 | 0.378 | 0.304 | 1 | 5 | Prkag2 | homeostatic |
| Ppp1r15b1 | 9.03E-05 | 0.2841951 | 0.716 | 0.66 | 1 | 5 | Ppp1r15b | homeostatic |
| Cyb5d2 | 9.78E-05 | 0.54671993 | 0.282 | 0.215 | 1 | 5 | Cyb5d2 | homeostatic |
| Adck2 | 0.00010077 | 0.67817199 | 0.326 | 0.247 | 1 | 5 | Adck2 | homeostatic |
| Fbxw9 | 0.00010659 | 0.29486242 | 0.299 | 0.255 | 1 | 5 | Fbxw9 | homeostatic |
| Rapgef11 | 0.00010826 | 0.29776392 | 0.761 | 0.745 | 1 | 5 | Rapgef1 | homeostatic |
| Psenen | 0.00010926 | 0.27406958 | 0.855 | 0.844 | 1 | 5 | Psenen | homeostatic |
| Apba1 | 0.00011877 | 0.51303928 | 0.376 | 0.309 | 1 | 5 | Apba1 | homeostatic |
| Vps45 | 0.00012217 | 0.42347339 | 0.305 | 0.259 | 1 | 5 | Vps45 | homeostatic |
| Rnf31 | 0.00012303 | 0.55434368 | 0.378 | 0.319 | 1 | 5 | Rnf31 | homeostatic |
| Asnsd11 | 0.00012333 | 0.4123009 | 0.59 | 0.536 | 1 | 5 | Asnsd1 | homeostatic |
| Mef2c1 | 0.00012889 | 0.43927108 | 0.369 | 0.274 | 1 | 5 | Mef2c | homeostatic |
| Tmem267 | 0.00012952 | 0.61518407 | 0.293 | 0.253 | 1 | 5 | Tmem267 | homeostatic |
| Nagk | 0.00013009 | 0.39990363 | 0.721 | 0.653 | 1 | 5 | Nagk | homeostatic |
| Tmem219 | 0.00013371 | 0.561663 | 0.44 | 0.366 | 1 | 5 | Tmem219 | homeostatic |
| Rexf1 | 0.00016105 | 0.25201622 | 0.426 | 0.364 | 1 | 5 | Rexf1 | homeostatic |
| Slc25a19 | 0.00016503 | 0.5061616 | 0.259 | 0.215 | 1 | 5 | Slc25a19 | homeostatic |
| Ercc5 | 0.00016662 | 0.54809618 | 0.284 | 0.23 | 1 | 5 | Ercc5 | homeostatic |
| Ppp2r5b | 0.00016982 | 0.33440484 | 0.585 | 0.542 | 1 | 5 | Ppp2r5b | homeostatic |
| Stim1 | 0.00018656 | 0.47492406 | 0.485 | 0.436 | 1 | 5 | Stim1 | homeostatic |
| Pdp11 | 0.00020261 | 0.51969113 | 0.294 | 0.232 | 1 | 5 | Pdp1 | homeostatic |
| Numb1 | 0.00021412 | 0.26334999 | 0.514 | 0.493 | 1 | 5 | Numb | homeostatic |
| Exoc3 | 0.00022084 | 0.35626291 | 0.504 | 0.472 | 1 | 5 | Exoc3 | homeostatic |
| Pik3r21 | 0.00022524 | 0.31543808 | 0.731 | 0.696 | 1 | 5 | Pik3r2 | homeostatic |
| Zfp4141 | 0.00022589 | 0.28439091 | 0.584 | 0.53 | 1 | 5 | Zfp414 | homeostatic |
| Dop1b1 | 0.00022659 | 0.44177626 | 0.343 | 0.27 | 1 | 5 | Dop1b | homeostatic |
| Atg2a1 | 0.00022896 | 0.35520108 | 0.515 | 0.475 | 1 | 5 | Atg2a | homeostatic |
| Ift122 | 0.00023694 | 0.5509071 | 0.253 | 0.201 | 1 | 5 | Ift122 | homeostatic |
| Cln81 | 0.00023816 | 0.54279018 | 0.278 | 0.221 | 1 | 5 | Cln8 | homeostatic |
| Zfp512b1 | 0.00025078 | 0.34334234 | 0.571 | 0.527 | 1 | 5 | Zfp512b | homeostatic |
| Rom1 | 0.00025799 | 0.58362946 | 0.307 | 0.258 | 1 | 5 | Rom1 | homeostatic |
| Armc10 | 0.00026737 | 0.42500624 | 0.299 | 0.244 | 1 | 5 | Armc10 | homeostatic |
| Sumf2 | 0.00027742 | 0.40319296 | 0.458 | 0.408 | 1 | 5 | Sumf2 | homeostatic |
| Lyrm2 | 0.0002804 | 0.57390528 | 0.362 | 0.302 | 1 | 5 | Lyrm2 | homeostatic |
| Abhd14a | 0.00028346 | 0.3913407 | 0.433 | 0.378 | 1 | 5 | Abhd14a | homeostatic |
| Traf5 | 0.00028857 | 0.57343718 | 0.256 | 0.216 | 1 | 5 | Traf5 | homeostatic |
| Irf21 | 0.00030121 | 0.35171603 | 0.452 | 0.405 | 1 | 5 | Irf2 | homeostatic |
| Acadvl | 0.00031033 | 0.27098939 | 0.632 | 0.59 | 1 | 5 | Acadvl | homeostatic |
| Aldh7a1 | 0.0003201 | 0.38948677 | 0.868 | 0.83 | 1 | 5 | Aldh7a1 | homeostatic |
| Gmppa | 0.00033398 | 0.29649638 | 0.694 | 0.644 | 1 | 5 | Gmppa | homeostatic |
| Msrb1 | 0.00034055 | 0.285194 | 0.755 | 0.723 | 1 | 5 | Msrb1 | homeostatic |
| Bloc1s5 | 0.0003413 | 0.4651861 | 0.294 | 0.245 | 1 | 5 | Bloc1s5 | homeostatic |
| Naa80 | 0.00034848 | 0.33226588 | 0.406 | 0.344 | 1 | 5 | Naa80 | homeostatic |
| Snap29 | 0.00034933 | 0.37992939 | 0.386 | 0.36 | 1 | 5 | Snap29 | homeostatic |
| St6galnac61 | 0.00036082 | 0.26602747 | 0.69 | 0.662 | 1 | 5 | St6galnac6 | homeostatic |
| Sidt2 | 0.00036989 | 0.37623983 | 0.877 | 0.838 | 1 | 5 | Sidt2 | homeostatic |
| Thra1 | 0.00037388 | 0.27630317 | 0.654 | 0.612 | 1 | 5 | Thra | homeostatic |
| Kitl1 | 0.00040272 | 0.47412333 | 0.274 | 0.214 | 1 | 5 | Kitl | homeostatic |
| Prkce1 | 0.00041602 | 0.34477136 | 0.332 | 0.281 | 1 | 5 | Prkce | homeostatic |
| R3hcc1l | 0.00042424 | 0.33082651 | 0.382 | 0.325 | 1 | 5 | R3hcc1l | homeostatic |
| Noct2 | 0.00043048 | 0.27837814 | 0.343 | 0.31 | 1 | 5 | Noct | homeostatic |
| Armc61 | 0.00044103 | 0.52927608 | 0.339 | 0.279 | 1 | 5 | Armc6 | homeostatic |
| Tctn31 | 0.00045652 | 0.57191114 | 0.359 | 0.297 | 1 | 5 | Tctn3 | homeostatic |
| Galnt111 | 0.00046189 | 0.34381422 | 0.437 | 0.399 | 1 | 5 | Galnt11 | homeostatic |
| Mapk7 | 0.00046642 | 0.54747654 | 0.278 | 0.237 | 1 | 5 | Mapk7 | homeostatic |
| Cyfip11 | 0.0004739 | 0.31855963 | 0.758 | 0.734 | 1 | 5 | Cyfip1 | homeostatic |
| Mafk | 0.00048194 | 0.42689398 | 0.416 | 0.354 | 1 | 5 | Mafk | homeostatic |
| Uggt11 | 0.00048507 | 0.32812468 | 0.825 | 0.794 | 1 | 5 | Uggt1 | homeostatic |
| Nudt7 | 0.00050792 | 0.48554232 | 0.31 | 0.257 | 1 | 5 | Nudt7 | homeostatic |
| Mcee1 | 0.00053407 | 0.33661579 | 0.268 | 0.222 | 1 | 5 | Mcee | homeostatic |
| Slc10a3 | 0.00053696 | 0.31472771 | 0.448 | 0.407 | 1 | 5 | Slc10a3 | homeostatic |
| Decr1 | 0.00057062 | 0.30801112 | 0.548 | 0.497 | 1 | 5 | Decr1 | homeostatic |
| Zfyve21 | 0.00058387 | 0.32235709 | 0.659 | 0.621 | 1 | 5 | Zfyve21 | homeostatic |
| Zfyve9 | 0.00058549 | 0.29182322 | 0.302 | 0.263 | 1 | 5 | Zfyve9 | homeostatic |
| Stxbp31 | 0.00059656 | 0.3682796 | 0.367 | 0.328 | 1 | 5 | Stxbp3 | homeostatic |
| Wdr7 | 0.00062645 | 0.30267275 | 0.467 | 0.416 | 1 | 5 | Wdr7 | homeostatic |
| Cds21 | 0.00062978 | 0.27763055 | 0.624 | 0.57 | 1 | 5 | Cds2 | homeostatic |
| Acvrl11 | 0.00064993 | 0.35630303 | 0.746 | 0.701 | 1 | 5 | Acvrl1 | homeostatic |
| Nt5c31 | 0.00065634 | 0.41188737 | 0.322 | 0.285 | 1 | 5 | Nt5c3 | homeostatic |
| Cndp2 | 0.00066145 | 0.38201291 | 0.702 | 0.638 | 1 | 5 | Cndp2 | homeostatic |
| Wdr6 | 0.00067131 | 0.41328402 | 0.435 | 0.369 | 1 | 5 | Wdr6 | homeostatic |
| Kif3b | 0.00070429 | 0.27336709 | 0.491 | 0.45 | 1 | 5 | Kif3b | homeostatic |
| Araf | 0.00071372 | 0.25754987 | 0.541 | 0.489 | 1 | 5 | Araf | homeostatic |
| Zxdc1 | 0.00073468 | 0.48966798 | 0.362 | 0.323 | 1 | 5 | Zxdc | homeostatic |
| Tepsin | 0.00075872 | 0.41819787 | 0.253 | 0.216 | 1 | 5 | Tepsin | homeostatic |
| Wdr41 | 0.0007651 | 0.47553228 | 0.459 | 0.396 | 1 | 5 | Wdr41 | homeostatic |
| Trim56 | 0.00077877 | 0.2696494 | 0.61 | 0.608 | 1 | 5 | Trim56 | homeostatic |
| Nit1 | 0.00079024 | 0.55971166 | 0.319 | 0.267 | 1 | 5 | Nit1 | homeostatic |
| Bcap29 | 0.00080467 | 0.35618487 | 0.272 | 0.23 | 1 | 5 | Bcap29 | homeostatic |
| Aldh3a21 | 0.00081014 | 0.42041356 | 0.484 | 0.425 | 1 | 5 | Aldh3a2 | homeostatic |
| Lgr4 | 0.00082481 | 0.35360999 | 0.388 | 0.336 | 1 | 5 | Lgr4 | homeostatic |
| Nedd4l1 | 0.00082502 | 0.38976083 | 0.322 | 0.283 | 1 | 5 | Nedd4l | homeostatic |
| Cyp2j6 | 0.00087202 | 0.52158049 | 0.287 | 0.231 | 1 | 5 | Cyp2j6 | homeostatic |
| Mpv17 | 0.0009015 | 0.28698189 | 0.638 | 0.618 | 1 | 5 | Mpv17 | homeostatic |
| Tmem1351 | 0.00091617 | 0.29737061 | 0.44 | 0.39 | 1 | 5 | Tmem135 | homeostatic |
| Gtf2h3 | 0.00091771 | 0.35108691 | 0.346 | 0.301 | 1 | 5 | Gtf2h3 | homeostatic |
| Tbc1d5 | 0.00093736 | 0.4040355 | 0.426 | 0.384 | 1 | 5 | Tbc1d5 | homeostatic |
| Vsig10 | 0.00097623 | 0.52996157 | 0.268 | 0.224 | 1 | 5 | Vsig10 | homeostatic |
| Cib1 | 0.00099497 | 0.29148995 | 0.514 | 0.463 | 1 | 5 | Cib1 | homeostatic |
| Cetn31 | 0.00100827 | 0.35221451 | 0.687 | 0.642 | 1 | 5 | Cetn3 | homeostatic |
| Trim24 | 0.00100892 | 0.25113426 | 0.411 | 0.37 | 1 | 5 | Trim24 | homeostatic |
| Slc41a3 | 0.00104033 | 0.38180642 | 0.344 | 0.315 | 1 | 5 | Slc41a3 | homeostatic |
| Glb1l | 0.00104056 | 0.39664754 | 0.383 | 0.328 | 1 | 5 | Glb1l | homeostatic |
| Nuak21 | 0.00109195 | 0.47787304 | 0.287 | 0.236 | 1 | 5 | Nuak2 | homeostatic |
| Auh1 | 0.00112857 | 0.29362917 | 0.664 | 0.599 | 1 | 5 | Auh | homeostatic |
| Pld2 | 0.00118495 | 0.31158899 | 0.387 | 0.346 | 1 | 5 | Pld2 | homeostatic |
| Trappc11 | 0.00127244 | 0.31405222 | 0.612 | 0.562 | 1 | 5 | Trappc1 | homeostatic |
| Slc25a20 | 0.00130063 | 0.35467455 | 0.576 | 0.549 | 1 | 5 | Slc25a20 | homeostatic |
| Mov101 | 0.00133944 | 0.33233816 | 0.354 | 0.304 | 1 | 5 | Mov10 | homeostatic |
| Mios | 0.00134879 | 0.32418803 | 0.426 | 0.384 | 1 | 5 | Mios | homeostatic |
| Rnf25 | 0.00141679 | 0.28185608 | 0.439 | 0.384 | 1 | 5 | Rnf25 | homeostatic |
| Rubcn1 | 0.00143623 | 0.278907 | 0.451 | 0.413 | 1 | 5 | Rubcn | homeostatic |
| Rusc11 | 0.00163837 | 0.306918 | 0.343 | 0.294 | 1 | 5 | Rusc1 | homeostatic |
| Htatip2 | 0.00167892 | 0.41817876 | 0.52 | 0.462 | 1 | 5 | Htatip2 | homeostatic |
| Ano10 | 0.00176048 | 0.30624836 | 0.695 | 0.659 | 1 | 5 | Ano10 | homeostatic |
| Asb32 | 0.00181334 | 0.42440884 | 0.329 | 0.292 | 1 | 5 | Asb3 | homeostatic |
| Manea1 | 0.00189941 | 0.41477743 | 0.355 | 0.31 | 1 | 5 | Manea | homeostatic |
| Kmt2c2 | 0.0019053 | 0.31678415 | 0.383 | 0.358 | 1 | 5 | Kmt2c | homeostatic |
| Ptdss1 | 0.0019097 | 0.29231869 | 0.758 | 0.739 | 1 | 5 | Ptdss1 | homeostatic |
| Gng10 | 0.00202366 | 0.26140454 | 0.761 | 0.749 | 1 | 5 | Gng10 | homeostatic |
| Anapc2 | 0.00205983 | 0.28136047 | 0.803 | 0.761 | 1 | 5 | Anapc2 | homeostatic |
| Dock71 | 0.00206217 | 0.25371042 | 0.782 | 0.758 | 1 | 5 | Dock7 | homeostatic |
| Cdk201 | 0.00208324 | 0.45890593 | 0.368 | 0.325 | 1 | 5 | Cdk20 | homeostatic |
| Tbk1 | 0.00209746 | 0.49746457 | 0.434 | 0.401 | 1 | 5 | Tbk1 | homeostatic |
| Spidr1 | 0.00211426 | 0.32310808 | 0.43 | 0.372 | 1 | 5 | Spidr | homeostatic |
| Wdsub11 | 0.00219892 | 0.39148044 | 0.303 | 0.249 | 1 | 5 | Wdsub1 | homeostatic |
| Ppox | 0.00220534 | 0.4161888 | 0.424 | 0.345 | 1 | 5 | Ppox | homeostatic |
| Synj1 | 0.0023781 | 0.44766534 | 0.332 | 0.287 | 1 | 5 | Synj1 | homeostatic |
| Cpt1c1 | 0.00240139 | 0.37621393 | 0.534 | 0.496 | 1 | 5 | Cpt1c | homeostatic |
| Scfd1 | 0.00244613 | 0.26471348 | 0.721 | 0.688 | 1 | 5 | Scfd1 | homeostatic |
| Isoc2b | 0.00248652 | 0.40152396 | 0.258 | 0.223 | 1 | 5 | Isoc2b | homeostatic |
| Tmem2371 | 0.00275821 | 0.57429821 | 0.277 | 0.227 | 1 | 5 | Tmem237 | homeostatic |
| Ctps21 | 0.00277222 | 0.29054903 | 0.666 | 0.628 | 1 | 5 | Ctps2 | homeostatic |
| St7 | 0.00279627 | 0.25474419 | 0.415 | 0.421 | 1 | 5 | St7 | homeostatic |
| Sspn | 0.00293709 | 0.50961932 | 0.383 | 0.322 | 1 | 5 | Sspn | homeostatic |
| Vps13b1 | 0.00294496 | 0.3180141 | 0.312 | 0.281 | 1 | 5 | Vps13b | homeostatic |
| Ngly11 | 0.00301022 | 0.31262216 | 0.51 | 0.459 | 1 | 5 | Ngly1 | homeostatic |
| Jpt11 | 0.00317397 | 0.27513028 | 0.811 | 0.743 | 1 | 5 | Jpt1 | homeostatic |
| Abcd41 | 0.00321707 | 0.43076347 | 0.256 | 0.213 | 1 | 5 | Abcd4 | homeostatic |
| Txndc15 | 0.00323631 | 0.2675546 | 0.74 | 0.718 | 1 | 5 | Txndc15 | homeostatic |
| C2cd51 | 0.00350103 | 0.64206299 | 0.291 | 0.23 | 1 | 5 | C2cd5 | homeostatic |
| Sdhaf4 | 0.00365019 | 0.3907919 | 0.325 | 0.29 | 1 | 5 | Sdhaf4 | homeostatic |
| Slc18b1 | 0.00370697 | 0.39867502 | 0.263 | 0.232 | 1 | 5 | Slc18b1 | homeostatic |
| Fkbp21 | 0.00383073 | 0.27020126 | 0.836 | 0.794 | 1 | 5 | Fkbp2 | homeostatic |
| Coq10a | 0.00390706 | 0.35679992 | 0.322 | 0.284 | 1 | 5 | Coq10a | homeostatic |
| Aifm2 | 0.00391856 | 0.34972865 | 0.36 | 0.318 | 1 | 5 | Aifm2 | homeostatic |
| Grin2d | 0.0040076 | 0.31300971 | 0.319 | 0.292 | 1 | 5 | Grin2d | homeostatic |
| Rtn21 | 0.00421232 | 0.25059655 | 0.269 | 0.252 | 1 | 5 | Rtn2 | homeostatic |
| Prorsd1 | 0.00429026 | 0.2906801 | 0.4 | 0.362 | 1 | 5 | Prorsd1 | homeostatic |
| Fitm2 | 0.00429844 | 0.29343924 | 0.279 | 0.238 | 1 | 5 | Fitm2 | homeostatic |

| | | | | | | | | |
|---|---|---|---|---|---|---|---|---|
| Rit1 | 0.00452712 | 0.32154097 | 0.661 | 0.596 | 1 | 5 | Rit1 | homeostatic |
| Magee1 | 0.00520118 | 0.37007755 | 0.437 | 0.402 | 1 | 5 | Magee1 | homeostatic |
| Ptk2b1 | 0.00568076 | 0.47228306 | 0.35 | 0.271 | 1 | 5 | Ptk2b | homeostatic |
| Stt3b1 | 0.00571917 | 0.28150689 | 0.897 | 0.878 | 1 | 5 | Stt3b | homeostatic |
| Atat1 | 0.00588538 | 0.3043208 | 0.277 | 0.248 | 1 | 5 | Atat1 | homeostatic |
| Parp41 | 0.00595745 | 0.25546782 | 0.424 | 0.372 | 1 | 5 | Parp4 | homeostatic |
| Wars1 | 0.00599912 | 0.27988308 | 0.523 | 0.492 | 1 | 5 | Wars | homeostatic |
| Gclm | 0.00603722 | 0.29163744 | 0.368 | 0.331 | 1 | 5 | Gclm | homeostatic |
| Hdac101 | 0.00612923 | 0.38883602 | 0.26 | 0.21 | 1 | 5 | Hdac10 | homeostatic |
| Lsm10 | 0.00631969 | 0.36812872 | 0.449 | 0.403 | 1 | 5 | Lsm10 | homeostatic |
| Ano8 | 0.00634731 | 0.30391687 | 0.355 | 0.314 | 1 | 5 | Ano8 | homeostatic |
| Smpd2 | 0.00636796 | 0.48329366 | 0.355 | 0.31 | 1 | 5 | Smpd2 | homeostatic |
| Hagh | 0.00655526 | 0.38663137 | 0.364 | 0.323 | 1 | 5 | Hagh | homeostatic |
| Dhx32 | 0.00667625 | 0.43641235 | 0.33 | 0.288 | 1 | 5 | Dhx32 | homeostatic |
| Magt11 | 0.00674841 | 0.25256299 | 0.799 | 0.753 | 1 | 5 | Magt1 | homeostatic |
| Acad11 | 0.0069625 | 0.46236713 | 0.289 | 0.255 | 1 | 5 | Acad11 | homeostatic |
| Cgrrf1 | 0.0070407 | 0.40262106 | 0.39 | 0.346 | 1 | 5 | Cgrrf1 | homeostatic |
| Nagpa | 0.00714092 | 0.32460384 | 0.387 | 0.358 | 1 | 5 | Nagpa | homeostatic |
| Tnfaip11 | 0.00734431 | 0.28122749 | 0.612 | 0.58 | 1 | 5 | Tnfaip1 | homeostatic |
| Cr1l | 0.00756859 | 0.44162447 | 0.376 | 0.328 | 1 | 5 | Cr1l | homeostatic |
| Slc22a51 | 0.00761006 | 0.51017841 | 0.251 | 0.219 | 1 | 5 | Slc22a5 | homeostatic |
| Tbl31 | 0.00796587 | 0.42385387 | 0.44 | 0.385 | 1 | 5 | Tbl3 | homeostatic |
| Dchs11 | 0.00798371 | 0.25681113 | 0.657 | 0.558 | 1 | 5 | Dchs1 | homeostatic |
| Tapbpl | 0.0080837 | 0.28961432 | 0.849 | 0.813 | 1 | 5 | Tapbp | homeostatic |
| Arl6 | 0.00813778 | 0.26915287 | 0.256 | 0.227 | 1 | 5 | Arl6 | homeostatic |
| Psmb81 | 0.00835651 | 0.40444145 | 0.59 | 0.521 | 1 | 5 | Psmb8 | homeostatic |
| Tacc2 | 0.00836718 | 0.29777085 | 0.397 | 0.372 | 1 | 5 | Tacc2 | homeostatic |
| Usp4 | 0.00890389 | 0.2529518 | 0.732 | 0.71 | 1 | 5 | Usp4 | homeostatic |
| Fbxo81 | 0.00936877 | 0.27662974 | 0.435 | 0.388 | 1 | 5 | Fbxo8 | homeostatic |
| Ghdc1 | 0.00974986 | 0.51243191 | 0.331 | 0.292 | 1 | 5 | Ghdc | homeostatic |
| Sp21 | 0.00985815 | 0.40478837 | 0.349 | 0.309 | 1 | 5 | Sp2 | homeostatic |
| Meox2 | 0 | 4.53895537 | 0.717 | 0.072 | 0 | 6 | Meox2 | intermediate |
| Sfrp41 | 0 | 3.19665923 | 0.788 | 0.184 | 0 | 6 | Sfrp4 | intermediate |
| Gas62 | 0 | 4.0121446 | 0.907 | 0.332 | 0 | 6 | Gas6 | intermediate |
| Tmeff22 | 0 | 2.51423237 | 0.894 | 0.321 | 0 | 6 | Tmeff2 | intermediate |
| Gdf10 | 0 | 5.71510117 | 0.614 | 0.051 | 0 | 6 | Gdf10 | intermediate |
| Cd341 | 0 | 1.60127598 | 0.894 | 0.356 | 0 | 6 | Cd34 | intermediate |
| Prelp2 | 0 | 1.69874393 | 0.949 | 0.421 | 0 | 6 | Prelp | intermediate |
| Cyp1b11 | 0 | 2.14113938 | 0.902 | 0.378 | 0 | 6 | Cyp1b1 | intermediate |
| Chrdl12 | 0 | 2.66741694 | 0.775 | 0.251 | 0 | 6 | Chrdl1 | intermediate |
| Ltbp42 | 0 | 3.77963522 | 0.806 | 0.288 | 0 | 6 | Ltbp4 | intermediate |
| Ccdc3 | 0 | 3.88559512 | 0.601 | 0.084 | 0 | 6 | Ccdc3 | intermediate |
| Clec3b1 | 0 | 1.64150703 | 0.646 | 0.131 | 0 | 6 | Clec3b | intermediate |
| Ccn52 | 0 | 1.63310583 | 0.939 | 0.433 | 0 | 6 | Ccn5 | intermediate |
| Mgp1 | 0 | 2.61951583 | 0.818 | 0.317 | 0 | 6 | Mgp | intermediate |
| Vcam13 | 0 | 2.17781099 | 0.826 | 0.326 | 0 | 6 | Vcam1 | intermediate |
| Tnfsf13b1 | 0 | 3.80595253 | 0.626 | 0.127 | 0 | 6 | Tnfsf13b | intermediate |
| Scn7a2 | 0 | 1.82897427 | 0.773 | 0.278 | 0 | 6 | Scn7a | intermediate |
| Tnxb1 | 0 | 1.02506569 | 0.773 | 0.279 | 0 | 6 | Tnxb | intermediate |
| Ly6c11 | 0 | 1.45873026 | 0.73 | 0.238 | 0 | 6 | Ly6c1 | intermediate |
| Trabd2b1 | 0 | 3.06298316 | 0.619 | 0.13 | 0 | 6 | Trabd2b | intermediate |
| Fhl11 | 0 | 2.70453693 | 0.79 | 0.303 | 0 | 6 | Fhl1 | intermediate |
| Bmper2 | 0 | 2.21119533 | 0.737 | 0.25 | 0 | 6 | Bmper | intermediate |
| Ces2g2 | 0 | 1.96945437 | 0.699 | 0.228 | 0 | 6 | Ces2g | intermediate |
| C4a2 | 0 | 1.55600322 | 0.836 | 0.368 | 0 | 6 | C4a | intermediate |
| Slc1a31 | 0 | 2.79260962 | 0.614 | 0.147 | 0 | 6 | Slc1a3 | intermediate |
| Ebf21 | 0 | 2.40849023 | 0.631 | 0.167 | 0 | 6 | Ebf2 | intermediate |
| Cilp1 | 0 | 3.04458989 | 0.707 | 0.245 | 0 | 6 | Cilp | intermediate |
| Spon21 | 0 | 2.05071519 | 0.689 | 0.231 | 0 | 6 | Spon2 | intermediate |
| Dapk12 | 0 | 1.99034898 | 0.705 | 0.249 | 0 | 6 | Dapk1 | intermediate |
| Tppp31 | 0 | 2.4212063 | 0.674 | 0.219 | 0 | 6 | Tppp3 | intermediate |
| Ly6a1 | 0 | 1.7443388 | 0.955 | 0.5 | 0 | 6 | Ly6a | intermediate |
| Il333 | 0 | 1.96407621 | 0.836 | 0.381 | 0 | 6 | Il33 | intermediate |
| Pltp2 | 0 | 2.40163604 | 0.747 | 0.297 | 0 | 6 | Pltp | intermediate |
| Steap42 | 0 | 1.84132654 | 0.745 | 0.295 | 0 | 6 | Steap4 | intermediate |
| Nfib3 | 0 | 2.21078673 | 0.864 | 0.417 | 0 | 6 | Nfib | intermediate |
| Sertad42 | 0 | 2.38043687 | 0.697 | 0.253 | 0 | 6 | Sertad4 | intermediate |
| Ccl192 | 0 | 3.57527514 | 0.606 | 0.162 | 0 | 6 | Ccl19 | intermediate |
| Fzd41 | 0 | 2.51337516 | 0.692 | 0.249 | 0 | 6 | Fzd4 | intermediate |
| Adgrd12 | 0 | 1.45000491 | 0.919 | 0.477 | 0 | 6 | Adgrd1 | intermediate |
| Itgbl12 | 0 | 3.07189932 | 0.659 | 0.224 | 0 | 6 | Itgbl1 | intermediate |
| Pcolce21 | 0 | 1.57807733 | 0.763 | 0.334 | 0 | 6 | Pcolce2 | intermediate |
| Zfp5212 | 0 | 2.07184798 | 0.788 | 0.36 | 0 | 6 | Zfp521 | intermediate |
| C4b2 | 0 | 1.59732105 | 0.955 | 0.53 | 0 | 6 | C4b | intermediate |
| Gabra31 | 0 | 2.36580982 | 0.568 | 0.145 | 0 | 6 | Gabra3 | intermediate |
| Ube2l61 | 0 | 1.82015726 | 0.808 | 0.386 | 0 | 6 | Ube2l6 | intermediate |
| Naalad22 | 0 | 2.05754654 | 0.659 | 0.244 | 0 | 6 | Naalad2 | intermediate |
| Entpd21 | 0 | 2.44176038 | 0.702 | 0.288 | 0 | 6 | Entpd2 | intermediate |
| Rtp41 | 0 | 2.1316484 | 0.586 | 0.173 | 0 | 6 | Rtp4 | intermediate |
| Osr11 | 0 | 1.77733858 | 0.669 | 0.259 | 0 | 6 | Osr1 | intermediate |
| Gstt12 | 0 | 2.1837352 | 0.591 | 0.182 | 0 | 6 | Gstt1 | intermediate |
| C32 | 0 | 1.44951146 | 0.927 | 0.523 | 0 | 6 | C3 | intermediate |
| Slfn52 | 0 | 1.55792388 | 0.929 | 0.532 | 0 | 6 | Slfn5 | intermediate |
| C22 | 0 | 1.98257994 | 0.859 | 0.468 | 0 | 6 | C2 | intermediate |
| Serpina3n2 | 0 | 1.00337461 | 0.965 | 0.575 | 0 | 6 | Serpina3n | intermediate |
| Podn2 | 0 | 2.14034447 | 0.937 | 0.548 | 0 | 6 | Podn | intermediate |
| Col14a12 | 0 | 1.4905877 | 0.922 | 0.541 | 0 | 6 | Col14a1 | intermediate |
| Pdgfrl1 | 0 | 2.17173467 | 0.737 | 0.357 | 0 | 6 | Pdgfrl | intermediate |
| Fgf72 | 0 | 1.23850064 | 0.854 | 0.477 | 0 | 6 | Fgf7 | intermediate |
| Ifi27l2a1 | 0 | 1.87167456 | 0.525 | 0.161 | 0 | 6 | Ifi27l2a | intermediate |
| Vit | 0 | 3.90713804 | 0.389 | 0.026 | 0 | 6 | Vit | intermediate |
| Bst22 | 0 | 1.80517627 | 0.904 | 0.545 | 0 | 6 | Bst2 | intermediate |
| Gas71 | 0 | 1.04349682 | 0.854 | 0.495 | 0 | 6 | Gas7 | intermediate |
| Sulf22 | 0 | 1.86522746 | 0.755 | 0.399 | 0 | 6 | Sulf2 | intermediate |
| Abcc92 | 0 | 2.04025899 | 0.566 | 0.215 | 0 | 6 | Abcc9 | intermediate |
| Clec11a1 | 0 | 2.228795 | 0.899 | 0.548 | 0 | 6 | Clec11a | intermediate |
| Cfi | 0 | 4.1733861 | 0.391 | 0.041 | 0 | 6 | Cfi | intermediate |
| Gbp22 | 0 | 1.77611393 | 0.601 | 0.255 | 0 | 6 | Gbp2 | intermediate |
| Ccn32 | 0 | 2.42917272 | 0.593 | 0.247 | 0 | 6 | Ccn3 | intermediate |
| Apod2 | 0 | 2.07002981 | 0.96 | 0.614 | 0 | 6 | Apod | intermediate |
| Isg151 | 0 | 2.37492016 | 0.619 | 0.275 | 0 | 6 | Isg15 | intermediate |
| Ntrk22 | 0 | 1.96972059 | 0.624 | 0.281 | 0 | 6 | Ntrk2 | intermediate |
| Scara51 | 0 | 1.86356211 | 0.434 | 0.092 | 0 | 6 | Scara5 | intermediate |
| Irf71 | 0 | 1.84249416 | 0.553 | 0.213 | 0 | 6 | Irf7 | intermediate |
| Cfb2 | 0 | 1.70383762 | 0.97 | 0.631 | 0 | 6 | Cfb | intermediate |
| F31 | 0 | 2.53379343 | 0.765 | 0.427 | 0 | 6 | F3 | intermediate |
| Ntn11 | 0 | 1.86633935 | 0.621 | 0.29 | 0 | 6 | Ntn1 | intermediate |
| Dpt2 | 0 | 1.23333025 | 0.972 | 0.643 | 0 | 6 | Dpt | intermediate |
| Eln2 | 0 | 2.45315534 | 0.848 | 0.532 | 0 | 6 | Eln | intermediate |
| Comp | 0 | 7.2845206 | 0.313 | 0.005 | 0 | 6 | Comp | intermediate |
| Thbs32 | 0 | 1.63529215 | 0.97 | 0.672 | 0 | 6 | Thbs3 | intermediate |
| Add32 | 0 | 1.51859515 | 0.846 | 0.549 | 0 | 6 | Add3 | intermediate |
| Dock9 | 0 | 2.85699256 | 0.371 | 0.075 | 0 | 6 | Dock9 | intermediate |
| C1qtnf31 | 0 | 2.14265972 | 0.604 | 0.313 | 0 | 6 | C1qtnf3 | intermediate |
| Ccn22 | 0 | 1.34344515 | 0.73 | 0.442 | 0 | 6 | Ccn2 | intermediate |
| Adamts52 | 0 | 0.96981595 | 0.811 | 0.524 | 0 | 6 | Adamts5 | intermediate |
| S100a42 | 0 | 1.12633449 | 0.808 | 0.524 | 0 | 6 | S100a4 | intermediate |
| Igfbp61 | 0 | 1.57819325 | 0.949 | 0.665 | 0 | 6 | Igfbp6 | intermediate |
| Srgn2 | 0 | 1.94030695 | 0.609 | 0.327 | 0 | 6 | Srgn | intermediate |
| Fmod | 0 | 3.51863423 | 0.396 | 0.116 | 0 | 6 | Fmod | intermediate |
| Prss231 | 0 | 0.98658346 | 0.914 | 0.635 | 0 | 6 | Prss23 | intermediate |
| Epha31 | 0 | 1.41565031 | 0.74 | 0.466 | 0 | 6 | Epha3 | intermediate |
| Rnf2132 | 0 | 1.42716527 | 0.722 | 0.449 | 0 | 6 | Rnf213 | intermediate |
| Ifit3 | 0 | 3.95065933 | 0.318 | 0.049 | 0 | 6 | Ifit3 | intermediate |
| Ptch21 | 0 | 2.35330058 | 0.49 | 0.225 | 0 | 6 | Ptch2 | intermediate |
| Igf11 | 0 | 1.43621847 | 0.876 | 0.617 | 0 | 6 | Igf1 | intermediate |
| Ndnf | 0 | 2.66683411 | 0.364 | 0.11 | 0 | 6 | Ndnf | intermediate |
| Fxyd62 | 0 | 1.23124601 | 0.884 | 0.632 | 0 | 6 | Fxyd6 | intermediate |
| Aqp1 | 0 | 2.2872985 | 0.379 | 0.131 | 0 | 6 | Aqp1 | intermediate |
| Alpl | 0 | 4.9289982 | 0.268 | 0.021 | 0 | 6 | Alpl | intermediate |
| Apoe | 0 | 2.50899791 | 0.828 | 0.582 | 0 | 6 | Apoe | intermediate |
| Arsi2 | 0 | 1.00967937 | 0.884 | 0.643 | 0 | 6 | Arsi | intermediate |
| Isyna11 | 0 | 1.61088344 | 0.705 | 0.464 | 0 | 6 | Isyna1 | intermediate |
| Lbp1 | 0 | 1.59011402 | 0.922 | 0.682 | 0 | 6 | Lbp | intermediate |
| Dpysl31 | 0 | 1.71890866 | 0.977 | 0.745 | 0 | 6 | Dpysl3 | intermediate |
| Ifit11 | 0 | 3.41230999 | 0.285 | 0.056 | 0 | 6 | Ifit1 | intermediate |
| Scn1b2 | 0 | 1.18879453 | 0.937 | 0.711 | 0 | 6 | Scn1b | intermediate |
| Lrrc322 | 0 | 0.97320086 | 0.914 | 0.694 | 0 | 6 | Lrrc32 | intermediate |
| Csrp23 | 0 | 1.32005738 | 0.891 | 0.672 | 0 | 6 | Csrp2 | intermediate |
| Col8a2 | 0 | 3.3310227 | 0.268 | 0.05 | 0 | 6 | Col8a2 | intermediate |
| Id31 | 0 | 0.76391047 | 0.861 | 0.653 | 0 | 6 | Id3 | intermediate |
| Olfml2b1 | 0 | 1.23153237 | 0.894 | 0.687 | 0 | 6 | Olfml2b | intermediate |
| Pam2 | 0 | 1.08192149 | 0.917 | 0.716 | 0 | 6 | Pam | intermediate |
| Ifi2052 | 0 | 0.82651858 | 0.828 | 0.628 | 0 | 6 | Ifi205 | intermediate |
| Heg11 | 0 | 1.12529463 | 0.975 | 0.775 | 0 | 6 | Heg1 | intermediate |
| Mfap42 | 0 | 1.08326097 | 0.801 | 0.605 | 0 | 6 | Mfap4 | intermediate |
| Selenop1 | 0 | 2.13123817 | 0.924 | 0.728 | 0 | 6 | Selenop | intermediate |
| Emb2 | 0 | 1.58496936 | 0.907 | 0.718 | 0 | 6 | Emb | intermediate |
| Tspan111 | 0 | 1.22477149 | 0.914 | 0.729 | 0 | 6 | Tspan11 | intermediate |
| Plac82 | 0 | 1.18466492 | 0.907 | 0.736 | 0 | 6 | Plac8 | intermediate |
| Cygb2 | 0 | 1.89733423 | 0.985 | 0.827 | 0 | 6 | Cygb | intermediate |
| Cxcl122 | 0 | 1.7251178 | 0.949 | 0.794 | 0 | 6 | Cxcl12 | intermediate |
| Cavin31 | 0 | 1.10498211 | 0.939 | 0.789 | 0 | 6 | Cavin3 | intermediate |
| Srpx1 | 0 | 1.46422757 | 0.914 | 0.765 | 0 | 6 | Srpx | intermediate |
| Igfbp51 | 0 | 1.57397933 | 0.348 | 0.201 | 0 | 6 | Igfbp5 | intermediate |
| Cxcl141 | 0 | 1.56130772 | 0.891 | 0.75 | 0 | 6 | Cxcl14 | intermediate |
| Sfrp11 | 0 | 1.71855109 | 0.843 | 0.705 | 0 | 6 | Sfrp1 | intermediate |
| Anpep2 | 0 | 1.07596458 | 0.904 | 0.767 | 0 | 6 | Anpep | intermediate |
| Aspn2 | 0 | 1.53634068 | 0.563 | 0.427 | 0 | 6 | Aspn | intermediate |
| Rbp12 | 0 | 1.02467787 | 0.952 | 0.817 | 0 | 6 | Rbp1 | intermediate |
| Mfap53 | 0 | 1.02201763 | 0.977 | 0.843 | 0 | 6 | Mfap5 | intermediate |
| Ltbp31 | 0 | 1.04356776 | 0.97 | 0.838 | 0 | 6 | Ltbp3 | intermediate |
| Ace2 | 0 | 0.94176461 | 0.856 | 0.735 | 0 | 6 | Ace | intermediate |
| Prnp2 | 0 | 1.00441783 | 0.952 | 0.837 | 0 | 6 | Prnp | intermediate |
| Plpp31 | 0 | 0.73200565 | 0.952 | 0.842 | 0 | 6 | Plpp3 | intermediate |
| S100a101 | 0 | 1.35842034 | 0.955 | 0.859 | 0 | 6 | S100a10 | intermediate |
| Sned11 | 0 | 1.3346631 | 0.856 | 0.761 | 0 | 6 | Sned1 | intermediate |
| Lrp61 | 0 | 1.04636541 | 0.934 | 0.843 | 0 | 6 | Lrp6 | intermediate |
| Igfbp41 | 0 | 2.18245957 | 0.96 | 0.87 | 0 | 6 | Igfbp4 | intermediate |
| Cavin11 | 0 | 0.92369049 | 0.985 | 0.898 | 0 | 6 | Cavin1 | intermediate |
| B2m2 | 0 | 0.90634476 | 0.929 | 0.844 | 0 | 6 | B2m | intermediate |
| C1ra2 | 0 | 0.80212795 | 0.962 | 0.88 | 0 | 6 | C1ra | intermediate |
| Lpl2 | 0 | 0.98889075 | 0.664 | 0.584 | 0 | 6 | Lpl | intermediate |
| C1s12 | 0 | 1.01498437 | 0.997 | 0.923 | 0 | 6 | C1s1 | intermediate |
| Il6st2 | 0 | 1.63883013 | 0.965 | 0.893 | 0 | 6 | Il6st | intermediate |
| Sod33 | 0 | 0.71584025 | 0.995 | 0.929 | 0 | 6 | Sod3 | intermediate |
| Lgals3bp2 | 0 | 0.97869227 | 0.977 | 0.912 | 0 | 6 | Lgals3bp | intermediate |
| Nbl12 | 0 | 1.0997156 | 0.98 | 0.923 | 0 | 6 | Nbl1 | intermediate |
| Sfrp22 | 0 | 0.66708383 | 0.785 | 0.841 | 0 | 6 | Sfrp2 | intermediate |
| Serpinb6a1 | 0 | 0.66772581 | 0.967 | 0.913 | 0 | 6 | Serpinb6a | intermediate |
| Nfix2 | 0 | 1.12487687 | 0.977 | 0.925 | 0 | 6 | Nfix | intermediate |
| Col4a11 | 0 | 0.60179088 | 0.934 | 0.893 | 0 | 6 | Col4a1 | intermediate |
| Crip12 | 0 | 1.08331222 | 0.96 | 0.919 | 0 | 6 | Crip1 | intermediate |
| Ahnak1 | 0 | 1.04346669 | 0.997 | 0.957 | 0 | 6 | Ahnak | intermediate |
| Cd2481 | 0 | 0.62444069 | 0.967 | 0.927 | 0 | 6 | Cd248 | intermediate |
| Col4a2 | 0 | 0.56106372 | 0.902 | 0.867 | 0 | 6 | Col4a2 | intermediate |
| Cyb5r31 | 0 | 0.64995599 | 0.977 | 0.943 | 0 | 6 | Cyb5r3 | intermediate |
| Serping12 | 0 | 1.20313289 | 0.995 | 0.963 | 0 | 6 | Serping1 | intermediate |
| Tgfbr21 | 0 | 0.5344553 | 0.995 | 0.963 | 0 | 6 | Tgfbr2 | intermediate |
| Fkbp92 | 0 | 0.5540623 | 0.995 | 0.964 | 0 | 6 | Fkbp9 | intermediate |
| Gsn1 | 0 | 1.29310506 | 0.997 | 0.967 | 0 | 6 | Gsn | intermediate |
| Grn1 | 0 | 0.82234678 | 0.992 | 0.962 | 0 | 6 | Grn | intermediate |
| Ly6e1 | 0 | 0.80213822 | 0.977 | 0.947 | 0 | 6 | Ly6e | intermediate |
| Sptbn11 | 0 | 0.73490019 | 0.985 | 0.956 | 0 | 6 | Sptbn1 | intermediate |
| Rnase42 | 0 | 0.54441835 | 0.972 | 0.946 | 0 | 6 | Rnase4 | intermediate |
| Pcolce3 | 0 | 0.72231813 | 1 | 0.983 | 0 | 6 | Pcolce | intermediate |
| Serpinf1 | 0 | 0.46390163 | 0.99 | 0.973 | 0 | 6 | Serpinf1 | intermediate |
| Timp31 | 0 | 1.29683699 | 0.689 | 0.672 | 0 | 6 | Timp3 | intermediate |
| Gpx32 | 0 | 0.40154104 | 0.965 | 0.951 | 0 | 6 | Gpx3 | intermediate |
| Fbn11 | 0 | 0.55015337 | 0.992 | 0.979 | 0 | 6 | Fbn1 | intermediate |
| Lmna1 | 0 | 0.7403261 | 0.992 | 0.979 | 0 | 6 | Lmna | intermediate |
| Fth12 | 0 | 1.11539105 | 0.997 | 0.984 | 0 | 6 | Fth1 | intermediate |
| Cst3 | 0 | 1.01281882 | 0.987 | 0.975 | 0 | 6 | Cst3 | intermediate |
| Anxa21 | 0 | 1.00521492 | 0.98 | 0.97 | 0 | 6 | Anxa2 | intermediate |
| Bgn1 | 0 | 0.37519294 | 1 | 0.99 | 0 | 6 | Bgn | intermediate |
| Tmsb4x1 | 0 | 0.69472287 | 0.992 | 0.986 | 0 | 6 | Tmsb4x | intermediate |
| Igfbp72 | 0 | 0.91022778 | 0.997 | 0.992 | 0 | 6 | Igfbp7 | intermediate |
| Col3a12 | 0 | 0.38259469 | 1 | 0.996 | 0 | 6 | Col3a1 | intermediate |
| Fstl11 | 0 | 0.56007239 | 0.995 | 0.991 | 0 | 6 | Fstl1 | intermediate |
| Mmp22 | 0 | 0.38815779 | 0.977 | 0.975 | 0 | 6 | Mmp2 | intermediate |
| Vim2 | 0 | 0.46046205 | 0.997 | 0.998 | 0 | 6 | Vim | intermediate |
| Thbd1 | 3.21E-293 | 1.26452826 | 0.566 | 0.397 | 6.21E-293 | 6 | Thbd | intermediate |
| Prg42 | 1.98E-293 | 0.60837722 | 0.707 | 0.302 | 3.82E-293 | 6 | Prg4 | intermediate |
| Chl12 | 1.25E-293 | 1.1036352 | 0.641 | 0.386 | 2.41E-293 | 6 | Chl1 | intermediate |
| Stat12 | 8.08E-293 | 1.30704038 | 0.765 | 0.548 | 1.56E-293 | 6 | Stat1 | intermediate |
| Iigp12 | 1.33E-293 | 1.87683241 | 0.472 | 0.16 | 2.57E-293 | 6 | Iigp1 | intermediate |
| Gli1 | 9.46E-293 | 3.0504873 | 0.369 | 0.065 | 1.83E-293 | 6 | Gli1 | intermediate |
| Efemp11 | 1.01E-293 | 0.90595754 | 0.71 | 0.324 | 1.95E-291 | 6 | Efemp1 | intermediate |
| Tspo2 | 1.24E-293 | 0.86077295 | 0.952 | 0.863 | 2.39E-291 | 6 | Tspo | intermediate |
| Fmo11 | 6.85E-293 | 2.42007243 | 0.525 | 0.114 | 1.32E-290 | 6 | Fmo1 | intermediate |
| Lum2 | 3.73E-293 | 0.41096942 | 0.949 | 0.844 | 7.20E-289 | 6 | Lum | intermediate |
| Jam22 | 6.27E-292 | 1.9451759 | 0.53 | 0.191 | 1.21E-287 | 6 | Jam2 | intermediate |
| Epdr11 | 9.15E-292 | 1.6342914 | 0.76 | 0.408 | 1.77E-287 | 6 | Epdr1 | intermediate |
| Adamtsl3 | 4.67E-290 | 3.18391395 | 0.409 | 0.06 | 9.03E-286 | 6 | Adamtsl3 | intermediate |
| Rassf21 | 7.02E-290 | 1.97744692 | 0.659 | 0.22 | 1.36E-285 | 6 | Rassf2 | intermediate |
| Ahr2 | 1.59E-288 | 1.59373356 | 0.659 | 0.325 | 3.08E-284 | 6 | Ahr | intermediate |
| Ccdc801 | 8.21E-288 | 0.40617851 | 0.972 | 0.956 | 1.59E-283 | 6 | Ccdc80 | intermediate |
| Gfra21 | 2.57E-287 | 1.97810033 | 0.48 | 0.151 | 4.97E-283 | 6 | Gfra2 | intermediate |
| Plxna41 | 6.24E-287 | 2.28809393 | 0.437 | 0.148 | 1.21E-282 | 6 | Plxna4 | intermediate |
| Anxa82 | 9.47E-286 | 2.1903796 | 0.472 | 0.132 | 1.83E-281 | 6 | Anxa8 | intermediate |
| Serpinb1a3 | 1.27E-284 | 1.62276279 | 0.654 | 0.237 | 2.46E-280 | 6 | Serpinb1a | intermediate |
| Tspan182 | 4.44E-278 | 1.95363114 | 0.54 | 0.196 | 8.58E-274 | 6 | Tspan18 | intermediate |
| Ildr21 | 1.15E-271 | 2.33099421 | 0.407 | 0.118 | 2.22E-267 | 6 | Ildr2 | intermediate |
| Capn22 | 2.73E-271 | 0.74748274 | 0.962 | 0.861 | 5.27E-267 | 6 | Capn2 | intermediate |
| Ogn1 | 3.93E-271 | 1.03946725 | 0.465 | 0.261 | 7.60E-267 | 6 | Ogn | intermediate |
| Nfasc3 | 2.55E-268 | 1.74213531 | 0.753 | 0.373 | 4.92E-264 | 6 | Nfasc | intermediate |
| Cltb1 | 3.31E-266 | 0.89394671 | 0.917 | 0.798 | 6.41E-262 | 6 | Cltb | intermediate |
| Gxylt21 | 2.32E-262 | 1.11435172 | 0.869 | 0.678 | 4.49E-258 | 6 | Gxylt2 | intermediate |
| Usp181 | 7.22E-262 | 2.40205966 | 0.323 | 0.094 | 1.40E-257 | 6 | Usp18 | intermediate |
| Irgm21 | 3.60E-261 | 1.66500828 | 0.609 | 0.284 | 6.96E-257 | 6 | Irgm2 | intermediate |
| Adcyap1r11 | 4.50E-258 | 1.88118308 | 0.422 | 0.171 | 8.70E-254 | 6 | Adcyap1r1 | intermediate |
| Oasl21 | 1.54E-255 | 1.45033238 | 0.657 | 0.345 | 2.98E-251 | 6 | Oasl2 | intermediate |
| Nr2f21 | 4.30E-255 | 1.46194026 | 0.669 | 0.392 | 8.31E-251 | 6 | Nr2f2 | intermediate |
| Svep12 | 9.45E-254 | 0.5421783 | 0.937 | 0.85 | 1.83E-249 | 6 | Svep1 | intermediate |
| Emp31 | 2.23E-251 | 0.75239903 | 0.942 | 0.847 | 4.32E-247 | 6 | Emp3 | intermediate |
| Plxdc21 | 5.48E-244 | 1.20101984 | 0.78 | 0.504 | 1.06E-239 | 6 | Plxdc2 | intermediate |
| Slc7a21 | 2.48E-241 | 2.21023112 | 0.505 | 0.181 | 4.80E-237 | 6 | Slc7a2 | intermediate |
| Ndn2 | 4.20E-238 | 1.04862095 | 0.889 | 0.655 | 8.12E-234 | 6 | Ndn | intermediate |
| Pid11 | 2.77E-236 | 1.94697127 | 0.444 | 0.199 | 5.35E-232 | 6 | Pid1 | intermediate |
| Slc43a32 | 6.01E-236 | 1.23269974 | 0.841 | 0.536 | 1.16E-231 | 6 | Slc43a3 | intermediate |
| Lamb23 | 2.16E-235 | 0.87171461 | 0.927 | 0.803 | 4.19E-231 | 6 | Lamb2 | intermediate |
| Acvr2a2 | 1.44E-232 | 1.70669113 | 0.715 | 0.357 | 2.79E-228 | 6 | Acvr2a | intermediate |
| Tmem176b1 | 4.01E-232 | 0.88301843 | 0.864 | 0.717 | 7.76E-228 | 6 | Tmem176b | intermediate |
| Plac9a1 | 4.38E-232 | 1.85178373 | 0.581 | 0.161 | 8.47E-228 | 6 | Plac9a | intermediate |
| Snx181 | 2.03E-231 | 1.0060719 | 0.856 | 0.741 | 3.92E-227 | 6 | Snx18 | intermediate |
| Irgm11 | 1.06E-226 | 1.52870996 | 0.591 | 0.351 | 2.06E-222 | 6 | Irgm1 | intermediate |
| Lgals32 | 1.90E-226 | 0.53745906 | 0.753 | 0.508 | 3.68E-222 | 6 | Lgals3 | intermediate |
| Gpx11 | 4.70E-225 | 0.55470567 | 0.962 | 0.914 | 9.08E-221 | 6 | Gpx1 | intermediate |
| Anxa51 | 2.74E-221 | 0.3435426 | 0.992 | 0.975 | 5.31E-217 | 6 | Anxa5 | intermediate |
| Phf11d1 | 7.56E-221 | 1.8172764 | 0.525 | 0.202 | 1.46E-216 | 6 | Phf11d | intermediate |
| Peg31 | 5.41E-219 | 1.52006038 | 0.46 | 0.326 | 1.05E-214 | 6 | Peg3 | intermediate |
| Kcnma11 | 2.29E-217 | 2.06859483 | 0.475 | 0.172 | 4.43E-213 | 6 | Kcnma1 | intermediate |
| Id4 | 5.78E-217 | 2.48121262 | 0.364 | 0.102 | 1.12E-212 | 6 | Id4 | intermediate |
| Samhd11 | 6.08E-214 | 1.17392953 | 0.811 | 0.546 | 1.17E-209 | 6 | Samhd1 | intermediate |
| Gbp31 | 4.59E-213 | 1.92172225 | 0.465 | 0.133 | 8.87E-209 | 6 | Gbp3 | intermediate |
| Xaf11 | 1.23E-211 | 1.72807516 | 0.609 | 0.232 | 2.38E-207 | 6 | Xaf1 | intermediate |
| Agmo | 6.65E-208 | 3.05260527 | 0.394 | 0.058 | 1.28E-203 | 6 | Agmo | intermediate |
| Tcea32 | 1.87E-207 | 2.52330218 | 0.497 | 0.121 | 3.62E-203 | 6 | Tcea3 | intermediate |
| Nod12 | 2.66E-207 | 1.26237525 | 0.848 | 0.544 | 5.14E-203 | 6 | Nod1 | intermediate |
| Bace21 | 1.66E-206 | 2.0177577 | 0.462 | 0.189 | 3.20E-202 | 6 | Bace2 | intermediate |
| Ifi271 | 2.19E-205 | 0.79838585 | 0.899 | 0.805 | 4.23E-201 | 6 | Ifi27 | intermediate |
| Ifitm31 | 6.62E-203 | 0.38458526 | 0.939 | 0.925 | 1.28E-198 | 6 | Ifitm3 | intermediate |
| Akap121 | 9.36E-203 | 1.59289437 | 0.641 | 0.409 | 1.81E-198 | 6 | Akap12 | intermediate |
| Glis21 | 5.17E-202 | 0.97698348 | 0.896 | 0.757 | 9.99E-198 | 6 | Glis2 | intermediate |
| Grb101 | 4.55E-201 | 0.86915119 | 0.891 | 0.74 | 8.80E-197 | 6 | Grb10 | intermediate |
| Nupr11 | 1.60E-200 | 0.72016847 | 0.904 | 0.808 | 3.09E-196 | 6 | Nupr1 | intermediate |
| Crip21 | 3.25E-197 | 1.69241532 | 0.525 | 0.195 | 6.29E-193 | 6 | Crip2 | intermediate |
| Gpnmb1 | 1.63E-195 | 1.63704891 | 0.447 | 0.213 | 3.15E-191 | 6 | Gpnmb | intermediate |
| Lgi22 | 2.75E-195 | 0.60033448 | 0.816 | 0.592 | 5.31E-191 | 6 | Lgi2 | intermediate |
| Rab3d2 | 4.91E-195 | 1.50384859 | 0.725 | 0.349 | 9.50E-191 | 6 | Rab3d | intermediate |
| Cpne82 | 5.00E-195 | 1.32509505 | 0.783 | 0.426 | 9.66E-191 | 6 | Cpne8 | intermediate |
| Man1a2 | 8.80E-195 | 0.71887802 | 0.955 | 0.852 | 1.70E-190 | 6 | Man1a | intermediate |
| Arap11 | 1.26E-190 | 0.81713886 | 0.881 | 0.813 | 2.44E-186 | 6 | Arap1 | intermediate |
| Tgfb31 | 1.57E-190 | 0.70931798 | 0.705 | 0.699 | 3.03E-186 | 6 | Tgfb3 | intermediate |
| Plod23 | 9.78E-190 | 0.72795874 | 0.816 | 0.645 | 1.89E-185 | 6 | Plod2 | intermediate |
| Oas21 | 3.11E-189 | 2.29375122 | 0.424 | 0.102 | 6.01E-185 | 6 | Oas2 | intermediate |
| Col28a12 | 1.24E-187 | 1.69081527 | 0.399 | 0.144 | 2.40E-183 | 6 | Col28a1 | intermediate |
| Ngef | 1.52E-187 | 3.2293817 | 0.273 | 0.034 | 2.94E-183 | 6 | Ngef | intermediate |
| Trim473 | 2.78E-187 | 1.87645941 | 0.573 | 0.234 | 5.37E-183 | 6 | Trim47 | intermediate |
| Rcn32 | 3.17E-187 | 0.27258617 | 0.995 | 0.98 | 6.12E-183 | 6 | Rcn3 | intermediate |
| Ablim11 | 6.79E-187 | 1.75166632 | 0.553 | 0.211 | 1.31E-182 | 6 | Ablim1 | intermediate |
| Steap31 | 2.39E-186 | 1.20477126 | 0.798 | 0.503 | 4.63E-182 | 6 | Steap3 | intermediate |
| Fgf182 | 9.04E-185 | 1.95226987 | 0.419 | 0.157 | 1.75E-180 | 6 | Fgf18 | intermediate |
| Sema3d1 | 1.47E-184 | 2.01850793 | 0.268 | 0.099 | 2.84E-180 | 6 | Sema3d | intermediate |
| Fxyd12 | 3.84E-184 | 0.95914785 | 0.891 | 0.624 | 7.43E-180 | 6 | Fxyd1 | intermediate |
| Tmem176a1 | 9.00E-183 | 0.78952488 | 0.795 | 0.676 | 1.74E-178 | 6 | Tmem176a | intermediate |
| Olfml12 | 2.39E-182 | 1.52335217 | 0.616 | 0.244 | 4.63E-178 | 6 | Olfml1 | intermediate |
| Atp2b41 | 7.56E-182 | 1.70197329 | 0.518 | 0.197 | 1.46E-177 | 6 | Atp2b4 | intermediate |
| Kcne41 | 8.70E-182 | 1.81862038 | 0.356 | 0.143 | 1.68E-177 | 6 | Kcne4 | intermediate |
| Cp2 | 2.56E-180 | 1.42565628 | 0.548 | 0.4 | 4.94E-176 | 6 | Cp | intermediate |
| Comt2 | 6.77E-180 | 0.84576622 | 0.927 | 0.735 | 1.31E-175 | 6 | Comt | intermediate |
| Tnfsf8 | 1.81E-179 | 2.46160633 | 0.275 | 0.064 | 3.50E-175 | 6 | Tnfsf8 | intermediate |
| Kctd122 | 1.93E-179 | 1.27278945 | 0.684 | 0.355 | 3.74E-175 | 6 | Kctd12 | intermediate |
| Gstm21 | 2.66E-178 | 0.65939037 | 0.924 | 0.852 | 5.15E-174 | 6 | Gstm2 | intermediate |
| Mgst12 | 1.10E-176 | 0.65833314 | 0.924 | 0.818 | 2.13E-172 | 6 | Mgst1 | intermediate |
| Epb41l21 | 5.03E-175 | 0.70773083 | 0.942 | 0.824 | 9.72E-171 | 6 | Epb41l2 | intermediate |
| Ifi471 | 1.49E-174 | 1.5937841 | 0.528 | 0.269 | 2.88E-170 | 6 | Ifi47 | intermediate |
| Hcfc1r12 | 5.29E-174 | 0.74845462 | 0.889 | 0.777 | 1.02E-169 | 6 | Hcfc1r1 | intermediate |
| Mxra71 | 7.21E-174 | 0.69678721 | 0.929 | 0.816 | 1.39E-169 | 6 | Mxra7 | intermediate |
| Smoc21 | 9.15E-174 | 0.79159894 | 0.573 | 0.476 | 1.77E-169 | 6 | Smoc2 | intermediate |
| Adamtsl12 | 1.26E-173 | 0.8350163 | 0.886 | 0.668 | 2.44E-169 | 6 | Adamtsl1 | intermediate |
| Synm | 5.59E-173 | 3.14923294 | 0.275 | 0.039 | 1.08E-168 | 6 | Synm | intermediate |
| Sparcl11 | 6.34E-173 | 0.56463065 | 0.543 | 0.243 | 1.23E-168 | 6 | Sparcl1 | intermediate |
| Cryab1 | 7.98E-173 | 1.01047402 | 0.639 | 0.329 | 1.54E-168 | 6 | Cryab | intermediate |
| Sptan12 | 4.47E-172 | 0.62384871 | 0.939 | 0.896 | 8.65E-168 | 6 | Sptan1 | intermediate |
| Cebpd2 | 1.51E-171 | 0.53692382 | 0.884 | 0.798 | 2.93E-167 | 6 | Cebpd | intermediate |
| Parp32 | 4.46E-171 | 0.9908319 | 0.904 | 0.65 | 8.62E-167 | 6 | Parp3 | intermediate |
| Asap32 | 2.50E-170 | 1.67180054 | 0.717 | 0.351 | 4.84E-166 | 6 | Asap3 | intermediate |
| H2-K12 | 3.84E-169 | 0.79763698 | 0.914 | 0.817 | 7.42E-165 | 6 | H2-K1 | intermediate |
| Itm2b | 5.28E-169 | 0.26976179 | 0.992 | 0.984 | 1.02E-164 | 6 | Itm2b | intermediate |
| Shisa52 | 6.37E-167 | 0.54673616 | 0.955 | 0.916 | 1.23E-162 | 6 | Shisa5 | intermediate |
| Tmtc1 | 8.53E-165 | 1.78165456 | 0.429 | 0.153 | 1.65E-160 | 6 | Tmtc1 | intermediate |
| Rasl11b1 | 5.45E-164 | 1.14433175 | 0.535 | 0.353 | 1.05E-159 | 6 | Rasl11b | intermediate |
| Serpinb9 | 2.56E-163 | 1.54831289 | 0.566 | 0.275 | 4.94E-159 | 6 | Serpinb9 | intermediate |
| Man2a12 | 8.30E-163 | 0.47591163 | 0.972 | 0.927 | 1.61E-158 | 6 | Man2a1 | intermediate |
| Ssc5d2 | 1.05E-162 | 1.00748654 | 0.828 | 0.501 | 2.02E-158 | 6 | Ssc5d | intermediate |
| Plpp12 | 1.58E-162 | 0.92848534 | 0.818 | 0.619 | 3.06E-158 | 6 | Plpp1 | intermediate |
| Gda3 | 9.60E-162 | 0.80636472 | 0.753 | 0.518 | 1.86E-157 | 6 | Gda | intermediate |
| Igtp1 | 2.28E-161 | 1.89888584 | 0.391 | 0.147 | 4.42E-157 | 6 | Igtp | intermediate |
| C1qtnf71 | 2.68E-161 | 2.13903343 | 0.424 | 0.111 | 5.17E-157 | 6 | C1qtnf7 | intermediate |
| Abi3bp1 | 9.53E-160 | 0.59301439 | 0.876 | 0.719 | 1.84E-155 | 6 | Abi3bp | intermediate |
| Adamts10 | 1.60E-159 | 1.13694527 | 0.846 | 0.658 | 3.09E-155 | 6 | Adamts10 | intermediate |
| Prrx21 | 4.42E-158 | 0.85171926 | 0.816 | 0.686 | 8.54E-154 | 6 | Prrx2 | intermediate |
| Scara32 | 8.10E-158 | 0.79533517 | 0.904 | 0.649 | 1.57E-153 | 6 | Scara3 | intermediate |
| Deptor | 1.90E-157 | 2.65519013 | 0.333 | 0.069 | 3.66E-153 | 6 | Deptor | intermediate |
| Cadm31 | 7.43E-155 | 0.91331893 | 0.566 | 0.271 | 1.44E-150 | 6 | Cadm3 | intermediate |
| Pkia | 8.25E-154 | 3.047662 | 0.288 | 0.05 | 1.59E-149 | 6 | Pkia | intermediate |
| Cd91 | 1.45E-153 | 0.4018532 | 0.955 | 0.9 | 2.81E-149 | 6 | Cd9 | intermediate |
| Itsn12 | 1.19E-152 | 0.89971784 | 0.823 | 0.646 | 2.31E-148 | 6 | Itsn1 | intermediate |
| Txnip2 | 4.68E-152 | 0.53751269 | 0.939 | 0.842 | 9.05E-148 | 6 | Txnip | intermediate |
| Ptprf | 4.66E-151 | 1.5970248 | 0.518 | 0.21 | 9.01E-147 | 6 | Ptprf | intermediate |
| Fgd5 | 3.58E-150 | 1.92816954 | 0.472 | 0.146 | 6.92E-146 | 6 | Fgd5 | intermediate |
| Lgals92 | 5.00E-149 | 0.79848735 | 0.856 | 0.674 | 9.67E-145 | 6 | Lgals9 | intermediate |
| Tmod21 | 5.87E-149 | 2.28429642 | 0.424 | 0.11 | 1.14E-144 | 6 | Tmod2 | intermediate |
| Ghr2 | 7.33E-149 | 1.34764558 | 0.679 | 0.421 | 1.42E-144 | 6 | Ghr | intermediate |
| Abca8a1 | 1.37E-148 | 1.68011919 | 0.346 | 0.121 | 2.65E-144 | 6 | Abca8a | intermediate |
| Tapbp2 | 6.68E-148 | 0.78040776 | 0.944 | 0.81 | 1.29E-143 | 6 | Tapbp | intermediate |
| Adamtsl41 | 2.06E-147 | 1.48352461 | 0.497 | 0.178 | 3.98E-143 | 6 | Adamtsl4 | intermediate |
| Crlf12 | 2.08E-147 | 0.84409647 | 0.485 | 0.394 | 4.03E-143 | 6 | Crlf1 | intermediate |
| Mustn11 | 6.30E-147 | 2.33408395 | 0.273 | 0.063 | 1.22E-142 | 6 | Mustn1 | intermediate |
| Flrt22 | 7.78E-147 | 0.73471951 | 0.869 | 0.643 | 1.50E-142 | 6 | Flrt2 | intermediate |
| Ifih11 | 1.31E-146 | 1.45249239 | 0.573 | 0.322 | 2.54E-142 | 6 | Ifih1 | intermediate |
| Fam102b1 | 2.88E-146 | 0.8564678 | 0.833 | 0.645 | 5.57E-142 | 6 | Fam102b | intermediate |
| Sep-81 | 3.06E-146 | 0.59104552 | 0.952 | 0.895 | 5.92E-142 | 6 | 8-Sep | intermediate |
| Tubb2a2 | 3.88E-146 | 0.72635319 | 0.881 | 0.711 | 7.49E-142 | 6 | Tubb2a | intermediate |
| Lgr41 | 5.32E-145 | 1.5968746 | 0.505 | 0.333 | 1.03E-140 | 6 | Lgr4 | intermediate |
| Anxa111 | 6.58E-145 | 0.96442565 | 0.859 | 0.642 | 1.27E-140 | 6 | Anxa11 | intermediate |
| Ccl73 | 1.52E-143 | 0.59051539 | 0.553 | 0.39 | 2.93E-139 | 6 | Ccl7 | intermediate |
| Sp1001 | 1.98E-143 | 1.67456221 | 0.561 | 0.219 | 3.83E-139 | 6 | Sp100 | intermediate |
| Gvin11 | 4.34E-142 | 1.82702742 | 0.396 | 0.115 | 8.39E-138 | 6 | Gvin1 | intermediate |
| Wnt9a1 | 8.97E-141 | 1.34173978 | 0.508 | 0.297 | 1.73E-136 | 6 | Wnt9a | intermediate |
| Ddost1 | 6.35E-139 | 0.38482528 | 0.96 | 0.95 | 1.23E-134 | 6 | Ddost | intermediate |
| F2r | 5.48E-138 | 0.53573595 | 0.773 | 0.699 | 1.06E-133 | 6 | F2r | intermediate |
| Col16a11 | 4.31E-137 | 0.36846027 | 0.98 | 0.962 | 8.33E-133 | 6 | Col16a1 | intermediate |
| Gpx72 | 1.08E-136 | 0.60800304 | 0.952 | 0.874 | 2.08E-132 | 6 | Gpx7 | intermediate |
| Tgm22 | 1.59E-136 | 0.72705617 | 0.657 | 0.47 | 3.08E-132 | 6 | Tgm2 | intermediate |
| Tor3a2 | 1.67E-136 | 1.49016487 | 0.497 | 0.234 | 3.23E-132 | 6 | Tor3a | intermediate |
| Adamts152 | 7.59E-136 | 0.69959016 | 0.636 | 0.423 | 1.47E-131 | 6 | Adamts15 | intermediate |
| Parp141 | 1.64E-135 | 1.46402767 | 0.513 | 0.285 | 3.16E-131 | 6 | Parp14 | intermediate |
| Pcdhgc32 | 1.19E-134 | 0.64401629 | 0.929 | 0.838 | 2.30E-130 | 6 | Pcdhgc3 | intermediate |
| Lhx8 | 2.42E-132 | 2.37260117 | 0.346 | 0.075 | 4.68E-128 | 6 | Lhx8 | intermediate |
| Ndrg22 | 3.44E-132 | 1.38401207 | 0.485 | 0.251 | 6.64E-128 | 6 | Ndrg2 | intermediate |
| Angptl43 | 9.27E-131 | 0.5116743 | 0.937 | 0.758 | 1.79E-126 | 6 | Angptl4 | intermediate |
| Nfia2 | 1.13E-130 | 0.91517689 | 0.833 | 0.585 | 2.18E-126 | 6 | Nfia | intermediate |
| Ecm2 | 4.19E-130 | 1.95564028 | 0.429 | 0.16 | 8.09E-126 | 6 | Ecm2 | intermediate |
| Scpep11 | 5.83E-129 | 0.40906476 | 0.96 | 0.945 | 1.13E-124 | 6 | Scpep1 | intermediate |
| Adh11 | 2.00E-128 | 1.90124886 | 0.255 | 0.058 | 3.87E-124 | 6 | Adh1 | intermediate |
| Hist1h2bc2 | 2.17E-127 | 0.59667793 | 0.823 | 0.651 | 4.20E-123 | 6 | Hist1h2bc | intermediate |
| Reep51 | 5.21E-127 | 0.47401269 | 0.965 | 0.922 | 1.01E-122 | 6 | Reep5 | intermediate |
| Mbnl1 | 6.27E-127 | 1.14983491 | 0.778 | 0.532 | 1.21E-122 | 6 | Mbnl1 | intermediate |
| Celf22 | 1.23E-126 | 0.9969004 | 0.775 | 0.493 | 2.38E-122 | 6 | Celf2 | intermediate |
| Ifi2032 | 1.62E-126 | 0.89648158 | 0.697 | 0.447 | 3.12E-122 | 6 | Ifi203 | intermediate |
| Plekha41 | 1.66E-126 | 1.60570803 | 0.568 | 0.234 | 3.20E-122 | 6 | Plekha4 | intermediate |
| Acp5 | 5.71E-125 | 1.85988943 | 0.472 | 0.174 | 1.10E-120 | 6 | Acp5 | intermediate |
| Cd1513 | 4.16E-124 | 0.8516919 | 0.803 | 0.635 | 8.04E-120 | 6 | Cd151 | intermediate |
| Clu3 | 4.69E-124 | 0.30797984 | 0.907 | 0.885 | 9.06E-120 | 6 | Clu | intermediate |
| Cd472 | 9.24E-123 | 0.30856547 | 0.949 | 0.916 | 1.79E-118 | 6 | Cd47 | intermediate |
| Pycard1 | 1.13E-122 | 1.61768797 | 0.538 | 0.193 | 2.18E-118 | 6 | Pycard | intermediate |
| Ccl112 | 9.27E-122 | 0.49768223 | 0.654 | 0.422 | 1.79E-117 | 6 | Ccl11 | intermediate |
| Tm4sf13 | 1.33E-121 | 0.60230686 | 0.886 | 0.726 | 2.57E-117 | 6 | Tm4sf1 | intermediate |
| Gnas | 1.38E-121 | 0.26173485 | 0.982 | 0.98 | 2.66E-117 | 6 | Gnas | intermediate |
| Tnfrsf1b2 | 3.21E-121 | 1.18046711 | 0.674 | 0.397 | 6.21E-117 | 6 | Tnfrsf1b | intermediate |
| Nid22 | 5.49E-121 | 1.02426664 | 0.621 | 0.443 | 1.06E-116 | 6 | Nid2 | intermediate |
| Plscr21 | 1.15E-120 | 1.65948489 | 0.354 | 0.109 | 2.23E-116 | 6 | Plscr2 | intermediate |
| Oas1a1 | 1.23E-120 | 1.87324265 | 0.396 | 0.125 | 2.38E-116 | 6 | Oas1a | intermediate |
| Map6 | 1.01E-118 | 1.06357765 | 0.747 | 0.506 | 1.95E-114 | 6 | Map6 | intermediate |
| Ctsz | 1.07E-118 | 0.39094934 | 0.982 | 0.958 | 2.07E-114 | 6 | Ctsz | intermediate |
| Ephb3 | 1.42E-118 | 1.09952257 | 0.687 | 0.5 | 2.74E-114 | 6 | Ephb3 | intermediate |
| Stat21 | 7.78E-118 | 1.4712774 | 0.566 | 0.272 | 1.50E-113 | 6 | Stat2 | intermediate |
| Bckdhb1 | 4.45E-117 | 1.52915043 | 0.52 | 0.251 | 8.60E-113 | 6 | Bckdhb | intermediate |
| Tshz22 | 4.69E-117 | 1.17965424 | 0.753 | 0.418 | 9.07E-113 | 6 | Tshz2 | intermediate |
| Tbrg12 | 5.61E-117 | 0.54054264 | 0.929 | 0.862 | 1.08E-112 | 6 | Tbrg1 | intermediate |
| Dpep12 | 4.54E-115 | 0.68037587 | 0.765 | 0.602 | 8.79E-111 | 6 | Dpep1 | intermediate |
| Masp12 | 4.74E-115 | 0.68608612 | 0.621 | 0.344 | 9.16E-111 | 6 | Masp1 | intermediate |
| Gata61 | 5.22E-115 | 1.74606084 | 0.432 | 0.172 | 1.01E-110 | 6 | Gata6 | intermediate |
| Cd812 | 1.90E-114 | 0.27781224 | 0.977 | 0.962 | 3.68E-110 | 6 | Cd81 | intermediate |
| Fgf101 | 2.98E-114 | 1.88361132 | 0.341 | 0.122 | 5.76E-110 | 6 | Fgf10 | intermediate |
| Nucb22 | 1.26E-113 | 0.76505774 | 0.851 | 0.68 | 2.44E-109 | 6 | Nucb2 | intermediate |
| Adcy71 | 1.38E-112 | 1.09688923 | 0.74 | 0.568 | 2.67E-108 | 6 | Adcy7 | intermediate |
| Ddx582 | 1.97E-112 | 1.18413635 | 0.601 | 0.356 | 3.81E-108 | 6 | Ddx58 | intermediate |
| Tiam13 | 4.28E-112 | 1.58735441 | 0.578 | 0.285 | 8.27E-108 | 6 | Tiam1 | intermediate |
| Nrp1 | 2.76E-111 | 0.75453282 | 0.843 | 0.717 | 5.33E-107 | 6 | Nrp1 | intermediate |
| Tmem254c1 | 3.11E-111 | 1.1753288 | 0.641 | 0.345 | 6.02E-107 | 6 | Tmem254c | intermediate |
| Nenf2 | 9.54E-111 | 0.46824019 | 0.965 | 0.913 | 1.84E-106 | 6 | Nenf | intermediate |
| Rab3il11 | 2.99E-110 | 0.79703619 | 0.806 | 0.548 | 5.78E-106 | 6 | Rab3il1 | intermediate |
| Samd5 | 3.58E-110 | 1.56853293 | 0.308 | 0.141 | 6.93E-106 | 6 | Samd5 | intermediate |
| Slpi2 | 3.85E-110 | 1.15294947 | 0.26 | 0.19 | 7.44E-106 | 6 | Slpi | intermediate |
| Mmp232 | 2.05E-109 | 0.56274423 | 0.934 | 0.841 | 3.96E-105 | 6 | Mmp23 | intermediate |
| Ski | 4.19E-109 | 0.37010262 | 0.972 | 0.95 | 8.11E-105 | 6 | Ski | intermediate |
| Adam151 | 2.89E-108 | 0.77790668 | 0.876 | 0.716 | 5.58E-104 | 6 | Adam15 | intermediate |
| Kng22 | 4.90E-108 | 1.35931615 | 0.457 | 0.226 | 9.47E-104 | 6 | Kng2 | intermediate |
| Slc1a52 | 5.25E-108 | 0.90576532 | 0.856 | 0.649 | 1.02E-103 | 6 | Slc1a5 | intermediate |
| Irf91 | 2.47E-107 | 1.20138663 | 0.672 | 0.379 | 4.77E-103 | 6 | Irf9 | intermediate |
| Pdlim12 | 8.04E-107 | 0.76817763 | 0.73 | 0.529 | 1.56E-102 | 6 | Pdlim1 | intermediate |
| Ahnak22 | 8.05E-107 | 0.65474236 | 0.836 | 0.695 | 1.56E-102 | 6 | Ahnak2 | intermediate |
| Lgi41 | 3.33E-106 | 1.42381769 | 0.545 | 0.236 | 6.44E-102 | 6 | Lgi4 | intermediate |
| Arhgap292 | 8.29E-106 | 0.79113486 | 0.798 | 0.585 | 1.60E-101 | 6 | Arhgap29 | intermediate |
| Slit31 | 1.20E-104 | 0.26317304 | 0.982 | 0.967 | 2.33E-100 | 6 | Slit3 | intermediate |
| Dtx41 | 1.71E-104 | 0.87417466 | 0.687 | 0.522 | 3.30E-100 | 6 | Dtx4 | intermediate |
| Ptch12 | 5.53E-104 | 1.54337811 | 0.379 | 0.217 | 1.07E-99 | 6 | Ptch1 | intermediate |
| Vcan2 | 2.68E-103 | 0.45176587 | 0.912 | 0.84 | 5.18E-99 | 6 | Vcan | intermediate |
| Zbp12 | 5.59E-103 | 0.75110316 | 0.662 | 0.5 | 1.08E-98 | 6 | Zbp1 | intermediate |
| Slfn82 | 6.16E-103 | 1.85718912 | 0.404 | 0.162 | 1.19E-98 | 6 | Slfn8 | intermediate |
| Tap11 | 7.11E-103 | 1.11544816 | 0.619 | 0.373 | 1.37E-98 | 6 | Tap1 | intermediate |
| Ndrg41 | 1.08E-102 | 0.99657543 | 0.669 | 0.433 | 2.09E-98 | 6 | Ndrg4 | intermediate |
| Anxa41 | 3.24E-102 | 0.73721641 | 0.879 | 0.701 | 6.27E-98 | 6 | Anxa4 | intermediate |
| Prrg32 | 4.92E-102 | 1.55309141 | 0.46 | 0.156 | 9.51E-98 | 6 | Prrg3 | intermediate |
| Ech11 | 1.76E-100 | 0.75243489 | 0.861 | 0.694 | 3.40E-96 | 6 | Ech1 | intermediate |
| Pnp2 | 6.82E-100 | 0.66090882 | 0.884 | 0.717 | 1.32E-95 | 6 | Pnp | intermediate |
| Gprc5b1 | 1.29E-98 | 1.03960805 | 0.573 | 0.36 | 2.49E-94 | 6 | Gprc5b | intermediate |
| Cpt1a2 | 8.70E-98 | 0.99842111 | 0.813 | 0.528 | 1.68E-93 | 6 | Cpt1a | intermediate |
| Prdx42 | 1.48E-97 | 0.57607738 | 0.866 | 0.796 | 2.87E-93 | 6 | Prdx4 | intermediate |
| Dclk11 | 2.90E-97 | 0.54313236 | 0.914 | 0.802 | 5.60E-93 | 6 | Dclk1 | intermediate |
| Sorcs21 | 3.04E-97 | 0.76440773 | 0.833 | 0.629 | 5.88E-93 | 6 | Sorcs2 | intermediate |
| Scarb21 | 4.03E-97 | 0.5529731 | 0.902 | 0.811 | 7.79E-93 | 6 | Scarb2 | intermediate |
| Tril2 | 4.92E-97 | 1.65914806 | 0.402 | 0.149 | 9.51E-93 | 6 | Tril | intermediate |
| Prr131 | 2.93E-96 | 0.75832919 | 0.826 | 0.653 | 5.67E-92 | 6 | Prr13 | intermediate |
| Nox42 | 5.14E-96 | 0.87919466 | 0.604 | 0.366 | 9.94E-92 | 6 | Nox4 | intermediate |
| Csf13 | 1.32E-95 | 0.61248707 | 0.869 | 0.566 | 2.55E-91 | 6 | Csf1 | intermediate |
| Cdon2 | 2.09E-95 | 0.86263261 | 0.861 | 0.669 | 4.04E-91 | 6 | Cdon | intermediate |
| Mxd41 | 2.50E-95 | 0.39832722 | 0.942 | 0.89 | 4.83E-91 | 6 | Mxd4 | intermediate |
| Slco3a11 | 3.91E-95 | 0.79632261 | 0.74 | 0.593 | 7.57E-91 | 6 | Slco3a1 | intermediate |
| Tnfsf181 | 7.59E-95 | 0.95024128 | 0.285 | 0.138 | 1.47E-90 | 6 | Tnfsf18 | intermediate |
| Lrrc75a1 | 5.97E-94 | 1.15657528 | 0.631 | 0.398 | 1.15E-89 | 6 | Lrrc75a | intermediate |
| Fbln12 | 4.16E-92 | 0.35041492 | 0.965 | 0.922 | 8.05E-88 | 6 | Fbln1 | intermediate |
| Zfpm21 | 2.98E-91 | 2.00799711 | 0.369 | 0.113 | 5.77E-87 | 6 | Zfpm2 | intermediate |
| Dag12 | 5.64E-91 | 0.50113171 | 0.919 | 0.854 | 1.09E-86 | 6 | Dag1 | intermediate |
| Klf23 | 1.02E-90 | 0.65623893 | 0.601 | 0.407 | 1.96E-86 | 6 | Klf2 | intermediate |
| Abca92 | 1.82E-90 | 1.50365253 | 0.513 | 0.216 | 3.52E-86 | 6 | Abca9 | intermediate |
| Phlda3 | 1.88E-90 | 0.90747932 | 0.672 | 0.477 | 3.63E-86 | 6 | Phlda3 | intermediate |
| Parm12 | 2.13E-90 | 1.28681507 | 0.351 | 0.217 | 4.11E-86 | 6 | Parm1 | intermediate |
| Wnt162 | 2.31E-90 | 1.08743676 | 0.278 | 0.133 | 4.47E-86 | 6 | Wnt16 | intermediate |

| | | | | | | | | |
|---|---|---|---|---|---|---|---|---|
| Tubb2b2 | 3.52E-90 | 0.83589334 | 0.722 | 0.485 | 6.81E-86 | 6 | Tubb2b | intermediate |
| Ifi351 | 4.39E-90 | 1.21970812 | 0.586 | 0.306 | 8.48E-86 | 6 | Ifi35 | intermediate |
| Pmp222 | 1.82E-89 | 0.43910301 | 0.818 | 0.727 | 3.51E-85 | 6 | Pmp22 | intermediate |
| Pdia3 | 1.92E-89 | 0.30841012 | 0.975 | 0.955 | 3.72E-85 | 6 | Pdia3 | intermediate |
| Galnt162 | 2.56E-89 | 0.48809275 | 0.629 | 0.366 | 4.94E-85 | 6 | Galnt16 | intermediate |
| Galnt21 | 1.81E-88 | 0.41349854 | 0.967 | 0.916 | 3.49E-84 | 6 | Galnt2 | intermediate |
| Fxyd52 | 2.28E-88 | 0.39878386 | 0.912 | 0.865 | 4.41E-84 | 6 | Fxyd5 | intermediate |
| Npdc12 | 7.61E-88 | 0.61808396 | 0.944 | 0.811 | 1.47E-83 | 6 | Npdc1 | intermediate |
| Dst2 | 8.46E-88 | 0.57316652 | 0.874 | 0.778 | 1.64E-83 | 6 | Dst | intermediate |
| Dnajc102 | 2.81E-87 | 0.47912067 | 0.912 | 0.869 | 5.44E-83 | 6 | Dnajc10 | intermediate |
| Cthrc13 | 3.59E-87 | 0.29570307 | 0.758 | 0.818 | 6.95E-83 | 6 | Cthrc1 | intermediate |
| Tmem432 | 5.24E-87 | 0.6435504 | 0.874 | 0.748 | 1.01E-82 | 6 | Tmem43 | intermediate |
| Rab302 | 1.73E-86 | 1.48494824 | 0.457 | 0.196 | 3.35E-82 | 6 | Rab30 | intermediate |
| Zfp36l22 | 3.22E-86 | 0.47161592 | 0.919 | 0.834 | 6.22E-82 | 6 | Zfp36l2 | intermediate |
| Wnt21 | 7.34E-85 | 0.87520281 | 0.333 | 0.186 | 1.42E-80 | 6 | Wnt2 | intermediate |
| Dbndd22 | 8.91E-85 | 1.04295801 | 0.52 | 0.343 | 1.72E-80 | 6 | Dbndd2 | intermediate |
| Glud11 | 2.09E-84 | 0.48799931 | 0.917 | 0.859 | 4.05E-80 | 6 | Glud1 | intermediate |
| Trim30a1 | 2.66E-84 | 1.55829854 | 0.359 | 0.144 | 5.14E-80 | 6 | Trim30a | intermediate |
| Itga112 | 7.82E-83 | 0.73938417 | 0.558 | 0.338 | 1.51E-78 | 6 | Itga11 | intermediate |
| Samd9l1 | 9.81E-83 | 1.51546743 | 0.467 | 0.194 | 1.90E-78 | 6 | Samd9l | intermediate |
| Zbtb7c | 1.17E-82 | 2.11545283 | 0.265 | 0.075 | 2.26E-78 | 6 | Zbtb7c | intermediate |
| Phf11b1 | 1.33E-82 | 1.84818221 | 0.25 | 0.074 | 2.57E-78 | 6 | Phf11b | intermediate |
| Grk52 | 2.49E-82 | 1.00956778 | 0.652 | 0.445 | 4.81E-78 | 6 | Grk5 | intermediate |
| Eng | 2.54E-82 | 0.49905658 | 0.629 | 0.517 | 4.91E-78 | 6 | Eng | intermediate |
| Lysmd22 | 3.30E-82 | 1.18252926 | 0.601 | 0.338 | 6.39E-78 | 6 | Lysmd2 | intermediate |
| DEPRECATED-l | 5.69E-82 | 0.767254 | 0.551 | 0.284 | 1.10E-77 | 6 | DEPRECATED-l | intermediate |
| Gjb5 | 1.65E-81 | 1.78315731 | 0.268 | 0.142 | 3.19E-77 | 6 | Gjb5 | intermediate |
| Pigt1 | 2.49E-81 | 0.58850798 | 0.876 | 0.789 | 4.81E-77 | 6 | Pigt | intermediate |
| Dap1 | 2.89E-81 | 0.36272759 | 0.924 | 0.892 | 5.58E-77 | 6 | Dap | intermediate |
| Lrig32 | 2.29E-80 | 1.29905447 | 0.492 | 0.265 | 4.43E-76 | 6 | Lrig3 | intermediate |
| Adamts3 | 2.59E-80 | 1.62023521 | 0.356 | 0.149 | 5.00E-76 | 6 | Adamts3 | intermediate |
| Clec2d2 | 3.40E-80 | 0.37103411 | 0.932 | 0.845 | 6.57E-76 | 6 | Clec2d | intermediate |
| Ifi2111 | 4.75E-80 | 0.59028102 | 0.763 | 0.642 | 9.18E-76 | 6 | Ifi211 | intermediate |
| Kcnt2 | 7.06E-80 | 2.20363206 | 0.253 | 0.064 | 1.37E-75 | 6 | Kcnt2 | intermediate |
| Agpat33 | 9.85E-80 | 0.82445617 | 0.737 | 0.492 | 1.91E-75 | 6 | Agpat3 | intermediate |
| Mar-32 | 1.59E-79 | 1.13354536 | 0.543 | 0.29 | 3.07E-75 | 6 | 3-Mar | intermediate |
| Myl12b1 | 3.59E-79 | 0.76030029 | 0.798 | 0.601 | 6.94E-75 | 6 | Myl12b | intermediate |
| Ramp21 | 7.48E-79 | 1.15433575 | 0.442 | 0.195 | 1.45E-74 | 6 | Ramp2 | intermediate |
| Tmem254a1 | 1.09E-78 | 1.26306526 | 0.583 | 0.281 | 2.10E-74 | 6 | Tmem254a | intermediate |
| Vill1 | 1.12E-78 | 1.46963732 | 0.384 | 0.17 | 2.16E-74 | 6 | Vill | intermediate |
| Wdr66 | 5.54E-78 | 1.95620264 | 0.333 | 0.099 | 1.07E-73 | 6 | Wdr66 | intermediate |
| Pqlc31 | 1.75E-77 | 1.20280195 | 0.487 | 0.256 | 3.39E-73 | 6 | Pqlc3 | intermediate |
| Sirpa2 | 2.77E-77 | 0.56420778 | 0.843 | 0.718 | 5.36E-73 | 6 | Sirpa | intermediate |
| Car5b | 1.27E-76 | 2.11957796 | 0.283 | 0.065 | 2.46E-72 | 6 | Car5b | intermediate |
| Gstt32 | 2.14E-76 | 1.4371611 | 0.457 | 0.181 | 4.14E-72 | 6 | Gstt3 | intermediate |
| Rab292 | 2.31E-76 | 1.06585873 | 0.601 | 0.355 | 4.46E-72 | 6 | Rab29 | intermediate |
| Dhx581 | 3.80E-76 | 1.66548853 | 0.293 | 0.089 | 7.34E-72 | 6 | Dhx58 | intermediate |
| Lamp1 | 1.14E-75 | 0.38274419 | 0.932 | 0.892 | 2.20E-71 | 6 | Lamp1 | intermediate |
| Idh11 | 2.88E-75 | 0.62315106 | 0.841 | 0.703 | 5.57E-71 | 6 | Idh1 | intermediate |
| Slc29a14 | 1.70E-74 | 1.0439426 | 0.548 | 0.296 | 3.30E-70 | 6 | Slc29a1 | intermediate |
| Psmb92 | 5.78E-74 | 1.35018006 | 0.404 | 0.204 | 1.12E-69 | 6 | Psmb9 | intermediate |
| Hexa1 | 6.31E-74 | 0.45204455 | 0.917 | 0.887 | 1.22E-69 | 6 | Hexa | intermediate |
| Mndal2 | 8.44E-74 | 1.26612788 | 0.495 | 0.263 | 1.63E-69 | 6 | Mndal | intermediate |
| Arhgef101 | 1.21E-73 | 1.15220704 | 0.598 | 0.34 | 2.33E-69 | 6 | Arhgef10 | intermediate |
| Kcnj81 | 1.36E-73 | 1.62342418 | 0.29 | 0.082 | 2.63E-69 | 6 | Kcnj8 | intermediate |
| Dpysl23 | 6.12E-73 | 0.42565035 | 0.879 | 0.77 | 1.18E-68 | 6 | Dpysl2 | intermediate |
| Lrrc173 | 7.83E-73 | 0.36592813 | 0.745 | 0.722 | 1.51E-68 | 6 | Lrrc17 | intermediate |
| Slc44a22 | 9.91E-73 | 0.56210877 | 0.874 | 0.801 | 1.92E-68 | 6 | Slc44a2 | intermediate |
| S100a131 | 2.93E-72 | 0.41261362 | 0.939 | 0.843 | 5.67E-68 | 6 | S100a13 | intermediate |
| Nav11 | 4.46E-72 | 0.4546881 | 0.934 | 0.853 | 8.63E-68 | 6 | Nav1 | intermediate |
| Atoh8 | 8.60E-72 | 1.86581779 | 0.27 | 0.11 | 1.66E-67 | 6 | Atoh8 | intermediate |
| Cyb5a2 | 1.23E-71 | 0.59524687 | 0.846 | 0.674 | 2.38E-67 | 6 | Cyb5a | intermediate |
| H2-D12 | 1.49E-71 | 1.07153116 | 0.604 | 0.365 | 2.89E-67 | 6 | H2-D1 | intermediate |
| Pkdcc3 | 2.01E-71 | 0.62319296 | 0.836 | 0.7 | 3.89E-67 | 6 | Pkdcc | intermediate |
| Gpx8 | 3.45E-71 | 0.36810461 | 0.952 | 0.898 | 6.66E-67 | 6 | Gpx8 | intermediate |
| Pcdh192 | 1.12E-70 | 0.9104781 | 0.669 | 0.437 | 2.17E-66 | 6 | Pcdh19 | intermediate |
| Naaa1 | 1.16E-70 | 0.92032073 | 0.646 | 0.402 | 2.25E-66 | 6 | Naaa | intermediate |
| Zbtb201 | 1.33E-70 | 0.86113531 | 0.71 | 0.49 | 2.57E-66 | 6 | Zbtb20 | intermediate |
| Vsir3 | 2.03E-70 | 0.81437676 | 0.692 | 0.448 | 3.92E-66 | 6 | Vsir | intermediate |
| Tor1aip11 | 2.11E-70 | 0.48133062 | 0.886 | 0.784 | 4.09E-66 | 6 | Tor1aip1 | intermediate |
| Zfp4231 | 2.47E-70 | 1.40690283 | 0.432 | 0.201 | 4.77E-66 | 6 | Zfp423 | intermediate |
| Ephb22 | 1.33E-69 | 0.72417171 | 0.798 | 0.607 | 2.58E-65 | 6 | Ephb2 | intermediate |
| Rpn22 | 2.98E-69 | 0.41288073 | 0.957 | 0.907 | 5.77E-65 | 6 | Rpn2 | intermediate |
| Gpc62 | 3.08E-69 | 0.80647059 | 0.662 | 0.528 | 5.96E-65 | 6 | Gpc6 | intermediate |
| Blvrb1 | 6.19E-69 | 0.90014579 | 0.667 | 0.466 | 1.20E-64 | 6 | Blvrb | intermediate |
| Nfic2 | 1.26E-68 | 0.43418796 | 0.912 | 0.841 | 2.44E-64 | 6 | Nfic | intermediate |
| Armh42 | 1.83E-68 | 1.11261111 | 0.593 | 0.294 | 3.54E-64 | 6 | Armh4 | intermediate |
| Tgfbi3 | 2.08E-68 | 0.61226029 | 0.679 | 0.483 | 4.03E-64 | 6 | Tgfbi | intermediate |
| Fat11 | 1.05E-67 | 0.42053725 | 0.927 | 0.868 | 2.02E-63 | 6 | Fat1 | intermediate |
| Plcb41 | 2.67E-67 | 1.59697547 | 0.328 | 0.105 | 5.17E-63 | 6 | Plcb4 | intermediate |
| Man2b22 | 3.02E-67 | 0.49835968 | 0.854 | 0.778 | 5.83E-63 | 6 | Man2b2 | intermediate |
| Rnf1501 | 1.04E-66 | 1.00654358 | 0.659 | 0.425 | 2.00E-62 | 6 | Rnf150 | intermediate |
| Gns2 | 1.18E-66 | 0.42331695 | 0.947 | 0.886 | 2.27E-62 | 6 | Gns | intermediate |
| Klhl131 | 1.63E-66 | 1.37395658 | 0.465 | 0.18 | 3.15E-62 | 6 | Klhl13 | intermediate |
| Cat1 | 1.93E-66 | 0.7229746 | 0.801 | 0.59 | 3.74E-62 | 6 | Cat | intermediate |
| Ap2a21 | 2.20E-66 | 0.44014521 | 0.934 | 0.854 | 4.25E-62 | 6 | Ap2a2 | intermediate |
| Icam11 | 2.53E-66 | 0.71958561 | 0.77 | 0.484 | 4.89E-62 | 6 | Icam1 | intermediate |
| Htra31 | 4.20E-66 | 0.25001547 | 0.912 | 0.832 | 8.12E-62 | 6 | Htra3 | intermediate |
| Lrrn4cl1 | 8.84E-66 | 0.50602973 | 0.629 | 0.375 | 1.71E-61 | 6 | Lrrn4cl | intermediate |
| Gbp71 | 1.13E-65 | 1.32037717 | 0.391 | 0.214 | 2.18E-61 | 6 | Gbp7 | intermediate |
| Eef2k1 | 1.16E-65 | 1.02142012 | 0.601 | 0.336 | 2.25E-61 | 6 | Eef2k | intermediate |
| Stx21 | 1.25E-65 | 0.70375577 | 0.778 | 0.594 | 2.41E-61 | 6 | Stx2 | intermediate |
| Pik3r3 | 1.88E-65 | 1.72124269 | 0.263 | 0.111 | 3.63E-61 | 6 | Pik3r3 | intermediate |
| Lama23 | 2.66E-65 | 0.4405846 | 0.806 | 0.664 | 5.15E-61 | 6 | Lama2 | intermediate |
| Ctsd | 3.76E-65 | 0.5616762 | 0.879 | 0.778 | 7.27E-61 | 6 | Ctsd | intermediate |
| Arl6ip52 | 9.08E-65 | 0.48165976 | 0.866 | 0.767 | 1.76E-60 | 6 | Arl6ip5 | intermediate |
| Iqgap2 | 9.82E-65 | 1.80973156 | 0.263 | 0.098 | 1.90E-60 | 6 | Iqgap2 | intermediate |
| Trafd11 | 9.87E-65 | 0.78091989 | 0.77 | 0.541 | 1.91E-60 | 6 | Trafd1 | intermediate |
| Hspa12a1 | 2.71E-64 | 1.22521375 | 0.404 | 0.184 | 5.25E-60 | 6 | Hspa12a | intermediate |
| Tcf7l22 | 5.04E-64 | 0.95835309 | 0.596 | 0.365 | 9.75E-60 | 6 | Tcf7l2 | intermediate |
| Cxcl162 | 6.22E-64 | 0.92525964 | 0.672 | 0.403 | 1.20E-59 | 6 | Cxcl16 | intermediate |
| Bicc1 | 7.19E-64 | 0.29221441 | 0.962 | 0.932 | 1.39E-59 | 6 | Bicc1 | intermediate |
| Syt111 | 7.87E-64 | 1.55864853 | 0.318 | 0.13 | 1.52E-59 | 6 | Syt11 | intermediate |
| Pik3ip11 | 1.26E-63 | 1.37579605 | 0.409 | 0.186 | 2.44E-59 | 6 | Pik3ip1 | intermediate |
| Vwa5a2 | 1.91E-63 | 0.45179518 | 0.866 | 0.729 | 3.69E-59 | 6 | Vwa5a | intermediate |
| Clip41 | 2.16E-63 | 1.36509908 | 0.505 | 0.183 | 4.17E-59 | 6 | Clip4 | intermediate |
| Vkorc11 | 2.28E-63 | 0.29290781 | 0.957 | 0.921 | 4.40E-59 | 6 | Vkorc1 | intermediate |
| Ids2 | 3.07E-63 | 0.72445826 | 0.697 | 0.493 | 5.93E-59 | 6 | Ids | intermediate |
| C1qtnf11 | 3.57E-63 | 0.38117188 | 0.846 | 0.742 | 6.91E-59 | 6 | C1qtnf1 | intermediate |
| Cdh131 | 5.58E-63 | 0.97069202 | 0.437 | 0.266 | 1.08E-58 | 6 | Cdh13 | intermediate |
| Eif2ak21 | 8.40E-63 | 0.79262004 | 0.578 | 0.419 | 1.62E-58 | 6 | Eif2ak2 | intermediate |
| Ism11 | 1.66E-62 | 1.42902901 | 0.331 | 0.162 | 3.21E-58 | 6 | Ism1 | intermediate |
| Cavin21 | 2.49E-62 | 1.6854053 | 0.278 | 0.091 | 4.81E-58 | 6 | Cavin2 | intermediate |
| Atp10a2 | 2.63E-62 | 0.90446442 | 0.614 | 0.397 | 5.09E-58 | 6 | Atp10a | intermediate |
| Parp91 | 2.65E-62 | 0.99640752 | 0.636 | 0.411 | 5.12E-58 | 6 | Parp9 | intermediate |
| Flot22 | 2.74E-62 | 0.57098536 | 0.854 | 0.722 | 5.31E-58 | 6 | Flot2 | intermediate |
| Txn12 | 4.58E-62 | 0.39380096 | 0.914 | 0.856 | 8.85E-58 | 6 | Txn1 | intermediate |
| Fuca1 | 6.09E-62 | 0.51860164 | 0.848 | 0.769 | 1.18E-57 | 6 | Fuca1 | intermediate |
| Man2b12 | 1.75E-61 | 0.66551372 | 0.801 | 0.671 | 3.39E-57 | 6 | Man2b1 | intermediate |
| Dmpk2 | 1.76E-61 | 1.06179023 | 0.5 | 0.255 | 3.40E-57 | 6 | Dmpk | intermediate |
| Rab321 | 2.40E-61 | 0.88292604 | 0.518 | 0.282 | 4.64E-57 | 6 | Rab32 | intermediate |
| Sigmar12 | 2.70E-61 | 0.43194265 | 0.912 | 0.813 | 5.21E-57 | 6 | Sigmar1 | intermediate |
| Ltbp12 | 4.44E-61 | 0.31563572 | 0.54 | 0.375 | 8.58E-57 | 6 | Ltbp1 | intermediate |
| Cmah1 | 4.66E-61 | 0.83723523 | 0.49 | 0.242 | 9.01E-57 | 6 | Cmah | intermediate |
| Nnmt2 | 4.96E-61 | 0.58766998 | 0.795 | 0.648 | 9.60E-57 | 6 | Nnmt | intermediate |
| Clic41 | 6.52E-61 | 0.26819348 | 0.932 | 0.919 | 1.26E-56 | 6 | Clic4 | intermediate |
| Rai21 | 7.06E-61 | 1.67212303 | 0.313 | 0.11 | 1.37E-56 | 6 | Rai2 | intermediate |
| Ldlrad42 | 1.34E-60 | 0.8332623 | 0.742 | 0.543 | 2.59E-56 | 6 | Ldlrad4 | intermediate |
| Adcy52 | 1.47E-60 | 1.55110541 | 0.351 | 0.158 | 2.83E-56 | 6 | Adcy5 | intermediate |
| Snapc51 | 2.69E-60 | 0.97316924 | 0.596 | 0.342 | 5.21E-56 | 6 | Snapc5 | intermediate |
| Chpf1 | 3.22E-60 | 0.43128974 | 0.972 | 0.888 | 6.23E-56 | 6 | Chpf | intermediate |
| Cap1 | 8.58E-60 | 0.2703071 | 0.96 | 0.934 | 1.66E-55 | 6 | Cap1 | intermediate |
| Hexb | 9.84E-60 | 0.51275651 | 0.859 | 0.75 | 1.90E-55 | 6 | Hexb | intermediate |
| Ccpg12 | 1.24E-59 | 0.49025919 | 0.864 | 0.733 | 2.40E-55 | 6 | Ccpg1 | intermediate |
| Sri2 | 5.09E-59 | 0.36654218 | 0.914 | 0.844 | 9.85E-55 | 6 | Sri | intermediate |
| Smpdl3a2 | 7.32E-59 | 1.03251565 | 0.452 | 0.238 | 1.42E-54 | 6 | Smpdl3a | intermediate |
| Pdcd42 | 2.48E-58 | 0.51740614 | 0.811 | 0.651 | 4.79E-54 | 6 | Pdcd4 | intermediate |
| Pycr11 | 2.53E-58 | 0.54353608 | 0.763 | 0.644 | 4.90E-54 | 6 | Pycr1 | intermediate |
| Psmb102 | 2.57E-58 | 0.83890682 | 0.631 | 0.411 | 4.97E-54 | 6 | Psmb10 | intermediate |
| Ctso2 | 2.76E-58 | 0.77698475 | 0.705 | 0.477 | 5.34E-54 | 6 | Ctso | intermediate |
| Sox92 | 3.37E-58 | 0.90595634 | 0.442 | 0.23 | 6.51E-54 | 6 | Sox9 | intermediate |
| Ifi2041 | 3.77E-58 | 0.52484222 | 0.72 | 0.534 | 7.29E-54 | 6 | Ifi204 | intermediate |
| Trim34a1 | 5.72E-58 | 1.75031821 | 0.295 | 0.106 | 1.11E-53 | 6 | Trim34a | intermediate |
| Ucp21 | 7.23E-58 | 1.14289141 | 0.265 | 0.188 | 1.40E-53 | 6 | Ucp2 | intermediate |
| Tmem106a2 | 1.61E-57 | 0.85443198 | 0.581 | 0.323 | 3.11E-53 | 6 | Tmem106a | intermediate |
| Esyt1 | 2.96E-57 | 0.42265181 | 0.914 | 0.829 | 5.73E-53 | 6 | Esyt1 | intermediate |
| Slc8a12 | 3.86E-57 | 0.74745977 | 0.737 | 0.514 | 7.46E-53 | 6 | Slc8a1 | intermediate |
| Plin31 | 8.18E-57 | 0.53674992 | 0.859 | 0.712 | 1.58E-52 | 6 | Plin3 | intermediate |
| Gabarap | 9.71E-57 | 0.30640392 | 0.927 | 0.89 | 1.88E-52 | 6 | Gabarap | intermediate |
| Nrbp22 | 1.69E-56 | 1.14931717 | 0.46 | 0.231 | 3.26E-52 | 6 | Nrbp2 | intermediate |
| Tlr31 | 1.82E-56 | 1.61775989 | 0.29 | 0.115 | 3.52E-52 | 6 | Tlr3 | intermediate |
| Aldh21 | 2.78E-56 | 0.32638025 | 0.929 | 0.884 | 5.38E-52 | 6 | Aldh2 | intermediate |
| Tmod31 | 5.47E-56 | 0.29755524 | 0.924 | 0.887 | 1.06E-51 | 6 | Tmod3 | intermediate |
| Palm1 | 2.24E-55 | 0.42320971 | 0.879 | 0.793 | 4.32E-51 | 6 | Palm | intermediate |
| Ddah22 | 3.88E-55 | 0.58305208 | 0.828 | 0.669 | 7.50E-51 | 6 | Ddah2 | intermediate |
| Castor1 | 4.13E-55 | 1.474208 | 0.381 | 0.178 | 7.98E-51 | 6 | Castor1 | intermediate |
| Dtx3l1 | 7.54E-55 | 0.84533645 | 0.588 | 0.392 | 1.46E-50 | 6 | Dtx3l | intermediate |
| Plscr3 | 8.92E-55 | 0.28173477 | 0.967 | 0.914 | 1.72E-50 | 6 | Plscr3 | intermediate |
| Rhou1 | 1.12E-54 | 0.61988192 | 0.866 | 0.694 | 2.16E-50 | 6 | Rhou | intermediate |
| St3gal42 | 1.56E-54 | 0.66010758 | 0.785 | 0.618 | 3.02E-50 | 6 | St3gal4 | intermediate |
| Psme12 | 5.77E-54 | 0.43007971 | 0.838 | 0.743 | 1.11E-49 | 6 | Psme1 | intermediate |
| Herc61 | 6.30E-54 | 1.39195186 | 0.26 | 0.128 | 1.22E-49 | 6 | Herc6 | intermediate |
| Smarca22 | 2.04E-53 | 0.70241959 | 0.745 | 0.599 | 3.94E-49 | 6 | Smarca2 | intermediate |
| Panx11 | 2.59E-53 | 0.98548277 | 0.591 | 0.359 | 5.01E-49 | 6 | Panx1 | intermediate |
| Mtmr111 | 6.49E-53 | 0.83817991 | 0.601 | 0.416 | 1.25E-48 | 6 | Mtmr11 | intermediate |
| Naglu2 | 7.66E-53 | 0.7713824 | 0.715 | 0.48 | 1.48E-48 | 6 | Naglu | intermediate |
| Ssr41 | 9.93E-53 | 0.27486618 | 0.922 | 0.889 | 1.92E-48 | 6 | Ssr4 | intermediate |
| Iqgap11 | 1.34E-52 | 0.25084735 | 0.975 | 0.937 | 2.59E-48 | 6 | Iqgap1 | intermediate |
| Has13 | 1.38E-52 | 0.87328217 | 0.346 | 0.252 | 2.66E-48 | 6 | Has1 | intermediate |
| Gpr39 | 4.55E-52 | 1.35245288 | 0.341 | 0.179 | 8.80E-48 | 6 | Gpr39 | intermediate |
| Agtr1a2 | 6.23E-52 | 1.062656 | 0.417 | 0.193 | 1.20E-47 | 6 | Agtr1a | intermediate |
| Art31 | 6.48E-52 | 1.01932583 | 0.301 | 0.179 | 1.25E-47 | 6 | Art3 | intermediate |
| Fcgrt1 | 3.52E-51 | 0.37475866 | 0.944 | 0.891 | 6.80E-47 | 6 | Fcgrt | intermediate |
| Fam114a11 | 3.77E-51 | 0.27185756 | 0.952 | 0.915 | 7.29E-47 | 6 | Fam114a1 | intermediate |
| Shox21 | 4.03E-51 | 0.76415988 | 0.631 | 0.403 | 7.80E-47 | 6 | Shox2 | intermediate |
| Dact12 | 4.07E-51 | 0.36059249 | 0.768 | 0.683 | 7.86E-47 | 6 | Dact1 | intermediate |
| Ptger31 | 4.44E-51 | 1.01311467 | 0.308 | 0.153 | 8.59E-47 | 6 | Ptger3 | intermediate |
| Hhipl12 | 5.91E-51 | 0.63084304 | 0.735 | 0.587 | 1.14E-46 | 6 | Hhipl1 | intermediate |
| Slfn21 | 6.68E-51 | 1.14594839 | 0.301 | 0.164 | 1.29E-46 | 6 | Slfn2 | intermediate |
| Parp121 | 6.79E-51 | 0.87224789 | 0.634 | 0.435 | 1.31E-46 | 6 | Parp12 | intermediate |
| Rsu11 | 6.87E-51 | 0.38710228 | 0.907 | 0.841 | 1.33E-46 | 6 | Rsu1 | intermediate |
| Dusp221 | 7.35E-51 | 1.34806315 | 0.361 | 0.134 | 1.42E-46 | 6 | Dusp22 | intermediate |
| Add11 | 8.09E-51 | 0.33346737 | 0.939 | 0.893 | 1.56E-46 | 6 | Add1 | intermediate |
| Hsd17b41 | 2.46E-50 | 0.57688006 | 0.773 | 0.613 | 4.75E-46 | 6 | Hsd17b4 | intermediate |
| Pcdh71 | 6.18E-50 | 0.64174043 | 0.747 | 0.595 | 1.20E-45 | 6 | Pcdh7 | intermediate |
| Ccl22 | 6.57E-50 | 0.58132521 | 0.298 | 0.219 | 1.27E-45 | 6 | Ccl2 | intermediate |
| Plce11 | 1.49E-49 | 1.12641583 | 0.369 | 0.166 | 2.88E-45 | 6 | Plce1 | intermediate |
| Acat12 | 3.29E-49 | 0.53541213 | 0.813 | 0.667 | 6.37E-45 | 6 | Acat1 | intermediate |
| Bok1 | 5.31E-49 | 1.12566199 | 0.442 | 0.235 | 1.03E-44 | 6 | Bok | intermediate |
| Hsd3b71 | 6.62E-49 | 0.56451928 | 0.639 | 0.501 | 1.28E-44 | 6 | Hsd3b7 | intermediate |
| Ltbr1 | 7.66E-49 | 0.42831126 | 0.886 | 0.787 | 1.48E-44 | 6 | Ltbr | intermediate |
| Lipa2 | 1.65E-48 | 0.47924174 | 0.813 | 0.687 | 3.19E-44 | 6 | Lipa | intermediate |
| Reep32 | 1.99E-48 | 0.3620038 | 0.848 | 0.774 | 3.85E-44 | 6 | Reep3 | intermediate |
| Cdk141 | 2.25E-48 | 0.65939238 | 0.758 | 0.602 | 4.35E-44 | 6 | Cdk14 | intermediate |
| Emilin22 | 3.34E-48 | 0.25144947 | 0.866 | 0.763 | 6.46E-44 | 6 | Emilin2 | intermediate |
| Capn12 | 5.50E-48 | 0.47527985 | 0.801 | 0.667 | 1.06E-43 | 6 | Capn1 | intermediate |
| Lrrk1 | 7.25E-48 | 0.76299535 | 0.727 | 0.529 | 1.40E-43 | 6 | Lrrk1 | intermediate |
| Apobec1 | 7.82E-48 | 1.5004462 | 0.298 | 0.108 | 1.51E-43 | 6 | Apobec1 | intermediate |
| Fas2 | 1.62E-47 | 1.18358249 | 0.477 | 0.233 | 3.14E-43 | 6 | Fas | intermediate |
| Fhl22 | 1.78E-47 | 0.68811947 | 0.548 | 0.446 | 3.44E-43 | 6 | Fhl2 | intermediate |
| Vav21 | 4.21E-47 | 1.12194573 | 0.52 | 0.314 | 8.15E-43 | 6 | Vav2 | intermediate |
| Arhgap212 | 6.06E-47 | 0.71995678 | 0.74 | 0.566 | 1.17E-42 | 6 | Arhgap21 | intermediate |
| Phldb12 | 8.71E-47 | 0.43745576 | 0.841 | 0.771 | 1.68E-42 | 6 | Phldb1 | intermediate |
| Il61 | 9.68E-47 | 0.88386436 | 0.26 | 0.146 | 1.87E-42 | 6 | Il6 | intermediate |
| Rin2 | 1.27E-46 | 0.57117519 | 0.692 | 0.516 | 2.46E-42 | 6 | Rin2 | intermediate |
| Lgmn2 | 1.30E-46 | 0.35197096 | 0.932 | 0.863 | 2.51E-42 | 6 | Lgmn | intermediate |
| Fam234a1 | 6.05E-46 | 0.66390948 | 0.803 | 0.608 | 1.17E-41 | 6 | Fam234a | intermediate |
| Cx3cl11 | 8.78E-46 | 1.12095296 | 0.298 | 0.178 | 1.70E-41 | 6 | Cx3cl1 | intermediate |
| Ppp1r22 | 1.02E-45 | 0.56686729 | 0.806 | 0.648 | 1.97E-41 | 6 | Ppp1r2 | intermediate |
| Pard3b | 1.27E-45 | 1.32961063 | 0.354 | 0.144 | 2.45E-41 | 6 | Pard3b | intermediate |
| Cdc42ep22 | 1.31E-45 | 0.84371763 | 0.591 | 0.367 | 2.53E-41 | 6 | Cdc42ep2 | intermediate |
| Znfx11 | 3.07E-45 | 1.05255353 | 0.465 | 0.269 | 5.93E-41 | 6 | Znfx1 | intermediate |
| Kctd171 | 7.17E-45 | 0.39964181 | 0.902 | 0.795 | 1.39E-40 | 6 | Kctd17 | intermediate |
| Flt3l1 | 4.11E-44 | 1.80800512 | 0.278 | 0.101 | 7.95E-40 | 6 | Flt3l | intermediate |
| Nsg13 | 5.17E-44 | 0.47521591 | 0.79 | 0.591 | 1.00E-39 | 6 | Nsg1 | intermediate |
| Pdlim52 | 8.58E-44 | 0.51761454 | 0.821 | 0.658 | 1.66E-39 | 6 | Pdlim5 | intermediate |
| Copz2 | 1.28E-43 | 0.26063484 | 0.939 | 0.899 | 2.47E-39 | 6 | Copz2 | intermediate |
| Helz21 | 4.75E-43 | 0.58866904 | 0.598 | 0.512 | 9.19E-39 | 6 | Helz2 | intermediate |
| Dctn2 | 1.43E-42 | 0.3173042 | 0.947 | 0.894 | 2.77E-38 | 6 | Dctn2 | intermediate |
| Xpnpep22 | 5.05E-42 | 0.9665417 | 0.273 | 0.147 | 9.76E-38 | 6 | Xpnpep2 | intermediate |
| Pygb1 | 5.55E-42 | 0.59216685 | 0.816 | 0.656 | 1.07E-37 | 6 | Pygb | intermediate |
| Paxx1 | 9.37E-42 | 1.12942664 | 0.487 | 0.247 | 1.81E-37 | 6 | Paxx | intermediate |
| Poglut31 | 1.37E-41 | 0.57044468 | 0.758 | 0.592 | 2.65E-37 | 6 | Poglut3 | intermediate |
| Cast1 | 1.48E-41 | 0.28747745 | 0.919 | 0.864 | 2.86E-37 | 6 | Cast | intermediate |
| Tmem1002 | 2.03E-41 | 0.55406025 | 0.674 | 0.444 | 3.92E-37 | 6 | Tmem100 | intermediate |
| Mn12 | 2.74E-41 | 1.2925103 | 0.419 | 0.223 | 5.30E-37 | 6 | Mn1 | intermediate |
| Mycbp22 | 3.45E-41 | 0.30827146 | 0.843 | 0.768 | 6.67E-37 | 6 | Mycbp2 | intermediate |
| Irak31 | 5.00E-41 | 0.68367161 | 0.606 | 0.401 | 9.66E-37 | 6 | Irak3 | intermediate |
| Nudt182 | 6.09E-41 | 0.89684586 | 0.505 | 0.305 | 1.18E-36 | 6 | Nudt18 | intermediate |
| Golga71 | 6.45E-41 | 0.27814121 | 0.869 | 0.851 | 1.25E-36 | 6 | Golga7 | intermediate |
| Nmi2 | 2.57E-40 | 1.14637611 | 0.381 | 0.182 | 4.96E-36 | 6 | Nmi | intermediate |
| Stk391 | 3.46E-40 | 0.88337719 | 0.566 | 0.365 | 6.69E-36 | 6 | Stk39 | intermediate |
| Ephx11 | 5.14E-40 | 1.01142437 | 0.419 | 0.257 | 9.94E-36 | 6 | Ephx1 | intermediate |
| Scp2 | 6.77E-40 | 0.61048486 | 0.687 | 0.516 | 1.31E-35 | 6 | Scp2 | intermediate |
| Tdrd71 | 6.93E-40 | 0.75252776 | 0.553 | 0.376 | 1.34E-35 | 6 | Tdrd7 | intermediate |
| Cgref12 | 9.27E-40 | 0.82593808 | 0.477 | 0.326 | 1.79E-35 | 6 | Cgref1 | intermediate |
| St3gal12 | 1.16E-39 | 0.58464824 | 0.725 | 0.575 | 2.25E-35 | 6 | St3gal1 | intermediate |
| Rdh102 | 1.35E-39 | 0.3132074 | 0.705 | 0.701 | 2.60E-35 | 6 | Rdh10 | intermediate |
| Fuca2 | 1.65E-39 | 1.27239714 | 0.323 | 0.164 | 3.20E-35 | 6 | Fuca2 | intermediate |
| Abhd14b1 | 2.02E-39 | 0.99357072 | 0.452 | 0.24 | 3.91E-35 | 6 | Abhd14b | intermediate |
| Maged2 | 3.61E-39 | 0.28363479 | 0.919 | 0.873 | 6.98E-35 | 6 | Maged2 | intermediate |
| Rexo21 | 3.96E-39 | 0.28772521 | 0.919 | 0.867 | 7.65E-35 | 6 | Rexo2 | intermediate |
| Stt3b2 | 4.66E-39 | 0.28784889 | 0.912 | 0.878 | 9.01E-35 | 6 | Stt3b | intermediate |
| Chrd2 | 4.91E-39 | 1.16315144 | 0.447 | 0.242 | 9.49E-35 | 6 | Chrd | intermediate |
| Eps83 | 7.79E-39 | 0.76207541 | 0.604 | 0.36 | 1.51E-34 | 6 | Eps8 | intermediate |
| Arsa1 | 7.97E-39 | 0.55116121 | 0.788 | 0.661 | 1.54E-34 | 6 | Arsa | intermediate |
| Ssh31 | 8.27E-39 | 1.02926541 | 0.47 | 0.242 | 1.60E-34 | 6 | Ssh3 | intermediate |
| Ank21 | 1.60E-38 | 0.61653863 | 0.626 | 0.471 | 3.09E-34 | 6 | Ank2 | intermediate |
| Tmbim12 | 2.68E-38 | 0.39725507 | 0.775 | 0.694 | 5.18E-34 | 6 | Tmbim1 | intermediate |
| Trim211 | 3.63E-38 | 1.16459384 | 0.326 | 0.18 | 7.01E-34 | 6 | Trim21 | intermediate |
| Ifi302 | 4.46E-38 | 0.60455375 | 0.646 | 0.447 | 8.62E-34 | 6 | Ifi30 | intermediate |
| Sqor2 | 5.45E-38 | 0.82662787 | 0.548 | 0.301 | 1.05E-33 | 6 | Sqor | intermediate |
| Galnt172 | 7.33E-38 | 0.53308374 | 0.667 | 0.519 | 1.42E-33 | 6 | Galnt17 | intermediate |
| Hsd17b111 | 8.63E-38 | 0.54964338 | 0.75 | 0.61 | 1.67E-33 | 6 | Hsd17b11 | intermediate |
| Grina3 | 9.09E-38 | 0.48689312 | 0.783 | 0.642 | 1.76E-33 | 6 | Grina | intermediate |
| Bscl21 | 1.46E-37 | 0.5115481 | 0.806 | 0.672 | 2.82E-33 | 6 | Bscl2 | intermediate |
| Gab21 | 1.98E-37 | 0.64749102 | 0.545 | 0.406 | 3.83E-33 | 6 | Gab2 | intermediate |
| Plscr41 | 2.63E-37 | 0.98260104 | 0.495 | 0.294 | 5.09E-33 | 6 | Plscr4 | intermediate |
| Naga | 8.12E-37 | 0.44352181 | 0.816 | 0.707 | 1.57E-32 | 6 | Naga | intermediate |
| Lynx11 | 9.77E-37 | 0.76169908 | 0.669 | 0.484 | 1.89E-32 | 6 | Lynx1 | intermediate |
| Capn52 | 1.24E-36 | 0.78932056 | 0.47 | 0.289 | 2.40E-32 | 6 | Capn5 | intermediate |
| Dgka1 | 1.26E-36 | 0.6757847 | 0.543 | 0.385 | 2.44E-32 | 6 | Dgka | intermediate |
| Park72 | 1.37E-36 | 0.37285744 | 0.869 | 0.78 | 2.65E-32 | 6 | Park7 | intermediate |
| Tmem140 | 2.13E-36 | 0.7609183 | 0.535 | 0.353 | 4.12E-32 | 6 | Tmem140 | intermediate |
| Erp291 | 2.35E-36 | 0.26270489 | 0.937 | 0.886 | 4.54E-32 | 6 | Erp29 | intermediate |
| Smarcd21 | 3.05E-36 | 0.63607986 | 0.73 | 0.547 | 5.91E-32 | 6 | Smarcd2 | intermediate |
| Dennd4a2 | 5.11E-36 | 0.45000488 | 0.629 | 0.567 | 9.87E-32 | 6 | Dennd4a | intermediate |
| Stac2 | 5.20E-36 | 1.17678518 | 0.253 | 0.137 | 1.00E-31 | 6 | Stac2 | intermediate |
| Hoxc101 | 5.66E-36 | 0.56397705 | 0.702 | 0.54 | 1.09E-31 | 6 | Hoxc10 | intermediate |
| Fkbp7 | 8.45E-36 | 0.29414766 | 0.859 | 0.828 | 1.63E-31 | 6 | Fkbp7 | intermediate |
| Gyg2 | 9.73E-36 | 0.76030398 | 0.538 | 0.374 | 1.88E-31 | 6 | Gyg | intermediate |
| Hspb82 | 1.22E-35 | 0.35009655 | 0.773 | 0.651 | 2.36E-31 | 6 | Hspb8 | intermediate |
| Atp9a | 1.26E-35 | 0.87717625 | 0.427 | 0.268 | 2.43E-31 | 6 | Atp9a | intermediate |
| Daxx1 | 1.41E-35 | 0.61382015 | 0.614 | 0.504 | 2.72E-31 | 6 | Daxx | intermediate |
| Sec14l11 | 2.12E-35 | 0.46121117 | 0.788 | 0.645 | 4.10E-31 | 6 | Sec14l1 | intermediate |
| Psmb82 | 4.46E-35 | 0.60368609 | 0.667 | 0.521 | 8.63E-31 | 6 | Psmb8 | intermediate |
| Map3k142 | 4.92E-35 | 0.795284 | 0.553 | 0.381 | 9.51E-31 | 6 | Map3k14 | intermediate |
| Fam49a | 8.39E-35 | 1.26314537 | 0.275 | 0.128 | 1.62E-30 | 6 | Fam49a | intermediate |
| Nt5c1 | 1.08E-34 | 0.43567468 | 0.727 | 0.605 | 2.10E-30 | 6 | Nt5c | intermediate |
| Iah11 | 1.25E-34 | 0.37235646 | 0.813 | 0.731 | 2.42E-30 | 6 | Iah1 | intermediate |
| Elovl52 | 1.89E-34 | 0.52663712 | 0.783 | 0.642 | 3.66E-30 | 6 | Elovl5 | intermediate |
| Cgnl11 | 2.48E-34 | 0.96845473 | 0.301 | 0.192 | 4.79E-30 | 6 | Cgnl1 | intermediate |
| Sppl2a1 | 2.74E-34 | 0.38883532 | 0.768 | 0.668 | 5.30E-30 | 6 | Sppl2a | intermediate |
| Praf21 | 2.93E-34 | 0.41388093 | 0.841 | 0.74 | 5.66E-30 | 6 | Praf2 | intermediate |
| Itpr12 | 3.16E-34 | 0.83283566 | 0.394 | 0.231 | 6.11E-30 | 6 | Itpr1 | intermediate |
| Dlc12 | 3.61E-34 | 0.37662706 | 0.795 | 0.655 | 6.98E-30 | 6 | Dlc1 | intermediate |
| Hoxd82 | 3.80E-34 | 0.62964916 | 0.576 | 0.399 | 7.35E-30 | 6 | Hoxd8 | intermediate |
| Plbd2 | 3.92E-34 | 0.32854962 | 0.891 | 0.801 | 7.57E-30 | 6 | Plbd2 | intermediate |
| Il3ra2 | 9.78E-34 | 0.25854765 | 0.912 | 0.836 | 1.89E-29 | 6 | Il3ra | intermediate |
| Phyh2 | 1.06E-33 | 0.85366227 | 0.528 | 0.302 | 2.05E-29 | 6 | Phyh | intermediate |
| Eml1 | 1.37E-33 | 0.59530663 | 0.684 | 0.561 | 2.64E-29 | 6 | Eml1 | intermediate |
| L3hypdh1 | 1.70E-33 | 0.62672136 | 0.619 | 0.461 | 3.29E-29 | 6 | L3hypdh | intermediate |
| Sgsm21 | 2.74E-33 | 0.98624348 | 0.472 | 0.264 | 5.30E-29 | 6 | Sgsm2 | intermediate |
| Mpp11 | 4.55E-33 | 0.69764638 | 0.548 | 0.385 | 8.80E-29 | 6 | Mpp1 | intermediate |
| Fam102a2 | 9.02E-33 | 0.49873775 | 0.662 | 0.53 | 1.74E-28 | 6 | Fam102a | intermediate |
| Rab40b1 | 9.56E-33 | 1.11845331 | 0.371 | 0.19 | 1.85E-28 | 6 | Rab40b | intermediate |
| Tsc22d31 | 1.42E-32 | 0.88785175 | 0.381 | 0.239 | 2.74E-28 | 6 | Tsc22d3 | intermediate |
| Sp1401 | 3.10E-32 | 1.365897 | 0.336 | 0.15 | 5.99E-28 | 6 | Sp140 | intermediate |
| Uap11 | 3.46E-32 | 0.35455432 | 0.848 | 0.748 | 6.69E-28 | 6 | Uap1 | intermediate |
| Ggct3 | 5.17E-32 | 0.98286356 | 0.371 | 0.169 | 9.99E-28 | 6 | Ggct | intermediate |
| Tcn22 | 5.87E-32 | 0.27613265 | 0.894 | 0.869 | 1.13E-27 | 6 | Tcn2 | intermediate |
| Kif13b1 | 7.59E-32 | 0.64701456 | 0.51 | 0.338 | 1.47E-27 | 6 | Kif13b | intermediate |
| Rgl11 | 8.82E-32 | 0.39440223 | 0.798 | 0.674 | 1.70E-27 | 6 | Rgl1 | intermediate |
| Castor21 | 9.31E-32 | 0.69626941 | 0.629 | 0.439 | 1.80E-27 | 6 | Castor2 | intermediate |
| Tbl1xr1 | 1.36E-31 | 0.50314404 | 0.727 | 0.594 | 2.64E-27 | 6 | Tbl1xr1 | intermediate |
| Gstm11 | 1.36E-31 | 0.9568474 | 0.313 | 0.138 | 2.64E-27 | 6 | Gstm1 | intermediate |
| Lmbr11 | 1.61E-31 | 0.85587064 | 0.477 | 0.289 | 3.12E-27 | 6 | Lmbr1 | intermediate |
| Camkk21 | 2.41E-31 | 0.72756056 | 0.606 | 0.441 | 4.67E-27 | 6 | Camkk2 | intermediate |
| Calhm21 | 3.76E-31 | 0.56278746 | 0.662 | 0.528 | 7.27E-27 | 6 | Calhm2 | intermediate |
| Aida2 | 5.06E-31 | 0.36337321 | 0.742 | 0.646 | 9.78E-27 | 6 | Aida | intermediate |
| Cpeb11 | 6.60E-31 | 1.00722825 | 0.326 | 0.176 | 1.28E-26 | 6 | Cpeb1 | intermediate |
| Ptgr12 | 6.80E-31 | 0.73912421 | 0.662 | 0.473 | 1.31E-26 | 6 | Ptgr1 | intermediate |
| Fgl23 | 7.02E-31 | 0.27427767 | 0.601 | 0.593 | 1.36E-26 | 6 | Fgl2 | intermediate |
| Jcad2 | 8.99E-31 | 0.68316312 | 0.598 | 0.45 | 1.74E-26 | 6 | Jcad | intermediate |
| Mrgprf2 | 1.27E-30 | 0.44797623 | 0.677 | 0.526 | 2.46E-26 | 6 | Mrgprf | intermediate |
| Acsl52 | 1.30E-30 | 0.44995529 | 0.74 | 0.619 | 2.51E-26 | 6 | Acsl5 | intermediate |
| Lrrc42 | 1.75E-30 | 0.73863017 | 0.629 | 0.419 | 3.38E-26 | 6 | Lrrc42 | intermediate |
| Mill21 | 2.24E-30 | 1.45579176 | 0.278 | 0.109 | 4.33E-26 | 6 | Mill2 | intermediate |
| Ebp2 | 3.22E-30 | 0.60482373 | 0.694 | 0.558 | 6.22E-26 | 6 | Ebp | intermediate |
| Foxo12 | 4.25E-30 | 0.65116002 | 0.548 | 0.384 | 8.22E-26 | 6 | Foxo1 | intermediate |
| Snx101 | 4.47E-30 | 1.06171683 | 0.379 | 0.193 | 8.64E-26 | 6 | Snx10 | intermediate |
| Slc8b11 | 5.45E-30 | 1.12231221 | 0.28 | 0.134 | 1.05E-25 | 6 | Slc8b1 | intermediate |
| 21-Mar | 5.70E-30 | 0.71408277 | 0.538 | 0.348 | 1.10E-25 | 6 | 2-Mar | intermediate |
| Trim252 | 6.53E-30 | 0.49320511 | 0.684 | 0.549 | 1.26E-25 | 6 | Trim25 | intermediate |
| Xpnpep1 | 6.99E-30 | 0.31385267 | 0.866 | 0.789 | 1.35E-25 | 6 | Xpnpep1 | intermediate |
| Loxl42 | 7.08E-30 | 0.59149415 | 0.492 | 0.321 | 1.37E-25 | 6 | Loxl4 | intermediate |
| Tlr41 | 7.58E-30 | 0.82760317 | 0.424 | 0.238 | 1.46E-25 | 6 | Tlr4 | intermediate |
| Sgcb | 7.92E-30 | 0.53259677 | 0.694 | 0.547 | 1.53E-25 | 6 | Sgcb | intermediate |
| Dync2li1 | 8.76E-30 | 1.1124575 | 0.323 | 0.143 | 1.69E-25 | 6 | Dync2li1 | intermediate |
| Pttg11 | 8.78E-30 | 0.63779591 | 0.598 | 0.435 | 1.70E-25 | 6 | Pttg1 | intermediate |
| Ackr31 | 9.93E-30 | 0.30927715 | 0.53 | 0.404 | 1.92E-25 | 6 | Ackr3 | intermediate |
| Erap11 | 1.02E-29 | 0.72800791 | 0.533 | 0.345 | 1.98E-25 | 6 | Erap1 | intermediate |
| Tmem2191 | 1.26E-29 | 0.72821282 | 0.558 | 0.364 | 2.43E-25 | 6 | Tmem219 | intermediate |
| Fmnl23 | 1.58E-29 | 0.62601433 | 0.641 | 0.419 | 3.05E-25 | 6 | Fmnl2 | intermediate |
| Lgals81 | 1.59E-29 | 0.47075594 | 0.72 | 0.561 | 3.08E-25 | 6 | Lgals8 | intermediate |
| Cab39l2 | 1.72E-29 | 0.88681679 | 0.384 | 0.242 | 3.32E-25 | 6 | Cab39l | intermediate |
| Arhgef251 | 1.93E-29 | 0.2833129 | 0.937 | 0.869 | 3.74E-25 | 6 | Arhgef25 | intermediate |
| Mpnd | 2.55E-29 | 0.47936809 | 0.773 | 0.609 | 4.93E-25 | 6 | Mpnd | intermediate |
| Sertad31 | 2.62E-29 | 0.70265087 | 0.422 | 0.28 | 5.06E-25 | 6 | Sertad3 | intermediate |
| Trim22 | 5.77E-29 | 0.72680743 | 0.508 | 0.328 | 1.11E-24 | 6 | Trim2 | intermediate |
| Ssbp21 | 6.27E-29 | 0.7085408 | 0.545 | 0.389 | 1.21E-24 | 6 | Ssbp2 | intermediate |
| Tent5a1 | 6.91E-29 | 0.46205053 | 0.755 | 0.623 | 1.34E-24 | 6 | Tent5a | intermediate |
| Psen22 | 7.59E-29 | 0.74435802 | 0.581 | 0.406 | 1.47E-24 | 6 | Psen2 | intermediate |
| Pcyox11 | 8.00E-29 | 0.31936933 | 0.879 | 0.785 | 1.55E-24 | 6 | Pcyox1 | intermediate |
| Tap21 | 8.15E-29 | 0.48817531 | 0.732 | 0.598 | 1.58E-24 | 6 | Tap2 | intermediate |
| Pigs2 | 1.01E-28 | 0.46026157 | 0.803 | 0.693 | 1.95E-24 | 6 | Pigs | intermediate |
| Suox1 | 1.15E-28 | 1.01048189 | 0.313 | 0.158 | 2.22E-24 | 6 | Suox | intermediate |
| Pdia52 | 1.29E-28 | 0.3164651 | 0.891 | 0.805 | 2.50E-24 | 6 | Pdia5 | intermediate |
| Tmem150a1 | 1.46E-28 | 0.45614005 | 0.722 | 0.586 | 2.83E-24 | 6 | Tmem150a | intermediate |
| Hoxc8 | 1.51E-28 | 0.3749416 | 0.768 | 0.665 | 2.91E-24 | 6 | Hoxc8 | intermediate |
| Mid2 | 1.56E-28 | 0.84815602 | 0.399 | 0.208 | 3.02E-24 | 6 | Mid2 | intermediate |
| Taldo11 | 1.72E-28 | 0.38176278 | 0.74 | 0.647 | 3.32E-24 | 6 | Taldo1 | intermediate |
| Fuz1 | 3.96E-28 | 0.89032305 | 0.381 | 0.21 | 7.66E-24 | 6 | Fuz | intermediate |
| Oxct12 | 4.50E-28 | 0.46723444 | 0.684 | 0.544 | 8.71E-24 | 6 | Oxct1 | intermediate |
| Abca21 | 5.15E-28 | 1.33310504 | 0.318 | 0.146 | 9.96E-24 | 6 | Abca2 | intermediate |
| Aff2 | 5.46E-28 | 1.12546146 | 0.273 | 0.152 | 1.06E-23 | 6 | Aff2 | intermediate |
| Trib22 | 5.52E-28 | 0.75118973 | 0.369 | 0.265 | 1.07E-23 | 6 | Trib2 | intermediate |
| Gpam1 | 5.56E-28 | 0.7387565 | 0.485 | 0.361 | 1.07E-23 | 6 | Gpam | intermediate |
| Stat62 | 6.78E-28 | 0.34535342 | 0.785 | 0.679 | 1.31E-23 | 6 | Stat6 | intermediate |
| Lmo4 | 7.55E-28 | 0.3542274 | 0.841 | 0.724 | 1.46E-23 | 6 | Lmo4 | intermediate |
| Gusb | 7.90E-28 | 0.39018764 | 0.785 | 0.669 | 1.53E-23 | 6 | Gusb | intermediate |
| Dact3 | 8.94E-28 | 0.46857749 | 0.654 | 0.508 | 1.73E-23 | 6 | Dact3 | intermediate |
| Sesn11 | 1.01E-27 | 1.01374083 | 0.364 | 0.187 | 1.96E-23 | 6 | Sesn1 | intermediate |
| Lpcat32 | 1.27E-27 | 0.41561396 | 0.747 | 0.654 | 2.45E-23 | 6 | Lpcat3 | intermediate |
| Jade12 | 1.31E-27 | 0.54040412 | 0.54 | 0.433 | 2.54E-23 | 6 | Jade1 | intermediate |
| Galnt13 | 1.40E-27 | 0.29066046 | 0.871 | 0.835 | 2.70E-23 | 6 | Galnt1 | intermediate |
| Thsd1 | 2.02E-27 | 1.32869437 | 0.255 | 0.105 | 3.90E-23 | 6 | Thsd1 | intermediate |
| Pitpna | 2.14E-27 | 0.25465107 | 0.899 | 0.832 | 4.13E-23 | 6 | Pitpna | intermediate |
| Yipf3 | 2.17E-27 | 0.31653965 | 0.914 | 0.849 | 4.20E-23 | 6 | Yipf3 | intermediate |
| Mef2c2 | 2.71E-27 | 0.66082347 | 0.447 | 0.274 | 5.25E-23 | 6 | Mef2c | intermediate |
| Boc | 3.04E-27 | 0.46881423 | 0.515 | 0.423 | 5.88E-23 | 6 | Boc | intermediate |
| Galk21 | 3.63E-27 | 0.49218774 | 0.636 | 0.517 | 7.02E-23 | 6 | Galk2 | intermediate |
| Rabgap1l1 | 4.59E-27 | 0.45902567 | 0.664 | 0.564 | 8.88E-23 | 6 | Rabgap1l | intermediate |
| Usp25 | 4.79E-27 | 0.36428906 | 0.785 | 0.72 | 9.25E-23 | 6 | Usp25 | intermediate |
| Adk2 | 5.41E-27 | 0.59390052 | 0.662 | 0.493 | 1.05E-22 | 6 | Adk | intermediate |
| C77080 | 5.57E-27 | 0.70878573 | 0.318 | 0.169 | 1.08E-22 | 6 | C77080 | intermediate |
| Setbp11 | 6.01E-27 | 0.5130795 | 0.712 | 0.575 | 1.16E-22 | 6 | Setbp1 | intermediate |
| Gclm1 | 8.68E-27 | 0.69067683 | 0.535 | 0.325 | 1.68E-22 | 6 | Gclm | intermediate |
| Myo1d1 | 1.07E-26 | 0.36676636 | 0.654 | 0.569 | 2.08E-22 | 6 | Myo1d | intermediate |
| Irs11 | 1.43E-26 | 0.30072906 | 0.737 | 0.695 | 2.77E-22 | 6 | Irs1 | intermediate |
| Zc2hc1a | 4.56E-26 | 0.61813941 | 0.551 | 0.402 | 8.81E-22 | 6 | Zc2hc1a | intermediate |
| Arhgap101 | 5.41E-26 | 0.51599904 | 0.682 | 0.516 | 1.05E-21 | 6 | Arhgap10 | intermediate |
| Tm2d21 | 6.42E-26 | 0.37920191 | 0.876 | 0.798 | 1.24E-21 | 6 | Tm2d2 | intermediate |
| Zdhhc81 | 8.64E-26 | 0.45952599 | 0.816 | 0.707 | 1.67E-21 | 6 | Zdhhc8 | intermediate |
| Sgce1 | 9.04E-26 | 0.35143082 | 0.823 | 0.714 | 1.75E-21 | 6 | Sgce | intermediate |
| Fkbp52 | 1.22E-25 | 0.54060746 | 0.659 | 0.514 | 2.36E-21 | 6 | Fkbp5 | intermediate |
| Dnpep1 | 1.25E-25 | 0.3782621 | 0.785 | 0.678 | 2.41E-21 | 6 | Dnpep | intermediate |
| Ptpn131 | 1.27E-25 | 0.73794273 | 0.621 | 0.443 | 2.46E-21 | 6 | Ptpn13 | intermediate |
| Hacd41 | 1.41E-25 | 0.58737145 | 0.465 | 0.346 | 2.72E-21 | 6 | Hacd4 | intermediate |
| Snap47 | 1.69E-25 | 0.53474981 | 0.646 | 0.536 | 3.27E-21 | 6 | Snap47 | intermediate |
| Pamr12 | 1.77E-25 | 0.4301786 | 0.283 | 0.175 | 3.41E-21 | 6 | Pamr1 | intermediate |
| Calhm5 | 1.99E-25 | 1.08331076 | 0.28 | 0.13 | 3.85E-21 | 6 | Calhm5 | intermediate |
| Tnfsf121 | 2.43E-25 | 0.37880074 | 0.864 | 0.757 | 4.70E-21 | 6 | Tnfsf12 | intermediate |
| Hectd3 | 2.45E-25 | 0.5993161 | 0.616 | 0.441 | 4.73E-21 | 6 | Hectd3 | intermediate |
| Hfe3 | 2.55E-25 | 0.97374149 | 0.351 | 0.207 | 4.93E-21 | 6 | Hfe | intermediate |
| Fam171a12 | 2.58E-25 | 0.57862727 | 0.689 | 0.529 | 4.98E-21 | 6 | Fam171a1 | intermediate |
| Clstn11 | 2.63E-25 | 0.30384968 | 0.856 | 0.756 | 5.09E-21 | 6 | Clstn1 | intermediate |
| Uba72 | 2.90E-25 | 0.61293095 | 0.551 | 0.385 | 5.61E-21 | 6 | Uba7 | intermediate |
| Pxk2 | 3.77E-25 | 0.38885383 | 0.798 | 0.701 | 7.29E-21 | 6 | Pxk | intermediate |
| Plagl13 | 5.06E-25 | 0.39193679 | 0.336 | 0.273 | 9.78E-21 | 6 | Plagl1 | intermediate |
| Htatip21 | 5.23E-25 | 0.46995557 | 0.576 | 0.462 | 1.01E-20 | 6 | Htatip2 | intermediate |
| Tbcb1 | 6.82E-25 | 0.28648842 | 0.864 | 0.79 | 1.32E-20 | 6 | Tbcb | intermediate |
| Ndufa41 | 1.04E-24 | 0.44065333 | 0.72 | 0.594 | 2.02E-20 | 6 | Ndufa4 | intermediate |
| Cd14 | 1.28E-24 | 0.47930752 | 0.26 | 0.177 | 2.47E-20 | 6 | Cd14 | intermediate |
| Syne32 | 1.30E-24 | 0.71055921 | 0.508 | 0.378 | 2.52E-20 | 6 | Syne3 | intermediate |
| Fes3 | 1.62E-24 | 0.57967961 | 0.553 | 0.375 | 3.12E-20 | 6 | Fes | intermediate |
| Ctnnbip1 | 1.72E-24 | 0.58187446 | 0.606 | 0.447 | 3.32E-20 | 6 | Ctnnbip1 | intermediate |
| Matn21 | 2.03E-24 | 0.48088926 | 0.758 | 0.665 | 3.93E-20 | 6 | Matn2 | intermediate |
| Hadh1 | 2.04E-24 | 0.41281192 | 0.667 | 0.529 | 3.94E-20 | 6 | Hadh | intermediate |
| Il18bp | 2.08E-24 | 0.8194905 | 0.29 | 0.139 | 4.02E-20 | 6 | Il18bp | intermediate |
| Pde8a2 | 2.12E-24 | 0.41527107 | 0.53 | 0.328 | 4.10E-20 | 6 | Pde8a | intermediate |
| Txndc9 | 2.29E-24 | 0.31198491 | 0.775 | 0.694 | 4.43E-20 | 6 | Txndc9 | intermediate |
| Mlkl2 | 4.16E-24 | 0.60191834 | 0.513 | 0.367 | 8.05E-20 | 6 | Mlkl | intermediate |
| Wwp11 | 4.21E-24 | 0.46888382 | 0.616 | 0.502 | 8.14E-20 | 6 | Wwp1 | intermediate |
| Hspb12 | 5.61E-24 | 0.3930728 | 0.538 | 0.387 | 1.08E-19 | 6 | Hspb1 | intermediate |
| Ptk2b2 | 5.90E-24 | 0.81198245 | 0.449 | 0.27 | 1.14E-19 | 6 | Ptk2b | intermediate |
| Apobec3 | 6.24E-24 | 0.65149766 | 0.588 | 0.433 | 1.21E-19 | 6 | Apobec3 | intermediate |
| Plaat31 | 6.60E-24 | 0.28431762 | 0.654 | 0.614 | 1.28E-19 | 6 | Plaat3 | intermediate |
| Bhlhe412 | 8.39E-24 | 0.51468031 | 0.407 | 0.299 | 1.62E-19 | 6 | Bhlhe41 | intermediate |
| Decr11 | 8.67E-24 | 0.46889078 | 0.631 | 0.495 | 1.68E-19 | 6 | Decr1 | intermediate |
| Arsb2 | 9.11E-24 | 0.45327886 | 0.775 | 0.628 | 1.76E-19 | 6 | Arsb | intermediate |
| Nedd91 | 1.44E-23 | 0.59260284 | 0.442 | 0.374 | 2.79E-19 | 6 | Nedd9 | intermediate |
| Fam129a2 | 1.59E-23 | 0.41323122 | 0.523 | 0.361 | 3.08E-19 | 6 | Fam129a | intermediate |
| Ccng11 | 1.82E-23 | 0.32975515 | 0.833 | 0.72 | 3.51E-19 | 6 | Ccng1 | intermediate |
| Uap1l11 | 2.06E-23 | 0.51600938 | 0.591 | 0.47 | 3.98E-19 | 6 | Uap1l1 | intermediate |
| Adar1 | 2.65E-23 | 0.42937118 | 0.586 | 0.463 | 5.13E-19 | 6 | Adar | intermediate |
| Gba | 3.00E-23 | 0.34165021 | 0.755 | 0.661 | 5.81E-19 | 6 | Gba | intermediate |
| Hoxc4 | 7.18E-23 | 0.91293196 | 0.328 | 0.167 | 1.39E-18 | 6 | Hoxc4 | intermediate |
| Ang2 | 9.42E-23 | 0.4781726 | 0.578 | 0.421 | 1.82E-18 | 6 | Ang | intermediate |
| Galnt101 | 1.53E-22 | 0.31345181 | 0.801 | 0.714 | 2.95E-18 | 6 | Galnt10 | intermediate |
| Abcd3 | 1.54E-22 | 0.55011 | 0.535 | 0.401 | 2.97E-18 | 6 | Abcd3 | intermediate |
| Bahcc13 | 2.19E-22 | 0.73770364 | 0.47 | 0.316 | 4.23E-18 | 6 | Bahcc1 | intermediate |
| Kidins2201 | 2.22E-22 | 0.38954675 | 0.712 | 0.615 | 4.30E-18 | 6 | Kidins220 | intermediate |
| Gne1 | 2.35E-22 | 0.7730636 | 0.419 | 0.243 | 4.55E-18 | 6 | Gne | intermediate |
| Gpd1l1 | 2.78E-22 | 0.85779977 | 0.386 | 0.206 | 5.37E-18 | 6 | Gpd1l | intermediate |
| Slc25a201 | 3.55E-22 | 0.348192 | 0.657 | 0.546 | 6.85E-18 | 6 | Slc25a20 | intermediate |
| Entpd52 | 3.74E-22 | 0.52938566 | 0.591 | 0.418 | 7.23E-18 | 6 | Entpd5 | intermediate |
| Gnptab | 6.67E-22 | 0.42129495 | 0.727 | 0.608 | 1.29E-17 | 6 | Gnptab | intermediate |
| Gnptg | 1.11E-21 | 0.58162606 | 0.636 | 0.458 | 2.15E-17 | 6 | Gnptg | intermediate |
| Dipk1a1 | 1.17E-21 | 1.02376715 | 0.27 | 0.133 | 2.26E-17 | 6 | Dipk1a | intermediate |
| Mcur1 | 1.19E-21 | 0.62181921 | 0.548 | 0.387 | 2.31E-17 | 6 | Mcur1 | intermediate |
| Ehbp11 | 1.57E-21 | 0.73475501 | 0.462 | 0.292 | 3.03E-17 | 6 | Ehbp1 | intermediate |
| Sash11 | 1.59E-21 | 0.29112663 | 0.836 | 0.766 | 3.07E-17 | 6 | Sash1 | intermediate |
| Radil1 | 1.83E-21 | 0.97609621 | 0.348 | 0.206 | 3.55E-17 | 6 | Radil | intermediate |
| Klf72 | 1.95E-21 | 0.58197736 | 0.583 | 0.428 | 3.78E-17 | 6 | Klf7 | intermediate |
| Cpt21 | 2.26E-21 | 0.61965811 | 0.588 | 0.421 | 4.37E-17 | 6 | Cpt2 | intermediate |
| Ttc30b | 2.48E-21 | 0.85034292 | 0.306 | 0.171 | 4.79E-17 | 6 | Ttc30b | intermediate |
| Ackr11 | 2.65E-21 | 0.63600296 | 0.27 | 0.12 | 5.13E-17 | 6 | Ackr1 | intermediate |
| Plgrkt1 | 2.90E-21 | 0.42730823 | 0.601 | 0.444 | 5.61E-17 | 6 | Plgrkt | intermediate |
| Cd551 | 3.60E-21 | 0.47448039 | 0.379 | 0.153 | 6.95E-17 | 6 | Cd55 | intermediate |
| Tle51 | 4.02E-21 | 0.26690802 | 0.866 | 0.773 | 7.77E-17 | 6 | Tle5 | intermediate |
| Scfd11 | 4.12E-21 | 0.32588019 | 0.773 | 0.688 | 7.98E-17 | 6 | Scfd1 | intermediate |
| Ide | 4.80E-21 | 0.29691384 | 0.808 | 0.707 | 9.28E-17 | 6 | Ide | intermediate |
| Coro2b1 | 5.18E-21 | 0.94780217 | 0.311 | 0.157 | 1.00E-16 | 6 | Coro2b | intermediate |
| Cyfip12 | 5.20E-21 | 0.25629619 | 0.821 | 0.732 | 1.01E-16 | 6 | Cyfip1 | intermediate |
| Gstm52 | 6.20E-21 | 0.66164662 | 0.439 | 0.304 | 1.20E-16 | 6 | Gstm5 | intermediate |
| Slc10a62 | 6.92E-21 | 0.65649171 | 0.402 | 0.232 | 1.34E-16 | 6 | Slc10a6 | intermediate |
| Mpdz2 | 7.81E-21 | 0.57244942 | 0.606 | 0.459 | 1.51E-16 | 6 | Mpdz | intermediate |
| Parp101 | 8.26E-21 | 0.89671306 | 0.311 | 0.2 | 1.60E-16 | 6 | Parp10 | intermediate |
| Txndc151 | 9.69E-21 | 0.29256056 | 0.788 | 0.716 | 1.87E-16 | 6 | Txndc15 | intermediate |
| Stx4a2 | 1.09E-20 | 0.28855028 | 0.848 | 0.733 | 2.10E-16 | 6 | Stx4a | intermediate |
| Smpd1 | 1.16E-20 | 0.42171603 | 0.737 | 0.653 | 2.24E-16 | 6 | Smpd1 | intermediate |

| | | | | | | | | |
|---|---|---|---|---|---|---|---|---|
| Etfbkmt1 | 1.26E-20 | 0.72937049 | 0.364 | 0.245 | 2.44E-16 | 6 | Etfbkmt | intermediate |
| Ccdc252 | 1.29E-20 | 0.42862131 | 0.631 | 0.489 | 2.50E-16 | 6 | Ccdc25 | intermediate |
| Edf1 | 1.31E-20 | 0.25401212 | 0.823 | 0.779 | 2.52E-16 | 6 | Edf1 | intermediate |
| Syne22 | 1.42E-20 | 0.7865264 | 0.444 | 0.246 | 2.75E-16 | 6 | Syne2 | intermediate |
| Zfp354c1 | 1.48E-20 | 0.79267014 | 0.475 | 0.325 | 2.87E-16 | 6 | Zfp354c | intermediate |
| Tep11 | 1.62E-20 | 0.47612123 | 0.551 | 0.397 | 3.13E-16 | 6 | Tep1 | intermediate |
| Tmem256 | 1.72E-20 | 0.28182548 | 0.881 | 0.801 | 3.32E-16 | 6 | Tmem256 | intermediate |
| Ksr12 | 1.84E-20 | 0.61162133 | 0.487 | 0.373 | 3.56E-16 | 6 | Ksr1 | intermediate |
| Parp111 | 1.93E-20 | 1.06036993 | 0.29 | 0.183 | 3.73E-16 | 6 | Parp11 | intermediate |
| Pir | 2.58E-20 | 1.11563382 | 0.253 | 0.114 | 4.98E-16 | 6 | Pir | intermediate |
| Fyn1 | 2.59E-20 | 0.31768812 | 0.672 | 0.561 | 5.01E-16 | 6 | Fyn | intermediate |
| Pcbp4 | 2.94E-20 | 0.40989331 | 0.674 | 0.559 | 5.68E-16 | 6 | Pcbp4 | intermediate |
| Insyn11 | 3.21E-20 | 0.39959867 | 0.621 | 0.548 | 6.21E-16 | 6 | Insyn1 | intermediate |
| Ppp3ca1 | 3.23E-20 | 0.26760028 | 0.803 | 0.718 | 6.25E-16 | 6 | Ppp3ca | intermediate |
| Dzip12 | 3.29E-20 | 0.62363852 | 0.503 | 0.344 | 6.36E-16 | 6 | Dzip1 | intermediate |
| Ccdc8 | 3.33E-20 | 0.29364822 | 0.753 | 0.643 | 6.44E-16 | 6 | Ccdc8 | intermediate |
| Prkca1 | 3.41E-20 | 0.41598205 | 0.444 | 0.342 | 6.58E-16 | 6 | Prkca | intermediate |
| Rgs19 | 4.44E-20 | 0.36523733 | 0.636 | 0.528 | 8.59E-16 | 6 | Rgs19 | intermediate |
| Ap1s2 | 4.62E-20 | 0.39554894 | 0.472 | 0.374 | 8.94E-16 | 6 | Ap1s2 | intermediate |
| Cetn32 | 5.04E-20 | 0.3194191 | 0.727 | 0.642 | 9.75E-16 | 6 | Cetn3 | intermediate |
| Crot1 | 5.21E-20 | 0.65551429 | 0.455 | 0.319 | 1.01E-15 | 6 | Crot | intermediate |
| Jtb | 6.00E-20 | 0.33120743 | 0.798 | 0.676 | 1.16E-15 | 6 | Jtb | intermediate |
| Hmcn12 | 6.01E-20 | 0.61120468 | 0.293 | 0.246 | 1.16E-15 | 6 | Hmcn1 | intermediate |
| Tmem106b2 | 6.76E-20 | 0.28893906 | 0.788 | 0.713 | 1.31E-15 | 6 | Tmem106b | intermediate |
| Plekhg11 | 7.07E-20 | 0.69869679 | 0.313 | 0.195 | 1.37E-15 | 6 | Plekhg1 | intermediate |
| Tprgl1 | 1.10E-19 | 0.39457375 | 0.753 | 0.596 | 2.13E-15 | 6 | Tprgl | intermediate |
| Map4k31 | 1.27E-19 | 0.32169823 | 0.667 | 0.567 | 2.45E-15 | 6 | Map4k3 | intermediate |
| Klf10 | 1.43E-19 | 0.5473738 | 0.583 | 0.404 | 2.76E-15 | 6 | Klf10 | intermediate |
| Slc48a11 | 1.43E-19 | 0.49512969 | 0.561 | 0.452 | 2.77E-15 | 6 | Slc48a1 | intermediate |
| Prcp1 | 1.79E-19 | 0.56981821 | 0.48 | 0.34 | 3.45E-15 | 6 | Prcp | intermediate |
| Aldh1a11 | 2.03E-19 | 0.47348322 | 0.316 | 0.125 | 3.92E-15 | 6 | Aldh1a1 | intermediate |
| Limk12 | 2.09E-19 | 0.27836146 | 0.73 | 0.664 | 4.05E-15 | 6 | Limk1 | intermediate |
| Plekhg31 | 2.13E-19 | 0.52012182 | 0.639 | 0.495 | 4.11E-15 | 6 | Plekhg3 | intermediate |
| Lrch11 | 2.20E-19 | 0.74048227 | 0.424 | 0.233 | 4.26E-15 | 6 | Lrch1 | intermediate |
| Slc31a21 | 2.30E-19 | 0.63703432 | 0.424 | 0.257 | 4.45E-15 | 6 | Slc31a2 | intermediate |
| Dock61 | 2.41E-19 | 0.65716678 | 0.462 | 0.307 | 4.66E-15 | 6 | Dock6 | intermediate |
| Tmc61 | 2.96E-19 | 0.57369088 | 0.503 | 0.358 | 5.73E-15 | 6 | Tmc6 | intermediate |
| Nagpa1 | 4.60E-19 | 0.60506528 | 0.525 | 0.353 | 8.89E-15 | 6 | Nagpa | intermediate |
| Tmem205 | 4.67E-19 | 0.77973528 | 0.328 | 0.188 | 9.02E-15 | 6 | Tmem205 | intermediate |
| Ophn1 | 5.05E-19 | 0.94341261 | 0.348 | 0.181 | 9.77E-15 | 6 | Ophn1 | intermediate |
| Ggcx | 5.28E-19 | 0.25886005 | 0.854 | 0.8 | 1.02E-14 | 6 | Ggcx | intermediate |
| Ift272 | 6.04E-19 | 0.45725239 | 0.662 | 0.517 | 1.17E-14 | 6 | Ift27 | intermediate |
| Spsb2 | 7.79E-19 | 0.733594 | 0.508 | 0.332 | 1.51E-14 | 6 | Spsb2 | intermediate |
| Neo12 | 8.11E-19 | 0.36651242 | 0.71 | 0.649 | 1.57E-14 | 6 | Neo1 | intermediate |
| Fam171a21 | 9.59E-19 | 0.52729642 | 0.482 | 0.347 | 1.85E-14 | 6 | Fam171a2 | intermediate |
| Gfra12 | 1.40E-18 | 0.42573842 | 0.523 | 0.343 | 2.70E-14 | 6 | Gfra1 | intermediate |
| Gulp11 | 1.43E-18 | 0.44310841 | 0.503 | 0.421 | 2.76E-14 | 6 | Gulp1 | intermediate |
| Tmem9b | 1.47E-18 | 0.3125919 | 0.732 | 0.657 | 2.84E-14 | 6 | Tmem9b | intermediate |
| Ece21 | 1.66E-18 | 0.90468768 | 0.328 | 0.204 | 3.20E-14 | 6 | Ece2 | intermediate |
| Fam53b | 1.81E-18 | 0.65072673 | 0.462 | 0.332 | 3.49E-14 | 6 | Fam53b | intermediate |
| Pccb1 | 1.86E-18 | 0.57066377 | 0.497 | 0.367 | 3.59E-14 | 6 | Pccb | intermediate |
| Zdhhc17 | 2.10E-18 | 0.47794627 | 0.563 | 0.419 | 4.06E-14 | 6 | Zdhhc1 | intermediate |
| Cdh22 | 2.11E-18 | 0.643222 | 0.359 | 0.267 | 4.07E-14 | 6 | Cdh2 | intermediate |
| Cyb5611 | 2.23E-18 | 0.53649286 | 0.528 | 0.375 | 4.31E-14 | 6 | Cyb561 | intermediate |
| Nck11 | 2.32E-18 | 0.46076683 | 0.659 | 0.482 | 4.49E-14 | 6 | Nck1 | intermediate |
| Taok31 | 2.43E-18 | 0.64662973 | 0.556 | 0.411 | 4.71E-14 | 6 | Taok3 | intermediate |
| Emc21 | 2.96E-18 | 0.3698816 | 0.697 | 0.574 | 5.73E-14 | 6 | Emc2 | intermediate |
| Fbxw171 | 3.04E-18 | 0.48882734 | 0.427 | 0.285 | 5.87E-14 | 6 | Fbxw17 | intermediate |
| Acyp22 | 3.67E-18 | 0.56924108 | 0.306 | 0.189 | 7.09E-14 | 6 | Acyp2 | intermediate |
| Rcor31 | 4.06E-18 | 0.48761746 | 0.583 | 0.415 | 7.84E-14 | 6 | Rcor3 | intermediate |
| Pmm11 | 4.34E-18 | 0.28930162 | 0.674 | 0.584 | 8.39E-14 | 6 | Pmm1 | intermediate |
| Npas21 | 5.37E-18 | 0.53213961 | 0.568 | 0.45 | 1.04E-13 | 6 | Npas2 | intermediate |
| Cib11 | 6.28E-18 | 0.50955953 | 0.609 | 0.461 | 1.21E-13 | 6 | Cib1 | intermediate |
| Tbx151 | 7.06E-18 | 0.30660166 | 0.783 | 0.675 | 1.37E-13 | 6 | Tbx15 | intermediate |
| Atp8b12 | 1.06E-17 | 0.3316362 | 0.727 | 0.613 | 2.04E-13 | 6 | Atp8b1 | intermediate |
| Chmp2b1 | 1.07E-17 | 0.39608864 | 0.705 | 0.591 | 2.06E-13 | 6 | Chmp2b | intermediate |
| Slc41a31 | 1.42E-17 | 0.64567941 | 0.434 | 0.312 | 2.74E-13 | 6 | Slc41a3 | intermediate |
| Stim11 | 1.57E-17 | 0.61688227 | 0.576 | 0.434 | 3.04E-13 | 6 | Stim1 | intermediate |
| Ggh1 | 1.64E-17 | 0.41876206 | 0.672 | 0.521 | 3.18E-13 | 6 | Ggh | intermediate |
| Cpne22 | 1.91E-17 | 0.74667238 | 0.452 | 0.283 | 3.69E-13 | 6 | Cpne2 | intermediate |
| Glrx2 | 1.91E-17 | 0.29632435 | 0.636 | 0.544 | 3.70E-13 | 6 | Glrx2 | intermediate |
| Hebp22 | 1.98E-17 | 0.51827849 | 0.48 | 0.377 | 3.83E-13 | 6 | Hebp2 | intermediate |
| Tradd | 2.19E-17 | 0.46848267 | 0.631 | 0.496 | 4.24E-13 | 6 | Tradd | intermediate |
| Uprt1 | 2.21E-17 | 0.5211362 | 0.588 | 0.461 | 4.28E-13 | 6 | Uprt | intermediate |
| Atp6v0a12 | 2.23E-17 | 0.5690954 | 0.444 | 0.274 | 4.31E-13 | 6 | Atp6v0a1 | intermediate |
| Aktip1 | 2.35E-17 | 0.59564876 | 0.558 | 0.389 | 4.55E-13 | 6 | Aktip | intermediate |
| Cbr32 | 2.39E-17 | 0.69620728 | 0.336 | 0.215 | 4.63E-13 | 6 | Cbr3 | intermediate |
| Uggt12 | 2.53E-17 | 0.2767071 | 0.871 | 0.793 | 4.90E-13 | 6 | Uggt1 | intermediate |
| Lst1 | 2.68E-17 | 1.00522794 | 0.263 | 0.132 | 5.18E-13 | 6 | Lst1 | intermediate |
| Sspn1 | 2.98E-17 | 0.70570523 | 0.48 | 0.32 | 5.77E-13 | 6 | Sspn | intermediate |
| Eif4e31 | 3.46E-17 | 0.55064288 | 0.712 | 0.552 | 6.69E-13 | 6 | Eif4e3 | intermediate |
| Osbpl1a2 | 3.53E-17 | 0.53812874 | 0.545 | 0.382 | 6.83E-13 | 6 | Osbpl1a | intermediate |
| Tcaf22 | 3.71E-17 | 0.66561391 | 0.338 | 0.215 | 7.18E-13 | 6 | Tcaf2 | intermediate |
| Glb1l1 | 3.72E-17 | 0.58622417 | 0.477 | 0.326 | 7.20E-13 | 6 | Glb1l | intermediate |
| Mgat4b1 | 4.39E-17 | 0.28520203 | 0.591 | 0.53 | 8.48E-13 | 6 | Mgat4b | intermediate |
| C1qtnf122 | 5.74E-17 | 0.32970933 | 0.677 | 0.553 | 1.11E-12 | 6 | C1qtnf12 | intermediate |
| Mpc2 | 5.75E-17 | 0.30159832 | 0.77 | 0.668 | 1.11E-12 | 6 | Mpc2 | intermediate |
| Syngr21 | 8.63E-17 | 0.33016092 | 0.596 | 0.513 | 1.67E-12 | 6 | Syngr2 | intermediate |
| Tcta | 9.58E-17 | 0.59142292 | 0.523 | 0.381 | 1.85E-12 | 6 | Tcta | intermediate |
| Pigk | 9.62E-17 | 0.31655914 | 0.725 | 0.648 | 1.86E-12 | 6 | Pigk | intermediate |
| Hgf1 | 9.95E-17 | 0.43698498 | 0.54 | 0.342 | 1.92E-12 | 6 | Hgf | intermediate |
| Vstm42 | 1.07E-16 | 0.37276391 | 0.619 | 0.495 | 2.06E-12 | 6 | Vstm4 | intermediate |
| Guk12 | 1.12E-16 | 0.31398233 | 0.707 | 0.601 | 2.16E-12 | 6 | Guk1 | intermediate |
| Sntb21 | 1.28E-16 | 0.26593237 | 0.72 | 0.634 | 2.48E-12 | 6 | Sntb2 | intermediate |
| Gcsh2 | 1.38E-16 | 0.32342747 | 0.71 | 0.598 | 2.66E-12 | 6 | Gcsh | intermediate |
| Ccdc1492 | 1.39E-16 | 0.87238112 | 0.275 | 0.157 | 2.70E-12 | 6 | Ccdc149 | intermediate |
| Flii1 | 1.53E-16 | 0.31043526 | 0.745 | 0.649 | 2.96E-12 | 6 | Flii | intermediate |
| Ostm1 | 1.85E-16 | 0.4189884 | 0.621 | 0.512 | 3.58E-12 | 6 | Ostm1 | intermediate |
| Slc4a32 | 1.98E-16 | 0.78890683 | 0.376 | 0.234 | 3.83E-12 | 6 | Slc4a3 | intermediate |
| Fam149a1 | 2.14E-16 | 0.42977925 | 0.619 | 0.485 | 4.14E-12 | 6 | Fam149a | intermediate |
| Ankfy11 | 2.28E-16 | 0.4747687 | 0.646 | 0.512 | 4.41E-12 | 6 | Ankfy1 | intermediate |
| Nub1 | 2.31E-16 | 0.40920345 | 0.619 | 0.5 | 4.46E-12 | 6 | Nub1 | intermediate |
| Ttc39c1 | 2.50E-16 | 0.56273389 | 0.444 | 0.306 | 4.84E-12 | 6 | Ttc39c | intermediate |
| Synj21 | 2.87E-16 | 0.36378743 | 0.561 | 0.442 | 5.56E-12 | 6 | Synj2 | intermediate |
| Gmds3 | 3.29E-16 | 0.45367049 | 0.553 | 0.404 | 6.36E-12 | 6 | Gmds | intermediate |
| Fut112 | 3.75E-16 | 0.31904988 | 0.677 | 0.568 | 7.25E-12 | 6 | Fut11 | intermediate |
| Pacs21 | 4.51E-16 | 0.6144229 | 0.364 | 0.235 | 8.72E-12 | 6 | Pacs2 | intermediate |
| Naprt1 | 4.73E-16 | 0.69928847 | 0.434 | 0.27 | 9.15E-12 | 6 | Naprt | intermediate |
| Cyp4v31 | 5.92E-16 | 0.65902239 | 0.46 | 0.307 | 1.14E-11 | 6 | Cyp4v3 | intermediate |
| Ldb2 | 6.54E-16 | 0.77938973 | 0.258 | 0.135 | 1.26E-11 | 6 | Ldb2 | intermediate |
| Aamdc1 | 7.92E-16 | 0.53034326 | 0.47 | 0.348 | 1.53E-11 | 6 | Aamdc | intermediate |
| Abhd16a1 | 8.15E-16 | 0.37906579 | 0.669 | 0.575 | 1.58E-11 | 6 | Abhd16a | intermediate |
| Xylt11 | 9.71E-16 | 0.46569256 | 0.545 | 0.408 | 1.88E-11 | 6 | Xylt1 | intermediate |
| Hadha | 1.01E-15 | 0.26102607 | 0.803 | 0.714 | 1.94E-11 | 6 | Hadha | intermediate |
| Mcc2 | 1.28E-15 | 0.50806401 | 0.556 | 0.38 | 2.48E-11 | 6 | Mcc | intermediate |
| Uhrf21 | 1.32E-15 | 0.3072769 | 0.753 | 0.642 | 2.54E-11 | 6 | Uhrf2 | intermediate |
| Siae1 | 1.35E-15 | 0.61376905 | 0.437 | 0.256 | 2.61E-11 | 6 | Siae | intermediate |
| Slc49a41 | 1.46E-15 | 0.37078686 | 0.71 | 0.614 | 2.81E-11 | 6 | Slc49a4 | intermediate |
| Mpst2 | 1.70E-15 | 0.48524703 | 0.495 | 0.372 | 3.28E-11 | 6 | Mpst | intermediate |
| Itprid2 | 1.72E-15 | 0.59927618 | 0.487 | 0.378 | 3.33E-11 | 6 | Itprid2 | intermediate |
| Mcub2 | 1.97E-15 | 0.51895144 | 0.518 | 0.352 | 3.81E-11 | 6 | Mcub | intermediate |
| Elf12 | 1.99E-15 | 0.28352431 | 0.694 | 0.572 | 3.84E-11 | 6 | Elf1 | intermediate |
| Rnf133 | 2.02E-15 | 0.30102357 | 0.679 | 0.565 | 3.91E-11 | 6 | Rnf13 | intermediate |
| Ccnd12 | 2.02E-15 | 0.55244286 | 0.295 | 0.199 | 3.91E-11 | 6 | Ccnd1 | intermediate |
| Plcd1 | 2.04E-15 | 0.46557173 | 0.588 | 0.421 | 3.95E-11 | 6 | Plcd1 | intermediate |
| Zfp46 | 2.15E-15 | 0.37869396 | 0.523 | 0.388 | 4.16E-11 | 6 | Zfp46 | intermediate |
| Nol31 | 2.27E-15 | 0.45996685 | 0.391 | 0.27 | 4.39E-11 | 6 | Nol3 | intermediate |
| Cers63 | 2.31E-15 | 0.3761393 | 0.626 | 0.499 | 4.46E-11 | 6 | Cers6 | intermediate |
| Daglb1 | 2.51E-15 | 0.52626906 | 0.376 | 0.228 | 4.86E-11 | 6 | Daglb | intermediate |
| Ccser21 | 2.94E-15 | 0.41807823 | 0.654 | 0.513 | 5.69E-11 | 6 | Ccser2 | intermediate |
| Camk1 | 4.04E-15 | 0.33616819 | 0.755 | 0.639 | 7.81E-11 | 6 | Camk1 | intermediate |
| Gmppa1 | 4.20E-15 | 0.29696383 | 0.747 | 0.644 | 8.12E-11 | 6 | Gmppa | intermediate |
| Prickle22 | 4.29E-15 | 0.44486751 | 0.508 | 0.36 | 8.30E-11 | 6 | Prickle2 | intermediate |
| Map1a | 4.60E-15 | 0.47163286 | 0.568 | 0.429 | 8.89E-11 | 6 | Map1a | intermediate |
| Rmnd5b | 5.11E-15 | 0.51702647 | 0.518 | 0.4 | 9.88E-11 | 6 | Rmnd5b | intermediate |
| Fbxo41 | 5.26E-15 | 0.39261516 | 0.503 | 0.378 | 1.02E-10 | 6 | Fbxo4 | intermediate |
| Lhpp2 | 5.99E-15 | 0.57773112 | 0.343 | 0.21 | 1.16E-10 | 6 | Lhpp | intermediate |
| Tiam2 | 6.12E-15 | 0.54075562 | 0.321 | 0.185 | 1.18E-10 | 6 | Tiam2 | intermediate |
| Tdrd31 | 6.44E-15 | 0.29118624 | 0.629 | 0.513 | 1.25E-10 | 6 | Tdrd3 | intermediate |
| Ift46 | 7.73E-15 | 0.31805114 | 0.715 | 0.62 | 1.50E-10 | 6 | Ift46 | intermediate |
| Pcmtd11 | 7.93E-15 | 0.51403605 | 0.495 | 0.349 | 1.53E-10 | 6 | Pcmtd1 | intermediate |
| Wdsub12 | 7.95E-15 | 0.74660025 | 0.376 | 0.248 | 1.54E-10 | 6 | Wdsub1 | intermediate |
| Zfhx32 | 9.02E-15 | 0.55268418 | 0.495 | 0.329 | 1.74E-10 | 6 | Zfhx3 | intermediate |
| Rit11 | 9.17E-15 | 0.26729994 | 0.674 | 0.599 | 1.77E-10 | 6 | Rit1 | intermediate |
| Zfp4142 | 9.47E-15 | 0.39623211 | 0.646 | 0.53 | 1.83E-10 | 6 | Zfp414 | intermediate |
| Cyp20a11 | 9.62E-15 | 0.26539212 | 0.659 | 0.603 | 1.86E-10 | 6 | Cyp20a1 | intermediate |
| Cd1091 | 9.75E-15 | 0.74076097 | 0.333 | 0.215 | 1.89E-10 | 6 | Cd109 | intermediate |
| Tns11 | 1.14E-14 | 0.46445459 | 0.601 | 0.473 | 2.21E-10 | 6 | Tns1 | intermediate |
| Cc2d2a2 | 1.17E-14 | 0.64800259 | 0.394 | 0.258 | 2.26E-10 | 6 | Cc2d2a | intermediate |
| Sgms22 | 1.63E-14 | 0.55133871 | 0.52 | 0.367 | 3.16E-10 | 6 | Sgms2 | intermediate |
| H2-T221 | 1.70E-14 | 0.53691016 | 0.47 | 0.34 | 3.29E-10 | 6 | H2-T22 | intermediate |
| Pgghg | 1.77E-14 | 0.36250141 | 0.631 | 0.489 | 3.42E-10 | 6 | Pgghg | intermediate |
| Trim12c | 1.80E-14 | 1.05536535 | 0.263 | 0.144 | 3.49E-10 | 6 | Trim12c | intermediate |
| Cdk2ap21 | 2.32E-14 | 0.33132593 | 0.611 | 0.472 | 4.49E-10 | 6 | Cdk2ap2 | intermediate |
| Stoml11 | 3.23E-14 | 0.46651693 | 0.545 | 0.408 | 6.24E-10 | 6 | Stoml1 | intermediate |
| Tango21 | 3.50E-14 | 0.59574075 | 0.472 | 0.323 | 6.77E-10 | 6 | Tango2 | intermediate |
| Gsdmd1 | 3.55E-14 | 0.36394295 | 0.73 | 0.613 | 6.86E-10 | 6 | Gsdmd | intermediate |
| Prkag22 | 3.58E-14 | 0.48187062 | 0.452 | 0.304 | 6.91E-10 | 6 | Prkag2 | intermediate |
| Abhd14a1 | 3.73E-14 | 0.5168965 | 0.52 | 0.377 | 7.21E-10 | 6 | Abhd14a | intermediate |
| Hspa4l | 3.81E-14 | 0.72122883 | 0.283 | 0.182 | 7.36E-10 | 6 | Hspa4l | intermediate |
| Rsrc12 | 4.02E-14 | 0.44023209 | 0.439 | 0.322 | 7.78E-10 | 6 | Rsrc1 | intermediate |
| Sfi11 | 4.32E-14 | 0.39833447 | 0.465 | 0.328 | 8.36E-10 | 6 | Sfi1 | intermediate |
| Bad1 | 4.88E-14 | 0.29547439 | 0.727 | 0.617 | 9.44E-10 | 6 | Bad | intermediate |
| Ltbp22 | 4.97E-14 | 0.34979893 | 0.47 | 0.425 | 9.60E-10 | 6 | Ltbp2 | intermediate |
| AW5519841 | 5.52E-14 | 0.26274003 | 0.503 | 0.412 | 1.07E-09 | 6 | AW551984 | intermediate |
| Mindy12 | 5.78E-14 | 0.40796229 | 0.662 | 0.523 | 1.12E-09 | 6 | Mindy1 | intermediate |
| Vps26a | 6.05E-14 | 0.39410998 | 0.667 | 0.521 | 1.17E-09 | 6 | Vps26a | intermediate |
| Wbp13 | 6.34E-14 | 0.3216891 | 0.578 | 0.469 | 1.22E-09 | 6 | Wbp1 | intermediate |
| Ubl3 | 6.36E-14 | 0.366723 | 0.768 | 0.676 | 1.23E-09 | 6 | Ubl3 | intermediate |
| Tmem160 | 6.72E-14 | 0.30215475 | 0.694 | 0.576 | 1.30E-09 | 6 | Tmem160 | intermediate |
| Nbas2 | 6.91E-14 | 0.27654838 | 0.745 | 0.647 | 1.34E-09 | 6 | Nbas | intermediate |
| Hebp13 | 7.17E-14 | 0.29125965 | 0.482 | 0.356 | 1.39E-09 | 6 | Hebp1 | intermediate |
| Hip1 | 7.90E-14 | 0.25473879 | 0.53 | 0.442 | 1.53E-09 | 6 | Hip1 | intermediate |
| Trp53inp12 | 7.94E-14 | 0.45883813 | 0.386 | 0.269 | 1.53E-09 | 6 | Trp53inp1 | intermediate |
| Edem21 | 7.98E-14 | 0.3731052 | 0.667 | 0.542 | 1.54E-09 | 6 | Edem2 | intermediate |
| Zbtb8a | 9.51E-14 | 0.61434633 | 0.402 | 0.267 | 1.84E-09 | 6 | Zbtb8a | intermediate |
| Bphl1 | 9.58E-14 | 0.6341554 | 0.404 | 0.285 | 1.85E-09 | 6 | Bphl | intermediate |
| Car133 | 9.69E-14 | 0.46450418 | 0.51 | 0.387 | 1.87E-09 | 6 | Car13 | intermediate |
| H2-Ke61 | 1.07E-13 | 0.3882196 | 0.394 | 0.285 | 2.07E-09 | 6 | H2-Ke6 | intermediate |
| Cnpy32 | 1.09E-13 | 0.34188314 | 0.72 | 0.579 | 2.11E-09 | 6 | Cnpy3 | intermediate |
| Emc91 | 1.11E-13 | 0.65040441 | 0.29 | 0.173 | 2.15E-09 | 6 | Emc9 | intermediate |
| Nudt71 | 1.14E-13 | 0.56870818 | 0.369 | 0.257 | 2.20E-09 | 6 | Nudt7 | intermediate |
| Ncoa72 | 1.14E-13 | 0.54161487 | 0.404 | 0.292 | 2.21E-09 | 6 | Ncoa7 | intermediate |
| Kif1b1 | 1.23E-13 | 0.26166371 | 0.667 | 0.544 | 2.39E-09 | 6 | Kif1b | intermediate |
| Rex1bd | 1.25E-13 | 0.35499593 | 0.679 | 0.541 | 2.42E-09 | 6 | Rex1bd | intermediate |
| D2hgdh | 1.26E-13 | 0.446621 | 0.409 | 0.284 | 2.43E-09 | 6 | D2hgdh | intermediate |
| Tnip1 | 1.27E-13 | 0.53214239 | 0.596 | 0.464 | 2.46E-09 | 6 | Tnip1 | intermediate |
| Prkce2 | 1.41E-13 | 0.60320413 | 0.369 | 0.282 | 2.72E-09 | 6 | Prkce | intermediate |
| Papss22 | 1.57E-13 | 0.33668053 | 0.72 | 0.622 | 3.04E-09 | 6 | Papss2 | intermediate |
| Ap1m11 | 1.61E-13 | 0.34594007 | 0.758 | 0.628 | 3.12E-09 | 6 | Ap1m1 | intermediate |
| Exoc41 | 2.05E-13 | 0.33927666 | 0.74 | 0.624 | 3.97E-09 | 6 | Exoc4 | intermediate |
| Ncoa3 | 2.06E-13 | 0.31500577 | 0.659 | 0.558 | 3.99E-09 | 6 | Ncoa3 | intermediate |
| Dnase1l12 | 2.25E-13 | 0.68256397 | 0.351 | 0.21 | 4.35E-09 | 6 | Dnase1l1 | intermediate |
| Gtf3a1 | 2.28E-13 | 0.37361205 | 0.646 | 0.52 | 4.41E-09 | 6 | Gtf3a | intermediate |
| Rbm432 | 2.36E-13 | 0.54676445 | 0.326 | 0.213 | 4.56E-09 | 6 | Rbm43 | intermediate |
| Lamtor4 | 2.39E-13 | 0.49213953 | 0.439 | 0.33 | 4.62E-09 | 6 | Lamtor4 | intermediate |
| Cryl12 | 2.54E-13 | 0.73070294 | 0.364 | 0.212 | 4.92E-09 | 6 | Cryl1 | intermediate |
| Sh3bp51 | 2.73E-13 | 0.40305223 | 0.535 | 0.433 | 5.28E-09 | 6 | Sh3bp5 | intermediate |
| Sumf21 | 2.74E-13 | 0.45386444 | 0.551 | 0.406 | 5.29E-09 | 6 | Sumf2 | intermediate |
| Acad91 | 2.78E-13 | 0.36737383 | 0.442 | 0.319 | 5.38E-09 | 6 | Acad9 | intermediate |
| Phka11 | 3.23E-13 | 0.86794515 | 0.318 | 0.181 | 6.24E-09 | 6 | Phka1 | intermediate |
| Tnfaip81 | 3.44E-13 | 0.53352104 | 0.348 | 0.227 | 6.64E-09 | 6 | Tnfaip8 | intermediate |
| Mpv171 | 3.91E-13 | 0.25800403 | 0.722 | 0.615 | 7.56E-09 | 6 | Mpv17 | intermediate |
| Dpy19l12 | 3.93E-13 | 0.28514857 | 0.793 | 0.671 | 7.59E-09 | 6 | Dpy19l1 | intermediate |
| Adam332 | 4.04E-13 | 0.5426095 | 0.321 | 0.205 | 7.81E-09 | 6 | Adam33 | intermediate |
| Gpsm22 | 4.51E-13 | 0.44427082 | 0.561 | 0.399 | 8.72E-09 | 6 | Gpsm2 | intermediate |
| Adcy9 | 4.61E-13 | 0.69695712 | 0.263 | 0.139 | 8.92E-09 | 6 | Adcy9 | intermediate |
| Stimate | 4.73E-13 | 0.68136328 | 0.485 | 0.354 | 9.15E-09 | 6 | Stimate | intermediate |
| Arntl3 | 5.26E-13 | 0.44740837 | 0.417 | 0.324 | 1.02E-08 | 6 | Arntl | intermediate |
| Czib | 5.47E-13 | 0.42078379 | 0.699 | 0.557 | 1.06E-08 | 6 | Czib | intermediate |
| Nfatc11 | 6.05E-13 | 0.31030572 | 0.485 | 0.405 | 1.17E-08 | 6 | Nfatc1 | intermediate |
| Pdk22 | 6.10E-13 | 0.64069844 | 0.409 | 0.282 | 1.18E-08 | 6 | Pdk2 | intermediate |
| Ino80c2 | 6.52E-13 | 0.3060485 | 0.487 | 0.404 | 1.26E-08 | 6 | Ino80c | intermediate |
| Cd99l21 | 6.56E-13 | 0.379747 | 0.588 | 0.485 | 1.27E-08 | 6 | Cd99l2 | intermediate |
| Ak51 | 7.38E-13 | 0.55683662 | 0.328 | 0.232 | 1.43E-08 | 6 | Ak5 | intermediate |
| Ttc7 | 7.75E-13 | 0.41159362 | 0.621 | 0.488 | 1.50E-08 | 6 | Ttc7 | intermediate |
| Med111 | 7.86E-13 | 0.44057237 | 0.503 | 0.369 | 1.52E-08 | 6 | Med11 | intermediate |
| Cldn102 | 9.64E-13 | 0.53054332 | 0.331 | 0.215 | 1.86E-08 | 6 | Cldn10 | intermediate |
| Anks3 | 9.65E-13 | 0.3455938 | 0.598 | 0.468 | 1.87E-08 | 6 | Anks3 | intermediate |
| Pdgfd1 | 1.06E-12 | 0.32295039 | 0.356 | 0.262 | 2.04E-08 | 6 | Pdgfd | intermediate |
| Il162 | 1.11E-12 | 0.50730172 | 0.303 | 0.191 | 2.14E-08 | 6 | Il16 | intermediate |
| Tmem175 | 1.16E-12 | 0.60608601 | 0.422 | 0.293 | 2.24E-08 | 6 | Tmem175 | intermediate |
| Pnpla2 | 1.56E-12 | 0.60487762 | 0.326 | 0.272 | 3.01E-08 | 6 | Pnpla2 | intermediate |
| Naaladl21 | 1.79E-12 | 1.03342844 | 0.258 | 0.141 | 3.47E-08 | 6 | Naaladl2 | intermediate |
| Nab11 | 1.92E-12 | 0.26434501 | 0.823 | 0.698 | 3.71E-08 | 6 | Nab1 | intermediate |
| Pyroxd21 | 2.00E-12 | 0.62612813 | 0.427 | 0.302 | 3.87E-08 | 6 | Pyroxd2 | intermediate |
| Zc3hav12 | 2.01E-12 | 0.32164991 | 0.596 | 0.53 | 3.88E-08 | 6 | Zc3hav1 | intermediate |
| Itpkb | 2.04E-12 | 0.39156798 | 0.47 | 0.344 | 3.94E-08 | 6 | Itpkb | intermediate |
| Pbxip12 | 2.21E-12 | 0.31176932 | 0.677 | 0.554 | 4.28E-08 | 6 | Pbxip1 | intermediate |
| Dpp71 | 2.26E-12 | 0.49831841 | 0.419 | 0.331 | 4.37E-08 | 6 | Dpp7 | intermediate |
| Abca71 | 2.56E-12 | 0.71334603 | 0.275 | 0.167 | 4.94E-08 | 6 | Abca7 | intermediate |
| Isg201 | 2.94E-12 | 0.25495103 | 0.54 | 0.477 | 5.68E-08 | 6 | Isg20 | intermediate |
| Tpcn11 | 3.09E-12 | 0.45580565 | 0.543 | 0.418 | 5.97E-08 | 6 | Tpcn1 | intermediate |
| Rasa3 | 3.11E-12 | 0.30159249 | 0.705 | 0.67 | 6.02E-08 | 6 | Rasa3 | intermediate |
| Diaph12 | 3.38E-12 | 0.3482208 | 0.583 | 0.448 | 6.53E-08 | 6 | Diaph1 | intermediate |
| Rab7b2 | 3.63E-12 | 0.42516239 | 0.439 | 0.307 | 7.01E-08 | 6 | Rab7b | intermediate |
| Cln82 | 3.89E-12 | 0.6535761 | 0.361 | 0.22 | 7.51E-08 | 6 | Cln8 | intermediate |
| Mta3 | 3.91E-12 | 0.26926339 | 0.659 | 0.516 | 7.55E-08 | 6 | Mta3 | intermediate |
| Ap3m21 | 4.18E-12 | 0.59964315 | 0.333 | 0.199 | 8.07E-08 | 6 | Ap3m2 | intermediate |
| Mitd11 | 4.39E-12 | 0.64129702 | 0.338 | 0.224 | 8.49E-08 | 6 | Mitd1 | intermediate |
| Runx1t11 | 4.40E-12 | 0.55972768 | 0.412 | 0.29 | 8.51E-08 | 6 | Runx1t1 | intermediate |
| Exoc31 | 4.46E-12 | 0.25342585 | 0.578 | 0.47 | 8.63E-08 | 6 | Exoc3 | intermediate |
| Adamtsl52 | 5.24E-12 | 0.35038954 | 0.548 | 0.496 | 1.01E-07 | 6 | Adamtsl5 | intermediate |
| Acad111 | 6.00E-12 | 0.55211191 | 0.336 | 0.255 | 1.16E-07 | 6 | Acad11 | intermediate |
| Plekhg51 | 6.58E-12 | 0.82722918 | 0.316 | 0.235 | 1.27E-07 | 6 | Plekhg5 | intermediate |
| Mapk71 | 7.78E-12 | 0.57857272 | 0.369 | 0.235 | 1.50E-07 | 6 | Mapk7 | intermediate |
| Acot11 | 1.14E-11 | 0.71464058 | 0.361 | 0.255 | 2.21E-07 | 6 | Acot11 | intermediate |
| Mfsd1 | 1.31E-11 | 0.38150633 | 0.523 | 0.415 | 2.53E-07 | 6 | Mfsd1 | intermediate |
| Lrba1 | 1.33E-11 | 0.7092573 | 0.295 | 0.179 | 2.56E-07 | 6 | Lrba | intermediate |
| Slc23a2 | 1.63E-11 | 0.49940888 | 0.462 | 0.328 | 3.15E-07 | 6 | Slc23a2 | intermediate |
| Nfkbie1 | 1.65E-11 | 0.62937693 | 0.273 | 0.201 | 3.20E-07 | 6 | Nfkbie | intermediate |
| Rnf1391 | 1.89E-11 | 0.42326 | 0.573 | 0.477 | 3.66E-07 | 6 | Rnf139 | intermediate |
| Rogdi1 | 2.00E-11 | 0.45258378 | 0.268 | 0.185 | 3.86E-07 | 6 | Rogdi | intermediate |
| Idua1 | 2.02E-11 | 0.4021858 | 0.399 | 0.27 | 3.90E-07 | 6 | Idua | intermediate |
| Evc3 | 2.06E-11 | 0.55270386 | 0.513 | 0.374 | 3.98E-07 | 6 | Evc | intermediate |
| Echdc21 | 2.21E-11 | 0.74061406 | 0.356 | 0.245 | 4.26E-07 | 6 | Echdc2 | intermediate |
| Prmt21 | 2.34E-11 | 0.38644873 | 0.376 | 0.265 | 4.53E-07 | 6 | Prmt2 | intermediate |
| Zfp451 | 2.56E-11 | 0.42416255 | 0.631 | 0.541 | 4.96E-07 | 6 | Zfp451 | intermediate |
| Apbb11 | 2.61E-11 | 0.45186718 | 0.417 | 0.283 | 5.05E-07 | 6 | Apbb1 | intermediate |
| Jmjd81 | 2.83E-11 | 0.38995706 | 0.497 | 0.4 | 5.47E-07 | 6 | Jmjd8 | intermediate |
| Slc2a131 | 2.94E-11 | 0.45807056 | 0.457 | 0.348 | 5.68E-07 | 6 | Slc2a13 | intermediate |
| Pdp12 | 3.30E-11 | 0.51388578 | 0.338 | 0.233 | 6.37E-07 | 6 | Pdp1 | intermediate |
| Ccdc28b | 3.48E-11 | 0.49527658 | 0.429 | 0.307 | 6.73E-07 | 6 | Ccdc28b | intermediate |
| Slc36a1 | 3.94E-11 | 0.3481579 | 0.518 | 0.398 | 7.62E-07 | 6 | Slc36a1 | intermediate |
| Gstm41 | 4.58E-11 | 0.69972946 | 0.278 | 0.173 | 8.85E-07 | 6 | Gstm4 | intermediate |
| Naa801 | 4.59E-11 | 0.42214124 | 0.475 | 0.343 | 8.88E-07 | 6 | Naa80 | intermediate |
| Sobp1 | 4.94E-11 | 0.41276352 | 0.386 | 0.276 | 9.56E-07 | 6 | Sobp | intermediate |
| Slc1a42 | 6.59E-11 | 0.32476891 | 0.424 | 0.348 | 1.27E-06 | 6 | Slc1a4 | intermediate |
| Dexi | 7.28E-11 | 0.31920864 | 0.636 | 0.543 | 1.41E-06 | 6 | Dexi | intermediate |
| Smarca11 | 7.77E-11 | 0.68439858 | 0.306 | 0.205 | 1.50E-06 | 6 | Smarca1 | intermediate |
| Hoxc91 | 8.72E-11 | 0.45763845 | 0.462 | 0.334 | 1.69E-06 | 6 | Hoxc9 | intermediate |
| C1galt11 | 8.78E-11 | 0.3167859 | 0.583 | 0.484 | 1.70E-06 | 6 | C1galt1 | intermediate |
| Nbeal1 | 9.94E-11 | 0.25131392 | 0.636 | 0.523 | 1.92E-06 | 6 | Nbeal1 | intermediate |
| Akr1b102 | 9.98E-11 | 0.43235307 | 0.472 | 0.348 | 1.93E-06 | 6 | Akr1b10 | intermediate |
| Myorg2 | 1.01E-10 | 0.5054391 | 0.402 | 0.289 | 1.95E-06 | 6 | Myorg | intermediate |
| Ubr1 | 1.08E-10 | 0.29713007 | 0.477 | 0.385 | 2.08E-06 | 6 | Ubr1 | intermediate |
| Dguok | 1.12E-10 | 0.27237804 | 0.52 | 0.42 | 2.16E-06 | 6 | Dguok | intermediate |
| Mapkapk3 | 1.14E-10 | 0.60828449 | 0.25 | 0.157 | 2.20E-06 | 6 | Mapkapk3 | intermediate |
| Ammecr11 | 1.14E-10 | 0.26483958 | 0.679 | 0.581 | 2.21E-06 | 6 | Ammecr1 | intermediate |
| Yipf1 | 1.19E-10 | 0.2648161 | 0.712 | 0.622 | 2.30E-06 | 6 | Yipf1 | intermediate |
| Sec16b | 1.42E-10 | 0.37995166 | 0.649 | 0.521 | 2.75E-06 | 6 | Sec16b | intermediate |
| Prr72 | 1.58E-10 | 0.44147084 | 0.308 | 0.213 | 3.06E-06 | 6 | Prr7 | intermediate |
| Pef1 | 1.65E-10 | 0.27115676 | 0.722 | 0.641 | 3.19E-06 | 6 | Pef1 | intermediate |
| Uri1 | 1.82E-10 | 0.26015029 | 0.614 | 0.527 | 3.53E-06 | 6 | Uri1 | intermediate |
| Tapbpl1 | 1.83E-10 | 0.4664449 | 0.394 | 0.287 | 3.54E-06 | 6 | Tapbpl | intermediate |
| Pdlim41 | 1.88E-10 | 0.3256372 | 0.636 | 0.604 | 3.64E-06 | 6 | Pdlim4 | intermediate |
| H2-M31 | 1.98E-10 | 0.42921569 | 0.5 | 0.377 | 3.82E-06 | 6 | H2-M3 | intermediate |
| Hs1bp31 | 2.04E-10 | 0.39337841 | 0.497 | 0.384 | 3.94E-06 | 6 | Hs1bp3 | intermediate |
| Serf1 | 2.04E-10 | 0.40213156 | 0.475 | 0.37 | 3.94E-06 | 6 | Serf1 | intermediate |
| Eif2ak32 | 2.15E-10 | 0.33303827 | 0.576 | 0.503 | 4.16E-06 | 6 | Eif2ak3 | intermediate |
| Mtto1 | 2.17E-10 | 0.48684484 | 0.455 | 0.32 | 4.20E-06 | 6 | Mtto1 | intermediate |
| Flcn | 2.37E-10 | 0.29953919 | 0.54 | 0.425 | 4.59E-06 | 6 | Flcn | intermediate |
| Oplah1 | 2.39E-10 | 0.66708324 | 0.25 | 0.146 | 4.62E-06 | 6 | Oplah | intermediate |
| Lyrm21 | 2.51E-10 | 0.41518145 | 0.396 | 0.303 | 4.86E-06 | 6 | Lyrm2 | intermediate |
| Zfhx41 | 2.59E-10 | 0.29282592 | 0.616 | 0.519 | 5.00E-06 | 6 | Zfhx4 | intermediate |
| Porcn2 | 2.97E-10 | 0.38204808 | 0.47 | 0.359 | 5.73E-06 | 6 | Porcn | intermediate |
| Nit21 | 3.08E-10 | 0.47038196 | 0.381 | 0.306 | 5.95E-06 | 6 | Nit2 | intermediate |
| Pcbp32 | 3.26E-10 | 0.62426627 | 0.258 | 0.177 | 6.30E-06 | 6 | Pcbp3 | intermediate |
| Pon3 | 3.44E-10 | 0.41775129 | 0.606 | 0.502 | 6.65E-06 | 6 | Pon3 | intermediate |
| B9d1 | 3.47E-10 | 0.44905387 | 0.371 | 0.267 | 6.72E-06 | 6 | B9d1 | intermediate |
| Slc10a31 | 3.58E-10 | 0.44118417 | 0.513 | 0.406 | 6.92E-06 | 6 | Slc10a3 | intermediate |
| Etfdh1 | 3.65E-10 | 0.33210051 | 0.571 | 0.456 | 7.06E-06 | 6 | Etfdh | intermediate |
| Eci1 | 4.01E-10 | 0.33165803 | 0.417 | 0.318 | 7.75E-06 | 6 | Eci1 | intermediate |
| Tctn1 | 4.40E-10 | 0.47459363 | 0.402 | 0.283 | 8.52E-06 | 6 | Tctn1 | intermediate |
| Lysmd4 | 4.83E-10 | 0.5578853 | 0.306 | 0.205 | 9.34E-06 | 6 | Lysmd4 | intermediate |
| Akr1e11 | 4.84E-10 | 0.53595483 | 0.336 | 0.225 | 9.36E-06 | 6 | Akr1e1 | intermediate |
| Ube2e22 | 4.89E-10 | 0.28228104 | 0.679 | 0.581 | 9.46E-06 | 6 | Ube2e2 | intermediate |
| Pdgfc2 | 5.65E-10 | 0.6305101 | 0.275 | 0.192 | 1.09E-05 | 6 | Pdgfc | intermediate |
| Commd6 | 6.36E-10 | 0.29282331 | 0.508 | 0.415 | 1.23E-05 | 6 | Commd6 | intermediate |
| Zfp260 | 6.56E-10 | 0.28090593 | 0.753 | 0.638 | 1.27E-05 | 6 | Zfp260 | intermediate |
| Slc25a282 | 6.65E-10 | 0.32137869 | 0.523 | 0.431 | 1.28E-05 | 6 | Slc25a28 | intermediate |
| Angpt11 | 6.71E-10 | 0.61955803 | 0.328 | 0.203 | 1.30E-05 | 6 | Angpt1 | intermediate |
| Tspan61 | 7.48E-10 | 0.26439482 | 0.644 | 0.572 | 1.45E-05 | 6 | Tspan6 | intermediate |
| Tmem50b1 | 7.58E-10 | 0.34219643 | 0.543 | 0.424 | 1.47E-05 | 6 | Tmem50b | intermediate |
| Traf21 | 8.25E-10 | 0.43295434 | 0.601 | 0.458 | 1.59E-05 | 6 | Traf2 | intermediate |
| Fam98a1 | 8.25E-10 | 0.28584896 | 0.677 | 0.6 | 1.60E-05 | 6 | Fam98a | intermediate |
| Dusp18 | 9.32E-10 | 0.55906849 | 0.275 | 0.181 | 1.80E-05 | 6 | Dusp18 | intermediate |
| Lipe | 9.53E-10 | 0.57355353 | 0.258 | 0.162 | 1.84E-05 | 6 | Lipe | intermediate |
| Lrrk21 | 9.53E-10 | 0.44542048 | 0.518 | 0.387 | 1.84E-05 | 6 | Lrrk2 | intermediate |
| Cox20 | 9.76E-10 | 0.32236057 | 0.47 | 0.353 | 1.89E-05 | 6 | Cox20 | intermediate |
| Ccdc90b1 | 9.88E-10 | 0.28559223 | 0.548 | 0.47 | 1.91E-05 | 6 | Ccdc90b | intermediate |
| Mpp62 | 1.01E-09 | 0.29308735 | 0.54 | 0.495 | 1.96E-05 | 6 | Mpp6 | intermediate |
| Pomk | 1.08E-09 | 0.53462121 | 0.298 | 0.188 | 2.08E-05 | 6 | Pomk | intermediate |
| Tmem2411 | 1.16E-09 | 0.45985512 | 0.371 | 0.251 | 2.23E-05 | 6 | Tmem241 | intermediate |
| Mtmr13 | 1.16E-09 | 0.54504341 | 0.29 | 0.198 | 2.25E-05 | 6 | Mtmr1 | intermediate |
| Cbx61 | 1.35E-09 | 0.30413949 | 0.649 | 0.529 | 2.61E-05 | 6 | Cbx6 | intermediate |
| Prdm51 | 1.41E-09 | 0.2879441 | 0.515 | 0.376 | 2.72E-05 | 6 | Prdm5 | intermediate |
| Alg11 | 1.43E-09 | 0.42843232 | 0.417 | 0.33 | 2.76E-05 | 6 | Alg1 | intermediate |
| Pigb | 1.51E-09 | 0.52077571 | 0.306 | 0.209 | 2.93E-05 | 6 | Pigb | intermediate |
| Nudt16l1 | 1.56E-09 | 0.2527629 | 0.641 | 0.532 | 3.01E-05 | 6 | Nudt16l1 | intermediate |
| Mpc1 | 1.62E-09 | 0.28101912 | 0.667 | 0.604 | 3.13E-05 | 6 | Mpc1 | intermediate |
| Synrg1 | 1.67E-09 | 0.38886353 | 0.311 | 0.211 | 3.23E-05 | 6 | Synrg | intermediate |
| Inpp12 | 1.87E-09 | 0.66749762 | 0.417 | 0.273 | 3.61E-05 | 6 | Inpp1 | intermediate |
| Kcnk62 | 1.93E-09 | 0.54372059 | 0.331 | 0.224 | 3.72E-05 | 6 | Kcnk6 | intermediate |
| Erlin13 | 2.08E-09 | 0.42094423 | 0.394 | 0.284 | 4.03E-05 | 6 | Erlin1 | intermediate |
| Map1b1 | 2.16E-09 | 0.27425172 | 0.609 | 0.508 | 4.18E-05 | 6 | Map1b | intermediate |
| Ano101 | 2.20E-09 | 0.26219326 | 0.763 | 0.658 | 4.25E-05 | 6 | Ano10 | intermediate |
| Hdgfl31 | 2.41E-09 | 0.38783593 | 0.472 | 0.344 | 4.65E-05 | 6 | Hdgfl3 | intermediate |
| Golph3l1 | 2.56E-09 | 0.37467364 | 0.492 | 0.364 | 4.96E-05 | 6 | Golph3l | intermediate |
| Stx161 | 2.93E-09 | 0.31248735 | 0.727 | 0.606 | 5.67E-05 | 6 | Stx16 | intermediate |
| Mcee2 | 3.01E-09 | 0.4415754 | 0.333 | 0.221 | 5.81E-05 | 6 | Mcee | intermediate |
| Synj11 | 3.46E-09 | 0.61941583 | 0.384 | 0.286 | 6.69E-05 | 6 | Synj1 | intermediate |
| Csf2ra3 | 3.47E-09 | 0.27950472 | 0.298 | 0.196 | 6.72E-05 | 6 | Csf2ra | intermediate |
| Tnfaip12 | 3.67E-09 | 0.26343643 | 0.679 | 0.579 | 7.10E-05 | 6 | Tnfaip1 | intermediate |
| Smad32 | 3.98E-09 | 0.25327222 | 0.742 | 0.631 | 7.70E-05 | 6 | Smad3 | intermediate |
| Vipas39 | 4.09E-09 | 0.31524393 | 0.556 | 0.43 | 7.92E-05 | 6 | Vipas39 | intermediate |
| Mtln | 4.10E-09 | 0.38193877 | 0.311 | 0.203 | 7.93E-05 | 6 | Mtln | intermediate |
| Rnaset2a | 4.30E-09 | 0.40089851 | 0.467 | 0.337 | 8.32E-05 | 6 | Rnaset2a | intermediate |
| Ankrd461 | 4.71E-09 | 0.29514436 | 0.621 | 0.478 | 9.11E-05 | 6 | Ankrd46 | intermediate |
| Csad | 4.82E-09 | 0.28236773 | 0.503 | 0.4 | 9.31E-05 | 6 | Csad | intermediate |
| Ncoa11 | 4.83E-09 | 0.31033732 | 0.348 | 0.242 | 9.35E-05 | 6 | Ncoa1 | intermediate |
| Nudt191 | 4.94E-09 | 0.47448925 | 0.467 | 0.35 | 9.56E-05 | 6 | Nudt19 | intermediate |
| Phkb1 | 5.02E-09 | 0.41878584 | 0.384 | 0.287 | 9.71E-05 | 6 | Phkb | intermediate |
| Adck21 | 5.11E-09 | 0.33989243 | 0.348 | 0.25 | 9.89E-05 | 6 | Adck2 | intermediate |
| Mov102 | 5.44E-09 | 0.46434024 | 0.412 | 0.304 | 0.00010522 | 6 | Mov10 | intermediate |
| Polb1 | 5.70E-09 | 0.25815446 | 0.624 | 0.53 | 0.00011024 | 6 | Polb | intermediate |
| Apba11 | 5.82E-09 | 0.36131518 | 0.414 | 0.31 | 0.00011251 | 6 | Apba1 | intermediate |
| Rab233 | 5.93E-09 | 0.32156499 | 0.621 | 0.499 | 0.0001146 | 6 | Rab23 | intermediate |
| St71 | 6.83E-09 | 0.2555321 | 0.48 | 0.418 | 0.000132 | 6 | St7 | intermediate |
| Ccdc93 | 7.17E-09 | 0.35248234 | 0.477 | 0.368 | 0.00013864 | 6 | Ccdc93 | intermediate |
| Wwp2 | 7.21E-09 | 0.41697117 | 0.485 | 0.354 | 0.00013934 | 6 | Wwp2 | intermediate |
| Fbxo25 | 7.41E-09 | 0.38821938 | 0.356 | 0.269 | 0.00014334 | 6 | Fbxo25 | intermediate |
| Hsdl21 | 7.71E-09 | 0.52641739 | 0.399 | 0.272 | 0.000149 | 6 | Hsdl2 | intermediate |
| Megf91 | 7.96E-09 | 0.37670965 | 0.323 | 0.207 | 0.00015386 | 6 | Megf9 | intermediate |
| Psme2b | 8.11E-09 | 0.26469681 | 0.508 | 0.416 | 0.00015677 | 6 | Psme2b | intermediate |
| Agtrap2 | 8.30E-09 | 0.29712119 | 0.442 | 0.34 | 0.0001604 | 6 | Agtrap | intermediate |
| Ccdc1571 | 8.38E-09 | 0.73920171 | 0.376 | 0.265 | 0.00016196 | 6 | Ccdc157 | intermediate |
| Tbc1d25 | 8.53E-09 | 0.55735438 | 0.27 | 0.171 | 0.00016485 | 6 | Tbc1d25 | intermediate |
| Mapkap1 | 8.56E-09 | 0.31807348 | 0.657 | 0.545 | 0.00016541 | 6 | Mapkap1 | intermediate |
| Mmut2 | 8.86E-09 | 0.34525501 | 0.306 | 0.224 | 0.00017129 | 6 | Mmut | intermediate |
| Gypc1 | 9.26E-09 | 0.38795464 | 0.568 | 0.487 | 0.00017903 | 6 | Gypc | intermediate |
| Med142 | 9.52E-09 | 0.29822908 | 0.485 | 0.404 | 0.00018405 | 6 | Med14 | intermediate |
| Krcc1 | 9.63E-09 | 0.25108234 | 0.649 | 0.567 | 0.00018609 | 6 | Krcc1 | intermediate |
| Trim46 | 9.66E-09 | 0.42422623 | 0.394 | 0.311 | 0.0001867 | 6 | Trim46 | intermediate |
| Neu1 | 9.80E-09 | 0.45649839 | 0.376 | 0.28 | 0.00018949 | 6 | Neu1 | intermediate |
| Ppil3 | 9.89E-09 | 0.42503874 | 0.404 | 0.304 | 0.00019118 | 6 | Ppil3 | intermediate |
| Galnt4 | 1.14E-08 | 0.33320618 | 0.538 | 0.415 | 0.00022135 | 6 | Galnt4 | intermediate |
| Pcdhga12 | 1.15E-08 | 0.41288794 | 0.275 | 0.236 | 0.00022315 | 6 | Pcdhga12 | intermediate |
| Plppr2 | 1.28E-08 | 0.26726897 | 0.548 | 0.478 | 0.00024657 | 6 | Plppr2 | intermediate |
| Ift1221 | 1.32E-08 | 0.43407282 | 0.311 | 0.2 | 0.00025497 | 6 | Ift122 | intermediate |
| Armc62 | 1.34E-08 | 0.41656321 | 0.399 | 0.279 | 0.00025999 | 6 | Armc6 | intermediate |
| D6Wsu163e | 1.41E-08 | 0.56751056 | 0.311 | 0.21 | 0.00027245 | 6 | D6Wsu163e | intermediate |
| Mppe11 | 1.48E-08 | 0.44487656 | 0.338 | 0.228 | 0.00028584 | 6 | Mppe1 | intermediate |
| Extl22 | 1.53E-08 | 0.36685399 | 0.462 | 0.362 | 0.00029501 | 6 | Extl2 | intermediate |
| Coro71 | 1.54E-08 | 0.36522269 | 0.396 | 0.319 | 0.00029839 | 6 | Coro7 | intermediate |
| Rasal21 | 1.56E-08 | 0.49106433 | 0.356 | 0.247 | 0.00030144 | 6 | Rasal2 | intermediate |
| Mief2 | 1.57E-08 | 0.58587047 | 0.288 | 0.196 | 0.00030355 | 6 | Mief2 | intermediate |
| Mical31 | 1.61E-08 | 0.42184728 | 0.442 | 0.35 | 0.00031139 | 6 | Mical3 | intermediate |
| Lonp21 | 1.65E-08 | 0.28558544 | 0.525 | 0.439 | 0.0003185 | 6 | Lonp2 | intermediate |
| Surf1 | 1.71E-08 | 0.42068788 | 0.568 | 0.434 | 0.00033092 | 6 | Surf1 | intermediate |
| Tmem41a | 1.78E-08 | 0.33230838 | 0.444 | 0.358 | 0.00034406 | 6 | Tmem41a | intermediate |
| Rmc11 | 1.81E-08 | 0.40266936 | 0.399 | 0.297 | 0.00034898 | 6 | Rmc1 | intermediate |
| Cfap2982 | 1.96E-08 | 0.41085375 | 0.351 | 0.24 | 0.00037853 | 6 | Cfap298 | intermediate |
| Cbr4 | 2.03E-08 | 0.64726072 | 0.384 | 0.256 | 0.00039307 | 6 | Cbr4 | intermediate |
| Hykk1 | 2.14E-08 | 0.50069563 | 0.293 | 0.19 | 0.00041365 | 6 | Hykk | intermediate |
| Fundc1 | 2.26E-08 | 0.25416846 | 0.725 | 0.613 | 0.00043705 | 6 | Fundc1 | intermediate |
| Arl4a1 | 2.48E-08 | 0.32369409 | 0.278 | 0.196 | 0.00047912 | 6 | Arl4a | intermediate |
| Kat2b | 2.48E-08 | 0.61194774 | 0.311 | 0.211 | 0.00047913 | 6 | Kat2b | intermediate |
| Rnf1672 | 2.52E-08 | 0.38336974 | 0.434 | 0.329 | 0.00048796 | 6 | Rnf167 | intermediate |
| Eef1akmt11 | 2.65E-08 | 0.30419689 | 0.51 | 0.415 | 0.00051258 | 6 | Eef1akmt1 | intermediate |
| Pml2 | 2.91E-08 | 0.29182035 | 0.641 | 0.551 | 0.0005624 | 6 | Pml | intermediate |
| Nqo22 | 2.98E-08 | 0.31651126 | 0.571 | 0.473 | 0.00057693 | 6 | Nqo2 | intermediate |
| Fam3a | 3.14E-08 | 0.30072132 | 0.518 | 0.43 | 0.00060655 | 6 | Fam3a | intermediate |
| Slc39a11 | 3.33E-08 | 0.27439274 | 0.644 | 0.539 | 0.00064428 | 6 | Slc39a11 | intermediate |
| Xpc | 3.72E-08 | 0.4897169 | 0.295 | 0.193 | 0.00071862 | 6 | Xpc | intermediate |
| Phyhd11 | 3.90E-08 | 0.69498936 | 0.283 | 0.191 | 0.0007547 | 6 | Phyhd1 | intermediate |
| Hspbap1 | 4.09E-08 | 0.58961982 | 0.293 | 0.208 | 0.0007908 | 6 | Hspbap1 | intermediate |
| Cyb561d21 | 4.09E-08 | 0.46555101 | 0.341 | 0.237 | 0.00079102 | 6 | Cyb561d2 | intermediate |
| Tgfbrap1 | 4.67E-08 | 0.25504581 | 0.414 | 0.302 | 0.00090317 | 6 | Tgfbrap1 | intermediate |
| Slc22a52 | 4.68E-08 | 0.60838357 | 0.323 | 0.218 | 0.00090567 | 6 | Slc22a5 | intermediate |
| Eno21 | 5.32E-08 | 0.28251879 | 0.331 | 0.244 | 0.00102831 | 6 | Eno2 | intermediate |
| Grin2d1 | 5.38E-08 | 0.52489926 | 0.399 | 0.29 | 0.00104005 | 6 | Grin2d | intermediate |
| Mfsd101 | 5.50E-08 | 0.25618778 | 0.722 | 0.636 | 0.00106295 | 6 | Mfsd10 | intermediate |
| Pgap2 | 5.64E-08 | 0.30816803 | 0.475 | 0.391 | 0.00109065 | 6 | Pgap2 | intermediate |
| Man1c1 | 5.71E-08 | 0.27567751 | 0.682 | 0.608 | 0.00110467 | 6 | Man1c1 | intermediate |
| Cyp2j61 | 5.79E-08 | 0.36690293 | 0.318 | 0.232 | 0.00111953 | 6 | Cyp2j6 | intermediate |
| Rcbtb21 | 6.29E-08 | 0.28085729 | 0.682 | 0.566 | 0.00121624 | 6 | Rcbtb2 | intermediate |
| Slc7a71 | 6.56E-08 | 0.49380345 | 0.303 | 0.224 | 0.00126808 | 6 | Slc7a7 | intermediate |
| Clec16a | 6.57E-08 | 0.55102654 | 0.313 | 0.23 | 0.00126939 | 6 | Clec16a | intermediate |
| Rfx51 | 6.60E-08 | 0.70236025 | 0.285 | 0.217 | 0.00127678 | 6 | Rfx5 | intermediate |
| Ap3m11 | 6.67E-08 | 0.37778828 | 0.331 | 0.241 | 0.00128963 | 6 | Ap3m1 | intermediate |
| Mras1 | 6.78E-08 | 0.45163981 | 0.419 | 0.343 | 0.00131047 | 6 | Mras | intermediate |
| Btd | 7.30E-08 | 0.32848291 | 0.51 | 0.401 | 0.00141173 | 6 | Btd | intermediate |
| Hdac102 | 7.33E-08 | 0.51849315 | 0.306 | 0.21 | 0.00141693 | 6 | Hdac10 | intermediate |
| Dcps1 | 7.59E-08 | 0.51064953 | 0.369 | 0.263 | 0.00146663 | 6 | Dcps | intermediate |
| Psph1 | 7.64E-08 | 0.38763188 | 0.366 | 0.264 | 0.00147801 | 6 | Psph | intermediate |
| Parn | 7.77E-08 | 0.33580308 | 0.497 | 0.387 | 0.00150248 | 6 | Parn | intermediate |
| Tcf7l11 | 8.07E-08 | 0.2902195 | 0.48 | 0.404 | 0.00156064 | 6 | Tcf7l1 | intermediate |
| Plcl1 | 8.22E-08 | 0.60672312 | 0.258 | 0.159 | 0.0015894 | 6 | Plcl1 | intermediate |
| Cry22 | 8.22E-08 | 0.27010238 | 0.424 | 0.332 | 0.00159006 | 6 | Cry2 | intermediate |
| Lztfl11 | 8.59E-08 | 0.26620284 | 0.354 | 0.268 | 0.00166123 | 6 | Lztfl1 | intermediate |
| Smim27 | 9.07E-08 | 0.52848259 | 0.278 | 0.205 | 0.00175288 | 6 | Smim27 | intermediate |
| Cyld1 | 9.83E-08 | 0.27744904 | 0.46 | 0.357 | 0.00190069 | 6 | Cyld | intermediate |
| Scfd2 | 1.14E-07 | 0.26083517 | 0.492 | 0.385 | 0.00220794 | 6 | Scfd2 | intermediate |
| Sec22a | 1.20E-07 | 0.31380026 | 0.503 | 0.395 | 0.0023145 | 6 | Sec22a | intermediate |
| Cr1l1 | 1.21E-07 | 0.35596354 | 0.412 | 0.328 | 0.00234476 | 6 | Cr1l | intermediate |
| Rusc12 | 1.31E-07 | 0.32723108 | 0.374 | 0.294 | 0.00253834 | 6 | Rusc1 | intermediate |
| Pter1 | 1.33E-07 | 0.54605101 | 0.29 | 0.187 | 0.00257259 | 6 | Pter | intermediate |
| Fbxw91 | 1.36E-07 | 0.29182546 | 0.346 | 0.254 | 0.00263372 | 6 | Fbxw9 | intermediate |
| Pi4kb | 1.37E-07 | 0.31631908 | 0.414 | 0.312 | 0.00265525 | 6 | Pi4kb | intermediate |
| Pcmtd22 | 1.54E-07 | 0.25038482 | 0.46 | 0.364 | 0.00297235 | 6 | Pcmtd2 | intermediate |
| Gpr272 | 1.59E-07 | 0.50296941 | 0.399 | 0.284 | 0.00307979 | 6 | Gpr27 | intermediate |
| Inpp5k | 1.61E-07 | 0.45901494 | 0.303 | 0.207 | 0.0031131 | 6 | Inpp5k | intermediate |
| Morc3 | 1.99E-07 | 0.27055343 | 0.472 | 0.405 | 0.00383803 | 6 | Morc3 | intermediate |
| Kif3a | 2.08E-07 | 0.30176691 | 0.399 | 0.291 | 0.00403 | 6 | Kif3a | intermediate |
| Ptpdc12 | 2.13E-07 | 0.42655249 | 0.341 | 0.263 | 0.00412608 | 6 | Ptpdc1 | intermediate |
| Rtkn1 | 2.35E-07 | 0.26759425 | 0.371 | 0.282 | 0.00453557 | 6 | Rtkn | intermediate |
| Poglut11 | 2.75E-07 | 0.28871405 | 0.523 | 0.408 | 0.00532487 | 6 | Poglut1 | intermediate |
| Scamp11 | 2.76E-07 | 0.31536427 | 0.452 | 0.367 | 0.00534091 | 6 | Scamp1 | intermediate |
| Vps361 | 3.09E-07 | 0.27652603 | 0.654 | 0.535 | 0.005975 | 6 | Vps36 | intermediate |
| Kiz1 | 3.21E-07 | 0.39603056 | 0.341 | 0.244 | 0.00620229 | 6 | Kiz | intermediate |
| Slc25a16 | 3.32E-07 | 0.28177328 | 0.495 | 0.387 | 0.0064278 | 6 | Slc25a16 | intermediate |
| Taz | 3.44E-07 | 0.45782664 | 0.586 | 0.475 | 0.006644 | 6 | Taz | intermediate |
| Mccc12 | 4.21E-07 | 0.4882751 | 0.371 | 0.261 | 0.0081308 | 6 | Mccc1 | intermediate |
| Ngly12 | 4.28E-07 | 0.28984961 | 0.561 | 0.459 | 0.00827045 | 6 | Ngly1 | intermediate |
| Eci22 | 4.47E-07 | 0.30947995 | 0.576 | 0.498 | 0.0086504 | 6 | Eci2 | intermediate |
| Pts2 | 4.74E-07 | 0.42067739 | 0.318 | 0.222 | 0.0091703 | 6 | Pts | intermediate |
| Sdr39u1 | 5.06E-07 | 0.62242472 | 0.295 | 0.207 | 0.00978852 | 6 | Sdr39u1 | intermediate |
| Tbce1 | 5.40E-07 | 0.33527408 | 0.46 | 0.355 | 0.01043232 | 6 | Tbce | intermediate |
| Ppp2r3a | 5.44E-07 | 0.49029338 | 0.369 | 0.273 | 0.01051196 | 6 | Ppp2r3a | intermediate |
| Nt5c32 | 5.73E-07 | 0.48899031 | 0.336 | 0.286 | 0.01107523 | 6 | Nt5c3 | intermediate |
| Wdr44 | 5.88E-07 | 0.51168811 | 0.27 | 0.188 | 0.01137178 | 6 | Wdr44 | intermediate |

| | | | | | | | | |
|---|---|---|---|---|---|---|---|---|
| Tmem106c2 | 5.97E-07 | 0.51197439 | 0.381 | 0.285 | 0.01154612 | 6 | Tmem106c | intermediate |
| Borcs51 | 6.02E-07 | 0.32015749 | 0.318 | 0.241 | 0.01164555 | 6 | Borcs5 | intermediate |
| Acad8 | 6.33E-07 | 0.36105547 | 0.503 | 0.409 | 0.01224212 | 6 | Acad8 | intermediate |
| C2cd52 | 7.02E-07 | 0.48744365 | 0.326 | 0.232 | 0.01356371 | 6 | C2cd5 | intermediate |
| Heatr31 | 7.38E-07 | 0.36158488 | 0.432 | 0.364 | 0.01427324 | 6 | Heatr3 | intermediate |
| Parp42 | 8.33E-07 | 0.25605605 | 0.48 | 0.372 | 0.01611067 | 6 | Parp4 | intermediate |
| Gamt2 | 1.03E-06 | 0.29373127 | 0.394 | 0.292 | 0.01989839 | 6 | Gamt | intermediate |
| Tbc1d51 | 1.03E-06 | 0.35556723 | 0.5 | 0.383 | 0.01993129 | 6 | Tbc1d5 | intermediate |
| Nsmaf1 | 1.04E-06 | 0.30066839 | 0.495 | 0.424 | 0.02013019 | 6 | Nsmaf | intermediate |
| Arrb12 | 1.05E-06 | 0.2812416 | 0.462 | 0.332 | 0.02022787 | 6 | Arrb1 | intermediate |
| Dapp1 | 1.06E-06 | 0.49481898 | 0.303 | 0.205 | 0.02043923 | 6 | Dapp1 | intermediate |
| Asah21 | 1.06E-06 | 0.27978021 | 0.331 | 0.238 | 0.02044993 | 6 | Asah2 | intermediate |
| Mthfsd | 1.07E-06 | 0.34056885 | 0.427 | 0.336 | 0.02077772 | 6 | Mthfsd | intermediate |
| Trim68 | 1.16E-06 | 0.51106449 | 0.265 | 0.187 | 0.02236862 | 6 | Trim68 | intermediate |
| Ints8 | 1.17E-06 | 0.37189533 | 0.447 | 0.373 | 0.02267832 | 6 | Ints8 | intermediate |
| Zbtb8os | 1.24E-06 | 0.27793334 | 0.404 | 0.331 | 0.02396901 | 6 | Zbtb8os | intermediate |
| Pla2r12 | 1.42E-06 | 0.29017974 | 0.462 | 0.394 | 0.02750471 | 6 | Pla2r1 | intermediate |
| Coq10a1 | 1.45E-06 | 0.55724606 | 0.379 | 0.283 | 0.02809769 | 6 | Coq10a | intermediate |
| Mon1a | 1.46E-06 | 0.36722271 | 0.356 | 0.269 | 0.02826291 | 6 | Mon1a | intermediate |
| Lin37 | 1.50E-06 | 0.6626488 | 0.323 | 0.23 | 0.02900756 | 6 | Lin37 | intermediate |
| Lpcat4 | 1.52E-06 | 0.37680025 | 0.318 | 0.235 | 0.02931772 | 6 | Lpcat4 | intermediate |
| Srgap12 | 1.52E-06 | 0.40215351 | 0.318 | 0.219 | 0.02938866 | 6 | Srgap1 | intermediate |
| Srr1 | 1.54E-06 | 0.37850834 | 0.288 | 0.219 | 0.02973223 | 6 | Srr | intermediate |
| Pld21 | 1.59E-06 | 0.25065833 | 0.439 | 0.346 | 0.03077987 | 6 | Pld2 | intermediate |
| Nit11 | 1.63E-06 | 0.36014873 | 0.356 | 0.268 | 0.03157054 | 6 | Nit1 | intermediate |
| Rom11 | 1.66E-06 | 0.2551347 | 0.333 | 0.259 | 0.03200925 | 6 | Rom1 | intermediate |
| Sept102 | 1.72E-06 | 0.28177361 | 0.563 | 0.428 | 0.0331689 | 6 | 10-Sep | intermediate |
| Thtpa | 1.77E-06 | 0.42001232 | 0.301 | 0.209 | 0.0342457 | 6 | Thtpa | intermediate |
| DEPRECATED-I | 1.77E-06 | 0.34533905 | 0.391 | 0.314 | 0.03429409 | 6 | DEPRECATED-I | intermediate |
| Dnmbp1 | 1.81E-06 | 0.35217823 | 0.495 | 0.353 | 0.03494849 | 6 | Dnmbp | intermediate |
| Zrsr1 | 1.89E-06 | 0.33772324 | 0.258 | 0.188 | 0.03659892 | 6 | Zrsr1 | intermediate |
| Magi31 | 1.98E-06 | 0.54267161 | 0.331 | 0.243 | 0.03823611 | 6 | Magi3 | intermediate |
| Fadd1 | 2.04E-06 | 0.38996034 | 0.482 | 0.376 | 0.03935586 | 6 | Fadd | intermediate |
| Bcs1l1 | 2.04E-06 | 0.33154051 | 0.351 | 0.267 | 0.03951404 | 6 | Bcs1l | intermediate |
| Trp53i131 | 2.05E-06 | 0.3296449 | 0.533 | 0.429 | 0.03971604 | 6 | Trp53i13 | intermediate |
| Sgk31 | 2.18E-06 | 0.64450857 | 0.268 | 0.183 | 0.04221685 | 6 | Sgk3 | intermediate |
| Smim112 | 2.20E-06 | 0.27468728 | 0.576 | 0.473 | 0.04253653 | 6 | Smim11 | intermediate |
| Wrb1 | 2.28E-06 | 0.45548772 | 0.303 | 0.216 | 0.04408089 | 6 | Wrb | intermediate |
| Armc9 | 2.55E-06 | 0.29277514 | 0.313 | 0.225 | 0.04923781 | 6 | Armc9 | intermediate |
| Gdap2 | 2.56E-06 | 0.27127518 | 0.515 | 0.416 | 0.04958427 | 6 | Gdap2 | intermediate |
| Ecsit | 2.66E-06 | 0.30717165 | 0.391 | 0.304 | 0.05142793 | 6 | Ecsit | intermediate |
| Zc3h10 | 3.42E-06 | 0.43482777 | 0.25 | 0.166 | 0.06614706 | 6 | Zc3h10 | intermediate |
| Dop1b2 | 3.42E-06 | 0.36939363 | 0.364 | 0.272 | 0.06617333 | 6 | Dop1b | intermediate |
| Tmem185a | 3.99E-06 | 0.29327249 | 0.364 | 0.279 | 0.07706367 | 6 | Tmem185a | intermediate |
| Armt11 | 4.12E-06 | 0.35685338 | 0.381 | 0.289 | 0.07964303 | 6 | Armt1 | intermediate |
| Serhl1 | 4.13E-06 | 0.27830159 | 0.452 | 0.351 | 0.07991585 | 6 | Serhl | intermediate |
| Ptgs11 | 4.34E-06 | 0.36234223 | 0.384 | 0.258 | 0.0838279 | 6 | Ptgs1 | intermediate |
| Jade21 | 4.39E-06 | 0.44896229 | 0.326 | 0.225 | 0.08483403 | 6 | Jade2 | intermediate |
| Hoxc51 | 4.46E-06 | 0.47659005 | 0.311 | 0.217 | 0.08625871 | 6 | Hoxc5 | intermediate |
| Nacc2 | 4.52E-06 | 0.44317244 | 0.407 | 0.297 | 0.0873336 | 6 | Nacc2 | intermediate |
| Cc2d1b | 4.59E-06 | 0.41492965 | 0.376 | 0.267 | 0.0881085 | 6 | Cc2d1b | intermediate |
| Xrcc51 | 4.60E-06 | 0.57470045 | 0.265 | 0.184 | 0.08890836 | 6 | Xrcc5 | intermediate |
| B4galt41 | 4.70E-06 | 0.46444456 | 0.348 | 0.255 | 0.09079997 | 6 | B4galt4 | intermediate |
| Ndufaf7 | 4.79E-06 | 0.34071417 | 0.369 | 0.285 | 0.09255372 | 6 | Ndufaf7 | intermediate |
| Slc25a451 | 4.87E-06 | 0.42391711 | 0.275 | 0.199 | 0.09422066 | 6 | Slc25a45 | intermediate |
| Ap5s1 | 5.03E-06 | 0.466549 | 0.275 | 0.189 | 0.0972423 | 6 | Ap5s1 | intermediate |
| Mrrf | 6.73E-06 | 0.28184528 | 0.422 | 0.323 | 0.13010048 | 6 | Mrrf | intermediate |
| Shf | 7.46E-06 | 0.35519738 | 0.356 | 0.28 | 0.14419915 | 6 | Shf | intermediate |
| Snx291 | 7.63E-06 | 0.30088396 | 0.556 | 0.468 | 0.14755312 | 6 | Snx29 | intermediate |
| Hibch | 7.74E-06 | 0.43538496 | 0.273 | 0.189 | 0.14966229 | 6 | Hibch | intermediate |
| Chfr | 7.99E-06 | 0.29209706 | 0.551 | 0.438 | 0.15454834 | 6 | Chfr | intermediate |
| Gstz1 | 8.00E-06 | 0.2911781 | 0.376 | 0.297 | 0.15474597 | 6 | Gstz1 | intermediate |
| Uchl32 | 8.02E-06 | 0.36235634 | 0.306 | 0.232 | 0.15514699 | 6 | Uchl3 | intermediate |
| Rpia2 | 8.04E-06 | 0.43182276 | 0.444 | 0.316 | 0.15551807 | 6 | Rpia | intermediate |
| Evc2 | 8.35E-06 | 0.33758802 | 0.679 | 0.551 | 0.16148174 | 6 | Evc2 | intermediate |
| Smim15 | 8.65E-06 | 0.25950954 | 0.598 | 0.534 | 0.16723843 | 6 | Smim15 | intermediate |
| Ssh21 | 1.17E-05 | 0.33049743 | 0.414 | 0.325 | 0.2264887 | 6 | Ssh2 | intermediate |
| Abcd42 | 1.22E-05 | 0.36763559 | 0.313 | 0.213 | 0.23670896 | 6 | Abcd4 | intermediate |
| Tmco6 | 1.23E-05 | 0.50113624 | 0.258 | 0.176 | 0.23849216 | 6 | Tmco6 | intermediate |
| Rnf141 | 1.31E-05 | 0.41121056 | 0.316 | 0.237 | 0.25360153 | 6 | Rnf141 | intermediate |
| Polr3k1 | 1.39E-05 | 0.29968551 | 0.639 | 0.548 | 0.26814767 | 6 | Polr3k | intermediate |
| Endod13 | 1.48E-05 | 0.31017092 | 0.346 | 0.295 | 0.28663262 | 6 | Endod1 | intermediate |
| Rangrf1 | 1.53E-05 | 0.26745672 | 0.386 | 0.319 | 0.29596193 | 6 | Rangrf | intermediate |
| Vps37c1 | 1.65E-05 | 0.29763872 | 0.437 | 0.364 | 0.31963768 | 6 | Vps37c | intermediate |
| Slc29a3 | 1.69E-05 | 0.33102459 | 0.447 | 0.369 | 0.32627438 | 6 | Slc29a3 | intermediate |
| Pecr | 1.95E-05 | 0.34352349 | 0.26 | 0.192 | 0.37672138 | 6 | Pecr | intermediate |
| Tpcn21 | 2.49E-05 | 0.28311637 | 0.434 | 0.351 | 0.48206382 | 6 | Tpcn2 | intermediate |
| Trmt2b1 | 2.62E-05 | 0.33139975 | 0.301 | 0.225 | 0.50714107 | 6 | Trmt2b | intermediate |
| Amacr1 | 2.66E-05 | 0.38786146 | 0.306 | 0.229 | 0.51464213 | 6 | Amacr | intermediate |
| Zfp267 | 2.78E-05 | 0.25398977 | 0.316 | 0.228 | 0.53703753 | 6 | Zfp267 | intermediate |
| Cgrrf11 | 2.78E-05 | 0.33148369 | 0.444 | 0.345 | 0.53772853 | 6 | Cgrrf1 | intermediate |
| Rubcn2 | 3.02E-05 | 0.30486228 | 0.497 | 0.412 | 0.58362746 | 6 | Rubcn | intermediate |
| Pnpt12 | 3.05E-05 | 0.34294289 | 0.384 | 0.3 | 0.58903969 | 6 | Pnpt1 | intermediate |
| Crbn1 | 3.18E-05 | 0.26874245 | 0.434 | 0.365 | 0.61480387 | 6 | Crbn | intermediate |
| Tsen21 | 3.27E-05 | 0.46343448 | 0.263 | 0.19 | 0.63207704 | 6 | Tsen2 | intermediate |
| Cfap97 | 3.57E-05 | 0.30917406 | 0.341 | 0.257 | 0.69041228 | 6 | Cfap97 | intermediate |
| Kdm1b | 3.64E-05 | 0.47808138 | 0.258 | 0.176 | 0.70332586 | 6 | Kdm1b | intermediate |
| Tmem1352 | 3.93E-05 | 0.30585854 | 0.447 | 0.391 | 0.76024023 | 6 | Tmem135 | intermediate |
| Dmap11 | 3.95E-05 | 0.32968622 | 0.328 | 0.25 | 0.7637827 | 6 | Dmap1 | intermediate |
| Lrrcc11 | 4.11E-05 | 0.33098845 | 0.341 | 0.24 | 0.79457526 | 6 | Lrrcc1 | intermediate |
| Dalrd3 | 4.19E-05 | 0.3559802 | 0.298 | 0.223 | 0.80963089 | 6 | Dalrd3 | intermediate |
| Krit11 | 4.29E-05 | 0.26329049 | 0.424 | 0.361 | 0.82988137 | 6 | Krit1 | intermediate |
| Galt2 | 4.40E-05 | 0.30047078 | 0.422 | 0.338 | 0.85065412 | 6 | Galt | intermediate |
| Bmyc | 4.65E-05 | 0.3986192 | 0.25 | 0.177 | 0.89906487 | 6 | Bmyc | intermediate |
| Ano81 | 5.07E-05 | 0.32372746 | 0.391 | 0.314 | 0.98022486 | 6 | Ano8 | intermediate |
| Cep781 | 5.32E-05 | 0.32149128 | 0.253 | 0.192 | 1 | 6 | Cep78 | intermediate |
| Rgs101 | 5.59E-05 | 0.27356602 | 0.424 | 0.361 | 1 | 6 | Rgs10 | intermediate |
| Rpap11 | 5.90E-05 | 0.28476267 | 0.295 | 0.229 | 1 | 6 | Rpap1 | intermediate |
| Armc101 | 5.92E-05 | 0.46510936 | 0.333 | 0.245 | 1 | 6 | Armc10 | intermediate |
| Pde6d1 | 6.24E-05 | 0.30198506 | 0.551 | 0.457 | 1 | 6 | Pde6d | intermediate |
| Vps53 | 6.34E-05 | 0.50457339 | 0.437 | 0.344 | 1 | 6 | Vps53 | intermediate |
| Smarcal1 | 6.35E-05 | 0.46645477 | 0.348 | 0.262 | 1 | 6 | Smarcal1 | intermediate |
| Hspa2 | 6.44E-05 | 0.28361711 | 0.298 | 0.213 | 1 | 6 | Hspa2 | intermediate |
| Sdhaf41 | 6.72E-05 | 0.32030148 | 0.384 | 0.289 | 1 | 6 | Sdhaf4 | intermediate |
| Smarcd3 | 6.75E-05 | 0.34884711 | 0.384 | 0.306 | 1 | 6 | Smarcd3 | intermediate |
| Nkiras1 | 6.84E-05 | 0.44826961 | 0.336 | 0.252 | 1 | 6 | Nkiras1 | intermediate |
| Lage3 | 7.16E-05 | 0.30874255 | 0.328 | 0.253 | 1 | 6 | Lage3 | intermediate |
| Elmod2 | 7.19E-05 | 0.28319667 | 0.417 | 0.346 | 1 | 6 | Elmod2 | intermediate |
| Zadh21 | 7.36E-05 | 0.26504394 | 0.641 | 0.534 | 1 | 6 | Zadh2 | intermediate |
| Hspb111 | 7.83E-05 | 0.38340122 | 0.394 | 0.313 | 1 | 6 | Hspb11 | intermediate |
| Thumpd1 | 8.03E-05 | 0.25341181 | 0.477 | 0.404 | 1 | 6 | Thumpd1 | intermediate |
| Afdn1 | 8.92E-05 | 0.32351611 | 0.452 | 0.365 | 1 | 6 | Afdn | intermediate |
| Kctd6 | 9.20E-05 | 0.27266387 | 0.333 | 0.267 | 1 | 6 | Kctd6 | intermediate |
| Pet100 | 9.44E-05 | 0.38171857 | 0.455 | 0.386 | 1 | 6 | Pet100 | intermediate |
| Xpr11 | 0.00010201 | 0.3497372 | 0.407 | 0.333 | 1 | 6 | Xpr1 | intermediate |
| Arl61 | 0.00010316 | 0.36179988 | 0.293 | 0.226 | 1 | 6 | Arl6 | intermediate |
| Kif7 | 0.00010492 | 0.33360893 | 0.28 | 0.19 | 1 | 6 | Kif7 | intermediate |
| Fbxl8 | 0.00010763 | 0.32611954 | 0.283 | 0.214 | 1 | 6 | Fbxl8 | intermediate |
| Tirap | 0.0001108 | 0.26924907 | 0.374 | 0.321 | 1 | 6 | Tirap | intermediate |
| Zbtb241 | 0.0001116 | 0.26075488 | 0.359 | 0.283 | 1 | 6 | Zbtb24 | intermediate |
| Decr21 | 0.00011203 | 0.34626677 | 0.417 | 0.343 | 1 | 6 | Decr2 | intermediate |
| Lrp12 | 0.0001199 | 0.34642548 | 0.485 | 0.412 | 1 | 6 | Lrp12 | intermediate |
| Taco11 | 0.00012668 | 0.38968228 | 0.27 | 0.209 | 1 | 6 | Taco1 | intermediate |
| Tmem120b | 0.00012735 | 0.49568348 | 0.26 | 0.183 | 1 | 6 | Tmem120b | intermediate |
| DEPRECATED-I | 0.00013106 | 0.29174099 | 0.308 | 0.248 | 1 | 6 | DEPRECATED-I | intermediate |
| Myo18a | 0.00014397 | 0.37110553 | 0.29 | 0.233 | 1 | 6 | Myo18a | intermediate |
| Map3k121 | 0.00014534 | 0.37283846 | 0.379 | 0.314 | 1 | 6 | Map3k12 | intermediate |
| Hspa12b1 | 0.00015082 | 0.28732467 | 0.273 | 0.199 | 1 | 6 | Hspa12b | intermediate |
| Stard91 | 0.0001526 | 0.46608152 | 0.273 | 0.192 | 1 | 6 | Stard9 | intermediate |
| Ptcd1 | 0.00015948 | 0.26134731 | 0.293 | 0.234 | 1 | 6 | Ptcd1 | intermediate |
| Eogt2 | 0.00016094 | 0.40790887 | 0.328 | 0.243 | 1 | 6 | Eogt | intermediate |
| Pxn1 | 0.00016683 | 0.26827853 | 0.482 | 0.426 | 1 | 6 | Pxn | intermediate |
| Hltf1 | 0.00017032 | 0.30909973 | 0.341 | 0.267 | 1 | 6 | Hltf | intermediate |
| Eef1aknmt | 0.00017175 | 0.45656088 | 0.29 | 0.221 | 1 | 6 | Eef1aknmt | intermediate |
| Fmc11 | 0.00017463 | 0.32654235 | 0.333 | 0.24 | 1 | 6 | Fmc1 | intermediate |
| Mpi1 | 0.00018453 | 0.3328462 | 0.407 | 0.328 | 1 | 6 | Mpi | intermediate |
| Bloc1s51 | 0.00018822 | 0.2648788 | 0.313 | 0.246 | 1 | 6 | Bloc1s5 | intermediate |
| Ptpn142 | 0.0001903 | 0.36133801 | 0.318 | 0.24 | 1 | 6 | Ptpn14 | intermediate |
| Blvra1 | 0.00019281 | 0.33672764 | 0.437 | 0.344 | 1 | 6 | Blvra | intermediate |
| Aldh3b11 | 0.00019463 | 0.53247964 | 0.308 | 0.229 | 1 | 6 | Aldh3b1 | intermediate |
| Senp7 | 0.00020232 | 0.25037893 | 0.394 | 0.325 | 1 | 6 | Senp7 | intermediate |
| Nme71 | 0.00020743 | 0.32548496 | 0.341 | 0.259 | 1 | 6 | Nme7 | intermediate |
| Iffo12 | 0.0002079 | 0.40068244 | 0.424 | 0.333 | 1 | 6 | Iffo1 | intermediate |
| Tstd31 | 0.00021106 | 0.25576244 | 0.265 | 0.189 | 1 | 6 | Tstd3 | intermediate |
| Slc19a1 | 0.00021115 | 0.29956048 | 0.283 | 0.231 | 1 | 6 | Slc19a1 | intermediate |
| Carmil11 | 0.00024162 | 0.48401162 | 0.381 | 0.303 | 1 | 6 | Carmil1 | intermediate |
| Dusp231 | 0.00024177 | 0.30731601 | 0.288 | 0.223 | 1 | 6 | Dusp23 | intermediate |
| Nat9 | 0.00024907 | 0.46183648 | 0.336 | 0.253 | 1 | 6 | Nat9 | intermediate |
| Casp91 | 0.00026003 | 0.27082426 | 0.311 | 0.227 | 1 | 6 | Casp9 | intermediate |
| Map3k4 | 0.00026384 | 0.41856735 | 0.253 | 0.2 | 1 | 6 | Map3k4 | intermediate |
| Cryzl21 | 0.0002887 | 0.28746753 | 0.321 | 0.26 | 1 | 6 | Cryzl2 | intermediate |
| Ints4 | 0.00033849 | 0.5970029 | 0.25 | 0.164 | 1 | 6 | Ints4 | intermediate |
| Tusc21 | 0.00035027 | 0.35425009 | 0.278 | 0.205 | 1 | 6 | Tusc2 | intermediate |
| Mst1r2 | 0.00035386 | 0.25489696 | 0.293 | 0.228 | 1 | 6 | Mst1r | intermediate |
| Slc45a4 | 0.00035555 | 0.25886568 | 0.457 | 0.386 | 1 | 6 | Slc45a4 | intermediate |
| Trappc2 | 0.00036691 | 0.27113423 | 0.255 | 0.202 | 1 | 6 | Trappc2 | intermediate |
| Fn3krp2 | 0.00037243 | 0.2664549 | 0.298 | 0.214 | 1 | 6 | Fn3krp | intermediate |
| Ints91 | 0.00037879 | 0.44923639 | 0.27 | 0.193 | 1 | 6 | Ints9 | intermediate |
| Pxmp4 | 0.00039876 | 0.30908401 | 0.253 | 0.194 | 1 | 6 | Pxmp4 | intermediate |
| Eef1akmt31 | 0.00048505 | 0.3608978 | 0.295 | 0.239 | 1 | 6 | Eef1akmt3 | intermediate |
| Dhx291 | 0.00048967 | 0.41766347 | 0.293 | 0.229 | 1 | 6 | Dhx29 | intermediate |
| Fhod12 | 0.00049199 | 0.45617822 | 0.359 | 0.284 | 1 | 6 | Fhod1 | intermediate |
| Zfp251 | 0.00051349 | 0.28390304 | 0.333 | 0.284 | 1 | 6 | Zfp251 | intermediate |
| Lancl21 | 0.00054813 | 0.29057266 | 0.346 | 0.281 | 1 | 6 | Lancl2 | intermediate |
| Trap12 | 0.00059217 | 0.25465881 | 0.601 | 0.498 | 1 | 6 | Trap1 | intermediate |
| Pias11 | 0.00064189 | 0.27096421 | 0.427 | 0.369 | 1 | 6 | Pias1 | intermediate |
| Ddr11 | 0.00065062 | 0.31149931 | 0.341 | 0.284 | 1 | 6 | Ddr1 | intermediate |
| Morc41 | 0.00070543 | 0.30324027 | 0.295 | 0.242 | 1 | 6 | Morc4 | intermediate |
| Klhl18 | 0.00070586 | 0.26800653 | 0.321 | 0.259 | 1 | 6 | Klhl18 | intermediate |
| Nif3l11 | 0.00072253 | 0.29706324 | 0.288 | 0.233 | 1 | 6 | Nif3l1 | intermediate |
| Pnpla6 | 0.0007237 | 0.55384683 | 0.369 | 0.276 | 1 | 6 | Pnpla6 | intermediate |
| Dhx34 | 0.00073103 | 0.35981334 | 0.253 | 0.19 | 1 | 6 | Dhx34 | intermediate |
| Prpf32 | 0.00075746 | 0.26777006 | 0.429 | 0.351 | 1 | 6 | Prpf3 | intermediate |
| Zdhhc212 | 0.00079096 | 0.26416423 | 0.295 | 0.218 | 1 | 6 | Zdhhc21 | intermediate |
| Nudt22 | 0.00079097 | 0.25722334 | 0.381 | 0.301 | 1 | 6 | Nudt2 | intermediate |
| Rsph3b | 0.00079485 | 0.33706392 | 0.253 | 0.194 | 1 | 6 | Rsph3b | intermediate |
| Vwa8 | 0.0008045 | 0.36773792 | 0.356 | 0.275 | 1 | 6 | Vwa8 | intermediate |
| Cby11 | 0.00081536 | 0.2831173 | 0.384 | 0.306 | 1 | 6 | Cby1 | intermediate |
| Hps5 | 0.00082138 | 0.29584571 | 0.356 | 0.271 | 1 | 6 | Hps5 | intermediate |
| Prmt9 | 0.00083169 | 0.48673371 | 0.255 | 0.192 | 1 | 6 | Prmt9 | intermediate |
| Akap1 | 0.00083662 | 0.32277085 | 0.303 | 0.241 | 1 | 6 | Akap1 | intermediate |
| Snta11 | 0.00091363 | 0.32298661 | 0.465 | 0.399 | 1 | 6 | Snta1 | intermediate |
| Mecr1 | 0.00096052 | 0.27270103 | 0.369 | 0.298 | 1 | 6 | Mecr | intermediate |
| Cpped1 | 0.00096283 | 0.43449614 | 0.293 | 0.226 | 1 | 6 | Cpped1 | intermediate |
| Abhd131 | 0.00096337 | 0.30598076 | 0.495 | 0.431 | 1 | 6 | Abhd13 | intermediate |
| Lrrc28 | 0.00117155 | 0.28149695 | 0.253 | 0.196 | 1 | 6 | Lrrc28 | intermediate |
| Spg11 | 0.00127139 | 0.25608175 | 0.394 | 0.353 | 1 | 6 | Spg11 | intermediate |
| C2cd21 | 0.00131792 | 0.30301764 | 0.253 | 0.194 | 1 | 6 | C2cd2 | intermediate |
| Ly962 | 0.00137769 | 0.28872988 | 0.301 | 0.246 | 1 | 6 | Ly96 | intermediate |
| Cetn21 | 0.00140688 | 0.27069308 | 0.402 | 0.334 | 1 | 6 | Cetn2 | intermediate |
| Cars21 | 0.00150216 | 0.28995457 | 0.374 | 0.298 | 1 | 6 | Cars2 | intermediate |
| Stx112 | 0.00159732 | 0.40960622 | 0.328 | 0.264 | 1 | 6 | Stx11 | intermediate |
| Erbb2 | 0.00175327 | 0.39789137 | 0.298 | 0.234 | 1 | 6 | Erbb2 | intermediate |
| Ift81 | 0.00188141 | 0.25112302 | 0.265 | 0.208 | 1 | 6 | Ift81 | intermediate |
| Txnrd21 | 0.00195224 | 0.26894882 | 0.326 | 0.27 | 1 | 6 | Txnrd2 | intermediate |
| Akip11 | 0.00195484 | 0.26085697 | 0.447 | 0.387 | 1 | 6 | Akip1 | intermediate |
| Tle22 | 0.00195626 | 0.26934387 | 0.298 | 0.249 | 1 | 6 | Tle2 | intermediate |
| Ptpn3 | 0.00206247 | 0.42678353 | 0.316 | 0.266 | 1 | 6 | Ptpn3 | intermediate |
| Dcaf171 | 0.0021052 | 0.3424097 | 0.268 | 0.213 | 1 | 6 | Dcaf17 | intermediate |
| Ttf11 | 0.00213042 | 0.43079058 | 0.308 | 0.237 | 1 | 6 | Ttf1 | intermediate |
| Gprasp1 | 0.00214096 | 0.29571035 | 0.263 | 0.203 | 1 | 6 | Gprasp1 | intermediate |
| Slc35e21 | 0.00233622 | 0.32767654 | 0.321 | 0.261 | 1 | 6 | Slc35e2 | intermediate |
| Utp14a1 | 0.0023639 | 0.41500703 | 0.346 | 0.287 | 1 | 6 | Utp14a | intermediate |
| Snapc3 | 0.00244747 | 0.27938801 | 0.442 | 0.376 | 1 | 6 | Snapc3 | intermediate |
| Alkbh3 | 0.00274445 | 0.26219236 | 0.404 | 0.325 | 1 | 6 | Alkbh3 | intermediate |
| Nphp1 | 0.00298191 | 0.4130114 | 0.288 | 0.224 | 1 | 6 | Nphp1 | intermediate |
| Rad51d | 0.00329832 | 0.349881 | 0.27 | 0.217 | 1 | 6 | Rad51d | intermediate |
| Zfp5121 | 0.00372981 | 0.3061408 | 0.28 | 0.214 | 1 | 6 | Zfp512 | intermediate |
| Ogfod2 | 0.00391338 | 0.31739847 | 0.285 | 0.211 | 1 | 6 | Ogfod2 | intermediate |
| Fzd52 | 0.00419976 | 0.3389173 | 0.321 | 0.263 | 1 | 6 | Fzd5 | intermediate |
| Tmem1071 | 0.00504691 | 0.28574542 | 0.268 | 0.212 | 1 | 6 | Tmem107 | intermediate |
| Bckdha | 0.0052518 | 0.26692628 | 0.321 | 0.268 | 1 | 6 | Bckdha | intermediate |
| Oxsm1 | 0.00525353 | 0.31877304 | 0.313 | 0.227 | 1 | 6 | Oxsm | intermediate |
| Gpatch2 | 0.00572242 | 0.30994825 | 0.348 | 0.291 | 1 | 6 | Gpatch2 | intermediate |
| Anks6 | 0.00615127 | 0.41902618 | 0.316 | 0.248 | 1 | 6 | Anks6 | intermediate |
| Nlrx11 | 0.00620947 | 0.30900683 | 0.356 | 0.29 | 1 | 6 | Nlrx1 | intermediate |
| Sgcd2 | 0.00631566 | 0.25580271 | 0.404 | 0.318 | 1 | 6 | Sgcd | intermediate |
| Herpud21 | 0.00715959 | 0.25883427 | 0.614 | 0.544 | 1 | 6 | Herpud2 | intermediate |
| Inpp5e1 | 0.0078001 | 0.30886739 | 0.275 | 0.213 | 1 | 6 | Inpp5e | intermediate |
| Fbxo46 | 0.00829093 | 0.40595261 | 0.255 | 0.203 | 1 | 6 | Fbxo46 | intermediate |
| Vps451 | 0.00839438 | 0.27849808 | 0.311 | 0.261 | 1 | 6 | Vps45 | intermediate |
| Tyk2 | 0.00929338 | 0.34679913 | 0.258 | 0.208 | 1 | 6 | Tyk2 | intermediate |
| Stk11ip | 0.00941232 | 0.29160602 | 0.341 | 0.283 | 1 | 6 | Stk11ip | intermediate |
| Tmem2671 | 0.00985621 | 0.33845405 | 0.29 | 0.255 | 1 | 6 | Tmem267 | intermediate |
| Wif1 | 0 | 6.31398534 | 0.675 | 0.033 | 0 | 7 | Wif1 | myCAF_3 |
| Egflam1 | 0 | 4.08182596 | 0.706 | 0.133 | 0 | 7 | Egflam | myCAF_3 |
| Alx4 | 0 | 7.42904071 | 0.562 | 0.016 | 0 | 7 | Alx4 | myCAF_3 |
| Lamc3 | 0 | 6.17918221 | 0.572 | 0.038 | 0 | 7 | Lamc3 | myCAF_3 |
| Runx3 | 0 | 5.3522302 | 0.557 | 0.048 | 0 | 7 | Runx3 | myCAF_3 |
| Syt13 | 0 | 6.11817875 | 0.51 | 0.024 | 0 | 7 | Syt13 | myCAF_3 |
| Nav2 | 0 | 5.64950019 | 0.51 | 0.025 | 0 | 7 | Nav2 | myCAF_3 |
| Ntn12 | 0 | 4.18775497 | 0.763 | 0.294 | 0 | 7 | Ntn1 | myCAF_3 |
| Dkk21 | 0 | 4.40724721 | 0.593 | 0.126 | 0 | 7 | Dkk2 | myCAF_3 |
| Cyp4b11 | 0 | 3.418089 | 0.655 | 0.202 | 0 | 7 | Cyp4b1 | myCAF_3 |
| Pianp | 0 | 5.33493864 | 0.474 | 0.034 | 0 | 7 | Pianp | myCAF_3 |
| Chchd101 | 0 | 4.15018828 | 0.603 | 0.185 | 0 | 7 | Chchd10 | myCAF_3 |
| Ednrb | 0 | 7.02259012 | 0.433 | 0.017 | 0 | 7 | Ednrb | myCAF_3 |
| Gpnmb2 | 0 | 3.23757172 | 0.608 | 0.215 | 0 | 7 | Gpnmb | myCAF_3 |
| Coch | 0 | 10.3796094 | 0.371 | 0.005 | 0 | 7 | Coch | myCAF_3 |
| Cyp26b11 | 0 | 2.91409533 | 0.856 | 0.499 | 0 | 7 | Cyp26b1 | myCAF_3 |
| Smoc22 | 0 | 2.6532119 | 0.825 | 0.473 | 0 | 7 | Smoc2 | myCAF_3 |
| Sema3g | 0 | 6.04161655 | 0.366 | 0.021 | 0 | 7 | Sema3g | myCAF_3 |
| Cpe1 | 0 | 2.73591296 | 0.562 | 0.231 | 0 | 7 | Cpe | myCAF_3 |
| Tnfrsf11b1 | 0 | 2.99336119 | 0.686 | 0.362 | 0 | 7 | Tnfrsf11b | myCAF_3 |
| Tnn1 | 0 | 2.66920977 | 0.552 | 0.261 | 0 | 7 | Tnn | myCAF_3 |
| Mmp111 | 0 | 3.60010286 | 0.876 | 0.589 | 0 | 7 | Mmp11 | myCAF_3 |
| Scd1 | 0 | 4.27670433 | 0.18 | 0.44 | 0 | 7 | Scd1 | myCAF_3 |
| Mmp132 | 0 | 1.25896387 | 0.691 | 0.431 | 0 | 7 | Mmp13 | myCAF_3 |
| Apod3 | 0 | 2.14661307 | 0.845 | 0.624 | 0 | 7 | Apod | myCAF_3 |
| S100a43 | 0 | 2.92103508 | 0.737 | 0.532 | 0 | 7 | S100a4 | myCAF_3 |
| Apoe1 | 0 | 3.74650106 | 0.763 | 0.588 | 0 | 7 | Apoe | myCAF_3 |
| Crispld21 | 0 | 2.15834998 | 0.881 | 0.751 | 0 | 7 | Crispld2 | myCAF_3 |
| Col7a12 | 0 | 1.18455328 | 0.711 | 0.598 | 0 | 7 | Col7a1 | myCAF_3 |
| F2r1 | 0 | 3.0829524 | 0.799 | 0.7 | 0 | 7 | F2r | myCAF_3 |
| Col4a21 | 0 | 1.86654819 | 0.933 | 0.867 | 0 | 7 | Col4a2 | myCAF_3 |
| Igfbp73 | 0 | 1.35983064 | 0.954 | 0.993 | 0 | 7 | Igfbp7 | myCAF_3 |
| Col4a12 | 0 | 1.442533 | 0.928 | 0.894 | 0 | 7 | Col4a1 | myCAF_3 |
| Vim3 | 0 | 0.42696236 | 0.985 | 0.999 | 0 | 7 | Vim | myCAF_3 |
| Lgals33 | 1.65E-238 | 2.53571786 | 0.789 | 0.512 | 3.18E-234 | 7 | Lgals3 | myCAF_3 |
| Slc6a17 | 5.37E-230 | 4.7554216 | 0.428 | 0.051 | 1.04E-225 | 7 | Slc6a17 | myCAF_3 |
| Bmp4 | 2.83E-229 | 5.50173824 | 0.402 | 0.036 | 5.47E-225 | 7 | Bmp4 | myCAF_3 |
| Ifitm12 | 6.92E-226 | 3.11158014 | 0.732 | 0.367 | 1.34E-221 | 7 | Ifitm1 | myCAF_3 |
| Trps11 | 1.24E-188 | 3.91355079 | 0.753 | 0.244 | 2.40E-184 | 7 | Trps1 | myCAF_3 |
| Itga8 | 2.53E-188 | 4.06220906 | 0.371 | 0.059 | 4.89E-184 | 7 | Itga8 | myCAF_3 |
| Fth13 | 1.89E-187 | 0.28948281 | 0.933 | 0.985 | 3.66E-183 | 7 | Fth1 | myCAF_3 |
| Tfap2c | 1.49E-182 | 7.23131479 | 0.299 | 0.007 | 2.88E-178 | 7 | Tfap2c | myCAF_3 |
| Col6a33 | 1.31E-176 | 0.31992911 | 0.871 | 0.991 | 2.54E-172 | 7 | Col6a3 | myCAF_3 |
| Junb3 | 8.30E-175 | 0.59740981 | 0.866 | 0.934 | 1.60E-170 | 7 | Junb | myCAF_3 |
| Kcnab2 | 2.10E-161 | 4.24221113 | 0.33 | 0.046 | 4.05E-157 | 7 | Kcnab2 | myCAF_3 |
| Bpgm2 | 5.28E-155 | 1.56877039 | 0.768 | 0.667 | 1.02E-150 | 7 | Bpgm | myCAF_3 |
| Zfp361 | 8.22E-152 | 0.63716832 | 0.711 | 0.827 | 1.59E-147 | 7 | Zfp36 | myCAF_3 |
| Mgst31 | 1.22E-132 | 2.29393315 | 0.603 | 0.397 | 2.37E-128 | 7 | Mgst3 | myCAF_3 |
| Kif1a | 8.23E-128 | 4.90814133 | 0.33 | 0.022 | 1.59E-123 | 7 | Kif1a | myCAF_3 |
| Tmem2042 | 4.24E-123 | 2.10965757 | 0.701 | 0.337 | 8.19E-119 | 7 | Tmem204 | myCAF_3 |
| Lmna2 | 3.57E-118 | 0.53211945 | 0.907 | 0.981 | 6.91E-114 | 7 | Lmna | myCAF_3 |
| Ier32 | 1.21E-117 | 0.36923479 | 0.727 | 0.8 | 2.35E-113 | 7 | Ier3 | myCAF_3 |
| C1qtnf13 | 1.29E-117 | 2.04975413 | 0.861 | 0.744 | 2.49E-113 | 7 | C1qtnf1 | myCAF_3 |
| Lama24 | 1.69E-115 | 1.74137826 | 0.768 | 0.668 | 3.28E-111 | 7 | Lama2 | myCAF_3 |
| Klhl33 | 1.50E-114 | 3.32006181 | 0.412 | 0.107 | 2.89E-110 | 7 | Klhl33 | myCAF_3 |
| Gnas1 | 1.83E-114 | 0.34294819 | 0.954 | 0.98 | 3.53E-110 | 7 | Gnas | myCAF_3 |
| Cdc42ep32 | 2.74E-111 | 2.50930544 | 0.557 | 0.226 | 5.30E-107 | 7 | Cdc42ep3 | myCAF_3 |
| Dusp13 | 8.26E-109 | 0.68639009 | 0.727 | 0.734 | 1.60E-104 | 7 | Dusp1 | myCAF_3 |
| Pappa | 8.91E-108 | 2.77243716 | 0.33 | 0.178 | 1.72E-103 | 7 | Pappa | myCAF_3 |
| Ets22 | 1.19E-107 | 2.64191261 | 0.763 | 0.565 | 2.31E-103 | 7 | Ets2 | myCAF_3 |
| Stc11 | 3.54E-100 | 2.94574956 | 0.258 | 0.121 | 6.84E-96 | 7 | Stc1 | myCAF_3 |
| Col11a1 | 4.67E-100 | 3.36496356 | 0.376 | 0.068 | 9.04E-96 | 7 | Col11a1 | myCAF_3 |
| Inhba3 | 5.94E-99 | 0.28151244 | 0.747 | 0.715 | 1.15E-94 | 7 | Inhba | myCAF_3 |
| Tfap2a | 7.61E-97 | 4.15940823 | 0.5 | 0.156 | 1.47E-92 | 7 | Tfap2a | myCAF_3 |
| Mmp93 | 9.08E-97 | 0.78484727 | 0.371 | 0.319 | 1.76E-92 | 7 | Mmp9 | myCAF_3 |
| S100a102 | 2.71E-95 | 0.25449794 | 0.814 | 0.864 | 5.24E-91 | 7 | S100a10 | myCAF_3 |
| Mtss21 | 8.26E-94 | 2.74457336 | 0.67 | 0.441 | 1.60E-89 | 7 | Mtss2 | myCAF_3 |
| Rhob2 | 2.57E-90 | 0.37080256 | 0.649 | 0.867 | 4.97E-86 | 7 | Rhob | myCAF_3 |
| Kazn2 | 3.08E-89 | 1.98479999 | 0.655 | 0.414 | 5.95E-85 | 7 | Kazn | myCAF_3 |
| Nr4a13 | 6.88E-84 | 1.16330264 | 0.603 | 0.634 | 1.33E-79 | 7 | Nr4a1 | myCAF_3 |
| Megf62 | 9.06E-84 | 1.89500963 | 0.345 | 0.199 | 1.75E-79 | 7 | Megf6 | myCAF_3 |
| Fam174b | 1.47E-83 | 3.04276228 | 0.428 | 0.132 | 2.85E-79 | 7 | Fam174b | myCAF_3 |
| Ier23 | 3.66E-83 | 0.46276374 | 0.675 | 0.801 | 7.07E-79 | 7 | Ier2 | myCAF_3 |
| Nkd21 | 1.22E-82 | 2.07375009 | 0.753 | 0.58 | 2.36E-78 | 7 | Nkd2 | myCAF_3 |
| Ubc2 | 2.08E-82 | 0.61691218 | 0.876 | 0.937 | 4.02E-78 | 7 | Ubc | myCAF_3 |
| Slc48a12 | 2.01E-75 | 2.3191855 | 0.67 | 0.452 | 3.89E-71 | 7 | Slc48a1 | myCAF_3 |
| Bcl11b | 7.50E-74 | 3.97567673 | 0.335 | 0.063 | 1.45E-69 | 7 | Bcl11b | myCAF_3 |
| Marcks1 | 1.12E-73 | 0.29324855 | 0.881 | 0.972 | 2.17E-69 | 7 | Marcks | myCAF_3 |
| Masp13 | 1.24E-73 | 1.76587815 | 0.588 | 0.351 | 2.40E-69 | 7 | Masp1 | myCAF_3 |
| Grem11 | 2.93E-73 | 1.16198593 | 0.479 | 0.383 | 5.66E-69 | 7 | Grem1 | myCAF_3 |
| Rarres11 | 1.76E-72 | 2.5631961 | 0.485 | 0.174 | 3.41E-68 | 7 | Rarres1 | myCAF_3 |
| Col23a11 | 1.08E-69 | 1.73768961 | 0.835 | 0.643 | 2.10E-65 | 7 | Col23a1 | myCAF_3 |
| Uba1 | 6.31E-68 | 0.25614562 | 0.856 | 0.948 | 1.22E-63 | 7 | Uba1 | myCAF_3 |
| Mgll1 | 3.12E-67 | 2.22559619 | 0.469 | 0.217 | 6.03E-63 | 7 | Mgll | myCAF_3 |
| Antxr11 | 2.87E-64 | 0.45505557 | 0.902 | 0.971 | 5.54E-60 | 7 | Antxr1 | myCAF_3 |
| Pmepa12 | 3.58E-64 | 0.27308159 | 0.871 | 0.969 | 6.93E-60 | 7 | Pmepa1 | myCAF_3 |
| Ptma1 | 5.05E-64 | 0.33343674 | 0.691 | 0.873 | 9.77E-60 | 7 | Ptma | myCAF_3 |
| Tagln1 | 1.11E-63 | 0.31533931 | 0.655 | 0.551 | 2.15E-59 | 7 | Tagln | myCAF_3 |
| Rabac1 | 1.68E-61 | 0.35494387 | 0.887 | 0.965 | 3.24E-57 | 7 | Rabac1 | myCAF_3 |
| Prdm13 | 3.98E-60 | 1.92586569 | 0.737 | 0.405 | 7.69E-56 | 7 | Prdm1 | myCAF_3 |
| Csrp12 | 2.53E-59 | 0.26264826 | 0.825 | 0.871 | 4.89E-55 | 7 | Csrp1 | myCAF_3 |
| Plxdc1 | 1.12E-58 | 3.23133343 | 0.284 | 0.073 | 2.17E-54 | 7 | Plxdc1 | myCAF_3 |
| Slc38a21 | 3.70E-56 | 0.80952395 | 0.83 | 0.919 | 7.16E-52 | 7 | Slc38a2 | myCAF_3 |
| Tspan42 | 8.83E-56 | 0.2519226 | 0.83 | 0.933 | 1.71E-51 | 7 | Tspan4 | myCAF_3 |
| Man1a3 | 1.10E-55 | 0.48626369 | 0.732 | 0.859 | 2.13E-51 | 7 | Man1a | myCAF_3 |
| Ramp1 | 2.34E-55 | 2.70046887 | 0.314 | 0.079 | 4.52E-51 | 7 | Ramp1 | myCAF_3 |
| Syncrip1 | 4.18E-55 | 0.25086859 | 0.716 | 0.911 | 8.08E-51 | 7 | Syncrip | myCAF_3 |
| Aqp11 | 4.99E-55 | 3.00397445 | 0.366 | 0.137 | 9.65E-51 | 7 | Aqp1 | myCAF_3 |
| Bmp31 | 2.41E-53 | 2.51948245 | 0.304 | 0.068 | 4.66E-49 | 7 | Bmp3 | myCAF_3 |
| Ctdsp21 | 8.17E-53 | 0.6388715 | 0.948 | 0.977 | 1.58E-48 | 7 | Ctdsp2 | myCAF_3 |
| Ahnak3 | 3.26E-51 | 0.53073831 | 0.907 | 0.96 | 6.31E-47 | 7 | Ahnak | myCAF_3 |
| Plin2 | 7.86E-51 | 1.74962247 | 0.809 | 0.722 | 1.52E-46 | 7 | Plin2 | myCAF_3 |
| Bcl2 | 7.84E-50 | 3.57053818 | 0.603 | 0.322 | 1.52E-45 | 7 | Bcl2 | myCAF_3 |
| Dazap11 | 2.81E-49 | 0.28230437 | 0.794 | 0.907 | 5.43E-45 | 7 | Dazap1 | myCAF_3 |
| Cyb5r32 | 2.83E-49 | 0.25800543 | 0.814 | 0.947 | 5.48E-45 | 7 | Cyb5r3 | myCAF_3 |
| Puf601 | 4.20E-49 | 0.25165045 | 0.753 | 0.908 | 8.12E-45 | 7 | Puf60 | myCAF_3 |
| Cst31 | 5.70E-49 | 1.26841789 | 0.912 | 0.977 | 1.10E-44 | 7 | Cst3 | myCAF_3 |
| Bhlhe402 | 8.09E-49 | 0.49892544 | 0.789 | 0.855 | 1.56E-44 | 7 | Bhlhe40 | myCAF_3 |
| Raly2 | 1.55E-48 | 0.41584265 | 0.804 | 0.917 | 3.00E-44 | 7 | Raly | myCAF_3 |
| Scara33 | 2.18E-48 | 0.31074736 | 0.608 | 0.661 | 4.21E-44 | 7 | Scara3 | myCAF_3 |
| Furin1 | 2.43E-48 | 0.36573119 | 0.881 | 0.947 | 4.70E-44 | 7 | Furin | myCAF_3 |
| Ddx3x | 2.48E-48 | 0.64063978 | 0.876 | 0.937 | 4.79E-44 | 7 | Ddx3x | myCAF_3 |
| Stac21 | 4.02E-47 | 3.5367679 | 0.412 | 0.136 | 7.77E-43 | 7 | Stac2 | myCAF_3 |
| Ctsz1 | 5.86E-47 | 0.33606508 | 0.845 | 0.962 | 1.13E-42 | 7 | Ctsz | myCAF_3 |
| Hspa1a2 | 8.93E-47 | 1.44746367 | 0.418 | 0.551 | 1.73E-42 | 7 | Hspa1a | myCAF_3 |
| Svep13 | 2.18E-46 | 0.31697488 | 0.778 | 0.855 | 4.21E-42 | 7 | Svep1 | myCAF_3 |
| Klf42 | 4.74E-46 | 0.47043287 | 0.495 | 0.457 | 9.16E-42 | 7 | Klf4 | myCAF_3 |
| DEPRECATED-I | 7.32E-46 | 1.88087212 | 0.521 | 0.29 | 1.41E-41 | 7 | DEPRECATED-I | myCAF_3 |
| Cdh132 | 7.59E-46 | 2.23726642 | 0.49 | 0.269 | 1.47E-41 | 7 | Cdh13 | myCAF_3 |
| Ifrd13 | 9.54E-45 | 0.2843942 | 0.629 | 0.75 | 1.84E-40 | 7 | Ifrd1 | myCAF_3 |
| Stat31 | 3.45E-44 | 0.3780216 | 0.789 | 0.914 | 6.68E-40 | 7 | Stat3 | myCAF_3 |
| Dnaja11 | 6.89E-44 | 0.29261177 | 0.5 | 0.772 | 1.33E-39 | 7 | Dnaja1 | myCAF_3 |
| Rbfox2 | 2.81E-43 | 0.38604911 | 0.753 | 0.899 | 5.43E-39 | 7 | Rbfox2 | myCAF_3 |
| Pum22 | 4.33E-43 | 0.26022739 | 0.634 | 0.845 | 8.37E-39 | 7 | Pum2 | myCAF_3 |
| Spry11 | 4.79E-43 | 1.76436438 | 0.747 | 0.581 | 9.27E-39 | 7 | Spry1 | myCAF_3 |
| Slc25a3 | 1.68E-42 | 0.44095044 | 0.835 | 0.931 | 3.25E-38 | 7 | Slc25a3 | myCAF_3 |
| Il343 | 4.18E-42 | 1.85034773 | 0.402 | 0.259 | 8.07E-38 | 7 | Il34 | myCAF_3 |
| Adap2 | 1.18E-41 | 2.47599485 | 0.325 | 0.085 | 2.28E-37 | 7 | Adap2 | myCAF_3 |
| Tnfrsf212 | 1.25E-41 | 1.26601202 | 0.438 | 0.353 | 2.41E-37 | 7 | Tnfrsf21 | myCAF_3 |
| Scpep12 | 1.59E-41 | 0.33659515 | 0.835 | 0.948 | 3.07E-37 | 7 | Scpep1 | myCAF_3 |
| Tnfaip21 | 2.06E-41 | 0.43012512 | 0.778 | 0.911 | 3.97E-37 | 7 | Tnfaip2 | myCAF_3 |
| Bcl3 | 2.60E-41 | 0.49059078 | 0.639 | 0.801 | 5.04E-37 | 7 | Bcl3 | myCAF_3 |
| Cilp2 | 5.10E-41 | 1.77804188 | 0.412 | 0.261 | 9.86E-37 | 7 | Cilp | myCAF_3 |
| Hnrnpul21 | 1.04E-40 | 0.40313578 | 0.758 | 0.899 | 2.02E-36 | 7 | Hnrnpul2 | myCAF_3 |
| Cd441 | 1.66E-40 | 0.34183195 | 0.763 | 0.88 | 3.21E-36 | 7 | Cd44 | myCAF_3 |
| Dync1h1 | 2.32E-40 | 0.35252743 | 0.794 | 0.913 | 4.48E-36 | 7 | Dync1h1 | myCAF_3 |
| Luc7l22 | 4.01E-40 | 0.51401015 | 0.768 | 0.902 | 7.76E-36 | 7 | Luc7l2 | myCAF_3 |
| Mmp32 | 1.06E-39 | 0.5634241 | 0.51 | 0.612 | 2.06E-35 | 7 | Mmp3 | myCAF_3 |
| Glis32 | 2.06E-39 | 0.5856499 | 0.804 | 0.888 | 3.99E-35 | 7 | Glis3 | myCAF_3 |
| Twist12 | 2.63E-39 | 0.5297961 | 0.84 | 0.907 | 5.08E-35 | 7 | Twist1 | myCAF_3 |
| Sec62 | 7.58E-39 | 0.33447641 | 0.789 | 0.904 | 1.46E-34 | 7 | Sec62 | myCAF_3 |
| Bag31 | 1.97E-38 | 0.8167211 | 0.572 | 0.826 | 3.81E-34 | 7 | Bag3 | myCAF_3 |
| Sh3pxd2b | 2.07E-38 | 0.4183231 | 0.856 | 0.961 | 4.01E-34 | 7 | Sh3pxd2b | myCAF_3 |
| Calm21 | 3.04E-38 | 0.32269121 | 0.691 | 0.841 | 5.88E-34 | 7 | Calm2 | myCAF_3 |
| Ndfip1 | 3.08E-38 | 0.38630448 | 0.861 | 0.957 | 5.96E-34 | 7 | Ndfip1 | myCAF_3 |
| Vax2 | 3.57E-38 | 2.97382114 | 0.376 | 0.1 | 6.90E-34 | 7 | Vax2 | myCAF_3 |
| Ctnna12 | 5.27E-38 | 0.33095448 | 0.778 | 0.896 | 1.02E-33 | 7 | Ctnna1 | myCAF_3 |
| Plec1 | 7.79E-38 | 0.59593592 | 0.933 | 0.971 | 1.51E-33 | 7 | Plec | myCAF_3 |
| Arrdc32 | 9.50E-38 | 3.23653659 | 0.588 | 0.372 | 1.84E-33 | 7 | Arrdc3 | myCAF_3 |
| Mcl11 | 1.12E-37 | 0.57694306 | 0.861 | 0.947 | 2.17E-33 | 7 | Mcl1 | myCAF_3 |
| Igfbp42 | 3.33E-37 | 0.62043312 | 0.866 | 0.873 | 6.44E-33 | 7 | Igfbp4 | myCAF_3 |
| Prrc2a1 | 5.00E-37 | 0.36070077 | 0.701 | 0.884 | 9.66E-33 | 7 | Prrc2a | myCAF_3 |
| Hspa1b2 | 6.25E-37 | 1.54296473 | 0.479 | 0.603 | 1.21E-32 | 7 | Hspa1b | myCAF_3 |
| Fam129b2 | 7.86E-37 | 0.42865572 | 0.789 | 0.883 | 1.52E-32 | 7 | Fam129b | myCAF_3 |
| Smad41 | 1.45E-36 | 0.30837497 | 0.732 | 0.897 | 2.80E-32 | 7 | Smad4 | myCAF_3 |
| Clta | 1.66E-36 | 0.44473691 | 0.856 | 0.966 | 3.20E-32 | 7 | Clta | myCAF_3 |
| Nsd32 | 4.13E-36 | 0.45960999 | 0.701 | 0.876 | 7.98E-32 | 7 | Nsd3 | myCAF_3 |
| Son1 | 6.72E-36 | 0.65216016 | 0.84 | 0.927 | 1.30E-31 | 7 | Son | myCAF_3 |
| Atf32 | 1.16E-35 | 0.32205548 | 0.139 | 0.319 | 2.24E-31 | 7 | Atf3 | myCAF_3 |
| Cd473 | 1.22E-35 | 0.5851873 | 0.851 | 0.919 | 2.36E-31 | 7 | Cd47 | myCAF_3 |
| Lpp2 | 2.68E-35 | 0.43398418 | 0.789 | 0.93 | 5.18E-31 | 7 | Lpp | myCAF_3 |
| Zcchc242 | 3.07E-35 | 0.29870752 | 0.814 | 0.939 | 5.94E-31 | 7 | Zcchc24 | myCAF_3 |
| Ddx171 | 4.59E-35 | 0.52529888 | 0.866 | 0.955 | 8.87E-31 | 7 | Ddx17 | myCAF_3 |
| Chtop1 | 2.45E-34 | 0.30402528 | 0.742 | 0.874 | 4.73E-30 | 7 | Chtop | myCAF_3 |
| Rab18 | 2.93E-34 | 0.41143769 | 0.799 | 0.937 | 5.66E-30 | 7 | Rab18 | myCAF_3 |
| Chd41 | 3.82E-34 | 0.36151161 | 0.691 | 0.819 | 7.38E-30 | 7 | Chd4 | myCAF_3 |
| Fgfr11 | 5.66E-34 | 1.04966281 | 0.99 | 0.978 | 1.09E-29 | 7 | Fgfr1 | myCAF_3 |
| Rbm31 | 1.17E-33 | 0.40678075 | 0.665 | 0.852 | 2.27E-29 | 7 | Rbm3 | myCAF_3 |
| Cblb2 | 1.57E-33 | 0.48028838 | 0.603 | 0.795 | 3.04E-29 | 7 | Cblb | myCAF_3 |
| Tmod32 | 2.11E-33 | 0.26939191 | 0.727 | 0.892 | 4.08E-29 | 7 | Tmod3 | myCAF_3 |
| Ptbp32 | 2.39E-33 | 0.44694499 | 0.753 | 0.874 | 4.62E-29 | 7 | Ptbp3 | myCAF_3 |
| Esyt21 | 2.56E-33 | 0.41536287 | 0.789 | 0.876 | 4.96E-29 | 7 | Esyt2 | myCAF_3 |
| Wsb11 | 4.43E-33 | 0.33570826 | 0.67 | 0.85 | 8.57E-29 | 7 | Wsb1 | myCAF_3 |
| Csnk1a1 | 4.83E-33 | 0.27922123 | 0.701 | 0.862 | 9.35E-29 | 7 | Csnk1a1 | myCAF_3 |
| Atxn2l1 | 7.10E-33 | 0.32738107 | 0.706 | 0.879 | 1.37E-28 | 7 | Atxn2l | myCAF_3 |
| Sfpq2 | 1.14E-32 | 0.60381244 | 0.716 | 0.86 | 2.20E-28 | 7 | Sfpq | myCAF_3 |
| Ctnnd12 | 1.32E-32 | 0.27770255 | 0.794 | 0.894 | 2.56E-28 | 7 | Ctnnd1 | myCAF_3 |
| Ptprf1 | 1.44E-32 | 2.75544899 | 0.536 | 0.217 | 2.79E-28 | 7 | Ptprf | myCAF_3 |
| Bmpr21 | 1.73E-32 | 0.25999438 | 0.773 | 0.898 | 3.35E-28 | 7 | Bmpr2 | myCAF_3 |
| Cbfb1 | 2.31E-32 | 0.49701371 | 0.83 | 0.93 | 4.46E-28 | 7 | Cbfb | myCAF_3 |
| Pnrc12 | 4.56E-32 | 0.73804294 | 0.778 | 0.877 | 8.81E-28 | 7 | Pnrc1 | myCAF_3 |
| Tspo3 | 4.67E-32 | 0.47582546 | 0.732 | 0.869 | 9.02E-28 | 7 | Tspo | myCAF_3 |
| Brd22 | 6.52E-32 | 0.60492132 | 0.577 | 0.814 | 1.26E-27 | 7 | Brd2 | myCAF_3 |
| U2af21 | 7.75E-32 | 0.26197038 | 0.701 | 0.854 | 1.50E-27 | 7 | U2af2 | myCAF_3 |
| Rbbp41 | 7.99E-32 | 0.28099518 | 0.634 | 0.803 | 1.55E-27 | 7 | Rbbp4 | myCAF_3 |
| Dynll21 | 8.92E-32 | 0.25056242 | 0.67 | 0.839 | 1.72E-27 | 7 | Dynll2 | myCAF_3 |
| Atrn11 | 1.19E-31 | 0.40314733 | 0.82 | 0.908 | 2.31E-27 | 7 | Atrn1 | myCAF_3 |
| Ddx211 | 2.16E-31 | 0.40360379 | 0.624 | 0.774 | 4.18E-27 | 7 | Ddx21 | myCAF_3 |
| Wdr261 | 3.19E-31 | 0.33440685 | 0.747 | 0.912 | 6.17E-27 | 7 | Wdr26 | myCAF_3 |
| Nfe2l11 | 4.53E-31 | 0.31334583 | 0.763 | 0.895 | 8.77E-27 | 7 | Nfe2l1 | myCAF_3 |
| Ogdh1 | 5.68E-31 | 0.3714238 | 0.732 | 0.877 | 1.10E-26 | 7 | Ogdh | myCAF_3 |
| Map7d1 | 7.75E-31 | 0.49925398 | 0.84 | 0.924 | 1.50E-26 | 7 | Map7d1 | myCAF_3 |
| Dlgap41 | 1.21E-30 | 0.35337805 | 0.747 | 0.906 | 2.33E-26 | 7 | Dlgap4 | myCAF_3 |
| Tns33 | 3.39E-30 | 1.8508713 | 0.84 | 0.664 | 6.55E-26 | 7 | Tns3 | myCAF_3 |
| Atp6v0c1 | 4.09E-30 | 0.3319868 | 0.768 | 0.869 | 7.91E-26 | 7 | Atp6v0c | myCAF_3 |
| Tmem176a2 | 7.70E-30 | 0.69548935 | 0.51 | 0.685 | 1.49E-25 | 7 | Tmem176a | myCAF_3 |
| Eif3g1 | 8.41E-30 | 0.37072331 | 0.655 | 0.811 | 1.63E-25 | 7 | Eif3g | myCAF_3 |
| Dock91 | 1.55E-29 | 3.42868843 | 0.32 | 0.082 | 2.99E-25 | 7 | Dock9 | myCAF_3 |
| Robo2 | 1.77E-29 | 2.80634408 | 0.289 | 0.084 | 3.42E-25 | 7 | Robo2 | myCAF_3 |
| Kdm2a1 | 2.11E-29 | 0.47332057 | 0.742 | 0.869 | 4.07E-25 | 7 | Kdm2a | myCAF_3 |
| Clec2d3 | 3.72E-29 | 0.33627938 | 0.768 | 0.85 | 7.18E-25 | 7 | Clec2d | myCAF_3 |
| Pth1r2 | 3.98E-29 | 0.50548559 | 0.66 | 0.79 | 7.69E-25 | 7 | Pth1r | myCAF_3 |
| Elf12 | 8.22E-29 | 0.27396932 | 0.557 | 0.76 | 1.59E-24 | 7 | Elf1 | myCAF_3 |
| Anxa72 | 8.75E-29 | 0.40302629 | 0.66 | 0.812 | 1.69E-24 | 7 | Anxa7 | myCAF_3 |
| Ddx61 | 1.06E-28 | 0.30538976 | 0.696 | 0.845 | 2.05E-24 | 7 | Ddx6 | myCAF_3 |
| Srsf23 | 1.24E-28 | 0.43231109 | 0.531 | 0.737 | 2.40E-24 | 7 | Srsf2 | myCAF_3 |
| Hnrnpab1 | 1.24E-28 | 0.35922249 | 0.634 | 0.756 | 2.40E-24 | 7 | Hnrnpab | myCAF_3 |
| Mme1 | 1.48E-28 | 2.37499645 | 0.356 | 0.102 | 2.86E-24 | 7 | Mme | myCAF_3 |
| Wbp1l2 | 1.77E-28 | 0.59246189 | 0.778 | 0.891 | 3.42E-24 | 7 | Wbp1l | myCAF_3 |
| Ptges31 | 2.09E-28 | 0.34509711 | 0.536 | 0.713 | 4.04E-24 | 7 | Ptges3 | myCAF_3 |
| Erf2 | 2.76E-28 | 0.50581806 | 0.562 | 0.732 | 5.34E-24 | 7 | Erf | myCAF_3 |
| Cavin12 | 3.25E-28 | 0.6547828 | 0.82 | 0.903 | 6.29E-24 | 7 | Cavin1 | myCAF_3 |
| Brd4 | 3.39E-28 | 0.60285326 | 0.763 | 0.884 | 6.56E-24 | 7 | Brd4 | myCAF_3 |
| Actn42 | 3.44E-28 | 0.5861374 | 0.856 | 0.951 | 6.66E-24 | 7 | Actn4 | myCAF_3 |
| Mprip | 3.55E-28 | 0.60393596 | 0.912 | 0.968 | 6.85E-24 | 7 | Mprip | myCAF_3 |
| Gas12 | 4.52E-28 | 0.66331998 | 0.82 | 0.856 | 8.74E-24 | 7 | Gas1 | myCAF_3 |
| Rnps11 | 6.01E-28 | 0.39747579 | 0.644 | 0.813 | 1.16E-23 | 7 | Rnps1 | myCAF_3 |
| Igsf82 | 6.47E-28 | 0.452432 | 0.541 | 0.757 | 1.25E-23 | 7 | Igsf8 | myCAF_3 |
| Gpx12 | 7.12E-28 | 1.12427641 | 0.856 | 0.917 | 1.38E-23 | 7 | Gpx1 | myCAF_3 |
| Hp1bp31 | 9.06E-28 | 0.69982684 | 0.814 | 0.887 | 1.75E-23 | 7 | Hp1bp3 | myCAF_3 |
| Tmem591 | 1.33E-27 | 0.27518175 | 0.768 | 0.917 | 2.57E-23 | 7 | Tmem59 | myCAF_3 |
| Dpysl24 | 1.41E-27 | 0.41818528 | 0.644 | 0.778 | 2.72E-23 | 7 | Dpysl2 | myCAF_3 |
| Rab211 | 1.42E-27 | 0.46406933 | 0.768 | 0.901 | 2.74E-23 | 7 | Rab21 | myCAF_3 |
| Ubald21 | 2.09E-27 | 0.43990125 | 0.665 | 0.806 | 4.05E-23 | 7 | Ubald2 | myCAF_3 |
| Sdc43 | 2.26E-27 | 0.91715121 | 0.696 | 0.767 | 4.36E-23 | 7 | Sdc4 | myCAF_3 |
| Sh3pxd2a | 2.29E-27 | 0.52117465 | 0.753 | 0.856 | 4.42E-23 | 7 | Sh3pxd2a | myCAF_3 |
| Pawr | 2.31E-27 | 2.35535545 | 0.309 | 0.106 | 4.46E-23 | 7 | Pawr | myCAF_3 |
| Cspg42 | 2.82E-27 | 1.78461932 | 0.68 | 0.365 | 5.46E-23 | 7 | Cspg4 | myCAF_3 |
| Wdr83os | 2.94E-27 | 0.30930035 | 0.624 | 0.835 | 5.69E-23 | 7 | Wdr83os | myCAF_3 |
| Smarcc2 | 2.97E-27 | 0.48413994 | 0.732 | 0.872 | 5.74E-23 | 7 | Smarcc2 | myCAF_3 |
| Pim12 | 4.39E-27 | 1.00315742 | 0.711 | 0.789 | 8.49E-23 | 7 | Pim1 | myCAF_3 |
| Serinc33 | 4.51E-27 | 0.43799862 | 0.856 | 0.957 | 8.73E-23 | 7 | Serinc3 | myCAF_3 |
| Map1lc3b2 | 4.82E-27 | 0.50089517 | 0.809 | 0.901 | 9.31E-23 | 7 | Map1lc3b | myCAF_3 |
| Cldnd1 | 7.41E-27 | 0.40359889 | 0.644 | 0.819 | 1.43E-22 | 7 | Cldnd1 | myCAF_3 |
| Ucn21 | 7.54E-27 | 1.22428007 | 0.567 | 0.375 | 1.46E-22 | 7 | Ucn2 | myCAF_3 |
| Far1 | 9.41E-27 | 0.2864344 | 0.603 | 0.814 | 1.82E-22 | 7 | Far1 | myCAF_3 |
| Lmo7 | 1.02E-26 | 2.4187067 | 0.284 | 0.091 | 1.97E-22 | 7 | Lmo7 | myCAF_3 |
| Bmpr1a1 | 1.05E-26 | 0.34252629 | 0.629 | 0.782 | 2.03E-22 | 7 | Bmpr1a | myCAF_3 |

| | | | | | | | | |
|---|---|---|---|---|---|---|---|---|
| Zfp36l23 | 1.19E-26 | 0.54781426 | 0.753 | 0.839 | 2.30E-22 | 7 | Zfp36l2 | myCAF_3 |
| Rnf1871 | 1.25E-26 | 0.35174488 | 0.742 | 0.908 | 2.43E-22 | 7 | Rnf187 | myCAF_3 |
| H2afv1 | 1.29E-26 | 0.51123083 | 0.603 | 0.74 | 2.50E-22 | 7 | H2afv | myCAF_3 |
| Midn | 1.65E-26 | 0.60764503 | 0.778 | 0.888 | 3.20E-22 | 7 | Midn | myCAF_3 |
| Sertad13 | 2.30E-26 | 0.77323691 | 0.474 | 0.65 | 4.44E-22 | 7 | Sertad1 | myCAF_3 |
| Pdgfra1 | 2.75E-26 | 0.63931573 | 0.876 | 0.946 | 5.32E-22 | 7 | Pdgfra | myCAF_3 |
| Dennd5a | 3.19E-26 | 0.30888901 | 0.68 | 0.852 | 6.17E-22 | 7 | Dennd5a | myCAF_3 |
| Cul1 | 3.60E-26 | 0.27647433 | 0.598 | 0.798 | 6.97E-22 | 7 | Cul1 | myCAF_3 |
| Tmem50a2 | 3.90E-26 | 0.42809257 | 0.851 | 0.93 | 7.54E-22 | 7 | Tmem50a | myCAF_3 |
| Fosl21 | 4.15E-26 | 0.6094797 | 0.799 | 0.87 | 8.03E-22 | 7 | Fosl2 | myCAF_3 |
| Sesn31 | 4.45E-26 | 1.72088698 | 0.644 | 0.526 | 8.61E-22 | 7 | Sesn3 | myCAF_3 |
| Nsd11 | 5.03E-26 | 0.33333546 | 0.696 | 0.866 | 9.73E-22 | 7 | Nsd1 | myCAF_3 |
| Eif4a22 | 5.86E-26 | 0.57571436 | 0.778 | 0.881 | 1.13E-21 | 7 | Eif4a2 | myCAF_3 |
| Ehmt2 | 7.53E-26 | 0.43969127 | 0.763 | 0.882 | 1.45E-21 | 7 | Ehmt2 | myCAF_3 |
| Sqstm12 | 8.98E-26 | 0.72642106 | 0.789 | 0.88 | 1.74E-21 | 7 | Sqstm1 | myCAF_3 |
| Nab21 | 1.30E-25 | 0.39906015 | 0.485 | 0.724 | 2.50E-21 | 7 | Nab2 | myCAF_3 |
| Kif5b1 | 1.69E-25 | 0.25582163 | 0.593 | 0.776 | 3.26E-21 | 7 | Kif5b | myCAF_3 |
| Tmem2561 | 1.79E-25 | 0.27490981 | 0.649 | 0.808 | 3.46E-21 | 7 | Tmem256 | myCAF_3 |
| Cdkn1b2 | 2.09E-25 | 0.25884441 | 0.582 | 0.775 | 4.04E-21 | 7 | Cdkn1b | myCAF_3 |
| Riok31 | 2.31E-25 | 0.30194018 | 0.593 | 0.773 | 4.47E-21 | 7 | Riok3 | myCAF_3 |
| Sash12 | 2.64E-25 | 0.31181114 | 0.531 | 0.774 | 5.10E-21 | 7 | Sash1 | myCAF_3 |
| Snai12 | 2.65E-25 | 0.80175425 | 0.747 | 0.806 | 5.12E-21 | 7 | Snai1 | myCAF_3 |
| Dnm11 | 2.68E-25 | 0.4526004 | 0.691 | 0.852 | 5.19E-21 | 7 | Dnm1 | myCAF_3 |
| Gng81 | 2.77E-25 | 3.13243884 | 0.366 | 0.165 | 5.36E-21 | 7 | Gng8 | myCAF_3 |
| Zmynd111 | 3.62E-25 | 0.3366833 | 0.649 | 0.841 | 7.01E-21 | 7 | Zmynd11 | myCAF_3 |
| Dpp41 | 3.65E-25 | 0.84579641 | 0.113 | 0.27 | 7.05E-21 | 7 | Dpp4 | myCAF_3 |
| Uqcrc12 | 3.85E-25 | 0.43541807 | 0.835 | 0.932 | 7.44E-21 | 7 | Uqcrc1 | myCAF_3 |
| Gsk3b1 | 4.08E-25 | 0.38449199 | 0.639 | 0.807 | 7.88E-21 | 7 | Gsk3b | myCAF_3 |
| Tnfrsf12a1 | 6.51E-25 | 0.29399536 | 0.686 | 0.82 | 1.26E-20 | 7 | Tnfrsf12a | myCAF_3 |
| Mapkapk22 | 8.46E-25 | 0.71759098 | 0.778 | 0.911 | 1.64E-20 | 7 | Mapkapk2 | myCAF_3 |
| Srebf21 | 8.54E-25 | 0.53858044 | 0.644 | 0.806 | 1.65E-20 | 7 | Srebf2 | myCAF_3 |
| Ier5l2 | 2.00E-24 | 0.3732912 | 0.557 | 0.735 | 3.86E-20 | 7 | Ier5l | myCAF_3 |
| Colec123 | 2.51E-24 | 0.34573205 | 0.804 | 0.914 | 4.85E-20 | 7 | Colec12 | myCAF_3 |
| Sel1l1 | 3.07E-24 | 0.341391 | 0.706 | 0.878 | 5.94E-20 | 7 | Sel1l | myCAF_3 |
| Kctd1 | 4.38E-24 | 3.68585654 | 0.268 | 0.104 | 8.47E-20 | 7 | Kctd1 | myCAF_3 |
| Pgrmc12 | 5.40E-24 | 0.4051423 | 0.686 | 0.836 | 1.04E-19 | 7 | Pgrmc1 | myCAF_3 |
| Stard5 | 6.87E-24 | 1.98671128 | 0.412 | 0.237 | 1.33E-19 | 7 | Stard5 | myCAF_3 |
| Sorbs1 | 7.53E-24 | 3.23342773 | 0.299 | 0.112 | 1.46E-19 | 7 | Sorbs1 | myCAF_3 |
| Sod22 | 9.60E-24 | 0.54597529 | 0.691 | 0.786 | 1.86E-19 | 7 | Sod2 | myCAF_3 |
| Epn1 | 1.01E-23 | 0.29747574 | 0.732 | 0.891 | 1.94E-19 | 7 | Epn1 | myCAF_3 |
| Eloa1 | 1.24E-23 | 0.33174615 | 0.598 | 0.801 | 2.40E-19 | 7 | Eloa | myCAF_3 |
| Trim351 | 1.50E-23 | 0.29865991 | 0.701 | 0.893 | 2.91E-19 | 7 | Trim35 | myCAF_3 |
| Tra2a2 | 1.65E-23 | 0.70270518 | 0.608 | 0.78 | 3.20E-19 | 7 | Tra2a | myCAF_3 |
| Med13l1 | 1.69E-23 | 0.81431306 | 0.763 | 0.867 | 3.28E-19 | 7 | Med13l | myCAF_3 |
| B2m3 | 1.73E-23 | 0.57776785 | 0.758 | 0.85 | 3.35E-19 | 7 | B2m | myCAF_3 |
| Tpbg1 | 1.88E-23 | 0.47136915 | 0.778 | 0.912 | 3.64E-19 | 7 | Tpbg | myCAF_3 |
| Rad211 | 2.03E-23 | 0.35927261 | 0.469 | 0.632 | 3.92E-19 | 7 | Rad21 | myCAF_3 |
| Appbp2 | 2.10E-23 | 0.25411287 | 0.464 | 0.721 | 4.06E-19 | 7 | Appbp2 | myCAF_3 |
| Cdkn1a3 | 2.26E-23 | 0.70072765 | 0.387 | 0.427 | 4.36E-19 | 7 | Cdkn1a | myCAF_3 |
| Ski1 | 2.28E-23 | 0.70358697 | 0.897 | 0.952 | 4.41E-19 | 7 | Ski | myCAF_3 |
| Grn2 | 2.40E-23 | 0.47091331 | 0.897 | 0.965 | 4.65E-19 | 7 | Grn | myCAF_3 |
| Mbtps11 | 2.52E-23 | 0.25726363 | 0.603 | 0.786 | 4.88E-19 | 7 | Mbtps1 | myCAF_3 |
| Ptp4a21 | 2.68E-23 | 0.33394966 | 0.448 | 0.692 | 5.18E-19 | 7 | Ptp4a2 | myCAF_3 |
| Golph3 | 3.71E-23 | 0.50947176 | 0.778 | 0.889 | 7.18E-19 | 7 | Golph3 | myCAF_3 |
| Slc3a21 | 3.78E-23 | 0.54899187 | 0.773 | 0.848 | 7.31E-19 | 7 | Slc3a2 | myCAF_3 |
| Rdh103 | 4.24E-23 | 0.42883256 | 0.613 | 0.703 | 8.19E-19 | 7 | Rdh10 | myCAF_3 |
| Mdh11 | 4.85E-23 | 0.32302392 | 0.624 | 0.794 | 9.38E-19 | 7 | Mdh1 | myCAF_3 |
| Insig12 | 5.15E-23 | 0.62201858 | 0.665 | 0.727 | 9.95E-19 | 7 | Insig1 | myCAF_3 |
| Gpat4 | 6.31E-23 | 0.31184057 | 0.624 | 0.806 | 1.22E-18 | 7 | Gpat4 | myCAF_3 |
| Dll1 | 6.37E-23 | 2.21864261 | 0.278 | 0.121 | 1.23E-18 | 7 | Dll1 | myCAF_3 |
| Dst3 | 6.39E-23 | 0.56839143 | 0.644 | 0.785 | 1.24E-18 | 7 | Dst | myCAF_3 |
| Hspb13 | 7.60E-23 | 2.59798537 | 0.608 | 0.388 | 1.47E-18 | 7 | Hspb1 | myCAF_3 |
| Tmem1401 | 7.61E-23 | 1.85700826 | 0.485 | 0.358 | 1.47E-18 | 7 | Tmem140 | myCAF_3 |
| Arhgef22 | 9.20E-23 | 0.49960081 | 0.639 | 0.793 | 1.78E-18 | 7 | Arhgef2 | myCAF_3 |
| Pls31 | 1.17E-22 | 0.42525607 | 0.696 | 0.832 | 2.26E-18 | 7 | Pls3 | myCAF_3 |
| Snx92 | 1.50E-22 | 0.3174793 | 0.784 | 0.921 | 2.89E-18 | 7 | Snx9 | myCAF_3 |
| Psmd15 | 1.61E-22 | 0.33282136 | 0.619 | 0.787 | 3.11E-18 | 7 | Psmd1 | myCAF_3 |
| Rnh11 | 1.86E-22 | 0.51751585 | 0.66 | 0.822 | 3.59E-18 | 7 | Rnh1 | myCAF_3 |
| Gaa2 | 2.07E-22 | 0.33194135 | 0.768 | 0.928 | 4.01E-18 | 7 | Gaa | myCAF_3 |
| Slc35b21 | 2.22E-22 | 0.27471344 | 0.603 | 0.808 | 4.29E-18 | 7 | Slc35b2 | myCAF_3 |
| Trip102 | 2.29E-22 | 0.2597975 | 0.464 | 0.71 | 4.43E-18 | 7 | Trip10 | myCAF_3 |
| Purb2 | 2.49E-22 | 0.47383601 | 0.665 | 0.807 | 4.82E-18 | 7 | Purb | myCAF_3 |
| Lrrc151 | 2.66E-22 | 0.43876574 | 0.763 | 0.759 | 5.15E-18 | 7 | Lrrc15 | myCAF_3 |
| Lnpep2 | 2.95E-22 | 0.38340722 | 0.655 | 0.85 | 5.70E-18 | 7 | Lnpep | myCAF_3 |
| Fubp11 | 3.12E-22 | 0.56274722 | 0.541 | 0.742 | 6.03E-18 | 7 | Fubp1 | myCAF_3 |
| Itgav2 | 3.22E-22 | 0.25423484 | 0.706 | 0.875 | 6.23E-18 | 7 | Itgav | myCAF_3 |
| Nfe2l21 | 3.39E-22 | 0.56500321 | 0.753 | 0.879 | 6.56E-18 | 7 | Nfe2l2 | myCAF_3 |
| Ndufs2 | 3.45E-22 | 0.31354987 | 0.737 | 0.898 | 6.67E-18 | 7 | Ndufs2 | myCAF_3 |
| Ampd32 | 4.98E-22 | 1.56473993 | 0.33 | 0.495 | 9.63E-18 | 7 | Ampd3 | myCAF_3 |
| Ilrun1 | 6.00E-22 | 0.34418017 | 0.593 | 0.797 | 1.16E-17 | 7 | Ilrun | myCAF_3 |
| Hnrnph21 | 8.94E-22 | 0.29313392 | 0.546 | 0.735 | 1.73E-17 | 7 | Hnrnph2 | myCAF_3 |
| Skp1a | 1.19E-21 | 0.41371714 | 0.593 | 0.81 | 2.30E-17 | 7 | Skp1a | myCAF_3 |
| Ranbp22 | 1.20E-21 | 0.64409604 | 0.593 | 0.747 | 2.32E-17 | 7 | Ranbp2 | myCAF_3 |
| Eftud21 | 1.34E-21 | 0.32431915 | 0.531 | 0.73 | 2.60E-17 | 7 | Eftud2 | myCAF_3 |
| Wdr821 | 1.35E-21 | 0.33577865 | 0.588 | 0.781 | 2.61E-17 | 7 | Wdr82 | myCAF_3 |
| Zfp36l13 | 1.41E-21 | 0.7919086 | 0.784 | 0.864 | 2.72E-17 | 7 | Zfp36l1 | myCAF_3 |
| Sep-82 | 1.42E-21 | 0.72921984 | 0.814 | 0.899 | 2.74E-17 | 7 | 8-Sep | myCAF_3 |
| Slk1 | 1.51E-21 | 0.495012 | 0.448 | 0.688 | 2.92E-17 | 7 | Slk | myCAF_3 |
| Cpsf61 | 1.57E-21 | 0.32468274 | 0.588 | 0.732 | 3.04E-17 | 7 | Cpsf6 | myCAF_3 |
| Adamts41 | 1.68E-21 | 0.57476242 | 0.804 | 0.909 | 3.26E-17 | 7 | Adamts4 | myCAF_3 |
| Gtf2i | 1.77E-21 | 0.34713434 | 0.67 | 0.832 | 3.42E-17 | 7 | Gtf2i | myCAF_3 |
| Rbm391 | 2.33E-21 | 0.39914067 | 0.593 | 0.775 | 4.50E-17 | 7 | Rbm39 | myCAF_3 |
| Cct21 | 2.78E-21 | 0.29556413 | 0.634 | 0.777 | 5.38E-17 | 7 | Cct2 | myCAF_3 |
| Actr1a1 | 2.94E-21 | 0.36809119 | 0.67 | 0.855 | 5.68E-17 | 7 | Actr1a | myCAF_3 |
| Pnpla21 | 3.17E-21 | 2.18608502 | 0.242 | 0.275 | 6.13E-17 | 7 | Pnpla2 | myCAF_3 |
| Kdm6b2 | 3.32E-21 | 0.32980665 | 0.582 | 0.711 | 6.43E-17 | 7 | Kdm6b | myCAF_3 |
| Itga113 | 3.51E-21 | 1.43487241 | 0.49 | 0.344 | 6.79E-17 | 7 | Itga11 | myCAF_3 |
| Ecm12 | 3.96E-21 | 0.25701615 | 0.763 | 0.959 | 7.67E-17 | 7 | Ecm1 | myCAF_3 |
| Gemin71 | 4.01E-21 | 0.33402296 | 0.521 | 0.736 | 7.76E-17 | 7 | Gemin7 | myCAF_3 |
| Utp111 | 4.29E-21 | 0.37351731 | 0.448 | 0.661 | 8.29E-17 | 7 | Utp11 | myCAF_3 |
| Huwe1 | 4.29E-21 | 0.48899123 | 0.706 | 0.854 | 8.30E-17 | 7 | Huwe1 | myCAF_3 |
| Slc6a61 | 4.56E-21 | 2.23280009 | 0.675 | 0.564 | 8.82E-17 | 7 | Slc6a6 | myCAF_3 |
| Adipor11 | 4.88E-21 | 0.67340906 | 0.881 | 0.933 | 9.44E-17 | 7 | Adipor1 | myCAF_3 |
| Cdc42bpb | 5.76E-21 | 0.34457421 | 0.603 | 0.806 | 1.11E-16 | 7 | Cdc42bpb | myCAF_3 |
| Ppp1r15b2 | 9.23E-21 | 0.26079468 | 0.402 | 0.67 | 1.78E-16 | 7 | Ppp1r15b | myCAF_3 |
| Dnajc14 | 1.01E-20 | 0.26078947 | 0.644 | 0.817 | 1.95E-16 | 7 | Dnajc14 | myCAF_3 |
| Ecpas1 | 1.07E-20 | 0.34359754 | 0.381 | 0.635 | 2.08E-16 | 7 | Ecpas | myCAF_3 |
| Rab351 | 1.38E-20 | 0.29742276 | 0.51 | 0.725 | 2.66E-16 | 7 | Rab35 | myCAF_3 |
| Ilf31 | 1.43E-20 | 0.34110523 | 0.309 | 0.6 | 2.77E-16 | 7 | Ilf3 | myCAF_3 |
| Lrp62 | 1.49E-20 | 0.72431279 | 0.701 | 0.85 | 2.88E-16 | 7 | Lrp6 | myCAF_3 |
| Lum21 | 1.70E-20 | 0.35462617 | 0.371 | 0.551 | 3.28E-16 | 7 | Lum2 | myCAF_3 |
| Gnai21 | 2.06E-20 | 0.29513969 | 0.778 | 0.917 | 3.99E-16 | 7 | Gnai2 | myCAF_3 |
| Srrm11 | 2.33E-20 | 0.35590891 | 0.459 | 0.69 | 4.50E-16 | 7 | Srrm1 | myCAF_3 |
| Ankrd17 | 2.39E-20 | 0.56130913 | 0.629 | 0.8 | 4.62E-16 | 7 | Ankrd17 | myCAF_3 |
| Tgfb12 | 2.42E-20 | 0.45235156 | 0.809 | 0.843 | 4.67E-16 | 7 | Tgfb1 | myCAF_3 |
| Bclaf11 | 3.14E-20 | 0.27623601 | 0.619 | 0.793 | 6.08E-16 | 7 | Bclaf1 | myCAF_3 |
| Psma32 | 3.55E-20 | 0.25503815 | 0.371 | 0.629 | 6.86E-16 | 7 | Psma3 | myCAF_3 |
| Irak1 | 3.97E-20 | 0.27208679 | 0.603 | 0.799 | 7.67E-16 | 7 | Irak1 | myCAF_3 |
| Ap2a1 | 4.02E-20 | 0.26931353 | 0.619 | 0.783 | 7.77E-16 | 7 | Ap2a1 | myCAF_3 |
| Ndufb61 | 4.06E-20 | 0.47730322 | 0.593 | 0.779 | 7.84E-16 | 7 | Ndufb6 | myCAF_3 |
| Arhgef13 | 4.84E-20 | 0.57960329 | 0.649 | 0.819 | 9.36E-16 | 7 | Arhgef1 | myCAF_3 |
| Plekhj1 | 5.07E-20 | 0.36467562 | 0.5 | 0.732 | 9.81E-16 | 7 | Plekhj1 | myCAF_3 |
| Srsf52 | 5.66E-20 | 0.93846657 | 0.871 | 0.92 | 1.09E-15 | 7 | Srsf5 | myCAF_3 |
| Mib11 | 5.77E-20 | 0.74888497 | 0.691 | 0.83 | 1.12E-15 | 7 | Mib1 | myCAF_3 |
| Sgpl12 | 6.10E-20 | 0.57883123 | 0.582 | 0.749 | 1.18E-15 | 7 | Sgpl1 | myCAF_3 |
| Anapc12 | 6.39E-20 | 0.34353249 | 0.536 | 0.727 | 1.24E-15 | 7 | Anapc1 | myCAF_3 |
| Wipf11 | 6.53E-20 | 0.31864113 | 0.727 | 0.858 | 1.26E-15 | 7 | Wipf1 | myCAF_3 |
| R3hdm2 | 6.92E-20 | 0.30237957 | 0.608 | 0.801 | 1.34E-15 | 7 | R3hdm2 | myCAF_3 |
| Slc27a12 | 7.03E-20 | 1.79591361 | 0.49 | 0.36 | 1.36E-15 | 7 | Slc27a1 | myCAF_3 |
| Ppp4r2 | 7.50E-20 | 0.4728809 | 0.639 | 0.8 | 1.45E-15 | 7 | Ppp4r2 | myCAF_3 |
| Rbm141 | 8.88E-20 | 0.44941605 | 0.526 | 0.738 | 1.72E-15 | 7 | Rbm14 | myCAF_3 |
| Ptar11 | 9.19E-20 | 0.46746766 | 0.67 | 0.812 | 1.78E-15 | 7 | Ptar1 | myCAF_3 |
| Hdac31 | 9.30E-20 | 0.28527983 | 0.515 | 0.715 | 1.80E-15 | 7 | Hdac3 | myCAF_3 |
| Smo1 | 1.13E-19 | 0.39201048 | 0.706 | 0.848 | 2.19E-15 | 7 | Smo | myCAF_3 |
| Tmem433 | 1.14E-19 | 0.48565041 | 0.624 | 0.756 | 2.19E-15 | 7 | Tmem43 | myCAF_3 |
| S1pr22 | 1.16E-19 | 0.36370932 | 0.557 | 0.765 | 2.25E-15 | 7 | S1pr2 | myCAF_3 |
| Crk1 | 1.22E-19 | 0.25152868 | 0.644 | 0.8 | 2.36E-15 | 7 | Crk | myCAF_3 |
| Srcap1 | 1.54E-19 | 0.60611095 | 0.603 | 0.796 | 2.99E-15 | 7 | Srcap | myCAF_3 |
| Rab102 | 1.54E-19 | 0.28591668 | 0.495 | 0.703 | 2.99E-15 | 7 | Rab10 | myCAF_3 |
| Fads11 | 2.19E-19 | 0.27469357 | 0.464 | 0.629 | 4.24E-15 | 7 | Fads1 | myCAF_3 |
| Ddx231 | 2.28E-19 | 0.39187234 | 0.649 | 0.781 | 4.40E-15 | 7 | Ddx23 | myCAF_3 |
| Rsrc21 | 2.52E-19 | 0.31031712 | 0.485 | 0.701 | 4.87E-15 | 7 | Rsrc2 | myCAF_3 |
| Prrc2b1 | 2.54E-19 | 0.47256011 | 0.737 | 0.847 | 4.92E-15 | 7 | Prrc2b | myCAF_3 |
| Cox14 | 2.76E-19 | 0.40542396 | 0.593 | 0.779 | 5.33E-15 | 7 | Cox14 | myCAF_3 |
| Prrx11 | 3.52E-19 | 0.64843948 | 0.902 | 0.966 | 6.80E-15 | 7 | Prrx1 | myCAF_3 |
| Wasf2 | 4.87E-19 | 0.31163889 | 0.613 | 0.796 | 9.42E-15 | 7 | Wasf2 | myCAF_3 |
| Tpd52l21 | 4.88E-19 | 0.25380042 | 0.665 | 0.862 | 9.43E-15 | 7 | Tpd52l2 | myCAF_3 |
| Mcmbp1 | 4.93E-19 | 0.3542002 | 0.49 | 0.687 | 9.54E-15 | 7 | Mcmbp | myCAF_3 |
| Arl8a | 5.47E-19 | 0.40746946 | 0.644 | 0.826 | 1.06E-14 | 7 | Arl8a | myCAF_3 |
| Smu11 | 5.59E-19 | 0.3045151 | 0.418 | 0.649 | 1.08E-14 | 7 | Smu1 | myCAF_3 |
| Poldip31 | 6.40E-19 | 0.47629716 | 0.613 | 0.792 | 1.24E-14 | 7 | Poldip3 | myCAF_3 |
| Cmtm31 | 6.76E-19 | 0.34965588 | 0.747 | 0.922 | 1.31E-14 | 7 | Cmtm3 | myCAF_3 |
| Ep400 | 7.00E-19 | 0.3230091 | 0.438 | 0.658 | 1.35E-14 | 7 | Ep400 | myCAF_3 |
| Csnk2a1 | 7.55E-19 | 0.47881394 | 0.546 | 0.73 | 1.46E-14 | 7 | Csnk2a1 | myCAF_3 |
| Usp41 | 9.13E-19 | 0.435848 | 0.505 | 0.717 | 1.77E-14 | 7 | Usp4 | myCAF_3 |
| Cops81 | 9.29E-19 | 0.36624323 | 0.536 | 0.73 | 1.80E-14 | 7 | Cops8 | myCAF_3 |
| Tmem259 | 1.05E-18 | 0.58929615 | 0.814 | 0.908 | 2.03E-14 | 7 | Tmem259 | myCAF_3 |
| Fam20a1 | 1.18E-18 | 1.08330192 | 0.727 | 0.798 | 2.27E-14 | 7 | Fam20a | myCAF_3 |
| Bag6 | 1.20E-18 | 0.32874729 | 0.649 | 0.821 | 2.32E-14 | 7 | Bag6 | myCAF_3 |
| Rgmb2 | 1.20E-18 | 0.2919972 | 0.474 | 0.706 | 2.32E-14 | 7 | Rgmb | myCAF_3 |
| Ndufa7 | 1.24E-18 | 0.40768875 | 0.572 | 0.748 | 2.39E-14 | 7 | Ndufa7 | myCAF_3 |
| Zmpste24 | 1.44E-18 | 0.30825119 | 0.598 | 0.791 | 2.79E-14 | 7 | Zmpste24 | myCAF_3 |
| Afg3l1 | 1.47E-18 | 0.27092001 | 0.515 | 0.717 | 2.84E-14 | 7 | Afg3l1 | myCAF_3 |
| Clic42 | 1.48E-18 | 0.54933595 | 0.799 | 0.922 | 2.86E-14 | 7 | Clic4 | myCAF_3 |
| Axl1 | 1.58E-18 | 0.61322246 | 0.881 | 0.967 | 3.06E-14 | 7 | Axl | myCAF_3 |
| Nr3c11 | 1.71E-18 | 0.76785208 | 0.505 | 0.7 | 3.31E-14 | 7 | Nr3c1 | myCAF_3 |
| Strn41 | 1.75E-18 | 0.27361917 | 0.593 | 0.784 | 3.39E-14 | 7 | Strn4 | myCAF_3 |
| Snx2 | 1.79E-18 | 0.28446551 | 0.515 | 0.715 | 3.47E-14 | 7 | Snx2 | myCAF_3 |
| Tmpo1 | 1.86E-18 | 0.28384763 | 0.155 | 0.329 | 3.59E-14 | 7 | Tmpo | myCAF_3 |
| Il17ra2 | 1.87E-18 | 0.45565375 | 0.546 | 0.67 | 3.61E-14 | 7 | Il17ra | myCAF_3 |
| Fam114a21 | 1.94E-18 | 0.25404285 | 0.428 | 0.645 | 3.75E-14 | 7 | Fam114a2 | myCAF_3 |
| Ddx39b1 | 1.97E-18 | 0.36542184 | 0.536 | 0.721 | 3.82E-14 | 7 | Ddx39b | myCAF_3 |
| Chmp5 | 2.11E-18 | 0.25743872 | 0.567 | 0.776 | 4.08E-14 | 7 | Chmp5 | myCAF_3 |
| Ebf13 | 2.12E-18 | 0.25071142 | 0.49 | 0.677 | 4.10E-14 | 7 | Ebf1 | myCAF_3 |
| Aamp1 | 2.25E-18 | 0.40389295 | 0.773 | 0.904 | 4.34E-14 | 7 | Aamp | myCAF_3 |
| Ssh32 | 2.35E-18 | 2.50969491 | 0.469 | 0.247 | 4.55E-14 | 7 | Ssh3 | myCAF_3 |
| Mex3c | 2.41E-18 | 0.47907283 | 0.552 | 0.732 | 4.66E-14 | 7 | Mex3c | myCAF_3 |
| Myh92 | 2.46E-18 | 0.49693317 | 0.84 | 0.93 | 4.75E-14 | 7 | Myh9 | myCAF_3 |
| Dync1li1 | 2.54E-18 | 0.52351658 | 0.541 | 0.75 | 4.92E-14 | 7 | Dync1li1 | myCAF_3 |
| Picalm2 | 2.74E-18 | 0.26224181 | 0.732 | 0.886 | 5.30E-14 | 7 | Picalm | myCAF_3 |
| Ndufv11 | 2.88E-18 | 0.46850098 | 0.629 | 0.795 | 5.57E-14 | 7 | Ndufv1 | myCAF_3 |
| Tmtc11 | 3.58E-18 | 2.35197959 | 0.34 | 0.161 | 6.93E-14 | 7 | Tmtc1 | myCAF_3 |
| Med131 | 3.82E-18 | 0.39318493 | 0.619 | 0.781 | 7.38E-14 | 7 | Med13 | myCAF_3 |
| Ppp1r15a3 | 4.06E-18 | 0.86654733 | 0.521 | 0.644 | 7.85E-14 | 7 | Ppp1r15a | myCAF_3 |
| Srrm21 | 4.30E-18 | 0.57999297 | 0.758 | 0.872 | 8.31E-14 | 7 | Srrm2 | myCAF_3 |
| Rab11b1 | 4.34E-18 | 0.45114493 | 0.593 | 0.802 | 8.39E-14 | 7 | Rab11b | myCAF_3 |
| Tmem176b2 | 4.35E-18 | 1.20211258 | 0.619 | 0.726 | 8.40E-14 | 7 | Tmem176b | myCAF_3 |
| Mxd42 | 4.50E-18 | 0.4202638 | 0.804 | 0.894 | 8.69E-14 | 7 | Mxd4 | myCAF_3 |
| Ncor22 | 4.58E-18 | 0.52055115 | 0.804 | 0.895 | 8.85E-14 | 7 | Ncor2 | myCAF_3 |
| Cebpd3 | 4.59E-18 | 0.99865523 | 0.722 | 0.804 | 8.87E-14 | 7 | Cebpd | myCAF_3 |
| Mllt61 | 4.79E-18 | 0.4192989 | 0.567 | 0.765 | 9.26E-14 | 7 | Mllt6 | myCAF_3 |
| Spag91 | 5.01E-18 | 0.37190612 | 0.619 | 0.784 | 9.69E-14 | 7 | Spag9 | myCAF_3 |
| Srf2 | 5.16E-18 | 0.44012965 | 0.66 | 0.755 | 9.98E-14 | 7 | Srf | myCAF_3 |
| Matr32 | 6.14E-18 | 0.30976904 | 0.474 | 0.677 | 1.19E-13 | 7 | Matr3 | myCAF_3 |
| Tbl1xr11 | 6.53E-18 | 0.29060693 | 0.387 | 0.604 | 1.26E-13 | 7 | Tbl1xr1 | myCAF_3 |
| Atp1b31 | 6.71E-18 | 0.38128981 | 0.598 | 0.745 | 1.30E-13 | 7 | Atp1b3 | myCAF_3 |
| Cab391 | 7.61E-18 | 0.40470601 | 0.644 | 0.808 | 1.47E-13 | 7 | Cab39 | myCAF_3 |
| Lrp101 | 7.91E-18 | 0.41777119 | 0.778 | 0.894 | 1.53E-13 | 7 | Lrp10 | myCAF_3 |
| Drg11 | 8.05E-18 | 0.35961058 | 0.567 | 0.767 | 1.56E-13 | 7 | Drg1 | myCAF_3 |
| Chmp3 | 8.94E-18 | 0.44920484 | 0.649 | 0.794 | 1.73E-13 | 7 | Chmp3 | myCAF_3 |
| Pxdc13 | 9.49E-18 | 1.29230386 | 0.603 | 0.545 | 1.83E-13 | 7 | Pxdc1 | myCAF_3 |
| Olfm11 | 9.59E-18 | 1.66479185 | 0.701 | 0.577 | 1.85E-13 | 7 | Olfm1 | myCAF_3 |
| Cdo11 | 1.08E-17 | 1.93712797 | 0.263 | 0.184 | 2.10E-13 | 7 | Cdo1 | myCAF_3 |
| Vps26b | 1.12E-17 | 0.27348875 | 0.588 | 0.778 | 2.16E-13 | 7 | Vps26b | myCAF_3 |
| Dbndd23 | 1.15E-17 | 1.64146502 | 0.526 | 0.347 | 2.23E-13 | 7 | Dbndd2 | myCAF_3 |
| Rabl61 | 1.17E-17 | 0.2678064 | 0.557 | 0.746 | 2.26E-13 | 7 | Rabl6 | myCAF_3 |
| Usp471 | 1.19E-17 | 0.60742604 | 0.577 | 0.755 | 2.30E-13 | 7 | Usp47 | myCAF_3 |
| Rbms31 | 1.29E-17 | 0.41438114 | 0.727 | 0.875 | 2.49E-13 | 7 | Rbms3 | myCAF_3 |
| Rarg1 | 1.56E-17 | 0.61554019 | 0.67 | 0.79 | 3.01E-13 | 7 | Rarg | myCAF_3 |
| Chmp1a | 1.56E-17 | 0.28340202 | 0.562 | 0.796 | 3.01E-13 | 7 | Chmp1a | myCAF_3 |
| Herpud11 | 1.73E-17 | 0.36895591 | 0.546 | 0.735 | 3.34E-13 | 7 | Herpud1 | myCAF_3 |
| Cs1 | 1.76E-17 | 0.25008651 | 0.381 | 0.599 | 3.41E-13 | 7 | Cs | myCAF_3 |
| Stat63 | 1.78E-17 | 0.25964793 | 0.505 | 0.687 | 3.44E-13 | 7 | Stat6 | myCAF_3 |
| Smg1 | 1.83E-17 | 0.2642045 | 0.438 | 0.672 | 3.55E-13 | 7 | Smg1 | myCAF_3 |
| Eif1a1 | 1.94E-17 | 0.56581482 | 0.665 | 0.801 | 3.75E-13 | 7 | Eif1a | myCAF_3 |
| Atp6v1g11 | 2.18E-17 | 0.39613708 | 0.727 | 0.861 | 4.21E-13 | 7 | Atp6v1g1 | myCAF_3 |
| Etnk11 | 2.24E-17 | 0.40327074 | 0.552 | 0.747 | 4.33E-13 | 7 | Etnk1 | myCAF_3 |
| Lpl3 | 2.61E-17 | 0.34100716 | 0.423 | 0.59 | 5.04E-13 | 7 | Lpl | myCAF_3 |
| Fbln51 | 2.74E-17 | 1.1454494 | 0.68 | 0.651 | 5.29E-13 | 7 | Fbln5 | myCAF_3 |
| Cox7a2l1 | 2.77E-17 | 0.2645091 | 0.608 | 0.798 | 5.36E-13 | 7 | Cox7a2l | myCAF_3 |
| Nudt31 | 2.90E-17 | 0.42012333 | 0.644 | 0.824 | 5.61E-13 | 7 | Nudt3 | myCAF_3 |
| Ppp1r9b | 2.95E-17 | 0.40750386 | 0.691 | 0.854 | 5.71E-13 | 7 | Ppp1r9b | myCAF_3 |
| Itga91 | 3.42E-17 | 0.47673585 | 0.284 | 0.524 | 6.62E-13 | 7 | Itga9 | myCAF_3 |
| Atxn7l3 | 3.59E-17 | 0.66751327 | 0.552 | 0.755 | 6.93E-13 | 7 | Atxn7l3 | myCAF_3 |
| Exoc5 | 3.69E-17 | 0.31036731 | 0.567 | 0.758 | 7.14E-13 | 7 | Exoc5 | myCAF_3 |
| Dstn1 | 3.73E-17 | 0.68679941 | 0.814 | 0.896 | 7.21E-13 | 7 | Dstn | myCAF_3 |
| Sympk1 | 3.78E-17 | 0.39549636 | 0.505 | 0.699 | 7.31E-13 | 7 | Sympk | myCAF_3 |
| Tsc22d1 | 3.79E-17 | 0.882035 | 0.763 | 0.892 | 7.33E-13 | 7 | Tsc22d1 | myCAF_3 |
| Ist11 | 3.81E-17 | 0.47847089 | 0.67 | 0.817 | 7.37E-13 | 7 | Ist1 | myCAF_3 |
| Pcdhgc33 | 4.01E-17 | 0.83234376 | 0.778 | 0.843 | 7.75E-13 | 7 | Pcdhgc3 | myCAF_3 |
| Raf1 | 4.26E-17 | 0.63550214 | 0.567 | 0.736 | 8.23E-13 | 7 | Raf1 | myCAF_3 |
| Rsu12 | 4.30E-17 | 0.35524537 | 0.696 | 0.846 | 8.32E-13 | 7 | Rsu1 | myCAF_3 |
| Emd1 | 4.62E-17 | 0.57232798 | 0.433 | 0.617 | 8.93E-13 | 7 | Emd | myCAF_3 |
| Hcfc11 | 4.64E-17 | 0.386087 | 0.515 | 0.687 | 8.98E-13 | 7 | Hcfc1 | myCAF_3 |
| Cinp1 | 4.66E-17 | 0.30790827 | 0.392 | 0.575 | 9.00E-13 | 7 | Cinp | myCAF_3 |
| M6pr2 | 4.71E-17 | 0.25288206 | 0.649 | 0.81 | 9.10E-13 | 7 | M6pr | myCAF_3 |
| Stk111 | 5.04E-17 | 0.36042655 | 0.567 | 0.768 | 9.75E-13 | 7 | Stk11 | myCAF_3 |
| Zbtb381 | 5.19E-17 | 0.81024735 | 0.531 | 0.7 | 1.00E-12 | 7 | Zbtb38 | myCAF_3 |
| Babam1 | 5.34E-17 | 0.32618292 | 0.515 | 0.736 | 1.03E-12 | 7 | Babam1 | myCAF_3 |
| Sema4b1 | 5.41E-17 | 0.37915687 | 0.273 | 0.538 | 1.05E-12 | 7 | Sema4b | myCAF_3 |
| Tshz23 | 5.81E-17 | 0.67038814 | 0.211 | 0.436 | 1.12E-12 | 7 | Tshz2 | myCAF_3 |
| Becn1 | 6.21E-17 | 0.31546116 | 0.541 | 0.747 | 1.20E-12 | 7 | Becn1 | myCAF_3 |
| Loxl43 | 6.98E-17 | 1.82855752 | 0.495 | 0.324 | 1.35E-12 | 7 | Loxl4 | myCAF_3 |
| Top2b1 | 8.07E-17 | 0.29491048 | 0.526 | 0.727 | 1.56E-12 | 7 | Top2b | myCAF_3 |
| Dync1li2 | 8.33E-17 | 0.71961081 | 0.747 | 0.835 | 1.61E-12 | 7 | Dync1li2 | myCAF_3 |
| Nktr2 | 8.82E-17 | 0.30060227 | 0.546 | 0.752 | 1.71E-12 | 7 | Nktr | myCAF_3 |
| Trappc6b | 9.33E-17 | 0.27972774 | 0.531 | 0.714 | 1.80E-12 | 7 | Trappc6b | myCAF_3 |
| Dctn4 | 9.87E-17 | 0.25234315 | 0.577 | 0.749 | 1.91E-12 | 7 | Dctn4 | myCAF_3 |
| Brwd12 | 1.04E-16 | 0.3684318 | 0.51 | 0.708 | 2.01E-12 | 7 | Brwd1 | myCAF_3 |
| Fam32a | 1.06E-16 | 0.26311464 | 0.588 | 0.778 | 2.05E-12 | 7 | Fam32a | myCAF_3 |
| Ddx24 | 1.14E-16 | 0.49771174 | 0.546 | 0.752 | 2.21E-12 | 7 | Ddx24 | myCAF_3 |
| Thoc71 | 1.18E-16 | 0.26940727 | 0.577 | 0.755 | 2.28E-12 | 7 | Thoc7 | myCAF_3 |
| Chp11 | 1.22E-16 | 0.48261207 | 0.727 | 0.857 | 2.36E-12 | 7 | Chp1 | myCAF_3 |
| Os9 | 1.23E-16 | 0.48339031 | 0.696 | 0.857 | 2.37E-12 | 7 | Os9 | myCAF_3 |
| Bckdk1 | 1.26E-16 | 0.31458026 | 0.443 | 0.633 | 2.44E-12 | 7 | Bckdk | myCAF_3 |
| Npdc13 | 1.29E-16 | 0.41063248 | 0.696 | 0.819 | 2.50E-12 | 7 | Npdc1 | myCAF_3 |
| Plekho23 | 1.44E-16 | 0.32124595 | 0.701 | 0.857 | 2.79E-12 | 7 | Plekho2 | myCAF_3 |
| Usp8 | 1.46E-16 | 0.27219165 | 0.552 | 0.737 | 2.82E-12 | 7 | Usp8 | myCAF_3 |
| Ilf21 | 1.66E-16 | 0.35302305 | 0.407 | 0.596 | 3.21E-12 | 7 | Ilf2 | myCAF_3 |
| Plpbp2 | 1.71E-16 | 0.26383413 | 0.351 | 0.56 | 3.30E-12 | 7 | Plpbp | myCAF_3 |
| DEPRECATED-l | 1.74E-16 | 0.32471513 | 0.577 | 0.718 | 3.37E-12 | 7 | DEPRECATED-l | myCAF_3 |
| Cnot6l2 | 1.77E-16 | 0.30342196 | 0.381 | 0.608 | 3.43E-12 | 7 | Cnot6l | myCAF_3 |
| Arf3 | 1.79E-16 | 0.41666721 | 0.711 | 0.867 | 3.45E-12 | 7 | Arf3 | myCAF_3 |
| Fgfr1op21 | 1.80E-16 | 0.32521006 | 0.649 | 0.815 | 3.48E-12 | 7 | Fgfr1op2 | myCAF_3 |
| Smarcb1 | 1.91E-16 | 0.4565505 | 0.531 | 0.737 | 3.69E-12 | 7 | Smarcb1 | myCAF_3 |
| Ptk22 | 2.00E-16 | 0.25382247 | 0.505 | 0.666 | 3.87E-12 | 7 | Ptk2 | myCAF_3 |
| Flot23 | 2.12E-16 | 0.27088944 | 0.521 | 0.732 | 4.10E-12 | 7 | Flot2 | myCAF_3 |
| Ap3d1 | 2.16E-16 | 0.28080478 | 0.593 | 0.795 | 4.17E-12 | 7 | Ap3d1 | myCAF_3 |
| Clu4 | 2.20E-16 | 0.57650645 | 0.866 | 0.887 | 4.25E-12 | 7 | Clu | myCAF_3 |
| Stx4a3 | 2.23E-16 | 0.43693316 | 0.552 | 0.742 | 4.31E-12 | 7 | Stx4a | myCAF_3 |
| Timm17a1 | 2.41E-16 | 0.34918024 | 0.526 | 0.7 | 4.65E-12 | 7 | Timm17a | myCAF_3 |
| Psap1 | 2.45E-16 | 0.52926901 | 0.84 | 0.952 | 4.75E-12 | 7 | Psap | myCAF_3 |
| Pdgfrb3 | 2.84E-16 | 0.52531525 | 0.866 | 0.946 | 5.49E-12 | 7 | Pdgfrb | myCAF_3 |
| Gsk3a1 | 2.94E-16 | 0.30037174 | 0.572 | 0.742 | 5.68E-12 | 7 | Gsk3a | myCAF_3 |
| Impa1 | 2.99E-16 | 0.34657226 | 0.412 | 0.639 | 5.78E-12 | 7 | Impa1 | myCAF_3 |
| Lix1l | 3.21E-16 | 0.69023822 | 0.851 | 0.939 | 6.21E-12 | 7 | Lix1l | myCAF_3 |
| Isyna12 | 3.23E-16 | 1.52754555 | 0.34 | 0.477 | 6.25E-12 | 7 | Isyna1 | myCAF_3 |
| Immt1 | 3.42E-16 | 0.36969762 | 0.479 | 0.639 | 6.62E-12 | 7 | Immt | myCAF_3 |
| Rgma2 | 3.58E-16 | 0.44309357 | 0.567 | 0.704 | 6.93E-12 | 7 | Rgma | myCAF_3 |
| Myo102 | 3.59E-16 | 1.50273638 | 0.67 | 0.548 | 6.93E-12 | 7 | Myo10 | myCAF_3 |
| Cebpa2 | 3.81E-16 | 2.26149955 | 0.68 | 0.63 | 7.36E-12 | 7 | Cebpa | myCAF_3 |
| Arglu11 | 4.04E-16 | 0.25740906 | 0.67 | 0.804 | 7.81E-12 | 7 | Arglu1 | myCAF_3 |
| Hibadh | 4.10E-16 | 0.39577961 | 0.526 | 0.71 | 7.93E-12 | 7 | Hibadh | myCAF_3 |
| Sh3d192 | 4.20E-16 | 0.25031022 | 0.608 | 0.79 | 8.11E-12 | 7 | Sh3d19 | myCAF_3 |
| Crebbp1 | 4.27E-16 | 0.5357309 | 0.485 | 0.71 | 8.26E-12 | 7 | Crebbp | myCAF_3 |
| Med15 | 4.71E-16 | 0.41205349 | 0.629 | 0.775 | 9.11E-12 | 7 | Med15 | myCAF_3 |
| Cyc11 | 4.77E-16 | 0.52208318 | 0.66 | 0.756 | 9.23E-12 | 7 | Cyc1 | myCAF_3 |
| Tmem1582 | 4.92E-16 | 1.8901373 | 0.392 | 0.247 | 9.51E-12 | 7 | Tmem158 | myCAF_3 |
| Naa151 | 5.09E-16 | 0.28314428 | 0.351 | 0.579 | 9.84E-12 | 7 | Naa15 | myCAF_3 |
| Keap1 | 5.24E-16 | 0.28267062 | 0.5 | 0.72 | 1.01E-11 | 7 | Keap1 | myCAF_3 |
| Naa201 | 5.26E-16 | 0.3200266 | 0.531 | 0.7 | 1.02E-11 | 7 | Naa20 | myCAF_3 |
| Rras1 | 6.25E-16 | 0.32377973 | 0.67 | 0.856 | 1.21E-11 | 7 | Rras | myCAF_3 |
| Aldh22 | 6.50E-16 | 0.39533505 | 0.773 | 0.888 | 1.26E-11 | 7 | Aldh2 | myCAF_3 |
| Atrx1 | 6.54E-16 | 0.54760592 | 0.515 | 0.696 | 1.27E-11 | 7 | Atrx | myCAF_3 |
| Csnk2b1 | 6.56E-16 | 0.39260269 | 0.469 | 0.642 | 1.27E-11 | 7 | Csnk2b | myCAF_3 |
| Mdfic2 | 6.65E-16 | 0.32037945 | 0.577 | 0.774 | 1.29E-11 | 7 | Mdfic | myCAF_3 |
| Srrt1 | 6.81E-16 | 0.67693279 | 0.479 | 0.639 | 1.32E-11 | 7 | Srrt | myCAF_3 |
| Foxp4 | 6.85E-16 | 0.46029596 | 0.454 | 0.672 | 1.32E-11 | 7 | Foxp4 | myCAF_3 |
| Pcf11 | 6.88E-16 | 0.60027225 | 0.51 | 0.68 | 1.33E-11 | 7 | Pcf11 | myCAF_3 |
| Mapk31 | 7.11E-16 | 0.38637326 | 0.526 | 0.711 | 1.38E-11 | 7 | Mapk3 | myCAF_3 |
| Mknk22 | 7.23E-16 | 0.33530651 | 0.433 | 0.655 | 1.40E-11 | 7 | Mknk2 | myCAF_3 |
| Zfp5323 | 7.35E-16 | 0.27448093 | 0.593 | 0.734 | 1.42E-11 | 7 | Zfp532 | myCAF_3 |
| Ing12 | 7.64E-16 | 0.43950165 | 0.5 | 0.673 | 1.48E-11 | 7 | Ing1 | myCAF_3 |
| Cdk41 | 7.87E-16 | 0.36237355 | 0.49 | 0.704 | 1.52E-11 | 7 | Cdk4 | myCAF_3 |
| Nipbl1 | 8.25E-16 | 0.45057663 | 0.387 | 0.618 | 1.60E-11 | 7 | Nipbl | myCAF_3 |
| Mon2 | 8.72E-16 | 0.36659457 | 0.552 | 0.734 | 1.69E-11 | 7 | Mon2 | myCAF_3 |
| Psen11 | 8.81E-16 | 0.34544788 | 0.521 | 0.684 | 1.70E-11 | 7 | Psen1 | myCAF_3 |
| Tut72 | 1.01E-15 | 0.53432362 | 0.593 | 0.764 | 1.96E-11 | 7 | Tut7 | myCAF_3 |
| Mef2d2 | 1.02E-15 | 0.54202026 | 0.644 | 0.788 | 1.98E-11 | 7 | Mef2d | myCAF_3 |
| Anapc16 | 1.06E-15 | 0.42464652 | 0.505 | 0.695 | 2.04E-11 | 7 | Anapc16 | myCAF_3 |
| Fzr11 | 1.08E-15 | 0.47158065 | 0.356 | 0.531 | 2.09E-11 | 7 | Fzr1 | myCAF_3 |
| Pcm12 | 1.20E-15 | 0.3675381 | 0.371 | 0.593 | 2.31E-11 | 7 | Pcm1 | myCAF_3 |
| Clmp1 | 1.24E-15 | 0.70595463 | 0.861 | 0.926 | 2.40E-11 | 7 | Clmp | myCAF_3 |
| Phc2 | 1.31E-15 | 0.46431696 | 0.675 | 0.787 | 2.53E-11 | 7 | Phc2 | myCAF_3 |
| Cic1 | 1.36E-15 | 0.27689275 | 0.459 | 0.687 | 2.62E-11 | 7 | Cic | myCAF_3 |
| Stau11 | 1.39E-15 | 0.46251213 | 0.593 | 0.753 | 2.69E-11 | 7 | Stau1 | myCAF_3 |
| Txndc11 | 1.41E-15 | 0.43367912 | 0.598 | 0.748 | 2.72E-11 | 7 | Txndc11 | myCAF_3 |
| Gpr1761 | 1.42E-15 | 0.34280772 | 0.598 | 0.695 | 2.74E-11 | 7 | Gpr176 | myCAF_3 |
| Tnrc18 | 1.48E-15 | 0.6265857 | 0.747 | 0.867 | 2.86E-11 | 7 | Tnrc18 | myCAF_3 |
| Rhbdf11 | 1.63E-15 | 0.2529624 | 0.577 | 0.773 | 3.15E-11 | 7 | Rhbdf1 | myCAF_3 |
| Glyr1 | 1.66E-15 | 0.52328794 | 0.582 | 0.767 | 3.20E-11 | 7 | Glyr1 | myCAF_3 |
| Cnih41 | 1.81E-15 | 0.25084626 | 0.381 | 0.572 | 3.50E-11 | 7 | Cnih4 | myCAF_3 |
| Clstn12 | 2.06E-15 | 0.33174156 | 0.593 | 0.763 | 3.99E-11 | 7 | Clstn1 | myCAF_3 |
| Sgrd1 | 2.30E-15 | 0.47418583 | 0.753 | 0.891 | 4.44E-11 | 7 | Sgrd1 | myCAF_3 |
| Fxyd13 | 2.32E-15 | 0.4648281 | 0.531 | 0.638 | 4.49E-11 | 7 | Fxyd1 | myCAF_3 |
| Gga1 | 2.47E-15 | 0.53224546 | 0.505 | 0.698 | 4.77E-11 | 7 | Gga1 | myCAF_3 |
| Fam102b2 | 2.55E-15 | 0.27195969 | 0.531 | 0.656 | 4.92E-11 | 7 | Fam102b | myCAF_3 |
| Ago22 | 2.68E-15 | 0.3984351 | 0.619 | 0.782 | 5.17E-11 | 7 | Ago2 | myCAF_3 |
| Hspa91 | 2.69E-15 | 0.3170414 | 0.49 | 0.649 | 5.19E-11 | 7 | Hspa9 | myCAF_3 |
| Rgs31 | 2.75E-15 | 0.28348926 | 0.5 | 0.691 | 5.32E-11 | 7 | Rgs3 | myCAF_3 |
| Ier52 | 2.77E-15 | 0.95664747 | 0.706 | 0.757 | 5.35E-11 | 7 | Ier5 | myCAF_3 |
| Mob1b | 2.89E-15 | 0.29884668 | 0.309 | 0.565 | 5.59E-11 | 7 | Mob1b | myCAF_3 |
| Ik1 | 3.78E-15 | 0.60637498 | 0.634 | 0.779 | 7.30E-11 | 7 | Ik | myCAF_3 |
| Pkn21 | 3.99E-15 | 0.38174704 | 0.381 | 0.592 | 7.71E-11 | 7 | Pkn2 | myCAF_3 |
| Snrnp200 | 4.23E-15 | 0.63344333 | 0.526 | 0.676 | 8.17E-11 | 7 | Snrnp200 | myCAF_3 |
| Sec24c1 | 4.24E-15 | 0.47229195 | 0.557 | 0.755 | 8.20E-11 | 7 | Sec24c | myCAF_3 |
| Usp251 | 4.25E-15 | 0.56070709 | 0.546 | 0.727 | 8.22E-11 | 7 | Usp25 | myCAF_3 |
| Ss182 | 4.50E-15 | 0.58475143 | 0.546 | 0.724 | 8.70E-11 | 7 | Ss18 | myCAF_3 |
| Bola21 | 4.56E-15 | 0.43239749 | 0.552 | 0.764 | 8.82E-11 | 7 | Bola2 | myCAF_3 |
| D8Ertd738e | 4.59E-15 | 0.2755302 | 0.613 | 0.8 | 8.87E-11 | 7 | D8Ertd738e | myCAF_3 |
| Leng82 | 4.90E-15 | 0.50006557 | 0.644 | 0.782 | 9.48E-11 | 7 | Leng8 | myCAF_3 |
| Lsm71 | 4.90E-15 | 0.34029083 | 0.469 | 0.698 | 9.48E-11 | 7 | Lsm7 | myCAF_3 |
| Txndc91 | 4.95E-15 | 0.36105731 | 0.521 | 0.701 | 9.58E-11 | 7 | Txndc9 | myCAF_3 |
| Arhgap351 | 5.01E-15 | 0.51475475 | 0.495 | 0.661 | 9.69E-11 | 7 | Arhgap35 | myCAF_3 |
| Cyb5612 | 5.31E-15 | 0.39694352 | 0.155 | 0.386 | 1.03E-10 | 7 | Cyb561 | myCAF_3 |
| Ppp1r12a1 | 5.59E-15 | 0.30881754 | 0.546 | 0.728 | 1.08E-10 | 7 | Ppp1r12a | myCAF_3 |
| Rad23a1 | 5.78E-15 | 0.32655759 | 0.546 | 0.701 | 1.12E-10 | 7 | Rad23a | myCAF_3 |
| Dyrk21 | 6.58E-15 | 0.55641512 | 0.423 | 0.614 | 1.27E-10 | 7 | Dyrk2 | myCAF_3 |
| Slc9a3r11 | 6.95E-15 | 0.30338063 | 0.309 | 0.473 | 1.34E-10 | 7 | Slc9a3r1 | myCAF_3 |
| Snx182 | 6.99E-15 | 0.2684918 | 0.577 | 0.75 | 1.35E-10 | 7 | Snx18 | myCAF_3 |
| Vbp11 | 7.52E-15 | 0.33016319 | 0.423 | 0.634 | 1.45E-10 | 7 | Vbp1 | myCAF_3 |
| Dnm21 | 7.86E-15 | 0.34813154 | 0.479 | 0.712 | 1.52E-10 | 7 | Dnm2 | myCAF_3 |
| Scaf1 | 8.80E-15 | 0.40591796 | 0.649 | 0.799 | 1.70E-10 | 7 | Scaf1 | myCAF_3 |
| Emp32 | 8.89E-15 | 0.64739387 | 0.768 | 0.852 | 1.72E-10 | 7 | Emp3 | myCAF_3 |
| Usp7 | 9.47E-15 | 0.47397167 | 0.552 | 0.725 | 1.83E-10 | 7 | Usp7 | myCAF_3 |
| Wac1 | 1.00E-14 | 0.2889232 | 0.361 | 0.592 | 1.94E-10 | 7 | Wac | myCAF_3 |
| Lzts21 | 1.08E-14 | 0.49055366 | 0.634 | 0.809 | 2.09E-10 | 7 | Lzts2 | myCAF_3 |
| Pde3a1 | 1.15E-14 | 2.13933369 | 0.443 | 0.319 | 2.21E-10 | 7 | Pde3a | myCAF_3 |
| Mal1 | 1.18E-14 | 0.50837846 | 0.613 | 0.763 | 2.29E-10 | 7 | Mal1 | myCAF_3 |
| Git21 | 1.20E-14 | 0.2599742 | 0.376 | 0.601 | 2.32E-10 | 7 | Git2 | myCAF_3 |
| Socs32 | 1.20E-14 | 1.18472482 | 0.722 | 0.795 | 2.33E-10 | 7 | Socs3 | myCAF_3 |
| Hmg20b1 | 1.26E-14 | 0.4604568 | 0.552 | 0.73 | 2.43E-10 | 7 | Hmg20b | myCAF_3 |
| Kpna41 | 1.26E-14 | 0.48902855 | 0.418 | 0.628 | 2.44E-10 | 7 | Kpna4 | myCAF_3 |
| Jak22 | 1.36E-14 | 0.79576945 | 0.753 | 0.814 | 2.63E-10 | 7 | Jak2 | myCAF_3 |
| Fbxo111 | 1.38E-14 | 0.52235834 | 0.464 | 0.671 | 2.67E-10 | 7 | Fbxo11 | myCAF_3 |
| Laptm4b1 | 1.41E-14 | 0.36414997 | 0.505 | 0.655 | 2.72E-10 | 7 | Laptm4b | myCAF_3 |
| Atp6ap1 | 1.42E-14 | 0.47836941 | 0.716 | 0.873 | 2.75E-10 | 7 | Atp6ap1 | myCAF_3 |
| Gdi1 | 1.42E-14 | 0.63307931 | 0.557 | 0.736 | 2.75E-10 | 7 | Gdi1 | myCAF_3 |
| Prkd31 | 1.46E-14 | 0.26157329 | 0.392 | 0.613 | 2.82E-10 | 7 | Prkd3 | myCAF_3 |
| Rbm251 | 1.46E-14 | 0.50308598 | 0.593 | 0.733 | 2.83E-10 | 7 | Rbm25 | myCAF_3 |
| Ccng22 | 1.52E-14 | 0.38568637 | 0.505 | 0.622 | 2.94E-10 | 7 | Ccng2 | myCAF_3 |
| Btbd12 | 1.66E-14 | 0.29378415 | 0.397 | 0.595 | 3.21E-10 | 7 | Btbd1 | myCAF_3 |
| Gatad2b | 1.79E-14 | 0.36207492 | 0.613 | 0.768 | 3.47E-10 | 7 | Gatad2b | myCAF_3 |
| Slc11a2 | 1.87E-14 | 0.25461186 | 0.366 | 0.591 | 3.62E-10 | 7 | Slc11a2 | myCAF_3 |
| Pank32 | 1.93E-14 | 0.32181974 | 0.464 | 0.633 | 3.73E-10 | 7 | Pank3 | myCAF_3 |
| Pacsin21 | 1.97E-14 | 0.57042098 | 0.381 | 0.595 | 3.80E-10 | 7 | Pacsin2 | myCAF_3 |
| Ppp2r2a | 2.05E-14 | 0.30933063 | 0.351 | 0.576 | 3.97E-10 | 7 | Ppp2r2a | myCAF_3 |
| Hat11 | 2.18E-14 | 0.43237003 | 0.201 | 0.333 | 4.22E-10 | 7 | Hat1 | myCAF_3 |
| Kdm5c | 2.27E-14 | 0.35831031 | 0.536 | 0.74 | 4.40E-10 | 7 | Kdm5c | myCAF_3 |
| Adamts121 | 2.29E-14 | 0.31646167 | 0.706 | 0.917 | 4.43E-10 | 7 | Adamts12 | myCAF_3 |
| Cdc341 | 2.43E-14 | 0.34347559 | 0.454 | 0.628 | 4.70E-10 | 7 | Cdc34 | myCAF_3 |
| Rere2 | 2.47E-14 | 0.40613661 | 0.572 | 0.742 | 4.78E-10 | 7 | Rere | myCAF_3 |
| Sbno11 | 2.68E-14 | 0.31384812 | 0.366 | 0.6 | 5.19E-10 | 7 | Sbno1 | myCAF_3 |
| Srsf111 | 2.75E-14 | 0.33414741 | 0.577 | 0.737 | 5.32E-10 | 7 | Srsf11 | myCAF_3 |
| Cdc271 | 2.85E-14 | 0.28240011 | 0.423 | 0.588 | 5.51E-10 | 7 | Cdc27 | myCAF_3 |
| G3bp11 | 3.09E-14 | 0.30965707 | 0.397 | 0.569 | 5.97E-10 | 7 | G3bp1 | myCAF_3 |
| Piezo11 | 3.19E-14 | 0.48814516 | 0.649 | 0.775 | 6.16E-10 | 7 | Piezo1 | myCAF_3 |
| Cdc42bpa2 | 3.23E-14 | 0.49432722 | 0.701 | 0.776 | 6.25E-10 | 7 | Cdc42bpa | myCAF_3 |
| Tmem1151 | 3.34E-14 | 0.32436651 | 0.459 | 0.688 | 6.46E-10 | 7 | Tmem115 | myCAF_3 |
| Ppp4r3a1 | 3.51E-14 | 0.38315206 | 0.278 | 0.517 | 6.79E-10 | 7 | Ppp4r3a | myCAF_3 |
| Smc31 | 3.58E-14 | 0.70939135 | 0.366 | 0.535 | 6.92E-10 | 7 | Smc3 | myCAF_3 |
| Etv61 | 3.65E-14 | 0.42713962 | 0.552 | 0.677 | 7.05E-10 | 7 | Etv6 | myCAF_3 |
| Sema4c | 3.72E-14 | 0.61301001 | 0.557 | 0.69 | 7.18E-10 | 7 | Sema4c | myCAF_3 |
| Oga2 | 3.77E-14 | 0.43422463 | 0.485 | 0.675 | 7.29E-10 | 7 | Oga | myCAF_3 |
| Pdcd43 | 3.92E-14 | 0.65603978 | 0.531 | 0.66 | 7.57E-10 | 7 | Pdcd4 | myCAF_3 |
| Ctsk2 | 3.94E-14 | 0.9208049 | 0.83 | 0.975 | 7.62E-10 | 7 | Ctsk | myCAF_3 |
| Nxf12 | 4.05E-14 | 0.57591834 | 0.531 | 0.696 | 7.84E-10 | 7 | Nxf1 | myCAF_3 |
| Ogfr | 4.10E-14 | 0.28207979 | 0.598 | 0.746 | 7.92E-10 | 7 | Ogfr | myCAF_3 |
| Spop | 4.22E-14 | 0.51850104 | 0.758 | 0.873 | 8.16E-10 | 7 | Spop | myCAF_3 |
| Limd12 | 4.39E-14 | 0.49835786 | 0.67 | 0.797 | 8.48E-10 | 7 | Limd1 | myCAF_3 |
| Cuta | 4.60E-14 | 0.35014729 | 0.613 | 0.776 | 8.89E-10 | 7 | Cuta | myCAF_3 |
| Prkar2b | 4.78E-14 | 1.77256513 | 0.289 | 0.143 | 9.24E-10 | 7 | Prkar2b | myCAF_3 |
| Ptbp21 | 4.79E-14 | 0.40535525 | 0.438 | 0.659 | 9.27E-10 | 7 | Ptbp2 | myCAF_3 |
| Mxi12 | 4.80E-14 | 0.29881703 | 0.345 | 0.514 | 9.28E-10 | 7 | Mxi1 | myCAF_3 |
| Slc4a22 | 5.04E-14 | 0.35965733 | 0.526 | 0.717 | 9.75E-10 | 7 | Slc4a2 | myCAF_3 |
| Stoml21 | 5.09E-14 | 0.34950455 | 0.438 | 0.611 | 9.85E-10 | 7 | Stoml2 | myCAF_3 |
| Xpo6 | 5.19E-14 | 0.56859228 | 0.531 | 0.719 | 1.00E-09 | 7 | Xpo6 | myCAF_3 |
| Mgrn12 | 5.26E-14 | 0.30368621 | 0.686 | 0.835 | 1.02E-09 | 7 | Mgrn1 | myCAF_3 |
| Srsf41 | 5.34E-14 | 0.31773869 | 0.505 | 0.683 | 1.03E-09 | 7 | Srsf4 | myCAF_3 |
| Hnrnpdl1 | 5.61E-14 | 0.51676673 | 0.381 | 0.554 | 1.08E-09 | 7 | Hnrnpdl | myCAF_3 |
| Tmem33 | 5.89E-14 | 0.3967132 | 0.469 | 0.649 | 1.14E-09 | 7 | Tmem33 | myCAF_3 |
| Man1a21 | 6.03E-14 | 0.54156561 | 0.557 | 0.702 | 1.17E-09 | 7 | Man1a2 | myCAF_3 |
| Spry22 | 7.34E-14 | 0.42893423 | 0.485 | 0.588 | 1.42E-09 | 7 | Spry2 | myCAF_3 |
| Tor1a | 7.96E-14 | 0.30516648 | 0.387 | 0.601 | 1.54E-09 | 7 | Tor1a | myCAF_3 |
| Psmd11 | 8.04E-14 | 0.62219901 | 0.588 | 0.73 | 1.55E-09 | 7 | Psmd11 | myCAF_3 |
| Aco1 | 8.28E-14 | 0.36598666 | 0.454 | 0.643 | 1.60E-09 | 7 | Aco1 | myCAF_3 |
| Tmem63b | 8.53E-14 | 0.47204717 | 0.361 | 0.579 | 1.65E-09 | 7 | Tmem63b | myCAF_3 |
| Yy12 | 9.55E-14 | 0.73555922 | 0.737 | 0.85 | 1.85E-09 | 7 | Yy1 | myCAF_3 |
| Birc31 | 9.93E-14 | 0.48406354 | 0.479 | 0.604 | 1.92E-09 | 7 | Birc3 | myCAF_3 |
| Chd21 | 1.00E-13 | 0.51123046 | 0.273 | 0.49 | 1.94E-09 | 7 | Chd2 | myCAF_3 |
| Ccdc501 | 1.01E-13 | 0.41125578 | 0.387 | 0.611 | 1.96E-09 | 7 | Ccdc50 | myCAF_3 |
| Psme4 | 1.05E-13 | 0.63112429 | 0.526 | 0.682 | 2.02E-09 | 7 | Psme4 | myCAF_3 |
| Adam171 | 1.11E-13 | 0.54884084 | 0.598 | 0.761 | 2.16E-09 | 7 | Adam17 | myCAF_3 |
| Nploc41 | 1.12E-13 | 0.31424134 | 0.557 | 0.702 | 2.16E-09 | 7 | Nploc4 | myCAF_3 |
| Cdk5rap3 | 1.16E-13 | 0.30104212 | 0.577 | 0.742 | 2.24E-09 | 7 | Cdk5rap3 | myCAF_3 |
| Fbxl172 | 1.16E-13 | 0.54391841 | 0.335 | 0.555 | 2.25E-09 | 7 | Fbxl17 | myCAF_3 |
| Clcn41 | 1.34E-13 | 0.38332416 | 0.51 | 0.737 | 2.60E-09 | 7 | Clcn4 | myCAF_3 |
| Atf71 | 1.36E-13 | 0.27599966 | 0.603 | 0.743 | 2.62E-09 | 7 | Atf7 | myCAF_3 |
| Sec23b | 1.36E-13 | 0.26077756 | 0.304 | 0.541 | 2.63E-09 | 7 | Sec23b | myCAF_3 |
| Cop11 | 1.43E-13 | 1.02635986 | 0.485 | 0.671 | 2.77E-09 | 7 | Cop1 | myCAF_3 |
| Sf3b41 | 1.48E-13 | 0.62808157 | 0.433 | 0.588 | 2.87E-09 | 7 | Sf3b4 | myCAF_3 |
| Ssbp22 | 1.52E-13 | 0.57739374 | 0.16 | 0.401 | 2.94E-09 | 7 | Ssbp2 | myCAF_3 |
| Cdk122 | 1.65E-13 | 0.25215164 | 0.309 | 0.54 | 3.18E-09 | 7 | Cdk12 | myCAF_3 |
| Arap12 | 1.88E-13 | 0.50384887 | 0.716 | 0.818 | 3.63E-09 | 7 | Arap1 | myCAF_3 |
| Rela | 1.96E-13 | 0.4375257 | 0.624 | 0.793 | 3.79E-09 | 7 | Rela | myCAF_3 |
| Dpp8 | 2.04E-13 | 0.3437918 | 0.526 | 0.739 | 3.94E-09 | 7 | Dpp8 | myCAF_3 |
| Coq10b | 2.07E-13 | 0.34967221 | 0.392 | 0.575 | 4.00E-09 | 7 | Coq10b | myCAF_3 |
| Tax1bp1 | 2.09E-13 | 0.33323901 | 0.624 | 0.769 | 4.04E-09 | 7 | Tax1bp1 | myCAF_3 |
| Fbxw111 | 2.14E-13 | 0.79721936 | 0.541 | 0.727 | 4.13E-09 | 7 | Fbxw11 | myCAF_3 |
| Dync1i21 | 2.17E-13 | 0.52855592 | 0.577 | 0.756 | 4.19E-09 | 7 | Dync1i2 | myCAF_3 |
| Setd71 | 2.24E-13 | 0.45234899 | 0.572 | 0.73 | 4.33E-09 | 7 | Setd7 | myCAF_3 |
| Kdm3a2 | 2.36E-13 | 0.6070441 | 0.314 | 0.537 | 4.56E-09 | 7 | Kdm3a | myCAF_3 |
| Mar-61 | 2.40E-13 | 0.4285964 | 0.515 | 0.719 | 4.64E-09 | 7 | 6-Mar | myCAF_3 |
| Cnpy41 | 2.41E-13 | 0.4151117 | 0.402 | 0.61 | 4.65E-09 | 7 | Cnpy4 | myCAF_3 |
| Dcaf81 | 2.52E-13 | 0.35761709 | 0.546 | 0.714 | 4.88E-09 | 7 | Dcaf8 | myCAF_3 |
| Taok21 | 2.64E-13 | 0.26024579 | 0.51 | 0.685 | 5.10E-09 | 7 | Taok2 | myCAF_3 |
| Setbp12 | 2.66E-13 | 0.63415451 | 0.392 | 0.585 | 5.14E-09 | 7 | Setbp1 | myCAF_3 |
| N4bp2l22 | 3.02E-13 | 0.67976705 | 0.361 | 0.564 | 5.84E-09 | 7 | N4bp2l2 | myCAF_3 |
| Kirrel | 3.07E-13 | 0.60752709 | 0.851 | 0.932 | 5.93E-09 | 7 | Kirrel | myCAF_3 |
| Flt11 | 3.14E-13 | 1.2667443 | 0.294 | 0.165 | 6.06E-09 | 7 | Flt1 | myCAF_3 |
| Ctdsp1 | 3.18E-13 | 0.51137559 | 0.747 | 0.899 | 6.15E-09 | 7 | Ctdsp1 | myCAF_3 |
| Tab2 | 3.26E-13 | 0.36776365 | 0.582 | 0.749 | 6.31E-09 | 7 | Tab2 | myCAF_3 |
| Smchd11 | 3.27E-13 | 0.75855112 | 0.284 | 0.477 | 6.32E-09 | 7 | Smchd1 | myCAF_3 |
| Spsb12 | 3.34E-13 | 0.45234492 | 0.485 | 0.586 | 6.46E-09 | 7 | Spsb1 | myCAF_3 |
| Ncaph21 | 3.57E-13 | 0.40676063 | 0.325 | 0.518 | 6.91E-09 | 7 | Ncaph2 | myCAF_3 |
| Acin11 | 3.62E-13 | 0.53645506 | 0.639 | 0.764 | 6.99E-09 | 7 | Acin1 | myCAF_3 |
| Jag12 | 3.62E-13 | 0.27789951 | 0.485 | 0.531 | 7.00E-09 | 7 | Jag1 | myCAF_3 |
| Tob13 | 3.67E-13 | 0.96747015 | 0.433 | 0.487 | 7.09E-09 | 7 | Tob1 | myCAF_3 |
| Atxn7l3b1 | 3.78E-13 | 0.38687254 | 0.701 | 0.833 | 7.31E-09 | 7 | Atxn7l3b | myCAF_3 |
| Stat5a2 | 3.90E-13 | 0.66443704 | 0.242 | 0.449 | 7.54E-09 | 7 | Stat5a | myCAF_3 |
| Nsmce4a1 | 3.93E-13 | 0.2513118 | 0.361 | 0.533 | 7.59E-09 | 7 | Nsmce4a | myCAF_3 |
| Dnajb6 | 3.99E-13 | 0.53692969 | 0.495 | 0.69 | 7.71E-09 | 7 | Dnajb6 | myCAF_3 |

| | | | | | | | | |
|---|---|---|---|---|---|---|---|---|
| Atl21 | 4.01E-13 | 0.39648256 | 0.237 | 0.48 | 7.76E-09 | 7 | Atl2 | myCAF_3 |
| Tlk21 | 4.11E-13 | 0.26010553 | 0.309 | 0.537 | 7.95E-09 | 7 | Tlk2 | myCAF_3 |
| Dcaf121 | 4.15E-13 | 0.25253575 | 0.428 | 0.646 | 8.02E-09 | 7 | Dcaf12 | myCAF_3 |
| Zfp911 | 4.30E-13 | 0.55896724 | 0.603 | 0.768 | 8.32E-09 | 7 | Zfp91 | myCAF_3 |
| Rbm261 | 4.42E-13 | 0.36007453 | 0.433 | 0.637 | 8.54E-09 | 7 | Rbm26 | myCAF_3 |
| Git11 | 4.44E-13 | 0.58197492 | 0.51 | 0.674 | 8.59E-09 | 7 | Git1 | myCAF_3 |
| Lars1 | 4.79E-13 | 0.59462445 | 0.423 | 0.607 | 9.26E-09 | 7 | Lars | myCAF_3 |
| Fam160b1 | 4.99E-13 | 0.45507987 | 0.531 | 0.706 | 9.65E-09 | 7 | Fam160b1 | myCAF_3 |
| Ltbp13 | 4.99E-13 | 1.83746697 | 0.289 | 0.384 | 9.66E-09 | 7 | Ltbp1 | myCAF_3 |
| Ubxn11 | 5.30E-13 | 0.38790382 | 0.619 | 0.808 | 1.02E-08 | 7 | Ubxn1 | myCAF_3 |
| Fcgrt2 | 5.45E-13 | 0.55026953 | 0.794 | 0.895 | 1.05E-08 | 7 | Fcgrt | myCAF_3 |
| Ubr2 | 6.05E-13 | 0.88784512 | 0.68 | 0.77 | 1.17E-08 | 7 | Ubr2 | myCAF_3 |
| Cd2bp21 | 6.32E-13 | 0.42247563 | 0.51 | 0.701 | 1.22E-08 | 7 | Cd2bp2 | myCAF_3 |
| Ralbp12 | 6.35E-13 | 0.45691663 | 0.572 | 0.735 | 1.23E-08 | 7 | Ralbp1 | myCAF_3 |
| Gpr1071 | 6.50E-13 | 0.39957149 | 0.572 | 0.72 | 1.26E-08 | 7 | Gpr107 | myCAF_3 |
| Smim151 | 6.66E-13 | 0.29068072 | 0.314 | 0.541 | 1.29E-08 | 7 | Smim15 | myCAF_3 |
| Ubr3 | 7.11E-13 | 1.0760875 | 0.572 | 0.722 | 1.38E-08 | 7 | Ubr3 | myCAF_3 |
| Tbpl12 | 7.21E-13 | 0.54036609 | 0.201 | 0.424 | 1.39E-08 | 7 | Tbpl1 | myCAF_3 |
| Rcbtb11 | 7.36E-13 | 0.59791228 | 0.351 | 0.541 | 1.42E-08 | 7 | Rcbtb1 | myCAF_3 |
| Fam104a | 7.40E-13 | 0.3245364 | 0.423 | 0.623 | 1.43E-08 | 7 | Fam104a | myCAF_3 |
| Elf13 | 7.73E-13 | 0.65460922 | 0.381 | 0.582 | 1.49E-08 | 7 | Elf1 | myCAF_3 |
| Mtmr9 | 8.48E-13 | 0.3809742 | 0.381 | 0.605 | 1.64E-08 | 7 | Mtmr9 | myCAF_3 |
| Rhog | 8.66E-13 | 0.26095003 | 0.49 | 0.671 | 1.68E-08 | 7 | Rhog | myCAF_3 |
| Kmt2d1 | 8.71E-13 | 0.50986606 | 0.526 | 0.74 | 1.68E-08 | 7 | Kmt2d | myCAF_3 |
| Numa11 | 8.96E-13 | 0.59144268 | 0.546 | 0.728 | 1.73E-08 | 7 | Numa1 | myCAF_3 |
| Ash1l1 | 9.77E-13 | 0.56830973 | 0.521 | 0.716 | 1.89E-08 | 7 | Ash1l | myCAF_3 |
| Irf11 | 9.94E-13 | 0.5648585 | 0.454 | 0.664 | 1.92E-08 | 7 | Irf1 | myCAF_3 |
| Khdrbs31 | 1.06E-12 | 0.31149616 | 0.278 | 0.428 | 2.04E-08 | 7 | Khdrbs3 | myCAF_3 |
| Klhdc10 | 1.06E-12 | 0.31748903 | 0.438 | 0.621 | 2.04E-08 | 7 | Klhdc10 | myCAF_3 |
| Arhgef121 | 1.06E-12 | 0.64808589 | 0.691 | 0.813 | 2.05E-08 | 7 | Arhgef12 | myCAF_3 |
| Gpr108 | 1.08E-12 | 0.30636971 | 0.412 | 0.587 | 2.08E-08 | 7 | Gpr108 | myCAF_3 |
| Tfe3 | 1.08E-12 | 0.26697557 | 0.459 | 0.653 | 2.09E-08 | 7 | Tfe3 | myCAF_3 |
| Dpagt12 | 1.13E-12 | 0.27356163 | 0.392 | 0.584 | 2.19E-08 | 7 | Dpagt1 | myCAF_3 |
| Pura2 | 1.18E-12 | 0.71374673 | 0.66 | 0.816 | 2.28E-08 | 7 | Pura | myCAF_3 |
| Ddx422 | 1.28E-12 | 0.69765471 | 0.402 | 0.593 | 2.48E-08 | 7 | Ddx42 | myCAF_3 |
| Ddrgk1 | 1.38E-12 | 0.49844892 | 0.546 | 0.718 | 2.66E-08 | 7 | Ddrgk1 | myCAF_3 |
| Mars1 | 1.43E-12 | 0.43139484 | 0.469 | 0.634 | 2.76E-08 | 7 | Mars | myCAF_3 |
| Rbbp61 | 1.44E-12 | 1.01172853 | 0.495 | 0.647 | 2.79E-08 | 7 | Rbbp6 | myCAF_3 |
| Cdc42ep11 | 1.48E-12 | 0.34014118 | 0.423 | 0.572 | 2.86E-08 | 7 | Cdc42ep1 | myCAF_3 |
| Prcc | 1.52E-12 | 0.64820474 | 0.505 | 0.715 | 2.95E-08 | 7 | Prcc | myCAF_3 |
| Tmcc1 | 1.57E-12 | 0.42772608 | 0.438 | 0.634 | 3.04E-08 | 7 | Tmcc1 | myCAF_3 |
| Tns22 | 1.76E-12 | 0.74050546 | 0.603 | 0.73 | 3.39E-08 | 7 | Tns2 | myCAF_3 |
| Glrx51 | 1.77E-12 | 0.29742458 | 0.474 | 0.641 | 3.42E-08 | 7 | Glrx5 | myCAF_3 |
| Ern11 | 1.81E-12 | 0.45997061 | 0.325 | 0.534 | 3.49E-08 | 7 | Ern1 | myCAF_3 |
| Ptk73 | 1.87E-12 | 0.47897578 | 0.835 | 0.82 | 3.61E-08 | 7 | Ptk7 | myCAF_3 |
| Gmps1 | 1.89E-12 | 0.42118851 | 0.356 | 0.538 | 3.66E-08 | 7 | Gmps | myCAF_3 |
| Fnbp13 | 1.92E-12 | 0.89771749 | 0.763 | 0.808 | 3.71E-08 | 7 | Fnbp1 | myCAF_3 |
| Ccnl2 | 1.97E-12 | 0.90306047 | 0.753 | 0.866 | 3.81E-08 | 7 | Ccnl2 | myCAF_3 |
| Irx12 | 2.04E-12 | 1.30512846 | 0.624 | 0.601 | 3.94E-08 | 7 | Irx1 | myCAF_3 |
| Nup1531 | 2.07E-12 | 0.35047763 | 0.34 | 0.529 | 4.00E-08 | 7 | Nup153 | myCAF_3 |
| Numbl | 2.11E-12 | 0.35828354 | 0.629 | 0.803 | 4.09E-08 | 7 | Numbl | myCAF_3 |
| Acap3 | 2.13E-12 | 0.25584197 | 0.454 | 0.633 | 4.11E-08 | 7 | Acap3 | myCAF_3 |
| Kat6a | 2.25E-12 | 0.4192594 | 0.423 | 0.634 | 4.36E-08 | 7 | Kat6a | myCAF_3 |
| B3gnt91 | 2.34E-12 | 0.31302621 | 0.691 | 0.887 | 4.52E-08 | 7 | B3gnt9 | myCAF_3 |
| Ccdc591 | 2.34E-12 | 0.68556229 | 0.371 | 0.557 | 4.53E-08 | 7 | Ccdc59 | myCAF_3 |
| Gtpbp41 | 2.37E-12 | 0.44663619 | 0.371 | 0.557 | 4.59E-08 | 7 | Gtpbp4 | myCAF_3 |
| Rab281 | 2.42E-12 | 0.36977601 | 0.552 | 0.697 | 4.67E-08 | 7 | Rab28 | myCAF_3 |
| Atp2c11 | 2.64E-12 | 0.39835917 | 0.598 | 0.761 | 5.10E-08 | 7 | Atp2c1 | myCAF_3 |
| Rab71 | 2.66E-12 | 0.36489029 | 0.644 | 0.82 | 5.15E-08 | 7 | Rab7 | myCAF_3 |
| Emc22 | 2.73E-12 | 0.3770856 | 0.392 | 0.583 | 5.27E-08 | 7 | Emc2 | myCAF_3 |
| Farsa1 | 2.77E-12 | 0.27433622 | 0.33 | 0.536 | 5.35E-08 | 7 | Farsa | myCAF_3 |
| Grk21 | 2.98E-12 | 0.32891264 | 0.376 | 0.589 | 5.75E-08 | 7 | Grk2 | myCAF_3 |
| Sf3a21 | 3.13E-12 | 0.67941392 | 0.448 | 0.587 | 6.04E-08 | 7 | Sf3a2 | myCAF_3 |
| Phf5a1 | 3.13E-12 | 0.30274516 | 0.376 | 0.57 | 6.06E-08 | 7 | Phf5a | myCAF_3 |
| Nrep2 | 3.23E-12 | 0.72789789 | 0.691 | 0.696 | 6.25E-08 | 7 | Nrep | myCAF_3 |
| Zfp384 | 3.23E-12 | 0.27471984 | 0.407 | 0.621 | 6.25E-08 | 7 | Zfp384 | myCAF_3 |
| Apcdd12 | 3.36E-12 | 2.66925838 | 0.557 | 0.47 | 6.49E-08 | 7 | Apcdd1 | myCAF_3 |
| Smad72 | 3.43E-12 | 0.30388705 | 0.49 | 0.632 | 6.63E-08 | 7 | Smad7 | myCAF_3 |
| Flii2 | 3.60E-12 | 0.50770893 | 0.505 | 0.656 | 6.96E-08 | 7 | Flii | myCAF_3 |
| Pid12 | 3.62E-12 | 1.45214768 | 0.299 | 0.207 | 7.00E-08 | 7 | Pid1 | myCAF_3 |
| Mapk141 | 3.81E-12 | 0.42553938 | 0.572 | 0.733 | 7.36E-08 | 7 | Mapk14 | myCAF_3 |
| Sbno21 | 3.84E-12 | 1.14256923 | 0.907 | 0.894 | 7.42E-08 | 7 | Sbno2 | myCAF_3 |
| Drg2 | 4.04E-12 | 0.46994216 | 0.433 | 0.617 | 7.81E-08 | 7 | Drg2 | myCAF_3 |
| Cdc73 | 4.36E-12 | 0.48110509 | 0.356 | 0.551 | 8.42E-08 | 7 | Cdc73 | myCAF_3 |
| Adamts71 | 4.40E-12 | 1.57156031 | 0.634 | 0.531 | 8.50E-08 | 7 | Adamts7 | myCAF_3 |
| Arhgap293 | 4.42E-12 | 0.52459465 | 0.521 | 0.595 | 8.55E-08 | 7 | Arhgap29 | myCAF_3 |
| Erbin1 | 4.59E-12 | 0.43043979 | 0.572 | 0.773 | 8.87E-08 | 7 | Erbin | myCAF_3 |
| Pitpnb1 | 4.62E-12 | 0.43329361 | 0.521 | 0.689 | 8.93E-08 | 7 | Pitpnb | myCAF_3 |
| Tpp21 | 4.70E-12 | 0.49069191 | 0.407 | 0.622 | 9.09E-08 | 7 | Tpp2 | myCAF_3 |
| Ehbp1l11 | 4.72E-12 | 0.47356456 | 0.515 | 0.686 | 9.12E-08 | 7 | Ehbp1l1 | myCAF_3 |
| Ccar11 | 4.80E-12 | 0.37908586 | 0.469 | 0.658 | 9.28E-08 | 7 | Ccar1 | myCAF_3 |
| Mamdc22 | 4.99E-12 | 1.39516647 | 0.371 | 0.295 | 9.64E-08 | 7 | Mamdc2 | myCAF_3 |
| Wdr331 | 5.03E-12 | 0.38827094 | 0.454 | 0.657 | 9.73E-08 | 7 | Wdr33 | myCAF_3 |
| Upf11 | 5.19E-12 | 0.50250748 | 0.567 | 0.725 | 1.00E-07 | 7 | Upf1 | myCAF_3 |
| Cacna1c3 | 5.62E-12 | 0.58651977 | 0.639 | 0.728 | 1.09E-07 | 7 | Cacna1c | myCAF_3 |
| Hoxc102 | 5.67E-12 | 0.38703457 | 0.443 | 0.549 | 1.10E-07 | 7 | Hoxc10 | myCAF_3 |
| Supt201 | 5.74E-12 | 0.64002491 | 0.5 | 0.671 | 1.11E-07 | 7 | Supt20 | myCAF_3 |
| Jarid22 | 6.16E-12 | 0.3560591 | 0.284 | 0.45 | 1.19E-07 | 7 | Jarid2 | myCAF_3 |
| Osbpl53 | 6.39E-12 | 0.29517106 | 0.567 | 0.708 | 1.24E-07 | 7 | Osbpl5 | myCAF_3 |
| Zmat2 | 6.54E-12 | 0.42098275 | 0.433 | 0.647 | 1.26E-07 | 7 | Zmat2 | myCAF_3 |
| Plrg11 | 6.64E-12 | 0.35213905 | 0.34 | 0.552 | 1.28E-07 | 7 | Plrg1 | myCAF_3 |
| Klhl221 | 6.73E-12 | 0.36816378 | 0.407 | 0.584 | 1.30E-07 | 7 | Klhl22 | myCAF_3 |
| Prkcd | 6.94E-12 | 0.36157993 | 0.345 | 0.519 | 1.34E-07 | 7 | Prkcd | myCAF_3 |
| Cherp1 | 7.12E-12 | 0.27582383 | 0.469 | 0.644 | 1.38E-07 | 7 | Cherp | myCAF_3 |
| Klhl213 | 7.41E-12 | 0.4986434 | 0.412 | 0.588 | 1.43E-07 | 7 | Klhl21 | myCAF_3 |
| Ubl31 | 7.57E-12 | 0.36849201 | 0.526 | 0.683 | 1.46E-07 | 7 | Ubl3 | myCAF_3 |
| Phf23 | 7.65E-12 | 0.27222139 | 0.531 | 0.716 | 1.48E-07 | 7 | Phf23 | myCAF_3 |
| Vamp71 | 7.72E-12 | 0.28524852 | 0.273 | 0.45 | 1.49E-07 | 7 | Vamp7 | myCAF_3 |
| Ginm11 | 8.07E-12 | 0.63518117 | 0.691 | 0.793 | 1.56E-07 | 7 | Ginm1 | myCAF_3 |
| Hmg20a | 8.08E-12 | 0.31715413 | 0.376 | 0.593 | 1.56E-07 | 7 | Hmg20a | myCAF_3 |
| Fstl32 | 8.23E-12 | 0.36148156 | 0.629 | 0.696 | 1.59E-07 | 7 | Fstl3 | myCAF_3 |
| Dennd4a3 | 8.24E-12 | 0.53847598 | 0.479 | 0.571 | 1.59E-07 | 7 | Dennd4a | myCAF_3 |
| Bcl6 | 8.53E-12 | 0.42866664 | 0.454 | 0.627 | 1.65E-07 | 7 | Bcl6 | myCAF_3 |
| Chd11 | 8.57E-12 | 0.50481199 | 0.433 | 0.594 | 1.66E-07 | 7 | Chd1 | myCAF_3 |
| Cd2ap | 8.59E-12 | 0.38426827 | 0.345 | 0.567 | 1.66E-07 | 7 | Cd2ap | myCAF_3 |
| Vezf1 | 8.67E-12 | 0.51069973 | 0.361 | 0.575 | 1.68E-07 | 7 | Vezf1 | myCAF_3 |
| Atp9b | 8.68E-12 | 0.31808293 | 0.423 | 0.633 | 1.68E-07 | 7 | Atp9b | myCAF_3 |
| Map3k31 | 8.73E-12 | 0.38233384 | 0.443 | 0.622 | 1.69E-07 | 7 | Map3k3 | myCAF_3 |
| Cbx42 | 8.78E-12 | 0.44622486 | 0.216 | 0.416 | 1.70E-07 | 7 | Cbx4 | myCAF_3 |
| Nubp11 | 8.88E-12 | 0.38047284 | 0.402 | 0.588 | 1.72E-07 | 7 | Nubp1 | myCAF_3 |
| Adcy62 | 8.93E-12 | 0.45007301 | 0.541 | 0.72 | 1.73E-07 | 7 | Adcy6 | myCAF_3 |
| Agps1 | 8.95E-12 | 0.89495832 | 0.268 | 0.469 | 1.73E-07 | 7 | Agps | myCAF_3 |
| Prpf38b1 | 9.08E-12 | 0.37304854 | 0.428 | 0.61 | 1.76E-07 | 7 | Prpf38b | myCAF_3 |
| Nop531 | 9.42E-12 | 0.34243423 | 0.552 | 0.733 | 1.82E-07 | 7 | Nop53 | myCAF_3 |
| Pakap3 | 9.71E-12 | 1.66427369 | 0.608 | 0.474 | 1.88E-07 | 7 | Pakap | myCAF_3 |
| Rsrp11 | 9.85E-12 | 0.87665996 | 0.82 | 0.899 | 1.90E-07 | 7 | Rsrp1 | myCAF_3 |
| Dnajc91 | 9.87E-12 | 0.6061267 | 0.309 | 0.352 | 1.91E-07 | 7 | Dnajc9 | myCAF_3 |
| Vmp12 | 9.89E-12 | 0.75470678 | 0.845 | 0.949 | 1.91E-07 | 7 | Vmp1 | myCAF_3 |
| Gtf3c51 | 1.03E-11 | 0.59799433 | 0.191 | 0.402 | 1.99E-07 | 7 | Gtf3c5 | myCAF_3 |
| Eif1ax1 | 1.04E-11 | 0.6667767 | 0.32 | 0.504 | 2.01E-07 | 7 | Eif1ax | myCAF_3 |
| Hdgfl21 | 1.05E-11 | 0.28535226 | 0.448 | 0.613 | 2.02E-07 | 7 | Hdgfl2 | myCAF_3 |
| Clk12 | 1.08E-11 | 1.14920794 | 0.763 | 0.816 | 2.08E-07 | 7 | Clk1 | myCAF_3 |
| Lig3 | 1.17E-11 | 0.36688401 | 0.351 | 0.533 | 2.26E-07 | 7 | Lig3 | myCAF_3 |
| Tufm1 | 1.19E-11 | 0.36385562 | 0.459 | 0.59 | 2.31E-07 | 7 | Tufm | myCAF_3 |
| Tent4a1 | 1.21E-11 | 0.26269526 | 0.371 | 0.541 | 2.34E-07 | 7 | Tent4a | myCAF_3 |
| Hgs1 | 1.22E-11 | 0.55165166 | 0.526 | 0.692 | 2.36E-07 | 7 | Hgs | myCAF_3 |
| Bcas21 | 1.22E-11 | 0.30933248 | 0.474 | 0.648 | 2.36E-07 | 7 | Bcas2 | myCAF_3 |
| Atf21 | 1.30E-11 | 0.26050402 | 0.438 | 0.655 | 2.51E-07 | 7 | Atf2 | myCAF_3 |
| Jdp2 | 1.31E-11 | 0.33578933 | 0.686 | 0.828 | 2.53E-07 | 7 | Jdp2 | myCAF_3 |
| Ak31 | 1.39E-11 | 0.49982739 | 0.701 | 0.786 | 2.68E-07 | 7 | Ak3 | myCAF_3 |
| Extl3 | 1.39E-11 | 0.68141992 | 0.773 | 0.867 | 2.68E-07 | 7 | Extl3 | myCAF_3 |
| Gas2l11 | 1.40E-11 | 0.38101985 | 0.49 | 0.668 | 2.70E-07 | 7 | Gas2l1 | myCAF_3 |
| Cfap201 | 1.42E-11 | 0.55505385 | 0.366 | 0.568 | 2.75E-07 | 7 | Cfap20 | myCAF_3 |
| Mlxip2 | 1.47E-11 | 0.59037621 | 0.531 | 0.733 | 2.84E-07 | 7 | Mlxip | myCAF_3 |
| Aggf1 | 1.50E-11 | 0.30892627 | 0.505 | 0.649 | 2.89E-07 | 7 | Aggf1 | myCAF_3 |
| Rock12 | 1.51E-11 | 0.70524229 | 0.448 | 0.639 | 2.92E-07 | 7 | Rock1 | myCAF_3 |
| Ankrd111 | 1.52E-11 | 0.52444153 | 0.593 | 0.755 | 2.94E-07 | 7 | Ankrd11 | myCAF_3 |
| Gna131 | 1.52E-11 | 0.52596922 | 0.696 | 0.815 | 2.94E-07 | 7 | Gna13 | myCAF_3 |
| Rprd22 | 1.52E-11 | 0.39983466 | 0.423 | 0.6 | 2.95E-07 | 7 | Rprd2 | myCAF_3 |
| Csnk1e | 1.58E-11 | 0.84177419 | 0.639 | 0.761 | 3.05E-07 | 7 | Csnk1e | myCAF_3 |
| Smarcd22 | 1.62E-11 | 0.35053083 | 0.387 | 0.558 | 3.14E-07 | 7 | Smarcd2 | myCAF_3 |
| Sh3bp52 | 1.70E-11 | 0.63481919 | 0.242 | 0.441 | 3.28E-07 | 7 | Sh3bp5 | myCAF_3 |
| Rnf7 | 1.72E-11 | 0.74880536 | 0.366 | 0.547 | 3.32E-07 | 7 | Rnf7 | myCAF_3 |
| Coa3 | 1.85E-11 | 0.32952883 | 0.428 | 0.605 | 3.57E-07 | 7 | Coa3 | myCAF_3 |
| Pgm2 | 1.85E-11 | 0.47245785 | 0.32 | 0.559 | 3.58E-07 | 7 | Pgm2 | myCAF_3 |
| Sik12 | 1.89E-11 | 0.97724495 | 0.418 | 0.448 | 3.65E-07 | 7 | Sik1 | myCAF_3 |
| Spg71 | 1.94E-11 | 0.60573621 | 0.505 | 0.674 | 3.75E-07 | 7 | Spg7 | myCAF_3 |
| Kbtbd22 | 1.98E-11 | 0.30104747 | 0.438 | 0.605 | 3.82E-07 | 7 | Kbtbd2 | myCAF_3 |
| Ctsd1 | 2.06E-11 | 0.31731295 | 0.613 | 0.786 | 3.99E-07 | 7 | Ctsd | myCAF_3 |
| Prpf6 | 2.07E-11 | 0.28888271 | 0.412 | 0.608 | 4.00E-07 | 7 | Prpf6 | myCAF_3 |
| Wdr131 | 2.13E-11 | 0.30442452 | 0.33 | 0.523 | 4.11E-07 | 7 | Wdr13 | myCAF_3 |
| Anxa42 | 2.19E-11 | 0.36594566 | 0.567 | 0.711 | 4.24E-07 | 7 | Anxa4 | myCAF_3 |
| Adh5 | 2.28E-11 | 0.48156813 | 0.572 | 0.726 | 4.41E-07 | 7 | Adh5 | myCAF_3 |
| Rnf11 | 2.36E-11 | 0.5076072 | 0.629 | 0.788 | 4.56E-07 | 7 | Rnf11 | myCAF_3 |
| Tapbp3 | 2.40E-11 | 0.6108885 | 0.655 | 0.819 | 4.64E-07 | 7 | Tapbp | myCAF_3 |
| Prr12 | 2.42E-11 | 0.34485134 | 0.485 | 0.687 | 4.68E-07 | 7 | Prr12 | myCAF_3 |
| Timm221 | 2.42E-11 | 0.30443497 | 0.387 | 0.565 | 4.69E-07 | 7 | Timm22 | myCAF_3 |
| Zc3hav13 | 2.47E-11 | 0.4945323 | 0.376 | 0.536 | 4.77E-07 | 7 | Zc3hav1 | myCAF_3 |
| Arfgap2 | 2.49E-11 | 0.65790849 | 0.485 | 0.631 | 4.82E-07 | 7 | Arfgap2 | myCAF_3 |
| Usp19 | 2.49E-11 | 0.59735016 | 0.552 | 0.715 | 4.82E-07 | 7 | Usp19 | myCAF_3 |
| Slc25a361 | 2.50E-11 | 0.30428389 | 0.34 | 0.552 | 4.84E-07 | 7 | Slc25a36 | myCAF_3 |
| Sppl31 | 2.56E-11 | 0.2579953 | 0.557 | 0.769 | 4.96E-07 | 7 | Sppl3 | myCAF_3 |
| Pex192 | 2.74E-11 | 0.4634185 | 0.515 | 0.66 | 5.30E-07 | 7 | Pex19 | myCAF_3 |
| Rabif | 2.78E-11 | 0.50361267 | 0.381 | 0.558 | 5.38E-07 | 7 | Rabif | myCAF_3 |
| Srpk11 | 2.80E-11 | 0.66885234 | 0.552 | 0.672 | 5.41E-07 | 7 | Srpk1 | myCAF_3 |
| Dpf2 | 2.94E-11 | 0.35707696 | 0.428 | 0.626 | 5.69E-07 | 7 | Dpf2 | myCAF_3 |
| Polr2b1 | 3.05E-11 | 0.47799822 | 0.356 | 0.518 | 5.90E-07 | 7 | Polr2b | myCAF_3 |
| Plac9a2 | 3.14E-11 | 1.89029241 | 0.371 | 0.175 | 6.08E-07 | 7 | Plac9a | myCAF_3 |
| Phldb13 | 3.32E-11 | 0.68309698 | 0.691 | 0.775 | 6.42E-07 | 7 | Phldb1 | myCAF_3 |
| Lysmd32 | 3.37E-11 | 0.28764901 | 0.258 | 0.465 | 6.52E-07 | 7 | Lysmd3 | myCAF_3 |
| Rabgap1l2 | 3.38E-11 | 0.26399758 | 0.418 | 0.571 | 6.53E-07 | 7 | Rabgap1l | myCAF_3 |
| Ubn11 | 3.39E-11 | 0.34081671 | 0.433 | 0.611 | 6.56E-07 | 7 | Ubn1 | myCAF_3 |
| Dlg52 | 3.43E-11 | 0.45462186 | 0.541 | 0.68 | 6.62E-07 | 7 | Dlg5 | myCAF_3 |
| Gatd3a2 | 3.88E-11 | 0.26204108 | 0.325 | 0.533 | 7.51E-07 | 7 | Gatd3a | myCAF_3 |
| Gpc41 | 3.95E-11 | 0.41650034 | 0.428 | 0.613 | 7.64E-07 | 7 | Gpc4 | myCAF_3 |
| Pef11 | 3.96E-11 | 0.30994076 | 0.49 | 0.648 | 7.65E-07 | 7 | Pef1 | myCAF_3 |
| Rara1 | 3.96E-11 | 0.3480444 | 0.469 | 0.633 | 7.66E-07 | 7 | Rara | myCAF_3 |
| Ergic3 | 4.02E-11 | 0.31316783 | 0.67 | 0.835 | 7.77E-07 | 7 | Ergic3 | myCAF_3 |
| Cmtm62 | 4.17E-11 | 0.56771303 | 0.521 | 0.7 | 8.07E-07 | 7 | Cmtm6 | myCAF_3 |
| Akap131 | 4.24E-11 | 0.6734345 | 0.742 | 0.852 | 8.21E-07 | 7 | Akap13 | myCAF_3 |
| Penk | 4.33E-11 | 1.80467729 | 0.268 | 0.134 | 8.38E-07 | 7 | Penk | myCAF_3 |
| Ldlr1 | 4.37E-11 | 0.62831418 | 0.495 | 0.617 | 8.45E-07 | 7 | Ldlr | myCAF_3 |
| Wdr51 | 4.43E-11 | 0.37334309 | 0.345 | 0.519 | 8.56E-07 | 7 | Wdr5 | myCAF_3 |
| Eapp1 | 4.55E-11 | 0.48710428 | 0.443 | 0.637 | 8.79E-07 | 7 | Eapp | myCAF_3 |
| Mark41 | 4.63E-11 | 0.29256452 | 0.433 | 0.633 | 8.95E-07 | 7 | Mark4 | myCAF_3 |
| Naga1 | 4.76E-11 | 0.39223922 | 0.546 | 0.715 | 9.20E-07 | 7 | Naga | myCAF_3 |
| Hipk1 | 4.87E-11 | 0.42413317 | 0.619 | 0.776 | 9.42E-07 | 7 | Hipk1 | myCAF_3 |
| Cdc42ep41 | 5.15E-11 | 0.61817065 | 0.567 | 0.706 | 9.97E-07 | 7 | Cdc42ep4 | myCAF_3 |
| Arid1a2 | 5.29E-11 | 0.83783021 | 0.392 | 0.582 | 1.02E-06 | 7 | Arid1a | myCAF_3 |
| Ncbp11 | 5.37E-11 | 0.37929006 | 0.304 | 0.503 | 1.04E-06 | 7 | Ncbp1 | myCAF_3 |
| Gramd1a1 | 5.72E-11 | 0.29295875 | 0.428 | 0.606 | 1.11E-06 | 7 | Gramd1a | myCAF_3 |
| Mtl2 | 5.76E-11 | 0.41953459 | 0.366 | 0.606 | 1.11E-06 | 7 | Mtl2 | myCAF_3 |
| Syf21 | 5.96E-11 | 0.70386525 | 0.691 | 0.794 | 1.15E-06 | 7 | Syf2 | myCAF_3 |
| Grk61 | 6.00E-11 | 0.38335019 | 0.412 | 0.55 | 1.16E-06 | 7 | Grk6 | myCAF_3 |
| Tnrc6c | 6.13E-11 | 0.39085825 | 0.448 | 0.651 | 1.19E-06 | 7 | Tnrc6c | myCAF_3 |
| Dpp3 | 6.15E-11 | 0.25692379 | 0.541 | 0.672 | 1.19E-06 | 7 | Dpp3 | myCAF_3 |
| Ahdc11 | 6.36E-11 | 0.96413311 | 0.567 | 0.673 | 1.23E-06 | 7 | Ahdc1 | myCAF_3 |
| Max1 | 6.40E-11 | 0.37839168 | 0.505 | 0.648 | 1.24E-06 | 7 | Max | myCAF_3 |
| Pspc11 | 6.55E-11 | 0.65956085 | 0.314 | 0.508 | 1.27E-06 | 7 | Pspc1 | myCAF_3 |
| Nudt16l11 | 6.56E-11 | 0.4703111 | 0.309 | 0.541 | 1.27E-06 | 7 | Nudt16l1 | myCAF_3 |
| Cltb2 | 6.62E-11 | 0.53006763 | 0.68 | 0.806 | 1.28E-06 | 7 | Cltb | myCAF_3 |
| Ireb2 | 6.64E-11 | 0.29593243 | 0.438 | 0.62 | 1.28E-06 | 7 | Ireb2 | myCAF_3 |
| H3f3a | 6.75E-11 | 0.65210326 | 0.809 | 0.903 | 1.31E-06 | 7 | H3f3a | myCAF_3 |
| Phf12 | 7.12E-11 | 0.44238407 | 0.552 | 0.692 | 1.38E-06 | 7 | Phf12 | myCAF_3 |
| Cstf2t1 | 7.23E-11 | 0.35144433 | 0.33 | 0.539 | 1.40E-06 | 7 | Cstf2t | myCAF_3 |
| Kmt2a1 | 7.34E-11 | 0.47486607 | 0.381 | 0.564 | 1.42E-06 | 7 | Kmt2a | myCAF_3 |
| Creb1 | 7.34E-11 | 0.44107027 | 0.387 | 0.579 | 1.42E-06 | 7 | Creb1 | myCAF_3 |
| Orc3 | 7.38E-11 | 0.4649653 | 0.206 | 0.411 | 1.43E-06 | 7 | Orc3 | myCAF_3 |
| Phb | 7.58E-11 | 0.38444091 | 0.289 | 0.498 | 1.47E-06 | 7 | Phb | myCAF_3 |
| Rae1 | 7.77E-11 | 0.39765148 | 0.433 | 0.598 | 1.50E-06 | 7 | Rae1 | myCAF_3 |
| Plbd21 | 7.92E-11 | 0.5459424 | 0.68 | 0.807 | 1.53E-06 | 7 | Plbd2 | myCAF_3 |
| Mdp1 | 7.96E-11 | 0.36087201 | 0.351 | 0.529 | 1.54E-06 | 7 | Mdp1 | myCAF_3 |
| Pip4k2b | 7.97E-11 | 0.36190449 | 0.423 | 0.62 | 1.54E-06 | 7 | Pip4k2b | myCAF_3 |
| Usp121 | 8.10E-11 | 0.37138792 | 0.361 | 0.526 | 1.57E-06 | 7 | Usp12 | myCAF_3 |
| Fbxl51 | 8.14E-11 | 0.36064054 | 0.361 | 0.526 | 1.57E-06 | 7 | Fbxl5 | myCAF_3 |
| Cacybp1 | 8.38E-11 | 0.42502097 | 0.371 | 0.525 | 1.62E-06 | 7 | Cacybp | myCAF_3 |
| Plppr32 | 8.99E-11 | 1.24182806 | 0.18 | 0.377 | 1.74E-06 | 7 | Plppr3 | myCAF_3 |
| Dchs12 | 9.16E-11 | 0.77188246 | 0.51 | 0.567 | 1.77E-06 | 7 | Dchs1 | myCAF_3 |
| Riok2 | 9.47E-11 | 0.36529454 | 0.227 | 0.439 | 1.83E-06 | 7 | Riok2 | myCAF_3 |
| Nrp22 | 9.61E-11 | 1.71081854 | 0.546 | 0.42 | 1.86E-06 | 7 | Nrp2 | myCAF_3 |
| Camsap2 | 1.00E-10 | 0.25711953 | 0.428 | 0.598 | 1.93E-06 | 7 | Camsap2 | myCAF_3 |
| Mfap3 | 1.05E-10 | 0.3958582 | 0.381 | 0.568 | 2.02E-06 | 7 | Mfap3 | myCAF_3 |
| Eif2b5 | 1.06E-10 | 0.44967893 | 0.49 | 0.628 | 2.04E-06 | 7 | Eif2b5 | myCAF_3 |
| Tubgcp41 | 1.07E-10 | 0.61119482 | 0.258 | 0.453 | 2.06E-06 | 7 | Tubgcp4 | myCAF_3 |
| Acaca1 | 1.08E-10 | 0.32375844 | 0.418 | 0.608 | 2.08E-06 | 7 | Acaca | myCAF_3 |
| Cxxc11 | 1.17E-10 | 0.4162855 | 0.448 | 0.621 | 2.27E-06 | 7 | Cxxc1 | myCAF_3 |
| Iws11 | 1.20E-10 | 0.67234099 | 0.33 | 0.498 | 2.32E-06 | 7 | Iws1 | myCAF_3 |
| Med143 | 1.21E-10 | 0.29703108 | 0.227 | 0.411 | 2.34E-06 | 7 | Med14 | myCAF_3 |
| Pbx22 | 1.22E-10 | 0.71854555 | 0.418 | 0.596 | 2.36E-06 | 7 | Pbx2 | myCAF_3 |
| Supt4a | 1.27E-10 | 0.45307619 | 0.562 | 0.739 | 2.46E-06 | 7 | Supt4a | myCAF_3 |
| Spr | 1.28E-10 | 0.55469393 | 0.464 | 0.644 | 2.48E-06 | 7 | Spr | myCAF_3 |
| Dyrk1a1 | 1.28E-10 | 0.44615759 | 0.366 | 0.549 | 2.48E-06 | 7 | Dyrk1a | myCAF_3 |
| Ncoa51 | 1.35E-10 | 0.59813201 | 0.284 | 0.48 | 2.61E-06 | 7 | Ncoa5 | myCAF_3 |
| Gnai12 | 1.38E-10 | 0.47015309 | 0.119 | 0.284 | 2.67E-06 | 7 | Gnai1 | myCAF_3 |
| Rxrb1 | 1.45E-10 | 0.47325409 | 0.443 | 0.623 | 2.81E-06 | 7 | Rxrb | myCAF_3 |
| Ece11 | 1.54E-10 | 0.42028207 | 0.655 | 0.811 | 2.97E-06 | 7 | Ece1 | myCAF_3 |
| Otulin1 | 1.54E-10 | 0.71132497 | 0.392 | 0.541 | 2.97E-06 | 7 | Otulin | myCAF_3 |
| Maf2 | 1.56E-10 | 0.91606754 | 0.794 | 0.83 | 3.02E-06 | 7 | Maf | myCAF_3 |
| Micos13 | 1.57E-10 | 0.33948316 | 0.567 | 0.767 | 3.03E-06 | 7 | Micos13 | myCAF_3 |
| Ppp2r5a2 | 1.59E-10 | 0.51510139 | 0.459 | 0.62 | 3.07E-06 | 7 | Ppp2r5a | myCAF_3 |
| Tsc22d4 | 1.63E-10 | 0.37608829 | 0.376 | 0.598 | 3.16E-06 | 7 | Tsc22d4 | myCAF_3 |
| Atp6v1a2 | 1.68E-10 | 0.39646975 | 0.552 | 0.724 | 3.25E-06 | 7 | Atp6v1a | myCAF_3 |
| Zfp4511 | 1.77E-10 | 0.70559722 | 0.361 | 0.548 | 3.42E-06 | 7 | Zfp451 | myCAF_3 |
| Cdk11b1 | 1.85E-10 | 0.56378996 | 0.443 | 0.621 | 3.57E-06 | 7 | Cdk11b | myCAF_3 |
| Fmnl24 | 1.86E-10 | 0.47509076 | 0.314 | 0.431 | 3.60E-06 | 7 | Fmnl2 | myCAF_3 |
| Bmp71 | 1.89E-10 | 3.16511216 | 0.335 | 0.131 | 3.65E-06 | 7 | Bmp7 | myCAF_3 |
| Cul5 | 1.89E-10 | 0.37528116 | 0.299 | 0.505 | 3.65E-06 | 7 | Cul5 | myCAF_3 |
| Glmp | 1.94E-10 | 0.4966109 | 0.789 | 0.903 | 3.75E-06 | 7 | Glmp | myCAF_3 |
| Cdk9 | 1.97E-10 | 0.72331904 | 0.629 | 0.765 | 3.80E-06 | 7 | Cdk9 | myCAF_3 |
| Zadh22 | 1.98E-10 | 0.25519509 | 0.366 | 0.542 | 3.83E-06 | 7 | Zadh2 | myCAF_3 |
| Mindy22 | 2.00E-10 | 0.53038943 | 0.237 | 0.448 | 3.87E-06 | 7 | Mindy2 | myCAF_3 |
| Pakap.11 | 2.01E-10 | 0.58128184 | 0.634 | 0.667 | 3.89E-06 | 7 | Pakap.1 | myCAF_3 |
| Srek11 | 2.04E-10 | 0.75930045 | 0.521 | 0.701 | 3.95E-06 | 7 | Srek1 | myCAF_3 |
| St131 | 2.12E-10 | 0.2848991 | 0.479 | 0.62 | 4.10E-06 | 7 | St13 | myCAF_3 |
| Tmem91 | 2.15E-10 | 0.61261071 | 0.598 | 0.747 | 4.16E-06 | 7 | Tmem9 | myCAF_3 |
| Tpp12 | 2.24E-10 | 0.41001739 | 0.634 | 0.778 | 4.34E-06 | 7 | Tpp1 | myCAF_3 |
| Apex11 | 2.27E-10 | 0.3146283 | 0.381 | 0.522 | 4.39E-06 | 7 | Apex1 | myCAF_3 |
| Zfp1311 | 2.28E-10 | 0.83684098 | 0.402 | 0.566 | 4.41E-06 | 7 | Zfp131 | myCAF_3 |
| Taldo12 | 2.36E-10 | 0.34390275 | 0.49 | 0.654 | 4.57E-06 | 7 | Taldo1 | myCAF_3 |
| Safb1 | 2.57E-10 | 0.40659593 | 0.49 | 0.602 | 4.96E-06 | 7 | Safb | myCAF_3 |
| Dnajc13 | 2.64E-10 | 0.5979318 | 0.521 | 0.673 | 5.11E-06 | 7 | Dnajc13 | myCAF_3 |
| Myo5a1 | 2.75E-10 | 0.41686559 | 0.5 | 0.68 | 5.31E-06 | 7 | Myo5a | myCAF_3 |
| Hsd3b72 | 2.80E-10 | 0.35400425 | 0.351 | 0.51 | 5.41E-06 | 7 | Hsd3b7 | myCAF_3 |
| Prkaa1 | 2.82E-10 | 0.60042294 | 0.32 | 0.511 | 5.46E-06 | 7 | Prkaa1 | myCAF_3 |
| Tasor2 | 2.86E-10 | 0.30534849 | 0.242 | 0.446 | 5.53E-06 | 7 | Tasor2 | myCAF_3 |
| Suds31 | 2.91E-10 | 0.65485519 | 0.649 | 0.773 | 5.62E-06 | 7 | Suds3 | myCAF_3 |
| Nmt11 | 2.94E-10 | 0.48293852 | 0.696 | 0.838 | 5.68E-06 | 7 | Nmt1 | myCAF_3 |
| Gli32 | 2.99E-10 | 0.44003796 | 0.433 | 0.594 | 5.78E-06 | 7 | Gli3 | myCAF_3 |
| Rnf1143 | 3.06E-10 | 0.29722461 | 0.278 | 0.473 | 5.92E-06 | 7 | Rnf114 | myCAF_3 |
| Nfkb21 | 3.07E-10 | 0.97839703 | 0.691 | 0.773 | 5.94E-06 | 7 | Nfkb2 | myCAF_3 |
| Synj2bp | 3.07E-10 | 0.67965278 | 0.495 | 0.653 | 5.94E-06 | 7 | Synj2bp | myCAF_3 |
| Ppdpf | 3.15E-10 | 0.56291507 | 0.505 | 0.663 | 6.10E-06 | 7 | Ppdpf | myCAF_3 |
| Supt6 | 3.27E-10 | 0.60851087 | 0.582 | 0.714 | 6.33E-06 | 7 | Supt6 | myCAF_3 |
| Exoc7 | 3.30E-10 | 0.57091541 | 0.392 | 0.591 | 6.37E-06 | 7 | Exoc7 | myCAF_3 |
| Kif1c1 | 3.32E-10 | 0.52343937 | 0.557 | 0.74 | 6.41E-06 | 7 | Kif1c | myCAF_3 |
| Kctd123 | 3.39E-10 | 0.93101622 | 0.227 | 0.372 | 6.56E-06 | 7 | Kctd12 | myCAF_3 |
| Pmm12 | 3.53E-10 | 0.29986133 | 0.402 | 0.591 | 6.82E-06 | 7 | Pmm1 | myCAF_3 |
| Smarce1 | 3.55E-10 | 0.47169767 | 0.567 | 0.702 | 6.87E-06 | 7 | Smarce1 | myCAF_3 |
| Fbxo322 | 3.56E-10 | 2.15299407 | 0.459 | 0.255 | 6.88E-06 | 7 | Fbxo32 | myCAF_3 |
| Pld31 | 3.79E-10 | 0.38184721 | 0.691 | 0.866 | 7.33E-06 | 7 | Pld3 | myCAF_3 |
| Phip1 | 3.80E-10 | 0.43789835 | 0.371 | 0.529 | 7.35E-06 | 7 | Phip | myCAF_3 |
| Eny21 | 3.84E-10 | 0.28070523 | 0.186 | 0.377 | 7.43E-06 | 7 | Eny2 | myCAF_3 |
| Tmem1341 | 4.04E-10 | 0.26750422 | 0.526 | 0.736 | 7.82E-06 | 7 | Tmem134 | myCAF_3 |
| Itm2c1 | 4.22E-10 | 0.55663029 | 0.881 | 0.973 | 8.15E-06 | 7 | Itm2c | myCAF_3 |
| Msi22 | 4.23E-10 | 0.36252585 | 0.376 | 0.601 | 8.17E-06 | 7 | Msi2 | myCAF_3 |
| Kank2 | 4.34E-10 | 0.77824179 | 0.773 | 0.866 | 8.39E-06 | 7 | Kank2 | myCAF_3 |
| Pphln11 | 4.35E-10 | 0.37798584 | 0.397 | 0.563 | 8.41E-06 | 7 | Pphln1 | myCAF_3 |
| Clk32 | 4.53E-10 | 0.30874622 | 0.335 | 0.537 | 8.76E-06 | 7 | Clk3 | myCAF_3 |
| Rbmx1 | 4.75E-10 | 0.67828995 | 0.247 | 0.388 | 9.18E-06 | 7 | Rbmx | myCAF_3 |
| Bok2 | 4.76E-10 | 2.03179768 | 0.423 | 0.24 | 9.20E-06 | 7 | Bok | myCAF_3 |
| Abhd17c2 | 4.82E-10 | 0.47985789 | 0.284 | 0.468 | 9.31E-06 | 7 | Abhd17c | myCAF_3 |
| Prps1l31 | 4.88E-10 | 0.2529636 | 0.299 | 0.462 | 9.44E-06 | 7 | Prps1l3 | myCAF_3 |
| Lin7c1 | 4.95E-10 | 0.53643835 | 0.505 | 0.644 | 9.56E-06 | 7 | Lin7c | myCAF_3 |
| Mtfr11 | 5.01E-10 | 0.57307605 | 0.356 | 0.514 | 9.69E-06 | 7 | Mtfr1 | myCAF_3 |
| Rpe | 5.05E-10 | 0.6553237 | 0.351 | 0.514 | 9.77E-06 | 7 | Rpe | myCAF_3 |
| Txndc152 | 5.23E-10 | 0.29603712 | 0.567 | 0.723 | 1.01E-05 | 7 | Txndc15 | myCAF_3 |
| Uchl51 | 5.38E-10 | 0.54517495 | 0.124 | 0.305 | 1.04E-05 | 7 | Uchl5 | myCAF_3 |
| Gna122 | 5.60E-10 | 0.31769838 | 0.665 | 0.848 | 1.08E-05 | 7 | Gna12 | myCAF_3 |
| Enoph11 | 5.80E-10 | 0.38283921 | 0.216 | 0.412 | 1.12E-05 | 7 | Enoph1 | myCAF_3 |
| Rbm281 | 5.95E-10 | 0.28280537 | 0.314 | 0.483 | 1.15E-05 | 7 | Rbm28 | myCAF_3 |
| Nol61 | 6.00E-10 | 0.33007263 | 0.304 | 0.512 | 1.16E-05 | 7 | Nol6 | myCAF_3 |
| Ppp1r37 | 6.08E-10 | 0.48287046 | 0.428 | 0.605 | 1.17E-05 | 7 | Ppp1r37 | myCAF_3 |
| Trp531 | 6.16E-10 | 0.66218375 | 0.603 | 0.683 | 1.19E-05 | 7 | Trp53 | myCAF_3 |
| Ccdc1861 | 6.17E-10 | 0.61078008 | 0.397 | 0.57 | 1.19E-05 | 7 | Ccdc186 | myCAF_3 |
| Gbp23 | 6.22E-10 | 0.35146828 | 0.165 | 0.272 | 1.20E-05 | 7 | Gbp2 | myCAF_3 |
| Pdzd81 | 6.27E-10 | 0.55354462 | 0.371 | 0.541 | 1.21E-05 | 7 | Pdzd8 | myCAF_3 |
| Trappc81 | 6.31E-10 | 0.43636859 | 0.34 | 0.522 | 1.22E-05 | 7 | Trappc8 | myCAF_3 |
| Zfp7401 | 6.67E-10 | 0.63613389 | 0.448 | 0.606 | 1.29E-05 | 7 | Zfp740 | myCAF_3 |
| Maml1 | 6.95E-10 | 0.84635036 | 0.505 | 0.665 | 1.34E-05 | 7 | Maml1 | myCAF_3 |
| Pdcd101 | 7.27E-10 | 0.51341406 | 0.376 | 0.539 | 1.41E-05 | 7 | Pdcd10 | myCAF_3 |
| Ttc39b | 7.33E-10 | 0.39476891 | 0.33 | 0.491 | 1.42E-05 | 7 | Ttc39b | myCAF_3 |
| Rybp | 7.36E-10 | 0.5962654 | 0.227 | 0.419 | 1.42E-05 | 7 | Rybp | myCAF_3 |
| Cyb5b1 | 7.44E-10 | 0.69828836 | 0.577 | 0.677 | 1.44E-05 | 7 | Cyb5b | myCAF_3 |
| Trpm71 | 7.47E-10 | 0.89649527 | 0.392 | 0.586 | 1.44E-05 | 7 | Trpm7 | myCAF_3 |
| Megf8 | 7.51E-10 | 0.49321273 | 0.505 | 0.712 | 1.45E-05 | 7 | Megf8 | myCAF_3 |
| Osbpl1a3 | 7.54E-10 | 0.42060553 | 0.216 | 0.392 | 1.46E-05 | 7 | Osbpl1a | myCAF_3 |
| Azin22 | 7.64E-10 | 0.57096178 | 0.577 | 0.694 | 1.48E-05 | 7 | Azin2 | myCAF_3 |
| Nr1h2 | 8.21E-10 | 0.48566224 | 0.407 | 0.604 | 1.59E-05 | 7 | Nr1h2 | myCAF_3 |
| Fibp1 | 8.27E-10 | 0.33545361 | 0.505 | 0.687 | 1.60E-05 | 7 | Fibp | myCAF_3 |
| Swap702 | 8.32E-10 | 0.33440446 | 0.361 | 0.522 | 1.61E-05 | 7 | Swap70 | myCAF_3 |
| Ppp6r1 | 8.39E-10 | 0.59448065 | 0.526 | 0.666 | 1.62E-05 | 7 | Ppp6r1 | myCAF_3 |
| Nf21 | 8.60E-10 | 0.37521616 | 0.51 | 0.693 | 1.66E-05 | 7 | Nf2 | myCAF_3 |
| Ppfibp11 | 8.62E-10 | 0.43651311 | 0.716 | 0.863 | 1.67E-05 | 7 | Ppfibp1 | myCAF_3 |
| Gpaa11 | 8.72E-10 | 0.56623557 | 0.335 | 0.525 | 1.69E-05 | 7 | Gpaa1 | myCAF_3 |
| Sumf1 | 8.83E-10 | 0.36296688 | 0.531 | 0.743 | 1.71E-05 | 7 | Sumf1 | myCAF_3 |
| Ap1ar1 | 8.86E-10 | 0.28091075 | 0.366 | 0.542 | 1.71E-05 | 7 | Ap1ar | myCAF_3 |
| Fam117b2 | 8.88E-10 | 0.65417813 | 0.5 | 0.583 | 1.72E-05 | 7 | Fam117b | myCAF_3 |
| Abl21 | 9.06E-10 | 0.43692403 | 0.387 | 0.566 | 1.75E-05 | 7 | Abl2 | myCAF_3 |
| Sike12 | 9.08E-10 | 0.32716116 | 0.531 | 0.698 | 1.76E-05 | 7 | Sike1 | myCAF_3 |
| Wwtr12 | 9.11E-10 | 0.59944362 | 0.784 | 0.891 | 1.76E-05 | 7 | Wwtr1 | myCAF_3 |
| Arih2 | 9.40E-10 | 0.54743017 | 0.495 | 0.629 | 1.82E-05 | 7 | Arih2 | myCAF_3 |
| Txnip3 | 9.41E-10 | 0.95276218 | 0.799 | 0.847 | 1.82E-05 | 7 | Txnip | myCAF_3 |
| Elf2 | 9.44E-10 | 0.58218739 | 0.454 | 0.64 | 1.83E-05 | 7 | Elf2 | myCAF_3 |
| Actl6a1 | 9.48E-10 | 0.67356987 | 0.325 | 0.484 | 1.83E-05 | 7 | Actl6a | myCAF_3 |
| Prkar2a1 | 9.49E-10 | 0.30029144 | 0.278 | 0.442 | 1.83E-05 | 7 | Prkar2a | myCAF_3 |
| Zhx11 | 9.58E-10 | 0.48429145 | 0.232 | 0.451 | 1.85E-05 | 7 | Zhx1 | myCAF_3 |
| Pank21 | 9.91E-10 | 0.53946181 | 0.412 | 0.58 | 1.92E-05 | 7 | Pank2 | myCAF_3 |
| Kifap31 | 1.02E-09 | 0.25507774 | 0.443 | 0.595 | 1.97E-05 | 7 | Kifap3 | myCAF_3 |
| Pelp11 | 1.05E-09 | 0.30338839 | 0.304 | 0.472 | 2.02E-05 | 7 | Pelp1 | myCAF_3 |
| Lrrc47 | 1.05E-09 | 0.7557951 | 0.448 | 0.61 | 2.03E-05 | 7 | Lrrc47 | myCAF_3 |
| Vapa2 | 1.05E-09 | 0.28636499 | 0.376 | 0.559 | 2.03E-05 | 7 | Vapa | myCAF_3 |
| Adamts22 | 1.10E-09 | 0.50003324 | 0.856 | 0.986 | 2.14E-05 | 7 | Adamts2 | myCAF_3 |
| Sp11 | 1.13E-09 | 0.61530209 | 0.366 | 0.53 | 2.19E-05 | 7 | Sp1 | myCAF_3 |
| Fnip11 | 1.14E-09 | 0.35519838 | 0.34 | 0.552 | 2.20E-05 | 7 | Fnip1 | myCAF_3 |
| Urm1 | 1.14E-09 | 0.40779754 | 0.407 | 0.596 | 2.21E-05 | 7 | Urm1 | myCAF_3 |
| Tbc1d141 | 1.16E-09 | 0.41978964 | 0.366 | 0.53 | 2.24E-05 | 7 | Tbc1d14 | myCAF_3 |
| Phrf11 | 1.17E-09 | 0.78580499 | 0.412 | 0.561 | 2.27E-05 | 7 | Phrf1 | myCAF_3 |
| Rapgef21 | 1.18E-09 | 0.31091227 | 0.335 | 0.514 | 2.28E-05 | 7 | Rapgef2 | myCAF_3 |
| Ddx10 | 1.29E-09 | 0.95593216 | 0.247 | 0.438 | 2.49E-05 | 7 | Ddx10 | myCAF_3 |
| Nfya2 | 1.29E-09 | 0.41138983 | 0.392 | 0.527 | 2.49E-05 | 7 | Nfya | myCAF_3 |
| Fdft11 | 1.30E-09 | 0.49082039 | 0.407 | 0.506 | 2.52E-05 | 7 | Fdft1 | myCAF_3 |
| Fam89b | 1.30E-09 | 0.54908396 | 0.515 | 0.677 | 2.52E-05 | 7 | Fam89b | myCAF_3 |
| Pnn1 | 1.31E-09 | 0.60638368 | 0.469 | 0.599 | 2.53E-05 | 7 | Pnn | myCAF_3 |
| Csrnp11 | 1.33E-09 | 0.85835072 | 0.371 | 0.516 | 2.58E-05 | 7 | Csrnp1 | myCAF_3 |
| Immp1l1 | 1.36E-09 | 0.43516981 | 0.242 | 0.417 | 2.63E-05 | 7 | Immp1l | myCAF_3 |
| Setd1a1 | 1.36E-09 | 0.72051104 | 0.515 | 0.65 | 2.63E-05 | 7 | Setd1a | myCAF_3 |
| Tmem87b1 | 1.36E-09 | 0.45545573 | 0.515 | 0.672 | 2.64E-05 | 7 | Tmem87b | myCAF_3 |
| Iqsec11 | 1.39E-09 | 1.10218384 | 0.567 | 0.695 | 2.69E-05 | 7 | Iqsec1 | myCAF_3 |
| Brms1l1 | 1.46E-09 | 0.50314965 | 0.345 | 0.509 | 2.83E-05 | 7 | Brms1l | myCAF_3 |
| Clip31 | 1.49E-09 | 0.877796 | 0.613 | 0.746 | 2.88E-05 | 7 | Clip3 | myCAF_3 |
| Ddx461 | 1.50E-09 | 0.45120332 | 0.474 | 0.624 | 2.90E-05 | 7 | Ddx46 | myCAF_3 |
| Lgals82 | 1.51E-09 | 0.32482892 | 0.418 | 0.571 | 2.92E-05 | 7 | Lgals8 | myCAF_3 |
| Xpo51 | 1.54E-09 | 0.37372535 | 0.278 | 0.493 | 2.97E-05 | 7 | Xpo5 | myCAF_3 |
| Itch | 1.55E-09 | 0.6243731 | 0.418 | 0.572 | 2.99E-05 | 7 | Itch | myCAF_3 |
| Rbm151 | 1.60E-09 | 0.49273724 | 0.335 | 0.504 | 3.10E-05 | 7 | Rbm15 | myCAF_3 |
| Rp91 | 1.61E-09 | 0.34497783 | 0.448 | 0.597 | 3.11E-05 | 7 | Rp9 | myCAF_3 |
| Tm7sf31 | 1.63E-09 | 0.395499 | 0.655 | 0.812 | 3.16E-05 | 7 | Tm7sf3 | myCAF_3 |
| Eri11 | 1.67E-09 | 0.64884637 | 0.222 | 0.387 | 3.24E-05 | 7 | Eri1 | myCAF_3 |
| Ppp6c | 1.69E-09 | 0.58748326 | 0.366 | 0.563 | 3.28E-05 | 7 | Ppp6c | myCAF_3 |
| En11 | 1.74E-09 | 0.3308771 | 0.448 | 0.594 | 3.37E-05 | 7 | En1 | myCAF_3 |
| Stn1 | 1.75E-09 | 0.82985277 | 0.175 | 0.36 | 3.38E-05 | 7 | Stn1 | myCAF_3 |
| Taf101 | 1.77E-09 | 0.31793131 | 0.278 | 0.447 | 3.43E-05 | 7 | Taf10 | myCAF_3 |
| Thada | 1.81E-09 | 0.71961361 | 0.18 | 0.365 | 3.51E-05 | 7 | Thada | myCAF_3 |
| Fnbp41 | 1.99E-09 | 0.48996133 | 0.284 | 0.472 | 3.84E-05 | 7 | Fnbp4 | myCAF_3 |
| Blvrb2 | 2.01E-09 | 0.4759012 | 0.299 | 0.478 | 3.88E-05 | 7 | Blvrb | myCAF_3 |
| Rab241 | 2.02E-09 | 0.50834904 | 0.536 | 0.69 | 3.91E-05 | 7 | Rab24 | myCAF_3 |
| Bud231 | 2.05E-09 | 1.02925417 | 0.33 | 0.481 | 3.96E-05 | 7 | Bud23 | myCAF_3 |
| Slc30a11 | 2.15E-09 | 0.40573488 | 0.289 | 0.451 | 4.16E-05 | 7 | Slc30a1 | myCAF_3 |
| Cnot111 | 2.15E-09 | 0.2759755 | 0.309 | 0.485 | 4.16E-05 | 7 | Cnot11 | myCAF_3 |
| Akap111 | 2.22E-09 | 0.46186107 | 0.423 | 0.591 | 4.30E-05 | 7 | Akap11 | myCAF_3 |
| Trip11 | 2.24E-09 | 0.4777779 | 0.469 | 0.616 | 4.33E-05 | 7 | Trip11 | myCAF_3 |
| Adgre53 | 2.25E-09 | 0.47874279 | 0.562 | 0.615 | 4.35E-05 | 7 | Adgre5 | myCAF_3 |
| Macf11 | 2.25E-09 | 0.52775576 | 0.495 | 0.66 | 4.35E-05 | 7 | Macf1 | myCAF_3 |
| Mau21 | 2.29E-09 | 0.66410912 | 0.448 | 0.618 | 4.42E-05 | 7 | Mau2 | myCAF_3 |
| Ranbp9 | 2.49E-09 | 0.3505578 | 0.345 | 0.529 | 4.81E-05 | 7 | Ranbp9 | myCAF_3 |
| B4galt3 | 2.49E-09 | 0.4643585 | 0.242 | 0.456 | 4.82E-05 | 7 | B4galt3 | myCAF_3 |
| Rgs102 | 2.50E-09 | 0.4600653 | 0.227 | 0.367 | 4.83E-05 | 7 | Rgs10 | myCAF_3 |
| Vps16 | 2.54E-09 | 0.54302129 | 0.505 | 0.657 | 4.92E-05 | 7 | Vps16 | myCAF_3 |
| Bri32 | 2.57E-09 | 0.56079329 | 0.742 | 0.863 | 4.97E-05 | 7 | Bri3 | myCAF_3 |
| Zhx2 | 2.59E-09 | 0.29890916 | 0.263 | 0.439 | 5.00E-05 | 7 | Zhx2 | myCAF_3 |
| Usf31 | 2.69E-09 | 0.39518971 | 0.222 | 0.42 | 5.20E-05 | 7 | Usf3 | myCAF_3 |
| Eif5b | 2.71E-09 | 0.25913712 | 0.387 | 0.555 | 5.23E-05 | 7 | Eif5b | myCAF_3 |
| Prr142 | 2.72E-09 | 0.31817902 | 0.407 | 0.577 | 5.26E-05 | 7 | Prr14 | myCAF_3 |
| Nrm1 | 2.74E-09 | 0.6953473 | 0.191 | 0.294 | 5.31E-05 | 7 | Nrm | myCAF_3 |
| BC0297222 | 2.75E-09 | 0.26097904 | 0.351 | 0.542 | 5.33E-05 | 7 | BC029722 | myCAF_3 |
| Larp71 | 2.76E-09 | 0.41611662 | 0.247 | 0.44 | 5.34E-05 | 7 | Larp7 | myCAF_3 |
| Atg2b | 2.77E-09 | 0.49406343 | 0.34 | 0.529 | 5.35E-05 | 7 | Atg2b | myCAF_3 |
| Gdap21 | 2.85E-09 | 0.37153848 | 0.232 | 0.424 | 5.52E-05 | 7 | Gdap2 | myCAF_3 |
| Zranb21 | 2.89E-09 | 0.3563229 | 0.309 | 0.487 | 5.58E-05 | 7 | Zranb2 | myCAF_3 |
| Sqle1 | 2.90E-09 | 0.86695092 | 0.351 | 0.392 | 5.61E-05 | 7 | Sqle | myCAF_3 |
| Nfil32 | 2.96E-09 | 1.03272987 | 0.577 | 0.652 | 5.72E-05 | 7 | Nfil3 | myCAF_3 |
| Fcf1 | 2.98E-09 | 0.33791 | 0.33 | 0.507 | 5.76E-05 | 7 | Fcf1 | myCAF_3 |
| Cep350 | 3.09E-09 | 0.45466034 | 0.345 | 0.547 | 5.97E-05 | 7 | Cep350 | myCAF_3 |
| Cnot72 | 3.10E-09 | 0.31658044 | 0.289 | 0.47 | 6.00E-05 | 7 | Cnot7 | myCAF_3 |
| Bag5 | 3.11E-09 | 0.6034519 | 0.428 | 0.577 | 6.02E-05 | 7 | Bag5 | myCAF_3 |
| Nbr12 | 3.27E-09 | 0.39790493 | 0.361 | 0.538 | 6.32E-05 | 7 | Nbr1 | myCAF_3 |
| Setd1b | 3.28E-09 | 0.77948046 | 0.438 | 0.61 | 6.34E-05 | 7 | Setd1b | myCAF_3 |
| Foxn2 | 3.32E-09 | 0.57952781 | 0.407 | 0.568 | 6.41E-05 | 7 | Foxn2 | myCAF_3 |
| Mark3 | 3.41E-09 | 0.28621315 | 0.459 | 0.626 | 6.59E-05 | 7 | Mark3 | myCAF_3 |
| Ccz11 | 3.41E-09 | 0.54214551 | 0.268 | 0.468 | 6.60E-05 | 7 | Ccz1 | myCAF_3 |
| Tmem120a | 3.44E-09 | 0.50654784 | 0.263 | 0.457 | 6.65E-05 | 7 | Tmem120a | myCAF_3 |
| Homer3 | 3.47E-09 | 0.44340685 | 0.469 | 0.635 | 6.71E-05 | 7 | Homer3 | myCAF_3 |
| Sbf21 | 3.49E-09 | 0.63893631 | 0.402 | 0.548 | 6.76E-05 | 7 | Sbf2 | myCAF_3 |
| Gak1 | 3.52E-09 | 0.39145827 | 0.392 | 0.564 | 6.80E-05 | 7 | Gak | myCAF_3 |
| Dcaf131 | 3.61E-09 | 0.4314163 | 0.376 | 0.528 | 6.98E-05 | 7 | Dcaf13 | myCAF_3 |
| Sart11 | 3.65E-09 | 0.46130825 | 0.438 | 0.588 | 7.06E-05 | 7 | Sart1 | myCAF_3 |
| Fbxl14 | 3.73E-09 | 0.53773549 | 0.376 | 0.552 | 7.22E-05 | 7 | Fbxl14 | myCAF_3 |
| Dhx361 | 3.79E-09 | 0.75534035 | 0.392 | 0.589 | 7.33E-05 | 7 | Dhx36 | myCAF_3 |
| Rnf14 | 3.86E-09 | 0.60815711 | 0.459 | 0.637 | 7.47E-05 | 7 | Rnf14 | myCAF_3 |
| Zdhhc82 | 4.06E-09 | 0.27913271 | 0.51 | 0.716 | 7.85E-05 | 7 | Zdhhc8 | myCAF_3 |
| Wdr48 | 4.17E-09 | 0.51932276 | 0.314 | 0.496 | 8.07E-05 | 7 | Wdr48 | myCAF_3 |
| Mier1 | 4.19E-09 | 0.90199489 | 0.557 | 0.724 | 8.11E-05 | 7 | Mier1 | myCAF_3 |
| Atg13 | 4.19E-09 | 0.61695381 | 0.294 | 0.481 | 8.11E-05 | 7 | Atg13 | myCAF_3 |
| Gapvd1 | 4.19E-09 | 0.63370055 | 0.448 | 0.591 | 8.11E-05 | 7 | Gapvd1 | myCAF_3 |
| Plaat32 | 4.32E-09 | 0.27243757 | 0.5 | 0.619 | 8.36E-05 | 7 | Plaat3 | myCAF_3 |
| Pcnx2 | 4.36E-09 | 0.4123488 | 0.464 | 0.634 | 8.43E-05 | 7 | Pcnx | myCAF_3 |
| Fam192a | 4.41E-09 | 0.58792588 | 0.325 | 0.498 | 8.52E-05 | 7 | Fam192a | myCAF_3 |
| Iars2 | 4.42E-09 | 0.37549829 | 0.423 | 0.593 | 8.54E-05 | 7 | Iars2 | myCAF_3 |
| Kalrn2 | 4.53E-09 | 0.69088551 | 0.433 | 0.502 | 8.76E-05 | 7 | Kalrn | myCAF_3 |
| Sirt6 | 4.61E-09 | 0.3325624 | 0.34 | 0.535 | 8.91E-05 | 7 | Sirt6 | myCAF_3 |
| Ppm1m1 | 4.62E-09 | 0.33839975 | 0.423 | 0.616 | 8.93E-05 | 7 | Ppm1m | myCAF_3 |
| Usp33 | 4.63E-09 | 0.41654138 | 0.423 | 0.57 | 8.94E-05 | 7 | Usp33 | myCAF_3 |
| Ltbp32 | 4.63E-09 | 0.58934556 | 0.696 | 0.847 | 8.95E-05 | 7 | Ltbp3 | myCAF_3 |
| Zfp5213 | 4.67E-09 | 0.41992872 | 0.237 | 0.381 | 9.04E-05 | 7 | Zfp521 | myCAF_3 |
| Snrpd21 | 4.72E-09 | 0.3488768 | 0.309 | 0.489 | 9.12E-05 | 7 | Snrpd2 | myCAF_3 |
| Wdtc11 | 4.75E-09 | 0.31153601 | 0.34 | 0.513 | 9.19E-05 | 7 | Wdtc1 | myCAF_3 |
| Rcl12 | 4.76E-09 | 1.07912537 | 0.196 | 0.375 | 9.19E-05 | 7 | Rcl1 | myCAF_3 |
| Aacs1 | 4.81E-09 | 0.50247853 | 0.356 | 0.522 | 9.30E-05 | 7 | Aacs | myCAF_3 |
| Rnf44 | 4.89E-09 | 0.33577435 | 0.593 | 0.782 | 9.45E-05 | 7 | Rnf44 | myCAF_3 |
| Sap1302 | 4.90E-09 | 0.58308594 | 0.351 | 0.529 | 9.48E-05 | 7 | Sap130 | myCAF_3 |
| Dazap21 | 5.22E-09 | 0.39878843 | 0.469 | 0.647 | 0.00010098 | 7 | Dazap2 | myCAF_3 |
| Plscr42 | 5.60E-09 | 1.19405097 | 0.129 | 0.306 | 0.0001082 | 7 | Plscr4 | myCAF_3 |
| Exosc31 | 5.60E-09 | 0.33766784 | 0.299 | 0.468 | 0.00010823 | 7 | Exosc3 | myCAF_3 |
| Mob41 | 5.61E-09 | 0.29201122 | 0.356 | 0.515 | 0.0001085 | 7 | Mob4 | myCAF_3 |
| Pepd | 5.61E-09 | 0.65059213 | 0.356 | 0.505 | 0.00010854 | 7 | Pepd | myCAF_3 |
| Pik3ca1 | 5.63E-09 | 0.72851889 | 0.459 | 0.628 | 0.00010887 | 7 | Pik3ca | myCAF_3 |
| Slc7a6os1 | 5.93E-09 | 0.56675914 | 0.268 | 0.452 | 0.00011468 | 7 | Slc7a6os | myCAF_3 |
| Impdh11 | 5.94E-09 | 0.29867439 | 0.433 | 0.607 | 0.00011488 | 7 | Impdh1 | myCAF_3 |
| Samd14 | 5.99E-09 | 0.25197489 | 0.247 | 0.445 | 0.00011574 | 7 | Samd14 | myCAF_3 |
| Kidins2202 | 6.03E-09 | 0.40142432 | 0.428 | 0.623 | 0.00011651 | 7 | Kidins220 | myCAF_3 |
| Ap3b11 | 6.06E-09 | 0.44498692 | 0.598 | 0.73 | 0.00011712 | 7 | Ap3b1 | myCAF_3 |
| Tcp11l23 | 6.07E-09 | 0.45052425 | 0.495 | 0.573 | 0.00011727 | 7 | Tcp11l2 | myCAF_3 |
| Dot1l2 | 6.16E-09 | 0.33265691 | 0.299 | 0.469 | 0.00011907 | 7 | Dot1l | myCAF_3 |
| Luc7l31 | 6.31E-09 | 0.7986059 | 0.345 | 0.524 | 0.00012203 | 7 | Luc7l3 | myCAF_3 |
| Abl11 | 6.45E-09 | 0.87560676 | 0.67 | 0.78 | 0.00012473 | 7 | Abl1 | myCAF_3 |
| Bcl9l | 6.53E-09 | 0.78922648 | 0.716 | 0.799 | 0.00012623 | 7 | Bcl9l | myCAF_3 |
| Gan2 | 6.60E-09 | 0.60505527 | 0.314 | 0.463 | 0.0001276 | 7 | Gan | myCAF_3 |
| Fbxw5 | 6.93E-09 | 0.3185754 | 0.351 | 0.546 | 0.00013408 | 7 | Fbxw5 | myCAF_3 |
| H2-K13 | 6.97E-09 | 1.36538331 | 0.789 | 0.822 | 0.0001347 | 7 | H2-K1 | myCAF_3 |
| Thoc11 | 7.15E-09 | 0.35107495 | 0.18 | 0.372 | 0.00013816 | 7 | Thoc1 | myCAF_3 |
| Thoc22 | 7.16E-09 | 0.52853144 | 0.314 | 0.493 | 0.00013847 | 7 | Thoc2 | myCAF_3 |
| Abhd52 | 7.18E-09 | 0.26403493 | 0.34 | 0.482 | 0.00013882 | 7 | Abhd5 | myCAF_3 |

| | | | | | | | | |
|---|---|---|---|---|---|---|---|---|
| Ripor32 | 7.21E-09 | 0.55617563 | 0.345 | 0.471 | 0.00013948 | 7 | Ripor3 | myCAF_3 |
| Herc2 | 7.31E-09 | 0.29747668 | 0.34 | 0.524 | 0.00014136 | 7 | Herc2 | myCAF_3 |
| Bicd21 | 7.36E-09 | 0.61971405 | 0.521 | 0.683 | 0.00014224 | 7 | Bicd2 | myCAF_3 |
| Cryab2 | 7.69E-09 | 1.1796042 | 0.459 | 0.339 | 0.00014868 | 7 | Cryab | myCAF_3 |
| Calhm22 | 7.89E-09 | 0.50992928 | 0.356 | 0.538 | 0.00015253 | 7 | Calhm2 | myCAF_3 |
| Rbbp5 | 8.27E-09 | 0.46271061 | 0.232 | 0.388 | 0.00015992 | 7 | Rbbp5 | myCAF_3 |
| Ktn11 | 8.55E-09 | 0.31650046 | 0.438 | 0.584 | 0.00016529 | 7 | Ktn1 | myCAF_3 |
| Dvl1 | 8.56E-09 | 0.83813554 | 0.448 | 0.587 | 0.00016547 | 7 | Dvl1 | myCAF_3 |
| Rab3il12 | 8.60E-09 | 0.42836898 | 0.485 | 0.56 | 0.00016629 | 7 | Rab3il1 | myCAF_3 |
| Mfn1 | 8.63E-09 | 0.32336128 | 0.407 | 0.564 | 0.00016678 | 7 | Mfn1 | myCAF_3 |
| Hectd1 | 8.97E-09 | 0.6333875 | 0.66 | 0.752 | 0.00017342 | 7 | Hectd1 | myCAF_3 |
| Prkd11 | 9.21E-09 | 0.59828263 | 0.139 | 0.288 | 0.00017801 | 7 | Prkd1 | myCAF_3 |
| Aagab | 9.26E-09 | 0.52467279 | 0.299 | 0.467 | 0.00017894 | 7 | Aagab | myCAF_3 |
| Opa31 | 9.42E-09 | 0.5468029 | 0.247 | 0.441 | 0.00018211 | 7 | Opa3 | myCAF_3 |
| Ift43 | 9.83E-09 | 0.4150808 | 0.49 | 0.644 | 0.00019002 | 7 | Ift43 | myCAF_3 |
| Rsf11 | 9.88E-09 | 0.71235811 | 0.289 | 0.464 | 0.00019103 | 7 | Rsf1 | myCAF_3 |
| Ctu21 | 9.89E-09 | 0.63000215 | 0.175 | 0.341 | 0.00019115 | 7 | Ctu2 | myCAF_3 |
| Wdr371 | 1.09E-08 | 0.43081197 | 0.216 | 0.386 | 0.00021064 | 7 | Wdr37 | myCAF_3 |
| Hipk2 | 1.09E-08 | 0.43372987 | 0.227 | 0.408 | 0.00021093 | 7 | Hipk2 | myCAF_3 |
| Map3k13 | 1.13E-08 | 0.38554991 | 0.149 | 0.336 | 0.00021754 | 7 | Map3k1 | myCAF_3 |
| Vps13c1 | 1.13E-08 | 0.84167824 | 0.381 | 0.518 | 0.00021926 | 7 | Vps13c | myCAF_3 |
| Pbrm11 | 1.13E-08 | 0.78583035 | 0.521 | 0.654 | 0.00021942 | 7 | Pbrm1 | myCAF_3 |
| Cars3 | 1.14E-08 | 0.55598464 | 0.407 | 0.563 | 0.00022101 | 7 | Cars | myCAF_3 |
| Prkci1 | 1.15E-08 | 0.525803 | 0.325 | 0.5 | 0.00022227 | 7 | Prkci | myCAF_3 |
| Ldb11 | 1.17E-08 | 1.00506985 | 0.753 | 0.807 | 0.00022603 | 7 | Ldb1 | myCAF_3 |
| Prpf311 | 1.17E-08 | 0.38555437 | 0.34 | 0.468 | 0.00022624 | 7 | Prpf31 | myCAF_3 |
| Socs6 | 1.17E-08 | 0.50262191 | 0.33 | 0.475 | 0.00022644 | 7 | Socs6 | myCAF_3 |
| Oxa1l | 1.19E-08 | 0.368434 | 0.412 | 0.6 | 0.00022924 | 7 | Oxa1l | myCAF_3 |
| Zbtb212 | 1.19E-08 | 0.56588015 | 0.227 | 0.407 | 0.00022926 | 7 | Zbtb21 | myCAF_3 |
| Tmem179b1 | 1.19E-08 | 0.25684854 | 0.289 | 0.454 | 0.00022953 | 7 | Tmem179b | myCAF_3 |
| Trappc3 | 1.19E-08 | 0.50848233 | 0.51 | 0.69 | 0.00023066 | 7 | Trappc3 | myCAF_3 |
| Akap91 | 1.20E-08 | 0.27306746 | 0.387 | 0.545 | 0.00023129 | 7 | Akap9 | myCAF_3 |
| Srebf11 | 1.23E-08 | 0.4920063 | 0.521 | 0.662 | 0.00023706 | 7 | Srebf1 | myCAF_3 |
| Fasn | 1.26E-08 | 0.65711942 | 0.639 | 0.716 | 0.00024325 | 7 | Fasn | myCAF_3 |
| Fxr22 | 1.27E-08 | 0.55509673 | 0.448 | 0.593 | 0.0002453 | 7 | Fxr2 | myCAF_3 |
| Ip6k11 | 1.30E-08 | 0.81006977 | 0.763 | 0.856 | 0.00025083 | 7 | Ip6k1 | myCAF_3 |
| Ahctf11 | 1.31E-08 | 0.53404609 | 0.366 | 0.527 | 0.00025352 | 7 | Ahctf1 | myCAF_3 |
| Ppp1r23 | 1.31E-08 | 0.41366399 | 0.51 | 0.658 | 0.00025424 | 7 | Ppp1r2 | myCAF_3 |
| Stat5b1 | 1.32E-08 | 0.5729439 | 0.335 | 0.493 | 0.00025575 | 7 | Stat5b | myCAF_3 |
| Prmt71 | 1.34E-08 | 0.42644439 | 0.17 | 0.342 | 0.0002591 | 7 | Prmt7 | myCAF_3 |
| Ppp1r12c2 | 1.35E-08 | 0.64300824 | 0.588 | 0.708 | 0.00026103 | 7 | Ppp1r12c | myCAF_3 |
| Ints1 | 1.35E-08 | 0.4023242 | 0.32 | 0.498 | 0.0002611 | 7 | Ints1 | myCAF_3 |
| Ankrd501 | 1.36E-08 | 0.40900262 | 0.423 | 0.56 | 0.00026382 | 7 | Ankrd50 | myCAF_3 |
| Plpp32 | 1.38E-08 | 0.71642555 | 0.799 | 0.848 | 0.00026755 | 7 | Plpp3 | myCAF_3 |
| Rai1 | 1.39E-08 | 0.71020775 | 0.531 | 0.675 | 0.00026853 | 7 | Rai1 | myCAF_3 |
| Wdr45b | 1.41E-08 | 0.32082647 | 0.454 | 0.613 | 0.00027256 | 7 | Wdr45b | myCAF_3 |
| Mpg2 | 1.43E-08 | 0.29346886 | 0.361 | 0.524 | 0.00027723 | 7 | Mpg | myCAF_3 |
| Borcs81 | 1.47E-08 | 0.57339517 | 0.479 | 0.622 | 0.00028454 | 7 | Borcs8 | myCAF_3 |
| Mfn2 | 1.53E-08 | 0.58364693 | 0.356 | 0.521 | 0.00029613 | 7 | Mfn2 | myCAF_3 |
| Fam193a | 1.55E-08 | 0.68557869 | 0.448 | 0.597 | 0.00029932 | 7 | Fam193a | myCAF_3 |
| Pan31 | 1.55E-08 | 0.41800093 | 0.304 | 0.472 | 0.00030051 | 7 | Pan3 | myCAF_3 |
| Mafb1 | 1.57E-08 | 0.9570004 | 0.629 | 0.665 | 0.00030298 | 7 | Mafb | myCAF_3 |
| Fkbp15 | 1.57E-08 | 0.33943316 | 0.418 | 0.58 | 0.00030324 | 7 | Fkbp15 | myCAF_3 |
| Bax | 1.64E-08 | 0.59661659 | 0.716 | 0.85 | 0.00031627 | 7 | Bax | myCAF_3 |
| Hoxa32 | 1.71E-08 | 0.43439875 | 0.119 | 0.268 | 0.00033063 | 7 | Hoxa3 | myCAF_3 |
| Znrf12 | 1.79E-08 | 0.85538361 | 0.557 | 0.695 | 0.00034539 | 7 | Znrf1 | myCAF_3 |
| Alyref4 | 1.81E-08 | 1.15998084 | 0.144 | 0.257 | 0.00034951 | 7 | Alyref | myCAF_3 |
| Sdad1 | 1.81E-08 | 0.35867751 | 0.232 | 0.401 | 0.00035028 | 7 | Sdad1 | myCAF_3 |
| Eed1 | 1.82E-08 | 1.44532855 | 0.206 | 0.355 | 0.00035154 | 7 | Eed | myCAF_3 |
| Sema3f | 1.84E-08 | 2.17545843 | 0.278 | 0.153 | 0.00035606 | 7 | Sema3f | myCAF_3 |
| Ccar2 | 1.85E-08 | 0.47000851 | 0.402 | 0.582 | 0.0003584 | 7 | Ccar2 | myCAF_3 |
| Vrk11 | 1.88E-08 | 0.43895895 | 0.149 | 0.264 | 0.00036373 | 7 | Vrk1 | myCAF_3 |
| Def8 | 1.89E-08 | 0.32070866 | 0.263 | 0.439 | 0.00036505 | 7 | Def8 | myCAF_3 |
| Ap1s21 | 1.89E-08 | 1.42383977 | 0.479 | 0.376 | 0.0003659 | 7 | Ap1s2 | myCAF_3 |
| Per12 | 1.93E-08 | 0.5266365 | 0.402 | 0.551 | 0.00037384 | 7 | Per1 | myCAF_3 |
| Ripor1 | 1.95E-08 | 0.52248053 | 0.433 | 0.574 | 0.00037681 | 7 | Ripor1 | myCAF_3 |
| Tinf21 | 1.96E-08 | 0.46322871 | 0.165 | 0.295 | 0.00037949 | 7 | Tinf2 | myCAF_3 |
| Sucla2 | 1.99E-08 | 0.29803264 | 0.469 | 0.625 | 0.00038487 | 7 | Sucla2 | myCAF_3 |
| Ap1b1 | 2.00E-08 | 0.45177446 | 0.387 | 0.531 | 0.0003865 | 7 | Ap1b1 | myCAF_3 |
| Tenm31 | 2.04E-08 | 0.55454336 | 0.443 | 0.503 | 0.00039483 | 7 | Tenm3 | myCAF_3 |
| Spryd3 | 2.09E-08 | 0.68292986 | 0.567 | 0.683 | 0.00040418 | 7 | Spryd3 | myCAF_3 |
| Paxip11 | 2.09E-08 | 0.43426283 | 0.284 | 0.414 | 0.00040481 | 7 | Paxip1 | myCAF_3 |
| Mtor1 | 2.10E-08 | 0.39712616 | 0.289 | 0.46 | 0.00040558 | 7 | Mtor | myCAF_3 |
| Ankrd281 | 2.12E-08 | 2.20479397 | 0.639 | 0.502 | 0.00041078 | 7 | Ankrd28 | myCAF_3 |
| Med25 | 2.20E-08 | 0.61129985 | 0.567 | 0.725 | 0.00042464 | 7 | Med25 | myCAF_3 |
| Smcr81 | 2.21E-08 | 0.61984191 | 0.387 | 0.535 | 0.00042684 | 7 | Smcr8 | myCAF_3 |
| Men11 | 2.21E-08 | 0.4460972 | 0.335 | 0.515 | 0.00042764 | 7 | Men1 | myCAF_3 |
| Gabpa1 | 2.26E-08 | 0.41029022 | 0.392 | 0.549 | 0.00043618 | 7 | Gabpa | myCAF_3 |
| Nfx1 | 2.27E-08 | 0.43066429 | 0.33 | 0.486 | 0.00043834 | 7 | Nfx1 | myCAF_3 |
| Myo1b2 | 2.27E-08 | 0.61160362 | 0.608 | 0.735 | 0.00043903 | 7 | Myo1b | myCAF_3 |
| Trp53i132 | 2.35E-08 | 0.34081917 | 0.247 | 0.437 | 0.0004553 | 7 | Trp53i13 | myCAF_3 |
| Rbm6 | 2.42E-08 | 0.83120872 | 0.464 | 0.6 | 0.00046834 | 7 | Rbm6 | myCAF_3 |
| Pex51 | 2.46E-08 | 0.72161846 | 0.5 | 0.657 | 0.0004761 | 7 | Pex5 | myCAF_3 |
| Rab91 | 2.48E-08 | 0.70396168 | 0.448 | 0.586 | 0.00047926 | 7 | Rab9 | myCAF_3 |
| Ebp3 | 2.50E-08 | 0.35498536 | 0.448 | 0.566 | 0.00048349 | 7 | Ebp | myCAF_3 |
| Tdrd72 | 2.56E-08 | 0.45164385 | 0.211 | 0.387 | 0.00049426 | 7 | Tdrd7 | myCAF_3 |
| Bop11 | 2.58E-08 | 0.44577498 | 0.232 | 0.384 | 0.00049809 | 7 | Bop1 | myCAF_3 |
| Nfatc12 | 2.67E-08 | 0.37906128 | 0.314 | 0.41 | 0.00051605 | 7 | Nfatc1 | myCAF_3 |
| St6galnac62 | 2.67E-08 | 0.50808196 | 0.505 | 0.668 | 0.00051704 | 7 | St6galnac6 | myCAF_3 |
| Odc11 | 2.68E-08 | 0.64865087 | 0.794 | 0.885 | 0.00051849 | 7 | Odc1 | myCAF_3 |
| Dhx33 | 2.69E-08 | 0.30436428 | 0.299 | 0.468 | 0.0005196 | 7 | Dhx33 | myCAF_3 |
| Oxr12 | 2.72E-08 | 0.66685313 | 0.309 | 0.474 | 0.00052602 | 7 | Oxr1 | myCAF_3 |
| Ndrg42 | 2.74E-08 | 0.38169903 | 0.33 | 0.445 | 0.00052983 | 7 | Ndrg4 | myCAF_3 |
| Agrn2 | 2.75E-08 | 0.63780581 | 0.464 | 0.605 | 0.00053117 | 7 | Agrn | myCAF_3 |
| Sh2b1 | 2.76E-08 | 0.41792611 | 0.371 | 0.549 | 0.00053361 | 7 | Sh2b1 | myCAF_3 |
| Dapk13 | 2.81E-08 | 0.89849721 | 0.175 | 0.27 | 0.00054268 | 7 | Dapk1 | myCAF_3 |
| Rprd1a | 2.90E-08 | 0.35218326 | 0.201 | 0.36 | 0.00056091 | 7 | Rprd1a | myCAF_3 |
| Stk38l | 2.91E-08 | 2.30182877 | 0.314 | 0.263 | 0.00056269 | 7 | Stk38l | myCAF_3 |
| Atp6v1e1 | 2.99E-08 | 0.49345149 | 0.603 | 0.752 | 0.00057818 | 7 | Atp6v1e1 | myCAF_3 |
| Fut82 | 3.03E-08 | 0.54032228 | 0.474 | 0.613 | 0.00058504 | 7 | Fut8 | myCAF_3 |
| Smad5 | 3.04E-08 | 0.95220387 | 0.526 | 0.648 | 0.0005871 | 7 | Smad5 | myCAF_3 |
| Ercc12 | 3.05E-08 | 0.25555756 | 0.376 | 0.54 | 0.0005899 | 7 | Ercc1 | myCAF_3 |
| Foxred11 | 3.07E-08 | 0.40378087 | 0.216 | 0.378 | 0.00059329 | 7 | Foxred1 | myCAF_3 |
| Zfp703 | 3.07E-08 | 0.96244923 | 0.608 | 0.673 | 0.00059338 | 7 | Zfp703 | myCAF_3 |
| Cog1 | 3.19E-08 | 0.3569205 | 0.356 | 0.549 | 0.00061702 | 7 | Cog1 | myCAF_3 |
| Timm441 | 3.19E-08 | 0.52134133 | 0.258 | 0.416 | 0.00061724 | 7 | Timm44 | myCAF_3 |
| Nsmf1 | 3.26E-08 | 0.25467759 | 0.536 | 0.702 | 0.0006303 | 7 | Nsmf | myCAF_3 |
| Ptcd31 | 3.39E-08 | 0.44951728 | 0.242 | 0.398 | 0.00065539 | 7 | Ptcd3 | myCAF_3 |
| Abcf21 | 3.48E-08 | 0.29016523 | 0.345 | 0.501 | 0.00067211 | 7 | Abcf2 | myCAF_3 |
| Parm13 | 3.50E-08 | 1.24666565 | 0.263 | 0.222 | 0.00067573 | 7 | Parm1 | myCAF_3 |
| Safb21 | 3.55E-08 | 0.41499807 | 0.387 | 0.548 | 0.0006865 | 7 | Safb2 | myCAF_3 |
| Zfp5921 | 3.55E-08 | 0.57382924 | 0.387 | 0.543 | 0.00068721 | 7 | Zfp592 | myCAF_3 |
| Wdfy31 | 3.57E-08 | 0.77821988 | 0.655 | 0.81 | 0.00069048 | 7 | Wdfy3 | myCAF_3 |
| Clock1 | 3.60E-08 | 0.59148355 | 0.304 | 0.458 | 0.00069518 | 7 | Clock | myCAF_3 |
| Rdh111 | 3.60E-08 | 0.79454665 | 0.17 | 0.352 | 0.00069679 | 7 | Rdh11 | myCAF_3 |
| Cdk8 | 3.65E-08 | 0.42427707 | 0.361 | 0.51 | 0.00070485 | 7 | Cdk8 | myCAF_3 |
| Trmt2a | 3.79E-08 | 0.50260719 | 0.289 | 0.438 | 0.0007335 | 7 | Trmt2a | myCAF_3 |
| Pop41 | 3.84E-08 | 0.28342579 | 0.258 | 0.411 | 0.00074276 | 7 | Pop4 | myCAF_3 |
| Aldh3a22 | 3.87E-08 | 0.35827488 | 0.253 | 0.433 | 0.00074838 | 7 | Aldh3a2 | myCAF_3 |
| Osbpl8 | 3.89E-08 | 0.66489465 | 0.392 | 0.578 | 0.00075175 | 7 | Osbpl8 | myCAF_3 |
| Ankib1 | 3.96E-08 | 0.49132972 | 0.309 | 0.462 | 0.00076594 | 7 | Ankib1 | myCAF_3 |
| Naa25 | 3.96E-08 | 0.59047407 | 0.201 | 0.37 | 0.00076646 | 7 | Naa25 | myCAF_3 |
| Dnmt3a1 | 4.05E-08 | 0.30370254 | 0.397 | 0.552 | 0.00078312 | 7 | Dnmt3a | myCAF_3 |
| Aktip2 | 4.06E-08 | 0.33374209 | 0.222 | 0.4 | 0.00078518 | 7 | Aktip | myCAF_3 |
| Ube4a1 | 4.09E-08 | 0.39144918 | 0.454 | 0.607 | 0.00079073 | 7 | Ube4a | myCAF_3 |
| Cd402 | 4.10E-08 | 1.20732449 | 0.284 | 0.177 | 0.00079209 | 7 | Cd40 | myCAF_3 |
| Gtf2b | 4.11E-08 | 0.42247509 | 0.495 | 0.633 | 0.00079411 | 7 | Gtf2b | myCAF_3 |
| Cuedc2 | 4.13E-08 | 0.5376681 | 0.51 | 0.677 | 0.00079763 | 7 | Cuedc2 | myCAF_3 |
| Bex32 | 4.14E-08 | 0.60513879 | 0.51 | 0.568 | 0.00080135 | 7 | Bex3 | myCAF_3 |
| Trabd | 4.15E-08 | 0.41595175 | 0.392 | 0.54 | 0.00080218 | 7 | Trabd | myCAF_3 |
| Lrch32 | 4.15E-08 | 0.78393355 | 0.433 | 0.6 | 0.00080332 | 7 | Lrch3 | myCAF_3 |
| Gzf11 | 4.16E-08 | 0.3367637 | 0.201 | 0.38 | 0.0008044 | 7 | Gzf1 | myCAF_3 |
| Fzd7 | 4.24E-08 | 0.70050921 | 0.531 | 0.655 | 0.00082073 | 7 | Fzd7 | myCAF_3 |
| Atf41 | 4.25E-08 | 0.84521886 | 0.454 | 0.574 | 0.00082245 | 7 | Atf4 | myCAF_3 |
| Zc2hc1a1 | 4.36E-08 | 0.41137592 | 0.237 | 0.412 | 0.00084225 | 7 | Zc2hc1a | myCAF_3 |
| Champ1 | 4.37E-08 | 0.41086102 | 0.273 | 0.446 | 0.00084413 | 7 | Champ1 | myCAF_3 |
| Arhgef172 | 4.37E-08 | 0.51799117 | 0.588 | 0.684 | 0.00084431 | 7 | Arhgef17 | myCAF_3 |
| Egln2 | 4.40E-08 | 0.3588518 | 0.495 | 0.675 | 0.00085127 | 7 | Egln2 | myCAF_3 |
| Itgb32 | 4.48E-08 | 0.28667498 | 0.582 | 0.68 | 0.00086573 | 7 | Itgb3 | myCAF_3 |
| Bap1 | 4.54E-08 | 0.31506697 | 0.32 | 0.496 | 0.00087823 | 7 | Bap1 | myCAF_3 |
| Spred22 | 4.58E-08 | 0.27980665 | 0.402 | 0.585 | 0.0008863 | 7 | Spred2 | myCAF_3 |
| Ddx47 | 4.62E-08 | 0.41275254 | 0.294 | 0.482 | 0.00089392 | 7 | Ddx47 | myCAF_3 |
| Baz1b1 | 4.65E-08 | 0.76685774 | 0.515 | 0.624 | 0.00089894 | 7 | Baz1b | myCAF_3 |
| Steap32 | 4.66E-08 | 0.39031014 | 0.412 | 0.518 | 0.00090187 | 7 | Steap3 | myCAF_3 |
| Ints71 | 4.70E-08 | 0.57521842 | 0.227 | 0.395 | 0.00090877 | 7 | Ints7 | myCAF_3 |
| Auts21 | 4.76E-08 | 0.92251151 | 0.433 | 0.555 | 0.00092086 | 7 | Auts2 | myCAF_3 |
| Igfbp32 | 4.85E-08 | 1.29714288 | 0.794 | 0.729 | 0.00093759 | 7 | Igfbp3 | myCAF_3 |
| Tex2642 | 4.96E-08 | 0.64976933 | 0.696 | 0.797 | 0.0009591 | 7 | Tex264 | myCAF_3 |
| Ddx561 | 5.03E-08 | 0.46079784 | 0.351 | 0.509 | 0.00097309 | 7 | Ddx56 | myCAF_3 |
| Cenpc11 | 5.12E-08 | 0.4279861 | 0.237 | 0.376 | 0.00099057 | 7 | Cenpc1 | myCAF_3 |
| Kdm5a1 | 5.18E-08 | 0.66403673 | 0.505 | 0.631 | 0.00100138 | 7 | Kdm5a | myCAF_3 |
| Zcchc81 | 5.21E-08 | 0.33599 | 0.258 | 0.408 | 0.00100695 | 7 | Zcchc8 | myCAF_3 |
| N4bp11 | 5.25E-08 | 0.48828567 | 0.505 | 0.621 | 0.0010157 | 7 | N4bp1 | myCAF_3 |
| Ambra1 | 5.45E-08 | 0.68272733 | 0.366 | 0.512 | 0.00105312 | 7 | Ambra1 | myCAF_3 |
| Zfp146 | 5.50E-08 | 0.25696275 | 0.258 | 0.44 | 0.00106429 | 7 | Zfp146 | myCAF_3 |
| Kansl11 | 5.51E-08 | 1.11622316 | 0.567 | 0.721 | 0.0010652 | 7 | Kansl1 | myCAF_3 |
| Gtpbp11 | 5.62E-08 | 0.6364662 | 0.335 | 0.489 | 0.0010858 | 7 | Gtpbp1 | myCAF_3 |
| Tstd2 | 5.64E-08 | 0.50236907 | 0.216 | 0.39 | 0.00109136 | 7 | Tstd2 | myCAF_3 |
| Fbrs1 | 5.74E-08 | 0.62425262 | 0.557 | 0.73 | 0.00110962 | 7 | Fbrs | myCAF_3 |
| Otud5 | 5.78E-08 | 0.51627363 | 0.335 | 0.502 | 0.0011173 | 7 | Otud5 | myCAF_3 |
| Napg | 5.79E-08 | 0.39783999 | 0.314 | 0.451 | 0.0011191 | 7 | Napg | myCAF_3 |
| Nt5c4 | 5.79E-08 | 0.63060647 | 0.49 | 0.612 | 0.00111987 | 7 | Nt5c | myCAF_3 |
| Fam204a1 | 5.80E-08 | 0.2799835 | 0.206 | 0.403 | 0.0011207 | 7 | Fam204a | myCAF_3 |
| Rhot11 | 5.80E-08 | 0.6709046 | 0.278 | 0.442 | 0.00112184 | 7 | Rhot1 | myCAF_3 |
| Lrrfip1 | 6.21E-08 | 0.47322155 | 0.613 | 0.709 | 0.00120146 | 7 | Lrrfip1 | myCAF_3 |
| Cyb5r11 | 6.37E-08 | 0.34790611 | 0.381 | 0.484 | 0.00123109 | 7 | Cyb5r1 | myCAF_3 |
| Pycr21 | 6.47E-08 | 0.54645264 | 0.345 | 0.48 | 0.00125015 | 7 | Pycr2 | myCAF_3 |
| Ctsa1 | 6.49E-08 | 0.58642037 | 0.804 | 0.915 | 0.00125463 | 7 | Ctsa | myCAF_3 |
| Adar2 | 6.53E-08 | 0.58646405 | 0.299 | 0.471 | 0.00126311 | 7 | Adar | myCAF_3 |
| Psmd4 | 6.71E-08 | 0.77478572 | 0.536 | 0.655 | 0.00129753 | 7 | Psmd4 | myCAF_3 |
| Arf21 | 6.74E-08 | 0.32131345 | 0.397 | 0.551 | 0.00130254 | 7 | Arf2 | myCAF_3 |
| Fam214a2 | 7.00E-08 | 1.29175607 | 0.438 | 0.392 | 0.00135257 | 7 | Fam214a | myCAF_3 |
| Rcc21 | 7.12E-08 | 0.80176582 | 0.593 | 0.68 | 0.00137576 | 7 | Rcc2 | myCAF_3 |
| Cln5 | 7.15E-08 | 0.59450448 | 0.567 | 0.705 | 0.00138258 | 7 | Cln5 | myCAF_3 |
| Smpd11 | 7.15E-08 | 0.26479263 | 0.5 | 0.66 | 0.00138327 | 7 | Smpd1 | myCAF_3 |
| Slc5a31 | 7.21E-08 | 2.2042128 | 0.325 | 0.349 | 0.00139365 | 7 | Slc5a3 | myCAF_3 |
| Plcg12 | 7.23E-08 | 0.72648845 | 0.557 | 0.668 | 0.00139823 | 7 | Plcg1 | myCAF_3 |
| Brd31 | 7.28E-08 | 0.7767521 | 0.376 | 0.507 | 0.00140842 | 7 | Brd3 | myCAF_3 |
| Ints111 | 7.43E-08 | 0.26742896 | 0.294 | 0.445 | 0.00143686 | 7 | Ints11 | myCAF_3 |
| Narf3 | 7.52E-08 | 0.50446699 | 0.289 | 0.454 | 0.00145464 | 7 | Narf | myCAF_3 |
| Pogz | 7.62E-08 | 0.30942612 | 0.268 | 0.427 | 0.00147264 | 7 | Pogz | myCAF_3 |
| Dtx31 | 7.63E-08 | 0.74421229 | 0.66 | 0.787 | 0.00147491 | 7 | Dtx3 | myCAF_3 |
| Hnrnph31 | 7.66E-08 | 0.56762043 | 0.309 | 0.465 | 0.00148031 | 7 | Hnrnph3 | myCAF_3 |
| Ncoa61 | 7.75E-08 | 0.43691774 | 0.412 | 0.561 | 0.00149905 | 7 | Ncoa6 | myCAF_3 |
| Phf31 | 7.78E-08 | 0.71704416 | 0.289 | 0.455 | 0.00150388 | 7 | Phf3 | myCAF_3 |
| Epc11 | 7.85E-08 | 0.80423553 | 0.521 | 0.659 | 0.00151711 | 7 | Epc1 | myCAF_3 |
| Rptor1 | 7.98E-08 | 0.56237366 | 0.299 | 0.48 | 0.0015435 | 7 | Rptor | myCAF_3 |
| Osbpl91 | 8.28E-08 | 0.7186852 | 0.572 | 0.67 | 0.00160019 | 7 | Osbpl9 | myCAF_3 |
| Trak11 | 8.31E-08 | 0.39336157 | 0.541 | 0.694 | 0.00160604 | 7 | Trak1 | myCAF_3 |
| Hmox13 | 8.36E-08 | 0.59836559 | 0.634 | 0.708 | 0.00161709 | 7 | Hmox1 | myCAF_3 |
| Auh2 | 8.62E-08 | 1.03252611 | 0.495 | 0.607 | 0.0016659 | 7 | Auh | myCAF_3 |
| Atf6b | 8.72E-08 | 0.36612619 | 0.371 | 0.5 | 0.00168536 | 7 | Atf6b | myCAF_3 |
| Manbal | 8.74E-08 | 0.88449191 | 0.531 | 0.639 | 0.00169038 | 7 | Manbal | myCAF_3 |
| Ddhd21 | 8.97E-08 | 0.3284601 | 0.284 | 0.477 | 0.0017344 | 7 | Ddhd2 | myCAF_3 |
| Zfp445 | 8.99E-08 | 0.59002368 | 0.454 | 0.582 | 0.00173749 | 7 | Zfp445 | myCAF_3 |
| Lmbrd11 | 8.99E-08 | 0.42428999 | 0.335 | 0.507 | 0.0017378 | 7 | Lmbrd1 | myCAF_3 |
| Mtmr31 | 9.50E-08 | 0.34859965 | 0.309 | 0.496 | 0.00183651 | 7 | Mtmr3 | myCAF_3 |
| Hp2 | 9.82E-08 | 1.16553913 | 0.263 | 0.278 | 0.00189853 | 7 | Hp | myCAF_3 |
| Rtf2 | 9.82E-08 | 0.51630893 | 0.407 | 0.566 | 0.00189874 | 7 | Rtf2 | myCAF_3 |
| Ints131 | 1.00E-07 | 0.29919479 | 0.263 | 0.41 | 0.00194213 | 7 | Ints13 | myCAF_3 |
| Aen | 1.01E-07 | 0.34877758 | 0.376 | 0.49 | 0.00195295 | 7 | Aen | myCAF_3 |
| Hexim12 | 1.05E-07 | 1.09145292 | 0.572 | 0.686 | 0.00203668 | 7 | Hexim1 | myCAF_3 |
| Kmt2e2 | 1.06E-07 | 0.70447574 | 0.515 | 0.647 | 0.00205269 | 7 | Kmt2e | myCAF_3 |
| Wdr461 | 1.07E-07 | 0.47743121 | 0.304 | 0.462 | 0.00206427 | 7 | Wdr46 | myCAF_3 |
| Tmem651 | 1.08E-07 | 0.49581242 | 0.598 | 0.738 | 0.00209241 | 7 | Tmem65 | myCAF_3 |
| Josd1 | 1.09E-07 | 0.39166724 | 0.423 | 0.587 | 0.00211009 | 7 | Josd1 | myCAF_3 |
| Acbd5 | 1.10E-07 | 0.9303168 | 0.314 | 0.458 | 0.00212926 | 7 | Acbd5 | myCAF_3 |
| Dlg11 | 1.11E-07 | 0.7192604 | 0.603 | 0.714 | 0.00214246 | 7 | Dlg1 | myCAF_3 |
| Ammecr1l1 | 1.12E-07 | 0.64222177 | 0.371 | 0.528 | 0.00215891 | 7 | Ammecr1l | myCAF_3 |
| Rnf401 | 1.15E-07 | 0.42416911 | 0.371 | 0.574 | 0.00223077 | 7 | Rnf40 | myCAF_3 |
| Aspscr1 | 1.18E-07 | 0.53992096 | 0.314 | 0.459 | 0.00228002 | 7 | Aspscr1 | myCAF_3 |
| Ephb31 | 1.21E-07 | 0.48238779 | 0.371 | 0.511 | 0.00234118 | 7 | Ephb3 | myCAF_3 |
| Hsph12 | 1.21E-07 | 0.52668813 | 0.237 | 0.421 | 0.00234513 | 7 | Hsph1 | myCAF_3 |
| Rabep1 | 1.24E-07 | 0.4963019 | 0.448 | 0.627 | 0.00239216 | 7 | Rabep1 | myCAF_3 |
| Aff1 | 1.26E-07 | 0.64122556 | 0.655 | 0.786 | 0.00243572 | 7 | Aff1 | myCAF_3 |
| Ttc41 | 1.26E-07 | 0.38911654 | 0.216 | 0.395 | 0.00243665 | 7 | Ttc4 | myCAF_3 |
| Rnf1301 | 1.28E-07 | 0.58286299 | 0.438 | 0.587 | 0.0024657 | 7 | Rnf130 | myCAF_3 |
| Bpnt11 | 1.31E-07 | 0.30247581 | 0.278 | 0.429 | 0.00253399 | 7 | Bpnt1 | myCAF_3 |
| Rnf34 | 1.35E-07 | 0.58965872 | 0.33 | 0.478 | 0.00260317 | 7 | Rnf34 | myCAF_3 |
| Pds5b2 | 1.39E-07 | 0.30109289 | 0.32 | 0.463 | 0.0026788 | 7 | Pds5b | myCAF_3 |
| Gjc12 | 1.39E-07 | 0.68062302 | 0.361 | 0.483 | 0.00269629 | 7 | Gjc1 | myCAF_3 |
| Ttl | 1.40E-07 | 0.57876543 | 0.304 | 0.466 | 0.00270773 | 7 | Ttl | myCAF_3 |
| Wbp4 | 1.40E-07 | 0.60637842 | 0.371 | 0.523 | 0.00271065 | 7 | Wbp4 | myCAF_3 |
| Pebp11 | 1.42E-07 | 0.66936771 | 0.706 | 0.828 | 0.002739 | 7 | Pebp1 | myCAF_3 |
| Trib11 | 1.45E-07 | 0.93712141 | 0.361 | 0.422 | 0.00279848 | 7 | Trib1 | myCAF_3 |
| Ebf41 | 1.46E-07 | 2.0213731 | 0.273 | 0.145 | 0.00282229 | 7 | Ebf4 | myCAF_3 |
| Rnaseh2a1 | 1.48E-07 | 0.80075 | 0.284 | 0.423 | 0.00286382 | 7 | Rnaseh2a | myCAF_3 |
| Tubgcp31 | 1.49E-07 | 0.78529013 | 0.304 | 0.43 | 0.00288079 | 7 | Tubgcp3 | myCAF_3 |
| Dip2c | 1.50E-07 | 0.7996636 | 0.309 | 0.442 | 0.00289922 | 7 | Dip2c | myCAF_3 |
| Pnisr1 | 1.57E-07 | 0.62396156 | 0.469 | 0.617 | 0.00302601 | 7 | Pnisr | myCAF_3 |
| Stag11 | 1.57E-07 | 0.31228065 | 0.253 | 0.419 | 0.00302807 | 7 | Stag1 | myCAF_3 |
| Gdpd52 | 1.57E-07 | 1.36631791 | 0.376 | 0.271 | 0.00304031 | 7 | Gdpd5 | myCAF_3 |
| Cpne23 | 1.59E-07 | 0.29563982 | 0.165 | 0.293 | 0.00306707 | 7 | Cpne2 | myCAF_3 |
| Rpp14 | 1.60E-07 | 0.31385641 | 0.196 | 0.351 | 0.00310164 | 7 | Rpp14 | myCAF_3 |
| Ntmt11 | 1.62E-07 | 0.62668086 | 0.325 | 0.482 | 0.0031254 | 7 | Ntmt1 | myCAF_3 |
| Slc25a22 | 1.64E-07 | 0.41462079 | 0.356 | 0.472 | 0.00317391 | 7 | Slc25a22 | myCAF_3 |
| Ylpm11 | 1.67E-07 | 0.52453862 | 0.381 | 0.54 | 0.00322453 | 7 | Ylpm1 | myCAF_3 |
| Hoxc61 | 1.70E-07 | 0.72690901 | 0.665 | 0.731 | 0.00329281 | 7 | Hoxc6 | myCAF_3 |
| Khdc41 | 1.73E-07 | 0.76491404 | 0.572 | 0.665 | 0.0033405 | 7 | Khdc4 | myCAF_3 |
| Matn22 | 1.73E-07 | 0.5420901 | 0.531 | 0.672 | 0.00335077 | 7 | Matn2 | myCAF_3 |
| Uri11 | 1.76E-07 | 0.66856513 | 0.387 | 0.533 | 0.00340153 | 7 | Uri1 | myCAF_3 |
| Sun12 | 1.77E-07 | 0.63639724 | 0.428 | 0.544 | 0.00342299 | 7 | Sun1 | myCAF_3 |
| Crebzf1 | 1.77E-07 | 0.3011524 | 0.366 | 0.528 | 0.0034239 | 7 | Crebzf | myCAF_3 |
| Atp13a1 | 1.79E-07 | 0.31319908 | 0.485 | 0.597 | 0.00345242 | 7 | Atp13a1 | myCAF_3 |
| Gpsm23 | 1.81E-07 | 0.31773009 | 0.32 | 0.408 | 0.00349213 | 7 | Gpsm2 | myCAF_3 |
| Npc1 | 1.83E-07 | 0.49370628 | 0.526 | 0.682 | 0.00352888 | 7 | Npc1 | myCAF_3 |
| Ntrk23 | 1.83E-07 | 1.01327109 | 0.33 | 0.295 | 0.00353527 | 7 | Ntrk2 | myCAF_3 |
| Phf132 | 1.84E-07 | 0.77166603 | 0.407 | 0.556 | 0.00356272 | 7 | Phf13 | myCAF_3 |
| Samhd12 | 1.88E-07 | 0.49938821 | 0.469 | 0.559 | 0.00364184 | 7 | Samhd1 | myCAF_3 |
| Dnajc4 | 1.89E-07 | 0.54024024 | 0.299 | 0.445 | 0.00365731 | 7 | Dnajc4 | myCAF_3 |
| Lactb1 | 1.95E-07 | 0.48753752 | 0.124 | 0.283 | 0.00376258 | 7 | Lactb | myCAF_3 |
| B3gat31 | 2.03E-07 | 0.26926938 | 0.247 | 0.446 | 0.00392586 | 7 | B3gat3 | myCAF_3 |
| Mtr1 | 2.03E-07 | 0.27271156 | 0.17 | 0.33 | 0.00392621 | 7 | Mtr | myCAF_3 |
| Mical32 | 2.04E-07 | 1.91128738 | 0.454 | 0.352 | 0.00393474 | 7 | Mical3 | myCAF_3 |
| Deaf1 | 2.07E-07 | 0.34224095 | 0.345 | 0.51 | 0.00400551 | 7 | Deaf1 | myCAF_3 |
| Eif3a1 | 2.08E-07 | 0.44945187 | 0.438 | 0.593 | 0.00402196 | 7 | Eif3a | myCAF_3 |
| Mapkbp12 | 2.12E-07 | 0.28105603 | 0.113 | 0.264 | 0.00409774 | 7 | Mapkbp1 | myCAF_3 |
| Ralgapb | 2.14E-07 | 0.37157772 | 0.381 | 0.547 | 0.00413651 | 7 | Ralgapb | myCAF_3 |
| Gnl3l | 2.16E-07 | 0.40745111 | 0.361 | 0.509 | 0.00417756 | 7 | Gnl3l | myCAF_3 |
| Plekhg32 | 2.17E-07 | 0.36105562 | 0.345 | 0.504 | 0.00419288 | 7 | Plekhg3 | myCAF_3 |
| Kdm7a1 | 2.17E-07 | 2.22569541 | 0.536 | 0.467 | 0.00420362 | 7 | Kdm7a | myCAF_3 |
| Pdhb1 | 2.18E-07 | 0.60592608 | 0.387 | 0.559 | 0.00421109 | 7 | Pdhb | myCAF_3 |
| Rcor32 | 2.21E-07 | 0.75868592 | 0.278 | 0.425 | 0.00426612 | 7 | Rcor3 | myCAF_3 |
| Scaf8 | 2.21E-07 | 0.76849052 | 0.299 | 0.485 | 0.00427524 | 7 | Scaf8 | myCAF_3 |
| Ralgapa1 | 2.21E-07 | 0.27139329 | 0.34 | 0.492 | 0.00427957 | 7 | Ralgapa1 | myCAF_3 |
| Meis12 | 2.37E-07 | 0.88338457 | 0.49 | 0.587 | 0.00458031 | 7 | Meis1 | myCAF_3 |
| Traf3ip22 | 2.37E-07 | 0.50678569 | 0.366 | 0.488 | 0.00458719 | 7 | Traf3ip2 | myCAF_3 |
| Ivd1 | 2.40E-07 | 0.8669576 | 0.232 | 0.398 | 0.00463749 | 7 | Ivd | myCAF_3 |
| Necap21 | 2.40E-07 | 0.41499049 | 0.5 | 0.668 | 0.00464192 | 7 | Necap2 | myCAF_3 |
| Fam76a | 2.45E-07 | 0.38276055 | 0.356 | 0.505 | 0.00473827 | 7 | Fam76a | myCAF_3 |
| Pitpnc12 | 2.49E-07 | 0.43877497 | 0.485 | 0.563 | 0.00480554 | 7 | Pitpnc1 | myCAF_3 |
| Rab341 | 2.50E-07 | 0.56632341 | 0.696 | 0.828 | 0.00483525 | 7 | Rab34 | myCAF_3 |
| Slc35c1 | 2.55E-07 | 0.34597783 | 0.237 | 0.403 | 0.00492772 | 7 | Slc35c1 | myCAF_3 |
| Znhit61 | 2.56E-07 | 0.48373798 | 0.165 | 0.303 | 0.00494396 | 7 | Znhit6 | myCAF_3 |
| Senp3 | 2.57E-07 | 0.58852413 | 0.407 | 0.568 | 0.00496572 | 7 | Senp3 | myCAF_3 |
| Gfm11 | 2.58E-07 | 0.84908492 | 0.34 | 0.468 | 0.00499629 | 7 | Gfm1 | myCAF_3 |
| Cebpg1 | 2.60E-07 | 1.1375422 | 0.536 | 0.638 | 0.00502767 | 7 | Cebpg | myCAF_3 |
| Cped12 | 2.62E-07 | 0.81943047 | 0.727 | 0.814 | 0.00507453 | 7 | Cped1 | myCAF_3 |
| Emsy1 | 2.68E-07 | 0.25007744 | 0.253 | 0.391 | 0.00518717 | 7 | Emsy | myCAF_3 |
| Gkap11 | 2.69E-07 | 0.33913967 | 0.263 | 0.403 | 0.00520475 | 7 | Gkap1 | myCAF_3 |
| Gab11 | 2.71E-07 | 0.45312329 | 0.263 | 0.406 | 0.00524747 | 7 | Gab1 | myCAF_3 |
| Cnot101 | 2.75E-07 | 0.64781073 | 0.309 | 0.456 | 0.00531478 | 7 | Cnot10 | myCAF_3 |
| Rbm341 | 2.78E-07 | 0.4533943 | 0.108 | 0.264 | 0.00536699 | 7 | Rbm34 | myCAF_3 |
| Klf31 | 2.80E-07 | 0.66855922 | 0.593 | 0.718 | 0.00541176 | 7 | Klf3 | myCAF_3 |
| Cnn21 | 2.80E-07 | 0.66138906 | 0.799 | 0.834 | 0.00541604 | 7 | Cnn2 | myCAF_3 |
| Akt31 | 2.82E-07 | 0.34467985 | 0.443 | 0.602 | 0.0054432 | 7 | Akt3 | myCAF_3 |
| Naa351 | 2.82E-07 | 0.35006864 | 0.191 | 0.354 | 0.00545892 | 7 | Naa35 | myCAF_3 |
| Zfp622 | 2.83E-07 | 0.51061283 | 0.253 | 0.404 | 0.00547896 | 7 | Zfp622 | myCAF_3 |
| Rpia3 | 2.84E-07 | 0.33976197 | 0.186 | 0.325 | 0.00549216 | 7 | Rpia | myCAF_3 |
| Brd11 | 2.87E-07 | 0.77870042 | 0.552 | 0.672 | 0.00554427 | 7 | Brd1 | myCAF_3 |
| Rc3h11 | 2.90E-07 | 0.59460654 | 0.495 | 0.657 | 0.00560469 | 7 | Rc3h1 | myCAF_3 |
| Rbck1 | 2.94E-07 | 0.44965674 | 0.433 | 0.568 | 0.00568023 | 7 | Rbck1 | myCAF_3 |
| Chm1 | 2.98E-07 | 0.46817765 | 0.278 | 0.434 | 0.00576684 | 7 | Chm | myCAF_3 |
| Samd11 | 3.05E-07 | 0.49505637 | 0.113 | 0.263 | 0.00589951 | 7 | Samd1 | myCAF_3 |
| Epm2aip12 | 3.07E-07 | 0.7746038 | 0.428 | 0.539 | 0.00593921 | 7 | Epm2aip1 | myCAF_3 |
| Zfp512b2 | 3.08E-07 | 0.4286324 | 0.387 | 0.534 | 0.00596074 | 7 | Zfp512b | myCAF_3 |
| Orai3 | 3.08E-07 | 0.54755717 | 0.361 | 0.532 | 0.00596194 | 7 | Orai3 | myCAF_3 |
| Irf2bpl1 | 3.09E-07 | 0.77741451 | 0.711 | 0.778 | 0.00597231 | 7 | Irf2bpl | myCAF_3 |
| Tbc1d13 | 3.09E-07 | 0.42129083 | 0.351 | 0.517 | 0.00597232 | 7 | Tbc1d13 | myCAF_3 |
| Gtf2e11 | 3.10E-07 | 0.5856121 | 0.144 | 0.298 | 0.00598475 | 7 | Gtf2e1 | myCAF_3 |
| Casc3 | 3.10E-07 | 0.71594171 | 0.448 | 0.592 | 0.00599702 | 7 | Casc3 | myCAF_3 |
| Dohh1 | 3.15E-07 | 0.35068791 | 0.284 | 0.448 | 0.00608298 | 7 | Dohh | myCAF_3 |
| Traf22 | 3.19E-07 | 0.71119025 | 0.314 | 0.467 | 0.00617531 | 7 | Traf2 | myCAF_3 |
| Cbx62 | 3.20E-07 | 0.39866819 | 0.407 | 0.536 | 0.00618309 | 7 | Cbx6 | myCAF_3 |
| Zfp2921 | 3.21E-07 | 0.82297841 | 0.387 | 0.542 | 0.00620953 | 7 | Zfp292 | myCAF_3 |
| Exosc11 | 3.27E-07 | 0.29545187 | 0.211 | 0.345 | 0.00631599 | 7 | Exosc1 | myCAF_3 |
| Ercc3 | 3.29E-07 | 0.80310815 | 0.366 | 0.525 | 0.0063543 | 7 | Ercc3 | myCAF_3 |
| Trim561 | 3.29E-07 | 0.69006506 | 0.485 | 0.611 | 0.0063593 | 7 | Trim56 | myCAF_3 |
| Klf16 | 3.31E-07 | 0.54576776 | 0.222 | 0.382 | 0.0063942 | 7 | Klf16 | myCAF_3 |
| Grem22 | 3.34E-07 | 0.53521445 | 0.737 | 0.818 | 0.00646303 | 7 | Grem2 | myCAF_3 |
| Ptpn91 | 3.46E-07 | 0.33348666 | 0.582 | 0.746 | 0.00669215 | 7 | Ptpn9 | myCAF_3 |
| Gusb1 | 3.51E-07 | 0.91908539 | 0.562 | 0.676 | 0.00678415 | 7 | Gusb | myCAF_3 |
| Diaph13 | 3.52E-07 | 0.49421476 | 0.299 | 0.457 | 0.00681521 | 7 | Diaph1 | myCAF_3 |
| Tm2d11 | 3.55E-07 | 0.36798239 | 0.201 | 0.365 | 0.00687304 | 7 | Tm2d1 | myCAF_3 |
| Ephb23 | 3.57E-07 | 0.36497003 | 0.541 | 0.616 | 0.00690439 | 7 | Ephb2 | myCAF_3 |
| Tada3 | 3.68E-07 | 0.51231703 | 0.16 | 0.334 | 0.00712091 | 7 | Tada3 | myCAF_3 |
| Dnlz1 | 3.73E-07 | 0.3427545 | 0.407 | 0.55 | 0.00721257 | 7 | Dnlz | myCAF_3 |
| Cnot4 | 3.78E-07 | 0.86240519 | 0.309 | 0.467 | 0.00730147 | 7 | Cnot4 | myCAF_3 |
| Cpt1a3 | 3.86E-07 | 0.38886452 | 0.443 | 0.541 | 0.00746587 | 7 | Cpt1a | myCAF_3 |
| Mrrf1 | 3.92E-07 | 0.409218 | 0.165 | 0.331 | 0.00758856 | 7 | Mrrf | myCAF_3 |
| Prkca2 | 3.93E-07 | 0.34428926 | 0.222 | 0.349 | 0.00759864 | 7 | Prkca | myCAF_3 |
| Ormdl31 | 3.95E-07 | 0.3496128 | 0.438 | 0.592 | 0.00763532 | 7 | Ormdl3 | myCAF_3 |
| Kdm3b1 | 3.97E-07 | 1.01395716 | 0.521 | 0.635 | 0.00768179 | 7 | Kdm3b | myCAF_3 |
| Sec61a2 | 4.05E-07 | 0.36039106 | 0.278 | 0.418 | 0.007826 | 7 | Sec61a2 | myCAF_3 |
| Qpct2 | 4.06E-07 | 1.2650197 | 0.515 | 0.36 | 0.00785871 | 7 | Qpct | myCAF_3 |
| Chordc1 | 4.07E-07 | 0.26199097 | 0.268 | 0.412 | 0.00787086 | 7 | Chordc1 | myCAF_3 |
| Ncoa73 | 4.12E-07 | 0.39782835 | 0.17 | 0.299 | 0.0079702 | 7 | Ncoa7 | myCAF_3 |
| Zfp326 | 4.22E-07 | 0.28251571 | 0.32 | 0.469 | 0.00815386 | 7 | Zfp326 | myCAF_3 |
| Mfge82 | 4.26E-07 | 0.8865519 | 0.851 | 0.949 | 0.00823585 | 7 | Mfge8 | myCAF_3 |
| Phf14 | 4.27E-07 | 0.50452943 | 0.371 | 0.523 | 0.00825725 | 7 | Phf14 | myCAF_3 |
| Morc31 | 4.27E-07 | 0.74910992 | 0.247 | 0.411 | 0.00825809 | 7 | Morc3 | myCAF_3 |
| Rbl2 | 4.43E-07 | 0.64124869 | 0.387 | 0.513 | 0.00855725 | 7 | Rbl2 | myCAF_3 |
| Atg4b | 4.44E-07 | 0.54428736 | 0.464 | 0.581 | 0.00857711 | 7 | Atg4b | myCAF_3 |
| Atg3 | 4.44E-07 | 0.37505704 | 0.387 | 0.544 | 0.00858113 | 7 | Atg3 | myCAF_3 |
| Mbnl11 | 4.45E-07 | 0.8448851 | 0.438 | 0.544 | 0.00859408 | 7 | Mbnl1 | myCAF_3 |
| Cchcr11 | 4.49E-07 | 0.58779098 | 0.144 | 0.288 | 0.0086854 | 7 | Cchcr1 | myCAF_3 |
| Tmem265 | 4.50E-07 | 0.51704413 | 0.216 | 0.372 | 0.00869201 | 7 | Tmem265 | myCAF_3 |
| Fntb2 | 4.53E-07 | 0.46244349 | 0.201 | 0.356 | 0.00875565 | 7 | Fntb | myCAF_3 |
| Acsl31 | 4.60E-07 | 0.36982178 | 0.201 | 0.336 | 0.0088883 | 7 | Acsl3 | myCAF_3 |
| Trim461 | 4.60E-07 | 0.44368933 | 0.175 | 0.317 | 0.00889422 | 7 | Trim46 | myCAF_3 |
| Frs21 | 4.64E-07 | 0.5839048 | 0.438 | 0.616 | 0.0089745 | 7 | Frs2 | myCAF_3 |
| Brox1 | 4.73E-07 | 0.31785036 | 0.253 | 0.438 | 0.00913786 | 7 | Brox | myCAF_3 |
| Akirin1 | 4.93E-07 | 0.31709268 | 0.242 | 0.385 | 0.00952632 | 7 | Akirin1 | myCAF_3 |
| Plekha51 | 5.10E-07 | 0.9727204 | 0.418 | 0.534 | 0.00986076 | 7 | Plekha5 | myCAF_3 |
| Jmy | 5.14E-07 | 0.77419527 | 0.572 | 0.678 | 0.00994709 | 7 | Jmy | myCAF_3 |
| Rpf11 | 5.19E-07 | 0.39327862 | 0.314 | 0.465 | 0.0100371 | 7 | Rpf1 | myCAF_3 |
| Ttc143 | 5.22E-07 | 0.72371198 | 0.381 | 0.505 | 0.01009909 | 7 | Ttc14 | myCAF_3 |
| Mcur11 | 5.23E-07 | 0.26780059 | 0.247 | 0.397 | 0.01010672 | 7 | Mcur1 | myCAF_3 |
| Mllt10 | 5.25E-07 | 0.52593348 | 0.273 | 0.42 | 0.010142 | 7 | Mllt10 | myCAF_3 |
| Kat72 | 5.25E-07 | 0.63394 | 0.387 | 0.512 | 0.01014215 | 7 | Kat7 | myCAF_3 |
| Tox41 | 5.27E-07 | 0.79918206 | 0.443 | 0.587 | 0.01018966 | 7 | Tox4 | myCAF_3 |
| Mkln11 | 5.48E-07 | 0.79601779 | 0.485 | 0.577 | 0.01059143 | 7 | Mkln1 | myCAF_3 |
| Rabgap11 | 5.71E-07 | 0.36684373 | 0.33 | 0.503 | 0.01103546 | 7 | Rabgap1 | myCAF_3 |
| Stx62 | 5.89E-07 | 0.49821721 | 0.598 | 0.709 | 0.01139711 | 7 | Stx6 | myCAF_3 |
| Dgkz1 | 5.93E-07 | 1.01254472 | 0.521 | 0.578 | 0.01147047 | 7 | Dgkz | myCAF_3 |
| Sh3rf31 | 5.94E-07 | 1.23339469 | 0.371 | 0.273 | 0.01148143 | 7 | Sh3rf3 | myCAF_3 |
| Crtc32 | 5.95E-07 | 0.93319013 | 0.397 | 0.54 | 0.01150244 | 7 | Crtc3 | myCAF_3 |
| Rpp25l1 | 5.96E-07 | 0.35718529 | 0.258 | 0.383 | 0.01152905 | 7 | Rpp25l | myCAF_3 |
| Ndrg23 | 6.05E-07 | 0.7104788 | 0.18 | 0.262 | 0.01170124 | 7 | Ndrg2 | myCAF_3 |
| Cstf21 | 6.21E-07 | 0.52488266 | 0.191 | 0.336 | 0.01201414 | 7 | Cstf2 | myCAF_3 |
| Dicer1 | 6.32E-07 | 0.8327766 | 0.371 | 0.479 | 0.0122204 | 7 | Dicer1 | myCAF_3 |
| Ncor1 | 6.37E-07 | 0.70683953 | 0.644 | 0.746 | 0.01231276 | 7 | Ncor1 | myCAF_3 |
| Klf101 | 6.39E-07 | 1.00183171 | 0.284 | 0.414 | 0.01234627 | 7 | Klf10 | myCAF_3 |
| Ifi303 | 6.47E-07 | 0.3950785 | 0.335 | 0.458 | 0.01250424 | 7 | Ifi30 | myCAF_3 |
| Usp21 | 6.47E-07 | 0.80402769 | 0.268 | 0.421 | 0.01251614 | 7 | Usp21 | myCAF_3 |
| Gtpbp21 | 6.80E-07 | 0.32538851 | 0.485 | 0.63 | 0.01314192 | 7 | Gtpbp2 | myCAF_3 |
| Apc | 6.80E-07 | 0.66521216 | 0.381 | 0.517 | 0.01314662 | 7 | Apc | myCAF_3 |
| Zw101 | 6.82E-07 | 0.55001726 | 0.211 | 0.344 | 0.01317753 | 7 | Zw10 | myCAF_3 |
| Parp43 | 6.83E-07 | 0.35566562 | 0.206 | 0.38 | 0.01319565 | 7 | Parp4 | myCAF_3 |
| Rnf1681 | 6.86E-07 | 0.49303871 | 0.227 | 0.361 | 0.01326412 | 7 | Rnf168 | myCAF_3 |
| Rgs191 | 6.89E-07 | 0.30070423 | 0.397 | 0.536 | 0.01331821 | 7 | Rgs19 | myCAF_3 |
| Bnip3l2 | 6.93E-07 | 0.29037881 | 0.299 | 0.415 | 0.0133892 | 7 | Bnip3l | myCAF_3 |
| Thap7 | 7.00E-07 | 0.52272856 | 0.361 | 0.509 | 0.01353802 | 7 | Thap7 | myCAF_3 |
| Hmbox1 | 7.01E-07 | 0.28568734 | 0.222 | 0.397 | 0.01355594 | 7 | Hmbox1 | myCAF_3 |
| Ints81 | 7.07E-07 | 0.68267959 | 0.227 | 0.379 | 0.01367106 | 7 | Ints8 | myCAF_3 |
| Ep3001 | 7.09E-07 | 1.04384081 | 0.495 | 0.626 | 0.01371482 | 7 | Ep300 | myCAF_3 |
| Tnrc6b1 | 7.19E-07 | 0.8728874 | 0.448 | 0.581 | 0.01390597 | 7 | Tnrc6b | myCAF_3 |
| Nudt72 | 7.22E-07 | 0.56085569 | 0.124 | 0.264 | 0.0139655 | 7 | Nudt7 | myCAF_3 |
| Olfml2b2 | 7.28E-07 | 0.3224932 | 0.536 | 0.699 | 0.01407809 | 7 | Olfml2b | myCAF_3 |
| Anapc7 | 7.29E-07 | 0.39194213 | 0.438 | 0.564 | 0.01409251 | 7 | Anapc7 | myCAF_3 |
| Zfp865 | 7.31E-07 | 0.7456387 | 0.361 | 0.498 | 0.01412462 | 7 | Zfp865 | myCAF_3 |
| Agap3 | 7.45E-07 | 0.77944777 | 0.655 | 0.739 | 0.01439693 | 7 | Agap3 | myCAF_3 |
| Lrrc421 | 7.45E-07 | 0.50427798 | 0.258 | 0.431 | 0.0143999 | 7 | Lrrc42 | myCAF_3 |
| Adam333 | 7.51E-07 | 1.14953336 | 0.351 | 0.206 | 0.01451892 | 7 | Adam33 | myCAF_3 |
| Nosip1 | 7.72E-07 | 0.73122411 | 0.407 | 0.545 | 0.01492257 | 7 | Nosip | myCAF_3 |
| Cpne1 | 7.98E-07 | 0.65784418 | 0.593 | 0.73 | 0.01542923 | 7 | Cpne1 | myCAF_3 |
| Zbtb112 | 8.01E-07 | 0.82979678 | 0.304 | 0.432 | 0.01547871 | 7 | Zbtb11 | myCAF_3 |
| Dus3l | 8.04E-07 | 0.63163598 | 0.366 | 0.541 | 0.01555018 | 7 | Dus3l | myCAF_3 |
| Slx41 | 8.10E-07 | 0.25990152 | 0.134 | 0.272 | 0.01566704 | 7 | Slx4 | myCAF_3 |
| Zfp4101 | 8.12E-07 | 0.66145867 | 0.34 | 0.493 | 0.01569602 | 7 | Zfp410 | myCAF_3 |
| Atp6v1d1 | 8.24E-07 | 0.27496873 | 0.464 | 0.61 | 0.01593894 | 7 | Atp6v1d | myCAF_3 |
| Chrac11 | 8.27E-07 | 0.60203186 | 0.325 | 0.477 | 0.01598825 | 7 | Chrac1 | myCAF_3 |
| Fbxw2 | 8.32E-07 | 0.38755948 | 0.438 | 0.588 | 0.0160899 | 7 | Fbxw2 | myCAF_3 |
| Rbfa | 8.39E-07 | 0.25017352 | 0.402 | 0.516 | 0.01622436 | 7 | Rbfa | myCAF_3 |
| Akap81 | 8.56E-07 | 1.20179303 | 0.428 | 0.568 | 0.01655414 | 7 | Akap8 | myCAF_3 |
| Ctnnbl1 | 8.59E-07 | 0.77019395 | 0.222 | 0.404 | 0.01660357 | 7 | Ctnnbl1 | myCAF_3 |
| Senp21 | 8.66E-07 | 0.77986463 | 0.258 | 0.423 | 0.01674471 | 7 | Senp2 | myCAF_3 |
| Pias12 | 8.66E-07 | 0.46376509 | 0.242 | 0.374 | 0.01674662 | 7 | Pias1 | myCAF_3 |
| Sgcb1 | 8.70E-07 | 0.27367295 | 0.412 | 0.557 | 0.0168187 | 7 | Sgcb | myCAF_3 |
| Acadvl1 | 8.71E-07 | 0.64399631 | 0.433 | 0.596 | 0.01683767 | 7 | Acadvl | myCAF_3 |
| Snx292 | 8.74E-07 | 0.7024009 | 0.351 | 0.474 | 0.0168942 | 7 | Snx29 | myCAF_3 |
| AI8371811 | 8.88E-07 | 0.44414174 | 0.521 | 0.66 | 0.01717701 | 7 | AI837181 | myCAF_3 |
| Bmt21 | 9.03E-07 | 0.83289434 | 0.356 | 0.477 | 0.01745401 | 7 | Bmt2 | myCAF_3 |
| Zdhhc4 | 9.08E-07 | 0.44466446 | 0.381 | 0.516 | 0.01755824 | 7 | Zdhhc4 | myCAF_3 |
| Ehmt11 | 9.18E-07 | 0.50752314 | 0.438 | 0.611 | 0.01775649 | 7 | Ehmt1 | myCAF_3 |
| Utp31 | 9.20E-07 | 0.31507336 | 0.335 | 0.472 | 0.01779564 | 7 | Utp3 | myCAF_3 |
| Map2k71 | 9.23E-07 | 0.56802543 | 0.469 | 0.605 | 0.01784059 | 7 | Map2k7 | myCAF_3 |
| Zbed31 | 9.25E-07 | 0.49314278 | 0.351 | 0.481 | 0.01788442 | 7 | Zbed3 | myCAF_3 |
| Wasl2 | 9.31E-07 | 0.50956701 | 0.521 | 0.664 | 0.01800241 | 7 | Wasl | myCAF_3 |
| Fbxo38 | 9.41E-07 | 0.4355952 | 0.438 | 0.565 | 0.01819379 | 7 | Fbxo38 | myCAF_3 |
| Uck1 | 9.42E-07 | 0.34832173 | 0.273 | 0.439 | 0.01820864 | 7 | Uck1 | myCAF_3 |
| a2 | 9.46E-07 | 1.87076541 | 0.469 | 0.31 | 0.01828828 | 7 | a | myCAF_3 |
| Abi21 | 9.47E-07 | 0.39707589 | 0.314 | 0.438 | 0.01831449 | 7 | Abi2 | myCAF_3 |
| Nepro1 | 9.54E-07 | 0.62292798 | 0.165 | 0.304 | 0.018438 | 7 | Nepro | myCAF_3 |
| Pim32 | 9.56E-07 | 0.70464431 | 0.345 | 0.42 | 0.01849097 | 7 | Pim3 | myCAF_3 |
| Tmem184c | 9.66E-07 | 0.47938441 | 0.242 | 0.401 | 0.0186739 | 7 | Tmem184c | myCAF_3 |
| Ddx271 | 9.99E-07 | 0.61308234 | 0.34 | 0.463 | 0.01930589 | 7 | Ddx27 | myCAF_3 |
| Dzip13 | 1.02E-06 | 0.36277646 | 0.206 | 0.354 | 0.0197236 | 7 | Dzip1 | myCAF_3 |
| Chmp7 | 1.03E-06 | 0.43251652 | 0.376 | 0.52 | 0.0199716 | 7 | Chmp7 | myCAF_3 |
| Slmap1 | 1.04E-06 | 0.66932091 | 0.469 | 0.589 | 0.02009709 | 7 | Slmap | myCAF_3 |
| Med62 | 1.05E-06 | 0.61369028 | 0.247 | 0.373 | 0.02031268 | 7 | Med6 | myCAF_3 |
| Strip11 | 1.06E-06 | 0.54190732 | 0.216 | 0.372 | 0.02043061 | 7 | Strip1 | myCAF_3 |
| Iffo21 | 1.06E-06 | 0.4157261 | 0.381 | 0.52 | 0.02045351 | 7 | Iffo2 | myCAF_3 |
| Trappc121 | 1.06E-06 | 0.53191366 | 0.273 | 0.444 | 0.02058054 | 7 | Trappc12 | myCAF_3 |
| Plpp22 | 1.07E-06 | 0.55945249 | 0.139 | 0.29 | 0.0206819 | 7 | Plpp2 | myCAF_3 |
| Gabarapl2 | 1.07E-06 | 0.6656099 | 0.552 | 0.679 | 0.02071675 | 7 | Gabarapl2 | myCAF_3 |
| Mpp12 | 1.09E-06 | 0.28053906 | 0.273 | 0.394 | 0.0211198 | 7 | Mpp1 | myCAF_3 |
| Sun22 | 1.11E-06 | 1.69158174 | 0.66 | 0.507 | 0.02154458 | 7 | Sun2 | myCAF_3 |
| Zbtb442 | 1.14E-06 | 0.51552685 | 0.371 | 0.513 | 0.02196428 | 7 | Zbtb44 | myCAF_3 |
| Etv51 | 1.16E-06 | 0.48513547 | 0.495 | 0.614 | 0.02240856 | 7 | Etv5 | myCAF_3 |
| Baz1a1 | 1.18E-06 | 0.65908755 | 0.335 | 0.475 | 0.02279777 | 7 | Baz1a | myCAF_3 |
| DEPRECATED-I | 1.19E-06 | 0.50509818 | 0.361 | 0.49 | 0.02294654 | 7 | DEPRECATED-I | myCAF_3 |
| Epg5 | 1.21E-06 | 0.33961607 | 0.232 | 0.365 | 0.02333965 | 7 | Epg5 | myCAF_3 |
| Wtip1 | 1.21E-06 | 0.43195263 | 0.582 | 0.734 | 0.02336086 | 7 | Wtip | myCAF_3 |
| Tmed52 | 1.22E-06 | 0.48882742 | 0.351 | 0.486 | 0.02357781 | 7 | Tmed5 | myCAF_3 |
| Cdyl1 | 1.22E-06 | 0.50969247 | 0.407 | 0.534 | 0.0236702 | 7 | Cdyl | myCAF_3 |
| Armc102 | 1.24E-06 | 0.286928 | 0.098 | 0.252 | 0.02391768 | 7 | Armc10 | myCAF_3 |
| Tbc1d22a | 1.24E-06 | 0.50014479 | 0.273 | 0.411 | 0.02400508 | 7 | Tbc1d22a | myCAF_3 |
| Uhmk11 | 1.24E-06 | 0.38669908 | 0.345 | 0.481 | 0.02403216 | 7 | Uhmk1 | myCAF_3 |
| D16Ertd472e | 1.25E-06 | 0.94735685 | 0.227 | 0.344 | 0.0241438 | 7 | D16Ertd472e | myCAF_3 |
| Dcaf6 | 1.25E-06 | 0.35725569 | 0.387 | 0.542 | 0.02419695 | 7 | Dcaf6 | myCAF_3 |
| Top3b | 1.26E-06 | 0.36616443 | 0.284 | 0.453 | 0.02426694 | 7 | Top3b | myCAF_3 |
| Rnf1461 | 1.26E-06 | 0.99921514 | 0.407 | 0.517 | 0.02429107 | 7 | Rnf146 | myCAF_3 |
| Mterf31 | 1.26E-06 | 0.38155292 | 0.232 | 0.383 | 0.02434927 | 7 | Mterf3 | myCAF_3 |
| Ttc172 | 1.27E-06 | 0.50858756 | 0.284 | 0.466 | 0.02453856 | 7 | Ttc17 | myCAF_3 |
| Mbtd11 | 1.28E-06 | 0.30817113 | 0.289 | 0.439 | 0.02472327 | 7 | Mbtd1 | myCAF_3 |
| Mre11a1 | 1.29E-06 | 0.70738579 | 0.134 | 0.276 | 0.0249946 | 7 | Mre11a | myCAF_3 |
| Dnajc21 | 1.29E-06 | 0.63234431 | 0.361 | 0.47 | 0.0250005 | 7 | Dnajc21 | myCAF_3 |
| Mark21 | 1.30E-06 | 0.65642228 | 0.351 | 0.483 | 0.02506465 | 7 | Mark2 | myCAF_3 |
| Naa30 | 1.31E-06 | 0.79607288 | 0.351 | 0.487 | 0.0253594 | 7 | Naa30 | myCAF_3 |
| Zfp6511 | 1.31E-06 | 0.45305658 | 0.464 | 0.605 | 0.0253765 | 7 | Zfp651 | myCAF_3 |
| Tia11 | 1.33E-06 | 0.50261719 | 0.309 | 0.445 | 0.02577119 | 7 | Tia1 | myCAF_3 |
| Dnttip21 | 1.34E-06 | 0.3577412 | 0.356 | 0.502 | 0.02588668 | 7 | Dnttip2 | myCAF_3 |
| Retsat1 | 1.34E-06 | 0.38686811 | 0.113 | 0.264 | 0.0259067 | 7 | Retsat | myCAF_3 |
| Eif4e32 | 1.35E-06 | 0.50191659 | 0.387 | 0.562 | 0.02610028 | 7 | Eif4e3 | myCAF_3 |
| Atp6v1f2 | 1.35E-06 | 0.43874898 | 0.567 | 0.711 | 0.02610555 | 7 | Atp6v1f | myCAF_3 |
| Elmsan1 | 1.35E-06 | 0.76110491 | 0.304 | 0.451 | 0.02612363 | 7 | Elmsan1 | myCAF_3 |
| Topors1 | 1.37E-06 | 0.74862372 | 0.335 | 0.473 | 0.02645124 | 7 | Topors | myCAF_3 |
| Pttg12 | 1.38E-06 | 1.04146464 | 0.284 | 0.445 | 0.02661101 | 7 | Pttg1 | myCAF_3 |
| Ddx54 | 1.38E-06 | 0.90790209 | 0.469 | 0.596 | 0.02666378 | 7 | Ddx54 | myCAF_3 |
| Coq9 | 1.42E-06 | 0.2811469 | 0.216 | 0.383 | 0.02751421 | 7 | Coq9 | myCAF_3 |
| Mlst81 | 1.43E-06 | 0.55977597 | 0.119 | 0.276 | 0.02764388 | 7 | Mlst8 | myCAF_3 |

| | | | | | | | | |
|---|---|---|---|---|---|---|---|---|
| Ankle2 | 1.45E-06 | 0.36572981 | 0.314 | 0.439 | 0.02794862 | 7 | Ankle2 | myCAF_3 |
| Oser11 | 1.45E-06 | 0.81302641 | 0.418 | 0.544 | 0.02795552 | 7 | Oser1 | myCAF_3 |
| Ric8b2 | 1.45E-06 | 0.70530483 | 0.345 | 0.496 | 0.02800668 | 7 | Ric8b | myCAF_3 |
| Mpnd1 | 1.49E-06 | 0.59736177 | 0.51 | 0.618 | 0.0288526 | 7 | Mpnd | myCAF_3 |
| Rad54l21 | 1.50E-06 | 0.50525681 | 0.371 | 0.499 | 0.02898212 | 7 | Rad54l2 | myCAF_3 |
| Cdc401 | 1.51E-06 | 0.980622 | 0.206 | 0.367 | 0.02909809 | 7 | Cdc40 | myCAF_3 |
| Rcc1l1 | 1.53E-06 | 0.31978053 | 0.232 | 0.359 | 0.02964569 | 7 | Rcc1l | myCAF_3 |
| Tank1 | 1.55E-06 | 0.64710833 | 0.273 | 0.446 | 0.03000644 | 7 | Tank | myCAF_3 |
| Bahcc14 | 1.56E-06 | 0.46567763 | 0.201 | 0.325 | 0.03008729 | 7 | Bahcc1 | myCAF_3 |
| Cpt1c2 | 1.56E-06 | 0.3097186 | 0.356 | 0.502 | 0.03022555 | 7 | Cpt1c | myCAF_3 |
| Zbtb12 | 1.58E-06 | 0.55635241 | 0.51 | 0.617 | 0.03051625 | 7 | Zbtb1 | myCAF_3 |
| Rreb11 | 1.64E-06 | 0.51492253 | 0.33 | 0.494 | 0.03169741 | 7 | Rreb1 | myCAF_3 |
| Sars1 | 1.65E-06 | 0.31972573 | 0.459 | 0.624 | 0.03199353 | 7 | Sars | myCAF_3 |
| Samd81 | 1.66E-06 | 0.41515627 | 0.263 | 0.415 | 0.03211101 | 7 | Samd8 | myCAF_3 |
| Mecp22 | 1.67E-06 | 0.40495184 | 0.356 | 0.502 | 0.03225417 | 7 | Mecp2 | myCAF_3 |
| Zcchc171 | 1.70E-06 | 0.58087864 | 0.258 | 0.396 | 0.03279539 | 7 | Zcchc17 | myCAF_3 |
| Cmklr12 | 1.70E-06 | 0.68351058 | 0.227 | 0.263 | 0.03293077 | 7 | Cmklr1 | myCAF_3 |
| Isg20l2 | 1.73E-06 | 0.70742801 | 0.268 | 0.411 | 0.03335565 | 7 | Isg20l2 | myCAF_3 |
| Zswim8 | 1.74E-06 | 0.65274822 | 0.696 | 0.797 | 0.03368329 | 7 | Zswim8 | myCAF_3 |
| Tomm34 | 1.75E-06 | 0.40735734 | 0.443 | 0.558 | 0.03386296 | 7 | Tomm34 | myCAF_3 |
| Gabarapl11 | 1.76E-06 | 0.79798069 | 0.737 | 0.817 | 0.03411245 | 7 | Gabarapl1 | myCAF_3 |
| Taf5l | 1.80E-06 | 0.30917558 | 0.284 | 0.411 | 0.03471314 | 7 | Taf5l | myCAF_3 |
| Rnf103 | 1.80E-06 | 0.61501688 | 0.469 | 0.572 | 0.03476062 | 7 | Rnf103 | myCAF_3 |
| Pex61 | 1.80E-06 | 0.52119855 | 0.325 | 0.488 | 0.03478041 | 7 | Pex6 | myCAF_3 |
| Gosr1 | 1.85E-06 | 0.79364963 | 0.392 | 0.535 | 0.03577473 | 7 | Gosr1 | myCAF_3 |
| Tpgs11 | 1.85E-06 | 0.43078465 | 0.464 | 0.613 | 0.03579116 | 7 | Tpgs1 | myCAF_3 |
| Azi21 | 1.86E-06 | 0.37231442 | 0.397 | 0.547 | 0.03589837 | 7 | Azi2 | myCAF_3 |
| Pgs1 | 1.87E-06 | 0.69505758 | 0.624 | 0.723 | 0.03612229 | 7 | Pgs1 | myCAF_3 |
| Rbm42 | 1.88E-06 | 0.89717428 | 0.505 | 0.646 | 0.03626592 | 7 | Rbm4 | myCAF_3 |
| Numb2 | 1.92E-06 | 0.53941229 | 0.34 | 0.498 | 0.03706243 | 7 | Numb | myCAF_3 |
| Ago3 | 1.92E-06 | 0.49903312 | 0.258 | 0.418 | 0.03719725 | 7 | Ago3 | myCAF_3 |
| Ssh11 | 1.95E-06 | 0.71821336 | 0.536 | 0.663 | 0.03773627 | 7 | Ssh1 | myCAF_3 |
| Lrpprc | 1.96E-06 | 0.42647788 | 0.186 | 0.328 | 0.037867 | 7 | Lrpprc | myCAF_3 |
| Trim161 | 1.98E-06 | 2.13771031 | 0.314 | 0.187 | 0.0382146 | 7 | Trim16 | myCAF_3 |
| Vamp4 | 2.01E-06 | 0.9255899 | 0.32 | 0.478 | 0.03887552 | 7 | Vamp4 | myCAF_3 |
| Nox43 | 2.06E-06 | 1.11116858 | 0.412 | 0.375 | 0.03981489 | 7 | Nox4 | myCAF_3 |
| Ptdss21 | 2.06E-06 | 0.5838411 | 0.258 | 0.413 | 0.03986608 | 7 | Ptdss2 | myCAF_3 |
| Frmd8 | 2.07E-06 | 0.5134652 | 0.448 | 0.591 | 0.04001773 | 7 | Frmd8 | myCAF_3 |
| Usp35 | 2.07E-06 | 0.57293191 | 0.253 | 0.417 | 0.04005669 | 7 | Usp3 | myCAF_3 |
| Mdm41 | 2.08E-06 | 0.6799572 | 0.397 | 0.526 | 0.04020524 | 7 | Mdm4 | myCAF_3 |
| Ppif1 | 2.10E-06 | 1.06881219 | 0.113 | 0.256 | 0.04062497 | 7 | Ppif | myCAF_3 |
| Ptpn11 | 2.12E-06 | 0.66973728 | 0.588 | 0.691 | 0.04091035 | 7 | Ptpn1 | myCAF_3 |
| Sumf22 | 2.13E-06 | 0.42748199 | 0.268 | 0.415 | 0.04109054 | 7 | Sumf2 | myCAF_3 |
| Adnp21 | 2.14E-06 | 0.51165997 | 0.345 | 0.493 | 0.04142528 | 7 | Adnp2 | myCAF_3 |
| Trmt111 | 2.22E-06 | 0.57395401 | 0.103 | 0.262 | 0.04287076 | 7 | Trmt11 | myCAF_3 |
| Ly963 | 2.24E-06 | 0.53379788 | 0.098 | 0.251 | 0.04321223 | 7 | Ly96 | myCAF_3 |
| Rint11 | 2.29E-06 | 0.25999285 | 0.237 | 0.384 | 0.0443441 | 7 | Rint1 | myCAF_3 |
| Sptlc1 | 2.30E-06 | 0.32477243 | 0.289 | 0.431 | 0.04456303 | 7 | Sptlc1 | myCAF_3 |
| Alas12 | 2.32E-06 | 0.34990134 | 0.325 | 0.446 | 0.04486644 | 7 | Alas1 | myCAF_3 |
| Dxo1 | 2.32E-06 | 0.45152 | 0.273 | 0.405 | 0.04489625 | 7 | Dxo | myCAF_3 |
| Ppig1 | 2.34E-06 | 0.41064721 | 0.376 | 0.526 | 0.04533358 | 7 | Ppig | myCAF_3 |
| Cstf11 | 2.36E-06 | 0.32422022 | 0.211 | 0.344 | 0.04558796 | 7 | Cstf1 | myCAF_3 |
| Noc4l1 | 2.36E-06 | 0.70676011 | 0.165 | 0.297 | 0.04570236 | 7 | Noc4l | myCAF_3 |
| Cpsf41 | 2.38E-06 | 0.40027341 | 0.232 | 0.376 | 0.04596193 | 7 | Cpsf4 | myCAF_3 |
| Dcun1d1 | 2.38E-06 | 0.42774604 | 0.211 | 0.364 | 0.04598137 | 7 | Dcun1d1 | myCAF_3 |
| Chmp61 | 2.41E-06 | 0.39506808 | 0.366 | 0.485 | 0.04653186 | 7 | Chmp6 | myCAF_3 |
| Pcnt1 | 2.43E-06 | 0.7799971 | 0.196 | 0.343 | 0.04697074 | 7 | Pcnt | myCAF_3 |
| Nadk22 | 2.47E-06 | 0.28203153 | 0.144 | 0.271 | 0.04769223 | 7 | Nadk2 | myCAF_3 |
| Gtf2h41 | 2.49E-06 | 0.76291944 | 0.232 | 0.371 | 0.04819538 | 7 | Gtf2h4 | myCAF_3 |
| Ubash3b2 | 2.50E-06 | 0.36357423 | 0.17 | 0.288 | 0.04836009 | 7 | Ubash3b | myCAF_3 |
| Rgp1 | 2.53E-06 | 0.40584447 | 0.33 | 0.466 | 0.0488584 | 7 | Rgp1 | myCAF_3 |
| Foxj2 | 2.54E-06 | 0.26848817 | 0.263 | 0.432 | 0.04913453 | 7 | Foxj2 | myCAF_3 |
| Letmd11 | 2.56E-06 | 0.25069291 | 0.242 | 0.373 | 0.04944414 | 7 | Letmd1 | myCAF_3 |
| Ap3s2 | 2.59E-06 | 0.25715914 | 0.139 | 0.277 | 0.05000586 | 7 | Ap3s2 | myCAF_3 |
| Nynrin1 | 2.59E-06 | 0.56526 | 0.572 | 0.677 | 0.05010599 | 7 | Nynrin | myCAF_3 |
| Tjp12 | 2.60E-06 | 0.81105715 | 0.541 | 0.621 | 0.05034617 | 7 | Tjp1 | myCAF_3 |
| Ccdc97 | 2.60E-06 | 0.99610536 | 0.423 | 0.532 | 0.05036314 | 7 | Ccdc97 | myCAF_3 |
| Cyth22 | 2.65E-06 | 0.68903267 | 0.613 | 0.733 | 0.0513227 | 7 | Cyth2 | myCAF_3 |
| Paf1 | 2.66E-06 | 0.3971793 | 0.335 | 0.461 | 0.0514065 | 7 | Paf1 | myCAF_3 |
| Pigm1 | 2.66E-06 | 0.54616092 | 0.304 | 0.439 | 0.05141043 | 7 | Pigm | myCAF_3 |
| Dkk32 | 2.69E-06 | 0.91077488 | 0.778 | 0.836 | 0.05193922 | 7 | Dkk3 | myCAF_3 |
| Ubp1 | 2.71E-06 | 0.81340539 | 0.649 | 0.743 | 0.05235924 | 7 | Ubp1 | myCAF_3 |
| Klhl71 | 2.74E-06 | 0.38168232 | 0.32 | 0.451 | 0.05293236 | 7 | Klhl7 | myCAF_3 |
| Actg21 | 2.74E-06 | 1.30026582 | 0.392 | 0.131 | 0.05301359 | 7 | Actg2 | myCAF_3 |
| Washc1 | 2.77E-06 | 0.38999762 | 0.459 | 0.594 | 0.05357785 | 7 | Washc1 | myCAF_3 |
| Parp21 | 2.77E-06 | 0.33489171 | 0.222 | 0.35 | 0.05359262 | 7 | Parp2 | myCAF_3 |
| Anks31 | 2.80E-06 | 0.27176227 | 0.34 | 0.476 | 0.05419032 | 7 | Anks3 | myCAF_3 |
| Gstp1 | 2.82E-06 | 0.63804106 | 0.68 | 0.805 | 0.05456551 | 7 | Gstp1 | myCAF_3 |
| Trappc101 | 2.88E-06 | 0.53437298 | 0.304 | 0.411 | 0.05566 | 7 | Trappc10 | myCAF_3 |
| Retreg2 | 2.91E-06 | 0.72584444 | 0.613 | 0.712 | 0.05616819 | 7 | Retreg2 | myCAF_3 |
| Tdp11 | 2.93E-06 | 0.45804712 | 0.165 | 0.289 | 0.0565999 | 7 | Tdp1 | myCAF_3 |
| Trmt12 | 2.94E-06 | 0.48693581 | 0.216 | 0.348 | 0.05689126 | 7 | Trmt1 | myCAF_3 |
| Ebag91 | 2.98E-06 | 0.34222135 | 0.18 | 0.322 | 0.05754042 | 7 | Ebag9 | myCAF_3 |
| Ints121 | 3.00E-06 | 0.28821727 | 0.16 | 0.303 | 0.05800536 | 7 | Ints12 | myCAF_3 |
| Zfp687 | 3.05E-06 | 0.85094714 | 0.242 | 0.375 | 0.05905409 | 7 | Zfp687 | myCAF_3 |
| Rnf220 | 3.06E-06 | 0.59013445 | 0.577 | 0.722 | 0.05924116 | 7 | Rnf220 | myCAF_3 |
| Rnaseh2c1 | 3.09E-06 | 0.69299399 | 0.495 | 0.558 | 0.05975035 | 7 | Rnaseh2c | myCAF_3 |
| Kat141 | 3.16E-06 | 0.6311303 | 0.345 | 0.487 | 0.06108125 | 7 | Kat14 | myCAF_3 |
| Wdr741 | 3.18E-06 | 1.03417968 | 0.299 | 0.426 | 0.06140042 | 7 | Wdr74 | myCAF_3 |
| Exosc72 | 3.18E-06 | 0.33126754 | 0.201 | 0.316 | 0.06145066 | 7 | Exosc7 | myCAF_3 |
| Atp7a2 | 3.22E-06 | 0.40454159 | 0.345 | 0.502 | 0.06228855 | 7 | Atp7a | myCAF_3 |
| Mypop | 3.27E-06 | 0.3688902 | 0.284 | 0.432 | 0.06312856 | 7 | Mypop | myCAF_3 |
| Tet32 | 3.27E-06 | 0.63410367 | 0.418 | 0.529 | 0.06319979 | 7 | Tet3 | myCAF_3 |
| Fam13b | 3.30E-06 | 0.76642051 | 0.289 | 0.422 | 0.06379515 | 7 | Fam13b | myCAF_3 |
| Zfand6 | 3.33E-06 | 0.5338609 | 0.371 | 0.493 | 0.06435298 | 7 | Zfand6 | myCAF_3 |
| Frg11 | 3.37E-06 | 0.71618649 | 0.356 | 0.467 | 0.06510243 | 7 | Frg1 | myCAF_3 |
| Tbx152 | 3.37E-06 | 0.85500045 | 0.521 | 0.683 | 0.06515784 | 7 | Tbx15 | myCAF_3 |
| Gtf2h11 | 3.42E-06 | 0.37978903 | 0.232 | 0.376 | 0.06620957 | 7 | Gtf2h1 | myCAF_3 |
| Med27 | 3.42E-06 | 0.33173436 | 0.253 | 0.376 | 0.06620966 | 7 | Med27 | myCAF_3 |
| Nudcd31 | 3.47E-06 | 0.75930194 | 0.588 | 0.694 | 0.06715204 | 7 | Nudcd3 | myCAF_3 |
| Rexlbd1 | 3.51E-06 | 0.42626479 | 0.418 | 0.549 | 0.06784092 | 7 | Rexlbd | myCAF_3 |
| Shf1 | 3.52E-06 | 0.4602263 | 0.155 | 0.285 | 0.06802832 | 7 | Shf | myCAF_3 |
| Pigu1 | 3.54E-06 | 0.42924517 | 0.309 | 0.439 | 0.06852743 | 7 | Pigu | myCAF_3 |
| Pym11 | 3.55E-06 | 0.82537487 | 0.17 | 0.311 | 0.06863391 | 7 | Pym1 | myCAF_3 |
| Arhgap422 | 3.56E-06 | 0.46030533 | 0.644 | 0.779 | 0.06888462 | 7 | Arhgap42 | myCAF_3 |
| Tut42 | 3.57E-06 | 0.25349183 | 0.211 | 0.395 | 0.06896559 | 7 | Tut4 | myCAF_3 |
| Zdhhc171 | 3.64E-06 | 0.25653169 | 0.242 | 0.376 | 0.07028463 | 7 | Zdhhc17 | myCAF_3 |
| Nt5c2 | 3.71E-06 | 0.78359533 | 0.392 | 0.527 | 0.07170527 | 7 | Nt5c2 | myCAF_3 |
| Vipas391 | 3.71E-06 | 0.48615843 | 0.294 | 0.438 | 0.07175052 | 7 | Vipas39 | myCAF_3 |
| Nck22 | 3.78E-06 | 0.40025765 | 0.294 | 0.436 | 0.07311153 | 7 | Nck2 | myCAF_3 |
| Timm101 | 3.81E-06 | 0.40264438 | 0.33 | 0.417 | 0.07363505 | 7 | Timm10 | myCAF_3 |
| Nsun4 | 3.83E-06 | 0.25630998 | 0.371 | 0.505 | 0.07403986 | 7 | Nsun4 | myCAF_3 |
| Mif4gd2 | 3.86E-06 | 0.36854775 | 0.294 | 0.422 | 0.07457597 | 7 | Mif4gd | myCAF_3 |
| Prpf33 | 3.87E-06 | 0.3397166 | 0.206 | 0.358 | 0.0747533 | 7 | Prpf3 | myCAF_3 |
| Heatr1 | 3.87E-06 | 0.40794003 | 0.175 | 0.3 | 0.07479446 | 7 | Heatr1 | myCAF_3 |
| Tfpt | 3.89E-06 | 0.65203968 | 0.407 | 0.541 | 0.07522611 | 7 | Tfpt | myCAF_3 |
| Pcdhga51 | 3.91E-06 | 0.8151489 | 0.201 | 0.291 | 0.0755473 | 7 | Pcdhga5 | myCAF_3 |
| Ccdc861 | 3.91E-06 | 0.98661061 | 0.227 | 0.355 | 0.07554752 | 7 | Ccdc86 | myCAF_3 |
| Smndc11 | 3.93E-06 | 0.39827242 | 0.371 | 0.481 | 0.0759318 | 7 | Smndc1 | myCAF_3 |
| Arhgap171 | 3.95E-06 | 0.55675092 | 0.345 | 0.472 | 0.07637631 | 7 | Arhgap17 | myCAF_3 |
| Zfp518b1 | 3.98E-06 | 0.29946803 | 0.186 | 0.324 | 0.07701142 | 7 | Zfp518b | myCAF_3 |
| Nr2c2 | 3.99E-06 | 0.52738459 | 0.376 | 0.516 | 0.07720728 | 7 | Nr2c2 | myCAF_3 |
| Foxo4 | 4.02E-06 | 0.60697197 | 0.345 | 0.495 | 0.07777177 | 7 | Foxo4 | myCAF_3 |
| Mga | 4.09E-06 | 0.51874334 | 0.32 | 0.478 | 0.07907158 | 7 | Mga | myCAF_3 |
| Irf31 | 4.13E-06 | 0.46458938 | 0.366 | 0.491 | 0.07985468 | 7 | Irf3 | myCAF_3 |
| Gtf3c1 | 4.24E-06 | 0.66352645 | 0.345 | 0.468 | 0.08196477 | 7 | Gtf3c1 | myCAF_3 |
| Paxbp11 | 4.24E-06 | 1.01946966 | 0.366 | 0.502 | 0.08198979 | 7 | Paxbp1 | myCAF_3 |
| Rnf215 | 4.28E-06 | 0.28014356 | 0.387 | 0.54 | 0.08273362 | 7 | Rnf215 | myCAF_3 |
| Pja1 | 4.33E-06 | 0.82314896 | 0.495 | 0.615 | 0.0837096 | 7 | Pja1 | myCAF_3 |
| Slc25a44 | 4.38E-06 | 0.3264056 | 0.211 | 0.344 | 0.08471334 | 7 | Slc25a44 | myCAF_3 |
| Htatip22 | 4.40E-06 | 0.59064911 | 0.351 | 0.469 | 0.0850825 | 7 | Htatip2 | myCAF_3 |
| Rnf2172 | 4.43E-06 | 0.67845159 | 0.32 | 0.439 | 0.08556865 | 7 | Rnf217 | myCAF_3 |
| D5Ertd579e | 4.43E-06 | 0.45361897 | 0.423 | 0.551 | 0.08557629 | 7 | D5Ertd579e | myCAF_3 |
| Thoc51 | 4.46E-06 | 0.49366214 | 0.201 | 0.331 | 0.08621953 | 7 | Thoc5 | myCAF_3 |
| Ncoa31 | 4.46E-06 | 0.93044724 | 0.459 | 0.564 | 0.08626787 | 7 | Ncoa3 | myCAF_3 |
| Tatdn2 | 4.61E-06 | 0.68273356 | 0.526 | 0.622 | 0.08912138 | 7 | Tatdn2 | myCAF_3 |
| Tead31 | 4.64E-06 | 0.68331975 | 0.557 | 0.672 | 0.08978651 | 7 | Tead3 | myCAF_3 |
| Bcorl1 | 4.68E-06 | 0.34437741 | 0.175 | 0.31 | 0.09043552 | 7 | Bcorl1 | myCAF_3 |
| Zranb12 | 4.72E-06 | 1.07964384 | 0.356 | 0.496 | 0.09129877 | 7 | Zranb1 | myCAF_3 |
| Atp2b42 | 4.79E-06 | 2.05068685 | 0.361 | 0.208 | 0.09257873 | 7 | Atp2b4 | myCAF_3 |
| Phactr4 | 4.85E-06 | 0.29636848 | 0.284 | 0.437 | 0.09383658 | 7 | Phactr4 | myCAF_3 |
| Acbd6 | 5.01E-06 | 0.67875102 | 0.407 | 0.554 | 0.09688211 | 7 | Acbd6 | myCAF_3 |
| Slc35f61 | 5.09E-06 | 0.40757031 | 0.366 | 0.485 | 0.09842435 | 7 | Slc35f6 | myCAF_3 |
| Ddhd11 | 5.11E-06 | 0.45357859 | 0.237 | 0.345 | 0.09888918 | 7 | Ddhd1 | myCAF_3 |
| Trp53inp21 | 5.15E-06 | 0.47137509 | 0.552 | 0.702 | 0.09961068 | 7 | Trp53inp2 | myCAF_3 |
| Sema5a1 | 5.16E-06 | 1.2777347 | 0.557 | 0.492 | 0.09969747 | 7 | Sema5a | myCAF_3 |
| Nol101 | 5.17E-06 | 0.46843517 | 0.17 | 0.308 | 0.10002887 | 7 | Nol10 | myCAF_3 |
| Agtrap3 | 5.22E-06 | 0.47034339 | 0.201 | 0.347 | 0.10095113 | 7 | Agtrap | myCAF_3 |
| Ube2e32 | 5.23E-06 | 0.49716501 | 0.479 | 0.58 | 0.10109195 | 7 | Ube2e3 | myCAF_3 |
| Ppil2 | 5.27E-06 | 0.41391124 | 0.407 | 0.556 | 0.10181157 | 7 | Ppil2 | myCAF_3 |
| Plekhf12 | 5.37E-06 | 0.32992953 | 0.428 | 0.524 | 0.10381552 | 7 | Plekhf1 | myCAF_3 |
| Vps37a2 | 5.39E-06 | 0.7549648 | 0.34 | 0.457 | 0.10412678 | 7 | Vps37a | myCAF_3 |
| Mcm91 | 5.47E-06 | 0.34225556 | 0.129 | 0.261 | 0.10582959 | 7 | Mcm9 | myCAF_3 |
| Ciao1 | 5.54E-06 | 0.56077228 | 0.351 | 0.49 | 0.1070661 | 7 | Ciao1 | myCAF_3 |
| Sorbs31 | 5.61E-06 | 0.77509209 | 0.361 | 0.487 | 0.10843846 | 7 | Sorbs3 | myCAF_3 |
| Kctd2 | 5.65E-06 | 0.31640077 | 0.304 | 0.461 | 0.10916247 | 7 | Kctd2 | myCAF_3 |
| Sertad21 | 5.69E-06 | 0.54697512 | 0.32 | 0.459 | 0.10996027 | 7 | Sertad2 | myCAF_3 |
| Usp42 | 5.75E-06 | 0.34837226 | 0.309 | 0.425 | 0.11124015 | 7 | Usp42 | myCAF_3 |
| Ikbkb1 | 5.79E-06 | 0.58477598 | 0.32 | 0.458 | 0.11188674 | 7 | Ikbkb | myCAF_3 |
| Pcdhgb22 | 5.80E-06 | 0.85417022 | 0.237 | 0.289 | 0.11206509 | 7 | Pcdhgb2 | myCAF_3 |
| Pold41 | 5.81E-06 | 0.42796435 | 0.443 | 0.609 | 0.11230101 | 7 | Pold4 | myCAF_3 |
| Taf61 | 5.84E-06 | 1.01707816 | 0.294 | 0.415 | 0.11284514 | 7 | Taf6 | myCAF_3 |
| Trak21 | 5.85E-06 | 0.69298514 | 0.567 | 0.672 | 0.11306353 | 7 | Trak2 | myCAF_3 |
| Ten11 | 5.85E-06 | 0.61814535 | 0.18 | 0.338 | 0.113082 | 7 | Ten1 | myCAF_3 |
| Naf11 | 5.95E-06 | 0.46664702 | 0.18 | 0.305 | 0.11495568 | 7 | Naf1 | myCAF_3 |
| Chuk | 6.05E-06 | 0.27939935 | 0.309 | 0.455 | 0.11705181 | 7 | Chuk | myCAF_3 |
| Chst14 | 6.09E-06 | 0.46304528 | 0.577 | 0.71 | 0.11769588 | 7 | Chst14 | myCAF_3 |
| Marf11 | 6.13E-06 | 0.67791217 | 0.294 | 0.423 | 0.11842148 | 7 | Marf1 | myCAF_3 |
| Rexf11 | 6.23E-06 | 0.47391991 | 0.268 | 0.371 | 0.12045241 | 7 | Rexf1 | myCAF_3 |
| Ppm1b | 6.23E-06 | 0.70154214 | 0.51 | 0.645 | 0.12052384 | 7 | Ppm1b | myCAF_3 |
| Alg3 | 6.25E-06 | 0.50739698 | 0.18 | 0.317 | 0.12087238 | 7 | Alg3 | myCAF_3 |
| Tmed8 | 6.29E-06 | 0.60911902 | 0.289 | 0.401 | 0.12170622 | 7 | Tmed8 | myCAF_3 |
| Med41 | 6.30E-06 | 1.05644849 | 0.253 | 0.371 | 0.12183319 | 7 | Med4 | myCAF_3 |
| Fkrp2 | 6.36E-06 | 0.58670085 | 0.459 | 0.576 | 0.12288265 | 7 | Fkrp | myCAF_3 |
| Kat5 | 6.45E-06 | 0.60965954 | 0.34 | 0.476 | 0.12474765 | 7 | Kat5 | myCAF_3 |
| Gid41 | 6.64E-06 | 0.37404809 | 0.299 | 0.412 | 0.12831135 | 7 | Gid4 | myCAF_3 |
| Aqr1 | 6.66E-06 | 0.43356523 | 0.314 | 0.439 | 0.12875481 | 7 | Aqr | myCAF_3 |
| Tmem41a1 | 6.71E-06 | 0.43671381 | 0.232 | 0.364 | 0.1297833 | 7 | Tmem41a | myCAF_3 |
| Gmeb12 | 6.74E-06 | 0.27798358 | 0.227 | 0.343 | 0.13034659 | 7 | Gmeb1 | myCAF_3 |
| Mycbp23 | 6.77E-06 | 0.70302706 | 0.675 | 0.774 | 0.13086272 | 7 | Mycbp2 | myCAF_3 |
| Rab3d3 | 6.82E-06 | 0.28193423 | 0.284 | 0.367 | 0.13182318 | 7 | Rab3d | myCAF_3 |
| Fam171a22 | 6.87E-06 | 0.29569707 | 0.232 | 0.355 | 0.13290934 | 7 | Fam171a2 | myCAF_3 |
| Adprhl2 | 6.89E-06 | 0.38798734 | 0.201 | 0.358 | 0.1331304 | 7 | Adprhl2 | myCAF_3 |
| Ints101 | 7.14E-06 | 0.52751403 | 0.191 | 0.323 | 0.13800426 | 7 | Ints10 | myCAF_3 |
| Dennd1a1 | 7.15E-06 | 0.349485 | 0.345 | 0.495 | 0.13831745 | 7 | Dennd1a | myCAF_3 |
| Ccdc822 | 7.35E-06 | 0.71138505 | 0.253 | 0.387 | 0.14211419 | 7 | Ccdc82 | myCAF_3 |
| Galnt112 | 7.43E-06 | 0.62856116 | 0.278 | 0.405 | 0.14359708 | 7 | Galnt11 | myCAF_3 |
| Naa401 | 7.47E-06 | 0.48423389 | 0.206 | 0.317 | 0.1444493 | 7 | Naa40 | myCAF_3 |
| Rab11fip52 | 7.58E-06 | 0.4435237 | 0.593 | 0.708 | 0.14655082 | 7 | Rab11fip5 | myCAF_3 |
| Mier3 | 7.61E-06 | 0.25190041 | 0.186 | 0.333 | 0.14714446 | 7 | Mier3 | myCAF_3 |
| Slc4a72 | 7.72E-06 | 0.40033841 | 0.149 | 0.268 | 0.14918525 | 7 | Slc4a7 | myCAF_3 |
| Adcy21 | 7.76E-06 | 0.37369011 | 0.304 | 0.412 | 0.15008773 | 7 | Adcy2 | myCAF_3 |
| Qser12 | 7.87E-06 | 0.57512904 | 0.191 | 0.319 | 0.15222253 | 7 | Qser1 | myCAF_3 |
| Hdgfl32 | 7.88E-06 | 0.84584806 | 0.247 | 0.352 | 0.15226919 | 7 | Hdgfl3 | myCAF_3 |
| Stk402 | 8.17E-06 | 0.63358295 | 0.541 | 0.659 | 0.15790448 | 7 | Stk40 | myCAF_3 |
| Dhodh1 | 8.54E-06 | 0.27994294 | 0.144 | 0.256 | 0.16513324 | 7 | Dhodh | myCAF_3 |
| Lcor | 8.79E-06 | 0.69969187 | 0.34 | 0.502 | 0.16996175 | 7 | Lcor | myCAF_3 |
| Tmbim13 | 8.98E-06 | 0.74964188 | 0.603 | 0.699 | 0.17366916 | 7 | Tmbim1 | myCAF_3 |
| Fmnl32 | 8.99E-06 | 0.65324841 | 0.691 | 0.739 | 0.17377181 | 7 | Fmnl3 | myCAF_3 |
| Pdlim13 | 9.07E-06 | 0.55902256 | 0.505 | 0.538 | 0.17545171 | 7 | Pdlim1 | myCAF_3 |
| Nid23 | 9.09E-06 | 0.37533072 | 0.443 | 0.451 | 0.17572103 | 7 | Nid2 | myCAF_3 |
| Vps72 | 9.21E-06 | 0.84126779 | 0.309 | 0.426 | 0.17811414 | 7 | Vps72 | myCAF_3 |
| Trim32 | 9.34E-06 | 0.63098306 | 0.469 | 0.561 | 0.18050326 | 7 | Trim3 | myCAF_3 |
| Lamtor1 | 9.36E-06 | 0.35355084 | 0.304 | 0.452 | 0.18101622 | 7 | Lamtor1 | myCAF_3 |
| Ptgir1 | 9.40E-06 | 2.32125041 | 0.474 | 0.331 | 0.18178647 | 7 | Ptgir | myCAF_3 |
| Eif4ebp31 | 9.45E-06 | 0.45818884 | 0.165 | 0.294 | 0.18277638 | 7 | Eif4ebp3 | myCAF_3 |
| Kif1bp | 9.61E-06 | 0.54848147 | 0.34 | 0.481 | 0.18574819 | 7 | Kif1bp | myCAF_3 |
| Hadh2 | 9.71E-06 | 0.64003803 | 0.381 | 0.538 | 0.18764989 | 7 | Hadh | myCAF_3 |
| Sfmbt11 | 9.95E-06 | 0.60376014 | 0.247 | 0.34 | 0.19232094 | 7 | Sfmbt1 | myCAF_3 |
| Trip4 | 9.97E-06 | 0.30835091 | 0.278 | 0.432 | 0.19267067 | 7 | Trip4 | myCAF_3 |
| Zbtb141 | 1.01E-05 | 0.25490565 | 0.186 | 0.301 | 0.19555288 | 7 | Zbtb14 | myCAF_3 |
| Vps391 | 1.03E-05 | 0.26297156 | 0.232 | 0.374 | 0.19896637 | 7 | Vps39 | myCAF_3 |
| Cbx11 | 1.03E-05 | 0.70073449 | 0.33 | 0.43 | 0.19937824 | 7 | Cbx1 | myCAF_3 |
| Ttll41 | 1.04E-05 | 0.4188818 | 0.165 | 0.284 | 0.20093436 | 7 | Ttll4 | myCAF_3 |
| Zbtb7a1 | 1.04E-05 | 0.92475569 | 0.67 | 0.735 | 0.20107239 | 7 | Zbtb7a | myCAF_3 |
| Haus41 | 1.04E-05 | 1.31692398 | 0.263 | 0.301 | 0.20122473 | 7 | Haus4 | myCAF_3 |
| Ciz11 | 1.05E-05 | 0.83983757 | 0.433 | 0.56 | 0.20261616 | 7 | Ciz1 | myCAF_3 |
| Brd71 | 1.06E-05 | 0.59049527 | 0.392 | 0.513 | 0.20487085 | 7 | Brd7 | myCAF_3 |
| Pias2 | 1.06E-05 | 0.63333089 | 0.433 | 0.573 | 0.20514592 | 7 | Pias2 | myCAF_3 |
| Polr2c1 | 1.06E-05 | 0.49351493 | 0.144 | 0.264 | 0.20565427 | 7 | Polr2c | myCAF_3 |
| Dusp42 | 1.07E-05 | 1.11320573 | 0.485 | 0.319 | 0.20733943 | 7 | Dusp4 | myCAF_3 |
| Klhdc41 | 1.08E-05 | 0.67324691 | 0.263 | 0.402 | 0.20967117 | 7 | Klhdc4 | myCAF_3 |
| Rsbn1l2 | 1.09E-05 | 0.25724482 | 0.201 | 0.33 | 0.21142781 | 7 | Rsbn1l | myCAF_3 |
| Xpo41 | 1.10E-05 | 0.66244228 | 0.222 | 0.326 | 0.21221264 | 7 | Xpo4 | myCAF_3 |
| Xrn11 | 1.12E-05 | 0.42803549 | 0.216 | 0.356 | 0.21632479 | 7 | Xrn1 | myCAF_3 |
| Secisbp2 | 1.12E-05 | 0.34523977 | 0.227 | 0.352 | 0.21726658 | 7 | Secisbp2 | myCAF_3 |
| Tpst2 | 1.13E-05 | 0.46135801 | 0.536 | 0.651 | 0.21752355 | 7 | Tpst2 | myCAF_3 |
| Notch22 | 1.14E-05 | 0.73900211 | 0.469 | 0.587 | 0.22009177 | 7 | Notch2 | myCAF_3 |
| Porcn3 | 1.15E-05 | 1.3144629 | 0.459 | 0.362 | 0.22140472 | 7 | Porcn | myCAF_3 |
| D17H6S53E | 1.15E-05 | 0.63509461 | 0.314 | 0.44 | 0.22281121 | 7 | D17H6S53E | myCAF_3 |
| Polr2i1 | 1.15E-05 | 0.82627101 | 0.258 | 0.368 | 0.22308611 | 7 | Polr2i | myCAF_3 |
| Washc2 | 1.16E-05 | 0.66970706 | 0.515 | 0.633 | 0.22383778 | 7 | Washc2 | myCAF_3 |
| Btaf1 | 1.17E-05 | 0.27207695 | 0.258 | 0.396 | 0.22583851 | 7 | Btaf1 | myCAF_3 |
| Vcpip11 | 1.17E-05 | 0.84511015 | 0.263 | 0.396 | 0.22583855 | 7 | Vcpip1 | myCAF_3 |
| Hira1 | 1.17E-05 | 0.43043524 | 0.284 | 0.408 | 0.22640249 | 7 | Hira | myCAF_3 |
| Irf2bp11 | 1.20E-05 | 0.51433564 | 0.356 | 0.483 | 0.23105551 | 7 | Irf2bp1 | myCAF_3 |
| Jam31 | 1.20E-05 | 0.56394984 | 0.526 | 0.621 | 0.23173798 | 7 | Jam3 | myCAF_3 |
| Cryzl12 | 1.20E-05 | 0.91936926 | 0.407 | 0.534 | 0.23228036 | 7 | Cryzl1 | myCAF_3 |
| Ube2q21 | 1.21E-05 | 0.79001419 | 0.356 | 0.505 | 0.23468798 | 7 | Ube2q2 | myCAF_3 |
| Aldh9a11 | 1.22E-05 | 0.49427349 | 0.515 | 0.652 | 0.23581235 | 7 | Aldh9a1 | myCAF_3 |
| Fbxo9 | 1.25E-05 | 0.38665663 | 0.294 | 0.421 | 0.24215826 | 7 | Fbxo9 | myCAF_3 |
| Pomt2 | 1.28E-05 | 0.55329828 | 0.278 | 0.397 | 0.24733542 | 7 | Pomt2 | myCAF_3 |
| Abhd132 | 1.29E-05 | 0.28402179 | 0.299 | 0.437 | 0.24880049 | 7 | Abhd13 | myCAF_3 |
| Ubxn81 | 1.30E-05 | 0.63389382 | 0.469 | 0.572 | 0.25064376 | 7 | Ubxn8 | myCAF_3 |
| Ubxn72 | 1.32E-05 | 0.9839314 | 0.567 | 0.645 | 0.25596541 | 7 | Ubxn7 | myCAF_3 |
| Ube2h2 | 1.35E-05 | 0.66457567 | 0.598 | 0.722 | 0.26123228 | 7 | Ube2h | myCAF_3 |
| Rit12 | 1.36E-05 | 0.27196517 | 0.469 | 0.605 | 0.26310763 | 7 | Rit1 | myCAF_3 |
| Nkiras2 | 1.37E-05 | 0.5140463 | 0.407 | 0.566 | 0.26576635 | 7 | Nkiras2 | myCAF_3 |
| Setx1 | 1.37E-05 | 0.46726367 | 0.242 | 0.369 | 0.2658098 | 7 | Setx | myCAF_3 |
| Ramac1 | 1.41E-05 | 0.81745245 | 0.299 | 0.429 | 0.27295728 | 7 | Ramac | myCAF_3 |
| Nsmce21 | 1.43E-05 | 0.75193382 | 0.433 | 0.532 | 0.27610511 | 7 | Nsmce2 | myCAF_3 |
| Rnf1502 | 1.43E-05 | 0.52032479 | 0.397 | 0.436 | 0.2763975 | 7 | Rnf150 | myCAF_3 |
| Dgcr61 | 1.45E-05 | 0.38486094 | 0.402 | 0.589 | 0.28013048 | 7 | Dgcr6 | myCAF_3 |
| Ndufaf3 | 1.45E-05 | 0.35781944 | 0.155 | 0.275 | 0.28026278 | 7 | Ndufaf3 | myCAF_3 |
| Cacfd1 | 1.47E-05 | 0.5990052 | 0.531 | 0.676 | 0.28479942 | 7 | Cacfd1 | myCAF_3 |
| Nubp2 | 1.50E-05 | 0.66302019 | 0.356 | 0.484 | 0.28986659 | 7 | Nubp2 | myCAF_3 |
| Urod | 1.51E-05 | 0.54433282 | 0.407 | 0.539 | 0.29251886 | 7 | Urod | myCAF_3 |
| Macrod1 | 1.54E-05 | 2.08797929 | 0.253 | 0.142 | 0.2969201 | 7 | Macrod1 | myCAF_3 |
| Itfg21 | 1.55E-05 | 0.71098505 | 0.216 | 0.359 | 0.29918084 | 7 | Itfg2 | myCAF_3 |
| Arid21 | 1.57E-05 | 0.79229297 | 0.526 | 0.634 | 0.30270674 | 7 | Arid2 | myCAF_3 |
| Mnat1 | 1.57E-05 | 0.48360777 | 0.294 | 0.412 | 0.30294698 | 7 | Mnat1 | myCAF_3 |
| Ppp2r1b1 | 1.57E-05 | 0.56425435 | 0.273 | 0.44 | 0.30340326 | 7 | Ppp2r1b | myCAF_3 |
| Fam220a.11 | 1.57E-05 | 0.4456864 | 0.402 | 0.511 | 0.30448985 | 7 | Fam220a.1 | myCAF_3 |
| Trafd12 | 1.58E-05 | 0.68382064 | 0.433 | 0.553 | 0.30557089 | 7 | Trafd1 | myCAF_3 |
| Tmem161a | 1.58E-05 | 0.33057062 | 0.361 | 0.525 | 0.30632688 | 7 | Tmem161a | myCAF_3 |
| Zfp236 | 1.59E-05 | 0.38021836 | 0.263 | 0.409 | 0.30645925 | 7 | Zfp236 | myCAF_3 |
| Rmnd5a1 | 1.60E-05 | 0.69403312 | 0.464 | 0.574 | 0.31013486 | 7 | Rmnd5a | myCAF_3 |
| Ppp4r11 | 1.61E-05 | 0.45236338 | 0.309 | 0.417 | 0.31053084 | 7 | Ppp4r1 | myCAF_3 |
| Clk2 | 1.61E-05 | 0.4846921 | 0.464 | 0.594 | 0.31131575 | 7 | Clk2 | myCAF_3 |
| Fbxl19 | 1.61E-05 | 0.50047788 | 0.232 | 0.368 | 0.31164989 | 7 | Fbxl19 | myCAF_3 |
| Uvssa | 1.63E-05 | 0.48552948 | 0.165 | 0.314 | 0.31423866 | 7 | Uvssa | myCAF_3 |
| Irf22 | 1.63E-05 | 0.92089916 | 0.299 | 0.411 | 0.31546462 | 7 | Irf2 | myCAF_3 |
| Hmgn31 | 1.66E-05 | 0.47498123 | 0.376 | 0.496 | 0.32012594 | 7 | Hmgn3 | myCAF_3 |
| Zfp5031 | 1.68E-05 | 1.01643431 | 0.423 | 0.503 | 0.32427274 | 7 | Zfp503 | myCAF_3 |
| Pex11b | 1.70E-05 | 0.30702247 | 0.242 | 0.37 | 0.32834741 | 7 | Pex11b | myCAF_3 |
| Dgkh | 1.70E-05 | 0.88812178 | 0.186 | 0.318 | 0.32895556 | 7 | Dgkh | myCAF_3 |
| Map3k143 | 1.71E-05 | 1.04384165 | 0.284 | 0.391 | 0.32984197 | 7 | Map3k14 | myCAF_3 |
| Ate11 | 1.71E-05 | 0.32318391 | 0.222 | 0.355 | 0.33152056 | 7 | Ate1 | myCAF_3 |
| Mcc3 | 1.72E-05 | 0.57907837 | 0.314 | 0.389 | 0.33225418 | 7 | Mcc | myCAF_3 |
| Nup2141 | 1.73E-05 | 0.40270593 | 0.278 | 0.402 | 0.33487943 | 7 | Nup214 | myCAF_3 |
| Lrrc451 | 1.76E-05 | 0.63078669 | 0.216 | 0.359 | 0.339863 | 7 | Lrrc45 | myCAF_3 |
| Zfp646 | 1.77E-05 | 0.98090633 | 0.139 | 0.268 | 0.34194345 | 7 | Zfp646 | myCAF_3 |
| Dcaf16 | 1.78E-05 | 0.43348264 | 0.211 | 0.324 | 0.34356918 | 7 | Dcaf1 | myCAF_3 |
| Urb21 | 1.78E-05 | 0.32371169 | 0.175 | 0.308 | 0.34390234 | 7 | Urb2 | myCAF_3 |
| Epn22 | 1.78E-05 | 0.56475084 | 0.366 | 0.476 | 0.3443381 | 7 | Epn2 | myCAF_3 |
| Clpb1 | 1.79E-05 | 0.32965135 | 0.175 | 0.295 | 0.34649136 | 7 | Clpb | myCAF_3 |
| Atg101 | 1.79E-05 | 0.88892785 | 0.428 | 0.504 | 0.34669635 | 7 | Atg101 | myCAF_3 |
| Rbsn | 1.81E-05 | 0.26883514 | 0.345 | 0.486 | 0.3504592 | 7 | Rbsn | myCAF_3 |
| Cdkal11 | 1.86E-05 | 0.58462429 | 0.206 | 0.334 | 0.3588437 | 7 | Cdkal1 | myCAF_3 |
| Fam53c | 1.87E-05 | 0.54977931 | 0.423 | 0.534 | 0.36136922 | 7 | Fam53c | myCAF_3 |
| Pla1a2 | 1.87E-05 | 0.4271592 | 0.691 | 0.884 | 0.36201656 | 7 | Pla1a | myCAF_3 |
| Ick | 1.88E-05 | 0.29280623 | 0.412 | 0.547 | 0.36396976 | 7 | Ick | myCAF_3 |
| Strn | 1.92E-05 | 0.33508323 | 0.366 | 0.521 | 0.37214727 | 7 | Strn | myCAF_3 |
| Rusc13 | 1.93E-05 | 0.27767648 | 0.18 | 0.3 | 0.37281752 | 7 | Rusc1 | myCAF_3 |
| Nob1 | 1.97E-05 | 1.15368903 | 0.201 | 0.324 | 0.38065689 | 7 | Nob1 | myCAF_3 |
| Zfp6182 | 1.97E-05 | 0.49447972 | 0.263 | 0.386 | 0.38102715 | 7 | Zfp618 | myCAF_3 |
| Gse11 | 1.97E-05 | 0.8506038 | 0.18 | 0.311 | 0.38152672 | 7 | Gse1 | myCAF_3 |
| Fbxo3 | 1.98E-05 | 0.72478392 | 0.392 | 0.499 | 0.38288999 | 7 | Fbxo3 | myCAF_3 |
| Agap12 | 1.98E-05 | 0.41111043 | 0.361 | 0.479 | 0.38326152 | 7 | Agap1 | myCAF_3 |
| Gcc11 | 1.99E-05 | 0.62903753 | 0.32 | 0.473 | 0.38536794 | 7 | Gcc1 | myCAF_3 |
| Snx4 | 2.00E-05 | 0.51755525 | 0.392 | 0.545 | 0.38696419 | 7 | Snx4 | myCAF_3 |
| Nr2c2ap | 2.00E-05 | 0.39235841 | 0.186 | 0.309 | 0.38722349 | 7 | Nr2c2ap | myCAF_3 |
| Zfp142 | 2.01E-05 | 0.53329249 | 0.242 | 0.355 | 0.3887047 | 7 | Zfp142 | myCAF_3 |
| Srbd11 | 2.04E-05 | 1.25782987 | 0.139 | 0.251 | 0.39353788 | 7 | Srbd1 | myCAF_3 |
| Nfs11 | 2.04E-05 | 0.71696764 | 0.201 | 0.328 | 0.39486531 | 7 | Nfs1 | myCAF_3 |
| Nfkbib | 2.05E-05 | 0.79706504 | 0.376 | 0.471 | 0.39556625 | 7 | Nfkbib | myCAF_3 |
| Rbm271 | 2.07E-05 | 0.85617533 | 0.314 | 0.432 | 0.39958656 | 7 | Rbm27 | myCAF_3 |
| Dhcr241 | 2.08E-05 | 0.29607232 | 0.253 | 0.336 | 0.40233175 | 7 | Dhcr24 | myCAF_3 |
| Tmem42 | 2.08E-05 | 0.38698388 | 0.165 | 0.29 | 0.40307045 | 7 | Tmem42 | myCAF_3 |
| Gtf3c3 | 2.17E-05 | 0.9252296 | 0.278 | 0.409 | 0.41961056 | 7 | Gtf3c3 | myCAF_3 |
| Dtx42 | 2.18E-05 | 0.43164031 | 0.459 | 0.531 | 0.42132428 | 7 | Dtx4 | myCAF_3 |
| Mfsd14a2 | 2.18E-05 | 0.32920058 | 0.299 | 0.439 | 0.42199733 | 7 | Mfsd14a | myCAF_3 |
| Pigq | 2.21E-05 | 0.5390259 | 0.381 | 0.504 | 0.42634147 | 7 | Pigq | myCAF_3 |
| Eif2b41 | 2.23E-05 | 0.37369994 | 0.309 | 0.414 | 0.43152498 | 7 | Eif2b4 | myCAF_3 |
| Alg21 | 2.26E-05 | 0.55027165 | 0.247 | 0.393 | 0.43652611 | 7 | Alg2 | myCAF_3 |
| Nr1d11 | 2.28E-05 | 0.82240755 | 0.459 | 0.492 | 0.44156885 | 7 | Nr1d1 | myCAF_3 |
| Vps531 | 2.29E-05 | 0.27872584 | 0.227 | 0.35 | 0.44305873 | 7 | Vps53 | myCAF_3 |
| Asf1a1 | 2.31E-05 | 0.78673055 | 0.149 | 0.29 | 0.44597147 | 7 | Asf1a | myCAF_3 |
| Dzip1l2 | 2.31E-05 | 0.3996463 | 0.356 | 0.487 | 0.44728489 | 7 | Dzip1l | myCAF_3 |
| Dis31 | 2.32E-05 | 0.51519752 | 0.16 | 0.268 | 0.4493944 | 7 | Dis3 | myCAF_3 |
| Spg111 | 2.35E-05 | 0.6193507 | 0.227 | 0.358 | 0.45391111 | 7 | Spg11 | myCAF_3 |
| Usp321 | 2.35E-05 | 0.87740217 | 0.345 | 0.476 | 0.45410944 | 7 | Usp32 | myCAF_3 |
| Ythdc11 | 2.38E-05 | 0.65740851 | 0.258 | 0.411 | 0.46056234 | 7 | Ythdc1 | myCAF_3 |
| Zfp334 | 2.39E-05 | 0.67873501 | 0.247 | 0.36 | 0.46191972 | 7 | Zfp334 | myCAF_3 |
| Evi51 | 2.39E-05 | 0.43818732 | 0.289 | 0.434 | 0.46200974 | 7 | Evi5 | myCAF_3 |
| Pdcd71 | 2.40E-05 | 0.54672051 | 0.211 | 0.321 | 0.46332358 | 7 | Pdcd7 | myCAF_3 |
| Epc21 | 2.42E-05 | 0.45218271 | 0.289 | 0.405 | 0.46733586 | 7 | Epc2 | myCAF_3 |
| Vps13d | 2.44E-05 | 0.89766046 | 0.485 | 0.561 | 0.47221748 | 7 | Vps13d | myCAF_3 |
| Tprgl2 | 2.46E-05 | 0.76824085 | 0.49 | 0.605 | 0.47489982 | 7 | Tprgl | myCAF_3 |
| Cep631 | 2.46E-05 | 0.6520178 | 0.18 | 0.311 | 0.47635743 | 7 | Cep63 | myCAF_3 |
| Rasl11b2 | 2.47E-05 | 2.04538434 | 0.469 | 0.358 | 0.47687651 | 7 | Rasl11b | myCAF_3 |
| Urgcp | 2.47E-05 | 0.54220006 | 0.289 | 0.42 | 0.47775133 | 7 | Urgcp | myCAF_3 |
| P2rx42 | 2.48E-05 | 0.25466364 | 0.51 | 0.637 | 0.47909341 | 7 | P2rx4 | myCAF_3 |
| Mtfr1l | 2.51E-05 | 0.53611364 | 0.423 | 0.534 | 0.48551258 | 7 | Mtfr1l | myCAF_3 |
| Yipf6 | 2.52E-05 | 0.53361306 | 0.258 | 0.387 | 0.48648594 | 7 | Yipf6 | myCAF_3 |
| Lrch41 | 2.52E-05 | 0.71021786 | 0.423 | 0.523 | 0.48815573 | 7 | Lrch4 | myCAF_3 |
| Cars22 | 2.53E-05 | 0.55528892 | 0.191 | 0.303 | 0.48940895 | 7 | Cars2 | myCAF_3 |
| Vamp21 | 2.54E-05 | 0.92206573 | 0.423 | 0.53 | 0.49111819 | 7 | Vamp2 | myCAF_3 |
| Helb1 | 2.56E-05 | 1.12947487 | 0.247 | 0.377 | 0.4947476 | 7 | Helb | myCAF_3 |
| Man2b23 | 2.60E-05 | 0.42732244 | 0.629 | 0.785 | 0.50358444 | 7 | Man2b2 | myCAF_3 |
| Hspbp11 | 2.65E-05 | 0.75917721 | 0.284 | 0.39 | 0.51316439 | 7 | Hspbp1 | myCAF_3 |
| Tbcc2 | 2.67E-05 | 0.34165533 | 0.299 | 0.436 | 0.51561648 | 7 | Tbcc | myCAF_3 |
| Fbxo42 | 2.67E-05 | 0.60758684 | 0.335 | 0.461 | 0.51600989 | 7 | Fbxo42 | myCAF_3 |
| Crif31 | 2.68E-05 | 0.44426366 | 0.227 | 0.341 | 0.51816401 | 7 | Crif3 | myCAF_3 |
| Acadm | 2.69E-05 | 0.43044156 | 0.196 | 0.313 | 0.51931307 | 7 | Acadm | myCAF_3 |
| Zkscan11 | 2.72E-05 | 0.95616522 | 0.201 | 0.329 | 0.5249403 | 7 | Zkscan1 | myCAF_3 |
| Ino80d1 | 2.75E-05 | 0.68950805 | 0.397 | 0.515 | 0.53119142 | 7 | Ino80d | myCAF_3 |
| Taz1 | 2.79E-05 | 0.94680978 | 0.376 | 0.482 | 0.54011087 | 7 | Taz | myCAF_3 |
| Cabin1 | 2.81E-05 | 0.45219813 | 0.361 | 0.484 | 0.54288378 | 7 | Cabin1 | myCAF_3 |
| Herc41 | 2.82E-05 | 1.0596931 | 0.314 | 0.445 | 0.54497829 | 7 | Herc4 | myCAF_3 |
| Man2b13 | 2.84E-05 | 0.34883935 | 0.546 | 0.679 | 0.54842335 | 7 | Man2b1 | myCAF_3 |
| Acox12 | 2.85E-05 | 0.74052073 | 0.438 | 0.557 | 0.55180561 | 7 | Acox1 | myCAF_3 |
| Prpf18 | 2.88E-05 | 0.98285085 | 0.366 | 0.466 | 0.5559592 | 7 | Prpf18 | myCAF_3 |
| Inpp4a | 2.91E-05 | 1.03876405 | 0.206 | 0.33 | 0.56223234 | 7 | Inpp4a | myCAF_3 |
| Pltp3 | 2.94E-05 | 1.49474785 | 0.387 | 0.314 | 0.56776723 | 7 | Pltp | myCAF_3 |
| Cdip12 | 2.94E-05 | 0.92556355 | 0.407 | 0.527 | 0.5678126 | 7 | Cdip1 | myCAF_3 |
| Phf20l11 | 2.99E-05 | 0.6372424 | 0.438 | 0.547 | 0.57853904 | 7 | Phf20l1 | myCAF_3 |
| Fbxo251 | 3.02E-05 | 0.28373759 | 0.155 | 0.275 | 0.58316855 | 7 | Fbxo25 | myCAF_3 |
| Dhx16 | 3.02E-05 | 0.42408017 | 0.376 | 0.492 | 0.58415163 | 7 | Dhx16 | myCAF_3 |
| Ghr3 | 3.02E-05 | 0.26525732 | 0.247 | 0.435 | 0.58442565 | 7 | Ghr | myCAF_3 |
| Gabbr12 | 3.02E-05 | 1.36547467 | 0.552 | 0.473 | 0.58481969 | 7 | Gabbr1 | myCAF_3 |
| Wdr411 | 3.04E-05 | 0.2749632 | 0.263 | 0.404 | 0.58690919 | 7 | Wdr41 | myCAF_3 |
| Syngap11 | 3.04E-05 | 0.37115667 | 0.397 | 0.531 | 0.58749089 | 7 | Syngap1 | myCAF_3 |
| Tmem132a2 | 3.07E-05 | 0.60461636 | 0.335 | 0.443 | 0.59439747 | 7 | Tmem132a | myCAF_3 |
| Inhbb1 | 3.09E-05 | 1.48461813 | 0.784 | 0.646 | 0.59744695 | 7 | Inhbb | myCAF_3 |
| Pithd11 | 3.14E-05 | 0.436795 | 0.268 | 0.39 | 0.6064898 | 7 | Pithd1 | myCAF_3 |
| Gtf2h21 | 3.15E-05 | 0.2858263 | 0.149 | 0.276 | 0.60983884 | 7 | Gtf2h2 | myCAF_3 |
| Heatr61 | 3.16E-05 | 0.34998914 | 0.216 | 0.352 | 0.61016113 | 7 | Heatr6 | myCAF_3 |
| Clasrp1 | 3.17E-05 | 0.70821593 | 0.335 | 0.495 | 0.61247834 | 7 | Clasrp | myCAF_3 |
| Rif11 | 3.18E-05 | 0.69713334 | 0.149 | 0.254 | 0.61540356 | 7 | Rif1 | myCAF_3 |
| Uqcc3 | 3.20E-05 | 0.59785552 | 0.418 | 0.547 | 0.61848362 | 7 | Uqcc3 | myCAF_3 |
| Efna51 | 3.21E-05 | 0.31561766 | 0.201 | 0.334 | 0.61975079 | 7 | Efna5 | myCAF_3 |
| Elp1 | 3.23E-05 | 0.81581565 | 0.304 | 0.448 | 0.62431446 | 7 | Elp1 | myCAF_3 |
| Adrm11 | 3.23E-05 | 0.67048091 | 0.33 | 0.463 | 0.62442821 | 7 | Adrm1 | myCAF_3 |
| Mecr2 | 3.24E-05 | 0.40095264 | 0.186 | 0.303 | 0.62577975 | 7 | Mecr | myCAF_3 |
| Pitpnm1 | 3.24E-05 | 2.28905894 | 0.376 | 0.27 | 0.62702534 | 7 | Pitpnm1 | myCAF_3 |
| Polr3e | 3.26E-05 | 0.35534725 | 0.273 | 0.387 | 0.63028918 | 7 | Polr3e | myCAF_3 |
| Ncoa22 | 3.29E-05 | 0.290915 | 0.299 | 0.414 | 0.63529515 | 7 | Ncoa2 | myCAF_3 |
| Casd11 | 3.30E-05 | 0.35063299 | 0.34 | 0.482 | 0.63762811 | 7 | Casd1 | myCAF_3 |
| Dvl3 | 3.32E-05 | 0.49848517 | 0.5 | 0.642 | 0.64218875 | 7 | Dvl3 | myCAF_3 |
| Eif2b2 | 3.36E-05 | 0.33908999 | 0.366 | 0.504 | 0.6505402 | 7 | Eif2b2 | myCAF_3 |
| Zfp461 | 3.42E-05 | 0.36730154 | 0.268 | 0.396 | 0.66063569 | 7 | Zfp46 | myCAF_3 |
| Rabep2 | 3.43E-05 | 0.46602099 | 0.191 | 0.31 | 0.66290489 | 7 | Rabep2 | myCAF_3 |
| Kdm2b1 | 3.43E-05 | 0.47776573 | 0.165 | 0.288 | 0.66296668 | 7 | Kdm2b | myCAF_3 |
| Ddx41 | 3.51E-05 | 0.54882297 | 0.423 | 0.563 | 0.67868808 | 7 | Ddx41 | myCAF_3 |
| Dtx3l2 | 3.52E-05 | 0.36119754 | 0.345 | 0.401 | 0.68032847 | 7 | Dtx3l | myCAF_3 |
| Ccnt22 | 3.52E-05 | 0.49456624 | 0.412 | 0.517 | 0.68060824 | 7 | Ccnt2 | myCAF_3 |
| Exoc11 | 3.54E-05 | 0.72930795 | 0.356 | 0.466 | 0.68382059 | 7 | Exoc1 | myCAF_3 |
| Exoc6b2 | 3.56E-05 | 1.05231706 | 0.438 | 0.546 | 0.68785395 | 7 | Exoc6b | myCAF_3 |
| Mex3a2 | 3.58E-05 | 0.38706623 | 0.144 | 0.255 | 0.69274058 | 7 | Mex3a | myCAF_3 |
| Tbck1 | 3.59E-05 | 0.3085013 | 0.186 | 0.313 | 0.69369385 | 7 | Tbck | myCAF_3 |
| Chkb | 3.63E-05 | 0.37933022 | 0.464 | 0.6 | 0.7010192 | 7 | Chkb | myCAF_3 |
| Cox201 | 3.63E-05 | 0.98454685 | 0.242 | 0.361 | 0.70239064 | 7 | Cox20 | myCAF_3 |
| Tarbp21 | 3.72E-05 | 0.46438296 | 0.191 | 0.313 | 0.71882737 | 7 | Tarbp2 | myCAF_3 |
| Phyh3 | 3.74E-05 | 0.38111318 | 0.211 | 0.314 | 0.72279356 | 7 | Phyh | myCAF_3 |
| Insr1 | 3.77E-05 | 0.85325765 | 0.418 | 0.569 | 0.72950045 | 7 | Insr | myCAF_3 |
| Nt5m | 3.78E-05 | 0.3058685 | 0.227 | 0.33 | 0.7306441 | 7 | Nt5m | myCAF_3 |
| Zbtb8a1 | 3.84E-05 | 0.31574731 | 0.175 | 0.274 | 0.74274215 | 7 | Zbtb8a | myCAF_3 |
| Lrrc8a1 | 3.89E-05 | 0.62080144 | 0.603 | 0.72 | 0.75124829 | 7 | Lrrc8a | myCAF_3 |
| Camta2 | 3.89E-05 | 0.72533217 | 0.335 | 0.461 | 0.75206444 | 7 | Camta2 | myCAF_3 |
| Rictor2 | 3.96E-05 | 0.55799633 | 0.381 | 0.505 | 0.76654047 | 7 | Rictor | myCAF_3 |
| Kxd11 | 4.02E-05 | 0.90179381 | 0.144 | 0.267 | 0.77768398 | 7 | Kxd1 | myCAF_3 |
| Dhx571 | 4.05E-05 | 0.47318886 | 0.175 | 0.297 | 0.78242106 | 7 | Dhx57 | myCAF_3 |
| Tmem2511 | 4.09E-05 | 0.30336943 | 0.17 | 0.298 | 0.79126404 | 7 | Tmem251 | myCAF_3 |
| Stx8 | 4.09E-05 | 0.60023018 | 0.469 | 0.594 | 0.79129174 | 7 | Stx8 | myCAF_3 |
| Sgta1 | 4.13E-05 | 0.47746522 | 0.242 | 0.384 | 0.79833529 | 7 | Sgta | myCAF_3 |
| Slu7 | 4.16E-05 | 0.28328476 | 0.335 | 0.442 | 0.80379636 | 7 | Slu7 | myCAF_3 |
| Znrd21 | 4.20E-05 | 0.4133697 | 0.247 | 0.37 | 0.81226329 | 7 | Znrd2 | myCAF_3 |
| Tshz31 | 4.24E-05 | 0.60445465 | 0.557 | 0.667 | 0.81974271 | 7 | Tshz3 | myCAF_3 |
| Arid4b2 | 4.27E-05 | 0.87581563 | 0.371 | 0.519 | 0.82486937 | 7 | Arid4b | myCAF_3 |
| Gyg3 | 4.28E-05 | 0.28572885 | 0.258 | 0.383 | 0.8269045 | 7 | Gyg | myCAF_3 |
| Heca1 | 4.30E-05 | 1.16197082 | 0.521 | 0.6 | 0.83082697 | 7 | Heca | myCAF_3 |
| Ift52 | 4.33E-05 | 0.53201195 | 0.356 | 0.501 | 0.83742962 | 7 | Ift52 | myCAF_3 |
| Ranbp10 | 4.34E-05 | 0.26482599 | 0.227 | 0.355 | 0.83843925 | 7 | Ranbp10 | myCAF_3 |
| Nrbf2 | 4.37E-05 | 0.78950779 | 0.443 | 0.562 | 0.84500776 | 7 | Nrbf2 | myCAF_3 |
| Slc35b3 | 4.39E-05 | 0.48184685 | 0.165 | 0.292 | 0.8492112 | 7 | Slc35b3 | myCAF_3 |
| Kif13a1 | 4.42E-05 | 0.32892985 | 0.325 | 0.464 | 0.85408318 | 7 | Kif13a | myCAF_3 |
| Fam76b1 | 4.42E-05 | 0.36396501 | 0.356 | 0.479 | 0.85479207 | 7 | Fam76b | myCAF_3 |
| Itga13 | 4.49E-05 | 0.32504643 | 0.577 | 0.631 | 0.86819068 | 7 | Itga1 | myCAF_3 |
| Zkscan171 | 4.50E-05 | 0.76329202 | 0.309 | 0.424 | 0.86910496 | 7 | Zkscan17 | myCAF_3 |
| Rbm5 | 4.51E-05 | 1.27830155 | 0.613 | 0.722 | 0.87168437 | 7 | Rbm5 | myCAF_3 |
| Ccnh1 | 4.51E-05 | 0.37836814 | 0.201 | 0.351 | 0.87219168 | 7 | Ccnh | myCAF_3 |
| Pdcd11 | 4.55E-05 | 0.6374218 | 0.34 | 0.455 | 0.8795635 | 7 | Pdcd11 | myCAF_3 |
| Foxo31 | 4.55E-05 | 1.19840917 | 0.433 | 0.533 | 0.88056738 | 7 | Foxo3 | myCAF_3 |
| Rilpl1 | 4.60E-05 | 0.36247237 | 0.304 | 0.461 | 0.88977146 | 7 | Rilpl1 | myCAF_3 |
| Ccdc71l | 4.61E-05 | 0.30586355 | 0.273 | 0.442 | 0.89189223 | 7 | Ccdc71l | myCAF_3 |
| Phf21a1 | 4.68E-05 | 0.80197058 | 0.485 | 0.593 | 0.90572437 | 7 | Phf21a | myCAF_3 |
| Isca21 | 4.69E-05 | 0.39676939 | 0.356 | 0.451 | 0.90630054 | 7 | Isca2 | myCAF_3 |
| Hmgb31 | 4.70E-05 | 0.70551707 | 0.335 | 0.38 | 0.90787127 | 7 | Hmgb3 | myCAF_3 |
| Rgs121 | 4.76E-05 | 0.40380477 | 0.366 | 0.488 | 0.92035011 | 7 | Rgs12 | myCAF_3 |
| Cc2d1a | 4.76E-05 | 1.18820703 | 0.314 | 0.42 | 0.92069128 | 7 | Cc2d1a | myCAF_3 |
| Commd4 | 4.76E-05 | 0.35284826 | 0.402 | 0.53 | 0.92103679 | 7 | Commd4 | myCAF_3 |
| Inpp5b | 4.77E-05 | 0.60557882 | 0.258 | 0.392 | 0.92204745 | 7 | Inpp5b | myCAF_3 |
| Lztfl12 | 4.78E-05 | 0.2721152 | 0.175 | 0.274 | 0.92454098 | 7 | Lztfl1 | myCAF_3 |
| Spindoc | 4.79E-05 | 0.52532777 | 0.418 | 0.513 | 0.9270631 | 7 | Spindoc | myCAF_3 |
| Hdac41 | 4.80E-05 | 0.73030887 | 0.495 | 0.593 | 0.92870602 | 7 | Hdac4 | myCAF_3 |
| Ddt2 | 4.81E-05 | 0.38666237 | 0.253 | 0.384 | 0.93026208 | 7 | Ddt | myCAF_3 |
| Slc25a122 | 4.84E-05 | 0.43949712 | 0.34 | 0.442 | 0.9362711 | 7 | Slc25a12 | myCAF_3 |
| Hmgb11 | 4.86E-05 | 0.76053936 | 0.448 | 0.551 | 0.93992671 | 7 | Hmgb1 | myCAF_3 |
| Atp6v1b21 | 4.87E-05 | 0.65045337 | 0.613 | 0.762 | 0.94149947 | 7 | Atp6v1b2 | myCAF_3 |
| Dpy19l41 | 4.89E-05 | 0.25471729 | 0.206 | 0.328 | 0.94470834 | 7 | Dpy19l4 | myCAF_3 |
| Ndufaf41 | 4.94E-05 | 0.75059828 | 0.149 | 0.261 | 0.95438524 | 7 | Ndufaf4 | myCAF_3 |
| Terf2ip | 4.97E-05 | 0.90812747 | 0.155 | 0.279 | 0.96160071 | 7 | Terf2ip | myCAF_3 |
| Arhgdib2 | 4.98E-05 | 0.80394424 | 0.639 | 0.615 | 0.96260155 | 7 | Arhgdib | myCAF_3 |
| Ghdc2 | 4.99E-05 | 0.30909788 | 0.175 | 0.297 | 0.96456969 | 7 | Ghdc | myCAF_3 |
| Kdm6a1 | 5.00E-05 | 1.23226213 | 0.314 | 0.431 | 0.96600253 | 7 | Kdm6a | myCAF_3 |
| Fbxo10 | 5.01E-05 | 0.35627969 | 0.232 | 0.36 | 0.96892885 | 7 | Fbxo10 | myCAF_3 |
| Bod1 | 5.02E-05 | 0.80417469 | 0.273 | 0.389 | 0.97118235 | 7 | Bod1 | myCAF_3 |
| Slc2a132 | 5.02E-05 | 1.59106014 | 0.247 | 0.354 | 0.97148134 | 7 | Slc2a13 | myCAF_3 |
| Znrf21 | 5.04E-05 | 0.53879963 | 0.284 | 0.416 | 0.97455536 | 7 | Znrf2 | myCAF_3 |
| Tap22 | 5.04E-05 | 0.82262706 | 0.515 | 0.605 | 0.97455868 | 7 | Tap2 | myCAF_3 |
| Scamp12 | 5.11E-05 | 0.37918997 | 0.211 | 0.374 | 0.98703272 | 7 | Scamp1 | myCAF_3 |
| Yes11 | 5.11E-05 | 1.07341882 | 0.33 | 0.43 | 0.98869289 | 7 | Yes1 | myCAF_3 |
| Sh3bp1 | 5.17E-05 | 1.15249685 | 0.273 | 0.396 | 0.99963959 | 7 | Sh3bp1 | myCAF_3 |
| Plcd11 | 5.19E-05 | 1.07513309 | 0.34 | 0.429 | 1 | 7 | Plcd1 | myCAF_3 |
| Gab22 | 5.19E-05 | 0.96459485 | 0.32 | 0.413 | 1 | 7 | Gab2 | myCAF_3 |
| Prmt31 | 5.25E-05 | 0.31610659 | 0.201 | 0.319 | 1 | 7 | Prmt3 | myCAF_3 |
| N6amt1 | 5.26E-05 | 0.45898763 | 0.201 | 0.366 | 1 | 7 | N6amt1 | myCAF_3 |
| Mtmr112 | 5.28E-05 | 0.60628377 | 0.309 | 0.426 | 1 | 7 | Mtmr11 | myCAF_3 |
| Kmt5b1 | 5.40E-05 | 0.94458088 | 0.531 | 0.599 | 1 | 7 | Kmt5b | myCAF_3 |
| Bsdc11 | 5.41E-05 | 0.5489958 | 0.356 | 0.502 | 1 | 7 | Bsdc1 | myCAF_3 |
| Scly | 5.44E-05 | 0.46611268 | 0.191 | 0.32 | 1 | 7 | Scly | myCAF_3 |

| | | | | | | | | |
|---|---|---|---|---|---|---|---|---|
| Tfeb | 5.49E-05 | 0.61143199 | 0.206 | 0.29 | 1 | 7 | Tfeb | myCAF_3 |
| Slc2a10 | 5.58E-05 | 0.5137097 | 0.216 | 0.356 | 1 | 7 | Slc2a10 | myCAF_3 |
| Mepce1 | 5.64E-05 | 0.72417756 | 0.356 | 0.43 | 1 | 7 | Mepce | myCAF_3 |
| Msh21 | 5.69E-05 | 0.39804054 | 0.18 | 0.273 | 1 | 7 | Msh2 | myCAF_3 |
| Slit32 | 5.72E-05 | 0.70487551 | 0.851 | 0.97 | 1 | 7 | Slit3 | myCAF_3 |
| Kif1b2 | 5.77E-05 | 0.96108346 | 0.479 | 0.551 | 1 | 7 | Kif1b | myCAF_3 |
| Nbdy1 | 5.78E-05 | 0.27819134 | 0.232 | 0.368 | 1 | 7 | Nbdy | myCAF_3 |
| Exoc2 | 5.79E-05 | 0.49778083 | 0.309 | 0.435 | 1 | 7 | Exoc2 | myCAF_3 |
| Tcf31 | 5.81E-05 | 0.6175318 | 0.433 | 0.553 | 1 | 7 | Tcf3 | myCAF_3 |
| Zfp275 | 5.81E-05 | 0.72511873 | 0.284 | 0.412 | 1 | 7 | Zfp275 | myCAF_3 |
| Ccdc1071 | 5.82E-05 | 0.3869297 | 0.361 | 0.464 | 1 | 7 | Ccdc107 | myCAF_3 |
| Fbxo331 | 5.84E-05 | 0.42372934 | 0.237 | 0.335 | 1 | 7 | Fbxo33 | myCAF_3 |
| Man2a22 | 5.86E-05 | 0.61156146 | 0.742 | 0.846 | 1 | 7 | Man2a2 | myCAF_3 |
| Pygo2 | 5.88E-05 | 0.47072896 | 0.443 | 0.567 | 1 | 7 | Pygo2 | myCAF_3 |
| DEPRECATED-I | 5.89E-05 | 0.40345363 | 0.253 | 0.373 | 1 | 7 | DEPRECATED-I | myCAF_3 |
| Lias | 5.91E-05 | 0.37599675 | 0.196 | 0.301 | 1 | 7 | Lias | myCAF_3 |
| Cdkn2aipnl1 | 5.91E-05 | 0.46186377 | 0.216 | 0.322 | 1 | 7 | Cdkn2aipnl | myCAF_3 |
| Terf21 | 5.94E-05 | 0.43342924 | 0.284 | 0.404 | 1 | 7 | Terf2 | myCAF_3 |
| Otud6b1 | 5.96E-05 | 0.38340939 | 0.273 | 0.387 | 1 | 7 | Otud6b | myCAF_3 |
| Pwwp3a1 | 5.99E-05 | 0.85496741 | 0.294 | 0.416 | 1 | 7 | Pwwp3a | myCAF_3 |
| Bmp2k | 5.99E-05 | 0.88225083 | 0.33 | 0.472 | 1 | 7 | Bmp2k | myCAF_3 |
| Cep85l1 | 6.02E-05 | 0.83923522 | 0.149 | 0.269 | 1 | 7 | Cep85l | myCAF_3 |
| Zfp2812 | 6.05E-05 | 0.90484579 | 0.526 | 0.607 | 1 | 7 | Zfp281 | myCAF_3 |
| Abcd1 | 6.09E-05 | 0.41013919 | 0.304 | 0.408 | 1 | 7 | Abcd1 | myCAF_3 |
| Mllt111 | 6.12E-05 | 0.26107487 | 0.144 | 0.264 | 1 | 7 | Mllt11 | myCAF_3 |
| Prr161 | 6.21E-05 | 1.76216354 | 0.588 | 0.419 | 1 | 7 | Prr16 | myCAF_3 |
| Rnf201 | 6.32E-05 | 0.67970196 | 0.387 | 0.517 | 1 | 7 | Rnf20 | myCAF_3 |
| Kmt2b | 6.43E-05 | 1.09631818 | 0.34 | 0.456 | 1 | 7 | Kmt2b | myCAF_3 |
| Cachd11 | 6.52E-05 | 0.86155607 | 0.314 | 0.471 | 1 | 7 | Cachd1 | myCAF_3 |
| Gng23 | 6.55E-05 | 1.33990271 | 0.448 | 0.318 | 1 | 7 | Gng2 | myCAF_3 |
| Mapk8ip31 | 6.59E-05 | 0.70589976 | 0.381 | 0.503 | 1 | 7 | Mapk8ip3 | myCAF_3 |
| Med16 | 6.60E-05 | 0.510818 | 0.314 | 0.442 | 1 | 7 | Med16 | myCAF_3 |
| Peli22 | 6.65E-05 | 0.28881434 | 0.351 | 0.493 | 1 | 7 | Peli2 | myCAF_3 |
| Itgbl13 | 6.67E-05 | 1.62981245 | 0.33 | 0.24 | 1 | 7 | Itgbl1 | myCAF_3 |
| Sirt2 | 6.76E-05 | 0.54885515 | 0.66 | 0.788 | 1 | 7 | Sirt2 | myCAF_3 |
| D2hgdh1 | 6.79E-05 | 0.53126079 | 0.18 | 0.292 | 1 | 7 | D2hgdh | myCAF_3 |
| Ift461 | 6.81E-05 | 0.45389129 | 0.515 | 0.626 | 1 | 7 | Ift46 | myCAF_3 |
| Micu31 | 6.84E-05 | 0.64883473 | 0.139 | 0.271 | 1 | 7 | Micu3 | myCAF_3 |
| Tmem472 | 6.86E-05 | 0.51132474 | 0.227 | 0.351 | 1 | 7 | Tmem47 | myCAF_3 |
| Ubap1 | 6.88E-05 | 0.36015448 | 0.33 | 0.43 | 1 | 7 | Ubap1 | myCAF_3 |
| Rnf2142 | 7.08E-05 | 0.57419072 | 0.376 | 0.517 | 1 | 7 | Rnf214 | myCAF_3 |
| Zfp5111 | 7.12E-05 | 0.34976271 | 0.247 | 0.355 | 1 | 7 | Zfp511 | myCAF_3 |
| Fbxl62 | 7.15E-05 | 0.34224879 | 0.325 | 0.449 | 1 | 7 | Fbxl6 | myCAF_3 |
| Parp33 | 7.23E-05 | 0.50764781 | 0.531 | 0.663 | 1 | 7 | Parp3 | myCAF_3 |
| Phc31 | 7.27E-05 | 0.55511475 | 0.464 | 0.546 | 1 | 7 | Phc3 | myCAF_3 |
| Zfp335 | 7.31E-05 | 0.65749187 | 0.294 | 0.387 | 1 | 7 | Zfp335 | myCAF_3 |
| Msmo11 | 7.34E-05 | 0.34601989 | 0.242 | 0.334 | 1 | 7 | Msmo1 | myCAF_3 |
| Smc51 | 7.38E-05 | 0.82868069 | 0.521 | 0.624 | 1 | 7 | Smc5 | myCAF_3 |
| Taf8 | 7.41E-05 | 0.26170149 | 0.258 | 0.384 | 1 | 7 | Taf8 | myCAF_3 |
| Mnt1 | 7.47E-05 | 0.27924834 | 0.258 | 0.364 | 1 | 7 | Mnt | myCAF_3 |
| Tsc1 | 7.48E-05 | 0.46724827 | 0.258 | 0.369 | 1 | 7 | Tsc1 | myCAF_3 |
| Lrig21 | 7.78E-05 | 0.55336739 | 0.284 | 0.386 | 1 | 7 | Lrig2 | myCAF_3 |
| Pink12 | 7.85E-05 | 1.45486611 | 0.742 | 0.718 | 1 | 7 | Pink1 | myCAF_3 |
| Otud7b | 7.96E-05 | 0.64271274 | 0.485 | 0.623 | 1 | 7 | Otud7b | myCAF_3 |
| Ccdc92 | 7.97E-05 | 0.93517274 | 0.521 | 0.623 | 1 | 7 | Ccdc9 | myCAF_3 |
| Ttc232 | 7.97E-05 | 0.64233618 | 0.206 | 0.322 | 1 | 7 | Ttc23 | myCAF_3 |
| Exoc8 | 7.98E-05 | 0.6462315 | 0.289 | 0.39 | 1 | 7 | Exoc8 | myCAF_3 |
| Zfp8712 | 8.14E-05 | 1.24420447 | 0.464 | 0.551 | 1 | 7 | Zfp871 | myCAF_3 |
| Mmgt2 | 8.18E-05 | 0.5847929 | 0.273 | 0.366 | 1 | 7 | Mmgt2 | myCAF_3 |
| Gem2 | 8.20E-05 | 2.28078796 | 0.495 | 0.526 | 1 | 7 | Gem | myCAF_3 |
| Ulk21 | 8.23E-05 | 0.52048963 | 0.485 | 0.635 | 1 | 7 | Ulk2 | myCAF_3 |
| Nat10 | 8.24E-05 | 0.38860358 | 0.211 | 0.324 | 1 | 7 | Nat10 | myCAF_3 |
| Togaram11 | 8.62E-05 | 0.56021711 | 0.299 | 0.449 | 1 | 7 | Togaram1 | myCAF_3 |
| Apba12 | 8.76E-05 | 0.31732911 | 0.211 | 0.317 | 1 | 7 | Apba1 | myCAF_3 |
| Mafg | 8.81E-05 | 0.89609506 | 0.536 | 0.65 | 1 | 7 | Mafg | myCAF_3 |
| Rab312 | 8.88E-05 | 0.93456464 | 0.789 | 0.857 | 1 | 7 | Rab31 | myCAF_3 |
| Zbtb202 | 8.92E-05 | 1.13387817 | 0.423 | 0.501 | 1 | 7 | Zbtb20 | myCAF_3 |
| Polr3gl1 | 8.94E-05 | 0.83839733 | 0.273 | 0.405 | 1 | 7 | Polr3gl | myCAF_3 |
| Bcor | 9.00E-05 | 0.94942321 | 0.34 | 0.437 | 1 | 7 | Bcor | myCAF_3 |
| Rnf1691 | 9.01E-05 | 0.54922205 | 0.33 | 0.444 | 1 | 7 | Rnf169 | myCAF_3 |
| Tex102 | 9.01E-05 | 0.37745926 | 0.345 | 0.429 | 1 | 7 | Tex10 | myCAF_3 |
| Cttnbp2nl1 | 9.07E-05 | 0.76737256 | 0.428 | 0.553 | 1 | 7 | Cttnbp2nl | myCAF_3 |
| Ccdc43 | 9.10E-05 | 0.36136931 | 0.201 | 0.335 | 1 | 7 | Ccdc43 | myCAF_3 |
| Appl22 | 9.12E-05 | 1.21865127 | 0.284 | 0.402 | 1 | 7 | Appl2 | myCAF_3 |
| Ogt2 | 9.12E-05 | 1.11156702 | 0.629 | 0.703 | 1 | 7 | Ogt | myCAF_3 |
| Rftn11 | 9.14E-05 | 0.56558124 | 0.66 | 0.792 | 1 | 7 | Rftn1 | myCAF_3 |
| Smim4 | 9.17E-05 | 0.47920368 | 0.304 | 0.429 | 1 | 7 | Smim4 | myCAF_3 |
| Pdcl | 9.37E-05 | 0.3415283 | 0.32 | 0.478 | 1 | 7 | Pdcl | myCAF_3 |
| Emc81 | 9.47E-05 | 0.49075679 | 0.237 | 0.339 | 1 | 7 | Emc8 | myCAF_3 |
| Mfsd11 | 9.49E-05 | 0.60522667 | 0.438 | 0.554 | 1 | 7 | Mfsd11 | myCAF_3 |
| Tmem39b1 | 9.52E-05 | 0.37977983 | 0.206 | 0.31 | 1 | 7 | Tmem39b | myCAF_3 |
| Tfip11 | 9.58E-05 | 0.7260359 | 0.371 | 0.498 | 1 | 7 | Tfip11 | myCAF_3 |
| Ppard1 | 9.64E-05 | 0.8910531 | 0.515 | 0.617 | 1 | 7 | Ppard | myCAF_3 |
| Myl91 | 9.81E-05 | 0.98085686 | 0.629 | 0.51 | 1 | 7 | Myl9 | myCAF_3 |
| Llgl1 | 9.88E-05 | 0.54038742 | 0.258 | 0.373 | 1 | 7 | Llgl1 | myCAF_3 |
| Pgghg1 | 0.00010047 | 1.08422429 | 0.376 | 0.498 | 1 | 7 | Pgghg | myCAF_3 |
| Ccpg13 | 0.00010293 | 0.7572126 | 0.629 | 0.741 | 1 | 7 | Ccpg1 | myCAF_3 |
| Fbf12 | 0.00010337 | 0.58501719 | 0.278 | 0.407 | 1 | 7 | Fbf1 | myCAF_3 |
| Katnbl12 | 0.00010358 | 0.53031624 | 0.284 | 0.398 | 1 | 7 | Katnbl1 | myCAF_3 |
| Cdk5 | 0.00010385 | 0.55641872 | 0.433 | 0.565 | 1 | 7 | Cdk5 | myCAF_3 |
| Pnpla81 | 0.00010449 | 0.72905037 | 0.433 | 0.577 | 1 | 7 | Pnpla8 | myCAF_3 |
| Xpa1 | 0.00010665 | 0.68804463 | 0.294 | 0.405 | 1 | 7 | Xpa | myCAF_3 |
| Lancl22 | 0.00010812 | 0.58107841 | 0.211 | 0.285 | 1 | 7 | Lancl2 | myCAF_3 |
| Ascc2 | 0.00010854 | 0.69564295 | 0.314 | 0.412 | 1 | 7 | Ascc2 | myCAF_3 |
| Pwwp2a2 | 0.00010861 | 0.67224316 | 0.412 | 0.487 | 1 | 7 | Pwwp2a | myCAF_3 |
| Vwa81 | 0.00010878 | 0.28610681 | 0.155 | 0.281 | 1 | 7 | Vwa8 | myCAF_3 |
| Vcl1 | 0.00011135 | 0.8020448 | 0.84 | 0.862 | 1 | 7 | Vcl | myCAF_3 |
| BC005624 | 0.00011303 | 0.50928788 | 0.34 | 0.474 | 1 | 7 | BC005624 | myCAF_3 |
| Nupr12 | 0.00011305 | 0.95350183 | 0.758 | 0.813 | 1 | 7 | Nupr1 | myCAF_3 |
| Klhl111 | 0.00011412 | 0.96476184 | 0.371 | 0.474 | 1 | 7 | Klhl11 | myCAF_3 |
| Lypla1 | 0.00011535 | 0.2992144 | 0.314 | 0.444 | 1 | 7 | Lypla1 | myCAF_3 |
| Slc15a4 | 0.00011548 | 0.58252677 | 0.268 | 0.375 | 1 | 7 | Slc15a4 | myCAF_3 |
| Gpbp1l11 | 0.00011729 | 0.79061482 | 0.397 | 0.509 | 1 | 7 | Gpbp1l1 | myCAF_3 |
| Rnpepl1 | 0.00011733 | 0.74449859 | 0.536 | 0.646 | 1 | 7 | Rnpepl1 | myCAF_3 |
| Armc81 | 0.00011798 | 0.56656793 | 0.201 | 0.295 | 1 | 7 | Armc8 | myCAF_3 |
| Errfi11 | 0.00011853 | 0.88213833 | 0.68 | 0.787 | 1 | 7 | Errfi1 | myCAF_3 |
| Arid5b2 | 0.00011996 | 1.08421765 | 0.66 | 0.705 | 1 | 7 | Arid5b | myCAF_3 |
| Ifi352 | 0.00012082 | 0.59421252 | 0.242 | 0.319 | 1 | 7 | Ifi35 | myCAF_3 |
| St7l1 | 0.00012159 | 0.30495424 | 0.165 | 0.275 | 1 | 7 | St7l | myCAF_3 |
| Wbp14 | 0.00012166 | 0.59828763 | 0.361 | 0.476 | 1 | 7 | Wbp1 | myCAF_3 |
| Lrp121 | 0.00012174 | 0.33787752 | 0.309 | 0.418 | 1 | 7 | Lrp12 | myCAF_3 |
| Ankhd11 | 0.00012176 | 0.36106473 | 0.196 | 0.308 | 1 | 7 | Ankhd1 | myCAF_3 |
| Myg11 | 0.00012212 | 0.3562129 | 0.268 | 0.378 | 1 | 7 | Myg1 | myCAF_3 |
| Mthfd11 | 0.00012311 | 0.32914214 | 0.201 | 0.292 | 1 | 7 | Mthfd1 | myCAF_3 |
| Arhgef102 | 0.00012321 | 0.261137 | 0.268 | 0.352 | 1 | 7 | Arhgef10 | myCAF_3 |
| Pheta21 | 0.00012351 | 0.54114329 | 0.294 | 0.425 | 1 | 7 | Pheta2 | myCAF_3 |
| Zmym3 | 0.00012369 | 0.28300471 | 0.222 | 0.346 | 1 | 7 | Zmym3 | myCAF_3 |
| Iqsec21 | 0.00012379 | 0.31518983 | 0.294 | 0.411 | 1 | 7 | Iqsec2 | myCAF_3 |
| Sppl2b | 0.00012381 | 0.48370661 | 0.361 | 0.472 | 1 | 7 | Sppl2b | myCAF_3 |
| Pikfyve | 0.00012381 | 0.36961866 | 0.165 | 0.309 | 1 | 7 | Pikfyve | myCAF_3 |
| Cds22 | 0.00012391 | 0.57701342 | 0.495 | 0.576 | 1 | 7 | Cds2 | myCAF_3 |
| Ift57 | 0.00012426 | 0.30601725 | 0.273 | 0.394 | 1 | 7 | Ift57 | myCAF_3 |
| Utp14a2 | 0.00012461 | 0.29648886 | 0.191 | 0.291 | 1 | 7 | Utp14a | myCAF_3 |
| Camkk22 | 0.00012627 | 0.82904594 | 0.345 | 0.45 | 1 | 7 | Camkk2 | myCAF_3 |
| Zfp1801 | 0.00012632 | 1.30834513 | 0.206 | 0.309 | 1 | 7 | Zfp180 | myCAF_3 |
| Acsf22 | 0.00012772 | 0.38675384 | 0.242 | 0.341 | 1 | 7 | Acsf2 | myCAF_3 |
| Gpatch111 | 0.00012803 | 0.59415839 | 0.247 | 0.363 | 1 | 7 | Gpatch11 | myCAF_3 |
| Osr12 | 0.0001293 | 0.44337433 | 0.309 | 0.276 | 1 | 7 | Osr1 | myCAF_3 |
| Adamts101 | 0.0001296 | 0.90718213 | 0.613 | 0.667 | 1 | 7 | Adamts10 | myCAF_3 |
| Bahd1 | 0.00013134 | 0.86083335 | 0.32 | 0.423 | 1 | 7 | Bahd1 | myCAF_3 |
| 5-Mar | 0.00013139 | 0.38121837 | 0.253 | 0.353 | 1 | 7 | 5-Mar | myCAF_3 |
| Pdk31 | 0.0001314 | 0.33229234 | 0.32 | 0.464 | 1 | 7 | Pdk3 | myCAF_3 |
| Shprh | 0.0001334 | 0.52299399 | 0.175 | 0.303 | 1 | 7 | Shprh | myCAF_3 |
| Cables21 | 0.00013678 | 0.26146241 | 0.175 | 0.311 | 1 | 7 | Cables2 | myCAF_3 |
| Xpr12 | 0.00013788 | 0.77183348 | 0.216 | 0.339 | 1 | 7 | Xpr1 | myCAF_3 |
| Pdzd41 | 0.00013977 | 0.33694427 | 0.237 | 0.335 | 1 | 7 | Pdzd4 | myCAF_3 |
| Sart32 | 0.00014007 | 0.65019894 | 0.216 | 0.324 | 1 | 7 | Sart3 | myCAF_3 |
| Sft2d1 | 0.00014137 | 0.35474685 | 0.216 | 0.333 | 1 | 7 | Sft2d1 | myCAF_3 |
| Gba1 | 0.00014299 | 0.48617749 | 0.557 | 0.667 | 1 | 7 | Gba | myCAF_3 |
| Apeh1 | 0.00014382 | 0.34712454 | 0.407 | 0.549 | 1 | 7 | Apeh | myCAF_3 |
| Ldlrad31 | 0.00014393 | 0.84934686 | 0.253 | 0.368 | 1 | 7 | Ldlrad3 | myCAF_3 |
| Cdc42se2 | 0.00014765 | 0.68266424 | 0.227 | 0.33 | 1 | 7 | Cdc42se2 | myCAF_3 |
| Mob2 | 0.00014821 | 0.44841735 | 0.407 | 0.514 | 1 | 7 | Mob2 | myCAF_3 |
| Map3k61 | 0.00014931 | 0.9129824 | 0.216 | 0.35 | 1 | 7 | Map3k6 | myCAF_3 |
| Phf8 | 0.00014937 | 1.03966937 | 0.34 | 0.441 | 1 | 7 | Phf8 | myCAF_3 |
| Wbp21 | 0.00015301 | 0.73745488 | 0.356 | 0.463 | 1 | 7 | Wbp2 | myCAF_3 |
| Il17rc | 0.00015456 | 0.4223334 | 0.268 | 0.379 | 1 | 7 | Il17rc | myCAF_3 |
| Lysmd23 | 0.00015607 | 0.53631343 | 0.268 | 0.35 | 1 | 7 | Lysmd2 | myCAF_3 |
| Rnf51 | 0.00015729 | 0.67512222 | 0.278 | 0.41 | 1 | 7 | Rnf5 | myCAF_3 |
| Lrrk22 | 0.00015832 | 0.83256719 | 0.253 | 0.396 | 1 | 7 | Lrrk2 | myCAF_3 |
| Mysm12 | 0.00016104 | 0.46323145 | 0.335 | 0.414 | 1 | 7 | Mysm1 | myCAF_3 |
| Stimate1 | 0.00016153 | 0.38500362 | 0.268 | 0.361 | 1 | 7 | Stimate | myCAF_3 |
| Parn1 | 0.00016205 | 0.57439639 | 0.289 | 0.394 | 1 | 7 | Parn | myCAF_3 |
| Vps541 | 0.00016271 | 0.74354734 | 0.325 | 0.442 | 1 | 7 | Vps54 | myCAF_3 |
| Nfat51 | 0.00016386 | 1.01875032 | 0.742 | 0.778 | 1 | 7 | Nfat5 | myCAF_3 |
| Itpr13 | 0.00016392 | 1.74177638 | 0.32 | 0.236 | 1 | 7 | Itpr1 | myCAF_3 |
| Spaca61 | 0.0001642 | 0.98193087 | 0.309 | 0.404 | 1 | 7 | Spaca6 | myCAF_3 |
| Fam174a | 0.00016437 | 0.50871588 | 0.33 | 0.465 | 1 | 7 | Fam174a | myCAF_3 |
| Fyco1 | 0.00016842 | 0.67610947 | 0.402 | 0.477 | 1 | 7 | Fyco1 | myCAF_3 |
| Kansl3 | 0.00017135 | 0.33032094 | 0.402 | 0.508 | 1 | 7 | Kansl3 | myCAF_3 |
| Uqcc1 | 0.00017255 | 0.50625779 | 0.186 | 0.296 | 1 | 7 | Uqcc1 | myCAF_3 |
| Thap11 | 0.00017419 | 0.59210986 | 0.376 | 0.467 | 1 | 7 | Thap11 | myCAF_3 |
| Zfand52 | 0.00017594 | 1.5675496 | 0.799 | 0.841 | 1 | 7 | Zfand5 | myCAF_3 |
| Rangrf2 | 0.00017947 | 0.63783921 | 0.242 | 0.323 | 1 | 7 | Rangrf | myCAF_3 |
| Tomm40l | 0.00018069 | 0.74748182 | 0.366 | 0.479 | 1 | 7 | Tomm40l | myCAF_3 |
| Fadd2 | 0.00018095 | 0.52845394 | 0.294 | 0.382 | 1 | 7 | Fadd | myCAF_3 |
| Pskh12 | 0.00018189 | 0.40069236 | 0.314 | 0.407 | 1 | 7 | Pskh1 | myCAF_3 |
| Klf92 | 0.00018618 | 1.69984607 | 0.881 | 0.854 | 1 | 7 | Klf9 | myCAF_3 |
| Mindy3 | 0.00018805 | 0.31446299 | 0.284 | 0.388 | 1 | 7 | Mindy3 | myCAF_3 |
| Efl1 | 0.00018872 | 0.56541491 | 0.206 | 0.324 | 1 | 7 | Efl1 | myCAF_3 |
| Depdc51 | 0.00019116 | 1.01916918 | 0.216 | 0.323 | 1 | 7 | Depdc5 | myCAF_3 |
| Myo61 | 0.0001912 | 1.84248166 | 0.335 | 0.237 | 1 | 7 | Myo6 | myCAF_3 |
| Lrrc75a2 | 0.00019285 | 0.56989805 | 0.33 | 0.41 | 1 | 7 | Lrrc75a | myCAF_3 |
| AW5498771 | 0.00019307 | 0.75674812 | 0.428 | 0.534 | 1 | 7 | AW549877 | myCAF_3 |
| Brpf3 | 0.00019337 | 1.00072234 | 0.284 | 0.398 | 1 | 7 | Brpf3 | myCAF_3 |
| Gpr1372 | 0.0001935 | 0.27295862 | 0.278 | 0.427 | 1 | 7 | Gpr137 | myCAF_3 |
| Rfx72 | 0.00019518 | 1.02124698 | 0.206 | 0.307 | 1 | 7 | Rfx7 | myCAF_3 |
| Slc35d1 | 0.00019667 | 0.73823844 | 0.186 | 0.285 | 1 | 7 | Slc35d1 | myCAF_3 |
| Ncbp31 | 0.00019731 | 0.6378618 | 0.387 | 0.479 | 1 | 7 | Ncbp3 | myCAF_3 |
| Ssh22 | 0.00019801 | 0.40098075 | 0.227 | 0.331 | 1 | 7 | Ssh2 | myCAF_3 |
| Katnb11 | 0.00019812 | 0.57916034 | 0.191 | 0.294 | 1 | 7 | Katnb1 | myCAF_3 |
| Ppox1 | 0.00020059 | 0.46690933 | 0.237 | 0.354 | 1 | 7 | Ppox | myCAF_3 |
| Fbxo28 | 0.00020311 | 0.69339953 | 0.351 | 0.461 | 1 | 7 | Fbxo28 | myCAF_3 |
| Dido11 | 0.00020521 | 0.87073944 | 0.268 | 0.384 | 1 | 7 | Dido1 | myCAF_3 |
| DEPRECATED-I | 0.00020732 | 0.49366116 | 0.155 | 0.253 | 1 | 7 | DEPRECATED-I | myCAF_3 |
| Alkbh31 | 0.00020913 | 0.37161231 | 0.227 | 0.33 | 1 | 7 | Alkbh3 | myCAF_3 |
| Slc25a161 | 0.00020928 | 0.37319097 | 0.289 | 0.393 | 1 | 7 | Slc25a16 | myCAF_3 |
| Ino80b | 0.00021034 | 0.6011569 | 0.335 | 0.446 | 1 | 7 | Ino80b | myCAF_3 |
| Luc7l1 | 0.00021035 | 0.67709709 | 0.247 | 0.38 | 1 | 7 | Luc7l | myCAF_3 |
| Med20 | 0.0002104 | 0.86968857 | 0.49 | 0.569 | 1 | 7 | Med20 | myCAF_3 |
| Mpp5 | 0.00021059 | 0.5268655 | 0.273 | 0.397 | 1 | 7 | Mpp5 | myCAF_3 |
| Ptges1 | 0.00021175 | 0.71020395 | 0.598 | 0.742 | 1 | 7 | Ptges | myCAF_3 |
| Adcy32 | 0.0002138 | 0.37361712 | 0.588 | 0.723 | 1 | 7 | Adcy3 | myCAF_3 |
| Borcs61 | 0.00021483 | 0.50525875 | 0.284 | 0.424 | 1 | 7 | Borcs6 | myCAF_3 |
| Nsmaf2 | 0.00021487 | 0.84509793 | 0.33 | 0.429 | 1 | 7 | Nsmaf | myCAF_3 |
| Ncdn | 0.00021947 | 0.48641402 | 0.387 | 0.51 | 1 | 7 | Ncdn | myCAF_3 |
| Prdm52 | 0.00021992 | 0.46584589 | 0.253 | 0.385 | 1 | 7 | Prdm5 | myCAF_3 |
| Mul1 | 0.00022048 | 0.37636779 | 0.289 | 0.387 | 1 | 7 | Mul1 | myCAF_3 |
| Crat1 | 0.0002209 | 0.6843437 | 0.258 | 0.386 | 1 | 7 | Crat | myCAF_3 |
| Apex21 | 0.00022239 | 0.42462747 | 0.201 | 0.324 | 1 | 7 | Apex2 | myCAF_3 |
| Telo2 | 0.00022256 | 0.37684701 | 0.16 | 0.256 | 1 | 7 | Telo2 | myCAF_3 |
| Atp11b | 0.00022647 | 0.47948873 | 0.206 | 0.318 | 1 | 7 | Atp11b | myCAF_3 |
| B3glct1 | 0.00022707 | 0.27299527 | 0.273 | 0.38 | 1 | 7 | B3glct | myCAF_3 |
| Sos21 | 0.00022963 | 1.13876262 | 0.206 | 0.311 | 1 | 7 | Sos2 | myCAF_3 |
| Atg16l1 | 0.00023131 | 0.77569526 | 0.273 | 0.401 | 1 | 7 | Atg16l1 | myCAF_3 |
| Nkapd1 | 0.0002338 | 0.46352182 | 0.263 | 0.379 | 1 | 7 | Nkapd1 | myCAF_3 |
| Vps51 | 0.00023463 | 0.44919777 | 0.34 | 0.468 | 1 | 7 | Vps51 | myCAF_3 |
| Aplp12 | 0.00023677 | 1.34575408 | 0.325 | 0.234 | 1 | 7 | Aplp1 | myCAF_3 |
| Mynn1 | 0.00023901 | 0.7303866 | 0.216 | 0.309 | 1 | 7 | Mynn | myCAF_3 |
| Cplane1 | 0.00023966 | 1.0932795 | 0.211 | 0.322 | 1 | 7 | Cplane1 | myCAF_3 |
| Samd41 | 0.00024048 | 0.71500348 | 0.608 | 0.683 | 1 | 7 | Samd4 | myCAF_3 |
| Gps21 | 0.00024111 | 0.78418149 | 0.289 | 0.383 | 1 | 7 | Gps2 | myCAF_3 |
| DEPRECATED-I | 0.00024579 | 0.40986821 | 0.18 | 0.275 | 1 | 7 | DEPRECATED-I | myCAF_3 |
| Ppp2r5b1 | 0.00024655 | 0.6688938 | 0.454 | 0.548 | 1 | 7 | Ppp2r5b | myCAF_3 |
| Arfrp11 | 0.0002483 | 0.91979242 | 0.253 | 0.386 | 1 | 7 | Arfrp1 | myCAF_3 |
| Nprl3 | 0.00024927 | 0.67970663 | 0.268 | 0.368 | 1 | 7 | Nprl3 | myCAF_3 |
| Nfatc41 | 0.00024968 | 1.15992288 | 0.804 | 0.823 | 1 | 7 | Nfatc4 | myCAF_3 |
| Sprtn | 0.00025078 | 0.31079461 | 0.191 | 0.305 | 1 | 7 | Sprtn | myCAF_3 |
| Scyl2 | 0.00025105 | 0.70995901 | 0.32 | 0.451 | 1 | 7 | Scyl2 | myCAF_3 |
| Dhx30 | 0.00025371 | 1.12133207 | 0.304 | 0.398 | 1 | 7 | Dhx30 | myCAF_3 |
| Smarcad1 | 0.0002541 | 0.77404369 | 0.469 | 0.553 | 1 | 7 | Smarcad1 | myCAF_3 |
| Mtch2 | 0.00025496 | 0.67328071 | 0.284 | 0.397 | 1 | 7 | Mtch2 | myCAF_3 |
| Rnf1851 | 0.00025625 | 1.05966956 | 0.289 | 0.384 | 1 | 7 | Rnf185 | myCAF_3 |
| Naa802 | 0.00025821 | 0.78127579 | 0.227 | 0.352 | 1 | 7 | Naa80 | myCAF_3 |
| Asb6 | 0.0002649 | 0.4100342 | 0.402 | 0.468 | 1 | 7 | Asb6 | myCAF_3 |
| Arhgef10l1 | 0.00026493 | 0.74640041 | 0.253 | 0.338 | 1 | 7 | Arhgef10l | myCAF_3 |
| Atrn | 0.00026672 | 0.54810458 | 0.433 | 0.539 | 1 | 7 | Atrn | myCAF_3 |
| Zfp6091 | 0.00027055 | 1.0928636 | 0.371 | 0.501 | 1 | 7 | Zfp609 | myCAF_3 |
| Mapkapk52 | 0.00027238 | 0.28200146 | 0.155 | 0.258 | 1 | 7 | Mapkapk5 | myCAF_3 |
| Nlrx12 | 0.00027305 | 0.31187904 | 0.18 | 0.295 | 1 | 7 | Nlrx1 | myCAF_3 |
| Rnd32 | 0.00027566 | 0.7869891 | 0.727 | 0.831 | 1 | 7 | Rnd3 | myCAF_3 |
| Dusp161 | 0.00027634 | 0.59517285 | 0.196 | 0.299 | 1 | 7 | Dusp16 | myCAF_3 |
| Smim26 | 0.00027801 | 0.60186985 | 0.16 | 0.25 | 1 | 7 | Smim26 | myCAF_3 |
| Crem | 0.00028004 | 1.15449445 | 0.273 | 0.343 | 1 | 7 | Crem | myCAF_3 |
| Rev11 | 0.0002822 | 0.66521805 | 0.17 | 0.274 | 1 | 7 | Rev1 | myCAF_3 |
| Fgfr1op1 | 0.00028374 | 0.43942015 | 0.201 | 0.289 | 1 | 7 | Fgfr1op | myCAF_3 |
| Asl | 0.00029595 | 0.64737862 | 0.196 | 0.291 | 1 | 7 | Asl | myCAF_3 |
| Cbl | 0.00030074 | 0.67483524 | 0.531 | 0.649 | 1 | 7 | Cbl | myCAF_3 |
| Agpat22 | 0.00030177 | 0.40674567 | 0.381 | 0.51 | 1 | 7 | Agpat2 | myCAF_3 |
| Pet1001 | 0.00030388 | 0.4193108 | 0.273 | 0.392 | 1 | 7 | Pet100 | myCAF_3 |
| Copg22 | 0.00030616 | 0.30303542 | 0.175 | 0.269 | 1 | 7 | Copg2 | myCAF_3 |
| Hif1an | 0.00030812 | 0.76704649 | 0.294 | 0.405 | 1 | 7 | Hif1an | myCAF_3 |
| Adipor2 | 0.00030846 | 0.62283018 | 0.33 | 0.484 | 1 | 7 | Adipor2 | myCAF_3 |
| Itsn2 | 0.00031143 | 0.52139129 | 0.299 | 0.443 | 1 | 7 | Itsn2 | myCAF_3 |
| Slc25a252 | 0.0003133 | 1.08400741 | 0.263 | 0.344 | 1 | 7 | Slc25a25 | myCAF_3 |
| Trp53rka | 0.00031339 | 0.48934706 | 0.155 | 0.261 | 1 | 7 | Trp53rka | myCAF_3 |
| Mpv17l21 | 0.00032053 | 0.4036713 | 0.32 | 0.448 | 1 | 7 | Mpv17l2 | myCAF_3 |
| Nsrp11 | 0.00032266 | 0.8154272 | 0.227 | 0.362 | 1 | 7 | Nsrp1 | myCAF_3 |
| Zfp6441 | 0.0003241 | 0.51781127 | 0.268 | 0.383 | 1 | 7 | Zfp644 | myCAF_3 |
| Utp252 | 0.00032527 | 0.7224363 | 0.18 | 0.279 | 1 | 7 | Utp25 | myCAF_3 |
| Zbtb25 | 0.0003258 | 0.67233413 | 0.196 | 0.289 | 1 | 7 | Zbtb25 | myCAF_3 |
| Mindy13 | 0.00033419 | 0.74318963 | 0.433 | 0.53 | 1 | 7 | Mindy1 | myCAF_3 |
| Cd741 | 0.00033561 | 0.50784722 | 0.5 | 0.532 | 1 | 7 | Cd74 | myCAF_3 |
| Med311 | 0.00033628 | 0.39804298 | 0.227 | 0.333 | 1 | 7 | Med31 | myCAF_3 |
| Patz13 | 0.00033891 | 0.81218657 | 0.392 | 0.458 | 1 | 7 | Patz1 | myCAF_3 |
| Ap5m1 | 0.00033962 | 0.6148865 | 0.216 | 0.301 | 1 | 7 | Ap5m1 | myCAF_3 |
| Vegfb1 | 0.00034039 | 0.59534483 | 0.361 | 0.465 | 1 | 7 | Vegfb | myCAF_3 |
| Zfp681 | 0.00034108 | 0.35014049 | 0.216 | 0.32 | 1 | 7 | Zfp68 | myCAF_3 |
| Mvk1 | 0.0003415 | 0.55443903 | 0.351 | 0.446 | 1 | 7 | Mvk | myCAF_3 |
| Dvl2 | 0.00034417 | 1.08854108 | 0.392 | 0.473 | 1 | 7 | Dvl2 | myCAF_3 |
| Ccdc322 | 0.00034485 | 0.29624911 | 0.144 | 0.261 | 1 | 7 | Ccdc32 | myCAF_3 |
| Cyth32 | 0.0003455 | 0.8105836 | 0.784 | 0.864 | 1 | 7 | Cyth3 | myCAF_3 |
| Zmym51 | 0.00034838 | 0.75010421 | 0.289 | 0.398 | 1 | 7 | Zmym5 | myCAF_3 |
| Cep104 | 0.00035063 | 0.43771748 | 0.263 | 0.369 | 1 | 7 | Cep104 | myCAF_3 |
| Katna11 | 0.00035101 | 0.93032982 | 0.16 | 0.273 | 1 | 7 | Katna1 | myCAF_3 |
| Zmiz2 | 0.00035535 | 0.55356097 | 0.268 | 0.356 | 1 | 7 | Zmiz2 | myCAF_3 |
| Tmem203 | 0.00035691 | 0.30994502 | 0.345 | 0.449 | 1 | 7 | Tmem203 | myCAF_3 |
| Mms191 | 0.0003586 | 0.29431818 | 0.227 | 0.336 | 1 | 7 | Mms19 | myCAF_3 |
| Ubxn2a | 0.00036188 | 0.84262158 | 0.227 | 0.352 | 1 | 7 | Ubxn2a | myCAF_3 |
| Crim11 | 0.00036396 | 0.84220864 | 0.278 | 0.344 | 1 | 7 | Crim1 | myCAF_3 |
| Pced1a1 | 0.00036575 | 0.77544207 | 0.34 | 0.458 | 1 | 7 | Pced1a | myCAF_3 |
| Zc3hc11 | 0.00036988 | 0.73526824 | 0.16 | 0.261 | 1 | 7 | Zc3hc1 | myCAF_3 |
| Pip4k2c2 | 0.00037035 | 0.58574425 | 0.454 | 0.548 | 1 | 7 | Pip4k2c | myCAF_3 |
| Rora2 | 0.00037441 | 0.52721129 | 0.567 | 0.641 | 1 | 7 | Rora | myCAF_3 |
| DEPRECATED-I | 0.00038003 | 0.6339226 | 0.49 | 0.623 | 1 | 7 | DEPRECATED-I | myCAF_3 |
| Eif2b31 | 0.00038155 | 1.10175437 | 0.18 | 0.301 | 1 | 7 | Eif2b3 | myCAF_3 |
| Etl42 | 0.00039629 | 1.71520373 | 0.299 | 0.193 | 1 | 7 | Etl4 | myCAF_3 |
| Grwd11 | 0.00039794 | 0.77577428 | 0.237 | 0.296 | 1 | 7 | Grwd1 | myCAF_3 |
| Zfp6552 | 0.00040146 | 0.50342195 | 0.438 | 0.542 | 1 | 7 | Zfp655 | myCAF_3 |
| Snx111 | 0.00041234 | 0.32392668 | 0.356 | 0.42 | 1 | 7 | Snx11 | myCAF_3 |
| Braf1 | 0.00041397 | 0.70651303 | 0.201 | 0.296 | 1 | 7 | Braf | myCAF_3 |
| Iffo13 | 0.00041423 | 0.68659466 | 0.237 | 0.339 | 1 | 7 | Iffo1 | myCAF_3 |
| Zc3h7b | 0.00041641 | 0.76543362 | 0.582 | 0.707 | 1 | 7 | Zc3h7b | myCAF_3 |
| Zbtb171 | 0.00042258 | 0.70129403 | 0.16 | 0.264 | 1 | 7 | Zbtb17 | myCAF_3 |
| Gpsm1 | 0.00042548 | 0.78656802 | 0.392 | 0.511 | 1 | 7 | Gpsm1 | myCAF_3 |
| Mylk1 | 0.00042729 | 0.87949809 | 0.428 | 0.336 | 1 | 7 | Mylk | myCAF_3 |
| Kat8 | 0.00042894 | 0.77352948 | 0.206 | 0.309 | 1 | 7 | Kat8 | myCAF_3 |
| Habp4 | 0.00042913 | 0.63372666 | 0.407 | 0.52 | 1 | 7 | Habp4 | myCAF_3 |
| Gstz11 | 0.00042936 | 0.44897721 | 0.191 | 0.302 | 1 | 7 | Gstz1 | myCAF_3 |
| Herc1 | 0.00043858 | 0.86624997 | 0.361 | 0.502 | 1 | 7 | Herc1 | myCAF_3 |
| Pccb2 | 0.00044638 | 0.37464689 | 0.278 | 0.374 | 1 | 7 | Pccb | myCAF_3 |
| Ubn21 | 0.00044957 | 0.8563532 | 0.381 | 0.485 | 1 | 7 | Ubn2 | myCAF_3 |
| Chfr1 | 0.00045404 | 0.79661846 | 0.335 | 0.445 | 1 | 7 | Chfr | myCAF_3 |
| D11Wsu47e1 | 0.0004563 | 0.3771097 | 0.186 | 0.289 | 1 | 7 | D11Wsu47e | myCAF_3 |
| Med17 | 0.0004606 | 0.79220197 | 0.144 | 0.257 | 1 | 7 | Med17 | myCAF_3 |
| Rmnd5b1 | 0.00046916 | 0.26055324 | 0.314 | 0.407 | 1 | 7 | Rmnd5b | myCAF_3 |
| Brf11 | 0.00047106 | 0.52169321 | 0.371 | 0.485 | 1 | 7 | Brf1 | myCAF_3 |
| Ldlrap12 | 0.00047548 | 0.48087436 | 0.469 | 0.57 | 1 | 7 | Ldlrap1 | myCAF_3 |
| Xab2 | 0.0004757 | 0.77547609 | 0.206 | 0.297 | 1 | 7 | Xab2 | myCAF_3 |
| Dennd6a1 | 0.00047699 | 0.62836089 | 0.32 | 0.412 | 1 | 7 | Dennd6a | myCAF_3 |
| Nelfcd1 | 0.00047808 | 1.50768781 | 0.242 | 0.337 | 1 | 7 | Nelfcd | myCAF_3 |
| Tmem1861 | 0.00047823 | 0.8376624 | 0.18 | 0.3 | 1 | 7 | Tmem186 | myCAF_3 |
| Rasa31 | 0.00047934 | 1.60936089 | 0.722 | 0.67 | 1 | 7 | Rasa3 | myCAF_3 |
| Chd8 | 0.00047999 | 0.88308455 | 0.464 | 0.559 | 1 | 7 | Chd8 | myCAF_3 |
| Psen23 | 0.00048182 | 0.84241663 | 0.309 | 0.415 | 1 | 7 | Psen2 | myCAF_3 |
| Fam222b | 0.00048691 | 0.60625543 | 0.201 | 0.314 | 1 | 7 | Fam222b | myCAF_3 |
| Tsr3 | 0.00048939 | 0.67871406 | 0.232 | 0.354 | 1 | 7 | Tsr3 | myCAF_3 |
| Impact1 | 0.00049019 | 0.64228579 | 0.521 | 0.602 | 1 | 7 | Impact | myCAF_3 |
| Wdr3 | 0.00049559 | 0.74026533 | 0.186 | 0.299 | 1 | 7 | Wdr3 | myCAF_3 |
| Rab6b2 | 0.00049608 | 0.44090461 | 0.186 | 0.276 | 1 | 7 | Rab6b | myCAF_3 |
| Adcy41 | 0.00049622 | 2.05375346 | 0.381 | 0.281 | 1 | 7 | Adcy4 | myCAF_3 |
| Pyroxd11 | 0.00049753 | 0.26086616 | 0.165 | 0.281 | 1 | 7 | Pyroxd1 | myCAF_3 |
| Rcan31 | 0.00049847 | 0.44091176 | 0.273 | 0.35 | 1 | 7 | Rcan3 | myCAF_3 |
| Pde5a2 | 0.00049954 | 0.5283798 | 0.423 | 0.462 | 1 | 7 | Pde5a | myCAF_3 |
| Hdac8 | 0.000501 | 0.503003 | 0.134 | 0.256 | 1 | 7 | Hdac8 | myCAF_3 |
| Snapin1 | 0.00050184 | 0.83061572 | 0.459 | 0.509 | 1 | 7 | Snapin | myCAF_3 |
| Fgl24 | 0.00051233 | 1.639157 | 0.485 | 0.595 | 1 | 7 | Fgl2 | myCAF_3 |
| Tjap11 | 0.00051973 | 0.67926192 | 0.526 | 0.638 | 1 | 7 | Tjap1 | myCAF_3 |
| Ctdspl1 | 0.00051994 | 2.03314102 | 0.371 | 0.28 | 1 | 7 | Ctdspl | myCAF_3 |
| Tbc1d10a1 | 0.000523 | 0.66396639 | 0.5 | 0.615 | 1 | 7 | Tbc1d10a | myCAF_3 |
| Vac141 | 0.00052578 | 0.90744385 | 0.284 | 0.358 | 1 | 7 | Vac14 | myCAF_3 |
| Guf11 | 0.00052856 | 0.2869873 | 0.165 | 0.265 | 1 | 7 | Guf1 | myCAF_3 |
| Zyg11b | 0.00053388 | 0.48410253 | 0.273 | 0.374 | 1 | 7 | Zyg11b | myCAF_3 |
| Smg9 | 0.00053424 | 0.70497073 | 0.258 | 0.377 | 1 | 7 | Smg9 | myCAF_3 |
| Ftsj11 | 0.00053577 | 0.73257986 | 0.361 | 0.454 | 1 | 7 | Ftsj1 | myCAF_3 |
| Zfp3191 | 0.00054171 | 0.63511373 | 0.216 | 0.324 | 1 | 7 | Zfp319 | myCAF_3 |
| Polm1 | 0.00055064 | 0.42745649 | 0.165 | 0.267 | 1 | 7 | Polm | myCAF_3 |
| Golga41 | 0.00055226 | 0.54158637 | 0.474 | 0.615 | 1 | 7 | Golga4 | myCAF_3 |
| Stat22 | 0.00055542 | 0.5822856 | 0.216 | 0.285 | 1 | 7 | Stat2 | myCAF_3 |
| Alg12 | 0.00055684 | 0.37808505 | 0.268 | 0.335 | 1 | 7 | Alg1 | myCAF_3 |
| Bhlhb91 | 0.00055901 | 0.27018789 | 0.247 | 0.315 | 1 | 7 | Bhlhb9 | myCAF_3 |
| Taf41 | 0.00056115 | 1.19817016 | 0.263 | 0.376 | 1 | 7 | Taf4 | myCAF_3 |
| Blvra2 | 0.00056412 | 0.3271905 | 0.258 | 0.35 | 1 | 7 | Blvra | myCAF_3 |
| Metrn1 | 0.00056547 | 0.46927282 | 0.191 | 0.272 | 1 | 7 | Metrn | myCAF_3 |
| Flywch11 | 0.00057325 | 1.04164216 | 0.443 | 0.543 | 1 | 7 | Flywch1 | myCAF_3 |
| Ptrh11 | 0.00057363 | 0.35797052 | 0.469 | 0.53 | 1 | 7 | Ptrh1 | myCAF_3 |
| Mavs | 0.00058432 | 0.38998424 | 0.196 | 0.282 | 1 | 7 | Mavs | myCAF_3 |
| Klc11 | 0.00058489 | 0.98465715 | 0.613 | 0.676 | 1 | 7 | Klc1 | myCAF_3 |
| Fbxw92 | 0.00058699 | 1.10046299 | 0.175 | 0.26 | 1 | 7 | Fbxw9 | myCAF_3 |
| Trp53inp13 | 0.00058866 | 1.76886232 | 0.34 | 0.272 | 1 | 7 | Trp53inp1 | myCAF_3 |
| Vps37b1 | 0.00060203 | 0.48714385 | 0.237 | 0.332 | 1 | 7 | Vps37b | myCAF_3 |
| Dop1a | 0.00060541 | 0.54456228 | 0.216 | 0.316 | 1 | 7 | Dop1a | myCAF_3 |
| Il1b1 | 0.00060788 | 0.97316993 | 0.294 | 0.387 | 1 | 7 | Il1b | myCAF_3 |
| Idh21 | 0.00061035 | 0.80392629 | 0.747 | 0.749 | 1 | 7 | Idh2 | myCAF_3 |
| Lsg11 | 0.00062092 | 0.65007408 | 0.289 | 0.375 | 1 | 7 | Lsg1 | myCAF_3 |
| Fbxo342 | 0.00062097 | 0.45707083 | 0.165 | 0.273 | 1 | 7 | Fbxo34 | myCAF_3 |
| Ecsit1 | 0.00062697 | 0.52025747 | 0.201 | 0.31 | 1 | 7 | Ecsit | myCAF_3 |
| Fbxo61 | 0.00062809 | 0.63146027 | 0.418 | 0.531 | 1 | 7 | Fbxo6 | myCAF_3 |
| Zfp280d2 | 0.00063 | 0.95347488 | 0.387 | 0.478 | 1 | 7 | Zfp280d | myCAF_3 |
| Ankra2 | 0.00063088 | 1.04645133 | 0.196 | 0.279 | 1 | 7 | Ankra2 | myCAF_3 |
| Dmap12 | 0.00063234 | 0.29287027 | 0.16 | 0.255 | 1 | 7 | Dmap1 | myCAF_3 |
| Syne23 | 0.00063437 | 0.88188465 | 0.191 | 0.255 | 1 | 7 | Syne2 | myCAF_3 |
| Snapc41 | 0.00063709 | 0.53612056 | 0.155 | 0.266 | 1 | 7 | Snapc4 | myCAF_3 |
| Cnnm41 | 0.00063906 | 0.69441413 | 0.299 | 0.382 | 1 | 7 | Cnnm4 | myCAF_3 |
| Zfp2511 | 0.00064745 | 0.35298189 | 0.18 | 0.289 | 1 | 7 | Zfp251 | myCAF_3 |
| Tjp22 | 0.00064822 | 0.49743382 | 0.18 | 0.282 | 1 | 7 | Tjp2 | myCAF_3 |
| Rbm12b21 | 0.00065913 | 0.52045993 | 0.206 | 0.307 | 1 | 7 | Rbm12b2 | myCAF_3 |
| Ndfip21 | 0.0006603 | 0.38050532 | 0.335 | 0.425 | 1 | 7 | Ndfip2 | myCAF_3 |
| Cep120 | 0.00066077 | 0.54729294 | 0.351 | 0.471 | 1 | 7 | Cep120 | myCAF_3 |
| Tubgcp61 | 0.00066213 | 0.6667997 | 0.16 | 0.268 | 1 | 7 | Tubgcp6 | myCAF_3 |
| Cdadc11 | 0.00066852 | 0.57274109 | 0.314 | 0.417 | 1 | 7 | Cdadc1 | myCAF_3 |
| Mier2 | 0.0006709 | 0.80444325 | 0.268 | 0.373 | 1 | 7 | Mier2 | myCAF_3 |
| Cog7 | 0.00067311 | 0.7264515 | 0.232 | 0.344 | 1 | 7 | Cog7 | myCAF_3 |
| Prpf39 | 0.00067612 | 0.7170769 | 0.557 | 0.612 | 1 | 7 | Prpf39 | myCAF_3 |
| Pcdhga9 | 0.00067645 | 0.49260296 | 0.186 | 0.258 | 1 | 7 | Pcdhga9 | myCAF_3 |
| Hdac52 | 0.0006845 | 1.10901594 | 0.644 | 0.683 | 1 | 7 | Hdac5 | myCAF_3 |
| Zc3h13 | 0.00068753 | 0.85980479 | 0.268 | 0.375 | 1 | 7 | Zc3h13 | myCAF_3 |
| Cc2d2a3 | 0.00068966 | 0.70453833 | 0.155 | 0.266 | 1 | 7 | Cc2d2a | myCAF_3 |
| Baz2a | 0.00069367 | 0.71096841 | 0.397 | 0.53 | 1 | 7 | Baz2a | myCAF_3 |
| Bptf | 0.00070181 | 0.9983942 | 0.613 | 0.681 | 1 | 7 | Bptf | myCAF_3 |
| Relb1 | 0.00071463 | 0.74304918 | 0.423 | 0.51 | 1 | 7 | Relb | myCAF_3 |
| Ccdc115 | 0.00071494 | 0.6189364 | 0.196 | 0.283 | 1 | 7 | Ccdc115 | myCAF_3 |
| Zgpat | 0.00071504 | 0.6945616 | 0.366 | 0.471 | 1 | 7 | Zgpat | myCAF_3 |
| Mgat51 | 0.00071592 | 0.77213988 | 0.299 | 0.396 | 1 | 7 | Mgat5 | myCAF_3 |
| Fhod13 | 0.00071641 | 1.50402316 | 0.366 | 0.285 | 1 | 7 | Fhod1 | myCAF_3 |
| Axl21 | 0.00072057 | 0.53861912 | 0.33 | 0.409 | 1 | 7 | Axl2 | myCAF_3 |
| Pmp223 | 0.00073019 | 0.89147297 | 0.753 | 0.73 | 1 | 7 | Pmp22 | myCAF_3 |
| Htra12 | 0.0007319 | 0.49146498 | 0.758 | 0.829 | 1 | 7 | Htra1 | myCAF_3 |
| Rxra1 | 0.00073263 | 0.83487183 | 0.557 | 0.662 | 1 | 7 | Rxra | myCAF_3 |
| Sde21 | 0.00073776 | 0.65039039 | 0.402 | 0.492 | 1 | 7 | Sde2 | myCAF_3 |
| Ralgds3 | 0.00074296 | 1.25527969 | 0.428 | 0.488 | 1 | 7 | Ralgds | myCAF_3 |
| Zfp62 | 0.00075002 | 0.36549668 | 0.227 | 0.334 | 1 | 7 | Zfp62 | myCAF_3 |
| Dock62 | 0.00075176 | 0.39562619 | 0.227 | 0.316 | 1 | 7 | Dock6 | myCAF_3 |
| Hbs1l | 0.00075751 | 1.35349594 | 0.247 | 0.351 | 1 | 7 | Hbs1l | myCAF_3 |
| Dtnb1 | 0.00076025 | 0.37346704 | 0.278 | 0.388 | 1 | 7 | Dtnb | myCAF_3 |
| Mcoln11 | 0.00077353 | 0.75422887 | 0.356 | 0.464 | 1 | 7 | Mcoln1 | myCAF_3 |
| Mgat33 | 0.00077641 | 0.41240747 | 0.34 | 0.429 | 1 | 7 | Mgat3 | myCAF_3 |
| Hps1 | 0.00078202 | 1.04583142 | 0.345 | 0.406 | 1 | 7 | Hps1 | myCAF_3 |
| Utp15 | 0.00078469 | 1.03828382 | 0.227 | 0.308 | 1 | 7 | Utp15 | myCAF_3 |
| Zfp2172 | 0.00078513 | 0.73398469 | 0.546 | 0.634 | 1 | 7 | Zfp217 | myCAF_3 |
| Inafm21 | 0.00078602 | 0.52055598 | 0.247 | 0.336 | 1 | 7 | Inafm2 | myCAF_3 |
| Mettl3 | 0.00078752 | 0.30566269 | 0.278 | 0.368 | 1 | 7 | Mettl3 | myCAF_3 |
| Lsm101 | 0.00079058 | 0.38699639 | 0.284 | 0.409 | 1 | 7 | Lsm10 | myCAF_3 |
| Fem1a | 0.00079574 | 0.89228038 | 0.206 | 0.292 | 1 | 7 | Fem1a | myCAF_3 |
| Vps26a1 | 0.00079697 | 0.52332151 | 0.412 | 0.529 | 1 | 7 | Vps26a | myCAF_3 |
| Spsb21 | 0.00079869 | 0.86277116 | 0.247 | 0.341 | 1 | 7 | Spsb2 | myCAF_3 |
| Golph3l2 | 0.00080707 | 0.85001327 | 0.289 | 0.371 | 1 | 7 | Golph3l | myCAF_3 |
| Camk11 | 0.00081 | 0.73119893 | 0.572 | 0.645 | 1 | 7 | Camk1 | myCAF_3 |
| Krr11 | 0.00081525 | 0.85704554 | 0.237 | 0.327 | 1 | 7 | Krr1 | myCAF_3 |
| Vps112 | 0.0008186 | 0.78482894 | 0.407 | 0.475 | 1 | 7 | Vps11 | myCAF_3 |
| Clcn7 | 0.00084186 | 0.50237878 | 0.381 | 0.504 | 1 | 7 | Clcn7 | myCAF_3 |
| Ascc11 | 0.00084363 | 0.73473121 | 0.206 | 0.305 | 1 | 7 | Ascc1 | myCAF_3 |
| Ppm1f | 0.00084741 | 0.41891433 | 0.268 | 0.364 | 1 | 7 | Ppm1f | myCAF_3 |
| Klhl181 | 0.00084773 | 0.55075028 | 0.16 | 0.264 | 1 | 7 | Klhl18 | myCAF_3 |
| Ccs1 | 0.00087048 | 0.43928965 | 0.278 | 0.391 | 1 | 7 | Ccs | myCAF_3 |
| Zfp6641 | 0.00087137 | 0.80586472 | 0.345 | 0.436 | 1 | 7 | Zfp664 | myCAF_3 |
| Ubl7 | 0.00087288 | 0.53875608 | 0.552 | 0.679 | 1 | 7 | Ubl7 | myCAF_3 |
| Sgk12 | 0.00087569 | 1.31478346 | 0.747 | 0.685 | 1 | 7 | Sgk1 | myCAF_3 |
| Akap101 | 0.00087931 | 1.06123663 | 0.284 | 0.387 | 1 | 7 | Akap10 | myCAF_3 |
| Dop1b3 | 0.00087992 | 0.91098799 | 0.175 | 0.278 | 1 | 7 | Dop1b | myCAF_3 |
| Ttc331 | 0.00088188 | 0.73127876 | 0.32 | 0.409 | 1 | 7 | Ttc33 | myCAF_3 |
| Btf3l42 | 0.00088324 | 0.75718928 | 0.211 | 0.316 | 1 | 7 | Btf3l4 | myCAF_3 |
| Zfp8301 | 0.00089545 | 0.26129015 | 0.196 | 0.29 | 1 | 7 | Zfp830 | myCAF_3 |
| Boc1 | 0.00091021 | 1.64989989 | 0.459 | 0.426 | 1 | 7 | Boc | myCAF_3 |
| Tmem181a | 0.00091074 | 0.88500568 | 0.32 | 0.439 | 1 | 7 | Tmem181a | myCAF_3 |
| Tapt11 | 0.00091508 | 0.55339642 | 0.258 | 0.398 | 1 | 7 | Tapt1 | myCAF_3 |
| Ccdc911 | 0.00092535 | 0.37347255 | 0.309 | 0.436 | 1 | 7 | Ccdc91 | myCAF_3 |
| Phlpp21 | 0.00092726 | 0.49722227 | 0.155 | 0.25 | 1 | 7 | Phlpp2 | myCAF_3 |
| Ninj12 | 0.00093009 | 0.71359537 | 0.582 | 0.623 | 1 | 7 | Ninj1 | myCAF_3 |
| Chst151 | 0.00093204 | 1.41158332 | 0.515 | 0.48 | 1 | 7 | Chst15 | myCAF_3 |
| Cdkn2aip | 0.00093432 | 0.8990161 | 0.216 | 0.315 | 1 | 7 | Cdkn2aip | myCAF_3 |
| Stk19 | 0.0009362 | 0.62987828 | 0.186 | 0.277 | 1 | 7 | Stk19 | myCAF_3 |
| Zfp282 | 0.00093636 | 1.20989145 | 0.18 | 0.296 | 1 | 7 | Zfp282 | myCAF_3 |
| Nsun51 | 0.00095418 | 0.45173982 | 0.18 | 0.275 | 1 | 7 | Nsun5 | myCAF_3 |
| Mtpap | 0.00096308 | 0.65544201 | 0.314 | 0.398 | 1 | 7 | Mtpap | myCAF_3 |
| Bloc1s52 | 0.00096622 | 0.56531758 | 0.144 | 0.251 | 1 | 7 | Bloc1s5 | myCAF_3 |
| Cflar1 | 0.00097017 | 0.62733908 | 0.49 | 0.618 | 1 | 7 | Cflar | myCAF_3 |
| Zfp407 | 0.00097215 | 0.51254467 | 0.232 | 0.335 | 1 | 7 | Zfp407 | myCAF_3 |
| Pcsk92 | 0.00098034 | 0.42440026 | 0.294 | 0.354 | 1 | 7 | Pcsk9 | myCAF_3 |
| Yod1 | 0.00098984 | 1.2559179 | 0.191 | 0.252 | 1 | 7 | Yod1 | myCAF_3 |
| Nrf11 | 0.00100791 | 1.26324812 | 0.227 | 0.3 | 1 | 7 | Nrf1 | myCAF_3 |
| Stim12 | 0.00101904 | 0.48511791 | 0.376 | 0.441 | 1 | 7 | Stim1 | myCAF_3 |
| Etfbkmt2 | 0.00102093 | 0.83063589 | 0.196 | 0.251 | 1 | 7 | Etfbkmt | myCAF_3 |
| Mon1a1 | 0.00102164 | 1.14788808 | 0.186 | 0.274 | 1 | 7 | Mon1a | myCAF_3 |
| Spsb3 | 0.00102249 | 0.88683049 | 0.459 | 0.548 | 1 | 7 | Spsb3 | myCAF_3 |
| Tmem223 | 0.00102391 | 0.60798441 | 0.33 | 0.422 | 1 | 7 | Tmem223 | myCAF_3 |
| Avpr1a1 | 0.00102534 | 1.12699529 | 0.588 | 0.644 | 1 | 7 | Avpr1a | myCAF_3 |
| Rb1cc1 | 0.00102653 | 0.97394344 | 0.443 | 0.534 | 1 | 7 | Rb1cc1 | myCAF_3 |
| Cacng71 | 0.00104148 | 1.06636292 | 0.216 | 0.25 | 1 | 7 | Cacng7 | myCAF_3 |
| Bicral1 | 0.00104324 | 0.67801211 | 0.253 | 0.341 | 1 | 7 | Bicral | myCAF_3 |
| Capn15 | 0.00104457 | 0.48136504 | 0.263 | 0.379 | 1 | 7 | Capn15 | myCAF_3 |
| Tceanc2 | 0.00104954 | 0.7102106 | 0.206 | 0.314 | 1 | 7 | Tceanc2 | myCAF_3 |
| Pigc | 0.00105006 | 0.91922075 | 0.464 | 0.538 | 1 | 7 | Pigc | myCAF_3 |
| Trim391 | 0.00105332 | 1.14439841 | 0.175 | 0.275 | 1 | 7 | Trim39 | myCAF_3 |
| Mphosph101 | 0.00108292 | 0.37349268 | 0.258 | 0.32 | 1 | 7 | Mphosph10 | myCAF_3 |
| Map3k10 | 0.00108956 | 0.36168285 | 0.289 | 0.405 | 1 | 7 | Map3k10 | myCAF_3 |
| Tent4b1 | 0.00109424 | 0.40604584 | 0.17 | 0.258 | 1 | 7 | Tent4b | myCAF_3 |
| Tspan141 | 0.00110266 | 0.78388458 | 0.294 | 0.406 | 1 | 7 | Tspan14 | myCAF_3 |
| Crtc2 | 0.00110761 | 0.82811353 | 0.454 | 0.511 | 1 | 7 | Crtc2 | myCAF_3 |
| E2f61 | 0.00111208 | 1.32388883 | 0.263 | 0.326 | 1 | 7 | E2f6 | myCAF_3 |
| Crip22 | 0.00112017 | 1.58917699 | 0.325 | 0.207 | 1 | 7 | Crip2 | myCAF_3 |
| Caap11 | 0.00112694 | 0.81371709 | 0.196 | 0.291 | 1 | 7 | Caap1 | myCAF_3 |
| Afdn2 | 0.00112771 | 0.63484255 | 0.289 | 0.37 | 1 | 7 | Afdn | myCAF_3 |
| Fam133b2 | 0.00113589 | 0.36724083 | 0.165 | 0.266 | 1 | 7 | Fam133b | myCAF_3 |
| Mtf21 | 0.00114072 | 0.5438926 | 0.175 | 0.284 | 1 | 7 | Mtf2 | myCAF_3 |
| Utp181 | 0.00114374 | 0.55875275 | 0.263 | 0.35 | 1 | 7 | Utp18 | myCAF_3 |
| Cactin1 | 0.00114509 | 0.27254869 | 0.175 | 0.273 | 1 | 7 | Cactin | myCAF_3 |
| Lrfn42 | 0.0011874 | 0.82630705 | 0.33 | 0.456 | 1 | 7 | Lrfn4 | myCAF_3 |
| Stx1a1 | 0.00119051 | 0.42041691 | 0.191 | 0.281 | 1 | 7 | Stx1a | myCAF_3 |
| Dcaf151 | 0.00119063 | 0.59779506 | 0.351 | 0.42 | 1 | 7 | Dcaf15 | myCAF_3 |
| Kansl21 | 0.00119983 | 0.7497132 | 0.686 | 0.81 | 1 | 7 | Kansl2 | myCAF_3 |
| Slc25a372 | 0.00120543 | 0.52454364 | 0.253 | 0.407 | 1 | 7 | Slc25a37 | myCAF_3 |

| | | | | | | | | |
|---|---|---|---|---|---|---|---|---|
| Eif2ak42 | 0.00120651 | 0.81241289 | 0.335 | 0.422 | 1 | 7 | Eif2ak4 | myCAF_3 |
| Sdc31 | 0.0012086 | 1.50155772 | 0.557 | 0.428 | 1 | 7 | Sdc3 | myCAF_3 |
| Bmyc1 | 0.00120937 | 1.44269664 | 0.263 | 0.178 | 1 | 7 | Bmyc | myCAF_3 |
| Atat11 | 0.00122498 | 0.83290795 | 0.155 | 0.252 | 1 | 7 | Atat1 | myCAF_3 |
| Map4k32 | 0.00122784 | 0.89771444 | 0.433 | 0.574 | 1 | 7 | Map4k3 | myCAF_3 |
| Gnl21 | 0.00122968 | 1.89663038 | 0.34 | 0.4 | 1 | 7 | Gnl2 | myCAF_3 |
| Phactr21 | 0.00123109 | 0.67016619 | 0.268 | 0.381 | 1 | 7 | Phactr2 | myCAF_3 |
| Mbd61 | 0.00125343 | 0.78796557 | 0.186 | 0.275 | 1 | 7 | Mbd6 | myCAF_3 |
| Dhx8 | 0.00125976 | 0.35227775 | 0.381 | 0.497 | 1 | 7 | Dhx8 | myCAF_3 |
| Ciapin11 | 0.00126815 | 0.38239044 | 0.304 | 0.392 | 1 | 7 | Ciapin1 | myCAF_3 |
| Angel2 | 0.0012687 | 0.60107673 | 0.211 | 0.319 | 1 | 7 | Angel2 | myCAF_3 |
| Cd99l22 | 0.00127047 | 0.51338562 | 0.361 | 0.492 | 1 | 7 | Cd99l2 | myCAF_3 |
| Gtpbp3 | 0.00127195 | 0.98803355 | 0.216 | 0.331 | 1 | 7 | Gtpbp3 | myCAF_3 |
| Pan22 | 0.00127583 | 0.59898067 | 0.242 | 0.317 | 1 | 7 | Pan2 | myCAF_3 |
| Uap1l12 | 0.00128212 | 1.08765342 | 0.443 | 0.476 | 1 | 7 | Uap1l1 | myCAF_3 |
| Kif3b1 | 0.00130523 | 0.33487604 | 0.32 | 0.456 | 1 | 7 | Kif3b | myCAF_3 |
| Nt5dc1 | 0.00133462 | 1.03289869 | 0.309 | 0.381 | 1 | 7 | Nt5dc1 | myCAF_3 |
| Zfyve11 | 0.00134957 | 0.79356275 | 0.433 | 0.561 | 1 | 7 | Zfyve1 | myCAF_3 |
| Pias41 | 0.00135456 | 0.55771764 | 0.18 | 0.28 | 1 | 7 | Pias4 | myCAF_3 |
| Tspyl11 | 0.00138026 | 0.82853273 | 0.557 | 0.635 | 1 | 7 | Tspyl1 | myCAF_3 |
| Arfgef21 | 0.00144236 | 0.90381386 | 0.237 | 0.333 | 1 | 7 | Arfgef2 | myCAF_3 |
| Dnajc24 | 0.00146933 | 0.85144365 | 0.258 | 0.353 | 1 | 7 | Dnajc24 | myCAF_3 |
| Zzef1 | 0.00146999 | 0.54614449 | 0.356 | 0.467 | 1 | 7 | Zzef1 | myCAF_3 |
| Pcnx3 | 0.00149727 | 0.75270246 | 0.443 | 0.55 | 1 | 7 | Pcnx3 | myCAF_3 |
| Ngrn1 | 0.00150087 | 0.47007646 | 0.175 | 0.253 | 1 | 7 | Ngrn | myCAF_3 |
| Wipi21 | 0.00151189 | 0.64832305 | 0.526 | 0.605 | 1 | 7 | Wipi2 | myCAF_3 |
| Pi4k2a1 | 0.00152337 | 1.1940658 | 0.495 | 0.575 | 1 | 7 | Pi4k2a | myCAF_3 |
| Fat12 | 0.0015537 | 0.61986097 | 0.758 | 0.873 | 1 | 7 | Fat1 | myCAF_3 |
| Vangl22 | 0.00155654 | 0.35953951 | 0.304 | 0.378 | 1 | 7 | Vangl2 | myCAF_3 |
| Zmat51 | 0.00155766 | 0.26567601 | 0.191 | 0.28 | 1 | 7 | Zmat5 | myCAF_3 |
| Wrn2 | 0.00156817 | 1.00559056 | 0.206 | 0.289 | 1 | 7 | Wrn | myCAF_3 |
| Plekho11 | 0.00157621 | 0.82002658 | 0.423 | 0.51 | 1 | 7 | Plekho1 | myCAF_3 |
| Coasy | 0.0015777 | 0.65713052 | 0.242 | 0.314 | 1 | 7 | Coasy | myCAF_3 |
| Pogk1 | 0.00158326 | 0.67600379 | 0.201 | 0.298 | 1 | 7 | Pogk | myCAF_3 |
| Agk | 0.00159428 | 0.2944053 | 0.232 | 0.323 | 1 | 7 | Agk | myCAF_3 |
| Gmeb21 | 0.0015943 | 0.82223169 | 0.258 | 0.346 | 1 | 7 | Gmeb2 | myCAF_3 |
| Smim19 | 0.00159686 | 0.61257532 | 0.34 | 0.432 | 1 | 7 | Smim19 | myCAF_3 |
| Mlycd1 | 0.0016009 | 0.34097399 | 0.237 | 0.32 | 1 | 7 | Mlycd | myCAF_3 |
| Lrrc201 | 0.00160841 | 0.8230009 | 0.165 | 0.26 | 1 | 7 | Lrrc20 | myCAF_3 |
| Nedd92 | 0.00161395 | 0.62406333 | 0.294 | 0.379 | 1 | 7 | Nedd9 | myCAF_3 |
| Slc35e22 | 0.00164893 | 0.77411947 | 0.191 | 0.265 | 1 | 7 | Slc35e2 | myCAF_3 |
| Dnajc16 | 0.00165188 | 0.56768986 | 0.222 | 0.307 | 1 | 7 | Dnajc16 | myCAF_3 |
| Lman2l1 | 0.00173994 | 0.61921008 | 0.242 | 0.332 | 1 | 7 | Lman2l | myCAF_3 |
| Bdp1 | 0.00175072 | 0.69801198 | 0.253 | 0.32 | 1 | 7 | Bdp1 | myCAF_3 |
| Slc4a1ap | 0.00175766 | 0.98798129 | 0.263 | 0.335 | 1 | 7 | Slc4a1ap | myCAF_3 |
| Pfn22 | 0.00176395 | 0.64951115 | 0.304 | 0.342 | 1 | 7 | Pfn2 | myCAF_3 |
| Ilkap | 0.00177093 | 0.5266084 | 0.247 | 0.336 | 1 | 7 | Ilkap | myCAF_3 |
| Fgf21 | 0.00177355 | 0.97025315 | 0.428 | 0.466 | 1 | 7 | Fgf2 | myCAF_3 |
| Xylt2 | 0.0017738 | 0.68142114 | 0.289 | 0.385 | 1 | 7 | Xylt2 | myCAF_3 |
| Prg43 | 0.00180126 | 0.50192425 | 0.052 | 0.324 | 1 | 7 | Prg4 | myCAF_3 |
| Dlg31 | 0.00180798 | 0.72204197 | 0.253 | 0.365 | 1 | 7 | Dlg3 | myCAF_3 |
| Kctd181 | 0.00181301 | 0.26324302 | 0.17 | 0.252 | 1 | 7 | Kctd18 | myCAF_3 |
| Mlip1 | 0.00185127 | 0.40761772 | 0.232 | 0.288 | 1 | 7 | Mlip | myCAF_3 |
| Pnkp1 | 0.0018628 | 0.59423617 | 0.258 | 0.322 | 1 | 7 | Pnkp | myCAF_3 |
| Spry43 | 0.00186646 | 0.40497417 | 0.351 | 0.434 | 1 | 7 | Spry4 | myCAF_3 |
| Cdc42bpg1 | 0.00188534 | 1.30819481 | 0.278 | 0.22 | 1 | 7 | Cdc42bpg | myCAF_3 |
| Slc22a171 | 0.00189126 | 0.4123467 | 0.284 | 0.404 | 1 | 7 | Slc22a17 | myCAF_3 |
| Tmem192 | 0.00191196 | 0.44070022 | 0.17 | 0.277 | 1 | 7 | Tmem192 | myCAF_3 |
| Trim26 | 0.00192739 | 0.83064336 | 0.258 | 0.337 | 1 | 7 | Trim2 | myCAF_3 |
| Sirt12 | 0.00193085 | 1.5421187 | 0.237 | 0.301 | 1 | 7 | Sirt1 | myCAF_3 |
| Ccnt11 | 0.00193208 | 0.57547957 | 0.34 | 0.443 | 1 | 7 | Ccnt1 | myCAF_3 |
| Traf62 | 0.00196323 | 0.43018681 | 0.253 | 0.342 | 1 | 7 | Traf6 | myCAF_3 |
| Vps26c1 | 0.00196368 | 0.34229318 | 0.387 | 0.496 | 1 | 7 | Vps26c | myCAF_3 |
| Atp13a21 | 0.00196641 | 0.6282086 | 0.206 | 0.273 | 1 | 7 | Atp13a2 | myCAF_3 |
| Zfp5231 | 0.00196743 | 0.60634267 | 0.34 | 0.447 | 1 | 7 | Zfp523 | myCAF_3 |
| Sipa1l22 | 0.00196901 | 0.68860893 | 0.247 | 0.326 | 1 | 7 | Sipa1l2 | myCAF_3 |
| Ublcp1 | 0.00198091 | 0.32199858 | 0.216 | 0.303 | 1 | 7 | Ublcp1 | myCAF_3 |
| Itpr22 | 0.0019965 | 0.57220333 | 0.263 | 0.357 | 1 | 7 | Itpr2 | myCAF_3 |
| Cnst | 0.00200627 | 0.47935217 | 0.222 | 0.307 | 1 | 7 | Cnst | myCAF_3 |
| Commd11 | 0.00201721 | 0.46817166 | 0.165 | 0.252 | 1 | 7 | Commd1 | myCAF_3 |
| Sco21 | 0.00203519 | 0.55666757 | 0.237 | 0.322 | 1 | 7 | Sco2 | myCAF_3 |
| Abhd14a2 | 0.00204347 | 0.36691834 | 0.309 | 0.384 | 1 | 7 | Abhd14a | myCAF_3 |
| Ndst1 | 0.00206683 | 0.89870922 | 0.737 | 0.812 | 1 | 7 | Ndst1 | myCAF_3 |
| DEPRECATED-I | 0.00207231 | 0.45556173 | 0.649 | 0.711 | 1 | 7 | DEPRECATED-I | myCAF_3 |
| Cspp11 | 0.00207385 | 0.51409288 | 0.247 | 0.313 | 1 | 7 | Cspp1 | myCAF_3 |
| Kat6b1 | 0.00213257 | 1.05977358 | 0.418 | 0.512 | 1 | 7 | Kat6b | myCAF_3 |
| Cyhr11 | 0.0021515 | 0.76596992 | 0.443 | 0.523 | 1 | 7 | Cyhr1 | myCAF_3 |
| Cldn12 | 0.00219265 | 0.57760956 | 0.289 | 0.378 | 1 | 7 | Cldn12 | myCAF_3 |
| Fbxo173 | 0.00219553 | 0.39839178 | 0.351 | 0.423 | 1 | 7 | Fbxo17 | myCAF_3 |
| Siah2 | 0.00220586 | 0.54610153 | 0.253 | 0.346 | 1 | 7 | Siah2 | myCAF_3 |
| Crbn2 | 0.00221208 | 0.27178316 | 0.263 | 0.37 | 1 | 7 | Crbn | myCAF_3 |
| Washc51 | 0.00225838 | 0.66073256 | 0.351 | 0.435 | 1 | 7 | Washc5 | myCAF_3 |
| Fcho21 | 0.00229094 | 0.76393786 | 0.237 | 0.321 | 1 | 7 | Fcho2 | myCAF_3 |
| Khnyn2 | 0.00230185 | 0.91095391 | 0.34 | 0.419 | 1 | 7 | Khnyn | myCAF_3 |
| Atg2a2 | 0.00230677 | 0.86470285 | 0.407 | 0.48 | 1 | 7 | Atg2a | myCAF_3 |
| Mink11 | 0.00231199 | 1.15488979 | 0.284 | 0.363 | 1 | 7 | Mink1 | myCAF_3 |
| Dnajc18 | 0.00234118 | 0.74595229 | 0.371 | 0.443 | 1 | 7 | Dnajc18 | myCAF_3 |
| Gna142 | 0.00234199 | 0.69094692 | 0.268 | 0.316 | 1 | 7 | Gna14 | myCAF_3 |
| Iscu | 0.00235057 | 0.42982431 | 0.289 | 0.403 | 1 | 7 | Iscu | myCAF_3 |
| Sipa1l11 | 0.00235123 | 1.00828164 | 0.381 | 0.46 | 1 | 7 | Sipa1l1 | myCAF_3 |
| Ccndbp11 | 0.00235225 | 0.45710916 | 0.397 | 0.502 | 1 | 7 | Ccndbp1 | myCAF_3 |
| Asb1 | 0.00237783 | 0.27896185 | 0.191 | 0.279 | 1 | 7 | Asb1 | myCAF_3 |
| Zfp821 | 0.00241083 | 0.45042398 | 0.273 | 0.364 | 1 | 7 | Zfp821 | myCAF_3 |
| H2-Ke62 | 0.00242632 | 0.86408837 | 0.216 | 0.291 | 1 | 7 | H2-Ke6 | myCAF_3 |
| Crnkl11 | 0.00245071 | 0.76244586 | 0.335 | 0.421 | 1 | 7 | Crnkl1 | myCAF_3 |
| Calcoco12 | 0.0024528 | 1.39442987 | 0.686 | 0.609 | 1 | 7 | Calcoco1 | myCAF_3 |
| Mbtps21 | 0.00246634 | 0.34392836 | 0.247 | 0.345 | 1 | 7 | Mbtps2 | myCAF_3 |
| Ap4e11 | 0.00251477 | 0.581045 | 0.216 | 0.29 | 1 | 7 | Ap4e1 | myCAF_3 |
| Rundc1 | 0.00253856 | 0.37614762 | 0.191 | 0.296 | 1 | 7 | Rundc1 | myCAF_3 |
| Rab11fip21 | 0.00254805 | 0.83347829 | 0.345 | 0.443 | 1 | 7 | Rab11fip2 | myCAF_3 |
| Slc12a62 | 0.00256953 | 0.87683019 | 0.268 | 0.343 | 1 | 7 | Slc12a6 | myCAF_3 |
| Hars2 | 0.00259789 | 1.24795894 | 0.258 | 0.324 | 1 | 7 | Hars2 | myCAF_3 |
| Mex3d1 | 0.00266052 | 0.87763563 | 0.598 | 0.664 | 1 | 7 | Mex3d | myCAF_3 |
| Pdcd2l1 | 0.00266363 | 0.64138322 | 0.18 | 0.277 | 1 | 7 | Pdcd2l | myCAF_3 |
| Hdac71 | 0.00268888 | 1.09700725 | 0.608 | 0.671 | 1 | 7 | Hdac7 | myCAF_3 |
| Klhl121 | 0.00269532 | 0.67205621 | 0.216 | 0.325 | 1 | 7 | Klhl12 | myCAF_3 |
| Ppil41 | 0.00270324 | 0.66060039 | 0.294 | 0.399 | 1 | 7 | Ppil4 | myCAF_3 |
| Proser11 | 0.00270667 | 0.54906054 | 0.186 | 0.262 | 1 | 7 | Proser1 | myCAF_3 |
| Klc41 | 0.00273575 | 0.56755424 | 0.227 | 0.323 | 1 | 7 | Klc4 | myCAF_3 |
| Zxdb1 | 0.00274306 | 0.25646848 | 0.186 | 0.265 | 1 | 7 | Zxdb | myCAF_3 |
| Hectd31 | 0.00275951 | 0.38561769 | 0.34 | 0.451 | 1 | 7 | Hectd3 | myCAF_3 |
| Hbp13 | 0.00277605 | 0.88301083 | 0.732 | 0.801 | 1 | 7 | Hbp1 | myCAF_3 |
| Ksr13 | 0.00278266 | 1.25520033 | 0.314 | 0.379 | 1 | 7 | Ksr1 | myCAF_3 |
| Rnf261 | 0.00281802 | 0.62142775 | 0.17 | 0.251 | 1 | 7 | Rnf26 | myCAF_3 |
| Coro72 | 0.00286638 | 0.47478703 | 0.232 | 0.324 | 1 | 7 | Coro7 | myCAF_3 |
| Mkrn1 | 0.00286826 | 0.6943626 | 0.232 | 0.32 | 1 | 7 | Mkrn1 | myCAF_3 |
| Clpx | 0.00287574 | 0.60170722 | 0.284 | 0.376 | 1 | 7 | Clpx | myCAF_3 |
| Zxdc2 | 0.00289057 | 0.63374963 | 0.232 | 0.328 | 1 | 7 | Zxdc | myCAF_3 |
| Rin21 | 0.0029119 | 1.51571255 | 0.552 | 0.523 | 1 | 7 | Rin2 | myCAF_3 |
| Slc35a51 | 0.0029169 | 1.16651455 | 0.325 | 0.393 | 1 | 7 | Slc35a5 | myCAF_3 |
| Lrwd11 | 0.00292463 | 0.31482882 | 0.258 | 0.354 | 1 | 7 | Lrwd1 | myCAF_3 |
| Dcaf111 | 0.00294581 | 0.33456804 | 0.34 | 0.42 | 1 | 7 | Dcaf11 | myCAF_3 |
| Arl22 | 0.00295697 | 0.45479213 | 0.469 | 0.557 | 1 | 7 | Arl2 | myCAF_3 |
| Aplp22 | 0.00296539 | 1.06057195 | 0.804 | 0.824 | 1 | 7 | Aplp2 | myCAF_3 |
| Clcf12 | 0.00298071 | 2.16063005 | 0.304 | 0.242 | 1 | 7 | Clcf1 | myCAF_3 |
| Atp11c3 | 0.00299895 | 1.11508527 | 0.232 | 0.307 | 1 | 7 | Atp11c | myCAF_3 |
| Axin1 | 0.00308126 | 0.45185469 | 0.314 | 0.477 | 1 | 7 | Axin1 | myCAF_3 |
| Fgfrl11 | 0.00312819 | 0.59098209 | 0.268 | 0.351 | 1 | 7 | Fgfrl1 | myCAF_3 |
| Fam216a1 | 0.00313111 | 0.72259551 | 0.273 | 0.335 | 1 | 7 | Fam216a | myCAF_3 |
| Jup1 | 0.00314248 | 0.7323864 | 0.242 | 0.262 | 1 | 7 | Jup | myCAF_3 |
| Nexn | 0.0031992 | 1.57619939 | 0.253 | 0.148 | 1 | 7 | Nexn | myCAF_3 |
| Mtt2 | 0.00321289 | 0.38831402 | 0.294 | 0.369 | 1 | 7 | Mtt2 | myCAF_3 |
| Mid1ip12 | 0.00321708 | 0.8111883 | 0.613 | 0.61 | 1 | 7 | Mid1ip1 | myCAF_3 |
| Skiv2l | 0.00323481 | 0.7164729 | 0.201 | 0.292 | 1 | 7 | Skiv2l | myCAF_3 |
| Ttc11 | 0.00333213 | 0.55630871 | 0.284 | 0.365 | 1 | 7 | Ttc1 | myCAF_3 |
| Plekhh32 | 0.00334707 | 0.61901631 | 0.201 | 0.267 | 1 | 7 | Plekhh3 | myCAF_3 |
| Ttbk21 | 0.00336888 | 0.91438716 | 0.191 | 0.274 | 1 | 7 | Ttbk2 | myCAF_3 |
| Pycrl1 | 0.00339616 | 0.31281993 | 0.186 | 0.286 | 1 | 7 | Pycrl | myCAF_3 |
| Gpatch2l | 0.00340104 | 0.42114005 | 0.222 | 0.328 | 1 | 7 | Gpatch2l | myCAF_3 |
| Nr1d21 | 0.0034314 | 0.8433646 | 0.397 | 0.463 | 1 | 7 | Nr1d2 | myCAF_3 |
| Wdr83 | 0.0035026 | 0.97173042 | 0.191 | 0.27 | 1 | 7 | Wdr83 | myCAF_3 |
| Tubgcp21 | 0.00350445 | 0.6550102 | 0.258 | 0.301 | 1 | 7 | Tubgcp2 | myCAF_3 |
| Scrib1 | 0.00350513 | 0.80987299 | 0.284 | 0.356 | 1 | 7 | Scrib | myCAF_3 |
| Pcbd21 | 0.00351063 | 0.49244848 | 0.242 | 0.316 | 1 | 7 | Pcbd2 | myCAF_3 |
| Tbcel | 0.00355409 | 0.63124628 | 0.412 | 0.47 | 1 | 7 | Tbcel | myCAF_3 |
| Tgfb32 | 0.00356219 | 0.84157185 | 0.691 | 0.699 | 1 | 7 | Tgfb3 | myCAF_3 |
| Stx17 | 0.00365505 | 0.69931733 | 0.335 | 0.431 | 1 | 7 | Stx17 | myCAF_3 |
| Casp81 | 0.00372063 | 0.40565158 | 0.33 | 0.433 | 1 | 7 | Casp8 | myCAF_3 |
| Nr2c1 | 0.00375762 | 0.84085294 | 0.278 | 0.34 | 1 | 7 | Nr2c1 | myCAF_3 |
| Mon1b | 0.0037781 | 0.6392689 | 0.237 | 0.314 | 1 | 7 | Mon1b | myCAF_3 |
| Mthfsd1 | 0.0038312 | 0.9858217 | 0.278 | 0.341 | 1 | 7 | Mthfsd | myCAF_3 |
| Pcdhga111 | 0.00383946 | 1.174457 | 0.17 | 0.258 | 1 | 7 | Pcdhga11 | myCAF_3 |
| Trim231 | 0.00385045 | 0.48377677 | 0.191 | 0.269 | 1 | 7 | Trim23 | myCAF_3 |
| DEPRECATED-I | 0.00385155 | 0.56702625 | 0.222 | 0.302 | 1 | 7 | DEPRECATED-I | myCAF_3 |
| Wdr71 | 0.00385214 | 0.44041508 | 0.34 | 0.422 | 1 | 7 | Wdr7 | myCAF_3 |
| Hoxc92 | 0.00386231 | 1.39248035 | 0.289 | 0.34 | 1 | 7 | Hoxc9 | myCAF_3 |
| Gcnt42 | 0.00387912 | 0.47269025 | 0.402 | 0.371 | 1 | 7 | Gcnt4 | myCAF_3 |
| Pnpla61 | 0.00389407 | 0.51779364 | 0.201 | 0.282 | 1 | 7 | Pnpla6 | myCAF_3 |
| Runx1t12 | 0.00390395 | 1.26032593 | 0.242 | 0.296 | 1 | 7 | Runx1t1 | myCAF_3 |
| Scyl3 | 0.00406854 | 0.58388704 | 0.222 | 0.286 | 1 | 7 | Scyl3 | myCAF_3 |
| Nfrkb | 0.00406901 | 1.35961427 | 0.206 | 0.284 | 1 | 7 | Nfrkb | myCAF_3 |
| Map61 | 0.00407517 | 0.7843252 | 0.474 | 0.517 | 1 | 7 | Map6 | myCAF_3 |
| H2-D13 | 0.00408328 | 1.66883678 | 0.423 | 0.374 | 1 | 7 | H2-D1 | myCAF_3 |
| Zbtb22 | 0.00419961 | 0.99339651 | 0.371 | 0.41 | 1 | 7 | Zbtb22 | myCAF_3 |
| Tbc1d52 | 0.00421759 | 0.46615994 | 0.299 | 0.389 | 1 | 7 | Tbc1d5 | myCAF_3 |
| Tcta1 | 0.0042249 | 1.20622833 | 0.273 | 0.389 | 1 | 7 | Tcta | myCAF_3 |
| Ptk2b3 | 0.0042504 | 0.79130501 | 0.227 | 0.278 | 1 | 7 | Ptk2b | myCAF_3 |
| Nckap5l1 | 0.00425519 | 0.71885445 | 0.196 | 0.28 | 1 | 7 | Nckap5l | myCAF_3 |
| Zdhhc121 | 0.00426111 | 0.95227974 | 0.149 | 0.254 | 1 | 7 | Zdhhc12 | myCAF_3 |
| Fam118a | 0.00433827 | 0.72190183 | 0.201 | 0.265 | 1 | 7 | Fam118a | myCAF_3 |
| Phc11 | 0.00435163 | 0.66335092 | 0.227 | 0.294 | 1 | 7 | Phc1 | myCAF_3 |
| Dmac2 | 0.00438324 | 0.6743813 | 0.186 | 0.265 | 1 | 7 | Dmac2 | myCAF_3 |
| Prepl | 0.00441797 | 0.77752506 | 0.16 | 0.263 | 1 | 7 | Prepl | myCAF_3 |
| Pcdhgc51 | 0.00443158 | 0.25427639 | 0.284 | 0.372 | 1 | 7 | Pcdhgc5 | myCAF_3 |
| Patl11 | 0.00444858 | 0.94732307 | 0.469 | 0.533 | 1 | 7 | Patl1 | myCAF_3 |
| Cpeb12 | 0.00446961 | 1.54808977 | 0.263 | 0.181 | 1 | 7 | Cpeb1 | myCAF_3 |
| Hsdl22 | 0.00452553 | 0.78434176 | 0.201 | 0.279 | 1 | 7 | Hsdl2 | myCAF_3 |
| Gorab1 | 0.00459712 | 0.8848487 | 0.253 | 0.332 | 1 | 7 | Gorab | myCAF_3 |
| Gclm2 | 0.00460623 | 0.48142374 | 0.247 | 0.336 | 1 | 7 | Gclm | myCAF_3 |
| Ankrd10 | 0.00460692 | 0.8589499 | 0.397 | 0.477 | 1 | 7 | Ankrd10 | myCAF_3 |
| Map2k31 | 0.0047116 | 0.53711349 | 0.619 | 0.71 | 1 | 7 | Map2k3 | myCAF_3 |
| Prkd2 | 0.00473598 | 0.61606916 | 0.175 | 0.266 | 1 | 7 | Prkd2 | myCAF_3 |
| Hoxd83 | 0.00476859 | 0.63075082 | 0.371 | 0.407 | 1 | 7 | Hoxd8 | myCAF_3 |
| Fam53a | 0.00478795 | 0.756183 | 0.273 | 0.353 | 1 | 7 | Fam53a | myCAF_3 |
| Prickle23 | 0.0048075 | 1.59780193 | 0.412 | 0.366 | 1 | 7 | Prickle2 | myCAF_3 |
| Foxk11 | 0.00481401 | 0.56528737 | 0.433 | 0.537 | 1 | 7 | Foxk1 | myCAF_3 |
| Slc2a8 | 0.00487148 | 1.05709981 | 0.17 | 0.251 | 1 | 7 | Slc2a8 | myCAF_3 |
| B3gnt21 | 0.00487312 | 0.84890429 | 0.335 | 0.403 | 1 | 7 | B3gnt2 | myCAF_3 |
| Gpatch21 | 0.00488145 | 0.8213676 | 0.201 | 0.296 | 1 | 7 | Gpatch2 | myCAF_3 |
| Lcmt1 | 0.00488833 | 0.76598744 | 0.227 | 0.311 | 1 | 7 | Lcmt1 | myCAF_3 |
| Zscan26 | 0.00490086 | 0.74884261 | 0.5 | 0.581 | 1 | 7 | Zscan26 | myCAF_3 |
| Fnbp1l2 | 0.00494447 | 0.25441134 | 0.253 | 0.299 | 1 | 7 | Fnbp1l | myCAF_3 |
| Smpd21 | 0.00497959 | 0.88003068 | 0.232 | 0.316 | 1 | 7 | Smpd2 | myCAF_3 |
| Rab40c | 0.00501953 | 1.08066321 | 0.381 | 0.454 | 1 | 7 | Rab40c | myCAF_3 |
| Uhrf1bp1 | 0.00503511 | 0.6231664 | 0.17 | 0.253 | 1 | 7 | Uhrf1bp1 | myCAF_3 |
| Maip11 | 0.00505236 | 0.37559869 | 0.186 | 0.272 | 1 | 7 | Maip1 | myCAF_3 |
| Rnf1252 | 0.00507478 | 1.90793718 | 0.33 | 0.32 | 1 | 7 | Rnf125 | myCAF_3 |
| Ppfia1 | 0.00510268 | 0.77130626 | 0.582 | 0.723 | 1 | 7 | Ppfia1 | myCAF_3 |
| Cc2d1b1 | 0.00514924 | 0.70836664 | 0.191 | 0.273 | 1 | 7 | Cc2d1b | myCAF_3 |
| Surf21 | 0.00518732 | 0.45090434 | 0.175 | 0.251 | 1 | 7 | Surf2 | myCAF_3 |
| Ppp6r2 | 0.00524647 | 0.68231952 | 0.381 | 0.467 | 1 | 7 | Ppp6r2 | myCAF_3 |
| Tbc1d161 | 0.00527062 | 1.31342344 | 0.356 | 0.288 | 1 | 7 | Tbc1d16 | myCAF_3 |
| Map111 | 0.00527269 | 0.68455896 | 0.247 | 0.331 | 1 | 7 | Map11 | myCAF_3 |
| Mrs2 | 0.00534428 | 0.67178895 | 0.314 | 0.359 | 1 | 7 | Mrs2 | myCAF_3 |
| Arid3a2 | 0.00538429 | 1.10777457 | 0.32 | 0.41 | 1 | 7 | Arid3a | myCAF_3 |
| Prune1 | 0.00539788 | 0.97542847 | 0.232 | 0.314 | 1 | 7 | Prune1 | myCAF_3 |
| Mapk8ip13 | 0.00550914 | 0.41432595 | 0.196 | 0.285 | 1 | 7 | Mapk8ip1 | myCAF_3 |
| Ccdc1171 | 0.00551348 | 1.00164912 | 0.268 | 0.345 | 1 | 7 | Ccdc117 | myCAF_3 |
| Icam12 | 0.00554134 | 1.19474339 | 0.541 | 0.495 | 1 | 7 | Icam1 | myCAF_3 |
| Ulmc11 | 0.00555911 | 0.68708115 | 0.201 | 0.274 | 1 | 7 | Ulmc1 | myCAF_3 |
| Atf6 | 0.00562931 | 0.5440909 | 0.485 | 0.626 | 1 | 7 | Atf6 | myCAF_3 |
| Armc5 | 0.00569653 | 0.90839824 | 0.196 | 0.279 | 1 | 7 | Armc5 | myCAF_3 |
| Zfpm11 | 0.00578039 | 0.36699538 | 0.289 | 0.366 | 1 | 7 | Zfpm1 | myCAF_3 |
| Sbf1 | 0.00580121 | 0.77105387 | 0.227 | 0.293 | 1 | 7 | Sbf1 | myCAF_3 |
| Rflnb1 | 0.0058015 | 0.48824328 | 0.438 | 0.486 | 1 | 7 | Rflnb | myCAF_3 |
| Pcmtd12 | 0.00580247 | 0.73465516 | 0.278 | 0.357 | 1 | 7 | Pcmtd1 | myCAF_3 |
| Etv11 | 0.0058439 | 1.56459874 | 0.268 | 0.186 | 1 | 7 | Etv1 | myCAF_3 |
| Grina4 | 0.00590971 | 1.02783985 | 0.608 | 0.649 | 1 | 7 | Grina | myCAF_3 |
| Mapre3 | 0.00611684 | 0.92087516 | 0.443 | 0.477 | 1 | 7 | Mapre3 | myCAF_3 |
| Tpcn22 | 0.0061946 | 0.99945171 | 0.258 | 0.357 | 1 | 7 | Tpcn2 | myCAF_3 |
| Wdr912 | 0.00622142 | 0.88532565 | 0.304 | 0.37 | 1 | 7 | Wdr91 | myCAF_3 |
| Cnppd1 | 0.00626466 | 0.77129377 | 0.603 | 0.671 | 1 | 7 | Cnppd1 | myCAF_3 |
| Kri11 | 0.00628534 | 0.70366905 | 0.33 | 0.428 | 1 | 7 | Kri1 | myCAF_3 |
| Snapc52 | 0.00638712 | 0.49242083 | 0.294 | 0.354 | 1 | 7 | Snapc5 | myCAF_3 |
| Vezt | 0.00645174 | 0.79150862 | 0.222 | 0.302 | 1 | 7 | Vezt | myCAF_3 |
| Rnf113a2 | 0.00646111 | 0.39194741 | 0.222 | 0.303 | 1 | 7 | Rnf113a2 | myCAF_3 |
| Aamdc2 | 0.00647848 | 0.34534106 | 0.258 | 0.355 | 1 | 7 | Aamdc | myCAF_3 |
| Inf21 | 0.00660023 | 0.6199497 | 0.675 | 0.74 | 1 | 7 | Inf2 | myCAF_3 |
| Fbxo311 | 0.00660655 | 0.97490702 | 0.268 | 0.376 | 1 | 7 | Fbxo31 | myCAF_3 |
| R3hcc1 | 0.00660656 | 0.39631747 | 0.299 | 0.38 | 1 | 7 | R3hcc1 | myCAF_3 |
| Rufy2 | 0.0066863 | 0.43742502 | 0.175 | 0.264 | 1 | 7 | Rufy2 | myCAF_3 |
| Ndst21 | 0.0067055 | 0.94862699 | 0.206 | 0.272 | 1 | 7 | Ndst2 | myCAF_3 |
| Pip5k1c | 0.00683654 | 1.48449209 | 0.649 | 0.645 | 1 | 7 | Pip5k1c | myCAF_3 |
| Dpp72 | 0.00688278 | 0.52739364 | 0.263 | 0.336 | 1 | 7 | Dpp7 | myCAF_3 |
| Cog41 | 0.00692852 | 0.72438419 | 0.309 | 0.377 | 1 | 7 | Cog4 | myCAF_3 |
| Secisbp2l | 0.00705394 | 0.86195463 | 0.541 | 0.612 | 1 | 7 | Secisbp2l | myCAF_3 |
| Pik3c3 | 0.00715044 | 1.26143506 | 0.33 | 0.413 | 1 | 7 | Pik3c3 | myCAF_3 |
| Ttc281 | 0.00715842 | 0.59524037 | 0.521 | 0.587 | 1 | 7 | Ttc28 | myCAF_3 |
| Pwp2 | 0.00716784 | 1.1746872 | 0.196 | 0.255 | 1 | 7 | Pwp2 | myCAF_3 |
| Usp6nl | 0.00720578 | 0.27898767 | 0.211 | 0.288 | 1 | 7 | Usp6nl | myCAF_3 |
| Bcl7b | 0.00731065 | 0.53516832 | 0.216 | 0.283 | 1 | 7 | Bcl7b | myCAF_3 |
| Baz2b1 | 0.00735086 | 1.02520429 | 0.237 | 0.324 | 1 | 7 | Baz2b | myCAF_3 |
| Ptgs12 | 0.00749685 | 1.05230146 | 0.227 | 0.264 | 1 | 7 | Ptgs1 | myCAF_3 |
| Snx131 | 0.00767309 | 0.4679609 | 0.345 | 0.446 | 1 | 7 | Snx13 | myCAF_3 |
| Mettl6 | 0.00769836 | 0.45939809 | 0.216 | 0.289 | 1 | 7 | Mettl6 | myCAF_3 |
| Zfp7152 | 0.00774236 | 0.76755404 | 0.191 | 0.283 | 1 | 7 | Zfp715 | myCAF_3 |
| Slc20a21 | 0.00775442 | 0.82778179 | 0.448 | 0.512 | 1 | 7 | Slc20a2 | myCAF_3 |
| Nrbp23 | 0.00781326 | 1.61908302 | 0.289 | 0.24 | 1 | 7 | Nrbp2 | myCAF_3 |
| Repin12 | 0.00783815 | 0.962033 | 0.222 | 0.336 | 1 | 7 | Repin1 | myCAF_3 |
| Enc11 | 0.00784212 | 0.55568892 | 0.67 | 0.808 | 1 | 7 | Enc1 | myCAF_3 |
| Cacnb31 | 0.00803973 | 0.66070301 | 0.691 | 0.787 | 1 | 7 | Cacnb3 | myCAF_3 |
| Tab1 | 0.00805332 | 0.40125778 | 0.222 | 0.297 | 1 | 7 | Tab1 | myCAF_3 |
| Wdr24 | 0.00817004 | 0.6266953 | 0.196 | 0.281 | 1 | 7 | Wdr24 | myCAF_3 |
| Zic12 | 0.00826344 | 0.79207053 | 0.273 | 0.274 | 1 | 7 | Zic1 | myCAF_3 |
| Ppp1r351 | 0.0083522 | 0.41785293 | 0.247 | 0.31 | 1 | 7 | Ppp1r35 | myCAF_3 |
| Brpf11 | 0.00851262 | 0.94666706 | 0.278 | 0.35 | 1 | 7 | Brpf1 | myCAF_3 |
| Atxn7 | 0.00853216 | 0.63609357 | 0.356 | 0.449 | 1 | 7 | Atxn7 | myCAF_3 |
| Stard31 | 0.00854676 | 0.39755452 | 0.381 | 0.48 | 1 | 7 | Stard3 | myCAF_3 |
| Plekhn13 | 0.00861543 | 0.76845142 | 0.345 | 0.411 | 1 | 7 | Plekhn1 | myCAF_3 |
| Nub11 | 0.00862451 | 1.01280589 | 0.469 | 0.506 | 1 | 7 | Nub1 | myCAF_3 |
| Strada | 0.00865622 | 0.72363864 | 0.211 | 0.291 | 1 | 7 | Strada | myCAF_3 |
| Phf21 | 0.0087059 | 0.99380746 | 0.572 | 0.636 | 1 | 7 | Phf2 | myCAF_3 |
| Stx113 | 0.00877216 | 0.32287261 | 0.201 | 0.268 | 1 | 7 | Stx11 | myCAF_3 |
| Tpcn12 | 0.00881026 | 0.85659336 | 0.325 | 0.425 | 1 | 7 | Tpcn1 | myCAF_3 |
| Gpr1611 | 0.00882326 | 0.64708284 | 0.222 | 0.299 | 1 | 7 | Gpr161 | myCAF_3 |
| Uspl11 | 0.00884095 | 0.66980666 | 0.309 | 0.341 | 1 | 7 | Uspl1 | myCAF_3 |
| Cpz1 | 0.00887204 | 0.78538789 | 0.526 | 0.632 | 1 | 7 | Cpz | myCAF_3 |
| Chd6 | 0.00899846 | 1.08991196 | 0.371 | 0.446 | 1 | 7 | Chd6 | myCAF_3 |
| Arl16 | 0.00908831 | 0.52143123 | 0.222 | 0.285 | 1 | 7 | Arl16 | myCAF_3 |
| Metap1d1 | 0.00925067 | 0.56817318 | 0.211 | 0.295 | 1 | 7 | Metap1d | myCAF_3 |
| Ehd41 | 0.00926108 | 0.47183753 | 0.253 | 0.343 | 1 | 7 | Ehd4 | myCAF_3 |
| Abr | 0.00946972 | 0.96174321 | 0.464 | 0.522 | 1 | 7 | Abr | myCAF_3 |
| Smarcd31 | 0.00948847 | 0.55871323 | 0.237 | 0.31 | 1 | 7 | Smarcd3 | myCAF_3 |
| Snrnp351 | 0.00951395 | 1.0191522 | 0.237 | 0.317 | 1 | 7 | Snrnp35 | myCAF_3 |
| Nudt81 | 0.00959275 | 0.69709019 | 0.242 | 0.323 | 1 | 7 | Nudt8 | myCAF_3 |
| Unc45a1 | 0.00960093 | 1.11477534 | 0.402 | 0.48 | 1 | 7 | Unc45a | myCAF_3 |
| Zfand2b1 | 0.00972592 | 1.01139478 | 0.325 | 0.384 | 1 | 7 | Zfand2b | myCAF_3 |
| Rasa21 | 0.00978894 | 0.75396535 | 0.216 | 0.323 | 1 | 7 | Rasa2 | myCAF_3 |
| Fam135a1 | 0.00979934 | 0.71886062 | 0.253 | 0.333 | 1 | 7 | Fam135a | myCAF_3 |
| Nbeal11 | 0.00983587 | 0.64301448 | 0.464 | 0.529 | 1 | 7 | Nbeal1 | myCAF_3 |
| Ankrd13b1 | 0.00995876 | 0.70904318 | 0.17 | 0.256 | 1 | 7 | Ankrd13b | myCAF_3 |

| | p_val | avg_log2FC | pct.1 | pct.2 | p_val_adj | cluster | gene |
|---|---|---|---|---|---|---|---|
| Itga9 | 0 | 1.73379495 | 0.766 | 0.284 | 0 | TAM.Vcam1 | Itga9 |
| Ptgs1 | 0 | 1.92400458 | 0.79 | 0.31 | 0 | TAM.Vcam1 | Ptgs1 |
| Ms4a7 | 0 | 1.5236787 | 0.881 | 0.403 | 0 | TAM.Vcam1 | Ms4a7 |
| Vcam1 | 0 | 2.83360998 | 0.599 | 0.13 | 0 | TAM.Vcam1 | Vcam1 |
| Cd81 | 0 | 1.56481677 | 0.897 | 0.436 | 0 | TAM.Vcam1 | Cd81 |
| Mmp13 | 0 | 1.6549011 | 0.649 | 0.229 | 0 | TAM.Vcam1 | Mmp13 |
| Mgl2 | 0 | 1.16503028 | 0.73 | 0.312 | 0 | TAM.Vcam1 | Mgl2 |
| Hpgds | 0 | 1.84732366 | 0.58 | 0.168 | 0 | TAM.Vcam1 | Hpgds |
| Fcrls | 0 | 1.36021936 | 0.725 | 0.32 | 0 | TAM.Vcam1 | Fcrls |
| Fcgrt | 0 | 0.71625032 | 0.87 | 0.467 | 0 | TAM.Vcam1 | Fcgrt |
| Hebp1 | 0 | 1.65178511 | 0.692 | 0.302 | 0 | TAM.Vcam1 | Hebp1 |
| Gas6 | 0 | 0.69200481 | 0.624 | 0.237 | 0 | TAM.Vcam1 | Gas6 |
| Frmd4b | 0 | 1.3399133 | 0.672 | 0.296 | 0 | TAM.Vcam1 | Frmd4b |
| Lair1 | 0 | 1.34486227 | 0.799 | 0.434 | 0 | TAM.Vcam1 | Lair1 |
| Npl | 0 | 1.39562912 | 0.574 | 0.211 | 0 | TAM.Vcam1 | Npl |
| Gbp8 | 0 | 1.71832423 | 0.565 | 0.204 | 0 | TAM.Vcam1 | Gbp8 |
| Maf | 0 | 0.6926768 | 0.868 | 0.507 | 0 | TAM.Vcam1 | Maf |
| C1qc | 0 | 1.2730199 | 0.984 | 0.632 | 0 | TAM.Vcam1 | C1qc |
| Tmem176a | 0 | 1.25323804 | 0.703 | 0.357 | 0 | TAM.Vcam1 | Tmem176a |
| Tmem176b | 0 | 1.08279205 | 0.8 | 0.455 | 0 | TAM.Vcam1 | Tmem176b |
| Stab1 | 0 | 1.03357896 | 0.781 | 0.436 | 0 | TAM.Vcam1 | Stab1 |
| Gna12 | 0 | 1.27473477 | 0.859 | 0.514 | 0 | TAM.Vcam1 | Gna12 |
| Abcc3 | 0 | 1.17737721 | 0.729 | 0.387 | 0 | TAM.Vcam1 | Abcc3 |
| Sash1 | 0 | 1.22120767 | 0.87 | 0.531 | 0 | TAM.Vcam1 | Sash1 |
| Pf4 | 0 | 0.94590879 | 0.833 | 0.495 | 0 | TAM.Vcam1 | Pf4 |
| Tlr12 | 0 | 3.66393645 | 0.381 | 0.046 | 0 | TAM.Vcam1 | Tlr12 |
| Cd72 | 0 | 1.31642334 | 0.561 | 0.227 | 0 | TAM.Vcam1 | Cd72 |
| Fcgr4 | 0 | 1.2014022 | 0.746 | 0.412 | 0 | TAM.Vcam1 | Fcgr4 |
| Fgd2 | 0 | 1.21537978 | 0.602 | 0.269 | 0 | TAM.Vcam1 | Fgd2 |
| Nxpe5 | 0 | 1.0697901 | 0.621 | 0.29 | 0 | TAM.Vcam1 | Nxpe5 |
| Cd93 | 0 | 1.26099768 | 0.672 | 0.342 | 0 | TAM.Vcam1 | Cd93 |
| Adgre1 | 0 | 0.87324819 | 0.835 | 0.505 | 0 | TAM.Vcam1 | Adgre1 |
| Lpcat2 | 0 | 1.29991191 | 0.72 | 0.393 | 0 | TAM.Vcam1 | Lpcat2 |
| Rgs1 | 0 | 1.29550546 | 0.825 | 0.499 | 0 | TAM.Vcam1 | Rgs1 |
| Pxd3 | 0 | 1.14007088 | 0.553 | 0.228 | 0 | TAM.Vcam1 | Pxd3 |
| Rgl1 | 0 | 0.94235748 | 0.667 | 0.344 | 0 | TAM.Vcam1 | Rgl1 |
| Sec14l1 | 0 | 1.11387166 | 0.765 | 0.442 | 0 | TAM.Vcam1 | Sec14l1 |
| Abcg3 | 0 | 1.44673903 | 0.582 | 0.26 | 0 | TAM.Vcam1 | Abcg3 |
| Rcbtb2 | 0 | 1.28987073 | 0.692 | 0.37 | 0 | TAM.Vcam1 | Rcbtb2 |
| Fam234b | 0 | 2.0431401 | 0.459 | 0.138 | 0 | TAM.Vcam1 | Fam234b |
| Pxdc1 | 0 | 1.11336922 | 0.674 | 0.356 | 0 | TAM.Vcam1 | Pxdc1 |
| C1qa | 0 | 1.18287305 | 0.995 | 0.677 | 0 | TAM.Vcam1 | C1qa |
| Pmepa1 | 0 | 1.16631525 | 0.841 | 0.524 | 0 | TAM.Vcam1 | Pmepa1 |
| Gatm | 0 | 1.13010888 | 0.65 | 0.334 | 0 | TAM.Vcam1 | Gatm |
| Rab3il1 | 0 | 0.97531258 | 0.932 | 0.619 | 0 | TAM.Vcam1 | Rab3il1 |
| Lag3 | 0 | 2.57168746 | 0.419 | 0.108 | 0 | TAM.Vcam1 | Lag3 |
| Tmem86a | 0 | 1.03196776 | 0.741 | 0.432 | 0 | TAM.Vcam1 | Tmem86a |
| Ccl12 | 0 | 1.6463407 | 0.486 | 0.178 | 0 | TAM.Vcam1 | Ccl12 |
| Pdgfc | 0 | 0.9595461 | 0.502 | 0.195 | 0 | TAM.Vcam1 | Pdgfc |
| Gdf15 | 0 | 1.62521443 | 0.526 | 0.221 | 0 | TAM.Vcam1 | Gdf15 |
| Acp5 | 0 | 0.65713255 | 0.923 | 0.618 | 0 | TAM.Vcam1 | Acp5 |
| Itga6 | 0 | 1.11026172 | 0.598 | 0.293 | 0 | TAM.Vcam1 | Itga6 |
| Ntpcr | 0 | 1.22520627 | 0.649 | 0.346 | 0 | TAM.Vcam1 | Ntpcr |
| Rhoh | 0 | 1.11201164 | 0.603 | 0.307 | 0 | TAM.Vcam1 | Rhoh |
| Atp8a1 | 0 | 0.96835595 | 0.71 | 0.414 | 0 | TAM.Vcam1 | Atp8a1 |
| Zmynd15 | 0 | 1.34506453 | 0.542 | 0.247 | 0 | TAM.Vcam1 | Zmynd15 |
| Scamp5 | 0 | 1.34640867 | 0.484 | 0.19 | 0 | TAM.Vcam1 | Scamp5 |
| Hfe | 0 | 0.77227795 | 0.735 | 0.444 | 0 | TAM.Vcam1 | Hfe |
| Lrp6 | 0 | 0.78441886 | 0.806 | 0.516 | 0 | TAM.Vcam1 | Lrp6 |
| Tgfbr2 | 0 | 0.94695878 | 0.839 | 0.55 | 0 | TAM.Vcam1 | Tgfbr2 |
| Selenop | 0 | 1.28582032 | 0.967 | 0.68 | 0 | TAM.Vcam1 | Selenop |
| Mertk | 0 | 0.83700129 | 0.803 | 0.517 | 0 | TAM.Vcam1 | Mertk |
| Ap1b1 | 0 | 1.01548643 | 0.865 | 0.58 | 0 | TAM.Vcam1 | Ap1b1 |
| Aoah | 0 | 0.72336006 | 0.67 | 0.385 | 0 | TAM.Vcam1 | Aoah |
| Dtx4 | 0 | 0.84558519 | 0.682 | 0.397 | 0 | TAM.Vcam1 | Dtx4 |
| Apobec1 | 0 | 0.80647302 | 0.71 | 0.425 | 0 | TAM.Vcam1 | Apobec1 |
| Camk1 | 0 | 0.87061806 | 0.763 | 0.479 | 0 | TAM.Vcam1 | Camk1 |
| Tmem37 | 0 | 0.84730856 | 0.645 | 0.363 | 0 | TAM.Vcam1 | Tmem37 |
| Dnase1l3 | 0 | 3.18009076 | 0.338 | 0.056 | 0 | TAM.Vcam1 | Dnase1l3 |
| C1qb | 0 | 0.98133866 | 0.997 | 0.719 | 0 | TAM.Vcam1 | C1qb |
| Ttyh2 | 0 | 1.31651517 | 0.52 | 0.242 | 0 | TAM.Vcam1 | Ttyh2 |
| Cd38 | 0 | 0.79883178 | 0.682 | 0.407 | 0 | TAM.Vcam1 | Cd38 |
| Cd63 | 0 | 0.87633488 | 0.973 | 0.698 | 0 | TAM.Vcam1 | Cd63 |
| Igfbp4 | 0 | 0.86838288 | 0.68 | 0.405 | 0 | TAM.Vcam1 | Igfbp4 |
| Timp2 | 0 | 0.33933284 | 0.784 | 0.509 | 0 | TAM.Vcam1 | Timp2 |
| Ly86 | 0 | 0.90486048 | 0.856 | 0.583 | 0 | TAM.Vcam1 | Ly86 |
| Ndst1 | 0 | 0.73933858 | 0.745 | 0.472 | 0 | TAM.Vcam1 | Ndst1 |
| Prune2 | 0 | 1.19695219 | 0.483 | 0.212 | 0 | TAM.Vcam1 | Prune2 |
| Mrc1 | 0 | 0.65214143 | 0.966 | 0.696 | 0 | TAM.Vcam1 | Mrc1 |
| Pacc1 | 0 | 1.32888503 | 0.517 | 0.248 | 0 | TAM.Vcam1 | Pacc1 |
| Etv5 | 0 | 0.92898237 | 0.669 | 0.402 | 0 | TAM.Vcam1 | Etv5 |
| Tifab | 0 | 0.93136196 | 0.706 | 0.44 | 0 | TAM.Vcam1 | Tifab |
| Axl | 0 | 0.83118825 | 0.917 | 0.652 | 0 | TAM.Vcam1 | Axl |
| Pld3 | 0 | 0.7924963 | 0.834 | 0.573 | 0 | TAM.Vcam1 | Pld3 |
| Ms4a4a | 0 | 0.69195413 | 0.919 | 0.66 | 0 | TAM.Vcam1 | Ms4a4a |
| Itgb5 | 0 | 1.05600259 | 0.911 | 0.654 | 0 | TAM.Vcam1 | Itgb5 |
| Snx6 | 0 | 0.79121293 | 0.707 | 0.45 | 0 | TAM.Vcam1 | Snx6 |
| Abi3 | 0 | 0.68797142 | 0.732 | 0.476 | 0 | TAM.Vcam1 | Abi3 |
| Itsn1 | 0 | 0.54904387 | 0.543 | 0.289 | 0 | TAM.Vcam1 | Itsn1 |
| Tnfaip8l2 | 0 | 0.75061629 | 0.724 | 0.47 | 0 | TAM.Vcam1 | Tnfaip8l2 |
| Mvb12b | 0 | 0.96562457 | 0.588 | 0.334 | 0 | TAM.Vcam1 | Mvb12b |
| P2ry6 | 0 | 0.84352619 | 0.856 | 0.605 | 0 | TAM.Vcam1 | P2ry6 |
| Sema4b | 0 | 0.92451119 | 0.569 | 0.318 | 0 | TAM.Vcam1 | Sema4b |
| Gbp7 | 0 | 1.00570752 | 0.528 | 0.278 | 0 | TAM.Vcam1 | Gbp7 |
| Dip2c | 0 | 0.9756274 | 0.547 | 0.298 | 0 | TAM.Vcam1 | Dip2c |
| Hsd17b4 | 0 | 0.90137618 | 0.663 | 0.414 | 0 | TAM.Vcam1 | Hsd17b4 |
| Dusp6 | 0 | 0.90479976 | 0.835 | 0.586 | 0 | TAM.Vcam1 | Dusp6 |
| Entpd1 | 0 | 0.85089177 | 0.833 | 0.584 | 0 | TAM.Vcam1 | Entpd1 |
| Tgfbr1 | 0 | 0.92089726 | 0.792 | 0.545 | 0 | TAM.Vcam1 | Tgfbr1 |
| Acp2 | 0 | 0.80417938 | 0.677 | 0.431 | 0 | TAM.Vcam1 | Acp2 |
| Pla2g15 | 0 | 0.68897638 | 0.855 | 0.609 | 0 | TAM.Vcam1 | Pla2g15 |
| Irf8 | 0 | 0.79214674 | 0.857 | 0.612 | 0 | TAM.Vcam1 | Irf8 |
| Aif1 | 0 | 0.99343228 | 0.965 | 0.72 | 0 | TAM.Vcam1 | Aif1 |
| Gpr65 | 0 | 0.71438363 | 0.597 | 0.353 | 0 | TAM.Vcam1 | Gpr65 |
| Abca9 | 0 | 0.78995905 | 0.51 | 0.267 | 0 | TAM.Vcam1 | Abca9 |
| Plau | 0 | 0.7983589 | 0.845 | 0.602 | 0 | TAM.Vcam1 | Plau |
| Fabp5 | 0 | 0.88457759 | 0.879 | 0.637 | 0 | TAM.Vcam1 | Fabp5 |
| Rnase4 | 0 | 0.54221738 | 0.606 | 0.364 | 0 | TAM.Vcam1 | Rnase4 |
| Slc9a9 | 0 | 0.53513627 | 0.508 | 0.267 | 0 | TAM.Vcam1 | Slc9a9 |
| Tent5c | 0 | 0.9701288 | 0.566 | 0.325 | 0 | TAM.Vcam1 | Tent5c |
| Ms4a6b | 0 | 0.46349816 | 0.655 | 0.415 | 0 | TAM.Vcam1 | Ms4a6b |
| Tspan13 | 0 | 1.07930279 | 0.509 | 0.269 | 0 | TAM.Vcam1 | Tspan13 |
| Rtcb | 0 | 0.87546608 | 0.789 | 0.55 | 0 | TAM.Vcam1 | Rtcb |
| Spint1 | 0 | 0.39924649 | 0.806 | 0.568 | 0 | TAM.Vcam1 | Spint1 |
| Mmp2 | 0 | 2.73540624 | 0.295 | 0.057 | 0 | TAM.Vcam1 | Mmp2 |
| Fads1 | 0 | 1.30951475 | 0.429 | 0.191 | 0 | TAM.Vcam1 | Fads1 |
| Snx29 | 0 | 0.52970321 | 0.671 | 0.434 | 0 | TAM.Vcam1 | Snx29 |
| P2ry13 | 0 | 1.5207291 | 0.393 | 0.156 | 0 | TAM.Vcam1 | P2ry13 |
| P2ry12 | 0 | 1.10410738 | 0.438 | 0.204 | 0 | TAM.Vcam1 | P2ry12 |
| Herpud1 | 0 | 0.87133764 | 0.618 | 0.384 | 0 | TAM.Vcam1 | Herpud1 |
| Rogdi | 0 | 0.80282284 | 0.541 | 0.309 | 0 | TAM.Vcam1 | Rogdi |
| Myo7a | 0 | 0.75149998 | 0.616 | 0.385 | 0 | TAM.Vcam1 | Myo7a |
| Tmem106a | 0 | 0.47430056 | 0.782 | 0.552 | 0 | TAM.Vcam1 | Tmem106a |
| Cd300a | 0 | 0.80664372 | 0.876 | 0.646 | 0 | TAM.Vcam1 | Cd300a |
| Gpx3 | 0 | 1.08210445 | 0.411 | 0.182 | 0 | TAM.Vcam1 | Gpx3 |
| Tmem273 | 0 | 1.24336138 | 0.41 | 0.181 | 0 | TAM.Vcam1 | Tmem273 |
| Tnfrsf11a | 0 | 0.57947255 | 0.659 | 0.431 | 0 | TAM.Vcam1 | Tnfrsf11a |
| Snx8 | 0 | 0.52187842 | 0.678 | 0.45 | 0 | TAM.Vcam1 | Snx8 |
| Pea15a | 0 | 0.66962697 | 0.71 | 0.483 | 0 | TAM.Vcam1 | Pea15a |
| Map11 | 0 | 0.76818367 | 0.659 | 0.433 | 0 | TAM.Vcam1 | Map11 |
| C3ar1 | 0 | 0.31913872 | 0.869 | 0.643 | 0 | TAM.Vcam1 | C3ar1 |
| Cd4 | 0 | 2.47458145 | 0.282 | 0.056 | 0 | TAM.Vcam1 | Cd4 |
| Fcgr1 | 0 | 0.32637513 | 0.809 | 0.584 | 0 | TAM.Vcam1 | Fcgr1 |
| Gaa | 0 | 0.71837687 | 0.641 | 0.416 | 0 | TAM.Vcam1 | Gaa |
| Cd83 | 0 | 0.65855453 | 0.735 | 0.51 | 0 | TAM.Vcam1 | Cd83 |
| Atp13a2 | 0 | 0.65287874 | 0.864 | 0.64 | 0 | TAM.Vcam1 | Atp13a2 |
| Man2b2 | 0 | 0.69126563 | 0.709 | 0.485 | 0 | TAM.Vcam1 | Man2b2 |
| Adap2 | 0 | 0.54427139 | 0.769 | 0.545 | 0 | TAM.Vcam1 | Adap2 |
| Prxl2b | 0 | 0.71414574 | 0.533 | 0.311 | 0 | TAM.Vcam1 | Prxl2b |
| Il16 | 0 | 1.135274 | 0.409 | 0.187 | 0 | TAM.Vcam1 | Il16 |
| Comt | 0 | 0.621388 | 0.723 | 0.502 | 0 | TAM.Vcam1 | Comt |
| Hmgcs1 | 0 | 0.94236907 | 0.562 | 0.342 | 0 | TAM.Vcam1 | Hmgcs1 |
| Cyp27a1 | 0 | 0.89776917 | 0.437 | 0.217 | 0 | TAM.Vcam1 | Cyp27a1 |
| Tbc1d10a | 0 | 0.89366136 | 0.545 | 0.327 | 0 | TAM.Vcam1 | Tbc1d10a |
| Abl1 | 0 | 0.71829222 | 0.618 | 0.4 | 0 | TAM.Vcam1 | Abl1 |
| Dhrs3 | 0 | 0.39696543 | 0.833 | 0.615 | 0 | TAM.Vcam1 | Dhrs3 |
| Tpp1 | 0 | 0.70854001 | 0.846 | 0.629 | 0 | TAM.Vcam1 | Tpp1 |
| Jaml | 0 | 0.69983359 | 0.577 | 0.36 | 0 | TAM.Vcam1 | Jaml |
| Lacc1 | 0 | 0.72176336 | 0.563 | 0.347 | 0 | TAM.Vcam1 | Lacc1 |
| Itgax | 0 | 0.29775156 | 0.787 | 0.572 | 0 | TAM.Vcam1 | Itgax |
| Vav2 | 0 | 0.88735035 | 0.483 | 0.269 | 0 | TAM.Vcam1 | Vav2 |
| Cd84 | 0 | 0.48714514 | 0.8 | 0.587 | 0 | TAM.Vcam1 | Cd84 |
| Evl | 0 | 0.59793787 | 0.773 | 0.56 | 0 | TAM.Vcam1 | Evl |
| Renbp | 0 | 0.55237925 | 0.72 | 0.507 | 0 | TAM.Vcam1 | Renbp |
| Gng2 | 0 | 0.58378973 | 0.682 | 0.47 | 0 | TAM.Vcam1 | Gng2 |
| Vipas39 | 0 | 0.78278408 | 0.527 | 0.315 | 0 | TAM.Vcam1 | Vipas39 |
| Dynll2 | 0 | 0.70206405 | 0.598 | 0.386 | 0 | TAM.Vcam1 | Dynll2 |
| Pdia4 | 0 | 0.62356414 | 0.799 | 0.589 | 0 | TAM.Vcam1 | Pdia4 |
| Cebpa | 0 | 0.73312199 | 0.937 | 0.727 | 0 | TAM.Vcam1 | Cebpa |
| Slc30a1 | 0 | 0.78722369 | 0.544 | 0.334 | 0 | TAM.Vcam1 | Slc30a1 |
| Man2a2 | 0 | 0.49725994 | 0.62 | 0.41 | 0 | TAM.Vcam1 | Man2a2 |
| Tbc1d9b | 0 | 0.61098745 | 0.609 | 0.399 | 0 | TAM.Vcam1 | Tbc1d9b |
| Mmp14 | 0 | 0.49211577 | 0.92 | 0.711 | 0 | TAM.Vcam1 | Mmp14 |
| P2rx7 | 0 | 0.78710246 | 0.513 | 0.304 | 0 | TAM.Vcam1 | P2rx7 |
| Spon1 | 0 | 1.24884782 | 0.369 | 0.16 | 0 | TAM.Vcam1 | Spon1 |
| Bcl2l1 | 0 | 0.67745242 | 0.617 | 0.41 | 0 | TAM.Vcam1 | Bcl2l1 |
| Rab11fip5 | 0 | 0.33459051 | 0.764 | 0.557 | 0 | TAM.Vcam1 | Rab11fip5 |
| Galc | 0 | 0.84038672 | 0.476 | 0.269 | 0 | TAM.Vcam1 | Galc |
| Daglb | 0 | 0.51536731 | 0.675 | 0.469 | 0 | TAM.Vcam1 | Daglb |
| Cd33 | 0 | 0.56312337 | 0.774 | 0.568 | 0 | TAM.Vcam1 | Cd33 |
| Cd300c2 | 0 | 0.58491583 | 0.779 | 0.573 | 0 | TAM.Vcam1 | Cd300c2 |
| Lilra5 | 0 | 1.9677907 | 0.292 | 0.086 | 0 | TAM.Vcam1 | Lilra5 |
| Adgrg5 | 0 | 1.29239131 | 0.35 | 0.144 | 0 | TAM.Vcam1 | Adgrg5 |
| Rin2 | 0 | 0.69143973 | 0.547 | 0.342 | 0 | TAM.Vcam1 | Rin2 |
| Mxd4 | 0 | 0.78889216 | 0.505 | 0.3 | 0 | TAM.Vcam1 | Mxd4 |
| Paox | 0 | 1.71106194 | 0.303 | 0.099 | 0 | TAM.Vcam1 | Paox |
| Lap3 | 0 | 0.91300557 | 0.479 | 0.275 | 0 | TAM.Vcam1 | Lap3 |
| Galnt1 | 0 | 0.60690713 | 0.702 | 0.498 | 0 | TAM.Vcam1 | Galnt1 |
| Slc29a3 | 0 | 0.61592367 | 0.644 | 0.441 | 0 | TAM.Vcam1 | Slc29a3 |
| Lpar6 | 0 | 0.81778584 | 0.499 | 0.296 | 0 | TAM.Vcam1 | Lpar6 |
| Emilin1 | 0 | 0.97459301 | 0.416 | 0.213 | 0 | TAM.Vcam1 | Emilin1 |
| Otulinl | 0 | 0.7025875 | 0.868 | 0.665 | 0 | TAM.Vcam1 | Otulinl |
| Lrrc25 | 0 | 0.66777405 | 0.88 | 0.678 | 0 | TAM.Vcam1 | Lrrc25 |
| P2rx4 | 0 | 0.52727743 | 0.8 | 0.599 | 0 | TAM.Vcam1 | P2rx4 |
| Ppt1 | 0 | 0.62655525 | 0.77 | 0.57 | 0 | TAM.Vcam1 | Ppt1 |
| Tlr1 | 0 | 0.98002129 | 0.402 | 0.203 | 0 | TAM.Vcam1 | Tlr1 |
| Cttnbp2nl | 0 | 0.47097196 | 0.703 | 0.504 | 0 | TAM.Vcam1 | Cttnbp2nl |
| Ccl4 | 0 | 0.9816422 | 0.604 | 0.405 | 0 | TAM.Vcam1 | Ccl4 |
| Abhd12 | 0 | 0.44452422 | 0.795 | 0.597 | 0 | TAM.Vcam1 | Abhd12 |
| Plxdc2 | 0 | 0.94228771 | 0.402 | 0.204 | 0 | TAM.Vcam1 | Plxdc2 |
| Coro7 | 0 | 0.56697721 | 0.633 | 0.435 | 0 | TAM.Vcam1 | Coro7 |
| Nceh1 | 0 | 0.67026475 | 0.553 | 0.356 | 0 | TAM.Vcam1 | Nceh1 |
| Snx5 | 0 | 0.63329848 | 0.713 | 0.516 | 0 | TAM.Vcam1 | Snx5 |
| Rgs10 | 0 | 0.58271604 | 0.83 | 0.633 | 0 | TAM.Vcam1 | Rgs10 |
| Mcfd2 | 0 | 0.57388694 | 0.751 | 0.554 | 0 | TAM.Vcam1 | Mcfd2 |
| Soga1 | 0 | 0.670246 | 0.537 | 0.341 | 0 | TAM.Vcam1 | Soga1 |
| Mef2c | 0 | 0.67604215 | 0.455 | 0.259 | 0 | TAM.Vcam1 | Mef2c |
| Ms4a14 | 0 | 1.59953479 | 0.292 | 0.096 | 0 | TAM.Vcam1 | Ms4a14 |
| Lgals3bp | 0 | 0.51019135 | 0.909 | 0.714 | 0 | TAM.Vcam1 | Lgals3bp |
| Pip4p1 | 0 | 0.49502681 | 0.664 | 0.469 | 0 | TAM.Vcam1 | Pip4p1 |
| Ttyh3 | 0 | 0.59474668 | 0.862 | 0.667 | 0 | TAM.Vcam1 | Ttyh3 |
| Tnfsf13 | 0 | 0.58825977 | 0.598 | 0.403 | 0 | TAM.Vcam1 | Tnfsf13 |
| DEPRECATED-I | 0 | 0.98400474 | 0.401 | 0.207 | 0 | TAM.Vcam1 | DEPRECATED-ENSMUSG00000002103 |
| Eef2k | 0 | 1.25131063 | 0.329 | 0.135 | 0 | TAM.Vcam1 | Eef2k |
| Mgat5 | 0 | 0.58051042 | 0.587 | 0.393 | 0 | TAM.Vcam1 | Mgat5 |
| Slc40a1 | 0 | 1.75463058 | 0.271 | 0.078 | 0 | TAM.Vcam1 | Slc40a1 |
| Igf1 | 0 | 0.87949466 | 0.358 | 0.165 | 0 | TAM.Vcam1 | Igf1 |
| Tbc1d22a | 0 | 0.48298755 | 0.738 | 0.545 | 0 | TAM.Vcam1 | Tbc1d22a |
| Txnip | 0 | 0.41248041 | 0.689 | 0.497 | 0 | TAM.Vcam1 | Txnip |
| Plekhm2 | 0 | 0.65088065 | 0.583 | 0.391 | 0 | TAM.Vcam1 | Plekhm2 |
| Marcks | 0 | 0.88613818 | 0.975 | 0.783 | 0 | TAM.Vcam1 | Marcks |
| Lst1 | 0 | 0.36900159 | 0.749 | 0.557 | 0 | TAM.Vcam1 | Lst1 |
| Klhl9 | 0 | 0.48647407 | 0.68 | 0.489 | 0 | TAM.Vcam1 | Klhl9 |
| Gns | 0 | 0.62029831 | 0.886 | 0.695 | 0 | TAM.Vcam1 | Gns |
| Lbh | 0 | 0.56925127 | 0.536 | 0.345 | 0 | TAM.Vcam1 | Lbh |
| Tor4a | 0 | 0.82317036 | 0.444 | 0.253 | 0 | TAM.Vcam1 | Tor4a |
| Ms4a6c | 0 | 0.26759773 | 0.697 | 0.506 | 0 | TAM.Vcam1 | Ms4a6c |
| Ehd4 | 0 | 0.51655373 | 0.803 | 0.613 | 0 | TAM.Vcam1 | Ehd4 |
| Zfp691 | 0 | 1.31927752 | 0.332 | 0.142 | 0 | TAM.Vcam1 | Zfp691 |
| Dnmt3a | 0 | 0.67805909 | 0.543 | 0.354 | 0 | TAM.Vcam1 | Dnmt3a |
| Ccdc102a | 0 | 1.22495002 | 0.333 | 0.144 | 0 | TAM.Vcam1 | Ccdc102a |
| Amdhd2 | 0 | 0.56411797 | 0.59 | 0.401 | 0 | TAM.Vcam1 | Amdhd2 |
| Itpripl2 | 0 | 0.51303073 | 0.717 | 0.531 | 0 | TAM.Vcam1 | Itpripl2 |
| Tanc2 | 0 | 1.29566071 | 0.317 | 0.132 | 0 | TAM.Vcam1 | Tanc2 |
| Cebpzos | 0 | 0.89825723 | 0.41 | 0.225 | 0 | TAM.Vcam1 | Cebpzos |
| Wdfy3 | 0 | 0.45188681 | 0.814 | 0.63 | 0 | TAM.Vcam1 | Wdfy3 |
| Trf | 0 | 0.51176024 | 0.954 | 0.77 | 0 | TAM.Vcam1 | Trf |
| Tagap | 0 | 0.76643957 | 0.386 | 0.203 | 0 | TAM.Vcam1 | Tagap |
| Inpp4b | 0 | 1.42584046 | 0.284 | 0.101 | 0 | TAM.Vcam1 | Inpp4b |
| Sdf2l1 | 0 | 0.52692795 | 0.707 | 0.524 | 0 | TAM.Vcam1 | Sdf2l1 |
| Limd2 | 0 | 0.55199585 | 0.535 | 0.353 | 0 | TAM.Vcam1 | Limd2 |
| Cerk | 0 | 0.54300862 | 0.621 | 0.439 | 0 | TAM.Vcam1 | Cerk |
| Slc7a7 | 0 | 0.77195855 | 0.434 | 0.253 | 0 | TAM.Vcam1 | Slc7a7 |
| Hes1 | 0 | 1.1466848 | 0.358 | 0.177 | 0 | TAM.Vcam1 | Hes1 |
| Cd22 | 0 | 1.82637565 | 0.265 | 0.085 | 0 | TAM.Vcam1 | Cd22 |
| Tmcc3 | 0 | 1.1022515 | 0.355 | 0.175 | 0 | TAM.Vcam1 | Tmcc3 |
| Ddhd1 | 0 | 0.61178992 | 0.657 | 0.478 | 0 | TAM.Vcam1 | Ddhd1 |
| Eid1 | 0 | 0.47289106 | 0.866 | 0.688 | 0 | TAM.Vcam1 | Eid1 |
| Slc43a2 | 0 | 0.49050525 | 0.764 | 0.586 | 0 | TAM.Vcam1 | Slc43a2 |
| Hgf | 0 | 1.09289784 | 0.326 | 0.148 | 0 | TAM.Vcam1 | Hgf |
| Abhd15 | 0 | 1.34490207 | 0.292 | 0.114 | 0 | TAM.Vcam1 | Abhd15 |
| Bmp2k | 0 | 0.37853963 | 0.761 | 0.584 | 0 | TAM.Vcam1 | Bmp2k |
| Nmt1 | 0 | 0.54944924 | 0.784 | 0.607 | 0 | TAM.Vcam1 | Nmt1 |
| Camk2d | 0 | 0.36076956 | 0.692 | 0.515 | 0 | TAM.Vcam1 | Camk2d |
| Icosl | 0 | 0.71585614 | 0.385 | 0.209 | 0 | TAM.Vcam1 | Icosl |
| Sirt2 | 0 | 0.41968188 | 0.738 | 0.562 | 0 | TAM.Vcam1 | Sirt2 |
| Rragc | 0 | 0.57108527 | 0.658 | 0.483 | 0 | TAM.Vcam1 | Rragc |
| Sh2d1b1 | 0 | 0.82761145 | 0.404 | 0.229 | 0 | TAM.Vcam1 | Sh2d1b1 |
| Cxcl16 | 0 | 0.86566932 | 0.975 | 0.8 | 0 | TAM.Vcam1 | Cxcl16 |
| Lgals9 | 0 | 0.41426374 | 0.728 | 0.554 | 0 | TAM.Vcam1 | Lgals9 |
| Dusp7 | 0 | 0.57640828 | 0.563 | 0.39 | 0 | TAM.Vcam1 | Dusp7 |
| Atp6v0a1 | 0 | 0.40555663 | 0.655 | 0.483 | 0 | TAM.Vcam1 | Atp6v0a1 |
| Ahr | 0 | 0.33594781 | 0.7 | 0.528 | 0 | TAM.Vcam1 | Ahr |
| Lgmn | 0 | 0.98987865 | 0.983 | 0.811 | 0 | TAM.Vcam1 | Lgmn |
| Lrp12 | 0 | 0.61895279 | 0.582 | 0.411 | 0 | TAM.Vcam1 | Lrp12 |
| Itm2c | 0 | 0.81678269 | 0.947 | 0.78 | 0 | TAM.Vcam1 | Itm2c |
| Rasgef1b | 0 | 0.45843246 | 0.718 | 0.552 | 0 | TAM.Vcam1 | Rasgef1b |
| Scpep1 | 0 | 0.31959791 | 0.871 | 0.706 | 0 | TAM.Vcam1 | Scpep1 |
| Gpr34 | 0 | 1.36286311 | 0.253 | 0.088 | 0 | TAM.Vcam1 | Gpr34 |
| Il18bp | 0 | 1.06080086 | 0.298 | 0.134 | 0 | TAM.Vcam1 | Il18bp |
| Hspa1b | 0 | 0.8724767 | 0.498 | 0.335 | 0 | TAM.Vcam1 | Hspa1b |
| Anxa3 | 0 | 0.37784571 | 0.911 | 0.749 | 0 | TAM.Vcam1 | Anxa3 |
| Asah1 | 0 | 0.53420543 | 0.819 | 0.657 | 0 | TAM.Vcam1 | Asah1 |
| Rhob | 0 | 0.48495438 | 0.895 | 0.734 | 0 | TAM.Vcam1 | Rhob |
| Mdfic | 0 | 0.40820914 | 0.653 | 0.493 | 0 | TAM.Vcam1 | Mdfic |
| Man2b1 | 0 | 0.75099275 | 0.927 | 0.768 | 0 | TAM.Vcam1 | Man2b1 |
| Syngr1 | 0 | 1.48413439 | 0.255 | 0.096 | 0 | TAM.Vcam1 | Syngr1 |
| Adcy7 | 0 | 0.37569881 | 0.714 | 0.556 | 0 | TAM.Vcam1 | Adcy7 |
| Stra6l | 0 | 1.14269195 | 0.296 | 0.139 | 0 | TAM.Vcam1 | Stra6l |
| Sox4 | 0 | 1.10885473 | 0.297 | 0.14 | 0 | TAM.Vcam1 | Sox4 |
| Ehmt2 | 0 | 0.35237559 | 0.759 | 0.603 | 0 | TAM.Vcam1 | Ehmt2 |
| Lrch4 | 0 | 0.3004686 | 0.808 | 0.653 | 0 | TAM.Vcam1 | Lrch4 |
| Adcy3 | 0 | 1.23015866 | 0.254 | 0.1 | 0 | TAM.Vcam1 | Adcy3 |
| Pdia3 | 0 | 0.44467085 | 0.792 | 0.639 | 0 | TAM.Vcam1 | Pdia3 |
| Nucb1 | 0 | 0.37977619 | 0.82 | 0.668 | 0 | TAM.Vcam1 | Nucb1 |
| Vav1 | 0 | 0.39501603 | 0.82 | 0.668 | 0 | TAM.Vcam1 | Vav1 |
| Cyp51 | 0 | 0.64548654 | 0.453 | 0.302 | 0 | TAM.Vcam1 | Cyp51 |
| Rab32 | 0 | 0.30769784 | 0.732 | 0.581 | 0 | TAM.Vcam1 | Rab32 |
| Arap1 | 0 | 0.34011928 | 0.848 | 0.698 | 0 | TAM.Vcam1 | Arap1 |
| Msr1 | 0 | 0.5374768 | 0.916 | 0.766 | 0 | TAM.Vcam1 | Msr1 |
| Fyb | 0 | 0.27476266 | 0.772 | 0.622 | 0 | TAM.Vcam1 | Fyb |
| Zfp622 | 0 | 0.4584603 | 0.6 | 0.45 | 0 | TAM.Vcam1 | Zfp622 |
| Glmp | 0 | 0.38575868 | 0.83 | 0.68 | 0 | TAM.Vcam1 | Glmp |
| Ccl3 | 0 | 0.55126315 | 0.694 | 0.544 | 0 | TAM.Vcam1 | Ccl3 |
| Klra2 | 0 | 0.42422995 | 0.459 | 0.31 | 0 | TAM.Vcam1 | Klra2 |
| Rilpl2 | 0 | 0.43909607 | 0.703 | 0.554 | 0 | TAM.Vcam1 | Rilpl2 |
| Neurl3 | 0 | 0.33981835 | 0.868 | 0.72 | 0 | TAM.Vcam1 | Neurl3 |
| C5ar1 | 0 | 0.38260429 | 0.907 | 0.759 | 0 | TAM.Vcam1 | C5ar1 |
| Hspa1a | 0 | 0.65197995 | 0.538 | 0.39 | 0 | TAM.Vcam1 | Hspa1a |
| Ptms | 0 | 0.70136245 | 0.978 | 0.831 | 0 | TAM.Vcam1 | Ptms |
| Gusb | 0 | 0.39571485 | 0.843 | 0.698 | 0 | TAM.Vcam1 | Gusb |
| H2-DMa | 0 | 0.36567476 | 0.845 | 0.701 | 0 | TAM.Vcam1 | H2-DMa |
| Mfsd1 | 0 | 0.39687043 | 0.828 | 0.686 | 0 | TAM.Vcam1 | Mfsd1 |
| Nr1h3 | 0 | 1.0647283 | 0.277 | 0.136 | 0 | TAM.Vcam1 | Nr1h3 |
| Dab2 | 0 | 0.69050992 | 0.97 | 0.829 | 0 | TAM.Vcam1 | Dab2 |
| Fnbp1 | 0 | 0.42009912 | 0.838 | 0.698 | 0 | TAM.Vcam1 | Fnbp1 |
| Pld4 | 0 | 0.27912937 | 0.908 | 0.768 | 0 | TAM.Vcam1 | Pld4 |
| Plekho1 | 0 | 0.37509721 | 0.877 | 0.738 | 0 | TAM.Vcam1 | Plekho1 |
| Hexa | 0 | 0.56848809 | 0.93 | 0.791 | 0 | TAM.Vcam1 | Hexa |
| Fermt3 | 0 | 0.39172816 | 0.866 | 0.728 | 0 | TAM.Vcam1 | Fermt3 |
| Insig1 | 0 | 0.52282825 | 0.742 | 0.606 | 0 | TAM.Vcam1 | Insig1 |
| Dgkz | 0 | 0.28265551 | 0.8 | 0.665 | 0 | TAM.Vcam1 | Dgkz |
| Rnf130 | 0 | 0.31628287 | 0.821 | 0.687 | 0 | TAM.Vcam1 | Rnf130 |
| Hexb | 0 | 0.41067127 | 0.756 | 0.626 | 0 | TAM.Vcam1 | Hexb |
| Snx2 | 0 | 0.27826115 | 0.834 | 0.704 | 0 | TAM.Vcam1 | Snx2 |
| Ccl7 | 0 | 0.42976853 | 0.336 | 0.208 | 0 | TAM.Vcam1 | Ccl7 |
| Tcn2 | 0 | 0.44920433 | 0.921 | 0.793 | 0 | TAM.Vcam1 | Tcn2 |
| Mafb | 0 | 0.48504382 | 0.986 | 0.859 | 0 | TAM.Vcam1 | Mafb |
| Calr | 0 | 0.32868457 | 0.837 | 0.712 | 0 | TAM.Vcam1 | Calr |
| Adipor1 | 0 | 0.43268374 | 0.894 | 0.774 | 0 | TAM.Vcam1 | Adipor1 |
| Lamp1 | 0 | 0.36465262 | 0.906 | 0.786 | 0 | TAM.Vcam1 | Lamp1 |
| Cxcr4 | 0 | 0.25248337 | 0.872 | 0.753 | 0 | TAM.Vcam1 | Cxcr4 |
| Irs2 | 0 | 0.33139679 | 0.796 | 0.677 | 0 | TAM.Vcam1 | Irs2 |
| Inpp5d | 0 | 0.26988306 | 0.869 | 0.751 | 0 | TAM.Vcam1 | Inpp5d |
| Cltc | 0 | 0.28935047 | 0.777 | 0.659 | 0 | TAM.Vcam1 | Cltc |
| Rab14 | 0 | 0.28313106 | 0.818 | 0.702 | 0 | TAM.Vcam1 | Rab14 |
| Arrb2 | 0 | 0.39792228 | 0.944 | 0.83 | 0 | TAM.Vcam1 | Arrb2 |
| Hnrnpll | 0 | 0.26980218 | 0.708 | 0.595 | 0 | TAM.Vcam1 | Hnrnpll |
| Nckap1l | 0 | 0.44848315 | 0.915 | 0.804 | 0 | TAM.Vcam1 | Nckap1l |
| Ciita | 0 | 0.45195779 | 0.476 | 0.365 | 0 | TAM.Vcam1 | Ciita |
| Atf3 | 0 | 0.28191065 | 0.79 | 0.681 | 0 | TAM.Vcam1 | Atf3 |
| Gdi2 | 0 | 0.27353968 | 0.831 | 0.722 | 0 | TAM.Vcam1 | Gdi2 |
| Apoe | 0 | 1.32956228 | 1 | 0.896 | 0 | TAM.Vcam1 | Apoe |
| Qk | 0 | 0.43392018 | 0.917 | 0.814 | 0 | TAM.Vcam1 | Qk |
| Rnasek | 0 | 0.27671502 | 0.856 | 0.753 | 0 | TAM.Vcam1 | Rnasek |
| Vsir | 0 | 0.48093398 | 0.93 | 0.828 | 0 | TAM.Vcam1 | Vsir |
| Unc93b1 | 0 | 0.59221453 | 0.96 | 0.858 | 0 | TAM.Vcam1 | Unc93b1 |
| Eef2 | 0 | 0.26661546 | 0.899 | 0.8 | 0 | TAM.Vcam1 | Eef2 |
| Mpeg1 | 0 | 0.75465876 | 0.979 | 0.881 | 0 | TAM.Vcam1 | Mpeg1 |
| Lyn | 0 | 0.32513304 | 0.894 | 0.802 | 0 | TAM.Vcam1 | Lyn |
| Serinc3 | 0 | 0.41902086 | 0.938 | 0.851 | 0 | TAM.Vcam1 | Serinc3 |
| Sat1 | 0 | 0.48185105 | 0.968 | 0.884 | 0 | TAM.Vcam1 | Sat1 |
| Ctsa | 0 | 0.61123282 | 0.975 | 0.895 | 0 | TAM.Vcam1 | Ctsa |
| Ctsh | 0 | 0.41848098 | 0.954 | 0.874 | 0 | TAM.Vcam1 | Ctsh |
| Fcgr3 | 0 | 0.36108889 | 0.963 | 0.888 | 0 | TAM.Vcam1 | Fcgr3 |
| Kctd12 | 0 | 0.45684095 | 0.989 | 0.918 | 0 | TAM.Vcam1 | Kctd12 |
| Mmp9 | 0 | 0.49942662 | 0.276 | 0.21 | 0 | TAM.Vcam1 | Mmp9 |
| Grn | 0 | 0.4817483 | 0.989 | 0.937 | 0 | TAM.Vcam1 | Grn |
| Ctsc | 0 | 0.62076683 | 0.996 | 0.949 | 0 | TAM.Vcam1 | Ctsc |
| Itm2b | 0 | 0.5530621 | 0.992 | 0.946 | 0 | TAM.Vcam1 | Itm2b |
| Sirpa | 0 | 0.34288352 | 0.968 | 0.922 | 0 | TAM.Vcam1 | Sirpa |
| Ctsb | 0 | 0.82860735 | 0.996 | 0.959 | 0 | TAM.Vcam1 | Ctsb |
| Clta | 0 | 0.36343999 | 0.989 | 0.956 | 0 | TAM.Vcam1 | Clta |
| Csf1r | 0 | 0.37295517 | 0.987 | 0.954 | 0 | TAM.Vcam1 | Csf1r |
| Cd74 | 0 | 0.39741306 | 0.967 | 0.935 | 0 | TAM.Vcam1 | Cd74 |
| Cotl1 | 0 | 0.27342519 | 0.989 | 0.959 | 0 | TAM.Vcam1 | Cotl1 |
| Ctsz | 0 | 0.47957134 | 0.996 | 0.967 | 0 | TAM.Vcam1 | Ctsz |
| Psap | 0 | 0.40316173 | 0.994 | 0.965 | 0 | TAM.Vcam1 | Psap |
| Ppfia4 | 6.03E-308 | 1.27277559 | 0.252 | 0.107 | 1.17E-303 | TAM.Vcam1 | Ppfia4 |
| Cmklr1 | 1.52E-307 | 0.33514351 | 0.751 | 0.586 | 2.94E-303 | TAM.Vcam1 | Cmklr1 |
| Cebpg | 2.06E-306 | 0.51560571 | 0.606 | 0.422 | 3.98E-302 | TAM.Vcam1 | Cebpg |
| Sgpl1 | 1.88E-305 | 0.30842511 | 0.798 | 0.668 | 3.64E-301 | TAM.Vcam1 | Sgpl1 |
| Nampt | 3.06E-305 | 0.30120591 | 0.767 | 0.624 | 5.92E-301 | TAM.Vcam1 | Nampt |
| Blnk | 1.35E-304 | 0.97907425 | 0.374 | 0.188 | 2.62E-300 | TAM.Vcam1 | Blnk |
| Flcn | 9.68E-304 | 0.36501717 | 0.666 | 0.507 | 1.87E-299 | TAM.Vcam1 | Flcn |
| Arhgap25 | 3.83E-303 | 0.71030536 | 0.457 | 0.273 | 7.40E-299 | TAM.Vcam1 | Arhgap25 |
| Man1c1 | 5.08E-303 | 0.51685654 | 0.568 | 0.35 | 9.82E-299 | TAM.Vcam1 | Man1c1 |
| Wdr81 | 1.64E-300 | 0.61484792 | 0.534 | 0.338 | 3.17E-296 | TAM.Vcam1 | Wdr81 |
| Zfp385a | 1.63E-298 | 0.47907512 | 0.625 | 0.451 | 3.15E-294 | TAM.Vcam1 | Zfp385a |
| Hsp90b1 | 2.80E-298 | 0.37303971 | 0.758 | 0.606 | 5.42E-294 | TAM.Vcam1 | Hsp90b1 |
| Pnpla7 | 1.12E-297 | 0.61581207 | 0.549 | 0.354 | 2.16E-293 | TAM.Vcam1 | Pnpla7 |
| Evi2a | 3.23E-295 | 0.29803108 | 0.716 | 0.569 | 6.24E-291 | TAM.Vcam1 | Evi2a |
| Fuca2 | 5.39E-295 | 0.47154421 | 0.644 | 0.473 | 1.04E-290 | TAM.Vcam1 | Fuca2 |
| Mknk1 | 5.54E-295 | 0.58359567 | 0.531 | 0.354 | 1.07E-290 | TAM.Vcam1 | Mknk1 |
| Usf1 | 9.66E-294 | 0.51360636 | 0.577 | 0.39 | 1.87E-289 | TAM.Vcam1 | Usf1 |
| Mfsd12 | 1.71E-291 | 0.50684915 | 0.654 | 0.463 | 3.31E-287 | TAM.Vcam1 | Mfsd12 |
| Manf | 2.13E-291 | 0.34384089 | 0.737 | 0.597 | 4.12E-287 | TAM.Vcam1 | Manf |
| Themis2 | 9.70E-291 | 0.28481264 | 0.616 | 0.439 | 1.88E-286 | TAM.Vcam1 | Themis2 |
| Mxra1 | 5.81E-289 | 0.87323258 | 0.384 | 0.223 | 1.12E-284 | TAM.Vcam1 | Mxra1 |
| Dcakd | 9.66E-289 | 1.14367932 | 0.281 | 0.127 | 1.87E-284 | TAM.Vcam1 | Dcakd |
| Amz1 | 8.97E-287 | 0.54958004 | 0.43 | 0.297 | 1.74E-282 | TAM.Vcam1 | Amz1 |
| Slamf9 | 4.26E-285 | 0.84751243 | 0.356 | 0.194 | 8.24E-281 | TAM.Vcam1 | Slamf9 |
| Apoc2 | 8.82E-285 | 0.40819614 | 0.588 | 0.388 | 1.71E-280 | TAM.Vcam1 | Apoc2 |
| Pecam1 | 1.44E-283 | 0.65374254 | 0.358 | 0.22 | 2.78E-279 | TAM.Vcam1 | Pecam1 |
| Ptpn18 | 1.72E-283 | 0.29531203 | 0.686 | 0.506 | 3.32E-279 | TAM.Vcam1 | Ptpn18 |
| Uap1l1 | 3.95E-283 | 0.52923621 | 0.635 | 0.444 | 7.63E-279 | TAM.Vcam1 | Uap1l1 |
| Cd300ld5 | 2.58E-282 | 0.53705868 | 0.457 | 0.225 | 4.99E-278 | TAM.Vcam1 | Cd300ld5 |
| Rxra | 6.52E-282 | 0.53973198 | 0.585 | 0.401 | 1.26E-277 | TAM.Vcam1 | Rxra |
| Fdps | 4.89E-281 | 0.68054982 | 0.414 | 0.267 | 9.46E-277 | TAM.Vcam1 | Fdps |
| Os9 | 6.11E-281 | 0.40312106 | 0.718 | 0.551 | 1.18E-276 | TAM.Vcam1 | Os9 |
| Ccdc93 | 3.81E-280 | 0.79279301 | 0.406 | 0.226 | 7.36E-276 | TAM.Vcam1 | Ccdc93 |
| Itgav | 1.24E-278 | 0.43862569 | 0.649 | 0.477 | 2.40E-274 | TAM.Vcam1 | Itgav |
| Ssh2 | 5.09E-277 | 0.38439108 | 0.545 | 0.393 | 9.83E-273 | TAM.Vcam1 | Ssh2 |
| Tnfrsf21 | 5.60E-277 | 0.42752373 | 0.589 | 0.406 | 1.08E-272 | TAM.Vcam1 | Tnfrsf21 |
| Slc7a8 | 1.96E-276 | 0.35417619 | 0.654 | 0.482 | 3.79E-272 | TAM.Vcam1 | Slc7a8 |
| Nek6 | 9.25E-275 | 0.49251751 | 0.632 | 0.447 | 1.79E-270 | TAM.Vcam1 | Nek6 |
| Dhcr7 | 1.73E-273 | 1.09269569 | 0.354 | 0.175 | 3.35E-269 | TAM.Vcam1 | Dhcr7 |
| Pdgfb | 3.21E-273 | 0.78425528 | 0.402 | 0.221 | 6.22E-269 | TAM.Vcam1 | Pdgfb |
| Sema4c | 2.35E-272 | 1.22561116 | 0.252 | 0.103 | 4.54E-268 | TAM.Vcam1 | Sema4c |
| Fkbp15 | 4.31E-271 | 0.39754154 | 0.633 | 0.462 | 8.34E-267 | TAM.Vcam1 | Fkbp15 |
| Ypel3 | 4.61E-271 | 0.31789873 | 0.644 | 0.48 | 8.91E-267 | TAM.Vcam1 | Ypel3 |
| Arhgap22 | 2.99E-270 | 0.97206112 | 0.295 | 0.14 | 5.77E-266 | TAM.Vcam1 | Arhgap22 |
| Tsc22d3 | 3.63E-270 | 0.38863888 | 0.547 | 0.377 | 7.01E-266 | TAM.Vcam1 | Tsc22d3 |
| Rap2b | 7.55E-270 | 0.33890166 | 0.639 | 0.466 | 1.46E-265 | TAM.Vcam1 | Rap2b |
| Aldh1b1 | 1.18E-269 | 0.98275076 | 0.304 | 0.148 | 2.28E-265 | TAM.Vcam1 | Aldh1b1 |
| Mink1 | 6.59E-268 | 0.59285601 | 0.561 | 0.389 | 1.27E-263 | TAM.Vcam1 | Mink1 |
| Mlf2 | 5.98E-267 | 0.31533653 | 0.787 | 0.652 | 1.16E-262 | TAM.Vcam1 | Mlf2 |
| Ctnnd1 | 1.04E-265 | 0.39400457 | 0.677 | 0.507 | 2.02E-261 | TAM.Vcam1 | Ctnnd1 |
| Rgl2 | 1.02E-264 | 0.41257619 | 0.616 | 0.439 | 1.97E-260 | TAM.Vcam1 | Rgl2 |
| Ncoa4 | 1.52E-264 | 0.44060361 | 0.672 | 0.499 | 2.94E-260 | TAM.Vcam1 | Ncoa4 |
| Cd300ld3 | 2.53E-262 | 0.74556604 | 0.331 | 0.166 | 4.90E-258 | TAM.Vcam1 | Cd300ld3 |
| Plbd2 | 1.26E-261 | 0.39306184 | 0.69 | 0.536 | 2.43E-257 | TAM.Vcam1 | Plbd2 |
| Ralgds | 2.09E-261 | 0.28309838 | 0.731 | 0.575 | 4.04E-257 | TAM.Vcam1 | Ralgds |
| Bmyc | 4.23E-261 | 0.86218521 | 0.366 | 0.198 | 8.18E-257 | TAM.Vcam1 | Bmyc |
| Aagab | 5.58E-261 | 0.48643903 | 0.562 | 0.382 | 1.08E-256 | TAM.Vcam1 | Aagab |
| Lpin1 | 1.16E-260 | 0.90541677 | 0.324 | 0.172 | 2.25E-256 | TAM.Vcam1 | Lpin1 |
| Rasgrp1 | 3.94E-260 | 0.51767556 | 0.524 | 0.349 | 7.62E-256 | TAM.Vcam1 | Rasgrp1 |
| Nfxl1 | 5.20E-259 | 0.59472822 | 0.41 | 0.232 | 1.01E-254 | TAM.Vcam1 | Nfxl1 |
| Smad3 | 2.02E-257 | 0.63973024 | 0.349 | 0.197 | 3.91E-253 | TAM.Vcam1 | Smad3 |
| Mtr | 1.64E-253 | 0.58897829 | 0.49 | 0.305 | 3.16E-249 | TAM.Vcam1 | Mtr |
| Spsb1 | 3.62E-253 | 0.7925775 | 0.339 | 0.191 | 7.00E-249 | TAM.Vcam1 | Spsb1 |
| Lmo2 | 4.09E-253 | 0.30231832 | 0.661 | 0.491 | 7.90E-249 | TAM.Vcam1 | Lmo2 |
| Slamf8 | 2.66E-252 | 0.63885742 | 0.425 | 0.263 | 5.14E-248 | TAM.Vcam1 | Slamf8 |
| Glul | 1.09E-249 | 0.27641361 | 0.705 | 0.556 | 2.10E-245 | TAM.Vcam1 | Glul |
| Sesn1 | 7.60E-249 | 0.80087249 | 0.326 | 0.166 | 1.47E-244 | TAM.Vcam1 | Sesn1 |
| Cant1 | 8.53E-249 | 0.54178825 | 0.552 | 0.378 | 1.65E-244 | TAM.Vcam1 | Cant1 |
| Prkacb | 2.75E-248 | 0.5868816 | 0.527 | 0.345 | 5.33E-244 | TAM.Vcam1 | Prkacb |
| Fdft1 | 4.30E-248 | 0.897776 | 0.331 | 0.186 | 8.31E-244 | TAM.Vcam1 | Fdft1 |
| Zfp90 | 5.70E-248 | 1.02845923 | 0.281 | 0.131 | 1.10E-243 | TAM.Vcam1 | Zfp90 |
| Htra3 | 3.58E-245 | 0.60757803 | 0.438 | 0.267 | 6.92E-241 | TAM.Vcam1 | Htra3 |
| Arl11 | 4.48E-244 | 0.66335283 | 0.449 | 0.277 | 8.65E-240 | TAM.Vcam1 | Arl11 |
| Mtus1 | 1.26E-243 | 0.65775808 | 0.345 | 0.187 | 2.43E-239 | TAM.Vcam1 | Mtus1 |
| Sla | 1.68E-242 | 0.25325677 | 0.695 | 0.537 | 3.24E-238 | TAM.Vcam1 | Sla |
| Pmvk | 7.61E-241 | 0.6597487 | 0.458 | 0.295 | 1.47E-236 | TAM.Vcam1 | Pmvk |
| Rasal3 | 1.65E-240 | 0.74347004 | 0.36 | 0.207 | 3.20E-236 | TAM.Vcam1 | Rasal3 |
| Agpat3 | 8.06E-239 | 0.67468576 | 0.42 | 0.238 | 1.56E-234 | TAM.Vcam1 | Agpat3 |
| Canx | 8.55E-235 | 0.34509668 | 0.744 | 0.6 | 1.65E-230 | TAM.Vcam1 | Canx |
| Npc1 | 1.00E-234 | 0.31498971 | 0.738 | 0.583 | 1.94E-230 | TAM.Vcam1 | Npc1 |
| Mmd | 4.64E-231 | 0.74766492 | 0.391 | 0.223 | 8.97E-227 | TAM.Vcam1 | Mmd |
| Nrp1 | 1.21E-230 | 0.55182623 | 0.499 | 0.295 | 2.34E-226 | TAM.Vcam1 | Nrp1 |
| Palld | 2.75E-230 | 0.72073832 | 0.401 | 0.236 | 5.31E-226 | TAM.Vcam1 | Palld |
| Hip1 | 1.11E-228 | 0.31856692 | 0.55 | 0.394 | 2.14E-224 | TAM.Vcam1 | Hip1 |
| Impact | 8.22E-228 | 0.31485963 | 0.594 | 0.462 | 1.59E-223 | TAM.Vcam1 | Impact |
| Frmd4a | 1.66E-226 | 0.51067604 | 0.5 | 0.33 | 3.21E-222 | TAM.Vcam1 | Frmd4a |
| Tnfaip8 | 3.07E-226 | 0.36163946 | 0.606 | 0.453 | 5.94E-222 | TAM.Vcam1 | Tnfaip8 |
| Wwp1 | 6.94E-225 | 0.4078736 | 0.459 | 0.276 | 1.34E-220 | TAM.Vcam1 | Wwp1 |
| Acox3 | 5.03E-224 | 0.46348955 | 0.58 | 0.409 | 9.72E-220 | TAM.Vcam1 | Acox3 |
| Tceal9 | 2.81E-223 | 0.56194157 | 0.495 | 0.33 | 5.44E-219 | TAM.Vcam1 | Tceal9 |
| Scimp | 3.52E-223 | 0.42939775 | 0.514 | 0.356 | 6.80E-219 | TAM.Vcam1 | Scimp |
| Med25 | 4.90E-222 | 0.43842631 | 0.591 | 0.425 | 9.48E-218 | TAM.Vcam1 | Med25 |
| Tmem50b | 2.33E-221 | 0.4933898 | 0.519 | 0.367 | 4.51E-217 | TAM.Vcam1 | Tmem50b |
| Ifngr2 | 1.26E-219 | 0.2663277 | 0.707 | 0.579 | 2.44E-215 | TAM.Vcam1 | Ifngr2 |
| Fblim1 | 2.98E-219 | 0.78175205 | 0.423 | 0.234 | 5.75E-215 | TAM.Vcam1 | Fblim1 |
| Rnf180 | 4.59E-219 | 0.47518975 | 0.504 | 0.332 | 8.88E-215 | TAM.Vcam1 | Rnf180 |
| Pisd | 3.27E-218 | 0.38398183 | 0.603 | 0.456 | 6.32E-214 | TAM.Vcam1 | Pisd |
| Stard4 | 1.71E-217 | 0.67093099 | 0.361 | 0.229 | 3.30E-213 | TAM.Vcam1 | Stard4 |
| Extl3 | 9.89E-217 | 0.39636139 | 0.544 | 0.39 | 1.91E-212 | TAM.Vcam1 | Extl3 |
| Mroh1 | 4.91E-216 | 0.43588519 | 0.585 | 0.414 | 9.50E-212 | TAM.Vcam1 | Mroh1 |
| Gtf2i | 3.41E-215 | 0.57524716 | 0.503 | 0.336 | 6.59E-211 | TAM.Vcam1 | Gtf2i |
| Plscr3 | 6.02E-215 | 0.50597648 | 0.496 | 0.336 | 1.16E-210 | TAM.Vcam1 | Plscr3 |
| Naga | 2.12E-214 | 0.35084826 | 0.69 | 0.527 | 4.09E-210 | TAM.Vcam1 | Naga |
| Echs1 | 3.12E-214 | 0.54788872 | 0.501 | 0.346 | 6.03E-210 | TAM.Vcam1 | Echs1 |
| Ggt5 | 1.80E-213 | 0.91778323 | 0.293 | 0.141 | 3.49E-209 | TAM.Vcam1 | Ggt5 |
| Saraf | 4.35E-213 | 0.2658731 | 0.721 | 0.592 | 8.41E-209 | TAM.Vcam1 | Saraf |
| Ulk2 | 7.90E-213 | 0.67904107 | 0.388 | 0.228 | 1.53E-208 | TAM.Vcam1 | Ulk2 |
| Tmem214 | 2.03E-212 | 0.63061662 | 0.423 | 0.271 | 3.92E-208 | TAM.Vcam1 | Tmem214 |
| Tspan32 | 2.88E-212 | 0.75780289 | 0.3 | 0.153 | 5.56E-208 | TAM.Vcam1 | Tspan32 |
| Cuta | 4.21E-212 | 0.35810679 | 0.659 | 0.51 | 8.14E-208 | TAM.Vcam1 | Cuta |
| Colgalt1 | 4.72E-212 | 0.30444677 | 0.786 | 0.633 | 9.13E-208 | TAM.Vcam1 | Colgalt1 |
| Hlx | 2.54E-211 | 0.35902884 | 0.618 | 0.457 | 4.90E-207 | TAM.Vcam1 | Hlx |
| Ggta1 | 6.49E-211 | 0.29044235 | 0.666 | 0.512 | 1.25E-206 | TAM.Vcam1 | Ggta1 |
| Tspan4 | 5.75E-209 | 0.66001327 | 0.484 | 0.305 | 1.11E-204 | TAM.Vcam1 | Tspan4 |
| Secisbp2l | 2.87E-208 | 0.39832767 | 0.591 | 0.438 | 5.55E-204 | TAM.Vcam1 | Secisbp2l |
| Txndc16 | 8.69E-208 | 0.80622449 | 0.331 | 0.183 | 1.68E-203 | TAM.Vcam1 | Txndc16 |
| Ap2s1 | 8.82E-208 | 0.26227597 | 0.751 | 0.625 | 1.70E-203 | TAM.Vcam1 | Ap2s1 |
| Bscl2 | 1.09E-207 | 0.30792231 | 0.683 | 0.539 | 2.10E-203 | TAM.Vcam1 | Bscl2 |
| Arhgef3 | 1.39E-207 | 0.43749197 | 0.431 | 0.256 | 2.68E-203 | TAM.Vcam1 | Arhgef3 |
| Chst14 | 1.60E-206 | 0.84877967 | 0.317 | 0.172 | 3.08E-202 | TAM.Vcam1 | Chst14 |
| Slc38a7 | 5.59E-206 | 0.7839364 | 0.374 | 0.217 | 1.08E-201 | TAM.Vcam1 | Slc38a7 |
| Kcnk6 | 6.91E-206 | 0.62310128 | 0.406 | 0.255 | 1.34E-201 | TAM.Vcam1 | Kcnk6 |
| H1f0 | 9.58E-206 | 0.36754378 | 0.538 | 0.391 | 1.85E-201 | TAM.Vcam1 | H1f0 |
| Fundc1 | 7.79E-205 | 0.39930407 | 0.611 | 0.444 | 1.51E-200 | TAM.Vcam1 | Fundc1 |
| Wdr41 | 3.11E-204 | 0.55012934 | 0.485 | 0.323 | 6.02E-200 | TAM.Vcam1 | Wdr41 |
| Gpr171 | 4.64E-204 | 0.30017365 | 0.49 | 0.342 | 8.98E-200 | TAM.Vcam1 | Gpr171 |
| Ncstn | 8.39E-204 | 0.30062632 | 0.724 | 0.584 | 1.62E-199 | TAM.Vcam1 | Ncstn |
| Leprot | 1.18E-203 | 0.40192051 | 0.661 | 0.507 | 2.28E-199 | TAM.Vcam1 | Leprot |
| Folr2 | 3.85E-203 | 0.34616584 | 0.456 | 0.234 | 7.44E-199 | TAM.Vcam1 | Folr2 |
| Jmy | 3.86E-203 | 0.28321308 | 0.481 | 0.358 | 7.45E-199 | TAM.Vcam1 | Jmy |
| Ncoa3 | 5.43E-202 | 0.25714398 | 0.668 | 0.533 | 1.05E-197 | TAM.Vcam1 | Ncoa3 |
| Slc13a3 | 6.67E-202 | 0.80176661 | 0.281 | 0.145 | 1.29E-197 | TAM.Vcam1 | Slc13a3 |
| Slc36a1 | 1.46E-201 | 0.3773128 | 0.578 | 0.41 | 2.82E-197 | TAM.Vcam1 | Slc36a1 |
| Tmem51 | 2.17E-200 | 0.42578376 | 0.507 | 0.352 | 4.20E-196 | TAM.Vcam1 | Tmem51 |
| Stx12 | 5.45E-197 | 0.27702632 | 0.662 | 0.532 | 1.05E-192 | TAM.Vcam1 | Stx12 |
| Snx9 | 8.40E-197 | 0.32438368 | 0.673 | 0.515 | 1.62E-192 | TAM.Vcam1 | Snx9 |
| Cmtm3 | 8.91E-197 | 0.32357968 | 0.702 | 0.545 | 1.72E-192 | TAM.Vcam1 | Cmtm3 |
| Trim8 | 3.53E-196 | 0.26571246 | 0.723 | 0.587 | 6.82E-192 | TAM.Vcam1 | Trim8 |
| Scarb2 | 9.36E-196 | 0.29774382 | 0.677 | 0.544 | 1.81E-191 | TAM.Vcam1 | Scarb2 |

| | | | | | | | |
|---|---|---|---|---|---|---|---|
| Vegfb | 1.76E-195 | 0.87601164 | 0.347 | 0.188 | 3.40E-191 | TAM.Vcam1 | Vegfb |
| Gpr157 | 2.53E-195 | 0.86491705 | 0.271 | 0.151 | 4.89E-191 | TAM.Vcam1 | Gpr157 |
| Vps18 | 3.77E-195 | 0.39154574 | 0.6 | 0.452 | 7.28E-191 | TAM.Vcam1 | Vps18 |
| Sptssa | 3.99E-195 | 0.47710366 | 0.567 | 0.401 | 7.72E-191 | TAM.Vcam1 | Sptssa |
| Hmgcr | 8.46E-195 | 0.39202149 | 0.547 | 0.427 | 1.64E-190 | TAM.Vcam1 | Hmgcr |
| Slc12a9 | 1.09E-193 | 0.6483908 | 0.338 | 0.198 | 2.10E-189 | TAM.Vcam1 | Slc12a9 |
| Mast3 | 2.80E-193 | 0.44918984 | 0.494 | 0.342 | 5.41E-189 | TAM.Vcam1 | Mast3 |
| C5ar2 | 5.26E-192 | 0.58692971 | 0.41 | 0.249 | 1.02E-187 | TAM.Vcam1 | C5ar2 |
| Sh3gl1 | 7.50E-192 | 0.36355505 | 0.569 | 0.411 | 1.45E-187 | TAM.Vcam1 | Sh3gl1 |
| Cdc42se2 | 1.21E-191 | 0.40483507 | 0.604 | 0.454 | 2.34E-187 | TAM.Vcam1 | Cdc42se2 |
| Src | 2.68E-191 | 0.35435369 | 0.712 | 0.532 | 5.19E-187 | TAM.Vcam1 | Src |
| Calhm2 | 5.06E-191 | 0.68185889 | 0.279 | 0.147 | 9.77E-187 | TAM.Vcam1 | Calhm2 |
| Ebi3 | 4.19E-189 | 0.78831641 | 0.346 | 0.193 | 8.10E-185 | TAM.Vcam1 | Ebi3 |
| Tbc1d14 | 6.54E-189 | 0.2830844 | 0.612 | 0.451 | 1.26E-184 | TAM.Vcam1 | Tbc1d14 |
| Slc39a7 | 8.36E-187 | 0.34991821 | 0.6 | 0.46 | 1.62E-182 | TAM.Vcam1 | Slc39a7 |
| Tmem268 | 8.56E-187 | 0.66637144 | 0.354 | 0.211 | 1.65E-182 | TAM.Vcam1 | Tmem268 |
| Plpbp | 1.44E-186 | 0.39282235 | 0.49 | 0.347 | 2.78E-182 | TAM.Vcam1 | Plpbp |
| Heatr1 | 9.73E-186 | 0.35907725 | 0.483 | 0.342 | 1.88E-181 | TAM.Vcam1 | Heatr1 |
| Slc45a4 | 4.23E-185 | 0.61642623 | 0.399 | 0.252 | 8.18E-181 | TAM.Vcam1 | Slc45a4 |
| Pik3cg | 7.85E-185 | 0.32402514 | 0.531 | 0.404 | 1.52E-180 | TAM.Vcam1 | Pik3cg |
| Vps41 | 1.13E-183 | 0.60418262 | 0.393 | 0.251 | 2.19E-179 | TAM.Vcam1 | Vps41 |
| Nucb2 | 1.32E-183 | 0.51963545 | 0.457 | 0.311 | 2.56E-179 | TAM.Vcam1 | Nucb2 |
| Eif1a | 2.57E-183 | 0.41079214 | 0.555 | 0.406 | 4.97E-179 | TAM.Vcam1 | Eif1a |
| Nagpa | 2.60E-183 | 0.45559999 | 0.532 | 0.382 | 5.03E-179 | TAM.Vcam1 | Nagpa |
| Batf3 | 5.99E-183 | 0.94406015 | 0.253 | 0.124 | 1.16E-178 | TAM.Vcam1 | Batf3 |
| Usp12 | 1.23E-181 | 0.40900867 | 0.47 | 0.333 | 2.38E-177 | TAM.Vcam1 | Usp12 |
| Ap3d1 | 2.52E-181 | 0.26544493 | 0.685 | 0.546 | 4.86E-177 | TAM.Vcam1 | Ap3d1 |
| Phyhd1 | 2.61E-181 | 0.42621614 | 0.421 | 0.21 | 5.05E-177 | TAM.Vcam1 | Phyhd1 |
| Slc38a10 | 8.79E-180 | 0.42715608 | 0.568 | 0.415 | 1.70E-175 | TAM.Vcam1 | Slc38a10 |
| Fam219a | 6.09E-179 | 0.75254079 | 0.344 | 0.194 | 1.18E-174 | TAM.Vcam1 | Fam219a |
| Lmbrd2 | 8.28E-179 | 0.70910024 | 0.331 | 0.196 | 1.60E-174 | TAM.Vcam1 | Lmbrd2 |
| Ptgr2 | 2.55E-178 | 0.57532537 | 0.445 | 0.291 | 4.94E-174 | TAM.Vcam1 | Ptgr2 |
| Kif13b | 1.17E-177 | 0.45372215 | 0.429 | 0.285 | 2.26E-173 | TAM.Vcam1 | Kif13b |
| Gbp3 | 7.83E-177 | 0.53700042 | 0.357 | 0.201 | 1.51E-172 | TAM.Vcam1 | Gbp3 |
| Tmem104 | 1.78E-176 | 0.31523034 | 0.582 | 0.425 | 3.44E-172 | TAM.Vcam1 | Tmem104 |
| Vps26a | 4.18E-176 | 0.25015071 | 0.693 | 0.569 | 8.09E-172 | TAM.Vcam1 | Vps26a |
| Tceal8 | 9.12E-176 | 0.51168354 | 0.446 | 0.298 | 1.76E-171 | TAM.Vcam1 | Tceal8 |
| Mgat4a | 1.26E-175 | 0.35862529 | 0.557 | 0.418 | 2.44E-171 | TAM.Vcam1 | Mgat4a |
| Hs6st1 | 3.76E-175 | 0.37492167 | 0.548 | 0.408 | 7.26E-171 | TAM.Vcam1 | Hs6st1 |
| Ddhd2 | 9.83E-175 | 0.73450906 | 0.321 | 0.192 | 1.90E-170 | TAM.Vcam1 | Ddhd2 |
| Washc2 | 8.01E-174 | 0.28964434 | 0.664 | 0.519 | 1.55E-169 | TAM.Vcam1 | Washc2 |
| Eva1b | 9.90E-174 | 0.3551838 | 0.53 | 0.386 | 1.91E-169 | TAM.Vcam1 | Eva1b |
| Pdxk | 4.68E-173 | 0.41126571 | 0.511 | 0.376 | 9.05E-169 | TAM.Vcam1 | Pdxk |
| Rcsd1 | 6.75E-173 | 0.35846423 | 0.478 | 0.321 | 1.31E-168 | TAM.Vcam1 | Rcsd1 |
| Lpcat1 | 1.04E-172 | 0.72693049 | 0.409 | 0.251 | 2.02E-168 | TAM.Vcam1 | Lpcat1 |
| Ccpg1 | 1.55E-172 | 0.50044505 | 0.476 | 0.318 | 3.00E-168 | TAM.Vcam1 | Ccpg1 |
| Naglu | 1.90E-172 | 0.53738441 | 0.502 | 0.344 | 3.67E-168 | TAM.Vcam1 | Naglu |
| Igsf8 | 7.74E-172 | 0.36247836 | 0.676 | 0.521 | 1.50E-167 | TAM.Vcam1 | Igsf8 |
| Lgals8 | 7.83E-172 | 0.34357381 | 0.543 | 0.4 | 1.51E-167 | TAM.Vcam1 | Lgals8 |
| Prkcb | 1.01E-171 | 0.39095723 | 0.63 | 0.424 | 1.96E-167 | TAM.Vcam1 | Prkcb |
| Slc12a7 | 1.15E-171 | 0.50750026 | 0.47 | 0.3 | 2.22E-167 | TAM.Vcam1 | Slc12a7 |
| Slc16a6 | 1.32E-171 | 0.40709318 | 0.46 | 0.299 | 2.55E-167 | TAM.Vcam1 | Slc16a6 |
| Ccni | 3.44E-171 | 0.2530532 | 0.719 | 0.595 | 6.65E-167 | TAM.Vcam1 | Ccni |
| Tmem250-ps | 1.43E-170 | 0.47950395 | 0.493 | 0.342 | 2.76E-166 | TAM.Vcam1 | Tmem250-ps |
| Rnasel | 3.88E-170 | 0.42439412 | 0.47 | 0.314 | 7.50E-166 | TAM.Vcam1 | Rnasel |
| Snx17 | 7.14E-170 | 0.27237221 | 0.653 | 0.532 | 1.38E-165 | TAM.Vcam1 | Snx17 |
| Ptov1 | 1.32E-168 | 0.39277327 | 0.543 | 0.397 | 2.55E-164 | TAM.Vcam1 | Ptov1 |
| Mospd2 | 2.17E-168 | 0.44631168 | 0.502 | 0.355 | 4.20E-164 | TAM.Vcam1 | Mospd2 |
| Cnpy2 | 2.49E-168 | 0.48602016 | 0.509 | 0.357 | 4.81E-164 | TAM.Vcam1 | Cnpy2 |
| Zscan26 | 5.52E-168 | 0.4734467 | 0.422 | 0.273 | 1.07E-163 | TAM.Vcam1 | Zscan26 |
| Lrp5 | 1.08E-167 | 0.35197329 | 0.504 | 0.356 | 2.08E-163 | TAM.Vcam1 | Lrp5 |
| Rnf13 | 4.24E-167 | 0.31459289 | 0.613 | 0.474 | 8.19E-163 | TAM.Vcam1 | Rnf13 |
| Slc35f6 | 6.29E-166 | 0.36928762 | 0.531 | 0.395 | 1.22E-161 | TAM.Vcam1 | Slc35f6 |
| Commd8 | 8.36E-166 | 0.31422146 | 0.591 | 0.455 | 1.62E-161 | TAM.Vcam1 | Commd8 |
| Scamp2 | 9.34E-166 | 0.29665447 | 0.606 | 0.465 | 1.81E-161 | TAM.Vcam1 | Scamp2 |
| Sdhc | 1.10E-165 | 0.28634086 | 0.71 | 0.58 | 2.13E-161 | TAM.Vcam1 | Sdhc |
| Jup | 1.22E-165 | 0.51242827 | 0.397 | 0.239 | 2.36E-161 | TAM.Vcam1 | Jup |
| Cd300ld | 4.66E-165 | 0.25941917 | 0.64 | 0.498 | 9.01E-161 | TAM.Vcam1 | Cd300ld |
| Zdhhc14 | 7.47E-165 | 0.4412094 | 0.457 | 0.281 | 1.44E-160 | TAM.Vcam1 | Zdhhc14 |
| Tmem173 | 3.68E-164 | 0.30432822 | 0.578 | 0.433 | 7.12E-160 | TAM.Vcam1 | Tmem173 |
| 3-Mar | 3.99E-164 | 0.57963811 | 0.35 | 0.246 | 7.71E-160 | TAM.Vcam1 | 3-Mar |
| Inppl1 | 4.48E-164 | 0.3790043 | 0.539 | 0.379 | 8.66E-160 | TAM.Vcam1 | Inppl1 |
| Heatr5a | 3.17E-163 | 0.48820907 | 0.456 | 0.303 | 6.14E-159 | TAM.Vcam1 | Heatr5a |
| Dclre1c | 4.09E-163 | 0.37125001 | 0.443 | 0.291 | 7.91E-159 | TAM.Vcam1 | Dclre1c |
| Usp30 | 4.94E-163 | 0.80717504 | 0.264 | 0.143 | 9.54E-159 | TAM.Vcam1 | Usp30 |
| Rap1gds1 | 4.94E-163 | 0.27600244 | 0.624 | 0.499 | 9.56E-159 | TAM.Vcam1 | Rap1gds1 |
| Mvd | 2.83E-162 | 0.76979682 | 0.283 | 0.163 | 5.47E-158 | TAM.Vcam1 | Mvd |
| Fasn | 4.45E-162 | 0.74698113 | 0.329 | 0.192 | 8.61E-158 | TAM.Vcam1 | Fasn |
| Limk1 | 1.80E-161 | 0.64431737 | 0.366 | 0.225 | 3.47E-157 | TAM.Vcam1 | Limk1 |
| Sipa1 | 8.13E-161 | 0.32250728 | 0.568 | 0.423 | 1.57E-156 | TAM.Vcam1 | Sipa1 |
| Akr1b10 | 1.91E-160 | 0.67309015 | 0.369 | 0.227 | 3.70E-156 | TAM.Vcam1 | Akr1b10 |
| Mfsd11 | 2.70E-159 | 0.56718313 | 0.408 | 0.263 | 5.22E-155 | TAM.Vcam1 | Mfsd11 |
| Idh2 | 1.23E-158 | 0.38368123 | 0.568 | 0.385 | 2.37E-154 | TAM.Vcam1 | Idh2 |
| Cox7a2l | 1.47E-158 | 0.25493691 | 0.689 | 0.564 | 2.84E-154 | TAM.Vcam1 | Cox7a2l |
| Zfp704 | 8.25E-158 | 0.66785766 | 0.327 | 0.182 | 1.59E-153 | TAM.Vcam1 | Zfp704 |
| Rnf150 | 1.26E-157 | 0.482738 | 0.355 | 0.203 | 2.43E-153 | TAM.Vcam1 | Rnf150 |
| Ltbr | 1.52E-157 | 0.25712677 | 0.649 | 0.511 | 2.93E-153 | TAM.Vcam1 | Ltbr |
| Dbndd2 | 3.88E-157 | 0.53280714 | 0.394 | 0.263 | 7.51E-153 | TAM.Vcam1 | Dbndd2 |
| Fam91a1 | 1.99E-155 | 0.34604921 | 0.56 | 0.427 | 3.85E-151 | TAM.Vcam1 | Fam91a1 |
| Slc35c2 | 7.09E-155 | 0.49571181 | 0.447 | 0.31 | 1.37E-150 | TAM.Vcam1 | Slc35c2 |
| Dnajc3 | 8.82E-154 | 0.3049986 | 0.67 | 0.53 | 1.70E-149 | TAM.Vcam1 | Dnajc3 |
| Adamtsl4 | 1.11E-153 | 0.4491202 | 0.411 | 0.278 | 2.14E-149 | TAM.Vcam1 | Adamtsl4 |
| Tm9sf2 | 3.21E-153 | 0.33862323 | 0.614 | 0.477 | 6.20E-149 | TAM.Vcam1 | Tm9sf2 |
| Socs6 | 7.34E-153 | 0.4280814 | 0.406 | 0.276 | 1.42E-148 | TAM.Vcam1 | Socs6 |
| Aco1 | 9.07E-153 | 0.48940827 | 0.484 | 0.33 | 1.75E-148 | TAM.Vcam1 | Aco1 |
| Cpq | 1.14E-152 | 0.86096351 | 0.263 | 0.137 | 2.21E-148 | TAM.Vcam1 | Cpq |
| Tlr7 | 1.10E-151 | 0.46656242 | 0.352 | 0.228 | 2.13E-147 | TAM.Vcam1 | Tlr7 |
| Wdr91 | 4.51E-151 | 0.40397727 | 0.472 | 0.321 | 8.73E-147 | TAM.Vcam1 | Wdr91 |
| Rmnd5a | 8.67E-151 | 0.39343291 | 0.544 | 0.408 | 1.68E-146 | TAM.Vcam1 | Rmnd5a |
| Chd3 | 1.52E-150 | 0.48927462 | 0.37 | 0.228 | 2.95E-146 | TAM.Vcam1 | Chd3 |
| Acvrl1 | 2.14E-150 | 0.44657054 | 0.433 | 0.292 | 4.14E-146 | TAM.Vcam1 | Acvrl1 |
| Klc4 | 7.64E-150 | 0.56879084 | 0.397 | 0.251 | 1.48E-145 | TAM.Vcam1 | Klc4 |
| Dok1 | 2.83E-148 | 0.52465618 | 0.382 | 0.241 | 5.47E-144 | TAM.Vcam1 | Dok1 |
| Ing4 | 3.83E-148 | 0.46949944 | 0.439 | 0.3 | 7.41E-144 | TAM.Vcam1 | Ing4 |
| Pink1 | 4.75E-148 | 0.37855243 | 0.496 | 0.352 | 9.19E-144 | TAM.Vcam1 | Pink1 |
| Twf1 | 2.19E-147 | 0.3836266 | 0.556 | 0.41 | 4.24E-143 | TAM.Vcam1 | Twf1 |
| Napa | 2.47E-147 | 0.25450731 | 0.678 | 0.548 | 4.78E-143 | TAM.Vcam1 | Napa |
| Washc1 | 6.58E-147 | 0.4214874 | 0.458 | 0.32 | 1.27E-142 | TAM.Vcam1 | Washc1 |
| Eif2b2 | 1.50E-146 | 0.39355655 | 0.469 | 0.331 | 2.90E-142 | TAM.Vcam1 | Eif2b2 |
| Nrip1 | 2.39E-146 | 0.51088498 | 0.342 | 0.219 | 4.61E-142 | TAM.Vcam1 | Nrip1 |
| Cln3 | 4.02E-146 | 0.30780377 | 0.546 | 0.399 | 7.78E-142 | TAM.Vcam1 | Cln3 |
| Dcaf8 | 2.01E-143 | 0.48742529 | 0.431 | 0.294 | 3.89E-139 | TAM.Vcam1 | Dcaf8 |
| Zfhx3 | 2.24E-143 | 0.43593323 | 0.456 | 0.319 | 4.32E-139 | TAM.Vcam1 | Zfhx3 |
| Mapre2 | 3.93E-143 | 0.5197735 | 0.441 | 0.301 | 7.59E-139 | TAM.Vcam1 | Mapre2 |
| Pald1 | 4.05E-143 | 0.45634197 | 0.481 | 0.327 | 7.83E-139 | TAM.Vcam1 | Pald1 |
| Ctns | 8.55E-143 | 0.57481115 | 0.377 | 0.244 | 1.65E-138 | TAM.Vcam1 | Ctns |
| Slc25a45 | 1.01E-142 | 0.45515866 | 0.393 | 0.263 | 1.95E-138 | TAM.Vcam1 | Slc25a45 |
| Dpp7 | 3.25E-142 | 0.55362107 | 0.399 | 0.254 | 6.28E-138 | TAM.Vcam1 | Dpp7 |
| Hs3st3b1 | 5.59E-142 | 0.39566648 | 0.38 | 0.258 | 1.08E-137 | TAM.Vcam1 | Hs3st3b1 |
| Pdgfa | 8.92E-142 | 0.5394537 | 0.358 | 0.233 | 1.72E-137 | TAM.Vcam1 | Pdgfa |
| Leprotl1 | 1.31E-141 | 0.32944898 | 0.584 | 0.445 | 2.53E-137 | TAM.Vcam1 | Leprotl1 |
| Trim47 | 1.23E-140 | 0.74811466 | 0.268 | 0.143 | 2.37E-136 | TAM.Vcam1 | Trim47 |
| Ndufc2 | 1.60E-140 | 0.25061094 | 0.721 | 0.584 | 3.09E-136 | TAM.Vcam1 | Ndufc2 |
| Zfp362 | 2.32E-140 | 0.55664974 | 0.367 | 0.228 | 4.49E-136 | TAM.Vcam1 | Zfp362 |
| Ocstamp | 2.70E-140 | 0.65838781 | 0.32 | 0.196 | 5.22E-136 | TAM.Vcam1 | Ocstamp |
| Usp21 | 3.11E-140 | 0.63960306 | 0.294 | 0.175 | 6.02E-136 | TAM.Vcam1 | Usp21 |
| Pdlim4 | 3.01E-139 | 0.56192285 | 0.402 | 0.25 | 5.81E-135 | TAM.Vcam1 | Pdlim4 |
| Mcrip1 | 6.05E-139 | 0.29462824 | 0.561 | 0.426 | 1.17E-134 | TAM.Vcam1 | Mcrip1 |
| Trem2 | 1.16E-137 | 0.33684629 | 0.549 | 0.403 | 2.25E-133 | TAM.Vcam1 | Trem2 |
| Selenon | 1.42E-135 | 0.38280414 | 0.473 | 0.342 | 2.75E-131 | TAM.Vcam1 | Selenon |
| Bex3 | 6.75E-135 | 0.63604607 | 0.307 | 0.177 | 1.31E-130 | TAM.Vcam1 | Bex3 |
| Dnajc13 | 6.91E-135 | 0.25031638 | 0.59 | 0.453 | 1.34E-130 | TAM.Vcam1 | Dnajc13 |
| Cyth3 | 9.99E-135 | 0.46636689 | 0.438 | 0.289 | 1.93E-130 | TAM.Vcam1 | Cyth3 |
| Rasa1 | 2.04E-134 | 0.48388498 | 0.376 | 0.247 | 3.95E-130 | TAM.Vcam1 | Rasa1 |
| Socs2 | 4.29E-134 | 0.53700996 | 0.437 | 0.288 | 8.30E-130 | TAM.Vcam1 | Socs2 |
| Snx11 | 7.97E-133 | 0.56209894 | 0.333 | 0.209 | 1.54E-128 | TAM.Vcam1 | Snx11 |
| Slc49a4 | 2.21E-132 | 0.25303426 | 0.552 | 0.423 | 4.27E-128 | TAM.Vcam1 | Slc49a4 |
| Acbd5 | 2.25E-131 | 0.46697265 | 0.419 | 0.288 | 4.34E-127 | TAM.Vcam1 | Acbd5 |
| Paip1 | 9.06E-131 | 0.40914649 | 0.466 | 0.337 | 1.75E-126 | TAM.Vcam1 | Paip1 |
| Lrmp | 9.20E-131 | 0.45061938 | 0.396 | 0.258 | 1.78E-126 | TAM.Vcam1 | Lrmp |
| Klhl18 | 1.36E-130 | 0.61631693 | 0.275 | 0.164 | 2.64E-126 | TAM.Vcam1 | Klhl18 |
| Pde1b | 1.45E-130 | 0.35520775 | 0.44 | 0.317 | 2.81E-126 | TAM.Vcam1 | Pde1b |
| Plcb2 | 2.86E-130 | 0.43364024 | 0.428 | 0.292 | 5.54E-126 | TAM.Vcam1 | Plcb2 |
| Gapvd1 | 4.87E-130 | 0.3087631 | 0.541 | 0.41 | 9.41E-126 | TAM.Vcam1 | Gapvd1 |
| Nudt16l1 | 1.14E-129 | 0.3605543 | 0.452 | 0.32 | 2.21E-125 | TAM.Vcam1 | Nudt16l1 |
| Wdr11 | 2.99E-129 | 0.63050855 | 0.269 | 0.16 | 5.78E-125 | TAM.Vcam1 | Wdr11 |
| Tlk1 | 4.34E-129 | 0.41801552 | 0.381 | 0.262 | 8.40E-125 | TAM.Vcam1 | Tlk1 |
| Letmd1 | 5.10E-129 | 0.66958799 | 0.27 | 0.155 | 9.85E-125 | TAM.Vcam1 | Letmd1 |
| Nenf | 5.55E-129 | 0.49106112 | 0.408 | 0.27 | 1.07E-124 | TAM.Vcam1 | Nenf |
| Clcn7 | 6.98E-129 | 0.31786702 | 0.597 | 0.444 | 1.35E-124 | TAM.Vcam1 | Clcn7 |
| Jak3 | 2.80E-128 | 0.34595858 | 0.429 | 0.297 | 5.41E-124 | TAM.Vcam1 | Jak3 |
| Mospd3 | 5.31E-128 | 0.38430649 | 0.454 | 0.332 | 1.03E-123 | TAM.Vcam1 | Mospd3 |
| Tmx1 | 7.44E-128 | 0.34710555 | 0.632 | 0.489 | 1.44E-123 | TAM.Vcam1 | Tmx1 |
| Tnfsf12 | 1.21E-127 | 0.2500633 | 0.572 | 0.439 | 2.34E-123 | TAM.Vcam1 | Tnfsf12 |
| Sgsh | 1.50E-127 | 0.6695459 | 0.273 | 0.162 | 2.90E-123 | TAM.Vcam1 | Sgsh |
| Dtx3 | 1.51E-127 | 0.59733372 | 0.323 | 0.199 | 2.91E-123 | TAM.Vcam1 | Dtx3 |
| Tk2 | 6.33E-127 | 0.59099531 | 0.3 | 0.18 | 1.22E-122 | TAM.Vcam1 | Tk2 |
| Slco2b1 | 8.58E-127 | 0.59135632 | 0.264 | 0.121 | 1.66E-122 | TAM.Vcam1 | Slco2b1 |
| Vkorc1 | 1.19E-126 | 0.67790575 | 0.346 | 0.213 | 2.30E-122 | TAM.Vcam1 | Vkorc1 |
| Brox | 2.28E-126 | 0.37330228 | 0.453 | 0.323 | 4.42E-122 | TAM.Vcam1 | Brox |
| Anpep | 2.49E-126 | 0.25907712 | 0.643 | 0.467 | 4.81E-122 | TAM.Vcam1 | Anpep |
| Pip4k2b | 3.23E-126 | 0.42452492 | 0.456 | 0.332 | 6.25E-122 | TAM.Vcam1 | Pip4k2b |
| Slc38a6 | 4.46E-126 | 0.35257094 | 0.515 | 0.351 | 8.61E-122 | TAM.Vcam1 | Slc38a6 |
| Trim41 | 5.21E-126 | 0.34833169 | 0.454 | 0.323 | 1.01E-121 | TAM.Vcam1 | Trim41 |
| Acox1 | 8.74E-126 | 0.3419811 | 0.509 | 0.38 | 1.69E-121 | TAM.Vcam1 | Acox1 |
| Zcchc24 | 1.14E-125 | 0.25306521 | 0.45 | 0.309 | 2.21E-121 | TAM.Vcam1 | Zcchc24 |
| Acss1 | 1.71E-125 | 0.53695975 | 0.279 | 0.176 | 3.31E-121 | TAM.Vcam1 | Acss1 |
| Phc2 | 1.83E-125 | 0.25426891 | 0.565 | 0.445 | 3.53E-121 | TAM.Vcam1 | Phc2 |
| Ubald1 | 2.56E-125 | 0.46537815 | 0.426 | 0.298 | 4.95E-121 | TAM.Vcam1 | Ubald1 |
| Nudt9 | 2.74E-124 | 0.38689562 | 0.499 | 0.36 | 5.31E-120 | TAM.Vcam1 | Nudt9 |
| Pcyox1 | 1.26E-123 | 0.38481317 | 0.517 | 0.353 | 2.44E-119 | TAM.Vcam1 | Pcyox1 |
| Rbfa | 2.07E-123 | 0.39854152 | 0.44 | 0.311 | 4.00E-119 | TAM.Vcam1 | Rbfa |
| Tecr | 4.00E-123 | 0.31107462 | 0.606 | 0.467 | 7.73E-119 | TAM.Vcam1 | Tecr |
| Dmac1 | 1.66E-122 | 0.48502141 | 0.361 | 0.241 | 3.21E-118 | TAM.Vcam1 | Dmac1 |
| Nnt | 3.36E-122 | 0.45900173 | 0.397 | 0.272 | 6.50E-118 | TAM.Vcam1 | Nnt |
| Hps3 | 3.93E-122 | 0.41590115 | 0.346 | 0.23 | 7.61E-118 | TAM.Vcam1 | Hps3 |
| Cib1 | 4.09E-122 | 0.34566861 | 0.433 | 0.321 | 7.91E-118 | TAM.Vcam1 | Cib1 |
| Arhgap27 | 2.78E-121 | 0.42445376 | 0.347 | 0.23 | 5.38E-117 | TAM.Vcam1 | Arhgap27 |
| Tfeb | 2.99E-121 | 0.41286965 | 0.408 | 0.283 | 5.79E-117 | TAM.Vcam1 | Tfeb |
| Rxrb | 4.57E-121 | 0.31595349 | 0.444 | 0.323 | 8.84E-117 | TAM.Vcam1 | Rxrb |
| Nrbf2 | 9.67E-121 | 0.45399964 | 0.36 | 0.237 | 1.87E-116 | TAM.Vcam1 | Nrbf2 |
| Ophn1 | 1.60E-120 | 0.54207308 | 0.265 | 0.146 | 3.09E-116 | TAM.Vcam1 | Ophn1 |
| Ifi27 | 9.12E-120 | 0.43879991 | 0.522 | 0.371 | 1.76E-115 | TAM.Vcam1 | Ifi27 |
| Pank3 | 1.67E-119 | 0.39254366 | 0.453 | 0.328 | 3.23E-115 | TAM.Vcam1 | Pank3 |
| Blvra | 1.86E-119 | 0.33324259 | 0.571 | 0.445 | 3.60E-115 | TAM.Vcam1 | Blvra |
| Arhgap18 | 2.59E-119 | 0.33980468 | 0.385 | 0.264 | 5.01E-115 | TAM.Vcam1 | Arhgap18 |
| B3gnt8 | 6.05E-119 | 0.47013725 | 0.333 | 0.216 | 1.17E-114 | TAM.Vcam1 | B3gnt8 |
| Fam3c | 8.07E-119 | 0.36557777 | 0.508 | 0.373 | 1.56E-114 | TAM.Vcam1 | Fam3c |
| Tbc1d13 | 1.03E-118 | 0.37766419 | 0.481 | 0.348 | 1.99E-114 | TAM.Vcam1 | Tbc1d13 |
| Txlna | 1.40E-118 | 0.31356511 | 0.471 | 0.344 | 2.71E-114 | TAM.Vcam1 | Txlna |
| Plagl2 | 2.21E-118 | 0.37929613 | 0.424 | 0.305 | 4.27E-114 | TAM.Vcam1 | Plagl2 |
| Vps8 | 8.24E-118 | 0.39111679 | 0.398 | 0.292 | 1.59E-113 | TAM.Vcam1 | Vps8 |
| Nod1 | 9.73E-118 | 0.31125349 | 0.317 | 0.205 | 1.88E-113 | TAM.Vcam1 | Nod1 |
| Rgmb | 2.85E-117 | 0.55574101 | 0.321 | 0.202 | 5.51E-113 | TAM.Vcam1 | Rgmb |
| Prkab1 | 1.52E-116 | 0.6805807 | 0.256 | 0.149 | 2.94E-112 | TAM.Vcam1 | Prkab1 |
| Med22 | 4.62E-116 | 0.55311183 | 0.303 | 0.197 | 8.94E-112 | TAM.Vcam1 | Med22 |
| Anks1 | 8.53E-116 | 0.2983089 | 0.448 | 0.336 | 1.65E-111 | TAM.Vcam1 | Anks1 |
| Hs1bp3 | 1.00E-115 | 0.59127609 | 0.304 | 0.189 | 1.94E-111 | TAM.Vcam1 | Hs1bp3 |
| Pcyt2 | 1.13E-115 | 0.61525129 | 0.288 | 0.179 | 2.18E-111 | TAM.Vcam1 | Pcyt2 |
| Anks3 | 1.27E-115 | 0.44338954 | 0.345 | 0.23 | 2.46E-111 | TAM.Vcam1 | Anks3 |
| Rnf145 | 1.52E-115 | 0.33809581 | 0.444 | 0.317 | 2.93E-111 | TAM.Vcam1 | Rnf145 |
| Pdcl3 | 3.70E-114 | 0.29415347 | 0.494 | 0.385 | 7.15E-110 | TAM.Vcam1 | Pdcl3 |
| Slc25a11 | 4.58E-114 | 0.29735179 | 0.566 | 0.437 | 8.86E-110 | TAM.Vcam1 | Slc25a11 |
| Pld2 | 1.08E-113 | 0.51646939 | 0.349 | 0.227 | 2.08E-109 | TAM.Vcam1 | Pld2 |
| Fam217b | 2.07E-112 | 0.61148314 | 0.253 | 0.153 | 4.01E-108 | TAM.Vcam1 | Fam217b |
| Slc30a5 | 4.87E-112 | 0.44238732 | 0.424 | 0.3 | 9.41E-108 | TAM.Vcam1 | Slc30a5 |
| Irak4 | 7.03E-112 | 0.28130009 | 0.5 | 0.379 | 1.36E-107 | TAM.Vcam1 | Irak4 |
| Ppp1r21 | 1.89E-111 | 0.29612147 | 0.451 | 0.329 | 3.65E-107 | TAM.Vcam1 | Ppp1r21 |
| Pigt | 3.03E-111 | 0.31472139 | 0.56 | 0.428 | 5.86E-107 | TAM.Vcam1 | Pigt |
| Fam53b | 1.16E-110 | 0.33323266 | 0.356 | 0.236 | 2.24E-106 | TAM.Vcam1 | Fam53b |
| Tbcc | 1.33E-110 | 0.5515615 | 0.287 | 0.179 | 2.57E-106 | TAM.Vcam1 | Tbcc |
| Tmem87a | 1.43E-110 | 0.49980282 | 0.328 | 0.219 | 2.76E-106 | TAM.Vcam1 | Tmem87a |
| Pik3r2 | 4.09E-110 | 0.38198562 | 0.407 | 0.281 | 7.90E-106 | TAM.Vcam1 | Pik3r2 |
| Vps11 | 4.30E-110 | 0.31608638 | 0.456 | 0.333 | 8.31E-106 | TAM.Vcam1 | Vps11 |
| Trim14 | 4.91E-110 | 0.31474899 | 0.393 | 0.274 | 9.50E-106 | TAM.Vcam1 | Trim14 |
| Cyb5r1 | 6.27E-110 | 0.42071629 | 0.367 | 0.252 | 1.21E-105 | TAM.Vcam1 | Cyb5r1 |
| Tns1 | 9.06E-110 | 0.59184629 | 0.295 | 0.165 | 1.75E-105 | TAM.Vcam1 | Tns1 |
| C2cd2l | 1.52E-109 | 0.38171751 | 0.42 | 0.297 | 2.94E-105 | TAM.Vcam1 | C2cd2l |
| Scn1b | 1.64E-109 | 0.32761862 | 0.358 | 0.222 | 3.17E-105 | TAM.Vcam1 | Scn1b |
| Wls | 2.93E-109 | 0.39171813 | 0.434 | 0.309 | 5.66E-105 | TAM.Vcam1 | Wls |
| Tex264 | 1.04E-108 | 0.37346265 | 0.487 | 0.347 | 2.02E-104 | TAM.Vcam1 | Tex264 |
| Slf2 | 1.36E-108 | 0.37689738 | 0.388 | 0.281 | 2.62E-104 | TAM.Vcam1 | Slf2 |
| Oxa1l | 1.74E-108 | 0.39726056 | 0.413 | 0.293 | 3.36E-104 | TAM.Vcam1 | Oxa1l |
| Snx30 | 3.70E-108 | 0.36149436 | 0.378 | 0.257 | 7.15E-104 | TAM.Vcam1 | Snx30 |
| Tle5 | 4.47E-108 | 0.28626256 | 0.617 | 0.476 | 8.65E-104 | TAM.Vcam1 | Tle5 |
| Atmin | 8.86E-108 | 0.40811115 | 0.372 | 0.25 | 1.71E-103 | TAM.Vcam1 | Atmin |
| Zfp513 | 1.64E-107 | 0.39271402 | 0.336 | 0.228 | 3.16E-103 | TAM.Vcam1 | Zfp513 |
| Cracr2b | 1.57E-106 | 0.63924021 | 0.309 | 0.191 | 3.03E-102 | TAM.Vcam1 | Cracr2b |
| Tpcn2 | 1.72E-106 | 0.34869446 | 0.474 | 0.336 | 3.32E-102 | TAM.Vcam1 | Tpcn2 |
| Nek7 | 2.44E-106 | 0.28097023 | 0.478 | 0.367 | 4.72E-102 | TAM.Vcam1 | Nek7 |
| Irf2bpl | 4.03E-106 | 0.42343577 | 0.278 | 0.174 | 7.78E-102 | TAM.Vcam1 | Irf2bpl |
| Sap25 | 4.72E-106 | 0.45229514 | 0.305 | 0.192 | 9.12E-102 | TAM.Vcam1 | Sap25 |
| Wbp2 | 1.18E-105 | 0.29746839 | 0.537 | 0.407 | 2.27E-101 | TAM.Vcam1 | Wbp2 |
| Bahd1 | 3.72E-105 | 0.39016403 | 0.369 | 0.255 | 7.20E-101 | TAM.Vcam1 | Bahd1 |
| Scly | 2.33E-104 | 0.56492095 | 0.267 | 0.161 | 4.50E-100 | TAM.Vcam1 | Scly |
| Ppm1h | 3.17E-104 | 0.38288692 | 0.378 | 0.264 | 6.13E-100 | TAM.Vcam1 | Ppm1h |
| Siglece | 3.36E-104 | 0.36406239 | 0.309 | 0.209 | 6.50E-100 | TAM.Vcam1 | Siglece |
| Ap5b1 | 4.01E-104 | 0.41288746 | 0.324 | 0.222 | 7.76E-100 | TAM.Vcam1 | Ap5b1 |
| Ap4m1 | 8.33E-104 | 0.51881489 | 0.278 | 0.176 | 1.61E-99 | TAM.Vcam1 | Ap4m1 |
| Slc25a1 | 6.33E-103 | 0.38004512 | 0.47 | 0.342 | 1.22E-98 | TAM.Vcam1 | Slc25a1 |
| Clasp2 | 7.28E-103 | 0.39648468 | 0.406 | 0.29 | 1.41E-98 | TAM.Vcam1 | Clasp2 |
| Pgpep1 | 1.82E-102 | 0.60359026 | 0.26 | 0.154 | 3.53E-98 | TAM.Vcam1 | Pgpep1 |
| Dock4 | 2.08E-102 | 0.31855572 | 0.356 | 0.245 | 4.03E-98 | TAM.Vcam1 | Dock4 |
| Ccnd1 | 2.21E-102 | 0.50705053 | 0.319 | 0.18 | 4.27E-98 | TAM.Vcam1 | Ccnd1 |
| Dnajc16 | 2.55E-102 | 0.50415818 | 0.307 | 0.199 | 4.93E-98 | TAM.Vcam1 | Dnajc16 |
| Arfgef2 | 7.38E-102 | 0.57797165 | 0.25 | 0.151 | 1.43E-97 | TAM.Vcam1 | Arfgef2 |
| Itpripl1 | 8.30E-101 | 0.52261636 | 0.257 | 0.161 | 1.60E-96 | TAM.Vcam1 | Itpripl1 |
| Tlr9 | 1.03E-100 | 0.39958208 | 0.271 | 0.157 | 2.00E-96 | TAM.Vcam1 | Tlr9 |
| Adi1 | 1.10E-100 | 0.61007523 | 0.29 | 0.18 | 2.12E-96 | TAM.Vcam1 | Adi1 |
| Mad1l1 | 1.13E-100 | 0.58926038 | 0.285 | 0.181 | 2.18E-96 | TAM.Vcam1 | Mad1l1 |
| Hic1 | 1.53E-100 | 0.37737359 | 0.259 | 0.165 | 2.95E-96 | TAM.Vcam1 | Hic1 |
| Dram2 | 2.17E-100 | 0.34448243 | 0.403 | 0.292 | 4.19E-96 | TAM.Vcam1 | Dram2 |
| Borcs6 | 6.64E-100 | 0.27527921 | 0.392 | 0.276 | 1.28E-95 | TAM.Vcam1 | Borcs6 |
| Gfpt1 | 8.88E-100 | 0.38421107 | 0.388 | 0.268 | 1.72E-95 | TAM.Vcam1 | Gfpt1 |
| Clcn5 | 1.84E-99 | 0.393154 | 0.386 | 0.271 | 3.56E-95 | TAM.Vcam1 | Clcn5 |
| Galk2 | 2.26E-99 | 0.36441459 | 0.437 | 0.321 | 4.37E-95 | TAM.Vcam1 | Galk2 |
| Rab7b | 2.37E-99 | 0.29109588 | 0.429 | 0.313 | 4.58E-95 | TAM.Vcam1 | Rab7b |
| Eif4ebp2 | 2.89E-99 | 0.25531058 | 0.554 | 0.436 | 5.59E-95 | TAM.Vcam1 | Eif4ebp2 |
| Gatd3a | 3.92E-99 | 0.48986966 | 0.311 | 0.207 | 7.57E-95 | TAM.Vcam1 | Gatd3a |
| Rab12 | 1.70E-98 | 0.32835066 | 0.454 | 0.339 | 3.29E-94 | TAM.Vcam1 | Rab12 |
| Tbc1d5 | 1.81E-98 | 0.26363558 | 0.445 | 0.334 | 3.49E-94 | TAM.Vcam1 | Tbc1d5 |
| Sash3 | 2.28E-98 | 0.34026892 | 0.405 | 0.293 | 4.41E-94 | TAM.Vcam1 | Sash3 |
| Ipo13 | 1.70E-97 | 0.46325591 | 0.38 | 0.258 | 3.28E-93 | TAM.Vcam1 | Ipo13 |
| Arhgap4 | 2.18E-97 | 0.30863374 | 0.371 | 0.258 | 4.22E-93 | TAM.Vcam1 | Arhgap4 |
| Atxn7l3b | 3.17E-97 | 0.25030419 | 0.488 | 0.362 | 6.13E-93 | TAM.Vcam1 | Atxn7l3b |
| Gga2 | 1.01E-96 | 0.70532745 | 0.314 | 0.186 | 1.96E-92 | TAM.Vcam1 | Gga2 |
| Arhgef6 | 1.01E-96 | 0.3076943 | 0.439 | 0.312 | 1.96E-92 | TAM.Vcam1 | Arhgef6 |
| Myc | 1.93E-96 | 0.33069202 | 0.38 | 0.251 | 3.74E-92 | TAM.Vcam1 | Myc |
| Acsf2 | 2.10E-96 | 0.5612994 | 0.251 | 0.152 | 4.05E-92 | TAM.Vcam1 | Acsf2 |
| Dmxl1 | 5.38E-96 | 0.35540357 | 0.393 | 0.282 | 1.04E-91 | TAM.Vcam1 | Dmxl1 |
| Haghl | 8.00E-96 | 0.53242369 | 0.253 | 0.159 | 1.55E-91 | TAM.Vcam1 | Haghl |
| Idua | 8.05E-96 | 0.47869383 | 0.359 | 0.242 | 1.56E-91 | TAM.Vcam1 | Idua |
| Zfyve26 | 1.27E-95 | 0.42143656 | 0.322 | 0.212 | 2.45E-91 | TAM.Vcam1 | Zfyve26 |
| Slc25a51 | 2.31E-95 | 0.28104692 | 0.519 | 0.398 | 4.46E-91 | TAM.Vcam1 | Slc25a51 |
| Atxn1 | 3.40E-95 | 0.40711032 | 0.366 | 0.257 | 6.58E-91 | TAM.Vcam1 | Atxn1 |
| Eri3 | 6.31E-95 | 0.58159596 | 0.288 | 0.187 | 1.22E-90 | TAM.Vcam1 | Eri3 |
| Kif3b | 7.54E-95 | 0.28622817 | 0.442 | 0.328 | 1.46E-90 | TAM.Vcam1 | Kif3b |
| Pkib | 9.97E-95 | 0.43350498 | 0.324 | 0.223 | 1.93E-90 | TAM.Vcam1 | Pkib |
| Pygb | 1.24E-94 | 0.34190024 | 0.418 | 0.297 | 2.40E-90 | TAM.Vcam1 | Pygb |
| Tmem229b | 1.40E-94 | 0.49674653 | 0.289 | 0.183 | 2.70E-90 | TAM.Vcam1 | Tmem229b |
| Apeh | 2.62E-94 | 0.37061764 | 0.368 | 0.256 | 5.07E-90 | TAM.Vcam1 | Apeh |
| Smim10l1 | 1.22E-93 | 0.46194325 | 0.361 | 0.246 | 2.36E-89 | TAM.Vcam1 | Smim10l1 |
| Usp36 | 1.93E-93 | 0.38313303 | 0.363 | 0.255 | 3.74E-89 | TAM.Vcam1 | Usp36 |
| Slc35a5 | 5.90E-93 | 0.39560672 | 0.366 | 0.261 | 1.14E-88 | TAM.Vcam1 | Slc35a5 |
| Vps39 | 2.49E-92 | 0.49925656 | 0.308 | 0.201 | 4.81E-88 | TAM.Vcam1 | Vps39 |
| Pkdcc | 6.66E-92 | 0.52627575 | 0.271 | 0.169 | 1.29E-87 | TAM.Vcam1 | Pkdcc |
| Cryl1 | 1.46E-91 | 0.41499972 | 0.41 | 0.287 | 2.82E-87 | TAM.Vcam1 | Cryl1 |
| Gnptg | 2.43E-91 | 0.49670845 | 0.304 | 0.197 | 4.70E-87 | TAM.Vcam1 | Gnptg |
| Eng | 2.30E-90 | 0.37202561 | 0.364 | 0.249 | 4.45E-86 | TAM.Vcam1 | Eng |
| Trp53inp2 | 2.78E-90 | 0.36039118 | 0.394 | 0.281 | 5.37E-86 | TAM.Vcam1 | Trp53inp2 |
| Rabl6 | 4.70E-90 | 0.33758579 | 0.429 | 0.318 | 9.09E-86 | TAM.Vcam1 | Rabl6 |
| Slc30a7 | 1.67E-89 | 0.32392356 | 0.392 | 0.29 | 3.23E-85 | TAM.Vcam1 | Slc30a7 |
| Mavs | 2.05E-89 | 0.54426626 | 0.273 | 0.174 | 3.97E-85 | TAM.Vcam1 | Mavs |
| Vps33a | 2.40E-89 | 0.45597817 | 0.281 | 0.186 | 4.63E-85 | TAM.Vcam1 | Vps33a |
| Dapk3 | 2.78E-89 | 0.33349766 | 0.401 | 0.295 | 5.37E-85 | TAM.Vcam1 | Dapk3 |
| Dvl2 | 1.36E-88 | 0.29220567 | 0.333 | 0.234 | 2.63E-84 | TAM.Vcam1 | Dvl2 |
| Dctn5 | 1.79E-88 | 0.29534414 | 0.392 | 0.285 | 3.47E-84 | TAM.Vcam1 | Dctn5 |
| Lpgat1 | 1.80E-88 | 0.28350713 | 0.443 | 0.329 | 3.47E-84 | TAM.Vcam1 | Lpgat1 |
| Fgd4 | 3.30E-88 | 0.26886742 | 0.325 | 0.219 | 6.38E-84 | TAM.Vcam1 | Fgd4 |
| Chd9 | 6.81E-88 | 0.32700547 | 0.35 | 0.243 | 1.32E-83 | TAM.Vcam1 | Chd9 |
| Arhgap39 | 9.29E-88 | 0.37553134 | 0.29 | 0.19 | 1.80E-83 | TAM.Vcam1 | Arhgap39 |
| Fadd | 1.39E-87 | 0.39884842 | 0.303 | 0.2 | 2.68E-83 | TAM.Vcam1 | Fadd |
| Asb8 | 1.78E-87 | 0.43430832 | 0.29 | 0.187 | 3.44E-83 | TAM.Vcam1 | Asb8 |
| Gba | 3.17E-87 | 0.25174641 | 0.542 | 0.436 | 6.13E-83 | TAM.Vcam1 | Gba |
| Helz | 3.90E-87 | 0.45058346 | 0.278 | 0.182 | 7.53E-83 | TAM.Vcam1 | Helz |
| Glb1 | 1.14E-86 | 0.36056772 | 0.455 | 0.315 | 2.21E-82 | TAM.Vcam1 | Glb1 |
| Qdpr | 1.58E-86 | 0.3292888 | 0.553 | 0.423 | 3.05E-82 | TAM.Vcam1 | Qdpr |
| Nbr1 | 2.09E-86 | 0.47290591 | 0.302 | 0.2 | 4.04E-82 | TAM.Vcam1 | Nbr1 |
| Tmed3 | 2.40E-86 | 0.26478932 | 0.534 | 0.422 | 4.65E-82 | TAM.Vcam1 | Tmed3 |
| Snx13 | 6.64E-86 | 0.35728096 | 0.353 | 0.244 | 1.28E-81 | TAM.Vcam1 | Snx13 |
| Fam173a | 7.98E-86 | 0.29991686 | 0.427 | 0.319 | 1.54E-81 | TAM.Vcam1 | Fam173a |
| Krt8 | 1.12E-85 | 0.31061934 | 0.333 | 0.232 | 2.16E-81 | TAM.Vcam1 | Krt8 |
| Iffo1 | 1.70E-85 | 0.32393472 | 0.267 | 0.175 | 3.28E-81 | TAM.Vcam1 | Iffo1 |
| Npepl1 | 2.73E-85 | 0.31837371 | 0.48 | 0.358 | 5.28E-81 | TAM.Vcam1 | Npepl1 |
| Oxct1 | 3.11E-85 | 0.42442233 | 0.374 | 0.252 | 6.00E-81 | TAM.Vcam1 | Oxct1 |
| Phactr2 | 3.82E-85 | 0.39481531 | 0.301 | 0.201 | 7.39E-81 | TAM.Vcam1 | Phactr2 |
| Vps52 | 4.45E-85 | 0.35003186 | 0.399 | 0.285 | 8.60E-81 | TAM.Vcam1 | Vps52 |
| Ldhb | 5.85E-85 | 0.58721448 | 0.251 | 0.155 | 1.13E-80 | TAM.Vcam1 | Ldhb |
| Mdh1 | 1.16E-84 | 0.27920372 | 0.587 | 0.449 | 2.24E-80 | TAM.Vcam1 | Mdh1 |
| Tmem106b | 1.54E-84 | 0.42741324 | 0.342 | 0.239 | 2.98E-80 | TAM.Vcam1 | Tmem106b |
| Trim35 | 3.62E-84 | 0.31547313 | 0.444 | 0.326 | 7.00E-80 | TAM.Vcam1 | Trim35 |
| Pggt1b | 4.89E-84 | 0.38993509 | 0.334 | 0.23 | 9.45E-80 | TAM.Vcam1 | Pggt1b |
| Patz1 | 5.22E-84 | 0.52938832 | 0.265 | 0.166 | 1.01E-79 | TAM.Vcam1 | Patz1 |
| Fmn1 | 1.04E-83 | 0.40484717 | 0.368 | 0.257 | 2.00E-79 | TAM.Vcam1 | Fmn1 |
| Plcb3 | 2.20E-83 | 0.35139618 | 0.358 | 0.256 | 4.25E-79 | TAM.Vcam1 | Plcb3 |
| Acads | 7.34E-83 | 0.40004042 | 0.398 | 0.285 | 1.42E-78 | TAM.Vcam1 | Acads |
| Il18 | 1.38E-82 | 0.30498488 | 0.256 | 0.16 | 2.67E-78 | TAM.Vcam1 | Il18 |
| Prr12 | 1.44E-82 | 0.36300749 | 0.37 | 0.259 | 2.77E-78 | TAM.Vcam1 | Prr12 |
| Smim15 | 1.65E-82 | 0.3319254 | 0.361 | 0.258 | 3.19E-78 | TAM.Vcam1 | Smim15 |
| Stx2 | 2.21E-82 | 0.37465241 | 0.379 | 0.266 | 4.27E-78 | TAM.Vcam1 | Stx2 |
| Zbtb38 | 3.27E-82 | 0.42802955 | 0.297 | 0.19 | 6.33E-78 | TAM.Vcam1 | Zbtb38 |
| Mms19 | 3.47E-82 | 0.46022209 | 0.289 | 0.191 | 6.71E-78 | TAM.Vcam1 | Mms19 |
| Rcan3 | 3.71E-82 | 0.48464602 | 0.298 | 0.198 | 7.17E-78 | TAM.Vcam1 | Rcan3 |
| Gng12 | 1.02E-81 | 0.26562364 | 0.395 | 0.288 | 1.97E-77 | TAM.Vcam1 | Gng12 |
| Tomm40l | 1.75E-81 | 0.39280253 | 0.352 | 0.247 | 3.38E-77 | TAM.Vcam1 | Tomm40l |
| Txndc15 | 5.17E-81 | 0.33730143 | 0.434 | 0.318 | 1.00E-76 | TAM.Vcam1 | Txndc15 |
| Mocs2 | 1.37E-80 | 0.31042096 | 0.426 | 0.306 | 2.64E-76 | TAM.Vcam1 | Mocs2 |
| Armcx3 | 2.02E-80 | 0.29502524 | 0.362 | 0.262 | 3.91E-76 | TAM.Vcam1 | Armcx3 |
| Gnpda1 | 2.96E-80 | 0.28745498 | 0.38 | 0.271 | 5.72E-76 | TAM.Vcam1 | Gnpda1 |
| Uqcc3 | 3.12E-80 | 0.35083268 | 0.342 | 0.248 | 6.02E-76 | TAM.Vcam1 | Uqcc3 |
| Swap70 | 4.07E-80 | 0.27415531 | 0.464 | 0.338 | 7.87E-76 | TAM.Vcam1 | Swap70 |
| Zfp740 | 4.44E-80 | 0.35336596 | 0.336 | 0.238 | 8.59E-76 | TAM.Vcam1 | Zfp740 |
| Akap8l | 6.35E-80 | 0.3794768 | 0.287 | 0.193 | 1.23E-75 | TAM.Vcam1 | Akap8l |
| Pilrb2 | 1.40E-79 | 0.25895777 | 0.407 | 0.288 | 2.71E-75 | TAM.Vcam1 | Pilrb2 |
| Relch | 2.74E-79 | 0.32043533 | 0.291 | 0.201 | 5.30E-75 | TAM.Vcam1 | Relch |
| Tesk1 | 6.01E-79 | 0.25169931 | 0.46 | 0.346 | 1.16E-74 | TAM.Vcam1 | Tesk1 |
| Sh3pxd2a | 7.47E-79 | 0.3746313 | 0.287 | 0.184 | 1.44E-74 | TAM.Vcam1 | Sh3pxd2a |
| Plaat3 | 3.01E-78 | 0.3995772 | 0.269 | 0.168 | 5.82E-74 | TAM.Vcam1 | Plaat3 |
| Fibp | 6.14E-78 | 0.27423192 | 0.437 | 0.337 | 1.19E-73 | TAM.Vcam1 | Fibp |
| Lsm10 | 8.16E-78 | 0.26698632 | 0.333 | 0.242 | 1.58E-73 | TAM.Vcam1 | Lsm10 |
| Gnl3 | 8.98E-78 | 0.30911812 | 0.412 | 0.31 | 1.74E-73 | TAM.Vcam1 | Gnl3 |
| Erlin2 | 1.24E-76 | 0.34789178 | 0.392 | 0.285 | 2.39E-72 | TAM.Vcam1 | Erlin2 |
| Ctso | 1.27E-76 | 0.33944159 | 0.384 | 0.275 | 2.45E-72 | TAM.Vcam1 | Ctso |
| Rufy3 | 3.36E-76 | 0.32879384 | 0.365 | 0.262 | 6.49E-72 | TAM.Vcam1 | Rufy3 |
| Wdr7 | 5.22E-76 | 0.31585407 | 0.344 | 0.245 | 1.01E-71 | TAM.Vcam1 | Wdr7 |
| Abtb1 | 1.89E-75 | 0.4811863 | 0.25 | 0.163 | 3.65E-71 | TAM.Vcam1 | Abtb1 |
| Fance | 2.15E-75 | 0.52514613 | 0.257 | 0.162 | 4.16E-71 | TAM.Vcam1 | Fance |
| Nfic | 6.66E-75 | 0.25041275 | 0.396 | 0.283 | 1.29E-70 | TAM.Vcam1 | Nfic |
| Scnm1 | 7.37E-75 | 0.40967321 | 0.257 | 0.172 | 1.43E-70 | TAM.Vcam1 | Scnm1 |
| Akt1s1 | 7.58E-75 | 0.33573916 | 0.333 | 0.233 | 1.47E-70 | TAM.Vcam1 | Akt1s1 |
| Ccdc47 | 7.91E-75 | 0.36100241 | 0.405 | 0.296 | 1.53E-70 | TAM.Vcam1 | Ccdc47 |
| Rab9 | 1.76E-74 | 0.34107619 | 0.33 | 0.238 | 3.40E-70 | TAM.Vcam1 | Rab9 |
| Clec4a2 | 1.97E-74 | 0.32797616 | 0.438 | 0.311 | 3.81E-70 | TAM.Vcam1 | Clec4a2 |
| Hist3h2a | 4.77E-74 | 0.34041374 | 0.267 | 0.175 | 9.23E-70 | TAM.Vcam1 | Hist3h2a |
| Mpv17 | 2.74E-73 | 0.43454179 | 0.312 | 0.216 | 5.29E-69 | TAM.Vcam1 | Mpv17 |
| Hes6 | 3.71E-73 | 0.40999596 | 0.304 | 0.21 | 7.18E-69 | TAM.Vcam1 | Hes6 |
| Slc11a2 | 1.47E-72 | 0.30932225 | 0.388 | 0.301 | 2.83E-68 | TAM.Vcam1 | Slc11a2 |
| Phf20 | 1.67E-72 | 0.35503263 | 0.323 | 0.223 | 3.23E-68 | TAM.Vcam1 | Phf20 |
| Rubcn | 1.36E-71 | 0.38672249 | 0.276 | 0.179 | 2.62E-67 | TAM.Vcam1 | Rubcn |
| Irf3 | 2.79E-71 | 0.42810767 | 0.304 | 0.207 | 5.39E-67 | TAM.Vcam1 | Irf3 |
| Atraid | 1.33E-70 | 0.29776612 | 0.408 | 0.313 | 2.56E-66 | TAM.Vcam1 | Atraid |
| Calcoco1 | 1.50E-70 | 0.3215039 | 0.294 | 0.199 | 2.89E-66 | TAM.Vcam1 | Calcoco1 |
| Tor1b | 1.62E-70 | 0.4563831 | 0.273 | 0.187 | 3.14E-66 | TAM.Vcam1 | Tor1b |
| Cd180 | 6.65E-70 | 0.3256063 | 0.252 | 0.167 | 1.29E-65 | TAM.Vcam1 | Cd180 |
| Arfgap1 | 7.37E-70 | 0.25067024 | 0.368 | 0.278 | 1.42E-65 | TAM.Vcam1 | Arfgap1 |
| Trim26 | 1.37E-69 | 0.26244815 | 0.33 | 0.238 | 2.65E-65 | TAM.Vcam1 | Trim26 |
| Hps5 | 4.34E-69 | 0.39988896 | 0.274 | 0.19 | 8.40E-65 | TAM.Vcam1 | Hps5 |
| Slc25a4 | 5.36E-69 | 0.34707241 | 0.431 | 0.313 | 1.04E-64 | TAM.Vcam1 | Slc25a4 |
| Washc4 | 7.71E-69 | 0.25507649 | 0.353 | 0.258 | 1.49E-64 | TAM.Vcam1 | Washc4 |
| Aim2 | 1.73E-68 | 0.2830786 | 0.305 | 0.22 | 3.35E-64 | TAM.Vcam1 | Aim2 |
| Ints3 | 2.16E-68 | 0.34642818 | 0.345 | 0.25 | 4.18E-64 | TAM.Vcam1 | Ints3 |
| Tex261 | 2.25E-68 | 0.2506484 | 0.517 | 0.411 | 4.35E-64 | TAM.Vcam1 | Tex261 |
| Ap5z1 | 3.59E-68 | 0.39487466 | 0.253 | 0.167 | 6.94E-64 | TAM.Vcam1 | Ap5z1 |
| Tmem168 | 3.63E-68 | 0.39731925 | 0.3 | 0.211 | 7.02E-64 | TAM.Vcam1 | Tmem168 |
| Cdk5rap3 | 4.50E-68 | 0.2943151 | 0.361 | 0.265 | 8.70E-64 | TAM.Vcam1 | Cdk5rap3 |
| Synrg | 4.79E-68 | 0.29388934 | 0.324 | 0.238 | 9.26E-64 | TAM.Vcam1 | Synrg |
| Rnf215 | 5.20E-68 | 0.3142151 | 0.31 | 0.222 | 1.00E-63 | TAM.Vcam1 | Rnf215 |
| Hectd3 | 1.31E-67 | 0.25074473 | 0.372 | 0.278 | 2.53E-63 | TAM.Vcam1 | Hectd3 |
| Stk11ip | 4.88E-67 | 0.38586848 | 0.279 | 0.185 | 9.44E-63 | TAM.Vcam1 | Stk11ip |
| Uhmk1 | 8.28E-67 | 0.26628578 | 0.459 | 0.36 | 1.60E-62 | TAM.Vcam1 | Uhmk1 |
| Selenos | 9.99E-67 | 0.2751438 | 0.419 | 0.323 | 1.93E-62 | TAM.Vcam1 | Selenos |
| Arl2 | 1.18E-66 | 0.42977232 | 0.274 | 0.182 | 2.28E-62 | TAM.Vcam1 | Arl2 |
| Kat2a | 2.58E-66 | 0.35372193 | 0.319 | 0.228 | 4.99E-62 | TAM.Vcam1 | Kat2a |
| Abhd16a | 8.34E-66 | 0.29513544 | 0.355 | 0.263 | 1.61E-61 | TAM.Vcam1 | Abhd16a |
| Stard5 | 1.82E-65 | 0.43211187 | 0.306 | 0.211 | 3.52E-61 | TAM.Vcam1 | Stard5 |
| Ltbp3 | 2.41E-65 | 0.33897185 | 0.427 | 0.303 | 4.67E-61 | TAM.Vcam1 | Ltbp3 |
| Ago1 | 1.25E-64 | 0.34295089 | 0.334 | 0.24 | 2.41E-60 | TAM.Vcam1 | Ago1 |
| Slc37a2 | 2.30E-64 | 0.30991004 | 0.497 | 0.356 | 4.45E-60 | TAM.Vcam1 | Slc37a2 |
| Dennd6a | 2.68E-64 | 0.38415541 | 0.271 | 0.187 | 5.17E-60 | TAM.Vcam1 | Dennd6a |
| Cdk16 | 7.26E-64 | 0.25157823 | 0.393 | 0.291 | 1.40E-59 | TAM.Vcam1 | Cdk16 |
| Herc2 | 2.82E-63 | 0.28545597 | 0.343 | 0.255 | 5.46E-59 | TAM.Vcam1 | Herc2 |
| Abhd4 | 3.73E-63 | 0.31723992 | 0.367 | 0.271 | 7.21E-59 | TAM.Vcam1 | Abhd4 |
| Ddx42 | 3.81E-63 | 0.31773161 | 0.348 | 0.255 | 7.37E-59 | TAM.Vcam1 | Ddx42 |
| Bmf | 9.04E-63 | 0.31029322 | 0.276 | 0.198 | 1.75E-58 | TAM.Vcam1 | Bmf |
| Adpgk | 1.28E-62 | 0.40821438 | 0.337 | 0.24 | 2.47E-58 | TAM.Vcam1 | Adpgk |
| Lrrc8a | 2.03E-62 | 0.34323631 | 0.312 | 0.217 | 3.93E-58 | TAM.Vcam1 | Lrrc8a |
| Fbxo3 | 3.66E-62 | 0.3300648 | 0.329 | 0.242 | 7.08E-58 | TAM.Vcam1 | Fbxo3 |
| Gpr107 | 7.32E-62 | 0.2676626 | 0.42 | 0.324 | 1.42E-57 | TAM.Vcam1 | Gpr107 |
| Srpk2 | 8.72E-62 | 0.37225021 | 0.266 | 0.179 | 1.69E-57 | TAM.Vcam1 | Srpk2 |
| Stard9 | 1.05E-61 | 0.25669631 | 0.295 | 0.213 | 2.03E-57 | TAM.Vcam1 | Stard9 |
| Hacd3 | 1.51E-61 | 0.45680131 | 0.282 | 0.191 | 2.92E-57 | TAM.Vcam1 | Hacd3 |
| Slc25a10 | 1.71E-61 | 0.44338413 | 0.329 | 0.218 | 3.30E-57 | TAM.Vcam1 | Slc25a10 |
| Pdcd2l | 2.01E-61 | 0.32542923 | 0.29 | 0.205 | 3.89E-57 | TAM.Vcam1 | Pdcd2l |
| Galns | 2.74E-61 | 0.3381367 | 0.323 | 0.234 | 5.31E-57 | TAM.Vcam1 | Galns |
| Exoc3 | 5.14E-61 | 0.34139011 | 0.303 | 0.215 | 9.94E-57 | TAM.Vcam1 | Exoc3 |
| Sft2d1 | 9.79E-61 | 0.35046588 | 0.325 | 0.233 | 1.89E-56 | TAM.Vcam1 | Sft2d1 |
| Tmem9b | 1.10E-60 | 0.28462803 | 0.412 | 0.324 | 2.12E-56 | TAM.Vcam1 | Tmem9b |
| Tspyl2 | 1.74E-60 | 0.27069637 | 0.268 | 0.191 | 3.37E-56 | TAM.Vcam1 | Tspyl2 |
| Ccdc22 | 2.01E-60 | 0.27537318 | 0.366 | 0.269 | 3.89E-56 | TAM.Vcam1 | Ccdc22 |
| Commd10 | 2.31E-60 | 0.30295333 | 0.379 | 0.281 | 4.46E-56 | TAM.Vcam1 | Commd10 |
| Spast | 2.98E-60 | 0.28867398 | 0.354 | 0.265 | 5.75E-56 | TAM.Vcam1 | Spast |
| AU040320 | 4.18E-60 | 0.31904139 | 0.285 | 0.203 | 8.07E-56 | TAM.Vcam1 | AU040320 |
| Gga3 | 6.27E-60 | 0.30928416 | 0.302 | 0.21 | 1.21E-55 | TAM.Vcam1 | Gga3 |
| Ppp1cc | 1.14E-59 | 0.27819087 | 0.45 | 0.349 | 2.21E-55 | TAM.Vcam1 | Ppp1cc |
| Topors | 3.08E-59 | 0.32600701 | 0.33 | 0.248 | 5.95E-55 | TAM.Vcam1 | Topors |
| Pycr2 | 5.75E-59 | 0.37001703 | 0.33 | 0.236 | 1.11E-54 | TAM.Vcam1 | Pycr2 |
| Tmem119 | 1.31E-58 | 0.25078472 | 0.412 | 0.28 | 2.54E-54 | TAM.Vcam1 | Tmem119 |
| Cops7a | 6.14E-58 | 0.30064814 | 0.359 | 0.269 | 1.19E-53 | TAM.Vcam1 | Cops7a |
| Hagh | 6.38E-58 | 0.32721857 | 0.352 | 0.259 | 1.23E-53 | TAM.Vcam1 | Hagh |
| Tmem19 | 7.67E-58 | 0.2749059 | 0.391 | 0.292 | 1.48E-53 | TAM.Vcam1 | Tmem19 |
| Kdm4a | 2.19E-57 | 0.27937458 | 0.32 | 0.236 | 4.24E-53 | TAM.Vcam1 | Kdm4a |
| Hspe1 | 5.94E-57 | 0.3745638 | 0.35 | 0.254 | 1.15E-52 | TAM.Vcam1 | Hspe1 |
| Ubfd1 | 7.97E-57 | 0.35570819 | 0.287 | 0.204 | 1.54E-52 | TAM.Vcam1 | Ubfd1 |
| Trappc12 | 8.76E-57 | 0.29629885 | 0.282 | 0.202 | 1.69E-52 | TAM.Vcam1 | Trappc12 |
| Bzw2 | 9.24E-57 | 0.3919684 | 0.307 | 0.214 | 1.79E-52 | TAM.Vcam1 | Bzw2 |
| Pon2 | 9.74E-57 | 0.25917045 | 0.516 | 0.392 | 1.88E-52 | TAM.Vcam1 | Pon2 |
| D10Wsu102e | 1.73E-56 | 0.34350251 | 0.282 | 0.209 | 3.35E-52 | TAM.Vcam1 | D10Wsu102e |
| Gatd1 | 5.33E-56 | 0.34443806 | 0.297 | 0.212 | 1.03E-51 | TAM.Vcam1 | Gatd1 |
| Manba | 9.33E-56 | 0.30544316 | 0.391 | 0.296 | 1.80E-51 | TAM.Vcam1 | Manba |
| Washc5 | 1.12E-55 | 0.2568516 | 0.351 | 0.265 | 2.17E-51 | TAM.Vcam1 | Washc5 |
| Cln6 | 1.52E-55 | 0.33261075 | 0.353 | 0.253 | 2.94E-51 | TAM.Vcam1 | Cln6 |
| Mfap3 | 7.36E-55 | 0.28878111 | 0.309 | 0.221 | 1.42E-50 | TAM.Vcam1 | Mfap3 |
| AI413582 | 9.35E-55 | 0.28500291 | 0.316 | 0.231 | 1.81E-50 | TAM.Vcam1 | AI413582 |
| Fkrp | 9.36E-55 | 0.25454945 | 0.352 | 0.269 | 1.81E-50 | TAM.Vcam1 | Fkrp |
| Eif2ak1 | 4.72E-54 | 0.26576605 | 0.378 | 0.286 | 9.13E-50 | TAM.Vcam1 | Eif2ak1 |
| Arl6ip1 | 5.44E-54 | 0.26526787 | 0.451 | 0.342 | 1.05E-49 | TAM.Vcam1 | Arl6ip1 |
| Ppox | 6.68E-54 | 0.29523473 | 0.275 | 0.198 | 1.29E-49 | TAM.Vcam1 | Ppox |
| Hps1 | 1.81E-53 | 0.25426778 | 0.344 | 0.26 | 3.50E-49 | TAM.Vcam1 | Hps1 |
| Dhcr24 | 2.94E-53 | 0.38565034 | 0.29 | 0.209 | 5.68E-49 | TAM.Vcam1 | Dhcr24 |
| Ccnd2 | 6.53E-53 | 0.2717269 | 0.338 | 0.214 | 1.26E-48 | TAM.Vcam1 | Ccnd2 |
| Nagk | 6.97E-53 | 0.36667764 | 0.368 | 0.264 | 1.35E-48 | TAM.Vcam1 | Nagk |
| Ptpn7 | 8.02E-53 | 0.32045621 | 0.334 | 0.252 | 1.55E-48 | TAM.Vcam1 | Ptpn7 |
| Orai3 | 2.03E-52 | 0.29781576 | 0.315 | 0.236 | 3.92E-48 | TAM.Vcam1 | Orai3 |
| BC029722 | 2.53E-52 | 0.27717353 | 0.302 | 0.223 | 4.89E-48 | TAM.Vcam1 | BC029722 |
| Umps | 3.76E-52 | 0.39948771 | 0.279 | 0.19 | 7.27E-48 | TAM.Vcam1 | Umps |
| Rbl2 | 3.39E-51 | 0.27559902 | 0.319 | 0.233 | 6.55E-47 | TAM.Vcam1 | Rbl2 |
| Cstf2t | 6.29E-51 | 0.36258325 | 0.267 | 0.185 | 1.22E-46 | TAM.Vcam1 | Cstf2t |
| Fundc2 | 6.46E-51 | 0.2522487 | 0.412 | 0.316 | 1.25E-46 | TAM.Vcam1 | Fundc2 |
| Exoc8 | 6.49E-51 | 0.28924596 | 0.25 | 0.175 | 1.25E-46 | TAM.Vcam1 | Exoc8 |
| Mbtps1 | 6.59E-51 | 0.25881023 | 0.333 | 0.247 | 1.27E-46 | TAM.Vcam1 | Mbtps1 |
| Klhl24 | 7.05E-51 | 0.25072007 | 0.361 | 0.27 | 1.36E-46 | TAM.Vcam1 | Klhl24 |
| Ptp4a3 | 7.32E-51 | 0.39449394 | 0.254 | 0.18 | 1.42E-46 | TAM.Vcam1 | Ptp4a3 |
| Creld1 | 1.41E-50 | 0.43345339 | 0.278 | 0.196 | 2.73E-46 | TAM.Vcam1 | Creld1 |
| Nipsnap2 | 2.57E-50 | 0.25528898 | 0.34 | 0.254 | 4.97E-46 | TAM.Vcam1 | Nipsnap2 |
| Tm7sf3 | 3.73E-50 | 0.27674209 | 0.322 | 0.237 | 7.21E-46 | TAM.Vcam1 | Tm7sf3 |
| Maged1 | 4.12E-50 | 0.45434715 | 0.262 | 0.171 | 7.97E-46 | TAM.Vcam1 | Maged1 |
| Ints14 | 6.79E-50 | 0.28748978 | 0.312 | 0.23 | 1.31E-45 | TAM.Vcam1 | Ints14 |
| Ncoa5 | 9.89E-50 | 0.2834218 | 0.27 | 0.195 | 1.91E-45 | TAM.Vcam1 | Ncoa5 |
| Tceanc2 | 1.30E-49 | 0.26378694 | 0.261 | 0.188 | 2.52E-45 | TAM.Vcam1 | Tceanc2 |
| Cryzl1 | 2.20E-49 | 0.2950818 | 0.276 | 0.201 | 4.26E-45 | TAM.Vcam1 | Cryzl1 |
| Ccar2 | 2.41E-49 | 0.31483982 | 0.271 | 0.195 | 4.65E-45 | TAM.Vcam1 | Ccar2 |
| Rab40c | 6.32E-49 | 0.28859735 | 0.255 | 0.184 | 1.22E-44 | TAM.Vcam1 | Rab40c |
| Pign | 1.30E-48 | 0.26357058 | 0.291 | 0.208 | 2.51E-44 | TAM.Vcam1 | Pign |
| Arfip2 | 1.74E-48 | 0.33239405 | 0.262 | 0.186 | 3.36E-44 | TAM.Vcam1 | Arfip2 |
| Sfmbt1 | 1.87E-48 | 0.32908334 | 0.332 | 0.239 | 3.62E-44 | TAM.Vcam1 | Sfmbt1 |
| Prpf19 | 1.02E-47 | 0.26922007 | 0.434 | 0.33 | 1.98E-43 | TAM.Vcam1 | Prpf19 |
| Efcab14 | 3.22E-47 | 0.33381528 | 0.263 | 0.191 | 6.22E-43 | TAM.Vcam1 | Efcab14 |
| Slc39a9 | 3.84E-47 | 0.25257591 | 0.35 | 0.266 | 7.43E-43 | TAM.Vcam1 | Slc39a9 |
| Erap1 | 4.45E-47 | 0.2594725 | 0.351 | 0.271 | 8.59E-43 | TAM.Vcam1 | Erap1 |
| Slc29a1 | 6.16E-47 | 0.33766427 | 0.412 | 0.304 | 1.19E-42 | TAM.Vcam1 | Slc29a1 |
| Thap7 | 2.41E-46 | 0.33755378 | 0.259 | 0.186 | 4.67E-42 | TAM.Vcam1 | Thap7 |
| Ppp5c | 2.45E-46 | 0.2844136 | 0.358 | 0.269 | 4.74E-42 | TAM.Vcam1 | Ppp5c |
| Arsa | 4.91E-46 | 0.29409287 | 0.325 | 0.24 | 9.49E-42 | TAM.Vcam1 | Arsa |
| Gnpat | 3.20E-45 | 0.26349839 | 0.336 | 0.254 | 6.18E-41 | TAM.Vcam1 | Gnpat |
| Sbf2 | 2.26E-44 | 0.25984484 | 0.331 | 0.243 | 4.38E-40 | TAM.Vcam1 | Sbf2 |
| Snapin | 2.28E-44 | 0.29095841 | 0.293 | 0.217 | 4.40E-40 | TAM.Vcam1 | Snapin |
| Fkbp5 | 6.32E-44 | 0.26856722 | 0.316 | 0.242 | 1.22E-39 | TAM.Vcam1 | Fkbp5 |
| Ext2 | 2.19E-43 | 0.25629277 | 0.378 | 0.288 | 4.23E-39 | TAM.Vcam1 | Ext2 |

| | | | | | | | |
|---|---|---|---|---|---|---|---|
| Rcc2 | 2.61E-43 | 0.25700941 | 0.45 | 0.341 | 5.05E-39 | TAM.Vcam1 | Rcc2 |
| Htra2 | 5.40E-43 | 0.32741994 | 0.318 | 0.228 | 1.04E-38 | TAM.Vcam1 | Htra2 |
| Mex3d | 5.55E-43 | 0.28595008 | 0.273 | 0.202 | 1.07E-38 | TAM.Vcam1 | Mex3d |
| Slc35b1 | 9.17E-43 | 0.26758623 | 0.297 | 0.223 | 1.77E-38 | TAM.Vcam1 | Slc35b1 |
| Trmt2a | 1.05E-42 | 0.27655996 | 0.32 | 0.235 | 2.03E-38 | TAM.Vcam1 | Trmt2a |
| Pja2 | 3.71E-42 | 0.26623239 | 0.286 | 0.208 | 7.18E-38 | TAM.Vcam1 | Pja2 |
| Plxnc1 | 3.37E-41 | 0.27938847 | 0.291 | 0.218 | 6.51E-37 | TAM.Vcam1 | Plxnc1 |
| Tvp23b | 1.41E-40 | 0.36401275 | 0.255 | 0.188 | 2.73E-36 | TAM.Vcam1 | Tvp23b |
| Myo9a | 2.44E-40 | 0.32026898 | 0.252 | 0.183 | 4.73E-36 | TAM.Vcam1 | Myo9a |
| Apex1 | 5.21E-40 | 0.34727214 | 0.29 | 0.206 | 1.01E-35 | TAM.Vcam1 | Apex1 |
| Ipo9 | 7.58E-40 | 0.27665539 | 0.356 | 0.264 | 1.47E-35 | TAM.Vcam1 | Ipo9 |
| Spg11 | 1.21E-39 | 0.25576079 | 0.252 | 0.191 | 2.34E-35 | TAM.Vcam1 | Spg11 |
| Dhdh | 1.46E-39 | 0.29675484 | 0.271 | 0.2 | 2.83E-35 | TAM.Vcam1 | Dhdh |
| Lman2l | 5.95E-39 | 0.26966858 | 0.264 | 0.195 | 1.15E-34 | TAM.Vcam1 | Lman2l |
| Dicer1 | 4.48E-37 | 0.27640521 | 0.261 | 0.189 | 8.67E-33 | TAM.Vcam1 | Dicer1 |
| Fubp3 | 8.36E-37 | 0.3073425 | 0.259 | 0.19 | 1.62E-32 | TAM.Vcam1 | Fubp3 |
| Slc50a1 | 1.90E-36 | 0.26993054 | 0.266 | 0.2 | 3.67E-32 | TAM.Vcam1 | Slc50a1 |
| Crat | 3.93E-36 | 0.25053384 | 0.324 | 0.244 | 7.59E-32 | TAM.Vcam1 | Crat |
| Cdkn2aipnl | 4.26E-36 | 0.36380794 | 0.27 | 0.19 | 8.23E-32 | TAM.Vcam1 | Cdkn2aipnl |
| Pithd1 | 1.80E-35 | 0.2716909 | 0.276 | 0.209 | 3.48E-31 | TAM.Vcam1 | Pithd1 |
| Galnt7 | 2.22E-35 | 0.25245825 | 0.323 | 0.251 | 4.30E-31 | TAM.Vcam1 | Galnt7 |
| Bckdk | 8.41E-35 | 0.28320219 | 0.321 | 0.239 | 1.63E-30 | TAM.Vcam1 | Bckdk |
| Hspd1 | 4.60E-34 | 0.29508883 | 0.391 | 0.298 | 8.89E-30 | TAM.Vcam1 | Hspd1 |
| Scap | 6.74E-34 | 0.28716908 | 0.273 | 0.203 | 1.30E-29 | TAM.Vcam1 | Scap |
| Ift22 | 2.63E-33 | 0.25279504 | 0.279 | 0.21 | 5.09E-29 | TAM.Vcam1 | Ift22 |
| Chordc1 | 2.57E-32 | 0.2900843 | 0.26 | 0.196 | 4.96E-28 | TAM.Vcam1 | Chordc1 |
| Tulp4 | 4.02E-32 | 0.30659094 | 0.253 | 0.185 | 7.78E-28 | TAM.Vcam1 | Tulp4 |
| Ilvbl | 4.23E-32 | 0.25794925 | 0.257 | 0.189 | 8.19E-28 | TAM.Vcam1 | Ilvbl |
| Acot13 | 1.31E-31 | 0.2642886 | 0.297 | 0.223 | 2.54E-27 | TAM.Vcam1 | Acot13 |
| Tmx2 | 1.88E-31 | 0.269274 | 0.284 | 0.216 | 3.63E-27 | TAM.Vcam1 | Tmx2 |
| Snx15 | 3.16E-27 | 0.26328279 | 0.257 | 0.199 | 6.12E-23 | TAM.Vcam1 | Snx15 |
| Pon3 | 1.56E-14 | 0.29857185 | 0.282 | 0.212 | 3.02E-10 | TAM.Vcam1 | Pon3 |
| Inhba | 0 | 3.21492977 | 0.768 | 0.151 | 0 | TAM.pro-infl. | Inhba |
| Dcstamp | 0 | 3.19478603 | 0.767 | 0.154 | 0 | TAM.pro-infl. | Dcstamp |
| Cxcl3 | 0 | 4.50051245 | 0.661 | 0.074 | 0 | TAM.pro-infl. | Cxcl3 |
| Gpc1 | 0 | 3.34424846 | 0.672 | 0.148 | 0 | TAM.pro-infl. | Gpc1 |
| Sdc1 | 0 | 2.41638398 | 0.787 | 0.27 | 0 | TAM.pro-infl. | Sdc1 |
| Slc7a11 | 0 | 2.28857718 | 0.917 | 0.411 | 0 | TAM.pro-infl. | Slc7a11 |
| Hvcn1 | 0 | 2.70232154 | 0.765 | 0.267 | 0 | TAM.pro-infl. | Hvcn1 |
| Mmp12 | 0 | 2.65255429 | 0.926 | 0.43 | 0 | TAM.pro-infl. | Mmp12 |
| Gclm | 0 | 2.73238177 | 0.722 | 0.232 | 0 | TAM.pro-infl. | Gclm |
| AA467197 | 0 | 1.64861331 | 0.832 | 0.348 | 0 | TAM.pro-infl. | AA467197 |
| Lpl | 0 | 3.56976905 | 0.598 | 0.123 | 0 | TAM.pro-infl. | Lpl |
| Ass1 | 0 | 2.35625625 | 0.778 | 0.317 | 0 | TAM.pro-infl. | Ass1 |
| Met | 0 | 3.10905219 | 0.564 | 0.104 | 0 | TAM.pro-infl. | Met |
| Procr | 0 | 3.82258148 | 0.535 | 0.082 | 0 | TAM.pro-infl. | Procr |
| Slpi | 0 | 2.6003606 | 0.571 | 0.118 | 0 | TAM.pro-infl. | Slpi |
| F7 | 0 | 2.14215605 | 0.63 | 0.191 | 0 | TAM.pro-infl. | F7 |
| Fpr2 | 0 | 2.68736963 | 0.538 | 0.111 | 0 | TAM.pro-infl. | Fpr2 |
| Srxn1 | 0 | 1.91347561 | 0.732 | 0.313 | 0 | TAM.pro-infl. | Srxn1 |
| F10 | 0 | 1.6116819 | 0.905 | 0.49 | 0 | TAM.pro-infl. | F10 |
| Il1a | 0 | 1.16697179 | 0.799 | 0.391 | 0 | TAM.pro-infl. | Il1a |
| Adam8 | 0 | 1.62012663 | 0.911 | 0.504 | 0 | TAM.pro-infl. | Adam8 |
| Abcc1 | 0 | 2.00751576 | 0.749 | 0.342 | 0 | TAM.pro-infl. | Abcc1 |
| Actn1 | 0 | 1.52012047 | 0.79 | 0.384 | 0 | TAM.pro-infl. | Actn1 |
| Matk | 0 | 1.61819342 | 0.692 | 0.296 | 0 | TAM.pro-infl. | Matk |
| Hilpda | 0 | 1.57669685 | 0.901 | 0.512 | 0 | TAM.pro-infl. | Hilpda |
| Rnf128 | 0 | 3.17750611 | 0.462 | 0.075 | 0 | TAM.pro-infl. | Rnf128 |
| Tnfrsf9 | 0 | 1.63366907 | 0.571 | 0.185 | 0 | TAM.pro-infl. | Tnfrsf9 |
| Dst | 0 | 1.8774417 | 0.623 | 0.241 | 0 | TAM.pro-infl. | Dst |
| Alas1 | 0 | 2.67205953 | 0.897 | 0.517 | 0 | TAM.pro-infl. | Alas1 |
| Fam20c | 0 | 1.48617593 | 0.825 | 0.445 | 0 | TAM.pro-infl. | Fam20c |
| Pdpn | 0 | 1.9272156 | 0.618 | 0.239 | 0 | TAM.pro-infl. | Pdpn |
| Bst1 | 0 | 1.60244534 | 0.693 | 0.315 | 0 | TAM.pro-infl. | Bst1 |
| Zc3h12c | 0 | 2.46415163 | 0.49 | 0.113 | 0 | TAM.pro-infl. | Zc3h12c |
| Ptgr1 | 0 | 3.13785129 | 0.598 | 0.229 | 0 | TAM.pro-infl. | Ptgr1 |
| Chst11 | 0 | 2.23268473 | 0.525 | 0.158 | 0 | TAM.pro-infl. | Chst11 |
| Blvrb | 0 | 1.69482427 | 0.862 | 0.497 | 0 | TAM.pro-infl. | Blvrb |
| Csf1 | 0 | 2.54852526 | 0.463 | 0.101 | 0 | TAM.pro-infl. | Csf1 |
| Plp2 | 0 | 1.2014361 | 0.675 | 0.314 | 0 | TAM.pro-infl. | Plp2 |
| Itgax1 | 0 | 1.39395178 | 0.912 | 0.551 | 0 | TAM.pro-infl. | Itgax |
| Rgcc | 0 | 1.14536095 | 0.656 | 0.296 | 0 | TAM.pro-infl. | Rgcc |
| Rhoc | 0 | 1.68364276 | 0.587 | 0.232 | 0 | TAM.pro-infl. | Rhoc |
| Slc6a8 | 0 | 1.7274618 | 0.587 | 0.233 | 0 | TAM.pro-infl. | Slc6a8 |
| Gstm1 | 0 | 2.38207027 | 0.482 | 0.131 | 0 | TAM.pro-infl. | Gstm1 |
| Clec4e | 0 | 1.28709307 | 0.72 | 0.369 | 0 | TAM.pro-infl. | Clec4e |
| Rab11fip1 | 0 | 1.13522878 | 0.846 | 0.497 | 0 | TAM.pro-infl. | Rab11fip1 |
| Mmp91 | 0 | 1.47650291 | 0.509 | 0.164 | 0 | TAM.pro-infl. | Mmp9 |
| Cyb5a | 0 | 1.833084 | 0.707 | 0.364 | 0 | TAM.pro-infl. | Cyb5a |
| Pla1a | 0 | 2.3054353 | 0.445 | 0.104 | 0 | TAM.pro-infl. | Pla1a |
| Arg2 | 0 | 1.19112393 | 0.622 | 0.281 | 0 | TAM.pro-infl. | Arg2 |
| Slc25a37 | 0 | 1.61990889 | 0.569 | 0.229 | 0 | TAM.pro-infl. | Slc25a37 |
| Vegfa | 0 | 0.8701172 | 0.918 | 0.58 | 0 | TAM.pro-infl. | Vegfa |
| Hp | 0 | 1.69067173 | 0.528 | 0.19 | 0 | TAM.pro-infl. | Hp |
| Clec4d | 0 | 1.47000962 | 0.932 | 0.595 | 0 | TAM.pro-infl. | Clec4d |
| Cd36 | 0 | 2.86274158 | 0.453 | 0.116 | 0 | TAM.pro-infl. | Cd36 |
| Lhfpl2 | 0 | 1.68472761 | 0.657 | 0.321 | 0 | TAM.pro-infl. | Lhfpl2 |
| Sgk1 | 0 | 1.66610767 | 0.901 | 0.569 | 0 | TAM.pro-infl. | Sgk1 |
| Sort1 | 0 | 2.20945492 | 0.433 | 0.106 | 0 | TAM.pro-infl. | Sort1 |
| Gclc | 0 | 1.89979688 | 0.547 | 0.223 | 0 | TAM.pro-infl. | Gclc |
| Clcf1 | 0 | 1.17098618 | 0.59 | 0.266 | 0 | TAM.pro-infl. | Clcf1 |
| Arg1 | 0 | 0.99128862 | 0.848 | 0.525 | 0 | TAM.pro-infl. | Arg1 |
| Vat1 | 0 | 1.25758604 | 0.867 | 0.544 | 0 | TAM.pro-infl. | Vat1 |
| Agap1 | 0 | 1.47781966 | 0.582 | 0.26 | 0 | TAM.pro-infl. | Agap1 |
| Best1 | 0 | 3.92420204 | 0.357 | 0.035 | 0 | TAM.pro-infl. | Best1 |
| Mcemp1 | 0 | 1.08073202 | 0.545 | 0.232 | 0 | TAM.pro-infl. | Mcemp1 |
| Itgal | 0 | 0.9932931 | 0.843 | 0.53 | 0 | TAM.pro-infl. | Itgal |
| Gsr | 0 | 1.24038062 | 0.925 | 0.613 | 0 | TAM.pro-infl. | Gsr |
| Stac2 | 0 | 2.19595627 | 0.411 | 0.1 | 0 | TAM.pro-infl. | Stac2 |
| Prdx1 | 0 | 2.7342541 | 0.897 | 0.587 | 0 | TAM.pro-infl. | Prdx1 |
| Abcc5 | 0 | 1.68065447 | 0.537 | 0.228 | 0 | TAM.pro-infl. | Abcc5 |
| Acot7 | 0 | 1.22694022 | 0.565 | 0.257 | 0 | TAM.pro-infl. | Acot7 |
| Ddit4 | 0 | 0.91711654 | 0.707 | 0.399 | 0 | TAM.pro-infl. | Ddit4 |
| Gfod1 | 0 | 1.59545777 | 0.451 | 0.146 | 0 | TAM.pro-infl. | Gfod1 |
| Fcrlb | 0 | 1.09677184 | 0.651 | 0.347 | 0 | TAM.pro-infl. | Fcrlb |
| Tmem165 | 0 | 1.35410485 | 0.698 | 0.394 | 0 | TAM.pro-infl. | Tmem165 |
| C3 | 0 | 0.76421236 | 0.83 | 0.527 | 0 | TAM.pro-infl. | C3 |
| Aldoc | 0 | 1.20104725 | 0.566 | 0.266 | 0 | TAM.pro-infl. | Aldoc |
| Cd274 | 0 | 0.84750444 | 0.728 | 0.429 | 0 | TAM.pro-infl. | Cd274 |
| Mmp19 | 0 | 1.6194783 | 0.856 | 0.557 | 0 | TAM.pro-infl. | Mmp19 |
| Ptprs | 0 | 1.20433249 | 0.685 | 0.387 | 0 | TAM.pro-infl. | Ptprs |
| Pik3cb | 0 | 1.2555384 | 0.606 | 0.308 | 0 | TAM.pro-infl. | Pik3cb |
| Dpep2 | 0 | 1.03611655 | 0.62 | 0.325 | 0 | TAM.pro-infl. | Dpep2 |
| Nrp2 | 0 | 0.97661634 | 0.69 | 0.396 | 0 | TAM.pro-infl. | Nrp2 |
| Smpdl3b | 0 | 2.09320203 | 0.388 | 0.095 | 0 | TAM.pro-infl. | Smpdl3b |
| Cd9 | 0 | 1.99663614 | 0.97 | 0.677 | 0 | TAM.pro-infl. | Cd9 |
| Acp51 | 0 | 0.9218534 | 0.916 | 0.625 | 0 | TAM.pro-infl. | Acp5 |
| Eid3 | 0 | 1.46809406 | 0.517 | 0.228 | 0 | TAM.pro-infl. | Eid3 |
| Lipa | 0 | 1.61269994 | 0.811 | 0.522 | 0 | TAM.pro-infl. | Lipa |
| Slc48a1 | 0 | 1.25116237 | 0.739 | 0.452 | 0 | TAM.pro-infl. | Slc48a1 |
| Igf1r | 0 | 1.34201743 | 0.544 | 0.258 | 0 | TAM.pro-infl. | Igf1r |
| Malt1 | 0 | 0.59615817 | 0.925 | 0.639 | 0 | TAM.pro-infl. | Malt1 |
| Iqsec1 | 0 | 1.22168481 | 0.652 | 0.367 | 0 | TAM.pro-infl. | Iqsec1 |
| Sgms2 | 0 | 1.50549876 | 0.42 | 0.136 | 0 | TAM.pro-infl. | Sgms2 |
| Sphk1 | 0 | 1.83209579 | 0.403 | 0.12 | 0 | TAM.pro-infl. | Sphk1 |
| Fpr1 | 0 | 3.67762881 | 0.316 | 0.033 | 0 | TAM.pro-infl. | Fpr1 |
| Psmb5 | 0 | 1.13250301 | 0.826 | 0.544 | 0 | TAM.pro-infl. | Psmb5 |
| Sema4d | 0 | 1.05903069 | 0.9 | 0.619 | 0 | TAM.pro-infl. | Sema4d |
| Atp1a3 | 0 | 1.4265248 | 0.474 | 0.193 | 0 | TAM.pro-infl. | Atp1a3 |
| Cd300lb | 0 | 1.14940758 | 0.576 | 0.299 | 0 | TAM.pro-infl. | Cd300lb |
| Capg | 0 | 1.33193054 | 0.924 | 0.648 | 0 | TAM.pro-infl. | Capg |
| Cxcl14 | 0 | 2.35822804 | 0.453 | 0.177 | 0 | TAM.pro-infl. | Cxcl14 |
| Anxa4 | 0 | 1.69775872 | 0.885 | 0.61 | 0 | TAM.pro-infl. | Anxa4 |
| Ece1 | 0 | 0.87941686 | 0.734 | 0.459 | 0 | TAM.pro-infl. | Ece1 |
| Cd381 | 0 | 0.9038654 | 0.686 | 0.412 | 0 | TAM.pro-infl. | Cd38 |
| Zdhhc18 | 0 | 1.31763002 | 0.613 | 0.339 | 0 | TAM.pro-infl. | Zdhhc18 |
| Gpr68 | 0 | 2.14918928 | 0.364 | 0.091 | 0 | TAM.pro-infl. | Gpr68 |
| Il7r | 0 | 1.84466234 | 0.381 | 0.11 | 0 | TAM.pro-infl. | Il7r |
| Carmil1 | 0 | 2.10505164 | 0.37 | 0.1 | 0 | TAM.pro-infl. | Carmil1 |
| Arl5a | 0 | 1.07860704 | 0.646 | 0.38 | 0 | TAM.pro-infl. | Arl5a |
| Map4k4 | 0 | 1.05937592 | 0.829 | 0.57 | 0 | TAM.pro-infl. | Map4k4 |
| Gsap | 0 | 0.93341531 | 0.728 | 0.469 | 0 | TAM.pro-infl. | Gsap |
| Gngt2 | 0 | 0.95108255 | 0.766 | 0.507 | 0 | TAM.pro-infl. | Gngt2 |
| Acap1 | 0 | 2.33827799 | 0.334 | 0.076 | 0 | TAM.pro-infl. | Acap1 |
| Plxna1 | 0 | 0.95113833 | 0.712 | 0.456 | 0 | TAM.pro-infl. | Plxna1 |
| Olr1 | 0 | 1.41025709 | 0.42 | 0.165 | 0 | TAM.pro-infl. | Olr1 |
| Kmt5a | 0 | 0.94194005 | 0.602 | 0.348 | 0 | TAM.pro-infl. | Kmt5a |
| Slc38a61 | 0 | 0.94148721 | 0.593 | 0.339 | 0 | TAM.pro-infl. | Slc38a6 |
| Il1rn | 0 | 1.13403434 | 0.957 | 0.704 | 0 | TAM.pro-infl. | Il1rn |
| Txn1 | 0 | 1.24902597 | 0.872 | 0.619 | 0 | TAM.pro-infl. | Txn1 |
| Dusp4 | 0 | 2.75166392 | 0.32 | 0.068 | 0 | TAM.pro-infl. | Dusp4 |
| Fnip2 | 0 | 0.93669216 | 0.714 | 0.462 | 0 | TAM.pro-infl. | Fnip2 |
| Myo1e | 0 | 0.78796288 | 0.704 | 0.452 | 0 | TAM.pro-infl. | Myo1e |
| Txnrd1 | 0 | 1.46194686 | 0.931 | 0.68 | 0 | TAM.pro-infl. | Txnrd1 |
| Foxp4 | 0 | 1.45818994 | 0.426 | 0.177 | 0 | TAM.pro-infl. | Foxp4 |
| Raf1 | 0 | 0.9721807 | 0.742 | 0.494 | 0 | TAM.pro-infl. | Raf1 |
| Lrrc32 | 0 | 3.88144998 | 0.282 | 0.034 | 0 | TAM.pro-infl. | Lrrc32 |
| Inf2 | 0 | 0.9764212 | 0.451 | 0.204 | 0 | TAM.pro-infl. | Inf2 |
| Fmnl2 | 0 | 1.08830395 | 0.482 | 0.235 | 0 | TAM.pro-infl. | Fmnl2 |
| Plec | 0 | 0.61630535 | 0.87 | 0.623 | 0 | TAM.pro-infl. | Plec |
| Alcam | 0 | 0.90691311 | 0.591 | 0.345 | 0 | TAM.pro-infl. | Alcam |
| Rusc2 | 0 | 1.34657997 | 0.427 | 0.182 | 0 | TAM.pro-infl. | Rusc2 |
| Sod2 | 0 | 1.49785823 | 0.851 | 0.606 | 0 | TAM.pro-infl. | Sod2 |
| Mindy1 | 0 | 1.0080059 | 0.535 | 0.291 | 0 | TAM.pro-infl. | Mindy1 |
| Dtx41 | 0 | 0.93883074 | 0.652 | 0.408 | 0 | TAM.pro-infl. | Dtx4 |
| Lonrf3 | 0 | 1.24894661 | 0.451 | 0.207 | 0 | TAM.pro-infl. | Lonrf3 |
| Src1 | 0 | 0.70449131 | 0.768 | 0.524 | 0 | TAM.pro-infl. | Src |
| Rnf217 | 0 | 1.01637376 | 0.481 | 0.237 | 0 | TAM.pro-infl. | Rnf217 |
| Acod1 | 0 | 2.14244886 | 0.321 | 0.078 | 0 | TAM.pro-infl. | Acod1 |
| Ahcyl1 | 0 | 0.99127573 | 0.729 | 0.487 | 0 | TAM.pro-infl. | Ahcyl1 |
| Tnfaip2 | 0 | 1.18128968 | 0.933 | 0.692 | 0 | TAM.pro-infl. | Tnfaip2 |
| Dusp5 | 0 | 0.69449676 | 0.924 | 0.684 | 0 | TAM.pro-infl. | Dusp5 |
| Asph | 0 | 0.80818407 | 0.649 | 0.41 | 0 | TAM.pro-infl. | Asph |
| Pdlim7 | 0 | 0.8065699 | 0.544 | 0.305 | 0 | TAM.pro-infl. | Pdlim7 |
| Tarm1 | 0 | 1.1111606 | 0.494 | 0.255 | 0 | TAM.pro-infl. | Tarm1 |
| Ppp2cb | 0 | 0.80478301 | 0.715 | 0.477 | 0 | TAM.pro-infl. | Ppp2cb |
| Gde1 | 0 | 0.87940678 | 0.7 | 0.462 | 0 | TAM.pro-infl. | Gde1 |
| Gpr84 | 0 | 1.21818913 | 0.4 | 0.163 | 0 | TAM.pro-infl. | Gpr84 |
| Hmox1 | 0 | 0.56494413 | 0.885 | 0.648 | 0 | TAM.pro-infl. | Hmox1 |
| Gabarapl1 | 0 | 0.83030159 | 0.698 | 0.461 | 0 | TAM.pro-infl. | Gabarapl1 |
| Cyp4v3 | 0 | 1.18253017 | 0.466 | 0.23 | 0 | TAM.pro-infl. | Cyp4v3 |
| Slc2a1 | 0 | 0.71437827 | 0.893 | 0.657 | 0 | TAM.pro-infl. | Slc2a1 |
| Clec5a | 0 | 1.04031924 | 0.593 | 0.357 | 0 | TAM.pro-infl. | Clec5a |
| Axl1 | 0 | 0.42780781 | 0.897 | 0.661 | 0 | TAM.pro-infl. | Axl |
| Tma16 | 0 | 1.82505186 | 0.353 | 0.117 | 0 | TAM.pro-infl. | Tma16 |
| Slc11a1 | 0 | 1.09938887 | 0.909 | 0.674 | 0 | TAM.pro-infl. | Slc11a1 |
| Ralgds1 | 0 | 0.81836157 | 0.799 | 0.565 | 0 | TAM.pro-infl. | Ralgds |
| Slc44a1 | 0 | 1.22671586 | 0.444 | 0.21 | 0 | TAM.pro-infl. | Slc44a1 |
| Jag1 | 0 | 1.55720785 | 0.361 | 0.127 | 0 | TAM.pro-infl. | Jag1 |
| Pdcd1lg2 | 0 | 1.17549164 | 0.433 | 0.199 | 0 | TAM.pro-infl. | Pdcd1lg2 |
| Dmxl2 | 0 | 0.72004823 | 0.565 | 0.331 | 0 | TAM.pro-infl. | Dmxl2 |
| Slc7a2 | 0 | 0.87175911 | 0.516 | 0.283 | 0 | TAM.pro-infl. | Slc7a2 |
| Ltbp31 | 0 | 1.00376343 | 0.52 | 0.287 | 0 | TAM.pro-infl. | Ltbp3 |
| Rbpms | 0 | 1.17395116 | 0.403 | 0.172 | 0 | TAM.pro-infl. | Rbpms |
| Tmem1191 | 0 | 0.79108431 | 0.496 | 0.265 | 0 | TAM.pro-infl. | Tmem119 |
| Dennd3 | 0 | 2.25378513 | 0.293 | 0.062 | 0 | TAM.pro-infl. | Dennd3 |
| Slc39a14 | 0 | 0.85914152 | 0.529 | 0.299 | 0 | TAM.pro-infl. | Slc39a14 |
| Aldh3b1 | 0 | 0.77611151 | 0.612 | 0.382 | 0 | TAM.pro-infl. | Aldh3b1 |
| Anxa1 | 0 | 0.69322702 | 0.74 | 0.51 | 0 | TAM.pro-infl. | Anxa1 |
| Fbxl5 | 0 | 1.29857104 | 0.554 | 0.325 | 0 | TAM.pro-infl. | Fbxl5 |
| Fam129b | 0 | 0.83140689 | 0.873 | 0.644 | 0 | TAM.pro-infl. | Fam129b |
| Myof | 0 | 1.09879569 | 0.862 | 0.634 | 0 | TAM.pro-infl. | Myof |
| Prkcb1 | 0 | 0.99246099 | 0.651 | 0.424 | 0 | TAM.pro-infl. | Prkcb |
| Isyna1 | 0 | 0.7383515 | 0.58 | 0.353 | 0 | TAM.pro-infl. | Isyna1 |
| Adora2a | 0 | 1.57741403 | 0.307 | 0.08 | 0 | TAM.pro-infl. | Adora2a |
| G6pdx | 0 | 0.91268466 | 0.589 | 0.362 | 0 | TAM.pro-infl. | G6pdx |
| Anpep1 | 0 | 0.94975302 | 0.688 | 0.461 | 0 | TAM.pro-infl. | Anpep |
| Abca1 | 0 | 0.77588767 | 0.916 | 0.69 | 0 | TAM.pro-infl. | Abca1 |
| Dram1 | 0 | 0.86680718 | 0.504 | 0.278 | 0 | TAM.pro-infl. | Dram1 |
| Gch1 | 0 | 0.62045696 | 0.617 | 0.391 | 0 | TAM.pro-infl. | Gch1 |
| Ak2 | 0 | 0.75833915 | 0.771 | 0.546 | 0 | TAM.pro-infl. | Ak2 |
| Pilra | 0 | 0.62977636 | 0.781 | 0.556 | 0 | TAM.pro-infl. | Pilra |
| Sema4b1 | 0 | 0.84351804 | 0.55 | 0.326 | 0 | TAM.pro-infl. | Sema4b |
| Cat | 0 | 1.07090238 | 0.692 | 0.468 | 0 | TAM.pro-infl. | Cat |
| Cd631 | 0 | 0.9805698 | 0.935 | 0.712 | 0 | TAM.pro-infl. | Cd63 |
| Il17ra | 0 | 0.51412593 | 0.642 | 0.419 | 0 | TAM.pro-infl. | Il17ra |
| Tlnrd1 | 0 | 0.81551178 | 0.652 | 0.429 | 0 | TAM.pro-infl. | Tlnrd1 |
| Dusp16 | 0 | 0.8760157 | 0.545 | 0.324 | 0 | TAM.pro-infl. | Dusp16 |
| Nt5e | 0 | 1.04698684 | 0.38 | 0.159 | 0 | TAM.pro-infl. | Nt5e |
| Mafg | 0 | 0.71289892 | 0.625 | 0.404 | 0 | TAM.pro-infl. | Mafg |
| Rai14 | 0 | 0.58659044 | 0.689 | 0.468 | 0 | TAM.pro-infl. | Rai14 |
| Prr13 | 0 | 0.77386493 | 0.672 | 0.452 | 0 | TAM.pro-infl. | Prr13 |
| Klhl21 | 0 | 1.03824855 | 0.47 | 0.25 | 0 | TAM.pro-infl. | Klhl21 |
| Nus1 | 0 | 0.76458352 | 0.718 | 0.498 | 0 | TAM.pro-infl. | Nus1 |
| Ric1 | 0 | 0.83206767 | 0.6 | 0.38 | 0 | TAM.pro-infl. | Ric1 |
| Plk3 | 0 | 0.78006434 | 0.787 | 0.568 | 0 | TAM.pro-infl. | Plk3 |
| Csf2rb2 | 0 | 0.75750729 | 0.805 | 0.586 | 0 | TAM.pro-infl. | Csf2rb2 |
| Cpd | 0 | 0.74650978 | 0.629 | 0.41 | 0 | TAM.pro-infl. | Cpd |
| Tnip1 | 0 | 0.83952149 | 0.61 | 0.391 | 0 | TAM.pro-infl. | Tnip1 |
| Wsb2 | 0 | 0.8489994 | 0.5 | 0.282 | 0 | TAM.pro-infl. | Wsb2 |
| Cyfip2 | 0 | 1.76581275 | 0.303 | 0.085 | 0 | TAM.pro-infl. | Cyfip2 |
| Dapk1 | 0 | 0.75520876 | 0.611 | 0.393 | 0 | TAM.pro-infl. | Dapk1 |
| Eaf1 | 0 | 0.79128393 | 0.566 | 0.348 | 0 | TAM.pro-infl. | Eaf1 |
| Card19 | 0 | 0.57187651 | 0.793 | 0.576 | 0 | TAM.pro-infl. | Card19 |
| Mmp141 | 0 | 0.81622048 | 0.93 | 0.713 | 0 | TAM.pro-infl. | Mmp14 |
| Mitf | 0 | 0.88058557 | 0.539 | 0.322 | 0 | TAM.pro-infl. | Mitf |
| Mllt6 | 0 | 0.62773941 | 0.683 | 0.466 | 0 | TAM.pro-infl. | Mllt6 |
| Map4k3 | 0 | 0.93915399 | 0.445 | 0.228 | 0 | TAM.pro-infl. | Map4k3 |
| Chpf2 | 0 | 0.88108135 | 0.571 | 0.354 | 0 | TAM.pro-infl. | Chpf2 |
| Pgam1 | 0 | 0.86991459 | 0.892 | 0.675 | 0 | TAM.pro-infl. | Pgam1 |
| Egr2 | 0 | 0.55737207 | 0.659 | 0.443 | 0 | TAM.pro-infl. | Egr2 |
| Slco4a1 | 0 | 3.37410002 | 0.255 | 0.039 | 0 | TAM.pro-infl. | Slco4a1 |
| Rassf3 | 0 | 0.73008025 | 0.581 | 0.365 | 0 | TAM.pro-infl. | Rassf3 |
| Ptgfrn | 0 | 0.73394909 | 0.544 | 0.329 | 0 | TAM.pro-infl. | Ptgfrn |
| Degs1 | 0 | 0.79221746 | 0.696 | 0.481 | 0 | TAM.pro-infl. | Degs1 |
| Osbpl8 | 0 | 0.70731017 | 0.743 | 0.528 | 0 | TAM.pro-infl. | Osbpl8 |
| Fem1b | 0 | 1.16099177 | 0.517 | 0.303 | 0 | TAM.pro-infl. | Fem1b |
| Ptpre | 0 | 0.7749 | 0.746 | 0.532 | 0 | TAM.pro-infl. | Ptpre |
| Slc16a3 | 0 | 0.52641331 | 0.806 | 0.594 | 0 | TAM.pro-infl. | Slc16a3 |
| Uap1l11 | 0 | 0.53437858 | 0.655 | 0.443 | 0 | TAM.pro-infl. | Uap1l1 |
| Tpm1 | 0 | 0.85796313 | 0.496 | 0.284 | 0 | TAM.pro-infl. | Tpm1 |
| Adssl1 | 0 | 0.61597723 | 0.75 | 0.54 | 0 | TAM.pro-infl. | Adssl1 |
| Grina | 0 | 1.84404642 | 0.946 | 0.736 | 0 | TAM.pro-infl. | Grina |
| Spp1 | 0 | 2.64231537 | 0.928 | 0.72 | 0 | TAM.pro-infl. | Spp1 |
| Trpv2 | 0 | 0.74096237 | 0.487 | 0.279 | 0 | TAM.pro-infl. | Trpv2 |
| Creg1 | 0 | 1.43898612 | 0.896 | 0.689 | 0 | TAM.pro-infl. | Creg1 |
| Prdx6 | 0 | 0.80960264 | 0.503 | 0.296 | 0 | TAM.pro-infl. | Prdx6 |
| Kcnn4 | 0 | 0.89164966 | 0.401 | 0.194 | 0 | TAM.pro-infl. | Kcnn4 |
| Abr | 0 | 0.86412796 | 0.89 | 0.683 | 0 | TAM.pro-infl. | Abr |
| Rffl | 0 | 0.80795611 | 0.473 | 0.267 | 0 | TAM.pro-infl. | Rffl |
| Tbc1d9 | 0 | 0.72957841 | 0.677 | 0.472 | 0 | TAM.pro-infl. | Tbc1d9 |
| Sntb2 | 0 | 0.97562227 | 0.404 | 0.199 | 0 | TAM.pro-infl. | Sntb2 |
| Nufip1 | 0 | 0.60282404 | 0.575 | 0.37 | 0 | TAM.pro-infl. | Nufip1 |
| Gnptab | 0 | 0.66574255 | 0.558 | 0.354 | 0 | TAM.pro-infl. | Gnptab |
| Chchd10 | 0 | 1.05917472 | 0.387 | 0.183 | 0 | TAM.pro-infl. | Chchd10 |
| Glipr1 | 0 | 1.11143838 | 0.403 | 0.199 | 0 | TAM.pro-infl. | Glipr1 |
| F11r | 0 | 0.49570798 | 0.733 | 0.529 | 0 | TAM.pro-infl. | F11r |
| Tatdn2 | 0 | 0.67510811 | 0.814 | 0.61 | 0 | TAM.pro-infl. | Tatdn2 |
| Pld31 | 0 | 0.61446711 | 0.79 | 0.587 | 0 | TAM.pro-infl. | Pld3 |
| Psmb6 | 0 | 0.66646525 | 0.613 | 0.41 | 0 | TAM.pro-infl. | Psmb6 |
| Cdadc1 | 0 | 1.71032181 | 0.324 | 0.121 | 0 | TAM.pro-infl. | Cdadc1 |
| Shc1 | 0 | 0.71772509 | 0.591 | 0.388 | 0 | TAM.pro-infl. | Shc1 |
| Anxa7 | 0 | 0.62735049 | 0.606 | 0.403 | 0 | TAM.pro-infl. | Anxa7 |
| Itga5 | 0 | 0.50870433 | 0.706 | 0.503 | 0 | TAM.pro-infl. | Itga5 |
| Sra1 | 0 | 0.74136303 | 0.733 | 0.53 | 0 | TAM.pro-infl. | Sra1 |
| Cxcl1 | 0 | 1.17704536 | 0.403 | 0.201 | 0 | TAM.pro-infl. | Cxcl1 |
| Bri3bp | 0 | 0.58828374 | 0.466 | 0.264 | 0 | TAM.pro-infl. | Bri3bp |
| Ampd2 | 0 | 0.99166999 | 0.428 | 0.226 | 0 | TAM.pro-infl. | Ampd2 |
| Ramp3 | 0 | 0.95568768 | 0.359 | 0.157 | 0 | TAM.pro-infl. | Ramp3 |
| Hcar2 | 0 | 0.72294884 | 0.556 | 0.355 | 0 | TAM.pro-infl. | Hcar2 |
| Clec4n | 0 | 1.55317326 | 0.961 | 0.76 | 0 | TAM.pro-infl. | Clec4n |
| Pgk1 | 0 | 0.64804815 | 0.754 | 0.553 | 0 | TAM.pro-infl. | Pgk1 |
| Cdk18 | 0 | 2.2590306 | 0.26 | 0.06 | 0 | TAM.pro-infl. | Cdk18 |
| Pcnx | 0 | 0.87523794 | 0.401 | 0.201 | 0 | TAM.pro-infl. | Pcnx |
| Lasp1 | 0 | 0.62564894 | 0.684 | 0.485 | 0 | TAM.pro-infl. | Lasp1 |
| Cln5 | 0 | 0.71778865 | 0.594 | 0.395 | 0 | TAM.pro-infl. | Cln5 |
| Piezo1 | 0 | 0.56083663 | 0.668 | 0.47 | 0 | TAM.pro-infl. | Piezo1 |
| Cxcl2 | 0 | 0.74796285 | 0.955 | 0.757 | 0 | TAM.pro-infl. | Cxcl2 |
| Pim3 | 0 | 0.42175881 | 0.724 | 0.526 | 0 | TAM.pro-infl. | Pim3 |
| Id2 | 0 | 0.53050615 | 0.847 | 0.65 | 0 | TAM.pro-infl. | Id2 |
| Parvg | 0 | 0.59918239 | 0.477 | 0.28 | 0 | TAM.pro-infl. | Parvg |
| Iah1 | 0 | 0.74822632 | 0.542 | 0.346 | 0 | TAM.pro-infl. | Iah1 |
| Sqor | 0 | 0.69255861 | 0.488 | 0.292 | 0 | TAM.pro-infl. | Sqor |
| Rassf8 | 0 | 0.92213113 | 0.499 | 0.303 | 0 | TAM.pro-infl. | Rassf8 |
| Pgap2 | 0 | 1.14623047 | 0.46 | 0.264 | 0 | TAM.pro-infl. | Pgap2 |
| Cds2 | 0 | 0.74460711 | 0.463 | 0.268 | 0 | TAM.pro-infl. | Cds2 |
| Tshz1 | 0 | 1.72575418 | 0.293 | 0.098 | 0 | TAM.pro-infl. | Tshz1 |
| Sema4a | 0 | 0.66864454 | 0.823 | 0.628 | 0 | TAM.pro-infl. | Sema4a |
| Cyb5r3 | 0 | 0.60444719 | 0.743 | 0.548 | 0 | TAM.pro-infl. | Cyb5r3 |
| Slc20a1 | 0 | 0.6450093 | 0.617 | 0.423 | 0 | TAM.pro-infl. | Slc20a1 |
| Metap1 | 0 | 0.95453266 | 0.424 | 0.23 | 0 | TAM.pro-infl. | Metap1 |
| Cmklr11 | 0 | 0.64031682 | 0.776 | 0.584 | 0 | TAM.pro-infl. | Cmklr1 |
| Neu1 | 0 | 0.54623666 | 0.679 | 0.489 | 0 | TAM.pro-infl. | Neu1 |
| Ldhb1 | 0 | 1.26553652 | 0.331 | 0.141 | 0 | TAM.pro-infl. | Ldhb |
| Gpd2 | 0 | 0.74747011 | 0.424 | 0.234 | 0 | TAM.pro-infl. | Gpd2 |
| Adgre5 | 0 | 0.35206313 | 0.743 | 0.553 | 0 | TAM.pro-infl. | Adgre5 |
| Pygl | 0 | 0.49904813 | 0.75 | 0.56 | 0 | TAM.pro-infl. | Pygl |
| Cd80 | 0 | 0.25059045 | 0.753 | 0.564 | 0 | TAM.pro-infl. | Cd80 |
| Chka | 0 | 0.71543829 | 0.515 | 0.326 | 0 | TAM.pro-infl. | Chka |
| Pfkp | 0 | 0.38459552 | 0.738 | 0.549 | 0 | TAM.pro-infl. | Pfkp |
| Ptpn12 | 0 | 0.63670272 | 0.793 | 0.605 | 0 | TAM.pro-infl. | Ptpn12 |
| Ikbkg | 0 | 1.05157729 | 0.403 | 0.215 | 0 | TAM.pro-infl. | Ikbkg |
| Gstp1 | 0 | 0.81307387 | 0.558 | 0.371 | 0 | TAM.pro-infl. | Gstp1 |
| Spn | 0 | 1.50053667 | 0.264 | 0.077 | 0 | TAM.pro-infl. | Spn |
| Gpalpp1 | 0 | 0.57352699 | 0.589 | 0.402 | 0 | TAM.pro-infl. | Gpalpp1 |
| Rhov | 0 | 0.54783473 | 0.334 | 0.148 | 0 | TAM.pro-infl. | Rhov |
| Lrrc8d | 0 | 0.71329064 | 0.527 | 0.341 | 0 | TAM.pro-infl. | Lrrc8d |
| Cav2 | 0 | 0.96558065 | 0.377 | 0.191 | 0 | TAM.pro-infl. | Cav2 |
| Irak3 | 0 | 0.9770265 | 0.357 | 0.171 | 0 | TAM.pro-infl. | Irak3 |
| Mtpn | 0 | 0.67828894 | 0.754 | 0.569 | 0 | TAM.pro-infl. | Mtpn |
| Cisd2 | 0 | 0.55650308 | 0.645 | 0.46 | 0 | TAM.pro-infl. | Cisd2 |
| Tmem43 | 0 | 0.55384234 | 0.527 | 0.342 | 0 | TAM.pro-infl. | Tmem43 |
| Samd8 | 0 | 0.65863177 | 0.51 | 0.325 | 0 | TAM.pro-infl. | Samd8 |
| Slc9a3r1 | 0 | 0.53258666 | 0.706 | 0.521 | 0 | TAM.pro-infl. | Slc9a3r1 |
| Rhbdf2 | 0 | 0.48224445 | 0.698 | 0.513 | 0 | TAM.pro-infl. | Rhbdf2 |
| Frrs1 | 0 | 0.63905181 | 0.507 | 0.323 | 0 | TAM.pro-infl. | Frrs1 |
| Htatip2 | 0 | 0.65855089 | 0.469 | 0.285 | 0 | TAM.pro-infl. | Htatip2 |
| Pgd | 0 | 1.46512866 | 0.922 | 0.739 | 0 | TAM.pro-infl. | Pgd |
| Fam71f2 | 0 | 1.56923018 | 0.277 | 0.094 | 0 | TAM.pro-infl. | Fam71f2 |
| Smox | 0 | 0.50523791 | 0.884 | 0.702 | 0 | TAM.pro-infl. | Smox |
| DEPRECATED-E | 0 | 0.59889149 | 0.529 | 0.347 | 0 | TAM.pro-infl. | DEPRECATED-ENSMUSG00000039782 |
| Aprt | 0 | 0.49299629 | 0.538 | 0.357 | 0 | TAM.pro-infl. | Aprt |
| Prdm1 | 0 | 0.43603338 | 0.543 | 0.362 | 0 | TAM.pro-infl. | Prdm1 |
| Blcap | 0 | 0.70096553 | 0.454 | 0.273 | 0 | TAM.pro-infl. | Blcap |
| Hnrnph3 | 0 | 1.18429958 | 0.356 | 0.175 | 0 | TAM.pro-infl. | Hnrnph3 |
| Phldb1 | 0 | 1.04443284 | 0.349 | 0.168 | 0 | TAM.pro-infl. | Phldb1 |
| Pcyt1a | 0 | 0.54255566 | 0.562 | 0.382 | 0 | TAM.pro-infl. | Pcyt1a |
| Stra6l1 | 0 | 1.29415424 | 0.318 | 0.138 | 0 | TAM.pro-infl. | Stra6l |
| Akr1b8 | 0 | 2.05954097 | 0.284 | 0.104 | 0 | TAM.pro-infl. | Akr1b8 |
| 31-Mar | 0 | 0.74970193 | 0.415 | 0.235 | 0 | TAM.pro-infl. | 3-Mar |
| Gadd45a | 0 | 0.71843588 | 0.408 | 0.228 | 0 | TAM.pro-infl. | Gadd45a |
| Ccl9 | 0 | 0.9772562 | 0.821 | 0.641 | 0 | TAM.pro-infl. | Ccl9 |
| Agap3 | 0 | 0.50922681 | 0.613 | 0.434 | 0 | TAM.pro-infl. | Agap3 |
| Runx2 | 0 | 0.82945835 | 0.347 | 0.168 | 0 | TAM.pro-infl. | Runx2 |
| Edem2 | 0 | 0.66899428 | 0.579 | 0.4 | 0 | TAM.pro-infl. | Edem2 |
| Mapkapk3 | 0 | 0.39313675 | 0.648 | 0.47 | 0 | TAM.pro-infl. | Mapkapk3 |
| H6pd | 0 | 0.62450221 | 0.5 | 0.322 | 0 | TAM.pro-infl. | H6pd |
| Pttg1ip | 0 | 0.56265542 | 0.74 | 0.562 | 0 | TAM.pro-infl. | Pttg1ip |
| Idh1 | 0 | 0.61934865 | 0.565 | 0.388 | 0 | TAM.pro-infl. | Idh1 |
| Limd1 | 0 | 0.53953343 | 0.587 | 0.411 | 0 | TAM.pro-infl. | Limd1 |
| Piwil2 | 0 | 1.42356458 | 0.29 | 0.115 | 0 | TAM.pro-infl. | Piwil2 |
| Rab7 | 0 | 0.78708922 | 0.838 | 0.663 | 0 | TAM.pro-infl. | Rab7 |
| Soat1 | 0 | 0.65253999 | 0.793 | 0.619 | 0 | TAM.pro-infl. | Soat1 |
| Tspan5 | 0 | 0.9723959 | 0.337 | 0.163 | 0 | TAM.pro-infl. | Tspan5 |
| Nfil3 | 0 | 0.32516182 | 0.863 | 0.69 | 0 | TAM.pro-infl. | Nfil3 |
| Psd4 | 0 | 0.47864795 | 0.589 | 0.416 | 0 | TAM.pro-infl. | Psd4 |
| Igsf81 | 0 | 0.44206826 | 0.693 | 0.52 | 0 | TAM.pro-infl. | Igsf8 |
| Tmem63a | 0 | 0.51925835 | 0.559 | 0.387 | 0 | TAM.pro-infl. | Tmem63a |
| Slc23a2 | 0 | 0.58072312 | 0.49 | 0.318 | 0 | TAM.pro-infl. | Slc23a2 |
| Retreg1 | 0 | 0.62657981 | 0.474 | 0.302 | 0 | TAM.pro-infl. | Retreg1 |
| Tfec | 0 | 0.54580464 | 0.536 | 0.365 | 0 | TAM.pro-infl. | Tfec |
| Cpeb2 | 0 | 0.59401056 | 0.58 | 0.409 | 0 | TAM.pro-infl. | Cpeb2 |
| Plxnc11 | 0 | 0.91633721 | 0.373 | 0.202 | 0 | TAM.pro-infl. | Plxnc1 |
| Tmem170b | 0 | 0.5269162 | 0.582 | 0.411 | 0 | TAM.pro-infl. | Tmem170b |
| Mpc1 | 0 | 0.60603205 | 0.588 | 0.417 | 0 | TAM.pro-infl. | Mpc1 |
| Irak2 | 0 | 0.31759218 | 0.731 | 0.56 | 0 | TAM.pro-infl. | Irak2 |
| Tmem120b | 0 | 1.34726056 | 0.28 | 0.11 | 0 | TAM.pro-infl. | Tmem120b |
| Ppt2 | 0 | 0.50845867 | 0.527 | 0.357 | 0 | TAM.pro-infl. | Ppt2 |
| Plekhn1 | 0 | 0.73091858 | 0.443 | 0.273 | 0 | TAM.pro-infl. | Plekhn1 |
| Tmbim1 | 0 | 0.55159628 | 0.696 | 0.526 | 0 | TAM.pro-infl. | Tmbim1 |
| Rilpl21 | 0 | 0.45852274 | 0.723 | 0.553 | 0 | TAM.pro-infl. | Rilpl2 |
| Uck2 | 0 | 0.25298093 | 0.863 | 0.694 | 0 | TAM.pro-infl. | Uck2 |
| Tmem189 | 0 | 0.63385156 | 0.901 | 0.732 | 0 | TAM.pro-infl. | Tmem189 |
| Ugdh | 0 | 0.63775463 | 0.531 | 0.362 | 0 | TAM.pro-infl. | Ugdh |
| Clcn71 | 0 | 0.50935227 | 0.613 | 0.444 | 0 | TAM.pro-infl. | Clcn7 |
| Med14 | 0 | 0.68314955 | 0.432 | 0.263 | 0 | TAM.pro-infl. | Med14 |
| Acsl4 | 0 | 0.56168842 | 0.6 | 0.431 | 0 | TAM.pro-infl. | Acsl4 |
| Gpr35 | 0 | 0.49253277 | 0.827 | 0.658 | 0 | TAM.pro-infl. | Gpr35 |
| Arrdc4 | 0 | 0.59598219 | 0.727 | 0.558 | 0 | TAM.pro-infl. | Arrdc4 |
| Irs21 | 0 | 0.52072751 | 0.839 | 0.67 | 0 | TAM.pro-infl. | Irs2 |
| Pfkl | 0 | 0.35626572 | 0.715 | 0.546 | 0 | TAM.pro-infl. | Pfkl |
| Ldlrap1 | 0 | 0.47063109 | 0.536 | 0.368 | 0 | TAM.pro-infl. | Ldlrap1 |
| Atp6v1e1 | 0 | 0.84387081 | 0.879 | 0.711 | 0 | TAM.pro-infl. | Atp6v1e1 |
| Arrb1 | 0 | 0.38963584 | 0.665 | 0.497 | 0 | TAM.pro-infl. | Arrb1 |
| Zadh2 | 0 | 1.14724177 | 0.321 | 0.154 | 0 | TAM.pro-infl. | Zadh2 |
| Atp6v1c1 | 0 | 0.48479365 | 0.701 | 0.534 | 0 | TAM.pro-infl. | Atp6v1c1 |
| Socs21 | 0 | 0.66496991 | 0.453 | 0.288 | 0 | TAM.pro-infl. | Socs2 |
| Plaur | 0 | 0.60776897 | 0.964 | 0.799 | 0 | TAM.pro-infl. | Plaur |
| Tgfbr11 | 0 | 0.52001287 | 0.727 | 0.563 | 0 | TAM.pro-infl. | Tgfbr1 |
| Emb | 0 | 0.64797388 | 0.883 | 0.719 | 0 | TAM.pro-infl. | Emb |
| Asap1 | 0 | 0.53329046 | 0.514 | 0.35 | 0 | TAM.pro-infl. | Asap1 |
| Tmem38b | 0 | 0.7971879 | 0.363 | 0.199 | 0 | TAM.pro-infl. | Tmem38b |
| Lrpap1 | 0 | 0.69389312 | 0.8 | 0.637 | 0 | TAM.pro-infl. | Lrpap1 |
| Ctsd | 0 | 0.75259495 | 0.883 | 0.72 | 0 | TAM.pro-infl. | Ctsd |
| Mfsd121 | 0 | 0.47135679 | 0.633 | 0.471 | 0 | TAM.pro-infl. | Mfsd12 |
| Got1 | 0 | 0.32513564 | 0.775 | 0.613 | 0 | TAM.pro-infl. | Got1 |
| Il2rg | 0 | 0.48084957 | 0.742 | 0.58 | 0 | TAM.pro-infl. | Il2rg |
| Slc25a25 | 0 | 0.70466857 | 0.388 | 0.226 | 0 | TAM.pro-infl. | Slc25a25 |
| Fkbp1a | 0 | 0.40962973 | 0.613 | 0.451 | 0 | TAM.pro-infl. | Fkbp1a |
| Ets1 | 0 | 1.35485192 | 0.273 | 0.112 | 0 | TAM.pro-infl. | Ets1 |
| Ergic1 | 0 | 0.5029872 | 0.534 | 0.373 | 0 | TAM.pro-infl. | Ergic1 |
| Glrx | 0 | 0.56277127 | 0.662 | 0.502 | 0 | TAM.pro-infl. | Glrx |
| Flna | 0 | 0.62265234 | 0.886 | 0.726 | 0 | TAM.pro-infl. | Flna |
| Katnbl1 | 0 | 0.88779576 | 0.344 | 0.184 | 0 | TAM.pro-infl. | Katnbl1 |
| Dgat1 | 0 | 0.76807523 | 0.353 | 0.193 | 0 | TAM.pro-infl. | Dgat1 |
| Pla2g7 | 0 | 1.47520455 | 0.955 | 0.795 | 0 | TAM.pro-infl. | Pla2g7 |
| Cd841 | 0 | 0.31213806 | 0.758 | 0.599 | 0 | TAM.pro-infl. | Cd84 |
| Sptan1 | 0 | 0.36182679 | 0.677 | 0.518 | 0 | TAM.pro-infl. | Sptan1 |
| Bhlhe40 | 0 | 0.36528945 | 0.935 | 0.776 | 0 | TAM.pro-infl. | Bhlhe40 |
| Sh3pxd2b | 0 | 0.56546842 | 0.874 | 0.715 | 0 | TAM.pro-infl. | Sh3pxd2b |
| Ptk2b | 0 | 0.3025148 | 0.735 | 0.576 | 0 | TAM.pro-infl. | Ptk2b |
| Tpm4 | 0 | 0.73171527 | 0.834 | 0.675 | 0 | TAM.pro-infl. | Tpm4 |
| Chp1 | 0 | 0.40945057 | 0.724 | 0.566 | 0 | TAM.pro-infl. | Chp1 |
| Cox17 | 0 | 0.46233337 | 0.724 | 0.566 | 0 | TAM.pro-infl. | Cox17 |
| Lpp | 0 | 0.3546323 | 0.711 | 0.553 | 0 | TAM.pro-infl. | Lpp |
| Ncoa41 | 0 | 0.55977382 | 0.662 | 0.505 | 0 | TAM.pro-infl. | Ncoa4 |
| Traf5 | 0 | 0.93161785 | 0.367 | 0.21 | 0 | TAM.pro-infl. | Traf5 |
| Myo1f | 0 | 0.59027604 | 0.822 | 0.665 | 0 | TAM.pro-infl. | Myo1f |
| S100a1 | 0 | 0.43713631 | 0.58 | 0.424 | 0 | TAM.pro-infl. | S100a1 |
| Arhgap31 | 0 | 0.39956926 | 0.623 | 0.467 | 0 | TAM.pro-infl. | Arhgap31 |
| Mertk1 | 0 | 0.27228158 | 0.699 | 0.543 | 0 | TAM.pro-infl. | Mertk |
| Tbca | 0 | 0.42295264 | 0.658 | 0.503 | 0 | TAM.pro-infl. | Tbca |
| Tnfrsf23 | 0 | 0.94103989 | 0.306 | 0.151 | 0 | TAM.pro-infl. | Tnfrsf23 |
| B4galt5 | 0 | 0.49261155 | 0.801 | 0.647 | 0 | TAM.pro-infl. | B4galt5 |
| Fbxo32 | 0 | 1.05320605 | 0.276 | 0.122 | 0 | TAM.pro-infl. | Fbxo32 |
| Npc11 | 0 | 0.37008937 | 0.74 | 0.586 | 0 | TAM.pro-infl. | Npc1 |
| Cpeb4 | 0 | 0.48608703 | 0.575 | 0.421 | 0 | TAM.pro-infl. | Cpeb4 |
| Ppp2r1a | 0 | 0.38310702 | 0.71 | 0.556 | 0 | TAM.pro-infl. | Ppp2r1a |
| Larp4b | 0 | 0.41051883 | 0.649 | 0.496 | 0 | TAM.pro-infl. | Larp4b |
| Apbb3 | 0 | 0.93151721 | 0.302 | 0.149 | 0 | TAM.pro-infl. | Apbb3 |
| Zc3h12a | 0 | 0.40832127 | 0.862 | 0.71 | 0 | TAM.pro-infl. | Zc3h12a |
| Rnf103 | 0 | 0.73435571 | 0.486 | 0.334 | 0 | TAM.pro-infl. | Rnf103 |
| Mapk3 | 0 | 0.42247482 | 0.693 | 0.542 | 0 | TAM.pro-infl. | Mapk3 |
| Stxbp2 | 0 | 0.39050902 | 0.685 | 0.535 | 0 | TAM.pro-infl. | Stxbp2 |
| Mdm2 | 0 | 0.36019263 | 0.714 | 0.564 | 0 | TAM.pro-infl. | Mdm2 |
| Anxa11 | 0 | 0.35759091 | 0.708 | 0.558 | 0 | TAM.pro-infl. | Anxa11 |
| Ets2 | 0 | 0.73249747 | 0.941 | 0.791 | 0 | TAM.pro-infl. | Ets2 |
| Hopx | 0 | 0.35449885 | 0.341 | 0.192 | 0 | TAM.pro-infl. | Hopx |
| Rtn3 | 0 | 0.49600817 | 0.804 | 0.655 | 0 | TAM.pro-infl. | Rtn3 |
| Fn1 | 0 | 0.7370312 | 0.963 | 0.815 | 0 | TAM.pro-infl. | Fn1 |
| Elf4e | 0 | 0.43058616 | 0.685 | 0.537 | 0 | TAM.pro-infl. | Elf4e |
| Atp6v0e | 0 | 0.56111491 | 0.796 | 0.649 | 0 | TAM.pro-infl. | Atp6v0e |
| Cndp2 | 0 | 0.48256057 | 0.796 | 0.649 | 0 | TAM.pro-infl. | Cndp2 |
| Plgrkt | 0 | 0.3653023 | 0.656 | 0.509 | 0 | TAM.pro-infl. | Plgrkt |
| Cybb | 0 | 0.90213866 | 0.856 | 0.709 | 0 | TAM.pro-infl. | Cybb |
| Cd82 | 0 | 0.35521183 | 0.624 | 0.478 | 0 | TAM.pro-infl. | Cd82 |
| Rpn1 | 0 | 0.38098741 | 0.743 | 0.597 | 0 | TAM.pro-infl. | Rpn1 |
| Abhd17c | 0 | 0.80769138 | 0.337 | 0.191 | 0 | TAM.pro-infl. | Abhd17c |
| Riok3 | 0 | 0.40175132 | 0.793 | 0.647 | 0 | TAM.pro-infl. | Riok3 |
| Myh9 | 0 | 0.4792527 | 0.797 | 0.652 | 0 | TAM.pro-infl. | Myh9 |
| Clic4 | 0 | 0.54737288 | 0.862 | 0.718 | 0 | TAM.pro-infl. | Clic4 |
| Colgalt11 | 0 | 0.38654984 | 0.781 | 0.637 | 0 | TAM.pro-infl. | Colgalt1 |
| Lrrc59 | 0 | 0.34267318 | 0.782 | 0.638 | 0 | TAM.pro-infl. | Lrrc59 |
| Hmgcr1 | 0 | 0.35531917 | 0.569 | 0.425 | 0 | TAM.pro-infl. | Hmgcr |
| Dhcr241 | 0 | 0.71177456 | 0.342 | 0.2 | 0 | TAM.pro-infl. | Dhcr24 |
| Gss | 0 | 1.15494201 | 0.277 | 0.136 | 0 | TAM.pro-infl. | Gss |
| Trim25 | 0 | 0.45606275 | 0.789 | 0.648 | 0 | TAM.pro-infl. | Trim25 |
| Prkar2b | 0 | 1.10566642 | 0.259 | 0.118 | 0 | TAM.pro-infl. | Prkar2b |
| Hexb1 | 0 | 0.28825159 | 0.768 | 0.627 | 0 | TAM.pro-infl. | Hexb |
| Hbegf | 0 | 0.62415887 | 0.401 | 0.26 | 0 | TAM.pro-infl. | Hbegf |
| Gpi1 | 0 | 0.42077732 | 0.864 | 0.725 | 0 | TAM.pro-infl. | Gpi1 |
| Csf2rb | 0 | 0.59241789 | 0.913 | 0.774 | 0 | TAM.pro-infl. | Csf2rb |
| Chd4 | 0 | 0.31547123 | 0.747 | 0.61 | 0 | TAM.pro-infl. | Chd4 |
| Aldoa | 0 | 0.84534399 | 0.942 | 0.805 | 0 | TAM.pro-infl. | Aldoa |
| Rabac1 | 0 | 0.30687569 | 0.793 | 0.658 | 0 | TAM.pro-infl. | Rabac1 |
| Igsf6 | 0 | 0.33125627 | 0.734 | 0.599 | 0 | TAM.pro-infl. | Igsf6 |
| Atp1b3 | 0 | 0.54984252 | 0.847 | 0.712 | 0 | TAM.pro-infl. | Atp1b3 |
| Cd300c21 | 0 | 0.2557154 | 0.723 | 0.588 | 0 | TAM.pro-infl. | Cd300c2 |
| Sem1 | 0 | 0.53510436 | 0.863 | 0.73 | 0 | TAM.pro-infl. | Sem1 |
| Bax | 0 | 0.45598778 | 0.892 | 0.759 | 0 | TAM.pro-infl. | Bax |
| Phlda1 | 0 | 0.26194574 | 0.578 | 0.445 | 0 | TAM.pro-infl. | Phlda1 |
| Nfkbid | 0 | 0.31270688 | 0.727 | 0.595 | 0 | TAM.pro-infl. | Nfkbid |
| Il1b | 0 | 0.30994722 | 0.963 | 0.832 | 0 | TAM.pro-infl. | Il1b |
| Taldo1 | 0 | 0.91614328 | 0.898 | 0.767 | 0 | TAM.pro-infl. | Taldo1 |
| Esyt1 | 0 | 0.27868419 | 0.721 | 0.591 | 0 | TAM.pro-infl. | Esyt1 |
| Dnajc14 | 0 | 0.37991724 | 0.618 | 0.488 | 0 | TAM.pro-infl. | Dnajc14 |
| Hivep3 | 0 | 0.5396833 | 0.486 | 0.357 | 0 | TAM.pro-infl. | Hivep3 |
| Rnh1 | 0 | 0.34770958 | 0.779 | 0.651 | 0 | TAM.pro-infl. | Rnh1 |
| Cers2 | 0 | 0.26421325 | 0.694 | 0.568 | 0 | TAM.pro-infl. | Cers2 |
| Macf1 | 0 | 0.27574833 | 0.701 | 0.575 | 0 | TAM.pro-infl. | Macf1 |
| Actn4 | 0 | 0.252416 | 0.798 | 0.672 | 0 | TAM.pro-infl. | Actn4 |
| Anxa31 | 0 | 0.35822996 | 0.883 | 0.758 | 0 | TAM.pro-infl. | Anxa3 |
| Nptn | 0 | 0.26835996 | 0.767 | 0.642 | 0 | TAM.pro-infl. | Nptn |
| Cd300ld1 | 0 | 0.26425588 | 0.628 | 0.503 | 0 | TAM.pro-infl. | Cd300ld |
| Cd14 | 0 | 0.81081427 | 0.981 | 0.856 | 0 | TAM.pro-infl. | Cd14 |
| Dstn | 0 | 0.30767214 | 0.706 | 0.582 | 0 | TAM.pro-infl. | Dstn |
| Atp6v1a | 0 | 0.31749912 | 0.84 | 0.717 | 0 | TAM.pro-infl. | Atp6v1a |
| Plod3 | 0 | 0.26402558 | 0.732 | 0.61 | 0 | TAM.pro-infl. | Plod3 |

| | | | | | | | |
|---|---|---|---|---|---|---|---|
| Nampt1 | 0 | 0.2674283 | 0.752 | 0.63 | 0 | TAM.pro-infl. | Nampt |
| Ampd3 | 0 | 0.85259636 | 0.264 | 0.144 | 0 | TAM.pro-infl. | Ampd3 |
| M6pr | 0 | 0.55950006 | 0.881 | 0.761 | 0 | TAM.pro-infl. | M6pr |
| Ctnnb1 | 0 | 0.39301031 | 0.795 | 0.675 | 0 | TAM.pro-infl. | Ctnnb1 |
| Sqstm1 | 0 | 0.38554304 | 0.873 | 0.753 | 0 | TAM.pro-infl. | Sqstm1 |
| Dusp3 | 0 | 0.36103489 | 0.845 | 0.725 | 0 | TAM.pro-infl. | Dusp3 |
| Tkt | 0 | 0.60025127 | 0.898 | 0.778 | 0 | TAM.pro-infl. | Tkt |
| Msrb1 | 0 | 0.60764243 | 0.915 | 0.795 | 0 | TAM.pro-infl. | Msrb1 |
| Sgpl11 | 0 | 0.26816915 | 0.791 | 0.672 | 0 | TAM.pro-infl. | Sgpl1 |
| Prdx2 | 0 | 0.27557502 | 0.709 | 0.591 | 0 | TAM.pro-infl. | Prdx2 |
| Mvp | 0 | 0.28141768 | 0.8 | 0.684 | 0 | TAM.pro-infl. | Mvp |
| Mxi1 | 0 | 0.32718515 | 0.688 | 0.573 | 0 | TAM.pro-infl. | Mxi1 |
| Surf4 | 0 | 0.2839413 | 0.718 | 0.605 | 0 | TAM.pro-infl. | Surf4 |
| Nfkb1 | 0 | 0.27569729 | 0.763 | 0.65 | 0 | TAM.pro-infl. | Nfkb1 |
| Tnfrsf1b | 0 | 0.58896851 | 0.886 | 0.774 | 0 | TAM.pro-infl. | Tnfrsf1b |
| Ywhag | 0 | 0.46793737 | 0.922 | 0.81 | 0 | TAM.pro-infl. | Ywhag |
| Vcp | 0 | 0.32827974 | 0.897 | 0.786 | 0 | TAM.pro-infl. | Vcp |
| Aldh2 | 0 | 0.2548127 | 0.817 | 0.707 | 0 | TAM.pro-infl. | Aldh2 |
| Gsn | 0 | 0.50291747 | 0.925 | 0.816 | 0 | TAM.pro-infl. | Gsn |
| Atp6v0b | 0 | 0.3918114 | 0.872 | 0.764 | 0 | TAM.pro-infl. | Atp6v0b |
| Atp6v1b2 | 0 | 0.39593349 | 0.89 | 0.782 | 0 | TAM.pro-infl. | Atp6v1b2 |
| Cyth1 | 0 | 0.33187558 | 0.859 | 0.752 | 0 | TAM.pro-infl. | Cyth1 |
| Csf2ra | 0 | 0.57083271 | 0.965 | 0.858 | 0 | TAM.pro-infl. | Csf2ra |
| Cd44 | 0 | 0.30851006 | 0.934 | 0.828 | 0 | TAM.pro-infl. | Cd44 |
| Tmem50a | 0 | 0.25788943 | 0.809 | 0.704 | 0 | TAM.pro-infl. | Tmem50a |
| Atp6ap2 | 0 | 0.29439801 | 0.853 | 0.748 | 0 | TAM.pro-infl. | Atp6ap2 |
| Anxa5 | 0 | 0.768654 | 0.947 | 0.844 | 0 | TAM.pro-infl. | Anxa5 |
| P4hb | 0 | 0.27602963 | 0.87 | 0.767 | 0 | TAM.pro-infl. | P4hb |
| Cfb | 0 | 0.28763102 | 0.482 | 0.38 | 0 | TAM.pro-infl. | Cfb |
| Adam17 | 0 | 0.26984015 | 0.782 | 0.681 | 0 | TAM.pro-infl. | Adam17 |
| Thbd | 0 | 0.7546982 | 0.288 | 0.189 | 0 | TAM.pro-infl. | Thbd |
| Myl12a | 0 | 0.26431014 | 0.852 | 0.756 | 0 | TAM.pro-infl. | Myl12a |
| Slc3a2 | 0 | 0.69912909 | 0.952 | 0.856 | 0 | TAM.pro-infl. | Slc3a2 |
| Cstb | 0 | 0.88678715 | 0.955 | 0.86 | 0 | TAM.pro-infl. | Cstb |
| Itgam | 0 | 0.31248659 | 0.885 | 0.791 | 0 | TAM.pro-infl. | Itgam |
| Pkm | 0 | 0.49495443 | 0.943 | 0.849 | 0 | TAM.pro-infl. | Pkm |
| Plxnd1 | 0 | 0.33315543 | 0.887 | 0.794 | 0 | TAM.pro-infl. | Plxnd1 |
| S100a11 | 0 | 0.84560126 | 0.946 | 0.854 | 0 | TAM.pro-infl. | S100a11 |
| Lgals3 | 0 | 1.12848424 | 0.983 | 0.891 | 0 | TAM.pro-infl. | Lgals3 |
| Plin2 | 0 | 0.62290135 | 0.939 | 0.848 | 0 | TAM.pro-infl. | Plin2 |
| Ptpn1 | 0 | 0.33315886 | 0.845 | 0.755 | 0 | TAM.pro-infl. | Ptpn1 |
| Slc39a1 | 0 | 0.32293935 | 0.898 | 0.81 | 0 | TAM.pro-infl. | Slc39a1 |
| Tcirg1 | 0 | 0.51810012 | 0.938 | 0.856 | 0 | TAM.pro-infl. | Tcirg1 |
| Prkcd | 0 | 0.43769619 | 0.941 | 0.861 | 0 | TAM.pro-infl. | Prkcd |
| Cd52 | 0 | 0.49777849 | 0.933 | 0.86 | 0 | TAM.pro-infl. | Cd52 |
| Ncf2 | 0 | 0.38396766 | 0.926 | 0.856 | 0 | TAM.pro-infl. | Ncf2 |
| Mpeg11 | 0 | 0.43367714 | 0.956 | 0.887 | 0 | TAM.pro-infl. | Mpeg1 |
| Slc15a3 | 0 | 0.62646721 | 0.979 | 0.912 | 0 | TAM.pro-infl. | Slc15a3 |
| Rnf149 | 0 | 0.71192393 | 0.978 | 0.915 | 0 | TAM.pro-infl. | Rnf149 |
| Lrp1 | 0 | 0.36033429 | 0.953 | 0.891 | 0 | TAM.pro-infl. | Lrp1 |
| Cd53 | 0 | 0.60477118 | 0.936 | 0.876 | 0 | TAM.pro-infl. | Cd53 |
| Fabp51 | 0 | 0.50110195 | 0.731 | 0.672 | 0 | TAM.pro-infl. | Fabp5 |
| Itgb2 | 0 | 1.35370184 | 0.976 | 0.917 | 0 | TAM.pro-infl. | Itgb2 |
| Prdx5 | 0 | 1.11706611 | 0.98 | 0.921 | 0 | TAM.pro-infl. | Prdx5 |
| Iqgap1 | 0 | 0.36251972 | 0.968 | 0.912 | 0 | TAM.pro-infl. | Iqgap1 |
| Atp1a1 | 0 | 0.50120974 | 0.936 | 0.883 | 0 | TAM.pro-infl. | Atp1a1 |
| Hcls1 | 0 | 0.55570357 | 0.947 | 0.895 | 0 | TAM.pro-infl. | Hcls1 |
| Ier3 | 0 | 0.31646086 | 0.961 | 0.911 | 0 | TAM.pro-infl. | Ier3 |
| Sh3bgrl3 | 0 | 0.55437272 | 0.945 | 0.896 | 0 | TAM.pro-infl. | Sh3bgrl3 |
| Syk | 0 | 0.35844776 | 0.944 | 0.898 | 0 | TAM.pro-infl. | Syk |
| Mfge8 | 0 | 1.32330223 | 0.283 | 0.24 | 0 | TAM.pro-infl. | Mfge8 |
| Atp6v0c | 0 | 0.27172027 | 0.936 | 0.9 | 0 | TAM.pro-infl. | Atp6v0c |
| Cyba | 0 | 0.64766406 | 0.961 | 0.926 | 0 | TAM.pro-infl. | Cyba |
| Lcp1 | 0 | 0.53621608 | 0.983 | 0.949 | 0 | TAM.pro-infl. | Lcp1 |
| Tnf | 0 | 0.33585485 | 0.643 | 0.612 | 0 | TAM.pro-infl. | Tnf |
| Akr1a1 | 0 | 0.47966978 | 0.96 | 0.933 | 0 | TAM.pro-infl. | Akr1a1 |
| Msn | 0 | 0.43290386 | 0.965 | 0.939 | 0 | TAM.pro-infl. | Msn |
| Cebpb | 0 | 0.41295137 | 0.995 | 0.974 | 0 | TAM.pro-infl. | Cebpb |
| Ctsz1 | 0 | 0.40160869 | 0.988 | 0.969 | 0 | TAM.pro-infl. | Ctsz |
| Lyz2 | 0 | 0.57479225 | 0.979 | 0.962 | 0 | TAM.pro-infl. | Lyz2 |
| Psap1 | 0 | 0.84478759 | 0.982 | 0.968 | 0 | TAM.pro-infl. | Psap |
| Fcer1g | 0 | 0.36347223 | 0.986 | 0.973 | 0 | TAM.pro-infl. | Fcer1g |
| Ucp2 | 0 | 0.62664774 | 0.985 | 0.981 | 0 | TAM.pro-infl. | Ucp2 |
| Fth1 | 0 | 1.381825 | 1 | 0.996 | 0 | TAM.pro-infl. | Fth1 |
| Ctss | 0 | 0.41512922 | 0.991 | 0.991 | 0 | TAM.pro-infl. | Ctss |
| Ncor2 | 1.17E-306 | 0.34668843 | 0.64 | 0.489 | 2.25E-302 | TAM.pro-infl. | Ncor2 |
| Kif1c | 1.31E-303 | 0.46388105 | 0.57 | 0.404 | 2.54E-299 | TAM.pro-infl. | Kif1c |
| Bola2 | 7.51E-302 | 0.26566794 | 0.654 | 0.514 | 1.45E-297 | TAM.pro-infl. | Bola2 |
| Pnpla2 | 6.16E-301 | 0.27597836 | 0.616 | 0.498 | 1.19E-296 | TAM.pro-infl. | Pnpla2 |
| BC028528 | 1.15E-296 | 0.55337093 | 0.404 | 0.245 | 2.22E-292 | TAM.pro-infl. | BC028528 |
| Pstpip2 | 7.12E-296 | 0.34358687 | 0.501 | 0.357 | 1.38E-291 | TAM.pro-infl. | Pstpip2 |
| Clk4 | 5.96E-295 | 0.47498374 | 0.58 | 0.416 | 1.15E-290 | TAM.pro-infl. | Clk4 |
| Ralgapa2 | 6.95E-292 | 1.00198706 | 0.298 | 0.148 | 1.34E-287 | TAM.pro-infl. | Ralgapa2 |
| Mocos | 1.33E-291 | 0.94556747 | 0.286 | 0.129 | 2.58E-287 | TAM.pro-infl. | Mocos |
| Fam89b | 4.98E-291 | 0.26601937 | 0.624 | 0.493 | 9.62E-287 | TAM.pro-infl. | Fam89b |
| Rab10 | 9.57E-291 | 0.43500806 | 0.537 | 0.394 | 1.85E-286 | TAM.pro-infl. | Rab10 |
| Pqlc3 | 2.55E-289 | 0.74527555 | 0.339 | 0.173 | 4.93E-285 | TAM.pro-infl. | Pqlc3 |
| Fam107b | 7.15E-289 | 0.26044522 | 0.561 | 0.434 | 1.38E-284 | TAM.pro-infl. | Fam107b |
| Dnajc5 | 6.64E-287 | 0.30941991 | 0.659 | 0.525 | 1.28E-282 | TAM.pro-infl. | Dnajc5 |
| Gga21 | 1.24E-285 | 0.83846435 | 0.324 | 0.186 | 2.39E-281 | TAM.pro-infl. | Gga2 |
| Ybx3 | 9.35E-284 | 0.4330207 | 0.575 | 0.398 | 1.81E-279 | TAM.pro-infl. | Ybx3 |
| Atp8b4 | 6.93E-283 | 0.59957891 | 0.352 | 0.19 | 1.34E-278 | TAM.pro-infl. | Atp8b4 |
| Cd302 | 1.87E-281 | 0.55856772 | 0.431 | 0.273 | 3.63E-277 | TAM.pro-infl. | Cd302 |
| Nufip2 | 3.52E-280 | 0.33035057 | 0.649 | 0.513 | 6.81E-276 | TAM.pro-infl. | Nufip2 |
| Slc41a2 | 3.68E-280 | 0.83238235 | 0.352 | 0.179 | 7.11E-276 | TAM.pro-infl. | Slc41a2 |
| Unc119 | 5.68E-280 | 0.79552153 | 0.316 | 0.164 | 1.10E-275 | TAM.pro-infl. | Unc119 |
| Cln8 | 1.15E-277 | 0.33378388 | 0.606 | 0.464 | 2.21E-273 | TAM.pro-infl. | Cln8 |
| Pik3r6 | 2.17E-275 | 0.67372948 | 0.41 | 0.248 | 4.20E-271 | TAM.pro-infl. | Pik3r6 |
| Tpd52 | 6.31E-272 | 0.3233737 | 0.599 | 0.445 | 1.22E-267 | TAM.pro-infl. | Tpd52 |
| Shtn1 | 6.43E-272 | 0.30185535 | 0.591 | 0.453 | 1.24E-267 | TAM.pro-infl. | Shtn1 |
| Galnt6 | 1.30E-269 | 0.65204372 | 0.484 | 0.31 | 2.51E-265 | TAM.pro-infl. | Galnt6 |
| Renbp1 | 1.47E-267 | 0.35324874 | 0.678 | 0.52 | 2.85E-263 | TAM.pro-infl. | Renbp |
| Psmd1 | 1.25E-266 | 0.436393 | 0.63 | 0.467 | 2.41E-262 | TAM.pro-infl. | Psmd1 |
| Psmd11 | 4.49E-266 | 0.43259017 | 0.588 | 0.429 | 8.69E-262 | TAM.pro-infl. | Psmd11 |
| Npepps | 8.13E-266 | 0.51335285 | 0.556 | 0.38 | 1.57E-261 | TAM.pro-infl. | Npepps |
| Mif | 4.13E-265 | 0.34499897 | 0.701 | 0.534 | 7.99E-261 | TAM.pro-infl. | Mif |
| Cd24a | 4.75E-264 | 0.33549782 | 0.34 | 0.245 | 9.19E-260 | TAM.pro-infl. | Cd24a |
| Cnppd1 | 1.24E-260 | 0.2796014 | 0.701 | 0.574 | 2.39E-256 | TAM.pro-infl. | Cnppd1 |
| Ago2 | 2.54E-259 | 0.28421086 | 0.665 | 0.526 | 4.90E-255 | TAM.pro-infl. | Ago2 |
| Micall1 | 2.51E-258 | 1.08862196 | 0.255 | 0.112 | 4.85E-254 | TAM.pro-infl. | Micall1 |
| Dnaja2 | 1.25E-257 | 0.27916466 | 0.693 | 0.569 | 2.43E-253 | TAM.pro-infl. | Dnaja2 |
| Cry1 | 4.90E-257 | 0.92497827 | 0.294 | 0.14 | 9.47E-253 | TAM.pro-infl. | Cry1 |
| Acsl1 | 2.53E-256 | 0.80720192 | 0.3 | 0.163 | 4.89E-252 | TAM.pro-infl. | Acsl1 |
| Ttc39c | 8.94E-256 | 0.87708769 | 0.298 | 0.162 | 1.73E-251 | TAM.pro-infl. | Ttc39c |
| Slc7a81 | 7.70E-255 | 0.41610631 | 0.635 | 0.489 | 1.49E-250 | TAM.pro-infl. | Slc7a8 |
| Hspa4l | 2.84E-254 | 0.76594561 | 0.35 | 0.215 | 5.50E-250 | TAM.pro-infl. | Hspa4l |
| Slc37a21 | 6.38E-253 | 0.46905642 | 0.516 | 0.355 | 1.23E-248 | TAM.pro-infl. | Slc37a2 |
| Sel1l | 7.53E-253 | 0.32136848 | 0.624 | 0.49 | 1.46E-248 | TAM.pro-infl. | Sel1l |
| Abracl | 1.06E-251 | 0.32723269 | 0.625 | 0.488 | 2.06E-247 | TAM.pro-infl. | Abracl |
| Agpat4 | 1.08E-251 | 0.31039903 | 0.537 | 0.377 | 2.10E-247 | TAM.pro-infl. | Agpat4 |
| Capn1 | 9.62E-251 | 0.41585757 | 0.529 | 0.371 | 1.86E-246 | TAM.pro-infl. | Capn1 |
| Cyth31 | 1.44E-250 | 0.54483488 | 0.431 | 0.293 | 2.78E-246 | TAM.pro-infl. | Cyth3 |
| Lin54 | 1.59E-250 | 0.49955803 | 0.414 | 0.264 | 3.07E-246 | TAM.pro-infl. | Lin54 |
| Mroh11 | 1.98E-250 | 0.40105926 | 0.575 | 0.42 | 3.83E-246 | TAM.pro-infl. | Mroh1 |
| Ezr | 2.61E-249 | 0.32103994 | 0.591 | 0.445 | 5.05E-245 | TAM.pro-infl. | Ezr |
| Pbxip1 | 8.95E-247 | 0.50276103 | 0.385 | 0.232 | 1.73E-242 | TAM.pro-infl. | Pbxip1 |
| Clip2 | 1.54E-246 | 0.48464713 | 0.516 | 0.35 | 2.97E-242 | TAM.pro-infl. | Clip2 |
| Flvcr1 | 1.14E-245 | 0.42620764 | 0.402 | 0.264 | 2.20E-241 | TAM.pro-infl. | Flvcr1 |
| Pitpnm1 | 1.54E-245 | 0.32271578 | 0.55 | 0.394 | 2.99E-241 | TAM.pro-infl. | Pitpnm1 |
| B3gnt7 | 2.03E-245 | 0.53324651 | 0.354 | 0.239 | 3.93E-241 | TAM.pro-infl. | B3gnt7 |
| Klf7 | 9.99E-243 | 0.50138797 | 0.469 | 0.321 | 1.93E-238 | TAM.pro-infl. | Klf7 |
| Vps181 | 3.79E-239 | 0.33570136 | 0.601 | 0.455 | 7.32E-235 | TAM.pro-infl. | Vps18 |
| B3gnt5 | 1.38E-234 | 0.5280454 | 0.277 | 0.134 | 2.67E-230 | TAM.pro-infl. | B3gnt5 |
| Lrp121 | 3.43E-234 | 0.38282388 | 0.553 | 0.42 | 6.64E-230 | TAM.pro-infl. | Lrp12 |
| Eprs | 1.29E-231 | 0.27116528 | 0.477 | 0.381 | 2.48E-227 | TAM.pro-infl. | Eprs |
| Abhd5 | 2.95E-230 | 0.66065956 | 0.342 | 0.192 | 5.71E-226 | TAM.pro-infl. | Abhd5 |
| Ikbke | 1.81E-228 | 0.49060441 | 0.554 | 0.393 | 3.50E-224 | TAM.pro-infl. | Ikbke |
| Dop1b | 4.34E-227 | 0.65887984 | 0.326 | 0.178 | 8.38E-223 | TAM.pro-infl. | Dop1b |
| Rmc1 | 5.29E-227 | 0.58894443 | 0.401 | 0.249 | 1.02E-222 | TAM.pro-infl. | Rmc1 |
| Dock5 | 1.70E-226 | 0.54421279 | 0.352 | 0.183 | 3.30E-222 | TAM.pro-infl. | Dock5 |
| Fam102b | 1.01E-225 | 0.40534069 | 0.461 | 0.345 | 1.96E-221 | TAM.pro-infl. | Fam102b |
| Snx12 | 1.42E-222 | 0.3619888 | 0.583 | 0.434 | 2.75E-218 | TAM.pro-infl. | Snx12 |
| Zcchc2 | 6.97E-222 | 0.31778364 | 0.475 | 0.338 | 1.35E-217 | TAM.pro-infl. | Zcchc2 |
| Txnl1 | 8.40E-222 | 0.3954737 | 0.561 | 0.415 | 1.62E-217 | TAM.pro-infl. | Txnl1 |
| Rapgef1 | 8.97E-222 | 0.34432815 | 0.536 | 0.394 | 1.73E-217 | TAM.pro-infl. | Rapgef1 |
| Tbc1d23 | 4.16E-221 | 0.77426553 | 0.311 | 0.169 | 8.05E-217 | TAM.pro-infl. | Tbc1d23 |
| Sgms1 | 1.93E-220 | 0.5723678 | 0.37 | 0.217 | 3.73E-216 | TAM.pro-infl. | Sgms1 |
| Spata13 | 2.27E-220 | 0.40810442 | 0.411 | 0.273 | 4.39E-216 | TAM.pro-infl. | Spata13 |
| Ckap4 | 4.44E-220 | 0.47487957 | 0.423 | 0.299 | 8.59E-216 | TAM.pro-infl. | Ckap4 |
| Comtd1 | 7.87E-220 | 0.83827755 | 0.324 | 0.173 | 1.52E-215 | TAM.pro-infl. | Comtd1 |
| Mrtfb | 2.77E-219 | 0.71509854 | 0.306 | 0.175 | 5.36E-215 | TAM.pro-infl. | Mrtfb |
| Mob3b | 1.09E-218 | 0.52008019 | 0.331 | 0.194 | 2.11E-214 | TAM.pro-infl. | Mob3b |
| Pla2g4a | 1.43E-218 | 0.39722556 | 0.505 | 0.367 | 2.76E-214 | TAM.pro-infl. | Pla2g4a |
| Atp6v1d | 5.48E-218 | 0.33094579 | 0.688 | 0.546 | 1.06E-213 | TAM.pro-infl. | Atp6v1d |
| Spryd7 | 1.36E-217 | 0.88421329 | 0.252 | 0.132 | 2.63E-213 | TAM.pro-infl. | Spryd7 |
| Dnase1l1 | 2.10E-217 | 0.28065541 | 0.563 | 0.429 | 4.06E-213 | TAM.pro-infl. | Dnase1l1 |
| Rbm47 | 1.44E-215 | 0.25378637 | 0.641 | 0.508 | 2.78E-211 | TAM.pro-infl. | Rbm47 |
| Slc1a5 | 2.82E-215 | 0.41400464 | 0.548 | 0.365 | 5.45E-211 | TAM.pro-infl. | Slc1a5 |
| Lpcat3 | 1.42E-214 | 0.54457709 | 0.426 | 0.275 | 2.75E-210 | TAM.pro-infl. | Lpcat3 |
| Dot1l | 1.45E-214 | 0.38786364 | 0.589 | 0.416 | 2.80E-210 | TAM.pro-infl. | Dot1l |
| Tpcn21 | 1.70E-214 | 0.46294227 | 0.473 | 0.339 | 3.29E-210 | TAM.pro-infl. | Tpcn2 |
| Nfe2l1 | 7.07E-214 | 0.25132801 | 0.65 | 0.523 | 1.37E-209 | TAM.pro-infl. | Nfe2l1 |
| Znrf1 | 5.42E-213 | 0.43977881 | 0.468 | 0.313 | 1.05E-208 | TAM.pro-infl. | Znrf1 |
| Gramd1b | 1.66E-212 | 0.61773225 | 0.35 | 0.224 | 3.21E-208 | TAM.pro-infl. | Gramd1b |
| Josd1 | 2.09E-211 | 0.75675498 | 0.336 | 0.195 | 4.04E-207 | TAM.pro-infl. | Josd1 |
| Ddhd11 | 1.07E-210 | 0.28835 | 0.614 | 0.49 | 2.07E-206 | TAM.pro-infl. | Ddhd1 |
| DEPRECATED-I | 2.24E-208 | 0.38925715 | 0.468 | 0.347 | 4.33E-204 | TAM.pro-infl. | DEPRECATED-ENSMUSG00000025025 |
| Fbxo22 | 7.47E-208 | 0.65397812 | 0.334 | 0.206 | 1.44E-203 | TAM.pro-infl. | Fbxo22 |
| Pef1 | 3.17E-207 | 0.6007024 | 0.351 | 0.21 | 6.12E-203 | TAM.pro-infl. | Pef1 |
| Cdc42se21 | 6.86E-207 | 0.27115536 | 0.583 | 0.461 | 1.33E-202 | TAM.pro-infl. | Cdc42se2 |
| Uhrf2 | 2.17E-206 | 0.44460389 | 0.469 | 0.358 | 4.20E-202 | TAM.pro-infl. | Uhrf2 |
| Tecpr1 | 4.95E-206 | 0.75869745 | 0.276 | 0.15 | 9.56E-202 | TAM.pro-infl. | Tecpr1 |
| Nedd9 | 1.68E-205 | 0.32839466 | 0.401 | 0.275 | 3.25E-201 | TAM.pro-infl. | Nedd9 |
| Fmn11 | 3.02E-205 | 0.59954864 | 0.398 | 0.253 | 5.85E-201 | TAM.pro-infl. | Fmn1 |
| Eif2ak4 | 1.12E-204 | 0.644259 | 0.372 | 0.228 | 2.17E-200 | TAM.pro-infl. | Eif2ak4 |
| Chst12 | 1.98E-202 | 0.45638599 | 0.397 | 0.264 | 3.82E-198 | TAM.pro-infl. | Chst12 |
| Acbd3 | 3.17E-202 | 0.37030822 | 0.517 | 0.379 | 6.12E-198 | TAM.pro-infl. | Acbd3 |
| Lyst | 1.83E-200 | 0.58500499 | 0.324 | 0.193 | 3.54E-196 | TAM.pro-infl. | Lyst |
| Cmas | 2.04E-200 | 0.39375326 | 0.464 | 0.323 | 3.94E-196 | TAM.pro-infl. | Cmas |
| Golph3l | 3.67E-200 | 0.60055422 | 0.349 | 0.229 | 7.10E-196 | TAM.pro-infl. | Golph3l |
| Lemd2 | 1.38E-199 | 0.25908688 | 0.601 | 0.473 | 2.68E-195 | TAM.pro-infl. | Lemd2 |
| Pi4k2b | 1.31E-198 | 0.85452085 | 0.25 | 0.121 | 2.54E-194 | TAM.pro-infl. | Pi4k2b |
| Elf6 | 1.53E-197 | 0.42559743 | 0.496 | 0.348 | 2.95E-193 | TAM.pro-infl. | Elf6 |
| Flrt3 | 2.75E-196 | 0.29375449 | 0.353 | 0.234 | 5.32E-192 | TAM.pro-infl. | Flrt3 |
| Gripap1 | 1.07E-195 | 0.41973774 | 0.462 | 0.321 | 2.07E-191 | TAM.pro-infl. | Gripap1 |
| Lrrc58 | 1.92E-195 | 0.59726997 | 0.385 | 0.237 | 3.71E-191 | TAM.pro-infl. | Lrrc58 |
| Ubxn8 | 2.02E-195 | 0.61910861 | 0.354 | 0.205 | 3.90E-191 | TAM.pro-infl. | Ubxn8 |
| Aebp2 | 4.03E-193 | 0.39425538 | 0.513 | 0.373 | 7.80E-189 | TAM.pro-infl. | Aebp2 |
| Spcs3 | 8.93E-193 | 0.37829016 | 0.541 | 0.408 | 1.73E-188 | TAM.pro-infl. | Spcs3 |
| Ercc6 | 3.83E-192 | 0.79726426 | 0.267 | 0.141 | 7.40E-188 | TAM.pro-infl. | Ercc6 |
| Suco | 7.85E-192 | 0.56277557 | 0.368 | 0.234 | 1.52E-187 | TAM.pro-infl. | Suco |
| Vamp4 | 1.09E-191 | 0.3870964 | 0.515 | 0.374 | 2.10E-187 | TAM.pro-infl. | Vamp4 |
| Lyz1 | 1.75E-190 | 0.26308039 | 0.446 | 0.283 | 3.39E-186 | TAM.pro-infl. | Lyz1 |
| Cyc1 | 3.12E-190 | 0.33246696 | 0.594 | 0.443 | 6.03E-186 | TAM.pro-infl. | Cyc1 |
| Denr | 6.47E-188 | 0.53816883 | 0.452 | 0.316 | 1.25E-183 | TAM.pro-infl. | Denr |
| Fzd7 | 8.89E-188 | 0.46658804 | 0.303 | 0.209 | 1.72E-183 | TAM.pro-infl. | Fzd7 |
| Mlxip | 1.89E-187 | 0.3603435 | 0.54 | 0.41 | 3.65E-183 | TAM.pro-infl. | Mlxip |
| Scd2 | 6.94E-186 | 0.6272219 | 0.33 | 0.198 | 1.34E-181 | TAM.pro-infl. | Scd2 |
| Qdpr1 | 1.56E-185 | 0.28996766 | 0.554 | 0.425 | 3.02E-181 | TAM.pro-infl. | Qdpr |
| Slc16a1 | 4.89E-185 | 0.55579258 | 0.302 | 0.187 | 9.45E-181 | TAM.pro-infl. | Slc16a1 |
| Psmd7 | 5.49E-185 | 0.29272241 | 0.599 | 0.458 | 1.06E-180 | TAM.pro-infl. | Psmd7 |
| Cdc34 | 1.70E-184 | 0.31072445 | 0.535 | 0.405 | 3.30E-180 | TAM.pro-infl. | Cdc34 |
| St6galnac6 | 5.82E-184 | 0.49040072 | 0.366 | 0.232 | 1.12E-179 | TAM.pro-infl. | St6galnac6 |
| Spen | 1.94E-183 | 0.31930463 | 0.499 | 0.358 | 3.74E-179 | TAM.pro-infl. | Spen |
| Agfg2 | 1.24E-182 | 0.67257901 | 0.303 | 0.173 | 2.40E-178 | TAM.pro-infl. | Agfg2 |
| Sbno1 | 3.84E-182 | 0.31465609 | 0.517 | 0.379 | 7.41E-178 | TAM.pro-infl. | Sbno1 |
| Slc36a4 | 2.49E-181 | 0.65516056 | 0.342 | 0.203 | 4.81E-177 | TAM.pro-infl. | Slc36a4 |
| Dedd2 | 6.57E-181 | 0.72914634 | 0.255 | 0.14 | 1.27E-176 | TAM.pro-infl. | Dedd2 |
| Acox11 | 1.05E-180 | 0.31047151 | 0.508 | 0.383 | 2.03E-176 | TAM.pro-infl. | Acox1 |
| Gtf2f2 | 1.19E-180 | 0.46237008 | 0.442 | 0.299 | 2.30E-176 | TAM.pro-infl. | Gtf2f2 |
| Grk5 | 2.01E-180 | 0.32210398 | 0.392 | 0.248 | 3.89E-176 | TAM.pro-infl. | Grk5 |
| Nploc4 | 2.91E-178 | 0.31934407 | 0.522 | 0.391 | 5.63E-174 | TAM.pro-infl. | Nploc4 |
| Mical1 | 1.35E-177 | 0.47196991 | 0.396 | 0.253 | 2.61E-173 | TAM.pro-infl. | Mical1 |
| Tapt1 | 1.99E-177 | 0.69004929 | 0.265 | 0.14 | 3.85E-173 | TAM.pro-infl. | Tapt1 |
| Stx6 | 4.76E-177 | 0.30116304 | 0.51 | 0.373 | 9.20E-173 | TAM.pro-infl. | Stx6 |
| Smurf1 | 7.20E-177 | 0.36506115 | 0.445 | 0.307 | 1.39E-172 | TAM.pro-infl. | Smurf1 |
| Gba1 | 8.01E-177 | 0.27431538 | 0.568 | 0.432 | 1.55E-172 | TAM.pro-infl. | Gba |
| Def6 | 1.42E-176 | 0.40575305 | 0.402 | 0.264 | 2.74E-172 | TAM.pro-infl. | Def6 |
| Vdac2 | 1.44E-176 | 0.25920913 | 0.551 | 0.431 | 2.79E-172 | TAM.pro-infl. | Vdac2 |
| Mink11 | 4.79E-175 | 0.26240152 | 0.522 | 0.4 | 9.26E-171 | TAM.pro-infl. | Mink1 |
| Mgat4b | 9.34E-175 | 0.45881706 | 0.435 | 0.294 | 1.81E-170 | TAM.pro-infl. | Mgat4b |
| Gnl1 | 1.51E-174 | 0.3053504 | 0.504 | 0.37 | 2.93E-170 | TAM.pro-infl. | Gnl1 |
| Csnk2b | 4.15E-174 | 0.33592571 | 0.512 | 0.377 | 8.03E-170 | TAM.pro-infl. | Csnk2b |
| Lcor | 3.13E-173 | 0.41780301 | 0.449 | 0.321 | 6.05E-169 | TAM.pro-infl. | Lcor |
| Tbc1d8 | 8.78E-173 | 0.43792037 | 0.411 | 0.266 | 1.70E-168 | TAM.pro-infl. | Tbc1d8 |
| Impa2 | 2.33E-172 | 0.52812576 | 0.332 | 0.206 | 4.51E-168 | TAM.pro-infl. | Impa2 |
| Slamf81 | 7.37E-172 | 0.35169466 | 0.392 | 0.273 | 1.42E-167 | TAM.pro-infl. | Slamf8 |
| Retreg2 | 1.20E-171 | 0.30619045 | 0.534 | 0.397 | 2.31E-167 | TAM.pro-infl. | Retreg2 |
| Ubap1 | 1.27E-171 | 0.44385398 | 0.419 | 0.283 | 2.45E-167 | TAM.pro-infl. | Ubap1 |
| Casp7 | 2.72E-171 | 0.44934482 | 0.369 | 0.249 | 5.25E-167 | TAM.pro-infl. | Casp7 |
| Snx10 | 2.17E-170 | 0.32708595 | 0.461 | 0.325 | 4.20E-166 | TAM.pro-infl. | Snx10 |
| Grpel1 | 2.75E-170 | 0.32399756 | 0.485 | 0.359 | 5.32E-166 | TAM.pro-infl. | Grpel1 |
| Elovl1 | 2.90E-170 | 0.28987768 | 0.493 | 0.362 | 5.61E-166 | TAM.pro-infl. | Elovl1 |
| Creb5 | 5.56E-169 | 0.44436998 | 0.495 | 0.335 | 1.08E-164 | TAM.pro-infl. | Creb5 |
| Egln3 | 1.30E-168 | 0.666046 | 0.317 | 0.186 | 2.52E-164 | TAM.pro-infl. | Egln3 |
| Slc25a28 | 2.57E-168 | 0.26318359 | 0.502 | 0.373 | 4.97E-164 | TAM.pro-infl. | Slc25a28 |
| Nck1 | 3.17E-168 | 0.59863096 | 0.34 | 0.212 | 6.12E-164 | TAM.pro-infl. | Nck1 |
| Clcn4 | 6.20E-168 | 0.29866145 | 0.455 | 0.356 | 1.20E-163 | TAM.pro-infl. | Clcn4 |
| Vps29 | 1.83E-166 | 0.25065968 | 0.604 | 0.482 | 3.53E-162 | TAM.pro-infl. | Vps29 |
| Socs1 | 2.42E-166 | 0.2993331 | 0.304 | 0.211 | 4.68E-162 | TAM.pro-infl. | Socs1 |
| Slc37a3 | 5.79E-166 | 0.57052933 | 0.273 | 0.153 | 1.12E-161 | TAM.pro-infl. | Slc37a3 |
| Rnf24 | 1.33E-165 | 0.72124981 | 0.253 | 0.128 | 2.58E-161 | TAM.pro-infl. | Rnf24 |
| Zbtb1 | 2.15E-165 | 0.29695309 | 0.451 | 0.337 | 4.16E-161 | TAM.pro-infl. | Zbtb1 |
| Pira2 | 1.63E-164 | 0.34537201 | 0.325 | 0.204 | 3.16E-160 | TAM.pro-infl. | Pira2 |
| Fam214b | 5.92E-164 | 0.40009257 | 0.44 | 0.308 | 1.14E-159 | TAM.pro-infl. | Fam214b |
| Luzp1 | 8.10E-164 | 0.50364782 | 0.406 | 0.271 | 1.57E-159 | TAM.pro-infl. | Luzp1 |
| Ccdc50 | 1.31E-163 | 0.31009257 | 0.53 | 0.413 | 2.53E-159 | TAM.pro-infl. | Ccdc50 |
| Plcb21 | 1.92E-163 | 0.38763599 | 0.412 | 0.298 | 3.70E-159 | TAM.pro-infl. | Plcb2 |
| Pilrb1 | 1.98E-163 | 0.29415159 | 0.429 | 0.299 | 3.82E-159 | TAM.pro-infl. | Pilrb1 |
| Hgsnat | 2.65E-162 | 0.3078235 | 0.377 | 0.274 | 5.13E-158 | TAM.pro-infl. | Hgsnat |
| Dtx31 | 2.84E-162 | 0.56354713 | 0.33 | 0.2 | 5.49E-158 | TAM.pro-infl. | Dtx3 |
| Zfand2a | 3.78E-162 | 0.27211686 | 0.606 | 0.434 | 7.30E-158 | TAM.pro-infl. | Zfand2a |
| Abtb2 | 7.79E-162 | 0.3695252 | 0.377 | 0.242 | 1.51E-157 | TAM.pro-infl. | Abtb2 |
| Naa20 | 3.05E-161 | 0.34790457 | 0.452 | 0.323 | 5.90E-157 | TAM.pro-infl. | Naa20 |
| Clip1 | 2.50E-160 | 0.33150305 | 0.44 | 0.313 | 4.83E-156 | TAM.pro-infl. | Clip1 |
| Gpr1571 | 3.25E-159 | 0.74692087 | 0.276 | 0.153 | 6.28E-155 | TAM.pro-infl. | Gpr157 |
| Slc27a4 | 1.36E-158 | 0.56557723 | 0.35 | 0.226 | 2.62E-154 | TAM.pro-infl. | Slc27a4 |
| Xylt1 | 5.72E-158 | 0.29811019 | 0.363 | 0.246 | 1.11E-153 | TAM.pro-infl. | Xylt1 |
| Agpat2 | 1.39E-157 | 0.64075335 | 0.252 | 0.136 | 2.68E-153 | TAM.pro-infl. | Agpat2 |
| Amfr | 1.44E-157 | 0.25564161 | 0.546 | 0.417 | 2.78E-153 | TAM.pro-infl. | Amfr |
| Ccdc88a | 2.21E-156 | 0.26453081 | 0.466 | 0.359 | 4.27E-152 | TAM.pro-infl. | Ccdc88a |
| Stk10 | 3.48E-156 | 0.25067894 | 0.471 | 0.333 | 6.73E-152 | TAM.pro-infl. | Stk10 |
| Gmfb | 5.82E-156 | 0.25471183 | 0.634 | 0.515 | 1.13E-151 | TAM.pro-infl. | Gmfb |
| Mfsd5 | 1.42E-155 | 0.26900107 | 0.502 | 0.376 | 2.74E-151 | TAM.pro-infl. | Mfsd5 |
| Alg9 | 2.46E-155 | 0.55247638 | 0.335 | 0.211 | 4.76E-151 | TAM.pro-infl. | Alg9 |
| Plin3 | 1.39E-154 | 0.27324327 | 0.498 | 0.382 | 2.69E-150 | TAM.pro-infl. | Plin3 |
| Ciapin1 | 5.62E-154 | 0.66250812 | 0.321 | 0.194 | 1.09E-149 | TAM.pro-infl. | Ciapin1 |
| Ccl24 | 3.93E-153 | 0.54561402 | 0.273 | 0.226 | 7.61E-149 | TAM.pro-infl. | Ccl24 |
| Ifi271 | 3.30E-152 | 0.28797454 | 0.501 | 0.378 | 6.37E-148 | TAM.pro-infl. | Ifi27 |
| Phf13 | 3.55E-152 | 0.59191129 | 0.305 | 0.185 | 6.87E-148 | TAM.pro-infl. | Phf13 |
| Parp10 | 4.74E-152 | 0.53067317 | 0.293 | 0.178 | 9.17E-148 | TAM.pro-infl. | Parp10 |
| Pde8a | 7.17E-152 | 0.45807658 | 0.398 | 0.267 | 1.39E-147 | TAM.pro-infl. | Pde8a |
| Ell | 1.00E-151 | 0.46379842 | 0.41 | 0.277 | 1.93E-147 | TAM.pro-infl. | Ell |
| Atp9b | 1.06E-151 | 0.25227152 | 0.512 | 0.396 | 2.06E-147 | TAM.pro-infl. | Atp9b |
| Ubash3b | 2.13E-151 | 0.33186322 | 0.452 | 0.334 | 4.12E-147 | TAM.pro-infl. | Ubash3b |
| Bnip3l | 7.74E-151 | 0.60279526 | 0.267 | 0.158 | 1.50E-146 | TAM.pro-infl. | Bnip3l |
| Fmnl3 | 2.21E-150 | 0.28083877 | 0.515 | 0.391 | 4.28E-146 | TAM.pro-infl. | Fmnl3 |
| Zbtb11 | 3.17E-149 | 0.34617173 | 0.461 | 0.34 | 6.14E-145 | TAM.pro-infl. | Zbtb11 |
| Gk | 5.37E-149 | 0.41701532 | 0.346 | 0.224 | 1.04E-144 | TAM.pro-infl. | Gk |
| Psma3 | 4.16E-148 | 0.29723361 | 0.499 | 0.372 | 8.03E-144 | TAM.pro-infl. | Psma3 |
| Psmd13 | 4.37E-148 | 0.25061156 | 0.561 | 0.432 | 8.44E-144 | TAM.pro-infl. | Psmd13 |
| Nsf | 2.21E-147 | 0.26566202 | 0.48 | 0.373 | 4.28E-143 | TAM.pro-infl. | Nsf |
| Arid3a | 2.54E-147 | 0.39330694 | 0.345 | 0.226 | 4.92E-143 | TAM.pro-infl. | Arid3a |
| Sephs2 | 3.10E-145 | 0.26195682 | 0.466 | 0.35 | 6.00E-141 | TAM.pro-infl. | Sephs2 |
| Srd5a3 | 1.56E-144 | 0.44083815 | 0.354 | 0.23 | 3.02E-140 | TAM.pro-infl. | Srd5a3 |
| Ppfibp1 | 2.10E-144 | 0.47532198 | 0.385 | 0.255 | 4.05E-140 | TAM.pro-infl. | Ppfibp1 |
| Pip4k2c | 5.04E-143 | 0.44955325 | 0.375 | 0.251 | 9.75E-139 | TAM.pro-infl. | Pip4k2c |
| Slc31a2 | 5.09E-143 | 0.30678669 | 0.477 | 0.348 | 9.85E-139 | TAM.pro-infl. | Slc31a2 |
| Arl1 | 6.24E-143 | 0.33446494 | 0.392 | 0.286 | 1.21E-138 | TAM.pro-infl. | Arl1 |
| Psmd10 | 3.37E-142 | 0.56319082 | 0.336 | 0.221 | 6.51E-138 | TAM.pro-infl. | Psmd10 |
| Pofut2 | 6.30E-142 | 0.34107558 | 0.405 | 0.278 | 1.22E-137 | TAM.pro-infl. | Pofut2 |
| Mkrn1 | 6.73E-142 | 0.32548468 | 0.425 | 0.304 | 1.30E-137 | TAM.pro-infl. | Mkrn1 |
| G6pc3 | 3.13E-141 | 0.52560626 | 0.346 | 0.223 | 6.06E-137 | TAM.pro-infl. | G6pc3 |
| Pqlc1 | 3.89E-141 | 0.28598904 | 0.44 | 0.319 | 7.51E-137 | TAM.pro-infl. | Pqlc1 |
| Klf16 | 2.19E-140 | 0.51200128 | 0.287 | 0.168 | 4.24E-136 | TAM.pro-infl. | Klf16 |
| Agtrap | 4.13E-138 | 0.50730733 | 0.318 | 0.195 | 7.98E-134 | TAM.pro-infl. | Agtrap |
| Ywhaz | 5.10E-138 | 0.36265825 | 0.399 | 0.273 | 9.86E-134 | TAM.pro-infl. | Ywhaz |
| Fam104a | 9.69E-137 | 0.41801372 | 0.376 | 0.261 | 1.87E-132 | TAM.pro-infl. | Fam104a |
| Pi4ka | 3.51E-136 | 0.28904445 | 0.469 | 0.358 | 6.78E-132 | TAM.pro-infl. | Pi4ka |
| Vps37c | 1.10E-135 | 0.54480968 | 0.329 | 0.214 | 2.12E-131 | TAM.pro-infl. | Vps37c |
| Slc39a11 | 1.59E-135 | 0.2742922 | 0.42 | 0.303 | 3.06E-131 | TAM.pro-infl. | Slc39a11 |
| Timm8b | 4.05E-135 | 0.36648203 | 0.43 | 0.311 | 7.83E-131 | TAM.pro-infl. | Timm8b |
| Cystm1 | 9.43E-135 | 0.38088894 | 0.292 | 0.191 | 1.82E-130 | TAM.pro-infl. | Cystm1 |
| Zfp608 | 7.26E-134 | 0.36461615 | 0.31 | 0.192 | 1.40E-129 | TAM.pro-infl. | Zfp608 |
| Plekhm3 | 1.93E-133 | 0.41236513 | 0.371 | 0.247 | 3.73E-129 | TAM.pro-infl. | Plekhm3 |
| Gzf1 | 6.66E-133 | 0.68401145 | 0.267 | 0.157 | 1.29E-128 | TAM.pro-infl. | Gzf1 |
| Mtm1 | 7.89E-133 | 0.49774644 | 0.274 | 0.165 | 1.53E-128 | TAM.pro-infl. | Mtm1 |
| Tjp2 | 1.80E-131 | 0.60449243 | 0.273 | 0.166 | 3.48E-127 | TAM.pro-infl. | Tjp2 |
| Pilrb21 | 9.73E-131 | 0.26215455 | 0.411 | 0.289 | 1.88E-126 | TAM.pro-infl. | Pilrb2 |
| Atn1 | 2.20E-130 | 0.29551004 | 0.468 | 0.35 | 4.26E-126 | TAM.pro-infl. | Atn1 |
| Rnf41 | 2.47E-130 | 0.39275645 | 0.357 | 0.246 | 4.78E-126 | TAM.pro-infl. | Rnf41 |
| Tubb4b | 6.49E-130 | 0.28292352 | 0.556 | 0.406 | 1.26E-125 | TAM.pro-infl. | Tubb4b |
| Ripk3 | 6.52E-130 | 0.30579992 | 0.419 | 0.311 | 1.26E-125 | TAM.pro-infl. | Ripk3 |
| Fbxo30 | 1.48E-129 | 0.36598274 | 0.365 | 0.252 | 2.86E-125 | TAM.pro-infl. | Fbxo30 |
| Tmc6 | 3.69E-129 | 0.38987339 | 0.286 | 0.178 | 7.14E-125 | TAM.pro-infl. | Tmc6 |
| Gpaa1 | 6.37E-129 | 0.46532363 | 0.356 | 0.237 | 1.23E-124 | TAM.pro-infl. | Gpaa1 |
| Ocstamp1 | 3.31E-128 | 0.41727816 | 0.3 | 0.202 | 6.40E-124 | TAM.pro-infl. | Ocstamp |
| Golt1b | 8.48E-128 | 0.44775078 | 0.41 | 0.286 | 1.64E-123 | TAM.pro-infl. | Golt1b |
| Galnt71 | 5.59E-127 | 0.44839011 | 0.363 | 0.244 | 1.08E-122 | TAM.pro-infl. | Galnt7 |
| Zfc3h1 | 8.52E-127 | 0.33891671 | 0.382 | 0.265 | 1.65E-122 | TAM.pro-infl. | Zfc3h1 |
| Abhd41 | 1.26E-126 | 0.31062762 | 0.383 | 0.27 | 2.44E-122 | TAM.pro-infl. | Abhd4 |
| Hdgf | 3.32E-126 | 0.29098642 | 0.531 | 0.397 | 6.41E-122 | TAM.pro-infl. | Hdgf |
| Cdk17 | 3.87E-126 | 0.28236048 | 0.397 | 0.279 | 7.49E-122 | TAM.pro-infl. | Cdk17 |
| Bcl2l2 | 1.76E-124 | 0.4983626 | 0.346 | 0.226 | 3.40E-120 | TAM.pro-infl. | Bcl2l2 |
| Nln | 7.95E-124 | 0.42359332 | 0.335 | 0.226 | 1.54E-119 | TAM.pro-infl. | Nln |
| Slc12a4 | 3.43E-123 | 0.34896929 | 0.328 | 0.219 | 6.64E-119 | TAM.pro-infl. | Slc12a4 |
| Pcmt1 | 1.11E-122 | 0.33484869 | 0.425 | 0.309 | 2.15E-118 | TAM.pro-infl. | Pcmt1 |
| Cyb5r11 | 1.15E-121 | 0.39658316 | 0.369 | 0.254 | 2.23E-117 | TAM.pro-infl. | Cyb5r1 |
| Pdgfb1 | 1.87E-121 | 0.37276105 | 0.342 | 0.237 | 3.62E-117 | TAM.pro-infl. | Pdgfb |
| Sap30l | 2.74E-121 | 0.35795004 | 0.365 | 0.261 | 5.30E-117 | TAM.pro-infl. | Sap30l |
| Trip10 | 4.99E-121 | 0.39588566 | 0.356 | 0.242 | 9.65E-117 | TAM.pro-infl. | Trip10 |
| Scamp3 | 1.39E-120 | 0.33556111 | 0.43 | 0.315 | 2.68E-116 | TAM.pro-infl. | Scamp3 |
| Arfgap3 | 1.63E-120 | 0.36643412 | 0.35 | 0.234 | 3.15E-116 | TAM.pro-infl. | Arfgap3 |
| Eef1aknmt | 3.57E-120 | 0.43966366 | 0.253 | 0.16 | 6.90E-116 | TAM.pro-infl. | Eef1aknmt |
| Galns1 | 7.78E-120 | 0.34258492 | 0.338 | 0.233 | 1.50E-115 | TAM.pro-infl. | Galns |
| Nudt4 | 1.06E-119 | 0.4375601 | 0.323 | 0.231 | 2.04E-115 | TAM.pro-infl. | Nudt4 |
| Wdfy1 | 2.94E-119 | 0.63946275 | 0.28 | 0.169 | 5.69E-115 | TAM.pro-infl. | Wdfy1 |
| Derl2 | 4.08E-119 | 0.28549257 | 0.46 | 0.341 | 7.89E-115 | TAM.pro-infl. | Derl2 |
| Arid5b | 7.43E-118 | 0.30394319 | 0.396 | 0.285 | 1.44E-113 | TAM.pro-infl. | Arid5b |
| Chd1 | 8.90E-118 | 0.25778823 | 0.5 | 0.394 | 1.72E-113 | TAM.pro-infl. | Chd1 |
| Ndufc1 | 1.13E-117 | 0.27062566 | 0.49 | 0.372 | 2.18E-113 | TAM.pro-infl. | Ndufc1 |
| Gan | 1.45E-117 | 0.43844818 | 0.268 | 0.179 | 2.81E-113 | TAM.pro-infl. | Gan |
| Aifm2 | 3.24E-117 | 0.45184701 | 0.274 | 0.166 | 6.27E-113 | TAM.pro-infl. | Aifm2 |
| Smg5 | 4.01E-117 | 0.37212806 | 0.443 | 0.313 | 7.76E-113 | TAM.pro-infl. | Smg5 |
| Hmg20b | 6.32E-117 | 0.26501811 | 0.452 | 0.333 | 1.22E-112 | TAM.pro-infl. | Hmg20b |
| Morf4l2 | 6.87E-116 | 0.25546722 | 0.46 | 0.346 | 1.33E-111 | TAM.pro-infl. | Morf4l2 |
| Acvr1b | 9.89E-116 | 0.42742518 | 0.31 | 0.197 | 1.91E-111 | TAM.pro-infl. | Acvr1b |
| Rgs3 | 2.42E-115 | 0.3761723 | 0.286 | 0.187 | 4.68E-111 | TAM.pro-infl. | Rgs3 |
| Trrap | 3.23E-115 | 0.30425259 | 0.411 | 0.308 | 6.25E-111 | TAM.pro-infl. | Trrap |
| Atg101 | 7.47E-115 | 0.2529931 | 0.425 | 0.311 | 1.44E-110 | TAM.pro-infl. | Atg101 |
| Homer3 | 1.79E-114 | 0.4907723 | 0.291 | 0.193 | 3.45E-110 | TAM.pro-infl. | Homer3 |
| Dpp71 | 2.47E-112 | 0.43315537 | 0.382 | 0.26 | 4.78E-108 | TAM.pro-infl. | Dpp7 |
| Senp2 | 2.77E-112 | 0.32254992 | 0.378 | 0.266 | 5.35E-108 | TAM.pro-infl. | Senp2 |
| Gas2l3 | 9.59E-112 | 0.37645641 | 0.392 | 0.268 | 1.85E-107 | TAM.pro-infl. | Gas2l3 |
| Golm1 | 7.80E-111 | 0.29873826 | 0.315 | 0.207 | 1.51E-106 | TAM.pro-infl. | Golm1 |
| Fbxo33 | 1.82E-110 | 0.31863183 | 0.331 | 0.234 | 3.51E-106 | TAM.pro-infl. | Fbxo33 |
| Zfp7041 | 2.97E-110 | 0.52198919 | 0.276 | 0.195 | 5.75E-106 | TAM.pro-infl. | Zfp704 |
| Pip5k1a | 3.98E-110 | 0.48717122 | 0.286 | 0.184 | 7.70E-106 | TAM.pro-infl. | Pip5k1a |
| Pdgfa1 | 2.20E-109 | 0.32786489 | 0.324 | 0.242 | 4.25E-105 | TAM.pro-infl. | Pdgfa |
| Entpd7 | 7.53E-108 | 0.25746581 | 0.432 | 0.319 | 1.46E-103 | TAM.pro-infl. | Entpd7 |
| Uso1 | 9.31E-108 | 0.28655934 | 0.441 | 0.323 | 1.80E-103 | TAM.pro-infl. | Uso1 |
| Slc25a33 | 7.39E-107 | 0.29611286 | 0.33 | 0.229 | 1.43E-102 | TAM.pro-infl. | Slc25a33 |
| Atp5g3 | 1.05E-105 | 0.2926091 | 0.461 | 0.349 | 2.04E-101 | TAM.pro-infl. | Atp5g3 |
| Tbrg1 | 1.27E-105 | 0.40452284 | 0.373 | 0.255 | 2.46E-101 | TAM.pro-infl. | Tbrg1 |
| Afg3l1 | 1.65E-104 | 0.28797682 | 0.448 | 0.329 | 3.20E-100 | TAM.pro-infl. | Afg3l1 |
| Ttyh21 | 2.17E-104 | 0.35760186 | 0.383 | 0.276 | 4.20E-100 | TAM.pro-infl. | Ttyh2 |
| Ipo131 | 4.28E-104 | 0.33957029 | 0.37 | 0.263 | 8.27E-100 | TAM.pro-infl. | Ipo13 |
| Slc25a24 | 2.03E-103 | 0.28855463 | 0.315 | 0.206 | 3.93E-99 | TAM.pro-infl. | Slc25a24 |
| Tm2d3 | 3.72E-103 | 0.46107238 | 0.285 | 0.187 | 7.20E-99 | TAM.pro-infl. | Tm2d3 |
| Ipmk | 1.27E-102 | 0.27020615 | 0.387 | 0.294 | 2.46E-98 | TAM.pro-infl. | Ipmk |
| Zpr1 | 1.48E-102 | 0.34215276 | 0.33 | 0.233 | 2.87E-98 | TAM.pro-infl. | Zpr1 |
| Zmpste24 | 2.79E-102 | 0.2876761 | 0.457 | 0.346 | 5.40E-98 | TAM.pro-infl. | Zmpste24 |
| Lsm14b | 4.28E-102 | 0.31286088 | 0.381 | 0.283 | 8.28E-98 | TAM.pro-infl. | Lsm14b |
| Stard51 | 3.07E-100 | 0.29291022 | 0.308 | 0.213 | 5.93E-96 | TAM.pro-infl. | Stard5 |
| Tmem135 | 2.34E-99 | 0.27629414 | 0.421 | 0.315 | 4.53E-95 | TAM.pro-infl. | Tmem135 |
| Pell2 | 7.31E-99 | 0.37864402 | 0.264 | 0.167 | 1.41E-94 | TAM.pro-infl. | Pell2 |
| Cers5 | 2.88E-98 | 0.27446156 | 0.427 | 0.315 | 5.56E-94 | TAM.pro-infl. | Cers5 |
| Zbtb21 | 4.44E-98 | 0.36660365 | 0.347 | 0.24 | 8.58E-94 | TAM.pro-infl. | Zbtb21 |
| Fbxo28 | 1.95E-97 | 0.28320441 | 0.363 | 0.261 | 3.77E-93 | TAM.pro-infl. | Fbxo28 |
| Tusc3 | 9.73E-97 | 0.38440448 | 0.296 | 0.197 | 1.88E-92 | TAM.pro-infl. | Tusc3 |
| Trim3 | 3.03E-96 | 0.29115439 | 0.297 | 0.204 | 5.86E-92 | TAM.pro-infl. | Trim3 |
| Guk1 | 5.30E-96 | 0.27475938 | 0.344 | 0.247 | 1.02E-91 | TAM.pro-infl. | Guk1 |
| Rbm4 | 3.86E-95 | 0.35105472 | 0.42 | 0.314 | 7.47E-91 | TAM.pro-infl. | Rbm4 |
| Dcaf13 | 4.34E-95 | 0.303798 | 0.387 | 0.281 | 8.39E-91 | TAM.pro-infl. | Dcaf13 |
| Ubxn6 | 6.42E-95 | 0.29311223 | 0.363 | 0.259 | 1.24E-90 | TAM.pro-infl. | Ubxn6 |
| Qsox1 | 6.76E-91 | 0.3345197 | 0.327 | 0.239 | 1.31E-86 | TAM.pro-infl. | Qsox1 |
| Uhrf1bp1l | 2.42E-90 | 0.34282697 | 0.333 | 0.228 | 4.68E-86 | TAM.pro-infl. | Uhrf1bp1l |
| Ppig | 1.58E-88 | 0.25718095 | 0.438 | 0.33 | 3.05E-84 | TAM.pro-infl. | Ppig |
| Tom1 | 3.18E-88 | 0.30363052 | 0.359 | 0.26 | 6.15E-84 | TAM.pro-infl. | Tom1 |
| Arhgap10 | 3.38E-88 | 0.46790685 | 0.29 | 0.185 | 6.53E-84 | TAM.pro-infl. | Arhgap10 |
| Washc51 | 1.84E-86 | 0.30496902 | 0.362 | 0.264 | 3.55E-82 | TAM.pro-infl. | Washc5 |
| Ptpn23 | 2.02E-86 | 0.26695112 | 0.437 | 0.332 | 3.90E-82 | TAM.pro-infl. | Ptpn23 |
| Timm23 | 9.98E-86 | 0.33358799 | 0.369 | 0.267 | 1.93E-81 | TAM.pro-infl. | Timm23 |
| Nagk1 | 2.07E-85 | 0.2737108 | 0.376 | 0.264 | 3.99E-81 | TAM.pro-infl. | Nagk |
| Fdft11 | 9.97E-85 | 0.28780192 | 0.281 | 0.199 | 1.93E-80 | TAM.pro-infl. | Fdft1 |
| Rad54l2 | 1.24E-83 | 0.44008262 | 0.273 | 0.182 | 2.40E-79 | TAM.pro-infl. | Rad54l2 |
| Yrdc | 2.41E-81 | 0.33984727 | 0.326 | 0.224 | 4.66E-77 | TAM.pro-infl. | Yrdc |
| Topors1 | 5.93E-80 | 0.28491668 | 0.337 | 0.248 | 1.15E-75 | TAM.pro-infl. | Topors |
| Pygo2 | 1.87E-79 | 0.31934785 | 0.307 | 0.21 | 3.62E-75 | TAM.pro-infl. | Pygo2 |
| Rnft1 | 3.07E-79 | 0.40040742 | 0.32 | 0.226 | 5.93E-75 | TAM.pro-infl. | Rnft1 |
| Rad9a | 7.71E-79 | 0.3339534 | 0.275 | 0.184 | 1.49E-74 | TAM.pro-infl. | Rad9a |
| Rnmt | 6.28E-78 | 0.37587108 | 0.273 | 0.184 | 1.21E-73 | TAM.pro-infl. | Rnmt |
| Tmco3 | 6.53E-78 | 0.45442778 | 0.251 | 0.171 | 1.26E-73 | TAM.pro-infl. | Tmco3 |
| Prkaa1 | 6.85E-78 | 0.28369232 | 0.323 | 0.234 | 1.32E-73 | TAM.pro-infl. | Prkaa1 |
| Emc4 | 9.23E-78 | 0.25165546 | 0.339 | 0.254 | 1.78E-73 | TAM.pro-infl. | Emc4 |
| Coq10a | 4.89E-76 | 0.33095201 | 0.264 | 0.184 | 9.46E-72 | TAM.pro-infl. | Coq10a |
| Rnf14 | 1.12E-75 | 0.44354445 | 0.283 | 0.188 | 2.17E-71 | TAM.pro-infl. | Rnf14 |
| Fbxo42 | 2.09E-75 | 0.26643002 | 0.345 | 0.253 | 4.04E-71 | TAM.pro-infl. | Fbxo42 |
| Otud4 | 2.94E-75 | 0.25242467 | 0.357 | 0.267 | 5.68E-71 | TAM.pro-infl. | Otud4 |
| Pbdc1 | 1.29E-74 | 0.40319155 | 0.289 | 0.197 | 2.50E-70 | TAM.pro-infl. | Pbdc1 |
| Fgd6 | 2.58E-74 | 0.31429449 | 0.269 | 0.174 | 4.99E-70 | TAM.pro-infl. | Fgd6 |
| Aldh3a2 | 7.58E-74 | 0.32148441 | 0.271 | 0.181 | 1.47E-69 | TAM.pro-infl. | Aldh3a2 |
| Impdh1 | 7.79E-74 | 0.37384564 | 0.272 | 0.18 | 1.51E-69 | TAM.pro-infl. | Impdh1 |
| Gskip | 1.05E-73 | 0.30716644 | 0.296 | 0.207 | 2.02E-69 | TAM.pro-infl. | Gskip |
| Synj2 | 1.43E-72 | 0.29793315 | 0.284 | 0.196 | 2.77E-68 | TAM.pro-infl. | Synj2 |
| Parp4 | 1.02E-71 | 0.28944201 | 0.302 | 0.21 | 1.98E-67 | TAM.pro-infl. | Parp4 |
| D17H6S53E | 3.26E-71 | 0.33950487 | 0.306 | 0.212 | 6.30E-67 | TAM.pro-infl. | D17H6S53E |
| Thap12 | 3.47E-71 | 0.34766612 | 0.304 | 0.221 | 6.71E-67 | TAM.pro-infl. | Thap12 |
| Rce1 | 2.03E-70 | 0.29709558 | 0.303 | 0.215 | 3.93E-66 | TAM.pro-infl. | Rce1 |
| Gbe1 | 6.37E-69 | 0.44283516 | 0.255 | 0.157 | 1.23E-64 | TAM.pro-infl. | Gbe1 |
| Arl3 | 1.11E-67 | 0.32995488 | 0.292 | 0.211 | 2.14E-63 | TAM.pro-infl. | Arl3 |
| Rnaseh2a | 1.56E-67 | 0.32120382 | 0.304 | 0.217 | 3.02E-63 | TAM.pro-infl. | Rnaseh2a |
| Fam76a | 2.49E-66 | 0.31891765 | 0.358 | 0.263 | 4.81E-62 | TAM.pro-infl. | Fam76a |
| Hmgxb3 | 2.57E-66 | 0.33911566 | 0.262 | 0.179 | 4.97E-62 | TAM.pro-infl. | Hmgxb3 |
| Mfsd10 | 2.78E-66 | 0.27298954 | 0.334 | 0.238 | 5.37E-62 | TAM.pro-infl. | Mfsd10 |
| Mnt | 3.55E-66 | 0.29662743 | 0.288 | 0.202 | 6.86E-62 | TAM.pro-infl. | Mnt |
| Tbcd | 8.15E-66 | 0.2791156 | 0.312 | 0.222 | 1.58E-61 | TAM.pro-infl. | Tbcd |
| Arpp19 | 8.19E-66 | 0.28522633 | 0.294 | 0.205 | 1.58E-61 | TAM.pro-infl. | Arpp19 |
| Zfp326 | 1.37E-65 | 0.29096573 | 0.313 | 0.228 | 2.64E-61 | TAM.pro-infl. | Zfp326 |
| Dhps | 4.38E-65 | 0.2860161 | 0.305 | 0.223 | 8.47E-61 | TAM.pro-infl. | Dhps |
| Gse1 | 7.88E-65 | 0.28800113 | 0.292 | 0.213 | 1.52E-60 | TAM.pro-infl. | Gse1 |
| Creld11 | 7.72E-64 | 0.36504243 | 0.283 | 0.196 | 1.49E-59 | TAM.pro-infl. | Creld1 |
| Gys1 | 4.19E-62 | 0.35096926 | 0.338 | 0.237 | 8.11E-58 | TAM.pro-infl. | Gys1 |
| Nomo1 | 1.27E-61 | 0.31379413 | 0.288 | 0.201 | 2.45E-57 | TAM.pro-infl. | Nomo1 |
| Slc38a71 | 6.21E-61 | 0.29161756 | 0.32 | 0.231 | 1.20E-56 | TAM.pro-infl. | Slc38a7 |
| Nmt2 | 1.05E-60 | 0.34377678 | 0.279 | 0.2 | 2.03E-56 | TAM.pro-infl. | Nmt2 |
| Pptc7 | 2.68E-59 | 0.25791214 | 0.267 | 0.192 | 5.19E-55 | TAM.pro-infl. | Pptc7 |
| Eif1ax | 2.99E-56 | 0.33956279 | 0.252 | 0.183 | 5.79E-52 | TAM.pro-infl. | Eif1ax |
| Tmem63b | 5.45E-54 | 0.28366274 | 0.252 | 0.18 | 1.05E-49 | TAM.pro-infl. | Tmem63b |
| Tspyl21 | 1.42E-53 | 0.31679193 | 0.269 | 0.192 | 2.74E-49 | TAM.pro-infl. | Tspyl2 |
| Fam45a | 2.45E-53 | 0.30641828 | 0.266 | 0.188 | 4.74E-49 | TAM.pro-infl. | Fam45a |
| Gpr137 | 3.25E-53 | 0.29017079 | 0.28 | 0.198 | 6.29E-49 | TAM.pro-infl. | Gpr137 |
| Ubxn2a | 3.59E-52 | 0.28124134 | 0.272 | 0.203 | 6.94E-48 | TAM.pro-infl. | Ubxn2a |
| Snx4 | 1.22E-51 | 0.25325112 | 0.296 | 0.22 | 2.36E-47 | TAM.pro-infl. | Snx4 |
| Ndufs5 | 7.09E-51 | 0.25739605 | 0.3 | 0.22 | 1.37E-46 | TAM.pro-infl. | Ndufs5 |
| Haus8 | 8.89E-49 | 0.25026708 | 0.293 | 0.22 | 1.72E-44 | TAM.pro-infl. | Haus8 |
| Pdcd7 | 5.65E-48 | 0.25916928 | 0.253 | 0.184 | 1.09E-43 | TAM.pro-infl. | Pdcd7 |
| Mast2 | 5.15E-47 | 0.25307791 | 0.295 | 0.213 | 9.95E-43 | TAM.pro-infl. | Mast2 |
| Napg | 1.34E-46 | 0.26277292 | 0.283 | 0.213 | 2.60E-42 | TAM.pro-infl. | Napg |
| Sms | 9.76E-40 | 0.28079427 | 0.257 | 0.18 | 1.89E-35 | TAM.pro-infl. | Sms |
| Il1r2 | 0 | 3.01755413 | 0.842 | 0.26 | 0 | TAM.Il1r2 | Il1r2 |
| Cdh1 | 0 | 2.22437067 | 0.529 | 0.164 | 0 | TAM.Il1r2 | Cdh1 |
| Cytip | 0 | 1.36841923 | 0.87 | 0.531 | 0 | TAM.Il1r2 | Cytip |
| Jaml1 | 0 | 1.25196217 | 0.691 | 0.354 | 0 | TAM.Il1r2 | Jaml |
| Nr4a2 | 0 | 1.48679464 | 0.832 | 0.497 | 0 | TAM.Il1r2 | Nr4a2 |
| Klrb1b | 0 | 2.37271582 | 0.48 | 0.15 | 0 | TAM.Il1r2 | Klrb1b |
| Coro2a | 0 | 1.39218553 | 0.691 | 0.372 | 0 | TAM.Il1r2 | Coro2a |
| Gpr1711 | 0 | 1.46328542 | 0.644 | 0.326 | 0 | TAM.Il1r2 | Gpr171 |
| Nr4a3 | 0 | 1.44959582 | 0.861 | 0.547 | 0 | TAM.Il1r2 | Nr4a3 |
| Spint11 | 0 | 1.02197807 | 0.883 | 0.57 | 0 | TAM.Il1r2 | Spint1 |
| Ccr2 | 0 | 0.74373655 | 0.684 | 0.376 | 0 | TAM.Il1r2 | Ccr2 |
| Ccl17 | 0 | 3.33259649 | 0.379 | 0.073 | 0 | TAM.Il1r2 | Ccl17 |
| Napsa | 0 | 1.32296637 | 0.535 | 0.234 | 0 | TAM.Il1r2 | Napsa |
| H2-DMb1 | 0 | 1.36458638 | 0.766 | 0.465 | 0 | TAM.Il1r2 | H2-DMb1 |
| Plet1 | 0 | 3.11298215 | 0.384 | 0.084 | 0 | TAM.Il1r2 | Plet1 |
| Ciita1 | 0 | 1.12220761 | 0.635 | 0.346 | 0 | TAM.Il1r2 | Ciita |
| Adgrg51 | 0 | 1.9206872 | 0.43 | 0.143 | 0 | TAM.Il1r2 | Adgrg5 |
| Malt11 | 0 | 1.1460855 | 0.926 | 0.649 | 0 | TAM.Il1r2 | Malt1 |
| D16Ertd472e | 0 | 1.70870674 | 0.52 | 0.247 | 0 | TAM.Il1r2 | D16Ertd472e |
| Ramp31 | 0 | 2.16735281 | 0.426 | 0.154 | 0 | TAM.Il1r2 | Ramp3 |
| Mgl21 | 0 | 1.38582725 | 0.624 | 0.353 | 0 | TAM.Il1r2 | Mgl2 |
| Cd801 | 0 | 1.23349124 | 0.829 | 0.559 | 0 | TAM.Il1r2 | Cd80 |
| Nrg1 | 0 | 1.29571571 | 0.611 | 0.342 | 0 | TAM.Il1r2 | Nrg1 |
| Bri3bp1 | 0 | 1.38992239 | 0.521 | 0.263 | 0 | TAM.Il1r2 | Bri3bp |
| Plbd1 | 0 | 1.19786831 | 0.902 | 0.647 | 0 | TAM.Il1r2 | Plbd1 |
| Dennd4a | 0 | 1.13587239 | 0.747 | 0.492 | 0 | TAM.Il1r2 | Dennd4a |
| Vrk1 | 0 | 1.42972602 | 0.571 | 0.317 | 0 | TAM.Il1r2 | Vrk1 |
| Naaa | 0 | 1.24761359 | 0.834 | 0.582 | 0 | TAM.Il1r2 | Naaa |
| Ptgs2 | 0 | 1.08520285 | 0.783 | 0.532 | 0 | TAM.Il1r2 | Ptgs2 |
| Ccnd21 | 0 | 1.66566155 | 0.451 | 0.203 | 0 | TAM.Il1r2 | Ccnd2 |
| Uck21 | 0 | 1.14690901 | 0.934 | 0.689 | 0 | TAM.Il1r2 | Uck2 |
| Rogdi1 | 0 | 1.22220468 | 0.563 | 0.319 | 0 | TAM.Il1r2 | Rogdi |
| Rin3 | 0 | 1.14463333 | 0.7 | 0.46 | 0 | TAM.Il1r2 | Rin3 |
| Ece11 | 0 | 0.95587112 | 0.71 | 0.473 | 0 | TAM.Il1r2 | Ece1 |
| Htra31 | 0 | 1.34262509 | 0.502 | 0.266 | 0 | TAM.Il1r2 | Htra3 |
| Hic11 | 0 | 1.90767855 | 0.385 | 0.15 | 0 | TAM.Il1r2 | Hic1 |
| Cd24a1 | 0 | 1.47723734 | 0.461 | 0.229 | 0 | TAM.Il1r2 | Cd24a |
| Cish | 0 | 1.14099419 | 0.623 | 0.391 | 0 | TAM.Il1r2 | Cish |

| | | | | | | | |
|---|---|---|---|---|---|---|---|
| Lta4h | 0 | 1.41465726 | 0.632 | 0.4 | 0 | TAM.Il1r2 | Lta4h |
| H2-DMa1 | 0 | 1.24221497 | 0.927 | 0.696 | 0 | TAM.Il1r2 | H2-DMa |
| Hs3st3b11 | 0 | 1.31968271 | 0.48 | 0.249 | 0 | TAM.Il1r2 | Hs3st3b1 |
| Crem | 0 | 1.30055628 | 0.85 | 0.62 | 0 | TAM.Il1r2 | Crem |
| Vdr | 0 | 1.21943311 | 0.576 | 0.348 | 0 | TAM.Il1r2 | Vdr |
| Trem1 | 0 | 0.94814053 | 0.544 | 0.32 | 0 | TAM.Il1r2 | Trem1 |
| Rab44 | 0 | 1.03908294 | 0.454 | 0.23 | 0 | TAM.Il1r2 | Rab44 |
| Actn11 | 0 | 1.10172451 | 0.642 | 0.423 | 0 | TAM.Il1r2 | Actn1 |
| Qpct | 0 | 2.60577565 | 0.301 | 0.082 | 0 | TAM.Il1r2 | Qpct |
| Dusp51 | 0 | 1.19172156 | 0.913 | 0.695 | 0 | TAM.Il1r2 | Dusp5 |
| Gng10 | 0 | 1.12723543 | 0.672 | 0.454 | 0 | TAM.Il1r2 | Gng10 |
| Htr7 | 0 | 1.55003497 | 0.365 | 0.147 | 0 | TAM.Il1r2 | Htr7 |
| H2-Eb1 | 0 | 1.3001374 | 0.974 | 0.756 | 0 | TAM.Il1r2 | H2-Eb1 |
| Vegfa1 | 0 | 0.74258656 | 0.825 | 0.607 | 0 | TAM.Il1r2 | Vegfa |
| S100a4 | 0 | 0.47398776 | 0.783 | 0.567 | 0 | TAM.Il1r2 | S100a4 |
| Hhex | 0 | 1.26064265 | 0.623 | 0.407 | 0 | TAM.Il1r2 | Hhex |
| Tes | 0 | 0.79449933 | 0.705 | 0.49 | 0 | TAM.Il1r2 | Tes |
| H2-Aa | 0 | 1.33061732 | 0.974 | 0.759 | 0 | TAM.Il1r2 | H2-Aa |
| Got11 | 0 | 1.1281501 | 0.825 | 0.611 | 0 | TAM.Il1r2 | Got1 |
| Gpr132 | 0 | 0.85798822 | 0.795 | 0.582 | 0 | TAM.Il1r2 | Gpr132 |
| Axl2 | 0 | 0.7841382 | 0.884 | 0.672 | 0 | TAM.Il1r2 | Axl |
| Maff | 0 | 0.87944621 | 0.826 | 0.616 | 0 | TAM.Il1r2 | Maff |
| Grasp | 0 | 1.65753595 | 0.375 | 0.166 | 0 | TAM.Il1r2 | Grasp |
| Mcm4 | 0 | 1.45655374 | 0.389 | 0.182 | 0 | TAM.Il1r2 | Mcm4 |
| Csrnp1 | 0 | 0.85941721 | 0.888 | 0.683 | 0 | TAM.Il1r2 | Csrnp1 |
| Smim3 | 0 | 1.10449922 | 0.524 | 0.321 | 0 | TAM.Il1r2 | Smim3 |
| Rel | 0 | 1.10622008 | 0.868 | 0.668 | 0 | TAM.Il1r2 | Rel |
| Jak2 | 0 | 0.82300498 | 0.688 | 0.488 | 0 | TAM.Il1r2 | Jak2 |
| Nfil31 | 0 | 0.84046265 | 0.89 | 0.692 | 0 | TAM.Il1r2 | Nfil3 |
| Pim31 | 0 | 0.84848552 | 0.73 | 0.532 | 0 | TAM.Il1r2 | Pim3 |
| Slc38a1 | 0 | 0.93367403 | 0.711 | 0.513 | 0 | TAM.Il1r2 | Slc38a1 |
| Antxr2 | 0 | 1.34037335 | 0.775 | 0.578 | 0 | TAM.Il1r2 | Antxr2 |
| Gcnt2 | 0 | 1.14955359 | 0.468 | 0.271 | 0 | TAM.Il1r2 | Gcnt2 |
| Isy1 | 0 | 1.01350151 | 0.707 | 0.51 | 0 | TAM.Il1r2 | Isy1 |
| Mycbp2 | 0 | 0.98771572 | 0.746 | 0.549 | 0 | TAM.Il1r2 | Mycbp2 |
| Fam117b | 0 | 1.240897 | 0.773 | 0.578 | 0 | TAM.Il1r2 | Fam117b |
| Runx3 | 0 | 0.80359887 | 0.899 | 0.704 | 0 | TAM.Il1r2 | Runx3 |
| Ifitm1 | 0 | 1.6579221 | 0.421 | 0.226 | 0 | TAM.Il1r2 | Ifitm1 |
| Jag2 | 0 | 1.70666273 | 0.322 | 0.129 | 0 | TAM.Il1r2 | Jag2 |
| Mdh2 | 0 | 1.41173332 | 0.585 | 0.394 | 0 | TAM.Il1r2 | Mdh2 |
| Kmo | 0 | 2.02696891 | 0.266 | 0.076 | 0 | TAM.Il1r2 | Kmo |
| S1pr3 | 0 | 2.09522044 | 0.265 | 0.075 | 0 | TAM.Il1r2 | S1pr3 |
| Enah | 0 | 1.94995077 | 0.277 | 0.09 | 0 | TAM.Il1r2 | Enah |
| Pde4b | 0 | 0.80789932 | 0.816 | 0.634 | 0 | TAM.Il1r2 | Pde4b |
| Spty2d1 | 0 | 1.19823232 | 0.74 | 0.56 | 0 | TAM.Il1r2 | Spty2d1 |
| Irf4 | 0 | 1.82884838 | 0.283 | 0.114 | 0 | TAM.Il1r2 | Irf4 |
| Crip1 | 0 | 0.57815536 | 0.906 | 0.741 | 0 | TAM.Il1r2 | Crip1 |
| Pxdc11 | 0 | 1.08226501 | 0.557 | 0.392 | 0 | TAM.Il1r2 | Pxdc1 |
| Tmem131 | 0 | 0.9939997 | 0.625 | 0.461 | 0 | TAM.Il1r2 | Tmem131 |
| Arl5b | 0 | 0.94179868 | 0.567 | 0.403 | 0 | TAM.Il1r2 | Arl5b |
| Nlrp3 | 0 | 0.75526385 | 0.861 | 0.699 | 0 | TAM.Il1r2 | Nlrp3 |
| Nr4a1 | 0 | 0.78823997 | 0.935 | 0.773 | 0 | TAM.Il1r2 | Nr4a1 |
| Bhlhe401 | 0 | 0.85126735 | 0.941 | 0.781 | 0 | TAM.Il1r2 | Bhlhe40 |
| Ell2 | 0 | 0.84820883 | 0.648 | 0.489 | 0 | TAM.Il1r2 | Ell2 |
| Gpr351 | 0 | 0.71937582 | 0.824 | 0.665 | 0 | TAM.Il1r2 | Gpr35 |
| Eif5a | 0 | 0.9218688 | 0.821 | 0.664 | 0 | TAM.Il1r2 | Eif5a |
| Eprs1 | 0 | 1.10255998 | 0.531 | 0.376 | 0 | TAM.Il1r2 | Eprs |
| Tiparp | 0 | 0.75022659 | 0.793 | 0.64 | 0 | TAM.Il1r2 | Tiparp |
| Il1b1 | 0 | 1.11212972 | 0.985 | 0.834 | 0 | TAM.Il1r2 | Il1b |
| H2-Ab1 | 0 | 1.37110575 | 0.982 | 0.831 | 0 | TAM.Il1r2 | H2-Ab1 |
| Cwc25 | 0 | 1.1129462 | 0.461 | 0.311 | 0 | TAM.Il1r2 | Cwc25 |
| Vps37b | 0 | 1.12367812 | 0.66 | 0.511 | 0 | TAM.Il1r2 | Vps37b |
| Kdm6b | 0 | 0.74020859 | 0.936 | 0.787 | 0 | TAM.Il1r2 | Kdm6b |
| Myc1 | 0 | 1.15065449 | 0.403 | 0.255 | 0 | TAM.Il1r2 | Myc |
| St8sia4 | 0 | 1.10273735 | 0.487 | 0.339 | 0 | TAM.Il1r2 | St8sia4 |
| Plaur1 | 0 | 0.78489546 | 0.953 | 0.807 | 0 | TAM.Il1r2 | Plaur |
| Cd86 | 0 | 0.81872309 | 0.837 | 0.694 | 0 | TAM.Il1r2 | Cd86 |
| Gadd45b | 0 | 0.6271267 | 0.923 | 0.781 | 0 | TAM.Il1r2 | Gadd45b |
| Pgf | 0 | 1.0894243 | 0.339 | 0.199 | 0 | TAM.Il1r2 | Pgf |
| Slc35b2 | 0 | 0.78408864 | 0.854 | 0.719 | 0 | TAM.Il1r2 | Slc35b2 |
| Fosb | 0 | 0.62011168 | 0.926 | 0.792 | 0 | TAM.Il1r2 | Fosb |
| Btg2 | 0 | 0.70743134 | 0.946 | 0.813 | 0 | TAM.Il1r2 | Btg2 |
| Ndel1 | 0 | 0.77145939 | 0.8 | 0.668 | 0 | TAM.Il1r2 | Ndel1 |
| Flrt31 | 0 | 1.01409092 | 0.368 | 0.237 | 0 | TAM.Il1r2 | Flrt3 |
| Odc1 | 0 | 0.88917346 | 0.788 | 0.657 | 0 | TAM.Il1r2 | Odc1 |
| Ccdc71l | 0 | 0.95334067 | 0.589 | 0.458 | 0 | TAM.Il1r2 | Ccdc71l |
| Csf2rb1 | 0 | 0.82805816 | 0.906 | 0.781 | 0 | TAM.Il1r2 | Csf2rb |
| Cdkn1a | 0 | 0.84017656 | 0.955 | 0.83 | 0 | TAM.Il1r2 | Cdkn1a |
| Rhoq | 0 | 0.84549243 | 0.661 | 0.537 | 0 | TAM.Il1r2 | Rhoq |
| Thbs1 | 0 | 0.49194077 | 0.785 | 0.671 | 0 | TAM.Il1r2 | Thbs1 |
| Ets21 | 0 | 0.57270161 | 0.914 | 0.801 | 0 | TAM.Il1r2 | Ets2 |
| Coro1a | 0 | 0.78577176 | 0.965 | 0.859 | 0 | TAM.Il1r2 | Coro1a |
| Fosl2 | 0 | 0.72019092 | 0.94 | 0.836 | 0 | TAM.Il1r2 | Fosl2 |
| Gsn1 | 0 | 0.789897 | 0.92 | 0.821 | 0 | TAM.Il1r2 | Gsn |
| Vasp | 0 | 0.7898709 | 0.956 | 0.864 | 0 | TAM.Il1r2 | Vasp |
| Pim1 | 0 | 0.6342715 | 0.97 | 0.88 | 0 | TAM.Il1r2 | Pim1 |
| S100a111 | 0 | 0.72870604 | 0.945 | 0.858 | 0 | TAM.Il1r2 | S100a11 |
| Gdi21 | 0 | 0.68872077 | 0.816 | 0.73 | 0 | TAM.Il1r2 | Gdi2 |
| Cd521 | 0 | 0.50733555 | 0.945 | 0.86 | 0 | TAM.Il1r2 | Cd52 |
| Cd141 | 0 | 0.53783341 | 0.949 | 0.866 | 0 | TAM.Il1r2 | Cd14 |
| Atox1 | 0 | 0.68552879 | 0.925 | 0.852 | 0 | TAM.Il1r2 | Atox1 |
| Cd741 | 0 | 1.05066159 | 0.995 | 0.933 | 0 | TAM.Il1r2 | Cd74 |
| Slc15a31 | 0 | 0.5465334 | 0.976 | 0.915 | 0 | TAM.Il1r2 | Slc15a3 |
| Plek | 0 | 0.58508119 | 0.98 | 0.932 | 0 | TAM.Il1r2 | Plek |
| Ccl241 | 0 | 0.86680534 | 0.273 | 0.228 | 0 | TAM.Il1r2 | Ccl24 |
| Junb | 0 | 0.42877711 | 0.99 | 0.952 | 0 | TAM.Il1r2 | Junb |
| Gm2a | 0 | 0.71359234 | 0.981 | 0.948 | 0 | TAM.Il1r2 | Gm2a |
| Cst3 | 0 | 0.8258775 | 0.988 | 0.973 | 0 | TAM.Il1r2 | Cst3 |
| Irf5 | 2.69E-307 | 0.56083729 | 0.895 | 0.817 | 5.20E-303 | TAM.Il1r2 | Irf5 |
| Syngr2 | 6.67E-299 | 0.84588892 | 0.727 | 0.592 | 1.29E-294 | TAM.Il1r2 | Syngr2 |
| Mxd1 | 5.66E-293 | 0.64542076 | 0.739 | 0.581 | 1.09E-288 | TAM.Il1r2 | Mxd1 |
| Basp1 | 4.21E-291 | 0.68581931 | 0.85 | 0.729 | 8.13E-287 | TAM.Il1r2 | Basp1 |
| Arih2 | 1.15E-280 | 0.86398713 | 0.684 | 0.52 | 2.23E-276 | TAM.Il1r2 | Arih2 |
| Lin541 | 1.20E-274 | 1.12830081 | 0.426 | 0.267 | 2.32E-270 | TAM.Il1r2 | Lin54 |
| Fam110a | 4.25E-272 | 0.9653018 | 0.429 | 0.294 | 8.22E-268 | TAM.Il1r2 | Fam110a |
| Per1 | 4.98E-272 | 0.66327358 | 0.819 | 0.683 | 9.63E-268 | TAM.Il1r2 | Per1 |
| Nfkbie | 1.23E-271 | 0.72038409 | 0.761 | 0.597 | 2.38E-267 | TAM.Il1r2 | Nfkbie |
| Tob2 | 4.71E-261 | 0.77960837 | 0.707 | 0.554 | 9.10E-257 | TAM.Il1r2 | Tob2 |
| Slc7a5 | 1.65E-259 | 1.14132759 | 0.437 | 0.245 | 3.19E-255 | TAM.Il1r2 | Slc7a5 |
| Pfkp1 | 1.54E-256 | 0.79477921 | 0.733 | 0.557 | 2.98E-252 | TAM.Il1r2 | Pfkp |
| Fgr | 3.23E-255 | 0.58193384 | 0.81 | 0.652 | 6.25E-251 | TAM.Il1r2 | Fgr |
| H2-DMb2 | 9.87E-253 | 1.78631575 | 0.264 | 0.101 | 1.91E-248 | TAM.Il1r2 | H2-DMb2 |
| Rab11fip11 | 2.30E-252 | 0.66510511 | 0.758 | 0.524 | 4.45E-248 | TAM.Il1r2 | Rab11fip1 |
| Homer1 | 2.10E-247 | 0.83151077 | 0.59 | 0.453 | 4.06E-243 | TAM.Il1r2 | Homer1 |
| Lmnb1 | 4.27E-247 | 0.88126668 | 0.649 | 0.396 | 8.26E-243 | TAM.Il1r2 | Lmnb1 |
| Chd7 | 5.65E-243 | 0.78681565 | 0.677 | 0.516 | 1.09E-238 | TAM.Il1r2 | Chd7 |
| Nuak2 | 4.31E-241 | 0.69342221 | 0.758 | 0.605 | 8.34E-237 | TAM.Il1r2 | Nuak2 |
| Rab43 | 4.58E-241 | 0.68490529 | 0.795 | 0.668 | 8.85E-237 | TAM.Il1r2 | Rab43 |
| Fam129a | 6.13E-239 | 0.82702217 | 0.494 | 0.337 | 1.19E-234 | TAM.Il1r2 | Fam129a |
| Bcl2l11 | 7.64E-238 | 0.57424573 | 0.793 | 0.664 | 1.48E-233 | TAM.Il1r2 | Bcl2l11 |
| Calm1 | 7.59E-235 | 0.54496961 | 0.908 | 0.852 | 1.47E-230 | TAM.Il1r2 | Calm1 |
| Lrrfip1 | 2.47E-232 | 0.54867785 | 0.861 | 0.746 | 4.78E-228 | TAM.Il1r2 | Lrrfip1 |
| Rnf2171 | 7.80E-232 | 1.0942828 | 0.409 | 0.258 | 1.51E-227 | TAM.Il1r2 | Rnf217 |
| Rac2 | 2.06E-230 | 0.59543453 | 0.906 | 0.818 | 3.98E-226 | TAM.Il1r2 | Rac2 |
| Pilra1 | 4.93E-226 | 0.74575401 | 0.757 | 0.568 | 9.53E-222 | TAM.Il1r2 | Pilra |
| Spag9 | 1.30E-225 | 0.67824183 | 0.788 | 0.669 | 2.51E-221 | TAM.Il1r2 | Spag9 |
| Sap18b | 9.82E-223 | 0.9582859 | 0.443 | 0.353 | 1.90E-218 | TAM.Il1r2 | Sap18b |
| Rabgef1 | 5.53E-216 | 0.72443519 | 0.654 | 0.53 | 1.07E-211 | TAM.Il1r2 | Rabgef1 |
| Impdh2 | 8.84E-216 | 0.95201105 | 0.544 | 0.383 | 1.71E-211 | TAM.Il1r2 | Impdh2 |
| Ahr1 | 7.15E-215 | 0.76642645 | 0.693 | 0.538 | 1.38E-210 | TAM.Il1r2 | Ahr |
| Cnbp | 8.00E-214 | 0.86866386 | 0.687 | 0.53 | 1.55E-209 | TAM.Il1r2 | Cnbp |
| Sorl1 | 9.20E-214 | 0.74605151 | 0.634 | 0.441 | 1.78E-209 | TAM.Il1r2 | Sorl1 |
| Xbp1 | 2.39E-209 | 0.55478198 | 0.871 | 0.792 | 4.62E-205 | TAM.Il1r2 | Xbp1 |
| Slc25a20 | 1.65E-207 | 0.92830714 | 0.464 | 0.322 | 3.18E-203 | TAM.Il1r2 | Slc25a20 |
| Dgat11 | 7.02E-207 | 1.28116185 | 0.346 | 0.2 | 1.36E-202 | TAM.Il1r2 | Dgat1 |
| Stk17b | 4.18E-201 | 0.68363754 | 0.675 | 0.551 | 8.07E-197 | TAM.Il1r2 | Stk17b |
| Fbl | 7.76E-201 | 1.01135596 | 0.555 | 0.376 | 1.50E-196 | TAM.Il1r2 | Fbl |
| Cxcl21 | 4.19E-200 | 0.30292573 | 0.868 | 0.778 | 8.09E-196 | TAM.Il1r2 | Cxcl2 |
| Rack1 | 4.62E-198 | 0.5317538 | 0.905 | 0.853 | 8.93E-194 | TAM.Il1r2 | Rack1 |
| Snrpf | 2.33E-196 | 0.85222037 | 0.649 | 0.489 | 4.51E-192 | TAM.Il1r2 | Snrpf |
| Fam107b1 | 4.74E-192 | 0.73190003 | 0.582 | 0.435 | 9.17E-188 | TAM.Il1r2 | Fam107b |
| Hnrnpll1 | 9.03E-190 | 0.73075732 | 0.738 | 0.596 | 1.75E-185 | TAM.Il1r2 | Hnrnpll |
| Fh1 | 4.62E-185 | 1.09818436 | 0.391 | 0.255 | 8.93E-181 | TAM.Il1r2 | Fh1 |
| AA4671971 | 5.19E-183 | 0.53522638 | 0.667 | 0.392 | 1.00E-178 | TAM.Il1r2 | AA467197 |
| Mcemp11 | 5.71E-183 | 0.74222489 | 0.454 | 0.257 | 1.10E-178 | TAM.Il1r2 | Mcemp1 |
| Emd | 6.03E-182 | 0.67199627 | 0.629 | 0.515 | 1.17E-177 | TAM.Il1r2 | Emd |
| Pygl1 | 1.09E-176 | 0.71668494 | 0.744 | 0.568 | 2.11E-172 | TAM.Il1r2 | Pygl |
| Neurl31 | 3.46E-176 | 0.59761099 | 0.844 | 0.732 | 6.69E-172 | TAM.Il1r2 | Neurl3 |
| Fmnl1 | 9.00E-172 | 0.46453758 | 0.768 | 0.641 | 1.74E-167 | TAM.Il1r2 | Fmnl1 |
| Olfm1 | 2.65E-170 | 0.66798622 | 0.695 | 0.557 | 5.12E-166 | TAM.Il1r2 | Olfm1 |
| Siglece1 | 3.62E-170 | 0.95694393 | 0.363 | 0.206 | 7.01E-166 | TAM.Il1r2 | Siglece |
| Pdcd1lg21 | 1.85E-167 | 1.07535241 | 0.364 | 0.219 | 3.59E-163 | TAM.Il1r2 | Pdcd1lg2 |
| Tkt1 | 1.22E-164 | 0.61205776 | 0.886 | 0.784 | 2.37E-160 | TAM.Il1r2 | Tkt |
| Hivep2 | 5.31E-163 | 1.00533638 | 0.395 | 0.247 | 1.03E-158 | TAM.Il1r2 | Hivep2 |
| Smchd1 | 1.47E-162 | 0.76571439 | 0.504 | 0.399 | 2.85E-158 | TAM.Il1r2 | Smchd1 |
| Med141 | 4.88E-161 | 1.02719978 | 0.412 | 0.272 | 9.43E-157 | TAM.Il1r2 | Med14 |
| Osgin2 | 9.32E-159 | 0.96232093 | 0.286 | 0.183 | 1.80E-154 | TAM.Il1r2 | Osgin2 |
| Slc38a2 | 1.40E-158 | 0.50789986 | 0.865 | 0.77 | 2.72E-154 | TAM.Il1r2 | Slc38a2 |
| H2afy | 2.90E-156 | 0.50452371 | 0.858 | 0.782 | 5.61E-152 | TAM.Il1r2 | H2afy |
| Nop58 | 1.52E-148 | 1.10178285 | 0.37 | 0.216 | 2.93E-144 | TAM.Il1r2 | Nop58 |
| Dot1l1 | 8.24E-148 | 0.76576778 | 0.578 | 0.424 | 1.59E-143 | TAM.Il1r2 | Dot1l |
| Parvg1 | 5.32E-147 | 0.97971359 | 0.463 | 0.289 | 1.03E-142 | TAM.Il1r2 | Parvg |
| B3gnt51 | 3.42E-145 | 1.01762422 | 0.275 | 0.14 | 6.60E-141 | TAM.Il1r2 | B3gnt5 |
| Hnrnpa1 | 4.93E-145 | 0.74331861 | 0.747 | 0.59 | 9.52E-141 | TAM.Il1r2 | Hnrnpa1 |
| Mapkapk31 | 2.70E-144 | 0.69619637 | 0.623 | 0.481 | 5.21E-140 | TAM.Il1r2 | Mapkapk3 |
| Ifrd1 | 4.63E-144 | 0.4264944 | 0.926 | 0.811 | 8.95E-140 | TAM.Il1r2 | Ifrd1 |
| Ddx21 | 6.40E-142 | 0.7074986 | 0.717 | 0.579 | 1.24E-137 | TAM.Il1r2 | Ddx21 |
| Prkcd1 | 7.49E-142 | 0.46375845 | 0.92 | 0.867 | 1.45E-137 | TAM.Il1r2 | Prkcd |
| Rasgrp11 | 4.66E-141 | 0.7656981 | 0.505 | 0.361 | 9.01E-137 | TAM.Il1r2 | Rasgrp1 |
| Dock10 | 3.40E-140 | 0.69810462 | 0.595 | 0.461 | 6.57E-136 | TAM.Il1r2 | Dock10 |
| Ybx31 | 3.79E-139 | 0.81613076 | 0.557 | 0.407 | 7.33E-135 | TAM.Il1r2 | Ybx3 |
| Itgax2 | 3.98E-136 | 0.66910436 | 0.857 | 0.573 | 7.70E-132 | TAM.Il1r2 | Itgax |
| Ppp1r3b | 5.48E-136 | 0.84016978 | 0.312 | 0.24 | 1.06E-131 | TAM.Il1r2 | Ppp1r3b |
| Gpd21 | 3.60E-135 | 0.94759467 | 0.376 | 0.249 | 6.97E-131 | TAM.Il1r2 | Gpd2 |
| Stk38l | 3.88E-133 | 0.8890484 | 0.402 | 0.268 | 7.51E-129 | TAM.Il1r2 | Stk38l |
| Cbfa2t3 | 7.52E-133 | 0.83374776 | 0.362 | 0.221 | 1.45E-128 | TAM.Il1r2 | Cbfa2t3 |
| St7 | 9.14E-132 | 0.93329475 | 0.41 | 0.272 | 1.77E-127 | TAM.Il1r2 | St7 |
| Ccdc102a1 | 1.02E-130 | 1.18553335 | 0.307 | 0.158 | 1.97E-126 | TAM.Il1r2 | Ccdc102a |
| Lsp1 | 3.53E-130 | 0.37710246 | 0.56 | 0.384 | 6.82E-126 | TAM.Il1r2 | Lsp1 |
| Snd1 | 8.11E-127 | 0.51553053 | 0.841 | 0.759 | 1.57E-122 | TAM.Il1r2 | Snd1 |
| Nup98 | 3.89E-126 | 0.71446636 | 0.566 | 0.446 | 7.53E-122 | TAM.Il1r2 | Nup98 |
| Nod2 | 1.75E-125 | 0.6509506 | 0.488 | 0.343 | 3.38E-121 | TAM.Il1r2 | Nod2 |
| B4galnt1 | 2.87E-125 | 0.72723174 | 0.649 | 0.502 | 5.55E-121 | TAM.Il1r2 | B4galnt1 |
| Bin2 | 2.44E-124 | 0.49485977 | 0.854 | 0.744 | 4.72E-120 | TAM.Il1r2 | Bin2 |
| Ywhah | 1.04E-123 | 0.57271252 | 0.725 | 0.649 | 2.02E-119 | TAM.Il1r2 | Ywhah |
| Slc1a51 | 1.11E-123 | 0.85063302 | 0.543 | 0.373 | 2.15E-119 | TAM.Il1r2 | Slc1a5 |
| Cdk2ap2 | 7.09E-122 | 0.6056426 | 0.657 | 0.576 | 1.37E-117 | TAM.Il1r2 | Cdk2ap2 |
| Dna2 | 1.22E-121 | 1.08996779 | 0.253 | 0.133 | 2.36E-117 | TAM.Il1r2 | Dna2 |
| Myo1g | 1.50E-121 | 0.53433321 | 0.697 | 0.588 | 2.90E-117 | TAM.Il1r2 | Myo1g |
| Bzw21 | 2.72E-121 | 1.05897146 | 0.346 | 0.213 | 5.26E-117 | TAM.Il1r2 | Bzw2 |
| Eef1akmt1 | 1.45E-120 | 1.24910834 | 0.269 | 0.161 | 2.80E-116 | TAM.Il1r2 | Eef1akmt |
| Nfkbiz | 3.94E-120 | 0.3835571 | 0.774 | 0.679 | 7.61E-116 | TAM.Il1r2 | Nfkbiz |
| Fam49b | 9.47E-120 | 0.3759696 | 0.857 | 0.809 | 1.83E-115 | TAM.Il1r2 | Fam49b |
| Cd300lf | 2.61E-119 | 0.61526837 | 0.73 | 0.555 | 5.06E-115 | TAM.Il1r2 | Cd300lf |
| Emb1 | 4.91E-119 | 0.49992418 | 0.858 | 0.729 | 9.49E-115 | TAM.Il1r2 | Emb |
| Eef1a1 | 5.29E-119 | 0.31302736 | 0.983 | 0.977 | 1.02E-114 | TAM.Il1r2 | Eef1a1 |
| Tspan131 | 4.11E-118 | 0.75367778 | 0.429 | 0.295 | 7.94E-114 | TAM.Il1r2 | Tspan13 |
| Aldoc1 | 1.08E-115 | 0.92453918 | 0.453 | 0.295 | 2.09E-111 | TAM.Il1r2 | Aldoc |
| Aldh18a1 | 6.57E-115 | 1.1801714 | 0.264 | 0.151 | 1.27E-110 | TAM.Il1r2 | Aldh18a1 |
| Rap2a | 2.65E-114 | 0.74920664 | 0.577 | 0.456 | 5.12E-110 | TAM.Il1r2 | Rap2a |
| Ccdc86 | 1.43E-113 | 0.78479894 | 0.595 | 0.447 | 2.76E-109 | TAM.Il1r2 | Ccdc86 |
| Picalm | 1.70E-113 | 0.44901397 | 0.924 | 0.872 | 3.29E-109 | TAM.Il1r2 | Picalm |
| Arf4 | 3.18E-113 | 0.5890532 | 0.661 | 0.584 | 6.14E-109 | TAM.Il1r2 | Arf4 |
| Sulf2 | 5.65E-113 | 0.57228672 | 0.738 | 0.601 | 1.09E-108 | TAM.Il1r2 | Sulf2 |
| Arih1 | 1.74E-112 | 0.58386484 | 0.668 | 0.584 | 3.37E-108 | TAM.Il1r2 | Arih1 |
| Tgfb1 | 2.22E-111 | 0.45651534 | 0.944 | 0.89 | 4.30E-107 | TAM.Il1r2 | Tgfb1 |
| Amz11 | 2.72E-111 | 0.77780219 | 0.464 | 0.299 | 5.26E-107 | TAM.Il1r2 | Amz1 |
| Lats2 | 3.51E-109 | 0.65197694 | 0.458 | 0.385 | 6.79E-105 | TAM.Il1r2 | Lats2 |
| Clcf11 | 6.07E-109 | 0.69532036 | 0.433 | 0.303 | 1.17E-104 | TAM.Il1r2 | Clcf1 |
| Gfod11 | 9.15E-108 | 0.99801191 | 0.317 | 0.178 | 1.77E-103 | TAM.Il1r2 | Gfod1 |
| Nufip11 | 1.44E-106 | 0.75138262 | 0.484 | 0.392 | 2.79E-102 | TAM.Il1r2 | Nufip1 |
| Ifngr1 | 6.11E-106 | 0.40603644 | 0.864 | 0.787 | 1.18E-101 | TAM.Il1r2 | Ifngr1 |
| Nt5e1 | 4.49E-105 | 0.92851051 | 0.293 | 0.181 | 8.68E-101 | TAM.Il1r2 | Nt5e |
| Psme1 | 1.94E-100 | 0.50647417 | 0.791 | 0.702 | 3.76E-96 | TAM.Il1r2 | Psme1 |
| F11r1 | 2.41E-100 | 0.59658519 | 0.713 | 0.539 | 4.66E-96 | TAM.Il1r2 | F11r |
| Tent5a | 1.23E-98 | 0.59906883 | 0.487 | 0.41 | 2.38E-94 | TAM.Il1r2 | Tent5a |
| Map4k1 | 1.75E-98 | 0.98908896 | 0.295 | 0.172 | 3.38E-94 | TAM.Il1r2 | Map4k1 |
| Nfe2l2 | 8.76E-98 | 0.42791567 | 0.925 | 0.863 | 1.69E-93 | TAM.Il1r2 | Nfe2l2 |
| Abtb21 | 8.11E-96 | 0.76028094 | 0.367 | 0.248 | 1.57E-91 | TAM.Il1r2 | Abtb2 |
| Slc25a3 | 2.83E-95 | 0.4963241 | 0.873 | 0.815 | 5.48E-91 | TAM.Il1r2 | Slc25a3 |
| Pvr | 2.14E-94 | 0.63158668 | 0.443 | 0.339 | 4.13E-90 | TAM.Il1r2 | Pvr |
| Med251 | 1.30E-93 | 0.71886927 | 0.559 | 0.44 | 2.52E-89 | TAM.Il1r2 | Med25 |
| Ralb | 1.47E-92 | 0.56434354 | 0.666 | 0.578 | 2.85E-88 | TAM.Il1r2 | Ralb |
| Trps1 | 3.32E-91 | 0.40049026 | 0.814 | 0.742 | 6.41E-87 | TAM.Il1r2 | Trps1 |
| Mybbp1a | 6.44E-91 | 0.83740635 | 0.497 | 0.355 | 1.24E-86 | TAM.Il1r2 | Mybbp1a |
| Cmtr1 | 2.89E-90 | 0.77307786 | 0.437 | 0.366 | 5.59E-86 | TAM.Il1r2 | Cmtr1 |
| Wnk1 | 3.70E-89 | 0.47594492 | 0.789 | 0.728 | 7.16E-85 | TAM.Il1r2 | Wnk1 |
| Bpnt1 | 4.29E-89 | 0.97477385 | 0.322 | 0.234 | 8.30E-85 | TAM.Il1r2 | Bpnt1 |
| Pilrb11 | 1.11E-88 | 0.77704812 | 0.435 | 0.303 | 2.14E-84 | TAM.Il1r2 | Pilrb1 |
| Atp13a3 | 1.93E-87 | 0.58595999 | 0.728 | 0.626 | 3.73E-83 | TAM.Il1r2 | Atp13a3 |
| Tmem123 | 2.15E-86 | 0.65397226 | 0.569 | 0.467 | 4.15E-82 | TAM.Il1r2 | Tmem123 |
| Bcl2a1b | 2.83E-86 | 0.52690593 | 0.836 | 0.721 | 5.46E-82 | TAM.Il1r2 | Bcl2a1b |
| Specc1l | 3.74E-86 | 0.66092331 | 0.539 | 0.447 | 7.23E-82 | TAM.Il1r2 | Specc1l |
| Gtpbp4 | 4.20E-86 | 0.77617646 | 0.479 | 0.357 | 8.12E-82 | TAM.Il1r2 | Gtpbp4 |
| Mthfr | 6.64E-85 | 0.6805944 | 0.4 | 0.317 | 1.28E-80 | TAM.Il1r2 | Mthfr |
| Cnnm4 | 1.11E-84 | 0.81743659 | 0.311 | 0.218 | 2.15E-80 | TAM.Il1r2 | Cnnm4 |
| Isyna11 | 1.46E-83 | 0.77245994 | 0.507 | 0.373 | 2.82E-79 | TAM.Il1r2 | Isyna1 |
| Runx1 | 4.86E-83 | 0.52770178 | 0.733 | 0.646 | 9.39E-79 | TAM.Il1r2 | Runx1 |
| St3gal4 | 7.83E-83 | 0.52045186 | 0.49 | 0.368 | 1.51E-78 | TAM.Il1r2 | St3gal4 |
| Dusp2 | 3.70E-82 | 0.53016105 | 0.393 | 0.306 | 7.15E-78 | TAM.Il1r2 | Dusp2 |
| Pip4k2a | 2.13E-81 | 0.6552471 | 0.518 | 0.406 | 4.12E-77 | TAM.Il1r2 | Pip4k2a |
| Btf3 | 1.94E-80 | 0.55590902 | 0.713 | 0.618 | 3.75E-76 | TAM.Il1r2 | Btf3 |
| Sec61b | 1.67E-79 | 0.68430711 | 0.473 | 0.373 | 3.22E-75 | TAM.Il1r2 | Sec61b |
| Pik3r1 | 5.59E-79 | 0.67450915 | 0.564 | 0.481 | 1.08E-74 | TAM.Il1r2 | Pik3r1 |
| Crtc3 | 1.81E-78 | 0.62954003 | 0.525 | 0.413 | 3.50E-74 | TAM.Il1r2 | Crtc3 |
| Usp38 | 2.70E-78 | 0.74765468 | 0.333 | 0.248 | 5.21E-74 | TAM.Il1r2 | Usp38 |
| Samsn1 | 1.44E-77 | 0.66916271 | 0.406 | 0.298 | 2.78E-73 | TAM.Il1r2 | Samsn1 |
| Il1rn1 | 5.75E-76 | 0.38340404 | 0.861 | 0.729 | 1.11E-71 | TAM.Il1r2 | Il1rn |
| Dusp161 | 1.29E-75 | 0.58593905 | 0.476 | 0.343 | 2.49E-71 | TAM.Il1r2 | Dusp16 |
| Hipk1 | 1.44E-75 | 0.49335877 | 0.713 | 0.643 | 2.78E-71 | TAM.Il1r2 | Hipk1 |
| Prkd3 | 8.24E-74 | 0.8275641 | 0.297 | 0.203 | 1.59E-69 | TAM.Il1r2 | Prkd3 |
| Pts | 1.40E-73 | 0.77336252 | 0.399 | 0.303 | 2.70E-69 | TAM.Il1r2 | Pts |
| Psmb8 | 3.24E-73 | 0.46541826 | 0.759 | 0.674 | 6.27E-69 | TAM.Il1r2 | Psmb8 |
| Kpna1 | 4.24E-73 | 0.67631004 | 0.418 | 0.34 | 8.20E-69 | TAM.Il1r2 | Kpna1 |
| Slc25a331 | 7.24E-73 | 0.8573454 | 0.334 | 0.232 | 1.40E-68 | TAM.Il1r2 | Slc25a33 |
| Plcb22 | 9.97E-72 | 0.78277307 | 0.425 | 0.3 | 1.93E-67 | TAM.Il1r2 | Plcb2 |
| Tmem185b | 2.06E-71 | 0.70281717 | 0.452 | 0.347 | 3.98E-67 | TAM.Il1r2 | Tmem185b |
| Exosc5 | 5.85E-71 | 0.8220239 | 0.392 | 0.284 | 1.13E-66 | TAM.Il1r2 | Exosc5 |
| Mad1l11 | 8.03E-71 | 0.8580015 | 0.281 | 0.187 | 1.55E-66 | TAM.Il1r2 | Mad1l1 |
| Klra21 | 1.34E-69 | 0.48504848 | 0.434 | 0.322 | 2.59E-65 | TAM.Il1r2 | Klra2 |
| Gar1 | 3.45E-69 | 0.92619709 | 0.317 | 0.208 | 6.66E-65 | TAM.Il1r2 | Gar1 |
| Emilin2 | 1.02E-68 | 0.29440911 | 0.865 | 0.786 | 1.98E-64 | TAM.Il1r2 | Emilin2 |
| Agpat41 | 3.53E-67 | 0.60955191 | 0.518 | 0.386 | 6.83E-63 | TAM.Il1r2 | Agpat4 |
| Gpr651 | 9.06E-66 | 0.57239176 | 0.452 | 0.39 | 1.75E-61 | TAM.Il1r2 | Gpr65 |
| Pak1 | 9.96E-66 | 0.72119919 | 0.393 | 0.289 | 1.93E-61 | TAM.Il1r2 | Pak1 |
| Elovl5 | 1.59E-65 | 0.69281923 | 0.359 | 0.27 | 3.07E-61 | TAM.Il1r2 | Elovl5 |
| Tsr1 | 1.61E-65 | 0.86243646 | 0.312 | 0.221 | 3.12E-61 | TAM.Il1r2 | Tsr1 |
| Coq10b | 1.38E-64 | 0.50601864 | 0.517 | 0.426 | 2.67E-60 | TAM.Il1r2 | Coq10b |
| Npm3 | 1.52E-64 | 0.89999235 | 0.268 | 0.175 | 2.93E-60 | TAM.Il1r2 | Npm3 |
| Nfkb11 | 9.22E-64 | 0.42729058 | 0.741 | 0.658 | 1.78E-59 | TAM.Il1r2 | Nfkb1 |
| Nbeal2 | 9.28E-64 | 0.71370019 | 0.285 | 0.196 | 1.79E-59 | TAM.Il1r2 | Nbeal2 |
| Ppm1m | 2.24E-63 | 0.62788115 | 0.51 | 0.397 | 4.34E-59 | TAM.Il1r2 | Ppm1m |
| Plk31 | 2.81E-63 | 0.47755219 | 0.701 | 0.589 | 5.44E-59 | TAM.Il1r2 | Plk3 |
| Eef1g | 1.66E-62 | 0.63425415 | 0.556 | 0.453 | 3.21E-58 | TAM.Il1r2 | Eef1g |
| Sirpb1c | 2.16E-62 | 0.57422288 | 0.465 | 0.35 | 4.18E-58 | TAM.Il1r2 | Sirpb1c |
| Eef1b2 | 3.42E-62 | 0.44176962 | 0.781 | 0.713 | 6.62E-58 | TAM.Il1r2 | Eef1b2 |
| Pilrb22 | 5.07E-62 | 0.77427258 | 0.406 | 0.295 | 9.81E-58 | TAM.Il1r2 | Pilrb2 |
| Tbc1d81 | 2.89E-61 | 0.74528549 | 0.39 | 0.274 | 5.58E-57 | TAM.Il1r2 | Tbc1d8 |
| Etv3 | 4.54E-61 | 0.59292262 | 0.516 | 0.451 | 8.77E-57 | TAM.Il1r2 | Etv3 |
| Trib1 | 2.29E-60 | 0.39940107 | 0.676 | 0.574 | 4.43E-56 | TAM.Il1r2 | Trib1 |
| Ocstamp2 | 8.80E-59 | 0.92421149 | 0.352 | 0.198 | 1.70E-54 | TAM.Il1r2 | Ocstamp |
| Slamf91 | 1.19E-58 | 0.77769695 | 0.313 | 0.21 | 2.30E-54 | TAM.Il1r2 | Slamf9 |
| Naca | 1.74E-58 | 0.46445002 | 0.646 | 0.563 | 3.36E-54 | TAM.Il1r2 | Naca |
| Smox1 | 1.60E-57 | 0.37018934 | 0.818 | 0.719 | 3.09E-53 | TAM.Il1r2 | Smox |
| Ltb4r1 | 4.90E-57 | 0.49693631 | 0.259 | 0.154 | 9.48E-53 | TAM.Il1r2 | Ltb4r1 |
| Snrpd3 | 5.35E-57 | 0.62660861 | 0.567 | 0.467 | 1.03E-52 | TAM.Il1r2 | Snrpd3 |
| Snx20 | 2.68E-56 | 0.43042858 | 0.745 | 0.679 | 5.19E-52 | TAM.Il1r2 | Snx20 |
| Hnrnpa2b1 | 3.40E-56 | 0.43199121 | 0.805 | 0.75 | 6.57E-52 | TAM.Il1r2 | Hnrnpa2b1 |
| Pmaip1 | 4.02E-56 | 0.5082873 | 0.489 | 0.411 | 7.77E-52 | TAM.Il1r2 | Pmaip1 |
| Nhp2 | 7.02E-55 | 0.70191385 | 0.515 | 0.392 | 1.36E-50 | TAM.Il1r2 | Nhp2 |
| Zbtb12 | 7.62E-55 | 0.64693339 | 0.406 | 0.348 | 1.47E-50 | TAM.Il1r2 | Zbtb1 |
| Tarm11 | 6.00E-54 | 0.61547138 | 0.425 | 0.275 | 1.16E-49 | TAM.Il1r2 | Tarm1 |
| Noc2l | 3.48E-52 | 0.62327178 | 0.561 | 0.452 | 6.74E-48 | TAM.Il1r2 | Noc2l |
| Snx18 | 4.04E-52 | 0.39390696 | 0.806 | 0.74 | 7.81E-48 | TAM.Il1r2 | Snx18 |
| S100a6 | 7.46E-52 | 0.26454602 | 0.833 | 0.762 | 1.44E-47 | TAM.Il1r2 | S100a6 |
| Havcr2 | 1.75E-51 | 0.37537804 | 0.457 | 0.362 | 3.38E-47 | TAM.Il1r2 | Havcr2 |
| Skil | 4.54E-51 | 0.37524081 | 0.836 | 0.761 | 8.79E-47 | TAM.Il1r2 | Skil |
| Cox171 | 7.27E-51 | 0.49725195 | 0.683 | 0.578 | 1.41E-46 | TAM.Il1r2 | Cox17 |
| Snrpa1 | 1.44E-50 | 0.76303208 | 0.348 | 0.252 | 2.79E-46 | TAM.Il1r2 | Snrpa1 |
| Rheb | 1.60E-50 | 0.55140423 | 0.628 | 0.543 | 3.09E-46 | TAM.Il1r2 | Rheb |
| Pqlc11 | 2.68E-50 | 0.69049001 | 0.415 | 0.327 | 5.18E-46 | TAM.Il1r2 | Pqlc1 |
| Nfkbid1 | 6.55E-50 | 0.45332812 | 0.711 | 0.603 | 1.27E-45 | TAM.Il1r2 | Nfkbid |
| Cdk11b | 1.38E-49 | 0.52739905 | 0.497 | 0.453 | 2.66E-45 | TAM.Il1r2 | Cdk11b |
| Pxn | 2.08E-49 | 0.49924442 | 0.602 | 0.532 | 4.03E-45 | TAM.Il1r2 | Pxn |
| Twf2 | 2.96E-49 | 0.52405857 | 0.596 | 0.496 | 5.71E-45 | TAM.Il1r2 | Twf2 |
| Sla1 | 8.20E-49 | 0.48085172 | 0.671 | 0.55 | 1.58E-44 | TAM.Il1r2 | Sla |
| Smad7 | 2.21E-48 | 0.49644451 | 0.436 | 0.36 | 4.27E-44 | TAM.Il1r2 | Smad7 |
| Stk40 | 3.41E-48 | 0.34908459 | 0.595 | 0.5 | 6.60E-44 | TAM.Il1r2 | Stk40 |
| Map3k14 | 5.86E-48 | 0.51716562 | 0.383 | 0.322 | 1.13E-43 | TAM.Il1r2 | Map3k14 |
| F101 | 9.73E-48 | 0.37933613 | 0.75 | 0.53 | 1.88E-43 | TAM.Il1r2 | F10 |
| Adora2b | 2.70E-47 | 0.49982973 | 0.291 | 0.216 | 5.21E-43 | TAM.Il1r2 | Adora2b |
| Klf4 | 2.99E-47 | 0.35746481 | 0.793 | 0.701 | 5.78E-43 | TAM.Il1r2 | Klf4 |
| Orai1 | 5.83E-47 | 0.46454475 | 0.569 | 0.489 | 1.13E-42 | TAM.Il1r2 | Orai1 |
| Tinf2 | 6.28E-47 | 0.64486532 | 0.293 | 0.226 | 1.21E-42 | TAM.Il1r2 | Tinf2 |
| Tnfrsf12a | 8.25E-47 | 0.49707098 | 0.489 | 0.407 | 1.59E-42 | TAM.Il1r2 | Tnfrsf12a |
| Rnf1801 | 1.16E-46 | 0.64712584 | 0.44 | 0.352 | 2.24E-42 | TAM.Il1r2 | Rnf180 |
| Zfp516 | 4.13E-46 | 0.3960886 | 0.701 | 0.636 | 7.98E-42 | TAM.Il1r2 | Zfp516 |
| Eif3i | 4.46E-46 | 0.63417524 | 0.463 | 0.37 | 8.62E-42 | TAM.Il1r2 | Eif3i |
| Purb | 6.91E-46 | 0.46206555 | 0.727 | 0.654 | 1.34E-41 | TAM.Il1r2 | Purb |
| Dapp1 | 1.68E-45 | 0.63328863 | 0.328 | 0.245 | 3.24E-41 | TAM.Il1r2 | Dapp1 |
| Fubp1 | 3.34E-45 | 0.57395603 | 0.535 | 0.434 | 6.47E-41 | TAM.Il1r2 | Fubp1 |
| Tfip11 | 3.94E-45 | 0.57389213 | 0.448 | 0.376 | 7.63E-41 | TAM.Il1r2 | Tfip11 |
| Epb41 | 2.17E-44 | 0.6008663 | 0.459 | 0.387 | 4.20E-40 | TAM.Il1r2 | Epb41 |
| Actg1 | 3.19E-44 | 0.28837514 | 0.956 | 0.936 | 6.16E-40 | TAM.Il1r2 | Actg1 |
| Tgif1 | 3.67E-44 | 0.35144889 | 0.846 | 0.762 | 7.09E-40 | TAM.Il1r2 | Tgif1 |
| Emg1 | 7.06E-44 | 0.59258488 | 0.545 | 0.444 | 1.36E-39 | TAM.Il1r2 | Emg1 |
| Lsm7 | 5.30E-43 | 0.67371136 | 0.395 | 0.301 | 1.02E-38 | TAM.Il1r2 | Lsm7 |
| Sik1 | 2.17E-42 | 0.42297146 | 0.818 | 0.69 | 4.20E-38 | TAM.Il1r2 | Sik1 |
| Pdk3 | 8.34E-42 | 0.7154205 | 0.293 | 0.213 | 1.61E-37 | TAM.Il1r2 | Pdk3 |
| Ago21 | 1.11E-41 | 0.48771057 | 0.631 | 0.537 | 2.14E-37 | TAM.Il1r2 | Ago2 |
| Ndufb1-ps | 3.94E-41 | 0.51730778 | 0.683 | 0.591 | 7.61E-37 | TAM.Il1r2 | Ndufb1-ps |
| Srsf7 | 4.61E-41 | 0.46381001 | 0.677 | 0.59 | 8.91E-37 | TAM.Il1r2 | Srsf7 |
| Wdr46 | 7.75E-41 | 0.81766189 | 0.329 | 0.238 | 1.50E-36 | TAM.Il1r2 | Wdr46 |
| Cct2 | 1.69E-40 | 0.64652433 | 0.563 | 0.455 | 3.26E-36 | TAM.Il1r2 | Cct2 |
| Atp5e | 2.10E-40 | 0.49646118 | 0.721 | 0.636 | 4.05E-36 | TAM.Il1r2 | Atp5e |
| Pik3cd | 7.39E-40 | 0.41374321 | 0.683 | 0.595 | 1.43E-35 | TAM.Il1r2 | Pik3cd |
| Man1b1 | 2.98E-39 | 0.62238534 | 0.373 | 0.32 | 5.76E-35 | TAM.Il1r2 | Man1b1 |
| Pld41 | 8.32E-39 | 0.3269136 | 0.89 | 0.779 | 1.61E-34 | TAM.Il1r2 | Pld4 |
| Ikbkb | 1.42E-38 | 0.42814322 | 0.579 | 0.503 | 2.74E-34 | TAM.Il1r2 | Ikbkb |
| Etfb | 2.76E-38 | 0.51917999 | 0.613 | 0.523 | 5.34E-34 | TAM.Il1r2 | Etfb |
| Appl1 | 3.79E-38 | 0.58364132 | 0.327 | 0.286 | 7.32E-34 | TAM.Il1r2 | Appl1 |
| Mthfd | 3.91E-37 | 0.6658566 | 0.266 | 0.205 | 7.55E-33 | TAM.Il1r2 | Mthfd |
| Scimp1 | 7.68E-37 | 0.52742159 | 0.48 | 0.37 | 1.48E-32 | TAM.Il1r2 | Scimp |
| Egr21 | 7.89E-37 | 0.54210997 | 0.598 | 0.461 | 1.52E-32 | TAM.Il1r2 | Egr2 |
| Eif2b21 | 1.10E-36 | 0.49405511 | 0.407 | 0.349 | 2.12E-32 | TAM.Il1r2 | Eif2b2 |
| Eml4 | 1.85E-36 | 0.49120082 | 0.467 | 0.384 | 3.59E-32 | TAM.Il1r2 | Eml4 |
| Ptp4a31 | 1.99E-36 | 0.74084642 | 0.264 | 0.182 | 3.84E-32 | TAM.Il1r2 | Ptp4a3 |
| Tubb6 | 2.35E-36 | 0.45393618 | 0.703 | 0.607 | 4.54E-32 | TAM.Il1r2 | Tubb6 |
| Edem1 | 3.44E-36 | 0.43784258 | 0.847 | 0.763 | 6.64E-32 | TAM.Il1r2 | Edem1 |
| Eif3g | 4.22E-36 | 0.54481509 | 0.568 | 0.474 | 8.16E-32 | TAM.Il1r2 | Eif3g |
| Sf3b1 | 8.57E-36 | 0.35824063 | 0.844 | 0.793 | 1.66E-31 | TAM.Il1r2 | Sf3b1 |
| Pa2g4 | 1.10E-35 | 0.78514232 | 0.332 | 0.241 | 2.12E-31 | TAM.Il1r2 | Pa2g4 |
| Serbp1 | 2.63E-35 | 0.53321221 | 0.622 | 0.527 | 5.09E-31 | TAM.Il1r2 | Serbp1 |
| Ccnd3 | 4.83E-35 | 0.36905927 | 0.372 | 0.292 | 9.33E-31 | TAM.Il1r2 | Ccnd3 |
| Ube2h | 5.26E-35 | 0.47849928 | 0.45 | 0.4 | 1.02E-30 | TAM.Il1r2 | Ube2h |
| Prex1 | 5.94E-35 | 0.41205858 | 0.823 | 0.75 | 1.15E-30 | TAM.Il1r2 | Prex1 |
| Slc8a1 | 1.92E-34 | 0.43359433 | 0.747 | 0.647 | 3.72E-30 | TAM.Il1r2 | Slc8a1 |
| Lsr | 2.32E-34 | 0.45142889 | 0.368 | 0.281 | 4.49E-30 | TAM.Il1r2 | Lsr |
| Ebna1bp2 | 2.61E-34 | 0.65819527 | 0.415 | 0.331 | 5.04E-30 | TAM.Il1r2 | Ebna1bp2 |
| Ptbp1 | 2.71E-34 | 0.43761764 | 0.8 | 0.727 | 5.24E-30 | TAM.Il1r2 | Ptbp1 |
| Yrdc1 | 4.22E-34 | 0.68368623 | 0.304 | 0.232 | 8.17E-30 | TAM.Il1r2 | Yrdc |
| Diaph2 | 4.44E-34 | 0.42518874 | 0.482 | 0.404 | 8.58E-30 | TAM.Il1r2 | Diaph2 |
| Lsm4 | 7.25E-34 | 0.55032703 | 0.508 | 0.424 | 1.40E-29 | TAM.Il1r2 | Lsm4 |
| Socs3 | 2.45E-33 | 0.26041899 | 0.837 | 0.775 | 4.74E-29 | TAM.Il1r2 | Socs3 |
| Ppm1g | 3.44E-33 | 0.59058635 | 0.554 | 0.461 | 6.65E-29 | TAM.Il1r2 | Ppm1g |
| Abce1 | 4.60E-33 | 0.74974771 | 0.295 | 0.214 | 8.90E-29 | TAM.Il1r2 | Abce1 |
| Prdm11 | 6.19E-33 | 0.50391816 | 0.476 | 0.379 | 1.20E-28 | TAM.Il1r2 | Prdm1 |
| Itprid2 | 6.41E-33 | 0.4174338 | 0.73 | 0.677 | 1.24E-28 | TAM.Il1r2 | Itprid2 |
| Fem1c | 9.43E-33 | 0.55726874 | 0.427 | 0.347 | 1.82E-28 | TAM.Il1r2 | Fem1c |
| Vill | 1.07E-32 | 0.48375371 | 0.31 | 0.249 | 2.07E-28 | TAM.Il1r2 | Vill |
| Gsto1 | 1.67E-32 | 0.50119394 | 0.73 | 0.635 | 3.23E-28 | TAM.Il1r2 | Gsto1 |
| Gcsh | 2.25E-32 | 0.62986949 | 0.375 | 0.31 | 4.34E-28 | TAM.Il1r2 | Gcsh |
| Cct8 | 2.33E-32 | 0.60062206 | 0.429 | 0.348 | 4.50E-28 | TAM.Il1r2 | Cct8 |
| Runx21 | 4.78E-32 | 0.60111572 | 0.265 | 0.188 | 9.24E-28 | TAM.Il1r2 | Runx2 |
| Bop1 | 1.01E-31 | 0.72365952 | 0.303 | 0.228 | 1.94E-27 | TAM.Il1r2 | Bop1 |
| Mpc11 | 2.89E-31 | 0.51977742 | 0.516 | 0.434 | 5.59E-27 | TAM.Il1r2 | Mpc1 |
| Ythdc1 | 3.10E-31 | 0.51246635 | 0.374 | 0.319 | 6.00E-27 | TAM.Il1r2 | Ythdc1 |
| Wdfy4 | 1.24E-30 | 0.53885584 | 0.434 | 0.337 | 2.41E-26 | TAM.Il1r2 | Wdfy4 |
| Sh2d1b11 | 2.30E-30 | 0.62890765 | 0.345 | 0.248 | 4.45E-26 | TAM.Il1r2 | Sh2d1b1 |
| Pmvk1 | 1.08E-29 | 0.60388693 | 0.38 | 0.316 | 2.08E-25 | TAM.Il1r2 | Pmvk |
| Atp5md | 1.12E-29 | 0.57904751 | 0.427 | 0.355 | 2.17E-25 | TAM.Il1r2 | Atp5md |
| Nop53 | 1.50E-29 | 0.40029227 | 0.595 | 0.537 | 2.91E-25 | TAM.Il1r2 | Nop53 |
| Snu13 | 1.60E-29 | 0.55644022 | 0.557 | 0.471 | 3.09E-25 | TAM.Il1r2 | Snu13 |
| Ptpn2 | 1.36E-28 | 0.55997159 | 0.409 | 0.337 | 2.64E-24 | TAM.Il1r2 | Ptpn2 |
| Nabp1 | 1.65E-28 | 0.3698618 | 0.435 | 0.379 | 3.18E-24 | TAM.Il1r2 | Nabp1 |
| Hif1a | 2.57E-28 | 0.38700175 | 0.641 | 0.563 | 4.96E-24 | TAM.Il1r2 | Hif1a |
| Cox7b | 2.62E-28 | 0.46793079 | 0.608 | 0.526 | 5.06E-24 | TAM.Il1r2 | Cox7b |
| Atp5d | 4.02E-28 | 0.46125156 | 0.784 | 0.7 | 7.78E-24 | TAM.Il1r2 | Atp5d |
| Btbd9 | 4.16E-28 | 0.47883139 | 0.383 | 0.319 | 8.05E-24 | TAM.Il1r2 | Btbd9 |
| Nsd3 | 4.31E-28 | 0.33650334 | 0.728 | 0.672 | 8.34E-24 | TAM.Il1r2 | Nsd3 |
| Stk11 | 6.64E-28 | 0.47166431 | 0.689 | 0.619 | 1.28E-23 | TAM.Il1r2 | Stk11 |
| Far1 | 9.19E-28 | 0.42198852 | 0.592 | 0.518 | 1.78E-23 | TAM.Il1r2 | Far1 |
| Cyp4f18 | 1.12E-27 | 0.44837245 | 0.529 | 0.422 | 2.16E-23 | TAM.Il1r2 | Cyp4f18 |
| Larp1 | 1.22E-27 | 0.50891076 | 0.595 | 0.515 | 2.37E-23 | TAM.Il1r2 | Larp1 |
| Elmsan1 | 2.20E-27 | 0.51936913 | 0.298 | 0.264 | 4.25E-23 | TAM.Il1r2 | Elmsan1 |
| Hivep31 | 2.72E-27 | 0.47802361 | 0.443 | 0.368 | 5.25E-23 | TAM.Il1r2 | Hivep3 |
| Pomp | 4.72E-27 | 0.43399882 | 0.632 | 0.557 | 9.13E-23 | TAM.Il1r2 | Pomp |
| Pop4 | 5.70E-27 | 0.7244904 | 0.284 | 0.22 | 1.10E-22 | TAM.Il1r2 | Pop4 |
| Zcchc21 | 1.12E-26 | 0.47457744 | 0.414 | 0.353 | 2.16E-22 | TAM.Il1r2 | Zcchc2 |
| Acly | 1.16E-26 | 0.39746389 | 0.846 | 0.778 | 2.25E-22 | TAM.Il1r2 | Acly |
| Rbm3 | 1.19E-26 | 0.43688857 | 0.68 | 0.604 | 2.30E-22 | TAM.Il1r2 | Rbm3 |
| 7-Mar | 1.53E-26 | 0.4429709 | 0.574 | 0.507 | 2.96E-22 | TAM.Il1r2 | 7-Mar |
| Mar-32 | 3.19E-26 | 0.6681131 | 0.336 | 0.254 | 6.16E-22 | TAM.Il1r2 | 3-Mar |
| Zbtb18 | 3.87E-26 | 0.46996981 | 0.392 | 0.333 | 7.49E-22 | TAM.Il1r2 | Zbtb18 |
| Lcp11 | 7.78E-26 | 0.33067862 | 0.975 | 0.951 | 1.50E-21 | TAM.Il1r2 | Lcp1 |
| Magoh | 7.84E-26 | 0.54150163 | 0.389 | 0.317 | 1.51E-21 | TAM.Il1r2 | Magoh |
| Eif3k | 1.43E-25 | 0.36160773 | 0.777 | 0.721 | 2.76E-21 | TAM.Il1r2 | Eif3k |
| Matk1 | 2.08E-25 | 0.57339936 | 0.482 | 0.344 | 4.03E-21 | TAM.Il1r2 | Matk |
| Osm | 2.11E-25 | 0.26854922 | 0.74 | 0.67 | 4.07E-21 | TAM.Il1r2 | Osm |
| Arpc2 | 2.14E-25 | 0.3431401 | 0.783 | 0.736 | 4.14E-21 | TAM.Il1r2 | Arpc2 |
| Olr11 | 2.52E-25 | 0.51667894 | 0.271 | 0.198 | 4.87E-21 | TAM.Il1r2 | Olr1 |
| Eif3b | 3.60E-25 | 0.55047488 | 0.533 | 0.444 | 6.96E-21 | TAM.Il1r2 | Eif3b |
| Eif4a1 | 5.17E-25 | 0.43397181 | 0.763 | 0.688 | 9.99E-21 | TAM.Il1r2 | Eif4a1 |
| Ola1 | 6.04E-25 | 0.54373015 | 0.396 | 0.325 | 1.17E-20 | TAM.Il1r2 | Ola1 |
| Deptor | 6.22E-25 | 0.5536159 | 0.288 | 0.229 | 1.20E-20 | TAM.Il1r2 | Deptor |
| Msl2 | 7.04E-25 | 0.36657827 | 0.557 | 0.505 | 1.36E-20 | TAM.Il1r2 | Msl2 |
| Fbxo34 | 7.74E-25 | 0.51028056 | 0.278 | 0.227 | 1.50E-20 | TAM.Il1r2 | Fbxo34 |
| Nme1 | 8.43E-25 | 0.58901554 | 0.451 | 0.367 | 1.63E-20 | TAM.Il1r2 | Nme1 |
| Pabpc1 | 9.74E-25 | 0.39919804 | 0.686 | 0.625 | 1.88E-20 | TAM.Il1r2 | Pabpc1 |
| Hnrnpl | 3.67E-24 | 0.46427834 | 0.698 | 0.616 | 7.10E-20 | TAM.Il1r2 | Hnrnpl |
| Zfp703 | 3.75E-24 | 0.36212689 | 0.864 | 0.811 | 7.26E-20 | TAM.Il1r2 | Zfp703 |
| Tbrg4 | 1.01E-23 | 0.58917971 | 0.39 | 0.31 | 1.96E-19 | TAM.Il1r2 | Tbrg4 |
| Tet3 | 1.51E-23 | 0.40006959 | 0.536 | 0.483 | 2.92E-19 | TAM.Il1r2 | Tet3 |
| Chsy1 | 1.99E-23 | 0.47572397 | 0.382 | 0.327 | 3.85E-19 | TAM.Il1r2 | Chsy1 |
| Slc25a5 | 2.56E-23 | 0.50165399 | 0.529 | 0.455 | 4.95E-19 | TAM.Il1r2 | Slc25a5 |
| Gramd4 | 2.60E-23 | 0.55844234 | 0.261 | 0.203 | 5.03E-19 | TAM.Il1r2 | Gramd4 |
| Fasn1 | 5.59E-23 | 0.73042887 | 0.279 | 0.208 | 1.08E-18 | TAM.Il1r2 | Fasn |
| Mbtd1 | 6.52E-23 | 0.5110909 | 0.331 | 0.272 | 1.26E-18 | TAM.Il1r2 | Mbtd1 |
| Sfxn1 | 8.10E-23 | 0.65812914 | 0.334 | 0.26 | 1.57E-18 | TAM.Il1r2 | Sfxn1 |
| Gk1 | 1.20E-22 | 0.57216956 | 0.292 | 0.237 | 2.32E-18 | TAM.Il1r2 | Gk |
| Eif4e1 | 1.46E-22 | 0.45266258 | 0.623 | 0.552 | 2.82E-18 | TAM.Il1r2 | Eif4e |
| Smdt1 | 1.74E-22 | 0.42257415 | 0.739 | 0.672 | 3.36E-18 | TAM.Il1r2 | Smdt1 |
| Ube2v2 | 9.92E-22 | 0.54364074 | 0.309 | 0.261 | 1.92E-17 | TAM.Il1r2 | Ube2v2 |
| Braf | 1.23E-21 | 0.62541438 | 0.263 | 0.207 | 2.38E-17 | TAM.Il1r2 | Braf |
| Uap1 | 1.39E-21 | 0.44370234 | 0.437 | 0.383 | 2.68E-17 | TAM.Il1r2 | Uap1 |
| Eif3f | 1.97E-21 | 0.33142219 | 0.635 | 0.588 | 3.80E-17 | TAM.Il1r2 | Eif3f |
| Cd244a | 2.08E-21 | 0.31435609 | 0.378 | 0.283 | 4.02E-17 | TAM.Il1r2 | Cd244a |
| Txn2 | 3.86E-21 | 0.51650799 | 0.471 | 0.398 | 7.47E-17 | TAM.Il1r2 | Txn2 |
| Heatr11 | 8.12E-21 | 0.45797506 | 0.423 | 0.359 | 1.57E-16 | TAM.Il1r2 | Heatr1 |
| Gosr2 | 1.05E-20 | 0.37102139 | 0.526 | 0.481 | 2.03E-16 | TAM.Il1r2 | Gosr2 |
| D8Ertd738e | 1.25E-20 | 0.35562678 | 0.666 | 0.608 | 2.42E-16 | TAM.Il1r2 | D8Ertd738e |
| Npm1 | 1.52E-20 | 0.44657447 | 0.769 | 0.682 | 2.95E-16 | TAM.Il1r2 | Npm1 |
| Myl12b | 1.53E-20 | 0.36390502 | 0.648 | 0.588 | 2.97E-16 | TAM.Il1r2 | Myl12b |
| Dhx15 | 1.75E-20 | 0.39533819 | 0.608 | 0.544 | 3.38E-16 | TAM.Il1r2 | Dhx15 |
| Dip2c1 | 2.58E-20 | 0.53444683 | 0.417 | 0.333 | 5.00E-16 | TAM.Il1r2 | Dip2c |
| Trp53 | 3.00E-20 | 0.46557754 | 0.488 | 0.411 | 5.80E-16 | TAM.Il1r2 | Trp53 |
| Cct6a | 3.47E-20 | 0.55937911 | 0.433 | 0.354 | 6.71E-16 | TAM.Il1r2 | Cct6a |
| Atf1 | 4.25E-20 | 0.44966568 | 0.585 | 0.521 | 8.21E-16 | TAM.Il1r2 | Atf1 |
| Siah2 | 4.90E-20 | 0.40076131 | 0.427 | 0.369 | 9.46E-16 | TAM.Il1r2 | Siah2 |
| Etf1 | 7.43E-20 | 0.44493001 | 0.467 | 0.403 | 1.44E-15 | TAM.Il1r2 | Etf1 |
| Psme3 | 9.55E-20 | 0.47856967 | 0.594 | 0.514 | 1.85E-15 | TAM.Il1r2 | Psme3 |
| Tra2a | 9.57E-20 | 0.38248341 | 0.645 | 0.583 | 1.85E-15 | TAM.Il1r2 | Tra2a |
| Myd88 | 1.31E-19 | 0.32927376 | 0.715 | 0.657 | 2.53E-15 | TAM.Il1r2 | Myd88 |
| Ddx6 | 1.94E-19 | 0.34569187 | 0.751 | 0.702 | 3.75E-15 | TAM.Il1r2 | Ddx6 |
| Glud1 | 2.25E-19 | 0.30222722 | 0.807 | 0.758 | 4.35E-15 | TAM.Il1r2 | Glud1 |
| Med29 | 2.56E-19 | 0.45243584 | 0.323 | 0.273 | 4.95E-15 | TAM.Il1r2 | Med29 |
| Prkar2a | 2.85E-19 | 0.63669104 | 0.253 | 0.198 | 5.51E-15 | TAM.Il1r2 | Prkar2a |
| Eva1b1 | 3.08E-19 | 0.46957535 | 0.474 | 0.403 | 5.96E-15 | TAM.Il1r2 | Eva1b |
| Midn | 3.45E-19 | 0.38055179 | 0.778 | 0.721 | 6.66E-15 | TAM.Il1r2 | Midn |
| Mthfs | 4.19E-19 | 0.450019 | 0.359 | 0.313 | 8.10E-15 | TAM.Il1r2 | Mthfs |
| Zfp6221 | 7.97E-19 | 0.43664303 | 0.524 | 0.471 | 1.54E-14 | TAM.Il1r2 | Zfp622 |
| Lsm12 | 1.05E-18 | 0.47011611 | 0.478 | 0.414 | 2.03E-14 | TAM.Il1r2 | Lsm12 |
| Prmt1 | 1.21E-18 | 0.68746568 | 0.306 | 0.235 | 2.34E-14 | TAM.Il1r2 | Prmt1 |
| Tmem14c | 2.84E-18 | 0.40946894 | 0.695 | 0.628 | 5.48E-14 | TAM.Il1r2 | Tmem14c |
| Tet2 | 4.68E-18 | 0.29404426 | 0.505 | 0.452 | 9.05E-14 | TAM.Il1r2 | Tet2 |
| Ssr4 | 5.59E-18 | 0.45468703 | 0.635 | 0.564 | 1.08E-13 | TAM.Il1r2 | Ssr4 |
| Pnpo | 6.16E-18 | 0.60975869 | 0.315 | 0.255 | 1.19E-13 | TAM.Il1r2 | Pnpo |
| Pecam11 | 9.63E-18 | 0.45110575 | 0.299 | 0.237 | 1.86E-13 | TAM.Il1r2 | Pecam1 |
| Syncrip | 1.03E-17 | 0.43300847 | 0.715 | 0.638 | 1.99E-13 | TAM.Il1r2 | Syncrip |
| Mogs | 1.35E-17 | 0.62410576 | 0.279 | 0.225 | 2.62E-13 | TAM.Il1r2 | Mogs |
| Hsp90ab1 | 1.72E-17 | 0.40006645 | 0.809 | 0.737 | 3.32E-13 | TAM.Il1r2 | Hsp90ab1 |
| Ncf4 | 1.99E-17 | 0.32240059 | 0.76 | 0.714 | 3.85E-13 | TAM.Il1r2 | Ncf4 |
| Sra11 | 2.51E-17 | 0.36150807 | 0.621 | 0.555 | 4.84E-13 | TAM.Il1r2 | Sra1 |
| Ube2j2 | 2.87E-17 | 0.51842567 | 0.39 | 0.329 | 5.55E-13 | TAM.Il1r2 | Ube2j2 |
| Sfpq | 3.35E-17 | 0.36382039 | 0.749 | 0.684 | 6.48E-13 | TAM.Il1r2 | Sfpq |
| Snrpg | 4.98E-17 | 0.51705815 | 0.334 | 0.272 | 9.63E-13 | TAM.Il1r2 | Snrpg |
| Ampd21 | 5.89E-17 | 0.58297752 | 0.313 | 0.252 | 1.14E-12 | TAM.Il1r2 | Ampd2 |
| Pprc1 | 6.62E-17 | 0.57746754 | 0.324 | 0.263 | 1.28E-12 | TAM.Il1r2 | Pprc1 |
| Actr3 | 7.91E-17 | 0.33459599 | 0.689 | 0.629 | 1.53E-12 | TAM.Il1r2 | Actr3 |
| Atic | 1.64E-16 | 0.59622239 | 0.438 | 0.349 | 3.17E-12 | TAM.Il1r2 | Atic |
| Swi | 1.71E-16 | 0.33768624 | 0.482 | 0.418 | 3.31E-12 | TAM.Il1r2 | Swi |
| U2af1 | 1.93E-16 | 0.54299428 | 0.351 | 0.291 | 3.74E-12 | TAM.Il1r2 | U2af1 |
| Nubp1 | 2.12E-16 | 0.40751431 | 0.487 | 0.421 | 4.10E-12 | TAM.Il1r2 | Nubp1 |
| Irf2bp2 | 2.24E-16 | 0.26739738 | 0.964 | 0.935 | 4.33E-12 | TAM.Il1r2 | Irf2bp2 |
| Kdm7a | 3.14E-16 | 0.27171792 | 0.725 | 0.685 | 6.07E-12 | TAM.Il1r2 | Kdm7a |
| Ifi30 | 3.63E-16 | 0.32560803 | 0.963 | 0.931 | 7.02E-12 | TAM.Il1r2 | Ifi30 |
| Cdkn1b | 4.99E-16 | 0.31946126 | 0.554 | 0.516 | 9.64E-12 | TAM.Il1r2 | Cdkn1b |
| Plcg2 | 4.99E-16 | 0.31918222 | 0.585 | 0.519 | 9.66E-12 | TAM.Il1r2 | Plcg2 |

| | | | | | | | |
|---|---|---|---|---|---|---|---|
| Ndrg1 | 6.20E-16 | 0.28375485 | 0.436 | 0.341 | 1.20E-11 | TAM.Il1r2 | Ndrg1 |
| Plxdc21 | 6.60E-16 | 0.55579018 | 0.276 | 0.235 | 1.28E-11 | TAM.Il1r2 | Plxdc2 |
| Sh3bgrl31 | 8.47E-16 | 0.33841259 | 0.941 | 0.898 | 1.64E-11 | TAM.Il1r2 | Sh3bgrl3 |
| Snrpd1 | 9.78E-16 | 0.52610344 | 0.411 | 0.33 | 1.89E-11 | TAM.Il1r2 | Snrpd1 |
| Tbc1d1 | 1.08E-15 | 0.46815433 | 0.417 | 0.361 | 2.10E-11 | TAM.Il1r2 | Tbc1d1 |
| Atp5g1 | 1.31E-15 | 0.57388711 | 0.4 | 0.336 | 2.54E-11 | TAM.Il1r2 | Atp5g1 |
| Bak1 | 1.89E-15 | 0.40592838 | 0.651 | 0.582 | 3.66E-11 | TAM.Il1r2 | Bak1 |
| Paf1 | 1.96E-15 | 0.44593064 | 0.386 | 0.341 | 3.80E-11 | TAM.Il1r2 | Paf1 |
| Ube2j1 | 1.98E-15 | 0.52666791 | 0.299 | 0.257 | 3.84E-11 | TAM.Il1r2 | Ube2j1 |
| Dnajb6 | 2.10E-15 | 0.41455283 | 0.59 | 0.535 | 4.06E-11 | TAM.Il1r2 | Dnajb6 |
| Tomm40 | 2.38E-15 | 0.54307826 | 0.454 | 0.378 | 4.60E-11 | TAM.Il1r2 | Tomm40 |
| Taf1d | 2.79E-15 | 0.44611938 | 0.484 | 0.428 | 5.39E-11 | TAM.Il1r2 | Taf1d |
| Med21 | 3.01E-15 | 0.53297925 | 0.253 | 0.226 | 5.82E-11 | TAM.Il1r2 | Med21 |
| Sf3b6 | 3.69E-15 | 0.36558858 | 0.52 | 0.469 | 7.13E-11 | TAM.Il1r2 | Sf3b6 |
| Mllt61 | 3.81E-15 | 0.39795633 | 0.568 | 0.493 | 7.36E-11 | TAM.Il1r2 | Mllt6 |
| Tcerg1 | 6.37E-15 | 0.52735437 | 0.299 | 0.243 | 1.23E-10 | TAM.Il1r2 | Tcerg1 |
| Gpbp1 | 9.06E-15 | 0.44328207 | 0.361 | 0.306 | 1.75E-10 | TAM.Il1r2 | Gpbp1 |
| Zmiz2 | 1.13E-14 | 0.4271621 | 0.584 | 0.505 | 2.18E-10 | TAM.Il1r2 | Zmiz2 |
| Taldo11 | 1.76E-14 | 0.35317031 | 0.868 | 0.777 | 3.40E-10 | TAM.Il1r2 | Taldo1 |
| Ppp2ca | 2.06E-14 | 0.27796781 | 0.745 | 0.705 | 3.99E-10 | TAM.Il1r2 | Ppp2ca |
| Lactb | 2.38E-14 | 0.4967206 | 0.333 | 0.284 | 4.60E-10 | TAM.Il1r2 | Lactb |
| Smarca5 | 3.10E-14 | 0.49775759 | 0.338 | 0.294 | 6.00E-10 | TAM.Il1r2 | Smarca5 |
| Igsf82 | 3.32E-14 | 0.43653396 | 0.622 | 0.538 | 6.43E-10 | TAM.Il1r2 | Igsf8 |
| Csk | 3.94E-14 | 0.32428896 | 0.659 | 0.61 | 7.62E-10 | TAM.Il1r2 | Csk |
| Socs22 | 4.30E-14 | 0.58648564 | 0.386 | 0.305 | 8.32E-10 | TAM.Il1r2 | Socs2 |
| Sipa1l3 | 4.32E-14 | 0.43240249 | 0.381 | 0.324 | 8.35E-10 | TAM.Il1r2 | Sipa1l3 |
| Nup153 | 4.68E-14 | 0.48000877 | 0.401 | 0.344 | 9.05E-10 | TAM.Il1r2 | Nup153 |
| Vopp1 | 5.37E-14 | 0.54975116 | 0.404 | 0.337 | 1.04E-09 | TAM.Il1r2 | Vopp1 |
| Baz1a | 6.05E-14 | 0.33747334 | 0.579 | 0.523 | 1.17E-09 | TAM.Il1r2 | Baz1a |
| Ripk31 | 6.27E-14 | 0.41279156 | 0.378 | 0.322 | 1.21E-09 | TAM.Il1r2 | Ripk3 |
| Wdr74 | 8.07E-14 | 0.60832587 | 0.254 | 0.204 | 1.56E-09 | TAM.Il1r2 | Wdr74 |
| Prdx3 | 8.31E-14 | 0.52545949 | 0.362 | 0.308 | 1.61E-09 | TAM.Il1r2 | Prdx3 |
| Scap1 | 1.02E-13 | 0.59851723 | 0.254 | 0.21 | 1.98E-09 | TAM.Il1r2 | Scap |
| Ppig1 | 1.03E-13 | 0.46441805 | 0.383 | 0.343 | 1.98E-09 | TAM.Il1r2 | Ppig |
| Hnrnpdl | 1.20E-13 | 0.41118411 | 0.331 | 0.281 | 2.31E-09 | TAM.Il1r2 | Hnrnpdl |
| Errfi1 | 1.27E-13 | 0.25548859 | 0.395 | 0.365 | 2.45E-09 | TAM.Il1r2 | Errfi1 |
| Park7 | 1.82E-13 | 0.41886417 | 0.661 | 0.579 | 3.52E-09 | TAM.Il1r2 | Park7 |
| Ctnnbip1 | 1.98E-13 | 0.46997192 | 0.295 | 0.249 | 3.82E-09 | TAM.Il1r2 | Ctnnbip1 |
| Elavl1 | 2.01E-13 | 0.36097824 | 0.584 | 0.532 | 3.88E-09 | TAM.Il1r2 | Elavl1 |
| Chd11 | 2.13E-13 | 0.37185601 | 0.459 | 0.405 | 4.11E-09 | TAM.Il1r2 | Chd1 |
| Gtpbp1 | 2.22E-13 | 0.42879738 | 0.326 | 0.278 | 4.29E-09 | TAM.Il1r2 | Gtpbp1 |
| Snrpe | 2.97E-13 | 0.50238772 | 0.357 | 0.296 | 5.74E-09 | TAM.Il1r2 | Snrpe |
| Lipe | 3.45E-13 | 0.32386109 | 0.32 | 0.28 | 6.66E-09 | TAM.Il1r2 | Lipe |
| Mark3 | 3.84E-13 | 0.37267453 | 0.34 | 0.298 | 7.43E-09 | TAM.Il1r2 | Mark3 |
| Ist1 | 4.46E-13 | 0.27742568 | 0.764 | 0.724 | 8.63E-09 | TAM.Il1r2 | Ist1 |
| Opa3 | 4.47E-13 | 0.50005796 | 0.292 | 0.253 | 8.65E-09 | TAM.Il1r2 | Opa3 |
| Plpp5 | 5.29E-13 | 0.46055092 | 0.365 | 0.311 | 1.02E-08 | TAM.Il1r2 | Plpp5 |
| Tmem2731 | 9.66E-13 | 0.56392942 | 0.278 | 0.214 | 1.87E-08 | TAM.Il1r2 | Tmem273 |
| Idh3a | 1.16E-12 | 0.55545953 | 0.368 | 0.294 | 2.25E-08 | TAM.Il1r2 | Idh3a |
| Zc3h12a1 | 1.33E-12 | 0.32590661 | 0.818 | 0.723 | 2.57E-08 | TAM.Il1r2 | Zc3h12a |
| Cyp511 | 1.33E-12 | 0.43269856 | 0.375 | 0.323 | 2.58E-08 | TAM.Il1r2 | Cyp51 |
| Grk51 | 1.59E-12 | 0.3418332 | 0.318 | 0.265 | 3.07E-08 | TAM.Il1r2 | Grk5 |
| Tcf7l2 | 1.75E-12 | 0.4566774 | 0.404 | 0.329 | 3.38E-08 | TAM.Il1r2 | Tcf7l2 |
| Rassf5 | 2.15E-12 | 0.30337115 | 0.522 | 0.462 | 4.16E-08 | TAM.Il1r2 | Rassf5 |
| Hnrnpd | 2.39E-12 | 0.35124119 | 0.562 | 0.509 | 4.62E-08 | TAM.Il1r2 | Hnrnpd |
| Emc6 | 3.10E-12 | 0.49804707 | 0.361 | 0.305 | 5.99E-08 | TAM.Il1r2 | Emc6 |
| Pald11 | 4.00E-12 | 0.48047848 | 0.419 | 0.346 | 7.74E-08 | TAM.Il1r2 | Pald1 |
| Ubash3b1 | 4.95E-12 | 0.43261727 | 0.409 | 0.345 | 9.56E-08 | TAM.Il1r2 | Ubash3b |
| Slc35c21 | 5.54E-12 | 0.49880274 | 0.389 | 0.327 | 1.07E-07 | TAM.Il1r2 | Slc35c2 |
| Rcc21 | 6.66E-12 | 0.51989916 | 0.424 | 0.352 | 1.29E-07 | TAM.Il1r2 | Rcc2 |
| Sdf2l11 | 9.19E-12 | 0.39895738 | 0.622 | 0.548 | 1.78E-07 | TAM.Il1r2 | Sdf2l1 |
| Dpy19l1 | 1.18E-11 | 0.43455419 | 0.268 | 0.219 | 2.29E-07 | TAM.Il1r2 | Dpy19l1 |
| Cox8a | 1.21E-11 | 0.41886597 | 0.755 | 0.689 | 2.34E-07 | TAM.Il1r2 | Cox8a |
| Rassf1 | 1.35E-11 | 0.32469945 | 0.534 | 0.485 | 2.61E-07 | TAM.Il1r2 | Rassf1 |
| Itprip | 1.43E-11 | 0.37547726 | 0.357 | 0.298 | 2.77E-07 | TAM.Il1r2 | Itprip |
| Jmy1 | 1.65E-11 | 0.45585437 | 0.402 | 0.377 | 3.18E-07 | TAM.Il1r2 | Jmy |
| Pdlim71 | 3.11E-11 | 0.37847875 | 0.396 | 0.338 | 6.02E-07 | TAM.Il1r2 | Pdlim7 |
| Eif3l | 3.39E-11 | 0.40879191 | 0.611 | 0.541 | 6.55E-07 | TAM.Il1r2 | Eif3l |
| Borcs8 | 3.98E-11 | 0.41372397 | 0.321 | 0.275 | 7.69E-07 | TAM.Il1r2 | Borcs8 |
| Lrrc8c | 3.98E-11 | 0.3555847 | 0.396 | 0.35 | 7.69E-07 | TAM.Il1r2 | Lrrc8c |
| Hnrnph1 | 5.05E-11 | 0.37086063 | 0.43 | 0.381 | 9.76E-07 | TAM.Il1r2 | Hnrnph1 |
| Tmem147 | 5.27E-11 | 0.51146617 | 0.33 | 0.283 | 1.02E-06 | TAM.Il1r2 | Tmem147 |
| Phb2 | 6.02E-11 | 0.49127491 | 0.468 | 0.397 | 1.16E-06 | TAM.Il1r2 | Phb2 |
| Lrp51 | 7.17E-11 | 0.33698453 | 0.436 | 0.376 | 1.39E-06 | TAM.Il1r2 | Lrp5 |
| Ccdc12 | 7.48E-11 | 0.27148318 | 0.513 | 0.477 | 1.45E-06 | TAM.Il1r2 | Ccdc12 |
| Sec24b | 7.62E-11 | 0.3751453 | 0.501 | 0.456 | 1.47E-06 | TAM.Il1r2 | Sec24b |
| Psmb10 | 1.09E-10 | 0.32949373 | 0.517 | 0.464 | 2.11E-06 | TAM.Il1r2 | Psmb10 |
| Noct | 1.18E-10 | 0.33686973 | 0.303 | 0.26 | 2.29E-06 | TAM.Il1r2 | Noct |
| Cysltr1 | 1.22E-10 | 0.34727098 | 0.276 | 0.234 | 2.36E-06 | TAM.Il1r2 | Cysltr1 |
| Akna | 1.36E-10 | 0.27512078 | 0.359 | 0.313 | 2.63E-06 | TAM.Il1r2 | Akna |
| Nudt21 | 1.54E-10 | 0.41223401 | 0.342 | 0.296 | 2.97E-06 | TAM.Il1r2 | Nudt21 |
| Cct5 | 1.56E-10 | 0.46415634 | 0.269 | 0.226 | 3.01E-06 | TAM.Il1r2 | Cct5 |
| Ltv1 | 1.93E-10 | 0.46281823 | 0.269 | 0.225 | 3.73E-06 | TAM.Il1r2 | Ltv1 |
| Scaf4 | 2.09E-10 | 0.39028385 | 0.35 | 0.315 | 4.04E-06 | TAM.Il1r2 | Scaf4 |
| Cpeb21 | 3.92E-10 | 0.35235843 | 0.482 | 0.431 | 7.57E-06 | TAM.Il1r2 | Cpeb2 |
| Dock51 | 4.25E-10 | 0.39630386 | 0.264 | 0.203 | 8.21E-06 | TAM.Il1r2 | Dock5 |
| Grk3 | 5.08E-10 | 0.37821217 | 0.351 | 0.29 | 9.83E-06 | TAM.Il1r2 | Grk3 |
| Cct7 | 5.46E-10 | 0.46329453 | 0.408 | 0.354 | 1.06E-05 | TAM.Il1r2 | Cct7 |
| Sowahc | 6.61E-10 | 0.31059363 | 0.569 | 0.504 | 1.28E-05 | TAM.Il1r2 | Sowahc |
| Med13l | 7.17E-10 | 0.34048494 | 0.541 | 0.479 | 1.39E-05 | TAM.Il1r2 | Med13l |
| Gnl31 | 7.75E-10 | 0.48290002 | 0.383 | 0.321 | 1.50E-05 | TAM.Il1r2 | Gnl3 |
| Tcp1 | 8.19E-10 | 0.45950868 | 0.519 | 0.456 | 1.58E-05 | TAM.Il1r2 | Tcp1 |
| Pum3 | 8.73E-10 | 0.45577094 | 0.34 | 0.291 | 1.69E-05 | TAM.Il1r2 | Pum3 |
| Wdr83os | 1.12E-09 | 0.39367226 | 0.64 | 0.579 | 2.17E-05 | TAM.Il1r2 | Wdr83os |
| Gart | 1.96E-09 | 0.53111574 | 0.333 | 0.273 | 3.79E-05 | TAM.Il1r2 | Gart |
| Lbr | 1.97E-09 | 0.33595482 | 0.349 | 0.273 | 3.82E-05 | TAM.Il1r2 | Lbr |
| Rai141 | 2.27E-09 | 0.39141993 | 0.591 | 0.492 | 4.38E-05 | TAM.Il1r2 | Rai14 |
| Ncl | 2.56E-09 | 0.36272124 | 0.629 | 0.576 | 4.95E-05 | TAM.Il1r2 | Ncl |
| Hspa4 | 2.59E-09 | 0.44977898 | 0.427 | 0.364 | 5.01E-05 | TAM.Il1r2 | Hspa4 |
| Sh3bp1 | 2.93E-09 | 0.26094315 | 0.577 | 0.524 | 5.66E-05 | TAM.Il1r2 | Sh3bp1 |
| Mical11 | 3.17E-09 | 0.39835817 | 0.318 | 0.27 | 6.14E-05 | TAM.Il1r2 | Mical1 |
| Acer3 | 4.19E-09 | 0.30869508 | 0.321 | 0.285 | 8.11E-05 | TAM.Il1r2 | Acer3 |
| Bloc1s2 | 4.47E-09 | 0.38321882 | 0.377 | 0.337 | 8.64E-05 | TAM.Il1r2 | Bloc1s2 |
| Ddx54 | 5.54E-09 | 0.37893691 | 0.449 | 0.405 | 0.00010705 | TAM.Il1r2 | Ddx54 |
| Ripk2 | 7.06E-09 | 0.31211915 | 0.277 | 0.234 | 0.00013649 | TAM.Il1r2 | Ripk2 |
| Polr1c | 7.12E-09 | 0.46318959 | 0.314 | 0.274 | 0.00013762 | TAM.Il1r2 | Polr1c |
| Aprt1 | 7.31E-09 | 0.4548845 | 0.445 | 0.378 | 0.00014126 | TAM.Il1r2 | Aprt |
| Bud23 | 7.67E-09 | 0.49273545 | 0.278 | 0.232 | 0.0001483 | TAM.Il1r2 | Bud23 |
| Atp5b | 1.05E-08 | 0.45659227 | 0.731 | 0.644 | 0.0002023 | TAM.Il1r2 | Atp5b |
| Enoph1 | 1.34E-08 | 0.56219501 | 0.256 | 0.218 | 0.00025816 | TAM.Il1r2 | Enoph1 |
| Josd11 | 1.82E-08 | 0.47896696 | 0.25 | 0.214 | 0.00035162 | TAM.Il1r2 | Josd1 |
| Set | 1.95E-08 | 0.51584493 | 0.394 | 0.33 | 0.00037789 | TAM.Il1r2 | Set |
| Atp5j2 | 1.96E-08 | 0.38158242 | 0.673 | 0.604 | 0.00037884 | TAM.Il1r2 | Atp5j2 |
| Acat1 | 1.97E-08 | 0.41097589 | 0.34 | 0.292 | 0.00038012 | TAM.Il1r2 | Acat1 |
| Mob3b1 | 1.99E-08 | 0.40547473 | 0.259 | 0.21 | 0.00038567 | TAM.Il1r2 | Mob3b |
| Timm231 | 2.20E-08 | 0.47778389 | 0.331 | 0.277 | 0.00042472 | TAM.Il1r2 | Timm23 |
| Prpf31 | 2.51E-08 | 0.47091567 | 0.37 | 0.322 | 0.00048508 | TAM.Il1r2 | Prpf31 |
| Eif3a | 2.64E-08 | 0.38973735 | 0.465 | 0.408 | 0.00051018 | TAM.Il1r2 | Eif3a |
| Snrpb | 2.92E-08 | 0.34960542 | 0.453 | 0.409 | 0.00056362 | TAM.Il1r2 | Snrpb |
| Pbxip11 | 3.50E-08 | 0.38924021 | 0.3 | 0.251 | 0.00067757 | TAM.Il1r2 | Pbxip1 |
| Nsa2 | 4.35E-08 | 0.37799455 | 0.272 | 0.244 | 0.00084012 | TAM.Il1r2 | Nsa2 |
| Eif3d | 4.61E-08 | 0.3823649 | 0.504 | 0.458 | 0.00089172 | TAM.Il1r2 | Eif3d |
| Inpp5b | 5.49E-08 | 0.42565713 | 0.306 | 0.265 | 0.00106158 | TAM.Il1r2 | Inpp5b |
| Ptger4 | 6.21E-08 | 0.26776342 | 0.365 | 0.369 | 0.00120063 | TAM.Il1r2 | Ptger4 |
| Stat3 | 6.23E-08 | 0.27923622 | 0.743 | 0.7 | 0.00120514 | TAM.Il1r2 | Stat3 |
| Casp6 | 6.33E-08 | 0.39488656 | 0.354 | 0.32 | 0.00122316 | TAM.Il1r2 | Casp6 |
| Lrrc591 | 8.66E-08 | 0.39242224 | 0.725 | 0.653 | 0.0016748 | TAM.Il1r2 | Lrrc59 |
| Srpk1 | 8.92E-08 | 0.40710999 | 0.441 | 0.397 | 0.001725 | TAM.Il1r2 | Srpk1 |
| Card9 | 1.16E-07 | 0.30294612 | 0.469 | 0.431 | 0.00224996 | TAM.Il1r2 | Card9 |
| Psmb2 | 1.20E-07 | 0.37391465 | 0.588 | 0.54 | 0.00231846 | TAM.Il1r2 | Psmb2 |
| Psma1 | 1.31E-07 | 0.33226305 | 0.569 | 0.523 | 0.00252546 | TAM.Il1r2 | Psma1 |
| Psmd14 | 1.31E-07 | 0.3812604 | 0.421 | 0.372 | 0.00253152 | TAM.Il1r2 | Psmd14 |
| Ndufb6 | 1.48E-07 | 0.48998736 | 0.572 | 0.491 | 0.00285524 | TAM.Il1r2 | Ndufb6 |
| Smndc1 | 1.68E-07 | 0.36061613 | 0.312 | 0.28 | 0.00323993 | TAM.Il1r2 | Smndc1 |
| Mrtfa | 1.81E-07 | 0.39478137 | 0.433 | 0.399 | 0.00350901 | TAM.Il1r2 | Mrtfa |
| Atp11b | 1.89E-07 | 0.32705154 | 0.288 | 0.255 | 0.00365703 | TAM.Il1r2 | Atp11b |
| Arl5c | 1.90E-07 | 0.27767719 | 0.331 | 0.33 | 0.00367095 | TAM.Il1r2 | Arl5c |
| Tmed5 | 1.96E-07 | 0.35132737 | 0.537 | 0.489 | 0.00379605 | TAM.Il1r2 | Tmed5 |
| Csf2ra1 | 2.51E-07 | 0.32573967 | 0.932 | 0.867 | 0.00486244 | TAM.Il1r2 | Csf2ra |
| Cluh | 2.59E-07 | 0.50708007 | 0.328 | 0.267 | 0.00501605 | TAM.Il1r2 | Cluh |
| Tbkbp1 | 2.95E-07 | 0.33047564 | 0.306 | 0.267 | 0.00570015 | TAM.Il1r2 | Tbkbp1 |
| Arf2 | 2.96E-07 | 0.30280124 | 0.394 | 0.375 | 0.00572444 | TAM.Il1r2 | Arf2 |
| Unc45a | 3.13E-07 | 0.39948682 | 0.253 | 0.221 | 0.00605758 | TAM.Il1r2 | Unc45a |
| Pak1ip1 | 3.39E-07 | 0.3865973 | 0.362 | 0.322 | 0.00655208 | TAM.Il1r2 | Pak1ip1 |
| Donson | 3.60E-07 | 0.43145655 | 0.291 | 0.252 | 0.00695835 | TAM.Il1r2 | Donson |
| Dok2 | 3.74E-07 | 0.38806694 | 0.534 | 0.482 | 0.00722331 | TAM.Il1r2 | Dok2 |
| Nsun2 | 3.87E-07 | 0.49394219 | 0.375 | 0.315 | 0.00748003 | TAM.Il1r2 | Nsun2 |
| Ddx39 | 4.37E-07 | 0.40695347 | 0.404 | 0.34 | 0.00845305 | TAM.Il1r2 | Ddx39 |
| Sf3b3 | 4.58E-07 | 0.3797257 | 0.447 | 0.398 | 0.00886038 | TAM.Il1r2 | Sf3b3 |
| Plekha1 | 5.11E-07 | 0.37151333 | 0.343 | 0.307 | 0.00987429 | TAM.Il1r2 | Plekha1 |
| Clasp21 | 6.22E-07 | 0.40354133 | 0.338 | 0.308 | 0.01203479 | TAM.Il1r2 | Clasp2 |
| Nrd1 | 6.25E-07 | 0.32657818 | 0.562 | 0.521 | 0.0120897 | TAM.Il1r2 | Nrd1 |
| Bccip | 7.56E-07 | 0.40768623 | 0.339 | 0.302 | 0.01460695 | TAM.Il1r2 | Bccip |
| Zc3h15 | 7.67E-07 | 0.39597806 | 0.316 | 0.28 | 0.014831 | TAM.Il1r2 | Zc3h15 |
| Cers6 | 7.81E-07 | 0.25996151 | 0.575 | 0.509 | 0.01509957 | TAM.Il1r2 | Cers6 |
| Ncbp1 | 9.18E-07 | 0.46972606 | 0.259 | 0.222 | 0.01773953 | TAM.Il1r2 | Ncbp1 |
| Irak21 | 1.03E-06 | 0.33864718 | 0.65 | 0.58 | 0.01984032 | TAM.Il1r2 | Irak2 |
| Txndc11 | 1.09E-06 | 0.39151281 | 0.407 | 0.371 | 0.02107448 | TAM.Il1r2 | Txndc11 |
| Abhd17a | 1.23E-06 | 0.36603996 | 0.449 | 0.4 | 0.02371717 | TAM.Il1r2 | Abhd17a |
| Cpsf6 | 1.42E-06 | 0.33927991 | 0.493 | 0.444 | 0.02752915 | TAM.Il1r2 | Cpsf6 |
| Ralgapa1 | 1.44E-06 | 0.29825418 | 0.272 | 0.244 | 0.0279063 | TAM.Il1r2 | Ralgapa1 |
| Zfp385a1 | 1.55E-06 | 0.33093785 | 0.528 | 0.476 | 0.02987327 | TAM.Il1r2 | Zfp385a |
| Cct3 | 1.73E-06 | 0.44959812 | 0.411 | 0.36 | 0.0335342 | TAM.Il1r2 | Cct3 |
| Pold4 | 1.93E-06 | 0.35801941 | 0.509 | 0.462 | 0.0372982 | TAM.Il1r2 | Pold4 |
| Dnajc15 | 2.23E-06 | 0.42420535 | 0.44 | 0.397 | 0.04316569 | TAM.Il1r2 | Dnajc15 |
| Psme2 | 2.25E-06 | 0.32447643 | 0.448 | 0.402 | 0.04355962 | TAM.Il1r2 | Psme2 |
| Ddt | 2.34E-06 | 0.48840836 | 0.31 | 0.276 | 0.04533316 | TAM.Il1r2 | Ddt |
| Arhgap9 | 2.62E-06 | 0.31212399 | 0.618 | 0.573 | 0.05058286 | TAM.Il1r2 | Arhgap9 |
| Eif5 | 2.70E-06 | 0.35702085 | 0.379 | 0.346 | 0.05225533 | TAM.Il1r2 | Eif5 |
| Agtpbp1 | 2.75E-06 | 0.2840323 | 0.274 | 0.247 | 0.05309924 | TAM.Il1r2 | Agtpbp1 |
| Vars | 2.90E-06 | 0.46483752 | 0.389 | 0.314 | 0.05601472 | TAM.Il1r2 | Vars |
| Hivep1 | 3.11E-06 | 0.3261414 | 0.448 | 0.409 | 0.06011064 | TAM.Il1r2 | Hivep1 |
| Tnfsf9 | 3.39E-06 | 0.27665984 | 0.379 | 0.318 | 0.06544754 | TAM.Il1r2 | Tnfsf9 |
| Prdx61 | 3.45E-06 | 0.34026414 | 0.381 | 0.323 | 0.06663685 | TAM.Il1r2 | Prdx6 |
| Ube2e1 | 4.02E-06 | 0.42235129 | 0.333 | 0.295 | 0.07765426 | TAM.Il1r2 | Ube2e1 |
| Kars | 4.03E-06 | 0.40867759 | 0.441 | 0.387 | 0.07789707 | TAM.Il1r2 | Kars |
| Tmem88 | 4.38E-06 | 0.37214377 | 0.385 | 0.317 | 0.08476604 | TAM.Il1r2 | Tmem88 |
| Psd41 | 4.75E-06 | 0.39758177 | 0.485 | 0.439 | 0.09185039 | TAM.Il1r2 | Psd4 |
| Tomm20 | 6.61E-06 | 0.38659499 | 0.404 | 0.358 | 0.12780636 | TAM.Il1r2 | Tomm20 |
| Stard7 | 7.14E-06 | 0.46601938 | 0.381 | 0.332 | 0.13806756 | TAM.Il1r2 | Stard7 |
| Nop10 | 7.35E-06 | 0.45950358 | 0.427 | 0.37 | 0.14219978 | TAM.Il1r2 | Nop10 |
| Lemd3 | 8.16E-06 | 0.35583628 | 0.302 | 0.279 | 0.15782412 | TAM.Il1r2 | Lemd3 |
| Mob3a | 8.28E-06 | 0.38218256 | 0.31 | 0.278 | 0.16016904 | TAM.Il1r2 | Mob3a |
| Tra2b | 8.40E-06 | 0.31372745 | 0.482 | 0.441 | 0.16240329 | TAM.Il1r2 | Tra2b |
| Nrbp1 | 8.46E-06 | 0.3162235 | 0.45 | 0.424 | 0.16353013 | TAM.Il1r2 | Nrbp1 |
| Sub1 | 8.74E-06 | 0.3270061 | 0.417 | 0.38 | 0.16900521 | TAM.Il1r2 | Sub1 |
| Hnrnpm | 9.05E-06 | 0.29876132 | 0.785 | 0.738 | 0.1748958 | TAM.Il1r2 | Hnrnpm |
| Arid5a | 9.09E-06 | 0.25569505 | 0.283 | 0.261 | 0.17572686 | TAM.Il1r2 | Arid5a |
| Hnrnpab | 9.48E-06 | 0.38246824 | 0.589 | 0.52 | 0.18327421 | TAM.Il1r2 | Hnrnpab |
| Trim27 | 9.90E-06 | 0.37976157 | 0.365 | 0.319 | 0.19149376 | TAM.Il1r2 | Trim27 |
| Psmb3 | 1.08E-05 | 0.41252915 | 0.266 | 0.231 | 0.20820072 | TAM.Il1r2 | Psmb3 |
| Srf | 1.32E-05 | 0.2989926 | 0.449 | 0.423 | 0.25525117 | TAM.Il1r2 | Srf |
| Cox5a | 1.42E-05 | 0.40981571 | 0.707 | 0.625 | 0.2744264 | TAM.Il1r2 | Cox5a |
| Nt5c | 1.51E-05 | 0.3188462 | 0.456 | 0.416 | 0.29243121 | TAM.Il1r2 | Nt5c |
| Tuba1b | 1.71E-05 | 0.30851974 | 0.659 | 0.581 | 0.33067911 | TAM.Il1r2 | Tuba1b |
| Nop56 | 1.72E-05 | 0.39429392 | 0.427 | 0.37 | 0.33259278 | TAM.Il1r2 | Nop56 |
| Slc39a111 | 1.91E-05 | 0.5234211 | 0.364 | 0.316 | 0.37022421 | TAM.Il1r2 | Slc39a11 |
| Src2 | 2.08E-05 | 0.26314455 | 0.637 | 0.554 | 0.40171286 | TAM.Il1r2 | Src |
| Bcap29 | 2.14E-05 | 0.55383764 | 0.26 | 0.22 | 0.41315358 | TAM.Il1r2 | Bcap29 |
| Eef1d | 2.28E-05 | 0.37422481 | 0.333 | 0.298 | 0.44125754 | TAM.Il1r2 | Eef1d |
| Tsg101 | 2.45E-05 | 0.30817712 | 0.335 | 0.302 | 0.47306346 | TAM.Il1r2 | Tsg101 |
| Ipo4 | 2.51E-05 | 0.44490687 | 0.305 | 0.27 | 0.48476351 | TAM.Il1r2 | Ipo4 |
| Taf10 | 2.58E-05 | 0.35838061 | 0.299 | 0.269 | 0.49895277 | TAM.Il1r2 | Taf10 |
| Tomm5 | 2.68E-05 | 0.39567981 | 0.408 | 0.357 | 0.51831789 | TAM.Il1r2 | Tomm5 |
| Srsf1 | 2.81E-05 | 0.3541743 | 0.558 | 0.497 | 0.54362082 | TAM.Il1r2 | Srsf1 |
| Atp5mpl | 2.88E-05 | 0.40703813 | 0.502 | 0.452 | 0.55638213 | TAM.Il1r2 | Atp5mpl |
| Gatm1 | 2.92E-05 | 0.25907088 | 0.468 | 0.38 | 0.56438259 | TAM.Il1r2 | Gatm |
| Mcts1 | 2.92E-05 | 0.36143362 | 0.335 | 0.303 | 0.56501794 | TAM.Il1r2 | Mcts1 |
| Prdx21 | 3.48E-05 | 0.38099571 | 0.657 | 0.604 | 0.67333278 | TAM.Il1r2 | Prdx2 |
| Zswim4 | 3.55E-05 | 0.25604759 | 0.441 | 0.398 | 0.68616823 | TAM.Il1r2 | Zswim4 |
| Tm6sf1 | 3.66E-05 | 0.25199302 | 0.698 | 0.662 | 0.7078624 | TAM.Il1r2 | Tm6sf1 |
| Atp8b41 | 3.69E-05 | 0.27265336 | 0.25 | 0.212 | 0.71256494 | TAM.Il1r2 | Atp8b4 |
| Prcp | 3.71E-05 | 0.32209586 | 0.553 | 0.514 | 0.71763665 | TAM.Il1r2 | Prcp |
| Slc35e1 | 3.83E-05 | 0.33069089 | 0.419 | 0.386 | 0.74139455 | TAM.Il1r2 | Slc35e1 |
| Zbtb111 | 4.05E-05 | 0.3036683 | 0.397 | 0.355 | 0.78208868 | TAM.Il1r2 | Zbtb11 |
| Polr1d | 4.42E-05 | 0.3200887 | 0.565 | 0.524 | 0.85431954 | TAM.Il1r2 | Polr1d |
| Prpf191 | 4.50E-05 | 0.3930838 | 0.394 | 0.342 | 0.87013593 | TAM.Il1r2 | Prpf19 |
| Ube2d3 | 4.97E-05 | 0.27775987 | 0.579 | 0.544 | 0.96081124 | TAM.Il1r2 | Ube2d3 |
| Ptges3 | 6.58E-05 | 0.39029387 | 0.421 | 0.37 | 1 | TAM.Il1r2 | Ptges3 |
| Hspa14 | 8.30E-05 | 0.40671846 | 0.307 | 0.269 | 1 | TAM.Il1r2 | Hspa14 |
| Top1 | 8.87E-05 | 0.30085246 | 0.596 | 0.553 | 1 | TAM.Il1r2 | Top1 |
| Khdc4 | 9.09E-05 | 0.2536514 | 0.42 | 0.398 | 1 | TAM.Il1r2 | Khdc4 |
| Eif3h | 9.85E-05 | 0.3199024 | 0.408 | 0.377 | 1 | TAM.Il1r2 | Eif3h |
| Cct4 | 0.00010773 | 0.38762719 | 0.415 | 0.367 | 1 | TAM.Il1r2 | Cct4 |
| Exoc6 | 0.00011238 | 0.33429932 | 0.252 | 0.227 | 1 | TAM.Il1r2 | Exoc6 |
| Pfdn6 | 0.00011405 | 0.36741592 | 0.39 | 0.346 | 1 | TAM.Il1r2 | Pfdn6 |
| Zfp330 | 0.00012487 | 0.37851825 | 0.262 | 0.235 | 1 | TAM.Il1r2 | Zfp330 |
| Eid31 | 0.00012783 | 0.37616376 | 0.33 | 0.269 | 1 | TAM.Il1r2 | Eid3 |
| Foxn3 | 0.00013078 | 0.29434788 | 0.376 | 0.349 | 1 | TAM.Il1r2 | Foxn3 |
| Cdyl | 0.00013402 | 0.30885529 | 0.306 | 0.279 | 1 | TAM.Il1r2 | Cdyl |
| Pole4 | 0.00013599 | 0.41309591 | 0.35 | 0.312 | 1 | TAM.Il1r2 | Pole4 |
| Ssb | 0.00013683 | 0.32430459 | 0.604 | 0.561 | 1 | TAM.Il1r2 | Ssb |
| Ndufa12 | 0.00014506 | 0.37226216 | 0.376 | 0.331 | 1 | TAM.Il1r2 | Ndufa12 |
| Ddx46 | 0.00017403 | 0.31072955 | 0.497 | 0.455 | 1 | TAM.Il1r2 | Ddx46 |
| Pabpc4 | 0.0001785 | 0.49637696 | 0.285 | 0.245 | 1 | TAM.Il1r2 | Pabpc4 |
| Cox6b1 | 0.00018528 | 0.25840043 | 0.708 | 0.665 | 1 | TAM.Il1r2 | Cox6b1 |
| Farsa | 0.00019545 | 0.40668324 | 0.328 | 0.292 | 1 | TAM.Il1r2 | Farsa |
| Atxn10 | 0.00019731 | 0.34671704 | 0.349 | 0.314 | 1 | TAM.Il1r2 | Atxn10 |
| Spata2 | 0.0002137 | 0.38398278 | 0.268 | 0.238 | 1 | TAM.Il1r2 | Spata2 |
| Chuk | 0.0002218 | 0.32485012 | 0.319 | 0.291 | 1 | TAM.Il1r2 | Chuk |
| Golm11 | 0.00022755 | 0.37455583 | 0.254 | 0.221 | 1 | TAM.Il1r2 | Golm1 |
| Pip5k1c | 0.00023127 | 0.32972679 | 0.825 | 0.767 | 1 | TAM.Il1r2 | Pip5k1c |
| Mfsd101 | 0.00024022 | 0.40103452 | 0.284 | 0.249 | 1 | TAM.Il1r2 | Mfsd10 |
| Snrnp40 | 0.0002452 | 0.43712648 | 0.271 | 0.233 | 1 | TAM.Il1r2 | Snrnp40 |
| Osgep | 0.00024602 | 0.40062659 | 0.331 | 0.291 | 1 | TAM.Il1r2 | Osgep |
| Huwe1 | 0.00025741 | 0.33500633 | 0.576 | 0.523 | 1 | TAM.Il1r2 | Huwe1 |
| Uggt1 | 0.00025752 | 0.34053528 | 0.544 | 0.499 | 1 | TAM.Il1r2 | Uggt1 |
| Apex11 | 0.00026566 | 0.54607859 | 0.254 | 0.217 | 1 | TAM.Il1r2 | Apex1 |
| Was | 0.00030308 | 0.2535377 | 0.664 | 0.629 | 1 | TAM.Il1r2 | Was |
| Ube2n | 0.00030769 | 0.2967621 | 0.277 | 0.255 | 1 | TAM.Il1r2 | Ube2n |
| Zfp609 | 0.00031078 | 0.36282164 | 0.315 | 0.286 | 1 | TAM.Il1r2 | Zfp609 |
| Arpc1b | 0.00034868 | 0.25344404 | 0.951 | 0.928 | 1 | TAM.Il1r2 | Arpc1b |
| Sec61g | 0.0003639 | 0.31370015 | 0.376 | 0.342 | 1 | TAM.Il1r2 | Sec61g |
| Eif2s1 | 0.00037569 | 0.36047821 | 0.417 | 0.376 | 1 | TAM.Il1r2 | Eif2s1 |
| Tmem160 | 0.00041133 | 0.3602198 | 0.439 | 0.405 | 1 | TAM.Il1r2 | Tmem160 |
| Gtf2f21 | 0.00041809 | 0.31175246 | 0.351 | 0.319 | 1 | TAM.Il1r2 | Gtf2f2 |
| Ergic11 | 0.00042309 | 0.25447026 | 0.414 | 0.398 | 1 | TAM.Il1r2 | Ergic1 |
| Etv6 | 0.00049293 | 0.28266538 | 0.596 | 0.562 | 1 | TAM.Il1r2 | Etv6 |
| Ndufb7 | 0.00050008 | 0.31064883 | 0.58 | 0.531 | 1 | TAM.Il1r2 | Ndufb7 |
| Pno1 | 0.0005123 | 0.42664731 | 0.25 | 0.22 | 1 | TAM.Il1r2 | Pno1 |
| Cdk5rap31 | 0.00052727 | 0.32947306 | 0.313 | 0.278 | 1 | TAM.Il1r2 | Cdk5rap3 |
| Idh3g | 0.00055031 | 0.34582236 | 0.533 | 0.479 | 1 | TAM.Il1r2 | Idh3g |
| Limk11 | 0.00068362 | 0.44197353 | 0.273 | 0.247 | 1 | TAM.Il1r2 | Limk1 |
| Mgat2 | 0.00082028 | 0.32069355 | 0.503 | 0.463 | 1 | TAM.Il1r2 | Mgat2 |
| Ndufaf8 | 0.0008312 | 0.35427252 | 0.301 | 0.273 | 1 | TAM.Il1r2 | Ndufaf8 |
| Ier31 | 0.00087289 | 0.25419725 | 0.939 | 0.917 | 1 | TAM.Il1r2 | Ier3 |
| Naa15 | 0.0009402 | 0.34069733 | 0.398 | 0.363 | 1 | TAM.Il1r2 | Naa15 |
| Eef21 | 0.00099255 | 0.28253229 | 0.856 | 0.812 | 1 | TAM.Il1r2 | Eef2 |
| Wdr18 | 0.00100635 | 0.40718743 | 0.274 | 0.243 | 1 | TAM.Il1r2 | Wdr18 |
| Thap121 | 0.00105421 | 0.41706122 | 0.25 | 0.232 | 1 | TAM.Il1r2 | Thap12 |
| Relt | 0.00114043 | 0.32446922 | 0.344 | 0.318 | 1 | TAM.Il1r2 | Relt |
| Ndst11 | 0.00115113 | 0.39399772 | 0.602 | 0.51 | 1 | TAM.Il1r2 | Ndst1 |
| Psma31 | 0.00117508 | 0.35105116 | 0.43 | 0.387 | 1 | TAM.Il1r2 | Psma3 |
| Pus1 | 0.00120856 | 0.39121531 | 0.28 | 0.255 | 1 | TAM.Il1r2 | Pus1 |
| Pitpnm11 | 0.0012132 | 0.25161725 | 0.446 | 0.417 | 1 | TAM.Il1r2 | Pitpnm1 |
| Rbck1 | 0.00125998 | 0.25852507 | 0.367 | 0.346 | 1 | TAM.Il1r2 | Rbck1 |
| Eif4a3 | 0.00136049 | 0.36617657 | 0.257 | 0.226 | 1 | TAM.Il1r2 | Eif4a3 |
| Psmg4 | 0.00148391 | 0.31940176 | 0.315 | 0.285 | 1 | TAM.Il1r2 | Psmg4 |
| Atp5f1 | 0.00161755 | 0.35465222 | 0.499 | 0.458 | 1 | TAM.Il1r2 | Atp5f1 |
| Pom121 | 0.00170494 | 0.30391409 | 0.298 | 0.271 | 1 | TAM.Il1r2 | Pom121 |
| Gpsm3 | 0.0017109 | 0.28516752 | 0.778 | 0.736 | 1 | TAM.Il1r2 | Gpsm3 |
| Ndufa8 | 0.00174867 | 0.31184672 | 0.666 | 0.611 | 1 | TAM.Il1r2 | Ndufa8 |
| Nudc | 0.00185512 | 0.26493068 | 0.536 | 0.504 | 1 | TAM.Il1r2 | Nudc |
| Rrn3 | 0.00185553 | 0.37319316 | 0.266 | 0.235 | 1 | TAM.Il1r2 | Rrn3 |
| Pdzd8 | 0.00227085 | 0.30294612 | 0.317 | 0.294 | 1 | TAM.Il1r2 | Pdzd8 |
| Pten | 0.00227452 | 0.28051325 | 0.639 | 0.609 | 1 | TAM.Il1r2 | Pten |
| Matr3 | 0.0024611 | 0.3402422 | 0.342 | 0.317 | 1 | TAM.Il1r2 | Matr3 |
| Qdpr2 | 0.00260208 | 0.28609507 | 0.48 | 0.441 | 1 | TAM.Il1r2 | Qdpr |
| Ranbp1 | 0.00288679 | 0.44628545 | 0.52 | 0.435 | 1 | TAM.Il1r2 | Ranbp1 |
| Dcaf131 | 0.00290202 | 0.30939539 | 0.319 | 0.296 | 1 | TAM.Il1r2 | Dcaf13 |
| Ddx49 | 0.00291254 | 0.33475946 | 0.283 | 0.255 | 1 | TAM.Il1r2 | Ddx49 |
| Polr2f | 0.00295028 | 0.36253791 | 0.402 | 0.367 | 1 | TAM.Il1r2 | Polr2f |
| Fcf1 | 0.00312226 | 0.30967638 | 0.326 | 0.296 | 1 | TAM.Il1r2 | Fcf1 |
| Cdk2ap1 | 0.00368415 | 0.35121414 | 0.334 | 0.296 | 1 | TAM.Il1r2 | Cdk2ap1 |
| Tomm6 | 0.00411053 | 0.38178562 | 0.518 | 0.477 | 1 | TAM.Il1r2 | Tomm6 |
| Btaf1 | 0.00415312 | 0.3004179 | 0.328 | 0.301 | 1 | TAM.Il1r2 | Btaf1 |
| Hspe11 | 0.00422981 | 0.38322037 | 0.302 | 0.267 | 1 | TAM.Il1r2 | Hspe1 |
| Iars | 0.00451563 | 0.36417149 | 0.264 | 0.233 | 1 | TAM.Il1r2 | Iars |
| Hnrnpul1 | 0.00451571 | 0.25412461 | 0.82 | 0.774 | 1 | TAM.Il1r2 | Hnrnpul1 |
| Rpn2 | 0.00453212 | 0.25433069 | 0.556 | 0.52 | 1 | TAM.Il1r2 | Rpn2 |
| Cox5b | 0.00488694 | 0.35274424 | 0.712 | 0.66 | 1 | TAM.Il1r2 | Cox5b |
| Asph1 | 0.00498762 | 0.42479555 | 0.493 | 0.443 | 1 | TAM.Il1r2 | Asph |
| Ruvbl2 | 0.00500564 | 0.29891549 | 0.252 | 0.223 | 1 | TAM.Il1r2 | Ruvbl2 |
| Prelid1 | 0.00540365 | 0.34907081 | 0.764 | 0.707 | 1 | TAM.Il1r2 | Prelid1 |
| Dazap1 | 0.00540931 | 0.30192981 | 0.65 | 0.596 | 1 | TAM.Il1r2 | Dazap1 |
| Dr1 | 0.00605197 | 0.31043996 | 0.413 | 0.384 | 1 | TAM.Il1r2 | Dr1 |
| Morc2a | 0.00609442 | 0.34851356 | 0.291 | 0.268 | 1 | TAM.Il1r2 | Morc2a |
| Snrpb2 | 0.006153 | 0.2570767 | 0.373 | 0.352 | 1 | TAM.Il1r2 | Snrpb2 |
| Srsf9 | 0.00624654 | 0.27725411 | 0.735 | 0.694 | 1 | TAM.Il1r2 | Srsf9 |
| Hax1 | 0.00677184 | 0.30302727 | 0.344 | 0.318 | 1 | TAM.Il1r2 | Hax1 |
| Cope | 0.00697378 | 0.2842527 | 0.641 | 0.596 | 1 | TAM.Il1r2 | Cope |
| Mia2 | 0.00732135 | 0.29320037 | 0.634 | 0.595 | 1 | TAM.Il1r2 | Mia2 |
| Cdc37 | 0.00749798 | 0.31204994 | 0.688 | 0.633 | 1 | TAM.Il1r2 | Cdc37 |
| Jdp2 | 0.00779498 | 0.34845095 | 0.594 | 0.572 | 1 | TAM.Il1r2 | Jdp2 |
| Tmem50b1 | 0.00892445 | 0.30959709 | 0.426 | 0.39 | 1 | TAM.Il1r2 | Tmem50b |
| Cetn3 | 0.00985036 | 0.35535247 | 0.387 | 0.355 | 1 | TAM.Il1r2 | Cetn3 |
| Irf7 | 0 | 2.83022937 | 0.62 | 0.211 | 0 | TAM.IFN-respc | Irf7 |
| Slfn5 | 0 | 2.4596313 | 0.583 | 0.234 | 0 | TAM.IFN-respc | Slfn5 |
| Ms4a4c | 0 | 2.88024093 | 0.457 | 0.109 | 0 | TAM.IFN-respc | Ms4a4c |
| Ccr21 | 0 | 1.53215558 | 0.71 | 0.378 | 0 | TAM.IFN-respc | Ccr2 |
| Oasl2 | 0 | 2.21605666 | 0.543 | 0.222 | 0 | TAM.IFN-respc | Oasl2 |
| Ly6a | 0 | 3.23045318 | 0.421 | 0.11 | 0 | TAM.IFN-respc | Ly6a |
| Rtp4 | 0 | 2.44967015 | 0.411 | 0.118 | 0 | TAM.IFN-respc | Rtp4 |
| Ifi211 | 0 | 2.23866261 | 0.498 | 0.218 | 0 | TAM.IFN-respc | Ifi211 |
| Clec4a1 | 0 | 1.70668588 | 0.601 | 0.326 | 0 | TAM.IFN-respc | Clec4a1 |
| Ms4a6b1 | 0 | 1.46100795 | 0.694 | 0.427 | 0 | TAM.IFN-respc | Ms4a6b |
| Zbp1 | 0 | 2.03793494 | 0.52 | 0.26 | 0 | TAM.IFN-respc | Zbp1 |
| Ifi27l2a | 0 | 1.2053549 | 0.628 | 0.368 | 0 | TAM.IFN-respc | Ifi27l2a |
| Oas2 | 0 | 1.73428694 | 0.414 | 0.156 | 0 | TAM.IFN-respc | Oas2 |
| Ifi204 | 0 | 2.13494855 | 0.528 | 0.277 | 0 | TAM.IFN-respc | Ifi204 |
| Rnf213 | 0 | 1.91108115 | 0.67 | 0.424 | 0 | TAM.IFN-respc | Rnf213 |
| Ifi209 | 0 | 2.2801943 | 0.484 | 0.238 | 0 | TAM.IFN-respc | Ifi209 |
| Fcgr11 | 0 | 1.51455759 | 0.84 | 0.596 | 0 | TAM.IFN-respc | Fcgr1 |
| Ifi213 | 0 | 3.32420113 | 0.273 | 0.042 | 0 | TAM.IFN-respc | Ifi213 |
| Parp14 | 0 | 1.87982689 | 0.569 | 0.353 | 0 | TAM.IFN-respc | Parp14 |
| Ifi203 | 0 | 1.98255243 | 0.451 | 0.236 | 0 | TAM.IFN-respc | Ifi203 |
| Slfn8 | 0 | 2.02316266 | 0.39 | 0.182 | 0 | TAM.IFN-respc | Slfn8 |
| Irgm1 | 0 | 2.07881609 | 0.409 | 0.201 | 0 | TAM.IFN-respc | Irgm1 |
| Oas3 | 0 | 2.5442768 | 0.275 | 0.068 | 0 | TAM.IFN-respc | Oas3 |
| Ifi205 | 0 | 1.78577727 | 0.428 | 0.222 | 0 | TAM.IFN-respc | Ifi205 |
| Ifitm3 | 0 | 1.2409871 | 0.697 | 0.492 | 0 | TAM.IFN-respc | Ifitm3 |
| Ifi47 | 0 | 2.27712637 | 0.383 | 0.181 | 0 | TAM.IFN-respc | Ifi47 |
| Ms4a6c1 | 0 | 1.10790676 | 0.717 | 0.518 | 0 | TAM.IFN-respc | Ms4a6c |
| Stat1 | 0 | 1.70904873 | 0.65 | 0.459 | 0 | TAM.IFN-respc | Stat1 |
| Isg15 | 0 | 2.20145625 | 0.521 | 0.331 | 0 | TAM.IFN-respc | Isg15 |
| Clec4a3 | 0 | 1.34568083 | 0.638 | 0.45 | 0 | TAM.IFN-respc | Clec4a3 |
| Xaf1 | 0 | 2.03298002 | 0.314 | 0.129 | 0 | TAM.IFN-respc | Xaf1 |
| Sp100 | 0 | 1.6132054 | 0.429 | 0.247 | 0 | TAM.IFN-respc | Sp100 |
| Tap1 | 0 | 1.40350419 | 0.6 | 0.418 | 0 | TAM.IFN-respc | Tap1 |
| Clec12a | 0 | 1.8210664 | 0.335 | 0.157 | 0 | TAM.IFN-respc | Clec12a |
| Id3 | 0 | 1.40017317 | 0.422 | 0.245 | 0 | TAM.IFN-respc | Id3 |
| Oas1a | 0 | 1.46756127 | 0.472 | 0.296 | 0 | TAM.IFN-respc | Oas1a |
| Dhx58 | 0 | 1.71570466 | 0.373 | 0.197 | 0 | TAM.IFN-respc | Dhx58 |
| Nlrc5 | 0 | 1.37544191 | 0.444 | 0.276 | 0 | TAM.IFN-respc | Nlrc5 |
| Ifi207 | 0 | 1.16824097 | 0.721 | 0.554 | 0 | TAM.IFN-respc | Ifi207 |
| Ddx58 | 0 | 1.98130682 | 0.312 | 0.146 | 0 | TAM.IFN-respc | Ddx58 |
| Thbs11 | 0 | 0.62167696 | 0.831 | 0.667 | 0 | TAM.IFN-respc | Thbs1 |
| Plac8 | 0 | 1.11617019 | 0.476 | 0.313 | 0 | TAM.IFN-respc | Plac8 |
| Stat2 | 0 | 1.63852399 | 0.356 | 0.202 | 0 | TAM.IFN-respc | Stat2 |
| Mndal | 0 | 1.88775647 | 0.288 | 0.136 | 0 | TAM.IFN-respc | Mndal |
| Marcksl1 | 0 | 0.97032365 | 0.837 | 0.686 | 0 | TAM.IFN-respc | Marcksl1 |
| Ly6e | 0 | 1.26579558 | 0.958 | 0.808 | 0 | TAM.IFN-respc | Ly6e |
| Herc6 | 0 | 2.00980554 | 0.299 | 0.15 | 0 | TAM.IFN-respc | Herc6 |
| Helz2 | 0 | 1.71881642 | 0.302 | 0.156 | 0 | TAM.IFN-respc | Helz2 |
| Cxcl10 | 0 | 1.89098259 | 0.336 | 0.194 | 0 | TAM.IFN-respc | Cxcl10 |
| C31 | 0 | 0.85677573 | 0.702 | 0.561 | 0 | TAM.IFN-respc | C3 |
| Fgl2 | 0 | 1.37753939 | 0.628 | 0.49 | 0 | TAM.IFN-respc | Fgl2 |
| Znfx1 | 0 | 1.66442615 | 0.391 | 0.254 | 0 | TAM.IFN-respc | Znfx1 |
| Gbp2 | 0 | 2.20122481 | 0.339 | 0.205 | 0 | TAM.IFN-respc | Gbp2 |
| Cd40 | 0 | 1.38154811 | 0.399 | 0.282 | 0 | TAM.IFN-respc | Cd40 |
| Gbp5 | 0 | 2.15923214 | 0.25 | 0.135 | 0 | TAM.IFN-respc | Gbp5 |
| Daxx | 0 | 1.4727597 | 0.424 | 0.314 | 0 | TAM.IFN-respc | Daxx |
| H2-Aa1 | 0 | 0.8041417 | 0.881 | 0.776 | 0 | TAM.IFN-respc | H2-Aa |
| H2-Eb11 | 0 | 0.75998089 | 0.872 | 0.774 | 0 | TAM.IFN-respc | H2-Eb1 |
| Lgmn1 | 0 | 0.34336741 | 0.926 | 0.831 | 0 | TAM.IFN-respc | Lgmn |
| Samhd1 | 0 | 0.81660706 | 0.826 | 0.747 | 0 | TAM.IFN-respc | Samhd1 |
| Fos | 0 | 1.08000533 | 0.964 | 0.901 | 0 | TAM.IFN-respc | Fos |
| Tgfbi | 0 | 0.73480854 | 0.979 | 0.942 | 0 | TAM.IFN-respc | Tgfbi |
| Zfp36 | 0 | 0.66164353 | 0.983 | 0.955 | 0 | TAM.IFN-respc | Zfp36 |
| Ctsc1 | 0 | 0.26468091 | 0.982 | 0.954 | 0 | TAM.IFN-respc | Ctsc |
| Itm2b1 | 0 | 0.31476014 | 0.979 | 0.951 | 0 | TAM.IFN-respc | Itm2b |
| Junb1 | 0 | 0.27307112 | 0.962 | 0.957 | 0 | TAM.IFN-respc | Junb |
| Irf1 | 1.52E-302 | 0.99371879 | 0.562 | 0.492 | 2.93E-298 | TAM.IFN-respc | Irf1 |
| Parp12 | 1.06E-294 | 1.49318974 | 0.426 | 0.282 | 2.06E-290 | TAM.IFN-respc | Parp12 |
| Slamf7 | 2.24E-280 | 1.66169646 | 0.266 | 0.144 | 4.33E-276 | TAM.IFN-respc | Slamf7 |
| Dtx3l | 1.00E-270 | 1.47923856 | 0.43 | 0.277 | 1.93E-266 | TAM.IFN-respc | Dtx3l |
| Tlr91 | 7.31E-269 | 1.74365898 | 0.32 | 0.158 | 1.41E-264 | TAM.IFN-respc | Tlr9 |
| Rrad | 8.78E-258 | 1.57711122 | 0.293 | 0.152 | 1.70E-253 | TAM.IFN-respc | Rrad |
| Ifih1 | 3.88E-249 | 1.20840811 | 0.483 | 0.336 | 7.49E-245 | TAM.IFN-respc | Ifih1 |
| Sirpa1 | 7.85E-243 | 0.29754791 | 0.954 | 0.928 | 1.52E-238 | TAM.IFN-respc | Sirpa |
| Gvin1 | 1.26E-237 | 1.37914726 | 0.39 | 0.241 | 2.44E-233 | TAM.IFN-respc | Gvin1 |
| Epsti1 | 1.21E-234 | 1.17298581 | 0.623 | 0.462 | 2.34E-230 | TAM.IFN-respc | Epsti1 |
| Egr1 | 7.59E-213 | 0.30369893 | 0.735 | 0.666 | 1.47E-208 | TAM.IFN-respc | Egr1 |
| Trim30d | 1.01E-211 | 1.68556854 | 0.265 | 0.126 | 1.95E-207 | TAM.IFN-respc | Trim30d |
| Bst2 | 1.52E-203 | 0.97921673 | 0.711 | 0.632 | 2.93E-199 | TAM.IFN-respc | Bst2 |
| Ube2l6 | 2.29E-203 | 1.60895228 | 0.272 | 0.16 | 4.43E-199 | TAM.IFN-respc | Ube2l6 |
| Lst11 | 3.58E-193 | 0.93966205 | 0.726 | 0.574 | 6.91E-189 | TAM.IFN-respc | Lst1 |
| Tpt1 | 8.44E-191 | 0.29559979 | 0.97 | 0.956 | 1.63E-186 | TAM.IFN-respc | Tpt1 |
| Arhgef10l | 4.31E-186 | 1.14133889 | 0.493 | 0.319 | 8.33E-182 | TAM.IFN-respc | Arhgef10l |
| Jarid2 | 2.07E-185 | 0.90695042 | 0.677 | 0.515 | 4.01E-181 | TAM.IFN-respc | Jarid2 |
| Cd522 | 2.60E-180 | 0.27378701 | 0.92 | 0.865 | 5.02E-176 | TAM.IFN-respc | Cd52 |
| Gbp9 | 1.76E-177 | 1.43299567 | 0.348 | 0.215 | 3.41E-173 | TAM.IFN-respc | Gbp9 |
| H2-Q7 | 1.55E-176 | 1.06714801 | 0.465 | 0.327 | 3.00E-172 | TAM.IFN-respc | H2-Q7 |
| Trim30a | 1.17E-172 | 1.29849357 | 0.396 | 0.266 | 2.27E-168 | TAM.IFN-respc | Trim30a |
| Cfb1 | 1.08E-165 | 0.95071558 | 0.466 | 0.388 | 2.08E-161 | TAM.IFN-respc | Cfb |
| Rasgrp4 | 3.99E-162 | 1.07634729 | 0.459 | 0.286 | 7.71E-158 | TAM.IFN-respc | Rasgrp4 |
| Hspa8 | 7.31E-161 | 0.29911701 | 0.957 | 0.934 | 1.41E-156 | TAM.IFN-respc | Hspa8 |
| Itgam1 | 7.75E-149 | 0.29117195 | 0.887 | 0.796 | 1.50E-144 | TAM.IFN-respc | Itgam |
| Cd721 | 1.59E-146 | 1.24215804 | 0.371 | 0.278 | 3.07E-142 | TAM.IFN-respc | Cd72 |
| Mxd11 | 1.80E-146 | 0.7161964 | 0.654 | 0.595 | 3.47E-142 | TAM.IFN-respc | Mxd1 |
| Birc3 | 5.69E-138 | 0.8997748 | 0.687 | 0.572 | 1.10E-133 | TAM.IFN-respc | Birc3 |
| Aif11 | 3.64E-136 | 0.81574011 | 0.911 | 0.745 | 7.04E-132 | TAM.IFN-respc | Aif1 |
| Pid1 | 8.93E-136 | 0.77651915 | 0.819 | 0.682 | 1.73E-131 | TAM.IFN-respc | Pid1 |
| Apobec3 | 4.02E-134 | 1.06369373 | 0.523 | 0.393 | 7.78E-130 | TAM.IFN-respc | Apobec3 |
| H2-Ab11 | 4.96E-132 | 0.55654497 | 0.902 | 0.844 | 9.58E-128 | TAM.IFN-respc | H2-Ab1 |
| Ccl41 | 2.79E-129 | 0.35977628 | 0.475 | 0.438 | 5.40E-125 | TAM.IFN-respc | Ccl4 |
| Npc2 | 3.07E-127 | 0.3467022 | 0.968 | 0.945 | 5.93E-123 | TAM.IFN-respc | Npc2 |
| Pnp | 1.30E-123 | 0.89332175 | 0.674 | 0.599 | 2.51E-119 | TAM.IFN-respc | Pnp |
| Eif2ak2 | 1.49E-121 | 1.32442194 | 0.352 | 0.24 | 2.87E-117 | TAM.IFN-respc | Eif2ak2 |
| Csf3r | 3.92E-120 | 0.90684718 | 0.53 | 0.385 | 7.58E-116 | TAM.IFN-respc | Csf3r |
| Plbd11 | 1.47E-117 | 0.68668781 | 0.818 | 0.663 | 2.84E-113 | TAM.IFN-respc | Plbd1 |
| Pml | 3.55E-114 | 1.45534362 | 0.252 | 0.16 | 6.87E-110 | TAM.IFN-respc | Pml |
| Cd300ld51 | 3.49E-109 | 1.07131461 | 0.398 | 0.25 | 6.74E-105 | TAM.IFN-respc | Cd300ld5 |
| Gbp31 | 5.31E-107 | 1.19504548 | 0.297 | 0.221 | 1.03E-102 | TAM.IFN-respc | Gbp3 |
| Parp9 | 2.69E-106 | 1.26031919 | 0.328 | 0.224 | 5.21E-102 | TAM.IFN-respc | Parp9 |
| Rab7b1 | 3.01E-106 | 1.18540214 | 0.436 | 0.321 | 5.81E-102 | TAM.IFN-respc | Rab7b |
| Fcgr41 | 2.74E-104 | 0.85857735 | 0.555 | 0.463 | 5.29E-100 | TAM.IFN-respc | Fcgr4 |
| Psmb9 | 2.70E-103 | 0.92678797 | 0.581 | 0.488 | 5.23E-99 | TAM.IFN-respc | Psmb9 |
| Rap1b | 1.09E-100 | 0.26031878 | 0.911 | 0.888 | 2.10E-96 | TAM.IFN-respc | Rap1b |
| Trafd1 | 3.22E-98 | 0.87648395 | 0.609 | 0.563 | 6.22E-94 | TAM.IFN-respc | Trafd1 |
| Ccl2 | 2.36E-97 | 0.25873322 | 0.313 | 0.245 | 4.56E-93 | TAM.IFN-respc | Ccl2 |
| Ptpro | 6.25E-96 | 1.07083108 | 0.409 | 0.293 | 1.21E-91 | TAM.IFN-respc | Ptpro |
| Plxnd11 | 2.44E-94 | 0.26666124 | 0.816 | 0.809 | 4.72E-90 | TAM.IFN-respc | Plxnd1 |
| Uba7 | 4.40E-94 | 1.14106404 | 0.322 | 0.225 | 8.50E-90 | TAM.IFN-respc | Uba7 |
| Spon11 | 1.49E-93 | 1.11379054 | 0.305 | 0.184 | 2.88E-89 | TAM.IFN-respc | Spon1 |
| Crybg1 | 2.52E-93 | 1.13192008 | 0.304 | 0.188 | 4.88E-89 | TAM.IFN-respc | Crybg1 |
| Traf1 | 1.27E-89 | 0.75892853 | 0.41 | 0.322 | 2.46E-85 | TAM.IFN-respc | Traf1 |
| Pik3ap1 | 1.63E-87 | 0.72724187 | 0.82 | 0.766 | 3.16E-83 | TAM.IFN-respc | Pik3ap1 |
| Arrb21 | 2.37E-86 | 0.31763048 | 0.874 | 0.848 | 4.58E-82 | TAM.IFN-respc | Arrb2 |
| Rab5c | 1.64E-85 | 0.33757839 | 0.87 | 0.834 | 3.17E-81 | TAM.IFN-respc | Rab5c |
| Casp4 | 8.29E-85 | 1.01832958 | 0.388 | 0.275 | 1.60E-80 | TAM.IFN-respc | Casp4 |
| Hspa1b1 | 6.93E-83 | 0.28532234 | 0.386 | 0.362 | 1.34E-78 | TAM.IFN-respc | Hspa1b |
| Itgb51 | 9.92E-83 | 0.3606078 | 0.761 | 0.694 | 1.92E-78 | TAM.IFN-respc | Itgb5 |
| Bcl6 | 7.51E-82 | 1.02434161 | 0.327 | 0.223 | 1.45E-77 | TAM.IFN-respc | Bcl6 |
| Lmna | 4.98E-78 | 0.40906659 | 0.746 | 0.724 | 9.62E-74 | TAM.IFN-respc | Lmna |
| Ubc | 9.91E-77 | 0.35365192 | 0.941 | 0.914 | 1.92E-72 | TAM.IFN-respc | Ubc |
| Naaa1 | 1.51E-76 | 0.67590452 | 0.671 | 0.609 | 2.92E-72 | TAM.IFN-respc | Naaa |
| Srgap2 | 2.63E-75 | 0.99960189 | 0.465 | 0.36 | 5.08E-71 | TAM.IFN-respc | Srgap2 |
| Ifrd11 | 2.97E-74 | 0.61920774 | 0.849 | 0.823 | 5.74E-70 | TAM.IFN-respc | Ifrd1 |
| Ms4a4a1 | 3.33E-73 | 0.71036461 | 0.853 | 0.688 | 6.45E-69 | TAM.IFN-respc | Ms4a4a |
| Adar | 3.50E-71 | 1.00296893 | 0.428 | 0.327 | 6.77E-67 | TAM.IFN-respc | Adar |
| Unc93b11 | 4.70E-71 | 0.38792904 | 0.924 | 0.871 | 9.08E-67 | TAM.IFN-respc | Unc93b1 |
| Rnase41 | 1.54E-70 | 0.25427948 | 0.48 | 0.399 | 2.98E-66 | TAM.IFN-respc | Rnase4 |
| Lamp2 | 3.32E-65 | 0.29642242 | 0.822 | 0.802 | 6.42E-61 | TAM.IFN-respc | Lamp2 |
| Gpr1321 | 1.82E-64 | 0.62903034 | 0.682 | 0.602 | 3.51E-60 | TAM.IFN-respc | Gpr132 |
| Tor3a | 2.98E-64 | 1.14566682 | 0.271 | 0.22 | 5.76E-60 | TAM.IFN-respc | Tor3a |
| Ccrl2 | 4.68E-64 | 0.29458899 | 0.681 | 0.677 | 9.04E-60 | TAM.IFN-respc | Ccrl2 |
| Mefv | 4.46E-60 | 0.78891595 | 0.389 | 0.264 | 8.61E-56 | TAM.IFN-respc | Mefv |
| Il10rb | 1.10E-59 | 0.29678176 | 0.75 | 0.737 | 2.13E-55 | TAM.IFN-respc | Il10rb |
| Vps37b1 | 1.03E-58 | 0.6768523 | 0.586 | 0.524 | 1.99E-54 | TAM.IFN-respc | Vps37b |
| Maf1 | 1.20E-58 | 0.31109556 | 0.682 | 0.559 | 2.32E-54 | TAM.IFN-respc | Maf |
| Zfp36l2 | 1.24E-58 | 0.63044267 | 0.88 | 0.835 | 2.39E-54 | TAM.IFN-respc | Zfp36l2 |
| Sh3glb1 | 1.04E-55 | 0.30982278 | 0.774 | 0.764 | 2.01E-51 | TAM.IFN-respc | Sh3glb1 |
| Rnf1301 | 5.52E-55 | 0.34395562 | 0.732 | 0.709 | 1.07E-50 | TAM.IFN-respc | Rnf130 |
| Rab3il11 | 8.05E-55 | 0.36328426 | 0.757 | 0.666 | 1.56E-50 | TAM.IFN-respc | Rab3il1 |
| Themis21 | 1.02E-54 | 0.7360391 | 0.512 | 0.466 | 1.98E-50 | TAM.IFN-respc | Themis2 |
| Parp8 | 1.46E-53 | 1.06127885 | 0.312 | 0.226 | 2.82E-49 | TAM.IFN-respc | Parp8 |
| Otulinl1 | 1.50E-53 | 0.35800124 | 0.732 | 0.699 | 2.90E-49 | TAM.IFN-respc | Otulinl |
| Cd831 | 6.30E-53 | 0.72422129 | 0.679 | 0.534 | 1.22E-48 | TAM.IFN-respc | Cd83 |
| S100a10 | 8.68E-53 | 0.31745761 | 0.826 | 0.769 | 1.68E-48 | TAM.IFN-respc | S100a10 |
| Jmy2 | 1.17E-52 | 0.81558002 | 0.423 | 0.375 | 2.27E-48 | TAM.IFN-respc | Jmy |
| Cltc1 | 1.92E-52 | 0.26037234 | 0.661 | 0.683 | 3.72E-48 | TAM.IFN-respc | Cltc |
| Lair11 | 7.47E-52 | 0.25179943 | 0.522 | 0.498 | 1.44E-47 | TAM.IFN-respc | Lair1 |
| Irf9 | 2.72E-51 | 0.87343791 | 0.407 | 0.317 | 5.25E-47 | TAM.IFN-respc | Irf9 |
| Adgre11 | 1.48E-50 | 0.81169072 | 0.713 | 0.546 | 2.87E-46 | TAM.IFN-respc | Adgre1 |
| Apobec11 | 7.20E-50 | 0.86303774 | 0.591 | 0.462 | 1.39E-45 | TAM.IFN-respc | Apobec1 |
| Itgav1 | 8.80E-50 | 0.32601281 | 0.5 | 0.51 | 1.70E-45 | TAM.IFN-respc | Itgav |
| Socs11 | 1.89E-48 | 0.93789405 | 0.279 | 0.219 | 3.65E-44 | TAM.IFN-respc | Socs1 |
| Rasa4 | 2.12E-48 | 0.96110355 | 0.442 | 0.349 | 4.09E-44 | TAM.IFN-respc | Rasa4 |
| Prex11 | 5.99E-48 | 0.28534431 | 0.767 | 0.759 | 1.16E-43 | TAM.IFN-respc | Prex1 |
| Abi1 | 1.40E-47 | 0.29920777 | 0.781 | 0.783 | 2.70E-43 | TAM.IFN-respc | Abi1 |
| Sp140 | 3.52E-46 | 1.04961919 | 0.26 | 0.202 | 6.80E-42 | TAM.IFN-respc | Sp140 |
| Nrg11 | 3.87E-46 | 0.57877064 | 0.432 | 0.372 | 7.48E-42 | TAM.IFN-respc | Nrg1 |
| Tdrd7 | 1.40E-45 | 1.00937644 | 0.334 | 0.253 | 2.70E-41 | TAM.IFN-respc | Tdrd7 |
| Ogfr | 4.18E-44 | 0.76947279 | 0.504 | 0.44 | 8.09E-40 | TAM.IFN-respc | Ogfr |
| Ifngr11 | 9.14E-43 | 0.64318548 | 0.844 | 0.791 | 1.77E-38 | TAM.IFN-respc | Ifngr1 |
| Aup1 | 2.04E-41 | 0.27921153 | 0.654 | 0.666 | 3.94E-37 | TAM.IFN-respc | Aup1 |
| Nampt2 | 2.87E-41 | 0.34553397 | 0.63 | 0.653 | 5.55E-37 | TAM.IFN-respc | Nampt |

| | | | | | | | |
|---|---|---|---|---|---|---|---|
| Evi2a1 | 1.85E-40 | 0.80432399 | 0.673 | 0.586 | 3.58E-36 | TAM.IFN-respc | Evi2a |
| Mef2d | 2.36E-40 | 0.25783504 | 0.62 | 0.646 | 4.57E-36 | TAM.IFN-respc | Mef2d |
| Irf81 | 3.03E-40 | 0.26062412 | 0.652 | 0.658 | 5.86E-36 | TAM.IFN-respc | Irf8 |
| Cd300ld2 | 7.48E-40 | 0.27020429 | 0.495 | 0.529 | 1.45E-35 | TAM.IFN-respc | Cd300ld |
| AB124611 | 1.72E-39 | 0.82341973 | 0.391 | 0.308 | 3.33E-35 | TAM.IFN-respc | AB124611 |
| Fosb1 | 1.96E-39 | 0.64969478 | 0.873 | 0.802 | 3.79E-35 | TAM.IFN-respc | Fosb |
| Trim251 | 2.57E-39 | 0.26239126 | 0.669 | 0.672 | 4.97E-35 | TAM.IFN-respc | Trim25 |
| Sik11 | 2.82E-39 | 0.65827221 | 0.76 | 0.701 | 5.44E-35 | TAM.IFN-respc | Sik1 |
| Ifi35 | 3.87E-39 | 0.85490494 | 0.315 | 0.262 | 7.48E-35 | TAM.IFN-respc | Ifi35 |
| Fxyd5 | 4.37E-39 | 0.38635686 | 0.856 | 0.815 | 8.44E-35 | TAM.IFN-respc | Fxyd5 |
| Klf61 | 1.62E-38 | 0.59920098 | 0.767 | 0.706 | 3.13E-34 | TAM.IFN-respc | Klf6 |
| Snx51 | 2.70E-38 | 0.32980593 | 0.556 | 0.552 | 5.23E-34 | TAM.IFN-respc | Snx5 |
| Tlr2 | 4.24E-38 | 0.57423149 | 0.799 | 0.724 | 8.19E-34 | TAM.IFN-respc | Tlr2 |
| Glul1 | 7.35E-38 | 0.38278465 | 0.603 | 0.581 | 1.42E-33 | TAM.IFN-respc | Glul |
| Cd861 | 9.80E-38 | 0.29052162 | 0.736 | 0.71 | 1.89E-33 | TAM.IFN-respc | Cd86 |
| Bmp2k1 | 2.66E-37 | 0.27077691 | 0.621 | 0.616 | 5.14E-33 | TAM.IFN-respc | Bmp2k |
| Osm1 | 3.28E-37 | 0.303163 | 0.7 | 0.677 | 6.34E-33 | TAM.IFN-respc | Osm |
| Tor1aip1 | 9.22E-36 | 0.70911633 | 0.726 | 0.682 | 1.78E-31 | TAM.IFN-respc | Tor1aip1 |
| Cd300ld31 | 3.99E-35 | 1.0302948 | 0.277 | 0.186 | 7.72E-31 | TAM.IFN-respc | Cd300ld3 |
| Rnf114 | 4.87E-35 | 0.81431459 | 0.422 | 0.356 | 9.41E-31 | TAM.IFN-respc | Rnf114 |
| Socs31 | 1.09E-33 | 0.2541565 | 0.801 | 0.781 | 2.10E-29 | TAM.IFN-respc | Socs3 |
| Txnip1 | 1.36E-33 | 0.30781837 | 0.537 | 0.532 | 2.63E-29 | TAM.IFN-respc | Txnip |
| Tagap1 | 2.32E-33 | 0.86554481 | 0.298 | 0.228 | 4.48E-29 | TAM.IFN-respc | Tagap |
| Fosl21 | 2.93E-33 | 0.31086528 | 0.842 | 0.852 | 5.66E-29 | TAM.IFN-respc | Fosl2 |
| Dek | 4.97E-33 | 0.3073539 | 0.504 | 0.507 | 9.61E-29 | TAM.IFN-respc | Dek |
| Ddx5 | 1.72E-32 | 0.30691364 | 0.703 | 0.692 | 3.33E-28 | TAM.IFN-respc | Ddx5 |
| Pip4p11 | 2.77E-32 | 0.26008331 | 0.488 | 0.508 | 5.36E-28 | TAM.IFN-respc | Pip4p1 |
| Arl4c | 6.02E-32 | 0.25252765 | 0.599 | 0.609 | 1.16E-27 | TAM.IFN-respc | Arl4c |
| Odc11 | 1.14E-31 | 0.40232072 | 0.638 | 0.681 | 2.21E-27 | TAM.IFN-respc | Odc1 |
| Ifitm2 | 3.08E-31 | 0.5119193 | 0.942 | 0.907 | 5.96E-27 | TAM.IFN-respc | Ifitm2 |
| Ppp1r9b | 3.21E-31 | 0.33009669 | 0.811 | 0.793 | 6.21E-27 | TAM.IFN-respc | Ppp1r9b |
| Metrnl | 4.15E-31 | 0.31900094 | 0.592 | 0.582 | 8.03E-27 | TAM.IFN-respc | Metrnl |
| Acp21 | 5.33E-31 | 0.26948086 | 0.446 | 0.481 | 1.03E-26 | TAM.IFN-respc | Acp2 |
| Hsd17b11 | 1.31E-30 | 0.26169554 | 0.543 | 0.571 | 2.53E-26 | TAM.IFN-respc | Hsd17b11 |
| Wasf2 | 2.14E-30 | 0.25593754 | 0.585 | 0.612 | 4.14E-26 | TAM.IFN-respc | Wasf2 |
| Kdm7a1 | 2.43E-29 | 0.2741555 | 0.685 | 0.691 | 4.70E-25 | TAM.IFN-respc | Kdm7a |
| Lfng | 1.26E-28 | 0.34465828 | 0.66 | 0.651 | 2.45E-24 | TAM.IFN-respc | Lfng |
| Spop | 1.82E-28 | 0.25430077 | 0.544 | 0.565 | 3.51E-24 | TAM.IFN-respc | Spop |
| Ascc3 | 4.90E-28 | 0.77484068 | 0.448 | 0.393 | 9.48E-24 | TAM.IFN-respc | Ascc3 |
| Clec2d | 1.02E-27 | 0.54453386 | 0.749 | 0.672 | 1.98E-23 | TAM.IFN-respc | Clec2d |
| Nfkbie1 | 1.23E-27 | 0.59628561 | 0.677 | 0.611 | 2.38E-23 | TAM.IFN-respc | Nfkbie |
| Dck | 5.07E-27 | 0.8702881 | 0.334 | 0.289 | 9.80E-23 | TAM.IFN-respc | Dck |
| Dynll1 | 7.13E-27 | 0.27069047 | 0.473 | 0.491 | 1.38E-22 | TAM.IFN-respc | Dynll1 |
| Rin21 | 1.59E-26 | 0.3196302 | 0.353 | 0.383 | 3.07E-22 | TAM.IFN-respc | Rin2 |
| Tent5c1 | 3.11E-26 | 0.25727735 | 0.348 | 0.373 | 6.01E-22 | TAM.IFN-respc | Tent5c |
| Was1 | 4.23E-26 | 0.29261031 | 0.63 | 0.634 | 8.17E-22 | TAM.IFN-respc | Was |
| Cwc251 | 1.80E-25 | 0.7158377 | 0.359 | 0.328 | 3.48E-21 | TAM.IFN-respc | Cwc25 |
| Il2rg1 | 1.98E-25 | 0.63505247 | 0.664 | 0.599 | 3.83E-21 | TAM.IFN-respc | Il2rg |
| Stk17b1 | 2.74E-25 | 0.25670558 | 0.558 | 0.569 | 5.30E-21 | TAM.IFN-respc | Stk17b |
| Ccr5 | 3.86E-25 | 0.6392164 | 0.865 | 0.78 | 7.47E-21 | TAM.IFN-respc | Ccr5 |
| H2afz | 5.03E-25 | 0.39444982 | 0.581 | 0.564 | 9.73E-21 | TAM.IFN-respc | H2afz |
| Mob1a | 6.75E-25 | 0.31747182 | 0.7 | 0.695 | 1.31E-20 | TAM.IFN-respc | Mob1a |
| Cited2 | 1.23E-24 | 0.38138877 | 0.558 | 0.508 | 2.38E-20 | TAM.IFN-respc | Cited2 |
| Dusp1 | 1.25E-24 | 0.56682166 | 0.931 | 0.888 | 2.41E-20 | TAM.IFN-respc | Dusp1 |
| Icam1 | 3.71E-24 | 0.27732088 | 0.6 | 0.625 | 7.17E-20 | TAM.IFN-respc | Icam1 |
| Oser1 | 3.79E-24 | 0.26390996 | 0.42 | 0.45 | 7.33E-20 | TAM.IFN-respc | Oser1 |
| Ncoa31 | 6.15E-24 | 0.26493404 | 0.538 | 0.561 | 1.19E-19 | TAM.IFN-respc | Ncoa3 |
| Atf31 | 9.06E-24 | 0.67415029 | 0.776 | 0.69 | 1.75E-19 | TAM.IFN-respc | Atf3 |
| Tmem1192 | 3.20E-23 | 0.98137977 | 0.36 | 0.296 | 6.19E-19 | TAM.IFN-respc | Tmem119 |
| Siglec1 | 3.44E-23 | 0.72258956 | 0.408 | 0.263 | 6.64E-19 | TAM.IFN-respc | Siglec1 |
| Ifngr21 | 4.37E-23 | 0.36310763 | 0.599 | 0.603 | 8.44E-19 | TAM.IFN-respc | Ifngr2 |
| Selplg | 5.85E-23 | 0.33425023 | 0.53 | 0.544 | 1.13E-18 | TAM.IFN-respc | Selplg |
| Impact1 | 1.04E-22 | 0.27996411 | 0.473 | 0.488 | 2.01E-18 | TAM.IFN-respc | Impact |
| Hmox2 | 2.03E-22 | 0.31915907 | 0.497 | 0.525 | 3.93E-18 | TAM.IFN-respc | Hmox2 |
| B2m | 6.16E-22 | 0.60974506 | 0.929 | 0.893 | 1.19E-17 | TAM.IFN-respc | B2m |
| Epb41l2 | 9.31E-22 | 0.29484233 | 0.604 | 0.608 | 1.80E-17 | TAM.IFN-respc | Epb41l2 |
| Pik3cg1 | 1.07E-21 | 0.29439134 | 0.405 | 0.431 | 2.08E-17 | TAM.IFN-respc | Pik3cg |
| Hspa1a1 | 1.57E-21 | 0.44614156 | 0.489 | 0.407 | 3.04E-17 | TAM.IFN-respc | Hspa1a |
| Ms4a6d | 1.89E-21 | 0.45277307 | 0.843 | 0.792 | 3.66E-17 | TAM.IFN-respc | Ms4a6d |
| Tmem176a1 | 2.04E-21 | 0.33046398 | 0.478 | 0.414 | 3.95E-17 | TAM.IFN-respc | Tmem176a |
| Nfam1 | 2.19E-21 | 0.28213451 | 0.458 | 0.477 | 4.24E-17 | TAM.IFN-respc | Nfam1 |
| Ndel11 | 4.31E-21 | 0.25284759 | 0.669 | 0.688 | 8.32E-17 | TAM.IFN-respc | Ndel1 |
| Rassf2 | 5.06E-21 | 0.67780125 | 0.635 | 0.533 | 9.79E-17 | TAM.IFN-respc | Rassf2 |
| Arl5b1 | 9.09E-21 | 0.65135229 | 0.461 | 0.42 | 1.76E-16 | TAM.IFN-respc | Arl5b |
| Cd931 | 1.89E-20 | 0.42593643 | 0.451 | 0.396 | 3.66E-16 | TAM.IFN-respc | Cd93 |
| Clk1 | 6.65E-20 | 0.31118855 | 0.716 | 0.714 | 1.29E-15 | TAM.IFN-respc | Clk1 |
| Lnpep | 7.38E-20 | 0.29000792 | 0.675 | 0.696 | 1.43E-15 | TAM.IFN-respc | Lnpep |
| Gng121 | 7.94E-20 | 0.25481988 | 0.284 | 0.311 | 1.54E-15 | TAM.IFN-respc | Gng12 |
| Pxn1 | 9.78E-20 | 0.26212518 | 0.52 | 0.544 | 1.89E-15 | TAM.IFN-respc | Pxn |
| Use1 | 1.33E-19 | 0.28749967 | 0.523 | 0.545 | 2.57E-15 | TAM.IFN-respc | Use1 |
| Extl31 | 2.01E-19 | 0.31114237 | 0.385 | 0.423 | 3.90E-15 | TAM.IFN-respc | Extl3 |
| Vav11 | 2.33E-19 | 0.3619033 | 0.711 | 0.694 | 4.50E-15 | TAM.IFN-respc | Vav1 |
| Kif3b1 | 2.85E-19 | 0.2842774 | 0.32 | 0.353 | 5.51E-15 | TAM.IFN-respc | Kif3b |
| Pim11 | 3.84E-19 | 0.31567132 | 0.912 | 0.89 | 7.43E-15 | TAM.IFN-respc | Pim1 |
| Grk2 | 4.04E-19 | 0.36150933 | 0.744 | 0.729 | 7.82E-15 | TAM.IFN-respc | Grk2 |
| Tmem1231 | 6.00E-19 | 0.30077455 | 0.461 | 0.483 | 1.16E-14 | TAM.IFN-respc | Tmem123 |
| Pik3cd1 | 1.18E-18 | 0.29740017 | 0.611 | 0.607 | 2.28E-14 | TAM.IFN-respc | Pik3cd |
| Smcr8 | 1.98E-18 | 0.30129509 | 0.457 | 0.491 | 3.82E-14 | TAM.IFN-respc | Smcr8 |
| Tsc22d31 | 2.02E-18 | 0.29100863 | 0.397 | 0.41 | 3.90E-14 | TAM.IFN-respc | Tsc22d3 |
| Nfkbid2 | 4.28E-18 | 0.34447099 | 0.661 | 0.612 | 8.28E-14 | TAM.IFN-respc | Nfkbid |
| Slc35b21 | 4.98E-18 | 0.4579989 | 0.737 | 0.737 | 9.64E-14 | TAM.IFN-respc | Slc35b2 |
| H2-Q6 | 5.80E-18 | 0.55835374 | 0.462 | 0.413 | 1.12E-13 | TAM.IFN-respc | H2-Q6 |
| Sgk3 | 6.37E-18 | 0.73298877 | 0.374 | 0.316 | 1.23E-13 | TAM.IFN-respc | Sgk3 |
| Sema4d1 | 6.43E-18 | 0.3656878 | 0.725 | 0.658 | 1.24E-13 | TAM.IFN-respc | Sema4d |
| Coro2a1 | 6.99E-18 | 0.27481931 | 0.418 | 0.415 | 1.35E-13 | TAM.IFN-respc | Coro2a |
| Ldlrap11 | 7.90E-18 | 0.27900473 | 0.366 | 0.4 | 1.53E-13 | TAM.IFN-respc | Ldlrap1 |
| Lsr1 | 1.46E-17 | 0.73785815 | 0.36 | 0.284 | 2.83E-13 | TAM.IFN-respc | Lsr |
| Frmd4a1 | 1.77E-17 | 0.86671084 | 0.414 | 0.355 | 3.42E-13 | TAM.IFN-respc | Frmd4a |
| Sh3bp2 | 2.11E-17 | 0.345068 | 0.578 | 0.584 | 4.08E-13 | TAM.IFN-respc | Sh3bp2 |
| Rab4b | 2.25E-17 | 0.32460745 | 0.511 | 0.529 | 4.34E-13 | TAM.IFN-respc | Rab4b |
| Ptafr | 6.74E-17 | 0.2775879 | 0.637 | 0.635 | 1.30E-12 | TAM.IFN-respc | Ptafr |
| Mbnl1 | 6.79E-17 | 0.37054696 | 0.503 | 0.512 | 1.31E-12 | TAM.IFN-respc | Mbnl1 |
| Tlr11 | 6.93E-17 | 0.91872884 | 0.292 | 0.232 | 1.34E-12 | TAM.IFN-respc | Tlr1 |
| Abcd1 | 7.09E-17 | 0.34482872 | 0.495 | 0.504 | 1.37E-12 | TAM.IFN-respc | Abcd1 |
| Pnrc2 | 8.53E-17 | 0.26418469 | 0.343 | 0.382 | 1.65E-12 | TAM.IFN-respc | Pnrc2 |
| Abca91 | 8.60E-17 | 0.25877315 | 0.312 | 0.312 | 1.66E-12 | TAM.IFN-respc | Abca9 |
| Plekho2 | 1.05E-16 | 0.34714709 | 0.854 | 0.829 | 2.04E-12 | TAM.IFN-respc | Plekho2 |
| Map3k8 | 1.66E-16 | 0.69187834 | 0.34 | 0.29 | 3.21E-12 | TAM.IFN-respc | Map3k8 |
| Tbl1xr1 | 1.89E-16 | 0.30251153 | 0.501 | 0.519 | 3.66E-12 | TAM.IFN-respc | Tbl1xr1 |
| Gas7 | 2.37E-16 | 0.32511005 | 0.587 | 0.582 | 4.58E-12 | TAM.IFN-respc | Gas7 |
| Rgs101 | 4.73E-16 | 0.44789723 | 0.705 | 0.664 | 9.15E-12 | TAM.IFN-respc | Rgs10 |
| Ptpn6 | 4.78E-16 | 0.39298787 | 0.764 | 0.75 | 9.25E-12 | TAM.IFN-respc | Ptpn6 |
| Dynlt1f | 4.89E-16 | 0.29508976 | 0.241 | 0.268 | 9.45E-12 | TAM.IFN-respc | Dynlt1f |
| Ell21 | 5.15E-16 | 0.25273126 | 0.484 | 0.514 | 9.96E-12 | TAM.IFN-respc | Ell2 |
| Ilf3 | 6.12E-16 | 0.2689092 | 0.304 | 0.334 | 1.18E-11 | TAM.IFN-respc | Ilf3 |
| Dazap2 | 8.32E-16 | 0.33522889 | 0.649 | 0.654 | 1.61E-11 | TAM.IFN-respc | Dazap2 |
| Cd821 | 1.18E-15 | 0.3972536 | 0.488 | 0.504 | 2.28E-11 | TAM.IFN-respc | Cd82 |
| Med1 | 1.32E-15 | 0.26544696 | 0.278 | 0.311 | 2.54E-11 | TAM.IFN-respc | Med1 |
| Pdxk1 | 2.18E-15 | 0.38933646 | 0.391 | 0.402 | 4.22E-11 | TAM.IFN-respc | Pdxk |
| Tnfsf91 | 2.89E-15 | 0.43997463 | 0.368 | 0.321 | 5.58E-11 | TAM.IFN-respc | Tnfsf9 |
| Rnf19b | 3.23E-15 | 0.40542707 | 0.72 | 0.699 | 6.25E-11 | TAM.IFN-respc | Rnf19b |
| Camk2d1 | 3.53E-15 | 0.32451716 | 0.552 | 0.548 | 6.83E-11 | TAM.IFN-respc | Camk2d |
| Tpst2 | 3.63E-15 | 0.38833767 | 0.492 | 0.495 | 7.01E-11 | TAM.IFN-respc | Tpst2 |
| Gbp71 | 3.69E-15 | 0.7900784 | 0.365 | 0.318 | 7.13E-11 | TAM.IFN-respc | Gbp7 |
| Rnf115 | 3.75E-15 | 0.25165021 | 0.357 | 0.397 | 7.26E-11 | TAM.IFN-respc | Rnf115 |
| Nfatc2 | 4.39E-15 | 0.79052846 | 0.387 | 0.316 | 8.49E-11 | TAM.IFN-respc | Nfatc2 |
| Nrros | 4.81E-15 | 0.63292537 | 0.811 | 0.726 | 9.29E-11 | TAM.IFN-respc | Nrros |
| Specc1l1 | 7.69E-15 | 0.25691843 | 0.445 | 0.461 | 1.49E-10 | TAM.IFN-respc | Specc1l |
| Homer11 | 9.40E-15 | 0.26388227 | 0.447 | 0.475 | 1.82E-10 | TAM.IFN-respc | Homer1 |
| Nek9 | 9.61E-15 | 0.27628416 | 0.272 | 0.314 | 1.86E-10 | TAM.IFN-respc | Nek9 |
| Serp1 | 1.01E-14 | 0.25859645 | 0.299 | 0.339 | 1.94E-10 | TAM.IFN-respc | Serp1 |
| Arhgap45 | 1.06E-14 | 0.27983729 | 0.65 | 0.659 | 2.04E-10 | TAM.IFN-respc | Arhgap45 |
| Nipbl | 1.42E-14 | 0.26827957 | 0.38 | 0.412 | 2.74E-10 | TAM.IFN-respc | Nipbl |
| Smg9 | 1.59E-14 | 0.26843004 | 0.285 | 0.325 | 3.07E-10 | TAM.IFN-respc | Smg9 |
| Plekhg2 | 1.83E-14 | 0.63696075 | 0.348 | 0.302 | 3.54E-10 | TAM.IFN-respc | Plekhg2 |
| Dok3 | 1.92E-14 | 0.31488747 | 0.465 | 0.482 | 3.72E-10 | TAM.IFN-respc | Dok3 |
| Myd881 | 2.19E-14 | 0.29385079 | 0.649 | 0.668 | 4.23E-10 | TAM.IFN-respc | Myd88 |
| Selenot | 2.68E-14 | 0.31714222 | 0.383 | 0.411 | 5.19E-10 | TAM.IFN-respc | Selenot |
| Adrb2 | 2.87E-14 | 0.27401376 | 0.25 | 0.262 | 5.56E-10 | TAM.IFN-respc | Adrb2 |
| Antxr21 | 3.47E-14 | 0.29060647 | 0.621 | 0.602 | 6.72E-10 | TAM.IFN-respc | Antxr2 |
| Nfkbiz1 | 3.79E-14 | 0.33977264 | 0.748 | 0.684 | 7.33E-10 | TAM.IFN-respc | Nfkbiz |
| Efhd2 | 4.22E-14 | 0.43498984 | 0.92 | 0.894 | 8.16E-10 | TAM.IFN-respc | Efhd2 |
| Rreb1 | 4.35E-14 | 0.34563179 | 0.542 | 0.553 | 8.41E-10 | TAM.IFN-respc | Rreb1 |
| Klf3 | 6.85E-14 | 0.26398452 | 0.5 | 0.514 | 1.32E-09 | TAM.IFN-respc | Klf3 |
| Mthfr1 | 7.38E-14 | 0.64535815 | 0.357 | 0.325 | 1.43E-09 | TAM.IFN-respc | Mthfr |
| Stard3nl | 8.47E-14 | 0.35602868 | 0.297 | 0.32 | 1.64E-09 | TAM.IFN-respc | Stard3nl |
| Actr31 | 1.17E-13 | 0.39549171 | 0.653 | 0.635 | 2.27E-09 | TAM.IFN-respc | Actr3 |
| Csrnp11 | 1.26E-13 | 0.25412984 | 0.705 | 0.712 | 2.43E-09 | TAM.IFN-respc | Csrnp1 |
| Cd47 | 1.28E-13 | 0.55985333 | 0.79 | 0.737 | 2.48E-09 | TAM.IFN-respc | Cd47 |
| Zfp36l1 | 1.56E-13 | 0.38589884 | 0.664 | 0.628 | 3.02E-09 | TAM.IFN-respc | Zfp36l1 |
| Elf2 | 1.89E-13 | 0.26582407 | 0.315 | 0.349 | 3.66E-09 | TAM.IFN-respc | Elf2 |
| Tns3 | 1.97E-13 | 0.36342071 | 0.479 | 0.467 | 3.81E-09 | TAM.IFN-respc | Tns3 |
| Rala | 1.99E-13 | 0.31732922 | 0.297 | 0.327 | 3.84E-09 | TAM.IFN-respc | Rala |
| Cmpk1 | 2.08E-13 | 0.31506797 | 0.462 | 0.475 | 4.03E-09 | TAM.IFN-respc | Cmpk1 |
| Tubb2a | 2.63E-13 | 0.41135012 | 0.407 | 0.422 | 5.09E-09 | TAM.IFN-respc | Tubb2a |
| Rap2b1 | 3.98E-13 | 0.36223087 | 0.5 | 0.498 | 7.69E-09 | TAM.IFN-respc | Rap2b |
| Nod11 | 4.32E-13 | 0.61651722 | 0.258 | 0.221 | 8.36E-09 | TAM.IFN-respc | Nod1 |
| Cd48 | 4.61E-13 | 0.38102824 | 0.411 | 0.421 | 8.91E-09 | TAM.IFN-respc | Cd48 |
| Ambra1 | 5.92E-13 | 0.26264222 | 0.274 | 0.312 | 1.14E-08 | TAM.IFN-respc | Ambra1 |
| Cd300c22 | 7.87E-13 | 0.5111844 | 0.66 | 0.604 | 1.52E-08 | TAM.IFN-respc | Cd300c2 |
| Creb51 | 8.19E-13 | 0.29583093 | 0.336 | 0.366 | 1.58E-08 | TAM.IFN-respc | Creb5 |
| Mapk6 | 8.53E-13 | 0.34504642 | 0.459 | 0.472 | 1.65E-08 | TAM.IFN-respc | Mapk6 |
| Ptpn181 | 1.06E-12 | 0.65848775 | 0.62 | 0.528 | 2.06E-08 | TAM.IFN-respc | Ptpn18 |
| Inpp5d1 | 1.81E-12 | 0.42338368 | 0.817 | 0.767 | 3.50E-08 | TAM.IFN-respc | Inpp5d |
| Tmub2 | 1.83E-12 | 0.25065731 | 0.243 | 0.278 | 3.54E-08 | TAM.IFN-respc | Tmub2 |
| Pstpip1 | 1.96E-12 | 0.43141928 | 0.545 | 0.526 | 3.78E-08 | TAM.IFN-respc | Pstpip1 |
| Elk3 | 2.50E-12 | 0.37067961 | 0.495 | 0.501 | 4.83E-08 | TAM.IFN-respc | Elk3 |
| Dse | 2.77E-12 | 0.72414971 | 0.421 | 0.351 | 5.35E-08 | TAM.IFN-respc | Dse |
| Pltp | 2.79E-12 | 0.35376881 | 0.743 | 0.596 | 5.39E-08 | TAM.IFN-respc | Pltp |
| Snx21 | 3.48E-12 | 0.47614361 | 0.777 | 0.721 | 6.72E-08 | TAM.IFN-respc | Snx2 |
| Usp25 | 5.17E-12 | 0.62445904 | 0.531 | 0.509 | 1.00E-07 | TAM.IFN-respc | Usp25 |
| Lat2 | 5.96E-12 | 0.45672716 | 0.55 | 0.527 | 1.15E-07 | TAM.IFN-respc | Lat2 |
| Ep400 | 6.09E-12 | 0.33272667 | 0.417 | 0.444 | 1.18E-07 | TAM.IFN-respc | Ep400 |
| Sin3a | 6.97E-12 | 0.25831454 | 0.271 | 0.297 | 1.35E-07 | TAM.IFN-respc | Sin3a |
| Tm6sf11 | 8.43E-12 | 0.40237889 | 0.695 | 0.663 | 1.63E-07 | TAM.IFN-respc | Tm6sf1 |
| Tent2 | 1.08E-11 | 0.34586697 | 0.338 | 0.366 | 2.08E-07 | TAM.IFN-respc | Tent2 |
| Epc1 | 1.20E-11 | 0.29482692 | 0.434 | 0.471 | 2.33E-07 | TAM.IFN-respc | Epc1 |
| Ppp2r2a | 1.71E-11 | 0.29892897 | 0.24 | 0.268 | 3.31E-07 | TAM.IFN-respc | Ppp2r2a |
| Fcho2 | 1.76E-11 | 0.25966446 | 0.359 | 0.382 | 3.41E-07 | TAM.IFN-respc | Fcho2 |
| Ids | 2.09E-11 | 0.35638594 | 0.371 | 0.389 | 4.04E-07 | TAM.IFN-respc | Ids |
| Ppp1r12c | 2.13E-11 | 0.29258608 | 0.624 | 0.629 | 4.11E-07 | TAM.IFN-respc | Ppp1r12c |
| Mex3c | 2.63E-11 | 0.32047921 | 0.362 | 0.39 | 5.08E-07 | TAM.IFN-respc | Mex3c |
| Tob21 | 2.80E-11 | 0.30471192 | 0.566 | 0.576 | 5.42E-07 | TAM.IFN-respc | Tob2 |
| Dclre1c1 | 3.12E-11 | 0.66043997 | 0.371 | 0.312 | 6.04E-07 | TAM.IFN-respc | Dclre1c |
| Arid1a | 3.19E-11 | 0.33831686 | 0.39 | 0.41 | 6.16E-07 | TAM.IFN-respc | Arid1a |
| Nsd31 | 3.20E-11 | 0.31730081 | 0.67 | 0.681 | 6.19E-07 | TAM.IFN-respc | Nsd3 |
| Nop531 | 4.37E-11 | 0.26848726 | 0.526 | 0.548 | 8.44E-07 | TAM.IFN-respc | Nop53 |
| Fbrsl1 | 4.48E-11 | 0.7116728 | 0.279 | 0.246 | 8.66E-07 | TAM.IFN-respc | Fbrsl1 |
| Snrpb21 | 4.74E-11 | 0.29226895 | 0.335 | 0.358 | 9.16E-07 | TAM.IFN-respc | Snrpb2 |
| Lacc11 | 5.63E-11 | 0.33450522 | 0.369 | 0.39 | 1.09E-06 | TAM.IFN-respc | Lacc1 |
| Elmo1 | 6.38E-11 | 0.68446169 | 0.494 | 0.431 | 1.23E-06 | TAM.IFN-respc | Elmo1 |
| Pip4k2a1 | 6.75E-11 | 0.2563817 | 0.406 | 0.423 | 1.30E-06 | TAM.IFN-respc | Pip4k2a |
| Dagla1 | 7.43E-11 | 0.42722385 | 0.523 | 0.505 | 1.44E-06 | TAM.IFN-respc | Dagla |
| Tfip111 | 7.76E-11 | 0.31048851 | 0.362 | 0.389 | 1.50E-06 | TAM.IFN-respc | Tfip11 |
| Chmp4b | 9.13E-11 | 0.38354878 | 0.563 | 0.575 | 1.77E-06 | TAM.IFN-respc | Chmp4b |
| Cmtm7 | 1.01E-10 | 0.35081871 | 0.611 | 0.611 | 1.95E-06 | TAM.IFN-respc | Cmtm7 |
| Epb411 | 1.02E-10 | 0.33237662 | 0.373 | 0.4 | 1.97E-06 | TAM.IFN-respc | Epb41 |
| Parp1 | 1.13E-10 | 0.46093406 | 0.4 | 0.396 | 2.19E-06 | TAM.IFN-respc | Parp1 |
| Lgals3bp1 | 1.26E-10 | 0.63380913 | 0.823 | 0.74 | 2.44E-06 | TAM.IFN-respc | Lgals3bp |
| Ddx24 | 1.57E-10 | 0.35180939 | 0.437 | 0.451 | 3.04E-06 | TAM.IFN-respc | Ddx24 |
| Cog5 | 1.73E-10 | 0.26504132 | 0.229 | 0.257 | 3.34E-06 | TAM.IFN-respc | Cog5 |
| Lcp2 | 2.00E-10 | 0.35343126 | 0.593 | 0.597 | 3.86E-06 | TAM.IFN-respc | Lcp2 |
| Zfp263 | 2.24E-10 | 0.26510829 | 0.268 | 0.295 | 4.33E-06 | TAM.IFN-respc | Zfp263 |
| Rab431 | 2.59E-10 | 0.31038624 | 0.683 | 0.686 | 5.00E-06 | TAM.IFN-respc | Rab43 |
| Susd3 | 2.65E-10 | 0.79029829 | 0.257 | 0.209 | 5.13E-06 | TAM.IFN-respc | Susd3 |
| Snap23 | 2.66E-10 | 0.31032419 | 0.261 | 0.285 | 5.15E-06 | TAM.IFN-respc | Snap23 |
| Il10ra | 2.85E-10 | 0.53891735 | 0.763 | 0.704 | 5.50E-06 | TAM.IFN-respc | Il10ra |
| Abcg1 | 2.88E-10 | 0.30714607 | 0.245 | 0.264 | 5.57E-06 | TAM.IFN-respc | Abcg1 |
| Igbp1 | 3.25E-10 | 0.32540794 | 0.341 | 0.361 | 6.28E-06 | TAM.IFN-respc | Igbp1 |
| Gsdmd | 3.26E-10 | 0.63060234 | 0.572 | 0.521 | 6.31E-06 | TAM.IFN-respc | Gsdmd |
| Ier5 | 3.39E-10 | 0.47431305 | 0.852 | 0.79 | 6.55E-06 | TAM.IFN-respc | Ier5 |
| Pld4 | 3.47E-10 | 0.40050328 | 0.878 | 0.782 | 6.71E-06 | TAM.IFN-respc | Pld4 |
| Foxj2 | 4.96E-10 | 0.27982566 | 0.241 | 0.276 | 9.58E-06 | TAM.IFN-respc | Foxj2 |
| Lin542 | 5.18E-10 | 0.27837506 | 0.264 | 0.292 | 1.00E-05 | TAM.IFN-respc | Lin54 |
| Tmem511 | 5.22E-10 | 0.35415946 | 0.379 | 0.381 | 1.01E-05 | TAM.IFN-respc | Tmem51 |
| Dock101 | 6.66E-10 | 0.41169077 | 0.49 | 0.478 | 1.29E-05 | TAM.IFN-respc | Dock10 |
| Rbfa1 | 7.25E-10 | 0.31215942 | 0.313 | 0.338 | 1.40E-05 | TAM.IFN-respc | Rbfa |
| Dnajc131 | 7.53E-10 | 0.33266516 | 0.472 | 0.479 | 1.46E-05 | TAM.IFN-respc | Dnajc13 |
| Capza2 | 1.29E-09 | 0.40958895 | 0.619 | 0.616 | 2.50E-05 | TAM.IFN-respc | Capza2 |
| Tle3 | 1.54E-09 | 0.28148025 | 0.472 | 0.501 | 2.98E-05 | TAM.IFN-respc | Tle3 |
| Eid11 | 1.59E-09 | 0.41455554 | 0.767 | 0.715 | 3.08E-05 | TAM.IFN-respc | Eid1 |
| Tmem128 | 1.88E-09 | 0.25992255 | 0.281 | 0.31 | 3.63E-05 | TAM.IFN-respc | Tmem128 |
| Plxnb2 | 2.04E-09 | 0.40042272 | 0.635 | 0.591 | 3.95E-05 | TAM.IFN-respc | Plxnb2 |
| Ppp4r2 | 2.06E-09 | 0.31528129 | 0.503 | 0.524 | 3.98E-05 | TAM.IFN-respc | Ppp4r2 |
| Arhgef61 | 2.12E-09 | 0.34017319 | 0.331 | 0.336 | 4.10E-05 | TAM.IFN-respc | Arhgef6 |
| Zcchc241 | 2.16E-09 | 0.31276397 | 0.331 | 0.336 | 4.17E-05 | TAM.IFN-respc | Zcchc24 |
| Dok11 | 2.93E-09 | 0.26276204 | 0.242 | 0.27 | 5.67E-05 | TAM.IFN-respc | Dok1 |
| Akap10 | 3.15E-09 | 0.32741419 | 0.225 | 0.254 | 6.09E-05 | TAM.IFN-respc | Akap10 |
| Rgl11 | 4.73E-09 | 0.69580561 | 0.473 | 0.394 | 9.14E-05 | TAM.IFN-respc | Rgl1 |
| H2-K1 | 5.41E-09 | 0.38996451 | 0.965 | 0.95 | 0.00010466 | TAM.IFN-respc | H2-K1 |
| Xrn2 | 5.42E-09 | 0.35043224 | 0.465 | 0.486 | 0.0001048 | TAM.IFN-respc | Xrn2 |
| Nsa21 | 5.66E-09 | 0.32077457 | 0.23 | 0.251 | 0.00010938 | TAM.IFN-respc | Nsa2 |
| Gps2 | 5.74E-09 | 0.31031248 | 0.25 | 0.274 | 0.00011098 | TAM.IFN-respc | Gps2 |
| Cnot2 | 5.88E-09 | 0.28508513 | 0.35 | 0.372 | 0.00011371 | TAM.IFN-respc | Cnot2 |
| Arid5b1 | 7.60E-09 | 0.32274023 | 0.289 | 0.306 | 0.00014698 | TAM.IFN-respc | Arid5b |
| Zeb2 | 7.61E-09 | 0.4413652 | 0.941 | 0.898 | 0.00014718 | TAM.IFN-respc | Zeb2 |
| Zfp7401 | 8.05E-09 | 0.31534234 | 0.232 | 0.259 | 0.00015569 | TAM.IFN-respc | Zfp740 |
| Tcof1 | 8.31E-09 | 0.28645454 | 0.278 | 0.312 | 0.00016073 | TAM.IFN-respc | Tcof1 |
| Kdm6a | 8.45E-09 | 0.29266391 | 0.315 | 0.335 | 0.00016336 | TAM.IFN-respc | Kdm6a |
| Tor1aip2 | 9.19E-09 | 0.3318263 | 0.474 | 0.523 | 0.00017768 | TAM.IFN-respc | Tor1aip2 |
| Lmnb11 | 1.03E-08 | 0.31671986 | 0.467 | 0.426 | 0.00019888 | TAM.IFN-respc | Lmnb1 |
| Erap11 | 1.28E-08 | 0.34293023 | 0.265 | 0.289 | 0.00024741 | TAM.IFN-respc | Erap1 |
| Skil1 | 1.42E-08 | 0.51303165 | 0.814 | 0.765 | 0.0002746 | TAM.IFN-respc | Skil |
| Shisa5 | 1.49E-08 | 0.37930435 | 0.823 | 0.802 | 0.00028781 | TAM.IFN-respc | Shisa5 |
| Rasgef1b1 | 1.69E-08 | 0.57825544 | 0.666 | 0.572 | 0.00032616 | TAM.IFN-respc | Rasgef1b |
| Hps31 | 1.70E-08 | 0.30853492 | 0.229 | 0.255 | 0.00032833 | TAM.IFN-respc | Hps3 |
| Rgs2 | 2.30E-08 | 0.50853012 | 0.642 | 0.609 | 0.00044386 | TAM.IFN-respc | Rgs2 |
| Dock2 | 2.31E-08 | 0.37581452 | 0.591 | 0.576 | 0.00044716 | TAM.IFN-respc | Dock2 |
| Sppl2a | 2.47E-08 | 0.36921291 | 0.589 | 0.603 | 0.00047665 | TAM.IFN-respc | Sppl2a |
| Cyfip1 | 2.57E-08 | 0.47336617 | 0.702 | 0.67 | 0.00049715 | TAM.IFN-respc | Cyfip1 |
| Setd1b | 3.11E-08 | 0.29352817 | 0.32 | 0.341 | 0.00060197 | TAM.IFN-respc | Setd1b |
| Selenow | 3.49E-08 | 0.40874716 | 0.662 | 0.667 | 0.00067435 | TAM.IFN-respc | Selenow |
| Ube2v21 | 4.41E-08 | 0.28238364 | 0.251 | 0.27 | 0.00085357 | TAM.IFN-respc | Ube2v2 |
| Rassf4 | 4.52E-08 | 0.43452332 | 0.631 | 0.606 | 0.00087323 | TAM.IFN-respc | Rassf4 |
| Cep170 | 4.85E-08 | 0.33668137 | 0.446 | 0.464 | 0.00093789 | TAM.IFN-respc | Cep170 |
| Mepce | 5.07E-08 | 0.29692707 | 0.301 | 0.327 | 0.00097995 | TAM.IFN-respc | Mepce |
| Crebbp | 5.94E-08 | 0.31696786 | 0.407 | 0.426 | 0.00114761 | TAM.IFN-respc | Crebbp |
| Fchsd2 | 6.04E-08 | 0.28556373 | 0.368 | 0.356 | 0.00116785 | TAM.IFN-respc | Fchsd2 |
| Tmem184b | 7.71E-08 | 0.37029191 | 0.387 | 0.427 | 0.00148995 | TAM.IFN-respc | Tmem184b |
| Tnrc6b | 7.91E-08 | 0.32126604 | 0.278 | 0.299 | 0.00152903 | TAM.IFN-respc | Tnrc6b |
| Gtpbp11 | 8.54E-08 | 0.33440008 | 0.265 | 0.287 | 0.0016517 | TAM.IFN-respc | Gtpbp1 |
| Etnk1 | 9.42E-08 | 0.61688103 | 0.39 | 0.401 | 0.00182213 | TAM.IFN-respc | Etnk1 |
| Rufy31 | 1.17E-07 | 0.27512184 | 0.253 | 0.285 | 0.00225885 | TAM.IFN-respc | Rufy3 |
| Klhl5 | 1.34E-07 | 0.39750747 | 0.283 | 0.3 | 0.00258859 | TAM.IFN-respc | Klhl5 |
| Arhgap271 | 1.95E-07 | 0.35719513 | 0.231 | 0.254 | 0.00377523 | TAM.IFN-respc | Arhgap27 |
| Ankrd44 | 2.12E-07 | 0.27110344 | 0.293 | 0.315 | 0.00409417 | TAM.IFN-respc | Ankrd44 |
| Hdac5 | 2.16E-07 | 0.28394982 | 0.423 | 0.444 | 0.0041669 | TAM.IFN-respc | Hdac5 |
| Dennd1a | 2.16E-07 | 0.71934014 | 0.419 | 0.368 | 0.00417867 | TAM.IFN-respc | Dennd1a |
| Slamf82 | 2.67E-07 | 0.57831373 | 0.304 | 0.291 | 0.00515267 | TAM.IFN-respc | Slamf8 |
| Max | 2.67E-07 | 0.35207085 | 0.388 | 0.416 | 0.00516557 | TAM.IFN-respc | Max |
| Vill1 | 2.94E-07 | 0.41712838 | 0.265 | 0.257 | 0.00568285 | TAM.IFN-respc | Vill |
| C3ar11 | 3.12E-07 | 0.65109729 | 0.8 | 0.669 | 0.00603465 | TAM.IFN-respc | C3ar1 |
| Lgals91 | 3.16E-07 | 0.50632212 | 0.583 | 0.587 | 0.00611467 | TAM.IFN-respc | Lgals9 |
| Sbno2 | 3.54E-07 | 0.30756039 | 0.773 | 0.759 | 0.0068423 | TAM.IFN-respc | Sbno2 |
| Ptger41 | 3.75E-07 | 0.56982129 | 0.397 | 0.365 | 0.00724942 | TAM.IFN-respc | Ptger4 |
| Nr4a11 | 4.19E-07 | 0.45912573 | 0.835 | 0.79 | 0.00810162 | TAM.IFN-respc | Nr4a1 |
| Pik3r5 | 4.24E-07 | 0.40675646 | 0.574 | 0.558 | 0.00819117 | TAM.IFN-respc | Pik3r5 |
| Lgals1 | 5.27E-07 | 0.39124842 | 0.271 | 0.273 | 0.01019761 | TAM.IFN-respc | Lgals1 |
| Atp6v0a2 | 5.36E-07 | 0.35006034 | 0.289 | 0.323 | 0.01037149 | TAM.IFN-respc | Atp6v0a2 |
| Thoc6 | 5.88E-07 | 0.26428375 | 0.249 | 0.268 | 0.01137281 | TAM.IFN-respc | Thoc6 |
| Serinc31 | 6.82E-07 | 0.44758695 | 0.91 | 0.861 | 0.01319118 | TAM.IFN-respc | Serinc3 |
| Cdk11b1 | 7.67E-07 | 0.353613 | 0.425 | 0.464 | 0.01483307 | TAM.IFN-respc | Cdk11b |
| Cd742 | 8.29E-07 | 0.50199334 | 0.974 | 0.937 | 0.01602234 | TAM.IFN-respc | Cd74 |
| Ppp1r15a | 8.29E-07 | 0.39653325 | 0.766 | 0.725 | 0.01602706 | TAM.IFN-respc | Ppp1r15a |
| Pirb | 9.14E-07 | 0.39052637 | 0.847 | 0.81 | 0.01766516 | TAM.IFN-respc | Pirb |
| Itga61 | 9.68E-07 | 0.83068883 | 0.417 | 0.34 | 0.01872274 | TAM.IFN-respc | Itga6 |
| Dock8 | 1.23E-06 | 0.68356255 | 0.522 | 0.468 | 0.02383227 | TAM.IFN-respc | Dock8 |
| Il21r | 1.34E-06 | 0.3986549 | 0.489 | 0.47 | 0.02582577 | TAM.IFN-respc | Il21r |
| Tnfsf14 | 1.34E-06 | 0.6790535 | 0.261 | 0.217 | 0.02585299 | TAM.IFN-respc | Tnfsf14 |
| Zfp5131 | 1.35E-06 | 0.32290047 | 0.23 | 0.25 | 0.02618973 | TAM.IFN-respc | Zfp513 |
| Nr4a21 | 1.50E-06 | 0.49332878 | 0.598 | 0.535 | 0.02892976 | TAM.IFN-respc | Nr4a2 |
| Casp8 | 1.68E-06 | 0.34476314 | 0.43 | 0.444 | 0.03248503 | TAM.IFN-respc | Casp8 |
| Tmem881 | 1.85E-06 | 0.57385406 | 0.361 | 0.321 | 0.03576228 | TAM.IFN-respc | Tmem88 |
| Zswim6 | 2.51E-06 | 0.41841211 | 0.438 | 0.441 | 0.04850976 | TAM.IFN-respc | Zswim6 |
| Rb1 | 2.62E-06 | 0.3363237 | 0.312 | 0.332 | 0.05063709 | TAM.IFN-respc | Rb1 |
| Psmb101 | 2.66E-06 | 0.53903256 | 0.486 | 0.469 | 0.051446 | TAM.IFN-respc | Psmb10 |
| Cpne2 | 3.16E-06 | 0.45166882 | 0.265 | 0.272 | 0.06109592 | TAM.IFN-respc | Cpne2 |
| Rybp | 4.13E-06 | 0.55050794 | 0.294 | 0.262 | 0.07985578 | TAM.IFN-respc | Rybp |
| Fam49b1 | 4.75E-06 | 0.48821893 | 0.853 | 0.81 | 0.0917424 | TAM.IFN-respc | Fam49b |
| Arhgap181 | 4.85E-06 | 0.40439532 | 0.277 | 0.287 | 0.09382477 | TAM.IFN-respc | Arhgap18 |
| Ncoa2 | 4.92E-06 | 0.28874883 | 0.262 | 0.282 | 0.09517782 | TAM.IFN-respc | Ncoa2 |
| Trim141 | 5.10E-06 | 0.66073311 | 0.309 | 0.294 | 0.09866469 | TAM.IFN-respc | Trim14 |
| Ccl121 | 5.60E-06 | 0.50952002 | 0.293 | 0.227 | 0.10825342 | TAM.IFN-respc | Ccl12 |
| Iscu | 5.96E-06 | 0.38964191 | 0.249 | 0.264 | 0.11523122 | TAM.IFN-respc | Iscu |
| Elmsan11 | 6.63E-06 | 0.63553019 | 0.296 | 0.265 | 0.12814992 | TAM.IFN-respc | Elmsan1 |
| Cept1 | 7.22E-06 | 0.34449475 | 0.366 | 0.393 | 0.13967922 | TAM.IFN-respc | Cept1 |
| Tle4 | 7.39E-06 | 0.32772419 | 0.282 | 0.298 | 0.1429278 | TAM.IFN-respc | Tle4 |
| Armcx31 | 8.18E-06 | 0.36198048 | 0.264 | 0.283 | 0.15821824 | TAM.IFN-respc | Armcx3 |
| Bcl3 | 8.53E-06 | 0.32059878 | 0.535 | 0.555 | 0.16500598 | TAM.IFN-respc | Bcl3 |
| Gna15 | 8.86E-06 | 0.39232374 | 0.24 | 0.254 | 0.17138728 | TAM.IFN-respc | Gna15 |
| Clec4a21 | 9.58E-06 | 0.77638393 | 0.373 | 0.329 | 0.18514974 | TAM.IFN-respc | Clec4a2 |
| Sowahc1 | 9.66E-06 | 0.3156419 | 0.512 | 0.513 | 0.18667041 | TAM.IFN-respc | Sowahc |
| Tap2 | 1.11E-05 | 0.52076279 | 0.526 | 0.523 | 0.2154396 | TAM.IFN-respc | Tap2 |
| Anks11 | 1.21E-05 | 0.34678296 | 0.341 | 0.359 | 0.23467996 | TAM.IFN-respc | Anks1 |
| Lifr | 1.38E-05 | 0.53537249 | 0.282 | 0.247 | 0.26596676 | TAM.IFN-respc | Lifr |
| Cflar | 1.52E-05 | 0.30176798 | 0.497 | 0.507 | 0.29315917 | TAM.IFN-respc | Cflar |
| Stx16 | 1.63E-05 | 0.4035705 | 0.354 | 0.37 | 0.31565712 | TAM.IFN-respc | Stx16 |
| Slc29a31 | 1.73E-05 | 0.43830809 | 0.492 | 0.477 | 0.33463627 | TAM.IFN-respc | Slc29a3 |
| Celf2 | 1.76E-05 | 0.33368986 | 0.445 | 0.446 | 0.34042487 | TAM.IFN-respc | Celf2 |
| Nup981 | 1.77E-05 | 0.29496365 | 0.446 | 0.464 | 0.34274282 | TAM.IFN-respc | Nup98 |
| Rnf34 | 1.78E-05 | 0.61062951 | 0.286 | 0.27 | 0.34472506 | TAM.IFN-respc | Rnf34 |
| Apaf1 | 2.12E-05 | 0.30081387 | 0.414 | 0.436 | 0.4107992 | TAM.IFN-respc | Apaf1 |
| Slc30a11 | 2.24E-05 | 0.60673636 | 0.382 | 0.371 | 0.4330694 | TAM.IFN-respc | Slc30a1 |
| Tifa | 2.49E-05 | 0.59328993 | 0.306 | 0.282 | 0.48194257 | TAM.IFN-respc | Tifa |
| Irf2 | 2.63E-05 | 0.55200566 | 0.403 | 0.371 | 0.50937784 | TAM.IFN-respc | Irf2 |
| Zfp710 | 2.69E-05 | 0.40190011 | 0.49 | 0.482 | 0.52104196 | TAM.IFN-respc | Zfp710 |
| Fli1 | 2.91E-05 | 0.44285553 | 0.524 | 0.502 | 0.56228554 | TAM.IFN-respc | Fli1 |
| Ep300 | 2.98E-05 | 0.35964114 | 0.513 | 0.509 | 0.575281 | TAM.IFN-respc | Ep300 |
| Tmem106a1 | 3.19E-05 | 0.5851229 | 0.666 | 0.585 | 0.61617058 | TAM.IFN-respc | Tmem106a |
| Rp2 | 4.30E-05 | 0.37788815 | 0.467 | 0.48 | 0.83089048 | TAM.IFN-respc | Rp2 |
| Gadd45b1 | 4.32E-05 | 0.49236653 | 0.843 | 0.795 | 0.83576242 | TAM.IFN-respc | Gadd45b |
| Psmb81 | 4.68E-05 | 0.54426698 | 0.695 | 0.685 | 0.90429963 | TAM.IFN-respc | Psmb8 |
| Gbp81 | 5.76E-05 | 0.26540962 | 0.262 | 0.272 | 1 | TAM.IFN-respc | Gbp8 |
| Pvr1 | 5.85E-05 | 0.52215069 | 0.365 | 0.351 | 1 | TAM.IFN-respc | Pvr |
| Zfp5161 | 6.57E-05 | 0.36801658 | 0.653 | 0.644 | 1 | TAM.IFN-respc | Zfp516 |
| Rap2c | 6.98E-05 | 0.62447114 | 0.367 | 0.345 | 1 | TAM.IFN-respc | Rap2c |
| Lgals81 | 7.44E-05 | 0.43552006 | 0.434 | 0.426 | 1 | TAM.IFN-respc | Lgals8 |
| Ap1s2 | 7.71E-05 | 0.27669704 | 0.305 | 0.306 | 1 | TAM.IFN-respc | Ap1s2 |
| Tppp3 | 8.27E-05 | 0.33437095 | 0.363 | 0.254 | 1 | TAM.IFN-respc | Tppp3 |
| Tbc1d141 | 9.63E-05 | 0.41927945 | 0.505 | 0.477 | 1 | TAM.IFN-respc | Tbc1d14 |
| Tiam1 | 0.00011514 | 0.31377522 | 0.267 | 0.213 | 1 | TAM.IFN-respc | Tiam1 |
| B4galt6 | 0.00011544 | 0.41785451 | 0.403 | 0.407 | 1 | TAM.IFN-respc | B4galt6 |
| Tlr71 | 0.00011785 | 0.4247115 | 0.247 | 0.251 | 1 | TAM.IFN-respc | Tlr7 |
| Bach1 | 0.00012088 | 0.49974017 | 0.796 | 0.738 | 1 | TAM.IFN-respc | Bach1 |
| Mycbp21 | 0.00013081 | 0.37925518 | 0.585 | 0.575 | 1 | TAM.IFN-respc | Mycbp2 |
| Mafk | 0.00013449 | 0.36086668 | 0.333 | 0.357 | 1 | TAM.IFN-respc | Mafk |
| Dock41 | 0.00013768 | 0.69137171 | 0.298 | 0.261 | 1 | TAM.IFN-respc | Dock4 |
| Pxk | 0.00014954 | 0.34129201 | 0.413 | 0.413 | 1 | TAM.IFN-respc | Pxk |
| Fnbp4 | 0.00017888 | 0.39610226 | 0.336 | 0.356 | 1 | TAM.IFN-respc | Fnbp4 |
| Psme21 | 0.00021134 | 0.60664798 | 0.419 | 0.406 | 1 | TAM.IFN-respc | Psme2 |
| Eng1 | 0.00021329 | 0.33178062 | 0.261 | 0.272 | 1 | TAM.IFN-respc | Eng |
| Tnfaip3 | 0.00022778 | 0.54506741 | 0.648 | 0.615 | 1 | TAM.IFN-respc | Tnfaip3 |
| Vwa5a | 0.00024893 | 0.4434956 | 0.503 | 0.492 | 1 | TAM.IFN-respc | Vwa5a |
| Eva1b2 | 0.00026302 | 0.38140028 | 0.399 | 0.414 | 1 | TAM.IFN-respc | Eva1b |
| Adap21 | 0.00026459 | 0.60119158 | 0.644 | 0.579 | 1 | TAM.IFN-respc | Adap2 |
| Pstpip21 | 0.00027967 | 0.30410704 | 0.366 | 0.383 | 1 | TAM.IFN-respc | Pstpip2 |
| Tbc1d51 | 0.00030485 | 0.42864299 | 0.352 | 0.355 | 1 | TAM.IFN-respc | Tbc1d5 |
| Vps54 | 0.00036903 | 0.3707872 | 0.289 | 0.308 | 1 | TAM.IFN-respc | Vps54 |
| Lyl1 | 0.0004154 | 0.42977728 | 0.254 | 0.21 | 1 | TAM.IFN-respc | Lyl1 |
| Camk1d | 0.00043713 | 0.64249001 | 0.519 | 0.462 | 1 | TAM.IFN-respc | Camk1d |
| Wsb1 | 0.00046089 | 0.40546829 | 0.822 | 0.805 | 1 | TAM.IFN-respc | Wsb1 |
| Med291 | 0.00047397 | 0.45577455 | 0.27 | 0.282 | 1 | TAM.IFN-respc | Med29 |
| Usp381 | 0.00048678 | 0.33645001 | 0.236 | 0.262 | 1 | TAM.IFN-respc | Usp38 |
| Lpxn | 0.00053402 | 0.64977816 | 0.38 | 0.34 | 1 | TAM.IFN-respc | Lpxn |
| Synj1 | 0.00053881 | 0.59453461 | 0.475 | 0.447 | 1 | TAM.IFN-respc | Synj1 |
| Madd | 0.00058337 | 0.50458749 | 0.354 | 0.326 | 1 | TAM.IFN-respc | Madd |
| Cers61 | 0.00075043 | 0.54185994 | 0.53 | 0.516 | 1 | TAM.IFN-respc | Cers6 |
| Dusp21 | 0.00087404 | 0.43140417 | 0.365 | 0.311 | 1 | TAM.IFN-respc | Dusp2 |
| Tet31 | 0.00097511 | 0.38677685 | 0.487 | 0.491 | 1 | TAM.IFN-respc | Tet3 |
| Ms4a71 | 0.00116725 | 0.57700141 | 0.566 | 0.481 | 1 | TAM.IFN-respc | Ms4a7 |
| Btg21 | 0.00117101 | 0.38853739 | 0.878 | 0.825 | 1 | TAM.IFN-respc | Btg2 |
| Rara | 0.00122067 | 0.43401334 | 0.582 | 0.542 | 1 | TAM.IFN-respc | Rara |
| Coq10b1 | 0.00124477 | 0.52708019 | 0.449 | 0.437 | 1 | TAM.IFN-respc | Coq10b |
| Gla | 0.00127585 | 0.47364628 | 0.545 | 0.537 | 1 | TAM.IFN-respc | Gla |
| Ifnar1 | 0.00150775 | 0.60166552 | 0.61 | 0.565 | 1 | TAM.IFN-respc | Ifnar1 |
| Crlf3 | 0.00171982 | 0.40173541 | 0.455 | 0.459 | 1 | TAM.IFN-respc | Crlf3 |
| Arl5c1 | 0.00182247 | 0.43672138 | 0.346 | 0.328 | 1 | TAM.IFN-respc | Arl5c |
| Aftph | 0.0018449 | 0.34228466 | 0.255 | 0.26 | 1 | TAM.IFN-respc | Aftph |
| Trim261 | 0.00229126 | 0.36176199 | 0.236 | 0.257 | 1 | TAM.IFN-respc | Trim26 |
| Krt80 | 0.00229862 | 0.39455194 | 0.278 | 0.224 | 1 | TAM.IFN-respc | Krt80 |
| Emp3 | 0.00235118 | 0.45209516 | 0.849 | 0.785 | 1 | TAM.IFN-respc | Emp3 |
| Peli1 | 0.00283796 | 0.36106533 | 0.454 | 0.47 | 1 | TAM.IFN-respc | Peli1 |
| Limd21 | 0.00297155 | 0.57823981 | 0.417 | 0.383 | 1 | TAM.IFN-respc | Limd2 |
| Klf2 | 0.00316641 | 0.43113191 | 0.437 | 0.362 | 1 | TAM.IFN-respc | Klf2 |
| Akna1 | 0.00324941 | 0.32508705 | 0.319 | 0.319 | 1 | TAM.IFN-respc | Akna |
| S100a41 | 0.00358372 | 0.39255579 | 0.771 | 0.573 | 1 | TAM.IFN-respc | S100a4 |
| Prkx | 0.00391776 | 0.35262221 | 0.308 | 0.331 | 1 | TAM.IFN-respc | Prkx |
| Pdlim41 | 0.0039599 | 0.46267604 | 0.282 | 0.277 | 1 | TAM.IFN-respc | Pdlim4 |
| Morc3 | 0.0047166 | 0.58945335 | 0.397 | 0.369 | 1 | TAM.IFN-respc | Morc3 |
| Ly861 | 0.00504281 | 0.50313213 | 0.718 | 0.622 | 1 | TAM.IFN-respc | Ly86 |
| Ifnar2 | 0.00538398 | 0.59054848 | 0.746 | 0.693 | 1 | TAM.IFN-respc | Ifnar2 |
| Dusp71 | 0.00610283 | 0.53666503 | 0.435 | 0.421 | 1 | TAM.IFN-respc | Dusp7 |
| H3f3b | 0.00626444 | 0.50551817 | 0.877 | 0.83 | 1 | TAM.IFN-respc | H3f3b |
| Jmjd1c | 0.00720517 | 0.53507808 | 0.641 | 0.614 | 1 | TAM.IFN-respc | Jmjd1c |
| Plcl2 | 0.00774222 | 0.56820468 | 0.389 | 0.358 | 1 | TAM.IFN-respc | Plcl2 |
| Il6ra | 0.00824804 | 0.33807409 | 0.507 | 0.492 | 1 | TAM.IFN-respc | Il6ra |
| Acer31 | 0.00828656 | 0.58532305 | 0.316 | 0.287 | 1 | TAM.IFN-respc | Acer3 |
| Zfp7031 | 0.00860913 | 0.39290808 | 0.825 | 0.817 | 1 | TAM.IFN-respc | Zfp703 |
| Psme2b | 0.00879302 | 0.59666178 | 0.373 | 0.359 | 1 | TAM.IFN-respc | Psme2b |
| Plac81 | 0 | 3.82473666 | 0.824 | 0.272 | 0 | monocytes | Plac8 |
| Chil3 | 0 | 4.44632996 | 0.591 | 0.095 | 0 | monocytes | Chil3 |
| Ly6c2 | 0 | 3.96727793 | 0.47 | 0.044 | 0 | monocytes | Ly6c2 |
| S100a42 | 0 | 1.94549121 | 0.969 | 0.551 | 0 | monocytes | S100a4 |
| Hp1 | 0 | 2.59044031 | 0.617 | 0.202 | 0 | monocytes | Hp |
| Ly6c1 | 0 | 3.81258132 | 0.419 | 0.04 | 0 | monocytes | Ly6c1 |
| Ms4a4c1 | 0 | 2.23685889 | 0.448 | 0.114 | 0 | monocytes | Ms4a4c |
| Rab441 | 0 | 2.41184035 | 0.554 | 0.224 | 0 | monocytes | Rab44 |
| Itgb7 | 0 | 3.051343 | 0.431 | 0.12 | 0 | monocytes | Itgb7 |
| Vcan | 0 | 2.89838974 | 0.39 | 0.09 | 0 | monocytes | Vcan |
| Clec4e1 | 0 | 1.40531381 | 0.694 | 0.395 | 0 | monocytes | Clec4e |
| Cytip1 | 0 | 1.5443167 | 0.835 | 0.546 | 0 | monocytes | Cytip |
| Ccr22 | 0 | 1.38679319 | 0.674 | 0.386 | 0 | monocytes | Ccr2 |
| Sell | 0 | 4.97523742 | 0.306 | 0.025 | 0 | monocytes | Sell |
| Tiam11 | 0 | 2.47500663 | 0.464 | 0.19 | 0 | monocytes | Tiam1 |
| Atp2b4 | 0 | 2.58110494 | 0.425 | 0.155 | 0 | monocytes | Atp2b4 |
| Ifitm6 | 0 | 4.75898233 | 0.289 | 0.031 | 0 | monocytes | Ifitm6 |
| Mefv1 | 0 | 2.12132031 | 0.503 | 0.252 | 0 | monocytes | Mefv |
| Gsr1 | 0 | 1.51843728 | 0.888 | 0.638 | 0 | monocytes | Gsr |
| Ifi27l2a1 | 0 | 1.50601463 | 0.621 | 0.372 | 0 | monocytes | Ifi27l2a |
| Slpi1 | 0 | 2.1810435 | 0.416 | 0.167 | 0 | monocytes | Slpi |
| Dgat2 | 0 | 3.15527986 | 0.368 | 0.121 | 0 | monocytes | Dgat2 |
| Cd244a1 | 0 | 2.29426205 | 0.514 | 0.269 | 0 | monocytes | Cd244a |
| Ltb4r11 | 0 | 2.20644082 | 0.383 | 0.142 | 0 | monocytes | Ltb4r1 |
| Glipr2 | 0 | 2.18686298 | 0.384 | 0.151 | 0 | monocytes | Glipr2 |
| Rhov1 | 0 | 2.63660016 | 0.384 | 0.154 | 0 | monocytes | Rhov |
| Nod21 | 0 | 1.77754729 | 0.567 | 0.338 | 0 | monocytes | Nod2 |
| Arhgap26 | 0 | 2.21038953 | 0.46 | 0.241 | 0 | monocytes | Arhgap26 |
| Jak21 | 0 | 1.74475794 | 0.711 | 0.492 | 0 | monocytes | Jak2 |
| Ifitm31 | 0 | 1.97849351 | 0.711 | 0.493 | 0 | monocytes | Ifitm3 |
| Havcr21 | 0 | 2.00986707 | 0.567 | 0.351 | 0 | monocytes | Havcr2 |
| Fmnl11 | 0 | 1.55576118 | 0.842 | 0.636 | 0 | monocytes | Fmnl1 |
| Rhou | 0 | 2.36787465 | 0.355 | 0.151 | 0 | monocytes | Rhou |
| Ifi2051 | 0 | 2.04809962 | 0.426 | 0.225 | 0 | monocytes | Ifi205 |
| Adora2b1 | 0 | 1.91724502 | 0.4 | 0.205 | 0 | monocytes | Adora2b |
| St3gal41 | 0 | 1.81747066 | 0.555 | 0.364 | 0 | monocytes | St3gal4 |
| B3gnt52 | 0 | 1.97559641 | 0.327 | 0.138 | 0 | monocytes | B3gnt5 |
| Jarid21 | 0 | 1.60853835 | 0.699 | 0.514 | 0 | monocytes | Jarid2 |
| Rara1 | 0 | 1.70259595 | 0.709 | 0.527 | 0 | monocytes | Rara |
| Tes1 | 0 | 1.55096556 | 0.668 | 0.502 | 0 | monocytes | Tes |
| S100a61 | 0 | 1.34402651 | 0.908 | 0.755 | 0 | monocytes | S100a6 |
| Emilin21 | 0 | 1.60145758 | 0.925 | 0.781 | 0 | monocytes | Emilin2 |
| Thbs12 | 0 | 1.79036222 | 0.8 | 0.673 | 0 | monocytes | Thbs1 |
| Osm2 | 0 | 1.31467174 | 0.791 | 0.666 | 0 | monocytes | Osm |
| Cd802 | 0 | 1.35396238 | 0.705 | 0.582 | 0 | monocytes | Cd80 |
| Cd441 | 0 | 1.59740526 | 0.942 | 0.834 | 0 | monocytes | Cd44 |
| Il1b2 | 0 | 1.23346639 | 0.951 | 0.843 | 0 | monocytes | Il1b |
| Samhd11 | 0 | 1.38974114 | 0.842 | 0.746 | 0 | monocytes | Samhd1 |
| Anxa2 | 0 | 1.44435694 | 0.942 | 0.847 | 0 | monocytes | Anxa2 |
| Aldh21 | 0 | 0.28007944 | 0.646 | 0.735 | 0 | monocytes | Aldh2 |
| Cxcl22 | 0 | 0.29042314 | 0.721 | 0.799 | 0 | monocytes | Cxcl2 |
| Il1rn2 | 0 | 0.30081022 | 0.682 | 0.755 | 0 | monocytes | Il1rn |
| Gna13 | 0 | 0.25762302 | 0.837 | 0.908 | 0 | monocytes | Gna13 |
| Ms4a4a2 | 0 | 0.34404825 | 0.77 | 0.701 | 0 | monocytes | Ms4a4a |
| Cd523 | 0 | 0.43162724 | 0.928 | 0.865 | 0 | monocytes | Cd52 |
| Cybb1 | 0 | 0.41804395 | 0.682 | 0.74 | 0 | monocytes | Cybb |
| Cd91 | 0 | 0.36262287 | 0.688 | 0.731 | 0 | monocytes | Cd9 |
| Irf2bp21 | 0 | 0.3401025 | 0.901 | 0.944 | 0 | monocytes | Irf2bp2 |
| Srgn | 0 | 1.57651656 | 0.981 | 0.942 | 0 | monocytes | Srgn |
| Calm11 | 0 | 0.27045152 | 0.825 | 0.864 | 0 | monocytes | Calm1 |
| Dusp12 | 0 | 0.3627579 | 0.922 | 0.89 | 0 | monocytes | Dusp1 |
| Itgb21 | 0 | 0.46567959 | 0.955 | 0.923 | 0 | monocytes | Itgb2 |
| Tgm2 | 0 | 1.72217067 | 0.836 | 0.809 | 0 | monocytes | Tgm2 |
| H3f3a | 0 | 0.36858448 | 0.863 | 0.889 | 0 | monocytes | H3f3a |
| Vim | 0 | 1.26294624 | 0.984 | 0.959 | 0 | monocytes | Vim |
| Zfp361 | 0 | 0.41005936 | 0.978 | 0.956 | 0 | monocytes | Zfp36 |
| Rap1b1 | 0 | 0.43868905 | 0.873 | 0.893 | 0 | monocytes | Rap1b |
| Btg22 | 0 | 0.29981201 | 0.848 | 0.829 | 0 | monocytes | Btg2 |
| Litaf | 0 | 0.35178004 | 0.91 | 0.929 | 0 | monocytes | Litaf |
| Capzb | 0 | 0.39014383 | 0.832 | 0.85 | 0 | monocytes | Capzb |
| H3f3b1 | 0 | 0.29799743 | 0.82 | 0.837 | 0 | monocytes | H3f3b |
| Rac21 | 0 | 0.40567467 | 0.845 | 0.828 | 0 | monocytes | Rac2 |
| Zc3h12a2 | 0 | 0.29979799 | 0.722 | 0.738 | 0 | monocytes | Zc3h12a |
| Tyrobp | 0 | 0.50005022 | 0.985 | 0.973 | 0 | monocytes | Tyrobp |
| Plek1 | 0 | 0.44193169 | 0.949 | 0.937 | 0 | monocytes | Plek |
| Tpt11 | 0 | 0.56533035 | 0.966 | 0.956 | 0 | monocytes | Tpt1 |
| Rnf1491 | 0 | 0.37347729 | 0.933 | 0.925 | 0 | monocytes | Rnf149 |
| Fau | 0 | 0.43479766 | 0.957 | 0.949 | 0 | monocytes | Fau |
| Spi1 | 0 | 0.46785019 | 0.857 | 0.862 | 0 | monocytes | Spi1 |
| Arpc1b1 | 0 | 0.42203979 | 0.936 | 0.931 | 0 | monocytes | Arpc1b |
| Laptm5 | 0 | 0.26745724 | 0.978 | 0.974 | 0 | monocytes | Laptm5 |
| Taldo12 | 0 | 0.35295203 | 0.792 | 0.789 | 0 | monocytes | Taldo1 |
| Gpx1 | 0 | 0.31124862 | 0.948 | 0.951 | 0 | monocytes | Gpx1 |
| Prdx51 | 0 | 0.48702135 | 0.934 | 0.931 | 0 | monocytes | Prdx5 |
| Cap1 | 0 | 0.40832803 | 0.882 | 0.883 | 0 | monocytes | Cap1 |
| Iqgap11 | 0 | 0.36369752 | 0.921 | 0.921 | 0 | monocytes | Iqgap1 |
| Adgre51 | 1.46E-306 | 1.61420254 | 0.751 | 0.565 | 2.82E-302 | monocytes | Adgre5 |
| Capg1 | 9.99E-304 | 0.35382886 | 0.743 | 0.689 | 1.93E-299 | monocytes | Capg |
| Pim12 | 6.56E-294 | 1.11718581 | 0.961 | 0.884 | 1.27E-289 | monocytes | Pim1 |

| | | | | | | | |
|---|---|---|---|---|---|---|---|
| Cdc42 | 7.64E-290 | 0.27612253 | 0.761 | 0.815 | 1.48E-285 | monocytes | Cdc42 |
| Slc2a11 | 1.29E-286 | 0.28181928 | 0.729 | 0.693 | 2.50E-282 | monocytes | Slc2a1 |
| Mcl1 | 4.71E-285 | 0.54017896 | 0.985 | 0.976 | 9.10E-281 | monocytes | Mcl1 |
| Zfp36l21 | 1.94E-280 | 0.43969386 | 0.775 | 0.848 | 3.76E-276 | monocytes | Zfp36l2 |
| Il17ra1 | 2.55E-270 | 1.77679855 | 0.634 | 0.435 | 4.93E-266 | monocytes | Il17ra |
| Inpp5d2 | 1.21E-269 | 0.27055086 | 0.708 | 0.781 | 2.34E-265 | monocytes | Inpp5d |
| Nr4a31 | 3.16E-268 | 0.30167094 | 0.621 | 0.587 | 6.12E-264 | monocytes | Nr4a3 |
| Furin | 1.79E-264 | 0.37779201 | 0.793 | 0.814 | 3.47E-260 | monocytes | Furin |
| Hcls11 | 5.31E-261 | 0.51421583 | 0.918 | 0.902 | 1.03E-256 | monocytes | Hcls1 |
| Mcemp12 | 2.21E-260 | 1.80299779 | 0.505 | 0.257 | 4.27E-256 | monocytes | Mcemp1 |
| Lsp11 | 4.15E-260 | 1.70080689 | 0.586 | 0.386 | 8.03E-256 | monocytes | Lsp1 |
| Cdh11 | 4.70E-260 | 1.59934402 | 0.351 | 0.197 | 9.09E-256 | monocytes | Cdh1 |
| Plaur2 | 5.09E-259 | 1.11508441 | 0.936 | 0.814 | 9.85E-255 | monocytes | Plaur |
| Apbb1ip | 1.01E-253 | 0.27310293 | 0.598 | 0.717 | 1.95E-249 | monocytes | Apbb1ip |
| Wdr26 | 1.04E-246 | 0.28216421 | 0.748 | 0.826 | 2.01E-242 | monocytes | Wdr26 |
| Stx11 | 2.33E-244 | 2.23619169 | 0.264 | 0.106 | 4.51E-240 | monocytes | Stx11 |
| Ddx3x | 7.43E-244 | 0.32117045 | 0.739 | 0.814 | 1.44E-239 | monocytes | Ddx3x |
| Adam81 | 1.35E-242 | 0.42173575 | 0.62 | 0.567 | 2.61E-238 | monocytes | Adam8 |
| Actr2 | 4.57E-240 | 0.26909273 | 0.674 | 0.745 | 8.84E-236 | monocytes | Actr2 |
| Abr1 | 4.66E-237 | 0.29572228 | 0.668 | 0.724 | 9.01E-233 | monocytes | Abr |
| Ptpn61 | 2.04E-235 | 0.3772377 | 0.692 | 0.759 | 3.95E-231 | monocytes | Ptpn6 |
| Arf1 | 4.86E-233 | 0.29492772 | 0.707 | 0.776 | 9.39E-229 | monocytes | Arf1 |
| Ccnd31 | 3.03E-232 | 1.92728613 | 0.466 | 0.283 | 5.85E-228 | monocytes | Ccnd3 |
| Slc44a2 | 3.68E-229 | 2.16268372 | 0.403 | 0.237 | 7.11E-225 | monocytes | Slc44a2 |
| Krt801 | 6.55E-227 | 1.95330821 | 0.381 | 0.212 | 1.27E-222 | monocytes | Krt80 |
| Spint12 | 5.34E-226 | 0.35307586 | 0.584 | 0.616 | 1.03E-221 | monocytes | Spint1 |
| Eif3k1 | 1.96E-225 | 0.30022698 | 0.653 | 0.738 | 3.79E-221 | monocytes | Eif3k |
| Acly1 | 5.42E-225 | 0.33947829 | 0.711 | 0.796 | 1.05E-220 | monocytes | Acly |
| Wsb11 | 2.07E-224 | 0.28940587 | 0.765 | 0.813 | 4.00E-220 | monocytes | Wsb1 |
| Atox11 | 9.80E-222 | 0.44391875 | 0.846 | 0.864 | 1.89E-217 | monocytes | Atox1 |
| Trem11 | 1.48E-221 | 1.51445082 | 0.529 | 0.329 | 2.86E-217 | monocytes | Trem1 |
| Ywhab | 4.61E-217 | 0.2763724 | 0.625 | 0.704 | 8.92E-213 | monocytes | Ywhab |
| Wipf1 | 5.88E-216 | 0.31935704 | 0.674 | 0.757 | 1.14E-211 | monocytes | Wipf1 |
| Rhoa | 1.61E-215 | 0.34798239 | 0.751 | 0.799 | 3.12E-211 | monocytes | Rhoa |
| Tnfaip31 | 3.29E-214 | 0.3525606 | 0.542 | 0.628 | 6.37E-210 | monocytes | Tnfaip3 |
| Psme11 | 2.36E-208 | 0.30440338 | 0.666 | 0.72 | 4.57E-204 | monocytes | Psme1 |
| Ptgs21 | 1.40E-207 | 0.99827053 | 0.657 | 0.556 | 2.71E-203 | monocytes | Ptgs2 |
| Tlr21 | 3.94E-204 | 1.13540108 | 0.826 | 0.722 | 7.61E-200 | monocytes | Tlr2 |
| Ppp1r15a1 | 2.22E-203 | 0.36178039 | 0.695 | 0.734 | 4.29E-199 | monocytes | Ppp1r15a |
| Tmsb10 | 6.33E-203 | 1.39180934 | 0.793 | 0.636 | 1.22E-198 | monocytes | Tmsb10 |
| Gnas | 1.39E-198 | 0.26648216 | 0.626 | 0.718 | 2.68E-194 | monocytes | Gnas |
| Myo1f1 | 1.66E-198 | 0.37014846 | 0.632 | 0.699 | 3.21E-194 | monocytes | Myo1f |
| Arpc5 | 5.61E-198 | 0.4996015 | 0.814 | 0.828 | 1.08E-193 | monocytes | Arpc5 |
| Lrp11 | 1.24E-197 | 0.58185907 | 0.933 | 0.897 | 2.40E-193 | monocytes | Lrp1 |
| Txnrd11 | 1.79E-196 | 0.45396618 | 0.757 | 0.718 | 3.47E-192 | monocytes | Txnrd1 |
| Tagln2 | 1.96E-196 | 0.65087355 | 0.907 | 0.901 | 3.79E-192 | monocytes | Tagln2 |
| Fes | 9.14E-194 | 0.33314164 | 0.693 | 0.774 | 1.77E-189 | monocytes | Fes |
| Abi31 | 3.30E-193 | 0.28652684 | 0.417 | 0.537 | 6.38E-189 | monocytes | Abi3 |
| Psmb82 | 1.86E-191 | 0.28370052 | 0.62 | 0.694 | 3.59E-187 | monocytes | Psmb8 |
| Pak2 | 2.39E-189 | 0.32296851 | 0.626 | 0.719 | 4.62E-185 | monocytes | Pak2 |
| Traf11 | 4.99E-188 | 1.46344032 | 0.457 | 0.317 | 9.65E-184 | monocytes | Traf1 |
| Pkm1 | 8.03E-188 | 0.52762447 | 0.867 | 0.865 | 1.55E-183 | monocytes | Pkm |
| Uba1 | 1.72E-187 | 0.36774907 | 0.683 | 0.755 | 3.33E-183 | monocytes | Uba1 |
| Irf11 | 3.22E-187 | 0.32737339 | 0.424 | 0.51 | 6.22E-183 | monocytes | Irf1 |
| Nfkbid3 | 6.11E-187 | 0.28504264 | 0.57 | 0.623 | 1.18E-182 | monocytes | Nfkbid |
| Plbd12 | 1.25E-186 | 0.4905627 | 0.725 | 0.676 | 2.42E-182 | monocytes | Plbd1 |
| Gpr1322 | 1.36E-185 | 1.24384039 | 0.769 | 0.592 | 2.63E-181 | monocytes | Gpr132 |
| Aplp2 | 1.47E-183 | 0.3082815 | 0.615 | 0.691 | 2.84E-179 | monocytes | Aplp2 |
| Cd862 | 4.07E-183 | 0.26830386 | 0.646 | 0.722 | 7.87E-179 | monocytes | Cd86 |
| AA4671972 | 1.26E-179 | 1.18368878 | 0.491 | 0.422 | 2.43E-175 | monocytes | AA467197 |
| Lfng1 | 5.37E-179 | 0.35472183 | 0.591 | 0.66 | 1.04E-174 | monocytes | Lfng |
| Pcbp1 | 4.15E-178 | 0.27658273 | 0.6 | 0.697 | 8.02E-174 | monocytes | Pcbp1 |
| Ets22 | 5.08E-177 | 0.47777388 | 0.798 | 0.819 | 9.82E-173 | monocytes | Ets2 |
| Tmod3 | 2.00E-175 | 0.26248169 | 0.628 | 0.732 | 3.86E-171 | monocytes | Tmod3 |
| Arpc4 | 3.53E-173 | 0.40341738 | 0.675 | 0.723 | 6.82E-169 | monocytes | Arpc4 |
| Clic1 | 1.61E-171 | 0.303619 | 0.635 | 0.713 | 3.12E-167 | monocytes | Clic1 |
| Ly6e1 | 2.83E-171 | 1.07434032 | 0.937 | 0.813 | 5.47E-167 | monocytes | Ly6e |
| Sema4d2 | 3.51E-171 | 0.45734449 | 0.678 | 0.665 | 6.79E-167 | monocytes | Sema4d |
| Ddx51 | 2.89E-169 | 0.2940891 | 0.62 | 0.702 | 5.58E-165 | monocytes | Ddx5 |
| Gda | 5.96E-169 | 0.52535009 | 0.727 | 0.75 | 1.15E-164 | monocytes | Gda |
| Eif5a1 | 1.70E-167 | 0.38179532 | 0.645 | 0.69 | 3.29E-163 | monocytes | Eif5a |
| S100a101 | 3.14E-167 | 1.13393431 | 0.869 | 0.765 | 6.06E-163 | monocytes | S100a10 |
| Gngt21 | 5.89E-167 | 0.4946478 | 0.479 | 0.56 | 1.14E-162 | monocytes | Gngt2 |
| Tppp31 | 9.82E-167 | 1.50591309 | 0.404 | 0.25 | 1.90E-162 | monocytes | Tppp3 |
| Fam49a | 2.24E-166 | 1.87135549 | 0.41 | 0.265 | 4.32E-162 | monocytes | Fam49a |
| Ppp1ca | 8.99E-166 | 0.25908503 | 0.542 | 0.636 | 1.74E-161 | monocytes | Ppp1ca |
| Rnf10 | 1.73E-165 | 0.33241754 | 0.611 | 0.702 | 3.35E-161 | monocytes | Rnf10 |
| Tgfbi1 | 3.97E-162 | 0.95627922 | 0.967 | 0.944 | 7.67E-158 | monocytes | Tgfbi |
| Slc11a11 | 9.18E-161 | 0.64847851 | 0.736 | 0.711 | 1.77E-156 | monocytes | Slc11a1 |
| Emb2 | 9.12E-159 | 0.51617399 | 0.754 | 0.746 | 1.76E-154 | monocytes | Emb |
| Capza21 | 3.03E-158 | 0.30861189 | 0.543 | 0.625 | 5.86E-154 | monocytes | Capza2 |
| Ciao2a | 3.12E-158 | 0.36216704 | 0.569 | 0.64 | 6.04E-154 | monocytes | Ciao2a |
| Dusp162 | 3.07E-157 | 1.49945584 | 0.465 | 0.348 | 5.94E-153 | monocytes | Dusp16 |
| Rab20 | 3.65E-157 | 0.42862331 | 0.688 | 0.739 | 7.06E-153 | monocytes | Rab20 |
| Cish1 | 4.19E-156 | 1.37655384 | 0.553 | 0.406 | 8.11E-152 | monocytes | Cish |
| Lrrfip11 | 2.42E-155 | 1.2462301 | 0.849 | 0.751 | 4.69E-151 | monocytes | Lrrfip1 |
| Degs11 | 4.41E-154 | 0.28819407 | 0.424 | 0.529 | 8.53E-150 | monocytes | Degs1 |
| Ddx61 | 2.98E-152 | 0.3360391 | 0.637 | 0.717 | 5.76E-148 | monocytes | Ddx6 |
| Copz1 | 7.41E-151 | 0.2566128 | 0.516 | 0.629 | 1.43E-146 | monocytes | Copz1 |
| Tgif11 | 3.47E-150 | 0.48611047 | 0.778 | 0.773 | 6.71E-146 | monocytes | Tgif1 |
| Smox2 | 4.41E-150 | 1.19705808 | 0.808 | 0.723 | 8.52E-146 | monocytes | Smox |
| Alox5ap | 2.86E-149 | 1.2220868 | 0.893 | 0.756 | 5.53E-145 | monocytes | Alox5ap |
| Cers21 | 4.23E-149 | 0.25623261 | 0.497 | 0.6 | 8.17E-145 | monocytes | Cers2 |
| Sh3bgrl | 9.27E-149 | 0.43051049 | 0.705 | 0.757 | 1.79E-144 | monocytes | Sh3bgrl |
| Zmiz1 | 7.83E-148 | 0.36533489 | 0.582 | 0.668 | 1.51E-143 | monocytes | Zmiz1 |
| Csf2rb3 | 4.88E-147 | 0.54790546 | 0.819 | 0.795 | 9.43E-143 | monocytes | Csf2rb |
| Kdm6b1 | 1.73E-146 | 1.10142455 | 0.913 | 0.794 | 3.34E-142 | monocytes | Kdm6b |
| Cggbp1 | 1.99E-146 | 0.26825528 | 0.54 | 0.644 | 3.84E-142 | monocytes | Cggbp1 |
| Ube2d2a | 8.67E-145 | 0.28419333 | 0.544 | 0.644 | 1.68E-140 | monocytes | Ube2d2a |
| Surf41 | 1.40E-144 | 0.34269872 | 0.553 | 0.633 | 2.70E-140 | monocytes | Surf4 |
| Sod21 | 4.84E-144 | 0.49950291 | 0.586 | 0.654 | 9.36E-140 | monocytes | Sod2 |
| Tram1 | 1.10E-143 | 0.43124993 | 0.623 | 0.685 | 2.13E-139 | monocytes | Tram1 |
| Pten1 | 6.65E-143 | 0.29802222 | 0.508 | 0.626 | 1.29E-138 | monocytes | Pten |
| Cdc42se1 | 5.31E-142 | 0.32769684 | 0.539 | 0.641 | 1.03E-137 | monocytes | Cdc42se1 |
| Runx31 | 2.05E-141 | 1.15667952 | 0.808 | 0.721 | 3.96E-137 | monocytes | Runx3 |
| Sem11 | 4.20E-141 | 0.31435628 | 0.691 | 0.76 | 8.13E-137 | monocytes | Sem1 |
| Apoc21 | 5.84E-141 | 0.27175193 | 0.296 | 0.44 | 1.13E-136 | monocytes | Apoc2 |
| Eif4g3 | 1.02E-140 | 0.34641307 | 0.579 | 0.684 | 1.98E-136 | monocytes | Eif4g3 |
| Pirb1 | 1.18E-139 | 0.5318346 | 0.806 | 0.815 | 2.28E-135 | monocytes | Pirb |
| Midn1 | 1.59E-139 | 0.44453724 | 0.693 | 0.733 | 3.07E-135 | monocytes | Midn |
| Picalm1 | 2.13E-139 | 0.53589929 | 0.874 | 0.88 | 4.12E-135 | monocytes | Picalm |
| Clec4d1 | 3.86E-139 | 0.52564236 | 0.71 | 0.645 | 7.47E-135 | monocytes | Clec4d |
| Ccr1 | 6.14E-137 | 0.43568282 | 0.634 | 0.7 | 1.19E-132 | monocytes | Ccr1 |
| Themis22 | 1.23E-134 | 0.26759605 | 0.369 | 0.484 | 2.37E-130 | monocytes | Themis2 |
| Sppl2a1 | 2.03E-132 | 0.35265668 | 0.528 | 0.61 | 3.92E-128 | monocytes | Sppl2a |
| Lims1 | 6.98E-132 | 0.31890167 | 0.493 | 0.596 | 1.35E-127 | monocytes | Lims1 |
| Ccnl1 | 8.24E-132 | 0.45029057 | 0.776 | 0.8 | 1.59E-127 | monocytes | Ccnl1 |
| Trem3 | 9.44E-132 | 1.95204951 | 0.274 | 0.135 | 1.83E-127 | monocytes | Trem3 |
| Btf31 | 2.82E-131 | 0.31896168 | 0.564 | 0.639 | 5.45E-127 | monocytes | Btf3 |
| Glud11 | 1.73E-130 | 0.56788537 | 0.736 | 0.768 | 3.34E-126 | monocytes | Glud1 |
| Arf41 | 1.97E-130 | 0.32605947 | 0.543 | 0.6 | 3.80E-126 | monocytes | Arf4 |
| Inhba1 | 3.72E-130 | 1.08316833 | 0.358 | 0.243 | 7.18E-126 | monocytes | Inhba |
| Fgr1 | 6.10E-130 | 1.32882604 | 0.828 | 0.655 | 1.18E-125 | monocytes | Fgr |
| Kif5b | 7.35E-130 | 0.26014226 | 0.417 | 0.538 | 1.42E-125 | monocytes | Kif5b |
| Ifngr12 | 9.02E-130 | 0.49672389 | 0.753 | 0.803 | 1.74E-125 | monocytes | Ifngr1 |
| Mtpn1 | 1.10E-129 | 0.41041225 | 0.538 | 0.608 | 2.13E-125 | monocytes | Mtpn |
| Pomp1 | 5.11E-128 | 0.27495772 | 0.486 | 0.577 | 9.87E-124 | monocytes | Pomp |
| Fbrs | 1.84E-127 | 0.26766884 | 0.55 | 0.664 | 3.56E-123 | monocytes | Fbrs |
| Snd11 | 8.44E-127 | 0.45197779 | 0.728 | 0.775 | 1.63E-122 | monocytes | Snd1 |
| Icam11 | 5.39E-126 | 0.50170956 | 0.595 | 0.625 | 1.04E-121 | monocytes | Icam1 |
| Myo1c | 5.64E-125 | 0.31337246 | 0.552 | 0.664 | 1.09E-120 | monocytes | Myo1c |
| Sertad1 | 8.49E-125 | 0.2725107 | 0.516 | 0.58 | 1.64E-120 | monocytes | Sertad1 |
| Hexim1 | 4.97E-124 | 0.27534503 | 0.525 | 0.601 | 9.61E-120 | monocytes | Hexim1 |
| Rab8a | 9.40E-124 | 0.30119469 | 0.48 | 0.585 | 1.82E-119 | monocytes | Rab8a |
| Pacsin2 | 1.52E-123 | 0.41111242 | 0.646 | 0.709 | 2.94E-119 | monocytes | Pacsin2 |
| Susd6 | 3.59E-122 | 0.3242811 | 0.562 | 0.663 | 6.94E-118 | monocytes | Susd6 |
| Srsf5 | 7.47E-122 | 0.28168798 | 0.599 | 0.695 | 1.44E-117 | monocytes | Srsf5 |
| Srp9 | 1.63E-121 | 0.29593897 | 0.509 | 0.614 | 3.16E-117 | monocytes | Srp9 |
| Ikzf1 | 5.90E-121 | 0.35389075 | 0.414 | 0.536 | 1.14E-116 | monocytes | Ikzf1 |
| Ptafr1 | 7.07E-120 | 1.22083457 | 0.705 | 0.627 | 1.37E-115 | monocytes | Ptafr |
| Tln1 | 1.37E-119 | 0.63308038 | 0.859 | 0.843 | 2.65E-115 | monocytes | Tln1 |
| Far11 | 5.36E-119 | 0.2511539 | 0.427 | 0.54 | 1.04E-114 | monocytes | Far1 |
| Antxr22 | 1.10E-118 | 0.26457394 | 0.547 | 0.612 | 2.12E-114 | monocytes | Antxr2 |
| Nfkbia | 1.75E-118 | 0.70736042 | 0.924 | 0.9 | 3.39E-114 | monocytes | Nfkbia |
| Rbm31 | 1.48E-117 | 0.25796044 | 0.534 | 0.624 | 2.86E-113 | monocytes | Rbm3 |
| Ptprj | 1.87E-117 | 0.43469156 | 0.563 | 0.647 | 3.61E-113 | monocytes | Ptprj |
| Bak11 | 3.75E-117 | 0.44973946 | 0.546 | 0.597 | 7.24E-113 | monocytes | Bak1 |
| H2afy1 | 2.28E-116 | 0.5579202 | 0.777 | 0.794 | 4.42E-112 | monocytes | H2afy |
| Helz21 | 2.31E-116 | 1.63988416 | 0.313 | 0.157 | 4.46E-112 | monocytes | Helz2 |
| Jmjd1c1 | 6.98E-116 | 0.33276402 | 0.531 | 0.628 | 1.35E-111 | monocytes | Jmjd1c |
| Slc38a21 | 1.22E-115 | 0.47553734 | 0.771 | 0.785 | 2.37E-111 | monocytes | Slc38a2 |
| Napsa1 | 1.84E-115 | 1.62722503 | 0.452 | 0.253 | 3.56E-111 | monocytes | Napsa |
| Smap2 | 4.37E-115 | 0.3813803 | 0.536 | 0.624 | 8.46E-111 | monocytes | Smap2 |
| Becn1 | 7.03E-115 | 0.28081091 | 0.455 | 0.574 | 1.36E-110 | monocytes | Becn1 |
| Lsm14a | 8.79E-115 | 0.27328288 | 0.452 | 0.571 | 1.70E-110 | monocytes | Lsm14a |
| Ralb1 | 9.94E-115 | 0.32833895 | 0.509 | 0.6 | 1.92E-110 | monocytes | Ralb |
| Actr32 | 1.64E-114 | 0.48525386 | 0.602 | 0.641 | 3.18E-110 | monocytes | Actr3 |
| Hipk11 | 4.75E-114 | 0.42311569 | 0.593 | 0.66 | 9.18E-110 | monocytes | Hipk1 |
| Pstpip11 | 6.71E-114 | 0.31493029 | 0.463 | 0.537 | 1.30E-109 | monocytes | Pstpip1 |
| Gnb2 | 9.43E-112 | 0.4871319 | 0.665 | 0.702 | 1.82E-107 | monocytes | Gnb2 |
| Erbin | 1.01E-110 | 0.27671214 | 0.441 | 0.56 | 1.96E-106 | monocytes | Erbin |
| Slc35b22 | 1.52E-110 | 0.39249396 | 0.691 | 0.743 | 2.94E-106 | monocytes | Slc35b2 |
| Uqcrfs1 | 6.38E-109 | 0.45136441 | 0.515 | 0.575 | 1.23E-104 | monocytes | Uqcrfs1 |
| Syk1 | 1.44E-108 | 0.60921164 | 0.907 | 0.906 | 2.78E-104 | monocytes | Syk |
| Rlim | 3.52E-108 | 0.35684648 | 0.544 | 0.634 | 6.81E-104 | monocytes | Rlim |
| Slc9a3r11 | 3.07E-107 | 0.30100767 | 0.465 | 0.563 | 5.93E-103 | monocytes | Slc9a3r1 |
| St6galnac4 | 3.24E-107 | 0.2578642 | 0.319 | 0.44 | 6.26E-103 | monocytes | St6galnac4 |
| Pum2 | 2.75E-106 | 0.39469358 | 0.557 | 0.634 | 5.31E-102 | monocytes | Pum2 |
| D8Ertd738e1 | 3.43E-106 | 0.28540674 | 0.521 | 0.627 | 6.63E-102 | monocytes | D8Ertd738e |
| F102 | 3.50E-106 | 1.1014394 | 0.807 | 0.53 | 6.76E-102 | monocytes | F10 |
| Ptpn121 | 2.50E-105 | 0.46675055 | 0.619 | 0.639 | 4.82E-101 | monocytes | Ptpn12 |
| Rab321 | 3.66E-105 | 0.52947914 | 0.565 | 0.614 | 7.07E-101 | monocytes | Rab32 |
| Pilra2 | 7.70E-105 | 0.42771566 | 0.537 | 0.601 | 1.49E-100 | monocytes | Pilra |
| Rgs19 | 1.05E-104 | 0.33135445 | 0.4 | 0.508 | 2.04E-100 | monocytes | Rgs19 |
| Stat6 | 1.37E-104 | 0.39120193 | 0.531 | 0.626 | 2.65E-100 | monocytes | Stat6 |
| Eif3f1 | 2.39E-104 | 0.47650392 | 0.527 | 0.603 | 4.63E-100 | monocytes | Eif3f |
| Arih11 | 3.78E-104 | 0.34316744 | 0.523 | 0.604 | 7.30E-100 | monocytes | Arih1 |
| Rac1 | 8.67E-104 | 0.30943723 | 0.424 | 0.526 | 1.68E-99 | monocytes | Rac1 |
| Itsn2 | 2.21E-103 | 0.25632116 | 0.396 | 0.515 | 4.27E-99 | monocytes | Itsn2 |
| Dock102 | 2.77E-103 | 0.25901055 | 0.395 | 0.49 | 5.36E-99 | monocytes | Dock10 |
| Tmed2 | 6.50E-103 | 0.29150618 | 0.417 | 0.516 | 1.26E-98 | monocytes | Tmed2 |
| Tmbim11 | 1.68E-102 | 0.40440832 | 0.436 | 0.569 | 3.24E-98 | monocytes | Tmbim1 |
| Cul3 | 2.50E-102 | 0.34782141 | 0.51 | 0.6 | 4.84E-98 | monocytes | Cul3 |
| Tor1aip11 | 2.88E-102 | 0.49145596 | 0.638 | 0.693 | 5.56E-98 | monocytes | Tor1aip1 |
| Glrx1 | 3.06E-102 | 0.48579319 | 0.465 | 0.537 | 5.92E-98 | monocytes | Glrx |
| Sod1 | 4.31E-102 | 0.2784665 | 0.376 | 0.488 | 8.33E-98 | monocytes | Sod1 |
| Preb | 5.72E-102 | 0.26719064 | 0.394 | 0.515 | 1.11E-97 | monocytes | Preb |
| Anxa111 | 7.73E-102 | 0.33321254 | 0.493 | 0.595 | 1.50E-97 | monocytes | Anxa11 |
| Rai142 | 1.95E-101 | 0.2991875 | 0.393 | 0.519 | 3.76E-97 | monocytes | Rai14 |
| Ilrun | 2.37E-101 | 0.34376109 | 0.425 | 0.532 | 4.58E-97 | monocytes | Ilrun |
| B4galt1 | 5.31E-101 | 0.46660619 | 0.653 | 0.751 | 1.03E-96 | monocytes | B4galt1 |
| Fndc3a | 1.12E-100 | 0.3422634 | 0.485 | 0.561 | 2.16E-96 | monocytes | Fndc3a |
| Cebpb1 | 9.52E-100 | 0.76661712 | 0.994 | 0.976 | 1.84E-95 | monocytes | Cebpb |
| Ccdc124 | 2.25E-99 | 0.26485038 | 0.375 | 0.488 | 4.36E-95 | monocytes | Ccdc124 |
| Syf2 | 1.22E-98 | 0.30933056 | 0.485 | 0.587 | 2.35E-94 | monocytes | Syf2 |
| Tpm41 | 1.24E-98 | 0.563706 | 0.7 | 0.702 | 2.39E-94 | monocytes | Tpm4 |
| Ap1g1 | 2.26E-98 | 0.28611481 | 0.415 | 0.526 | 4.38E-94 | monocytes | Ap1g1 |
| Myl6 | 5.88E-98 | 0.28756159 | 0.427 | 0.533 | 1.14E-93 | monocytes | Myl6 |
| Plod31 | 6.93E-98 | 0.49864189 | 0.571 | 0.638 | 1.34E-93 | monocytes | Plod3 |
| Mbd2 | 8.78E-98 | 0.473031 | 0.724 | 0.772 | 1.70E-93 | monocytes | Mbd2 |
| Anxa6 | 2.02E-97 | 0.39882147 | 0.421 | 0.527 | 3.90E-93 | monocytes | Anxa6 |
| Tor1a | 2.01E-96 | 0.26421565 | 0.329 | 0.445 | 3.89E-92 | monocytes | Tor1a |
| Prrc2c | 3.19E-96 | 0.39872225 | 0.503 | 0.591 | 6.16E-92 | monocytes | Prrc2c |
| Dynll11 | 4.79E-96 | 0.25705669 | 0.4 | 0.5 | 9.27E-92 | monocytes | Dynll1 |
| Supt4a | 1.89E-95 | 0.25663794 | 0.375 | 0.5 | 3.65E-91 | monocytes | Supt4a |
| Cope1 | 1.95E-95 | 0.39538436 | 0.535 | 0.61 | 3.76E-91 | monocytes | Cope |
| Tpst21 | 2.22E-95 | 0.30882277 | 0.398 | 0.506 | 4.30E-91 | monocytes | Tpst2 |
| Twf21 | 1.07E-94 | 0.34106131 | 0.434 | 0.519 | 2.07E-90 | monocytes | Twf2 |
| Stk38 | 1.85E-94 | 0.33176682 | 0.349 | 0.494 | 3.57E-90 | monocytes | Stk38 |
| Vps37b2 | 7.15E-94 | 0.36105329 | 0.51 | 0.534 | 1.38E-89 | monocytes | Vps37b |
| Mef2a | 6.67E-93 | 0.37114584 | 0.545 | 0.615 | 1.29E-88 | monocytes | Mef2a |
| Dazap21 | 8.82E-93 | 0.45477519 | 0.594 | 0.661 | 1.70E-88 | monocytes | Dazap2 |
| Gsk3a | 1.91E-92 | 0.25229691 | 0.337 | 0.467 | 3.70E-88 | monocytes | Gsk3a |
| Lilr4b | 6.31E-92 | 0.42506052 | 0.641 | 0.67 | 1.22E-87 | monocytes | Lilr4b |
| Dhx151 | 1.90E-91 | 0.35042559 | 0.482 | 0.562 | 3.67E-87 | monocytes | Dhx15 |
| Hopx1 | 3.18E-91 | 1.67609765 | 0.359 | 0.2 | 6.16E-87 | monocytes | Hopx |
| Sik12 | 4.71E-91 | 0.48674399 | 0.698 | 0.709 | 9.11E-87 | monocytes | Sik1 |
| Ndufb1-ps1 | 1.39E-90 | 0.434571 | 0.552 | 0.61 | 2.69E-86 | monocytes | Ndufb1-ps |
| Zbtb7a | 5.31E-90 | 0.33143055 | 0.464 | 0.569 | 1.03E-85 | monocytes | Zbtb7a |
| Dennd5a | 1.96E-89 | 0.45690571 | 0.617 | 0.67 | 3.80E-85 | monocytes | Dennd5a |
| Sgms21 | 2.07E-89 | 1.55274977 | 0.285 | 0.172 | 4.00E-85 | monocytes | Sgms2 |
| Ankrd11 | 3.34E-89 | 0.26066196 | 0.477 | 0.581 | 6.46E-85 | monocytes | Ankrd11 |
| Ythdf3 | 6.01E-89 | 0.47117834 | 0.618 | 0.684 | 1.16E-84 | monocytes | Ythdf3 |
| Ppp1r18 | 6.57E-89 | 0.50834767 | 0.658 | 0.704 | 1.27E-84 | monocytes | Ppp1r18 |
| Ube2f | 8.34E-89 | 0.34035922 | 0.486 | 0.581 | 1.61E-84 | monocytes | Ube2f |
| Maff1 | 3.84E-88 | 0.43663486 | 0.63 | 0.646 | 7.43E-84 | monocytes | Maff |
| Sdc11 | 5.06E-88 | 0.57159403 | 0.352 | 0.358 | 9.78E-84 | monocytes | Sdc1 |
| Arhgef1 | 6.97E-88 | 0.46027911 | 0.577 | 0.665 | 1.35E-83 | monocytes | Arhgef1 |
| Arf5 | 1.39E-87 | 0.55022939 | 0.727 | 0.754 | 2.69E-83 | monocytes | Arf5 |
| Rbm42 | 1.72E-87 | 0.39476916 | 0.512 | 0.6 | 3.32E-83 | monocytes | Rbm42 |
| Ehbp1l1 | 8.47E-87 | 0.45152342 | 0.634 | 0.698 | 1.64E-82 | monocytes | Ehbp1l1 |
| Tmem164 | 1.59E-86 | 0.3297025 | 0.383 | 0.487 | 3.07E-82 | monocytes | Tmem164 |
| Mvp1 | 3.69E-86 | 0.48763312 | 0.656 | 0.709 | 7.13E-82 | monocytes | Mvp |
| Rassf11 | 6.37E-86 | 0.27756143 | 0.397 | 0.504 | 1.23E-81 | monocytes | Rassf1 |
| Stk401 | 8.13E-86 | 1.32768506 | 0.602 | 0.502 | 1.57E-81 | monocytes | Stk40 |
| Selenot1 | 1.21E-85 | 0.30649746 | 0.328 | 0.417 | 2.34E-81 | monocytes | Selenot |
| Ms4a6b2 | 1.24E-85 | 0.42820525 | 0.465 | 0.459 | 2.40E-81 | monocytes | Ms4a6b |
| Pycard | 2.70E-85 | 0.55331231 | 0.628 | 0.667 | 5.22E-81 | monocytes | Pycard |
| Ptbp3 | 5.49E-85 | 0.56723921 | 0.796 | 0.815 | 1.06E-80 | monocytes | Ptbp3 |
| Kcna3 | 1.43E-84 | 1.62417936 | 0.291 | 0.178 | 2.77E-80 | monocytes | Kcna3 |
| Ncf21 | 3.96E-84 | 0.74365575 | 0.899 | 0.864 | 7.66E-80 | monocytes | Ncf2 |
| Copb1 | 2.13E-83 | 0.30281675 | 0.378 | 0.475 | 4.11E-79 | monocytes | Copb1 |
| Psmb102 | 2.81E-83 | 0.29307769 | 0.385 | 0.481 | 5.43E-79 | monocytes | Psmb10 |
| Prr131 | 2.83E-83 | 0.55171694 | 0.421 | 0.497 | 5.47E-79 | monocytes | Prr13 |
| Mtmr6 | 3.01E-83 | 0.33782119 | 0.468 | 0.56 | 5.81E-79 | monocytes | Mtmr6 |
| Pik3cg2 | 5.47E-83 | 0.29403575 | 0.34 | 0.438 | 1.06E-78 | monocytes | Pik3cg |
| Ago22 | 1.57E-82 | 0.37119566 | 0.489 | 0.557 | 3.04E-78 | monocytes | Ago2 |
| Sptlc2 | 4.01E-82 | 0.2533333 | 0.339 | 0.452 | 7.75E-78 | monocytes | Sptlc2 |
| Uck22 | 8.61E-82 | 1.02710734 | 0.813 | 0.711 | 1.67E-77 | monocytes | Uck2 |
| Nlrp31 | 1.53E-81 | 0.52325135 | 0.751 | 0.717 | 2.95E-77 | monocytes | Nlrp3 |
| Nrd11 | 4.26E-81 | 0.35707867 | 0.454 | 0.535 | 8.24E-77 | monocytes | Nrd1 |
| Tns31 | 1.07E-80 | 0.39444642 | 0.414 | 0.475 | 2.08E-76 | monocytes | Tns3 |
| Arpc21 | 1.53E-80 | 0.63508909 | 0.763 | 0.74 | 2.96E-76 | monocytes | Arpc2 |
| Rbbp6 | 2.16E-80 | 0.27703092 | 0.41 | 0.51 | 4.18E-76 | monocytes | Rbbp6 |
| Cyp512 | 2.41E-80 | 0.29863866 | 0.26 | 0.338 | 4.66E-76 | monocytes | Cyp51 |
| Prpf40a | 3.21E-80 | 0.26865657 | 0.368 | 0.474 | 6.20E-76 | monocytes | Prpf40a |
| Tab2 | 7.52E-80 | 0.41843609 | 0.447 | 0.523 | 1.45E-75 | monocytes | Tab2 |
| Scand1 | 8.52E-80 | 0.30286457 | 0.449 | 0.556 | 1.65E-75 | monocytes | Scand1 |
| Tspan14 | 1.01E-79 | 0.45042795 | 0.457 | 0.533 | 1.96E-75 | monocytes | Tspan14 |
| Zfp106 | 1.28E-79 | 0.37400019 | 0.396 | 0.492 | 2.47E-75 | monocytes | Zfp106 |
| Stom | 1.68E-79 | 0.41914672 | 0.31 | 0.42 | 3.24E-75 | monocytes | Stom |
| Lnpep1 | 2.35E-78 | 0.51955357 | 0.656 | 0.698 | 4.54E-74 | monocytes | Lnpep |
| Dnajc10 | 2.47E-78 | 0.30410973 | 0.318 | 0.419 | 4.78E-74 | monocytes | Dnajc10 |
| Kmt2d | 2.92E-78 | 0.27880314 | 0.488 | 0.588 | 5.65E-74 | monocytes | Kmt2d |
| Gsdmd1 | 1.89E-77 | 0.34094965 | 0.454 | 0.536 | 3.66E-73 | monocytes | Gsdmd |
| Edem11 | 3.50E-77 | 0.58499731 | 0.775 | 0.774 | 6.76E-73 | monocytes | Edem1 |
| S100a13 | 4.13E-77 | 0.32132126 | 0.437 | 0.517 | 7.98E-73 | monocytes | S100a13 |
| Rhog | 1.59E-76 | 0.60323693 | 0.761 | 0.767 | 3.07E-72 | monocytes | Rhog |
| Srrm1 | 1.83E-76 | 0.35947556 | 0.443 | 0.529 | 3.54E-72 | monocytes | Srrm1 |
| Myh91 | 2.53E-76 | 0.53481243 | 0.646 | 0.68 | 4.89E-72 | monocytes | Myh9 |
| Clk11 | 2.85E-76 | 0.45238104 | 0.676 | 0.719 | 5.51E-72 | monocytes | Clk1 |
| Snrpb1 | 3.27E-76 | 0.30239915 | 0.337 | 0.425 | 6.32E-72 | monocytes | Snrpb |
| Mapkapk2 | 3.34E-76 | 0.63640722 | 0.843 | 0.84 | 6.45E-72 | monocytes | Mapkapk2 |
| Csnk1e | 5.31E-76 | 0.26443775 | 0.34 | 0.442 | 1.03E-71 | monocytes | Csnk1e |
| Elmo11 | 1.68E-75 | 0.28273183 | 0.364 | 0.447 | 3.25E-71 | monocytes | Elmo1 |
| Kcmf1 | 3.33E-75 | 0.2922571 | 0.38 | 0.473 | 6.44E-71 | monocytes | Kcmf1 |
| Cnot1 | 3.79E-75 | 0.25479892 | 0.322 | 0.432 | 7.33E-71 | monocytes | Cnot1 |
| Ass11 | 4.75E-75 | 1.11583291 | 0.556 | 0.375 | 9.18E-71 | monocytes | Ass1 |
| Tsc22d2 | 2.12E-74 | 0.26059256 | 0.355 | 0.442 | 4.09E-70 | monocytes | Tsc22d2 |
| Slc1a52 | 2.56E-74 | 0.2613639 | 0.33 | 0.404 | 4.94E-70 | monocytes | Slc1a5 |
| Tbl1xr11 | 4.55E-74 | 0.38521467 | 0.439 | 0.526 | 8.80E-70 | monocytes | Tbl1xr1 |
| Tox4 | 5.43E-74 | 0.26752149 | 0.326 | 0.436 | 1.05E-69 | monocytes | Tox4 |
| Nab1 | 7.45E-74 | 1.23524172 | 0.623 | 0.52 | 1.44E-69 | monocytes | Nab1 |
| Ep3001 | 9.41E-74 | 0.33287674 | 0.414 | 0.521 | 1.82E-69 | monocytes | Ep300 |
| Gpr108 | 1.19E-73 | 0.26756247 | 0.264 | 0.377 | 2.31E-69 | monocytes | Gpr108 |
| Flii | 1.82E-73 | 0.36145289 | 0.397 | 0.499 | 3.52E-69 | monocytes | Flii |
| Cd2741 | 2.39E-73 | 0.47810411 | 0.407 | 0.489 | 4.63E-69 | monocytes | Cd274 |
| Cmtm71 | 2.63E-73 | 0.44590015 | 0.554 | 0.618 | 5.08E-69 | monocytes | Cmtm7 |
| Gatad2b | 8.08E-73 | 0.29268882 | 0.341 | 0.446 | 1.56E-68 | monocytes | Gatad2b |
| Rap2a1 | 1.91E-72 | 0.37218565 | 0.397 | 0.482 | 3.70E-68 | monocytes | Rap2a |
| Arhgap451 | 3.53E-72 | 0.49311502 | 0.575 | 0.668 | 6.82E-68 | monocytes | Arhgap45 |
| Mycbp22 | 5.87E-72 | 0.30077236 | 0.484 | 0.588 | 1.13E-67 | monocytes | Mycbp2 |
| Esyt11 | 6.87E-72 | 0.54577554 | 0.587 | 0.616 | 1.33E-67 | monocytes | Esyt1 |
| Rab6a | 1.31E-71 | 0.29790264 | 0.335 | 0.424 | 2.53E-67 | monocytes | Rab6a |
| Zfr | 3.21E-71 | 0.2622225 | 0.287 | 0.392 | 6.20E-67 | monocytes | Zfr |
| Card191 | 4.10E-71 | 0.53341075 | 0.569 | 0.618 | 7.92E-67 | monocytes | Card19 |
| Med15 | 5.24E-71 | 0.45259826 | 0.55 | 0.62 | 1.01E-66 | monocytes | Med15 |
| Adam10 | 1.26E-70 | 0.51247104 | 0.523 | 0.585 | 2.44E-66 | monocytes | Adam10 |
| Kmt5a1 | 1.48E-70 | 0.30066425 | 0.323 | 0.399 | 2.86E-66 | monocytes | Kmt5a |
| Tmem63a1 | 2.04E-70 | 0.26525511 | 0.327 | 0.427 | 3.94E-66 | monocytes | Tmem63a |
| Kdelr2 | 2.13E-70 | 0.37879068 | 0.352 | 0.437 | 4.12E-66 | monocytes | Kdelr2 |
| Ctbp2 | 3.52E-70 | 0.37588609 | 0.385 | 0.476 | 6.80E-66 | monocytes | Ctbp2 |
| Api5 | 6.72E-70 | 0.38200873 | 0.372 | 0.456 | 1.30E-65 | monocytes | Api5 |
| Fgd3 | 7.33E-70 | 0.25195185 | 0.298 | 0.407 | 1.42E-65 | monocytes | Fgd3 |
| Sra12 | 8.84E-70 | 0.42286279 | 0.506 | 0.571 | 1.71E-65 | monocytes | Sra1 |
| Usp7 | 1.12E-69 | 0.32923237 | 0.384 | 0.48 | 2.17E-65 | monocytes | Usp7 |
| Pid11 | 1.75E-69 | 0.46784743 | 0.7 | 0.699 | 3.38E-65 | monocytes | Pid1 |
| Ldlrap12 | 4.02E-69 | 0.27918765 | 0.3 | 0.408 | 7.77E-65 | monocytes | Ldlrap1 |
| Rnf111 | 4.31E-69 | 0.26243676 | 0.342 | 0.445 | 8.33E-65 | monocytes | Rnf111 |
| Psme22 | 3.95E-68 | 0.30548358 | 0.332 | 0.417 | 7.65E-64 | monocytes | Psme2 |
| Tmem167 | 5.11E-68 | 0.30413554 | 0.308 | 0.395 | 9.88E-64 | monocytes | Tmem167 |
| Usp121 | 6.37E-68 | 0.3057879 | 0.264 | 0.37 | 1.23E-63 | monocytes | Usp12 |
| Ncf41 | 7.19E-68 | 0.54215456 | 0.692 | 0.724 | 1.39E-63 | monocytes | Ncf4 |
| Ppp2cb1 | 1.13E-67 | 0.48564217 | 0.474 | 0.523 | 2.19E-63 | monocytes | Ppp2cb |
| Trappc1 | 3.18E-67 | 0.29771168 | 0.269 | 0.362 | 6.14E-63 | monocytes | Trappc1 |
| Birc6 | 6.94E-67 | 0.38660389 | 0.406 | 0.492 | 1.34E-62 | monocytes | Birc6 |
| Clint1 | 1.33E-66 | 0.45724326 | 0.385 | 0.481 | 2.57E-62 | monocytes | Clint1 |
| Akirin1 | 1.79E-66 | 0.2594687 | 0.251 | 0.343 | 3.45E-62 | monocytes | Akirin1 |
| Fis1 | 2.27E-66 | 0.55954421 | 0.626 | 0.663 | 4.39E-62 | monocytes | Fis1 |
| Itch | 2.41E-66 | 0.31756538 | 0.292 | 0.387 | 4.65E-62 | monocytes | Itch |
| Reep3 | 2.80E-66 | 0.42978803 | 0.434 | 0.519 | 5.40E-62 | monocytes | Reep3 |
| Aip | 4.59E-66 | 0.41902208 | 0.399 | 0.484 | 8.87E-62 | monocytes | Aip |
| Hivep11 | 5.77E-66 | 0.27249793 | 0.318 | 0.426 | 1.12E-61 | monocytes | Hivep1 |
| Ssr3 | 7.31E-66 | 0.37917048 | 0.421 | 0.508 | 1.41E-61 | monocytes | Ssr3 |
| Nr4a12 | 1.73E-65 | 0.98327246 | 0.919 | 0.78 | 3.34E-61 | monocytes | Nr4a1 |
| Ccnl2 | 2.38E-65 | 0.34817029 | 0.477 | 0.559 | 4.59E-61 | monocytes | Ccnl2 |
| Ethe1 | 3.27E-65 | 0.29105073 | 0.227 | 0.328 | 6.32E-61 | monocytes | Ethe1 |
| Per11 | 5.05E-65 | 0.46581427 | 0.696 | 0.702 | 9.77E-61 | monocytes | Per1 |
| Rbm39 | 6.16E-65 | 0.38205879 | 0.448 | 0.543 | 1.19E-60 | monocytes | Rbm39 |
| Pnrc21 | 7.45E-65 | 0.27639995 | 0.289 | 0.388 | 1.44E-60 | monocytes | Pnrc2 |
| Ube2k | 1.02E-64 | 0.34319392 | 0.338 | 0.431 | 1.96E-60 | monocytes | Ube2k |
| Was2 | 1.67E-64 | 0.55755535 | 0.593 | 0.639 | 3.23E-60 | monocytes | Was |
| Eps8 | 2.03E-64 | 0.3966198 | 0.382 | 0.469 | 3.93E-60 | monocytes | Eps8 |
| Tnf1 | 4.05E-64 | 0.64290852 | 0.537 | 0.627 | 7.84E-60 | monocytes | Tnf |
| Stat31 | 5.55E-64 | 0.56621208 | 0.665 | 0.711 | 1.07E-59 | monocytes | Stat3 |
| Smchd11 | 6.85E-64 | 0.37824766 | 0.339 | 0.422 | 1.33E-59 | monocytes | Smchd1 |
| Pafah1b1 | 6.92E-64 | 0.33874378 | 0.386 | 0.483 | 1.34E-59 | monocytes | Pafah1b1 |
| Ssh21 | 8.64E-64 | 0.38974587 | 0.358 | 0.429 | 1.67E-59 | monocytes | Ssh2 |
| Sf3b11 | 1.47E-63 | 0.60423273 | 0.798 | 0.8 | 2.84E-59 | monocytes | Sf3b1 |
| Cmpk11 | 1.69E-63 | 0.37479396 | 0.395 | 0.484 | 3.27E-59 | monocytes | Cmpk1 |
| Lrp10 | 2.80E-63 | 0.61623324 | 0.693 | 0.717 | 5.41E-59 | monocytes | Lrp10 |
| Smad4 | 7.47E-63 | 0.41425665 | 0.418 | 0.513 | 1.44E-58 | monocytes | Smad4 |
| Csnk2b1 | 7.66E-63 | 0.35275957 | 0.317 | 0.41 | 1.48E-58 | monocytes | Csnk2b |
| Slc7a111 | 8.99E-63 | 0.63075384 | 0.556 | 0.489 | 1.74E-58 | monocytes | Slc7a11 |
| Chd71 | 1.16E-62 | 1.10876278 | 0.624 | 0.527 | 2.24E-58 | monocytes | Chd7 |
| Arid1a1 | 3.16E-62 | 0.30807414 | 0.316 | 0.419 | 6.11E-58 | monocytes | Arid1a |
| Wnk11 | 3.38E-62 | 0.57620557 | 0.691 | 0.742 | 6.54E-58 | monocytes | Wnk1 |
| Arhgap30 | 3.48E-62 | 0.5353194 | 0.602 | 0.647 | 6.73E-58 | monocytes | Arhgap30 |
| Lsm121 | 4.92E-62 | 0.32741303 | 0.343 | 0.433 | 9.52E-58 | monocytes | Lsm12 |
| Rap1a | 6.45E-62 | 0.56281685 | 0.637 | 0.69 | 1.25E-57 | monocytes | Rap1a |
| Pcf11 | 7.47E-62 | 0.44608504 | 0.635 | 0.674 | 1.44E-57 | monocytes | Pcf11 |
| Arid1b | 7.49E-62 | 0.27160163 | 0.284 | 0.391 | 1.45E-57 | monocytes | Arid1b |
| Ubr5 | 7.57E-62 | 0.28938822 | 0.335 | 0.433 | 1.46E-57 | monocytes | Ubr5 |
| Smc1a | 1.31E-61 | 0.2986438 | 0.241 | 0.332 | 2.53E-57 | monocytes | Smc1a |
| Mboat7 | 1.78E-61 | 0.39704085 | 0.365 | 0.47 | 3.43E-57 | monocytes | Mboat7 |
| Traf7 | 2.48E-61 | 0.25723375 | 0.287 | 0.391 | 4.79E-57 | monocytes | Traf7 |
| Gorasp2 | 3.02E-61 | 0.37288802 | 0.356 | 0.438 | 5.84E-57 | monocytes | Gorasp2 |
| Nfkbiz2 | 3.25E-61 | 0.59438517 | 0.669 | 0.694 | 6.28E-57 | monocytes | Nfkbiz |
| Sowahc2 | 3.92E-61 | 0.48509963 | 0.506 | 0.514 | 7.57E-57 | monocytes | Sowahc |
| Fam117b1 | 3.98E-61 | 0.48064246 | 0.593 | 0.606 | 7.70E-57 | monocytes | Fam117b |
| Tnfrsf1b1 | 5.75E-61 | 0.77218524 | 0.797 | 0.793 | 1.11E-56 | monocytes | Tnfrsf1b |
| Atp8b42 | 1.03E-60 | 1.4914203 | 0.335 | 0.203 | 1.99E-56 | monocytes | Atp8b4 |
| 11-Sep | 1.04E-60 | 0.35394468 | 0.302 | 0.363 | 2.00E-56 | monocytes | 11-Sep |
| Nfatc1 | 1.56E-60 | 0.46690673 | 0.399 | 0.465 | 3.03E-56 | monocytes | Nfatc1 |
| Fam49b2 | 2.17E-60 | 1.07200873 | 0.87 | 0.809 | 4.19E-56 | monocytes | Fam49b |
| Rsrc2 | 2.52E-60 | 0.37765336 | 0.404 | 0.486 | 4.88E-56 | monocytes | Rsrc2 |
| Ppp6r3 | 3.22E-60 | 0.35758083 | 0.34 | 0.428 | 6.22E-56 | monocytes | Ppp6r3 |
| Arl5b2 | 4.19E-60 | 0.28099802 | 0.378 | 0.431 | 8.10E-56 | monocytes | Arl5b |
| Tmem431 | 4.21E-60 | 0.3333887 | 0.295 | 0.383 | 8.15E-56 | monocytes | Tmem43 |
| Cd531 | 4.67E-60 | 0.74864097 | 0.909 | 0.883 | 9.04E-56 | monocytes | Cd53 |
| Snrpb22 | 5.20E-60 | 0.27151177 | 0.273 | 0.365 | 1.01E-55 | monocytes | Snrpb2 |
| Csnk2a2 | 8.60E-60 | 0.26391928 | 0.249 | 0.349 | 1.66E-55 | monocytes | Csnk2a2 |
| Clec4a31 | 9.64E-60 | 0.50631391 | 0.414 | 0.48 | 1.86E-55 | monocytes | Clec4a3 |
| Agfg1 | 1.07E-59 | 0.32578618 | 0.307 | 0.398 | 2.06E-55 | monocytes | Agfg1 |
| Cbl | 1.47E-59 | 0.53000931 | 0.575 | 0.629 | 2.84E-55 | monocytes | Cbl |
| Atp7a | 2.04E-59 | 0.45546069 | 0.369 | 0.45 | 3.94E-55 | monocytes | Atp7a |
| Ccdc71l1 | 2.06E-59 | 0.97371056 | 0.5 | 0.473 | 3.98E-55 | monocytes | Ccdc71l |
| Lmnb12 | 2.29E-59 | 1.16735152 | 0.56 | 0.415 | 4.42E-55 | monocytes | Lmnb1 |
| Nup1531 | 2.37E-59 | 0.27954722 | 0.269 | 0.362 | 4.58E-55 | monocytes | Nup153 |
| St3gal1 | 2.54E-59 | 1.23821412 | 0.645 | 0.539 | 4.92E-55 | monocytes | St3gal1 |
| Crk | 3.39E-59 | 0.3789822 | 0.439 | 0.53 | 6.56E-55 | monocytes | Crk |
| Myl12a1 | 3.70E-59 | 0.63239333 | 0.759 | 0.773 | 7.15E-55 | monocytes | Myl12a |
| Snf8 | 4.37E-59 | 0.3511333 | 0.397 | 0.487 | 8.45E-55 | monocytes | Snf8 |
| Coro2a2 | 6.47E-59 | 0.37238918 | 0.389 | 0.419 | 1.25E-54 | monocytes | Coro2a |
| Gch11 | 6.57E-59 | 1.24945745 | 0.545 | 0.415 | 1.27E-54 | monocytes | Gch1 |
| Nkiras2 | 2.02E-58 | 0.38695119 | 0.293 | 0.385 | 3.90E-54 | monocytes | Nkiras2 |
| Adprh | 2.81E-58 | 0.56169022 | 0.599 | 0.65 | 5.43E-54 | monocytes | Adprh |
| Lilrb4a | 4.05E-58 | 0.70659311 | 0.909 | 0.878 | 7.83E-54 | monocytes | Lilrb4a |
| Tmcc1 | 4.67E-58 | 0.42229621 | 0.314 | 0.426 | 9.03E-54 | monocytes | Tmcc1 |
| Sqor1 | 4.99E-58 | 0.25179693 | 0.238 | 0.336 | 9.65E-54 | monocytes | Sqor |
| Arnt | 5.23E-58 | 0.30446206 | 0.358 | 0.462 | 1.01E-53 | monocytes | Arnt |
| Tut7 | 8.49E-58 | 0.44627259 | 0.624 | 0.668 | 1.64E-53 | monocytes | Tut7 |
| Dusp11 | 2.03E-57 | 0.34099518 | 0.337 | 0.425 | 3.93E-53 | monocytes | Dusp11 |
| Ikbke1 | 2.71E-57 | 0.27299856 | 0.329 | 0.432 | 5.23E-53 | monocytes | Ikbke |
| Usp47 | 3.00E-57 | 0.28904547 | 0.235 | 0.332 | 5.81E-53 | monocytes | Usp47 |
| Taok1 | 3.54E-57 | 0.3197247 | 0.365 | 0.455 | 6.84E-53 | monocytes | Taok1 |
| Epc11 | 4.53E-57 | 0.33060124 | 0.378 | 0.477 | 8.76E-53 | monocytes | Epc1 |
| Chd2 | 8.41E-57 | 0.32895128 | 0.374 | 0.455 | 1.63E-52 | monocytes | Chd2 |
| Cmas1 | 8.58E-57 | 0.29734991 | 0.264 | 0.357 | 1.66E-52 | monocytes | Cmas |
| Wdr33 | 9.20E-57 | 0.25781182 | 0.273 | 0.365 | 1.78E-52 | monocytes | Wdr33 |
| Ptprc | 1.54E-56 | 0.75139075 | 0.802 | 0.799 | 2.97E-52 | monocytes | Ptprc |
| Ppp3cb | 1.63E-56 | 0.2513627 | 0.293 | 0.39 | 3.16E-52 | monocytes | Ppp3cb |
| Prdm2 | 1.71E-56 | 0.32028109 | 0.313 | 0.396 | 3.30E-52 | monocytes | Prdm2 |
| Nop532 | 1.94E-56 | 0.44836442 | 0.477 | 0.553 | 3.75E-52 | monocytes | Nop53 |
| Ubp1 | 2.20E-56 | 0.35558191 | 0.393 | 0.478 | 4.26E-52 | monocytes | Ubp1 |
| Tmem512 | 3.85E-56 | 0.48802813 | 0.316 | 0.389 | 7.44E-52 | monocytes | Tmem51 |
| Bloc1s21 | 3.98E-56 | 0.3002745 | 0.263 | 0.353 | 7.70E-52 | monocytes | Bloc1s2 |
| Baz2a | 4.90E-56 | 0.30777211 | 0.31 | 0.407 | 9.47E-52 | monocytes | Baz2a |
| Pds5a | 9.20E-56 | 0.36585569 | 0.341 | 0.421 | 1.78E-51 | monocytes | Pds5a |
| Rnf6 | 9.25E-56 | 0.25818802 | 0.275 | 0.374 | 1.79E-51 | monocytes | Rnf6 |
| Pgm2 | 1.12E-55 | 0.2632391 | 0.264 | 0.359 | 2.16E-51 | monocytes | Pgm2 |
| Cnih4 | 1.96E-55 | 0.32098986 | 0.274 | 0.357 | 3.80E-51 | monocytes | Cnih4 |
| Eif3h1 | 2.71E-55 | 0.38136735 | 0.308 | 0.39 | 5.24E-51 | monocytes | Eif3h |
| Tnfrsf1a | 4.08E-55 | 0.71623491 | 0.833 | 0.832 | 7.89E-51 | monocytes | Tnfrsf1a |
| Klhl51 | 6.27E-55 | 0.30854146 | 0.217 | 0.307 | 1.21E-50 | monocytes | Klhl5 |
| Iqgap2 | 6.59E-55 | 0.32792164 | 0.214 | 0.302 | 1.27E-50 | monocytes | Iqgap2 |
| Zdhhc5 | 6.83E-55 | 0.44874883 | 0.407 | 0.491 | 1.32E-50 | monocytes | Zdhhc5 |
| Tnrc18 | 7.36E-55 | 0.36786229 | 0.452 | 0.539 | 1.42E-50 | monocytes | Tnrc18 |
| Psmd111 | 8.79E-55 | 0.40897724 | 0.398 | 0.463 | 1.70E-50 | monocytes | Psmd11 |
| Tcf7l21 | 9.39E-55 | 1.32867046 | 0.39 | 0.333 | 1.82E-50 | monocytes | Tcf7l2 |
| Map2k1 | 2.51E-54 | 0.52387448 | 0.437 | 0.507 | 4.85E-50 | monocytes | Map2k1 |
| Smg1 | 4.20E-54 | 0.35305477 | 0.376 | 0.464 | 8.12E-50 | monocytes | Smg1 |
| Tmem179b | 4.93E-54 | 0.26774981 | 0.254 | 0.344 | 9.52E-50 | monocytes | Tmem179b |
| Rap1gap2 | 8.22E-54 | 1.67886161 | 0.266 | 0.188 | 1.59E-49 | monocytes | Rap1gap2 |
| Psen1 | 1.04E-53 | 0.46511674 | 0.377 | 0.449 | 2.01E-49 | monocytes | Psen1 |
| Atxn7l1 | 1.58E-53 | 0.33638243 | 0.331 | 0.424 | 3.06E-49 | monocytes | Atxn7l1 |
| Cpeb22 | 1.99E-53 | 0.32084337 | 0.37 | 0.446 | 3.85E-49 | monocytes | Cpeb2 |
| Pitpna | 2.26E-53 | 0.67829868 | 0.887 | 0.88 | 4.37E-49 | monocytes | Pitpna |
| Zfp131 | 2.43E-53 | 0.28287051 | 0.364 | 0.447 | 4.69E-49 | monocytes | Zfp131 |
| Acap2 | 2.49E-53 | 0.42818682 | 0.394 | 0.469 | 4.81E-49 | monocytes | Acap2 |
| Ppp4c | 4.18E-53 | 0.4935838 | 0.481 | 0.545 | 8.07E-49 | monocytes | Ppp4c |
| Clec4a11 | 7.24E-53 | 0.429602 | 0.32 | 0.364 | 1.40E-48 | monocytes | Clec4a1 |
| Atxn7l3 | 9.24E-53 | 0.30516744 | 0.315 | 0.411 | 1.79E-48 | monocytes | Atxn7l3 |
| Atp2c1 | 1.62E-52 | 0.52974522 | 0.432 | 0.515 | 3.12E-48 | monocytes | Atp2c1 |
| Naip2 | 3.54E-52 | 1.31696549 | 0.508 | 0.436 | 6.85E-48 | monocytes | Naip2 |
| Hdac1 | 6.87E-52 | 0.31039099 | 0.278 | 0.356 | 1.33E-47 | monocytes | Hdac1 |
| Rela | 9.80E-52 | 0.47942136 | 0.497 | 0.568 | 1.90E-47 | monocytes | Rela |
| Grk21 | 1.02E-51 | 0.63496296 | 0.712 | 0.734 | 1.98E-47 | monocytes | Grk2 |
| Brd4 | 1.56E-51 | 0.56276308 | 0.65 | 0.685 | 3.02E-47 | monocytes | Brd4 |
| Klra22 | 1.76E-51 | 1.37738658 | 0.453 | 0.323 | 3.41E-47 | monocytes | Klra2 |
| Nr4a22 | 2.89E-51 | 0.47229671 | 0.548 | 0.542 | 5.59E-47 | monocytes | Nr4a2 |
| Srebf2 | 2.91E-51 | 0.52390913 | 0.476 | 0.533 | 5.63E-47 | monocytes | Srebf2 |
| Kdm5c | 3.17E-51 | 0.29976892 | 0.318 | 0.408 | 6.13E-47 | monocytes | Kdm5c |
| Tax1bp1 | 3.37E-51 | 0.47395356 | 0.513 | 0.575 | 6.51E-47 | monocytes | Tax1bp1 |
| Cdk2ap11 | 4.76E-51 | 0.26418684 | 0.223 | 0.311 | 9.20E-47 | monocytes | Cdk2ap1 |
| Pdpk1 | 5.58E-51 | 0.25665541 | 0.271 | 0.364 | 1.08E-46 | monocytes | Pdpk1 |
| Pip4k2a2 | 6.60E-51 | 0.4013422 | 0.345 | 0.431 | 1.28E-46 | monocytes | Pip4k2a |
| Gosr21 | 9.74E-51 | 0.48629419 | 0.429 | 0.494 | 1.88E-46 | monocytes | Gosr2 |
| Camk2d2 | 1.28E-50 | 0.54901181 | 0.508 | 0.553 | 2.47E-46 | monocytes | Camk2d |
| Chsy11 | 1.28E-50 | 0.31878727 | 0.264 | 0.343 | 2.47E-46 | monocytes | Chsy1 |
| Asap11 | 1.31E-50 | 0.46066848 | 0.309 | 0.386 | 2.53E-46 | monocytes | Asap1 |
| Adssl11 | 1.42E-50 | 0.61933909 | 0.601 | 0.572 | 2.74E-46 | monocytes | Adssl1 |
| Fxyd51 | 6.44E-50 | 0.73612588 | 0.862 | 0.815 | 1.25E-45 | monocytes | Fxyd5 |
| Dhrs1 | 6.62E-50 | 0.40042449 | 0.313 | 0.402 | 1.28E-45 | monocytes | Dhrs1 |
| Hax11 | 8.70E-50 | 0.2565168 | 0.238 | 0.332 | 1.68E-45 | monocytes | Hax1 |
| Tcof11 | 9.23E-50 | 0.36114314 | 0.226 | 0.318 | 1.79E-45 | monocytes | Tcof1 |
| Pdlim5 | 1.13E-49 | 0.44399291 | 0.432 | 0.501 | 2.18E-45 | monocytes | Pdlim5 |
| Pdcd10 | 1.20E-49 | 0.28590948 | 0.283 | 0.371 | 2.32E-45 | monocytes | Pdcd10 |
| Lrch1 | 1.22E-49 | 0.27834396 | 0.283 | 0.374 | 2.36E-45 | monocytes | Lrch1 |
| Ubap11 | 1.41E-49 | 0.27389587 | 0.23 | 0.315 | 2.72E-45 | monocytes | Ubap1 |
| Tmem251 | 1.70E-49 | 0.30593703 | 0.259 | 0.343 | 3.29E-45 | monocytes | Tmem251 |
| Crybg11 | 1.94E-49 | 1.42674001 | 0.281 | 0.192 | 3.75E-45 | monocytes | Crybg1 |
| Map3k81 | 2.18E-49 | 0.37235155 | 0.218 | 0.306 | 4.21E-45 | monocytes | Map3k8 |
| Hivep32 | 2.41E-49 | 0.30847871 | 0.276 | 0.391 | 4.66E-45 | monocytes | Hivep3 |
| Elovl51 | 4.44E-49 | 0.39529471 | 0.22 | 0.29 | 8.58E-45 | monocytes | Elovl5 |
| Scimp2 | 7.40E-49 | 0.3988231 | 0.318 | 0.393 | 1.43E-44 | monocytes | Scimp |
| Ppdpf | 7.60E-49 | 0.26567593 | 0.194 | 0.275 | 1.47E-44 | monocytes | Ppdpf |
| Ranbp9 | 1.44E-48 | 0.29313672 | 0.257 | 0.347 | 2.79E-44 | monocytes | Ranbp9 |
| Hpse | 1.56E-48 | 0.34376425 | 0.244 | 0.275 | 3.02E-44 | monocytes | Hpse |
| Sec61a1 | 1.74E-48 | 0.42800984 | 0.328 | 0.407 | 3.36E-44 | monocytes | Sec61a1 |
| Tlr13 | 1.78E-48 | 0.46317259 | 0.257 | 0.311 | 3.45E-44 | monocytes | Tlr13 |
| Ier32 | 2.71E-48 | 0.72437001 | 0.933 | 0.918 | 5.23E-44 | monocytes | Ier3 |
| Cib11 | 3.01E-48 | 0.4100548 | 0.28 | 0.349 | 5.82E-44 | monocytes | Cib1 |
| Entr1 | 3.97E-48 | 0.31182378 | 0.249 | 0.34 | 7.67E-44 | monocytes | Entr1 |
| Clk3 | 4.65E-48 | 0.29815326 | 0.294 | 0.394 | 8.99E-44 | monocytes | Clk3 |
| Mex3c1 | 5.82E-48 | 0.32433814 | 0.315 | 0.396 | 1.12E-43 | monocytes | Mex3c |
| Slc35e11 | 6.73E-48 | 0.46999714 | 0.328 | 0.398 | 1.30E-43 | monocytes | Slc35e1 |
| Slc12a6 | 7.25E-48 | 0.38740626 | 0.308 | 0.399 | 1.40E-43 | monocytes | Slc12a6 |
| Mob3a1 | 9.04E-48 | 0.34700923 | 0.204 | 0.292 | 1.75E-43 | monocytes | Mob3a |
| Man2a1 | 9.40E-48 | 0.63406954 | 0.658 | 0.715 | 1.82E-43 | monocytes | Man2a1 |
| Cybc1 | 1.18E-47 | 0.40245543 | 0.326 | 0.399 | 2.27E-43 | monocytes | Cybc1 |
| Amz12 | 1.96E-47 | 0.48058943 | 0.285 | 0.326 | 3.78E-43 | monocytes | Amz1 |
| Skil2 | 3.38E-47 | 0.56115602 | 0.767 | 0.772 | 6.53E-43 | monocytes | Skil |
| Heatr12 | 4.90E-47 | 0.42023248 | 0.313 | 0.375 | 9.47E-43 | monocytes | Heatr1 |
| Cnot3 | 5.26E-47 | 0.26068212 | 0.25 | 0.335 | 1.02E-42 | monocytes | Cnot3 |
| Erp44 | 6.11E-47 | 0.34742327 | 0.251 | 0.329 | 1.18E-42 | monocytes | Erp44 |
| Rhbdf21 | 6.56E-47 | 0.54027433 | 0.496 | 0.55 | 1.27E-42 | monocytes | Rhbdf2 |
| Hectd1 | 6.80E-47 | 0.341492 | 0.278 | 0.369 | 1.31E-42 | monocytes | Hectd1 |

| | | | | | | | |
|---|---|---|---|---|---|---|---|
| Vbp1 | 7.09E-47 | 0.35419126 | 0.27 | 0.355 | 1.37E-42 | monocytes | Vbp1 |
| Ggh | 8.13E-47 | 0.33282655 | 0.251 | 0.331 | 1.57E-42 | monocytes | Ggh |
| Zswim61 | 1.07E-46 | 0.41167343 | 0.382 | 0.448 | 2.06E-42 | monocytes | Zswim6 |
| Khdc41 | 1.32E-46 | 0.31253407 | 0.323 | 0.41 | 2.55E-42 | monocytes | Khdc4 |
| Nabp11 | 1.49E-46 | 0.43305454 | 0.342 | 0.392 | 2.89E-42 | monocytes | Nabp1 |
| Ncoa6 | 1.73E-46 | 0.26801844 | 0.24 | 0.332 | 3.34E-42 | monocytes | Ncoa6 |
| Serp11 | 1.86E-46 | 0.30528223 | 0.265 | 0.342 | 3.60E-42 | monocytes | Serp1 |
| Bin3 | 2.07E-46 | 0.41000679 | 0.295 | 0.38 | 4.00E-42 | monocytes | Bin3 |
| Tspo | 2.43E-46 | 0.64420606 | 0.837 | 0.821 | 4.70E-42 | monocytes | Tspo |
| Map4k41 | 5.31E-46 | 0.57343975 | 0.585 | 0.617 | 1.03E-41 | monocytes | Map4k4 |
| Adar1 | 5.86E-46 | 0.28225836 | 0.259 | 0.349 | 1.13E-41 | monocytes | Adar |
| Fcf11 | 6.26E-46 | 0.26613218 | 0.224 | 0.31 | 1.21E-41 | monocytes | Fcf1 |
| Diaph1 | 6.44E-46 | 1.15774074 | 0.744 | 0.643 | 1.25E-41 | monocytes | Diaph1 |
| Add3 | 9.36E-46 | 1.58336311 | 0.292 | 0.256 | 1.81E-41 | monocytes | Add3 |
| DEPRECATED-I | 9.49E-46 | 0.3270117 | 0.322 | 0.385 | 1.83E-41 | monocytes | DEPRECATED-ENSMUSG00000039782 |
| Nup62 | 2.76E-45 | 0.26328608 | 0.18 | 0.259 | 5.33E-41 | monocytes | Nup62 |
| Dhx8 | 3.23E-45 | 0.28598707 | 0.222 | 0.314 | 6.24E-41 | monocytes | Dhx8 |
| Arhgap272 | 3.56E-45 | 0.36633534 | 0.186 | 0.26 | 6.87E-41 | monocytes | Arhgap27 |
| Slc9a1 | 4.03E-45 | 0.35526187 | 0.251 | 0.335 | 7.80E-41 | monocytes | Slc9a1 |
| Capza1 | 5.37E-45 | 0.45433055 | 0.351 | 0.409 | 1.04E-40 | monocytes | Capza1 |
| Bmpr2 | 6.16E-45 | 0.44574197 | 0.331 | 0.417 | 1.19E-40 | monocytes | Bmpr2 |
| Strn3 | 8.42E-45 | 0.32137105 | 0.254 | 0.339 | 1.63E-40 | monocytes | Strn3 |
| Fndc3b | 8.56E-45 | 0.50731747 | 0.485 | 0.54 | 1.66E-40 | monocytes | Fndc3b |
| Alcam1 | 8.89E-45 | 0.42938651 | 0.355 | 0.39 | 1.72E-40 | monocytes | Alcam |
| Setd2 | 1.08E-44 | 0.33516074 | 0.264 | 0.35 | 2.09E-40 | monocytes | Setd2 |
| Hnrnpd1 | 1.29E-44 | 0.46381264 | 0.464 | 0.522 | 2.49E-40 | monocytes | Hnrnpd |
| Ino80d | 1.49E-44 | 0.29547768 | 0.236 | 0.324 | 2.88E-40 | monocytes | Ino80d |
| Psd42 | 1.52E-44 | 0.44428988 | 0.381 | 0.453 | 2.94E-40 | monocytes | Psd4 |
| Slain2 | 1.93E-44 | 0.29308575 | 0.194 | 0.278 | 3.72E-40 | monocytes | Slain2 |
| Vps13d | 2.07E-44 | 0.37089338 | 0.276 | 0.361 | 4.01E-40 | monocytes | Vps13d |
| Mxd12 | 2.23E-44 | 0.98553011 | 0.686 | 0.592 | 4.31E-40 | monocytes | Mxd1 |
| Irf21 | 2.48E-44 | 0.35993359 | 0.293 | 0.385 | 4.80E-40 | monocytes | Irf2 |
| Gng101 | 2.73E-44 | 0.54910209 | 0.479 | 0.484 | 5.28E-40 | monocytes | Gng10 |
| Slc25a371 | 3.69E-44 | 0.29283138 | 0.239 | 0.293 | 7.13E-40 | monocytes | Slc25a37 |
| Kctd10 | 3.81E-44 | 0.35767109 | 0.305 | 0.388 | 7.36E-40 | monocytes | Kctd10 |
| Slc25a12 | 5.71E-44 | 0.32466233 | 0.193 | 0.271 | 1.10E-39 | monocytes | Slc25a12 |
| Nfatc3 | 7.19E-44 | 0.28147561 | 0.197 | 0.276 | 1.39E-39 | monocytes | Nfatc3 |
| Med13 | 8.56E-44 | 0.44800125 | 0.49 | 0.554 | 1.65E-39 | monocytes | Med13 |
| Rpp21 | 9.14E-44 | 0.26058611 | 0.179 | 0.268 | 1.77E-39 | monocytes | Rpp21 |
| Slc8b1 | 1.05E-43 | 0.56158837 | 0.425 | 0.496 | 2.04E-39 | monocytes | Slc8b1 |
| Limd11 | 1.08E-43 | 0.47064298 | 0.382 | 0.448 | 2.09E-39 | monocytes | Limd1 |
| Exoc2 | 1.14E-43 | 0.30238764 | 0.226 | 0.313 | 2.21E-39 | monocytes | Exoc2 |
| Ctps2 | 1.26E-43 | 0.38934079 | 0.271 | 0.348 | 2.43E-39 | monocytes | Ctps2 |
| Akap13 | 1.65E-43 | 0.5188906 | 0.504 | 0.581 | 3.19E-39 | monocytes | Akap13 |
| Clk41 | 1.69E-43 | 0.41087365 | 0.384 | 0.451 | 3.27E-39 | monocytes | Clk4 |
| Rock1 | 2.22E-43 | 0.4556316 | 0.401 | 0.475 | 4.30E-39 | monocytes | Rock1 |
| Dda1 | 2.26E-43 | 0.31745341 | 0.236 | 0.322 | 4.37E-39 | monocytes | Dda1 |
| Trak2 | 2.44E-43 | 0.26149632 | 0.231 | 0.311 | 4.72E-39 | monocytes | Trak2 |
| Shkbp1 | 2.52E-43 | 0.31347586 | 0.223 | 0.307 | 4.87E-39 | monocytes | Shkbp1 |
| Pvr2 | 2.81E-43 | 1.11982781 | 0.43 | 0.343 | 5.42E-39 | monocytes | Pvr |
| Vasp1 | 3.27E-43 | 0.69382959 | 0.886 | 0.876 | 6.33E-39 | monocytes | Vasp |
| Bri3bp2 | 3.30E-43 | 0.44312994 | 0.258 | 0.303 | 6.38E-39 | monocytes | Bri3bp |
| Sfxn3 | 4.36E-43 | 0.50018704 | 0.418 | 0.473 | 8.43E-39 | monocytes | Sfxn3 |
| Ugcg | 4.70E-43 | 0.4280016 | 0.354 | 0.431 | 9.09E-39 | monocytes | Ugcg |
| Card91 | 5.56E-43 | 0.40880898 | 0.363 | 0.445 | 1.08E-38 | monocytes | Card9 |
| Wapl | 5.68E-43 | 0.43624318 | 0.326 | 0.395 | 1.10E-38 | monocytes | Wapl |
| Dock21 | 5.71E-43 | 0.68937197 | 0.556 | 0.58 | 1.10E-38 | monocytes | Dock2 |
| Gcsh1 | 1.35E-42 | 0.37082825 | 0.227 | 0.33 | 2.61E-38 | monocytes | Gcsh |
| Kat6a | 1.46E-42 | 0.39136645 | 0.289 | 0.374 | 2.82E-38 | monocytes | Kat6a |
| Casp81 | 2.07E-42 | 0.42333659 | 0.376 | 0.45 | 4.00E-38 | monocytes | Casp8 |
| Sap30l1 | 6.04E-42 | 0.29672728 | 0.206 | 0.288 | 1.17E-37 | monocytes | Sap30l |
| Paf11 | 7.85E-42 | 0.36345415 | 0.279 | 0.355 | 1.52E-37 | monocytes | Paf1 |
| Ifi2071 | 8.61E-42 | 0.558266 | 0.556 | 0.576 | 1.67E-37 | monocytes | Ifi207 |
| Lypla2 | 1.10E-41 | 0.38360028 | 0.291 | 0.37 | 2.12E-37 | monocytes | Lypla2 |
| Elf1 | 1.12E-41 | 0.5528474 | 0.557 | 0.593 | 2.17E-37 | monocytes | Elf1 |
| Hnrnpc | 1.22E-41 | 0.39219345 | 0.257 | 0.33 | 2.36E-37 | monocytes | Hnrnpc |
| Sf3b61 | 1.33E-41 | 0.53148858 | 0.431 | 0.482 | 2.57E-37 | monocytes | Sf3b6 |
| Pitpnm12 | 1.60E-41 | 0.43479374 | 0.354 | 0.429 | 3.09E-37 | monocytes | Pitpnm1 |
| Ptk2b1 | 1.86E-41 | 0.69809379 | 0.591 | 0.604 | 3.60E-37 | monocytes | Ptk2b |
| Setd5 | 2.46E-41 | 0.46913651 | 0.403 | 0.477 | 4.75E-37 | monocytes | Setd5 |
| Setd1b1 | 3.66E-41 | 0.32844221 | 0.264 | 0.348 | 7.07E-37 | monocytes | Setd1b |
| Crtc2 | 5.85E-41 | 0.2767585 | 0.245 | 0.332 | 1.13E-36 | monocytes | Crtc2 |
| Myl12b1 | 6.92E-41 | 0.62684537 | 0.581 | 0.598 | 1.34E-36 | monocytes | Myl12b |
| Atf7ip | 9.84E-41 | 0.4316196 | 0.322 | 0.387 | 1.90E-36 | monocytes | Atf7ip |
| Cop1 | 1.01E-40 | 0.32337534 | 0.236 | 0.313 | 1.95E-36 | monocytes | Cop1 |
| Uso11 | 1.23E-40 | 0.36234347 | 0.277 | 0.35 | 2.38E-36 | monocytes | Uso1 |
| Prrc1 | 1.73E-40 | 0.32300853 | 0.214 | 0.293 | 3.35E-36 | monocytes | Prrc1 |
| Kdm3a | 1.79E-40 | 0.30987009 | 0.256 | 0.334 | 3.46E-36 | monocytes | Kdm3a |
| Phf20l1 | 2.00E-40 | 0.293235 | 0.245 | 0.332 | 3.87E-36 | monocytes | Phf20l1 |
| Ubn2 | 2.34E-40 | 0.25709156 | 0.187 | 0.275 | 4.52E-36 | monocytes | Ubn2 |
| Rnf1141 | 3.01E-40 | 0.37240104 | 0.299 | 0.372 | 5.81E-36 | monocytes | Rnf114 |
| Med12 | 3.02E-40 | 0.32732774 | 0.237 | 0.316 | 5.84E-36 | monocytes | Med1 |
| Azin1 | 4.08E-40 | 0.45853537 | 0.351 | 0.413 | 7.88E-36 | monocytes | Azin1 |
| Ints6 | 4.44E-40 | 0.31177922 | 0.28 | 0.335 | 8.59E-36 | monocytes | Ints6 |
| Btk | 6.43E-40 | 0.32998375 | 0.201 | 0.275 | 1.24E-35 | monocytes | Btk |
| Dyrk1a | 1.10E-39 | 0.33876253 | 0.285 | 0.353 | 2.13E-35 | monocytes | Dyrk1a |
| Nfkb12 | 1.31E-39 | 1.08835681 | 0.724 | 0.663 | 2.54E-35 | monocytes | Nfkb1 |
| Ogfr1 | 1.40E-39 | 0.49682641 | 0.386 | 0.455 | 2.70E-35 | monocytes | Ogfr |
| Nsa22 | 2.40E-39 | 0.25217791 | 0.179 | 0.257 | 4.63E-35 | monocytes | Nsa2 |
| Epb41l21 | 2.50E-39 | 0.54541051 | 0.546 | 0.615 | 4.84E-35 | monocytes | Epb41l2 |
| Tgfb11 | 3.51E-39 | 1.05301833 | 0.934 | 0.893 | 6.79E-35 | monocytes | Tgfb1 |
| Dock81 | 4.00E-39 | 0.50775225 | 0.411 | 0.482 | 7.74E-35 | monocytes | Dock8 |
| BC0285281 | 5.25E-39 | 0.45876146 | 0.204 | 0.28 | 1.01E-34 | monocytes | BC028528 |
| Klhl2 | 5.92E-39 | 1.49495549 | 0.263 | 0.18 | 1.14E-34 | monocytes | Klhl2 |
| Sp3 | 6.45E-39 | 0.43436714 | 0.309 | 0.381 | 1.25E-34 | monocytes | Sp3 |
| Nedd91 | 6.62E-39 | 1.3857114 | 0.383 | 0.286 | 1.28E-34 | monocytes | Nedd9 |
| Pilrb23 | 6.72E-39 | 0.42457375 | 0.247 | 0.318 | 1.30E-34 | monocytes | Pilrb2 |
| Mocs1 | 6.85E-39 | 0.28594728 | 0.185 | 0.269 | 1.32E-34 | monocytes | Mocs1 |
| Pet100 | 6.86E-39 | 0.29687957 | 0.258 | 0.336 | 1.33E-34 | monocytes | Pet100 |
| Igsf61 | 7.47E-39 | 0.76543083 | 0.626 | 0.621 | 1.44E-34 | monocytes | Igsf6 |
| Arid2 | 7.87E-39 | 0.29653454 | 0.253 | 0.338 | 1.52E-34 | monocytes | Arid2 |
| Mob1b | 1.22E-38 | 0.25145299 | 0.216 | 0.295 | 2.35E-34 | monocytes | Mob1b |
| Psma7 | 1.29E-38 | 0.46907651 | 0.27 | 0.327 | 2.49E-34 | monocytes | Psma7 |
| Ppp4r21 | 1.50E-38 | 0.50262517 | 0.471 | 0.527 | 2.89E-34 | monocytes | Ppp4r2 |
| Cpsf7 | 2.46E-38 | 0.37086519 | 0.329 | 0.41 | 4.75E-34 | monocytes | Cpsf7 |
| Zfp5162 | 2.84E-38 | 0.51582128 | 0.609 | 0.649 | 5.49E-34 | monocytes | Zfp516 |
| Trpv21 | 4.24E-38 | 0.37276136 | 0.255 | 0.321 | 8.21E-34 | monocytes | Trpv2 |
| Elf4e3 | 4.97E-38 | 0.33782856 | 0.208 | 0.288 | 9.61E-34 | monocytes | Elf4e3 |
| Tet21 | 5.83E-38 | 1.21878231 | 0.544 | 0.449 | 1.13E-33 | monocytes | Tet2 |
| Trim56 | 8.81E-38 | 0.26006685 | 0.179 | 0.259 | 1.70E-33 | monocytes | Trim56 |
| Hnrnph2 | 1.18E-37 | 0.50421398 | 0.455 | 0.514 | 2.28E-33 | monocytes | Hnrnph2 |
| Vrk11 | 1.26E-37 | 0.45059624 | 0.314 | 0.357 | 2.43E-33 | monocytes | Vrk1 |
| Rras | 1.26E-37 | 0.47023152 | 0.22 | 0.3 | 2.43E-33 | monocytes | Rras |
| Sema4a1 | 1.60E-37 | 0.62289042 | 0.696 | 0.656 | 3.09E-33 | monocytes | Sema4a |
| Gyg | 1.78E-37 | 0.69419532 | 0.353 | 0.415 | 3.44E-33 | monocytes | Gyg |
| Ifi2111 | 1.82E-37 | 1.17986758 | 0.344 | 0.241 | 3.53E-33 | monocytes | Ifi211 |
| Dlg1 | 3.25E-37 | 0.28128142 | 0.213 | 0.293 | 6.29E-33 | monocytes | Dlg1 |
| Taf101 | 3.32E-37 | 0.29362713 | 0.208 | 0.281 | 6.42E-33 | monocytes | Taf10 |
| Atf11 | 3.83E-37 | 0.5356625 | 0.486 | 0.535 | 7.41E-33 | monocytes | Atf1 |
| Heca | 7.52E-37 | 0.43374541 | 0.348 | 0.422 | 1.45E-32 | monocytes | Heca |
| Ccnk | 7.87E-37 | 0.37960597 | 0.235 | 0.31 | 1.52E-32 | monocytes | Ccnk |
| Klf31 | 1.51E-36 | 0.59726944 | 0.461 | 0.519 | 2.92E-32 | monocytes | Klf3 |
| Myd882 | 2.06E-36 | 0.66440467 | 0.661 | 0.666 | 3.99E-32 | monocytes | Myd88 |
| Rchy1 | 2.35E-36 | 0.29285888 | 0.194 | 0.264 | 4.55E-32 | monocytes | Rchy1 |
| Mark4 | 2.58E-36 | 0.4003373 | 0.225 | 0.308 | 5.00E-32 | monocytes | Mark4 |
| Lamtor4 | 3.04E-36 | 0.37388513 | 0.201 | 0.276 | 5.87E-32 | monocytes | Lamtor4 |
| Nfkbib | 5.45E-36 | 0.4640766 | 0.458 | 0.515 | 1.05E-31 | monocytes | Nfkbib |
| Osbp | 6.06E-36 | 0.40915917 | 0.249 | 0.318 | 1.17E-31 | monocytes | Osbp |
| Cdk13 | 7.15E-36 | 0.41378472 | 0.332 | 0.402 | 1.38E-31 | monocytes | Cdk13 |
| Foxj21 | 1.27E-35 | 0.30120521 | 0.202 | 0.28 | 2.46E-31 | monocytes | Foxj2 |
| Vhl | 1.59E-35 | 0.313016 | 0.194 | 0.27 | 3.08E-31 | monocytes | Vhl |
| Stat5b | 1.71E-35 | 0.48535824 | 0.251 | 0.318 | 3.30E-31 | monocytes | Stat5b |
| Tyk2 | 1.81E-35 | 0.34365171 | 0.178 | 0.256 | 3.50E-31 | monocytes | Tyk2 |
| Sde2 | 2.17E-35 | 0.49689454 | 0.406 | 0.465 | 4.19E-31 | monocytes | Sde2 |
| Arl6ip5 | 2.33E-35 | 0.65942819 | 0.541 | 0.57 | 4.50E-31 | monocytes | Arl6ip5 |
| Nfkb2 | 2.39E-35 | 0.60975366 | 0.723 | 0.738 | 4.62E-31 | monocytes | Nfkb2 |
| Ero1lb | 2.73E-35 | 0.33194762 | 0.258 | 0.332 | 5.29E-31 | monocytes | Ero1lb |
| Blcap1 | 2.75E-35 | 0.32758433 | 0.247 | 0.31 | 5.33E-31 | monocytes | Blcap |
| Chmp2a | 2.87E-35 | 0.37930675 | 0.291 | 0.367 | 5.55E-31 | monocytes | Chmp2a |
| Git1 | 3.40E-35 | 0.34976893 | 0.239 | 0.319 | 6.57E-31 | monocytes | Git1 |
| Pygl2 | 3.66E-35 | 0.6052004 | 0.594 | 0.591 | 7.07E-31 | monocytes | Pygl |
| Dapp11 | 5.27E-35 | 0.34973864 | 0.196 | 0.264 | 1.02E-30 | monocytes | Dapp1 |
| Atf6 | 5.77E-35 | 0.5174613 | 0.389 | 0.451 | 1.12E-30 | monocytes | Atf6 |
| Frg1 | 6.69E-35 | 0.32301288 | 0.197 | 0.269 | 1.29E-30 | monocytes | Frg1 |
| Abtb22 | 7.83E-35 | 1.1523652 | 0.348 | 0.254 | 1.51E-30 | monocytes | Abtb2 |
| Axin1 | 8.90E-35 | 0.3464973 | 0.178 | 0.254 | 1.72E-30 | monocytes | Axin1 |
| Ppp2ca1 | 9.41E-35 | 0.65212211 | 0.693 | 0.713 | 1.82E-30 | monocytes | Ppp2ca |
| Zfand3 | 9.56E-35 | 0.63535307 | 0.663 | 0.691 | 1.85E-30 | monocytes | Zfand3 |
| Cisd21 | 1.08E-34 | 0.59095401 | 0.467 | 0.494 | 2.08E-30 | monocytes | Cisd2 |
| Adap1 | 1.24E-34 | 0.56862202 | 0.325 | 0.387 | 2.40E-30 | monocytes | Adap1 |
| Mark2 | 1.30E-34 | 0.36809652 | 0.28 | 0.36 | 2.52E-30 | monocytes | Mark2 |
| Plekha11 | 1.31E-34 | 0.32975914 | 0.239 | 0.321 | 2.53E-30 | monocytes | Plekha1 |
| Fnbp41 | 1.91E-34 | 0.33870287 | 0.288 | 0.362 | 3.69E-30 | monocytes | Fnbp4 |
| Rprd2 | 1.95E-34 | 0.37119116 | 0.263 | 0.341 | 3.76E-30 | monocytes | Rprd2 |
| Spen1 | 1.98E-34 | 0.4538844 | 0.325 | 0.388 | 3.82E-30 | monocytes | Spen |
| Grk6 | 2.10E-34 | 0.51474663 | 0.25 | 0.323 | 4.05E-30 | monocytes | Grk6 |
| Elmo2 | 2.31E-34 | 0.45609612 | 0.193 | 0.273 | 4.47E-30 | monocytes | Elmo2 |
| Ppp2r2a1 | 4.12E-34 | 0.31220568 | 0.199 | 0.272 | 7.96E-30 | monocytes | Ppp2r2a |
| Sorl11 | 5.38E-34 | 1.22367952 | 0.573 | 0.455 | 1.04E-29 | monocytes | Sorl1 |
| Sarnp | 6.24E-34 | 0.37912275 | 0.214 | 0.284 | 1.21E-29 | monocytes | Sarnp |
| Prr14 | 6.76E-34 | 0.38000433 | 0.241 | 0.316 | 1.31E-29 | monocytes | Prr14 |
| Birc31 | 1.07E-33 | 0.60479836 | 0.574 | 0.587 | 2.06E-29 | monocytes | Birc3 |
| Prkaa11 | 1.12E-33 | 0.34043124 | 0.184 | 0.257 | 2.16E-29 | monocytes | Prkaa1 |
| Selplg1 | 1.33E-33 | 0.70502134 | 0.515 | 0.546 | 2.58E-29 | monocytes | Selplg |
| Crip11 | 1.35E-33 | 0.85173673 | 0.865 | 0.751 | 2.61E-29 | monocytes | Crip1 |
| Bcl10 | 1.41E-33 | 0.47772388 | 0.393 | 0.459 | 2.73E-29 | monocytes | Bcl10 |
| Ppig2 | 1.57E-33 | 0.36824668 | 0.288 | 0.355 | 3.03E-29 | monocytes | Ppig |
| Ythdf1 | 2.16E-33 | 0.41111731 | 0.254 | 0.326 | 4.18E-29 | monocytes | Ythdf1 |
| Tatdn21 | 2.39E-33 | 0.65199743 | 0.612 | 0.649 | 4.63E-29 | monocytes | Tatdn2 |
| Pik3c2a | 3.05E-33 | 0.35027205 | 0.236 | 0.305 | 5.90E-29 | monocytes | Pik3c2a |
| Ube2d31 | 5.13E-33 | 0.59556518 | 0.519 | 0.553 | 9.93E-29 | monocytes | Ube2d3 |
| Madd1 | 6.12E-33 | 1.14156794 | 0.397 | 0.321 | 1.18E-28 | monocytes | Madd |
| Pde8a1 | 6.73E-33 | 0.40644274 | 0.232 | 0.296 | 1.30E-28 | monocytes | Pde8a |
| Plekhg21 | 6.92E-33 | 0.32512026 | 0.286 | 0.311 | 1.34E-28 | monocytes | Plekhg2 |
| Apba3 | 7.09E-33 | 0.32727631 | 0.226 | 0.3 | 1.37E-28 | monocytes | Apba3 |
| Wac | 7.72E-33 | 0.38928756 | 0.273 | 0.338 | 1.49E-28 | monocytes | Wac |
| Ccdc88a1 | 7.94E-33 | 0.5366593 | 0.339 | 0.382 | 1.53E-28 | monocytes | Ccdc88a |
| Itgal1 | 8.68E-33 | 1.35644876 | 0.734 | 0.565 | 1.68E-28 | monocytes | Itgal |
| Hdac51 | 1.07E-32 | 0.4281419 | 0.377 | 0.449 | 2.08E-28 | monocytes | Hdac5 |
| Dhx40 | 1.11E-32 | 0.40872214 | 0.251 | 0.294 | 2.15E-28 | monocytes | Dhx40 |
| Mfsd14b | 1.68E-32 | 0.58141198 | 0.335 | 0.391 | 3.24E-28 | monocytes | Mfsd14b |
| Ppp1r10 | 1.69E-32 | 0.57830413 | 0.485 | 0.482 | 3.27E-28 | monocytes | Ppp1r10 |
| Ier2 | 2.27E-32 | 0.6760729 | 0.722 | 0.659 | 4.38E-28 | monocytes | Ier2 |
| Bsdc1 | 2.28E-32 | 0.33026192 | 0.208 | 0.277 | 4.42E-28 | monocytes | Bsdc1 |
| Vapa | 2.62E-32 | 0.43841202 | 0.231 | 0.299 | 5.07E-28 | monocytes | Vapa |
| Pkn2 | 4.46E-32 | 0.26554249 | 0.209 | 0.282 | 8.62E-28 | monocytes | Pkn2 |
| Ugp2 | 5.12E-32 | 0.38163011 | 0.219 | 0.284 | 9.91E-28 | monocytes | Ugp2 |
| Samd4b | 9.13E-32 | 0.39545906 | 0.266 | 0.336 | 1.77E-27 | monocytes | Samd4b |
| Lcp21 | 1.09E-31 | 0.74679652 | 0.586 | 0.598 | 2.10E-27 | monocytes | Lcp2 |
| Stk24 | 1.26E-31 | 0.68156055 | 0.522 | 0.545 | 2.44E-27 | monocytes | Stk24 |
| Tspan132 | 1.27E-31 | 0.62762106 | 0.332 | 0.311 | 2.45E-27 | monocytes | Tspan13 |
| Pfkp2 | 1.36E-31 | 0.56168723 | 0.605 | 0.578 | 2.62E-27 | monocytes | Pfkp |
| Ldlr | 1.73E-31 | 0.72830685 | 0.482 | 0.49 | 3.34E-27 | monocytes | Ldlr |
| Etf11 | 1.94E-31 | 0.51752905 | 0.366 | 0.418 | 3.75E-27 | monocytes | Etf1 |
| Sub11 | 2.34E-31 | 0.51428095 | 0.329 | 0.392 | 4.52E-27 | monocytes | Sub1 |
| Sidt2 | 2.96E-31 | 0.607575 | 0.515 | 0.549 | 5.73E-27 | monocytes | Sidt2 |
| Scaf41 | 3.18E-31 | 0.39666194 | 0.26 | 0.327 | 6.15E-27 | monocytes | Scaf4 |
| Ahnak | 3.83E-31 | 0.96834548 | 0.865 | 0.737 | 7.40E-27 | monocytes | Ahnak |
| Nfam11 | 5.24E-31 | 0.65397844 | 0.428 | 0.481 | 1.01E-26 | monocytes | Nfam1 |
| Chic2 | 5.64E-31 | 0.32932613 | 0.197 | 0.264 | 1.09E-26 | monocytes | Chic2 |
| Kansl3 | 5.65E-31 | 0.29822911 | 0.192 | 0.263 | 1.09E-26 | monocytes | Kansl3 |
| Ccnt1 | 6.42E-31 | 0.37973207 | 0.241 | 0.308 | 1.24E-26 | monocytes | Ccnt1 |
| Zbtb211 | 9.84E-31 | 0.2739278 | 0.191 | 0.266 | 1.90E-26 | monocytes | Zbtb21 |
| Rnh11 | 1.02E-30 | 0.76159309 | 0.648 | 0.676 | 1.98E-26 | monocytes | Rnh1 |
| Oasl21 | 1.30E-30 | 0.33923527 | 0.25 | 0.262 | 2.51E-26 | monocytes | Oasl2 |
| Dhx16 | 2.57E-30 | 0.38120749 | 0.238 | 0.305 | 4.98E-26 | monocytes | Dhx16 |
| Chp11 | 2.59E-30 | 0.64008614 | 0.564 | 0.596 | 5.01E-26 | monocytes | Chp1 |
| Rgcc1 | 3.65E-30 | 0.51131414 | 0.4 | 0.351 | 7.06E-26 | monocytes | Rgcc |
| Fam53c | 3.76E-30 | 0.29250034 | 0.189 | 0.254 | 7.26E-26 | monocytes | Fam53c |
| Lrrfip2 | 4.56E-30 | 0.5081121 | 0.32 | 0.374 | 8.82E-26 | monocytes | Lrrfip2 |
| Id31 | 5.18E-30 | 1.11732849 | 0.33 | 0.259 | 1.00E-25 | monocytes | Id3 |
| Stk4 | 6.18E-30 | 0.45294203 | 0.259 | 0.323 | 1.20E-25 | monocytes | Stk4 |
| Usp34 | 6.29E-30 | 0.31061143 | 0.22 | 0.286 | 1.22E-25 | monocytes | Usp34 |
| Usb1 | 8.55E-30 | 0.36178343 | 0.234 | 0.3 | 1.65E-25 | monocytes | Usb1 |
| Kras | 8.55E-30 | 0.55306605 | 0.429 | 0.491 | 1.65E-25 | monocytes | Kras |
| Slc25a30 | 9.22E-30 | 0.40363343 | 0.201 | 0.264 | 1.78E-25 | monocytes | Slc25a30 |
| Emd1 | 9.27E-30 | 0.60224147 | 0.527 | 0.531 | 1.79E-25 | monocytes | Emd |
| Lyn1 | 1.18E-29 | 0.82168464 | 0.852 | 0.815 | 2.29E-25 | monocytes | Lyn |
| Dok31 | 1.63E-29 | 0.62685726 | 0.438 | 0.485 | 3.16E-25 | monocytes | Dok3 |
| Phf2 | 1.74E-29 | 0.34223411 | 0.18 | 0.251 | 3.37E-25 | monocytes | Phf2 |
| Arfip1 | 1.87E-29 | 0.39271373 | 0.203 | 0.278 | 3.62E-25 | monocytes | Arfip1 |
| Ccdc97 | 2.50E-29 | 0.49909613 | 0.258 | 0.323 | 4.83E-25 | monocytes | Ccdc97 |
| Rbm27 | 3.10E-29 | 0.28264891 | 0.183 | 0.256 | 5.99E-25 | monocytes | Rbm27 |
| Ank | 3.48E-29 | 0.34335267 | 0.207 | 0.259 | 6.72E-25 | monocytes | Ank |
| Chmp4b1 | 4.03E-29 | 0.6252528 | 0.541 | 0.578 | 7.79E-25 | monocytes | Chmp4b |
| Pak1ip11 | 5.71E-29 | 0.53673248 | 0.269 | 0.335 | 1.10E-24 | monocytes | Pak1ip1 |
| Def61 | 1.85E-28 | 0.41991149 | 0.229 | 0.294 | 3.58E-24 | monocytes | Def6 |
| Rnf2172 | 2.36E-28 | 0.38880492 | 0.257 | 0.281 | 4.57E-24 | monocytes | Rnf217 |
| Thoc2 | 2.43E-28 | 0.35086549 | 0.196 | 0.264 | 4.70E-24 | monocytes | Thoc2 |
| Tnfrsf12a1 | 2.62E-28 | 0.42969756 | 0.373 | 0.424 | 5.06E-24 | monocytes | Tnfrsf12a |
| Relt1 | 3.06E-28 | 0.45303349 | 0.265 | 0.329 | 5.92E-24 | monocytes | Relt |
| Notch1 | 3.42E-28 | 0.4785501 | 0.287 | 0.358 | 6.62E-24 | monocytes | Notch1 |
| Tmem248 | 3.47E-28 | 0.43341535 | 0.224 | 0.296 | 6.70E-24 | monocytes | Tmem248 |
| Snap231 | 5.77E-28 | 0.40520139 | 0.227 | 0.289 | 1.12E-23 | monocytes | Snap23 |
| Errfi11 | 6.42E-28 | 1.08157578 | 0.407 | 0.364 | 1.24E-23 | monocytes | Errfi1 |
| Noct1 | 6.89E-28 | 1.21804523 | 0.336 | 0.257 | 1.33E-23 | monocytes | Noct |
| Ube2v22 | 7.95E-28 | 0.32147809 | 0.214 | 0.274 | 1.54E-23 | monocytes | Ube2v2 |
| Nsf1 | 8.15E-28 | 0.48173972 | 0.335 | 0.398 | 1.58E-23 | monocytes | Nsf |
| N4bp1 | 1.20E-27 | 1.03599176 | 0.676 | 0.633 | 2.32E-23 | monocytes | N4bp1 |
| Elk4 | 1.23E-27 | 0.32859825 | 0.229 | 0.301 | 2.38E-23 | monocytes | Elk4 |
| Ambra11 | 1.44E-27 | 0.40090278 | 0.245 | 0.315 | 2.78E-23 | monocytes | Ambra1 |
| Rel1 | 1.51E-27 | 0.63452869 | 0.718 | 0.692 | 2.91E-23 | monocytes | Rel |
| Arid4b | 1.59E-27 | 0.38005261 | 0.241 | 0.31 | 3.08E-23 | monocytes | Arid4b |
| Cdk19 | 1.70E-27 | 0.48115404 | 0.213 | 0.273 | 3.28E-23 | monocytes | Cdk19 |
| Stk17b2 | 1.74E-27 | 0.65369251 | 0.552 | 0.57 | 3.36E-23 | monocytes | Stk17b |
| Smad71 | 1.80E-27 | 0.55793627 | 0.341 | 0.374 | 3.48E-23 | monocytes | Smad7 |
| Bcl2l111 | 1.84E-27 | 0.55599356 | 0.663 | 0.684 | 3.56E-23 | monocytes | Bcl2l11 |
| Ifi2091 | 1.84E-27 | 1.14156525 | 0.35 | 0.258 | 3.56E-23 | monocytes | Ifi209 |
| Sestd1 | 2.47E-27 | 0.37702441 | 0.289 | 0.35 | 4.77E-23 | monocytes | Sestd1 |
| Clec5a1 | 2.60E-27 | 1.07944124 | 0.406 | 0.396 | 5.03E-23 | monocytes | Clec5a |
| Parp141 | 2.66E-27 | 0.45721003 | 0.361 | 0.381 | 5.14E-23 | monocytes | Parp14 |
| Smpdl3a | 2.93E-27 | 0.97011875 | 0.399 | 0.438 | 5.66E-23 | monocytes | Smpdl3a |
| Ppp1r12a | 4.07E-27 | 0.56419358 | 0.441 | 0.489 | 7.88E-23 | monocytes | Ppp1r12a |
| Nsd2 | 4.84E-27 | 0.50295908 | 0.38 | 0.398 | 9.37E-23 | monocytes | Nsd2 |
| Dock52 | 6.44E-27 | 1.342043 | 0.284 | 0.203 | 1.25E-22 | monocytes | Dock5 |
| Nfkbie2 | 7.55E-27 | 0.52754309 | 0.607 | 0.621 | 1.46E-22 | monocytes | Nfkbie |
| Chuk1 | 9.16E-27 | 0.38701434 | 0.232 | 0.303 | 1.77E-22 | monocytes | Chuk |
| Eif2b22 | 9.38E-27 | 0.5626883 | 0.306 | 0.363 | 1.81E-22 | monocytes | Eif2b2 |
| Ptpre1 | 1.10E-26 | 0.70727628 | 0.575 | 0.567 | 2.12E-22 | monocytes | Ptpre |
| Chfr | 1.25E-26 | 0.35202278 | 0.207 | 0.273 | 2.42E-22 | monocytes | Chfr |
| Flot2 | 1.86E-26 | 0.68693007 | 0.286 | 0.344 | 3.60E-22 | monocytes | Flot2 |
| Cdk12 | 2.49E-26 | 0.42621442 | 0.289 | 0.356 | 4.82E-22 | monocytes | Cdk12 |
| Smim31 | 4.27E-26 | 0.47869709 | 0.32 | 0.352 | 8.26E-22 | monocytes | Smim3 |
| Slc23a21 | 5.25E-26 | 0.4791744 | 0.293 | 0.354 | 1.02E-21 | monocytes | Slc23a2 |
| Med142 | 6.67E-26 | 0.43529367 | 0.257 | 0.296 | 1.29E-21 | monocytes | Med14 |
| Gxylt1 | 9.18E-26 | 0.38336763 | 0.218 | 0.284 | 1.78E-21 | monocytes | Gxylt1 |
| Flna1 | 1.08E-25 | 0.83666917 | 0.827 | 0.744 | 2.08E-21 | monocytes | Flna |
| Ptpn21 | 1.57E-25 | 0.50958649 | 0.303 | 0.353 | 3.04E-21 | monocytes | Ptpn2 |
| Tap11 | 2.11E-25 | 0.50461468 | 0.412 | 0.443 | 4.08E-21 | monocytes | Tap1 |
| Bach11 | 2.26E-25 | 0.74912029 | 0.729 | 0.747 | 4.36E-21 | monocytes | Bach1 |
| Tbc1d11 | 2.47E-25 | 0.5442074 | 0.311 | 0.376 | 4.77E-21 | monocytes | Tbc1d1 |
| Sec23b | 3.77E-25 | 0.44126728 | 0.232 | 0.297 | 7.29E-21 | monocytes | Sec23b |
| Mindy3 | 5.06E-25 | 0.409764 | 0.203 | 0.26 | 9.78E-21 | monocytes | Mindy3 |
| Abhd2 | 6.13E-25 | 0.39886537 | 0.226 | 0.29 | 1.19E-20 | monocytes | Abhd2 |
| Zc3h7a | 8.96E-25 | 0.58707315 | 0.446 | 0.5 | 1.73E-20 | monocytes | Zc3h7a |
| Bbip1 | 1.37E-24 | 0.42888523 | 0.226 | 0.285 | 2.66E-20 | monocytes | Bbip1 |
| Appbp2 | 1.58E-24 | 0.4387758 | 0.273 | 0.337 | 3.05E-20 | monocytes | Appbp2 |
| Mapk61 | 1.75E-24 | 0.55743701 | 0.414 | 0.477 | 3.38E-20 | monocytes | Mapk6 |
| Crebbp1 | 1.79E-24 | 0.51664521 | 0.373 | 0.43 | 3.46E-20 | monocytes | Crebbp |
| Ripk32 | 3.06E-24 | 0.48444148 | 0.283 | 0.335 | 5.91E-20 | monocytes | Ripk3 |
| Zranb1 | 6.87E-24 | 0.26243282 | 0.19 | 0.251 | 1.33E-19 | monocytes | Zranb1 |
| Lmo4 | 9.90E-24 | 0.65203994 | 0.295 | 0.331 | 1.91E-19 | monocytes | Lmo4 |
| Magoh1 | 1.42E-23 | 0.50038373 | 0.284 | 0.332 | 2.74E-19 | monocytes | Magoh |
| Hnrnpdl1 | 1.43E-23 | 0.34948947 | 0.238 | 0.294 | 2.77E-19 | monocytes | Hnrnpdl |
| Nipbl1 | 1.45E-23 | 0.60750325 | 0.367 | 0.413 | 2.80E-19 | monocytes | Nipbl |
| Srf1 | 1.53E-23 | 0.56418867 | 0.368 | 0.434 | 2.95E-19 | monocytes | Srf |
| Smndc11 | 2.11E-23 | 0.4782966 | 0.242 | 0.29 | 4.09E-19 | monocytes | Smndc1 |
| Ube2a | 2.24E-23 | 0.56495974 | 0.391 | 0.433 | 4.33E-19 | monocytes | Ube2a |
| Ece12 | 2.40E-23 | 0.6975699 | 0.492 | 0.508 | 4.64E-19 | monocytes | Ece1 |
| Pofut21 | 3.73E-23 | 0.44291496 | 0.249 | 0.305 | 7.21E-19 | monocytes | Pofut2 |
| Nin | 3.82E-23 | 0.61355526 | 0.332 | 0.379 | 7.38E-19 | monocytes | Nin |
| Pbxip12 | 4.08E-23 | 0.49660903 | 0.212 | 0.263 | 7.88E-19 | monocytes | Pbxip1 |
| Rabgef11 | 5.73E-23 | 0.53462064 | 0.542 | 0.547 | 1.11E-18 | monocytes | Rabgef1 |
| Sertad2 | 6.47E-23 | 0.50666468 | 0.249 | 0.304 | 1.25E-18 | monocytes | Sertad2 |
| Mar-71 | 6.64E-23 | 0.59917214 | 0.484 | 0.52 | 1.28E-18 | monocytes | 7-Mar |
| Msn1 | 1.02E-22 | 0.87353836 | 0.969 | 0.94 | 1.96E-18 | monocytes | Msn |
| Kmt2c | 1.33E-22 | 0.42487164 | 0.193 | 0.253 | 2.57E-18 | monocytes | Kmt2c |
| Cript | 1.54E-22 | 0.50871908 | 0.243 | 0.301 | 2.97E-18 | monocytes | Cript |
| Nuak21 | 3.04E-22 | 0.68605693 | 0.647 | 0.624 | 5.88E-18 | monocytes | Nuak2 |
| Prkag1 | 3.34E-22 | 0.50577406 | 0.222 | 0.27 | 6.45E-18 | monocytes | Prkag1 |
| Adipor2 | 3.68E-22 | 0.58088436 | 0.296 | 0.345 | 7.11E-18 | monocytes | Adipor2 |
| Plekhg3 | 3.95E-22 | 1.17792817 | 0.342 | 0.282 | 7.63E-18 | monocytes | Plekhg3 |
| Kdm7a2 | 4.11E-22 | 0.71113139 | 0.685 | 0.691 | 7.94E-18 | monocytes | Kdm7a |
| Apobr | 4.20E-22 | 0.46846763 | 0.218 | 0.273 | 8.13E-18 | monocytes | Apobr |
| Tmem1281 | 4.50E-22 | 0.4636162 | 0.256 | 0.313 | 8.70E-18 | monocytes | Tmem128 |
| Zfp800 | 4.85E-22 | 0.44295879 | 0.229 | 0.284 | 9.38E-18 | monocytes | Zfp800 |
| Arhgdib | 5.51E-22 | 0.83109973 | 0.841 | 0.813 | 1.06E-17 | monocytes | Arhgdib |
| Ccdc121 | 8.41E-22 | 0.63975946 | 0.455 | 0.485 | 1.63E-17 | monocytes | Ccdc12 |
| Cux1 | 1.14E-21 | 0.66611359 | 0.688 | 0.698 | 2.20E-17 | monocytes | Cux1 |
| Tle31 | 1.31E-21 | 0.7494278 | 0.423 | 0.506 | 2.53E-17 | monocytes | Tle3 |
| Tle41 | 1.32E-21 | 0.4458727 | 0.251 | 0.301 | 2.54E-17 | monocytes | Tle4 |
| Uap11 | 1.44E-21 | 0.56383644 | 0.35 | 0.395 | 2.79E-17 | monocytes | Uap1 |
| Mthfs1 | 1.46E-21 | 0.45406632 | 0.271 | 0.326 | 2.82E-17 | monocytes | Mthfs |
| Pgs1 | 2.69E-21 | 0.6473455 | 0.519 | 0.546 | 5.21E-17 | monocytes | Pgs1 |
| Auh | 2.84E-21 | 0.54361591 | 0.274 | 0.333 | 5.49E-17 | monocytes | Auh |
| Xylt11 | 3.59E-21 | 1.21329465 | 0.336 | 0.257 | 6.93E-17 | monocytes | Xylt1 |
| Ppp4r3b | 5.05E-21 | 0.50817203 | 0.25 | 0.304 | 9.77E-17 | monocytes | Ppp4r3b |
| Flot1 | 5.60E-21 | 1.04172015 | 0.485 | 0.42 | 1.08E-16 | monocytes | Flot1 |
| Rap2c1 | 5.97E-21 | 0.5801015 | 0.311 | 0.352 | 1.15E-16 | monocytes | Rap2c |
| Crtc31 | 6.17E-21 | 1.0629344 | 0.48 | 0.422 | 1.19E-16 | monocytes | Crtc3 |
| Sav1 | 6.51E-21 | 0.61593513 | 0.48 | 0.524 | 1.26E-16 | monocytes | Sav1 |
| Sec61b1 | 6.56E-21 | 0.62810652 | 0.366 | 0.389 | 1.27E-16 | monocytes | Sec61b |
| Ap1s21 | 8.81E-21 | 0.53823078 | 0.271 | 0.311 | 1.70E-16 | monocytes | Ap1s2 |
| H2-Q71 | 9.87E-21 | 0.51469545 | 0.315 | 0.347 | 1.91E-16 | monocytes | H2-Q7 |
| Btbd91 | 1.07E-20 | 0.41288678 | 0.276 | 0.334 | 2.07E-16 | monocytes | Btbd9 |
| Bcl31 | 1.07E-20 | 1.12515991 | 0.591 | 0.548 | 2.07E-16 | monocytes | Bcl3 |
| Ppp1r3b1 | 1.26E-20 | 1.138949 | 0.254 | 0.249 | 2.43E-16 | monocytes | Ppp1r3b |
| Kdm2a | 1.39E-20 | 0.62570559 | 0.714 | 0.733 | 2.68E-16 | monocytes | Kdm2a |
| Ppard | 1.42E-20 | 0.5577398 | 0.275 | 0.316 | 2.75E-16 | monocytes | Ppard |
| Cmtr11 | 1.46E-20 | 0.46240928 | 0.328 | 0.382 | 2.82E-16 | monocytes | Cmtr1 |
| Usp32 | 1.47E-20 | 0.48399216 | 0.247 | 0.305 | 2.85E-16 | monocytes | Usp32 |
| Rhoq1 | 1.93E-20 | 0.56739909 | 0.536 | 0.556 | 3.74E-16 | monocytes | Rhoq |
| Prkcd2 | 2.15E-20 | 0.78119788 | 0.881 | 0.873 | 4.15E-16 | monocytes | Prkcd |
| Arid3a1 | 2.40E-20 | 0.44626456 | 0.204 | 0.252 | 4.64E-16 | monocytes | Arid3a |
| Xbp11 | 2.92E-20 | 1.04700172 | 0.849 | 0.797 | 5.65E-16 | monocytes | Xbp1 |
| Sys1 | 5.22E-20 | 0.63977484 | 0.364 | 0.407 | 1.01E-15 | monocytes | Sys1 |
| H13 | 1.58E-19 | 0.70805289 | 0.538 | 0.562 | 3.05E-15 | monocytes | H13 |
| Gtf2b | 3.11E-19 | 0.605429 | 0.347 | 0.395 | 6.01E-15 | monocytes | Gtf2b |
| Cd142 | 5.73E-19 | 0.76421302 | 0.911 | 0.873 | 1.11E-14 | monocytes | Cd14 |
| Eed | 7.21E-19 | 0.4888878 | 0.242 | 0.295 | 1.39E-14 | monocytes | Eed |
| Ube2j11 | 9.38E-19 | 0.44121133 | 0.216 | 0.268 | 1.81E-14 | monocytes | Ube2j1 |
| Slfn2 | 1.07E-18 | 0.79103179 | 0.885 | 0.852 | 2.07E-14 | monocytes | Slfn2 |
| Tifa1 | 1.11E-18 | 0.51349349 | 0.239 | 0.291 | 2.15E-14 | monocytes | Tifa |
| Creb1 | 1.30E-18 | 0.58346967 | 0.354 | 0.405 | 2.51E-14 | monocytes | Creb1 |
| Hivep21 | 2.53E-18 | 0.35187418 | 0.221 | 0.273 | 4.89E-14 | monocytes | Hivep2 |
| Tmed51 | 2.91E-18 | 0.65899529 | 0.462 | 0.5 | 5.62E-14 | monocytes | Tmed5 |
| Nupr1 | 2.94E-18 | 0.62042946 | 0.407 | 0.388 | 5.68E-14 | monocytes | Nupr1 |
| Usp251 | 3.29E-18 | 0.63125648 | 0.479 | 0.515 | 6.37E-14 | monocytes | Usp25 |
| Siah21 | 4.07E-18 | 0.66375452 | 0.342 | 0.381 | 7.86E-14 | monocytes | Siah2 |
| Ern1 | 4.49E-18 | 0.61987617 | 0.476 | 0.499 | 8.68E-14 | monocytes | Ern1 |
| Ppp1r2 | 5.08E-18 | 0.64467525 | 0.265 | 0.311 | 9.83E-14 | monocytes | Ppp1r2 |
| Mta3 | 5.20E-18 | 0.54526292 | 0.308 | 0.352 | 1.01E-13 | monocytes | Mta3 |
| Msrb11 | 6.09E-18 | 0.97147096 | 0.874 | 0.808 | 1.18E-13 | monocytes | Msrb1 |
| Lipe1 | 6.44E-18 | 0.57831687 | 0.253 | 0.289 | 1.25E-13 | monocytes | Lipe |
| Tnip11 | 6.76E-18 | 1.17415057 | 0.454 | 0.425 | 1.31E-13 | monocytes | Tnip1 |
| Fam110a1 | 7.52E-18 | 0.9870882 | 0.362 | 0.306 | 1.45E-13 | monocytes | Fam110a |
| Trim30a1 | 8.69E-18 | 0.55120735 | 0.231 | 0.288 | 1.68E-13 | monocytes | Trim30a |
| Cyp4f181 | 9.37E-18 | 1.14272734 | 0.53 | 0.425 | 1.81E-13 | monocytes | Cyp4f18 |
| Carhsp1 | 9.51E-18 | 0.60289181 | 0.337 | 0.403 | 1.84E-13 | monocytes | Carhsp1 |
| Uhrf1bp1l1 | 1.05E-17 | 0.47790021 | 0.208 | 0.25 | 2.03E-13 | monocytes | Uhrf1bp1l |
| Ezr1 | 1.32E-17 | 0.68301649 | 0.444 | 0.473 | 2.56E-13 | monocytes | Ezr |
| Herc4 | 1.44E-17 | 0.61803785 | 0.262 | 0.315 | 2.78E-13 | monocytes | Herc4 |
| Med11 | 1.79E-17 | 0.42359126 | 0.241 | 0.288 | 3.46E-13 | monocytes | Med11 |
| Lst12 | 1.94E-17 | 0.75063527 | 0.624 | 0.588 | 3.75E-13 | monocytes | Lst1 |
| Gvin11 | 3.24E-17 | 0.5585694 | 0.216 | 0.265 | 6.27E-13 | monocytes | Gvin1 |
| Ralgapa11 | 4.82E-17 | 0.54648911 | 0.205 | 0.253 | 9.31E-13 | monocytes | Ralgapa1 |
| Itgam2 | 4.96E-17 | 0.81756687 | 0.897 | 0.796 | 9.59E-13 | monocytes | Itgam |
| Pilrb12 | 6.68E-17 | 0.6532901 | 0.297 | 0.324 | 1.29E-12 | monocytes | Pilrb1 |
| Daxx1 | 7.59E-17 | 0.63580754 | 0.29 | 0.332 | 1.47E-12 | monocytes | Daxx |
| AB1246111 | 9.86E-17 | 1.2179732 | 0.373 | 0.311 | 1.91E-12 | monocytes | AB124611 |
| Gtf2e2 | 1.01E-16 | 0.53116071 | 0.207 | 0.255 | 1.95E-12 | monocytes | Gtf2e2 |
| Stag2 | 1.71E-16 | 0.57748908 | 0.294 | 0.341 | 3.30E-12 | monocytes | Stag2 |
| Lemd31 | 2.49E-16 | 0.52643646 | 0.24 | 0.287 | 4.81E-12 | monocytes | Lemd3 |
| Usp382 | 3.24E-16 | 0.50814831 | 0.221 | 0.264 | 6.26E-12 | monocytes | Usp38 |
| Cd300lf1 | 3.32E-16 | 0.7071281 | 0.586 | 0.578 | 6.42E-12 | monocytes | Cd300lf |
| Socs32 | 3.49E-16 | 1.00066924 | 0.858 | 0.775 | 6.74E-12 | monocytes | Socs3 |
| Elk31 | 5.06E-16 | 0.63798385 | 0.466 | 0.505 | 9.79E-12 | monocytes | Elk3 |
| Ptpro1 | 6.41E-16 | 0.6279384 | 0.26 | 0.312 | 1.24E-11 | monocytes | Ptpro |
| Clcf12 | 7.52E-16 | 0.56825035 | 0.301 | 0.323 | 1.45E-11 | monocytes | Clcf1 |
| Tbc1d2b | 9.26E-16 | 1.08184474 | 0.488 | 0.428 | 1.79E-11 | monocytes | Tbc1d2b |
| Gpr352 | 9.53E-16 | 0.98388542 | 0.778 | 0.676 | 1.84E-11 | monocytes | Gpr35 |
| Uvrag | 1.05E-15 | 0.70044799 | 0.45 | 0.487 | 2.04E-11 | monocytes | Uvrag |
| Arfgef1 | 1.25E-15 | 0.57384934 | 0.284 | 0.33 | 2.42E-11 | monocytes | Arfgef1 |
| Cd471 | 1.32E-15 | 0.74538439 | 0.75 | 0.743 | 2.55E-11 | monocytes | Cd47 |
| Tmem185b1 | 1.37E-15 | 0.56755488 | 0.319 | 0.367 | 2.66E-11 | monocytes | Tmem185b |
| Cd24a2 | 1.67E-15 | 0.55853453 | 0.28 | 0.258 | 3.24E-11 | monocytes | Cd24a |
| Prkx1 | 2.10E-15 | 0.64843668 | 0.284 | 0.334 | 4.06E-11 | monocytes | Prkx |
| Rapgef11 | 2.23E-15 | 0.67533897 | 0.395 | 0.421 | 4.31E-11 | monocytes | Rapgef1 |
| Zyx | 4.67E-15 | 0.85658838 | 0.78 | 0.757 | 9.02E-11 | monocytes | Zyx |
| Arhgef10l1 | 4.73E-15 | 1.09315871 | 0.395 | 0.334 | 9.14E-11 | monocytes | Arhgef10l |
| Plcg21 | 4.96E-15 | 0.72953952 | 0.506 | 0.531 | 9.59E-11 | monocytes | Plcg2 |
| Il6st | 5.93E-15 | 0.70013759 | 0.322 | 0.361 | 1.15E-10 | monocytes | Il6st |
| Gsap1 | 6.15E-15 | 0.68595534 | 0.501 | 0.514 | 1.19E-10 | monocytes | Gsap |
| Rasa3 | 9.99E-15 | 0.81026633 | 0.371 | 0.41 | 1.93E-10 | monocytes | Rasa3 |
| Brpf1 | 1.12E-14 | 0.59028567 | 0.383 | 0.422 | 2.16E-10 | monocytes | Brpf1 |
| Chd12 | 1.22E-14 | 0.57046196 | 0.377 | 0.416 | 2.35E-10 | monocytes | Chd1 |
| C32 | 1.66E-14 | 1.01360693 | 0.803 | 0.551 | 3.21E-10 | monocytes | C3 |
| Rnf341 | 1.83E-14 | 0.53586904 | 0.231 | 0.277 | 3.55E-10 | monocytes | Rnf34 |
| Zfp6091 | 2.86E-14 | 0.49553302 | 0.244 | 0.295 | 5.53E-10 | monocytes | Zfp609 |
| Mtmr3 | 3.17E-14 | 0.6684149 | 0.321 | 0.363 | 6.13E-10 | monocytes | Mtmr3 |
| Cpne1 | 5.33E-14 | 0.60202586 | 0.264 | 0.304 | 1.03E-09 | monocytes | Cpne1 |
| Map2k3 | 1.18E-13 | 0.78717883 | 0.644 | 0.649 | 2.28E-09 | monocytes | Map2k3 |
| Cldnd1 | 1.44E-13 | 0.6379132 | 0.367 | 0.409 | 2.79E-09 | monocytes | Cldnd1 |
| Baz1a1 | 1.47E-13 | 0.68511294 | 0.517 | 0.532 | 2.84E-09 | monocytes | Baz1a |
| Zc3hav1 | 1.94E-13 | 0.66640326 | 0.441 | 0.458 | 3.74E-09 | monocytes | Zc3hav1 |
| Rnf2131 | 2.26E-13 | 0.64183702 | 0.452 | 0.454 | 4.38E-09 | monocytes | Rnf213 |
| Lats21 | 3.37E-13 | 0.60428998 | 0.387 | 0.396 | 6.52E-09 | monocytes | Lats2 |
| Hbegf1 | 4.14E-13 | 0.67216577 | 0.272 | 0.285 | 8.00E-09 | monocytes | Hbegf |
| Tdrd71 | 4.19E-13 | 0.53291081 | 0.229 | 0.267 | 8.10E-09 | monocytes | Tdrd7 |
| Csf3r1 | 4.37E-13 | 1.07917858 | 0.468 | 0.394 | 8.45E-09 | monocytes | Csf3r |
| Zfp592 | 4.43E-13 | 0.60027347 | 0.266 | 0.305 | 8.57E-09 | monocytes | Zfp592 |
| Btg1 | 5.65E-13 | 0.81010492 | 0.893 | 0.856 | 1.09E-08 | monocytes | Btg1 |
| Arg21 | 6.41E-13 | 1.01097915 | 0.408 | 0.33 | 1.24E-08 | monocytes | Arg2 |
| Ascc31 | 1.59E-12 | 0.55557587 | 0.368 | 0.404 | 3.08E-08 | monocytes | Ascc3 |
| Shisa51 | 1.84E-12 | 0.80808415 | 0.829 | 0.802 | 3.56E-08 | monocytes | Shisa5 |
| Cast | 1.84E-12 | 0.74159536 | 0.495 | 0.498 | 3.56E-08 | monocytes | Cast |
| Spata131 | 1.96E-12 | 1.21870024 | 0.336 | 0.291 | 3.80E-08 | monocytes | Spata13 |
| Dusp22 | 2.44E-12 | 1.05931217 | 0.382 | 0.31 | 4.71E-08 | monocytes | Dusp2 |
| Abca7 | 2.80E-12 | 1.24105681 | 0.326 | 0.272 | 5.41E-08 | monocytes | Abca7 |
| Sec24b1 | 3.07E-12 | 0.70954904 | 0.429 | 0.466 | 5.93E-08 | monocytes | Sec24b |
| Dram11 | 3.11E-12 | 0.63144959 | 0.297 | 0.319 | 6.02E-08 | monocytes | Dram1 |
| Il6ra1 | 3.12E-12 | 1.0922943 | 0.547 | 0.487 | 6.03E-08 | monocytes | Il6ra |
| Malt12 | 3.25E-12 | 0.87084761 | 0.732 | 0.682 | 6.28E-08 | monocytes | Malt1 |
| Il10ra1 | 3.53E-12 | 0.76753389 | 0.752 | 0.706 | 6.82E-08 | monocytes | Il10ra |
| Atp1a31 | 4.75E-12 | 1.03842145 | 0.253 | 0.239 | 9.19E-08 | monocytes | Atp1a3 |
| Zfp6081 | 6.89E-12 | 1.12943447 | 0.25 | 0.208 | 1.33E-07 | monocytes | Zfp608 |
| Zfp2631 | 8.44E-12 | 0.61072568 | 0.254 | 0.296 | 1.63E-07 | monocytes | Zfp263 |
| Pdlim72 | 8.81E-12 | 0.57456992 | 0.335 | 0.347 | 1.70E-07 | monocytes | Pdlim7 |
| Fbxo11 | 1.04E-11 | 0.67232914 | 0.472 | 0.496 | 2.00E-07 | monocytes | Fbxo11 |
| Nrg12 | 1.10E-11 | 0.47893049 | 0.428 | 0.373 | 2.12E-07 | monocytes | Nrg1 |
| Plec1 | 1.47E-11 | 0.74707848 | 0.701 | 0.66 | 2.85E-07 | monocytes | Plec |
| Irf91 | 1.62E-11 | 0.70294668 | 0.294 | 0.332 | 3.13E-07 | monocytes | Irf9 |
| Actg11 | 1.81E-11 | 0.91181754 | 0.96 | 0.936 | 3.49E-07 | monocytes | Actg1 |
| Snx201 | 2.10E-11 | 0.81212354 | 0.696 | 0.687 | 4.07E-07 | monocytes | Snx20 |
| Slfn51 | 2.72E-11 | 0.96179438 | 0.342 | 0.269 | 5.25E-07 | monocytes | Slfn5 |
| Grk31 | 2.86E-11 | 0.7048964 | 0.263 | 0.303 | 5.54E-07 | monocytes | Grk3 |
| Mllt62 | 3.29E-11 | 0.64713861 | 0.461 | 0.508 | 6.35E-07 | monocytes | Mllt6 |
| Pdzd81 | 3.95E-11 | 0.64927548 | 0.267 | 0.3 | 7.64E-07 | monocytes | Pdzd8 |
| Rybp1 | 6.51E-11 | 1.03202357 | 0.311 | 0.26 | 1.26E-06 | monocytes | Rybp |
| Acer32 | 6.71E-11 | 0.64414335 | 0.263 | 0.294 | 1.30E-06 | monocytes | Acer3 |
| Ppp1r15b | 6.78E-11 | 0.81499759 | 0.412 | 0.438 | 1.31E-06 | monocytes | Ppp1r15b |
| Cdyl1 | 9.33E-11 | 0.60757971 | 0.253 | 0.286 | 1.80E-06 | monocytes | Cdyl |
| Prdx62 | 1.39E-10 | 0.69488671 | 0.332 | 0.331 | 2.69E-06 | monocytes | Prdx6 |
| Agpat42 | 3.10E-10 | 1.03307544 | 0.465 | 0.396 | 6.00E-06 | monocytes | Agpat4 |
| Trib11 | 3.17E-10 | 0.78661006 | 0.634 | 0.582 | 6.13E-06 | monocytes | Trib1 |
| Gmfg | 4.00E-10 | 0.72245973 | 0.406 | 0.417 | 7.73E-06 | monocytes | Gmfg |
| Cox172 | 4.16E-10 | 0.95443042 | 0.627 | 0.588 | 8.04E-06 | monocytes | Cox17 |
| Csgalnact2 | 5.27E-10 | 0.74705989 | 0.249 | 0.291 | 1.02E-05 | monocytes | Csgalnact2 |
| Ptpn11 | 6.82E-10 | 1.07899857 | 0.819 | 0.764 | 1.32E-05 | monocytes | Ptpn1 |
| Dhx38 | 8.11E-10 | 0.72041627 | 0.522 | 0.536 | 1.57E-05 | monocytes | Dhx38 |
| Map3k141 | 9.37E-10 | 0.65320551 | 0.302 | 0.334 | 1.81E-05 | monocytes | Map3k14 |
| Mical12 | 9.73E-10 | 0.75967946 | 0.267 | 0.278 | 1.88E-05 | monocytes | Mical1 |
| Slc16a31 | 9.82E-10 | 0.96319649 | 0.716 | 0.619 | 1.90E-05 | monocytes | Slc16a3 |
| Atp11a | 1.39E-09 | 0.62971392 | 0.348 | 0.382 | 2.68E-05 | monocytes | Atp11a |
| Vps541 | 1.39E-09 | 0.61548477 | 0.275 | 0.31 | 2.69E-05 | monocytes | Vps54 |
| Fam129a1 | 1.41E-09 | 0.76723451 | 0.345 | 0.36 | 2.72E-05 | monocytes | Fam129a |
| Ifi2041 | 1.54E-09 | 0.99560925 | 0.369 | 0.299 | 2.98E-05 | monocytes | Ifi204 |
| Rassf51 | 1.62E-09 | 0.84158292 | 0.46 | 0.471 | 3.14E-05 | monocytes | Rassf5 |
| Elf4 | 2.00E-09 | 0.72895025 | 0.365 | 0.386 | 3.86E-05 | monocytes | Elf4 |
| Ywhaz1 | 2.14E-09 | 0.69474144 | 0.274 | 0.296 | 4.14E-05 | monocytes | Ywhaz |
| Kmt2e | 2.30E-09 | 0.68533328 | 0.371 | 0.383 | 4.44E-05 | monocytes | Kmt2e |
| Plekhf2 | 3.19E-09 | 0.71371501 | 0.28 | 0.321 | 6.17E-05 | monocytes | Plekhf2 |
| Hs3st3b12 | 3.26E-09 | 0.57260756 | 0.255 | 0.284 | 6.30E-05 | monocytes | Hs3st3b1 |
| Msl21 | 3.52E-09 | 0.71176491 | 0.497 | 0.514 | 6.80E-05 | monocytes | Msl2 |
| Ndel12 | 3.99E-09 | 0.87612408 | 0.728 | 0.681 | 7.72E-05 | monocytes | Ndel1 |
| Sec61g1 | 4.28E-09 | 0.60802693 | 0.318 | 0.35 | 8.27E-05 | monocytes | Sec61g |
| Mcub | 4.91E-09 | 0.72112274 | 0.322 | 0.333 | 9.50E-05 | monocytes | Mcub |
| Por | 5.23E-09 | 0.7368348 | 0.346 | 0.369 | 0.00010112 | monocytes | Por |
| D16Ertd472e | 5.78E-09 | 0.9675965 | 0.347 | 0.277 | 0.00011168 | monocytes | D16Ertd472e |
| Myo1g1 | 6.01E-09 | 0.84039395 | 0.617 | 0.601 | 0.00011617 | monocytes | Myo1g |
| Cflar1 | 7.38E-09 | 0.67989454 | 0.479 | 0.509 | 0.00014277 | monocytes | Cflar |
| Gcnt21 | 8.33E-09 | 1.12042415 | 0.377 | 0.288 | 0.00016099 | monocytes | Gcnt2 |
| Fam129b1 | 9.94E-09 | 1.02294351 | 0.754 | 0.674 | 0.00019223 | monocytes | Fam129b |
| Ifitm11 | 1.19E-08 | 0.59925569 | 0.299 | 0.247 | 0.00023071 | monocytes | Ifitm1 |
| Ist11 | 1.52E-08 | 0.75187097 | 0.737 | 0.729 | 0.00029423 | monocytes | Ist1 |
| Nfat5 | 1.90E-08 | 0.90477309 | 0.46 | 0.434 | 0.00036704 | monocytes | Nfat5 |
| Rnf38 | 2.06E-08 | 0.67558814 | 0.339 | 0.366 | 0.00039847 | monocytes | Rnf38 |
| Gk2 | 2.62E-08 | 1.05285834 | 0.282 | 0.24 | 0.00050692 | monocytes | Gk |
| Prdm12 | 5.37E-08 | 1.01416803 | 0.438 | 0.387 | 0.00103792 | monocytes | Prdm1 |
| Casp41 | 5.72E-08 | 0.65369815 | 0.268 | 0.291 | 0.00110634 | monocytes | Casp4 |
| Snx181 | 7.12E-08 | 0.77520573 | 0.774 | 0.746 | 0.00137589 | monocytes | Snx18 |
| Gpd22 | 7.82E-08 | 0.58916801 | 0.253 | 0.268 | 0.00151213 | monocytes | Gpd2 |
| Fam107b2 | 8.97E-08 | 0.98287248 | 0.496 | 0.45 | 0.00173507 | monocytes | Fam107b |
| Gpcpd1 | 9.58E-08 | 0.92181995 | 0.355 | 0.414 | 0.00185198 | monocytes | Gpcpd1 |
| Foxn31 | 1.30E-07 | 0.74696333 | 0.324 | 0.357 | 0.00251621 | monocytes | Foxn3 |
| Zbp11 | 1.34E-07 | 0.91661008 | 0.365 | 0.283 | 0.00259828 | monocytes | Zbp1 |
| Arid5a1 | 1.52E-07 | 1.07810876 | 0.298 | 0.26 | 0.00293093 | monocytes | Arid5a |
| Gpbp11 | 2.33E-07 | 0.65469622 | 0.299 | 0.316 | 0.00450587 | monocytes | Gpbp1 |

| | | | | | | | |
|---|---|---|---|---|---|---|---|
| Flrt32 | 3.31E-07 | 0.88959241 | 0.261 | 0.254 | 0.00640696 | monocytes | Flrt3 |
| Atp11b1 | 3.35E-07 | 1.01159291 | 0.278 | 0.257 | 0.00646836 | monocytes | Atp11b |
| Lrrc8c1 | 4.13E-07 | 0.81660091 | 0.356 | 0.356 | 0.00797643 | monocytes | Lrrc8c |
| Nup982 | 4.52E-07 | 0.66064436 | 0.45 | 0.464 | 0.00873479 | monocytes | Nup98 |
| Irf71 | 4.74E-07 | 0.71587057 | 0.363 | 0.248 | 0.00915711 | monocytes | Irf7 |
| Pcyt1a1 | 4.80E-07 | 0.74119755 | 0.353 | 0.42 | 0.00928666 | monocytes | Pcyt1a |
| Tmem1311 | 5.07E-07 | 0.80163249 | 0.477 | 0.485 | 0.00979422 | monocytes | Tmem131 |
| Slk | 5.94E-07 | 1.05522716 | 0.482 | 0.436 | 0.01148432 | monocytes | Slk |
| Vamp41 | 6.12E-07 | 0.70880934 | 0.391 | 0.399 | 0.01183611 | monocytes | Vamp4 |
| Tarm12 | 7.17E-07 | 1.08207595 | 0.378 | 0.285 | 0.01385888 | monocytes | Tarm1 |
| Bin21 | 7.46E-07 | 0.92769484 | 0.793 | 0.755 | 0.01441882 | monocytes | Bin2 |
| Scarb1 | 8.67E-07 | 0.92944957 | 0.362 | 0.375 | 0.01676007 | monocytes | Scarb1 |
| Cbfa2t31 | 9.79E-07 | 1.13789341 | 0.276 | 0.236 | 0.01892574 | monocytes | Cbfa2t3 |
| Dennd4a1 | 9.86E-07 | 0.61321198 | 0.55 | 0.524 | 0.01906052 | monocytes | Dennd4a |
| Vill2 | 1.03E-06 | 0.72478412 | 0.256 | 0.258 | 0.01985074 | monocytes | Vill |
| Dock42 | 1.81E-06 | 0.64176713 | 0.253 | 0.267 | 0.03495541 | monocytes | Dock4 |
| Akna2 | 3.45E-06 | 1.10617772 | 0.362 | 0.314 | 0.06676046 | monocytes | Akna |
| Mbp | 5.12E-06 | 0.92610192 | 0.286 | 0.327 | 0.09904946 | monocytes | Mbp |
| Ifi2031 | 6.83E-06 | 0.89803455 | 0.304 | 0.257 | 0.13207731 | monocytes | Ifi203 |
| Ankrd441 | 8.26E-06 | 0.72872474 | 0.286 | 0.316 | 0.15960398 | monocytes | Ankrd44 |
| Il1a1 | 9.07E-06 | 0.8425962 | 0.378 | 0.47 | 0.1752922 | monocytes | Il1a |
| Olr12 | 9.32E-06 | 0.93480755 | 0.275 | 0.2 | 0.18015492 | monocytes | Olr1 |
| Foxn2 | 1.24E-05 | 0.76473696 | 0.469 | 0.479 | 0.24045397 | monocytes | Foxn2 |
| Maml1 | 1.37E-05 | 0.68305357 | 0.276 | 0.3 | 0.2639317 | monocytes | Maml1 |
| Coro1a1 | 2.20E-05 | 0.99159831 | 0.966 | 0.862 | 0.42453814 | monocytes | Coro1a |
| Svil1 | 2.25E-05 | 0.80808726 | 0.426 | 0.427 | 0.43467179 | monocytes | Svil |
| Irf51 | 2.34E-05 | 0.86393631 | 0.869 | 0.822 | 0.45266609 | monocytes | Irf5 |
| Tpd521 | 3.93E-05 | 0.81699337 | 0.468 | 0.472 | 0.75961528 | monocytes | Tpd52 |
| Cdkn1a1 | 4.23E-05 | 0.75660642 | 0.91 | 0.839 | 0.81784731 | monocytes | Cdkn1a |
| Arih21 | 4.37E-05 | 0.67371437 | 0.542 | 0.542 | 0.8449176 | monocytes | Arih2 |
| Foxp1 | 5.89E-05 | 0.7441376 | 0.324 | 0.345 | 1 | monocytes | Foxp1 |
| Abhd17b | 0.00011281 | 0.7429866 | 0.241 | 0.276 | 1 | monocytes | Abhd17b |
| Samsn11 | 0.00012597 | 1.14097489 | 0.349 | 0.309 | 1 | monocytes | Samsn1 |
| Cdk2ap21 | 0.0001457 | 0.83667449 | 0.574 | 0.589 | 1 | monocytes | Cdk2ap2 |
| Stk101 | 0.00017876 | 0.96210602 | 0.337 | 0.359 | 1 | monocytes | Stk10 |
| Cnn2 | 0.00020464 | 0.92266007 | 0.578 | 0.547 | 1 | monocytes | Cnn2 |
| Hck | 0.00023485 | 1.02236285 | 0.736 | 0.665 | 1 | monocytes | Hck |
| Chst111 | 0.00026624 | 1.11288901 | 0.301 | 0.21 | 1 | monocytes | Chst11 |
| Traf6 | 0.00029352 | 0.72320809 | 0.311 | 0.331 | 1 | monocytes | Traf6 |
| Grk52 | 0.00041077 | 1.08720729 | 0.31 | 0.268 | 1 | monocytes | Grk5 |
| Irak22 | 0.00043051 | 0.74296087 | 0.595 | 0.588 | 1 | monocytes | Irak2 |
| Dmxl21 | 0.00043974 | 0.86998231 | 0.365 | 0.371 | 1 | monocytes | Dmxl2 |
| Fosl22 | 0.00052356 | 0.85590993 | 0.912 | 0.843 | 1 | monocytes | Fosl2 |
| Inf21 | 0.00054576 | 0.78254053 | 0.26 | 0.244 | 1 | monocytes | Inf2 |
| Ifitm21 | 0.00057048 | 0.86242913 | 0.921 | 0.91 | 1 | monocytes | Ifitm2 |
| St71 | 0.00120055 | 0.69241715 | 0.26 | 0.295 | 1 | monocytes | St7 |
| Sik3 | 0.00124046 | 1.01568742 | 0.309 | 0.289 | 1 | monocytes | Sik3 |
| Donson1 | 0.00129625 | 0.66134804 | 0.249 | 0.259 | 1 | monocytes | Donson |
| Milr1 | 0.00131932 | 0.74736808 | 0.355 | 0.361 | 1 | monocytes | Milr1 |
| Cxcr41 | 0.00152865 | 0.88310098 | 0.813 | 0.771 | 1 | monocytes | Cxcr4 |
| Runx11 | 0.00187195 | 0.89998106 | 0.683 | 0.655 | 1 | monocytes | Runx1 |
| Diaph21 | 0.00316578 | 0.71231598 | 0.415 | 0.415 | 1 | monocytes | Diaph2 |
| Rasgrp41 | 0.00350826 | 0.8498522 | 0.342 | 0.303 | 1 | monocytes | Rasgrp4 |
| Cmip | 0.0040385 | 0.942086 | 0.773 | 0.743 | 1 | monocytes | Cmip |
| Zcchc22 | 0.00410958 | 0.70029512 | 0.353 | 0.362 | 1 | monocytes | Zcchc2 |
| Ms4a6c2 | 0.00517204 | 0.82586151 | 0.586 | 0.536 | 1 | monocytes | Ms4a6c |
| Eml41 | 0.00544498 | 0.90936627 | 0.404 | 0.394 | 1 | monocytes | Eml4 |
| Met1 | 0.00604109 | 1.08402356 | 0.314 | 0.166 | 1 | monocytes | Met |
| Slc25a201 | 0.00717837 | 0.97726277 | 0.37 | 0.338 | 1 | monocytes | Slc25a20 |
| Baz2b | 0.00742545 | 0.74041033 | 0.292 | 0.303 | 1 | monocytes | Baz2b |
| Gadd45a1 | 0.00983953 | 0.79361008 | 0.248 | 0.259 | 1 | monocytes | Gadd45a |
| Cbr2 | 0 | 4.52648376 | 0.81 | 0.094 | 0 | TAM.Cd209 | Cbr2 |
| Ccl8 | 0 | 4.48279924 | 0.837 | 0.149 | 0 | TAM.Cd209 | Ccl8 |
| C4b | 0 | 4.55628954 | 0.821 | 0.134 | 0 | TAM.Cd209 | C4b |
| Gas61 | 0 | 3.39560533 | 0.887 | 0.249 | 0 | TAM.Cd209 | Gas6 |
| Pmp22 | 0 | 3.38181165 | 0.762 | 0.129 | 0 | TAM.Cd209 | Pmp22 |
| Fcna | 0 | 3.98853404 | 0.773 | 0.166 | 0 | TAM.Cd209 | Fcna |
| Rcn3 | 0 | 3.77732047 | 0.726 | 0.119 | 0 | TAM.Cd209 | Rcn3 |
| Folr21 | 0 | 3.36799227 | 0.821 | 0.218 | 0 | TAM.Cd209 | Folr2 |
| F13a1 | 0 | 3.07759155 | 0.934 | 0.351 | 0 | TAM.Cd209 | F13a1 |
| Dnm1 | 0 | 5.31864795 | 0.592 | 0.036 | 0 | TAM.Cd209 | Dnm1 |
| C4a | 0 | 4.52420108 | 0.613 | 0.061 | 0 | TAM.Cd209 | C4a |
| Tmem8 | 0 | 3.63761655 | 0.713 | 0.169 | 0 | TAM.Cd209 | Tmem8 |
| Lrg1 | 0 | 3.79456907 | 0.619 | 0.08 | 0 | TAM.Cd209 | Lrg1 |
| Siglec11 | 0 | 3.18736566 | 0.763 | 0.23 | 0 | TAM.Cd209 | Siglec1 |
| Fcrls1 | 0 | 2.07575 | 0.871 | 0.345 | 0 | TAM.Cd209 | Fcrls |
| Fgfr1 | 0 | 3.98411511 | 0.601 | 0.077 | 0 | TAM.Cd209 | Fgfr1 |
| Phyhd11 | 0 | 3.05544246 | 0.721 | 0.2 | 0 | TAM.Cd209 | Phyhd1 |
| Cd209f | 0 | 7.05938713 | 0.53 | 0.015 | 0 | TAM.Cd209 | Cd209f |
| Stard8 | 0 | 3.25693897 | 0.71 | 0.198 | 0 | TAM.Cd209 | Stard8 |
| Cd163 | 0 | 5.09019119 | 0.57 | 0.06 | 0 | TAM.Cd209 | Cd163 |
| Lyve1 | 0 | 5.88283817 | 0.526 | 0.024 | 0 | TAM.Cd209 | Lyve1 |
| Slco2b11 | 0 | 3.43191771 | 0.59 | 0.101 | 0 | TAM.Cd209 | Slco2b1 |
| Npl1 | 0 | 2.5483224 | 0.715 | 0.233 | 0 | TAM.Cd209 | Npl |
| Chp2 | 0 | 4.94934467 | 0.516 | 0.038 | 0 | TAM.Cd209 | Chp2 |
| Serpinb6a | 0 | 2.71590297 | 0.763 | 0.291 | 0 | TAM.Cd209 | Serpinb6a |
| Scn1b1 | 0 | 2.83659296 | 0.665 | 0.204 | 0 | TAM.Cd209 | Scn1b |
| Wwp11 | 0 | 2.68906537 | 0.726 | 0.266 | 0 | TAM.Cd209 | Wwp1 |
| Fcgrt1 | 0 | 2.96963606 | 0.955 | 0.499 | 0 | TAM.Cd209 | Fcgrt |
| Pdgfc1 | 0 | 2.69047763 | 0.661 | 0.209 | 0 | TAM.Cd209 | Pdgfc |
| Maf2 | 0 | 2.48497335 | 0.98 | 0.532 | 0 | TAM.Cd209 | Maf |
| Aldh7a1 | 0 | 3.27970168 | 0.573 | 0.128 | 0 | TAM.Cd209 | Aldh7a1 |
| Ccl71 | 0 | 2.40863828 | 0.63 | 0.19 | 0 | TAM.Cd209 | Ccl7 |
| Slc9a91 | 0 | 2.47854505 | 0.708 | 0.27 | 0 | TAM.Cd209 | Slc9a9 |
| Itsn11 | 0 | 2.48144548 | 0.725 | 0.295 | 0 | TAM.Cd209 | Itsn1 |
| Rnase42 | 0 | 2.47248502 | 0.792 | 0.368 | 0 | TAM.Cd209 | Rnase4 |
| Cd209g | 0 | 7.40321231 | 0.432 | 0.009 | 0 | TAM.Cd209 | Cd209g |
| Ctla2b | 0 | 3.28724566 | 0.509 | 0.101 | 0 | TAM.Cd209 | Ctla2b |
| Cfh | 0 | 2.87931886 | 0.665 | 0.258 | 0 | TAM.Cd209 | Cfh |
| Pltp1 | 0 | 3.06046984 | 0.98 | 0.576 | 0 | TAM.Cd209 | Pltp |
| Fchsd21 | 0 | 2.37721152 | 0.723 | 0.319 | 0 | TAM.Cd209 | Fchsd2 |
| Kank2 | 0 | 3.75755123 | 0.467 | 0.069 | 0 | TAM.Cd209 | Kank2 |
| Igf11 | 0 | 2.30923231 | 0.561 | 0.164 | 0 | TAM.Cd209 | Igf1 |
| Timp21 | 0 | 2.63590105 | 0.916 | 0.523 | 0 | TAM.Cd209 | Timp2 |
| Clec10a | 0 | 3.36279348 | 0.578 | 0.187 | 0 | TAM.Cd209 | Clec10a |
| Wfdc17 | 0 | 3.13891998 | 0.851 | 0.466 | 0 | TAM.Cd209 | Wfdc17 |
| Stab11 | 0 | 2.24164531 | 0.849 | 0.464 | 0 | TAM.Cd209 | Stab1 |
| Napa51 | 0 | 2.18078693 | 0.699 | 0.315 | 0 | TAM.Cd209 | Napa5 |
| S1pr1 | 0 | 2.81662995 | 0.495 | 0.115 | 0 | TAM.Cd209 | S1pr1 |
| Cd300ld52 | 0 | 2.57501728 | 0.608 | 0.232 | 0 | TAM.Cd209 | Cd300ld5 |
| Gprc5c | 0 | 2.81747525 | 0.488 | 0.116 | 0 | TAM.Cd209 | Gprc5c |
| Cd37 | 0 | 2.0730888 | 0.656 | 0.294 | 0 | TAM.Cd209 | Cd37 |
| Hist1h2bc | 0 | 1.91459089 | 0.683 | 0.326 | 0 | TAM.Cd209 | Hist1h2bc |
| Tmem140 | 0 | 2.73245626 | 0.477 | 0.122 | 0 | TAM.Cd209 | Tmem140 |
| Ms4a72 | 0 | 1.1823999 | 0.812 | 0.458 | 0 | TAM.Cd209 | Ms4a7 |
| Alox5 | 0 | 2.41861119 | 0.483 | 0.13 | 0 | TAM.Cd209 | Alox5 |
| Rnf1501 | 0 | 2.21880078 | 0.547 | 0.199 | 0 | TAM.Cd209 | Rnf150 |
| Ednrb | 0 | 3.42422084 | 0.406 | 0.062 | 0 | TAM.Cd209 | Ednrb |
| Arhgef31 | 0 | 2.07707669 | 0.595 | 0.256 | 0 | TAM.Cd209 | Arhgef3 |
| C6 | 0 | 4.52088494 | 0.371 | 0.032 | 0 | TAM.Cd209 | C6 |
| Sgpp1 | 0 | 2.05564969 | 0.681 | 0.347 | 0 | TAM.Cd209 | Sgpp1 |
| Igfbp41 | 0 | 1.63139509 | 0.758 | 0.425 | 0 | TAM.Cd209 | Igfbp4 |
| Bin1 | 0 | 2.0179528 | 0.723 | 0.39 | 0 | TAM.Cd209 | Bin1 |
| Entpd3 | 0 | 5.27803249 | 0.348 | 0.018 | 0 | TAM.Cd209 | Entpd3 |
| Abca92 | 0 | 1.83782431 | 0.61 | 0.281 | 0 | TAM.Cd209 | Abca9 |
| Cela1 | 0 | 2.99960327 | 0.403 | 0.076 | 0 | TAM.Cd209 | Cela1 |
| Nrp11 | 0 | 2.00786657 | 0.628 | 0.302 | 0 | TAM.Cd209 | Nrp1 |
| Cacna1d | 0 | 1.9625047 | 0.558 | 0.233 | 0 | TAM.Cd209 | Cacna1d |
| Fam43a | 0 | 2.57391356 | 0.447 | 0.122 | 0 | TAM.Cd209 | Fam43a |
| Adgrg1 | 0 | 3.96014368 | 0.364 | 0.04 | 0 | TAM.Cd209 | Adgrg1 |
| Man1c11 | 0 | 1.97195999 | 0.681 | 0.36 | 0 | TAM.Cd209 | Man1c1 |
| Gpt2 | 0 | 2.92470271 | 0.417 | 0.1 | 0 | TAM.Cd209 | Gpt2 |
| Lyl11 | 0 | 2.08472376 | 0.501 | 0.185 | 0 | TAM.Cd209 | Lyl1 |
| Ang | 0 | 3.29878004 | 0.374 | 0.06 | 0 | TAM.Cd209 | Ang |
| mt-Nd6 | 0 | 2.22297759 | 0.921 | 0.608 | 0 | TAM.Cd209 | mt-Nd6 |
| Etv1 | 0 | 3.09295955 | 0.387 | 0.074 | 0 | TAM.Cd209 | Etv1 |
| Zdhhc141 | 0 | 1.82228899 | 0.594 | 0.284 | 0 | TAM.Cd209 | Zdhhc14 |
| Nupr11 | 0 | 1.7334749 | 0.67 | 0.361 | 0 | TAM.Cd209 | Nupr1 |
| Pf41 | 0 | 0.84922697 | 0.837 | 0.529 | 0 | TAM.Cd209 | Pf4 |
| Gja1 | 0 | 3.23471725 | 0.374 | 0.067 | 0 | TAM.Cd209 | Gja1 |
| Fxyd2 | 0 | 3.24351202 | 0.371 | 0.065 | 0 | TAM.Cd209 | Fxyd2 |
| Txnip2 | 0 | 1.45855894 | 0.806 | 0.504 | 0 | TAM.Cd209 | Txnip |
| Adcy31 | 0 | 2.21922758 | 0.401 | 0.1 | 0 | TAM.Cd209 | Adcy3 |
| Plekhg5 | 0 | 4.33886862 | 0.348 | 0.049 | 0 | TAM.Cd209 | Plekhg5 |
| Hpse1 | 0 | 1.97095477 | 0.541 | 0.244 | 0 | TAM.Cd209 | Hpse |
| Xdh | 0 | 1.66003868 | 0.773 | 0.476 | 0 | TAM.Cd209 | Xdh |
| Sesn11 | 0 | 2.03092682 | 0.464 | 0.168 | 0 | TAM.Cd209 | Sesn1 |
| Epb41l1 | 0 | 2.27790819 | 0.434 | 0.138 | 0 | TAM.Cd209 | Epb41l1 |
| Reps2 | 0 | 3.89036945 | 0.33 | 0.035 | 0 | TAM.Cd209 | Reps2 |
| Rab11fip51 | 0 | 1.9363998 | 0.862 | 0.568 | 0 | TAM.Cd209 | Rab11fip5 |
| Aoah1 | 0 | 1.70835244 | 0.704 | 0.41 | 0 | TAM.Cd209 | Aoah |
| Ccl122 | 0 | 1.43173232 | 0.5 | 0.207 | 0 | TAM.Cd209 | Ccl12 |
| A4galt | 0 | 5.14980441 | 0.309 | 0.017 | 0 | TAM.Cd209 | A4galt |
| Saa3 | 0 | 4.35929202 | 0.358 | 0.068 | 0 | TAM.Cd209 | Saa3 |
| C1qtnf1 | 0 | 4.83785971 | 0.308 | 0.019 | 0 | TAM.Cd209 | C1qtnf1 |
| Oas21 | 0 | 2.03868064 | 0.443 | 0.16 | 0 | TAM.Cd209 | Oas2 |
| Lrp61 | 0 | 1.88296847 | 0.825 | 0.543 | 0 | TAM.Cd209 | Lrp6 |
| Kif3a | 0 | 2.79865623 | 0.373 | 0.092 | 0 | TAM.Cd209 | Kif3a |
| Ophn11 | 0 | 2.14942303 | 0.42 | 0.142 | 0 | TAM.Cd209 | Ophn1 |
| Prxl2b1 | 0 | 1.65533393 | 0.599 | 0.326 | 0 | TAM.Cd209 | Prxl2b |
| C1qa1 | 0 | 1.18601468 | 0.982 | 0.71 | 0 | TAM.Cd209 | C1qa |
| 8-Sep | 0 | 1.56881666 | 0.646 | 0.374 | 0 | TAM.Cd209 | 8-Sep |
| Ccl21 | 0 | 1.65579973 | 0.5 | 0.228 | 0 | TAM.Cd209 | Ccl2 |
| Snx291 | 0 | 1.65592923 | 0.724 | 0.452 | 0 | TAM.Cd209 | Snx29 |
| Tns11 | 0 | 2.21037392 | 0.435 | 0.164 | 0 | TAM.Cd209 | Tns1 |
| Padi4 | 0 | 3.91935157 | 0.308 | 0.038 | 0 | TAM.Cd209 | Padi4 |
| C1qc1 | 0 | 1.37365643 | 0.941 | 0.671 | 0 | TAM.Cd209 | C1qc |
| Tspan3 | 0 | 1.61003446 | 0.713 | 0.444 | 0 | TAM.Cd209 | Tspan3 |
| Gpx31 | 0 | 2.03787765 | 0.467 | 0.199 | 0 | TAM.Cd209 | Gpx3 |
| Agpat31 | 0 | 1.73404928 | 0.513 | 0.247 | 0 | TAM.Cd209 | Agpat3 |
| Cfb2 | 0 | 1.22158928 | 0.637 | 0.372 | 0 | TAM.Cd209 | Cfb |
| Cd300ld32 | 0 | 1.91946251 | 0.435 | 0.172 | 0 | TAM.Cd209 | Cd300ld3 |
| Gpr146 | 0 | 1.70708025 | 0.509 | 0.248 | 0 | TAM.Cd209 | Gpr146 |
| Fez2 | 0 | 1.65868313 | 0.573 | 0.312 | 0 | TAM.Cd209 | Fez2 |
| Plod1 | 0 | 1.5629766 | 0.62 | 0.36 | 0 | TAM.Cd209 | Plod1 |
| Hpgds1 | 0 | 1.59060299 | 0.48 | 0.22 | 0 | TAM.Cd209 | Hpgds |
| Cp | 0 | 3.38008713 | 0.299 | 0.04 | 0 | TAM.Cd209 | Cp |
| Cd209d | 0 | 5.62740115 | 0.272 | 0.013 | 0 | TAM.Cd209 | Cd209d |
| Ddx60 | 0 | 3.1417209 | 0.316 | 0.058 | 0 | TAM.Cd209 | Ddx60 |
| Selenop1 | 0 | 2.11784908 | 0.966 | 0.709 | 0 | TAM.Cd209 | Selenop |
| Mgst1 | 0 | 1.52192614 | 0.54 | 0.284 | 0 | TAM.Cd209 | Mgst1 |
| Sorbs3 | 0 | 2.78610178 | 0.327 | 0.071 | 0 | TAM.Cd209 | Sorbs3 |
| Appl2 | 0 | 1.55337466 | 0.59 | 0.334 | 0 | TAM.Cd209 | Appl2 |
| Pros1 | 0 | 2.01381464 | 0.466 | 0.213 | 0 | TAM.Cd209 | Pros1 |
| Tcf4 | 0 | 1.53143611 | 0.55 | 0.298 | 0 | TAM.Cd209 | Tcf4 |
| Ccl6 | 0 | 1.62782712 | 0.835 | 0.583 | 0 | TAM.Cd209 | Ccl6 |
| Slc28a2 | 0 | 2.64709274 | 0.318 | 0.067 | 0 | TAM.Cd209 | Slc28a2 |
| Eps15 | 0 | 1.80640036 | 0.714 | 0.463 | 0 | TAM.Cd209 | Eps15 |
| Nfatc21 | 0 | 1.42317564 | 0.551 | 0.301 | 0 | TAM.Cd209 | Nfatc2 |
| Mrc11 | 0 | 1.83670648 | 0.972 | 0.722 | 0 | TAM.Cd209 | Mrc1 |
| C1qb1 | 0 | 1.1950375 | 0.995 | 0.747 | 0 | TAM.Cd209 | C1qb |
| Hyal1 | 0 | 2.40674898 | 0.325 | 0.078 | 0 | TAM.Cd209 | Hyal1 |
| Adgre12 | 0 | 1.29214256 | 0.79 | 0.543 | 0 | TAM.Cd209 | Adgre1 |
| Scamp51 | 0 | 1.76951203 | 0.467 | 0.222 | 0 | TAM.Cd209 | Scamp5 |
| Man1a | 0 | 1.5288458 | 0.828 | 0.588 | 0 | TAM.Cd209 | Man1a |
| Rab3il12 | 0 | 1.19594387 | 0.894 | 0.654 | 0 | TAM.Cd209 | Rab3il1 |
| Dhrs31 | 0 | 1.57924828 | 0.87 | 0.633 | 0 | TAM.Cd209 | Dhrs3 |
| Mfsd6 | 0 | 2.15690173 | 0.317 | 0.081 | 0 | TAM.Cd209 | Mfsd6 |
| Mpi | 0 | 2.33561812 | 0.37 | 0.136 | 0 | TAM.Cd209 | Mpi |
| Iqgap21 | 0 | 1.33121634 | 0.504 | 0.27 | 0 | TAM.Cd209 | Iqgap2 |
| Inpp4a | 0 | 1.52985176 | 0.534 | 0.301 | 0 | TAM.Cd209 | Inpp4a |
| Sh3gl11 | 0 | 1.46367909 | 0.65 | 0.418 | 0 | TAM.Cd209 | Sh3gl1 |
| Pcyox11 | 0 | 1.46164439 | 0.594 | 0.362 | 0 | TAM.Cd209 | Pcyox1 |
| Tsku | 0 | 2.16270918 | 0.355 | 0.123 | 0 | TAM.Cd209 | Tsku |
| Ifi27l2a2 | 0 | 1.48403829 | 0.609 | 0.377 | 0 | TAM.Cd209 | Ifi27l2a |
| Hfe1 | 0 | 1.34799437 | 0.708 | 0.476 | 0 | TAM.Cd209 | Hfe |
| Tmem86a1 | 0 | 1.41606967 | 0.696 | 0.468 | 0 | TAM.Cd209 | Tmem86a |
| Ang | 0 | 2.22959544 | 0.331 | 0.104 | 0 | TAM.Cd209 | Ang |
| Prune21 | 0 | 1.47565174 | 0.466 | 0.241 | 0 | TAM.Cd209 | Prune2 |
| Xylt2 | 0 | 1.75992622 | 0.432 | 0.208 | 0 | TAM.Cd209 | Xylt2 |
| Cebpd | 0 | 1.25149552 | 0.744 | 0.52 | 0 | TAM.Cd209 | Cebpd |
| Zcchc242 | 0 | 1.48288432 | 0.537 | 0.314 | 0 | TAM.Cd209 | Zcchc24 |
| Osbpl11 | 0 | 1.40884639 | 0.503 | 0.282 | 0 | TAM.Cd209 | Osbpl11 |
| Pigz | 0 | 2.45124773 | 0.292 | 0.072 | 0 | TAM.Cd209 | Pigz |
| Irf72 | 0 | 0.85181102 | 0.458 | 0.24 | 0 | TAM.Cd209 | Irf7 |
| Mcoln2 | 0 | 2.2382871 | 0.313 | 0.095 | 0 | TAM.Cd209 | Mcoln2 |
| Snx61 | 0 | 1.44606218 | 0.695 | 0.477 | 0 | TAM.Cd209 | Snx6 |
| Ap2a2 | 0 | 1.56178013 | 0.798 | 0.581 | 0 | TAM.Cd209 | Ap2a2 |
| Snx81 | 0 | 1.36060243 | 0.688 | 0.471 | 0 | TAM.Cd209 | Snx8 |
| Rgl12 | 0 | 1.07457478 | 0.598 | 0.383 | 0 | TAM.Cd209 | Rgl1 |
| Lmo21 | 0 | 1.38487022 | 0.717 | 0.502 | 0 | TAM.Cd209 | Lmo2 |
| Trem21 | 0 | 1.33415523 | 0.624 | 0.41 | 0 | TAM.Cd209 | Trem2 |
| Tpcn1 | 0 | 1.24375179 | 0.518 | 0.306 | 0 | TAM.Cd209 | Tpcn1 |
| Rnf141 | 0 | 1.56577115 | 0.42 | 0.208 | 0 | TAM.Cd209 | Rnf141 |
| Naa80 | 0 | 1.59433443 | 0.407 | 0.195 | 0 | TAM.Cd209 | Naa80 |
| Trpv4 | 0 | 2.30979549 | 0.322 | 0.112 | 0 | TAM.Cd209 | Trpv4 |
| Eef2k1 | 0 | 1.77081775 | 0.361 | 0.151 | 0 | TAM.Cd209 | Eef2k |
| Tmem371 | 0 | 1.06694148 | 0.605 | 0.396 | 0 | TAM.Cd209 | Tmem37 |
| Ypel31 | 0 | 1.24868166 | 0.699 | 0.491 | 0 | TAM.Cd209 | Ypel3 |
| Gpr341 | 0 | 2.14584121 | 0.306 | 0.099 | 0 | TAM.Cd209 | Gpr34 |
| B3galnt1 | 0 | 2.67032277 | 0.265 | 0.06 | 0 | TAM.Cd209 | B3galnt1 |
| Mtss1 | 0 | 1.3247673 | 0.743 | 0.538 | 0 | TAM.Cd209 | Mtss1 |
| Dclre1c2 | 0 | 1.37738892 | 0.504 | 0.3 | 0 | TAM.Cd209 | Dclre1c |
| Ap1s22 | 0 | 1.45037053 | 0.491 | 0.287 | 0 | TAM.Cd209 | Ap1s2 |
| Abcd4 | 0 | 2.23844123 | 0.306 | 0.105 | 0 | TAM.Cd209 | Abcd4 |
| Tbc1d142 | 0 | 1.24721139 | 0.662 | 0.462 | 0 | TAM.Cd209 | Tbc1d14 |
| Nacc2 | 0 | 1.7821958 | 0.34 | 0.143 | 0 | TAM.Cd209 | Nacc2 |
| Slc12a71 | 0 | 1.31235788 | 0.51 | 0.313 | 0 | TAM.Cd209 | Slc12a7 |
| Pde2a | 0 | 1.64191502 | 0.387 | 0.192 | 0 | TAM.Cd209 | Pde2a |
| Dennd4c | 0 | 1.52883175 | 0.433 | 0.239 | 0 | TAM.Cd209 | Dennd4c |
| Tgfbr21 | 0 | 1.33868713 | 0.778 | 0.585 | 0 | TAM.Cd209 | Tgfbr2 |
| Hist3h2a1 | 0 | 1.55631681 | 0.367 | 0.174 | 0 | TAM.Cd209 | Hist3h2a |
| Tmem106a2 | 0 | 1.29649 | 0.768 | 0.577 | 0 | TAM.Cd209 | Tmem106a |
| Stom1 | 0 | 1.17207717 | 0.581 | 0.39 | 0 | TAM.Cd209 | Stom |
| Rnasel1 | 0 | 1.29738348 | 0.515 | 0.325 | 0 | TAM.Cd209 | Rnasel |
| Ltc4s | 0 | 1.932104 | 0.425 | 0.235 | 0 | TAM.Cd209 | Ltc4s |
| Hmox11 | 0 | 1.14663307 | 0.86 | 0.67 | 0 | TAM.Cd209 | Hmox1 |
| P2ry121 | 0 | 1.31879404 | 0.418 | 0.229 | 0 | TAM.Cd209 | P2ry12 |
| Serpinb8 | 0 | 1.35984605 | 0.335 | 0.151 | 0 | TAM.Cd209 | Serpinb8 |
| Cemip2 | 0 | 1.41815288 | 0.432 | 0.248 | 0 | TAM.Cd209 | Cemip2 |
| Camk11 | 0 | 1.32163478 | 0.699 | 0.515 | 0 | TAM.Cd209 | Camk1 |
| St6galnac41 | 0 | 1.16886098 | 0.592 | 0.41 | 0 | TAM.Cd209 | St6galnac4 |
| Dse1 | 0 | 1.13112178 | 0.521 | 0.343 | 0 | TAM.Cd209 | Dse |
| Slc13a31 | 0 | 1.84861385 | 0.33 | 0.153 | 0 | TAM.Cd209 | Slc13a3 |
| Sec14l11 | 0 | 1.06298386 | 0.661 | 0.485 | 0 | TAM.Cd209 | Sec14l1 |
| Ninj1 | 0 | 1.67121977 | 0.867 | 0.691 | 0 | TAM.Cd209 | Ninj1 |
| Glb11 | 0 | 1.21016499 | 0.5 | 0.324 | 0 | TAM.Cd209 | Glb1 |
| Eps81 | 0 | 1.02697464 | 0.618 | 0.443 | 0 | TAM.Cd209 | Eps8 |
| Hspa1a2 | 0 | 0.54586358 | 0.576 | 0.401 | 0 | TAM.Cd209 | Hspa1a |
| Plxnb21 | 0 | 1.08309937 | 0.753 | 0.58 | 0 | TAM.Cd209 | Plxnb2 |
| H2-T24 | 0 | 1.70493962 | 0.261 | 0.088 | 0 | TAM.Cd209 | H2-T24 |
| Tnfsf141 | 0 | 1.02903737 | 0.379 | 0.206 | 0 | TAM.Cd209 | Tnfsf14 |
| Ap1b11 | 0 | 1.07243178 | 0.789 | 0.617 | 0 | TAM.Cd209 | Ap1b1 |
| Glul2 | 0 | 1.21153175 | 0.739 | 0.568 | 0 | TAM.Cd209 | Glul |
| Il18bp1 | 0 | 1.59181682 | 0.319 | 0.148 | 0 | TAM.Cd209 | Il18bp |
| Pde7b | 0 | 1.90425182 | 0.269 | 0.098 | 0 | TAM.Cd209 | Pde7b |
| Rassf21 | 0 | 1.22096253 | 0.7 | 0.529 | 0 | TAM.Cd209 | Rassf2 |
| Clec2d1 | 0 | 1.2459633 | 0.836 | 0.665 | 0 | TAM.Cd209 | Clec2d |
| Pid12 | 0 | 1.0063813 | 0.852 | 0.683 | 0 | TAM.Cd209 | Pid1 |
| Oasl22 | 0 | 1.0725784 | 0.413 | 0.245 | 0 | TAM.Cd209 | Oasl2 |
| Rab31 | 0 | 1.17601875 | 0.741 | 0.576 | 0 | TAM.Cd209 | Rab31 |
| Neurl2 | 0 | 2.11091901 | 0.264 | 0.099 | 0 | TAM.Cd209 | Neurl2 |
| Emp1 | 0 | 0.96513102 | 0.707 | 0.543 | 0 | TAM.Cd209 | Emp1 |
| Myo5a | 0 | 1.21362926 | 0.762 | 0.6 | 0 | TAM.Cd209 | Myo5a |
| Selenbp1 | 0 | 1.92481883 | 0.251 | 0.09 | 0 | TAM.Cd209 | Selenbp1 |
| Wdfy31 | 0 | 1.17678464 | 0.809 | 0.649 | 0 | TAM.Cd209 | Wdfy3 |
| Tmem1041 | 0 | 1.03449154 | 0.598 | 0.439 | 0 | TAM.Cd209 | Tmem104 |
| Rtp41 | 0 | 1.21042369 | 0.297 | 0.138 | 0 | TAM.Cd209 | Rtp4 |
| Plaat31 | 0 | 1.35023621 | 0.33 | 0.172 | 0 | TAM.Cd209 | Plaat3 |
| C3ar12 | 0 | 0.99359089 | 0.828 | 0.67 | 0 | TAM.Cd209 | C3ar1 |
| Hsbp1 | 0 | 1.57813989 | 0.807 | 0.65 | 0 | TAM.Cd209 | Hsbp1 |
| Ly6a1 | 0 | 1.55856298 | 0.288 | 0.133 | 0 | TAM.Cd209 | Ly6a |
| Pfkfb3 | 0 | 1.06430127 | 0.523 | 0.37 | 0 | TAM.Cd209 | Pfkfb3 |
| Ccl91 | 0 | 0.78425061 | 0.81 | 0.657 | 0 | TAM.Cd209 | Ccl9 |
| Abca11 | 0 | 0.8776495 | 0.864 | 0.714 | 0 | TAM.Cd209 | Abca1 |
| Bmp2k2 | 0 | 1.18350282 | 0.75 | 0.602 | 0 | TAM.Cd209 | Bmp2k |
| Ms4a4a3 | 0 | 0.84205932 | 0.84 | 0.694 | 0 | TAM.Cd209 | Ms4a4a |
| Twf11 | 0 | 1.0693621 | 0.567 | 0.423 | 0 | TAM.Cd209 | Twf1 |
| Atp13a21 | 0 | 1.10780844 | 0.81 | 0.668 | 0 | TAM.Cd209 | Atp13a2 |
| Scamp21 | 0 | 1.08349555 | 0.617 | 0.478 | 0 | TAM.Cd209 | Scamp2 |
| Sdc3 | 0 | 1.27369974 | 0.893 | 0.755 | 0 | TAM.Cd209 | Sdc3 |
| Mgat1 | 0 | 1.02906659 | 0.61 | 0.473 | 0 | TAM.Cd209 | Mgat1 |
| Ms4a6b3 | 0 | 0.67562257 | 0.583 | 0.447 | 0 | TAM.Cd209 | Ms4a6b |
| Ctsd1 | 0 | 1.28569989 | 0.869 | 0.735 | 0 | TAM.Cd209 | Ctsd |
| Klhl91 | 0 | 0.96633334 | 0.645 | 0.512 | 0 | TAM.Cd209 | Klhl9 |
| Trf1 | 0 | 1.39610548 | 0.924 | 0.791 | 0 | TAM.Cd209 | Trf |
| Ecm1 | 0 | 1.05559552 | 0.83 | 0.699 | 0 | TAM.Cd209 | Ecm1 |
| Nisch | 0 | 1.11163631 | 0.819 | 0.688 | 0 | TAM.Cd209 | Nisch |
| Pla2g151 | 0 | 1.06870883 | 0.772 | 0.642 | 0 | TAM.Cd209 | Pla2g15 |
| Marcks1 | 0 | 1.08193471 | 0.935 | 0.806 | 0 | TAM.Cd209 | Marcks |
| Abhd121 | 0 | 1.05675908 | 0.748 | 0.621 | 0 | TAM.Cd209 | Abhd12 |
| Atp6v0a11 | 0 | 0.94769526 | 0.629 | 0.503 | 0 | TAM.Cd209 | Atp6v0a1 |
| Ctsl | 0 | 1.13733323 | 0.87 | 0.745 | 0 | TAM.Cd209 | Ctsl |
| Osbpl9 | 0 | 1.20577071 | 0.677 | 0.553 | 0 | TAM.Cd209 | Osbpl9 |
| Comt1 | 0 | 0.98061951 | 0.653 | 0.531 | 0 | TAM.Cd209 | Comt |
| Lgmn2 | 0 | 1.3154527 | 0.949 | 0.831 | 0 | TAM.Cd209 | Lgmn |
| Tbxas1 | 0 | 0.95865 | 0.818 | 0.705 | 0 | TAM.Cd209 | Tbxas1 |
| Slc43a21 | 0 | 0.84889867 | 0.718 | 0.609 | 0 | TAM.Cd209 | Slc43a2 |
| P2rx41 | 0 | 0.95122902 | 0.733 | 0.626 | 0 | TAM.Cd209 | P2rx4 |
| Aldh22 | 0 | 1.08710609 | 0.815 | 0.716 | 0 | TAM.Cd209 | Aldh2 |
| Adam15 | 0 | 0.88305414 | 0.755 | 0.657 | 0 | TAM.Cd209 | Adam15 |
| Ehd41 | 0 | 0.93287812 | 0.736 | 0.639 | 0 | TAM.Cd209 | Ehd4 |
| Bst21 | 0 | 1.12324916 | 0.728 | 0.632 | 0 | TAM.Cd209 | Bst2 |
| Lgals3bp2 | 0 | 0.96082785 | 0.836 | 0.741 | 0 | TAM.Cd209 | Lgals3bp |
| Apoe1 | 0 | 1.68648726 | 0.999 | 0.907 | 0 | TAM.Cd209 | Apoe |
| Dab21 | 0 | 0.91357049 | 0.937 | 0.846 | 0 | TAM.Cd209 | Dab2 |
| Necap2 | 0 | 0.98407229 | 0.755 | 0.664 | 0 | TAM.Cd209 | Necap2 |
| Tpp11 | 0 | 0.87920741 | 0.751 | 0.661 | 0 | TAM.Cd209 | Tpp1 |
| Mafb1 | 0 | 0.49057087 | 0.963 | 0.874 | 0 | TAM.Cd209 | Mafb |
| Snx22 | 0 | 1.07615748 | 0.807 | 0.72 | 0 | TAM.Cd209 | Snx2 |
| Sulf21 | 0 | 0.85560176 | 0.698 | 0.612 | 0 | TAM.Cd209 | Sulf2 |
| Ncf1 | 0 | 0.82572878 | 0.838 | 0.757 | 0 | TAM.Cd209 | Ncf1 |
| Scpep11 | 0 | 0.88951411 | 0.801 | 0.729 | 0 | TAM.Cd209 | Scpep1 |
| Ly6e2 | 0 | 0.90537353 | 0.891 | 0.819 | 0 | TAM.Cd209 | Ly6e |
| Scarb21 | 0 | 0.83448218 | 0.634 | 0.562 | 0 | TAM.Cd209 | Scarb2 |
| Serinc32 | 0 | 1.22892937 | 0.931 | 0.86 | 0 | TAM.Cd209 | Serinc3 |
| Dbnl | 0 | 0.78833338 | 0.711 | 0.641 | 0 | TAM.Cd209 | Dbnl |
| Aplp21 | 0 | 0.86852401 | 0.745 | 0.676 | 0 | TAM.Cd209 | Aplp2 |
| Gns1 | 0 | 0.84807272 | 0.789 | 0.724 | 0 | TAM.Cd209 | Gns |
| B2m1 | 0 | 0.30043505 | 0.841 | 0.903 | 0 | TAM.Cd209 | B2m |
| Adam9 | 0 | 1.04950507 | 0.799 | 0.737 | 0 | TAM.Cd209 | Adam9 |
| Fndc3a1 | 0 | 0.74218157 | 0.608 | 0.547 | 0 | TAM.Cd209 | Fndc3a |
| Il4ra | 0 | 1.48603494 | 0.935 | 0.876 | 0 | TAM.Cd209 | Il4ra |
| Tcn21 | 0 | 1.17536706 | 0.861 | 0.812 | 0 | TAM.Cd209 | Tcn2 |
| Ndfip1 | 0 | 0.91755082 | 0.754 | 0.706 | 0 | TAM.Cd209 | Ndfip1 |
| H2-D1 | 0 | 0.78056124 | 0.717 | 0.673 | 0 | TAM.Cd209 | H2-D1 |
| Grn1 | 0 | 1.41553621 | 0.984 | 0.943 | 0 | TAM.Cd209 | Grn |
| Itm2b2 | 0 | 1.77024668 | 0.991 | 0.951 | 0 | TAM.Cd209 | Itm2b |
| Jund | 0 | 0.54160744 | 0.9 | 0.937 | 0 | TAM.Cd209 | Jund |
| Cd68 | 0 | 0.47488527 | 0.853 | 0.889 | 0 | TAM.Cd209 | Cd68 |
| Cfp | 0 | 0.67277622 | 0.932 | 0.898 | 0 | TAM.Cd209 | Cfp |
| Lamp11 | 0 | 1.03026452 | 0.839 | 0.805 | 0 | TAM.Cd209 | Lamp1 |
| Fcgr31 | 0 | 0.31835001 | 0.872 | 0.905 | 0 | TAM.Cd209 | Fcgr3 |
| Jun | 0 | 0.97112946 | 0.886 | 0.853 | 0 | TAM.Cd209 | Jun |
| Qk1 | 0 | 0.90483409 | 0.86 | 0.83 | 0 | TAM.Cd209 | Qk |
| Ms4a6d1 | 0 | 0.4743135 | 0.774 | 0.801 | 0 | TAM.Cd209 | Ms4a6d |
| Ctsb1 | 0 | 1.25281761 | 0.989 | 0.963 | 0 | TAM.Cd209 | Ctsb |
| Egr11 | 0 | 0.70911048 | 0.694 | 0.673 | 0 | TAM.Cd209 | Egr1 |
| Clta1 | 0 | 0.55805996 | 0.951 | 0.963 | 0 | TAM.Cd209 | Clta |
| Ap2m1 | 0 | 0.77789787 | 0.766 | 0.757 | 0 | TAM.Cd209 | Ap2m1 |
| App | 0 | 0.91934726 | 0.962 | 0.955 | 0 | TAM.Cd209 | App |
| Lamp21 | 0 | 0.65613568 | 0.811 | 0.804 | 0 | TAM.Cd209 | Lamp2 |
| Itm2c1 | 0 | 0.71352892 | 0.815 | 0.811 | 0 | TAM.Cd209 | Itm2c |
| Hexa1 | 0 | 0.70323302 | 0.82 | 0.817 | 0 | TAM.Cd209 | Hexa |
| H2-K11 | 0 | 0.89032446 | 0.954 | 0.952 | 0 | TAM.Cd209 | H2-K1 |
| Fcgr2b | 0 | 0.49212786 | 0.958 | 0.959 | 0 | TAM.Cd209 | Fcgr2b |
| Ifitm22 | 0 | 0.53365576 | 0.91 | 0.911 | 0 | TAM.Cd209 | Ifitm2 |
| Plk2 | 5.80E-308 | 0.81587949 | 0.544 | 0.396 | 1.12E-303 | TAM.Cd209 | Plk2 |
| Ctsh1 | 1.10E-307 | 0.36216695 | 0.862 | 0.892 | 2.12E-303 | TAM.Cd209 | Ctsh |
| Smpdl3a1 | 1.75E-307 | 0.67047102 | 0.521 | 0.425 | 3.38E-303 | TAM.Cd209 | Smpdl3a |
| Mt2 | 8.51E-307 | 0.78380955 | 0.539 | 0.318 | 1.65E-302 | TAM.Cd209 | Mt2 |
| Slc16a61 | 1.21E-305 | 1.36387272 | 0.499 | 0.312 | 2.33E-301 | TAM.Cd209 | Slc16a6 |
| Adap22 | 1.63E-304 | 0.95038224 | 0.726 | 0.572 | 3.14E-300 | TAM.Cd209 | Adap2 |
| Pink11 | 1.60E-302 | 1.05414297 | 0.528 | 0.363 | 3.08E-298 | TAM.Cd209 | Pink1 |
| Rnf1302 | 7.44E-301 | 0.6465722 | 0.737 | 0.71 | 1.44E-296 | TAM.Cd209 | Rnf130 |
| Sirt21 | 2.36E-300 | 0.81239169 | 0.671 | 0.587 | 4.57E-296 | TAM.Cd209 | Sirt2 |
| Susd31 | 5.16E-300 | 1.44634582 | 0.395 | 0.196 | 9.97E-296 | TAM.Cd209 | Susd3 |
| Fkbp151 | 8.59E-299 | 0.96451142 | 0.608 | 0.482 | 1.66E-294 | TAM.Cd209 | Fkbp15 |
| Gda1 | 1.61E-298 | 0.63474473 | 0.801 | 0.742 | 3.11E-294 | TAM.Cd209 | Gda |
| Gnpda11 | 2.75E-298 | 1.44493077 | 0.443 | 0.275 | 5.32E-294 | TAM.Cd209 | Gnpda1 |
| Nfix | 7.56E-297 | 1.37906764 | 0.424 | 0.228 | 1.46E-292 | TAM.Cd209 | Nfix |
| Lifr1 | 1.10E-294 | 1.57591186 | 0.4 | 0.235 | 2.12E-290 | TAM.Cd209 | Lifr |
| Tnfsf121 | 2.74E-294 | 1.02598738 | 0.615 | 0.448 | 5.30E-290 | TAM.Cd209 | Tnfsf12 |
| Mocs21 | 4.38E-293 | 1.24962446 | 0.477 | 0.312 | 8.47E-289 | TAM.Cd209 | Mocs2 |
| Rxra1 | 8.28E-292 | 1.09169795 | 0.587 | 0.419 | 1.60E-287 | TAM.Cd209 | Rxra |
| Fam234a | 1.86E-287 | 1.22796116 | 0.55 | 0.369 | 3.60E-283 | TAM.Cd209 | Fam234a |
| Snx27 | 1.69E-285 | 1.3107673 | 0.437 | 0.264 | 3.26E-281 | TAM.Cd209 | Snx27 |
| Ifi471 | 1.09E-282 | 0.92463609 | 0.327 | 0.193 | 2.10E-278 | TAM.Cd209 | Ifi47 |
| Rin22 | 2.78E-281 | 1.05176388 | 0.523 | 0.365 | 5.37E-277 | TAM.Cd209 | Rin2 |
| Zfyve21 | 3.57E-279 | 1.89114167 | 0.266 | 0.103 | 6.90E-275 | TAM.Cd209 | Zfyve21 |
| Smim20 | 9.90E-277 | 1.16537271 | 0.435 | 0.272 | 1.92E-272 | TAM.Cd209 | Smim20 |
| Prkacb1 | 2.44E-276 | 1.07935529 | 0.52 | 0.364 | 4.71E-272 | TAM.Cd209 | Prkacb |
| Ly9 | 1.09E-273 | 1.29751056 | 0.35 | 0.165 | 2.10E-269 | TAM.Cd209 | Ly9 |
| P4ha1 | 1.34E-272 | 0.70499923 | 0.623 | 0.554 | 2.59E-268 | TAM.Cd209 | P4ha1 |
| Snx91 | 3.86E-271 | 0.94808669 | 0.656 | 0.532 | 7.45E-267 | TAM.Cd209 | Snx9 |
| Ptpn182 | 1.08E-267 | 0.91387782 | 0.654 | 0.527 | 2.09E-263 | TAM.Cd209 | Ptpn18 |
| Kif13b1 | 1.67E-267 | 1.26280562 | 0.484 | 0.293 | 3.23E-263 | TAM.Cd209 | Kif13b |
| Frmd4b1 | 1.48E-265 | 1.20021581 | 0.554 | 0.346 | 2.87E-261 | TAM.Cd209 | Frmd4b |
| Gna121 | 4.64E-265 | 0.86606468 | 0.712 | 0.564 | 8.98E-261 | TAM.Cd209 | Gna12 |
| Pon31 | 1.13E-264 | 1.36698202 | 0.38 | 0.208 | 2.19E-260 | TAM.Cd209 | Pon3 |
| Agtrap1 | 1.55E-263 | 1.4938694 | 0.374 | 0.199 | 2.99E-259 | TAM.Cd209 | Agtrap |
| Ctsf | 1.74E-259 | 1.77159871 | 0.259 | 0.094 | 3.36E-255 | TAM.Cd209 | Ctsf |
| Pon21 | 5.49E-257 | 0.92021356 | 0.534 | 0.403 | 1.06E-252 | TAM.Cd209 | Pon2 |
| Zfp652 | 2.37E-255 | 1.06452436 | 0.505 | 0.356 | 4.59E-251 | TAM.Cd209 | Zfp652 |
| Arhgap12 | 2.29E-253 | 1.41926854 | 0.347 | 0.173 | 4.43E-249 | TAM.Cd209 | Arhgap12 |
| Anxa61 | 2.92E-252 | 0.87066767 | 0.601 | 0.506 | 5.64E-248 | TAM.Cd209 | Anxa6 |
| Npc21 | 1.79E-251 | 0.35355163 | 0.924 | 0.95 | 3.46E-247 | TAM.Cd209 | Npc2 |
| Rap2b2 | 3.04E-251 | 0.99791189 | 0.633 | 0.484 | 5.87E-247 | TAM.Cd209 | Rap2b |
| Gadd45g | 1.88E-248 | 0.55981593 | 0.446 | 0.393 | 3.63E-244 | TAM.Cd209 | Gadd45g |
| Dnajc132 | 2.05E-248 | 1.11093031 | 0.61 | 0.465 | 3.96E-244 | TAM.Cd209 | Dnajc13 |
| Ubtd1 | 1.75E-247 | 0.99015103 | 0.565 | 0.432 | 3.38E-243 | TAM.Cd209 | Ubtd1 |
| Galc1 | 3.63E-243 | 1.20363596 | 0.474 | 0.29 | 7.03E-239 | TAM.Cd209 | Galc |
| Gcnt1 | 3.96E-241 | 1.04186716 | 0.328 | 0.193 | 7.65E-237 | TAM.Cd209 | Gcnt1 |
| Alox5ap1 | 9.05E-241 | 0.27897432 | 0.802 | 0.768 | 1.75E-236 | TAM.Cd209 | Alox5ap |
| Nfxl1 | 1.17E-240 | 1.42117805 | 0.427 | 0.248 | 2.27E-236 | TAM.Cd209 | Nfxl1 |
| Cd811 | 1.06E-238 | 0.66577947 | 0.749 | 0.498 | 2.05E-234 | TAM.Cd209 | Cd81 |
| Sbf21 | 5.92E-237 | 1.45103637 | 0.433 | 0.241 | 1.15E-232 | TAM.Cd209 | Sbf2 |
| Vegfb1 | 9.67E-234 | 1.42309755 | 0.389 | 0.2 | 1.87E-229 | TAM.Cd209 | Vegfb |
| P2ry61 | 2.73E-233 | 0.64787504 | 0.727 | 0.643 | 5.28E-229 | TAM.Cd209 | P2ry6 |
| Rnf1451 | 4.81E-232 | 1.14458562 | 0.48 | 0.326 | 9.31E-228 | TAM.Cd209 | Rnf145 |
| Cd3021 | 1.31E-228 | 1.166225 | 0.43 | 0.286 | 2.53E-224 | TAM.Cd209 | Cd302 |
| Mef2a1 | 1.34E-227 | 0.70752187 | 0.672 | 0.601 | 2.58E-223 | TAM.Cd209 | Mef2a |
| C5ar11 | 4.50E-227 | 0.58869045 | 0.861 | 0.779 | 8.71E-223 | TAM.Cd209 | C5ar1 |
| Egln1 | 4.09E-226 | 0.67178277 | 0.59 | 0.555 | 7.91E-222 | TAM.Cd209 | Egln1 |
| Cmtm31 | 1.03E-224 | 0.79156831 | 0.663 | 0.565 | 1.99E-220 | TAM.Cd209 | Cmtm3 |
| Bnip3 | 1.94E-223 | 1.3635591 | 0.256 | 0.13 | 3.75E-219 | TAM.Cd209 | Bnip3 |
| Tspan321 | 1.67E-219 | 1.71158496 | 0.343 | 0.163 | 3.23E-215 | TAM.Cd209 | Tspan32 |
| Tnfrsf211 | 4.38E-219 | 0.83626155 | 0.557 | 0.428 | 8.47E-215 | TAM.Cd209 | Tnfrsf21 |
| Fam219a1 | 5.37E-219 | 1.35577126 | 0.377 | 0.205 | 1.04E-214 | TAM.Cd209 | Fam219a |
| Tnfaip8l21 | 6.93E-217 | 0.89652558 | 0.638 | 0.505 | 1.34E-212 | TAM.Cd209 | Tnfaip8l2 |
| Gas71 | 1.55E-215 | 0.94440797 | 0.715 | 0.569 | 3.00E-211 | TAM.Cd209 | Gas7 |
| Plbd21 | 3.70E-215 | 0.78644752 | 0.646 | 0.556 | 7.15E-211 | TAM.Cd209 | Plbd2 |
| Il161 | 6.71E-215 | 1.25078763 | 0.395 | 0.211 | 1.30E-210 | TAM.Cd209 | Il16 |
| Tlr8 | 1.01E-214 | 1.13968036 | 0.366 | 0.214 | 1.96E-210 | TAM.Cd209 | Tlr8 |
| Slc17a5 | 9.09E-214 | 1.43970512 | 0.27 | 0.123 | 1.76E-209 | TAM.Cd209 | Slc17a5 |
| Crtap | 6.27E-213 | 1.18355187 | 0.419 | 0.258 | 1.21E-208 | TAM.Cd209 | Crtap |
| Cebpa1 | 3.33E-208 | 0.73852519 | 0.895 | 0.753 | 6.45E-204 | TAM.Cd209 | Cebpa |
| Hacd4 | 2.06E-207 | 0.83871131 | 0.487 | 0.376 | 3.97E-203 | TAM.Cd209 | Hacd4 |
| Ccpg11 | 6.66E-204 | 0.93993047 | 0.474 | 0.334 | 1.29E-199 | TAM.Cd209 | Ccpg1 |
| Klf21 | 4.74E-203 | 1.03905179 | 0.535 | 0.354 | 9.16E-199 | TAM.Cd209 | Klf2 |
| Il10rb1 | 4.46E-202 | 0.51217678 | 0.743 | 0.738 | 8.62E-198 | TAM.Cd209 | Il10rb |
| Snx131 | 3.01E-201 | 1.22233001 | 0.416 | 0.248 | 5.82E-197 | TAM.Cd209 | Snx13 |
| Snx301 | 1.27E-199 | 1.27913147 | 0.435 | 0.263 | 2.45E-195 | TAM.Cd209 | Snx30 |
| Lyz21 | 2.06E-199 | 0.34569432 | 0.989 | 0.962 | 3.98E-195 | TAM.Cd209 | Lyz2 |
| Rcsd11 | 2.08E-199 | 1.0459063 | 0.508 | 0.334 | 4.02E-195 | TAM.Cd209 | Rcsd1 |
| Clec4a12 | 3.27E-199 | 0.68184275 | 0.495 | 0.345 | 6.32E-195 | TAM.Cd209 | Clec4a1 |
| Ip6k1 | 2.81E-197 | 1.06175994 | 0.615 | 0.472 | 5.43E-193 | TAM.Cd209 | Ip6k1 |
| Sh3bp5 | 9.64E-197 | 1.3782368 | 0.272 | 0.119 | 1.86E-192 | TAM.Cd209 | Sh3bp5 |
| Csf1r1 | 1.14E-196 | 0.3869823 | 0.952 | 0.961 | 2.20E-192 | TAM.Cd209 | Csf1r |
| Stat11 | 1.56E-195 | 0.38768926 | 0.474 | 0.483 | 3.01E-191 | TAM.Cd209 | Stat1 |
| Stat5b1 | 2.63E-195 | 0.91045581 | 0.416 | 0.3 | 5.08E-191 | TAM.Cd209 | Stat5b |
| Secisbp2l1 | 1.19E-191 | 0.85017077 | 0.565 | 0.456 | 2.31E-187 | TAM.Cd209 | Secisbp2l |
| Tspo1 | 1.45E-190 | 0.46783351 | 0.807 | 0.824 | 2.80E-186 | TAM.Cd209 | Tspo |
| Lancl1 | 6.02E-190 | 1.53375938 | 0.265 | 0.13 | 1.16E-185 | TAM.Cd209 | Lancl1 |
| Tk21 | 4.72E-189 | 1.29150223 | 0.343 | 0.188 | 9.13E-185 | TAM.Cd209 | Tk2 |
| Rubcn1 | 5.51E-189 | 1.37867278 | 0.337 | 0.183 | 1.07E-184 | TAM.Cd209 | Rubcn |
| Cpq1 | 4.07E-188 | 1.51477627 | 0.295 | 0.146 | 7.87E-184 | TAM.Cd209 | Cpq |
| Nav1 | 8.60E-186 | 1.17820045 | 0.425 | 0.247 | 1.66E-181 | TAM.Cd209 | Nav1 |
| Pepd | 6.38E-185 | 1.13321197 | 0.59 | 0.471 | 1.23E-180 | TAM.Cd209 | Pepd |
| Kifc3 | 9.56E-185 | 1.37007562 | 0.312 | 0.157 | 1.85E-180 | TAM.Cd209 | Kifc3 |
| Mtus11 | 2.04E-184 | 1.25020476 | 0.349 | 0.202 | 3.94E-180 | TAM.Cd209 | Mtus1 |
| Sgsh1 | 6.78E-184 | 1.4677749 | 0.319 | 0.168 | 1.31E-179 | TAM.Cd209 | Sgsh |
| Rras1 | 1.11E-182 | 1.01026782 | 0.427 | 0.277 | 2.14E-178 | TAM.Cd209 | Rras |
| Stx121 | 2.71E-181 | 0.68926046 | 0.613 | 0.55 | 5.23E-177 | TAM.Cd209 | Stx12 |
| Mms191 | 5.46E-181 | 1.2471515 | 0.347 | 0.195 | 1.06E-176 | TAM.Cd209 | Mms19 |
| Rab11a | 1.49E-180 | 0.87120986 | 0.514 | 0.423 | 2.89E-176 | TAM.Cd209 | Rab11a |
| Sord | 6.46E-178 | 1.38601948 | 0.314 | 0.156 | 1.25E-173 | TAM.Cd209 | Sord |
| Fcho21 | 1.18E-177 | 0.95675948 | 0.492 | 0.368 | 2.29E-173 | TAM.Cd209 | Fcho2 |
| Vamp3 | 3.62E-176 | 0.6367978 | 0.518 | 0.47 | 6.99E-172 | TAM.Cd209 | Vamp3 |
| Dhx581 | 5.43E-174 | 0.97609354 | 0.32 | 0.207 | 1.05E-169 | TAM.Cd209 | Dhx58 |
| Fli11 | 3.12E-170 | 0.70003877 | 0.589 | 0.496 | 6.04E-166 | TAM.Cd209 | Fli1 |
| Plekhb2 | 1.26E-169 | 0.71346239 | 0.672 | 0.602 | 2.44E-165 | TAM.Cd209 | Plekhb2 |
| Psd31 | 3.89E-168 | 1.37648515 | 0.494 | 0.267 | 7.53E-164 | TAM.Cd209 | Psd3 |
| Creg11 | 1.82E-167 | 0.47357875 | 0.744 | 0.722 | 3.52E-163 | TAM.Cd209 | Creg1 |
| Ampd31 | 9.09E-167 | 1.25687884 | 0.329 | 0.147 | 1.76E-162 | TAM.Cd209 | Ampd3 |
| Wdr911 | 1.32E-166 | 1.1133035 | 0.512 | 0.332 | 2.56E-162 | TAM.Cd209 | Wdr91 |
| Tgfbrap1 | 1.58E-166 | 1.49377645 | 0.365 | 0.212 | 3.05E-162 | TAM.Cd209 | Tgfbrap1 |
| Slc27a1 | 7.70E-166 | 1.09560123 | 0.591 | 0.413 | 1.49E-161 | TAM.Cd209 | Slc27a1 |
| Ms4a141 | 8.57E-166 | 1.37285082 | 0.265 | 0.118 | 1.66E-161 | TAM.Cd209 | Ms4a14 |
| Irgm11 | 1.02E-165 | 0.62469429 | 0.316 | 0.217 | 1.98E-161 | TAM.Cd209 | Irgm1 |
| Tbc1d9b1 | 9.24E-165 | 0.93225629 | 0.539 | 0.428 | 1.79E-160 | TAM.Cd209 | Tbc1d9b |
| Col3a1 | 9.43E-165 | 1.21191261 | 0.485 | 0.363 | 1.82E-160 | TAM.Cd209 | Col3a1 |
| Ly96 | 2.04E-164 | 1.29300867 | 0.303 | 0.163 | 3.95E-160 | TAM.Cd209 | Ly96 |
| Fam53b1 | 3.18E-163 | 1.20017012 | 0.42 | 0.241 | 6.16E-159 | TAM.Cd209 | Fam53b |
| 1-Mar | 2.28E-162 | 0.75899958 | 0.553 | 0.453 | 4.41E-158 | TAM.Cd209 | 1-Mar |
| Ifi2072 | 2.63E-161 | 0.40022418 | 0.623 | 0.569 | 5.09E-157 | TAM.Cd209 | Ifi207 |
| Oxct11 | 2.64E-161 | 1.09052604 | 0.42 | 0.259 | 5.10E-157 | TAM.Cd209 | Oxct1 |
| Pcmtd1 | 3.77E-161 | 1.23316164 | 0.285 | 0.157 | 7.29E-157 | TAM.Cd209 | Pcmtd1 |
| Pitpnc1 | 4.63E-161 | 1.11946907 | 0.452 | 0.297 | 8.95E-157 | TAM.Cd209 | Pitpnc1 |
| St6gal1 | 1.68E-160 | 0.94344626 | 0.436 | 0.29 | 3.25E-156 | TAM.Cd209 | St6gal1 |
| Slc8b11 | 2.31E-160 | 0.78083066 | 0.6 | 0.476 | 4.47E-156 | TAM.Cd209 | Slc8b1 |
| Cd842 | 4.47E-160 | 0.75055663 | 0.726 | 0.616 | 8.63E-156 | TAM.Cd209 | Cd84 |
| Kcnk13 | 3.37E-159 | 1.34523694 | 0.295 | 0.155 | 6.51E-155 | TAM.Cd209 | Kcnk13 |
| Snx3 | 5.32E-158 | 0.5623755 | 0.794 | 0.798 | 1.03E-153 | TAM.Cd209 | Snx3 |
| Lrrc8a1 | 6.39E-157 | 1.19318818 | 0.369 | 0.221 | 1.24E-152 | TAM.Cd209 | Lrrc8a |
| Pld32 | 2.68E-156 | 0.62485933 | 0.719 | 0.611 | 5.18E-152 | TAM.Cd209 | Pld3 |
| Cmtm6 | 1.44E-155 | 0.62611514 | 0.699 | 0.65 | 2.79E-151 | TAM.Cd209 | Cmtm6 |
| Ulk21 | 3.08E-155 | 1.17904571 | 0.383 | 0.245 | 5.96E-151 | TAM.Cd209 | Ulk2 |

| | | | | | | | |
|---|---|---|---|---|---|---|---|
| Smpd1 | 1.20E-153 | 1.41887874 | 0.292 | 0.141 | 2.32E-149 | TAM.Cd209 | Smpd1 |
| Galnt2 | 2.12E-153 | 0.87961884 | 0.633 | 0.503 | 4.09E-149 | TAM.Cd209 | Galnt2 |
| Anp32a | 2.08E-152 | 0.61616281 | 0.751 | 0.713 | 4.03E-148 | TAM.Cd209 | Anp32a |
| Mid1ip1 | 1.22E-150 | 1.27395255 | 0.408 | 0.227 | 2.37E-146 | TAM.Cd209 | Mid1ip1 |
| Laptm4a | 8.56E-150 | 0.61205881 | 0.515 | 0.468 | 1.66E-145 | TAM.Cd209 | Laptm4a |
| Rab1a | 6.24E-149 | 0.42005666 | 0.797 | 0.835 | 1.21E-144 | TAM.Cd209 | Rab1a |
| Adrb21 | 1.04E-148 | 0.93217452 | 0.374 | 0.249 | 2.01E-144 | TAM.Cd209 | Adrb2 |
| Unc93b12 | 4.53E-148 | 0.35293987 | 0.865 | 0.878 | 8.76E-144 | TAM.Cd209 | Unc93b1 |
| Mfsd111 | 4.78E-148 | 1.03942144 | 0.416 | 0.277 | 9.25E-144 | TAM.Cd209 | Mfsd11 |
| B3gnt2 | 9.18E-148 | 1.04272801 | 0.372 | 0.238 | 1.77E-143 | TAM.Cd209 | B3gnt2 |
| DEPRECATED-I | 1.55E-147 | 1.06658726 | 0.43 | 0.282 | 3.00E-143 | TAM.Cd209 | DEPRECATED-ENSMUSG00000032667 |
| Tpra1 | 4.81E-146 | 1.26998647 | 0.301 | 0.167 | 9.29E-142 | TAM.Cd209 | Tpra1 |
| Arrdc3 | 2.81E-145 | 0.96976187 | 0.386 | 0.256 | 5.43E-141 | TAM.Cd209 | Arrdc3 |
| Man2b21 | 4.92E-145 | 0.65801819 | 0.592 | 0.52 | 9.51E-141 | TAM.Cd209 | Man2b2 |
| Pml1 | 7.37E-145 | 1.32270282 | 0.296 | 0.158 | 1.42E-140 | TAM.Cd209 | Pml |
| Ago3 | 3.42E-143 | 1.22219702 | 0.418 | 0.272 | 6.61E-139 | TAM.Cd209 | Ago3 |
| Ctso1 | 3.99E-142 | 0.96377448 | 0.406 | 0.284 | 7.71E-138 | TAM.Cd209 | Ctso |
| Tut71 | 7.79E-142 | 0.67197673 | 0.699 | 0.66 | 1.51E-137 | TAM.Cd209 | Tut7 |
| Rasgrp42 | 3.59E-139 | 1.10460889 | 0.455 | 0.292 | 6.93E-135 | TAM.Cd209 | Rasgrp4 |
| Tlr72 | 9.21E-139 | 0.81754557 | 0.346 | 0.241 | 1.78E-134 | TAM.Cd209 | Tlr7 |
| Sft2d11 | 1.72E-138 | 0.9688565 | 0.357 | 0.239 | 3.32E-134 | TAM.Cd209 | Sft2d1 |
| Serpine1 | 1.14E-137 | 0.5589654 | 0.31 | 0.199 | 2.20E-133 | TAM.Cd209 | Serpine1 |
| Mef2c1 | 1.25E-136 | 0.99521471 | 0.43 | 0.282 | 2.41E-132 | TAM.Cd209 | Mef2c |
| Plcl21 | 2.49E-136 | 0.95647182 | 0.482 | 0.349 | 4.81E-132 | TAM.Cd209 | Plcl2 |
| Tsc22d32 | 2.51E-136 | 0.80297801 | 0.55 | 0.394 | 4.85E-132 | TAM.Cd209 | Tsc22d3 |
| Spire1 | 5.85E-136 | 1.09152351 | 0.302 | 0.179 | 1.13E-131 | TAM.Cd209 | Spire1 |
| Tuba1a | 6.84E-134 | 0.93959444 | 0.457 | 0.329 | 1.32E-129 | TAM.Cd209 | Tuba1a |
| Tex2641 | 8.56E-133 | 0.94294479 | 0.491 | 0.361 | 1.66E-128 | TAM.Cd209 | Tex264 |
| Zbp12 | 2.05E-132 | 0.61317503 | 0.371 | 0.283 | 3.97E-128 | TAM.Cd209 | Zbp1 |
| Gltp | 7.86E-131 | 0.63932882 | 0.69 | 0.646 | 1.52E-126 | TAM.Cd209 | Gltp |
| Tmem87b | 8.13E-131 | 0.89922219 | 0.505 | 0.372 | 1.57E-126 | TAM.Cd209 | Tmem87b |
| Slc35f61 | 2.89E-130 | 0.78089256 | 0.524 | 0.41 | 5.60E-126 | TAM.Cd209 | Slc35f6 |
| Bst11 | 8.94E-130 | 0.67695315 | 0.52 | 0.364 | 1.73E-125 | TAM.Cd209 | Bst1 |
| Nagk2 | 1.70E-129 | 1.08581062 | 0.398 | 0.271 | 3.29E-125 | TAM.Cd209 | Nagk |
| Dtnbp1 | 2.03E-129 | 0.77525322 | 0.597 | 0.503 | 3.93E-125 | TAM.Cd209 | Dtnbp1 |
| Il3ra | 2.06E-128 | 1.02217111 | 0.392 | 0.261 | 3.98E-124 | TAM.Cd209 | Il3ra |
| Hmox21 | 6.03E-128 | 0.67603208 | 0.582 | 0.516 | 1.17E-123 | TAM.Cd209 | Hmox2 |
| Marcksl11 | 3.07E-127 | 0.41474585 | 0.764 | 0.698 | 5.94E-123 | TAM.Cd209 | Marcksl1 |
| Hip11 | 3.99E-127 | 0.81622907 | 0.524 | 0.413 | 7.71E-123 | TAM.Cd209 | Hip1 |
| Maged11 | 1.63E-125 | 1.13801766 | 0.307 | 0.175 | 3.14E-121 | TAM.Cd209 | Maged1 |
| Zbtb4 | 2.01E-125 | 1.38664662 | 0.27 | 0.144 | 3.88E-121 | TAM.Cd209 | Zbtb4 |
| Rasa11 | 7.27E-125 | 1.14372008 | 0.389 | 0.259 | 1.41E-120 | TAM.Cd209 | Rasa1 |
| Dusp72 | 2.49E-124 | 0.71499912 | 0.526 | 0.412 | 4.81E-120 | TAM.Cd209 | Dusp7 |
| Tmem250-ps1 | 1.47E-123 | 0.96237377 | 0.487 | 0.358 | 2.84E-119 | TAM.Cd209 | Tmem250-ps |
| Col1a1 | 6.10E-123 | 1.06601502 | 0.603 | 0.492 | 1.18E-118 | TAM.Cd209 | Col1a1 |
| Arpc3 | 2.60E-122 | 0.38029928 | 0.721 | 0.77 | 5.03E-118 | TAM.Cd209 | Arpc3 |
| Col1a2 | 3.31E-122 | 1.26329183 | 0.429 | 0.302 | 6.39E-118 | TAM.Cd209 | Col1a2 |
| Tspan41 | 1.25E-120 | 0.94773313 | 0.46 | 0.325 | 2.42E-116 | TAM.Cd209 | Tspan4 |
| Zbtb381 | 1.93E-120 | 1.22470268 | 0.322 | 0.198 | 3.72E-116 | TAM.Cd209 | Zbtb38 |
| Vezt | 7.64E-120 | 1.3634262 | 0.253 | 0.133 | 1.48E-115 | TAM.Cd209 | Vezt |
| Tmem141 | 2.96E-119 | 1.3100339 | 0.272 | 0.153 | 5.72E-115 | TAM.Cd209 | Tmem141 |
| Nlrc51 | 9.56E-119 | 0.94705175 | 0.411 | 0.285 | 1.85E-114 | TAM.Cd209 | Nlrc5 |
| Aftph1 | 9.90E-119 | 0.98108775 | 0.356 | 0.249 | 1.91E-114 | TAM.Cd209 | Aftph |
| Stxbp3 | 1.07E-118 | 1.25314496 | 0.408 | 0.269 | 2.07E-114 | TAM.Cd209 | Stxbp3 |
| Stx21 | 6.98E-118 | 1.05351953 | 0.397 | 0.275 | 1.35E-113 | TAM.Cd209 | Stx2 |
| Slfn52 | 1.46E-117 | 0.4375239 | 0.391 | 0.265 | 2.82E-113 | TAM.Cd209 | Slfn5 |
| Pacs1 | 1.63E-117 | 0.9550706 | 0.303 | 0.193 | 3.16E-113 | TAM.Cd209 | Pacs1 |
| Calcoco11 | 1.66E-117 | 1.27816453 | 0.342 | 0.204 | 3.21E-113 | TAM.Cd209 | Calcoco1 |
| Fbxw4 | 3.70E-117 | 1.14801878 | 0.254 | 0.137 | 7.15E-113 | TAM.Cd209 | Fbxw4 |
| Hspa1b2 | 9.50E-117 | 0.29431002 | 0.48 | 0.353 | 1.84E-112 | TAM.Cd209 | Hspa1b |
| Idh21 | 1.30E-116 | 1.05899492 | 0.554 | 0.405 | 2.52E-112 | TAM.Cd209 | Idh2 |
| Atxn11 | 2.12E-116 | 0.88812947 | 0.392 | 0.266 | 4.11E-112 | TAM.Cd209 | Atxn1 |
| Zfp36l11 | 4.18E-116 | 0.44829099 | 0.661 | 0.63 | 8.09E-112 | TAM.Cd209 | Zfp36l1 |
| Ddx581 | 8.42E-116 | 0.8753201 | 0.26 | 0.157 | 1.63E-111 | TAM.Cd209 | Ddx58 |
| Nfic1 | 4.66E-115 | 1.10084511 | 0.433 | 0.29 | 9.01E-111 | TAM.Cd209 | Nfic |
| Lima1 | 1.62E-114 | 1.1050151 | 0.278 | 0.165 | 3.13E-110 | TAM.Cd209 | Lima1 |
| Clu | 2.70E-113 | 0.81898735 | 0.82 | 0.747 | 5.21E-109 | TAM.Cd209 | Clu |
| Sparc | 3.73E-113 | 1.2556629 | 0.367 | 0.242 | 7.21E-109 | TAM.Cd209 | Sparc |
| Phf201 | 7.36E-113 | 1.06879669 | 0.353 | 0.23 | 1.42E-108 | TAM.Cd209 | Phf20 |
| 8-Mar | 8.49E-113 | 1.17594408 | 0.279 | 0.167 | 1.64E-108 | TAM.Cd209 | 8-Mar |
| Rab5c1 | 2.60E-112 | 0.32017697 | 0.79 | 0.843 | 5.03E-108 | TAM.Cd209 | Rab5c |
| Nr3c1 | 5.05E-112 | 0.85471902 | 0.461 | 0.348 | 9.77E-108 | TAM.Cd209 | Nr3c1 |
| Arl21 | 4.87E-110 | 1.15697308 | 0.308 | 0.188 | 9.41E-106 | TAM.Cd209 | Arl2 |
| Mctp1 | 1.25E-108 | 1.49405807 | 0.253 | 0.124 | 2.41E-104 | TAM.Cd209 | Mctp1 |
| Cystm11 | 3.38E-108 | 1.04382519 | 0.306 | 0.198 | 6.54E-104 | TAM.Cd209 | Cystm1 |
| Ttc33 | 6.38E-108 | 1.33394375 | 0.297 | 0.17 | 1.23E-103 | TAM.Cd209 | Ttc33 |
| Fgd61 | 4.93E-107 | 1.29138576 | 0.316 | 0.177 | 9.53E-103 | TAM.Cd209 | Fgd6 |
| Sash11 | 5.48E-106 | 0.58372713 | 0.712 | 0.581 | 1.06E-101 | TAM.Cd209 | Sash1 |
| Lgals11 | 6.40E-106 | 0.84087404 | 0.366 | 0.263 | 1.24E-101 | TAM.Cd209 | Lgals1 |
| Eid12 | 3.03E-105 | 0.53868207 | 0.775 | 0.715 | 5.86E-101 | TAM.Cd209 | Eid1 |
| Ssh3 | 4.99E-103 | 1.03750866 | 0.432 | 0.295 | 9.65E-99 | TAM.Cd209 | Ssh3 |
| Crebrf | 1.21E-102 | 0.88379512 | 0.335 | 0.223 | 2.34E-98 | TAM.Cd209 | Crebrf |
| Plekhm1 | 1.24E-102 | 1.11385802 | 0.321 | 0.208 | 2.40E-98 | TAM.Cd209 | Plekhm1 |
| Mpp1 | 1.57E-101 | 0.71365375 | 0.462 | 0.374 | 3.04E-97 | TAM.Cd209 | Mpp1 |
| Adi11 | 2.36E-101 | 1.08387991 | 0.306 | 0.19 | 4.57E-97 | TAM.Cd209 | Adi1 |
| Slc48a11 | 4.27E-101 | 0.6251384 | 0.575 | 0.492 | 8.26E-97 | TAM.Cd209 | Slc48a1 |
| Lbh1 | 5.31E-101 | 0.656209 | 0.466 | 0.372 | 1.03E-96 | TAM.Cd209 | Lbh |
| Sft2d2 | 6.07E-100 | 0.85301791 | 0.455 | 0.341 | 1.17E-95 | TAM.Cd209 | Sft2d2 |
| Ssh22 | 2.97E-99 | 0.59684502 | 0.5 | 0.413 | 5.74E-95 | TAM.Cd209 | Ssh2 |
| B4galt61 | 3.34E-99 | 0.73821796 | 0.504 | 0.396 | 6.45E-95 | TAM.Cd209 | B4galt6 |
| Cdk20 | 1.58E-98 | 1.44183184 | 0.253 | 0.136 | 3.06E-94 | TAM.Cd209 | Cdk20 |
| Rnf131 | 2.79E-98 | 0.69573304 | 0.56 | 0.494 | 5.39E-94 | TAM.Cd209 | Rnf13 |
| Cysltr11 | 3.32E-98 | 0.9334386 | 0.339 | 0.23 | 6.41E-94 | TAM.Cd209 | Cysltr1 |
| Fam3c1 | 1.67E-97 | 0.67689395 | 0.464 | 0.391 | 3.22E-93 | TAM.Cd209 | Fam3c |
| Lpin11 | 1.73E-97 | 1.00247634 | 0.308 | 0.189 | 3.34E-93 | TAM.Cd209 | Lpin1 |
| Stxbp5 | 1.29E-96 | 1.1136307 | 0.262 | 0.149 | 2.50E-92 | TAM.Cd209 | Stxbp5 |
| Pygb1 | 4.27E-96 | 0.97914654 | 0.419 | 0.309 | 8.25E-92 | TAM.Cd209 | Pygb |
| Idua1 | 4.34E-96 | 1.05979723 | 0.369 | 0.253 | 8.40E-92 | TAM.Cd209 | Idua |
| P2rx71 | 1.23E-95 | 0.86564525 | 0.432 | 0.334 | 2.37E-91 | TAM.Cd209 | P2rx7 |
| Sypl | 2.63E-95 | 0.52162308 | 0.584 | 0.542 | 5.09E-91 | TAM.Cd209 | Sypl |
| Hs1bp31 | 2.68E-95 | 1.18152125 | 0.327 | 0.198 | 5.19E-91 | TAM.Cd209 | Hs1bp3 |
| Ptgr21 | 2.82E-95 | 0.8717618 | 0.426 | 0.308 | 5.46E-91 | TAM.Cd209 | Ptgr2 |
| Galnt11 | 8.09E-95 | 0.56505149 | 0.591 | 0.531 | 1.56E-90 | TAM.Cd209 | Galnt1 |
| Pgm2l1 | 3.43E-94 | 1.09600497 | 0.256 | 0.155 | 6.64E-90 | TAM.Cd209 | Pgm2l1 |
| Gng122 | 1.31E-93 | 0.81672494 | 0.415 | 0.296 | 2.53E-89 | TAM.Cd209 | Gng12 |
| Dram21 | 1.93E-93 | 0.73388223 | 0.385 | 0.305 | 3.72E-89 | TAM.Cd209 | Dram2 |
| Cenpb | 8.42E-93 | 0.73953254 | 0.663 | 0.584 | 1.63E-88 | TAM.Cd209 | Cenpb |
| Rbpj | 1.22E-92 | 0.43582219 | 0.687 | 0.681 | 2.36E-88 | TAM.Cd209 | Rbpj |
| Sgpl12 | 2.37E-92 | 0.47310723 | 0.706 | 0.691 | 4.58E-88 | TAM.Cd209 | Sgpl1 |
| Rbms2 | 2.38E-92 | 0.91957979 | 0.328 | 0.236 | 4.60E-88 | TAM.Cd209 | Rbms2 |
| Rgmb1 | 5.87E-92 | 0.94652996 | 0.332 | 0.212 | 1.14E-87 | TAM.Cd209 | Rgmb |
| Tmem106b1 | 2.24E-91 | 0.90766684 | 0.359 | 0.248 | 4.33E-87 | TAM.Cd209 | Tmem106b |
| Vat11 | 1.05E-90 | 0.63697404 | 0.718 | 0.586 | 2.03E-86 | TAM.Cd209 | Vat1 |
| Dennd1a1 | 2.58E-90 | 0.7404144 | 0.471 | 0.364 | 4.98E-86 | TAM.Cd209 | Dennd1a |
| Pgpep11 | 3.04E-90 | 1.25334805 | 0.283 | 0.162 | 5.87E-86 | TAM.Cd209 | Pgpep1 |
| Ap2s11 | 6.45E-89 | 0.65546021 | 0.692 | 0.644 | 1.25E-84 | TAM.Cd209 | Ap2s1 |
| Mon2 | 1.28E-88 | 1.1752655 | 0.491 | 0.369 | 2.47E-84 | TAM.Cd209 | Mon2 |
| Lgals92 | 1.52E-88 | 0.381195 | 0.6 | 0.585 | 2.94E-84 | TAM.Cd209 | Lgals9 |
| Csf3r2 | 2.75E-88 | 0.553106 | 0.482 | 0.394 | 5.31E-84 | TAM.Cd209 | Csf3r |
| Mfsd13 | 1.04E-87 | 0.43592807 | 0.71 | 0.713 | 2.01E-83 | TAM.Cd209 | Mfsd1 |
| Dhrs7 | 2.09E-87 | 0.80660974 | 0.314 | 0.213 | 4.03E-83 | TAM.Cd209 | Dhrs7 |
| DEPRECATED-I | 1.84E-86 | 0.82462746 | 0.34 | 0.233 | 3.56E-82 | TAM.Cd209 | DEPRECATED-ENSMUSG00000002103 |
| Dmxl11 | 2.69E-86 | 0.92883441 | 0.401 | 0.292 | 5.20E-82 | TAM.Cd209 | Dmxl1 |
| Txndc161 | 1.18E-85 | 1.07435647 | 0.312 | 0.2 | 2.28E-81 | TAM.Cd209 | Txndc16 |
| Nucb11 | 1.55E-85 | 0.56168842 | 0.728 | 0.693 | 2.99E-81 | TAM.Cd209 | Nucb1 |
| AW549877 | 2.63E-85 | 1.23500797 | 0.295 | 0.178 | 5.09E-81 | TAM.Cd209 | AW549877 |
| Il181 | 2.69E-85 | 0.81939148 | 0.268 | 0.168 | 5.20E-81 | TAM.Cd209 | Il18 |
| Metrnl1 | 5.70E-85 | 0.46554908 | 0.652 | 0.576 | 1.10E-80 | TAM.Cd209 | Metrnl |
| Ccdc931 | 5.84E-85 | 0.92942434 | 0.36 | 0.249 | 1.13E-80 | TAM.Cd209 | Ccdc93 |
| Zfp901 | 1.97E-84 | 1.05561344 | 0.252 | 0.149 | 3.80E-80 | TAM.Cd209 | Zfp90 |
| Tlr5 | 1.99E-84 | 0.80365805 | 0.264 | 0.161 | 3.84E-80 | TAM.Cd209 | Tlr5 |
| Rab121 | 3.11E-84 | 0.69834253 | 0.44 | 0.352 | 6.02E-80 | TAM.Cd209 | Rab12 |
| Oxr1 | 5.38E-84 | 0.7557235 | 0.385 | 0.294 | 1.04E-79 | TAM.Cd209 | Oxr1 |
| Kctd2 | 1.02E-83 | 0.99009544 | 0.305 | 0.199 | 1.97E-79 | TAM.Cd209 | Kctd2 |
| Klhl241 | 4.23E-83 | 0.78949597 | 0.374 | 0.278 | 8.17E-79 | TAM.Cd209 | Klhl24 |
| Dag1 | 7.65E-83 | 0.76763826 | 0.338 | 0.238 | 1.48E-78 | TAM.Cd209 | Dag1 |
| Triap1 | 7.76E-83 | 0.93908451 | 0.279 | 0.18 | 1.50E-78 | TAM.Cd209 | Triap1 |
| Lrrc251 | 9.43E-83 | 0.51599512 | 0.746 | 0.712 | 1.82E-78 | TAM.Cd209 | Lrrc25 |
| Agfg11 | 1.56E-82 | 0.65892267 | 0.438 | 0.382 | 3.02E-78 | TAM.Cd209 | Agfg1 |
| Rpain | 9.54E-82 | 1.2101922 | 0.291 | 0.183 | 1.85E-77 | TAM.Cd209 | Rpain |
| Ggt51 | 3.94E-81 | 1.40956965 | 0.264 | 0.159 | 7.61E-77 | TAM.Cd209 | Ggt5 |
| Trafd11 | 5.43E-81 | 0.58080922 | 0.601 | 0.565 | 1.05E-76 | TAM.Cd209 | Trafd1 |
| Ctsa1 | 7.02E-81 | 0.29661739 | 0.872 | 0.914 | 1.36E-76 | TAM.Cd209 | Ctsa |
| Cebpzos1 | 1.36E-80 | 0.83340634 | 0.35 | 0.25 | 2.62E-76 | TAM.Cd209 | Cebpzos |
| Slc35f5 | 1.47E-80 | 0.82270044 | 0.269 | 0.182 | 2.85E-76 | TAM.Cd209 | Slc35f5 |
| Gdf151 | 3.08E-80 | 0.47621268 | 0.348 | 0.27 | 5.96E-76 | TAM.Cd209 | Gdf15 |
| Nrros1 | 3.67E-80 | 0.40072975 | 0.755 | 0.735 | 7.09E-76 | TAM.Cd209 | Nrros |
| Pnpla8 | 6.63E-80 | 0.71439633 | 0.341 | 0.248 | 1.28E-75 | TAM.Cd209 | Pnpla8 |
| Tbc1d22a1 | 1.38E-79 | 0.80863992 | 0.663 | 0.573 | 2.68E-75 | TAM.Cd209 | Tbc1d22a |
| Akr1b101 | 1.40E-79 | 0.90214293 | 0.344 | 0.244 | 2.71E-75 | TAM.Cd209 | Akr1b10 |
| Hgsnat1 | 3.92E-79 | 0.75208836 | 0.3 | 0.29 | 7.57E-75 | TAM.Cd209 | Hgsnat |
| Fut8 | 4.13E-79 | 1.00597721 | 0.258 | 0.162 | 7.98E-75 | TAM.Cd209 | Fut8 |
| Blnk1 | 2.68E-78 | 1.0237193 | 0.334 | 0.211 | 5.19E-74 | TAM.Cd209 | Blnk |
| Fam162a | 2.98E-78 | 0.50670554 | 0.416 | 0.375 | 5.77E-74 | TAM.Cd209 | Fam162a |
| Fads11 | 3.41E-78 | 0.92663836 | 0.335 | 0.225 | 6.59E-74 | TAM.Cd209 | Fads1 |
| Usp361 | 6.76E-78 | 0.92913406 | 0.374 | 0.264 | 1.31E-73 | TAM.Cd209 | Usp36 |
| Epn1 | 1.66E-77 | 0.50158111 | 0.812 | 0.814 | 3.21E-73 | TAM.Cd209 | Epn1 |
| mt-Cytb | 5.30E-76 | 0.36584845 | 0.659 | 0.708 | 1.02E-71 | TAM.Cd209 | mt-Cytb |
| Naglu1 | 4.22E-75 | 0.76873906 | 0.472 | 0.363 | 8.16E-71 | TAM.Cd209 | Naglu |
| Fbxw7 | 9.35E-75 | 1.01219199 | 0.256 | 0.166 | 1.81E-70 | TAM.Cd209 | Fbxw7 |
| Bgn | 3.85E-74 | 1.24641537 | 0.306 | 0.21 | 7.45E-70 | TAM.Cd209 | Bgn |
| Slc31a1 | 5.61E-74 | 0.53050671 | 0.345 | 0.285 | 1.09E-69 | TAM.Cd209 | Slc31a1 |
| Ogt | 8.77E-74 | 0.92340332 | 0.546 | 0.46 | 1.70E-69 | TAM.Cd209 | Ogt |
| Cracr2b1 | 1.66E-73 | 1.09982749 | 0.313 | 0.203 | 3.21E-69 | TAM.Cd209 | Cracr2b |
| Themis23 | 2.98E-73 | 0.58288248 | 0.551 | 0.463 | 5.77E-69 | TAM.Cd209 | Themis2 |
| Dtx3l1 | 1.38E-72 | 0.51545309 | 0.356 | 0.289 | 2.66E-68 | TAM.Cd209 | Dtx3l |
| Tpd52l2 | 2.17E-72 | 0.64243792 | 0.516 | 0.44 | 4.20E-68 | TAM.Cd209 | Tpd52l2 |
| Rgs31 | 2.39E-72 | 0.84474433 | 0.277 | 0.196 | 4.61E-68 | TAM.Cd209 | Rgs3 |
| Wls1 | 3.27E-72 | 0.7011093 | 0.419 | 0.323 | 6.31E-68 | TAM.Cd209 | Wls |
| Sil1 | 5.67E-72 | 1.13463574 | 0.265 | 0.162 | 1.10E-67 | TAM.Cd209 | Sil1 |
| Mcfd21 | 6.57E-72 | 0.50191138 | 0.631 | 0.586 | 1.27E-67 | TAM.Cd209 | Mcfd2 |
| Nrp21 | 1.43E-71 | 0.65216734 | 0.567 | 0.433 | 2.76E-67 | TAM.Cd209 | Nrp2 |
| Stard3nl1 | 7.99E-71 | 0.70452007 | 0.385 | 0.31 | 1.55E-66 | TAM.Cd209 | Stard3nl |
| B4gat1 | 9.93E-71 | 1.00821424 | 0.27 | 0.167 | 1.92E-66 | TAM.Cd209 | B4gat1 |
| Arid5b2 | 1.26E-69 | 0.60995994 | 0.38 | 0.296 | 2.44E-65 | TAM.Cd209 | Arid5b |
| Il21r1 | 2.38E-68 | 0.46073491 | 0.557 | 0.464 | 4.61E-64 | TAM.Cd209 | Il21r |
| Mfhas1 | 1.10E-67 | 0.81600653 | 0.438 | 0.322 | 2.13E-63 | TAM.Cd209 | Mfhas1 |
| Pfkl1 | 1.97E-67 | 0.45598282 | 0.581 | 0.574 | 3.81E-63 | TAM.Cd209 | Pfkl |
| Tmem59 | 2.05E-67 | 0.49871542 | 0.544 | 0.512 | 3.97E-63 | TAM.Cd209 | Tmem59 |
| Foxp11 | 2.42E-67 | 0.75254692 | 0.442 | 0.332 | 4.68E-63 | TAM.Cd209 | Foxp1 |
| Ccl31 | 2.82E-67 | 0.27209177 | 0.594 | 0.569 | 5.45E-63 | TAM.Cd209 | Ccl3 |
| Parp3 | 5.30E-67 | 1.10575745 | 0.259 | 0.162 | 1.02E-62 | TAM.Cd209 | Parp3 |
| Ypel5 | 5.36E-66 | 0.46743425 | 0.51 | 0.462 | 1.04E-61 | TAM.Cd209 | Ypel5 |
| Nod12 | 9.60E-66 | 0.63644499 | 0.306 | 0.217 | 1.86E-61 | TAM.Cd209 | Nod1 |
| Jak31 | 1.33E-65 | 0.93759842 | 0.417 | 0.312 | 2.58E-61 | TAM.Cd209 | Jak3 |
| Sh3glb11 | 2.09E-65 | 0.35046731 | 0.742 | 0.767 | 4.05E-61 | TAM.Cd209 | Sh3glb1 |
| Cln31 | 4.81E-65 | 0.56673313 | 0.485 | 0.42 | 9.30E-61 | TAM.Cd209 | Cln3 |
| Pgm1 | 6.21E-65 | 0.59811794 | 0.356 | 0.281 | 1.20E-60 | TAM.Cd209 | Pgm1 |
| Ier5l | 9.43E-65 | 1.12540987 | 0.274 | 0.152 | 1.82E-60 | TAM.Cd209 | Ier5l |
| Micu1 | 1.42E-64 | 0.71452602 | 0.318 | 0.244 | 2.75E-60 | TAM.Cd209 | Micu1 |
| Rnpep | 1.56E-64 | 0.40149107 | 0.731 | 0.735 | 3.02E-60 | TAM.Cd209 | Rnpep |
| Ing41 | 1.76E-63 | 0.88308758 | 0.424 | 0.316 | 3.40E-59 | TAM.Cd209 | Ing4 |
| Akap7 | 5.50E-63 | 0.99493579 | 0.27 | 0.197 | 1.06E-58 | TAM.Cd209 | Akap7 |
| Slc38a62 | 4.18E-62 | 1.01433304 | 0.522 | 0.367 | 8.09E-58 | TAM.Cd209 | Slc38a6 |
| Fam126a | 4.72E-62 | 0.86836642 | 0.268 | 0.191 | 9.12E-58 | TAM.Cd209 | Fam126a |
| Hk3 | 5.54E-62 | 0.68520317 | 0.388 | 0.3 | 1.07E-57 | TAM.Cd209 | Hk3 |
| Grcc10 | 8.65E-62 | 0.46392891 | 0.624 | 0.603 | 1.67E-57 | TAM.Cd209 | Grcc10 |
| Stat21 | 9.20E-62 | 0.77704911 | 0.288 | 0.214 | 1.78E-57 | TAM.Cd209 | Stat2 |
| Dlgap4 | 1.29E-61 | 0.59839495 | 0.572 | 0.509 | 2.49E-57 | TAM.Cd209 | Dlgap4 |
| Nbr11 | 3.19E-61 | 0.87036354 | 0.308 | 0.21 | 6.17E-57 | TAM.Cd209 | Nbr1 |
| mt-Nd4l | 8.41E-61 | 0.43484991 | 0.647 | 0.693 | 1.63E-56 | TAM.Cd209 | mt-Nd4l |
| Tpi1 | 1.48E-59 | 0.27639395 | 0.625 | 0.667 | 2.86E-55 | TAM.Cd209 | Tpi1 |
| Manbal | 1.54E-59 | 0.70803003 | 0.426 | 0.345 | 2.98E-55 | TAM.Cd209 | Manbal |
| Sfmbt11 | 4.29E-59 | 0.85444477 | 0.358 | 0.245 | 8.29E-55 | TAM.Cd209 | Sfmbt1 |
| Mtmr12 | 9.29E-59 | 0.96299241 | 0.255 | 0.164 | 1.80E-54 | TAM.Cd209 | Mtmr12 |
| Tsen34 | 1.78E-58 | 0.98507579 | 0.337 | 0.235 | 3.45E-54 | TAM.Cd209 | Tsen34 |
| Ms4a6c3 | 4.36E-58 | 0.33230656 | 0.59 | 0.537 | 8.42E-54 | TAM.Cd209 | Ms4a6c |
| Snx52 | 4.85E-58 | 0.47564629 | 0.566 | 0.551 | 9.38E-54 | TAM.Cd209 | Snx5 |
| Map1lc3b | 4.89E-58 | 0.36682794 | 0.653 | 0.678 | 9.45E-54 | TAM.Cd209 | Map1lc3b |
| Gng21 | 2.09E-57 | 0.56321091 | 0.569 | 0.503 | 4.05E-53 | TAM.Cd209 | Gng2 |
| Card192 | 3.45E-57 | 0.38818683 | 0.642 | 0.61 | 6.67E-53 | TAM.Cd209 | Card19 |
| Rit1 | 4.98E-57 | 0.71357616 | 0.389 | 0.305 | 9.63E-53 | TAM.Cd209 | Rit1 |
| Uhrf1bp1l2 | 5.08E-57 | 0.66431493 | 0.32 | 0.238 | 9.82E-53 | TAM.Cd209 | Uhrf1bp1l |
| Nrbf21 | 3.19E-56 | 0.83615809 | 0.344 | 0.251 | 6.16E-52 | TAM.Cd209 | Nrbf2 |
| Map1lc3a | 3.27E-56 | 0.63490602 | 0.307 | 0.232 | 6.33E-52 | TAM.Cd209 | Map1lc3a |
| Snap29 | 5.08E-56 | 0.80886006 | 0.359 | 0.288 | 9.82E-52 | TAM.Cd209 | Snap29 |
| Wdfy2 | 1.43E-55 | 0.89727887 | 0.544 | 0.433 | 2.76E-51 | TAM.Cd209 | Wdfy2 |
| Ptpro2 | 8.36E-55 | 0.5210144 | 0.388 | 0.298 | 1.62E-50 | TAM.Cd209 | Ptpro |
| Acp22 | 1.15E-54 | 0.46536255 | 0.534 | 0.47 | 2.22E-50 | TAM.Cd209 | Acp2 |
| Tmcc11 | 2.16E-54 | 0.68497373 | 0.483 | 0.406 | 4.17E-50 | TAM.Cd209 | Tmcc1 |
| Iffo11 | 5.47E-54 | 1.15499089 | 0.284 | 0.183 | 1.06E-49 | TAM.Cd209 | Iffo1 |
| Nt5dc2 | 5.76E-54 | 0.63229718 | 0.294 | 0.22 | 1.11E-49 | TAM.Cd209 | Nt5dc2 |
| Smim10l11 | 1.55E-53 | 0.93945661 | 0.356 | 0.258 | 3.00E-49 | TAM.Cd209 | Smim10l1 |
| Gramd1b1 | 5.13E-53 | 0.76895745 | 0.324 | 0.237 | 9.91E-49 | TAM.Cd209 | Gramd1b |
| Sp1001 | 5.94E-53 | 0.52242573 | 0.335 | 0.262 | 1.15E-48 | TAM.Cd209 | Sp100 |
| Gramd1a | 1.61E-52 | 0.67793138 | 0.509 | 0.43 | 3.11E-48 | TAM.Cd209 | Gramd1a |
| Cant11 | 1.89E-52 | 0.747638 | 0.493 | 0.401 | 3.65E-48 | TAM.Cd209 | Cant1 |
| Mfngd | 3.33E-52 | 0.70054338 | 0.437 | 0.365 | 6.44E-48 | TAM.Cd209 | Mfngd |
| Fundc11 | 3.61E-52 | 0.52083815 | 0.523 | 0.47 | 6.98E-48 | TAM.Cd209 | Fundc1 |
| Dcaf81 | 5.83E-52 | 0.65747995 | 0.391 | 0.312 | 1.13E-47 | TAM.Cd209 | Dcaf8 |
| Cerk1 | 6.40E-52 | 0.70530965 | 0.553 | 0.465 | 1.24E-47 | TAM.Cd209 | Cerk |
| Tor4a1 | 1.90E-51 | 0.74441645 | 0.382 | 0.279 | 3.67E-47 | TAM.Cd209 | Tor4a |
| Pip4p12 | 2.91E-51 | 0.53230209 | 0.558 | 0.5 | 5.63E-47 | TAM.Cd209 | Pip4p1 |
| Ulk1 | 3.74E-51 | 0.66833708 | 0.365 | 0.29 | 7.22E-47 | TAM.Cd209 | Ulk1 |
| Wdr811 | 5.41E-51 | 0.76859484 | 0.452 | 0.366 | 1.05E-46 | TAM.Cd209 | Wdr81 |
| Mxd41 | 2.29E-50 | 0.73818514 | 0.422 | 0.329 | 4.43E-46 | TAM.Cd209 | Mxd4 |
| Cpne21 | 2.94E-50 | 0.74549256 | 0.347 | 0.264 | 5.68E-46 | TAM.Cd209 | Cpne2 |
| Srgap21 | 4.85E-50 | 0.76266252 | 0.441 | 0.365 | 9.37E-46 | TAM.Cd209 | Srgap2 |
| Lamtor1 | 5.10E-50 | 0.99674888 | 0.384 | 0.28 | 9.85E-46 | TAM.Cd209 | Lamtor1 |
| Uba71 | 6.35E-50 | 0.98329895 | 0.319 | 0.228 | 1.23E-45 | TAM.Cd209 | Uba7 |
| Vps26c | 1.90E-49 | 0.76930667 | 0.343 | 0.264 | 3.68E-45 | TAM.Cd209 | Vps26c |
| Acadl | 2.38E-49 | 0.6820774 | 0.386 | 0.32 | 4.61E-45 | TAM.Cd209 | Acadl |
| Arap11 | 6.97E-49 | 0.5709044 | 0.764 | 0.722 | 1.35E-44 | TAM.Cd209 | Arap1 |
| Gbp72 | 1.14E-48 | 0.41435587 | 0.358 | 0.32 | 2.21E-44 | TAM.Cd209 | Gbp7 |
| Trim262 | 1.21E-48 | 0.85565302 | 0.336 | 0.246 | 2.33E-44 | TAM.Cd209 | Trim26 |
| Cd200r1 | 1.41E-48 | 0.85716338 | 0.381 | 0.274 | 2.72E-44 | TAM.Cd209 | Cd200r1 |
| Cpne3 | 2.17E-48 | 0.56800312 | 0.363 | 0.307 | 4.19E-44 | TAM.Cd209 | Cpne3 |
| Slc31a21 | 3.71E-48 | 0.50208792 | 0.434 | 0.363 | 7.18E-44 | TAM.Cd209 | Slc31a2 |
| Hp1bp3 | 6.99E-48 | 0.61896868 | 0.564 | 0.487 | 1.35E-43 | TAM.Cd209 | Hp1bp3 |
| Slc45a41 | 7.79E-48 | 0.77037518 | 0.353 | 0.272 | 1.51E-43 | TAM.Cd209 | Slc45a4 |
| Slc25a511 | 1.15E-47 | 0.56952971 | 0.482 | 0.414 | 2.23E-43 | TAM.Cd209 | Slc25a51 |
| Gdap2 | 1.35E-47 | 0.71282007 | 0.293 | 0.221 | 2.62E-43 | TAM.Cd209 | Gdap2 |
| Bri3 | 1.64E-47 | 0.41610452 | 0.754 | 0.748 | 3.17E-43 | TAM.Cd209 | Bri3 |
| Scly1 | 3.28E-47 | 1.04375348 | 0.261 | 0.173 | 6.34E-43 | TAM.Cd209 | Scly |
| Pnpla71 | 9.75E-47 | 0.64487448 | 0.457 | 0.383 | 1.89E-42 | TAM.Cd209 | Pnpla7 |
| Uqcc31 | 1.57E-46 | 0.79136944 | 0.353 | 0.256 | 3.04E-42 | TAM.Cd209 | Uqcc3 |
| Rsrp1 | 3.34E-46 | 0.32368362 | 0.765 | 0.824 | 6.46E-42 | TAM.Cd209 | Rsrp1 |
| Arhgap452 | 4.23E-46 | 0.60681292 | 0.703 | 0.653 | 8.18E-42 | TAM.Cd209 | Arhgap45 |
| Chst141 | 7.26E-46 | 0.71859394 | 0.27 | 0.192 | 1.40E-41 | TAM.Cd209 | Chst14 |
| Hmg20a | 8.81E-46 | 0.83595997 | 0.318 | 0.226 | 1.70E-41 | TAM.Cd209 | Hmg20a |
| Ube2q2 | 1.41E-45 | 0.59568969 | 0.337 | 0.265 | 2.73E-41 | TAM.Cd209 | Ube2q2 |
| Zeb21 | 2.02E-45 | 0.31540992 | 0.889 | 0.904 | 3.90E-41 | TAM.Cd209 | Zeb2 |
| Trim30a2 | 2.87E-45 | 0.50662051 | 0.338 | 0.276 | 5.54E-41 | TAM.Cd209 | Trim30a |
| Tmem50a1 | 3.21E-45 | 0.32430455 | 0.694 | 0.725 | 6.21E-41 | TAM.Cd209 | Tmem50a |
| Crlf2 | 4.16E-45 | 0.7209181 | 0.627 | 0.555 | 8.04E-41 | TAM.Cd209 | Crlf2 |
| Vps391 | 5.43E-45 | 0.81224976 | 0.289 | 0.213 | 1.05E-40 | TAM.Cd209 | Vps39 |
| Mylip | 9.91E-45 | 0.64362301 | 0.396 | 0.304 | 1.92E-40 | TAM.Cd209 | Mylip |
| Kif3b2 | 2.57E-44 | 0.57854585 | 0.409 | 0.343 | 4.98E-40 | TAM.Cd209 | Kif3b |
| Apobec31 | 2.83E-44 | 0.70598023 | 0.468 | 0.402 | 5.48E-40 | TAM.Cd209 | Apobec3 |
| Lgals82 | 3.01E-44 | 0.40771058 | 0.469 | 0.422 | 5.81E-40 | TAM.Cd209 | Lgals8 |
| Asb81 | 3.92E-44 | 0.95604436 | 0.285 | 0.198 | 7.57E-40 | TAM.Cd209 | Asb8 |
| Rgs102 | 4.52E-44 | 0.45244425 | 0.702 | 0.666 | 8.73E-40 | TAM.Cd209 | Rgs10 |
| Rinl | 6.44E-44 | 0.8619693 | 0.31 | 0.224 | 1.25E-39 | TAM.Cd209 | Rinl |
| Mpv171 | 8.87E-44 | 0.84128079 | 0.299 | 0.227 | 1.72E-39 | TAM.Cd209 | Mpv17 |
| Scamp1 | 1.26E-43 | 0.49538549 | 0.345 | 0.287 | 2.44E-39 | TAM.Cd209 | Scamp1 |
| Cd300ld4 | 3.41E-43 | 0.47844058 | 0.55 | 0.522 | 6.58E-39 | TAM.Cd209 | Cd300ld |
| Slc7a82 | 6.55E-43 | 0.46998604 | 0.592 | 0.506 | 1.27E-38 | TAM.Cd209 | Slc7a8 |
| Nenf1 | 7.66E-43 | 0.75213583 | 0.372 | 0.287 | 1.48E-38 | TAM.Cd209 | Nenf |
| Exoc7 | 1.08E-42 | 0.69029948 | 0.345 | 0.281 | 2.08E-38 | TAM.Cd209 | Exoc7 |
| Dctn51 | 1.26E-42 | 0.94035341 | 0.387 | 0.297 | 2.43E-38 | TAM.Cd209 | Dctn5 |
| mt-Nd1 | 7.46E-42 | 0.49075071 | 0.536 | 0.592 | 1.44E-37 | TAM.Cd209 | mt-Nd1 |
| Plin21 | 1.23E-41 | 0.28565941 | 0.866 | 0.863 | 2.37E-37 | TAM.Cd209 | Plin2 |
| Ptger42 | 1.83E-41 | 0.46794164 | 0.46 | 0.359 | 3.53E-37 | TAM.Cd209 | Ptger4 |
| Dstn1 | 2.13E-41 | 0.45451593 | 0.631 | 0.6 | 4.12E-37 | TAM.Cd209 | Dstn |
| Phyh | 2.27E-41 | 0.62103113 | 0.285 | 0.219 | 4.38E-37 | TAM.Cd209 | Phyh |
| Pura | 4.15E-41 | 0.71861878 | 0.47 | 0.397 | 8.03E-37 | TAM.Cd209 | Pura |
| Nudt91 | 4.54E-41 | 0.64777747 | 0.458 | 0.379 | 8.78E-37 | TAM.Cd209 | Nudt9 |
| Parp142 | 9.92E-41 | 0.39124126 | 0.435 | 0.373 | 1.92E-36 | TAM.Cd209 | Parp14 |
| Marveld1 | 1.07E-40 | 1.20813938 | 0.326 | 0.24 | 2.06E-36 | TAM.Cd209 | Marveld1 |
| Npc12 | 2.00E-40 | 0.53663745 | 0.648 | 0.608 | 3.86E-36 | TAM.Cd209 | Npc1 |
| Bsg | 2.51E-40 | 0.30126035 | 0.501 | 0.501 | 4.86E-36 | TAM.Cd209 | Bsg |
| Galk21 | 2.62E-40 | 0.67755012 | 0.392 | 0.337 | 5.07E-36 | TAM.Cd209 | Galk2 |
| Mcoln1 | 2.73E-40 | 0.8974062 | 0.369 | 0.284 | 5.27E-36 | TAM.Cd209 | Mcoln1 |
| Klf32 | 3.94E-40 | 0.48585531 | 0.571 | 0.507 | 7.61E-36 | TAM.Cd209 | Klf3 |
| Gnptg1 | 4.27E-40 | 0.78537257 | 0.277 | 0.21 | 8.26E-36 | TAM.Cd209 | Gnptg |
| Lmbrd1 | 6.26E-40 | 0.44947525 | 0.476 | 0.442 | 1.21E-35 | TAM.Cd209 | Lmbrd1 |
| Gtf2i1 | 7.39E-40 | 0.76372884 | 0.446 | 0.359 | 1.43E-35 | TAM.Cd209 | Gtf2i |
| Amdhd21 | 1.54E-39 | 0.5902626 | 0.489 | 0.43 | 2.98E-35 | TAM.Cd209 | Amdhd2 |
| Mindy2 | 1.58E-39 | 0.53183801 | 0.415 | 0.359 | 3.05E-35 | TAM.Cd209 | Mindy2 |
| Cnpy21 | 2.29E-39 | 0.58580828 | 0.436 | 0.38 | 4.43E-35 | TAM.Cd209 | Cnpy2 |
| Zfp217 | 2.48E-39 | 0.6270934 | 0.399 | 0.315 | 4.80E-35 | TAM.Cd209 | Zfp217 |
| Naa60 | 2.78E-39 | 0.64562869 | 0.47 | 0.426 | 5.38E-35 | TAM.Cd209 | Naa60 |
| Slfn81 | 3.36E-39 | 0.41636477 | 0.261 | 0.202 | 6.50E-35 | TAM.Cd209 | Slfn8 |
| Znfx11 | 5.71E-39 | 0.54511789 | 0.318 | 0.265 | 1.10E-34 | TAM.Cd209 | Znfx1 |
| Clcn51 | 9.52E-39 | 0.73780902 | 0.364 | 0.285 | 1.84E-34 | TAM.Cd209 | Clcn5 |
| Inpp5d3 | 2.06E-38 | 0.28983464 | 0.748 | 0.776 | 3.98E-34 | TAM.Cd209 | Inpp5d |
| Pttg1ip1 | 2.94E-38 | 0.42166381 | 0.613 | 0.59 | 5.68E-34 | TAM.Cd209 | Pttg1ip |
| Pde3b | 3.57E-38 | 0.59990398 | 0.396 | 0.319 | 6.90E-34 | TAM.Cd209 | Pde3b |
| Rere | 4.33E-38 | 0.81594431 | 0.404 | 0.312 | 8.37E-34 | TAM.Cd209 | Rere |
| Atraid1 | 9.78E-38 | 0.52364779 | 0.387 | 0.324 | 1.89E-33 | TAM.Cd209 | Atraid |
| Ralbp1 | 1.69E-37 | 0.62683414 | 0.449 | 0.4 | 3.26E-33 | TAM.Cd209 | Ralbp1 |
| Washc41 | 2.60E-37 | 0.7047442 | 0.34 | 0.269 | 5.02E-33 | TAM.Cd209 | Washc4 |
| mt-Atp8 | 5.82E-37 | 0.54269594 | 0.78 | 0.779 | 1.13E-32 | TAM.Cd209 | mt-Atp8 |
| Peak1 | 6.57E-37 | 0.74848654 | 0.254 | 0.182 | 1.27E-32 | TAM.Cd209 | Peak1 |
| Os91 | 8.21E-37 | 0.54217186 | 0.621 | 0.578 | 1.59E-32 | TAM.Cd209 | Os9 |
| Zscan261 | 1.23E-36 | 0.86362469 | 0.379 | 0.292 | 2.37E-32 | TAM.Cd209 | Zscan26 |
| Osbpl1a | 1.30E-36 | 0.91624464 | 0.254 | 0.177 | 2.52E-32 | TAM.Cd209 | Osbpl1a |
| Rab29 | 2.43E-36 | 0.64884963 | 0.263 | 0.199 | 4.69E-32 | TAM.Cd209 | Rab29 |
| Sumf1 | 3.17E-36 | 0.45370174 | 0.497 | 0.471 | 6.13E-32 | TAM.Cd209 | Sumf1 |
| Tspyl1 | 3.76E-36 | 0.76551049 | 0.385 | 0.31 | 7.28E-32 | TAM.Cd209 | Tspyl1 |
| Glmp1 | 4.93E-36 | 0.43007785 | 0.72 | 0.707 | 9.52E-32 | TAM.Cd209 | Glmp |
| C2cd2l1 | 4.99E-36 | 0.73111352 | 0.387 | 0.313 | 9.64E-32 | TAM.Cd209 | C2cd2l |
| Atxn7l3b1 | 5.78E-36 | 0.60906273 | 0.457 | 0.378 | 1.12E-31 | TAM.Cd209 | Atxn7l3b |
| Alkbh5 | 6.03E-36 | 0.36972998 | 0.58 | 0.583 | 1.17E-31 | TAM.Cd209 | Alkbh5 |
| Slc35a51 | 8.00E-36 | 0.58984435 | 0.334 | 0.275 | 1.55E-31 | TAM.Cd209 | Slc35a5 |
| Akt1s11 | 8.11E-36 | 0.93803958 | 0.327 | 0.243 | 1.57E-31 | TAM.Cd209 | Akt1s1 |
| Mesd | 9.56E-36 | 0.69060403 | 0.328 | 0.257 | 1.85E-31 | TAM.Cd209 | Mesd |
| Vps33a1 | 1.11E-35 | 0.64317067 | 0.255 | 0.198 | 2.15E-31 | TAM.Cd209 | Vps33a |
| Zfp992 | 1.21E-35 | 0.46881235 | 0.32 | 0.271 | 2.33E-31 | TAM.Cd209 | Zfp992 |
| Zfp3621 | 1.93E-35 | 0.75984692 | 0.326 | 0.246 | 3.73E-31 | TAM.Cd209 | Zfp362 |
| Cd151 | 3.36E-35 | 0.7667959 | 0.337 | 0.259 | 6.50E-31 | TAM.Cd209 | Cd151 |
| Parp91 | 4.71E-35 | 0.46287221 | 0.278 | 0.232 | 9.10E-31 | TAM.Cd209 | Parp9 |
| Akr1b3 | 8.18E-35 | 0.47861161 | 0.428 | 0.399 | 1.58E-30 | TAM.Cd209 | Akr1b3 |
| Reep31 | 8.83E-35 | 0.33974299 | 0.518 | 0.508 | 1.71E-30 | TAM.Cd209 | Reep3 |
| Ntpcr1 | 9.70E-35 | 0.58486085 | 0.469 | 0.395 | 1.88E-30 | TAM.Cd209 | Ntpcr |
| Asah11 | 1.32E-34 | 0.3211025 | 0.666 | 0.69 | 2.55E-30 | TAM.Cd209 | Asah1 |
| Bscl21 | 1.61E-34 | 0.37419749 | 0.582 | 0.564 | 3.12E-30 | TAM.Cd209 | Bscl2 |
| Pja21 | 1.95E-34 | 0.73007566 | 0.283 | 0.217 | 3.78E-30 | TAM.Cd209 | Pja2 |
| Arhgap101 | 3.08E-34 | 0.78667564 | 0.284 | 0.194 | 5.95E-30 | TAM.Cd209 | Arhgap10 |
| Eif2ak21 | 3.42E-34 | 0.53278308 | 0.305 | 0.248 | 6.61E-30 | TAM.Cd209 | Eif2ak2 |
| Gaa1 | 3.77E-34 | 0.58251989 | 0.527 | 0.45 | 7.28E-30 | TAM.Cd209 | Gaa |
| Tmem2141 | 4.10E-34 | 0.68520583 | 0.358 | 0.293 | 7.93E-30 | TAM.Cd209 | Tmem214 |
| Rab322 | 4.18E-34 | 0.27582452 | 0.604 | 0.609 | 8.08E-30 | TAM.Cd209 | Rab32 |
| Sptssa1 | 5.92E-34 | 0.50496237 | 0.478 | 0.427 | 1.14E-29 | TAM.Cd209 | Sptssa |
| Pirb2 | 7.66E-34 | 0.28986258 | 0.784 | 0.817 | 1.48E-29 | TAM.Cd209 | Pirb |
| Snapin1 | 9.97E-34 | 0.65844894 | 0.286 | 0.225 | 1.93E-29 | TAM.Cd209 | Snapin |
| Pea15a1 | 1.58E-33 | 0.3764381 | 0.549 | 0.523 | 3.05E-29 | TAM.Cd209 | Pea15a |
| Snx111 | 2.55E-33 | 0.72604386 | 0.296 | 0.225 | 4.94E-29 | TAM.Cd209 | Snx11 |
| Nek61 | 2.88E-33 | 0.52252727 | 0.548 | 0.474 | 5.58E-29 | TAM.Cd209 | Nek6 |
| Aagab1 | 2.91E-33 | 0.69158556 | 0.483 | 0.408 | 5.62E-29 | TAM.Cd209 | Aagab |
| Blvrb1 | 4.08E-33 | 0.58346692 | 0.7 | 0.544 | 7.89E-29 | TAM.Cd209 | Blvrb |
| Rabac11 | 6.07E-33 | 0.34909479 | 0.686 | 0.68 | 1.17E-28 | TAM.Cd209 | Rabac1 |
| Tmem65 | 6.76E-33 | 0.4109159 | 0.506 | 0.449 | 1.31E-28 | TAM.Cd209 | Tmem65 |
| Cept11 | 1.08E-32 | 0.54437687 | 0.425 | 0.386 | 2.09E-28 | TAM.Cd209 | Cept1 |
| Vps13c | 2.53E-32 | 0.79370327 | 0.357 | 0.284 | 4.89E-28 | TAM.Cd209 | Vps13c |
| Ddrgk1 | 3.15E-32 | 0.55297193 | 0.545 | 0.49 | 6.09E-28 | TAM.Cd209 | Ddrgk1 |
| Arhgap182 | 5.06E-32 | 0.76403364 | 0.369 | 0.278 | 9.78E-28 | TAM.Cd209 | Arhgap18 |
| Rhob1 | 5.23E-32 | 0.32591986 | 0.791 | 0.761 | 1.01E-27 | TAM.Cd209 | Rhob |
| Wdr411 | 6.20E-32 | 0.7560918 | 0.416 | 0.347 | 1.20E-27 | TAM.Cd209 | Wdr41 |
| Gas2l1 | 2.00E-31 | 0.74135316 | 0.429 | 0.346 | 3.86E-27 | TAM.Cd209 | Gas2l1 |
| Cpt1a | 2.16E-31 | 0.54116979 | 0.435 | 0.391 | 4.17E-27 | TAM.Cd209 | Cpt1a |
| Flcn1 | 2.45E-31 | 0.58895354 | 0.6 | 0.53 | 4.74E-27 | TAM.Cd209 | Flcn |
| Abcc31 | 4.75E-31 | 0.50055854 | 0.525 | 0.443 | 9.18E-27 | TAM.Cd209 | Abcc3 |
| Phactr21 | 5.45E-31 | 0.66023663 | 0.272 | 0.214 | 1.05E-26 | TAM.Cd209 | Phactr2 |
| Pcbd2 | 7.62E-31 | 0.67395316 | 0.265 | 0.2 | 1.47E-26 | TAM.Cd209 | Pcbd2 |
| Tbc1d2b1 | 1.06E-30 | 0.50313058 | 0.476 | 0.431 | 2.05E-26 | TAM.Cd209 | Tbc1d2b |
| Sephs21 | 1.09E-30 | 0.46330141 | 0.431 | 0.363 | 2.11E-26 | TAM.Cd209 | Sephs2 |
| Akap8l1 | 1.99E-30 | 1.01383945 | 0.275 | 0.204 | 3.85E-26 | TAM.Cd209 | Akap8l |
| Idh11 | 2.24E-30 | 0.49058198 | 0.467 | 0.412 | 4.33E-26 | TAM.Cd209 | Idh1 |
| Washc11 | 4.28E-30 | 0.68731842 | 0.405 | 0.339 | 8.28E-26 | TAM.Cd209 | Washc1 |
| Cebpg1 | 7.47E-30 | 0.4831073 | 0.518 | 0.449 | 1.44E-25 | TAM.Cd209 | Cebpg |
| Fam174a | 1.57E-29 | 0.66693478 | 0.458 | 0.397 | 3.03E-25 | TAM.Cd209 | Fam174a |
| Cln61 | 1.69E-29 | 0.53044936 | 0.327 | 0.266 | 3.27E-25 | TAM.Cd209 | Cln6 |
| Ski | 1.79E-29 | 0.38438875 | 0.816 | 0.819 | 3.46E-25 | TAM.Cd209 | Ski |
| Cyp27a11 | 2.02E-29 | 0.86888505 | 0.324 | 0.251 | 3.90E-25 | TAM.Cd209 | Cyp27a1 |
| Smim151 | 2.15E-29 | 0.698841 | 0.335 | 0.271 | 4.16E-25 | TAM.Cd209 | Smim15 |
| Gbp91 | 2.17E-29 | 0.33413943 | 0.276 | 0.226 | 4.20E-25 | TAM.Cd209 | Gbp9 |
| Phax | 4.63E-29 | 0.65474932 | 0.298 | 0.238 | 8.95E-25 | TAM.Cd209 | Phax |
| Zfp292 | 6.01E-29 | 0.61033406 | 0.396 | 0.34 | 1.16E-24 | TAM.Cd209 | Zfp292 |
| Fblim11 | 6.13E-29 | 0.73962055 | 0.361 | 0.259 | 1.18E-24 | TAM.Cd209 | Fblim1 |
| Mier1 | 4.06E-28 | 0.49508545 | 0.487 | 0.435 | 7.86E-24 | TAM.Cd209 | Mier1 |
| Rb11 | 8.14E-28 | 0.61468624 | 0.379 | 0.325 | 1.57E-23 | TAM.Cd209 | Rb1 |
| Sphk2 | 9.40E-28 | 0.62749725 | 0.385 | 0.318 | 1.82E-23 | TAM.Cd209 | Sphk2 |
| Rell1 | 9.63E-28 | 0.50605923 | 0.334 | 0.267 | 1.86E-23 | TAM.Cd209 | Rell1 |
| Pxk1 | 1.31E-27 | 0.35721052 | 0.447 | 0.409 | 2.53E-23 | TAM.Cd209 | Pxk |
| Ccdc471 | 3.45E-27 | 0.49011553 | 0.356 | 0.312 | 6.67E-23 | TAM.Cd209 | Ccdc47 |
| D5Ertd579e | 4.67E-27 | 0.66248742 | 0.366 | 0.303 | 9.02E-23 | TAM.Cd209 | D5Ertd579e |
| Rbl21 | 5.68E-27 | 0.77641636 | 0.303 | 0.243 | 1.10E-22 | TAM.Cd209 | Rbl2 |
| Trappc6a | 7.16E-27 | 0.63218592 | 0.379 | 0.313 | 1.38E-22 | TAM.Cd209 | Trappc6a |
| Vipas391 | 1.01E-26 | 0.59744858 | 0.41 | 0.348 | 1.96E-22 | TAM.Cd209 | Vipas39 |
| Slc16a10 | 1.24E-26 | 0.63427546 | 0.355 | 0.287 | 2.39E-22 | TAM.Cd209 | Slc16a10 |
| Plin31 | 2.09E-26 | 0.49359547 | 0.448 | 0.396 | 4.05E-22 | TAM.Cd209 | Plin3 |
| Arhgef62 | 2.62E-26 | 0.56025386 | 0.387 | 0.33 | 5.06E-22 | TAM.Cd209 | Arhgef6 |
| Gab1 | 8.90E-26 | 0.38179704 | 0.398 | 0.323 | 1.72E-21 | TAM.Cd209 | Gab1 |
| Tmem2681 | 1.03E-25 | 0.65233319 | 0.29 | 0.232 | 1.99E-21 | TAM.Cd209 | Tmem268 |
| Akt2 | 1.83E-25 | 0.49674096 | 0.36 | 0.322 | 3.53E-21 | TAM.Cd209 | Akt2 |
| Rab91 | 1.87E-25 | 0.51190997 | 0.308 | 0.249 | 3.62E-21 | TAM.Cd209 | Rab9 |
| Atg4c | 1.94E-25 | 0.6285563 | 0.294 | 0.238 | 3.74E-21 | TAM.Cd209 | Atg4c |
| Smim4 | 2.42E-25 | 0.65298499 | 0.304 | 0.247 | 4.67E-21 | TAM.Cd209 | Smim4 |
| Ptpra | 2.67E-25 | 0.33161307 | 0.469 | 0.443 | 5.16E-21 | TAM.Cd209 | Ptpra |
| Rgl21 | 3.06E-25 | 0.72962347 | 0.543 | 0.464 | 5.91E-21 | TAM.Cd209 | Rgl2 |
| Pttg1 | 3.41E-25 | 0.51958592 | 0.3 | 0.25 | 6.60E-21 | TAM.Cd209 | Pttg1 |
| Spata6 | 6.52E-25 | 0.84991928 | 0.26 | 0.197 | 1.26E-20 | TAM.Cd209 | Spata6 |
| Cirbp | 8.90E-25 | 0.59623464 | 0.361 | 0.301 | 1.72E-20 | TAM.Cd209 | Cirbp |
| Rbfa2 | 9.79E-25 | 0.63047901 | 0.385 | 0.33 | 1.89E-20 | TAM.Cd209 | Rbfa |
| Ipmk1 | 1.26E-24 | 0.42853184 | 0.36 | 0.305 | 2.44E-20 | TAM.Cd209 | Ipmk |
| AI837181 | 1.39E-24 | 0.68505401 | 0.253 | 0.193 | 2.69E-20 | TAM.Cd209 | AI837181 |
| Egln2 | 1.49E-24 | 0.66556785 | 0.406 | 0.341 | 2.88E-20 | TAM.Cd209 | Egln2 |
| Dhdh1 | 2.09E-24 | 0.71200377 | 0.272 | 0.207 | 4.03E-20 | TAM.Cd209 | Dhdh |
| Cd481 | 2.16E-24 | 0.39116726 | 0.455 | 0.416 | 4.17E-20 | TAM.Cd209 | Cd48 |
| Trim411 | 2.35E-24 | 0.6012442 | 0.407 | 0.341 | 4.54E-20 | TAM.Cd209 | Trim41 |
| Cttnbp2nl1 | 2.76E-24 | 0.48512122 | 0.566 | 0.538 | 5.33E-20 | TAM.Cd209 | Cttnbp2nl |
| Nosip | 2.84E-24 | 0.52073304 | 0.306 | 0.255 | 5.49E-20 | TAM.Cd209 | Nosip |
| Prxl2c | 3.40E-24 | 0.72604012 | 0.256 | 0.196 | 6.57E-20 | TAM.Cd209 | Prxl2c |
| AI4135821 | 4.67E-24 | 0.50772753 | 0.287 | 0.242 | 9.03E-20 | TAM.Cd209 | AI413582 |
| Atp8a11 | 4.90E-24 | 0.5995847 | 0.53 | 0.462 | 9.47E-20 | TAM.Cd209 | Atp8a1 |
| Cryzl11 | 5.04E-24 | 0.7014082 | 0.264 | 0.209 | 9.74E-20 | TAM.Cd209 | Cryzl1 |
| Txndc151 | 7.54E-24 | 0.43573503 | 0.368 | 0.336 | 1.46E-19 | TAM.Cd209 | Txndc15 |
| Chmp1b | 7.84E-24 | 0.55024015 | 0.471 | 0.424 | 1.52E-19 | TAM.Cd209 | Chmp1b |
| Cab39 | 1.09E-23 | 0.39118717 | 0.659 | 0.646 | 2.12E-19 | TAM.Cd209 | Cab39 |
| Krt81 | 1.31E-23 | 0.83719607 | 0.306 | 0.245 | 2.53E-19 | TAM.Cd209 | Krt8 |
| Gnai2 | 1.46E-23 | 0.25133751 | 0.885 | 0.904 | 2.83E-19 | TAM.Cd209 | Gnai2 |
| Tlr4 | 3.26E-23 | 0.36965706 | 0.448 | 0.403 | 6.31E-19 | TAM.Cd209 | Tlr4 |
| Hbp1 | 3.76E-23 | 0.42373318 | 0.492 | 0.459 | 7.26E-19 | TAM.Cd209 | Hbp1 |
| Apc | 4.83E-23 | 0.61429805 | 0.337 | 0.282 | 9.34E-19 | TAM.Cd209 | Apc |
| Crat1 | 4.87E-23 | 0.62445939 | 0.309 | 0.253 | 9.42E-19 | TAM.Cd209 | Crat |
| Klf9 | 6.31E-23 | 0.65413373 | 0.267 | 0.211 | 1.22E-18 | TAM.Cd209 | Klf9 |
| Gna151 | 7.00E-23 | 0.62303891 | 0.308 | 0.247 | 1.35E-18 | TAM.Cd209 | Gna15 |
| Eng2 | 7.07E-23 | 0.63696763 | 0.327 | 0.264 | 1.37E-18 | TAM.Cd209 | Eng |
| Renbp2 | 1.01E-22 | 0.52136844 | 0.603 | 0.541 | 1.95E-18 | TAM.Cd209 | Renbp |
| mt-Co3 | 1.17E-22 | 0.54250087 | 0.736 | 0.746 | 2.26E-18 | TAM.Cd209 | mt-Co3 |
| Arl111 | 1.20E-22 | 0.55308333 | 0.374 | 0.302 | 2.33E-18 | TAM.Cd209 | Arl11 |
| Akap101 | 1.44E-22 | 0.69724332 | 0.306 | 0.244 | 2.79E-18 | TAM.Cd209 | Akap10 |
| H6pd1 | 1.46E-22 | 0.52795306 | 0.403 | 0.346 | 2.82E-18 | TAM.Cd209 | H6pd |
| Col4a3bp | 2.10E-22 | 0.50135872 | 0.458 | 0.402 | 4.07E-18 | TAM.Cd209 | Col4a3bp |
| Hagh1 | 2.64E-22 | 0.63748127 | 0.335 | 0.27 | 5.10E-18 | TAM.Cd209 | Hagh |
| Slc7a71 | 2.70E-22 | 0.49268941 | 0.325 | 0.283 | 5.22E-18 | TAM.Cd209 | Slc7a7 |
| Tppp32 | 3.21E-22 | 0.25676399 | 0.239 | 0.27 | 6.20E-18 | TAM.Cd209 | Tppp3 |
| Socs61 | 8.80E-22 | 0.46170405 | 0.353 | 0.294 | 1.70E-17 | TAM.Cd209 | Socs6 |
| Eri31 | 1.14E-21 | 0.76284355 | 0.258 | 0.201 | 2.20E-17 | TAM.Cd209 | Eri3 |
| Bet1 | 1.45E-21 | 0.62331016 | 0.276 | 0.225 | 2.81E-17 | TAM.Cd209 | Bet1 |
| Ppp1r12c1 | 1.52E-21 | 0.59393453 | 0.665 | 0.625 | 2.94E-17 | TAM.Cd209 | Ppp1r12c |
| Dpp72 | 3.65E-21 | 0.53600607 | 0.325 | 0.276 | 7.06E-17 | TAM.Cd209 | Dpp7 |
| Ift221 | 3.85E-21 | 0.6214528 | 0.27 | 0.218 | 7.44E-17 | TAM.Cd209 | Ift22 |
| Lmbrd21 | 4.72E-21 | 0.59185385 | 0.278 | 0.215 | 9.13E-17 | TAM.Cd209 | Lmbrd2 |
| mt-Co2 | 4.80E-21 | 0.62299015 | 0.893 | 0.88 | 9.27E-17 | TAM.Cd209 | mt-Co2 |
| Chmp5 | 6.64E-21 | 0.29671374 | 0.535 | 0.535 | 1.28E-16 | TAM.Cd209 | Chmp5 |
| Fam102b1 | 7.11E-21 | 0.37697699 | 0.41 | 0.36 | 1.37E-16 | TAM.Cd209 | Fam102b |
| Vav21 | 1.53E-20 | 0.57254631 | 0.365 | 0.303 | 2.95E-16 | TAM.Cd209 | Vav2 |
| Tm9sf21 | 4.07E-20 | 0.3982499 | 0.514 | 0.501 | 7.87E-16 | TAM.Cd209 | Tm9sf2 |
| Clock | 5.22E-20 | 0.56933485 | 0.251 | 0.2 | 1.01E-15 | TAM.Cd209 | Clock |
| Map1s | 7.43E-20 | 0.7426826 | 0.351 | 0.29 | 1.44E-15 | TAM.Cd209 | Map1s |
| Gga31 | 8.05E-20 | 0.69521504 | 0.276 | 0.222 | 1.56E-15 | TAM.Cd209 | Gga3 |
| Nek71 | 8.28E-20 | 0.42968741 | 0.428 | 0.384 | 1.60E-15 | TAM.Cd209 | Nek7 |
| Morc31 | 8.30E-20 | 0.51055917 | 0.406 | 0.369 | 1.60E-15 | TAM.Cd209 | Morc3 |
| Acot131 | 1.03E-19 | 0.60924307 | 0.284 | 0.231 | 1.99E-15 | TAM.Cd209 | Acot13 |
| Etnk11 | 1.51E-19 | 0.44747073 | 0.432 | 0.396 | 2.92E-15 | TAM.Cd209 | Etnk1 |
| Itpkb | 2.59E-19 | 0.60854892 | 0.429 | 0.375 | 5.00E-15 | TAM.Cd209 | Itpkb |
| Rbbp8 | 3.07E-19 | 0.55159926 | 0.278 | 0.23 | 5.93E-15 | TAM.Cd209 | Rbbp8 |
| H2-M3 | 3.09E-19 | 0.47678387 | 0.441 | 0.408 | 5.98E-15 | TAM.Cd209 | H2-M3 |
| Homer31 | 3.68E-19 | 0.74131707 | 0.268 | 0.203 | 7.12E-15 | TAM.Cd209 | Homer3 |
| Arl8a | 4.30E-19 | 0.36793274 | 0.666 | 0.671 | 8.32E-15 | TAM.Cd209 | Arl8a |
| Hmbox1 | 4.55E-19 | 0.714737 | 0.307 | 0.254 | 8.79E-15 | TAM.Cd209 | Hmbox1 |
| Ppp3ca | 4.75E-19 | 0.54985967 | 0.481 | 0.434 | 9.19E-15 | TAM.Cd209 | Ppp3ca |
| Vps411 | 4.76E-19 | 0.59598875 | 0.321 | 0.273 | 9.21E-15 | TAM.Cd209 | Vps41 |
| Nkiras21 | 4.92E-19 | 0.49274992 | 0.413 | 0.371 | 9.50E-15 | TAM.Cd209 | Nkiras2 |
| Retreg3 | 5.73E-19 | 0.42365975 | 0.4 | 0.353 | 1.11E-14 | TAM.Cd209 | Retreg3 |
| Ifi2032 | 6.78E-19 | 0.28803233 | 0.314 | 0.257 | 1.31E-14 | TAM.Cd209 | Ifi203 |
| Zhx1 | 1.03E-18 | 0.55306192 | 0.271 | 0.222 | 1.99E-14 | TAM.Cd209 | Zhx1 |
| Zfyve1 | 1.08E-18 | 0.57571201 | 0.268 | 0.211 | 2.09E-14 | TAM.Cd209 | Zfyve1 |
| Use11 | 1.66E-18 | 0.31715939 | 0.553 | 0.541 | 3.21E-14 | TAM.Cd209 | Use1 |
| Nudt16l11 | 1.68E-18 | 0.59682609 | 0.396 | 0.339 | 3.24E-14 | TAM.Cd209 | Nudt16l1 |
| Ccng2 | 1.84E-18 | 0.40080445 | 0.393 | 0.348 | 3.57E-14 | TAM.Cd209 | Ccng2 |
| Chmp6 | 2.21E-18 | 0.54252914 | 0.275 | 0.223 | 4.26E-14 | TAM.Cd209 | Chmp6 |
| Retreg21 | 2.22E-18 | 0.55494753 | 0.469 | 0.415 | 4.29E-14 | TAM.Cd209 | Retreg2 |
| Insr | 2.35E-18 | 0.78401165 | 0.266 | 0.208 | 4.54E-14 | TAM.Cd209 | Insr |
| Tec | 2.63E-18 | 0.38471486 | 0.386 | 0.353 | 5.09E-14 | TAM.Cd209 | Tec |
| Dnajb9 | 3.64E-18 | 0.3191495 | 0.385 | 0.376 | 7.04E-14 | TAM.Cd209 | Dnajb9 |
| Shb | 4.13E-18 | 0.86950014 | 0.29 | 0.237 | 7.99E-14 | TAM.Cd209 | Shb |
| Rnf181 | 4.46E-18 | 0.45902947 | 0.308 | 0.269 | 8.63E-14 | TAM.Cd209 | Rnf181 |
| Wdr71 | 5.54E-18 | 0.77165253 | 0.311 | 0.258 | 1.07E-13 | TAM.Cd209 | Wdr7 |
| Chmp3 | 6.39E-18 | 0.47653075 | 0.508 | 0.475 | 1.24E-13 | TAM.Cd209 | Chmp3 |
| Prr121 | 1.06E-17 | 0.58016687 | 0.333 | 0.274 | 2.05E-13 | TAM.Cd209 | Prr12 |
| Vkorc11 | 1.24E-17 | 0.60105416 | 0.293 | 0.232 | 2.40E-13 | TAM.Cd209 | Vkorc1 |
| Wbp21 | 1.34E-17 | 0.42836114 | 0.464 | 0.427 | 2.60E-13 | TAM.Cd209 | Wbp2 |
| Borcs61 | 1.38E-17 | 0.6700266 | 0.35 | 0.292 | 2.68E-13 | TAM.Cd209 | Borcs6 |
| Numa1 | 2.59E-17 | 0.51649116 | 0.431 | 0.376 | 5.02E-13 | TAM.Cd209 | Numa1 |
| C5ar21 | 3.71E-17 | 0.35176517 | 0.325 | 0.274 | 7.18E-13 | TAM.Cd209 | C5ar2 |

| | | | | | | | |
|---|---|---|---|---|---|---|---|
| Rab3gap1 | 4.00E-17 | 0.56720735 | 0.284 | 0.233 | 7.73E-13 | TAM.Cd209 | Rab3gap1 |
| Ncoa42 | 4.10E-17 | 0.30892341 | 0.547 | 0.53 | 7.92E-13 | TAM.Cd209 | Ncoa4 |
| Vti1b | 4.18E-17 | 0.43133425 | 0.385 | 0.348 | 8.08E-13 | TAM.Cd209 | Vti1b |
| Parp121 | 4.70E-17 | 0.36764765 | 0.321 | 0.297 | 9.09E-13 | TAM.Cd209 | Parp12 |
| Pdlim42 | 5.92E-17 | 0.52239101 | 0.326 | 0.273 | 1.15E-12 | TAM.Cd209 | Pdlim4 |
| Abhd42 | 6.27E-17 | 0.55272208 | 0.336 | 0.284 | 1.21E-12 | TAM.Cd209 | Abhd4 |
| Txndc17 | 6.37E-17 | 0.45313912 | 0.636 | 0.621 | 1.23E-12 | TAM.Cd209 | Txndc17 |
| Keap1 | 1.15E-16 | 0.66420176 | 0.372 | 0.312 | 2.22E-12 | TAM.Cd209 | Keap1 |
| Ppp1cc1 | 1.56E-16 | 0.56532927 | 0.411 | 0.363 | 3.02E-12 | TAM.Cd209 | Ppp1cc |
| Slc25a36 | 1.67E-16 | 0.60946575 | 0.343 | 0.298 | 3.23E-12 | TAM.Cd209 | Slc25a36 |
| Fis11 | 1.75E-16 | 0.2716974 | 0.642 | 0.661 | 3.38E-12 | TAM.Cd209 | Fis1 |
| Galk1 | 3.17E-16 | 0.64016558 | 0.292 | 0.239 | 6.12E-12 | TAM.Cd209 | Galk1 |
| Trp53inp21 | 3.93E-16 | 0.57789301 | 0.35 | 0.297 | 7.59E-12 | TAM.Cd209 | Trp53inp2 |
| Ppp1r211 | 4.65E-16 | 0.62601048 | 0.401 | 0.346 | 8.99E-12 | TAM.Cd209 | Ppp1r21 |
| Pnisr | 5.56E-16 | 0.84311997 | 0.389 | 0.34 | 1.08E-11 | TAM.Cd209 | Pnisr |
| Serinc1 | 6.36E-16 | 0.27483488 | 0.525 | 0.529 | 1.23E-11 | TAM.Cd209 | Serinc1 |
| Fam214b1 | 1.20E-15 | 0.46040917 | 0.38 | 0.325 | 2.31E-11 | TAM.Cd209 | Fam214b |
| Ago11 | 1.36E-15 | 0.58756297 | 0.297 | 0.253 | 2.62E-11 | TAM.Cd209 | Ago1 |
| Hsd3b7 | 1.37E-15 | 0.5986258 | 0.347 | 0.288 | 2.64E-11 | TAM.Cd209 | Hsd3b7 |
| Rnf44 | 1.90E-15 | 0.33819909 | 0.572 | 0.557 | 3.67E-11 | TAM.Cd209 | Rnf44 |
| Mertk2 | 3.28E-15 | 0.48002308 | 0.658 | 0.56 | 6.33E-11 | TAM.Cd209 | Mertk |
| Dnajb1 | 4.48E-15 | 0.28192252 | 0.376 | 0.4 | 8.67E-11 | TAM.Cd209 | Dnajb1 |
| Lpin2 | 5.04E-15 | 0.43878126 | 0.347 | 0.312 | 9.74E-11 | TAM.Cd209 | Lpin2 |
| Vps81 | 6.56E-15 | 0.61893392 | 0.353 | 0.307 | 1.27E-10 | TAM.Cd209 | Vps8 |
| Saraf1 | 7.09E-15 | 0.31667085 | 0.622 | 0.615 | 1.37E-10 | TAM.Cd209 | Saraf |
| Washc3 | 7.62E-15 | 0.50194361 | 0.3 | 0.259 | 1.47E-10 | TAM.Cd209 | Washc3 |
| Naxd | 8.67E-15 | 0.49390686 | 0.264 | 0.218 | 1.68E-10 | TAM.Cd209 | Naxd |
| Lyz11 | 1.03E-14 | 0.63181519 | 0.354 | 0.306 | 1.99E-10 | TAM.Cd209 | Lyz1 |
| Pdrg1 | 1.28E-14 | 0.66551026 | 0.277 | 0.222 | 2.47E-10 | TAM.Cd209 | Pdrg1 |
| Erlin21 | 1.35E-14 | 0.43379985 | 0.334 | 0.302 | 2.62E-10 | TAM.Cd209 | Erlin2 |
| Swap701 | 1.45E-14 | 0.4624983 | 0.405 | 0.357 | 2.81E-10 | TAM.Cd209 | Swap70 |
| Sbds | 1.70E-14 | 0.3221777 | 0.42 | 0.4 | 3.28E-10 | TAM.Cd209 | Sbds |
| Blvra1 | 1.78E-14 | 0.35932136 | 0.479 | 0.467 | 3.44E-10 | TAM.Cd209 | Blvra |
| Man2a21 | 1.96E-14 | 0.59703403 | 0.502 | 0.443 | 3.78E-10 | TAM.Cd209 | Man2a2 |
| Rab24 | 2.23E-14 | 0.46084101 | 0.523 | 0.489 | 4.32E-10 | TAM.Cd209 | Rab24 |
| Creld12 | 2.40E-14 | 0.60877592 | 0.256 | 0.206 | 4.64E-10 | TAM.Cd209 | Creld1 |
| Commd101 | 3.25E-14 | 0.51043153 | 0.333 | 0.296 | 6.28E-10 | TAM.Cd209 | Commd10 |
| Gga1 | 4.72E-14 | 0.34632042 | 0.567 | 0.563 | 9.12E-10 | TAM.Cd209 | Gga1 |
| Vamp8 | 4.90E-14 | 0.25975459 | 0.778 | 0.814 | 9.47E-10 | TAM.Cd209 | Vamp8 |
| Mettl9 | 5.73E-14 | 0.37780726 | 0.373 | 0.334 | 1.11E-09 | TAM.Cd209 | Mettl9 |
| mt-Atp6 | 6.74E-14 | 0.61105186 | 0.797 | 0.775 | 1.30E-09 | TAM.Cd209 | mt-Atp6 |
| Ext1 | 7.39E-14 | 0.40107359 | 0.487 | 0.453 | 1.43E-09 | TAM.Cd209 | Ext1 |
| Rufy32 | 1.45E-13 | 0.53774577 | 0.321 | 0.277 | 2.81E-09 | TAM.Cd209 | Rufy3 |
| Rnf187 | 1.92E-13 | 0.4402331 | 0.668 | 0.642 | 3.72E-09 | TAM.Cd209 | Rnf187 |
| Hs2st1 | 1.97E-13 | 0.42594165 | 0.289 | 0.248 | 3.81E-09 | TAM.Cd209 | Hs2st1 |
| Atf7 | 2.61E-13 | 0.45856417 | 0.359 | 0.321 | 5.05E-09 | TAM.Cd209 | Atf7 |
| Pggt1b1 | 3.59E-13 | 0.56918542 | 0.29 | 0.245 | 6.94E-09 | TAM.Cd209 | Pggt1b |
| Snap232 | 4.11E-13 | 0.29412723 | 0.305 | 0.28 | 7.95E-09 | TAM.Cd209 | Snap23 |
| Slc25a101 | 4.18E-13 | 0.72388461 | 0.301 | 0.232 | 8.07E-09 | TAM.Cd209 | Slc25a10 |
| Pgrmc2 | 5.38E-13 | 0.49674093 | 0.335 | 0.287 | 1.04E-08 | TAM.Cd209 | Pgrmc2 |
| Tbc1d131 | 7.40E-13 | 0.46008342 | 0.402 | 0.369 | 1.43E-08 | TAM.Cd209 | Tbc1d13 |
| Camkk2 | 8.94E-13 | 0.57088537 | 0.259 | 0.217 | 1.73E-08 | TAM.Cd209 | Camkk2 |
| Wbp1l | 9.79E-13 | 0.34437716 | 0.5 | 0.495 | 1.89E-08 | TAM.Cd209 | Wbp1l |
| Usp24 | 1.01E-12 | 0.69805005 | 0.315 | 0.269 | 1.94E-08 | TAM.Cd209 | Usp24 |
| Ddx17 | 1.20E-12 | 0.26894238 | 0.7 | 0.743 | 2.32E-08 | TAM.Cd209 | Ddx17 |
| Rab18 | 1.28E-12 | 0.30026905 | 0.569 | 0.578 | 2.48E-08 | TAM.Cd209 | Rab18 |
| Adcy71 | 1.52E-12 | 0.25526577 | 0.555 | 0.588 | 2.94E-08 | TAM.Cd209 | Adcy7 |
| Stard52 | 2.00E-12 | 0.47428882 | 0.269 | 0.224 | 3.86E-08 | TAM.Cd209 | Stard5 |
| Trappc3 | 2.03E-12 | 0.37876861 | 0.447 | 0.42 | 3.92E-08 | TAM.Cd209 | Trappc3 |
| Ech1 | 2.52E-12 | 0.39608091 | 0.422 | 0.395 | 4.88E-08 | TAM.Cd209 | Ech1 |
| Fuca1 | 2.55E-12 | 0.28110516 | 0.504 | 0.5 | 4.94E-08 | TAM.Cd209 | Fuca1 |
| Ano6 | 3.44E-12 | 0.36224331 | 0.428 | 0.397 | 6.66E-08 | TAM.Cd209 | Ano6 |
| Tnfaip81 | 3.85E-12 | 0.27507797 | 0.491 | 0.48 | 7.44E-08 | TAM.Cd209 | Tnfaip8 |
| Atp11c | 4.05E-12 | 0.53411411 | 0.288 | 0.251 | 7.83E-08 | TAM.Cd209 | Atp11c |
| Fadd1 | 4.34E-12 | 0.70312333 | 0.261 | 0.215 | 8.39E-08 | TAM.Cd209 | Fadd |
| Map3k82 | 4.48E-12 | 0.4559912 | 0.345 | 0.291 | 8.67E-08 | TAM.Cd209 | Map3k8 |
| Add1 | 4.63E-12 | 0.40383832 | 0.523 | 0.503 | 8.94E-08 | TAM.Cd209 | Add1 |
| Fam173a1 | 6.19E-12 | 0.65943489 | 0.385 | 0.334 | 1.20E-07 | TAM.Cd209 | Fam173a |
| Stim1 | 7.69E-12 | 0.44409076 | 0.303 | 0.265 | 1.49E-07 | TAM.Cd209 | Stim1 |
| Rnf5 | 8.39E-12 | 0.49834783 | 0.304 | 0.267 | 1.62E-07 | TAM.Cd209 | Rnf5 |
| Chd91 | 8.85E-12 | 0.43894489 | 0.304 | 0.258 | 1.71E-07 | TAM.Cd209 | Chd9 |
| Cyth32 | 1.04E-11 | 0.44651957 | 0.358 | 0.312 | 2.01E-07 | TAM.Cd209 | Cyth3 |
| Tmem9b1 | 1.14E-11 | 0.36429912 | 0.362 | 0.338 | 2.20E-07 | TAM.Cd209 | Tmem9b |
| Ctnnd11 | 1.31E-11 | 0.41949002 | 0.569 | 0.535 | 2.53E-07 | TAM.Cd209 | Ctnnd1 |
| Golgb1 | 1.35E-11 | 0.62436799 | 0.313 | 0.265 | 2.61E-07 | TAM.Cd209 | Golgb1 |
| H2-Ke6 | 1.77E-11 | 0.49318275 | 0.27 | 0.235 | 3.42E-07 | TAM.Cd209 | H2-Ke6 |
| Ccdc6 | 2.08E-11 | 0.57133542 | 0.366 | 0.322 | 4.02E-07 | TAM.Cd209 | Ccdc6 |
| Tm6sf12 | 3.73E-11 | 0.27323952 | 0.656 | 0.668 | 7.21E-07 | TAM.Cd209 | Tm6sf1 |
| Chfr1 | 3.81E-11 | 0.36418316 | 0.229 | 0.27 | 7.37E-07 | TAM.Cd209 | Chfr |
| Gmfb1 | 4.17E-11 | 0.33133774 | 0.542 | 0.534 | 8.07E-07 | TAM.Cd209 | Gmfb |
| Pcm1 | 4.45E-11 | 0.54905941 | 0.281 | 0.244 | 8.60E-07 | TAM.Cd209 | Pcm1 |
| Rabep1 | 5.76E-11 | 0.4499439 | 0.561 | 0.532 | 1.11E-06 | TAM.Cd209 | Rabep1 |
| Tm4sf1 | 5.97E-11 | 0.82974679 | 0.327 | 0.269 | 1.15E-06 | TAM.Cd209 | Tm4sf1 |
| Abcc51 | 6.29E-11 | 0.47934255 | 0.314 | 0.276 | 1.22E-06 | TAM.Cd209 | Abcc5 |
| Stard91 | 6.84E-11 | 0.679788 | 0.264 | 0.224 | 1.32E-06 | TAM.Cd209 | Stard9 |
| Rnf169 | 8.22E-11 | 0.43504226 | 0.352 | 0.316 | 1.59E-06 | TAM.Cd209 | Rnf169 |
| Surf1 | 8.87E-11 | 0.5341458 | 0.251 | 0.206 | 1.72E-06 | TAM.Cd209 | Surf1 |
| Ppm1h1 | 1.08E-10 | 0.4979272 | 0.312 | 0.283 | 2.08E-06 | TAM.Cd209 | Ppm1h |
| Slc23a22 | 1.10E-10 | 0.32462371 | 0.379 | 0.344 | 2.13E-06 | TAM.Cd209 | Slc23a2 |
| Klc1 | 1.20E-10 | 0.69287554 | 0.355 | 0.315 | 2.32E-06 | TAM.Cd209 | Klc1 |
| Ak3 | 1.34E-10 | 0.42832077 | 0.318 | 0.273 | 2.60E-06 | TAM.Cd209 | Ak3 |
| Exoc4 | 1.39E-10 | 0.69539287 | 0.422 | 0.377 | 2.68E-06 | TAM.Cd209 | Exoc4 |
| Sipa1l2 | 2.33E-10 | 0.47963315 | 0.258 | 0.219 | 4.51E-06 | TAM.Cd209 | Sipa1l2 |
| Taz | 2.36E-10 | 0.43023229 | 0.327 | 0.297 | 4.57E-06 | TAM.Cd209 | Taz |
| Mdfic1 | 2.58E-10 | 0.34311637 | 0.536 | 0.521 | 4.98E-06 | TAM.Cd209 | Mdfic |
| Pkig | 2.59E-10 | 0.57249114 | 0.35 | 0.301 | 5.01E-06 | TAM.Cd209 | Pkig |
| Stk25 | 2.69E-10 | 0.54610279 | 0.356 | 0.323 | 5.21E-06 | TAM.Cd209 | Stk25 |
| Tprgl | 2.86E-10 | 0.41696801 | 0.327 | 0.297 | 5.54E-06 | TAM.Cd209 | Tprgl |
| Znrf11 | 2.92E-10 | 0.39451163 | 0.374 | 0.336 | 5.65E-06 | TAM.Cd209 | Znrf1 |
| Eif4a2 | 3.29E-10 | 0.39181903 | 0.65 | 0.638 | 6.36E-06 | TAM.Cd209 | Eif4a2 |
| Daglb2 | 4.64E-10 | 0.37645252 | 0.539 | 0.504 | 8.97E-06 | TAM.Cd209 | Daglb |
| Coa5 | 4.93E-10 | 0.39870366 | 0.492 | 0.482 | 9.53E-06 | TAM.Cd209 | Coa5 |
| Fbxo221 | 5.14E-10 | 0.44970026 | 0.257 | 0.224 | 9.93E-06 | TAM.Cd209 | Fbxo22 |
| Atg4b | 5.49E-10 | 0.63715321 | 0.282 | 0.242 | 1.06E-05 | TAM.Cd209 | Atg4b |
| Ino80b | 5.63E-10 | 0.5843971 | 0.304 | 0.265 | 1.09E-05 | TAM.Cd209 | Ino80b |
| Phf21a | 5.88E-10 | 0.29766354 | 0.292 | 0.263 | 1.14E-05 | TAM.Cd209 | Phf21a |
| Htatip21 | 8.21E-10 | 0.28724041 | 0.332 | 0.315 | 1.59E-05 | TAM.Cd209 | Htatip2 |
| Trim561 | 8.84E-10 | 0.51821928 | 0.289 | 0.247 | 1.71E-05 | TAM.Cd209 | Trim56 |
| Eea1 | 9.06E-10 | 0.34920572 | 0.367 | 0.334 | 1.75E-05 | TAM.Cd209 | Eea1 |
| Ppox1 | 9.86E-10 | 0.76543689 | 0.251 | 0.208 | 1.91E-05 | TAM.Cd209 | Ppox |
| Axxl2 | 9.95E-10 | 0.48568839 | 0.311 | 0.267 | 1.92E-05 | TAM.Cd209 | Axxl2 |
| Nipa2 | 1.63E-09 | 0.27212521 | 0.428 | 0.429 | 3.14E-05 | TAM.Cd209 | Nipa2 |
| Sdf2 | 1.71E-09 | 0.4083174 | 0.424 | 0.394 | 3.31E-05 | TAM.Cd209 | Sdf2 |
| Ocrl | 2.83E-09 | 0.57196827 | 0.291 | 0.25 | 5.47E-05 | TAM.Cd209 | Ocrl |
| BC0297221 | 2.86E-09 | 0.57865139 | 0.272 | 0.234 | 5.52E-05 | TAM.Cd209 | BC029722 |
| Chrac1 | 2.87E-09 | 0.4480087 | 0.261 | 0.231 | 5.56E-05 | TAM.Cd209 | Chrac1 |
| Slc36a11 | 3.58E-09 | 0.66252465 | 0.498 | 0.435 | 6.92E-05 | TAM.Cd209 | Slc36a1 |
| Morf4l11 | 3.88E-09 | 0.30562703 | 0.707 | 0.73 | 7.50E-05 | TAM.Cd209 | Morf4l1 |
| S100a12 | 4.13E-09 | 0.38583349 | 0.48 | 0.447 | 7.98E-05 | TAM.Cd209 | S100a1 |
| 6-Mar | 4.44E-09 | 0.42765534 | 0.477 | 0.457 | 8.58E-05 | TAM.Cd209 | 6-Mar |
| Oaz2 | 5.82E-09 | 0.39182223 | 0.326 | 0.295 | 0.00011249 | TAM.Cd209 | Oaz2 |
| Elf21 | 6.06E-09 | 0.33507423 | 0.37 | 0.342 | 0.00011721 | TAM.Cd209 | Elf2 |
| Pik3c3 | 6.47E-09 | 0.55559822 | 0.261 | 0.23 | 0.000125 | TAM.Cd209 | Pik3c3 |
| Hectd31 | 7.09E-09 | 0.52737007 | 0.335 | 0.291 | 0.00013707 | TAM.Cd209 | Hectd3 |
| Gmfg1 | 7.35E-09 | 0.25914091 | 0.422 | 0.416 | 0.00014214 | TAM.Cd209 | Gmfg |
| Fgd41 | 8.07E-09 | 0.60046892 | 0.28 | 0.234 | 0.00015602 | TAM.Cd209 | Fgd4 |
| Ctps21 | 8.21E-09 | 0.48790032 | 0.37 | 0.336 | 0.00015878 | TAM.Cd209 | Ctps2 |
| Nceh11 | 9.89E-09 | 0.31025315 | 0.429 | 0.389 | 0.00019127 | TAM.Cd209 | Nceh1 |
| Cd382 | 9.95E-09 | 0.29233924 | 0.506 | 0.453 | 0.00019243 | TAM.Cd209 | Cd38 |
| Slc41a21 | 1.03E-08 | 0.54850107 | 0.264 | 0.202 | 0.00019918 | TAM.Cd209 | Slc41a2 |
| Nck11 | 1.08E-08 | 0.40066256 | 0.27 | 0.23 | 0.00020924 | TAM.Cd209 | Nck1 |
| Tifa2 | 1.24E-08 | 0.33854344 | 0.313 | 0.282 | 0.00023921 | TAM.Cd209 | Tifa |
| Gigyf1 | 1.44E-08 | 0.70419649 | 0.259 | 0.216 | 0.00027929 | TAM.Cd209 | Gigyf1 |
| Paip11 | 1.62E-08 | 0.30533932 | 0.374 | 0.359 | 0.00031243 | TAM.Cd209 | Paip1 |
| R3hdm4 | 2.04E-08 | 0.37301178 | 0.49 | 0.474 | 0.00039492 | TAM.Cd209 | R3hdm4 |
| Fam193b | 2.12E-08 | 0.73345021 | 0.348 | 0.311 | 0.00041074 | TAM.Cd209 | Fam193b |
| Zmynd11 | 2.16E-08 | 0.51142969 | 0.44 | 0.407 | 0.000417 | TAM.Cd209 | Zmynd11 |
| Dnajc31 | 2.35E-08 | 0.34259905 | 0.563 | 0.555 | 0.0004543 | TAM.Cd209 | Dnajc3 |
| Zmynd8 | 2.72E-08 | 0.45394382 | 0.279 | 0.247 | 0.00052667 | TAM.Cd209 | Zmynd8 |
| mt-Nd4 | 2.89E-08 | 0.5570812 | 0.379 | 0.414 | 0.00055946 | TAM.Cd209 | mt-Nd4 |
| Rreb11 | 3.29E-08 | 0.29160519 | 0.561 | 0.551 | 0.00063518 | TAM.Cd209 | Rreb1 |
| Fech | 4.08E-08 | 0.37125919 | 0.265 | 0.224 | 0.00078846 | TAM.Cd209 | Fech |
| Ppm1a | 4.58E-08 | 0.27284339 | 0.502 | 0.499 | 0.00088499 | TAM.Cd209 | Ppm1a |
| Selenow1 | 4.77E-08 | 0.29725046 | 0.65 | 0.668 | 0.00092158 | TAM.Cd209 | Selenow |
| Arid3a2 | 5.18E-08 | 0.51522613 | 0.282 | 0.243 | 0.0010017 | TAM.Cd209 | Arid3a |
| Cox14 | 5.22E-08 | 0.29002396 | 0.543 | 0.536 | 0.00100916 | TAM.Cd209 | Cox14 |
| Vgll4 | 5.63E-08 | 0.26448126 | 0.434 | 0.406 | 0.00108824 | TAM.Cd209 | Vgll4 |
| Mmd1 | 6.18E-08 | 0.33228531 | 0.279 | 0.251 | 0.00119399 | TAM.Cd209 | Mmd |
| Smim14 | 7.02E-08 | 0.3495641 | 0.427 | 0.404 | 0.00135679 | TAM.Cd209 | Smim14 |
| Cltc2 | 1.05E-07 | 0.29665525 | 0.648 | 0.684 | 0.00203434 | TAM.Cd209 | Cltc |
| Arl6ip11 | 1.08E-07 | 0.27679555 | 0.377 | 0.36 | 0.00209566 | TAM.Cd209 | Arl6ip1 |
| Gapvd11 | 1.38E-07 | 0.49312575 | 0.453 | 0.433 | 0.00266627 | TAM.Cd209 | Gapvd1 |
| Nipsnap21 | 1.52E-07 | 0.46757294 | 0.298 | 0.267 | 0.00293603 | TAM.Cd209 | Nipsnap2 |
| Selenof | 1.61E-07 | 0.27712315 | 0.313 | 0.296 | 0.0031219 | TAM.Cd209 | Selenof |
| Atl3 | 1.77E-07 | 0.38508278 | 0.536 | 0.52 | 0.00342004 | TAM.Cd209 | Atl3 |
| Dctn6 | 1.83E-07 | 0.31491884 | 0.277 | 0.254 | 0.00353104 | TAM.Cd209 | Dctn6 |
| Mocs11 | 1.84E-07 | 0.62193089 | 0.289 | 0.257 | 0.00356186 | TAM.Cd209 | Mocs1 |
| Add31 | 1.85E-07 | 0.25445058 | 0.297 | 0.256 | 0.00358186 | TAM.Cd209 | Add3 |
| Trim142 | 1.85E-07 | 0.60775709 | 0.327 | 0.292 | 0.00358467 | TAM.Cd209 | Trim14 |
| Ncbp2 | 2.19E-07 | 0.56759178 | 0.323 | 0.289 | 0.00423401 | TAM.Cd209 | Ncbp2 |
| Chkb | 2.27E-07 | 0.58326452 | 0.391 | 0.354 | 0.00439399 | TAM.Cd209 | Chkb |
| Pdk1 | 2.38E-07 | 0.31539905 | 0.335 | 0.31 | 0.00460487 | TAM.Cd209 | Pdk1 |
| Tmed4 | 2.88E-07 | 0.4222058 | 0.385 | 0.359 | 0.00556879 | TAM.Cd209 | Tmed4 |
| Dpp8 | 3.19E-07 | 0.43300668 | 0.458 | 0.432 | 0.00615944 | TAM.Cd209 | Dpp8 |
| Tnfsf131 | 3.20E-07 | 0.39897841 | 0.465 | 0.437 | 0.00619337 | TAM.Cd209 | Tnfsf13 |
| Herpud2 | 3.22E-07 | 0.31774083 | 0.25 | 0.227 | 0.00623111 | TAM.Cd209 | Herpud2 |
| Ift52 | 3.24E-07 | 0.49991984 | 0.27 | 0.237 | 0.00627041 | TAM.Cd209 | Ift52 |
| Ergic3 | 3.44E-07 | 0.51766743 | 0.418 | 0.389 | 0.00665885 | TAM.Cd209 | Ergic3 |
| Slc38a72 | 3.72E-07 | 0.40772269 | 0.274 | 0.243 | 0.00719719 | TAM.Cd209 | Slc38a7 |
| Cuedc2 | 3.80E-07 | 0.4006292 | 0.491 | 0.474 | 0.00734048 | TAM.Cd209 | Cuedc2 |
| Txndc12 | 3.93E-07 | 0.39582422 | 0.261 | 0.239 | 0.0076062 | TAM.Cd209 | Txndc12 |
| Mospd21 | 4.03E-07 | 0.32584407 | 0.4 | 0.38 | 0.00779611 | TAM.Cd209 | Mospd2 |
| Fam114a2 | 5.18E-07 | 0.2609029 | 0.277 | 0.258 | 0.01002419 | TAM.Cd209 | Fam114a2 |
| Celf21 | 5.91E-07 | 0.68213725 | 0.485 | 0.442 | 0.01142878 | TAM.Cd209 | Celf2 |
| Ppil2 | 6.15E-07 | 0.4011019 | 0.317 | 0.299 | 0.01189122 | TAM.Cd209 | Ppil2 |
| Dynlt1f1 | 6.17E-07 | 0.2916479 | 0.287 | 0.262 | 0.01192576 | TAM.Cd209 | Dynlt1f |
| Usp48 | 6.23E-07 | 0.44180284 | 0.388 | 0.364 | 0.01205064 | TAM.Cd209 | Usp48 |
| Szrd1 | 6.42E-07 | 0.3044106 | 0.608 | 0.626 | 0.01240484 | TAM.Cd209 | Szrd1 |
| H2afv | 7.14E-07 | 0.54606439 | 0.371 | 0.298 | 0.01381241 | TAM.Cd209 | H2afv |
| Plxna11 | 7.42E-07 | 0.52241564 | 0.558 | 0.493 | 0.01434764 | TAM.Cd209 | Plxna1 |
| Lsm101 | 7.43E-07 | 0.38288464 | 0.284 | 0.256 | 0.01436039 | TAM.Cd209 | Lsm10 |
| Klhl52 | 7.45E-07 | 0.26608302 | 0.316 | 0.296 | 0.01440258 | TAM.Cd209 | Klhl5 |
| Dync1li2 | 7.66E-07 | 0.489931 | 0.408 | 0.376 | 0.01481141 | TAM.Cd209 | Dync1li2 |
| Tceal91 | 8.12E-07 | 0.40868369 | 0.401 | 0.356 | 0.01569325 | TAM.Cd209 | Tceal9 |
| Ccdc115 | 8.32E-07 | 0.44425463 | 0.338 | 0.301 | 0.01607916 | TAM.Cd209 | Ccdc115 |
| Lrp52 | 9.87E-07 | 0.51337632 | 0.42 | 0.38 | 0.01908488 | TAM.Cd209 | Lrp5 |
| Ap5b11 | 1.01E-06 | 0.51772295 | 0.272 | 0.239 | 0.01953842 | TAM.Cd209 | Ap5b1 |
| Krcc1 | 1.03E-06 | 0.30103305 | 0.304 | 0.288 | 0.01998733 | TAM.Cd209 | Krcc1 |
| Cd2ap | 1.05E-06 | 0.30738627 | 0.387 | 0.352 | 0.0202932 | TAM.Cd209 | Cd2ap |
| Orai31 | 1.06E-06 | 0.36497953 | 0.272 | 0.248 | 0.02049812 | TAM.Cd209 | Orai3 |
| Phc3 | 1.11E-06 | 0.52198574 | 0.321 | 0.293 | 0.02144242 | TAM.Cd209 | Phc3 |
| Tceal81 | 1.23E-06 | 0.47386621 | 0.355 | 0.322 | 0.02371402 | TAM.Cd209 | Tceal8 |
| Usp8 | 1.28E-06 | 0.30024296 | 0.532 | 0.527 | 0.02480142 | TAM.Cd209 | Usp8 |
| Plekhm21 | 1.34E-06 | 0.37555664 | 0.461 | 0.423 | 0.02589662 | TAM.Cd209 | Plekhm2 |
| Map3k142 | 1.42E-06 | 0.41058148 | 0.363 | 0.327 | 0.02749865 | TAM.Cd209 | Map3k14 |
| Selenok | 1.69E-06 | 0.4094714 | 0.471 | 0.444 | 0.03264023 | TAM.Cd209 | Selenok |
| Pi4k2a | 1.69E-06 | 0.32545312 | 0.452 | 0.428 | 0.03276626 | TAM.Cd209 | Pi4k2a |
| Dennd1c | 1.71E-06 | 0.65468995 | 0.282 | 0.248 | 0.03297505 | TAM.Cd209 | Dennd1c |
| Hes61 | 1.80E-06 | 0.49011162 | 0.256 | 0.224 | 0.03483906 | TAM.Cd209 | Hes6 |
| Nudcd3 | 1.91E-06 | 0.53712568 | 0.39 | 0.365 | 0.03694808 | TAM.Cd209 | Nudcd3 |
| Myo7a1 | 1.92E-06 | 0.4988475 | 0.466 | 0.424 | 0.03703419 | TAM.Cd209 | Myo7a |
| Mapk8ip3 | 1.98E-06 | 0.48702132 | 0.315 | 0.293 | 0.03836035 | TAM.Cd209 | Mapk8ip3 |
| Fyb1 | 2.07E-06 | 0.26734962 | 0.66 | 0.648 | 0.04004473 | TAM.Cd209 | Fyb |
| Kcnab2 | 2.13E-06 | 0.29874061 | 0.464 | 0.444 | 0.04127535 | TAM.Cd209 | Kcnab2 |
| Scaf11 | 2.18E-06 | 0.38314375 | 0.441 | 0.424 | 0.0421788 | TAM.Cd209 | Scaf11 |
| Sptlc1 | 2.64E-06 | 0.39918718 | 0.269 | 0.245 | 0.05107221 | TAM.Cd209 | Sptlc1 |
| Pex5 | 2.67E-06 | 0.37696249 | 0.28 | 0.256 | 0.051614 | TAM.Cd209 | Pex5 |
| Slc25a451 | 2.91E-06 | 0.44524181 | 0.316 | 0.284 | 0.0561982 | TAM.Cd209 | Slc25a45 |
| Sigmar1 | 2.94E-06 | 0.34684053 | 0.351 | 0.331 | 0.05693061 | TAM.Cd209 | Sigmar1 |
| Pias1 | 3.35E-06 | 0.4071339 | 0.34 | 0.32 | 0.06480875 | TAM.Cd209 | Pias1 |
| Tm9sf4 | 3.36E-06 | 0.36854017 | 0.263 | 0.239 | 0.06503956 | TAM.Cd209 | Tm9sf4 |
| Stx161 | 3.88E-06 | 0.4708454 | 0.398 | 0.365 | 0.07500639 | TAM.Cd209 | Stx16 |
| Akap8 | 4.93E-06 | 0.3703834 | 0.381 | 0.399 | 0.09529448 | TAM.Cd209 | Akap8 |
| Snw1 | 4.97E-06 | 0.39269933 | 0.313 | 0.293 | 0.09603716 | TAM.Cd209 | Snw1 |
| Atg5 | 5.50E-06 | 0.4085569 | 0.327 | 0.306 | 0.10642258 | TAM.Cd209 | Atg5 |
| Prpf6 | 6.20E-06 | 0.42322779 | 0.287 | 0.265 | 0.11978732 | TAM.Cd209 | Prpf6 |
| Bad | 6.30E-06 | 0.48312212 | 0.362 | 0.335 | 0.12177347 | TAM.Cd209 | Bad |
| Ptov11 | 6.72E-06 | 0.48824629 | 0.453 | 0.421 | 0.12994145 | TAM.Cd209 | Ptov1 |
| Ubxn1 | 7.25E-06 | 0.38625307 | 0.626 | 0.632 | 0.14017811 | TAM.Cd209 | Ubxn1 |
| Inppl11 | 8.02E-06 | 0.35702611 | 0.411 | 0.409 | 0.15512099 | TAM.Cd209 | Inppl1 |
| Arhgef10l2 | 8.34E-06 | 0.33844254 | 0.384 | 0.336 | 0.16127762 | TAM.Cd209 | Arhgef10l |
| Exoc31 | 9.63E-06 | 0.39956633 | 0.25 | 0.229 | 0.18628103 | TAM.Cd209 | Exoc3 |
| Rnf7 | 1.01E-05 | 0.255894 | 0.27 | 0.254 | 0.19579205 | TAM.Cd209 | Rnf7 |
| Bmt2 | 1.02E-05 | 0.39146587 | 0.365 | 0.347 | 0.19779611 | TAM.Cd209 | Bmt2 |
| Gdi1 | 1.15E-05 | 0.58232078 | 0.469 | 0.431 | 0.22202341 | TAM.Cd209 | Gdi1 |
| Ccng1 | 1.21E-05 | 0.27504019 | 0.424 | 0.413 | 0.2339122 | TAM.Cd209 | Ccng1 |
| Mxi1 | 1.27E-05 | 0.31910724 | 0.504 | 0.496 | 0.24487657 | TAM.Cd209 | Mxi1 |
| Cnpy3 | 1.29E-05 | 0.48875825 | 0.536 | 0.502 | 0.24844846 | TAM.Cd209 | Cnpy3 |
| Nt5c2 | 1.61E-05 | 0.41026965 | 0.349 | 0.325 | 0.31175928 | TAM.Cd209 | Nt5c2 |
| Herc1 | 1.69E-05 | 0.54831248 | 0.349 | 0.317 | 0.32679191 | TAM.Cd209 | Herc1 |
| Unc50 | 1.82E-05 | 0.29656513 | 0.259 | 0.248 | 0.35200115 | TAM.Cd209 | Unc50 |
| Cep1701 | 2.05E-05 | 0.40898557 | 0.472 | 0.461 | 0.39547917 | TAM.Cd209 | Cep170 |
| Ttc7b | 2.91E-05 | 0.26002642 | 0.235 | 0.256 | 0.56189164 | TAM.Cd209 | Ttc7b |
| Brf1 | 3.16E-05 | 0.28824796 | 0.264 | 0.289 | 0.61075557 | TAM.Cd209 | Brf1 |
| Sdf4 | 3.50E-05 | 0.2793344 | 0.634 | 0.656 | 0.67626391 | TAM.Cd209 | Sdf4 |
| Ndufv3 | 3.92E-05 | 0.38869491 | 0.616 | 0.612 | 0.75757813 | TAM.Cd209 | Ndufv3 |
| Tpcn22 | 4.35E-05 | 0.28758835 | 0.367 | 0.361 | 0.84117427 | TAM.Cd209 | Tpcn2 |
| Arhgef7 | 4.58E-05 | 0.4342827 | 0.377 | 0.367 | 0.88599079 | TAM.Cd209 | Arhgef7 |
| Mcur1 | 4.86E-05 | 0.31231586 | 0.362 | 0.354 | 0.93919968 | TAM.Cd209 | Mcur1 |
| Cul4a | 4.94E-05 | 0.45215261 | 0.351 | 0.323 | 0.95577882 | TAM.Cd209 | Cul4a |
| Eapp | 4.97E-05 | 0.3545395 | 0.359 | 0.345 | 0.96008976 | TAM.Cd209 | Eapp |
| Ube2g1 | 5.13E-05 | 0.27452356 | 0.335 | 0.328 | 0.99212133 | TAM.Cd209 | Ube2g1 |
| Hook3 | 5.20E-05 | 0.45353048 | 0.329 | 0.301 | 1 | TAM.Cd209 | Hook3 |
| Ssu72 | 6.81E-05 | 0.33961018 | 0.529 | 0.53 | 1 | TAM.Cd209 | Ssu72 |
| Fbh1 | 8.20E-05 | 0.54011966 | 0.318 | 0.285 | 1 | TAM.Cd209 | Fbh1 |
| Herc41 | 8.20E-05 | 0.31923526 | 0.318 | 0.308 | 1 | TAM.Cd209 | Herc4 |
| Dpm3 | 8.79E-05 | 0.39680941 | 0.335 | 0.313 | 1 | TAM.Cd209 | Dpm3 |
| Pcnx3 | 9.14E-05 | 0.30455984 | 0.259 | 0.289 | 1 | TAM.Cd209 | Pcnx3 |
| Ctdsp1 | 9.25E-05 | 0.29328405 | 0.594 | 0.587 | 1 | TAM.Cd209 | Ctdsp1 |
| Tmem167b | 0.00010136 | 0.28723503 | 0.308 | 0.295 | 1 | TAM.Cd209 | Tmem167b |
| Trim44 | 0.00010396 | 0.25864143 | 0.34 | 0.361 | 1 | TAM.Cd209 | Trim44 |
| Dnase1l11 | 0.00011016 | 0.32362944 | 0.477 | 0.449 | 1 | TAM.Cd209 | Dnase1l1 |
| Dbi | 0.00011833 | 0.27668845 | 0.471 | 0.476 | 1 | TAM.Cd209 | Dbi |
| Rnpepl1 | 0.00013713 | 0.44144863 | 0.479 | 0.46 | 1 | TAM.Cd209 | Rnpepl1 |
| Tpr | 0.0001732 | 0.28019119 | 0.438 | 0.437 | 1 | TAM.Cd209 | Tpr |
| Mrfap1 | 0.00017683 | 0.43463902 | 0.626 | 0.613 | 1 | TAM.Cd209 | Mrfap1 |
| Dynlt3 | 0.00022736 | 0.31109375 | 0.343 | 0.325 | 1 | TAM.Cd209 | Dynlt3 |
| Plscr31 | 0.00022862 | 0.39728733 | 0.378 | 0.364 | 1 | TAM.Cd209 | Plscr3 |
| Mapk9 | 0.00024289 | 0.28291608 | 0.264 | 0.254 | 1 | TAM.Cd209 | Mapk9 |
| 2-Mar | 0.00026831 | 0.38787546 | 0.277 | 0.251 | 1 | TAM.Cd209 | 2-Mar |
| Ddx421 | 0.00026903 | 0.36808245 | 0.287 | 0.27 | 1 | TAM.Cd209 | Ddx42 |
| Znhit1 | 0.00033152 | 0.26058069 | 0.473 | 0.467 | 1 | TAM.Cd209 | Znhit1 |
| Ubqln2 | 0.00034035 | 0.36966337 | 0.364 | 0.35 | 1 | TAM.Cd209 | Ubqln2 |
| Stx4a | 0.00034856 | 0.39561788 | 0.585 | 0.574 | 1 | TAM.Cd209 | Stx4a |
| Mindy11 | 0.00036416 | 0.28601695 | 0.361 | 0.329 | 1 | TAM.Cd209 | Mindy1 |
| Leng8 | 0.00039096 | 0.75973262 | 0.527 | 0.508 | 1 | TAM.Cd209 | Leng8 |
| Clcn72 | 0.00041069 | 0.35707321 | 0.501 | 0.47 | 1 | TAM.Cd209 | Clcn7 |
| Golga2 | 0.0005217 | 0.35851949 | 0.35 | 0.333 | 1 | TAM.Cd209 | Golga2 |
| Rassf41 | 0.00053424 | 0.26785388 | 0.614 | 0.609 | 1 | TAM.Cd209 | Rassf4 |
| Dctn1 | 0.00069107 | 0.26299948 | 0.464 | 0.453 | 1 | TAM.Cd209 | Dctn1 |
| Dynll21 | 0.0007826 | 0.38006331 | 0.482 | 0.419 | 1 | TAM.Cd209 | Dynll2 |
| Myo18a | 0.00091859 | 0.46834298 | 0.344 | 0.319 | 1 | TAM.Cd209 | Myo18a |
| Tmem115 | 0.00104658 | 0.42619178 | 0.28 | 0.256 | 1 | TAM.Cd209 | Tmem115 |
| Heca1 | 0.00108804 | 0.30628489 | 0.43 | 0.413 | 1 | TAM.Cd209 | Heca |
| Tep1 | 0.00118163 | 0.40682608 | 0.324 | 0.332 | 1 | TAM.Cd209 | Tep1 |
| Zdhhc20 | 0.00138366 | 0.25753844 | 0.327 | 0.317 | 1 | TAM.Cd209 | Zdhhc20 |
| Ssbp4 | 0.00155581 | 0.48730823 | 0.554 | 0.536 | 1 | TAM.Cd209 | Ssbp4 |
| Ccm2 | 0.00170229 | 0.31883846 | 0.353 | 0.341 | 1 | TAM.Cd209 | Ccm2 |
| Prdx4 | 0.00173357 | 0.29665381 | 0.305 | 0.279 | 1 | TAM.Cd209 | Prdx4 |
| Mroh12 | 0.00178189 | 0.27339344 | 0.438 | 0.447 | 1 | TAM.Cd209 | Mroh1 |
| Tmem87a1 | 0.00185671 | 0.37369728 | 0.257 | 0.237 | 1 | TAM.Cd209 | Tmem87a |
| Dym | 0.00200747 | 0.25262823 | 0.274 | 0.286 | 1 | TAM.Cd209 | Dym |
| Mkln1 | 0.00237403 | 0.38435065 | 0.366 | 0.357 | 1 | TAM.Cd209 | Mkln1 |
| Cxxc1 | 0.00241868 | 0.30293922 | 0.285 | 0.301 | 1 | TAM.Cd209 | Cxxc1 |
| Klc41 | 0.00247781 | 0.52960035 | 0.302 | 0.276 | 1 | TAM.Cd209 | Klc4 |
| Fnta | 0.00259563 | 0.50262185 | 0.337 | 0.312 | 1 | TAM.Cd209 | Fnta |
| Ldb1 | 0.00273737 | 0.44608752 | 0.485 | 0.469 | 1 | TAM.Cd209 | Ldb1 |
| Tmem242 | 0.00285242 | 0.30912638 | 0.322 | 0.302 | 1 | TAM.Cd209 | Tmem242 |
| Hadh | 0.00309596 | 0.41109202 | 0.298 | 0.271 | 1 | TAM.Cd209 | Hadh |
| Dusp31 | 0.00344319 | 0.26172061 | 0.74 | 0.746 | 1 | TAM.Cd209 | Dusp3 |
| Cox16 | 0.00376531 | 0.27383008 | 0.244 | 0.258 | 1 | TAM.Cd209 | Cox16 |
| Phf10 | 0.00376557 | 0.3682899 | 0.288 | 0.276 | 1 | TAM.Cd209 | Phf10 |
| Dock7 | 0.00390138 | 0.43269842 | 0.452 | 0.432 | 1 | TAM.Cd209 | Dock7 |
| Tmx11 | 0.00406904 | 0.36829727 | 0.528 | 0.514 | 1 | TAM.Cd209 | Tmx1 |
| Sh3glb2 | 0.00416227 | 0.43744829 | 0.282 | 0.259 | 1 | TAM.Cd209 | Sh3glb2 |
| Lrch41 | 0.00446842 | 0.35782571 | 0.678 | 0.682 | 1 | TAM.Cd209 | Lrch4 |
| Rock2 | 0.0050791 | 0.28203104 | 0.345 | 0.334 | 1 | TAM.Cd209 | Rock2 |
| Anapc4 | 0.00521796 | 0.32477102 | 0.253 | 0.231 | 1 | TAM.Cd209 | Anapc4 |
| Ankrd40 | 0.00535072 | 0.28060932 | 0.464 | 0.454 | 1 | TAM.Cd209 | Ankrd40 |
| Tnrc6b1 | 0.00565239 | 0.28348264 | 0.304 | 0.295 | 1 | TAM.Cd209 | Tnrc6b |
| Cltb | 0.00618539 | 0.41745408 | 0.44 | 0.411 | 1 | TAM.Cd209 | Cltb |
| Rabl61 | 0.00716006 | 0.26553606 | 0.349 | 0.337 | 1 | TAM.Cd209 | Rabl6 |
| Ubap2l | 0.00746632 | 0.25038363 | 0.543 | 0.556 | 1 | TAM.Cd209 | Ubap2l |
| Gtpbp2 | 0.00792216 | 0.33840285 | 0.411 | 0.396 | 1 | TAM.Cd209 | Gtpbp2 |
| Nr1h2 | 0.00819641 | 0.44396024 | 0.384 | 0.365 | 1 | TAM.Cd209 | Nr1h2 |
| Zwint | 0.00838009 | 0.32887112 | 0.29 | 0.272 | 1 | TAM.Cd209 | Zwint |
| Rnf2151 | 0.00925777 | 0.39865015 | 0.251 | 0.237 | 1 | TAM.Cd209 | Rnf215 |
| Foxo3 | 0.00978875 | 0.6513311 | 0.393 | 0.369 | 1 | TAM.Cd209 | Foxo3 |
| Slc7a21 | 0 | 2.82720094 | 0.689 | 0.285 | 0 | TAM.Ndrg1 | Slc7a2 |
| Ndrg11 | 0 | 2.41472894 | 0.719 | 0.316 | 0 | TAM.Ndrg1 | Ndrg1 |
| Egln31 | 0 | 2.62082944 | 0.571 | 0.171 | 0 | TAM.Ndrg1 | Egln3 |
| Mt21 | 0 | 2.79014733 | 0.694 | 0.302 | 0 | TAM.Ndrg1 | Mt2 |
| Rgcc2 | 0 | 1.49467755 | 0.674 | 0.324 | 0 | TAM.Ndrg1 | Rgcc |
| Arg11 | 0 | 2.19601971 | 0.892 | 0.547 | 0 | TAM.Ndrg1 | Arg1 |
| Phlda11 | 0 | 1.68669408 | 0.77 | 0.436 | 0 | TAM.Ndrg1 | Phlda1 |
| Cd812 | 0 | 0.77689893 | 0.812 | 0.491 | 0 | TAM.Ndrg1 | Cd81 |
| Fam162a1 | 0 | 1.62305282 | 0.662 | 0.35 | 0 | TAM.Ndrg1 | Fam162a |
| Serpine11 | 0 | 3.25098952 | 0.485 | 0.181 | 0 | TAM.Ndrg1 | Serpine1 |
| Appl21 | 0 | 1.54162418 | 0.628 | 0.33 | 0 | TAM.Ndrg1 | Appl2 |
| Tnfsf92 | 0 | 1.65917959 | 0.597 | 0.299 | 0 | TAM.Ndrg1 | Tnfsf9 |
| Tnfrsf91 | 0 | 2.35848743 | 0.517 | 0.223 | 0 | TAM.Ndrg1 | Tnfrsf9 |
| P4ha2 | 0 | 3.39811036 | 0.33 | 0.043 | 0 | TAM.Ndrg1 | P4ha2 |
| Klf22 | 0 | 1.32636989 | 0.629 | 0.345 | 0 | TAM.Ndrg1 | Klf2 |
| Emp11 | 0 | 1.38835132 | 0.816 | 0.532 | 0 | TAM.Ndrg1 | Emp1 |
| Gys11 | 0 | 1.65507208 | 0.509 | 0.227 | 0 | TAM.Ndrg1 | Gys1 |
| Ccl22 | 0 | 1.5662898 | 0.508 | 0.227 | 0 | TAM.Ndrg1 | Ccl2 |
| Ero1l | 0 | 2.25451003 | 0.759 | 0.48 | 0 | TAM.Ndrg1 | Ero1l |
| Plxdc22 | 0 | 1.66444923 | 0.491 | 0.214 | 0 | TAM.Ndrg1 | Plxdc2 |
| Rhoc1 | 0 | 1.32111623 | 0.541 | 0.266 | 0 | TAM.Ndrg1 | Rhoc |
| Alox51 | 0 | 1.85728338 | 0.407 | 0.138 | 0 | TAM.Ndrg1 | Alox5 |
| Baiap2 | 0 | 1.36266501 | 0.663 | 0.396 | 0 | TAM.Ndrg1 | Baiap2 |
| Bnip31 | 0 | 2.04444405 | 0.381 | 0.117 | 0 | TAM.Ndrg1 | Bnip3 |
| Cited21 | 0 | 1.54142214 | 0.753 | 0.489 | 0 | TAM.Ndrg1 | Cited2 |
| Egln11 | 0 | 1.42827802 | 0.796 | 0.534 | 0 | TAM.Ndrg1 | Egln1 |
| Prelid2 | 0 | 2.31857793 | 0.35 | 0.088 | 0 | TAM.Ndrg1 | Prelid2 |
| Slc5a3 | 0 | 1.68589909 | 0.537 | 0.276 | 0 | TAM.Ndrg1 | Slc5a3 |
| Tob1 | 0 | 1.30910158 | 0.695 | 0.434 | 0 | TAM.Ndrg1 | Tob1 |
| Cxcl11 | 0 | 1.82267521 | 0.471 | 0.211 | 0 | TAM.Ndrg1 | Cxcl1 |
| Pdpn1 | 0 | 1.15673045 | 0.538 | 0.279 | 0 | TAM.Ndrg1 | Pdpn |
| Slc16a32 | 0 | 0.9979848 | 0.862 | 0.605 | 0 | TAM.Ndrg1 | Slc16a3 |
| Hilpda1 | 0 | 1.13261697 | 0.81 | 0.554 | 0 | TAM.Ndrg1 | Hilpda |
| Egr12 | 0 | 1.85105983 | 0.905 | 0.651 | 0 | TAM.Ndrg1 | Egr1 |
| Pgm11 | 0 | 1.34529068 | 0.515 | 0.265 | 0 | TAM.Ndrg1 | Pgm1 |
| Jup1 | 0 | 1.5258636 | 0.491 | 0.245 | 0 | TAM.Ndrg1 | Jup |
| Mgarp | 0 | 3.04979731 | 0.286 | 0.04 | 0 | TAM.Ndrg1 | Mgarp |
| Adrb22 | 0 | 1.35114184 | 0.481 | 0.238 | 0 | TAM.Ndrg1 | Adrb2 |
| Vegfa2 | 0 | 0.92680331 | 0.857 | 0.615 | 0 | TAM.Ndrg1 | Vegfa |
| Ankrd37 | 0 | 1.8268352 | 0.361 | 0.123 | 0 | TAM.Ndrg1 | Ankrd37 |
| Plk21 | 0 | 1.52679843 | 0.625 | 0.387 | 0 | TAM.Ndrg1 | Plk2 |
| Hspa1a3 | 0 | 0.93889975 | 0.633 | 0.395 | 0 | TAM.Ndrg1 | Hspa1a |
| Hist1h2bc1 | 0 | 0.99559803 | 0.575 | 0.338 | 0 | TAM.Ndrg1 | Hist1h2bc |
| Tpi11 | 0 | 1.18195851 | 0.876 | 0.641 | 0 | TAM.Ndrg1 | Tpi1 |
| Cacna1d1 | 0 | 1.37251469 | 0.477 | 0.242 | 0 | TAM.Ndrg1 | Cacna1d |
| Lsmem1 | 0 | 3.10986077 | 0.276 | 0.045 | 0 | TAM.Ndrg1 | Lsmem1 |
| Stac21 | 0 | 1.56283879 | 0.362 | 0.131 | 0 | TAM.Ndrg1 | Stac2 |
| Selenbp11 | 0 | 2.40110027 | 0.31 | 0.084 | 0 | TAM.Ndrg1 | Selenbp1 |
| Slc41a22 | 0 | 1.4407465 | 0.413 | 0.187 | 0 | TAM.Ndrg1 | Slc41a2 |
| Polg2 | 0 | 1.68029465 | 0.384 | 0.159 | 0 | TAM.Ndrg1 | Polg2 |
| Sgms11 | 0 | 1.29734338 | 0.447 | 0.222 | 0 | TAM.Ndrg1 | Sgms1 |
| Mfhas11 | 0 | 1.04795067 | 0.536 | 0.312 | 0 | TAM.Ndrg1 | Mfhas1 |
| Mtss11 | 0 | 0.91106876 | 0.759 | 0.536 | 0 | TAM.Ndrg1 | Mtss1 |
| Hk1 | 0 | 1.08915937 | 0.657 | 0.435 | 0 | TAM.Ndrg1 | Hk1 |
| Galnt61 | 0 | 1.10808298 | 0.539 | 0.318 | 0 | TAM.Ndrg1 | Galnt6 |
| Ddit41 | 0 | 0.69887098 | 0.65 | 0.43 | 0 | TAM.Ndrg1 | Ddit4 |
| Afg3l2 | 0 | 1.10949955 | 0.628 | 0.409 | 0 | TAM.Ndrg1 | Afg3l2 |
| Krt802 | 0 | 1.13922151 | 0.429 | 0.21 | 0 | TAM.Ndrg1 | Krt80 |
| Mthfd1l | 0 | 1.54003578 | 0.381 | 0.164 | 0 | TAM.Ndrg1 | Mthfd1l |
| Pfkp3 | 0 | 0.90062691 | 0.778 | 0.561 | 0 | TAM.Ndrg1 | Pfkp |
| Cdc42ep4 | 0 | 1.18406954 | 0.456 | 0.24 | 0 | TAM.Ndrg1 | Cdc42ep4 |
| Pfkl2 | 0 | 0.85601737 | 0.77 | 0.554 | 0 | TAM.Ndrg1 | Pfkl |
| Vcl | 0 | 1.25669132 | 0.454 | 0.239 | 0 | TAM.Ndrg1 | Vcl |
| Hmox12 | 0 | 0.89305237 | 0.883 | 0.668 | 0 | TAM.Ndrg1 | Hmox1 |
| Impdh11 | 0 | 1.5791491 | 0.389 | 0.176 | 0 | TAM.Ndrg1 | Impdh1 |
| Adam82 | 0 | 0.74507693 | 0.765 | 0.553 | 0 | TAM.Ndrg1 | Adam8 |
| Tnfaip32 | 0 | 0.96305914 | 0.81 | 0.599 | 0 | TAM.Ndrg1 | Tnfaip3 |
| C3ar13 | 0 | 0.76587698 | 0.876 | 0.665 | 0 | TAM.Ndrg1 | C3ar1 |
| F13a11 | 0 | 0.76422529 | 0.597 | 0.386 | 0 | TAM.Ndrg1 | F13a1 |
| Errfi12 | 0 | 1.30213074 | 0.557 | 0.35 | 0 | TAM.Ndrg1 | Errfi1 |
| Mif1 | 0 | 0.93494906 | 0.75 | 0.543 | 0 | TAM.Ndrg1 | Mif |
| Dock71 | 0 | 1.02787335 | 0.621 | 0.415 | 0 | TAM.Ndrg1 | Dock7 |
| Tnip3 | 0 | 1.84816969 | 0.332 | 0.126 | 0 | TAM.Ndrg1 | Tnip3 |
| Tnf2 | 0 | 0.6941298 | 0.801 | 0.598 | 0 | TAM.Ndrg1 | Tnf |
| Slc2a12 | 0 | 1.40590583 | 0.881 | 0.678 | 0 | TAM.Ndrg1 | Slc2a1 |
| P4ha11 | 0 | 0.92477146 | 0.744 | 0.541 | 0 | TAM.Ndrg1 | P4ha1 |
| Id21 | 0 | 0.9804653 | 0.867 | 0.664 | 0 | TAM.Ndrg1 | Id2 |
| Pitpnc11 | 0 | 0.99139359 | 0.494 | 0.293 | 0 | TAM.Ndrg1 | Pitpnc1 |
| Plau1 | 0 | 1.08445536 | 0.829 | 0.629 | 0 | TAM.Ndrg1 | Plau |
| Dnajb91 | 0 | 1.18564573 | 0.556 | 0.358 | 0 | TAM.Ndrg1 | Dnajb9 |
| Pgk11 | 0 | 1.12339354 | 0.767 | 0.569 | 0 | TAM.Ndrg1 | Pgk1 |
| Ripk21 | 0 | 1.26025744 | 0.419 | 0.221 | 0 | TAM.Ndrg1 | Ripk2 |
| Higd1a | 0 | 1.03896703 | 0.666 | 0.469 | 0 | TAM.Ndrg1 | Higd1a |
| Ccl61 | 0 | 0.96050243 | 0.786 | 0.589 | 0 | TAM.Ndrg1 | Ccl6 |
| Sept111 | 0 | 1.07326402 | 0.535 | 0.338 | 0 | TAM.Ndrg1 | 11-Sep |
| Zfp395 | 0 | 1.62763574 | 0.32 | 0.123 | 0 | TAM.Ndrg1 | Zfp395 |
| Nabp12 | 0 | 0.87942519 | 0.564 | 0.368 | 0 | TAM.Ndrg1 | Nabp1 |
| Bsg1 | 0 | 1.07802291 | 0.677 | 0.482 | 0 | TAM.Ndrg1 | Bsg |
| Pltp2 | 0 | 0.35625445 | 0.789 | 0.596 | 0 | TAM.Ndrg1 | Pltp |
| Rgs11 | 0 | 0.71409873 | 0.734 | 0.541 | 0 | TAM.Ndrg1 | Rgs1 |
| Tcp11l2 | 0 | 0.84673837 | 0.571 | 0.378 | 0 | TAM.Ndrg1 | Tcp11l2 |
| Lmna1 | 0 | 1.08439297 | 0.901 | 0.709 | 0 | TAM.Ndrg1 | Lmna |
| Metrnl2 | 0 | 0.82911026 | 0.757 | 0.566 | 0 | TAM.Ndrg1 | Metrnl |
| Nfkbiz3 | 0 | 0.9131892 | 0.865 | 0.674 | 0 | TAM.Ndrg1 | Nfkbiz |
| Fam71f21 | 0 | 1.85071138 | 0.297 | 0.107 | 0 | TAM.Ndrg1 | Fam71f2 |
| Sertad11 | 0 | 0.86182614 | 0.746 | 0.556 | 0 | TAM.Ndrg1 | Sertad1 |
| Hspa1b3 | 0 | 0.81469356 | 0.532 | 0.348 | 0 | TAM.Ndrg1 | Hspa1b |
| Ahnak1 | 0 | 1.14181937 | 0.916 | 0.733 | 0 | TAM.Ndrg1 | Ahnak |
| Il6 | 0 | 2.08672438 | 0.286 | 0.103 | 0 | TAM.Ndrg1 | Il6 |
| Palld1 | 0 | 1.00392378 | 0.432 | 0.249 | 0 | TAM.Ndrg1 | Palld |
| Egr22 | 0 | 0.71566365 | 0.645 | 0.462 | 0 | TAM.Ndrg1 | Egr2 |
| Arrb11 | 0 | 0.76192865 | 0.691 | 0.508 | 0 | TAM.Ndrg1 | Arrb1 |
| Lsp12 | 0 | 0.76813599 | 0.574 | 0.391 | 0 | TAM.Ndrg1 | Lsp1 |
| Trib12 | 0 | 0.84036993 | 0.753 | 0.571 | 0 | TAM.Ndrg1 | Trib1 |
| Gadd45a2 | 0 | 1.3455462 | 0.423 | 0.241 | 0 | TAM.Ndrg1 | Gadd45a |
| Atf32 | 0 | 0.76663976 | 0.865 | 0.684 | 0 | TAM.Ndrg1 | Atf3 |
| Gbe11 | 0 | 1.64051266 | 0.337 | 0.157 | 0 | TAM.Ndrg1 | Gbe1 |
| Ccl92 | 0 | 0.78652396 | 0.834 | 0.654 | 0 | TAM.Ndrg1 | Ccl9 |
| Vdac1 | 0 | 0.99288811 | 0.541 | 0.362 | 0 | TAM.Ndrg1 | Vdac1 |
| Mdm21 | 0 | 1.18199267 | 0.75 | 0.572 | 0 | TAM.Ndrg1 | Mdm2 |
| Pmp221 | 0 | 0.78669157 | 0.35 | 0.172 | 0 | TAM.Ndrg1 | Pmp22 |
| Capg2 | 0 | 0.41377793 | 0.856 | 0.678 | 0 | TAM.Ndrg1 | Capg |
| Tppp33 | 0 | 2.02658751 | 0.426 | 0.25 | 0 | TAM.Ndrg1 | Tppp3 |
| Fcrlb1 | 0 | 0.80741355 | 0.556 | 0.382 | 0 | TAM.Ndrg1 | Fcrlb |
| Egr3 | 0 | 2.01595729 | 0.269 | 0.095 | 0 | TAM.Ndrg1 | Egr3 |
| Lrg11 | 0 | 1.20487164 | 0.289 | 0.115 | 0 | TAM.Ndrg1 | Lrg1 |
| Sdc31 | 0 | 0.85115043 | 0.923 | 0.752 | 0 | TAM.Ndrg1 | Sdc3 |
| Hk2 | 0 | 0.8135106 | 0.812 | 0.641 | 0 | TAM.Ndrg1 | Hk2 |
| Eif4ebp1 | 0 | 0.79253556 | 0.645 | 0.474 | 0 | TAM.Ndrg1 | Eif4ebp1 |
| Rassf81 | 0 | 1.07903507 | 0.49 | 0.32 | 0 | TAM.Ndrg1 | Rassf8 |
| Ecm11 | 0 | 1.02008819 | 0.862 | 0.696 | 0 | TAM.Ndrg1 | Ecm1 |
| Gadd45g1 | 0 | 0.80807907 | 0.548 | 0.382 | 0 | TAM.Ndrg1 | Gadd45g |
| Impa21 | 0 | 1.09203774 | 0.378 | 0.212 | 0 | TAM.Ndrg1 | Impa2 |
| Timp1 | 0 | 1.37604008 | 0.331 | 0.165 | 0 | TAM.Ndrg1 | Timp1 |
| Pgam11 | 0 | 0.76644813 | 0.861 | 0.696 | 0 | TAM.Ndrg1 | Pgam1 |
| Klf62 | 0 | 0.80516622 | 0.862 | 0.698 | 0 | TAM.Ndrg1 | Klf6 |
| Thbs13 | 0 | 0.35420852 | 0.833 | 0.672 | 0 | TAM.Ndrg1 | Thbs1 |
| Ier21 | 0 | 0.81736253 | 0.811 | 0.651 | 0 | TAM.Ndrg1 | Ier2 |
| Sowahc3 | 0 | 0.66857739 | 0.658 | 0.498 | 0 | TAM.Ndrg1 | Sowahc |
| Fbxo321 | 0 | 1.88361953 | 0.292 | 0.133 | 0 | TAM.Ndrg1 | Fbxo32 |
| Cbx4 | 0 | 0.99286483 | 0.509 | 0.351 | 0 | TAM.Ndrg1 | Cbx4 |
| Cd832 | 0 | 0.47365552 | 0.695 | 0.537 | 0 | TAM.Ndrg1 | Cd83 |
| Anxa12 | 0 | 0.85779857 | 0.692 | 0.534 | 0 | TAM.Ndrg1 | Anxa1 |
| Cxcl23 | 0 | 0.94513443 | 0.932 | 0.776 | 0 | TAM.Ndrg1 | Cxcl2 |
| Slc20a11 | 0 | 0.73284389 | 0.596 | 0.441 | 0 | TAM.Ndrg1 | Slc20a1 |
| Pdlim1 | 0 | 1.36651758 | 0.264 | 0.113 | 0 | TAM.Ndrg1 | Pdlim1 |
| Trem12 | 0 | 0.7178442 | 0.488 | 0.337 | 0 | TAM.Ndrg1 | Trem1 |
| Brd2 | 0 | 0.78541112 | 0.796 | 0.646 | 0 | TAM.Ndrg1 | Brd2 |
| Jag11 | 0 | 1.27011793 | 0.303 | 0.153 | 0 | TAM.Ndrg1 | Jag1 |
| Fn11 | 0 | 1.90971931 | 0.974 | 0.826 | 0 | TAM.Ndrg1 | Fn1 |
| Mt1 | 0 | 1.2358335 | 0.936 | 0.788 | 0 | TAM.Ndrg1 | Mt1 |
| Osbpl81 | 0 | 0.79771917 | 0.697 | 0.551 | 0 | TAM.Ndrg1 | Osbpl8 |
| Lsr2 | 0 | 0.82203531 | 0.425 | 0.28 | 0 | TAM.Ndrg1 | Lsr |
| Rbpj1 | 0 | 0.77583677 | 0.813 | 0.668 | 0 | TAM.Ndrg1 | Rbpj |
| Hopx2 | 0 | 1.37879464 | 0.346 | 0.204 | 0 | TAM.Ndrg1 | Hopx |
| Ltb4r12 | 0 | 0.9988749 | 0.295 | 0.155 | 0 | TAM.Ndrg1 | Ltb4r1 |
| Hbegf2 | 0 | 0.94756554 | 0.41 | 0.27 | 0 | TAM.Ndrg1 | Hbegf |
| Adssl12 | 0 | 0.59915312 | 0.702 | 0.562 | 0 | TAM.Ndrg1 | Adssl1 |
| Tmsb101 | 0 | 0.54914317 | 0.778 | 0.64 | 0 | TAM.Ndrg1 | Tmsb10 |
| Lilr4b1 | 0 | 0.52999027 | 0.792 | 0.654 | 0 | TAM.Ndrg1 | Lilr4b |
| Basp11 | 0 | 0.72142684 | 0.87 | 0.733 | 0 | TAM.Ndrg1 | Basp1 |
| Fndc3a2 | 0 | 0.70461993 | 0.676 | 0.54 | 0 | TAM.Ndrg1 | Fndc3a |
| Ccrl21 | 0 | 0.51987292 | 0.799 | 0.665 | 0 | TAM.Ndrg1 | Ccrl2 |
| Il1a2 | 0 | 0.96587042 | 0.582 | 0.448 | 0 | TAM.Ndrg1 | Il1a |
| Gpi11 | 0 | 1.08228839 | 0.866 | 0.736 | 0 | TAM.Ndrg1 | Gpi1 |
| Sqstm11 | 0 | 0.70960391 | 0.891 | 0.761 | 0 | TAM.Ndrg1 | Sqstm1 |
| Clec4d2 | 0 | 0.29059187 | 0.767 | 0.64 | 0 | TAM.Ndrg1 | Clec4d |
| Ier51 | 0 | 0.80626872 | 0.911 | 0.786 | 0 | TAM.Ndrg1 | Ier5 |
| Ctsl1 | 0 | 1.06551469 | 0.87 | 0.745 | 0 | TAM.Ndrg1 | Ctsl |
| Ccl72 | 0 | 0.78959326 | 0.345 | 0.22 | 0 | TAM.Ndrg1 | Ccl7 |
| Furin1 | 0 | 1.05981915 | 0.924 | 0.8 | 0 | TAM.Ndrg1 | Furin |
| Sgk11 | 0 | 0.62799904 | 0.738 | 0.614 | 0 | TAM.Ndrg1 | Sgk1 |
| Nlrp32 | 0 | 0.61892695 | 0.833 | 0.709 | 0 | TAM.Ndrg1 | Nlrp3 |
| Mmp191 | 0 | 0.57405517 | 0.719 | 0.596 | 0 | TAM.Ndrg1 | Mmp19 |
| Sdc4 | 0 | 0.59298866 | 0.816 | 0.694 | 0 | TAM.Ndrg1 | Sdc4 |
| Klf6 | 0 | 0.96522714 | 0.938 | 0.816 | 0 | TAM.Ndrg1 | Klf6 |
| Ppp1r15a2 | 0 | 0.54729465 | 0.839 | 0.719 | 0 | TAM.Ndrg1 | Ppp1r15a |
| Csrnp12 | 0 | 0.33069321 | 0.82 | 0.7 | 0 | TAM.Ndrg1 | Csrnp1 |
| Abca12 | 0 | 0.47920955 | 0.835 | 0.717 | 0 | TAM.Ndrg1 | Abca1 |
| Gadd45b2 | 0 | 0.32943531 | 0.908 | 0.79 | 0 | TAM.Ndrg1 | Gadd45b |
| Hexim11 | 0 | 0.68916521 | 0.698 | 0.581 | 0 | TAM.Ndrg1 | Hexim1 |
| Itgam3 | 0 | 0.6811058 | 0.91 | 0.796 | 0 | TAM.Ndrg1 | Itgam |
| Dhx401 | 0 | 1.26182038 | 0.393 | 0.279 | 0 | TAM.Ndrg1 | Dhx40 |
| Fosb2 | 0 | 0.7972275 | 0.913 | 0.8 | 0 | TAM.Ndrg1 | Fosb |
| Bcl2l112 | 0 | 0.4953703 | 0.783 | 0.671 | 0 | TAM.Ndrg1 | Bcl2l11 |
| Anxa21 | 0 | 1.10682196 | 0.959 | 0.847 | 0 | TAM.Ndrg1 | Anxa2 |
| Bhlhe402 | 0 | 0.31129289 | 0.903 | 0.793 | 0 | TAM.Ndrg1 | Bhlhe40 |
| Cdkn1a2 | 0 | 0.42597535 | 0.946 | 0.837 | 0 | TAM.Ndrg1 | Cdkn1a |
| Gls | 0 | 0.68169305 | 0.717 | 0.609 | 0 | TAM.Ndrg1 | Gls |
| Aplp22 | 0 | 0.54880178 | 0.779 | 0.672 | 0 | TAM.Ndrg1 | Aplp2 |

| | | | | | | | |
|---|---|---|---|---|---|---|---|
| Ldha | 0 | 0.86696734 | 0.937 | 0.831 | 0 | TAM.Ndrg1 | Ldha |
| Ehd1 | 0 | 0.5156697 | 0.798 | 0.694 | 0 | TAM.Ndrg1 | Ehd1 |
| Capn2 | 0 | 0.55087087 | 0.702 | 0.599 | 0 | TAM.Ndrg1 | Capn2 |
| Zfp36l12 | 0 | 0.58784572 | 0.726 | 0.623 | 0 | TAM.Ndrg1 | Zfp36l1 |
| Tmem1891 | 0 | 0.55876739 | 0.853 | 0.751 | 0 | TAM.Ndrg1 | Tmem189 |
| Rhob2 | 0 | 0.6424096 | 0.857 | 0.755 | 0 | TAM.Ndrg1 | Rhob |
| Aldoa1 | 0 | 0.70872622 | 0.92 | 0.819 | 0 | TAM.Ndrg1 | Aldoa |
| Ifrd12 | 0 | 0.51826548 | 0.915 | 0.817 | 0 | TAM.Ndrg1 | Ifrd1 |
| Marcksl12 | 0 | 0.65200613 | 0.79 | 0.695 | 0 | TAM.Ndrg1 | Marcksl1 |
| Tbxas11 | 0 | 0.47201871 | 0.801 | 0.707 | 0 | TAM.Ndrg1 | Tbxas1 |
| Btg23 | 0 | 0.60195396 | 0.914 | 0.822 | 0 | TAM.Ndrg1 | Btg2 |
| Wsb12 | 0 | 0.58066028 | 0.89 | 0.799 | 0 | TAM.Ndrg1 | Wsb1 |
| Tgif12 | 0 | 0.32564919 | 0.855 | 0.765 | 0 | TAM.Ndrg1 | Tgif1 |
| S100a102 | 0 | 0.68721487 | 0.854 | 0.768 | 0 | TAM.Ndrg1 | S100a10 |
| Zyx1 | 0 | 0.47412849 | 0.838 | 0.752 | 0 | TAM.Ndrg1 | Zyx |
| Socs33 | 0 | 0.31086887 | 0.862 | 0.776 | 0 | TAM.Ndrg1 | Socs3 |
| Pnrc1 | 0 | 0.68795 | 0.916 | 0.831 | 0 | TAM.Ndrg1 | Pnrc1 |
| Pgf1 | 0 | 1.36444703 | 0.295 | 0.21 | 0 | TAM.Ndrg1 | Pgf |
| Gda2 | 0 | 0.48217055 | 0.824 | 0.739 | 0 | TAM.Ndrg1 | Gda |
| Ccl32 | 0 | 0.46641465 | 0.647 | 0.564 | 0 | TAM.Ndrg1 | Ccl3 |
| Pkm2 | 0 | 0.60495281 | 0.94 | 0.857 | 0 | TAM.Ndrg1 | Pkm |
| Cd143 | 0 | 0.289414 | 0.952 | 0.869 | 0 | TAM.Ndrg1 | Cd14 |
| Arrdc41 | 0 | 0.50262219 | 0.661 | 0.579 | 0 | TAM.Ndrg1 | Arrdc4 |
| Mmp121 | 0 | 0.99001191 | 0.588 | 0.506 | 0 | TAM.Ndrg1 | Mmp12 |
| Mknk2 | 0 | 0.55663701 | 0.868 | 0.786 | 0 | TAM.Ndrg1 | Mknk2 |
| Emilin22 | 0 | 0.30678395 | 0.871 | 0.789 | 0 | TAM.Ndrg1 | Emilin2 |
| 1-Jun | 0 | 0.63596244 | 0.929 | 0.848 | 0 | TAM.Ndrg1 | Jun |
| Ptgs22 | 0 | 0.34474222 | 0.639 | 0.559 | 0 | TAM.Ndrg1 | Ptgs2 |
| Dusp13 | 0 | 0.92721457 | 0.966 | 0.886 | 0 | TAM.Ndrg1 | Dusp1 |
| Btg11 | 0 | 0.58229995 | 0.932 | 0.853 | 0 | TAM.Ndrg1 | Btg1 |
| Lgals31 | 0 | 0.75308757 | 0.975 | 0.9 | 0 | TAM.Ndrg1 | Lgals3 |
| Rnf19b1 | 0 | 0.50414347 | 0.767 | 0.695 | 0 | TAM.Ndrg1 | Rnf19b |
| P4hb1 | 0 | 0.50882589 | 0.848 | 0.777 | 0 | TAM.Ndrg1 | P4hb |
| Nfkbia1 | 0 | 0.78758992 | 0.967 | 0.896 | 0 | TAM.Ndrg1 | Nfkbia |
| Crip12 | 0 | 1.15685698 | 0.828 | 0.757 | 0 | TAM.Ndrg1 | Crip1 |
| Pld13 | 0 | 0.2890691 | 0.762 | 0.692 | 0 | TAM.Ndrg1 | Pld1 |
| Myadm | 0 | 0.46023757 | 0.723 | 0.655 | 0 | TAM.Ndrg1 | Myadm |
| Lilrb4a1 | 0 | 0.48825458 | 0.942 | 0.875 | 0 | TAM.Ndrg1 | Lilrb4a |
| Fxyd52 | 0 | 0.5359777 | 0.88 | 0.814 | 0 | TAM.Ndrg1 | Fxyd5 |
| H3f3b2 | 0 | 0.49109 | 0.894 | 0.829 | 0 | TAM.Ndrg1 | H3f3b |
| Acly2 | 0 | 0.53429356 | 0.846 | 0.781 | 0 | TAM.Ndrg1 | Acly |
| Mmp131 | 0 | 1.51458996 | 0.362 | 0.301 | 0 | TAM.Ndrg1 | Mmp13 |
| Ncf11 | 0 | 0.41958842 | 0.819 | 0.759 | 0 | TAM.Ndrg1 | Ncf1 |
| Lyz12 | 0 | 1.4250173 | 0.364 | 0.305 | 0 | TAM.Ndrg1 | Lyz1 |
| Lrp13 | 0 | 0.30586164 | 0.952 | 0.896 | 0 | TAM.Ndrg1 | Lrp1 |
| Jund1 | 0 | 0.73779161 | 0.983 | 0.928 | 0 | TAM.Ndrg1 | Jund |
| Fos1 | 0 | 0.60556626 | 0.957 | 0.903 | 0 | TAM.Ndrg1 | Fos |
| Ier33 | 0 | 0.42934277 | 0.969 | 0.915 | 0 | TAM.Ndrg1 | Ier3 |
| Ubc1 | 0 | 0.55487114 | 0.961 | 0.913 | 0 | TAM.Ndrg1 | Ubc |
| Iqgap12 | 0 | 0.47536794 | 0.963 | 0.917 | 0 | TAM.Ndrg1 | Iqgap1 |
| Slfn21 | 0 | 0.34337254 | 0.896 | 0.851 | 0 | TAM.Ndrg1 | Slfn2 |
| Zeb22 | 0 | 0.29414611 | 0.938 | 0.899 | 0 | TAM.Ndrg1 | Zeb2 |
| Trf2 | 0 | 0.77302476 | 0.839 | 0.8 | 0 | TAM.Ndrg1 | Trf |
| Srgn1 | 0 | 0.8464347 | 0.981 | 0.943 | 0 | TAM.Ndrg1 | Srgn |
| Vim1 | 0 | 0.92652746 | 0.995 | 0.959 | 0 | TAM.Ndrg1 | Vim |
| Kctd121 | 0 | 0.65139038 | 0.961 | 0.928 | 0 | TAM.Ndrg1 | Kctd12 |
| Junb2 | 0 | 0.38624741 | 0.986 | 0.954 | 0 | TAM.Ndrg1 | Junb |
| Tagln21 | 0 | 0.2773153 | 0.929 | 0.899 | 0 | TAM.Ndrg1 | Tagln2 |
| Zfp363 | 0 | 0.30104487 | 0.984 | 0.956 | 0 | TAM.Ndrg1 | Zfp36 |
| Plek2 | 0 | 0.439306 | 0.964 | 0.936 | 0 | TAM.Ndrg1 | Plek |
| App1 | 0 | 0.39199265 | 0.976 | 0.953 | 0 | TAM.Ndrg1 | App |
| Actg12 | 0 | 0.26182658 | 0.954 | 0.937 | 0 | TAM.Ndrg1 | Actg1 |
| Mcl11 | 0 | 0.55437091 | 0.991 | 0.975 | 0 | TAM.Ndrg1 | Mcl1 |
| Gpx11 | 0 | 0.27711228 | 0.96 | 0.95 | 0 | TAM.Ndrg1 | Gpx1 |
| Cxcl141 | 4.65E-308 | 0.81395682 | 0.359 | 0.21 | 8.99E-304 | TAM.Ndrg1 | Cxcl14 |
| Pdk11 | 1.10E-306 | 0.9554665 | 0.494 | 0.294 | 2.13E-302 | TAM.Ndrg1 | Pdk1 |
| Ccng21 | 2.82E-306 | 0.86194383 | 0.507 | 0.337 | 5.45E-302 | TAM.Ndrg1 | Ccng2 |
| Ephx1 | 6.13E-306 | 1.38867062 | 0.297 | 0.143 | 1.19E-301 | TAM.Ndrg1 | Ephx1 |
| Nt5e2 | 1.92E-303 | 0.91790578 | 0.295 | 0.186 | 3.71E-299 | TAM.Ndrg1 | Nt5e |
| Nfkbid4 | 2.31E-302 | 0.42694976 | 0.71 | 0.608 | 4.47E-298 | TAM.Ndrg1 | Nfkbid |
| Fzd71 | 6.97E-302 | 1.13768866 | 0.348 | 0.212 | 1.35E-297 | TAM.Ndrg1 | Fzd7 |
| Akap71 | 2.70E-300 | 1.15853561 | 0.369 | 0.187 | 5.23E-296 | TAM.Ndrg1 | Akap7 |
| Fzd5 | 1.42E-296 | 1.29047243 | 0.357 | 0.197 | 2.75E-292 | TAM.Ndrg1 | Fzd5 |
| Ccnl11 | 1.41E-295 | 0.36871838 | 0.876 | 0.789 | 2.73E-291 | TAM.Ndrg1 | Ccnl1 |
| Glipr21 | 1.46E-294 | 0.81100291 | 0.293 | 0.165 | 2.82E-290 | TAM.Ndrg1 | Glipr2 |
| Pfkfb31 | 2.80E-292 | 0.83948351 | 0.52 | 0.371 | 5.42E-288 | TAM.Ndrg1 | Pfkfb3 |
| Id32 | 6.30E-288 | 0.55600895 | 0.39 | 0.254 | 1.22E-283 | TAM.Ndrg1 | Id3 |
| Plekhg22 | 7.79E-284 | 1.05798014 | 0.397 | 0.299 | 1.51E-279 | TAM.Ndrg1 | Plekhg2 |
| Sra13 | 1.74E-283 | 0.55898114 | 0.668 | 0.553 | 3.36E-279 | TAM.Ndrg1 | Sra1 |
| Clk12 | 4.66E-283 | 0.41662235 | 0.801 | 0.705 | 9.00E-279 | TAM.Ndrg1 | Clk1 |
| Dusp32 | 1.73E-281 | 0.39913603 | 0.831 | 0.737 | 3.35E-277 | TAM.Ndrg1 | Dusp3 |
| Tiparp1 | 4.72E-281 | 0.3702975 | 0.753 | 0.651 | 9.12E-277 | TAM.Ndrg1 | Tiparp |
| Fam129b2 | 2.56E-279 | 0.34530568 | 0.798 | 0.671 | 4.96E-275 | TAM.Ndrg1 | Fam129b |
| Dhrs32 | 1.70E-275 | 0.43949989 | 0.808 | 0.639 | 3.29E-271 | TAM.Ndrg1 | Dhrs3 |
| Rnd3 | 2.70E-275 | 0.95743542 | 0.428 | 0.268 | 5.23E-271 | TAM.Ndrg1 | Rnd3 |
| Plp21 | 1.77E-274 | 0.80893126 | 0.539 | 0.358 | 3.41E-270 | TAM.Ndrg1 | Plp2 |
| Npepps1 | 1.61E-270 | 0.79424776 | 0.549 | 0.395 | 3.11E-266 | TAM.Ndrg1 | Npepps |
| Nop2 | 3.40E-268 | 1.17789215 | 0.367 | 0.262 | 6.58E-264 | TAM.Ndrg1 | Nop2 |
| Trappc6a1 | 6.78E-268 | 0.92770801 | 0.484 | 0.302 | 1.31E-263 | TAM.Ndrg1 | Trappc6a |
| Kcnn41 | 1.15E-267 | 1.17665168 | 0.392 | 0.212 | 2.23E-263 | TAM.Ndrg1 | Kcnn4 |
| Kif21b | 1.29E-265 | 1.08466969 | 0.409 | 0.245 | 2.50E-261 | TAM.Ndrg1 | Kif21b |
| Ltc4s1 | 1.76E-265 | 0.84181076 | 0.325 | 0.245 | 3.39E-261 | TAM.Ndrg1 | Ltc4s |
| Itgb1 | 1.17E-258 | 0.37020606 | 0.843 | 0.776 | 2.26E-254 | TAM.Ndrg1 | Itgb1 |
| Irf12 | 4.93E-258 | 0.43706102 | 0.619 | 0.489 | 9.54E-254 | TAM.Ndrg1 | Irf1 |
| Ier5l1 | 1.84E-256 | 1.29907218 | 0.288 | 0.15 | 3.56E-252 | TAM.Ndrg1 | Ier5l |
| Slc6a81 | 2.45E-255 | 0.96740496 | 0.455 | 0.276 | 4.74E-251 | TAM.Ndrg1 | Slc6a8 |
| Kcnab21 | 3.44E-255 | 0.72231165 | 0.583 | 0.432 | 6.66E-251 | TAM.Ndrg1 | Kcnab2 |
| Epb41l11 | 8.71E-254 | 1.10610749 | 0.336 | 0.148 | 1.68E-249 | TAM.Ndrg1 | Epb41l1 |
| Cd2742 | 3.83E-251 | 0.53595294 | 0.597 | 0.468 | 7.41E-247 | TAM.Ndrg1 | Cd274 |
| Sat11 | 2.52E-249 | 0.31106425 | 0.917 | 0.897 | 4.88E-245 | TAM.Ndrg1 | Sat1 |
| Mcub1 | 2.92E-248 | 0.73163103 | 0.437 | 0.321 | 5.65E-244 | TAM.Ndrg1 | Mcub |
| Osgin21 | 4.17E-248 | 0.95757768 | 0.319 | 0.184 | 8.07E-244 | TAM.Ndrg1 | Osgin2 |
| Dpep21 | 7.35E-248 | 0.71414793 | 0.513 | 0.36 | 1.42E-243 | TAM.Ndrg1 | Dpep2 |
| Xdh1 | 8.48E-246 | 0.4924996 | 0.672 | 0.487 | 1.64E-241 | TAM.Ndrg1 | Xdh |
| Xylt12 | 2.65E-244 | 0.82747574 | 0.429 | 0.249 | 5.13E-240 | TAM.Ndrg1 | Xylt1 |
| Cd200r11 | 2.35E-243 | 1.01810469 | 0.417 | 0.27 | 4.54E-239 | TAM.Ndrg1 | Cd200r1 |
| Zdhhc142 | 1.30E-235 | 0.82542542 | 0.497 | 0.294 | 2.51E-231 | TAM.Ndrg1 | Zdhhc14 |
| Fgfr11 | 2.75E-229 | 0.982678 | 0.282 | 0.11 | 5.32E-225 | TAM.Ndrg1 | Fgfr1 |
| Maged12 | 1.30E-228 | 1.05984809 | 0.307 | 0.175 | 2.50E-224 | TAM.Ndrg1 | Maged1 |
| B3gnt71 | 1.48E-227 | 0.90006344 | 0.39 | 0.245 | 2.86E-223 | TAM.Ndrg1 | B3gnt7 |
| Rtn4 | 7.51E-227 | 0.33657716 | 0.81 | 0.751 | 1.45E-222 | TAM.Ndrg1 | Rtn4 |
| Mat2a | 3.88E-226 | 0.54874438 | 0.778 | 0.666 | 7.50E-222 | TAM.Ndrg1 | Mat2a |
| Pygl3 | 3.52E-224 | 0.36373124 | 0.702 | 0.58 | 6.80E-220 | TAM.Ndrg1 | Pygl |
| Add32 | 3.82E-221 | 0.75335729 | 0.381 | 0.248 | 7.39E-217 | TAM.Ndrg1 | Add3 |
| Rgs21 | 1.37E-220 | 0.42178414 | 0.684 | 0.606 | 2.64E-216 | TAM.Ndrg1 | Rgs2 |
| Arrdc31 | 1.46E-216 | 0.80845254 | 0.379 | 0.257 | 2.81E-212 | TAM.Ndrg1 | Arrdc3 |
| Galk11 | 2.28E-215 | 0.92993581 | 0.382 | 0.23 | 4.41E-211 | TAM.Ndrg1 | Galk1 |
| Tuba1c | 1.09E-214 | 0.37799671 | 0.816 | 0.723 | 2.11E-210 | TAM.Ndrg1 | Tuba1c |
| Vat12 | 5.88E-212 | 0.46905385 | 0.784 | 0.579 | 1.14E-207 | TAM.Ndrg1 | Vat1 |
| Dnajb11 | 8.54E-211 | 0.83045877 | 0.526 | 0.384 | 1.65E-206 | TAM.Ndrg1 | Dnajb1 |
| Tmem154 | 1.32E-210 | 1.04836465 | 0.277 | 0.159 | 2.56E-206 | TAM.Ndrg1 | Tmem154 |
| Stom2 | 6.31E-208 | 0.57311141 | 0.56 | 0.392 | 1.22E-203 | TAM.Ndrg1 | Stom |
| Zfp532 | 1.03E-205 | 1.51012559 | 0.264 | 0.108 | 2.00E-201 | TAM.Ndrg1 | Zfp532 |
| Chka1 | 2.00E-203 | 0.83915433 | 0.463 | 0.347 | 3.86E-199 | TAM.Ndrg1 | Chka |
| Chd41 | 9.77E-203 | 0.5237998 | 0.721 | 0.625 | 1.89E-198 | TAM.Ndrg1 | Chd4 |
| Ccl42 | 8.20E-202 | 0.35662381 | 0.514 | 0.435 | 1.59E-197 | TAM.Ndrg1 | Ccl4 |
| Sep-81 | 7.85E-200 | 0.68034614 | 0.57 | 0.382 | 1.52E-195 | TAM.Ndrg1 | 8-Sep |
| Hsp90aa1 | 3.08E-199 | 0.59135512 | 0.705 | 0.589 | 5.96E-195 | TAM.Ndrg1 | Hsp90aa1 |
| Samsn12 | 2.82E-198 | 0.58393229 | 0.449 | 0.299 | 5.46E-194 | TAM.Ndrg1 | Samsn1 |
| Zfand2a1 | 1.27E-197 | 0.69600493 | 0.611 | 0.447 | 2.46E-193 | TAM.Ndrg1 | Zfand2a |
| Olr13 | 6.05E-193 | 0.40693756 | 0.263 | 0.202 | 1.17E-188 | TAM.Ndrg1 | Olr1 |
| Net1 | 3.07E-192 | 0.81634218 | 0.336 | 0.215 | 5.94E-188 | TAM.Ndrg1 | Net1 |
| Tmem230 | 1.65E-191 | 0.94139176 | 0.41 | 0.28 | 3.19E-187 | TAM.Ndrg1 | Tmem230 |
| Tubb61 | 9.71E-190 | 0.47023965 | 0.733 | 0.609 | 1.88E-185 | TAM.Ndrg1 | Tubb6 |
| Cflar2 | 2.84E-189 | 0.46682965 | 0.625 | 0.493 | 5.49E-185 | TAM.Ndrg1 | Cflar |
| Ppp1r3b2 | 2.23E-188 | 0.60712262 | 0.359 | 0.238 | 4.30E-184 | TAM.Ndrg1 | Ppp1r3b |
| Tmem1193 | 3.64E-185 | 0.67908106 | 0.418 | 0.292 | 7.05E-181 | TAM.Ndrg1 | Tmem119 |
| Prr5l | 1.99E-184 | 1.02210432 | 0.279 | 0.131 | 3.85E-180 | TAM.Ndrg1 | Prr5l |
| Pttg1ip2 | 1.23E-182 | 0.46829962 | 0.71 | 0.58 | 2.39E-178 | TAM.Ndrg1 | Pttg1ip |
| Hspa5 | 5.58E-182 | 0.4060563 | 0.771 | 0.716 | 1.08E-177 | TAM.Ndrg1 | Hspa5 |
| Asph2 | 6.61E-181 | 0.52992803 | 0.611 | 0.434 | 1.28E-176 | TAM.Ndrg1 | Asph |
| Itga51 | 2.67E-177 | 0.4458136 | 0.66 | 0.525 | 5.16E-173 | TAM.Ndrg1 | Itga5 |
| Gch12 | 2.96E-177 | 0.36558467 | 0.53 | 0.419 | 5.72E-173 | TAM.Ndrg1 | Gch1 |
| Fam20c1 | 5.52E-176 | 0.59149728 | 0.648 | 0.495 | 1.07E-171 | TAM.Ndrg1 | Fam20c |
| Cgnl1 | 8.02E-174 | 1.13447937 | 0.265 | 0.122 | 1.55E-169 | TAM.Ndrg1 | Cgnl1 |
| Mthfs2 | 5.14E-172 | 0.70557866 | 0.442 | 0.307 | 9.94E-168 | TAM.Ndrg1 | Mthfs |
| Lrp122 | 3.05E-168 | 0.62290232 | 0.572 | 0.429 | 5.90E-164 | TAM.Ndrg1 | Lrp12 |
| Rbms21 | 1.98E-165 | 0.89291374 | 0.387 | 0.23 | 3.82E-161 | TAM.Ndrg1 | Rbms2 |
| Zfand5 | 6.77E-164 | 0.31846566 | 0.848 | 0.783 | 1.31E-159 | TAM.Ndrg1 | Zfand5 |
| Tnfrsf231 | 1.91E-162 | 0.97416827 | 0.278 | 0.167 | 3.70E-158 | TAM.Ndrg1 | Tnfrsf23 |
| Card193 | 8.95E-161 | 0.31172788 | 0.717 | 0.602 | 1.73E-156 | TAM.Ndrg1 | Card19 |
| Ppp1r101 | 8.65E-160 | 0.50973488 | 0.571 | 0.473 | 1.67E-155 | TAM.Ndrg1 | Ppp1r10 |
| Rsrp11 | 2.82E-159 | 0.3167354 | 0.867 | 0.814 | 5.45E-155 | TAM.Ndrg1 | Rsrp1 |
| Maff2 | 3.92E-159 | 0.26834711 | 0.773 | 0.631 | 7.59E-155 | TAM.Ndrg1 | Maff |
| Tuba1a1 | 5.21E-159 | 0.60902709 | 0.436 | 0.332 | 1.01E-154 | TAM.Ndrg1 | Tuba1a |
| Ell1 | 1.90E-158 | 0.70443504 | 0.429 | 0.286 | 3.67E-154 | TAM.Ndrg1 | Ell |
| Psd32 | 1.99E-158 | 0.73308254 | 0.422 | 0.275 | 3.84E-154 | TAM.Ndrg1 | Psd3 |
| Dnaja1 | 4.32E-158 | 0.69800577 | 0.523 | 0.4 | 8.35E-154 | TAM.Ndrg1 | Dnaja1 |
| Lonrf31 | 3.41E-157 | 0.86153227 | 0.363 | 0.236 | 6.60E-153 | TAM.Ndrg1 | Lonrf3 |
| Rai143 | 7.08E-155 | 0.46329567 | 0.631 | 0.493 | 1.37E-150 | TAM.Ndrg1 | Rai14 |
| Icam12 | 4.34E-154 | 0.27438104 | 0.628 | 0.621 | 8.39E-150 | TAM.Ndrg1 | Icam1 |
| Mfge81 | 1.32E-153 | 0.68311491 | 0.296 | 0.243 | 2.55E-149 | TAM.Ndrg1 | Mfge8 |
| Alcam2 | 1.79E-153 | 0.54571786 | 0.522 | 0.372 | 3.45E-149 | TAM.Ndrg1 | Alcam |
| Mctp11 | 1.96E-151 | 0.98911865 | 0.252 | 0.124 | 3.79E-147 | TAM.Ndrg1 | Mctp1 |
| Rarg | 3.22E-150 | 0.8841255 | 0.265 | 0.141 | 6.23E-146 | TAM.Ndrg1 | Rarg |
| Cd371 | 6.05E-150 | 0.5487654 | 0.484 | 0.312 | 1.17E-145 | TAM.Ndrg1 | Cd37 |
| H2-Q61 | 4.26E-149 | 0.47373901 | 0.512 | 0.409 | 8.24E-145 | TAM.Ndrg1 | H2-Q6 |
| Grhpr | 4.43E-149 | 1.07921966 | 0.273 | 0.139 | 8.56E-145 | TAM.Ndrg1 | Grhpr |
| Elf11 | 8.05E-149 | 0.41299218 | 0.683 | 0.58 | 1.56E-144 | TAM.Ndrg1 | Elf1 |
| F71 | 4.75E-147 | 0.71492118 | 0.431 | 0.248 | 9.19E-143 | TAM.Ndrg1 | F7 |
| Adam151 | 6.81E-144 | 0.34237254 | 0.72 | 0.661 | 1.32E-139 | TAM.Ndrg1 | Adam15 |
| Acvrl11 | 5.81E-143 | 0.67483534 | 0.428 | 0.307 | 1.12E-138 | TAM.Ndrg1 | Acvrl1 |
| Bmt21 | 1.21E-142 | 0.65349375 | 0.476 | 0.336 | 2.33E-138 | TAM.Ndrg1 | Bmt2 |
| Stk241 | 1.57E-142 | 0.41060944 | 0.642 | 0.532 | 3.04E-138 | TAM.Ndrg1 | Stk24 |
| Homer12 | 2.24E-142 | 0.38772296 | 0.563 | 0.462 | 4.33E-138 | TAM.Ndrg1 | Homer1 |
| Ptgfrn1 | 1.86E-141 | 0.59505657 | 0.483 | 0.353 | 3.60E-137 | TAM.Ndrg1 | Ptgfrn |
| Tnip2 | 1.16E-140 | 0.53808399 | 0.61 | 0.466 | 2.25E-136 | TAM.Ndrg1 | Tnip2 |
| Cass4 | 1.54E-140 | 0.89570745 | 0.303 | 0.201 | 2.98E-136 | TAM.Ndrg1 | Cass4 |
| Pafah1b3 | 1.72E-139 | 0.85798681 | 0.32 | 0.188 | 3.33E-135 | TAM.Ndrg1 | Pafah1b3 |
| Sptan11 | 5.39E-139 | 0.50451764 | 0.657 | 0.534 | 1.04E-134 | TAM.Ndrg1 | Sptan1 |
| Cast1 | 1.10E-138 | 0.39915827 | 0.586 | 0.489 | 2.12E-134 | TAM.Ndrg1 | Cast |
| Rabac12 | 1.79E-138 | 0.31108194 | 0.762 | 0.672 | 3.45E-134 | TAM.Ndrg1 | Rabac1 |
| Plec2 | 2.38E-138 | 0.29653822 | 0.751 | 0.656 | 4.60E-134 | TAM.Ndrg1 | Plec |
| Tmem882 | 1.31E-137 | 0.51863738 | 0.426 | 0.316 | 2.53E-133 | TAM.Ndrg1 | Tmem88 |
| Diaph11 | 2.60E-137 | 0.30191028 | 0.742 | 0.645 | 5.02E-133 | TAM.Ndrg1 | Diaph1 |
| H2-Q72 | 1.43E-135 | 0.44730529 | 0.46 | 0.331 | 2.76E-131 | TAM.Ndrg1 | H2-Q7 |
| Ddx52 | 9.94E-135 | 0.2599336 | 0.739 | 0.689 | 1.92E-130 | TAM.Ndrg1 | Ddx5 |
| Cblb | 7.54E-134 | 0.76059039 | 0.323 | 0.213 | 1.46E-129 | TAM.Ndrg1 | Cblb |
| Tiam12 | 1.70E-133 | 0.51591213 | 0.335 | 0.208 | 3.29E-129 | TAM.Ndrg1 | Tiam1 |
| Rusc21 | 1.14E-132 | 0.84632921 | 0.341 | 0.211 | 2.21E-128 | TAM.Ndrg1 | Rusc2 |
| Alkbh51 | 2.69E-131 | 0.41502534 | 0.671 | 0.573 | 5.20E-127 | TAM.Ndrg1 | Alkbh5 |
| Pde3b1 | 7.28E-131 | 0.58695139 | 0.464 | 0.312 | 1.41E-126 | TAM.Ndrg1 | Pde3b |
| Aifm21 | 1.95E-130 | 0.92228867 | 0.31 | 0.171 | 3.77E-126 | TAM.Ndrg1 | Aifm2 |
| Eps82 | 7.03E-130 | 0.42830459 | 0.583 | 0.447 | 1.36E-125 | TAM.Ndrg1 | Eps8 |
| Ndufv31 | 2.11E-128 | 0.40809765 | 0.71 | 0.602 | 4.08E-124 | TAM.Ndrg1 | Ndufv3 |
| Pstpip12 | 4.86E-128 | 0.42305748 | 0.64 | 0.517 | 9.40E-124 | TAM.Ndrg1 | Pstpip1 |
| Itpk1 | 3.63E-127 | 0.61249258 | 0.438 | 0.327 | 7.01E-123 | TAM.Ndrg1 | Itpk1 |
| Fnip21 | 3.84E-126 | 0.49543936 | 0.56 | 0.499 | 7.42E-122 | TAM.Ndrg1 | Fnip2 |
| Atp8b43 | 6.73E-126 | 0.61536358 | 0.338 | 0.205 | 1.30E-121 | TAM.Ndrg1 | Atp8b4 |
| Tsc22d1 | 1.06E-125 | 0.64468899 | 0.463 | 0.346 | 2.06E-121 | TAM.Ndrg1 | Tsc22d1 |
| Rcan1 | 1.36E-125 | 0.62061951 | 0.292 | 0.222 | 2.63E-121 | TAM.Ndrg1 | Rcan1 |
| Irs22 | 3.67E-125 | 0.36802321 | 0.807 | 0.688 | 7.10E-121 | TAM.Ndrg1 | Irs2 |
| Cxcl101 | 4.52E-125 | 0.49130259 | 0.312 | 0.201 | 8.74E-121 | TAM.Ndrg1 | Cxcl10 |
| Hcar21 | 2.47E-124 | 0.52143042 | 0.482 | 0.38 | 4.77E-120 | TAM.Ndrg1 | Hcar2 |
| Serpinb6a1 | 4.28E-124 | 0.47844635 | 0.555 | 0.313 | 8.27E-120 | TAM.Ndrg1 | Serpinb6a |
| Raph1 | 7.81E-123 | 0.81722391 | 0.28 | 0.156 | 1.51E-118 | TAM.Ndrg1 | Raph1 |
| Ogt1 | 3.90E-122 | 0.44056144 | 0.571 | 0.458 | 7.53E-118 | TAM.Ndrg1 | Ogt |
| Clic41 | 9.64E-122 | 0.26634438 | 0.812 | 0.735 | 1.86E-117 | TAM.Ndrg1 | Clic4 |
| Sap30 | 1.21E-121 | 0.59921066 | 0.402 | 0.344 | 2.34E-117 | TAM.Ndrg1 | Sap30 |
| Socs12 | 3.36E-120 | 0.8258535 | 0.356 | 0.213 | 6.50E-116 | TAM.Ndrg1 | Socs1 |
| Plod11 | 4.54E-119 | 0.56454855 | 0.499 | 0.372 | 8.78E-115 | TAM.Ndrg1 | Plod1 |
| Tubb4b1 | 1.12E-118 | 0.67840786 | 0.572 | 0.416 | 2.16E-114 | TAM.Ndrg1 | Tubb4b |
| Sgpp11 | 2.10E-118 | 0.47723964 | 0.477 | 0.368 | 4.06E-114 | TAM.Ndrg1 | Sgpp1 |
| Tmem372 | 9.26E-118 | 0.50510184 | 0.549 | 0.402 | 1.79E-113 | TAM.Ndrg1 | Tmem37 |
| Ugcg1 | 1.89E-117 | 0.39540851 | 0.515 | 0.413 | 3.65E-113 | TAM.Ndrg1 | Ugcg |
| Nufip21 | 2.50E-116 | 0.4916035 | 0.645 | 0.525 | 4.83E-112 | TAM.Ndrg1 | Nufip2 |
| Ints61 | 4.37E-114 | 0.65023058 | 0.419 | 0.32 | 8.45E-110 | TAM.Ndrg1 | Ints6 |
| Cep1702 | 3.34E-113 | 0.41533322 | 0.584 | 0.45 | 6.45E-109 | TAM.Ndrg1 | Cep170 |
| Nupr12 | 5.49E-113 | 0.43331394 | 0.504 | 0.379 | 1.06E-108 | TAM.Ndrg1 | Nupr1 |
| Lamc1 | 7.98E-113 | 0.95397623 | 0.26 | 0.143 | 1.54E-108 | TAM.Ndrg1 | Lamc1 |
| Zswim62 | 2.40E-111 | 0.42128687 | 0.532 | 0.431 | 4.63E-107 | TAM.Ndrg1 | Zswim6 |
| Hipk2 | 3.63E-111 | 0.44959369 | 0.573 | 0.474 | 7.03E-107 | TAM.Ndrg1 | Hipk2 |
| Rapgef2 | 1.37E-110 | 0.60292641 | 0.422 | 0.319 | 2.65E-106 | TAM.Ndrg1 | Rapgef2 |
| Trim252 | 5.17E-110 | 0.25685346 | 0.735 | 0.665 | 1.00E-105 | TAM.Ndrg1 | Trim25 |
| Flrt33 | 1.78E-109 | 0.31968978 | 0.32 | 0.248 | 3.43E-105 | TAM.Ndrg1 | Flrt3 |
| Srsf2 | 2.00E-109 | 0.53660996 | 0.671 | 0.574 | 3.87E-105 | TAM.Ndrg1 | Srsf2 |
| Gfpt11 | 4.44E-109 | 0.62349358 | 0.383 | 0.28 | 8.59E-105 | TAM.Ndrg1 | Gfpt1 |
| Pdlim73 | 5.07E-108 | 0.49072664 | 0.465 | 0.333 | 9.81E-104 | TAM.Ndrg1 | Pdlim7 |
| Ahcyl2 | 5.68E-108 | 0.52547289 | 0.488 | 0.381 | 1.10E-103 | TAM.Ndrg1 | Ahcyl2 |
| Cyb5r31 | 3.76E-107 | 0.3675697 | 0.676 | 0.571 | 7.28E-103 | TAM.Ndrg1 | Cyb5r3 |
| Cd24a3 | 6.93E-107 | 0.40587686 | 0.34 | 0.253 | 1.34E-102 | TAM.Ndrg1 | Cd24a |
| Nfix1 | 1.64E-106 | 0.5529142 | 0.364 | 0.234 | 3.17E-102 | TAM.Ndrg1 | Nfix |
| Casp42 | 3.69E-106 | 0.5443393 | 0.401 | 0.277 | 7.14E-102 | TAM.Ndrg1 | Casp4 |
| Aff1 | 1.17E-104 | 0.2711646 | 0.691 | 0.609 | 2.26E-100 | TAM.Ndrg1 | Aff1 |
| Akap131 | 1.81E-104 | 0.33821415 | 0.663 | 0.563 | 3.51E-100 | TAM.Ndrg1 | Akap13 |
| Ptprs1 | 1.83E-104 | 0.43837835 | 0.54 | 0.427 | 3.53E-100 | TAM.Ndrg1 | Ptprs |
| Dag11 | 5.23E-102 | 0.72780453 | 0.366 | 0.235 | 1.01E-97 | TAM.Ndrg1 | Dag1 |
| Map2k11 | 1.30E-101 | 0.35370719 | 0.581 | 0.491 | 2.51E-97 | TAM.Ndrg1 | Map2k1 |
| Foxp12 | 2.75E-101 | 0.44654172 | 0.446 | 0.332 | 5.32E-97 | TAM.Ndrg1 | Foxp1 |
| Sfxn31 | 5.88E-101 | 0.32756958 | 0.565 | 0.456 | 1.14E-96 | TAM.Ndrg1 | Sfxn3 |
| Prdm21 | 1.06E-100 | 0.51299586 | 0.499 | 0.375 | 2.05E-96 | TAM.Ndrg1 | Prdm2 |
| Ubash3b2 | 4.02E-100 | 0.42899278 | 0.447 | 0.344 | 7.77E-96 | TAM.Ndrg1 | Ubash3b |
| Eaf11 | 9.69E-100 | 0.48991351 | 0.466 | 0.377 | 1.87E-95 | TAM.Ndrg1 | Eaf1 |
| Pcnx1 | 3.62E-98 | 0.62066407 | 0.312 | 0.227 | 7.00E-94 | TAM.Ndrg1 | Pcnx |
| Zmiz11 | 1.61E-97 | 0.26742281 | 0.717 | 0.653 | 3.11E-93 | TAM.Ndrg1 | Zmiz1 |
| Adss | 8.99E-97 | 0.54027696 | 0.469 | 0.371 | 1.74E-92 | TAM.Ndrg1 | Adss |
| Insig11 | 4.63E-96 | 0.29844459 | 0.712 | 0.623 | 8.96E-92 | TAM.Ndrg1 | Insig1 |
| Dpysl2 | 1.84E-94 | 0.35090317 | 0.589 | 0.525 | 3.56E-90 | TAM.Ndrg1 | Dpysl2 |
| Sirt1 | 1.32E-92 | 0.80514951 | 0.302 | 0.207 | 2.56E-88 | TAM.Ndrg1 | Sirt1 |
| Jmjd1c2 | 1.47E-92 | 0.27631068 | 0.688 | 0.61 | 2.85E-88 | TAM.Ndrg1 | Jmjd1c |
| Camk2g | 2.34E-91 | 0.63741346 | 0.35 | 0.242 | 4.53E-87 | TAM.Ndrg1 | Camk2g |
| Sbf22 | 1.20E-90 | 0.67697355 | 0.39 | 0.245 | 2.33E-86 | TAM.Ndrg1 | Sbf2 |
| Inpp4a1 | 5.89E-90 | 0.44963563 | 0.427 | 0.313 | 1.14E-85 | TAM.Ndrg1 | Inpp4a |
| Aldoc2 | 1.23E-89 | 0.50456317 | 0.424 | 0.306 | 2.39E-85 | TAM.Ndrg1 | Aldoc |
| Ppme1 | 3.16E-89 | 0.61031033 | 0.341 | 0.237 | 6.10E-85 | TAM.Ndrg1 | Ppme1 |
| Lrrfip21 | 7.00E-89 | 0.4937394 | 0.471 | 0.358 | 1.35E-84 | TAM.Ndrg1 | Lrrfip2 |
| Anpep2 | 4.71E-88 | 0.33377365 | 0.591 | 0.49 | 9.10E-84 | TAM.Ndrg1 | Anpep |
| Cd843 | 7.71E-88 | 0.2649267 | 0.725 | 0.616 | 1.49E-83 | TAM.Ndrg1 | Cd84 |
| Rnf1411 | 8.88E-88 | 0.60958511 | 0.318 | 0.219 | 1.72E-83 | TAM.Ndrg1 | Rnf141 |
| Atp7a1 | 1.32E-87 | 0.40409821 | 0.523 | 0.433 | 2.55E-83 | TAM.Ndrg1 | Atp7a |
| Nav11 | 2.66E-87 | 0.53715829 | 0.359 | 0.254 | 5.13E-83 | TAM.Ndrg1 | Nav1 |
| Gas2l11 | 4.08E-87 | 0.47524195 | 0.464 | 0.342 | 7.88E-83 | TAM.Ndrg1 | Gas2l1 |
| Mthfd11 | 4.50E-87 | 0.53764887 | 0.295 | 0.205 | 8.69E-83 | TAM.Ndrg1 | Mthfd1 |
| Dapk11 | 9.18E-87 | 0.43287603 | 0.512 | 0.421 | 1.77E-82 | TAM.Ndrg1 | Dapk1 |
| Sh3kbp1 | 5.44E-86 | 0.43280246 | 0.448 | 0.352 | 1.05E-81 | TAM.Ndrg1 | Sh3kbp1 |
| Tec1 | 6.25E-85 | 0.56322431 | 0.454 | 0.346 | 1.21E-80 | TAM.Ndrg1 | Tec |
| Ttc39c1 | 1.27E-84 | 0.66095304 | 0.264 | 0.176 | 2.45E-80 | TAM.Ndrg1 | Ttc39c |
| Macf11 | 7.99E-84 | 0.33723194 | 0.671 | 0.589 | 1.55E-79 | TAM.Ndrg1 | Macf1 |
| Tmem87b1 | 1.81E-83 | 0.49620761 | 0.486 | 0.374 | 3.50E-79 | TAM.Ndrg1 | Tmem87b |
| Arhgap261 | 4.58E-83 | 0.32019299 | 0.346 | 0.257 | 8.85E-79 | TAM.Ndrg1 | Arhgap26 |
| Ccnyl1 | 1.37E-82 | 0.67313782 | 0.254 | 0.166 | 2.66E-78 | TAM.Ndrg1 | Ccnyl1 |
| Srxn11 | 1.50E-82 | 0.45652132 | 0.477 | 0.374 | 2.91E-78 | TAM.Ndrg1 | Srxn1 |
| Snx1 | 1.58E-81 | 0.2815692 | 0.635 | 0.557 | 3.05E-77 | TAM.Ndrg1 | Snx1 |
| Tlr41 | 1.31E-80 | 0.33232676 | 0.507 | 0.397 | 2.53E-76 | TAM.Ndrg1 | Tlr4 |
| Dok21 | 5.50E-80 | 0.34382623 | 0.535 | 0.484 | 1.06E-75 | TAM.Ndrg1 | Dok2 |
| Fndc3b1 | 5.65E-80 | 0.30479251 | 0.614 | 0.526 | 1.09E-75 | TAM.Ndrg1 | Fndc3b |
| Fam102b2 | 8.07E-80 | 0.49825473 | 0.444 | 0.357 | 1.56E-75 | TAM.Ndrg1 | Fam102b |
| Zfp2921 | 1.54E-79 | 0.4646166 | 0.45 | 0.335 | 2.98E-75 | TAM.Ndrg1 | Zfp292 |
| Grk53 | 6.20E-79 | 0.48804485 | 0.354 | 0.264 | 1.20E-74 | TAM.Ndrg1 | Grk5 |
| Rras2 | 4.23E-78 | 0.40248486 | 0.389 | 0.281 | 8.18E-74 | TAM.Ndrg1 | Rras |
| C5ar22 | 6.13E-77 | 0.5285167 | 0.332 | 0.273 | 1.19E-72 | TAM.Ndrg1 | C5ar2 |
| Apbb31 | 1.84E-76 | 0.65577918 | 0.258 | 0.166 | 3.55E-72 | TAM.Ndrg1 | Apbb3 |
| Itpkb1 | 2.25E-76 | 0.40591605 | 0.469 | 0.371 | 4.36E-72 | TAM.Ndrg1 | Itpkb |
| Psmd8 | 4.61E-76 | 0.39599242 | 0.535 | 0.445 | 8.90E-72 | TAM.Ndrg1 | Psmd8 |
| Klhl211 | 4.89E-76 | 0.75791329 | 0.373 | 0.278 | 9.46E-72 | TAM.Ndrg1 | Klhl21 |
| Gas72 | 8.25E-76 | 0.27196324 | 0.65 | 0.576 | 1.59E-71 | TAM.Ndrg1 | Gas7 |
| Pi4k2a1 | 2.16E-75 | 0.47229031 | 0.537 | 0.419 | 4.18E-71 | TAM.Ndrg1 | Pi4k2a |
| Sipa1l1 | 3.04E-75 | 0.63223483 | 0.25 | 0.152 | 5.88E-71 | TAM.Ndrg1 | Sipa1l1 |
| Arhgap102 | 3.93E-75 | 0.84011617 | 0.308 | 0.192 | 7.60E-71 | TAM.Ndrg1 | Arhgap10 |
| Bnip3l1 | 7.41E-74 | 0.77551294 | 0.267 | 0.167 | 1.43E-69 | TAM.Ndrg1 | Bnip3l |
| Ddx461 | 7.57E-74 | 0.35066806 | 0.534 | 0.453 | 1.46E-69 | TAM.Ndrg1 | Ddx46 |
| Rnf19a | 2.83E-73 | 0.60429268 | 0.323 | 0.222 | 5.47E-69 | TAM.Ndrg1 | Rnf19a |
| Dlgap41 | 4.05E-73 | 0.33273907 | 0.601 | 0.506 | 7.84E-69 | TAM.Ndrg1 | Dlgap4 |
| Man2a22 | 6.74E-73 | 0.44572386 | 0.508 | 0.443 | 1.30E-68 | TAM.Ndrg1 | Man2a2 |
| Kmt2e1 | 1.56E-72 | 0.37916413 | 0.465 | 0.373 | 3.01E-68 | TAM.Ndrg1 | Kmt2e |
| Kdm4b | 2.58E-72 | 0.64159747 | 0.285 | 0.179 | 4.99E-68 | TAM.Ndrg1 | Kdm4b |
| Pik3r51 | 3.44E-72 | 0.28013723 | 0.619 | 0.554 | 6.65E-68 | TAM.Ndrg1 | Pik3r5 |
| Naa801 | 4.83E-72 | 0.6715865 | 0.297 | 0.206 | 9.34E-68 | TAM.Ndrg1 | Naa80 |
| Rbpms1 | 5.66E-72 | 0.47956644 | 0.297 | 0.202 | 1.09E-67 | TAM.Ndrg1 | Rbpms |
| Atf4 | 6.06E-72 | 0.46285174 | 0.506 | 0.394 | 1.17E-67 | TAM.Ndrg1 | Atf4 |
| Rab6a1 | 8.68E-72 | 0.30799673 | 0.495 | 0.406 | 1.68E-67 | TAM.Ndrg1 | Rab6a |
| Iqgap3 | 1.37E-71 | 0.98293114 | 0.263 | 0.128 | 2.66E-67 | TAM.Ndrg1 | Iqgap3 |
| Polr2a | 1.63E-70 | 0.36090656 | 0.63 | 0.544 | 3.15E-66 | TAM.Ndrg1 | Polr2a |
| St6gal11 | 2.42E-70 | 0.43303716 | 0.404 | 0.293 | 4.69E-66 | TAM.Ndrg1 | St6gal1 |
| DEPRECATED-I | 7.65E-70 | 0.28481778 | 0.418 | 0.362 | 1.48E-65 | TAM.Ndrg1 | DEPRECATED-ENSMUSG00000025025 |
| Cd1511 | 8.59E-70 | 0.57179107 | 0.378 | 0.255 | 1.66E-65 | TAM.Ndrg1 | Cd151 |
| Kdm3a1 | 1.06E-69 | 0.51856646 | 0.421 | 0.315 | 2.05E-65 | TAM.Ndrg1 | Kdm3a |
| Foxo1 | 1.67E-69 | 0.60929262 | 0.294 | 0.199 | 3.23E-65 | TAM.Ndrg1 | Foxo1 |
| Kat6b | 2.17E-69 | 0.69947623 | 0.264 | 0.166 | 4.20E-65 | TAM.Ndrg1 | Kat6b |
| Trio | 4.44E-69 | 0.37187194 | 0.563 | 0.457 | 8.59E-65 | TAM.Ndrg1 | Trio |
| Hmgcl | 7.27E-69 | 0.26965813 | 0.638 | 0.551 | 1.41E-64 | TAM.Ndrg1 | Hmgcl |
| Clcn3 | 1.08E-68 | 0.71564435 | 0.273 | 0.182 | 2.08E-64 | TAM.Ndrg1 | Clcn3 |
| Lpp1 | 1.65E-68 | 0.29543342 | 0.669 | 0.571 | 3.19E-64 | TAM.Ndrg1 | Lpp |
| Peak11 | 2.10E-68 | 0.70127419 | 0.286 | 0.179 | 4.07E-64 | TAM.Ndrg1 | Peak1 |
| Golph3l1 | 2.20E-68 | 0.51961897 | 0.326 | 0.241 | 4.25E-64 | TAM.Ndrg1 | Golph3l |
| R3hdm1 | 6.07E-68 | 0.52149417 | 0.32 | 0.241 | 1.17E-63 | TAM.Ndrg1 | R3hdm1 |
| Phf21a1 | 6.53E-68 | 0.51578741 | 0.357 | 0.256 | 1.26E-63 | TAM.Ndrg1 | Phf21a |
| Ddit3 | 6.53E-68 | 0.70825543 | 0.283 | 0.209 | 1.26E-63 | TAM.Ndrg1 | Ddit3 |
| Pgp | 9.90E-68 | 0.53315321 | 0.317 | 0.232 | 1.91E-63 | TAM.Ndrg1 | Pgp |
| Rftn1 | 1.01E-67 | 0.38826709 | 0.594 | 0.489 | 1.96E-63 | TAM.Ndrg1 | Rftn1 |
| Kcnk131 | 1.29E-67 | 0.79664948 | 0.256 | 0.159 | 2.49E-63 | TAM.Ndrg1 | Kcnk13 |
| Me2 | 8.08E-67 | 0.5362488 | 0.256 | 0.186 | 1.56E-62 | TAM.Ndrg1 | Me2 |
| Ltbp32 | 8.17E-67 | 0.51774735 | 0.438 | 0.315 | 1.58E-62 | TAM.Ndrg1 | Ltbp3 |
| Dynll12 | 3.39E-66 | 0.32759791 | 0.559 | 0.482 | 6.56E-62 | TAM.Ndrg1 | Dynll1 |
| Bag3 | 1.34E-65 | 0.4360251 | 0.469 | 0.372 | 2.58E-61 | TAM.Ndrg1 | Bag3 |
| Sptbn1 | 3.58E-65 | 0.41130847 | 0.403 | 0.326 | 6.91E-61 | TAM.Ndrg1 | Sptbn1 |
| Tnfsf142 | 8.75E-65 | 0.41036667 | 0.305 | 0.214 | 1.69E-60 | TAM.Ndrg1 | Tnfsf14 |
| Eea11 | 9.35E-65 | 0.45913184 | 0.433 | 0.327 | 1.81E-60 | TAM.Ndrg1 | Eea1 |
| Slc7a83 | 3.02E-64 | 0.26170104 | 0.592 | 0.506 | 5.84E-60 | TAM.Ndrg1 | Slc7a8 |
| Fam114a21 | 5.18E-64 | 0.51333533 | 0.344 | 0.251 | 1.00E-59 | TAM.Ndrg1 | Fam114a2 |
| Uap12 | 8.90E-64 | 0.32749547 | 0.453 | 0.384 | 1.72E-59 | TAM.Ndrg1 | Uap1 |
| Dot1l2 | 2.45E-63 | 0.25202159 | 0.517 | 0.438 | 4.73E-59 | TAM.Ndrg1 | Dot1l |
| Pcgf5 | 4.79E-63 | 0.48433264 | 0.303 | 0.22 | 9.27E-59 | TAM.Ndrg1 | Pcgf5 |
| Hsd3b71 | 1.49E-62 | 0.56790908 | 0.394 | 0.283 | 2.89E-58 | TAM.Ndrg1 | Hsd3b7 |
| Esyt2 | 2.01E-62 | 0.41484429 | 0.481 | 0.407 | 3.90E-58 | TAM.Ndrg1 | Esyt2 |
| Fnip1 | 2.46E-62 | 0.43890107 | 0.503 | 0.419 | 4.75E-58 | TAM.Ndrg1 | Fnip1 |
| Lemd21 | 5.61E-62 | 0.3376511 | 0.583 | 0.486 | 1.09E-57 | TAM.Ndrg1 | Lemd2 |
| Fbxo301 | 1.01E-61 | 0.53548851 | 0.348 | 0.263 | 1.94E-57 | TAM.Ndrg1 | Fbxo30 |
| Ifih11 | 1.24E-61 | 0.37464512 | 0.356 | 0.354 | 2.40E-57 | TAM.Ndrg1 | Ifih1 |
| Mindy21 | 2.31E-61 | 0.42018842 | 0.456 | 0.355 | 4.47E-57 | TAM.Ndrg1 | Mindy2 |
| Sertad21 | 3.64E-61 | 0.40851908 | 0.389 | 0.289 | 7.04E-57 | TAM.Ndrg1 | Sertad2 |
| Exoc5 | 6.39E-61 | 0.28835945 | 0.573 | 0.497 | 1.23E-56 | TAM.Ndrg1 | Exoc5 |
| Tmem1042 | 2.22E-59 | 0.3152394 | 0.551 | 0.444 | 4.30E-55 | TAM.Ndrg1 | Tmem104 |
| Slc27a11 | 4.28E-59 | 0.40356986 | 0.536 | 0.419 | 8.27E-55 | TAM.Ndrg1 | Slc27a1 |
| Abl2 | 4.47E-59 | 0.45950781 | 0.53 | 0.435 | 8.64E-55 | TAM.Ndrg1 | Abl2 |
| Scd21 | 4.97E-59 | 0.5386088 | 0.273 | 0.215 | 9.61E-55 | TAM.Ndrg1 | Scd2 |
| Eng3 | 5.36E-59 | 0.45747373 | 0.352 | 0.262 | 1.04E-54 | TAM.Ndrg1 | Eng |
| Pde1b1 | 8.95E-59 | 0.46716028 | 0.433 | 0.33 | 1.73E-54 | TAM.Ndrg1 | Pde1b |
| St8sia41 | 5.84E-58 | 0.27894982 | 0.451 | 0.35 | 1.13E-53 | TAM.Ndrg1 | St8sia4 |
| Blcap2 | 1.45E-57 | 0.30930139 | 0.378 | 0.295 | 2.81E-53 | TAM.Ndrg1 | Blcap |
| Nfe2l11 | 1.94E-57 | 0.32512925 | 0.623 | 0.536 | 3.76E-53 | TAM.Ndrg1 | Nfe2l1 |
| Pdlim51 | 7.20E-57 | 0.25245106 | 0.57 | 0.485 | 1.39E-52 | TAM.Ndrg1 | Pdlim5 |
| Slc16a101 | 2.34E-56 | 0.61614733 | 0.361 | 0.287 | 4.52E-52 | TAM.Ndrg1 | Slc16a10 |
| Tmx4 | 4.32E-56 | 0.52182297 | 0.309 | 0.225 | 8.36E-52 | TAM.Ndrg1 | Tmx4 |
| Fgd62 | 1.37E-54 | 0.66206908 | 0.288 | 0.18 | 2.65E-50 | TAM.Ndrg1 | Fgd6 |
| Spidr | 3.28E-54 | 0.60990946 | 0.251 | 0.161 | 6.34E-50 | TAM.Ndrg1 | Spidr |
| Tgoln1 | 4.70E-54 | 0.27027022 | 0.606 | 0.54 | 9.09E-50 | TAM.Ndrg1 | Tgoln1 |
| Ttc39b | 4.78E-54 | 0.34125667 | 0.413 | 0.327 | 9.24E-50 | TAM.Ndrg1 | Ttc39b |
| Tank | 5.63E-54 | 0.27968515 | 0.552 | 0.473 | 1.09E-49 | TAM.Ndrg1 | Tank |
| Parvb | 6.31E-54 | 0.45860289 | 0.355 | 0.263 | 1.22E-49 | TAM.Ndrg1 | Parvb |
| Cerk2 | 2.18E-53 | 0.31805353 | 0.569 | 0.463 | 4.21E-49 | TAM.Ndrg1 | Cerk |
| Klf71 | 1.39E-52 | 0.32835071 | 0.436 | 0.337 | 2.69E-48 | TAM.Ndrg1 | Klf7 |
| Fam53c1 | 2.05E-52 | 0.47854337 | 0.323 | 0.239 | 3.96E-48 | TAM.Ndrg1 | Fam53c |
| Creb52 | 5.08E-52 | 0.44984209 | 0.453 | 0.353 | 9.83E-48 | TAM.Ndrg1 | Creb5 |
| Arfgap31 | 6.36E-52 | 0.43806824 | 0.326 | 0.246 | 1.23E-47 | TAM.Ndrg1 | Arfgap3 |
| Phldb11 | 7.30E-52 | 0.480565 | 0.276 | 0.19 | 1.41E-47 | TAM.Ndrg1 | Phldb1 |
| Lyst1 | 9.79E-52 | 0.45758915 | 0.282 | 0.209 | 1.89E-47 | TAM.Ndrg1 | Lyst |
| Jak32 | 1.05E-51 | 0.33853207 | 0.407 | 0.313 | 2.02E-47 | TAM.Ndrg1 | Jak3 |
| Rab3gap2 | 2.18E-51 | 0.42236051 | 0.393 | 0.305 | 4.21E-47 | TAM.Ndrg1 | Rab3gap2 |
| Tmem432 | 2.62E-51 | 0.37282058 | 0.449 | 0.366 | 5.06E-47 | TAM.Ndrg1 | Tmem43 |
| Gas2l31 | 2.83E-50 | 0.53939219 | 0.369 | 0.281 | 5.48E-46 | TAM.Ndrg1 | Gas2l3 |
| Jmjd6 | 1.08E-49 | 0.41835844 | 0.409 | 0.322 | 2.09E-45 | TAM.Ndrg1 | Jmjd6 |
| Rbm38 | 1.28E-49 | 0.50298654 | 0.362 | 0.303 | 2.48E-45 | TAM.Ndrg1 | Rbm38 |
| Atg4c1 | 2.19E-49 | 0.47371931 | 0.32 | 0.236 | 4.23E-45 | TAM.Ndrg1 | Atg4c |
| Cpeb41 | 3.11E-49 | 0.41999563 | 0.52 | 0.44 | 6.01E-45 | TAM.Ndrg1 | Cpeb4 |
| Pcna | 2.61E-48 | 0.39381405 | 0.711 | 0.6 | 5.05E-44 | TAM.Ndrg1 | Pcna |
| Slc30a9 | 2.95E-48 | 0.4369277 | 0.357 | 0.273 | 5.70E-44 | TAM.Ndrg1 | Slc30a9 |
| Plekhn11 | 2.78E-47 | 0.33623856 | 0.373 | 0.294 | 5.38E-43 | TAM.Ndrg1 | Plekhn1 |
| Plekha2 | 1.57E-46 | 0.27655059 | 0.509 | 0.428 | 3.04E-42 | TAM.Ndrg1 | Plekha2 |
| Fem1b1 | 4.66E-46 | 0.30260092 | 0.383 | 0.335 | 9.01E-42 | TAM.Ndrg1 | Fem1b |
| Srsf3 | 7.03E-46 | 0.43487994 | 0.361 | 0.285 | 1.36E-41 | TAM.Ndrg1 | Srsf3 |
| Gab2 | 1.25E-45 | 0.26182201 | 0.576 | 0.506 | 2.42E-41 | TAM.Ndrg1 | Gab2 |
| Azin11 | 2.46E-45 | 0.31384077 | 0.487 | 0.398 | 4.76E-41 | TAM.Ndrg1 | Azin1 |
| Camkk21 | 2.98E-45 | 0.32565608 | 0.273 | 0.216 | 5.77E-41 | TAM.Ndrg1 | Camkk2 |
| Synj21 | 4.03E-45 | 0.51415687 | 0.287 | 0.203 | 7.79E-41 | TAM.Ndrg1 | Synj2 |
| Syf21 | 4.11E-45 | 0.30654692 | 0.626 | 0.571 | 7.95E-41 | TAM.Ndrg1 | Syf2 |
| Rexf1 | 1.87E-44 | 0.33221062 | 0.338 | 0.287 | 3.61E-40 | TAM.Ndrg1 | Rexf1 |
| Abhd51 | 2.05E-44 | 0.57464364 | 0.281 | 0.21 | 3.96E-40 | TAM.Ndrg1 | Abhd5 |
| Vps13c1 | 6.50E-44 | 0.40907149 | 0.369 | 0.283 | 1.26E-39 | TAM.Ndrg1 | Vps13c |
| Hcfc1r1 | 6.61E-44 | 0.26120575 | 0.503 | 0.436 | 1.28E-39 | TAM.Ndrg1 | Hcfc1r1 |
| Klf91 | 9.33E-44 | 0.42021514 | 0.298 | 0.208 | 1.80E-39 | TAM.Ndrg1 | Klf9 |
| Inf22 | 1.35E-43 | 0.36309789 | 0.337 | 0.236 | 2.61E-39 | TAM.Ndrg1 | Inf2 |
| Ern11 | 1.86E-43 | 0.25280647 | 0.585 | 0.488 | 3.59E-39 | TAM.Ndrg1 | Ern1 |
| Arl31 | 8.27E-43 | 0.41012478 | 0.297 | 0.217 | 1.60E-38 | TAM.Ndrg1 | Arl3 |
| Pon22 | 1.79E-42 | 0.26388104 | 0.483 | 0.408 | 3.46E-38 | TAM.Ndrg1 | Pon2 |
| Mex3c2 | 3.33E-42 | 0.27414922 | 0.438 | 0.382 | 6.44E-38 | TAM.Ndrg1 | Mex3c |
| Orai2 | 5.39E-42 | 0.3306705 | 0.285 | 0.239 | 1.04E-37 | TAM.Ndrg1 | Orai2 |
| Idh12 | 5.50E-42 | 0.27425687 | 0.486 | 0.41 | 1.06E-37 | TAM.Ndrg1 | Idh1 |
| Mapk8ip31 | 1.11E-41 | 0.38993035 | 0.372 | 0.287 | 2.14E-37 | TAM.Ndrg1 | Mapk8ip3 |
| Tsc22d33 | 1.54E-41 | 0.28785175 | 0.48 | 0.401 | 2.97E-37 | TAM.Ndrg1 | Tsc22d3 |
| Pgrmc1 | 3.88E-41 | 0.43029281 | 0.372 | 0.293 | 7.51E-37 | TAM.Ndrg1 | Pgrmc1 |
| Fmnl21 | 4.44E-41 | 0.43367341 | 0.334 | 0.271 | 8.59E-37 | TAM.Ndrg1 | Fmnl2 |
| Mepce1 | 7.83E-41 | 0.41429077 | 0.394 | 0.317 | 1.51E-36 | TAM.Ndrg1 | Mepce |
| Notch11 | 1.79E-40 | 0.2952023 | 0.429 | 0.342 | 3.46E-36 | TAM.Ndrg1 | Notch1 |
| Zfp1311 | 2.31E-40 | 0.26289933 | 0.51 | 0.431 | 4.46E-36 | TAM.Ndrg1 | Zfp131 |
| H1f01 | 3.31E-40 | 0.30450312 | 0.455 | 0.415 | 6.41E-36 | TAM.Ndrg1 | H1f0 |
| Rnf126 | 3.82E-39 | 0.39139453 | 0.417 | 0.333 | 7.39E-35 | TAM.Ndrg1 | Rnf126 |
| Fam214b2 | 5.00E-39 | 0.26732715 | 0.413 | 0.322 | 9.67E-35 | TAM.Ndrg1 | Fam214b |
| Akt3 | 1.14E-38 | 0.3520386 | 0.253 | 0.183 | 2.21E-34 | TAM.Ndrg1 | Akt3 |
| Tmem159 | 1.23E-38 | 0.57779594 | 0.257 | 0.179 | 2.38E-34 | TAM.Ndrg1 | Tmem159 |
| Phactr4 | 3.12E-38 | 0.45093437 | 0.291 | 0.218 | 6.03E-34 | TAM.Ndrg1 | Phactr4 |
| Mettl91 | 1.11E-37 | 0.27647065 | 0.409 | 0.33 | 2.15E-33 | TAM.Ndrg1 | Mettl9 |
| Elmo21 | 8.05E-36 | 0.29647889 | 0.327 | 0.257 | 1.56E-31 | TAM.Ndrg1 | Elmo2 |
| Dhdh2 | 1.93E-35 | 0.42161052 | 0.284 | 0.206 | 3.74E-31 | TAM.Ndrg1 | Dhdh |
| Hk31 | 4.79E-35 | 0.35872045 | 0.366 | 0.302 | 9.25E-31 | TAM.Ndrg1 | Hk3 |
| Iffo2 | 6.07E-35 | 0.54900921 | 0.25 | 0.177 | 1.17E-30 | TAM.Ndrg1 | Iffo2 |
| Tex2642 | 6.96E-35 | 0.34448837 | 0.462 | 0.364 | 1.35E-30 | TAM.Ndrg1 | Tex264 |
| Tmem63b1 | 8.84E-35 | 0.36762236 | 0.258 | 0.185 | 1.71E-30 | TAM.Ndrg1 | Tmem63b |
| Ep4001 | 1.66E-34 | 0.30459597 | 0.501 | 0.435 | 3.20E-30 | TAM.Ndrg1 | Ep400 |
| Ank1 | 2.78E-34 | 0.41463696 | 0.309 | 0.248 | 5.38E-30 | TAM.Ndrg1 | Ank |
| Ipmk2 | 3.03E-34 | 0.29922362 | 0.364 | 0.304 | 5.85E-30 | TAM.Ndrg1 | Ipmk |
| Ugp21 | 3.97E-34 | 0.33289979 | 0.334 | 0.271 | 7.68E-30 | TAM.Ndrg1 | Ugp2 |
| Wdr912 | 2.59E-33 | 0.40166627 | 0.431 | 0.34 | 5.00E-29 | TAM.Ndrg1 | Wdr91 |
| Tubb2a1 | 3.14E-33 | 0.30619551 | 0.494 | 0.413 | 6.06E-29 | TAM.Ndrg1 | Tubb2a |
| Morf4l21 | 3.34E-33 | 0.29910311 | 0.424 | 0.359 | 6.46E-29 | TAM.Ndrg1 | Morf4l2 |
| Cdc42bpb | 5.89E-33 | 0.26539991 | 0.448 | 0.382 | 1.14E-28 | TAM.Ndrg1 | Cdc42bpb |
| Slc25a241 | 7.52E-33 | 0.36396556 | 0.281 | 0.219 | 1.45E-28 | TAM.Ndrg1 | Slc25a24 |
| Gse11 | 7.63E-33 | 0.4797371 | 0.295 | 0.219 | 1.48E-28 | TAM.Ndrg1 | Gse1 |
| Spire11 | 1.15E-32 | 0.55703336 | 0.268 | 0.183 | 2.23E-28 | TAM.Ndrg1 | Spire1 |
| Cds21 | 3.01E-32 | 0.33491655 | 0.375 | 0.293 | 5.81E-28 | TAM.Ndrg1 | Cds2 |
| Klf10 | 3.23E-32 | 0.33412563 | 0.437 | 0.38 | 6.25E-28 | TAM.Ndrg1 | Klf10 |
| Socs62 | 3.60E-32 | 0.3920803 | 0.34 | 0.296 | 6.96E-28 | TAM.Ndrg1 | Socs6 |
| Nr3c11 | 1.35E-30 | 0.26637851 | 0.415 | 0.353 | 2.61E-26 | TAM.Ndrg1 | Nr3c1 |
| Creld13 | 1.59E-30 | 0.46186912 | 0.281 | 0.204 | 3.07E-26 | TAM.Ndrg1 | Creld1 |
| Sema6b | 2.55E-30 | 0.41421636 | 0.257 | 0.184 | 4.92E-26 | TAM.Ndrg1 | Sema6b |
| Khnyn | 1.04E-29 | 0.41137694 | 0.267 | 0.206 | 2.01E-25 | TAM.Ndrg1 | Khnyn |
| Hbp11 | 1.52E-29 | 0.30809211 | 0.532 | 0.455 | 2.93E-25 | TAM.Ndrg1 | Hbp1 |
| Katnbl11 | 2.57E-29 | 0.43157154 | 0.262 | 0.206 | 4.97E-25 | TAM.Ndrg1 | Katnbl1 |
| Trip101 | 5.49E-29 | 0.36732692 | 0.336 | 0.254 | 1.06E-24 | TAM.Ndrg1 | Trip10 |
| Fam168a | 6.07E-29 | 0.29537862 | 0.374 | 0.292 | 1.17E-24 | TAM.Ndrg1 | Fam168a |
| Tusc31 | 8.20E-29 | 0.42823727 | 0.274 | 0.207 | 1.59E-24 | TAM.Ndrg1 | Tusc3 |
| Tmem170b1 | 2.90E-28 | 0.29320015 | 0.507 | 0.433 | 5.60E-24 | TAM.Ndrg1 | Tmem170b |
| Luzp11 | 3.62E-28 | 0.38551856 | 0.363 | 0.287 | 7.01E-24 | TAM.Ndrg1 | Luzp1 |
| Plin32 | 4.43E-28 | 0.33713788 | 0.481 | 0.393 | 8.57E-24 | TAM.Ndrg1 | Plin3 |
| Akap9 | 5.27E-28 | 0.42034462 | 0.268 | 0.202 | 1.02E-23 | TAM.Ndrg1 | Akap9 |
| Sigmar11 | 5.87E-28 | 0.30643833 | 0.394 | 0.327 | 1.14E-23 | TAM.Ndrg1 | Sigmar1 |
| Slbp | 9.39E-28 | 0.32595545 | 0.501 | 0.414 | 1.82E-23 | TAM.Ndrg1 | Slbp |
| Fblim12 | 9.68E-28 | 0.41514794 | 0.333 | 0.262 | 1.87E-23 | TAM.Ndrg1 | Fblim1 |
| Ube2g11 | 1.38E-27 | 0.26056212 | 0.386 | 0.323 | 2.66E-23 | TAM.Ndrg1 | Ube2g1 |
| S100a14 | 1.77E-27 | 0.26420201 | 0.512 | 0.444 | 3.43E-23 | TAM.Ndrg1 | S100a1 |
| Eepd1 | 1.92E-27 | 0.2945899 | 0.432 | 0.366 | 3.72E-23 | TAM.Ndrg1 | Eepd1 |
| Oxct12 | 2.49E-27 | 0.38447746 | 0.36 | 0.266 | 4.81E-23 | TAM.Ndrg1 | Oxct1 |
| Slc25a301 | 5.32E-27 | 0.32239885 | 0.307 | 0.252 | 1.03E-22 | TAM.Ndrg1 | Slc25a30 |
| Plekhm31 | 1.42E-26 | 0.32815725 | 0.336 | 0.261 | 2.74E-22 | TAM.Ndrg1 | Plekhm3 |
| Slc16a11 | 1.55E-26 | 0.4211284 | 0.269 | 0.2 | 2.99E-22 | TAM.Ndrg1 | Slc16a1 |
| Mitf1 | 4.24E-26 | 0.33449949 | 0.441 | 0.35 | 8.20E-22 | TAM.Ndrg1 | Mitf |
| Katna1 | 6.10E-26 | 0.35891221 | 0.268 | 0.207 | 1.18E-21 | TAM.Ndrg1 | Katna1 |
| Clip21 | 8.72E-26 | 0.28672588 | 0.428 | 0.373 | 1.69E-21 | TAM.Ndrg1 | Clip2 |
| Spata61 | 1.23E-25 | 0.38749703 | 0.259 | 0.198 | 2.37E-21 | TAM.Ndrg1 | Spata6 |
| Agap11 | 1.54E-25 | 0.3310159 | 0.392 | 0.307 | 2.99E-21 | TAM.Ndrg1 | Agap1 |
| Atg12 | 2.23E-25 | 0.3822772 | 0.274 | 0.221 | 4.31E-21 | TAM.Ndrg1 | Atg12 |
| Rlf | 5.92E-25 | 0.35999819 | 0.278 | 0.225 | 1.14E-20 | TAM.Ndrg1 | Rlf |
| Vps13d1 | 8.31E-25 | 0.26153047 | 0.42 | 0.344 | 1.61E-20 | TAM.Ndrg1 | Vps13d |
| Rmc11 | 1.34E-24 | 0.32517701 | 0.337 | 0.268 | 2.59E-20 | TAM.Ndrg1 | Rmc1 |
| Lrrc8a2 | 1.35E-24 | 0.41284517 | 0.296 | 0.228 | 2.62E-20 | TAM.Ndrg1 | Lrrc8a |
| Inppl12 | 1.59E-24 | 0.30265759 | 0.491 | 0.4 | 3.08E-20 | TAM.Ndrg1 | Inppl1 |
| Wipi2 | 2.18E-24 | 0.39486081 | 0.297 | 0.244 | 4.22E-20 | TAM.Ndrg1 | Wipi2 |
| Suco1 | 4.80E-24 | 0.28533564 | 0.305 | 0.252 | 9.27E-20 | TAM.Ndrg1 | Suco |
| Vps37a | 6.81E-24 | 0.31553438 | 0.307 | 0.25 | 1.32E-19 | TAM.Ndrg1 | Vps37a |
| Kif13b2 | 5.23E-23 | 0.29685594 | 0.374 | 0.305 | 1.01E-18 | TAM.Ndrg1 | Kif13b |
| Rell11 | 9.65E-23 | 0.2888511 | 0.338 | 0.267 | 1.87E-18 | TAM.Ndrg1 | Rell1 |
| Soga11 | 1.69E-22 | 0.36979154 | 0.452 | 0.37 | 3.27E-18 | TAM.Ndrg1 | Soga1 |
| Sars | 3.56E-22 | 0.32247306 | 0.345 | 0.28 | 6.88E-18 | TAM.Ndrg1 | Sars |
| Ifi272 | 3.85E-22 | 0.26194505 | 0.477 | 0.391 | 7.45E-18 | TAM.Ndrg1 | Ifi27 |
| Slc7a51 | 1.94E-21 | 0.40780401 | 0.318 | 0.266 | 3.74E-17 | TAM.Ndrg1 | Slc7a5 |
| Ing1 | 2.42E-21 | 0.31456067 | 0.344 | 0.276 | 4.68E-17 | TAM.Ndrg1 | Ing1 |
| B3gnt21 | 3.45E-21 | 0.28657429 | 0.298 | 0.246 | 6.66E-17 | TAM.Ndrg1 | B3gnt2 |
| Lpin12 | 4.48E-21 | 0.41761296 | 0.258 | 0.194 | 8.65E-17 | TAM.Ndrg1 | Lpin1 |
| Klhl242 | 5.30E-21 | 0.28955771 | 0.347 | 0.281 | 1.02E-16 | TAM.Ndrg1 | Klhl24 |
| Ulk11 | 1.04E-20 | 0.31128631 | 0.351 | 0.292 | 2.01E-16 | TAM.Ndrg1 | Ulk1 |
| Hspa4l1 | 1.15E-20 | 0.26777232 | 0.285 | 0.233 | 2.22E-16 | TAM.Ndrg1 | Hspa4l |
| Pdcd4 | 1.80E-20 | 0.33252755 | 0.294 | 0.236 | 3.48E-16 | TAM.Ndrg1 | Pdcd4 |
| DEPRECATED-I | 2.68E-20 | 0.27983229 | 0.346 | 0.29 | 5.19E-16 | TAM.Ndrg1 | DEPRECATED-ENSMUSG00000032667 |
| Fam98c | 1.44E-19 | 0.28076387 | 0.329 | 0.262 | 2.79E-15 | TAM.Ndrg1 | Fam98c |
| Dnmt3a1 | 2.65E-19 | 0.25259487 | 0.439 | 0.384 | 5.12E-15 | TAM.Ndrg1 | Dnmt3a |
| Narf | 4.12E-19 | 0.26520232 | 0.274 | 0.221 | 7.97E-15 | TAM.Ndrg1 | Narf |
| Necap1 | 4.29E-19 | 0.32727238 | 0.252 | 0.206 | 8.28E-15 | TAM.Ndrg1 | Necap1 |
| Crat2 | 6.22E-18 | 0.27943952 | 0.305 | 0.254 | 1.20E-13 | TAM.Ndrg1 | Crat |

| | | | | | | | |
|---|---|---|---|---|---|---|---|
| Pdgfa2 | 6.65E-18 | 0.27648029 | 0.303 | 0.251 | 1.28E-13 | TAM.Ndrg1 | Pdgfa |
| Hdac4 | 1.08E-17 | 0.29960278 | 0.339 | 0.28 | 2.09E-13 | TAM.Ndrg1 | Hdac4 |
| Erf | 2.23E-17 | 0.27472565 | 0.421 | 0.369 | 4.32E-13 | TAM.Ndrg1 | Erf |
| Aim21 | 3.16E-17 | 0.26554785 | 0.283 | 0.231 | 6.11E-13 | TAM.Ndrg1 | Aim2 |
| Gramd41 | 6.84E-17 | 0.26133312 | 0.259 | 0.206 | 1.32E-12 | TAM.Ndrg1 | Gramd4 |
| Dennd4c1 | 9.63E-17 | 0.28135356 | 0.318 | 0.251 | 1.86E-12 | TAM.Ndrg1 | Dennd4c |
| Mylip1 | 2.99E-16 | 0.30161708 | 0.342 | 0.31 | 5.79E-12 | TAM.Ndrg1 | Mylip |
| Acadsb | 3.61E-16 | 0.29261022 | 0.257 | 0.211 | 6.98E-12 | TAM.Ndrg1 | Acadsb |
| Blnk2 | 3.51E-15 | 0.34069167 | 0.276 | 0.217 | 6.78E-11 | TAM.Ndrg1 | Blnk |
| Crebrf1 | 3.54E-15 | 0.30484388 | 0.283 | 0.229 | 6.84E-11 | TAM.Ndrg1 | Crebrf |
| Marveld11 | 9.42E-15 | 0.36479334 | 0.288 | 0.244 | 1.82E-10 | TAM.Ndrg1 | Marveld1 |
| Abhd17c1 | 9.57E-14 | 0.29023428 | 0.254 | 0.212 | 1.85E-09 | TAM.Ndrg1 | Abhd17c |
| Galnt72 | 1.29E-13 | 0.25262444 | 0.308 | 0.259 | 2.49E-09 | TAM.Ndrg1 | Galnt7 |
| Slc44a11 | 1.24E-12 | 0.26177781 | 0.306 | 0.244 | 2.40E-08 | TAM.Ndrg1 | Slc44a1 |
| D10Wsu102e | 2.14E-12 | 0.36374249 | 0.252 | 0.22 | 4.13E-08 | TAM.Ndrg1 | D10Wsu102e |
| Rest | 3.04E-11 | 0.26652689 | 0.282 | 0.237 | 5.89E-07 | TAM.Ndrg1 | Rest |
| Tmem41b | 1.33E-10 | 0.25114596 | 0.257 | 0.218 | 2.57E-06 | TAM.Ndrg1 | Tmem41b |
| Chd31 | 2.26E-10 | 0.2753368 | 0.3 | 0.25 | 4.37E-06 | TAM.Ndrg1 | Chd3 |
| Nop561 | 2.93E-08 | 0.26345982 | 0.426 | 0.373 | 0.00056652 | TAM.Ndrg1 | Nop56 |
| Topors2 | 1.33E-07 | 0.27849269 | 0.288 | 0.261 | 0.00257167 | TAM.Ndrg1 | Topors |
| Flvcr11 | 0.00011823 | 0.28946493 | 0.325 | 0.284 | 1 | TAM.Ndrg1 | Flvcr1 |
| Stmn1 | 0 | 4.57175697 | 0.858 | 0.135 | 0 | TAM.prolif | Stmn1 |
| Top2a | 0 | 6.09148583 | 0.661 | 0.039 | 0 | TAM.prolif | Top2a |
| Mcm7 | 0 | 4.01622721 | 0.693 | 0.081 | 0 | TAM.prolif | Mcm7 |
| Mcm5 | 0 | 4.35116824 | 0.654 | 0.052 | 0 | TAM.prolif | Mcm5 |
| Tacc3 | 0 | 4.31842791 | 0.66 | 0.059 | 0 | TAM.prolif | Tacc3 |
| Tk1 | 0 | 5.30782101 | 0.632 | 0.031 | 0 | TAM.prolif | Tk1 |
| Ncapd2 | 0 | 4.62767282 | 0.653 | 0.057 | 0 | TAM.prolif | Ncapd2 |
| Rrm1 | 0 | 3.57512976 | 0.706 | 0.121 | 0 | TAM.prolif | Rrm1 |
| Uhrf1 | 0 | 5.32292039 | 0.598 | 0.026 | 0 | TAM.prolif | Uhrf1 |
| Mcm6 | 0 | 3.81682235 | 0.678 | 0.112 | 0 | TAM.prolif | Mcm6 |
| Mki67 | 0 | 3.90748497 | 0.669 | 0.11 | 0 | TAM.prolif | Mki67 |
| Cdca3 | 0 | 5.73557848 | 0.586 | 0.029 | 0 | TAM.prolif | Cdca3 |
| Mcm3 | 0 | 3.03075136 | 0.743 | 0.192 | 0 | TAM.prolif | Mcm3 |
| Cdk1 | 0 | 4.40152931 | 0.597 | 0.058 | 0 | TAM.prolif | Cdk1 |
| Mcm2 | 0 | 3.73148672 | 0.623 | 0.087 | 0 | TAM.prolif | Mcm2 |
| Dnajc9 | 0 | 2.55843324 | 0.665 | 0.131 | 0 | TAM.prolif | Dnajc9 |
| Racgap1 | 0 | 4.98445996 | 0.56 | 0.029 | 0 | TAM.prolif | Racgap1 |
| Fen1 | 0 | 2.76237626 | 0.693 | 0.165 | 0 | TAM.prolif | Fen1 |
| Lig1 | 0 | 3.3982043 | 0.604 | 0.089 | 0 | TAM.prolif | Lig1 |
| Birc5 | 0 | 5.4308785 | 0.537 | 0.023 | 0 | TAM.prolif | Birc5 |
| Tpx2 | 0 | 5.3114804 | 0.52 | 0.019 | 0 | TAM.prolif | Tpx2 |
| Rrm2 | 0 | 5.57236356 | 0.532 | 0.033 | 0 | TAM.prolif | Rrm2 |
| Ccna2 | 0 | 5.82060042 | 0.51 | 0.018 | 0 | TAM.prolif | Ccna2 |
| Pclaf | 0 | 4.90143835 | 0.524 | 0.035 | 0 | TAM.prolif | Pclaf |
| Pmf1 | 0 | 2.46897831 | 0.631 | 0.144 | 0 | TAM.prolif | Pmf1 |
| Smc2 | 0 | 4.12425653 | 0.53 | 0.044 | 0 | TAM.prolif | Smc2 |
| Hirip3 | 0 | 3.20926409 | 0.557 | 0.071 | 0 | TAM.prolif | Hirip3 |
| Tmpo | 0 | 2.38993502 | 0.627 | 0.151 | 0 | TAM.prolif | Tmpo |
| Chaf1a | 0 | 3.9131911 | 0.515 | 0.041 | 0 | TAM.prolif | Chaf1a |
| Pola1 | 0 | 2.97471793 | 0.56 | 0.088 | 0 | TAM.prolif | Pola1 |
| Asf1b | 0 | 4.7385429 | 0.492 | 0.025 | 0 | TAM.prolif | Asf1b |
| Rangap1 | 0 | 2.06933618 | 0.703 | 0.243 | 0 | TAM.prolif | Rangap1 |
| Pold1 | 0 | 2.8169413 | 0.54 | 0.085 | 0 | TAM.prolif | Pold1 |
| Vars1 | 0 | 1.68465848 | 0.748 | 0.296 | 0 | TAM.prolif | Vars |
| Selenoh | 0 | 2.49795423 | 0.563 | 0.111 | 0 | TAM.prolif | Selenoh |
| Sae1 | 0 | 1.89397755 | 0.75 | 0.301 | 0 | TAM.prolif | Sae1 |
| Cbx5 | 0 | 2.63489933 | 0.558 | 0.112 | 0 | TAM.prolif | Cbx5 |
| Atad2 | 0 | 3.16650119 | 0.527 | 0.082 | 0 | TAM.prolif | Atad2 |
| Lmnb13 | 0 | 1.56958172 | 0.848 | 0.403 | 0 | TAM.prolif | Lmnb1 |
| Gmnn | 0 | 2.99398053 | 0.537 | 0.093 | 0 | TAM.prolif | Gmnn |
| Haus4 | 0 | 2.9021116 | 0.531 | 0.088 | 0 | TAM.prolif | Haus4 |
| Dtymk | 0 | 2.06678601 | 0.64 | 0.197 | 0 | TAM.prolif | Dtymk |
| Knstrn | 0 | 4.63302021 | 0.469 | 0.026 | 0 | TAM.prolif | Knstrn |
| Topbp1 | 0 | 2.36890401 | 0.554 | 0.114 | 0 | TAM.prolif | Topbp1 |
| Dnmt1 | 0 | 2.11411696 | 0.713 | 0.275 | 0 | TAM.prolif | Dnmt1 |
| Hist1h2ap | 0 | 5.70470382 | 0.503 | 0.066 | 0 | TAM.prolif | Hist1h2ap |
| Mcm41 | 0 | 2.08236551 | 0.62 | 0.184 | 0 | TAM.prolif | Mcm4 |
| H2afv1 | 0 | 2.0407318 | 0.713 | 0.277 | 0 | TAM.prolif | H2afv |
| Nrm | 0 | 2.46503641 | 0.536 | 0.102 | 0 | TAM.prolif | Nrm |
| Cdca8 | 0 | 4.70572823 | 0.46 | 0.027 | 0 | TAM.prolif | Cdca8 |
| Spc24 | 0 | 6.08465112 | 0.436 | 0.007 | 0 | TAM.prolif | Spc24 |
| Mgl22 | 0 | 1.54486587 | 0.791 | 0.363 | 0 | TAM.prolif | Mgl2 |
| Gins2 | 0 | 3.07930546 | 0.49 | 0.063 | 0 | TAM.prolif | Gins2 |
| Pkmyt1 | 0 | 5.19529397 | 0.44 | 0.014 | 0 | TAM.prolif | Pkmyt1 |
| Rpa1 | 0 | 2.08360984 | 0.62 | 0.196 | 0 | TAM.prolif | Rpa1 |
| Anln | 0 | 4.94909254 | 0.443 | 0.02 | 0 | TAM.prolif | Anln |
| Hist1h1e | 0 | 3.69770513 | 0.573 | 0.15 | 0 | TAM.prolif | Hist1h1e |
| Tipin | 0 | 2.9564525 | 0.504 | 0.083 | 0 | TAM.prolif | Tipin |
| Dut | 0 | 3.6210577 | 0.468 | 0.051 | 0 | TAM.prolif | Dut |
| Incenp | 0 | 2.71040425 | 0.541 | 0.125 | 0 | TAM.prolif | Incenp |
| Rbbp7 | 0 | 1.69568588 | 0.776 | 0.361 | 0 | TAM.prolif | Rbbp7 |
| Kifc1 | 0 | 5.33831575 | 0.427 | 0.013 | 0 | TAM.prolif | Kifc1 |
| Lsm2 | 0 | 1.91421116 | 0.599 | 0.185 | 0 | TAM.prolif | Lsm2 |
| Nucks1 | 0 | 1.99482855 | 0.621 | 0.209 | 0 | TAM.prolif | Nucks1 |
| Ncaph | 0 | 4.22539654 | 0.44 | 0.028 | 0 | TAM.prolif | Ncaph |
| Dlgap5 | 0 | 6.13328643 | 0.418 | 0.007 | 0 | TAM.prolif | Dlgap5 |
| Lbr1 | 0 | 1.55713593 | 0.668 | 0.258 | 0 | TAM.prolif | Lbr |
| Shmt2 | 0 | 1.82112862 | 0.638 | 0.228 | 0 | TAM.prolif | Shmt2 |
| Psip1 | 0 | 2.21184295 | 0.532 | 0.123 | 0 | TAM.prolif | Psip1 |
| Nup210 | 0 | 3.12331023 | 0.464 | 0.056 | 0 | TAM.prolif | Nup210 |
| Hjurp | 0 | 2.09455259 | 0.592 | 0.184 | 0 | TAM.prolif | Hjurp |
| Ezh2 | 0 | 2.55835879 | 0.495 | 0.09 | 0 | TAM.prolif | Ezh2 |
| Hist1h1b | 0 | 5.89358328 | 0.437 | 0.033 | 0 | TAM.prolif | Hist1h1b |
| Hells | 0 | 3.78112896 | 0.443 | 0.041 | 0 | TAM.prolif | Hells |
| Bub1b | 0 | 2.8577916 | 0.481 | 0.08 | 0 | TAM.prolif | Bub1b |
| Ndc1 | 0 | 2.85829562 | 0.472 | 0.072 | 0 | TAM.prolif | Ndc1 |
| Mad2l1 | 0 | 4.18965057 | 0.427 | 0.027 | 0 | TAM.prolif | Mad2l1 |
| Ccnb2 | 0 | 4.19788453 | 0.44 | 0.041 | 0 | TAM.prolif | Ccnb2 |
| Ube2c | 0 | 5.2311427 | 0.423 | 0.024 | 0 | TAM.prolif | Ube2c |
| Exosc8 | 0 | 1.68652195 | 0.603 | 0.205 | 0 | TAM.prolif | Exosc8 |
| Smc4 | 0 | 2.30828309 | 0.547 | 0.153 | 0 | TAM.prolif | Smc4 |
| Slc25a102 | 0 | 1.70011147 | 0.605 | 0.214 | 0 | TAM.prolif | Slc25a10 |
| H2afx | 0 | 2.40667806 | 0.613 | 0.224 | 0 | TAM.prolif | H2afx |
| Hist1h2bn | 0 | 5.59352433 | 0.429 | 0.04 | 0 | TAM.prolif | Hist1h2bn |
| Tfdp1 | 0 | 1.6295074 | 0.713 | 0.324 | 0 | TAM.prolif | Tfdp1 |
| Tyms | 0 | 3.61704971 | 0.431 | 0.043 | 0 | TAM.prolif | Tyms |
| Syce2 | 0 | 2.83752525 | 0.461 | 0.073 | 0 | TAM.prolif | Syce2 |
| Csrp1 | 0 | 1.6507629 | 0.597 | 0.212 | 0 | TAM.prolif | Csrp1 |
| Ppil1 | 0 | 2.13670928 | 0.509 | 0.124 | 0 | TAM.prolif | Ppil1 |
| Gatm2 | 0 | 1.47428383 | 0.752 | 0.369 | 0 | TAM.prolif | Gatm |
| Kif11 | 0 | 6.49303838 | 0.389 | 0.006 | 0 | TAM.prolif | Kif11 |
| Nasp | 0 | 1.43336129 | 0.692 | 0.31 | 0 | TAM.prolif | Nasp |
| Usp1 | 0 | 2.0479244 | 0.509 | 0.129 | 0 | TAM.prolif | Usp1 |
| Rfc2 | 0 | 1.58520294 | 0.656 | 0.276 | 0 | TAM.prolif | Rfc2 |
| Jpt2 | 0 | 1.88248371 | 0.54 | 0.16 | 0 | TAM.prolif | Jpt2 |
| Anp32e | 0 | 1.73809903 | 0.854 | 0.475 | 0 | TAM.prolif | Anp32e |
| Pole | 0 | 5.26925116 | 0.389 | 0.01 | 0 | TAM.prolif | Pole |
| Rbl1 | 0 | 2.38737419 | 0.469 | 0.095 | 0 | TAM.prolif | Rbl1 |
| Kif15 | 0 | 6.03867927 | 0.38 | 0.006 | 0 | TAM.prolif | Kif15 |
| Prr11 | 0 | 5.92371329 | 0.383 | 0.009 | 0 | TAM.prolif | Prr11 |
| Nusap1 | 0 | 6.11113929 | 0.381 | 0.008 | 0 | TAM.prolif | Nusap1 |
| Cks1b | 0 | 3.44801567 | 0.409 | 0.038 | 0 | TAM.prolif | Cks1b |
| Prc1 | 0 | 5.69810549 | 0.382 | 0.011 | 0 | TAM.prolif | Prc1 |
| Cdt1 | 0 | 1.88363226 | 0.577 | 0.207 | 0 | TAM.prolif | Cdt1 |
| Dynll22 | 0 | 1.35802253 | 0.771 | 0.402 | 0 | TAM.prolif | Dynll2 |
| Nup85 | 0 | 1.99041049 | 0.486 | 0.118 | 0 | TAM.prolif | Nup85 |
| Rcc1 | 0 | 1.88009124 | 0.524 | 0.157 | 0 | TAM.prolif | Rcc1 |
| Hist1h2ae | 0 | 5.99663857 | 0.394 | 0.028 | 0 | TAM.prolif | Hist1h2ae |
| Ranbp11 | 0 | 1.61576869 | 0.789 | 0.424 | 0 | TAM.prolif | Ranbp1 |
| Dek1 | 0 | 1.61949961 | 0.849 | 0.484 | 0 | TAM.prolif | Dek |
| Gins1 | 0 | 3.3115031 | 0.402 | 0.038 | 0 | TAM.prolif | Gins1 |
| Kif23 | 0 | 4.19494149 | 0.39 | 0.026 | 0 | TAM.prolif | Kif23 |
| Rcc22 | 0 | 1.16195933 | 0.703 | 0.339 | 0 | TAM.prolif | Rcc2 |
| Cdkn2c | 0 | 3.88100344 | 0.401 | 0.038 | 0 | TAM.prolif | Cdkn2c |
| Rad21 | 0 | 1.56770139 | 0.668 | 0.306 | 0 | TAM.prolif | Rad21 |
| Cip2a | 0 | 4.13624663 | 0.387 | 0.025 | 0 | TAM.prolif | Cip2a |
| Rnaseh2c | 0 | 1.14770638 | 0.673 | 0.312 | 0 | TAM.prolif | Rnaseh2c |
| Ect2 | 0 | 4.49889511 | 0.381 | 0.02 | 0 | TAM.prolif | Ect2 |
| Rpa2 | 0 | 2.96783423 | 0.413 | 0.052 | 0 | TAM.prolif | Rpa2 |
| Cryl11 | 0 | 1.22858411 | 0.647 | 0.287 | 0 | TAM.prolif | Cryl1 |
| Wdhd1 | 0 | 3.38459307 | 0.397 | 0.038 | 0 | TAM.prolif | Wdhd1 |
| Tubg1 | 0 | 2.28508599 | 0.46 | 0.101 | 0 | TAM.prolif | Tubg1 |
| Hat1 | 0 | 1.47447147 | 0.627 | 0.269 | 0 | TAM.prolif | Hat1 |
| Kif20a | 0 | 5.15835254 | 0.372 | 0.014 | 0 | TAM.prolif | Kif20a |
| Ccnd11 | 0 | 1.75557321 | 0.541 | 0.184 | 0 | TAM.prolif | Ccnd1 |
| Vrk12 | 0 | 1.21107643 | 0.687 | 0.33 | 0 | TAM.prolif | Vrk1 |
| Snrpd11 | 0 | 1.3308695 | 0.676 | 0.319 | 0 | TAM.prolif | Snrpd1 |
| Fanca | 0 | 4.3458944 | 0.374 | 0.017 | 0 | TAM.prolif | Fanca |
| Aurkb | 0 | 4.78099815 | 0.372 | 0.015 | 0 | TAM.prolif | Aurkb |
| Idh22 | 0 | 1.20137839 | 0.752 | 0.397 | 0 | TAM.prolif | Idh2 |
| Rad54l | 0 | 5.39466513 | 0.363 | 0.009 | 0 | TAM.prolif | Rad54l |
| Aaas | 0 | 2.17824639 | 0.439 | 0.086 | 0 | TAM.prolif | Aaas |
| Cmc2 | 0 | 2.13813402 | 0.456 | 0.105 | 0 | TAM.prolif | Cmc2 |
| Ncapg | 0 | 6.09846389 | 0.356 | 0.005 | 0 | TAM.prolif | Ncapg |
| Arhgap11a | 0 | 2.07928475 | 0.463 | 0.114 | 0 | TAM.prolif | Arhgap11a |
| Cdkn2aipnl1 | 0 | 1.46418329 | 0.532 | 0.183 | 0 | TAM.prolif | Cdkn2aipnl |
| Prdx41 | 0 | 1.43166442 | 0.608 | 0.26 | 0 | TAM.prolif | Prdx4 |
| Bex31 | 0 | 1.53377486 | 0.527 | 0.18 | 0 | TAM.prolif | Bex3 |
| Cit | 0 | 5.73314929 | 0.353 | 0.006 | 0 | TAM.prolif | Cit |
| Tuba1b1 | 0 | 1.95680768 | 0.916 | 0.57 | 0 | TAM.prolif | Tuba1b |
| Anp32b | 0 | 1.45289288 | 0.544 | 0.198 | 0 | TAM.prolif | Anp32b |
| Bckdk1 | 0 | 1.37037971 | 0.578 | 0.232 | 0 | TAM.prolif | Bckdk |
| Tcf19 | 0 | 4.75538155 | 0.363 | 0.017 | 0 | TAM.prolif | Tcf19 |
| Aasdt1 | 0 | 1.5186702 | 0.525 | 0.18 | 0 | TAM.prolif | Aasdt1 |
| Mcm10 | 0 | 5.55146514 | 0.351 | 0.006 | 0 | TAM.prolif | Mcm10 |
| Nudc1 | 0 | 1.29066604 | 0.831 | 0.487 | 0 | TAM.prolif | Nudc |
| Idh3a1 | 0 | 1.28190054 | 0.625 | 0.283 | 0 | TAM.prolif | Idh3a |
| Rad51 | 0 | 6.28289868 | 0.346 | 0.004 | 0 | TAM.prolif | Rad51 |
| Cdc20 | 0 | 4.88975073 | 0.358 | 0.016 | 0 | TAM.prolif | Cdc20 |
| Fignl1 | 0 | 5.32524397 | 0.351 | 0.009 | 0 | TAM.prolif | Fignl1 |
| Hmmr | 0 | 5.86276397 | 0.349 | 0.007 | 0 | TAM.prolif | Hmmr |
| Plk1 | 0 | 5.6562608 | 0.351 | 0.01 | 0 | TAM.prolif | Plk1 |
| Smc1a1 | 0 | 1.21496872 | 0.641 | 0.301 | 0 | TAM.prolif | Smc1a |
| Pold2 | 0 | 2.41671388 | 0.419 | 0.079 | 0 | TAM.prolif | Pold2 |
| Ssrp1 | 0 | 1.45375663 | 0.554 | 0.215 | 0 | TAM.prolif | Ssrp1 |
| Set1 | 0 | 1.18782636 | 0.657 | 0.318 | 0 | TAM.prolif | Set |
| Hpf1 | 0 | 1.43646094 | 0.528 | 0.189 | 0 | TAM.prolif | Hpf1 |
| Ipo91 | 0 | 1.21782586 | 0.599 | 0.26 | 0 | TAM.prolif | Ipo9 |
| Srsf12 | 0 | 1.10720838 | 0.822 | 0.484 | 0 | TAM.prolif | Srsf1 |
| Dhfr | 0 | 3.833301 | 0.364 | 0.026 | 0 | TAM.prolif | Dhfr |
| Nsd21 | 0 | 1.1811882 | 0.712 | 0.375 | 0 | TAM.prolif | Nsd2 |
| Cep128 | 0 | 2.47271291 | 0.407 | 0.07 | 0 | TAM.prolif | Cep128 |
| Pa2g41 | 0 | 1.3849773 | 0.569 | 0.233 | 0 | TAM.prolif | Pa2g4 |
| Suv39h1 | 0 | 2.02864948 | 0.426 | 0.09 | 0 | TAM.prolif | Suv39h1 |
| Clspn | 0 | 6.25304313 | 0.339 | 0.004 | 0 | TAM.prolif | Clspn |
| Timm50 | 0 | 1.25629476 | 0.572 | 0.237 | 0 | TAM.prolif | Timm50 |
| Hist1h1a | 0 | 5.36860663 | 0.355 | 0.021 | 0 | TAM.prolif | Hist1h1a |
| Iqgap31 | 0 | 2.41530944 | 0.453 | 0.12 | 0 | TAM.prolif | Iqgap3 |
| Cdk4 | 0 | 1.34860233 | 0.535 | 0.203 | 0 | TAM.prolif | Cdk4 |
| Cenpm | 0 | 5.33029347 | 0.34 | 0.008 | 0 | TAM.prolif | Cenpm |
| Mybl2 | 0 | 5.31458017 | 0.342 | 0.011 | 0 | TAM.prolif | Mybl2 |
| Ndc80 | 0 | 5.23025163 | 0.34 | 0.009 | 0 | TAM.prolif | Ndc80 |
| Mis18a | 0 | 1.43953177 | 0.501 | 0.17 | 0 | TAM.prolif | Mis18a |
| Gart1 | 0 | 1.28283258 | 0.592 | 0.261 | 0 | TAM.prolif | Gart |
| Rfc5 | 0 | 2.57757774 | 0.386 | 0.056 | 0 | TAM.prolif | Rfc5 |
| Pola2 | 0 | 1.85288895 | 0.442 | 0.112 | 0 | TAM.prolif | Pola2 |
| Nek2 | 0 | 5.88388022 | 0.335 | 0.006 | 0 | TAM.prolif | Nek2 |
| E2f8 | 0 | 3.28246583 | 0.37 | 0.041 | 0 | TAM.prolif | E2f8 |
| Spag5 | 0 | 6.20106675 | 0.334 | 0.005 | 0 | TAM.prolif | Spag5 |
| Cep55 | 0 | 5.58289979 | 0.335 | 0.007 | 0 | TAM.prolif | Cep55 |
| Fh11 | 0 | 1.03737821 | 0.581 | 0.253 | 0 | TAM.prolif | Fh1 |
| Cse1l | 0 | 1.29552267 | 0.564 | 0.236 | 0 | TAM.prolif | Cse1l |
| Nhp21 | 0 | 1.08360488 | 0.717 | 0.389 | 0 | TAM.prolif | Nhp2 |
| Apobec32 | 0 | 1.01091541 | 0.716 | 0.388 | 0 | TAM.prolif | Apobec3 |
| Cbx3 | 0 | 1.71004064 | 0.481 | 0.154 | 0 | TAM.prolif | Cbx3 |
| Ckap2l | 0 | 1.60111438 | 0.406 | 0.08 | 0 | TAM.prolif | Ckap2l |
| Xpo1 | 0 | 1.15552877 | 0.665 | 0.34 | 0 | TAM.prolif | Xpo1 |
| Fasn2 | 0 | 1.3682674 | 0.522 | 0.197 | 0 | TAM.prolif | Fasn |
| Tmem109 | 0 | 1.09861166 | 0.668 | 0.343 | 0 | TAM.prolif | Tmem109 |
| Parp11 | 0 | 1.1346364 | 0.702 | 0.377 | 0 | TAM.prolif | Parp1 |
| Supt16 | 0 | 1.02360227 | 0.701 | 0.376 | 0 | TAM.prolif | Supt16 |
| Ncapd3 | 0 | 2.36517207 | 0.4 | 0.076 | 0 | TAM.prolif | Ncapd3 |
| Cenpx | 0 | 0.92319907 | 0.665 | 0.341 | 0 | TAM.prolif | Cenpx |
| Ube2s | 0 | 1.42446934 | 0.68 | 0.358 | 0 | TAM.prolif | Ube2s |
| Ddx391 | 0 | 1.0522626 | 0.651 | 0.329 | 0 | TAM.prolif | Ddx39 |
| Nop581 | 0 | 1.27513455 | 0.538 | 0.217 | 0 | TAM.prolif | Nop58 |
| Cenpe | 0 | 5.56303338 | 0.328 | 0.007 | 0 | TAM.prolif | Cenpe |
| Nudt5 | 0 | 1.34837082 | 0.508 | 0.188 | 0 | TAM.prolif | Nudt5 |
| Rogdi2 | 0 | 1.19299434 | 0.652 | 0.332 | 0 | TAM.prolif | Rogdi |
| Impdh21 | 0 | 0.8750562 | 0.705 | 0.385 | 0 | TAM.prolif | Impdh2 |
| Slamf92 | 0 | 1.35459605 | 0.524 | 0.205 | 0 | TAM.prolif | Slamf9 |
| Spc25 | 0 | 4.81938778 | 0.331 | 0.012 | 0 | TAM.prolif | Spc25 |
| Dck1 | 0 | 1.09420332 | 0.594 | 0.275 | 0 | TAM.prolif | Dck |
| Itga62 | 0 | 0.87842783 | 0.648 | 0.33 | 0 | TAM.prolif | Itga6 |
| Ubap2 | 0 | 1.10114509 | 0.566 | 0.248 | 0 | TAM.prolif | Ubap2 |
| Ckap5 | 0 | 1.5259266 | 0.56 | 0.243 | 0 | TAM.prolif | Ckap5 |
| Stip1 | 0 | 1.03408413 | 0.655 | 0.338 | 0 | TAM.prolif | Stip1 |
| Kpnb1 | 0 | 1.21962631 | 0.587 | 0.27 | 0 | TAM.prolif | Kpnb1 |
| Snrpa11 | 0 | 1.09217296 | 0.561 | 0.245 | 0 | TAM.prolif | Snrpa1 |
| Dpy30 | 0 | 1.28566948 | 0.56 | 0.244 | 0 | TAM.prolif | Dpy30 |
| Atic1 | 0 | 1.09800705 | 0.658 | 0.342 | 0 | TAM.prolif | Atic |
| Fam111a | 0 | 1.34132099 | 0.486 | 0.17 | 0 | TAM.prolif | Fam111a |
| Fgd21 | 0 | 0.82499077 | 0.626 | 0.311 | 0 | TAM.prolif | Fgd2 |
| Kif2c | 0 | 6.63036688 | 0.318 | 0.004 | 0 | TAM.prolif | Kif2c |
| Prpf311 | 0 | 0.93487832 | 0.623 | 0.309 | 0 | TAM.prolif | Prpf31 |
| Ncapg2 | 0 | 1.72103597 | 0.45 | 0.136 | 0 | TAM.prolif | Ncapg2 |
| Espl1 | 0 | 3.0930209 | 0.367 | 0.053 | 0 | TAM.prolif | Espl1 |
| Nde1 | 0 | 1.61045425 | 0.472 | 0.158 | 0 | TAM.prolif | Nde1 |
| Lsm41 | 0 | 0.87895087 | 0.729 | 0.416 | 0 | TAM.prolif | Lsm4 |
| Cdc45 | 0 | 4.32719158 | 0.327 | 0.014 | 0 | TAM.prolif | Cdc45 |
| Sfxn11 | 0 | 1.1332125 | 0.564 | 0.251 | 0 | TAM.prolif | Sfxn1 |
| Atpif1 | 0 | 0.9673157 | 0.766 | 0.454 | 0 | TAM.prolif | Atpif1 |
| Exosc51 | 0 | 0.96347096 | 0.591 | 0.28 | 0 | TAM.prolif | Exosc5 |
| Maz | 0 | 0.9675791 | 0.781 | 0.471 | 0 | TAM.prolif | Maz |
| Haus81 | 0 | 1.20840024 | 0.523 | 0.213 | 0 | TAM.prolif | Haus8 |
| Eif1ad | 0 | 1.36533809 | 0.518 | 0.208 | 0 | TAM.prolif | Eif1ad |
| Smc3 | 0 | 1.23597255 | 0.511 | 0.201 | 0 | TAM.prolif | Smc3 |
| Ccnd22 | 0 | 1.23698625 | 0.527 | 0.218 | 0 | TAM.prolif | Ccnd2 |
| Nsmce4a | 0 | 1.18180161 | 0.548 | 0.239 | 0 | TAM.prolif | Nsmce4a |
| Chtf18 | 0 | 4.51213277 | 0.322 | 0.013 | 0 | TAM.prolif | Chtf18 |
| Nap1l1 | 0 | 1.41620898 | 0.522 | 0.214 | 0 | TAM.prolif | Nap1l1 |
| Prpf192 | 0 | 1.01872439 | 0.638 | 0.33 | 0 | TAM.prolif | Prpf19 |
| Nup155 | 0 | 1.41112713 | 0.458 | 0.151 | 0 | TAM.prolif | Nup155 |
| Stard71 | 0 | 0.85838519 | 0.626 | 0.319 | 0 | TAM.prolif | Stard7 |
| Cdk2 | 0 | 1.36603035 | 0.507 | 0.2 | 0 | TAM.prolif | Cdk2 |
| Anapc11 | 0 | 1.07322262 | 0.753 | 0.446 | 0 | TAM.prolif | Anapc11 |
| Mtr1 | 0 | 0.9173549 | 0.627 | 0.32 | 0 | TAM.prolif | Mtr |
| Nubp11 | 0 | 0.83825595 | 0.718 | 0.411 | 0 | TAM.prolif | Nubp1 |
| Hdgf1 | 0 | 0.88996943 | 0.708 | 0.401 | 0 | TAM.prolif | Hdgf |
| Naa40 | 0 | 1.31554799 | 0.52 | 0.214 | 0 | TAM.prolif | Naa40 |
| Aurka | 0 | 3.71560429 | 0.335 | 0.029 | 0 | TAM.prolif | Aurka |
| Kif4 | 0 | 5.55193149 | 0.312 | 0.006 | 0 | TAM.prolif | Kif4 |
| Hspd11 | 0 | 1.12115614 | 0.601 | 0.296 | 0 | TAM.prolif | Hspd1 |
| Apex12 | 0 | 1.2536695 | 0.508 | 0.203 | 0 | TAM.prolif | Apex1 |
| Ncaph2 | 0 | 1.25183772 | 0.507 | 0.202 | 0 | TAM.prolif | Ncaph2 |
| Tcp11 | 0 | 0.85520209 | 0.751 | 0.446 | 0 | TAM.prolif | Tcp1 |
| Stub1 | 0 | 0.88469339 | 0.792 | 0.488 | 0 | TAM.prolif | Stub1 |
| Umps1 | 0 | 1.27330989 | 0.492 | 0.188 | 0 | TAM.prolif | Umps |
| Gar11 | 0 | 1.18588839 | 0.507 | 0.204 | 0 | TAM.prolif | Gar1 |
| Srsf4 | 0 | 0.8092658 | 0.656 | 0.353 | 0 | TAM.prolif | Srsf4 |
| Melk | 0 | 5.7235156 | 0.309 | 0.006 | 0 | TAM.prolif | Melk |
| Brip1 | 0 | 3.96886077 | 0.321 | 0.018 | 0 | TAM.prolif | Brip1 |
| Bub1 | 0 | 5.32592897 | 0.309 | 0.007 | 0 | TAM.prolif | Bub1 |
| Paics | 0 | 1.37922178 | 0.477 | 0.175 | 0 | TAM.prolif | Paics |
| Hint1 | 0 | 1.02946939 | 0.629 | 0.327 | 0 | TAM.prolif | Hint1 |
| Hspa141 | 0 | 1.01317904 | 0.557 | 0.256 | 0 | TAM.prolif | Hspa14 |
| Prmt11 | 0 | 1.06462468 | 0.528 | 0.227 | 0 | TAM.prolif | Prmt1 |
| Gins4 | 0 | 1.35854304 | 0.464 | 0.163 | 0 | TAM.prolif | Gins4 |
| Zfp367 | 0 | 2.58808222 | 0.367 | 0.066 | 0 | TAM.prolif | Zfp367 |
| Rbbp4 | 0 | 1.2138267 | 0.75 | 0.45 | 0 | TAM.prolif | Rbbp4 |
| Sdf2l12 | 0 | 0.9803992 | 0.839 | 0.539 | 0 | TAM.prolif | Sdf2l1 |
| Mpp6 | 0 | 1.10872944 | 0.544 | 0.245 | 0 | TAM.prolif | Mpp6 |
| Rfwd3 | 0 | 1.63458059 | 0.427 | 0.128 | 0 | TAM.prolif | Rfwd3 |
| Nt5dc21 | 0 | 1.15015363 | 0.507 | 0.209 | 0 | TAM.prolif | Nt5dc2 |
| Cep192 | 0 | 1.36387491 | 0.456 | 0.158 | 0 | TAM.prolif | Cep192 |
| Baz1b | 0 | 1.00290377 | 0.612 | 0.314 | 0 | TAM.prolif | Baz1b |
| Cd722 | 0 | 0.90546302 | 0.568 | 0.27 | 0 | TAM.prolif | Cd72 |
| Ptgs11 | 0 | 1.32146334 | 0.677 | 0.38 | 0 | TAM.prolif | Ptgs1 |
| Smarcc1 | 0 | 1.0314632 | 0.584 | 0.287 | 0 | TAM.prolif | Smarcc1 |
| Nup107 | 0 | 1.55320886 | 0.408 | 0.111 | 0 | TAM.prolif | Nup107 |
| Dynlt1f2 | 0 | 1.02095504 | 0.542 | 0.246 | 0 | TAM.prolif | Dynlt1f |
| Tubb4b2 | 0 | 1.18569507 | 0.709 | 0.413 | 0 | TAM.prolif | Tubb4b |
| Lsm3 | 0 | 1.2897328 | 0.471 | 0.175 | 0 | TAM.prolif | Lsm3 |
| Dctpp1 | 0 | 2.14858522 | 0.381 | 0.085 | 0 | TAM.prolif | Dctpp1 |
| Tmem176a2 | 0 | 0.78456326 | 0.699 | 0.403 | 0 | TAM.prolif | Tmem176a |
| Ilf2 | 0 | 1.29072026 | 0.463 | 0.168 | 0 | TAM.prolif | Ilf2 |
| Rnaseh2b | 0 | 1.96416279 | 0.389 | 0.094 | 0 | TAM.prolif | Rnaseh2b |
| Eftud2 | 0 | 0.8742792 | 0.697 | 0.402 | 0 | TAM.prolif | Eftud2 |
| Ssna1 | 0 | 0.90163547 | 0.661 | 0.367 | 0 | TAM.prolif | Ssna1 |
| Sf3b31 | 0 | 0.80974086 | 0.68 | 0.386 | 0 | TAM.prolif | Sf3b3 |
| Snrpe1 | 0 | 0.96823528 | 0.58 | 0.286 | 0 | TAM.prolif | Snrpe |
| Nans | 0 | 1.05377027 | 0.584 | 0.29 | 0 | TAM.prolif | Nans |
| Gtse1 | 0 | 4.67672541 | 0.305 | 0.011 | 0 | TAM.prolif | Gtse1 |
| Tmem176b1 | 0 | 0.781057 | 0.794 | 0.501 | 0 | TAM.prolif | Tmem176b |
| Uba2 | 0 | 0.9587309 | 0.659 | 0.366 | 0 | TAM.prolif | Uba2 |
| Numa11 | 0 | 0.86678982 | 0.656 | 0.363 | 0 | TAM.prolif | Numa1 |
| Snrpf1 | 0 | 0.82796053 | 0.786 | 0.493 | 0 | TAM.prolif | Snrpf |
| Bub3 | 0 | 1.09366161 | 0.578 | 0.285 | 0 | TAM.prolif | Bub3 |
| Cdca5 | 0 | 4.58780328 | 0.307 | 0.014 | 0 | TAM.prolif | Cdca5 |
| Rdx | 0 | 0.94952489 | 0.566 | 0.273 | 0 | TAM.prolif | Rdx |
| Cltb1 | 0 | 0.87287201 | 0.688 | 0.395 | 0 | TAM.prolif | Cltb |
| Myef2 | 0 | 1.47096207 | 0.445 | 0.153 | 0 | TAM.prolif | Myef2 |
| Cct6a1 | 0 | 0.9128151 | 0.639 | 0.347 | 0 | TAM.prolif | Cct6a |
| Cnot9 | 0 | 1.16207319 | 0.495 | 0.203 | 0 | TAM.prolif | Cnot9 |
| Foxm1 | 0 | 5.35645955 | 0.299 | 0.007 | 0 | TAM.prolif | Foxm1 |
| Cad | 0 | 1.72347522 | 0.396 | 0.105 | 0 | TAM.prolif | Cad |
| Tomm401 | 0 | 0.84646583 | 0.661 | 0.37 | 0 | TAM.prolif | Tomm40 |
| Rad51ap1 | 0 | 6.41845071 | 0.294 | 0.003 | 0 | TAM.prolif | Rad51ap1 |
| Prim1 | 0 | 3.7057075 | 0.311 | 0.02 | 0 | TAM.prolif | Prim1 |
| Paxip1 | 0 | 1.33425063 | 0.458 | 0.168 | 0 | TAM.prolif | Paxip1 |
| Pygb2 | 0 | 0.79635538 | 0.591 | 0.301 | 0 | TAM.prolif | Pygb |
| Alg8 | 0 | 1.77393587 | 0.392 | 0.103 | 0 | TAM.prolif | Alg8 |
| Thoc7 | 0 | 0.87129618 | 0.562 | 0.273 | 0 | TAM.prolif | Thoc7 |
| Psmb91 | 0 | 0.82359163 | 0.771 | 0.482 | 0 | TAM.prolif | Psmb9 |
| Cdc123 | 0 | 0.90338057 | 0.575 | 0.286 | 0 | TAM.prolif | Cdc123 |
| E2f1 | 0 | 2.10708259 | 0.391 | 0.103 | 0 | TAM.prolif | E2f1 |
| Fam234b1 | 0 | 1.25281816 | 0.467 | 0.179 | 0 | TAM.prolif | Fam234b |
| Trim28 | 0 | 1.06039884 | 0.512 | 0.224 | 0 | TAM.prolif | Trim28 |
| Ipo5 | 0 | 1.13668657 | 0.634 | 0.346 | 0 | TAM.prolif | Ipo5 |
| Nuf2 | 0 | 5.70808948 | 0.294 | 0.006 | 0 | TAM.prolif | Nuf2 |
| Hist1h3a | 0 | 3.77385841 | 0.315 | 0.027 | 0 | TAM.prolif | Hist1h3a |
| Cdca2 | 0 | 6.09325033 | 0.292 | 0.004 | 0 | TAM.prolif | Cdca2 |
| Dbf4 | 0 | 2.12224873 | 0.371 | 0.084 | 0 | TAM.prolif | Dbf4 |
| Slc29a11 | 0 | 1.04705063 | 0.593 | 0.306 | 0 | TAM.prolif | Slc29a1 |
| Tceal92 | 0 | 0.80238125 | 0.63 | 0.343 | 0 | TAM.prolif | Tceal9 |
| Adgrg52 | 0 | 1.29065331 | 0.451 | 0.165 | 0 | TAM.prolif | Adgrg5 |
| Reep4 | 0 | 1.17829749 | 0.53 | 0.244 | 0 | TAM.prolif | Reep4 |
| Sephs1 | 0 | 1.17621851 | 0.481 | 0.195 | 0 | TAM.prolif | Sephs1 |
| Cdk2ap12 | 0 | 0.80544001 | 0.569 | 0.283 | 0 | TAM.prolif | Cdk2ap1 |
| Uchl5 | 0 | 1.07814311 | 0.503 | 0.218 | 0 | TAM.prolif | Uchl5 |
| Neil3 | 0 | 7.07104271 | 0.287 | 0.002 | 0 | TAM.prolif | Neil3 |
| Nedd1 | 0 | 1.42684508 | 0.421 | 0.136 | 0 | TAM.prolif | Nedd1 |
| Rnps1 | 0 | 0.79962904 | 0.741 | 0.456 | 0 | TAM.prolif | Rnps1 |
| Eri1 | 0 | 1.11142677 | 0.495 | 0.211 | 0 | TAM.prolif | Eri1 |
| Pcnt | 0 | 1.03715618 | 0.463 | 0.179 | 0 | TAM.prolif | Pcnt |
| Cenpa | 0 | 1.83243042 | 0.419 | 0.135 | 0 | TAM.prolif | Cenpa |
| BC030867 | 0 | 5.10686125 | 0.294 | 0.01 | 0 | TAM.prolif | BC030867 |
| Gspt1 | 0 | 0.84307972 | 0.623 | 0.339 | 0 | TAM.prolif | Gspt1 |
| Hnrnpab1 | 0 | 0.89854671 | 0.795 | 0.512 | 0 | TAM.prolif | Hnrnpab |
| Mms22l | 0 | 3.32440988 | 0.313 | 0.031 | 0 | TAM.prolif | Mms22l |
| Snrnp401 | 0 | 0.97638657 | 0.503 | 0.221 | 0 | TAM.prolif | Snrnp40 |
| Wbp11 | 0 | 0.70875324 | 0.631 | 0.349 | 0 | TAM.prolif | Wbp11 |
| Ntpcr2 | 0 | 0.87400015 | 0.666 | 0.384 | 0 | TAM.prolif | Ntpcr |
| Snrpa | 0 | 0.77120927 | 0.593 | 0.311 | 0 | TAM.prolif | Snrpa |
| Mdh11 | 0 | 0.86109212 | 0.739 | 0.457 | 0 | TAM.prolif | Mdh1 |
| Slbp1 | 0 | 1.00532071 | 0.686 | 0.405 | 0 | TAM.prolif | Slbp |
| Sf3b5 | 0 | 0.75090906 | 0.65 | 0.369 | 0 | TAM.prolif | Sf3b5 |
| H2afz1 | 0 | 1.30356104 | 0.829 | 0.548 | 0 | TAM.prolif | H2afz |
| Tsn | 0 | 0.76198266 | 0.648 | 0.368 | 0 | TAM.prolif | Tsn |
| Nop101 | 0 | 0.76815158 | 0.64 | 0.36 | 0 | TAM.prolif | Nop10 |
| Lsm8 | 0 | 1.05018188 | 0.508 | 0.228 | 0 | TAM.prolif | Lsm8 |
| G3bp1 | 0 | 0.73949617 | 0.638 | 0.358 | 0 | TAM.prolif | G3bp1 |
| Smc6 | 0 | 1.04616795 | 0.517 | 0.237 | 0 | TAM.prolif | Smc6 |
| Nelfe | 0 | 1.03153072 | 0.504 | 0.224 | 0 | TAM.prolif | Nelfe |
| Pkig1 | 0 | 0.83986297 | 0.568 | 0.288 | 0 | TAM.prolif | Pkig |
| Shcbp1 | 0 | 5.92838083 | 0.284 | 0.004 | 0 | TAM.prolif | Shcbp1 |
| Srm | 0 | 1.38718812 | 0.44 | 0.161 | 0 | TAM.prolif | Srm |
| Emg11 | 0 | 0.66690886 | 0.72 | 0.441 | 0 | TAM.prolif | Emg1 |
| Cd813 | 0 | 0.9526294 | 0.782 | 0.504 | 0 | TAM.prolif | Cd81 |
| Pbk | 0 | 6.06678809 | 0.282 | 0.004 | 0 | TAM.prolif | Pbk |
| Nono | 0 | 0.96421168 | 0.834 | 0.556 | 0 | TAM.prolif | Nono |
| Fbl1 | 0 | 0.60419972 | 0.661 | 0.384 | 0 | TAM.prolif | Fbl |
| Trp531 | 0 | 0.73635554 | 0.681 | 0.404 | 0 | TAM.prolif | Trp53 |
| Kif18b | 0 | 6.40878932 | 0.28 | 0.003 | 0 | TAM.prolif | Kif18b |
| Hmgb3 | 0 | 2.16234364 | 0.343 | 0.066 | 0 | TAM.prolif | Hmgb3 |
| Cd2bp2 | 0 | 0.76856283 | 0.592 | 0.315 | 0 | TAM.prolif | Cd2bp2 |
| Cdca4 | 0 | 1.0632518 | 0.477 | 0.2 | 0 | TAM.prolif | Cdca4 |
| Lyar | 0 | 1.15137381 | 0.446 | 0.17 | 0 | TAM.prolif | Lyar |
| Ulbp1 | 0 | 2.94401132 | 0.31 | 0.034 | 0 | TAM.prolif | Ulbp1 |
| Tle51 | 0 | 0.77278664 | 0.761 | 0.485 | 0 | TAM.prolif | Tle5 |
| Sumo3 | 0 | 0.65589541 | 0.699 | 0.423 | 0 | TAM.prolif | Sumo3 |
| Ybx32 | 0 | 0.83240045 | 0.685 | 0.41 | 0 | TAM.prolif | Ybx3 |
| Ptpa | 0 | 0.77007029 | 0.665 | 0.39 | 0 | TAM.prolif | Ptpa |
| Plk4 | 0 | 2.854502 | 0.31 | 0.035 | 0 | TAM.prolif | Plk4 |
| Fancd2 | 0 | 5.58425711 | 0.28 | 0.005 | 0 | TAM.prolif | Fancd2 |
| Mars | 0 | 1.04166515 | 0.488 | 0.213 | 0 | TAM.prolif | Mars |
| Kif22 | 0 | 5.13507833 | 0.283 | 0.008 | 0 | TAM.prolif | Kif22 |
| Slc25a41 | 0 | 0.80840842 | 0.593 | 0.318 | 0 | TAM.prolif | Slc25a4 |
| Snu131 | 0 | 0.82148582 | 0.74 | 0.465 | 0 | TAM.prolif | Snu13 |
| Nmral1 | 0 | 1.97147403 | 0.354 | 0.079 | 0 | TAM.prolif | Nmral1 |
| Ahsa1 | 0 | 0.9178504 | 0.835 | 0.56 | 0 | TAM.prolif | Ahsa1 |
| Orc6 | 0 | 2.39072261 | 0.33 | 0.056 | 0 | TAM.prolif | Orc6 |
| Ppp2r5d | 0 | 1.03501651 | 0.481 | 0.207 | 0 | TAM.prolif | Ppp2r5d |
| Banf1 | 0 | 1.17614981 | 0.443 | 0.169 | 0 | TAM.prolif | Banf1 |
| Arhgap19 | 0 | 3.44797431 | 0.304 | 0.03 | 0 | TAM.prolif | Arhgap19 |
| Hars | 0 | 0.82020166 | 0.585 | 0.311 | 0 | TAM.prolif | Hars |
| Hsd17b41 | 0 | 0.82485297 | 0.717 | 0.443 | 0 | TAM.prolif | Hsd17b4 |
| Srrt | 0 | 0.80723947 | 0.634 | 0.361 | 0 | TAM.prolif | Srrt |
| Akr1b31 | 0 | 0.72187475 | 0.658 | 0.385 | 0 | TAM.prolif | Akr1b3 |
| Dcp2 | 0 | 0.68569237 | 0.537 | 0.264 | 0 | TAM.prolif | Dcp2 |
| Cct21 | 0 | 0.66590835 | 0.725 | 0.452 | 0 | TAM.prolif | Cct2 |
| Hnrnpa11 | 0 | 1.1361356 | 0.867 | 0.595 | 0 | TAM.prolif | Hnrnpa1 |
| Smchd12 | 0 | 0.63248891 | 0.668 | 0.396 | 0 | TAM.prolif | Smchd1 |
| Slc43a3 | 0 | 4.4172427 | 0.283 | 0.011 | 0 | TAM.prolif | Slc43a3 |
| Ppm1g1 | 0 | 0.87199326 | 0.729 | 0.457 | 0 | TAM.prolif | Ppm1g |
| Nup621 | 0 | 0.91028285 | 0.505 | 0.234 | 0 | TAM.prolif | Nup62 |
| Pimreg | 0 | 6.04170187 | 0.276 | 0.005 | 0 | TAM.prolif | Pimreg |
| Srsf10 | 0 | 0.75427169 | 0.608 | 0.337 | 0 | TAM.prolif | Srsf10 |
| Aars | 0 | 0.71173906 | 0.603 | 0.333 | 0 | TAM.prolif | Aars |
| Sf3a1 | 0 | 0.71022683 | 0.589 | 0.319 | 0 | TAM.prolif | Sf3a1 |
| Ndufb11 | 0 | 0.69729189 | 0.716 | 0.446 | 0 | TAM.prolif | Ndufb11 |
| Sptssa2 | 0 | 0.69440327 | 0.684 | 0.415 | 0 | TAM.prolif | Sptssa |
| Mvb12b1 | 0 | 0.69488299 | 0.633 | 0.364 | 0 | TAM.prolif | Mvb12b |
| Ppat | 0 | 1.53440839 | 0.385 | 0.116 | 0 | TAM.prolif | Ppat |
| Drg1 | 0 | 0.7507282 | 0.659 | 0.39 | 0 | TAM.prolif | Drg1 |
| Paqr4 | 0 | 2.21227787 | 0.33 | 0.061 | 0 | TAM.prolif | Paqr4 |
| Slc35a4 | 0 | 0.66447164 | 0.676 | 0.408 | 0 | TAM.prolif | Slc35a4 |
| Prim2 | 0 | 1.97947134 | 0.339 | 0.071 | 0 | TAM.prolif | Prim2 |
| Sh2d1b12 | 0 | 0.98955542 | 0.513 | 0.245 | 0 | TAM.prolif | Sh2d1b1 |
| Snrpb3 | 0 | 0.68240266 | 0.666 | 0.398 | 0 | TAM.prolif | Snrpb |
| Chchd3 | 0 | 0.6694957 | 0.642 | 0.374 | 0 | TAM.prolif | Chchd3 |
| Mbd3 | 0 | 0.840139 | 0.808 | 0.54 | 0 | TAM.prolif | Mbd3 |
| Eif2s11 | 0 | 0.7450764 | 0.633 | 0.365 | 0 | TAM.prolif | Eif2s1 |
| Diaph3 | 0 | 4.70873516 | 0.278 | 0.01 | 0 | TAM.prolif | Diaph3 |
| Abcg31 | 0 | 0.75431539 | 0.571 | 0.303 | 0 | TAM.prolif | Abcg3 |
| Cetn31 | 0 | 0.72926812 | 0.611 | 0.343 | 0 | TAM.prolif | Cetn3 |
| Serbp11 | 0 | 0.87264149 | 0.791 | 0.524 | 0 | TAM.prolif | Serbp1 |
| Rad18 | 0 | 2.62526002 | 0.309 | 0.042 | 0 | TAM.prolif | Rad18 |
| Kars1 | 0 | 0.67622738 | 0.645 | 0.378 | 0 | TAM.prolif | Kars |
| Poc1a | 0 | 3.01710183 | 0.302 | 0.035 | 0 | TAM.prolif | Poc1a |
| Mybbp1a1 | 0 | 0.7917307 | 0.625 | 0.358 | 0 | TAM.prolif | Mybbp1a |
| Suz12 | 0 | 1.06511178 | 0.492 | 0.225 | 0 | TAM.prolif | Suz12 |
| Phf5a | 0 | 0.80680831 | 0.511 | 0.244 | 0 | TAM.prolif | Phf5a |
| Haus1 | 0 | 2.09566953 | 0.333 | 0.066 | 0 | TAM.prolif | Haus1 |
| Pdap1 | 0 | 0.68520393 | 0.719 | 0.452 | 0 | TAM.prolif | Pdap1 |
| Pnn | 0 | 0.62537572 | 0.582 | 0.315 | 0 | TAM.prolif | Pnn |
| Cct31 | 0 | 0.75627718 | 0.617 | 0.351 | 0 | TAM.prolif | Cct3 |
| Vkorc12 | 0 | 0.96899105 | 0.487 | 0.221 | 0 | TAM.prolif | Vkorc1 |
| Pfas | 0 | 1.69778988 | 0.367 | 0.101 | 0 | TAM.prolif | Pfas |
| Oat | 0 | 0.68543173 | 0.587 | 0.321 | 0 | TAM.prolif | Oat |
| Mtap | 0 | 1.1039871 | 0.461 | 0.196 | 0 | TAM.prolif | Mtap |
| Snrpg1 | 0 | 0.8892066 | 0.529 | 0.264 | 0 | TAM.prolif | Snrpg |
| Etfb1 | 0 | 0.6861689 | 0.784 | 0.519 | 0 | TAM.prolif | Etfb |
| Cks2 | 0 | 1.24195401 | 0.407 | 0.142 | 0 | TAM.prolif | Cks2 |
| Brd8 | 0 | 1.09177296 | 0.507 | 0.242 | 0 | TAM.prolif | Brd8 |
| Snrpd31 | 0 | 0.71507123 | 0.729 | 0.464 | 0 | TAM.prolif | Snrpd3 |
| Lsm71 | 0 | 0.75476935 | 0.563 | 0.298 | 0 | TAM.prolif | Lsm7 |
| Grk61 | 0 | 0.76571466 | 0.563 | 0.298 | 0 | TAM.prolif | Grk6 |
| Got2 | 0 | 0.69531717 | 0.752 | 0.488 | 0 | TAM.prolif | Got2 |
| Hfe2 | 0 | 0.5135277 | 0.745 | 0.481 | 0 | TAM.prolif | Hfe |
| Naa50 | 0 | 0.89705896 | 0.806 | 0.542 | 0 | TAM.prolif | Naa50 |
| Khsrp | 0 | 0.7854272 | 0.812 | 0.548 | 0 | TAM.prolif | Khsrp |
| Ddx39b | 0 | 0.66866259 | 0.714 | 0.45 | 0 | TAM.prolif | Ddx39b |
| Fundc21 | 0 | 0.72020349 | 0.581 | 0.317 | 0 | TAM.prolif | Fundc2 |
| Knl1 | 0 | 5.97692647 | 0.267 | 0.004 | 0 | TAM.prolif | Knl1 |
| Ddx27 | 0 | 0.89024212 | 0.503 | 0.24 | 0 | TAM.prolif | Ddx27 |
| Sf3a3 | 0 | 0.91639079 | 0.487 | 0.224 | 0 | TAM.prolif | Sf3a3 |
| Cct71 | 0 | 0.76922995 | 0.608 | 0.345 | 0 | TAM.prolif | Cct7 |
| Ctc1 | 0 | 1.42241117 | 0.377 | 0.114 | 0 | TAM.prolif | Ctc1 |
| Hist1h1d | 0 | 4.7117027 | 0.277 | 0.014 | 0 | TAM.prolif | Hist1h1d |
| Rfc4 | 0 | 2.80678779 | 0.297 | 0.034 | 0 | TAM.prolif | Rfc4 |
| Thrap3 | 0 | 0.65383467 | 0.59 | 0.327 | 0 | TAM.prolif | Thrap3 |
| Eif3b1 | 0 | 0.61053224 | 0.703 | 0.44 | 0 | TAM.prolif | Eif3b |
| Hcfc1 | 0 | 0.74800604 | 0.587 | 0.324 | 0 | TAM.prolif | Hcfc1 |
| Atp5g31 | 0 | 0.69264696 | 0.614 | 0.352 | 0 | TAM.prolif | Atp5g3 |
| Hnrnpr | 0 | 0.92699467 | 0.503 | 0.241 | 0 | TAM.prolif | Hnrnpr |
| Mogs1 | 0 | 0.89766328 | 0.478 | 0.216 | 0 | TAM.prolif | Mogs |
| Cgas | 0 | 1.3221438 | 0.372 | 0.11 | 0 | TAM.prolif | Cgas |
| Ddx1 | 0 | 0.75358464 | 0.531 | 0.269 | 0 | TAM.prolif | Ddx1 |
| Lpcat11 | 0 | 0.87286316 | 0.526 | 0.264 | 0 | TAM.prolif | Lpcat1 |
| Dkc1 | 0 | 1.55551595 | 0.379 | 0.117 | 0 | TAM.prolif | Dkc1 |
| Ppp1r7 | 0 | 0.92338697 | 0.5 | 0.239 | 0 | TAM.prolif | Ppp1r7 |
| Cenpf | 0 | 5.43051988 | 0.267 | 0.006 | 0 | TAM.prolif | Cenpf |
| Actl6a | 0 | 0.97844202 | 0.477 | 0.216 | 0 | TAM.prolif | Actl6a |
| Zdhhc3 | 0 | 0.60855701 | 0.641 | 0.38 | 0 | TAM.prolif | Zdhhc3 |
| Mlec | 0 | 0.83965491 | 0.815 | 0.554 | 0 | TAM.prolif | Mlec |
| Cacybp | 0 | 0.87060989 | 0.534 | 0.274 | 0 | TAM.prolif | Cacybp |
| Prelid3b | 0 | 0.95933976 | 0.496 | 0.236 | 0 | TAM.prolif | Prelid3b |
| Ppp1r8 | 0 | 0.93394409 | 0.443 | 0.183 | 0 | TAM.prolif | Ppp1r8 |
| Ppp5c1 | 0 | 0.7936948 | 0.53 | 0.27 | 0 | TAM.prolif | Ppp5c |
| Trmt2a1 | 0 | 0.91528604 | 0.495 | 0.235 | 0 | TAM.prolif | Trmt2a |
| Ung | 0 | 3.84309085 | 0.284 | 0.024 | 0 | TAM.prolif | Ung |
| Pfdn61 | 0 | 0.63527444 | 0.596 | 0.336 | 0 | TAM.prolif | Pfdn6 |
| Eef1g1 | 0 | 0.84164574 | 0.711 | 0.451 | 0 | TAM.prolif | Eef1g |
| Psmd141 | 0 | 0.57354991 | 0.622 | 0.363 | 0 | TAM.prolif | Psmd14 |
| Rfc3 | 0 | 2.0786002 | 0.326 | 0.067 | 0 | TAM.prolif | Rfc3 |
| Cln62 | 0 | 0.81679767 | 0.515 | 0.256 | 0 | TAM.prolif | Cln6 |

| | | | | | | | |
|---|---|---|---|---|---|---|---|
| Lmnb2 | 0 | 2.50835911 | 0.309 | 0.05 | 0 | TAM.prolif | Lmnb2 |
| Igfbp42 | 0 | 0.71419679 | 0.699 | 0.44 | 0 | TAM.prolif | Igfbp4 |
| Odf2 | 0 | 0.8941393 | 0.552 | 0.294 | 0 | TAM.prolif | Odf2 |
| Acat11 | 0 | 0.66880784 | 0.541 | 0.283 | 0 | TAM.prolif | Acat1 |
| Pelp1 | 0 | 1.01277129 | 0.434 | 0.176 | 0 | TAM.prolif | Pelp1 |
| Nme11 | 0 | 0.55914315 | 0.62 | 0.362 | 0 | TAM.prolif | Nme1 |
| Commd1 | 0 | 0.69787782 | 0.676 | 0.419 | 0 | TAM.prolif | Commd1 |
| Eef1d1 | 0 | 0.73539724 | 0.544 | 0.287 | 0 | TAM.prolif | Eef1d |
| Tcof12 | 0 | 0.65910308 | 0.549 | 0.292 | 0 | TAM.prolif | Tcof1 |
| Rnaseh2a1 | 0 | 0.85695368 | 0.473 | 0.216 | 0 | TAM.prolif | Rnaseh2a |
| Zwilch | 0 | 4.57613262 | 0.265 | 0.008 | 0 | TAM.prolif | Zwilch |
| Rufy1 | 0 | 0.7911113 | 0.517 | 0.26 | 0 | TAM.prolif | Rufy1 |
| Tmx12 | 0 | 0.65465356 | 0.756 | 0.499 | 0 | TAM.prolif | Tmx1 |
| Snrpc | 0 | 0.61113265 | 0.635 | 0.378 | 0 | TAM.prolif | Snrpc |
| Nol7 | 0 | 0.75997511 | 0.566 | 0.309 | 0 | TAM.prolif | Nol7 |
| Ccdc34 | 0 | 2.29901909 | 0.311 | 0.055 | 0 | TAM.prolif | Ccdc34 |
| Npepl11 | 0 | 0.66129498 | 0.621 | 0.365 | 0 | TAM.prolif | Npepl1 |
| Micos10 | 0 | 0.6350307 | 0.726 | 0.47 | 0 | TAM.prolif | Micos10 |
| Ckap2 | 0 | 4.29700805 | 0.267 | 0.011 | 0 | TAM.prolif | Ckap2 |
| Lmf2 | 0 | 1.2266099 | 0.417 | 0.161 | 0 | TAM.prolif | Lmf2 |
| Nop562 | 0 | 0.75394703 | 0.617 | 0.362 | 0 | TAM.prolif | Nop56 |
| Hp1bp31 | 0 | 0.77596387 | 0.733 | 0.478 | 0 | TAM.prolif | Hp1bp3 |
| Ruvbl21 | 0 | 0.97224831 | 0.466 | 0.211 | 0 | TAM.prolif | Ruvbl2 |
| Cnot6 | 0 | 0.77519947 | 0.527 | 0.272 | 0 | TAM.prolif | Cnot6 |
| Hmgn1 | 0 | 1.73924101 | 0.347 | 0.092 | 0 | TAM.prolif | Hmgn1 |
| Commd81 | 0 | 0.60185663 | 0.72 | 0.465 | 0 | TAM.prolif | Commd8 |
| Eif3i1 | 0 | 0.56557269 | 0.621 | 0.367 | 0 | TAM.prolif | Eif3i |
| Nup133 | 0 | 1.46813196 | 0.365 | 0.111 | 0 | TAM.prolif | Nup133 |
| Kpna2 | 0 | 1.63264394 | 0.407 | 0.153 | 0 | TAM.prolif | Kpna2 |
| Cenph | 0 | 5.00735237 | 0.261 | 0.007 | 0 | TAM.prolif | Cenph |
| Tubb5 | 0 | 2.10456817 | 0.942 | 0.688 | 0 | TAM.prolif | Tubb5 |
| Ebna1bp21 | 0 | 0.69944358 | 0.581 | 0.327 | 0 | TAM.prolif | Ebna1bp2 |
| Tardbp | 0 | 0.63043115 | 0.692 | 0.438 | 0 | TAM.prolif | Tardbp |
| Uqcr10 | 0 | 0.72536451 | 0.554 | 0.301 | 0 | TAM.prolif | Uqcr10 |
| Tbrg41 | 0 | 0.63873281 | 0.558 | 0.305 | 0 | TAM.prolif | Tbrg4 |
| Tspan133 | 0 | 0.94450024 | 0.551 | 0.298 | 0 | TAM.prolif | Tspan13 |
| Exosc10 | 0 | 0.75477925 | 0.488 | 0.235 | 0 | TAM.prolif | Exosc10 |
| Anapc5 | 0 | 1.27981844 | 0.39 | 0.137 | 0 | TAM.prolif | Anapc5 |
| Tcerg11 | 0 | 0.90616106 | 0.488 | 0.235 | 0 | TAM.prolif | Tcerg1 |
| Prps2 | 0 | 1.34812769 | 0.38 | 0.127 | 0 | TAM.prolif | Prps2 |
| Atp5g11 | 0 | 0.62981405 | 0.582 | 0.329 | 0 | TAM.prolif | Atp5g1 |
| Micos13 | 0 | 0.54546276 | 0.688 | 0.435 | 0 | TAM.prolif | Micos13 |
| Usf11 | 0 | 0.53942496 | 0.661 | 0.409 | 0 | TAM.prolif | Usf1 |
| Fmr1 | 0 | 0.68833148 | 0.553 | 0.301 | 0 | TAM.prolif | Fmr1 |
| Cep250 | 0 | 1.57803974 | 0.346 | 0.094 | 0 | TAM.prolif | Cep250 |
| Ash2l | 0 | 0.67916277 | 0.509 | 0.257 | 0 | TAM.prolif | Ash2l |
| Gadd45gip1 | 0 | 0.84882372 | 0.494 | 0.242 | 0 | TAM.prolif | Gadd45gip1 |
| Bzw22 | 0 | 0.93164301 | 0.468 | 0.216 | 0 | TAM.prolif | Bzw2 |
| Myc2 | 0 | 0.82439787 | 0.512 | 0.26 | 0 | TAM.prolif | Myc |
| Gps1 | 0 | 0.75880782 | 0.58 | 0.328 | 0 | TAM.prolif | Gps1 |
| Dars | 0 | 0.57196922 | 0.687 | 0.436 | 0 | TAM.prolif | Dars |
| Trim271 | 0 | 0.70066992 | 0.561 | 0.31 | 0 | TAM.prolif | Trim27 |
| Ndufc11 | 0 | 0.58687956 | 0.627 | 0.376 | 0 | TAM.prolif | Ndufc1 |
| Pds5a1 | 0 | 0.62634178 | 0.648 | 0.397 | 0 | TAM.prolif | Pds5a |
| Cnih1 | 0 | 0.69508692 | 0.574 | 0.323 | 0 | TAM.prolif | Cnih1 |
| Pole41 | 0 | 0.64621523 | 0.551 | 0.301 | 0 | TAM.prolif | Pole4 |
| U2af11 | 0 | 0.62839804 | 0.534 | 0.284 | 0 | TAM.prolif | U2af1 |
| Ctcf | 0 | 0.87191813 | 0.492 | 0.242 | 0 | TAM.prolif | Ctcf |
| Thop1 | 0 | 1.3285148 | 0.391 | 0.141 | 0 | TAM.prolif | Thop1 |
| Fkbp3 | 0 | 1.17302186 | 0.399 | 0.149 | 0 | TAM.prolif | Fkbp3 |
| Adsl | 0 | 1.01571408 | 0.433 | 0.183 | 0 | TAM.prolif | Adsl |
| Eif4g1 | 0 | 0.5833848 | 0.635 | 0.385 | 0 | TAM.prolif | Eif4g1 |
| Metrn | 0 | 2.02186348 | 0.312 | 0.062 | 0 | TAM.prolif | Metrn |
| Nudt41 | 0 | 0.77447705 | 0.481 | 0.231 | 0 | TAM.prolif | Nudt4 |
| Cyb5b | 0 | 0.65223327 | 0.54 | 0.291 | 0 | TAM.prolif | Cyb5b |
| Cpsf61 | 0 | 0.50617491 | 0.684 | 0.435 | 0 | TAM.prolif | Cpsf6 |
| Nsun21 | 0 | 0.83581809 | 0.557 | 0.308 | 0 | TAM.prolif | Nsun2 |
| Mthfd2 | 0 | 1.17889527 | 0.402 | 0.153 | 0 | TAM.prolif | Mthfd2 |
| Hspe12 | 0 | 0.80418403 | 0.506 | 0.257 | 0 | TAM.prolif | Hspe1 |
| Bard1 | 0 | 4.41955675 | 0.258 | 0.009 | 0 | TAM.prolif | Bard1 |
| Ptma | 0 | 1.2501796 | 0.913 | 0.664 | 0 | TAM.prolif | Ptma |
| Anapc1 | 0 | 0.67823098 | 0.517 | 0.268 | 0 | TAM.prolif | Anapc1 |
| Pih1d1 | 0 | 0.97845686 | 0.445 | 0.196 | 0 | TAM.prolif | Pih1d1 |
| Dazap11 | 0 | 0.77499616 | 0.837 | 0.588 | 0 | TAM.prolif | Dazap1 |
| Ska2 | 0 | 3.90753224 | 0.264 | 0.015 | 0 | TAM.prolif | Ska2 |
| Asb2 | 0 | 2.09030825 | 0.316 | 0.067 | 0 | TAM.prolif | Asb2 |
| Nubp2 | 0 | 0.58330114 | 0.612 | 0.363 | 0 | TAM.prolif | Nubp2 |
| Nudt3 | 0 | 0.52925477 | 0.743 | 0.494 | 0 | TAM.prolif | Nudt3 |
| Fkbp51 | 0 | 0.76917511 | 0.489 | 0.24 | 0 | TAM.prolif | Fkbp5 |
| Ubl4a | 0 | 1.01273507 | 0.445 | 0.196 | 0 | TAM.prolif | Ubl4a |
| Acsl5 | 0 | 0.71412433 | 0.706 | 0.457 | 0 | TAM.prolif | Acsl5 |
| Mcmbp | 0 | 0.63741465 | 0.681 | 0.433 | 0 | TAM.prolif | Mcmbp |
| Cdca7 | 0 | 3.65634412 | 0.266 | 0.018 | 0 | TAM.prolif | Cdca7 |
| Tmem245 | 0 | 1.05097427 | 0.402 | 0.154 | 0 | TAM.prolif | Tmem245 |
| Trim37 | 0 | 1.64291359 | 0.335 | 0.087 | 0 | TAM.prolif | Trim37 |
| Ndufb61 | 0 | 0.5738858 | 0.735 | 0.487 | 0 | TAM.prolif | Ndufb6 |
| Txlna1 | 0 | 0.61332138 | 0.6 | 0.352 | 0 | TAM.prolif | Txlna |
| Recql | 0 | 1.40614429 | 0.363 | 0.115 | 0 | TAM.prolif | Recql |
| Ptges31 | 0 | 0.78778193 | 0.61 | 0.362 | 0 | TAM.prolif | Ptges3 |
| Mthfd1 | 0 | 1.37301961 | 0.371 | 0.123 | 0 | TAM.prolif | Mthfd1 |
| Eif3l1 | 0 | 0.72705886 | 0.783 | 0.535 | 0 | TAM.prolif | Eif3l |
| Lars | 0 | 0.71817562 | 0.511 | 0.263 | 0 | TAM.prolif | Lars |
| Pgp1 | 0 | 0.82112665 | 0.472 | 0.224 | 0 | TAM.prolif | Pgp |
| Qdpr3 | 0 | 0.61427779 | 0.678 | 0.431 | 0 | TAM.prolif | Qdpr |
| Sipa11 | 0 | 0.53125983 | 0.681 | 0.434 | 0 | TAM.prolif | Sipa1 |
| Tlr121 | 0 | 1.57347697 | 0.34 | 0.093 | 0 | TAM.prolif | Tlr12 |
| Rfc1 | 0 | 1.23937917 | 0.398 | 0.151 | 0 | TAM.prolif | Rfc1 |
| Atad5 | 0 | 2.88696881 | 0.274 | 0.027 | 0 | TAM.prolif | Atad5 |
| Prkacb2 | 0 | 0.53565272 | 0.61 | 0.363 | 0 | TAM.prolif | Prkacb |
| Hnrnpd2 | 0 | 0.63773255 | 0.748 | 0.501 | 0 | TAM.prolif | Hnrnpd |
| Ndufa5 | 0 | 0.79104265 | 0.504 | 0.257 | 0 | TAM.prolif | Ndufa5 |
| Wdr181 | 0 | 0.91502192 | 0.479 | 0.232 | 0 | TAM.prolif | Wdr18 |
| Golt1b1 | 0 | 0.69922431 | 0.538 | 0.292 | 0 | TAM.prolif | Golt1b |
| Mast21 | 0 | 0.91439329 | 0.458 | 0.212 | 0 | TAM.prolif | Mast2 |
| Phb21 | 0 | 0.6113285 | 0.637 | 0.391 | 0 | TAM.prolif | Phb2 |
| Lair12 | 0 | 0.67952339 | 0.732 | 0.486 | 0 | TAM.prolif | Lair1 |
| Hprt | 0 | 0.68081391 | 0.732 | 0.486 | 0 | TAM.prolif | Hprt |
| Nup205 | 0 | 1.34116037 | 0.346 | 0.1 | 0 | TAM.prolif | Nup205 |
| Eif2b23 | 0 | 0.53997649 | 0.588 | 0.342 | 0 | TAM.prolif | Eif2b2 |
| Dip2c2 | 0 | 0.6334658 | 0.575 | 0.329 | 0 | TAM.prolif | Dip2c |
| Ndufs8 | 0 | 0.62980818 | 0.714 | 0.468 | 0 | TAM.prolif | Ndufs8 |
| Rbm17 | 0 | 0.63153689 | 0.68 | 0.435 | 0 | TAM.prolif | Rbm17 |
| Cinp | 0 | 0.88645682 | 0.464 | 0.219 | 0 | TAM.prolif | Cinp |
| Med30 | 0 | 0.92058297 | 0.468 | 0.223 | 0 | TAM.prolif | Med30 |
| Fzr1 | 0 | 1.05884954 | 0.493 | 0.248 | 0 | TAM.prolif | Fzr1 |
| Nagpa1 | 0 | 0.58333282 | 0.64 | 0.395 | 0 | TAM.prolif | Nagpa |
| Sf3a2 | 0 | 0.64339956 | 0.575 | 0.33 | 0 | TAM.prolif | Sf3a2 |
| Fubp11 | 0 | 0.46088663 | 0.677 | 0.433 | 0 | TAM.prolif | Fubp1 |
| Atp5a1 | 0 | 0.55110276 | 0.687 | 0.443 | 0 | TAM.prolif | Atp5a1 |
| Tmx21 | 0 | 0.88616659 | 0.457 | 0.213 | 0 | TAM.prolif | Tmx2 |
| Ddx56 | 0 | 0.86583628 | 0.454 | 0.21 | 0 | TAM.prolif | Ddx56 |
| Tmem2732 | 0 | 0.91870119 | 0.452 | 0.208 | 0 | TAM.prolif | Tmem273 |
| Tecr1 | 0 | 0.63532291 | 0.722 | 0.478 | 0 | TAM.prolif | Tecr |
| Slc7a52 | 0 | 0.74609221 | 0.5 | 0.256 | 0 | TAM.prolif | Slc7a5 |
| Med9 | 0 | 0.98015446 | 0.423 | 0.179 | 0 | TAM.prolif | Med9 |
| Parp2 | 0 | 0.81226643 | 0.447 | 0.203 | 0 | TAM.prolif | Parp2 |
| Pck2 | 0 | 1.39598869 | 0.367 | 0.123 | 0 | TAM.prolif | Pck2 |
| Smpd4 | 0 | 1.10170437 | 0.381 | 0.137 | 0 | TAM.prolif | Smpd4 |
| Idh3g1 | 0 | 0.56889255 | 0.715 | 0.471 | 0 | TAM.prolif | Idh3g |
| Spdl1 | 0 | 2.50994486 | 0.284 | 0.04 | 0 | TAM.prolif | Spdl1 |
| Afg3l11 | 0 | 0.55036478 | 0.578 | 0.334 | 0 | TAM.prolif | Afg3l1 |
| Arhgef63 | 0 | 0.61461731 | 0.564 | 0.32 | 0 | TAM.prolif | Arhgef6 |
| Ptpn71 | 0 | 0.9067095 | 0.495 | 0.252 | 0 | TAM.prolif | Ptpn7 |
| Frmd4b2 | 0 | 0.52106881 | 0.594 | 0.351 | 0 | TAM.prolif | Frmd4b |
| Cluh1 | 0 | 0.83439286 | 0.503 | 0.26 | 0 | TAM.prolif | Cluh |
| Tars | 0 | 0.7046113 | 0.523 | 0.28 | 0 | TAM.prolif | Tars |
| Usp14 | 0 | 0.73287882 | 0.515 | 0.272 | 0 | TAM.prolif | Usp14 |
| Atp5f11 | 0 | 0.59428736 | 0.691 | 0.448 | 0 | TAM.prolif | Atp5f1 |
| Naa38 | 0 | 0.58310837 | 0.573 | 0.33 | 0 | TAM.prolif | Naa38 |
| Atp5o | 0 | 0.72098468 | 0.702 | 0.459 | 0 | TAM.prolif | Atp5o |
| Prep | 0 | 0.63397397 | 0.556 | 0.314 | 0 | TAM.prolif | Prep |
| Osgep1 | 0 | 0.72427187 | 0.524 | 0.282 | 0 | TAM.prolif | Osgep |
| Ints7 | 0 | 1.3027776 | 0.373 | 0.131 | 0 | TAM.prolif | Ints7 |
| Dnajc101 | 0 | 0.60938261 | 0.635 | 0.393 | 0 | TAM.prolif | Dnajc10 |
| Alad | 0 | 1.39439575 | 0.353 | 0.111 | 0 | TAM.prolif | Alad |
| Cherp | 0 | 0.65073238 | 0.572 | 0.33 | 0 | TAM.prolif | Cherp |
| Hnrnpa0 | 0 | 0.44101 | 0.693 | 0.451 | 0 | TAM.prolif | Hnrnpa0 |
| Ide | 0 | 0.74125388 | 0.573 | 0.331 | 0 | TAM.prolif | Ide |
| Sec13 | 0 | 0.53947876 | 0.674 | 0.433 | 0 | TAM.prolif | Sec13 |
| Slc30a51 | 0 | 0.67538247 | 0.549 | 0.308 | 0 | TAM.prolif | Slc30a5 |
| Wdr76 | 0 | 2.96146806 | 0.272 | 0.031 | 0 | TAM.prolif | Wdr76 |
| Sass6 | 0 | 1.43032681 | 0.335 | 0.094 | 0 | TAM.prolif | Sass6 |
| Cdca7l | 0 | 2.87869283 | 0.273 | 0.032 | 0 | TAM.prolif | Cdca7l |
| Ndufaf2 | 0 | 1.01833365 | 0.392 | 0.151 | 0 | TAM.prolif | Ndufaf2 |
| Adpgk1 | 0 | 0.75790754 | 0.484 | 0.243 | 0 | TAM.prolif | Adpgk |
| Creld2 | 0 | 0.745041 | 0.489 | 0.248 | 0 | TAM.prolif | Creld2 |
| Ankrd52 | 0 | 0.4980566 | 0.564 | 0.323 | 0 | TAM.prolif | Ankrd52 |
| Magt1 | 0 | 0.44357465 | 0.705 | 0.464 | 0 | TAM.prolif | Magt1 |
| Rif1 | 0 | 1.36832224 | 0.342 | 0.102 | 0 | TAM.prolif | Rif1 |
| Magoh2 | 0 | 0.57757101 | 0.552 | 0.312 | 0 | TAM.prolif | Magoh |
| Ranbp3 | 0 | 0.58422073 | 0.554 | 0.314 | 0 | TAM.prolif | Ranbp3 |
| Bora | 0 | 2.93965962 | 0.275 | 0.035 | 0 | TAM.prolif | Bora |
| Npl2 | 0 | 0.74936113 | 0.503 | 0.263 | 0 | TAM.prolif | Npl |
| Gle1 | 0 | 1.34933945 | 0.349 | 0.109 | 0 | TAM.prolif | Gle1 |
| Ssb1 | 0 | 0.60110704 | 0.792 | 0.552 | 0 | TAM.prolif | Ssb |
| Cyc11 | 0 | 0.61157331 | 0.694 | 0.454 | 0 | TAM.prolif | Cyc1 |
| Ndufa10 | 0 | 0.5606097 | 0.661 | 0.422 | 0 | TAM.prolif | Ndufa10 |
| Kdelr21 | 0 | 0.57528072 | 0.652 | 0.413 | 0 | TAM.prolif | Kdelr2 |
| Pak11 | 0 | 0.70972085 | 0.528 | 0.289 | 0 | TAM.prolif | Pak1 |
| Psmd131 | 0 | 0.54523919 | 0.678 | 0.439 | 0 | TAM.prolif | Psmd13 |
| Rpn21 | 0 | 0.64414417 | 0.749 | 0.51 | 0 | TAM.prolif | Rpn2 |
| Park71 | 0 | 0.66126436 | 0.814 | 0.575 | 0 | TAM.prolif | Park7 |
| Gbp82 | 0 | 0.58061659 | 0.495 | 0.256 | 0 | TAM.prolif | Gbp8 |
| Arl6ip4 | 0 | 0.74017574 | 0.492 | 0.253 | 0 | TAM.prolif | Arl6ip4 |
| Gtf3a | 0 | 0.89360071 | 0.442 | 0.203 | 0 | TAM.prolif | Gtf3a |
| Psme31 | 0 | 0.61230328 | 0.749 | 0.51 | 0 | TAM.prolif | Psme3 |
| Comt2 | 0 | 0.50218723 | 0.767 | 0.528 | 0 | TAM.prolif | Comt |
| Srpk11 | 0 | 0.48476674 | 0.627 | 0.388 | 0 | TAM.prolif | Srpk1 |
| Ndufv2 | 0 | 0.59876285 | 0.617 | 0.378 | 0 | TAM.prolif | Ndufv2 |
| Rnf1261 | 0 | 0.55194397 | 0.565 | 0.326 | 0 | TAM.prolif | Rnf126 |
| Acadl1 | 0 | 0.64397374 | 0.549 | 0.311 | 0 | TAM.prolif | Acadl |
| Nudt211 | 0 | 0.70340473 | 0.525 | 0.287 | 0 | TAM.prolif | Nudt21 |
| Ncbp11 | 0 | 0.82443139 | 0.45 | 0.212 | 0 | TAM.prolif | Ncbp1 |
| Dhx9 | 0 | 0.69313037 | 0.743 | 0.505 | 0 | TAM.prolif | Dhx9 |
| Nup188 | 0 | 0.89683561 | 0.415 | 0.177 | 0 | TAM.prolif | Nup188 |
| Psmc4 | 0 | 0.56876533 | 0.637 | 0.399 | 0 | TAM.prolif | Psmc4 |
| Lrp53 | 0 | 0.45753991 | 0.607 | 0.369 | 0 | TAM.prolif | Lrp5 |
| Ilf31 | 0 | 0.60414922 | 0.553 | 0.316 | 0 | TAM.prolif | Ilf3 |
| Larp7 | 0 | 0.91320761 | 0.401 | 0.164 | 0 | TAM.prolif | Larp7 |
| Igsf83 | 0 | 0.40794953 | 0.772 | 0.535 | 0 | TAM.prolif | Igsf8 |
| Mdh21 | 0 | 0.89025206 | 0.642 | 0.405 | 0 | TAM.prolif | Mdh2 |
| Snrnp25 | 0 | 1.17074192 | 0.374 | 0.137 | 0 | TAM.prolif | Snrnp25 |
| Smarce1 | 0 | 0.4592578 | 0.608 | 0.371 | 0 | TAM.prolif | Smarce1 |
| Nap1l4 | 0 | 0.59386612 | 0.573 | 0.336 | 0 | TAM.prolif | Nap1l4 |
| Tm9sf1 | 0 | 0.59637224 | 0.578 | 0.341 | 0 | TAM.prolif | Tm9sf1 |
| Fkbp2 | 0 | 0.6048252 | 0.802 | 0.566 | 0 | TAM.prolif | Fkbp2 |
| Imp4 | 0 | 0.56988599 | 0.554 | 0.318 | 0 | TAM.prolif | Imp4 |
| Ndufv1 | 0 | 0.52929581 | 0.649 | 0.413 | 0 | TAM.prolif | Ndufv1 |
| Pcna1 | 0 | 1.27493209 | 0.832 | 0.596 | 0 | TAM.prolif | Pcna |
| Arl6ip12 | 0 | 0.88294877 | 0.583 | 0.347 | 0 | TAM.prolif | Arl6ip1 |
| Zmiz21 | 0 | 0.56952364 | 0.737 | 0.501 | 0 | TAM.prolif | Zmiz2 |
| Nudcd2 | 0 | 1.16815834 | 0.376 | 0.14 | 0 | TAM.prolif | Nudcd2 |
| Prpf8 | 0 | 0.64400463 | 0.709 | 0.473 | 0 | TAM.prolif | Prpf8 |
| Cops5 | 0 | 0.50258395 | 0.569 | 0.333 | 0 | TAM.prolif | Cops5 |
| Rad23b | 0 | 0.47000937 | 0.664 | 0.429 | 0 | TAM.prolif | Rad23b |
| Nme2 | 0 | 0.50357474 | 0.649 | 0.414 | 0 | TAM.prolif | Nme2 |
| Htra21 | 0 | 0.73707307 | 0.465 | 0.23 | 0 | TAM.prolif | Htra2 |
| Fkbp4 | 0 | 1.15710877 | 0.391 | 0.156 | 0 | TAM.prolif | Fkbp4 |
| Siglece2 | 0 | 0.78805185 | 0.448 | 0.213 | 0 | TAM.prolif | Siglece |
| Rhoh1 | 0 | 0.5817607 | 0.582 | 0.347 | 0 | TAM.prolif | Rhoh |
| Gtf2i2 | 0 | 0.53208693 | 0.587 | 0.352 | 0 | TAM.prolif | Gtf2i |
| Camkk1 | 0 | 1.34299929 | 0.361 | 0.126 | 0 | TAM.prolif | Camkk1 |
| Limd22 | 0 | 0.57878492 | 0.607 | 0.372 | 0 | TAM.prolif | Limd2 |
| Nop16 | 0 | 1.00426793 | 0.413 | 0.178 | 0 | TAM.prolif | Nop16 |
| Polr2f1 | 0 | 0.611657 | 0.592 | 0.357 | 0 | TAM.prolif | Polr2f |
| Pebp1 | 0 | 0.62316045 | 0.551 | 0.317 | 0 | TAM.prolif | Pebp1 |
| Mta1 | 0 | 0.6208603 | 0.54 | 0.306 | 0 | TAM.prolif | Mta1 |
| Ccdc102a2 | 0 | 0.91235665 | 0.398 | 0.164 | 0 | TAM.prolif | Ccdc102a |
| Tsr11 | 0 | 0.76730784 | 0.453 | 0.219 | 0 | TAM.prolif | Tsr1 |
| Ubr7 | 0 | 1.07439497 | 0.396 | 0.162 | 0 | TAM.prolif | Ubr7 |
| Iars1 | 0 | 0.7834595 | 0.457 | 0.223 | 0 | TAM.prolif | Iars |
| Nolc1 | 0 | 1.15388964 | 0.393 | 0.159 | 0 | TAM.prolif | Nolc1 |
| Nutf2-ps1 | 0 | 1.0307791 | 0.376 | 0.142 | 0 | TAM.prolif | Nutf2-ps1 |
| Bola21 | 0 | 0.58404798 | 0.757 | 0.523 | 0 | TAM.prolif | Bola2 |
| Prkar2a1 | 0 | 0.97588916 | 0.425 | 0.191 | 0 | TAM.prolif | Prkar2a |
| Pank2 | 0 | 0.64155887 | 0.52 | 0.287 | 0 | TAM.prolif | Pank2 |
| Rbm15 | 0 | 0.66334663 | 0.54 | 0.307 | 0 | TAM.prolif | Rbm15 |
| Med252 | 0 | 0.50057956 | 0.675 | 0.442 | 0 | TAM.prolif | Med25 |
| Babam1 | 0 | 0.54048831 | 0.662 | 0.429 | 0 | TAM.prolif | Babam1 |
| Eif3d1 | 0 | 0.52165178 | 0.683 | 0.45 | 0 | TAM.prolif | Eif3d |
| Slc25a39 | 0 | 0.575183 | 0.748 | 0.515 | 0 | TAM.prolif | Slc25a39 |
| Nup50 | 0 | 0.70722469 | 0.475 | 0.242 | 0 | TAM.prolif | Nup50 |
| Msh2 | 0 | 1.80024116 | 0.317 | 0.084 | 0 | TAM.prolif | Msh2 |
| Cox7b1 | 0 | 0.48353568 | 0.756 | 0.523 | 0 | TAM.prolif | Cox7b |
| Bcas2 | 0 | 0.60183127 | 0.554 | 0.322 | 0 | TAM.prolif | Bcas2 |
| Rad23a | 0 | 0.50197127 | 0.639 | 0.407 | 0 | TAM.prolif | Rad23a |
| Polr2e | 0 | 0.4899208 | 0.665 | 0.433 | 0 | TAM.prolif | Polr2e |
| Txndc5 | 0 | 0.503158 | 0.67 | 0.438 | 0 | TAM.prolif | Txndc5 |
| Psmd2 | 0 | 0.46886765 | 0.683 | 0.451 | 0 | TAM.prolif | Psmd2 |
| Rpa3 | 0 | 1.51235451 | 0.321 | 0.089 | 0 | TAM.prolif | Rpa3 |
| C1qbp | 0 | 1.17754306 | 0.393 | 0.161 | 0 | TAM.prolif | C1qbp |
| Tmco1 | 0 | 0.63318355 | 0.591 | 0.359 | 0 | TAM.prolif | Tmco1 |
| Ssr2 | 0 | 0.51183677 | 0.767 | 0.535 | 0 | TAM.prolif | Ssr2 |
| Cops6 | 0 | 0.53276316 | 0.738 | 0.506 | 0 | TAM.prolif | Cops6 |
| Ppia | 0 | 0.85481451 | 0.843 | 0.611 | 0 | TAM.prolif | Ppia |
| Utp6 | 0 | 0.70986323 | 0.5 | 0.268 | 0 | TAM.prolif | Utp6 |
| Gpr652 | 0 | 0.48459841 | 0.616 | 0.384 | 0 | TAM.prolif | Gpr65 |
| Cebpzos2 | 0 | 0.80387696 | 0.477 | 0.245 | 0 | TAM.prolif | Cebpzos |
| Galnt12 | 0 | 0.52930956 | 0.754 | 0.522 | 0 | TAM.prolif | Galnt1 |
| Utp11 | 0 | 0.6204249 | 0.529 | 0.298 | 0 | TAM.prolif | Utp11 |
| Tmed31 | 0 | 0.52043624 | 0.66 | 0.429 | 0 | TAM.prolif | Tmed3 |
| Cdk5rap2 | 0 | 1.52304916 | 0.317 | 0.086 | 0 | TAM.prolif | Cdk5rap2 |
| Ndufab1 | 0 | 0.83041794 | 0.452 | 0.221 | 0 | TAM.prolif | Ndufab1 |
| Psmc2 | 0 | 0.51913477 | 0.631 | 0.4 | 0 | TAM.prolif | Psmc2 |
| Suds3 | 0 | 0.55297787 | 0.631 | 0.4 | 0 | TAM.prolif | Suds3 |
| Hebp11 | 0 | 0.70438184 | 0.591 | 0.36 | 0 | TAM.prolif | Hebp1 |
| Ube2m | 0 | 0.55574395 | 0.722 | 0.491 | 0 | TAM.prolif | Ube2m |
| Vac14 | 0 | 0.45689973 | 0.623 | 0.392 | 0 | TAM.prolif | Vac14 |
| Wdr82 | 0 | 0.51020613 | 0.624 | 0.393 | 0 | TAM.prolif | Wdr82 |
| Metap2 | 0 | 0.56822035 | 0.697 | 0.466 | 0 | TAM.prolif | Metap2 |
| Atp5k | 0 | 0.52736079 | 0.658 | 0.428 | 0 | TAM.prolif | Atp5k |
| Snrnp200 | 0 | 0.49562679 | 0.628 | 0.398 | 0 | TAM.prolif | Snrnp200 |
| Tmco4 | 0 | 1.19023758 | 0.356 | 0.126 | 0 | TAM.prolif | Tmco4 |
| Akip1 | 0 | 1.40582037 | 0.344 | 0.114 | 0 | TAM.prolif | Akip1 |
| Ssbp41 | 0 | 0.41603894 | 0.753 | 0.523 | 0 | TAM.prolif | Ssbp4 |
| Timm13 | 0 | 0.59999011 | 0.792 | 0.562 | 0 | TAM.prolif | Timm13 |
| Blmh | 0 | 0.7215961 | 0.482 | 0.252 | 0 | TAM.prolif | Blmh |
| Ndufb10 | 0 | 0.57772516 | 0.773 | 0.543 | 0 | TAM.prolif | Ndufb10 |
| Cuta1 | 0 | 0.52119299 | 0.753 | 0.523 | 0 | TAM.prolif | Cuta |
| Sf3b2 | 0 | 0.68419272 | 0.807 | 0.578 | 0 | TAM.prolif | Sf3b2 |
| Pole3 | 0 | 1.38593961 | 0.332 | 0.103 | 0 | TAM.prolif | Pole3 |
| Gtf2h5 | 0 | 0.4930419 | 0.646 | 0.417 | 0 | TAM.prolif | Gtf2h5 |
| Caprin1 | 0 | 0.53214832 | 0.649 | 0.421 | 0 | TAM.prolif | Caprin1 |
| Bcap291 | 0 | 0.80560134 | 0.439 | 0.211 | 0 | TAM.prolif | Bcap29 |
| Pdcl31 | 0 | 0.56143998 | 0.619 | 0.391 | 0 | TAM.prolif | Pdcl3 |
| Ufc1 | 0 | 0.47224512 | 0.734 | 0.506 | 0 | TAM.prolif | Ufc1 |
| Fdps1 | 0 | 0.93306902 | 0.508 | 0.28 | 0 | TAM.prolif | Fdps |
| Haus6 | 0 | 2.03329484 | 0.284 | 0.056 | 0 | TAM.prolif | Haus6 |
| Nup37 | 0 | 1.82286381 | 0.297 | 0.069 | 0 | TAM.prolif | Nup37 |
| Hspa41 | 0 | 0.56818034 | 0.586 | 0.358 | 0 | TAM.prolif | Hspa4 |
| Cstf2 | 0 | 1.11477565 | 0.365 | 0.137 | 0 | TAM.prolif | Cstf2 |
| Cand1 | 0 | 0.53335855 | 0.567 | 0.339 | 0 | TAM.prolif | Cand1 |
| Ak21 | 0 | 0.55707806 | 0.797 | 0.57 | 0 | TAM.prolif | Ak2 |
| E2f2 | 0 | 1.72743778 | 0.274 | 0.047 | 0 | TAM.prolif | E2f2 |
| Tubgcp2 | 0 | 1.41595197 | 0.325 | 0.098 | 0 | TAM.prolif | Tubgcp2 |
| Fads12 | 0 | 0.89841368 | 0.448 | 0.221 | 0 | TAM.prolif | Fads1 |
| Elavl11 | 0 | 0.49677147 | 0.752 | 0.525 | 0 | TAM.prolif | Elavl1 |
| Etfa | 0 | 0.57137904 | 0.712 | 0.485 | 0 | TAM.prolif | Etfa |
| Cox7a2 | 0 | 0.59515848 | 0.575 | 0.348 | 0 | TAM.prolif | Cox7a2 |
| Sec11c | 0 | 0.55586097 | 0.691 | 0.464 | 0 | TAM.prolif | Sec11c |
| Pigt1 | 0 | 0.47016166 | 0.664 | 0.438 | 0 | TAM.prolif | Pigt |
| Tap12 | 0 | 0.35686438 | 0.652 | 0.426 | 0 | TAM.prolif | Tap1 |
| Tinf21 | 0 | 0.57557219 | 0.447 | 0.221 | 0 | TAM.prolif | Tinf2 |
| Rad50 | 0 | 1.15444239 | 0.362 | 0.136 | 0 | TAM.prolif | Rad50 |
| Tmem97 | 0 | 1.21491489 | 0.366 | 0.14 | 0 | TAM.prolif | Tmem97 |
| Gna122 | 0 | 0.58044476 | 0.789 | 0.564 | 0 | TAM.prolif | Gna12 |
| Mmd2 | 0 | 0.73559706 | 0.465 | 0.24 | 0 | TAM.prolif | Mmd |
| Psmd15 | 0 | 0.40872653 | 0.705 | 0.48 | 0 | TAM.prolif | Psmd1 |
| Adss1 | 0 | 0.41143368 | 0.592 | 0.367 | 0 | TAM.prolif | Adss |
| Cnot12 | 0 | 0.46760656 | 0.631 | 0.406 | 0 | TAM.prolif | Cnot1 |
| Ncl1 | 0 | 0.77942229 | 0.793 | 0.569 | 0 | TAM.prolif | Ncl |
| Cdc25b | 0 | 2.65649405 | 0.279 | 0.055 | 0 | TAM.prolif | Cdc25b |
| Batf31 | 0 | 1.19269925 | 0.358 | 0.134 | 0 | TAM.prolif | Batf3 |
| Ppp1cc2 | 0 | 0.53179961 | 0.578 | 0.354 | 0 | TAM.prolif | Ppp1cc |
| Far12 | 0 | 0.4511559 | 0.738 | 0.514 | 0 | TAM.prolif | Far1 |
| Jaml2 | 0 | 0.48875333 | 0.61 | 0.386 | 0 | TAM.prolif | Jaml |
| Ciao2a1 | 0 | 0.67659732 | 0.843 | 0.619 | 0 | TAM.prolif | Ciao2a |
| Ndufs7 | 0 | 0.39435769 | 0.703 | 0.479 | 0 | TAM.prolif | Ndufs7 |
| Abi32 | 0 | 0.36685078 | 0.734 | 0.51 | 0 | TAM.prolif | Abi3 |
| Psmc1 | 0 | 0.56053847 | 0.581 | 0.357 | 0 | TAM.prolif | Psmc1 |
| Etv51 | 0 | 0.53995543 | 0.661 | 0.438 | 0 | TAM.prolif | Etv5 |
| Lnpk | 0 | 1.0726745 | 0.364 | 0.141 | 0 | TAM.prolif | Lnpk |
| Tagap2 | 0 | 0.80637673 | 0.446 | 0.223 | 0 | TAM.prolif | Tagap |
| Samm50 | 0 | 0.54665294 | 0.619 | 0.396 | 0 | TAM.prolif | Samm50 |
| Aurkaip1 | 0 | 0.45670733 | 0.648 | 0.426 | 0 | TAM.prolif | Aurkaip1 |
| Blm | 0 | 2.53584662 | 0.256 | 0.034 | 0 | TAM.prolif | Blm |
| Pea15a2 | 0 | 0.50279929 | 0.734 | 0.512 | 0 | TAM.prolif | Pea15a |
| Mbnl11 | 0 | 0.43363226 | 0.719 | 0.497 | 0 | TAM.prolif | Mbnl1 |
| Ap1s1 | 0 | 0.59663769 | 0.701 | 0.479 | 0 | TAM.prolif | Ap1s1 |
| Sephs22 | 0 | 0.47261395 | 0.578 | 0.356 | 0 | TAM.prolif | Sephs2 |
| Gatad2a | 0 | 0.3786587 | 0.677 | 0.456 | 0 | TAM.prolif | Gatad2a |
| Nup1532 | 0 | 0.59702766 | 0.559 | 0.338 | 0 | TAM.prolif | Nup153 |
| Wdfy41 | 0 | 0.88398847 | 0.558 | 0.337 | 0 | TAM.prolif | Wdfy4 |
| Chaf1b | 0 | 2.15329059 | 0.275 | 0.054 | 0 | TAM.prolif | Chaf1b |
| Sac3d1 | 0 | 1.21577259 | 0.336 | 0.115 | 0 | TAM.prolif | Sac3d1 |
| Fcrls2 | 0 | 0.84474321 | 0.602 | 0.381 | 0 | TAM.prolif | Fcrls |
| Psmb21 | 0 | 0.5411588 | 0.754 | 0.533 | 0 | TAM.prolif | Psmb2 |
| Ncln | 0 | 0.43528508 | 0.702 | 0.481 | 0 | TAM.prolif | Ncln |
| Tomm7 | 0 | 0.46749561 | 0.689 | 0.468 | 0 | TAM.prolif | Tomm7 |
| Rtcb1 | 0 | 0.53183072 | 0.801 | 0.581 | 0 | TAM.prolif | Rtcb |
| Arl6ip6 | 0 | 0.69509246 | 0.462 | 0.242 | 0 | TAM.prolif | Arl6ip6 |
| Ywhaq | 0 | 0.77436548 | 0.463 | 0.243 | 0 | TAM.prolif | Ywhaq |
| Fuca21 | 0 | 0.5529646 | 0.711 | 0.491 | 0 | TAM.prolif | Fuca2 |
| Cnpy31 | 0 | 0.34934167 | 0.711 | 0.491 | 0 | TAM.prolif | Cnpy3 |
| Casp71 | 0 | 0.79443154 | 0.475 | 0.255 | 0 | TAM.prolif | Casp7 |
| Grwd1 | 0 | 1.2827435 | 0.324 | 0.105 | 0 | TAM.prolif | Grwd1 |
| Uqcr11 | 0 | 0.4701071 | 0.663 | 0.444 | 0 | TAM.prolif | Uqcr11 |
| Matr31 | 0 | 0.50735308 | 0.525 | 0.306 | 0 | TAM.prolif | Matr3 |
| Rpn11 | 0 | 0.5606713 | 0.827 | 0.608 | 0 | TAM.prolif | Rpn1 |
| Txn21 | 0 | 0.49049391 | 0.613 | 0.394 | 0 | TAM.prolif | Txn2 |
| Cct81 | 0 | 0.60755793 | 0.564 | 0.345 | 0 | TAM.prolif | Cct8 |
| Cbfb | 0 | 0.99840501 | 0.937 | 0.719 | 0 | TAM.prolif | Cbfb |
| Bak12 | 0 | 0.57059439 | 0.796 | 0.578 | 0 | TAM.prolif | Bak1 |
| Sod11 | 0 | 0.60433883 | 0.68 | 0.462 | 0 | TAM.prolif | Sod1 |
| Nup160 | 0 | 1.28389564 | 0.319 | 0.101 | 0 | TAM.prolif | Nup160 |
| Pole2 | 0 | 2.34083969 | 0.257 | 0.039 | 0 | TAM.prolif | Pole2 |
| Ttf2 | 0 | 1.73494141 | 0.294 | 0.076 | 0 | TAM.prolif | Ttf2 |
| Rexo2 | 0 | 0.41775794 | 0.689 | 0.471 | 0 | TAM.prolif | Rexo2 |
| DEPRECATED-I | 0 | 1.64383554 | 0.293 | 0.075 | 0 | TAM.prolif | DEPRECATED-ENSMUSG00000038759 |
| Eif4ebp21 | 0 | 0.36761572 | 0.662 | 0.445 | 0 | TAM.prolif | Eif4ebp2 |
| Syncrip1 | 0 | 0.57851939 | 0.852 | 0.635 | 0 | TAM.prolif | Syncrip |
| Itga91 | 0 | 0.57979122 | 0.577 | 0.36 | 0 | TAM.prolif | Itga9 |
| Dna21 | 0 | 0.8721461 | 0.353 | 0.136 | 0 | TAM.prolif | Dna2 |
| Srsf71 | 0 | 0.49724649 | 0.805 | 0.589 | 0 | TAM.prolif | Srsf7 |
| Mprip | 0 | 0.42190605 | 0.634 | 0.418 | 0 | TAM.prolif | Mprip |
| E2f4 | 0 | 0.44126476 | 0.708 | 0.492 | 0 | TAM.prolif | E2f4 |
| Cox5a1 | 0 | 0.73826062 | 0.839 | 0.623 | 0 | TAM.prolif | Cox5a |
| Psmc3 | 0 | 0.60834386 | 0.785 | 0.57 | 0 | TAM.prolif | Psmc3 |
| Tpr1 | 0 | 0.47287449 | 0.638 | 0.423 | 0 | TAM.prolif | Tpr |
| Cox6a1 | 0 | 0.3644708 | 0.647 | 0.432 | 0 | TAM.prolif | Cox6a1 |
| Herpud11 | 0 | 0.51421358 | 0.629 | 0.414 | 0 | TAM.prolif | Herpud1 |
| Mpdu1 | 0 | 0.40543679 | 0.643 | 0.428 | 0 | TAM.prolif | Mpdu1 |
| Tubb2a2 | 0 | 0.43933849 | 0.622 | 0.407 | 0 | TAM.prolif | Tubb2a |
| Ccl123 | 0 | 0.91491491 | 0.437 | 0.222 | 0 | TAM.prolif | Ccl12 |
| Ndufb9 | 0 | 0.37280307 | 0.702 | 0.487 | 0 | TAM.prolif | Ndufb9 |
| Ehmt21 | 0 | 0.50943268 | 0.834 | 0.619 | 0 | TAM.prolif | Ehmt2 |
| Ganab | 0 | 0.39783579 | 0.72 | 0.505 | 0 | TAM.prolif | Ganab |
| Psmb7 | 0 | 0.43383554 | 0.628 | 0.414 | 0 | TAM.prolif | Psmb7 |
| Hdac11 | 0 | 0.53508226 | 0.548 | 0.334 | 0 | TAM.prolif | Hdac1 |
| Pcmt11 | 0 | 0.55747196 | 0.53 | 0.316 | 0 | TAM.prolif | Pcmt1 |
| Psat1 | 0 | 2.02085072 | 0.261 | 0.047 | 0 | TAM.prolif | Psat1 |
| Ifrd2 | 0 | 1.48507196 | 0.306 | 0.092 | 0 | TAM.prolif | Ifrd2 |
| Uggt11 | 0 | 0.36197193 | 0.706 | 0.492 | 0 | TAM.prolif | Uggt1 |
| Ndufb71 | 0 | 0.45466042 | 0.739 | 0.525 | 0 | TAM.prolif | Ndufb7 |
| Manf1 | 0 | 0.60472975 | 0.824 | 0.61 | 0 | TAM.prolif | Manf |
| Cct41 | 0 | 0.48180002 | 0.574 | 0.36 | 0 | TAM.prolif | Cct4 |
| Ddx211 | 0 | 0.53314429 | 0.797 | 0.584 | 0 | TAM.prolif | Ddx21 |
| Mmp132 | 0 | 0.40285371 | 0.506 | 0.293 | 0 | TAM.prolif | Mmp13 |
| Hic12 | 0 | 0.96204288 | 0.382 | 0.169 | 0 | TAM.prolif | Hic1 |
| Ptpn183 | 0 | 0.30728478 | 0.739 | 0.526 | 0 | TAM.prolif | Ptpn18 |
| Gnl32 | 0 | 0.56124279 | 0.528 | 0.316 | 0 | TAM.prolif | Gnl3 |
| Hpgds2 | 0 | 0.69218161 | 0.443 | 0.231 | 0 | TAM.prolif | Hpgds |
| Psd33 | 0 | 0.48653532 | 0.487 | 0.275 | 0 | TAM.prolif | Psd3 |
| Heatr13 | 0 | 0.45241244 | 0.567 | 0.355 | 0 | TAM.prolif | Heatr1 |
| Hmgcs11 | 0 | 0.61710993 | 0.582 | 0.37 | 0 | TAM.prolif | Hmgcs1 |
| Sla2 | 0 | 0.43187988 | 0.765 | 0.553 | 0 | TAM.prolif | Sla |
| Blvra2 | 0 | 0.40566734 | 0.666 | 0.455 | 0 | TAM.prolif | Blvra |
| Nxt1 | 0 | 1.25808844 | 0.315 | 0.104 | 0 | TAM.prolif | Nxt1 |
| Sun2 | 0 | 0.66050734 | 0.638 | 0.427 | 0 | TAM.prolif | Sun2 |
| Vcam11 | 0 | 0.86308568 | 0.415 | 0.204 | 0 | TAM.prolif | Vcam1 |
| Chtop | 0 | 0.38439713 | 0.762 | 0.551 | 0 | TAM.prolif | Chtop |
| Hnrnpl1 | 0 | 0.60650049 | 0.825 | 0.614 | 0 | TAM.prolif | Hnrnpl |
| Stt3b | 0 | 0.47581677 | 0.726 | 0.515 | 0 | TAM.prolif | Stt3b |
| Mtch1 | 0 | 0.51476106 | 0.817 | 0.606 | 0 | TAM.prolif | Mtch1 |
| Scaf111 | 0 | 0.42935251 | 0.623 | 0.413 | 0 | TAM.prolif | Scaf11 |
| Msh6 | 0 | 1.7856652 | 0.276 | 0.066 | 0 | TAM.prolif | Msh6 |
| Slc25a52 | 0 | 0.62654013 | 0.662 | 0.452 | 0 | TAM.prolif | Slc25a5 |
| Hsd17b12 | 0 | 0.48982618 | 0.587 | 0.377 | 0 | TAM.prolif | Hsd17b12 |
| Tma7 | 0 | 0.48952433 | 0.734 | 0.524 | 0 | TAM.prolif | Tma7 |
| 2-Sep | 0 | 0.4934968 | 0.775 | 0.566 | 0 | TAM.prolif | 2-Sep |
| Ctbp1 | 0 | 0.46950657 | 0.799 | 0.59 | 0 | TAM.prolif | Ctbp1 |
| Atp5j21 | 0 | 0.55389076 | 0.809 | 0.6 | 0 | TAM.prolif | Atp5j2 |
| Bclaf1 | 0 | 0.36599443 | 0.672 | 0.463 | 0 | TAM.prolif | Bclaf1 |
| Psmc6 | 0 | 0.33147362 | 0.665 | 0.456 | 0 | TAM.prolif | Psmc6 |
| Cox5b1 | 0 | 0.56860439 | 0.863 | 0.654 | 0 | TAM.prolif | Cox5b |
| Actr1b | 0 | 0.43971573 | 0.585 | 0.376 | 0 | TAM.prolif | Actr1b |
| Gtf3c2 | 0 | 0.41303095 | 0.596 | 0.387 | 0 | TAM.prolif | Gtf3c2 |
| Slc1a53 | 0 | 0.36837045 | 0.592 | 0.383 | 0 | TAM.prolif | Slc1a5 |
| Tmem1471 | 0 | 0.57419659 | 0.485 | 0.276 | 0 | TAM.prolif | Tmem147 |
| Idh3b | 0 | 0.43215385 | 0.687 | 0.479 | 0 | TAM.prolif | Idh3b |
| Mrfap11 | 0 | 0.49630433 | 0.81 | 0.602 | 0 | TAM.prolif | Mrfap1 |
| Zw10 | 0 | 1.33422183 | 0.298 | 0.09 | 0 | TAM.prolif | Zw10 |
| Ppm1m1 | 0 | 0.51040658 | 0.608 | 0.4 | 0 | TAM.prolif | Ppm1m |
| Nin1 | 0 | 0.37078851 | 0.569 | 0.361 | 0 | TAM.prolif | Nin |
| Cox7c | 0 | 0.46337288 | 0.798 | 0.591 | 0 | TAM.prolif | Cox7c |
| Sumo2 | 0 | 0.74651022 | 0.465 | 0.258 | 0 | TAM.prolif | Sumo2 |
| Pik3cg3 | 0 | 0.41463588 | 0.622 | 0.415 | 0 | TAM.prolif | Pik3cg |
| Ndufa81 | 0 | 0.51424767 | 0.813 | 0.606 | 0 | TAM.prolif | Ndufa8 |
| Calm3 | 0 | 0.73669645 | 0.878 | 0.671 | 0 | TAM.prolif | Calm3 |
| Actn41 | 0 | 0.71352723 | 0.887 | 0.68 | 0 | TAM.prolif | Actn4 |
| Cenpt | 0 | 1.46672257 | 0.284 | 0.077 | 0 | TAM.prolif | Cenpt |
| Ikzf11 | 0 | 0.37347478 | 0.717 | 0.51 | 0 | TAM.prolif | Ikzf1 |
| Ly862 | 0 | 0.59410474 | 0.828 | 0.621 | 0 | TAM.prolif | Ly86 |
| Nfya | 0 | 0.38571717 | 0.611 | 0.404 | 0 | TAM.prolif | Nfya |
| Ndst12 | 0 | 0.39253034 | 0.716 | 0.509 | 0 | TAM.prolif | Ndst1 |
| Psmd71 | 0 | 0.3663933 | 0.675 | 0.469 | 0 | TAM.prolif | Psmd7 |
| Tnfaip82 | 0 | 0.40371137 | 0.674 | 0.468 | 0 | TAM.prolif | Tnfaip8 |
| Aldh9a1 | 0 | 0.3278085 | 0.716 | 0.51 | 0 | TAM.prolif | Aldh9a1 |
| Ap1b12 | 0 | 0.48641499 | 0.826 | 0.62 | 0 | TAM.prolif | Ap1b1 |
| Prrc2a | 0 | 0.46391154 | 0.787 | 0.582 | 0 | TAM.prolif | Prrc2a |
| Ndufb8 | 0 | 0.51429566 | 0.79 | 0.585 | 0 | TAM.prolif | Ndufb8 |
| H1f02 | 0 | 0.67065202 | 0.611 | 0.406 | 0 | TAM.prolif | H1f0 |
| Cenpb1 | 0 | 0.39760063 | 0.783 | 0.579 | 0 | TAM.prolif | Cenpb |
| Puf60 | 0 | 0.45872264 | 0.794 | 0.59 | 0 | TAM.prolif | Puf60 |
| Gstcd | 0 | 1.5799246 | 0.271 | 0.067 | 0 | TAM.prolif | Gstcd |
| Chek2 | 0 | 1.74662898 | 0.275 | 0.071 | 0 | TAM.prolif | Chek2 |
| Mgat21 | 0 | 0.40823877 | 0.66 | 0.456 | 0 | TAM.prolif | Mgat2 |
| Jade1 | 0 | 1.33126196 | 0.296 | 0.092 | 0 | TAM.prolif | Jade1 |
| Zmynd19 | 0 | 1.59423382 | 0.288 | 0.084 | 0 | TAM.prolif | Zmynd19 |
| Rnf4 | 0 | 0.43657802 | 0.748 | 0.544 | 0 | TAM.prolif | Rnf4 |
| Abracl1 | 0 | 0.34072561 | 0.702 | 0.498 | 0 | TAM.prolif | Abracl |
| B4galnt11 | 0 | 0.39002415 | 0.713 | 0.509 | 0 | TAM.prolif | B4galnt1 |
| Psmc5 | 0 | 0.46801893 | 0.739 | 0.535 | 0 | TAM.prolif | Psmc5 |
| Huwe11 | 0 | 0.32137419 | 0.721 | 0.517 | 0 | TAM.prolif | Huwe1 |
| Sdhb | 0 | 0.45672223 | 0.774 | 0.571 | 0 | TAM.prolif | Sdhb |
| Snrnp70 | 0 | 0.47073582 | 0.81 | 0.607 | 0 | TAM.prolif | Snrnp70 |
| Dock103 | 0 | 0.3715546 | 0.67 | 0.467 | 0 | TAM.prolif | Dock10 |
| Cenpo | 0 | 1.51724617 | 0.276 | 0.073 | 0 | TAM.prolif | Cenpo |
| Srrm11 | 0 | 0.34497611 | 0.71 | 0.507 | 0 | TAM.prolif | Srrm1 |
| Hsp90b11 | 0 | 0.56704688 | 0.824 | 0.621 | 0 | TAM.prolif | Hsp90b1 |
| Naca1 | 0 | 0.45628896 | 0.765 | 0.562 | 0 | TAM.prolif | Naca |
| Npm11 | 0 | 0.79506591 | 0.885 | 0.682 | 0 | TAM.prolif | Npm1 |
| Rnf1871 | 0 | 0.45796915 | 0.835 | 0.632 | 0 | TAM.prolif | Rnf187 |
| Dpysl21 | 0 | 0.40110038 | 0.721 | 0.518 | 0 | TAM.prolif | Dpysl2 |
| Rbm32 | 0 | 0.47897646 | 0.804 | 0.602 | 0 | TAM.prolif | Rbm3 |
| Cntln | 0 | 1.82905715 | 0.257 | 0.055 | 0 | TAM.prolif | Cntln |
| Ctps | 0 | 1.90782833 | 0.257 | 0.055 | 0 | TAM.prolif | Ctps |
| Cnbp1 | 0 | 0.38373917 | 0.741 | 0.539 | 0 | TAM.prolif | Cnbp |
| Larp11 | 0 | 0.34566947 | 0.715 | 0.513 | 0 | TAM.prolif | Larp1 |
| Mapre1 | 0 | 0.53301443 | 0.809 | 0.608 | 0 | TAM.prolif | Mapre1 |
| Ndufc21 | 0 | 0.49857109 | 0.798 | 0.597 | 0 | TAM.prolif | Ndufc2 |
| Med28 | 0 | 0.43467599 | 0.621 | 0.42 | 0 | TAM.prolif | Med28 |
| Mydgf | 0 | 0.40577834 | 0.725 | 0.524 | 0 | TAM.prolif | Mydgf |
| Prpf40a1 | 0 | 0.3565963 | 0.65 | 0.45 | 0 | TAM.prolif | Prpf40a |
| Adh5 | 0 | 0.42723329 | 0.632 | 0.432 | 0 | TAM.prolif | Adh5 |
| Tial1 | 0 | 0.29371304 | 0.642 | 0.442 | 0 | TAM.prolif | Tial1 |
| Sdhd | 0 | 0.42972344 | 0.758 | 0.558 | 0 | TAM.prolif | Sdhd |
| Ogdh | 0 | 0.49482297 | 0.857 | 0.657 | 0 | TAM.prolif | Ogdh |
| Dlst | 0 | 0.35022304 | 0.718 | 0.518 | 0 | TAM.prolif | Dlst |
| Spcs31 | 0 | 0.43103299 | 0.617 | 0.418 | 0 | TAM.prolif | Spcs3 |
| Pomp2 | 0 | 0.39430312 | 0.754 | 0.555 | 0 | TAM.prolif | Pomp |
| Galnt21 | 0 | 0.31338972 | 0.702 | 0.503 | 0 | TAM.prolif | Galnt2 |
| Twf22 | 0 | 0.39966414 | 0.696 | 0.497 | 0 | TAM.prolif | Twf2 |
| Tm9sf22 | 0 | 0.37911044 | 0.689 | 0.49 | 0 | TAM.prolif | Tm9sf2 |
| Dnajb111 | 0 | 0.3855043 | 0.76 | 0.561 | 0 | TAM.prolif | Dnajb11 |
| Tnfrsf11a1 | 0 | 0.35475449 | 0.659 | 0.461 | 0 | TAM.prolif | Tnfrsf11a |
| Taf1d1 | 0 | 0.28275811 | 0.622 | 0.424 | 0 | TAM.prolif | Taf1d |
| Hnrnpf | 0 | 0.60605461 | 0.826 | 0.628 | 0 | TAM.prolif | Hnrnpf |
| Psma11 | 0 | 0.41405683 | 0.715 | 0.517 | 0 | TAM.prolif | Psma1 |
| Cdc371 | 0 | 0.54784885 | 0.826 | 0.628 | 0 | TAM.prolif | Cdc37 |
| Tmem1671 | 0 | 0.48833782 | 0.571 | 0.373 | 0 | TAM.prolif | Tmem167 |
| Snx62 | 0 | 0.39223674 | 0.682 | 0.485 | 0 | TAM.prolif | Snx6 |
| Smarcc2 | 0 | 0.37203744 | 0.769 | 0.573 | 0 | TAM.prolif | Smarcc2 |
| Lpcat21 | 0 | 0.45341255 | 0.638 | 0.442 | 0 | TAM.prolif | Lpcat2 |
| Srsf21 | 0 | 0.25878223 | 0.766 | 0.571 | 0 | TAM.prolif | Srsf2 |
| Tubb62 | 0 | 0.42848791 | 0.803 | 0.608 | 0 | TAM.prolif | Tubb6 |
| Sf3b62 | 0 | 0.40632505 | 0.659 | 0.464 | 0 | TAM.prolif | Sf3b6 |
| Elmo12 | 0 | 0.30657018 | 0.621 | 0.426 | 0 | TAM.prolif | Elmo1 |

| | | | | | | | |
|---|---|---|---|---|---|---|---|
| Khdrbs1 | 0 | 0.42025267 | 0.753 | 0.558 | 0 | TAM.prolif | Khdrbs1 |
| Dhx152 | 0 | 0.3844428 | 0.736 | 0.541 | 0 | TAM.prolif | Dhx15 |
| U2af2 | 0 | 0.50950153 | 0.847 | 0.652 | 0 | TAM.prolif | U2af2 |
| Ywhae | 0 | 0.53847801 | 0.866 | 0.671 | 0 | TAM.prolif | Ywhae |
| Smarca4 | 0 | 0.40139188 | 0.771 | 0.577 | 0 | TAM.prolif | Smarca4 |
| Map4k11 | 0 | 0.91807957 | 0.37 | 0.176 | 0 | TAM.prolif | Map4k1 |
| Tra2b1 | 0 | 0.29404865 | 0.629 | 0.435 | 0 | TAM.prolif | Tra2b |
| Ap2s12 | 0 | 0.37186695 | 0.83 | 0.636 | 0 | TAM.prolif | Ap2s1 |
| Dcaf7 | 0 | 0.30141968 | 0.694 | 0.5 | 0 | TAM.prolif | Dcaf7 |
| Pik3cd2 | 0 | 0.34331102 | 0.788 | 0.595 | 0 | TAM.prolif | Pik3cd |
| Pdia41 | 0 | 0.49144131 | 0.809 | 0.616 | 0 | TAM.prolif | Pdia4 |
| Eif4b | 0 | 0.31327801 | 0.802 | 0.609 | 0 | TAM.prolif | Eif4b |
| Rer1 | 0 | 0.35664129 | 0.647 | 0.454 | 0 | TAM.prolif | Rer1 |
| Rp21 | 0 | 0.31540748 | 0.66 | 0.467 | 0 | TAM.prolif | Rp2 |
| Bag1 | 0 | 0.50336495 | 0.748 | 0.555 | 0 | TAM.prolif | Bag1 |
| Sdha | 0 | 0.61196931 | 0.842 | 0.649 | 0 | TAM.prolif | Sdha |
| Acin1 | 0 | 0.3200791 | 0.722 | 0.529 | 0 | TAM.prolif | Acin1 |
| Tmem256 | 0 | 0.34920745 | 0.759 | 0.567 | 0 | TAM.prolif | Tmem256 |
| Rac11 | 0 | 0.31485967 | 0.695 | 0.503 | 0 | TAM.prolif | Rac1 |
| Lrrc592 | 0 | 0.40505997 | 0.843 | 0.651 | 0 | TAM.prolif | Lrrc59 |
| Ivns1abp | 0 | 0.52001918 | 0.807 | 0.616 | 0 | TAM.prolif | Ivns1abp |
| Ndufs2 | 0 | 0.43076654 | 0.794 | 0.603 | 0 | TAM.prolif | Ndufs2 |
| Sec14l12 | 0 | 0.45288913 | 0.681 | 0.49 | 0 | TAM.prolif | Sec14l1 |
| Rhno1 | 0 | 1.56912392 | 0.266 | 0.075 | 0 | TAM.prolif | Rhno1 |
| Rgs12 | 0 | 0.60124437 | 0.738 | 0.547 | 0 | TAM.prolif | Rgs1 |
| Coro2a3 | 0 | 0.6572456 | 0.595 | 0.404 | 0 | TAM.prolif | Coro2a |
| Uqcrc2 | 0 | 0.37715231 | 0.569 | 0.378 | 0 | TAM.prolif | Uqcrc2 |
| Eif3g1 | 0 | 0.36751296 | 0.665 | 0.475 | 0 | TAM.prolif | Eif3g |
| Atp5b1 | 0 | 0.61965751 | 0.835 | 0.645 | 0 | TAM.prolif | Atp5b |
| Nmt11 | 0 | 0.4293635 | 0.818 | 0.628 | 0 | TAM.prolif | Nmt1 |
| Sfr1 | 0 | 0.26942789 | 0.734 | 0.544 | 0 | TAM.prolif | Sfr1 |
| Cdkn2d | 0 | 1.1316754 | 0.309 | 0.12 | 0 | TAM.prolif | Cdkn2d |
| Tmem14c1 | 0 | 0.45447315 | 0.814 | 0.625 | 0 | TAM.prolif | Tmem14c |
| Irak1 | 0 | 0.28614999 | 0.696 | 0.507 | 0 | TAM.prolif | Irak1 |
| Rin31 | 0 | 0.40488207 | 0.669 | 0.481 | 0 | TAM.prolif | Rin3 |
| Spon12 | 0 | 0.74626138 | 0.375 | 0.187 | 0 | TAM.prolif | Spon1 |
| Trmt112 | 0 | 0.35985712 | 0.72 | 0.532 | 0 | TAM.prolif | Trmt112 |
| Ssr41 | 0 | 0.43864249 | 0.75 | 0.562 | 0 | TAM.prolif | Ssr4 |
| Sdhc1 | 0 | 0.45540985 | 0.779 | 0.592 | 0 | TAM.prolif | Sdhc |
| Psma4 | 0 | 0.41022341 | 0.773 | 0.586 | 0 | TAM.prolif | Psma4 |
| Dnase1l31 | 0 | 1.21508461 | 0.284 | 0.097 | 0 | TAM.prolif | Dnase1l3 |
| Dnajc32 | 0 | 0.28865698 | 0.731 | 0.544 | 0 | TAM.prolif | Dnajc3 |
| Rab3il13 | 0 | 0.45358219 | 0.852 | 0.665 | 0 | TAM.prolif | Rab3il1 |
| Ppp6r1 | 0 | 0.35289754 | 0.757 | 0.571 | 0 | TAM.prolif | Ppp6r1 |
| Ch25h | 0 | 1.57575419 | 0.264 | 0.078 | 0 | TAM.prolif | Ch25h |
| Sh3bp11 | 0 | 0.32644337 | 0.706 | 0.52 | 0 | TAM.prolif | Sh3bp1 |
| Ankrd13a | 0 | 0.46914213 | 0.818 | 0.633 | 0 | TAM.prolif | Ankrd13a |
| Wdr83os1 | 0 | 0.32054861 | 0.76 | 0.576 | 0 | TAM.prolif | Wdr83os |
| Tgfbr22 | 0 | 0.29071514 | 0.776 | 0.592 | 0 | TAM.prolif | Tgfbr2 |
| Hsp90aa11 | 0 | 0.41511988 | 0.772 | 0.588 | 0 | TAM.prolif | Hsp90aa1 |
| mt-Nd5 | 0 | 0.33237589 | 0.759 | 0.575 | 0 | TAM.prolif | mt-Nd5 |
| Api51 | 0 | 0.30841188 | 0.619 | 0.435 | 0 | TAM.prolif | Api5 |
| Lta4h1 | 0 | 0.53836316 | 0.604 | 0.42 | 0 | TAM.prolif | Lta4h |
| Ubqln1 | 0 | 0.36990929 | 0.705 | 0.521 | 0 | TAM.prolif | Ubqln1 |
| Cwc15 | 0 | 0.36218979 | 0.774 | 0.591 | 0 | TAM.prolif | Cwc15 |
| Rtn1 | 0 | 1.60603603 | 0.259 | 0.076 | 0 | TAM.prolif | Rtn1 |
| Lamtor2 | 0 | 0.29531186 | 0.714 | 0.531 | 0 | TAM.prolif | Lamtor2 |
| Smdt11 | 0 | 0.54353919 | 0.853 | 0.67 | 0 | TAM.prolif | Smdt1 |
| Psmb83 | 0 | 0.50716396 | 0.858 | 0.675 | 0 | TAM.prolif | Psmb8 |
| Zfp91 | 0 | 0.35281738 | 0.698 | 0.515 | 0 | TAM.prolif | Zfp91 |
| Clptm1 | 0 | 0.374431 | 0.796 | 0.614 | 0 | TAM.prolif | Clptm1 |
| Uqcrfs11 | 0 | 0.39930662 | 0.739 | 0.557 | 0 | TAM.prolif | Uqcrfs1 |
| Plau2 | 0 | 0.3298887 | 0.818 | 0.636 | 0 | TAM.prolif | Plau |
| Pdia6 | 0 | 0.41445553 | 0.795 | 0.614 | 0 | TAM.prolif | Pdia6 |
| Ppp2r1a1 | 0 | 0.30182333 | 0.752 | 0.571 | 0 | TAM.prolif | Ppp2r1a |
| Elovl52 | 0 | 0.53267373 | 0.452 | 0.271 | 0 | TAM.prolif | Elovl5 |
| Hnrnpu | 0 | 0.50444688 | 0.84 | 0.659 | 0 | TAM.prolif | Hnrnpu |
| Rgs103 | 0 | 0.378856 | 0.839 | 0.658 | 0 | TAM.prolif | Rgs10 |
| Coro1c | 0 | 0.35098203 | 0.651 | 0.471 | 0 | TAM.prolif | Coro1c |
| Ttyh31 | 0 | 0.49717014 | 0.872 | 0.692 | 0 | TAM.prolif | Ttyh3 |
| Sash12 | 0 | 0.49743453 | 0.762 | 0.582 | 0 | TAM.prolif | Sash1 |
| Oaz1 | 0 | 0.29776861 | 0.796 | 0.616 | 0 | TAM.prolif | Oaz1 |
| Vdac21 | 0 | 0.39216116 | 0.62 | 0.44 | 0 | TAM.prolif | Vdac2 |
| Stk111 | 0 | 0.28398207 | 0.796 | 0.617 | 0 | TAM.prolif | Stk11 |
| Noc2l1 | 0 | 0.30042716 | 0.635 | 0.456 | 0 | TAM.prolif | Noc2l |
| Psma6 | 0 | 0.31022338 | 0.755 | 0.577 | 0 | TAM.prolif | Psma6 |
| Was3 | 0 | 0.3557177 | 0.801 | 0.623 | 0 | TAM.prolif | Was |
| Parvg2 | 0 | 0.48435871 | 0.48 | 0.302 | 0 | TAM.prolif | Parvg |
| Srsf6 | 0 | 0.35575178 | 0.849 | 0.671 | 0 | TAM.prolif | Srsf6 |
| Ppp4c1 | 0 | 0.25051756 | 0.705 | 0.527 | 0 | TAM.prolif | Ppp4c |
| Arf6 | 0 | 0.44121036 | 0.826 | 0.648 | 0 | TAM.prolif | Arf6 |
| Ssr1 | 0 | 0.58516038 | 0.84 | 0.662 | 0 | TAM.prolif | Ssr1 |
| Vps35 | 0 | 0.36192261 | 0.604 | 0.427 | 0 | TAM.prolif | Vps35 |
| Rap1gds11 | 0 | 0.27014772 | 0.688 | 0.511 | 0 | TAM.prolif | Rap1gds1 |
| Camk12 | 0 | 0.30645647 | 0.698 | 0.521 | 0 | TAM.prolif | Camk1 |
| Tifab1 | 0 | 0.33882176 | 0.655 | 0.479 | 0 | TAM.prolif | Tifab |
| Ndufa3 | 0 | 0.43305589 | 0.846 | 0.67 | 0 | TAM.prolif | Ndufa3 |
| Cox6b11 | 0 | 0.36901223 | 0.836 | 0.66 | 0 | TAM.prolif | Cox6b1 |
| Colgalt12 | 0 | 0.44154109 | 0.826 | 0.651 | 0 | TAM.prolif | Colgalt1 |
| Lims11 | 0 | 0.29875333 | 0.748 | 0.574 | 0 | TAM.prolif | Lims1 |
| Nfatc11 | 0 | 0.3159535 | 0.621 | 0.447 | 0 | TAM.prolif | Nfatc1 |
| Ddost | 0 | 0.61093083 | 0.833 | 0.659 | 0 | TAM.prolif | Ddost |
| Fus | 0 | 0.45052609 | 0.834 | 0.66 | 0 | TAM.prolif | Fus |
| Raly | 0 | 0.65564344 | 0.863 | 0.69 | 0 | TAM.prolif | Raly |
| Arhgap91 | 0 | 0.37014051 | 0.742 | 0.569 | 0 | TAM.prolif | Arhgap9 |
| C1qc2 | 0 | 0.6127014 | 0.859 | 0.686 | 0 | TAM.prolif | C1qc |
| Arl8b | 0 | 0.29889261 | 0.721 | 0.548 | 0 | TAM.prolif | Arl8b |
| Arcn1 | 0 | 0.28122246 | 0.737 | 0.565 | 0 | TAM.prolif | Arcn1 |
| Anp32a1 | 0 | 0.42567386 | 0.877 | 0.706 | 0 | TAM.prolif | Anp32a |
| Ranbp2 | 0 | 0.31226496 | 0.74 | 0.569 | 0 | TAM.prolif | Ranbp2 |
| Ppp1ca1 | 0 | 0.42260321 | 0.786 | 0.615 | 0 | TAM.prolif | Ppp1ca |
| Pcbp11 | 0 | 0.39673197 | 0.846 | 0.675 | 0 | TAM.prolif | Pcbp1 |
| Erp29 | 0 | 0.59362383 | 0.9 | 0.73 | 0 | TAM.prolif | Erp29 |
| Csk1 | 0 | 0.28570983 | 0.776 | 0.606 | 0 | TAM.prolif | Csk |
| Ubap2l1 | 0 | 0.29815217 | 0.714 | 0.544 | 0 | TAM.prolif | Ubap2l |
| Elob | 0 | 0.36393876 | 0.818 | 0.648 | 0 | TAM.prolif | Elob |
| Srp91 | 0 | 0.28031788 | 0.761 | 0.592 | 0 | TAM.prolif | Srp9 |
| 7-Sep | 0 | 0.33115037 | 0.763 | 0.594 | 0 | TAM.prolif | 7-Sep |
| Pdia31 | 0 | 0.48910672 | 0.826 | 0.657 | 0 | TAM.prolif | Pdia3 |
| Spint13 | 0 | 0.41980288 | 0.77 | 0.602 | 0 | TAM.prolif | Spint1 |
| Srsf91 | 0 | 0.3855711 | 0.857 | 0.689 | 0 | TAM.prolif | Srsf9 |
| Ndufb1-ps2 | 0 | 0.27838115 | 0.761 | 0.593 | 0 | TAM.prolif | Ndufb1-ps |
| Atl31 | 0 | 0.44100049 | 0.679 | 0.511 | 0 | TAM.prolif | Atl3 |
| Mlf21 | 0 | 0.39177721 | 0.834 | 0.666 | 0 | TAM.prolif | Mlf2 |
| Atp5e1 | 0 | 0.30309283 | 0.804 | 0.637 | 0 | TAM.prolif | Atp5e |
| Lman2 | 0 | 0.34246901 | 0.81 | 0.643 | 0 | TAM.prolif | Lman2 |
| Cpd1 | 0 | 0.32918698 | 0.604 | 0.437 | 0 | TAM.prolif | Cpd |
| Ccni1 | 0 | 0.2817207 | 0.773 | 0.607 | 0 | TAM.prolif | Ccni |
| Cox8a1 | 0 | 0.3932811 | 0.854 | 0.688 | 0 | TAM.prolif | Cox8a |
| Lrch42 | 0 | 0.271041 | 0.837 | 0.671 | 0 | TAM.prolif | Lrch4 |
| Hnrnpul2 | 0 | 0.4229865 | 0.823 | 0.657 | 0 | TAM.prolif | Hnrnpul2 |
| Gng102 | 0 | 0.4425652 | 0.638 | 0.473 | 0 | TAM.prolif | Gng10 |
| Pmepa11 | 0 | 0.3451515 | 0.737 | 0.573 | 0 | TAM.prolif | Pmepa1 |
| Jpt1 | 0 | 0.83960176 | 0.897 | 0.734 | 0 | TAM.prolif | Jpt1 |
| Eif5a2 | 0 | 0.42083583 | 0.838 | 0.675 | 0 | TAM.prolif | Eif5a |
| Gusb1 | 0 | 0.51517794 | 0.876 | 0.715 | 0 | TAM.prolif | Gusb |
| Cox4i1 | 0 | 0.28314241 | 0.766 | 0.605 | 0 | TAM.prolif | Cox4i1 |
| Stab12 | 0 | 0.30943495 | 0.651 | 0.49 | 0 | TAM.prolif | Stab1 |
| Ywhab1 | 0 | 0.33131423 | 0.846 | 0.685 | 0 | TAM.prolif | Ywhab |
| Ubald2 | 0 | 0.40390632 | 0.693 | 0.532 | 0 | TAM.prolif | Ubald2 |
| Sfpq1 | 0 | 0.27413397 | 0.843 | 0.683 | 0 | TAM.prolif | Sfpq |
| Rab1b | 0 | 0.33943351 | 0.847 | 0.687 | 0 | TAM.prolif | Rab1b |
| Ywhah1 | 0 | 0.54535355 | 0.809 | 0.65 | 0 | TAM.prolif | Ywhah |
| Pcbp2 | 0 | 0.26090314 | 0.678 | 0.519 | 0 | TAM.prolif | Pcbp2 |
| Drap1 | 0 | 0.34710576 | 0.808 | 0.649 | 0 | TAM.prolif | Drap1 |
| Uqcrc1 | 0 | 0.45797595 | 0.871 | 0.713 | 0 | TAM.prolif | Uqcrc1 |
| Axl3 | 0 | 0.42169996 | 0.848 | 0.691 | 0 | TAM.prolif | Axl |
| Ciita2 | 0 | 0.40707694 | 0.533 | 0.376 | 0 | TAM.prolif | Ciita |
| Clic11 | 0 | 0.40092924 | 0.852 | 0.695 | 0 | TAM.prolif | Clic1 |
| Cnn21 | 0 | 0.29588284 | 0.698 | 0.541 | 0 | TAM.prolif | Cnn2 |
| Capza22 | 0 | 0.25146241 | 0.762 | 0.606 | 0 | TAM.prolif | Capza2 |
| Dok22 | 0 | 0.25401181 | 0.635 | 0.479 | 0 | TAM.prolif | Dok2 |
| mt-Nd4l1 | 0 | 0.38668227 | 0.835 | 0.679 | 0 | TAM.prolif | mt-Nd4l |
| Tgfbr12 | 0 | 0.31105838 | 0.736 | 0.581 | 0 | TAM.prolif | Tgfbr1 |
| Reep5 | 0 | 0.35360807 | 0.846 | 0.691 | 0 | TAM.prolif | Reep5 |
| mt-Cytb1 | 0 | 0.44076128 | 0.849 | 0.694 | 0 | TAM.prolif | mt-Cytb |
| Ybx1 | 0 | 0.80883305 | 0.943 | 0.788 | 0 | TAM.prolif | Ybx1 |
| Eid13 | 0 | 0.25519542 | 0.865 | 0.711 | 0 | TAM.prolif | Eid1 |
| C1qa2 | 0 | 0.56527713 | 0.879 | 0.726 | 0 | TAM.prolif | C1qa |
| mt-Nd11 | 0 | 0.49088816 | 0.729 | 0.577 | 0 | TAM.prolif | mt-Nd1 |
| Actn12 | 0 | 0.49580162 | 0.595 | 0.443 | 0 | TAM.prolif | Actn1 |
| Papola | 0 | 0.28088386 | 0.813 | 0.661 | 0 | TAM.prolif | Papola |
| Myd883 | 0 | 0.34292335 | 0.807 | 0.656 | 0 | TAM.prolif | Myd88 |
| Atp5d1 | 0 | 0.3462482 | 0.853 | 0.702 | 0 | TAM.prolif | Atp5d |
| Btf32 | 0 | 0.32241287 | 0.773 | 0.622 | 0 | TAM.prolif | Btf3 |
| H2-DMb21 | 0 | 1.25544492 | 0.265 | 0.114 | 0 | TAM.prolif | H2-DMb2 |
| Hexb2 | 0 | 0.32765706 | 0.791 | 0.641 | 0 | TAM.prolif | Hexb |
| Pabpc11 | 0 | 0.32463093 | 0.774 | 0.624 | 0 | TAM.prolif | Pabpc1 |
| Ndufa2 | 0 | 0.28781692 | 0.854 | 0.705 | 0 | TAM.prolif | Ndufa2 |
| Eef1b21 | 0 | 0.53899836 | 0.861 | 0.713 | 0 | TAM.prolif | Eef1b2 |
| Fnbp11 | 0 | 0.26343262 | 0.863 | 0.715 | 0 | TAM.prolif | Fnbp1 |
| Fyn | 0 | 0.75459332 | 0.326 | 0.178 | 0 | TAM.prolif | Fyn |
| Psme12 | 0 | 0.31212317 | 0.853 | 0.705 | 0 | TAM.prolif | Psme1 |
| Ptbp11 | 0 | 0.42726919 | 0.875 | 0.728 | 0 | TAM.prolif | Ptbp1 |
| Nedd8 | 0 | 0.40737115 | 0.876 | 0.73 | 0 | TAM.prolif | Nedd8 |
| Hnrnpm1 | 0 | 0.41837666 | 0.881 | 0.735 | 0 | TAM.prolif | Hnrnpm |
| Hnrnpul11 | 0 | 0.75511892 | 0.916 | 0.771 | 0 | TAM.prolif | Hnrnpul1 |
| Otulinl2 | 0 | 0.28477404 | 0.837 | 0.694 | 0 | TAM.prolif | Otulinl |
| Eif3k2 | 0 | 0.29122865 | 0.862 | 0.72 | 0 | TAM.prolif | Eif3k |
| Canx1 | 0 | 0.26235896 | 0.759 | 0.617 | 0 | TAM.prolif | Canx |
| Bzw1 | 0 | 0.26873536 | 0.759 | 0.619 | 0 | TAM.prolif | Bzw1 |
| Aif12 | 0 | 0.68895589 | 0.897 | 0.757 | 0 | TAM.prolif | Aif1 |
| H2afy2 | 0 | 0.65167176 | 0.923 | 0.784 | 0 | TAM.prolif | H2afy |
| Cxcl102 | 0 | 0.31590145 | 0.341 | 0.203 | 0 | TAM.prolif | Cxcl10 |
| Actr33 | 0 | 0.3156979 | 0.766 | 0.629 | 0 | TAM.prolif | Actr3 |
| Cops9 | 0 | 0.32123637 | 0.845 | 0.709 | 0 | TAM.prolif | Cops9 |
| Hnrnpk | 0 | 0.25190265 | 0.817 | 0.681 | 0 | TAM.prolif | Hnrnpk |
| C1qb2 | 0 | 0.46477457 | 0.896 | 0.763 | 0 | TAM.prolif | C1qb |
| Hsp90ab11 | 0 | 0.44132904 | 0.871 | 0.739 | 0 | TAM.prolif | Hsp90ab1 |
| Snx23 | 0 | 0.42290121 | 0.852 | 0.72 | 0 | TAM.prolif | Snx2 |
| Plekho11 | 0 | 0.30166856 | 0.885 | 0.756 | 0 | TAM.prolif | Plekho1 |
| Hnrnpa2b11 | 0 | 0.5854564 | 0.876 | 0.749 | 0 | TAM.prolif | Hnrnpa2b1 |
| mt-Co1 | 0 | 0.3109402 | 0.858 | 0.731 | 0 | TAM.prolif | mt-Co1 |
| Tuba1c1 | 0 | 0.57514605 | 0.85 | 0.724 | 0 | TAM.prolif | Tuba1c |
| Fermt31 | 0 | 0.25998746 | 0.872 | 0.746 | 0 | TAM.prolif | Fermt3 |
| Ptms1 | 0 | 0.6401574 | 0.975 | 0.851 | 0 | TAM.prolif | Ptms |
| Trir | 0 | 0.28125986 | 0.855 | 0.732 | 0 | TAM.prolif | Trir |
| Rab141 | 0 | 0.25991445 | 0.838 | 0.716 | 0 | TAM.prolif | Rab14 |
| Calr1 | 0 | 0.28237827 | 0.85 | 0.728 | 0 | TAM.prolif | Calr |
| Arpc41 | 0 | 0.29210907 | 0.832 | 0.711 | 0 | TAM.prolif | Arpc4 |
| Dusp61 | 0 | 0.27309458 | 0.745 | 0.624 | 0 | TAM.prolif | Dusp6 |
| Marcks2 | 0 | 0.41650572 | 0.931 | 0.811 | 0 | TAM.prolif | Marcks |
| Ccl242 | 0 | 0.40384438 | 0.346 | 0.226 | 0 | TAM.prolif | Ccl24 |
| Rnpep1 | 0 | 0.30918584 | 0.844 | 0.727 | 0 | TAM.prolif | Rnpep |
| Bin22 | 0 | 0.25529521 | 0.868 | 0.752 | 0 | TAM.prolif | Bin2 |
| mt-Atp81 | 0 | 0.39031897 | 0.888 | 0.772 | 0 | TAM.prolif | mt-Atp8 |
| Gdi22 | 0 | 0.30407969 | 0.848 | 0.735 | 0 | TAM.prolif | Gdi2 |
| Cd164 | 0 | 0.3163317 | 0.891 | 0.779 | 0 | TAM.prolif | Cd164 |
| Vcp1 | 0 | 0.31377197 | 0.909 | 0.798 | 0 | TAM.prolif | Vcp |
| Glud12 | 0 | 0.27077149 | 0.869 | 0.758 | 0 | TAM.prolif | Glud1 |
| H2-DMa2 | 0 | 0.25664242 | 0.832 | 0.721 | 0 | TAM.prolif | H2-DMa |
| Mfsd14 | 0 | 0.27099229 | 0.815 | 0.706 | 0 | TAM.prolif | Mfsd1 |
| Snx31 | 0 | 0.28683759 | 0.898 | 0.791 | 0 | TAM.prolif | Snx3 |
| Wdr1 | 0 | 0.29139694 | 0.887 | 0.781 | 0 | TAM.prolif | Wdr1 |
| Eef22 | 0 | 0.3379327 | 0.916 | 0.812 | 0 | TAM.prolif | Eef2 |
| Akt1 | 0 | 0.2675251 | 0.899 | 0.795 | 0 | TAM.prolif | Akt1 |
| Tkt2 | 0 | 0.32969321 | 0.895 | 0.792 | 0 | TAM.prolif | Tkt |
| mt-Atp61 | 0 | 0.35882547 | 0.872 | 0.771 | 0 | TAM.prolif | mt-Atp6 |
| Epn11 | 0 | 0.26066532 | 0.907 | 0.808 | 0 | TAM.prolif | Epn1 |
| Eif4g2 | 0 | 0.33955506 | 0.854 | 0.756 | 0 | TAM.prolif | Eif4g2 |
| Man2b11 | 0 | 0.28752953 | 0.887 | 0.791 | 0 | TAM.prolif | Man2b1 |
| Arhgdia | 0 | 0.44680665 | 0.946 | 0.854 | 0 | TAM.prolif | Arhgdia |
| Serinc33 | 0 | 0.57591522 | 0.951 | 0.862 | 0 | TAM.prolif | Serinc3 |
| Rac22 | 0 | 0.44179866 | 0.912 | 0.824 | 0 | TAM.prolif | Rac2 |
| Hspa1b4 | 0 | 0.32250791 | 0.448 | 0.36 | 0 | TAM.prolif | Hspa1b |
| Rack11 | 0 | 0.50375602 | 0.939 | 0.855 | 0 | TAM.prolif | Rack1 |
| Tagln22 | 0 | 0.62069278 | 0.973 | 0.897 | 0 | TAM.prolif | Tagln2 |
| Itgb52 | 0 | 0.28265169 | 0.765 | 0.698 | 0 | TAM.prolif | Itgb5 |
| Ifitm12 | 0 | 0.82044332 | 0.296 | 0.249 | 0 | TAM.prolif | Ifitm1 |
| Pfn1 | 0 | 0.47428432 | 0.986 | 0.945 | 0 | TAM.prolif | Pfn1 |
| Ifi301 | 0 | 0.43071326 | 0.972 | 0.933 | 0 | TAM.prolif | Ifi30 |
| Msr11 | 0 | 0.39081507 | 0.812 | 0.792 | 0 | TAM.prolif | Msr1 |
| Clta2 | 0 | 0.27896403 | 0.98 | 0.961 | 0 | TAM.prolif | Clta |
| Cotl11 | 0 | 0.37626479 | 0.98 | 0.963 | 0 | TAM.prolif | Cotl1 |
| Cfl1 | 0 | 0.44175465 | 0.97 | 0.956 | 0 | TAM.prolif | Cfl1 |
| Cst31 | 0 | 0.44894742 | 0.982 | 0.974 | 0 | TAM.prolif | Cst3 |
| Tmsb4x | 0 | 0.44553006 | 1 | 0.998 | 0 | TAM.prolif | Tmsb4x |
| Smarca51 | 4.76E-307 | 0.47700415 | 0.489 | 0.287 | 9.21E-303 | TAM.prolif | Smarca5 |
| Icosl1 | 9.65E-307 | 0.55084485 | 0.443 | 0.229 | 1.87E-302 | TAM.prolif | Icosl |
| Eif4h | 9.70E-307 | 0.46258689 | 0.602 | 0.369 | 1.87E-302 | TAM.prolif | Eif4h |
| Cntrl | 1.26E-306 | 0.63064964 | 0.479 | 0.238 | 2.43E-302 | TAM.prolif | Cntrl |
| Pycr21 | 1.68E-306 | 0.73504917 | 0.488 | 0.238 | 3.25E-302 | TAM.prolif | Pycr2 |
| Timm8a1 | 2.34E-306 | 1.17157671 | 0.307 | 0.107 | 4.53E-302 | TAM.prolif | Timm8a1 |
| Pold3 | 3.51E-306 | 0.9784706 | 0.346 | 0.134 | 6.78E-302 | TAM.prolif | Pold3 |
| Ddx19a | 6.73E-306 | 0.66252612 | 0.486 | 0.261 | 1.30E-301 | TAM.prolif | Ddx19a |
| Trappc11 | 1.21E-305 | 0.49331992 | 0.549 | 0.338 | 2.34E-301 | TAM.prolif | Trappc1 |
| Cdc23 | 1.59E-305 | 1.0760016 | 0.351 | 0.131 | 3.07E-301 | TAM.prolif | Cdc23 |
| Dnajc8 | 3.23E-305 | 0.52049858 | 0.57 | 0.358 | 6.24E-301 | TAM.prolif | Dnajc8 |
| Polr1d1 | 5.24E-305 | 0.2737407 | 0.695 | 0.519 | 1.01E-300 | TAM.prolif | Polr1d |
| Ccnd32 | 5.68E-305 | 0.2683891 | 0.439 | 0.294 | 1.10E-300 | TAM.prolif | Ccnd3 |
| Palld2 | 9.58E-305 | 0.71475329 | 0.449 | 0.255 | 1.85E-300 | TAM.prolif | Palld |
| Phc21 | 7.54E-304 | 0.35796691 | 0.663 | 0.454 | 1.46E-299 | TAM.prolif | Phc2 |
| Pip4k2a3 | 8.29E-304 | 0.30057678 | 0.604 | 0.409 | 1.60E-299 | TAM.prolif | Pip4k2a |
| Hdgfl2 | 1.60E-303 | 0.72520786 | 0.491 | 0.246 | 3.10E-299 | TAM.prolif | Hdgfl2 |
| Wdr5 | 2.15E-303 | 0.79732321 | 0.436 | 0.213 | 4.16E-299 | TAM.prolif | Wdr5 |
| Dpy19l11 | 2.84E-302 | 0.70466593 | 0.436 | 0.212 | 5.50E-298 | TAM.prolif | Dpy19l1 |
| Psma5 | 7.50E-302 | 0.35756999 | 0.65 | 0.447 | 1.45E-297 | TAM.prolif | Psma5 |
| Erh | 5.92E-301 | 1.19687314 | 0.328 | 0.111 | 1.14E-296 | TAM.prolif | Erh |
| Ino80e | 6.33E-301 | 0.61050375 | 0.524 | 0.282 | 1.22E-296 | TAM.prolif | Ino80e |
| Dcaf15 | 7.18E-301 | 0.72906591 | 0.472 | 0.23 | 1.39E-296 | TAM.prolif | Dcaf15 |
| Prmt5 | 9.38E-301 | 1.07615539 | 0.365 | 0.142 | 1.81E-296 | TAM.prolif | Prmt5 |
| Celf22 | 1.48E-300 | 0.31888375 | 0.648 | 0.433 | 2.87E-296 | TAM.prolif | Celf2 |
| Ccdc122 | 2.79E-300 | 0.31572651 | 0.655 | 0.47 | 5.39E-296 | TAM.prolif | Ccdc12 |
| Zmat2 | 6.92E-300 | 0.43432259 | 0.624 | 0.401 | 1.34E-295 | TAM.prolif | Zmat2 |
| Dnmt3a2 | 1.31E-299 | 0.4175354 | 0.582 | 0.377 | 2.53E-295 | TAM.prolif | Dnmt3a |
| Oser11 | 1.49E-299 | 0.29815491 | 0.619 | 0.434 | 2.88E-295 | TAM.prolif | Oser1 |
| Skp2 | 2.76E-299 | 1.71958491 | 0.25 | 0.062 | 5.34E-295 | TAM.prolif | Skp2 |
| Aimp2 | 1.61E-297 | 1.03518701 | 0.354 | 0.144 | 3.11E-293 | TAM.prolif | Aimp2 |
| Dcps | 1.78E-297 | 1.11765301 | 0.346 | 0.122 | 3.44E-293 | TAM.prolif | Dcps |
| Fam3c2 | 2.61E-297 | 0.413213 | 0.599 | 0.385 | 5.06E-293 | TAM.prolif | Fam3c |
| Plp22 | 2.93E-297 | 0.47724032 | 0.484 | 0.368 | 5.66E-293 | TAM.prolif | Plp2 |
| Srebf21 | 3.82E-297 | 0.26072246 | 0.668 | 0.517 | 7.38E-293 | TAM.prolif | Srebf2 |
| Pitrm1 | 5.48E-297 | 0.75440187 | 0.436 | 0.205 | 1.06E-292 | TAM.prolif | Pitrm1 |
| Fam120a | 9.78E-297 | 0.41972816 | 0.606 | 0.407 | 1.89E-292 | TAM.prolif | Fam120a |
| Wdr77 | 4.57E-296 | 0.8592452 | 0.399 | 0.175 | 8.83E-292 | TAM.prolif | Wdr77 |
| Upf1 | 5.98E-296 | 0.28322741 | 0.687 | 0.474 | 1.16E-291 | TAM.prolif | Upf1 |
| Pafah1b2 | 1.15E-295 | 0.35379763 | 0.635 | 0.446 | 2.23E-291 | TAM.prolif | Pafah1b2 |
| Psmd12 | 3.21E-295 | 0.39560015 | 0.587 | 0.383 | 6.20E-291 | TAM.prolif | Psmd12 |
| Immt | 4.46E-295 | 0.47005979 | 0.56 | 0.345 | 8.63E-291 | TAM.prolif | Immt |
| Ift27 | 5.52E-295 | 1.35751128 | 0.298 | 0.091 | 1.07E-290 | TAM.prolif | Ift27 |
| Pabpc41 | 1.42E-294 | 0.68235741 | 0.459 | 0.237 | 2.74E-290 | TAM.prolif | Pabpc4 |
| Atxn101 | 2.12E-294 | 0.55796857 | 0.518 | 0.305 | 4.11E-290 | TAM.prolif | Atxn10 |
| Hdac3 | 3.11E-294 | 0.55839449 | 0.509 | 0.272 | 6.01E-290 | TAM.prolif | Hdac3 |
| Naa151 | 5.07E-294 | 0.36273429 | 0.557 | 0.355 | 9.81E-290 | TAM.prolif | Naa15 |
| Dpp9 | 7.85E-294 | 0.3831061 | 0.65 | 0.43 | 1.52E-289 | TAM.prolif | Dpp9 |
| Dgcr8 | 1.01E-293 | 1.30752765 | 0.3 | 0.095 | 1.96E-289 | TAM.prolif | Dgcr8 |
| Rgs191 | 1.06E-293 | 0.28927817 | 0.682 | 0.484 | 2.06E-289 | TAM.prolif | Rgs19 |
| Pgls | 1.70E-293 | 0.27698765 | 0.663 | 0.475 | 3.29E-289 | TAM.prolif | Pgls |
| Ppp3cb1 | 2.04E-293 | 0.40121998 | 0.572 | 0.366 | 3.95E-289 | TAM.prolif | Ppp3cb |
| Emc1 | 2.07E-293 | 0.42292899 | 0.594 | 0.375 | 4.01E-289 | TAM.prolif | Emc1 |
| Wdr461 | 5.50E-293 | 0.66053926 | 0.463 | 0.237 | 1.06E-288 | TAM.prolif | Wdr46 |
| Daglb3 | 7.32E-293 | 0.2563118 | 0.697 | 0.495 | 1.41E-288 | TAM.prolif | Daglb |
| Polr3k | 7.47E-293 | 0.78193267 | 0.449 | 0.217 | 1.44E-288 | TAM.prolif | Polr3k |
| Acp1 | 8.94E-293 | 0.63940832 | 0.49 | 0.256 | 1.73E-288 | TAM.prolif | Acp1 |
| Copg1 | 1.24E-292 | 0.41126209 | 0.618 | 0.401 | 2.39E-288 | TAM.prolif | Copg1 |
| Ola11 | 1.69E-292 | 0.44203243 | 0.53 | 0.322 | 3.26E-288 | TAM.prolif | Ola1 |
| Yars | 1.73E-292 | 1.04715174 | 0.36 | 0.142 | 3.34E-288 | TAM.prolif | Yars |
| Pds5b | 2.01E-292 | 1.0619745 | 0.324 | 0.117 | 3.89E-288 | TAM.prolif | Pds5b |
| Kdm3b | 5.41E-292 | 0.32957469 | 0.656 | 0.435 | 1.05E-287 | TAM.prolif | Kdm3b |
| Sf3b4 | 9.11E-292 | 0.65329124 | 0.512 | 0.276 | 1.76E-287 | TAM.prolif | Sf3b4 |
| DEPRECATED-I | 1.62E-291 | 0.44533416 | 0.597 | 0.38 | 3.12E-287 | TAM.prolif | DEPRECATED-ENSMUSG00000017221 |
| Gpaa11 | 3.32E-291 | 0.7578965 | 0.486 | 0.242 | 6.42E-287 | TAM.prolif | Gpaa1 |
| Nnt1 | 3.51E-291 | 0.64268549 | 0.514 | 0.281 | 6.79E-287 | TAM.prolif | Nnt |
| Gtpbp41 | 4.52E-291 | 0.42280899 | 0.573 | 0.361 | 8.74E-287 | TAM.prolif | Gtpbp4 |
| Psmd6 | 4.79E-291 | 0.45867061 | 0.577 | 0.357 | 9.27E-287 | TAM.prolif | Psmd6 |
| Ddx241 | 5.85E-291 | 0.29612474 | 0.627 | 0.437 | 1.13E-286 | TAM.prolif | Ddx24 |
| Ctdspl2 | 9.55E-291 | 0.81641901 | 0.394 | 0.175 | 1.85E-286 | TAM.prolif | Ctdspl2 |
| Gpr183 | 1.44E-290 | 0.63347961 | 0.364 | 0.174 | 2.78E-286 | TAM.prolif | Gpr183 |
| Tmem229b1 | 3.26E-290 | 0.8739725 | 0.418 | 0.189 | 6.30E-286 | TAM.prolif | Tmem229b |
| Lman1 | 8.85E-290 | 0.45938113 | 0.624 | 0.395 | 1.71E-285 | TAM.prolif | Lman1 |
| Cd2ap1 | 1.01E-289 | 0.42611014 | 0.559 | 0.342 | 1.95E-285 | TAM.prolif | Cd2ap |
| Abce11 | 3.98E-289 | 0.79206183 | 0.421 | 0.212 | 7.70E-285 | TAM.prolif | Abce1 |
| Atp5l | 4.34E-289 | 0.30980477 | 0.682 | 0.484 | 8.40E-285 | TAM.prolif | Atp5l |
| Cs | 8.05E-289 | 0.57002725 | 0.51 | 0.284 | 1.56E-284 | TAM.prolif | Cs |
| Ccdc1241 | 8.81E-289 | 0.33144587 | 0.664 | 0.464 | 1.70E-284 | TAM.prolif | Ccdc124 |
| Swi5 | 3.49E-288 | 0.55474588 | 0.574 | 0.336 | 6.75E-284 | TAM.prolif | Swi5 |
| Ccdc861 | 8.29E-288 | 0.28757151 | 0.647 | 0.455 | 1.60E-283 | TAM.prolif | Ccdc86 |
| Hyou1 | 8.49E-288 | 0.28540093 | 0.628 | 0.437 | 1.64E-283 | TAM.prolif | Hyou1 |
| Ubtf | 1.88E-287 | 0.30563421 | 0.549 | 0.382 | 3.63E-283 | TAM.prolif | Ubtf |
| G3bp2 | 5.24E-287 | 0.31412779 | 0.677 | 0.483 | 1.01E-282 | TAM.prolif | G3bp2 |
| Mad1l12 | 1.22E-286 | 0.87030507 | 0.427 | 0.185 | 2.37E-282 | TAM.prolif | Mad1l1 |
| Sms1 | 2.76E-286 | 0.7592976 | 0.366 | 0.182 | 5.34E-282 | TAM.prolif | Sms |
| Ddx23 | 3.58E-286 | 0.25900781 | 0.709 | 0.526 | 6.93E-282 | TAM.prolif | Ddx23 |
| Eif3c | 4.10E-286 | 0.34731313 | 0.622 | 0.423 | 7.94E-282 | TAM.prolif | Eif3c |
| Swap702 | 4.86E-286 | 0.43823523 | 0.576 | 0.347 | 9.40E-282 | TAM.prolif | Swap70 |
| Ppp1r14b | 8.53E-286 | 0.58580905 | 0.538 | 0.331 | 1.65E-281 | TAM.prolif | Ppp1r14b |
| Asna1 | 1.11E-285 | 0.33629484 | 0.671 | 0.477 | 2.15E-281 | TAM.prolif | Asna1 |
| Tufm | 1.84E-285 | 0.63765387 | 0.489 | 0.268 | 3.56E-281 | TAM.prolif | Tufm |
| Hmgcr2 | 4.19E-285 | 0.32700297 | 0.607 | 0.439 | 8.09E-281 | TAM.prolif | Hmgcr |
| Dnajc2 | 4.53E-285 | 0.93570813 | 0.358 | 0.151 | 8.77E-281 | TAM.prolif | Dnajc2 |
| Mfsd102 | 5.07E-285 | 0.67480412 | 0.468 | 0.24 | 9.80E-281 | TAM.prolif | Mfsd10 |
| Xrcc1 | 1.39E-284 | 0.87157706 | 0.372 | 0.15 | 2.70E-280 | TAM.prolif | Xrcc1 |
| Asnsd1 | 1.19E-283 | 0.62623303 | 0.496 | 0.274 | 2.30E-279 | TAM.prolif | Asnsd1 |
| Cuedc21 | 2.08E-283 | 0.34432071 | 0.667 | 0.463 | 4.02E-279 | TAM.prolif | Cuedc2 |
| Gbp73 | 3.10E-283 | 0.53548446 | 0.531 | 0.31 | 5.99E-279 | TAM.prolif | Gbp7 |
| Txnl11 | 4.01E-283 | 0.38770954 | 0.651 | 0.426 | 7.75E-279 | TAM.prolif | Txnl1 |
| Gmps | 1.03E-282 | 0.70817258 | 0.461 | 0.229 | 2.00E-278 | TAM.prolif | Gmps |
| Pcyox12 | 1.17E-282 | 0.56157037 | 0.579 | 0.371 | 2.26E-278 | TAM.prolif | Pcyox1 |
| Abl11 | 2.77E-282 | 0.39659848 | 0.648 | 0.427 | 5.35E-278 | TAM.prolif | Abl1 |
| Tbc1d10b | 3.81E-282 | 0.43215055 | 0.644 | 0.429 | 7.37E-278 | TAM.prolif | Tbc1d10b |
| Coq7 | 4.73E-282 | 1.26808784 | 0.308 | 0.105 | 9.15E-278 | TAM.prolif | Coq7 |
| Lmo41 | 7.61E-282 | 0.40208565 | 0.525 | 0.314 | 1.47E-277 | TAM.prolif | Lmo4 |
| Fbxl14 | 1.23E-281 | 0.57454176 | 0.518 | 0.283 | 2.38E-277 | TAM.prolif | Fbxl14 |
| Impa1 | 4.50E-281 | 0.63462362 | 0.551 | 0.317 | 8.71E-277 | TAM.prolif | Impa1 |
| Atrx | 7.64E-281 | 0.34168802 | 0.604 | 0.403 | 1.48E-276 | TAM.prolif | Atrx |
| Itpripl11 | 1.13E-280 | 0.8324051 | 0.383 | 0.165 | 2.18E-276 | TAM.prolif | Itpripl1 |
| Fundc12 | 1.56E-280 | 0.3881846 | 0.67 | 0.462 | 3.01E-276 | TAM.prolif | Fundc1 |
| Smyd2 | 1.59E-280 | 1.24188842 | 0.305 | 0.113 | 3.07E-276 | TAM.prolif | Smyd2 |
| Usp5 | 3.23E-280 | 0.51860175 | 0.586 | 0.362 | 6.25E-276 | TAM.prolif | Usp5 |
| Ints11 | 3.25E-280 | 0.88784333 | 0.389 | 0.165 | 6.28E-276 | TAM.prolif | Ints11 |
| Vbp11 | 4.02E-280 | 0.4286624 | 0.535 | 0.334 | 7.76E-276 | TAM.prolif | Vbp1 |
| Mef2c2 | 7.50E-280 | 0.48370809 | 0.506 | 0.282 | 1.45E-275 | TAM.prolif | Mef2c |
| Kmt5a2 | 7.94E-280 | 0.34584392 | 0.589 | 0.378 | 1.53E-275 | TAM.prolif | Kmt5a |
| Glod4 | 1.11E-279 | 0.63237188 | 0.501 | 0.273 | 2.15E-275 | TAM.prolif | Glod4 |
| Atp5mpl1 | 4.33E-279 | 0.32108133 | 0.644 | 0.446 | 8.36E-275 | TAM.prolif | Atp5mpl |
| Vegfb2 | 4.36E-279 | 0.7908607 | 0.426 | 0.204 | 8.43E-275 | TAM.prolif | Vegfb |
| Mdc1 | 4.51E-279 | 1.11215851 | 0.324 | 0.114 | 8.72E-275 | TAM.prolif | Mdc1 |
| Nup93 | 4.91E-279 | 0.94832367 | 0.385 | 0.16 | 9.49E-275 | TAM.prolif | Nup93 |
| Tomm51 | 5.55E-279 | 0.42892367 | 0.546 | 0.352 | 1.07E-274 | TAM.prolif | Tomm5 |
| Taf6 | 1.08E-278 | 1.00599789 | 0.366 | 0.145 | 2.09E-274 | TAM.prolif | Taf6 |
| Lrmp1 | 2.55E-278 | 0.56823655 | 0.502 | 0.269 | 4.92E-274 | TAM.prolif | Lrmp |
| Ube2e3 | 4.54E-278 | 0.67617758 | 0.467 | 0.236 | 8.78E-274 | TAM.prolif | Ube2e3 |
| Map4 | 5.78E-278 | 0.34993093 | 0.675 | 0.48 | 1.12E-273 | TAM.prolif | Map4 |
| Get4 | 1.20E-277 | 0.60300058 | 0.497 | 0.272 | 2.32E-273 | TAM.prolif | Get4 |
| Rabl62 | 1.28E-277 | 0.6430199 | 0.555 | 0.324 | 2.48E-273 | TAM.prolif | Rabl6 |
| Akirin2 | 1.42E-277 | 0.41081582 | 0.541 | 0.353 | 2.75E-273 | TAM.prolif | Akirin2 |
| Hmbs | 2.70E-277 | 0.88034793 | 0.4 | 0.182 | 5.22E-273 | TAM.prolif | Hmbs |
| Kat14 | 1.42E-276 | 0.63537974 | 0.501 | 0.263 | 2.75E-272 | TAM.prolif | Kat14 |
| Rars | 2.92E-276 | 0.50952342 | 0.533 | 0.315 | 5.64E-272 | TAM.prolif | Rars |
| Vps36 | 1.04E-275 | 0.63871642 | 0.481 | 0.257 | 2.01E-271 | TAM.prolif | Vps36 |
| Reps1 | 1.10E-275 | 0.50815975 | 0.502 | 0.286 | 2.13E-271 | TAM.prolif | Reps1 |
| Mapk1ip1l | 1.55E-275 | 0.2856713 | 0.669 | 0.464 | 3.00E-271 | TAM.prolif | Mapk1ip1l |
| Rnf51 | 2.10E-275 | 0.72959708 | 0.491 | 0.256 | 4.05E-271 | TAM.prolif | Rnf5 |
| Tbl3 | 8.03E-275 | 0.78774396 | 0.398 | 0.183 | 1.55E-270 | TAM.prolif | Tbl3 |
| Sumo1 | 1.40E-274 | 0.36597255 | 0.59 | 0.394 | 2.71E-270 | TAM.prolif | Sumo1 |
| Mre11a | 1.50E-274 | 1.07800996 | 0.297 | 0.103 | 2.89E-270 | TAM.prolif | Mre11a |
| Gtf2a1 | 1.60E-274 | 0.38634308 | 0.641 | 0.435 | 3.10E-270 | TAM.prolif | Gtf2a1 |
| Hmgn5 | 2.40E-274 | 1.0128709 | 0.334 | 0.125 | 4.65E-270 | TAM.prolif | Hmgn5 |
| Brix1 | 2.79E-274 | 0.80878091 | 0.433 | 0.193 | 5.40E-270 | TAM.prolif | Brix1 |
| Prps1l3 | 3.83E-274 | 0.98192069 | 0.342 | 0.136 | 7.41E-270 | TAM.prolif | Prps1l3 |
| Tbc1d9b2 | 4.35E-274 | 0.32717458 | 0.628 | 0.425 | 8.41E-270 | TAM.prolif | Tbc1d9b |
| Donson2 | 7.19E-274 | 0.46601812 | 0.447 | 0.245 | 1.39E-269 | TAM.prolif | Donson |
| Cep295 | 2.04E-273 | 1.2348875 | 0.278 | 0.089 | 3.94E-269 | TAM.prolif | Cep295 |
| Arpc5l | 2.43E-273 | 0.70181945 | 0.457 | 0.239 | 4.69E-269 | TAM.prolif | Arpc5l |
| Dele1 | 3.65E-273 | 1.03037213 | 0.349 | 0.134 | 7.06E-269 | TAM.prolif | Dele1 |
| Trap1 | 8.79E-273 | 0.57917005 | 0.445 | 0.249 | 1.70E-268 | TAM.prolif | Trap1 |
| Pum31 | 1.20E-272 | 0.52975563 | 0.504 | 0.284 | 2.31E-268 | TAM.prolif | Pum3 |
| Sema6b1 | 1.36E-272 | 0.70243423 | 0.371 | 0.179 | 2.63E-268 | TAM.prolif | Sema6b |
| Slc35e12 | 3.30E-272 | 0.35930343 | 0.585 | 0.378 | 6.38E-268 | TAM.prolif | Slc35e1 |
| Zc3h4 | 5.26E-272 | 0.45032635 | 0.573 | 0.356 | 1.02E-267 | TAM.prolif | Zc3h4 |
| Mlip | 7.01E-272 | 1.2398661 | 0.271 | 0.105 | 1.35E-267 | TAM.prolif | Mlip |
| Smu1 | 8.19E-272 | 0.41196104 | 0.568 | 0.364 | 1.58E-267 | TAM.prolif | Smu1 |
| Ndufa4 | 8.80E-272 | 0.61748583 | 0.414 | 0.215 | 1.70E-267 | TAM.prolif | Ndufa4 |
| Smarcd2 | 9.49E-272 | 0.4985326 | 0.574 | 0.344 | 1.83E-267 | TAM.prolif | Smarcd2 |
| Heatr5a1 | 1.32E-271 | 0.48408016 | 0.538 | 0.318 | 2.56E-267 | TAM.prolif | Heatr5a |
| Psmg2 | 1.73E-271 | 1.23978482 | 0.291 | 0.097 | 3.34E-267 | TAM.prolif | Psmg2 |
| Sfmbt12 | 2.54E-271 | 0.6135627 | 0.459 | 0.243 | 4.90E-267 | TAM.prolif | Sfmbt1 |
| Arsb | 2.66E-271 | 1.03199636 | 0.35 | 0.115 | 5.15E-267 | TAM.prolif | Arsb |
| Anapc13 | 2.88E-271 | 0.40650761 | 0.626 | 0.413 | 5.57E-267 | TAM.prolif | Anapc13 |
| Zfp385a2 | 3.17E-271 | 0.34575854 | 0.661 | 0.472 | 6.12E-267 | TAM.prolif | Zfp385a |
| Ergic12 | 8.06E-271 | 0.42058689 | 0.604 | 0.387 | 1.56E-266 | TAM.prolif | Ergic1 |
| Exosc7 | 1.14E-270 | 0.99568222 | 0.358 | 0.147 | 2.20E-266 | TAM.prolif | Exosc7 |
| Tdp1 | 2.10E-270 | 1.49419917 | 0.258 | 0.069 | 4.06E-266 | TAM.prolif | Tdp1 |
| Dctn11 | 6.15E-270 | 0.37554284 | 0.642 | 0.442 | 1.19E-265 | TAM.prolif | Dctn1 |
| Snrpd2 | 7.26E-270 | 0.83685208 | 0.412 | 0.188 | 1.40E-265 | TAM.prolif | Snrpd2 |
| Naa201 | 8.82E-270 | 0.37484449 | 0.528 | 0.332 | 1.71E-265 | TAM.prolif | Naa20 |
| Rbm14 | 1.29E-269 | 0.35676772 | 0.601 | 0.391 | 2.50E-265 | TAM.prolif | Rbm14 |
| Pom1211 | 1.61E-269 | 0.51842121 | 0.479 | 0.261 | 3.11E-265 | TAM.prolif | Pom121 |
| Snx171 | 2.69E-269 | 0.29275684 | 0.729 | 0.543 | 5.20E-265 | TAM.prolif | Snx17 |
| Farsb | 8.98E-269 | 0.77370032 | 0.399 | 0.193 | 1.74E-264 | TAM.prolif | Farsb |
| Gng22 | 2.22E-268 | 0.31056738 | 0.699 | 0.496 | 4.29E-264 | TAM.prolif | Gng2 |
| Abhd17a1 | 1.22E-267 | 0.4084748 | 0.607 | 0.394 | 2.35E-263 | TAM.prolif | Abhd17a |
| Fam91a11 | 2.88E-267 | 0.32055359 | 0.638 | 0.439 | 5.57E-263 | TAM.prolif | Fam91a1 |
| Rabggtb | 3.39E-267 | 0.35837067 | 0.564 | 0.368 | 6.55E-263 | TAM.prolif | Rabggtb |
| Srcap | 3.54E-267 | 0.26614716 | 0.682 | 0.478 | 6.85E-263 | TAM.prolif | Srcap |
| Psmd3 | 1.54E-266 | 0.43830804 | 0.572 | 0.353 | 2.97E-262 | TAM.prolif | Psmd3 |
| Aip1 | 3.04E-266 | 0.25217643 | 0.653 | 0.463 | 5.88E-262 | TAM.prolif | Aip |
| Cdk191 | 3.38E-266 | 0.54884279 | 0.452 | 0.254 | 6.53E-262 | TAM.prolif | Cdk19 |
| Eif4a31 | 7.65E-266 | 0.66784618 | 0.434 | 0.217 | 1.48E-261 | TAM.prolif | Eif4a3 |
| Tmem209 | 1.14E-265 | 1.09820779 | 0.327 | 0.12 | 2.20E-261 | TAM.prolif | Tmem209 |
| Trmt6 | 1.52E-265 | 0.96371547 | 0.356 | 0.145 | 2.93E-261 | TAM.prolif | Trmt6 |
| Ptov12 | 2.18E-265 | 0.3616649 | 0.624 | 0.411 | 4.21E-261 | TAM.prolif | Ptov1 |
| Gbp32 | 2.69E-265 | 0.56741569 | 0.445 | 0.216 | 5.20E-261 | TAM.prolif | Gbp3 |
| Cd24a4 | 4.96E-265 | 0.33103847 | 0.306 | 0.258 | 9.59E-261 | TAM.prolif | Cd24a |
| Eif2b5 | 1.99E-264 | 0.54837834 | 0.539 | 0.302 | 3.85E-260 | TAM.prolif | Eif2b5 |
| Me21 | 5.89E-264 | 0.72419101 | 0.381 | 0.18 | 1.14E-259 | TAM.prolif | Me2 |
| Frmd8 | 1.82E-263 | 0.58433723 | 0.553 | 0.327 | 3.53E-259 | TAM.prolif | Frmd8 |
| Uqcc2 | 2.24E-263 | 0.59282287 | 0.49 | 0.28 | 4.33E-259 | TAM.prolif | Uqcc2 |
| Hdac2 | 3.73E-263 | 0.88865949 | 0.364 | 0.151 | 7.22E-259 | TAM.prolif | Hdac2 |
| Abcc32 | 2.41E-262 | 0.26781619 | 0.596 | 0.441 | 4.66E-258 | TAM.prolif | Abcc3 |
| Stard41 | 3.29E-262 | 0.63095513 | 0.442 | 0.241 | 6.35E-258 | TAM.prolif | Stard4 |
| Flnb | 6.98E-262 | 0.78411019 | 0.312 | 0.144 | 1.35E-257 | TAM.prolif | Flnb |
| Rnf220 | 7.20E-261 | 0.41796987 | 0.625 | 0.407 | 1.39E-256 | TAM.prolif | Rnf220 |
| Ocstamp3 | 1.09E-260 | 0.52361141 | 0.386 | 0.208 | 2.11E-256 | TAM.prolif | Ocstamp |
| Ebp | 8.45E-260 | 0.57329777 | 0.538 | 0.309 | 1.63E-255 | TAM.prolif | Ebp |
| Pbrm1 | 9.00E-260 | 0.43022998 | 0.535 | 0.335 | 1.74E-255 | TAM.prolif | Pbrm1 |
| Slc38a101 | 3.76E-259 | 0.36281305 | 0.651 | 0.429 | 7.27E-255 | TAM.prolif | Slc38a10 |
| Psme23 | 6.53E-259 | 0.35025648 | 0.594 | 0.396 | 1.26E-254 | TAM.prolif | Psme2 |
| Srbd1 | 7.54E-259 | 1.26454816 | 0.257 | 0.08 | 1.46E-254 | TAM.prolif | Srbd1 |
| Myo7a2 | 1.84E-258 | 0.32595381 | 0.604 | 0.416 | 3.55E-254 | TAM.prolif | Myo7a |
| Nsmce1 | 2.16E-258 | 1.15994829 | 0.275 | 0.094 | 4.18E-254 | TAM.prolif | Nsmce1 |
| Zc3h7b | 2.27E-258 | 0.8915881 | 0.389 | 0.169 | 4.39E-254 | TAM.prolif | Zc3h7b |
| Faf1 | 2.61E-258 | 0.42065214 | 0.48 | 0.285 | 5.05E-254 | TAM.prolif | Faf1 |
| Pacc11 | 2.72E-258 | 0.56311914 | 0.507 | 0.284 | 5.26E-254 | TAM.prolif | Pacc1 |
| Srsf31 | 8.11E-257 | 0.40811945 | 0.478 | 0.28 | 1.57E-252 | TAM.prolif | Srsf3 |
| Ostc | 1.12E-256 | 0.37844486 | 0.641 | 0.438 | 2.17E-252 | TAM.prolif | Ostc |
| Rnf168 | 7.30E-256 | 0.92986236 | 0.323 | 0.128 | 1.41E-251 | TAM.prolif | Rnf168 |
| Cct51 | 8.83E-256 | 0.71211522 | 0.431 | 0.219 | 1.71E-251 | TAM.prolif | Cct5 |
| Nifk | 1.36E-255 | 0.84652453 | 0.376 | 0.177 | 2.63E-251 | TAM.prolif | Nifk |
| E2f6 | 1.55E-255 | 1.24142383 | 0.26 | 0.086 | 2.99E-251 | TAM.prolif | E2f6 |
| Ndufa121 | 1.83E-255 | 0.45106739 | 0.534 | 0.324 | 3.54E-251 | TAM.prolif | Ndufa12 |
| Pdhb | 5.50E-255 | 0.39749602 | 0.514 | 0.323 | 1.06E-250 | TAM.prolif | Pdhb |
| Htra32 | 7.80E-255 | 0.45214192 | 0.497 | 0.286 | 1.51E-250 | TAM.prolif | Htra3 |
| Orc2 | 1.02E-254 | 1.24524504 | 0.253 | 0.081 | 1.98E-250 | TAM.prolif | Orc2 |
| Larp4 | 1.03E-254 | 0.32171222 | 0.596 | 0.399 | 1.99E-250 | TAM.prolif | Larp4 |
| Slc25a111 | 1.25E-254 | 0.35901857 | 0.655 | 0.448 | 2.42E-250 | TAM.prolif | Slc25a11 |
| Setd3 | 6.01E-254 | 0.35708436 | 0.638 | 0.428 | 1.16E-249 | TAM.prolif | Setd3 |
| Lsm6 | 7.40E-254 | 1.07036511 | 0.326 | 0.117 | 1.43E-249 | TAM.prolif | Lsm6 |
| Scly2 | 1.17E-253 | 0.85109596 | 0.378 | 0.168 | 2.27E-249 | TAM.prolif | Scly |
| Fadd2 | 1.18E-253 | 0.70377059 | 0.421 | 0.206 | 2.28E-249 | TAM.prolif | Fadd |
| Oxct13 | 3.05E-253 | 0.51890405 | 0.438 | 0.264 | 5.90E-249 | TAM.prolif | Oxct1 |
| Uqcrb | 3.93E-253 | 0.45322189 | 0.556 | 0.35 | 7.60E-249 | TAM.prolif | Uqcrb |
| Arpp191 | 1.20E-252 | 0.67510836 | 0.417 | 0.207 | 2.32E-248 | TAM.prolif | Arpp19 |
| Kcnab22 | 1.99E-252 | 0.29820628 | 0.611 | 0.435 | 3.84E-248 | TAM.prolif | Kcnab2 |
| Cfdp1 | 2.91E-252 | 0.46735989 | 0.529 | 0.339 | 5.62E-248 | TAM.prolif | Cfdp1 |
| Bag6 | 3.55E-252 | 0.26660176 | 0.676 | 0.474 | 6.86E-248 | TAM.prolif | Bag6 |
| Eif1a1 | 1.03E-251 | 0.32010277 | 0.618 | 0.421 | 2.00E-247 | TAM.prolif | Eif1a |
| Emc10 | 1.94E-251 | 0.27278941 | 0.691 | 0.491 | 3.76E-247 | TAM.prolif | Emc10 |
| Akap81 | 2.91E-251 | 0.38706011 | 0.6 | 0.384 | 5.62E-247 | TAM.prolif | Akap8 |
| Trim351 | 3.58E-251 | 0.47215297 | 0.55 | 0.335 | 6.93E-247 | TAM.prolif | Trim35 |
| Cnot31 | 7.57E-251 | 0.43010873 | 0.528 | 0.312 | 1.46E-246 | TAM.prolif | Cnot3 |
| Erg28 | 1.71E-250 | 0.85611704 | 0.37 | 0.16 | 3.31E-246 | TAM.prolif | Erg28 |
| Polr2b | 1.87E-250 | 0.67661996 | 0.402 | 0.198 | 3.62E-246 | TAM.prolif | Polr2b |
| Dbi1 | 3.48E-250 | 0.28517214 | 0.652 | 0.464 | 6.72E-246 | TAM.prolif | Dbi |
| Dhrs4 | 1.69E-249 | 0.86454924 | 0.402 | 0.184 | 3.26E-245 | TAM.prolif | Dhrs4 |
| Gatad1 | 1.76E-249 | 0.5109648 | 0.51 | 0.305 | 3.41E-245 | TAM.prolif | Gatad1 |
| Erap12 | 2.05E-249 | 0.44744959 | 0.477 | 0.273 | 3.96E-245 | TAM.prolif | Erap1 |
| Polr3h | 4.44E-249 | 1.30682425 | 0.251 | 0.071 | 8.58E-245 | TAM.prolif | Polr3h |
| Sema4b2 | 7.91E-249 | 0.34678858 | 0.523 | 0.354 | 1.53E-244 | TAM.prolif | Sema4b |
| Timm17a | 8.62E-249 | 0.45781741 | 0.523 | 0.315 | 1.67E-244 | TAM.prolif | Timm17a |
| Nacc1 | 9.44E-249 | 0.44128458 | 0.546 | 0.337 | 1.83E-244 | TAM.prolif | Nacc1 |
| Lap31 | 9.95E-249 | 0.501932 | 0.504 | 0.3 | 1.92E-244 | TAM.prolif | Lap3 |
| Prdx31 | 5.26E-248 | 0.4948267 | 0.519 | 0.302 | 1.02E-243 | TAM.prolif | Prdx3 |
| Eif3a1 | 7.65E-248 | 0.32146171 | 0.598 | 0.404 | 1.48E-243 | TAM.prolif | Eif3a |
| Kat2a1 | 9.33E-248 | 0.61530391 | 0.453 | 0.231 | 1.80E-243 | TAM.prolif | Kat2a |
| Prkcsh | 1.64E-247 | 0.2804078 | 0.625 | 0.438 | 3.17E-243 | TAM.prolif | Prkcsh |
| Nop21 | 1.74E-247 | 0.4945243 | 0.49 | 0.258 | 3.36E-243 | TAM.prolif | Nop2 |
| Sash31 | 3.60E-247 | 0.48199986 | 0.511 | 0.3 | 6.97E-243 | TAM.prolif | Sash3 |
| Aebp21 | 4.90E-247 | 0.31354497 | 0.575 | 0.385 | 9.48E-243 | TAM.prolif | Aebp2 |
| Ankrd401 | 5.20E-247 | 0.34455602 | 0.642 | 0.443 | 1.01E-242 | TAM.prolif | Ankrd40 |
| Strap | 3.20E-246 | 0.6598869 | 0.427 | 0.222 | 6.18E-242 | TAM.prolif | Strap |
| Isyna12 | 6.26E-246 | 0.35022176 | 0.564 | 0.38 | 1.21E-241 | TAM.prolif | Isyna1 |
| Pald12 | 6.59E-246 | 0.4534472 | 0.552 | 0.343 | 1.27E-241 | TAM.prolif | Pald1 |
| Rrs1 | 1.91E-245 | 0.83063904 | 0.359 | 0.165 | 3.70E-241 | TAM.prolif | Rrs1 |
| Npm31 | 2.68E-245 | 0.71116604 | 0.379 | 0.175 | 5.18E-241 | TAM.prolif | Npm3 |
| Ncbp3 | 2.88E-245 | 0.58772037 | 0.451 | 0.239 | 5.58E-241 | TAM.prolif | Ncbp3 |
| Scarb11 | 7.26E-245 | 0.26252457 | 0.574 | 0.36 | 1.40E-240 | TAM.prolif | Scarb1 |
| Stoml2 | 1.07E-244 | 0.5547474 | 0.455 | 0.244 | 2.07E-240 | TAM.prolif | Stoml2 |

| | | | | | | | |
|---|---|---|---|---|---|---|---|
| Foxk2 | 1.28E-243 | 0.60889779 | 0.447 | 0.235 | 2.47E-239 | TAM.prolif | Foxk2 |
| Agpat5 | 1.71E-243 | 0.52206033 | 0.45 | 0.251 | 3.31E-239 | TAM.prolif | Agpat5 |
| Atad3a | 1.84E-243 | 0.90203527 | 0.362 | 0.165 | 3.57E-239 | TAM.prolif | Atad3a |
| Inppl13 | 3.16E-243 | 0.35451456 | 0.597 | 0.396 | 6.10E-239 | TAM.prolif | Inppl1 |
| P2ry122 | 5.36E-243 | 0.51587687 | 0.432 | 0.235 | 1.04E-238 | TAM.prolif | P2ry12 |
| Rbck11 | 1.52E-242 | 0.42744945 | 0.559 | 0.335 | 2.94E-238 | TAM.prolif | Rbck1 |
| Tssc4 | 1.62E-242 | 0.85169351 | 0.356 | 0.146 | 3.13E-238 | TAM.prolif | Tssc4 |
| Dnajc7 | 1.90E-242 | 0.29599111 | 0.606 | 0.413 | 3.67E-238 | TAM.prolif | Dnajc7 |
| Dcun1d5 | 3.20E-242 | 0.32796163 | 0.546 | 0.365 | 6.18E-238 | TAM.prolif | Dcun1d5 |
| Vdac11 | 3.28E-242 | 0.45355558 | 0.57 | 0.366 | 6.34E-238 | TAM.prolif | Vdac1 |
| Uchl3 | 3.09E-241 | 1.18532278 | 0.28 | 0.09 | 5.97E-237 | TAM.prolif | Uchl3 |
| Seh1l | 4.67E-241 | 0.41106055 | 0.59 | 0.382 | 9.03E-237 | TAM.prolif | Seh1l |
| Dnm1l | 4.93E-241 | 0.43031115 | 0.545 | 0.33 | 9.54E-237 | TAM.prolif | Dnm1l |
| Timm232 | 5.14E-241 | 0.5782965 | 0.486 | 0.271 | 9.94E-237 | TAM.prolif | Timm23 |
| Haus7 | 1.68E-240 | 1.23443781 | 0.275 | 0.087 | 3.25E-236 | TAM.prolif | Haus7 |
| Psmb1 | 3.45E-240 | 0.34373454 | 0.574 | 0.379 | 6.68E-236 | TAM.prolif | Psmb1 |
| Spout1 | 3.61E-240 | 0.96864724 | 0.361 | 0.147 | 6.97E-236 | TAM.prolif | Spout1 |
| Zc3h18 | 3.69E-240 | 0.52249075 | 0.465 | 0.258 | 7.13E-236 | TAM.prolif | Zc3h18 |
| Myg1 | 1.54E-239 | 0.93320827 | 0.343 | 0.139 | 2.97E-235 | TAM.prolif | Myg1 |
| Slirp | 1.90E-239 | 0.55838207 | 0.499 | 0.288 | 3.67E-235 | TAM.prolif | Slirp |
| Sys11 | 2.02E-239 | 0.31018041 | 0.587 | 0.39 | 3.91E-235 | TAM.prolif | Sys1 |
| Zmpste241 | 3.16E-239 | 0.4135463 | 0.555 | 0.352 | 6.10E-235 | TAM.prolif | Zmpste24 |
| Slc7a1 | 4.59E-239 | 0.93799447 | 0.261 | 0.093 | 8.88E-235 | TAM.prolif | Slc7a1 |
| Grpel11 | 7.24E-239 | 0.41839977 | 0.569 | 0.368 | 1.40E-234 | TAM.prolif | Grpel1 |
| Dpp3 | 7.32E-239 | 0.3264957 | 0.597 | 0.403 | 1.41E-234 | TAM.prolif | Dpp3 |
| Gtpbp12 | 1.59E-238 | 0.43875528 | 0.473 | 0.272 | 3.08E-234 | TAM.prolif | Gtpbp1 |
| Prune22 | 2.97E-238 | 0.59942273 | 0.438 | 0.25 | 5.73E-234 | TAM.prolif | Prune2 |
| Tmem50b2 | 3.42E-238 | 0.28077061 | 0.562 | 0.384 | 6.62E-234 | TAM.prolif | Tmem50b |
| Pop41 | 5.75E-238 | 0.62043156 | 0.418 | 0.216 | 1.11E-233 | TAM.prolif | Pop4 |
| Kmt5c | 3.19E-237 | 0.43486528 | 0.545 | 0.334 | 6.16E-233 | TAM.prolif | Kmt5c |
| Gsk3a1 | 5.64E-237 | 0.28702354 | 0.645 | 0.44 | 1.09E-232 | TAM.prolif | Gsk3a |
| Cpsf3 | 1.21E-236 | 0.45910739 | 0.479 | 0.278 | 2.34E-232 | TAM.prolif | Cpsf3 |
| Atp5j | 2.35E-236 | 0.52093739 | 0.513 | 0.308 | 4.55E-232 | TAM.prolif | Atp5j |
| Gfm1 | 5.14E-236 | 0.59108994 | 0.41 | 0.212 | 9.93E-232 | TAM.prolif | Gfm1 |
| Prpf38a | 1.24E-235 | 0.64141887 | 0.436 | 0.223 | 2.39E-231 | TAM.prolif | Prpf38a |
| Ppp1cb | 1.43E-235 | 0.26163667 | 0.67 | 0.482 | 2.77E-231 | TAM.prolif | Ppp1cb |
| Psmb61 | 1.76E-235 | 0.26096 | 0.614 | 0.433 | 3.40E-231 | TAM.prolif | Psmb6 |
| Cdc341 | 2.66E-235 | 0.32097104 | 0.615 | 0.415 | 5.14E-231 | TAM.prolif | Cdc34 |
| Ppa1 | 1.39E-234 | 0.8012153 | 0.353 | 0.162 | 2.68E-230 | TAM.prolif | Ppa1 |
| Cisd1 | 2.54E-234 | 0.97250662 | 0.332 | 0.141 | 4.90E-230 | TAM.prolif | Cisd1 |
| mt-Nd3 | 4.32E-234 | 0.42231607 | 0.499 | 0.332 | 8.34E-230 | TAM.prolif | mt-Nd3 |
| Nol6 | 5.31E-234 | 0.80000189 | 0.401 | 0.197 | 1.03E-229 | TAM.prolif | Nol6 |
| Utp3 | 6.54E-234 | 0.62718666 | 0.45 | 0.237 | 1.26E-229 | TAM.prolif | Utp3 |
| P2ry131 | 6.84E-234 | 0.66682769 | 0.367 | 0.189 | 1.32E-229 | TAM.prolif | P2ry13 |
| Ptpn111 | 9.72E-234 | 0.38307713 | 0.592 | 0.375 | 1.88E-229 | TAM.prolif | Ptpn11 |
| Agps | 1.14E-233 | 0.25233091 | 0.581 | 0.4 | 2.20E-229 | TAM.prolif | Agps |
| Lrwd1 | 1.38E-233 | 0.75526447 | 0.413 | 0.202 | 2.68E-229 | TAM.prolif | Lrwd1 |
| Hspa9 | 2.06E-233 | 0.48016888 | 0.509 | 0.313 | 3.98E-229 | TAM.prolif | Hspa9 |
| Bmyc1 | 2.13E-233 | 0.68052055 | 0.426 | 0.216 | 4.11E-229 | TAM.prolif | Bmyc |
| Qars | 3.88E-233 | 0.33938067 | 0.601 | 0.401 | 7.51E-229 | TAM.prolif | Qars |
| Bcat2 | 4.79E-233 | 0.45787339 | 0.563 | 0.351 | 9.25E-229 | TAM.prolif | Bcat2 |
| Brd7 | 1.01E-232 | 0.45283649 | 0.523 | 0.32 | 1.94E-228 | TAM.prolif | Brd7 |
| Rbmxl1 | 1.06E-232 | 0.80550004 | 0.372 | 0.169 | 2.06E-228 | TAM.prolif | Rbmxl1 |
| Cyp513 | 1.21E-232 | 0.37046836 | 0.48 | 0.32 | 2.34E-228 | TAM.prolif | Cyp51 |
| Acot71 | 3.03E-232 | 0.43068173 | 0.47 | 0.298 | 5.86E-228 | TAM.prolif | Acot7 |
| Ndufs3 | 1.08E-231 | 0.39077195 | 0.554 | 0.353 | 2.10E-227 | TAM.prolif | Ndufs3 |
| Taf1 | 1.42E-231 | 0.52611757 | 0.434 | 0.228 | 2.74E-227 | TAM.prolif | Taf1 |
| Cnpy22 | 4.43E-231 | 0.39741209 | 0.58 | 0.372 | 8.57E-227 | TAM.prolif | Cnpy2 |
| Pmvk2 | 5.19E-231 | 0.46917544 | 0.505 | 0.313 | 1.00E-226 | TAM.prolif | Pmvk |
| Cpsf4 | 7.38E-231 | 0.64553845 | 0.441 | 0.229 | 1.43E-226 | TAM.prolif | Cpsf4 |
| Ncbp21 | 7.75E-231 | 0.5140083 | 0.497 | 0.279 | 1.50E-226 | TAM.prolif | Ncbp2 |
| Kpna3 | 1.13E-230 | 0.67139498 | 0.394 | 0.207 | 2.18E-226 | TAM.prolif | Kpna3 |
| U2surp | 1.29E-230 | 0.43138597 | 0.523 | 0.307 | 2.50E-226 | TAM.prolif | U2surp |
| Plekhj1 | 1.54E-230 | 0.25267237 | 0.683 | 0.492 | 2.99E-226 | TAM.prolif | Plekhj1 |
| Ppp3r1 | 2.00E-230 | 0.28871153 | 0.607 | 0.446 | 3.86E-226 | TAM.prolif | Ppp3r1 |
| Dnaja3 | 4.76E-230 | 0.65189681 | 0.431 | 0.223 | 9.20E-226 | TAM.prolif | Dnaja3 |
| Vdac3 | 1.42E-229 | 0.62606102 | 0.453 | 0.237 | 2.75E-225 | TAM.prolif | Vdac3 |
| Bicd2 | 1.46E-229 | 0.40618983 | 0.57 | 0.371 | 2.82E-225 | TAM.prolif | Bicd2 |
| Clcc1 | 2.84E-229 | 0.76132096 | 0.396 | 0.19 | 5.49E-225 | TAM.prolif | Clcc1 |
| Arhgap183 | 3.94E-229 | 0.4931295 | 0.468 | 0.274 | 7.62E-225 | TAM.prolif | Arhgap18 |
| Spr | 1.17E-228 | 0.52782862 | 0.493 | 0.285 | 2.25E-224 | TAM.prolif | Spr |
| Gpat4 | 4.06E-228 | 0.3320682 | 0.553 | 0.346 | 7.85E-224 | TAM.prolif | Gpat4 |
| B4galt62 | 6.36E-228 | 0.30963365 | 0.568 | 0.396 | 1.23E-223 | TAM.prolif | B4galt6 |
| Wdr36 | 7.95E-228 | 0.56448272 | 0.413 | 0.219 | 1.54E-223 | TAM.prolif | Wdr36 |
| Ube2g2 | 8.35E-228 | 0.36505424 | 0.613 | 0.401 | 1.61E-223 | TAM.prolif | Ube2g2 |
| Ttll12 | 2.89E-227 | 1.12540437 | 0.285 | 0.093 | 5.58E-223 | TAM.prolif | Ttll12 |
| Tomm22 | 6.95E-227 | 0.27864509 | 0.579 | 0.414 | 1.34E-222 | TAM.prolif | Tomm22 |
| Oard1 | 1.01E-226 | 0.64234492 | 0.401 | 0.195 | 1.95E-222 | TAM.prolif | Oard1 |
| Soga12 | 1.07E-226 | 0.41055199 | 0.577 | 0.364 | 2.07E-222 | TAM.prolif | Soga1 |
| Clec4a22 | 1.52E-226 | 0.32861174 | 0.514 | 0.322 | 2.94E-222 | TAM.prolif | Clec4a2 |
| Hadhb | 1.72E-226 | 0.3092808 | 0.563 | 0.376 | 3.33E-222 | TAM.prolif | Hadhb |
| Chchd1 | 1.81E-226 | 0.61763542 | 0.414 | 0.203 | 3.50E-222 | TAM.prolif | Chchd1 |
| Bccip1 | 3.40E-226 | 0.47127397 | 0.498 | 0.295 | 6.58E-222 | TAM.prolif | Bccip |
| Dpagt1 | 3.71E-226 | 0.67067409 | 0.435 | 0.234 | 7.18E-222 | TAM.prolif | Dpagt1 |
| Tex2611 | 4.53E-226 | 0.26075237 | 0.604 | 0.419 | 8.76E-222 | TAM.prolif | Tex261 |
| Ube2e11 | 5.40E-226 | 0.4828524 | 0.481 | 0.288 | 1.04E-221 | TAM.prolif | Ube2e1 |
| Vps16 | 1.87E-225 | 0.43941342 | 0.462 | 0.258 | 3.61E-221 | TAM.prolif | Vps16 |
| Noc4l | 2.58E-225 | 0.88492849 | 0.338 | 0.14 | 4.98E-221 | TAM.prolif | Noc4l |
| Dld | 3.23E-225 | 0.33236834 | 0.518 | 0.336 | 6.24E-221 | TAM.prolif | Dld |
| Tep11 | 3.91E-225 | 0.40410196 | 0.517 | 0.319 | 7.56E-221 | TAM.prolif | Tep1 |
| Cpsf2 | 6.00E-225 | 0.5861107 | 0.442 | 0.239 | 1.16E-220 | TAM.prolif | Cpsf2 |
| Kat7 | 9.78E-225 | 0.53016337 | 0.451 | 0.247 | 1.89E-220 | TAM.prolif | Kat7 |
| Pcnp | 3.72E-224 | 0.42615966 | 0.529 | 0.321 | 7.19E-220 | TAM.prolif | Pcnp |
| Adnp | 3.13E-223 | 0.30898053 | 0.576 | 0.386 | 6.06E-219 | TAM.prolif | Adnp |
| Exoc41 | 1.98E-222 | 0.32398219 | 0.565 | 0.369 | 3.83E-218 | TAM.prolif | Exoc4 |
| Patz11 | 2.55E-222 | 0.70545152 | 0.369 | 0.172 | 4.93E-218 | TAM.prolif | Patz1 |
| Aagab2 | 3.93E-222 | 0.42166183 | 0.624 | 0.402 | 7.59E-218 | TAM.prolif | Aagab |
| Rpf1 | 8.31E-222 | 0.5717583 | 0.466 | 0.253 | 1.61E-217 | TAM.prolif | Rpf1 |
| Ahctf1 | 5.87E-221 | 0.50659059 | 0.432 | 0.244 | 1.14E-216 | TAM.prolif | Ahctf1 |
| Prkd31 | 7.05E-221 | 0.50551412 | 0.374 | 0.206 | 1.36E-216 | TAM.prolif | Prkd3 |
| Inpp5b1 | 9.56E-221 | 0.46105209 | 0.46 | 0.258 | 1.85E-216 | TAM.prolif | Inpp5b |
| Ints1 | 1.08E-220 | 0.6013968 | 0.486 | 0.262 | 2.08E-216 | TAM.prolif | Ints1 |
| Ndufb5 | 1.12E-220 | 0.3822159 | 0.519 | 0.317 | 2.16E-216 | TAM.prolif | Ndufb5 |
| Rpia | 1.43E-220 | 0.5846046 | 0.379 | 0.189 | 2.77E-216 | TAM.prolif | Rpia |
| mt-Nd41 | 1.78E-220 | 0.29349203 | 0.552 | 0.401 | 3.45E-216 | TAM.prolif | mt-Nd4 |
| Emilin11 | 4.69E-220 | 0.63933683 | 0.435 | 0.238 | 9.07E-216 | TAM.prolif | Emilin1 |
| Scamp52 | 6.51E-220 | 0.58757383 | 0.437 | 0.232 | 1.26E-215 | TAM.prolif | Scamp5 |
| Tmem2421 | 1.12E-219 | 0.52663566 | 0.504 | 0.291 | 2.17E-215 | TAM.prolif | Tmem242 |
| Eml3 | 1.94E-219 | 0.48816138 | 0.462 | 0.257 | 3.75E-215 | TAM.prolif | Eml3 |
| Nvl | 2.70E-219 | 0.62524266 | 0.431 | 0.214 | 5.23E-215 | TAM.prolif | Nvl |
| Usp39 | 3.90E-219 | 0.69509869 | 0.367 | 0.168 | 7.55E-215 | TAM.prolif | Usp39 |
| Cnih41 | 6.42E-219 | 0.28277208 | 0.502 | 0.338 | 1.24E-214 | TAM.prolif | Cnih4 |
| Elof1 | 7.40E-219 | 0.71867748 | 0.409 | 0.2 | 1.43E-214 | TAM.prolif | Elof1 |
| Iws1 | 1.10E-218 | 0.56432198 | 0.377 | 0.188 | 2.12E-214 | TAM.prolif | Iws1 |
| Abcf1 | 1.99E-218 | 0.38929929 | 0.543 | 0.33 | 3.85E-214 | TAM.prolif | Abcf1 |
| Echs11 | 1.99E-218 | 0.37208123 | 0.55 | 0.363 | 3.86E-214 | TAM.prolif | Echs1 |
| Nol11 | 2.23E-218 | 0.87756562 | 0.331 | 0.142 | 4.31E-214 | TAM.prolif | Nol11 |
| Tomm70a | 3.11E-218 | 0.55551287 | 0.381 | 0.208 | 6.02E-214 | TAM.prolif | Tomm70a |
| Smc5 | 3.26E-218 | 0.62100852 | 0.418 | 0.221 | 6.30E-214 | TAM.prolif | Smc5 |
| Usp122 | 3.98E-217 | 0.30727908 | 0.549 | 0.346 | 7.69E-213 | TAM.prolif | Usp12 |
| Gnpat1 | 4.63E-217 | 0.48383052 | 0.457 | 0.257 | 8.96E-213 | TAM.prolif | Gnpat |
| Ccdc221 | 6.11E-217 | 0.5810783 | 0.491 | 0.274 | 1.18E-212 | TAM.prolif | Ccdc22 |
| C2cd5 | 6.55E-217 | 0.92966398 | 0.306 | 0.123 | 1.27E-212 | TAM.prolif | C2cd5 |
| Atp13a1 | 8.52E-217 | 0.3688453 | 0.533 | 0.323 | 1.65E-212 | TAM.prolif | Atp13a1 |
| Naa10 | 8.74E-217 | 0.69836107 | 0.371 | 0.172 | 1.69E-212 | TAM.prolif | Naa10 |
| Sart1 | 1.20E-216 | 0.30607939 | 0.566 | 0.367 | 2.32E-212 | TAM.prolif | Sart1 |
| Gpd1l | 1.93E-216 | 0.90246728 | 0.364 | 0.164 | 3.74E-212 | TAM.prolif | Gpd1l |
| Map3k3 | 2.68E-216 | 0.30226549 | 0.591 | 0.395 | 5.18E-212 | TAM.prolif | Map3k3 |
| Gpx32 | 3.80E-216 | 0.51012746 | 0.377 | 0.215 | 7.36E-212 | TAM.prolif | Gpx3 |
| Cops7a1 | 6.53E-216 | 0.54613267 | 0.484 | 0.273 | 1.26E-211 | TAM.prolif | Cops7a |
| Trappc5 | 1.20E-215 | 0.5005528 | 0.439 | 0.231 | 2.31E-211 | TAM.prolif | Trappc5 |
| Anapc15 | 2.14E-215 | 1.0930393 | 0.284 | 0.099 | 4.13E-211 | TAM.prolif | Anapc15 |
| Ptp4a32 | 3.75E-215 | 0.73128808 | 0.378 | 0.181 | 7.25E-211 | TAM.prolif | Ptp4a3 |
| Grsf1 | 5.12E-215 | 0.26880193 | 0.557 | 0.366 | 9.91E-211 | TAM.prolif | Grsf1 |
| Gid8 | 5.26E-215 | 0.37883502 | 0.521 | 0.324 | 1.02E-210 | TAM.prolif | Gid8 |
| Tdp2 | 1.64E-214 | 0.9596749 | 0.315 | 0.115 | 3.17E-210 | TAM.prolif | Tdp2 |
| Smarcb1 | 3.01E-214 | 0.4773136 | 0.484 | 0.274 | 5.83E-210 | TAM.prolif | Smarcb1 |
| Abcf2 | 3.33E-214 | 0.80309287 | 0.413 | 0.211 | 6.43E-210 | TAM.prolif | Abcf2 |
| Eif2b1 | 3.52E-214 | 0.78759413 | 0.34 | 0.158 | 6.81E-210 | TAM.prolif | Eif2b1 |
| Fam173a2 | 5.63E-214 | 0.43658987 | 0.541 | 0.326 | 1.09E-209 | TAM.prolif | Fam173a |
| Ppp4r3a | 8.01E-214 | 0.30694994 | 0.5 | 0.305 | 1.55E-209 | TAM.prolif | Ppp4r3a |
| Ppan | 8.98E-214 | 0.71167712 | 0.347 | 0.163 | 1.74E-209 | TAM.prolif | Ppan |
| Wdr412 | 9.26E-214 | 0.41420302 | 0.551 | 0.34 | 1.79E-209 | TAM.prolif | Wdr41 |
| Ap2a1 | 1.33E-213 | 0.3435106 | 0.577 | 0.377 | 2.57E-209 | TAM.prolif | Ap2a1 |
| Ap1ar | 1.72E-213 | 0.58815311 | 0.447 | 0.237 | 3.33E-209 | TAM.prolif | Ap1ar |
| Rpp30 | 2.17E-213 | 1.13089731 | 0.264 | 0.094 | 4.19E-209 | TAM.prolif | Rpp30 |
| Hadha | 4.04E-213 | 0.29637318 | 0.603 | 0.418 | 7.82E-209 | TAM.prolif | Hadha |
| Thoc3 | 7.78E-213 | 0.91283372 | 0.3 | 0.12 | 1.50E-208 | TAM.prolif | Thoc3 |
| Tomm34 | 9.67E-213 | 0.36480592 | 0.551 | 0.349 | 1.87E-208 | TAM.prolif | Tomm34 |
| Kdm2b | 5.31E-212 | 0.66543623 | 0.421 | 0.207 | 1.03E-207 | TAM.prolif | Kdm2b |
| Aldh18a11 | 6.41E-212 | 0.78090999 | 0.344 | 0.155 | 1.24E-207 | TAM.prolif | Aldh18a1 |
| Pfdn2 | 1.24E-211 | 0.34679246 | 0.541 | 0.354 | 2.40E-207 | TAM.prolif | Pfdn2 |
| Rae1 | 1.36E-211 | 0.48118771 | 0.465 | 0.255 | 2.63E-207 | TAM.prolif | Rae1 |
| Ints10 | 1.85E-211 | 1.06543397 | 0.294 | 0.106 | 3.58E-207 | TAM.prolif | Ints10 |
| Gfer | 2.13E-211 | 0.81146705 | 0.383 | 0.177 | 4.11E-207 | TAM.prolif | Gfer |
| Sec61a11 | 2.49E-211 | 0.30760517 | 0.583 | 0.386 | 4.82E-207 | TAM.prolif | Sec61a1 |
| Pbdc11 | 4.68E-211 | 0.71458549 | 0.404 | 0.2 | 9.06E-207 | TAM.prolif | Pbdc1 |
| Utp20 | 4.77E-211 | 0.87707466 | 0.296 | 0.122 | 9.23E-207 | TAM.prolif | Utp20 |
| Nek72 | 5.56E-211 | 0.28277062 | 0.564 | 0.376 | 1.07E-206 | TAM.prolif | Nek7 |
| Prcc | 6.49E-211 | 0.35809134 | 0.568 | 0.361 | 1.26E-206 | TAM.prolif | Prcc |
| Tcea1 | 1.07E-210 | 0.30250233 | 0.524 | 0.349 | 2.06E-206 | TAM.prolif | Tcea1 |
| Prmt7 | 1.60E-210 | 1.04711196 | 0.305 | 0.123 | 3.09E-206 | TAM.prolif | Prmt7 |
| Zwint1 | 3.15E-210 | 0.50667124 | 0.473 | 0.26 | 6.10E-206 | TAM.prolif | Zwint |
| Coro71 | 3.18E-210 | 0.28791365 | 0.657 | 0.459 | 6.16E-206 | TAM.prolif | Coro7 |
| Pnrc22 | 4.82E-210 | 0.32573662 | 0.557 | 0.365 | 9.31E-206 | TAM.prolif | Pnrc2 |
| Pgm2l11 | 7.24E-210 | 0.73307327 | 0.336 | 0.153 | 1.40E-205 | TAM.prolif | Pgm2l1 |
| Poc1b | 7.27E-210 | 0.90265142 | 0.285 | 0.109 | 1.41E-205 | TAM.prolif | Poc1b |
| Farsa1 | 1.20E-209 | 0.41226885 | 0.487 | 0.284 | 2.32E-205 | TAM.prolif | Farsa |
| Exosc9 | 1.58E-209 | 0.66339995 | 0.372 | 0.178 | 3.05E-205 | TAM.prolif | Exosc9 |
| Zc3h151 | 2.24E-209 | 0.5041573 | 0.468 | 0.273 | 4.34E-205 | TAM.prolif | Zc3h15 |
| Pip4k2b1 | 7.70E-209 | 0.39270133 | 0.555 | 0.342 | 1.49E-204 | TAM.prolif | Pip4k2b |
| Med8 | 1.02E-208 | 0.49508225 | 0.472 | 0.279 | 1.98E-204 | TAM.prolif | Med8 |
| Ndufv1 | 1.13E-208 | 0.52617351 | 0.432 | 0.226 | 2.18E-204 | TAM.prolif | Ndufv1 |
| Nfatc31 | 1.14E-208 | 0.43980026 | 0.442 | 0.256 | 2.21E-204 | TAM.prolif | Nfatc3 |
| Denr1 | 1.61E-208 | 0.34106296 | 0.513 | 0.327 | 3.11E-204 | TAM.prolif | Denr |
| Ubqln21 | 2.66E-208 | 0.43075046 | 0.535 | 0.339 | 5.15E-204 | TAM.prolif | Ubqln2 |
| Lamtor11 | 3.47E-208 | 0.58219912 | 0.495 | 0.277 | 6.71E-204 | TAM.prolif | Lamtor1 |
| Nup214 | 3.48E-208 | 0.54878946 | 0.424 | 0.23 | 6.73E-204 | TAM.prolif | Nup214 |
| Arhgap41 | 3.66E-208 | 0.47141791 | 0.472 | 0.266 | 7.07E-204 | TAM.prolif | Arhgap4 |
| Cd1801 | 6.08E-208 | 0.58812877 | 0.362 | 0.17 | 1.18E-203 | TAM.prolif | Cd180 |
| Drg2 | 6.43E-208 | 0.60291339 | 0.431 | 0.228 | 1.24E-203 | TAM.prolif | Drg2 |
| Zcchc8 | 7.28E-208 | 0.50870748 | 0.423 | 0.229 | 1.41E-203 | TAM.prolif | Zcchc8 |
| Senp1 | 9.97E-208 | 0.68693502 | 0.333 | 0.147 | 1.93E-203 | TAM.prolif | Senp1 |
| Hypk | 3.62E-207 | 0.46470599 | 0.515 | 0.3 | 7.00E-203 | TAM.prolif | Hypk |
| Arfgap2 | 4.70E-207 | 0.35401085 | 0.56 | 0.35 | 9.08E-203 | TAM.prolif | Arfgap2 |
| Naxe | 1.04E-206 | 0.5267751 | 0.436 | 0.248 | 2.02E-202 | TAM.prolif | Naxe |
| Clasp22 | 1.29E-206 | 0.34716627 | 0.49 | 0.3 | 2.50E-202 | TAM.prolif | Clasp2 |
| Prkag11 | 1.71E-206 | 0.42870526 | 0.441 | 0.253 | 3.31E-202 | TAM.prolif | Prkag1 |
| Emc61 | 1.93E-206 | 0.46041303 | 0.502 | 0.3 | 3.74E-202 | TAM.prolif | Emc6 |
| Lag31 | 2.32E-206 | 0.67647556 | 0.313 | 0.156 | 4.49E-202 | TAM.prolif | Lag3 |
| Ss18 | 2.56E-206 | 0.27933043 | 0.55 | 0.37 | 4.95E-202 | TAM.prolif | Ss18 |
| Tyw1 | 2.79E-206 | 0.80747095 | 0.376 | 0.164 | 5.39E-202 | TAM.prolif | Tyw1 |
| Ftsj3 | 3.11E-206 | 0.90066905 | 0.336 | 0.147 | 6.02E-202 | TAM.prolif | Ftsj3 |
| Psma2 | 3.81E-206 | 0.30869627 | 0.536 | 0.353 | 7.37E-202 | TAM.prolif | Psma2 |
| Srp72 | 5.31E-206 | 0.25921741 | 0.617 | 0.431 | 1.03E-201 | TAM.prolif | Srp72 |
| Msl3 | 1.02E-205 | 0.82770082 | 0.335 | 0.144 | 1.97E-201 | TAM.prolif | Msl3 |
| Bloc1s22 | 1.20E-205 | 0.42001438 | 0.495 | 0.333 | 2.31E-201 | TAM.prolif | Bloc1s2 |
| Samd1 | 2.41E-205 | 0.75740158 | 0.328 | 0.149 | 4.65E-201 | TAM.prolif | Samd1 |
| Ppie | 2.68E-205 | 0.66814155 | 0.395 | 0.191 | 5.18E-201 | TAM.prolif | Ppie |
| Luc7l3 | 2.83E-205 | 0.32571851 | 0.581 | 0.383 | 5.47E-201 | TAM.prolif | Luc7l3 |
| Cep57 | 4.81E-205 | 1.02836691 | 0.273 | 0.101 | 9.31E-201 | TAM.prolif | Cep57 |
| Arhgef40 | 5.69E-205 | 0.66937331 | 0.292 | 0.138 | 1.10E-200 | TAM.prolif | Arhgef40 |
| Pign1 | 6.18E-205 | 0.52961571 | 0.408 | 0.211 | 1.19E-200 | TAM.prolif | Pign |
| Atg3 | 1.16E-204 | 0.28007699 | 0.543 | 0.36 | 2.24E-200 | TAM.prolif | Atg3 |
| Rbbp5 | 1.38E-204 | 0.69935349 | 0.386 | 0.19 | 2.67E-200 | TAM.prolif | Rbbp5 |
| L3mbtl2 | 1.48E-204 | 1.21835678 | 0.264 | 0.091 | 2.86E-200 | TAM.prolif | L3mbtl2 |
| Fhl3 | 2.14E-204 | 0.45268752 | 0.492 | 0.297 | 4.15E-200 | TAM.prolif | Fhl3 |
| Ttc9c | 2.56E-204 | 0.66129476 | 0.422 | 0.218 | 4.96E-200 | TAM.prolif | Ttc9c |
| Mri1 | 4.02E-204 | 0.8923864 | 0.321 | 0.141 | 7.78E-200 | TAM.prolif | Mri1 |
| Hnrnph11 | 1.01E-203 | 0.28152138 | 0.57 | 0.375 | 1.95E-199 | TAM.prolif | Hnrnph1 |
| Ptp4a2 | 1.07E-203 | 0.26415913 | 0.587 | 0.399 | 2.07E-199 | TAM.prolif | Ptp4a2 |
| Srsf11 | 2.11E-203 | 0.25570721 | 0.624 | 0.429 | 4.08E-199 | TAM.prolif | Srsf11 |
| Dpm31 | 3.78E-203 | 0.4385167 | 0.506 | 0.302 | 7.30E-199 | TAM.prolif | Dpm3 |
| Pgam5 | 3.82E-203 | 0.66224365 | 0.395 | 0.194 | 7.38E-199 | TAM.prolif | Pgam5 |
| Ints31 | 5.90E-203 | 0.52281721 | 0.469 | 0.254 | 1.14E-198 | TAM.prolif | Ints3 |
| Cpne22 | 7.08E-203 | 0.52981254 | 0.449 | 0.26 | 1.37E-198 | TAM.prolif | Cpne2 |
| Coa3 | 8.35E-203 | 0.58595033 | 0.473 | 0.271 | 1.61E-198 | TAM.prolif | Coa3 |
| Commd102 | 1.28E-202 | 0.54689157 | 0.49 | 0.287 | 2.47E-198 | TAM.prolif | Commd10 |
| Mtrex | 2.33E-202 | 0.58705436 | 0.396 | 0.204 | 4.51E-198 | TAM.prolif | Mtrex |
| Tubgcp3 | 3.44E-202 | 0.53470321 | 0.381 | 0.196 | 6.65E-198 | TAM.prolif | Tubgcp3 |
| Nfyb | 9.91E-202 | 0.65643313 | 0.33 | 0.153 | 1.92E-197 | TAM.prolif | Nfyb |
| Itpa | 1.35E-201 | 0.55131371 | 0.396 | 0.202 | 2.62E-197 | TAM.prolif | Itpa |
| Commd4 | 1.37E-201 | 0.320085 | 0.571 | 0.375 | 2.65E-197 | TAM.prolif | Commd4 |
| Ddx491 | 1.65E-201 | 0.53787214 | 0.449 | 0.246 | 3.19E-197 | TAM.prolif | Ddx49 |
| Pspc1 | 2.32E-201 | 0.27213217 | 0.516 | 0.328 | 4.48E-197 | TAM.prolif | Pspc1 |
| Twsg1 | 2.73E-201 | 0.87115309 | 0.334 | 0.149 | 5.28E-197 | TAM.prolif | Twsg1 |
| Wdr43 | 9.02E-201 | 0.45657478 | 0.345 | 0.178 | 1.74E-196 | TAM.prolif | Wdr43 |
| Prkaca | 3.06E-200 | 0.3018488 | 0.591 | 0.41 | 5.92E-196 | TAM.prolif | Prkaca |
| Hspbp1 | 4.62E-200 | 0.6577231 | 0.403 | 0.207 | 8.94E-196 | TAM.prolif | Hspbp1 |
| Uhrf21 | 5.26E-200 | 0.25646396 | 0.546 | 0.366 | 1.02E-195 | TAM.prolif | Uhrf2 |
| Treml4 | 1.07E-199 | 0.45595403 | 0.331 | 0.16 | 2.07E-195 | TAM.prolif | Treml4 |
| Dapp12 | 1.24E-199 | 0.3559498 | 0.412 | 0.246 | 2.40E-195 | TAM.prolif | Dapp1 |
| Fcho1 | 1.60E-199 | 1.0953831 | 0.254 | 0.101 | 3.09E-195 | TAM.prolif | Fcho1 |
| Smarcad1 | 3.65E-199 | 0.50046788 | 0.454 | 0.24 | 7.05E-195 | TAM.prolif | Smarcad1 |
| Ecd | 6.39E-199 | 0.62332637 | 0.388 | 0.199 | 1.24E-194 | TAM.prolif | Ecd |
| Enoph11 | 6.89E-199 | 0.61873153 | 0.418 | 0.21 | 1.33E-194 | TAM.prolif | Enoph1 |
| Smg51 | 8.89E-199 | 0.30422007 | 0.53 | 0.322 | 1.72E-194 | TAM.prolif | Smg5 |
| Skp1a | 2.83E-198 | 0.33400707 | 0.588 | 0.395 | 5.48E-194 | TAM.prolif | Skp1a |
| Uqcrq | 5.18E-198 | 0.44389009 | 0.481 | 0.284 | 1.00E-193 | TAM.prolif | Uqcrq |
| mt-Nd2 | 6.35E-198 | 0.36061197 | 0.513 | 0.326 | 1.23E-193 | TAM.prolif | mt-Nd2 |
| Fam98b | 7.07E-198 | 0.63331904 | 0.374 | 0.185 | 1.37E-193 | TAM.prolif | Fam98b |
| Trim471 | 7.63E-198 | 0.71885573 | 0.335 | 0.155 | 1.47E-193 | TAM.prolif | Trim47 |
| Apmap | 8.97E-198 | 0.48638876 | 0.436 | 0.242 | 1.73E-193 | TAM.prolif | Apmap |
| Pcbd21 | 1.55E-197 | 0.76055717 | 0.389 | 0.194 | 2.99E-193 | TAM.prolif | Pcbd2 |
| Spg7 | 2.24E-197 | 0.34807352 | 0.54 | 0.346 | 4.33E-193 | TAM.prolif | Spg7 |
| Pus11 | 2.42E-197 | 0.4985094 | 0.431 | 0.247 | 4.68E-193 | TAM.prolif | Pus1 |
| Asxl1 | 3.33E-197 | 0.45030163 | 0.459 | 0.263 | 6.44E-193 | TAM.prolif | Asxl1 |
| Ccar1 | 4.60E-197 | 0.26962504 | 0.542 | 0.346 | 8.90E-193 | TAM.prolif | Ccar1 |
| Plxdc23 | 5.12E-197 | 0.36856407 | 0.378 | 0.231 | 9.91E-193 | TAM.prolif | Plxdc2 |
| Pdzd11 | 1.29E-196 | 0.69980613 | 0.349 | 0.164 | 2.50E-192 | TAM.prolif | Pdzd11 |
| Ptcd3 | 4.88E-196 | 0.84309942 | 0.29 | 0.117 | 9.44E-192 | TAM.prolif | Ptcd3 |
| Psme2b1 | 9.66E-196 | 0.32803811 | 0.55 | 0.348 | 1.87E-191 | TAM.prolif | Psme2b |
| Keap11 | 1.11E-195 | 0.42452489 | 0.518 | 0.304 | 2.15E-191 | TAM.prolif | Keap1 |
| Acaca | 1.83E-195 | 0.61435969 | 0.409 | 0.207 | 3.54E-191 | TAM.prolif | Acaca |
| Oxa1l1 | 2.41E-195 | 0.43306822 | 0.504 | 0.303 | 4.65E-191 | TAM.prolif | Oxa1l |
| Bud231 | 3.30E-195 | 0.60214473 | 0.433 | 0.225 | 6.38E-191 | TAM.prolif | Bud23 |
| Timm22 | 5.63E-195 | 0.63166385 | 0.39 | 0.199 | 1.09E-190 | TAM.prolif | Timm22 |
| Nfic2 | 1.28E-194 | 0.36629186 | 0.468 | 0.293 | 2.48E-190 | TAM.prolif | Nfic |
| Cops3 | 1.75E-194 | 0.86359484 | 0.312 | 0.129 | 3.39E-190 | TAM.prolif | Cops3 |
| Pithd11 | 3.48E-194 | 0.66512987 | 0.409 | 0.209 | 6.73E-190 | TAM.prolif | Pithd1 |
| Mospd31 | 5.04E-194 | 0.37018813 | 0.535 | 0.342 | 9.74E-190 | TAM.prolif | Mospd3 |
| Pmpca | 3.68E-193 | 0.51882645 | 0.44 | 0.259 | 7.11E-189 | TAM.prolif | Pmpca |
| Ubqln4 | 9.41E-193 | 0.59757304 | 0.412 | 0.224 | 1.82E-188 | TAM.prolif | Ubqln4 |
| Tsen341 | 1.12E-192 | 0.59668183 | 0.429 | 0.232 | 2.17E-188 | TAM.prolif | Tsen34 |
| Tfg | 1.19E-192 | 0.29787861 | 0.589 | 0.408 | 2.30E-188 | TAM.prolif | Tfg |
| Cdc27 | 2.66E-192 | 0.63854682 | 0.352 | 0.173 | 5.15E-188 | TAM.prolif | Cdc27 |
| Fip1l1 | 4.13E-192 | 0.45219878 | 0.457 | 0.262 | 7.98E-188 | TAM.prolif | Fip1l1 |
| Srp19 | 1.09E-191 | 0.41158124 | 0.507 | 0.316 | 2.12E-187 | TAM.prolif | Srp19 |
| Plaa | 1.15E-191 | 0.39060229 | 0.54 | 0.339 | 2.22E-187 | TAM.prolif | Plaa |
| Gipc1 | 1.21E-191 | 0.57397957 | 0.438 | 0.236 | 2.34E-187 | TAM.prolif | Gipc1 |
| Mtmr4 | 1.59E-191 | 1.0334038 | 0.275 | 0.109 | 3.07E-187 | TAM.prolif | Mtmr4 |
| Rrn31 | 1.84E-191 | 0.64450592 | 0.413 | 0.228 | 3.57E-187 | TAM.prolif | Rrn3 |
| Nfxl12 | 2.13E-191 | 0.35409014 | 0.424 | 0.254 | 4.11E-187 | TAM.prolif | Nfxl1 |
| Sptbn11 | 3.08E-191 | 0.36302718 | 0.511 | 0.321 | 5.95E-187 | TAM.prolif | Sptbn1 |
| Top2b | 3.87E-191 | 0.36617076 | 0.488 | 0.295 | 7.49E-187 | TAM.prolif | Top2b |
| Cib12 | 4.44E-191 | 0.31304651 | 0.497 | 0.331 | 8.58E-187 | TAM.prolif | Cib1 |
| Ddx18 | 7.02E-191 | 0.78029548 | 0.331 | 0.149 | 1.36E-186 | TAM.prolif | Ddx18 |
| Zfp3622 | 1.03E-190 | 0.53950793 | 0.45 | 0.241 | 2.00E-186 | TAM.prolif | Zfp362 |
| Tsr2 | 1.51E-190 | 1.10722264 | 0.274 | 0.101 | 2.92E-186 | TAM.prolif | Tsr2 |
| Dr11 | 2.36E-190 | 0.28868366 | 0.559 | 0.376 | 4.57E-186 | TAM.prolif | Dr1 |
| Acy1 | 4.95E-190 | 1.0984148 | 0.254 | 0.096 | 9.57E-186 | TAM.prolif | Acy1 |
| Abhd151 | 5.33E-190 | 0.84262595 | 0.317 | 0.135 | 1.03E-185 | TAM.prolif | Abhd15 |
| Tsfm | 5.76E-190 | 0.77986534 | 0.31 | 0.14 | 1.11E-185 | TAM.prolif | Tsfm |
| Zfp706 | 7.81E-190 | 0.58802787 | 0.42 | 0.221 | 1.51E-185 | TAM.prolif | Zfp706 |
| Nomo11 | 1.54E-189 | 0.52952252 | 0.398 | 0.204 | 2.97E-185 | TAM.prolif | Nomo1 |
| Eed1 | 1.64E-189 | 0.29150488 | 0.461 | 0.278 | 3.18E-185 | TAM.prolif | Eed |
| Cdk6 | 1.73E-189 | 0.37517789 | 0.376 | 0.221 | 3.35E-185 | TAM.prolif | Cdk6 |
| Mzt1 | 3.44E-189 | 0.81275189 | 0.339 | 0.169 | 6.65E-185 | TAM.prolif | Mzt1 |
| Usp37 | 6.42E-189 | 0.70852037 | 0.348 | 0.159 | 1.24E-184 | TAM.prolif | Usp37 |
| Pcif1 | 1.25E-188 | 0.43151948 | 0.443 | 0.25 | 2.41E-184 | TAM.prolif | Pcif1 |
| Rinl1 | 2.72E-188 | 0.62492577 | 0.422 | 0.22 | 5.26E-184 | TAM.prolif | Rinl |
| Ruvbl1 | 3.51E-188 | 0.89658303 | 0.321 | 0.131 | 6.79E-184 | TAM.prolif | Ruvbl1 |
| Derl21 | 5.58E-188 | 0.37525461 | 0.544 | 0.349 | 1.08E-183 | TAM.prolif | Derl2 |
| Thyn1 | 6.65E-188 | 1.05562746 | 0.299 | 0.121 | 1.29E-183 | TAM.prolif | Thyn1 |
| Taf102 | 6.67E-188 | 0.46920903 | 0.454 | 0.261 | 1.29E-183 | TAM.prolif | Taf10 |
| Nudt92 | 8.45E-188 | 0.33691334 | 0.577 | 0.373 | 1.63E-183 | TAM.prolif | Nudt9 |
| Otud41 | 1.10E-187 | 0.4485897 | 0.457 | 0.27 | 2.13E-183 | TAM.prolif | Otud4 |
| Gmpr2 | 1.58E-187 | 0.5779707 | 0.444 | 0.255 | 3.06E-183 | TAM.prolif | Gmpr2 |
| Cars | 5.76E-187 | 0.71034941 | 0.363 | 0.18 | 1.11E-182 | TAM.prolif | Cars |
| Cxxc11 | 1.19E-186 | 0.35630795 | 0.489 | 0.287 | 2.30E-182 | TAM.prolif | Cxxc1 |
| Xpo5 | 1.22E-186 | 0.87496579 | 0.29 | 0.124 | 2.36E-182 | TAM.prolif | Xpo5 |
| Sart3 | 2.33E-186 | 0.76689982 | 0.35 | 0.16 | 4.50E-182 | TAM.prolif | Sart3 |
| Ifi273 | 2.35E-186 | 0.2764378 | 0.548 | 0.389 | 4.55E-182 | TAM.prolif | Ifi27 |
| Usp10 | 4.43E-186 | 0.78575835 | 0.291 | 0.119 | 8.57E-182 | TAM.prolif | Usp10 |
| Klhdc4 | 5.82E-186 | 0.46834264 | 0.415 | 0.22 | 1.13E-181 | TAM.prolif | Klhdc4 |
| Il162 | 6.36E-186 | 0.48523746 | 0.412 | 0.216 | 1.23E-181 | TAM.prolif | Il16 |
| Setd1a | 8.76E-186 | 0.3310567 | 0.531 | 0.333 | 1.69E-181 | TAM.prolif | Setd1a |
| Slc35a3 | 1.13E-185 | 0.45768149 | 0.448 | 0.258 | 2.19E-181 | TAM.prolif | Slc35a3 |
| Pak1ip12 | 1.72E-185 | 0.29020628 | 0.493 | 0.317 | 3.33E-181 | TAM.prolif | Pak1ip1 |
| Aida | 2.03E-185 | 0.59255807 | 0.347 | 0.173 | 3.93E-181 | TAM.prolif | Aida |
| Rcbtb1 | 2.06E-185 | 0.77604732 | 0.336 | 0.159 | 3.99E-181 | TAM.prolif | Rcbtb1 |
| Cox6c | 2.81E-185 | 0.33466824 | 0.531 | 0.328 | 5.43E-181 | TAM.prolif | Cox6c |
| Urm1 | 3.86E-185 | 0.52097411 | 0.464 | 0.257 | 7.47E-181 | TAM.prolif | Urm1 |
| Pphln1 | 1.36E-184 | 0.3808336 | 0.487 | 0.291 | 2.63E-180 | TAM.prolif | Pphln1 |
| Nabp2 | 1.78E-184 | 0.64181481 | 0.393 | 0.192 | 3.43E-180 | TAM.prolif | Nabp2 |
| Ube2l3 | 2.41E-184 | 0.27426763 | 0.553 | 0.374 | 4.67E-180 | TAM.prolif | Ube2l3 |
| Vrk2 | 2.67E-184 | 0.53823777 | 0.349 | 0.172 | 5.16E-180 | TAM.prolif | Vrk2 |
| Arhgap273 | 6.00E-184 | 0.38416857 | 0.428 | 0.24 | 1.16E-179 | TAM.prolif | Arhgap27 |
| Rbmx | 9.70E-184 | 0.80597063 | 0.289 | 0.118 | 1.88E-179 | TAM.prolif | Rbmx |
| Pycrl | 2.92E-183 | 0.88348521 | 0.266 | 0.105 | 5.65E-179 | TAM.prolif | Pycrl |
| Ccar21 | 3.22E-183 | 0.5672702 | 0.384 | 0.197 | 6.23E-179 | TAM.prolif | Ccar2 |
| Tuba1a2 | 4.15E-183 | 0.3585177 | 0.49 | 0.332 | 8.03E-179 | TAM.prolif | Tuba1a |
| Slc7a72 | 6.95E-183 | 0.41728608 | 0.455 | 0.276 | 1.34E-178 | TAM.prolif | Slc7a7 |
| Hras | 8.16E-183 | 0.49057365 | 0.419 | 0.227 | 1.58E-178 | TAM.prolif | Hras |
| Emc2 | 9.13E-183 | 0.28003165 | 0.546 | 0.349 | 1.77E-178 | TAM.prolif | Emc2 |
| Osbpl1a1 | 1.14E-182 | 0.64845129 | 0.345 | 0.173 | 2.20E-178 | TAM.prolif | Osbpl1a |
| Ccdc82 | 1.37E-182 | 0.63457011 | 0.345 | 0.166 | 2.65E-178 | TAM.prolif | Ccdc82 |
| Hps32 | 1.63E-182 | 0.34390994 | 0.427 | 0.24 | 3.14E-178 | TAM.prolif | Hps3 |
| Ptpn22 | 1.77E-182 | 0.69430973 | 0.272 | 0.123 | 3.42E-178 | TAM.prolif | Ptpn22 |
| Phf101 | 2.36E-182 | 0.50103388 | 0.454 | 0.265 | 4.56E-178 | TAM.prolif | Phf10 |
| Ptpmt1 | 2.88E-182 | 0.42057681 | 0.447 | 0.262 | 5.58E-178 | TAM.prolif | Ptpmt1 |
| Sin3a1 | 3.09E-182 | 0.35461821 | 0.451 | 0.283 | 5.98E-178 | TAM.prolif | Sin3a |
| Tfam | 3.17E-182 | 0.78265276 | 0.322 | 0.133 | 6.13E-178 | TAM.prolif | Tfam |
| Nrip11 | 3.31E-182 | 0.41479341 | 0.413 | 0.231 | 6.40E-178 | TAM.prolif | Nrip1 |
| Plscr1 | 5.35E-182 | 0.65727087 | 0.272 | 0.151 | 1.04E-177 | TAM.prolif | Plscr1 |
| Nudt19 | 7.80E-182 | 0.98835825 | 0.286 | 0.121 | 1.51E-177 | TAM.prolif | Nudt19 |
| Pin1 | 9.23E-182 | 0.73659284 | 0.334 | 0.153 | 1.78E-177 | TAM.prolif | Pin1 |
| Ap1m1 | 1.29E-181 | 0.32879275 | 0.463 | 0.282 | 2.50E-177 | TAM.prolif | Ap1m1 |
| Sgf29 | 1.39E-181 | 0.90361103 | 0.257 | 0.094 | 2.70E-177 | TAM.prolif | Sgf29 |
| Med4 | 2.91E-181 | 0.66800717 | 0.314 | 0.136 | 5.62E-177 | TAM.prolif | Med4 |
| Gemin7 | 5.23E-181 | 0.37114438 | 0.509 | 0.317 | 1.01E-176 | TAM.prolif | Gemin7 |
| Dcakd1 | 5.64E-181 | 0.70934993 | 0.318 | 0.145 | 1.09E-176 | TAM.prolif | Dcakd |
| Naglu2 | 6.90E-181 | 0.3138128 | 0.541 | 0.362 | 1.33E-176 | TAM.prolif | Naglu |
| Cstf1 | 1.02E-180 | 0.76159705 | 0.318 | 0.139 | 1.98E-176 | TAM.prolif | Cstf1 |
| Apip | 2.31E-180 | 0.86370425 | 0.315 | 0.139 | 4.47E-176 | TAM.prolif | Apip |
| Tpst1 | 2.34E-180 | 0.60851328 | 0.35 | 0.166 | 4.53E-176 | TAM.prolif | Tpst1 |
| Ogfrl1 | 4.06E-180 | 0.45422395 | 0.394 | 0.202 | 7.85E-176 | TAM.prolif | Ogfrl1 |
| Lpgat11 | 4.88E-180 | 0.30395874 | 0.53 | 0.339 | 9.43E-176 | TAM.prolif | Lpgat1 |
| Exosc3 | 1.47E-179 | 0.52658218 | 0.403 | 0.218 | 2.84E-175 | TAM.prolif | Exosc3 |
| Csnk2b2 | 2.47E-179 | 0.25167649 | 0.566 | 0.389 | 4.78E-175 | TAM.prolif | Csnk2b |
| Cnot10 | 3.47E-179 | 0.6378537 | 0.358 | 0.169 | 6.71E-175 | TAM.prolif | Cnot10 |
| Rcan31 | 4.63E-179 | 0.69609354 | 0.396 | 0.204 | 8.96E-175 | TAM.prolif | Rcan3 |
| Tmem250-ps2 | 5.01E-179 | 0.32858536 | 0.543 | 0.359 | 9.70E-175 | TAM.prolif | Tmem250-ps |
| Ints141 | 5.72E-179 | 0.46637396 | 0.408 | 0.235 | 1.11E-174 | TAM.prolif | Ints14 |
| Katnb1 | 1.28E-178 | 1.03283851 | 0.251 | 0.092 | 2.47E-174 | TAM.prolif | Katnb1 |
| Nucb21 | 2.19E-178 | 0.26821831 | 0.486 | 0.328 | 4.23E-174 | TAM.prolif | Nucb2 |
| Tsg1011 | 2.59E-178 | 0.39651666 | 0.479 | 0.295 | 5.01E-174 | TAM.prolif | Tsg101 |
| Taok2 | 2.77E-178 | 0.29842171 | 0.5 | 0.316 | 5.36E-174 | TAM.prolif | Taok2 |
| Pitpnb | 3.17E-178 | 0.40161632 | 0.465 | 0.268 | 6.12E-174 | TAM.prolif | Pitpnb |
| Nol12 | 3.46E-178 | 0.77238338 | 0.367 | 0.176 | 6.69E-174 | TAM.prolif | Nol12 |
| Mbtps11 | 6.66E-178 | 0.44945647 | 0.442 | 0.251 | 1.29E-173 | TAM.prolif | Mbtps1 |
| Pmpcb | 7.88E-178 | 0.41049454 | 0.431 | 0.251 | 1.52E-173 | TAM.prolif | Pmpcb |
| Phb | 1.02E-177 | 0.62012256 | 0.387 | 0.201 | 1.96E-173 | TAM.prolif | Phb |
| Fbxw8 | 1.07E-177 | 0.38962816 | 0.509 | 0.315 | 2.06E-173 | TAM.prolif | Fbxw8 |
| Twf12 | 1.29E-177 | 0.25850737 | 0.617 | 0.425 | 2.50E-173 | TAM.prolif | Twf1 |
| Meaf6 | 1.41E-177 | 0.45808423 | 0.401 | 0.222 | 2.72E-173 | TAM.prolif | Meaf6 |
| Ppp1r212 | 4.38E-177 | 0.31723465 | 0.528 | 0.34 | 8.46E-173 | TAM.prolif | Ppp1r21 |
| Eef1e1 | 9.21E-177 | 0.59780208 | 0.305 | 0.154 | 1.78E-172 | TAM.prolif | Eef1e1 |
| Ciapin11 | 1.09E-176 | 0.46932893 | 0.392 | 0.203 | 2.11E-172 | TAM.prolif | Ciapin1 |
| Dxo | 1.37E-176 | 0.53085072 | 0.391 | 0.202 | 2.64E-172 | TAM.prolif | Dxo |
| Hs2st11 | 1.58E-176 | 0.40610812 | 0.415 | 0.241 | 3.05E-172 | TAM.prolif | Hs2st1 |
| Clns1a | 1.78E-176 | 0.65863876 | 0.381 | 0.19 | 3.43E-172 | TAM.prolif | Clns1a |
| Tnpo2 | 1.93E-176 | 0.28898036 | 0.529 | 0.348 | 3.72E-172 | TAM.prolif | Tnpo2 |
| Scrib | 2.56E-176 | 0.69857522 | 0.372 | 0.179 | 4.95E-172 | TAM.prolif | Scrib |
| Ermp1 | 4.55E-176 | 0.6473946 | 0.326 | 0.149 | 8.80E-172 | TAM.prolif | Ermp1 |
| Snrpb23 | 4.93E-176 | 0.25924748 | 0.511 | 0.345 | 9.54E-172 | TAM.prolif | Snrpb2 |
| C2cd3 | 5.23E-176 | 0.98639056 | 0.259 | 0.092 | 1.01E-171 | TAM.prolif | C2cd3 |
| Tomm40l1 | 6.16E-176 | 0.43203082 | 0.454 | 0.254 | 1.19E-171 | TAM.prolif | Tomm40l |
| Mid1ip11 | 1.80E-175 | 0.44193349 | 0.402 | 0.234 | 3.47E-171 | TAM.prolif | Mid1ip1 |
| Cep83 | 2.85E-175 | 0.48544391 | 0.369 | 0.192 | 5.52E-171 | TAM.prolif | Cep83 |
| Gtf2a2 | 3.06E-175 | 0.46101837 | 0.366 | 0.194 | 5.92E-171 | TAM.prolif | Gtf2a2 |
| Abcb7 | 5.35E-175 | 0.45730295 | 0.373 | 0.198 | 1.03E-170 | TAM.prolif | Abcb7 |
| Dnttip2 | 1.31E-174 | 0.45286486 | 0.435 | 0.246 | 2.53E-170 | TAM.prolif | Dnttip2 |
| Lrrc8b | 1.35E-174 | 0.71812806 | 0.298 | 0.136 | 2.62E-170 | TAM.prolif | Lrrc8b |
| Htatsf1 | 1.47E-174 | 0.75024561 | 0.338 | 0.167 | 2.84E-170 | TAM.prolif | Htatsf1 |
| Xpo7 | 1.80E-174 | 0.53443081 | 0.409 | 0.223 | 3.48E-170 | TAM.prolif | Xpo7 |
| Sucla2 | 1.83E-174 | 0.50389898 | 0.429 | 0.243 | 3.54E-170 | TAM.prolif | Sucla2 |
| Akr7a5 | 2.13E-174 | 0.68505999 | 0.346 | 0.175 | 4.11E-170 | TAM.prolif | Akr7a5 |
| Pdha1 | 2.16E-174 | 0.27813261 | 0.544 | 0.361 | 4.17E-170 | TAM.prolif | Pdha1 |
| Gabpa | 2.29E-174 | 0.4053432 | 0.419 | 0.246 | 4.42E-170 | TAM.prolif | Gabpa |
| Dus1l | 2.68E-174 | 0.66785074 | 0.336 | 0.163 | 5.19E-170 | TAM.prolif | Dus1l |
| Mllt1 | 3.70E-174 | 0.67895851 | 0.327 | 0.15 | 7.16E-170 | TAM.prolif | Mllt1 |
| Klrb1b1 | 7.05E-174 | 0.32443148 | 0.31 | 0.187 | 1.36E-169 | TAM.prolif | Klrb1b |
| Fam76b | 7.76E-174 | 0.44464761 | 0.447 | 0.259 | 1.50E-169 | TAM.prolif | Fam76b |
| Wdr741 | 8.54E-174 | 0.57881988 | 0.393 | 0.199 | 1.65E-169 | TAM.prolif | Wdr74 |
| Glrx5 | 3.00E-173 | 0.56925007 | 0.414 | 0.225 | 5.80E-169 | TAM.prolif | Glrx5 |
| Nudt16l12 | 4.08E-173 | 0.29267297 | 0.53 | 0.332 | 7.88E-169 | TAM.prolif | Nudt16l1 |
| Rbbp81 | 4.93E-173 | 0.39078126 | 0.408 | 0.223 | 9.53E-169 | TAM.prolif | Rbbp8 |
| Timm10 | 6.60E-173 | 0.59072845 | 0.357 | 0.178 | 1.28E-168 | TAM.prolif | Timm10 |
| Casp3 | 8.31E-173 | 0.38618997 | 0.442 | 0.257 | 1.61E-168 | TAM.prolif | Casp3 |
| Rtca | 9.03E-173 | 0.63262063 | 0.367 | 0.182 | 1.75E-168 | TAM.prolif | Rtca |
| Zcchc17 | 9.09E-173 | 0.70584307 | 0.342 | 0.153 | 1.76E-168 | TAM.prolif | Zcchc17 |
| Rprd1b | 1.28E-172 | 0.64625128 | 0.309 | 0.137 | 2.47E-168 | TAM.prolif | Rprd1b |
| Cnot11 | 2.14E-172 | 0.6079037 | 0.351 | 0.171 | 4.14E-168 | TAM.prolif | Cnot11 |
| Emc3 | 4.33E-172 | 0.38539593 | 0.432 | 0.263 | 8.36E-168 | TAM.prolif | Emc3 |
| Tex10 | 6.08E-172 | 0.42169668 | 0.375 | 0.204 | 1.18E-167 | TAM.prolif | Tex10 |
| Rbm22 | 9.68E-172 | 0.34625122 | 0.447 | 0.277 | 1.87E-167 | TAM.prolif | Rbm22 |
| Map1s1 | 1.43E-171 | 0.42970174 | 0.495 | 0.283 | 2.77E-167 | TAM.prolif | Map1s |
| Ndufa11 | 1.92E-171 | 0.39101079 | 0.491 | 0.302 | 3.72E-167 | TAM.prolif | Ndufa11 |
| Nup88 | 1.97E-171 | 0.55809412 | 0.395 | 0.207 | 3.80E-167 | TAM.prolif | Nup88 |
| Mpv17l2 | 2.46E-171 | 0.40798141 | 0.477 | 0.29 | 4.76E-167 | TAM.prolif | Mpv17l2 |
| Fdft12 | 3.04E-171 | 0.55133046 | 0.371 | 0.202 | 5.88E-167 | TAM.prolif | Fdft1 |
| Aldh16a1 | 3.23E-171 | 0.73848637 | 0.371 | 0.184 | 6.25E-167 | TAM.prolif | Aldh16a1 |
| Kansl2 | 5.37E-171 | 0.32486991 | 0.498 | 0.309 | 1.04E-166 | TAM.prolif | Kansl2 |
| Carm1 | 9.58E-171 | 0.51137302 | 0.409 | 0.225 | 1.85E-166 | TAM.prolif | Carm1 |
| Eif2s2 | 1.15E-170 | 0.51634152 | 0.342 | 0.184 | 2.22E-166 | TAM.prolif | Eif2s2 |
| Dhcr71 | 1.45E-170 | 0.65547324 | 0.388 | 0.196 | 2.79E-166 | TAM.prolif | Dhcr7 |
| Eif2a | 1.59E-170 | 0.2844016 | 0.477 | 0.302 | 3.07E-166 | TAM.prolif | Eif2a |
| Usp211 | 2.90E-170 | 0.63915913 | 0.37 | 0.185 | 5.61E-166 | TAM.prolif | Usp21 |
| Nf2 | 6.12E-170 | 0.29049934 | 0.514 | 0.327 | 1.18E-165 | TAM.prolif | Nf2 |
| Phip | 1.35E-169 | 0.27542491 | 0.521 | 0.327 | 2.60E-165 | TAM.prolif | Phip |
| Wls2 | 1.94E-169 | 0.31920646 | 0.493 | 0.321 | 3.75E-165 | TAM.prolif | Wls |
| Bop11 | 2.34E-169 | 0.46620615 | 0.414 | 0.227 | 4.53E-165 | TAM.prolif | Bop1 |
| Cars2 | 2.54E-169 | 0.75725574 | 0.304 | 0.138 | 4.91E-165 | TAM.prolif | Cars2 |
| Ccdc25 | 5.18E-169 | 0.62491721 | 0.346 | 0.167 | 1.00E-164 | TAM.prolif | Ccdc25 |
| Ap1g2 | 5.54E-169 | 0.5222138 | 0.399 | 0.211 | 1.07E-164 | TAM.prolif | Ap1g2 |
| Aimp1 | 9.33E-169 | 0.45304539 | 0.428 | 0.251 | 1.80E-164 | TAM.prolif | Aimp1 |
| Gas2l32 | 1.00E-168 | 0.32358986 | 0.423 | 0.28 | 1.94E-164 | TAM.prolif | Gas2l3 |
| Elovl11 | 1.38E-168 | 0.26150912 | 0.544 | 0.374 | 2.66E-164 | TAM.prolif | Elovl1 |
| Arl2bp | 1.93E-168 | 0.70100245 | 0.325 | 0.148 | 3.73E-164 | TAM.prolif | Arl2bp |
| Yif1b | 2.11E-168 | 0.54530972 | 0.446 | 0.253 | 4.08E-164 | TAM.prolif | Yif1b |
| Phrf1 | 2.60E-168 | 0.31159708 | 0.45 | 0.274 | 5.02E-164 | TAM.prolif | Phrf1 |
| Bcl7c | 3.18E-168 | 0.55288284 | 0.401 | 0.212 | 6.14E-164 | TAM.prolif | Bcl7c |
| Tnpo3 | 5.25E-168 | 0.32016885 | 0.488 | 0.314 | 1.02E-163 | TAM.prolif | Tnpo3 |
| C1d | 6.96E-168 | 0.50272607 | 0.409 | 0.225 | 1.35E-163 | TAM.prolif | C1d |
| Aco2 | 1.60E-167 | 0.45998698 | 0.414 | 0.232 | 3.10E-163 | TAM.prolif | Aco2 |
| Gcn1 | 2.83E-167 | 0.3314181 | 0.466 | 0.282 | 5.47E-163 | TAM.prolif | Gcn1 |
| Pigq | 4.23E-167 | 0.63622917 | 0.373 | 0.192 | 8.17E-163 | TAM.prolif | Pigq |
| Tspan42 | 5.51E-167 | 0.29230139 | 0.485 | 0.328 | 1.07E-162 | TAM.prolif | Tspan4 |
| Tmem106c | 6.93E-167 | 0.88187315 | 0.324 | 0.147 | 1.34E-162 | TAM.prolif | Tmem106c |
| Thumpd1 | 9.29E-167 | 0.68707983 | 0.306 | 0.139 | 1.80E-162 | TAM.prolif | Thumpd1 |
| Ivd | 9.91E-167 | 0.69322313 | 0.326 | 0.147 | 1.92E-162 | TAM.prolif | Ivd |
| Polr2m | 1.26E-166 | 0.74070537 | 0.318 | 0.139 | 2.45E-162 | TAM.prolif | Polr2m |
| Cpsf1 | 1.35E-166 | 0.52137237 | 0.429 | 0.236 | 2.60E-162 | TAM.prolif | Cpsf1 |
| Mpst | 1.63E-166 | 0.77593948 | 0.331 | 0.154 | 3.15E-162 | TAM.prolif | Mpst |
| Ibtk | 1.94E-166 | 0.50038909 | 0.362 | 0.2 | 3.75E-162 | TAM.prolif | Ibtk |
| Dohh | 5.01E-166 | 0.46425994 | 0.425 | 0.238 | 9.69E-162 | TAM.prolif | Dohh |
| Gatd3a1 | 5.71E-166 | 0.58162734 | 0.407 | 0.214 | 1.10E-161 | TAM.prolif | Gatd3a |
| Trappc121 | 1.27E-165 | 0.5671373 | 0.386 | 0.206 | 2.46E-161 | TAM.prolif | Trappc12 |
| Zc3h14 | 1.62E-165 | 0.38864781 | 0.423 | 0.249 | 3.13E-161 | TAM.prolif | Zc3h14 |
| Elp5 | 1.72E-165 | 0.32696663 | 0.507 | 0.311 | 3.32E-161 | TAM.prolif | Elp5 |
| Ing11 | 1.87E-165 | 0.41517896 | 0.455 | 0.271 | 3.61E-161 | TAM.prolif | Ing1 |
| Nfatc22 | 2.25E-165 | 0.28302139 | 0.485 | 0.314 | 4.36E-161 | TAM.prolif | Nfatc2 |
| Txnl4a | 2.38E-165 | 0.44066217 | 0.377 | 0.211 | 4.61E-161 | TAM.prolif | Txnl4a |
| D6Wsu163e | 5.34E-165 | 0.81582538 | 0.294 | 0.127 | 1.03E-160 | TAM.prolif | D6Wsu163e |
| Sema4c1 | 3.60E-164 | 0.83491768 | 0.284 | 0.121 | 6.97E-160 | TAM.prolif | Sema4c |
| Stx22 | 3.73E-164 | 0.42926116 | 0.465 | 0.275 | 7.21E-160 | TAM.prolif | Stx2 |
| Nenf2 | 3.88E-164 | 0.45397099 | 0.47 | 0.284 | 7.50E-160 | TAM.prolif | Nenf |
| Trmt2b | 4.55E-164 | 0.97914751 | 0.259 | 0.096 | 8.80E-160 | TAM.prolif | Trmt2b |
| Wars | 6.83E-164 | 0.39604293 | 0.442 | 0.258 | 1.32E-159 | TAM.prolif | Wars |
| Sap30bp | 9.75E-164 | 0.4358493 | 0.392 | 0.209 | 1.88E-159 | TAM.prolif | Sap30bp |
| Rex1bd | 1.12E-163 | 0.38332655 | 0.468 | 0.272 | 2.16E-159 | TAM.prolif | Rex1bd |
| Pqbp1 | 3.04E-163 | 0.29731284 | 0.491 | 0.296 | 5.88E-159 | TAM.prolif | Pqbp1 |
| Taf12 | 3.49E-163 | 0.55079978 | 0.376 | 0.198 | 6.76E-159 | TAM.prolif | Taf12 |
| Rtraf | 5.81E-163 | 0.43999953 | 0.431 | 0.25 | 1.12E-158 | TAM.prolif | Rtraf |
| Sigmar12 | 6.64E-163 | 0.28477164 | 0.509 | 0.322 | 1.28E-158 | TAM.prolif | Sigmar1 |
| Nqo2 | 1.62E-162 | 0.73772729 | 0.311 | 0.148 | 3.13E-158 | TAM.prolif | Nqo2 |
| Cdc5l | 1.89E-162 | 0.38349089 | 0.453 | 0.274 | 3.65E-158 | TAM.prolif | Cdc5l |
| Ak6 | 4.37E-162 | 0.62220069 | 0.359 | 0.178 | 8.45E-158 | TAM.prolif | Ak6 |
| Pno11 | 5.25E-162 | 0.41363577 | 0.386 | 0.214 | 1.02E-157 | TAM.prolif | Pno1 |
| Polr2j | 7.10E-162 | 0.6146299 | 0.353 | 0.186 | 1.37E-157 | TAM.prolif | Polr2j |
| Cops2 | 7.41E-162 | 0.35676869 | 0.454 | 0.271 | 1.43E-157 | TAM.prolif | Cops2 |
| Phka2 | 1.38E-161 | 0.52564422 | 0.34 | 0.17 | 2.67E-157 | TAM.prolif | Phka2 |
| Mcts11 | 1.46E-161 | 0.25953269 | 0.459 | 0.297 | 2.82E-157 | TAM.prolif | Mcts1 |
| Arhgap1 | 1.79E-161 | 0.29758385 | 0.512 | 0.329 | 3.46E-157 | TAM.prolif | Arhgap1 |
| Sap130 | 2.73E-161 | 0.346214 | 0.429 | 0.25 | 5.28E-157 | TAM.prolif | Sap130 |
| Sqle | 3.17E-161 | 0.6868933 | 0.282 | 0.146 | 6.13E-157 | TAM.prolif | Sqle |
| Terf2 | 4.24E-161 | 0.65126365 | 0.315 | 0.151 | 8.21E-157 | TAM.prolif | Terf2 |
| Krtcap2 | 5.11E-161 | 0.45908412 | 0.383 | 0.208 | 9.88E-157 | TAM.prolif | Krtcap2 |
| Pprc11 | 7.27E-161 | 0.306616 | 0.433 | 0.261 | 1.41E-156 | TAM.prolif | Pprc1 |
| Fryl | 2.54E-160 | 0.31072459 | 0.413 | 0.253 | 4.91E-156 | TAM.prolif | Fryl |
| Mical13 | 2.65E-160 | 0.33372339 | 0.433 | 0.266 | 5.13E-156 | TAM.prolif | Mical1 |
| Dap3 | 3.54E-160 | 0.3341429 | 0.469 | 0.288 | 6.84E-156 | TAM.prolif | Dap3 |
| Khk | 3.76E-160 | 0.57656719 | 0.323 | 0.161 | 7.28E-156 | TAM.prolif | Khk |
| Lilra51 | 7.19E-160 | 0.84190531 | 0.26 | 0.116 | 1.39E-155 | TAM.prolif | Lilra5 |
| Rap1gap21 | 7.46E-160 | 0.28708034 | 0.352 | 0.186 | 1.44E-155 | TAM.prolif | Rap1gap2 |
| Tsta3 | 3.65E-159 | 0.55292328 | 0.427 | 0.239 | 7.06E-155 | TAM.prolif | Tsta3 |
| Gmds | 4.64E-159 | 0.74812301 | 0.278 | 0.109 | 8.96E-155 | TAM.prolif | Gmds |

| | | | | | | | |
|---|---|---|---|---|---|---|---|
| Gpn1 | 4.96E-159 | 0.70640554 | 0.299 | 0.131 | 9.59E-155 | TAM.prolif | Gpn1 |
| Zbtb8os | 5.45E-159 | 0.54806321 | 0.339 | 0.167 | 1.05E-154 | TAM.prolif | Zbtb8os |
| Hira | 5.54E-159 | 0.68637347 | 0.341 | 0.158 | 1.07E-154 | TAM.prolif | Hira |
| Krt82 | 6.48E-159 | 0.29914061 | 0.374 | 0.243 | 1.25E-154 | TAM.prolif | Krt8 |
| Chchd4 | 6.66E-159 | 0.74934892 | 0.273 | 0.121 | 1.29E-154 | TAM.prolif | Chchd4 |
| Klhdc10 | 1.05E-158 | 0.42556865 | 0.434 | 0.252 | 2.03E-154 | TAM.prolif | Klhdc10 |
| Cbx1 | 1.46E-158 | 0.88708579 | 0.278 | 0.114 | 2.81E-154 | TAM.prolif | Cbx1 |
| Pmm2 | 2.71E-158 | 0.30693794 | 0.477 | 0.301 | 5.23E-154 | TAM.prolif | Pmm2 |
| Dhdds | 3.02E-158 | 0.39820393 | 0.422 | 0.24 | 5.84E-154 | TAM.prolif | Dhdds |
| Commd3 | 3.73E-158 | 0.50247557 | 0.466 | 0.274 | 7.21E-154 | TAM.prolif | Commd3 |
| Rpp211 | 9.30E-158 | 0.30173976 | 0.425 | 0.248 | 1.80E-153 | TAM.prolif | Rpp21 |
| Nelfcd | 1.23E-157 | 0.83347299 | 0.275 | 0.115 | 2.38E-153 | TAM.prolif | Nelfcd |
| Elpr1 | 2.86E-157 | 0.48218444 | 0.368 | 0.206 | 5.53E-153 | TAM.prolif | Elpr1 |
| Smyd5 | 3.22E-157 | 0.81194666 | 0.276 | 0.129 | 6.22E-153 | TAM.prolif | Smyd5 |
| Dnajc11 | 3.76E-157 | 0.48428102 | 0.355 | 0.186 | 7.26E-153 | TAM.prolif | Dnajc11 |
| Zranb2 | 5.80E-157 | 0.56852559 | 0.353 | 0.176 | 1.12E-152 | TAM.prolif | Zranb2 |
| Arl22 | 7.55E-157 | 0.62528089 | 0.372 | 0.188 | 1.46E-152 | TAM.prolif | Arl2 |
| Paox1 | 1.75E-156 | 0.82651346 | 0.288 | 0.127 | 3.38E-152 | TAM.prolif | Paox |
| Spag7 | 1.76E-156 | 0.30439748 | 0.47 | 0.286 | 3.41E-152 | TAM.prolif | Spag7 |
| St13 | 2.23E-156 | 0.34203125 | 0.494 | 0.326 | 4.31E-152 | TAM.prolif | St13 |
| Cirbp1 | 3.34E-156 | 0.33481611 | 0.481 | 0.296 | 6.45E-152 | TAM.prolif | Cirbp |
| Net11 | 5.22E-156 | 0.55525468 | 0.394 | 0.215 | 1.01E-151 | TAM.prolif | Net1 |
| Dlat | 5.27E-156 | 0.43736748 | 0.395 | 0.234 | 1.02E-151 | TAM.prolif | Dlat |
| Lonp1 | 5.84E-156 | 0.25847211 | 0.508 | 0.322 | 1.13E-151 | TAM.prolif | Lonp1 |
| Snw11 | 8.96E-156 | 0.30441443 | 0.455 | 0.284 | 1.73E-151 | TAM.prolif | Snw1 |
| Mettl16 | 1.10E-155 | 0.81991209 | 0.285 | 0.12 | 2.13E-151 | TAM.prolif | Mettl16 |
| Plrg1 | 1.67E-155 | 0.6251519 | 0.361 | 0.189 | 3.22E-151 | TAM.prolif | Plrg1 |
| Rasal31 | 2.40E-155 | 0.40928281 | 0.395 | 0.225 | 4.64E-151 | TAM.prolif | Rasal3 |
| Fdx2 | 2.53E-155 | 0.37389666 | 0.441 | 0.266 | 4.90E-151 | TAM.prolif | Fdx2 |
| Pop5 | 2.62E-155 | 0.79988106 | 0.28 | 0.119 | 5.07E-151 | TAM.prolif | Pop5 |
| Xpnpep1 | 5.21E-155 | 0.31303557 | 0.506 | 0.324 | 1.01E-150 | TAM.prolif | Xpnpep1 |
| Pik3r21 | 5.99E-155 | 0.33133742 | 0.489 | 0.292 | 1.16E-150 | TAM.prolif | Pik3r2 |
| Taf4 | 9.52E-155 | 0.41483439 | 0.441 | 0.251 | 1.84E-150 | TAM.prolif | Taf4 |
| Srp68 | 1.77E-154 | 0.28933822 | 0.512 | 0.328 | 3.43E-150 | TAM.prolif | Srp68 |
| Hltf | 2.09E-154 | 0.77676899 | 0.295 | 0.129 | 4.03E-150 | TAM.prolif | Hltf |
| Traf3ip3 | 2.41E-154 | 0.71169488 | 0.306 | 0.137 | 4.66E-150 | TAM.prolif | Traf3ip3 |
| Ube2n1 | 2.66E-154 | 0.3682316 | 0.416 | 0.248 | 5.15E-150 | TAM.prolif | Ube2n |
| Zfyve19 | 4.34E-154 | 0.70115144 | 0.302 | 0.136 | 8.39E-150 | TAM.prolif | Zfyve19 |
| Tbrg11 | 4.48E-154 | 0.44713165 | 0.443 | 0.264 | 8.67E-150 | TAM.prolif | Tbrg1 |
| Wrnip1 | 8.42E-154 | 0.49852176 | 0.342 | 0.179 | 1.63E-149 | TAM.prolif | Wrnip1 |
| Cdk5rap32 | 1.12E-153 | 0.34850374 | 0.459 | 0.271 | 2.16E-149 | TAM.prolif | Cdk5rap3 |
| Ndufaf81 | 1.63E-153 | 0.39273936 | 0.444 | 0.266 | 3.15E-149 | TAM.prolif | Ndufaf8 |
| Ltv11 | 1.67E-153 | 0.49164048 | 0.397 | 0.22 | 3.22E-149 | TAM.prolif | Ltv1 |
| Ext21 | 1.96E-153 | 0.33235524 | 0.477 | 0.293 | 3.78E-149 | TAM.prolif | Ext2 |
| Brd9 | 2.65E-153 | 0.59540557 | 0.312 | 0.149 | 5.12E-149 | TAM.prolif | Brd9 |
| Clec12a1 | 3.09E-153 | 0.37736083 | 0.278 | 0.172 | 5.97E-149 | TAM.prolif | Clec12a |
| Nsfl1c | 4.52E-153 | 0.4242412 | 0.466 | 0.274 | 8.74E-149 | TAM.prolif | Nsfl1c |
| Sympk | 5.33E-153 | 0.26953971 | 0.493 | 0.32 | 1.03E-148 | TAM.prolif | Sympk |
| Foxred1 | 6.61E-153 | 0.48698146 | 0.345 | 0.173 | 1.28E-148 | TAM.prolif | Foxred1 |
| Papss1 | 7.01E-153 | 0.33529835 | 0.469 | 0.287 | 1.36E-148 | TAM.prolif | Papss1 |
| Ino80c | 9.89E-153 | 0.4987011 | 0.364 | 0.191 | 1.91E-148 | TAM.prolif | Ino80c |
| Psmd9 | 1.34E-152 | 0.50433693 | 0.373 | 0.194 | 2.59E-148 | TAM.prolif | Psmd9 |
| Clasp1 | 1.45E-152 | 0.33397975 | 0.414 | 0.241 | 2.80E-148 | TAM.prolif | Clasp1 |
| Bcl2l12 | 2.10E-152 | 0.89995641 | 0.271 | 0.113 | 4.07E-148 | TAM.prolif | Bcl2l12 |
| Ndufs6 | 6.15E-152 | 0.46117693 | 0.386 | 0.222 | 1.19E-147 | TAM.prolif | Ndufs6 |
| Polr2g | 8.41E-152 | 0.268465 | 0.469 | 0.304 | 1.63E-147 | TAM.prolif | Polr2g |
| Fam207a | 9.54E-152 | 0.38189842 | 0.431 | 0.248 | 1.84E-147 | TAM.prolif | Fam207a |
| Mesd1 | 1.41E-151 | 0.47045083 | 0.431 | 0.253 | 2.73E-147 | TAM.prolif | Mesd |
| Vkorc1l1 | 1.85E-151 | 0.51659746 | 0.322 | 0.161 | 3.58E-147 | TAM.prolif | Vkorc1l1 |
| Ube3c | 3.04E-151 | 0.42351057 | 0.405 | 0.236 | 5.88E-147 | TAM.prolif | Ube3c |
| Pfdn1 | 3.61E-151 | 0.44653543 | 0.408 | 0.228 | 6.98E-147 | TAM.prolif | Pfdn1 |
| Fnta1 | 3.75E-151 | 0.28896467 | 0.473 | 0.304 | 7.24E-147 | TAM.prolif | Fnta |
| Ripor1 | 3.92E-151 | 0.60629226 | 0.343 | 0.177 | 7.58E-147 | TAM.prolif | Ripor1 |
| Trmt1 | 5.18E-151 | 0.69599502 | 0.308 | 0.145 | 1.00E-146 | TAM.prolif | Trmt1 |
| Dhcr242 | 5.22E-151 | 0.56297 | 0.369 | 0.214 | 1.01E-146 | TAM.prolif | Dhcr24 |
| Tesk11 | 8.17E-151 | 0.30022836 | 0.536 | 0.356 | 1.58E-146 | TAM.prolif | Tesk1 |
| Ube4b | 8.68E-151 | 0.27991526 | 0.493 | 0.316 | 1.68E-146 | TAM.prolif | Ube4b |
| Socs23 | 1.29E-150 | 0.26335062 | 0.479 | 0.305 | 2.49E-146 | TAM.prolif | Socs2 |
| Zdhhc16 | 2.50E-150 | 0.63414744 | 0.33 | 0.155 | 4.84E-146 | TAM.prolif | Zdhhc16 |
| Cenpc1 | 3.94E-150 | 0.58465675 | 0.305 | 0.142 | 7.61E-146 | TAM.prolif | Cenpc1 |
| Nfyc | 4.42E-150 | 0.39866576 | 0.39 | 0.212 | 8.54E-146 | TAM.prolif | Nfyc |
| Ddb1 | 4.89E-150 | 0.5439639 | 0.375 | 0.198 | 9.44E-146 | TAM.prolif | Ddb1 |
| Polr1a | 5.42E-150 | 0.60413111 | 0.285 | 0.131 | 1.05E-145 | TAM.prolif | Polr1a |
| Tcf3 | 9.37E-150 | 0.70062597 | 0.329 | 0.158 | 1.81E-145 | TAM.prolif | Tcf3 |
| Gars | 1.06E-149 | 0.7794484 | 0.29 | 0.131 | 2.06E-145 | TAM.prolif | Gars |
| Tbc1d52 | 1.32E-149 | 0.25450785 | 0.526 | 0.343 | 2.56E-145 | TAM.prolif | Tbc1d5 |
| Prpf18 | 1.68E-149 | 0.42685484 | 0.408 | 0.239 | 3.25E-145 | TAM.prolif | Prpf18 |
| Ptar1 | 1.80E-149 | 0.56497386 | 0.342 | 0.169 | 3.49E-145 | TAM.prolif | Ptar1 |
| Sarnp1 | 2.43E-149 | 0.33420713 | 0.434 | 0.266 | 4.70E-145 | TAM.prolif | Sarnp |
| Zfp3301 | 2.72E-149 | 0.38874685 | 0.404 | 0.228 | 5.25E-145 | TAM.prolif | Zfp330 |
| Ccdc107 | 3.02E-149 | 0.53253667 | 0.326 | 0.169 | 5.84E-145 | TAM.prolif | Ccdc107 |
| Rad9a1 | 3.38E-149 | 0.48236935 | 0.366 | 0.189 | 6.54E-145 | TAM.prolif | Rad9a |
| Nop9 | 7.87E-149 | 0.70792728 | 0.287 | 0.124 | 1.52E-144 | TAM.prolif | Nop9 |
| Rbx1 | 8.43E-149 | 0.35939724 | 0.445 | 0.272 | 1.63E-144 | TAM.prolif | Rbx1 |
| Bola3 | 2.64E-148 | 0.6767794 | 0.3 | 0.138 | 5.11E-144 | TAM.prolif | Bola3 |
| Slc25a17 | 3.12E-148 | 0.34326144 | 0.422 | 0.243 | 6.04E-144 | TAM.prolif | Slc25a17 |
| Snrnp35 | 3.76E-148 | 0.67704603 | 0.327 | 0.162 | 7.26E-144 | TAM.prolif | Snrnp35 |
| Zdhhc6 | 4.66E-148 | 0.53539616 | 0.366 | 0.188 | 9.01E-144 | TAM.prolif | Zdhhc6 |
| Hikeshi | 5.09E-148 | 0.58676429 | 0.37 | 0.189 | 9.83E-144 | TAM.prolif | Hikeshi |
| Ndufs51 | 5.60E-148 | 0.46337494 | 0.393 | 0.223 | 1.08E-143 | TAM.prolif | Ndufs5 |
| Tiprl | 6.91E-148 | 0.27498468 | 0.484 | 0.311 | 1.34E-143 | TAM.prolif | Tiprl |
| Tars2 | 7.28E-148 | 0.67909173 | 0.293 | 0.126 | 1.41E-143 | TAM.prolif | Tars2 |
| Suclg1 | 2.75E-147 | 0.70529748 | 0.338 | 0.163 | 5.31E-143 | TAM.prolif | Suclg1 |
| Gga22 | 3.66E-147 | 0.59749442 | 0.362 | 0.199 | 7.08E-143 | TAM.prolif | Gga2 |
| Prr122 | 4.60E-147 | 0.29735222 | 0.445 | 0.269 | 8.90E-143 | TAM.prolif | Prr12 |
| Ppfia41 | 4.82E-147 | 0.7890034 | 0.289 | 0.124 | 9.32E-143 | TAM.prolif | Ppfia4 |
| Dmac11 | 1.02E-146 | 0.42307298 | 0.426 | 0.252 | 1.98E-142 | TAM.prolif | Dmac1 |
| Thap122 | 1.03E-146 | 0.41506201 | 0.404 | 0.224 | 1.99E-142 | TAM.prolif | Thap12 |
| Smim152 | 1.07E-146 | 0.38120704 | 0.442 | 0.266 | 2.08E-142 | TAM.prolif | Smim15 |
| Dicer11 | 1.30E-146 | 0.46883097 | 0.359 | 0.192 | 2.52E-142 | TAM.prolif | Dicer1 |
| Hnrnpdl2 | 1.40E-146 | 0.27055137 | 0.45 | 0.277 | 2.71E-142 | TAM.prolif | Hnrnpdl |
| Prmt3 | 2.04E-146 | 0.80574516 | 0.254 | 0.105 | 3.95E-142 | TAM.prolif | Prmt3 |
| Utp18 | 2.09E-146 | 0.43613223 | 0.371 | 0.206 | 4.05E-142 | TAM.prolif | Utp18 |
| Bpnt11 | 3.34E-146 | 0.3929144 | 0.407 | 0.235 | 6.45E-142 | TAM.prolif | Bpnt1 |
| Slc25a19 | 3.39E-146 | 0.32734222 | 0.407 | 0.241 | 6.55E-142 | TAM.prolif | Slc25a19 |
| Uck1 | 5.05E-146 | 0.64392194 | 0.319 | 0.158 | 9.76E-142 | TAM.prolif | Uck1 |
| Gtf3c6 | 7.99E-146 | 0.37227991 | 0.409 | 0.239 | 1.55E-141 | TAM.prolif | Gtf3c6 |
| Srp14 | 8.48E-146 | 0.25053408 | 0.584 | 0.413 | 1.64E-141 | TAM.prolif | Srp14 |
| Cdk161 | 1.71E-145 | 0.33304052 | 0.488 | 0.298 | 3.30E-141 | TAM.prolif | Cdk16 |
| Ccnh | 3.70E-145 | 0.59813762 | 0.3 | 0.138 | 7.15E-141 | TAM.prolif | Ccnh |
| Uhmk11 | 6.65E-145 | 0.28005146 | 0.539 | 0.367 | 1.29E-140 | TAM.prolif | Uhmk1 |
| Pdcd11 | 9.77E-145 | 0.43557317 | 0.344 | 0.175 | 1.89E-140 | TAM.prolif | Pdcd11 |
| Adi12 | 1.14E-144 | 0.55813948 | 0.361 | 0.19 | 2.20E-140 | TAM.prolif | Adi1 |
| Lrpprc | 1.33E-144 | 0.60509264 | 0.339 | 0.173 | 2.56E-140 | TAM.prolif | Lrpprc |
| Irf2bp1 | 1.35E-144 | 0.64862806 | 0.307 | 0.146 | 2.60E-140 | TAM.prolif | Irf2bp1 |
| Xpo4 | 1.45E-144 | 0.88293258 | 0.251 | 0.097 | 2.80E-140 | TAM.prolif | Xpo4 |
| Glrx3 | 1.52E-144 | 0.43039887 | 0.416 | 0.238 | 2.94E-140 | TAM.prolif | Glrx3 |
| Pfdn4 | 1.76E-144 | 0.68385589 | 0.289 | 0.144 | 3.40E-140 | TAM.prolif | Pfdn4 |
| Crybg3 | 2.44E-144 | 0.44913708 | 0.385 | 0.206 | 4.71E-140 | TAM.prolif | Crybg3 |
| Pwp1 | 3.02E-144 | 0.58272344 | 0.289 | 0.136 | 5.83E-140 | TAM.prolif | Pwp1 |
| Glo1 | 4.47E-144 | 0.36070776 | 0.456 | 0.282 | 8.64E-140 | TAM.prolif | Glo1 |
| Clpp | 4.52E-144 | 0.75240087 | 0.276 | 0.128 | 8.74E-140 | TAM.prolif | Clpp |
| Utp15 | 5.47E-144 | 0.52021905 | 0.352 | 0.183 | 1.06E-139 | TAM.prolif | Utp15 |
| Vrk3 | 6.69E-144 | 0.34421565 | 0.496 | 0.308 | 1.29E-139 | TAM.prolif | Vrk3 |
| Upf3a | 7.36E-144 | 0.41229497 | 0.402 | 0.223 | 1.42E-139 | TAM.prolif | Upf3a |
| Pcm11 | 1.18E-143 | 0.36300208 | 0.415 | 0.237 | 2.29E-139 | TAM.prolif | Pcm1 |
| Bcl2l13 | 1.27E-143 | 0.50871942 | 0.34 | 0.173 | 2.46E-139 | TAM.prolif | Bcl2l13 |
| Lzic | 1.58E-143 | 0.66633113 | 0.318 | 0.162 | 3.06E-139 | TAM.prolif | Lzic |
| Iars2 | 1.74E-143 | 0.38824248 | 0.423 | 0.243 | 3.37E-139 | TAM.prolif | Iars2 |
| Gnl3l | 3.80E-143 | 0.62589067 | 0.314 | 0.145 | 7.35E-139 | TAM.prolif | Gnl3l |
| Rnf40 | 6.98E-143 | 0.31817669 | 0.484 | 0.312 | 1.35E-138 | TAM.prolif | Rnf40 |
| Mtch2 | 1.38E-142 | 0.65112672 | 0.306 | 0.155 | 2.67E-138 | TAM.prolif | Mtch2 |
| Kctd5 | 1.50E-142 | 0.48552437 | 0.381 | 0.209 | 2.90E-138 | TAM.prolif | Kctd5 |
| Pdcd2 | 1.60E-142 | 0.78769444 | 0.275 | 0.125 | 3.10E-138 | TAM.prolif | Pdcd2 |
| Ado | 2.14E-142 | 0.49351582 | 0.388 | 0.213 | 4.14E-138 | TAM.prolif | Ado |
| Rnf1412 | 2.82E-142 | 0.52944116 | 0.375 | 0.219 | 5.45E-138 | TAM.prolif | Rnf141 |
| Tmem2682 | 3.47E-142 | 0.43845331 | 0.392 | 0.227 | 6.70E-138 | TAM.prolif | Tmem268 |
| Bcl2 | 3.56E-142 | 0.61070539 | 0.281 | 0.136 | 6.88E-138 | TAM.prolif | Bcl2 |
| Limk12 | 4.72E-142 | 0.43174364 | 0.417 | 0.24 | 9.13E-138 | TAM.prolif | Limk1 |
| Paip12 | 6.32E-142 | 0.2597882 | 0.505 | 0.351 | 1.22E-137 | TAM.prolif | Paip1 |
| Ctnnbl1 | 7.50E-142 | 0.43606793 | 0.4 | 0.228 | 1.45E-137 | TAM.prolif | Ctnnbl1 |
| Tgs1 | 7.83E-142 | 0.65911898 | 0.302 | 0.142 | 1.51E-137 | TAM.prolif | Tgs1 |
| Alg10b | 9.20E-142 | 0.46482881 | 0.418 | 0.246 | 1.78E-137 | TAM.prolif | Alg10b |
| Vps521 | 1.03E-141 | 0.39022935 | 0.479 | 0.295 | 2.00E-137 | TAM.prolif | Vps52 |
| Fto | 1.05E-141 | 0.26481867 | 0.467 | 0.288 | 2.04E-137 | TAM.prolif | Fto |
| Gtf3c5 | 1.11E-141 | 0.76028798 | 0.262 | 0.118 | 2.15E-137 | TAM.prolif | Gtf3c5 |
| Ufsp2 | 1.42E-141 | 0.26894152 | 0.46 | 0.291 | 2.74E-137 | TAM.prolif | Ufsp2 |
| Rmi1 | 3.34E-141 | 0.71338755 | 0.27 | 0.121 | 6.46E-137 | TAM.prolif | Rmi1 |
| Setdb1 | 4.13E-141 | 0.37574403 | 0.416 | 0.241 | 7.98E-137 | TAM.prolif | Setdb1 |
| Pdk31 | 6.51E-141 | 0.39799746 | 0.379 | 0.214 | 1.26E-136 | TAM.prolif | Pdk3 |
| Ndufs4 | 8.68E-141 | 0.32320855 | 0.455 | 0.286 | 1.68E-136 | TAM.prolif | Ndufs4 |
| Ciao1 | 8.93E-141 | 0.47152894 | 0.388 | 0.218 | 1.73E-136 | TAM.prolif | Ciao1 |
| Gng123 | 9.48E-141 | 0.28195408 | 0.435 | 0.299 | 1.83E-136 | TAM.prolif | Gng12 |
| Ngdn | 1.04E-140 | 0.35811831 | 0.408 | 0.237 | 2.01E-136 | TAM.prolif | Ngdn |
| Cstf3 | 1.28E-140 | 0.43342704 | 0.34 | 0.174 | 2.47E-136 | TAM.prolif | Cstf3 |
| Opa1 | 2.32E-140 | 0.4566294 | 0.405 | 0.236 | 4.48E-136 | TAM.prolif | Opa1 |
| Hadh1 | 2.39E-140 | 0.3395481 | 0.417 | 0.264 | 4.62E-136 | TAM.prolif | Hadh |
| Vav22 | 3.52E-140 | 0.35737279 | 0.473 | 0.298 | 6.81E-136 | TAM.prolif | Vav2 |
| Dhrs7b | 3.76E-140 | 0.58047527 | 0.316 | 0.158 | 7.27E-136 | TAM.prolif | Dhrs7b |
| Traf2 | 7.00E-140 | 0.34971934 | 0.399 | 0.227 | 1.35E-135 | TAM.prolif | Traf2 |
| Slc25a452 | 9.47E-140 | 0.31305879 | 0.46 | 0.276 | 1.83E-135 | TAM.prolif | Slc25a45 |
| Tbl1x | 1.36E-139 | 0.38393514 | 0.381 | 0.217 | 2.62E-135 | TAM.prolif | Tbl1x |
| Gtf2h2 | 1.92E-139 | 0.50224669 | 0.312 | 0.172 | 3.71E-135 | TAM.prolif | Gtf2h2 |
| Ppil21 | 2.86E-139 | 0.27276049 | 0.463 | 0.29 | 5.53E-135 | TAM.prolif | Ppil2 |
| Emc8 | 3.03E-139 | 0.68789258 | 0.293 | 0.133 | 5.85E-135 | TAM.prolif | Emc8 |
| Adprhl2 | 7.44E-139 | 0.73067809 | 0.271 | 0.122 | 1.44E-134 | TAM.prolif | Adprhl2 |
| Poldip2 | 8.28E-139 | 0.84499705 | 0.282 | 0.122 | 1.60E-134 | TAM.prolif | Poldip2 |
| Blnk3 | 1.73E-138 | 0.60106317 | 0.37 | 0.213 | 3.35E-134 | TAM.prolif | Blnk |
| Spidr1 | 2.85E-138 | 0.53888428 | 0.32 | 0.16 | 5.52E-134 | TAM.prolif | Spidr |
| Sec23b1 | 4.72E-138 | 0.26658432 | 0.456 | 0.279 | 9.12E-134 | TAM.prolif | Sec23b |
| Nip7 | 5.10E-138 | 0.57031227 | 0.322 | 0.163 | 9.87E-134 | TAM.prolif | Nip7 |
| Smim11 | 1.07E-137 | 0.60299283 | 0.326 | 0.161 | 2.07E-133 | TAM.prolif | Smim11 |
| Eloc | 1.43E-137 | 0.34480965 | 0.394 | 0.236 | 2.77E-133 | TAM.prolif | Eloc |
| Slc39a91 | 2.21E-137 | 0.41442201 | 0.457 | 0.27 | 4.27E-133 | TAM.prolif | Slc39a9 |
| Zfp414 | 2.46E-137 | 0.39035166 | 0.352 | 0.195 | 4.75E-133 | TAM.prolif | Zfp414 |
| Hmgb1 | 2.61E-137 | 0.50994362 | 0.304 | 0.165 | 5.05E-133 | TAM.prolif | Hmgb1 |
| Cops8 | 2.92E-137 | 0.35304566 | 0.475 | 0.302 | 5.65E-133 | TAM.prolif | Cops8 |
| Uxs1 | 3.29E-137 | 0.50577653 | 0.359 | 0.195 | 6.36E-133 | TAM.prolif | Uxs1 |
| Nfx1 | 1.55E-136 | 0.33035149 | 0.379 | 0.211 | 3.00E-132 | TAM.prolif | Nfx1 |
| Stag1 | 1.57E-136 | 0.44387409 | 0.311 | 0.157 | 3.04E-132 | TAM.prolif | Stag1 |
| Cox15 | 1.80E-136 | 0.56801504 | 0.33 | 0.168 | 3.47E-132 | TAM.prolif | Cox15 |
| Gabpb2 | 1.96E-136 | 0.37901603 | 0.362 | 0.201 | 3.79E-132 | TAM.prolif | Gabpb2 |
| Rfng | 2.00E-136 | 0.65563287 | 0.284 | 0.138 | 3.87E-132 | TAM.prolif | Rfng |
| Nol9 | 4.17E-136 | 0.63317318 | 0.284 | 0.137 | 8.07E-132 | TAM.prolif | Nol9 |
| Cdc16 | 4.60E-136 | 0.66441078 | 0.279 | 0.13 | 8.90E-132 | TAM.prolif | Cdc16 |
| Maged13 | 4.68E-136 | 0.54601184 | 0.318 | 0.179 | 9.05E-132 | TAM.prolif | Maged1 |
| Mvk | 3.57E-135 | 0.72785585 | 0.314 | 0.155 | 6.91E-131 | TAM.prolif | Mvk |
| Pigu | 9.97E-135 | 0.45221102 | 0.367 | 0.194 | 1.93E-130 | TAM.prolif | Pigu |
| Polr1c1 | 1.13E-134 | 0.33457483 | 0.438 | 0.269 | 2.18E-130 | TAM.prolif | Polr1c |
| Arhgap21 | 1.66E-134 | 0.42889789 | 0.303 | 0.158 | 3.20E-130 | TAM.prolif | Arhgap21 |
| Phf21 | 1.77E-134 | 0.3597043 | 0.406 | 0.233 | 3.42E-130 | TAM.prolif | Phf2 |
| Bysl | 2.36E-134 | 0.69558907 | 0.266 | 0.116 | 4.56E-130 | TAM.prolif | Bysl |
| Scd22 | 4.06E-134 | 0.42938135 | 0.358 | 0.211 | 7.85E-130 | TAM.prolif | Scd2 |
| Yeats4 | 8.41E-134 | 0.4560524 | 0.34 | 0.185 | 1.63E-129 | TAM.prolif | Yeats4 |
| Exosc4 | 8.51E-134 | 0.6144952 | 0.334 | 0.168 | 1.65E-129 | TAM.prolif | Exosc4 |
| Elf3a | 1.21E-133 | 0.27103 | 0.422 | 0.266 | 2.34E-129 | TAM.prolif | Elf3a |
| Slc35b4 | 1.49E-133 | 0.41883435 | 0.372 | 0.209 | 2.87E-129 | TAM.prolif | Slc35b4 |
| Ncdn | 2.08E-133 | 0.5501086 | 0.338 | 0.172 | 4.01E-129 | TAM.prolif | Ncdn |
| Sp2 | 2.42E-133 | 0.43264072 | 0.344 | 0.182 | 4.69E-129 | TAM.prolif | Sp2 |
| Cnot8 | 2.49E-133 | 0.27338968 | 0.496 | 0.326 | 4.81E-129 | TAM.prolif | Cnot8 |
| Parl | 2.72E-133 | 0.36680098 | 0.366 | 0.219 | 5.27E-129 | TAM.prolif | Parl |
| Armc1 | 2.82E-133 | 0.51566353 | 0.331 | 0.175 | 5.45E-129 | TAM.prolif | Armc1 |
| Prpf3 | 6.20E-133 | 0.51101473 | 0.349 | 0.178 | 1.20E-128 | TAM.prolif | Prpf3 |
| Pkib1 | 7.27E-133 | 0.4362175 | 0.37 | 0.233 | 1.41E-128 | TAM.prolif | Pkib |
| Ssbp3 | 8.63E-133 | 0.68032707 | 0.256 | 0.116 | 1.67E-128 | TAM.prolif | Ssbp3 |
| Vcpkmt | 1.00E-132 | 0.89169972 | 0.272 | 0.12 | 1.93E-128 | TAM.prolif | Vcpkmt |
| Kri1 | 1.01E-132 | 0.38823672 | 0.417 | 0.239 | 1.96E-128 | TAM.prolif | Kri1 |
| Serhl | 1.21E-132 | 0.74086913 | 0.275 | 0.121 | 2.34E-128 | TAM.prolif | Serhl |
| Elp1 | 1.79E-132 | 0.61200356 | 0.295 | 0.143 | 3.45E-128 | TAM.prolif | Elp1 |
| Bnip1 | 4.58E-132 | 0.91628329 | 0.25 | 0.104 | 8.86E-128 | TAM.prolif | Bnip1 |
| Aven | 4.76E-132 | 0.57620529 | 0.274 | 0.131 | 9.21E-128 | TAM.prolif | Aven |
| Lrrc47 | 1.50E-131 | 0.28307907 | 0.496 | 0.331 | 2.90E-127 | TAM.prolif | Lrrc47 |
| Exoc61 | 2.44E-131 | 0.32044889 | 0.378 | 0.22 | 4.71E-127 | TAM.prolif | Exoc6 |
| Otud6b | 2.97E-131 | 0.6255646 | 0.329 | 0.169 | 5.75E-127 | TAM.prolif | Otud6b |
| Mtx2 | 4.62E-131 | 0.39775092 | 0.392 | 0.225 | 8.94E-127 | TAM.prolif | Mtx2 |
| Rbm19 | 6.57E-131 | 0.50380551 | 0.31 | 0.159 | 1.27E-126 | TAM.prolif | Rbm19 |
| Hsph1 | 9.14E-131 | 0.53789644 | 0.282 | 0.153 | 1.77E-126 | TAM.prolif | Hsph1 |
| Tada1 | 1.06E-130 | 0.36543478 | 0.32 | 0.168 | 2.05E-126 | TAM.prolif | Tada1 |
| Isoc1 | 1.20E-130 | 0.58708876 | 0.339 | 0.169 | 2.31E-126 | TAM.prolif | Isoc1 |
| Klhdc2 | 4.31E-130 | 0.44953133 | 0.342 | 0.183 | 8.33E-126 | TAM.prolif | Klhdc2 |
| Nmd3 | 6.99E-130 | 0.48525712 | 0.317 | 0.17 | 1.35E-125 | TAM.prolif | Nmd3 |
| Mtln | 9.53E-130 | 0.43608424 | 0.389 | 0.225 | 1.84E-125 | TAM.prolif | Mtln |
| Scap2 | 1.59E-129 | 0.47769649 | 0.378 | 0.205 | 3.07E-125 | TAM.prolif | Scap |
| Tlr14 | 1.59E-129 | 0.32524011 | 0.385 | 0.23 | 3.08E-125 | TAM.prolif | Tlr1 |
| Gtf2h4 | 2.92E-129 | 0.71447983 | 0.268 | 0.122 | 5.64E-125 | TAM.prolif | Gtf2h4 |
| Tmem87a2 | 5.69E-129 | 0.34480163 | 0.395 | 0.229 | 1.10E-124 | TAM.prolif | Tmem87a |
| Ddx422 | 5.92E-129 | 0.25971917 | 0.436 | 0.261 | 1.14E-124 | TAM.prolif | Ddx42 |
| Lactb1 | 6.31E-129 | 0.28352771 | 0.439 | 0.281 | 1.22E-124 | TAM.prolif | Lactb |
| Ppdpf1 | 7.92E-129 | 0.28611245 | 0.39 | 0.258 | 1.53E-124 | TAM.prolif | Ppdpf |
| Znrd2 | 9.97E-129 | 0.57804277 | 0.268 | 0.12 | 1.93E-124 | TAM.prolif | Znrd2 |
| Pofut1 | 1.21E-128 | 0.44019437 | 0.36 | 0.189 | 2.34E-124 | TAM.prolif | Pofut1 |
| Pgrmc21 | 2.12E-128 | 0.3236364 | 0.44 | 0.281 | 4.09E-124 | TAM.prolif | Pgrmc2 |
| Ylpm1 | 2.23E-128 | 0.38883156 | 0.386 | 0.218 | 4.30E-124 | TAM.prolif | Ylpm1 |
| Tmem2481 | 4.44E-128 | 0.28895972 | 0.453 | 0.278 | 8.58E-124 | TAM.prolif | Tmem248 |
| Mycbp | 7.20E-128 | 0.67519275 | 0.268 | 0.124 | 1.39E-123 | TAM.prolif | Mycbp |
| Memo1 | 8.06E-128 | 0.42119624 | 0.347 | 0.199 | 1.56E-123 | TAM.prolif | Memo1 |
| Golim4 | 1.05E-127 | 0.43729497 | 0.352 | 0.188 | 2.04E-123 | TAM.prolif | Golim4 |
| Ercc3 | 1.94E-127 | 0.34752991 | 0.457 | 0.281 | 3.76E-123 | TAM.prolif | Ercc3 |
| Champ1 | 2.05E-127 | 0.64564778 | 0.254 | 0.112 | 3.96E-123 | TAM.prolif | Champ1 |
| Taf6l | 7.38E-127 | 0.66224127 | 0.257 | 0.112 | 1.43E-122 | TAM.prolif | Taf6l |
| Mphosph10 | 8.70E-127 | 0.4876739 | 0.318 | 0.171 | 1.68E-122 | TAM.prolif | Mphosph10 |
| Cfap20 | 1.23E-126 | 0.50110364 | 0.297 | 0.155 | 2.37E-122 | TAM.prolif | Cfap20 |
| Acbd6 | 1.24E-126 | 0.39949775 | 0.427 | 0.25 | 2.39E-122 | TAM.prolif | Acbd6 |
| Ccz1 | 2.71E-126 | 0.41803163 | 0.389 | 0.236 | 5.24E-122 | TAM.prolif | Ccz1 |
| Cox20 | 2.78E-126 | 0.56051623 | 0.319 | 0.156 | 5.38E-122 | TAM.prolif | Cox20 |
| Man2c1 | 4.26E-126 | 0.29111586 | 0.491 | 0.316 | 8.23E-122 | TAM.prolif | Man2c1 |
| Mpnd | 4.90E-126 | 0.28856594 | 0.459 | 0.288 | 9.47E-122 | TAM.prolif | Mpnd |
| Clasrp | 5.67E-126 | 0.41466258 | 0.386 | 0.219 | 1.10E-121 | TAM.prolif | Clasrp |
| Eci2 | 5.96E-126 | 0.45304715 | 0.356 | 0.204 | 1.15E-121 | TAM.prolif | Eci2 |
| Ppp2r1b | 7.84E-126 | 0.51888059 | 0.37 | 0.203 | 1.52E-121 | TAM.prolif | Ppp2r1b |
| Ost4 | 9.82E-126 | 0.27882343 | 0.46 | 0.302 | 1.90E-121 | TAM.prolif | Ost4 |
| Las1l | 1.36E-125 | 0.55894275 | 0.34 | 0.175 | 2.63E-121 | TAM.prolif | Las1l |
| Commd9 | 1.53E-125 | 0.67512299 | 0.3 | 0.143 | 2.96E-121 | TAM.prolif | Commd9 |
| Ebi31 | 1.95E-125 | 0.4476903 | 0.368 | 0.212 | 3.76E-121 | TAM.prolif | Ebi3 |
| Fam98a | 2.13E-125 | 0.57274717 | 0.256 | 0.116 | 4.12E-121 | TAM.prolif | Fam98a |
| Mtf2 | 2.47E-125 | 0.56807066 | 0.28 | 0.138 | 4.78E-121 | TAM.prolif | Mtf2 |
| Ndufa9 | 2.50E-125 | 0.66210716 | 0.284 | 0.139 | 4.83E-121 | TAM.prolif | Ndufa9 |
| Pcyt21 | 2.95E-125 | 0.4821152 | 0.341 | 0.19 | 5.71E-121 | TAM.prolif | Pcyt2 |
| Opa31 | 3.10E-125 | 0.28664349 | 0.408 | 0.249 | 5.99E-121 | TAM.prolif | Opa3 |
| Desi1 | 3.64E-125 | 0.43179683 | 0.352 | 0.193 | 7.03E-121 | TAM.prolif | Desi1 |
| Slc25a121 | 8.95E-125 | 0.28248175 | 0.408 | 0.253 | 1.73E-120 | TAM.prolif | Slc25a12 |
| Dgcr2 | 9.17E-125 | 0.28082126 | 0.443 | 0.276 | 1.77E-120 | TAM.prolif | Dgcr2 |
| Psmg1 | 1.13E-124 | 0.5321718 | 0.273 | 0.136 | 2.18E-120 | TAM.prolif | Psmg1 |
| Brd3 | 1.53E-124 | 0.70893773 | 0.265 | 0.126 | 2.95E-120 | TAM.prolif | Brd3 |
| Erlin1 | 1.96E-124 | 0.44672134 | 0.361 | 0.195 | 3.78E-120 | TAM.prolif | Erlin1 |
| Spast1 | 3.37E-124 | 0.26383029 | 0.424 | 0.272 | 6.52E-120 | TAM.prolif | Spast |
| Nup54 | 3.64E-124 | 0.4551907 | 0.29 | 0.144 | 7.03E-120 | TAM.prolif | Nup54 |
| Mtmr9 | 4.86E-124 | 0.44035433 | 0.422 | 0.247 | 9.40E-120 | TAM.prolif | Mtmr9 |
| Rnf1811 | 1.52E-123 | 0.37847173 | 0.427 | 0.263 | 2.94E-119 | TAM.prolif | Rnf181 |
| Ppm1d | 2.01E-123 | 0.48100637 | 0.28 | 0.138 | 3.88E-119 | TAM.prolif | Ppm1d |
| Psmg3 | 4.15E-123 | 0.62454711 | 0.279 | 0.131 | 8.01E-119 | TAM.prolif | Psmg3 |
| Zfp6911 | 6.50E-123 | 0.49307608 | 0.3 | 0.169 | 1.26E-118 | TAM.prolif | Zfp691 |
| Trappc13 | 1.29E-122 | 0.68976232 | 0.279 | 0.135 | 2.50E-118 | TAM.prolif | Trappc13 |
| Fam192a | 1.53E-122 | 0.25324254 | 0.42 | 0.255 | 2.96E-118 | TAM.prolif | Fam192a |
| Calhm21 | 1.65E-122 | 0.43048128 | 0.325 | 0.162 | 3.19E-118 | TAM.prolif | Calhm2 |
| Psmd5 | 2.18E-122 | 0.52139673 | 0.326 | 0.164 | 4.21E-118 | TAM.prolif | Psmd5 |
| Crebzf | 3.15E-122 | 0.30106141 | 0.429 | 0.253 | 6.08E-118 | TAM.prolif | Crebzf |
| Ipo41 | 5.02E-122 | 0.28010148 | 0.418 | 0.265 | 9.71E-118 | TAM.prolif | Ipo4 |
| Hsd17b10 | 5.42E-122 | 0.33967329 | 0.424 | 0.262 | 1.05E-117 | TAM.prolif | Hsd17b10 |
| Rexo4 | 8.69E-122 | 0.44435572 | 0.376 | 0.203 | 1.68E-117 | TAM.prolif | Rexo4 |
| Anapc16 | 1.01E-121 | 0.25418864 | 0.481 | 0.318 | 1.95E-117 | TAM.prolif | Anapc16 |
| Mecr | 1.05E-121 | 0.52460303 | 0.266 | 0.121 | 2.03E-117 | TAM.prolif | Mecr |
| Tbcd1 | 1.07E-121 | 0.40602322 | 0.383 | 0.227 | 2.08E-117 | TAM.prolif | Tbcd |
| Gbf1 | 1.57E-121 | 0.29402548 | 0.371 | 0.223 | 3.03E-117 | TAM.prolif | Gbf1 |
| Ptgr22 | 2.05E-121 | 0.26289293 | 0.482 | 0.309 | 3.97E-117 | TAM.prolif | Ptgr2 |
| Tmem70 | 8.09E-121 | 0.61282248 | 0.278 | 0.133 | 1.56E-116 | TAM.prolif | Tmem70 |
| Eif4enif1 | 1.35E-120 | 0.5770594 | 0.273 | 0.126 | 2.62E-116 | TAM.prolif | Eif4enif1 |
| Ift521 | 1.90E-120 | 0.40882079 | 0.389 | 0.23 | 3.67E-116 | TAM.prolif | Ift52 |
| Kpna6 | 1.91E-120 | 0.33964807 | 0.413 | 0.253 | 3.69E-116 | TAM.prolif | Kpna6 |
| Cmkl1 | 1.93E-120 | 0.45391055 | 0.374 | 0.209 | 3.73E-116 | TAM.prolif | Cmkl1 |
| Ptcd2 | 2.19E-120 | 0.56022222 | 0.26 | 0.119 | 4.23E-116 | TAM.prolif | Ptcd2 |
| Heatr3 | 2.37E-120 | 0.57095243 | 0.289 | 0.138 | 4.58E-116 | TAM.prolif | Heatr3 |
| Mthfd1l1 | 2.78E-120 | 0.44916063 | 0.337 | 0.174 | 5.37E-116 | TAM.prolif | Mthfd1l |
| Med292 | 3.82E-120 | 0.271903 | 0.432 | 0.27 | 7.38E-116 | TAM.prolif | Med29 |
| Sdad1 | 4.89E-120 | 0.42834086 | 0.341 | 0.181 | 9.45E-116 | TAM.prolif | Sdad1 |
| Topors3 | 8.15E-120 | 0.3045768 | 0.414 | 0.253 | 1.58E-115 | TAM.prolif | Topors |
| Ppfibp11 | 8.89E-120 | 0.34495635 | 0.42 | 0.268 | 1.72E-115 | TAM.prolif | Ppfibp1 |
| Urod | 1.05E-119 | 0.46697445 | 0.353 | 0.196 | 2.03E-115 | TAM.prolif | Urod |
| Nudt2 | 1.50E-119 | 0.61967524 | 0.267 | 0.129 | 2.89E-115 | TAM.prolif | Nudt2 |
| Tm7sf31 | 2.88E-119 | 0.28298682 | 0.401 | 0.243 | 5.56E-115 | TAM.prolif | Tm7sf3 |
| Taf2 | 3.05E-119 | 0.47319576 | 0.318 | 0.169 | 5.89E-115 | TAM.prolif | Taf2 |
| Aqr | 4.74E-119 | 0.36236228 | 0.378 | 0.213 | 9.17E-115 | TAM.prolif | Aqr |
| Elac2 | 6.48E-119 | 0.61053815 | 0.279 | 0.133 | 1.25E-114 | TAM.prolif | Elac2 |
| Rfk | 6.82E-119 | 0.30239228 | 0.447 | 0.297 | 1.32E-114 | TAM.prolif | Rfk |
| Med24 | 7.02E-119 | 0.4720803 | 0.312 | 0.161 | 1.36E-114 | TAM.prolif | Med24 |
| Ddx47 | 7.04E-119 | 0.29144554 | 0.399 | 0.249 | 1.36E-114 | TAM.prolif | Ddx47 |
| Tent4a | 8.06E-119 | 0.37178587 | 0.355 | 0.208 | 1.56E-114 | TAM.prolif | Tent4a |
| Nr2c2ap | 8.83E-119 | 0.28636953 | 0.399 | 0.243 | 1.71E-114 | TAM.prolif | Nr2c2ap |
| Imp3 | 1.04E-118 | 0.40629031 | 0.351 | 0.193 | 2.01E-114 | TAM.prolif | Imp3 |
| Fgfr1op | 1.27E-118 | 0.53264966 | 0.271 | 0.136 | 2.45E-114 | TAM.prolif | Fgfr1op |
| Nmt21 | 1.97E-118 | 0.40551122 | 0.356 | 0.204 | 3.80E-114 | TAM.prolif | Nmt2 |
| Thoc5 | 2.70E-118 | 0.53455349 | 0.32 | 0.168 | 5.23E-114 | TAM.prolif | Thoc5 |
| Depdc5 | 2.84E-118 | 0.48100479 | 0.307 | 0.162 | 5.50E-114 | TAM.prolif | Depdc5 |
| Srprb | 2.89E-118 | 0.37174583 | 0.363 | 0.2 | 5.59E-114 | TAM.prolif | Srprb |
| Eif2s3x | 4.07E-118 | 0.57280542 | 0.273 | 0.131 | 7.86E-114 | TAM.prolif | Eif2s3x |
| Plekha3 | 5.64E-118 | 0.25869178 | 0.436 | 0.275 | 1.09E-113 | TAM.prolif | Plekha3 |
| Ncoa51 | 8.56E-118 | 0.39835887 | 0.356 | 0.199 | 1.65E-113 | TAM.prolif | Ncoa5 |
| Tor4a2 | 8.89E-118 | 0.26875092 | 0.44 | 0.279 | 1.72E-113 | TAM.prolif | Tor4a |
| Slc35b11 | 2.37E-117 | 0.29513742 | 0.39 | 0.227 | 4.58E-113 | TAM.prolif | Slc35b1 |
| Nsmaf | 2.75E-117 | 0.38920963 | 0.318 | 0.18 | 5.31E-113 | TAM.prolif | Nsmaf |
| Luc7l | 3.11E-117 | 0.36926837 | 0.352 | 0.19 | 6.01E-113 | TAM.prolif | Luc7l |
| Lsg1 | 3.88E-117 | 0.4685672 | 0.32 | 0.168 | 7.49E-113 | TAM.prolif | Lsg1 |
| Orc3 | 4.66E-117 | 0.52600498 | 0.287 | 0.149 | 9.02E-113 | TAM.prolif | Orc3 |
| Psmf1 | 6.73E-117 | 0.37642481 | 0.368 | 0.208 | 1.30E-112 | TAM.prolif | Psmf1 |
| Spata5 | 8.28E-117 | 0.36545363 | 0.331 | 0.187 | 1.60E-112 | TAM.prolif | Spata5 |
| Drosha | 1.05E-116 | 0.68687749 | 0.253 | 0.115 | 2.04E-112 | TAM.prolif | Drosha |
| Anapc41 | 1.17E-116 | 0.33871037 | 0.377 | 0.224 | 2.26E-112 | TAM.prolif | Anapc4 |
| Mfn2 | 1.21E-116 | 0.45264509 | 0.344 | 0.194 | 2.35E-112 | TAM.prolif | Mfn2 |
| Aifm1 | 1.26E-116 | 0.64007075 | 0.263 | 0.119 | 2.44E-112 | TAM.prolif | Aifm1 |
| Ipo132 | 1.47E-116 | 0.36050666 | 0.438 | 0.27 | 2.83E-112 | TAM.prolif | Ipo13 |
| Kcnn42 | 1.91E-116 | 0.30506886 | 0.301 | 0.224 | 3.70E-112 | TAM.prolif | Kcnn4 |
| Smim201 | 3.27E-116 | 0.25884142 | 0.445 | 0.277 | 6.32E-112 | TAM.prolif | Smim20 |
| Ttc13 | 3.30E-116 | 0.37113205 | 0.333 | 0.187 | 6.37E-112 | TAM.prolif | Ttc13 |
| Spsb3 | 4.10E-116 | 0.32650128 | 0.374 | 0.218 | 7.93E-112 | TAM.prolif | Spsb3 |
| Tmem94 | 4.98E-116 | 0.46406065 | 0.325 | 0.167 | 9.63E-112 | TAM.prolif | Tmem94 |
| Tmem161a | 9.55E-116 | 0.50258763 | 0.317 | 0.158 | 1.85E-111 | TAM.prolif | Tmem161a |
| Dis3l | 3.67E-115 | 0.68411565 | 0.266 | 0.125 | 7.10E-111 | TAM.prolif | Dis3l |
| Hccs | 4.46E-115 | 0.30295095 | 0.373 | 0.223 | 8.63E-111 | TAM.prolif | Hccs |
| Crtap1 | 5.14E-115 | 0.31679954 | 0.417 | 0.264 | 9.94E-111 | TAM.prolif | Crtap |
| Cul4b | 5.33E-115 | 0.36911533 | 0.36 | 0.209 | 1.03E-110 | TAM.prolif | Cul4b |
| Med221 | 9.37E-115 | 0.47715649 | 0.356 | 0.207 | 1.81E-110 | TAM.prolif | Med22 |
| Nosip1 | 1.51E-114 | 0.38072886 | 0.409 | 0.25 | 2.93E-110 | TAM.prolif | Nosip |
| Uri1 | 1.81E-114 | 0.47608759 | 0.342 | 0.19 | 3.50E-110 | TAM.prolif | Uri1 |
| Slc13a32 | 2.39E-114 | 0.46772468 | 0.271 | 0.163 | 4.61E-110 | TAM.prolif | Slc13a3 |
| Mfsd112 | 3.46E-114 | 0.29666816 | 0.432 | 0.28 | 6.69E-110 | TAM.prolif | Mfsd11 |
| Virma | 9.66E-114 | 0.50184393 | 0.302 | 0.152 | 1.87E-109 | TAM.prolif | Virma |
| Ubfd11 | 1.29E-113 | 0.43907995 | 0.37 | 0.209 | 2.49E-109 | TAM.prolif | Ubfd1 |
| Thap71 | 1.81E-113 | 0.52516862 | 0.353 | 0.189 | 3.51E-109 | TAM.prolif | Thap7 |
| Akr1b102 | 3.69E-113 | 0.34370306 | 0.41 | 0.243 | 7.14E-109 | TAM.prolif | Akr1b10 |
| Irf31 | 5.30E-113 | 0.28961761 | 0.376 | 0.215 | 1.02E-108 | TAM.prolif | Irf3 |
| Mtus12 | 6.04E-113 | 0.26381161 | 0.364 | 0.206 | 1.17E-108 | TAM.prolif | Mtus1 |
| Thada | 7.19E-113 | 0.34308267 | 0.289 | 0.15 | 1.39E-108 | TAM.prolif | Thada |
| Nelfb | 7.48E-113 | 0.26426767 | 0.438 | 0.273 | 1.45E-108 | TAM.prolif | Nelfb |
| Pex19 | 1.60E-112 | 0.36019834 | 0.395 | 0.242 | 3.10E-108 | TAM.prolif | Pex19 |
| Fopnl | 1.67E-112 | 0.80142279 | 0.251 | 0.116 | 3.22E-108 | TAM.prolif | Fopnl |
| Cnep1r1 | 1.70E-112 | 0.36958799 | 0.358 | 0.199 | 3.28E-108 | TAM.prolif | Cnep1r1 |
| Ddhd21 | 1.85E-112 | 0.34027732 | 0.361 | 0.206 | 3.59E-108 | TAM.prolif | Ddhd2 |
| Elp3 | 3.02E-112 | 0.69324746 | 0.253 | 0.111 | 5.84E-108 | TAM.prolif | Elp3 |
| Cc2d1a | 3.22E-112 | 0.33561284 | 0.374 | 0.214 | 6.23E-108 | TAM.prolif | Cc2d1a |
| Mto1 | 3.40E-112 | 0.50742684 | 0.276 | 0.134 | 6.58E-108 | TAM.prolif | Mto1 |
| Scfd1 | 3.97E-112 | 0.33247703 | 0.354 | 0.21 | 7.67E-108 | TAM.prolif | Scfd1 |
| Tnfaip1 | 7.20E-112 | 0.48937354 | 0.286 | 0.146 | 1.39E-107 | TAM.prolif | Tnfaip1 |
| Gtf3c4 | 7.62E-112 | 0.56852296 | 0.275 | 0.138 | 1.47E-107 | TAM.prolif | Gtf3c4 |
| Mmadhc | 1.21E-111 | 0.32986485 | 0.377 | 0.228 | 2.34E-107 | TAM.prolif | Mmadhc |
| Mak16 | 1.84E-111 | 0.37104596 | 0.326 | 0.186 | 3.55E-107 | TAM.prolif | Mak16 |
| Ppil4 | 2.70E-111 | 0.32580678 | 0.356 | 0.192 | 5.22E-107 | TAM.prolif | Ppil4 |
| Dhx30 | 2.77E-111 | 0.47036455 | 0.284 | 0.145 | 5.36E-107 | TAM.prolif | Dhx30 |
| Ccdc117 | 4.89E-111 | 0.3791128 | 0.328 | 0.18 | 9.46E-107 | TAM.prolif | Ccdc117 |
| Tysnd1 | 1.30E-110 | 0.63129261 | 0.292 | 0.146 | 2.52E-106 | TAM.prolif | Tysnd1 |
| Nom1 | 1.60E-110 | 0.43402569 | 0.294 | 0.155 | 3.10E-106 | TAM.prolif | Nom1 |
| Minpp1 | 4.86E-110 | 0.42235374 | 0.306 | 0.162 | 9.40E-106 | TAM.prolif | Minpp1 |
| Spryd3 | 5.01E-110 | 0.27277645 | 0.447 | 0.28 | 9.69E-106 | TAM.prolif | Spryd3 |
| Zdhhc4 | 1.02E-109 | 0.41707939 | 0.365 | 0.208 | 1.97E-105 | TAM.prolif | Zdhhc4 |
| Eif3e | 1.57E-109 | 0.37267641 | 0.338 | 0.204 | 3.04E-105 | TAM.prolif | Eif3e |
| Sh3pxd2a1 | 1.74E-109 | 0.31876039 | 0.343 | 0.194 | 3.36E-105 | TAM.prolif | Sh3pxd2a |
| Ing5 | 4.85E-109 | 0.49089187 | 0.261 | 0.127 | 9.38E-105 | TAM.prolif | Ing5 |
| Actr10 | 6.06E-109 | 0.27195732 | 0.43 | 0.274 | 1.17E-104 | TAM.prolif | Actr10 |
| Stk16 | 1.26E-108 | 0.27748956 | 0.377 | 0.223 | 2.45E-104 | TAM.prolif | Stk16 |
| D10Wsu102e | 4.93E-108 | 0.27242796 | 0.366 | 0.213 | 9.54E-104 | TAM.prolif | D10Wsu102e |
| Galnt10 | 5.61E-108 | 0.26093772 | 0.37 | 0.218 | 1.08E-103 | TAM.prolif | Galnt10 |
| Psmb31 | 6.25E-108 | 0.35221669 | 0.373 | 0.227 | 1.21E-103 | TAM.prolif | Psmb3 |
| Pkp4 | 1.46E-107 | 0.67401186 | 0.252 | 0.118 | 2.82E-103 | TAM.prolif | Pkp4 |
| Eng4 | 1.46E-107 | 0.26683356 | 0.399 | 0.262 | 2.82E-103 | TAM.prolif | Eng |
| Prr14l | 1.48E-107 | 0.27399806 | 0.38 | 0.229 | 2.86E-103 | TAM.prolif | Prr14l |
| Exoc32 | 1.54E-107 | 0.3710573 | 0.372 | 0.222 | 2.97E-103 | TAM.prolif | Exoc3 |
| Oxsr1 | 1.55E-107 | 0.43135969 | 0.374 | 0.22 | 3.00E-103 | TAM.prolif | Oxsr1 |
| Dus3l | 3.00E-107 | 0.29155337 | 0.409 | 0.252 | 5.81E-103 | TAM.prolif | Dus3l |
| Triobp | 4.05E-107 | 0.34106432 | 0.394 | 0.234 | 7.84E-103 | TAM.prolif | Triobp |
| Sfxn5 | 7.40E-107 | 0.50402306 | 0.272 | 0.142 | 1.43E-102 | TAM.prolif | Sfxn5 |
| Riok2 | 9.93E-107 | 0.41986303 | 0.323 | 0.175 | 1.92E-102 | TAM.prolif | Riok2 |
| Bad1 | 1.09E-106 | 0.25728271 | 0.481 | 0.328 | 2.10E-102 | TAM.prolif | Bad |
| Eri32 | 1.29E-106 | 0.30936719 | 0.348 | 0.197 | 2.49E-102 | TAM.prolif | Eri3 |
| Hacd31 | 1.99E-106 | 0.52175668 | 0.342 | 0.199 | 3.86E-102 | TAM.prolif | Hacd3 |
| Helb | 2.13E-106 | 0.43829664 | 0.305 | 0.166 | 4.13E-102 | TAM.prolif | Helb |
| Wdr55 | 2.35E-106 | 0.54810933 | 0.297 | 0.156 | 4.54E-102 | TAM.prolif | Wdr55 |
| Ift46 | 6.34E-106 | 0.43123604 | 0.323 | 0.178 | 1.23E-101 | TAM.prolif | Ift46 |
| Galk12 | 6.58E-106 | 0.3795597 | 0.399 | 0.234 | 1.27E-101 | TAM.prolif | Galk1 |
| Rce11 | 1.51E-105 | 0.29670316 | 0.371 | 0.22 | 2.92E-101 | TAM.prolif | Rce1 |
| Galns2 | 1.55E-105 | 0.30737791 | 0.395 | 0.241 | 3.00E-101 | TAM.prolif | Galns |
| Pnpt1 | 2.04E-105 | 0.38830345 | 0.27 | 0.134 | 3.95E-101 | TAM.prolif | Pnpt1 |
| Klhl181 | 2.08E-105 | 0.44677352 | 0.324 | 0.175 | 4.03E-101 | TAM.prolif | Klhl18 |
| Trnau1ap | 4.81E-105 | 0.30402567 | 0.361 | 0.21 | 9.30E-101 | TAM.prolif | Trnau1ap |
| Chmp61 | 5.60E-105 | 0.30940964 | 0.366 | 0.219 | 1.08E-100 | TAM.prolif | Chmp6 |
| Nae1 | 2.20E-104 | 0.60240259 | 0.265 | 0.135 | 4.25E-100 | TAM.prolif | Nae1 |
| Dexi1 | 2.31E-104 | 0.54954286 | 0.273 | 0.133 | 4.47E-100 | TAM.prolif | Dexi1 |
| Cactin | 2.87E-104 | 0.71653854 | 0.268 | 0.128 | 5.55E-100 | TAM.prolif | Cactin |
| Ccdc59 | 4.10E-104 | 0.27461309 | 0.402 | 0.237 | 7.93E-100 | TAM.prolif | Ccdc59 |
| Dars2 | 4.81E-104 | 0.48180687 | 0.256 | 0.108 | 9.30E-100 | TAM.prolif | Dars2 |
| Pml2 | 5.46E-104 | 0.32585952 | 0.298 | 0.163 | 1.06E-99 | TAM.prolif | Pml |
| BC003965 | 6.92E-104 | 0.67427091 | 0.282 | 0.133 | 1.34E-99 | TAM.prolif | BC003965 |
| Polr2d | 1.54E-103 | 0.58666275 | 0.267 | 0.13 | 2.98E-99 | TAM.prolif | Polr2d |
| Cstf2t1 | 1.77E-103 | 0.31794139 | 0.34 | 0.191 | 3.42E-99 | TAM.prolif | Cstf2t |
| Rtf1 | 1.93E-103 | 0.37871717 | 0.349 | 0.2 | 3.73E-99 | TAM.prolif | Rtf1 |
| Sh3glb21 | 2.16E-103 | 0.28310062 | 0.405 | 0.252 | 4.18E-99 | TAM.prolif | Sh3glb2 |
| Gpn3 | 5.37E-103 | 0.44562926 | 0.319 | 0.166 | 1.04E-98 | TAM.prolif | Gpn3 |
| Tmem1681 | 5.84E-103 | 0.27739174 | 0.355 | 0.219 | 1.13E-98 | TAM.prolif | Tmem168 |
| Cops4 | 9.08E-103 | 0.29018944 | 0.349 | 0.204 | 1.76E-98 | TAM.prolif | Cops4 |
| Dera | 9.23E-103 | 0.27508301 | 0.342 | 0.195 | 1.78E-98 | TAM.prolif | Dera |
| Rbm45 | 1.20E-102 | 0.39725589 | 0.317 | 0.173 | 2.32E-98 | TAM.prolif | Rbm45 |
| Wdr48 | 2.04E-102 | 0.29000771 | 0.325 | 0.188 | 3.95E-98 | TAM.prolif | Wdr48 |
| Ppip5k2 | 2.31E-102 | 0.34132602 | 0.319 | 0.182 | 4.47E-98 | TAM.prolif | Ppip5k2 |
| Pym1 | 2.34E-102 | 0.51007019 | 0.275 | 0.136 | 4.53E-98 | TAM.prolif | Pym1 |
| Eif1ax1 | 2.47E-102 | 0.36031328 | 0.322 | 0.186 | 4.78E-98 | TAM.prolif | Eif1ax |
| Dnlz | 4.75E-102 | 0.59784992 | 0.27 | 0.135 | 9.19E-98 | TAM.prolif | Dnlz |
| Rnf71 | 4.92E-102 | 0.28575428 | 0.402 | 0.246 | 9.51E-98 | TAM.prolif | Rnf7 |
| Ttyh22 | 5.44E-102 | 0.30649993 | 0.427 | 0.285 | 1.05E-97 | TAM.prolif | Ttyh2 |
| Zfp410 | 8.74E-102 | 0.4269616 | 0.321 | 0.17 | 1.69E-97 | TAM.prolif | Zfp410 |
| Tceanc21 | 1.00E-101 | 0.31966 | 0.341 | 0.192 | 1.93E-97 | TAM.prolif | Tceanc2 |
| Ndufb2 | 1.02E-101 | 0.46960052 | 0.309 | 0.166 | 1.97E-97 | TAM.prolif | Ndufb2 |
| Ctr9 | 1.06E-101 | 0.57098097 | 0.299 | 0.153 | 2.05E-97 | TAM.prolif | Ctr9 |
| Unk | 1.25E-101 | 0.40927065 | 0.284 | 0.147 | 2.42E-97 | TAM.prolif | Unk |
| Akap11 | 1.92E-101 | 0.45458688 | 0.281 | 0.149 | 3.72E-97 | TAM.prolif | Akap11 |
| Mphosph6 | 2.40E-101 | 0.62401523 | 0.3 | 0.157 | 4.64E-97 | TAM.prolif | Mphosph6 |
| Krr1 | 2.77E-101 | 0.41823897 | 0.28 | 0.143 | 5.36E-97 | TAM.prolif | Krr1 |
| Jup2 | 5.51E-101 | 0.27088796 | 0.394 | 0.26 | 1.07E-96 | TAM.prolif | Jup |
| Pnpo1 | 6.70E-101 | 0.3125446 | 0.404 | 0.254 | 1.30E-96 | TAM.prolif | Pnpo |
| Zfp593 | 7.66E-101 | 0.55421028 | 0.25 | 0.12 | 1.48E-96 | TAM.prolif | Zfp593 |
| Kctd20 | 9.90E-101 | 0.31197507 | 0.362 | 0.209 | 1.91E-96 | TAM.prolif | Kctd20 |
| Cnpy4 | 1.09E-100 | 0.3606621 | 0.3 | 0.16 | 2.11E-96 | TAM.prolif | Cnpy4 |
| Nop14 | 2.11E-100 | 0.46082956 | 0.259 | 0.129 | 4.08E-96 | TAM.prolif | Nop14 |
| Etfdh | 2.12E-100 | 0.36557557 | 0.336 | 0.194 | 4.10E-96 | TAM.prolif | Etfdh |
| Chordc11 | 4.33E-100 | 0.33884688 | 0.338 | 0.199 | 8.37E-96 | TAM.prolif | Chordc1 |
| Eef1akmt1 | 4.36E-100 | 0.48117913 | 0.303 | 0.16 | 8.43E-96 | TAM.prolif | Eef1akmt1 |
| Apeh1 | 4.65E-100 | 0.26792242 | 0.414 | 0.268 | 9.00E-96 | TAM.prolif | Apeh |
| Aen | 9.01E-100 | 0.40937316 | 0.27 | 0.139 | 1.74E-95 | TAM.prolif | Aen |
| Slc7a6 | 1.03E-99 | 0.3894779 | 0.258 | 0.132 | 1.99E-95 | TAM.prolif | Slc7a6 |
| Mlycd | 1.09E-99 | 0.46473801 | 0.281 | 0.144 | 2.10E-95 | TAM.prolif | Mlycd |
| Ewsr1 | 1.99E-99 | 0.29138086 | 0.338 | 0.191 | 3.85E-95 | TAM.prolif | Ewsr1 |
| Taf11 | 2.00E-99 | 0.47864847 | 0.342 | 0.191 | 3.86E-95 | TAM.prolif | Taf11 |
| Zfp3261 | 2.71E-99 | 0.25086373 | 0.371 | 0.233 | 5.24E-95 | TAM.prolif | Zfp326 |
| Zfp280d | 6.01E-99 | 0.60140061 | 0.259 | 0.124 | 1.16E-94 | TAM.prolif | Zfp280d |
| Ago12 | 7.81E-99 | 0.34478159 | 0.403 | 0.247 | 1.51E-94 | TAM.prolif | Ago1 |
| Slc50a11 | 1.52E-98 | 0.33187271 | 0.342 | 0.204 | 2.94E-94 | TAM.prolif | Slc50a1 |
| Xxylt1 | 3.37E-98 | 0.29148742 | 0.283 | 0.156 | 6.51E-94 | TAM.prolif | Xxylt1 |
| Coq2 | 3.82E-98 | 0.2906965 | 0.302 | 0.168 | 7.38E-94 | TAM.prolif | Coq2 |
| Arhgap221 | 6.01E-98 | 0.39238178 | 0.299 | 0.16 | 1.16E-93 | TAM.prolif | Arhgap22 |
| Utp4 | 2.17E-97 | 0.31997198 | 0.307 | 0.179 | 4.19E-93 | TAM.prolif | Utp4 |
| Dhx33 | 3.16E-97 | 0.35233558 | 0.316 | 0.179 | 6.11E-93 | TAM.prolif | Dhx33 |
| Pde6d | 4.83E-97 | 0.50818202 | 0.257 | 0.129 | 9.34E-93 | TAM.prolif | Pde6d |
| Ccdc43 | 5.31E-97 | 0.41252498 | 0.317 | 0.18 | 1.03E-92 | TAM.prolif | Ccdc43 |
| Scfd2 | 5.72E-97 | 0.49072592 | 0.277 | 0.138 | 1.11E-92 | TAM.prolif | Scfd2 |
| Wbp1 | 9.38E-97 | 0.47238318 | 0.312 | 0.17 | 1.81E-92 | TAM.prolif | Wbp1 |
| Letm1 | 1.02E-96 | 0.33367903 | 0.335 | 0.187 | 1.98E-92 | TAM.prolif | Letm1 |
| Rwdd1 | 2.56E-96 | 0.4624932 | 0.289 | 0.159 | 4.96E-92 | TAM.prolif | Rwdd1 |
| Xpot | 4.29E-96 | 0.50286851 | 0.29 | 0.157 | 8.29E-92 | TAM.prolif | Xpot |
| Cd276 | 4.68E-96 | 0.30162527 | 0.275 | 0.166 | 9.05E-92 | TAM.prolif | Cd276 |
| Fam122a | 7.96E-96 | 0.40544598 | 0.284 | 0.155 | 1.54E-91 | TAM.prolif | Fam122a |
| Slc39a6 | 5.73E-95 | 0.35957574 | 0.277 | 0.148 | 1.11E-90 | TAM.prolif | Slc39a6 |
| Mzt2 | 5.81E-95 | 0.52555753 | 0.252 | 0.122 | 1.12E-90 | TAM.prolif | Mzt2 |
| Znhit2 | 6.21E-95 | 0.48337363 | 0.26 | 0.131 | 1.20E-90 | TAM.prolif | Znhit2 |
| Fech1 | 7.88E-95 | 0.34103005 | 0.355 | 0.22 | 1.52E-90 | TAM.prolif | Fech |
| Ttc37 | 8.50E-95 | 0.37633333 | 0.279 | 0.145 | 1.64E-90 | TAM.prolif | Ttc37 |
| Pus10 | 1.23E-94 | 0.36362876 | 0.278 | 0.149 | 2.37E-90 | TAM.prolif | Pus10 |
| Ssbp1 | 1.36E-94 | 0.36185534 | 0.344 | 0.207 | 2.64E-90 | TAM.prolif | Ssbp1 |
| Leo1 | 1.38E-94 | 0.5707914 | 0.258 | 0.128 | 2.67E-90 | TAM.prolif | Leo1 |
| Polr3e | 1.98E-94 | 0.51341843 | 0.266 | 0.135 | 3.83E-90 | TAM.prolif | Polr3e |
| Rgs32 | 4.07E-94 | 0.33432971 | 0.311 | 0.197 | 7.87E-90 | TAM.prolif | Rgs3 |
| Ipo8 | 4.96E-94 | 0.31140717 | 0.366 | 0.218 | 9.59E-90 | TAM.prolif | Ipo8 |
| Rpp25l | 5.36E-94 | 0.61389326 | 0.264 | 0.127 | 1.04E-89 | TAM.prolif | Rpp25l |
| Lrrc45 | 5.55E-94 | 0.5689105 | 0.25 | 0.116 | 1.07E-89 | TAM.prolif | Lrrc45 |
| Aak1 | 1.06E-93 | 0.40680925 | 0.312 | 0.172 | 2.04E-89 | TAM.prolif | Aak1 |
| Aspscr1 | 1.19E-93 | 0.26564326 | 0.393 | 0.231 | 2.30E-89 | TAM.prolif | Aspscr1 |
| Ctu2 | 1.25E-93 | 0.36592518 | 0.287 | 0.152 | 2.41E-89 | TAM.prolif | Ctu2 |
| Hps51 | 1.28E-93 | 0.34986473 | 0.343 | 0.197 | 2.47E-89 | TAM.prolif | Hps5 |
| Thap11 | 1.65E-93 | 0.36992684 | 0.343 | 0.194 | 3.19E-89 | TAM.prolif | Thap11 |
| Palm | 1.78E-93 | 0.35913995 | 0.293 | 0.173 | 3.43E-89 | TAM.prolif | Palm |
| Mtx1 | 2.06E-93 | 0.6788038 | 0.25 | 0.123 | 3.99E-89 | TAM.prolif | Mtx1 |
| Gabpb1 | 2.25E-93 | 0.38530724 | 0.262 | 0.141 | 4.35E-89 | TAM.prolif | Gabpb1 |

| | | | | | | | |
|---|---|---|---|---|---|---|---|
| Aim22 | 3.13E-93 | 0.31598272 | 0.363 | 0.227 | 6.06E-89 | TAM.prolif | Aim2 |
| Dok12 | 3.33E-93 | 0.29768801 | 0.409 | 0.258 | 6.43E-89 | TAM.prolif | Dok1 |
| Ormdl2 | 4.92E-93 | 0.49950303 | 0.281 | 0.146 | 9.51E-89 | TAM.prolif | Ormdl2 |
| Nat10 | 5.38E-93 | 0.61370154 | 0.261 | 0.129 | 1.04E-88 | TAM.prolif | Nat10 |
| Fam172a | 5.48E-93 | 0.25098038 | 0.351 | 0.208 | 1.06E-88 | TAM.prolif | Fam172a |
| Cep63 | 5.91E-93 | 0.29978626 | 0.296 | 0.163 | 1.14E-88 | TAM.prolif | Cep63 |
| Nsmce3 | 6.22E-93 | 0.49989871 | 0.312 | 0.17 | 1.20E-88 | TAM.prolif | Nsmce3 |
| Commd2 | 1.59E-92 | 0.41102705 | 0.317 | 0.179 | 3.07E-88 | TAM.prolif | Commd2 |
| Nol10 | 1.97E-92 | 0.37911569 | 0.25 | 0.123 | 3.80E-88 | TAM.prolif | Nol10 |
| Nelfa | 2.22E-92 | 0.29706016 | 0.314 | 0.179 | 4.30E-88 | TAM.prolif | Nelfa |
| Uxt | 8.65E-92 | 0.43073299 | 0.266 | 0.141 | 1.67E-87 | TAM.prolif | Uxt |
| Tmem126a | 1.03E-91 | 0.45102719 | 0.284 | 0.159 | 1.99E-87 | TAM.prolif | Tmem126a |
| Nedd4l | 1.13E-91 | 0.36996602 | 0.314 | 0.178 | 2.18E-87 | TAM.prolif | Nedd4l |
| Rbm34 | 2.20E-91 | 0.52623642 | 0.259 | 0.133 | 4.26E-87 | TAM.prolif | Rbm34 |
| Ttc1 | 2.23E-91 | 0.26247826 | 0.404 | 0.26 | 4.32E-87 | TAM.prolif | Ttc1 |
| Fance1 | 3.51E-91 | 0.39147505 | 0.316 | 0.171 | 6.79E-87 | TAM.prolif | Fance |
| Dhx36 | 4.10E-91 | 0.25810014 | 0.381 | 0.236 | 7.92E-87 | TAM.prolif | Dhx36 |
| Bet11 | 5.20E-91 | 0.31763842 | 0.367 | 0.221 | 1.01E-86 | TAM.prolif | Bet1 |
| Tor1b1 | 6.74E-91 | 0.35003848 | 0.334 | 0.195 | 1.30E-86 | TAM.prolif | Tor1b |
| Pdrg11 | 1.19E-90 | 0.28314967 | 0.356 | 0.219 | 2.30E-86 | TAM.prolif | Pdrg1 |
| Gpatch11 | 1.28E-90 | 0.53765333 | 0.273 | 0.138 | 2.48E-86 | TAM.prolif | Gpatch11 |
| Naa25 | 2.23E-90 | 0.30042694 | 0.301 | 0.172 | 4.30E-86 | TAM.prolif | Naa25 |
| Mief1 | 2.37E-90 | 0.41795341 | 0.286 | 0.149 | 4.58E-86 | TAM.prolif | Mief1 |
| Comtd11 | 3.24E-90 | 0.30814296 | 0.316 | 0.191 | 6.27E-86 | TAM.prolif | Comtd1 |
| Cnnm3 | 3.99E-90 | 0.37969249 | 0.282 | 0.152 | 7.71E-86 | TAM.prolif | Cnnm3 |
| Cnp | 5.33E-90 | 0.37160957 | 0.316 | 0.177 | 1.03E-85 | TAM.prolif | Cnp |
| G6pc31 | 7.38E-90 | 0.26903867 | 0.394 | 0.234 | 1.43E-85 | TAM.prolif | G6pc3 |
| Mtmr1 | 8.53E-90 | 0.47193571 | 0.266 | 0.14 | 1.65E-85 | TAM.prolif | Mtmr1 |
| Acad8 | 8.68E-90 | 0.53971687 | 0.262 | 0.135 | 1.68E-85 | TAM.prolif | Acad8 |
| Mapkap1 | 2.27E-89 | 0.27021219 | 0.377 | 0.235 | 4.39E-85 | TAM.prolif | Mapkap1 |
| Bms1 | 4.66E-89 | 0.53719143 | 0.27 | 0.137 | 9.01E-85 | TAM.prolif | Bms1 |
| Msmo11 | 7.63E-89 | 0.28082153 | 0.367 | 0.245 | 1.48E-84 | TAM.prolif | Msmo1 |
| Tmem222 | 8.31E-89 | 0.26237701 | 0.348 | 0.202 | 1.61E-84 | TAM.prolif | Tmem222 |
| Ddx521 | 9.13E-89 | 0.34284988 | 0.283 | 0.151 | 1.76E-84 | TAM.prolif | Ddx52 |
| Micu11 | 1.12E-88 | 0.29102614 | 0.392 | 0.242 | 2.16E-84 | TAM.prolif | Micu1 |
| Klhl6 | 1.57E-88 | 0.45933354 | 0.252 | 0.132 | 3.03E-84 | TAM.prolif | Klhl6 |
| Gon4l | 1.87E-88 | 0.30562717 | 0.312 | 0.172 | 3.62E-84 | TAM.prolif | Gon4l |
| Polr2h | 1.91E-88 | 0.36139622 | 0.273 | 0.152 | 3.69E-84 | TAM.prolif | Polr2h |
| Lancl11 | 2.38E-88 | 0.49252131 | 0.273 | 0.134 | 4.61E-84 | TAM.prolif | Lancl1 |
| Nfu1 | 3.00E-88 | 0.59099795 | 0.287 | 0.15 | 5.80E-84 | TAM.prolif | Nfu1 |
| Mavs1 | 1.03E-87 | 0.3351536 | 0.315 | 0.184 | 1.99E-83 | TAM.prolif | Mavs |
| Alkbh1 | 1.04E-87 | 0.33423873 | 0.283 | 0.153 | 2.02E-83 | TAM.prolif | Alkbh1 |
| Ikzf5 | 1.47E-87 | 0.28572687 | 0.305 | 0.169 | 2.85E-83 | TAM.prolif | Ikzf5 |
| Lcmt1 | 1.68E-87 | 0.486024 | 0.304 | 0.17 | 3.25E-83 | TAM.prolif | Lcmt1 |
| Aldh1b11 | 1.82E-87 | 0.40085228 | 0.305 | 0.169 | 3.53E-83 | TAM.prolif | Aldh1b1 |
| Asb82 | 1.98E-87 | 0.2892307 | 0.339 | 0.198 | 3.83E-83 | TAM.prolif | Asb8 |
| Smarca2 | 2.08E-87 | 0.37111674 | 0.302 | 0.174 | 4.02E-83 | TAM.prolif | Smarca2 |
| Trim65 | 2.90E-87 | 0.56670766 | 0.251 | 0.126 | 5.61E-83 | TAM.prolif | Trim65 |
| Men1 | 3.71E-87 | 0.44138123 | 0.279 | 0.151 | 7.18E-83 | TAM.prolif | Men1 |
| Pex2 | 5.58E-87 | 0.25093785 | 0.324 | 0.196 | 1.08E-82 | TAM.prolif | Pex2 |
| Gna152 | 8.94E-87 | 0.31389608 | 0.382 | 0.244 | 1.73E-82 | TAM.prolif | Gna15 |
| Mbd1 | 9.06E-87 | 0.31639777 | 0.339 | 0.2 | 1.75E-82 | TAM.prolif | Mbd1 |
| Fam217b1 | 9.75E-87 | 0.41850994 | 0.283 | 0.164 | 1.89E-82 | TAM.prolif | Fam217b |
| Cc2d1b | 1.87E-86 | 0.44413121 | 0.303 | 0.162 | 3.61E-82 | TAM.prolif | Cc2d1b |
| Sugp2 | 3.68E-86 | 0.4243977 | 0.279 | 0.148 | 7.12E-82 | TAM.prolif | Sugp2 |
| Tusc32 | 4.61E-86 | 0.38182588 | 0.34 | 0.205 | 8.91E-82 | TAM.prolif | Tusc3 |
| Zfyve261 | 6.45E-86 | 0.2779856 | 0.362 | 0.224 | 1.25E-81 | TAM.prolif | Zfyve26 |
| Nrf1 | 8.66E-86 | 0.2632226 | 0.255 | 0.13 | 1.67E-81 | TAM.prolif | Nrf1 |
| Ap4m11 | 1.46E-85 | 0.30812655 | 0.321 | 0.187 | 2.83E-81 | TAM.prolif | Ap4m1 |
| Dag12 | 1.60E-85 | 0.31813605 | 0.376 | 0.239 | 3.09E-81 | TAM.prolif | Dag1 |
| Cyp20a1 | 1.84E-85 | 0.48734247 | 0.272 | 0.155 | 3.56E-81 | TAM.prolif | Cyp20a1 |
| Smim12 | 2.42E-85 | 0.26270957 | 0.321 | 0.197 | 4.68E-81 | TAM.prolif | Smim12 |
| Slc10a7 | 3.24E-85 | 0.29753962 | 0.261 | 0.146 | 6.26E-81 | TAM.prolif | Slc10a7 |
| Sc5d | 6.29E-85 | 0.52033147 | 0.251 | 0.147 | 1.22E-80 | TAM.prolif | Sc5d |
| Mms192 | 7.20E-85 | 0.26163054 | 0.339 | 0.201 | 1.39E-80 | TAM.prolif | Mms19 |
| Prkab11 | 1.05E-84 | 0.30163365 | 0.283 | 0.161 | 2.02E-80 | TAM.prolif | Prkab1 |
| Gnptg2 | 1.26E-84 | 0.31861269 | 0.346 | 0.208 | 2.44E-80 | TAM.prolif | Gnptg |
| Fam117a | 1.75E-84 | 0.41356636 | 0.268 | 0.147 | 3.39E-80 | TAM.prolif | Fam117a |
| Twistnb | 3.14E-84 | 0.31607123 | 0.267 | 0.15 | 6.06E-80 | TAM.prolif | Twistnb |
| Gnl2 | 3.67E-84 | 0.27178293 | 0.3 | 0.173 | 7.09E-80 | TAM.prolif | Gnl2 |
| Pde4dip | 6.34E-84 | 0.32692945 | 0.277 | 0.162 | 1.23E-79 | TAM.prolif | Pde4dip |
| Agfg21 | 1.26E-83 | 0.38090709 | 0.316 | 0.187 | 2.43E-79 | TAM.prolif | Agfg2 |
| Rgmb2 | 1.30E-83 | 0.3338701 | 0.344 | 0.216 | 2.52E-79 | TAM.prolif | Rgmb |
| Pank4 | 1.33E-83 | 0.33750298 | 0.259 | 0.136 | 2.57E-79 | TAM.prolif | Pank4 |
| Cyb561a3 | 3.01E-83 | 0.4135508 | 0.289 | 0.172 | 5.83E-79 | TAM.prolif | Cyb561a3 |
| Pctp | 4.30E-83 | 0.51534295 | 0.292 | 0.155 | 8.31E-79 | TAM.prolif | Pctp |
| Myo9a1 | 5.71E-83 | 0.31255557 | 0.313 | 0.188 | 1.10E-78 | TAM.prolif | Myo9a |
| Immp1l | 6.96E-83 | 0.4981378 | 0.267 | 0.142 | 1.34E-78 | TAM.prolif | Immp1l |
| Ntan1 | 7.79E-83 | 0.37350007 | 0.297 | 0.167 | 1.51E-78 | TAM.prolif | Ntan1 |
| Aatf | 1.17E-82 | 0.39368343 | 0.256 | 0.138 | 2.26E-78 | TAM.prolif | Aatf |
| Ankrd10 | 1.30E-82 | 0.32784666 | 0.304 | 0.179 | 2.51E-78 | TAM.prolif | Ankrd10 |
| Mtpap | 1.66E-82 | 0.46585491 | 0.288 | 0.163 | 3.21E-78 | TAM.prolif | Mtpap |
| Prpsap2 | 2.82E-82 | 0.34119156 | 0.268 | 0.141 | 5.45E-78 | TAM.prolif | Prpsap2 |
| Ei24 | 3.00E-82 | 0.39718086 | 0.307 | 0.176 | 5.81E-78 | TAM.prolif | Ei24 |
| Bap1 | 5.07E-82 | 0.31354953 | 0.311 | 0.187 | 9.81E-78 | TAM.prolif | Bap1 |
| Vps13b | 7.61E-82 | 0.29201023 | 0.293 | 0.168 | 1.47E-77 | TAM.prolif | Vps13b |
| Eif2b4 | 8.08E-82 | 0.36980295 | 0.261 | 0.139 | 1.56E-77 | TAM.prolif | Eif2b4 |
| Hint2 | 2.20E-81 | 0.50246711 | 0.254 | 0.134 | 4.26E-77 | TAM.prolif | Hint2 |
| Wdr3 | 3.02E-81 | 0.25528853 | 0.314 | 0.185 | 5.83E-77 | TAM.prolif | Wdr3 |
| Rab28 | 3.68E-81 | 0.26516809 | 0.383 | 0.235 | 7.11E-77 | TAM.prolif | Rab28 |
| Med17 | 1.08E-80 | 0.45812999 | 0.277 | 0.148 | 2.09E-76 | TAM.prolif | Med17 |
| Romo1 | 2.11E-80 | 0.30346504 | 0.373 | 0.238 | 4.08E-76 | TAM.prolif | Romo1 |
| Tsnax | 2.41E-80 | 0.36008996 | 0.281 | 0.157 | 4.65E-76 | TAM.prolif | Tsnax |
| Rmdn3 | 8.72E-80 | 0.54293766 | 0.266 | 0.137 | 1.69E-75 | TAM.prolif | Rmdn3 |
| Pdcd5 | 9.58E-80 | 0.37475035 | 0.29 | 0.161 | 1.85E-75 | TAM.prolif | Pdcd5 |
| Alg5 | 9.66E-80 | 0.29123467 | 0.311 | 0.177 | 1.87E-75 | TAM.prolif | Alg5 |
| Aacs | 1.37E-79 | 0.38744814 | 0.298 | 0.175 | 2.64E-75 | TAM.prolif | Aacs |
| Eif3m | 1.55E-79 | 0.39537068 | 0.273 | 0.149 | 2.99E-75 | TAM.prolif | Eif3m |
| Pdpr | 2.05E-79 | 0.43677773 | 0.266 | 0.139 | 3.96E-75 | TAM.prolif | Pdpr |
| Rnf138 | 2.66E-79 | 0.26976382 | 0.328 | 0.201 | 5.14E-75 | TAM.prolif | Rnf138 |
| Polr2i | 3.46E-79 | 0.4382815 | 0.266 | 0.147 | 6.68E-75 | TAM.prolif | Polr2i |
| Gatd11 | 8.53E-79 | 0.26650516 | 0.352 | 0.219 | 1.65E-74 | TAM.prolif | Gatd1 |
| Nsrp1 | 1.18E-78 | 0.26372075 | 0.256 | 0.139 | 2.29E-74 | TAM.prolif | Nsrp1 |
| Pdhx | 1.99E-78 | 0.40768504 | 0.254 | 0.133 | 3.85E-74 | TAM.prolif | Pdhx |
| Triap11 | 3.12E-78 | 0.36393773 | 0.311 | 0.181 | 6.03E-74 | TAM.prolif | Triap1 |
| Uba6 | 4.26E-78 | 0.33088716 | 0.263 | 0.144 | 8.24E-74 | TAM.prolif | Uba6 |
| Wdr61 | 9.91E-78 | 0.28211121 | 0.284 | 0.163 | 1.92E-73 | TAM.prolif | Wdr61 |
| Ppp2r2d | 1.08E-77 | 0.37080363 | 0.251 | 0.142 | 2.09E-73 | TAM.prolif | Ppp2r2d |
| Cul2 | 1.00E-76 | 0.33682652 | 0.272 | 0.158 | 1.94E-72 | TAM.prolif | Cul2 |
| Rwdd4a | 1.29E-76 | 0.41570475 | 0.265 | 0.147 | 2.49E-72 | TAM.prolif | Rwdd4a |
| Mrs2 | 1.61E-76 | 0.43453912 | 0.294 | 0.161 | 3.12E-72 | TAM.prolif | Mrs2 |
| Washc31 | 2.51E-76 | 0.26068995 | 0.392 | 0.255 | 4.86E-72 | TAM.prolif | Washc3 |
| Yif1a | 3.18E-76 | 0.2757032 | 0.321 | 0.192 | 6.15E-72 | TAM.prolif | Yif1a |
| Icmt | 4.35E-76 | 0.33524027 | 0.335 | 0.196 | 8.42E-72 | TAM.prolif | Icmt |
| Abhd11 | 5.12E-76 | 0.30338958 | 0.309 | 0.189 | 9.90E-72 | TAM.prolif | Abhd11 |
| Bloc1s6 | 5.65E-76 | 0.54978408 | 0.287 | 0.154 | 1.09E-71 | TAM.prolif | Bloc1s6 |
| Ppp2r3c | 6.19E-76 | 0.4189105 | 0.256 | 0.15 | 1.20E-71 | TAM.prolif | Ppp2r3c |
| Cetn2 | 7.32E-76 | 0.4066043 | 0.272 | 0.148 | 1.42E-71 | TAM.prolif | Cetn2 |
| Plaat32 | 7.37E-76 | 0.32912737 | 0.293 | 0.18 | 1.42E-71 | TAM.prolif | Plaat3 |
| Fiz1 | 1.04E-75 | 0.29252039 | 0.289 | 0.166 | 2.01E-71 | TAM.prolif | Fiz1 |
| Pygo21 | 1.09E-75 | 0.25607902 | 0.355 | 0.218 | 2.10E-71 | TAM.prolif | Pygo2 |
| Eef2k2 | 2.17E-75 | 0.3073973 | 0.293 | 0.163 | 4.19E-71 | TAM.prolif | Eef2k |
| Vps33a2 | 9.45E-75 | 0.30532932 | 0.321 | 0.196 | 1.83E-70 | TAM.prolif | Vps33a |
| Tada3 | 1.46E-74 | 0.51527506 | 0.268 | 0.145 | 2.83E-70 | TAM.prolif | Tada3 |
| Naxd1 | 5.92E-74 | 0.35830745 | 0.346 | 0.214 | 1.14E-69 | TAM.prolif | Naxd |
| Dph3 | 1.27E-73 | 0.2937258 | 0.273 | 0.159 | 2.45E-69 | TAM.prolif | Dph3 |
| Eef1akmt4 | 3.67E-73 | 0.32380112 | 0.285 | 0.163 | 7.09E-69 | TAM.prolif | Eef1akmt4 |
| Tbcc1 | 5.81E-73 | 0.26251491 | 0.32 | 0.191 | 1.12E-68 | TAM.prolif | Tbcc |
| Snx19 | 1.29E-72 | 0.26731943 | 0.299 | 0.173 | 2.50E-68 | TAM.prolif | Snx19 |
| Abhd17c2 | 1.64E-72 | 0.3270032 | 0.336 | 0.208 | 3.18E-68 | TAM.prolif | Abhd17c |
| Nfs1 | 2.11E-72 | 0.34780222 | 0.263 | 0.143 | 4.08E-68 | TAM.prolif | Nfs1 |
| Lsm1 | 2.39E-72 | 0.39337182 | 0.254 | 0.138 | 4.62E-68 | TAM.prolif | Lsm1 |
| Bag5 | 3.01E-72 | 0.34145876 | 0.252 | 0.148 | 5.82E-68 | TAM.prolif | Bag5 |
| Smarcal1 | 3.13E-72 | 0.44540437 | 0.271 | 0.148 | 6.04E-68 | TAM.prolif | Smarcal1 |
| Ciao3 | 3.33E-72 | 0.25345854 | 0.263 | 0.143 | 6.44E-68 | TAM.prolif | Ciao3 |
| Dgcr6 | 5.03E-72 | 0.35255289 | 0.293 | 0.17 | 9.73E-68 | TAM.prolif | Dgcr6 |
| Pip4p2 | 8.93E-72 | 0.41119888 | 0.285 | 0.159 | 1.73E-67 | TAM.prolif | Pip4p2 |
| Panx1 | 1.03E-71 | 0.37363177 | 0.264 | 0.143 | 1.99E-67 | TAM.prolif | Panx1 |
| Prkci | 1.10E-71 | 0.3397945 | 0.257 | 0.141 | 2.12E-67 | TAM.prolif | Prkci |
| Zfp687 | 1.23E-71 | 0.30416363 | 0.267 | 0.147 | 2.39E-67 | TAM.prolif | Zfp687 |
| Phkg2 | 5.22E-71 | 0.25806585 | 0.28 | 0.16 | 1.01E-66 | TAM.prolif | Phkg2 |
| BC0297222 | 6.95E-71 | 0.29252246 | 0.365 | 0.23 | 1.34E-66 | TAM.prolif | BC029722 |
| Med16 | 8.38E-71 | 0.31504603 | 0.294 | 0.177 | 1.62E-66 | TAM.prolif | Med16 |
| Taf3 | 1.15E-69 | 0.41214106 | 0.252 | 0.13 | 2.23E-65 | TAM.prolif | Taf3 |
| Wdr111 | 1.63E-69 | 0.32695629 | 0.289 | 0.173 | 3.15E-65 | TAM.prolif | Wdr11 |
| Tns4 | 3.65E-69 | 0.33041375 | 0.274 | 0.157 | 7.06E-65 | TAM.prolif | Tns4 |
| Smad1 | 3.72E-69 | 0.47021607 | 0.281 | 0.165 | 7.19E-65 | TAM.prolif | Smad1 |
| B4gat11 | 7.01E-69 | 0.3644986 | 0.289 | 0.169 | 1.35E-64 | TAM.prolif | B4gat1 |
| Helz1 | 7.45E-69 | 0.27016846 | 0.323 | 0.192 | 1.44E-64 | TAM.prolif | Helz |
| Mettl3 | 1.05E-68 | 0.27082156 | 0.265 | 0.148 | 2.04E-64 | TAM.prolif | Mettl3 |
| Ggt52 | 1.37E-68 | 0.25701818 | 0.281 | 0.162 | 2.65E-64 | TAM.prolif | Ggt5 |
| Vps4a | 1.62E-68 | 0.25296955 | 0.318 | 0.194 | 3.13E-64 | TAM.prolif | Vps4a |
| Cdc26 | 1.64E-68 | 0.30594472 | 0.271 | 0.153 | 3.16E-64 | TAM.prolif | Cdc26 |
| Setx | 2.89E-68 | 0.27225799 | 0.275 | 0.153 | 5.58E-64 | TAM.prolif | Setx |
| Ttc4 | 3.12E-68 | 0.33016707 | 0.262 | 0.139 | 6.04E-64 | TAM.prolif | Ttc4 |
| Svbp | 4.48E-68 | 0.39255623 | 0.253 | 0.151 | 8.67E-64 | TAM.prolif | Svbp |
| Nkapd1 | 4.80E-68 | 0.36389705 | 0.259 | 0.148 | 9.28E-64 | TAM.prolif | Nkapd1 |
| Ankrd27 | 3.06E-67 | 0.31819978 | 0.259 | 0.154 | 5.92E-63 | TAM.prolif | Ankrd27 |
| Msra | 6.39E-67 | 0.26667469 | 0.281 | 0.17 | 1.24E-62 | TAM.prolif | Msra |
| Eif2ak3 | 1.02E-66 | 0.25504527 | 0.298 | 0.178 | 1.97E-62 | TAM.prolif | Eif2ak3 |
| Bloc1s1 | 1.77E-66 | 0.26882085 | 0.324 | 0.196 | 3.42E-62 | TAM.prolif | Bloc1s1 |
| Ppp1r35 | 1.92E-66 | 0.37988034 | 0.275 | 0.157 | 3.72E-62 | TAM.prolif | Ppp1r35 |
| Bod1l | 4.43E-66 | 0.30580096 | 0.29 | 0.17 | 8.56E-62 | TAM.prolif | Bod1l |
| Sord1 | 6.19E-66 | 0.32583233 | 0.28 | 0.164 | 1.20E-61 | TAM.prolif | Sord |
| Smim19 | 2.12E-65 | 0.34586926 | 0.255 | 0.142 | 4.09E-61 | TAM.prolif | Smim19 |
| Fig4 | 5.12E-65 | 0.25020675 | 0.27 | 0.153 | 9.90E-61 | TAM.prolif | Fig4 |
| Fbxo7 | 6.16E-65 | 0.28167916 | 0.27 | 0.16 | 1.19E-60 | TAM.prolif | Fbxo7 |
| Gtf3c1 | 1.33E-64 | 0.31968177 | 0.255 | 0.142 | 2.58E-60 | TAM.prolif | Gtf3c1 |
| Eci1 | 3.08E-64 | 0.26783067 | 0.262 | 0.155 | 5.96E-60 | TAM.prolif | Eci1 |
| Yaf2 | 3.98E-64 | 0.34091258 | 0.259 | 0.15 | 7.70E-60 | TAM.prolif | Yaf2 |
| Tstd2 | 5.40E-63 | 0.33669807 | 0.264 | 0.148 | 1.04E-58 | TAM.prolif | Tstd2 |
| Ate1 | 8.08E-63 | 0.32478895 | 0.274 | 0.162 | 1.56E-58 | TAM.prolif | Ate1 |
| Kif2a | 1.80E-62 | 0.30908628 | 0.251 | 0.141 | 3.48E-58 | TAM.prolif | Kif2a |
| Kif1bp | 4.75E-62 | 0.28931452 | 0.306 | 0.179 | 9.18E-58 | TAM.prolif | Kif1bp |
| Srek1ip1 | 1.31E-61 | 0.50204417 | 0.26 | 0.144 | 2.53E-57 | TAM.prolif | Srek1ip1 |
| Rft1 | 6.77E-61 | 0.30843107 | 0.252 | 0.144 | 1.31E-56 | TAM.prolif | Rft1 |
| Faap20 | 3.98E-60 | 0.28014065 | 0.308 | 0.192 | 7.70E-56 | TAM.prolif | Faap20 |
| Skiv2l | 6.43E-59 | 0.2746838 | 0.258 | 0.144 | 1.24E-54 | TAM.prolif | Skiv2l |
| Zadh21 | 4.66E-58 | 0.30877793 | 0.303 | 0.174 | 9.02E-54 | TAM.prolif | Zadh2 |
| Zfpl1 | 2.80E-57 | 0.30702665 | 0.267 | 0.155 | 5.42E-53 | TAM.prolif | Zfpl1 |
| Sugp1 | 3.15E-57 | 0.28650333 | 0.278 | 0.161 | 6.10E-53 | TAM.prolif | Sugp1 |
| Capn15 | 1.19E-56 | 0.31596903 | 0.261 | 0.147 | 2.31E-52 | TAM.prolif | Capn15 |
| Ppp1r16a | 1.87E-56 | 0.31379115 | 0.254 | 0.151 | 3.61E-52 | TAM.prolif | Ppp1r16a |
| Hspa13 | 4.28E-55 | 0.2561743 | 0.252 | 0.151 | 8.28E-51 | TAM.prolif | Hspa13 |
| Def8 | 9.06E-55 | 0.29850382 | 0.271 | 0.167 | 1.75E-50 | TAM.prolif | Def8 |
| Arfgef21 | 6.79E-54 | 0.25416217 | 0.274 | 0.162 | 1.31E-49 | TAM.prolif | Arfgef2 |
| Cog7 | 9.11E-54 | 0.34826589 | 0.28 | 0.167 | 1.76E-49 | TAM.prolif | Cog7 |
| Wiz | 3.02E-53 | 0.279298 | 0.267 | 0.157 | 5.84E-49 | TAM.prolif | Wiz |
| Tango2 | 1.84E-48 | 0.30523967 | 0.258 | 0.155 | 3.57E-44 | TAM.prolif | Tango2 |
| Casp9 | 1.08E-47 | 0.27176235 | 0.27 | 0.166 | 2.08E-43 | TAM.prolif | Casp9 |
| Aga | 5.10E-42 | 0.34653004 | 0.26 | 0.16 | 9.87E-38 | TAM.prolif | Aga |
| Serpinb6a2 | 0 | 1.67957855 | 0.805 | 0.324 | 0 | TAM.Gdf15 | Serpinb6a |
| Gdf152 | 0 | 2.67140771 | 0.745 | 0.266 | 0 | TAM.Gdf15 | Gdf15 |
| Pmp222 | 0 | 2.21600595 | 0.653 | 0.177 | 0 | TAM.Gdf15 | Pmp22 |
| Gclm1 | 0 | 2.48319566 | 0.757 | 0.303 | 0 | TAM.Gdf15 | Gclm |
| Cd361 | 0 | 2.62025073 | 0.608 | 0.162 | 0 | TAM.Gdf15 | Cd36 |
| Slc6a82 | 0 | 1.80938213 | 0.705 | 0.282 | 0 | TAM.Gdf15 | Slc6a8 |
| Slc48a12 | 0 | 2.28792273 | 0.873 | 0.491 | 0 | TAM.Gdf15 | Slc48a1 |
| Sep-82 | 0 | 1.49984766 | 0.769 | 0.39 | 0 | TAM.Gdf15 | 8-Sep |
| Hist1h2bc2 | 0 | 1.72309918 | 0.72 | 0.351 | 0 | TAM.Gdf15 | Hist1h2bc |
| S1pr11 | 0 | 2.05033526 | 0.503 | 0.142 | 0 | TAM.Gdf15 | S1pr1 |
| Pf42 | 0 | 1.92995786 | 0.905 | 0.549 | 0 | TAM.Gdf15 | Pf4 |
| Srxn12 | 0 | 1.68923172 | 0.728 | 0.376 | 0 | TAM.Gdf15 | Srxn1 |
| Akr1b81 | 0 | 2.78884737 | 0.473 | 0.126 | 0 | TAM.Gdf15 | Akr1b8 |
| Pdpn2 | 0 | 1.48321769 | 0.641 | 0.295 | 0 | TAM.Gdf15 | Pdpn |
| Ednrb1 | 0 | 2.58169554 | 0.428 | 0.086 | 0 | TAM.Gdf15 | Ednrb |
| Vat13 | 0 | 1.34320192 | 0.929 | 0.59 | 0 | TAM.Gdf15 | Vat1 |
| Adam83 | 0 | 1.3657481 | 0.903 | 0.565 | 0 | TAM.Gdf15 | Adam8 |
| Dnmt3l | 0 | 3.60996602 | 0.377 | 0.041 | 0 | TAM.Gdf15 | Dnmt3l |
| Htatip22 | 0 | 1.70105402 | 0.643 | 0.308 | 0 | TAM.Gdf15 | Htatip2 |
| Abcc11 | 0 | 1.3763502 | 0.738 | 0.403 | 0 | TAM.Gdf15 | Abcc1 |
| Tspan31 | 0 | 1.28968968 | 0.792 | 0.461 | 0 | TAM.Gdf15 | Tspan3 |
| Blvrb2 | 0 | 1.24987525 | 0.881 | 0.551 | 0 | TAM.Gdf15 | Blvrb |
| Ampd32 | 0 | 2.54901883 | 0.487 | 0.157 | 0 | TAM.Gdf15 | Ampd3 |
| Cgnl11 | 0 | 1.97816365 | 0.45 | 0.127 | 0 | TAM.Gdf15 | Cgnl1 |
| Plk22 | 0 | 1.79238244 | 0.721 | 0.402 | 0 | TAM.Gdf15 | Plk2 |
| Mindy12 | 0 | 1.39078351 | 0.643 | 0.325 | 0 | TAM.Gdf15 | Mindy1 |
| Ccl23 | 0 | 1.49773862 | 0.563 | 0.246 | 0 | TAM.Gdf15 | Ccl2 |
| Timp22 | 0 | 0.73283225 | 0.869 | 0.552 | 0 | TAM.Gdf15 | Timp2 |
| Cd814 | 0 | 0.67414257 | 0.829 | 0.514 | 0 | TAM.Gdf15 | Cd81 |
| Emp12 | 0 | 0.98702826 | 0.863 | 0.551 | 0 | TAM.Gdf15 | Emp1 |
| Xdh2 | 0 | 1.19571392 | 0.807 | 0.497 | 0 | TAM.Gdf15 | Xdh |
| Cd200r12 | 0 | 1.35766704 | 0.586 | 0.276 | 0 | TAM.Gdf15 | Cd200r1 |
| Mgst11 | 0 | 1.3081722 | 0.603 | 0.301 | 0 | TAM.Gdf15 | Mgst1 |
| Cyb5a1 | 0 | 1.63535819 | 0.717 | 0.415 | 0 | TAM.Gdf15 | Cyb5a |
| Fnip22 | 0 | 1.18118864 | 0.798 | 0.498 | 0 | TAM.Gdf15 | Fnip2 |
| Lipa1 | 0 | 1.45993178 | 0.85 | 0.564 | 0 | TAM.Gdf15 | Lipa |
| Gabarapl11 | 0 | 1.13250935 | 0.778 | 0.494 | 0 | TAM.Gdf15 | Gabarapl1 |
| Plxna12 | 0 | 1.04058702 | 0.773 | 0.492 | 0 | TAM.Gdf15 | Plxna1 |
| Atf5 | 0 | 2.80199178 | 0.38 | 0.099 | 0 | TAM.Gdf15 | Atf5 |
| Prdx11 | 0 | 1.82052119 | 0.908 | 0.633 | 0 | TAM.Gdf15 | Prdx1 |
| Clec4d3 | 0 | 1.43894578 | 0.92 | 0.646 | 0 | TAM.Gdf15 | Clec4d |
| Slc40a11 | 0 | 2.92002453 | 0.378 | 0.107 | 0 | TAM.Gdf15 | Slc40a1 |
| Serpine12 | 0 | 1.47843109 | 0.474 | 0.203 | 0 | TAM.Gdf15 | Serpine1 |
| Mcoln21 | 0 | 2.28992637 | 0.379 | 0.109 | 0 | TAM.Gdf15 | Mcoln2 |
| Hmox13 | 0 | 3.45215372 | 0.948 | 0.682 | 0 | TAM.Gdf15 | Hmox1 |
| Mt22 | 0 | 1.1094449 | 0.595 | 0.333 | 0 | TAM.Gdf15 | Mt2 |
| Bst12 | 0 | 0.96223407 | 0.633 | 0.372 | 0 | TAM.Gdf15 | Bst1 |
| Rnf1281 | 0 | 1.78921219 | 0.395 | 0.134 | 0 | TAM.Gdf15 | Rnf128 |
| Metrnl3 | 0 | 1.15059584 | 0.837 | 0.577 | 0 | TAM.Gdf15 | Metrnl |
| Gadd45g2 | 0 | 0.98209152 | 0.645 | 0.392 | 0 | TAM.Gdf15 | Gadd45g |
| Pld33 | 0 | 1.05153139 | 0.864 | 0.615 | 0 | TAM.Gdf15 | Pld3 |
| C3ar14 | 0 | 0.90150874 | 0.924 | 0.679 | 0 | TAM.Gdf15 | C3ar1 |
| Ccl62 | 0 | 1.6183611 | 0.841 | 0.601 | 0 | TAM.Gdf15 | Ccl6 |
| Cat1 | 0 | 1.63024886 | 0.738 | 0.5 | 0 | TAM.Gdf15 | Cat |
| Ccl73 | 0 | 1.01000772 | 0.455 | 0.226 | 0 | TAM.Gdf15 | Ccl7 |
| Card194 | 0 | 0.93916442 | 0.833 | 0.608 | 0 | TAM.Gdf15 | Card19 |
| Cd632 | 0 | 0.66673885 | 0.967 | 0.744 | 0 | TAM.Gdf15 | Cd63 |
| Fam129b3 | 0 | 0.83682899 | 0.898 | 0.678 | 0 | TAM.Gdf15 | Fam129b |
| Lmna2 | 0 | 0.8566214 | 0.941 | 0.722 | 0 | TAM.Gdf15 | Lmna |
| Ninj11 | 0 | 0.94593037 | 0.916 | 0.703 | 0 | TAM.Gdf15 | Ninj1 |
| Slc11a12 | 0 | 1.05668651 | 0.914 | 0.709 | 0 | TAM.Gdf15 | Slc11a1 |
| Pdia42 | 0 | 1.11770013 | 0.828 | 0.623 | 0 | TAM.Gdf15 | Pdia4 |
| Anxa41 | 0 | 0.67010388 | 0.857 | 0.652 | 0 | TAM.Gdf15 | Anxa4 |
| Plec3 | 0 | 0.74999821 | 0.865 | 0.66 | 0 | TAM.Gdf15 | Plec |
| Creg12 | 0 | 1.66417203 | 0.923 | 0.719 | 0 | TAM.Gdf15 | Creg1 |
| Psmb51 | 0 | 0.92233991 | 0.79 | 0.587 | 0 | TAM.Gdf15 | Psmb5 |
| Txn11 | 0 | 0.73347946 | 0.86 | 0.657 | 0 | TAM.Gdf15 | Txn1 |
| Ecm12 | 0 | 1.00701109 | 0.907 | 0.707 | 0 | TAM.Gdf15 | Ecm1 |
| Wfdc171 | 0 | 0.58939765 | 0.695 | 0.497 | 0 | TAM.Gdf15 | Wfdc17 |
| Ccl33 | 0 | 0.82585529 | 0.763 | 0.567 | 0 | TAM.Gdf15 | Ccl3 |
| Anpep3 | 0 | 0.73467396 | 0.688 | 0.495 | 0 | TAM.Gdf15 | Anpep |
| Ccl93 | 0 | 0.93183537 | 0.86 | 0.667 | 0 | TAM.Gdf15 | Ccl9 |
| Ctsd2 | 0 | 1.36740494 | 0.934 | 0.743 | 0 | TAM.Gdf15 | Ctsd |
| Sqstm12 | 0 | 1.9169266 | 0.959 | 0.769 | 0 | TAM.Gdf15 | Sqstm1 |
| Capn21 | 0 | 0.77950014 | 0.791 | 0.604 | 0 | TAM.Gdf15 | Capn2 |
| Cndp21 | 0 | 0.73266193 | 0.856 | 0.669 | 0 | TAM.Gdf15 | Cndp2 |
| Cd844 | 0 | 0.674549 | 0.799 | 0.622 | 0 | TAM.Gdf15 | Cd84 |
| Mmp192 | 0 | 0.53345623 | 0.781 | 0.604 | 0 | TAM.Gdf15 | Mmp19 |
| Lilr4b2 | 0 | 0.99636505 | 0.837 | 0.663 | 0 | TAM.Gdf15 | Lilr4b |
| Arg12 | 0 | 0.74778968 | 0.747 | 0.575 | 0 | TAM.Gdf15 | Arg1 |
| Txnrd12 | 0 | 0.67589827 | 0.886 | 0.718 | 0 | TAM.Gdf15 | Txnrd1 |
| Cebpa2 | 0 | 0.68265333 | 0.929 | 0.762 | 0 | TAM.Gdf15 | Cebpa |
| Aldh23 | 0 | 0.85180992 | 0.881 | 0.721 | 0 | TAM.Gdf15 | Aldh2 |
| Scpep12 | 0 | 0.71762551 | 0.891 | 0.732 | 0 | TAM.Gdf15 | Scpep1 |
| Atf33 | 0 | 0.43275479 | 0.849 | 0.697 | 0 | TAM.Gdf15 | Atf3 |
| Ctsl2 | 0 | 0.80056929 | 0.903 | 0.753 | 0 | TAM.Gdf15 | Ctsl |
| Hilpda2 | 0 | 0.32124744 | 0.723 | 0.574 | 0 | TAM.Gdf15 | Hilpda |
| Pgd1 | 0 | 0.97877175 | 0.912 | 0.767 | 0 | TAM.Gdf15 | Pgd |
| C5ar12 | 0 | 0.66973002 | 0.92 | 0.783 | 0 | TAM.Gdf15 | C5ar1 |
| Ywhag1 | 0 | 0.91424866 | 0.96 | 0.826 | 0 | TAM.Gdf15 | Ywhag |
| Fabp52 | 0 | 1.09014499 | 0.808 | 0.679 | 0 | TAM.Gdf15 | Fabp5 |
| Id22 | 0 | 0.4620022 | 0.809 | 0.68 | 0 | TAM.Gdf15 | Id2 |
| Vcp2 | 0 | 0.66890241 | 0.929 | 0.802 | 0 | TAM.Gdf15 | Vcp |
| Clec4n1 | 0 | 0.68774181 | 0.913 | 0.791 | 0 | TAM.Gdf15 | Clec4n |
| Plin22 | 0 | 1.34315919 | 0.974 | 0.86 | 0 | TAM.Gdf15 | Plin2 |
| Ap2m11 | 0 | 0.65825422 | 0.865 | 0.756 | 0 | TAM.Gdf15 | Ap2m1 |
| Dab22 | 0 | 0.36857937 | 0.945 | 0.853 | 0 | TAM.Gdf15 | Dab2 |
| Alox5ap2 | 0 | 0.33989115 | 0.859 | 0.769 | 0 | TAM.Gdf15 | Alox5ap |
| Mt11 | 0 | 0.51155973 | 0.887 | 0.8 | 0 | TAM.Gdf15 | Mt1 |
| Lgals32 | 0 | 1.18045005 | 0.992 | 0.905 | 0 | TAM.Gdf15 | Lgals3 |
| 2-Jun | 0 | 0.43910373 | 0.94 | 0.854 | 0 | TAM.Gdf15 | Jun |
| Cstb1 | 0 | 0.90085042 | 0.96 | 0.874 | 0 | TAM.Gdf15 | Cstb |
| Ms4a6d2 | 0 | 0.46049563 | 0.878 | 0.796 | 0 | TAM.Gdf15 | Ms4a6d |
| Anxa51 | 0 | 0.50429633 | 0.938 | 0.859 | 0 | TAM.Gdf15 | Anxa5 |
| Msr12 | 0 | 0.48289561 | 0.869 | 0.792 | 0 | TAM.Gdf15 | Msr1 |
| Pla2g71 | 0 | 0.26914804 | 0.894 | 0.82 | 0 | TAM.Gdf15 | Pla2g7 |
| Mafb2 | 0 | 0.26313894 | 0.953 | 0.881 | 0 | TAM.Gdf15 | Mafb |
| Lamp12 | 0 | 0.38994402 | 0.879 | 0.807 | 0 | TAM.Gdf15 | Lamp1 |
| Lilrb4a2 | 0 | 0.84854663 | 0.952 | 0.88 | 0 | TAM.Gdf15 | Lilrb4a |
| Slc3a21 | 0 | 0.54899083 | 0.942 | 0.871 | 0 | TAM.Gdf15 | Slc3a2 |
| Pkm3 | 0 | 0.36740012 | 0.931 | 0.863 | 0 | TAM.Gdf15 | Pkm |
| Taldo13 | 0 | 0.34129276 | 0.851 | 0.788 | 0 | TAM.Gdf15 | Taldo1 |
| Dusp14 | 0 | 0.44794575 | 0.953 | 0.892 | 0 | TAM.Gdf15 | Dusp1 |
| Iqgap13 | 0 | 0.6268069 | 0.974 | 0.92 | 0 | TAM.Gdf15 | Iqgap1 |
| Lrp14 | 0 | 0.25256367 | 0.945 | 0.9 | 0 | TAM.Gdf15 | Lrp1 |
| Cd681 | 0 | 0.47688899 | 0.928 | 0.884 | 0 | TAM.Gdf15 | Cd68 |
| Ctsa2 | 0 | 0.39887787 | 0.951 | 0.909 | 0 | TAM.Gdf15 | Ctsa |
| Ubc2 | 0 | 0.48705568 | 0.957 | 0.917 | 0 | TAM.Gdf15 | Ubc |
| Akr1a11 | 0 | 0.59980466 | 0.974 | 0.937 | 0 | TAM.Gdf15 | Akr1a1 |
| Sdcbp | 0 | 0.43738556 | 0.965 | 0.932 | 0 | TAM.Gdf15 | Sdcbp |
| Cd532 | 0 | 0.30253533 | 0.917 | 0.885 | 0 | TAM.Gdf15 | Cd53 |
| Grn2 | 0 | 0.43935433 | 0.977 | 0.946 | 0 | TAM.Gdf15 | Grn |
| Lyz22 | 0 | 0.39635862 | 0.935 | 0.966 | 0 | TAM.Gdf15 | Lyz2 |
| Ctsb2 | 0 | 0.81597614 | 0.992 | 0.965 | 0 | TAM.Gdf15 | Ctsb |
| Cebpb2 | 0 | 0.26064825 | 0.994 | 0.977 | 0 | TAM.Gdf15 | Cebpb |
| App2 | 0 | 0.4035822 | 0.971 | 0.955 | 0 | TAM.Gdf15 | App |
| Ctsz2 | 0 | 0.29845355 | 0.983 | 0.972 | 0 | TAM.Gdf15 | Ctsz |
| Psap2 | 0 | 0.54854166 | 0.977 | 0.97 | 0 | TAM.Gdf15 | Psap |
| Fth11 | 0 | 0.9992468 | 0.999 | 0.996 | 0 | TAM.Gdf15 | Fth1 |
| Atp6v1b21 | 3.53E-307 | 0.60848953 | 0.916 | 0.798 | 6.82E-303 | TAM.Gdf15 | Atp6v1b2 |
| Emp31 | 1.39E-298 | 0.29207197 | 0.891 | 0.79 | 2.68E-294 | TAM.Gdf15 | Emp3 |
| Slc6a6 | 3.34E-298 | 0.59633001 | 0.851 | 0.736 | 6.45E-294 | TAM.Gdf15 | Slc6a6 |
| Atp6v1a1 | 3.52E-296 | 0.71310891 | 0.897 | 0.733 | 6.81E-292 | TAM.Gdf15 | Atp6v1a |
| Klf61 | 6.99E-290 | 0.49928686 | 0.926 | 0.825 | 1.35E-285 | TAM.Gdf15 | Klf6 |
| Slc43a22 | 1.05E-284 | 0.75192881 | 0.819 | 0.614 | 2.03E-280 | TAM.Gdf15 | Slc43a2 |
| Arl8a1 | 2.85E-283 | 0.82730646 | 0.856 | 0.666 | 5.50E-279 | TAM.Gdf15 | Arl8a |
| Chpf21 | 3.11E-282 | 1.18472485 | 0.671 | 0.384 | 6.01E-278 | TAM.Gdf15 | Chpf2 |
| Dhrs33 | 1.12E-281 | 0.6313884 | 0.858 | 0.65 | 2.17E-277 | TAM.Gdf15 | Dhrs3 |
| Capg3 | 2.26E-281 | 0.46883406 | 0.887 | 0.69 | 4.37E-277 | TAM.Gdf15 | Capg |
| Jund2 | 8.16E-281 | 0.34335972 | 0.974 | 0.932 | 1.58E-276 | TAM.Gdf15 | Jund |
| Ankrd33b | 8.19E-280 | 2.1748771 | 0.393 | 0.11 | 1.58E-275 | TAM.Gdf15 | Ankrd33b |
| Msrb12 | 1.79E-277 | 0.32052582 | 0.891 | 0.813 | 3.46E-273 | TAM.Gdf15 | Msrb1 |
| Gclc1 | 3.65E-274 | 1.28264589 | 0.54 | 0.271 | 7.06E-270 | TAM.Gdf15 | Gclc |
| Zfand2a2 | 1.77E-271 | 1.00829808 | 0.743 | 0.456 | 3.42E-267 | TAM.Gdf15 | Zfand2a |
| Cacna1d2 | 1.56E-264 | 0.89346386 | 0.519 | 0.258 | 3.02E-260 | TAM.Gdf15 | Cacna1d |
| Rnh12 | 2.11E-264 | 0.6285804 | 0.826 | 0.669 | 4.07E-260 | TAM.Gdf15 | Rnh1 |
| Lyz13 | 1.43E-262 | 0.85153833 | 0.453 | 0.307 | 2.77E-258 | TAM.Gdf15 | Lyz1 |
| Aldoa2 | 9.61E-261 | 0.34447989 | 0.921 | 0.826 | 1.86E-256 | TAM.Gdf15 | Aldoa |
| Ahnak3 | 1.62E-258 | 0.30973303 | 0.924 | 0.746 | 3.14E-254 | TAM.Gdf15 | Ahnak |
| Gns2 | 1.67E-252 | 0.4834487 | 0.877 | 0.727 | 3.22E-248 | TAM.Gdf15 | Gns |
| Hyal11 | 1.82E-252 | 2.28192184 | 0.333 | 0.096 | 3.52E-248 | TAM.Gdf15 | Hyal1 |
| Cd1512 | 8.47E-250 | 1.3722373 | 0.547 | 0.259 | 1.64E-245 | TAM.Gdf15 | Cd151 |
| Smpdl3a2 | 6.47E-247 | 0.70042911 | 0.641 | 0.429 | 1.25E-242 | TAM.Gdf15 | Smpdl3a |
| Prr132 | 2.57E-245 | 0.88469121 | 0.7 | 0.484 | 4.97E-241 | TAM.Gdf15 | Prr13 |
| Osbpl82 | 1.74E-244 | 1.00281285 | 0.741 | 0.56 | 3.36E-240 | TAM.Gdf15 | Osbpl8 |
| Slc27a12 | 3.81E-244 | 1.01489395 | 0.716 | 0.423 | 7.38E-240 | TAM.Gdf15 | Slc27a1 |
| Riok31 | 1.49E-242 | 0.73958397 | 0.83 | 0.668 | 2.88E-238 | TAM.Gdf15 | Riok3 |
| Adamtsl5 | 3.89E-241 | 2.67940993 | 0.288 | 0.056 | 7.53E-237 | TAM.Gdf15 | Adamtsl5 |
| Ly91 | 2.17E-239 | 1.48966749 | 0.42 | 0.176 | 4.20E-235 | TAM.Gdf15 | Ly9 |
| Nceh12 | 2.77E-239 | 1.09238252 | 0.649 | 0.386 | 5.36E-235 | TAM.Gdf15 | Nceh1 |
| Rtn41 | 3.49E-237 | 0.57344392 | 0.872 | 0.754 | 6.76E-233 | TAM.Gdf15 | Rtn4 |
| Mfhas12 | 2.66E-235 | 1.09225204 | 0.629 | 0.326 | 5.15E-231 | TAM.Gdf15 | Mfhas1 |
| Tbxas12 | 1.49E-233 | 0.3706162 | 0.821 | 0.713 | 2.88E-229 | TAM.Gdf15 | Tbxas1 |
| Alcam3 | 1.56E-231 | 1.01158811 | 0.631 | 0.38 | 3.01E-227 | TAM.Gdf15 | Alcam |
| Tmem431 | 4.46E-228 | 1.16438363 | 0.652 | 0.367 | 8.61E-224 | TAM.Gdf15 | Tmem43 |
| Cd300lb1 | 3.44E-227 | 1.08632498 | 0.65 | 0.338 | 6.65E-223 | TAM.Gdf15 | Cd300lb |
| Tpp12 | 1.39E-223 | 0.58707241 | 0.837 | 0.665 | 2.68E-219 | TAM.Gdf15 | Tpp1 |
| Por1 | 8.32E-223 | 1.03226724 | 0.614 | 0.36 | 1.61E-218 | TAM.Gdf15 | Por |
| Tmem1892 | 1.17E-222 | 0.52308688 | 0.89 | 0.758 | 2.27E-218 | TAM.Gdf15 | Tmem189 |
| F72 | 1.59E-219 | 0.66235296 | 0.402 | 0.262 | 3.08E-215 | TAM.Gdf15 | F7 |
| Gda3 | 2.73E-219 | 0.26353101 | 0.821 | 0.745 | 5.28E-215 | TAM.Gdf15 | Gda |
| Myo5a1 | 7.04E-219 | 0.67566104 | 0.82 | 0.61 | 1.36E-214 | TAM.Gdf15 | Myo5a |
| Igf12 | 1.17E-211 | 1.41913396 | 0.431 | 0.196 | 2.26E-207 | TAM.Gdf15 | Igf1 |
| Lrp8 | 3.41E-207 | 2.03261371 | 0.265 | 0.064 | 6.60E-203 | TAM.Gdf15 | Lrp8 |
| Nrp22 | 4.30E-205 | 0.8204754 | 0.714 | 0.439 | 8.31E-201 | TAM.Gdf15 | Nrp2 |
| Atp6v0b1 | 1.04E-203 | 0.4667448 | 0.865 | 0.78 | 2.02E-199 | TAM.Gdf15 | Atp6v0b |
| Fam219a2 | 2.17E-200 | 1.5225236 | 0.49 | 0.215 | 4.19E-196 | TAM.Gdf15 | Fam219a |
| Dnase1l12 | 2.80E-200 | 1.08998852 | 0.693 | 0.445 | 5.42E-196 | TAM.Gdf15 | Dnase1l1 |
| Igf2r | 1.26E-199 | 1.63287952 | 0.374 | 0.113 | 2.44E-195 | TAM.Gdf15 | Igf2r |
| Anxa32 | 3.59E-199 | 0.36162566 | 0.894 | 0.776 | 6.95E-195 | TAM.Gdf15 | Anxa3 |
| Snx92 | 2.07E-198 | 0.67359205 | 0.729 | 0.54 | 4.00E-194 | TAM.Gdf15 | Snx9 |
| Cxcl12 | 2.76E-198 | 0.49731681 | 0.324 | 0.233 | 5.33E-194 | TAM.Gdf15 | Cxcl1 |
| Cers22 | 4.43E-198 | 0.66698841 | 0.777 | 0.584 | 8.56E-194 | TAM.Gdf15 | Cers2 |
| Nus11 | 8.32E-197 | 0.86472146 | 0.76 | 0.53 | 1.61E-192 | TAM.Gdf15 | Nus1 |
| Eps83 | 2.52E-196 | 0.67901046 | 0.666 | 0.455 | 4.86E-192 | TAM.Gdf15 | Eps8 |
| Pitpnc12 | 3.44E-196 | 1.14015069 | 0.584 | 0.305 | 6.66E-192 | TAM.Gdf15 | Pitpnc1 |
| Npc13 | 1.46E-194 | 0.64517107 | 0.806 | 0.607 | 2.82E-190 | TAM.Gdf15 | Npc1 |
| Naa802 | 2.62E-194 | 1.74326068 | 0.463 | 0.209 | 5.07E-190 | TAM.Gdf15 | Naa80 |
| Clcn73 | 1.16E-193 | 1.06383727 | 0.727 | 0.466 | 2.24E-189 | TAM.Gdf15 | Clcn7 |
| Zdhhc181 | 1.23E-191 | 1.20216537 | 0.69 | 0.377 | 2.37E-187 | TAM.Gdf15 | Zdhhc18 |
| Atp6ap1 | 2.27E-189 | 0.33989033 | 0.867 | 0.799 | 4.38E-185 | TAM.Gdf15 | Atp6ap1 |
| Ahnak2 | 8.64E-189 | 2.1019811 | 0.299 | 0.085 | 1.67E-184 | TAM.Gdf15 | Ahnak2 |
| Dusp33 | 5.29E-188 | 0.402582 | 0.862 | 0.743 | 1.02E-183 | TAM.Gdf15 | Dusp3 |
| Ypel51 | 1.45E-187 | 0.8411072 | 0.662 | 0.461 | 2.80E-183 | TAM.Gdf15 | Ypel5 |
| Tcirg11 | 4.37E-187 | 0.34284659 | 0.927 | 0.868 | 8.46E-183 | TAM.Gdf15 | Tcirg1 |
| Rab11fip52 | 1.06E-186 | 0.57543086 | 0.831 | 0.59 | 2.04E-182 | TAM.Gdf15 | Rab11fip5 |
| Rxra2 | 1.68E-186 | 0.83322389 | 0.699 | 0.428 | 3.24E-182 | TAM.Gdf15 | Rxra |
| Bax1 | 2.71E-186 | 0.4579467 | 0.881 | 0.779 | 5.24E-182 | TAM.Gdf15 | Bax |
| Gaa2 | 7.32E-186 | 0.82290891 | 0.688 | 0.452 | 1.41E-181 | TAM.Gdf15 | Gaa |
| Rabac13 | 1.06E-185 | 0.59689021 | 0.845 | 0.677 | 2.05E-181 | TAM.Gdf15 | Rabac1 |
| Arhgap103 | 3.00E-185 | 1.52265862 | 0.498 | 0.196 | 5.80E-181 | TAM.Gdf15 | Arhgap10 |
| Tnfaip21 | 1.82E-184 | 0.27495482 | 0.855 | 0.73 | 3.51E-180 | TAM.Gdf15 | Tnfaip2 |
| Lhfpl21 | 4.17E-184 | 0.8191627 | 0.624 | 0.372 | 8.07E-180 | TAM.Gdf15 | Lhfpl2 |
| Ddit31 | 3.35E-181 | 1.75567209 | 0.399 | 0.212 | 6.47E-177 | TAM.Gdf15 | Ddit3 |
| Epb41l12 | 4.68E-181 | 1.47246208 | 0.444 | 0.159 | 9.05E-177 | TAM.Gdf15 | Epb41l1 |
| Slc35f62 | 1.21E-180 | 0.75330012 | 0.63 | 0.415 | 2.34E-176 | TAM.Gdf15 | Slc35f6 |
| Anxa71 | 2.63E-179 | 0.90109467 | 0.637 | 0.432 | 5.08E-175 | TAM.Gdf15 | Anxa7 |
| Nfe2l12 | 1.84E-178 | 0.91082195 | 0.758 | 0.539 | 3.56E-174 | TAM.Gdf15 | Nfe2l1 |
| Lamp22 | 7.86E-178 | 0.28521162 | 0.866 | 0.803 | 1.52E-173 | TAM.Gdf15 | Lamp2 |
| Ephx11 | 1.08E-175 | 1.28783588 | 0.347 | 0.153 | 2.08E-171 | TAM.Gdf15 | Ephx1 |
| Rhob3 | 9.98E-175 | 0.46198756 | 0.871 | 0.762 | 1.93E-170 | TAM.Gdf15 | Rhob |
| Folr22 | 1.77E-173 | 0.6187577 | 0.608 | 0.267 | 3.42E-169 | TAM.Gdf15 | Folr2 |
| Tmem373 | 6.37E-170 | 0.54142764 | 0.6 | 0.411 | 1.23E-165 | TAM.Gdf15 | Tmem37 |
| Mfsd122 | 1.47E-169 | 0.8930474 | 0.707 | 0.494 | 2.85E-165 | TAM.Gdf15 | Mfsd12 |
| Cyb5r32 | 7.56E-167 | 0.58957703 | 0.75 | 0.577 | 1.46E-162 | TAM.Gdf15 | Cyb5r3 |
| Lrp123 | 1.44E-165 | 0.81509682 | 0.656 | 0.437 | 2.79E-161 | TAM.Gdf15 | Lrp12 |
| Cd200r4 | 1.81E-164 | 1.77274274 | 0.283 | 0.093 | 3.50E-160 | TAM.Gdf15 | Cd200r4 |
| Trpv41 | 2.25E-164 | 1.52510925 | 0.364 | 0.126 | 4.35E-160 | TAM.Gdf15 | Trpv4 |
| Trem22 | 7.74E-164 | 0.77782409 | 0.671 | 0.424 | 1.50E-159 | TAM.Gdf15 | Trem2 |
| Adap23 | 2.28E-163 | 0.54102713 | 0.794 | 0.581 | 4.40E-159 | TAM.Gdf15 | Adap2 |
| Wdr913 | 6.92E-163 | 1.11127367 | 0.638 | 0.342 | 1.34E-158 | TAM.Gdf15 | Wdr91 |
| Psmd16 | 2.24E-161 | 0.8452195 | 0.699 | 0.489 | 4.33E-157 | TAM.Gdf15 | Psmd1 |
| Psmd72 | 3.81E-161 | 0.90995057 | 0.721 | 0.475 | 7.37E-157 | TAM.Gdf15 | Psmd7 |
| Ptgr11 | 1.43E-160 | 0.70713792 | 0.43 | 0.288 | 2.77E-156 | TAM.Gdf15 | Ptgr1 |
| Pon32 | 5.51E-160 | 1.1793059 | 0.463 | 0.219 | 1.07E-155 | TAM.Gdf15 | Pon3 |
| Gltp1 | 1.13E-156 | 0.4326208 | 0.767 | 0.647 | 2.18E-152 | TAM.Gdf15 | Gltp |
| Cds22 | 1.14E-156 | 0.92261623 | 0.52 | 0.295 | 2.19E-152 | TAM.Gdf15 | Cds2 |
| Mcfd22 | 1.34E-155 | 0.60317259 | 0.772 | 0.586 | 2.59E-151 | TAM.Gdf15 | Mcfd2 |
| Aars1 | 4.35E-155 | 1.17309614 | 0.55 | 0.344 | 8.40E-151 | TAM.Gdf15 | Aars |
| Rhoc2 | 1.55E-151 | 1.14905675 | 0.56 | 0.285 | 2.99E-147 | TAM.Gdf15 | Rhoc |
| Arap12 | 1.24E-150 | 0.52040222 | 0.869 | 0.723 | 2.39E-146 | TAM.Gdf15 | Arap1 |
| Tmem1651 | 1.57E-149 | 0.71401402 | 0.65 | 0.44 | 3.03E-145 | TAM.Gdf15 | Tmem165 |
| Dtnbp11 | 3.42E-149 | 0.65288938 | 0.719 | 0.507 | 6.61E-145 | TAM.Gdf15 | Dtnbp1 |
| Tnfsf143 | 4.46E-149 | 0.68686266 | 0.424 | 0.218 | 8.62E-145 | TAM.Gdf15 | Tnfsf14 |
| Rit11 | 7.15E-149 | 1.09608462 | 0.565 | 0.307 | 1.38E-144 | TAM.Gdf15 | Rit1 |
| Stac22 | 4.67E-148 | 1.29174414 | 0.376 | 0.147 | 9.02E-144 | TAM.Gdf15 | Stac2 |
| Tnfsf122 | 3.98E-147 | 0.62831805 | 0.68 | 0.458 | 7.70E-143 | TAM.Gdf15 | Tnfsf12 |
| Ikbkg1 | 2.02E-145 | 1.28721283 | 0.456 | 0.241 | 3.91E-141 | TAM.Gdf15 | Ikbkg |
| Mertk3 | 7.45E-145 | 0.41183393 | 0.714 | 0.566 | 1.44E-140 | TAM.Gdf15 | Mertk |
| Adam152 | 1.16E-144 | 0.34381003 | 0.787 | 0.663 | 2.25E-140 | TAM.Gdf15 | Adam15 |
| Atp6v1h | 1.57E-144 | 0.67269464 | 0.665 | 0.495 | 3.03E-140 | TAM.Gdf15 | Atp6v1h |
| Fbxo302 | 2.59E-144 | 0.93361081 | 0.448 | 0.267 | 5.01E-140 | TAM.Gdf15 | Fbxo30 |
| Gla1 | 4.43E-143 | 0.50614107 | 0.726 | 0.533 | 8.56E-139 | TAM.Gdf15 | Gla |
| Rab1a1 | 1.88E-142 | 0.25505456 | 0.887 | 0.83 | 3.64E-138 | TAM.Gdf15 | Rab1a |
| Tma161 | 1.37E-141 | 1.51977366 | 0.362 | 0.152 | 2.66E-137 | TAM.Gdf15 | Tma16 |
| G6pdx1 | 6.71E-141 | 0.64869698 | 0.537 | 0.397 | 1.30E-136 | TAM.Gdf15 | G6pdx |
| Mtss12 | 1.49E-140 | 0.7799263 | 0.784 | 0.551 | 2.88E-136 | TAM.Gdf15 | Mtss1 |
| Sash13 | 3.36E-140 | 0.50874796 | 0.806 | 0.588 | 6.50E-136 | TAM.Gdf15 | Sash1 |
| Abhd122 | 4.03E-139 | 0.37775454 | 0.756 | 0.63 | 7.78E-135 | TAM.Gdf15 | Abhd12 |
| Lamc11 | 1.43E-137 | 1.64126964 | 0.381 | 0.148 | 2.76E-133 | TAM.Gdf15 | Lamc1 |
| Rusc22 | 4.81E-137 | 1.35859213 | 0.479 | 0.217 | 9.30E-133 | TAM.Gdf15 | Rusc2 |
| Kcnab23 | 8.58E-137 | 0.58847261 | 0.621 | 0.442 | 1.66E-132 | TAM.Gdf15 | Kcnab2 |
| Pdlim43 | 9.60E-137 | 1.02159604 | 0.493 | 0.273 | 1.86E-132 | TAM.Gdf15 | Pdlim4 |
| Atp6v1g1 | 1.64E-135 | 0.31822237 | 0.845 | 0.754 | 3.18E-131 | TAM.Gdf15 | Atp6v1g1 |
| Rexo21 | 1.82E-135 | 0.67845703 | 0.683 | 0.479 | 3.51E-131 | TAM.Gdf15 | Rexo2 |
| Impact2 | 4.38E-134 | 0.62421288 | 0.677 | 0.481 | 8.46E-130 | TAM.Gdf15 | Impact |
| Slc41a23 | 6.92E-134 | 1.21590329 | 0.451 | 0.202 | 1.34E-129 | TAM.Gdf15 | Slc41a2 |
| Pnrc11 | 7.60E-134 | 0.36567309 | 0.916 | 0.837 | 1.47E-129 | TAM.Gdf15 | Pnrc1 |
| Osgin1 | 1.05E-133 | 1.75259445 | 0.312 | 0.144 | 2.03E-129 | TAM.Gdf15 | Osgin1 |
| Palld3 | 2.80E-133 | 0.92019648 | 0.47 | 0.262 | 5.41E-129 | TAM.Gdf15 | Palld |
| Sgpl13 | 3.23E-133 | 0.38912903 | 0.819 | 0.689 | 6.25E-129 | TAM.Gdf15 | Sgpl1 |
| Tlr131 | 4.09E-133 | 0.83776953 | 0.483 | 0.301 | 7.92E-129 | TAM.Gdf15 | Tlr13 |
| Atp6v0a12 | 5.98E-132 | 0.46365747 | 0.671 | 0.511 | 1.16E-127 | TAM.Gdf15 | Atp6v0a1 |
| Shtn11 | 1.49E-131 | 0.56121131 | 0.641 | 0.472 | 2.87E-127 | TAM.Gdf15 | Shtn1 |
| Nek62 | 1.99E-131 | 0.60731205 | 0.658 | 0.477 | 3.86E-127 | TAM.Gdf15 | Nek6 |
| Rbm471 | 4.52E-131 | 0.73851318 | 0.74 | 0.525 | 8.74E-127 | TAM.Gdf15 | Rbm47 |
| Kcnk132 | 1.59E-130 | 1.31343349 | 0.408 | 0.162 | 3.07E-126 | TAM.Gdf15 | Kcnk13 |
| Mafg1 | 1.78E-129 | 0.91650304 | 0.652 | 0.436 | 3.44E-125 | TAM.Gdf15 | Mafg |
| Gtpbp21 | 6.31E-129 | 1.16907039 | 0.563 | 0.394 | 1.22E-124 | TAM.Gdf15 | Gtpbp2 |
| Rab181 | 3.62E-128 | 0.58949677 | 0.729 | 0.573 | 7.00E-124 | TAM.Gdf15 | Rab18 |
| P2rx72 | 3.95E-128 | 1.02906888 | 0.501 | 0.339 | 7.63E-124 | TAM.Gdf15 | P2rx7 |
| Dennd4c2 | 1.20E-127 | 1.01543776 | 0.499 | 0.251 | 2.32E-123 | TAM.Gdf15 | Dennd4c |
| Bri31 | 3.51E-127 | 0.29090343 | 0.841 | 0.747 | 6.78E-123 | TAM.Gdf15 | Bri3 |
| Gstm11 | 3.89E-127 | 1.13656035 | 0.41 | 0.185 | 7.53E-123 | TAM.Gdf15 | Gstm1 |
| Jag12 | 1.13E-125 | 1.32935835 | 0.362 | 0.162 | 2.18E-121 | TAM.Gdf15 | Jag1 |
| P2rx42 | 1.90E-125 | 0.46625427 | 0.805 | 0.632 | 3.68E-121 | TAM.Gdf15 | P2rx4 |
| Slc8b12 | 2.82E-125 | 0.73452552 | 0.716 | 0.482 | 5.45E-121 | TAM.Gdf15 | Slc8b1 |
| Il10rb2 | 3.52E-124 | 0.28959343 | 0.838 | 0.736 | 6.80E-120 | TAM.Gdf15 | Il10rb |
| Abcc33 | 5.23E-124 | 0.578842 | 0.63 | 0.446 | 1.01E-119 | TAM.Gdf15 | Abcc3 |
| Zwint2 | 6.35E-124 | 1.30785911 | 0.467 | 0.269 | 1.23E-119 | TAM.Gdf15 | Zwint |
| Degs12 | 1.02E-123 | 0.40442924 | 0.657 | 0.514 | 1.97E-119 | TAM.Gdf15 | Degs1 |
| Atp2a2 | 4.11E-123 | 0.34316793 | 0.826 | 0.717 | 7.94E-119 | TAM.Gdf15 | Atp2a2 |
| Atp6v1e11 | 8.06E-123 | 0.27095929 | 0.821 | 0.737 | 1.56E-118 | TAM.Gdf15 | Atp6v1e1 |
| Tpcn23 | 9.32E-123 | 0.87054308 | 0.599 | 0.356 | 1.80E-118 | TAM.Gdf15 | Tpcn2 |
| Rnasek1 | 1.07E-122 | 0.29504521 | 0.856 | 0.77 | 2.07E-118 | TAM.Gdf15 | Rnasek |
| Lonrf32 | 1.27E-122 | 1.12019385 | 0.451 | 0.243 | 2.46E-118 | TAM.Gdf15 | Lonrf3 |
| Bcar3 | 5.66E-122 | 1.45569819 | 0.374 | 0.146 | 1.09E-117 | TAM.Gdf15 | Bcar3 |
| Serinc11 | 1.16E-121 | 0.52061925 | 0.691 | 0.524 | 2.23E-117 | TAM.Gdf15 | Serinc1 |
| Tuba1c2 | 1.24E-121 | 0.35184467 | 0.828 | 0.729 | 2.39E-117 | TAM.Gdf15 | Tuba1c |
| Kpna4 | 1.29E-121 | 0.52282083 | 0.7 | 0.568 | 2.49E-117 | TAM.Gdf15 | Kpna4 |
| Mvp2 | 1.98E-121 | 0.45987451 | 0.822 | 0.701 | 3.82E-117 | TAM.Gdf15 | Mvp |
| Prdx63 | 2.19E-121 | 0.62505809 | 0.488 | 0.327 | 4.24E-117 | TAM.Gdf15 | Prdx6 |
| Fbxo322 | 1.24E-119 | 1.17021746 | 0.287 | 0.144 | 2.40E-115 | TAM.Gdf15 | Fbxo32 |
| Homer32 | 3.24E-119 | 1.17851867 | 0.423 | 0.204 | 6.26E-115 | TAM.Gdf15 | Homer3 |
| Cab391 | 3.98E-118 | 0.58791305 | 0.803 | 0.644 | 7.69E-114 | TAM.Gdf15 | Cab39 |
| Tnfrsf26 | 8.36E-118 | 1.60371637 | 0.31 | 0.116 | 1.62E-113 | TAM.Gdf15 | Tnfrsf26 |
| Alox52 | 9.68E-118 | 0.82807731 | 0.365 | 0.158 | 1.87E-113 | TAM.Gdf15 | Alox5 |

| | | | | | | | |
|---|---|---|---|---|---|---|---|
| Pon23 | 2.58E-117 | 0.62360509 | 0.594 | 0.411 | 5.00E-113 | TAM.Gdf15 | Pon2 |
| Trim253 | 4.55E-115 | 0.3416425 | 0.745 | 0.67 | 8.79E-111 | TAM.Gdf15 | Trim25 |
| Tmem1043 | 7.82E-115 | 0.60286853 | 0.663 | 0.449 | 1.51E-110 | TAM.Gdf15 | Tmem104 |
| Fam214b3 | 1.16E-114 | 1.09918377 | 0.552 | 0.325 | 2.24E-110 | TAM.Gdf15 | Fam214b |
| Renbp3 | 1.45E-113 | 0.56493258 | 0.745 | 0.541 | 2.80E-109 | TAM.Gdf15 | Renbp |
| Slc39a71 | 3.34E-113 | 0.69790713 | 0.657 | 0.482 | 6.46E-109 | TAM.Gdf15 | Slc39a7 |
| Fbxl51 | 7.45E-113 | 0.64072001 | 0.528 | 0.36 | 1.44E-108 | TAM.Gdf15 | Fbxl5 |
| Soat11 | 8.89E-113 | 0.36715003 | 0.758 | 0.645 | 1.72E-108 | TAM.Gdf15 | Soat1 |
| Rbpj2 | 1.08E-112 | 0.30151683 | 0.776 | 0.68 | 2.10E-108 | TAM.Gdf15 | Rbpj |
| Ptger2 | 1.47E-112 | 1.33535219 | 0.27 | 0.098 | 2.84E-108 | TAM.Gdf15 | Ptger2 |
| Ifi274 | 1.53E-112 | 0.70704672 | 0.621 | 0.393 | 2.95E-108 | TAM.Gdf15 | Ifi27 |
| Camk13 | 1.72E-112 | 0.37100118 | 0.693 | 0.528 | 3.33E-108 | TAM.Gdf15 | Camk1 |
| Tmbim12 | 9.42E-112 | 0.52189638 | 0.716 | 0.551 | 1.82E-107 | TAM.Gdf15 | Tmbim1 |
| Zmiz12 | 3.81E-111 | 0.49942569 | 0.8 | 0.655 | 7.37E-107 | TAM.Gdf15 | Zmiz1 |
| Fgfr12 | 3.35E-110 | 0.97975104 | 0.372 | 0.12 | 6.48E-106 | TAM.Gdf15 | Fgfr1 |
| Tmem651 | 4.73E-109 | 0.57843221 | 0.653 | 0.45 | 9.15E-105 | TAM.Gdf15 | Tmem65 |
| Slc23a23 | 5.36E-109 | 0.91068744 | 0.558 | 0.342 | 1.04E-104 | TAM.Gdf15 | Slc23a2 |
| Ubap12 | 6.20E-109 | 1.05018972 | 0.499 | 0.301 | 1.20E-104 | TAM.Gdf15 | Ubap1 |
| Abca13 | 1.81E-108 | 0.28122771 | 0.89 | 0.724 | 3.50E-104 | TAM.Gdf15 | Abca1 |
| Fblim13 | 4.08E-108 | 1.14900678 | 0.523 | 0.263 | 7.90E-104 | TAM.Gdf15 | Fblim1 |
| Uap1l12 | 2.23E-107 | 0.61913718 | 0.685 | 0.474 | 4.30E-103 | TAM.Gdf15 | Uap1l1 |
| Plekhm22 | 6.91E-107 | 0.79451243 | 0.613 | 0.422 | 1.34E-102 | TAM.Gdf15 | Plekhm2 |
| Klhl212 | 7.33E-107 | 0.83775644 | 0.495 | 0.282 | 1.42E-102 | TAM.Gdf15 | Klhl21 |
| Eea12 | 8.88E-107 | 0.97840915 | 0.55 | 0.332 | 1.72E-102 | TAM.Gdf15 | Eea1 |
| Eaf12 | 1.24E-106 | 0.83097598 | 0.588 | 0.38 | 2.40E-102 | TAM.Gdf15 | Eaf1 |
| Myof1 | 5.94E-106 | 0.33879822 | 0.816 | 0.669 | 1.15E-101 | TAM.Gdf15 | Myof |
| Tatdn22 | 1.48E-105 | 0.31029244 | 0.742 | 0.642 | 2.86E-101 | TAM.Gdf15 | Tatdn2 |
| Pde2a1 | 3.18E-105 | 0.90660763 | 0.442 | 0.205 | 6.15E-101 | TAM.Gdf15 | Pde2a |
| Tex2643 | 7.37E-105 | 0.71913892 | 0.593 | 0.368 | 1.43E-100 | TAM.Gdf15 | Tex264 |
| Surf42 | 9.99E-105 | 0.37633677 | 0.726 | 0.622 | 1.93E-100 | TAM.Gdf15 | Surf4 |
| Nisch1 | 4.37E-104 | 0.40323154 | 0.85 | 0.697 | 8.46E-100 | TAM.Gdf15 | Nisch |
| Ftl1-ps1 | 7.34E-104 | 1.51875655 | 0.258 | 0.09 | 1.42E-99 | TAM.Gdf15 | Ftl1-ps1 |
| Slc7a84 | 8.79E-104 | 0.42471735 | 0.699 | 0.509 | 1.70E-99 | TAM.Gdf15 | Slc7a8 |
| Sh3bp21 | 1.57E-103 | 0.4292172 | 0.742 | 0.579 | 3.03E-99 | TAM.Gdf15 | Sh3bp2 |
| Fam20c2 | 1.72E-103 | 0.35244759 | 0.697 | 0.505 | 3.32E-99 | TAM.Gdf15 | Fam20c |
| H2-D11 | 5.10E-103 | 0.30874606 | 0.77 | 0.675 | 9.86E-99 | TAM.Gdf15 | H2-D1 |
| Rap2b3 | 2.11E-102 | 0.50143164 | 0.699 | 0.493 | 4.07E-98 | TAM.Gdf15 | Rap2b |
| Ralgds2 | 3.15E-102 | 0.37679529 | 0.723 | 0.601 | 6.09E-98 | TAM.Gdf15 | Ralgds |
| Plekhb21 | 6.08E-102 | 0.39776388 | 0.759 | 0.605 | 1.18E-97 | TAM.Gdf15 | Plekhb2 |
| Mitf2 | 2.06E-101 | 0.83093943 | 0.594 | 0.352 | 3.97E-97 | TAM.Gdf15 | Mitf |
| Twf13 | 4.57E-101 | 0.63276284 | 0.62 | 0.432 | 8.84E-97 | TAM.Gdf15 | Twf1 |
| Tuba1a3 | 2.80E-100 | 0.79511582 | 0.534 | 0.337 | 5.41E-96 | TAM.Gdf15 | Tuba1a |
| Sh3gl12 | 6.93E-100 | 0.6167983 | 0.658 | 0.435 | 1.34E-95 | TAM.Gdf15 | Sh3gl1 |
| Atp6ap21 | 8.03E-100 | 0.2867216 | 0.857 | 0.764 | 1.55E-95 | TAM.Gdf15 | Atp6ap2 |
| C2cd2l2 | 1.30E-99 | 0.6810901 | 0.52 | 0.315 | 2.51E-95 | TAM.Gdf15 | C2cd2l |
| Tfec1 | 2.27E-99 | 0.70029197 | 0.597 | 0.389 | 4.38E-95 | TAM.Gdf15 | Tfec |
| Gnai3 | 8.54E-99 | 0.41450979 | 0.774 | 0.639 | 1.65E-94 | TAM.Gdf15 | Gnai3 |
| Tmem106a3 | 9.39E-99 | 0.40689267 | 0.777 | 0.59 | 1.82E-94 | TAM.Gdf15 | Tmem106a |
| Sgms12 | 2.31E-98 | 1.00844413 | 0.453 | 0.238 | 4.47E-94 | TAM.Gdf15 | Sgms1 |
| Pi4k2a2 | 1.08E-97 | 0.7887128 | 0.648 | 0.425 | 2.09E-93 | TAM.Gdf15 | Pi4k2a |
| Idh13 | 2.14E-97 | 0.46336368 | 0.572 | 0.414 | 4.14E-93 | TAM.Gdf15 | Idh1 |
| Cd300ld6 | 6.23E-97 | 0.41833821 | 0.697 | 0.52 | 1.20E-92 | TAM.Gdf15 | Cd300ld |
| Rragc1 | 7.25E-97 | 0.48744052 | 0.68 | 0.511 | 1.40E-92 | TAM.Gdf15 | Rragc |
| Rab5if | 8.32E-97 | 0.32476429 | 0.734 | 0.609 | 1.61E-92 | TAM.Gdf15 | Rab5if |
| Psmd112 | 4.53E-96 | 0.63969626 | 0.603 | 0.452 | 8.75E-92 | TAM.Gdf15 | Psmd11 |
| Brd21 | 1.03E-95 | 0.41569727 | 0.762 | 0.657 | 1.98E-91 | TAM.Gdf15 | Brd2 |
| Gadd45a3 | 1.32E-95 | 0.90903554 | 0.448 | 0.254 | 2.54E-91 | TAM.Gdf15 | Gadd45a |
| Gas62 | 1.75E-95 | 0.41890366 | 0.593 | 0.302 | 3.38E-91 | TAM.Gdf15 | Gas6 |
| Raf11 | 3.68E-95 | 0.60150209 | 0.714 | 0.531 | 7.11E-91 | TAM.Gdf15 | Raf1 |
| Tcp11l1 | 5.10E-95 | 1.19612602 | 0.324 | 0.142 | 9.86E-91 | TAM.Gdf15 | Tcp11l1 |
| Pstpip13 | 7.20E-95 | 0.46411823 | 0.703 | 0.524 | 1.39E-90 | TAM.Gdf15 | Pstpip1 |
| Fndc3a3 | 8.56E-95 | 0.49683242 | 0.72 | 0.549 | 1.66E-90 | TAM.Gdf15 | Fndc3a |
| Psma51 | 8.85E-95 | 0.58627608 | 0.585 | 0.457 | 1.71E-90 | TAM.Gdf15 | Psma5 |
| Psmd51 | 1.00E-94 | 1.14232394 | 0.352 | 0.17 | 1.94E-90 | TAM.Gdf15 | Psmd5 |
| Hspa51 | 1.89E-94 | 0.25707651 | 0.783 | 0.719 | 3.65E-90 | TAM.Gdf15 | Hspa5 |
| Fcna1 | 3.80E-94 | 0.46962001 | 0.537 | 0.216 | 7.35E-90 | TAM.Gdf15 | Fcna |
| Psmc61 | 9.77E-94 | 0.70032893 | 0.634 | 0.465 | 1.89E-89 | TAM.Gdf15 | Psmc6 |
| Plin33 | 1.54E-92 | 0.75258815 | 0.601 | 0.396 | 2.98E-88 | TAM.Gdf15 | Plin3 |
| Ap1s11 | 2.70E-92 | 0.38733955 | 0.612 | 0.49 | 5.22E-88 | TAM.Gdf15 | Ap1s1 |
| Dync1h1 | 3.37E-92 | 0.45154792 | 0.765 | 0.633 | 6.52E-88 | TAM.Gdf15 | Dync1h1 |
| Slc25a372 | 1.02E-91 | 0.90866177 | 0.52 | 0.281 | 1.97E-87 | TAM.Gdf15 | Slc25a37 |
| Psmc31 | 2.99E-91 | 0.43072091 | 0.717 | 0.58 | 5.77E-87 | TAM.Gdf15 | Psmc3 |
| Dock72 | 3.04E-91 | 0.7663521 | 0.616 | 0.429 | 5.89E-87 | TAM.Gdf15 | Dock7 |
| Plekhm11 | 4.56E-91 | 1.05399573 | 0.397 | 0.214 | 8.82E-87 | TAM.Gdf15 | Plekhm1 |
| Cd372 | 5.62E-91 | 0.73503832 | 0.559 | 0.322 | 1.09E-86 | TAM.Gdf15 | Cd37 |
| Plekhn12 | 1.19E-90 | 0.91227686 | 0.484 | 0.297 | 2.30E-86 | TAM.Gdf15 | Plekhn1 |
| Snx14 | 1.60E-90 | 0.48160713 | 0.716 | 0.561 | 3.09E-86 | TAM.Gdf15 | Snx1 |
| Ogdh1 | 2.01E-90 | 0.28009305 | 0.772 | 0.666 | 3.88E-86 | TAM.Gdf15 | Ogdh |
| Lrrfip22 | 6.08E-90 | 0.7260417 | 0.564 | 0.364 | 1.18E-85 | TAM.Gdf15 | Lrrfip2 |
| Hk32 | 1.49E-89 | 0.91041812 | 0.507 | 0.303 | 2.88E-85 | TAM.Gdf15 | Hk3 |
| Elf4g21 | 2.05E-89 | 0.25854867 | 0.852 | 0.759 | 3.97E-85 | TAM.Gdf15 | Eif4g2 |
| Psmd121 | 2.10E-89 | 0.7858977 | 0.558 | 0.392 | 4.06E-85 | TAM.Gdf15 | Psmd12 |
| Sort11 | 8.31E-89 | 0.73518595 | 0.276 | 0.159 | 1.61E-84 | TAM.Gdf15 | Sort1 |
| Cebpg2 | 1.54E-88 | 0.76676475 | 0.65 | 0.451 | 2.97E-84 | TAM.Gdf15 | Cebpg |
| Vcl1 | 3.77E-88 | 0.6157916 | 0.413 | 0.255 | 7.30E-84 | TAM.Gdf15 | Vcl |
| Calm2 | 7.30E-88 | 0.33790219 | 0.648 | 0.519 | 1.41E-83 | TAM.Gdf15 | Calm2 |
| Psmc21 | 6.70E-87 | 0.69941112 | 0.591 | 0.41 | 1.30E-82 | TAM.Gdf15 | Psmc2 |
| Mbnl2 | 6.97E-87 | 0.40873723 | 0.755 | 0.631 | 1.35E-82 | TAM.Gdf15 | Mbnl2 |
| Atp6v1c11 | 1.19E-86 | 0.42091399 | 0.702 | 0.558 | 2.30E-82 | TAM.Gdf15 | Atp6v1c1 |
| Pdxk2 | 1.55E-86 | 0.37254198 | 0.535 | 0.397 | 3.01E-82 | TAM.Gdf15 | Pdxk |
| Nploc41 | 1.64E-86 | 0.79473959 | 0.588 | 0.409 | 3.18E-82 | TAM.Gdf15 | Nploc4 |
| Cyth11 | 2.67E-86 | 0.28640809 | 0.873 | 0.767 | 5.16E-82 | TAM.Gdf15 | Cyth1 |
| Emc7 | 4.20E-86 | 0.51116651 | 0.644 | 0.48 | 8.12E-82 | TAM.Gdf15 | Emc7 |
| Galnt22 | 9.56E-86 | 0.45351897 | 0.713 | 0.51 | 1.85E-81 | TAM.Gdf15 | Galnt2 |
| H6pd2 | 9.27E-85 | 0.60825756 | 0.541 | 0.347 | 1.79E-80 | TAM.Gdf15 | H6pd |
| Psmc51 | 1.31E-84 | 0.45730884 | 0.715 | 0.544 | 2.54E-80 | TAM.Gdf15 | Psmc5 |
| Rnf132 | 2.36E-84 | 0.45807125 | 0.659 | 0.496 | 4.56E-80 | TAM.Gdf15 | Rnf13 |
| Fam234a1 | 2.43E-84 | 0.58510132 | 0.592 | 0.381 | 4.70E-80 | TAM.Gdf15 | Fam234a |
| Cltb2 | 1.03E-83 | 0.53241761 | 0.595 | 0.409 | 1.99E-79 | TAM.Gdf15 | Cltb |
| Stom3 | 1.51E-83 | 0.56676498 | 0.591 | 0.404 | 2.92E-79 | TAM.Gdf15 | Stom |
| Impdh12 | 1.93E-83 | 1.03252103 | 0.39 | 0.191 | 3.73E-79 | TAM.Gdf15 | Impdh1 |
| Psmc11 | 2.78E-83 | 0.70189776 | 0.55 | 0.367 | 5.37E-79 | TAM.Gdf15 | Psmc1 |
| Rab311 | 2.31E-82 | 0.2590185 | 0.733 | 0.588 | 4.47E-78 | TAM.Gdf15 | Rab31 |
| Ptpra1 | 4.01E-82 | 0.75551025 | 0.622 | 0.441 | 7.75E-78 | TAM.Gdf15 | Ptpra |
| Gstp11 | 4.71E-82 | 0.61212871 | 0.527 | 0.399 | 9.10E-78 | TAM.Gdf15 | Gstp1 |
| Lemd22 | 5.59E-82 | 0.73824197 | 0.686 | 0.49 | 1.08E-77 | TAM.Gdf15 | Lemd2 |
| Bag31 | 5.75E-82 | 0.62838264 | 0.566 | 0.377 | 1.11E-77 | TAM.Gdf15 | Bag3 |
| Pef11 | 6.16E-82 | 0.83912066 | 0.412 | 0.229 | 1.19E-77 | TAM.Gdf15 | Pef1 |
| Stxbp51 | 2.81E-81 | 1.12717055 | 0.329 | 0.155 | 5.44E-77 | TAM.Gdf15 | Stxbp5 |
| Morf4l11 | 3.58E-81 | 0.32023445 | 0.827 | 0.725 | 6.92E-77 | TAM.Gdf15 | Morf4l1 |
| Ubxn4 | 5.15E-81 | 0.59048616 | 0.648 | 0.482 | 9.97E-77 | TAM.Gdf15 | Ubxn4 |
| Cln81 | 5.52E-81 | 0.50137289 | 0.65 | 0.484 | 1.07E-76 | TAM.Gdf15 | Cln8 |
| Gss1 | 5.73E-81 | 0.97375343 | 0.293 | 0.156 | 1.11E-76 | TAM.Gdf15 | Gss |
| Fmn12 | 1.07E-80 | 0.89357508 | 0.497 | 0.272 | 2.06E-76 | TAM.Gdf15 | Fmn1 |
| Pdgfc2 | 1.36E-80 | 0.66517284 | 0.488 | 0.246 | 2.64E-76 | TAM.Gdf15 | Pdgfc |
| Txnl12 | 2.40E-80 | 0.61386717 | 0.592 | 0.436 | 4.64E-76 | TAM.Gdf15 | Txnl1 |
| Gpc11 | 4.66E-80 | 0.65428758 | 0.428 | 0.232 | 9.02E-76 | TAM.Gdf15 | Gpc1 |
| Plbd22 | 6.82E-80 | 0.37953376 | 0.697 | 0.561 | 1.32E-75 | TAM.Gdf15 | Plbd2 |
| Mdm22 | 1.38E-79 | 0.57503717 | 0.731 | 0.586 | 2.66E-75 | TAM.Gdf15 | Mdm2 |
| Pttg1ip3 | 3.37E-79 | 0.34156954 | 0.73 | 0.589 | 6.52E-75 | TAM.Gdf15 | Pttg1ip |
| Ubtd11 | 5.24E-79 | 0.5887973 | 0.628 | 0.44 | 1.01E-74 | TAM.Gdf15 | Ubtd1 |
| Tmem87b2 | 5.59E-79 | 0.734982 | 0.592 | 0.379 | 1.08E-74 | TAM.Gdf15 | Tmem87b |
| Ppp1r14b1 | 1.30E-78 | 0.63253802 | 0.526 | 0.339 | 2.52E-74 | TAM.Gdf15 | Ppp1r14b |
| Tmem1541 | 1.33E-78 | 0.4946921 | 0.284 | 0.167 | 2.57E-74 | TAM.Gdf15 | Tmem154 |
| Dync1i2 | 1.59E-78 | 0.55348432 | 0.651 | 0.478 | 3.08E-74 | TAM.Gdf15 | Dync1i2 |
| Sh2b2 | 1.62E-78 | 0.39896154 | 0.678 | 0.522 | 3.13E-74 | TAM.Gdf15 | Sh2b2 |
| Arhgef12 | 2.05E-78 | 1.40202473 | 0.357 | 0.15 | 3.95E-74 | TAM.Gdf15 | Arhgef12 |
| Akap72 | 5.33E-78 | 0.75765557 | 0.376 | 0.2 | 1.03E-73 | TAM.Gdf15 | Akap7 |
| Zfp7042 | 5.56E-78 | 1.1803147 | 0.408 | 0.204 | 1.08E-73 | TAM.Gdf15 | Zfp704 |
| Ptgfrn2 | 7.67E-78 | 0.35990119 | 0.505 | 0.362 | 1.48E-73 | TAM.Gdf15 | Ptgfrn |
| Cav21 | 2.38E-77 | 0.86382995 | 0.402 | 0.218 | 4.61E-73 | TAM.Gdf15 | Cav2 |
| Mocs22 | 3.95E-77 | 0.67540712 | 0.519 | 0.323 | 7.63E-73 | TAM.Gdf15 | Mocs2 |
| Mpp11 | 1.08E-76 | 0.51623665 | 0.562 | 0.378 | 2.08E-72 | TAM.Gdf15 | Mpp1 |
| Lpar1 | 1.57E-76 | 1.43767563 | 0.25 | 0.085 | 3.03E-72 | TAM.Gdf15 | Lpar1 |
| Mrtfb1 | 3.32E-75 | 0.8953525 | 0.373 | 0.193 | 6.42E-71 | TAM.Gdf15 | Mrtfb |
| Bin11 | 3.45E-75 | 0.53680642 | 0.652 | 0.416 | 6.67E-71 | TAM.Gdf15 | Bin1 |
| Dapk12 | 6.03E-75 | 0.40209255 | 0.581 | 0.426 | 1.17E-70 | TAM.Gdf15 | Dapk1 |
| Hmox22 | 6.21E-74 | 0.30771087 | 0.645 | 0.519 | 1.20E-69 | TAM.Gdf15 | Hmox2 |
| Cpeb42 | 2.84E-73 | 0.64234121 | 0.635 | 0.443 | 5.49E-69 | TAM.Gdf15 | Cpeb4 |
| Atf41 | 4.52E-73 | 0.8543605 | 0.595 | 0.4 | 8.74E-69 | TAM.Gdf15 | Atf4 |
| Ctps22 | 4.68E-73 | 0.56591699 | 0.503 | 0.335 | 9.04E-69 | TAM.Gdf15 | Ctps2 |
| Sod12 | 6.48E-73 | 0.43239162 | 0.612 | 0.472 | 1.25E-68 | TAM.Gdf15 | Sod1 |
| Cmtm32 | 8.60E-73 | 0.43042646 | 0.74 | 0.57 | 1.66E-68 | TAM.Gdf15 | Cmtm3 |
| Abcd41 | 1.18E-72 | 1.18359901 | 0.287 | 0.12 | 2.29E-68 | TAM.Gdf15 | Abcd4 |
| Mdfic2 | 3.58E-72 | 0.46695344 | 0.68 | 0.519 | 6.93E-68 | TAM.Gdf15 | Mdfic |
| Zdhhc143 | 3.62E-72 | 0.61773077 | 0.53 | 0.308 | 7.00E-68 | TAM.Gdf15 | Zdhhc14 |
| Agfg12 | 3.67E-72 | 0.65775802 | 0.542 | 0.384 | 7.09E-68 | TAM.Gdf15 | Agfg1 |
| Dpp73 | 1.06E-71 | 0.83625208 | 0.448 | 0.277 | 2.05E-67 | TAM.Gdf15 | Dpp7 |
| Naglu3 | 1.38E-71 | 0.56792681 | 0.544 | 0.369 | 2.66E-67 | TAM.Gdf15 | Naglu |
| Zbtb7a1 | 4.69E-71 | 0.47669407 | 0.703 | 0.554 | 9.08E-67 | TAM.Gdf15 | Zbtb7a |
| Cenpb2 | 4.74E-71 | 0.3589225 | 0.763 | 0.587 | 9.17E-67 | TAM.Gdf15 | Cenpb |
| Mgat4b1 | 1.27E-70 | 0.75469045 | 0.498 | 0.314 | 2.46E-66 | TAM.Gdf15 | Mgat4b |
| Acbd31 | 1.37E-70 | 0.75111956 | 0.564 | 0.398 | 2.65E-66 | TAM.Gdf15 | Acbd3 |
| Pdgfb2 | 1.59E-70 | 1.01865063 | 0.374 | 0.251 | 3.08E-66 | TAM.Gdf15 | Pdgfb |
| Nav12 | 2.32E-70 | 0.69929632 | 0.451 | 0.259 | 4.48E-66 | TAM.Gdf15 | Nav1 |
| Slc35e4 | 2.53E-70 | 1.11703399 | 0.309 | 0.141 | 4.88E-66 | TAM.Gdf15 | Slc35e4 |
| Gde11 | 8.96E-70 | 0.39202496 | 0.644 | 0.498 | 1.73E-65 | TAM.Gdf15 | Gde1 |
| Oga | 1.29E-69 | 0.65006202 | 0.634 | 0.429 | 2.49E-65 | TAM.Gdf15 | Oga |
| Tsku1 | 1.40E-69 | 0.97252745 | 0.316 | 0.14 | 2.71E-65 | TAM.Gdf15 | Tsku |
| Azin12 | 1.87E-68 | 0.55775739 | 0.565 | 0.403 | 3.61E-64 | TAM.Gdf15 | Azin1 |
| Clip22 | 1.96E-68 | 0.69340112 | 0.551 | 0.374 | 3.79E-64 | TAM.Gdf15 | Clip2 |
| Eepd11 | 2.03E-68 | 0.45905868 | 0.523 | 0.368 | 3.93E-64 | TAM.Gdf15 | Eepd1 |
| Fmnl22 | 2.30E-68 | 0.67921416 | 0.445 | 0.273 | 4.45E-64 | TAM.Gdf15 | Fmnl2 |
| Sbf23 | 4.74E-68 | 0.90061065 | 0.485 | 0.253 | 9.17E-64 | TAM.Gdf15 | Sbf2 |
| Cd2743 | 7.06E-68 | 0.2669474 | 0.601 | 0.477 | 1.36E-63 | TAM.Gdf15 | Cd274 |
| Tpd52l21 | 8.94E-68 | 0.56261345 | 0.624 | 0.443 | 1.73E-63 | TAM.Gdf15 | Tpd52l2 |
| Mcoln11 | 1.40E-67 | 0.60672594 | 0.452 | 0.288 | 2.70E-63 | TAM.Gdf15 | Mcoln1 |
| Gsk3b | 2.19E-67 | 0.53973882 | 0.634 | 0.473 | 4.23E-63 | TAM.Gdf15 | Gsk3b |
| Camk2d3 | 2.33E-67 | 0.31058316 | 0.681 | 0.545 | 4.51E-63 | TAM.Gdf15 | Camk2d |
| Sfxn32 | 6.51E-67 | 0.34656987 | 0.614 | 0.463 | 1.26E-62 | TAM.Gdf15 | Sfxn3 |
| Tlr42 | 6.88E-67 | 0.45699243 | 0.566 | 0.404 | 1.33E-62 | TAM.Gdf15 | Tlr4 |
| Dbi2 | 9.19E-67 | 0.25515508 | 0.559 | 0.473 | 1.78E-62 | TAM.Gdf15 | Dbi |
| Fkbp21 | 1.10E-66 | 0.29869995 | 0.71 | 0.578 | 2.13E-62 | TAM.Gdf15 | Fkbp2 |
| Slc30a12 | 1.55E-66 | 0.37935692 | 0.507 | 0.369 | 2.99E-62 | TAM.Gdf15 | Slc30a1 |
| Arl12 | 1.59E-66 | 0.55937161 | 0.437 | 0.3 | 3.07E-62 | TAM.Gdf15 | Arl1 |
| Map7d1 | 1.89E-66 | 0.26603167 | 0.829 | 0.751 | 3.66E-62 | TAM.Gdf15 | Map7d1 |
| Kif13b3 | 2.22E-66 | 0.73516194 | 0.503 | 0.307 | 4.29E-62 | TAM.Gdf15 | Kif13b |
| Samd81 | 3.29E-66 | 0.62530749 | 0.523 | 0.352 | 6.37E-62 | TAM.Gdf15 | Samd8 |
| Kif3a1 | 3.96E-66 | 1.37019215 | 0.28 | 0.114 | 7.66E-62 | TAM.Gdf15 | Kif3a |
| Tmed9 | 1.90E-65 | 0.26399313 | 0.722 | 0.619 | 3.68E-61 | TAM.Gdf15 | Tmed9 |
| Thbd1 | 3.82E-65 | 0.34597043 | 0.331 | 0.203 | 7.39E-61 | TAM.Gdf15 | Thbd |
| Fndc3b2 | 4.16E-65 | 0.43638261 | 0.679 | 0.53 | 8.05E-61 | TAM.Gdf15 | Fndc3b |
| Lat21 | 4.40E-65 | 0.32757499 | 0.627 | 0.527 | 8.51E-61 | TAM.Gdf15 | Lat2 |
| Dlgap42 | 4.77E-65 | 0.43619578 | 0.68 | 0.511 | 9.23E-61 | TAM.Gdf15 | Dlgap4 |
| Kcmf11 | 5.33E-65 | 0.62818063 | 0.622 | 0.459 | 1.03E-60 | TAM.Gdf15 | Kcmf1 |
| Slc25a512 | 9.23E-65 | 0.50354655 | 0.579 | 0.416 | 1.78E-60 | TAM.Gdf15 | Slc25a51 |
| Ano61 | 1.46E-64 | 0.51792766 | 0.573 | 0.395 | 2.82E-60 | TAM.Gdf15 | Ano6 |
| Tanc21 | 2.63E-64 | 1.29786991 | 0.34 | 0.162 | 5.09E-60 | TAM.Gdf15 | Tanc2 |
| Psmd21 | 3.29E-64 | 0.56123615 | 0.62 | 0.461 | 6.36E-60 | TAM.Gdf15 | Psmd2 |
| Cdk14 | 4.03E-64 | 1.13658647 | 0.28 | 0.118 | 7.78E-60 | TAM.Gdf15 | Cdk14 |
| Ube2q21 | 5.16E-64 | 0.75758636 | 0.442 | 0.268 | 9.98E-60 | TAM.Gdf15 | Ube2q2 |
| Zswim8 | 5.61E-64 | 0.69531726 | 0.63 | 0.436 | 1.08E-59 | TAM.Gdf15 | Zswim8 |
| Plxnc12 | 8.62E-64 | 0.86722274 | 0.4 | 0.227 | 1.67E-59 | TAM.Gdf15 | Plxnc1 |
| Sirt22 | 1.48E-63 | 0.30630184 | 0.716 | 0.592 | 2.86E-59 | TAM.Gdf15 | Sirt2 |
| Gbe12 | 1.54E-63 | 1.11341639 | 0.35 | 0.169 | 2.98E-59 | TAM.Gdf15 | Gbe1 |
| Syngr11 | 2.13E-63 | 1.29968063 | 0.27 | 0.122 | 4.12E-59 | TAM.Gdf15 | Syngr1 |
| Hspa91 | 2.13E-63 | 0.80355961 | 0.5 | 0.321 | 4.13E-59 | TAM.Gdf15 | Hspa9 |
| Nup622 | 2.49E-63 | 0.8978814 | 0.416 | 0.246 | 4.82E-59 | TAM.Gdf15 | Nup62 |
| Tlnrd11 | 3.40E-63 | 0.60883888 | 0.672 | 0.461 | 6.57E-59 | TAM.Gdf15 | Tlnrd1 |
| Tubb2a3 | 8.00E-63 | 0.41234739 | 0.552 | 0.417 | 1.55E-58 | TAM.Gdf15 | Tubb2a |
| Tbcel | 1.45E-62 | 0.96733223 | 0.321 | 0.181 | 2.80E-58 | TAM.Gdf15 | Tbcel |
| Leng81 | 1.57E-62 | 0.59929906 | 0.652 | 0.507 | 3.04E-58 | TAM.Gdf15 | Leng8 |
| Baiap21 | 2.69E-62 | 0.46757025 | 0.558 | 0.418 | 5.20E-58 | TAM.Gdf15 | Baiap2 |
| Ptpre2 | 3.95E-62 | 0.27881957 | 0.684 | 0.565 | 7.63E-58 | TAM.Gdf15 | Ptpre |
| Klc11 | 4.33E-62 | 0.83876469 | 0.523 | 0.314 | 8.37E-58 | TAM.Gdf15 | Klc1 |
| Ipo133 | 7.15E-62 | 0.81422371 | 0.451 | 0.277 | 1.38E-57 | TAM.Gdf15 | Ipo13 |
| Washc21 | 1.96E-61 | 0.32907984 | 0.678 | 0.543 | 3.79E-57 | TAM.Gdf15 | Washc2 |
| Rabep11 | 2.29E-61 | 0.50131971 | 0.697 | 0.53 | 4.43E-57 | TAM.Gdf15 | Rabep1 |
| DEPRECATED-I | 3.47E-61 | 0.6523393 | 0.564 | 0.389 | 6.71E-57 | TAM.Gdf15 | DEPRECATED-ENSMUSG00000017221 |
| Mctp12 | 3.53E-61 | 0.79724115 | 0.287 | 0.132 | 6.82E-57 | TAM.Gdf15 | Mctp1 |
| Gas2l12 | 6.96E-61 | 0.5582952 | 0.544 | 0.349 | 1.35E-56 | TAM.Gdf15 | Gas2l1 |
| AW5498771 | 1.48E-60 | 1.10032567 | 0.358 | 0.185 | 2.86E-56 | TAM.Gdf15 | AW549877 |
| Lpp2 | 1.99E-60 | 0.3686315 | 0.728 | 0.576 | 3.84E-56 | TAM.Gdf15 | Lpp |
| Hint11 | 5.50E-60 | 0.44700472 | 0.485 | 0.343 | 1.06E-55 | TAM.Gdf15 | Hint1 |
| Peak12 | 5.70E-60 | 1.04360882 | 0.363 | 0.185 | 1.10E-55 | TAM.Gdf15 | Peak1 |
| Lmbrd11 | 1.37E-59 | 0.4664226 | 0.597 | 0.441 | 2.65E-55 | TAM.Gdf15 | Lmbrd1 |
| Psmd81 | 1.69E-59 | 0.58591041 | 0.598 | 0.45 | 3.26E-55 | TAM.Gdf15 | Psmd8 |
| Tmem1401 | 3.00E-59 | 1.10060033 | 0.316 | 0.152 | 5.80E-55 | TAM.Gdf15 | Tmem140 |
| DEPRECATED-I | 3.08E-59 | 0.64503824 | 0.455 | 0.292 | 5.95E-55 | TAM.Gdf15 | DEPRECATED-ENSMUSG00000032667 |
| Ttc39c2 | 3.54E-59 | 0.65362598 | 0.322 | 0.181 | 6.85E-55 | TAM.Gdf15 | Ttc39c |
| Oxct14 | 1.95E-58 | 0.55578345 | 0.457 | 0.27 | 3.78E-54 | TAM.Gdf15 | Oxct1 |
| Rmc12 | 2.72E-58 | 0.59278665 | 0.436 | 0.27 | 5.25E-54 | TAM.Gdf15 | Rmc1 |
| Vezt1 | 6.42E-58 | 1.13317251 | 0.293 | 0.141 | 1.24E-53 | TAM.Gdf15 | Vezt |
| Ckap41 | 6.68E-58 | 0.4959421 | 0.487 | 0.316 | 1.29E-53 | TAM.Gdf15 | Ckap4 |
| Slc5a31 | 8.12E-58 | 0.45300761 | 0.42 | 0.297 | 1.57E-53 | TAM.Gdf15 | Slc5a3 |
| Ankrd50 | 9.81E-58 | 0.9123815 | 0.331 | 0.168 | 1.90E-53 | TAM.Gdf15 | Ankrd50 |
| Ppfibp12 | 1.20E-57 | 0.57246227 | 0.451 | 0.273 | 2.33E-53 | TAM.Gdf15 | Ppfibp1 |
| Gpcpd11 | 2.34E-57 | 0.35393844 | 0.534 | 0.405 | 4.52E-53 | TAM.Gdf15 | Gpcpd1 |
| Saraf2 | 3.03E-57 | 0.2632683 | 0.721 | 0.613 | 5.86E-53 | TAM.Gdf15 | Saraf |
| Gtf2a11 | 3.10E-57 | 0.52558302 | 0.599 | 0.444 | 6.00E-53 | TAM.Gdf15 | Gtf2a1 |
| Lgals83 | 3.84E-57 | 0.45650684 | 0.584 | 0.423 | 7.42E-53 | TAM.Gdf15 | Lgals8 |
| Arrdc42 | 7.41E-57 | 0.27340809 | 0.714 | 0.583 | 1.43E-52 | TAM.Gdf15 | Arrdc4 |
| Pkp41 | 8.28E-57 | 1.11541022 | 0.274 | 0.123 | 1.60E-52 | TAM.Gdf15 | Pkp4 |
| Fhl31 | 9.83E-57 | 0.64462364 | 0.473 | 0.305 | 1.90E-52 | TAM.Gdf15 | Fhl3 |
| Sema4b3 | 2.71E-56 | 0.53462212 | 0.528 | 0.36 | 5.24E-52 | TAM.Gdf15 | Sema4b |
| Nacc21 | 4.28E-56 | 0.94325754 | 0.334 | 0.157 | 8.27E-52 | TAM.Gdf15 | Nacc2 |
| Rheb1 | 8.06E-56 | 0.2802513 | 0.672 | 0.552 | 1.56E-51 | TAM.Gdf15 | Rheb |
| Naa501 | 1.12E-55 | 0.41562475 | 0.706 | 0.555 | 2.16E-51 | TAM.Gdf15 | Naa50 |
| Tmem184b1 | 2.13E-55 | 0.49443271 | 0.57 | 0.419 | 4.12E-51 | TAM.Gdf15 | Tmem184b |
| Lgals12 | 2.72E-55 | 0.53423921 | 0.434 | 0.269 | 5.26E-51 | TAM.Gdf15 | Lgals1 |
| Akt1s12 | 5.23E-55 | 0.82933123 | 0.437 | 0.247 | 1.01E-50 | TAM.Gdf15 | Akt1s1 |
| Dnajc151 | 7.21E-55 | 0.3972387 | 0.54 | 0.4 | 1.39E-50 | TAM.Gdf15 | Dnajc15 |
| Sars1 | 7.55E-55 | 0.9121174 | 0.456 | 0.282 | 1.46E-50 | TAM.Gdf15 | Sars |
| Snx302 | 9.41E-55 | 0.58034639 | 0.447 | 0.275 | 1.82E-50 | TAM.Gdf15 | Snx30 |
| Nkiras22 | 1.03E-54 | 0.44521579 | 0.529 | 0.372 | 2.00E-50 | TAM.Gdf15 | Nkiras2 |
| Hsph11 | 1.42E-54 | 0.74142212 | 0.303 | 0.158 | 2.74E-50 | TAM.Gdf15 | Hsph1 |
| Slc25a242 | 1.80E-54 | 0.64385721 | 0.372 | 0.221 | 3.47E-50 | TAM.Gdf15 | Slc25a24 |
| Nupr13 | 1.89E-54 | 0.46906033 | 0.553 | 0.386 | 3.66E-50 | TAM.Gdf15 | Nupr1 |
| Crat3 | 2.19E-54 | 0.64580128 | 0.4 | 0.255 | 4.23E-50 | TAM.Gdf15 | Crat |
| Mgat11 | 4.12E-54 | 0.31583641 | 0.629 | 0.482 | 7.96E-50 | TAM.Gdf15 | Mgat1 |
| Katnbl12 | 4.82E-54 | 1.01162872 | 0.377 | 0.207 | 9.31E-50 | TAM.Gdf15 | Katnbl1 |
| Psmd4 | 5.60E-54 | 0.59894204 | 0.533 | 0.386 | 1.08E-49 | TAM.Gdf15 | Psmd4 |
| Tspan17 | 6.97E-54 | 1.20877032 | 0.253 | 0.107 | 1.35E-49 | TAM.Gdf15 | Tspan17 |
| Coro1c1 | 1.07E-53 | 0.41691773 | 0.602 | 0.479 | 2.06E-49 | TAM.Gdf15 | Coro1c |
| Galnt62 | 1.13E-53 | 0.57348531 | 0.52 | 0.335 | 2.19E-49 | TAM.Gdf15 | Galnt6 |
| Rab5a | 1.20E-53 | 0.43928676 | 0.647 | 0.531 | 2.32E-49 | TAM.Gdf15 | Rab5a |
| Hyal2 | 3.86E-53 | 1.17745729 | 0.258 | 0.11 | 7.47E-49 | TAM.Gdf15 | Hyal2 |
| Hgs | 4.04E-53 | 0.61150452 | 0.57 | 0.388 | 7.81E-49 | TAM.Gdf15 | Hgs |
| Col4a3bp1 | 4.16E-53 | 0.57531502 | 0.567 | 0.404 | 8.03E-49 | TAM.Gdf15 | Col4a3bp |
| Pla2g4a1 | 5.34E-53 | 0.48531529 | 0.522 | 0.387 | 1.03E-48 | TAM.Gdf15 | Pla2g4a |
| Rab3gap21 | 9.45E-53 | 0.75640104 | 0.484 | 0.309 | 1.83E-48 | TAM.Gdf15 | Rab3gap2 |
| Adcy32 | 1.16E-52 | 1.16127672 | 0.262 | 0.125 | 2.24E-48 | TAM.Gdf15 | Adcy3 |
| Snx121 | 1.31E-52 | 0.38733982 | 0.591 | 0.456 | 2.54E-48 | TAM.Gdf15 | Snx12 |
| Kif1c1 | 1.44E-52 | 0.48642135 | 0.579 | 0.429 | 2.79E-48 | TAM.Gdf15 | Kif1c |
| Psmb62 | 1.94E-52 | 0.46888171 | 0.572 | 0.441 | 3.76E-48 | TAM.Gdf15 | Psmb6 |
| Derl22 | 3.26E-52 | 0.57400341 | 0.527 | 0.357 | 6.29E-48 | TAM.Gdf15 | Derl2 |
| Mfsd113 | 4.26E-52 | 0.665984 | 0.466 | 0.286 | 8.25E-48 | TAM.Gdf15 | Mfsd11 |
| Znrf12 | 7.05E-52 | 0.78215587 | 0.5 | 0.336 | 1.36E-47 | TAM.Gdf15 | Znrf1 |
| Kif1b | 7.40E-52 | 0.74154654 | 0.426 | 0.269 | 1.43E-47 | TAM.Gdf15 | Kif1b |
| Lamtor3 | 8.53E-52 | 0.33987101 | 0.678 | 0.536 | 1.65E-47 | TAM.Gdf15 | Lamtor3 |
| Il6st1 | 1.18E-51 | 0.40456428 | 0.484 | 0.354 | 2.27E-47 | TAM.Gdf15 | Il6st |
| Rassf82 | 1.18E-51 | 0.78304026 | 0.523 | 0.331 | 2.28E-47 | TAM.Gdf15 | Rassf8 |
| Tubb4b3 | 1.60E-51 | 0.50944978 | 0.595 | 0.427 | 3.09E-47 | TAM.Gdf15 | Tubb4b |
| Glipr11 | 3.43E-51 | 0.83590982 | 0.384 | 0.229 | 6.64E-47 | TAM.Gdf15 | Glipr1 |
| Stam2 | 5.36E-51 | 0.77367052 | 0.407 | 0.25 | 1.04E-46 | TAM.Gdf15 | Stam2 |
| Slc12a72 | 5.37E-51 | 0.42633926 | 0.494 | 0.328 | 1.04E-46 | TAM.Gdf15 | Slc12a7 |
| Golga4 | 5.61E-51 | 0.60962192 | 0.509 | 0.352 | 1.09E-46 | TAM.Gdf15 | Golga4 |
| Ankrd402 | 6.01E-51 | 0.39669152 | 0.606 | 0.452 | 1.16E-46 | TAM.Gdf15 | Ankrd40 |
| Nr1d2 | 7.89E-51 | 1.06178953 | 0.265 | 0.132 | 1.53E-46 | TAM.Gdf15 | Nr1d2 |
| Tspan43 | 1.14E-50 | 0.49786981 | 0.506 | 0.334 | 2.21E-46 | TAM.Gdf15 | Tspan4 |
| Hsd3b72 | 2.32E-50 | 0.80787864 | 0.448 | 0.289 | 4.49E-46 | TAM.Gdf15 | Hsd3b7 |
| Atg7 | 2.60E-50 | 0.5520777 | 0.509 | 0.352 | 5.02E-46 | TAM.Gdf15 | Atg7 |
| Piezo11 | 3.90E-50 | 0.44294629 | 0.65 | 0.5 | 7.54E-46 | TAM.Gdf15 | Piezo1 |
| Trappc6b | 4.36E-50 | 0.48097353 | 0.507 | 0.337 | 8.43E-46 | TAM.Gdf15 | Trappc6b |
| Tec2 | 1.44E-49 | 0.40604636 | 0.501 | 0.353 | 2.79E-45 | TAM.Gdf15 | Tec |
| Psmd31 | 1.70E-49 | 0.57015327 | 0.501 | 0.364 | 3.30E-45 | TAM.Gdf15 | Psmd3 |
| Adss2 | 1.86E-49 | 0.62110317 | 0.52 | 0.377 | 3.60E-45 | TAM.Gdf15 | Adss |
| Rab7b2 | 2.49E-49 | 0.40919938 | 0.483 | 0.331 | 4.81E-45 | TAM.Gdf15 | Rab7b |
| Dync1li21 | 2.98E-49 | 0.73637991 | 0.53 | 0.376 | 5.77E-45 | TAM.Gdf15 | Dync1li2 |
| Cd932 | 4.56E-49 | 0.28128748 | 0.541 | 0.399 | 8.82E-45 | TAM.Gdf15 | Cd93 |
| Fez21 | 7.53E-49 | 0.51003112 | 0.517 | 0.332 | 1.46E-44 | TAM.Gdf15 | Fez2 |
| Derl1 | 1.17E-48 | 0.25341229 | 0.694 | 0.605 | 2.26E-44 | TAM.Gdf15 | Derl1 |
| Rnf1151 | 1.55E-48 | 0.46733524 | 0.54 | 0.389 | 2.99E-44 | TAM.Gdf15 | Rnf115 |
| Casp31 | 2.75E-48 | 0.83377035 | 0.399 | 0.265 | 5.32E-44 | TAM.Gdf15 | Casp3 |
| Psmd142 | 3.66E-48 | 0.53671235 | 0.512 | 0.375 | 7.07E-44 | TAM.Gdf15 | Psmd14 |
| Cacul1 | 5.03E-48 | 0.54407661 | 0.527 | 0.358 | 9.72E-44 | TAM.Gdf15 | Cacul1 |
| Phf23 | 7.39E-48 | 0.29078446 | 0.641 | 0.538 | 1.43E-43 | TAM.Gdf15 | Phf23 |
| Steap3 | 9.89E-48 | 0.64724602 | 0.264 | 0.134 | 1.91E-43 | TAM.Gdf15 | Steap3 |
| Golt1b2 | 1.10E-47 | 0.64306926 | 0.463 | 0.303 | 2.12E-43 | TAM.Gdf15 | Golt1b |
| Prkar2b1 | 1.28E-47 | 1.0328792 | 0.28 | 0.139 | 2.48E-43 | TAM.Gdf15 | Prkar2b |
| Slc37a22 | 1.59E-47 | 0.44442351 | 0.551 | 0.378 | 3.07E-43 | TAM.Gdf15 | Slc37a2 |
| Ctnnd12 | 2.14E-47 | 0.40123515 | 0.703 | 0.535 | 4.15E-43 | TAM.Gdf15 | Ctnnd1 |
| Kifc31 | 2.20E-47 | 0.67681592 | 0.327 | 0.167 | 4.26E-43 | TAM.Gdf15 | Kifc3 |
| Apobec12 | 3.54E-47 | 0.29619312 | 0.61 | 0.474 | 6.85E-43 | TAM.Gdf15 | Apobec1 |
| Mink12 | 4.59E-47 | 0.3484035 | 0.572 | 0.417 | 8.87E-43 | TAM.Gdf15 | Mink1 |
| Rufy33 | 4.85E-47 | 0.81169034 | 0.45 | 0.277 | 9.38E-43 | TAM.Gdf15 | Rufy3 |
| Clec10a1 | 5.21E-47 | 0.61068105 | 0.322 | 0.222 | 1.01E-42 | TAM.Gdf15 | Clec10a |
| Stx122 | 7.44E-47 | 0.28665591 | 0.692 | 0.553 | 1.44E-42 | TAM.Gdf15 | Stx12 |
| Tgoln11 | 7.96E-47 | 0.2727494 | 0.677 | 0.543 | 1.54E-42 | TAM.Gdf15 | Tgoln1 |
| Ube2m1 | 1.35E-46 | 0.39851596 | 0.648 | 0.502 | 2.62E-42 | TAM.Gdf15 | Ube2m |
| Snx271 | 1.49E-46 | 0.66911942 | 0.451 | 0.277 | 2.89E-42 | TAM.Gdf15 | Snx27 |
| Dnajb4 | 1.55E-46 | 0.92462255 | 0.25 | 0.139 | 2.99E-42 | TAM.Gdf15 | Dnajb4 |
| Rab2a | 2.69E-46 | 0.27652274 | 0.68 | 0.56 | 5.20E-42 | TAM.Gdf15 | Rab2a |
| Laptm4a1 | 2.88E-46 | 0.2666246 | 0.587 | 0.47 | 5.56E-42 | TAM.Gdf15 | Laptm4a |
| Lpxn1 | 4.08E-46 | 0.62861356 | 0.516 | 0.34 | 7.90E-42 | TAM.Gdf15 | Lpxn |
| Mroh13 | 5.44E-46 | 0.47641509 | 0.59 | 0.442 | 1.05E-41 | TAM.Gdf15 | Mroh1 |
| Pik3r52 | 6.39E-46 | 0.40657837 | 0.686 | 0.557 | 1.24E-41 | TAM.Gdf15 | Pik3r5 |
| Nfic3 | 6.63E-46 | 0.56545592 | 0.464 | 0.3 | 1.28E-41 | TAM.Gdf15 | Nfic |
| Dnajb92 | 7.84E-46 | 0.38103683 | 0.501 | 0.373 | 1.52E-41 | TAM.Gdf15 | Dnajb9 |
| Selenos1 | 8.19E-46 | 0.56836782 | 0.492 | 0.337 | 1.58E-41 | TAM.Gdf15 | Selenos |
| Sephs23 | 9.25E-46 | 0.48104324 | 0.524 | 0.366 | 1.79E-41 | TAM.Gdf15 | Sephs2 |
| Mocos1 | 1.13E-45 | 0.89409508 | 0.286 | 0.152 | 2.18E-41 | TAM.Gdf15 | Mocos |
| Gnaq | 1.19E-45 | 0.25064459 | 0.724 | 0.612 | 2.31E-41 | TAM.Gdf15 | Gnaq |
| Nagk3 | 1.45E-45 | 0.60020561 | 0.451 | 0.279 | 2.80E-41 | TAM.Gdf15 | Nagk |
| Tshz11 | 1.55E-45 | 1.21105666 | 0.262 | 0.127 | 2.99E-41 | TAM.Gdf15 | Tshz1 |
| Tmem106b2 | 2.21E-45 | 0.56576624 | 0.393 | 0.255 | 4.27E-41 | TAM.Gdf15 | Tmem106b |
| Tom1l2 | 2.79E-45 | 1.04772668 | 0.255 | 0.132 | 5.39E-41 | TAM.Gdf15 | Tom1l2 |
| Rad23b1 | 2.83E-45 | 0.37889955 | 0.569 | 0.441 | 5.46E-41 | TAM.Gdf15 | Rad23b |
| Klhl92 | 3.05E-45 | 0.41893463 | 0.7 | 0.52 | 5.90E-41 | TAM.Gdf15 | Klhl9 |
| Slc49a41 | 3.06E-45 | 0.32231696 | 0.588 | 0.443 | 5.92E-41 | TAM.Gdf15 | Slc49a4 |
| Psma32 | 3.13E-45 | 0.30335919 | 0.502 | 0.391 | 6.04E-41 | TAM.Gdf15 | Psma3 |
| Rock21 | 7.31E-45 | 0.59911025 | 0.493 | 0.331 | 1.41E-40 | TAM.Gdf15 | Rock2 |
| Bicd21 | 1.07E-44 | 0.44172994 | 0.531 | 0.38 | 2.07E-40 | TAM.Gdf15 | Bicd2 |
| Exoc6b | 1.11E-44 | 0.82191262 | 0.327 | 0.186 | 2.15E-40 | TAM.Gdf15 | Exoc6b |
| Sntb21 | 1.38E-44 | 0.94977836 | 0.367 | 0.23 | 2.66E-40 | TAM.Gdf15 | Sntb2 |
| Car13 | 1.52E-44 | 0.66971538 | 0.295 | 0.153 | 2.94E-40 | TAM.Gdf15 | Car13 |
| Rab101 | 2.20E-44 | 0.39952787 | 0.573 | 0.414 | 4.25E-40 | TAM.Gdf15 | Rab10 |
| Chst121 | 3.57E-44 | 0.36746713 | 0.43 | 0.283 | 6.89E-40 | TAM.Gdf15 | Chst12 |
| Sept112 | 4.45E-44 | 0.28268394 | 0.503 | 0.353 | 8.60E-40 | TAM.Gdf15 | 11-Sep |
| Mia3 | 4.91E-44 | 0.34846775 | 0.603 | 0.47 | 9.49E-40 | TAM.Gdf15 | Mia3 |
| Creld14 | 6.60E-44 | 0.75305886 | 0.352 | 0.208 | 1.28E-39 | TAM.Gdf15 | Creld1 |
| Gmfb2 | 1.02E-43 | 0.31648455 | 0.679 | 0.531 | 1.97E-39 | TAM.Gdf15 | Gmfb |
| Ghitm | 1.08E-43 | 0.43520222 | 0.549 | 0.422 | 2.10E-39 | TAM.Gdf15 | Ghitm |
| Cdc42bpb1 | 1.56E-43 | 0.47782859 | 0.533 | 0.384 | 3.02E-39 | TAM.Gdf15 | Cdc42bpb |
| Ell3 | 1.79E-43 | 0.65390423 | 0.462 | 0.296 | 3.45E-39 | TAM.Gdf15 | Ell |
| Snx82 | 2.13E-43 | 0.31251181 | 0.65 | 0.488 | 4.11E-39 | TAM.Gdf15 | Snx8 |
| Acadl2 | 4.03E-43 | 0.5440909 | 0.477 | 0.322 | 7.78E-39 | TAM.Gdf15 | Acadl |
| Selenok1 | 4.97E-43 | 0.34033479 | 0.585 | 0.444 | 9.60E-39 | TAM.Gdf15 | Selenok |
| Ubxn11 | 5.23E-43 | 0.25480361 | 0.747 | 0.629 | 1.01E-38 | TAM.Gdf15 | Ubxn1 |
| Ptges32 | 5.34E-43 | 0.39960233 | 0.519 | 0.374 | 1.03E-38 | TAM.Gdf15 | Ptges3 |
| Cln51 | 5.52E-43 | 0.36009275 | 0.576 | 0.425 | 1.07E-38 | TAM.Gdf15 | Cln5 |
| Cttn | 6.09E-43 | 0.84291759 | 0.266 | 0.134 | 1.18E-38 | TAM.Gdf15 | Cttn |
| Tbc1d20 | 1.71E-42 | 0.35977507 | 0.524 | 0.395 | 3.31E-38 | TAM.Gdf15 | Tbc1d20 |
| Pip4p21 | 1.91E-42 | 0.81717072 | 0.291 | 0.164 | 3.69E-38 | TAM.Gdf15 | Pip4p2 |
| Snx292 | 1.95E-42 | 0.4267138 | 0.649 | 0.473 | 3.77E-38 | TAM.Gdf15 | Snx29 |
| Hdgf2 | 3.51E-42 | 0.49375945 | 0.579 | 0.416 | 6.78E-38 | TAM.Gdf15 | Hdgf |
| Pafah1b21 | 4.00E-42 | 0.28330846 | 0.59 | 0.454 | 7.73E-38 | TAM.Gdf15 | Pafah1b2 |
| Polg21 | 4.24E-42 | 0.81446615 | 0.297 | 0.177 | 8.20E-38 | TAM.Gdf15 | Polg2 |
| Pnpla81 | 4.32E-42 | 0.69919481 | 0.394 | 0.253 | 8.35E-38 | TAM.Gdf15 | Pnpla8 |
| Flcn2 | 5.26E-42 | 0.32699157 | 0.653 | 0.533 | 1.02E-37 | TAM.Gdf15 | Flcn |
| Eif1a2 | 5.37E-42 | 0.36524688 | 0.571 | 0.43 | 1.04E-37 | TAM.Gdf15 | Eif1a |
| Wdtc1 | 5.59E-42 | 0.89728219 | 0.299 | 0.172 | 1.08E-37 | TAM.Gdf15 | Wdtc1 |
| Ufd1 | 1.09E-41 | 0.48235534 | 0.428 | 0.299 | 2.10E-37 | TAM.Gdf15 | Ufd1 |
| Ubr4 | 1.12E-41 | 0.34975605 | 0.564 | 0.467 | 2.16E-37 | TAM.Gdf15 | Ubr4 |
| Rab6a2 | 1.19E-41 | 0.40991141 | 0.528 | 0.411 | 2.31E-37 | TAM.Gdf15 | Rab6a |
| BC031181 | 1.29E-41 | 0.32741356 | 0.473 | 0.369 | 2.50E-37 | TAM.Gdf15 | BC031181 |
| Slc38a73 | 1.71E-41 | 0.76179173 | 0.409 | 0.242 | 3.30E-37 | TAM.Gdf15 | Slc38a7 |
| Parp31 | 3.10E-41 | 0.75566606 | 0.317 | 0.168 | 5.99E-37 | TAM.Gdf15 | Parp3 |
| Pip4p13 | 4.23E-41 | 0.27421372 | 0.634 | 0.502 | 8.18E-37 | TAM.Gdf15 | Pip4p1 |
| Psmc41 | 4.41E-41 | 0.38195929 | 0.542 | 0.41 | 8.52E-37 | TAM.Gdf15 | Psmc4 |
| Inppl14 | 4.48E-41 | 0.48136927 | 0.56 | 0.405 | 8.66E-37 | TAM.Gdf15 | Inppl1 |
| Npepps2 | 4.88E-41 | 0.5782071 | 0.579 | 0.405 | 9.44E-37 | TAM.Gdf15 | Npepps |
| Fbxo222 | 6.53E-41 | 0.80986218 | 0.37 | 0.224 | 1.26E-36 | TAM.Gdf15 | Fbxo22 |
| Tbk1 | 8.80E-41 | 0.43690471 | 0.578 | 0.431 | 1.70E-36 | TAM.Gdf15 | Tbk1 |
| Itga52 | 9.17E-41 | 0.27728905 | 0.633 | 0.535 | 1.77E-36 | TAM.Gdf15 | Itga5 |
| Stard53 | 1.19E-40 | 0.76731806 | 0.393 | 0.225 | 2.29E-36 | TAM.Gdf15 | Stard5 |
| Ppt21 | 1.47E-40 | 0.35470391 | 0.531 | 0.382 | 2.84E-36 | TAM.Gdf15 | Ppt2 |
| Ide1 | 1.71E-40 | 0.45334225 | 0.483 | 0.343 | 3.31E-36 | TAM.Gdf15 | Ide |
| Gas2l33 | 2.15E-40 | 0.53877378 | 0.452 | 0.285 | 4.17E-36 | TAM.Gdf15 | Gas2l3 |
| Cln32 | 2.46E-40 | 0.34676633 | 0.588 | 0.422 | 4.75E-36 | TAM.Gdf15 | Cln3 |
| Aopep | 2.58E-40 | 0.99110241 | 0.26 | 0.143 | 4.99E-36 | TAM.Gdf15 | Aopep |
| B4galt7 | 2.71E-40 | 0.99866128 | 0.259 | 0.133 | 5.25E-36 | TAM.Gdf15 | B4galt7 |
| Aldh3b11 | 3.57E-40 | 0.29115875 | 0.543 | 0.418 | 6.91E-36 | TAM.Gdf15 | Aldh3b1 |
| Hgsnat2 | 5.55E-40 | 0.30979095 | 0.348 | 0.29 | 1.07E-35 | TAM.Gdf15 | Hgsnat |
| B3gnt22 | 6.32E-40 | 0.61283059 | 0.38 | 0.248 | 1.22E-35 | TAM.Gdf15 | B3gnt2 |
| Oser12 | 9.10E-40 | 0.51019255 | 0.579 | 0.443 | 1.76E-35 | TAM.Gdf15 | Oser1 |
| Kansl21 | 1.10E-39 | 0.6702906 | 0.455 | 0.317 | 2.13E-35 | TAM.Gdf15 | Kansl2 |
| Cerk3 | 1.11E-39 | 0.25664781 | 0.595 | 0.47 | 2.14E-35 | TAM.Gdf15 | Cerk |
| Memo11 | 1.74E-39 | 0.77411055 | 0.341 | 0.205 | 3.36E-35 | TAM.Gdf15 | Memo1 |
| Mphosph101 | 1.96E-39 | 0.78997405 | 0.313 | 0.177 | 3.79E-35 | TAM.Gdf15 | Mphosph10 |
| Tmem81 | 2.23E-39 | 0.43480277 | 0.394 | 0.216 | 4.31E-35 | TAM.Gdf15 | Tmem8 |
| Fam45a1 | 2.60E-39 | 0.66095971 | 0.348 | 0.198 | 5.04E-35 | TAM.Gdf15 | Fam45a |
| Akr1b32 | 3.57E-39 | 0.31326637 | 0.521 | 0.399 | 6.91E-35 | TAM.Gdf15 | Akr1b3 |
| Xylt13 | 5.16E-39 | 0.71112928 | 0.41 | 0.262 | 9.97E-35 | TAM.Gdf15 | Xylt1 |
| Usp51 | 5.46E-39 | 0.54798104 | 0.507 | 0.373 | 1.05E-34 | TAM.Gdf15 | Usp5 |
| Notch2 | 5.56E-39 | 0.29598525 | 0.651 | 0.552 | 1.08E-34 | TAM.Gdf15 | Notch2 |
| Chd32 | 5.90E-39 | 0.54438767 | 0.401 | 0.251 | 1.14E-34 | TAM.Gdf15 | Chd3 |
| Abhd43 | 1.47E-38 | 0.53409754 | 0.42 | 0.286 | 2.85E-34 | TAM.Gdf15 | Abhd4 |
| Blnk4 | 1.70E-38 | 0.60761026 | 0.377 | 0.219 | 3.28E-34 | TAM.Gdf15 | Blnk |
| Crtap2 | 2.36E-38 | 0.4040074 | 0.409 | 0.27 | 4.55E-34 | TAM.Gdf15 | Crtap |
| St6gal12 | 2.43E-38 | 0.37252461 | 0.441 | 0.3 | 4.69E-34 | TAM.Gdf15 | St6gal1 |
| Sigmar13 | 2.69E-38 | 0.43059227 | 0.464 | 0.33 | 5.21E-34 | TAM.Gdf15 | Sigmar1 |
| Mfsd51 | 3.39E-38 | 0.42915061 | 0.542 | 0.394 | 6.55E-34 | TAM.Gdf15 | Mfsd5 |
| Psd34 | 3.69E-38 | 0.57502041 | 0.424 | 0.285 | 7.12E-34 | TAM.Gdf15 | Psd3 |
| Crlf31 | 4.01E-38 | 0.31396495 | 0.58 | 0.455 | 7.75E-34 | TAM.Gdf15 | Crlf3 |
| Fgd63 | 4.73E-38 | 0.98468459 | 0.336 | 0.186 | 9.14E-34 | TAM.Gdf15 | Fgd6 |
| Magt11 | 4.88E-38 | 0.25755717 | 0.588 | 0.476 | 9.43E-34 | TAM.Gdf15 | Magt1 |
| Tfeb1 | 5.22E-38 | 0.41110084 | 0.455 | 0.303 | 1.01E-33 | TAM.Gdf15 | Tfeb |
| Fam50a | 5.26E-38 | 0.32108471 | 0.563 | 0.424 | 1.02E-33 | TAM.Gdf15 | Fam50a |
| Iqgap22 | 5.81E-38 | 0.62206069 | 0.424 | 0.289 | 1.12E-33 | TAM.Gdf15 | Iqgap2 |
| Cdc342 | 6.18E-38 | 0.41427861 | 0.573 | 0.424 | 1.19E-33 | TAM.Gdf15 | Cdc34 |
| Tns32 | 6.75E-38 | 0.3571048 | 0.631 | 0.464 | 1.31E-33 | TAM.Gdf15 | Tns3 |
| Pafah1b11 | 9.12E-38 | 0.37051402 | 0.59 | 0.469 | 1.76E-33 | TAM.Gdf15 | Pafah1b1 |
| Vav3 | 1.22E-37 | 0.6486199 | 0.371 | 0.224 | 2.36E-33 | TAM.Gdf15 | Vav3 |
| Dennd1a2 | 1.68E-37 | 0.36855729 | 0.513 | 0.37 | 3.25E-33 | TAM.Gdf15 | Dennd1a |
| Arhgap121 | 1.99E-37 | 0.83166301 | 0.329 | 0.186 | 3.85E-33 | TAM.Gdf15 | Arhgap12 |
| Atg51 | 2.03E-37 | 0.33660079 | 0.43 | 0.305 | 3.92E-33 | TAM.Gdf15 | Atg5 |
| Tbc1d231 | 2.18E-37 | 0.83852227 | 0.317 | 0.19 | 4.21E-33 | TAM.Gdf15 | Tbc1d23 |
| Pdlim11 | 2.46E-37 | 0.46219149 | 0.251 | 0.124 | 4.75E-33 | TAM.Gdf15 | Pdlim1 |
| Trappc4 | 2.67E-37 | 0.44711182 | 0.565 | 0.423 | 5.16E-33 | TAM.Gdf15 | Trappc4 |
| Sec24d | 2.90E-37 | 0.73154071 | 0.281 | 0.166 | 5.61E-33 | TAM.Gdf15 | Sec24d |
| Smcr81 | 3.22E-37 | 0.38832186 | 0.649 | 0.483 | 6.23E-33 | TAM.Gdf15 | Smcr8 |
| Tmem63b2 | 3.51E-37 | 0.76285657 | 0.333 | 0.189 | 6.78E-33 | TAM.Gdf15 | Tmem63b |
| Morf4l22 | 4.84E-37 | 0.42179441 | 0.478 | 0.362 | 9.35E-33 | TAM.Gdf15 | Morf4l2 |
| Taok11 | 5.77E-37 | 0.2868763 | 0.566 | 0.443 | 1.12E-32 | TAM.Gdf15 | Taok1 |
| Gnl33 | 6.69E-37 | 0.42659858 | 0.449 | 0.326 | 1.29E-32 | TAM.Gdf15 | Gnl3 |
| Nudt93 | 7.24E-37 | 0.33556856 | 0.529 | 0.382 | 1.40E-32 | TAM.Gdf15 | Nudt9 |
| Med151 | 9.82E-37 | 0.27764541 | 0.707 | 0.61 | 1.90E-32 | TAM.Gdf15 | Med15 |
| Numa12 | 1.09E-36 | 0.41943556 | 0.533 | 0.377 | 2.10E-32 | TAM.Gdf15 | Numa1 |
| Plaa1 | 1.42E-36 | 0.57607914 | 0.488 | 0.348 | 2.75E-32 | TAM.Gdf15 | Plaa |
| Ttyh23 | 2.34E-36 | 0.56090878 | 0.449 | 0.29 | 4.52E-32 | TAM.Gdf15 | Ttyh2 |
| Blcap3 | 3.05E-36 | 0.60752702 | 0.43 | 0.3 | 5.89E-32 | TAM.Gdf15 | Blcap |
| Srsf51 | 3.08E-36 | 0.28877803 | 0.777 | 0.682 | 5.95E-32 | TAM.Gdf15 | Srsf5 |
| S1pr2 | 3.19E-36 | 0.5559651 | 0.316 | 0.187 | 6.16E-32 | TAM.Gdf15 | S1pr2 |
| Tnpo21 | 4.29E-36 | 0.45341042 | 0.485 | 0.357 | 8.29E-32 | TAM.Gdf15 | Tnpo2 |
| Idua2 | 6.85E-36 | 0.5238941 | 0.392 | 0.261 | 1.32E-31 | TAM.Gdf15 | Idua |
| Pced1b | 1.13E-35 | 0.98625937 | 0.251 | 0.117 | 2.19E-31 | TAM.Gdf15 | Pced1b |
| Scyl1 | 1.23E-35 | 0.3553719 | 0.626 | 0.492 | 2.38E-31 | TAM.Gdf15 | Scyl1 |
| Wbp22 | 2.68E-35 | 0.30168922 | 0.576 | 0.427 | 5.19E-31 | TAM.Gdf15 | Wbp2 |
| Sfmbt13 | 3.09E-35 | 0.62808105 | 0.394 | 0.253 | 5.97E-31 | TAM.Gdf15 | Sfmbt1 |
| Pde3b2 | 3.31E-35 | 0.48250498 | 0.465 | 0.322 | 6.40E-31 | TAM.Gdf15 | Pde3b |
| Spg20 | 5.26E-35 | 0.87044545 | 0.265 | 0.143 | 1.02E-30 | TAM.Gdf15 | Spg20 |
| Dnajc51 | 6.74E-35 | 0.28803109 | 0.671 | 0.545 | 1.30E-30 | TAM.Gdf15 | Dnajc5 |
| Keap12 | 7.75E-35 | 0.38699512 | 0.448 | 0.314 | 1.50E-30 | TAM.Gdf15 | Keap1 |
| Osbpl1a2 | 8.95E-35 | 0.64460661 | 0.319 | 0.181 | 1.73E-30 | TAM.Gdf15 | Osbpl1a |
| Snx112 | 1.10E-34 | 0.58622096 | 0.365 | 0.228 | 2.13E-30 | TAM.Gdf15 | Snx11 |
| Uhrf1bp1l3 | 1.67E-34 | 0.59756052 | 0.358 | 0.243 | 3.23E-30 | TAM.Gdf15 | Uhrf1bp1l |
| Cops51 | 1.90E-34 | 0.405454 | 0.476 | 0.344 | 3.68E-30 | TAM.Gdf15 | Cops5 |
| Ccdc472 | 1.95E-34 | 0.46240145 | 0.444 | 0.313 | 3.77E-30 | TAM.Gdf15 | Ccdc47 |
| Ogt2 | 2.01E-34 | 0.34015628 | 0.594 | 0.465 | 3.89E-30 | TAM.Gdf15 | Ogt |
| Gnptg3 | 2.03E-34 | 0.60693611 | 0.35 | 0.213 | 3.92E-30 | TAM.Gdf15 | Gnptg |
| Ufm1 | 2.13E-34 | 0.35001163 | 0.486 | 0.365 | 4.12E-30 | TAM.Gdf15 | Ufm1 |
| Tns12 | 2.60E-34 | 0.59307976 | 0.329 | 0.186 | 5.02E-30 | TAM.Gdf15 | Tns1 |
| Smim7 | 2.62E-34 | 0.26308628 | 0.484 | 0.358 | 5.07E-30 | TAM.Gdf15 | Smim7 |
| Lrrc8a3 | 2.87E-34 | 0.62611894 | 0.372 | 0.231 | 5.54E-30 | TAM.Gdf15 | Lrrc8a |
| Larp41 | 3.09E-34 | 0.42357002 | 0.55 | 0.407 | 5.97E-30 | TAM.Gdf15 | Larp4 |
| Rab122 | 4.19E-34 | 0.36994125 | 0.493 | 0.357 | 8.10E-30 | TAM.Gdf15 | Rab12 |
| Frrs11 | 5.30E-34 | 0.43671786 | 0.49 | 0.35 | 1.02E-29 | TAM.Gdf15 | Frrs1 |
| Tbc1d15 | 5.59E-34 | 0.49848209 | 0.523 | 0.394 | 1.08E-29 | TAM.Gdf15 | Tbc1d15 |
| Tom11 | 5.62E-34 | 0.57091766 | 0.423 | 0.273 | 1.09E-29 | TAM.Gdf15 | Tom1 |
| Spire12 | 6.05E-34 | 0.67609289 | 0.319 | 0.187 | 1.17E-29 | TAM.Gdf15 | Spire1 |
| Rassf12 | 7.54E-34 | 0.37003904 | 0.564 | 0.49 | 1.46E-29 | TAM.Gdf15 | Rassf1 |
| Rex1bd1 | 9.58E-34 | 0.31398953 | 0.402 | 0.281 | 1.85E-29 | TAM.Gdf15 | Rex1bd |
| Hist3h2a2 | 1.01E-33 | 0.69939527 | 0.307 | 0.189 | 1.95E-29 | TAM.Gdf15 | Hist3h2a |
| Pnpla21 | 1.25E-33 | 0.36174968 | 0.633 | 0.515 | 2.42E-29 | TAM.Gdf15 | Pnpla2 |
| Ccdc9 | 1.33E-33 | 0.48299869 | 0.567 | 0.434 | 2.57E-29 | TAM.Gdf15 | Ccdc9 |
| Mid1ip12 | 2.43E-33 | 0.641217 | 0.4 | 0.241 | 4.69E-29 | TAM.Gdf15 | Mid1ip1 |
| Comtd12 | 2.53E-33 | 0.5188981 | 0.33 | 0.196 | 4.90E-29 | TAM.Gdf15 | Comtd1 |
| Nras | 2.70E-33 | 0.30935237 | 0.612 | 0.477 | 5.21E-29 | TAM.Gdf15 | Nras |
| Ptpmt11 | 3.01E-33 | 0.54008135 | 0.407 | 0.27 | 5.82E-29 | TAM.Gdf15 | Ptpmt1 |
| Spen2 | 3.07E-33 | 0.43161754 | 0.517 | 0.378 | 5.95E-29 | TAM.Gdf15 | Spen |
| Dnajb12 | 3.55E-33 | 0.46766469 | 0.57 | 0.403 | 6.85E-29 | TAM.Gdf15 | Dnajb12 |
| Gab21 | 5.22E-33 | 0.36743887 | 0.645 | 0.509 | 1.01E-28 | TAM.Gdf15 | Gab2 |
| Tmem2142 | 7.13E-33 | 0.44375909 | 0.43 | 0.296 | 1.38E-28 | TAM.Gdf15 | Tmem214 |
| Itpripl21 | 7.31E-33 | 0.30307042 | 0.697 | 0.562 | 1.41E-28 | TAM.Gdf15 | Itpripl2 |
| Lactb2 | 8.52E-33 | 0.80795478 | 0.27 | 0.149 | 1.65E-28 | TAM.Gdf15 | Lactb2 |
| Heatr5a2 | 9.20E-33 | 0.43174728 | 0.473 | 0.328 | 1.78E-28 | TAM.Gdf15 | Heatr5a |
| Ppm1a1 | 9.31E-33 | 0.37633208 | 0.641 | 0.495 | 1.80E-28 | TAM.Gdf15 | Ppm1a |
| Ggnbp2 | 1.01E-32 | 0.35514098 | 0.478 | 0.36 | 1.96E-28 | TAM.Gdf15 | Ggnbp2 |
| Tmem41b1 | 1.34E-32 | 0.66672764 | 0.333 | 0.219 | 2.60E-28 | TAM.Gdf15 | Tmem41b |
| Arfgap32 | 1.40E-32 | 0.67509894 | 0.386 | 0.25 | 2.71E-28 | TAM.Gdf15 | Arfgap3 |
| B4galt63 | 1.80E-32 | 0.26535804 | 0.531 | 0.403 | 3.48E-28 | TAM.Gdf15 | B4galt6 |
| Nsfl1c1 | 2.55E-32 | 0.5699266 | 0.415 | 0.283 | 4.92E-28 | TAM.Gdf15 | Nsfl1c |
| Mocs12 | 2.64E-32 | 0.38403497 | 0.384 | 0.257 | 5.10E-28 | TAM.Gdf15 | Mocs1 |
| Etfrf1 | 3.08E-32 | 0.75998053 | 0.271 | 0.15 | 5.96E-28 | TAM.Gdf15 | Etfrf1 |
| Zfp911 | 3.19E-32 | 0.29949321 | 0.633 | 0.523 | 6.16E-28 | TAM.Gdf15 | Zfp91 |
| Nktr | 4.20E-32 | 0.372605 | 0.556 | 0.44 | 8.12E-28 | TAM.Gdf15 | Nktr |
| Afg3l21 | 5.53E-32 | 0.43391343 | 0.569 | 0.426 | 1.07E-27 | TAM.Gdf15 | Afg3l2 |

| | | | | | | | |
|---|---|---|---|---|---|---|---|
| Abl21 | 6.03E-32 | 0.36336687 | 0.583 | 0.441 | 1.17E-27 | TAM.Gdf15 | Abl2 |
| Bahd11 | 7.09E-32 | 0.64930089 | 0.426 | 0.272 | 1.37E-27 | TAM.Gdf15 | Bahd1 |
| Klc42 | 7.53E-32 | 0.46544431 | 0.414 | 0.275 | 1.46E-27 | TAM.Gdf15 | Klc4 |
| Esyt21 | 1.04E-31 | 0.2876237 | 0.513 | 0.411 | 2.01E-27 | TAM.Gdf15 | Esyt2 |
| Ube2g21 | 1.44E-31 | 0.3922828 | 0.55 | 0.411 | 2.79E-27 | TAM.Gdf15 | Ube2g2 |
| Ube2g12 | 1.53E-31 | 0.56605999 | 0.476 | 0.325 | 2.96E-27 | TAM.Gdf15 | Ube2g1 |
| Atg121 | 2.56E-31 | 0.53689899 | 0.338 | 0.223 | 4.94E-27 | TAM.Gdf15 | Atg12 |
| Rnf11 | 2.96E-31 | 0.440797 | 0.421 | 0.289 | 5.72E-27 | TAM.Gdf15 | Rnf11 |
| Pdgfa3 | 2.96E-31 | 0.47962765 | 0.394 | 0.253 | 5.72E-27 | TAM.Gdf15 | Pdgfa |
| Cryzl12 | 3.19E-31 | 0.52330627 | 0.34 | 0.211 | 6.16E-27 | TAM.Gdf15 | Cryzl1 |
| Ergic13 | 3.22E-31 | 0.26552718 | 0.521 | 0.397 | 6.23E-27 | TAM.Gdf15 | Ergic1 |
| Vti1b1 | 3.28E-31 | 0.25541431 | 0.462 | 0.349 | 6.34E-27 | TAM.Gdf15 | Vti1b |
| Slc11a21 | 3.47E-31 | 0.4632838 | 0.458 | 0.314 | 6.70E-27 | TAM.Gdf15 | Slc11a2 |
| Adrm1 | 3.94E-31 | 0.66088417 | 0.363 | 0.224 | 7.62E-27 | TAM.Gdf15 | Adrm1 |
| Stard3nl2 | 5.66E-31 | 0.37788825 | 0.436 | 0.314 | 1.10E-26 | TAM.Gdf15 | Stard3nl |
| Pctp1 | 8.13E-31 | 0.55024794 | 0.265 | 0.161 | 1.57E-26 | TAM.Gdf15 | Pctp |
| Rras3 | 1.54E-30 | 0.329482 | 0.409 | 0.288 | 2.98E-26 | TAM.Gdf15 | Rras |
| Vegfb3 | 1.79E-30 | 0.68797433 | 0.364 | 0.214 | 3.46E-26 | TAM.Gdf15 | Vegfb |
| Dop1b1 | 2.32E-30 | 0.56895467 | 0.329 | 0.2 | 4.49E-26 | TAM.Gdf15 | Dop1b |
| Cmas2 | 2.79E-30 | 0.29817049 | 0.458 | 0.344 | 5.40E-26 | TAM.Gdf15 | Cmas |
| Rcn2 | 3.18E-30 | 0.31183036 | 0.434 | 0.301 | 6.15E-26 | TAM.Gdf15 | Rcn2 |
| Herpud21 | 3.70E-30 | 0.44106182 | 0.341 | 0.227 | 7.15E-26 | TAM.Gdf15 | Herpud2 |
| Mesd2 | 4.51E-30 | 0.38921942 | 0.386 | 0.261 | 8.71E-26 | TAM.Gdf15 | Mesd |
| Psmb71 | 5.13E-30 | 0.34651694 | 0.556 | 0.424 | 9.93E-26 | TAM.Gdf15 | Psmb7 |
| Npepl12 | 5.44E-30 | 0.45093655 | 0.53 | 0.377 | 1.05E-25 | TAM.Gdf15 | Npepl1 |
| Yars1 | 7.21E-30 | 0.72048649 | 0.278 | 0.152 | 1.39E-25 | TAM.Gdf15 | Yars |
| Trim472 | 7.60E-30 | 0.75649175 | 0.301 | 0.163 | 1.47E-25 | TAM.Gdf15 | Trim47 |
| Scamp53 | 9.20E-30 | 0.41907553 | 0.403 | 0.241 | 1.78E-25 | TAM.Gdf15 | Scamp5 |
| Specc1 | 9.77E-30 | 0.58891837 | 0.338 | 0.216 | 1.89E-25 | TAM.Gdf15 | Specc1 |
| Ubr3 | 1.27E-29 | 0.45761296 | 0.495 | 0.377 | 2.46E-25 | TAM.Gdf15 | Ubr3 |
| Tbc1d10b1 | 1.30E-29 | 0.40571471 | 0.541 | 0.44 | 2.51E-25 | TAM.Gdf15 | Tbc1d10b |
| Srpk21 | 1.44E-29 | 0.58862523 | 0.306 | 0.192 | 2.78E-25 | TAM.Gdf15 | Srpk2 |
| Mthfd21 | 1.45E-29 | 0.88622126 | 0.273 | 0.166 | 2.80E-25 | TAM.Gdf15 | Mthfd2 |
| Lcor1 | 1.75E-29 | 0.53050471 | 0.459 | 0.34 | 3.38E-25 | TAM.Gdf15 | Lcor |
| Nars | 1.76E-29 | 0.30985162 | 0.477 | 0.36 | 3.39E-25 | TAM.Gdf15 | Nars |
| Agap31 | 1.94E-29 | 0.32509996 | 0.601 | 0.461 | 3.74E-25 | TAM.Gdf15 | Agap3 |
| Abcc52 | 2.23E-29 | 0.51784752 | 0.457 | 0.276 | 4.31E-25 | TAM.Gdf15 | Abcc5 |
| Smg11 | 2.29E-29 | 0.3355414 | 0.56 | 0.452 | 4.43E-25 | TAM.Gdf15 | Smg1 |
| Lrrc8d1 | 2.76E-29 | 0.26434375 | 0.476 | 0.37 | 5.34E-25 | TAM.Gdf15 | Lrrc8d |
| Sec62 | 2.83E-29 | 0.40305817 | 0.599 | 0.474 | 5.47E-25 | TAM.Gdf15 | Sec62 |
| Cyp20a11 | 2.88E-29 | 0.49494012 | 0.265 | 0.16 | 5.57E-25 | TAM.Gdf15 | Cyp20a1 |
| Pde8a2 | 3.87E-29 | 0.47302691 | 0.421 | 0.286 | 7.48E-25 | TAM.Gdf15 | Pde8a |
| Elf22 | 4.17E-29 | 0.40035157 | 0.472 | 0.341 | 8.07E-25 | TAM.Gdf15 | Elf2 |
| Hdac41 | 6.06E-29 | 0.26141291 | 0.378 | 0.283 | 1.17E-24 | TAM.Gdf15 | Hdac4 |
| Tulp41 | 6.08E-29 | 0.62313458 | 0.315 | 0.195 | 1.18E-24 | TAM.Gdf15 | Tulp4 |
| Rnf62 | 6.40E-29 | 0.26409334 | 0.649 | 0.554 | 1.24E-24 | TAM.Gdf15 | Rnf6 |
| Cpeb23 | 8.58E-29 | 0.26409146 | 0.555 | 0.435 | 1.66E-24 | TAM.Gdf15 | Cpeb2 |
| Tmcc12 | 9.25E-29 | 0.25476996 | 0.534 | 0.411 | 1.79E-24 | TAM.Gdf15 | Tmcc1 |
| Uso12 | 9.95E-29 | 0.33031738 | 0.445 | 0.34 | 1.92E-24 | TAM.Gdf15 | Uso1 |
| Dpagt11 | 1.32E-28 | 0.46446938 | 0.358 | 0.243 | 2.55E-24 | TAM.Gdf15 | Dpagt1 |
| Mpv172 | 1.57E-28 | 0.40964365 | 0.363 | 0.231 | 3.04E-24 | TAM.Gdf15 | Mpv17 |
| Ppp4r1 | 1.94E-28 | 0.55052819 | 0.33 | 0.208 | 3.74E-24 | TAM.Gdf15 | Ppp4r1 |
| DEPRECATED-I | 2.02E-28 | 0.40583481 | 0.359 | 0.24 | 3.90E-24 | TAM.Gdf15 | DEPRECATED-ENSMUSG00000002103 |
| Mindy22 | 2.06E-28 | 0.27625959 | 0.471 | 0.362 | 3.99E-24 | TAM.Gdf15 | Mindy2 |
| Commd31 | 2.20E-28 | 0.36366522 | 0.401 | 0.283 | 4.26E-24 | TAM.Gdf15 | Commd3 |
| Zfp655 | 2.63E-28 | 0.60644464 | 0.435 | 0.308 | 5.08E-24 | TAM.Gdf15 | Zfp655 |
| Snx132 | 2.95E-28 | 0.55898338 | 0.392 | 0.261 | 5.71E-24 | TAM.Gdf15 | Snx13 |
| Cct32 | 3.29E-28 | 0.28155506 | 0.476 | 0.365 | 6.37E-24 | TAM.Gdf15 | Cct3 |
| Slamf71 | 3.34E-28 | 0.6711956 | 0.271 | 0.156 | 6.45E-24 | TAM.Gdf15 | Slamf7 |
| Nucb22 | 3.73E-28 | 0.27209139 | 0.428 | 0.336 | 7.21E-24 | TAM.Gdf15 | Nucb2 |
| Elf51 | 3.97E-28 | 0.33437616 | 0.457 | 0.348 | 7.68E-24 | TAM.Gdf15 | Elf5 |
| Prpf39 | 4.01E-28 | 0.50080682 | 0.485 | 0.382 | 7.75E-24 | TAM.Gdf15 | Prpf39 |
| Tank1 | 5.06E-28 | 0.36391482 | 0.601 | 0.477 | 9.79E-24 | TAM.Gdf15 | Tank |
| Prkaa12 | 6.00E-28 | 0.4347088 | 0.371 | 0.247 | 1.16E-23 | TAM.Gdf15 | Prkaa1 |
| Ptpn231 | 8.18E-28 | 0.45518887 | 0.46 | 0.347 | 1.58E-23 | TAM.Gdf15 | Ptpn23 |
| Cyb5r12 | 1.03E-27 | 0.7190486 | 0.379 | 0.271 | 1.99E-23 | TAM.Gdf15 | Cyb5r1 |
| Atf12 | 1.15E-27 | 0.27598816 | 0.648 | 0.527 | 2.23E-23 | TAM.Gdf15 | Atf1 |
| Faf11 | 1.31E-27 | 0.43897273 | 0.423 | 0.294 | 2.53E-23 | TAM.Gdf15 | Faf1 |
| Ttc331 | 1.31E-27 | 0.55449684 | 0.298 | 0.179 | 2.54E-23 | TAM.Gdf15 | Ttc33 |
| Sec24a | 1.38E-27 | 0.34010473 | 0.465 | 0.35 | 2.66E-23 | TAM.Gdf15 | Sec24a |
| Serpinb81 | 1.69E-27 | 0.54160802 | 0.285 | 0.165 | 3.26E-23 | TAM.Gdf15 | Serpinb8 |
| Ubald11 | 1.83E-27 | 0.33683335 | 0.447 | 0.319 | 3.53E-23 | TAM.Gdf15 | Ubald1 |
| Kctd6 | 2.53E-27 | 0.7257563 | 0.271 | 0.156 | 4.90E-23 | TAM.Gdf15 | Kctd6 |
| Slc31a22 | 2.54E-27 | 0.46003557 | 0.502 | 0.366 | 4.90E-23 | TAM.Gdf15 | Slc31a2 |
| Pno12 | 2.54E-27 | 0.58948721 | 0.338 | 0.222 | 4.92E-23 | TAM.Gdf15 | Pno1 |
| Scamp11 | 2.70E-27 | 0.38073614 | 0.399 | 0.289 | 5.22E-23 | TAM.Gdf15 | Scamp1 |
| Tbce | 2.73E-27 | 0.81855893 | 0.257 | 0.148 | 5.28E-23 | TAM.Gdf15 | Tbce |
| Plod12 | 3.49E-27 | 0.26212747 | 0.537 | 0.38 | 6.75E-23 | TAM.Gdf15 | Plod1 |
| Pdlim74 | 4.13E-27 | 0.7142156 | 0.476 | 0.343 | 7.99E-23 | TAM.Gdf15 | Pdlim7 |
| Ykt6 | 5.71E-27 | 0.25576622 | 0.533 | 0.429 | 1.10E-22 | TAM.Gdf15 | Ykt6 |
| Xylt21 | 6.54E-27 | 0.48008936 | 0.36 | 0.226 | 1.26E-22 | TAM.Gdf15 | Xylt2 |
| D5Ertd579e1 | 7.49E-27 | 0.41098879 | 0.419 | 0.306 | 1.45E-22 | TAM.Gdf15 | D5Ertd579e |
| Smg52 | 8.02E-27 | 0.43535074 | 0.462 | 0.332 | 1.55E-22 | TAM.Gdf15 | Smg5 |
| Tpra11 | 1.07E-26 | 0.57747883 | 0.297 | 0.177 | 2.07E-22 | TAM.Gdf15 | Tpra1 |
| Ccdc932 | 1.22E-26 | 0.37099704 | 0.377 | 0.256 | 2.37E-22 | TAM.Gdf15 | Ccdc93 |
| Flvcr12 | 1.26E-26 | 0.37568199 | 0.416 | 0.284 | 2.44E-22 | TAM.Gdf15 | Flvcr1 |
| Ecpas | 1.42E-26 | 0.34707923 | 0.484 | 0.374 | 2.75E-22 | TAM.Gdf15 | Ecpas |
| C5ar23 | 1.46E-26 | 0.49734439 | 0.371 | 0.277 | 2.83E-22 | TAM.Gdf15 | C5ar2 |
| Tor1aip21 | 1.54E-26 | 0.30639841 | 0.626 | 0.515 | 2.98E-22 | TAM.Gdf15 | Tor1aip2 |
| Bet12 | 1.77E-26 | 0.50326115 | 0.35 | 0.227 | 3.43E-22 | TAM.Gdf15 | Bet1 |
| Zbtb7b | 2.02E-26 | 0.32233047 | 0.494 | 0.387 | 3.91E-22 | TAM.Gdf15 | Zbtb7b |
| Pik3c2a1 | 2.32E-26 | 0.57914634 | 0.416 | 0.295 | 4.49E-22 | TAM.Gdf15 | Pik3c2a |
| Sike1 | 2.74E-26 | 0.38877382 | 0.413 | 0.28 | 5.30E-22 | TAM.Gdf15 | Sike1 |
| Crk1 | 2.90E-26 | 0.26962856 | 0.636 | 0.517 | 5.60E-22 | TAM.Gdf15 | Crk |
| Chmp7 | 3.42E-26 | 0.73353549 | 0.323 | 0.199 | 6.61E-22 | TAM.Gdf15 | Chmp7 |
| Alg1 | 3.51E-26 | 0.49821984 | 0.387 | 0.254 | 6.79E-22 | TAM.Gdf15 | Alg1 |
| Tollip | 5.29E-26 | 0.41739084 | 0.501 | 0.387 | 1.02E-21 | TAM.Gdf15 | Tollip |
| Lima11 | 5.36E-26 | 0.49264215 | 0.283 | 0.173 | 1.04E-21 | TAM.Gdf15 | Lima1 |
| Ly961 | 5.54E-26 | 0.6547099 | 0.295 | 0.173 | 1.07E-21 | TAM.Gdf15 | Ly96 |
| Rhbdd1 | 6.06E-26 | 0.57483285 | 0.323 | 0.204 | 1.17E-21 | TAM.Gdf15 | Rhbdd1 |
| Git11 | 6.36E-26 | 0.46744554 | 0.428 | 0.307 | 1.23E-21 | TAM.Gdf15 | Git1 |
| Rab22a | 6.44E-26 | 0.28945539 | 0.507 | 0.394 | 1.25E-21 | TAM.Gdf15 | Rab22a |
| Samd4b1 | 7.46E-26 | 0.40410401 | 0.444 | 0.325 | 1.44E-21 | TAM.Gdf15 | Samd4b |
| Golga5 | 9.25E-26 | 0.55941327 | 0.313 | 0.203 | 1.79E-21 | TAM.Gdf15 | Golga5 |
| Lmbrd22 | 9.75E-26 | 0.43609093 | 0.319 | 0.218 | 1.88E-21 | TAM.Gdf15 | Lmbrd2 |
| Tmem2301 | 1.11E-25 | 0.3544174 | 0.406 | 0.289 | 2.15E-21 | TAM.Gdf15 | Tmem230 |
| Inf23 | 1.19E-25 | 0.45502295 | 0.369 | 0.242 | 2.30E-21 | TAM.Gdf15 | Inf2 |
| Tmbim4 | 1.34E-25 | 0.29580394 | 0.522 | 0.404 | 2.60E-21 | TAM.Gdf15 | Tmbim4 |
| Psmd61 | 1.46E-25 | 0.27539658 | 0.483 | 0.367 | 2.83E-21 | TAM.Gdf15 | Psmd6 |
| Ssh31 | 1.70E-25 | 0.3442349 | 0.445 | 0.305 | 3.28E-21 | TAM.Gdf15 | Ssh3 |
| Senp5 | 2.20E-25 | 0.46359249 | 0.348 | 0.255 | 4.25E-21 | TAM.Gdf15 | Senp5 |
| Prkcsh1 | 3.08E-25 | 0.27573455 | 0.57 | 0.447 | 5.95E-21 | TAM.Gdf15 | Prkcsh |
| Mospd22 | 3.64E-25 | 0.30848892 | 0.503 | 0.379 | 7.04E-21 | TAM.Gdf15 | Mospd2 |
| Rell12 | 4.22E-25 | 0.4250326 | 0.39 | 0.271 | 8.17E-21 | TAM.Gdf15 | Rell1 |
| Vac141 | 4.80E-25 | 0.27824838 | 0.522 | 0.403 | 9.28E-21 | TAM.Gdf15 | Vac14 |
| Amdhd22 | 4.87E-25 | 0.30663023 | 0.563 | 0.433 | 9.42E-21 | TAM.Gdf15 | Amdhd2 |
| Psen2 | 6.33E-25 | 0.33472191 | 0.491 | 0.355 | 1.22E-20 | TAM.Gdf15 | Psen2 |
| Unc501 | 6.55E-25 | 0.41753636 | 0.358 | 0.247 | 1.27E-20 | TAM.Gdf15 | Unc50 |
| Ahctf11 | 7.26E-25 | 0.42295182 | 0.357 | 0.253 | 1.40E-20 | TAM.Gdf15 | Ahctf1 |
| Chmp1b1 | 1.12E-24 | 0.30553915 | 0.541 | 0.426 | 2.16E-20 | TAM.Gdf15 | Chmp1b |
| Slc13a33 | 1.22E-24 | 0.63835729 | 0.263 | 0.168 | 2.36E-20 | TAM.Gdf15 | Slc13a3 |
| Rnf1502 | 1.41E-24 | 0.30638371 | 0.367 | 0.228 | 2.72E-20 | TAM.Gdf15 | Rnf150 |
| Ltv12 | 1.74E-24 | 0.40381162 | 0.333 | 0.229 | 3.36E-20 | TAM.Gdf15 | Ltv1 |
| Parvb1 | 1.83E-24 | 0.4677589 | 0.38 | 0.269 | 3.54E-20 | TAM.Gdf15 | Parvb |
| Lyst2 | 1.96E-24 | 0.46593531 | 0.305 | 0.213 | 3.79E-20 | TAM.Gdf15 | Lyst |
| Dynll23 | 2.70E-24 | 0.25731175 | 0.558 | 0.422 | 5.22E-20 | TAM.Gdf15 | Dynll2 |
| Tmem167b1 | 2.81E-24 | 0.43129353 | 0.41 | 0.294 | 5.43E-20 | TAM.Gdf15 | Tmem167b |
| Sec63 | 3.17E-24 | 0.34597924 | 0.422 | 0.311 | 6.13E-20 | TAM.Gdf15 | Sec63 |
| Rars1 | 3.37E-24 | 0.34771111 | 0.43 | 0.326 | 6.51E-20 | TAM.Gdf15 | Rars |
| Cars1 | 3.54E-24 | 0.65987387 | 0.288 | 0.189 | 6.84E-20 | TAM.Gdf15 | Cars |
| Riok21 | 4.83E-24 | 0.62387417 | 0.286 | 0.181 | 9.34E-20 | TAM.Gdf15 | Riok2 |
| DEPRECATED-I | 5.36E-24 | 0.29020673 | 0.472 | 0.376 | 1.04E-19 | TAM.Gdf15 | DEPRECATED-ENSMUSG00000039782 |
| Smpd11 | 6.62E-24 | 0.47202686 | 0.257 | 0.153 | 1.28E-19 | TAM.Gdf15 | Smpd1 |
| Uba3 | 7.04E-24 | 0.31475715 | 0.336 | 0.235 | 1.36E-19 | TAM.Gdf15 | Uba3 |
| Tsc41 | 7.40E-24 | 0.61600219 | 0.264 | 0.157 | 1.43E-19 | TAM.Gdf15 | Tsc4 |
| Soga13 | 7.61E-24 | 0.430073 | 0.507 | 0.374 | 1.47E-19 | TAM.Gdf15 | Soga1 |
| Uba5 | 9.38E-24 | 0.43038098 | 0.312 | 0.195 | 1.81E-19 | TAM.Gdf15 | Uba5 |
| Def81 | 1.25E-23 | 0.36047916 | 0.265 | 0.171 | 2.41E-19 | TAM.Gdf15 | Def8 |
| Wwp2 | 1.31E-23 | 0.31116289 | 0.497 | 0.387 | 2.53E-19 | TAM.Gdf15 | Wwp2 |
| Uprt | 1.33E-23 | 0.55304877 | 0.265 | 0.159 | 2.57E-19 | TAM.Gdf15 | Uprt |
| Snx151 | 1.55E-23 | 0.52363685 | 0.309 | 0.207 | 3.00E-19 | TAM.Gdf15 | Snx15 |
| Cyth2 | 1.56E-23 | 0.39560352 | 0.499 | 0.392 | 3.01E-19 | TAM.Gdf15 | Cyth2 |
| Utp111 | 1.63E-23 | 0.42956801 | 0.44 | 0.309 | 3.16E-19 | TAM.Gdf15 | Utp11 |
| Armcx32 | 1.65E-23 | 0.40057502 | 0.369 | 0.278 | 3.19E-19 | TAM.Gdf15 | Armcx3 |
| Sec22b | 1.81E-23 | 0.42571733 | 0.402 | 0.298 | 3.50E-19 | TAM.Gdf15 | Sec22b |
| Akirin11 | 2.27E-23 | 0.26808719 | 0.422 | 0.33 | 4.39E-19 | TAM.Gdf15 | Akirin1 |
| Plpp2 | 2.91E-23 | 0.51537511 | 0.297 | 0.172 | 5.62E-19 | TAM.Gdf15 | Plpp2 |
| Nr1h31 | 3.09E-23 | 0.56301171 | 0.251 | 0.16 | 5.98E-19 | TAM.Gdf15 | Nr1h3 |
| Cnih42 | 3.10E-23 | 0.27975407 | 0.453 | 0.346 | 5.99E-19 | TAM.Gdf15 | Cnih4 |
| Golgb11 | 3.21E-23 | 0.52317756 | 0.394 | 0.266 | 6.21E-19 | TAM.Gdf15 | Golgb1 |
| Sel1l1 | 4.13E-23 | 0.25032207 | 0.613 | 0.51 | 7.99E-19 | TAM.Gdf15 | Sel1l |
| Fdx21 | 4.21E-23 | 0.35879627 | 0.403 | 0.273 | 8.14E-19 | TAM.Gdf15 | Fdx2 |
| Sbds1 | 4.43E-23 | 0.2734215 | 0.502 | 0.4 | 8.56E-19 | TAM.Gdf15 | Sbds |
| Klc2 | 4.73E-23 | 0.7921452 | 0.253 | 0.15 | 9.15E-19 | TAM.Gdf15 | Klc2 |
| Tmem1151 | 5.83E-23 | 0.43211505 | 0.369 | 0.256 | 1.13E-18 | TAM.Gdf15 | Tmem115 |
| Vapb | 6.01E-23 | 0.2904858 | 0.549 | 0.426 | 1.16E-18 | TAM.Gdf15 | Vapb |
| Prkx2 | 6.83E-23 | 0.26390512 | 0.427 | 0.326 | 1.32E-18 | TAM.Gdf15 | Prkx |
| Lars1 | 8.39E-23 | 0.37621371 | 0.395 | 0.275 | 1.62E-18 | TAM.Gdf15 | Lars |
| Slc45a42 | 8.85E-23 | 0.41242891 | 0.412 | 0.276 | 1.71E-18 | TAM.Gdf15 | Slc45a4 |
| Tnrc6a | 1.03E-22 | 0.49221963 | 0.386 | 0.263 | 1.99E-18 | TAM.Gdf15 | Tnrc6a |
| Napg1 | 1.05E-22 | 0.64796192 | 0.342 | 0.222 | 2.03E-18 | TAM.Gdf15 | Napg |
| Kbtbd2 | 1.29E-22 | 0.40662823 | 0.45 | 0.342 | 2.49E-18 | TAM.Gdf15 | Kbtbd2 |
| Xpot1 | 1.53E-22 | 0.80178118 | 0.265 | 0.162 | 2.96E-18 | TAM.Gdf15 | Xpot |
| Eif2s21 | 2.05E-22 | 0.41661363 | 0.288 | 0.191 | 3.96E-18 | TAM.Gdf15 | Eif2s2 |
| Sqor2 | 2.06E-22 | 0.26877142 | 0.413 | 0.323 | 3.98E-18 | TAM.Gdf15 | Sqor |
| Cnppd11 | 2.41E-22 | 0.26210348 | 0.686 | 0.593 | 4.66E-18 | TAM.Gdf15 | Cnppd1 |
| Ubr2 | 2.41E-22 | 0.27627412 | 0.557 | 0.451 | 4.66E-18 | TAM.Gdf15 | Ubr2 |
| Emc31 | 2.74E-22 | 0.40061187 | 0.392 | 0.271 | 5.29E-18 | TAM.Gdf15 | Emc3 |
| Trappc10 | 2.91E-22 | 0.41649341 | 0.37 | 0.276 | 5.62E-18 | TAM.Gdf15 | Trappc10 |
| Zgpat | 3.10E-22 | 0.46249284 | 0.355 | 0.239 | 5.98E-18 | TAM.Gdf15 | Zgpat |
| Hax12 | 3.81E-22 | 0.39049916 | 0.423 | 0.319 | 7.36E-18 | TAM.Gdf15 | Hax1 |
| Ubxn61 | 4.49E-22 | 0.42221359 | 0.383 | 0.274 | 8.68E-18 | TAM.Gdf15 | Ubxn6 |
| Fam102b3 | 4.95E-22 | 0.25919033 | 0.464 | 0.362 | 9.57E-18 | TAM.Gdf15 | Fam102b |
| Mtm11 | 5.42E-22 | 0.35579581 | 0.283 | 0.181 | 1.05E-17 | TAM.Gdf15 | Mtm1 |
| Atxn12 | 5.50E-22 | 0.40856634 | 0.378 | 0.275 | 1.06E-17 | TAM.Gdf15 | Atxn1 |
| Slmap | 5.87E-22 | 0.4505068 | 0.38 | 0.276 | 1.13E-17 | TAM.Gdf15 | Slmap |
| Hagh2 | 5.97E-22 | 0.26939372 | 0.394 | 0.273 | 1.15E-17 | TAM.Gdf15 | Hagh |
| Slc4a2 | 6.11E-22 | 0.61359827 | 0.447 | 0.3 | 1.18E-17 | TAM.Gdf15 | Slc4a2 |
| Rabggtb1 | 6.32E-22 | 0.32006151 | 0.503 | 0.377 | 1.22E-17 | TAM.Gdf15 | Rabggtb |
| Ttc11 | 6.68E-22 | 0.35238707 | 0.36 | 0.267 | 1.29E-17 | TAM.Gdf15 | Ttc1 |
| Dnmt3a3 | 6.77E-22 | 0.3056103 | 0.52 | 0.386 | 1.31E-17 | TAM.Gdf15 | Dnmt3a |
| Vps26c1 | 7.08E-22 | 0.36532242 | 0.371 | 0.269 | 1.37E-17 | TAM.Gdf15 | Vps26c |
| Ube3c1 | 7.86E-22 | 0.38010105 | 0.351 | 0.244 | 1.52E-17 | TAM.Gdf15 | Ube3c |
| Casp1 | 8.70E-22 | 0.41266544 | 0.336 | 0.233 | 1.68E-17 | TAM.Gdf15 | Casp1 |
| Krcc11 | 1.09E-21 | 0.31127223 | 0.377 | 0.288 | 2.11E-17 | TAM.Gdf15 | Krcc1 |
| Manbal1 | 1.11E-21 | 0.28424085 | 0.48 | 0.35 | 2.16E-17 | TAM.Gdf15 | Manbal |
| Cyth33 | 1.45E-21 | 0.25385089 | 0.431 | 0.313 | 2.79E-17 | TAM.Gdf15 | Cyth3 |
| Dtx2 | 1.69E-21 | 0.42783435 | 0.309 | 0.201 | 3.26E-17 | TAM.Gdf15 | Dtx2 |
| Fgd31 | 1.87E-21 | 0.2544572 | 0.486 | 0.392 | 3.61E-17 | TAM.Gdf15 | Fgd3 |
| Nr2c2ap1 | 1.99E-21 | 0.38552136 | 0.351 | 0.25 | 3.84E-17 | TAM.Gdf15 | Nr2c2ap |
| Tmem250-ps1 | 1.99E-21 | 0.26030321 | 0.499 | 0.367 | 3.84E-17 | TAM.Gdf15 | Tmem250-ps |
| Nup931 | 2.03E-21 | 0.48236029 | 0.28 | 0.171 | 3.92E-17 | TAM.Gdf15 | Nup93 |
| Cacybp1 | 2.80E-21 | 0.25052707 | 0.403 | 0.288 | 5.41E-17 | TAM.Gdf15 | Cacybp |
| Atxn3 | 3.64E-21 | 0.32367683 | 0.384 | 0.268 | 7.03E-17 | TAM.Gdf15 | Atxn3 |
| Ascc2 | 3.68E-21 | 0.38186611 | 0.378 | 0.267 | 7.12E-17 | TAM.Gdf15 | Ascc2 |
| Zfp2632 | 5.86E-21 | 0.37436196 | 0.377 | 0.289 | 1.13E-16 | TAM.Gdf15 | Zfp263 |
| Ubxn81 | 6.15E-21 | 0.33228244 | 0.326 | 0.228 | 1.19E-16 | TAM.Gdf15 | Ubxn8 |
| Snx101 | 6.48E-21 | 0.28793611 | 0.453 | 0.346 | 1.25E-16 | TAM.Gdf15 | Snx10 |
| Stam | 7.16E-21 | 0.7801662 | 0.286 | 0.186 | 1.38E-16 | TAM.Gdf15 | Stam |
| Acsl3 | 9.67E-21 | 0.41747413 | 0.26 | 0.172 | 1.87E-16 | TAM.Gdf15 | Acsl3 |
| Eef1akmt41 | 9.81E-21 | 0.53808746 | 0.273 | 0.168 | 1.90E-16 | TAM.Gdf15 | Eef1akmt4 |
| Ube2z | 1.14E-20 | 0.33843019 | 0.595 | 0.481 | 2.20E-16 | TAM.Gdf15 | Ube2z |
| Fam126a1 | 1.47E-20 | 0.32989857 | 0.288 | 0.196 | 2.84E-16 | TAM.Gdf15 | Fam126a |
| Emc41 | 1.51E-20 | 0.29022543 | 0.374 | 0.266 | 2.92E-16 | TAM.Gdf15 | Emc4 |
| Gps11 | 1.56E-20 | 0.29661737 | 0.449 | 0.341 | 3.01E-16 | TAM.Gdf15 | Gps1 |
| Pak1ip13 | 1.64E-20 | 0.28942243 | 0.43 | 0.325 | 3.17E-16 | TAM.Gdf15 | Pak1ip1 |
| Slc35b41 | 1.83E-20 | 0.27579049 | 0.307 | 0.217 | 3.54E-16 | TAM.Gdf15 | Slc35b4 |
| Timm101 | 1.91E-20 | 0.58967825 | 0.303 | 0.187 | 3.69E-16 | TAM.Gdf15 | Timm10 |
| Crbn | 2.08E-20 | 0.37754311 | 0.258 | 0.163 | 4.02E-16 | TAM.Gdf15 | Crbn |
| Anapc2 | 2.49E-20 | 0.34848749 | 0.531 | 0.41 | 4.82E-16 | TAM.Gdf15 | Anapc2 |
| Zfp9921 | 2.68E-20 | 0.30722049 | 0.387 | 0.273 | 5.17E-16 | TAM.Gdf15 | Zfp992 |
| Vps37a1 | 3.26E-20 | 0.4652013 | 0.356 | 0.253 | 6.30E-16 | TAM.Gdf15 | Vps37a |
| Rnf1812 | 3.39E-20 | 0.42790598 | 0.37 | 0.271 | 6.56E-16 | TAM.Gdf15 | Rnf181 |
| Ube4b1 | 4.11E-20 | 0.39598166 | 0.438 | 0.324 | 7.94E-16 | TAM.Gdf15 | Ube4b |
| Tmco31 | 4.59E-20 | 0.35320472 | 0.277 | 0.182 | 8.88E-16 | TAM.Gdf15 | Tmco3 |
| Rbm28 | 4.67E-20 | 0.48279459 | 0.343 | 0.249 | 9.03E-16 | TAM.Gdf15 | Rbm28 |
| Spryd31 | 5.05E-20 | 0.38454231 | 0.394 | 0.288 | 9.76E-16 | TAM.Gdf15 | Spryd3 |
| Nop22 | 5.59E-20 | 0.35118139 | 0.378 | 0.27 | 1.08E-15 | TAM.Gdf15 | Nop2 |
| Klf92 | 5.88E-20 | 0.57276456 | 0.323 | 0.214 | 1.14E-15 | TAM.Gdf15 | Klf9 |
| Aco11 | 6.02E-20 | 0.28156504 | 0.473 | 0.356 | 1.16E-15 | TAM.Gdf15 | Aco1 |
| Adi13 | 6.13E-20 | 0.57865919 | 0.303 | 0.198 | 1.19E-15 | TAM.Gdf15 | Adi1 |
| Asxl11 | 6.92E-20 | 0.42765849 | 0.374 | 0.273 | 1.34E-15 | TAM.Gdf15 | Asxl1 |
| Spsb2 | 6.93E-20 | 0.71597923 | 0.279 | 0.171 | 1.34E-15 | TAM.Gdf15 | Spsb2 |
| Golga21 | 7.00E-20 | 0.25702028 | 0.459 | 0.331 | 1.35E-15 | TAM.Gdf15 | Golga2 |
| Yipf5 | 7.62E-20 | 0.4413707 | 0.351 | 0.256 | 1.47E-15 | TAM.Gdf15 | Yipf5 |
| Pgpep12 | 7.89E-20 | 0.52561704 | 0.286 | 0.171 | 1.53E-15 | TAM.Gdf15 | Pgpep1 |
| Arhgap222 | 8.34E-20 | 0.68851584 | 0.27 | 0.167 | 1.61E-15 | TAM.Gdf15 | Arhgap22 |
| Nbr12 | 8.89E-20 | 0.59085018 | 0.33 | 0.216 | 1.72E-15 | TAM.Gdf15 | Nbr1 |
| Arid5b3 | 9.67E-20 | 0.31531474 | 0.398 | 0.301 | 1.87E-15 | TAM.Gdf15 | Arid5b |
| Slc25a122 | 9.86E-20 | 0.25663923 | 0.359 | 0.26 | 1.91E-15 | TAM.Gdf15 | Slc25a12 |
| Ppp2r5b | 9.94E-20 | 0.67778178 | 0.258 | 0.158 | 1.92E-15 | TAM.Gdf15 | Ppp2r5b |
| Immt1 | 1.01E-19 | 0.25264303 | 0.466 | 0.355 | 1.95E-15 | TAM.Gdf15 | Immt |
| Slc25a44 | 1.10E-19 | 0.35370899 | 0.259 | 0.167 | 2.14E-15 | TAM.Gdf15 | Slc25a44 |
| Myo9a2 | 1.20E-19 | 0.40729073 | 0.307 | 0.193 | 2.32E-15 | TAM.Gdf15 | Myo9a |
| Pcmt12 | 1.61E-19 | 0.27869179 | 0.427 | 0.326 | 3.11E-15 | TAM.Gdf15 | Pcmt1 |
| Snap291 | 1.85E-19 | 0.27733251 | 0.387 | 0.293 | 3.58E-15 | TAM.Gdf15 | Snap29 |
| Acvr1 | 2.05E-19 | 0.54786901 | 0.271 | 0.169 | 3.97E-15 | TAM.Gdf15 | Acvr1 |
| Sec31a | 2.05E-19 | 0.29152381 | 0.453 | 0.343 | 3.97E-15 | TAM.Gdf15 | Sec31a |
| Zadh22 | 2.16E-19 | 0.5363832 | 0.281 | 0.18 | 4.17E-15 | TAM.Gdf15 | Zadh2 |
| Mafk1 | 2.83E-19 | 0.46220573 | 0.469 | 0.351 | 5.47E-15 | TAM.Gdf15 | Mafk |
| Asb6 | 3.27E-19 | 0.37710108 | 0.308 | 0.204 | 6.33E-15 | TAM.Gdf15 | Asb6 |
| Foxj3 | 3.44E-19 | 0.39389282 | 0.281 | 0.172 | 6.65E-15 | TAM.Gdf15 | Foxj3 |
| B3gat3 | 3.60E-19 | 0.48704532 | 0.295 | 0.193 | 6.96E-15 | TAM.Gdf15 | B3gat3 |
| Stx3 | 3.70E-19 | 0.51263598 | 0.266 | 0.177 | 7.16E-15 | TAM.Gdf15 | Stx3 |
| Htra22 | 4.60E-19 | 0.46083543 | 0.342 | 0.242 | 8.90E-15 | TAM.Gdf15 | Htra2 |
| Mast22 | 5.48E-19 | 0.74515917 | 0.335 | 0.225 | 1.06E-14 | TAM.Gdf15 | Mast2 |
| Plekhm32 | 6.38E-19 | 0.37038908 | 0.376 | 0.266 | 1.23E-14 | TAM.Gdf15 | Plekhm3 |
| Mars1 | 7.05E-19 | 0.43716211 | 0.341 | 0.227 | 1.36E-14 | TAM.Gdf15 | Mars |
| Uhrf22 | 7.81E-19 | 0.29198012 | 0.48 | 0.375 | 1.51E-14 | TAM.Gdf15 | Uhrf2 |
| Ago31 | 1.09E-18 | 0.30085083 | 0.401 | 0.283 | 2.10E-14 | TAM.Gdf15 | Ago3 |
| Arhgap211 | 1.18E-18 | 0.52732943 | 0.256 | 0.164 | 2.29E-14 | TAM.Gdf15 | Arhgap21 |
| Caml | 1.20E-18 | 0.65993573 | 0.278 | 0.172 | 2.31E-14 | TAM.Gdf15 | Caml |
| Sptlc11 | 1.41E-18 | 0.34484851 | 0.343 | 0.245 | 2.72E-14 | TAM.Gdf15 | Sptlc1 |
| Tmem2221 | 1.69E-18 | 0.27560197 | 0.302 | 0.208 | 3.27E-14 | TAM.Gdf15 | Tmem222 |
| Hmg20b1 | 1.80E-18 | 0.36041033 | 0.458 | 0.35 | 3.48E-14 | TAM.Gdf15 | Hmg20b |
| Hook31 | 1.86E-18 | 0.25763563 | 0.408 | 0.301 | 3.60E-14 | TAM.Gdf15 | Hook3 |
| Zfyve27 | 1.89E-18 | 0.492196 | 0.358 | 0.255 | 3.66E-14 | TAM.Gdf15 | Zfyve27 |
| Ip6k11 | 1.95E-18 | 0.28169586 | 0.598 | 0.483 | 3.76E-14 | TAM.Gdf15 | Ip6k1 |
| Ttc7b1 | 2.07E-18 | 0.4568784 | 0.344 | 0.252 | 4.00E-14 | TAM.Gdf15 | Ttc7b |
| Nabp13 | 2.12E-18 | 0.25120171 | 0.472 | 0.384 | 4.11E-14 | TAM.Gdf15 | Nabp1 |
| Pwwp2b | 2.23E-18 | 0.31808378 | 0.298 | 0.191 | 4.31E-14 | TAM.Gdf15 | Pwwp2b |
| Pdpk11 | 2.36E-18 | 0.28511079 | 0.457 | 0.351 | 4.56E-14 | TAM.Gdf15 | Pdpk1 |
| Acot132 | 2.68E-18 | 0.44848879 | 0.349 | 0.233 | 5.18E-14 | TAM.Gdf15 | Acot13 |
| Hspa131 | 3.09E-18 | 0.50391443 | 0.252 | 0.155 | 5.98E-14 | TAM.Gdf15 | Hspa13 |
| Pros11 | 3.34E-18 | 0.25811992 | 0.337 | 0.235 | 6.45E-14 | TAM.Gdf15 | Pros1 |
| Ing42 | 3.58E-18 | 0.40954713 | 0.444 | 0.323 | 6.92E-14 | TAM.Gdf15 | Ing4 |
| Drg11 | 3.95E-18 | 0.25703176 | 0.507 | 0.404 | 7.64E-14 | TAM.Gdf15 | Drg1 |
| Mapk91 | 4.11E-18 | 0.34854404 | 0.358 | 0.252 | 7.94E-14 | TAM.Gdf15 | Mapk9 |
| Vapa1 | 4.26E-18 | 0.34648248 | 0.381 | 0.289 | 8.23E-14 | TAM.Gdf15 | Vapa |
| Pja22 | 4.70E-18 | 0.36174649 | 0.323 | 0.22 | 9.09E-14 | TAM.Gdf15 | Pja2 |
| Gspt11 | 4.95E-18 | 0.29918632 | 0.459 | 0.354 | 9.57E-14 | TAM.Gdf15 | Gspt1 |
| Galc2 | 5.85E-18 | 0.43883665 | 0.436 | 0.304 | 1.13E-13 | TAM.Gdf15 | Galc |
| Tesk12 | 7.00E-18 | 0.3525435 | 0.477 | 0.365 | 1.35E-13 | TAM.Gdf15 | Tesk1 |
| Ndfip2 | 7.14E-18 | 0.45301457 | 0.302 | 0.199 | 1.38E-13 | TAM.Gdf15 | Ndfip2 |
| Eif4ebp3 | 7.20E-18 | 0.51816998 | 0.343 | 0.234 | 1.39E-13 | TAM.Gdf15 | Eif4ebp3 |
| Nlk | 7.32E-18 | 0.344258 | 0.258 | 0.168 | 1.42E-13 | TAM.Gdf15 | Nlk |
| Dhrs71 | 8.33E-18 | 0.42057142 | 0.315 | 0.22 | 1.61E-13 | TAM.Gdf15 | Dhrs7 |
| Dnaja31 | 9.31E-18 | 0.55246804 | 0.336 | 0.233 | 1.80E-13 | TAM.Gdf15 | Dnaja3 |
| Tcea11 | 9.86E-18 | 0.26348588 | 0.465 | 0.357 | 1.91E-13 | TAM.Gdf15 | Tcea1 |
| Rexf11 | 1.36E-17 | 0.29543577 | 0.367 | 0.29 | 2.63E-13 | TAM.Gdf15 | Rexf1 |
| Atg13 | 1.39E-17 | 0.56480713 | 0.272 | 0.18 | 2.68E-13 | TAM.Gdf15 | Atg13 |
| Sec23b2 | 1.41E-17 | 0.26793065 | 0.385 | 0.288 | 2.72E-13 | TAM.Gdf15 | Sec23b |
| Phyhd12 | 1.49E-17 | 0.54028497 | 0.428 | 0.245 | 2.89E-13 | TAM.Gdf15 | Phyhd1 |
| Bcl2l21 | 1.51E-17 | 0.29173069 | 0.327 | 0.244 | 2.92E-13 | TAM.Gdf15 | Bcl2l2 |
| Bmi1 | 1.59E-17 | 0.57526418 | 0.263 | 0.166 | 3.07E-13 | TAM.Gdf15 | Bmi1 |
| Zbtb212 | 1.59E-17 | 0.41994156 | 0.343 | 0.256 | 3.08E-13 | TAM.Gdf15 | Zbtb21 |
| Egln21 | 1.98E-17 | 0.37720371 | 0.46 | 0.344 | 3.82E-13 | TAM.Gdf15 | Egln2 |
| Gigyf11 | 2.08E-17 | 0.44618512 | 0.315 | 0.218 | 4.03E-13 | TAM.Gdf15 | Gigyf1 |
| Naa251 | 2.31E-17 | 0.62219032 | 0.269 | 0.178 | 4.47E-13 | TAM.Gdf15 | Naa25 |
| Aldh7a11 | 2.53E-17 | 0.37409572 | 0.292 | 0.167 | 4.90E-13 | TAM.Gdf15 | Aldh7a1 |
| Atg2a | 2.58E-17 | 0.26164814 | 0.543 | 0.46 | 4.98E-13 | TAM.Gdf15 | Atg2a |
| Ppp2r2a2 | 2.64E-17 | 0.31445187 | 0.364 | 0.262 | 5.10E-13 | TAM.Gdf15 | Ppp2r2a |
| Ppp1r22 | 2.82E-17 | 0.27079752 | 0.4 | 0.304 | 5.46E-13 | TAM.Gdf15 | Ppp1r2 |
| Slc16a102 | 2.84E-17 | 0.34225021 | 0.415 | 0.291 | 5.49E-13 | TAM.Gdf15 | Slc16a10 |
| Ier5l2 | 2.84E-17 | 0.39546455 | 0.262 | 0.161 | 5.50E-13 | TAM.Gdf15 | Ier5l |
| Scaf112 | 3.53E-17 | 0.28151253 | 0.506 | 0.424 | 6.83E-13 | TAM.Gdf15 | Scaf11 |
| Ngly1 | 3.67E-17 | 0.32081647 | 0.278 | 0.19 | 7.10E-13 | TAM.Gdf15 | Ngly1 |
| Srprb1 | 3.69E-17 | 0.39923134 | 0.297 | 0.208 | 7.13E-13 | TAM.Gdf15 | Srprb |
| Micu12 | 3.77E-17 | 0.28035614 | 0.349 | 0.248 | 7.29E-13 | TAM.Gdf15 | Micu1 |
| Syne3 | 4.06E-17 | 0.42985003 | 0.25 | 0.155 | 7.85E-13 | TAM.Gdf15 | Syne3 |
| Tbrg12 | 4.08E-17 | 0.29031975 | 0.353 | 0.273 | 7.88E-13 | TAM.Gdf15 | Tbrg1 |
| Trim412 | 4.26E-17 | 0.33470499 | 0.458 | 0.345 | 8.25E-13 | TAM.Gdf15 | Trim41 |
| Washc52 | 4.31E-17 | 0.29719659 | 0.378 | 0.278 | 8.32E-13 | TAM.Gdf15 | Washc5 |
| St131 | 4.81E-17 | 0.28219223 | 0.43 | 0.334 | 9.29E-13 | TAM.Gdf15 | St13 |
| Pnisr1 | 6.60E-17 | 0.34402707 | 0.433 | 0.343 | 1.28E-12 | TAM.Gdf15 | Pnisr |
| Slc12a41 | 6.63E-17 | 0.26876557 | 0.326 | 0.235 | 1.28E-12 | TAM.Gdf15 | Slc12a4 |
| Kidins220 | 7.67E-17 | 0.27996533 | 0.497 | 0.39 | 1.48E-12 | TAM.Gdf15 | Kidins220 |
| Srm1 | 8.55E-17 | 0.53575883 | 0.277 | 0.176 | 1.65E-12 | TAM.Gdf15 | Srm |
| Stra6l2 | 1.00E-16 | 0.44346848 | 0.269 | 0.166 | 1.94E-12 | TAM.Gdf15 | Stra6l |
| Myo18a1 | 1.02E-16 | 0.30685579 | 0.433 | 0.318 | 1.96E-12 | TAM.Gdf15 | Myo18a |
| Creb3 | 1.23E-16 | 0.25481578 | 0.344 | 0.249 | 2.38E-12 | TAM.Gdf15 | Creb3 |
| Pprc12 | 1.32E-16 | 0.39579519 | 0.379 | 0.269 | 2.56E-12 | TAM.Gdf15 | Pprc1 |
| Il3ra1 | 1.34E-16 | 0.38937403 | 0.384 | 0.271 | 2.59E-12 | TAM.Gdf15 | Il3ra |
| Cystm12 | 1.61E-16 | 0.31469625 | 0.286 | 0.206 | 3.11E-12 | TAM.Gdf15 | Cystm1 |
| Osbpl111 | 1.62E-16 | 0.35409956 | 0.405 | 0.3 | 3.13E-12 | TAM.Gdf15 | Osbpl11 |
| Mtln1 | 1.83E-16 | 0.40666605 | 0.324 | 0.233 | 3.54E-12 | TAM.Gdf15 | Mtln |
| Vps392 | 1.99E-16 | 0.40434361 | 0.319 | 0.218 | 3.86E-12 | TAM.Gdf15 | Vps39 |
| Sco2 | 2.04E-16 | 0.34142185 | 0.251 | 0.16 | 3.95E-12 | TAM.Gdf15 | Sco2 |
| Pgrmc22 | 2.12E-16 | 0.2690172 | 0.393 | 0.289 | 4.09E-12 | TAM.Gdf15 | Pgrmc2 |
| Dag13 | 2.30E-16 | 0.37934045 | 0.35 | 0.245 | 4.44E-12 | TAM.Gdf15 | Dag1 |
| Zdhhc8 | 3.28E-16 | 0.50586373 | 0.29 | 0.194 | 6.34E-12 | TAM.Gdf15 | Zdhhc8 |
| Cnep1r11 | 3.43E-16 | 0.37893825 | 0.302 | 0.206 | 6.63E-12 | TAM.Gdf15 | Cnep1r1 |
| Mga | 3.71E-16 | 0.50210967 | 0.279 | 0.186 | 7.17E-12 | TAM.Gdf15 | Mga |
| Pbdc12 | 3.91E-16 | 0.42034241 | 0.307 | 0.21 | 7.57E-12 | TAM.Gdf15 | Pbdc1 |
| Cdk61 | 3.94E-16 | 0.63644702 | 0.315 | 0.229 | 7.63E-12 | TAM.Gdf15 | Cdk6 |
| Fadd3 | 4.00E-16 | 0.45500413 | 0.319 | 0.216 | 7.73E-12 | TAM.Gdf15 | Fadd |
| Rab40c1 | 4.28E-16 | 0.46062729 | 0.287 | 0.195 | 8.27E-12 | TAM.Gdf15 | Rab40c |
| Lpin13 | 4.46E-16 | 0.41069674 | 0.281 | 0.198 | 8.62E-12 | TAM.Gdf15 | Lpin1 |
| Numb | 5.52E-16 | 0.5858489 | 0.331 | 0.235 | 1.07E-11 | TAM.Gdf15 | Numb |
| Dnpep | 6.21E-16 | 0.3492602 | 0.299 | 0.215 | 1.20E-11 | TAM.Gdf15 | Dnpep |
| Ccdc61 | 6.36E-16 | 0.29305765 | 0.437 | 0.323 | 1.23E-11 | TAM.Gdf15 | Ccdc6 |
| Scaf1 | 6.51E-16 | 0.30595894 | 0.51 | 0.391 | 1.26E-11 | TAM.Gdf15 | Scaf1 |
| Washc42 | 7.34E-16 | 0.41549807 | 0.376 | 0.273 | 1.42E-11 | TAM.Gdf15 | Washc4 |
| Aifm22 | 8.41E-16 | 0.25858656 | 0.267 | 0.182 | 1.63E-11 | TAM.Gdf15 | Aifm2 |
| Ccdc186 | 8.48E-16 | 0.58139421 | 0.28 | 0.184 | 1.64E-11 | TAM.Gdf15 | Ccdc186 |
| Casc3 | 8.73E-16 | 0.34637235 | 0.431 | 0.336 | 1.69E-11 | TAM.Gdf15 | Casc3 |
| Tmx22 | 8.89E-16 | 0.26982827 | 0.322 | 0.226 | 1.72E-11 | TAM.Gdf15 | Tmx2 |
| Vps37c1 | 9.43E-16 | 0.50476202 | 0.32 | 0.231 | 1.82E-11 | TAM.Gdf15 | Vps37c |
| Trabd | 9.49E-16 | 0.38108487 | 0.373 | 0.272 | 1.84E-11 | TAM.Gdf15 | Trabd |
| Mob2 | 1.04E-15 | 0.47158177 | 0.33 | 0.228 | 2.02E-11 | TAM.Gdf15 | Mob2 |
| Atxn2 | 1.13E-15 | 0.31099175 | 0.334 | 0.229 | 2.19E-11 | TAM.Gdf15 | Atxn2 |
| Tsc2 | 1.36E-15 | 0.27590346 | 0.351 | 0.258 | 2.62E-11 | TAM.Gdf15 | Tsc2 |
| Exosc41 | 1.55E-15 | 0.34828833 | 0.265 | 0.176 | 3.00E-11 | TAM.Gdf15 | Exosc4 |
| Fech2 | 1.64E-15 | 0.25395508 | 0.31 | 0.226 | 3.16E-11 | TAM.Gdf15 | Fech |
| Pde1b2 | 1.69E-15 | 0.40038153 | 0.441 | 0.337 | 3.27E-11 | TAM.Gdf15 | Pde1b |
| Cog71 | 1.75E-15 | 0.35628378 | 0.258 | 0.172 | 3.39E-11 | TAM.Gdf15 | Cog7 |
| Patl1 | 1.77E-15 | 0.28646423 | 0.536 | 0.433 | 3.42E-11 | TAM.Gdf15 | Patl1 |
| Trappc9 | 2.00E-15 | 0.30478677 | 0.26 | 0.176 | 3.86E-11 | TAM.Gdf15 | Trappc9 |
| Hs1bp32 | 2.06E-15 | 0.39464311 | 0.307 | 0.208 | 3.98E-11 | TAM.Gdf15 | Hs1bp3 |
| Gzf11 | 2.23E-15 | 0.36757676 | 0.258 | 0.174 | 4.32E-11 | TAM.Gdf15 | Gzf1 |
| Mark41 | 2.24E-15 | 0.28563933 | 0.387 | 0.297 | 4.34E-11 | TAM.Gdf15 | Mark4 |
| Gfm11 | 2.27E-15 | 0.3467651 | 0.312 | 0.222 | 4.38E-11 | TAM.Gdf15 | Gfm1 |
| Ktn1 | 2.46E-15 | 0.47136478 | 0.35 | 0.234 | 4.76E-11 | TAM.Gdf15 | Ktn1 |
| Srp54b | 2.97E-15 | 0.45403844 | 0.308 | 0.223 | 5.74E-11 | TAM.Gdf15 | Srp54b |
| Mar-81 | 3.46E-15 | 0.56623429 | 0.266 | 0.175 | 6.69E-11 | TAM.Gdf15 | 8-Mar |
| Aspscr11 | 3.52E-15 | 0.27205361 | 0.335 | 0.239 | 6.81E-11 | TAM.Gdf15 | Aspscr1 |
| Asxl21 | 3.99E-15 | 0.26149283 | 0.377 | 0.269 | 7.72E-11 | TAM.Gdf15 | Asxl2 |
| Nmd31 | 4.37E-15 | 0.49247824 | 0.255 | 0.177 | 8.46E-11 | TAM.Gdf15 | Nmd3 |
| Gipc11 | 4.46E-15 | 0.27805124 | 0.328 | 0.247 | 8.62E-11 | TAM.Gdf15 | Gipc1 |
| Ate11 | 4.53E-15 | 0.41406001 | 0.25 | 0.167 | 8.75E-11 | TAM.Gdf15 | Ate1 |
| Rabgap1 | 4.67E-15 | 0.37184669 | 0.259 | 0.172 | 9.02E-11 | TAM.Gdf15 | Rabgap1 |
| Cdkn2aipnl2 | 5.12E-15 | 0.46683623 | 0.295 | 0.202 | 9.90E-11 | TAM.Gdf15 | Cdkn2aipnl |
| Cops7a2 | 5.13E-15 | 0.39490053 | 0.399 | 0.283 | 9.92E-11 | TAM.Gdf15 | Cops7a |
| Trip102 | 5.22E-15 | 0.32988643 | 0.329 | 0.26 | 1.01E-10 | TAM.Gdf15 | Trip10 |
| Calcoco12 | 5.28E-15 | 0.55029239 | 0.316 | 0.214 | 1.02E-10 | TAM.Gdf15 | Calcoco1 |
| Tubgcp4 | 6.12E-15 | 0.55078146 | 0.265 | 0.18 | 1.18E-10 | TAM.Gdf15 | Tubgcp4 |
| Faf2 | 7.12E-15 | 0.2693802 | 0.298 | 0.205 | 1.38E-10 | TAM.Gdf15 | Faf2 |
| Timm17a1 | 1.07E-14 | 0.28466886 | 0.431 | 0.325 | 2.07E-10 | TAM.Gdf15 | Timm17a |
| Fkrp1 | 1.08E-14 | 0.27908742 | 0.387 | 0.282 | 2.09E-10 | TAM.Gdf15 | Fkrp |
| Tcf12 | 1.33E-14 | 0.2845113 | 0.381 | 0.278 | 2.56E-10 | TAM.Gdf15 | Tcf12 |
| Ralgapb | 1.69E-14 | 0.32277857 | 0.356 | 0.264 | 3.26E-10 | TAM.Gdf15 | Ralgapb |
| Ubqln41 | 1.96E-14 | 0.25969087 | 0.327 | 0.234 | 3.79E-10 | TAM.Gdf15 | Ubqln4 |
| Crebrf2 | 2.03E-14 | 0.30189325 | 0.306 | 0.232 | 3.92E-10 | TAM.Gdf15 | Crebrf |
| Exoc1 | 2.53E-14 | 0.2894636 | 0.405 | 0.3 | 4.90E-10 | TAM.Gdf15 | Exoc1 |
| Gpr1071 | 2.76E-14 | 0.26718873 | 0.437 | 0.34 | 5.34E-10 | TAM.Gdf15 | Gpr107 |
| Nmt22 | 2.94E-14 | 0.47471532 | 0.302 | 0.211 | 5.68E-10 | TAM.Gdf15 | Nmt2 |
| Fbxl17 | 3.02E-14 | 0.44390503 | 0.279 | 0.204 | 5.84E-10 | TAM.Gdf15 | Fbxl17 |
| Pld21 | 3.33E-14 | 0.41333431 | 0.347 | 0.248 | 6.45E-10 | TAM.Gdf15 | Pld2 |
| Scaf8 | 3.38E-14 | 0.4139864 | 0.303 | 0.218 | 6.53E-10 | TAM.Gdf15 | Scaf8 |
| Togaram1 | 4.82E-14 | 0.37778921 | 0.333 | 0.248 | 9.33E-10 | TAM.Gdf15 | Togaram1 |
| Vps50 | 7.63E-14 | 0.30339678 | 0.264 | 0.174 | 1.48E-09 | TAM.Gdf15 | Vps50 |
| Trnau1ap1 | 8.84E-14 | 0.29268354 | 0.314 | 0.217 | 1.71E-09 | TAM.Gdf15 | Trnau1ap |
| Ercc31 | 8.90E-14 | 0.32136621 | 0.391 | 0.289 | 1.72E-09 | TAM.Gdf15 | Ercc3 |
| Gtf2h1 | 9.07E-14 | 0.30998681 | 0.314 | 0.223 | 1.75E-09 | TAM.Gdf15 | Gtf2h1 |
| Kdm4a1 | 9.87E-14 | 0.36015321 | 0.347 | 0.249 | 1.91E-09 | TAM.Gdf15 | Kdm4a |
| Kctd21 | 1.00E-13 | 0.40579376 | 0.301 | 0.207 | 1.94E-09 | TAM.Gdf15 | Kctd2 |
| Brd71 | 1.04E-13 | 0.292699 | 0.433 | 0.33 | 2.01E-09 | TAM.Gdf15 | Brd7 |
| Zranb21 | 1.07E-13 | 0.54731579 | 0.27 | 0.185 | 2.06E-09 | TAM.Gdf15 | Zranb2 |
| Agtrap2 | 1.14E-13 | 0.36122998 | 0.317 | 0.213 | 2.21E-09 | TAM.Gdf15 | Agtrap |
| Chmp62 | 1.35E-13 | 0.36594321 | 0.314 | 0.226 | 2.61E-09 | TAM.Gdf15 | Chmp6 |
| Ino80d1 | 1.40E-13 | 0.27648719 | 0.424 | 0.312 | 2.72E-09 | TAM.Gdf15 | Ino80d |
| Atad3a1 | 1.48E-13 | 0.55616197 | 0.278 | 0.175 | 2.86E-09 | TAM.Gdf15 | Atad3a |
| Pex14 | 1.49E-13 | 0.35733189 | 0.314 | 0.233 | 2.88E-09 | TAM.Gdf15 | Pex14 |
| Sestd11 | 1.53E-13 | 0.30452434 | 0.43 | 0.341 | 2.96E-09 | TAM.Gdf15 | Sestd1 |
| Bsdc11 | 1.59E-13 | 0.28373686 | 0.335 | 0.268 | 3.07E-09 | TAM.Gdf15 | Bsdc1 |
| Cklf | 1.75E-13 | 0.54047002 | 0.279 | 0.193 | 3.38E-09 | TAM.Gdf15 | Cklf |
| Clcn31 | 1.77E-13 | 0.40933073 | 0.27 | 0.188 | 3.43E-09 | TAM.Gdf15 | Clcn3 |
| Rhot1 | 2.00E-13 | 0.39998461 | 0.308 | 0.222 | 3.87E-09 | TAM.Gdf15 | Rhot1 |
| Kmt2a | 2.35E-13 | 0.25027915 | 0.326 | 0.229 | 4.55E-09 | TAM.Gdf15 | Kmt2a |
| Trappc111 | 2.95E-13 | 0.35703145 | 0.33 | 0.23 | 5.70E-09 | TAM.Gdf15 | Trappc11 |
| Sirt11 | 3.11E-13 | 0.59546587 | 0.297 | 0.214 | 6.01E-09 | TAM.Gdf15 | Sirt1 |
| Fam76a1 | 3.40E-13 | 0.29465095 | 0.369 | 0.277 | 6.58E-09 | TAM.Gdf15 | Fam76a |
| Appl22 | 3.65E-13 | 0.30327184 | 0.52 | 0.354 | 7.06E-09 | TAM.Gdf15 | Appl2 |
| Nln1 | 3.77E-13 | 0.28154607 | 0.323 | 0.243 | 7.30E-09 | TAM.Gdf15 | Nln |
| Phf202 | 3.89E-13 | 0.3551596 | 0.329 | 0.239 | 7.53E-09 | TAM.Gdf15 | Phf20 |
| Abhd17c3 | 4.52E-13 | 0.32988593 | 0.281 | 0.214 | 8.75E-09 | TAM.Gdf15 | Abhd17c |
| Anapc42 | 4.65E-13 | 0.27761592 | 0.309 | 0.231 | 9.00E-09 | TAM.Gdf15 | Anapc4 |
| Gnl21 | 4.67E-13 | 0.35761038 | 0.252 | 0.179 | 9.02E-09 | TAM.Gdf15 | Gnl2 |
| Cluh2 | 4.93E-13 | 0.27891616 | 0.372 | 0.273 | 9.54E-09 | TAM.Gdf15 | Cluh |
| Wipi21 | 5.87E-13 | 0.28735362 | 0.327 | 0.247 | 1.13E-08 | TAM.Gdf15 | Wipi2 |
| Tbcc2 | 6.36E-13 | 0.26378725 | 0.271 | 0.198 | 1.23E-08 | TAM.Gdf15 | Tbcc |
| Alg51 | 6.91E-13 | 0.28326981 | 0.256 | 0.183 | 1.34E-08 | TAM.Gdf15 | Alg5 |
| Tspan51 | 6.93E-13 | 0.28638651 | 0.274 | 0.191 | 1.34E-08 | TAM.Gdf15 | Tspan5 |
| Agpat32 | 7.33E-13 | 0.27310538 | 0.355 | 0.27 | 1.42E-08 | TAM.Gdf15 | Agpat3 |
| Rce12 | 8.17E-13 | 0.27883462 | 0.314 | 0.228 | 1.58E-08 | TAM.Gdf15 | Rce1 |
| Prkag2 | 8.60E-13 | 0.30283931 | 0.269 | 0.182 | 1.66E-08 | TAM.Gdf15 | Prkag2 |
| Tecpr11 | 9.28E-13 | 0.46793996 | 0.253 | 0.17 | 1.79E-08 | TAM.Gdf15 | Tecpr1 |
| Reps11 | 9.32E-13 | 0.31923659 | 0.386 | 0.297 | 1.80E-08 | TAM.Gdf15 | Reps1 |
| Pdrg12 | 9.59E-13 | 0.32447501 | 0.303 | 0.226 | 1.85E-08 | TAM.Gdf15 | Pdrg1 |
| Slc35f51 | 1.05E-12 | 0.40272135 | 0.281 | 0.188 | 2.04E-08 | TAM.Gdf15 | Slc35f5 |
| Abcg11 | 1.42E-12 | 0.26409441 | 0.326 | 0.26 | 2.74E-08 | TAM.Gdf15 | Abcg1 |
| Rxylt1 | 1.61E-12 | 0.27459231 | 0.3 | 0.223 | 3.10E-08 | TAM.Gdf15 | Rxylt1 |
| Necap11 | 2.12E-12 | 0.43716423 | 0.298 | 0.208 | 4.09E-08 | TAM.Gdf15 | Necap1 |
| Pacs2 | 2.41E-12 | 0.38700897 | 0.276 | 0.189 | 4.65E-08 | TAM.Gdf15 | Pacs2 |
| Gng5 | 4.00E-12 | 0.27886335 | 0.317 | 0.235 | 7.73E-08 | TAM.Gdf15 | Gng5 |
| Gramd1b2 | 4.06E-12 | 0.28039299 | 0.334 | 0.243 | 7.86E-08 | TAM.Gdf15 | Gramd1b |
| Sms2 | 4.20E-12 | 0.45929667 | 0.271 | 0.191 | 8.12E-08 | TAM.Gdf15 | Sms |
| Mtfr1l | 4.60E-12 | 0.29227087 | 0.31 | 0.229 | 8.89E-08 | TAM.Gdf15 | Mtfr1l |
| Nbeal1 | 4.95E-12 | 0.53620236 | 0.265 | 0.181 | 9.56E-08 | TAM.Gdf15 | Nbeal1 |
| Tnfrsf232 | 4.95E-12 | 0.42786394 | 0.263 | 0.175 | 9.57E-08 | TAM.Gdf15 | Tnfrsf23 |
| Rb1cc1 | 5.82E-12 | 0.39646481 | 0.267 | 0.204 | 1.12E-07 | TAM.Gdf15 | Rb1cc1 |
| Dennd6a1 | 6.28E-12 | 0.43758464 | 0.276 | 0.201 | 1.21E-07 | TAM.Gdf15 | Dennd6a |
| Dnajc21 | 7.92E-12 | 0.46919265 | 0.264 | 0.186 | 1.53E-07 | TAM.Gdf15 | Dnajc21 |
| Ppp1r37 | 8.00E-12 | 0.59193923 | 0.271 | 0.174 | 1.55E-07 | TAM.Gdf15 | Ppp1r37 |
| Yif1a1 | 1.03E-11 | 0.3387146 | 0.265 | 0.198 | 2.00E-07 | TAM.Gdf15 | Yif1a |
| Arl23 | 1.31E-11 | 0.40241642 | 0.272 | 0.197 | 2.52E-07 | TAM.Gdf15 | Arl2 |
| Arpc5l1 | 1.32E-11 | 0.26538464 | 0.338 | 0.251 | 2.56E-07 | TAM.Gdf15 | Arpc5l |
| Ubfd12 | 1.55E-11 | 0.43169614 | 0.301 | 0.217 | 3.00E-07 | TAM.Gdf15 | Ubfd1 |
| Krtcap21 | 1.61E-11 | 0.40668288 | 0.29 | 0.217 | 3.11E-07 | TAM.Gdf15 | Krtcap2 |
| Mfn1 | 1.66E-11 | 0.292433 | 0.317 | 0.234 | 3.21E-07 | TAM.Gdf15 | Mfn1 |
| Prrc11 | 1.82E-11 | 0.26329774 | 0.37 | 0.283 | 3.51E-07 | TAM.Gdf15 | Prrc1 |
| Tk22 | 1.86E-11 | 0.3257608 | 0.281 | 0.201 | 3.59E-07 | TAM.Gdf15 | Tk2 |
| Morc2a1 | 2.42E-11 | 0.45214047 | 0.335 | 0.269 | 4.67E-07 | TAM.Gdf15 | Morc2a |
| Trap11 | 2.49E-11 | 0.2833094 | 0.359 | 0.259 | 4.82E-07 | TAM.Gdf15 | Trap1 |
| Stk251 | 2.57E-11 | 0.33896507 | 0.421 | 0.324 | 4.96E-07 | TAM.Gdf15 | Stk25 |
| Dhcr72 | 2.70E-11 | 0.26826993 | 0.28 | 0.206 | 5.23E-07 | TAM.Gdf15 | Dhcr7 |
| Ncdn1 | 3.14E-11 | 0.41959452 | 0.264 | 0.181 | 6.06E-07 | TAM.Gdf15 | Ncdn |
| Golga3 | 3.17E-11 | 0.30641776 | 0.286 | 0.209 | 6.13E-07 | TAM.Gdf15 | Golga3 |
| Pip5k1a1 | 3.26E-11 | 0.31724257 | 0.274 | 0.2 | 6.30E-07 | TAM.Gdf15 | Pip5k1a |
| Arl32 | 3.79E-11 | 0.25170304 | 0.298 | 0.223 | 7.32E-07 | TAM.Gdf15 | Arl3 |
| Ncoa7 | 4.08E-11 | 0.44236214 | 0.288 | 0.197 | 7.88E-07 | TAM.Gdf15 | Ncoa7 |
| Mzt11 | 4.93E-11 | 0.42323489 | 0.255 | 0.178 | 9.54E-07 | TAM.Gdf15 | Mzt1 |
| Rrs11 | 5.39E-11 | 0.2596861 | 0.258 | 0.175 | 1.04E-06 | TAM.Gdf15 | Rrs1 |
| Usp3 | 5.45E-11 | 0.32854337 | 0.374 | 0.291 | 1.05E-06 | TAM.Gdf15 | Usp3 |
| Triap12 | 5.56E-11 | 0.272174 | 0.262 | 0.187 | 1.08E-06 | TAM.Gdf15 | Triap1 |
| Tlk2 | 5.76E-11 | 0.31274516 | 0.334 | 0.257 | 1.11E-06 | TAM.Gdf15 | Tlk2 |
| Maco1 | 6.56E-11 | 0.35011499 | 0.321 | 0.244 | 1.27E-06 | TAM.Gdf15 | Maco1 |
| Rab92 | 7.24E-11 | 0.32588382 | 0.328 | 0.253 | 1.40E-06 | TAM.Gdf15 | Rab9 |
| Tfpt | 8.89E-11 | 0.35328522 | 0.33 | 0.236 | 1.72E-06 | TAM.Gdf15 | Tfpt |
| Lsg11 | 9.24E-11 | 0.30571609 | 0.262 | 0.176 | 1.79E-06 | TAM.Gdf15 | Lsg1 |
| Arsa1 | 9.88E-11 | 0.25046199 | 0.32 | 0.254 | 1.91E-06 | TAM.Gdf15 | Arsa |
| Taz1 | 1.04E-10 | 0.30250538 | 0.391 | 0.297 | 2.01E-06 | TAM.Gdf15 | Taz |
| Lztr1 | 1.07E-10 | 0.27345649 | 0.251 | 0.186 | 2.07E-06 | TAM.Gdf15 | Lztr1 |
| Akap91 | 1.11E-10 | 0.33431013 | 0.288 | 0.206 | 2.15E-06 | TAM.Gdf15 | Akap9 |
| Tsr12 | 1.21E-10 | 0.33370732 | 0.305 | 0.231 | 2.34E-06 | TAM.Gdf15 | Tsr1 |
| Gosr1 | 1.32E-10 | 0.28644039 | 0.279 | 0.208 | 2.55E-06 | TAM.Gdf15 | Gosr1 |
| Klhl243 | 1.42E-10 | 0.26980106 | 0.376 | 0.285 | 2.75E-06 | TAM.Gdf15 | Klhl24 |
| Chchd101 | 1.61E-10 | 0.37748762 | 0.333 | 0.215 | 3.11E-06 | TAM.Gdf15 | Chchd10 |
| Procr1 | 1.79E-10 | 0.79046085 | 0.314 | 0.154 | 3.46E-06 | TAM.Gdf15 | Procr |
| Dvl1 | 2.03E-10 | 0.46144698 | 0.309 | 0.245 | 3.92E-06 | TAM.Gdf15 | Dvl1 |
| Wbp4 | 2.75E-10 | 0.27681289 | 0.328 | 0.263 | 5.31E-06 | TAM.Gdf15 | Wbp4 |
| Frs2 | 3.02E-10 | 0.25693517 | 0.29 | 0.21 | 5.85E-06 | TAM.Gdf15 | Frs2 |
| Hook2 | 4.05E-10 | 0.35150306 | 0.25 | 0.172 | 7.83E-06 | TAM.Gdf15 | Hook2 |
| AU0403201 | 4.43E-10 | 0.39705676 | 0.293 | 0.217 | 8.56E-06 | TAM.Gdf15 | AU040320 |
| Mex3d1 | 4.83E-10 | 0.37891865 | 0.292 | 0.213 | 9.33E-06 | TAM.Gdf15 | Mex3d |
| Slc44a12 | 6.13E-10 | 0.28320789 | 0.345 | 0.247 | 1.19E-05 | TAM.Gdf15 | Slc44a1 |
| Ccdc591 | 6.90E-10 | 0.30418327 | 0.319 | 0.246 | 1.33E-05 | TAM.Gdf15 | Ccdc59 |
| Arid3a3 | 7.01E-10 | 0.26969458 | 0.331 | 0.244 | 1.36E-05 | TAM.Gdf15 | Arid3a |
| Copl11 | 7.56E-10 | 0.28436961 | 0.384 | 0.303 | 1.46E-05 | TAM.Gdf15 | Copl1 |
| Plekhj11 | 9.18E-10 | 0.26335312 | 0.565 | 0.502 | 1.77E-05 | TAM.Gdf15 | Plekhj1 |
| Slc27a41 | 1.07E-09 | 0.26917904 | 0.308 | 0.245 | 2.06E-05 | TAM.Gdf15 | Slc27a4 |
| Scfd11 | 1.58E-09 | 0.26978114 | 0.278 | 0.217 | 3.06E-05 | TAM.Gdf15 | Scfd1 |
| Hmg20a1 | 1.92E-09 | 0.2797148 | 0.312 | 0.233 | 3.71E-05 | TAM.Gdf15 | Hmg20a |
| Dcaf10 | 2.22E-09 | 0.37704286 | 0.273 | 0.209 | 4.29E-05 | TAM.Gdf15 | Dcaf10 |
| Ezh1 | 2.36E-09 | 0.40694416 | 0.252 | 0.176 | 4.57E-05 | TAM.Gdf15 | Ezh1 |
| Ufl1 | 2.70E-09 | 0.3073362 | 0.253 | 0.185 | 5.21E-05 | TAM.Gdf15 | Ufl1 |
| Hras1 | 2.79E-09 | 0.26907315 | 0.308 | 0.237 | 5.40E-05 | TAM.Gdf15 | Hras |
| Zbtb382 | 4.78E-09 | 0.3012364 | 0.283 | 0.208 | 9.25E-05 | TAM.Gdf15 | Zbtb38 |

| | | | | | | | |
|---|---|---|---|---|---|---|---|
| Naa30 | 5.25E-09 | 0.32889052 | 0.326 | 0.254 | 0.00010152 | TAM.Gdf15 | Naa30 |
| Mbd11 | 5.96E-09 | 0.25423142 | 0.285 | 0.207 | 0.00011528 | TAM.Gdf15 | Mbd1 |
| Ciao11 | 7.40E-09 | 0.28682547 | 0.298 | 0.227 | 0.00014304 | TAM.Gdf15 | Ciao1 |
| Trim31 | 8.81E-09 | 0.37879528 | 0.287 | 0.218 | 0.0001704 | TAM.Gdf15 | Trim3 |
| Senp21 | 9.10E-09 | 0.25768112 | 0.358 | 0.283 | 0.00017588 | TAM.Gdf15 | Senp2 |
| Rnf146 | 1.01E-08 | 0.28835834 | 0.294 | 0.219 | 0.00019449 | TAM.Gdf15 | Rnf146 |
| Ap5b12 | 1.08E-08 | 0.25279601 | 0.315 | 0.241 | 0.0002095 | TAM.Gdf15 | Ap5b1 |
| Wdr742 | 1.46E-08 | 0.31657893 | 0.284 | 0.209 | 0.00028151 | TAM.Gdf15 | Wdr74 |
| Chst142 | 2.37E-08 | 0.2715432 | 0.285 | 0.197 | 0.0004588 | TAM.Gdf15 | Chst14 |
| Arfrp1 | 2.49E-08 | 0.30334509 | 0.286 | 0.222 | 0.00048084 | TAM.Gdf15 | Arfrp1 |
| AI8371811 | 2.56E-08 | 0.37690449 | 0.271 | 0.197 | 0.00049516 | TAM.Gdf15 | AI837181 |
| Wdr481 | 3.25E-08 | 0.33199184 | 0.252 | 0.195 | 0.00062889 | TAM.Gdf15 | Wdr48 |
| Rnmt1 | 3.54E-08 | 0.27074365 | 0.263 | 0.197 | 0.00068501 | TAM.Gdf15 | Rnmt |
| Setd1a1 | 5.51E-08 | 0.31139412 | 0.424 | 0.343 | 0.0010655 | TAM.Gdf15 | Setd1a |
| Eya3 | 1.71E-07 | 0.25969929 | 0.26 | 0.196 | 0.0033084 | TAM.Gdf15 | Eya3 |
| Parp41 | 2.34E-07 | 0.28634589 | 0.286 | 0.224 | 0.00452428 | TAM.Gdf15 | Parp4 |
| Trim562 | 2.43E-07 | 0.38390299 | 0.322 | 0.249 | 0.00470332 | TAM.Gdf15 | Trim56 |
| Gga32 | 2.49E-07 | 0.30540207 | 0.299 | 0.225 | 0.00481644 | TAM.Gdf15 | Gga3 |
| Desi2 | 2.52E-07 | 0.27380888 | 0.334 | 0.261 | 0.00486982 | TAM.Gdf15 | Desi2 |
| Hmgxb31 | 2.62E-07 | 0.3160264 | 0.256 | 0.192 | 0.00506005 | TAM.Gdf15 | Hmgxb3 |
| Luc7l1 | 3.22E-07 | 0.25885841 | 0.264 | 0.198 | 0.00622991 | TAM.Gdf15 | Luc7l |
| Tasor | 4.57E-07 | 0.2795457 | 0.262 | 0.202 | 0.00883653 | TAM.Gdf15 | Tasor |
| Lats1 | 5.22E-07 | 0.32654263 | 0.256 | 0.195 | 0.01009372 | TAM.Gdf15 | Lats1 |
| Phactr22 | 9.03E-07 | 0.31068569 | 0.29 | 0.217 | 0.01745808 | TAM.Gdf15 | Phactr2 |
| Fam13b | 1.65E-06 | 0.30356628 | 0.303 | 0.238 | 0.03186752 | TAM.Gdf15 | Fam13b |
| Isca2 | 3.91E-06 | 0.28164981 | 0.299 | 0.234 | 0.0755787 | TAM.Gdf15 | Isca2 |
| Psmf11 | 7.24E-06 | 0.28107762 | 0.28 | 0.217 | 0.13999427 | TAM.Gdf15 | Psmf1 |
| Rbm41 | 1.33E-05 | 0.27646812 | 0.39 | 0.331 | 0.25703182 | TAM.Gdf15 | Rbm4 |
| Axin11 | 1.62E-05 | 0.27183 | 0.297 | 0.244 | 0.3132946 | TAM.Gdf15 | Axin1 |
| Kansl31 | 0.00012266 | 0.26442216 | 0.287 | 0.255 | 1 | TAM.Gdf15 | Kansl3 |
| Cyp2s1 | 0 | 7.49691552 | 0.941 | 0.017 | 0 | osteoclast.like | Cyp2s1 |
| Ctsk | 0 | 9.05357571 | 1 | 0.113 | 0 | osteoclast.like | Ctsk |
| Slc9b2 | 0 | 9.98647897 | 0.878 | 0.003 | 0 | osteoclast.like | Slc9b2 |
| Atp6v0d2 | 0 | 5.22409033 | 0.946 | 0.093 | 0 | osteoclast.like | Atp6v0d2 |
| Coq8a | 0 | 6.41556786 | 0.883 | 0.031 | 0 | osteoclast.like | Coq8a |
| Myo10 | 0 | 4.64938156 | 0.878 | 0.069 | 0 | osteoclast.like | Myo10 |
| Myo1b | 0 | 4.84554355 | 0.873 | 0.071 | 0 | osteoclast.like | Myo1b |
| Plekhf1 | 0 | 4.63206634 | 0.873 | 0.082 | 0 | osteoclast.like | Plekhf1 |
| Rufy4 | 0 | 9.42229422 | 0.785 | 0.002 | 0 | osteoclast.like | Rufy4 |
| Spns2 | 0 | 6.3949827 | 0.79 | 0.015 | 0 | osteoclast.like | Spns2 |
| Igf2r1 | 0 | 3.57840543 | 0.888 | 0.115 | 0 | osteoclast.like | Igf2r |
| Mmp92 | 0 | 4.78411684 | 0.985 | 0.218 | 0 | osteoclast.like | Mmp9 |
| Fam102a | 0 | 5.30803531 | 0.834 | 0.069 | 0 | osteoclast.like | Fam102a |
| Nav2 | 0 | 5.71562906 | 0.785 | 0.029 | 0 | osteoclast.like | Nav2 |
| Chchd102 | 0 | 4.17836513 | 0.961 | 0.213 | 0 | osteoclast.like | Chchd10 |
| C77080 | 0 | 5.28655852 | 0.78 | 0.034 | 0 | osteoclast.like | C77080 |
| Fnbp1l | 0 | 3.72639759 | 0.873 | 0.133 | 0 | osteoclast.like | Fnbp1l |
| Car2 | 0 | 5.81174285 | 0.785 | 0.049 | 0 | osteoclast.like | Car2 |
| Adcy33 | 0 | 3.94414156 | 0.854 | 0.124 | 0 | osteoclast.like | Adcy3 |
| Arsb1 | 0 | 6.02923442 | 0.854 | 0.125 | 0 | osteoclast.like | Arsb |
| Dusp41 | 0 | 3.51944867 | 0.824 | 0.106 | 0 | osteoclast.like | Dusp4 |
| Ppargc1b | 0 | 3.89721201 | 0.815 | 0.1 | 0 | osteoclast.like | Ppargc1b |
| Scin | 0 | 6.6776516 | 0.702 | 0.012 | 0 | osteoclast.like | Scin |
| Twsg11 | 0 | 3.27302346 | 0.844 | 0.156 | 0 | osteoclast.like | Twsg1 |
| Serinc5 | 0 | 3.50104191 | 0.79 | 0.106 | 0 | osteoclast.like | Serinc5 |
| Myo1d | 0 | 6.8091094 | 0.702 | 0.018 | 0 | osteoclast.like | Myo1d |
| Gask1b | 0 | 5.12494928 | 0.741 | 0.058 | 0 | osteoclast.like | Gask1b |
| Slc4a21 | 0 | 3.41244962 | 0.98 | 0.3 | 0 | osteoclast.like | Slc4a2 |
| Hmgb31 | 0 | 3.50676169 | 0.756 | 0.08 | 0 | osteoclast.like | Hmgb3 |
| Clec12a2 | 0 | 2.73673026 | 0.844 | 0.175 | 0 | osteoclast.like | Clec12a |
| Cd109 | 0 | 7.25993034 | 0.678 | 0.009 | 0 | osteoclast.like | Cd109 |
| Slc6a83 | 0 | 2.52559614 | 0.956 | 0.289 | 0 | osteoclast.like | Slc6a8 |
| Kbtbd11 | 0 | 4.33605995 | 0.771 | 0.111 | 0 | osteoclast.like | Kbtbd11 |
| Lpl1 | 0 | 1.52418694 | 0.859 | 0.2 | 0 | osteoclast.like | Lpl |
| Tfrc | 0 | 4.16524184 | 0.732 | 0.076 | 0 | osteoclast.like | Tfrc |
| Serpinb82 | 0 | 2.46366975 | 0.82 | 0.165 | 0 | osteoclast.like | Serpinb8 |
| Plxna4 | 0 | 4.3506954 | 0.698 | 0.044 | 0 | osteoclast.like | Plxna4 |
| Cttn1 | 0 | 3.38492784 | 0.785 | 0.133 | 0 | osteoclast.like | Cttn |
| Xpr1 | 0 | 3.36629877 | 0.951 | 0.302 | 0 | osteoclast.like | Xpr1 |
| Kcnn43 | 0 | 2.04681193 | 0.873 | 0.225 | 0 | osteoclast.like | Kcnn4 |
| Hsd17b101 | 0 | 3.10420363 | 0.912 | 0.268 | 0 | osteoclast.like | Hsd17b10 |
| Siglec15 | 0 | 7.42433618 | 0.644 | 0.005 | 0 | osteoclast.like | Siglec15 |
| Mid1ip13 | 0 | 2.82778691 | 0.878 | 0.241 | 0 | osteoclast.like | Mid1ip1 |
| Ndufa41 | 0 | 2.87596667 | 0.859 | 0.223 | 0 | osteoclast.like | Ndufa4 |
| Msi2 | 0 | 3.35514905 | 0.756 | 0.12 | 0 | osteoclast.like | Msi2 |
| Spry1 | 0 | 3.84105923 | 0.732 | 0.098 | 0 | osteoclast.like | Spry1 |
| Itgb3 | 0 | 6.38465973 | 0.649 | 0.017 | 0 | osteoclast.like | Itgb3 |
| Nudt22 | 0 | 4.06065989 | 0.717 | 0.087 | 0 | osteoclast.like | Nudt22 |
| Ckb | 0 | 4.86972368 | 0.732 | 0.103 | 0 | osteoclast.like | Ckb |
| Inf24 | 0 | 2.29913888 | 0.868 | 0.242 | 0 | osteoclast.like | Inf2 |
| Sms3 | 0 | 2.73418979 | 0.815 | 0.189 | 0 | osteoclast.like | Sms |
| Ermp11 | 0 | 3.19225111 | 0.78 | 0.156 | 0 | osteoclast.like | Ermp1 |
| Dock53 | 0 | 2.5902845 | 0.829 | 0.208 | 0 | osteoclast.like | Dock5 |
| Pde4dip1 | 0 | 2.50065862 | 0.785 | 0.165 | 0 | osteoclast.like | Pde4dip |
| Kazn | 0 | 6.02333634 | 0.634 | 0.016 | 0 | osteoclast.like | Kazn |
| Ocstamp4 | 0 | 3.58790583 | 0.829 | 0.215 | 0 | osteoclast.like | Ocstamp |
| Tsc22d11 | 0 | 3.64115199 | 0.966 | 0.353 | 0 | osteoclast.like | Tsc22d1 |
| Oxct15 | 0 | 2.46684184 | 0.883 | 0.271 | 0 | osteoclast.like | Oxct1 |
| Cd200 | 0 | 6.0147532 | 0.624 | 0.013 | 0 | osteoclast.like | Cd200 |
| Sh3pxd2a2 | 0 | 2.51050956 | 0.81 | 0.2 | 0 | osteoclast.like | Sh3pxd2a |
| Creb3l2 | 0 | 2.97879675 | 0.751 | 0.141 | 0 | osteoclast.like | Creb3l2 |
| Aco21 | 0 | 2.55822296 | 0.849 | 0.24 | 0 | osteoclast.like | Aco2 |
| Akap6 | 0 | 8.59389016 | 0.61 | 0.002 | 0 | osteoclast.like | Akap6 |
| Akip11 | 0 | 3.20353501 | 0.732 | 0.125 | 0 | osteoclast.like | Akip1 |
| Hsd3b73 | 0 | 2.64624215 | 0.893 | 0.29 | 0 | osteoclast.like | Hsd3b7 |
| Oaf | 0 | 3.9475203 | 0.663 | 0.06 | 0 | osteoclast.like | Oaf |
| Sdc12 | 0 | 2.15784629 | 0.956 | 0.354 | 0 | osteoclast.like | Sdc1 |
| Grk32 | 0 | 2.33650221 | 0.893 | 0.295 | 0 | osteoclast.like | Grk3 |
| Ptpn221 | 0 | 2.69804256 | 0.722 | 0.129 | 0 | osteoclast.like | Ptpn22 |
| Fblim14 | 0 | 2.53833108 | 0.859 | 0.266 | 0 | osteoclast.like | Fblim1 |
| Pkn3 | 0 | 4.2273917 | 0.629 | 0.037 | 0 | osteoclast.like | Pkn3 |
| Ankrd13b | 0 | 4.3658663 | 0.62 | 0.031 | 0 | osteoclast.like | Ankrd13b |
| Slc1a54 | 0 | 2.60565026 | 0.98 | 0.393 | 0 | osteoclast.like | Slc1a5 |
| Specc11 | 0 | 2.41428507 | 0.8 | 0.216 | 0 | osteoclast.like | Specc1 |
| Slc39a13 | 0 | 2.68491595 | 0.741 | 0.158 | 0 | osteoclast.like | Slc39a13 |
| Cluh3 | 0 | 2.62613281 | 0.854 | 0.272 | 0 | osteoclast.like | Cluh |
| Pcdhgc3 | 0 | 4.41204649 | 0.61 | 0.028 | 0 | osteoclast.like | Pcdhgc3 |
| Pheta1 | 0 | 4.36308152 | 0.639 | 0.06 | 0 | osteoclast.like | Pheta1 |
| Uqcr101 | 0 | 2.57150203 | 0.893 | 0.314 | 0 | osteoclast.like | Uqcr10 |
| Idh3a2 | 0 | 2.77914474 | 0.878 | 0.301 | 0 | osteoclast.like | Idh3a |
| Tpm11 | 0 | 2.00533632 | 0.893 | 0.317 | 0 | osteoclast.like | Tpm1 |
| Pgpep13 | 0 | 2.64252771 | 0.746 | 0.17 | 0 | osteoclast.like | Pgpep1 |
| Agap12 | 0 | 2.41749937 | 0.883 | 0.311 | 0 | osteoclast.like | Agap1 |
| Fhl32 | 0 | 2.72231156 | 0.878 | 0.306 | 0 | osteoclast.like | Fhl3 |
| Slc37a23 | 0 | 3.68612647 | 0.951 | 0.379 | 0 | osteoclast.like | Slc37a2 |
| Slc25a42 | 0 | 2.21931585 | 0.902 | 0.332 | 0 | osteoclast.like | Slc25a4 |
| Slc44a13 | 0 | 2.34997383 | 0.815 | 0.246 | 0 | osteoclast.like | Slc44a1 |
| Arhgap104 | 0 | 2.07078021 | 0.766 | 0.2 | 0 | osteoclast.like | Arhgap10 |
| Plp23 | 0 | 1.96680176 | 0.937 | 0.372 | 0 | osteoclast.like | Plp2 |
| Col27a1 | 0 | 5.00593407 | 0.59 | 0.026 | 0 | osteoclast.like | Col27a1 |
| Cav22 | 0 | 2.12529097 | 0.78 | 0.219 | 0 | osteoclast.like | Cav2 |
| Plekhg51 | 0 | 2.68142047 | 0.634 | 0.074 | 0 | osteoclast.like | Plekhg5 |
| Lnx2 | 0 | 3.02472507 | 0.654 | 0.095 | 0 | osteoclast.like | Lnx2 |
| Letm11 | 0 | 2.29341023 | 0.751 | 0.193 | 0 | osteoclast.like | Letm1 |
| Srgap3 | 0 | 4.39412558 | 0.59 | 0.032 | 0 | osteoclast.like | Srgap3 |
| Lpcat12 | 0 | 2.18206431 | 0.834 | 0.277 | 0 | osteoclast.like | Lpcat1 |
| Mst1r | 0 | 7.60160096 | 0.561 | 0.005 | 0 | osteoclast.like | Mst1r |
| Tmem26 | 0 | 4.24830046 | 0.595 | 0.044 | 0 | osteoclast.like | Tmem26 |
| Hagh3 | 0 | 2.2802936 | 0.824 | 0.273 | 0 | osteoclast.like | Hagh |
| Gpr1831 | 0 | 2.398443 | 0.732 | 0.182 | 0 | osteoclast.like | Gpr183 |
| Ank2 | 0 | 2.38002735 | 0.8 | 0.25 | 0 | osteoclast.like | Ank |
| Dcstamp1 | 0 | 1.06598201 | 0.805 | 0.255 | 0 | osteoclast.like | Dcstamp |
| Crtap3 | 0 | 2.45975029 | 0.82 | 0.27 | 0 | osteoclast.like | Crtap |
| Gnptab1 | 0 | 2.79413285 | 0.932 | 0.385 | 0 | osteoclast.like | Gnptab |
| Rassf83 | 0 | 2.26715714 | 0.878 | 0.333 | 0 | osteoclast.like | Rassf8 |
| Nfatc12 | 0 | 3.85533323 | 1 | 0.455 | 0 | osteoclast.like | Nfatc1 |
| Gpr681 | 0 | 2.37059202 | 0.678 | 0.134 | 0 | osteoclast.like | Gpr68 |
| Acaa2 | 0 | 2.29889296 | 0.863 | 0.319 | 0 | osteoclast.like | Acaa2 |
| Matk2 | 0 | 1.66582526 | 0.902 | 0.36 | 0 | osteoclast.like | Matk |
| Pop42 | 0 | 2.54926322 | 0.766 | 0.225 | 0 | osteoclast.like | Pop4 |
| Slc16a7 | 0 | 2.6307064 | 0.678 | 0.137 | 0 | osteoclast.like | Slc16a7 |
| Plekhn13 | 0 | 1.87674528 | 0.839 | 0.298 | 0 | osteoclast.like | Plekhn1 |
| Tufm1 | 0 | 2.01410779 | 0.82 | 0.279 | 0 | osteoclast.like | Tufm |
| Pdhb1 | 0 | 2.357061 | 0.873 | 0.332 | 0 | osteoclast.like | Pdhb |
| Mcrip2 | 0 | 2.88432807 | 0.644 | 0.104 | 0 | osteoclast.like | Mcrip2 |
| Nav13 | 0 | 1.65609956 | 0.8 | 0.261 | 0 | osteoclast.like | Nav1 |
| Nsd22 | 0 | 2.57523635 | 0.932 | 0.393 | 0 | osteoclast.like | Nsd2 |
| Bcar31 | 0 | 2.72497738 | 0.688 | 0.149 | 0 | osteoclast.like | Bcar3 |
| Syne31 | 0 | 2.05583374 | 0.693 | 0.154 | 0 | osteoclast.like | Syne3 |
| Prdm13 | 0 | 2.19598185 | 0.927 | 0.39 | 0 | osteoclast.like | Prdm1 |
| Git12 | 0 | 1.86635906 | 0.844 | 0.307 | 0 | osteoclast.like | Git1 |
| Naa803 | 0 | 1.97411707 | 0.746 | 0.212 | 0 | osteoclast.like | Naa80 |
| Mtx21 | 0 | 2.13196863 | 0.766 | 0.233 | 0 | osteoclast.like | Mtx2 |
| Acat12 | 0 | 1.9663622 | 0.829 | 0.296 | 0 | osteoclast.like | Acat1 |
| Glrx51 | 0 | 2.00860548 | 0.766 | 0.234 | 0 | osteoclast.like | Glrx5 |
| Cd2761 | 0 | 2.32110191 | 0.702 | 0.17 | 0 | osteoclast.like | Cd276 |
| Fchsd22 | 0 | 1.79428851 | 0.883 | 0.354 | 0 | osteoclast.like | Fchsd2 |
| Nrp23 | 0 | 3.08593249 | 0.971 | 0.443 | 0 | osteoclast.like | Nrp2 |
| Etfrf11 | 0 | 2.53165637 | 0.678 | 0.15 | 0 | osteoclast.like | Etfrf1 |
| Ablim1 | 0 | 5.02084227 | 0.546 | 0.018 | 0 | osteoclast.like | Ablim1 |
| Pabpc42 | 0 | 2.06376588 | 0.771 | 0.248 | 0 | osteoclast.like | Pabpc4 |
| Pik3cb1 | 0 | 1.69641143 | 0.878 | 0.356 | 0 | osteoclast.like | Pik3cb |
| Actn13 | 0 | 2.31894755 | 0.971 | 0.449 | 0 | osteoclast.like | Actn1 |
| Net12 | 0 | 2.07215379 | 0.746 | 0.224 | 0 | osteoclast.like | Net1 |
| Atp5g12 | 0 | 2.15960787 | 0.863 | 0.342 | 0 | osteoclast.like | Atp5g1 |
| Dmxl12 | 0 | 2.31051662 | 0.82 | 0.299 | 0 | osteoclast.like | Dmxl1 |
| Dlat1 | 0 | 2.05708105 | 0.761 | 0.241 | 0 | osteoclast.like | Dlat |
| Tango21 | 0 | 2.45282141 | 0.678 | 0.159 | 0 | osteoclast.like | Tango2 |
| Ndufab11 | 0 | 2.13204534 | 0.751 | 0.233 | 0 | osteoclast.like | Ndufab1 |
| Fam110b | 0 | 3.42651638 | 0.58 | 0.064 | 0 | osteoclast.like | Fam110b |
| Sipa1l31 | 0 | 1.83229951 | 0.844 | 0.328 | 0 | osteoclast.like | Sipa1l3 |
| Parvb2 | 0 | 1.83938128 | 0.785 | 0.269 | 0 | osteoclast.like | Parvb |
| Chchd31 | 0 | 1.8522121 | 0.902 | 0.387 | 0 | osteoclast.like | Chchd3 |
| Fkbp11 | 0 | 4.71253353 | 0.541 | 0.026 | 0 | osteoclast.like | Fkbp11 |
| Atp5g32 | 0 | 2.28912696 | 0.878 | 0.365 | 0 | osteoclast.like | Atp5g3 |
| Nsun22 | 0 | 2.25465671 | 0.834 | 0.321 | 0 | osteoclast.like | Nsun2 |
| Hs2st12 | 0 | 2.10790165 | 0.761 | 0.249 | 0 | osteoclast.like | Hs2st1 |
| Idh23 | 0 | 1.96087231 | 0.927 | 0.416 | 0 | osteoclast.like | Idh2 |
| Abcb4 | 0 | 5.97385602 | 0.517 | 0.009 | 0 | osteoclast.like | Abcb4 |
| Wdr914 | 0 | 1.99715389 | 0.854 | 0.346 | 0 | osteoclast.like | Wdr91 |
| Hspa92 | 0 | 1.82921155 | 0.829 | 0.323 | 0 | osteoclast.like | Hspa9 |
| Ext22 | 0 | 2.22386025 | 0.805 | 0.301 | 0 | osteoclast.like | Ext2 |
| Swap703 | 0 | 1.78832713 | 0.863 | 0.359 | 0 | osteoclast.like | Swap70 |
| Hspd12 | 0 | 2.17872063 | 0.815 | 0.312 | 0 | osteoclast.like | Hspd1 |
| Ext11 | 0 | 2.59691211 | 0.956 | 0.453 | 0 | osteoclast.like | Ext1 |
| Oscar | 0 | 10.5080262 | 0.502 | 0.001 | 0 | osteoclast.like | Oscar |
| Uqcr111 | 0 | 2.33520944 | 0.956 | 0.455 | 0 | osteoclast.like | Uqcr11 |
| Mdh22 | 0 | 2.39868639 | 0.917 | 0.417 | 0 | osteoclast.like | Mdh2 |
| Slc27a13 | 0 | 1.7653317 | 0.927 | 0.427 | 0 | osteoclast.like | Slc27a1 |
| Tns33 | 0 | 2.65235471 | 0.966 | 0.466 | 0 | osteoclast.like | Tns3 |
| Mitf3 | 0 | 2.0473474 | 0.854 | 0.355 | 0 | osteoclast.like | Mitf |
| Ybx33 | 0 | 1.75700102 | 0.922 | 0.425 | 0 | osteoclast.like | Ybx3 |
| Ndufb51 | 0 | 1.98213217 | 0.824 | 0.327 | 0 | osteoclast.like | Ndufb5 |
| Oxr11 | 0 | 1.78080019 | 0.795 | 0.3 | 0 | osteoclast.like | Oxr1 |
| Tnfrsf11a2 | 0 | 2.14876763 | 0.966 | 0.471 | 0 | osteoclast.like | Tnfrsf11a |
| Ndufv11 | 0 | 1.94174122 | 0.917 | 0.425 | 0 | osteoclast.like | Ndufv1 |
| Sema7a | 0 | 3.388595 | 0.522 | 0.033 | 0 | osteoclast.like | Sema7a |
| Mdh12 | 0 | 2.58511573 | 0.961 | 0.472 | 0 | osteoclast.like | Mdh1 |
| Map4k31 | 0 | 1.93520569 | 0.751 | 0.262 | 0 | osteoclast.like | Map4k3 |
| Tspan44 | 0 | 2.0322792 | 0.824 | 0.335 | 0 | osteoclast.like | Tspan4 |
| Ndufa101 | 0 | 2.2587833 | 0.922 | 0.434 | 0 | osteoclast.like | Ndufa10 |
| Stard72 | 0 | 1.90775183 | 0.824 | 0.336 | 0 | osteoclast.like | Stard7 |
| Ndufs31 | 0 | 1.80857837 | 0.849 | 0.362 | 0 | osteoclast.like | Ndufs3 |
| Cox6c1 | 0 | 1.65591157 | 0.824 | 0.337 | 0 | osteoclast.like | Cox6c |
| Utp112 | 0 | 1.81938899 | 0.795 | 0.309 | 0 | osteoclast.like | Utp11 |
| Bbs9 | 0 | 3.40862533 | 0.537 | 0.051 | 0 | osteoclast.like | Bbs9 |
| Uqcrq1 | 0 | 2.04005624 | 0.78 | 0.294 | 0 | osteoclast.like | Uqcrq |
| Tmem652 | 0 | 2.701091 | 0.937 | 0.452 | 0 | osteoclast.like | Tmem65 |
| Sept113 | 0 | 1.66766911 | 0.839 | 0.354 | 0 | osteoclast.like | 11-Sep |
| Ifi275 | 0 | 2.32365498 | 0.878 | 0.396 | 0 | osteoclast.like | Ifi27 |
| Itga2 | 0 | 4.40625634 | 0.502 | 0.02 | 0 | osteoclast.like | Itga2 |
| Glb12 | 0 | 2.3877912 | 0.82 | 0.338 | 0 | osteoclast.like | Glb1 |
| Cyc12 | 0 | 2.73039672 | 0.946 | 0.466 | 0 | osteoclast.like | Cyc1 |
| Cox7a21 | 0 | 1.77428253 | 0.839 | 0.36 | 0 | osteoclast.like | Cox7a2 |
| Frrs12 | 0 | 1.7409736 | 0.829 | 0.351 | 0 | osteoclast.like | Frrs1 |
| Cblb1 | 0 | 2.00586505 | 0.698 | 0.221 | 0 | osteoclast.like | Cblb |
| Uqcrc21 | 0 | 2.43226488 | 0.863 | 0.387 | 0 | osteoclast.like | Uqcrc2 |
| Ndufc12 | 0 | 1.9981874 | 0.863 | 0.389 | 0 | osteoclast.like | Ndufc1 |
| Phb22 | 0 | 1.82463173 | 0.878 | 0.404 | 0 | osteoclast.like | Phb2 |
| F2r | 0 | 2.58935216 | 0.58 | 0.107 | 0 | osteoclast.like | F2r |
| Ndufv21 | 0 | 2.19282563 | 0.863 | 0.391 | 0 | osteoclast.like | Ndufv2 |
| Samm501 | 0 | 1.91843349 | 0.878 | 0.407 | 0 | osteoclast.like | Samm50 |
| Ndufs11 | 0 | 1.94323526 | 0.707 | 0.236 | 0 | osteoclast.like | Ndufs1 |
| Atp5md1 | 0 | 1.77571163 | 0.834 | 0.363 | 0 | osteoclast.like | Atp5md |
| Tbc1d2b2 | 0 | 1.95999139 | 0.902 | 0.432 | 0 | osteoclast.like | Tbc1d2b |
| Ccdc88a2 | 0 | 1.68268084 | 0.844 | 0.374 | 0 | osteoclast.like | Ccdc88a |
| Micos131 | 0 | 1.84334881 | 0.917 | 0.448 | 0 | osteoclast.like | Micos13 |
| Pstpip14 | 0 | 3.01045036 | 0.995 | 0.526 | 0 | osteoclast.like | Pstpip1 |
| Mt3 | 0 | 8.38852044 | 0.473 | 0.005 | 0 | osteoclast.like | Mt3 |
| Itgav2 | 0 | 2.75832503 | 0.971 | 0.506 | 0 | osteoclast.like | Itgav |
| Rai144 | 0 | 2.2681866 | 0.966 | 0.503 | 0 | osteoclast.like | Rai14 |
| Atp5k1 | 0 | 1.94340881 | 0.902 | 0.44 | 0 | osteoclast.like | Atp5k |
| Uap1l13 | 0 | 1.70118288 | 0.937 | 0.476 | 0 | osteoclast.like | Uap1l1 |
| Dld1 | 0 | 1.85883822 | 0.805 | 0.344 | 0 | osteoclast.like | Dld |
| Tmem204 | 0 | 6.92613361 | 0.463 | 0.003 | 0 | osteoclast.like | Tmem204 |
| Cited22 | 0 | 2.11297459 | 0.971 | 0.511 | 0 | osteoclast.like | Cited2 |
| Cox6a11 | 0 | 1.64461888 | 0.902 | 0.443 | 0 | osteoclast.like | Cox6a1 |
| Abl22 | 0 | 1.91140184 | 0.898 | 0.441 | 0 | osteoclast.like | Abl2 |
| St6galnac42 | 0 | 1.85272025 | 0.878 | 0.424 | 0 | osteoclast.like | St6galnac4 |
| Acap21 | 0 | 1.91461166 | 0.912 | 0.458 | 0 | osteoclast.like | Acap2 |
| Ddt1 | 0 | 1.81573514 | 0.732 | 0.278 | 0 | osteoclast.like | Ddt |
| Ndufb91 | 0 | 2.0928252 | 0.951 | 0.498 | 0 | osteoclast.like | Ndufb9 |
| Dnmt3a4 | 0 | 1.88180702 | 0.839 | 0.387 | 0 | osteoclast.like | Dnmt3a |
| Atp5o1 | 0 | 2.20366816 | 0.922 | 0.471 | 0 | osteoclast.like | Atp5o |
| Tomm402 | 0 | 1.68212945 | 0.834 | 0.385 | 0 | osteoclast.like | Tomm40 |
| Mfsd123 | 0 | 1.89289223 | 0.941 | 0.496 | 0 | osteoclast.like | Mfsd12 |
| Fam234a2 | 0 | 1.607341 | 0.829 | 0.384 | 0 | osteoclast.like | Fam234a |
| Camk2a | 0 | 4.26644528 | 0.473 | 0.028 | 0 | osteoclast.like | Camk2a |
| Pdha11 | 0 | 1.89861606 | 0.815 | 0.37 | 0 | osteoclast.like | Pdha1 |
| Shtn12 | 0 | 1.64728668 | 0.917 | 0.474 | 0 | osteoclast.like | Shtn1 |
| Idh3b1 | 0 | 2.17967171 | 0.932 | 0.49 | 0 | osteoclast.like | Idh3b |
| Ldhd | 0 | 6.43343472 | 0.454 | 0.012 | 0 | osteoclast.like | Ldhd |
| Ndufs71 | 0 | 1.89662964 | 0.932 | 0.491 | 0 | osteoclast.like | Ndufs7 |
| Sec11c1 | 0 | 1.59475004 | 0.917 | 0.476 | 0 | osteoclast.like | Sec11c |
| Ndufb62 | 0 | 1.77305751 | 0.941 | 0.5 | 0 | osteoclast.like | Ndufb6 |
| Piezo12 | 0 | 1.59377346 | 0.941 | 0.501 | 0 | osteoclast.like | Piezo1 |
| DEPRECATED-I | 0 | 4.21493741 | 0.468 | 0.029 | 0 | osteoclast.like | DEPRECATED-ENSMUSG00000037907 |
| Txn22 | 0 | 1.64427212 | 0.844 | 0.405 | 0 | osteoclast.like | Txn2 |
| Slc45a3 | 0 | 6.70776516 | 0.439 | 0.005 | 0 | osteoclast.like | Slc45a3 |
| Idh3g2 | 0 | 2.04817367 | 0.917 | 0.484 | 0 | osteoclast.like | Idh3g |
| Kidins2201 | 0 | 1.89985105 | 0.824 | 0.391 | 0 | osteoclast.like | Kidins220 |
| Atp5f12 | 0 | 1.60982289 | 0.893 | 0.461 | 0 | osteoclast.like | Atp5f1 |
| Adgra3 | 0 | 5.04406187 | 0.444 | 0.013 | 0 | osteoclast.like | Adgra3 |
| St5 | 0 | 5.84173645 | 0.439 | 0.008 | 0 | osteoclast.like | St5 |
| Nop102 | 0 | 1.87599998 | 0.805 | 0.375 | 0 | osteoclast.like | Nop10 |
| Klhl12 | 0 | 4.0074724 | 0.454 | 0.024 | 0 | osteoclast.like | Klhl12 |
| Hadhb1 | 0 | 1.72999791 | 0.815 | 0.385 | 0 | osteoclast.like | Hadhb |
| Jdp21 | 0 | 3.83835005 | 1 | 0.572 | 0 | osteoclast.like | Jdp2 |
| Tmem267 | 0 | 5.02811569 | 0.459 | 0.031 | 0 | osteoclast.like | Tmem267 |
| Steap4 | 0 | 7.41002863 | 0.429 | 0.004 | 0 | osteoclast.like | Steap4 |
| Ccr3 | 0 | 5.67285745 | 0.434 | 0.01 | 0 | osteoclast.like | Ccr3 |
| Atp5a11 | 0 | 1.76641689 | 0.878 | 0.456 | 0 | osteoclast.like | Atp5a1 |
| Epn2 | 0 | 4.20019451 | 0.439 | 0.019 | 0 | osteoclast.like | Epn2 |
| Micos101 | 0 | 1.71360106 | 0.902 | 0.483 | 0 | osteoclast.like | Micos10 |
| Tsku2 | 0 | 2.59177188 | 0.561 | 0.142 | 0 | osteoclast.like | Tsku |
| Slc25a391 | 0 | 2.25489948 | 0.946 | 0.527 | 0 | osteoclast.like | Slc25a39 |
| Tomm52 | 0 | 1.71560886 | 0.78 | 0.362 | 0 | osteoclast.like | Tomm5 |
| Dap | 0 | 2.19897118 | 0.878 | 0.462 | 0 | osteoclast.like | Dap |
| Eif4ebp11 | 0 | 1.47386334 | 0.902 | 0.487 | 0 | osteoclast.like | Eif4ebp1 |
| Mpc2 | 0 | 1.80211452 | 0.756 | 0.341 | 0 | osteoclast.like | Mpc2 |
| Gja11 | 0 | 2.23527356 | 0.507 | 0.093 | 0 | osteoclast.like | Gja1 |
| Zfp462 | 0 | 6.48574029 | 0.415 | 0.003 | 0 | osteoclast.like | Zfp462 |
| Etfa1 | 0 | 1.66238705 | 0.907 | 0.497 | 0 | osteoclast.like | Etfa |
| Anxa13 | 0 | 1.85483501 | 0.956 | 0.546 | 0 | osteoclast.like | Anxa1 |
| Pter | 0 | 3.65747658 | 0.449 | 0.04 | 0 | osteoclast.like | Pter |
| Src3 | 0 | 1.97793743 | 0.971 | 0.563 | 0 | osteoclast.like | Src |
| Atp6v1d1 | 0 | 2.54475107 | 0.976 | 0.568 | 0 | osteoclast.like | Atp6v1d |
| Atp5mpl2 | 0 | 1.47999181 | 0.863 | 0.456 | 0 | osteoclast.like | Atp5mpl |
| Agap32 | 0 | 1.38047213 | 0.868 | 0.462 | 0 | osteoclast.like | Agap3 |
| Naa502 | 0 | 1.93861618 | 0.961 | 0.556 | 0 | osteoclast.like | Naa50 |
| Cox7b2 | 0 | 1.61504732 | 0.937 | 0.535 | 0 | osteoclast.like | Cox7b |
| S100a43 | 0 | 2.4922344 | 0.995 | 0.594 | 0 | osteoclast.like | S100a4 |
| Polr1d2 | 0 | 1.43653266 | 0.927 | 0.527 | 0 | osteoclast.like | Polr1d |
| Sod13 | 0 | 1.51976332 | 0.873 | 0.473 | 0 | osteoclast.like | Sod1 |
| Atp5l1 | 0 | 1.63691433 | 0.893 | 0.494 | 0 | osteoclast.like | Atp5l |
| Ntn1 | 0 | 7.68711649 | 0.4 | 0.003 | 0 | osteoclast.like | Ntn1 |
| Ndst13 | 0 | 1.19454996 | 0.917 | 0.52 | 0 | osteoclast.like | Ndst1 |
| Lat | 0 | 6.00006147 | 0.4 | 0.006 | 0 | osteoclast.like | Lat |
| Myo1e1 | 0 | 1.35637441 | 0.883 | 0.492 | 0 | osteoclast.like | Myo1e |
| Dlst1 | 0 | 1.48380696 | 0.917 | 0.528 | 0 | osteoclast.like | Dlst |
| Cadm1 | 0 | 3.03315032 | 0.454 | 0.066 | 0 | osteoclast.like | Cadm1 |
| Slc25a53 | 0 | 1.69095019 | 0.849 | 0.463 | 0 | osteoclast.like | Slc25a5 |
| Atp6v1c12 | 0 | 2.052521 | 0.946 | 0.56 | 0 | osteoclast.like | Atp6v1c1 |
| Ak22 | 0 | 2.09636114 | 0.966 | 0.581 | 0 | osteoclast.like | Ak2 |
| Uqcrfs12 | 0 | 2.1921634 | 0.951 | 0.566 | 0 | osteoclast.like | Uqcrfs1 |
| Sdhb1 | 0 | 2.21327938 | 0.966 | 0.582 | 0 | osteoclast.like | Sdhb |
| Gtf2a12 | 0 | 1.40231611 | 0.829 | 0.446 | 0 | osteoclast.like | Gtf2a1 |
| Snx102 | 0 | 1.88409045 | 0.727 | 0.346 | 0 | osteoclast.like | Snx10 |
| Ssc5d | 0 | 6.13617931 | 0.385 | 0.004 | 0 | osteoclast.like | Ssc5d |
| Lasp11 | 0 | 1.6793798 | 0.898 | 0.517 | 0 | osteoclast.like | Lasp1 |
| Vegfc | 0 | 5.43890841 | 0.385 | 0.007 | 0 | osteoclast.like | Vegfc |
| Ndufb101 | 0 | 2.00283098 | 0.927 | 0.555 | 0 | osteoclast.like | Ndufb10 |
| St8sia6 | 0 | 5.64999265 | 0.376 | 0.009 | 0 | osteoclast.like | St8sia6 |
| Sdhd1 | 0 | 1.85583491 | 0.932 | 0.568 | 0 | osteoclast.like | Sdhd |
| Map2k2 | 0 | 1.2255232 | 0.937 | 0.573 | 0 | osteoclast.like | Map2k2 |
| Itpripl22 | 0 | 1.26501582 | 0.927 | 0.564 | 0 | osteoclast.like | Itpripl2 |
| Antxr23 | 0 | 1.25172507 | 0.966 | 0.603 | 0 | osteoclast.like | Antxr2 |
| Plxnb22 | 0 | 1.53040672 | 0.956 | 0.594 | 0 | osteoclast.like | Plxnb2 |
| Pgls1 | 0 | 1.31559445 | 0.844 | 0.485 | 0 | osteoclast.like | Pgls |
| Nkain1 | 0 | 4.86765172 | 0.366 | 0.008 | 0 | osteoclast.like | Nkain1 |
| Olt3 | 0 | 7.00762921 | 0.361 | 0.003 | 0 | osteoclast.like | Olt3 |
| Ubald21 | 0 | 1.23933491 | 0.898 | 0.54 | 0 | osteoclast.like | Ubald2 |
| Arl8b1 | 0 | 1.50355324 | 0.912 | 0.556 | 0 | osteoclast.like | Arl8b |
| Cox7c1 | 0 | 1.54765772 | 0.956 | 0.601 | 0 | osteoclast.like | Cox7c |
| Ncoa43 | 0 | 1.46588996 | 0.883 | 0.529 | 0 | osteoclast.like | Ncoa4 |
| Cox141 | 0 | 1.25887847 | 0.888 | 0.535 | 0 | osteoclast.like | Cox14 |
| Ephx2 | 0 | 6.87412384 | 0.356 | 0.004 | 0 | osteoclast.like | Ephx2 |
| Spred1 | 0 | 2.07110881 | 0.966 | 0.615 | 0 | osteoclast.like | Spred1 |
| Rabep12 | 0 | 1.33996494 | 0.883 | 0.532 | 0 | osteoclast.like | Rabep1 |
| Clcn74 | 0 | 1.51134428 | 0.82 | 0.47 | 0 | osteoclast.like | Clcn7 |
| Park72 | 0 | 1.55806348 | 0.937 | 0.588 | 0 | osteoclast.like | Park7 |
| Ndufs21 | 0 | 1.92382565 | 0.961 | 0.613 | 0 | osteoclast.like | Ndufs2 |
| Rnd1 | 0 | 2.79519207 | 0.444 | 0.096 | 0 | osteoclast.like | Rnd1 |
| Atp5j22 | 0 | 1.82596971 | 0.956 | 0.611 | 0 | osteoclast.like | Atp5j2 |
| Ndufb81 | 0 | 1.55132345 | 0.941 | 0.596 | 0 | osteoclast.like | Ndufb8 |
| Sdhc2 | 0 | 1.9882649 | 0.946 | 0.602 | 0 | osteoclast.like | Sdhc |
| Myo5a2 | 0 | 1.49296989 | 0.956 | 0.613 | 0 | osteoclast.like | Myo5a |
| Atp6v1h1 | 0 | 1.36368249 | 0.839 | 0.497 | 0 | osteoclast.like | Atp6v1h |
| Adgre52 | 0 | 1.14203201 | 0.922 | 0.583 | 0 | osteoclast.like | Adgre5 |
| Ndufc22 | 0 | 1.62379415 | 0.946 | 0.607 | 0 | osteoclast.like | Ndufc2 |
| Emp13 | 0 | 0.69361704 | 0.893 | 0.557 | 0 | osteoclast.like | Emp1 |
| Adssl13 | 0 | 0.99773294 | 0.907 | 0.573 | 0 | osteoclast.like | Adssl1 |
| Cox5a2 | 0 | 1.65380158 | 0.966 | 0.634 | 0 | osteoclast.like | Cox5a |
| Prss46 | 0 | 5.9668765 | 0.337 | 0.006 | 0 | osteoclast.like | Prss46 |
| Ndufa82 | 0 | 1.29319331 | 0.946 | 0.617 | 0 | osteoclast.like | Ndufa8 |
| Acp52 | 0 | 4.68266303 | 1 | 0.672 | 0 | osteoclast.like | Acp5 |
| Map4k42 | 0 | 1.11711761 | 0.937 | 0.611 | 0 | osteoclast.like | Map4k4 |
| Ivns1abp1 | 0 | 1.60213094 | 0.951 | 0.626 | 0 | osteoclast.like | Ivns1abp |
| Timm131 | 0 | 1.40898329 | 0.898 | 0.575 | 0 | osteoclast.like | Timm13 |
| Cox7a2l1 | 0 | 1.42149977 | 0.907 | 0.585 | 0 | osteoclast.like | Cox7a2l |
| Sdha1 | 0 | 2.50956691 | 0.98 | 0.659 | 0 | osteoclast.like | Sdha |
| Cox4i11 | 0 | 1.75149891 | 0.932 | 0.613 | 0 | osteoclast.like | Cox4i1 |
| Tgfbr13 | 0 | 1.1438568 | 0.907 | 0.589 | 0 | osteoclast.like | Tgfbr1 |
| Sema4d3 | 0 | 1.64911446 | 0.976 | 0.664 | 0 | osteoclast.like | Sema4d |
| Atp5b2 | 0 | 2.12025297 | 0.966 | 0.655 | 0 | osteoclast.like | Atp5b |
| Ndufa1 | 0 | 1.38429856 | 0.898 | 0.588 | 0 | osteoclast.like | Ndufa1 |
| Hspa1b5 | 0 | 0.54143903 | 0.673 | 0.364 | 0 | osteoclast.like | Hspa1b |
| Tubb63 | 0 | 1.04599973 | 0.927 | 0.618 | 0 | osteoclast.like | Tubb6 |
| Map2k31 | 0 | 1.34541868 | 0.956 | 0.647 | 0 | osteoclast.like | Map2k3 |
| Rab5if1 | 0 | 1.27382221 | 0.917 | 0.61 | 0 | osteoclast.like | Rab5if |
| Lims12 | 0 | 1.11206773 | 0.888 | 0.583 | 0 | osteoclast.like | Lims1 |
| Tap21 | 0 | 1.26984778 | 0.824 | 0.522 | 0 | osteoclast.like | Tap2 |
| Clec2d2 | 0 | 1.19222393 | 0.98 | 0.679 | 0 | osteoclast.like | Clec2d |
| 9-Sep | 0 | 0.98046965 | 0.966 | 0.665 | 0 | osteoclast.like | 9-Sep |
| Ndufa6 | 0 | 1.35296107 | 0.917 | 0.617 | 0 | osteoclast.like | Ndufa6 |
| Atp5e2 | 0 | 1.2680749 | 0.946 | 0.646 | 0 | osteoclast.like | Atp5e |
| Atp6v0e1 | 0 | 2.09259212 | 0.971 | 0.672 | 0 | osteoclast.like | Atp6v0e |
| Dstn2 | 0 | 1.05747927 | 0.898 | 0.601 | 0 | osteoclast.like | Dstn |
| Capg4 | 0 | 1.62505035 | 0.99 | 0.693 | 0 | osteoclast.like | Capg |
| Cox5b2 | 0 | 1.66862555 | 0.961 | 0.665 | 0 | osteoclast.like | Cox5b |
| Sin3b | 0 | 1.70036531 | 0.966 | 0.671 | 0 | osteoclast.like | Sin3b |
| Myof2 | 0 | 1.21344772 | 0.966 | 0.671 | 0 | osteoclast.like | Myof |
| Plec4 | 0 | 1.57725416 | 0.956 | 0.663 | 0 | osteoclast.like | Plec |
| Gsr2 | 0 | 0.67784832 | 0.956 | 0.664 | 0 | osteoclast.like | Gsr |
| Lfng2 | 0 | 0.89947503 | 0.941 | 0.651 | 0 | osteoclast.like | Lfng |
| Ogdh2 | 0 | 1.35550591 | 0.956 | 0.667 | 0 | osteoclast.like | Ogdh |
| Hexb3 | 0 | 1.74883982 | 0.937 | 0.649 | 0 | osteoclast.like | Hexb |
| Uchl1 | 0 | 5.09698209 | 0.293 | 0.009 | 0 | osteoclast.like | Uchl1 |
| Txn12 | 0 | 1.73494442 | 0.941 | 0.66 | 0 | osteoclast.like | Txn1 |
| Pabpc12 | 0 | 1.20950401 | 0.912 | 0.632 | 0 | osteoclast.like | Pabpc1 |
| Gsto11 | 0 | 0.96724552 | 0.927 | 0.647 | 0 | osteoclast.like | Gsto1 |
| Ptpn122 | 0 | 1.10281209 | 0.912 | 0.635 | 0 | osteoclast.like | Ptpn12 |
| Lmna3 | 0 | 2.09639764 | 1 | 0.725 | 0 | osteoclast.like | Lmna |
| Atp5d2 | 0 | 1.62050461 | 0.985 | 0.71 | 0 | osteoclast.like | Atp5d |
| Cox6b12 | 0 | 1.48200748 | 0.941 | 0.669 | 0 | osteoclast.like | Cox6b1 |
| Actn42 | 0 | 0.97397959 | 0.956 | 0.691 | 0 | osteoclast.like | Actn4 |
| Slc18a1 | 0 | 6.91185979 | 0.268 | 0.003 | 0 | osteoclast.like | Slc18a1 |
| Cux11 | 0 | 1.0737377 | 0.956 | 0.696 | 0 | osteoclast.like | Cux1 |
| Uqcrc11 | 0 | 2.06071288 | 0.98 | 0.721 | 0 | osteoclast.like | Uqcrc1 |
| Ppia1 | 0 | 1.10537575 | 0.883 | 0.624 | 0 | osteoclast.like | Ppia |
| Npm12 | 0 | 1.13196688 | 0.951 | 0.693 | 0 | osteoclast.like | Npm1 |
| Ndufa13 | 0 | 1.35133546 | 0.941 | 0.683 | 0 | osteoclast.like | Ndufa13 |
| Ndufa31 | 0 | 1.26417161 | 0.937 | 0.68 | 0 | osteoclast.like | Ndufa3 |
| Ptpre3 | 0 | 1.21652193 | 0.824 | 0.567 | 0 | osteoclast.like | Ptpre |
| Fam129b4 | 0 | 0.686893 | 0.932 | 0.682 | 0 | osteoclast.like | Fam129b |
| Atp6v1a2 | 0 | 2.44904605 | 0.985 | 0.736 | 0 | osteoclast.like | Atp6v1a |
| Cox8a2 | 0 | 1.37267906 | 0.946 | 0.697 | 0 | osteoclast.like | Cox8a |
| Rnf19b2 | 0 | 1.56693051 | 0.946 | 0.7 | 0 | osteoclast.like | Rnf19b |
| Cd633 | 0 | 0.78408793 | 0.99 | 0.748 | 0 | osteoclast.like | Cd63 |
| Atp6v1g11 | 0 | 2.37385724 | 0.995 | 0.754 | 0 | osteoclast.like | Atp6v1g1 |
| Mmp142 | 0 | 1.51175682 | 0.99 | 0.749 | 0 | osteoclast.like | Mmp14 |
| Ndufa21 | 0 | 0.92644231 | 0.951 | 0.713 | 0 | osteoclast.like | Ndufa2 |
| Ttyh32 | 0 | 0.86535368 | 0.932 | 0.702 | 0 | osteoclast.like | Ttyh3 |
| Mvp3 | 0 | 1.30500385 | 0.932 | 0.702 | 0 | osteoclast.like | Mvp |
| Ccl94 | 0 | 0.80660382 | 0.898 | 0.67 | 0 | osteoclast.like | Ccl9 |
| Gns3 | 0 | 0.93261029 | 0.956 | 0.729 | 0 | osteoclast.like | Gns |
| Erp291 | 0 | 1.01126256 | 0.961 | 0.739 | 0 | osteoclast.like | Erp29 |
| Eef1b22 | 0 | 1.22640683 | 0.941 | 0.721 | 0 | osteoclast.like | Eef1b2 |
| Atp6v1e12 | 0 | 1.73162262 | 0.956 | 0.738 | 0 | osteoclast.like | Atp6v1e1 |
| Atp6v1f | 0 | 1.24211943 | 0.907 | 0.695 | 0 | osteoclast.like | Atp6v1f |
| S100a103 | 0 | 1.51381011 | 0.98 | 0.775 | 0 | osteoclast.like | S100a10 |
| Calr2 | 0 | 0.86577749 | 0.932 | 0.734 | 0 | osteoclast.like | Calr |
| Sh3glb12 | 0 | 0.86913113 | 0.961 | 0.764 | 0 | osteoclast.like | Sh3glb1 |
| Atp6v1b22 | 0 | 3.0298612 | 0.995 | 0.799 | 0 | osteoclast.like | Atp6v1b2 |
| Atp6v0b2 | 0 | 1.83467381 | 0.976 | 0.781 | 0 | osteoclast.like | Atp6v0b |
| Rnasek2 | 0 | 1.2840776 | 0.966 | 0.771 | 0 | osteoclast.like | Rnasek |
| Hsp90ab12 | 0 | 0.84863731 | 0.941 | 0.746 | 0 | osteoclast.like | Hsp90ab1 |
| Edem12 | 0 | 1.08528627 | 0.966 | 0.773 | 0 | osteoclast.like | Edem1 |
| Cops91 | 0 | 0.82203187 | 0.907 | 0.717 | 0 | osteoclast.like | Cops9 |
| Clic42 | 0 | 0.63484073 | 0.927 | 0.741 | 0 | osteoclast.like | Clic4 |
| Atp6ap22 | 0 | 1.32054766 | 0.951 | 0.765 | 0 | osteoclast.like | Atp6ap2 |
| Snd12 | 0 | 0.79301093 | 0.951 | 0.769 | 0 | osteoclast.like | Snd1 |
| Arf11 | 0 | 0.98159776 | 0.946 | 0.767 | 0 | osteoclast.like | Arf1 |
| Bhlhe403 | 0 | 0.37820743 | 0.976 | 0.802 | 0 | osteoclast.like | Bhlhe40 |
| Zfand51 | 0 | 0.79514909 | 0.961 | 0.788 | 0 | osteoclast.like | Zfand5 |
| Hexa2 | 0 | 1.51284017 | 0.98 | 0.816 | 0 | osteoclast.like | Hexa |
| Ybx11 | 0 | 0.83956355 | 0.961 | 0.797 | 0 | osteoclast.like | Ybx1 |
| Fxyd53 | 0 | 1.0044751 | 0.98 | 0.819 | 0 | osteoclast.like | Fxyd5 |
| Chchd2 | 0 | 1.1848169 | 0.961 | 0.803 | 0 | osteoclast.like | Chchd2 |
| Aldoa3 | 0 | 0.55830079 | 0.985 | 0.828 | 0 | osteoclast.like | Aldoa |
| Abi11 | 0 | 0.78012539 | 0.937 | 0.781 | 0 | osteoclast.like | Abi1 |
| Furin2 | 0 | 0.74359644 | 0.961 | 0.811 | 0 | osteoclast.like | Furin |
| Slc39a12 | 0 | 1.11110549 | 0.971 | 0.824 | 0 | osteoclast.like | Slc39a1 |
| Plxnd12 | 0 | 1.20163002 | 0.956 | 0.809 | 0 | osteoclast.like | Plxnd1 |
| M6pr1 | 0 | 0.76960798 | 0.927 | 0.781 | 0 | osteoclast.like | M6pr |
| Emp32 | 0 | 0.5054608 | 0.937 | 0.792 | 0 | osteoclast.like | Emp3 |
| Eef23 | 0 | 1.08407737 | 0.961 | 0.817 | 0 | osteoclast.like | Eef2 |
| Slc25a31 | 0 | 1.17854061 | 0.966 | 0.822 | 0 | osteoclast.like | Slc25a3 |
| Anxa22 | 0 | 0.56900505 | 0.99 | 0.857 | 0 | osteoclast.like | Anxa2 |
| Rack12 | 0 | 1.04882656 | 0.99 | 0.86 | 0 | osteoclast.like | Rack1 |
| Tln11 | 0 | 0.51039126 | 0.966 | 0.844 | 0 | osteoclast.like | Tln1 |
| Fosl23 | 0 | 0.58779733 | 0.971 | 0.85 | 0 | osteoclast.like | Fosl2 |
| Tcirg12 | 0 | 1.77331103 | 0.99 | 0.869 | 0 | osteoclast.like | Tcirg1 |
| Spi11 | 0 | 0.82046447 | 0.976 | 0.861 | 0 | osteoclast.like | Spi1 |
| Cstb2 | 0 | 1.14080752 | 0.985 | 0.876 | 0 | osteoclast.like | Cstb |
| Cd682 | 0 | 1.30624025 | 0.99 | 0.885 | 0 | osteoclast.like | Cd68 |
| Rac23 | 0 | 0.95433551 | 0.922 | 0.829 | 0 | osteoclast.like | Rac2 |
| Atp6v0c1 | 0 | 1.55501779 | 0.995 | 0.906 | 0 | osteoclast.like | Atp6v0c |
| Sh3bgrl32 | 0 | 1.6471619 | 0.99 | 0.904 | 0 | osteoclast.like | Sh3bgrl3 |
| Ctsa3 | 0 | 0.64244124 | 0.976 | 0.91 | 0 | osteoclast.like | Ctsa |
| Gna131 | 0 | 0.36063727 | 0.966 | 0.9 | 0 | osteoclast.like | Gna13 |
| Ifitm23 | 0 | 0.37206001 | 0.971 | 0.911 | 0 | osteoclast.like | Ifitm2 |
| Npc22 | 0 | 1.16743503 | 1 | 0.948 | 0 | osteoclast.like | Npc2 |
| Msn2 | 0 | 0.36770065 | 0.995 | 0.943 | 0 | osteoclast.like | Msn |
| Gpx12 | 0 | 1.09665283 | 0.995 | 0.951 | 0 | osteoclast.like | Gpx1 |
| Tpt12 | 0 | 0.37809649 | 1 | 0.957 | 0 | osteoclast.like | Tpt1 |
| Akr1a12 | 0 | 0.50995127 | 0.98 | 0.937 | 0 | osteoclast.like | Akr1a1 |
| Ifi302 | 0 | 0.97705364 | 0.976 | 0.935 | 0 | osteoclast.like | Ifi30 |
| Vim2 | 0 | 2.06843582 | 1 | 0.962 | 0 | osteoclast.like | Vim |
| Fau1 | 0 | 0.36352603 | 0.985 | 0.949 | 0 | osteoclast.like | Fau |
| Csf1r2 | 0 | 0.32789627 | 0.99 | 0.96 | 0 | osteoclast.like | Csf1r |
| Eef1a11 | 0 | 0.90297353 | 1 | 0.977 | 0 | osteoclast.like | Eef1a1 |
| Tyrobp1 | 0 | 0.53450317 | 0.995 | 0.974 | 0 | osteoclast.like | Tyrobp |
| Cst32 | 0 | 0.68889123 | 0.995 | 0.975 | 0 | osteoclast.like | Cst3 |
| Lcp22 | 1.19E-307 | 0.93983569 | 0.878 | 0.595 | 2.31E-303 | osteoclast.like | Lcp2 |
| Hnrnpa12 | 6.74E-306 | 1.11065248 | 0.863 | 0.61 | 1.30E-301 | osteoclast.like | Hnrnpa1 |
| Osbpl3 | 1.08E-305 | 3.00115954 | 0.58 | 0.075 | 2.10E-301 | osteoclast.like | Osbpl3 |
| Hyal12 | 1.56E-305 | 2.40114405 | 0.585 | 0.099 | 3.01E-301 | osteoclast.like | Hyal1 |
| Eif3l2 | 5.86E-305 | 1.18379169 | 0.902 | 0.549 | 1.13E-300 | osteoclast.like | Eif3l |
| Hyal21 | 7.28E-305 | 2.62535235 | 0.605 | 0.11 | 1.41E-300 | osteoclast.like | Hyal2 |
| Eef1g2 | 8.31E-304 | 1.32933199 | 0.829 | 0.465 | 1.61E-299 | osteoclast.like | Eef1g |
| Uqcrb1 | 3.97E-303 | 1.67213128 | 0.81 | 0.36 | 7.67E-299 | osteoclast.like | Uqcrb |
| Btbd11 | 8.92E-303 | 4.9072643 | 0.356 | 0.009 | 1.72E-298 | osteoclast.like | Btbd11 |
| Clec4a23 | 8.63E-302 | 1.47189402 | 0.722 | 0.332 | 1.67E-297 | osteoclast.like | Clec4a2 |
| Eif3f2 | 1.15E-301 | 1.09024687 | 0.878 | 0.593 | 2.22E-297 | osteoclast.like | Eif3f |
| Klhl244 | 6.60E-301 | 1.58584639 | 0.737 | 0.285 | 1.28E-296 | osteoclast.like | Klhl24 |
| Mllt63 | 7.64E-301 | 1.20373218 | 0.883 | 0.501 | 1.48E-296 | osteoclast.like | Mllt6 |
| Nenf3 | 1.56E-300 | 1.76987582 | 0.81 | 0.292 | 3.01E-296 | osteoclast.like | Nenf |
| Vdac12 | 1.69E-299 | 1.51959318 | 0.8 | 0.376 | 3.27E-295 | osteoclast.like | Vdac1 |
| Gsdmd2 | 1.02E-298 | 1.19143652 | 0.893 | 0.525 | 1.97E-294 | osteoclast.like | Gsdmd |
| Iqsec11 | 3.16E-298 | 1.39287213 | 0.834 | 0.413 | 6.12E-294 | osteoclast.like | Iqsec1 |
| Itgb71 | 2.58E-296 | 1.49032315 | 0.566 | 0.151 | 4.99E-292 | osteoclast.like | Itgb7 |
| Rasa41 | 1.12E-294 | 1.47068427 | 0.849 | 0.357 | 2.16E-290 | osteoclast.like | Rasa4 |
| Immt2 | 2.28E-294 | 1.55125038 | 0.8 | 0.355 | 4.41E-290 | osteoclast.like | Immt |
| Ndufa122 | 2.48E-292 | 1.62925997 | 0.795 | 0.334 | 4.80E-288 | osteoclast.like | Ndufa12 |
| Syne1 | 6.90E-292 | 3.67928397 | 0.405 | 0.033 | 1.33E-287 | osteoclast.like | Syne1 |
| Fam53b2 | 6.45E-291 | 1.64798432 | 0.761 | 0.255 | 1.25E-286 | osteoclast.like | Fam53b |
| Pcdh7 | 9.72E-290 | 4.13398656 | 0.293 | 0.014 | 1.88E-285 | osteoclast.like | Pcdh7 |

| | | | | | | | |
|---|---|---|---|---|---|---|---|
| Smpd12 | 2.65E-288 | 2.41795517 | 0.644 | 0.152 | 5.12E-284 | osteoclast.like | Smpd1 |
| Vwa8 | 2.72E-288 | 2.63560479 | 0.595 | 0.112 | 5.27E-284 | osteoclast.like | Vwa8 |
| Hibadh | 1.26E-287 | 1.55069336 | 0.834 | 0.369 | 2.43E-283 | osteoclast.like | Hibadh |
| Atpif11 | 1.46E-286 | 1.4361989 | 0.878 | 0.471 | 2.83E-282 | osteoclast.like | Atpif1 |
| Ndufa51 | 6.06E-286 | 1.85669627 | 0.776 | 0.269 | 1.17E-281 | osteoclast.like | Ndufa5 |
| Ak1 | 2.61E-285 | 3.72918874 | 0.337 | 0.029 | 5.05E-281 | osteoclast.like | Ak1 |
| Rab38 | 3.02E-285 | 4.34656145 | 0.361 | 0.02 | 5.84E-281 | osteoclast.like | Rab38 |
| Rtn4r | 8.73E-283 | 7.0263844 | 0.307 | 0.002 | 1.69E-278 | osteoclast.like | Rtn4r |
| St18 | 1.62E-282 | 3.98704597 | 0.468 | 0.028 | 3.13E-278 | osteoclast.like | St18 |
| Dars1 | 2.75E-282 | 1.33162536 | 0.854 | 0.45 | 5.32E-278 | osteoclast.like | Dars |
| Dynlrb1 | 8.60E-282 | 0.98840934 | 0.912 | 0.616 | 1.66E-277 | osteoclast.like | Dynlrb1 |
| Ktn11 | 1.27E-281 | 1.89985061 | 0.717 | 0.234 | 2.46E-277 | osteoclast.like | Ktn1 |
| Prkch | 3.50E-279 | 1.64409473 | 0.776 | 0.287 | 6.76E-275 | osteoclast.like | Prkch |
| Oxa1l2 | 3.94E-279 | 1.63175805 | 0.815 | 0.313 | 7.62E-275 | osteoclast.like | Oxa1l |
| Lgalsl | 2.99E-278 | 4.366213 | 0.41 | 0.019 | 5.78E-274 | osteoclast.like | Lgalsl |
| Psmd101 | 3.36E-277 | 1.89475175 | 0.693 | 0.238 | 6.49E-273 | osteoclast.like | Psmd10 |
| Ckap42 | 7.80E-277 | 1.51729283 | 0.785 | 0.317 | 1.51E-272 | osteoclast.like | Ckap4 |
| Srp681 | 1.75E-276 | 1.60029021 | 0.854 | 0.336 | 3.39E-272 | osteoclast.like | Srp68 |
| Ndufb111 | 8.56E-276 | 1.20423306 | 0.815 | 0.46 | 1.65E-271 | osteoclast.like | Ndufb11 |
| Rapgef12 | 3.10E-275 | 1.26865679 | 0.834 | 0.416 | 5.99E-271 | osteoclast.like | Rapgef1 |
| Dyrk2 | 3.43E-272 | 1.61281392 | 0.815 | 0.325 | 6.63E-268 | osteoclast.like | Dyrk2 |
| Cyb5r33 | 1.12E-271 | 1.10225844 | 0.912 | 0.579 | 2.17E-267 | osteoclast.like | Cyb5r3 |
| Pwwp2b1 | 1.14E-270 | 1.93493802 | 0.702 | 0.191 | 2.20E-266 | osteoclast.like | Pwwp2b |
| Sbsn | 1.21E-270 | 3.73393609 | 0.371 | 0.025 | 2.35E-266 | osteoclast.like | Sbsn |
| Ik | 2.54E-270 | 0.85891347 | 0.917 | 0.672 | 4.92E-266 | osteoclast.like | Ik |
| Apbb1 | 6.22E-270 | 5.65460003 | 0.302 | 0.004 | 1.20E-265 | osteoclast.like | Apbb1 |
| Use12 | 1.94E-269 | 1.04698917 | 0.834 | 0.541 | 3.74E-265 | osteoclast.like | Use1 |
| Psmd73 | 2.98E-268 | 1.34181731 | 0.859 | 0.479 | 5.75E-264 | osteoclast.like | Psmd7 |
| Zfp827 | 9.72E-268 | 4.72620004 | 0.307 | 0.008 | 1.88E-263 | osteoclast.like | Zfp827 |
| Ciao2a2 | 7.27E-267 | 0.90644665 | 0.883 | 0.631 | 1.41E-262 | osteoclast.like | Ciao2a |
| Fundc22 | 1.25E-266 | 1.56626053 | 0.78 | 0.331 | 2.42E-262 | osteoclast.like | Fundc2 |
| Rilpl22 | 3.11E-266 | 0.78954961 | 0.883 | 0.58 | 6.01E-262 | osteoclast.like | Rilpl2 |
| Ankrd28 | 3.72E-266 | 2.68293043 | 0.585 | 0.078 | 7.20E-262 | osteoclast.like | Ankrd28 |
| Ppp2r3a | 1.12E-265 | 2.979635 | 0.517 | 0.053 | 2.16E-261 | osteoclast.like | Ppp2r3a |
| Tuba1c3 | 1.41E-265 | 0.5954727 | 0.902 | 0.73 | 2.73E-261 | osteoclast.like | Tuba1c |
| Sft2d21 | 1.03E-264 | 1.43265425 | 0.82 | 0.349 | 2.00E-260 | osteoclast.like | Sft2d2 |
| Cacnb2 | 1.07E-264 | 5.70170938 | 0.283 | 0.004 | 2.07E-260 | osteoclast.like | Cacnb2 |
| Fkbp9 | 9.18E-264 | 2.62429526 | 0.507 | 0.074 | 1.77E-259 | osteoclast.like | Fkbp9 |
| Nhp22 | 2.45E-263 | 1.49539814 | 0.824 | 0.407 | 4.74E-259 | osteoclast.like | Nhp2 |
| C1qbp1 | 5.24E-263 | 2.34587231 | 0.659 | 0.173 | 1.01E-258 | osteoclast.like | C1qbp |
| Acsl11 | 6.27E-263 | 2.10301681 | 0.717 | 0.183 | 1.21E-258 | osteoclast.like | Acsl1 |
| Fastk | 6.34E-263 | 1.74800458 | 0.761 | 0.27 | 1.23E-258 | osteoclast.like | Fastk |
| Cct33 | 2.21E-262 | 1.34833269 | 0.761 | 0.365 | 4.27E-258 | osteoclast.like | Cct3 |
| Tbrg13 | 7.87E-262 | 1.78495125 | 0.751 | 0.272 | 1.52E-257 | osteoclast.like | Tbrg1 |
| Tmed32 | 1.01E-261 | 1.24562549 | 0.761 | 0.441 | 1.96E-257 | osteoclast.like | Tmed3 |
| Agrn | 1.42E-261 | 2.12869276 | 0.649 | 0.148 | 2.74E-257 | osteoclast.like | Agrn |
| Atp6ap11 | 2.34E-261 | 0.56862151 | 0.951 | 0.799 | 4.53E-257 | osteoclast.like | Atp6ap1 |
| Ndufb72 | 3.31E-260 | 1.08025368 | 0.893 | 0.536 | 6.39E-256 | osteoclast.like | Ndufb7 |
| Vdac22 | 5.63E-260 | 1.22875276 | 0.805 | 0.449 | 1.09E-255 | osteoclast.like | Vdac2 |
| Atp5j1 | 2.56E-259 | 1.55986413 | 0.761 | 0.318 | 4.95E-255 | osteoclast.like | Atp5j |
| Fahd1 | 9.74E-259 | 3.1527464 | 0.522 | 0.05 | 1.88E-254 | osteoclast.like | Fahd1 |
| Ranbp12 | 1.97E-255 | 1.32598719 | 0.805 | 0.444 | 3.82E-251 | osteoclast.like | Ranbp1 |
| Prdx32 | 7.90E-255 | 1.68571598 | 0.79 | 0.312 | 1.53E-250 | osteoclast.like | Prdx3 |
| Timp23 | 1.37E-254 | 0.73217693 | 0.932 | 0.558 | 2.65E-250 | osteoclast.like | Timp2 |
| Pdgfb3 | 4.19E-252 | 1.74848652 | 0.717 | 0.252 | 8.10E-248 | osteoclast.like | Pdgfb |
| Naca2 | 2.95E-250 | 1.03060956 | 0.912 | 0.572 | 5.71E-246 | osteoclast.like | Naca |
| Mdfic3 | 1.93E-248 | 1.06923529 | 0.888 | 0.52 | 3.74E-244 | osteoclast.like | Mdfic |
| Lif | 2.81E-248 | 1.86036032 | 0.561 | 0.092 | 5.43E-244 | osteoclast.like | Lif |
| Coq71 | 2.17E-246 | 2.5870321 | 0.566 | 0.115 | 4.20E-242 | osteoclast.like | Coq7 |
| Sgpl14 | 2.68E-246 | 0.72476742 | 0.907 | 0.691 | 5.18E-242 | osteoclast.like | Sgpl1 |
| Tpp13 | 4.36E-246 | 0.76384742 | 0.902 | 0.668 | 8.43E-242 | osteoclast.like | Tpp1 |
| Cant12 | 1.44E-245 | 1.33189088 | 0.829 | 0.407 | 2.79E-241 | osteoclast.like | Cant1 |
| Nipsnap22 | 4.78E-245 | 1.7208082 | 0.707 | 0.267 | 9.24E-241 | osteoclast.like | Nipsnap2 |
| Atp2b41 | 5.80E-245 | 1.44109433 | 0.634 | 0.182 | 1.12E-240 | osteoclast.like | Atp2b4 |
| Aagab3 | 2.42E-244 | 1.32647718 | 0.805 | 0.413 | 4.68E-240 | osteoclast.like | Aagab |
| Sptssa3 | 4.10E-244 | 1.30535878 | 0.824 | 0.43 | 7.94E-240 | osteoclast.like | Sptssa |
| Rcc23 | 5.35E-244 | 1.56616296 | 0.761 | 0.359 | 1.03E-239 | osteoclast.like | Rcc2 |
| Mapk62 | 1.68E-242 | 1.11140239 | 0.829 | 0.468 | 3.26E-238 | osteoclast.like | Mapk6 |
| Cs1 | 5.10E-241 | 1.66222886 | 0.756 | 0.295 | 9.87E-237 | osteoclast.like | Cs |
| Las1l1 | 1.57E-240 | 1.96736343 | 0.673 | 0.183 | 3.03E-236 | osteoclast.like | Las1l |
| Vcl2 | 3.00E-240 | 1.58820424 | 0.766 | 0.256 | 5.79E-236 | osteoclast.like | Vcl |
| Htra23 | 3.91E-240 | 1.68456631 | 0.756 | 0.242 | 7.55E-236 | osteoclast.like | Htra2 |
| Dnaja32 | 6.16E-240 | 1.87510785 | 0.732 | 0.233 | 1.19E-235 | osteoclast.like | Dnaja3 |
| Acadm | 3.91E-239 | 1.68216154 | 0.673 | 0.243 | 7.56E-235 | osteoclast.like | Acadm |
| Layn | 4.70E-239 | 3.32546036 | 0.415 | 0.036 | 9.08E-235 | osteoclast.like | Layn |
| Dok32 | 5.61E-239 | 1.04977744 | 0.844 | 0.478 | 1.09E-234 | osteoclast.like | Dok3 |
| Clybl | 4.23E-238 | 2.38495151 | 0.624 | 0.11 | 8.19E-234 | osteoclast.like | Clybl |
| Mgst3 | 6.09E-238 | 1.97227036 | 0.62 | 0.18 | 1.18E-233 | osteoclast.like | Mgst3 |
| Abhd17c4 | 1.64E-236 | 1.85394763 | 0.649 | 0.213 | 3.16E-232 | osteoclast.like | Abhd17c |
| Tm4sf19 | 9.65E-236 | 2.41468121 | 0.454 | 0.085 | 1.87E-231 | osteoclast.like | Tm4sf19 |
| Abcb8 | 1.31E-235 | 2.69692898 | 0.546 | 0.081 | 2.53E-231 | osteoclast.like | Abcb8 |
| Ppp1r12b | 2.06E-233 | 2.59814977 | 0.6 | 0.093 | 3.97E-229 | osteoclast.like | Ppp1r12b |
| Mfn21 | 7.25E-233 | 1.96998085 | 0.698 | 0.2 | 1.40E-228 | osteoclast.like | Mfn2 |
| Padi2 | 1.05E-231 | 2.19957296 | 0.507 | 0.133 | 2.03E-227 | osteoclast.like | Padi2 |
| Eci11 | 1.37E-231 | 2.02713129 | 0.615 | 0.159 | 2.66E-227 | osteoclast.like | Eci1 |
| Plin34 | 2.84E-230 | 1.25459552 | 0.8 | 0.399 | 5.50E-226 | osteoclast.like | Plin3 |
| Edil3 | 7.44E-230 | 3.36136369 | 0.298 | 0.028 | 1.44E-225 | osteoclast.like | Edil3 |
| Trap12 | 1.22E-229 | 1.78089848 | 0.727 | 0.259 | 2.36E-225 | osteoclast.like | Trap1 |
| Zmiz13 | 2.13E-229 | 0.78578841 | 0.912 | 0.657 | 4.11E-225 | osteoclast.like | Zmiz1 |
| Ppp1r14b2 | 4.77E-229 | 1.60953991 | 0.785 | 0.341 | 9.23E-225 | osteoclast.like | Ppp1r14b |
| Retsat | 1.01E-228 | 2.61798423 | 0.537 | 0.099 | 1.95E-224 | osteoclast.like | Retsat |
| Elovl12 | 1.82E-228 | 1.38208992 | 0.766 | 0.382 | 3.52E-224 | osteoclast.like | Elovl1 |
| Repin1 | 1.58E-225 | 2.50972741 | 0.576 | 0.096 | 3.06E-221 | osteoclast.like | Repin1 |
| Cdk181 | 3.65E-225 | 2.40569028 | 0.576 | 0.091 | 7.06E-221 | osteoclast.like | Cdk18 |
| Gtf2i3 | 5.42E-225 | 1.41163332 | 0.829 | 0.364 | 1.05E-220 | osteoclast.like | Gtf2i |
| G3bp11 | 2.07E-224 | 1.40146658 | 0.849 | 0.372 | 4.00E-220 | osteoclast.like | G3bp1 |
| Timm17a2 | 3.63E-224 | 1.57186958 | 0.78 | 0.325 | 7.03E-220 | osteoclast.like | Timm17a |
| Cyp4v31 | 7.32E-223 | 1.55641727 | 0.663 | 0.267 | 1.42E-218 | osteoclast.like | Cyp4v3 |
| Gstt1 | 2.30E-222 | 3.45324922 | 0.283 | 0.021 | 4.45E-218 | osteoclast.like | Gstt1 |
| Fhod1 | 3.95E-222 | 1.3274254 | 0.785 | 0.331 | 7.64E-218 | osteoclast.like | Fhod1 |
| Spint2 | 6.19E-222 | 2.99279156 | 0.39 | 0.042 | 1.20E-217 | osteoclast.like | Spint2 |
| Zc3h3 | 6.22E-222 | 2.12861893 | 0.649 | 0.146 | 1.20E-217 | osteoclast.like | Zc3h3 |
| Suclg11 | 1.16E-221 | 2.10892084 | 0.649 | 0.171 | 2.24E-217 | osteoclast.like | Suclg1 |
| Gga23 | 1.92E-221 | 1.98896891 | 0.688 | 0.207 | 3.71E-217 | osteoclast.like | Gga2 |
| Arhgef71 | 7.16E-221 | 1.34877582 | 0.771 | 0.366 | 1.38E-216 | osteoclast.like | Arhgef7 |
| Efnb1 | 1.16E-220 | 2.65848611 | 0.488 | 0.059 | 2.25E-216 | osteoclast.like | Efnb1 |
| Rgs104 | 1.62E-220 | 0.66139381 | 0.912 | 0.668 | 3.13E-216 | osteoclast.like | Rgs10 |
| Eif4b1 | 5.41E-220 | 0.93807463 | 0.927 | 0.619 | 1.05E-215 | osteoclast.like | Eif4b |
| Atad3a2 | 7.34E-220 | 2.03392566 | 0.644 | 0.175 | 1.42E-215 | osteoclast.like | Atad3a |
| Gucd1 | 2.59E-219 | 1.97360347 | 0.546 | 0.173 | 5.00E-215 | osteoclast.like | Gucd1 |
| Btf33 | 1.05E-217 | 0.82052102 | 0.868 | 0.63 | 2.02E-213 | osteoclast.like | Btf3 |
| Gprin3 | 1.08E-217 | 7.14430694 | 0.278 | 0.001 | 2.08E-213 | osteoclast.like | Gprin3 |
| Psma61 | 1.64E-217 | 0.9300804 | 0.849 | 0.586 | 3.17E-213 | osteoclast.like | Psma6 |
| Rabggtb2 | 7.53E-217 | 1.3105651 | 0.805 | 0.378 | 1.46E-212 | osteoclast.like | Rabggtb |
| Cnih43 | 1.33E-216 | 1.27401432 | 0.737 | 0.346 | 2.56E-212 | osteoclast.like | Cnih4 |
| Ssr42 | 4.76E-216 | 0.92113946 | 0.883 | 0.572 | 9.21E-212 | osteoclast.like | Ssr4 |
| Spr1 | 6.29E-215 | 1.49145285 | 0.732 | 0.296 | 1.22E-210 | osteoclast.like | Spr |
| Pls3 | 1.83E-214 | 2.07287032 | 0.424 | 0.11 | 3.53E-210 | osteoclast.like | Pls3 |
| Pik3r53 | 2.58E-213 | 0.9287755 | 0.844 | 0.559 | 5.00E-209 | osteoclast.like | Pik3r5 |
| Prdx22 | 8.83E-213 | 0.80304621 | 0.854 | 0.61 | 1.71E-208 | osteoclast.like | Prdx2 |
| Eif4g22 | 5.33E-211 | 0.58742002 | 0.898 | 0.761 | 1.03E-206 | osteoclast.like | Eif4g2 |
| Coro1c2 | 3.31E-209 | 1.07555911 | 0.863 | 0.48 | 6.41E-205 | osteoclast.like | Coro1c |
| Gnsf11 | 3.70E-209 | 1.26946265 | 0.79 | 0.375 | 7.15E-205 | osteoclast.like | Gnsf1 |
| Nsmaf1 | 2.17E-208 | 1.80058862 | 0.693 | 0.185 | 4.20E-204 | osteoclast.like | Nsmaf |
| Cpeb43 | 6.77E-208 | 1.11157164 | 0.81 | 0.445 | 1.31E-203 | osteoclast.like | Cpeb4 |
| H2afz2 | 9.81E-207 | 0.85011893 | 0.829 | 0.564 | 1.90E-202 | osteoclast.like | H2afz |
| Ttc7 | 1.10E-206 | 0.83463459 | 0.912 | 0.598 | 2.13E-202 | osteoclast.like | Ttc7 |
| Dazap12 | 6.12E-206 | 0.85485661 | 0.907 | 0.602 | 1.18E-201 | osteoclast.like | Dazap1 |
| Iars3 | 6.34E-206 | 1.67169872 | 0.707 | 0.235 | 1.23E-201 | osteoclast.like | Iars |
| Cmtr12 | 3.39E-205 | 1.1590865 | 0.81 | 0.373 | 6.55E-201 | osteoclast.like | Cmtr1 |
| Ndufb1-ps3 | 6.32E-205 | 0.87981975 | 0.893 | 0.602 | 1.22E-200 | osteoclast.like | Ndufb1-ps |
| Ap3d11 | 8.07E-205 | 0.77974963 | 0.844 | 0.571 | 1.56E-200 | osteoclast.like | Ap3d1 |
| Rhou1 | 2.26E-204 | 1.5068228 | 0.605 | 0.17 | 4.38E-200 | osteoclast.like | Rhou |
| Gfm12 | 8.32E-204 | 1.75710818 | 0.668 | 0.221 | 1.61E-199 | osteoclast.like | Gfm1 |
| Fryl1 | 1.03E-203 | 1.64118607 | 0.717 | 0.26 | 2.00E-199 | osteoclast.like | Fryl |
| Grpel12 | 3.40E-203 | 1.35679525 | 0.824 | 0.378 | 6.57E-199 | osteoclast.like | Grpel1 |
| Comtd13 | 1.60E-202 | 1.93261163 | 0.678 | 0.196 | 3.10E-198 | osteoclast.like | Comtd1 |
| Ndufs52 | 1.71E-202 | 1.78321339 | 0.683 | 0.231 | 3.31E-198 | osteoclast.like | Ndufs5 |
| Aebp22 | 4.61E-202 | 1.3599624 | 0.815 | 0.394 | 8.90E-198 | osteoclast.like | Aebp2 |
| Cisd11 | 7.97E-202 | 2.15142457 | 0.629 | 0.15 | 1.54E-197 | osteoclast.like | Cisd1 |
| Tnip12 | 1.17E-201 | 1.02863117 | 0.849 | 0.425 | 2.26E-197 | osteoclast.like | Tnip1 |
| Pip5k1c1 | 1.80E-201 | 0.52162059 | 0.932 | 0.774 | 3.49E-197 | osteoclast.like | Pip5k1c |
| Lrpprc1 | 3.33E-201 | 2.10144414 | 0.624 | 0.18 | 6.44E-197 | osteoclast.like | Lrpprc |
| Pmepa12 | 7.29E-201 | 0.75968312 | 0.898 | 0.581 | 1.41E-196 | osteoclast.like | Pmepa1 |
| Vamp81 | 3.79E-200 | 0.44964309 | 0.937 | 0.81 | 7.32E-196 | osteoclast.like | Vamp8 |
| Cct6a2 | 1.56E-199 | 1.34281144 | 0.741 | 0.363 | 3.02E-195 | osteoclast.like | Cct6a |
| Slc5a32 | 1.57E-199 | 1.12068393 | 0.678 | 0.298 | 3.04E-195 | osteoclast.like | Slc5a3 |
| Sik13 | 1.83E-199 | 0.41891389 | 0.834 | 0.707 | 3.54E-195 | osteoclast.like | Sik1 |
| Ncl2 | 5.05E-199 | 0.94752186 | 0.859 | 0.582 | 9.76E-195 | osteoclast.like | Ncl |
| Smim202 | 4.43E-198 | 1.48671143 | 0.732 | 0.285 | 8.56E-194 | osteoclast.like | Smim20 |
| Usp6nl | 5.91E-198 | 1.96229589 | 0.649 | 0.147 | 1.14E-193 | osteoclast.like | Usp6nl |
| Eml2 | 1.10E-197 | 2.35292159 | 0.541 | 0.099 | 2.14E-193 | osteoclast.like | Eml2 |
| Slc29a12 | 1.99E-197 | 1.34950031 | 0.698 | 0.321 | 3.85E-193 | osteoclast.like | Slc29a1 |
| Arcn11 | 9.19E-197 | 0.81120203 | 0.824 | 0.574 | 1.78E-192 | osteoclast.like | Arcn1 |
| Atxn102 | 4.62E-196 | 1.53680881 | 0.737 | 0.316 | 8.94E-192 | osteoclast.like | Atxn10 |
| Serbp12 | 7.89E-196 | 0.9899976 | 0.844 | 0.538 | 1.53E-191 | osteoclast.like | Serbp1 |
| Atf13 | 3.82E-195 | 0.88731283 | 0.79 | 0.528 | 7.39E-191 | osteoclast.like | Atf1 |
| Agfg22 | 4.45E-195 | 1.82383333 | 0.644 | 0.192 | 8.60E-191 | osteoclast.like | Agfg2 |
| Cdt11 | 9.36E-195 | 1.7568235 | 0.712 | 0.227 | 1.81E-190 | osteoclast.like | Cdt1 |
| Krba1 | 1.85E-194 | 4.58770657 | 0.288 | 0.011 | 3.57E-190 | osteoclast.like | Krba1 |
| Ube2a1 | 6.62E-194 | 1.06827418 | 0.815 | 0.426 | 1.28E-189 | osteoclast.like | Ube2a |
| Wipf11 | 9.23E-194 | 0.59212643 | 0.932 | 0.747 | 1.79E-189 | osteoclast.like | Wipf1 |
| Rreb12 | 1.21E-193 | 0.88982413 | 0.854 | 0.55 | 2.34E-189 | osteoclast.like | Rreb1 |
| Vcp3 | 2.24E-193 | 0.42693174 | 0.927 | 0.804 | 4.34E-189 | osteoclast.like | Vcp |
| Stard3 | 2.30E-193 | 1.32662645 | 0.829 | 0.382 | 4.44E-189 | osteoclast.like | Stard3 |
| Ostc1 | 2.38E-193 | 1.05038107 | 0.81 | 0.449 | 4.61E-189 | osteoclast.like | Ostc |
| Bcar1 | 3.04E-193 | 3.31509499 | 0.327 | 0.025 | 5.88E-189 | osteoclast.like | Bcar1 |
| Nup371 | 6.39E-193 | 2.61355836 | 0.527 | 0.08 | 1.24E-188 | osteoclast.like | Nup37 |
| Naglu4 | 2.75E-192 | 1.36211482 | 0.79 | 0.371 | 5.31E-188 | osteoclast.like | Naglu |
| Naa202 | 1.76E-191 | 1.24651464 | 0.727 | 0.342 | 3.41E-187 | osteoclast.like | Naa20 |
| Ppp2ca2 | 5.97E-191 | 0.57711683 | 0.927 | 0.71 | 1.15E-186 | osteoclast.like | Ppp2ca |
| Arpin | 3.88E-190 | 2.37679447 | 0.502 | 0.088 | 7.49E-186 | osteoclast.like | Arpin |
| Phb1 | 5.85E-190 | 1.72759252 | 0.654 | 0.21 | 1.13E-185 | osteoclast.like | Phb |
| Epas1 | 1.50E-189 | 2.56396637 | 0.405 | 0.058 | 2.90E-185 | osteoclast.like | Epas1 |
| Dennd2a | 6.41E-189 | 2.66717592 | 0.551 | 0.09 | 1.24E-184 | osteoclast.like | Dennd2a |
| Hspe13 | 1.59E-188 | 1.46068431 | 0.712 | 0.27 | 3.07E-184 | osteoclast.like | Hspe1 |
| Acads1 | 2.37E-188 | 1.43932833 | 0.722 | 0.303 | 4.59E-184 | osteoclast.like | Acads |
| Lix1l | 7.98E-188 | 2.09264669 | 0.546 | 0.119 | 1.54E-183 | osteoclast.like | Lix1l |
| Galm | 2.14E-187 | 2.0345131 | 0.561 | 0.148 | 4.14E-183 | osteoclast.like | Galm |
| Agpat43 | 2.90E-187 | 1.0845137 | 0.839 | 0.401 | 5.60E-183 | osteoclast.like | Agpat4 |
| Sh3kbp11 | 3.60E-187 | 1.21846868 | 0.761 | 0.359 | 6.97E-183 | osteoclast.like | Sh3kbp1 |
| Sep-71 | 4.19E-187 | 0.7391617 | 0.844 | 0.603 | 8.11E-183 | osteoclast.like | 7-Sep |
| Tmem35b | 1.42E-186 | 2.50380019 | 0.507 | 0.097 | 2.74E-182 | osteoclast.like | Tmem35b |
| Pde8a3 | 2.32E-186 | 1.31097953 | 0.737 | 0.286 | 4.49E-182 | osteoclast.like | Pde8a |
| Schip1 | 1.94E-185 | 3.13012713 | 0.327 | 0.027 | 3.76E-181 | osteoclast.like | Schip1 |
| Esrra | 2.29E-185 | 1.15713609 | 0.834 | 0.404 | 4.43E-181 | osteoclast.like | Esrra |
| Polr2f2 | 9.22E-185 | 1.3437478 | 0.771 | 0.37 | 1.78E-180 | osteoclast.like | Polr2f |
| Ap1s12 | 1.72E-184 | 1.01399599 | 0.839 | 0.491 | 3.32E-180 | osteoclast.like | Ap1s1 |
| Timm44 | 1.10E-183 | 1.68779251 | 0.678 | 0.211 | 2.12E-179 | osteoclast.like | Timm44 |
| Glud13 | 1.93E-183 | 0.5063636 | 0.946 | 0.764 | 3.73E-179 | osteoclast.like | Glud1 |
| Slc25a112 | 3.41E-183 | 1.09280803 | 0.829 | 0.458 | 6.60E-179 | osteoclast.like | Slc25a11 |
| Ptges2 | 8.45E-183 | 2.24417049 | 0.551 | 0.114 | 1.63E-178 | osteoclast.like | Ptges2 |
| Cryl12 | 9.93E-182 | 1.3711162 | 0.751 | 0.307 | 1.92E-177 | osteoclast.like | Cryl1 |
| Arl6ip51 | 1.96E-181 | 0.883233 | 0.868 | 0.565 | 3.80E-177 | osteoclast.like | Arl6ip5 |
| Ipo51 | 2.61E-181 | 1.33074245 | 0.815 | 0.361 | 5.04E-177 | osteoclast.like | Ipo5 |
| Nmnat3 | 2.63E-181 | 2.70145036 | 0.478 | 0.07 | 5.08E-177 | osteoclast.like | Nmnat3 |
| Ctbp21 | 5.79E-181 | 1.07111839 | 0.854 | 0.464 | 1.12E-176 | osteoclast.like | Ctbp2 |
| Chst112 | 2.79E-180 | 1.29770265 | 0.722 | 0.217 | 5.39E-176 | osteoclast.like | Chst11 |
| Gfpt12 | 2.90E-180 | 1.25120652 | 0.678 | 0.288 | 5.61E-176 | osteoclast.like | Gfpt1 |
| Fam122a1 | 3.52E-180 | 1.96107837 | 0.62 | 0.16 | 6.80E-176 | osteoclast.like | Fam122a |
| Rnf241 | 4.38E-180 | 1.9343096 | 0.585 | 0.146 | 8.47E-176 | osteoclast.like | Rnf24 |
| Rmnd1 | 4.49E-180 | 2.24693179 | 0.502 | 0.095 | 8.68E-176 | osteoclast.like | Rmnd1 |
| Orai21 | 4.72E-180 | 1.49767024 | 0.707 | 0.241 | 9.12E-176 | osteoclast.like | Orai2 |
| Tle52 | 8.73E-180 | 1.044937 | 0.902 | 0.5 | 1.69E-175 | osteoclast.like | Tle5 |
| Actr1b1 | 9.60E-180 | 1.26169289 | 0.805 | 0.387 | 1.86E-175 | osteoclast.like | Actr1b |
| Tbc1d10b2 | 1.04E-179 | 0.97421711 | 0.824 | 0.44 | 2.01E-175 | osteoclast.like | Tbc1d10b |
| Cd99l2 | 1.23E-179 | 2.37355085 | 0.493 | 0.079 | 2.37E-175 | osteoclast.like | Cd99l2 |
| Timm102 | 2.27E-178 | 1.79492979 | 0.649 | 0.187 | 4.38E-174 | osteoclast.like | Timm10 |
| Pak12 | 3.87E-178 | 1.30826769 | 0.732 | 0.301 | 7.49E-174 | osteoclast.like | Pak1 |
| Pdia32 | 8.05E-178 | 0.6383656 | 0.893 | 0.666 | 1.56E-173 | osteoclast.like | Pdia3 |
| Apbb2 | 1.84E-177 | 2.76810968 | 0.454 | 0.058 | 3.56E-173 | osteoclast.like | Apbb2 |
| Aldh4a1 | 1.85E-177 | 2.23555996 | 0.551 | 0.115 | 3.59E-173 | osteoclast.like | Aldh4a1 |
| Hspa1a4 | 5.86E-177 | 0.26896674 | 0.629 | 0.416 | 1.13E-172 | osteoclast.like | Hspa1a |
| Stambpl1 | 5.92E-177 | 1.96485242 | 0.556 | 0.139 | 1.15E-172 | osteoclast.like | Stambpl1 |
| Hk11 | 1.91E-176 | 0.82048105 | 0.795 | 0.454 | 3.69E-172 | osteoclast.like | Hk1 |
| Ptgr23 | 2.59E-176 | 1.2249949 | 0.722 | 0.317 | 5.00E-172 | osteoclast.like | Ptgr2 |
| Slc38a63 | 8.87E-176 | 1.0791722 | 0.766 | 0.379 | 1.72E-171 | osteoclast.like | Slc38a6 |
| Ssr21 | 1.06E-175 | 0.90365398 | 0.883 | 0.547 | 2.04E-171 | osteoclast.like | Ssr2 |
| Tgfbr23 | 5.73E-175 | 0.52174965 | 0.829 | 0.602 | 1.11E-170 | osteoclast.like | Tgfbr2 |
| Tiam2 | 6.00E-175 | 2.99913795 | 0.312 | 0.035 | 1.16E-170 | osteoclast.like | Tiam2 |
| Pdgfa4 | 6.30E-175 | 1.32752616 | 0.605 | 0.254 | 1.22E-170 | osteoclast.like | Pdgfa |
| Ociad1 | 1.27E-174 | 1.31268794 | 0.81 | 0.369 | 2.45E-170 | osteoclast.like | Ociad1 |
| Hacd41 | 1.72E-174 | 1.09981807 | 0.712 | 0.384 | 3.32E-170 | osteoclast.like | Hacd4 |
| Aebp1 | 1.09E-173 | 2.88394624 | 0.38 | 0.034 | 2.11E-169 | osteoclast.like | Aebp1 |
| Ubxn12 | 3.12E-173 | 0.84960142 | 0.917 | 0.63 | 6.03E-169 | osteoclast.like | Ubxn1 |
| Ube2f1 | 4.98E-173 | 0.83140902 | 0.863 | 0.569 | 9.63E-169 | osteoclast.like | Ube2f |
| Ddx212 | 6.22E-173 | 0.73550218 | 0.863 | 0.596 | 1.20E-168 | osteoclast.like | Ddx21 |
| Dtx42 | 7.60E-173 | 1.06027677 | 0.81 | 0.447 | 1.47E-168 | osteoclast.like | Dtx4 |
| Coa51 | 3.64E-171 | 0.91812193 | 0.79 | 0.481 | 7.04E-167 | osteoclast.like | Coa5 |
| Cox10 | 1.14E-170 | 2.08723105 | 0.546 | 0.118 | 2.20E-166 | osteoclast.like | Cox10 |
| Raph11 | 1.31E-170 | 1.66862361 | 0.566 | 0.165 | 2.53E-166 | osteoclast.like | Raph1 |
| Ywhag2 | 1.41E-170 | 0.43281698 | 0.966 | 0.828 | 2.72E-166 | osteoclast.like | Ywhag |
| Ywhae1 | 1.49E-170 | 0.65406859 | 0.893 | 0.682 | 2.87E-166 | osteoclast.like | Ywhae |
| Hmgn3 | 2.42E-170 | 4.00422257 | 0.259 | 0.016 | 4.69E-166 | osteoclast.like | Hmgn3 |
| Anxa42 | 5.34E-170 | 0.60669314 | 0.912 | 0.655 | 1.03E-165 | osteoclast.like | Anxa4 |
| Htra1 | 2.60E-169 | 3.61559035 | 0.312 | 0.017 | 5.03E-165 | osteoclast.like | Htra1 |
| Cmas3 | 4.08E-169 | 1.24719926 | 0.771 | 0.344 | 7.89E-165 | osteoclast.like | Cmas |
| Got21 | 1.50E-168 | 0.914964 | 0.815 | 0.503 | 2.91E-164 | osteoclast.like | Got2 |
| Otud7b | 3.73E-168 | 3.06587289 | 0.366 | 0.035 | 7.20E-164 | osteoclast.like | Otud7b |
| Pkig2 | 1.00E-167 | 1.43076068 | 0.702 | 0.303 | 1.94E-163 | osteoclast.like | Pkig |
| Sbno21 | 1.22E-167 | 0.29271659 | 0.902 | 0.76 | 2.35E-163 | osteoclast.like | Sbno2 |
| Pex10 | 2.42E-167 | 2.70466267 | 0.405 | 0.054 | 4.68E-163 | osteoclast.like | Pex10 |
| Fh12 | 2.78E-167 | 1.30061528 | 0.712 | 0.271 | 5.38E-163 | osteoclast.like | Fh1 |
| Zbtb381 | 3.85E-167 | 1.56785292 | 0.659 | 0.207 | 7.44E-163 | osteoclast.like | Zbtb38 |
| Flvcr13 | 1.65E-166 | 1.48080117 | 0.732 | 0.285 | 3.19E-162 | osteoclast.like | Flvcr1 |
| Psmb52 | 5.35E-166 | 0.74341314 | 0.917 | 0.59 | 1.04E-161 | osteoclast.like | Psmb5 |
| Rilpl1 | 1.47E-165 | 2.38456929 | 0.42 | 0.064 | 2.84E-161 | osteoclast.like | Rilpl1 |
| Hsp90aa12 | 5.85E-165 | 0.57960308 | 0.839 | 0.598 | 1.13E-160 | osteoclast.like | Hsp90aa1 |
| mt-Cytb2 | 1.76E-164 | 0.46651427 | 0.888 | 0.702 | 3.40E-160 | osteoclast.like | mt-Cytb |
| Rnf63 | 6.80E-164 | 0.75747467 | 0.815 | 0.555 | 1.31E-159 | osteoclast.like | Rnf6 |
| Rdh13 | 1.13E-163 | 2.50951228 | 0.454 | 0.073 | 2.19E-159 | osteoclast.like | Rdh13 |
| Coq9 | 2.22E-163 | 2.09577839 | 0.629 | 0.137 | 4.30E-159 | osteoclast.like | Coq9 |
| Rpn12 | 6.71E-162 | 0.64924105 | 0.863 | 0.62 | 1.30E-157 | osteoclast.like | Rpn1 |
| Ndufv32 | 1.27E-161 | 0.68000517 | 0.888 | 0.611 | 2.46E-157 | osteoclast.like | Ndufv3 |
| Coa31 | 1.35E-161 | 1.54401343 | 0.732 | 0.281 | 2.61E-157 | osteoclast.like | Coa3 |
| Slc35a41 | 4.43E-161 | 1.0155391 | 0.785 | 0.422 | 8.56E-157 | osteoclast.like | Slc35a4 |
| Snrpd32 | 1.56E-160 | 1.01209081 | 0.805 | 0.479 | 3.02E-156 | osteoclast.like | Snrpd3 |
| Slc12a73 | 7.34E-160 | 1.10703241 | 0.732 | 0.329 | 1.42E-155 | osteoclast.like | Slc12a7 |
| Ctnna1 | 9.26E-160 | 0.68835319 | 0.839 | 0.573 | 1.79E-155 | osteoclast.like | Ctnna1 |
| Entpd71 | 1.12E-159 | 1.22496017 | 0.717 | 0.336 | 2.17E-155 | osteoclast.like | Entpd7 |
| Heatr5a3 | 2.02E-159 | 1.22798075 | 0.732 | 0.329 | 3.91E-155 | osteoclast.like | Heatr5a |
| Xylb | 4.93E-159 | 3.17140546 | 0.366 | 0.031 | 9.54E-155 | osteoclast.like | Xylb |
| Smad72 | 6.02E-159 | 0.96565459 | 0.712 | 0.369 | 1.16E-154 | osteoclast.like | Smad7 |
| Oaz11 | 2.39E-157 | 0.82166182 | 0.927 | 0.626 | 4.62E-153 | osteoclast.like | Oaz1 |
| Gsap2 | 2.07E-156 | 0.80823896 | 0.82 | 0.511 | 4.00E-152 | osteoclast.like | Gsap |
| Marveld12 | 4.72E-156 | 1.5427873 | 0.673 | 0.245 | 9.13E-152 | osteoclast.like | Marveld1 |
| Tomm70a1 | 8.82E-156 | 1.6597853 | 0.6 | 0.217 | 1.70E-151 | osteoclast.like | Tomm70a |
| Adss3 | 2.11E-154 | 1.12784412 | 0.766 | 0.378 | 4.08E-150 | osteoclast.like | Adss |
| Wdr261 | 6.49E-154 | 0.28545749 | 0.941 | 0.816 | 1.25E-149 | osteoclast.like | Wdr26 |
| Tbrg42 | 7.38E-154 | 1.22119848 | 0.722 | 0.319 | 1.43E-149 | osteoclast.like | Tbrg4 |
| Cdk62 | 8.30E-154 | 1.52024928 | 0.702 | 0.228 | 1.61E-149 | osteoclast.like | Cdk6 |
| Gusb2 | 8.46E-154 | 0.49970204 | 0.941 | 0.724 | 1.64E-149 | osteoclast.like | Gusb |
| Eif2s12 | 1.27E-153 | 1.19554228 | 0.78 | 0.38 | 2.45E-149 | osteoclast.like | Eif2s1 |
| Nub1 | 1.91E-153 | 1.3383939 | 0.688 | 0.329 | 3.69E-149 | osteoclast.like | Nub1 |
| Fignl2 | 1.33E-152 | 2.66788596 | 0.38 | 0.052 | 2.56E-148 | osteoclast.like | Fignl2 |
| Vps393 | 2.46E-152 | 1.55118427 | 0.629 | 0.218 | 4.75E-148 | osteoclast.like | Vps39 |
| Fkbp41 | 3.25E-152 | 1.76033624 | 0.576 | 0.169 | 6.29E-148 | osteoclast.like | Fkbp4 |
| St3gal11 | 4.22E-152 | 0.52630551 | 0.82 | 0.549 | 8.17E-148 | osteoclast.like | St3gal1 |
| Npepl13 | 4.50E-152 | 1.13498457 | 0.746 | 0.378 | 8.69E-148 | osteoclast.like | Npepl1 |
| Uqcc1 | 7.48E-152 | 2.15197432 | 0.551 | 0.115 | 1.45E-147 | osteoclast.like | Uqcc1 |
| Hccs1 | 1.25E-151 | 1.51392904 | 0.605 | 0.23 | 2.42E-147 | osteoclast.like | Hccs |
| Flii1 | 2.08E-151 | 0.86828917 | 0.839 | 0.485 | 4.03E-147 | osteoclast.like | Flii |
| Ppfibp13 | 2.45E-151 | 1.38781391 | 0.683 | 0.275 | 4.73E-147 | osteoclast.like | Ppfibp1 |
| Cd59a | 4.81E-151 | 3.82442339 | 0.288 | 0.018 | 9.30E-147 | osteoclast.like | Cd59a |
| Sipa1l21 | 5.01E-151 | 1.41985568 | 0.58 | 0.221 | 9.68E-147 | osteoclast.like | Sipa1l2 |
| Uqcc21 | 5.15E-151 | 1.34642421 | 0.688 | 0.291 | 9.95E-147 | osteoclast.like | Uqcc2 |
| Ghitm1 | 6.64E-151 | 0.91450889 | 0.771 | 0.423 | 1.28E-146 | osteoclast.like | Ghitm |
| Tomm221 | 1.02E-150 | 0.97454298 | 0.751 | 0.423 | 1.97E-146 | osteoclast.like | Tomm22 |
| Eif4a21 | 1.26E-150 | 0.59586578 | 0.873 | 0.638 | 2.43E-146 | osteoclast.like | Eif4a2 |
| Pnkd | 1.91E-150 | 1.88487336 | 0.537 | 0.141 | 3.70E-146 | osteoclast.like | Pnkd |
| Cox151 | 4.54E-150 | 1.62319507 | 0.61 | 0.176 | 8.77E-146 | osteoclast.like | Cox15 |
| Ndufb21 | 1.40E-149 | 1.74616738 | 0.58 | 0.173 | 2.70E-145 | osteoclast.like | Ndufb2 |
| Gnb4 | 1.43E-149 | 3.80376993 | 0.259 | 0.014 | 2.76E-145 | osteoclast.like | Gnb4 |
| Pxk2 | 5.02E-149 | 0.88309459 | 0.741 | 0.411 | 9.71E-145 | osteoclast.like | Pxk |
| Hnrnpa2b12 | 5.87E-149 | 0.42771111 | 0.898 | 0.756 | 1.14E-144 | osteoclast.like | Hnrnpa2b1 |
| Dync1li1 | 1.04E-148 | 0.76530491 | 0.81 | 0.518 | 2.01E-144 | osteoclast.like | Dync1li1 |
| Cst7 | 1.38E-148 | 2.97608522 | 0.371 | 0.037 | 2.67E-144 | osteoclast.like | Cst7 |
| Hnrnpm2 | 2.47E-148 | 0.42502881 | 0.912 | 0.743 | 4.78E-144 | osteoclast.like | Hnrnpm |
| Rnpep2 | 1.08E-147 | 0.48633351 | 0.912 | 0.733 | 2.08E-143 | osteoclast.like | Rnpep |
| Tmem41a | 1.42E-147 | 2.6876746 | 0.376 | 0.053 | 2.75E-143 | osteoclast.like | Tmem41a |
| Hadha1 | 1.91E-147 | 1.02199016 | 0.771 | 0.428 | 3.70E-143 | osteoclast.like | Hadha |
| Pbx1 | 2.21E-147 | 1.52434177 | 0.507 | 0.129 | 4.27E-143 | osteoclast.like | Pbx1 |
| Zfp7101 | 3.79E-147 | 0.80500101 | 0.81 | 0.481 | 7.32E-143 | osteoclast.like | Zfp710 |
| Tcf121 | 5.67E-147 | 1.34204723 | 0.712 | 0.278 | 1.10E-142 | osteoclast.like | Tcf12 |
| Ndfip11 | 1.17E-146 | 0.39541555 | 0.907 | 0.71 | 2.27E-142 | osteoclast.like | Ndfip1 |
| Angptl2 | 1.25E-146 | 2.25071047 | 0.288 | 0.056 | 2.42E-142 | osteoclast.like | Angptl2 |
| Tcp12 | 1.55E-146 | 0.92767378 | 0.795 | 0.463 | 3.00E-142 | osteoclast.like | Tcp1 |
| Ech11 | 2.50E-146 | 1.03080606 | 0.722 | 0.396 | 4.84E-142 | osteoclast.like | Ech1 |
| Acy11 | 4.25E-146 | 2.26757522 | 0.507 | 0.104 | 8.23E-142 | osteoclast.like | Acy1 |
| Pdcd41 | 4.42E-146 | 1.30340167 | 0.702 | 0.239 | 8.54E-142 | osteoclast.like | Pdcd4 |
| Vegfb4 | 9.07E-146 | 1.49841823 | 0.649 | 0.215 | 1.75E-141 | osteoclast.like | Vegfb |
| Etfdh1 | 9.50E-146 | 1.66316332 | 0.61 | 0.2 | 1.84E-141 | osteoclast.like | Etfdh |
| Ssna11 | 1.25E-145 | 1.18424222 | 0.78 | 0.383 | 2.42E-141 | osteoclast.like | Ssna1 |
| Psmb72 | 2.14E-145 | 1.00126415 | 0.776 | 0.425 | 4.13E-141 | osteoclast.like | Psmb7 |
| Gadd45gip11 | 2.63E-145 | 1.39693737 | 0.649 | 0.255 | 5.08E-141 | osteoclast.like | Gadd45gip1 |
| Midn2 | 1.46E-144 | 0.27449999 | 0.898 | 0.728 | 2.83E-140 | osteoclast.like | Midn |
| Rasal2 | 1.58E-144 | 2.92553498 | 0.405 | 0.044 | 3.06E-140 | osteoclast.like | Rasal2 |
| H1f03 | 3.75E-144 | 0.67700685 | 0.683 | 0.417 | 7.26E-140 | osteoclast.like | H1f0 |
| Cav1 | 4.93E-144 | 1.60378058 | 0.351 | 0.08 | 9.52E-140 | osteoclast.like | Cav1 |
| Jarid22 | 8.85E-144 | 0.46302318 | 0.78 | 0.533 | 1.71E-139 | osteoclast.like | Jarid2 |
| Hnmt | 9.19E-144 | 2.36020942 | 0.405 | 0.08 | 1.78E-139 | osteoclast.like | Hnmt |
| Clptm1l | 9.27E-144 | 0.80919588 | 0.883 | 0.578 | 1.79E-139 | osteoclast.like | Clptm1l |
| Ap1g21 | 1.03E-143 | 1.46646032 | 0.634 | 0.221 | 2.00E-139 | osteoclast.like | Ap1g2 |
| Eci21 | 1.14E-143 | 1.50532661 | 0.61 | 0.211 | 2.20E-139 | osteoclast.like | Eci2 |
| Maea | 1.65E-143 | 0.85328047 | 0.829 | 0.493 | 3.18E-139 | osteoclast.like | Maea |
| Mtdh | 4.49E-142 | 1.08593257 | 0.849 | 0.411 | 8.68E-138 | osteoclast.like | Mtdh |
| Vti1b2 | 4.62E-142 | 1.16047797 | 0.712 | 0.349 | 8.93E-138 | osteoclast.like | Vti1b |
| Ndufs41 | 6.37E-142 | 1.29488725 | 0.707 | 0.294 | 1.23E-137 | osteoclast.like | Ndufs4 |
| Znrf13 | 8.40E-142 | 1.05460062 | 0.712 | 0.337 | 1.62E-137 | osteoclast.like | Znrf1 |
| Fkbp22 | 9.52E-142 | 0.72888014 | 0.829 | 0.579 | 1.84E-137 | osteoclast.like | Fkbp2 |
| Ccr11 | 1.44E-141 | 0.34350743 | 0.883 | 0.692 | 2.79E-137 | osteoclast.like | Ccr1 |
| Ppp2r1a2 | 1.45E-141 | 0.70374674 | 0.863 | 0.58 | 2.80E-137 | osteoclast.like | Ppp2r1a |
| Sh3bgrl1 | 2.21E-141 | 0.32490576 | 0.868 | 0.75 | 4.27E-137 | osteoclast.like | Sh3bgrl |
| Rad23b2 | 2.47E-141 | 1.07194375 | 0.771 | 0.442 | 4.78E-137 | osteoclast.like | Rad23b |
| Anxa72 | 2.56E-141 | 0.83233273 | 0.805 | 0.435 | 4.96E-137 | osteoclast.like | Anxa7 |
| Ttyh24 | 3.80E-141 | 1.28693068 | 0.673 | 0.292 | 7.34E-137 | osteoclast.like | Ttyh2 |
| Calcrl | 4.59E-141 | 2.0938171 | 0.395 | 0.094 | 8.88E-137 | osteoclast.like | Calcrl |
| Vwa5a1 | 9.30E-141 | 0.77180362 | 0.751 | 0.492 | 1.80E-136 | osteoclast.like | Vwa5a |
| Ptpmt12 | 1.27E-140 | 1.43216744 | 0.683 | 0.271 | 2.46E-136 | osteoclast.like | Ptpmt1 |
| Rhof | 1.46E-140 | 2.63870816 | 0.38 | 0.054 | 2.82E-136 | osteoclast.like | Rhof |
| Sec31a1 | 1.63E-140 | 1.18483742 | 0.751 | 0.343 | 3.15E-136 | osteoclast.like | Sec31a |
| Ndufs81 | 5.97E-140 | 0.86822796 | 0.795 | 0.482 | 1.16E-135 | osteoclast.like | Ndufs8 |
| Arhgap311 | 6.18E-140 | 0.79329217 | 0.805 | 0.492 | 1.20E-135 | osteoclast.like | Arhgap31 |
| Ifrd21 | 6.21E-139 | 2.08905174 | 0.478 | 0.103 | 1.20E-134 | osteoclast.like | Ifrd2 |
| Nedd81 | 6.63E-139 | 0.50345822 | 0.902 | 0.738 | 1.28E-134 | osteoclast.like | Nedd8 |
| Gstt2 | 7.51E-139 | 2.29669748 | 0.39 | 0.079 | 1.45E-134 | osteoclast.like | Gstt2 |
| Mprip1 | 9.35E-139 | 0.92470649 | 0.78 | 0.43 | 1.81E-134 | osteoclast.like | Mprip |
| Hdgf3 | 1.12E-138 | 1.03687373 | 0.761 | 0.418 | 2.17E-134 | osteoclast.like | Hdgf |
| Zc3h7b1 | 1.50E-138 | 1.98359287 | 0.634 | 0.18 | 2.90E-134 | osteoclast.like | Zc3h7b |
| Timm233 | 2.01E-138 | 1.43790625 | 0.761 | 0.282 | 3.89E-134 | osteoclast.like | Timm23 |
| Ide2 | 2.23E-138 | 1.21385606 | 0.741 | 0.344 | 4.32E-134 | osteoclast.like | Ide |
| Hipk3 | 5.68E-138 | 0.84818662 | 0.824 | 0.489 | 1.10E-133 | osteoclast.like | Hipk3 |
| Apex13 | 1.06E-137 | 1.68830898 | 0.615 | 0.22 | 2.05E-133 | osteoclast.like | Apex1 |
| Qars1 | 1.79E-137 | 1.04905444 | 0.776 | 0.411 | 3.46E-133 | osteoclast.like | Qars |
| Dnajc152 | 2.00E-137 | 1.16618475 | 0.79 | 0.401 | 3.87E-133 | osteoclast.like | Dnajc15 |
| Supt3 | 2.22E-137 | 2.33751731 | 0.473 | 0.081 | 4.28E-133 | osteoclast.like | Supt3 |
| Pon33 | 3.25E-137 | 1.50418861 | 0.639 | 0.222 | 6.28E-133 | osteoclast.like | Pon3 |
| Psmb22 | 5.01E-137 | 0.72550929 | 0.805 | 0.545 | 9.68E-133 | osteoclast.like | Psmb2 |
| Dnajb14 | 9.71E-137 | 1.17977816 | 0.727 | 0.325 | 1.88E-132 | osteoclast.like | Dnajb14 |
| Rab123 | 2.50E-136 | 1.09466414 | 0.737 | 0.358 | 4.84E-132 | osteoclast.like | Rab12 |
| Lrrc471 | 2.93E-136 | 1.13248809 | 0.712 | 0.339 | 5.67E-132 | osteoclast.like | Lrrc47 |
| Zfp1061 | 3.99E-136 | 0.7526304 | 0.776 | 0.48 | 7.71E-132 | osteoclast.like | Zfp106 |
| Tomm201 | 1.15E-135 | 1.08904895 | 0.722 | 0.362 | 2.22E-131 | osteoclast.like | Tomm20 |
| Nus12 | 1.34E-135 | 0.8491831 | 0.907 | 0.533 | 2.59E-131 | osteoclast.like | Nus1 |
| Tab21 | 1.58E-135 | 0.76562149 | 0.8 | 0.513 | 3.05E-131 | osteoclast.like | Tab2 |
| Phka21 | 1.63E-135 | 1.55254102 | 0.576 | 0.178 | 3.15E-131 | osteoclast.like | Phka2 |
| Pdxk3 | 1.00E-134 | 1.00369985 | 0.79 | 0.399 | 1.94E-130 | osteoclast.like | Pdxk |
| Tbcc3 | 1.07E-134 | 1.66810547 | 0.595 | 0.197 | 2.06E-130 | osteoclast.like | Tbcc |
| Tmem701 | 1.28E-134 | 1.92785025 | 0.532 | 0.14 | 2.48E-130 | osteoclast.like | Tmem70 |
| Tecr2 | 2.33E-134 | 0.83327835 | 0.829 | 0.491 | 4.50E-130 | osteoclast.like | Tecr |
| Ubqln11 | 2.70E-134 | 0.80404926 | 0.824 | 0.531 | 5.23E-130 | osteoclast.like | Ubqln1 |
| Txnip3 | 4.03E-134 | 0.52630829 | 0.732 | 0.531 | 7.79E-130 | osteoclast.like | Txnip |
| Lrp101 | 6.66E-134 | 0.43390861 | 0.917 | 0.713 | 1.29E-129 | osteoclast.like | Lrp10 |
| Kdelr1 | 9.02E-134 | 0.7715444 | 0.795 | 0.511 | 1.74E-129 | osteoclast.like | Kdelr1 |
| Ptma1 | 1.40E-133 | 0.59144536 | 0.873 | 0.679 | 2.71E-129 | osteoclast.like | Ptma |
| Ctdsp2 | 1.53E-133 | 0.5232462 | 0.902 | 0.672 | 2.96E-129 | osteoclast.like | Ctdsp2 |
| Eif3g2 | 2.02E-133 | 0.79514601 | 0.795 | 0.485 | 3.90E-129 | osteoclast.like | Eif3g |
| Snu132 | 5.76E-133 | 0.80599089 | 0.78 | 0.481 | 1.11E-128 | osteoclast.like | Snu13 |
| Manba1 | 2.42E-132 | 1.09189169 | 0.678 | 0.312 | 4.67E-128 | osteoclast.like | Manba |
| Stip11 | 2.44E-132 | 1.12544437 | 0.673 | 0.356 | 4.71E-128 | osteoclast.like | Stip1 |
| Cstf2t2 | 3.19E-132 | 1.67698525 | 0.61 | 0.198 | 6.16E-128 | osteoclast.like | Cstf2t |
| Huwe12 | 1.55E-131 | 0.74956451 | 0.834 | 0.528 | 3.00E-127 | osteoclast.like | Huwe1 |
| Arhgef2 | 1.83E-131 | 0.47390544 | 0.907 | 0.681 | 3.54E-127 | osteoclast.like | Arhgef2 |
| Ecsit | 3.16E-130 | 2.05761017 | 0.493 | 0.096 | 6.11E-126 | osteoclast.like | Ecsit |
| Irak11 | 3.89E-130 | 0.78566015 | 0.824 | 0.517 | 7.51E-126 | osteoclast.like | Irak1 |
| Bzw23 | 3.92E-130 | 1.47441756 | 0.654 | 0.229 | 7.57E-126 | osteoclast.like | Bzw2 |
| Papola1 | 4.75E-130 | 0.56382099 | 0.883 | 0.669 | 9.19E-126 | osteoclast.like | Papola |
| Trim352 | 6.08E-130 | 1.10024093 | 0.746 | 0.346 | 1.18E-125 | osteoclast.like | Trim35 |
| Tjp21 | 7.20E-130 | 1.56247434 | 0.576 | 0.181 | 1.39E-125 | osteoclast.like | Tjp2 |
| Manf2 | 1.52E-129 | 0.51292807 | 0.863 | 0.622 | 2.95E-125 | osteoclast.like | Manf |
| Ctnnb11 | 3.53E-129 | 0.41408751 | 0.902 | 0.694 | 6.83E-125 | osteoclast.like | Ctnnb1 |
| Scarb12 | 6.59E-129 | 0.87175204 | 0.707 | 0.372 | 1.27E-124 | osteoclast.like | Scarb1 |
| Tmem192 | 8.19E-129 | 1.25069545 | 0.654 | 0.249 | 1.58E-124 | osteoclast.like | Tmem192 |
| B4galt11 | 9.10E-129 | 0.37327079 | 0.941 | 0.739 | 1.76E-124 | osteoclast.like | B4galt1 |
| Nbr13 | 2.97E-128 | 1.48733285 | 0.605 | 0.217 | 5.74E-124 | osteoclast.like | Nbr1 |
| Atl32 | 3.15E-128 | 0.75592777 | 0.844 | 0.52 | 6.10E-124 | osteoclast.like | Atl3 |
| Nudc2 | 8.33E-128 | 0.91115156 | 0.849 | 0.506 | 1.61E-123 | osteoclast.like | Nudc |
| Sf3b32 | 1.14E-127 | 0.99977911 | 0.732 | 0.403 | 2.21E-123 | osteoclast.like | Sf3b3 |
| Bola31 | 1.66E-127 | 1.84810283 | 0.527 | 0.146 | 3.20E-123 | osteoclast.like | Bola3 |
| Eif2b11 | 2.30E-127 | 1.75275641 | 0.6 | 0.167 | 4.45E-123 | osteoclast.like | Eif2b1 |
| Canx2 | 2.66E-127 | 0.65209911 | 0.863 | 0.625 | 5.15E-123 | osteoclast.like | Canx |
| Dtx21 | 6.34E-127 | 1.56666988 | 0.644 | 0.201 | 1.23E-122 | osteoclast.like | Dtx2 |
| Mul1 | 7.48E-127 | 1.74123917 | 0.556 | 0.146 | 1.45E-122 | osteoclast.like | Mul1 |
| Anpep4 | 9.19E-127 | 0.76629807 | 0.82 | 0.498 | 1.78E-122 | osteoclast.like | Anpep |
| Slc25a513 | 9.43E-127 | 0.87698625 | 0.766 | 0.418 | 1.82E-122 | osteoclast.like | Slc25a51 |
| Zmpste242 | 1.11E-126 | 0.97710442 | 0.727 | 0.363 | 2.14E-122 | osteoclast.like | Zmpste24 |
| Nudt13 | 1.33E-126 | 2.17324171 | 0.444 | 0.069 | 2.57E-122 | osteoclast.like | Nudt13 |
| Swi51 | 1.58E-126 | 1.17836089 | 0.737 | 0.348 | 3.06E-122 | osteoclast.like | Swi5 |
| Stk11ip1 | 2.12E-126 | 1.50188781 | 0.585 | 0.2 | 4.09E-122 | osteoclast.like | Stk11ip |
| Traf12 | 4.20E-126 | 0.74009287 | 0.698 | 0.331 | 8.12E-122 | osteoclast.like | Traf1 |
| Bcat21 | 9.45E-126 | 1.10539658 | 0.78 | 0.362 | 1.83E-121 | osteoclast.like | Bcat2 |
| Wdr83os2 | 1.55E-125 | 0.64289191 | 0.868 | 0.586 | 3.00E-121 | osteoclast.like | Wdr83os |
| Trip12 | 1.57E-125 | 0.4727426 | 0.859 | 0.656 | 3.03E-121 | osteoclast.like | Trip12 |
| Myc3 | 7.54E-125 | 1.24984629 | 0.571 | 0.273 | 1.46E-120 | osteoclast.like | Myc |
| Tmco11 | 9.33E-125 | 0.95908645 | 0.717 | 0.371 | 1.80E-120 | osteoclast.like | Tmco1 |
| Dgkz1 | 9.52E-125 | 0.51312565 | 0.937 | 0.688 | 1.84E-120 | osteoclast.like | Dgkz |
| Fdxr | 1.29E-124 | 2.07244039 | 0.429 | 0.085 | 2.50E-120 | osteoclast.like | Fdxr |
| Pmpcb1 | 1.65E-124 | 1.28369851 | 0.624 | 0.26 | 3.20E-120 | osteoclast.like | Pmpcb |
| Poldip21 | 2.13E-124 | 1.82847236 | 0.527 | 0.129 | 4.12E-120 | osteoclast.like | Poldip2 |
| Hist1h2bc3 | 3.52E-124 | 0.67361131 | 0.693 | 0.358 | 6.81E-120 | osteoclast.like | Hist1h2bc |
| Cd822 | 4.59E-124 | 0.71472335 | 0.771 | 0.501 | 8.87E-120 | osteoclast.like | Cd82 |
| Mars2 | 5.03E-124 | 1.46136631 | 0.654 | 0.227 | 9.73E-120 | osteoclast.like | Mars |
| Flad1 | 5.78E-124 | 1.58509543 | 0.541 | 0.157 | 1.12E-119 | osteoclast.like | Flad1 |
| Tomm71 | 5.87E-124 | 0.87388436 | 0.834 | 0.48 | 1.14E-119 | osteoclast.like | Tomm7 |
| Eif3d2 | 1.15E-123 | 0.87249953 | 0.8 | 0.463 | 2.23E-119 | osteoclast.like | Eif3d |
| Traf61 | 1.57E-123 | 0.96954442 | 0.678 | 0.327 | 3.04E-119 | osteoclast.like | Traf6 |
| Anapc131 | 2.23E-123 | 0.90304239 | 0.829 | 0.423 | 4.32E-119 | osteoclast.like | Anapc13 |
| Ak31 | 4.45E-123 | 1.13531722 | 0.654 | 0.275 | 8.61E-119 | osteoclast.like | Ak3 |
| 21-Sep | 5.69E-123 | 0.61958588 | 0.844 | 0.577 | 1.10E-118 | osteoclast.like | 2-Sep |
| Cdc372 | 6.74E-123 | 0.6478622 | 0.888 | 0.639 | 1.30E-118 | osteoclast.like | Cdc37 |
| Psmd17 | 9.36E-123 | 0.76310591 | 0.78 | 0.493 | 1.81E-118 | osteoclast.like | Psmd1 |
| Larp42 | 1.25E-122 | 0.89947931 | 0.727 | 0.409 | 2.41E-118 | osteoclast.like | Larp4 |
| Gdi11 | 1.51E-122 | 0.79313129 | 0.698 | 0.433 | 2.93E-118 | osteoclast.like | Gdi1 |
| Klhdc21 | 1.52E-122 | 1.60449314 | 0.58 | 0.19 | 2.94E-118 | osteoclast.like | Klhdc2 |
| Med152 | 2.47E-122 | 0.59476401 | 0.878 | 0.611 | 4.78E-118 | osteoclast.like | Med15 |
| Enc1 | 3.96E-122 | 1.41551635 | 0.444 | 0.159 | 7.67E-118 | osteoclast.like | Enc1 |
| Prmt12 | 4.14E-122 | 1.1995067 | 0.644 | 0.243 | 8.00E-118 | osteoclast.like | Prmt1 |
| Cct22 | 5.25E-122 | 0.71600824 | 0.785 | 0.468 | 1.01E-117 | osteoclast.like | Cct2 |
| Edem3 | 6.33E-122 | 1.27288437 | 0.62 | 0.249 | 1.22E-117 | osteoclast.like | Edem3 |
| Eri33 | 6.48E-122 | 1.62108652 | 0.59 | 0.204 | 1.25E-117 | osteoclast.like | Eri3 |
| Iars21 | 1.81E-121 | 1.33673917 | 0.668 | 0.252 | 3.50E-117 | osteoclast.like | Iars2 |
| Pnpla22 | 5.05E-121 | 0.77107635 | 0.839 | 0.516 | 9.76E-117 | osteoclast.like | Pnpla2 |
| Ykt61 | 5.36E-121 | 0.97051616 | 0.746 | 0.43 | 1.04E-116 | osteoclast.like | Ykt6 |
| Acadvl | 5.52E-121 | 1.23146535 | 0.654 | 0.257 | 1.07E-116 | osteoclast.like | Acadvl |
| Uba31 | 8.60E-121 | 1.32568542 | 0.634 | 0.235 | 1.66E-116 | osteoclast.like | Uba3 |
| Tmem250-ps4 | 1.34E-120 | 0.93810108 | 0.707 | 0.368 | 2.60E-116 | osteoclast.like | Tmem250-ps |
| Slc13a34 | 1.93E-120 | 1.39622676 | 0.59 | 0.168 | 3.73E-116 | osteoclast.like | Slc13a3 |
| Cast2 | 3.45E-120 | 0.77261166 | 0.829 | 0.496 | 6.67E-116 | osteoclast.like | Cast |
| Bckdk2 | 4.46E-120 | 1.51921026 | 0.688 | 0.251 | 8.62E-116 | osteoclast.like | Bckdk |
| Ndufa111 | 4.55E-120 | 1.05854835 | 0.698 | 0.311 | 8.79E-116 | osteoclast.like | Ndufa11 |
| Polr2e1 | 6.85E-120 | 0.92012399 | 0.829 | 0.445 | 1.32E-115 | osteoclast.like | Polr2e |
| Slc46a1 | 1.00E-119 | 2.45954575 | 0.42 | 0.068 | 1.94E-115 | osteoclast.like | Slc46a1 |
| Gnas1 | 1.27E-119 | 0.4485831 | 0.888 | 0.707 | 2.46E-115 | osteoclast.like | Gnas |
| Umps2 | 1.50E-119 | 1.37849002 | 0.571 | 0.204 | 2.90E-115 | osteoclast.like | Umps |
| Nono1 | 1.55E-119 | 0.59889585 | 0.844 | 0.571 | 2.99E-115 | osteoclast.like | Nono |
| Mybbp1a2 | 2.73E-119 | 1.04675552 | 0.683 | 0.373 | 5.28E-115 | osteoclast.like | Mybbp1a |
| Agpat51 | 3.29E-119 | 1.33938064 | 0.659 | 0.261 | 6.36E-115 | osteoclast.like | Agpat5 |
| Dot1l3 | 3.49E-119 | 0.79505506 | 0.8 | 0.443 | 6.75E-115 | osteoclast.like | Dot1l |
| Cmtm4 | 3.88E-119 | 1.85761996 | 0.4 | 0.104 | 7.50E-115 | osteoclast.like | Cmtm4 |
| Ndufs61 | 4.34E-119 | 1.30949828 | 0.615 | 0.23 | 8.39E-115 | osteoclast.like | Ndufs6 |
| Srp191 | 5.32E-119 | 1.11413816 | 0.688 | 0.326 | 1.03E-114 | osteoclast.like | Srp19 |
| Sinhcaf | 5.80E-119 | 2.513744 | 0.42 | 0.065 | 1.12E-114 | osteoclast.like | Sinhcaf |
| Ppa11 | 6.02E-119 | 1.70350974 | 0.507 | 0.172 | 1.16E-114 | osteoclast.like | Ppa1 |
| Mical3 | 2.36E-118 | 2.54975548 | 0.356 | 0.046 | 4.56E-114 | osteoclast.like | Mical3 |
| Zeb1 | 3.52E-118 | 2.46054478 | 0.327 | 0.049 | 6.80E-114 | osteoclast.like | Zeb1 |
| Pcgf51 | 3.82E-118 | 1.21216819 | 0.571 | 0.226 | 7.38E-114 | osteoclast.like | Pcgf5 |
| Abi33 | 9.96E-118 | 0.54303164 | 0.854 | 0.522 | 1.93E-113 | osteoclast.like | Abi3 |
| Epn12 | 1.49E-117 | 0.28214452 | 0.927 | 0.813 | 2.88E-113 | osteoclast.like | Epn1 |
| Glce | 1.75E-117 | 2.11249991 | 0.424 | 0.075 | 3.38E-113 | osteoclast.like | Glce |
| Gaa3 | 2.23E-117 | 0.75060514 | 0.751 | 0.456 | 4.30E-113 | osteoclast.like | Gaa |
| mt-Nd51 | 2.74E-117 | 0.46046997 | 0.795 | 0.585 | 5.30E-113 | osteoclast.like | mt-Nd5 |
| Tank2 | 3.04E-117 | 0.83350144 | 0.795 | 0.478 | 5.88E-113 | osteoclast.like | Tank |
| Ube2g13 | 3.28E-117 | 1.12057196 | 0.722 | 0.326 | 6.33E-113 | osteoclast.like | Ube2g1 |
| Elavl12 | 7.39E-117 | 0.67555333 | 0.795 | 0.538 | 1.43E-112 | osteoclast.like | Elavl1 |
| Endog | 9.73E-117 | 1.63869197 | 0.541 | 0.142 | 1.88E-112 | osteoclast.like | Endog |
| Ptcd31 | 1.43E-116 | 1.68302713 | 0.473 | 0.126 | 2.77E-112 | osteoclast.like | Ptcd3 |
| Sacm1l | 1.99E-116 | 1.21899519 | 0.712 | 0.326 | 3.85E-112 | osteoclast.like | Sacm1l |
| Eaf13 | 2.64E-116 | 1.08320204 | 0.8 | 0.383 | 5.11E-112 | osteoclast.like | Eaf1 |
| Csnk1a1 | 2.78E-116 | 0.84168351 | 0.824 | 0.508 | 5.37E-112 | osteoclast.like | Csnk1a1 |
| Nptn1 | 3.91E-116 | 0.47640765 | 0.878 | 0.662 | 7.57E-112 | osteoclast.like | Nptn |
| Hdlbp | 4.00E-116 | 0.36673515 | 0.878 | 0.666 | 7.73E-112 | osteoclast.like | Hdlbp |
| Tmem1472 | 4.69E-116 | 1.14633347 | 0.668 | 0.287 | 9.06E-112 | osteoclast.like | Tmem147 |
| Trmt10c | 1.43E-115 | 1.91508743 | 0.42 | 0.107 | 2.76E-111 | osteoclast.like | Trmt10c |
| Ppp2r3c1 | 1.45E-115 | 1.55103286 | 0.551 | 0.154 | 2.80E-111 | osteoclast.like | Ppp2r3c |
| Traf3ip2 | 1.50E-115 | 2.60977014 | 0.322 | 0.034 | 2.90E-111 | osteoclast.like | Traf3ip2 |
| Phldb12 | 1.88E-115 | 1.23535986 | 0.571 | 0.196 | 3.64E-111 | osteoclast.like | Phldb1 |
| Bag32 | 4.43E-115 | 0.93198228 | 0.717 | 0.379 | 8.56E-111 | osteoclast.like | Bag3 |
| Dnaja21 | 5.14E-115 | 0.54426943 | 0.829 | 0.589 | 9.94E-111 | osteoclast.like | Dnaja2 |

| | | | | | | | |
|---|---|---|---|---|---|---|---|
| Mcm61 | 5.68E-115 | 1.63386099 | 0.517 | 0.145 | 1.10E-110 | osteoclast.like | Mcm6 |
| Dnm1l1 | 5.75E-115 | 1.02745331 | 0.727 | 0.341 | 1.11E-110 | osteoclast.like | Dnm1l |
| Prrc12 | 6.19E-115 | 1.08377594 | 0.634 | 0.283 | 1.20E-110 | osteoclast.like | Prrc1 |
| Nol71 | 6.34E-115 | 1.06496766 | 0.717 | 0.323 | 1.23E-110 | osteoclast.like | Nol7 |
| Ciao31 | 1.36E-114 | 1.7649235 | 0.522 | 0.148 | 2.63E-110 | osteoclast.like | Ciao3 |
| Ndufa7 | 2.16E-114 | 0.87269968 | 0.766 | 0.444 | 4.18E-110 | osteoclast.like | Ndufa7 |
| Sac3d11 | 4.56E-114 | 1.87455495 | 0.449 | 0.126 | 8.81E-110 | osteoclast.like | Sac3d1 |
| Rab312 | 2.50E-113 | 0.54688051 | 0.868 | 0.59 | 4.83E-109 | osteoclast.like | Rab31 |
| Metap21 | 3.60E-113 | 0.78270187 | 0.761 | 0.479 | 6.96E-109 | osteoclast.like | Metap2 |
| Kars2 | 3.77E-113 | 0.9419904 | 0.751 | 0.392 | 7.28E-109 | osteoclast.like | Kars |
| Pi4k2a3 | 4.43E-113 | 0.79140696 | 0.776 | 0.428 | 8.57E-109 | osteoclast.like | Pi4k2a |
| Lonp11 | 6.99E-113 | 1.10679633 | 0.761 | 0.331 | 1.35E-108 | osteoclast.like | Lonp1 |
| Akr1b82 | 1.59E-112 | 1.4171428 | 0.571 | 0.132 | 3.07E-108 | osteoclast.like | Akr1b8 |
| Etfb2 | 1.71E-112 | 0.73859962 | 0.844 | 0.533 | 3.31E-108 | osteoclast.like | Etfb |
| Plpp1 | 1.73E-112 | 3.09607367 | 0.278 | 0.025 | 3.34E-108 | osteoclast.like | Plpp1 |
| Atxn21 | 1.96E-112 | 1.62580602 | 0.595 | 0.23 | 3.79E-108 | osteoclast.like | Atxn2 |
| Prep1 | 2.13E-112 | 1.12015813 | 0.663 | 0.327 | 4.11E-108 | osteoclast.like | Prep |
| Ctsf1 | 2.29E-112 | 2.13667894 | 0.41 | 0.108 | 4.42E-108 | osteoclast.like | Ctsf |
| C1d1 | 2.74E-112 | 1.29592341 | 0.605 | 0.234 | 5.30E-108 | osteoclast.like | C1d |
| Set2 | 2.95E-112 | 0.99951759 | 0.659 | 0.337 | 5.70E-108 | osteoclast.like | Set |
| Cd373 | 4.55E-112 | 0.94207194 | 0.766 | 0.325 | 8.79E-108 | osteoclast.like | Cd37 |
| Ppif | 4.79E-112 | 1.88543975 | 0.449 | 0.093 | 9.26E-108 | osteoclast.like | Ppif |
| Plod32 | 5.71E-112 | 0.44753472 | 0.868 | 0.629 | 1.10E-107 | osteoclast.like | Plod3 |
| Gnb1l | 6.34E-112 | 2.95602536 | 0.298 | 0.029 | 1.23E-107 | osteoclast.like | Gnb1l |
| Cfdp11 | 7.85E-112 | 1.03355816 | 0.732 | 0.348 | 1.52E-107 | osteoclast.like | Cfdp1 |
| Pa2g42 | 2.10E-111 | 1.36408332 | 0.615 | 0.251 | 4.05E-107 | osteoclast.like | Pa2g4 |
| Opa11 | 2.32E-111 | 1.41623736 | 0.62 | 0.244 | 4.48E-107 | osteoclast.like | Opa1 |
| Wdfy21 | 2.69E-111 | 0.70516307 | 0.78 | 0.442 | 5.21E-107 | osteoclast.like | Wdfy2 |
| Ahctf12 | 3.30E-111 | 1.34303808 | 0.654 | 0.254 | 6.37E-107 | osteoclast.like | Ahctf1 |
| Chpf | 3.51E-111 | 2.41324977 | 0.39 | 0.061 | 6.79E-107 | osteoclast.like | Chpf |
| Pcyt1a2 | 8.07E-111 | 0.74600513 | 0.746 | 0.411 | 1.56E-106 | osteoclast.like | Pcyt1a |
| Golga7 | 9.58E-111 | 0.6253731 | 0.815 | 0.535 | 1.85E-106 | osteoclast.like | Golga7 |
| Emc62 | 1.26E-110 | 1.17891166 | 0.698 | 0.31 | 2.43E-106 | osteoclast.like | Emc6 |
| Rnf1872 | 1.69E-110 | 0.49556983 | 0.893 | 0.643 | 3.26E-106 | osteoclast.like | Rnf187 |
| Nme21 | 1.95E-110 | 0.92769691 | 0.756 | 0.427 | 3.76E-106 | osteoclast.like | Nme2 |
| Noc2l2 | 2.73E-110 | 0.85360806 | 0.751 | 0.465 | 5.28E-106 | osteoclast.like | Noc2l |
| Gss2 | 3.89E-110 | 1.44373447 | 0.546 | 0.157 | 7.52E-106 | osteoclast.like | Gss |
| Tmem191 | 1.11E-109 | 1.10040734 | 0.693 | 0.308 | 2.15E-105 | osteoclast.like | Tmem19 |
| Mpdu11 | 1.16E-109 | 0.80910343 | 0.776 | 0.439 | 2.25E-105 | osteoclast.like | Mpdu1 |
| Nadk | 1.52E-109 | 0.46432561 | 0.82 | 0.579 | 2.93E-105 | osteoclast.like | Nadk |
| Fgd64 | 2.09E-109 | 1.43188214 | 0.585 | 0.188 | 4.04E-105 | osteoclast.like | Fgd6 |
| Eif5a3 | 2.30E-109 | 0.43975981 | 0.898 | 0.684 | 4.44E-105 | osteoclast.like | Eif5a |
| Ldhb2 | 3.07E-109 | 1.32072216 | 0.483 | 0.171 | 5.94E-105 | osteoclast.like | Ldhb |
| Lsm42 | 4.74E-109 | 0.82227976 | 0.771 | 0.434 | 9.17E-105 | osteoclast.like | Lsm4 |
| Akap1 | 6.65E-109 | 2.77616667 | 0.293 | 0.044 | 1.28E-104 | osteoclast.like | Akap1 |
| U2af21 | 6.86E-109 | 0.47339177 | 0.878 | 0.663 | 1.33E-104 | osteoclast.like | U2af2 |
| Kdm3a2 | 9.13E-109 | 0.92371113 | 0.595 | 0.324 | 1.77E-104 | osteoclast.like | Kdm3a |
| Mpp5 | 9.91E-109 | 1.59255735 | 0.502 | 0.146 | 1.92E-104 | osteoclast.like | Mpp5 |
| Lsm72 | 1.02E-108 | 1.05757793 | 0.639 | 0.312 | 1.97E-104 | osteoclast.like | Lsm7 |
| Syncrip2 | 1.51E-108 | 0.49763159 | 0.883 | 0.647 | 2.91E-104 | osteoclast.like | Syncrip |
| Abracl2 | 3.05E-108 | 0.61705479 | 0.766 | 0.509 | 5.90E-104 | osteoclast.like | Abracl |
| Kdelr22 | 3.18E-108 | 0.7995718 | 0.722 | 0.426 | 6.15E-104 | osteoclast.like | Kdelr2 |
| Copa | 4.09E-108 | 0.45389576 | 0.898 | 0.691 | 7.90E-104 | osteoclast.like | Copa |
| Cyfip11 | 4.31E-108 | 0.34436349 | 0.834 | 0.673 | 8.34E-104 | osteoclast.like | Cyfip1 |
| Aup11 | 4.48E-108 | 0.44059074 | 0.873 | 0.663 | 8.66E-104 | osteoclast.like | Aup1 |
| Tmem1351 | 5.61E-108 | 0.95477166 | 0.654 | 0.331 | 1.08E-103 | osteoclast.like | Tmem135 |
| Slc11a22 | 1.45E-107 | 0.98147312 | 0.634 | 0.315 | 2.80E-103 | osteoclast.like | Slc11a2 |
| Sucla21 | 1.72E-107 | 1.32001386 | 0.624 | 0.252 | 3.33E-103 | osteoclast.like | Sucla2 |
| Emc42 | 1.92E-107 | 1.10129866 | 0.629 | 0.266 | 3.72E-103 | osteoclast.like | Emc4 |
| Sh3gl13 | 1.94E-107 | 0.7669083 | 0.829 | 0.438 | 3.75E-103 | osteoclast.like | Sh3gl1 |
| Bmf1 | 3.42E-107 | 1.30429146 | 0.571 | 0.21 | 6.62E-103 | osteoclast.like | Bmf |
| Sap301 | 3.50E-107 | 0.87473402 | 0.668 | 0.348 | 6.76E-103 | osteoclast.like | Sap30 |
| Exoc6b1 | 4.12E-107 | 1.53891151 | 0.58 | 0.187 | 7.97E-103 | osteoclast.like | Exoc6b |
| Eif3k3 | 6.37E-107 | 0.42468329 | 0.898 | 0.728 | 1.23E-102 | osteoclast.like | Eif3k |
| Cpt2 | 8.96E-107 | 1.95704744 | 0.459 | 0.102 | 1.73E-102 | osteoclast.like | Cpt2 |
| Qdpr4 | 1.05E-106 | 0.92831909 | 0.829 | 0.444 | 2.03E-102 | osteoclast.like | Qdpr |
| Pja1 | 1.55E-106 | 1.51600389 | 0.561 | 0.167 | 3.01E-102 | osteoclast.like | Pja1 |
| Tsfm1 | 1.62E-106 | 1.80288013 | 0.532 | 0.148 | 3.14E-102 | osteoclast.like | Tsfm |
| Trmt61a | 2.17E-106 | 2.26719262 | 0.41 | 0.077 | 4.20E-102 | osteoclast.like | Trmt61a |
| Fcf12 | 3.37E-106 | 0.99777775 | 0.698 | 0.298 | 6.52E-102 | osteoclast.like | Fcf1 |
| Stxbp31 | 5.29E-106 | 1.03092548 | 0.712 | 0.28 | 1.02E-101 | osteoclast.like | Stxbp3 |
| Prpf81 | 3.19E-105 | 0.65845901 | 0.751 | 0.486 | 6.17E-101 | osteoclast.like | Prpf8 |
| Cacybp2 | 1.26E-104 | 1.31152344 | 0.634 | 0.288 | 2.44E-100 | osteoclast.like | Cacybp |
| Bag11 | 1.43E-104 | 0.69993973 | 0.849 | 0.566 | 2.76E-100 | osteoclast.like | Bag1 |
| Ahsa11 | 2.06E-104 | 0.64101798 | 0.844 | 0.576 | 3.98E-100 | osteoclast.like | Ahsa1 |
| Klhl93 | 2.66E-104 | 0.55411588 | 0.824 | 0.523 | 5.13E-100 | osteoclast.like | Klhl9 |
| Cbr4 | 3.30E-104 | 1.97255235 | 0.454 | 0.093 | 6.39E-100 | osteoclast.like | Cbr4 |
| Aifm23 | 3.93E-104 | 1.37414743 | 0.561 | 0.182 | 7.59E-100 | osteoclast.like | Aifm2 |
| Afg3l22 | 5.31E-104 | 0.82229445 | 0.766 | 0.427 | 1.03E-99 | osteoclast.like | Afg3l2 |
| Eif4g11 | 1.49E-103 | 0.9312924 | 0.761 | 0.399 | 2.89E-99 | osteoclast.like | Eif4g1 |
| Galnt23 | 2.05E-103 | 0.76106417 | 0.863 | 0.513 | 3.97E-99 | osteoclast.like | Galnt2 |
| Prkcsh2 | 2.27E-103 | 0.76640894 | 0.795 | 0.448 | 4.39E-99 | osteoclast.like | Prkcsh |
| Kif13a | 2.70E-103 | 0.97131733 | 0.659 | 0.311 | 5.22E-99 | osteoclast.like | Kif13a |
| Aco12 | 2.87E-103 | 0.86170874 | 0.712 | 0.357 | 5.55E-99 | osteoclast.like | Aco1 |
| Tm9sf3 | 3.17E-103 | 0.2934872 | 0.868 | 0.712 | 6.13E-99 | osteoclast.like | Tm9sf3 |
| Tbkbp11 | 3.42E-103 | 1.09260895 | 0.663 | 0.27 | 6.61E-99 | osteoclast.like | Tbkbp1 |
| Psmd22 | 5.27E-103 | 0.72379527 | 0.815 | 0.463 | 1.02E-98 | osteoclast.like | Psmd2 |
| Hacd1 | 7.52E-103 | 2.45415931 | 0.332 | 0.048 | 1.45E-98 | osteoclast.like | Hacd1 |
| Mtap1 | 1.16E-102 | 1.31461297 | 0.551 | 0.211 | 2.25E-98 | osteoclast.like | Mtap |
| Cct42 | 1.29E-102 | 0.92614746 | 0.707 | 0.372 | 2.50E-98 | osteoclast.like | Cct4 |
| Cnpy23 | 1.57E-102 | 0.8645682 | 0.732 | 0.383 | 3.04E-98 | osteoclast.like | Cnpy2 |
| Mlec1 | 3.86E-102 | 0.59052709 | 0.844 | 0.569 | 7.45E-98 | osteoclast.like | Mlec |
| Bcap31 | 5.82E-102 | 0.66594876 | 0.839 | 0.531 | 1.13E-97 | osteoclast.like | Bcap31 |
| Drg12 | 7.16E-102 | 0.96530565 | 0.776 | 0.405 | 1.38E-97 | osteoclast.like | Drg1 |
| Gabarapl2 | 1.21E-101 | 0.74527598 | 0.712 | 0.393 | 2.34E-97 | osteoclast.like | Gabarapl2 |
| Ppm1h2 | 1.29E-101 | 0.94439162 | 0.6 | 0.284 | 2.49E-97 | osteoclast.like | Ppm1h |
| Srm2 | 2.47E-101 | 1.49786932 | 0.478 | 0.176 | 4.78E-97 | osteoclast.like | Srm |
| Actr34 | 2.72E-101 | 0.32000834 | 0.824 | 0.636 | 5.26E-97 | osteoclast.like | Actr3 |
| Hbp12 | 3.27E-101 | 0.71694768 | 0.766 | 0.46 | 6.32E-97 | osteoclast.like | Hbp1 |
| Spry2 | 4.15E-101 | 2.09968932 | 0.385 | 0.059 | 8.03E-97 | osteoclast.like | Spry2 |
| Noa1 | 6.60E-101 | 2.18217208 | 0.41 | 0.07 | 1.28E-96 | osteoclast.like | Noa1 |
| Klhl213 | 7.19E-101 | 0.96882797 | 0.654 | 0.285 | 1.39E-96 | osteoclast.like | Klhl21 |
| Rbbp71 | 8.66E-101 | 0.8928867 | 0.766 | 0.384 | 1.67E-96 | osteoclast.like | Rbbp7 |
| Bcl2l131 | 9.30E-101 | 1.29612817 | 0.551 | 0.181 | 1.80E-96 | osteoclast.like | Bcl2l13 |
| Bnip2 | 1.01E-100 | 0.58792707 | 0.844 | 0.563 | 1.96E-96 | osteoclast.like | Bnip2 |
| Itpr3 | 9.78E-100 | 2.17409142 | 0.395 | 0.069 | 1.89E-95 | osteoclast.like | Itpr3 |
| Galnt63 | 1.36E-99 | 0.9377669 | 0.624 | 0.337 | 2.63E-95 | osteoclast.like | Galnt6 |
| Frmd4a2 | 1.38E-99 | 0.80827507 | 0.693 | 0.36 | 2.67E-95 | osteoclast.like | Frmd4a |
| Acot133 | 1.57E-99 | 1.21931117 | 0.561 | 0.234 | 3.05E-95 | osteoclast.like | Acot13 |
| Ifngr22 | 1.73E-99 | 0.41000049 | 0.824 | 0.601 | 3.35E-95 | osteoclast.like | Ifngr2 |
| Rae11 | 2.12E-99 | 1.16963033 | 0.659 | 0.266 | 4.10E-95 | osteoclast.like | Rae1 |
| Emc21 | 3.00E-99 | 0.98425869 | 0.707 | 0.359 | 5.80E-95 | osteoclast.like | Emc2 |
| Golm12 | 3.27E-99 | 1.23471693 | 0.541 | 0.223 | 6.32E-95 | osteoclast.like | Golm1 |
| Dnph1 | 4.26E-99 | 3.08449346 | 0.263 | 0.031 | 8.23E-95 | osteoclast.like | Dnph1 |
| Ndufaf4 | 7.83E-99 | 1.8017472 | 0.41 | 0.092 | 1.51E-94 | osteoclast.like | Ndufaf4 |
| Cops61 | 8.44E-99 | 0.62453501 | 0.8 | 0.519 | 1.63E-94 | osteoclast.like | Cops6 |
| Rnf142 | 1.00E-98 | 1.32270382 | 0.576 | 0.202 | 1.94E-94 | osteoclast.like | Rnf14 |
| Elp31 | 1.71E-98 | 1.75648168 | 0.502 | 0.118 | 3.31E-94 | osteoclast.like | Elp3 |
| Zfp2633 | 3.07E-98 | 0.90469705 | 0.624 | 0.289 | 5.94E-94 | osteoclast.like | Zfp263 |
| Macf12 | 4.18E-98 | 0.42163576 | 0.824 | 0.595 | 8.07E-94 | osteoclast.like | Macf1 |
| Wdr821 | 5.65E-98 | 0.87345238 | 0.751 | 0.405 | 1.09E-93 | osteoclast.like | Wdr82 |
| Sike11 | 5.81E-98 | 0.97619599 | 0.629 | 0.281 | 1.12E-93 | osteoclast.like | Sike1 |
| Tomm61 | 7.26E-98 | 0.63683481 | 0.771 | 0.481 | 1.40E-93 | osteoclast.like | Tomm6 |
| Nifk1 | 7.63E-98 | 1.47393564 | 0.517 | 0.188 | 1.48E-93 | osteoclast.like | Nifk |
| Naa381 | 1.72E-97 | 0.95076288 | 0.732 | 0.343 | 3.34E-93 | osteoclast.like | Naa38 |
| Pofut22 | 2.81E-97 | 1.07744193 | 0.693 | 0.297 | 5.44E-93 | osteoclast.like | Pofut2 |
| Prkag21 | 3.40E-97 | 1.2171275 | 0.532 | 0.182 | 6.57E-93 | osteoclast.like | Prkag2 |
| Ndufaf5 | 4.08E-97 | 2.21744188 | 0.4 | 0.067 | 7.88E-93 | osteoclast.like | Ndufaf5 |
| Agk | 4.46E-97 | 1.99347967 | 0.41 | 0.069 | 8.62E-93 | osteoclast.like | Agk |
| Inppl15 | 4.54E-97 | 0.82590983 | 0.741 | 0.407 | 8.78E-93 | osteoclast.like | Inppl1 |
| Slc25a43 | 5.42E-97 | 2.60530229 | 0.307 | 0.035 | 1.05E-92 | osteoclast.like | Slc25a43 |
| Mydgf1 | 6.02E-97 | 0.67047566 | 0.849 | 0.534 | 1.16E-92 | osteoclast.like | Mydgf |
| Insl6 | 9.41E-97 | 2.43311378 | 0.312 | 0.051 | 1.82E-92 | osteoclast.like | Insl6 |
| Smim121 | 1.16E-96 | 1.23903478 | 0.6 | 0.202 | 2.25E-92 | osteoclast.like | Smim12 |
| Lman21 | 2.79E-96 | 0.43899475 | 0.868 | 0.652 | 5.39E-92 | osteoclast.like | Lman2 |
| N6amt1 | 5.31E-96 | 2.09775212 | 0.41 | 0.082 | 1.03E-91 | osteoclast.like | N6amt1 |
| Ostm1 | 9.31E-96 | 0.92310771 | 0.673 | 0.38 | 1.80E-91 | osteoclast.like | Ostm1 |
| Cpne23 | 1.50E-95 | 1.1520255 | 0.634 | 0.269 | 2.89E-91 | osteoclast.like | Cpne2 |
| Rere1 | 2.22E-95 | 1.12878944 | 0.683 | 0.319 | 4.28E-91 | osteoclast.like | Rere |
| Abcb71 | 2.57E-95 | 1.31984682 | 0.566 | 0.206 | 4.96E-91 | osteoclast.like | Abcb7 |
| Nyap2 | 3.97E-95 | 10.5170017 | 0.263 | 0 | 7.67E-91 | osteoclast.like | Nyap2 |
| Rtn31 | 4.21E-95 | 0.35815 | 0.863 | 0.679 | 8.13E-91 | osteoclast.like | Rtn3 |
| Rnps11 | 4.31E-95 | 0.66779351 | 0.805 | 0.471 | 8.34E-91 | osteoclast.like | Rnps1 |
| Kmt5c1 | 5.01E-95 | 0.84668589 | 0.717 | 0.345 | 9.69E-91 | osteoclast.like | Kmt5c |
| Psmd122 | 6.72E-95 | 0.84251843 | 0.727 | 0.394 | 1.30E-90 | osteoclast.like | Psmd12 |
| Twf23 | 6.89E-95 | 0.60030584 | 0.795 | 0.508 | 1.33E-90 | osteoclast.like | Twf2 |
| Faim | 7.31E-95 | 0.83497784 | 0.668 | 0.33 | 1.41E-90 | osteoclast.like | Faim |
| Mff | 1.39E-94 | 1.05837175 | 0.639 | 0.287 | 2.69E-90 | osteoclast.like | Mff |
| Slc35e41 | 1.40E-94 | 1.32472911 | 0.449 | 0.143 | 2.70E-90 | osteoclast.like | Slc35e4 |
| Rhno11 | 1.79E-94 | 2.18457228 | 0.424 | 0.085 | 3.45E-90 | osteoclast.like | Rhno1 |
| Spcs1 | 3.57E-94 | 0.48101938 | 0.82 | 0.615 | 6.91E-90 | osteoclast.like | Spcs1 |
| Ddost1 | 4.80E-94 | 0.4680017 | 0.878 | 0.669 | 9.29E-90 | osteoclast.like | Ddost |
| Smarca41 | 5.95E-94 | 0.47393083 | 0.834 | 0.587 | 1.15E-89 | osteoclast.like | Smarca4 |
| Ap2s13 | 7.45E-94 | 0.36590902 | 0.873 | 0.647 | 1.44E-89 | osteoclast.like | Ap2s1 |
| Mbtps12 | 7.47E-94 | 1.15311732 | 0.668 | 0.26 | 1.44E-89 | osteoclast.like | Mbtps1 |
| Tmem60 | 9.20E-94 | 1.89329148 | 0.42 | 0.093 | 1.78E-89 | osteoclast.like | Tmem60 |
| Sfxn12 | 1.17E-93 | 1.13942275 | 0.6 | 0.268 | 2.26E-89 | osteoclast.like | Sfxn1 |
| Drap11 | 1.35E-93 | 0.4755698 | 0.873 | 0.658 | 2.60E-89 | osteoclast.like | Drap1 |
| Slc20a2 | 3.40E-93 | 2.00175598 | 0.42 | 0.087 | 6.56E-89 | osteoclast.like | Slc20a2 |
| Golph3 | 3.89E-93 | 0.56466643 | 0.82 | 0.558 | 7.52E-89 | osteoclast.like | Golph3 |
| Smyd21 | 8.04E-93 | 1.6216467 | 0.454 | 0.123 | 1.56E-88 | osteoclast.like | Smyd2 |
| Cul31 | 8.41E-93 | 0.45533083 | 0.824 | 0.589 | 1.63E-88 | osteoclast.like | Cul3 |
| Prex12 | 8.99E-93 | 0.27979865 | 0.927 | 0.759 | 1.74E-88 | osteoclast.like | Prex1 |
| Atxn31 | 1.93E-92 | 1.07032898 | 0.634 | 0.269 | 3.73E-88 | osteoclast.like | Atxn3 |
| Lrwd11 | 1.94E-92 | 1.49646326 | 0.62 | 0.213 | 3.75E-88 | osteoclast.like | Lrwd1 |
| Msantd3 | 6.17E-92 | 2.37232824 | 0.302 | 0.034 | 1.19E-87 | osteoclast.like | Msantd3 |
| Sf3a11 | 6.71E-92 | 1.06794723 | 0.707 | 0.333 | 1.30E-87 | osteoclast.like | Sf3a1 |
| Plekhm12 | 7.81E-92 | 1.14633122 | 0.566 | 0.216 | 1.51E-87 | osteoclast.like | Plekhm1 |
| Ift43 | 2.24E-91 | 2.36405209 | 0.439 | 0.093 | 4.32E-87 | osteoclast.like | Ift43 |
| Eif61 | 2.56E-91 | 0.95686213 | 0.722 | 0.371 | 4.94E-87 | osteoclast.like | Eif6 |
| Dap31 | 2.91E-91 | 1.06809873 | 0.659 | 0.297 | 5.63E-87 | osteoclast.like | Dap3 |
| Phtf2 | 3.83E-91 | 1.80528227 | 0.424 | 0.093 | 7.40E-87 | osteoclast.like | Phtf2 |
| Pkn1 | 4.28E-91 | 0.26588172 | 0.946 | 0.751 | 8.27E-87 | osteoclast.like | Pkn1 |
| Timm221 | 1.46E-90 | 1.28630045 | 0.561 | 0.209 | 2.83E-86 | osteoclast.like | Timm22 |
| Eif3h2 | 3.16E-90 | 0.81987231 | 0.722 | 0.379 | 6.11E-86 | osteoclast.like | Eif3h |
| Cct82 | 4.31E-90 | 0.82364539 | 0.659 | 0.357 | 8.34E-86 | osteoclast.like | Cct8 |
| Stt3b1 | 8.00E-90 | 0.60346693 | 0.81 | 0.526 | 1.55E-85 | osteoclast.like | Stt3b |
| Sgsh2 | 1.29E-89 | 1.30425333 | 0.6 | 0.18 | 2.50E-85 | osteoclast.like | Sgsh |
| Spcs32 | 1.33E-89 | 0.80952823 | 0.766 | 0.428 | 2.57E-85 | osteoclast.like | Spcs3 |
| St3gal2 | 1.85E-89 | 2.10821261 | 0.327 | 0.056 | 3.58E-85 | osteoclast.like | St3gal2 |
| Fnip23 | 2.18E-89 | 0.50992452 | 0.815 | 0.503 | 4.21E-85 | osteoclast.like | Fnip2 |
| Smc1a2 | 2.38E-89 | 1.0359251 | 0.702 | 0.32 | 4.61E-85 | osteoclast.like | Smc1a |
| Zswim41 | 2.55E-89 | 0.62458022 | 0.741 | 0.402 | 4.93E-85 | osteoclast.like | Zswim4 |
| Prr15 | 2.63E-89 | 1.79055282 | 0.332 | 0.091 | 5.08E-85 | osteoclast.like | Prr15 |
| Mpnd1 | 3.01E-89 | 1.10569106 | 0.702 | 0.296 | 5.83E-85 | osteoclast.like | Mpnd |
| Adh51 | 3.87E-89 | 0.70640933 | 0.737 | 0.442 | 7.48E-85 | osteoclast.like | Adh5 |
| Ubtf1 | 4.41E-89 | 0.6831806 | 0.698 | 0.39 | 8.53E-85 | osteoclast.like | Ubtf |
| Ube2q1 | 4.42E-89 | 0.50063766 | 0.81 | 0.567 | 8.55E-85 | osteoclast.like | Ube2q1 |
| Gtf2e21 | 7.94E-89 | 1.13435446 | 0.644 | 0.247 | 1.54E-84 | osteoclast.like | Gtf2e2 |
| Tars21 | 9.12E-89 | 1.5219276 | 0.502 | 0.134 | 1.76E-84 | osteoclast.like | Tars2 |
| Hnrnpl2 | 9.92E-89 | 0.38650802 | 0.8 | 0.626 | 1.92E-84 | osteoclast.like | Hnrnpl |
| Cryzl2 | 1.15E-88 | 1.94646991 | 0.366 | 0.076 | 2.23E-84 | osteoclast.like | Cryzl2 |
| Plxna13 | 1.29E-88 | 0.61396494 | 0.805 | 0.497 | 2.49E-84 | osteoclast.like | Plxna1 |
| Ubl7 | 1.40E-88 | 0.90961827 | 0.707 | 0.344 | 2.71E-84 | osteoclast.like | Ubl7 |
| Sec131 | 1.65E-88 | 0.588624 | 0.712 | 0.446 | 3.20E-84 | osteoclast.like | Sec13 |
| Mbd31 | 1.69E-88 | 0.59929337 | 0.834 | 0.555 | 3.27E-84 | osteoclast.like | Mbd3 |
| Mmachc | 1.88E-88 | 2.19350693 | 0.371 | 0.06 | 3.63E-84 | osteoclast.like | Mmachc |
| Dglucy | 1.95E-88 | 1.26612168 | 0.488 | 0.177 | 3.76E-84 | osteoclast.like | Dglucy |
| Cnih11 | 2.01E-88 | 0.98928792 | 0.688 | 0.337 | 3.89E-84 | osteoclast.like | Cnih1 |
| Ndufa91 | 2.01E-88 | 1.70970206 | 0.498 | 0.146 | 3.89E-84 | osteoclast.like | Ndufa9 |
| Ipo42 | 2.42E-88 | 1.08449388 | 0.629 | 0.273 | 4.67E-84 | osteoclast.like | Ipo4 |
| Dnajc111 | 2.65E-88 | 1.26679579 | 0.576 | 0.195 | 5.13E-84 | osteoclast.like | Dnajc11 |
| Zpr11 | 2.70E-88 | 1.03273464 | 0.605 | 0.247 | 5.22E-84 | osteoclast.like | Zpr1 |
| Rab71 | 4.98E-88 | 0.32154747 | 0.868 | 0.692 | 9.63E-84 | osteoclast.like | Rab7 |
| Fdx22 | 5.83E-88 | 1.20789991 | 0.639 | 0.274 | 1.13E-83 | osteoclast.like | Fdx2 |
| Ndufaf1 | 5.97E-88 | 1.96484756 | 0.351 | 0.067 | 1.16E-83 | osteoclast.like | Ndufaf1 |
| Tomm40l2 | 9.79E-88 | 1.13411035 | 0.668 | 0.264 | 1.89E-83 | osteoclast.like | Tomm40l |
| Tmem127 | 1.11E-87 | 0.83298168 | 0.683 | 0.376 | 2.14E-83 | osteoclast.like | Tmem127 |
| Rdx1 | 1.86E-87 | 1.14930196 | 0.673 | 0.289 | 3.59E-83 | osteoclast.like | Rdx |
| Psme32 | 2.39E-87 | 0.67469681 | 0.815 | 0.523 | 4.62E-83 | osteoclast.like | Psme3 |
| Atic2 | 2.96E-87 | 0.91941191 | 0.654 | 0.36 | 5.72E-83 | osteoclast.like | Atic |
| Abca2 | 3.28E-87 | 1.38481213 | 0.468 | 0.148 | 6.34E-83 | osteoclast.like | Abca2 |
| Nr1d1 | 3.53E-87 | 1.99862541 | 0.351 | 0.053 | 6.82E-83 | osteoclast.like | Nr1d1 |
| Impdh22 | 6.34E-87 | 0.66745606 | 0.673 | 0.403 | 1.23E-82 | osteoclast.like | Impdh2 |
| Nars1 | 7.04E-87 | 0.90751675 | 0.668 | 0.361 | 1.36E-82 | osteoclast.like | Nars |
| Mgat4b2 | 7.61E-87 | 0.87370849 | 0.629 | 0.316 | 1.47E-82 | osteoclast.like | Mgat4b |
| Iscu1 | 1.47E-86 | 1.13753355 | 0.624 | 0.26 | 2.84E-82 | osteoclast.like | Iscu |
| Nadk2 | 1.79E-86 | 1.8068313 | 0.449 | 0.108 | 3.46E-82 | osteoclast.like | Nadk2 |
| Tmem258 | 2.97E-86 | 0.59972246 | 0.805 | 0.499 | 5.73E-82 | osteoclast.like | Tmem258 |
| Polr2a1 | 3.06E-86 | 0.45647797 | 0.79 | 0.551 | 5.91E-82 | osteoclast.like | Polr2a |
| Puf601 | 3.32E-86 | 0.44479631 | 0.859 | 0.601 | 6.41E-82 | osteoclast.like | Puf60 |
| Arhgef11 | 3.72E-86 | 0.30677899 | 0.863 | 0.654 | 7.19E-82 | osteoclast.like | Arhgef1 |
| Ptgfrn3 | 5.09E-86 | 0.76947091 | 0.654 | 0.364 | 9.85E-82 | osteoclast.like | Ptgfrn |
| Cers23 | 5.22E-86 | 0.50114921 | 0.824 | 0.588 | 1.01E-81 | osteoclast.like | Cers2 |
| Ankrd403 | 7.02E-86 | 0.80985894 | 0.8 | 0.453 | 1.36E-81 | osteoclast.like | Ankrd40 |
| Eef1d2 | 1.25E-85 | 1.04377771 | 0.649 | 0.301 | 2.42E-81 | osteoclast.like | Eef1d |
| Abcc53 | 1.43E-85 | 0.96493201 | 0.683 | 0.278 | 2.77E-81 | osteoclast.like | Abcc5 |
| Lman11 | 2.28E-85 | 0.83206307 | 0.756 | 0.407 | 4.42E-81 | osteoclast.like | Lman1 |
| Aarsd11 | 2.70E-85 | 1.39151265 | 0.576 | 0.199 | 5.21E-81 | osteoclast.like | Aarsd1 |
| Lrrc61 | 3.72E-85 | 1.70156117 | 0.405 | 0.106 | 7.20E-81 | osteoclast.like | Lrrc61 |
| Usp52 | 4.13E-85 | 0.78836195 | 0.702 | 0.374 | 7.99E-81 | osteoclast.like | Usp5 |
| Nsd1 | 4.24E-85 | 0.47902951 | 0.8 | 0.579 | 8.20E-81 | osteoclast.like | Nsd1 |
| Ubr51 | 7.53E-85 | 0.73522283 | 0.727 | 0.421 | 1.46E-80 | osteoclast.like | Ubr5 |
| Hsp90b12 | 8.87E-85 | 0.32734605 | 0.844 | 0.633 | 1.71E-80 | osteoclast.like | Hsp90b1 |
| Hdac21 | 9.99E-85 | 1.47268738 | 0.512 | 0.162 | 1.93E-80 | osteoclast.like | Hdac2 |
| Mad1l13 | 1.88E-84 | 1.31244186 | 0.498 | 0.198 | 3.63E-80 | osteoclast.like | Mad1l1 |
| Abhd111 | 2.54E-84 | 1.30354013 | 0.576 | 0.194 | 4.90E-80 | osteoclast.like | Abhd11 |
| Fam210a | 2.62E-84 | 1.40619743 | 0.488 | 0.149 | 5.06E-80 | osteoclast.like | Fam210a |
| Fdx1 | 2.88E-84 | 1.93399795 | 0.405 | 0.089 | 5.56E-80 | osteoclast.like | Fdx1 |
| Gdpd5 | 2.96E-84 | 1.97219753 | 0.385 | 0.068 | 5.73E-80 | osteoclast.like | Gdpd5 |
| Clptm11 | 3.41E-84 | 0.38943966 | 0.839 | 0.624 | 6.58E-80 | osteoclast.like | Clptm1 |
| Rab3gap22 | 4.82E-84 | 0.91763837 | 0.659 | 0.312 | 9.32E-80 | osteoclast.like | Rab3gap2 |
| Shmt21 | 5.19E-84 | 1.0941504 | 0.502 | 0.252 | 1.00E-79 | osteoclast.like | Shmt2 |
| Hars1 | 5.45E-84 | 0.96612656 | 0.673 | 0.326 | 1.05E-79 | osteoclast.like | Hars |
| Stxbp52 | 6.13E-84 | 1.45831239 | 0.512 | 0.158 | 1.19E-79 | osteoclast.like | Stxbp5 |
| Pomp3 | 6.28E-84 | 0.46489475 | 0.815 | 0.565 | 1.21E-79 | osteoclast.like | Pomp |
| Srp721 | 6.40E-84 | 0.63049384 | 0.702 | 0.441 | 1.24E-79 | osteoclast.like | Srp72 |
| H6pd3 | 1.43E-83 | 0.75858886 | 0.668 | 0.35 | 2.77E-79 | osteoclast.like | H6pd |
| Cct72 | 1.70E-83 | 0.81937592 | 0.668 | 0.36 | 3.29E-79 | osteoclast.like | Cct7 |
| Slc25a13 | 2.19E-83 | 2.15624861 | 0.41 | 0.081 | 4.23E-79 | osteoclast.like | Slc25a13 |
| Pabpn1 | 2.25E-83 | 0.43714771 | 0.82 | 0.611 | 4.35E-79 | osteoclast.like | Pabpn1 |
| Pepd1 | 2.46E-83 | 0.53074114 | 0.78 | 0.48 | 4.75E-79 | osteoclast.like | Pepd |
| Grwd11 | 3.51E-83 | 1.65721728 | 0.41 | 0.117 | 6.78E-79 | osteoclast.like | Grwd1 |
| Npc14 | 3.76E-83 | 0.36327269 | 0.815 | 0.611 | 7.28E-79 | osteoclast.like | Npc1 |
| Cpsf41 | 3.86E-83 | 1.03268804 | 0.595 | 0.24 | 7.47E-79 | osteoclast.like | Cpsf4 |
| Ssb2 | 7.40E-83 | 0.47288499 | 0.78 | 0.566 | 1.43E-78 | osteoclast.like | Ssb |
| Eif2b51 | 9.78E-83 | 0.88778868 | 0.624 | 0.315 | 1.89E-78 | osteoclast.like | Eif2b5 |
| Pigt2 | 1.14E-82 | 0.63013094 | 0.732 | 0.45 | 2.21E-78 | osteoclast.like | Pigt |
| Amacr | 1.80E-82 | 3.04896743 | 0.263 | 0.029 | 3.49E-78 | osteoclast.like | Amacr |
| Copb2 | 2.58E-82 | 0.62791843 | 0.732 | 0.455 | 4.99E-78 | osteoclast.like | Copb2 |
| Usp141 | 3.69E-82 | 1.06487621 | 0.624 | 0.285 | 7.14E-78 | osteoclast.like | Usp14 |
| Bclaf11 | 5.06E-82 | 0.68019337 | 0.8 | 0.474 | 9.79E-78 | osteoclast.like | Bclaf1 |
| Dynll13 | 5.36E-82 | 0.57561362 | 0.746 | 0.488 | 1.04E-77 | osteoclast.like | Dynll1 |
| Scp2 | 5.44E-82 | 0.78124523 | 0.707 | 0.371 | 1.05E-77 | osteoclast.like | Scp2 |
| Lactb3 | 7.39E-82 | 0.91677092 | 0.629 | 0.289 | 1.43E-77 | osteoclast.like | Lactb |
| Farsb1 | 8.77E-82 | 1.12157042 | 0.473 | 0.204 | 1.70E-77 | osteoclast.like | Farsb |
| Pqlc31 | 9.77E-82 | 1.11097525 | 0.488 | 0.199 | 1.89E-77 | osteoclast.like | Pqlc3 |
| Mesd3 | 1.21E-81 | 1.19423376 | 0.615 | 0.262 | 2.34E-77 | osteoclast.like | Mesd |
| Pdk12 | 1.77E-81 | 0.80085162 | 0.58 | 0.311 | 3.42E-77 | osteoclast.like | Pdk1 |
| Mtfr1 | 2.23E-81 | 1.41029309 | 0.498 | 0.171 | 4.31E-77 | osteoclast.like | Mtfr1 |
| Guf1 | 2.35E-81 | 2.25475815 | 0.346 | 0.07 | 4.55E-77 | osteoclast.like | Guf1 |
| Xxylt11 | 2.61E-81 | 1.40526945 | 0.502 | 0.162 | 5.06E-77 | osteoclast.like | Xxylt1 |
| Nudt8 | 2.85E-81 | 1.72440233 | 0.488 | 0.136 | 5.52E-77 | osteoclast.like | Nudt8 |
| Rpp14 | 2.99E-81 | 1.87992221 | 0.38 | 0.1 | 5.77E-77 | osteoclast.like | Rpp14 |
| Rgs192 | 3.72E-81 | 0.53805516 | 0.785 | 0.494 | 7.19E-77 | osteoclast.like | Rgs19 |
| Pgam51 | 4.27E-81 | 1.33846451 | 0.566 | 0.204 | 8.26E-77 | osteoclast.like | Pgam5 |
| Nrbf22 | 5.84E-81 | 0.99561879 | 0.615 | 0.258 | 1.13E-76 | osteoclast.like | Nrbf2 |
| Cope2 | 8.28E-81 | 0.46742505 | 0.82 | 0.601 | 1.60E-76 | osteoclast.like | Cope |
| Mbnl21 | 8.55E-81 | 0.39747575 | 0.859 | 0.632 | 1.65E-76 | osteoclast.like | Mbnl2 |
| Timm8a11 | 8.69E-81 | 1.77088119 | 0.454 | 0.117 | 1.68E-76 | osteoclast.like | Timm8a1 |
| Ssr31 | 9.66E-81 | 0.55540931 | 0.751 | 0.497 | 1.87E-76 | osteoclast.like | Ssr3 |
| S100a15 | 2.01E-80 | 0.69392582 | 0.717 | 0.449 | 3.89E-76 | osteoclast.like | S100a1 |
| Tent21 | 2.37E-80 | 0.86649484 | 0.722 | 0.361 | 4.58E-76 | osteoclast.like | Tent2 |
| Wdr771 | 4.01E-80 | 1.30771399 | 0.532 | 0.187 | 7.74E-76 | osteoclast.like | Wdr77 |
| Eif3i2 | 4.35E-80 | 0.90707893 | 0.717 | 0.381 | 8.41E-76 | osteoclast.like | Eif3i |
| Agpat1 | 4.53E-80 | 0.97764041 | 0.62 | 0.299 | 8.75E-76 | osteoclast.like | Agpat1 |
| Zfp451 | 4.86E-80 | 1.06212027 | 0.566 | 0.255 | 9.39E-76 | osteoclast.like | Zfp451 |
| Acad10 | 8.50E-80 | 2.75828678 | 0.278 | 0.031 | 1.64E-75 | osteoclast.like | Acad10 |
| Serhl1 | 8.89E-80 | 1.4277499 | 0.42 | 0.129 | 1.72E-75 | osteoclast.like | Serhl |
| Cmc21 | 9.16E-80 | 1.6666676 | 0.488 | 0.124 | 1.77E-75 | osteoclast.like | Cmc2 |
| Ctbp11 | 1.01E-79 | 0.42036 | 0.849 | 0.602 | 1.94E-75 | osteoclast.like | Ctbp1 |
| Pcbp21 | 1.01E-79 | 0.5943103 | 0.776 | 0.527 | 1.95E-75 | osteoclast.like | Pcbp2 |
| Chpt1 | 1.04E-79 | 1.08591557 | 0.498 | 0.194 | 2.02E-75 | osteoclast.like | Chpt1 |
| Sgsm3 | 1.31E-79 | 1.59844084 | 0.356 | 0.095 | 2.54E-75 | osteoclast.like | Sgsm3 |
| Ahnak21 | 2.35E-79 | 1.78571436 | 0.385 | 0.088 | 4.55E-75 | osteoclast.like | Ahnak2 |
| Coq5 | 2.54E-79 | 1.45041092 | 0.42 | 0.124 | 4.91E-75 | osteoclast.like | Coq5 |
| Bola22 | 3.73E-79 | 0.60009823 | 0.805 | 0.536 | 7.21E-75 | osteoclast.like | Bola2 |
| Znhit11 | 3.84E-79 | 0.65211486 | 0.741 | 0.466 | 7.42E-75 | osteoclast.like | Znhit1 |
| Naxe1 | 4.75E-79 | 1.02568913 | 0.576 | 0.258 | 9.18E-75 | osteoclast.like | Naxe |
| Zfp7061 | 6.39E-79 | 1.1622842 | 0.61 | 0.231 | 1.24E-74 | osteoclast.like | Zfp706 |
| Tinrd12 | 7.26E-79 | 0.56250193 | 0.776 | 0.465 | 1.40E-74 | osteoclast.like | Tinrd1 |
| Pum32 | 7.66E-79 | 0.96621797 | 0.659 | 0.296 | 1.48E-74 | osteoclast.like | Pum3 |
| Sh2b3 | 8.19E-79 | 0.53944588 | 0.78 | 0.493 | 1.58E-74 | osteoclast.like | Sh2b3 |
| Mrfap12 | 8.63E-79 | 0.42876284 | 0.863 | 0.613 | 1.67E-74 | osteoclast.like | Mrfap1 |
| Tardbp1 | 9.70E-79 | 0.66687618 | 0.717 | 0.453 | 1.88E-74 | osteoclast.like | Tardbp |
| Dnajc33 | 1.30E-78 | 0.47604858 | 0.795 | 0.554 | 2.51E-74 | osteoclast.like | Dnajc3 |
| Kmt5a3 | 1.81E-78 | 0.72568293 | 0.722 | 0.389 | 3.49E-74 | osteoclast.like | Kmt5a |
| Hnrnpf1 | 2.40E-78 | 0.40100404 | 0.844 | 0.639 | 4.64E-74 | osteoclast.like | Hnrnpf |
| St6gal13 | 2.47E-78 | 0.96106144 | 0.629 | 0.302 | 4.77E-74 | osteoclast.like | St6gal1 |
| Tax1bp3 | 2.64E-78 | 0.61078433 | 0.727 | 0.423 | 5.10E-74 | osteoclast.like | Tax1bp3 |
| Trappc112 | 2.68E-78 | 1.12885389 | 0.605 | 0.23 | 5.18E-74 | osteoclast.like | Trappc11 |
| Uhrf23 | 2.92E-78 | 0.80934247 | 0.683 | 0.375 | 5.65E-74 | osteoclast.like | Uhrf2 |
| Mtx11 | 3.02E-78 | 1.4639393 | 0.478 | 0.128 | 5.84E-74 | osteoclast.like | Mtx1 |
| Vasn | 3.47E-78 | 1.70752623 | 0.283 | 0.065 | 6.70E-74 | osteoclast.like | Vasn |
| Trim81 | 3.50E-78 | 0.31766043 | 0.859 | 0.611 | 6.77E-74 | osteoclast.like | Trim8 |
| Birc2 | 3.56E-78 | 1.58970984 | 0.356 | 0.11 | 6.88E-74 | osteoclast.like | Birc2 |
| Rxrb1 | 3.56E-78 | 0.8494824 | 0.717 | 0.343 | 6.89E-74 | osteoclast.like | Rxrb |
| Yipf4 | 4.08E-78 | 0.89690272 | 0.585 | 0.258 | 7.89E-74 | osteoclast.like | Yipf4 |
| Nfu11 | 4.27E-78 | 1.4964467 | 0.517 | 0.156 | 8.25E-74 | osteoclast.like | Nfu1 |
| Zfp467 | 5.71E-78 | 1.35216138 | 0.454 | 0.142 | 1.10E-73 | osteoclast.like | Zfp467 |
| Ivd1 | 6.26E-78 | 1.59260368 | 0.522 | 0.156 | 1.21E-73 | osteoclast.like | Ivd |
| Ndufb4 | 7.64E-78 | 1.43835584 | 0.415 | 0.128 | 1.48E-73 | osteoclast.like | Ndufb4 |
| Spag71 | 7.84E-78 | 0.76592559 | 0.61 | 0.296 | 1.52E-73 | osteoclast.like | Spag7 |
| Ppa2 | 8.23E-78 | 1.51426323 | 0.41 | 0.105 | 1.59E-73 | osteoclast.like | Ppa2 |
| 2-Mar | 9.00E-78 | 0.59349428 | 0.776 | 0.474 | 1.74E-73 | osteoclast.like | 2-Mar |
| Lamtor21 | 1.02E-77 | 0.45120278 | 0.771 | 0.541 | 1.98E-73 | osteoclast.like | Lamtor2 |
| Taco1 | 1.24E-77 | 1.87443594 | 0.361 | 0.07 | 2.40E-73 | osteoclast.like | Taco1 |
| Pdcl | 1.70E-77 | 1.25515343 | 0.566 | 0.221 | 3.29E-73 | osteoclast.like | Pdcl |
| Maz1 | 1.74E-77 | 0.61255839 | 0.79 | 0.488 | 3.36E-73 | osteoclast.like | Maz |
| Foxred11 | 2.06E-77 | 1.25166139 | 0.522 | 0.182 | 3.98E-73 | osteoclast.like | Foxred1 |
| Cpped1 | 2.08E-77 | 1.68522629 | 0.405 | 0.088 | 4.02E-73 | osteoclast.like | Cpped1 |
| Coq3 | 2.50E-77 | 1.94295688 | 0.361 | 0.076 | 4.83E-73 | osteoclast.like | Coq3 |
| Fkbp31 | 2.66E-77 | 1.48298772 | 0.498 | 0.163 | 5.14E-73 | osteoclast.like | Fkbp3 |
| Gmpr | 2.68E-77 | 2.92610384 | 0.263 | 0.036 | 5.18E-73 | osteoclast.like | Gmpr |
| Pigq1 | 3.63E-77 | 1.14097462 | 0.541 | 0.201 | 7.02E-73 | osteoclast.like | Pigq |
| Ube2v1 | 4.13E-77 | 0.40237935 | 0.863 | 0.638 | 7.99E-73 | osteoclast.like | Ube2v1 |
| Emg12 | 4.47E-77 | 0.70986833 | 0.751 | 0.456 | 8.64E-73 | osteoclast.like | Emg1 |
| Cttnbp2nl2 | 4.81E-77 | 0.40887619 | 0.8 | 0.539 | 9.30E-73 | osteoclast.like | Cttnbp2nl |
| Riok32 | 5.58E-77 | 0.26716326 | 0.883 | 0.67 | 1.08E-72 | osteoclast.like | Riok3 |
| Plcb23 | 5.88E-77 | 0.74995132 | 0.605 | 0.316 | 1.14E-72 | osteoclast.like | Plcb2 |
| Bcap292 | 8.11E-77 | 1.20852509 | 0.551 | 0.223 | 1.57E-72 | osteoclast.like | Bcap29 |
| Cep571 | 8.39E-77 | 1.43777189 | 0.376 | 0.11 | 1.62E-72 | osteoclast.like | Cep57 |
| Klc43 | 9.28E-77 | 1.11098604 | 0.629 | 0.276 | 1.79E-72 | osteoclast.like | Klc4 |
| Kpna61 | 1.30E-76 | 1.04114221 | 0.605 | 0.261 | 2.51E-72 | osteoclast.like | Kpna6 |
| Chmp1b2 | 1.35E-76 | 0.71949451 | 0.707 | 0.427 | 2.61E-72 | osteoclast.like | Chmp1b |
| Elob1 | 1.60E-76 | 0.40626268 | 0.868 | 0.657 | 3.09E-72 | osteoclast.like | Elob |
| Parl1 | 1.79E-76 | 1.01072916 | 0.546 | 0.226 | 3.45E-72 | osteoclast.like | Parl |
| Ndufb3 | 2.12E-76 | 1.83137378 | 0.4 | 0.094 | 4.10E-72 | osteoclast.like | Ndufb3 |
| Adsl1 | 2.33E-76 | 1.19135431 | 0.502 | 0.196 | 4.50E-72 | osteoclast.like | Adsl |
| Larp12 | 2.77E-76 | 0.49450842 | 0.776 | 0.524 | 5.35E-72 | osteoclast.like | Larp1 |
| Tra2a1 | 5.03E-76 | 0.40673672 | 0.854 | 0.59 | 9.73E-72 | osteoclast.like | Tra2a |
| Sh3rf1 | 5.66E-76 | 1.92066717 | 0.371 | 0.075 | 1.09E-71 | osteoclast.like | Sh3rf1 |
| Hadh2 | 5.66E-76 | 1.07137964 | 0.624 | 0.272 | 1.09E-71 | osteoclast.like | Hadh |
| Xpot2 | 5.74E-76 | 1.15714911 | 0.473 | 0.163 | 1.11E-71 | osteoclast.like | Xpot |
| Dnajb13 | 7.04E-76 | 0.5089122 | 0.683 | 0.396 | 1.36E-71 | osteoclast.like | Dnajb1 |
| Hax13 | 9.06E-76 | 0.80758886 | 0.629 | 0.32 | 1.75E-71 | osteoclast.like | Hax1 |
| Tpk1 | 1.03E-75 | 1.85737149 | 0.356 | 0.071 | 2.00E-71 | osteoclast.like | Tpk1 |
| Abcf11 | 1.10E-75 | 0.83132247 | 0.668 | 0.341 | 2.13E-71 | osteoclast.like | Abcf1 |
| Larp4b1 | 1.54E-75 | 0.65581486 | 0.81 | 0.52 | 2.99E-71 | osteoclast.like | Larp4b |
| Ano10 | 2.14E-75 | 1.21681885 | 0.434 | 0.147 | 4.13E-71 | osteoclast.like | Ano10 |
| Snrpf2 | 2.19E-75 | 0.53566956 | 0.751 | 0.51 | 4.23E-71 | osteoclast.like | Snrpf |
| Nf21 | 3.20E-75 | 0.89141578 | 0.663 | 0.336 | 6.18E-71 | osteoclast.like | Nf2 |
| Cracr2b2 | 4.50E-75 | 1.06587683 | 0.537 | 0.211 | 8.71E-71 | osteoclast.like | Cracr2b |
| Tbc1d24 | 5.39E-75 | 1.45867103 | 0.463 | 0.137 | 1.04E-70 | osteoclast.like | Tbc1d24 |
| Ubap2l2 | 7.63E-75 | 0.47115174 | 0.79 | 0.553 | 1.48E-70 | osteoclast.like | Ubap2l |
| Zfp568 | 8.07E-75 | 1.92558911 | 0.366 | 0.091 | 1.56E-70 | osteoclast.like | Zfp568 |
| Gcn11 | 8.70E-75 | 0.76528739 | 0.605 | 0.292 | 1.68E-70 | osteoclast.like | Gcn1 |
| Hnrnpc1 | 9.18E-75 | 0.8254825 | 0.605 | 0.32 | 1.77E-70 | osteoclast.like | Hnrnpc |
| Morf4l23 | 9.96E-75 | 0.83076159 | 0.649 | 0.364 | 1.93E-70 | osteoclast.like | Morf4l2 |
| Sri | 1.07E-74 | 0.30492016 | 0.79 | 0.611 | 2.06E-70 | osteoclast.like | Sri |
| Pdap11 | 1.34E-74 | 0.56696617 | 0.732 | 0.467 | 2.60E-70 | osteoclast.like | Pdap1 |
| Cdk162 | 1.59E-74 | 0.83875365 | 0.659 | 0.308 | 3.07E-70 | osteoclast.like | Cdk16 |
| Akirin21 | 2.23E-74 | 0.70904743 | 0.668 | 0.363 | 4.31E-70 | osteoclast.like | Akirin2 |
| Psma52 | 2.93E-74 | 0.63746218 | 0.732 | 0.458 | 5.67E-70 | osteoclast.like | Psma5 |
| Snx293 | 4.40E-74 | 0.35349679 | 0.717 | 0.476 | 8.51E-70 | osteoclast.like | Snx29 |
| Wdfy11 | 5.93E-74 | 1.06572785 | 0.498 | 0.186 | 1.15E-69 | osteoclast.like | Wdfy1 |
| Khsrp1 | 6.22E-74 | 0.5028793 | 0.854 | 0.563 | 1.20E-69 | osteoclast.like | Khsrp |
| Pdpr1 | 1.17E-73 | 1.31309633 | 0.463 | 0.145 | 2.26E-69 | osteoclast.like | Pdpr |
| Nubp21 | 1.19E-73 | 0.75465817 | 0.727 | 0.376 | 2.31E-69 | osteoclast.like | Nubp2 |
| Hint12 | 1.22E-73 | 0.87067142 | 0.605 | 0.345 | 2.36E-69 | osteoclast.like | Hint1 |
| Maged14 | 1.48E-73 | 1.27026342 | 0.478 | 0.186 | 2.85E-69 | osteoclast.like | Maged1 |
| Dynlt31 | 1.48E-73 | 0.82085062 | 0.629 | 0.325 | 2.86E-69 | osteoclast.like | Dynlt3 |
| Csde1 | 1.52E-73 | 0.39659798 | 0.805 | 0.585 | 2.94E-69 | osteoclast.like | Csde1 |
| Atp2c11 | 2.37E-73 | 0.50094918 | 0.78 | 0.504 | 4.58E-69 | osteoclast.like | Atp2c1 |
| Aph1a | 7.22E-73 | 0.35611923 | 0.849 | 0.609 | 1.40E-68 | osteoclast.like | Aph1a |
| Ssbp31 | 8.46E-73 | 1.29803214 | 0.439 | 0.122 | 1.64E-68 | osteoclast.like | Ssbp3 |
| Cpsf62 | 8.67E-73 | 0.53974307 | 0.722 | 0.449 | 1.68E-68 | osteoclast.like | Cpsf6 |
| Dync1h11 | 1.16E-72 | 0.33502895 | 0.849 | 0.635 | 2.24E-68 | osteoclast.like | Dync1h1 |
| Kdm5c1 | 1.24E-72 | 0.68013356 | 0.693 | 0.396 | 2.41E-68 | osteoclast.like | Kdm5c |
| Phip1 | 1.52E-72 | 0.71862491 | 0.688 | 0.337 | 2.94E-68 | osteoclast.like | Phip |
| Mthfd114 | 1.75E-72 | 0.96270974 | 0.62 | 0.288 | 3.38E-68 | osteoclast.like | Mthfd11 |
| Dkc11 | 2.16E-72 | 1.38804043 | 0.376 | 0.132 | 4.17E-68 | osteoclast.like | Dkc1 |
| Ube2m2 | 2.52E-72 | 0.49671587 | 0.761 | 0.504 | 4.87E-68 | osteoclast.like | Ube2m |
| Timm501 | 2.64E-72 | 1.24794 | 0.605 | 0.256 | 5.11E-68 | osteoclast.like | Timm50 |
| Arsa2 | 2.70E-72 | 0.9516544 | 0.595 | 0.254 | 5.22E-68 | osteoclast.like | Arsa |
| Crot | 2.88E-72 | 1.47077213 | 0.454 | 0.135 | 5.57E-68 | osteoclast.like | Crot |
| Hdhd2 | 3.45E-72 | 1.38082769 | 0.454 | 0.138 | 6.67E-68 | osteoclast.like | Hdhd2 |
| Casp61 | 4.89E-72 | 0.76541319 | 0.659 | 0.323 | 9.45E-68 | osteoclast.like | Casp6 |
| Cyth21 | 5.70E-72 | 0.60981355 | 0.683 | 0.393 | 1.10E-67 | osteoclast.like | Cyth2 |
| Farsa2 | 6.17E-72 | 0.83496212 | 0.634 | 0.295 | 1.19E-67 | osteoclast.like | Farsa |
| Rab34 | 7.38E-72 | 1.82572429 | 0.327 | 0.066 | 1.43E-67 | osteoclast.like | Rab34 |
| Dpf2 | 7.64E-72 | 0.62515043 | 0.644 | 0.365 | 1.48E-67 | osteoclast.like | Dpf2 |
| Tspyl11 | 9.04E-72 | 0.82544576 | 0.693 | 0.315 | 1.75E-67 | osteoclast.like | Tspyl1 |
| Fbh11 | 9.46E-72 | 0.97370557 | 0.624 | 0.286 | 1.83E-67 | osteoclast.like | Fbh1 |
| Sppl3 | 1.76E-71 | 0.824424 | 0.702 | 0.343 | 3.41E-67 | osteoclast.like | Sppl3 |
| Adgrl2 | 1.89E-71 | 2.18191095 | 0.322 | 0.055 | 3.65E-67 | osteoclast.like | Adgrl2 |
| Tcp11l21 | 1.96E-71 | 0.45167052 | 0.702 | 0.395 | 3.80E-67 | osteoclast.like | Tcp11l2 |
| Prodh | 2.40E-71 | 2.01944325 | 0.356 | 0.072 | 4.65E-67 | osteoclast.like | Prodh |
| Cpsf31 | 4.10E-71 | 0.8941332 | 0.595 | 0.289 | 7.92E-67 | osteoclast.like | Cpsf3 |
| Tpd52l22 | 4.24E-71 | 0.59029909 | 0.737 | 0.446 | 8.19E-67 | osteoclast.like | Tpd52l2 |
| Atg31 | 5.99E-71 | 0.74205177 | 0.727 | 0.37 | 1.16E-66 | osteoclast.like | Atg3 |
| Med10 | 6.87E-71 | 0.82370613 | 0.639 | 0.332 | 1.33E-66 | osteoclast.like | Med10 |
| Gtpbp42 | 9.39E-71 | 0.63540314 | 0.659 | 0.372 | 1.82E-66 | osteoclast.like | Gtpbp4 |
| Rcan11 | 1.25E-70 | 0.88404689 | 0.527 | 0.227 | 2.41E-66 | osteoclast.like | Rcan1 |
| Txnl13 | 1.42E-70 | 0.5940593 | 0.751 | 0.438 | 2.75E-66 | osteoclast.like | Txnl1 |
| Lrch11 | 1.81E-70 | 0.6745159 | 0.707 | 0.362 | 3.49E-66 | osteoclast.like | Lrch1 |
| Sumo31 | 2.54E-70 | 0.67244352 | 0.756 | 0.439 | 4.91E-66 | osteoclast.like | Sumo3 |
| Chchd41 | 3.05E-70 | 1.56415627 | 0.429 | 0.129 | 5.90E-66 | osteoclast.like | Chchd4 |
| Cdk5rap21 | 3.30E-70 | 1.6442141 | 0.405 | 0.099 | 6.39E-66 | osteoclast.like | Cdk5rap2 |
| Fam214b4 | 3.51E-70 | 0.68393208 | 0.62 | 0.329 | 6.80E-66 | osteoclast.like | Fam214b |
| Aamp | 3.61E-70 | 0.32876415 | 0.878 | 0.649 | 6.99E-66 | osteoclast.like | Aamp |
| Prpf38b | 3.66E-70 | 0.66588907 | 0.785 | 0.477 | 7.08E-66 | osteoclast.like | Prpf38b |
| Pex191 | 3.70E-70 | 0.96224282 | 0.58 | 0.25 | 7.14E-66 | osteoclast.like | Pex19 |
| Atf61 | 3.70E-70 | 0.46462912 | 0.722 | 0.443 | 7.16E-66 | osteoclast.like | Atf6 |
| Dtymk1 | 3.72E-70 | 1.150211 | 0.59 | 0.222 | 7.19E-66 | osteoclast.like | Dtymk |
| Rab1b1 | 6.33E-70 | 0.31743928 | 0.922 | 0.696 | 1.22E-65 | osteoclast.like | Rab1b |
| Kat141 | 8.68E-70 | 0.93204568 | 0.605 | 0.275 | 1.68E-65 | osteoclast.like | Kat14 |
| Capn22 | 8.96E-70 | 0.25154655 | 0.746 | 0.607 | 1.73E-65 | osteoclast.like | Capn2 |
| Klhl61 | 1.44E-69 | 1.31659554 | 0.468 | 0.138 | 2.79E-65 | osteoclast.like | Klhl6 |
| Sbno11 | 1.51E-69 | 0.52785501 | 0.702 | 0.401 | 2.92E-65 | osteoclast.like | Sbno1 |
| Lmtk2 | 1.63E-69 | 2.22623376 | 0.288 | 0.046 | 3.15E-65 | osteoclast.like | Lmtk2 |
| Nop23 | 1.64E-69 | 0.66278859 | 0.517 | 0.271 | 3.18E-65 | osteoclast.like | Nop2 |
| Mov10 | 1.85E-69 | 1.59555756 | 0.385 | 0.091 | 3.58E-65 | osteoclast.like | Mov10 |
| Phf102 | 2.08E-69 | 0.82344985 | 0.59 | 0.275 | 4.02E-65 | osteoclast.like | Phf10 |
| Sec24c | 2.33E-69 | 0.63768919 | 0.732 | 0.447 | 4.51E-65 | osteoclast.like | Sec24c |
| Fech3 | 2.90E-69 | 1.07567113 | 0.541 | 0.226 | 5.60E-65 | osteoclast.like | Fech |
| Sfxn33 | 3.20E-69 | 0.55377685 | 0.756 | 0.465 | 6.19E-65 | osteoclast.like | Sfxn3 |
| Tmed1 | 5.58E-69 | 1.85381428 | 0.366 | 0.081 | 1.08E-64 | osteoclast.like | Tmed1 |
| Surf6 | 5.96E-69 | 1.57614794 | 0.39 | 0.101 | 1.15E-64 | osteoclast.like | Surf6 |
| Timmdc1 | 6.52E-69 | 1.43287328 | 0.424 | 0.122 | 1.26E-64 | osteoclast.like | Timmdc1 |
| Brd72 | 7.86E-69 | 0.68846425 | 0.639 | 0.331 | 1.52E-64 | osteoclast.like | Brd7 |
| Dpy301 | 9.17E-69 | 1.11489155 | 0.605 | 0.262 | 1.77E-64 | osteoclast.like | Dpy30 |
| Urod1 | 9.95E-69 | 1.16523501 | 0.561 | 0.203 | 1.92E-64 | osteoclast.like | Urod |
| Tmx23 | 1.16E-68 | 1.15609941 | 0.527 | 0.226 | 2.23E-64 | osteoclast.like | Tmx2 |
| Tars1 | 1.18E-68 | 0.91575607 | 0.571 | 0.294 | 2.28E-64 | osteoclast.like | Tars |
| Pxn2 | 1.25E-68 | 0.29540405 | 0.761 | 0.54 | 2.41E-64 | osteoclast.like | Pxn |
| Usp81 | 1.71E-68 | 0.39490179 | 0.785 | 0.526 | 3.31E-64 | osteoclast.like | Usp8 |
| Galt | 2.34E-68 | 1.45350907 | 0.463 | 0.137 | 4.53E-64 | osteoclast.like | Galt |
| Espl11 | 3.29E-68 | 2.01696596 | 0.351 | 0.071 | 6.37E-64 | osteoclast.like | Espl1 |
| Tulp42 | 3.81E-68 | 0.90954647 | 0.478 | 0.196 | 7.36E-64 | osteoclast.like | Tulp4 |
| Aurkaip11 | 4.48E-68 | 0.6492514 | 0.737 | 0.438 | 8.67E-64 | osteoclast.like | Aurkaip1 |
| Trappc14 | 4.82E-68 | 0.67905263 | 0.62 | 0.35 | 9.32E-64 | osteoclast.like | Trappc1 |
| Fra10ac1 | 6.90E-68 | 1.58529651 | 0.366 | 0.088 | 1.33E-63 | osteoclast.like | Fra10ac1 |
| Eif3c1 | 7.02E-68 | 0.73813949 | 0.771 | 0.434 | 1.36E-63 | osteoclast.like | Eif3c |
| Rlim1 | 7.88E-68 | 0.25758148 | 0.854 | 0.622 | 1.52E-63 | osteoclast.like | Rlim |
| Rpn22 | 8.62E-68 | 0.50254108 | 0.776 | 0.524 | 1.67E-63 | osteoclast.like | Rpn2 |
| Tmed10 | 8.77E-68 | 0.52534898 | 0.732 | 0.478 | 1.70E-63 | osteoclast.like | Tmed10 |
| Ssr11 | 1.22E-67 | 0.3487831 | 0.868 | 0.672 | 2.36E-63 | osteoclast.like | Ssr1 |
| Pptc71 | 1.33E-67 | 1.06469398 | 0.546 | 0.202 | 2.58E-63 | osteoclast.like | Pptc7 |
| B4galt71 | 1.73E-67 | 1.47794049 | 0.405 | 0.135 | 3.34E-63 | osteoclast.like | B4galt7 |
| Sars2 | 1.90E-67 | 1.61841604 | 0.371 | 0.084 | 3.67E-63 | osteoclast.like | Sars2 |
| Fmc1 | 2.49E-67 | 1.33130888 | 0.39 | 0.106 | 4.81E-63 | osteoclast.like | Fmc1 |
| Yif1a2 | 2.50E-67 | 1.15245483 | 0.546 | 0.198 | 4.83E-63 | osteoclast.like | Yif1a |
| Spg71 | 2.59E-67 | 0.71336906 | 0.717 | 0.356 | 5.01E-63 | osteoclast.like | Spg7 |
| Gipc12 | 2.75E-67 | 0.98534681 | 0.585 | 0.247 | 5.32E-63 | osteoclast.like | Gipc1 |
| Ube3a | 3.16E-67 | 0.67575865 | 0.668 | 0.391 | 6.11E-63 | osteoclast.like | Ube3a |
| Dnajb121 | 3.46E-67 | 0.69074936 | 0.688 | 0.405 | 6.69E-63 | osteoclast.like | Dnajb12 |
| Srsf13 | 3.59E-67 | 0.63666055 | 0.8 | 0.503 | 6.95E-63 | osteoclast.like | Srsf1 |
| Atp5c1 | 5.49E-67 | 1.59805668 | 0.415 | 0.11 | 1.06E-62 | osteoclast.like | Atp5c1 |
| Mocs23 | 5.83E-67 | 0.87935229 | 0.688 | 0.326 | 1.13E-62 | osteoclast.like | Mocs2 |
| Mecp2 | 6.72E-67 | 1.22534092 | 0.585 | 0.2 | 1.30E-62 | osteoclast.like | Mecp2 |
| Dhx91 | 7.03E-67 | 0.44760984 | 0.722 | 0.518 | 1.36E-62 | osteoclast.like | Dhx9 |
| Tor4a3 | 8.78E-67 | 0.79159544 | 0.624 | 0.287 | 1.70E-62 | osteoclast.like | Tor4a |
| Fam136a | 1.49E-66 | 1.68542054 | 0.444 | 0.114 | 2.89E-62 | osteoclast.like | Fam136a |
| Lpp3 | 2.04E-66 | 0.43076145 | 0.8 | 0.578 | 3.95E-62 | osteoclast.like | Lpp |
| Plod13 | 2.04E-66 | 0.7548115 | 0.751 | 0.382 | 3.95E-62 | osteoclast.like | Plod1 |
| Selenof1 | 2.28E-66 | 0.76668029 | 0.561 | 0.296 | 4.40E-62 | osteoclast.like | Selenof |
| Ly92 | 2.63E-66 | 1.07429973 | 0.532 | 0.18 | 5.08E-62 | osteoclast.like | Ly9 |
| Txndc9 | 3.02E-66 | 0.60706348 | 0.732 | 0.42 | 5.84E-62 | osteoclast.like | Txndc9 |
| Gpbp1l1 | 3.16E-66 | 0.77194956 | 0.698 | 0.348 | 6.11E-62 | osteoclast.like | Gpbp1l1 |
| Skp1a1 | 3.64E-66 | 0.65869675 | 0.702 | 0.405 | 7.04E-62 | osteoclast.like | Skp1a |
| Atxn7l3b2 | 5.61E-66 | 0.50210634 | 0.673 | 0.383 | 1.09E-61 | osteoclast.like | Atxn7l3b |
| Dok13 | 7.31E-66 | 0.86174345 | 0.58 | 0.265 | 1.41E-61 | osteoclast.like | Dok1 |
| Armc11 | 9.19E-66 | 1.33155637 | 0.507 | 0.183 | 1.78E-61 | osteoclast.like | Armc1 |
| Sf3b51 | 9.67E-66 | 0.70144954 | 0.678 | 0.384 | 1.87E-61 | osteoclast.like | Sf3b5 |
| Ap1g11 | 1.68E-65 | 0.2701009 | 0.756 | 0.512 | 3.26E-61 | osteoclast.like | Ap1g1 |
| Stoml21 | 1.75E-65 | 1.01301537 | 0.6 | 0.255 | 3.38E-61 | osteoclast.like | Stoml2 |
| Eif4e2 | 1.87E-65 | 0.59940792 | 0.746 | 0.462 | 3.61E-61 | osteoclast.like | Eif4e2 |

| | | | | | | | |
|---|---|---|---|---|---|---|---|
| Mycbp1 | 1.99E-65 | 1.20682917 | 0.41 | 0.132 | 3.84E-61 | osteoclast.like | Mycbp |
| Birc61 | 2.16E-65 | 0.47633413 | 0.771 | 0.481 | 4.18E-61 | osteoclast.like | Birc6 |
| Eef1e11 | 2.40E-65 | 1.36723057 | 0.488 | 0.161 | 4.64E-61 | osteoclast.like | Eef1e1 |
| Mief2 | 4.25E-65 | 1.64205172 | 0.361 | 0.08 | 8.22E-61 | osteoclast.like | Mief2 |
| St8sia42 | 5.19E-65 | 0.4910256 | 0.6 | 0.358 | 1.00E-60 | osteoclast.like | St8sia4 |
| Nap1l41 | 5.31E-65 | 0.8610068 | 0.712 | 0.349 | 1.03E-60 | osteoclast.like | Nap1l4 |
| Pfkfb2 | 5.58E-65 | 1.3749396 | 0.376 | 0.118 | 1.08E-60 | osteoclast.like | Pfkfb2 |
| Yars3 | 5.73E-65 | 1.59631948 | 0.459 | 0.154 | 1.11E-60 | osteoclast.like | Yars |
| Pgm21 | 5.82E-65 | 0.69121852 | 0.634 | 0.347 | 1.12E-60 | osteoclast.like | Pgm2 |
| Ywhah2 | 6.76E-65 | 0.29555136 | 0.873 | 0.658 | 1.31E-60 | osteoclast.like | Ywhah |
| Steap31 | 7.68E-65 | 1.48489607 | 0.488 | 0.135 | 1.49E-60 | osteoclast.like | Steap3 |
| Mcfd23 | 8.85E-65 | 0.38551722 | 0.824 | 0.589 | 1.71E-60 | osteoclast.like | Mcfd2 |
| Ss18l2 | 1.02E-64 | 1.60870759 | 0.346 | 0.089 | 1.98E-60 | osteoclast.like | Ss18l2 |
| Glrx31 | 1.42E-64 | 1.091438 | 0.58 | 0.247 | 2.75E-60 | osteoclast.like | Glrx3 |
| Akr1c13 | 1.44E-64 | 1.54055916 | 0.366 | 0.106 | 2.78E-60 | osteoclast.like | Akr1c13 |
| Ddx242 | 1.73E-64 | 0.42687126 | 0.702 | 0.448 | 3.35E-60 | osteoclast.like | Ddx24 |
| Mmut | 1.87E-64 | 1.80900846 | 0.366 | 0.085 | 3.62E-60 | osteoclast.like | Mmut |
| Ngly11 | 2.47E-64 | 1.03319333 | 0.488 | 0.19 | 4.78E-60 | osteoclast.like | Ngly1 |
| Eif2a1 | 2.53E-64 | 0.77193345 | 0.6 | 0.311 | 4.89E-60 | osteoclast.like | Eif2a |
| Add11 | 3.07E-64 | 0.47420668 | 0.776 | 0.503 | 5.93E-60 | osteoclast.like | Add1 |
| Poglut3 | 3.62E-64 | 1.86220086 | 0.341 | 0.069 | 6.99E-60 | osteoclast.like | Poglut3 |
| Psen21 | 3.72E-64 | 0.694995 | 0.693 | 0.357 | 7.20E-60 | osteoclast.like | Psen2 |
| Psmb63 | 4.29E-64 | 0.63588696 | 0.766 | 0.443 | 8.29E-60 | osteoclast.like | Psmb6 |
| Prkaca1 | 4.48E-64 | 0.58231705 | 0.673 | 0.42 | 8.66E-60 | osteoclast.like | Prkaca |
| Dbndd21 | 4.88E-64 | 0.8447945 | 0.62 | 0.285 | 9.43E-60 | osteoclast.like | Dbndd2 |
| Tbca1 | 6.13E-64 | 0.47229492 | 0.8 | 0.528 | 1.18E-59 | osteoclast.like | Tbca |
| Eif3a2 | 7.02E-64 | 0.56012634 | 0.727 | 0.414 | 1.36E-59 | osteoclast.like | Eif3a |
| Snap233 | 7.61E-64 | 0.74198668 | 0.561 | 0.281 | 1.47E-59 | osteoclast.like | Snap23 |
| Cd1513 | 8.31E-64 | 0.99313417 | 0.59 | 0.264 | 1.61E-59 | osteoclast.like | Cd151 |
| Sox41 | 9.34E-64 | 1.21958694 | 0.444 | 0.168 | 1.81E-59 | osteoclast.like | Sox4 |
| Pcbd22 | 1.12E-63 | 0.91471028 | 0.517 | 0.205 | 2.17E-59 | osteoclast.like | Pcbd2 |
| Abcb6 | 1.26E-63 | 2.33106082 | 0.288 | 0.045 | 2.43E-59 | osteoclast.like | Abcb6 |
| Ppdpf2 | 1.30E-63 | 0.80649233 | 0.59 | 0.264 | 2.51E-59 | osteoclast.like | Ppdpf |
| Glyr1 | 1.50E-63 | 0.4311184 | 0.81 | 0.549 | 2.91E-59 | osteoclast.like | Glyr1 |
| Impdh13 | 1.76E-63 | 1.21647286 | 0.517 | 0.194 | 3.40E-59 | osteoclast.like | Impdh1 |
| Wdr182 | 1.83E-63 | 1.0347102 | 0.556 | 0.245 | 3.54E-59 | osteoclast.like | Wdr18 |
| Pmpca1 | 1.87E-63 | 0.85373048 | 0.556 | 0.268 | 3.61E-59 | osteoclast.like | Pmpca |
| Nkiras23 | 2.31E-63 | 0.53522118 | 0.702 | 0.373 | 4.47E-59 | osteoclast.like | Nkiras2 |
| Pgrmc11 | 3.05E-63 | 0.68247101 | 0.59 | 0.299 | 5.90E-59 | osteoclast.like | Pgrmc1 |
| Uqcrh | 3.09E-63 | 2.27905856 | 0.288 | 0.043 | 5.97E-59 | osteoclast.like | Uqcrh |
| Snrnp2001 | 3.48E-63 | 0.60718486 | 0.717 | 0.41 | 6.73E-59 | osteoclast.like | Snrnp200 |
| Dhrs7b1 | 3.57E-63 | 1.09501454 | 0.454 | 0.166 | 6.89E-59 | osteoclast.like | Dhrs7b |
| Slco4a11 | 3.62E-63 | 1.34806886 | 0.356 | 0.074 | 7.00E-59 | osteoclast.like | Slco4a1 |
| Baiap22 | 3.92E-63 | 0.48121164 | 0.668 | 0.42 | 7.57E-59 | osteoclast.like | Baiap2 |
| Trmt5 | 4.47E-63 | 2.0434483 | 0.312 | 0.055 | 8.65E-59 | osteoclast.like | Trmt5 |
| Cwc151 | 4.70E-63 | 0.32723724 | 0.839 | 0.601 | 9.09E-59 | osteoclast.like | Cwc15 |
| Taz2 | 5.48E-63 | 0.98140102 | 0.615 | 0.298 | 1.06E-58 | osteoclast.like | Taz |
| Ankrd17 | 7.74E-63 | 0.37964538 | 0.732 | 0.492 | 1.50E-58 | osteoclast.like | Ankrd17 |
| Snrpa2 | 8.04E-63 | 0.72737974 | 0.58 | 0.327 | 1.55E-58 | osteoclast.like | Snrpa |
| Supv3l1 | 8.63E-63 | 1.48102715 | 0.4 | 0.103 | 1.67E-58 | osteoclast.like | Supv3l1 |
| Anp32e1 | 8.94E-63 | 0.4478281 | 0.741 | 0.497 | 1.73E-58 | osteoclast.like | Anp32e |
| Dnajc71 | 9.82E-63 | 0.46928629 | 0.688 | 0.423 | 1.90E-58 | osteoclast.like | Dnajc7 |
| Uba51 | 1.04E-62 | 1.0601526 | 0.502 | 0.196 | 2.00E-58 | osteoclast.like | Uba5 |
| Trim473 | 1.12E-62 | 1.2055005 | 0.498 | 0.165 | 2.16E-58 | osteoclast.like | Trim47 |
| Eif1b | 1.16E-62 | 0.83744106 | 0.683 | 0.333 | 2.24E-58 | osteoclast.like | Eif1b |
| Acad11 | 1.55E-62 | 1.51665432 | 0.337 | 0.092 | 3.00E-58 | osteoclast.like | Acad11 |
| Gnpat2 | 1.70E-62 | 0.93043337 | 0.59 | 0.268 | 3.29E-58 | osteoclast.like | Gnpat |
| Ddx561 | 1.89E-62 | 0.96849614 | 0.546 | 0.223 | 3.66E-58 | osteoclast.like | Ddx56 |
| Oxsm | 1.94E-62 | 1.68921507 | 0.332 | 0.072 | 3.76E-58 | osteoclast.like | Oxsm |
| Ube2g22 | 2.01E-62 | 0.66738991 | 0.722 | 0.413 | 3.89E-58 | osteoclast.like | Ube2g2 |
| Hmg20b2 | 2.11E-62 | 0.59349089 | 0.634 | 0.351 | 4.09E-58 | osteoclast.like | Hmg20b |
| Mapk1 | 2.21E-62 | 0.3977246 | 0.79 | 0.557 | 4.28E-58 | osteoclast.like | Mapk1 |
| Uchl31 | 2.36E-62 | 1.71794833 | 0.41 | 0.1 | 4.57E-58 | osteoclast.like | Uchl3 |
| Wdr361 | 2.53E-62 | 1.08199986 | 0.551 | 0.229 | 4.89E-58 | osteoclast.like | Wdr36 |
| Csnk2a21 | 2.61E-62 | 0.68505723 | 0.62 | 0.336 | 5.04E-58 | osteoclast.like | Csnk2a2 |
| Lpcat4 | 2.84E-62 | 1.56860445 | 0.366 | 0.082 | 5.50E-58 | osteoclast.like | Lpcat4 |
| Psmc32 | 3.55E-62 | 0.36570622 | 0.776 | 0.582 | 6.86E-58 | osteoclast.like | Psmc3 |
| Btk1 | 3.98E-62 | 0.96018795 | 0.6 | 0.265 | 7.69E-58 | osteoclast.like | Btk |
| Ebna1bp22 | 4.62E-62 | 0.67965158 | 0.624 | 0.341 | 8.93E-58 | osteoclast.like | Ebna1bp2 |
| Rab2a1 | 4.90E-62 | 0.38673267 | 0.82 | 0.561 | 9.48E-58 | osteoclast.like | Rab2a |
| Glo11 | 5.66E-62 | 1.11304821 | 0.654 | 0.29 | 1.09E-57 | osteoclast.like | Glo1 |
| Ubr21 | 6.15E-62 | 0.35354434 | 0.688 | 0.452 | 1.19E-57 | osteoclast.like | Ubr2 |
| Macrod1 | 7.51E-62 | 1.7974189 | 0.346 | 0.075 | 1.45E-57 | osteoclast.like | Macrod1 |
| Tmem38b1 | 8.69E-62 | 0.88656617 | 0.556 | 0.224 | 1.68E-57 | osteoclast.like | Tmem38b |
| Denr2 | 1.78E-61 | 0.93360838 | 0.678 | 0.337 | 3.45E-57 | osteoclast.like | Denr |
| Msh21 | 2.02E-61 | 1.48960937 | 0.356 | 0.097 | 3.90E-57 | osteoclast.like | Msh2 |
| Cstf21 | 2.67E-61 | 1.51956862 | 0.473 | 0.149 | 5.16E-57 | osteoclast.like | Cstf2 |
| Tmed21 | 3.09E-61 | 0.48721232 | 0.741 | 0.504 | 5.98E-57 | osteoclast.like | Tmed2 |
| Tsc22d34 | 3.36E-61 | 0.46627516 | 0.649 | 0.407 | 6.49E-57 | osteoclast.like | Tsc22d3 |
| Gtf3c51 | 3.63E-61 | 1.43507214 | 0.42 | 0.125 | 7.01E-57 | osteoclast.like | Gtf3c5 |
| Ptcd21 | 3.78E-61 | 1.78745961 | 0.424 | 0.126 | 7.32E-57 | osteoclast.like | Ptcd2 |
| Rai1 | 4.00E-61 | 1.09456569 | 0.502 | 0.209 | 7.73E-57 | osteoclast.like | Rai1 |
| Trmt2b1 | 4.53E-61 | 1.61037323 | 0.371 | 0.105 | 8.77E-57 | osteoclast.like | Trmt2b |
| Snrpe2 | 4.83E-61 | 0.89917857 | 0.624 | 0.303 | 9.33E-57 | osteoclast.like | Snrpe |
| Khdrbs11 | 4.96E-61 | 0.35707979 | 0.776 | 0.569 | 9.59E-57 | osteoclast.like | Khdrbs1 |
| Herpud22 | 4.98E-61 | 0.92996283 | 0.512 | 0.228 | 9.63E-57 | osteoclast.like | Herpud2 |
| Tmed91 | 5.14E-61 | 0.3730636 | 0.82 | 0.621 | 9.93E-57 | osteoclast.like | Tmed9 |
| Msl31 | 7.00E-61 | 1.39675181 | 0.493 | 0.154 | 1.35E-56 | osteoclast.like | Msl3 |
| Pgghg | 1.02E-60 | 1.54080146 | 0.361 | 0.09 | 1.98E-56 | osteoclast.like | Pgghg |
| Crybg31 | 1.13E-60 | 0.8758196 | 0.473 | 0.216 | 2.19E-56 | osteoclast.like | Crybg3 |
| Zmiz22 | 1.20E-60 | 0.46486367 | 0.771 | 0.515 | 2.31E-56 | osteoclast.like | Zmiz2 |
| Tmem11 | 1.68E-60 | 1.0047925 | 0.629 | 0.272 | 3.24E-56 | osteoclast.like | Tmem11 |
| Ccdc1221 | 1.79E-60 | 1.9861612 | 0.263 | 0.044 | 3.47E-56 | osteoclast.like | Ccdc122 |
| Magt12 | 2.29E-60 | 0.43930933 | 0.746 | 0.477 | 4.43E-56 | osteoclast.like | Magt1 |
| Arap3 | 2.42E-60 | 1.5184776 | 0.429 | 0.123 | 4.68E-56 | osteoclast.like | Arap3 |
| Rbfox2 | 2.79E-60 | 1.91507114 | 0.254 | 0.045 | 5.40E-56 | osteoclast.like | Rbfox2 |
| Mettl1 | 3.03E-60 | 1.60924667 | 0.337 | 0.089 | 5.86E-56 | osteoclast.like | Mettl1 |
| Suclg2 | 3.58E-60 | 1.63682503 | 0.385 | 0.088 | 6.92E-56 | osteoclast.like | Suclg2 |
| Gtf2h51 | 4.96E-60 | 0.60350496 | 0.732 | 0.43 | 9.59E-56 | osteoclast.like | Gtf2h5 |
| Trp53inp1 | 5.20E-60 | 1.37726121 | 0.312 | 0.111 | 1.01E-55 | osteoclast.like | Trp53inp1 |
| Pnrc23 | 5.90E-60 | 0.5148335 | 0.668 | 0.376 | 1.14E-55 | osteoclast.like | Pnrc2 |
| Krt7 | 1.01E-59 | 1.07880577 | 0.322 | 0.103 | 1.96E-55 | osteoclast.like | Krt7 |
| Fzr11 | 1.04E-59 | 0.69432434 | 0.512 | 0.261 | 2.02E-55 | osteoclast.like | Fzr1 |
| Gmfb3 | 1.06E-59 | 0.38401062 | 0.82 | 0.533 | 2.05E-55 | osteoclast.like | Gmfb |
| Dgka | 1.30E-59 | 1.06033791 | 0.41 | 0.15 | 2.52E-55 | osteoclast.like | Dgka |
| Eif2b3 | 1.38E-59 | 1.68750689 | 0.337 | 0.089 | 2.67E-55 | osteoclast.like | Eif2b3 |
| Acadl3 | 1.45E-59 | 0.65602908 | 0.629 | 0.324 | 2.80E-55 | osteoclast.like | Acadl |
| C9orf72 | 1.58E-59 | 1.07662227 | 0.463 | 0.175 | 3.06E-55 | osteoclast.like | C9orf72 |
| Arhgap184 | 1.76E-59 | 0.65663385 | 0.595 | 0.284 | 3.40E-55 | osteoclast.like | Arhgap18 |
| Akr1b33 | 2.61E-59 | 0.59702275 | 0.693 | 0.4 | 5.05E-55 | osteoclast.like | Akr1b3 |
| Zfp207 | 4.22E-59 | 0.31102473 | 0.766 | 0.54 | 8.16E-55 | osteoclast.like | Zfp207 |
| Bcr | 4.84E-59 | 1.03330188 | 0.385 | 0.122 | 9.36E-55 | osteoclast.like | Bcr |
| Wdr72 | 5.21E-59 | 1.00098257 | 0.634 | 0.261 | 1.01E-54 | osteoclast.like | Wdr7 |
| Igbp11 | 5.38E-59 | 0.71642949 | 0.688 | 0.357 | 1.04E-54 | osteoclast.like | Igbp1 |
| Htt | 5.41E-59 | 0.49929296 | 0.6 | 0.352 | 1.05E-54 | osteoclast.like | Htt |
| Ciao12 | 7.35E-59 | 1.12061324 | 0.585 | 0.226 | 1.42E-54 | osteoclast.like | Ciao1 |
| Rnf441 | 9.26E-59 | 0.39750312 | 0.78 | 0.557 | 1.79E-54 | osteoclast.like | Rnf44 |
| Prpf193 | 1.04E-58 | 0.72024387 | 0.649 | 0.348 | 2.02E-54 | osteoclast.like | Prpf19 |
| Rala1 | 1.13E-58 | 0.67074305 | 0.629 | 0.321 | 2.19E-54 | osteoclast.like | Rala |
| Mtss13 | 1.25E-58 | 0.32119729 | 0.815 | 0.555 | 2.41E-54 | osteoclast.like | Mtss1 |
| Dag14 | 1.25E-58 | 1.01016673 | 0.58 | 0.246 | 2.41E-54 | osteoclast.like | Dag1 |
| Msh3 | 1.38E-58 | 1.34666282 | 0.395 | 0.101 | 2.67E-54 | osteoclast.like | Msh3 |
| Polr1c2 | 1.46E-58 | 0.91230791 | 0.561 | 0.278 | 2.83E-54 | osteoclast.like | Polr1c |
| Rbm472 | 1.49E-58 | 0.32551349 | 0.805 | 0.529 | 2.87E-54 | osteoclast.like | Rbm47 |
| Golim41 | 1.64E-58 | 1.09626785 | 0.507 | 0.196 | 3.17E-54 | osteoclast.like | Golim4 |
| Plekhj12 | 1.75E-58 | 0.41355129 | 0.761 | 0.502 | 3.37E-54 | osteoclast.like | Plekhj1 |
| Fkbp1a1 | 1.91E-58 | 0.51548921 | 0.785 | 0.477 | 3.68E-54 | osteoclast.like | Fkbp1a |
| Coa4 | 3.49E-58 | 1.23141496 | 0.424 | 0.129 | 6.75E-54 | osteoclast.like | Coa4 |
| Gnpnat1 | 3.74E-58 | 1.52311459 | 0.322 | 0.083 | 7.23E-54 | osteoclast.like | Gnpnat1 |
| Hmgn11 | 4.56E-58 | 1.83422656 | 0.395 | 0.106 | 8.81E-54 | osteoclast.like | Hmgn1 |
| Hint3 | 4.59E-58 | 1.22649903 | 0.42 | 0.127 | 8.87E-54 | osteoclast.like | Hint3 |
| Psma12 | 5.49E-58 | 0.41865358 | 0.746 | 0.528 | 1.06E-53 | osteoclast.like | Psma1 |
| Cpsf21 | 5.78E-58 | 0.74205123 | 0.546 | 0.25 | 1.12E-53 | osteoclast.like | Cpsf2 |
| Cep85 | 5.80E-58 | 1.45435323 | 0.385 | 0.105 | 1.12E-53 | osteoclast.like | Cep85 |
| Ebag9 | 5.92E-58 | 1.11995084 | 0.415 | 0.141 | 1.14E-53 | osteoclast.like | Ebag9 |
| Ano62 | 6.20E-58 | 0.57155378 | 0.663 | 0.398 | 1.20E-53 | osteoclast.like | Ano6 |
| Fbxo35 | 6.20E-58 | 0.86353766 | 0.61 | 0.256 | 1.20E-53 | osteoclast.like | Fbxo3 |
| Gorasp21 | 6.39E-58 | 0.55114538 | 0.712 | 0.428 | 1.24E-53 | osteoclast.like | Gorasp2 |
| Dlgap43 | 6.90E-58 | 0.31094818 | 0.756 | 0.513 | 1.33E-53 | osteoclast.like | Dlgap4 |
| Rnd31 | 7.52E-58 | 0.61343341 | 0.615 | 0.281 | 1.45E-53 | osteoclast.like | Rnd3 |
| Pebp11 | 8.66E-58 | 0.87389694 | 0.624 | 0.33 | 1.68E-53 | osteoclast.like | Pebp1 |
| Cuedc22 | 8.98E-58 | 0.44822075 | 0.737 | 0.475 | 1.74E-53 | osteoclast.like | Cuedc2 |
| Fbl2 | 1.15E-57 | 0.42690578 | 0.668 | 0.399 | 2.21E-53 | osteoclast.like | Fbl |
| Arel1 | 1.52E-57 | 0.92886003 | 0.502 | 0.215 | 2.95E-53 | osteoclast.like | Arel1 |
| Gnl34 | 1.62E-57 | 0.74856568 | 0.571 | 0.328 | 3.14E-53 | osteoclast.like | Gnl3 |
| Vars2 | 1.70E-57 | 0.75476353 | 0.585 | 0.323 | 3.29E-53 | osteoclast.like | Vars |
| Oma1 | 1.78E-57 | 1.47257704 | 0.332 | 0.092 | 3.45E-53 | osteoclast.like | Oma1 |
| Alg10b1 | 1.79E-57 | 1.02274744 | 0.58 | 0.255 | 3.47E-53 | osteoclast.like | Alg10b |
| Pla2g12a | 3.27E-57 | 1.5833862 | 0.395 | 0.09 | 6.32E-53 | osteoclast.like | Pla2g12a |
| Arhgef121 | 3.62E-57 | 1.19172268 | 0.449 | 0.153 | 6.99E-53 | osteoclast.like | Arhgef12 |
| Cln82 | 4.34E-57 | 0.41970521 | 0.741 | 0.487 | 8.40E-53 | osteoclast.like | Cln8 |
| Mta11 | 4.40E-57 | 0.81475341 | 0.659 | 0.319 | 8.50E-53 | osteoclast.like | Mta1 |
| Srsf61 | 4.43E-57 | 0.25923328 | 0.863 | 0.681 | 8.57E-53 | osteoclast.like | Srsf6 |
| Gatad2a1 | 4.56E-57 | 0.4761397 | 0.727 | 0.469 | 8.82E-53 | osteoclast.like | Gatad2a |
| Rangrf | 4.85E-57 | 1.98970577 | 0.298 | 0.067 | 9.37E-53 | osteoclast.like | Rangrf |
| Zfp768 | 5.55E-57 | 1.54913857 | 0.322 | 0.088 | 1.07E-52 | osteoclast.like | Zfp768 |
| Rnf2201 | 6.18E-57 | 0.54909948 | 0.727 | 0.419 | 1.19E-52 | osteoclast.like | Rnf220 |
| Prcc1 | 6.28E-57 | 0.58572911 | 0.688 | 0.372 | 1.21E-52 | osteoclast.like | Prcc |
| Banf11 | 6.36E-57 | 1.2269723 | 0.498 | 0.185 | 1.23E-52 | osteoclast.like | Banf1 |
| Ntmt1 | 6.70E-57 | 1.39400944 | 0.385 | 0.118 | 1.30E-52 | osteoclast.like | Ntmt1 |
| Elof11 | 8.57E-57 | 1.09180469 | 0.59 | 0.211 | 1.66E-52 | osteoclast.like | Elof1 |
| Rrs12 | 1.26E-56 | 1.26467688 | 0.483 | 0.175 | 2.44E-52 | osteoclast.like | Rrs1 |
| Cep70 | 1.29E-56 | 1.54082177 | 0.317 | 0.084 | 2.49E-52 | osteoclast.like | Cep70 |
| Cul1 | 1.30E-56 | 0.37653865 | 0.702 | 0.483 | 2.52E-52 | osteoclast.like | Cul1 |
| Mapkap11 | 1.60E-56 | 1.02188104 | 0.498 | 0.243 | 3.09E-52 | osteoclast.like | Mapkap1 |
| Glod41 | 1.67E-56 | 0.97096162 | 0.605 | 0.285 | 3.22E-52 | osteoclast.like | Glod4 |
| Prxl2c1 | 2.03E-56 | 1.02589629 | 0.488 | 0.2 | 3.93E-52 | osteoclast.like | Prxl2c |
| Ints101 | 3.14E-56 | 1.39038476 | 0.385 | 0.116 | 6.07E-52 | osteoclast.like | Ints10 |
| Cenpx1 | 3.29E-56 | 0.61512658 | 0.698 | 0.359 | 6.37E-52 | osteoclast.like | Cenpx |
| Gar12 | 3.50E-56 | 0.98910022 | 0.522 | 0.221 | 6.76E-52 | osteoclast.like | Gar1 |
| Nop533 | 3.50E-56 | 0.26405161 | 0.8 | 0.544 | 6.78E-52 | osteoclast.like | Nop53 |
| Afg1l | 3.93E-56 | 2.01420424 | 0.283 | 0.052 | 7.59E-52 | osteoclast.like | Afg1l |
| Erbin1 | 4.65E-56 | 0.31629305 | 0.8 | 0.546 | 8.98E-52 | osteoclast.like | Erbin |
| Tmem2511 | 5.39E-56 | 0.5532108 | 0.634 | 0.332 | 1.04E-51 | osteoclast.like | Tmem251 |
| Nprl3 | 5.95E-56 | 1.57298798 | 0.405 | 0.11 | 1.15E-51 | osteoclast.like | Nprl3 |
| Taf1d2 | 6.58E-56 | 0.41947394 | 0.668 | 0.435 | 1.27E-51 | osteoclast.like | Taf1d |
| Mcrip11 | 8.20E-56 | 0.51272416 | 0.727 | 0.449 | 1.59E-51 | osteoclast.like | Mcrip1 |
| Asb83 | 8.21E-56 | 0.97843599 | 0.473 | 0.205 | 1.59E-51 | osteoclast.like | Asb8 |
| Tmem2422 | 1.27E-55 | 0.7321493 | 0.59 | 0.303 | 2.46E-51 | osteoclast.like | Tmem242 |
| Hnrnpab2 | 1.40E-55 | 0.30039295 | 0.771 | 0.528 | 2.71E-51 | osteoclast.like | Hnrnpab |
| Mat2b | 1.85E-55 | 0.81281457 | 0.532 | 0.239 | 3.57E-51 | osteoclast.like | Mat2b |
| Ccz11 | 1.92E-55 | 1.01209491 | 0.556 | 0.243 | 3.71E-51 | osteoclast.like | Ccz1 |
| Hspa4l2 | 1.98E-55 | 0.6906755 | 0.537 | 0.236 | 3.83E-51 | osteoclast.like | Hspa4l |
| Spire13 | 2.05E-55 | 1.08284666 | 0.512 | 0.189 | 3.96E-51 | osteoclast.like | Spire1 |
| Ddx39b1 | 2.12E-55 | 0.5025969 | 0.727 | 0.465 | 4.10E-51 | osteoclast.like | Ddx39b |
| Wrb | 3.16E-55 | 2.07425899 | 0.263 | 0.046 | 6.11E-51 | osteoclast.like | Wrb |
| Fbxo4 | 4.17E-55 | 1.2878354 | 0.312 | 0.076 | 8.06E-51 | osteoclast.like | Fbxo4 |
| Lag32 | 4.82E-55 | 0.71996168 | 0.395 | 0.164 | 9.32E-51 | osteoclast.like | Lag3 |
| Znrf2 | 5.06E-55 | 0.27020934 | 0.707 | 0.517 | 9.78E-51 | osteoclast.like | Znrf2 |
| Ganab1 | 5.66E-55 | 0.36832943 | 0.756 | 0.517 | 1.09E-50 | osteoclast.like | Ganab |
| Spred2 | 6.37E-55 | 0.90219068 | 0.507 | 0.209 | 1.23E-50 | osteoclast.like | Spred2 |
| Ppp2r2a3 | 6.43E-55 | 0.66729932 | 0.537 | 0.263 | 1.24E-50 | osteoclast.like | Ppp2r2a |
| Fam32a | 7.22E-55 | 0.33914212 | 0.756 | 0.509 | 1.40E-50 | osteoclast.like | Fam32a |
| Plekhb22 | 8.19E-55 | 0.25361085 | 0.824 | 0.607 | 1.58E-50 | osteoclast.like | Plekhb2 |
| Coq21 | 8.68E-55 | 0.90825664 | 0.488 | 0.174 | 1.68E-50 | osteoclast.like | Coq2 |
| Mink13 | 8.98E-55 | 0.59204889 | 0.746 | 0.419 | 1.74E-50 | osteoclast.like | Mink1 |
| Ssh1 | 1.02E-54 | 0.71152247 | 0.683 | 0.324 | 1.98E-50 | osteoclast.like | Ssh1 |
| Atp9b1 | 1.05E-54 | 0.56456229 | 0.678 | 0.414 | 2.02E-50 | osteoclast.like | Atp9b |
| Usp19 | 1.22E-54 | 0.64102474 | 0.707 | 0.386 | 2.35E-50 | osteoclast.like | Usp19 |
| Nudcd1 | 1.42E-54 | 1.8484786 | 0.273 | 0.058 | 2.74E-50 | osteoclast.like | Nudcd1 |
| Fbxo323 | 1.51E-54 | 1.0736537 | 0.463 | 0.146 | 2.91E-50 | osteoclast.like | Fbxo32 |
| Telo2 | 1.57E-54 | 1.64777563 | 0.327 | 0.069 | 3.04E-50 | osteoclast.like | Telo2 |
| Ttll121 | 1.60E-54 | 1.33992354 | 0.351 | 0.103 | 3.10E-50 | osteoclast.like | Ttll12 |
| Rab182 | 1.71E-54 | 0.27666129 | 0.824 | 0.575 | 3.30E-50 | osteoclast.like | Rab18 |
| Nolc11 | 1.83E-54 | 1.15440224 | 0.444 | 0.172 | 3.53E-50 | osteoclast.like | Nolc1 |
| Zfp1312 | 1.87E-54 | 0.43652428 | 0.644 | 0.437 | 3.62E-50 | osteoclast.like | Zfp131 |
| Pfdn5 | 2.04E-54 | 0.27592956 | 0.776 | 0.549 | 3.95E-50 | osteoclast.like | Pfdn5 |
| Rnf1461 | 2.35E-54 | 0.61801899 | 0.463 | 0.22 | 4.53E-50 | osteoclast.like | Rnf146 |
| Gars1 | 2.45E-54 | 1.19774461 | 0.405 | 0.139 | 4.75E-50 | osteoclast.like | Gars |
| Tmem143 | 2.62E-54 | 1.84666058 | 0.312 | 0.066 | 5.06E-50 | osteoclast.like | Tmem143 |
| Mtch21 | 3.29E-54 | 1.26552826 | 0.488 | 0.162 | 6.37E-50 | osteoclast.like | Mtch2 |
| Ddx41 | 3.49E-54 | 0.77889316 | 0.624 | 0.31 | 6.76E-50 | osteoclast.like | Ddx41 |
| Usp33 | 4.10E-54 | 0.65396403 | 0.522 | 0.244 | 7.93E-50 | osteoclast.like | Usp33 |
| Cdc5l1 | 4.30E-54 | 0.70895575 | 0.576 | 0.283 | 8.31E-50 | osteoclast.like | Cdc5l |
| Ube2l31 | 4.34E-54 | 0.5208372 | 0.624 | 0.384 | 8.40E-50 | osteoclast.like | Ube2l3 |
| Slc30a13 | 5.05E-54 | 0.47496989 | 0.663 | 0.371 | 9.77E-50 | osteoclast.like | Slc30a1 |
| Emc32 | 8.62E-54 | 0.70932244 | 0.527 | 0.272 | 1.67E-49 | osteoclast.like | Emc3 |
| Itga53 | 9.05E-54 | 0.28536267 | 0.815 | 0.536 | 1.75E-49 | osteoclast.like | Itga5 |
| Ranbp10 | 9.45E-54 | 1.14652198 | 0.522 | 0.174 | 1.83E-49 | osteoclast.like | Ranbp10 |
| Cacfd1 | 1.00E-53 | 1.31396356 | 0.361 | 0.116 | 1.94E-49 | osteoclast.like | Cacfd1 |
| BC0039651 | 1.09E-53 | 1.14601667 | 0.444 | 0.14 | 2.11E-49 | osteoclast.like | BC003965 |
| Ufsp21 | 1.11E-53 | 0.76715474 | 0.585 | 0.3 | 2.14E-49 | osteoclast.like | Ufsp2 |
| Fam192a1 | 1.15E-53 | 0.84716039 | 0.6 | 0.263 | 2.23E-49 | osteoclast.like | Fam192a |
| Ufc11 | 1.17E-53 | 0.44072254 | 0.79 | 0.518 | 2.26E-49 | osteoclast.like | Ufc1 |
| Supt4a1 | 1.26E-53 | 0.42265639 | 0.717 | 0.485 | 2.44E-49 | osteoclast.like | Supt4a |
| Chmp31 | 1.38E-53 | 0.52215643 | 0.761 | 0.477 | 2.66E-49 | osteoclast.like | Chmp3 |
| Snrpd21 | 1.51E-53 | 1.04342135 | 0.527 | 0.2 | 2.93E-49 | osteoclast.like | Snrpd2 |
| Chmp51 | 1.91E-53 | 0.39620401 | 0.78 | 0.534 | 3.70E-49 | osteoclast.like | Chmp5 |
| Slc35a31 | 1.93E-53 | 0.8627805 | 0.576 | 0.268 | 3.73E-49 | osteoclast.like | Slc35a3 |
| Agl | 1.96E-53 | 1.19842015 | 0.439 | 0.13 | 3.80E-49 | osteoclast.like | Agl |
| Cuta2 | 1.99E-53 | 0.41178859 | 0.78 | 0.536 | 3.84E-49 | osteoclast.like | Cuta |
| Psmd62 | 2.06E-53 | 0.55480966 | 0.629 | 0.369 | 3.98E-49 | osteoclast.like | Psmd6 |
| Acsf3 | 2.20E-53 | 1.28367363 | 0.376 | 0.1 | 4.26E-49 | osteoclast.like | Acsf3 |
| Pus12 | 2.78E-53 | 0.88038562 | 0.571 | 0.257 | 5.38E-49 | osteoclast.like | Pus1 |
| Eif4h1 | 2.84E-53 | 0.65238193 | 0.683 | 0.382 | 5.48E-49 | osteoclast.like | Eif4h |
| Ei241 | 5.01E-53 | 1.11579858 | 0.512 | 0.182 | 9.68E-49 | osteoclast.like | Ei24 |
| Tmem161b | 5.04E-53 | 1.41189269 | 0.385 | 0.108 | 9.75E-49 | osteoclast.like | Tmem161b |
| Gatd3a2 | 5.23E-53 | 0.91843316 | 0.537 | 0.224 | 1.01E-48 | osteoclast.like | Gatd3a |
| Impa11 | 5.94E-53 | 0.74967649 | 0.644 | 0.33 | 1.15E-48 | osteoclast.like | Impa1 |
| Ap1m11 | 6.81E-53 | 0.78384982 | 0.59 | 0.292 | 1.32E-48 | osteoclast.like | Ap1m1 |
| Rnf52 | 6.94E-53 | 0.95699052 | 0.59 | 0.268 | 1.34E-48 | osteoclast.like | Rnf5 |
| Ube4b2 | 7.69E-53 | 0.54688121 | 0.605 | 0.325 | 1.49E-48 | osteoclast.like | Ube4b |
| Abraxas2 | 8.47E-53 | 0.99230955 | 0.502 | 0.187 | 1.64E-48 | osteoclast.like | Abraxas2 |
| Slc39a112 | 8.77E-53 | 0.67241222 | 0.668 | 0.321 | 1.70E-48 | osteoclast.like | Slc39a11 |
| Dus3l1 | 9.13E-53 | 0.64722735 | 0.58 | 0.259 | 1.76E-48 | osteoclast.like | Dus3l |
| Usp252 | 9.61E-53 | 0.25694963 | 0.771 | 0.51 | 1.86E-48 | osteoclast.like | Usp25 |
| Prmt51 | 1.05E-52 | 1.48415158 | 0.439 | 0.155 | 2.04E-48 | osteoclast.like | Prmt5 |
| Prps1l31 | 1.35E-52 | 1.09109539 | 0.405 | 0.147 | 2.61E-48 | osteoclast.like | Prps1l3 |
| Psmd113 | 1.45E-52 | 0.39125515 | 0.707 | 0.454 | 2.81E-48 | osteoclast.like | Psmd11 |
| Bag61 | 1.61E-52 | 0.4022038 | 0.756 | 0.485 | 3.12E-48 | osteoclast.like | Bag6 |
| Hspa42 | 1.64E-52 | 0.8119071 | 0.659 | 0.371 | 3.18E-48 | osteoclast.like | Hspa4 |
| Chchd7 | 1.98E-52 | 1.07597261 | 0.41 | 0.155 | 3.83E-48 | osteoclast.like | Chchd7 |
| Pnpla72 | 2.96E-52 | 0.61628224 | 0.688 | 0.389 | 5.73E-48 | osteoclast.like | Pnpla7 |
| Gemin5 | 3.00E-52 | 1.49518406 | 0.341 | 0.092 | 5.80E-48 | osteoclast.like | Gemin5 |
| Mpp12 | 3.31E-52 | 0.31128189 | 0.507 | 0.381 | 6.40E-48 | osteoclast.like | Mpp1 |
| Acvr11 | 3.65E-52 | 0.96924016 | 0.459 | 0.17 | 7.05E-48 | osteoclast.like | Acvr1 |
| Zzz3 | 3.72E-52 | 0.66444921 | 0.585 | 0.312 | 7.19E-48 | osteoclast.like | Zzz3 |
| Pigk | 3.79E-52 | 0.74596176 | 0.507 | 0.237 | 7.33E-48 | osteoclast.like | Pigk |
| Ppp1cb1 | 3.86E-52 | 0.34282354 | 0.761 | 0.492 | 7.46E-48 | osteoclast.like | Ppp1cb |
| Cdk41 | 3.89E-52 | 0.94099554 | 0.483 | 0.222 | 7.52E-48 | osteoclast.like | Cdk4 |
| Psmd32 | 4.08E-52 | 0.76165013 | 0.659 | 0.365 | 7.88E-48 | osteoclast.like | Psmd3 |
| Dctn2 | 4.88E-52 | 0.32898498 | 0.854 | 0.617 | 9.44E-48 | osteoclast.like | Dctn2 |
| Hgh1 | 7.00E-52 | 1.64172732 | 0.288 | 0.066 | 1.35E-47 | osteoclast.like | Hgh1 |
| Trappc31 | 7.09E-52 | 0.51289845 | 0.707 | 0.421 | 1.37E-47 | osteoclast.like | Trappc3 |
| D6Wsu163e1 | 7.26E-52 | 1.28815562 | 0.376 | 0.136 | 1.40E-47 | osteoclast.like | D6Wsu163e |
| Tmbim41 | 7.39E-52 | 0.43124409 | 0.683 | 0.405 | 1.43E-47 | osteoclast.like | Tmbim4 |
| Hps11 | 8.15E-52 | 1.09374388 | 0.566 | 0.274 | 1.58E-47 | osteoclast.like | Hps1 |
| Rhbdf1 | 8.98E-52 | 2.16291911 | 0.278 | 0.049 | 1.74E-47 | osteoclast.like | Rhbdf1 |
| Higd2a | 9.95E-52 | 0.50608011 | 0.668 | 0.394 | 1.92E-47 | osteoclast.like | Higd2a |
| Mbp1 | 1.28E-51 | 0.63834274 | 0.605 | 0.321 | 2.47E-47 | osteoclast.like | Mbp |
| Cars3 | 1.39E-51 | 1.02773775 | 0.468 | 0.19 | 2.70E-47 | osteoclast.like | Cars |
| Prkci1 | 1.46E-51 | 1.18575082 | 0.434 | 0.146 | 2.83E-47 | osteoclast.like | Prkci |
| Ppp2r5c | 1.58E-51 | 0.28684379 | 0.805 | 0.547 | 3.05E-47 | osteoclast.like | Ppp2r5c |
| Erlin22 | 1.68E-51 | 0.75008471 | 0.624 | 0.303 | 3.24E-47 | osteoclast.like | Erlin2 |
| Cnbp2 | 1.84E-51 | 0.27496186 | 0.761 | 0.551 | 3.56E-47 | osteoclast.like | Cnbp |
| Psmc62 | 1.84E-51 | 0.46061496 | 0.732 | 0.468 | 3.57E-47 | osteoclast.like | Psmc6 |
| Dusp73 | 1.93E-51 | 0.401711 | 0.698 | 0.421 | 3.73E-47 | osteoclast.like | Dusp7 |
| Ccar11 | 2.02E-51 | 0.59562338 | 0.6 | 0.357 | 3.90E-47 | osteoclast.like | Ccar1 |
| Rufy11 | 2.25E-51 | 0.70013844 | 0.566 | 0.274 | 4.36E-47 | osteoclast.like | Rufy1 |
| Col15a1 | 2.64E-51 | 1.33228533 | 0.307 | 0.091 | 5.10E-47 | osteoclast.like | Col15a1 |
| Osbp1 | 2.77E-51 | 0.55838801 | 0.576 | 0.309 | 5.35E-47 | osteoclast.like | Osbp |
| Gatb | 3.61E-51 | 1.48595738 | 0.312 | 0.082 | 6.99E-47 | osteoclast.like | Gatb |
| Clpx | 4.11E-51 | 1.28560376 | 0.4 | 0.139 | 7.95E-47 | osteoclast.like | Clpx |
| Krcc12 | 5.10E-51 | 0.6799772 | 0.541 | 0.288 | 9.86E-47 | osteoclast.like | Krcc1 |
| Plekha21 | 5.23E-51 | 0.50034384 | 0.659 | 0.434 | 1.01E-46 | osteoclast.like | Plekha2 |
| Rad211 | 5.34E-51 | 0.67488108 | 0.644 | 0.327 | 1.03E-46 | osteoclast.like | Rad21 |
| Nop563 | 5.83E-51 | 0.54139314 | 0.644 | 0.376 | 1.13E-46 | osteoclast.like | Nop56 |
| Akt1s13 | 7.41E-51 | 0.88124645 | 0.556 | 0.25 | 1.43E-46 | osteoclast.like | Akt1s1 |
| Pfdn41 | 7.41E-51 | 1.15320284 | 0.429 | 0.151 | 1.43E-46 | osteoclast.like | Pfdn4 |
| Mthfd22 | 8.71E-51 | 0.84956701 | 0.332 | 0.167 | 1.68E-46 | osteoclast.like | Mthfd2 |
| Ccdc91 | 9.27E-51 | 1.40497616 | 0.366 | 0.096 | 1.79E-46 | osteoclast.like | Ccdc91 |
| Calu | 1.03E-50 | 1.75885205 | 0.366 | 0.094 | 1.99E-46 | osteoclast.like | Calu |
| Rgl22 | 1.09E-50 | 0.41459734 | 0.722 | 0.47 | 2.11E-46 | osteoclast.like | Rgl2 |
| Ten1 | 1.12E-50 | 0.81361031 | 0.6 | 0.267 | 2.16E-46 | osteoclast.like | Ten1 |
| Gspt12 | 1.18E-50 | 0.62936255 | 0.615 | 0.355 | 2.27E-46 | osteoclast.like | Gspt1 |
| Tfg1 | 1.40E-50 | 0.52748514 | 0.688 | 0.418 | 2.70E-46 | osteoclast.like | Tfg |
| Pbxip13 | 1.43E-50 | 0.62767826 | 0.517 | 0.256 | 2.77E-46 | osteoclast.like | Pbxip1 |
| Synj2bp | 1.48E-50 | 0.77066005 | 0.595 | 0.307 | 2.86E-46 | osteoclast.like | Synj2bp |
| Usp4 | 1.55E-50 | 0.51676716 | 0.698 | 0.422 | 3.00E-46 | osteoclast.like | Usp4 |
| Naxd2 | 1.93E-50 | 0.85615104 | 0.541 | 0.22 | 3.74E-46 | osteoclast.like | Naxd |
| Cops7a3 | 2.01E-50 | 0.70247255 | 0.576 | 0.284 | 3.89E-46 | osteoclast.like | Cops7a |
| Deptor1 | 2.21E-50 | 0.71280384 | 0.571 | 0.236 | 4.28E-46 | osteoclast.like | Deptor |
| Cops52 | 2.24E-50 | 0.49981081 | 0.634 | 0.346 | 4.34E-46 | osteoclast.like | Cops5 |
| Wdr45 | 2.32E-50 | 1.10577444 | 0.459 | 0.165 | 4.48E-46 | osteoclast.like | Wdr45 |
| Ano8 | 2.36E-50 | 1.38972029 | 0.38 | 0.115 | 4.56E-46 | osteoclast.like | Ano8 |
| Med293 | 2.40E-50 | 0.66644238 | 0.561 | 0.279 | 4.64E-46 | osteoclast.like | Med29 |
| Smim191 | 2.96E-50 | 1.04781645 | 0.4 | 0.148 | 5.72E-46 | osteoclast.like | Smim19 |
| Rabl63 | 3.18E-50 | 0.70158359 | 0.644 | 0.337 | 6.15E-46 | osteoclast.like | Rabl6 |
| Itfg1 | 3.83E-50 | 0.62924366 | 0.595 | 0.31 | 7.40E-46 | osteoclast.like | Itfg1 |
| Nr2f6 | 4.42E-50 | 1.32833384 | 0.459 | 0.168 | 8.55E-46 | osteoclast.like | Nr2f6 |
| Irf2bp11 | 7.84E-50 | 0.91283849 | 0.424 | 0.155 | 1.52E-45 | osteoclast.like | Irf2bp1 |
| Wipi22 | 8.25E-50 | 0.8291763 | 0.532 | 0.247 | 1.59E-45 | osteoclast.like | Wipi2 |
| Fibp1 | 8.96E-50 | 0.61493583 | 0.639 | 0.354 | 1.73E-45 | osteoclast.like | Fibp |
| Coa6 | 9.07E-50 | 1.38445075 | 0.341 | 0.115 | 1.75E-45 | osteoclast.like | Coa6 |
| Ccny | 9.15E-50 | 0.58216884 | 0.683 | 0.386 | 1.77E-45 | osteoclast.like | Ccny |
| Csrp11 | 1.14E-49 | 0.69898232 | 0.502 | 0.234 | 2.20E-45 | osteoclast.like | Csrp1 |
| Paip2 | 1.19E-49 | 0.31437758 | 0.727 | 0.513 | 2.30E-45 | osteoclast.like | Paip2 |
| Vps522 | 1.20E-49 | 0.58827921 | 0.59 | 0.304 | 2.31E-45 | osteoclast.like | Vps52 |
| Lamtor31 | 1.47E-49 | 0.32304186 | 0.78 | 0.538 | 2.84E-45 | osteoclast.like | Lamtor3 |
| Dus1l1 | 1.83E-49 | 1.03773761 | 0.468 | 0.172 | 3.55E-45 | osteoclast.like | Dus1l |
| Kif1c2 | 2.26E-49 | 0.44648236 | 0.698 | 0.431 | 4.37E-45 | osteoclast.like | Kif1c |
| Mak161 | 2.41E-49 | 1.11849198 | 0.498 | 0.193 | 4.67E-45 | osteoclast.like | Mak16 |
| Slc26a2 | 2.51E-49 | 0.63426067 | 0.473 | 0.244 | 4.85E-45 | osteoclast.like | Slc26a2 |
| Cmc1 | 2.57E-49 | 0.3490768 | 0.61 | 0.356 | 4.97E-45 | osteoclast.like | Cmc1 |
| Wbp1l1 | 2.60E-49 | 0.3457902 | 0.712 | 0.494 | 5.03E-45 | osteoclast.like | Wbp1l |
| Commd11 | 2.64E-49 | 0.41330395 | 0.698 | 0.433 | 5.11E-45 | osteoclast.like | Commd1 |
| Thop11 | 2.79E-49 | 1.20644662 | 0.39 | 0.155 | 5.40E-45 | osteoclast.like | Thop1 |
| Fam120a1 | 2.85E-49 | 0.45711547 | 0.673 | 0.418 | 5.51E-45 | osteoclast.like | Fam120a |
| Ola12 | 3.80E-49 | 0.67151868 | 0.576 | 0.334 | 7.35E-45 | osteoclast.like | Ola1 |
| Spast2 | 4.68E-49 | 0.70796959 | 0.571 | 0.28 | 9.05E-45 | osteoclast.like | Spast |
| Sf3a21 | 4.90E-49 | 0.62264088 | 0.634 | 0.343 | 9.47E-45 | osteoclast.like | Sf3a2 |
| Fam160a2 | 6.69E-49 | 1.33397859 | 0.322 | 0.073 | 1.29E-44 | osteoclast.like | Fam160a2 |
| Cisd3 | 7.14E-49 | 0.8257695 | 0.488 | 0.213 | 1.38E-44 | osteoclast.like | Cisd3 |
| Tbc1d232 | 7.16E-49 | 0.90304868 | 0.488 | 0.191 | 1.38E-44 | osteoclast.like | Tbc1d23 |
| Gmds1 | 8.01E-49 | 1.2557179 | 0.371 | 0.118 | 1.55E-44 | osteoclast.like | Gmds |
| Dcaf1 | 1.35E-48 | 1.03162265 | 0.4 | 0.145 | 2.60E-44 | osteoclast.like | Dcaf1 |
| Mapre21 | 1.62E-48 | 0.77494236 | 0.629 | 0.325 | 3.13E-44 | osteoclast.like | Mapre2 |
| Prpf312 | 1.73E-48 | 0.66312255 | 0.61 | 0.327 | 3.35E-44 | osteoclast.like | Prpf31 |
| Trmt1121 | 1.76E-48 | 0.30129178 | 0.746 | 0.542 | 3.41E-44 | osteoclast.like | Trmt112 |
| Cdk8 | 1.91E-48 | 0.66377905 | 0.571 | 0.302 | 3.69E-44 | osteoclast.like | Cdk8 |
| Wdr611 | 2.04E-48 | 0.86639827 | 0.42 | 0.169 | 3.95E-44 | osteoclast.like | Wdr61 |
| Hdac31 | 3.02E-48 | 0.7068739 | 0.58 | 0.285 | 5.84E-44 | osteoclast.like | Hdac3 |
| Fbxo421 | 3.16E-48 | 0.88231602 | 0.566 | 0.267 | 6.10E-44 | osteoclast.like | Fbxo42 |
| Tusc2 | 3.28E-48 | 0.97326558 | 0.444 | 0.153 | 6.35E-44 | osteoclast.like | Tusc2 |
| DEPRECATED-I | 3.29E-48 | 1.42702326 | 0.38 | 0.095 | 6.37E-44 | osteoclast.like | DEPRECATED-ENSMUSG00000050608 |
| Tesk13 | 3.38E-48 | 0.58945188 | 0.678 | 0.366 | 6.53E-44 | osteoclast.like | Tesk1 |
| Fem1a | 3.54E-48 | 1.21492988 | 0.395 | 0.135 | 6.85E-44 | osteoclast.like | Fem1a |
| Rpf11 | 3.75E-48 | 0.63880531 | 0.561 | 0.265 | 7.26E-44 | osteoclast.like | Rpf1 |
| Eif3e1 | 4.72E-48 | 0.84541941 | 0.483 | 0.21 | 9.12E-44 | osteoclast.like | Eif3e |
| Dvl11 | 4.87E-48 | 0.74249578 | 0.541 | 0.245 | 9.41E-44 | osteoclast.like | Dvl1 |
| Pmf11 | 5.55E-48 | 1.21298282 | 0.468 | 0.173 | 1.07E-43 | osteoclast.like | Pmf1 |
| Pip4p22 | 5.56E-48 | 1.16922313 | 0.449 | 0.165 | 1.08E-43 | osteoclast.like | Pip4p2 |
| Impad1 | 5.80E-48 | 0.63285996 | 0.585 | 0.324 | 1.12E-43 | osteoclast.like | Impad1 |
| Smim26 | 6.62E-48 | 1.28762221 | 0.366 | 0.106 | 1.28E-43 | osteoclast.like | Smim26 |
| Cyld | 6.62E-48 | 0.45188097 | 0.595 | 0.321 | 1.28E-43 | osteoclast.like | Cyld |
| Ppie1 | 6.94E-48 | 1.05046545 | 0.493 | 0.202 | 1.34E-43 | osteoclast.like | Ppie |
| Nomo12 | 7.33E-48 | 0.97466044 | 0.517 | 0.214 | 1.42E-43 | osteoclast.like | Nomo1 |
| Phf12 | 7.65E-48 | 0.51172928 | 0.707 | 0.409 | 1.48E-43 | osteoclast.like | Phf12 |
| Ccdc62 | 9.44E-48 | 0.61045186 | 0.62 | 0.324 | 1.82E-43 | osteoclast.like | Ccdc6 |
| Tmem1411 | 1.06E-47 | 0.92923605 | 0.424 | 0.163 | 2.05E-43 | osteoclast.like | Tmem141 |
| Pex16 | 1.11E-47 | 1.54897425 | 0.346 | 0.101 | 2.15E-43 | osteoclast.like | Pex16 |
| N4bp2 | 1.19E-47 | 1.18264623 | 0.332 | 0.091 | 2.31E-43 | osteoclast.like | N4bp2 |
| Thg1l | 1.28E-47 | 1.71104021 | 0.322 | 0.077 | 2.48E-43 | osteoclast.like | Thg1l |
| Isoc2b | 1.81E-47 | 1.40228245 | 0.307 | 0.08 | 3.49E-43 | osteoclast.like | Isoc2b |
| Chrac11 | 2.41E-47 | 0.78300543 | 0.537 | 0.232 | 4.67E-43 | osteoclast.like | Chrac1 |
| Sqor3 | 2.42E-47 | 0.62978299 | 0.517 | 0.324 | 4.67E-43 | osteoclast.like | Sqor |
| Selenot2 | 3.31E-47 | 0.31940487 | 0.61 | 0.406 | 6.41E-43 | osteoclast.like | Selenot |
| Psmg11 | 3.73E-47 | 1.16276382 | 0.385 | 0.143 | 7.20E-43 | osteoclast.like | Psmg1 |
| Vapb1 | 3.76E-47 | 0.52262928 | 0.702 | 0.427 | 7.27E-43 | osteoclast.like | Vapb |
| Abcc12 | 4.15E-47 | 0.46618088 | 0.79 | 0.409 | 8.02E-43 | osteoclast.like | Abcc1 |
| Cox201 | 4.51E-47 | 1.20356822 | 0.459 | 0.164 | 8.72E-43 | osteoclast.like | Cox20 |
| Tmem199 | 4.71E-47 | 1.34467001 | 0.41 | 0.137 | 9.12E-43 | osteoclast.like | Tmem199 |
| Trmt1l | 5.07E-47 | 0.81517784 | 0.439 | 0.168 | 9.81E-43 | osteoclast.like | Trmt1l |
| Aggf1 | 6.14E-47 | 0.54640429 | 0.566 | 0.317 | 1.19E-42 | osteoclast.like | Aggf1 |
| Pdk2 | 6.73E-47 | 1.55247854 | 0.351 | 0.091 | 1.30E-42 | osteoclast.like | Pdk2 |
| Cdv3 | 6.90E-47 | 0.48968829 | 0.693 | 0.426 | 1.33E-42 | osteoclast.like | Cdv3 |
| Caprin11 | 7.07E-47 | 0.57466344 | 0.673 | 0.434 | 1.37E-42 | osteoclast.like | Caprin1 |
| Cog4 | 8.50E-47 | 1.1741286 | 0.39 | 0.114 | 1.64E-42 | osteoclast.like | Cog4 |
| AU0403202 | 9.16E-47 | 0.78026767 | 0.459 | 0.217 | 1.77E-42 | osteoclast.like | AU040320 |
| Micall11 | 9.81E-47 | 1.0077043 | 0.371 | 0.135 | 1.90E-42 | osteoclast.like | Micall1 |
| Trub1 | 1.10E-46 | 1.87979144 | 0.317 | 0.067 | 2.13E-42 | osteoclast.like | Trub1 |
| Cul4a1 | 1.24E-46 | 0.70705683 | 0.615 | 0.324 | 2.40E-42 | osteoclast.like | Cul4a |
| Fubp12 | 1.26E-46 | 0.38229061 | 0.624 | 0.447 | 2.44E-42 | osteoclast.like | Fubp1 |
| Ptges33 | 1.34E-46 | 0.59968891 | 0.58 | 0.376 | 2.60E-42 | osteoclast.like | Ptges3 |
| Zdhhc51 | 1.65E-46 | 0.36584757 | 0.702 | 0.48 | 3.18E-42 | osteoclast.like | Zdhhc5 |
| Trmt11 | 1.69E-46 | 1.2496916 | 0.449 | 0.154 | 3.27E-42 | osteoclast.like | Trmt1 |
| Eif3b2 | 1.91E-46 | 0.35209222 | 0.737 | 0.455 | 3.69E-42 | osteoclast.like | Eif3b |
| Nubp12 | 2.15E-46 | 0.47676143 | 0.683 | 0.428 | 4.15E-42 | osteoclast.like | Nubp1 |
| Ttc28 | 2.49E-46 | 2.14514948 | 0.283 | 0.048 | 4.81E-42 | osteoclast.like | Ttc28 |
| Zfp912 | 2.51E-46 | 0.27622583 | 0.766 | 0.525 | 4.85E-42 | osteoclast.like | Zfp91 |
| Fnta2 | 2.66E-46 | 0.55675616 | 0.615 | 0.313 | 5.13E-42 | osteoclast.like | Fnta |
| Slc39a72 | 2.76E-46 | 0.44110244 | 0.756 | 0.484 | 5.33E-42 | osteoclast.like | Slc39a7 |
| Map2k7 | 2.76E-46 | 0.58248175 | 0.561 | 0.294 | 5.33E-42 | osteoclast.like | Map2k7 |
| Prmt71 | 2.90E-46 | 1.06630579 | 0.371 | 0.133 | 5.61E-42 | osteoclast.like | Prmt7 |
| Prpf4b | 2.97E-46 | 0.33880584 | 0.644 | 0.402 | 5.74E-42 | osteoclast.like | Prpf4b |
| Rasa31 | 3.21E-46 | 0.37248463 | 0.615 | 0.405 | 6.21E-42 | osteoclast.like | Rasa3 |
| Dmac2 | 3.60E-46 | 1.0556483 | 0.366 | 0.139 | 6.95E-42 | osteoclast.like | Dmac2 |
| Tnrc181 | 4.06E-46 | 0.36766845 | 0.771 | 0.528 | 7.84E-42 | osteoclast.like | Tnrc18 |
| Tm9sf11 | 4.07E-46 | 0.54010142 | 0.668 | 0.354 | 7.86E-42 | osteoclast.like | Tm9sf1 |
| Hs6st11 | 4.62E-46 | 0.43576868 | 0.673 | 0.432 | 8.94E-42 | osteoclast.like | Hs6st1 |
| Ptpra2 | 4.93E-46 | 0.3426918 | 0.678 | 0.444 | 9.52E-42 | osteoclast.like | Ptpra |
| Foxp13 | 5.02E-46 | 0.46729121 | 0.585 | 0.341 | 9.70E-42 | osteoclast.like | Foxp1 |
| Tiprl1 | 5.03E-46 | 0.53784747 | 0.6 | 0.32 | 9.72E-42 | osteoclast.like | Tiprl |
| Wdr431 | 5.32E-46 | 1.11305002 | 0.415 | 0.187 | 1.03E-41 | osteoclast.like | Wdr43 |
| Mcat | 5.72E-46 | 1.6831289 | 0.263 | 0.069 | 1.11E-41 | osteoclast.like | Mcat |
| Srp141 | 6.26E-46 | 0.35914923 | 0.659 | 0.422 | 1.21E-41 | osteoclast.like | Srp14 |
| Fam241a | 6.91E-46 | 0.72778283 | 0.395 | 0.176 | 1.34E-41 | osteoclast.like | Fam241a |
| Ireb2 | 7.49E-46 | 0.76130612 | 0.532 | 0.253 | 1.45E-41 | osteoclast.like | Ireb2 |
| Kctd101 | 7.49E-46 | 0.47712983 | 0.595 | 0.378 | 1.45E-41 | osteoclast.like | Kctd10 |
| Ppan1 | 8.11E-46 | 0.86176821 | 0.429 | 0.173 | 1.57E-41 | osteoclast.like | Ppan |
| Nop582 | 8.30E-46 | 0.81984556 | 0.424 | 0.236 | 1.60E-41 | osteoclast.like | Nop58 |
| Trim143 | 8.49E-46 | 0.46442721 | 0.527 | 0.294 | 1.64E-41 | osteoclast.like | Trim14 |
| Lsm21 | 9.09E-46 | 1.01177077 | 0.512 | 0.208 | 1.76E-41 | osteoclast.like | Lsm2 |
| Cad1 | 9.41E-46 | 1.15045846 | 0.317 | 0.122 | 1.82E-41 | osteoclast.like | Cad |
| Sec61b2 | 9.47E-46 | 0.44400404 | 0.624 | 0.385 | 1.83E-41 | osteoclast.like | Sec61b |
| Nucks11 | 9.94E-46 | 0.96151799 | 0.498 | 0.233 | 1.92E-41 | osteoclast.like | Nucks1 |
| Rest1 | 1.03E-45 | 0.74132397 | 0.556 | 0.239 | 2.00E-41 | osteoclast.like | Rest |
| Ddx11 | 1.13E-45 | 0.66219123 | 0.512 | 0.284 | 2.19E-41 | osteoclast.like | Ddx1 |
| Sympk1 | 1.14E-45 | 0.53281199 | 0.595 | 0.33 | 2.20E-41 | osteoclast.like | Sympk |
| Nos1ap | 1.21E-45 | 1.71919118 | 0.293 | 0.066 | 2.35E-41 | osteoclast.like | Nos1ap |
| Uba21 | 1.26E-45 | 0.51801846 | 0.629 | 0.382 | 2.44E-41 | osteoclast.like | Uba2 |
| Gtf2b1 | 1.28E-45 | 0.39812456 | 0.649 | 0.389 | 2.48E-41 | osteoclast.like | Gtf2b |
| Snrpc1 | 1.29E-45 | 0.54813808 | 0.639 | 0.393 | 2.50E-41 | osteoclast.like | Snrpc |
| Yif1b1 | 1.38E-45 | 0.73849124 | 0.537 | 0.263 | 2.68E-41 | osteoclast.like | Yif1b |
| Tmem2451 | 1.50E-45 | 1.05364684 | 0.424 | 0.167 | 2.89E-41 | osteoclast.like | Tmem245 |
| Fbxw5 | 2.41E-45 | 0.7553514 | 0.546 | 0.241 | 4.65E-41 | osteoclast.like | Fbxw5 |
| Cct52 | 2.48E-45 | 0.93941502 | 0.493 | 0.231 | 4.80E-41 | osteoclast.like | Cct5 |
| Dut1 | 2.75E-45 | 1.38631612 | 0.356 | 0.075 | 5.31E-41 | osteoclast.like | Dut |
| Ubqln22 | 2.80E-45 | 0.57020042 | 0.624 | 0.349 | 5.41E-41 | osteoclast.like | Ubqln2 |
| Gtf3a1 | 2.93E-45 | 0.93909795 | 0.493 | 0.216 | 5.66E-41 | osteoclast.like | Gtf3a |
| Thoc71 | 3.08E-45 | 0.5985519 | 0.576 | 0.289 | 5.95E-41 | osteoclast.like | Thoc7 |
| Adnp1 | 3.12E-45 | 0.45131732 | 0.663 | 0.397 | 6.04E-41 | osteoclast.like | Adnp |
| Bsdc12 | 3.17E-45 | 0.8083207 | 0.546 | 0.268 | 6.12E-41 | osteoclast.like | Bsdc1 |
| Pam | 3.22E-45 | 1.12991541 | 0.337 | 0.107 | 6.23E-41 | osteoclast.like | Pam |
| Rer11 | 3.31E-45 | 0.3160846 | 0.678 | 0.464 | 6.40E-41 | osteoclast.like | Rer1 |
| Plaa2 | 3.32E-45 | 0.56011725 | 0.629 | 0.35 | 6.41E-41 | osteoclast.like | Plaa |
| Kti12 | 3.51E-45 | 1.252548 | 0.361 | 0.119 | 6.79E-41 | osteoclast.like | Kti12 |
| Vkorc1l11 | 3.67E-45 | 0.83161789 | 0.405 | 0.169 | 7.10E-41 | osteoclast.like | Vkorc1l1 |
| Vdac31 | 3.93E-45 | 0.81129402 | 0.527 | 0.249 | 7.60E-41 | osteoclast.like | Vdac3 |
| Cemip21 | 4.34E-45 | 0.56954947 | 0.546 | 0.264 | 8.39E-41 | osteoclast.like | Cemip2 |
| Rnmt2 | 4.64E-45 | 0.9340744 | 0.498 | 0.197 | 8.98E-41 | osteoclast.like | Rnmt |
| Bcs1l | 4.93E-45 | 1.70651407 | 0.293 | 0.067 | 9.54E-41 | osteoclast.like | Bcs1l |
| Tfpt1 | 4.94E-45 | 0.6550889 | 0.522 | 0.236 | 9.56E-41 | osteoclast.like | Tfpt |
| Serp12 | 5.23E-45 | 0.43741002 | 0.595 | 0.332 | 1.01E-40 | osteoclast.like | Serp1 |
| Npm32 | 5.54E-45 | 1.007948 | 0.434 | 0.187 | 1.07E-40 | osteoclast.like | Npm3 |
| Elf23 | 5.75E-45 | 0.46965337 | 0.595 | 0.343 | 1.11E-40 | osteoclast.like | Elf2 |
| Psmc52 | 7.80E-45 | 0.27784946 | 0.741 | 0.547 | 1.51E-40 | osteoclast.like | Psmc5 |
| Polr2m1 | 7.88E-45 | 1.29355976 | 0.41 | 0.149 | 1.52E-40 | osteoclast.like | Polr2m |
| Txndc51 | 7.95E-45 | 0.36208668 | 0.693 | 0.451 | 1.54E-40 | osteoclast.like | Txndc5 |
| Mtpap1 | 8.00E-45 | 0.83368971 | 0.444 | 0.169 | 1.55E-40 | osteoclast.like | Mtpap |
| Mtg1 | 9.13E-45 | 1.67076175 | 0.317 | 0.083 | 1.77E-40 | osteoclast.like | Mtg1 |
| Slirp1 | 9.45E-45 | 0.74661437 | 0.556 | 0.3 | 1.83E-40 | osteoclast.like | Slirp |
| Apool | 1.05E-44 | 1.1055451 | 0.371 | 0.126 | 2.03E-40 | osteoclast.like | Apool |
| Atg10 | 1.18E-44 | 1.36333723 | 0.332 | 0.097 | 2.28E-40 | osteoclast.like | Atg10 |
| Srsf41 | 1.46E-44 | 0.35825865 | 0.624 | 0.371 | 2.82E-40 | osteoclast.like | Srsf4 |
| Sil11 | 1.50E-44 | 1.10289419 | 0.454 | 0.17 | 2.90E-40 | osteoclast.like | Sil1 |
| Psmc22 | 1.50E-44 | 0.55117067 | 0.678 | 0.413 | 2.91E-40 | osteoclast.like | Psmc2 |
| Tbc1d151 | 1.58E-44 | 0.38659527 | 0.639 | 0.396 | 3.05E-40 | osteoclast.like | Tbc1d15 |
| Phyhd13 | 1.62E-44 | 0.56636161 | 0.654 | 0.247 | 3.13E-40 | osteoclast.like | Phyhd1 |
| Ccdc92 | 1.84E-44 | 0.41053667 | 0.741 | 0.435 | 3.56E-40 | osteoclast.like | Ccdc9 |
| Zbtb44 | 2.03E-44 | 0.71226752 | 0.507 | 0.248 | 3.93E-40 | osteoclast.like | Zbtb44 |
| Ciapin12 | 2.11E-44 | 0.94794483 | 0.463 | 0.214 | 4.08E-40 | osteoclast.like | Ciapin1 |
| Sephs11 | 2.29E-44 | 0.99280667 | 0.517 | 0.211 | 4.44E-40 | osteoclast.like | Sephs1 |
| Uso13 | 2.44E-44 | 0.7228512 | 0.639 | 0.341 | 4.72E-40 | osteoclast.like | Uso1 |
| Mtmr11 | 2.83E-44 | 1.14386228 | 0.42 | 0.146 | 5.48E-40 | osteoclast.like | Mtmr1 |
| Stx162 | 3.36E-44 | 0.56592579 | 0.683 | 0.366 | 6.50E-40 | osteoclast.like | Stx16 |
| Ppp1r12a1 | 3.55E-44 | 0.3198887 | 0.746 | 0.482 | 6.87E-40 | osteoclast.like | Ppp1r12a |
| Exosc91 | 3.57E-44 | 1.03477839 | 0.449 | 0.188 | 6.90E-40 | osteoclast.like | Exosc9 |
| Cops81 | 3.61E-44 | 0.73071926 | 0.605 | 0.311 | 6.97E-40 | osteoclast.like | Cops8 |
| Nr1d21 | 3.89E-44 | 1.09038982 | 0.415 | 0.134 | 7.52E-40 | osteoclast.like | Nr1d2 |
| Zfp146 | 4.32E-44 | 1.32686782 | 0.312 | 0.095 | 8.35E-40 | osteoclast.like | Zfp146 |
| Pgm3 | 4.40E-44 | 1.24209495 | 0.341 | 0.102 | 8.51E-40 | osteoclast.like | Pgm3 |
| Snx122 | 4.48E-44 | 0.33944946 | 0.693 | 0.458 | 8.66E-40 | osteoclast.like | Snx12 |
| Kremen1 | 4.82E-44 | 0.92675516 | 0.439 | 0.176 | 9.32E-40 | osteoclast.like | Kremen1 |
| Ruvbl22 | 5.58E-44 | 0.94292505 | 0.468 | 0.225 | 1.08E-39 | osteoclast.like | Ruvbl2 |
| Mcee | 5.98E-44 | 1.1052483 | 0.376 | 0.123 | 1.16E-39 | osteoclast.like | Mcee |
| Timm21 | 6.35E-44 | 1.54035593 | 0.307 | 0.069 | 1.23E-39 | osteoclast.like | Timm21 |
| Slk1 | 6.47E-44 | 0.28825916 | 0.702 | 0.44 | 1.25E-39 | osteoclast.like | Slk |
| Kansl22 | 7.88E-44 | 0.63087043 | 0.6 | 0.319 | 1.52E-39 | osteoclast.like | Kansl2 |
| Babam11 | 9.98E-44 | 0.40048475 | 0.683 | 0.443 | 1.93E-39 | osteoclast.like | Babam1 |
| Kctd9 | 1.01E-43 | 1.29456308 | 0.371 | 0.102 | 1.95E-39 | osteoclast.like | Kctd9 |
| Ppil22 | 1.05E-43 | 0.67128811 | 0.6 | 0.299 | 2.03E-39 | osteoclast.like | Ppil2 |
| Abce12 | 1.05E-43 | 0.91257767 | 0.478 | 0.224 | 2.04E-39 | osteoclast.like | Abce1 |
| Trak21 | 1.17E-43 | 0.69235527 | 0.615 | 0.3 | 2.26E-39 | osteoclast.like | Trak2 |
| Ddb11 | 1.24E-43 | 0.88767346 | 0.463 | 0.208 | 2.40E-39 | osteoclast.like | Ddb1 |
| Rcc11 | 1.35E-43 | 1.00107576 | 0.444 | 0.179 | 2.61E-39 | osteoclast.like | Rcc1 |
| Vkorc13 | 1.36E-43 | 0.65038066 | 0.493 | 0.236 | 2.63E-39 | osteoclast.like | Vkorc1 |
| Ift522 | 1.41E-43 | 0.72238751 | 0.532 | 0.238 | 2.72E-39 | osteoclast.like | Ift52 |

| | | | | | | | |
|---|---|---|---|---|---|---|---|
| Chd6 | 1.57E-43 | 1.104214 | 0.459 | 0.183 | 3.03E-39 | osteoclast.like | Chd6 |
| Tmem87a3 | 1.58E-43 | 0.61445934 | 0.478 | 0.238 | 3.06E-39 | osteoclast.like | Tmem87a |
| Aars2 | 1.59E-43 | 0.64124764 | 0.576 | 0.348 | 3.08E-39 | osteoclast.like | Aars |
| Anapc12 | 1.60E-43 | 0.52905891 | 0.551 | 0.282 | 3.10E-39 | osteoclast.like | Anapc1 |
| Mecr1 | 1.83E-43 | 1.24610031 | 0.4 | 0.129 | 3.54E-39 | osteoclast.like | Mecr |
| Zfp281 | 1.88E-43 | 0.9388466 | 0.42 | 0.153 | 3.64E-39 | osteoclast.like | Zfp281 |
| Gpsm2 | 1.89E-43 | 1.68912834 | 0.263 | 0.051 | 3.65E-39 | osteoclast.like | Gpsm2 |
| H2-Ke61 | 1.94E-43 | 0.85314576 | 0.502 | 0.237 | 3.74E-39 | osteoclast.like | H2-Ke6 |
| Bet13 | 1.95E-43 | 0.82782263 | 0.532 | 0.228 | 3.77E-39 | osteoclast.like | Bet1 |
| Smg53 | 2.16E-43 | 0.59908926 | 0.639 | 0.333 | 4.17E-39 | osteoclast.like | Smg5 |
| Dnaja11 | 2.40E-43 | 0.41225911 | 0.668 | 0.41 | 4.65E-39 | osteoclast.like | Dnaja1 |
| Cdc161 | 2.41E-43 | 1.25266858 | 0.38 | 0.137 | 4.65E-39 | osteoclast.like | Cdc16 |
| Imp41 | 2.47E-43 | 0.66957307 | 0.58 | 0.331 | 4.78E-39 | osteoclast.like | Imp4 |
| Dele11 | 2.79E-43 | 0.96900561 | 0.385 | 0.146 | 5.40E-39 | osteoclast.like | Dele1 |
| Txndc121 | 3.15E-43 | 0.70987761 | 0.522 | 0.239 | 6.09E-39 | osteoclast.like | Txndc12 |
| Pdhx1 | 3.20E-43 | 0.98682304 | 0.376 | 0.139 | 6.19E-39 | osteoclast.like | Pdhx |
| Romo11 | 3.46E-43 | 0.81051531 | 0.512 | 0.245 | 6.69E-39 | osteoclast.like | Romo1 |
| Magoh3 | 3.51E-43 | 0.50130396 | 0.546 | 0.326 | 6.79E-39 | osteoclast.like | Magoh |
| Txndc111 | 4.19E-43 | 0.40207985 | 0.659 | 0.375 | 8.10E-39 | osteoclast.like | Txndc11 |
| Chchd11 | 4.38E-43 | 0.9332754 | 0.478 | 0.214 | 8.47E-39 | osteoclast.like | Chchd1 |
| Elf1 | 4.43E-43 | 0.41811961 | 0.683 | 0.423 | 8.56E-39 | osteoclast.like | Elf1 |
| Gcsh2 | 5.37E-43 | 0.46312846 | 0.62 | 0.317 | 1.04E-38 | osteoclast.like | Gcsh |
| Sco21 | 5.40E-43 | 1.28011099 | 0.439 | 0.161 | 1.04E-38 | osteoclast.like | Sco2 |
| Snrpa12 | 5.49E-43 | 0.73097486 | 0.517 | 0.263 | 1.06E-38 | osteoclast.like | Snrpa1 |
| Mtln2 | 5.57E-43 | 0.82396372 | 0.537 | 0.234 | 1.08E-38 | osteoclast.like | Mtln |
| Pdcd6 | 6.19E-43 | 0.37045327 | 0.668 | 0.451 | 1.20E-38 | osteoclast.like | Pdcd6 |
| Lamtor5 | 8.51E-43 | 0.67437423 | 0.537 | 0.255 | 1.65E-38 | osteoclast.like | Lamtor5 |
| Nup2141 | 1.01E-42 | 0.6670096 | 0.522 | 0.24 | 1.94E-38 | osteoclast.like | Nup214 |
| Slc25a243 | 1.14E-42 | 0.86289447 | 0.493 | 0.223 | 2.21E-38 | osteoclast.like | Slc25a24 |
| Trp532 | 1.24E-42 | 0.53465679 | 0.663 | 0.42 | 2.39E-38 | osteoclast.like | Trp53 |
| Imp31 | 1.43E-42 | 0.76921428 | 0.488 | 0.201 | 2.77E-38 | osteoclast.like | Imp3 |
| Haus41 | 1.49E-42 | 1.17222966 | 0.4 | 0.114 | 2.88E-38 | osteoclast.like | Haus4 |
| Mien1 | 1.61E-42 | 0.39920517 | 0.629 | 0.366 | 3.11E-38 | osteoclast.like | Mien1 |
| Mbtps2 | 1.67E-42 | 0.94375763 | 0.42 | 0.154 | 3.22E-38 | osteoclast.like | Mbtps2 |
| Rxylt11 | 1.68E-42 | 0.87369345 | 0.507 | 0.223 | 3.24E-38 | osteoclast.like | Rxylt1 |
| Eif1ad1 | 1.73E-42 | 0.94925941 | 0.502 | 0.226 | 3.34E-38 | osteoclast.like | Eif1ad |
| Vps182 | 1.73E-42 | 0.38172598 | 0.732 | 0.478 | 3.35E-38 | osteoclast.like | Vps18 |
| Guk11 | 1.79E-42 | 0.80627408 | 0.546 | 0.262 | 3.47E-38 | osteoclast.like | Guk1 |
| Mthfd11 | 1.85E-42 | 1.12393815 | 0.4 | 0.137 | 3.58E-38 | osteoclast.like | Mthfd1 |
| Pdcl32 | 1.87E-42 | 0.54041142 | 0.644 | 0.404 | 3.61E-38 | osteoclast.like | Pdcl3 |
| Rab22a1 | 1.93E-42 | 0.44377183 | 0.683 | 0.395 | 3.73E-38 | osteoclast.like | Rab22a |
| Sbds2 | 1.99E-42 | 0.3365699 | 0.654 | 0.401 | 3.84E-38 | osteoclast.like | Sbds |
| Rap2c2 | 2.04E-42 | 0.57448715 | 0.615 | 0.346 | 3.94E-38 | osteoclast.like | Rap2c |
| Idua3 | 2.25E-42 | 0.73255396 | 0.556 | 0.262 | 4.36E-38 | osteoclast.like | Idua |
| Rbm141 | 2.37E-42 | 0.46550879 | 0.688 | 0.402 | 4.58E-38 | osteoclast.like | Rbm14 |
| Dync1i21 | 2.52E-42 | 0.32140721 | 0.693 | 0.481 | 4.87E-38 | osteoclast.like | Dync1i2 |
| Tbl31 | 3.32E-42 | 0.80500258 | 0.429 | 0.195 | 6.42E-38 | osteoclast.like | Tbl3 |
| Afg3l12 | 3.38E-42 | 0.46161683 | 0.644 | 0.348 | 6.53E-38 | osteoclast.like | Afg3l1 |
| Psmb32 | 3.42E-42 | 0.65380408 | 0.454 | 0.235 | 6.60E-38 | osteoclast.like | Psmb3 |
| DEPRECATED-I | 3.72E-42 | 0.49835571 | 0.668 | 0.392 | 7.20E-38 | osteoclast.like | DEPRECATED-ENSMUSG00000017221 |
| Ccdc1242 | 3.75E-42 | 0.38000017 | 0.712 | 0.475 | 7.26E-38 | osteoclast.like | Ccdc124 |
| Znrd21 | 4.11E-42 | 1.1260254 | 0.376 | 0.128 | 7.94E-38 | osteoclast.like | Znrd2 |
| Stk381 | 4.18E-42 | 0.26485264 | 0.707 | 0.476 | 8.08E-38 | osteoclast.like | Stk38 |
| Tsn1 | 4.51E-42 | 0.50839217 | 0.62 | 0.384 | 8.73E-38 | osteoclast.like | Tsn |
| Smyd51 | 4.91E-42 | 1.1225931 | 0.366 | 0.137 | 9.50E-38 | osteoclast.like | Smyd5 |
| Rcor1 | 5.26E-42 | 0.49898528 | 0.649 | 0.369 | 1.02E-37 | osteoclast.like | Rcor1 |
| Ercc32 | 5.72E-42 | 0.52906568 | 0.546 | 0.29 | 1.11E-37 | osteoclast.like | Ercc3 |
| Psme24 | 5.83E-42 | 0.40524994 | 0.683 | 0.406 | 1.13E-37 | osteoclast.like | Psme2 |
| AW5498772 | 5.87E-42 | 0.75741395 | 0.468 | 0.188 | 1.13E-37 | osteoclast.like | AW549877 |
| Atraid2 | 5.99E-42 | 0.59480013 | 0.585 | 0.329 | 1.16E-37 | osteoclast.like | Atraid |
| Slc30a52 | 6.51E-42 | 0.53911174 | 0.551 | 0.322 | 1.26E-37 | osteoclast.like | Slc30a5 |
| Nudt51 | 7.97E-42 | 0.99870502 | 0.507 | 0.206 | 1.54E-37 | osteoclast.like | Nudt5 |
| Kbtbd21 | 9.52E-42 | 0.43639458 | 0.61 | 0.343 | 1.84E-37 | osteoclast.like | Kbtbd2 |
| Exosc42 | 9.54E-42 | 0.95598441 | 0.439 | 0.176 | 1.85E-37 | osteoclast.like | Exosc4 |
| Gsk3a2 | 9.85E-42 | 0.41990253 | 0.673 | 0.451 | 1.91E-37 | osteoclast.like | Gsk3a |
| Znfx12 | 9.96E-42 | 0.5245661 | 0.615 | 0.268 | 1.93E-37 | osteoclast.like | Znfx1 |
| Rexf12 | 1.05E-41 | 0.50954687 | 0.561 | 0.29 | 2.03E-37 | osteoclast.like | Rexf1 |
| Zfand1 | 1.18E-41 | 1.82062352 | 0.288 | 0.066 | 2.28E-37 | osteoclast.like | Zfand1 |
| Aven1 | 1.19E-41 | 1.16086233 | 0.371 | 0.138 | 2.31E-37 | osteoclast.like | Aven |
| Sar1a | 1.51E-41 | 0.31607595 | 0.707 | 0.489 | 2.91E-37 | osteoclast.like | Sar1a |
| Nsf2 | 1.52E-41 | 0.41523998 | 0.654 | 0.389 | 2.95E-37 | osteoclast.like | Nsf |
| Prag1 | 1.74E-41 | 1.19404027 | 0.322 | 0.083 | 3.36E-37 | osteoclast.like | Prag1 |
| Ciz1 | 1.76E-41 | 0.5708325 | 0.649 | 0.335 | 3.40E-37 | osteoclast.like | Ciz1 |
| Gtf3c41 | 2.07E-41 | 1.07014284 | 0.385 | 0.145 | 4.00E-37 | osteoclast.like | Gtf3c4 |
| Ppm1a2 | 2.55E-41 | 0.26003235 | 0.727 | 0.498 | 4.92E-37 | osteoclast.like | Ppm1a |
| Cog6 | 2.92E-41 | 1.17290008 | 0.41 | 0.15 | 5.65E-37 | osteoclast.like | Cog6 |
| Immp1l1 | 3.13E-41 | 0.89452737 | 0.415 | 0.148 | 6.06E-37 | osteoclast.like | Immp1l |
| Sco1 | 3.26E-41 | 1.68746676 | 0.254 | 0.056 | 6.30E-37 | osteoclast.like | Sco1 |
| Eif2b41 | 3.35E-41 | 1.07336494 | 0.42 | 0.145 | 6.47E-37 | osteoclast.like | Eif2b4 |
| Acbd32 | 3.36E-41 | 0.32722919 | 0.663 | 0.401 | 6.49E-37 | osteoclast.like | Acbd3 |
| Trappc2l | 3.44E-41 | 0.27883366 | 0.673 | 0.459 | 6.65E-37 | osteoclast.like | Trappc2l |
| Fam98b1 | 3.58E-41 | 0.90865915 | 0.463 | 0.195 | 6.92E-37 | osteoclast.like | Fam98b |
| Med81 | 3.63E-41 | 0.6944391 | 0.546 | 0.289 | 7.01E-37 | osteoclast.like | Med8 |
| Eif4ebp22 | 3.75E-41 | 0.44427492 | 0.702 | 0.457 | 7.25E-37 | osteoclast.like | Eif4ebp2 |
| Kdm3b1 | 3.75E-41 | 0.28402622 | 0.712 | 0.447 | 7.26E-37 | osteoclast.like | Kdm3b |
| Herc3 | 3.75E-41 | 1.41470303 | 0.38 | 0.113 | 7.26E-37 | osteoclast.like | Herc3 |
| Tmem106c1 | 4.10E-41 | 1.19842975 | 0.444 | 0.157 | 7.92E-37 | osteoclast.like | Tmem106c |
| Smu11 | 4.46E-41 | 0.4743449 | 0.615 | 0.375 | 8.62E-37 | osteoclast.like | Smu1 |
| Rbbp41 | 4.58E-41 | 0.4249133 | 0.707 | 0.467 | 8.85E-37 | osteoclast.like | Rbbp4 |
| Exosc52 | 5.63E-41 | 0.75090202 | 0.576 | 0.297 | 1.09E-36 | osteoclast.like | Exosc5 |
| Nfam12 | 5.81E-41 | 0.25069924 | 0.693 | 0.474 | 1.12E-36 | osteoclast.like | Nfam1 |
| Pfdn21 | 5.82E-41 | 0.60212951 | 0.624 | 0.364 | 1.12E-36 | osteoclast.like | Pfdn2 |
| Tbc1d53 | 6.42E-41 | 0.37476986 | 0.624 | 0.353 | 1.24E-36 | osteoclast.like | Tbc1d5 |
| Mief11 | 7.20E-41 | 1.12572753 | 0.424 | 0.156 | 1.39E-36 | osteoclast.like | Mief1 |
| Slc35f63 | 8.16E-41 | 0.41802531 | 0.673 | 0.419 | 1.58E-36 | osteoclast.like | Slc35f6 |
| Lsm14b1 | 8.30E-41 | 0.58633139 | 0.58 | 0.298 | 1.60E-36 | osteoclast.like | Lsm14b |
| Zadh23 | 9.71E-41 | 0.87365482 | 0.449 | 0.18 | 1.88E-36 | osteoclast.like | Zadh2 |
| Cops21 | 1.05E-40 | 0.71497839 | 0.566 | 0.281 | 2.03E-36 | osteoclast.like | Cops2 |
| Srrt1 | 1.06E-40 | 0.49246702 | 0.649 | 0.377 | 2.06E-36 | osteoclast.like | Srrt |
| Hspbp11 | 1.17E-40 | 0.78812574 | 0.483 | 0.218 | 2.26E-36 | osteoclast.like | Hspbp1 |
| Fars2 | 1.32E-40 | 0.64964352 | 0.493 | 0.224 | 2.55E-36 | osteoclast.like | Fars2 |
| Rtn4ip1 | 1.33E-40 | 1.60950557 | 0.263 | 0.058 | 2.57E-36 | osteoclast.like | Rtn4ip1 |
| Cnot13 | 1.62E-40 | 0.3671425 | 0.698 | 0.418 | 3.14E-36 | osteoclast.like | Cnot1 |
| Sec61a12 | 1.74E-40 | 0.43996452 | 0.654 | 0.397 | 3.37E-36 | osteoclast.like | Sec61a1 |
| Srsf101 | 2.23E-40 | 0.47535383 | 0.6 | 0.352 | 4.31E-36 | osteoclast.like | Srsf10 |
| Dpp31 | 2.37E-40 | 0.37256983 | 0.683 | 0.414 | 4.58E-36 | osteoclast.like | Dpp3 |
| Snrpg2 | 2.37E-40 | 0.72215727 | 0.532 | 0.279 | 4.58E-36 | osteoclast.like | Snrpg |
| Faf12 | 2.46E-40 | 0.71895635 | 0.6 | 0.296 | 4.75E-36 | osteoclast.like | Faf1 |
| Tarbp2 | 2.51E-40 | 1.2070216 | 0.4 | 0.12 | 4.85E-36 | osteoclast.like | Tarbp2 |
| Pds5a2 | 2.53E-40 | 0.3172562 | 0.639 | 0.411 | 4.90E-36 | osteoclast.like | Pds5a |
| Atad1 | 2.71E-40 | 0.88843926 | 0.488 | 0.204 | 5.25E-36 | osteoclast.like | Atad1 |
| Eif1ax2 | 3.09E-40 | 0.94418418 | 0.439 | 0.193 | 5.96E-36 | osteoclast.like | Eif1ax |
| Fam111a1 | 3.14E-40 | 0.75228064 | 0.434 | 0.189 | 6.07E-36 | osteoclast.like | Fam111a |
| Micu2 | 3.45E-40 | 0.60242403 | 0.59 | 0.307 | 6.66E-36 | osteoclast.like | Micu2 |
| Ilf21 | 3.57E-40 | 0.87934664 | 0.415 | 0.185 | 6.91E-36 | osteoclast.like | Ilf2 |
| Daam1 | 3.70E-40 | 1.22673216 | 0.371 | 0.133 | 7.16E-36 | osteoclast.like | Daam1 |
| Smim71 | 4.38E-40 | 0.4800189 | 0.615 | 0.36 | 8.47E-36 | osteoclast.like | Smim7 |
| Tacc2 | 4.83E-40 | 1.44773781 | 0.263 | 0.065 | 9.33E-36 | osteoclast.like | Tacc2 |
| Ddx492 | 5.59E-40 | 0.6469721 | 0.522 | 0.257 | 1.08E-35 | osteoclast.like | Ddx49 |
| Trak1 | 6.71E-40 | 0.30789711 | 0.727 | 0.506 | 1.30E-35 | osteoclast.like | Trak1 |
| Aimp11 | 7.27E-40 | 0.77269425 | 0.522 | 0.261 | 1.40E-35 | osteoclast.like | Aimp1 |
| Ndufaf82 | 7.94E-40 | 0.66093879 | 0.502 | 0.276 | 1.53E-35 | osteoclast.like | Ndufaf8 |
| Cops31 | 9.21E-40 | 1.14459704 | 0.39 | 0.139 | 1.78E-35 | osteoclast.like | Cops3 |
| Slc16a103 | 1.06E-39 | 0.42957147 | 0.62 | 0.292 | 2.05E-35 | osteoclast.like | Slc16a10 |
| Orc31 | 1.19E-39 | 0.90938764 | 0.38 | 0.157 | 2.31E-35 | osteoclast.like | Orc3 |
| Ipo11 | 1.34E-39 | 1.36074026 | 0.327 | 0.091 | 2.58E-35 | osteoclast.like | Ipo11 |
| Bloc1s61 | 1.36E-39 | 0.8468613 | 0.424 | 0.161 | 2.64E-35 | osteoclast.like | Bloc1s6 |
| Suds31 | 1.57E-39 | 0.35676194 | 0.663 | 0.413 | 3.03E-35 | osteoclast.like | Suds3 |
| Tmem241 | 1.59E-39 | 1.14038336 | 0.327 | 0.117 | 3.07E-35 | osteoclast.like | Tmem241 |
| Apeh2 | 1.62E-39 | 0.64102326 | 0.571 | 0.275 | 3.14E-35 | osteoclast.like | Apeh |
| Acsl31 | 1.64E-39 | 0.79669651 | 0.376 | 0.173 | 3.17E-35 | osteoclast.like | Acsl3 |
| Dcaf12 | 1.65E-39 | 0.40855418 | 0.644 | 0.389 | 3.20E-35 | osteoclast.like | Dcaf12 |
| Map2k5 | 1.87E-39 | 1.28149409 | 0.351 | 0.111 | 3.61E-35 | osteoclast.like | Map2k5 |
| Faf21 | 1.89E-39 | 0.73866619 | 0.449 | 0.206 | 3.66E-35 | osteoclast.like | Faf2 |
| Llgl1 | 1.93E-39 | 1.2237572 | 0.317 | 0.092 | 3.72E-35 | osteoclast.like | Llgl1 |
| Mipep | 2.07E-39 | 1.32506972 | 0.307 | 0.086 | 4.01E-35 | osteoclast.like | Mipep |
| Dnase1l13 | 2.34E-39 | 0.35827157 | 0.683 | 0.45 | 4.53E-35 | osteoclast.like | Dnase1l1 |
| Nop161 | 2.36E-39 | 0.87975436 | 0.385 | 0.191 | 4.57E-35 | osteoclast.like | Nop16 |
| Spata132 | 2.40E-39 | 0.61377433 | 0.634 | 0.294 | 4.65E-35 | osteoclast.like | Spata13 |
| Psmd82 | 3.00E-39 | 0.2875566 | 0.693 | 0.452 | 5.79E-35 | osteoclast.like | Psmd8 |
| Zfp2171 | 3.08E-39 | 0.45066048 | 0.585 | 0.322 | 5.96E-35 | osteoclast.like | Zfp217 |
| Zfp6551 | 3.30E-39 | 0.69067828 | 0.58 | 0.309 | 6.38E-35 | osteoclast.like | Zfp655 |
| Efl1 | 3.66E-39 | 1.14588889 | 0.371 | 0.131 | 7.08E-35 | osteoclast.like | Efl1 |
| Scamp31 | 3.89E-39 | 0.48734486 | 0.605 | 0.333 | 7.52E-35 | osteoclast.like | Scamp3 |
| Xpnpep11 | 3.95E-39 | 0.51973335 | 0.595 | 0.334 | 7.63E-35 | osteoclast.like | Xpnpep1 |
| Tle42 | 3.99E-39 | 0.45985491 | 0.517 | 0.294 | 7.72E-35 | osteoclast.like | Tle4 |
| Gatad11 | 4.15E-39 | 0.43470742 | 0.532 | 0.317 | 8.02E-35 | osteoclast.like | Gatad1 |
| Cirbp2 | 4.61E-39 | 0.65322783 | 0.59 | 0.305 | 8.91E-35 | osteoclast.like | Cirbp |
| Ap2a11 | 4.68E-39 | 0.4340883 | 0.634 | 0.388 | 9.05E-35 | osteoclast.like | Ap2a1 |
| Smurf11 | 4.99E-39 | 0.52171219 | 0.58 | 0.329 | 9.64E-35 | osteoclast.like | Smurf1 |
| Slc7a12 | 5.27E-39 | 1.22364569 | 0.293 | 0.102 | 1.02E-34 | osteoclast.like | Slc7a1 |
| St132 | 5.34E-39 | 0.51854141 | 0.6 | 0.335 | 1.03E-34 | osteoclast.like | St13 |
| Scaf12 | 5.52E-39 | 0.39077145 | 0.668 | 0.393 | 1.07E-34 | osteoclast.like | Scaf1 |
| Csnk2a1 | 5.56E-39 | 0.50768733 | 0.61 | 0.361 | 1.07E-34 | osteoclast.like | Csnk2a1 |
| Dhx35 | 5.77E-39 | 1.12223115 | 0.307 | 0.089 | 1.11E-34 | osteoclast.like | Dhx35 |
| Actl6a1 | 5.81E-39 | 0.7198833 | 0.468 | 0.231 | 1.12E-34 | osteoclast.like | Actl6a |
| Zfp8001 | 5.83E-39 | 0.75684087 | 0.551 | 0.276 | 1.13E-34 | osteoclast.like | Zfp800 |
| Myo18a2 | 5.95E-39 | 0.64960733 | 0.6 | 0.32 | 1.15E-34 | osteoclast.like | Myo18a |
| Mmadhc1 | 6.13E-39 | 0.99375452 | 0.541 | 0.236 | 1.18E-34 | osteoclast.like | Mmadhc |
| Ash2l1 | 6.45E-39 | 0.54482253 | 0.512 | 0.272 | 1.25E-34 | osteoclast.like | Ash2l |
| Phf5a1 | 7.49E-39 | 0.86481739 | 0.527 | 0.259 | 1.45E-34 | osteoclast.like | Phf5a |
| Kcmf12 | 8.20E-39 | 0.27782158 | 0.702 | 0.461 | 1.59E-34 | osteoclast.like | Kcmf1 |
| Zfp335 | 9.17E-39 | 0.83993215 | 0.459 | 0.204 | 1.77E-34 | osteoclast.like | Zfp335 |
| Arhgef64 | 9.68E-39 | 0.49732383 | 0.551 | 0.334 | 1.87E-34 | osteoclast.like | Arhgef6 |
| Fmnl31 | 9.82E-39 | 0.38581249 | 0.698 | 0.411 | 1.90E-34 | osteoclast.like | Fmnl3 |
| Ecd1 | 1.23E-38 | 0.94548493 | 0.473 | 0.209 | 2.38E-34 | osteoclast.like | Ecd |
| Cldnd11 | 1.24E-38 | 0.28227897 | 0.649 | 0.403 | 2.40E-34 | osteoclast.like | Cldnd1 |
| Polr2b1 | 1.27E-38 | 0.89299491 | 0.478 | 0.21 | 2.45E-34 | osteoclast.like | Polr2b |
| Prelid3b1 | 1.28E-38 | 0.5839776 | 0.512 | 0.25 | 2.47E-34 | osteoclast.like | Prelid3b |
| Fgd32 | 1.52E-38 | 0.3922071 | 0.634 | 0.393 | 2.94E-34 | osteoclast.like | Fgd3 |
| Dctn12 | 1.54E-38 | 0.27749839 | 0.702 | 0.453 | 2.97E-34 | osteoclast.like | Dctn1 |
| Emc101 | 1.63E-38 | 0.41560722 | 0.776 | 0.502 | 3.15E-34 | osteoclast.like | Emc10 |
| Ino80e1 | 1.75E-38 | 0.58986747 | 0.585 | 0.295 | 3.37E-34 | osteoclast.like | Ino80e |
| Pdcd21 | 1.88E-38 | 1.09737625 | 0.38 | 0.133 | 3.63E-34 | osteoclast.like | Pdcd2 |
| Phax1 | 1.96E-38 | 0.70701395 | 0.527 | 0.242 | 3.78E-34 | osteoclast.like | Phax |
| Gtf2f22 | 2.03E-38 | 0.4690165 | 0.566 | 0.322 | 3.92E-34 | osteoclast.like | Gtf2f2 |
| Zdhhc21 | 2.08E-38 | 0.83221206 | 0.454 | 0.18 | 4.03E-34 | osteoclast.like | Zdhhc21 |
| U2surp1 | 2.24E-38 | 0.53258892 | 0.571 | 0.319 | 4.33E-34 | osteoclast.like | U2surp |
| Pias11 | 2.48E-38 | 0.43640652 | 0.576 | 0.321 | 4.80E-34 | osteoclast.like | Pias1 |
| Ndufaf21 | 2.56E-38 | 1.25923325 | 0.449 | 0.164 | 4.96E-34 | osteoclast.like | Ndufaf2 |
| Rbm221 | 2.73E-38 | 0.59704933 | 0.556 | 0.286 | 5.29E-34 | osteoclast.like | Rbm22 |
| Gcc1 | 2.80E-38 | 0.90558799 | 0.38 | 0.144 | 5.42E-34 | osteoclast.like | Gcc1 |
| Traf21 | 3.17E-38 | 0.73625364 | 0.468 | 0.237 | 6.13E-34 | osteoclast.like | Traf2 |
| Cse1l1 | 3.66E-38 | 0.70334693 | 0.502 | 0.255 | 7.07E-34 | osteoclast.like | Cse1l |
| Lyar1 | 3.79E-38 | 1.0136987 | 0.449 | 0.186 | 7.33E-34 | osteoclast.like | Lyar |
| Tmem9b2 | 3.93E-38 | 0.53371009 | 0.615 | 0.338 | 7.60E-34 | osteoclast.like | Tmem9b |
| Srp54b1 | 4.04E-38 | 0.73826946 | 0.493 | 0.223 | 7.82E-34 | osteoclast.like | Srp54b |
| Senp22 | 4.46E-38 | 0.6271691 | 0.58 | 0.283 | 8.62E-34 | osteoclast.like | Senp2 |
| Pdzd111 | 4.48E-38 | 0.98774268 | 0.424 | 0.174 | 8.65E-34 | osteoclast.like | Pdzd11 |
| Ctnnbl11 | 4.77E-38 | 0.75789885 | 0.502 | 0.237 | 9.22E-34 | osteoclast.like | Ctnnbl1 |
| Aimp21 | 5.18E-38 | 1.02929284 | 0.41 | 0.155 | 1.00E-33 | osteoclast.like | Aimp2 |
| Nek91 | 5.27E-38 | 0.51942835 | 0.6 | 0.307 | 1.02E-33 | osteoclast.like | Nek9 |
| Nutf2-ps11 | 5.65E-38 | 1.07917521 | 0.351 | 0.155 | 1.09E-33 | osteoclast.like | Nutf2-ps1 |
| Pfdn11 | 5.96E-38 | 0.61725953 | 0.478 | 0.238 | 1.15E-33 | osteoclast.like | Pfdn1 |
| Tprn | 6.01E-38 | 1.48194708 | 0.341 | 0.107 | 1.16E-33 | osteoclast.like | Tprn |
| Gmip | 6.69E-38 | 0.43446559 | 0.527 | 0.31 | 1.29E-33 | osteoclast.like | Gmip |
| Upf11 | 6.74E-38 | 0.40307475 | 0.766 | 0.485 | 1.30E-33 | osteoclast.like | Upf1 |
| Plekha31 | 6.91E-38 | 0.6275336 | 0.532 | 0.284 | 1.34E-33 | osteoclast.like | Plekha3 |
| Cdk5 | 6.96E-38 | 0.68555774 | 0.522 | 0.25 | 1.35E-33 | osteoclast.like | Cdk5 |
| Chaf1a1 | 7.29E-38 | 1.69379565 | 0.263 | 0.07 | 1.41E-33 | osteoclast.like | Chaf1a |
| Exoc33 | 7.39E-38 | 0.64035421 | 0.478 | 0.23 | 1.43E-33 | osteoclast.like | Exoc3 |
| Numa13 | 7.52E-38 | 0.45280841 | 0.659 | 0.38 | 1.45E-33 | osteoclast.like | Numa1 |
| Hacd32 | 7.55E-38 | 0.91898904 | 0.478 | 0.206 | 1.46E-33 | osteoclast.like | Hacd3 |
| Pelo | 7.59E-38 | 0.95260993 | 0.337 | 0.124 | 1.47E-33 | osteoclast.like | Pelo |
| Zfp639 | 7.61E-38 | 0.93043472 | 0.366 | 0.136 | 1.47E-33 | osteoclast.like | Zfp639 |
| Dcun1d51 | 8.14E-38 | 0.59105266 | 0.629 | 0.375 | 1.57E-33 | osteoclast.like | Dcun1d5 |
| Ccdc90b | 8.69E-38 | 1.26238049 | 0.366 | 0.114 | 1.68E-33 | osteoclast.like | Ccdc90b |
| Nfyc1 | 9.81E-38 | 0.63810696 | 0.459 | 0.222 | 1.90E-33 | osteoclast.like | Nfyc |
| Sirt6 | 1.04E-37 | 1.02561073 | 0.429 | 0.174 | 2.02E-33 | osteoclast.like | Sirt6 |
| Rbm171 | 1.11E-37 | 0.31282874 | 0.678 | 0.449 | 2.15E-33 | osteoclast.like | Rbm17 |
| Map1lc3a1 | 1.18E-37 | 0.60377124 | 0.459 | 0.238 | 2.28E-33 | osteoclast.like | Map1lc3a |
| Foxo11 | 1.23E-37 | 0.6913089 | 0.478 | 0.206 | 2.38E-33 | osteoclast.like | Foxo1 |
| Golt1b3 | 1.30E-37 | 0.51997108 | 0.541 | 0.306 | 2.51E-33 | osteoclast.like | Golt1b |
| Commd32 | 1.31E-37 | 0.69697663 | 0.59 | 0.285 | 2.53E-33 | osteoclast.like | Commd3 |
| Alg91 | 1.42E-37 | 0.78215161 | 0.522 | 0.23 | 2.74E-33 | osteoclast.like | Alg9 |
| Echdc1 | 1.46E-37 | 1.25461704 | 0.322 | 0.096 | 2.83E-33 | osteoclast.like | Echdc1 |
| Med301 | 1.52E-37 | 0.66784576 | 0.468 | 0.237 | 2.95E-33 | osteoclast.like | Med30 |
| Zmat21 | 1.85E-37 | 0.45709422 | 0.649 | 0.413 | 3.57E-33 | osteoclast.like | Zmat2 |
| Chchd5 | 2.10E-37 | 1.54236087 | 0.327 | 0.107 | 4.07E-33 | osteoclast.like | Chchd5 |
| Smg91 | 2.12E-37 | 0.51460741 | 0.595 | 0.318 | 4.10E-33 | osteoclast.like | Smg9 |
| Polr2g1 | 2.50E-37 | 0.51910335 | 0.556 | 0.313 | 4.83E-33 | osteoclast.like | Polr2g |
| Mlycd1 | 2.67E-37 | 0.90638137 | 0.434 | 0.151 | 5.16E-33 | osteoclast.like | Mlycd |
| Hint21 | 2.99E-37 | 1.05534346 | 0.385 | 0.14 | 5.77E-33 | osteoclast.like | Hint2 |
| Larp1b | 2.99E-37 | 1.08427746 | 0.371 | 0.147 | 5.79E-33 | osteoclast.like | Larp1b |
| Rars3 | 3.23E-37 | 0.49146818 | 0.541 | 0.327 | 6.24E-33 | osteoclast.like | Rars |
| Rab93 | 3.38E-37 | 0.6255316 | 0.507 | 0.253 | 6.54E-33 | osteoclast.like | Rab9 |
| Snapc2 | 3.52E-37 | 1.56831878 | 0.298 | 0.078 | 6.81E-33 | osteoclast.like | Snapc2 |
| Ccdc58 | 3.54E-37 | 1.08456841 | 0.38 | 0.142 | 6.84E-33 | osteoclast.like | Ccdc58 |
| Mcts12 | 4.58E-37 | 0.51297644 | 0.493 | 0.306 | 8.86E-33 | osteoclast.like | Mcts1 |
| Rbm451 | 4.69E-37 | 1.00131485 | 0.42 | 0.181 | 9.06E-33 | osteoclast.like | Rbm45 |
| Gpr55 | 5.41E-37 | 1.11610192 | 0.259 | 0.073 | 1.05E-32 | osteoclast.like | Gpr55 |
| Gosr11 | 5.94E-37 | 0.68139349 | 0.473 | 0.208 | 1.15E-32 | osteoclast.like | Gosr1 |
| Cntrob | 6.13E-37 | 1.86371317 | 0.293 | 0.066 | 1.19E-32 | osteoclast.like | Cntrob |
| Zfc3h11 | 6.39E-37 | 0.58239252 | 0.498 | 0.284 | 1.23E-32 | osteoclast.like | Zfc3h1 |
| Tsr13 | 6.75E-37 | 0.70503746 | 0.483 | 0.232 | 1.30E-32 | osteoclast.like | Tsr1 |
| Rars2 | 6.84E-37 | 1.54414684 | 0.293 | 0.085 | 1.32E-32 | osteoclast.like | Rars2 |
| Eea13 | 7.30E-37 | 0.44922544 | 0.654 | 0.335 | 1.41E-32 | osteoclast.like | Eea1 |
| Commd21 | 7.71E-37 | 0.81084665 | 0.444 | 0.186 | 1.49E-32 | osteoclast.like | Commd2 |
| Ube3c2 | 8.17E-37 | 0.75871219 | 0.527 | 0.245 | 1.58E-32 | osteoclast.like | Ube3c |
| Supt6 | 9.06E-37 | 0.34028171 | 0.659 | 0.403 | 1.75E-32 | osteoclast.like | Supt6 |
| Eef1akmt42 | 9.11E-37 | 0.86786338 | 0.424 | 0.169 | 1.76E-32 | osteoclast.like | Eef1akmt4 |
| Hnrnpr1 | 9.78E-37 | 0.57448183 | 0.468 | 0.256 | 1.89E-32 | osteoclast.like | Hnrnpr |
| Stub11 | 1.01E-36 | 0.28385159 | 0.732 | 0.505 | 1.94E-32 | osteoclast.like | Stub1 |
| Mmgt2 | 1.04E-36 | 1.55198928 | 0.273 | 0.076 | 2.00E-32 | osteoclast.like | Mmgt2 |
| Htatsf11 | 1.07E-36 | 0.77024641 | 0.415 | 0.176 | 2.07E-32 | osteoclast.like | Htatsf1 |
| Drg21 | 1.09E-36 | 0.72702859 | 0.522 | 0.239 | 2.10E-32 | osteoclast.like | Drg2 |
| Bcl9l | 1.23E-36 | 1.19393349 | 0.361 | 0.124 | 2.37E-32 | osteoclast.like | Bcl9l |
| Ccar22 | 1.23E-36 | 0.86133326 | 0.439 | 0.208 | 2.38E-32 | osteoclast.like | Ccar2 |
| Ccng11 | 1.25E-36 | 0.34458061 | 0.663 | 0.413 | 2.41E-32 | osteoclast.like | Ccng1 |
| Kxd1 | 1.31E-36 | 0.75122976 | 0.454 | 0.186 | 2.54E-32 | osteoclast.like | Kxd1 |
| Bid | 1.34E-36 | 0.86879037 | 0.351 | 0.131 | 2.60E-32 | osteoclast.like | Bid |
| Bap11 | 1.39E-36 | 1.08388487 | 0.444 | 0.193 | 2.69E-32 | osteoclast.like | Bap1 |
| Fmnl23 | 1.40E-36 | 0.54042116 | 0.517 | 0.276 | 2.70E-32 | osteoclast.like | Fmnl2 |
| Parp13 | 1.41E-36 | 0.37767555 | 0.649 | 0.395 | 2.72E-32 | osteoclast.like | Parp1 |
| Abhd17a2 | 1.54E-36 | 0.46192161 | 0.649 | 0.405 | 2.98E-32 | osteoclast.like | Abhd17a |
| C1qtnf12 | 1.56E-36 | 1.3332829 | 0.283 | 0.083 | 3.01E-32 | osteoclast.like | C1qtnf12 |
| Dusp111 | 1.62E-36 | 0.31274032 | 0.634 | 0.414 | 3.13E-32 | osteoclast.like | Dusp11 |
| Copz2 | 1.79E-36 | 1.73646338 | 0.283 | 0.068 | 3.46E-32 | osteoclast.like | Copz2 |
| Nagk4 | 1.98E-36 | 0.48751721 | 0.527 | 0.282 | 3.82E-32 | osteoclast.like | Nagk |
| Mlxip1 | 2.07E-36 | 0.30307412 | 0.693 | 0.43 | 4.01E-32 | osteoclast.like | Mlxip |
| Pex21 | 2.11E-36 | 0.68330577 | 0.449 | 0.202 | 4.08E-32 | osteoclast.like | Pex2 |
| Psmd52 | 2.11E-36 | 0.93924448 | 0.41 | 0.173 | 4.08E-32 | osteoclast.like | Psmd5 |
| Supt161 | 2.18E-36 | 0.37840031 | 0.62 | 0.395 | 4.21E-32 | osteoclast.like | Supt16 |
| Trappc6b1 | 2.18E-36 | 0.36028599 | 0.59 | 0.339 | 4.22E-32 | osteoclast.like | Trappc6b |
| Cog72 | 2.47E-36 | 0.89246046 | 0.424 | 0.173 | 4.78E-32 | osteoclast.like | Cog7 |
| Glg1 | 2.52E-36 | 0.27205963 | 0.771 | 0.523 | 4.87E-32 | osteoclast.like | Glg1 |
| Spen3 | 3.13E-36 | 0.41257924 | 0.615 | 0.38 | 6.04E-32 | osteoclast.like | Spen |
| Pip4k2b2 | 3.13E-36 | 0.53581341 | 0.668 | 0.353 | 6.05E-32 | osteoclast.like | Pip4k2b |
| Dapk31 | 3.21E-36 | 0.40212087 | 0.527 | 0.314 | 6.20E-32 | osteoclast.like | Dapk3 |
| Fuom | 3.21E-36 | 1.54258363 | 0.278 | 0.072 | 6.20E-32 | osteoclast.like | Fuom |
| Pml3 | 3.55E-36 | 0.6057091 | 0.424 | 0.17 | 6.86E-32 | osteoclast.like | Pml |
| Arl13 | 4.09E-36 | 0.5649686 | 0.541 | 0.302 | 7.92E-32 | osteoclast.like | Arl1 |
| Psmc42 | 4.19E-36 | 0.51387466 | 0.624 | 0.412 | 8.10E-32 | osteoclast.like | Psmc4 |
| Cdc42ep41 | 4.27E-36 | 0.46260613 | 0.551 | 0.259 | 8.25E-32 | osteoclast.like | Cdc42ep4 |
| Lypla1 | 4.58E-36 | 0.77221728 | 0.571 | 0.258 | 8.85E-32 | osteoclast.like | Lypla1 |
| Yars2 | 4.74E-36 | 1.19153378 | 0.288 | 0.094 | 9.16E-32 | osteoclast.like | Yars2 |
| Polr1a1 | 4.86E-36 | 0.9566316 | 0.395 | 0.139 | 9.41E-32 | osteoclast.like | Polr1a |
| Anapc111 | 5.08E-36 | 0.29641858 | 0.644 | 0.464 | 9.83E-32 | osteoclast.like | Anapc11 |
| Eftud21 | 5.11E-36 | 0.441255 | 0.644 | 0.419 | 9.89E-32 | osteoclast.like | Eftud2 |
| Bin31 | 5.23E-36 | 0.48093843 | 0.649 | 0.369 | 1.01E-31 | osteoclast.like | Bin3 |
| Sh3glb22 | 5.59E-36 | 0.66106831 | 0.541 | 0.259 | 1.08E-31 | osteoclast.like | Sh3glb2 |
| Uqcc32 | 5.97E-36 | 0.65330625 | 0.527 | 0.264 | 1.15E-31 | osteoclast.like | Uqcc3 |
| Arpp192 | 6.14E-36 | 0.85963124 | 0.488 | 0.219 | 1.19E-31 | osteoclast.like | Arpp19 |
| Hbegf3 | 6.61E-36 | 0.33922317 | 0.507 | 0.282 | 1.28E-31 | osteoclast.like | Hbegf |
| Emc71 | 6.94E-36 | 0.29153813 | 0.712 | 0.483 | 1.34E-31 | osteoclast.like | Emc7 |
| BC0311811 | 7.53E-36 | 0.35049505 | 0.624 | 0.37 | 1.46E-31 | osteoclast.like | BC031181 |
| Dnajc1 | 7.75E-36 | 0.88434781 | 0.371 | 0.145 | 1.50E-31 | osteoclast.like | Dnajc1 |
| Erc1 | 9.15E-36 | 1.20319947 | 0.268 | 0.075 | 1.77E-31 | osteoclast.like | Erc1 |
| Lsm81 | 1.07E-35 | 0.54299953 | 0.478 | 0.244 | 2.08E-31 | osteoclast.like | Lsm8 |
| Hgsnat3 | 1.11E-35 | 0.51149477 | 0.6 | 0.289 | 2.15E-31 | osteoclast.like | Hgsnat |
| Eny2 | 1.13E-35 | 1.24070991 | 0.283 | 0.092 | 2.18E-31 | osteoclast.like | Eny2 |
| Ubap21 | 1.14E-35 | 0.8093833 | 0.556 | 0.266 | 2.21E-31 | osteoclast.like | Ubap2 |
| Cetn32 | 1.19E-35 | 0.28324129 | 0.585 | 0.358 | 2.31E-31 | osteoclast.like | Cetn3 |
| Dera1 | 1.26E-35 | 0.68256358 | 0.473 | 0.203 | 2.43E-31 | osteoclast.like | Dera |
| Mgat22 | 1.27E-35 | 0.4119155 | 0.668 | 0.467 | 2.46E-31 | osteoclast.like | Mgat2 |
| Ier3ip1 | 1.29E-35 | 0.29724572 | 0.668 | 0.41 | 2.49E-31 | osteoclast.like | Ier3ip1 |
| Snrpd12 | 1.33E-35 | 0.59128284 | 0.585 | 0.339 | 2.56E-31 | osteoclast.like | Snrpd1 |
| Pex141 | 1.44E-35 | 0.51970294 | 0.473 | 0.233 | 2.79E-31 | osteoclast.like | Pex14 |
| Fpgs | 1.58E-35 | 1.52503983 | 0.263 | 0.078 | 3.06E-31 | osteoclast.like | Fpgs |
| Cand11 | 1.60E-35 | 0.3050928 | 0.546 | 0.352 | 3.09E-31 | osteoclast.like | Cand1 |
| Rnf1262 | 1.64E-35 | 0.52220511 | 0.644 | 0.339 | 3.18E-31 | osteoclast.like | Rnf126 |
| Aldh1b12 | 1.67E-35 | 0.91464779 | 0.395 | 0.176 | 3.23E-31 | osteoclast.like | Aldh1b1 |
| R3hdm11 | 1.83E-35 | 0.71305047 | 0.532 | 0.247 | 3.54E-31 | osteoclast.like | R3hdm1 |
| Ergic31 | 1.88E-35 | 0.40950022 | 0.629 | 0.391 | 3.64E-31 | osteoclast.like | Ergic3 |
| Msto1 | 1.89E-35 | 1.16914795 | 0.361 | 0.123 | 3.65E-31 | osteoclast.like | Msto1 |
| Dcaf82 | 2.19E-35 | 0.53020789 | 0.522 | 0.318 | 4.23E-31 | osteoclast.like | Dcaf8 |
| Tomm341 | 2.21E-35 | 0.523481 | 0.624 | 0.36 | 4.27E-31 | osteoclast.like | Tomm34 |
| Mfn11 | 2.25E-35 | 0.66837342 | 0.517 | 0.234 | 4.35E-31 | osteoclast.like | Mfn1 |
| Slc25a202 | 2.31E-35 | 0.25846871 | 0.556 | 0.34 | 4.46E-31 | osteoclast.like | Slc25a20 |
| Prr123 | 2.53E-35 | 0.53316464 | 0.522 | 0.278 | 4.88E-31 | osteoclast.like | Prr12 |
| Dgkd | 2.56E-35 | 0.32307078 | 0.649 | 0.375 | 4.95E-31 | osteoclast.like | Dgkd |
| Tdg | 2.66E-35 | 1.13073315 | 0.351 | 0.12 | 5.15E-31 | osteoclast.like | Tdg |
| Commd6 | 2.73E-35 | 0.59879582 | 0.439 | 0.217 | 5.28E-31 | osteoclast.like | Commd6 |
| Ftsj31 | 2.77E-35 | 0.81051575 | 0.39 | 0.157 | 5.36E-31 | osteoclast.like | Ftsj3 |
| Mtfr1l1 | 3.07E-35 | 0.63091847 | 0.507 | 0.229 | 5.94E-31 | osteoclast.like | Mtfr1l |
| Copl12 | 3.12E-35 | 0.44891849 | 0.551 | 0.304 | 6.03E-31 | osteoclast.like | Copl1 |
| Mpp7 | 3.20E-35 | 1.39681424 | 0.288 | 0.069 | 6.19E-31 | osteoclast.like | Mpp7 |
| Copg2 | 3.23E-35 | 0.63290257 | 0.434 | 0.2 | 6.24E-31 | osteoclast.like | Copg2 |
| Cyb5r4 | 3.34E-35 | 0.25147352 | 0.683 | 0.449 | 6.45E-31 | osteoclast.like | Cyb5r4 |
| Pcca | 3.45E-35 | 0.66249757 | 0.434 | 0.181 | 6.66E-31 | osteoclast.like | Pcca |
| Bop12 | 3.81E-35 | 0.77716256 | 0.478 | 0.237 | 7.36E-31 | osteoclast.like | Bop1 |
| Trappc122 | 3.81E-35 | 0.83854787 | 0.498 | 0.215 | 7.37E-31 | osteoclast.like | Trappc12 |
| Ccnt2 | 3.82E-35 | 0.4669338 | 0.541 | 0.285 | 7.38E-31 | osteoclast.like | Ccnt2 |
| Acsl51 | 3.98E-35 | 0.30762267 | 0.678 | 0.471 | 7.70E-31 | osteoclast.like | Acsl5 |
| Ssbp11 | 4.52E-35 | 0.70949337 | 0.468 | 0.214 | 8.73E-31 | osteoclast.like | Ssbp1 |
| Nudt23 | 4.65E-35 | 1.02620497 | 0.371 | 0.137 | 8.98E-31 | osteoclast.like | Nudt2 |
| Hcfc11 | 5.63E-35 | 0.42033496 | 0.566 | 0.339 | 1.09E-30 | osteoclast.like | Hcfc1 |
| Nde11 | 6.77E-35 | 0.7550336 | 0.429 | 0.176 | 1.31E-30 | osteoclast.like | Nde1 |
| Ppp3cb2 | 6.78E-35 | 0.38804517 | 0.615 | 0.378 | 1.31E-30 | osteoclast.like | Ppp3cb |
| Smap1 | 7.32E-35 | 0.7629962 | 0.522 | 0.235 | 1.42E-30 | osteoclast.like | Smap1 |
| Plrg11 | 8.16E-35 | 0.86667339 | 0.459 | 0.199 | 1.58E-30 | osteoclast.like | Plrg1 |
| Mcrs1 | 8.80E-35 | 0.98463244 | 0.361 | 0.134 | 1.70E-30 | osteoclast.like | Mcrs1 |
| Ranbp31 | 9.14E-35 | 0.40491057 | 0.595 | 0.327 | 1.77E-30 | osteoclast.like | Ranbp3 |
| Letmd11 | 9.87E-35 | 0.80006616 | 0.415 | 0.175 | 1.91E-30 | osteoclast.like | Letmd1 |
| Rbbp61 | 1.02E-34 | 0.31109594 | 0.727 | 0.498 | 1.98E-30 | osteoclast.like | Rbbp6 |
| Avpi1 | 1.05E-34 | 1.18577007 | 0.356 | 0.111 | 2.03E-30 | osteoclast.like | Avpi1 |
| Tmem208 | 1.22E-34 | 0.3958429 | 0.537 | 0.309 | 2.37E-30 | osteoclast.like | Tmem208 |
| Naa152 | 1.23E-34 | 0.34399588 | 0.59 | 0.367 | 2.39E-30 | osteoclast.like | Naa15 |
| Strn31 | 1.30E-34 | 0.44089072 | 0.522 | 0.329 | 2.52E-30 | osteoclast.like | Strn3 |
| Irf32 | 1.37E-34 | 0.87547411 | 0.498 | 0.223 | 2.66E-30 | osteoclast.like | Irf3 |
| Thra | 1.38E-34 | 1.66783865 | 0.273 | 0.066 | 2.67E-30 | osteoclast.like | Thra |
| Setdb11 | 1.41E-34 | 0.60552623 | 0.493 | 0.251 | 2.73E-30 | osteoclast.like | Setdb1 |
| Esf1 | 1.54E-34 | 1.07636507 | 0.337 | 0.116 | 2.98E-30 | osteoclast.like | Esf1 |
| Api52 | 1.59E-34 | 0.28247347 | 0.693 | 0.445 | 3.08E-30 | osteoclast.like | Api5 |
| Nfyb1 | 1.61E-34 | 0.75915191 | 0.395 | 0.163 | 3.12E-30 | osteoclast.like | Nfyb |
| Nit2 | 1.67E-34 | 1.06375546 | 0.341 | 0.127 | 3.22E-30 | osteoclast.like | Nit2 |
| Polr3k1 | 1.73E-34 | 0.70921115 | 0.493 | 0.23 | 3.34E-30 | osteoclast.like | Polr3k |
| Dpm32 | 1.75E-34 | 0.39530971 | 0.541 | 0.314 | 3.38E-30 | osteoclast.like | Dpm3 |
| Orc61 | 1.75E-34 | 1.50397986 | 0.278 | 0.072 | 3.38E-30 | osteoclast.like | Orc6 |
| Psmb11 | 1.78E-34 | 0.40916706 | 0.615 | 0.39 | 3.43E-30 | osteoclast.like | Psmb1 |
| Copg11 | 1.83E-34 | 0.32843023 | 0.649 | 0.413 | 3.54E-30 | osteoclast.like | Copg1 |
| Sec11a | 2.01E-34 | 0.28513174 | 0.712 | 0.466 | 3.89E-30 | osteoclast.like | Sec11a |
| Bpnt12 | 2.03E-34 | 0.46112495 | 0.498 | 0.244 | 3.93E-30 | osteoclast.like | Bpnt1 |
| Atr | 2.08E-34 | 1.47135369 | 0.263 | 0.073 | 4.03E-30 | osteoclast.like | Atr |
| Pitpnb1 | 2.44E-34 | 0.46171491 | 0.566 | 0.278 | 4.71E-30 | osteoclast.like | Pitpnb |
| Desi21 | 2.90E-34 | 0.610418 | 0.537 | 0.261 | 5.61E-30 | osteoclast.like | Desi2 |
| Psma21 | 2.95E-34 | 0.36978375 | 0.59 | 0.363 | 5.71E-30 | osteoclast.like | Psma2 |
| Vipas392 | 3.04E-34 | 0.41144574 | 0.615 | 0.352 | 5.88E-30 | osteoclast.like | Vipas39 |
| Nqo21 | 3.16E-34 | 1.05621399 | 0.439 | 0.157 | 6.10E-30 | osteoclast.like | Nqo2 |
| Nsfl1c2 | 3.55E-34 | 0.57419105 | 0.551 | 0.284 | 6.87E-30 | osteoclast.like | Nsfl1c |
| Gid81 | 3.60E-34 | 0.42997313 | 0.615 | 0.334 | 6.96E-30 | osteoclast.like | Gid8 |
| Nploc42 | 3.70E-34 | 0.31238791 | 0.639 | 0.412 | 7.15E-30 | osteoclast.like | Nploc4 |
| Pwp2 | 3.94E-34 | 1.49421446 | 0.317 | 0.105 | 7.63E-30 | osteoclast.like | Pwp2 |
| Zswim7 | 5.60E-34 | 1.2829112 | 0.273 | 0.072 | 1.08E-29 | osteoclast.like | Zswim7 |
| Aldh9a11 | 5.75E-34 | 0.25098782 | 0.741 | 0.521 | 1.11E-29 | osteoclast.like | Aldh9a1 |
| Stoml1 | 5.85E-34 | 0.87877759 | 0.356 | 0.118 | 1.13E-29 | osteoclast.like | Stoml1 |
| Pik3ca | 5.87E-34 | 0.53724431 | 0.454 | 0.225 | 1.13E-29 | osteoclast.like | Pik3ca |
| Ppm1g2 | 6.24E-34 | 0.3085999 | 0.678 | 0.472 | 1.21E-29 | osteoclast.like | Ppm1g |
| Ubxn82 | 6.33E-34 | 0.63533178 | 0.522 | 0.229 | 1.22E-29 | osteoclast.like | Ubxn8 |
| Pcmt13 | 6.37E-34 | 0.55348732 | 0.629 | 0.327 | 1.23E-29 | osteoclast.like | Pcmt1 |
| Ripor11 | 7.03E-34 | 0.6552668 | 0.454 | 0.186 | 1.36E-29 | osteoclast.like | Ripor1 |
| Otud6b1 | 8.02E-34 | 0.92581504 | 0.405 | 0.178 | 1.55E-29 | osteoclast.like | Otud6b |
| Mpv17l21 | 9.17E-34 | 0.41950199 | 0.556 | 0.3 | 1.77E-29 | osteoclast.like | Mpv17l2 |
| Exosc71 | 9.28E-34 | 0.81430212 | 0.415 | 0.159 | 1.80E-29 | osteoclast.like | Exosc7 |
| Snrnp27 | 9.65E-34 | 0.74547232 | 0.434 | 0.215 | 1.87E-29 | osteoclast.like | Snrnp27 |
| Acp11 | 1.08E-33 | 0.60178608 | 0.512 | 0.269 | 2.09E-29 | osteoclast.like | Acp1 |
| Cgnl12 | 1.09E-33 | 0.85098578 | 0.39 | 0.134 | 2.10E-29 | osteoclast.like | Cgnl1 |
| Rbmx1 | 1.30E-33 | 0.98498185 | 0.346 | 0.128 | 2.52E-29 | osteoclast.like | Rbmx |
| Mtrf1l | 1.33E-33 | 1.51541069 | 0.273 | 0.073 | 2.58E-29 | osteoclast.like | Mtrf1l |
| Wsb21 | 1.38E-33 | 0.51268658 | 0.571 | 0.317 | 2.67E-29 | osteoclast.like | Wsb2 |
| Gfm2 | 1.39E-33 | 0.9529602 | 0.317 | 0.116 | 2.68E-29 | osteoclast.like | Gfm2 |
| Ccdc1071 | 1.40E-33 | 0.7787594 | 0.41 | 0.177 | 2.71E-29 | osteoclast.like | Ccdc107 |
| Usp20 | 1.55E-33 | 1.52484364 | 0.259 | 0.06 | 3.01E-29 | osteoclast.like | Usp20 |
| Fbxw81 | 1.60E-33 | 0.47601221 | 0.59 | 0.325 | 3.09E-29 | osteoclast.like | Fbxw8 |
| Tor2a | 1.62E-33 | 0.88582837 | 0.415 | 0.172 | 3.12E-29 | osteoclast.like | Tor2a |
| Sdhaf1 | 1.81E-33 | 0.82894222 | 0.341 | 0.128 | 3.49E-29 | osteoclast.like | Sdhaf1 |
| Smim10l12 | 1.92E-33 | 0.57910813 | 0.546 | 0.266 | 3.72E-29 | osteoclast.like | Smim10l1 |
| Celf1 | 2.28E-33 | 0.51408024 | 0.517 | 0.235 | 4.40E-29 | osteoclast.like | Celf1 |
| Akap82 | 2.28E-33 | 0.3099696 | 0.644 | 0.396 | 4.42E-29 | osteoclast.like | Akap8 |
| Ddx392 | 2.51E-33 | 0.40198741 | 0.522 | 0.348 | 4.86E-29 | osteoclast.like | Ddx39 |
| Decr1 | 2.66E-33 | 1.32732169 | 0.317 | 0.099 | 5.14E-29 | osteoclast.like | Decr1 |
| Exoc71 | 2.67E-33 | 0.36758058 | 0.532 | 0.285 | 5.17E-29 | osteoclast.like | Exoc7 |
| Scfd12 | 2.74E-33 | 0.43666364 | 0.434 | 0.218 | 5.30E-29 | osteoclast.like | Scfd1 |
| Lsm31 | 2.83E-33 | 0.81835867 | 0.439 | 0.192 | 5.46E-29 | osteoclast.like | Lsm3 |
| Mplkip | 3.59E-33 | 0.9944502 | 0.317 | 0.105 | 6.95E-29 | osteoclast.like | Mplkip |
| Itfg2 | 3.86E-33 | 1.56326228 | 0.254 | 0.064 | 7.46E-29 | osteoclast.like | Itfg2 |
| Tmem120a | 3.97E-33 | 0.99695578 | 0.356 | 0.118 | 7.67E-29 | osteoclast.like | Tmem120a |
| Gatd12 | 4.01E-33 | 0.68731632 | 0.483 | 0.226 | 7.75E-29 | osteoclast.like | Gatd1 |
| Slc25a171 | 4.92E-33 | 0.54398134 | 0.473 | 0.253 | 9.52E-29 | osteoclast.like | Slc25a17 |
| Mto11 | 5.46E-33 | 1.01796363 | 0.351 | 0.141 | 1.05E-28 | osteoclast.like | Mto1 |
| Trip103 | 5.53E-33 | 0.60034616 | 0.488 | 0.26 | 1.07E-28 | osteoclast.like | Trip10 |
| Bccip2 | 5.82E-33 | 0.37611807 | 0.527 | 0.306 | 1.13E-28 | osteoclast.like | Bccip |
| Tceal82 | 6.22E-33 | 0.41886196 | 0.527 | 0.324 | 1.20E-28 | osteoclast.like | Tceal8 |
| Isca21 | 6.63E-33 | 0.68253197 | 0.473 | 0.234 | 1.28E-28 | osteoclast.like | Isca2 |
| Mphosph61 | 6.90E-33 | 0.87462042 | 0.376 | 0.164 | 1.33E-28 | osteoclast.like | Mphosph6 |
| Topors4 | 7.68E-33 | 0.5356916 | 0.532 | 0.262 | 1.48E-28 | osteoclast.like | Topors |
| Alad1 | 8.71E-33 | 1.10234458 | 0.332 | 0.125 | 1.68E-28 | osteoclast.like | Alad |
| Prorp | 9.91E-33 | 1.53153413 | 0.273 | 0.063 | 1.92E-28 | osteoclast.like | Prorp |
| Mras | 1.22E-32 | 1.13784797 | 0.302 | 0.106 | 2.36E-28 | osteoclast.like | Mras |
| Ttc27 | 1.26E-32 | 1.29258394 | 0.278 | 0.082 | 2.44E-28 | osteoclast.like | Ttc27 |
| Slc25a46 | 1.27E-32 | 0.90544535 | 0.346 | 0.13 | 2.45E-28 | osteoclast.like | Slc25a46 |
| Spopl | 1.44E-32 | 0.53744089 | 0.42 | 0.204 | 2.78E-28 | osteoclast.like | Spopl |
| Ublcp1 | 1.62E-32 | 1.22637249 | 0.254 | 0.071 | 3.12E-28 | osteoclast.like | Ublcp1 |
| Phkb | 1.71E-32 | 0.7839669 | 0.356 | 0.148 | 3.31E-28 | osteoclast.like | Phkb |
| Secisbp2 | 1.86E-32 | 0.62975432 | 0.459 | 0.214 | 3.60E-28 | osteoclast.like | Secisbp2 |
| Lmf1 | 1.99E-32 | 0.53238551 | 0.488 | 0.234 | 3.85E-28 | osteoclast.like | Lmf1 |
| Eif2d | 2.53E-32 | 1.36336208 | 0.288 | 0.093 | 4.89E-28 | osteoclast.like | Eif2d |
| Pfdn62 | 2.58E-32 | 0.49214227 | 0.605 | 0.351 | 4.99E-28 | osteoclast.like | Pfdn6 |
| Creb31 | 2.61E-32 | 0.51668655 | 0.498 | 0.25 | 5.04E-28 | osteoclast.like | Creb3 |
| Ascc1 | 2.65E-32 | 0.91692562 | 0.385 | 0.143 | 5.12E-28 | osteoclast.like | Ascc1 |
| Phyh1 | 2.67E-32 | 0.3564407 | 0.39 | 0.224 | 5.16E-28 | osteoclast.like | Phyh |
| Gga33 | 3.06E-32 | 0.61142086 | 0.478 | 0.226 | 5.92E-28 | osteoclast.like | Gga3 |
| Wdr331 | 3.10E-32 | 0.2847518 | 0.585 | 0.354 | 6.00E-28 | osteoclast.like | Wdr33 |
| Nudcd21 | 3.48E-32 | 0.85863427 | 0.4 | 0.153 | 6.73E-28 | osteoclast.like | Nudcd2 |
| Dcaf11 | 3.96E-32 | 0.38360652 | 0.522 | 0.301 | 7.65E-28 | osteoclast.like | Dcaf11 |
| Naa101 | 4.32E-32 | 0.8481534 | 0.405 | 0.183 | 8.36E-28 | osteoclast.like | Naa10 |
| Fam98a1 | 4.63E-32 | 1.13818166 | 0.346 | 0.123 | 8.95E-28 | osteoclast.like | Fam98a |
| Gle11 | 4.73E-32 | 1.03189724 | 0.337 | 0.123 | 9.15E-28 | osteoclast.like | Gle1 |
| Drosha1 | 4.76E-32 | 1.08288826 | 0.356 | 0.122 | 9.20E-28 | osteoclast.like | Drosha |
| Smad5 | 4.85E-32 | 0.86903481 | 0.42 | 0.147 | 9.38E-28 | osteoclast.like | Smad5 |
| Utp61 | 4.92E-32 | 0.52222735 | 0.532 | 0.281 | 9.51E-28 | osteoclast.like | Utp6 |
| Mau2 | 5.27E-32 | 0.27976583 | 0.624 | 0.381 | 1.02E-27 | osteoclast.like | Mau2 |
| Bcas21 | 5.42E-32 | 0.28394809 | 0.58 | 0.335 | 1.05E-27 | osteoclast.like | Bcas2 |
| Brix11 | 5.66E-32 | 0.7384024 | 0.449 | 0.206 | 1.09E-27 | osteoclast.like | Brix1 |
| Pitrm11 | 6.05E-32 | 0.76679109 | 0.424 | 0.218 | 1.17E-27 | osteoclast.like | Pitrm1 |
| Pex6 | 6.63E-32 | 0.69208612 | 0.459 | 0.203 | 1.28E-27 | osteoclast.like | Pex6 |
| Mob21 | 8.23E-32 | 0.76902537 | 0.527 | 0.229 | 1.59E-27 | osteoclast.like | Mob2 |
| Actr101 | 8.34E-32 | 0.44756435 | 0.512 | 0.282 | 1.61E-27 | osteoclast.like | Actr10 |
| Nr2c2ap2 | 8.48E-32 | 0.76682205 | 0.546 | 0.251 | 1.64E-27 | osteoclast.like | Nr2c2ap |
| Echs12 | 8.56E-32 | 0.44983779 | 0.615 | 0.373 | 1.66E-27 | osteoclast.like | Echs1 |
| Kdm5a | 8.75E-32 | 0.4243686 | 0.478 | 0.277 | 1.69E-27 | osteoclast.like | Kdm5a |
| Strap1 | 8.79E-32 | 0.59931885 | 0.478 | 0.234 | 1.70E-27 | osteoclast.like | Strap |
| Atp13a11 | 1.01E-31 | 0.34248921 | 0.59 | 0.335 | 1.95E-27 | osteoclast.like | Atp13a1 |
| Uxs11 | 1.02E-31 | 0.62978471 | 0.444 | 0.204 | 1.97E-27 | osteoclast.like | Uxs1 |
| Akr1e1 | 1.04E-31 | 1.51310999 | 0.273 | 0.083 | 2.00E-27 | osteoclast.like | Akr1e1 |
| Setd1a2 | 1.08E-31 | 0.44385552 | 0.615 | 0.344 | 2.09E-27 | osteoclast.like | Setd1a |
| Tsta31 | 1.47E-31 | 0.54526133 | 0.468 | 0.249 | 2.84E-27 | osteoclast.like | Tsta3 |
| Mettl4 | 1.48E-31 | 1.56996252 | 0.298 | 0.078 | 2.87E-27 | osteoclast.like | Mettl4 |
| Asnsd11 | 1.51E-31 | 0.55099634 | 0.541 | 0.287 | 2.93E-27 | osteoclast.like | Asnsd1 |
| Senp6 | 1.67E-31 | 0.54274607 | 0.483 | 0.24 | 3.23E-27 | osteoclast.like | Senp6 |
| Ppip5k1 | 2.11E-31 | 1.04659306 | 0.259 | 0.079 | 4.08E-27 | osteoclast.like | Ppip5k1 |
| Txnl4a1 | 2.43E-31 | 0.84260566 | 0.502 | 0.22 | 4.70E-27 | osteoclast.like | Txnl4a |
| Setd1b2 | 2.47E-31 | 0.28342686 | 0.58 | 0.337 | 4.78E-27 | osteoclast.like | Setd1b |
| Kifc32 | 2.53E-31 | 0.50288874 | 0.395 | 0.17 | 4.90E-27 | osteoclast.like | Kifc3 |
| Hypk1 | 2.54E-31 | 0.51558206 | 0.595 | 0.312 | 4.92E-27 | osteoclast.like | Hypk |
| Nfs11 | 2.70E-31 | 1.10364382 | 0.38 | 0.149 | 5.22E-27 | osteoclast.like | Nfs1 |
| Ammecr1 | 2.85E-31 | 0.98866969 | 0.346 | 0.122 | 5.51E-27 | osteoclast.like | Ammecr1 |
| Mccc2 | 2.88E-31 | 1.53229935 | 0.278 | 0.082 | 5.57E-27 | osteoclast.like | Mccc2 |
| Acad9 | 2.99E-31 | 1.01967264 | 0.307 | 0.098 | 5.78E-27 | osteoclast.like | Acad9 |
| Ints32 | 3.16E-31 | 0.45647933 | 0.498 | 0.266 | 6.11E-27 | osteoclast.like | Ints3 |
| Ralgapa12 | 3.98E-31 | 0.52032911 | 0.512 | 0.246 | 7.69E-27 | osteoclast.like | Ralgapa1 |
| Crybg12 | 4.07E-31 | 0.3537992 | 0.41 | 0.201 | 7.87E-27 | osteoclast.like | Crybg1 |
| Numb1 | 4.59E-31 | 0.81557576 | 0.502 | 0.236 | 8.87E-27 | osteoclast.like | Numb |
| Dnpep1 | 5.46E-31 | 0.85040984 | 0.459 | 0.216 | 1.06E-26 | osteoclast.like | Dnpep |
| Xrcc6 | 5.80E-31 | 1.52955109 | 0.268 | 0.072 | 1.12E-26 | osteoclast.like | Xrcc6 |
| Blcap4 | 5.98E-31 | 0.30436539 | 0.561 | 0.302 | 1.16E-26 | osteoclast.like | Blcap |
| Ubxn7 | 6.28E-31 | 0.53797863 | 0.493 | 0.237 | 1.21E-26 | osteoclast.like | Ubxn7 |
| Trappc51 | 6.48E-31 | 0.56723601 | 0.473 | 0.242 | 1.25E-26 | osteoclast.like | Trappc5 |
| Vps542 | 6.88E-31 | 0.39806133 | 0.532 | 0.305 | 1.33E-26 | osteoclast.like | Vps54 |
| Ercc61 | 6.93E-31 | 0.66941763 | 0.395 | 0.161 | 1.34E-26 | osteoclast.like | Ercc6 |
| Me22 | 7.17E-31 | 0.60243672 | 0.449 | 0.191 | 1.39E-26 | osteoclast.like | Me2 |
| Acox12 | 7.35E-31 | 0.35090129 | 0.683 | 0.402 | 1.42E-26 | osteoclast.like | Acox1 |
| Dohh1 | 7.67E-31 | 0.55852453 | 0.507 | 0.248 | 1.48E-26 | osteoclast.like | Dohh |
| Cul5 | 7.74E-31 | 0.52670784 | 0.41 | 0.187 | 1.50E-26 | osteoclast.like | Cul5 |
| Snf81 | 7.81E-31 | 0.32717651 | 0.732 | 0.475 | 1.51E-26 | osteoclast.like | Snf8 |
| Btbd2 | 8.24E-31 | 0.91205599 | 0.385 | 0.167 | 1.59E-26 | osteoclast.like | Btbd2 |
| Inpp5a | 8.44E-31 | 0.51731836 | 0.385 | 0.172 | 1.63E-26 | osteoclast.like | Inpp5a |
| Gtpbp3 | 8.98E-31 | 0.92998577 | 0.312 | 0.114 | 1.74E-26 | osteoclast.like | Gtpbp3 |
| Tmem184b2 | 9.53E-31 | 0.32551994 | 0.678 | 0.421 | 1.84E-26 | osteoclast.like | Tmem184b |
| Sec24d1 | 1.06E-30 | 0.89632334 | 0.41 | 0.167 | 2.04E-26 | osteoclast.like | Sec24d |
| Stk161 | 1.16E-30 | 0.58018505 | 0.532 | 0.231 | 2.24E-26 | osteoclast.like | Stk16 |
| Osbpl2 | 1.21E-30 | 0.84586713 | 0.332 | 0.129 | 2.35E-26 | osteoclast.like | Osbpl2 |
| Txlna2 | 1.26E-30 | 0.33271858 | 0.585 | 0.366 | 2.43E-26 | osteoclast.like | Txlna |
| Josd12 | 1.30E-30 | 0.73222051 | 0.454 | 0.217 | 2.52E-26 | osteoclast.like | Josd1 |
| Enox2 | 1.50E-30 | 1.03970388 | 0.332 | 0.12 | 2.90E-26 | osteoclast.like | Enox2 |
| Metap11 | 2.08E-30 | 0.43011419 | 0.493 | 0.261 | 4.03E-26 | osteoclast.like | Metap1 |
| Dhx37 | 2.10E-30 | 0.93198828 | 0.341 | 0.116 | 4.06E-26 | osteoclast.like | Dhx37 |
| Trappc8 | 2.14E-30 | 0.39270939 | 0.595 | 0.327 | 4.14E-26 | osteoclast.like | Trappc8 |

| | | | | | | | |
|---|---|---|---|---|---|---|---|
| Snw12 | 2.24E-30 | 0.46739424 | 0.561 | 0.293 | 4.33E-26 | osteoclast.like | Snw1 |
| E2f61 | 2.41E-30 | 1.17158659 | 0.317 | 0.095 | 4.67E-26 | osteoclast.like | E2f6 |
| Crbn1 | 2.80E-30 | 0.86850783 | 0.395 | 0.164 | 5.41E-26 | osteoclast.like | Crbn |
| Gart2 | 2.88E-30 | 0.47116686 | 0.498 | 0.28 | 5.56E-26 | osteoclast.like | Gart |
| Orai32 | 2.95E-30 | 0.44814398 | 0.454 | 0.25 | 5.70E-26 | osteoclast.like | Orai3 |
| Aldh16a11 | 3.06E-30 | 0.73171235 | 0.434 | 0.194 | 5.91E-26 | osteoclast.like | Aldh16a1 |
| Timm9 | 3.20E-30 | 1.26219113 | 0.273 | 0.085 | 6.18E-26 | osteoclast.like | Timm9 |
| Sh3bgrl2 | 3.21E-30 | 1.23487531 | 0.273 | 0.067 | 6.21E-26 | osteoclast.like | Sh3bgrl2 |
| Ppp5c2 | 3.34E-30 | 0.5954205 | 0.532 | 0.284 | 6.46E-26 | osteoclast.like | Ppp5c |
| Psmg31 | 3.43E-30 | 1.08116969 | 0.351 | 0.139 | 6.63E-26 | osteoclast.like | Psmg3 |
| Gzf12 | 3.46E-30 | 0.79424132 | 0.41 | 0.174 | 6.68E-26 | osteoclast.like | Gzf1 |
| Nudt16l13 | 3.57E-30 | 0.3141501 | 0.6 | 0.343 | 6.89E-26 | osteoclast.like | Nudt16l1 |
| Ppp1r71 | 3.57E-30 | 0.55430787 | 0.493 | 0.254 | 6.90E-26 | osteoclast.like | Ppp1r7 |
| Rchy11 | 3.78E-30 | 0.29410803 | 0.454 | 0.255 | 7.31E-26 | osteoclast.like | Rchy1 |
| Sphk21 | 4.06E-30 | 0.33328993 | 0.546 | 0.323 | 7.86E-26 | osteoclast.like | Sphk2 |
| Prkra | 4.13E-30 | 1.11104737 | 0.307 | 0.118 | 7.99E-26 | osteoclast.like | Prkra |
| Yme1l11 | 4.21E-30 | 0.27089558 | 0.561 | 0.344 | 8.15E-26 | osteoclast.like | Yme1l1 |
| Tfam1 | 4.33E-30 | 0.76418304 | 0.341 | 0.144 | 8.37E-26 | osteoclast.like | Tfam |
| Washc32 | 4.37E-30 | 0.47504348 | 0.502 | 0.262 | 8.45E-26 | osteoclast.like | Washc3 |
| Zfp787 | 5.15E-30 | 0.6749218 | 0.38 | 0.148 | 9.95E-26 | osteoclast.like | Zfp787 |
| Usp22 | 5.36E-30 | 0.34815719 | 0.551 | 0.308 | 1.04E-25 | osteoclast.like | Usp22 |
| Polr2j1 | 5.43E-30 | 0.5667729 | 0.415 | 0.195 | 1.05E-25 | osteoclast.like | Polr2j |
| Tex101 | 5.53E-30 | 0.58433285 | 0.424 | 0.214 | 1.07E-25 | osteoclast.like | Tex10 |
| Agfg13 | 5.64E-30 | 0.3715293 | 0.644 | 0.386 | 1.09E-25 | osteoclast.like | Agfg1 |
| Cherp1 | 5.97E-30 | 0.47771987 | 0.595 | 0.344 | 1.15E-25 | osteoclast.like | Cherp |
| Lars2 | 6.41E-30 | 0.48219592 | 0.502 | 0.277 | 1.24E-25 | osteoclast.like | Lars |
| Smpd41 | 6.48E-30 | 0.94650498 | 0.395 | 0.151 | 1.25E-25 | osteoclast.like | Smpd4 |
| Bag51 | 6.92E-30 | 0.87183994 | 0.395 | 0.153 | 1.34E-25 | osteoclast.like | Bag5 |
| Cdc231 | 6.95E-30 | 0.85125083 | 0.351 | 0.144 | 1.34E-25 | osteoclast.like | Cdc23 |
| D5Ertd579e2 | 6.97E-30 | 0.35330832 | 0.546 | 0.307 | 1.35E-25 | osteoclast.like | D5Ertd579e |
| Cdc42ep2 | 7.00E-30 | 1.29838401 | 0.298 | 0.087 | 1.35E-25 | osteoclast.like | Cdc42ep2 |
| Hspb11 | 7.82E-30 | 1.14620671 | 0.293 | 0.093 | 1.51E-25 | osteoclast.like | Hspb11 |
| Usp341 | 8.75E-30 | 0.55567518 | 0.512 | 0.277 | 1.69E-25 | osteoclast.like | Usp34 |
| Galk22 | 9.04E-30 | 0.36196015 | 0.6 | 0.341 | 1.75E-25 | osteoclast.like | Galk2 |
| Jtb | 9.28E-30 | 0.461053 | 0.58 | 0.334 | 1.79E-25 | osteoclast.like | Jtb |
| Sae11 | 9.40E-30 | 0.66239407 | 0.571 | 0.328 | 1.82E-25 | osteoclast.like | Sae1 |
| Nob1 | 9.63E-30 | 1.08992164 | 0.302 | 0.111 | 1.86E-25 | osteoclast.like | Nob1 |
| Prmt31 | 9.91E-30 | 1.28128121 | 0.346 | 0.113 | 1.92E-25 | osteoclast.like | Prmt3 |
| Champ11 | 1.01E-29 | 1.09429696 | 0.332 | 0.12 | 1.96E-25 | osteoclast.like | Champ1 |
| Ikbkg2 | 1.11E-29 | 0.61463337 | 0.527 | 0.245 | 2.15E-25 | osteoclast.like | Ikbkg |
| Usp383 | 1.18E-29 | 0.41230019 | 0.541 | 0.258 | 2.28E-25 | osteoclast.like | Usp38 |
| Ost41 | 1.23E-29 | 0.46604275 | 0.537 | 0.311 | 2.38E-25 | osteoclast.like | Ost4 |
| Rtraf1 | 1.23E-29 | 0.56520671 | 0.463 | 0.26 | 2.38E-25 | osteoclast.like | Rtraf |
| Dffa | 1.37E-29 | 0.95664102 | 0.341 | 0.124 | 2.64E-25 | osteoclast.like | Dffa |
| Fmn13 | 1.46E-29 | 0.58548729 | 0.502 | 0.276 | 2.83E-25 | osteoclast.like | Fmn1 |
| Afdn | 1.47E-29 | 1.06165139 | 0.376 | 0.144 | 2.84E-25 | osteoclast.like | Afdn |
| Nol111 | 1.59E-29 | 0.84398297 | 0.356 | 0.153 | 3.07E-25 | osteoclast.like | Nol11 |
| Ltv13 | 1.59E-29 | 0.64135199 | 0.488 | 0.23 | 3.07E-25 | osteoclast.like | Ltv1 |
| Wdsub1 | 1.64E-29 | 1.03358537 | 0.317 | 0.112 | 3.17E-25 | osteoclast.like | Wdsub1 |
| Heatr31 | 1.71E-29 | 0.84219699 | 0.351 | 0.146 | 3.31E-25 | osteoclast.like | Heatr3 |
| Lin7c | 1.81E-29 | 0.2645286 | 0.673 | 0.466 | 3.51E-25 | osteoclast.like | Lin7c |
| Tnpo31 | 1.93E-29 | 0.41122009 | 0.571 | 0.323 | 3.73E-25 | osteoclast.like | Tnpo3 |
| Trim441 | 2.01E-29 | 0.46692094 | 0.59 | 0.358 | 3.89E-25 | osteoclast.like | Trim44 |
| Gtf3c61 | 2.27E-29 | 0.73609798 | 0.507 | 0.248 | 4.39E-25 | osteoclast.like | Gtf3c6 |
| Nsa23 | 2.36E-29 | 0.65489069 | 0.454 | 0.247 | 4.55E-25 | osteoclast.like | Nsa2 |
| Pccb | 2.42E-29 | 1.06015159 | 0.337 | 0.118 | 4.68E-25 | osteoclast.like | Pccb |
| Trpv22 | 2.45E-29 | 0.25612648 | 0.532 | 0.313 | 4.73E-25 | osteoclast.like | Trpv2 |
| Cnot81 | 2.67E-29 | 0.33747647 | 0.576 | 0.336 | 5.16E-25 | osteoclast.like | Cnot8 |
| Mri11 | 2.72E-29 | 0.69864888 | 0.395 | 0.15 | 5.25E-25 | osteoclast.like | Mri1 |
| Clpb | 2.84E-29 | 1.13355807 | 0.322 | 0.112 | 5.49E-25 | osteoclast.like | Clpb |
| Tpr2 | 2.84E-29 | 0.30986676 | 0.615 | 0.436 | 5.50E-25 | osteoclast.like | Tpr |
| Phf14 | 2.86E-29 | 0.29034277 | 0.478 | 0.284 | 5.53E-25 | osteoclast.like | Phf14 |
| Bloc1s5 | 2.89E-29 | 0.8502825 | 0.317 | 0.102 | 5.59E-25 | osteoclast.like | Bloc1s5 |
| Ptpn9 | 2.95E-29 | 0.56358519 | 0.488 | 0.234 | 5.71E-25 | osteoclast.like | Ptpn9 |
| Smarcc11 | 3.05E-29 | 0.56070482 | 0.551 | 0.304 | 5.91E-25 | osteoclast.like | Smarcc1 |
| Atf2 | 3.22E-29 | 0.389622 | 0.585 | 0.328 | 6.23E-25 | osteoclast.like | Atf2 |
| Golga41 | 3.46E-29 | 0.32128161 | 0.58 | 0.354 | 6.70E-25 | osteoclast.like | Golga4 |
| Utp201 | 3.70E-29 | 1.03765885 | 0.337 | 0.132 | 7.16E-25 | osteoclast.like | Utp20 |
| Nipsnap3b | 4.37E-29 | 0.70838141 | 0.551 | 0.276 | 8.44E-25 | osteoclast.like | Nipsnap3b |
| Vbp12 | 4.46E-29 | 0.30481902 | 0.595 | 0.345 | 8.62E-25 | osteoclast.like | Vbp1 |
| Hectd32 | 4.62E-29 | 0.4456156 | 0.541 | 0.294 | 8.93E-25 | osteoclast.like | Hectd3 |
| Arfgap21 | 4.70E-29 | 0.40825568 | 0.595 | 0.362 | 9.09E-25 | osteoclast.like | Arfgap2 |
| Nae11 | 5.06E-29 | 0.78136136 | 0.356 | 0.142 | 9.78E-25 | osteoclast.like | Nae1 |
| Ube2s1 | 5.49E-29 | 0.37621818 | 0.654 | 0.376 | 1.06E-24 | osteoclast.like | Ube2s |
| Ston2 | 5.82E-29 | 0.57806554 | 0.307 | 0.144 | 1.13E-24 | osteoclast.like | Ston2 |
| Usp471 | 5.91E-29 | 0.35147987 | 0.541 | 0.32 | 1.14E-24 | osteoclast.like | Usp47 |
| mt-Nd21 | 5.92E-29 | 0.25035595 | 0.541 | 0.336 | 1.15E-24 | osteoclast.like | mt-Nd2 |
| Pmm1 | 6.57E-29 | 1.24671628 | 0.307 | 0.097 | 1.27E-24 | osteoclast.like | Pmm1 |
| Dynlt1f3 | 6.81E-29 | 0.68813896 | 0.502 | 0.263 | 1.32E-24 | osteoclast.like | Dynlt1f |
| Lyrm2 | 6.85E-29 | 0.9968949 | 0.327 | 0.105 | 1.32E-24 | osteoclast.like | Lyrm2 |
| Tcf20 | 6.91E-29 | 0.39557127 | 0.541 | 0.308 | 1.34E-24 | osteoclast.like | Tcf20 |
| Wdr13 | 7.15E-29 | 0.54096763 | 0.488 | 0.264 | 1.38E-24 | osteoclast.like | Wdr13 |
| Fbxo6 | 7.40E-29 | 0.48466414 | 0.502 | 0.276 | 1.43E-24 | osteoclast.like | Fbxo6 |
| Ube2e12 | 7.74E-29 | 0.55037563 | 0.537 | 0.299 | 1.50E-24 | osteoclast.like | Ube2e1 |
| Parn | 7.79E-29 | 1.1504168 | 0.327 | 0.114 | 1.51E-24 | osteoclast.like | Parn |
| Xpo11 | 7.81E-29 | 0.39175444 | 0.595 | 0.359 | 1.51E-24 | osteoclast.like | Xpo1 |
| Gtf2a21 | 8.43E-29 | 0.66558249 | 0.454 | 0.203 | 1.63E-24 | osteoclast.like | Gtf2a2 |
| Bckdha | 8.72E-29 | 1.03809184 | 0.351 | 0.122 | 1.69E-24 | osteoclast.like | Bckdha |
| Fbxl20 | 8.96E-29 | 0.58321804 | 0.459 | 0.235 | 1.73E-24 | osteoclast.like | Fbxl20 |
| Cetn21 | 9.77E-29 | 0.9927606 | 0.376 | 0.155 | 1.89E-24 | osteoclast.like | Cetn2 |
| Nfatc32 | 1.05E-28 | 0.58875838 | 0.498 | 0.266 | 2.03E-24 | osteoclast.like | Nfatc3 |
| Bcor | 1.21E-28 | 0.37543523 | 0.376 | 0.212 | 2.35E-24 | osteoclast.like | Bcor |
| Psmd132 | 1.30E-28 | 0.27073592 | 0.698 | 0.452 | 2.52E-24 | osteoclast.like | Psmd13 |
| Dgcr61 | 1.34E-28 | 0.88942031 | 0.42 | 0.176 | 2.59E-24 | osteoclast.like | Dgcr6 |
| Smarcd21 | 1.35E-28 | 0.34874708 | 0.629 | 0.356 | 2.61E-24 | osteoclast.like | Smarcd2 |
| Taf103 | 1.38E-28 | 0.4563029 | 0.507 | 0.271 | 2.67E-24 | osteoclast.like | Taf10 |
| Vapa2 | 1.43E-28 | 0.46784263 | 0.512 | 0.29 | 2.76E-24 | osteoclast.like | Vapa |
| Pbrm11 | 1.47E-28 | 0.34019309 | 0.585 | 0.346 | 2.83E-24 | osteoclast.like | Pbrm1 |
| Stam1 | 1.59E-28 | 0.71715464 | 0.41 | 0.187 | 3.07E-24 | osteoclast.like | Stam |
| Zfp3623 | 1.82E-28 | 0.65160979 | 0.493 | 0.253 | 3.52E-24 | osteoclast.like | Zfp362 |
| Brms1 | 1.85E-28 | 0.34820771 | 0.561 | 0.358 | 3.57E-24 | osteoclast.like | Brms1 |
| Ehmt1 | 1.85E-28 | 0.36555863 | 0.493 | 0.264 | 3.58E-24 | osteoclast.like | Ehmt1 |
| Slc19a1 | 1.89E-28 | 1.3235805 | 0.254 | 0.083 | 3.65E-24 | osteoclast.like | Slc19a1 |
| Nt5c3 | 1.91E-28 | 0.85934901 | 0.4 | 0.138 | 3.70E-24 | osteoclast.like | Nt5c3 |
| Usp391 | 2.03E-28 | 0.67777726 | 0.395 | 0.179 | 3.93E-24 | osteoclast.like | Usp39 |
| Cox161 | 2.14E-28 | 0.65625848 | 0.532 | 0.255 | 4.13E-24 | osteoclast.like | Cox16 |
| Fiz11 | 2.18E-28 | 0.65965674 | 0.39 | 0.172 | 4.22E-24 | osteoclast.like | Fiz1 |
| Trim281 | 2.33E-28 | 0.48086479 | 0.424 | 0.241 | 4.51E-24 | osteoclast.like | Trim28 |
| Fam173a3 | 2.45E-28 | 0.47548675 | 0.605 | 0.337 | 4.73E-24 | osteoclast.like | Fam173a |
| Bloc1s11 | 2.48E-28 | 0.76290303 | 0.449 | 0.202 | 4.80E-24 | osteoclast.like | Bloc1s1 |
| Gmnn1 | 2.49E-28 | 1.13123029 | 0.337 | 0.12 | 4.82E-24 | osteoclast.like | Gmnn |
| Fip1l11 | 2.73E-28 | 0.52082286 | 0.566 | 0.272 | 5.29E-24 | osteoclast.like | Fip1l1 |
| Trmt2a2 | 2.82E-28 | 0.56660753 | 0.517 | 0.249 | 5.46E-24 | osteoclast.like | Trmt2a |
| Exosc31 | 2.83E-28 | 0.39330702 | 0.468 | 0.228 | 5.47E-24 | osteoclast.like | Exosc3 |
| Anapc7 | 3.16E-28 | 0.58160273 | 0.473 | 0.238 | 6.11E-24 | osteoclast.like | Anapc7 |
| Urm11 | 3.30E-28 | 0.75587706 | 0.527 | 0.268 | 6.38E-24 | osteoclast.like | Urm1 |
| Blmh1 | 3.83E-28 | 0.58665823 | 0.498 | 0.265 | 7.40E-24 | osteoclast.like | Blmh |
| Rab3gap11 | 3.93E-28 | 0.79251469 | 0.493 | 0.236 | 7.59E-24 | osteoclast.like | Rab3gap1 |
| Acot2 | 4.10E-28 | 1.10600308 | 0.298 | 0.092 | 7.92E-24 | osteoclast.like | Acot2 |
| Dtd2 | 4.10E-28 | 1.07240447 | 0.298 | 0.108 | 7.93E-24 | osteoclast.like | Dtd2 |
| Prpf32 | 4.30E-28 | 0.57084969 | 0.444 | 0.187 | 8.32E-24 | osteoclast.like | Prpf3 |
| Cnot91 | 4.99E-28 | 0.82537008 | 0.488 | 0.219 | 9.64E-24 | osteoclast.like | Cnot9 |
| Pym11 | 5.14E-28 | 0.78115688 | 0.366 | 0.144 | 9.94E-24 | osteoclast.like | Pym1 |
| Tti2 | 5.48E-28 | 1.08540162 | 0.293 | 0.096 | 1.06E-23 | osteoclast.like | Tti2 |
| Ncbp12 | 5.92E-28 | 0.66080912 | 0.42 | 0.226 | 1.14E-23 | osteoclast.like | Ncbp1 |
| Nup932 | 6.10E-28 | 0.67022241 | 0.39 | 0.172 | 1.18E-23 | osteoclast.like | Nup93 |
| Lrrc14 | 6.62E-28 | 0.93245644 | 0.288 | 0.107 | 1.28E-23 | osteoclast.like | Lrrc14 |
| Ring1 | 6.81E-28 | 0.51601988 | 0.473 | 0.247 | 1.32E-23 | osteoclast.like | Ring1 |
| Pop51 | 7.39E-28 | 0.79150712 | 0.317 | 0.128 | 1.43E-23 | osteoclast.like | Pop5 |
| Hpf11 | 7.52E-28 | 0.6156087 | 0.449 | 0.209 | 1.45E-23 | osteoclast.like | Hpf1 |
| Gpnmb | 7.68E-28 | 0.81838866 | 0.322 | 0.09 | 1.49E-23 | osteoclast.like | Gpnmb |
| Trnt1 | 8.05E-28 | 0.97778952 | 0.371 | 0.141 | 1.56E-23 | osteoclast.like | Trnt1 |
| Stx5a | 9.90E-28 | 0.26012634 | 0.624 | 0.352 | 1.91E-23 | osteoclast.like | Stx5a |
| Adrm11 | 1.04E-27 | 0.6490682 | 0.459 | 0.226 | 2.00E-23 | osteoclast.like | Adrm1 |
| Tmed41 | 1.04E-27 | 0.28900181 | 0.576 | 0.36 | 2.01E-23 | osteoclast.like | Tmed4 |
| Gmppa | 1.13E-27 | 0.58462596 | 0.493 | 0.245 | 2.19E-23 | osteoclast.like | Gmppa |
| Clpp1 | 1.17E-27 | 0.95582948 | 0.346 | 0.136 | 2.27E-23 | osteoclast.like | Clpp |
| Gtdc1 | 1.23E-27 | 0.76969366 | 0.302 | 0.123 | 2.38E-23 | osteoclast.like | Gtdc1 |
| Wdr51 | 1.24E-27 | 0.90178627 | 0.488 | 0.225 | 2.39E-23 | osteoclast.like | Wdr5 |
| Krtcap22 | 1.29E-27 | 0.51136845 | 0.449 | 0.217 | 2.50E-23 | osteoclast.like | Krtcap2 |
| Helz3 | 1.40E-27 | 0.58751854 | 0.415 | 0.199 | 2.71E-23 | osteoclast.like | Helz |
| Coq10a1 | 1.46E-27 | 0.5528849 | 0.429 | 0.196 | 2.82E-23 | osteoclast.like | Coq10a |
| Aifm11 | 1.47E-27 | 0.81730277 | 0.322 | 0.127 | 2.85E-23 | osteoclast.like | Aifm1 |
| Dcaf101 | 1.54E-27 | 0.65074725 | 0.463 | 0.209 | 2.98E-23 | osteoclast.like | Dcaf10 |
| Itpkc | 1.57E-27 | 0.55733427 | 0.346 | 0.134 | 3.04E-23 | osteoclast.like | Itpkc |
| Ap1ar1 | 1.60E-27 | 0.53968562 | 0.507 | 0.248 | 3.08E-23 | osteoclast.like | Ap1ar |
| Xpo51 | 1.67E-27 | 1.06279143 | 0.341 | 0.133 | 3.23E-23 | osteoclast.like | Xpo5 |
| Cfap298 | 1.74E-27 | 0.96887605 | 0.278 | 0.102 | 3.37E-23 | osteoclast.like | Cfap298 |
| Eif2s3x1 | 1.76E-27 | 0.99891443 | 0.351 | 0.138 | 3.40E-23 | osteoclast.like | Eif2s3x |
| Zc3h152 | 2.03E-27 | 0.41900147 | 0.522 | 0.283 | 3.92E-23 | osteoclast.like | Zc3h15 |
| Ube3b | 2.05E-27 | 0.82138215 | 0.434 | 0.189 | 3.96E-23 | osteoclast.like | Ube3b |
| Zmym3 | 2.06E-27 | 0.82053846 | 0.341 | 0.125 | 3.98E-23 | osteoclast.like | Zmym3 |
| Phrf11 | 2.19E-27 | 0.39682237 | 0.527 | 0.283 | 4.22E-23 | osteoclast.like | Phrf1 |
| Pik3r22 | 2.27E-27 | 0.5509879 | 0.532 | 0.303 | 4.39E-23 | osteoclast.like | Pik3r2 |
| Nmd32 | 2.72E-27 | 0.87266246 | 0.395 | 0.178 | 5.25E-23 | osteoclast.like | Nmd3 |
| Dnajc81 | 2.76E-27 | 0.29938651 | 0.615 | 0.369 | 5.33E-23 | osteoclast.like | Dnajc8 |
| Sgta | 2.79E-27 | 0.71254562 | 0.337 | 0.165 | 5.40E-23 | osteoclast.like | Sgta |
| Ino80b1 | 2.81E-27 | 0.52336541 | 0.507 | 0.267 | 5.44E-23 | osteoclast.like | Ino80b |
| Mettl92 | 2.81E-27 | 0.30544202 | 0.585 | 0.336 | 5.44E-23 | osteoclast.like | Mettl9 |
| Nme12 | 2.86E-27 | 0.34081816 | 0.566 | 0.377 | 5.53E-23 | osteoclast.like | Nme1 |
| Mlkl | 2.92E-27 | 0.87886018 | 0.322 | 0.122 | 5.64E-23 | osteoclast.like | Mlkl |
| Kmt2c1 | 2.92E-27 | 0.56883091 | 0.463 | 0.245 | 5.64E-23 | osteoclast.like | Kmt2c |
| Cd2bp21 | 3.32E-27 | 0.46018971 | 0.6 | 0.331 | 6.42E-23 | osteoclast.like | Cd2bp2 |
| Kdm5b | 3.36E-27 | 0.64662642 | 0.351 | 0.142 | 6.50E-23 | osteoclast.like | Kdm5b |
| Thyn11 | 3.38E-27 | 0.90652654 | 0.302 | 0.131 | 6.53E-23 | osteoclast.like | Thyn1 |
| Rabgap1l | 3.38E-27 | 0.76398031 | 0.268 | 0.117 | 6.54E-23 | osteoclast.like | Rabgap1l |
| Hsf2 | 3.43E-27 | 1.25770061 | 0.298 | 0.094 | 6.63E-23 | osteoclast.like | Hsf2 |
| Tmem106b3 | 3.79E-27 | 0.44398999 | 0.454 | 0.257 | 7.34E-23 | osteoclast.like | Tmem106b |
| Vps72 | 3.92E-27 | 0.69537581 | 0.468 | 0.198 | 7.59E-23 | osteoclast.like | Vps72 |
| Mogs2 | 4.29E-27 | 0.437523 | 0.415 | 0.231 | 8.30E-23 | osteoclast.like | Mogs |
| Pdrg13 | 4.35E-27 | 0.58484791 | 0.405 | 0.226 | 8.41E-23 | osteoclast.like | Pdrg1 |
| Nol91 | 4.38E-27 | 0.72656088 | 0.341 | 0.145 | 8.48E-23 | osteoclast.like | Nol9 |
| Farp2 | 4.59E-27 | 1.36695396 | 0.259 | 0.072 | 8.87E-23 | osteoclast.like | Farp2 |
| Prepl | 5.39E-27 | 1.26346738 | 0.278 | 0.08 | 1.04E-22 | osteoclast.like | Prepl |
| Tmem205 | 5.45E-27 | 1.00022751 | 0.302 | 0.109 | 1.05E-22 | osteoclast.like | Tmem205 |
| Pik3c31 | 5.50E-27 | 0.49413557 | 0.468 | 0.231 | 1.06E-22 | osteoclast.like | Pik3c3 |
| Pogk | 7.02E-27 | 0.75240797 | 0.288 | 0.105 | 1.36E-22 | osteoclast.like | Pogk |
| Zc3h141 | 7.11E-27 | 0.53738212 | 0.478 | 0.258 | 1.38E-22 | osteoclast.like | Zc3h14 |
| Mapk8 | 7.29E-27 | 1.06728451 | 0.288 | 0.103 | 1.41E-22 | osteoclast.like | Mapk8 |
| Zfp3302 | 7.39E-27 | 0.63144597 | 0.463 | 0.238 | 1.43E-22 | osteoclast.like | Zfp330 |
| Dph5 | 7.47E-27 | 0.81637866 | 0.317 | 0.128 | 1.44E-22 | osteoclast.like | Dph5 |
| Acbd61 | 7.52E-27 | 0.70449501 | 0.488 | 0.26 | 1.45E-22 | osteoclast.like | Acbd6 |
| Eif2ak11 | 7.57E-27 | 0.415036 | 0.571 | 0.302 | 1.46E-22 | osteoclast.like | Eif2ak1 |
| Aopep1 | 8.78E-27 | 0.48807971 | 0.346 | 0.144 | 1.70E-22 | osteoclast.like | Aopep |
| Zfand6 | 8.98E-27 | 0.48861748 | 0.493 | 0.291 | 1.74E-22 | osteoclast.like | Zfand6 |
| Gemin71 | 9.51E-27 | 0.39866205 | 0.566 | 0.328 | 1.84E-22 | osteoclast.like | Gemin7 |
| Coa7 | 9.70E-27 | 1.22791225 | 0.298 | 0.09 | 1.87E-22 | osteoclast.like | Coa7 |
| Metap1d | 9.98E-27 | 0.65593852 | 0.332 | 0.123 | 1.93E-22 | osteoclast.like | Metap1d |
| Eif2ak31 | 1.07E-26 | 0.49974962 | 0.415 | 0.184 | 2.06E-22 | osteoclast.like | Eif2ak3 |
| Gba2 | 1.08E-26 | 0.98548973 | 0.283 | 0.096 | 2.08E-22 | osteoclast.like | Gba2 |
| Ankrd49 | 1.14E-26 | 1.22281654 | 0.254 | 0.074 | 2.21E-22 | osteoclast.like | Ankrd49 |
| Inpp5b2 | 1.46E-26 | 0.46051573 | 0.424 | 0.27 | 2.83E-22 | osteoclast.like | Inpp5b |
| Slc17a51 | 1.55E-26 | 0.74221287 | 0.327 | 0.136 | 3.00E-22 | osteoclast.like | Slc17a5 |
| Ap5s1 | 1.59E-26 | 0.70285931 | 0.38 | 0.147 | 3.07E-22 | osteoclast.like | Ap5s1 |
| Wdr112 | 1.61E-26 | 0.61421982 | 0.371 | 0.179 | 3.10E-22 | osteoclast.like | Wdr11 |
| Rnf2152 | 1.70E-26 | 0.42013568 | 0.439 | 0.237 | 3.28E-22 | osteoclast.like | Rnf215 |
| Pigc | 1.85E-26 | 0.65524659 | 0.356 | 0.134 | 3.58E-22 | osteoclast.like | Pigc |
| Fam204a | 2.05E-26 | 0.58827863 | 0.39 | 0.173 | 3.96E-22 | osteoclast.like | Fam204a |
| Dcaf132 | 2.10E-26 | 0.51971397 | 0.527 | 0.298 | 4.05E-22 | osteoclast.like | Dcaf13 |
| Sars3 | 2.10E-26 | 0.35547154 | 0.517 | 0.285 | 4.06E-22 | osteoclast.like | Sars |
| Rabif | 2.19E-26 | 0.64635824 | 0.356 | 0.15 | 4.23E-22 | osteoclast.like | Rabif |
| Dhx361 | 2.43E-26 | 0.48940482 | 0.429 | 0.244 | 4.70E-22 | osteoclast.like | Dhx36 |
| Ralgapb1 | 2.67E-26 | 0.42374053 | 0.498 | 0.265 | 5.15E-22 | osteoclast.like | Ralgapb |
| Selenos2 | 2.82E-26 | 0.27865829 | 0.571 | 0.339 | 5.45E-22 | osteoclast.like | Selenos |
| Alg81 | 2.82E-26 | 1.03849928 | 0.346 | 0.12 | 5.46E-22 | osteoclast.like | Alg8 |
| Oaz21 | 2.82E-26 | 0.4433701 | 0.537 | 0.296 | 5.46E-22 | osteoclast.like | Oaz2 |
| Ctbs | 3.11E-26 | 0.41795054 | 0.376 | 0.2 | 6.01E-22 | osteoclast.like | Ctbs |
| Tspan171 | 3.39E-26 | 0.79134073 | 0.298 | 0.109 | 6.55E-22 | osteoclast.like | Tspan17 |
| Zfp638 | 3.99E-26 | 0.41154186 | 0.463 | 0.229 | 7.71E-22 | osteoclast.like | Zfp638 |
| Neurl21 | 4.22E-26 | 0.81369285 | 0.278 | 0.114 | 8.16E-22 | osteoclast.like | Neurl2 |
| Srek1ip11 | 4.42E-26 | 0.81168712 | 0.385 | 0.15 | 8.55E-22 | osteoclast.like | Srek1ip1 |
| Lrrc41 | 4.51E-26 | 0.39570972 | 0.546 | 0.313 | 8.72E-22 | osteoclast.like | Lrrc41 |
| Hikeshi1 | 4.55E-26 | 0.83510464 | 0.424 | 0.199 | 8.80E-22 | osteoclast.like | Hikeshi |
| Pfas1 | 4.57E-26 | 0.92393509 | 0.322 | 0.116 | 8.84E-22 | osteoclast.like | Pfas |
| Cdc261 | 4.73E-26 | 0.67138154 | 0.371 | 0.159 | 9.14E-22 | osteoclast.like | Cdc26 |
| Dctn3 | 5.62E-26 | 0.28704392 | 0.644 | 0.38 | 1.09E-21 | osteoclast.like | Dctn3 |
| Nagpa2 | 5.78E-26 | 0.33647613 | 0.624 | 0.409 | 1.12E-21 | osteoclast.like | Nagpa |
| Sec631 | 6.18E-26 | 0.32818175 | 0.532 | 0.312 | 1.20E-21 | osteoclast.like | Sec63 |
| Fubp31 | 6.48E-26 | 0.71530178 | 0.424 | 0.201 | 1.25E-21 | osteoclast.like | Fubp3 |
| Slc25a16 | 6.53E-26 | 0.94951715 | 0.346 | 0.128 | 1.26E-21 | osteoclast.like | Slc25a16 |
| Trim272 | 6.56E-26 | 0.41204897 | 0.576 | 0.324 | 1.27E-21 | osteoclast.like | Trim27 |
| Pphln11 | 7.13E-26 | 0.31403478 | 0.537 | 0.302 | 1.38E-21 | osteoclast.like | Pphln1 |
| Ccser2 | 7.63E-26 | 1.05117309 | 0.278 | 0.101 | 1.47E-21 | osteoclast.like | Ccser2 |
| Nol61 | 8.87E-26 | 0.74073139 | 0.4 | 0.208 | 1.71E-21 | osteoclast.like | Nol6 |
| Esyt22 | 9.08E-26 | 0.2926435 | 0.644 | 0.412 | 1.76E-21 | osteoclast.like | Esyt2 |
| Rrn32 | 9.19E-26 | 0.63537078 | 0.483 | 0.238 | 1.78E-21 | osteoclast.like | Rrn3 |
| Ccdc163 | 1.04E-25 | 1.03637962 | 0.327 | 0.099 | 2.01E-21 | osteoclast.like | Ccdc163 |
| Cdk5rap33 | 1.05E-25 | 0.41920833 | 0.532 | 0.282 | 2.02E-21 | osteoclast.like | Cdk5rap3 |
| Ube2j21 | 1.10E-25 | 0.3055591 | 0.541 | 0.336 | 2.12E-21 | osteoclast.like | Ube2j2 |
| Jkamp | 1.25E-25 | 0.60169453 | 0.395 | 0.188 | 2.42E-21 | osteoclast.like | Jkamp |
| Rbmxl1 | 1.30E-25 | 0.66695185 | 0.405 | 0.18 | 2.51E-21 | osteoclast.like | Rbmxl1 |
| Abhd21 | 1.32E-25 | 0.29328524 | 0.541 | 0.281 | 2.55E-21 | osteoclast.like | Abhd2 |
| Usf3 | 1.42E-25 | 0.33816033 | 0.424 | 0.226 | 2.74E-21 | osteoclast.like | Usf3 |
| Ndfip21 | 1.43E-25 | 0.56350057 | 0.39 | 0.201 | 2.77E-21 | osteoclast.like | Ndfip2 |
| Ccdc592 | 1.52E-25 | 0.43912668 | 0.498 | 0.246 | 2.93E-21 | osteoclast.like | Ccdc59 |
| Arfgap11 | 1.65E-25 | 0.28416051 | 0.502 | 0.294 | 3.20E-21 | osteoclast.like | Arfgap1 |
| Pold21 | 2.06E-25 | 1.23387678 | 0.327 | 0.099 | 3.99E-21 | osteoclast.like | Pold2 |
| Arfip21 | 2.07E-25 | 0.55037034 | 0.415 | 0.199 | 4.00E-21 | osteoclast.like | Arfip2 |
| Elp2 | 2.32E-25 | 1.04678507 | 0.312 | 0.121 | 4.48E-21 | osteoclast.like | Elp2 |
| Jagn1 | 2.51E-25 | 0.74002471 | 0.298 | 0.135 | 4.85E-21 | osteoclast.like | Jagn1 |
| Rell13 | 2.54E-25 | 0.35212242 | 0.463 | 0.273 | 4.91E-21 | osteoclast.like | Rell1 |
| Snrnp351 | 2.54E-25 | 0.50901411 | 0.395 | 0.171 | 4.91E-21 | osteoclast.like | Snrnp35 |
| Akr1b103 | 2.61E-25 | 0.47400504 | 0.478 | 0.252 | 5.05E-21 | osteoclast.like | Akr1b10 |
| G6pc32 | 2.79E-25 | 0.57064067 | 0.488 | 0.243 | 5.39E-21 | osteoclast.like | G6pc3 |
| Smad11 | 2.80E-25 | 0.72218154 | 0.38 | 0.171 | 5.42E-21 | osteoclast.like | Smad1 |
| Pop7 | 2.86E-25 | 0.9663807 | 0.263 | 0.087 | 5.52E-21 | osteoclast.like | Pop7 |
| Hras2 | 2.95E-25 | 0.54593302 | 0.502 | 0.237 | 5.71E-21 | osteoclast.like | Hras |
| Taf21 | 2.95E-25 | 0.54849304 | 0.39 | 0.177 | 5.71E-21 | osteoclast.like | Taf2 |
| Wdr743 | 2.97E-25 | 0.49085547 | 0.429 | 0.209 | 5.74E-21 | osteoclast.like | Wdr74 |
| Met2 | 2.98E-25 | 0.36825051 | 0.537 | 0.18 | 5.76E-21 | osteoclast.like | Met |
| Kif16b | 3.32E-25 | 0.80778837 | 0.298 | 0.12 | 6.42E-21 | osteoclast.like | Kif16b |
| Syvn1 | 3.79E-25 | 0.60882569 | 0.483 | 0.272 | 7.32E-21 | osteoclast.like | Syvn1 |
| Ddx181 | 3.91E-25 | 0.79004605 | 0.395 | 0.159 | 7.57E-21 | osteoclast.like | Ddx18 |
| Nelfcd1 | 4.09E-25 | 0.96101727 | 0.332 | 0.124 | 7.90E-21 | osteoclast.like | Nelfcd |
| Abcf21 | 4.25E-25 | 0.83009316 | 0.415 | 0.223 | 8.22E-21 | osteoclast.like | Abcf2 |
| Rab3ip | 4.60E-25 | 0.76801798 | 0.337 | 0.129 | 8.89E-21 | osteoclast.like | Rab3ip |
| Pcyox13 | 5.08E-25 | 0.25436561 | 0.634 | 0.382 | 9.81E-21 | osteoclast.like | Pcyox1 |
| Abcd3 | 5.41E-25 | 0.78452903 | 0.317 | 0.124 | 1.05E-20 | osteoclast.like | Abcd3 |
| Tysnd11 | 5.51E-25 | 0.52664029 | 0.361 | 0.154 | 1.06E-20 | osteoclast.like | Tysnd1 |
| Desi11 | 5.74E-25 | 0.46885787 | 0.405 | 0.202 | 1.11E-20 | osteoclast.like | Desi1 |
| Rab40c2 | 6.12E-25 | 0.73497621 | 0.434 | 0.196 | 1.18E-20 | osteoclast.like | Rab40c |
| Pno13 | 6.24E-25 | 0.60378208 | 0.434 | 0.223 | 1.21E-20 | osteoclast.like | Pno1 |
| Tdrd3 | 6.28E-25 | 1.01406936 | 0.351 | 0.126 | 1.21E-20 | osteoclast.like | Tdrd3 |
| Wdr31 | 6.42E-25 | 0.71787489 | 0.38 | 0.191 | 1.24E-20 | osteoclast.like | Wdr3 |
| Ints142 | 7.29E-25 | 0.48603721 | 0.478 | 0.244 | 1.41E-20 | osteoclast.like | Ints14 |
| Churc1 | 7.36E-25 | 0.84525485 | 0.307 | 0.115 | 1.42E-20 | osteoclast.like | Churc1 |
| Bola1 | 7.53E-25 | 1.22798432 | 0.259 | 0.103 | 1.46E-20 | osteoclast.like | Bola1 |
| Mzt12 | 7.60E-25 | 0.75787439 | 0.361 | 0.179 | 1.47E-20 | osteoclast.like | Mzt1 |
| Ints13 | 7.72E-25 | 0.41548853 | 0.483 | 0.274 | 1.49E-20 | osteoclast.like | Ints1 |
| Nectin1 | 7.74E-25 | 0.7486832 | 0.307 | 0.118 | 1.50E-20 | osteoclast.like | Nectin1 |
| Smarcd1 | 9.29E-25 | 0.951457 | 0.327 | 0.126 | 1.80E-20 | osteoclast.like | Smarcd1 |
| Cyb5r13 | 9.43E-25 | 0.31545503 | 0.527 | 0.272 | 1.82E-20 | osteoclast.like | Cyb5r1 |
| Sdhaf4 | 9.54E-25 | 0.93704097 | 0.283 | 0.105 | 1.84E-20 | osteoclast.like | Sdhaf4 |
| Ate12 | 1.11E-24 | 0.64693195 | 0.376 | 0.167 | 2.15E-20 | osteoclast.like | Ate1 |
| Ppip5k21 | 1.17E-24 | 0.65456657 | 0.395 | 0.19 | 2.27E-20 | osteoclast.like | Ppip5k2 |
| Pbdc13 | 1.21E-24 | 0.63629404 | 0.444 | 0.211 | 2.33E-20 | osteoclast.like | Pbdc1 |
| Larp71 | 1.21E-24 | 0.73241519 | 0.415 | 0.177 | 2.34E-20 | osteoclast.like | Larp7 |
| Tmem39b | 1.22E-24 | 0.82090772 | 0.298 | 0.137 | 2.35E-20 | osteoclast.like | Tmem39b |
| Fbxo331 | 1.22E-24 | 0.38104884 | 0.444 | 0.249 | 2.36E-20 | osteoclast.like | Fbxo33 |
| Foxk21 | 1.26E-24 | 0.49206711 | 0.459 | 0.247 | 2.43E-20 | osteoclast.like | Foxk2 |
| Stard54 | 1.44E-24 | 0.55401532 | 0.415 | 0.228 | 2.79E-20 | osteoclast.like | Stard5 |
| Thap4 | 1.45E-24 | 0.74762606 | 0.307 | 0.131 | 2.80E-20 | osteoclast.like | Thap4 |
| Shoc2 | 1.51E-24 | 0.49731771 | 0.502 | 0.236 | 2.92E-20 | osteoclast.like | Shoc2 |
| Micu13 | 1.59E-24 | 0.54138153 | 0.444 | 0.25 | 3.08E-20 | osteoclast.like | Micu1 |
| Eya31 | 1.59E-24 | 0.64487587 | 0.41 | 0.197 | 3.08E-20 | osteoclast.like | Eya3 |
| Rfxap | 1.69E-24 | 0.69865834 | 0.41 | 0.21 | 3.26E-20 | osteoclast.like | Rfxap |
| Srpr | 1.79E-24 | 0.35132609 | 0.571 | 0.314 | 3.46E-20 | osteoclast.like | Srpr |
| Sarnp2 | 1.79E-24 | 0.35708666 | 0.517 | 0.275 | 3.46E-20 | osteoclast.like | Sarnp |
| Usp101 | 1.89E-24 | 0.93948152 | 0.288 | 0.128 | 3.65E-20 | osteoclast.like | Usp10 |
| Nudt42 | 1.99E-24 | 0.37144243 | 0.459 | 0.245 | 3.85E-20 | osteoclast.like | Nudt4 |
| Thap2 | 2.07E-24 | 0.84979908 | 0.395 | 0.164 | 4.01E-20 | osteoclast.like | Thap2 |
| Alkbh3 | 2.27E-24 | 1.00872283 | 0.298 | 0.121 | 4.38E-20 | osteoclast.like | Alkbh3 |
| Slc39a10 | 2.49E-24 | 0.75335112 | 0.302 | 0.114 | 4.82E-20 | osteoclast.like | Slc39a10 |
| Trp53bp1 | 2.73E-24 | 0.8188408 | 0.298 | 0.122 | 5.27E-20 | osteoclast.like | Trp53bp1 |
| Slc38a102 | 2.75E-24 | 0.25998157 | 0.649 | 0.442 | 5.32E-20 | osteoclast.like | Slc38a10 |
| Nom11 | 3.28E-24 | 0.60690155 | 0.371 | 0.162 | 6.34E-20 | osteoclast.like | Nom1 |
| Flnb1 | 3.44E-24 | 0.59696916 | 0.361 | 0.153 | 6.65E-20 | osteoclast.like | Flnb |
| Eef1akmt2 | 3.48E-24 | 0.60996824 | 0.341 | 0.133 | 6.73E-20 | osteoclast.like | Eef1akmt2 |
| Tmem175 | 3.53E-24 | 1.16812652 | 0.273 | 0.098 | 6.83E-20 | osteoclast.like | Tmem175 |
| Chd33 | 3.63E-24 | 0.42085142 | 0.42 | 0.254 | 7.02E-20 | osteoclast.like | Chd3 |
| Armh3 | 3.95E-24 | 0.71917274 | 0.263 | 0.094 | 7.64E-20 | osteoclast.like | Armh3 |
| Pdcd111 | 4.08E-24 | 0.96550205 | 0.376 | 0.184 | 7.88E-20 | osteoclast.like | Pdcd11 |
| Jade11 | 4.85E-24 | 0.97403059 | 0.302 | 0.103 | 9.38E-20 | osteoclast.like | Jade1 |
| Selenoi | 5.01E-24 | 0.73108219 | 0.341 | 0.141 | 9.68E-20 | osteoclast.like | Selenoi |
| Pdcd71 | 5.39E-24 | 0.36879509 | 0.395 | 0.194 | 1.04E-19 | osteoclast.like | Pdcd7 |
| Mast31 | 6.92E-24 | 0.3238955 | 0.61 | 0.368 | 1.34E-19 | osteoclast.like | Mast3 |
| Gse12 | 7.01E-24 | 0.3854846 | 0.468 | 0.225 | 1.36E-19 | osteoclast.like | Gse1 |
| Nabp21 | 7.15E-24 | 0.6625142 | 0.454 | 0.203 | 1.38E-19 | osteoclast.like | Nabp2 |
| Elmod2 | 7.25E-24 | 0.84341196 | 0.283 | 0.102 | 1.40E-19 | osteoclast.like | Elmod2 |
| Rwdd4a1 | 7.45E-24 | 0.79794505 | 0.346 | 0.153 | 1.44E-19 | osteoclast.like | Rwdd4a |
| Nsun5 | 7.67E-24 | 0.84057977 | 0.268 | 0.093 | 1.48E-19 | osteoclast.like | Nsun5 |
| Nhlrc2 | 7.68E-24 | 0.56701789 | 0.405 | 0.213 | 1.49E-19 | osteoclast.like | Nhlrc2 |
| Nedd11 | 8.08E-24 | 0.73835457 | 0.366 | 0.152 | 1.56E-19 | osteoclast.like | Nedd1 |
| Rnf401 | 8.18E-24 | 0.28393864 | 0.566 | 0.322 | 1.58E-19 | osteoclast.like | Rnf40 |
| Ttc7b2 | 8.28E-24 | 0.43982451 | 0.483 | 0.253 | 1.60E-19 | osteoclast.like | Ttc7b |
| Tmem2222 | 8.43E-24 | 0.65691815 | 0.449 | 0.209 | 1.63E-19 | osteoclast.like | Tmem222 |
| Cyb561d2 | 8.64E-24 | 0.776058 | 0.341 | 0.139 | 1.67E-19 | osteoclast.like | Cyb561d2 |
| Casp72 | 9.26E-24 | 0.41053393 | 0.488 | 0.268 | 1.79E-19 | osteoclast.like | Casp7 |
| Thap123 | 9.40E-24 | 0.40278311 | 0.468 | 0.233 | 1.82E-19 | osteoclast.like | Thap12 |
| Rpp212 | 9.62E-24 | 0.31396005 | 0.468 | 0.257 | 1.86E-19 | osteoclast.like | Rpp21 |
| Alg11 | 1.00E-23 | 0.52653927 | 0.502 | 0.255 | 1.94E-19 | osteoclast.like | Alg1 |
| Galns3 | 1.02E-23 | 0.43805046 | 0.463 | 0.249 | 1.97E-19 | osteoclast.like | Galns |
| Rp9 | 1.09E-23 | 0.42009505 | 0.537 | 0.325 | 2.10E-19 | osteoclast.like | Rp9 |
| Ubr1 | 1.17E-23 | 0.76088254 | 0.312 | 0.122 | 2.27E-19 | osteoclast.like | Ubr1 |
| Manbal2 | 1.27E-23 | 0.25365052 | 0.576 | 0.351 | 2.46E-19 | osteoclast.like | Manbal |
| St6galnac61 | 1.37E-23 | 0.43411254 | 0.478 | 0.253 | 2.65E-19 | osteoclast.like | St6galnac6 |
| Ndc11 | 1.38E-23 | 1.11597816 | 0.254 | 0.096 | 2.66E-19 | osteoclast.like | Ndc1 |
| Golgb12 | 1.61E-23 | 0.39394231 | 0.502 | 0.268 | 3.11E-19 | osteoclast.like | Golgb1 |
| Kcnab24 | 1.75E-23 | 0.32663483 | 0.702 | 0.445 | 3.37E-19 | osteoclast.like | Kcnab2 |
| Scamp54 | 1.88E-23 | 0.43377194 | 0.493 | 0.243 | 3.64E-19 | osteoclast.like | Scamp5 |
| Scfd21 | 1.92E-23 | 0.87152628 | 0.361 | 0.145 | 3.72E-19 | osteoclast.like | Scfd2 |
| Ccdc431 | 2.02E-23 | 0.45525993 | 0.376 | 0.187 | 3.91E-19 | osteoclast.like | Ccdc43 |
| Ezh11 | 2.22E-23 | 0.41349995 | 0.341 | 0.177 | 4.29E-19 | osteoclast.like | Ezh1 |
| Mfge82 | 2.25E-23 | 0.33814305 | 0.473 | 0.246 | 4.35E-19 | osteoclast.like | Mfge8 |
| Vps111 | 2.29E-23 | 0.30560331 | 0.541 | 0.355 | 4.42E-19 | osteoclast.like | Vps11 |
| D17H6S53E1 | 2.37E-23 | 0.44970946 | 0.429 | 0.226 | 4.57E-19 | osteoclast.like | D17H6S53E |
| Lrrc42 | 2.71E-23 | 0.76886838 | 0.346 | 0.144 | 5.25E-19 | osteoclast.like | Lrrc42 |
| Golga51 | 2.73E-23 | 0.61187379 | 0.439 | 0.204 | 5.29E-19 | osteoclast.like | Golga5 |
| Mrgbp | 2.86E-23 | 0.99140336 | 0.278 | 0.086 | 5.53E-19 | osteoclast.like | Mrgbp |
| Ube4a | 2.92E-23 | 0.44220164 | 0.537 | 0.3 | 5.64E-19 | osteoclast.like | Ube4a |
| Dpm2 | 3.34E-23 | 0.42125196 | 0.468 | 0.245 | 6.45E-19 | osteoclast.like | Dpm2 |
| Sik31 | 3.56E-23 | 0.32559424 | 0.527 | 0.289 | 6.88E-19 | osteoclast.like | Sik3 |
| Mtor | 3.77E-23 | 0.62117051 | 0.434 | 0.236 | 7.28E-19 | osteoclast.like | Mtor |
| Med18 | 4.00E-23 | 0.28026584 | 0.502 | 0.306 | 7.73E-19 | osteoclast.like | Med1 |
| Tspan52 | 4.12E-23 | 0.43169901 | 0.42 | 0.191 | 7.97E-19 | osteoclast.like | Tspan5 |
| Wdr462 | 4.30E-23 | 0.40613442 | 0.463 | 0.249 | 8.32E-19 | osteoclast.like | Wdr46 |
| Pomt1 | 4.47E-23 | 1.3195929 | 0.254 | 0.08 | 8.64E-19 | osteoclast.like | Pomt1 |
| Rwdd11 | 4.71E-23 | 0.71897691 | 0.41 | 0.165 | 9.11E-19 | osteoclast.like | Rwdd1 |
| Kctd51 | 4.72E-23 | 0.41949451 | 0.42 | 0.218 | 9.13E-19 | osteoclast.like | Kctd5 |
| Kctd61 | 4.92E-23 | 0.6000867 | 0.356 | 0.157 | 9.51E-19 | osteoclast.like | Kctd6 |
| Nans1 | 5.08E-23 | 0.51007076 | 0.551 | 0.307 | 9.81E-19 | osteoclast.like | Nans |
| Klhdc101 | 5.10E-23 | 0.41547708 | 0.483 | 0.262 | 9.85E-19 | osteoclast.like | Klhdc10 |
| Txndc152 | 5.12E-23 | 0.30212433 | 0.527 | 0.338 | 9.90E-19 | osteoclast.like | Txndc15 |
| Zfp7043 | 5.82E-23 | 0.72120791 | 0.502 | 0.207 | 1.13E-18 | osteoclast.like | Zfp704 |
| Carm11 | 5.86E-23 | 0.45591939 | 0.429 | 0.236 | 1.13E-18 | osteoclast.like | Carm1 |
| Adi14 | 6.04E-23 | 0.60058907 | 0.449 | 0.199 | 1.17E-18 | osteoclast.like | Adi1 |
| Kansl32 | 6.45E-23 | 0.41048409 | 0.473 | 0.254 | 1.25E-18 | osteoclast.like | Kansl3 |
| Spg111 | 6.89E-23 | 0.50628264 | 0.42 | 0.201 | 1.33E-18 | osteoclast.like | Spg11 |
| Lhpp | 7.04E-23 | 0.98379658 | 0.293 | 0.116 | 1.36E-18 | osteoclast.like | Lhpp |
| Zfyve262 | 7.27E-23 | 0.36617084 | 0.454 | 0.231 | 1.41E-18 | osteoclast.like | Zfyve26 |
| Wdr482 | 7.33E-23 | 0.34723059 | 0.38 | 0.196 | 1.42E-18 | osteoclast.like | Wdr48 |
| Ift140 | 7.40E-23 | 0.8061228 | 0.273 | 0.103 | 1.43E-18 | osteoclast.like | Ift140 |
| Slain21 | 7.63E-23 | 0.26152811 | 0.459 | 0.268 | 1.48E-18 | osteoclast.like | Slain2 |
| Yod1 | 7.67E-23 | 0.76576813 | 0.268 | 0.108 | 1.48E-18 | osteoclast.like | Yod1 |
| Ubfd13 | 8.22E-23 | 0.67758882 | 0.449 | 0.218 | 1.59E-18 | osteoclast.like | Ubfd1 |
| Rnf112 | 8.90E-23 | 0.43330034 | 0.498 | 0.291 | 1.72E-18 | osteoclast.like | Rnf11 |
| Lonp2 | 9.10E-23 | 0.54713571 | 0.351 | 0.163 | 1.76E-18 | osteoclast.like | Lonp2 |
| Dnmt11 | 9.34E-23 | 0.39459256 | 0.522 | 0.301 | 1.81E-18 | osteoclast.like | Dnmt1 |
| Ccdc71 | 9.52E-23 | 0.7359424 | 0.332 | 0.139 | 1.84E-18 | osteoclast.like | Ccdc71 |
| Pycrl1 | 9.83E-23 | 0.846132 | 0.263 | 0.114 | 1.90E-18 | osteoclast.like | Pycrl |
| Papss2 | 9.93E-23 | 0.34259351 | 0.268 | 0.141 | 1.92E-18 | osteoclast.like | Papss2 |
| Spg201 | 1.00E-22 | 0.92763956 | 0.366 | 0.144 | 1.94E-18 | osteoclast.like | Spg20 |
| Cdk2ap13 | 1.05E-22 | 0.38510502 | 0.522 | 0.3 | 2.02E-18 | osteoclast.like | Cdk2ap1 |
| Gmpr21 | 1.07E-22 | 0.47336574 | 0.478 | 0.265 | 2.06E-18 | osteoclast.like | Gmpr2 |
| Trim263 | 1.07E-22 | 0.27707994 | 0.424 | 0.254 | 2.06E-18 | osteoclast.like | Trim26 |
| Unc502 | 1.16E-22 | 0.36481148 | 0.468 | 0.248 | 2.24E-18 | osteoclast.like | Unc50 |
| Eif5b | 1.16E-22 | 0.28608256 | 0.512 | 0.326 | 2.25E-18 | osteoclast.like | Eif5b |
| Adck1 | 1.22E-22 | 0.88738028 | 0.356 | 0.139 | 2.36E-18 | osteoclast.like | Adck1 |
| Gtf3c11 | 1.32E-22 | 0.8404346 | 0.356 | 0.148 | 2.54E-18 | osteoclast.like | Gtf3c1 |
| Rbm191 | 1.35E-22 | 0.74264388 | 0.385 | 0.167 | 2.60E-18 | osteoclast.like | Rbm19 |
| Eef2k3 | 1.35E-22 | 0.54228732 | 0.424 | 0.17 | 2.61E-18 | osteoclast.like | Eef2k |
| Ppp4r3a1 | 1.45E-22 | 0.3714928 | 0.541 | 0.316 | 2.79E-18 | osteoclast.like | Ppp4r3a |
| Iws11 | 1.62E-22 | 0.52592979 | 0.39 | 0.199 | 3.13E-18 | osteoclast.like | Iws1 |
| Ncor1 | 1.65E-22 | 0.32477384 | 0.659 | 0.424 | 3.20E-18 | osteoclast.like | Ncor1 |
| Med111 | 1.80E-22 | 0.38786292 | 0.512 | 0.282 | 3.47E-18 | osteoclast.like | Med11 |
| Golga22 | 1.86E-22 | 0.26130705 | 0.532 | 0.333 | 3.60E-18 | osteoclast.like | Golga2 |
| Psme4 | 2.39E-22 | 0.27382768 | 0.576 | 0.349 | 4.62E-18 | osteoclast.like | Psme4 |
| Tmem1152 | 2.43E-22 | 0.48314662 | 0.468 | 0.257 | 4.71E-18 | osteoclast.like | Tmem115 |
| Hist3h2a3 | 2.53E-22 | 0.39469257 | 0.38 | 0.191 | 4.89E-18 | osteoclast.like | Hist3h2a |
| Krr11 | 2.57E-22 | 0.71325025 | 0.332 | 0.15 | 4.98E-18 | osteoclast.like | Krr1 |
| Uxt1 | 2.75E-22 | 0.59519381 | 0.327 | 0.147 | 5.32E-18 | osteoclast.like | Uxt |
| Tle1 | 3.11E-22 | 0.60491065 | 0.351 | 0.135 | 6.02E-18 | osteoclast.like | Tle1 |
| Smim41 | 3.16E-22 | 0.4081949 | 0.483 | 0.251 | 6.10E-18 | osteoclast.like | Smim4 |
| Rgs33 | 3.22E-22 | 0.39031399 | 0.444 | 0.203 | 6.22E-18 | osteoclast.like | Rgs3 |
| Ammecr1l | 3.27E-22 | 0.28560021 | 0.434 | 0.236 | 6.32E-18 | osteoclast.like | Ammecr1l |
| Fbxl4 | 3.64E-22 | 0.8792367 | 0.254 | 0.08 | 7.03E-18 | osteoclast.like | Fbxl4 |
| Fam98c1 | 3.75E-22 | 0.37414925 | 0.502 | 0.267 | 7.24E-18 | osteoclast.like | Fam98c |
| Rasa2 | 3.79E-22 | 0.85073112 | 0.322 | 0.118 | 7.32E-18 | osteoclast.like | Rasa2 |
| Pgrmc23 | 4.07E-22 | 0.52825392 | 0.488 | 0.29 | 7.86E-18 | osteoclast.like | Pgrmc2 |
| Anapc51 | 4.14E-22 | 0.75633157 | 0.327 | 0.152 | 8.00E-18 | osteoclast.like | Anapc5 |
| Ruvbl11 | 4.15E-22 | 0.73796128 | 0.332 | 0.142 | 8.03E-18 | osteoclast.like | Ruvbl1 |
| Hmbs1 | 4.54E-22 | 0.64568603 | 0.434 | 0.194 | 8.78E-18 | osteoclast.like | Hmbs |
| Btf3l4 | 4.60E-22 | 0.71011302 | 0.273 | 0.109 | 8.89E-18 | osteoclast.like | Btf3l4 |
| Ppid | 4.99E-22 | 1.0533011 | 0.273 | 0.094 | 9.65E-18 | osteoclast.like | Ppid |
| Smarcad11 | 5.26E-22 | 0.63208122 | 0.478 | 0.252 | 1.02E-17 | osteoclast.like | Smarcad1 |
| Cbx31 | 5.73E-22 | 0.56321965 | 0.346 | 0.173 | 1.11E-17 | osteoclast.like | Cbx3 |
| Minpp11 | 5.91E-22 | 0.66473001 | 0.376 | 0.17 | 1.14E-17 | osteoclast.like | Minpp1 |
| Dvl3 | 6.08E-22 | 0.86436137 | 0.376 | 0.163 | 1.18E-17 | osteoclast.like | Dvl3 |
| Rab102 | 6.76E-22 | 0.27261067 | 0.615 | 0.417 | 1.31E-17 | osteoclast.like | Rab10 |
| Dhx81 | 6.84E-22 | 0.28701939 | 0.498 | 0.303 | 1.32E-17 | osteoclast.like | Dhx8 |
| Calcoco13 | 7.13E-22 | 0.5371059 | 0.454 | 0.216 | 1.38E-17 | osteoclast.like | Calcoco1 |
| Tmem183a | 8.07E-22 | 0.50959278 | 0.561 | 0.314 | 1.56E-17 | osteoclast.like | Tmem183a |
| Wdr4 | 8.11E-22 | 1.06642764 | 0.327 | 0.11 | 1.57E-17 | osteoclast.like | Wdr4 |
| Noc4l1 | 8.67E-22 | 0.82483941 | 0.337 | 0.151 | 1.68E-17 | osteoclast.like | Noc4l |
| Exosc81 | 8.74E-22 | 0.37083479 | 0.424 | 0.229 | 1.69E-17 | osteoclast.like | Exosc8 |
| Cnot111 | 8.77E-22 | 0.67185471 | 0.371 | 0.181 | 1.70E-17 | osteoclast.like | Cnot11 |
| Dph31 | 8.81E-22 | 0.52280402 | 0.356 | 0.165 | 1.70E-17 | osteoclast.like | Dph3 |
| Traf51 | 9.00E-22 | 0.27347925 | 0.444 | 0.235 | 1.74E-17 | osteoclast.like | Traf5 |
| Ints12 | 9.02E-22 | 0.46500995 | 0.498 | 0.239 | 1.74E-17 | osteoclast.like | Ints12 |
| Txnrd2 | 9.43E-22 | 0.89090319 | 0.41 | 0.19 | 1.82E-17 | osteoclast.like | Txnrd2 |
| Yipf3 | 9.87E-22 | 0.32241432 | 0.629 | 0.402 | 1.91E-17 | osteoclast.like | Yipf3 |
| Aig1 | 1.08E-21 | 0.56272386 | 0.39 | 0.189 | 2.08E-17 | osteoclast.like | Aig1 |
| Vps45 | 1.14E-21 | 0.57104095 | 0.337 | 0.144 | 2.20E-17 | osteoclast.like | Vps45 |
| Alg3 | 1.19E-21 | 1.06226606 | 0.254 | 0.094 | 2.30E-17 | osteoclast.like | Alg3 |
| Hars2 | 1.34E-21 | 0.85734435 | 0.312 | 0.133 | 2.58E-17 | osteoclast.like | Hars2 |
| Ptar11 | 1.84E-21 | 0.34958866 | 0.38 | 0.178 | 3.55E-17 | osteoclast.like | Ptar1 |
| Irf2bpl1 | 1.89E-21 | 0.34207169 | 0.415 | 0.192 | 3.65E-17 | osteoclast.like | Irf2bpl |
| Rfk1 | 1.94E-21 | 0.32158653 | 0.517 | 0.305 | 3.75E-17 | osteoclast.like | Rfk |
| Ddx271 | 1.96E-21 | 0.6087554 | 0.493 | 0.255 | 3.79E-17 | osteoclast.like | Ddx27 |
| Wars1 | 2.01E-21 | 0.25947273 | 0.439 | 0.268 | 3.89E-17 | osteoclast.like | Wars |
| Apopt1 | 2.14E-21 | 0.93165229 | 0.293 | 0.112 | 4.13E-17 | osteoclast.like | Apopt1 |
| Pih1d11 | 2.26E-21 | 0.66779584 | 0.434 | 0.21 | 4.36E-17 | osteoclast.like | Pih1d1 |
| Gdap21 | 2.36E-21 | 0.29767133 | 0.444 | 0.227 | 4.55E-17 | osteoclast.like | Gdap2 |
| Tada11 | 2.59E-21 | 0.49986922 | 0.341 | 0.177 | 5.01E-17 | osteoclast.like | Tada1 |
| Cdk7 | 2.78E-21 | 0.72879141 | 0.366 | 0.156 | 5.38E-17 | osteoclast.like | Cdk7 |
| Taf1b | 3.02E-21 | 0.75948339 | 0.312 | 0.135 | 5.83E-17 | osteoclast.like | Taf1b |
| Mtif2 | 3.60E-21 | 0.86154243 | 0.307 | 0.105 | 6.96E-17 | osteoclast.like | Mtif2 |
| Mnat1 | 3.62E-21 | 0.58523495 | 0.293 | 0.125 | 7.00E-17 | osteoclast.like | Mnat1 |
| Golph3l2 | 3.64E-21 | 0.4321973 | 0.498 | 0.247 | 7.04E-17 | osteoclast.like | Golph3l |
| Ppm1f | 3.92E-21 | 1.18227239 | 0.263 | 0.087 | 7.58E-17 | osteoclast.like | Ppm1f |
| Lipo3 | 4.00E-21 | 0.85454862 | 0.263 | 0.098 | 7.74E-17 | osteoclast.like | Lipo3 |
| Rbbp51 | 4.13E-21 | 0.57487843 | 0.41 | 0.201 | 7.98E-17 | osteoclast.like | Rbbp5 |
| Rnf1813 | 4.40E-21 | 0.26444397 | 0.478 | 0.272 | 8.51E-17 | osteoclast.like | Rnf181 |
| Fkrp2 | 4.71E-21 | 0.40362595 | 0.541 | 0.283 | 9.11E-17 | osteoclast.like | Fkrp |
| Ywhaz2 | 4.75E-21 | 0.34608113 | 0.468 | 0.293 | 9.19E-17 | osteoclast.like | Ywhaz |
| Hbs1l | 5.07E-21 | 0.62463841 | 0.39 | 0.157 | 9.80E-17 | osteoclast.like | Hbs1l |
| Wipi1 | 5.25E-21 | 0.91396326 | 0.254 | 0.103 | 1.02E-16 | osteoclast.like | Wipi1 |
| Nelfa1 | 5.47E-21 | 0.58484466 | 0.385 | 0.186 | 1.06E-16 | osteoclast.like | Nelfa |
| Rcan32 | 5.62E-21 | 0.59132597 | 0.439 | 0.215 | 1.09E-16 | osteoclast.like | Rcan3 |
| Armt1 | 5.62E-21 | 0.81926622 | 0.268 | 0.103 | 1.09E-16 | osteoclast.like | Armt1 |
| Kri11 | 5.71E-21 | 0.33550536 | 0.468 | 0.249 | 1.10E-16 | osteoclast.like | Kri1 |
| Chid1 | 5.74E-21 | 0.55985852 | 0.38 | 0.183 | 1.11E-16 | osteoclast.like | Chid1 |
| Zfp598 | 6.35E-21 | 0.38066162 | 0.566 | 0.346 | 1.23E-16 | osteoclast.like | Zfp598 |
| Trappc101 | 6.86E-21 | 0.50056929 | 0.483 | 0.277 | 1.33E-16 | osteoclast.like | Trappc10 |
| Zfp52 | 6.89E-21 | 0.57555083 | 0.288 | 0.128 | 1.33E-16 | osteoclast.like | Zfp52 |
| Axl | 7.09E-21 | 0.32876571 | 0.527 | 0.336 | 1.37E-16 | osteoclast.like | Axl |
| Tmtc3 | 7.17E-21 | 0.68996438 | 0.273 | 0.117 | 1.39E-16 | osteoclast.like | Tmtc3 |
| Gid4 | 7.50E-21 | 0.79703517 | 0.278 | 0.119 | 1.45E-16 | osteoclast.like | Gid4 |
| Tbc1d16 | 7.80E-21 | 0.87874415 | 0.288 | 0.115 | 1.51E-16 | osteoclast.like | Tbc1d16 |
| Lzic1 | 7.80E-21 | 0.60273444 | 0.371 | 0.171 | 1.51E-16 | osteoclast.like | Lzic |
| Pom1212 | 8.10E-21 | 0.41045623 | 0.502 | 0.274 | 1.57E-16 | osteoclast.like | Pom121 |
| Yipf51 | 8.56E-21 | 0.61119127 | 0.488 | 0.257 | 1.66E-16 | osteoclast.like | Yipf5 |
| Gnl3l1 | 8.92E-21 | 0.70564 | 0.366 | 0.154 | 1.73E-16 | osteoclast.like | Gnl3l |
| Cdk131 | 9.01E-21 | 0.26487994 | 0.605 | 0.393 | 1.74E-16 | osteoclast.like | Cdk13 |
| Sumo21 | 9.13E-21 | 0.32582161 | 0.463 | 0.27 | 1.77E-16 | osteoclast.like | Sumo2 |
| Ankrd101 | 9.33E-21 | 0.50907808 | 0.42 | 0.186 | 1.80E-16 | osteoclast.like | Ankrd10 |
| Urgcp | 9.47E-21 | 0.40831278 | 0.341 | 0.152 | 1.83E-16 | osteoclast.like | Urgcp |
| Maco11 | 9.73E-21 | 0.42529509 | 0.444 | 0.245 | 1.88E-16 | osteoclast.like | Maco1 |
| Tvp23b1 | 1.06E-20 | 0.51123141 | 0.434 | 0.199 | 2.05E-16 | osteoclast.like | Tvp23b |
| Mospd32 | 1.06E-20 | 0.27142483 | 0.58 | 0.353 | 2.05E-16 | osteoclast.like | Mospd3 |
| Vopp11 | 1.18E-20 | 0.29311536 | 0.498 | 0.345 | 2.28E-16 | osteoclast.like | Vopp1 |
| Utp151 | 1.20E-20 | 0.53377237 | 0.371 | 0.192 | 2.32E-16 | osteoclast.like | Utp15 |
| Smg6 | 1.21E-20 | 0.37708185 | 0.463 | 0.257 | 2.34E-16 | osteoclast.like | Smg6 |
| Memo12 | 1.28E-20 | 0.5056709 | 0.444 | 0.207 | 2.47E-16 | osteoclast.like | Memo1 |
| Patz12 | 1.33E-20 | 0.6528748 | 0.41 | 0.183 | 2.58E-16 | osteoclast.like | Patz1 |
| Gnl22 | 1.45E-20 | 0.55187606 | 0.361 | 0.18 | 2.80E-16 | osteoclast.like | Gnl2 |
| Top2b1 | 1.54E-20 | 0.46792631 | 0.488 | 0.306 | 2.98E-16 | osteoclast.like | Top2b |
| Pigyl | 1.62E-20 | 0.90685861 | 0.268 | 0.1 | 3.14E-16 | osteoclast.like | Pigyl |
| Def82 | 1.66E-20 | 0.69019529 | 0.317 | 0.172 | 3.21E-16 | osteoclast.like | Def8 |
| Clns1a1 | 1.99E-20 | 0.50548682 | 0.385 | 0.2 | 3.84E-16 | osteoclast.like | Clns1a |
| Sdhaf2 | 2.03E-20 | 0.41569816 | 0.502 | 0.312 | 3.93E-16 | osteoclast.like | Sdhaf2 |
| Nvl1 | 2.18E-20 | 0.54201844 | 0.439 | 0.226 | 4.22E-16 | osteoclast.like | Nvl |
| Sfmbt14 | 2.38E-20 | 0.28202557 | 0.459 | 0.255 | 4.60E-16 | osteoclast.like | Sfmbt1 |
| Zfp445 | 2.48E-20 | 0.3557781 | 0.488 | 0.284 | 4.80E-16 | osteoclast.like | Zfp445 |
| Wdtc11 | 2.66E-20 | 0.460099 | 0.385 | 0.174 | 5.15E-16 | osteoclast.like | Wdtc1 |
| Psmc12 | 2.73E-20 | 0.33140609 | 0.502 | 0.37 | 5.27E-16 | osteoclast.like | Psmc1 |
| Bbip11 | 3.27E-20 | 0.25812551 | 0.463 | 0.278 | 6.32E-16 | osteoclast.like | Bbip1 |
| Cnot32 | 3.32E-20 | 0.26405106 | 0.527 | 0.325 | 6.41E-16 | osteoclast.like | Cnot3 |

| | | | | | | | |
|---|---|---|---|---|---|---|---|
| Eif2s22 | 3.34E-20 | 0.49682628 | 0.376 | 0.192 | 6.46E-16 | osteoclast.like | Eif2s2 |
| Zkscan3 | 3.61E-20 | 0.61100718 | 0.312 | 0.151 | 6.99E-16 | osteoclast.like | Zkscan3 |
| Megf8 | 3.99E-20 | 0.65234069 | 0.366 | 0.152 | 7.72E-16 | osteoclast.like | Megf8 |
| Capn7 | 4.01E-20 | 0.50904086 | 0.41 | 0.224 | 7.75E-16 | osteoclast.like | Capn7 |
| Pnpt11 | 4.13E-20 | 0.61467994 | 0.366 | 0.141 | 7.98E-16 | osteoclast.like | Pnpt1 |
| Paics1 | 4.30E-20 | 0.43772374 | 0.415 | 0.193 | 8.31E-16 | osteoclast.like | Paics |
| Elac21 | 4.31E-20 | 0.67406044 | 0.312 | 0.141 | 8.34E-16 | osteoclast.like | Elac2 |
| Alg52 | 4.42E-20 | 0.64469361 | 0.429 | 0.184 | 8.55E-16 | osteoclast.like | Alg5 |
| Hscb | 4.59E-20 | 0.87961662 | 0.317 | 0.123 | 8.87E-16 | osteoclast.like | Hscb |
| Ints71 | 4.79E-20 | 0.67732106 | 0.288 | 0.145 | 9.26E-16 | osteoclast.like | Ints7 |
| Cog1 | 4.84E-20 | 0.43295316 | 0.41 | 0.227 | 9.35E-16 | osteoclast.like | Cog1 |
| Ubl4a1 | 4.85E-20 | 0.43771481 | 0.444 | 0.21 | 9.38E-16 | osteoclast.like | Ubl4a |
| Mtg2 | 4.93E-20 | 1.2549283 | 0.254 | 0.09 | 9.53E-16 | osteoclast.like | Mtg2 |
| Galnt73 | 5.04E-20 | 0.51316022 | 0.468 | 0.263 | 9.74E-16 | osteoclast.like | Galnt7 |
| Wrnip11 | 5.15E-20 | 0.50874409 | 0.395 | 0.188 | 9.96E-16 | osteoclast.like | Wrnip1 |
| Apip1 | 5.26E-20 | 0.64524849 | 0.341 | 0.149 | 1.02E-15 | osteoclast.like | Apip |
| Coasy | 5.35E-20 | 0.82788821 | 0.332 | 0.131 | 1.04E-15 | osteoclast.like | Coasy |
| Rnf72 | 5.67E-20 | 0.51141558 | 0.449 | 0.254 | 1.10E-15 | osteoclast.like | Rnf7 |
| Aatf1 | 6.30E-20 | 0.5763771 | 0.317 | 0.144 | 1.22E-15 | osteoclast.like | Aatf |
| Fam149b | 6.79E-20 | 0.69941727 | 0.302 | 0.103 | 1.31E-15 | osteoclast.like | Fam149b |
| Tm2d31 | 7.03E-20 | 0.64480325 | 0.41 | 0.202 | 1.36E-15 | osteoclast.like | Tm2d3 |
| Chd1l | 7.23E-20 | 0.83918435 | 0.283 | 0.104 | 1.40E-15 | osteoclast.like | Chd1l |
| Ddx10 | 7.30E-20 | 0.32382442 | 0.434 | 0.245 | 1.41E-15 | osteoclast.like | Ddx10 |
| Tsnax1 | 7.56E-20 | 0.48223283 | 0.346 | 0.164 | 1.46E-15 | osteoclast.like | Tsnax |
| Gbf11 | 7.92E-20 | 0.56641984 | 0.463 | 0.23 | 1.53E-15 | osteoclast.like | Gbf1 |
| Polr3e1 | 8.69E-20 | 0.87646638 | 0.307 | 0.142 | 1.68E-15 | osteoclast.like | Polr3e |
| Hmces | 8.79E-20 | 0.47092353 | 0.332 | 0.143 | 1.70E-15 | osteoclast.like | Hmces |
| Hddc2 | 9.48E-20 | 1.09744634 | 0.263 | 0.085 | 1.83E-15 | osteoclast.like | Hddc2 |
| Sema6b2 | 9.55E-20 | 0.31686426 | 0.39 | 0.189 | 1.85E-15 | osteoclast.like | Sema6b |
| Btbd1 | 1.08E-19 | 0.27261181 | 0.39 | 0.225 | 2.08E-15 | osteoclast.like | Btbd1 |
| Tmem126a1 | 1.13E-19 | 0.66919536 | 0.366 | 0.166 | 2.19E-15 | osteoclast.like | Tmem126a |
| Bmyc2 | 1.21E-19 | 0.34604213 | 0.385 | 0.228 | 2.35E-15 | osteoclast.like | Bmyc |
| Nsmce31 | 1.23E-19 | 0.50612607 | 0.4 | 0.178 | 2.38E-15 | osteoclast.like | Nsmce3 |
| Eif3m1 | 1.25E-19 | 0.55142874 | 0.327 | 0.156 | 2.42E-15 | osteoclast.like | Eif3m |
| Gng51 | 1.27E-19 | 0.44556237 | 0.38 | 0.236 | 2.46E-15 | osteoclast.like | Gng5 |
| Nip71 | 1.28E-19 | 0.5411273 | 0.366 | 0.172 | 2.47E-15 | osteoclast.like | Nip7 |
| Mpst1 | 1.28E-19 | 0.82133146 | 0.4 | 0.164 | 2.47E-15 | osteoclast.like | Mpst |
| Fbxl6 | 1.29E-19 | 0.50700664 | 0.39 | 0.205 | 2.49E-15 | osteoclast.like | Fbxl6 |
| Mettl14 | 1.54E-19 | 1.14144538 | 0.273 | 0.104 | 2.98E-15 | osteoclast.like | Mettl14 |
| Top3b | 1.58E-19 | 0.38464944 | 0.327 | 0.17 | 3.05E-15 | osteoclast.like | Top3b |
| Vps4a1 | 1.76E-19 | 0.55803654 | 0.41 | 0.201 | 3.40E-15 | osteoclast.like | Vps4a |
| Thumpd11 | 1.89E-19 | 0.72423947 | 0.322 | 0.149 | 3.65E-15 | osteoclast.like | Thumpd1 |
| Abhd17b1 | 2.11E-19 | 0.3523936 | 0.488 | 0.271 | 4.07E-15 | osteoclast.like | Abhd17b |
| Leo11 | 2.24E-19 | 0.87332354 | 0.307 | 0.135 | 4.34E-15 | osteoclast.like | Leo1 |
| Gtf3c3 | 2.33E-19 | 0.97643505 | 0.307 | 0.13 | 4.51E-15 | osteoclast.like | Gtf3c3 |
| Lsm61 | 2.40E-19 | 0.74331189 | 0.288 | 0.129 | 4.65E-15 | osteoclast.like | Lsm6 |
| mt-Nd31 | 2.43E-19 | 0.28644978 | 0.517 | 0.341 | 4.70E-15 | osteoclast.like | mt-Nd3 |
| Rfc21 | 2.47E-19 | 0.34439807 | 0.488 | 0.298 | 4.77E-15 | osteoclast.like | Rfc2 |
| Erlec1 | 2.51E-19 | 0.78212661 | 0.317 | 0.148 | 4.86E-15 | osteoclast.like | Erlec1 |
| Zdhhc61 | 2.57E-19 | 0.61570709 | 0.4 | 0.198 | 4.96E-15 | osteoclast.like | Zdhhc6 |
| Lig3 | 2.61E-19 | 0.98444015 | 0.312 | 0.133 | 5.04E-15 | osteoclast.like | Lig3 |
| Akap102 | 2.64E-19 | 0.36181377 | 0.473 | 0.249 | 5.11E-15 | osteoclast.like | Akap10 |
| Thada1 | 2.75E-19 | 0.64032879 | 0.356 | 0.157 | 5.31E-15 | osteoclast.like | Thada |
| Usp31 | 2.79E-19 | 0.27408274 | 0.493 | 0.292 | 5.38E-15 | osteoclast.like | Usp3 |
| Mettl161 | 2.84E-19 | 0.87581944 | 0.302 | 0.129 | 5.50E-15 | osteoclast.like | Mettl16 |
| Ccdc473 | 2.89E-19 | 0.28737516 | 0.473 | 0.315 | 5.59E-15 | osteoclast.like | Ccdc47 |
| Prpsap21 | 2.89E-19 | 0.57387879 | 0.302 | 0.148 | 5.59E-15 | osteoclast.like | Prpsap2 |
| Cbfa2t2 | 2.93E-19 | 0.54037078 | 0.273 | 0.112 | 5.66E-15 | osteoclast.like | Cbfa2t2 |
| Ppp1r16a1 | 2.97E-19 | 0.67378194 | 0.351 | 0.156 | 5.75E-15 | osteoclast.like | Ppp1r16a |
| Rgp1 | 2.99E-19 | 0.53077772 | 0.293 | 0.143 | 5.79E-15 | osteoclast.like | Rgp1 |
| Tm7sf32 | 3.00E-19 | 0.32350532 | 0.434 | 0.252 | 5.79E-15 | osteoclast.like | Tm7sf3 |
| Tubgcp41 | 3.13E-19 | 0.43604134 | 0.322 | 0.181 | 6.04E-15 | osteoclast.like | Tubgcp4 |
| Ndufaf3 | 3.25E-19 | 0.60887905 | 0.337 | 0.154 | 6.29E-15 | osteoclast.like | Ndufaf3 |
| Nt5dc1 | 3.31E-19 | 0.54718976 | 0.41 | 0.203 | 6.40E-15 | osteoclast.like | Nt5dc1 |
| Thoc51 | 3.86E-19 | 0.71838936 | 0.39 | 0.177 | 7.45E-15 | osteoclast.like | Thoc5 |
| Mtrex1 | 3.88E-19 | 0.3436313 | 0.4 | 0.215 | 7.51E-15 | osteoclast.like | Mtrex |
| Tmem223 | 3.90E-19 | 0.52115272 | 0.385 | 0.163 | 7.54E-15 | osteoclast.like | Tmem223 |
| Usp40 | 4.04E-19 | 0.91504567 | 0.273 | 0.1 | 7.82E-15 | osteoclast.like | Usp40 |
| Slc7a6os | 4.04E-19 | 0.40762024 | 0.463 | 0.278 | 7.82E-15 | osteoclast.like | Slc7a6os |
| Yipf1 | 4.20E-19 | 0.35670199 | 0.556 | 0.324 | 8.12E-15 | osteoclast.like | Yipf1 |
| Snx191 | 4.28E-19 | 0.43638578 | 0.405 | 0.18 | 8.28E-15 | osteoclast.like | Snx19 |
| Pcmtd2 | 4.52E-19 | 0.77117333 | 0.327 | 0.134 | 8.74E-15 | osteoclast.like | Pcmtd2 |
| Ascc21 | 4.84E-19 | 0.4069541 | 0.493 | 0.268 | 9.36E-15 | osteoclast.like | Ascc2 |
| Nr2c2 | 4.91E-19 | 0.58514582 | 0.351 | 0.177 | 9.49E-15 | osteoclast.like | Nr2c2 |
| Exosc101 | 4.99E-19 | 0.29503559 | 0.42 | 0.25 | 9.64E-15 | osteoclast.like | Exosc10 |
| Aga1 | 5.19E-19 | 0.65021825 | 0.376 | 0.165 | 1.00E-14 | osteoclast.like | Aga |
| Sh2b1 | 5.69E-19 | 0.4008198 | 0.4 | 0.216 | 1.10E-14 | osteoclast.like | Sh2b1 |
| Pex7 | 5.84E-19 | 0.6893845 | 0.312 | 0.12 | 1.13E-14 | osteoclast.like | Pex7 |
| Nat101 | 5.87E-19 | 0.90573181 | 0.293 | 0.136 | 1.13E-14 | osteoclast.like | Nat10 |
| Dop1b2 | 6.10E-19 | 0.62930322 | 0.415 | 0.202 | 1.18E-14 | osteoclast.like | Dop1b |
| Zfp5132 | 6.14E-19 | 0.36103756 | 0.468 | 0.247 | 1.19E-14 | osteoclast.like | Zfp513 |
| Edc3 | 6.26E-19 | 0.47167707 | 0.351 | 0.168 | 1.21E-14 | osteoclast.like | Edc3 |
| Prpf61 | 6.26E-19 | 0.39154615 | 0.488 | 0.265 | 1.21E-14 | osteoclast.like | Prpf6 |
| Mrs21 | 6.50E-19 | 0.73664009 | 0.371 | 0.168 | 1.26E-14 | osteoclast.like | Mrs2 |
| Hat11 | 6.57E-19 | 0.32251821 | 0.502 | 0.29 | 1.27E-14 | osteoclast.like | Hat1 |
| Riok22 | 6.97E-19 | 0.49671019 | 0.395 | 0.182 | 1.35E-14 | osteoclast.like | Riok2 |
| Ranbp91 | 7.09E-19 | 0.25031558 | 0.502 | 0.336 | 1.37E-14 | osteoclast.like | Ranbp9 |
| Lmbrd23 | 7.25E-19 | 0.45622683 | 0.415 | 0.22 | 1.40E-14 | osteoclast.like | Lmbrd2 |
| Foxk1 | 7.65E-19 | 0.2785903 | 0.415 | 0.203 | 1.48E-14 | osteoclast.like | Foxk1 |
| Gabpa1 | 8.13E-19 | 0.38610999 | 0.459 | 0.255 | 1.57E-14 | osteoclast.like | Gabpa |
| Ptdss2 | 8.18E-19 | 0.88538432 | 0.259 | 0.096 | 1.58E-14 | osteoclast.like | Ptdss2 |
| Acad81 | 8.28E-19 | 0.5745692 | 0.322 | 0.142 | 1.60E-14 | osteoclast.like | Acad8 |
| Klhl7 | 9.79E-19 | 0.5876697 | 0.312 | 0.145 | 1.89E-14 | osteoclast.like | Klhl7 |
| Fbxo8 | 9.95E-19 | 0.64812425 | 0.259 | 0.11 | 1.92E-14 | osteoclast.like | Fbxo8 |
| Sord2 | 1.03E-18 | 0.26739044 | 0.278 | 0.171 | 2.00E-14 | osteoclast.like | Sord |
| Chordc12 | 1.06E-18 | 0.48523019 | 0.424 | 0.207 | 2.06E-14 | osteoclast.like | Chordc1 |
| Sec22b1 | 1.06E-18 | 0.26433288 | 0.522 | 0.299 | 2.06E-14 | osteoclast.like | Sec22b |
| Otud5 | 1.11E-18 | 0.26915835 | 0.488 | 0.315 | 2.15E-14 | osteoclast.like | Otud5 |
| Xpo41 | 1.13E-18 | 0.91377967 | 0.254 | 0.106 | 2.18E-14 | osteoclast.like | Xpo4 |
| Zfp6092 | 1.20E-18 | 0.28048467 | 0.463 | 0.289 | 2.32E-14 | osteoclast.like | Zfp609 |
| Relch1 | 1.21E-18 | 0.47539713 | 0.439 | 0.216 | 2.35E-14 | osteoclast.like | Relch |
| Blzf1 | 1.36E-18 | 0.60585539 | 0.273 | 0.109 | 2.62E-14 | osteoclast.like | Blzf1 |
| Arl6ip61 | 1.41E-18 | 0.41270126 | 0.459 | 0.255 | 2.72E-14 | osteoclast.like | Arl6ip6 |
| Katnbl13 | 1.46E-18 | 0.4223703 | 0.449 | 0.21 | 2.82E-14 | osteoclast.like | Katnbl1 |
| Erh1 | 1.50E-18 | 0.80925701 | 0.302 | 0.123 | 2.91E-14 | osteoclast.like | Erh |
| Gtf2h41 | 1.60E-18 | 0.78515786 | 0.298 | 0.13 | 3.10E-14 | osteoclast.like | Gtf2h4 |
| Isoc11 | 1.71E-18 | 0.76131694 | 0.39 | 0.179 | 3.30E-14 | osteoclast.like | Isoc1 |
| Gatc | 1.78E-18 | 0.6562274 | 0.293 | 0.117 | 3.44E-14 | osteoclast.like | Gatc |
| Pnpo2 | 2.22E-18 | 0.29152947 | 0.439 | 0.262 | 4.30E-14 | osteoclast.like | Pnpo |
| Ctr91 | 2.24E-18 | 0.76717886 | 0.332 | 0.161 | 4.32E-14 | osteoclast.like | Ctr9 |
| Wdr24 | 2.77E-18 | 0.73889341 | 0.283 | 0.101 | 5.35E-14 | osteoclast.like | Wdr24 |
| Pus101 | 2.95E-18 | 0.53915035 | 0.351 | 0.156 | 5.70E-14 | osteoclast.like | Pus10 |
| Amz2 | 3.18E-18 | 0.69721489 | 0.317 | 0.138 | 6.16E-14 | osteoclast.like | Amz2 |
| Med241 | 3.36E-18 | 0.44472441 | 0.327 | 0.169 | 6.50E-14 | osteoclast.like | Med24 |
| Crat4 | 3.51E-18 | 0.28191252 | 0.459 | 0.258 | 6.79E-14 | osteoclast.like | Crat |
| Csib | 3.82E-18 | 0.54386544 | 0.332 | 0.157 | 7.38E-14 | osteoclast.like | Csib |
| Mvb12a | 3.86E-18 | 0.26570584 | 0.478 | 0.27 | 7.46E-14 | osteoclast.like | Mvb12a |
| Arl6ip41 | 3.98E-18 | 0.34284905 | 0.454 | 0.267 | 7.69E-14 | osteoclast.like | Arl6ip4 |
| Naa252 | 4.10E-18 | 0.51697954 | 0.327 | 0.179 | 7.93E-14 | osteoclast.like | Naa25 |
| Ap5z11 | 4.14E-18 | 0.55653556 | 0.337 | 0.182 | 8.01E-14 | osteoclast.like | Ap5z1 |
| Tsc1 | 4.54E-18 | 0.61629684 | 0.351 | 0.161 | 8.78E-14 | osteoclast.like | Tsc1 |
| Sf3a31 | 5.16E-18 | 0.39057278 | 0.434 | 0.239 | 9.97E-14 | osteoclast.like | Sf3a3 |
| Aspscr12 | 5.36E-18 | 0.32635078 | 0.463 | 0.24 | 1.04E-13 | osteoclast.like | Aspscr1 |
| Rexo41 | 5.45E-18 | 0.34614885 | 0.38 | 0.213 | 1.05E-13 | osteoclast.like | Rexo4 |
| Vta1 | 5.49E-18 | 0.42113052 | 0.341 | 0.16 | 1.06E-13 | osteoclast.like | Vta1 |
| Nudt212 | 5.92E-18 | 0.33595362 | 0.507 | 0.301 | 1.15E-13 | osteoclast.like | Nudt21 |
| Sntb22 | 6.09E-18 | 0.33180595 | 0.439 | 0.232 | 1.18E-13 | osteoclast.like | Sntb2 |
| Zfp142 | 6.17E-18 | 0.67948021 | 0.254 | 0.091 | 1.19E-13 | osteoclast.like | Zfp142 |
| Tmem971 | 6.23E-18 | 0.58276873 | 0.317 | 0.153 | 1.20E-13 | osteoclast.like | Tmem97 |
| Cpsf11 | 6.81E-18 | 0.43132741 | 0.424 | 0.247 | 1.32E-13 | osteoclast.like | Cpsf1 |
| Clasrp1 | 7.03E-18 | 0.36390794 | 0.405 | 0.228 | 1.36E-13 | osteoclast.like | Clasrp |
| Cbx11 | 7.83E-18 | 0.78511954 | 0.293 | 0.123 | 1.51E-13 | osteoclast.like | Cbx1 |
| N4bp2l2 | 8.09E-18 | 0.3881249 | 0.522 | 0.291 | 1.56E-13 | osteoclast.like | N4bp2l2 |
| Zfp628 | 8.81E-18 | 0.68710706 | 0.346 | 0.147 | 1.70E-13 | osteoclast.like | Zfp628 |
| Sf3b41 | 9.03E-18 | 0.47852662 | 0.488 | 0.289 | 1.75E-13 | osteoclast.like | Sf3b4 |
| Slc31a11 | 9.09E-18 | 0.41283973 | 0.459 | 0.29 | 1.76E-13 | osteoclast.like | Slc31a1 |
| Fam104a1 | 9.95E-18 | 0.29295854 | 0.473 | 0.279 | 1.92E-13 | osteoclast.like | Fam104a |
| Snrk | 1.00E-17 | 0.78965644 | 0.283 | 0.119 | 1.93E-13 | osteoclast.like | Snrk |
| Bphl | 1.06E-17 | 0.83150342 | 0.302 | 0.116 | 2.04E-13 | osteoclast.like | Bphl |
| Coq8b | 1.11E-17 | 0.42465183 | 0.371 | 0.164 | 2.15E-13 | osteoclast.like | Coq8b |
| Tbc1d17 | 1.26E-17 | 0.53823859 | 0.366 | 0.166 | 2.43E-13 | osteoclast.like | Tbc1d17 |
| Ufm11 | 1.33E-17 | 0.28202016 | 0.58 | 0.366 | 2.57E-13 | osteoclast.like | Ufm1 |
| Ankra2 | 1.42E-17 | 0.92058734 | 0.259 | 0.096 | 2.75E-13 | osteoclast.like | Ankra2 |
| B4gat12 | 1.46E-17 | 0.50310692 | 0.317 | 0.176 | 2.83E-13 | osteoclast.like | B4gat1 |
| Pdcd51 | 1.61E-17 | 0.68857895 | 0.346 | 0.168 | 3.12E-13 | osteoclast.like | Pdcd5 |
| U2af12 | 1.67E-17 | 0.29645461 | 0.498 | 0.298 | 3.23E-13 | osteoclast.like | U2af1 |
| Pspc11 | 1.72E-17 | 0.32444639 | 0.498 | 0.339 | 3.33E-13 | osteoclast.like | Pspc1 |
| Alkbh6 | 1.76E-17 | 0.62852019 | 0.351 | 0.173 | 3.39E-13 | osteoclast.like | Alkbh6 |
| Mapk1ip1 | 1.85E-17 | 0.92656287 | 0.259 | 0.095 | 3.57E-13 | osteoclast.like | Mapk1ip1 |
| Fut81 | 1.88E-17 | 0.30856978 | 0.312 | 0.171 | 3.64E-13 | osteoclast.like | Fut8 |
| Dhx301 | 1.88E-17 | 0.70455098 | 0.346 | 0.152 | 3.64E-13 | osteoclast.like | Dhx30 |
| Aqr1 | 1.94E-17 | 0.3523719 | 0.39 | 0.222 | 3.76E-13 | osteoclast.like | Aqr |
| Twistnb1 | 1.99E-17 | 0.64730466 | 0.337 | 0.156 | 3.85E-13 | osteoclast.like | Twistnb |
| Impa22 | 2.12E-17 | 0.31314504 | 0.434 | 0.226 | 4.09E-13 | osteoclast.like | Impa2 |
| Polr2k | 2.23E-17 | 0.39011029 | 0.283 | 0.129 | 4.30E-13 | osteoclast.like | Polr2k |
| Gpn11 | 2.33E-17 | 0.52099823 | 0.302 | 0.14 | 4.50E-13 | osteoclast.like | Gpn1 |
| Xpo71 | 2.35E-17 | 0.41170962 | 0.444 | 0.233 | 4.54E-13 | osteoclast.like | Xpo7 |
| Myg11 | 2.44E-17 | 0.53496859 | 0.332 | 0.151 | 4.72E-13 | osteoclast.like | Myg1 |
| Dedd21 | 2.44E-17 | 0.44973719 | 0.298 | 0.159 | 4.72E-13 | osteoclast.like | Dedd2 |
| Dnttip21 | 2.54E-17 | 0.46176651 | 0.454 | 0.257 | 4.91E-13 | osteoclast.like | Dnttip2 |
| Hsf1 | 2.59E-17 | 0.40508291 | 0.4 | 0.203 | 5.01E-13 | osteoclast.like | Hsf1 |
| Agpat21 | 2.60E-17 | 0.44093981 | 0.322 | 0.155 | 5.02E-13 | osteoclast.like | Agpat2 |
| Ndrg3 | 2.80E-17 | 0.70221834 | 0.307 | 0.139 | 5.42E-13 | osteoclast.like | Ndrg3 |
| Mad2l1bp | 3.04E-17 | 0.65857718 | 0.288 | 0.136 | 5.87E-13 | osteoclast.like | Mad2l1bp |
| Slc25a103 | 3.17E-17 | 0.35185307 | 0.434 | 0.237 | 6.12E-13 | osteoclast.like | Slc25a10 |
| Ap3s2 | 3.30E-17 | 0.77774666 | 0.278 | 0.123 | 6.38E-13 | osteoclast.like | Ap3s2 |
| Smurf2 | 3.66E-17 | 0.40340876 | 0.4 | 0.225 | 7.07E-13 | osteoclast.like | Smurf2 |
| Eloc1 | 3.75E-17 | 0.45847099 | 0.415 | 0.245 | 7.26E-13 | osteoclast.like | Eloc |
| Apmap1 | 3.79E-17 | 0.3853321 | 0.468 | 0.253 | 7.32E-13 | osteoclast.like | Apmap |
| Xab2 | 3.92E-17 | 0.43044991 | 0.366 | 0.19 | 7.58E-13 | osteoclast.like | Xab2 |
| Rhbdd11 | 4.37E-17 | 0.34933549 | 0.41 | 0.206 | 8.45E-13 | osteoclast.like | Rhbdd1 |
| Tef | 4.79E-17 | 0.83752333 | 0.278 | 0.094 | 9.26E-13 | osteoclast.like | Tef |
| Tmem2091 | 5.14E-17 | 0.46865777 | 0.278 | 0.132 | 9.94E-13 | osteoclast.like | Tmem209 |
| Wdr551 | 5.17E-17 | 0.46739888 | 0.356 | 0.164 | 1.00E-12 | osteoclast.like | Wdr55 |
| Cog3 | 5.20E-17 | 0.43495995 | 0.327 | 0.154 | 1.01E-12 | osteoclast.like | Cog3 |
| Cul21 | 6.06E-17 | 0.46272075 | 0.312 | 0.164 | 1.17E-12 | osteoclast.like | Cul2 |
| E2f3 | 6.07E-17 | 0.77228064 | 0.268 | 0.112 | 1.17E-12 | osteoclast.like | E2f3 |
| Tceanc22 | 6.36E-17 | 0.3991152 | 0.395 | 0.2 | 1.23E-12 | osteoclast.like | Tceanc2 |
| Dnajc25 | 6.80E-17 | 0.59550033 | 0.283 | 0.123 | 1.31E-12 | osteoclast.like | Dnajc25 |
| Abhd13 | 7.56E-17 | 0.49812728 | 0.356 | 0.182 | 1.46E-12 | osteoclast.like | Abhd13 |
| Gmps1 | 7.59E-17 | 0.35117401 | 0.439 | 0.242 | 1.47E-12 | osteoclast.like | Gmps |
| Arfgef22 | 8.16E-17 | 0.44762709 | 0.361 | 0.168 | 1.58E-12 | osteoclast.like | Arfgef2 |
| DEPRECATED-l | 8.25E-17 | 0.56213439 | 0.351 | 0.184 | 1.59E-12 | osteoclast.like | DEPRECATED-ENSMUSG00000033323 |
| Cog2 | 8.34E-17 | 0.36417231 | 0.351 | 0.183 | 1.61E-12 | osteoclast.like | Cog2 |
| Mdm4 | 9.21E-17 | 0.33130698 | 0.429 | 0.258 | 1.78E-12 | osteoclast.like | Mdm4 |
| Ergic2 | 1.00E-16 | 0.41133595 | 0.351 | 0.175 | 1.93E-12 | osteoclast.like | Ergic2 |
| Rhot2 | 1.03E-16 | 0.77831016 | 0.283 | 0.117 | 1.98E-12 | osteoclast.like | Rhot2 |
| Fam114a22 | 1.16E-16 | 0.35394921 | 0.454 | 0.259 | 2.24E-12 | osteoclast.like | Fam114a2 |
| Gon4l1 | 1.21E-16 | 0.44750866 | 0.361 | 0.18 | 2.35E-12 | osteoclast.like | Gon4l |
| Uri11 | 1.25E-16 | 0.48504553 | 0.39 | 0.199 | 2.41E-12 | osteoclast.like | Uri1 |
| Cyp20a12 | 1.27E-16 | 0.52423055 | 0.337 | 0.161 | 2.46E-12 | osteoclast.like | Cyp20a1 |
| Kank21 | 1.29E-16 | 0.54795633 | 0.293 | 0.105 | 2.49E-12 | osteoclast.like | Kank2 |
| Spsb31 | 1.30E-16 | 0.43720524 | 0.429 | 0.227 | 2.51E-12 | osteoclast.like | Spsb3 |
| Fadd4 | 1.34E-16 | 0.32166695 | 0.439 | 0.218 | 2.58E-12 | osteoclast.like | Fadd |
| Trp53rka | 1.34E-16 | 0.68976973 | 0.273 | 0.11 | 2.59E-12 | osteoclast.like | Trp53rka |
| Psmd91 | 1.71E-16 | 0.42252905 | 0.39 | 0.204 | 3.30E-12 | osteoclast.like | Psmd9 |
| Mocos2 | 2.01E-16 | 0.41234679 | 0.332 | 0.154 | 3.88E-12 | osteoclast.like | Mocos |
| Nsdhl | 2.37E-16 | 0.63651042 | 0.288 | 0.125 | 4.58E-12 | osteoclast.like | Nsdhl |
| Actr8 | 2.55E-16 | 1.08259823 | 0.302 | 0.121 | 4.93E-12 | osteoclast.like | Actr8 |
| Prpsap1 | 2.62E-16 | 0.35154396 | 0.454 | 0.279 | 5.06E-12 | osteoclast.like | Prpsap1 |
| Ttc332 | 2.68E-16 | 0.51353701 | 0.385 | 0.181 | 5.17E-12 | osteoclast.like | Ttc33 |
| Ccdc28b | 2.99E-16 | 0.77108563 | 0.268 | 0.126 | 5.78E-12 | osteoclast.like | Ccdc28b |
| Rhot11 | 3.06E-16 | 0.35841142 | 0.439 | 0.223 | 5.91E-12 | osteoclast.like | Rhot1 |
| Sfi1 | 3.09E-16 | 0.41244206 | 0.337 | 0.159 | 5.97E-12 | osteoclast.like | Sfi1 |
| Tctex1d2 | 3.90E-16 | 0.45120346 | 0.337 | 0.163 | 7.54E-12 | osteoclast.like | Tctex1d2 |
| Acp6 | 4.04E-16 | 0.5729085 | 0.302 | 0.127 | 7.81E-12 | osteoclast.like | Acp6 |
| Rif11 | 4.23E-16 | 0.79994748 | 0.293 | 0.116 | 8.17E-12 | osteoclast.like | Rif1 |
| Synj22 | 4.27E-16 | 0.39911347 | 0.366 | 0.21 | 8.27E-12 | osteoclast.like | Synj2 |
| Cnnm31 | 4.50E-16 | 0.41611149 | 0.337 | 0.159 | 8.69E-12 | osteoclast.like | Cnnm3 |
| Mphosph102 | 4.81E-16 | 0.53256539 | 0.332 | 0.18 | 9.30E-12 | osteoclast.like | Mphosph10 |
| Rbx11 | 4.86E-16 | 0.36478134 | 0.463 | 0.282 | 9.39E-12 | osteoclast.like | Rbx1 |
| Vps361 | 5.05E-16 | 0.26888374 | 0.459 | 0.27 | 9.76E-12 | osteoclast.like | Vps36 |
| Med6 | 5.05E-16 | 0.4628068 | 0.322 | 0.147 | 9.76E-12 | osteoclast.like | Med6 |
| Dpm1 | 5.44E-16 | 0.38653575 | 0.434 | 0.263 | 1.05E-11 | osteoclast.like | Dpm1 |
| Haghl1 | 5.53E-16 | 0.47552364 | 0.371 | 0.176 | 1.07E-11 | osteoclast.like | Haghl |
| Cdip1 | 5.94E-16 | 0.3487517 | 0.385 | 0.214 | 1.15E-11 | osteoclast.like | Cdip1 |
| Rnf41 | 6.63E-16 | 0.29199011 | 0.468 | 0.264 | 1.28E-11 | osteoclast.like | Rnf41 |
| Cars21 | 8.02E-16 | 0.78300724 | 0.327 | 0.148 | 1.55E-11 | osteoclast.like | Cars2 |
| Bnip11 | 8.11E-16 | 0.53660963 | 0.263 | 0.113 | 1.57E-11 | osteoclast.like | Bnip1 |
| Fam193b1 | 8.11E-16 | 0.29564204 | 0.512 | 0.313 | 1.57E-11 | osteoclast.like | Fam193b |
| Pskh1 | 8.15E-16 | 0.6611562 | 0.273 | 0.104 | 1.58E-11 | osteoclast.like | Pskh1 |
| Thap111 | 8.25E-16 | 0.40456281 | 0.4 | 0.202 | 1.60E-11 | osteoclast.like | Thap11 |
| Wbp12 | 8.83E-16 | 0.45751388 | 0.361 | 0.178 | 1.71E-11 | osteoclast.like | Wbp1 |
| Thoc61 | 9.62E-16 | 0.26233099 | 0.439 | 0.265 | 1.86E-11 | osteoclast.like | Thoc6 |
| Tbcd2 | 1.03E-15 | 0.33825245 | 0.444 | 0.236 | 1.99E-11 | osteoclast.like | Tbcd |
| Med91 | 1.05E-15 | 0.44457216 | 0.405 | 0.193 | 2.03E-11 | osteoclast.like | Med9 |
| Isg20l2 | 1.07E-15 | 0.50651605 | 0.312 | 0.151 | 2.08E-11 | osteoclast.like | Isg20l2 |
| Zc3h13 | 1.08E-15 | 0.34059978 | 0.351 | 0.181 | 2.10E-11 | osteoclast.like | Zc3h13 |
| DEPRECATED-l | 1.20E-15 | 0.59846313 | 0.283 | 0.12 | 2.32E-11 | osteoclast.like | DEPRECATED-ENSMUSG00000056493 |
| Agpat33 | 1.34E-15 | 0.29955013 | 0.459 | 0.271 | 2.59E-11 | osteoclast.like | Agpat3 |
| Orc4 | 1.36E-15 | 0.3168495 | 0.444 | 0.261 | 2.63E-11 | osteoclast.like | Orc4 |
| Sptlc12 | 1.39E-15 | 0.40079332 | 0.429 | 0.246 | 2.69E-11 | osteoclast.like | Sptlc1 |
| Tada2b | 1.41E-15 | 0.47449658 | 0.293 | 0.14 | 2.73E-11 | osteoclast.like | Tada2b |
| Bud232 | 1.48E-15 | 0.2685059 | 0.41 | 0.237 | 2.86E-11 | osteoclast.like | Bud23 |
| Upf2 | 1.65E-15 | 0.49339584 | 0.351 | 0.159 | 3.18E-11 | osteoclast.like | Upf2 |
| Tsr3 | 1.73E-15 | 0.49163257 | 0.288 | 0.128 | 3.34E-11 | osteoclast.like | Tsr3 |
| Gcdh | 1.75E-15 | 0.61065512 | 0.254 | 0.103 | 3.37E-11 | osteoclast.like | Gcdh |
| Dis3l1 | 2.11E-15 | 0.35416277 | 0.254 | 0.133 | 4.08E-11 | osteoclast.like | Dis3l |
| Mtmr91 | 2.17E-15 | 0.41942325 | 0.483 | 0.256 | 4.19E-11 | osteoclast.like | Mtmr9 |
| Zranb22 | 2.19E-15 | 0.44723516 | 0.337 | 0.186 | 4.24E-11 | osteoclast.like | Zranb2 |
| Cdadc11 | 2.34E-15 | 0.48769897 | 0.322 | 0.154 | 4.52E-11 | osteoclast.like | Cdadc1 |
| Jpt21 | 2.41E-15 | 0.56713791 | 0.346 | 0.183 | 4.67E-11 | osteoclast.like | Jpt2 |
| Dnttip1 | 2.48E-15 | 0.32377364 | 0.4 | 0.22 | 4.80E-11 | osteoclast.like | Dnttip1 |
| Unk1 | 2.72E-15 | 0.38156119 | 0.293 | 0.155 | 5.26E-11 | osteoclast.like | Unk |
| Epg5 | 2.80E-15 | 0.53717314 | 0.307 | 0.144 | 5.40E-11 | osteoclast.like | Epg5 |
| Bub31 | 3.08E-15 | 0.25568328 | 0.468 | 0.303 | 5.95E-11 | osteoclast.like | Bub3 |
| Cnot101 | 3.14E-15 | 0.42593173 | 0.376 | 0.18 | 6.07E-11 | osteoclast.like | Cnot10 |
| Rbl22 | 3.31E-15 | 0.33052369 | 0.41 | 0.248 | 6.40E-11 | osteoclast.like | Rbl2 |
| Kif1bp1 | 3.46E-15 | 0.2813874 | 0.361 | 0.186 | 6.70E-11 | osteoclast.like | Kif1bp |
| Tcf31 | 3.51E-15 | 0.52073679 | 0.351 | 0.168 | 6.79E-11 | osteoclast.like | Tcf3 |
| Glrx2 | 3.65E-15 | 0.32154512 | 0.42 | 0.234 | 7.05E-11 | osteoclast.like | Glrx2 |
| Ewsr11 | 3.94E-15 | 0.46045268 | 0.38 | 0.199 | 7.61E-11 | osteoclast.like | Ewsr1 |
| Ufl11 | 4.14E-15 | 0.40697278 | 0.376 | 0.186 | 8.00E-11 | osteoclast.like | Ufl1 |
| Nit1 | 4.39E-15 | 0.45484051 | 0.302 | 0.162 | 8.48E-11 | osteoclast.like | Nit1 |
| Zbtb43 | 4.60E-15 | 0.49992579 | 0.317 | 0.153 | 8.89E-11 | osteoclast.like | Zbtb43 |
| Zfp24 | 4.64E-15 | 0.41484935 | 0.278 | 0.134 | 8.98E-11 | osteoclast.like | Zfp24 |
| Eps15l1 | 4.82E-15 | 0.55205032 | 0.307 | 0.142 | 9.31E-11 | osteoclast.like | Eps15l1 |
| Napg2 | 5.00E-15 | 0.36668188 | 0.415 | 0.224 | 9.67E-11 | osteoclast.like | Napg |
| Cstf11 | 5.17E-15 | 0.41586182 | 0.332 | 0.149 | 1.00E-10 | osteoclast.like | Cstf1 |
| Hmgb11 | 5.20E-15 | 0.33575354 | 0.351 | 0.172 | 1.00E-10 | osteoclast.like | Hmgb1 |
| Cask | 5.58E-15 | 0.52559469 | 0.278 | 0.118 | 1.08E-10 | osteoclast.like | Cask |
| DEPRECATED-l | 5.63E-15 | 0.40854234 | 0.405 | 0.241 | 1.09E-10 | osteoclast.like | DEPRECATED-ENSMUSG00000022521 |
| Psmg21 | 5.65E-15 | 0.6579052 | 0.259 | 0.108 | 1.09E-10 | osteoclast.like | Psmg2 |
| Spout11 | 5.67E-15 | 0.32199001 | 0.322 | 0.159 | 1.10E-10 | osteoclast.like | Spout1 |
| Cmkl11 | 5.67E-15 | 0.37616055 | 0.376 | 0.218 | 1.10E-10 | osteoclast.like | Cmkl1 |
| Tsen342 | 6.00E-15 | 0.38438819 | 0.473 | 0.243 | 1.16E-10 | osteoclast.like | Tsen34 |
| Arhgap24 | 6.53E-15 | 0.40014523 | 0.283 | 0.134 | 1.26E-10 | osteoclast.like | Arhgap24 |
| Ift222 | 6.70E-15 | 0.30436702 | 0.395 | 0.222 | 1.30E-10 | osteoclast.like | Ift22 |
| Tmem161a1 | 7.10E-15 | 0.41305374 | 0.317 | 0.167 | 1.37E-10 | osteoclast.like | Tmem161a |
| Mdn1 | 7.13E-15 | 0.85609292 | 0.268 | 0.133 | 1.38E-10 | osteoclast.like | Mdn1 |
| Anapc43 | 7.50E-15 | 0.26955759 | 0.41 | 0.232 | 1.45E-10 | osteoclast.like | Anapc4 |
| Rnf123 | 7.54E-15 | 0.38034763 | 0.351 | 0.179 | 1.46E-10 | osteoclast.like | Rnf123 |
| Ubqln42 | 8.01E-15 | 0.335607 | 0.424 | 0.235 | 1.55E-10 | osteoclast.like | Ubqln4 |
| Dcaf6 | 8.84E-15 | 0.50854331 | 0.288 | 0.12 | 1.71E-10 | osteoclast.like | Dcaf6 |
| Nosip2 | 9.09E-15 | 0.28637564 | 0.434 | 0.259 | 1.76E-10 | osteoclast.like | Nosip |
| Miip1 | 9.45E-15 | 0.65070601 | 0.293 | 0.114 | 1.83E-10 | osteoclast.like | Miip |
| Ipo7 | 9.85E-15 | 0.65031041 | 0.259 | 0.111 | 1.91E-10 | osteoclast.like | Ipo7 |
| Aen1 | 1.06E-14 | 0.44326736 | 0.307 | 0.146 | 2.04E-10 | osteoclast.like | Aen |
| Tmem39a | 1.08E-14 | 0.59266204 | 0.283 | 0.127 | 2.08E-10 | osteoclast.like | Tmem39a |
| Kdm1a | 1.12E-14 | 0.35420933 | 0.327 | 0.177 | 2.17E-10 | osteoclast.like | Kdm1a |
| Wiz1 | 1.16E-14 | 0.29868324 | 0.327 | 0.163 | 2.24E-10 | osteoclast.like | Wiz |
| Fam222b | 1.20E-14 | 0.45782678 | 0.268 | 0.125 | 2.32E-10 | osteoclast.like | Fam222b |
| Sppl2b | 1.25E-14 | 0.44818088 | 0.337 | 0.164 | 2.42E-10 | osteoclast.like | Sppl2b |
| Dph6 | 1.27E-14 | 0.9525683 | 0.273 | 0.107 | 2.46E-10 | osteoclast.like | Dph6 |
| Lsg12 | 1.38E-14 | 0.34616646 | 0.332 | 0.177 | 2.66E-10 | osteoclast.like | Lsg1 |
| Fam53a | 1.43E-14 | 0.47109623 | 0.263 | 0.12 | 2.76E-10 | osteoclast.like | Fam53a |
| Unc13d | 1.63E-14 | 0.40731084 | 0.288 | 0.148 | 3.15E-10 | osteoclast.like | Unc13d |
| Eif4enif11 | 1.64E-14 | 0.50004231 | 0.278 | 0.134 | 3.18E-10 | osteoclast.like | Eif4enif1 |
| Vma21 | 1.84E-14 | 0.77665271 | 0.288 | 0.128 | 3.55E-10 | osteoclast.like | Vma21 |
| Ube2d1 | 1.89E-14 | 0.42220087 | 0.405 | 0.215 | 3.65E-10 | osteoclast.like | Ube2d1 |
| Cwf19l2 | 1.95E-14 | 0.46710338 | 0.332 | 0.15 | 3.78E-10 | osteoclast.like | Cwf19l2 |
| Snrnp402 | 1.98E-14 | 0.35121553 | 0.424 | 0.237 | 3.82E-10 | osteoclast.like | Snrnp40 |
| Fam172a1 | 2.03E-14 | 0.4647749 | 0.39 | 0.215 | 3.92E-10 | osteoclast.like | Fam172a |
| Zcrb1 | 2.09E-14 | 0.33441026 | 0.463 | 0.267 | 4.04E-10 | osteoclast.like | Zcrb1 |
| Nfrkb | 2.15E-14 | 0.75962724 | 0.298 | 0.124 | 4.15E-10 | osteoclast.like | Nfrkb |
| Psip11 | 2.25E-14 | 0.32538638 | 0.322 | 0.148 | 4.35E-10 | osteoclast.like | Psip1 |
| Eif4a32 | 2.46E-14 | 0.52665819 | 0.39 | 0.23 | 4.75E-10 | osteoclast.like | Eif4a3 |
| Gpatch2l | 2.50E-14 | 0.30104746 | 0.395 | 0.228 | 4.83E-10 | osteoclast.like | Gpatch2l |
| Ikzf51 | 2.58E-14 | 0.36566857 | 0.327 | 0.176 | 4.98E-10 | osteoclast.like | Ikzf5 |
| Rcor3 | 2.75E-14 | 0.65542089 | 0.307 | 0.141 | 5.31E-10 | osteoclast.like | Rcor3 |
| Foxo4 | 2.88E-14 | 0.4373828 | 0.312 | 0.158 | 5.57E-10 | osteoclast.like | Foxo4 |
| Nap1l11 | 2.96E-14 | 0.43243384 | 0.415 | 0.232 | 5.72E-10 | osteoclast.like | Nap1l1 |
| Aptx | 3.43E-14 | 0.66383442 | 0.273 | 0.108 | 6.63E-10 | osteoclast.like | Aptx |
| Bysl1 | 3.45E-14 | 0.43011583 | 0.254 | 0.124 | 6.68E-10 | osteoclast.like | Bysl |
| Rbm341 | 3.46E-14 | 0.44913715 | 0.259 | 0.14 | 6.69E-10 | osteoclast.like | Rbm34 |
| Hmgn51 | 3.53E-14 | 0.56642692 | 0.302 | 0.137 | 6.82E-10 | osteoclast.like | Hmgn5 |
| Ppp4r11 | 3.65E-14 | 0.33982804 | 0.361 | 0.21 | 7.06E-10 | osteoclast.like | Ppp4r1 |
| Slc35a1 | 3.74E-14 | 0.5863226 | 0.298 | 0.133 | 7.23E-10 | osteoclast.like | Slc35a1 |
| Polr3gl | 4.13E-14 | 0.68763361 | 0.263 | 0.107 | 7.99E-10 | osteoclast.like | Polr3gl |
| St7l | 4.41E-14 | 0.88783202 | 0.254 | 0.104 | 8.52E-10 | osteoclast.like | St7l |
| Ngdn1 | 4.67E-14 | 0.2662819 | 0.434 | 0.247 | 9.03E-10 | osteoclast.like | Ngdn |
| Nrm1 | 4.78E-14 | 0.49662819 | 0.293 | 0.128 | 9.23E-10 | osteoclast.like | Nrm |
| Arl24 | 4.94E-14 | 0.384231 | 0.405 | 0.198 | 9.54E-10 | osteoclast.like | Arl2 |
| Gpatch2 | 5.51E-14 | 0.52833121 | 0.307 | 0.152 | 1.06E-09 | osteoclast.like | Gpatch2 |
| Med27 | 5.52E-14 | 0.31532628 | 0.332 | 0.168 | 1.07E-09 | osteoclast.like | Med27 |
| Nup623 | 6.31E-14 | 0.25001681 | 0.405 | 0.25 | 1.22E-09 | osteoclast.like | Nup62 |
| Spns1 | 6.55E-14 | 0.49076285 | 0.283 | 0.143 | 1.27E-09 | osteoclast.like | Spns1 |
| Pigu1 | 6.66E-14 | 0.58084159 | 0.376 | 0.204 | 1.29E-09 | osteoclast.like | Pigu |
| Atpaf2 | 7.15E-14 | 0.54991592 | 0.268 | 0.119 | 1.38E-09 | osteoclast.like | Atpaf2 |
| Ftsj1 | 7.53E-14 | 0.68335332 | 0.259 | 0.109 | 1.46E-09 | osteoclast.like | Ftsj1 |
| Zmat5 | 7.83E-14 | 0.30647355 | 0.346 | 0.175 | 1.51E-09 | osteoclast.like | Zmat5 |
| Brd31 | 7.95E-14 | 0.7624356 | 0.268 | 0.134 | 1.54E-09 | osteoclast.like | Brd3 |
| Ly962 | 8.15E-14 | 0.30204446 | 0.322 | 0.175 | 1.58E-09 | osteoclast.like | Ly96 |
| Fbxo31 | 1.04E-13 | 0.4895336 | 0.298 | 0.136 | 2.02E-09 | osteoclast.like | Fbxo31 |
| Ift461 | 1.06E-13 | 0.338635 | 0.332 | 0.186 | 2.05E-09 | osteoclast.like | Ift46 |
| Pelp11 | 1.11E-13 | 0.33683653 | 0.371 | 0.191 | 2.15E-09 | osteoclast.like | Pelp1 |
| Zmym4 | 1.14E-13 | 0.53678338 | 0.254 | 0.127 | 2.20E-09 | osteoclast.like | Zmym4 |
| Armc10 | 1.17E-13 | 0.67408145 | 0.263 | 0.116 | 2.26E-09 | osteoclast.like | Armc10 |
| Mettl31 | 1.20E-13 | 0.36728694 | 0.341 | 0.154 | 2.32E-09 | osteoclast.like | Mettl3 |
| Ddx19a1 | 1.26E-13 | 0.27634297 | 0.454 | 0.274 | 2.44E-09 | osteoclast.like | Ddx19a |
| Nup1881 | 1.30E-13 | 0.3742319 | 0.346 | 0.191 | 2.51E-09 | osteoclast.like | Nup188 |
| Aktip | 1.31E-13 | 0.6477813 | 0.288 | 0.139 | 2.52E-09 | osteoclast.like | Aktip |
| Psmf12 | 1.31E-13 | 0.45353784 | 0.38 | 0.217 | 2.54E-09 | osteoclast.like | Psmf1 |
| Tmem2683 | 1.33E-13 | 0.27805735 | 0.415 | 0.236 | 2.57E-09 | osteoclast.like | Tmem268 |
| Ints9 | 1.35E-13 | 0.7310359 | 0.263 | 0.126 | 2.62E-09 | osteoclast.like | Ints9 |
| Trim651 | 1.36E-13 | 0.42049268 | 0.273 | 0.133 | 2.63E-09 | osteoclast.like | Trim65 |
| Tbce1 | 1.47E-13 | 0.46899498 | 0.312 | 0.15 | 2.84E-09 | osteoclast.like | Tbce |
| Zc3h12c1 | 1.53E-13 | 0.31497611 | 0.415 | 0.175 | 2.95E-09 | osteoclast.like | Zc3h12c |
| Kpnb11 | 1.60E-13 | 0.25334242 | 0.463 | 0.288 | 3.09E-09 | osteoclast.like | Kpnb1 |
| Utp41 | 1.64E-13 | 0.39875328 | 0.322 | 0.186 | 3.16E-09 | osteoclast.like | Utp4 |
| Sap30bp1 | 1.84E-13 | 0.29738094 | 0.38 | 0.219 | 3.57E-09 | osteoclast.like | Sap30bp |
| Pde12 | 1.85E-13 | 0.66871676 | 0.259 | 0.107 | 3.58E-09 | osteoclast.like | Pde12 |
| Scaf81 | 1.90E-13 | 0.3151618 | 0.405 | 0.219 | 3.67E-09 | osteoclast.like | Scaf8 |
| Parg | 2.23E-13 | 0.31326513 | 0.346 | 0.191 | 4.31E-09 | osteoclast.like | Parg |
| Dhps1 | 2.39E-13 | 0.39640453 | 0.366 | 0.236 | 4.62E-09 | osteoclast.like | Dhps |
| Pcm12 | 2.82E-13 | 0.38168459 | 0.463 | 0.246 | 5.46E-09 | osteoclast.like | Pcm1 |
| Blnk5 | 3.05E-13 | 0.29656982 | 0.371 | 0.222 | 5.91E-09 | osteoclast.like | Blnk |
| D2hgdh | 3.09E-13 | 0.83387713 | 0.283 | 0.111 | 5.97E-09 | osteoclast.like | D2hgdh |
| Srpk22 | 3.36E-13 | 0.39061214 | 0.371 | 0.194 | 6.49E-09 | osteoclast.like | Srpk2 |
| Fig41 | 3.61E-13 | 0.38764828 | 0.322 | 0.16 | 6.99E-09 | osteoclast.like | Fig4 |
| Ak61 | 3.77E-13 | 0.55472449 | 0.361 | 0.188 | 7.29E-09 | osteoclast.like | Ak6 |
| Znhit21 | 3.95E-13 | 0.56137969 | 0.298 | 0.138 | 7.64E-09 | osteoclast.like | Znhit2 |
| Zdhhc41 | 4.02E-13 | 0.33882571 | 0.41 | 0.217 | 7.76E-09 | osteoclast.like | Zdhhc4 |
| Kat71 | 4.70E-13 | 0.2777895 | 0.449 | 0.258 | 9.08E-09 | osteoclast.like | Kat7 |
| Phf3 | 4.93E-13 | 0.48236558 | 0.371 | 0.213 | 9.54E-09 | osteoclast.like | Phf3 |
| Car131 | 4.98E-13 | 0.40017413 | 0.346 | 0.156 | 9.63E-09 | osteoclast.like | Car13 |
| Mcm71 | 5.06E-13 | 0.56657833 | 0.263 | 0.118 | 9.78E-09 | osteoclast.like | Mcm7 |
| Selenoh1 | 5.30E-13 | 0.5513868 | 0.283 | 0.138 | 1.02E-08 | osteoclast.like | Selenoh |
| Cables2 | 5.90E-13 | 0.60842634 | 0.283 | 0.113 | 1.14E-08 | osteoclast.like | Cables2 |
| Ift20 | 5.92E-13 | 0.31176801 | 0.429 | 0.261 | 1.15E-08 | osteoclast.like | Ift20 |
| Dgkq | 5.99E-13 | 0.68698467 | 0.307 | 0.137 | 1.16E-08 | osteoclast.like | Dgkq |
| Ncdn2 | 6.11E-13 | 0.32967308 | 0.322 | 0.182 | 1.18E-08 | osteoclast.like | Ncdn |
| Slc33a1 | 6.48E-13 | 0.25237433 | 0.371 | 0.205 | 1.25E-08 | osteoclast.like | Slc33a1 |
| Ipo81 | 6.74E-13 | 0.3667175 | 0.376 | 0.226 | 1.30E-08 | osteoclast.like | Ipo8 |
| Gpkow | 6.80E-13 | 0.33720776 | 0.327 | 0.155 | 1.31E-08 | osteoclast.like | Gpkow |
| Ddx50 | 7.91E-13 | 0.32501197 | 0.439 | 0.252 | 1.53E-08 | osteoclast.like | Ddx50 |
| Zfp574 | 7.97E-13 | 0.78843718 | 0.273 | 0.122 | 1.54E-08 | osteoclast.like | Zfp574 |
| Nsmce4a1 | 8.52E-13 | 0.44963977 | 0.41 | 0.258 | 1.65E-08 | osteoclast.like | Nsmce4a |
| Dnajc161 | 8.71E-13 | 0.47636943 | 0.385 | 0.218 | 1.68E-08 | osteoclast.like | Dnajc16 |
| Arl14ep | 8.91E-13 | 0.30825532 | 0.288 | 0.138 | 1.72E-08 | osteoclast.like | Arl14ep |
| Lactb21 | 8.95E-13 | 0.37193909 | 0.312 | 0.151 | 1.73E-08 | osteoclast.like | Lactb2 |
| Fam120b | 1.08E-12 | 0.48090984 | 0.293 | 0.145 | 2.08E-08 | osteoclast.like | Fam120b |
| Rnf25 | 1.08E-12 | 0.58458522 | 0.259 | 0.119 | 2.09E-08 | osteoclast.like | Rnf25 |
| Brpf3 | 1.18E-12 | 0.26946158 | 0.263 | 0.138 | 2.28E-08 | osteoclast.like | Brpf3 |
| Insig2 | 1.20E-12 | 0.63057003 | 0.254 | 0.131 | 2.31E-08 | osteoclast.like | Insig2 |
| Mavs2 | 1.20E-12 | 0.39645666 | 0.337 | 0.191 | 2.33E-08 | osteoclast.like | Mavs |
| Adpgk2 | 1.25E-12 | 0.3229795 | 0.434 | 0.257 | 2.42E-08 | osteoclast.like | Adpgk |
| Dnajc30 | 1.32E-12 | 0.64618321 | 0.254 | 0.113 | 2.54E-08 | osteoclast.like | Dnajc30 |
| Angel2 | 1.40E-12 | 0.46461448 | 0.268 | 0.124 | 2.70E-08 | osteoclast.like | Angel2 |
| Lats11 | 1.41E-12 | 0.35524392 | 0.337 | 0.195 | 2.72E-08 | osteoclast.like | Lats1 |
| Cyhr1 | 1.46E-12 | 0.38381733 | 0.341 | 0.189 | 2.83E-08 | osteoclast.like | Cyhr1 |
| Ttc41 | 1.78E-12 | 0.53202626 | 0.263 | 0.146 | 3.43E-08 | osteoclast.like | Ttc4 |
| Cox19 | 1.78E-12 | 0.35480872 | 0.307 | 0.148 | 3.43E-08 | osteoclast.like | Cox19 |
| Dnajc22 | 1.78E-12 | 0.75138787 | 0.302 | 0.163 | 3.44E-08 | osteoclast.like | Dnajc2 |
| Ciao2b | 1.86E-12 | 0.33794045 | 0.298 | 0.152 | 3.60E-08 | osteoclast.like | Ciao2b |
| Rft11 | 1.89E-12 | 0.49243521 | 0.307 | 0.149 | 3.65E-08 | osteoclast.like | Rft1 |
| Gnptg4 | 1.99E-12 | 0.54706224 | 0.405 | 0.216 | 3.85E-08 | osteoclast.like | Gnptg |
| Pwwp3a | 2.04E-12 | 0.49956952 | 0.268 | 0.108 | 3.94E-08 | osteoclast.like | Pwwp3a |
| Rnf121 | 2.07E-12 | 0.40569909 | 0.317 | 0.16 | 4.00E-08 | osteoclast.like | Rnf121 |
| Nup501 | 2.31E-12 | 0.31851336 | 0.439 | 0.256 | 4.46E-08 | osteoclast.like | Nup50 |
| Prpf38a1 | 2.31E-12 | 0.33854626 | 0.41 | 0.235 | 4.47E-08 | osteoclast.like | Prpf38a |
| Tmem260 | 2.58E-12 | 0.66590916 | 0.263 | 0.12 | 4.99E-08 | osteoclast.like | Tmem260 |
| Lsm11 | 2.58E-12 | 0.65833319 | 0.283 | 0.144 | 4.99E-08 | osteoclast.like | Lsm1 |
| Bmi11 | 2.82E-12 | 0.32685184 | 0.327 | 0.168 | 5.45E-08 | osteoclast.like | Bmi1 |
| Mcoln12 | 2.83E-12 | 0.33112834 | 0.424 | 0.291 | 5.48E-08 | osteoclast.like | Mcoln1 |
| Siae | 2.84E-12 | 0.41748768 | 0.317 | 0.163 | 5.50E-08 | osteoclast.like | Siae |
| Ssrp11 | 3.55E-12 | 0.42971774 | 0.463 | 0.234 | 6.87E-08 | osteoclast.like | Ssrp1 |
| Trmt61 | 3.66E-12 | 0.57943286 | 0.283 | 0.158 | 7.08E-08 | osteoclast.like | Trmt6 |
| Zkscan17 | 3.69E-12 | 0.4424891 | 0.254 | 0.124 | 7.14E-08 | osteoclast.like | Zkscan17 |
| Kctd201 | 4.05E-12 | 0.27142367 | 0.38 | 0.218 | 7.83E-08 | osteoclast.like | Kctd20 |
| Zfp664 | 4.18E-12 | 0.61842432 | 0.268 | 0.127 | 8.08E-08 | osteoclast.like | Zfp664 |
| Ttc9c1 | 4.31E-12 | 0.28098402 | 0.376 | 0.23 | 8.33E-08 | osteoclast.like | Ttc9c |
| Ppp2r2d1 | 4.40E-12 | 0.54041437 | 0.293 | 0.148 | 8.50E-08 | osteoclast.like | Ppp2r2d |
| Srfbp1 | 5.04E-12 | 0.63041922 | 0.268 | 0.119 | 9.74E-08 | osteoclast.like | Srfbp1 |
| Klc21 | 5.43E-12 | 0.63317139 | 0.302 | 0.152 | 1.05E-07 | osteoclast.like | Klc2 |
| Smarcal11 | 5.44E-12 | 0.42456518 | 0.302 | 0.154 | 1.05E-07 | osteoclast.like | Smarcal1 |
| Xrcc4 | 5.95E-12 | 0.51712821 | 0.322 | 0.15 | 1.15E-07 | osteoclast.like | Xrcc4 |
| Aldh18a12 | 6.76E-12 | 0.3127888 | 0.307 | 0.166 | 1.31E-07 | osteoclast.like | Aldh18a1 |
| Ankrd501 | 6.97E-12 | 0.47445381 | 0.317 | 0.172 | 1.35E-07 | osteoclast.like | Ankrd50 |
| Ramac | 8.39E-12 | 0.40124471 | 0.341 | 0.208 | 1.62E-07 | osteoclast.like | Ramac |
| Dusp221 | 9.13E-12 | 0.34395156 | 0.317 | 0.184 | 1.76E-07 | osteoclast.like | Dusp22 |
| Slc41a24 | 9.62E-12 | 0.26024697 | 0.38 | 0.207 | 1.86E-07 | osteoclast.like | Slc41a2 |
| Slc10a71 | 1.11E-11 | 0.27307906 | 0.278 | 0.152 | 2.14E-07 | osteoclast.like | Slc10a7 |
| Zbtb14 | 1.32E-11 | 0.46743648 | 0.263 | 0.114 | 2.55E-07 | osteoclast.like | Zbtb14 |
| Faap201 | 1.33E-11 | 0.2818728 | 0.376 | 0.198 | 2.57E-07 | osteoclast.like | Faap20 |
| Cfap201 | 1.42E-11 | 0.55359143 | 0.317 | 0.163 | 2.75E-07 | osteoclast.like | Cfap20 |
| Rnf167 | 1.93E-11 | 0.34150982 | 0.317 | 0.171 | 3.72E-07 | osteoclast.like | Rnf167 |
| Decr2 | 2.03E-11 | 0.43067246 | 0.283 | 0.128 | 3.92E-07 | osteoclast.like | Decr2 |
| Trappc131 | 2.06E-11 | 0.41806025 | 0.298 | 0.143 | 3.98E-07 | osteoclast.like | Trappc13 |
| Pwp11 | 2.29E-11 | 0.40785801 | 0.283 | 0.144 | 4.44E-07 | osteoclast.like | Pwp1 |
| Rabgap11 | 2.33E-11 | 0.41286275 | 0.312 | 0.173 | 4.50E-07 | osteoclast.like | Rabgap1 |
| Ago13 | 2.51E-11 | 0.35017887 | 0.434 | 0.256 | 4.86E-07 | osteoclast.like | Ago1 |
| Arl15 | 2.69E-11 | 0.25664875 | 0.298 | 0.155 | 5.19E-07 | osteoclast.like | Arl15 |
| Rsbn1l | 2.74E-11 | 0.25284152 | 0.337 | 0.217 | 5.30E-07 | osteoclast.like | Rsbn1l |
| Nup851 | 2.88E-11 | 0.47480301 | 0.268 | 0.14 | 5.56E-07 | osteoclast.like | Nup85 |
| Chmp63 | 2.90E-11 | 0.25711058 | 0.395 | 0.227 | 5.61E-07 | osteoclast.like | Chmp6 |
| Gpn31 | 3.48E-11 | 0.29935118 | 0.322 | 0.175 | 6.73E-07 | osteoclast.like | Gpn3 |
| Stard92 | 3.51E-11 | 0.29704263 | 0.366 | 0.227 | 6.79E-07 | osteoclast.like | Stard9 |
| Nmt23 | 4.02E-11 | 0.4417945 | 0.366 | 0.212 | 7.77E-07 | osteoclast.like | Nmt2 |
| Ldah | 4.36E-11 | 0.25670421 | 0.298 | 0.173 | 8.42E-07 | osteoclast.like | Ldah |
| Ttc371 | 4.57E-11 | 0.29940631 | 0.307 | 0.152 | 8.84E-07 | osteoclast.like | Ttc37 |
| Vezf1 | 4.84E-11 | 0.34887488 | 0.288 | 0.145 | 9.36E-07 | osteoclast.like | Vezf1 |
| Xrcc11 | 5.12E-11 | 0.25883822 | 0.302 | 0.163 | 9.91E-07 | osteoclast.like | Xrcc1 |
| Ccnh1 | 5.18E-11 | 0.30941626 | 0.273 | 0.147 | 1.00E-06 | osteoclast.like | Ccnh |
| Ap4m12 | 5.20E-11 | 0.46837624 | 0.366 | 0.194 | 1.00E-06 | osteoclast.like | Ap4m1 |
| Sec23a | 5.21E-11 | 0.51912659 | 0.307 | 0.159 | 1.01E-06 | osteoclast.like | Sec23a |
| Uckl1 | 5.37E-11 | 0.27975604 | 0.415 | 0.23 | 1.04E-06 | osteoclast.like | Uckl1 |
| Armc8 | 5.41E-11 | 0.39285644 | 0.298 | 0.143 | 1.05E-06 | osteoclast.like | Armc8 |
| Yipf6 | 5.94E-11 | 0.54216295 | 0.259 | 0.134 | 1.15E-06 | osteoclast.like | Yipf6 |
| Tm2d1 | 7.33E-11 | 0.51919523 | 0.273 | 0.143 | 1.42E-06 | osteoclast.like | Tm2d1 |
| Tbcel1 | 8.04E-11 | 0.28045318 | 0.341 | 0.183 | 1.55E-06 | osteoclast.like | Tbcel |
| Med41 | 9.53E-11 | 0.49554993 | 0.263 | 0.146 | 1.84E-06 | osteoclast.like | Med4 |
| Rarg1 | 1.13E-10 | 0.46856944 | 0.307 | 0.152 | 2.19E-06 | osteoclast.like | Rarg |
| Pin11 | 1.41E-10 | 0.27965073 | 0.283 | 0.164 | 2.72E-06 | osteoclast.like | Pin1 |
| Lamc12 | 1.42E-10 | 0.30051397 | 0.307 | 0.153 | 2.74E-06 | osteoclast.like | Lamc1 |
| Pcgf3 | 1.42E-10 | 0.60417499 | 0.278 | 0.143 | 2.75E-06 | osteoclast.like | Pcgf3 |
| Lancl12 | 1.82E-10 | 0.44603867 | 0.298 | 0.142 | 3.53E-06 | osteoclast.like | Lancl1 |
| Rusc1 | 1.93E-10 | 0.42601742 | 0.278 | 0.146 | 3.73E-06 | osteoclast.like | Rusc1 |
| Eef1akmt11 | 2.10E-10 | 0.46286786 | 0.307 | 0.168 | 4.06E-06 | osteoclast.like | Eef1akmt1 |
| Ppat1 | 2.18E-10 | 0.66542765 | 0.283 | 0.132 | 4.21E-06 | osteoclast.like | Ppat |
| Rpp25l1 | 2.30E-10 | 0.34805441 | 0.278 | 0.134 | 4.44E-06 | osteoclast.like | Rpp25l |
| Zc3hc1 | 2.42E-10 | 0.31911041 | 0.259 | 0.115 | 4.68E-06 | osteoclast.like | Zc3hc1 |
| Zfpl11 | 2.52E-10 | 0.52574727 | 0.278 | 0.162 | 4.88E-06 | osteoclast.like | Zfpl1 |
| Rngtt | 2.74E-10 | 0.27511179 | 0.259 | 0.133 | 5.31E-06 | osteoclast.like | Rngtt |
| Hes62 | 2.77E-10 | 0.37768934 | 0.415 | 0.226 | 5.36E-06 | osteoclast.like | Hes6 |
| Nbdy | 2.85E-10 | 0.49679997 | 0.268 | 0.135 | 5.52E-06 | osteoclast.like | Nbdy |
| Tada31 | 3.03E-10 | 0.25791057 | 0.298 | 0.152 | 5.85E-06 | osteoclast.like | Tada3 |
| Wwc2 | 3.65E-10 | 0.30936226 | 0.283 | 0.146 | 7.06E-06 | osteoclast.like | Wwc2 |
| Lcmt11 | 3.79E-10 | 0.40176932 | 0.341 | 0.178 | 7.32E-06 | osteoclast.like | Lcmt1 |
| Anp32b1 | 4.23E-10 | 0.38698423 | 0.351 | 0.218 | 8.17E-06 | osteoclast.like | Anp32b |
| Zyg11b | 4.66E-10 | 0.3640718 | 0.288 | 0.144 | 9.02E-06 | osteoclast.like | Zyg11b |
| Thap72 | 4.97E-10 | 0.46115001 | 0.317 | 0.199 | 9.61E-06 | osteoclast.like | Thap7 |
| Ints111 | 5.43E-10 | 0.48157765 | 0.337 | 0.178 | 1.05E-05 | osteoclast.like | Ints11 |
| Fbxo9 | 6.38E-10 | 0.51165565 | 0.307 | 0.151 | 1.23E-05 | osteoclast.like | Fbxo9 |
| Stmn11 | 6.59E-10 | 0.34037345 | 0.317 | 0.179 | 1.27E-05 | osteoclast.like | Stmn1 |
| Urb2 | 7.36E-10 | 0.56673585 | 0.254 | 0.115 | 1.42E-05 | osteoclast.like | Urb2 |
| Bnip3l2 | 7.84E-10 | 0.33695372 | 0.317 | 0.176 | 1.52E-05 | osteoclast.like | Bnip3l |
| Polr3d | 8.09E-10 | 0.5319303 | 0.273 | 0.124 | 1.56E-05 | osteoclast.like | Polr3d |
| Wwox | 8.44E-10 | 0.38864042 | 0.263 | 0.131 | 1.63E-05 | osteoclast.like | Wwox |
| Zfyve191 | 9.18E-10 | 0.34474207 | 0.283 | 0.145 | 1.77E-05 | osteoclast.like | Zfyve19 |
| Zfp280d1 | 1.01E-09 | 0.44477148 | 0.283 | 0.131 | 1.96E-05 | osteoclast.like | Zfp280d |
| Cdkn2aipnl3 | 1.26E-09 | 0.27154931 | 0.356 | 0.204 | 2.43E-05 | osteoclast.like | Cdkn2aipnl |
| Ap3m1 | 1.27E-09 | 0.32079423 | 0.254 | 0.12 | 2.46E-05 | osteoclast.like | Ap3m1 |
| Thoc31 | 1.31E-09 | 0.26662695 | 0.259 | 0.13 | 2.53E-05 | osteoclast.like | Thoc3 |
| Kctd13 | 1.39E-09 | 0.36826368 | 0.259 | 0.154 | 2.70E-05 | osteoclast.like | Kctd13 |
| Slc4a1ap | 1.89E-09 | 0.41339467 | 0.293 | 0.143 | 3.65E-05 | osteoclast.like | Slc4a1ap |
| Sos2 | 1.95E-09 | 0.36630756 | 0.332 | 0.197 | 3.78E-05 | osteoclast.like | Sos2 |
| Polr2i1 | 2.07E-09 | 0.2526258 | 0.307 | 0.154 | 4.00E-05 | osteoclast.like | Polr2i |
| Dcps1 | 2.24E-09 | 0.28643664 | 0.254 | 0.135 | 4.34E-05 | osteoclast.like | Dcps |
| Nepro | 2.29E-09 | 0.32593327 | 0.254 | 0.131 | 4.42E-05 | osteoclast.like | Nepro |
| Arhgap11a1 | 2.40E-09 | 0.5276064 | 0.278 | 0.134 | 4.64E-05 | osteoclast.like | Arhgap11a |
| Deaf11 | 2.50E-09 | 0.32088804 | 0.254 | 0.141 | 4.83E-05 | osteoclast.like | Deaf1 |
| Acap3 | 2.75E-09 | 0.31597858 | 0.278 | 0.143 | 5.31E-05 | osteoclast.like | Acap3 |
| Avl9 | 2.86E-09 | 0.28493854 | 0.273 | 0.152 | 5.53E-05 | osteoclast.like | Avl9 |
| Mllt11 | 3.10E-09 | 0.30871421 | 0.293 | 0.16 | 6.00E-05 | osteoclast.like | Mllt1 |
| Polg | 3.16E-09 | 0.29060704 | 0.278 | 0.151 | 6.11E-05 | osteoclast.like | Polg |
| Luc7l2 | 3.43E-09 | 0.28632552 | 0.337 | 0.199 | 6.64E-05 | osteoclast.like | Luc7l |
| Fam160b1 | 4.65E-09 | 0.31936831 | 0.385 | 0.221 | 8.99E-05 | osteoclast.like | Fam160b1 |

| | | | | | | |
|---|---|---|---|---|---|---|
| Tmem941 | 4.91E-09 | 0.29911034 | 0.302 | 0.176 | 9.49E-05 | osteoclast.like:Tmem94 |
| Sart31 | 5.08E-09 | 0.38160412 | 0.327 | 0.171 | 9.82E-05 | osteoclast.like:Sart3 |
| Tmem120b1 | 5.16E-09 | 0.25646871 | 0.259 | 0.138 | 9.98E-05 | osteoclast.like:Tmem120b |
| Trim33 | 5.31E-09 | 0.38520425 | 0.307 | 0.168 | 0.00010267 | osteoclast.like:Trim33 |
| Crcp | 5.84E-09 | 0.35288909 | 0.356 | 0.199 | 0.00011284 | osteoclast.like:Crcp |
| Ttc131 | 6.49E-09 | 0.30277374 | 0.356 | 0.195 | 0.0001255 | osteoclast.like:Ttc13 |
| Elp11 | 6.93E-09 | 0.6439756 | 0.293 | 0.152 | 0.00013408 | osteoclast.like:Elp1 |
| Mettl23 | 7.56E-09 | 0.40377904 | 0.254 | 0.127 | 0.00014626 | osteoclast.like:Mettl23 |
| Rictor | 8.02E-09 | 0.25412919 | 0.351 | 0.201 | 0.00015506 | osteoclast.like:Rictor |
| Dicer12 | 9.49E-09 | 0.35458156 | 0.366 | 0.201 | 0.00018353 | osteoclast.like:Dicer1 |
| B3gat31 | 1.13E-08 | 0.27591338 | 0.317 | 0.195 | 0.00021871 | osteoclast.like:B3gat3 |
| Tbpl1 | 1.26E-08 | 0.34910904 | 0.302 | 0.193 | 0.00024299 | osteoclast.like:Tbpl1 |
| Rprd1b1 | 1.45E-08 | 0.33671763 | 0.273 | 0.147 | 0.00028075 | osteoclast.like:Rprd1b |
| Aida1 | 1.78E-08 | 0.34516215 | 0.312 | 0.183 | 0.00034453 | osteoclast.like:Aida |
| Surf2 | 2.07E-08 | 0.35438929 | 0.273 | 0.145 | 0.00039934 | osteoclast.like:Surf2 |
| Zfp5931 | 2.42E-08 | 0.45152147 | 0.263 | 0.127 | 0.00046764 | osteoclast.like:Zfp593 |
| Abcf3 | 3.58E-08 | 0.30408615 | 0.302 | 0.176 | 0.00069277 | osteoclast.like:Abcf3 |
| Bcl7b | 4.32E-08 | 0.29628515 | 0.254 | 0.129 | 0.00083488 | osteoclast.like:Bcl7b |
| Tstd21 | 4.40E-08 | 0.41511357 | 0.283 | 0.154 | 0.0008516 | osteoclast.like:Tstd2 |
| Fbxl171 | 5.31E-08 | 0.26904795 | 0.332 | 0.205 | 0.00102716 | osteoclast.like:Fbxl17 |
| Nbeal11 | 8.75E-08 | 0.25134275 | 0.298 | 0.182 | 0.00169165 | osteoclast.like:Nbeal1 |
| Prkar2a2 | 9.41E-08 | 0.2556124 | 0.351 | 0.205 | 0.0018193 | osteoclast.like:Prkar2a |
| Ccnc | 1.03E-07 | 0.31360634 | 0.283 | 0.168 | 0.00198804 | osteoclast.like:Ccnc |
| Zfp64 | 1.19E-07 | 0.36878417 | 0.254 | 0.131 | 0.00229481 | osteoclast.like:Zfp64 |
| Ppil11 | 1.51E-07 | 0.26595233 | 0.283 | 0.147 | 0.00292169 | osteoclast.like:Ppil1 |
| Rad54l21 | 1.71E-07 | 0.26114651 | 0.327 | 0.196 | 0.00329647 | osteoclast.like:Rad54l2 |
| Caml1 | 1.75E-07 | 0.26761765 | 0.273 | 0.174 | 0.00337874 | osteoclast.like:Caml |
| Nsun4 | 1.97E-07 | 0.363457 | 0.322 | 0.177 | 0.00381423 | osteoclast.like:Nsun4 |
| Map4k5 | 2.02E-07 | 0.46209217 | 0.259 | 0.14 | 0.00390101 | osteoclast.like:Map4k5 |
| Tmco32 | 2.09E-07 | 0.31052713 | 0.312 | 0.183 | 0.00404933 | osteoclast.like:Tmco3 |
| Cep631 | 2.42E-07 | 0.2559541 | 0.293 | 0.17 | 0.00467664 | osteoclast.like:Cep63 |
| Rmdn31 | 2.93E-07 | 0.30814682 | 0.268 | 0.144 | 0.00566459 | osteoclast.like:Rmdn3 |
| C1galt1 | 3.08E-07 | 0.25511139 | 0.268 | 0.158 | 0.00595833 | osteoclast.like:C1galt1 |
| Polr2h1 | 3.60E-07 | 0.3248605 | 0.259 | 0.159 | 0.00695175 | osteoclast.like:Polr2h |
| Smc31 | 5.80E-07 | 0.32583493 | 0.346 | 0.219 | 0.01121442 | osteoclast.like:Smc3 |
| Pias4 | 6.28E-07 | 0.40125839 | 0.273 | 0.158 | 0.01214279 | osteoclast.like:Pias4 |
| Pank41 | 6.41E-07 | 0.39202101 | 0.259 | 0.143 | 0.01239524 | osteoclast.like:Pank4 |
| Amd1 | 1.80E-06 | 0.30215199 | 0.263 | 0.15 | 0.03482783 | osteoclast.like:Amd1 |
| Ptp4a33 | 6.13E-06 | 0.29555581 | 0.283 | 0.193 | 0.11850538 | osteoclast.like:Ptp4a3 |
| Sugp11 | 1.30E-05 | 0.25384179 | 0.283 | 0.167 | 0.2506814 | osteoclast.like:Sugp1 |

| | p_val | avg_log2FC | pct.1 | pct.2 | p_val_adj | cluster | gene | annotation_1 |
|---|---|---|---|---|---|---|---|---|
| Osm | 0 | 2.34515945 | 0.697 | 0.214 | 0 | 0 | Osm | neutrophil.1 |
| Slfn4 | 0 | 1.59098409 | 0.719 | 0.284 | 0 | 0 | Slfn4 | neutrophil.1 |
| Lmnb1 | 0 | 1.53807698 | 0.8 | 0.383 | 0 | 0 | Lmnb1 | neutrophil.1 |
| Lrg1 | 0 | 1.95914912 | 0.625 | 0.211 | 0 | 0 | Lrg1 | neutrophil.1 |
| Sell | 0 | 1.43975737 | 0.637 | 0.244 | 0 | 0 | Sell | neutrophil.1 |
| S100a9 | 0 | 1.97020283 | 0.911 | 0.523 | 0 | 0 | S100a9 | neutrophil.1 |
| Entpd1 | 0 | 1.52853467 | 0.799 | 0.412 | 0 | 0 | Entpd1 | neutrophil.1 |
| Smox | 0 | 1.43874925 | 0.772 | 0.406 | 0 | 0 | Smox | neutrophil.1 |
| Adam8 | 0 | 1.45109859 | 0.576 | 0.213 | 0 | 0 | Adam8 | neutrophil.1 |
| Hdc | 0 | 1.38687564 | 0.911 | 0.549 | 0 | 0 | Hdc | neutrophil.1 |
| Lilr4b | 0 | 1.30446653 | 0.783 | 0.422 | 0 | 0 | Lilr4b | neutrophil.1 |
| Slc15a3 | 0 | 1.49678251 | 0.924 | 0.586 | 0 | 0 | Slc15a3 | neutrophil.1 |
| Ccr1 | 0 | 1.36418128 | 0.754 | 0.416 | 0 | 0 | Ccr1 | neutrophil.1 |
| DEPRECATED-l | 0 | 1.55505712 | 0.618 | 0.28 | 0 | 0 | DEPRECATED- | neutrophil.1 |
| Ptgs2 | 0 | 0.97686839 | 0.895 | 0.56 | 0 | 0 | Ptgs2 | neutrophil.1 |
| Slc16a3 | 0 | 1.34907615 | 0.842 | 0.512 | 0 | 0 | Slc16a3 | neutrophil.1 |
| Trib1 | 0 | 1.20075831 | 0.793 | 0.466 | 0 | 0 | Trib1 | neutrophil.1 |
| Ccl6 | 0 | 1.81305902 | 0.462 | 0.136 | 0 | 0 | Ccl6 | neutrophil.1 |
| S100a8 | 0 | 1.50671743 | 0.87 | 0.546 | 0 | 0 | S100a8 | neutrophil.1 |
| Homer1 | 0 | 1.1890144 | 0.604 | 0.284 | 0 | 0 | Homer1 | neutrophil.1 |
| Arg2 | 0 | 1.28236279 | 0.69 | 0.374 | 0 | 0 | Arg2 | neutrophil.1 |
| Ndel1 | 0 | 1.05586883 | 0.745 | 0.434 | 0 | 0 | Ndel1 | neutrophil.1 |
| Nsd3 | 0 | 1.31925693 | 0.674 | 0.364 | 0 | 0 | Nsd3 | neutrophil.1 |
| Acod1 | 0 | 0.95034656 | 0.867 | 0.564 | 0 | 0 | Acod1 | neutrophil.1 |
| Trem1 | 0 | 1.43851498 | 0.822 | 0.519 | 0 | 0 | Trem1 | neutrophil.1 |
| Rabgef1 | 0 | 1.06649872 | 0.672 | 0.372 | 0 | 0 | Rabgef1 | neutrophil.1 |
| Hcar2 | 0 | 1.09833502 | 0.897 | 0.604 | 0 | 0 | Hcar2 | neutrophil.1 |
| Vps37b | 0 | 1.28199745 | 0.671 | 0.38 | 0 | 0 | Vps37b | neutrophil.1 |
| Gda | 0 | 1.46677705 | 0.497 | 0.208 | 0 | 0 | Gda | neutrophil.1 |
| Gk | 0 | 2.40134547 | 0.377 | 0.091 | 0 | 0 | Gk | neutrophil.1 |
| Nfil3 | 0 | 1.21522021 | 0.574 | 0.288 | 0 | 0 | Nfil3 | neutrophil.1 |
| Jaml | 0 | 2.06970398 | 0.407 | 0.122 | 0 | 0 | Jaml | neutrophil.1 |
| Cd300ld | 0 | 1.26688439 | 0.569 | 0.287 | 0 | 0 | Cd300ld | neutrophil.1 |
| Nr4a3 | 0 | 0.97028457 | 0.65 | 0.371 | 0 | 0 | Nr4a3 | neutrophil.1 |
| Jdp2 | 0 | 1.03756315 | 0.656 | 0.379 | 0 | 0 | Jdp2 | neutrophil.1 |
| Ppp1r3b | 0 | 1.50631635 | 0.502 | 0.226 | 0 | 0 | Ppp1r3b | neutrophil.1 |
| G0s2 | 0 | 1.25344339 | 0.802 | 0.527 | 0 | 0 | G0s2 | neutrophil.1 |
| Ell2 | 0 | 1.06278599 | 0.567 | 0.292 | 0 | 0 | Ell2 | neutrophil.1 |
| Cd14 | 0 | 1.32866816 | 0.921 | 0.647 | 0 | 0 | Cd14 | neutrophil.1 |
| Clec4d | 0 | 1.19778699 | 0.957 | 0.683 | 0 | 0 | Clec4d | neutrophil.1 |
| Fgl2 | 0 | 0.98820513 | 0.702 | 0.429 | 0 | 0 | Fgl2 | neutrophil.1 |
| Csrnp1 | 0 | 0.85712811 | 0.917 | 0.645 | 0 | 0 | Csrnp1 | neutrophil.1 |
| Nlrp3 | 0 | 1.08841556 | 0.924 | 0.655 | 0 | 0 | Nlrp3 | neutrophil.1 |
| Wfdc17 | 0 | 1.90544611 | 0.413 | 0.144 | 0 | 0 | Wfdc17 | neutrophil.1 |
| Zyx | 0 | 1.05522104 | 0.703 | 0.434 | 0 | 0 | Zyx | neutrophil.1 |
| Cxcr2 | 0 | 0.83825802 | 0.703 | 0.436 | 0 | 0 | Cxcr2 | neutrophil.1 |
| Nfkbiz | 0 | 0.88817063 | 0.887 | 0.623 | 0 | 0 | Nfkbiz | neutrophil.1 |
| Pde4b | 0 | 1.13696497 | 0.655 | 0.392 | 0 | 0 | Pde4b | neutrophil.1 |
| Samsn1 | 0 | 0.92242054 | 0.753 | 0.49 | 0 | 0 | Samsn1 | neutrophil.1 |
| S100a11 | 0 | 1.10073606 | 0.887 | 0.627 | 0 | 0 | S100a11 | neutrophil.1 |
| Ankrd33b | 0 | 0.82074541 | 0.508 | 0.248 | 0 | 0 | Ankrd33b | neutrophil.1 |
| Ifitm1 | 0 | 0.94964595 | 0.836 | 0.577 | 0 | 0 | Ifitm1 | neutrophil.1 |
| C5ar1 | 0 | 0.68397867 | 0.844 | 0.587 | 0 | 0 | C5ar1 | neutrophil.1 |
| Grina | 0 | 0.92255292 | 0.882 | 0.626 | 0 | 0 | Grina | neutrophil.1 |
| Selenon | 0 | 1.48035875 | 0.414 | 0.159 | 0 | 0 | Selenon | neutrophil.1 |
| Socs3 | 0 | 0.9555212 | 0.755 | 0.502 | 0 | 0 | Socs3 | neutrophil.1 |
| Retnlg | 0 | 2.30304389 | 0.308 | 0.055 | 0 | 0 | Retnlg | neutrophil.1 |
| Fosl2 | 0 | 0.71932896 | 0.881 | 0.633 | 0 | 0 | Fosl2 | neutrophil.1 |
| Rasip1 | 0 | 1.88678532 | 0.363 | 0.116 | 0 | 0 | Rasip1 | neutrophil.1 |
| Mcemp1 | 0 | 1.68625034 | 0.387 | 0.14 | 0 | 0 | Mcemp1 | neutrophil.1 |
| Hp | 0 | 0.76618124 | 0.667 | 0.422 | 0 | 0 | Hp | neutrophil.1 |
| Cxcl3 | 0 | 2.25412575 | 0.358 | 0.115 | 0 | 0 | Cxcl3 | neutrophil.1 |
| Antxr2 | 0 | 0.46142026 | 0.737 | 0.496 | 0 | 0 | Antxr2 | neutrophil.1 |
| Tgoln1 | 0 | 0.97113915 | 0.562 | 0.323 | 0 | 0 | Tgoln1 | neutrophil.1 |
| Lilrb4a | 0 | 0.91359329 | 0.925 | 0.687 | 0 | 0 | Lilrb4a | neutrophil.1 |
| Egr1 | 0 | 0.5644223 | 0.935 | 0.7 | 0 | 0 | Egr1 | neutrophil.1 |
| Dgat2 | 0 | 1.6069804 | 0.373 | 0.141 | 0 | 0 | Dgat2 | neutrophil.1 |
| Cd33 | 0 | 0.79033069 | 0.716 | 0.484 | 0 | 0 | Cd33 | neutrophil.1 |
| Themis2 | 0 | 0.70285659 | 0.684 | 0.453 | 0 | 0 | Themis2 | neutrophil.1 |
| Thbs1 | 0 | 1.61035917 | 0.477 | 0.248 | 0 | 0 | Thbs1 | neutrophil.1 |
| Actg1 | 0 | 1.07460693 | 0.875 | 0.648 | 0 | 0 | Actg1 | neutrophil.1 |
| Dgat1 | 0 | 1.56175639 | 0.37 | 0.144 | 0 | 0 | Dgat1 | neutrophil.1 |
| Trim30b | 0 | 1.20998776 | 0.408 | 0.182 | 0 | 0 | Trim30b | neutrophil.1 |
| Rab44 | 0 | 0.71214813 | 0.685 | 0.46 | 0 | 0 | Rab44 | neutrophil.1 |
| Slc7a11 | 0 | 0.39816507 | 0.84 | 0.615 | 0 | 0 | Slc7a11 | neutrophil.1 |
| Il1rap | 0 | 1.33168972 | 0.4 | 0.177 | 0 | 0 | Il1rap | neutrophil.1 |
| Fos | 0 | 0.45142964 | 0.737 | 0.514 | 0 | 0 | Fos | neutrophil.1 |
| Plaur | 0 | 1.52859469 | 0.93 | 0.711 | 0 | 0 | Plaur | neutrophil.1 |
| Atf3 | 0 | 0.86782322 | 0.561 | 0.344 | 0 | 0 | Atf3 | neutrophil.1 |
| Tes | 0 | 0.92627627 | 0.549 | 0.332 | 0 | 0 | Tes | neutrophil.1 |
| Kctd12 | 0 | 0.42637608 | 0.804 | 0.589 | 0 | 0 | Kctd12 | neutrophil.1 |
| Il1r2 | 0 | 0.38540992 | 0.914 | 0.7 | 0 | 0 | Il1r2 | neutrophil.1 |
| Chd7 | 0 | 1.01615536 | 0.479 | 0.266 | 0 | 0 | Chd7 | neutrophil.1 |
| Mxd1 | 0 | 0.68704912 | 0.921 | 0.71 | 0 | 0 | Mxd1 | neutrophil.1 |
| Glrx | 0 | 1.76678667 | 0.34 | 0.13 | 0 | 0 | Glrx | neutrophil.1 |
| Rab8b | 0 | 0.62473953 | 0.607 | 0.397 | 0 | 0 | Rab8b | neutrophil.1 |
| Clec4e | 0 | 0.31486863 | 0.78 | 0.571 | 0 | 0 | Clec4e | neutrophil.1 |
| Rdh12 | 0 | 3.06428575 | 0.25 | 0.041 | 0 | 0 | Rdh12 | neutrophil.1 |
| Ifitm2 | 0 | 0.77800848 | 0.855 | 0.646 | 0 | 0 | Ifitm2 | neutrophil.1 |
| Sp140 | 0 | 1.58337244 | 0.328 | 0.121 | 0 | 0 | Sp140 | neutrophil.1 |
| Lsp1 | 0 | 1.15308046 | 0.451 | 0.245 | 0 | 0 | Lsp1 | neutrophil.1 |
| Per1 | 0 | 0.6580927 | 0.569 | 0.369 | 0 | 0 | Per1 | neutrophil.1 |
| Rnasel | 0 | 2.31663658 | 0.261 | 0.063 | 0 | 0 | Rnasel | neutrophil.1 |
| Il10ra | 0 | 0.70911403 | 0.579 | 0.382 | 0 | 0 | Il10ra | neutrophil.1 |
| Slc2a3 | 0 | 1.55210479 | 0.304 | 0.108 | 0 | 0 | Slc2a3 | neutrophil.1 |
| Ifrd1 | 0 | 0.6462694 | 0.696 | 0.5 | 0 | 0 | Ifrd1 | neutrophil.1 |
| Tnfrsf1a | 0 | 0.58213647 | 0.705 | 0.51 | 0 | 0 | Tnfrsf1a | neutrophil.1 |
| Ddx6 | 0 | 0.93612068 | 0.567 | 0.373 | 0 | 0 | Ddx6 | neutrophil.1 |
| Ets2 | 0 | 0.80577066 | 0.952 | 0.759 | 0 | 0 | Ets2 | neutrophil.1 |
| Fam107b | 0 | 1.2699523 | 0.416 | 0.225 | 0 | 0 | Fam107b | neutrophil.1 |
| Eif5a | 0 | 0.92013147 | 0.453 | 0.264 | 0 | 0 | Eif5a | neutrophil.1 |
| Gpr35 | 0 | 0.84509127 | 0.477 | 0.291 | 0 | 0 | Gpr35 | neutrophil.1 |
| Pi16 | 0 | 1.81640773 | 0.264 | 0.078 | 0 | 0 | Pi16 | neutrophil.1 |
| Cass4 | 0 | 0.95579778 | 0.543 | 0.358 | 0 | 0 | Cass4 | neutrophil.1 |
| Rgcc | 0 | 0.8450082 | 0.443 | 0.258 | 0 | 0 | Rgcc | neutrophil.1 |
| Gadd45a | 0 | 0.95556067 | 0.425 | 0.241 | 0 | 0 | Gadd45a | neutrophil.1 |
| Tiparp | 0 | 0.90234306 | 0.409 | 0.225 | 0 | 0 | Tiparp | neutrophil.1 |
| Tlr2 | 0 | 1.02737815 | 0.416 | 0.232 | 0 | 0 | Tlr2 | neutrophil.1 |
| Tgm1 | 0 | 2.05061014 | 0.265 | 0.082 | 0 | 0 | Tgm1 | neutrophil.1 |
| Cd44 | 0 | 0.53330983 | 0.876 | 0.694 | 0 | 0 | Cd44 | neutrophil.1 |
| Zswim4 | 0 | 0.7951884 | 0.45 | 0.272 | 0 | 0 | Zswim4 | neutrophil.1 |
| Tinf2 | 0 | 1.37239045 | 0.318 | 0.142 | 0 | 0 | Tinf2 | neutrophil.1 |
| Sorl1 | 0 | 0.50296796 | 0.735 | 0.559 | 0 | 0 | Sorl1 | neutrophil.1 |
| Dusp1 | 0 | 0.92818961 | 0.987 | 0.811 | 0 | 0 | Dusp1 | neutrophil.1 |
| Sra1 | 0 | 1.20029371 | 0.403 | 0.228 | 0 | 0 | Sra1 | neutrophil.1 |
| Rhoh | 0 | 1.11284661 | 0.378 | 0.203 | 0 | 0 | Rhoh | neutrophil.1 |
| Alox5ap | 0 | 0.50651184 | 0.585 | 0.411 | 0 | 0 | Alox5ap | neutrophil.1 |
| Btg1 | 0 | 0.52708574 | 0.817 | 0.643 | 0 | 0 | Btg1 | neutrophil.1 |
| Stx11 | 0 | 1.20193868 | 0.363 | 0.192 | 0 | 0 | Stx11 | neutrophil.1 |
| Litaf | 0 | 0.59295088 | 0.927 | 0.758 | 0 | 0 | Litaf | neutrophil.1 |
| DEPRECATED-l | 0 | 0.55093931 | 0.508 | 0.339 | 0 | 0 | DEPRECATED- | neutrophil.1 |
| Rab7 | 0 | 0.71357706 | 0.573 | 0.404 | 0 | 0 | Rab7 | neutrophil.1 |
| Ppp1r15a | 0 | 0.44190401 | 0.814 | 0.645 | 0 | 0 | Ppp1r15a | neutrophil.1 |
| Cpeb2 | 0 | 0.50931544 | 0.549 | 0.381 | 0 | 0 | Cpeb2 | neutrophil.1 |
| Arih2 | 0 | 0.56216847 | 0.528 | 0.361 | 0 | 0 | Arih2 | neutrophil.1 |
| Tpd52 | 0 | 0.91449088 | 0.381 | 0.214 | 0 | 0 | Tpd52 | neutrophil.1 |
| Emilin2 | 0 | 0.42652856 | 0.731 | 0.565 | 0 | 0 | Emilin2 | neutrophil.1 |
| Nfkbid | 0 | 0.33684299 | 0.789 | 0.626 | 0 | 0 | Nfkbid | neutrophil.1 |
| Hcls1 | 0 | 0.41622595 | 0.644 | 0.481 | 0 | 0 | Hcls1 | neutrophil.1 |
| Kdm6b | 0 | 0.50620139 | 0.911 | 0.749 | 0 | 0 | Kdm6b | neutrophil.1 |
| Ripor2 | 0 | 0.76687694 | 0.343 | 0.182 | 0 | 0 | Ripor2 | neutrophil.1 |
| Cd53 | 0 | 0.32397821 | 0.818 | 0.658 | 0 | 0 | Cd53 | neutrophil.1 |
| Ifitm3 | 0 | 1.27675886 | 0.265 | 0.106 | 0 | 0 | Ifitm3 | neutrophil.1 |
| Btg2 | 0 | 0.62582603 | 0.965 | 0.806 | 0 | 0 | Btg2 | neutrophil.1 |
| Il18rap | 0 | 0.97189852 | 0.34 | 0.182 | 0 | 0 | Il18rap | neutrophil.1 |
| Ier3 | 0 | 0.69394898 | 0.957 | 0.801 | 0 | 0 | Ier3 | neutrophil.1 |
| Tlr4 | 0 | 0.90721242 | 0.356 | 0.205 | 0 | 0 | Tlr4 | neutrophil.1 |
| Fcgr3 | 0 | 0.59992614 | 0.603 | 0.455 | 0 | 0 | Fcgr3 | neutrophil.1 |
| Gpcpd1 | 0 | 0.90800454 | 0.319 | 0.176 | 0 | 0 | Gpcpd1 | neutrophil.1 |
| Sgms2 | 0 | 0.77728284 | 0.324 | 0.181 | 0 | 0 | Sgms2 | neutrophil.1 |
| Pglyrp1 | 0 | 0.75102021 | 0.341 | 0.201 | 0 | 0 | Pglyrp1 | neutrophil.1 |
| Trem3 | 0 | 0.59435665 | 0.508 | 0.368 | 0 | 0 | Trem3 | neutrophil.1 |
| Rnf149 | 0 | 0.58882451 | 0.906 | 0.772 | 0 | 0 | Rnf149 | neutrophil.1 |
| Tgm2 | 0 | 0.67993253 | 0.834 | 0.7 | 0 | 0 | Tgm2 | neutrophil.1 |
| Cyth1 | 0 | 0.94743738 | 0.282 | 0.152 | 0 | 0 | Cyth1 | neutrophil.1 |
| Dhx38 | 0 | 0.74742698 | 0.356 | 0.231 | 0 | 0 | Dhx38 | neutrophil.1 |
| Cd300lf | 0 | 0.3029448 | 0.823 | 0.699 | 0 | 0 | Cd300lf | neutrophil.1 |
| Cxcl2 | 0 | 0.80175334 | 0.982 | 0.862 | 0 | 0 | Cxcl2 | neutrophil.1 |
| Pik3ap1 | 0 | 0.35748651 | 0.749 | 0.64 | 0 | 0 | Pik3ap1 | neutrophil.1 |
| Zfp36 | 0 | 0.48838853 | 0.964 | 0.858 | 0 | 0 | Zfp36 | neutrophil.1 |
| Plek | 0 | 0.72757101 | 0.988 | 0.882 | 0 | 0 | Plek | neutrophil.1 |
| Ccl4 | 0 | 0.95826814 | 0.43 | 0.327 | 0 | 0 | Ccl4 | neutrophil.1 |
| Gadd45b | 0 | 0.27421878 | 0.852 | 0.762 | 0 | 0 | Gadd45b | neutrophil.1 |
| Xbp1 | 0 | 0.40839568 | 0.555 | 0.466 | 0 | 0 | Xbp1 | neutrophil.1 |
| Il1b | 0 | 0.7435488 | 0.999 | 0.91 | 0 | 0 | Il1b | neutrophil.1 |
| Pim1 | 0 | 0.25670307 | 0.958 | 0.87 | 0 | 0 | Pim1 | neutrophil.1 |
| Ccl3 | 0 | 0.37595189 | 0.671 | 0.589 | 0 | 0 | Ccl3 | neutrophil.1 |
| Junb | 0 | 0.42600032 | 0.989 | 0.921 | 0 | 0 | Junb | neutrophil.1 |
| Mcl1 | 0 | 0.37182769 | 0.987 | 0.923 | 0 | 0 | Mcl1 | neutrophil.1 |
| Fth1 | 0 | 0.31808396 | 0.98 | 0.923 | 0 | 0 | Fth1 | neutrophil.1 |
| Tmsb4x | 0 | 0.28126652 | 0.967 | 0.911 | 0 | 0 | Tmsb4x | neutrophil.1 |
| Srgn | 0 | 0.89761484 | 0.997 | 0.942 | 0 | 0 | Srgn | neutrophil.1 |
| Abtb2 | 1.34E-304 | 0.71013679 | 0.397 | 0.246 | 2.60E-300 | 0 | Abtb2 | neutrophil.1 |
| Isg15 | 5.90E-301 | 0.6565569 | 0.391 | 0.224 | 1.14E-296 | 0 | Isg15 | neutrophil.1 |
| Fxyd5 | 1.26E-297 | 0.41219026 | 0.605 | 0.462 | 2.44E-293 | 0 | Fxyd5 | neutrophil.1 |
| Arih1 | 3.13E-296 | 0.61731357 | 0.445 | 0.302 | 6.05E-292 | 0 | Arih1 | neutrophil.1 |
| Stk17b | 3.14E-294 | 0.58726136 | 0.446 | 0.295 | 6.07E-290 | 0 | Stk17b | neutrophil.1 |
| Arl5b | 6.27E-279 | 0.89802262 | 0.285 | 0.153 | 1.21E-274 | 0 | Arl5b | neutrophil.1 |
| Ccpg1 | 6.58E-274 | 0.73674688 | 0.295 | 0.17 | 1.27E-269 | 0 | Ccpg1 | neutrophil.1 |
| Pnrc1 | 7.42E-273 | 0.26417111 | 0.789 | 0.65 | 1.43E-268 | 0 | Pnrc1 | neutrophil.1 |
| Plk3 | 3.60E-270 | 0.39037014 | 0.581 | 0.45 | 6.96E-266 | 0 | Plk3 | neutrophil.1 |
| Tet2 | 5.05E-270 | 0.81650056 | 0.324 | 0.189 | 9.77E-266 | 0 | Tet2 | neutrophil.1 |
| Cfp | 3.23E-265 | 0.64046456 | 0.391 | 0.242 | 6.24E-261 | 0 | Cfp | neutrophil.1 |
| Lamp2 | 1.25E-264 | 0.47602314 | 0.444 | 0.335 | 2.42E-260 | 0 | Lamp2 | neutrophil.1 |
| Cux1 | 1.57E-263 | 0.67725439 | 0.365 | 0.219 | 3.03E-259 | 0 | Cux1 | neutrophil.1 |
| Egln3 | 5.85E-260 | 0.36158998 | 0.474 | 0.379 | 1.13E-255 | 0 | Egln3 | neutrophil.1 |
| Rab20 | 3.46E-258 | 0.29866349 | 0.615 | 0.502 | 6.69E-254 | 0 | Rab20 | neutrophil.1 |
| Nabp1 | 2.17E-254 | 0.35868756 | 0.523 | 0.364 | 4.20E-250 | 0 | Nabp1 | neutrophil.1 |
| R3hdm4 | 1.63E-252 | 0.86023982 | 0.251 | 0.149 | 3.15E-248 | 0 | R3hdm4 | neutrophil.1 |
| Rnf11 | 8.87E-249 | 0.82032256 | 0.294 | 0.16 | 1.71E-244 | 0 | Rnf11 | neutrophil.1 |
| Fmnl1 | 5.56E-248 | 0.38390077 | 0.583 | 0.418 | 1.08E-243 | 0 | Fmnl1 | neutrophil.1 |
| Coq10b | 1.95E-247 | 0.79491174 | 0.325 | 0.188 | 3.77E-243 | 0 | Coq10b | neutrophil.1 |
| 7-Mar | 2.41E-242 | 0.55322685 | 0.461 | 0.326 | 4.66E-238 | 0 | 7-Mar | neutrophil.1 |
| Selplg | 4.76E-241 | 0.4556934 | 0.505 | 0.372 | 9.21E-237 | 0 | Selplg | neutrophil.1 |
| Ist1 | 6.23E-234 | 0.52663911 | 0.396 | 0.271 | 1.21E-229 | 0 | Ist1 | neutrophil.1 |
| Dmxl2 | 2.58E-231 | 0.76058606 | 0.3 | 0.177 | 4.99E-227 | 0 | Dmxl2 | neutrophil.1 |
| Zc3h12a | 5.32E-229 | 0.28924792 | 0.583 | 0.476 | 1.03E-224 | 0 | Zc3h12a | neutrophil.1 |
| Slc2a1 | 2.70E-226 | 0.29434653 | 0.576 | 0.458 | 5.22E-222 | 0 | Slc2a1 | neutrophil.1 |
| App | 1.77E-219 | 0.41289502 | 0.577 | 0.441 | 3.43E-215 | 0 | App | neutrophil.1 |
| Myadm | 3.78E-218 | 0.468165 | 0.42 | 0.305 | 7.31E-214 | 0 | Myadm | neutrophil.1 |
| Picalm | 2.69E-213 | 0.38393768 | 0.571 | 0.426 | 5.21E-209 | 0 | Picalm | neutrophil.1 |
| Map2k3 | 3.25E-208 | 0.42095641 | 0.456 | 0.323 | 6.29E-204 | 0 | Map2k3 | neutrophil.1 |
| Sde2 | 1.84E-197 | 0.60830292 | 0.316 | 0.206 | 3.55E-193 | 0 | Sde2 | neutrophil.1 |
| Carhsp1 | 3.13E-189 | 0.62746627 | 0.329 | 0.21 | 6.06E-185 | 0 | Carhsp1 | neutrophil.1 |
| Nudt4 | 5.40E-183 | 0.59579157 | 0.342 | 0.222 | 1.04E-178 | 0 | Nudt4 | neutrophil.1 |
| Pirb | 4.03E-182 | 0.38657571 | 0.456 | 0.334 | 7.79E-178 | 0 | Pirb | neutrophil.1 |
| Snx20 | 7.46E-179 | 0.27690094 | 0.649 | 0.532 | 1.44E-174 | 0 | Snx20 | neutrophil.1 |
| Cish | 5.93E-178 | 0.3792751 | 0.345 | 0.256 | 1.15E-173 | 0 | Cish | neutrophil.1 |
| Tnrc18 | 1.12E-177 | 0.55720148 | 0.312 | 0.202 | 2.17E-173 | 0 | Tnrc18 | neutrophil.1 |
| Atg2a | 4.13E-175 | 0.44205801 | 0.335 | 0.224 | 7.98E-171 | 0 | Atg2a | neutrophil.1 |
| Btbd9 | 3.89E-174 | 0.61581771 | 0.28 | 0.167 | 7.53E-170 | 0 | Btbd9 | neutrophil.1 |
| Tgfbi | 3.53E-172 | 0.28393826 | 0.467 | 0.329 | 6.83E-168 | 0 | Tgfbi | neutrophil.1 |
| Tcn2 | 1.33E-170 | 0.47352505 | 0.359 | 0.252 | 2.57E-166 | 0 | Tcn2 | neutrophil.1 |
| Fpr2 | 6.16E-169 | 0.35663598 | 0.442 | 0.314 | 1.19E-164 | 0 | Fpr2 | neutrophil.1 |
| Fbxl5 | 9.33E-160 | 0.5456871 | 0.254 | 0.179 | 1.80E-155 | 0 | Fbxl5 | neutrophil.1 |
| Flot1 | 5.80E-157 | 0.57142838 | 0.283 | 0.174 | 1.12E-152 | 0 | Flot1 | neutrophil.1 |
| Lrrfip1 | 2.32E-152 | 0.28817972 | 0.439 | 0.331 | 4.49E-148 | 0 | Lrrfip1 | neutrophil.1 |
| Spag9 | 2.43E-152 | 0.37571983 | 0.506 | 0.381 | 4.69E-148 | 0 | Spag9 | neutrophil.1 |
| Rab31 | 1.78E-150 | 0.59076053 | 0.272 | 0.177 | 3.45E-146 | 0 | Rab31 | neutrophil.1 |
| Tut7 | 2.04E-149 | 0.25221446 | 0.538 | 0.431 | 3.94E-145 | 0 | Tut7 | neutrophil.1 |
| Riok3 | 1.42E-148 | 0.40478171 | 0.377 | 0.273 | 2.74E-144 | 0 | Riok3 | neutrophil.1 |
| Notch2 | 2.16E-145 | 0.34696771 | 0.39 | 0.277 | 4.18E-141 | 0 | Notch2 | neutrophil.1 |
| Arpc3 | 1.71E-143 | 0.50402688 | 0.353 | 0.242 | 3.31E-139 | 0 | Arpc3 | neutrophil.1 |
| Tmem88 | 7.17E-143 | 0.47324094 | 0.355 | 0.238 | 1.39E-138 | 0 | Tmem88 | neutrophil.1 |
| Zfp36l1 | 2.02E-142 | 0.37953427 | 0.371 | 0.288 | 3.90E-138 | 0 | Zfp36l1 | neutrophil.1 |
| Sbno2 | 1.27E-141 | 0.33518404 | 0.479 | 0.385 | 2.45E-137 | 0 | Sbno2 | neutrophil.1 |
| Crem | 6.63E-138 | 0.50843062 | 0.317 | 0.185 | 1.28E-133 | 0 | Crem | neutrophil.1 |
| Cd84 | 1.72E-135 | 0.3915753 | 0.26 | 0.164 | 3.32E-131 | 0 | Cd84 | neutrophil.1 |
| Wnk1 | 1.75E-133 | 0.3605217 | 0.438 | 0.357 | 3.39E-129 | 0 | Wnk1 | neutrophil.1 |
| Anxa11 | 3.46E-128 | 0.37854662 | 0.396 | 0.288 | 6.70E-124 | 0 | Anxa11 | neutrophil.1 |
| Ttc7 | 7.78E-128 | 0.52477876 | 0.282 | 0.188 | 1.50E-123 | 0 | Ttc7 | neutrophil.1 |
| Arid5a | 9.91E-128 | 0.60616346 | 0.255 | 0.163 | 1.92E-123 | 0 | Arid5a | neutrophil.1 |
| Emb | 1.53E-126 | 0.57284725 | 0.27 | 0.174 | 2.96E-122 | 0 | Emb | neutrophil.1 |
| Rbms1 | 6.52E-125 | 0.41500061 | 0.375 | 0.272 | 1.26E-120 | 0 | Rbms1 | neutrophil.1 |
| Itprip | 1.22E-123 | 0.50030575 | 0.278 | 0.182 | 2.36E-119 | 0 | Itprip | neutrophil.1 |
| Cox17 | 1.10E-122 | 0.30499421 | 0.451 | 0.36 | 2.12E-118 | 0 | Cox17 | neutrophil.1 |
| Il13ra1 | 2.66E-121 | 0.43654491 | 0.368 | 0.267 | 5.15E-117 | 0 | Il13ra1 | neutrophil.1 |
| Snx18 | 2.55E-120 | 0.32512258 | 0.365 | 0.279 | 4.94E-116 | 0 | Snx18 | neutrophil.1 |
| Ube2b | 4.76E-118 | 0.4765362 | 0.321 | 0.225 | 9.19E-114 | 0 | Ube2b | neutrophil.1 |
| Tra2a | 2.80E-116 | 0.29128529 | 0.385 | 0.302 | 5.41E-112 | 0 | Tra2a | neutrophil.1 |
| Gla | 1.27E-106 | 0.42791886 | 0.326 | 0.226 | 2.45E-102 | 0 | Gla | neutrophil.1 |
| Fosb | 1.72E-106 | 0.26763104 | 0.385 | 0.264 | 3.33E-102 | 0 | Fosb | neutrophil.1 |
| Egr3 | 2.53E-103 | 0.36404353 | 0.275 | 0.183 | 4.89E-99 | 0 | Egr3 | neutrophil.1 |
| Rbpj | 7.58E-102 | 0.33217285 | 0.298 | 0.228 | 1.47E-97 | 0 | Rbpj | neutrophil.1 |
| Preb | 9.39E-96 | 0.46226736 | 0.259 | 0.174 | 1.82E-91 | 0 | Preb | neutrophil.1 |
| Nfkb1 | 9.73E-94 | 0.3198967 | 0.26 | 0.201 | 1.88E-89 | 0 | Nfkb1 | neutrophil.1 |
| Igf1r | 1.52E-93 | 0.4449764 | 0.27 | 0.187 | 2.95E-89 | 0 | Igf1r | neutrophil.1 |
| Cd244a | 6.67E-93 | 0.31583958 | 0.268 | 0.188 | 1.29E-88 | 0 | Cd244a | neutrophil.1 |
| Lbr | 1.12E-90 | 0.40595257 | 0.27 | 0.196 | 2.17E-86 | 0 | Lbr | neutrophil.1 |
| Kdm7a | 1.89E-90 | 0.40939691 | 0.265 | 0.186 | 3.66E-86 | 0 | Kdm7a | neutrophil.1 |
| Eif1 | 4.10E-90 | 0.29806645 | 0.434 | 0.33 | 7.93E-86 | 0 | Eif1 | neutrophil.1 |
| Sik1 | 1.19E-79 | 0.27892721 | 0.394 | 0.296 | 2.30E-75 | 0 | Sik1 | neutrophil.1 |
| Chp1 | 1.63E-75 | 0.38952093 | 0.281 | 0.207 | 3.15E-71 | 0 | Chp1 | neutrophil.1 |
| Slc25a3 | 3.78E-74 | 0.3321006 | 0.311 | 0.223 | 7.30E-70 | 0 | Slc25a3 | neutrophil.1 |
| Xylt1 | 3.78E-68 | 0.35789028 | 0.317 | 0.225 | 7.31E-64 | 0 | Xylt1 | neutrophil.1 |
| Tnfrsf23 | 1.61E-63 | 0.33875608 | 0.277 | 0.213 | 3.11E-59 | 0 | Tnfrsf23 | neutrophil.1 |
| Tpm4 | 1.04E-62 | 0.2542812 | 0.348 | 0.274 | 2.02E-58 | 0 | Tpm4 | neutrophil.1 |
| Nup98 | 3.16E-55 | 0.29118659 | 0.289 | 0.215 | 6.10E-51 | 0 | Nup98 | neutrophil.1 |
| Emd | 6.38E-49 | 0.25168413 | 0.348 | 0.244 | 1.23E-44 | 0 | Emd | neutrophil.1 |
| Gmfg | 6.24E-47 | 0.28746368 | 0.263 | 0.196 | 1.21E-42 | 0 | Gmfg | neutrophil.1 |
| Ffar2 | 0 | 1.57978472 | 0.498 | 0.199 | 0 | 1 | Ffar2 | neutrophil.2 |
| Cd300a | 0 | 1.11052542 | 0.713 | 0.428 | 0 | 1 | Cd300a | neutrophil.2 |
| Gpx1 | 0 | 1.05002025 | 0.699 | 0.43 | 0 | 1 | Gpx1 | neutrophil.2 |
| Cst3 | 0 | 1.05077911 | 0.795 | 0.547 | 0 | 1 | Cst3 | neutrophil.2 |
| Havcr2 | 0 | 1.79306435 | 0.365 | 0.118 | 0 | 1 | Havcr2 | neutrophil.2 |
| Txnip | 0 | 1.00698819 | 0.67 | 0.43 | 0 | 1 | Txnip | neutrophil.2 |
| Gpr84 | 0 | 1.40222498 | 0.461 | 0.226 | 0 | 1 | Gpr84 | neutrophil.2 |
| Ikbke | 0 | 0.95340021 | 0.474 | 0.239 | 0 | 1 | Ikbke | neutrophil.2 |
| Itgax | 0 | 1.07836427 | 0.515 | 0.288 | 0 | 1 | Itgax | neutrophil.2 |
| Sh3bp1 | 0 | 1.51237416 | 0.363 | 0.142 | 0 | 1 | Sh3bp1 | neutrophil.2 |
| Rac2 | 0 | 0.8644502 | 0.889 | 0.67 | 0 | 1 | Rac2 | neutrophil.2 |
| Adgre5 | 0 | 0.75638355 | 0.84 | 0.626 | 0 | 1 | Adgre5 | neutrophil.2 |
| Il1a | 0 | 0.80218473 | 0.724 | 0.513 | 0 | 1 | Il1a | neutrophil.2 |
| Mmp9 | 0 | 1.01170507 | 0.831 | 0.622 | 0 | 1 | Mmp9 | neutrophil.2 |
| Fgr | 0 | 0.71927095 | 0.755 | 0.558 | 0 | 1 | Fgr | neutrophil.2 |
| Cxcr4 | 0 | 0.80187259 | 0.733 | 0.54 | 0 | 1 | Cxcr4 | neutrophil.2 |
| Rgs3 | 0 | 1.05541225 | 0.454 | 0.263 | 0 | 1 | Rgs3 | neutrophil.2 |
| Laptm5 | 0 | 0.69406626 | 0.858 | 0.668 | 0 | 1 | Laptm5 | neutrophil.2 |
| Cdk2ap2 | 0 | 0.9037866 | 0.601 | 0.417 | 0 | 1 | Cdk2ap2 | neutrophil.2 |
| Gngt2 | 0 | 1.19561818 | 0.353 | 0.169 | 0 | 1 | Gngt2 | neutrophil.2 |
| Atox1 | 0 | 0.89929786 | 0.469 | 0.288 | 0 | 1 | Atox1 | neutrophil.2 |
| Lst1 | 0 | 0.74877159 | 0.58 | 0.399 | 0 | 1 | Lst1 | neutrophil.2 |
| Plin2 | 0 | 0.75500102 | 0.496 | 0.319 | 0 | 1 | Plin2 | neutrophil.2 |
| Icam1 | 0 | 0.65008832 | 0.775 | 0.602 | 0 | 1 | Icam1 | neutrophil.2 |
| Naaa | 0 | 1.34727776 | 0.316 | 0.148 | 0 | 1 | Naaa | neutrophil.2 |
| B4galt1 | 0 | 1.09232791 | 0.388 | 0.222 | 0 | 1 | B4galt1 | neutrophil.2 |
| Dusp2 | 0 | 0.65795496 | 0.516 | 0.351 | 0 | 1 | Dusp2 | neutrophil.2 |
| Clec5a | 0 | 1.15685427 | 0.385 | 0.222 | 0 | 1 | Clec5a | neutrophil.2 |
| Scimp | 0 | 1.45345803 | 0.283 | 0.124 | 0 | 1 | Scimp | neutrophil.2 |
| Hic1 | 0 | 1.19863308 | 0.256 | 0.105 | 0 | 1 | Hic1 | neutrophil.2 |
| Csf3r | 0 | 0.46236458 | 0.963 | 0.831 | 0 | 1 | Csf3r | neutrophil.2 |
| Gsr | 0 | 0.47062932 | 0.907 | 0.79 | 0 | 1 | Gsr | neutrophil.2 |
| Tnf | 0 | 0.6640095 | 0.653 | 0.575 | 0 | 1 | Tnf | neutrophil.2 |
| Il23a | 0 | 0.81225645 | 0.287 | 0.212 | 0 | 1 | Il23a | neutrophil.2 |
| Tgfbi | 5.14E-305 | 0.79616426 | 0.47 | 0.334 | 9.93E-301 | 1 | Tgfbi | neutrophil.2 |
| Ptpn6 | 2.57E-304 | 0.75605889 | 0.618 | 0.425 | 4.97E-300 | 1 | Ptpn6 | neutrophil.2 |
| Traf1 | 1.05E-300 | 0.68923789 | 0.419 | 0.281 | 2.04E-296 | 1 | Traf1 | neutrophil.2 |
| Camkk1 | 2.99E-288 | 1.07436101 | 0.325 | 0.164 | 5.78E-284 | 1 | Camkk1 | neutrophil.2 |
| Abi3 | 1.01E-283 | 1.17789022 | 0.28 | 0.134 | 1.96E-279 | 1 | Abi3 | neutrophil.2 |
| Ndst1 | 1.87E-281 | 1.05557002 | 0.339 | 0.181 | 3.61E-277 | 1 | Ndst1 | neutrophil.2 |
| Coro2a | 6.48E-281 | 1.04745279 | 0.296 | 0.154 | 1.25E-276 | 1 | Coro2a | neutrophil.2 |
| Sla | 3.81E-279 | 1.02894411 | 0.286 | 0.143 | 7.36E-275 | 1 | Sla | neutrophil.2 |
| Otulinl | 5.05E-277 | 1.30720141 | 0.264 | 0.12 | 9.76E-273 | 1 | Otulinl | neutrophil.2 |
| Ppp1r9b | 2.91E-273 | 0.92298005 | 0.415 | 0.242 | 5.62E-269 | 1 | Ppp1r9b | neutrophil.2 |
| Ptk2b | 1.31E-269 | 0.68010202 | 0.578 | 0.391 | 2.53E-265 | 1 | Ptk2b | neutrophil.2 |
| Tsc22d3 | 4.64E-266 | 0.97438779 | 0.284 | 0.155 | 8.96E-262 | 1 | Tsc22d3 | neutrophil.2 |
| Arrb2 | 2.22E-262 | 0.7848129 | 0.53 | 0.346 | 4.30E-258 | 1 | Arrb2 | neutrophil.2 |
| Ncf1 | 4.02E-260 | 0.32490738 | 0.736 | 0.605 | 7.77E-256 | 1 | Ncf1 | neutrophil.2 |
| Oasl2 | 2.64E-251 | 0.66360157 | 0.469 | 0.354 | 5.11E-247 | 1 | Oasl2 | neutrophil.2 |
| St3gal1 | 2.18E-247 | 0.93080993 | 0.401 | 0.24 | 4.22E-243 | 1 | St3gal1 | neutrophil.2 |
| Arf2 | 2.91E-246 | 0.93515073 | 0.359 | 0.201 | 5.62E-242 | 1 | Arf2 | neutrophil.2 |
| Mllt6 | 2.30E-245 | 0.84405314 | 0.386 | 0.218 | 4.44E-241 | 1 | Mllt6 | neutrophil.2 |
| Atp6v0a1 | 8.69E-244 | 0.8995755 | 0.384 | 0.235 | 1.68E-239 | 1 | Atp6v0a1 | neutrophil.2 |
| Tgfbr1 | 9.09E-241 | 0.9850193 | 0.324 | 0.184 | 1.76E-236 | 1 | Tgfbr1 | neutrophil.2 |
| Sh3bgrl3 | 3.16E-233 | 0.73554713 | 0.54 | 0.378 | 6.12E-229 | 1 | Sh3bgrl3 | neutrophil.2 |
| P2ry13 | 1.31E-231 | 1.15017262 | 0.277 | 0.152 | 2.53E-227 | 1 | P2ry13 | neutrophil.2 |
| Fcgr2b | 1.02E-228 | 0.81581459 | 0.347 | 0.226 | 1.98E-224 | 1 | Fcgr2b | neutrophil.2 |
| Ninj1 | 6.86E-224 | 0.54057574 | 0.697 | 0.524 | 1.33E-219 | 1 | Ninj1 | neutrophil.2 |
| Vasp | 3.48E-207 | 0.31901457 | 0.912 | 0.801 | 6.72E-203 | 1 | Vasp | neutrophil.2 |
| C3 | 1.87E-206 | 0.46648902 | 0.645 | 0.462 | 3.61E-202 | 1 | C3 | neutrophil.2 |
| Irf1 | 4.67E-204 | 0.58654271 | 0.468 | 0.318 | 9.03E-200 | 1 | Irf1 | neutrophil.2 |
| Psd4 | 5.62E-203 | 0.70441748 | 0.374 | 0.232 | 1.09E-198 | 1 | Psd4 | neutrophil.2 |
| Fam49b | 2.50E-201 | 0.70616967 | 0.536 | 0.369 | 4.84E-197 | 1 | Fam49b | neutrophil.2 |
| Arpc1b | 1.85E-198 | 0.52102349 | 0.655 | 0.498 | 3.57E-194 | 1 | Arpc1b | neutrophil.2 |
| Cd80 | 6.79E-194 | 0.51151881 | 0.681 | 0.52 | 1.31E-189 | 1 | Cd80 | neutrophil.2 |
| Cd74 | 3.16E-191 | 0.46749279 | 0.475 | 0.355 | 6.11E-187 | 1 | Cd74 | neutrophil.2 |
| Pilra | 3.24E-191 | 0.71378566 | 0.416 | 0.276 | 6.26E-187 | 1 | Pilra | neutrophil.2 |
| Siglecf | 4.30E-187 | 0.79487965 | 0.321 | 0.182 | 8.32E-183 | 1 | Siglecf | neutrophil.2 |
| Was | 7.86E-184 | 1.06157705 | 0.286 | 0.157 | 1.52E-179 | 1 | Was | neutrophil.2 |
| Ttyh3 | 4.78E-180 | 0.68842491 | 0.459 | 0.32 | 9.23E-176 | 1 | Ttyh3 | neutrophil.2 |
| Atp1a1 | 6.03E-177 | 0.63602183 | 0.472 | 0.318 | 1.17E-172 | 1 | Atp1a1 | neutrophil.2 |
| Fcer1g | 1.14E-176 | 0.38899053 | 0.913 | 0.8 | 2.21E-172 | 1 | Fcer1g | neutrophil.2 |
| Hipk1 | 3.80E-175 | 0.91543215 | 0.347 | 0.208 | 7.35E-171 | 1 | Hipk1 | neutrophil.2 |
| Pfn1 | 2.42E-172 | 0.51221712 | 0.675 | 0.521 | 4.69E-168 | 1 | Pfn1 | neutrophil.2 |
| Vamp8 | 3.11E-170 | 0.73623166 | 0.419 | 0.279 | 6.02E-166 | 1 | Vamp8 | neutrophil.2 |
| Slc38a2 | 3.15E-169 | 0.50975008 | 0.578 | 0.444 | 6.09E-165 | 1 | Slc38a2 | neutrophil.2 |
| Treml4 | 7.16E-167 | 0.65595717 | 0.445 | 0.288 | 1.38E-162 | 1 | Treml4 | neutrophil.2 |
| Entpd3 | 7.47E-167 | 1.01078506 | 0.273 | 0.156 | 1.44E-162 | 1 | Entpd3 | neutrophil.2 |
| Lcp1 | 1.09E-164 | 0.41420706 | 0.883 | 0.747 | 2.12E-160 | 1 | Lcp1 | neutrophil.2 |
| Reep5 | 3.02E-163 | 0.97089355 | 0.257 | 0.141 | 5.84E-159 | 1 | Reep5 | neutrophil.2 |
| Gsn | 6.39E-163 | 0.47760675 | 0.639 | 0.488 | 1.24E-158 | 1 | Gsn | neutrophil.2 |
| Irf2bp2 | 7.43E-162 | 0.43319612 | 0.591 | 0.439 | 1.44E-157 | 1 | Irf2bp2 | neutrophil.2 |
| Calm3 | 1.94E-159 | 0.80695409 | 0.343 | 0.214 | 3.76E-155 | 1 | Calm3 | neutrophil.2 |
| Myd88 | 2.26E-159 | 0.63190637 | 0.535 | 0.389 | 4.37E-155 | 1 | Myd88 | neutrophil.2 |
| Vsir | 7.96E-157 | 0.58760107 | 0.568 | 0.425 | 1.54E-152 | 1 | Vsir | neutrophil.2 |
| Clec4n | 4.15E-154 | 0.45657326 | 0.73 | 0.549 | 8.03E-150 | 1 | Clec4n | neutrophil.2 |
| Pip5k1c | 1.57E-147 | 0.77564882 | 0.354 | 0.224 | 3.04E-143 | 1 | Pip5k1c | neutrophil.2 |
| Msn | 4.40E-147 | 0.3924949 | 0.88 | 0.744 | 8.51E-143 | 1 | Msn | neutrophil.2 |
| Gm2a | 1.29E-146 | 0.4942048 | 0.554 | 0.426 | 2.49E-142 | 1 | Gm2a | neutrophil.2 |
| Pkn1 | 1.35E-144 | 0.60822705 | 0.412 | 0.278 | 2.61E-140 | 1 | Pkn1 | neutrophil.2 |
| Susd6 | 2.27E-142 | 0.57990748 | 0.514 | 0.384 | 4.38E-138 | 1 | Susd6 | neutrophil.2 |
| Csf1r | 2.31E-141 | 0.71512914 | 0.372 | 0.243 | 4.47E-137 | 1 | Csf1r | neutrophil.2 |
| Prex1 | 1.98E-139 | 0.64600667 | 0.384 | 0.266 | 3.82E-135 | 1 | Prex1 | neutrophil.2 |
| Cotl1 | 1.51E-138 | 0.35670411 | 0.734 | 0.593 | 2.92E-134 | 1 | Cotl1 | neutrophil.2 |
| Ctsa | 2.53E-137 | 0.75486722 | 0.336 | 0.216 | 4.90E-133 | 1 | Ctsa | neutrophil.2 |
| Cbfb | 9.10E-137 | 0.72685573 | 0.267 | 0.156 | 1.76E-132 | 1 | Cbfb | neutrophil.2 |
| Ncf4 | 3.55E-136 | 0.56485721 | 0.472 | 0.346 | 6.86E-132 | 1 | Ncf4 | neutrophil.2 |
| Hnrnpul1 | 1.74E-135 | 0.66979838 | 0.407 | 0.272 | 3.37E-131 | 1 | Hnrnpul1 | neutrophil.2 |
| Coro1a | 2.78E-135 | 0.43941909 | 0.721 | 0.59 | 5.37E-131 | 1 | Coro1a | neutrophil.2 |
| Syk | 3.84E-135 | 0.3405988 | 0.801 | 0.688 | 7.42E-131 | 1 | Syk | neutrophil.2 |
| Ctdsp2 | 4.65E-132 | 0.72777839 | 0.299 | 0.187 | 8.99E-128 | 1 | Ctdsp2 | neutrophil.2 |
| Grk2 | 6.31E-130 | 0.57484694 | 0.544 | 0.4 | 1.22E-125 | 1 | Grk2 | neutrophil.2 |
| Atf1 | 2.59E-129 | 0.59711605 | 0.35 | 0.238 | 5.02E-125 | 1 | Atf1 | neutrophil.2 |
| Xbp1 | 9.30E-128 | 0.50773431 | 0.599 | 0.46 | 1.80E-123 | 1 | Xbp1 | neutrophil.2 |
| Rabac1 | 1.74E-125 | 0.5893223 | 0.413 | 0.293 | 3.37E-121 | 1 | Rabac1 | neutrophil.2 |
| Brk1 | 1.76E-125 | 0.67964726 | 0.365 | 0.248 | 3.41E-121 | 1 | Brk1 | neutrophil.2 |
| Gnai2 | 8.19E-125 | 0.38104037 | 0.54 | 0.412 | 1.58E-120 | 1 | Gnai2 | neutrophil.2 |
| Adprh | 8.82E-125 | 0.58232534 | 0.506 | 0.381 | 1.70E-120 | 1 | Adprh | neutrophil.2 |
| Hspa8 | 3.31E-115 | 0.42805039 | 0.527 | 0.398 | 6.40E-111 | 1 | Hspa8 | neutrophil.2 |
| Mob1a | 4.06E-114 | 0.71562231 | 0.355 | 0.233 | 7.84E-110 | 1 | Mob1a | neutrophil.2 |
| Rassf5 | 9.93E-114 | 0.74483746 | 0.285 | 0.183 | 1.92E-109 | 1 | Rassf5 | neutrophil.2 |
| Ralgds | 2.82E-112 | 0.6375026 | 0.298 | 0.192 | 5.45E-108 | 1 | Ralgds | neutrophil.2 |
| Tcirg1 | 1.17E-111 | 0.64132652 | 0.309 | 0.196 | 2.27E-107 | 1 | Tcirg1 | neutrophil.2 |
| Slc39a1 | 1.45E-110 | 0.5604787 | 0.347 | 0.233 | 2.80E-106 | 1 | Slc39a1 | neutrophil.2 |
| Irf5 | 3.40E-110 | 0.48652304 | 0.327 | 0.228 | 6.58E-106 | 1 | Irf5 | neutrophil.2 |
| Nod2 | 1.44E-109 | 0.70881842 | 0.272 | 0.179 | 2.79E-105 | 1 | Nod2 | neutrophil.2 |
| Bin2 | 4.26E-108 | 0.53673749 | 0.366 | 0.264 | 8.24E-104 | 1 | Bin2 | neutrophil.2 |
| Csk | 1.43E-107 | 0.77943771 | 0.262 | 0.165 | 2.76E-103 | 1 | Csk | neutrophil.2 |
| Rin3 | 5.25E-105 | 0.6269955 | 0.354 | 0.238 | 1.02E-100 | 1 | Rin3 | neutrophil.2 |
| Zfp36l2 | 9.37E-105 | 0.41693447 | 0.427 | 0.322 | 1.81E-100 | 1 | Zfp36l2 | neutrophil.2 |
| Fam168b | 1.53E-104 | 0.76668174 | 0.26 | 0.16 | 2.96E-100 | 1 | Fam168b | neutrophil.2 |
| Spi1 | 2.42E-104 | 0.50064015 | 0.622 | 0.489 | 4.67E-100 | 1 | Spi1 | neutrophil.2 |
| Actn1 | 1.11E-103 | 0.62120232 | 0.404 | 0.288 | 2.15E-99 | 1 | Actn1 | neutrophil.2 |
| Man2b1 | 5.97E-103 | 0.58858116 | 0.275 | 0.174 | 1.15E-98 | 1 | Man2b1 | neutrophil.2 |
| Pik3cd | 9.97E-101 | 0.56839357 | 0.314 | 0.213 | 1.93E-96 | 1 | Pik3cd | neutrophil.2 |
| Ppp2cb | 3.51E-100 | 0.69316889 | 0.257 | 0.164 | 6.79E-96 | 1 | Ppp2cb | neutrophil.2 |
| Ltb | 4.20E-100 | 0.60267526 | 0.377 | 0.252 | 8.13E-96 | 1 | Ltb | neutrophil.2 |
| Jak2 | 5.83E-100 | 0.51281518 | 0.603 | 0.462 | 1.13E-95 | 1 | Jak2 | neutrophil.2 |
| Tln1 | 1.74E-99 | 0.4800829 | 0.475 | 0.363 | 3.35E-95 | 1 | Tln1 | neutrophil.2 |
| Cmtm6 | 9.08E-99 | 0.56057389 | 0.537 | 0.397 | 1.76E-94 | 1 | Cmtm6 | neutrophil.2 |
| Bax | 2.26E-98 | 0.63573385 | 0.253 | 0.166 | 4.37E-94 | 1 | Bax | neutrophil.2 |
| Mpeg1 | 5.69E-98 | 0.42925244 | 0.554 | 0.374 | 1.10E-93 | 1 | Mpeg1 | neutrophil.2 |
| Atp6v1g1 | 2.13E-96 | 0.58541526 | 0.439 | 0.328 | 4.12E-92 | 1 | Atp6v1g1 | neutrophil.2 |
| Arhgdib | 2.79E-95 | 0.58468238 | 0.412 | 0.296 | 5.39E-91 | 1 | Arhgdib | neutrophil.2 |
| Atrnl1 | 4.88E-95 | 0.71093316 | 0.331 | 0.228 | 9.44E-91 | 1 | Atrnl1 | neutrophil.2 |
| Cyba | 7.66E-95 | 0.45507631 | 0.533 | 0.412 | 1.48E-90 | 1 | Cyba | neutrophil.2 |
| Lrch4 | 8.88E-95 | 0.56576189 | 0.267 | 0.17 | 1.72E-90 | 1 | Lrch4 | neutrophil.2 |
| Grn | 1.86E-93 | 0.59100712 | 0.264 | 0.173 | 3.59E-89 | 1 | Grn | neutrophil.2 |
| Bcl2a1b | 3.12E-93 | 0.38980645 | 0.652 | 0.538 | 6.03E-89 | 1 | Bcl2a1b | neutrophil.2 |
| Rhoa | 4.22E-93 | 0.55396975 | 0.398 | 0.288 | 8.17E-89 | 1 | Rhoa | neutrophil.2 |
| Capg | 2.69E-92 | 0.58615416 | 0.3 | 0.201 | 5.19E-88 | 1 | Capg | neutrophil.2 |
| Clta | 3.21E-92 | 0.42481975 | 0.452 | 0.336 | 6.21E-88 | 1 | Clta | neutrophil.2 |
| Igsf6 | 8.47E-92 | 0.4878666 | 0.462 | 0.352 | 1.64E-87 | 1 | Igsf6 | neutrophil.2 |
| Vav1 | 3.81E-91 | 0.5559139 | 0.452 | 0.339 | 7.36E-87 | 1 | Vav1 | neutrophil.2 |
| Il1rn | 6.20E-91 | 0.25576319 | 0.922 | 0.824 | 1.20E-86 | 1 | Il1rn | neutrophil.2 |
| B2m | 1.18E-90 | 0.45800265 | 0.684 | 0.555 | 2.29E-86 | 1 | B2m | neutrophil.2 |
| Pcbp1 | 8.34E-90 | 0.73027068 | 0.257 | 0.16 | 1.61E-85 | 1 | Pcbp1 | neutrophil.2 |
| Akt1 | 3.92E-89 | 0.50352147 | 0.407 | 0.299 | 7.59E-85 | 1 | Akt1 | neutrophil.2 |
| Myo1f | 2.08E-88 | 0.50991059 | 0.441 | 0.329 | 4.02E-84 | 1 | Myo1f | neutrophil.2 |
| Nfam1 | 4.82E-88 | 0.63821261 | 0.4 | 0.29 | 9.32E-84 | 1 | Nfam1 | neutrophil.2 |
| Cd164 | 1.06E-87 | 0.53442611 | 0.391 | 0.288 | 2.05E-83 | 1 | Cd164 | neutrophil.2 |
| Stx7 | 2.20E-87 | 0.68289421 | 0.295 | 0.198 | 4.25E-83 | 1 | Stx7 | neutrophil.2 |
| Lrp10 | 4.23E-87 | 0.54879658 | 0.437 | 0.317 | 8.17E-83 | 1 | Lrp10 | neutrophil.2 |
| Mvp | 2.54E-86 | 0.66901322 | 0.299 | 0.205 | 4.91E-82 | 1 | Mvp | neutrophil.2 |
| Jpt1 | 2.93E-85 | 0.32348153 | 0.308 | 0.213 | 5.66E-81 | 1 | Jpt1 | neutrophil.2 |
| Arhgef2 | 5.88E-84 | 0.55472507 | 0.303 | 0.207 | 1.14E-79 | 1 | Arhgef2 | neutrophil.2 |
| Usf2 | 1.01E-83 | 0.56255339 | 0.275 | 0.184 | 1.96E-79 | 1 | Usf2 | neutrophil.2 |
| Atp6ap1 | 4.10E-83 | 0.59242307 | 0.373 | 0.274 | 7.92E-79 | 1 | Atp6ap1 | neutrophil.2 |
| Tnip1 | 2.87E-82 | 0.55127656 | 0.331 | 0.227 | 5.54E-78 | 1 | Tnip1 | neutrophil.2 |
| AA467197 | 1.77E-81 | 0.53065484 | 0.398 | 0.296 | 3.43E-77 | 1 | AA467197 | neutrophil.2 |
| Ptprc | 7.85E-80 | 0.43526416 | 0.62 | 0.51 | 1.52E-75 | 1 | Ptprc | neutrophil.2 |
| Siglece | 6.73E-79 | 0.51868413 | 0.534 | 0.413 | 1.30E-74 | 1 | Siglece | neutrophil.2 |
| Trir | 2.48E-77 | 0.41768798 | 0.36 | 0.264 | 4.80E-73 | 1 | Trir | neutrophil.2 |
| Sirpa | 5.10E-77 | 0.42506037 | 0.557 | 0.442 | 9.86E-73 | 1 | Sirpa | neutrophil.2 |
| Cdkn1b | 1.27E-76 | 0.60975317 | 0.291 | 0.205 | 2.46E-72 | 1 | Cdkn1b | neutrophil.2 |
| Yy1 | 4.78E-74 | 0.37802397 | 0.293 | 0.203 | 9.25E-70 | 1 | Yy1 | neutrophil.2 |
| Shisa5 | 4.63E-73 | 0.60727055 | 0.355 | 0.257 | 8.96E-69 | 1 | Shisa5 | neutrophil.2 |
| Dgkz | 4.03E-71 | 0.45811239 | 0.282 | 0.196 | 7.79E-67 | 1 | Dgkz | neutrophil.2 |
| Gsk3a | 9.78E-69 | 0.54995947 | 0.288 | 0.202 | 1.89E-64 | 1 | Gsk3a | neutrophil.2 |
| Rab5c | 9.86E-69 | 0.58584026 | 0.38 | 0.275 | 1.91E-64 | 1 | Rab5c | neutrophil.2 |
| Runx1 | 2.64E-68 | 0.62127948 | 0.253 | 0.174 | 5.11E-64 | 1 | Runx1 | neutrophil.2 |
| Cox8a | 9.90E-68 | 0.57999189 | 0.289 | 0.202 | 1.91E-63 | 1 | Cox8a | neutrophil.2 |
| Skap2 | 1.52E-67 | 0.42789957 | 0.456 | 0.354 | 2.93E-63 | 1 | Skap2 | neutrophil.2 |
| Mthfs | 8.04E-67 | 0.523681 | 0.327 | 0.26 | 1.55E-62 | 1 | Mthfs | neutrophil.2 |
| Csf2rb2 | 1.68E-66 | 0.56651006 | 0.259 | 0.182 | 3.26E-62 | 1 | Csf2rb2 | neutrophil.2 |
| Arf6 | 2.36E-66 | 0.59006821 | 0.301 | 0.214 | 4.57E-62 | 1 | Arf6 | neutrophil.2 |
| Arhgap9 | 1.21E-65 | 0.64296994 | 0.276 | 0.189 | 2.34E-61 | 1 | Arhgap9 | neutrophil.2 |
| Pik3r5 | 5.23E-64 | 0.5786831 | 0.288 | 0.2 | 1.01E-59 | 1 | Pik3r5 | neutrophil.2 |
| Tapbp | 1.01E-63 | 0.42463703 | 0.45 | 0.34 | 1.96E-59 | 1 | Tapbp | neutrophil.2 |
| Ptpre | 1.26E-63 | 0.29595077 | 0.365 | 0.285 | 2.44E-59 | 1 | Ptpre | neutrophil.2 |
| Son | 2.67E-63 | 0.37798477 | 0.486 | 0.381 | 5.16E-59 | 1 | Son | neutrophil.2 |
| Gpr132 | 4.11E-63 | 0.45688667 | 0.393 | 0.304 | 7.95E-59 | 1 | Gpr132 | neutrophil.2 |
| Psenen | 2.51E-62 | 0.58564151 | 0.291 | 0.213 | 4.86E-58 | 1 | Psenen | neutrophil.2 |
| Itgb1 | 2.61E-62 | 0.46667787 | 0.33 | 0.233 | 5.05E-58 | 1 | Itgb1 | neutrophil.2 |
| Ddx17 | 3.94E-62 | 0.32267315 | 0.366 | 0.277 | 7.61E-58 | 1 | Ddx17 | neutrophil.2 |
| Arf1 | 5.34E-62 | 0.49608152 | 0.364 | 0.276 | 1.03E-57 | 1 | Arf1 | neutrophil.2 |
| Pak2 | 1.88E-61 | 0.49122902 | 0.394 | 0.298 | 3.63E-57 | 1 | Pak2 | neutrophil.2 |
| Prkcd | 4.02E-61 | 0.38872023 | 0.467 | 0.373 | 7.76E-57 | 1 | Prkcd | neutrophil.2 |
| Ctss | 4.67E-61 | 0.46878975 | 0.362 | 0.26 | 9.03E-57 | 1 | Ctss | neutrophil.2 |
| Uck2 | 8.86E-61 | 0.60146367 | 0.253 | 0.169 | 1.71E-56 | 1 | Uck2 | neutrophil.2 |
| Dusp16 | 1.01E-60 | 0.3379046 | 0.535 | 0.456 | 1.95E-56 | 1 | Dusp16 | neutrophil.2 |
| Fes | 3.19E-60 | 0.41860416 | 0.334 | 0.241 | 6.17E-56 | 1 | Fes | neutrophil.2 |
| Cggbp1 | 5.43E-60 | 0.52551308 | 0.297 | 0.213 | 1.05E-55 | 1 | Cggbp1 | neutrophil.2 |
| Atp6v1b2 | 2.20E-59 | 0.47050991 | 0.364 | 0.267 | 4.26E-55 | 1 | Atp6v1b2 | neutrophil.2 |
| Rtn3 | 2.34E-59 | 0.49592051 | 0.319 | 0.239 | 4.53E-55 | 1 | Rtn3 | neutrophil.2 |
| Ski | 2.03E-58 | 0.28436413 | 0.504 | 0.397 | 3.92E-54 | 1 | Ski | neutrophil.2 |
| Gnb2 | 2.71E-57 | 0.58922321 | 0.32 | 0.236 | 5.24E-53 | 1 | Gnb2 | neutrophil.2 |
| Cxcr2 | 2.99E-57 | 0.45187858 | 0.58 | 0.475 | 5.79E-53 | 1 | Cxcr2 | neutrophil.2 |
| Resf1 | 6.88E-57 | 0.42719321 | 0.446 | 0.379 | 1.33E-52 | 1 | Resf1 | neutrophil.2 |
| Dok3 | 1.20E-55 | 0.58490528 | 0.254 | 0.175 | 2.33E-51 | 1 | Dok3 | neutrophil.2 |
| Hnrnpm | 1.37E-54 | 0.38835952 | 0.278 | 0.2 | 2.66E-50 | 1 | Hnrnpm | neutrophil.2 |
| Map7d1 | 1.27E-53 | 0.38868201 | 0.369 | 0.277 | 2.46E-49 | 1 | Map7d1 | neutrophil.2 |
| Elob | 1.02E-52 | 0.61358776 | 0.253 | 0.174 | 1.98E-48 | 1 | Elob | neutrophil.2 |
| Bach1 | 7.87E-52 | 0.52283563 | 0.3 | 0.213 | 1.52E-47 | 1 | Bach1 | neutrophil.2 |
| Abr | 3.24E-51 | 0.3902922 | 0.51 | 0.399 | 6.27E-47 | 1 | Abr | neutrophil.2 |
| Lfng | 9.94E-51 | 0.38770271 | 0.485 | 0.371 | 1.92E-46 | 1 | Lfng | neutrophil.2 |
| Tmbim6 | 1.39E-50 | 0.4540398 | 0.518 | 0.412 | 2.68E-46 | 1 | Tmbim6 | neutrophil.2 |
| Cdc42 | 2.29E-50 | 0.39473797 | 0.39 | 0.301 | 4.42E-46 | 1 | Cdc42 | neutrophil.2 |
| Alox5 | 3.68E-50 | 0.41136661 | 0.26 | 0.187 | 7.12E-46 | 1 | Alox5 | neutrophil.2 |
| Ypel3 | 4.20E-50 | 0.45179448 | 0.344 | 0.271 | 8.13E-46 | 1 | Ypel3 | neutrophil.2 |
| Arhgap30 | 4.35E-50 | 0.41399876 | 0.291 | 0.217 | 8.42E-46 | 1 | Arhgap30 | neutrophil.2 |
| Atp6v0c | 1.44E-49 | 0.34611553 | 0.724 | 0.596 | 2.78E-45 | 1 | Atp6v0c | neutrophil.2 |
| Col4a3bp | 3.83E-49 | 0.43697577 | 0.276 | 0.202 | 7.40E-45 | 1 | Col4a3bp | neutrophil.2 |
| Akr1a1 | 4.62E-49 | 0.49812207 | 0.266 | 0.183 | 8.93E-45 | 1 | Akr1a1 | neutrophil.2 |
| Actn4 | 3.30E-48 | 0.39635692 | 0.284 | 0.199 | 6.38E-44 | 1 | Actn4 | neutrophil.2 |
| Il4ra | 7.66E-48 | 0.37407138 | 0.606 | 0.476 | 1.48E-43 | 1 | Il4ra | neutrophil.2 |
| Cish | 2.02E-47 | 0.4620758 | 0.339 | 0.261 | 3.90E-43 | 1 | Cish | neutrophil.2 |
| Gnb1 | 3.28E-47 | 0.40239005 | 0.315 | 0.235 | 6.35E-43 | 1 | Gnb1 | neutrophil.2 |
| Stk24 | 5.61E-47 | 0.50549756 | 0.273 | 0.198 | 1.08E-42 | 1 | Stk24 | neutrophil.2 |
| Il17ra | 1.22E-46 | 0.53293267 | 0.312 | 0.234 | 2.35E-42 | 1 | Il17ra | neutrophil.2 |
| Grb2 | 3.21E-46 | 0.5349718 | 0.318 | 0.24 | 6.20E-42 | 1 | Grb2 | neutrophil.2 |
| Slc11a1 | 1.19E-45 | 0.29147218 | 0.676 | 0.562 | 2.30E-41 | 1 | Slc11a1 | neutrophil.2 |
| Ythdf3 | 1.72E-45 | 0.44779374 | 0.301 | 0.228 | 3.32E-41 | 1 | Ythdf3 | neutrophil.2 |
| Ppp1r18 | 2.80E-45 | 0.39150102 | 0.436 | 0.349 | 5.41E-41 | 1 | Ppp1r18 | neutrophil.2 |
| Snx3 | 1.17E-44 | 0.4647354 | 0.308 | 0.229 | 2.26E-40 | 1 | Snx3 | neutrophil.2 |
| Actr2 | 4.61E-44 | 0.52941425 | 0.335 | 0.255 | 8.91E-40 | 1 | Actr2 | neutrophil.2 |
| Ly6e | 1.01E-43 | 0.39169224 | 0.464 | 0.349 | 1.95E-39 | 1 | Ly6e | neutrophil.2 |
| Cers6 | 5.91E-43 | 0.38229389 | 0.36 | 0.275 | 1.14E-38 | 1 | Cers6 | neutrophil.2 |
| Edem1 | 1.30E-42 | 0.44991679 | 0.254 | 0.187 | 2.52E-38 | 1 | Edem1 | neutrophil.2 |
| Plp2 | 2.05E-42 | 0.45289999 | 0.274 | 0.208 | 3.97E-38 | 1 | Plp2 | neutrophil.2 |
| Arf3 | 2.16E-41 | 0.43909953 | 0.262 | 0.198 | 4.18E-37 | 1 | Arf3 | neutrophil.2 |
| Itm2b | 8.07E-41 | 0.30858908 | 0.508 | 0.41 | 1.56E-36 | 1 | Itm2b | neutrophil.2 |
| Pgs1 | 2.59E-40 | 0.48528354 | 0.282 | 0.211 | 5.02E-36 | 1 | Pgs1 | neutrophil.2 |
| Hnrnpu | 4.69E-40 | 0.25761228 | 0.255 | 0.185 | 9.06E-36 | 1 | Hnrnpu | neutrophil.2 |
| Adam17 | 1.31E-39 | 0.44592962 | 0.311 | 0.234 | 2.54E-35 | 1 | Adam17 | neutrophil.2 |
| Cbl | 3.19E-39 | 0.52940713 | 0.251 | 0.187 | 6.17E-35 | 1 | Cbl | neutrophil.2 |
| Ccnl2 | 2.90E-37 | 0.35289373 | 0.332 | 0.259 | 5.60E-33 | 1 | Ccnl2 | neutrophil.2 |

| | p_val | avg_log2FC | pct.1 | pct.2 | p_val_adj | cluster | gene |
|---|---|---|---|---|---|---|---|
| Cd209a | 0 | 4.17899267 | 0.689 | 0.099 | 0 | cDC.Cd209 | Cd209a |
| Csf1r | 0 | 2.0178987 | 0.764 | 0.245 | 0 | cDC.Cd209 | Csf1r |
| Tgfbi | 0 | 2.36562078 | 0.783 | 0.297 | 0 | cDC.Cd209 | Tgfbi |
| Fam129b | 0 | 2.74740584 | 0.661 | 0.202 | 0 | cDC.Cd209 | Fam129b |
| Ms4a4c | 0 | 4.27303856 | 0.518 | 0.06 | 0 | cDC.Cd209 | Ms4a4c |
| Emilin2 | 0 | 2.46458153 | 0.722 | 0.264 | 0 | cDC.Cd209 | Emilin2 |
| Olfm1 | 0 | 2.00745043 | 0.799 | 0.362 | 0 | cDC.Cd209 | Olfm1 |
| S100a4 | 0 | 1.988331 | 0.971 | 0.537 | 0 | cDC.Cd209 | S100a4 |
| Ahnak | 0 | 1.92950272 | 0.783 | 0.363 | 0 | cDC.Cd209 | Ahnak |
| Emp3 | 0 | 1.88925189 | 0.752 | 0.336 | 0 | cDC.Cd209 | Emp3 |
| Ccr5 | 0 | 1.7778093 | 0.725 | 0.327 | 0 | cDC.Cd209 | Ccr5 |
| Fcgr2b | 0 | 2.07305795 | 0.691 | 0.295 | 0 | cDC.Cd209 | Fcgr2b |
| Clec10a | 0 | 2.47655565 | 0.579 | 0.184 | 0 | cDC.Cd209 | Clec10a |
| Cfp | 0 | 1.34843649 | 0.93 | 0.545 | 0 | cDC.Cd209 | Cfp |
| Plaur | 0 | 1.72548464 | 0.66 | 0.279 | 0 | cDC.Cd209 | Plaur |
| Fcer1g | 0 | 1.6253912 | 0.955 | 0.584 | 0 | cDC.Cd209 | Fcer1g |
| Fgr | 0 | 1.32532525 | 0.862 | 0.491 | 0 | cDC.Cd209 | Fgr |
| Dab2 | 0 | 1.73282277 | 0.745 | 0.378 | 0 | cDC.Cd209 | Dab2 |
| Ccr2 | 0 | 3.69986839 | 0.417 | 0.055 | 0 | cDC.Cd209 | Ccr2 |
| S100a10 | 0 | 1.55454373 | 0.893 | 0.532 | 0 | cDC.Cd209 | S100a10 |
| Ifitm3 | 0 | 1.77212049 | 0.839 | 0.478 | 0 | cDC.Cd209 | Ifitm3 |
| Shisa5 | 0 | 1.78509603 | 0.838 | 0.48 | 0 | cDC.Cd209 | Shisa5 |
| Tlr2 | 0 | 2.28334898 | 0.506 | 0.151 | 0 | cDC.Cd209 | Tlr2 |
| Ms4a6d | 0 | 2.0888517 | 0.615 | 0.264 | 0 | cDC.Cd209 | Ms4a6d |
| Ifi27l2a | 0 | 2.82365919 | 0.453 | 0.114 | 0 | cDC.Cd209 | Ifi27l2a |
| Cd53 | 0 | 1.26053173 | 0.913 | 0.576 | 0 | cDC.Cd209 | Cd53 |
| Cdh1 | 0 | 1.29747926 | 0.685 | 0.353 | 0 | cDC.Cd209 | Cdh1 |
| Klrd1 | 0 | 2.01589631 | 0.529 | 0.218 | 0 | cDC.Cd209 | Klrd1 |
| Fosl2 | 0 | 1.04911337 | 0.87 | 0.56 | 0 | cDC.Cd209 | Fosl2 |
| Zfp36l2 | 0 | 1.12204128 | 0.838 | 0.541 | 0 | cDC.Cd209 | Zfp36l2 |
| Fxyd5 | 0 | 1.43277822 | 0.852 | 0.573 | 0 | cDC.Cd209 | Fxyd5 |
| Osm | 0 | 1.66511497 | 0.499 | 0.232 | 0 | cDC.Cd209 | Osm |
| Il1r2 | 0 | 0.7229865 | 0.739 | 0.493 | 0 | cDC.Cd209 | Il1r2 |
| S100a6 | 0 | 1.23464677 | 0.954 | 0.717 | 0 | cDC.Cd209 | S100a6 |
| AA467197 | 0 | 1.41340758 | 0.718 | 0.486 | 0 | cDC.Cd209 | AA467197 |
| Pitpna | 0 | 1.092535 | 0.905 | 0.679 | 0 | cDC.Cd209 | Pitpna |
| Ifitm2 | 0 | 1.14272502 | 0.974 | 0.757 | 0 | cDC.Cd209 | Ifitm2 |
| Ly6a | 0 | 1.84580548 | 0.306 | 0.098 | 0 | cDC.Cd209 | Ly6a |
| Cd7 | 0 | 2.31535432 | 0.334 | 0.127 | 0 | cDC.Cd209 | Cd7 |
| Malt1 | 0 | 0.95127894 | 0.842 | 0.653 | 0 | cDC.Cd209 | Malt1 |
| Tyrobp | 0 | 0.78192697 | 0.968 | 0.782 | 0 | cDC.Cd209 | Tyrobp |
| Il1b | 0 | 0.95850663 | 0.906 | 0.742 | 0 | cDC.Cd209 | Il1b |
| Prkcd | 0 | 0.96035942 | 0.897 | 0.741 | 0 | cDC.Cd209 | Prkcd |
| H2-Ab1 | 0 | 0.61835908 | 0.964 | 0.823 | 0 | cDC.Cd209 | H2-Ab1 |
| Bhlhe40 | 0 | 0.70132001 | 0.954 | 0.815 | 0 | cDC.Cd209 | Bhlhe40 |
| Laptm5 | 0 | 0.81144852 | 0.981 | 0.848 | 0 | cDC.Cd209 | Laptm5 |
| Msn | 0 | 0.87392037 | 0.978 | 0.846 | 0 | cDC.Cd209 | Msn |
| Pim1 | 0 | 0.77157257 | 0.972 | 0.846 | 0 | cDC.Cd209 | Pim1 |
| H2-Eb1 | 0 | 0.53036025 | 0.955 | 0.857 | 0 | cDC.Cd209 | H2-Eb1 |
| Ctss | 0 | 0.71871676 | 0.988 | 0.892 | 0 | cDC.Cd209 | Ctss |
| H2-Aa | 0 | 0.75200784 | 0.956 | 0.862 | 0 | cDC.Cd209 | H2-Aa |
| Vim | 0 | 1.265324 | 0.988 | 0.911 | 0 | cDC.Cd209 | Vim |
| Cd74 | 0 | 0.55540181 | 0.992 | 0.963 | 0 | cDC.Cd209 | Cd74 |
| Itgam | 2.96E-303 | 1.86941245 | 0.602 | 0.258 | 5.71E-299 | cDC.Cd209 | Itgam |
| Plec | 2.53E-296 | 1.93970834 | 0.615 | 0.275 | 4.90E-292 | cDC.Cd209 | Plec |
| Capg | 1.95E-293 | 1.6509617 | 0.549 | 0.218 | 3.78E-289 | cDC.Cd209 | Capg |
| Irf2bp2 | 3.33E-293 | 0.81619496 | 0.937 | 0.73 | 6.44E-289 | cDC.Cd209 | Irf2bp2 |
| Cdkn1a | 5.95E-286 | 0.62337445 | 0.883 | 0.668 | 1.15E-281 | cDC.Cd209 | Cdkn1a |
| Ccl17 | 9.31E-283 | 0.39950571 | 0.524 | 0.492 | 1.80E-278 | cDC.Cd209 | Ccl17 |
| Socs3 | 4.30E-280 | 1.13196872 | 0.808 | 0.519 | 8.31E-276 | cDC.Cd209 | Socs3 |
| Cd44 | 2.69E-278 | 1.23663065 | 0.789 | 0.512 | 5.20E-274 | cDC.Cd209 | Cd44 |
| Cyth4 | 2.03E-277 | 1.10516888 | 0.864 | 0.533 | 3.93E-273 | cDC.Cd209 | Cyth4 |
| Vrk1 | 6.17E-277 | 1.24977474 | 0.815 | 0.486 | 1.19E-272 | cDC.Cd209 | Vrk1 |
| Klrb1b | 4.97E-268 | 1.17486503 | 0.599 | 0.395 | 9.61E-264 | cDC.Cd209 | Klrb1b |
| Lrp1 | 9.16E-267 | 1.94352387 | 0.522 | 0.176 | 1.77E-262 | cDC.Cd209 | Lrp1 |
| Rin3 | 2.13E-265 | 1.18074415 | 0.904 | 0.543 | 4.13E-261 | cDC.Cd209 | Rin3 |
| Ngfr | 6.98E-258 | 2.75410407 | 0.418 | 0.111 | 1.35E-253 | cDC.Cd209 | Ngfr |
| Pid1 | 9.00E-250 | 2.4927078 | 0.48 | 0.153 | 1.74E-245 | cDC.Cd209 | Pid1 |
| Tppp3 | 1.05E-249 | 2.46387375 | 0.421 | 0.197 | 2.04E-245 | cDC.Cd209 | Tppp3 |
| Alox5ap | 6.85E-248 | 1.01608411 | 0.878 | 0.541 | 1.32E-243 | cDC.Cd209 | Alox5ap |
| Ank | 2.99E-243 | 2.3475354 | 0.522 | 0.165 | 5.79E-239 | cDC.Cd209 | Ank |
| Sirpa | 9.84E-240 | 1.12511329 | 0.858 | 0.484 | 1.90E-235 | cDC.Cd209 | Sirpa |
| Nrros | 5.05E-239 | 1.27932576 | 0.822 | 0.528 | 9.75E-235 | cDC.Cd209 | Nrros |
| Bcl3 | 2.13E-231 | 1.25565181 | 0.766 | 0.524 | 4.11E-227 | cDC.Cd209 | Bcl3 |
| Arhgdib | 8.09E-230 | 0.84408549 | 0.875 | 0.645 | 1.56E-225 | cDC.Cd209 | Arhgdib |
| Cd209c | 5.35E-229 | 4.11214612 | 0.343 | 0.029 | 1.03E-224 | cDC.Cd209 | Cd209c |
| Timp1 | 3.00E-224 | 2.09133599 | 0.325 | 0.104 | 5.81E-220 | cDC.Cd209 | Timp1 |
| Cish | 6.63E-224 | 1.28383537 | 0.61 | 0.375 | 1.28E-219 | cDC.Cd209 | Cish |
| Plac8 | 3.54E-221 | 0.95666828 | 0.621 | 0.424 | 6.85E-217 | cDC.Cd209 | Plac8 |
| Lilrb4a | 3.30E-220 | 1.00344497 | 0.787 | 0.476 | 6.38E-216 | cDC.Cd209 | Lilrb4a |
| Cd14 | 3.70E-219 | 0.91021481 | 0.456 | 0.264 | 7.15E-215 | cDC.Cd209 | Cd14 |
| Mrc1 | 5.69E-217 | 1.96265784 | 0.342 | 0.102 | 1.10E-212 | cDC.Cd209 | Mrc1 |
| Nr4a1 | 4.42E-209 | 0.78218495 | 0.881 | 0.632 | 8.54E-205 | cDC.Cd209 | Nr4a1 |
| Wfdc17 | 1.10E-204 | 1.60431438 | 0.622 | 0.278 | 2.13E-200 | cDC.Cd209 | Wfdc17 |
| Rara | 3.35E-204 | 1.42927785 | 0.703 | 0.403 | 6.47E-200 | cDC.Cd209 | Rara |
| Dok2 | 1.22E-202 | 2.74537331 | 0.339 | 0.068 | 2.35E-198 | cDC.Cd209 | Dok2 |
| H2-DMa | 1.55E-200 | 0.77051879 | 0.915 | 0.622 | 2.99E-196 | cDC.Cd209 | H2-DMa |
| Tgm2 | 1.22E-197 | 0.90640565 | 0.734 | 0.498 | 2.36E-193 | cDC.Cd209 | Tgm2 |
| Ehd4 | 3.13E-191 | 1.62191869 | 0.61 | 0.267 | 6.05E-187 | cDC.Cd209 | Ehd4 |
| Fcgrt | 2.10E-190 | 1.47849158 | 0.705 | 0.404 | 4.06E-186 | cDC.Cd209 | Fcgrt |
| Fosb | 4.91E-188 | 0.6493625 | 0.768 | 0.608 | 9.49E-184 | cDC.Cd209 | Fosb |
| Ms4a6c | 1.10E-186 | 2.22941234 | 0.497 | 0.182 | 2.13E-182 | cDC.Cd209 | Ms4a6c |
| Ltb4r1 | 4.78E-186 | 3.58081697 | 0.319 | 0.044 | 9.23E-182 | cDC.Cd209 | Ltb4r1 |
| St3gal4 | 1.33E-185 | 1.23364498 | 0.764 | 0.462 | 2.57E-181 | cDC.Cd209 | St3gal4 |
| Slfn5 | 2.27E-184 | 1.52077592 | 0.46 | 0.225 | 4.39E-180 | cDC.Cd209 | Slfn5 |
| Isg15 | 7.88E-183 | 1.29600774 | 0.362 | 0.229 | 1.52E-178 | cDC.Cd209 | Isg15 |
| Slc15a3 | 9.68E-183 | 0.57480216 | 0.82 | 0.569 | 1.87E-178 | cDC.Cd209 | Slc15a3 |
| Pycard | 8.26E-180 | 1.274399 | 0.681 | 0.419 | 1.60E-175 | cDC.Cd209 | Pycard |
| Cd300a | 2.03E-179 | 1.10252802 | 0.82 | 0.487 | 3.92E-175 | cDC.Cd209 | Cd300a |
| Bach1 | 3.16E-179 | 1.25156909 | 0.722 | 0.446 | 6.11E-175 | cDC.Cd209 | Bach1 |
| Ccnd3 | 2.21E-178 | 1.11722994 | 0.8 | 0.5 | 4.27E-174 | cDC.Cd209 | Ccnd3 |
| Fh1 | 1.76E-175 | 1.50778096 | 0.545 | 0.28 | 3.40E-171 | cDC.Cd209 | Fh1 |
| Cdk2ap2 | 5.03E-175 | 1.22046265 | 0.654 | 0.433 | 9.72E-171 | cDC.Cd209 | Cdk2ap2 |
| Pgf | 6.18E-172 | 1.3735585 | 0.329 | 0.159 | 1.19E-167 | cDC.Cd209 | Pgf |
| Fcgr1 | 1.42E-171 | 2.62065717 | 0.358 | 0.088 | 2.74E-167 | cDC.Cd209 | Fcgr1 |
| Irf7 | 1.15E-169 | 1.85322293 | 0.319 | 0.109 | 2.22E-165 | cDC.Cd209 | Irf7 |
| Emp1 | 1.26E-168 | 1.69078003 | 0.421 | 0.2 | 2.43E-164 | cDC.Cd209 | Emp1 |
| Jun | 1.54E-167 | 0.81964968 | 0.828 | 0.609 | 2.97E-163 | cDC.Cd209 | Jun |
| H13 | 2.06E-166 | 1.11619048 | 0.686 | 0.366 | 3.98E-162 | cDC.Cd209 | H13 |
| Padi2 | 5.06E-166 | 2.35778999 | 0.357 | 0.076 | 9.78E-162 | cDC.Cd209 | Padi2 |
| Mgl2 | 3.98E-162 | 0.46028497 | 0.566 | 0.48 | 7.70E-158 | cDC.Cd209 | Mgl2 |
| H2-DMb1 | 7.14E-160 | 1.2020947 | 0.737 | 0.437 | 1.38E-155 | cDC.Cd209 | H2-DMb1 |
| Anxa2 | 6.78E-159 | 0.47018058 | 0.932 | 0.797 | 1.31E-154 | cDC.Cd209 | Anxa2 |
| Vps37b | 8.09E-157 | 0.86758569 | 0.683 | 0.453 | 1.56E-152 | cDC.Cd209 | Vps37b |
| Ly6e | 1.19E-154 | 0.59256155 | 0.912 | 0.604 | 2.30E-150 | cDC.Cd209 | Ly6e |
| Rab3il1 | 2.70E-154 | 2.15228851 | 0.466 | 0.157 | 5.21E-150 | cDC.Cd209 | Rab3il1 |
| Sbno2 | 4.76E-152 | 0.86041807 | 0.862 | 0.658 | 9.20E-148 | cDC.Cd209 | Sbno2 |
| Cxcr4 | 8.53E-152 | 0.84873813 | 0.677 | 0.487 | 1.65E-147 | cDC.Cd209 | Cxcr4 |
| Picalm | 4.51E-148 | 0.7947029 | 0.894 | 0.682 | 8.72E-144 | cDC.Cd209 | Picalm |
| Tpt1 | 1.33E-147 | 0.43704539 | 0.967 | 0.894 | 2.57E-143 | cDC.Cd209 | Tpt1 |
| Zeb2 | 2.09E-147 | 1.26051768 | 0.612 | 0.277 | 4.03E-143 | cDC.Cd209 | Zeb2 |
| Lat2 | 2.91E-147 | 1.7095175 | 0.477 | 0.197 | 5.62E-143 | cDC.Cd209 | Lat2 |
| B4galt1 | 1.10E-145 | 1.1031665 | 0.64 | 0.431 | 2.13E-141 | cDC.Cd209 | B4galt1 |
| Ms4a4a | 1.14E-145 | 2.27396962 | 0.379 | 0.08 | 2.21E-141 | cDC.Cd209 | Ms4a4a |
| Mcemp1 | 1.36E-144 | 1.30593817 | 0.457 | 0.242 | 2.62E-140 | cDC.Cd209 | Mcemp1 |
| Fam49b | 1.52E-143 | 0.70187198 | 0.909 | 0.739 | 2.94E-139 | cDC.Cd209 | Fam49b |
| Ifngr1 | 1.82E-143 | 0.83584175 | 0.867 | 0.618 | 3.52E-139 | cDC.Cd209 | Ifngr1 |
| Emb | 2.58E-143 | 1.15392335 | 0.655 | 0.364 | 4.98E-139 | cDC.Cd209 | Emb |
| Il6ra | 2.83E-142 | 1.79406708 | 0.529 | 0.226 | 5.48E-138 | cDC.Cd209 | Il6ra |
| Oas1a | 1.17E-141 | 2.30925204 | 0.372 | 0.128 | 2.26E-137 | cDC.Cd209 | Oas1a |
| Ms4a6b | 1.34E-141 | 2.55166728 | 0.375 | 0.115 | 2.58E-137 | cDC.Cd209 | Ms4a6b |
| Ifitm1 | 7.01E-141 | 0.45040525 | 0.802 | 0.562 | 1.35E-136 | cDC.Cd209 | Ifitm1 |
| Kdm6b | 2.81E-140 | 0.66164985 | 0.884 | 0.734 | 5.42E-136 | cDC.Cd209 | Kdm6b |
| Lyn | 5.13E-139 | 0.97521066 | 0.856 | 0.603 | 9.91E-135 | cDC.Cd209 | Lyn |
| Csrnp1 | 7.93E-139 | 0.80643601 | 0.835 | 0.632 | 1.53E-134 | cDC.Cd209 | Csrnp1 |
| Tgfb1 | 5.56E-137 | 0.86150558 | 0.838 | 0.643 | 1.07E-132 | cDC.Cd209 | Tgfb1 |
| Fmnl1 | 2.71E-135 | 0.62906721 | 0.889 | 0.733 | 5.25E-131 | cDC.Cd209 | Fmnl1 |
| Il17ra | 1.12E-133 | 1.87881325 | 0.439 | 0.18 | 2.17E-129 | cDC.Cd209 | Il17ra |
| Gpr35 | 5.79E-133 | 1.17254332 | 0.709 | 0.43 | 1.12E-128 | cDC.Cd209 | Gpr35 |
| Ybx3 | 5.81E-132 | 1.14750996 | 0.63 | 0.375 | 1.12E-127 | cDC.Cd209 | Ybx3 |
| Uck2 | 2.96E-131 | 0.66685425 | 0.855 | 0.663 | 5.71E-127 | cDC.Cd209 | Uck2 |
| Pnp | 7.52E-130 | 1.1791835 | 0.627 | 0.399 | 1.45E-125 | cDC.Cd209 | Pnp |
| Il2rg | 1.45E-128 | 0.8750633 | 0.755 | 0.5 | 2.80E-124 | cDC.Cd209 | Il2rg |
| Sulf2 | 5.74E-127 | 1.19593025 | 0.565 | 0.339 | 1.11E-122 | cDC.Cd209 | Sulf2 |
| Ubl3 | 9.71E-127 | 0.75644204 | 0.851 | 0.642 | 1.88E-122 | cDC.Cd209 | Ubl3 |
| Ifi30 | 6.62E-124 | 0.43255384 | 0.985 | 0.895 | 1.28E-119 | cDC.Cd209 | Ifi30 |
| Zfp36 | 8.49E-123 | 0.44018029 | 0.971 | 0.884 | 1.64E-118 | cDC.Cd209 | Zfp36 |
| Matk | 1.02E-122 | 1.78438148 | 0.373 | 0.127 | 1.97E-118 | cDC.Cd209 | Matk |
| Plxnb2 | 4.76E-121 | 1.30670658 | 0.607 | 0.303 | 9.20E-117 | cDC.Cd209 | Plxnb2 |
| Tnfsf9 | 5.54E-120 | 1.28658591 | 0.446 | 0.249 | 1.07E-115 | cDC.Cd209 | Tnfsf9 |
| Vdr | 5.21E-118 | 0.86974377 | 0.624 | 0.379 | 1.01E-113 | cDC.Cd209 | Vdr |
| Icosl | 9.71E-118 | 1.09664622 | 0.543 | 0.316 | 1.88E-113 | cDC.Cd209 | Icosl |
| Slfn2 | 1.14E-116 | 0.99852912 | 0.704 | 0.47 | 2.20E-112 | cDC.Cd209 | Slfn2 |
| Rhoq | 7.87E-116 | 0.77301267 | 0.599 | 0.428 | 1.52E-111 | cDC.Cd209 | Rhoq |
| Ifi203 | 4.81E-115 | 1.89925809 | 0.305 | 0.129 | 9.30E-111 | cDC.Cd209 | Ifi203 |
| Ier2 | 5.27E-114 | 0.97546617 | 0.785 | 0.533 | 1.02E-109 | cDC.Cd209 | Ier2 |
| Syk | 2.60E-113 | 0.54261978 | 0.898 | 0.749 | 5.03E-109 | cDC.Cd209 | Syk |
| Sell | 3.50E-113 | 1.00379184 | 0.268 | 0.075 | 6.76E-109 | cDC.Cd209 | Sell |
| Runx3 | 3.96E-113 | 0.59681714 | 0.858 | 0.725 | 7.66E-109 | cDC.Cd209 | Runx3 |
| Ikzf1 | 6.54E-113 | 0.92994575 | 0.738 | 0.513 | 1.26E-108 | cDC.Cd209 | Ikzf1 |
| Plek | 4.04E-111 | 0.48377354 | 0.96 | 0.809 | 7.81E-107 | cDC.Cd209 | Plek |
| Ifi207 | 1.11E-109 | 1.666464 | 0.37 | 0.173 | 2.15E-105 | cDC.Cd209 | Ifi207 |
| Pxdc1 | 1.16E-109 | 0.84354337 | 0.474 | 0.313 | 2.23E-105 | cDC.Cd209 | Pxdc1 |
| Atf3 | 1.56E-107 | 0.898646 | 0.584 | 0.439 | 3.01E-103 | cDC.Cd209 | Atf3 |
| Rnf213 | 1.32E-106 | 1.19116125 | 0.472 | 0.292 | 2.56E-102 | cDC.Cd209 | Rnf213 |
| Spint1 | 7.98E-106 | 0.81216357 | 0.727 | 0.512 | 1.54E-101 | cDC.Cd209 | Spint1 |
| Hivep2 | 1.19E-104 | 1.36660342 | 0.564 | 0.299 | 2.31E-100 | cDC.Cd209 | Hivep2 |
| Eif5a | 1.99E-104 | 0.78803516 | 0.824 | 0.62 | 3.84E-100 | cDC.Cd209 | Eif5a |
| Kmo | 3.22E-102 | 0.82578423 | 0.778 | 0.48 | 6.22E-98 | cDC.Cd209 | Kmo |
| Ass1 | 1.07E-101 | 0.5971232 | 0.66 | 0.413 | 2.07E-97 | cDC.Cd209 | Ass1 |
| Runx1 | 2.46E-101 | 1.01393833 | 0.77 | 0.472 | 4.76E-97 | cDC.Cd209 | Runx1 |
| Lgals3bp | 2.93E-101 | 1.07289004 | 0.627 | 0.386 | 5.67E-97 | cDC.Cd209 | Lgals3bp |
| Map2k3 | 2.59E-100 | 0.99485312 | 0.674 | 0.435 | 5.01E-96 | cDC.Cd209 | Map2k3 |
| Trappc5 | 6.05E-100 | 1.26239277 | 0.427 | 0.219 | 1.17E-95 | cDC.Cd209 | Trappc5 |
| Gcsh | 1.64E-99 | 1.45969957 | 0.356 | 0.163 | 3.18E-95 | cDC.Cd209 | Gcsh |
| Ddit4 | 1.34E-97 | 0.94347697 | 0.453 | 0.242 | 2.59E-93 | cDC.Cd209 | Ddit4 |
| Cux1 | 1.89E-97 | 0.85188323 | 0.719 | 0.505 | 3.65E-93 | cDC.Cd209 | Cux1 |
| Itgb7 | 8.89E-97 | 1.01552109 | 0.726 | 0.434 | 1.72E-92 | cDC.Cd209 | Itgb7 |
| Eng | 2.56E-96 | 1.25801315 | 0.378 | 0.142 | 4.94E-92 | cDC.Cd209 | Eng |
| Jarid2 | 3.39E-95 | 1.04740464 | 0.734 | 0.484 | 6.55E-91 | cDC.Cd209 | Jarid2 |
| Ctsa | 1.30E-94 | 0.78345367 | 0.85 | 0.578 | 2.52E-90 | cDC.Cd209 | Ctsa |
| Tiam1 | 4.60E-94 | 1.44477126 | 0.485 | 0.265 | 8.90E-90 | cDC.Cd209 | Tiam1 |
| Capn2 | 2.40E-93 | 0.98585231 | 0.627 | 0.419 | 4.64E-89 | cDC.Cd209 | Capn2 |
| Adam15 | 2.45E-93 | 1.46485746 | 0.432 | 0.206 | 4.73E-89 | cDC.Cd209 | Adam15 |
| Sdc4 | 6.57E-93 | 0.95851875 | 0.434 | 0.256 | 1.27E-88 | cDC.Cd209 | Sdc4 |
| Ccnd1 | 7.97E-93 | 0.91416936 | 0.563 | 0.34 | 1.54E-88 | cDC.Cd209 | Ccnd1 |
| Gpx1 | 1.92E-91 | 0.36146748 | 0.968 | 0.886 | 3.70E-87 | cDC.Cd209 | Gpx1 |
| Mcub | 2.61E-91 | 1.79253028 | 0.317 | 0.116 | 5.06E-87 | cDC.Cd209 | Mcub |
| Ccl9 | 3.44E-91 | 1.86118627 | 0.305 | 0.097 | 6.64E-87 | cDC.Cd209 | Ccl9 |
| Nfkb1 | 1.26E-87 | 0.56832315 | 0.885 | 0.739 | 2.44E-83 | cDC.Cd209 | Nfkb1 |
| Mapkapk3 | 1.39E-87 | 1.01272736 | 0.531 | 0.345 | 2.69E-83 | cDC.Cd209 | Mapkapk3 |
| Arl6ip5 | 1.45E-87 | 1.05055282 | 0.655 | 0.41 | 2.80E-83 | cDC.Cd209 | Arl6ip5 |
| Klrk1 | 1.66E-86 | 0.95580767 | 0.473 | 0.291 | 3.21E-82 | cDC.Cd209 | Klrk1 |
| Cebpb | 3.29E-85 | 0.77158741 | 0.637 | 0.41 | 6.37E-81 | cDC.Cd209 | Cebpb |
| Slk | 6.92E-85 | 1.17677308 | 0.579 | 0.353 | 1.34E-80 | cDC.Cd209 | Slk |
| Srgn | 1.25E-83 | 0.42004094 | 0.972 | 0.88 | 2.42E-79 | cDC.Cd209 | Srgn |
| Ptpro | 1.58E-83 | 1.88873197 | 0.284 | 0.106 | 3.06E-79 | cDC.Cd209 | Ptpro |
| Arhgef10 | 3.67E-83 | 1.4605675 | 0.329 | 0.144 | 7.09E-79 | cDC.Cd209 | Arhgef10 |
| Fyn | 1.38E-81 | 0.50826936 | 0.81 | 0.657 | 2.66E-77 | cDC.Cd209 | Fyn |
| Axl | 8.75E-81 | 0.92030687 | 0.326 | 0.204 | 1.69E-76 | cDC.Cd209 | Axl |
| Spi1 | 9.70E-81 | 0.62345235 | 0.905 | 0.725 | 1.88E-76 | cDC.Cd209 | Spi1 |
| Arl8b | 1.56E-80 | 0.80988625 | 0.727 | 0.523 | 3.01E-76 | cDC.Cd209 | Arl8b |
| Lta4h | 4.83E-80 | 0.69787829 | 0.721 | 0.508 | 9.33E-76 | cDC.Cd209 | Lta4h |
| Fos | 5.74E-80 | 0.60479177 | 0.815 | 0.633 | 1.11E-75 | cDC.Cd209 | Fos |
| Msrb1 | 1.29E-79 | 0.74826407 | 0.684 | 0.477 | 2.49E-75 | cDC.Cd209 | Msrb1 |
| Prnp | 4.29E-79 | 0.87362616 | 0.276 | 0.175 | 8.29E-75 | cDC.Cd209 | Prnp |
| B4galt5 | 6.39E-78 | 0.79525559 | 0.675 | 0.487 | 1.24E-73 | cDC.Cd209 | B4galt5 |
| Gvin1 | 1.44E-77 | 1.49846098 | 0.385 | 0.183 | 2.79E-73 | cDC.Cd209 | Gvin1 |
| Grap2 | 3.22E-77 | 1.56176046 | 0.436 | 0.194 | 6.23E-73 | cDC.Cd209 | Grap2 |
| Nfkbid | 4.51E-77 | 0.82081487 | 0.633 | 0.401 | 8.73E-73 | cDC.Cd209 | Nfkbid |
| Adssl1 | 3.34E-76 | 1.12711671 | 0.397 | 0.204 | 6.46E-72 | cDC.Cd209 | Adssl1 |
| Cd68 | 3.85E-76 | 0.87026326 | 0.746 | 0.485 | 7.44E-72 | cDC.Cd209 | Cd68 |
| Edem1 | 6.14E-75 | 0.48975927 | 0.884 | 0.753 | 1.19E-70 | cDC.Cd209 | Edem1 |
| Tgif1 | 2.40E-74 | 0.64756521 | 0.699 | 0.517 | 4.63E-70 | cDC.Cd209 | Tgif1 |
| Samhd1 | 7.38E-74 | 0.31009258 | 0.838 | 0.753 | 1.43E-69 | cDC.Cd209 | Samhd1 |
| Hif1a | 1.52E-73 | 0.96322479 | 0.55 | 0.343 | 2.93E-69 | cDC.Cd209 | Hif1a |
| Tmem176b | 3.53E-73 | 0.6080463 | 0.628 | 0.419 | 6.83E-69 | cDC.Cd209 | Tmem176b |
| Nfkbiz | 4.05E-72 | 0.71365015 | 0.633 | 0.469 | 7.84E-68 | cDC.Cd209 | Nfkbiz |
| Klf6 | 1.74E-71 | 0.60965065 | 0.593 | 0.449 | 3.36E-67 | cDC.Cd209 | Klf6 |
| Med13l | 3.72E-71 | 0.88206497 | 0.708 | 0.509 | 7.18E-67 | cDC.Cd209 | Med13l |
| Ciita | 9.24E-71 | 0.6933724 | 0.823 | 0.547 | 1.79E-66 | cDC.Cd209 | Ciita |
| Cebpa | 1.25E-70 | 1.23197799 | 0.483 | 0.237 | 2.42E-66 | cDC.Cd209 | Cebpa |
| Irgm1 | 2.91E-70 | 1.31354858 | 0.27 | 0.141 | 5.62E-66 | cDC.Cd209 | Irgm1 |
| Elmsan1 | 5.99E-70 | 1.1196617 | 0.498 | 0.311 | 1.16E-65 | cDC.Cd209 | Elmsan1 |
| Fem1c | 4.35E-68 | 1.34269223 | 0.472 | 0.271 | 8.41E-64 | cDC.Cd209 | Fem1c |
| Junb | 2.18E-67 | 0.33917893 | 0.967 | 0.861 | 4.22E-63 | cDC.Cd209 | Junb |
| Tatdn2 | 4.29E-67 | 0.97057836 | 0.615 | 0.432 | 8.29E-63 | cDC.Cd209 | Tatdn2 |
| Tnfrsf1a | 6.82E-67 | 0.75710016 | 0.784 | 0.567 | 1.32E-62 | cDC.Cd209 | Tnfrsf1a |
| Galnt6 | 4.36E-66 | 1.4726965 | 0.376 | 0.177 | 8.43E-62 | cDC.Cd209 | Galnt6 |
| Nfkbie | 5.52E-65 | 0.69876968 | 0.699 | 0.506 | 1.07E-60 | cDC.Cd209 | Nfkbie |
| Nlrp3 | 1.31E-63 | 0.72786808 | 0.6 | 0.376 | 2.53E-59 | cDC.Cd209 | Nlrp3 |
| Slc7a5 | 8.47E-63 | 0.72403528 | 0.605 | 0.429 | 1.64E-58 | cDC.Cd209 | Slc7a5 |
| Mefv | 1.86E-62 | 1.46305913 | 0.326 | 0.161 | 3.60E-58 | cDC.Cd209 | Mefv |
| Snx18 | 1.98E-62 | 0.58632904 | 0.651 | 0.45 | 3.83E-58 | cDC.Cd209 | Snx18 |
| Syngr2 | 2.43E-62 | 0.27078857 | 0.904 | 0.821 | 4.69E-58 | cDC.Cd209 | Syngr2 |
| Arl5b | 2.50E-62 | 0.96166568 | 0.441 | 0.288 | 4.84E-58 | cDC.Cd209 | Arl5b |
| Dpysl2 | 3.98E-62 | 0.88308342 | 0.694 | 0.464 | 7.69E-58 | cDC.Cd209 | Dpysl2 |
| Rassf4 | 5.93E-62 | 0.71757271 | 0.559 | 0.37 | 1.15E-57 | cDC.Cd209 | Rassf4 |
| Adora2b | 8.35E-62 | 1.77807936 | 0.289 | 0.118 | 1.61E-57 | cDC.Cd209 | Adora2b |
| Oasl2 | 1.46E-61 | 1.17190208 | 0.315 | 0.163 | 2.82E-57 | cDC.Cd209 | Oasl2 |
| Ndel1 | 1.68E-61 | 0.66169956 | 0.679 | 0.529 | 3.24E-57 | cDC.Cd209 | Ndel1 |
| Stk10 | 4.69E-61 | 1.33748555 | 0.431 | 0.222 | 9.08E-57 | cDC.Cd209 | Stk10 |
| Cebpd | 1.55E-60 | 1.0578909 | 0.398 | 0.225 | 2.99E-56 | cDC.Cd209 | Cebpd |
| Man2a2 | 3.74E-60 | 1.07935412 | 0.436 | 0.219 | 7.23E-56 | cDC.Cd209 | Man2a2 |
| Ddr1 | 4.26E-60 | 0.93999621 | 0.513 | 0.299 | 8.24E-56 | cDC.Cd209 | Ddr1 |
| Alcam | 7.05E-60 | 0.79449852 | 0.614 | 0.395 | 1.36E-55 | cDC.Cd209 | Alcam |
| Tiparp | 1.00E-59 | 0.80770728 | 0.573 | 0.372 | 1.94E-55 | cDC.Cd209 | Tiparp |
| Stk40 | 3.15E-59 | 0.80022694 | 0.672 | 0.468 | 6.09E-55 | cDC.Cd209 | Stk40 |
| Il10ra | 7.78E-59 | 0.66253855 | 0.699 | 0.526 | 1.50E-54 | cDC.Cd209 | Il10ra |
| Smox | 1.30E-58 | 0.72953768 | 0.445 | 0.306 | 2.51E-54 | cDC.Cd209 | Smox |
| Krt80 | 2.33E-58 | 0.91201277 | 0.471 | 0.302 | 4.51E-54 | cDC.Cd209 | Krt80 |
| Atp6v0c | 3.02E-58 | 0.57770694 | 0.831 | 0.647 | 5.83E-54 | cDC.Cd209 | Atp6v0c |
| Ywhah | 3.99E-58 | 0.40795969 | 0.786 | 0.689 | 7.71E-54 | cDC.Cd209 | Ywhah |
| Sdcbp | 6.79E-58 | 0.6093902 | 0.84 | 0.64 | 1.31E-53 | cDC.Cd209 | Sdcbp |
| Sp100 | 1.07E-57 | 1.06267159 | 0.432 | 0.27 | 2.07E-53 | cDC.Cd209 | Sp100 |
| Crtc3 | 2.93E-57 | 0.82007036 | 0.586 | 0.391 | 5.66E-53 | cDC.Cd209 | Crtc3 |
| Casp6 | 8.50E-57 | 0.77018004 | 0.565 | 0.405 | 1.64E-52 | cDC.Cd209 | Casp6 |
| Def6 | 9.03E-57 | 1.40138981 | 0.415 | 0.206 | 1.74E-52 | cDC.Cd209 | Def6 |
| Per1 | 9.29E-57 | 0.6076556 | 0.72 | 0.546 | 1.80E-52 | cDC.Cd209 | Per1 |
| Arhgap26 | 9.43E-57 | 0.78015505 | 0.518 | 0.326 | 1.82E-52 | cDC.Cd209 | Arhgap26 |
| Nod2 | 2.26E-56 | 0.71752794 | 0.485 | 0.344 | 4.37E-52 | cDC.Cd209 | Nod2 |
| Rarg | 3.77E-56 | 1.54809017 | 0.25 | 0.084 | 7.28E-52 | cDC.Cd209 | Rarg |
| Tnfrsf12a | 4.46E-56 | 0.78593647 | 0.374 | 0.21 | 8.63E-52 | cDC.Cd209 | Tnfrsf12a |
| Clec4n | 8.48E-56 | 0.60360245 | 0.559 | 0.353 | 1.64E-51 | cDC.Cd209 | Clec4n |
| Rnh1 | 4.15E-53 | 0.6351857 | 0.773 | 0.635 | 8.02E-49 | cDC.Cd209 | Rnh1 |
| Sidt2 | 1.01E-52 | 0.87546847 | 0.483 | 0.295 | 1.95E-48 | cDC.Cd209 | Sidt2 |
| Dusp3 | 3.58E-52 | 0.78623624 | 0.659 | 0.471 | 6.93E-48 | cDC.Cd209 | Dusp3 |
| Dusp1 | 3.66E-52 | 0.49392735 | 0.814 | 0.706 | 7.08E-48 | cDC.Cd209 | Dusp1 |
| Sfxn3 | 7.30E-52 | 1.0102557 | 0.443 | 0.262 | 1.41E-47 | cDC.Cd209 | Sfxn3 |
| Gdi2 | 1.38E-51 | 0.40128062 | 0.807 | 0.696 | 2.67E-47 | cDC.Cd209 | Gdi2 |
| Ly86 | 1.43E-51 | 0.83431031 | 0.662 | 0.476 | 2.76E-47 | cDC.Cd209 | Ly86 |
| Gramd4 | 5.62E-51 | 0.88593637 | 0.547 | 0.363 | 1.09E-46 | cDC.Cd209 | Gramd4 |
| Dennd4a | 2.77E-50 | 0.46711758 | 0.743 | 0.602 | 5.36E-46 | cDC.Cd209 | Dennd4a |
| Myo1e | 3.82E-50 | 1.24939165 | 0.282 | 0.128 | 7.39E-46 | cDC.Cd209 | Myo1e |
| Kdm7a | 4.37E-50 | 0.78938061 | 0.598 | 0.404 | 8.46E-46 | cDC.Cd209 | Kdm7a |
| Rasgef1b | 1.41E-49 | 0.94763587 | 0.49 | 0.302 | 2.73E-45 | cDC.Cd209 | Rasgef1b |
| Cldnd1 | 2.49E-49 | 0.87975207 | 0.575 | 0.32 | 4.82E-45 | cDC.Cd209 | Cldnd1 |
| Gpr146 | 1.11E-48 | 1.31744729 | 0.298 | 0.139 | 2.14E-44 | cDC.Cd209 | Gpr146 |
| Slc29a3 | 2.40E-48 | 0.79816127 | 0.414 | 0.243 | 4.64E-44 | cDC.Cd209 | Slc29a3 |
| Slc2a1 | 3.24E-48 | 0.70009378 | 0.623 | 0.428 | 6.25E-44 | cDC.Cd209 | Slc2a1 |
| Parp14 | 4.56E-48 | 0.88053139 | 0.379 | 0.251 | 8.83E-44 | cDC.Cd209 | Parp14 |
| Ncf2 | 1.10E-47 | 0.66138825 | 0.693 | 0.488 | 2.12E-43 | cDC.Cd209 | Ncf2 |
| St3gal1 | 1.15E-47 | 0.79755785 | 0.547 | 0.385 | 2.22E-43 | cDC.Cd209 | St3gal1 |
| Orai1 | 1.50E-47 | 0.71044279 | 0.715 | 0.508 | 2.89E-43 | cDC.Cd209 | Orai1 |
| Esyt1 | 1.92E-47 | 0.5768336 | 0.81 | 0.618 | 3.71E-43 | cDC.Cd209 | Esyt1 |
| Epha2 | 5.24E-47 | 0.85625441 | 0.363 | 0.215 | 1.01E-42 | cDC.Cd209 | Epha2 |
| Pim3 | 8.80E-47 | 0.79072979 | 0.536 | 0.344 | 1.70E-42 | cDC.Cd209 | Pim3 |
| Ptafr | 1.47E-46 | 0.70319759 | 0.484 | 0.353 | 2.84E-42 | cDC.Cd209 | Ptafr |
| Agpat4 | 1.76E-46 | 0.74890729 | 0.583 | 0.379 | 3.40E-42 | cDC.Cd209 | Agpat4 |
| Lamp2 | 2.88E-46 | 0.72023325 | 0.686 | 0.501 | 5.57E-42 | cDC.Cd209 | Lamp2 |
| Pirb | 3.98E-46 | 0.51298339 | 0.805 | 0.658 | 7.70E-42 | cDC.Cd209 | Pirb |
| Ccnl1 | 4.10E-46 | 0.50070816 | 0.777 | 0.642 | 7.92E-42 | cDC.Cd209 | Ccnl1 |
| Midn | 3.73E-45 | 0.58465846 | 0.752 | 0.573 | 7.21E-41 | cDC.Cd209 | Midn |
| Hilpda | 4.60E-45 | 0.49143101 | 0.433 | 0.33 | 8.90E-41 | cDC.Cd209 | Hilpda |
| Ccdc71l | 1.24E-44 | 0.46509834 | 0.36 | 0.267 | 2.40E-40 | cDC.Cd209 | Ccdc71l |
| Evi2a | 2.58E-44 | 0.81389322 | 0.614 | 0.415 | 4.99E-40 | cDC.Cd209 | Evi2a |
| Trem1 | 2.60E-44 | 0.97948862 | 0.269 | 0.146 | 5.03E-40 | cDC.Cd209 | Trem1 |
| Cd164 | 3.28E-44 | 0.46966041 | 0.768 | 0.562 | 6.34E-40 | cDC.Cd209 | Cd164 |
| Ptgs2 | 2.28E-43 | 0.42887871 | 0.384 | 0.285 | 4.41E-39 | cDC.Cd209 | Ptgs2 |
| Gpr132 | 3.01E-43 | 0.32317917 | 0.814 | 0.727 | 5.83E-39 | cDC.Cd209 | Gpr132 |
| Xbp1 | 6.83E-43 | 0.45278307 | 0.806 | 0.668 | 1.32E-38 | cDC.Cd209 | Xbp1 |
| Slc44a2 | 2.40E-42 | 0.69285665 | 0.662 | 0.418 | 4.64E-38 | cDC.Cd209 | Slc44a2 |
| Vsir | 3.72E-42 | 0.97185553 | 0.501 | 0.319 | 7.19E-38 | cDC.Cd209 | Vsir |
| Lmna | 3.77E-42 | 0.88875407 | 0.326 | 0.232 | 7.29E-38 | cDC.Cd209 | Lmna |
| Tagap | 4.62E-42 | 1.11777374 | 0.333 | 0.192 | 8.93E-38 | cDC.Cd209 | Tagap |
| Bcl2 | 5.17E-42 | 1.2085188 | 0.273 | 0.126 | 1.00E-37 | cDC.Cd209 | Bcl2 |
| Nfe2l2 | 8.71E-42 | 0.44996756 | 0.803 | 0.664 | 1.68E-37 | cDC.Cd209 | Nfe2l2 |
| Ptpn6 | 2.51E-41 | 0.56952493 | 0.784 | 0.595 | 4.86E-37 | cDC.Cd209 | Ptpn6 |
| Lgals9 | 2.68E-41 | 0.97359695 | 0.41 | 0.234 | 5.18E-37 | cDC.Cd209 | Lgals9 |
| Acvr2a | 4.86E-41 | 1.2252968 | 0.264 | 0.133 | 9.40E-37 | cDC.Cd209 | Acvr2a |
| Hnrnpll | 6.52E-41 | 0.57196557 | 0.631 | 0.484 | 1.26E-36 | cDC.Cd209 | Hnrnpll |
| Zbtb18 | 6.90E-41 | 0.29726369 | 0.662 | 0.553 | 1.33E-36 | cDC.Cd209 | Zbtb18 |
| Aplp2 | 8.16E-41 | 0.40577788 | 0.577 | 0.431 | 1.58E-36 | cDC.Cd209 | Aplp2 |
| Rtl5 | 1.05E-40 | 1.05846927 | 0.306 | 0.17 | 2.02E-36 | cDC.Cd209 | Rtl5 |
| Dock4 | 1.31E-40 | 1.2467827 | 0.294 | 0.14 | 2.53E-36 | cDC.Cd209 | Dock4 |
| Pepd | 1.79E-40 | 0.9113067 | 0.496 | 0.336 | 3.47E-36 | cDC.Cd209 | Pepd |
| Rap1a | 3.30E-40 | 0.64913026 | 0.678 | 0.503 | 6.38E-36 | cDC.Cd209 | Rap1a |
| Tmem176a | 5.42E-40 | 0.42259646 | 0.539 | 0.373 | 1.05E-35 | cDC.Cd209 | Tmem176a |
| Bcl11a | 5.91E-40 | 1.0111579 | 0.255 | 0.141 | 1.14E-35 | cDC.Cd209 | Bcl11a |
| Coro1a | 8.74E-40 | 0.35761676 | 0.983 | 0.883 | 1.69E-35 | cDC.Cd209 | Coro1a |
| Fcho2 | 9.61E-40 | 0.6290703 | 0.453 | 0.3 | 1.86E-35 | cDC.Cd209 | Fcho2 |
| Klf13 | 1.32E-38 | 0.81983499 | 0.503 | 0.33 | 2.54E-34 | cDC.Cd209 | Klf13 |
| Flna | 1.55E-38 | 0.4128395 | 0.848 | 0.674 | 3.01E-34 | cDC.Cd209 | Flna |
| Atp1a1 | 4.01E-38 | 0.44853304 | 0.905 | 0.755 | 7.75E-34 | cDC.Cd209 | Atp1a1 |
| Nfam1 | 8.96E-38 | 0.9487405 | 0.451 | 0.291 | 1.73E-33 | cDC.Cd209 | Nfam1 |
| Atp6ap2 | 1.93E-37 | 0.65936812 | 0.727 | 0.555 | 3.74E-33 | cDC.Cd209 | Atp6ap2 |
| Ptbp3 | 4.08E-37 | 0.45307011 | 0.847 | 0.706 | 7.90E-33 | cDC.Cd209 | Ptbp3 |
| Bin2 | 6.34E-37 | 0.61762751 | 0.822 | 0.581 | 1.23E-32 | cDC.Cd209 | Bin2 |
| Nfkbia | 7.22E-37 | 0.3189594 | 0.817 | 0.699 | 1.40E-32 | cDC.Cd209 | Nfkbia |
| Ucp2 | 1.02E-36 | 0.25239979 | 0.983 | 0.9 | 1.98E-32 | cDC.Cd209 | Ucp2 |
| Ier3 | 1.11E-36 | 0.57590331 | 0.573 | 0.405 | 2.15E-32 | cDC.Cd209 | Ier3 |
| Cdkn2d | 1.30E-36 | 1.21306305 | 0.339 | 0.19 | 2.51E-32 | cDC.Cd209 | Cdkn2d |
| Siah2 | 2.18E-36 | 1.09353302 | 0.416 | 0.249 | 4.21E-32 | cDC.Cd209 | Siah2 |
| H2afy | 3.88E-36 | 0.37824004 | 0.946 | 0.827 | 7.51E-32 | cDC.Cd209 | H2afy |
| Rassf5 | 5.07E-36 | 0.63709673 | 0.692 | 0.501 | 9.81E-32 | cDC.Cd209 | Rassf5 |
| Lcp2 | 5.90E-36 | 0.98709614 | 0.49 | 0.305 | 1.14E-31 | cDC.Cd209 | Lcp2 |
| D16Ertd472e | 8.44E-36 | 0.66537341 | 0.407 | 0.302 | 1.63E-31 | cDC.Cd209 | D16Ertd472e |
| Chd7 | 1.01E-35 | 0.47955832 | 0.701 | 0.585 | 1.96E-31 | cDC.Cd209 | Chd7 |
| Hs3st3b1 | 1.33E-35 | 0.90508158 | 0.31 | 0.175 | 2.56E-31 | cDC.Cd209 | Hs3st3b1 |
| Ski | 1.37E-35 | 0.62315965 | 0.587 | 0.42 | 2.65E-31 | cDC.Cd209 | Ski |
| Card19 | 3.73E-35 | 0.82233105 | 0.483 | 0.332 | 7.22E-31 | cDC.Cd209 | Card19 |
| Ugcg | 1.48E-34 | 1.02114692 | 0.315 | 0.194 | 2.85E-30 | cDC.Cd209 | Ugcg |
| Kbtbd11 | 2.42E-34 | 0.72331696 | 0.312 | 0.213 | 4.68E-30 | cDC.Cd209 | Kbtbd11 |
| Egr1 | 2.74E-34 | 0.65672402 | 0.46 | 0.344 | 5.30E-30 | cDC.Cd209 | Egr1 |
| Csf2rb2 | 4.61E-34 | 0.320707 | 0.642 | 0.462 | 8.91E-30 | cDC.Cd209 | Csf2rb2 |
| Mboat7 | 6.11E-34 | 0.92292677 | 0.469 | 0.294 | 1.18E-29 | cDC.Cd209 | Mboat7 |
| Ndufb1-ps | 7.14E-34 | 0.48561285 | 0.705 | 0.568 | 1.38E-29 | cDC.Cd209 | Ndufb1-ps |
| 11-Sep | 8.63E-34 | 1.0592459 | 0.344 | 0.198 | 1.67E-29 | cDC.Cd209 | 11-Sep |
| Maff | 1.34E-33 | 0.52292129 | 0.635 | 0.466 | 2.58E-29 | cDC.Cd209 | Maff |
| Chsy1 | 1.88E-33 | 0.97819124 | 0.407 | 0.243 | 3.64E-29 | cDC.Cd209 | Chsy1 |
| Ssh2 | 1.95E-33 | 1.03442868 | 0.414 | 0.259 | 3.76E-29 | cDC.Cd209 | Ssh2 |
| Eml4 | 2.23E-33 | 0.78041758 | 0.504 | 0.352 | 4.31E-29 | cDC.Cd209 | Eml4 |
| Sipa1l3 | 4.10E-33 | 0.64098919 | 0.526 | 0.355 | 7.93E-29 | cDC.Cd209 | Sipa1l3 |
| Bst2 | 5.71E-33 | 0.29838433 | 0.653 | 0.457 | 1.10E-28 | cDC.Cd209 | Bst2 |
| Ifnar2 | 6.69E-33 | 0.56196121 | 0.614 | 0.449 | 1.29E-28 | cDC.Cd209 | Ifnar2 |
| Rab11fip5 | 1.13E-32 | 1.47255725 | 0.288 | 0.113 | 2.19E-28 | cDC.Cd209 | Rab11fip5 |
| Cd48 | 1.23E-32 | 0.72333595 | 0.562 | 0.417 | 2.38E-28 | cDC.Cd209 | Cd48 |
| Irak2 | 1.50E-32 | 0.76414078 | 0.466 | 0.334 | 2.90E-28 | cDC.Cd209 | Irak2 |
| Lfng | 1.61E-32 | 0.58431598 | 0.534 | 0.397 | 3.12E-28 | cDC.Cd209 | Lfng |
| Slc39a14 | 1.92E-32 | 0.78353559 | 0.28 | 0.151 | 3.72E-28 | cDC.Cd209 | Slc39a14 |
| Prcp | 2.24E-32 | 0.52474088 | 0.674 | 0.553 | 4.34E-28 | cDC.Cd209 | Prcp |
| Lrp12 | 2.76E-32 | 1.42208151 | 0.253 | 0.127 | 5.34E-28 | cDC.Cd209 | Lrp12 |
| Sh3bp2 | 1.17E-31 | 0.99369996 | 0.353 | 0.212 | 2.26E-27 | cDC.Cd209 | Sh3bp2 |
| Irf2 | 1.45E-31 | 0.79147344 | 0.424 | 0.275 | 2.80E-27 | cDC.Cd209 | Irf2 |
| Rnf149 | 2.35E-31 | 0.46570089 | 0.752 | 0.582 | 4.55E-27 | cDC.Cd209 | Rnf149 |
| Dhx58 | 2.96E-31 | 1.1533009 | 0.255 | 0.147 | 5.72E-27 | cDC.Cd209 | Dhx58 |
| H3f3b | 3.10E-31 | 0.43992433 | 0.822 | 0.682 | 6.00E-27 | cDC.Cd209 | H3f3b |
| Trps1 | 3.84E-31 | 0.50552317 | 0.699 | 0.518 | 7.42E-27 | cDC.Cd209 | Trps1 |
| Slc14a1 | 3.99E-31 | 1.11976085 | 0.257 | 0.154 | 7.71E-27 | cDC.Cd209 | Slc14a1 |
| Pstpip1 | 4.07E-31 | 0.86664911 | 0.434 | 0.274 | 7.86E-27 | cDC.Cd209 | Pstpip1 |
| Afdn | 8.03E-31 | 0.82886989 | 0.469 | 0.316 | 1.55E-26 | cDC.Cd209 | Afdn |
| Sec24b | 8.49E-31 | 0.66329692 | 0.554 | 0.402 | 1.64E-26 | cDC.Cd209 | Sec24b |
| Ptpn1 | 1.15E-30 | 0.44948468 | 0.774 | 0.656 | 2.22E-26 | cDC.Cd209 | Ptpn1 |
| Tmem189 | 1.21E-30 | 0.59827101 | 0.492 | 0.376 | 2.34E-26 | cDC.Cd209 | Tmem189 |
| Zc3h12a | 2.47E-30 | 0.4697678 | 0.576 | 0.469 | 4.77E-26 | cDC.Cd209 | Zc3h12a |
| Eif4g3 | 2.98E-30 | 0.69072073 | 0.763 | 0.539 | 5.76E-26 | cDC.Cd209 | Eif4g3 |
| Zcchc2 | 3.65E-30 | 1.00582076 | 0.302 | 0.18 | 7.06E-26 | cDC.Cd209 | Zcchc2 |
| Il4ra | 4.32E-30 | 0.34879107 | 0.861 | 0.727 | 8.35E-26 | cDC.Cd209 | Il4ra |
| Sik1 | 2.19E-29 | 0.44137624 | 0.706 | 0.53 | 4.23E-25 | cDC.Cd209 | Sik1 |
| Acer3 | 2.77E-29 | 1.01306626 | 0.334 | 0.193 | 5.37E-25 | cDC.Cd209 | Acer3 |
| Foxp1 | 2.83E-29 | 0.55691466 | 0.517 | 0.372 | 5.47E-25 | cDC.Cd209 | Foxp1 |
| Htra3 | 1.44E-28 | 0.83198227 | 0.31 | 0.183 | 2.79E-24 | cDC.Cd209 | Htra3 |
| Ptk2b | 1.54E-28 | 0.41151278 | 0.671 | 0.524 | 2.98E-24 | cDC.Cd209 | Ptk2b |
| Plcg2 | 1.78E-28 | 0.69757229 | 0.642 | 0.453 | 3.45E-24 | cDC.Cd209 | Plcg2 |
| Metrnl | 3.08E-28 | 0.8120562 | 0.436 | 0.296 | 5.96E-24 | cDC.Cd209 | Metrnl |
| Atf6 | 4.01E-28 | 0.79197083 | 0.515 | 0.349 | 7.76E-24 | cDC.Cd209 | Atf6 |
| Pfkp | 7.24E-28 | 0.3893989 | 0.823 | 0.663 | 1.40E-23 | cDC.Cd209 | Pfkp |
| Nsd3 | 8.49E-28 | 0.42850859 | 0.775 | 0.649 | 1.64E-23 | cDC.Cd209 | Nsd3 |
| Gabarapl1 | 1.20E-27 | 0.63141924 | 0.534 | 0.422 | 2.31E-23 | cDC.Cd209 | Gabarapl1 |
| Il13ra1 | 1.34E-27 | 0.72252209 | 0.466 | 0.322 | 2.58E-23 | cDC.Cd209 | Il13ra1 |
| Mfng | 1.49E-27 | 1.16453428 | 0.294 | 0.161 | 2.89E-23 | cDC.Cd209 | Mfng |
| Limd2 | 2.75E-27 | 0.56239901 | 0.596 | 0.469 | 5.32E-23 | cDC.Cd209 | Limd2 |
| Mef2c | 2.96E-27 | 1.14618657 | 0.3 | 0.154 | 5.72E-23 | cDC.Cd209 | Mef2c |
| Diaph1 | 3.29E-27 | 0.47352996 | 0.71 | 0.56 | 6.36E-23 | cDC.Cd209 | Diaph1 |
| Ist1 | 3.32E-27 | 0.35791387 | 0.738 | 0.605 | 6.42E-23 | cDC.Cd209 | Ist1 |
| Treml2 | 3.49E-27 | 0.83785596 | 0.323 | 0.193 | 6.75E-23 | cDC.Cd209 | Treml2 |
| Slc30a4 | 3.53E-27 | 0.71929184 | 0.433 | 0.318 | 6.82E-23 | cDC.Cd209 | Slc30a4 |
| Pdia6 | 3.71E-27 | 0.6276117 | 0.633 | 0.452 | 7.17E-23 | cDC.Cd209 | Pdia6 |
| Prkch | 3.95E-27 | 0.89783214 | 0.354 | 0.216 | 7.64E-23 | cDC.Cd209 | Prkch |
| Ppp1r9b | 7.63E-27 | 0.35151063 | 0.784 | 0.634 | 1.47E-22 | cDC.Cd209 | Ppp1r9b |
| Sec61b | 1.21E-26 | 0.56213305 | 0.569 | 0.428 | 2.34E-22 | cDC.Cd209 | Sec61b |
| Plxna1 | 1.37E-26 | 0.79161314 | 0.338 | 0.231 | 2.64E-22 | cDC.Cd209 | Plxna1 |
| Abl2 | 1.96E-26 | 0.89666295 | 0.379 | 0.24 | 3.79E-22 | cDC.Cd209 | Abl2 |
| Nme2 | 2.01E-26 | 0.82756389 | 0.493 | 0.326 | 3.88E-22 | cDC.Cd209 | Nme2 |
| Rac2 | 2.16E-26 | 0.3521159 | 0.906 | 0.661 | 4.17E-22 | cDC.Cd209 | Rac2 |
| Itga5 | 3.59E-26 | 0.69525256 | 0.459 | 0.318 | 6.95E-22 | cDC.Cd209 | Itga5 |
| Fermt3 | 4.31E-26 | 0.46408795 | 0.733 | 0.555 | 8.33E-22 | cDC.Cd209 | Fermt3 |
| Diaph2 | 5.94E-26 | 0.59569684 | 0.446 | 0.304 | 1.15E-21 | cDC.Cd209 | Diaph2 |
| Tkt | 9.57E-26 | 0.47875263 | 0.844 | 0.678 | 1.85E-21 | cDC.Cd209 | Tkt |
| Prkx | 1.13E-25 | 0.78532958 | 0.413 | 0.291 | 2.18E-21 | cDC.Cd209 | Prkx |
| Hsd17b10 | 1.26E-25 | 0.74871644 | 0.363 | 0.258 | 2.44E-21 | cDC.Cd209 | Hsd17b10 |
| Hdac5 | 1.33E-25 | 0.66287947 | 0.303 | 0.189 | 2.57E-21 | cDC.Cd209 | Hdac5 |
| Lrrfip2 | 1.49E-25 | 0.6679315 | 0.441 | 0.312 | 2.87E-21 | cDC.Cd209 | Lrrfip2 |
| Cybb | 2.16E-25 | 0.80420536 | 0.492 | 0.29 | 4.17E-21 | cDC.Cd209 | Cybb |
| Ssr1 | 2.30E-25 | 0.50836356 | 0.731 | 0.562 | 4.45E-21 | cDC.Cd209 | Ssr1 |
| Atp13a2 | 2.47E-25 | 0.91878021 | 0.303 | 0.169 | 4.77E-21 | cDC.Cd209 | Atp13a2 |
| Reep5 | 3.43E-25 | 0.51768984 | 0.734 | 0.568 | 6.64E-21 | cDC.Cd209 | Reep5 |
| Casp8 | 4.43E-25 | 0.6267626 | 0.468 | 0.339 | 8.56E-21 | cDC.Cd209 | Casp8 |
| Cnnm4 | 4.80E-25 | 0.62317582 | 0.317 | 0.205 | 9.28E-21 | cDC.Cd209 | Cnnm4 |
| Crlf2 | 5.11E-25 | 0.58907531 | 0.559 | 0.407 | 9.89E-21 | cDC.Cd209 | Crlf2 |
| Atf1 | 6.75E-25 | 0.55703112 | 0.561 | 0.413 | 1.31E-20 | cDC.Cd209 | Atf1 |
| Syncrip | 8.54E-25 | 0.4591561 | 0.674 | 0.532 | 1.65E-20 | cDC.Cd209 | Syncrip |
| Tmem185b | 1.12E-24 | 0.831675 | 0.403 | 0.265 | 2.16E-20 | cDC.Cd209 | Tmem185b |
| Ube2h | 3.58E-24 | 0.41465462 | 0.596 | 0.476 | 6.92E-20 | cDC.Cd209 | Ube2h |
| Zc3hav1 | 4.42E-24 | 0.54707108 | 0.577 | 0.463 | 8.55E-20 | cDC.Cd209 | Zc3hav1 |
| Mapk14 | 4.91E-24 | 0.62393542 | 0.65 | 0.451 | 9.50E-20 | cDC.Cd209 | Mapk14 |
| Sertad2 | 5.68E-24 | 0.66893533 | 0.347 | 0.247 | 1.10E-19 | cDC.Cd209 | Sertad2 |
| Trib1 | 6.29E-24 | 0.60567626 | 0.612 | 0.423 | 1.22E-19 | cDC.Cd209 | Trib1 |
| Acot7 | 6.69E-24 | 0.3940111 | 0.637 | 0.515 | 1.29E-19 | cDC.Cd209 | Acot7 |
| Lats2 | 7.99E-24 | 0.69091386 | 0.432 | 0.301 | 1.55E-19 | cDC.Cd209 | Lats2 |
| Rnf130 | 1.22E-23 | 0.71745053 | 0.621 | 0.446 | 2.36E-19 | cDC.Cd209 | Rnf130 |
| Plbd1 | 1.39E-23 | 0.43015712 | 0.947 | 0.635 | 2.69E-19 | cDC.Cd209 | Plbd1 |
| Elmo2 | 2.45E-23 | 0.98565437 | 0.383 | 0.25 | 4.74E-19 | cDC.Cd209 | Elmo2 |
| Tmem198b | 3.75E-23 | 0.76836387 | 0.256 | 0.154 | 7.26E-19 | cDC.Cd209 | Tmem198b |
| Prdx6 | 3.92E-23 | 0.94526439 | 0.34 | 0.218 | 7.57E-19 | cDC.Cd209 | Prdx6 |
| Plod3 | 3.99E-23 | 0.89244013 | 0.352 | 0.232 | 7.71E-19 | cDC.Cd209 | Plod3 |
| Spty2d1 | 6.04E-23 | 0.28606724 | 0.55 | 0.489 | 1.17E-18 | cDC.Cd209 | Spty2d1 |
| Enah | 9.21E-23 | 0.67026717 | 0.356 | 0.253 | 1.78E-18 | cDC.Cd209 | Enah |
| Milr1 | 1.80E-22 | 1.23009031 | 0.267 | 0.148 | 3.47E-18 | cDC.Cd209 | Milr1 |
| Heatr1 | 1.81E-22 | 1.05678686 | 0.324 | 0.192 | 3.51E-18 | cDC.Cd209 | Heatr1 |
| Tagln2 | 1.89E-22 | 0.27686512 | 0.965 | 0.862 | 3.66E-18 | cDC.Cd209 | Tagln2 |
| Degs1 | 1.97E-22 | 0.60288661 | 0.515 | 0.395 | 3.80E-18 | cDC.Cd209 | Degs1 |
| Selenop | 2.01E-22 | 0.53334608 | 0.333 | 0.158 | 3.89E-18 | cDC.Cd209 | Selenop |
| Ptprc | 2.43E-22 | 0.41883133 | 0.847 | 0.671 | 4.70E-18 | cDC.Cd209 | Ptprc |
| Hnrnpa1 | 3.60E-22 | 0.40212247 | 0.779 | 0.611 | 6.97E-18 | cDC.Cd209 | Hnrnpa1 |
| Parp12 | 4.68E-22 | 0.66901828 | 0.289 | 0.202 | 9.05E-18 | cDC.Cd209 | Parp12 |
| Scimp | 5.15E-22 | 0.60809401 | 0.379 | 0.27 | 9.96E-18 | cDC.Cd209 | Scimp |
| P2ry6 | 6.37E-22 | 1.01496858 | 0.321 | 0.183 | 1.23E-17 | cDC.Cd209 | P2ry6 |
| Acly | 7.01E-22 | 0.41293525 | 0.756 | 0.636 | 1.36E-17 | cDC.Cd209 | Acly |
| Acat1 | 7.22E-22 | 0.61753298 | 0.399 | 0.273 | 1.40E-17 | cDC.Cd209 | Acat1 |
| Ifrd1 | 8.95E-22 | 0.28385248 | 0.777 | 0.644 | 1.73E-17 | cDC.Cd209 | Ifrd1 |
| Capza2 | 1.01E-21 | 0.46412922 | 0.609 | 0.478 | 1.95E-17 | cDC.Cd209 | Capza2 |
| Txnrd1 | 1.31E-21 | 0.39450943 | 0.752 | 0.59 | 2.53E-17 | cDC.Cd209 | Txnrd1 |
| Tmem65 | 1.53E-21 | 0.76090864 | 0.3 | 0.181 | 2.97E-17 | cDC.Cd209 | Tmem65 |
| Sf3b6 | 1.98E-21 | 0.51640321 | 0.581 | 0.443 | 3.83E-17 | cDC.Cd209 | Sf3b6 |
| Wdr46 | 2.67E-21 | 0.88825194 | 0.285 | 0.187 | 5.16E-17 | cDC.Cd209 | Wdr46 |
| Noc2l | 2.97E-21 | 0.64733652 | 0.498 | 0.354 | 5.75E-17 | cDC.Cd209 | Noc2l |
| Snx1 | 3.50E-21 | 0.45985641 | 0.556 | 0.451 | 6.78E-17 | cDC.Cd209 | Snx1 |
| Tspan14 | 4.03E-21 | 0.8529503 | 0.404 | 0.269 | 7.78E-17 | cDC.Cd209 | Tspan14 |
| Rexo2 | 4.74E-21 | 0.4427774 | 0.529 | 0.387 | 9.16E-17 | cDC.Cd209 | Rexo2 |
| Gadd45g | 5.12E-21 | 0.6055886 | 0.311 | 0.212 | 9.90E-17 | cDC.Cd209 | Gadd45g |
| Kcnab2 | 5.77E-21 | 0.7729217 | 0.381 | 0.258 | 1.12E-16 | cDC.Cd209 | Kcnab2 |
| Lrrfip1 | 5.86E-21 | 0.28593215 | 0.842 | 0.723 | 1.13E-16 | cDC.Cd209 | Lrrfip1 |
| Ankle2 | 6.32E-21 | 0.81002807 | 0.317 | 0.207 | 1.22E-16 | cDC.Cd209 | Ankle2 |
| Wdr83os | 6.71E-21 | 0.49815085 | 0.611 | 0.468 | 1.30E-16 | cDC.Cd209 | Wdr83os |
| Sgk3 | 9.02E-21 | 0.53832404 | 0.46 | 0.336 | 1.74E-16 | cDC.Cd209 | Sgk3 |
| App | 1.33E-20 | 0.30428658 | 0.856 | 0.715 | 2.58E-16 | cDC.Cd209 | App |
| Tent5a | 1.52E-20 | 0.75863302 | 0.364 | 0.234 | 2.94E-16 | cDC.Cd209 | Tent5a |
| Smad7 | 1.55E-20 | 0.52834236 | 0.482 | 0.369 | 3.00E-16 | cDC.Cd209 | Smad7 |
| Tmc6 | 2.04E-20 | 0.66587507 | 0.321 | 0.224 | 3.94E-16 | cDC.Cd209 | Tmc6 |
| Cers2 | 2.12E-20 | 0.41367281 | 0.49 | 0.375 | 4.10E-16 | cDC.Cd209 | Cers2 |
| Tm6sf1 | 3.03E-20 | 0.5139307 | 0.582 | 0.405 | 5.85E-16 | cDC.Cd209 | Tm6sf1 |
| Nop58 | 4.05E-20 | 0.84612916 | 0.4 | 0.24 | 7.83E-16 | cDC.Cd209 | Nop58 |
| Bzw2 | 5.99E-20 | 0.64124957 | 0.352 | 0.26 | 1.16E-15 | cDC.Cd209 | Bzw2 |
| Irf4 | 6.30E-20 | 0.62525209 | 0.416 | 0.284 | 1.22E-15 | cDC.Cd209 | Irf4 |
| Wsb1 | 6.92E-20 | 0.56035807 | 0.66 | 0.52 | 1.34E-15 | cDC.Cd209 | Wsb1 |
| Dgkd | 9.40E-20 | 0.6964911 | 0.405 | 0.295 | 1.82E-15 | cDC.Cd209 | Dgkd |
| Dock2 | 1.13E-19 | 0.29919945 | 0.706 | 0.571 | 2.19E-15 | cDC.Cd209 | Dock2 |
| Flot1 | 1.28E-19 | 0.52229372 | 0.562 | 0.418 | 2.48E-15 | cDC.Cd209 | Flot1 |
| Smdt1 | 1.36E-19 | 0.40368779 | 0.775 | 0.627 | 2.63E-15 | cDC.Cd209 | Smdt1 |
| Ndufa6 | 1.45E-19 | 0.31576252 | 0.748 | 0.63 | 2.80E-15 | cDC.Cd209 | Ndufa6 |
| Tor1aip1 | 1.54E-19 | 0.45836078 | 0.706 | 0.566 | 2.97E-15 | cDC.Cd209 | Tor1aip1 |
| Fam117b | 1.54E-19 | 0.34248396 | 0.665 | 0.555 | 2.97E-15 | cDC.Cd209 | Fam117b |
| Rhbdf2 | 1.64E-19 | 0.56681075 | 0.363 | 0.25 | 3.17E-15 | cDC.Cd209 | Rhbdf2 |
| Sorl1 | 2.00E-19 | 0.48167442 | 0.469 | 0.33 | 3.86E-15 | cDC.Cd209 | Sorl1 |
| Ppard | 2.56E-19 | 0.75350299 | 0.296 | 0.185 | 4.96E-15 | cDC.Cd209 | Ppard |
| Sde2 | 2.71E-19 | 0.80260205 | 0.355 | 0.238 | 5.25E-15 | cDC.Cd209 | Sde2 |

| | | | | | | | |
|---|---|---|---|---|---|---|---|
| Zfand5 | 3.16E-19 | 0.5049704 | 0.754 | 0.602 | 6.10E-15 | cDC.Cd209 | Zfand5 |
| Nadk | 3.48E-19 | 0.61514494 | 0.61 | 0.437 | 6.73E-15 | cDC.Cd209 | Nadk |
| D8Ertd738e | 4.39E-19 | 0.35383191 | 0.636 | 0.524 | 8.50E-15 | cDC.Cd209 | D8Ertd738e |
| H2-DMb2 | 5.38E-19 | 0.51749684 | 0.476 | 0.372 | 1.04E-14 | cDC.Cd209 | H2-DMb2 |
| Tmem147 | 5.68E-19 | 0.84348967 | 0.33 | 0.213 | 1.10E-14 | cDC.Cd209 | Tmem147 |
| Hip1 | 6.48E-19 | 0.86725106 | 0.369 | 0.238 | 1.25E-14 | cDC.Cd209 | Hip1 |
| Osgep | 6.84E-19 | 0.69996082 | 0.41 | 0.284 | 1.32E-14 | cDC.Cd209 | Osgep |
| Slc35b2 | 6.88E-19 | 0.33081398 | 0.667 | 0.543 | 1.33E-14 | cDC.Cd209 | Slc35b2 |
| Ccr1 | 9.21E-19 | 0.69714005 | 0.381 | 0.239 | 1.78E-14 | cDC.Cd209 | Ccr1 |
| Ddx21 | 1.01E-18 | 0.43941947 | 0.652 | 0.502 | 1.96E-14 | cDC.Cd209 | Ddx21 |
| Emg1 | 1.23E-18 | 0.55651073 | 0.561 | 0.412 | 2.38E-14 | cDC.Cd209 | Emg1 |
| Was | 1.46E-18 | 0.5144152 | 0.674 | 0.512 | 2.83E-14 | cDC.Cd209 | Was |
| Fes | 1.55E-18 | 0.34676123 | 0.766 | 0.632 | 2.99E-14 | cDC.Cd209 | Fes |
| Fcgr3 | 1.95E-18 | 0.52962696 | 0.425 | 0.363 | 3.77E-14 | cDC.Cd209 | Fcgr3 |
| Rwdd1 | 2.44E-18 | 0.82911309 | 0.268 | 0.189 | 4.73E-14 | cDC.Cd209 | Rwdd1 |
| Zfp703 | 2.76E-18 | 0.31072434 | 0.499 | 0.409 | 5.34E-14 | cDC.Cd209 | Zfp703 |
| Cerk | 3.51E-18 | 0.82133402 | 0.294 | 0.173 | 6.79E-14 | cDC.Cd209 | Cerk |
| Uhrf1bp1l | 3.62E-18 | 0.94346599 | 0.276 | 0.176 | 6.99E-14 | cDC.Cd209 | Uhrf1bp1l |
| Sh3pxd2b | 4.11E-18 | 0.31191847 | 0.689 | 0.586 | 7.94E-14 | cDC.Cd209 | Sh3pxd2b |
| Acaa2 | 4.58E-18 | 0.82258452 | 0.285 | 0.186 | 8.86E-14 | cDC.Cd209 | Acaa2 |
| Apex1 | 5.09E-18 | 0.91136964 | 0.314 | 0.2 | 9.83E-14 | cDC.Cd209 | Apex1 |
| Ube2d3 | 7.60E-18 | 0.50113569 | 0.572 | 0.454 | 1.47E-13 | cDC.Cd209 | Ube2d3 |
| Wdr26 | 8.10E-18 | 0.35490423 | 0.777 | 0.645 | 1.57E-13 | cDC.Cd209 | Wdr26 |
| Atp6ap1 | 9.16E-18 | 0.5162388 | 0.669 | 0.512 | 1.77E-13 | cDC.Cd209 | Atp6ap1 |
| Tacc1 | 1.38E-17 | 0.48273309 | 0.624 | 0.486 | 2.67E-13 | cDC.Cd209 | Tacc1 |
| Tubb6 | 1.79E-17 | 0.43587371 | 0.487 | 0.387 | 3.46E-13 | cDC.Cd209 | Tubb6 |
| Camk2g | 1.79E-17 | 0.67741106 | 0.413 | 0.299 | 3.47E-13 | cDC.Cd209 | Camk2g |
| Cd244a | 2.08E-17 | 0.71477298 | 0.54 | 0.355 | 4.02E-13 | cDC.Cd209 | Cd244a |
| Adam10 | 3.82E-17 | 0.66188204 | 0.428 | 0.303 | 7.39E-13 | cDC.Cd209 | Adam10 |
| Ascc3 | 4.20E-17 | 0.51722979 | 0.403 | 0.29 | 8.12E-13 | cDC.Cd209 | Ascc3 |
| Dapk1 | 4.34E-17 | 0.52902643 | 0.588 | 0.449 | 8.40E-13 | cDC.Cd209 | Dapk1 |
| Purb | 4.62E-17 | 0.26571674 | 0.786 | 0.703 | 8.94E-13 | cDC.Cd209 | Purb |
| Mycbp2 | 4.67E-17 | 0.3903864 | 0.6 | 0.477 | 9.03E-13 | cDC.Cd209 | Mycbp2 |
| Svil | 6.50E-17 | 0.69609687 | 0.254 | 0.168 | 1.26E-12 | cDC.Cd209 | Svil |
| Tram1 | 6.57E-17 | 0.53139053 | 0.649 | 0.502 | 1.27E-12 | cDC.Cd209 | Tram1 |
| Trpv2 | 7.74E-17 | 0.53720412 | 0.437 | 0.309 | 1.50E-12 | cDC.Cd209 | Trpv2 |
| Snrpd3 | 9.97E-17 | 0.46131019 | 0.603 | 0.463 | 1.93E-12 | cDC.Cd209 | Snrpd3 |
| Rbms1 | 1.09E-16 | 0.46013841 | 0.565 | 0.454 | 2.11E-12 | cDC.Cd209 | Rbms1 |
| Madd | 1.13E-16 | 0.51004021 | 0.471 | 0.365 | 2.19E-12 | cDC.Cd209 | Madd |
| Sdhb | 1.57E-16 | 0.48843867 | 0.559 | 0.417 | 3.04E-12 | cDC.Cd209 | Sdhb |
| Ifih1 | 1.65E-16 | 0.64024027 | 0.307 | 0.223 | 3.18E-12 | cDC.Cd209 | Ifih1 |
| Htr5a | 1.74E-16 | 0.75879551 | 0.321 | 0.215 | 3.36E-12 | cDC.Cd209 | Htr5a |
| Ttc7 | 2.14E-16 | 0.41040402 | 0.727 | 0.577 | 4.13E-12 | cDC.Cd209 | Ttc7 |
| Sdhc | 2.70E-16 | 0.47060103 | 0.61 | 0.472 | 5.22E-12 | cDC.Cd209 | Sdhc |
| Furin | 3.43E-16 | 0.44112592 | 0.553 | 0.478 | 6.64E-12 | cDC.Cd209 | Furin |
| Tmed5 | 4.43E-16 | 0.6380898 | 0.488 | 0.359 | 8.57E-12 | cDC.Cd209 | Tmed5 |
| Oxtc | 4.80E-16 | 0.51163566 | 0.474 | 0.346 | 9.29E-12 | cDC.Cd209 | Oxtc |
| Tet2 | 5.27E-16 | 0.64322957 | 0.511 | 0.385 | 1.02E-11 | cDC.Cd209 | Tet2 |
| Cysltr1 | 7.47E-16 | 0.69952705 | 0.275 | 0.185 | 1.44E-11 | cDC.Cd209 | Cysltr1 |
| Ctbp2 | 9.17E-16 | 0.59735036 | 0.374 | 0.265 | 1.77E-11 | cDC.Cd209 | Ctbp2 |
| Cd300lf | 1.22E-15 | 0.45722071 | 0.475 | 0.343 | 2.35E-11 | cDC.Cd209 | Cd300lf |
| Tmem109 | 1.69E-15 | 0.63519485 | 0.445 | 0.315 | 3.27E-11 | cDC.Cd209 | Tmem109 |
| Fuca2 | 1.81E-15 | 0.67294914 | 0.444 | 0.302 | 3.50E-11 | cDC.Cd209 | Fuca2 |
| Fam3c | 1.85E-15 | 0.56808883 | 0.319 | 0.219 | 3.58E-11 | cDC.Cd209 | Fam3c |
| Sap30 | 2.12E-15 | 0.94229723 | 0.296 | 0.188 | 4.10E-11 | cDC.Cd209 | Sap30 |
| Hipk1 | 2.95E-15 | 0.39915262 | 0.727 | 0.578 | 5.71E-11 | cDC.Cd209 | Hipk1 |
| Npm3 | 3.52E-15 | 0.79144434 | 0.289 | 0.194 | 6.80E-11 | cDC.Cd209 | Npm3 |
| Map7d1 | 3.54E-15 | 0.3236524 | 0.724 | 0.579 | 6.85E-11 | cDC.Cd209 | Map7d1 |
| Bag1 | 4.44E-15 | 0.41456038 | 0.547 | 0.424 | 8.58E-11 | cDC.Cd209 | Bag1 |
| Psenen | 4.96E-15 | 0.39412991 | 0.643 | 0.514 | 9.60E-11 | cDC.Cd209 | Psenen |
| Lrrc59 | 5.00E-15 | 0.35882697 | 0.653 | 0.55 | 9.67E-11 | cDC.Cd209 | Lrrc59 |
| Sh3bgrl | 5.48E-15 | 0.25683674 | 0.766 | 0.649 | 1.06E-10 | cDC.Cd209 | Sh3bgrl |
| Ech1 | 5.66E-15 | 0.45405695 | 0.453 | 0.341 | 1.09E-10 | cDC.Cd209 | Ech1 |
| Nsf | 7.10E-15 | 0.51260269 | 0.392 | 0.289 | 1.37E-10 | cDC.Cd209 | Nsf |
| Twf2 | 7.88E-15 | 0.41606026 | 0.643 | 0.501 | 1.52E-10 | cDC.Cd209 | Twf2 |
| Tep1 | 9.91E-15 | 0.52076014 | 0.515 | 0.376 | 1.92E-10 | cDC.Cd209 | Tep1 |
| Tmed3 | 1.26E-14 | 0.49626097 | 0.436 | 0.326 | 2.43E-10 | cDC.Cd209 | Tmed3 |
| Rxra | 1.28E-14 | 1.12699811 | 0.251 | 0.148 | 2.47E-10 | cDC.Cd209 | Rxra |
| Gmfg | 1.49E-14 | 0.69198208 | 0.374 | 0.244 | 2.88E-10 | cDC.Cd209 | Gmfg |
| Sap18b | 1.57E-14 | 0.42175048 | 0.33 | 0.287 | 3.04E-10 | cDC.Cd209 | Sap18b |
| Ciao2a | 1.78E-14 | 0.30242482 | 0.637 | 0.523 | 3.45E-10 | cDC.Cd209 | Ciao2a |
| Nab1 | 2.13E-14 | 0.45786356 | 0.623 | 0.499 | 4.12E-10 | cDC.Cd209 | Nab1 |
| Aldh9a1 | 2.55E-14 | 0.44757633 | 0.578 | 0.441 | 4.93E-10 | cDC.Cd209 | Aldh9a1 |
| Nsd2 | 2.98E-14 | 0.46788033 | 0.446 | 0.335 | 5.77E-10 | cDC.Cd209 | Nsd2 |
| Trmt112 | 3.04E-14 | 0.45401264 | 0.58 | 0.455 | 5.88E-10 | cDC.Cd209 | Trmt112 |
| Fndc3b | 3.11E-14 | 0.55882473 | 0.314 | 0.225 | 6.01E-10 | cDC.Cd209 | Fndc3b |
| Abhd17b | 3.51E-14 | 0.4485998 | 0.373 | 0.267 | 6.79E-10 | cDC.Cd209 | Abhd17b |
| Trak1 | 3.68E-14 | 0.50526865 | 0.493 | 0.379 | 7.11E-10 | cDC.Cd209 | Trak1 |
| Capns1 | 3.91E-14 | 0.38850414 | 0.615 | 0.492 | 7.55E-10 | cDC.Cd209 | Capns1 |
| Sft2d2 | 5.23E-14 | 0.52271921 | 0.478 | 0.369 | 1.01E-09 | cDC.Cd209 | Sft2d2 |
| Pgs1 | 5.60E-14 | 0.49715263 | 0.598 | 0.461 | 1.08E-09 | cDC.Cd209 | Pgs1 |
| Eif4a1 | 5.61E-14 | 0.41721851 | 0.69 | 0.565 | 1.08E-09 | cDC.Cd209 | Eif4a1 |
| Card9 | 5.74E-14 | 0.33238242 | 0.422 | 0.329 | 1.11E-09 | cDC.Cd209 | Card9 |
| Clic1 | 6.40E-14 | 0.46995248 | 0.755 | 0.587 | 1.24E-09 | cDC.Cd209 | Clic1 |
| Cdc42ep4 | 6.61E-14 | 0.71174872 | 0.252 | 0.173 | 1.28E-09 | cDC.Cd209 | Cdc42ep4 |
| Tmed10 | 1.06E-13 | 0.60510905 | 0.427 | 0.3 | 2.06E-09 | cDC.Cd209 | Tmed10 |
| Hcls1 | 1.35E-13 | 0.31365726 | 0.934 | 0.735 | 2.61E-09 | cDC.Cd209 | Hcls1 |
| Hs6st1 | 1.70E-13 | 0.57092347 | 0.501 | 0.392 | 3.28E-09 | cDC.Cd209 | Hs6st1 |
| Fli1 | 1.72E-13 | 0.72350165 | 0.412 | 0.287 | 3.32E-09 | cDC.Cd209 | Fli1 |
| Tsc22d2 | 2.01E-13 | 0.53750656 | 0.376 | 0.289 | 3.90E-09 | cDC.Cd209 | Tsc22d2 |
| Glud1 | 2.51E-13 | 0.28157848 | 0.813 | 0.698 | 4.85E-09 | cDC.Cd209 | Glud1 |
| Ap2a2 | 2.68E-13 | 0.56982114 | 0.557 | 0.426 | 5.19E-09 | cDC.Cd209 | Ap2a2 |
| Spag9 | 2.92E-13 | 0.27484399 | 0.793 | 0.686 | 5.65E-09 | cDC.Cd209 | Spag9 |
| Ube2f | 3.08E-13 | 0.33773861 | 0.64 | 0.534 | 5.96E-09 | cDC.Cd209 | Ube2f |
| Cltc | 3.32E-13 | 0.44949328 | 0.612 | 0.495 | 6.42E-09 | cDC.Cd209 | Cltc |
| Txn2 | 3.33E-13 | 0.55736124 | 0.487 | 0.363 | 6.45E-09 | cDC.Cd209 | Txn2 |
| Atrnl1 | 3.73E-13 | 0.32983849 | 0.538 | 0.408 | 7.22E-09 | cDC.Cd209 | Atrnl1 |
| Idh3a | 3.74E-13 | 0.68595435 | 0.368 | 0.252 | 7.22E-09 | cDC.Cd209 | Idh3a |
| Hsd17b12 | 3.79E-13 | 0.71388315 | 0.367 | 0.257 | 7.33E-09 | cDC.Cd209 | Hsd17b12 |
| Npc1 | 3.94E-13 | 0.28146537 | 0.395 | 0.26 | 7.62E-09 | cDC.Cd209 | Npc1 |
| Atp8b2 | 4.49E-13 | 0.58103971 | 0.342 | 0.248 | 8.69E-09 | cDC.Cd209 | Atp8b2 |
| Chmp4b | 5.02E-13 | 0.41268712 | 0.595 | 0.494 | 9.70E-09 | cDC.Cd209 | Chmp4b |
| Ube2a | 6.30E-13 | 0.57792518 | 0.357 | 0.255 | 1.22E-08 | cDC.Cd209 | Ube2a |
| Slc25a30 | 6.40E-13 | 0.61818544 | 0.301 | 0.219 | 1.24E-08 | cDC.Cd209 | Slc25a30 |
| Trim30a | 6.56E-13 | 0.68684205 | 0.351 | 0.255 | 1.27E-08 | cDC.Cd209 | Trim30a |
| Sla | 7.70E-13 | 0.54279712 | 0.7 | 0.467 | 1.49E-08 | cDC.Cd209 | Sla |
| Ncln | 8.91E-13 | 0.62919909 | 0.423 | 0.306 | 1.72E-08 | cDC.Cd209 | Ncln |
| Fbl | 1.06E-12 | 0.38894974 | 0.547 | 0.403 | 2.04E-08 | cDC.Cd209 | Fbl |
| Tcof1 | 1.12E-12 | 0.70914118 | 0.342 | 0.23 | 2.16E-08 | cDC.Cd209 | Tcof1 |
| Slc15a4 | 1.24E-12 | 0.31958907 | 0.432 | 0.344 | 2.40E-08 | cDC.Cd209 | Slc15a4 |
| Ppp2ca | 1.33E-12 | 0.39284369 | 0.751 | 0.637 | 2.58E-08 | cDC.Cd209 | Ppp2ca |
| Rasa3 | 1.38E-12 | 0.72892877 | 0.278 | 0.191 | 2.67E-08 | cDC.Cd209 | Rasa3 |
| Psmd8 | 1.50E-12 | 0.48634557 | 0.526 | 0.394 | 2.89E-08 | cDC.Cd209 | Psmd8 |
| Smim3 | 1.59E-12 | 0.38845914 | 0.443 | 0.33 | 3.07E-08 | cDC.Cd209 | Smim3 |
| Zfp516 | 1.75E-12 | 0.25091064 | 0.752 | 0.654 | 3.37E-08 | cDC.Cd209 | Zfp516 |
| B4galt3 | 1.98E-12 | 0.63057432 | 0.342 | 0.253 | 3.83E-08 | cDC.Cd209 | B4galt3 |
| Daxx | 1.98E-12 | 0.44105598 | 0.335 | 0.275 | 3.84E-08 | cDC.Cd209 | Daxx |
| Uqcrc1 | 2.28E-12 | 0.3369389 | 0.767 | 0.644 | 4.41E-08 | cDC.Cd209 | Uqcrc1 |
| Arpc4 | 2.40E-12 | 0.27958145 | 0.799 | 0.681 | 4.65E-08 | cDC.Cd209 | Arpc4 |
| Dynll1 | 3.85E-12 | 0.43141348 | 0.464 | 0.353 | 7.44E-08 | cDC.Cd209 | Dynll1 |
| Rap2c | 4.20E-12 | 0.45142591 | 0.308 | 0.218 | 8.11E-08 | cDC.Cd209 | Rap2c |
| Hdlbp | 4.92E-12 | 0.37090257 | 0.648 | 0.531 | 9.51E-08 | cDC.Cd209 | Hdlbp |
| Nufip1 | 6.07E-12 | 0.56058837 | 0.277 | 0.202 | 1.17E-07 | cDC.Cd209 | Nufip1 |
| Gch1 | 6.33E-12 | 0.6496288 | 0.427 | 0.299 | 1.22E-07 | cDC.Cd209 | Gch1 |
| Rnpep | 6.51E-12 | 0.45408869 | 0.54 | 0.426 | 1.26E-07 | cDC.Cd209 | Rnpep |
| Selenot | 6.54E-12 | 0.51381942 | 0.443 | 0.333 | 1.26E-07 | cDC.Cd209 | Selenot |
| Gpr183 | 7.14E-12 | 0.54558207 | 0.402 | 0.324 | 1.38E-07 | cDC.Cd209 | Gpr183 |
| Pdha1 | 7.46E-12 | 0.57879851 | 0.342 | 0.26 | 1.44E-07 | cDC.Cd209 | Pdha1 |
| Ssr4 | 7.60E-12 | 0.37929742 | 0.609 | 0.497 | 1.47E-07 | cDC.Cd209 | Ssr4 |
| Hspa1a | 7.85E-12 | 0.26719113 | 0.385 | 0.352 | 1.52E-07 | cDC.Cd209 | Hspa1a |
| Pcbp1 | 8.20E-12 | 0.31610148 | 0.748 | 0.622 | 1.58E-07 | cDC.Cd209 | Pcbp1 |
| Otulinl | 1.01E-11 | 0.56458874 | 0.684 | 0.484 | 1.95E-07 | cDC.Cd209 | Otulinl |
| Ezr | 1.08E-11 | 0.43203517 | 0.626 | 0.528 | 2.09E-07 | cDC.Cd209 | Ezr |
| Tmed9 | 1.11E-11 | 0.39761641 | 0.58 | 0.458 | 2.14E-07 | cDC.Cd209 | Tmed9 |
| Il6st | 1.12E-11 | 0.33315371 | 0.458 | 0.368 | 2.17E-07 | cDC.Cd209 | Il6st |
| Zdhhc5 | 1.27E-11 | 0.63713075 | 0.432 | 0.324 | 2.45E-07 | cDC.Cd209 | Zdhhc5 |
| Emc6 | 1.37E-11 | 0.49096159 | 0.405 | 0.301 | 2.65E-07 | cDC.Cd209 | Emc6 |
| Arid5a | 1.38E-11 | 0.4918228 | 0.385 | 0.292 | 2.67E-07 | cDC.Cd209 | Arid5a |
| Sypl | 1.57E-11 | 0.57539425 | 0.32 | 0.228 | 3.03E-07 | cDC.Cd209 | Sypl |
| Lilr4b | 1.99E-11 | 0.47399433 | 0.563 | 0.392 | 3.85E-07 | cDC.Cd209 | Lilr4b |
| Dlst | 2.02E-11 | 0.46571508 | 0.547 | 0.431 | 3.91E-07 | cDC.Cd209 | Dlst |
| Mta3 | 2.31E-11 | 0.44756646 | 0.495 | 0.39 | 4.46E-07 | cDC.Cd209 | Mta3 |
| Ptpn12 | 2.33E-11 | 0.41697965 | 0.542 | 0.426 | 4.51E-07 | cDC.Cd209 | Ptpn12 |
| Snx8 | 2.86E-11 | 0.49026439 | 0.386 | 0.301 | 5.53E-07 | cDC.Cd209 | Snx8 |
| Bzw1 | 3.33E-11 | 0.32025175 | 0.665 | 0.548 | 6.45E-07 | cDC.Cd209 | Bzw1 |
| Snap23 | 3.38E-11 | 0.5324497 | 0.379 | 0.291 | 6.53E-07 | cDC.Cd209 | Snap23 |
| Vamp3 | 4.12E-11 | 0.58098991 | 0.404 | 0.301 | 7.97E-07 | cDC.Cd209 | Vamp3 |
| Pak1ip1 | 4.45E-11 | 0.53214413 | 0.447 | 0.331 | 8.60E-07 | cDC.Cd209 | Pak1ip1 |
| Tmed7 | 5.21E-11 | 0.38989691 | 0.621 | 0.502 | 1.01E-06 | cDC.Cd209 | Tmed7 |
| Ipo4 | 6.84E-11 | 0.64382914 | 0.258 | 0.171 | 1.32E-06 | cDC.Cd209 | Ipo4 |
| Irf9 | 8.47E-11 | 0.48804897 | 0.403 | 0.318 | 1.64E-06 | cDC.Cd209 | Irf9 |
| Psmd11 | 8.51E-11 | 0.44952602 | 0.499 | 0.396 | 1.64E-06 | cDC.Cd209 | Psmd11 |
| Hspa5 | 8.59E-11 | 0.45451982 | 0.695 | 0.548 | 1.66E-06 | cDC.Cd209 | Hspa5 |
| Lifr | 8.66E-11 | 0.41990062 | 0.361 | 0.241 | 1.67E-06 | cDC.Cd209 | Lifr |
| Snrpf | 8.70E-11 | 0.42766465 | 0.639 | 0.5 | 1.68E-06 | cDC.Cd209 | Snrpf |
| Atp13a3 | 8.86E-11 | 0.43061707 | 0.637 | 0.524 | 1.71E-06 | cDC.Cd209 | Atp13a3 |
| Plk2 | 8.87E-11 | 0.4141395 | 0.256 | 0.182 | 1.72E-06 | cDC.Cd209 | Plk2 |
| Tspan31 | 9.34E-11 | 0.48174887 | 0.453 | 0.361 | 1.81E-06 | cDC.Cd209 | Tspan31 |
| Rab24 | 1.03E-10 | 0.46324166 | 0.479 | 0.374 | 1.98E-06 | cDC.Cd209 | Rab24 |
| Itgb2 | 1.14E-10 | 0.46864033 | 0.887 | 0.619 | 2.19E-06 | cDC.Cd209 | Itgb2 |
| Kras | 1.23E-10 | 0.36746779 | 0.503 | 0.413 | 2.38E-06 | cDC.Cd209 | Kras |
| Ddt | 1.29E-10 | 0.34429302 | 0.336 | 0.316 | 2.50E-06 | cDC.Cd209 | Ddt |
| Gprc5c | 1.30E-10 | 0.51719039 | 0.371 | 0.328 | 2.52E-06 | cDC.Cd209 | Gprc5c |
| Rfk | 1.34E-10 | 0.5455519 | 0.333 | 0.262 | 2.60E-06 | cDC.Cd209 | Rfk |
| Tcf4 | 1.46E-10 | 0.33199421 | 0.36 | 0.261 | 2.83E-06 | cDC.Cd209 | Tcf4 |
| Msl2 | 1.54E-10 | 0.37114393 | 0.495 | 0.414 | 2.98E-06 | cDC.Cd209 | Msl2 |
| Coro1c | 1.55E-10 | 0.51549168 | 0.53 | 0.405 | 3.01E-06 | cDC.Cd209 | Coro1c |
| Ifi209 | 1.58E-10 | 0.31018251 | 0.308 | 0.253 | 3.05E-06 | cDC.Cd209 | Ifi209 |
| Mdh2 | 1.67E-10 | 0.32696508 | 0.642 | 0.522 | 3.23E-06 | cDC.Cd209 | Mdh2 |
| Akr7a5 | 1.86E-10 | 0.53046308 | 0.316 | 0.229 | 3.59E-06 | cDC.Cd209 | Akr7a5 |
| Sptbn1 | 2.03E-10 | 0.63503888 | 0.324 | 0.226 | 3.92E-06 | cDC.Cd209 | Sptbn1 |
| Dad1 | 2.16E-10 | 0.28922653 | 0.739 | 0.597 | 4.17E-06 | cDC.Cd209 | Dad1 |
| Sec11a | 2.23E-10 | 0.37775336 | 0.476 | 0.382 | 4.32E-06 | cDC.Cd209 | Sec11a |
| Ppp2r5a | 2.29E-10 | 0.45047238 | 0.559 | 0.46 | 4.42E-06 | cDC.Cd209 | Ppp2r5a |
| Qsox1 | 2.35E-10 | 0.40521599 | 0.287 | 0.228 | 4.53E-06 | cDC.Cd209 | Qsox1 |
| Mcur1 | 2.72E-10 | 0.46484605 | 0.365 | 0.274 | 5.25E-06 | cDC.Cd209 | Mcur1 |
| Napsa | 3.15E-10 | 0.40988947 | 0.791 | 0.571 | 6.08E-06 | cDC.Cd209 | Napsa |
| Etfb | 3.16E-10 | 0.40236861 | 0.542 | 0.413 | 6.11E-06 | cDC.Cd209 | Etfb |
| Ltbr | 3.28E-10 | 0.6299955 | 0.42 | 0.313 | 6.35E-06 | cDC.Cd209 | Ltbr |
| Prrc2c | 3.41E-10 | 0.36040592 | 0.637 | 0.526 | 6.60E-06 | cDC.Cd209 | Prrc2c |
| Ndufb8 | 3.57E-10 | 0.30480131 | 0.622 | 0.507 | 6.90E-06 | cDC.Cd209 | Ndufb8 |
| Eif4e | 3.59E-10 | 0.41352049 | 0.496 | 0.385 | 6.94E-06 | cDC.Cd209 | Eif4e |
| Scarb1 | 3.90E-10 | 0.78454415 | 0.338 | 0.238 | 7.54E-06 | cDC.Cd209 | Scarb1 |
| Csnk1e | 4.27E-10 | 0.42534458 | 0.397 | 0.31 | 8.25E-06 | cDC.Cd209 | Csnk1e |
| Mapkapk2 | 4.61E-10 | 0.27538597 | 0.773 | 0.66 | 8.92E-06 | cDC.Cd209 | Mapkapk2 |
| Naa60 | 4.77E-10 | 0.58676346 | 0.398 | 0.303 | 9.22E-06 | cDC.Cd209 | Naa60 |
| Nfatc1 | 5.32E-10 | 0.50546828 | 0.394 | 0.311 | 1.03E-05 | cDC.Cd209 | Nfatc1 |
| Maml1 | 5.64E-10 | 0.65362006 | 0.334 | 0.249 | 1.09E-05 | cDC.Cd209 | Maml1 |
| Eif1ax | 5.73E-10 | 0.63106785 | 0.257 | 0.179 | 1.11E-05 | cDC.Cd209 | Eif1ax |
| Fubp1 | 5.98E-10 | 0.27271845 | 0.515 | 0.413 | 1.16E-05 | cDC.Cd209 | Fubp1 |
| Stom | 6.52E-10 | 0.79472381 | 0.252 | 0.186 | 1.26E-05 | cDC.Cd209 | Stom |
| Emc1 | 7.01E-10 | 0.60465849 | 0.314 | 0.223 | 1.36E-05 | cDC.Cd209 | Emc1 |
| Timm10b | 7.14E-10 | 0.41297686 | 0.446 | 0.354 | 1.38E-05 | cDC.Cd209 | Timm10b |
| Abr | 8.01E-10 | 0.40794375 | 0.62 | 0.486 | 1.55E-05 | cDC.Cd209 | Abr |
| Stx7 | 8.47E-10 | 0.29043645 | 0.768 | 0.644 | 1.64E-05 | cDC.Cd209 | Stx7 |
| Mfsd1 | 8.61E-10 | 0.38301513 | 0.456 | 0.336 | 1.66E-05 | cDC.Cd209 | Mfsd1 |
| Atp6v0d1 | 1.02E-09 | 0.29690516 | 0.575 | 0.492 | 1.98E-05 | cDC.Cd209 | Atp6v0d1 |
| Slc1a5 | 1.04E-09 | 0.46401377 | 0.528 | 0.401 | 2.01E-05 | cDC.Cd209 | Slc1a5 |
| Myd88 | 1.11E-09 | 0.38726247 | 0.744 | 0.558 | 2.14E-05 | cDC.Cd209 | Myd88 |
| Zswim4 | 1.12E-09 | 0.39370202 | 0.34 | 0.272 | 2.16E-05 | cDC.Cd209 | Zswim4 |
| Rraga | 1.13E-09 | 0.55048578 | 0.321 | 0.234 | 2.19E-05 | cDC.Cd209 | Rraga |
| Ddx10 | 1.39E-09 | 0.61453152 | 0.257 | 0.179 | 2.68E-05 | cDC.Cd209 | Ddx10 |
| Ranbp2 | 1.46E-09 | 0.29693236 | 0.649 | 0.53 | 2.82E-05 | cDC.Cd209 | Ranbp2 |
| Cct8 | 1.64E-09 | 0.52466215 | 0.451 | 0.345 | 3.17E-05 | cDC.Cd209 | Cct8 |
| Naa25 | 1.68E-09 | 0.72149246 | 0.256 | 0.179 | 3.24E-05 | cDC.Cd209 | Naa25 |
| Macf1 | 1.82E-09 | 0.5044832 | 0.515 | 0.383 | 3.53E-05 | cDC.Cd209 | Macf1 |
| Cpt1a | 1.87E-09 | 0.4901033 | 0.283 | 0.209 | 3.61E-05 | cDC.Cd209 | Cpt1a |
| Cuedc2 | 1.96E-09 | 0.32143098 | 0.44 | 0.347 | 3.79E-05 | cDC.Cd209 | Cuedc2 |
| Prpf31 | 1.98E-09 | 0.54734385 | 0.38 | 0.303 | 3.83E-05 | cDC.Cd209 | Prpf31 |
| Nmi | 2.01E-09 | 0.56775911 | 0.268 | 0.204 | 3.88E-05 | cDC.Cd209 | Nmi |
| Fam107b | 2.12E-09 | 0.33986195 | 0.53 | 0.393 | 4.10E-05 | cDC.Cd209 | Fam107b |
| N4bp1 | 2.37E-09 | 0.27640143 | 0.538 | 0.459 | 4.58E-05 | cDC.Cd209 | N4bp1 |
| Tmem243 | 2.77E-09 | 0.3271106 | 0.345 | 0.267 | 5.35E-05 | cDC.Cd209 | Tmem243 |
| Mybbp1a | 3.29E-09 | 0.40199896 | 0.441 | 0.343 | 6.37E-05 | cDC.Cd209 | Mybbp1a |
| Atxn3 | 3.34E-09 | 0.54840353 | 0.261 | 0.191 | 6.46E-05 | cDC.Cd209 | Atxn3 |
| Eif3k | 3.61E-09 | 0.26214329 | 0.815 | 0.687 | 6.97E-05 | cDC.Cd209 | Eif3k |
| Rnf13 | 3.67E-09 | 0.48557105 | 0.421 | 0.317 | 7.09E-05 | cDC.Cd209 | Rnf13 |
| Ffar2 | 3.67E-09 | 0.51062864 | 0.339 | 0.232 | 7.10E-05 | cDC.Cd209 | Ffar2 |
| Zfp609 | 3.95E-09 | 0.50730822 | 0.353 | 0.269 | 7.63E-05 | cDC.Cd209 | Zfp609 |
| Parp1 | 3.96E-09 | 0.56136403 | 0.427 | 0.325 | 7.66E-05 | cDC.Cd209 | Parp1 |
| Actr2 | 5.13E-09 | 0.25485124 | 0.8 | 0.685 | 9.92E-05 | cDC.Cd209 | Actr2 |
| Plscr3 | 5.24E-09 | 0.58817194 | 0.335 | 0.248 | 0.00010136 | cDC.Cd209 | Plscr3 |
| Ctnnd2 | 5.54E-09 | 0.44752576 | 0.44 | 0.285 | 0.00010712 | cDC.Cd209 | Ctnnd2 |
| Gde1 | 6.30E-09 | 0.52172777 | 0.252 | 0.188 | 0.00012179 | cDC.Cd209 | Gde1 |
| Tmem50a | 8.30E-09 | 0.46177916 | 0.584 | 0.46 | 0.00016042 | cDC.Cd209 | Tmem50a |
| Lsr | 8.70E-09 | 0.36323557 | 0.384 | 0.323 | 0.00016811 | cDC.Cd209 | Lsr |
| Ptpre | 8.87E-09 | 0.50902704 | 0.461 | 0.354 | 0.00017148 | cDC.Cd209 | Ptpre |
| Anp32e | 9.42E-09 | 0.38382083 | 0.704 | 0.531 | 0.00018209 | cDC.Cd209 | Anp32e |
| Cd72 | 9.95E-09 | 0.35065705 | 0.27 | 0.233 | 0.00019237 | cDC.Cd209 | Cd72 |
| Tfe3 | 1.03E-08 | 0.38853218 | 0.371 | 0.293 | 0.0001996 | cDC.Cd209 | Tfe3 |
| Lgals1 | 1.11E-08 | 0.55076189 | 0.259 | 0.18 | 0.00021538 | cDC.Cd209 | Lgals1 |
| Ndufs8 | 1.14E-08 | 0.36010611 | 0.516 | 0.413 | 0.00022028 | cDC.Cd209 | Ndufs8 |
| Atg7 | 1.18E-08 | 0.47383338 | 0.264 | 0.197 | 0.00022831 | cDC.Cd209 | Atg7 |
| Hmgcl | 1.20E-08 | 0.30003608 | 0.542 | 0.434 | 0.00023124 | cDC.Cd209 | Hmgcl |
| Rnf166 | 1.21E-08 | 0.52856618 | 0.417 | 0.325 | 0.00023467 | cDC.Cd209 | Rnf166 |
| Snx2 | 1.37E-08 | 0.60819345 | 0.515 | 0.375 | 0.00026402 | cDC.Cd209 | Snx2 |
| Tbc1d2b | 1.44E-08 | 0.59931808 | 0.301 | 0.228 | 0.00027759 | cDC.Cd209 | Tbc1d2b |
| Tob2 | 1.50E-08 | 0.30987673 | 0.583 | 0.482 | 0.00028945 | cDC.Cd209 | Tob2 |
| Arih1 | 1.60E-08 | 0.29057805 | 0.678 | 0.575 | 0.00030874 | cDC.Cd209 | Arih1 |
| Rock1 | 1.61E-08 | 0.41501569 | 0.478 | 0.401 | 0.00031157 | cDC.Cd209 | Rock1 |
| Mta2 | 1.74E-08 | 0.36898535 | 0.669 | 0.549 | 0.00033611 | cDC.Cd209 | Mta2 |
| Serp1 | 1.74E-08 | 0.49833815 | 0.293 | 0.207 | 0.00033625 | cDC.Cd209 | Serp1 |
| P2ry10 | 2.72E-08 | 0.2828919 | 0.643 | 0.56 | 0.00052684 | cDC.Cd209 | P2ry10 |
| Ncor2 | 2.98E-08 | 0.26056736 | 0.388 | 0.306 | 0.00057595 | cDC.Cd209 | Ncor2 |
| Hnrnpl | 3.11E-08 | 0.34375867 | 0.695 | 0.586 | 0.00060178 | cDC.Cd209 | Hnrnpl |
| Osbpl9 | 3.19E-08 | 0.26681775 | 0.579 | 0.481 | 0.00061761 | cDC.Cd209 | Osbpl9 |
| Eif2a | 3.32E-08 | 0.45635811 | 0.321 | 0.236 | 0.00064258 | cDC.Cd209 | Eif2a |
| Med15 | 3.36E-08 | 0.30624196 | 0.506 | 0.43 | 0.00065035 | cDC.Cd209 | Med15 |
| Matr3 | 3.65E-08 | 0.37096212 | 0.394 | 0.302 | 0.0007063 | cDC.Cd209 | Matr3 |
| Selenof | 3.79E-08 | 0.60680966 | 0.262 | 0.189 | 0.00073337 | cDC.Cd209 | Selenof |
| Surf4 | 3.94E-08 | 0.3687202 | 0.624 | 0.504 | 0.00076233 | cDC.Cd209 | Surf4 |
| Tmem173 | 4.39E-08 | 0.40391475 | 0.54 | 0.449 | 0.00084952 | cDC.Cd209 | Tmem173 |
| Tfip11 | 4.97E-08 | 0.27725185 | 0.333 | 0.286 | 0.00096003 | cDC.Cd209 | Tfip11 |
| Nab2 | 5.66E-08 | 0.4196537 | 0.252 | 0.206 | 0.00109442 | cDC.Cd209 | Nab2 |
| Nrd1 | 6.29E-08 | 0.30438368 | 0.555 | 0.461 | 0.00121531 | cDC.Cd209 | Nrd1 |
| Hyou1 | 7.21E-08 | 0.41659984 | 0.465 | 0.354 | 0.00139343 | cDC.Cd209 | Hyou1 |
| Nol6 | 7.26E-08 | 0.44850642 | 0.271 | 0.208 | 0.00140307 | cDC.Cd209 | Nol6 |
| Larp1 | 7.61E-08 | 0.27403174 | 0.584 | 0.49 | 0.0014705 | cDC.Cd209 | Larp1 |
| Pet100 | 7.69E-08 | 0.43181428 | 0.367 | 0.295 | 0.00148718 | cDC.Cd209 | Pet100 |
| Zmiz1 | 8.33E-08 | 0.29230319 | 0.616 | 0.516 | 0.00160962 | cDC.Cd209 | Zmiz1 |
| Copb1 | 8.41E-08 | 0.41758008 | 0.48 | 0.393 | 0.00162686 | cDC.Cd209 | Copb1 |
| Mex3c | 8.52E-08 | 0.35973448 | 0.369 | 0.298 | 0.00164682 | cDC.Cd209 | Mex3c |
| Cope | 9.02E-08 | 0.27090976 | 0.64 | 0.555 | 0.00174384 | cDC.Cd209 | Cope |
| Slc12a6 | 1.00E-07 | 0.46098696 | 0.284 | 0.206 | 0.00193347 | cDC.Cd209 | Slc12a6 |
| Abce1 | 1.06E-07 | 0.58064959 | 0.262 | 0.191 | 0.00204734 | cDC.Cd209 | Abce1 |
| Ube2g2 | 1.08E-07 | 0.4236265 | 0.466 | 0.37 | 0.00208415 | cDC.Cd209 | Ube2g2 |
| Atp5o | 1.14E-07 | 0.39695302 | 0.494 | 0.398 | 0.00219693 | cDC.Cd209 | Atp5o |
| Magoh | 1.23E-07 | 0.51796797 | 0.451 | 0.331 | 0.00237053 | cDC.Cd209 | Magoh |
| Bccip | 1.44E-07 | 0.56231705 | 0.37 | 0.271 | 0.00278429 | cDC.Cd209 | Bccip |
| Rassf1 | 1.45E-07 | 0.36928009 | 0.487 | 0.397 | 0.00280685 | cDC.Cd209 | Rassf1 |
| Rybp | 1.48E-07 | 0.60537182 | 0.254 | 0.184 | 0.00286756 | cDC.Cd209 | Rybp |
| Dna2 | 1.60E-07 | 0.64223731 | 0.337 | 0.243 | 0.00309493 | cDC.Cd209 | Dna2 |
| Dbi | 1.64E-07 | 0.42354642 | 0.379 | 0.285 | 0.00317851 | cDC.Cd209 | Dbi |
| Tmed2 | 1.76E-07 | 0.35971639 | 0.504 | 0.389 | 0.00339399 | cDC.Cd209 | Tmed2 |
| Zfp263 | 1.84E-07 | 0.35064421 | 0.319 | 0.265 | 0.00356487 | cDC.Cd209 | Zfp263 |
| Utp18 | 1.92E-07 | 0.4923172 | 0.301 | 0.229 | 0.00371948 | cDC.Cd209 | Utp18 |
| Rreb1 | 1.94E-07 | 0.48638139 | 0.319 | 0.238 | 0.00375576 | cDC.Cd209 | Rreb1 |
| Pea15a | 2.07E-07 | 0.36296143 | 0.432 | 0.365 | 0.00399899 | cDC.Cd209 | Pea15a |
| Prkag1 | 2.12E-07 | 0.52274467 | 0.275 | 0.202 | 0.00410522 | cDC.Cd209 | Prkag1 |
| Chn2 | 2.44E-07 | 0.61209054 | 0.269 | 0.191 | 0.00470848 | cDC.Cd209 | Chn2 |
| Tbxas1 | 2.85E-07 | 0.49738828 | 0.396 | 0.31 | 0.00551248 | cDC.Cd209 | Tbxas1 |
| Slc35b1 | 2.88E-07 | 0.51949205 | 0.271 | 0.201 | 0.0055697 | cDC.Cd209 | Slc35b1 |
| Dyrk2 | 2.97E-07 | 0.40269827 | 0.298 | 0.237 | 0.00574503 | cDC.Cd209 | Dyrk2 |
| Lbr | 2.99E-07 | 0.68765186 | 0.464 | 0.319 | 0.00578104 | cDC.Cd209 | Lbr |
| Htt | 3.29E-07 | 0.42755522 | 0.37 | 0.294 | 0.00635339 | cDC.Cd209 | Htt |
| Plekha3 | 3.31E-07 | 0.30377977 | 0.276 | 0.232 | 0.00639289 | cDC.Cd209 | Plekha3 |
| Atic | 3.63E-07 | 0.53200489 | 0.352 | 0.279 | 0.00702723 | cDC.Cd209 | Atic |
| Btk | 3.64E-07 | 0.36873801 | 0.289 | 0.223 | 0.00704668 | cDC.Cd209 | Btk |
| Arl1 | 3.77E-07 | 0.42755774 | 0.295 | 0.245 | 0.00729623 | cDC.Cd209 | Arl1 |
| Sarnp | 3.79E-07 | 0.56200438 | 0.349 | 0.265 | 0.00733047 | cDC.Cd209 | Sarnp |
| Lsm7 | 4.05E-07 | 0.41496109 | 0.416 | 0.324 | 0.00782876 | cDC.Cd209 | Lsm7 |
| Mydgf | 4.33E-07 | 0.37357524 | 0.511 | 0.389 | 0.00836366 | cDC.Cd209 | Mydgf |
| Rpn1 | 4.35E-07 | 0.55917145 | 0.611 | 0.433 | 0.00840911 | cDC.Cd209 | Rpn1 |
| Atp5mpl | 4.38E-07 | 0.3491762 | 0.446 | 0.356 | 0.0084669 | cDC.Cd209 | Atp5mpl |
| Jaml | 4.69E-07 | 0.31619853 | 0.696 | 0.532 | 0.00905805 | cDC.Cd209 | Jaml |
| Slc39a11 | 4.82E-07 | 0.50818893 | 0.277 | 0.211 | 0.00931744 | cDC.Cd209 | Slc39a11 |
| Lsm12 | 5.02E-07 | 0.2629754 | 0.46 | 0.372 | 0.00971042 | cDC.Cd209 | Lsm12 |
| Naa15 | 5.48E-07 | 0.44803317 | 0.431 | 0.331 | 0.01059711 | cDC.Cd209 | Naa15 |
| Med29 | 5.54E-07 | 0.47510123 | 0.306 | 0.247 | 0.01070434 | cDC.Cd209 | Med29 |
| Slc43a2 | 5.65E-07 | 0.68842841 | 0.319 | 0.21 | 0.01092333 | cDC.Cd209 | Slc43a2 |
| 2-Mar | 5.69E-07 | 0.37818533 | 0.455 | 0.367 | 0.01099275 | cDC.Cd209 | 2-Mar |
| Rab1a | 5.73E-07 | 0.33854585 | 0.707 | 0.587 | 0.01107688 | cDC.Cd209 | Rab1a |
| Usp38 | 5.77E-07 | 0.34679521 | 0.329 | 0.277 | 0.01116522 | cDC.Cd209 | Usp38 |
| Hnrnpk | 6.69E-07 | 0.32926367 | 0.734 | 0.615 | 0.01293869 | cDC.Cd209 | Hnrnpk |
| Eef1g | 6.89E-07 | 0.36742639 | 0.616 | 0.505 | 0.01333013 | cDC.Cd209 | Eef1g |
| Gart | 7.48E-07 | 0.55301625 | 0.31 | 0.22 | 0.01446218 | cDC.Cd209 | Gart |
| Hk1 | 7.53E-07 | 0.37621482 | 0.473 | 0.394 | 0.01455162 | cDC.Cd209 | Hk1 |
| Mapk6 | 7.69E-07 | 0.45259741 | 0.337 | 0.267 | 0.01486201 | cDC.Cd209 | Mapk6 |
| Senp6 | 7.80E-07 | 0.40965178 | 0.358 | 0.289 | 0.01508667 | cDC.Cd209 | Senp6 |
| Tln1 | 7.85E-07 | 0.25629797 | 0.894 | 0.767 | 0.01517302 | cDC.Cd209 | Tln1 |
| Zfp106 | 8.80E-07 | 0.35793037 | 0.637 | 0.522 | 0.01701139 | cDC.Cd209 | Zfp106 |
| Zbtb11 | 1.05E-06 | 0.53015547 | 0.347 | 0.271 | 0.02027953 | cDC.Cd209 | Zbtb11 |
| Dcaf12 | 1.09E-06 | 0.45185868 | 0.439 | 0.35 | 0.021041 | cDC.Cd209 | Dcaf12 |
| Tmem256 | 1.10E-06 | 0.39813056 | 0.568 | 0.461 | 0.02130741 | cDC.Cd209 | Tmem256 |
| Ube2v2 | 1.11E-06 | 0.46202825 | 0.269 | 0.222 | 0.02149652 | cDC.Cd209 | Ube2v2 |
| Lman2 | 1.40E-06 | 0.33769704 | 0.644 | 0.518 | 0.02711123 | cDC.Cd209 | Lman2 |
| Trak2 | 1.45E-06 | 0.4880226 | 0.294 | 0.233 | 0.02794866 | cDC.Cd209 | Trak2 |
| Pxk | 1.51E-06 | 0.3796492 | 0.299 | 0.218 | 0.02912144 | cDC.Cd209 | Pxk |
| Snrpb2 | 1.57E-06 | 0.38326746 | 0.363 | 0.288 | 0.03029047 | cDC.Cd209 | Snrpb2 |
| Arf2 | 1.65E-06 | 0.49185665 | 0.34 | 0.278 | 0.03190281 | cDC.Cd209 | Arf2 |
| Calr | 1.71E-06 | 0.3587867 | 0.708 | 0.569 | 0.0330781 | cDC.Cd209 | Calr |
| Lyz2 | 1.76E-06 | 0.62189272 | 0.395 | 0.249 | 0.03409419 | cDC.Cd209 | Lyz2 |
| Reps1 | 1.82E-06 | 0.40999589 | 0.351 | 0.285 | 0.03523884 | cDC.Cd209 | Reps1 |
| Serinc1 | 2.65E-06 | 0.34049856 | 0.451 | 0.38 | 0.05116285 | cDC.Cd209 | Serinc1 |
| Ppp3ca | 2.68E-06 | 0.58055423 | 0.344 | 0.258 | 0.05185382 | cDC.Cd209 | Ppp3ca |
| Arcn1 | 2.76E-06 | 0.2527066 | 0.602 | 0.514 | 0.05337553 | cDC.Cd209 | Arcn1 |
| Metap2 | 3.04E-06 | 0.3803107 | 0.442 | 0.365 | 0.05879548 | cDC.Cd209 | Metap2 |
| Lrpap1 | 3.07E-06 | 0.34407596 | 0.346 | 0.291 | 0.05936511 | cDC.Cd209 | Lrpap1 |
| Wdr18 | 3.12E-06 | 0.46062099 | 0.285 | 0.218 | 0.06033869 | cDC.Cd209 | Wdr18 |
| Samm50 | 3.21E-06 | 0.36675661 | 0.416 | 0.336 | 0.06202258 | cDC.Cd209 | Samm50 |
| Cep170 | 3.26E-06 | 0.26597227 | 0.501 | 0.434 | 0.06302156 | cDC.Cd209 | Cep170 |
| Sogal | 3.66E-06 | 0.26750978 | 0.326 | 0.255 | 0.07069012 | cDC.Cd209 | Sogal |
| Cast | 3.78E-06 | 0.34313479 | 0.458 | 0.389 | 0.07316376 | cDC.Cd209 | Cast |
| Pgm2 | 3.95E-06 | 0.38322388 | 0.331 | 0.268 | 0.0764343 | cDC.Cd209 | Pgm2 |
| Iqgap2 | 4.07E-06 | 0.66048372 | 0.423 | 0.32 | 0.07877975 | cDC.Cd209 | Iqgap2 |
| Tomm20 | 4.25E-06 | 0.32151607 | 0.375 | 0.303 | 0.08207999 | cDC.Cd209 | Tomm20 |
| Srsf3 | 4.50E-06 | 0.43151628 | 0.375 | 0.305 | 0.08704084 | cDC.Cd209 | Srsf3 |
| Ebp | 4.74E-06 | 0.46023507 | 0.282 | 0.218 | 0.09156857 | cDC.Cd209 | Ebp |
| Farsb | 4.91E-06 | 0.54222405 | 0.254 | 0.2 | 0.09494281 | cDC.Cd209 | Farsb |
| Sar1b | 5.24E-06 | 0.43582461 | 0.349 | 0.287 | 0.10124258 | cDC.Cd209 | Sar1b |
| Cops6 | 5.28E-06 | 0.26445685 | 0.54 | 0.444 | 0.10208588 | cDC.Cd209 | Cops6 |
| Extl3 | 5.32E-06 | 0.38785991 | 0.34 | 0.259 | 0.10288684 | cDC.Cd209 | Extl3 |
| Tmem179b | 5.34E-06 | 0.39538178 | 0.344 | 0.272 | 0.10328047 | cDC.Cd209 | Tmem179b |
| Emc10 | 5.66E-06 | 0.41521605 | 0.515 | 0.409 | 0.10937454 | cDC.Cd209 | Emc10 |
| Ndufc2 | 5.71E-06 | 0.29410307 | 0.6 | 0.489 | 0.11030381 | cDC.Cd209 | Ndufc2 |
| Zfp385a | 5.82E-06 | 0.41122489 | 0.482 | 0.371 | 0.11247638 | cDC.Cd209 | Zfp385a |
| Cmas | 5.89E-06 | 0.43086931 | 0.385 | 0.323 | 0.11389101 | cDC.Cd209 | Cmas |
| Gnb2 | 6.02E-06 | 0.33614932 | 0.768 | 0.636 | 0.1163916 | cDC.Cd209 | Gnb2 |
| Azin1 | 6.03E-06 | 0.46082642 | 0.356 | 0.283 | 0.11651784 | cDC.Cd209 | Azin1 |
| Atp7a | 6.13E-06 | 0.42112316 | 0.357 | 0.284 | 0.11859032 | cDC.Cd209 | Atp7a |
| Ubtd1 | 6.28E-06 | 0.39121421 | 0.315 | 0.241 | 0.12140232 | cDC.Cd209 | Ubtd1 |
| Acap2 | 6.33E-06 | 0.309817 | 0.415 | 0.367 | 0.12244536 | cDC.Cd209 | Acap2 |
| Glg1 | 7.18E-06 | 0.33955412 | 0.496 | 0.407 | 0.1387459 | cDC.Cd209 | Glg1 |
| Bscl2 | 7.67E-06 | 0.39720764 | 0.413 | 0.336 | 0.14825804 | cDC.Cd209 | Bscl2 |
| Vcl | 8.19E-06 | 0.29412366 | 0.319 | 0.266 | 0.15841291 | cDC.Cd209 | Vcl |
| Sppl2a | 8.77E-06 | 0.30414581 | 0.589 | 0.487 | 0.16964126 | cDC.Cd209 | Sppl2a |
| Pa2g4 | 1.00E-05 | 0.57962484 | 0.313 | 0.232 | 0.19326201 | cDC.Cd209 | Pa2g4 |
| Usp22 | 1.07E-05 | 0.44188563 | 0.386 | 0.305 | 0.20742826 | cDC.Cd209 | Usp22 |
| Ube2d2a | 1.08E-05 | 0.2679966 | 0.671 | 0.587 | 0.20793018 | cDC.Cd209 | Ube2d2a |
| Nagpa | 1.08E-05 | 0.57020162 | 0.31 | 0.244 | 0.2087517 | cDC.Cd209 | Nagpa |
| Impdh2 | 1.10E-05 | 0.2592375 | 0.456 | 0.363 | 0.21333398 | cDC.Cd209 | Impdh2 |
| Grpel1 | 1.13E-05 | 0.33688756 | 0.424 | 0.351 | 0.21762989 | cDC.Cd209 | Grpel1 |
| Smarca5 | 1.16E-05 | 0.40334647 | 0.31 | 0.256 | 0.22413683 | cDC.Cd209 | Smarca5 |
| Tet3 | 1.30E-05 | 0.3130072 | 0.458 | 0.382 | 0.25073501 | cDC.Cd209 | Tet3 |
| Psma7 | 1.45E-05 | 0.32751378 | 0.418 | 0.338 | 0.28108797 | cDC.Cd209 | Psma7 |
| Pqbp1 | 1.57E-05 | 0.31743707 | 0.362 | 0.303 | 0.30447571 | cDC.Cd209 | Pqbp1 |
| Tmem88 | 1.64E-05 | 0.35803178 | 0.303 | 0.232 | 0.31745226 | cDC.Cd209 | Tmem88 |
| Trp53 | 1.66E-05 | 0.37814674 | 0.458 | 0.375 | 0.3200228 | cDC.Cd209 | Trp53 |
| Rheb | 1.72E-05 | 0.28268903 | 0.589 | 0.498 | 0.33232701 | cDC.Cd209 | Rheb |
| Hspa4 | 1.77E-05 | 0.39741536 | 0.407 | 0.334 | 0.34137049 | cDC.Cd209 | Hspa4 |
| Sptlc2 | 1.87E-05 | 0.44585769 | 0.338 | 0.281 | 0.36163969 | cDC.Cd209 | Sptlc2 |
| Vav1 | 2.04E-05 | 0.43540086 | 0.582 | 0.47 | 0.39440535 | cDC.Cd209 | Vav1 |
| Tnfaip8l2 | 2.16E-05 | 0.47950217 | 0.391 | 0.287 | 0.41757916 | cDC.Cd209 | Tnfaip8l2 |
| Adar | 2.16E-05 | 0.3206751 | 0.404 | 0.343 | 0.41810755 | cDC.Cd209 | Adar |
| Chtop | 2.22E-05 | 0.29416882 | 0.568 | 0.482 | 0.42830482 | cDC.Cd209 | Chtop |
| Nfatc2 | 2.23E-05 | 0.37326422 | 0.505 | 0.374 | 0.43210719 | cDC.Cd209 | Nfatc2 |
| Gtf2h5 | 2.35E-05 | 0.34277673 | 0.414 | 0.344 | 0.45482598 | cDC.Cd209 | Gtf2h5 |
| Txnip | 2.44E-05 | 0.4168016 | 0.335 | 0.287 | 0.47132089 | cDC.Cd209 | Txnip |
| Rnf38 | 2.47E-05 | 0.34342574 | 0.399 | 0.33 | 0.47796377 | cDC.Cd209 | Rnf38 |
| Mthfs | 2.67E-05 | 0.26680239 | 0.352 | 0.283 | 0.51707902 | cDC.Cd209 | Mthfs |
| Sigmar1 | 2.76E-05 | 0.52063818 | 0.307 | 0.242 | 0.53265617 | cDC.Cd209 | Sigmar1 |
| Canx | 3.32E-05 | 0.31536361 | 0.57 | 0.48 | 0.64167048 | cDC.Cd209 | Canx |
| Map11 | 3.39E-05 | 0.42938468 | 0.309 | 0.246 | 0.65462538 | cDC.Cd209 | Map11 |
| Nop10 | 3.51E-05 | 0.43559589 | 0.383 | 0.305 | 0.67854269 | cDC.Cd209 | Nop10 |
| Tmem184b | 3.53E-05 | 0.28847638 | 0.425 | 0.357 | 0.68173994 | cDC.Cd209 | Tmem184b |
| Hnrnpdl | 3.86E-05 | 0.37021242 | 0.358 | 0.295 | 0.74695841 | cDC.Cd209 | Hnrnpdl |
| Mtmr3 | 3.90E-05 | 0.42940779 | 0.314 | 0.253 | 0.75369275 | cDC.Cd209 | Mtmr3 |
| Ube2j1 | 3.98E-05 | 0.3166433 | 0.285 | 0.24 | 0.76946257 | cDC.Cd209 | Ube2j1 |
| Psmb2 | 4.23E-05 | 0.33304877 | 0.593 | 0.506 | 0.81808061 | cDC.Cd209 | Psmb2 |
| Anxa1 | 4.27E-05 | 0.5251201 | 0.489 | 0.386 | 0.82522579 | cDC.Cd209 | Anxa1 |
| Pno1 | 4.37E-05 | 0.37144133 | 0.275 | 0.214 | 0.84440248 | cDC.Cd209 | Pno1 |
| Ndufv1 | 4.43E-05 | 0.37010024 | 0.444 | 0.361 | 0.85631818 | cDC.Cd209 | Ndufv1 |
| Ndufs3 | 4.53E-05 | 0.31614214 | 0.348 | 0.288 | 0.87526095 | cDC.Cd209 | Ndufs3 |
| Tcerg1 | 4.64E-05 | 0.36558513 | 0.328 | 0.27 | 0.8979504 | cDC.Cd209 | Tcerg1 |
| Gstp1 | 4.86E-05 | 0.34255869 | 0.368 | 0.289 | 0.94052108 | cDC.Cd209 | Gstp1 |
| Dhcr24 | 4.97E-05 | 0.48432731 | 0.316 | 0.26 | 0.96048892 | cDC.Cd209 | Dhcr24 |
| Gtf2f2 | 5.34E-05 | 0.27592243 | 0.356 | 0.293 | 1 | cDC.Cd209 | Gtf2f2 |
| Akr1b3 | 5.52E-05 | 0.36719597 | 0.399 | 0.313 | 1 | cDC.Cd209 | Akr1b3 |
| Arhgap9 | 5.53E-05 | 0.29259905 | 0.601 | 0.488 | 1 | cDC.Cd209 | Arhgap9 |
| Mapk1 | 5.57E-05 | 0.26408005 | 0.621 | 0.537 | 1 | cDC.Cd209 | Mapk1 |
| Ssbp4 | 5.60E-05 | 0.45021332 | 0.474 | 0.345 | 1 | cDC.Cd209 | Ssbp4 |
| Ids | 5.68E-05 | 0.27998211 | 0.448 | 0.368 | 1 | cDC.Cd209 | Ids |
| Gda | 5.79E-05 | 0.29184204 | 0.371 | 0.312 | 1 | cDC.Cd209 | Gda |
| Bcap31 | 5.92E-05 | 0.37093807 | 0.458 | 0.387 | 1 | cDC.Cd209 | Bcap31 |
| Sec61g | 6.71E-05 | 0.26502055 | 0.389 | 0.329 | 1 | cDC.Cd209 | Sec61g |
| Lmo2 | 7.34E-05 | 0.3779768 | 0.453 | 0.351 | 1 | cDC.Cd209 | Lmo2 |
| Exoc6 | 7.53E-05 | 0.426739 | 0.265 | 0.211 | 1 | cDC.Cd209 | Exoc6 |
| Homer1 | 7.59E-05 | 0.25310718 | 0.37 | 0.307 | 1 | cDC.Cd209 | Homer1 |
| Ppm1g | 7.90E-05 | 0.26349522 | 0.538 | 0.458 | 1 | cDC.Cd209 | Ppm1g |
| Slc35e1 | 7.92E-05 | 0.40507753 | 0.469 | 0.406 | 1 | cDC.Cd209 | Slc35e1 |
| Ppig | 7.93E-05 | 0.43857489 | 0.328 | 0.26 | 1 | cDC.Cd209 | Ppig |
| Cd300c2 | 8.85E-05 | 0.34668028 | 0.264 | 0.214 | 1 | cDC.Cd209 | Cd300c2 |
| Flot2 | 9.52E-05 | 0.27631416 | 0.324 | 0.268 | 1 | cDC.Cd209 | Flot2 |
| Gpr108 | 9.77E-05 | 0.42110822 | 0.32 | 0.251 | 1 | cDC.Cd209 | Gpr108 |
| Amz1 | 0.00010042 | 0.4611 | 0.471 | 0.341 | 1 | cDC.Cd209 | Amz1 |
| Setd1b | 0.00010261 | 0.32662978 | 0.344 | 0.281 | 1 | cDC.Cd209 | Setd1b |
| Ythdf1 | 0.00010276 | 0.49520047 | 0.323 | 0.257 | 1 | cDC.Cd209 | Ythdf1 |
| Gba | 0.00010736 | 0.41486292 | 0.365 | 0.321 | 1 | cDC.Cd209 | Gba |
| Cog5 | 0.00010882 | 0.25087587 | 0.326 | 0.275 | 1 | cDC.Cd209 | Cog5 |
| Gng10 | 0.00011223 | 0.2523732 | 0.837 | 0.618 | 1 | cDC.Cd209 | Gng10 |
| Utp6 | 0.00011421 | 0.35846632 | 0.28 | 0.227 | 1 | cDC.Cd209 | Utp6 |
| Rnf141 | 0.00011462 | 0.48924957 | 0.275 | 0.218 | 1 | cDC.Cd209 | Rnf141 |
| Morc3 | 0.00011471 | 0.45193729 | 0.35 | 0.278 | 1 | cDC.Cd209 | Morc3 |
| Slc35a4 | 0.00011515 | 0.30994223 | 0.411 | 0.336 | 1 | cDC.Cd209 | Slc35a4 |
| Reep3 | 0.0001253 | 0.38182171 | 0.425 | 0.343 | 1 | cDC.Cd209 | Reep3 |
| Pira2 | 0.00012845 | 0.42365731 | 0.263 | 0.21 | 1 | cDC.Cd209 | Pira2 |
| Lman1 | 0.00013265 | 0.36075514 | 0.315 | 0.254 | 1 | cDC.Cd209 | Lman1 |
| Tbkbp1 | 0.00014711 | 0.27811193 | 0.257 | 0.195 | 1 | cDC.Cd209 | Tbkbp1 |
| Mob3b | 0.00015094 | 0.46596331 | 0.317 | 0.248 | 1 | cDC.Cd209 | Mob3b |
| Rap1gds1 | 0.00015715 | 0.2877635 | 0.473 | 0.387 | 1 | cDC.Cd209 | Rap1gds1 |
| Pigt | 0.00015954 | 0.42622624 | 0.421 | 0.348 | 1 | cDC.Cd209 | Pigt |
| Cdc73 | 0.00016298 | 0.30579534 | 0.297 | 0.244 | 1 | cDC.Cd209 | Cdc73 |
| Donson | 0.0001662 | 0.35657793 | 0.342 | 0.281 | 1 | cDC.Cd209 | Donson |
| Mafk | 0.00017308 | 0.34927241 | 0.299 | 0.243 | 1 | cDC.Cd209 | Mafk |
| Slamf9 | 0.00017476 | 0.38809157 | 0.267 | 0.208 | 1 | cDC.Cd209 | Slamf9 |
| Srf | 0.00017646 | 0.30293474 | 0.54 | 0.468 | 1 | cDC.Cd209 | Srf |
| Lrmp | 0.00018896 | 0.42655108 | 0.279 | 0.21 | 1 | cDC.Cd209 | Lrmp |
| Atp1b3 | 0.0001909 | 0.27153988 | 0.497 | 0.423 | 1 | cDC.Cd209 | Atp1b3 |
| Pspc1 | 0.00020092 | 0.33605909 | 0.35 | 0.275 | 1 | cDC.Cd209 | Pspc1 |
| Nup153 | 0.00021193 | 0.41858007 | 0.529 | 0.434 | 1 | cDC.Cd209 | Nup153 |
| Irak4 | 0.00021694 | 0.39470058 | 0.436 | 0.365 | 1 | cDC.Cd209 | Irak4 |
| Fkbp2 | 0.00022807 | 0.39254334 | 0.453 | 0.352 | 1 | cDC.Cd209 | Fkbp2 |
| Setdb1 | 0.00023713 | 0.33511346 | 0.28 | 0.232 | 1 | cDC.Cd209 | Setdb1 |
| Naa50 | 0.00025112 | 0.31217707 | 0.519 | 0.41 | 1 | cDC.Cd209 | Naa50 |
| Bop1 | 0.00025564 | 0.31283369 | 0.261 | 0.209 | 1 | cDC.Cd209 | Bop1 |
| Timm17a | 0.00027723 | 0.36591383 | 0.313 | 0.252 | 1 | cDC.Cd209 | Timm17a |
| Gripap1 | 0.00028638 | 0.32597796 | 0.315 | 0.255 | 1 | cDC.Cd209 | Gripap1 |
| Sec23b | 0.00028713 | 0.36273687 | 0.318 | 0.249 | 1 | cDC.Cd209 | Sec23b |
| Ncbp1 | 0.00029132 | 0.40492042 | 0.267 | 0.219 | 1 | cDC.Cd209 | Ncbp1 |
| Ddx24 | 0.00029138 | 0.33908471 | 0.428 | 0.365 | 1 | cDC.Cd209 | Ddx24 |
| Rhoh | 0.00029695 | 0.30792457 | 0.277 | 0.217 | 1 | cDC.Cd209 | Rhoh |
| Pik3cd | 0.00032074 | 0.31603883 | 0.672 | 0.522 | 1 | cDC.Cd209 | Pik3cd |
| Elp5 | 0.00032588 | 0.326501 | 0.397 | 0.347 | 1 | cDC.Cd209 | Elp5 |
| Tle3 | 0.00033518 | 0.36565652 | 0.706 | 0.615 | 1 | cDC.Cd209 | Tle3 |
| Rexf1 | 0.00034387 | 0.34403601 | 0.253 | 0.21 | 1 | cDC.Cd209 | Rexf1 |
| Fbxo34 | 0.00035633 | 0.41184775 | 0.257 | 0.204 | 1 | cDC.Cd209 | Fbxo34 |
| Dhx38 | 0.00036186 | 0.25842216 | 0.499 | 0.432 | 1 | cDC.Cd209 | Dhx38 |
| Usb1 | 0.00036756 | 0.378467 | 0.252 | 0.208 | 1 | cDC.Cd209 | Usb1 |
| Atp5g1 | 0.00037539 | 0.31374902 | 0.373 | 0.301 | 1 | cDC.Cd209 | Atp5g1 |
| Slc25a5 | 0.00038706 | 0.39477429 | 0.43 | 0.339 | 1 | cDC.Cd209 | Slc25a5 |
| Notch2 | 0.00042616 | 0.3986703 | 0.559 | 0.458 | 1 | cDC.Cd209 | Notch2 |
| Mef2d | 0.00043615 | 0.33744485 | 0.606 | 0.5 | 1 | cDC.Cd209 | Mef2d |
| Psmd6 | 0.00043897 | 0.36702631 | 0.402 | 0.337 | 1 | cDC.Cd209 | Psmd6 |
| Rictor | 0.00047506 | 0.40887183 | 0.265 | 0.222 | 1 | cDC.Cd209 | Rictor |
| Dhx40 | 0.00048091 | 0.57043856 | 0.312 | 0.212 | 1 | cDC.Cd209 | Dhx40 |
| Ankfy1 | 0.00050939 | 0.38954419 | 0.324 | 0.258 | 1 | cDC.Cd209 | Ankfy1 |
| Ncstn | 0.00053085 | 0.30407792 | 0.504 | 0.42 | 1 | cDC.Cd209 | Ncstn |
| Rnd3 | 0.00053501 | 0.36425082 | 0.284 | 0.244 | 1 | cDC.Cd209 | Rnd3 |
| Dars | 0.00055929 | 0.33132311 | 0.495 | 0.397 | 1 | cDC.Cd209 | Dars |
| Dpm1 | 0.00056949 | 0.29556111 | 0.252 | 0.213 | 1 | cDC.Cd209 | Dpm1 |
| G6pdx | 0.00057641 | 0.49217717 | 0.289 | 0.225 | 1 | cDC.Cd209 | G6pdx |
| Utp11 | 0.0005783 | 0.46267531 | 0.334 | 0.273 | 1 | cDC.Cd209 | Utp11 |
| Nme1 | 0.00060816 | 0.39938007 | 0.336 | 0.272 | 1 | cDC.Cd209 | Nme1 |
| Bmpr2 | 0.00062641 | 0.38202193 | 0.308 | 0.25 | 1 | cDC.Cd209 | Bmpr2 |
| Mri1 | 0.00062932 | 0.49931627 | 0.261 | 0.206 | 1 | cDC.Cd209 | Mri1 |
| Phf8 | 0.00063369 | 0.33469323 | 0.29 | 0.237 | 1 | cDC.Cd209 | Phf8 |
| Dyrk1a | 0.00063433 | 0.35958184 | 0.416 | 0.342 | 1 | cDC.Cd209 | Dyrk1a |
| Morf4l2 | 0.00063561 | 0.29743388 | 0.351 | 0.296 | 1 | cDC.Cd209 | Morf4l2 |
| Bud23 | 0.00065179 | 0.25056751 | 0.273 | 0.219 | 1 | cDC.Cd209 | Bud23 |
| Stag2 | 0.00068769 | 0.31660891 | 0.382 | 0.31 | 1 | cDC.Cd209 | Stag2 |
| Uggt1 | 0.00069019 | 0.2762728 | 0.568 | 0.485 | 1 | cDC.Cd209 | Uggt1 |
| Gnl3 | 0.0006949 | 0.40035798 | 0.259 | 0.201 | 1 | cDC.Cd209 | Gnl3 |
| Vamp4 | 0.00070149 | 0.31327937 | 0.337 | 0.275 | 1 | cDC.Cd209 | Vamp4 |
| Ccdc47 | 0.00072126 | 0.64919248 | 0.271 | 0.219 | 1 | cDC.Cd209 | Ccdc47 |
| Kdelr2 | 0.00086767 | 0.37458386 | 0.364 | 0.297 | 1 | cDC.Cd209 | Kdelr2 |
| Pds5a | 0.00087051 | 0.36659359 | 0.464 | 0.384 | 1 | cDC.Cd209 | Pds5a |
| Tmco1 | 0.00092632 | 0.34602606 | 0.345 | 0.271 | 1 | cDC.Cd209 | Tmco1 |
| Adgrg5 | 0.0009575 | 0.29630115 | 0.621 | 0.476 | 1 | cDC.Cd209 | Adgrg5 |

| | | | | | | |
|---|---|---|---|---|---|---|
| Chst12 | 0.00102402 | 0.38707044 | 0.262 | 0.215 | 1 | cDC.Cd209 Chst12 |
| Sec24a | 0.00106705 | 0.38081442 | 0.256 | 0.204 | 1 | cDC.Cd209 Sec24a |
| Pip4k2b | 0.00107515 | 0.30526575 | 0.35 | 0.292 | 1 | cDC.Cd209 Pip4k2b |
| Rpn2 | 0.00121903 | 0.4110764 | 0.483 | 0.382 | 1 | cDC.Cd209 Rpn2 |
| Hdac4 | 0.00125217 | 0.42885321 | 0.265 | 0.216 | 1 | cDC.Cd209 Hdac4 |
| Bcl10 | 0.00127234 | 0.35446196 | 0.36 | 0.311 | 1 | cDC.Cd209 Bcl10 |
| Sf3b3 | 0.00133295 | 0.29650762 | 0.486 | 0.401 | 1 | cDC.Cd209 Sf3b3 |
| Cnot6l | 0.00141185 | 0.29668307 | 0.502 | 0.42 | 1 | cDC.Cd209 Cnot6l |
| Strn3 | 0.00141621 | 0.27486845 | 0.321 | 0.264 | 1 | cDC.Cd209 Strn3 |
| Camk2d | 0.00148948 | 0.30241753 | 0.259 | 0.204 | 1 | cDC.Cd209 Camk2d |
| Eif3l | 0.00150615 | 0.27956139 | 0.615 | 0.51 | 1 | cDC.Cd209 Eif3l |
| Ndufaf8 | 0.00156691 | 0.41263147 | 0.296 | 0.239 | 1 | cDC.Cd209 Ndufaf8 |
| Gla | 0.00156786 | 0.32412304 | 0.253 | 0.212 | 1 | cDC.Cd209 Gla |
| Vapa | 0.00173453 | 0.33819771 | 0.251 | 0.211 | 1 | cDC.Cd209 Vapa |
| Stard3 | 0.00186264 | 0.45575729 | 0.269 | 0.219 | 1 | cDC.Cd209 Stard3 |
| Dnajb11 | 0.00205434 | 0.2784473 | 0.517 | 0.452 | 1 | cDC.Cd209 Dnajb11 |
| Plcb2 | 0.00233524 | 0.32994494 | 0.481 | 0.358 | 1 | cDC.Cd209 Plcb2 |
| Mcfd2 | 0.00238718 | 0.40476413 | 0.31 | 0.254 | 1 | cDC.Cd209 Mcfd2 |
| Api5 | 0.0024328 | 0.28776726 | 0.515 | 0.438 | 1 | cDC.Cd209 Api5 |
| Cept1 | 0.00243468 | 0.27544463 | 0.307 | 0.275 | 1 | cDC.Cd209 Cept1 |
| Ncl | 0.00247464 | 0.27321469 | 0.582 | 0.485 | 1 | cDC.Cd209 Ncl |
| Pdia4 | 0.0024842 | 0.37728118 | 0.46 | 0.376 | 1 | cDC.Cd209 Pdia4 |
| Pprc1 | 0.00264027 | 0.41761887 | 0.252 | 0.201 | 1 | cDC.Cd209 Pprc1 |
| Comt | 0.00264233 | 0.25583023 | 0.337 | 0.291 | 1 | cDC.Cd209 Comt |
| Sar1a | 0.00265232 | 0.3005512 | 0.52 | 0.462 | 1 | cDC.Cd209 Sar1a |
| Isyna1 | 0.00274539 | 0.2887599 | 0.506 | 0.402 | 1 | cDC.Cd209 Isyna1 |
| Lypla1 | 0.00285878 | 0.336931 | 0.259 | 0.222 | 1 | cDC.Cd209 Lypla1 |
| C1d | 0.00302237 | 0.375415 | 0.252 | 0.207 | 1 | cDC.Cd209 C1d |
| Ppp2r5d | 0.00312888 | 0.28757266 | 0.261 | 0.223 | 1 | cDC.Cd209 Ppp2r5d |
| Arid1a | 0.00335038 | 0.30616504 | 0.441 | 0.386 | 1 | cDC.Cd209 Arid1a |
| Arhgap27 | 0.00337251 | 0.25721173 | 0.326 | 0.275 | 1 | cDC.Cd209 Arhgap27 |
| Tbrg4 | 0.00354972 | 0.27914877 | 0.349 | 0.299 | 1 | cDC.Cd209 Tbrg4 |
| Cdipt | 0.00362275 | 0.26565528 | 0.331 | 0.279 | 1 | cDC.Cd209 Cdipt |
| Stoml2 | 0.00373011 | 0.35836553 | 0.275 | 0.232 | 1 | cDC.Cd209 Stoml2 |
| Psmd3 | 0.00373279 | 0.2696245 | 0.389 | 0.334 | 1 | cDC.Cd209 Psmd3 |
| Hars | 0.00398223 | 0.43433904 | 0.325 | 0.265 | 1 | cDC.Cd209 Hars |
| Cltb | 0.00422027 | 0.29090356 | 0.492 | 0.404 | 1 | cDC.Cd209 Cltb |
| Smim7 | 0.00434042 | 0.30772136 | 0.302 | 0.251 | 1 | cDC.Cd209 Smim7 |
| Plk3 | 0.00436355 | 0.35589975 | 0.271 | 0.22 | 1 | cDC.Cd209 Plk3 |
| Tmem59 | 0.00447763 | 0.25416262 | 0.453 | 0.391 | 1 | cDC.Cd209 Tmem59 |
| Atp2a2 | 0.00473272 | 0.30606508 | 0.572 | 0.47 | 1 | cDC.Cd209 Atp2a2 |
| Hint1 | 0.00518691 | 0.32342836 | 0.342 | 0.285 | 1 | cDC.Cd209 Hint1 |
| Pigs | 0.00558351 | 0.26606682 | 0.349 | 0.302 | 1 | cDC.Cd209 Pigs |
| Arnt | 0.00580926 | 0.26986928 | 0.416 | 0.357 | 1 | cDC.Cd209 Arnt |
| Sp3 | 0.00592511 | 0.29123371 | 0.33 | 0.281 | 1 | cDC.Cd209 Sp3 |
| Gatad2a | 0.0061777 | 0.25355514 | 0.542 | 0.463 | 1 | cDC.Cd209 Gatad2a |
| Rabl6 | 0.00643286 | 0.27164224 | 0.324 | 0.284 | 1 | cDC.Cd209 Rabl6 |
| Ibtk | 0.00649039 | 0.29262344 | 0.255 | 0.208 | 1 | cDC.Cd209 Ibtk |
| Lmnb1 | 0.0065188 | 0.32379258 | 0.714 | 0.575 | 1 | cDC.Cd209 Lmnb1 |
| Mgat2 | 0.00669837 | 0.28818144 | 0.448 | 0.386 | 1 | cDC.Cd209 Mgat2 |
| Anapc16 | 0.00677196 | 0.34492463 | 0.326 | 0.28 | 1 | cDC.Cd209 Anapc16 |
| Ncbp3 | 0.00685784 | 0.25793748 | 0.306 | 0.26 | 1 | cDC.Cd209 Ncbp3 |
| Crebbp | 0.00700998 | 0.25993451 | 0.439 | 0.38 | 1 | cDC.Cd209 Crebbp |
| Ubald2 | 0.00703767 | 0.26455478 | 0.671 | 0.565 | 1 | cDC.Cd209 Ubald2 |
| Nucb1 | 0.00720976 | 0.30194543 | 0.533 | 0.438 | 1 | cDC.Cd209 Nucb1 |
| Sf3b4 | 0.00750467 | 0.32793598 | 0.338 | 0.262 | 1 | cDC.Cd209 Sf3b4 |
| Smndc1 | 0.00761791 | 0.25977321 | 0.343 | 0.289 | 1 | cDC.Cd209 Smndc1 |
| Hk2 | 0.00771449 | 0.315247 | 0.564 | 0.444 | 1 | cDC.Cd209 Hk2 |
| Vps36 | 0.00777912 | 0.25411949 | 0.289 | 0.25 | 1 | cDC.Cd209 Vps36 |
| Cnih4 | 0.00786802 | 0.31896809 | 0.279 | 0.218 | 1 | cDC.Cd209 Cnih4 |
| Adcy7 | 0.00806758 | 0.27886813 | 0.48 | 0.387 | 1 | cDC.Cd209 Adcy7 |
| Dnaja1 | 0.00838667 | 0.32228787 | 0.487 | 0.413 | 1 | cDC.Cd209 Dnaja1 |
| Tomm40 | 0.00856173 | 0.27138102 | 0.377 | 0.321 | 1 | cDC.Cd209 Tomm40 |
| Fkbp15 | 0.0087984 | 0.29039364 | 0.458 | 0.41 | 1 | cDC.Cd209 Fkbp15 |
| Prep | 0.00881603 | 0.26914491 | 0.315 | 0.261 | 1 | cDC.Cd209 Prep |
| Tob1 | 0.00901 | 0.31810872 | 0.337 | 0.295 | 1 | cDC.Cd209 Tob1 |
| Cpsf3 | 0.00932282 | 0.32811303 | 0.328 | 0.278 | 1 | cDC.Cd209 Cpsf3 |
| Dpm3 | 0.00952332 | 0.28688261 | 0.271 | 0.224 | 1 | cDC.Cd209 Dpm3 |
| Fscn1 | 0 | 5.60400552 | 0.884 | 0.066 | 0 | cDC.migratory Fscn1 |
| Cacnb3 | 0 | 5.25429003 | 0.921 | 0.108 | 0 | cDC.migratory Cacnb3 |
| Ccl5 | 0 | 4.94644332 | 0.793 | 0.077 | 0 | cDC.migratory Ccl5 |
| Mreg | 0 | 4.67031338 | 0.803 | 0.088 | 0 | cDC.migratory Mreg |
| Ccr7 | 0 | 3.56285579 | 0.917 | 0.233 | 0 | cDC.migratory Ccr7 |
| Adcy6 | 0 | 4.98670924 | 0.702 | 0.067 | 0 | cDC.migratory Adcy6 |
| Anxa3 | 0 | 4.14930804 | 0.842 | 0.226 | 0 | cDC.migratory Anxa3 |
| Ccl22 | 0 | 3.71964127 | 0.811 | 0.22 | 0 | cDC.migratory Ccl22 |
| Il4i1 | 0 | 2.64036753 | 0.708 | 0.143 | 0 | cDC.migratory Il4i1 |
| Strip2 | 0 | 4.31055589 | 0.654 | 0.101 | 0 | cDC.migratory Strip2 |
| Mmp25 | 0 | 3.50033464 | 0.807 | 0.257 | 0 | cDC.migratory Mmp25 |
| Slc4a8 | 0 | 5.69666616 | 0.581 | 0.041 | 0 | cDC.migratory Slc4a8 |
| Tnfrsf9 | 0 | 2.80066507 | 0.674 | 0.145 | 0 | cDC.migratory Tnfrsf9 |
| Rabgap1l | 0 | 3.2624731 | 0.756 | 0.23 | 0 | cDC.migratory Rabgap1l |
| Adora2a | 0 | 3.81873104 | 0.54 | 0.075 | 0 | cDC.migratory Adora2a |
| Il15ra | 0 | 4.97021566 | 0.529 | 0.07 | 0 | cDC.migratory Il15ra |
| Ly75 | 0 | 3.5004456 | 0.699 | 0.242 | 0 | cDC.migratory Ly75 |
| Cd63 | 0 | 2.55004902 | 0.738 | 0.283 | 0 | cDC.migratory Cd63 |
| Syn3 | 0 | 5.18695909 | 0.494 | 0.043 | 0 | cDC.migratory Syn3 |
| Cblb | 0 | 3.47342466 | 0.722 | 0.276 | 0 | cDC.migratory Cblb |
| Serpinb9 | 0 | 3.38300134 | 0.693 | 0.261 | 0 | cDC.migratory Serpinb9 |
| Stat4 | 0 | 3.26966901 | 0.778 | 0.349 | 0 | cDC.migratory Stat4 |
| Poglut1 | 0 | 3.64734817 | 0.566 | 0.139 | 0 | cDC.migratory Poglut1 |
| Cd200 | 0 | 4.22929928 | 0.489 | 0.068 | 0 | cDC.migratory Cd200 |
| Serpinb6b | 0 | 3.80823191 | 0.487 | 0.082 | 0 | cDC.migratory Serpinb6b |
| Marcksl1 | 0 | 2.44931853 | 0.838 | 0.44 | 0 | cDC.migratory Marcksl1 |
| H2-Q6 | 0 | 3.05977164 | 0.839 | 0.443 | 0 | cDC.migratory H2-Q6 |
| Samsn1 | 0 | 2.85690817 | 0.818 | 0.431 | 0 | cDC.migratory Samsn1 |
| Fam49a | 0 | 2.69913914 | 0.675 | 0.291 | 0 | cDC.migratory Fam49a |
| Ktn1 | 0 | 3.11646609 | 0.694 | 0.31 | 0 | cDC.migratory Ktn1 |
| Gypc | 0 | 3.95103349 | 0.458 | 0.077 | 0 | cDC.migratory Gypc |
| Foxp4 | 0 | 2.53316292 | 0.627 | 0.248 | 0 | cDC.migratory Foxp4 |
| Arl5c | 0 | 2.51558035 | 0.621 | 0.244 | 0 | cDC.migratory Arl5c |
| Socs2 | 0 | 3.1338374 | 0.572 | 0.195 | 0 | cDC.migratory Socs2 |
| Relb | 0 | 2.68263585 | 0.824 | 0.452 | 0 | cDC.migratory Relb |
| Rap2b | 0 | 3.0190443 | 0.667 | 0.296 | 0 | cDC.migratory Rap2b |
| Cmc2 | 0 | 3.57839519 | 0.501 | 0.13 | 0 | cDC.migratory Cmc2 |
| Eno3 | 0 | 4.0481723 | 0.456 | 0.091 | 0 | cDC.migratory Eno3 |
| Zmynd15 | 0 | 3.65892611 | 0.482 | 0.122 | 0 | cDC.migratory Zmynd15 |
| Itga4 | 0 | 2.68516984 | 0.771 | 0.412 | 0 | cDC.migratory Itga4 |
| Plxnc1 | 0 | 3.07088504 | 0.535 | 0.179 | 0 | cDC.migratory Plxnc1 |
| Pcgf5 | 0 | 2.87870563 | 0.645 | 0.29 | 0 | cDC.migratory Pcgf5 |
| Net1 | 0 | 3.21261533 | 0.759 | 0.408 | 0 | cDC.migratory Net1 |
| Mex3b | 0 | 3.87069434 | 0.428 | 0.084 | 0 | cDC.migratory Mex3b |
| Ssh1 | 0 | 2.79811332 | 0.714 | 0.373 | 0 | cDC.migratory Ssh1 |
| Pfkfb3 | 0 | 2.92571991 | 0.664 | 0.328 | 0 | cDC.migratory Pfkfb3 |
| Rhoc | 0 | 2.98997589 | 0.506 | 0.172 | 0 | cDC.migratory Rhoc |
| Mab21l3 | 0 | 4.48219351 | 0.369 | 0.037 | 0 | cDC.migratory Mab21l3 |
| Adgrg6 | 0 | 3.02107158 | 0.508 | 0.176 | 0 | cDC.migratory Adgrg6 |
| Fabp5 | 0 | 2.5710764 | 0.792 | 0.462 | 0 | cDC.migratory Fabp5 |
| Il2ra | 0 | 4.49269545 | 0.379 | 0.051 | 0 | cDC.migratory Il2ra |
| Scin | 0 | 4.11522711 | 0.385 | 0.076 | 0 | cDC.migratory Scin |
| H2-Q7 | 0 | 2.43705574 | 0.675 | 0.368 | 0 | cDC.migratory H2-Q7 |
| Sema7a | 0 | 2.30213364 | 0.724 | 0.42 | 0 | cDC.migratory Sema7a |
| Aebp2 | 0 | 2.34574871 | 0.778 | 0.475 | 0 | cDC.migratory Aebp2 |
| Ramp3 | 0 | 2.00151499 | 0.814 | 0.516 | 0 | cDC.migratory Ramp3 |
| Car2 | 0 | 3.72147521 | 0.4 | 0.103 | 0 | cDC.migratory Car2 |
| Tbc1d8 | 0 | 1.87154365 | 0.805 | 0.519 | 0 | cDC.migratory Tbc1d8 |
| Uap1 | 0 | 2.95846562 | 0.708 | 0.422 | 0 | cDC.migratory Uap1 |
| Tmem39a | 0 | 2.78612295 | 0.509 | 0.228 | 0 | cDC.migratory Tmem39a |
| Id2 | 0 | 1.8709997 | 0.861 | 0.586 | 0 | cDC.migratory Id2 |
| Map4k4 | 0 | 2.35698906 | 0.822 | 0.564 | 0 | cDC.migratory Map4k4 |
| Tmem123 | 0 | 2.75536252 | 0.84 | 0.585 | 0 | cDC.migratory Tmem123 |
| Glipr2 | 0 | 2.6645048 | 0.649 | 0.401 | 0 | cDC.migratory Glipr2 |
| Swap70 | 0 | 2.43631171 | 0.693 | 0.448 | 0 | cDC.migratory Swap70 |
| Rassf2 | 0 | 2.3533137 | 0.775 | 0.532 | 0 | cDC.migratory Rassf2 |
| Cd40 | 0 | 2.40711951 | 0.568 | 0.351 | 0 | cDC.migratory Cd40 |
| Helz2 | 0 | 2.46276774 | 0.514 | 0.301 | 0 | cDC.migratory Helz2 |
| Gyg | 0 | 2.49723567 | 0.711 | 0.516 | 0 | cDC.migratory Gyg |
| Tspan3 | 0 | 2.43797474 | 0.506 | 0.314 | 0 | cDC.migratory Tspan3 |
| Traf1 | 0 | 2.13403084 | 0.887 | 0.702 | 0 | cDC.migratory Traf1 |
| Rogdi | 0 | 1.97057654 | 0.873 | 0.688 | 0 | cDC.migratory Rogdi |
| Marcks | 0 | 1.97147593 | 0.903 | 0.757 | 0 | cDC.migratory Marcks |
| Csrp1 | 0 | 1.96370061 | 0.652 | 0.518 | 0 | cDC.migratory Csrp1 |
| Myo1g | 0 | 2.09078905 | 0.906 | 0.803 | 0 | cDC.migratory Myo1g |
| Clic4 | 0 | 1.81819396 | 0.802 | 0.7 | 0 | cDC.migratory Clic4 |
| Nfkb2 | 0 | 1.6011765 | 0.844 | 0.759 | 0 | cDC.migratory Nfkb2 |
| Stat3 | 0 | 1.73328008 | 0.804 | 0.741 | 0 | cDC.migratory Stat3 |
| Clu | 0 | 1.54520289 | 0.747 | 0.719 | 0 | cDC.migratory Clu |
| Spp1 | 0 | 1.20392126 | 0.583 | 0.608 | 0 | cDC.migratory Spp1 |
| Crip1 | 0 | 1.39219263 | 0.907 | 0.913 | 0 | cDC.migratory Crip1 |
| Calm1 | 0 | 1.7115229 | 0.888 | 0.885 | 0 | cDC.migratory Calm1 |
| Epsti1 | 1.26E-302 | 2.30286824 | 0.594 | 0.498 | 2.44E-298 | cDC.migratory Epsti1 |
| Adap1 | 1.65E-299 | 2.57437103 | 0.669 | 0.344 | 3.18E-295 | cDC.migratory Adap1 |
| Ogfrl1 | 8.52E-291 | 2.73423888 | 0.571 | 0.269 | 1.65E-286 | cDC.migratory Ogfrl1 |
| Txndc17 | 1.85E-286 | 2.12456228 | 0.66 | 0.576 | 3.57E-282 | cDC.migratory Txndc17 |
| Nudt17 | 7.45E-286 | 5.40063519 | 0.421 | 0.024 | 1.44E-281 | cDC.migratory Nudt17 |
| Tbc1d4 | 2.82E-283 | 3.31321777 | 0.469 | 0.116 | 5.46E-279 | cDC.migratory Tbc1d4 |
| Slc22a23 | 1.39E-280 | 4.42981836 | 0.395 | 0.048 | 2.70E-276 | cDC.migratory Slc22a23 |
| Rab8b | 3.70E-279 | 1.68295755 | 0.815 | 0.738 | 7.16E-275 | cDC.migratory Rab8b |
| Tmcc3 | 1.11E-271 | 3.45093618 | 0.474 | 0.111 | 2.14E-267 | cDC.migratory Tmcc3 |
| Rftn1 | 4.41E-270 | 2.13128147 | 0.655 | 0.328 | 8.53E-266 | cDC.migratory Rftn1 |
| Rcsd1 | 5.07E-264 | 2.27855199 | 0.522 | 0.337 | 9.80E-260 | cDC.migratory Rcsd1 |
| Gga2 | 1.03E-259 | 2.8917287 | 0.461 | 0.152 | 1.99E-255 | cDC.migratory Gga2 |
| Arhgef40 | 5.55E-253 | 2.21140256 | 0.66 | 0.446 | 1.07E-248 | cDC.migratory Arhgef40 |
| Rassf3 | 1.47E-245 | 2.49366114 | 0.573 | 0.263 | 2.84E-241 | cDC.migratory Rassf3 |
| Ankrd33b | 6.36E-245 | 4.49520934 | 0.429 | 0.054 | 1.23E-240 | cDC.migratory Ankrd33b |
| Lad1 | 1.70E-240 | 4.15660741 | 0.282 | 0.039 | 3.29E-236 | cDC.migratory Lad1 |
| Atp11a | 5.55E-237 | 2.79147221 | 0.587 | 0.254 | 1.07E-232 | cDC.migratory Atp11a |
| Vwa5a | 4.98E-236 | 1.99175875 | 0.605 | 0.477 | 9.63E-232 | cDC.migratory Vwa5a |
| Klf6 | 1.36E-234 | 1.46776706 | 0.804 | 0.699 | 2.62E-230 | cDC.migratory Klf6 |
| Kif21b | 1.80E-225 | 2.30523524 | 0.635 | 0.346 | 3.47E-221 | cDC.migratory Kif21b |
| Specc1 | 4.88E-223 | 3.09067017 | 0.403 | 0.134 | 9.43E-219 | cDC.migratory Specc1 |
| Mfhas1 | 4.47E-222 | 2.34372891 | 0.579 | 0.247 | 8.64E-218 | cDC.migratory Mfhas1 |
| Synpo2 | 1.07E-219 | 5.61566991 | 0.317 | 0.016 | 2.06E-215 | cDC.migratory Synpo2 |
| Ikzf4 | 3.62E-218 | 3.74126428 | 0.374 | 0.064 | 7.00E-214 | cDC.migratory Ikzf4 |
| Nlrc5 | 1.02E-214 | 2.27070509 | 0.568 | 0.297 | 1.98E-210 | cDC.migratory Nlrc5 |
| Lsp1 | 1.80E-210 | 1.1335323 | 0.862 | 0.913 | 3.47E-206 | cDC.migratory Lsp1 |
| Nostrin | 1.26E-209 | 3.31253696 | 0.412 | 0.105 | 2.44E-205 | cDC.migratory Nostrin |
| Casc1 | 2.64E-203 | 5.24599353 | 0.257 | 0.02 | 5.10E-199 | cDC.migratory Casc1 |
| Tuba1a | 2.17E-201 | 1.80685844 | 0.702 | 0.582 | 4.19E-197 | cDC.migratory Tuba1a |
| Gbp5 | 3.28E-201 | 2.74662445 | 0.453 | 0.159 | 6.34E-197 | cDC.migratory Gbp5 |
| Pcyt1a | 1.28E-200 | 2.15890351 | 0.565 | 0.317 | 2.48E-196 | cDC.migratory Pcyt1a |
| Pdlim4 | 1.35E-198 | 2.99065538 | 0.401 | 0.132 | 2.61E-194 | cDC.migratory Pdlim4 |
| Tmem150c | 1.03E-197 | 5.48349968 | 0.352 | 0.017 | 2.00E-193 | cDC.migratory Tmem150c |
| Tmtc2 | 1.24E-197 | 3.57566409 | 0.371 | 0.068 | 2.39E-193 | cDC.migratory Tmtc2 |
| Hsf2 | 2.41E-197 | 3.54735546 | 0.377 | 0.088 | 4.66E-193 | cDC.migratory Hsf2 |
| Arl5a | 2.69E-197 | 2.14056951 | 0.577 | 0.351 | 5.21E-193 | cDC.migratory Arl5a |
| Arc | 1.88E-193 | 4.88059877 | 0.328 | 0.025 | 3.63E-189 | cDC.migratory Arc |
| Basp1 | 1.01E-191 | 1.38299211 | 0.884 | 0.805 | 1.95E-187 | cDC.migratory Basp1 |
| Akap5 | 3.56E-191 | 2.6974264 | 0.332 | 0.143 | 6.89E-187 | cDC.migratory Akap5 |
| Cytip | 7.02E-190 | 1.1569281 | 0.906 | 0.907 | 1.36E-185 | cDC.migratory Cytip |
| Pkib | 3.83E-188 | 1.89640118 | 0.609 | 0.489 | 7.40E-184 | cDC.migratory Pkib |
| Tpm1 | 3.93E-187 | 2.63828343 | 0.518 | 0.268 | 7.60E-183 | cDC.migratory Tpm1 |
| N4bp2l1 | 3.54E-185 | 3.00308273 | 0.397 | 0.128 | 6.85E-181 | cDC.migratory N4bp2l1 |
| Gucd1 | 2.14E-183 | 2.32964458 | 0.5 | 0.22 | 4.15E-179 | cDC.migratory Gucd1 |
| Tmem131 | 1.33E-181 | 1.49996563 | 0.708 | 0.642 | 2.56E-177 | cDC.migratory Tmem131 |
| Laptm4b | 7.58E-181 | 2.45325905 | 0.363 | 0.167 | 1.47E-176 | cDC.migratory Laptm4b |
| Pik3r1 | 4.47E-180 | 1.64275414 | 0.735 | 0.602 | 8.65E-176 | cDC.migratory Pik3r1 |
| Birc2 | 4.84E-180 | 2.6100337 | 0.471 | 0.187 | 9.36E-176 | cDC.migratory Birc2 |
| Pakap.1 | 6.08E-180 | 5.19935704 | 0.314 | 0.021 | 1.17E-175 | cDC.migratory Pakap.1 |
| Npr1 | 8.07E-175 | 6.07713399 | 0.267 | 0.011 | 1.56E-170 | cDC.migratory Npr1 |
| Papss2 | 8.49E-175 | 3.94282788 | 0.315 | 0.049 | 1.64E-170 | cDC.migratory Papss2 |
| Castor2 | 1.89E-173 | 2.47212491 | 0.422 | 0.168 | 3.65E-169 | cDC.migratory Castor2 |
| Hopx | 1.67E-168 | 2.4864469 | 0.339 | 0.167 | 3.22E-164 | cDC.migratory Hopx |
| Inf2 | 7.40E-167 | 3.25903021 | 0.434 | 0.124 | 1.43E-162 | cDC.migratory Inf2 |
| Psme2 | 4.84E-163 | 1.73204732 | 0.629 | 0.537 | 9.36E-159 | cDC.migratory Psme2 |
| Sh3bp4 | 3.04E-162 | 2.730763 | 0.404 | 0.123 | 5.88E-158 | cDC.migratory Sh3bp4 |
| Tnfrsf11a | 3.37E-162 | 2.48217407 | 0.572 | 0.284 | 6.52E-158 | cDC.migratory Tnfrsf11a |
| Zfp36l1 | 3.91E-162 | 1.51087617 | 0.671 | 0.518 | 7.57E-158 | cDC.migratory Zfp36l1 |
| Nfkbib | 2.06E-159 | 1.70304621 | 0.662 | 0.538 | 3.97E-155 | cDC.migratory Nfkbib |
| Vsig10 | 6.58E-159 | 3.63683332 | 0.316 | 0.057 | 1.27E-154 | cDC.migratory Vsig10 |
| Cd274 | 1.24E-158 | 1.65156537 | 0.763 | 0.535 | 2.40E-154 | cDC.migratory Cd274 |
| Lrrk1 | 1.26E-158 | 1.92187719 | 0.686 | 0.5 | 2.44E-154 | cDC.migratory Lrrk1 |
| Ccser2 | 1.53E-158 | 2.89397311 | 0.395 | 0.134 | 2.95E-154 | cDC.migratory Ccser2 |
| Hmgn3 | 2.01E-155 | 3.5550611 | 0.288 | 0.068 | 3.88E-151 | cDC.migratory Hmgn3 |
| Vopp1 | 3.07E-152 | 2.09480173 | 0.522 | 0.328 | 5.93E-148 | cDC.migratory Vopp1 |
| Mrtfa | 7.06E-149 | 1.54765926 | 0.743 | 0.66 | 1.37E-144 | cDC.migratory Mrtfa |
| Ncoa7 | 9.09E-149 | 1.78702682 | 0.541 | 0.309 | 1.76E-144 | cDC.migratory Ncoa7 |
| Nck2 | 1.56E-148 | 2.94421454 | 0.377 | 0.104 | 3.03E-144 | cDC.migratory Nck2 |
| Fau | 1.54E-145 | 0.28269868 | 0.881 | 0.97 | 2.97E-141 | cDC.migratory Fau |
| Aldh1a2 | 5.09E-144 | 2.29205051 | 0.272 | 0.136 | 9.84E-140 | cDC.migratory Aldh1a2 |
| Mllt6 | 6.27E-143 | 1.53033409 | 0.733 | 0.6 | 1.21E-138 | cDC.migratory Mllt6 |
| Serinc5 | 1.82E-142 | 2.7220584 | 0.362 | 0.096 | 3.53E-138 | cDC.migratory Serinc5 |
| Stap2 | 2.13E-142 | 3.83690841 | 0.284 | 0.041 | 4.12E-138 | cDC.migratory Stap2 |
| Plaat3 | 1.09E-140 | 2.65665159 | 0.396 | 0.213 | 2.11E-136 | cDC.migratory Plaat3 |
| Tnfrsf4 | 1.49E-139 | 5.95615385 | 0.267 | 0.012 | 2.87E-135 | cDC.migratory Tnfrsf4 |
| Pfn1 | 1.62E-139 | 0.28144562 | 0.881 | 0.97 | 3.12E-135 | cDC.migratory Pfn1 |
| Map3k1 | 4.47E-138 | 1.63021434 | 0.719 | 0.595 | 8.64E-134 | cDC.migratory Map3k1 |
| Rras2 | 5.04E-138 | 2.05350661 | 0.486 | 0.258 | 9.75E-134 | cDC.migratory Rras2 |
| Dnajc12 | 7.79E-137 | 4.07110314 | 0.274 | 0.044 | 1.51E-132 | cDC.migratory Dnajc12 |
| Stk4 | 8.98E-137 | 1.84610017 | 0.591 | 0.43 | 1.74E-132 | cDC.migratory Stk4 |
| Stxbp6 | 1.90E-135 | 2.59336235 | 0.282 | 0.088 | 3.67E-131 | cDC.migratory Stxbp6 |
| Gnai3 | 1.03E-133 | 1.49456144 | 0.688 | 0.617 | 1.99E-129 | cDC.migratory Gnai3 |
| Fas | 5.79E-131 | 3.45912372 | 0.281 | 0.059 | 1.12E-126 | cDC.migratory Fas |
| Fnbp1l | 2.98E-130 | 2.06504923 | 0.546 | 0.29 | 5.76E-126 | cDC.migratory Fnbp1l |
| Arap2 | 1.38E-128 | 1.65114422 | 0.422 | 0.288 | 2.66E-124 | cDC.migratory Arap2 |
| Cx3cl1 | 3.41E-128 | 2.54006217 | 0.343 | 0.119 | 6.58E-124 | cDC.migratory Cx3cl1 |
| Nudt9 | 1.97E-127 | 2.01836274 | 0.462 | 0.301 | 3.82E-123 | cDC.migratory Nudt9 |
| Ceacam1 | 4.53E-127 | 2.8105297 | 0.27 | 0.104 | 8.76E-123 | cDC.migratory Ceacam1 |
| Plekhm2 | 7.63E-126 | 2.11123331 | 0.539 | 0.314 | 1.48E-121 | cDC.migratory Plekhm2 |
| Rnf19b | 2.15E-125 | 1.563576 | 0.671 | 0.514 | 4.15E-121 | cDC.migratory Rnf19b |
| Col27a1 | 3.46E-125 | 3.08361257 | 0.258 | 0.06 | 6.69E-121 | cDC.migratory Col27a1 |
| Ccdc6 | 3.49E-125 | 2.13057095 | 0.599 | 0.353 | 6.75E-121 | cDC.migratory Ccdc6 |
| Stx11 | 9.11E-123 | 2.19273036 | 0.444 | 0.224 | 1.76E-118 | cDC.migratory Stx11 |
| Lima1 | 2.33E-120 | 2.59920634 | 0.319 | 0.118 | 4.50E-116 | cDC.migratory Lima1 |
| Arpin | 4.46E-120 | 2.34772412 | 0.388 | 0.117 | 8.61E-116 | cDC.migratory Arpin |
| Actg1 | 1.39E-117 | 0.333275 | 0.869 | 0.963 | 2.68E-113 | cDC.migratory Actg1 |
| Map3k14 | 1.08E-116 | 1.45900396 | 0.797 | 0.668 | 2.09E-112 | cDC.migratory Map3k14 |
| Ppdpf | 1.45E-115 | 1.53719463 | 0.558 | 0.427 | 2.81E-111 | cDC.migratory Ppdpf |
| Tbc1d15 | 2.79E-115 | 1.77253749 | 0.556 | 0.391 | 5.40E-111 | cDC.migratory Tbc1d15 |
| mt-Co2 | 3.27E-114 | 0.55084431 | 0.747 | 0.838 | 6.32E-110 | cDC.migratory mt-Co2 |
| Cnn2 | 7.11E-114 | 1.13256287 | 0.813 | 0.85 | 1.38E-109 | cDC.migratory Cnn2 |
| Nrp2 | 1.67E-111 | 2.39870751 | 0.382 | 0.14 | 3.24E-107 | cDC.migratory Nrp2 |
| Clec2d | 5.00E-111 | 1.2692324 | 0.757 | 0.684 | 9.67E-107 | cDC.migratory Clec2d |
| Gnb4 | 7.74E-110 | 1.9895302 | 0.454 | 0.275 | 1.50E-105 | cDC.migratory Gnb4 |
| Bmf | 3.48E-108 | 2.58795529 | 0.363 | 0.153 | 6.72E-104 | cDC.migratory Bmf |
| Lactb | 7.85E-107 | 1.9519272 | 0.554 | 0.401 | 1.52E-102 | cDC.migratory Lactb |
| Bmp2k | 2.71E-105 | 1.51985965 | 0.696 | 0.637 | 5.23E-101 | cDC.migratory Bmp2k |
| Rasa2 | 3.16E-105 | 2.57908385 | 0.343 | 0.137 | 6.12E-101 | cDC.migratory Rasa2 |
| Tmem131l | 3.93E-104 | 1.81073513 | 0.534 | 0.369 | 7.61E-100 | cDC.migratory Tmem131l |
| Arhgap31 | 2.49E-103 | 1.85220074 | 0.502 | 0.348 | 4.81E-99 | cDC.migratory Arhgap31 |
| Psme2b | 9.54E-103 | 1.73489164 | 0.6 | 0.459 | 1.84E-98 | cDC.migratory Psme2b |
| Adam23 | 1.97E-102 | 1.65299096 | 0.577 | 0.373 | 3.81E-98 | cDC.migratory Adam23 |
| Actn1 | 3.95E-102 | 0.41316108 | 0.778 | 0.752 | 7.63E-98 | cDC.migratory Actn1 |
| Fam32a | 2.96E-101 | 1.41623525 | 0.598 | 0.529 | 5.71E-97 | cDC.migratory Fam32a |
| Sinhcaf | 5.96E-101 | 2.22998321 | 0.406 | 0.202 | 1.15E-96 | cDC.migratory Sinhcaf |
| Tmem8 | 8.35E-101 | 1.78908745 | 0.514 | 0.337 | 1.61E-96 | cDC.migratory Tmem8 |
| Cdc42ep3 | 8.48E-101 | 2.79505605 | 0.252 | 0.086 | 1.64E-96 | cDC.migratory Cdc42ep3 |
| Tspan33 | 1.55E-100 | 1.94189762 | 0.494 | 0.281 | 3.00E-96 | cDC.migratory Tspan33 |
| Mical3 | 1.12E-99 | 2.46509171 | 0.372 | 0.171 | 2.16E-95 | cDC.migratory Mical3 |
| Flrt3 | 1.39E-99 | 1.60588758 | 0.507 | 0.302 | 2.69E-95 | cDC.migratory Flrt3 |
| Rgmb | 2.08E-99 | 2.62385412 | 0.317 | 0.12 | 4.02E-95 | cDC.migratory Rgmb |
| Ncf1 | 2.55E-99 | 1.33137823 | 0.8 | 0.73 | 4.94E-95 | cDC.migratory Ncf1 |
| Spsb1 | 3.16E-99 | 2.3237409 | 0.431 | 0.21 | 6.11E-95 | cDC.migratory Spsb1 |
| Atmin | 5.16E-99 | 1.77884225 | 0.454 | 0.298 | 9.98E-95 | cDC.migratory Atmin |
| Mmd | 1.25E-98 | 2.24549869 | 0.358 | 0.167 | 2.41E-94 | cDC.migratory Mmd |
| Etv3 | 1.25E-97 | 1.3968594 | 0.798 | 0.667 | 2.41E-93 | cDC.migratory Etv3 |
| Tank | 1.02E-96 | 1.93172353 | 0.552 | 0.368 | 1.97E-92 | cDC.migratory Tank |
| Ube2l6 | 1.05E-96 | 2.33051844 | 0.331 | 0.15 | 2.03E-92 | cDC.migratory Ube2l6 |
| Nfat5 | 2.47E-96 | 1.85486009 | 0.542 | 0.36 | 4.77E-92 | cDC.migratory Nfat5 |
| Spred1 | 1.88E-95 | 1.86401216 | 0.627 | 0.438 | 3.64E-91 | cDC.migratory Spred1 |
| Abtb2 | 4.83E-94 | 2.38977747 | 0.355 | 0.141 | 9.34E-90 | cDC.migratory Abtb2 |
| Ric1 | 9.89E-92 | 1.91148634 | 0.541 | 0.373 | 1.91E-87 | cDC.migratory Ric1 |
| Stat5a | 4.41E-91 | 1.34226574 | 0.5 | 0.393 | 8.53E-87 | cDC.migratory Stat5a |
| Nrip1 | 8.50E-90 | 1.84138139 | 0.441 | 0.296 | 1.64E-85 | cDC.migratory Nrip1 |
| Rab21 | 1.32E-89 | 1.34300477 | 0.79 | 0.729 | 2.54E-85 | cDC.migratory Rab21 |
| Iffo2 | 1.64E-89 | 2.48731162 | 0.335 | 0.145 | 3.17E-85 | cDC.migratory Iffo2 |
| Gramd1b | 6.38E-89 | 2.23094284 | 0.432 | 0.268 | 1.23E-84 | cDC.migratory Gramd1b |
| Irf1 | 6.53E-88 | 1.50729536 | 0.607 | 0.414 | 1.26E-83 | cDC.migratory Irf1 |
| Rgs1 | 8.53E-88 | 1.1021898 | 0.713 | 0.555 | 1.65E-83 | cDC.migratory Rgs1 |
| Rnf115 | 1.55E-87 | 1.87828297 | 0.458 | 0.326 | 3.00E-83 | cDC.migratory Rnf115 |
| Htra2 | 6.14E-87 | 2.18088895 | 0.445 | 0.233 | 1.19E-82 | cDC.migratory Htra2 |
| Csf2rb | 7.35E-87 | 0.33675747 | 0.764 | 0.84 | 1.42E-82 | cDC.migratory Csf2rb |
| Bcl2l1 | 8.60E-87 | 1.7663225 | 0.336 | 0.292 | 1.66E-82 | cDC.migratory Bcl2l1 |
| Anxa4 | 1.11E-86 | 1.63303375 | 0.665 | 0.484 | 2.15E-82 | cDC.migratory Anxa4 |
| Zc3h7b | 1.04E-85 | 2.03813087 | 0.409 | 0.238 | 2.00E-81 | cDC.migratory Zc3h7b |
| Iscu | 3.50E-85 | 1.99827641 | 0.428 | 0.258 | 6.76E-81 | cDC.migratory Iscu |
| Procr | 6.03E-85 | 1.85814385 | 0.353 | 0.168 | 1.17E-80 | cDC.migratory Procr |
| Gadd45b | 6.16E-82 | 1.09313509 | 0.824 | 0.601 | 1.19E-77 | cDC.migratory Gadd45b |
| Ptpn3 | 3.85E-81 | 2.32367558 | 0.329 | 0.153 | 7.44E-77 | cDC.migratory Ptpn3 |
| Pdcd1lg2 | 9.22E-81 | 1.27982718 | 0.627 | 0.492 | 1.78E-76 | cDC.migratory Pdcd1lg2 |
| Washc1 | 7.40E-79 | 1.77375302 | 0.464 | 0.309 | 1.43E-74 | cDC.migratory Washc1 |
| Mxi1 | 2.01E-78 | 1.76216629 | 0.557 | 0.358 | 3.89E-74 | cDC.migratory Mxi1 |
| Ctnna1 | 3.86E-77 | 1.30357746 | 0.712 | 0.714 | 7.47E-73 | cDC.migratory Ctnna1 |
| Sema4c | 3.80E-76 | 2.42137543 | 0.315 | 0.143 | 7.35E-72 | cDC.migratory Sema4c |
| Nans | 1.59E-75 | 1.75934118 | 0.421 | 0.327 | 3.08E-71 | cDC.migratory Nans |
| Atp6v0a1 | 1.84E-75 | 1.77182566 | 0.439 | 0.279 | 3.56E-71 | cDC.migratory Atp6v0a1 |
| Tmem63b | 3.37E-75 | 2.3850676 | 0.295 | 0.118 | 6.52E-71 | cDC.migratory Tmem63b |
| Spr | 3.63E-75 | 1.80410818 | 0.483 | 0.293 | 7.03E-71 | cDC.migratory Spr |
| mt-Atp8 | 4.78E-75 | 0.3926405 | 0.592 | 0.704 | 9.25E-71 | cDC.migratory mt-Atp8 |
| Atxn7l1 | 8.46E-74 | 1.59056661 | 0.597 | 0.478 | 1.63E-69 | cDC.migratory Atxn7l1 |
| Zc3h12c | 4.12E-73 | 2.38380606 | 0.26 | 0.098 | 7.96E-69 | cDC.migratory Zc3h12c |
| mt-Co1 | 2.53E-71 | 0.30704827 | 0.535 | 0.676 | 4.89E-67 | cDC.migratory mt-Co1 |
| Nfe2l1 | 8.02E-71 | 1.47304026 | 0.582 | 0.479 | 1.55E-66 | cDC.migratory Nfe2l1 |
| St3gal3 | 1.80E-69 | 2.07788196 | 0.369 | 0.186 | 3.48E-65 | cDC.migratory St3gal3 |
| mt-Atp6 | 2.03E-68 | 0.56917631 | 0.608 | 0.715 | 3.92E-64 | cDC.migratory mt-Atp6 |
| H2-K1 | 1.07E-67 | 1.32543932 | 0.925 | 0.902 | 2.07E-63 | cDC.migratory H2-K1 |
| Rnaset2a | 2.24E-67 | 1.81884726 | 0.344 | 0.242 | 4.33E-63 | cDC.migratory Rnaset2a |
| Plekhg2 | 4.17E-66 | 1.75775987 | 0.462 | 0.29 | 8.06E-62 | cDC.migratory Plekhg2 |
| Spint2 | 1.07E-65 | 1.56697931 | 0.543 | 0.392 | 2.08E-61 | cDC.migratory Spint2 |
| Dok1 | 1.69E-65 | 1.89690483 | 0.42 | 0.262 | 3.27E-61 | cDC.migratory Dok1 |
| H2afz | 5.15E-65 | 0.27920798 | 0.562 | 0.703 | 9.96E-61 | cDC.migratory H2afz |
| Nhlrc2 | 1.02E-64 | 2.13539802 | 0.392 | 0.231 | 1.97E-60 | cDC.migratory Nhlrc2 |
| Smurf1 | 1.08E-64 | 1.79169429 | 0.487 | 0.302 | 2.10E-60 | cDC.migratory Smurf1 |
| Sbf2 | 6.01E-64 | 2.46828364 | 0.281 | 0.115 | 1.16E-59 | cDC.migratory Sbf2 |
| Zfand6 | 2.24E-63 | 1.77006677 | 0.436 | 0.313 | 4.32E-59 | cDC.migratory Zfand6 |
| Phlpp1 | 1.65E-62 | 1.74271217 | 0.513 | 0.357 | 3.18E-58 | cDC.migratory Phlpp1 |
| Sat1 | 1.79E-62 | 0.25821352 | 0.584 | 0.709 | 3.46E-58 | cDC.migratory Sat1 |
| Adam11 | 3.20E-62 | 2.1382802 | 0.25 | 0.072 | 6.18E-58 | cDC.migratory Adam11 |
| Pnpla8 | 1.01E-61 | 1.97276391 | 0.361 | 0.196 | 1.96E-57 | cDC.migratory Pnpla8 |
| Tnfrsf18 | 5.02E-61 | 1.69689568 | 0.348 | 0.243 | 9.71E-57 | cDC.migratory Tnfrsf18 |
| Polr3c | 2.54E-60 | 1.87850501 | 0.394 | 0.24 | 4.91E-56 | cDC.migratory Polr3c |
| H2-Q4 | 4.85E-60 | 2.38545516 | 0.308 | 0.157 | 9.38E-56 | cDC.migratory H2-Q4 |
| Dstn | 1.69E-59 | 1.54395946 | 0.55 | 0.462 | 3.28E-55 | cDC.migratory Dstn |
| Cd83 | 2.14E-59 | 0.89783998 | 0.775 | 0.697 | 4.13E-55 | cDC.migratory Cd83 |
| Plxnd1 | 2.48E-59 | 1.43011849 | 0.693 | 0.591 | 4.79E-55 | cDC.migratory Plxnd1 |
| Auh | 7.42E-59 | 1.99063861 | 0.506 | 0.349 | 1.43E-54 | cDC.migratory Auh |
| Dusp5 | 8.24E-59 | 0.49854065 | 0.728 | 0.749 | 1.59E-54 | cDC.migratory Dusp5 |
| Casp3 | 3.08E-58 | 1.85201413 | 0.401 | 0.264 | 5.96E-54 | cDC.migratory Casp3 |
| Psmg4 | 7.23E-58 | 1.77201755 | 0.454 | 0.327 | 1.40E-53 | cDC.migratory Psmg4 |
| Rnf157 | 1.80E-57 | 2.5424807 | 0.268 | 0.126 | 3.48E-53 | cDC.migratory Rnf157 |
| Epb41 | 2.56E-57 | 1.55342714 | 0.576 | 0.487 | 4.95E-53 | cDC.migratory Epb41 |
| Flvcr1 | 3.17E-55 | 2.18110052 | 0.383 | 0.209 | 6.13E-51 | cDC.migratory Flvcr1 |
| Son | 3.63E-55 | 0.31689182 | 0.627 | 0.772 | 7.02E-51 | cDC.migratory Son |
| Ostf1 | 1.23E-54 | 1.17388559 | 0.716 | 0.738 | 2.38E-50 | cDC.migratory Ostf1 |
| Phldb1 | 5.28E-54 | 2.37362114 | 0.266 | 0.101 | 1.02E-49 | cDC.migratory Phldb1 |
| Zbtb46 | 8.00E-54 | 1.37659898 | 0.559 | 0.435 | 1.55E-49 | cDC.migratory Zbtb46 |
| Cep85l | 9.68E-54 | 1.68940434 | 0.348 | 0.239 | 1.87E-49 | cDC.migratory Cep85l |
| Traf3 | 1.33E-53 | 1.87774077 | 0.391 | 0.22 | 2.58E-49 | cDC.migratory Traf3 |
| Ddhd1 | 1.44E-53 | 1.7275175 | 0.473 | 0.329 | 2.77E-49 | cDC.migratory Ddhd1 |
| Ccdc88a | 4.07E-53 | 1.32341386 | 0.622 | 0.535 | 7.87E-49 | cDC.migratory Ccdc88a |
| mt-Nd4l | 9.35E-53 | 0.26460959 | 0.442 | 0.591 | 1.81E-48 | cDC.migratory mt-Nd4l |
| Tapbpl | 4.36E-52 | 1.57916348 | 0.387 | 0.263 | 8.44E-48 | cDC.migratory Tapbpl |
| Il7r | 6.22E-52 | 1.59460219 | 0.447 | 0.256 | 1.20E-47 | cDC.migratory Il7r |
| Ptger4 | 2.75E-51 | 1.0558929 | 0.681 | 0.597 | 5.32E-47 | cDC.migratory Ptger4 |
| Ccng2 | 2.84E-51 | 1.84454603 | 0.384 | 0.236 | 5.50E-47 | cDC.migratory Ccng2 |
| Pnkd | 6.70E-51 | 2.46333373 | 0.255 | 0.139 | 1.30E-46 | cDC.migratory Pnkd |
| Dnajc10 | 7.90E-51 | 1.45700043 | 0.476 | 0.385 | 1.53E-46 | cDC.migratory Dnajc10 |
| Dnajb2 | 1.29E-50 | 2.52829055 | 0.251 | 0.101 | 2.48E-46 | cDC.migratory Dnajb2 |
| Ccl171 | 9.95E-50 | 0.45107712 | 0.583 | 0.475 | 1.92E-45 | cDC.migratory Ccl17 |
| Mthfd2 | 1.34E-49 | 1.64320903 | 0.444 | 0.35 | 2.59E-45 | cDC.migratory Mthfd2 |
| Tspo | 2.25E-49 | 0.36453522 | 0.754 | 0.886 | 4.34E-45 | cDC.migratory Tspo |
| Gtpbp2 | 6.91E-49 | 1.76616408 | 0.474 | 0.334 | 1.34E-44 | cDC.migratory Gtpbp2 |
| Rsrp1 | 8.83E-49 | 0.57107133 | 0.603 | 0.716 | 1.71E-44 | cDC.migratory Rsrp1 |
| Gpc1 | 1.21E-48 | 1.93526498 | 0.384 | 0.246 | 2.34E-44 | cDC.migratory Gpc1 |
| Samhd11 | 1.06E-47 | 0.54886315 | 0.735 | 0.794 | 2.05E-43 | cDC.migratory Samhd1 |
| Atl3 | 1.13E-47 | 1.28139042 | 0.578 | 0.53 | 2.18E-43 | cDC.migratory Atl3 |
| Cers6 | 1.76E-47 | 1.405095 | 0.587 | 0.522 | 3.41E-43 | cDC.migratory Cers6 |
| Dynll2 | 2.77E-47 | 1.39930159 | 0.612 | 0.504 | 5.36E-43 | cDC.migratory Dynll2 |
| Rab11fip1 | 1.38E-46 | 0.26295292 | 0.59 | 0.674 | 2.66E-42 | cDC.migratory Rab11fip1 |
| Tcf25 | 1.73E-46 | 0.3252644 | 0.608 | 0.751 | 3.35E-42 | cDC.migratory Tcf25 |
| Cmtm6 | 3.15E-46 | 0.29443167 | 0.533 | 0.693 | 6.09E-42 | cDC.migratory Cmtm6 |
| Polb | 3.46E-46 | 1.56398542 | 0.417 | 0.328 | 6.69E-42 | cDC.migratory Polb |
| Luc7l2 | 6.42E-46 | 0.27888637 | 0.507 | 0.689 | 1.24E-41 | cDC.migratory Luc7l2 |
| Hivep1 | 8.10E-46 | 1.50287393 | 0.595 | 0.489 | 1.57E-41 | cDC.migratory Hivep1 |
| Fam129a | 2.14E-45 | 1.18465562 | 0.583 | 0.527 | 4.13E-41 | cDC.migratory Fam129a |
| Crebl2 | 3.26E-45 | 2.28404032 | 0.264 | 0.122 | 6.30E-41 | cDC.migratory Crebl2 |
| mt-Co3 | 4.69E-44 | 0.51132457 | 0.557 | 0.669 | 9.07E-40 | cDC.migratory mt-Co3 |
| Tmem19 | 4.96E-44 | 1.71886099 | 0.371 | 0.241 | 9.58E-40 | cDC.migratory Tmem19 |
| Sfpq | 5.28E-44 | 0.29284002 | 0.549 | 0.702 | 1.02E-39 | cDC.migratory Sfpq |
| Nsun4 | 1.28E-43 | 1.69279568 | 0.343 | 0.205 | 2.47E-39 | cDC.migratory Nsun4 |
| Clip2 | 1.71E-43 | 1.80119557 | 0.456 | 0.309 | 3.30E-39 | cDC.migratory Clip2 |
| DEPRECATED-I | 2.47E-43 | 1.92966596 | 0.348 | 0.2 | 4.77E-39 | cDC.migratory DEPRECATED-ENSMUSG00000025025 |
| Tmbim4 | 2.48E-43 | 1.73848132 | 0.343 | 0.295 | 4.79E-39 | cDC.migratory Tmbim4 |
| Cstb | 2.88E-43 | 0.28622159 | 0.493 | 0.599 | 5.57E-39 | cDC.migratory Cstb |
| Herc6 | 3.31E-43 | 2.12048314 | 0.3 | 0.158 | 6.39E-39 | cDC.migratory Herc6 |
| Lgmn | 7.72E-43 | 0.36413296 | 0.317 | 0.305 | 1.49E-38 | cDC.migratory Lgmn |
| Il21r | 1.69E-42 | 1.2113466 | 0.693 | 0.567 | 3.26E-38 | cDC.migratory Il21r |
| Thap12 | 2.70E-42 | 1.66230415 | 0.31 | 0.222 | 5.22E-38 | cDC.migratory Thap12 |
| Skil | 1.07E-41 | 0.32971566 | 0.609 | 0.696 | 2.07E-37 | cDC.migratory Skil |
| Calm3 | 1.10E-41 | 0.29067932 | 0.594 | 0.74 | 2.13E-37 | cDC.migratory Calm3 |
| Sepsecs | 1.11E-41 | 1.86290188 | 0.267 | 0.135 | 2.14E-37 | cDC.migratory Sepsecs |
| Icam1 | 3.26E-41 | 1.01304059 | 0.664 | 0.563 | 6.30E-37 | cDC.migratory Icam1 |
| Kcnk6 | 7.61E-41 | 1.57515088 | 0.482 | 0.353 | 1.47E-36 | cDC.migratory Kcnk6 |
| Adam19 | 1.52E-40 | 1.51359141 | 0.389 | 0.247 | 2.93E-36 | cDC.migratory Adam19 |
| Srsf5 | 5.24E-40 | 0.26143969 | 0.513 | 0.684 | 1.01E-35 | cDC.migratory Srsf5 |
| Ifi2091 | 1.22E-39 | 1.32190077 | 0.372 | 0.236 | 2.37E-35 | cDC.migratory Ifi209 |
| Rras | 1.27E-39 | 1.65949088 | 0.314 | 0.196 | 2.45E-35 | cDC.migratory Rras |
| Dennd5a | 4.17E-39 | 1.07025392 | 0.675 | 0.668 | 8.07E-35 | cDC.migratory Dennd5a |
| Sap25 | 4.78E-39 | 2.04078133 | 0.302 | 0.163 | 9.24E-35 | cDC.migratory Sap25 |
| Dlgap4 | 8.53E-39 | 1.36294215 | 0.603 | 0.523 | 1.65E-34 | cDC.migratory Dlgap4 |
| Rgs12 | 1.01E-38 | 1.6087942 | 0.454 | 0.29 | 1.95E-34 | cDC.migratory Rgs12 |
| Irf8 | 1.61E-38 | 0.28456716 | 0.737 | 0.555 | 3.11E-34 | cDC.migratory Irf8 |
| Vgll4 | 5.31E-38 | 1.51777266 | 0.481 | 0.378 | 1.03E-33 | cDC.migratory Vgll4 |
| Zbtb38 | 7.66E-38 | 1.82476099 | 0.252 | 0.154 | 1.48E-33 | cDC.migratory Zbtb38 |
| Triobp | 9.09E-38 | 1.71255158 | 0.37 | 0.251 | 1.76E-33 | cDC.migratory Triobp |
| Adprh | 1.21E-37 | 0.27246729 | 0.553 | 0.693 | 2.34E-33 | cDC.migratory Adprh |
| Itprid2 | 6.39E-37 | 0.35230877 | 0.446 | 0.586 | 1.24E-32 | cDC.migratory Itprid2 |
| Palld | 7.77E-37 | 1.682093 | 0.363 | 0.211 | 1.50E-32 | cDC.migratory Palld |
| Gdi1 | 8.00E-37 | 1.54412451 | 0.568 | 0.484 | 1.55E-32 | cDC.migratory Gdi1 |
| Marveld1 | 8.51E-37 | 1.72224157 | 0.375 | 0.242 | 1.65E-32 | cDC.migratory Marveld1 |
| Micall1 | 1.17E-36 | 1.96511591 | 0.27 | 0.149 | 2.26E-32 | cDC.migratory Micall1 |
| Nub1 | 1.94E-36 | 1.34396731 | 0.501 | 0.418 | 3.76E-32 | cDC.migratory Nub1 |
| Asl | 3.64E-36 | 1.51209992 | 0.466 | 0.346 | 7.04E-32 | cDC.migratory Asl |
| Sec11c | 8.36E-36 | 0.32870123 | 0.368 | 0.559 | 1.62E-31 | cDC.migratory Sec11c |
| Sik3 | 1.17E-35 | 1.52165484 | 0.412 | 0.304 | 2.26E-31 | cDC.migratory Sik3 |
| Bcl2a1b | 1.28E-35 | 1.08277508 | 0.757 | 0.661 | 2.47E-31 | cDC.migratory Bcl2a1b |
| Batf3 | 1.50E-35 | 1.49619521 | 0.617 | 0.448 | 2.90E-31 | cDC.migratory Batf3 |
| Bax | 1.60E-35 | 0.41905488 | 0.592 | 0.726 | 3.09E-31 | cDC.migratory Bax |
| Dip2b | 1.66E-35 | 1.62673263 | 0.479 | 0.355 | 3.22E-31 | cDC.migratory Dip2b |
| Vti1a | 1.86E-35 | 1.55405685 | 0.478 | 0.414 | 3.59E-31 | cDC.migratory Vti1a |
| Stat1 | 3.67E-35 | 1.22409189 | 0.59 | 0.509 | 7.10E-31 | cDC.migratory Stat1 |
| Nfkbia1 | 4.41E-35 | 0.51689378 | 0.798 | 0.715 | 8.53E-31 | cDC.migratory Nfkbia |
| Prmp1 | 6.60E-35 | 1.29822906 | 0.312 | 0.171 | 1.28E-30 | cDC.migratory Prmp |
| Slc25a3 | 9.89E-35 | 0.29284524 | 0.656 | 0.812 | 1.91E-30 | cDC.migratory Slc25a3 |
| Dnajc1 | 1.18E-34 | 2.14599366 | 0.255 | 0.154 | 2.29E-30 | cDC.migratory Dnajc1 |
| Zfp598 | 1.66E-34 | 1.48393793 | 0.435 | 0.35 | 3.21E-30 | cDC.migratory Zfp598 |
| Gfpt1 | 2.02E-34 | 1.27946899 | 0.498 | 0.412 | 3.90E-30 | cDC.migratory Gfpt1 |
| Nudt3 | 3.58E-34 | 0.44197384 | 0.378 | 0.534 | 6.93E-30 | cDC.migratory Nudt3 |
| Bin1 | 4.72E-34 | 0.26879879 | 0.45 | 0.583 | 9.12E-30 | cDC.migratory Bin1 |
| Slc38a2 | 7.66E-34 | 0.40302498 | 0.638 | 0.743 | 1.48E-29 | cDC.migratory Slc38a2 |
| Cxcl16 | 8.77E-34 | 1.13292324 | 0.847 | 0.689 | 1.70E-29 | cDC.migratory Cxcl16 |
| Vps39 | 1.49E-33 | 1.7175311 | 0.308 | 0.193 | 2.88E-29 | cDC.migratory Vps39 |
| Dclre1c | 4.13E-33 | 1.68068595 | 0.33 | 0.205 | 7.98E-29 | cDC.migratory Dclre1c |
| Cdc37 | 4.66E-33 | 0.41793845 | 0.573 | 0.689 | 9.00E-29 | cDC.migratory Cdc37 |
| Zfp217 | 7.41E-33 | 1.36035062 | 0.525 | 0.412 | 1.43E-28 | cDC.migratory Zfp217 |
| Gbp8 | 1.05E-32 | 2.2765265 | 0.27 | 0.141 | 2.04E-28 | cDC.migratory Gbp8 |
| Tnip3 | 2.45E-32 | 1.35287114 | 0.464 | 0.286 | 4.74E-28 | cDC.migratory Tnip3 |
| Grk3 | 4.82E-32 | 1.39373281 | 0.581 | 0.488 | 9.33E-28 | cDC.migratory Grk3 |
| Fbrsl1 | 6.16E-32 | 1.71588674 | 0.436 | 0.314 | 1.19E-27 | cDC.migratory Fbrsl1 |
| Filip1l | 1.17E-31 | 1.71689937 | 0.305 | 0.18 | 2.27E-27 | cDC.migratory Filip1l |
| B2m | 1.72E-31 | 0.96764655 | 0.789 | 0.831 | 3.33E-27 | cDC.migratory B2m |
| Ikbkb | 2.90E-31 | 1.18200048 | 0.692 | 0.679 | 5.61E-27 | cDC.migratory Ikbkb |
| Ndrg1 | 3.19E-31 | 0.99215102 | 0.599 | 0.535 | 6.17E-27 | cDC.migratory Ndrg1 |
| Trim24 | 3.30E-31 | 1.70797653 | 0.256 | 0.169 | 6.39E-27 | cDC.migratory Trim24 |
| H2-M3 | 6.80E-31 | 0.31062476 | 0.303 | 0.453 | 1.31E-26 | cDC.migratory H2-M3 |
| Dph5 | 1.39E-30 | 2.15497696 | 0.274 | 0.152 | 2.69E-26 | cDC.migratory Dph5 |
| Alkbh6 | 2.88E-30 | 1.54006353 | 0.338 | 0.229 | 5.58E-26 | cDC.migratory Alkbh6 |
| Gbp2 | 3.50E-30 | 1.66729525 | 0.39 | 0.212 | 6.77E-26 | cDC.migratory Gbp2 |
| Tnip2 | 1.38E-29 | 0.25605836 | 0.447 | 0.574 | 2.66E-25 | cDC.migratory Tnip2 |
| Ppp1r16a | 1.74E-29 | 1.88451775 | 0.27 | 0.157 | 3.36E-25 | cDC.migratory Ppp1r16a |
| Tecr | 2.01E-29 | 0.29829187 | 0.371 | 0.512 | 3.89E-25 | cDC.migratory Tecr |
| Ttyh2 | 2.19E-29 | 2.19177341 | 0.259 | 0.137 | 4.24E-25 | cDC.migratory Ttyh2 |
| Yy1 | 2.51E-29 | 0.51548663 | 0.561 | 0.692 | 4.86E-25 | cDC.migratory Yy1 |
| Tap1 | 3.83E-29 | 0.36647782 | 0.642 | 0.725 | 7.40E-25 | cDC.migratory Tap1 |
| Gtpbp1 | 4.40E-29 | 1.44845805 | 0.432 | 0.356 | 8.51E-25 | cDC.migratory Gtpbp1 |
| Chka | 7.59E-29 | 2.06821335 | 0.289 | 0.178 | 1.47E-24 | cDC.migratory Chka |
| mt-Nd5 | 8.46E-29 | 0.34642437 | 0.377 | 0.501 | 1.64E-24 | cDC.migratory mt-Nd5 |
| Sh3glb1 | 8.49E-29 | 0.34168122 | 0.578 | 0.696 | 1.64E-24 | cDC.migratory Sh3glb1 |
| Etnk1 | 9.62E-29 | 1.25244794 | 0.571 | 0.527 | 1.86E-24 | cDC.migratory Etnk1 |
| Srsf11 | 1.11E-28 | 0.26122395 | 0.343 | 0.496 | 2.14E-24 | cDC.migratory Srsf11 |
| Brk1 | 1.56E-28 | 0.3997892 | 0.647 | 0.787 | 3.02E-24 | cDC.migratory Brk1 |
| Stk24 | 1.98E-28 | 0.30824172 | 0.553 | 0.681 | 3.83E-24 | cDC.migratory Stk24 |
| Mif4gd | 2.40E-28 | 1.25424759 | 0.531 | 0.464 | 4.64E-24 | cDC.migratory Mif4gd |
| Srsf7 | 2.98E-28 | 0.61756118 | 0.619 | 0.703 | 5.76E-24 | cDC.migratory Srsf7 |
| Gnptab | 3.49E-28 | 1.40820846 | 0.377 | 0.335 | 6.75E-24 | cDC.migratory Gnptab |
| Rbm39 | 3.85E-28 | 0.31581601 | 0.408 | 0.567 | 7.45E-24 | cDC.migratory Rbm39 |
| Zeb1 | 3.92E-28 | 1.68919908 | 0.301 | 0.185 | 7.58E-24 | cDC.migratory Zeb1 |
| Zmiz2 | 5.05E-28 | 0.31624357 | 0.623 | 0.74 | 9.76E-24 | cDC.migratory Zmiz2 |
| Spg11 | 6.98E-28 | 1.73348171 | 0.302 | 0.191 | 1.35E-23 | cDC.migratory Spg11 |
| Tmem120b | 8.87E-28 | 1.69859828 | 0.289 | 0.176 | 1.71E-23 | cDC.migratory Tmem120b |
| Tmod3 | 1.32E-27 | 0.27307587 | 0.594 | 0.734 | 2.55E-23 | cDC.migratory Tmod3 |
| Adam8 | 2.30E-27 | 1.18455419 | 0.601 | 0.531 | 4.46E-23 | cDC.migratory Adam8 |
| Zfc3h1 | 4.77E-27 | 1.59281905 | 0.349 | 0.258 | 9.22E-23 | cDC.migratory Zfc3h1 |
| Ptms | 6.42E-27 | 0.59571946 | 0.873 | 0.907 | 1.24E-22 | cDC.migratory Ptms |
| Tubb5 | 7.33E-27 | 0.64323887 | 0.654 | 0.691 | 1.42E-22 | cDC.migratory Tubb5 |
| Prr5l | 7.70E-27 | 1.64821524 | 0.356 | 0.233 | 1.49E-22 | cDC.migratory Prr5l |
| Eif4a2 | 8.87E-27 | 0.41016664 | 0.572 | 0.679 | 1.72E-22 | cDC.migratory Eif4a2 |
| Cnpy3 | 9.99E-27 | 0.40282667 | 0.39 | 0.526 | 1.93E-22 | cDC.migratory Cnpy3 |
| Stxbp3 | 1.60E-26 | 1.93227207 | 0.361 | 0.223 | 3.10E-22 | cDC.migratory Stxbp3 |
| Tmsb4x | 2.35E-26 | 0.65750928 | 0.989 | 0.999 | 4.55E-22 | cDC.migratory Tmsb4x |
| Sin3b | 3.26E-26 | 0.30957249 | 0.557 | 0.697 | 6.30E-22 | cDC.migratory Sin3b |
| Cflar | 8.59E-26 | 1.38407802 | 0.454 | 0.345 | 1.66E-21 | cDC.migratory Cflar |
| Git2 | 1.15E-25 | 0.29960538 | 0.411 | 0.546 | 2.22E-21 | cDC.migratory Git2 |
| Cyb5r3 | 1.33E-25 | 0.25367081 | 0.403 | 0.548 | 2.58E-21 | cDC.migratory Cyb5r3 |
| Plekhj1 | 2.16E-25 | 0.27950255 | 0.413 | 0.526 | 4.17E-21 | cDC.migratory Plekhj1 |
| Mtmr4 | 2.34E-25 | 1.20664014 | 0.487 | 0.422 | 4.52E-21 | cDC.migratory Mtmr4 |
| Pus10 | 3.48E-25 | 1.85990575 | 0.258 | 0.171 | 6.73E-21 | cDC.migratory Pus10 |
| Traf2 | 3.53E-25 | 1.58975773 | 0.352 | 0.258 | 6.83E-21 | cDC.migratory Traf2 |

| | | | | | | | |
|---|---|---|---|---|---|---|---|
| Phc2 | 3.53E-25 | 0.34244471 | 0.399 | 0.505 | 6.83E-21 | cDC.migratory | Phc2 |
| Becn1 | 7.11E-25 | 0.43180678 | 0.45 | 0.578 | 1.37E-20 | cDC.migratory | Becn1 |
| Irak1 | 7.92E-25 | 0.32354122 | 0.441 | 0.581 | 1.53E-20 | cDC.migratory | Irak1 |
| Kdm2a | 8.81E-25 | 0.43888355 | 0.588 | 0.705 | 1.70E-20 | cDC.migratory | Kdm2a |
| Mapre1 | 1.31E-24 | 0.30803286 | 0.539 | 0.663 | 2.53E-20 | cDC.migratory | Mapre1 |
| Rab9 | 1.42E-24 | 1.62921448 | 0.252 | 0.155 | 2.75E-20 | cDC.migratory | Rab9 |
| Luc7l3 | 1.57E-24 | 0.36989651 | 0.286 | 0.427 | 3.04E-20 | cDC.migratory | Luc7l3 |
| Myo1c | 1.86E-24 | 1.22721155 | 0.639 | 0.58 | 3.59E-20 | cDC.migratory | Myo1c |
| Sdhaf1 | 3.03E-24 | 1.62233511 | 0.267 | 0.159 | 5.86E-20 | cDC.migratory | Sdhaf1 |
| Ttyh3 | 3.12E-24 | 1.0309803 | 0.671 | 0.717 | 6.02E-20 | cDC.migratory | Ttyh3 |
| Pbxip1 | 4.32E-24 | 1.63098104 | 0.393 | 0.281 | 8.35E-20 | cDC.migratory | Pbxip1 |
| Sfi1 | 4.93E-24 | 1.92980897 | 0.312 | 0.213 | 9.54E-20 | cDC.migratory | Sfi1 |
| Ap1g2 | 1.13E-23 | 1.46592135 | 0.388 | 0.28 | 2.18E-19 | cDC.migratory | Ap1g2 |
| Map4 | 1.16E-23 | 1.25358077 | 0.544 | 0.492 | 2.24E-19 | cDC.migratory | Map4 |
| Nabp1 | 1.17E-23 | 0.43383851 | 0.395 | 0.483 | 2.27E-19 | cDC.migratory | Nabp1 |
| Nuak2 | 1.33E-23 | 1.03348214 | 0.736 | 0.726 | 2.56E-19 | cDC.migratory | Nuak2 |
| Srsf10 | 1.65E-23 | 0.30300285 | 0.286 | 0.417 | 3.19E-19 | cDC.migratory | Srsf10 |
| Slc33a1 | 1.87E-23 | 1.3453908 | 0.395 | 0.327 | 3.62E-19 | cDC.migratory | Slc33a1 |
| Cyld | 2.06E-23 | 0.31115062 | 0.381 | 0.509 | 3.98E-19 | cDC.migratory | Cyld |
| Prr14 | 4.36E-23 | 1.33778061 | 0.374 | 0.29 | 8.44E-19 | cDC.migratory | Prr14 |
| Stk38 | 4.64E-23 | 0.35705791 | 0.35 | 0.473 | 8.97E-19 | cDC.migratory | Stk38 |
| Dapp1 | 6.13E-23 | 0.265587 | 0.493 | 0.585 | 1.18E-18 | cDC.migratory | Dapp1 |
| Dnm1l | 7.84E-23 | 1.46378406 | 0.432 | 0.343 | 1.52E-18 | cDC.migratory | Dnm1l |
| Rapgef2 | 1.20E-22 | 1.35994202 | 0.323 | 0.255 | 2.32E-18 | cDC.migratory | Rapgef2 |
| Zfp366 | 1.95E-22 | 0.25139316 | 0.455 | 0.49 | 3.76E-18 | cDC.migratory | Zfp366 |
| Khnyn | 2.52E-22 | 1.70967134 | 0.331 | 0.22 | 4.87E-18 | cDC.migratory | Khnyn |
| Upf1 | 2.67E-22 | 0.32288761 | 0.357 | 0.486 | 5.16E-18 | cDC.migratory | Upf1 |
| Sema4a | 2.77E-22 | 1.13817689 | 0.658 | 0.581 | 5.35E-18 | cDC.migratory | Sema4a |
| Tubb2a | 4.33E-22 | 1.39155293 | 0.354 | 0.275 | 8.38E-18 | cDC.migratory | Tubb2a |
| Dpp4 | 4.76E-22 | 0.36640893 | 0.425 | 0.53 | 9.20E-18 | cDC.migratory | Dpp4 |
| Nelfe | 4.92E-22 | 0.29911629 | 0.202 | 0.298 | 9.52E-18 | cDC.migratory | Nelfe |
| Rhof | 6.19E-22 | 1.2443466 | 0.328 | 0.258 | 1.20E-17 | cDC.migratory | Rhof |
| Flt3 | 1.28E-21 | 1.31621533 | 0.441 | 0.318 | 2.48E-17 | cDC.migratory | Flt3 |
| Rubcn | 2.24E-21 | 1.69093752 | 0.253 | 0.16 | 4.32E-17 | cDC.migratory | Rubcn |
| Nop53 | 2.75E-21 | 0.49767967 | 0.506 | 0.619 | 5.31E-17 | cDC.migratory | Nop53 |
| Supt20 | 3.85E-21 | 0.55673187 | 0.391 | 0.509 | 7.44E-17 | cDC.migratory | Supt20 |
| Chchd2 | 4.00E-21 | 0.50758058 | 0.682 | 0.801 | 7.73E-17 | cDC.migratory | Chchd2 |
| Sub1 | 6.01E-21 | 0.26235359 | 0.491 | 0.595 | 1.16E-16 | cDC.migratory | Sub1 |
| Stub1 | 7.47E-21 | 0.41336766 | 0.364 | 0.472 | 1.44E-16 | cDC.migratory | Stub1 |
| Arrdc4 | 7.82E-21 | 0.37217077 | 0.321 | 0.428 | 1.51E-16 | cDC.migratory | Arrdc4 |
| Thoc7 | 7.86E-21 | 0.36113875 | 0.223 | 0.343 | 1.52E-16 | cDC.migratory | Thoc7 |
| Trim35 | 8.62E-21 | 1.19998781 | 0.62 | 0.461 | 1.67E-16 | cDC.migratory | Trim35 |
| Zmynd11 | 8.76E-21 | 0.36144834 | 0.352 | 0.467 | 1.69E-16 | cDC.migratory | Zmynd11 |
| Srrm2 | 9.09E-21 | 0.62889465 | 0.553 | 0.652 | 1.76E-16 | cDC.migratory | Srrm2 |
| Gcnt2 | 1.48E-20 | 0.2863596 | 0.4 | 0.497 | 2.86E-16 | cDC.migratory | Gcnt2 |
| Mob1a | 1.59E-20 | 0.44420461 | 0.616 | 0.739 | 3.08E-16 | cDC.migratory | Mob1a |
| Foxn2 | 1.61E-20 | 1.24214716 | 0.51 | 0.442 | 3.12E-16 | cDC.migratory | Foxn2 |
| R3hdm2 | 1.64E-20 | 0.30109102 | 0.399 | 0.52 | 3.17E-16 | cDC.migratory | R3hdm2 |
| Snd1 | 1.73E-20 | 0.56704589 | 0.738 | 0.842 | 3.34E-16 | cDC.migratory | Snd1 |
| Cxcr41 | 2.19E-20 | 0.26112176 | 0.528 | 0.551 | 4.24E-16 | cDC.migratory | Cxcr4 |
| Crybg1 | 3.10E-20 | 1.19308344 | 0.388 | 0.303 | 5.99E-16 | cDC.migratory | Crybg1 |
| Rbm5 | 3.11E-20 | 0.33186518 | 0.38 | 0.497 | 6.00E-16 | cDC.migratory | Rbm5 |
| Lrch4 | 3.52E-20 | 1.2270099 | 0.678 | 0.64 | 6.81E-16 | cDC.migratory | Lrch4 |
| Tagln21 | 3.81E-20 | 0.63349817 | 0.835 | 0.914 | 7.36E-16 | cDC.migratory | Tagln2 |
| Rab35 | 3.90E-20 | 0.25798925 | 0.355 | 0.478 | 7.54E-16 | cDC.migratory | Rab35 |
| 2-Sep | 6.13E-20 | 0.40843013 | 0.487 | 0.595 | 1.18E-15 | cDC.migratory | 2-Sep |
| Rbm42 | 6.42E-20 | 0.29848297 | 0.517 | 0.662 | 1.24E-15 | cDC.migratory | Rbm42 |
| Smarcc2 | 7.21E-20 | 0.53729364 | 0.467 | 0.586 | 1.39E-15 | cDC.migratory | Smarcc2 |
| Map2k2 | 9.68E-20 | 0.4140022 | 0.418 | 0.547 | 1.87E-15 | cDC.migratory | Map2k2 |
| Plekha1 | 1.08E-19 | 0.25580196 | 0.362 | 0.472 | 2.08E-15 | cDC.migratory | Plekha1 |
| Bmt2 | 1.17E-19 | 1.57464037 | 0.384 | 0.293 | 2.26E-15 | cDC.migratory | Bmt2 |
| Galnt12 | 1.43E-19 | 1.46207294 | 0.317 | 0.233 | 2.77E-15 | cDC.migratory | Galnt12 |
| Nup98 | 1.57E-19 | 0.32253886 | 0.45 | 0.543 | 3.03E-15 | cDC.migratory | Nup98 |
| Slc25a10 | 1.62E-19 | 1.70831281 | 0.304 | 0.183 | 3.13E-15 | cDC.migratory | Slc25a10 |
| Aamp | 1.82E-19 | 0.37557986 | 0.55 | 0.678 | 3.51E-15 | cDC.migratory | Aamp |
| Clasp1 | 1.98E-19 | 1.5735119 | 0.397 | 0.301 | 3.82E-15 | cDC.migratory | Clasp1 |
| Trip12 | 2.27E-19 | 0.30542794 | 0.554 | 0.694 | 4.39E-15 | cDC.migratory | Trip12 |
| Ip6k1 | 2.39E-19 | 0.25679595 | 0.3 | 0.429 | 4.63E-15 | cDC.migratory | Ip6k1 |
| Setd3 | 3.18E-19 | 0.30475392 | 0.322 | 0.446 | 6.14E-15 | cDC.migratory | Setd3 |
| Aup1 | 3.38E-19 | 0.46924542 | 0.495 | 0.608 | 6.54E-15 | cDC.migratory | Aup1 |
| Trip10 | 3.66E-19 | 1.52172503 | 0.332 | 0.231 | 7.07E-15 | cDC.migratory | Trip10 |
| Kdm4a | 3.80E-19 | 1.44069561 | 0.376 | 0.286 | 7.34E-15 | cDC.migratory | Kdm4a |
| Map2k7 | 4.05E-19 | 0.28250426 | 0.162 | 0.274 | 7.84E-15 | cDC.migratory | Map2k7 |
| Ccdc50 | 4.56E-19 | 1.29880696 | 0.5 | 0.456 | 8.82E-15 | cDC.migratory | Ccdc50 |
| Setd7 | 5.47E-19 | 0.40847202 | 0.255 | 0.377 | 1.06E-14 | cDC.migratory | Setd7 |
| Cyfip1 | 6.36E-19 | 1.05792538 | 0.579 | 0.56 | 1.23E-14 | cDC.migratory | Cyfip1 |
| Ccdc71l1 | 7.37E-19 | 0.57406723 | 0.253 | 0.31 | 1.43E-14 | cDC.migratory | Ccdc71l |
| Srpk1 | 7.48E-19 | 0.34380087 | 0.352 | 0.465 | 1.45E-14 | cDC.migratory | Srpk1 |
| Anxa21 | 1.46E-18 | 0.6925299 | 0.814 | 0.847 | 2.82E-14 | cDC.migratory | Anxa2 |
| F11r | 1.47E-18 | 1.26795373 | 0.428 | 0.328 | 2.83E-14 | cDC.migratory | F11r |
| Ankrd13c | 1.56E-18 | 0.25847279 | 0.233 | 0.352 | 3.01E-14 | cDC.migratory | Ankrd13c |
| Zfp622 | 1.79E-18 | 0.27055697 | 0.266 | 0.361 | 3.46E-14 | cDC.migratory | Zfp622 |
| Gnb1 | 2.41E-18 | 0.42470987 | 0.664 | 0.785 | 4.66E-14 | cDC.migratory | Gnb1 |
| St6galnac6 | 2.68E-18 | 1.4079034 | 0.281 | 0.183 | 5.18E-14 | cDC.migratory | St6galnac6 |
| Ss18 | 2.87E-18 | 0.32842827 | 0.228 | 0.36 | 5.55E-14 | cDC.migratory | Ss18 |
| Rtn4 | 3.11E-18 | 0.39081057 | 0.614 | 0.735 | 6.01E-14 | cDC.migratory | Rtn4 |
| Ccnl2 | 3.93E-18 | 0.40950327 | 0.423 | 0.525 | 7.61E-14 | cDC.migratory | Ccnl2 |
| Btbd9 | 3.99E-18 | 0.44098881 | 0.278 | 0.384 | 7.72E-14 | cDC.migratory | Btbd9 |
| Rac1 | 5.29E-18 | 0.39378577 | 0.398 | 0.525 | 1.02E-13 | cDC.migratory | Rac1 |
| Tut4 | 8.28E-18 | 0.25467164 | 0.176 | 0.286 | 1.60E-13 | cDC.migratory | Tut4 |
| Smarce1 | 8.39E-18 | 1.15438598 | 0.491 | 0.467 | 1.62E-13 | cDC.migratory | Smarce1 |
| Chd1 | 8.60E-18 | 0.28746103 | 0.355 | 0.468 | 1.66E-13 | cDC.migratory | Chd1 |
| Apaf1 | 9.26E-18 | 1.19932994 | 0.568 | 0.536 | 1.79E-13 | cDC.migratory | Apaf1 |
| Rab14 | 1.43E-17 | 0.48128886 | 0.561 | 0.672 | 2.76E-13 | cDC.migratory | Rab14 |
| Itm2c | 1.43E-17 | 0.51742386 | 0.738 | 0.851 | 2.76E-13 | cDC.migratory | Itm2c |
| Tbcd | 1.52E-17 | 0.30008059 | 0.185 | 0.292 | 2.93E-13 | cDC.migratory | Tbcd |
| Arfrp1 | 1.60E-17 | 1.54069946 | 0.321 | 0.241 | 3.09E-13 | cDC.migratory | Arfrp1 |
| Slc30a7 | 1.61E-17 | 0.25804835 | 0.212 | 0.313 | 3.11E-13 | cDC.migratory | Slc30a7 |
| Slc12a7 | 1.64E-17 | 0.40980735 | 0.142 | 0.252 | 3.17E-13 | cDC.migratory | Slc12a7 |
| Rufy3 | 1.67E-17 | 1.6108607 | 0.253 | 0.156 | 3.23E-13 | cDC.migratory | Rufy3 |
| Cep350 | 1.87E-17 | 1.24476784 | 0.465 | 0.388 | 3.61E-13 | cDC.migratory | Cep350 |
| Osbpl8 | 1.89E-17 | 1.18594848 | 0.489 | 0.433 | 3.66E-13 | cDC.migratory | Osbpl8 |
| Cox7a2l | 2.11E-17 | 0.29409828 | 0.545 | 0.657 | 4.08E-13 | cDC.migratory | Cox7a2l |
| Dync1h1 | 2.24E-17 | 0.44860558 | 0.527 | 0.644 | 4.33E-13 | cDC.migratory | Dync1h1 |
| Ncoa4 | 2.61E-17 | 1.04495899 | 0.449 | 0.42 | 5.05E-13 | cDC.migratory | Ncoa4 |
| Hvcn1 | 3.31E-17 | 1.4603525 | 0.259 | 0.139 | 6.40E-13 | cDC.migratory | Hvcn1 |
| Myh9 | 4.23E-17 | 1.04699692 | 0.7 | 0.724 | 8.17E-13 | cDC.migratory | Myh9 |
| Rabep1 | 4.40E-17 | 0.43429349 | 0.363 | 0.484 | 8.51E-13 | cDC.migratory | Rabep1 |
| Fhod1 | 4.85E-17 | 1.39456095 | 0.316 | 0.246 | 9.38E-13 | cDC.migratory | Fhod1 |
| Atxn7l3b | 8.04E-17 | 0.33588743 | 0.268 | 0.383 | 1.55E-12 | cDC.migratory | Atxn7l3b |
| Mxd1 | 9.13E-17 | 1.17435391 | 0.697 | 0.586 | 1.76E-12 | cDC.migratory | Mxd1 |
| Actr1a | 9.36E-17 | 0.26604578 | 0.431 | 0.558 | 1.81E-12 | cDC.migratory | Actr1a |
| Ago1 | 9.63E-17 | 1.52601283 | 0.307 | 0.222 | 1.86E-12 | cDC.migratory | Ago1 |
| Lpcat1 | 9.93E-17 | 1.30716128 | 0.367 | 0.27 | 1.92E-12 | cDC.migratory | Lpcat1 |
| Acadl | 1.24E-16 | 0.30043627 | 0.331 | 0.41 | 2.41E-12 | cDC.migratory | Acadl |
| Cmc1 | 1.60E-16 | 0.29969595 | 0.242 | 0.354 | 3.10E-12 | cDC.migratory | Cmc1 |
| Traf6 | 1.67E-16 | 1.14684861 | 0.43 | 0.36 | 3.22E-12 | cDC.migratory | Traf6 |
| Phf21a | 1.70E-16 | 1.65844501 | 0.288 | 0.21 | 3.29E-12 | cDC.migratory | Phf21a |
| Uba7 | 1.76E-16 | 1.3822982 | 0.428 | 0.365 | 3.39E-12 | cDC.migratory | Uba7 |
| Ep400 | 1.84E-16 | 0.26210272 | 0.351 | 0.455 | 3.56E-12 | cDC.migratory | Ep400 |
| Ncoa2 | 2.14E-16 | 1.34569812 | 0.37 | 0.306 | 4.13E-12 | cDC.migratory | Ncoa2 |
| Sdccag8 | 2.91E-16 | 0.48382543 | 0.261 | 0.363 | 5.62E-12 | cDC.migratory | Sdccag8 |
| Cdk17 | 3.08E-16 | 1.53095873 | 0.317 | 0.218 | 5.95E-12 | cDC.migratory | Cdk17 |
| Ap2b1 | 3.30E-16 | 0.30671248 | 0.439 | 0.569 | 6.39E-12 | cDC.migratory | Ap2b1 |
| Ctcf | 3.62E-16 | 0.28823237 | 0.2 | 0.299 | 6.99E-12 | cDC.migratory | Ctcf |
| Fbxo11 | 4.02E-16 | 1.12924334 | 0.527 | 0.462 | 7.78E-12 | cDC.migratory | Fbxo11 |
| Ctnnb1 | 4.17E-16 | 0.40647465 | 0.471 | 0.588 | 8.06E-12 | cDC.migratory | Ctnnb1 |
| Vdac1 | 4.49E-16 | 0.27559983 | 0.258 | 0.339 | 8.68E-12 | cDC.migratory | Vdac1 |
| Trafd1 | 5.01E-16 | 1.31135844 | 0.621 | 0.551 | 9.68E-12 | cDC.migratory | Trafd1 |
| Vps26a | 5.24E-16 | 0.31448094 | 0.332 | 0.436 | 1.01E-11 | cDC.migratory | Vps26a |
| Paf1 | 5.33E-16 | 0.40371108 | 0.272 | 0.389 | 1.03E-11 | cDC.migratory | Paf1 |
| Apeh | 5.38E-16 | 0.29484092 | 0.22 | 0.319 | 1.04E-11 | cDC.migratory | Apeh |
| Sp140 | 6.37E-16 | 1.30180501 | 0.344 | 0.287 | 1.23E-11 | cDC.migratory | Sp140 |
| Cul1 | 6.68E-16 | 0.43195747 | 0.381 | 0.478 | 1.29E-11 | cDC.migratory | Cul1 |
| Itpripl2 | 8.01E-16 | 0.41249998 | 0.406 | 0.501 | 1.55E-11 | cDC.migratory | Itpripl2 |
| Mtl1 | 8.15E-16 | 0.27387686 | 0.34 | 0.455 | 1.58E-11 | cDC.migratory | Mtl1 |
| Ak2 | 8.29E-16 | 0.43595923 | 0.432 | 0.523 | 1.60E-11 | cDC.migratory | Ak2 |
| Rbck1 | 8.48E-16 | 0.35942017 | 0.338 | 0.448 | 1.64E-11 | cDC.migratory | Rbck1 |
| Setd2 | 8.57E-16 | 0.33825499 | 0.292 | 0.392 | 1.66E-11 | cDC.migratory | Setd2 |
| Ppm1a | 9.01E-16 | 0.32322953 | 0.308 | 0.421 | 1.74E-11 | cDC.migratory | Ppm1a |
| Nampt | 9.36E-16 | 1.04670355 | 0.448 | 0.435 | 1.81E-11 | cDC.migratory | Nampt |
| Rnf6 | 1.08E-15 | 0.32111557 | 0.256 | 0.368 | 2.09E-11 | cDC.migratory | Rnf6 |
| Scaf11 | 1.16E-15 | 0.26284074 | 0.259 | 0.375 | 2.24E-11 | cDC.migratory | Scaf11 |
| Selplg | 1.27E-15 | 0.74546238 | 0.687 | 0.721 | 2.46E-11 | cDC.migratory | Selplg |
| Apc | 1.30E-15 | 0.25972043 | 0.156 | 0.25 | 2.51E-11 | cDC.migratory | Apc |
| Golga4 | 1.33E-15 | 0.28785719 | 0.204 | 0.309 | 2.56E-11 | cDC.migratory | Golga4 |
| Kmt2e | 1.34E-15 | 0.32392169 | 0.272 | 0.372 | 2.59E-11 | cDC.migratory | Kmt2e |
| Zc3h4 | 1.39E-15 | 0.38947151 | 0.276 | 0.385 | 2.69E-11 | cDC.migratory | Zc3h4 |
| Arid4a | 1.64E-15 | 0.41420471 | 0.224 | 0.321 | 3.16E-11 | cDC.migratory | Arid4a |
| Ripk2 | 1.71E-15 | 1.25719557 | 0.295 | 0.243 | 3.30E-11 | cDC.migratory | Ripk2 |
| Zbtb181 | 1.79E-15 | 1.04387389 | 0.635 | 0.571 | 3.46E-11 | cDC.migratory | Zbtb18 |
| Pcbp2 | 2.00E-15 | 0.38448553 | 0.41 | 0.545 | 3.86E-11 | cDC.migratory | Pcbp2 |
| Nrbf2 | 2.09E-15 | 1.63600402 | 0.263 | 0.187 | 4.05E-11 | cDC.migratory | Nrbf2 |
| Gsto1 | 2.12E-15 | 0.96751114 | 0.564 | 0.627 | 4.09E-11 | cDC.migratory | Gsto1 |
| Ubp1 | 2.16E-15 | 0.42547959 | 0.385 | 0.497 | 4.17E-11 | cDC.migratory | Ubp1 |
| Agap3 | 2.66E-15 | 0.26639305 | 0.259 | 0.349 | 5.15E-11 | cDC.migratory | Agap3 |
| Eif1a | 2.86E-15 | 0.26306728 | 0.266 | 0.395 | 5.52E-11 | cDC.migratory | Eif1a |
| Galnt1 | 3.45E-15 | 0.34379113 | 0.424 | 0.554 | 6.66E-11 | cDC.migratory | Galnt1 |
| Sav1 | 3.69E-15 | 1.19366618 | 0.482 | 0.441 | 7.14E-11 | cDC.migratory | Sav1 |
| Fip1l1 | 4.11E-15 | 0.36201645 | 0.186 | 0.29 | 7.95E-11 | cDC.migratory | Fip1l1 |
| Mcrip1 | 4.79E-15 | 0.28914085 | 0.314 | 0.427 | 9.27E-11 | cDC.migratory | Mcrip1 |
| Orc4 | 5.20E-15 | 0.42124597 | 0.197 | 0.284 | 1.01E-10 | cDC.migratory | Orc4 |
| Mt2 | 5.75E-15 | 1.17139867 | 0.278 | 0.184 | 1.11E-10 | cDC.migratory | Mt2 |
| Vbp1 | 6.04E-15 | 0.3422393 | 0.278 | 0.38 | 1.17E-10 | cDC.migratory | Vbp1 |
| Sri | 6.47E-15 | 0.96902341 | 0.708 | 0.712 | 1.25E-10 | cDC.migratory | Sri |
| Abi1 | 7.29E-15 | 0.49292237 | 0.606 | 0.713 | 1.41E-10 | cDC.migratory | Abi1 |
| Tgoln1 | 7.49E-15 | 0.28019628 | 0.329 | 0.442 | 1.45E-10 | cDC.migratory | Tgoln1 |
| Maea | 8.22E-15 | 0.42015718 | 0.401 | 0.505 | 1.59E-10 | cDC.migratory | Maea |
| Med13 | 1.04E-14 | 0.35233848 | 0.423 | 0.546 | 2.00E-10 | cDC.migratory | Med13 |
| Map3k3 | 1.11E-14 | 0.25515766 | 0.323 | 0.431 | 2.15E-10 | cDC.migratory | Map3k3 |
| Mpc1 | 1.27E-14 | 0.51440982 | 0.489 | 0.591 | 2.45E-10 | cDC.migratory | Mpc1 |
| Lcor | 1.33E-14 | 0.43346372 | 0.23 | 0.341 | 2.57E-10 | cDC.migratory | Lcor |
| Kmt2d | 1.39E-14 | 0.47534939 | 0.489 | 0.586 | 2.69E-10 | cDC.migratory | Kmt2d |
| Rbbp4 | 1.49E-14 | 0.37058455 | 0.397 | 0.5 | 2.88E-10 | cDC.migratory | Rbbp4 |
| Cmip | 1.55E-14 | 0.39763637 | 0.656 | 0.765 | 2.99E-10 | cDC.migratory | Cmip |
| Kpna3 | 1.94E-14 | 1.64887884 | 0.263 | 0.194 | 3.75E-10 | cDC.migratory | Kpna3 |
| Lamp1 | 1.95E-14 | 1.30016267 | 0.542 | 0.438 | 3.77E-10 | cDC.migratory | Lamp1 |
| Pde4b | 2.20E-14 | 0.58524491 | 0.655 | 0.706 | 4.26E-10 | cDC.migratory | Pde4b |
| Safb | 2.35E-14 | 0.25949491 | 0.228 | 0.324 | 4.55E-10 | cDC.migratory | Safb |
| Rbm33 | 2.43E-14 | 0.47562667 | 0.248 | 0.351 | 4.70E-10 | cDC.migratory | Rbm33 |
| Atp2b1 | 2.66E-14 | 0.52208599 | 0.392 | 0.517 | 5.14E-10 | cDC.migratory | Atp2b1 |
| Tbc1d10b | 2.72E-14 | 0.26160301 | 0.349 | 0.46 | 5.26E-10 | cDC.migratory | Tbc1d10b |
| Ahcyl1 | 3.26E-14 | 0.36272063 | 0.344 | 0.448 | 6.29E-10 | cDC.migratory | Ahcyl1 |
| Pisd | 3.35E-14 | 0.43246921 | 0.272 | 0.36 | 6.48E-10 | cDC.migratory | Pisd |
| Fhl3 | 3.45E-14 | 0.27618195 | 0.297 | 0.375 | 6.67E-10 | cDC.migratory | Fhl3 |
| Fxr2 | 3.98E-14 | 0.31059755 | 0.229 | 0.316 | 7.69E-10 | cDC.migratory | Fxr2 |
| Dcaf8 | 4.67E-14 | 0.25271826 | 0.193 | 0.294 | 9.03E-10 | cDC.migratory | Dcaf8 |
| Thoc2 | 5.38E-14 | 0.35478726 | 0.178 | 0.279 | 1.04E-09 | cDC.migratory | Thoc2 |
| Ampd2 | 5.53E-14 | 1.40326547 | 0.3 | 0.235 | 1.07E-09 | cDC.migratory | Ampd2 |
| Ube2e3 | 5.54E-14 | 0.26586532 | 0.253 | 0.332 | 1.07E-09 | cDC.migratory | Ube2e3 |
| Nras | 5.64E-14 | 0.26522379 | 0.298 | 0.422 | 1.09E-09 | cDC.migratory | Nras |
| Agtpbp1 | 5.84E-14 | 0.40461978 | 0.185 | 0.267 | 1.13E-09 | cDC.migratory | Agtpbp1 |
| Esrra | 6.66E-14 | 0.26741 | 0.289 | 0.395 | 1.29E-09 | cDC.migratory | Esrra |
| Vamp2 | 6.88E-14 | 0.29363773 | 0.163 | 0.258 | 1.33E-09 | cDC.migratory | Vamp2 |
| Creb5 | 7.00E-14 | 0.2819685 | 0.215 | 0.289 | 1.35E-09 | cDC.migratory | Creb5 |
| Exoc4 | 8.96E-14 | 0.75587966 | 0.28 | 0.374 | 1.73E-09 | cDC.migratory | Exoc4 |
| Arrdc1 | 9.22E-14 | 1.56871367 | 0.417 | 0.333 | 1.78E-09 | cDC.migratory | Arrdc1 |
| Ssu72 | 9.63E-14 | 0.28210195 | 0.435 | 0.533 | 1.86E-09 | cDC.migratory | Ssu72 |
| Dvl1 | 9.73E-14 | 1.49319762 | 0.312 | 0.241 | 1.88E-09 | cDC.migratory | Dvl1 |
| Tnfaip8 | 1.07E-13 | 0.43551165 | 0.575 | 0.669 | 2.07E-09 | cDC.migratory | Tnfaip8 |
| Gramd1a | 1.17E-13 | 0.55800719 | 0.314 | 0.4 | 2.26E-09 | cDC.migratory | Gramd1a |
| Tap2 | 1.20E-13 | 0.54762839 | 0.635 | 0.698 | 2.31E-09 | cDC.migratory | Tap2 |
| mt-Nd4 | 1.24E-13 | 0.29044713 | 0.231 | 0.313 | 2.40E-09 | cDC.migratory | mt-Nd4 |
| Flii | 1.33E-13 | 0.34133759 | 0.384 | 0.483 | 2.56E-09 | cDC.migratory | Flii |
| Mta1 | 1.34E-13 | 0.42073823 | 0.252 | 0.358 | 2.59E-09 | cDC.migratory | Mta1 |
| Ccdc59 | 1.44E-13 | 0.32306313 | 0.16 | 0.255 | 2.78E-09 | cDC.migratory | Ccdc59 |
| Plscr1 | 1.60E-13 | 0.45913993 | 0.434 | 0.483 | 3.09E-09 | cDC.migratory | Plscr1 |
| Prr12 | 1.74E-13 | 1.50056935 | 0.322 | 0.241 | 3.36E-09 | cDC.migratory | Prr12 |
| P2rx4 | 1.92E-13 | 0.54070947 | 0.343 | 0.418 | 3.71E-09 | cDC.migratory | P2rx4 |
| Psmb8 | 2.29E-13 | 0.56182702 | 0.763 | 0.865 | 4.42E-09 | cDC.migratory | Psmb8 |
| Med11 | 2.31E-13 | 0.28480201 | 0.233 | 0.334 | 4.46E-09 | cDC.migratory | Med11 |
| Ppp1cb | 3.50E-13 | 0.26100354 | 0.388 | 0.517 | 6.77E-09 | cDC.migratory | Ppp1cb |
| Elf2 | 4.26E-13 | 0.29589538 | 0.208 | 0.29 | 8.24E-09 | cDC.migratory | Elf2 |
| Gmps | 6.04E-13 | 0.32179725 | 0.175 | 0.267 | 1.17E-08 | cDC.migratory | Gmps |
| Pabpn1 | 8.47E-13 | 0.63057854 | 0.561 | 0.649 | 1.64E-08 | cDC.migratory | Pabpn1 |
| Sertad1 | 8.57E-13 | 0.32673334 | 0.24 | 0.309 | 1.66E-08 | cDC.migratory | Sertad1 |
| Mfsd14b | 9.50E-13 | 0.26353231 | 0.3 | 0.41 | 1.84E-08 | cDC.migratory | Mfsd14b |
| mt-Nd3 | 1.05E-12 | 0.37765535 | 0.205 | 0.292 | 2.03E-08 | cDC.migratory | mt-Nd3 |
| Atxn7l3 | 1.08E-12 | 0.38664682 | 0.308 | 0.405 | 2.09E-08 | cDC.migratory | Atxn7l3 |
| Dync1i2 | 1.11E-12 | 0.27316976 | 0.299 | 0.412 | 2.15E-08 | cDC.migratory | Dync1i2 |
| Wars | 1.15E-12 | 0.29495373 | 0.197 | 0.28 | 2.23E-08 | cDC.migratory | Wars |
| Hsd3b7 | 1.32E-12 | 0.44293188 | 0.219 | 0.285 | 2.55E-08 | cDC.migratory | Hsd3b7 |
| Tax1bp1 | 1.36E-12 | 0.53722794 | 0.483 | 0.583 | 2.62E-08 | cDC.migratory | Tax1bp1 |
| Uvrag | 1.45E-12 | 1.09879061 | 0.674 | 0.667 | 2.80E-08 | cDC.migratory | Uvrag |
| Tbc1d1 | 2.01E-12 | 1.11530416 | 0.483 | 0.449 | 3.88E-08 | cDC.migratory | Tbc1d1 |
| Elf4 | 2.05E-12 | 0.38014122 | 0.336 | 0.421 | 3.96E-08 | cDC.migratory | Elf4 |
| Akap9 | 2.42E-12 | 1.42074404 | 0.317 | 0.261 | 4.68E-08 | cDC.migratory | Akap9 |
| Aagab | 2.49E-12 | 0.41241634 | 0.274 | 0.363 | 4.81E-08 | cDC.migratory | Aagab |
| Dnajc5 | 2.52E-12 | 0.51405892 | 0.414 | 0.497 | 4.86E-08 | cDC.migratory | Dnajc5 |
| Lnx2 | 2.72E-12 | 1.19784035 | 0.303 | 0.234 | 5.27E-08 | cDC.migratory | Lnx2 |
| Psd4 | 2.99E-12 | 0.38044719 | 0.414 | 0.518 | 5.78E-08 | cDC.migratory | Psd4 |
| Coa3 | 3.04E-12 | 0.27842374 | 0.177 | 0.268 | 5.87E-08 | cDC.migratory | Coa3 |
| Pla2g7 | 3.06E-12 | 0.76428941 | 0.251 | 0.24 | 5.92E-08 | cDC.migratory | Pla2g7 |
| Pmaip1 | 3.08E-12 | 0.46248057 | 0.533 | 0.56 | 5.96E-08 | cDC.migratory | Pmaip1 |
| Tmbim1 | 3.26E-12 | 0.34754949 | 0.35 | 0.442 | 6.31E-08 | cDC.migratory | Tmbim1 |
| Cnot2 | 3.32E-12 | 0.38064209 | 0.314 | 0.402 | 6.43E-08 | cDC.migratory | Cnot2 |
| Mink1 | 3.74E-12 | 1.20302593 | 0.517 | 0.48 | 7.24E-08 | cDC.migratory | Mink1 |
| Slc20a1 | 3.79E-12 | 0.57619647 | 0.265 | 0.297 | 7.33E-08 | cDC.migratory | Slc20a1 |
| Plgrkt | 4.52E-12 | 0.4375774 | 0.372 | 0.461 | 8.73E-08 | cDC.migratory | Plgrkt |
| Stx8 | 5.86E-12 | 1.42619304 | 0.433 | 0.355 | 1.13E-07 | cDC.migratory | Stx8 |
| Flna1 | 6.30E-12 | 0.41221936 | 0.622 | 0.763 | 1.22E-07 | cDC.migratory | Flna |
| Dock7 | 6.32E-12 | 0.56183724 | 0.248 | 0.324 | 1.22E-07 | cDC.migratory | Dock7 |
| Lrch1 | 6.85E-12 | 0.53337076 | 0.284 | 0.395 | 1.32E-07 | cDC.migratory | Lrch1 |
| Chmp1a | 7.00E-12 | 0.28559399 | 0.37 | 0.462 | 1.35E-07 | cDC.migratory | Chmp1a |
| Kmt5a | 7.69E-12 | 0.71894157 | 0.373 | 0.467 | 1.49E-07 | cDC.migratory | Kmt5a |
| Tbc1d9b | 7.74E-12 | 0.3378437 | 0.305 | 0.399 | 1.50E-07 | cDC.migratory | Tbc1d9b |
| Nmt1 | 8.19E-12 | 0.46086959 | 0.529 | 0.63 | 1.58E-07 | cDC.migratory | Nmt1 |
| Ppp4r2 | 8.31E-12 | 1.02938884 | 0.513 | 0.511 | 1.61E-07 | cDC.migratory | Ppp4r2 |
| Ints3 | 8.53E-12 | 1.14484454 | 0.305 | 0.276 | 1.65E-07 | cDC.migratory | Ints3 |
| Fmr1 | 9.74E-12 | 0.32957984 | 0.2 | 0.281 | 1.88E-07 | cDC.migratory | Fmr1 |
| Phf20l1 | 1.04E-11 | 0.33946326 | 0.185 | 0.27 | 2.01E-07 | cDC.migratory | Phf20l1 |
| Nek6 | 1.10E-11 | 1.36805213 | 0.38 | 0.317 | 2.12E-07 | cDC.migratory | Nek6 |
| Vdr1 | 1.56E-11 | 0.35024197 | 0.404 | 0.472 | 3.03E-07 | cDC.migratory | Vdr |
| Med151 | 1.63E-11 | 0.26042917 | 0.377 | 0.479 | 3.15E-07 | cDC.migratory | Med15 |
| Acbd3 | 1.85E-11 | 0.402049 | 0.283 | 0.384 | 3.58E-07 | cDC.migratory | Acbd3 |
| Avpi1 | 1.98E-11 | 1.05079443 | 0.354 | 0.289 | 3.84E-07 | cDC.migratory | Avpi1 |
| Ccdc9 | 2.35E-11 | 0.43796926 | 0.305 | 0.393 | 4.54E-07 | cDC.migratory | Ccdc9 |
| Mbp | 2.36E-11 | 1.14005286 | 0.497 | 0.45 | 4.56E-07 | cDC.migratory | Mbp |
| Pitpnm1 | 2.38E-11 | 0.55754956 | 0.333 | 0.416 | 4.61E-07 | cDC.migratory | Pitpnm1 |
| Tbc1d10a | 2.48E-11 | 1.40446542 | 0.323 | 0.243 | 4.79E-07 | cDC.migratory | Tbc1d10a |
| Rexo4 | 2.61E-11 | 0.27706303 | 0.179 | 0.269 | 5.05E-07 | cDC.migratory | Rexo4 |
| Smg1 | 2.64E-11 | 0.40935927 | 0.404 | 0.495 | 5.10E-07 | cDC.migratory | Smg1 |
| Sppl2a1 | 2.72E-11 | 0.3138591 | 0.443 | 0.544 | 5.26E-07 | cDC.migratory | Sppl2a |
| Brd8 | 2.92E-11 | 0.47144243 | 0.206 | 0.271 | 5.64E-07 | cDC.migratory | Brd8 |
| Nvl | 2.99E-11 | 0.47521206 | 0.187 | 0.268 | 5.78E-07 | cDC.migratory | Nvl |
| Chd8 | 3.38E-11 | 0.387921 | 0.237 | 0.328 | 6.53E-07 | cDC.migratory | Chd8 |
| Rai1 | 3.48E-11 | 0.37765685 | 0.191 | 0.269 | 6.73E-07 | cDC.migratory | Rai1 |
| Zgpat | 4.24E-11 | 1.52764564 | 0.274 | 0.223 | 8.20E-07 | cDC.migratory | Zgpat |
| Frmd8 | 4.47E-11 | 0.256277 | 0.248 | 0.35 | 8.65E-07 | cDC.migratory | Frmd8 |
| Gosr2 | 4.86E-11 | 0.36278616 | 0.558 | 0.673 | 9.39E-07 | cDC.migratory | Gosr2 |
| Mthfs1 | 4.94E-11 | 0.36297481 | 0.265 | 0.317 | 9.54E-07 | cDC.migratory | Mthfs |
| Atf7ip | 6.34E-11 | 0.27719448 | 0.274 | 0.351 | 1.23E-06 | cDC.migratory | Atf7ip |
| Smcr8 | 7.24E-11 | 0.28255674 | 0.232 | 0.307 | 1.40E-06 | cDC.migratory | Smcr8 |
| Pdcd10 | 7.28E-11 | 0.28530395 | 0.3 | 0.371 | 1.41E-06 | cDC.migratory | Pdcd10 |
| Icosl1 | 7.47E-11 | 0.33173135 | 0.284 | 0.421 | 1.44E-06 | cDC.migratory | Icosl |
| Otulin | 8.35E-11 | 0.47786389 | 0.333 | 0.441 | 1.61E-06 | cDC.migratory | Otulin |
| Ppil2 | 8.60E-11 | 0.40252086 | 0.244 | 0.321 | 1.66E-06 | cDC.migratory | Ppil2 |
| Soga11 | 8.69E-11 | 0.35325 | 0.217 | 0.297 | 1.68E-06 | cDC.migratory | Soga1 |
| Fastk | 9.22E-11 | 0.30781781 | 0.167 | 0.25 | 1.78E-06 | cDC.migratory | Fastk |
| Foxn3 | 1.09E-10 | 1.15572643 | 0.388 | 0.342 | 2.10E-06 | cDC.migratory | Foxn3 |
| Gtf2a1 | 1.22E-10 | 1.11917029 | 0.482 | 0.443 | 2.36E-06 | cDC.migratory | Gtf2a1 |
| Gnas | 1.24E-10 | 0.43312223 | 0.645 | 0.761 | 2.40E-06 | cDC.migratory | Gnas |
| Nsmce2 | 1.32E-10 | 0.40994709 | 0.188 | 0.269 | 2.56E-06 | cDC.migratory | Nsmce2 |
| Ap3b1 | 1.48E-10 | 1.22112752 | 0.517 | 0.549 | 2.86E-06 | cDC.migratory | Ap3b1 |
| Snx10 | 1.60E-10 | 0.34509528 | 0.179 | 0.258 | 3.09E-06 | cDC.migratory | Snx10 |
| Hmgcs1 | 1.70E-10 | 0.28355729 | 0.395 | 0.46 | 3.29E-06 | cDC.migratory | Hmgcs1 |
| Cnppd1 | 1.79E-10 | 0.3134759 | 0.362 | 0.46 | 3.46E-06 | cDC.migratory | Cnppd1 |
| Clk3 | 1.82E-10 | 0.40816901 | 0.256 | 0.345 | 3.52E-06 | cDC.migratory | Clk3 |
| Mgat4a | 2.08E-10 | 0.33321475 | 0.236 | 0.308 | 4.03E-06 | cDC.migratory | Mgat4a |
| Clk2 | 2.24E-10 | 0.40277092 | 0.232 | 0.32 | 4.32E-06 | cDC.migratory | Clk2 |
| Nr4a3 | 2.32E-10 | 0.62697501 | 0.722 | 0.683 | 4.48E-06 | cDC.migratory | Nr4a3 |
| Usp19 | 2.32E-10 | 0.27210317 | 0.3 | 0.392 | 4.48E-06 | cDC.migratory | Usp19 |
| Sdf4 | 2.34E-10 | 0.57023066 | 0.547 | 0.65 | 4.52E-06 | cDC.migratory | Sdf4 |
| Pgap2 | 2.38E-10 | 1.24736109 | 0.378 | 0.318 | 4.61E-06 | cDC.migratory | Pgap2 |
| Eapp | 2.48E-10 | 0.39097242 | 0.288 | 0.368 | 4.80E-06 | cDC.migratory | Eapp |
| Pnpla2 | 2.61E-10 | 0.47615463 | 0.377 | 0.458 | 5.05E-06 | cDC.migratory | Pnpla2 |
| Coa5 | 2.66E-10 | 0.53046147 | 0.382 | 0.471 | 5.14E-06 | cDC.migratory | Coa5 |
| Txn1 | 2.71E-10 | 1.19760463 | 0.526 | 0.496 | 5.25E-06 | cDC.migratory | Txn1 |
| Vps35l | 3.13E-10 | 0.32894459 | 0.27 | 0.367 | 6.04E-06 | cDC.migratory | Vps35l |
| Utp3 | 3.34E-10 | 0.38813488 | 0.179 | 0.262 | 6.45E-06 | cDC.migratory | Utp3 |
| Zfand3 | 3.42E-10 | 0.38637524 | 0.561 | 0.663 | 6.61E-06 | cDC.migratory | Zfand3 |
| Gtpbp4 | 3.68E-10 | 0.32083976 | 0.38 | 0.515 | 7.11E-06 | cDC.migratory | Gtpbp4 |
| Mtor | 3.78E-10 | 1.3060848 | 0.361 | 0.325 | 7.32E-06 | cDC.migratory | Mtor |
| Taok2 | 3.95E-10 | 0.75302139 | 0.228 | 0.311 | 7.64E-06 | cDC.migratory | Taok2 |
| Rptor | 4.09E-10 | 1.22205911 | 0.33 | 0.271 | 7.91E-06 | cDC.migratory | Rptor |
| Rbm17 | 4.38E-10 | 0.63604842 | 0.487 | 0.561 | 8.46E-06 | cDC.migratory | Rbm17 |
| Setd1a | 4.51E-10 | 0.42311494 | 0.241 | 0.334 | 8.72E-06 | cDC.migratory | Setd1a |
| Zswim8 | 4.55E-10 | 0.48021252 | 0.329 | 0.411 | 8.79E-06 | cDC.migratory | Zswim8 |
| Nfil3 | 4.56E-10 | 0.70299199 | 0.617 | 0.618 | 8.82E-06 | cDC.migratory | Nfil3 |
| Gpbp1l1 | 4.85E-10 | 0.30730995 | 0.289 | 0.377 | 9.37E-06 | cDC.migratory | Gpbp1l1 |
| Capn1 | 5.23E-10 | 0.54896659 | 0.325 | 0.403 | 1.01E-05 | cDC.migratory | Capn1 |
| Mprip | 5.84E-10 | 0.57662641 | 0.336 | 0.435 | 1.13E-05 | cDC.migratory | Mprip |
| Scarb2 | 6.40E-10 | 0.28902079 | 0.311 | 0.394 | 1.24E-05 | cDC.migratory | Scarb2 |
| Fbxo6 | 6.63E-10 | 0.26681128 | 0.215 | 0.302 | 1.28E-05 | cDC.migratory | Fbxo6 |
| Arfgap1 | 6.95E-10 | 0.38967933 | 0.195 | 0.265 | 1.34E-05 | cDC.migratory | Arfgap1 |
| N4bp2l2 | 7.47E-10 | 0.34866864 | 0.233 | 0.308 | 1.44E-05 | cDC.migratory | N4bp2l2 |
| Dda1 | 7.82E-10 | 0.29598411 | 0.226 | 0.313 | 1.51E-05 | cDC.migratory | Dda1 |
| Nedd4l | 7.91E-10 | 1.22155671 | 0.309 | 0.267 | 1.53E-05 | cDC.migratory | Nedd4l |
| Ift52 | 8.34E-10 | 0.4459893 | 0.228 | 0.305 | 1.61E-05 | cDC.migratory | Ift52 |
| Tfg | 8.50E-10 | 0.33556088 | 0.325 | 0.423 | 1.64E-05 | cDC.migratory | Tfg |
| Mier1 | 8.80E-10 | 0.43458143 | 0.353 | 0.423 | 1.70E-05 | cDC.migratory | Mier1 |
| Mtmr14 | 9.02E-10 | 1.23179673 | 0.27 | 0.217 | 1.74E-05 | cDC.migratory | Mtmr14 |
| Psmb10 | 9.86E-10 | 0.4912661 | 0.432 | 0.524 | 1.91E-05 | cDC.migratory | Psmb10 |
| Csf2rb21 | 9.99E-10 | 0.38808378 | 0.491 | 0.526 | 1.93E-05 | cDC.migratory | Csf2rb2 |
| Heca | 1.04E-09 | 0.47045357 | 0.266 | 0.35 | 2.00E-05 | cDC.migratory | Heca |
| Crebzf | 1.14E-09 | 0.78358722 | 0.23 | 0.303 | 2.21E-05 | cDC.migratory | Crebzf |
| Fbrs | 1.18E-09 | 0.37982885 | 0.551 | 0.672 | 2.28E-05 | cDC.migratory | Fbrs |
| Bbx | 1.19E-09 | 0.34915046 | 0.175 | 0.263 | 2.30E-05 | cDC.migratory | Bbx |
| Rtraf | 1.25E-09 | 0.37444525 | 0.237 | 0.307 | 2.42E-05 | cDC.migratory | Rtraf |
| Jak2 | 1.36E-09 | 0.8713597 | 0.882 | 0.909 | 2.64E-05 | cDC.migratory | Jak2 |
| Prpf6 | 1.47E-09 | 0.511285 | 0.206 | 0.284 | 2.84E-05 | cDC.migratory | Prpf6 |
| Mtmr6 | 1.51E-09 | 0.49200443 | 0.439 | 0.525 | 2.93E-05 | cDC.migratory | Mtmr6 |
| Tbrg1 | 1.51E-09 | 0.46177825 | 0.373 | 0.435 | 2.93E-05 | cDC.migratory | Tbrg1 |
| Cpne1 | 1.59E-09 | 0.34376949 | 0.294 | 0.379 | 3.08E-05 | cDC.migratory | Cpne1 |
| Myo9b | 1.79E-09 | 0.45580507 | 0.27 | 0.354 | 3.46E-05 | cDC.migratory | Myo9b |
| Mysm1 | 1.81E-09 | 0.27556283 | 0.215 | 0.278 | 3.51E-05 | cDC.migratory | Mysm1 |
| Ccnd2 | 2.07E-09 | 0.42379012 | 0.527 | 0.664 | 3.99E-05 | cDC.migratory | Ccnd2 |
| Nsfl1c | 2.18E-09 | 0.64300902 | 0.241 | 0.321 | 4.22E-05 | cDC.migratory | Nsfl1c |
| Gpd2 | 2.20E-09 | 1.19830553 | 0.455 | 0.417 | 4.26E-05 | cDC.migratory | Gpd2 |
| Dnajb12 | 2.23E-09 | 0.41628108 | 0.261 | 0.339 | 4.31E-05 | cDC.migratory | Dnajb12 |
| Ubxn1 | 2.34E-09 | 0.39136256 | 0.526 | 0.618 | 4.52E-05 | cDC.migratory | Ubxn1 |
| Lpin2 | 2.54E-09 | 1.63875902 | 0.288 | 0.21 | 4.91E-05 | cDC.migratory | Lpin2 |
| Glyr1 | 2.58E-09 | 0.48881851 | 0.48 | 0.571 | 4.99E-05 | cDC.migratory | Glyr1 |
| Ubash3b | 2.59E-09 | 0.56838276 | 0.24 | 0.314 | 5.01E-05 | cDC.migratory | Ubash3b |
| Mkln1 | 2.76E-09 | 0.3649095 | 0.254 | 0.331 | 5.33E-05 | cDC.migratory | Mkln1 |
| Zswim41 | 2.81E-09 | 0.33384839 | 0.243 | 0.31 | 5.44E-05 | cDC.migratory | Zswim4 |
| Tnrc18 | 2.95E-09 | 0.52650347 | 0.429 | 0.506 | 5.71E-05 | cDC.migratory | Tnrc18 |
| Meaf6 | 3.36E-09 | 0.64710529 | 0.29 | 0.341 | 6.49E-05 | cDC.migratory | Meaf6 |
| Ppp1r15a | 3.72E-09 | 0.56594808 | 0.593 | 0.629 | 7.19E-05 | cDC.migratory | Ppp1r15a |
| Hp1bp3 | 4.33E-09 | 0.41182785 | 0.387 | 0.475 | 8.37E-05 | cDC.migratory | Hp1bp3 |
| Gpr107 | 4.53E-09 | 0.31427815 | 0.199 | 0.269 | 8.76E-05 | cDC.migratory | Gpr107 |
| Zfp638 | 4.89E-09 | 0.29861092 | 0.197 | 0.286 | 9.46E-05 | cDC.migratory | Zfp638 |
| Cirbp | 4.98E-09 | 1.18302312 | 0.414 | 0.359 | 9.63E-05 | cDC.migratory | Cirbp |
| Appl1 | 5.14E-09 | 1.13852031 | 0.449 | 0.406 | 9.94E-05 | cDC.migratory | Appl1 |
| Fibp | 5.17E-09 | 0.53990831 | 0.302 | 0.371 | 9.99E-05 | cDC.migratory | Fibp |
| Tdrd7 | 5.77E-09 | 1.09419088 | 0.269 | 0.229 | 0.00011157 | cDC.migratory | Tdrd7 |
| Usp34 | 6.27E-09 | 0.32569332 | 0.219 | 0.299 | 0.00012127 | cDC.migratory | Usp34 |
| Epc1 | 6.54E-09 | 0.51235715 | 0.386 | 0.438 | 0.00012649 | cDC.migratory | Epc1 |
| Prex1 | 6.76E-09 | 0.67919179 | 0.785 | 0.848 | 0.00013076 | cDC.migratory | Prex1 |
| Socs1 | 7.06E-09 | 1.37448924 | 0.381 | 0.259 | 0.00013649 | cDC.migratory | Socs1 |
| Ube4a | 7.99E-09 | 0.25126016 | 0.199 | 0.266 | 0.00015451 | cDC.migratory | Ube4a |
| Tnrc6c | 8.03E-09 | 0.45644761 | 0.222 | 0.298 | 0.00015524 | cDC.migratory | Tnrc6c |
| Cdk12 | 8.37E-09 | 0.59993996 | 0.323 | 0.389 | 0.00016181 | cDC.migratory | Cdk12 |
| Brms1 | 9.13E-09 | 0.29530686 | 0.267 | 0.36 | 0.00017654 | cDC.migratory | Brms1 |
| Baz2b | 9.32E-09 | 0.52819301 | 0.305 | 0.364 | 0.00018012 | cDC.migratory | Baz2b |
| Tpd52l2 | 1.05E-08 | 0.4556602 | 0.287 | 0.384 | 0.00020313 | cDC.migratory | Tpd52l2 |
| Gatad1 | 1.07E-08 | 0.2986923 | 0.221 | 0.297 | 0.00020594 | cDC.migratory | Gatad1 |
| Ino80b | 1.09E-08 | 0.28687839 | 0.191 | 0.263 | 0.00021153 | cDC.migratory | Ino80b |
| Cdk5 | 1.22E-08 | 0.4641428 | 0.177 | 0.251 | 0.00023558 | cDC.migratory | Cdk5 |
| Akap13 | 1.27E-08 | 0.90228416 | 0.693 | 0.72 | 0.00024527 | cDC.migratory | Akap13 |
| Cdyl | 1.43E-08 | 0.38899323 | 0.291 | 0.379 | 0.00027626 | cDC.migratory | Cdyl |
| Rasa4 | 1.56E-08 | 0.42317815 | 0.279 | 0.367 | 0.00030116 | cDC.migratory | Rasa4 |
| Riok3 | 1.59E-08 | 0.5132642 | 0.584 | 0.687 | 0.00030817 | cDC.migratory | Riok3 |
| Ing1 | 1.67E-08 | 0.36203748 | 0.208 | 0.281 | 0.00032208 | cDC.migratory | Ing1 |
| Syap1 | 1.73E-08 | 0.36154081 | 0.246 | 0.321 | 0.00033518 | cDC.migratory | Syap1 |
| Gpalpp1 | 1.82E-08 | 0.30501342 | 0.195 | 0.265 | 0.00035112 | cDC.migratory | Gpalpp1 |
| Nxf1 | 1.82E-08 | 1.14518984 | 0.515 | 0.496 | 0.00035256 | cDC.migratory | Nxf1 |
| Cox17 | 1.92E-08 | 0.4989087 | 0.627 | 0.715 | 0.00037181 | cDC.migratory | Cox17 |
| Med20 | 1.93E-08 | 0.45277713 | 0.204 | 0.272 | 0.00037317 | cDC.migratory | Med20 |
| Bptf | 2.23E-08 | 0.33005395 | 0.281 | 0.362 | 0.0004309 | cDC.migratory | Bptf |
| Ciz1 | 2.29E-08 | 0.45913737 | 0.239 | 0.299 | 0.00044264 | cDC.migratory | Ciz1 |
| Tsta3 | 2.30E-08 | 0.29414494 | 0.228 | 0.299 | 0.00044376 | cDC.migratory | Tsta3 |
| Cpne2 | 3.10E-08 | 1.14949463 | 0.449 | 0.412 | 0.00060022 | cDC.migratory | Cpne2 |
| Tmx3 | 3.29E-08 | 0.53886405 | 0.307 | 0.4 | 0.00063632 | cDC.migratory | Tmx3 |
| Sdf2 | 3.30E-08 | 0.3559161 | 0.3 | 0.378 | 0.00063709 | cDC.migratory | Sdf2 |
| Trappc6b | 3.42E-08 | 0.37493325 | 0.247 | 0.327 | 0.00066216 | cDC.migratory | Trappc6b |
| Lmbr1l | 3.45E-08 | 1.17941283 | 0.284 | 0.23 | 0.00066717 | cDC.migratory | Lmbr1l |
| Fam104a | 3.50E-08 | 0.39093856 | 0.214 | 0.288 | 0.0006774 | cDC.migratory | Fam104a |
| Vcp | 3.55E-08 | 0.60615248 | 0.723 | 0.834 | 0.00068677 | cDC.migratory | Vcp |
| Stard7 | 3.56E-08 | 1.17568065 | 0.416 | 0.363 | 0.00068766 | cDC.migratory | Stard7 |
| Psen2 | 3.64E-08 | 1.43687441 | 0.275 | 0.227 | 0.00070454 | cDC.migratory | Psen2 |
| Akt3 | 3.70E-08 | 1.15609276 | 0.387 | 0.35 | 0.00071487 | cDC.migratory | Akt3 |
| Fam98c | 3.75E-08 | 0.43597293 | 0.322 | 0.41 | 0.00072433 | cDC.migratory | Fam98c |
| Appbp2 | 4.26E-08 | 0.32719145 | 0.215 | 0.285 | 0.0008246 | cDC.migratory | Appbp2 |
| Ilrun | 4.39E-08 | 1.05563482 | 0.494 | 0.485 | 0.00084838 | cDC.migratory | Ilrun |
| Pwwp2a | 4.49E-08 | 0.36763196 | 0.197 | 0.265 | 0.00086903 | cDC.migratory | Pwwp2a |
| Hook3 | 4.50E-08 | 0.41827983 | 0.208 | 0.268 | 0.00086944 | cDC.migratory | Hook3 |
| Abhd16a | 4.50E-08 | 0.48632357 | 0.191 | 0.268 | 0.00087039 | cDC.migratory | Abhd16a |
| Cand1 | 4.85E-08 | 0.43252908 | 0.356 | 0.43 | 0.00093774 | cDC.migratory | Cand1 |
| Ccni | 4.87E-08 | 0.97790276 | 0.517 | 0.524 | 0.00094223 | cDC.migratory | Ccni |
| Tollip | 5.16E-08 | 0.32740529 | 0.219 | 0.286 | 0.00099791 | cDC.migratory | Tollip |
| Golgb1 | 5.53E-08 | 0.58617848 | 0.215 | 0.271 | 0.00106893 | cDC.migratory | Golgb1 |
| Ehbp1l1 | 5.89E-08 | 0.65070665 | 0.673 | 0.759 | 0.0011386 | cDC.migratory | Ehbp1l1 |
| Ambra1 | 6.38E-08 | 0.37627161 | 0.29 | 0.369 | 0.00123406 | cDC.migratory | Ambra1 |
| Scpep1 | 7.32E-08 | 0.88933074 | 0.611 | 0.63 | 0.00141435 | cDC.migratory | Scpep1 |
| Stk38l | 8.30E-08 | 0.50444334 | 0.391 | 0.454 | 0.00160494 | cDC.migratory | Stk38l |
| Fam192a | 8.41E-08 | 0.46690779 | 0.22 | 0.278 | 0.00162615 | cDC.migratory | Fam192a |
| Kdm3b | 8.50E-08 | 0.63380999 | 0.483 | 0.559 | 0.00164349 | cDC.migratory | Kdm3b |
| Rbm14 | 8.51E-08 | 0.63375214 | 0.376 | 0.452 | 0.00164586 | cDC.migratory | Rbm14 |
| Pmvk | 8.77E-08 | 1.09252323 | 0.496 | 0.469 | 0.00169469 | cDC.migratory | Pmvk |
| Srcap | 8.79E-08 | 0.7093907 | 0.362 | 0.433 | 0.00170011 | cDC.migratory | Srcap |
| Add1 | 8.81E-08 | 1.14201918 | 0.508 | 0.476 | 0.00170262 | cDC.migratory | Add1 |
| Mbtd1 | 9.00E-08 | 0.36137858 | 0.348 | 0.435 | 0.0017391 | cDC.migratory | Mbtd1 |
| Sik2 | 9.44E-08 | 0.2886725 | 0.224 | 0.288 | 0.00182537 | cDC.migratory | Sik2 |
| Suz12 | 9.84E-08 | 0.46355621 | 0.238 | 0.299 | 0.0019017 | cDC.migratory | Suz12 |
| Atf7 | 1.05E-07 | 0.36292722 | 0.179 | 0.255 | 0.00202809 | cDC.migratory | Atf7 |
| Pfdn5 | 1.12E-07 | 0.5469155 | 0.531 | 0.609 | 0.00217286 | cDC.migratory | Pfdn5 |
| Sbno1 | 1.20E-07 | 1.07976291 | 0.447 | 0.398 | 0.00231282 | cDC.migratory | Sbno1 |
| Ugdh | 1.44E-07 | 0.43289742 | 0.241 | 0.331 | 0.00278658 | cDC.migratory | Ugdh |
| Rala | 1.45E-07 | 0.41963734 | 0.365 | 0.436 | 0.00279781 | cDC.migratory | Rala |
| St8sia4 | 1.50E-07 | 0.87658016 | 0.652 | 0.582 | 0.00290255 | cDC.migratory | St8sia4 |
| Plcl2 | 1.57E-07 | 1.14346525 | 0.336 | 0.313 | 0.00302918 | cDC.migratory | Plcl2 |
| Polr2e | 1.67E-07 | 0.43345529 | 0.421 | 0.511 | 0.00322073 | cDC.migratory | Polr2e |
| Atp9b | 1.77E-07 | 0.54756978 | 0.284 | 0.35 | 0.00342244 | cDC.migratory | Atp9b |
| Phrf1 | 1.82E-07 | 0.53685954 | 0.23 | 0.292 | 0.00351309 | cDC.migratory | Phrf1 |
| Zfr | 1.89E-07 | 0.38795591 | 0.281 | 0.349 | 0.00364578 | cDC.migratory | Zfr |
| Nfkb11 | 1.90E-07 | 0.61152229 | 0.722 | 0.805 | 0.00367745 | cDC.migratory | Nfkb1 |
| Vps51 | 2.06E-07 | 1.19942938 | 0.388 | 0.349 | 0.00398404 | cDC.migratory | Vps51 |
| Rbl2 | 2.26E-07 | 1.24811093 | 0.258 | 0.207 | 0.00436755 | cDC.migratory | Rbl2 |
| Prr13 | 2.72E-07 | 0.78403085 | 0.469 | 0.454 | 0.00525334 | cDC.migratory | Prr13 |
| Snx6 | 3.00E-07 | 0.41483924 | 0.214 | 0.276 | 0.0058015 | cDC.migratory | Snx6 |
| Safb2 | 3.01E-07 | 0.70421513 | 0.435 | 0.504 | 0.00582697 | cDC.migratory | Safb2 |
| Mepce | 3.05E-07 | 0.51268818 | 0.193 | 0.255 | 0.00589736 | cDC.migratory | Mepce |
| Itgav | 3.05E-07 | 0.52696659 | 0.205 | 0.291 | 0.0058984 | cDC.migratory | Itgav |
| Sun2 | 3.12E-07 | 0.59182805 | 0.358 | 0.405 | 0.00602459 | cDC.migratory | Sun2 |
| Ddx42 | 3.17E-07 | 0.36063736 | 0.239 | 0.3 | 0.00612431 | cDC.migratory | Ddx42 |
| Sh2b3 | 3.26E-07 | 0.63004608 | 0.37 | 0.482 | 0.00631094 | cDC.migratory | Sh2b3 |
| Pank2 | 3.30E-07 | 0.57371068 | 0.35 | 0.42 | 0.00638824 | cDC.migratory | Pank2 |
| Cop1 | 3.42E-07 | 0.28337786 | 0.292 | 0.355 | 0.00660843 | cDC.migratory | Cop1 |
| Gga3 | 3.71E-07 | 1.25219534 | 0.259 | 0.228 | 0.0071778 | cDC.migratory | Gga3 |
| Cat | 3.83E-07 | 0.60196851 | 0.384 | 0.475 | 0.00741089 | cDC.migratory | Cat |
| Tmem2431 | 3.93E-07 | 0.42315782 | 0.239 | 0.309 | 0.00759065 | cDC.migratory | Tmem243 |
| Fbxw11 | 4.49E-07 | 0.97552359 | 0.486 | 0.471 | 0.00867184 | cDC.migratory | Fbxw11 |
| Myo18a | 4.78E-07 | 0.61152725 | 0.273 | 0.345 | 0.00924176 | cDC.migratory | Myo18a |
| Tor1aip2 | 4.80E-07 | 1.03189111 | 0.53 | 0.541 | 0.00927249 | cDC.migratory | Tor1aip2 |
| Tmem127 | 4.92E-07 | 0.96011218 | 0.359 | 0.318 | 0.00951172 | cDC.migratory | Tmem127 |
| Igbp1 | 5.08E-07 | 0.47181811 | 0.267 | 0.336 | 0.00982256 | cDC.migratory | Igbp1 |
| Kmt5c | 5.32E-07 | 0.75522369 | 0.285 | 0.341 | 0.0102852 | cDC.migratory | Kmt5c |
| Vdac3 | 5.37E-07 | 0.39556298 | 0.195 | 0.252 | 0.01038897 | cDC.migratory | Vdac3 |
| Traf7 | 5.68E-07 | 0.4725587 | 0.318 | 0.395 | 0.01097538 | cDC.migratory | Traf7 |
| Fxr1 | 5.70E-07 | 0.45595666 | 0.404 | 0.499 | 0.01101418 | cDC.migratory | Fxr1 |
| [illegible] | [illegible] | [illegible] | [illegible] | [illegible] | [illegible] | [illegible] | [illegible] |

| | | | | | | | |
|---|---|---|---|---|---|---|---|
| Spin1 | 2.45E-06 | 0.25885861 | 0.189 | 0.265 | 0.04727881 | cDC.migratory | Spin1 |
| Znfx1 | 2.48E-06 | 1.04835721 | 0.284 | 0.243 | 0.04796135 | cDC.migratory | Znfx1 |
| Mfsd6 | 2.55E-06 | 0.5188989 | 0.228 | 0.289 | 0.04934394 | cDC.migratory | Mfsd6 |
| Tpm4 | 2.77E-06 | 0.58478452 | 0.699 | 0.792 | 0.05353352 | cDC.migratory | Tpm4 |
| Nploc4 | 2.86E-06 | 0.47221798 | 0.347 | 0.414 | 0.05537667 | cDC.migratory | Nploc4 |
| Arl8a | 2.89E-06 | 0.59897591 | 0.458 | 0.523 | 0.05590171 | cDC.migratory | Arl8a |
| Tomm34 | 3.37E-06 | 0.6853263 | 0.344 | 0.393 | 0.06509569 | cDC.migratory | Tomm34 |
| Gsap | 3.52E-06 | 0.67922516 | 0.341 | 0.405 | 0.06796964 | cDC.migratory | Gsap |
| Ercc3 | 3.53E-06 | 0.61600763 | 0.246 | 0.3 | 0.06819936 | cDC.migratory | Ercc3 |
| Paxbp1 | 3.53E-06 | 1.40300055 | 0.456 | 0.398 | 0.06821735 | cDC.migratory | Paxbp1 |
| Fmnl2 | 3.56E-06 | 0.53207218 | 0.29 | 0.34 | 0.06886194 | cDC.migratory | Fmnl2 |
| Map3k11 | 3.69E-06 | 0.51160841 | 0.307 | 0.378 | 0.07129974 | cDC.migratory | Map3k11 |
| Ubr2 | 3.81E-06 | 0.62657441 | 0.344 | 0.411 | 0.07360624 | cDC.migratory | Ubr2 |
| Dhrs3 | 4.16E-06 | 0.63691841 | 0.237 | 0.258 | 0.08052361 | cDC.migratory | Dhrs3 |
| Nt5c2 | 4.18E-06 | 0.52315448 | 0.304 | 0.369 | 0.08075438 | cDC.migratory | Nt5c2 |
| Anxa7 | 4.67E-06 | 0.40315446 | 0.419 | 0.523 | 0.09030859 | cDC.migratory | Anxa7 |
| Hmg20b | 5.13E-06 | 1.2174584 | 0.362 | 0.334 | 0.09920522 | cDC.migratory | Hmg20b |
| Zfp362 | 5.17E-06 | 0.94896141 | 0.338 | 0.322 | 0.0999583 | cDC.migratory | Zfp362 |
| Tmem183a | 5.88E-06 | 0.52846054 | 0.243 | 0.31 | 0.11366171 | cDC.migratory | Tmem183a |
| Crk | 7.26E-06 | 0.51147243 | 0.359 | 0.438 | 0.14045116 | cDC.migratory | Crk |
| Pptc7 | 7.49E-06 | 0.33949426 | 0.207 | 0.262 | 0.14485903 | cDC.migratory | Pptc7 |
| Dennd3 | 7.88E-06 | 1.08536908 | 0.348 | 0.307 | 0.15229067 | cDC.migratory | Dennd3 |
| Dctn4 | 8.01E-06 | 0.60039994 | 0.358 | 0.421 | 0.15495555 | cDC.migratory | Dctn4 |
| Vps4b | 9.05E-06 | 0.59673504 | 0.224 | 0.271 | 0.17505639 | cDC.migratory | Vps4b |
| H1f0 | 9.20E-06 | 0.51066092 | 0.337 | 0.37 | 0.17781062 | cDC.migratory | H1f0 |
| Klhl5 | 9.22E-06 | 0.99291824 | 0.349 | 0.331 | 0.17830361 | cDC.migratory | Klhl5 |
| Arhgap22 | 9.53E-06 | 1.21570628 | 0.308 | 0.309 | 0.18427207 | cDC.migratory | Arhgap22 |
| Ak3 | 9.81E-06 | 0.38764236 | 0.201 | 0.264 | 0.18976125 | cDC.migratory | Ak3 |
| Plekhb2 | 9.88E-06 | 1.04894001 | 0.569 | 0.534 | 0.19102415 | cDC.migratory | Plekhb2 |
| Mdm2 | 9.89E-06 | 0.64003338 | 0.341 | 0.402 | 0.19130313 | cDC.migratory | Mdm2 |
| Ttc14 | 1.00E-05 | 0.57549683 | 0.198 | 0.251 | 0.19333751 | cDC.migratory | Ttc14 |
| Parp8 | 1.03E-05 | 0.51254322 | 0.252 | 0.325 | 0.19983383 | cDC.migratory | Parp8 |
| Usf1 | 1.15E-05 | 0.64730461 | 0.386 | 0.438 | 0.22284781 | cDC.migratory | Usf1 |
| Mff | 1.28E-05 | 0.27826975 | 0.189 | 0.258 | 0.24671366 | cDC.migratory | Mff |
| Gpbp1 | 1.41E-05 | 0.98062453 | 0.417 | 0.427 | 0.27326034 | cDC.migratory | Gpbp1 |
| Abhd2 | 1.52E-05 | 1.18343567 | 0.4 | 0.414 | 0.29304503 | cDC.migratory | Abhd2 |
| Btg1 | 1.58E-05 | 0.78114535 | 0.742 | 0.749 | 0.30615394 | cDC.migratory | Btg1 |
| Mapk3 | 1.62E-05 | 0.98008228 | 0.425 | 0.434 | 0.31280952 | cDC.migratory | Mapk3 |
| Arrb2 | 1.69E-05 | 0.8042899 | 0.77 | 0.786 | 0.32755786 | cDC.migratory | Arrb2 |
| Dock8 | 1.78E-05 | 0.61041555 | 0.453 | 0.562 | 0.34414143 | cDC.migratory | Dock8 |
| Map2k1 | 1.82E-05 | 0.91196869 | 0.432 | 0.446 | 0.3510857 | cDC.migratory | Map2k1 |
| Ppfibp1 | 1.88E-05 | 0.61645359 | 0.223 | 0.278 | 0.36388818 | cDC.migratory | Ppfibp1 |
| B4galnt1 | 1.91E-05 | 0.58079654 | 0.492 | 0.575 | 0.36834664 | cDC.migratory | B4galnt1 |
| Suco | 1.96E-05 | 0.48473335 | 0.219 | 0.267 | 0.37857409 | cDC.migratory | Suco |
| Ggnbp2 | 2.01E-05 | 0.56535356 | 0.237 | 0.277 | 0.38848346 | cDC.migratory | Ggnbp2 |
| Cdk19 | 2.12E-05 | 0.42764584 | 0.208 | 0.257 | 0.40897852 | cDC.migratory | Cdk19 |
| Eif4e2 | 2.15E-05 | 0.58594969 | 0.396 | 0.483 | 0.416425 | cDC.migratory | Eif4e2 |
| Tbc1d20 | 2.36E-05 | 0.33946943 | 0.266 | 0.336 | 0.45660483 | cDC.migratory | Tbc1d20 |
| Parp9 | 2.47E-05 | 0.28851247 | 0.227 | 0.284 | 0.47702137 | cDC.migratory | Parp9 |
| Lemd3 | 2.72E-05 | 0.44378841 | 0.254 | 0.319 | 0.52511449 | cDC.migratory | Lemd3 |
| Atad2b | 2.87E-05 | 0.75839052 | 0.193 | 0.251 | 0.55528188 | cDC.migratory | Atad2b |
| Aftph | 2.91E-05 | 1.02563716 | 0.325 | 0.3 | 0.5626356 | cDC.migratory | Aftph |
| Kmt2b | 2.99E-05 | 0.44668094 | 0.242 | 0.3 | 0.57897373 | cDC.migratory | Kmt2b |
| Ubqln2 | 3.56E-05 | 0.52044653 | 0.266 | 0.33 | 0.68781489 | cDC.migratory | Ubqln2 |
| Cntrl | 3.89E-05 | 0.82236912 | 0.355 | 0.404 | 0.75147502 | cDC.migratory | Cntrl |
| Prpf39 | 4.07E-05 | 0.68184009 | 0.329 | 0.382 | 0.78701254 | cDC.migratory | Prpf39 |
| Ddx49 | 4.66E-05 | 0.60906952 | 0.245 | 0.324 | 0.90157755 | cDC.migratory | Ddx49 |
| Sugt1 | 4.74E-05 | 0.97118819 | 0.377 | 0.362 | 0.9163661 | cDC.migratory | Sugt1 |
| Stk3 | 5.03E-05 | 0.57056312 | 0.218 | 0.273 | 0.97313569 | cDC.migratory | Stk3 |
| Hbp1 | 5.18E-05 | 1.05874656 | 0.347 | 0.311 | 1 | cDC.migratory | Hbp1 |
| Rab22a | 5.19E-05 | 1.21284065 | 0.374 | 0.342 | 1 | cDC.migratory | Rab22a |
| Rel | 5.25E-05 | 0.86411352 | 0.82 | 0.79 | 1 | cDC.migratory | Rel |
| Stx12 | 5.53E-05 | 0.64908078 | 0.436 | 0.508 | 1 | cDC.migratory | Stx12 |
| Dyrk21 | 6.08E-05 | 0.36131027 | 0.21 | 0.271 | 1 | cDC.migratory | Dyrk2 |
| Ino80d | 6.19E-05 | 1.18114456 | 0.336 | 0.307 | 1 | cDC.migratory | Ino80d |
| Ptpn2 | 6.21E-05 | 0.42676644 | 0.316 | 0.394 | 1 | cDC.migratory | Ptpn2 |
| Nisch | 6.37E-05 | 0.93958484 | 0.523 | 0.587 | 1 | cDC.migratory | Nisch |
| Csnk1d | 6.50E-05 | 0.57724576 | 0.528 | 0.618 | 1 | cDC.migratory | Csnk1d |
| Tnpo2 | 6.69E-05 | 0.56060598 | 0.259 | 0.309 | 1 | cDC.migratory | Tnpo2 |
| Pik3r5 | 6.87E-05 | 0.77887897 | 0.624 | 0.691 | 1 | cDC.migratory | Pik3r5 |
| Ier5 | 7.05E-05 | 0.84058213 | 0.767 | 0.767 | 1 | cDC.migratory | Ier5 |
| Mdm4 | 7.09E-05 | 1.301035 | 0.321 | 0.278 | 1 | cDC.migratory | Mdm4 |
| Pan3 | 7.36E-05 | 0.59703251 | 0.309 | 0.364 | 1 | cDC.migratory | Pan3 |
| Ldlr | 8.21E-05 | 0.49474129 | 0.491 | 0.515 | 1 | cDC.migratory | Ldlr |
| Slc25a4 | 8.50E-05 | 0.43697057 | 0.332 | 0.403 | 1 | cDC.migratory | Slc25a4 |
| Cbx4 | 8.64E-05 | 0.46482173 | 0.201 | 0.253 | 1 | cDC.migratory | Cbx4 |
| Tnpo1 | 8.68E-05 | 0.69429867 | 0.414 | 0.487 | 1 | cDC.migratory | Tnpo1 |
| Vti1b | 9.05E-05 | 0.3885389 | 0.274 | 0.326 | 1 | cDC.migratory | Vti1b |
| Lipe | 9.68E-05 | 0.6000857 | 0.414 | 0.491 | 1 | cDC.migratory | Lipe |
| Shb | 9.78E-05 | 1.16844834 | 0.323 | 0.281 | 1 | cDC.migratory | Shb |
| Etv6 | 9.89E-05 | 0.76404601 | 0.641 | 0.674 | 1 | cDC.migratory | Etv6 |
| Trrap | 9.99E-05 | 0.61725181 | 0.218 | 0.256 | 1 | cDC.migratory | Trrap |
| Patl1 | 0.00010106 | 0.56596694 | 0.278 | 0.329 | 1 | cDC.migratory | Patl1 |
| Ptpn23 | 0.00010412 | 0.33323274 | 0.223 | 0.282 | 1 | cDC.migratory | Ptpn23 |
| Wdr91 | 0.00010762 | 0.48684344 | 0.18 | 0.256 | 1 | cDC.migratory | Wdr91 |
| Arid1b | 0.00010798 | 0.55591925 | 0.369 | 0.437 | 1 | cDC.migratory | Arid1b |
| Chp1 | 0.00010845 | 0.83070596 | 0.559 | 0.603 | 1 | cDC.migratory | Chp1 |
| Msi2 | 0.00011119 | 1.09872728 | 0.257 | 0.238 | 1 | cDC.migratory | Msi2 |
| Smap2 | 0.00011321 | 0.52144452 | 0.514 | 0.6 | 1 | cDC.migratory | Smap2 |
| Slc41a1 | 0.00011593 | 0.94207244 | 0.361 | 0.315 | 1 | cDC.migratory | Slc41a1 |
| Slc25a51 | 0.00012602 | 0.32508323 | 0.27 | 0.33 | 1 | cDC.migratory | Slc25a51 |
| Eps15 | 0.00012737 | 1.04743329 | 0.483 | 0.449 | 1 | cDC.migratory | Eps15 |
| Tmcc1 | 0.00013084 | 1.00394218 | 0.351 | 0.327 | 1 | cDC.migratory | Tmcc1 |
| Pcnx3 | 0.00013276 | 0.6259599 | 0.209 | 0.262 | 1 | cDC.migratory | Pcnx3 |
| Clip1 | 0.00014092 | 1.18220871 | 0.306 | 0.263 | 1 | cDC.migratory | Clip1 |
| Samd4b | 0.00015341 | 0.48860627 | 0.281 | 0.33 | 1 | cDC.migratory | Samd4b |
| Itgb1 | 0.00016028 | 0.73027278 | 0.69 | 0.766 | 1 | cDC.migratory | Itgb1 |
| Atxn7 | 0.00016225 | 1.03640512 | 0.268 | 0.248 | 1 | cDC.migratory | Atxn7 |
| Leng8 | 0.00017667 | 1.27809551 | 0.525 | 0.487 | 1 | cDC.migratory | Leng8 |
| Ncor21 | 0.00018308 | 0.75643525 | 0.294 | 0.344 | 1 | cDC.migratory | Ncor2 |
| Rsu1 | 0.00018734 | 0.64644328 | 0.528 | 0.607 | 1 | cDC.migratory | Rsu1 |
| Agpat1 | 0.00018784 | 0.50048164 | 0.284 | 0.334 | 1 | cDC.migratory | Agpat1 |
| Atp6v1b2 | 0.00019523 | 0.68591444 | 0.503 | 0.582 | 1 | cDC.migratory | Atp6v1b2 |
| Piezo1 | 0.00020703 | 0.55755989 | 0.34 | 0.416 | 1 | cDC.migratory | Piezo1 |
| Use1 | 0.00020811 | 0.67224774 | 0.479 | 0.555 | 1 | cDC.migratory | Use1 |
| Ppm1b | 0.00021071 | 0.67705058 | 0.33 | 0.38 | 1 | cDC.migratory | Ppm1b |
| Vps13d | 0.00023608 | 0.57426405 | 0.204 | 0.257 | 1 | cDC.migratory | Vps13d |
| Peli1 | 0.00023791 | 0.8760009 | 0.441 | 0.42 | 1 | cDC.migratory | Peli1 |
| Mef2a | 0.00024796 | 0.99704624 | 0.419 | 0.398 | 1 | cDC.migratory | Mef2a |
| Tgfbr1 | 0.00025145 | 0.64837447 | 0.26 | 0.272 | 1 | cDC.migratory | Tgfbr1 |
| Ypel5 | 0.00025655 | 0.34390378 | 0.29 | 0.343 | 1 | cDC.migratory | Ypel5 |
| Herc2 | 0.00026013 | 0.63941374 | 0.215 | 0.267 | 1 | cDC.migratory | Herc2 |
| Usp3 | 0.00028731 | 0.41437638 | 0.226 | 0.276 | 1 | cDC.migratory | Usp3 |
| Parp3 | 0.00028862 | 1.21773398 | 0.258 | 0.238 | 1 | cDC.migratory | Parp3 |
| Mlf2 | 0.00029295 | 0.62571506 | 0.506 | 0.638 | 1 | cDC.migratory | Mlf2 |
| Zfp800 | 0.00029546 | 0.58659505 | 0.259 | 0.306 | 1 | cDC.migratory | Zfp800 |
| Mafg | 0.00030424 | 0.81464629 | 0.2 | 0.251 | 1 | cDC.migratory | Mafg |
| Zcrb1 | 0.00030628 | 0.49029628 | 0.22 | 0.264 | 1 | cDC.migratory | Zcrb1 |
| Faf1 | 0.00031424 | 0.84064243 | 0.269 | 0.319 | 1 | cDC.migratory | Faf1 |
| Cd47 | 0.00031837 | 0.62573043 | 0.759 | 0.817 | 1 | cDC.migratory | Cd47 |
| Ensa | 0.00032326 | 0.32158994 | 0.323 | 0.385 | 1 | cDC.migratory | Ensa |
| Chst11 | 0.00033304 | 0.46405499 | 0.324 | 0.359 | 1 | cDC.migratory | Chst11 |
| Mtmr12 | 0.00033365 | 0.95016494 | 0.27 | 0.242 | 1 | cDC.migratory | Mtmr12 |
| Ppp1r12a | 0.00033684 | 0.95789241 | 0.547 | 0.566 | 1 | cDC.migratory | Ppp1r12a |
| Wbp2 | 0.0003397 | 0.59452487 | 0.284 | 0.349 | 1 | cDC.migratory | Wbp2 |
| Nrbp1 | 0.00034693 | 0.83309165 | 0.432 | 0.458 | 1 | cDC.migratory | Nrbp1 |
| Vav3 | 0.0003538 | 0.38407587 | 0.229 | 0.282 | 1 | cDC.migratory | Vav3 |
| Gpr65 | 0.00035727 | 1.27079371 | 0.353 | 0.266 | 1 | cDC.migratory | Gpr65 |
| Ptov1 | 0.00040596 | 0.59312455 | 0.334 | 0.383 | 1 | cDC.migratory | Ptov1 |
| Ttc7b | 0.00043821 | 1.34885623 | 0.325 | 0.289 | 1 | cDC.migratory | Ttc7b |
| Numa1 | 0.00046362 | 0.71803223 | 0.321 | 0.35 | 1 | cDC.migratory | Numa1 |
| Epb41l2 | 0.00046735 | 0.98603603 | 0.502 | 0.453 | 1 | cDC.migratory | Epb41l2 |
| Pkn2 | 0.00048149 | 0.52467931 | 0.207 | 0.255 | 1 | cDC.migratory | Pkn2 |
| Camta2 | 0.00053678 | 0.72161332 | 0.538 | 0.582 | 1 | cDC.migratory | Camta2 |
| Stat5b | 0.00056632 | 0.87205456 | 0.356 | 0.358 | 1 | cDC.migratory | Stat5b |
| Ggta1 | 0.00060618 | 0.912483 | 0.623 | 0.6 | 1 | cDC.migratory | Ggta1 |
| Kmt2a | 0.00061392 | 1.3352803 | 0.287 | 0.243 | 1 | cDC.migratory | Kmt2a |
| Med10 | 0.00062608 | 0.52586314 | 0.259 | 0.308 | 1 | cDC.migratory | Med10 |
| Tnfrsf1b | 0.00074073 | 0.94955237 | 0.643 | 0.59 | 1 | cDC.migratory | Tnfrsf1b |
| Chfr | 0.00074323 | 0.71416971 | 0.226 | 0.281 | 1 | cDC.migratory | Chfr |
| Slc6a6 | 0.00077292 | 0.62526206 | 0.57 | 0.646 | 1 | cDC.migratory | Slc6a6 |
| Phip | 0.00077431 | 0.63915343 | 0.285 | 0.335 | 1 | cDC.migratory | Phip |
| Vhl | 0.0007798 | 1.01032686 | 0.311 | 0.294 | 1 | cDC.migratory | Vhl |
| Mbtps1 | 0.00078822 | 0.57309762 | 0.268 | 0.317 | 1 | cDC.migratory | Mbtps1 |
| Mea1 | 0.00091824 | 0.39463785 | 0.27 | 0.341 | 1 | cDC.migratory | Mea1 |
| Ube2r2 | 0.00092519 | 0.7980762 | 0.476 | 0.529 | 1 | cDC.migratory | Ube2r2 |
| Sumo2 | 0.00097939 | 0.46266166 | 0.285 | 0.358 | 1 | cDC.migratory | Sumo2 |
| Cd86 | 0.00101631 | 0.63857167 | 0.648 | 0.701 | 1 | cDC.migratory | Cd86 |
| Tbl1x | 0.00110207 | 0.99637349 | 0.318 | 0.278 | 1 | cDC.migratory | Tbl1x |
| Mapre2 | 0.00114124 | 1.19741123 | 0.255 | 0.234 | 1 | cDC.migratory | Mapre2 |
| Casc3 | 0.00126752 | 0.67198958 | 0.391 | 0.448 | 1 | cDC.migratory | Casc3 |
| Tuba1c | 0.00136808 | 0.6568318 | 0.515 | 0.562 | 1 | cDC.migratory | Tuba1c |
| Sirt7 | 0.00137621 | 0.8437 | 0.322 | 0.367 | 1 | cDC.migratory | Sirt7 |
| Pias1 | 0.00141073 | 0.63494033 | 0.245 | 0.305 | 1 | cDC.migratory | Pias1 |
| Lpp | 0.00146231 | 0.99658311 | 0.472 | 0.466 | 1 | cDC.migratory | Lpp |
| Rsbn1l | 0.00151119 | 0.61270919 | 0.229 | 0.278 | 1 | cDC.migratory | Rsbn1l |
| Arhgap30 | 0.00152341 | 0.698288 | 0.678 | 0.761 | 1 | cDC.migratory | Arhgap30 |
| Lsm14b | 0.00155814 | 0.64516133 | 0.229 | 0.272 | 1 | cDC.migratory | Lsm14b |
| Plagl2 | 0.00165196 | 0.96469822 | 0.28 | 0.255 | 1 | cDC.migratory | Plagl2 |
| Sirt2 | 0.00166977 | 0.45195903 | 0.344 | 0.433 | 1 | cDC.migratory | Sirt2 |
| Mark4 | 0.00170436 | 0.47430872 | 0.217 | 0.262 | 1 | cDC.migratory | Mark4 |
| Sqstm1 | 0.00180383 | 0.63377822 | 0.589 | 0.645 | 1 | cDC.migratory | Sqstm1 |
| Grasp | 0.00184718 | 0.88718666 | 0.557 | 0.49 | 1 | cDC.migratory | Grasp |
| Stx6 | 0.00187007 | 0.82977615 | 0.33 | 0.318 | 1 | cDC.migratory | Stx6 |
| Zfp451 | 0.00202492 | 0.68413872 | 0.204 | 0.25 | 1 | cDC.migratory | Zfp451 |
| Mapkap1 | 0.00205687 | 0.53201799 | 0.215 | 0.267 | 1 | cDC.migratory | Mapkap1 |
| Spag7 | 0.00230419 | 0.73183702 | 0.284 | 0.331 | 1 | cDC.migratory | Spag7 |
| Clasp2 | 0.0023139 | 1.11817407 | 0.346 | 0.349 | 1 | cDC.migratory | Clasp2 |
| Sel1l | 0.00239164 | 0.82189694 | 0.409 | 0.445 | 1 | cDC.migratory | Sel1l |
| Rabac1 | 0.00248076 | 0.62343196 | 0.501 | 0.557 | 1 | cDC.migratory | Rabac1 |
| Stat6 | 0.00255268 | 0.44317992 | 0.514 | 0.606 | 1 | cDC.migratory | Stat6 |
| Clptm1 | 0.00259839 | 0.56121145 | 0.453 | 0.532 | 1 | cDC.migratory | Clptm1 |
| Selenok | 0.00267253 | 0.56457771 | 0.301 | 0.361 | 1 | cDC.migratory | Selenok |
| Emd | 0.00271687 | 0.57887714 | 0.567 | 0.648 | 1 | cDC.migratory | Emd |
| Slc4a2 | 0.00293682 | 1.00810949 | 0.263 | 0.248 | 1 | cDC.migratory | Slc4a2 |
| Fam76b | 0.00308918 | 0.81398858 | 0.206 | 0.251 | 1 | cDC.migratory | Fam76b |
| Arid4b | 0.00397687 | 0.74978025 | 0.303 | 0.35 | 1 | cDC.migratory | Arid4b |
| Tmem176a1 | 0.00432515 | 0.88295185 | 0.501 | 0.399 | 1 | cDC.migratory | Tmem176a |
| Rapgef6 | 0.00439593 | 0.43485029 | 0.24 | 0.301 | 1 | cDC.migratory | Rapgef6 |
| Vps16 | 0.00471163 | 0.48190128 | 0.261 | 0.312 | 1 | cDC.migratory | Vps16 |
| Rab12 | 0.00478424 | 1.20824489 | 0.271 | 0.238 | 1 | cDC.migratory | Rab12 |
| Bahd1 | 0.00521057 | 0.94443186 | 0.298 | 0.267 | 1 | cDC.migratory | Bahd1 |
| Ilk | 0.00527102 | 0.53688972 | 0.49 | 0.565 | 1 | cDC.migratory | Ilk |
| Mdfic | 0.00553804 | 0.68010406 | 0.253 | 0.284 | 1 | cDC.migratory | Mdfic |
| Apobec3 | 0.00560842 | 0.97791938 | 0.677 | 0.665 | 1 | cDC.migratory | Apobec3 |
| Yme1l1 | 0.00561577 | 1.0154347 | 0.336 | 0.34 | 1 | cDC.migratory | Yme1l1 |
| Ecpas | 0.00562321 | 0.91597019 | 0.303 | 0.349 | 1 | cDC.migratory | Ecpas |
| Nfya | 0.00635554 | 0.58509761 | 0.45 | 0.483 | 1 | cDC.migratory | Nfya |
| Rela | 0.00653785 | 0.5179274 | 0.493 | 0.585 | 1 | cDC.migratory | Rela |
| Arl4c | 0.00739691 | 0.60101713 | 0.43 | 0.441 | 1 | cDC.migratory | Arl4c |
| Myl6 | 0.00770801 | 0.79367374 | 0.569 | 0.61 | 1 | cDC.migratory | Myl6 |
| Acsf2 | 0.00777207 | 0.59394014 | 0.22 | 0.255 | 1 | cDC.migratory | Acsf2 |
| Dnajb6 | 0.00831298 | 0.91063188 | 0.552 | 0.568 | 1 | cDC.migratory | Dnajb6 |
| Zc3h7a | 0.00872641 | 0.61213346 | 0.461 | 0.533 | 1 | cDC.migratory | Zc3h7a |
| Smad4 | 0.00996038 | 0.68448631 | 0.583 | 0.641 | 1 | cDC.migratory | Smad4 |
| Itgae | 0 | 1.50340024 | 0.872 | 0.255 | 0 | cDC2.Mgl2 | Itgae |
| Pilra | 0 | 2.93354335 | 0.74 | 0.143 | 0 | cDC2.Mgl2 | Pilra |
| Siglece | 0 | 3.21133366 | 0.711 | 0.134 | 0 | cDC2.Mgl2 | Siglece |
| Ppp1r14a | 0 | 2.61902951 | 0.701 | 0.153 | 0 | cDC2.Mgl2 | Ppp1r14a |
| Plet1 | 0 | 1.4833668 | 0.966 | 0.419 | 0 | cDC2.Mgl2 | Plet1 |
| Cdh17 | 0 | 3.84240898 | 0.606 | 0.074 | 0 | cDC2.Mgl2 | Cdh17 |
| Mmp12 | 0 | 2.03816774 | 0.703 | 0.181 | 0 | cDC2.Mgl2 | Mmp12 |
| Ppp1r1a | 0 | 2.47613763 | 0.644 | 0.132 | 0 | cDC2.Mgl2 | Ppp1r1a |
| Ffar21 | 0 | 2.17514649 | 0.67 | 0.17 | 0 | cDC2.Mgl2 | Ffar2 |
| Cdh11 | 0 | 1.31902456 | 0.86 | 0.361 | 0 | cDC2.Mgl2 | Cdh1 |
| Dcstamp | 0 | 2.08397957 | 0.657 | 0.158 | 0 | cDC2.Mgl2 | Dcstamp |
| Ctnnd21 | 0 | 1.99811429 | 0.735 | 0.239 | 0 | cDC2.Mgl2 | Ctnnd2 |
| Clec4n1 | 0 | 1.53140918 | 0.818 | 0.323 | 0 | cDC2.Mgl2 | Clec4n |
| Mgl21 | 0 | 1.99713462 | 0.906 | 0.413 | 0 | cDC2.Mgl2 | Mgl2 |
| Qpct | 0 | 1.88354904 | 0.688 | 0.197 | 0 | cDC2.Mgl2 | Qpct |
| Atp2a3 | 0 | 2.22792171 | 0.668 | 0.183 | 0 | cDC2.Mgl2 | Atp2a3 |
| H2-Oa | 0 | 1.9879072 | 0.762 | 0.29 | 0 | cDC2.Mgl2 | H2-Oa |
| Krt801 | 0 | 1.71294806 | 0.735 | 0.265 | 0 | cDC2.Mgl2 | Krt80 |
| Cd33 | 0 | 1.95170758 | 0.641 | 0.173 | 0 | cDC2.Mgl2 | Cd33 |
| Cd24a | 0 | 1.5301457 | 0.934 | 0.469 | 0 | cDC2.Mgl2 | Cd24a |
| Cd101 | 0 | 4.55815763 | 0.489 | 0.031 | 0 | cDC2.Mgl2 | Cd101 |
| H2-DMb21 | 0 | 1.66316948 | 0.765 | 0.319 | 0 | cDC2.Mgl2 | H2-DMb2 |
| Ndst1 | 0 | 1.69469922 | 0.714 | 0.27 | 0 | cDC2.Mgl2 | Ndst1 |
| Hic1 | 0 | 1.57837037 | 0.653 | 0.218 | 0 | cDC2.Mgl2 | Hic1 |
| Il1r21 | 0 | 1.7415849 | 0.92 | 0.487 | 0 | cDC2.Mgl2 | Il1r2 |
| Tgm21 | 0 | 1.29370254 | 0.92 | 0.489 | 0 | cDC2.Mgl2 | Tgm2 |
| Camkk1 | 0 | 2.02998387 | 0.615 | 0.187 | 0 | cDC2.Mgl2 | Camkk1 |
| B4galt6 | 0 | 1.82691956 | 0.746 | 0.319 | 0 | cDC2.Mgl2 | B4galt6 |
| Cyp4f16 | 0 | 2.31989576 | 0.559 | 0.146 | 0 | cDC2.Mgl2 | Cyp4f16 |
| Gem | 0 | 2.12511704 | 0.603 | 0.19 | 0 | cDC2.Mgl2 | Gem |
| Pygl | 0 | 1.58859151 | 0.711 | 0.301 | 0 | cDC2.Mgl2 | Pygl |
| Emb1 | 0 | 1.23952836 | 0.783 | 0.377 | 0 | cDC2.Mgl2 | Emb |
| Immp2l | 0 | 2.06449459 | 0.561 | 0.156 | 0 | cDC2.Mgl2 | Immp2l |
| Pltp | 0 | 1.47842125 | 0.738 | 0.335 | 0 | cDC2.Mgl2 | Pltp |
| Epcam | 0 | 3.12331792 | 0.469 | 0.066 | 0 | cDC2.Mgl2 | Epcam |
| Mdh21 | 0 | 1.88459243 | 0.881 | 0.483 | 0 | cDC2.Mgl2 | Mdh2 |
| Ifitm11 | 0 | 1.40482319 | 0.957 | 0.561 | 0 | cDC2.Mgl2 | Ifitm1 |
| Wdfy4 | 0 | 0.68927158 | 0.906 | 0.512 | 0 | cDC2.Mgl2 | Wdfy4 |
| Mcm4 | 0 | 1.72443803 | 0.621 | 0.232 | 0 | cDC2.Mgl2 | Mcm4 |
| Hilpda1 | 0 | 1.47518391 | 0.675 | 0.288 | 0 | cDC2.Mgl2 | Hilpda |
| Themis2 | 0 | 1.39553138 | 0.855 | 0.475 | 0 | cDC2.Mgl2 | Themis2 |
| Tcea3 | 0 | 1.85503138 | 0.548 | 0.168 | 0 | cDC2.Mgl2 | Tcea3 |
| Nlrp31 | 0 | 1.4037686 | 0.746 | 0.375 | 0 | cDC2.Mgl2 | Nlrp3 |
| Mt1 | 0 | 1.81921322 | 0.904 | 0.535 | 0 | cDC2.Mgl2 | Mt1 |
| Pak1 | 0 | 1.23973901 | 0.735 | 0.37 | 0 | cDC2.Mgl2 | Pak1 |
| Lilr4b1 | 0 | 1.27418591 | 0.737 | 0.376 | 0 | cDC2.Mgl2 | Lilr4b |
| Ccrl2 | 0 | 1.1742333 | 0.684 | 0.325 | 0 | cDC2.Mgl2 | Ccrl2 |
| Fgl2 | 0 | 1.39846336 | 0.909 | 0.551 | 0 | cDC2.Mgl2 | Fgl2 |
| Jdp2 | 0 | 1.61763594 | 0.705 | 0.347 | 0 | cDC2.Mgl2 | Jdp2 |
| Hexb | 0 | 1.36803582 | 0.756 | 0.401 | 0 | cDC2.Mgl2 | Hexb |
| Itgax | 0 | 1.45116154 | 0.908 | 0.554 | 0 | cDC2.Mgl2 | Itgax |
| Cd300a1 | 0 | 0.97958376 | 0.876 | 0.522 | 0 | cDC2.Mgl2 | Cd300a |
| Spint11 | 0 | 0.95852044 | 0.861 | 0.512 | 0 | cDC2.Mgl2 | Spint1 |
| Lta4h1 | 0 | 1.28353675 | 0.856 | 0.508 | 0 | cDC2.Mgl2 | Lta4h |
| Cfp1 | 0 | 1.13280342 | 0.944 | 0.598 | 0 | cDC2.Mgl2 | Cfp |
| Sema4d | 0 | 1.0250166 | 0.847 | 0.505 | 0 | cDC2.Mgl2 | Sema4d |
| Neurl3 | 0 | 1.44917706 | 0.903 | 0.566 | 0 | cDC2.Mgl2 | Neurl3 |
| Odc1 | 0 | 1.37434666 | 0.904 | 0.568 | 0 | cDC2.Mgl2 | Odc1 |
| Arhgap91 | 0 | 1.10651367 | 0.789 | 0.461 | 0 | cDC2.Mgl2 | Arhgap9 |
| Napsa1 | 0 | 1.5129054 | 0.897 | 0.578 | 0 | cDC2.Mgl2 | Napsa |
| Tiparp1 | 0 | 1.17866837 | 0.693 | 0.374 | 0 | cDC2.Mgl2 | Tiparp |
| Cd1641 | 0 | 1.06162422 | 0.88 | 0.566 | 0 | cDC2.Mgl2 | Cd164 |
| Nabp11 | 0 | 1.01278092 | 0.714 | 0.402 | 0 | cDC2.Mgl2 | Nabp1 |
| Irs2 | 0 | 1.1790635 | 0.715 | 0.406 | 0 | cDC2.Mgl2 | Irs2 |
| Rin31 | 0 | 0.74249566 | 0.904 | 0.596 | 0 | cDC2.Mgl2 | Rin3 |
| Alox5ap1 | 0 | 0.55098062 | 0.892 | 0.587 | 0 | cDC2.Mgl2 | Alox5ap |
| Rhoq1 | 0 | 1.00651295 | 0.725 | 0.424 | 0 | cDC2.Mgl2 | Rhoq |
| Ly6e1 | 0 | 0.48919123 | 0.938 | 0.644 | 0 | cDC2.Mgl2 | Ly6e |
| Fosb1 | 0 | 1.19443245 | 0.892 | 0.603 | 0 | cDC2.Mgl2 | Fosb |
| Man2b1 | 0 | 1.02481033 | 0.869 | 0.584 | 0 | cDC2.Mgl2 | Man2b1 |
| Fosl21 | 0 | 0.87639045 | 0.883 | 0.602 | 0 | cDC2.Mgl2 | Fosl2 |
| Bin21 | 0 | 0.82673987 | 0.884 | 0.603 | 0 | cDC2.Mgl2 | Bin2 |
| Cxcl2 | 0 | 0.99561535 | 0.592 | 0.311 | 0 | cDC2.Mgl2 | Cxcl2 |
| Rgs11 | 0 | 0.74308148 | 0.816 | 0.542 | 0 | cDC2.Mgl2 | Rgs1 |
| Slc35b21 | 0 | 0.97877863 | 0.8 | 0.53 | 0 | cDC2.Mgl2 | Slc35b2 |
| Ctsc | 0 | 0.963658 | 0.897 | 0.631 | 0 | cDC2.Mgl2 | Ctsc |
| H2-DMa1 | 0 | 0.76204027 | 0.926 | 0.663 | 0 | cDC2.Mgl2 | H2-DMa |
| Itgb21 | 0 | 0.65969279 | 0.914 | 0.652 | 0 | cDC2.Mgl2 | Itgb2 |
| Gsn | 0 | 1.83712654 | 0.983 | 0.729 | 0 | cDC2.Mgl2 | Gsn |
| Rac21 | 0 | 1.08891889 | 0.942 | 0.689 | 0 | cDC2.Mgl2 | Rac2 |
| Coro2a | 0 | 0.8805512 | 0.857 | 0.611 | 0 | cDC2.Mgl2 | Coro2a |
| Gng101 | 0 | 0.72161027 | 0.884 | 0.639 | 0 | cDC2.Mgl2 | Gng10 |
| Nr4a2 | 0 | 0.84372808 | 0.791 | 0.557 | 0 | cDC2.Mgl2 | Nr4a2 |
| Ptgs21 | 0 | 1.07868725 | 0.503 | 0.271 | 0 | cDC2.Mgl2 | Ptgs2 |
| Uck21 | 0 | 0.74953856 | 0.91 | 0.678 | 0 | cDC2.Mgl2 | Uck2 |
| Actn11 | 0 | 0.45618924 | 0.941 | 0.715 | 0 | cDC2.Mgl2 | Actn1 |
| Stk17b | 0 | 0.70258623 | 0.837 | 0.617 | 0 | cDC2.Mgl2 | Stk17b |
| Ifrd11 | 0 | 0.74933229 | 0.859 | 0.644 | 0 | cDC2.Mgl2 | Ifrd1 |
| Gm2a | 0 | 0.86523537 | 0.982 | 0.769 | 0 | cDC2.Mgl2 | Gm2a |
| Hcls11 | 0 | 0.79600871 | 0.965 | 0.757 | 0 | cDC2.Mgl2 | Hcls1 |
| Crem | 0 | 0.8130058 | 0.868 | 0.669 | 0 | cDC2.Mgl2 | Crem |
| App1 | 0 | 0.77748142 | 0.912 | 0.722 | 0 | cDC2.Mgl2 | App |
| Nr4a11 | 0 | 0.41621673 | 0.863 | 0.673 | 0 | cDC2.Mgl2 | Nr4a1 |
| Tspan13 | 0 | 0.61132408 | 0.885 | 0.696 | 0 | cDC2.Mgl2 | Tspan13 |
| Ldha | 0 | 0.71684286 | 0.918 | 0.741 | 0 | cDC2.Mgl2 | Ldha |
| Btg2 | 0 | 0.6698192 | 0.914 | 0.752 | 0 | cDC2.Mgl2 | Btg2 |
| H2-Ab11 | 0 | 0.71090066 | 0.997 | 0.836 | 0 | cDC2.Mgl2 | H2-Ab1 |
| Il1b1 | 0 | 0.52665392 | 0.921 | 0.762 | 0 | cDC2.Mgl2 | Il1b |
| H2-Eb11 | 0 | 0.72312178 | 0.999 | 0.861 | 0 | cDC2.Mgl2 | H2-Eb1 |
| Tyrobp1 | 0 | 0.49949804 | 0.949 | 0.814 | 0 | cDC2.Mgl2 | Tyrobp |
| H2-Aa1 | 0 | 0.54458091 | 0.996 | 0.867 | 0 | cDC2.Mgl2 | H2-Aa |
| Cotl1 | 0 | 0.69594805 | 0.974 | 0.858 | 0 | cDC2.Mgl2 | Cotl1 |
| Vasp | 0 | 0.66269974 | 0.969 | 0.858 | 0 | cDC2.Mgl2 | Vasp |
| Cd52 | 0 | 0.43703002 | 0.944 | 0.841 | 0 | cDC2.Mgl2 | Cd52 |
| Plek1 | 0 | 0.47329353 | 0.937 | 0.837 | 0 | cDC2.Mgl2 | Plek |
| Junb1 | 0 | 0.57846391 | 0.95 | 0.881 | 0 | cDC2.Mgl2 | Junb |
| Ctss1 | 0 | 0.38866274 | 0.97 | 0.911 | 0 | cDC2.Mgl2 | Ctss |
| Fth1 | 0 | 0.35928878 | 0.991 | 0.95 | 0 | cDC2.Mgl2 | Fth1 |
| Cd741 | 0 | 0.58780011 | 0.998 | 0.966 | 0 | cDC2.Mgl2 | Cd74 |
| Cst3 | 0 | 0.35198425 | 1 | 0.983 | 0 | cDC2.Mgl2 | Cst3 |
| Myadm | 7.09E-307 | 0.56197802 | 0.93 | 0.761 | 1.37E-302 | cDC2.Mgl2 | Myadm |
| Tln11 | 5.96E-306 | 0.5180979 | 0.901 | 0.784 | 1.15E-301 | cDC2.Mgl2 | Tln1 |
| Kdm6b1 | 9.71E-302 | 0.49561059 | 0.899 | 0.753 | 1.88E-297 | cDC2.Mgl2 | Kdm6b |
| Rrad | 3.35E-300 | 1.07046127 | 0.449 | 0.284 | 6.48E-296 | cDC2.Mgl2 | Rrad |
| Lilrb4a1 | 4.31E-298 | 0.8387586 | 0.844 | 0.508 | 8.34E-294 | cDC2.Mgl2 | Lilrb4a |
| Dck | 4.70E-293 | 1.53604784 | 0.656 | 0.299 | 9.08E-289 | cDC2.Mgl2 | Dck |
| Hipk11 | 4.30E-292 | 0.82158777 | 0.822 | 0.577 | 8.32E-288 | cDC2.Mgl2 | Hipk1 |
| Igflr1 | 1.77E-291 | 2.89329107 | 0.439 | 0.074 | 3.43E-287 | cDC2.Mgl2 | Igflr1 |
| Rpa1 | 2.04E-289 | 1.5469353 | 0.649 | 0.277 | 3.94E-285 | cDC2.Mgl2 | Rpa1 |
| Ccr11 | 9.46E-286 | 1.7342033 | 0.61 | 0.207 | 1.83E-281 | cDC2.Mgl2 | Ccr1 |
| Actr1b | 1.18E-284 | 1.16098394 | 0.728 | 0.393 | 2.28E-280 | cDC2.Mgl2 | Actr1b |
| Fgr1 | 3.02E-284 | 0.6978358 | 0.859 | 0.547 | 5.85E-280 | cDC2.Mgl2 | Fgr |
| Tm6sf11 | 2.12E-283 | 1.16091926 | 0.757 | 0.39 | 4.09E-279 | cDC2.Mgl2 | Tm6sf1 |
| Card11 | 8.71E-281 | 1.70391044 | 0.624 | 0.197 | 1.68E-276 | cDC2.Mgl2 | Card11 |
| Htra31 | 4.09E-278 | 2.05770719 | 0.56 | 0.143 | 7.90E-274 | cDC2.Mgl2 | Htra3 |
| Nup210 | 2.36E-276 | 0.89304343 | 0.765 | 0.45 | 4.56E-272 | cDC2.Mgl2 | Nup210 |
| Slc16a3 | 1.04E-275 | 1.17311079 | 0.71 | 0.372 | 2.02E-271 | cDC2.Mgl2 | Slc16a3 |
| Aldoc | 5.88E-273 | 1.85007255 | 0.536 | 0.219 | 1.14E-268 | cDC2.Mgl2 | Aldoc |
| Kmo1 | 9.66E-271 | 0.69676429 | 0.836 | 0.51 | 1.87E-266 | cDC2.Mgl2 | Kmo |
| Ifngr11 | 2.75E-269 | 0.5813679 | 0.871 | 0.654 | 5.31E-265 | cDC2.Mgl2 | Ifngr1 |
| Snx20 | 2.67E-268 | 0.57635637 | 0.888 | 0.716 | 5.17E-264 | cDC2.Mgl2 | Snx20 |
| Irf41 | 1.13E-267 | 1.14872325 | 0.551 | 0.272 | 2.18E-263 | cDC2.Mgl2 | Irf4 |
| Irf2bp21 | 3.03E-264 | 0.30809013 | 0.929 | 0.762 | 5.86E-260 | cDC2.Mgl2 | Irf2bp2 |
| Myd881 | 8.05E-264 | 0.82549483 | 0.82 | 0.568 | 1.56E-259 | cDC2.Mgl2 | Myd88 |
| Cd244a1 | 1.15E-263 | 1.18966943 | 0.72 | 0.34 | 2.22E-259 | cDC2.Mgl2 | Cd244a |
| Ptpn61 | 2.92E-263 | 0.64610045 | 0.847 | 0.608 | 5.64E-259 | cDC2.Mgl2 | Ptpn6 |
| Slamf91 | 3.21E-263 | 1.80796883 | 0.543 | 0.152 | 6.21E-259 | cDC2.Mgl2 | Slamf9 |
| Klri1 | 1.28E-261 | 1.48625597 | 0.614 | 0.232 | 2.47E-257 | cDC2.Mgl2 | Klri1 |
| Klf41 | 1.12E-258 | 0.79138469 | 0.719 | 0.441 | 2.16E-254 | cDC2.Mgl2 | Klf4 |
| Ndufa8 | 5.86E-257 | 1.06057326 | 0.785 | 0.508 | 1.13E-252 | cDC2.Mgl2 | Ndufa8 |
| Elovl5 | 2.14E-256 | 1.01884694 | 0.775 | 0.482 | 4.14E-252 | cDC2.Mgl2 | Elovl5 |
| Maff1 | 9.94E-254 | 0.92928187 | 0.771 | 0.459 | 1.92E-249 | cDC2.Mgl2 | Maff |
| Arpc1b | 1.23E-252 | 0.35925551 | 0.943 | 0.867 | 2.38E-248 | cDC2.Mgl2 | Arpc1b |
| Tpcn1 | 5.08E-251 | 1.70172562 | 0.574 | 0.201 | 9.83E-247 | cDC2.Mgl2 | Tpcn1 |
| Ccnd21 | 9.14E-250 | 0.59413761 | 0.862 | 0.575 | 1.77E-245 | cDC2.Mgl2 | Ccnd2 |
| Arih11 | 4.46E-247 | 0.70286737 | 0.766 | 0.569 | 8.62E-243 | cDC2.Mgl2 | Arih1 |
| Runx11 | 1.75E-245 | 0.80258374 | 0.782 | 0.513 | 3.39E-241 | cDC2.Mgl2 | Runx1 |
| Treml21 | 3.22E-245 | 1.76881848 | 0.535 | 0.162 | 6.22E-241 | cDC2.Mgl2 | Treml2 |
| Fgd3 | 1.20E-243 | 1.40618588 | 0.639 | 0.312 | 2.33E-239 | cDC2.Mgl2 | Fgd3 |
| Zfand51 | 1.64E-241 | 0.64818409 | 0.801 | 0.613 | 3.17E-237 | cDC2.Mgl2 | Zfand5 |
| Rell1 | 1.27E-239 | 1.15661713 | 0.614 | 0.281 | 2.46E-235 | cDC2.Mgl2 | Rell1 |
| Kif1a | 3.39E-238 | 1.56096362 | 0.53 | 0.191 | 6.55E-234 | cDC2.Mgl2 | Kif1a |
| Dock10 | 1.69E-235 | 0.65848664 | 0.825 | 0.615 | 3.27E-231 | cDC2.Mgl2 | Dock10 |
| Havcr2 | 1.30E-234 | 1.12600358 | 0.71 | 0.336 | 2.51E-230 | cDC2.Mgl2 | Havcr2 |
| Mthfr | 3.01E-233 | 1.46793207 | 0.553 | 0.275 | 5.82E-229 | cDC2.Mgl2 | Mthfr |
| Nfkbiz1 | 1.59E-231 | 0.58590495 | 0.688 | 0.48 | 3.08E-227 | cDC2.Mgl2 | Nfkbiz |
| Hnrnpll1 | 1.00E-228 | 0.96565234 | 0.754 | 0.477 | 1.94E-224 | cDC2.Mgl2 | Hnrnpll |
| Adgrg51 | 3.76E-228 | 0.88050843 | 0.78 | 0.46 | 7.26E-224 | cDC2.Mgl2 | Adgrg5 |
| Rtn1 | 3.98E-228 | 1.43455644 | 0.682 | 0.277 | 7.70E-224 | cDC2.Mgl2 | Rtn1 |
| Cd721 | 2.51E-227 | 1.58692656 | 0.535 | 0.176 | 4.86E-223 | cDC2.Mgl2 | Cd72 |
| Fam110a | 2.62E-225 | 1.02203295 | 0.586 | 0.344 | 5.06E-221 | cDC2.Mgl2 | Fam110a |
| Colgalt1 | 1.24E-221 | 0.70260743 | 0.817 | 0.573 | 2.40E-217 | cDC2.Mgl2 | Colgalt1 |
| Taldo1 | 1.40E-220 | 0.51208392 | 0.904 | 0.742 | 2.70E-216 | cDC2.Mgl2 | Taldo1 |
| Mcm7 | 9.10E-220 | 2.09730271 | 0.308 | 0.101 | 1.76E-215 | cDC2.Mgl2 | Mcm7 |
| Cad | 1.04E-218 | 2.41346604 | 0.458 | 0.105 | 2.00E-214 | cDC2.Mgl2 | Cad |
| Myo1f | 1.21E-218 | 1.06661031 | 0.756 | 0.408 | 2.34E-214 | cDC2.Mgl2 | Myo1f |
| Ctsa1 | 2.52E-218 | 0.63091672 | 0.853 | 0.617 | 4.88E-214 | cDC2.Mgl2 | Ctsa |
| Med25 | 2.57E-214 | 1.382395 | 0.679 | 0.345 | 4.97E-210 | cDC2.Mgl2 | Med25 |
| Vps37b1 | 8.25E-214 | 0.63324793 | 0.704 | 0.481 | 1.59E-209 | cDC2.Mgl2 | Vps37b |
| Plscr11 | 3.26E-213 | 0.76751265 | 0.687 | 0.42 | 6.31E-209 | cDC2.Mgl2 | Plscr1 |
| Prelid1 | 4.50E-213 | 0.82508847 | 0.862 | 0.615 | 8.70E-209 | cDC2.Mgl2 | Prelid1 |
| Csf2rb1 | 5.13E-213 | 0.25543222 | 0.88 | 0.807 | 9.91E-209 | cDC2.Mgl2 | Csf2rb |
| Acap1 | 1.00E-210 | 1.35852877 | 0.628 | 0.253 | 1.94E-206 | cDC2.Mgl2 | Acap1 |
| Cwc25 | 1.11E-210 | 1.28095982 | 0.483 | 0.247 | 2.15E-206 | cDC2.Mgl2 | Cwc25 |
| Rilpl2 | 3.13E-210 | 0.75226164 | 0.714 | 0.446 | 6.05E-206 | cDC2.Mgl2 | Rilpl2 |
| Mcm3 | 9.25E-210 | 1.52153198 | 0.391 | 0.191 | 1.79E-205 | cDC2.Mgl2 | Mcm3 |
| Pkm | 2.58E-209 | 0.58932574 | 0.865 | 0.686 | 4.98E-205 | cDC2.Mgl2 | Pkm |
| Tkt1 | 2.14E-208 | 0.54638845 | 0.875 | 0.695 | 4.13E-204 | cDC2.Mgl2 | Tkt |
| Eprs | 3.92E-206 | 1.13636224 | 0.614 | 0.342 | 7.57E-202 | cDC2.Mgl2 | Eprs |
| Slc38a1 | 1.47E-205 | 0.75872911 | 0.795 | 0.517 | 2.84E-201 | cDC2.Mgl2 | Slc38a1 |
| Ahr | 6.51E-205 | 0.78637106 | 0.818 | 0.532 | 1.26E-200 | cDC2.Mgl2 | Ahr |
| Grk6 | 7.05E-204 | 0.84994626 | 0.669 | 0.389 | 1.36E-199 | cDC2.Mgl2 | Grk6 |
| Grasp1 | 1.14E-203 | 0.49586352 | 0.668 | 0.469 | 2.20E-199 | cDC2.Mgl2 | Grasp |
| Zfp36l21 | 1.76E-203 | 0.42385141 | 0.828 | 0.587 | 3.39E-199 | cDC2.Mgl2 | Zfp36l2 |
| Mcm2 | 1.99E-203 | 1.61888744 | 0.32 | 0.124 | 3.85E-199 | cDC2.Mgl2 | Mcm2 |
| Erp29 | 4.65E-203 | 0.58418848 | 0.855 | 0.656 | 8.99E-199 | cDC2.Mgl2 | Erp29 |
| Tnfrsf1a1 | 9.00E-203 | 0.65562978 | 0.833 | 0.588 | 1.74E-198 | cDC2.Mgl2 | Tnfrsf1a |
| Hk21 | 9.90E-202 | 0.95043107 | 0.732 | 0.422 | 1.91E-197 | cDC2.Mgl2 | Hk2 |
| Pcna | 5.19E-201 | 0.8667032 | 0.673 | 0.443 | 1.00E-196 | cDC2.Mgl2 | Pcna |
| Chn21 | 1.58E-200 | 1.77763954 | 0.504 | 0.148 | 3.06E-196 | cDC2.Mgl2 | Chn2 |
| Jaml1 | 2.72E-200 | 0.6606687 | 0.829 | 0.525 | 5.26E-196 | cDC2.Mgl2 | Jaml |
| Adra2a | 4.02E-200 | 2.85423026 | 0.333 | 0.05 | 7.77E-196 | cDC2.Mgl2 | Adra2a |
| Eif2b2 | 1.85E-199 | 1.05508861 | 0.615 | 0.366 | 3.58E-195 | cDC2.Mgl2 | Eif2b2 |
| Pik3ap1 | 2.02E-198 | 0.58880241 | 0.825 | 0.562 | 3.90E-194 | cDC2.Mgl2 | Pik3ap1 |
| Dok3 | 3.06E-198 | 2.40597609 | 0.428 | 0.094 | 5.92E-194 | cDC2.Mgl2 | Dok3 |
| Cyth41 | 3.20E-198 | 0.50973955 | 0.865 | 0.582 | 6.19E-194 | cDC2.Mgl2 | Cyth4 |
| Efhd2 | 1.02E-197 | 0.34916857 | 0.94 | 0.818 | 1.98E-193 | cDC2.Mgl2 | Efhd2 |
| Pilrb1 | 2.21E-197 | 2.00378546 | 0.51 | 0.14 | 4.27E-193 | cDC2.Mgl2 | Pilrb1 |
| Atf31 | 3.71E-197 | 0.56388815 | 0.644 | 0.446 | 7.17E-193 | cDC2.Mgl2 | Atf3 |
| Arl8b1 | 1.22E-196 | 0.69781054 | 0.775 | 0.542 | 2.36E-192 | cDC2.Mgl2 | Arl8b |
| Sh2d1b1 | 1.91E-191 | 1.83488312 | 0.488 | 0.126 | 3.69E-187 | cDC2.Mgl2 | Sh2d1b1 |
| Hexa | 1.00E-190 | 0.91474368 | 0.813 | 0.516 | 1.94E-186 | cDC2.Mgl2 | Hexa |
| Cyp51 | 7.75E-190 | 0.82102665 | 0.639 | 0.384 | 1.50E-185 | cDC2.Mgl2 | Cyp51 |
| Cd531 | 2.49E-189 | 0.38869177 | 0.873 | 0.635 | 4.80E-185 | cDC2.Mgl2 | Cd53 |
| Ptprc1 | 2.31E-188 | 0.41015055 | 0.863 | 0.693 | 4.46E-184 | cDC2.Mgl2 | Ptprc |
| Mcm6 | 2.56E-188 | 1.28987248 | 0.352 | 0.17 | 4.94E-184 | cDC2.Mgl2 | Mcm6 |
| Hsd17b4 | 3.23E-188 | 1.00386168 | 0.688 | 0.411 | 6.25E-184 | cDC2.Mgl2 | Hsd17b4 |
| Ndrg11 | 1.12E-187 | 0.55615963 | 0.699 | 0.516 | 2.16E-183 | cDC2.Mgl2 | Ndrg1 |
| Cap1 | 6.91E-187 | 0.30417406 | 0.924 | 0.833 | 1.34E-182 | cDC2.Mgl2 | Cap1 |
| Tmem14c | 7.30E-186 | 0.76265165 | 0.763 | 0.53 | 1.41E-181 | cDC2.Mgl2 | Tmem14c |
| Rack1 | 3.57E-185 | 0.31120469 | 0.934 | 0.844 | 6.91E-181 | cDC2.Mgl2 | Rack1 |
| Eml3 | 9.62E-185 | 1.10694772 | 0.618 | 0.313 | 1.86E-180 | cDC2.Mgl2 | Eml3 |
| Nedd9 | 1.22E-184 | 1.27121689 | 0.53 | 0.258 | 2.35E-180 | cDC2.Mgl2 | Nedd9 |
| Eps8 | 4.83E-184 | 0.95598994 | 0.698 | 0.393 | 9.34E-180 | cDC2.Mgl2 | Eps8 |
| Cd69 | 4.21E-183 | 1.2606509 | 0.368 | 0.15 | 8.15E-179 | cDC2.Mgl2 | Cd69 |
| Fasn | 4.85E-182 | 1.16732061 | 0.63 | 0.292 | 9.38E-178 | cDC2.Mgl2 | Fasn |
| Picalm1 | 2.12E-181 | 0.46722223 | 0.891 | 0.714 | 4.09E-177 | cDC2.Mgl2 | Picalm |
| Mtss1 | 9.30E-179 | 0.91699589 | 0.737 | 0.409 | 1.80E-174 | cDC2.Mgl2 | Mtss1 |
| Dpysl21 | 1.62E-178 | 0.68535124 | 0.773 | 0.48 | 3.13E-174 | cDC2.Mgl2 | Dpysl2 |
| Tpi1 | 1.86E-178 | 0.98735554 | 0.717 | 0.423 | 3.59E-174 | cDC2.Mgl2 | Tpi1 |
| H2-Ob | 5.10E-178 | 1.22860964 | 0.604 | 0.264 | 9.86E-174 | cDC2.Mgl2 | H2-Ob |
| Slc7a51 | 5.47E-178 | 0.59507085 | 0.722 | 0.428 | 1.06E-173 | cDC2.Mgl2 | Slc7a5 |
| Slc8a2 | 2.09E-177 | 5.06456312 | 0.301 | 0.013 | 4.04E-173 | cDC2.Mgl2 | Slc8a2 |
| Scimp1 | 5.49E-177 | 1.35293071 | 0.565 | 0.243 | 1.06E-172 | cDC2.Mgl2 | Scimp |
| Por | 9.17E-177 | 1.6099322 | 0.51 | 0.171 | 1.77E-172 | cDC2.Mgl2 | Por |
| Ncf21 | 2.19E-175 | 0.702439 | 0.784 | 0.497 | 4.24E-171 | cDC2.Mgl2 | Ncf2 |
| Ap1s2 | 3.03E-175 | 1.24263397 | 0.551 | 0.256 | 5.86E-171 | cDC2.Mgl2 | Ap1s2 |
| Ltb | 3.26E-175 | 1.64040768 | 0.469 | 0.134 | 6.30E-171 | cDC2.Mgl2 | Ltb |
| Prkcb | 1.68E-174 | 1.50717201 | 0.514 | 0.164 | 3.25E-170 | cDC2.Mgl2 | Prkcb |
| Slc44a21 | 1.98E-174 | 0.48164943 | 0.746 | 0.434 | 3.83E-170 | cDC2.Mgl2 | Slc44a2 |
| Insig1 | 4.32E-174 | 0.68318952 | 0.685 | 0.458 | 8.36E-170 | cDC2.Mgl2 | Insig1 |
| Jup | 1.36E-173 | 0.79169055 | 0.657 | 0.35 | 2.63E-169 | cDC2.Mgl2 | Jup |
| Sdf2l1 | 7.50E-172 | 0.81350222 | 0.734 | 0.436 | 1.45E-167 | cDC2.Mgl2 | Sdf2l1 |
| Homer11 | 4.48E-171 | 1.13938759 | 0.535 | 0.277 | 8.66E-167 | cDC2.Mgl2 | Homer1 |
| Gpi1 | 1.76E-170 | 0.65473769 | 0.833 | 0.63 | 3.40E-166 | cDC2.Mgl2 | Gpi1 |
| Sh3pxd2b1 | 1.39E-169 | 0.60719213 | 0.801 | 0.575 | 2.68E-165 | cDC2.Mgl2 | Sh3pxd2b |
| Pilrb2 | 2.65E-168 | 2.40922843 | 0.411 | 0.087 | 5.12E-164 | cDC2.Mgl2 | Pilrb2 |
| Acox3 | 3.98E-167 | 0.88735411 | 0.727 | 0.436 | 7.70E-163 | cDC2.Mgl2 | Acox3 |
| Arhgdib1 | 1.02E-166 | 0.34599384 | 0.873 | 0.679 | 1.97E-162 | cDC2.Mgl2 | Arhgdib |
| Cystm1 | 1.14E-166 | 0.88198853 | 0.673 | 0.362 | 2.21E-162 | cDC2.Mgl2 | Cystm1 |
| Lifr1 | 2.34E-166 | 0.9771827 | 0.578 | 0.208 | 4.53E-162 | cDC2.Mgl2 | Lifr |
| Ifitm6 | 8.75E-165 | 0.91886662 | 0.502 | 0.197 | 1.69E-160 | cDC2.Mgl2 | Ifitm6 |
| Lmo21 | 1.20E-164 | 1.04732108 | 0.628 | 0.325 | 2.31E-160 | cDC2.Mgl2 | Lmo2 |
| Abi3 | 7.47E-164 | 0.9465529 | 0.679 | 0.35 | 1.44E-159 | cDC2.Mgl2 | Abi3 |
| Srebf2 | 6.72E-162 | 0.45559941 | 0.871 | 0.708 | 1.30E-157 | cDC2.Mgl2 | Srebf2 |
| Spi11 | 9.08E-162 | 0.35614148 | 0.904 | 0.751 | 1.76E-157 | cDC2.Mgl2 | Spi1 |
| Dennd4a1 | 1.64E-160 | 0.40259104 | 0.785 | 0.613 | 3.16E-156 | cDC2.Mgl2 | Dennd4a |
| Smdt11 | 2.48E-160 | 0.5868462 | 0.823 | 0.638 | 4.79E-156 | cDC2.Mgl2 | Smdt1 |
| Tnfaip3 | 4.04E-160 | 0.38821667 | 0.549 | 0.458 | 7.81E-156 | cDC2.Mgl2 | Tnfaip3 |
| Mif | 4.30E-160 | 1.0275107 | 0.667 | 0.383 | 8.32E-156 | cDC2.Mgl2 | Mif |
| Fgfr1 | 5.27E-160 | 1.40694142 | 0.48 | 0.168 | 1.02E-155 | cDC2.Mgl2 | Fgfr1 |
| Sash1 | 7.66E-160 | 2.20334069 | 0.426 | 0.114 | 1.48E-155 | cDC2.Mgl2 | Sash1 |
| Siglecf | 1.69E-158 | 3.05424006 | 0.253 | 0.035 | 3.26E-154 | cDC2.Mgl2 | Siglecf |
| Tep11 | 1.34E-157 | 0.96605501 | 0.669 | 0.361 | 2.60E-153 | cDC2.Mgl2 | Tep1 |
| Ywhah1 | 2.36E-157 | 0.36492627 | 0.825 | 0.695 | 4.57E-153 | cDC2.Mgl2 | Ywhah |
| Ddr11 | 2.44E-157 | 0.79683653 | 0.619 | 0.305 | 4.73E-153 | cDC2.Mgl2 | Ddr1 |
| Ccnd31 | 5.85E-157 | 0.33017486 | 0.777 | 0.55 | 1.13E-152 | cDC2.Mgl2 | Ccnd3 |
| Stk32c | 1.15E-156 | 1.87104873 | 0.467 | 0.141 | 2.23E-152 | cDC2.Mgl2 | Stk32c |
| Epn1 | 1.52E-155 | 0.56843328 | 0.856 | 0.672 | 2.93E-151 | cDC2.Mgl2 | Epn1 |
| Slc2a11 | 9.59E-154 | 0.7710505 | 0.698 | 0.439 | 1.85E-149 | cDC2.Mgl2 | Slc2a1 |
| Pygb | 1.04E-153 | 1.07470693 | 0.607 | 0.322 | 2.00E-149 | cDC2.Mgl2 | Pygb |
| Csf2ra | 3.98E-152 | 0.4753381 | 0.881 | 0.711 | 7.69E-148 | cDC2.Mgl2 | Csf2ra |
| Pim31 | 5.44E-152 | 0.8762081 | 0.627 | 0.351 | 1.05E-147 | cDC2.Mgl2 | Pim3 |
| B4galt51 | 1.20E-151 | 0.69559919 | 0.738 | 0.5 | 2.33E-147 | cDC2.Mgl2 | B4galt5 |
| Ddost | 9.48E-151 | 0.56531919 | 0.783 | 0.581 | 1.83E-146 | cDC2.Mgl2 | Ddost |
| Hmgcr | 1.58E-149 | 0.57149379 | 0.706 | 0.51 | 3.06E-145 | cDC2.Mgl2 | Hmgcr |
| Mcm5 | 1.75E-149 | 1.74561131 | 0.28 | 0.098 | 3.38E-145 | cDC2.Mgl2 | Mcm5 |
| Gpd1l | 2.41E-149 | 1.84567185 | 0.441 | 0.146 | 4.66E-145 | cDC2.Mgl2 | Gpd1l |
| Vrk11 | 3.15E-149 | 0.41460856 | 0.778 | 0.544 | 6.09E-145 | cDC2.Mgl2 | Vrk1 |
| Antxr2 | 5.86E-149 | 0.58796439 | 0.684 | 0.538 | 1.13E-144 | cDC2.Mgl2 | Antxr2 |
| Tm2d2 | 7.03E-149 | 0.99632418 | 0.659 | 0.385 | 1.36E-144 | cDC2.Mgl2 | Tm2d2 |
| Arrb21 | 1.54E-148 | 0.28077602 | 0.885 | 0.758 | 2.99E-144 | cDC2.Mgl2 | Arrb2 |
| Pram1 | 2.80E-148 | 2.13609769 | 0.413 | 0.114 | 5.42E-144 | cDC2.Mgl2 | Pram1 |
| Scd2 | 6.79E-148 | 1.32753544 | 0.526 | 0.226 | 1.31E-143 | cDC2.Mgl2 | Scd2 |
| Adcy71 | 6.83E-148 | 0.95431536 | 0.668 | 0.356 | 1.32E-143 | cDC2.Mgl2 | Adcy7 |
| Pik3cd1 | 1.18E-147 | 0.63334622 | 0.792 | 0.516 | 2.28E-143 | cDC2.Mgl2 | Pik3cd |
| Ptk2b1 | 8.11E-147 | 0.62840981 | 0.74 | 0.529 | 1.57E-142 | cDC2.Mgl2 | Ptk2b |
| Ndel11 | 9.57E-147 | 0.56111445 | 0.739 | 0.537 | 1.85E-142 | cDC2.Mgl2 | Ndel1 |
| Slc3a2 | 9.14E-146 | 0.35973351 | 0.848 | 0.712 | 1.77E-141 | cDC2.Mgl2 | Slc3a2 |
| Exosc5 | 5.30E-145 | 1.04930131 | 0.594 | 0.317 | 1.03E-140 | cDC2.Mgl2 | Exosc5 |
| Ptpn121 | 1.81E-144 | 0.95611034 | 0.688 | 0.408 | 3.50E-140 | cDC2.Mgl2 | Ptpn12 |
| Pgam1 | 2.71E-144 | 0.90800881 | 0.741 | 0.457 | 5.25E-140 | cDC2.Mgl2 | Pgam1 |
| Siglecg | 2.87E-144 | 0.88479814 | 0.548 | 0.265 | 5.55E-140 | cDC2.Mgl2 | Siglecg |
| Ptpn18 | 4.09E-143 | 0.91414926 | 0.707 | 0.364 | 7.91E-139 | cDC2.Mgl2 | Ptpn18 |
| Fhl31 | 7.44E-143 | 0.95825948 | 0.593 | 0.3 | 1.44E-138 | cDC2.Mgl2 | Fhl3 |
| Stx71 | 8.85E-143 | 0.42971209 | 0.821 | 0.65 | 1.71E-138 | cDC2.Mgl2 | Stx7 |
| Pmvk1 | 1.54E-141 | 0.48974011 | 0.64 | 0.438 | 2.97E-137 | cDC2.Mgl2 | Pmvk |
| Pfkp1 | 2.91E-139 | 0.37811544 | 0.885 | 0.672 | 5.63E-135 | cDC2.Mgl2 | Pfkp |
| Nfe2l21 | 2.10E-138 | 0.35936544 | 0.845 | 0.675 | 4.07E-134 | cDC2.Mgl2 | Nfe2l2 |
| Enah1 | 8.56E-138 | 1.17067168 | 0.529 | 0.228 | 1.65E-133 | cDC2.Mgl2 | Enah |
| Gnai2 | 1.06E-137 | 0.26211516 | 0.94 | 0.853 | 2.05E-133 | cDC2.Mgl2 | Gnai2 |
| Ssbp41 | 1.91E-137 | 1.04622911 | 0.638 | 0.326 | 3.70E-133 | cDC2.Mgl2 | Ssbp4 |
| Gpr4 | 1.21E-136 | 1.52739828 | 0.366 | 0.127 | 2.35E-132 | cDC2.Mgl2 | Gpr4 |

| | | | | | | | |
|---|---|---|---|---|---|---|---|
| Hck | 5.91E-135 | 0.52997956 | 0.784 | 0.555 | 1.14E-130 | cDC2.Mgl2 | Hck |
| Rab1b | 9.43E-135 | 0.49112197 | 0.787 | 0.608 | 1.82E-130 | cDC2.Mgl2 | Rab1b |
| Aldoa | 1.01E-134 | 0.48537325 | 0.83 | 0.663 | 1.96E-130 | cDC2.Mgl2 | Aldoa |
| Arid5a1 | 1.66E-134 | 1.11352335 | 0.567 | 0.263 | 3.20E-130 | cDC2.Mgl2 | Arid5a |
| Snx181 | 6.07E-133 | 0.51432598 | 0.724 | 0.463 | 1.17E-128 | cDC2.Mgl2 | Snx18 |
| Tent5a1 | 1.71E-132 | 1.25217389 | 0.502 | 0.22 | 3.31E-128 | cDC2.Mgl2 | Tent5a |
| Tmem50b | 5.76E-132 | 0.90041576 | 0.619 | 0.353 | 1.11E-127 | cDC2.Mgl2 | Tmem50b |
| Jag2 | 8.64E-132 | 1.32893996 | 0.515 | 0.232 | 1.67E-127 | cDC2.Mgl2 | Jag2 |
| Limd21 | 7.18E-131 | 0.48816588 | 0.68 | 0.468 | 1.39E-126 | cDC2.Mgl2 | Limd2 |
| Slc16a6 | 8.92E-131 | 1.45833625 | 0.443 | 0.17 | 1.73E-126 | cDC2.Mgl2 | Slc16a6 |
| Asb2 | 1.05E-129 | 0.74940072 | 0.638 | 0.215 | 2.02E-125 | cDC2.Mgl2 | Asb2 |
| Osgin2 | 3.70E-127 | 1.00215053 | 0.491 | 0.254 | 7.16E-123 | cDC2.Mgl2 | Osgin2 |
| Rab241 | 8.66E-126 | 0.87141159 | 0.633 | 0.353 | 1.68E-121 | cDC2.Mgl2 | Rab24 |
| Slc25a33 | 9.22E-126 | 1.41683261 | 0.443 | 0.169 | 1.78E-121 | cDC2.Mgl2 | Slc25a33 |
| Pgf1 | 2.06E-125 | 0.98387623 | 0.376 | 0.173 | 3.97E-121 | cDC2.Mgl2 | Pgf |
| Khk | 6.18E-125 | 1.34159108 | 0.482 | 0.197 | 1.19E-120 | cDC2.Mgl2 | Khk |
| Tnfaip81 | 1.53E-124 | 0.34502882 | 0.784 | 0.613 | 2.95E-120 | cDC2.Mgl2 | Tnfaip8 |
| Mycbp21 | 2.29E-124 | 0.54334135 | 0.703 | 0.471 | 4.43E-120 | cDC2.Mgl2 | Mycbp2 |
| Slc15a31 | 8.25E-124 | 0.41533747 | 0.871 | 0.594 | 1.59E-119 | cDC2.Mgl2 | Slc15a3 |
| Otulinl1 | 2.71E-123 | 0.57081618 | 0.759 | 0.495 | 5.23E-119 | cDC2.Mgl2 | Otulinl |
| St6gal1 | 5.58E-123 | 1.26263574 | 0.538 | 0.25 | 1.08E-118 | cDC2.Mgl2 | St6gal1 |
| Zbtb1 | 8.35E-123 | 0.78662253 | 0.506 | 0.321 | 1.61E-118 | cDC2.Mgl2 | Zbtb1 |
| Fndc5 | 2.36E-122 | 2.06402094 | 0.327 | 0.064 | 4.56E-118 | cDC2.Mgl2 | Fndc5 |
| Parvg | 5.20E-122 | 0.82369495 | 0.63 | 0.338 | 1.01E-117 | cDC2.Mgl2 | Parvg |
| Ubald21 | 1.14E-119 | 0.47802944 | 0.761 | 0.56 | 2.20E-115 | cDC2.Mgl2 | Ubald2 |
| Dab2ip | 1.96E-119 | 2.34729577 | 0.298 | 0.054 | 3.79E-115 | cDC2.Mgl2 | Dab2ip |
| Jund | 2.17E-119 | 0.2608976 | 0.92 | 0.789 | 4.19E-115 | cDC2.Mgl2 | Jund |
| Pdk1 | 3.53E-119 | 1.67856785 | 0.421 | 0.155 | 6.83E-115 | cDC2.Mgl2 | Pdk1 |
| Ppp1r3b | 3.65E-119 | 1.90474063 | 0.309 | 0.092 | 7.05E-115 | cDC2.Mgl2 | Ppp1r3b |
| Trp53i11 | 1.29E-118 | 0.90948583 | 0.328 | 0.118 | 2.49E-114 | cDC2.Mgl2 | Trp53i11 |
| Per11 | 1.83E-118 | 0.42122136 | 0.767 | 0.56 | 3.54E-114 | cDC2.Mgl2 | Per1 |
| Dbnl | 6.55E-118 | 0.30330163 | 0.886 | 0.746 | 1.27E-113 | cDC2.Mgl2 | Dbnl |
| Lmnb11 | 1.84E-117 | 0.52105214 | 0.802 | 0.575 | 3.55E-113 | cDC2.Mgl2 | Lmnb1 |
| Isy1 | 2.00E-117 | 0.61323998 | 0.746 | 0.526 | 3.86E-113 | cDC2.Mgl2 | Isy1 |
| Cd681 | 1.04E-116 | 0.54980298 | 0.777 | 0.516 | 2.01E-112 | cDC2.Mgl2 | Cd68 |
| Acsl5 | 2.77E-114 | 0.74936536 | 0.648 | 0.421 | 5.35E-110 | cDC2.Mgl2 | Acsl5 |
| Csk | 3.97E-114 | 0.5361053 | 0.734 | 0.53 | 7.68E-110 | cDC2.Mgl2 | Csk |
| Klk8 | 1.41E-113 | 2.28358201 | 0.273 | 0.062 | 2.72E-109 | cDC2.Mgl2 | Klk8 |
| D16Ertd472e1 | 1.14E-112 | 0.83892646 | 0.536 | 0.288 | 2.21E-108 | cDC2.Mgl2 | D16Ertd472e |
| Dgkz | 1.85E-112 | 0.99060043 | 0.596 | 0.308 | 3.58E-108 | cDC2.Mgl2 | Dgkz |
| Nsd31 | 1.95E-112 | 0.32092459 | 0.799 | 0.662 | 3.76E-108 | cDC2.Mgl2 | Nsd3 |
| Sema6b | 4.09E-112 | 1.9224535 | 0.337 | 0.101 | 7.91E-108 | cDC2.Mgl2 | Sema6b |
| Slc52a3 | 1.03E-111 | 1.67250639 | 0.351 | 0.117 | 1.99E-107 | cDC2.Mgl2 | Slc52a3 |
| Phc21 | 2.07E-111 | 0.60104891 | 0.651 | 0.438 | 4.00E-107 | cDC2.Mgl2 | Phc2 |
| Ivns1abp | 2.75E-111 | 0.25059679 | 0.802 | 0.639 | 5.33E-107 | cDC2.Mgl2 | Ivns1abp |
| Gpr55 | 6.04E-111 | 0.87443682 | 0.529 | 0.269 | 1.17E-106 | cDC2.Mgl2 | Gpr55 |
| Litaf | 7.44E-111 | 0.27235902 | 0.872 | 0.755 | 1.44E-106 | cDC2.Mgl2 | Litaf |
| Atp13a31 | 9.28E-111 | 0.56328716 | 0.74 | 0.517 | 1.79E-106 | cDC2.Mgl2 | Atp13a3 |
| Ly861 | 2.07E-110 | 0.42137808 | 0.726 | 0.488 | 3.99E-106 | cDC2.Mgl2 | Ly86 |
| Anpep | 3.05E-110 | 0.60897049 | 0.754 | 0.481 | 5.89E-106 | cDC2.Mgl2 | Anpep |
| Ncf4 | 3.62E-110 | 0.7831359 | 0.702 | 0.403 | 7.00E-106 | cDC2.Mgl2 | Ncf4 |
| Spty2d11 | 1.20E-109 | 0.59480265 | 0.645 | 0.475 | 2.32E-105 | cDC2.Mgl2 | Spty2d1 |
| Pnpo | 2.91E-109 | 1.20743538 | 0.467 | 0.2 | 5.62E-105 | cDC2.Mgl2 | Pnpo |
| Twf21 | 2.96E-109 | 0.5898571 | 0.733 | 0.501 | 5.72E-105 | cDC2.Mgl2 | Twf2 |
| Gfod1 | 3.13E-109 | 1.10866726 | 0.48 | 0.221 | 6.06E-105 | cDC2.Mgl2 | Gfod1 |
| Coq10b | 5.56E-109 | 0.56059962 | 0.653 | 0.446 | 1.08E-104 | cDC2.Mgl2 | Coq10b |
| Plcb21 | 1.82E-108 | 0.76463845 | 0.637 | 0.34 | 3.53E-104 | cDC2.Mgl2 | Plcb2 |
| Anxa6 | 2.01E-108 | 0.51080356 | 0.81 | 0.527 | 3.90E-104 | cDC2.Mgl2 | Anxa6 |
| Rabgef1 | 3.37E-108 | 0.67265084 | 0.578 | 0.368 | 6.52E-104 | cDC2.Mgl2 | Rabgef1 |
| Itgam1 | 2.39E-107 | 0.56982337 | 0.601 | 0.309 | 4.63E-103 | cDC2.Mgl2 | Itgam |
| Stap1 | 6.70E-107 | 1.45545019 | 0.366 | 0.13 | 1.30E-102 | cDC2.Mgl2 | Stap1 |
| Atrnl11 | 1.38E-105 | 0.48143319 | 0.611 | 0.41 | 2.67E-101 | cDC2.Mgl2 | Atrnl1 |
| Tbc1d9 | 2.10E-105 | 0.46707529 | 0.784 | 0.53 | 4.06E-101 | cDC2.Mgl2 | Tbc1d9 |
| Cnot6l1 | 2.69E-105 | 0.63862088 | 0.612 | 0.406 | 5.19E-101 | cDC2.Mgl2 | Cnot6l |
| Cysltr11 | 3.96E-105 | 1.44779101 | 0.424 | 0.163 | 7.65E-101 | cDC2.Mgl2 | Cysltr1 |
| Sesn2 | 1.13E-104 | 0.80439816 | 0.561 | 0.298 | 2.18E-100 | cDC2.Mgl2 | Sesn2 |
| Ppm1j | 1.17E-104 | 1.71469631 | 0.32 | 0.098 | 2.27E-100 | cDC2.Mgl2 | Ppm1j |
| Eif4g31 | 2.79E-104 | 0.48015433 | 0.787 | 0.567 | 5.39E-100 | cDC2.Mgl2 | Eif4g3 |
| Ccdc88b | 2.03E-103 | 0.29346027 | 0.798 | 0.621 | 3.92E-99 | cDC2.Mgl2 | Ccdc88b |
| Stk241 | 2.94E-103 | 0.35314481 | 0.782 | 0.617 | 5.68E-99 | cDC2.Mgl2 | Stk24 |
| Ell2 | 1.53E-102 | 0.60061922 | 0.59 | 0.365 | 2.96E-98 | cDC2.Mgl2 | Ell2 |
| Ccr6 | 2.27E-102 | 1.19399539 | 0.282 | 0.093 | 4.39E-98 | cDC2.Mgl2 | Ccr6 |
| Akr7a51 | 3.20E-102 | 1.3070017 | 0.468 | 0.206 | 6.19E-98 | cDC2.Mgl2 | Akr7a5 |
| Pld4 | 4.84E-102 | 0.26575062 | 0.846 | 0.605 | 9.36E-98 | cDC2.Mgl2 | Pld4 |
| Rrm1 | 5.11E-102 | 1.26121626 | 0.303 | 0.135 | 9.88E-98 | cDC2.Mgl2 | Rrm1 |
| Il18rap | 1.28E-101 | 1.39334813 | 0.378 | 0.129 | 2.48E-97 | cDC2.Mgl2 | Il18rap |
| Soat1 | 2.22E-101 | 0.94999166 | 0.618 | 0.364 | 4.29E-97 | cDC2.Mgl2 | Soat1 |
| Ptpn11 | 2.60E-101 | 0.31073435 | 0.835 | 0.659 | 5.03E-97 | cDC2.Mgl2 | Ptpn1 |
| Vars | 3.78E-101 | 0.86468788 | 0.625 | 0.361 | 7.30E-97 | cDC2.Mgl2 | Vars |
| Slc38a9 | 1.64E-100 | 1.0535956 | 0.479 | 0.222 | 3.17E-96 | cDC2.Mgl2 | Slc38a9 |
| Rasgef1b1 | 2.06E-100 | 0.66965824 | 0.51 | 0.325 | 3.98E-96 | cDC2.Mgl2 | Rasgef1b |
| Ocstamp | 9.43E-100 | 1.38688028 | 0.389 | 0.149 | 1.82E-95 | cDC2.Mgl2 | Ocstamp |
| Nfkbie1 | 3.37E-99 | 0.42055434 | 0.729 | 0.528 | 6.52E-95 | cDC2.Mgl2 | Nfkbie |
| Mlec | 5.26E-99 | 0.48010899 | 0.753 | 0.492 | 1.02E-94 | cDC2.Mgl2 | Mlec |
| Dnmt3a | 1.02E-98 | 0.87100048 | 0.538 | 0.303 | 1.98E-94 | cDC2.Mgl2 | Dnmt3a |
| Kit | 1.17E-98 | 0.5152212 | 0.623 | 0.37 | 2.25E-94 | cDC2.Mgl2 | Kit |
| Iqgap21 | 3.97E-98 | 0.70017227 | 0.576 | 0.299 | 7.68E-94 | cDC2.Mgl2 | Iqgap2 |
| Fermt31 | 3.97E-98 | 0.46218863 | 0.783 | 0.569 | 7.68E-94 | cDC2.Mgl2 | Fermt3 |
| Ciita1 | 1.24E-97 | 0.25805902 | 0.848 | 0.582 | 2.40E-93 | cDC2.Mgl2 | Ciita |
| Hfe | 1.20E-96 | 0.68138544 | 0.724 | 0.404 | 2.33E-92 | cDC2.Mgl2 | Hfe |
| Ddx211 | 1.30E-96 | 0.44868536 | 0.703 | 0.512 | 2.50E-92 | cDC2.Mgl2 | Ddx21 |
| P4hb | 1.82E-96 | 0.64794004 | 0.785 | 0.544 | 3.51E-92 | cDC2.Mgl2 | P4hb |
| Hs3st3b11 | 6.04E-96 | 1.36809727 | 0.426 | 0.168 | 1.17E-91 | cDC2.Mgl2 | Hs3st3b1 |
| Xbp11 | 6.84E-96 | 0.33863199 | 0.846 | 0.679 | 1.32E-91 | cDC2.Mgl2 | Xbp1 |
| Ldlr1 | 7.88E-96 | 0.33927944 | 0.656 | 0.474 | 1.52E-91 | cDC2.Mgl2 | Ldlr |
| Lgals91 | 1.28E-95 | 0.9933879 | 0.518 | 0.235 | 2.48E-91 | cDC2.Mgl2 | Lgals9 |
| Lrrc8c | 1.46E-95 | 0.33594553 | 0.684 | 0.527 | 2.82E-91 | cDC2.Mgl2 | Lrrc8c |
| Atp6v0c1 | 1.61E-95 | 0.37293831 | 0.84 | 0.672 | 3.10E-91 | cDC2.Mgl2 | Atp6v0c |
| Rnf1301 | 1.88E-95 | 0.4848484 | 0.664 | 0.462 | 3.63E-91 | cDC2.Mgl2 | Rnf130 |
| Hmgcl1 | 2.52E-95 | 0.86549835 | 0.679 | 0.417 | 4.87E-91 | cDC2.Mgl2 | Hmgcl |
| Nab11 | 4.53E-95 | 0.46900313 | 0.687 | 0.502 | 8.76E-91 | cDC2.Mgl2 | Nab1 |
| Impact | 8.19E-95 | 0.96172311 | 0.46 | 0.234 | 1.58E-90 | cDC2.Mgl2 | Impact |
| Sla1 | 1.00E-94 | 0.42012035 | 0.756 | 0.488 | 1.93E-90 | cDC2.Mgl2 | Sla |
| Mvb12b | 1.43E-94 | 1.1311071 | 0.435 | 0.169 | 2.77E-90 | cDC2.Mgl2 | Mvb12b |
| Plekha5 | 2.61E-94 | 0.82598995 | 0.546 | 0.268 | 5.05E-90 | cDC2.Mgl2 | Plekha5 |
| Ndufa61 | 3.19E-94 | 0.33595478 | 0.788 | 0.638 | 6.16E-90 | cDC2.Mgl2 | Ndufa6 |
| Isyna11 | 4.74E-94 | 0.76403577 | 0.642 | 0.386 | 9.16E-90 | cDC2.Mgl2 | Isyna1 |
| Ppp1r10 | 5.97E-94 | 0.61672775 | 0.591 | 0.375 | 1.15E-89 | cDC2.Mgl2 | Ppp1r10 |
| Sdcbp1 | 2.30E-93 | 0.40052966 | 0.84 | 0.67 | 4.45E-89 | cDC2.Mgl2 | Sdcbp |
| Sfxn1 | 4.34E-93 | 0.90950998 | 0.477 | 0.237 | 8.39E-89 | cDC2.Mgl2 | Sfxn1 |
| Amz11 | 4.38E-93 | 0.85264644 | 0.595 | 0.331 | 8.47E-89 | cDC2.Mgl2 | Amz1 |
| Tnfaip2 | 9.86E-93 | 0.58800901 | 0.59 | 0.387 | 1.91E-88 | cDC2.Mgl2 | Tnfaip2 |
| Agpat41 | 1.67E-91 | 0.69469856 | 0.633 | 0.397 | 3.22E-87 | cDC2.Mgl2 | Agpat4 |
| Slc39a6 | 2.04E-91 | 1.41624902 | 0.363 | 0.139 | 3.95E-87 | cDC2.Mgl2 | Slc39a6 |
| Mia2 | 2.90E-91 | 0.53152095 | 0.717 | 0.521 | 5.61E-87 | cDC2.Mgl2 | Mia2 |
| Plau | 4.37E-91 | 1.45453767 | 0.3 | 0.129 | 8.44E-87 | cDC2.Mgl2 | Plau |
| Lbh | 6.73E-91 | 0.84118827 | 0.543 | 0.297 | 1.30E-86 | cDC2.Mgl2 | Lbh |
| Lims1 | 1.07E-90 | 0.40593429 | 0.788 | 0.587 | 2.07E-86 | cDC2.Mgl2 | Lims1 |
| Cd82 | 1.23E-90 | 0.39575299 | 0.786 | 0.517 | 2.37E-86 | cDC2.Mgl2 | Cd82 |
| Pip5k1c | 1.37E-90 | 0.25334197 | 0.849 | 0.696 | 2.64E-86 | cDC2.Mgl2 | Pip5k1c |
| Elmo1 | 2.91E-90 | 0.38603261 | 0.659 | 0.465 | 5.63E-86 | cDC2.Mgl2 | Elmo1 |
| Etv5 | 3.15E-90 | 1.33747148 | 0.413 | 0.173 | 6.10E-86 | cDC2.Mgl2 | Etv5 |
| Milr11 | 3.21E-90 | 1.32024297 | 0.383 | 0.138 | 6.21E-86 | cDC2.Mgl2 | Milr1 |
| Ppm1m | 3.25E-90 | 0.47994301 | 0.82 | 0.568 | 6.29E-86 | cDC2.Mgl2 | Ppm1m |
| Ggt5 | 3.69E-90 | 1.62742701 | 0.353 | 0.115 | 7.13E-86 | cDC2.Mgl2 | Ggt5 |
| Gcnt1 | 1.05E-89 | 1.52233332 | 0.326 | 0.12 | 2.04E-85 | cDC2.Mgl2 | Gcnt1 |
| Prpf18 | 3.28E-89 | 0.76233571 | 0.515 | 0.29 | 6.34E-85 | cDC2.Mgl2 | Prpf18 |
| Lbr1 | 1.07E-88 | 0.58606347 | 0.564 | 0.316 | 2.07E-84 | cDC2.Mgl2 | Lbr |
| Rgs121 | 1.47E-88 | 0.47256346 | 0.547 | 0.281 | 2.84E-84 | cDC2.Mgl2 | Rgs12 |
| Sorl11 | 1.57E-88 | 0.61100748 | 0.578 | 0.325 | 3.03E-84 | cDC2.Mgl2 | Sorl1 |
| Gabbr1 | 8.44E-88 | 1.10152275 | 0.373 | 0.141 | 1.63E-83 | cDC2.Mgl2 | Gabbr1 |
| Eci2 | 9.83E-88 | 0.9388551 | 0.486 | 0.255 | 1.90E-83 | cDC2.Mgl2 | Eci2 |
| Eef2k | 1.72E-87 | 1.28715189 | 0.363 | 0.139 | 3.33E-83 | cDC2.Mgl2 | Eef2k |
| Aldh2 | 1.86E-87 | 0.4803255 | 0.708 | 0.518 | 3.59E-83 | cDC2.Mgl2 | Aldh2 |
| Cox7a2l1 | 3.69E-87 | 0.42269839 | 0.779 | 0.594 | 7.13E-83 | cDC2.Mgl2 | Cox7a2l |
| Zc3h12a1 | 6.34E-87 | 0.42611787 | 0.669 | 0.463 | 1.23E-82 | cDC2.Mgl2 | Zc3h12a |
| Rnf1491 | 1.02E-86 | 0.34892572 | 0.799 | 0.596 | 1.97E-82 | cDC2.Mgl2 | Rnf149 |
| Sgpp1 | 1.22E-86 | 0.79537293 | 0.556 | 0.312 | 2.37E-82 | cDC2.Mgl2 | Sgpp1 |
| Ccdc12 | 6.76E-86 | 0.37882256 | 0.755 | 0.568 | 1.31E-81 | cDC2.Mgl2 | Ccdc12 |
| Nasp | 7.22E-86 | 0.9086438 | 0.393 | 0.227 | 1.40E-81 | cDC2.Mgl2 | Nasp |
| Nop581 | 8.22E-86 | 0.80426875 | 0.458 | 0.25 | 1.59E-81 | cDC2.Mgl2 | Nop58 |
| Dtx4 | 1.26E-85 | 1.49759614 | 0.297 | 0.108 | 2.43E-81 | cDC2.Mgl2 | Dtx4 |
| Etfa | 1.51E-85 | 0.64668552 | 0.695 | 0.48 | 2.91E-81 | cDC2.Mgl2 | Etfa |
| Midn1 | 2.98E-85 | 0.29812828 | 0.754 | 0.599 | 5.76E-81 | cDC2.Mgl2 | Midn |
| Palm | 9.25E-85 | 0.69354118 | 0.513 | 0.264 | 1.79E-80 | cDC2.Mgl2 | Palm |
| Tnip31 | 1.06E-84 | 0.43200797 | 0.466 | 0.299 | 2.05E-80 | cDC2.Mgl2 | Tnip3 |
| P2ry101 | 1.36E-84 | 0.27884694 | 0.695 | 0.56 | 2.62E-80 | cDC2.Mgl2 | P2ry10 |
| Rassf51 | 2.72E-84 | 0.39090328 | 0.722 | 0.522 | 5.26E-80 | cDC2.Mgl2 | Rassf5 |
| Prelid2 | 2.97E-84 | 1.51834594 | 0.324 | 0.118 | 5.74E-80 | cDC2.Mgl2 | Prelid2 |
| Atp6v1a | 9.30E-84 | 0.67389397 | 0.695 | 0.461 | 1.80E-79 | cDC2.Mgl2 | Atp6v1a |
| Pla2g4a | 1.88E-83 | 1.4608186 | 0.343 | 0.115 | 3.64E-79 | cDC2.Mgl2 | Pla2g4a |
| Rgs2 | 1.92E-83 | 0.52497326 | 0.6 | 0.421 | 3.72E-79 | cDC2.Mgl2 | Rgs2 |
| Dpp41 | 4.03E-83 | 0.37429804 | 0.671 | 0.464 | 7.79E-79 | cDC2.Mgl2 | Dpp4 |
| Rbm47 | 7.12E-83 | 0.69218011 | 0.638 | 0.401 | 1.38E-78 | cDC2.Mgl2 | Rbm47 |
| Jmjd1c | 8.24E-83 | 0.41677242 | 0.688 | 0.509 | 1.59E-78 | cDC2.Mgl2 | Jmjd1c |
| Rfc2 | 1.04E-82 | 0.62275103 | 0.445 | 0.271 | 2.01E-78 | cDC2.Mgl2 | Rfc2 |
| Lcp21 | 3.06E-82 | 0.73703211 | 0.565 | 0.314 | 5.92E-78 | cDC2.Mgl2 | Lcp2 |
| Chd71 | 3.10E-82 | 0.29704336 | 0.784 | 0.583 | 5.99E-78 | cDC2.Mgl2 | Chd7 |
| Was1 | 4.89E-82 | 0.42293929 | 0.739 | 0.52 | 9.46E-78 | cDC2.Mgl2 | Was |
| Macf11 | 1.11E-81 | 0.46312596 | 0.597 | 0.383 | 2.15E-77 | cDC2.Mgl2 | Macf1 |
| Ptma | 1.69E-81 | 0.27181939 | 0.795 | 0.652 | 3.27E-77 | cDC2.Mgl2 | Ptma |
| Tnip21 | 6.61E-81 | 0.43914058 | 0.699 | 0.505 | 1.28E-76 | cDC2.Mgl2 | Tnip2 |
| Entpd1 | 8.97E-81 | 0.96650671 | 0.481 | 0.236 | 1.73E-76 | cDC2.Mgl2 | Entpd1 |
| Gpsm3 | 1.78E-80 | 0.29216907 | 0.849 | 0.672 | 3.44E-76 | cDC2.Mgl2 | Gpsm3 |
| Smpdl3a | 1.84E-80 | 0.69661689 | 0.535 | 0.304 | 3.56E-76 | cDC2.Mgl2 | Smpdl3a |
| Tspan141 | 1.91E-80 | 0.8114564 | 0.478 | 0.272 | 3.69E-76 | cDC2.Mgl2 | Tspan14 |
| Tnks | 4.68E-80 | 0.78436459 | 0.492 | 0.281 | 9.04E-76 | cDC2.Mgl2 | Tnks |
| Mta31 | 8.31E-80 | 0.55332116 | 0.604 | 0.38 | 1.61E-75 | cDC2.Mgl2 | Mta3 |
| Smad71 | 1.41E-79 | 0.4870313 | 0.535 | 0.373 | 2.73E-75 | cDC2.Mgl2 | Smad7 |
| Myc | 1.77E-79 | 1.12456163 | 0.333 | 0.159 | 3.43E-75 | cDC2.Mgl2 | Myc |
| Plpp5 | 2.29E-79 | 1.07107145 | 0.481 | 0.231 | 4.44E-75 | cDC2.Mgl2 | Plpp5 |
| Cops9 | 2.41E-79 | 0.38642274 | 0.809 | 0.642 | 4.65E-75 | cDC2.Mgl2 | Cops9 |
| Mical1 | 2.67E-79 | 0.59539349 | 0.699 | 0.434 | 5.16E-75 | cDC2.Mgl2 | Mical1 |
| Hmgcs11 | 5.14E-79 | 0.26370971 | 0.583 | 0.411 | 9.93E-75 | cDC2.Mgl2 | Hmgcs1 |
| Anxa11 | 6.77E-79 | 0.53167492 | 0.708 | 0.523 | 1.31E-74 | cDC2.Mgl2 | Anxa11 |
| Fes1 | 1.19E-78 | 0.30612702 | 0.811 | 0.641 | 2.29E-74 | cDC2.Mgl2 | Fes |
| Map111 | 1.34E-77 | 0.96809406 | 0.453 | 0.222 | 2.58E-73 | cDC2.Mgl2 | Map11 |
| Ptbp1 | 3.16E-77 | 0.27826576 | 0.812 | 0.654 | 6.10E-73 | cDC2.Mgl2 | Ptbp1 |
| Ssr11 | 3.48E-77 | 0.3183482 | 0.76 | 0.58 | 6.73E-73 | cDC2.Mgl2 | Ssr1 |
| Trem11 | 5.06E-77 | 1.09365859 | 0.373 | 0.14 | 9.79E-73 | cDC2.Mgl2 | Trem1 |
| Ago2 | 1.86E-76 | 0.38316379 | 0.67 | 0.479 | 3.59E-72 | cDC2.Mgl2 | Ago2 |
| Trps11 | 2.49E-76 | 0.26244076 | 0.714 | 0.541 | 4.82E-72 | cDC2.Mgl2 | Trps1 |
| Sphk2 | 3.75E-76 | 1.00854244 | 0.447 | 0.251 | 7.25E-72 | cDC2.Mgl2 | Sphk2 |
| Eipr1 | 9.08E-76 | 0.49871921 | 0.57 | 0.363 | 1.76E-71 | cDC2.Mgl2 | Eipr1 |
| Tob21 | 9.91E-76 | 0.43661878 | 0.666 | 0.478 | 1.92E-71 | cDC2.Mgl2 | Tob2 |
| Pip4k2a | 3.64E-75 | 0.25903783 | 0.769 | 0.611 | 7.05E-71 | cDC2.Mgl2 | Pip4k2a |
| Dhx401 | 5.13E-75 | 0.72006622 | 0.343 | 0.219 | 9.93E-71 | cDC2.Mgl2 | Dhx40 |
| Anp32e1 | 6.00E-75 | 0.47789017 | 0.756 | 0.544 | 1.16E-70 | cDC2.Mgl2 | Anp32e |
| Drap1 | 7.43E-75 | 0.32988356 | 0.746 | 0.583 | 1.44E-70 | cDC2.Mgl2 | Drap1 |
| Dnaja11 | 1.18E-74 | 0.5336895 | 0.586 | 0.401 | 2.28E-70 | cDC2.Mgl2 | Dnaja1 |
| Prkar2a | 1.37E-74 | 0.68245445 | 0.582 | 0.332 | 2.65E-70 | cDC2.Mgl2 | Prkar2a |
| Srgap3 | 2.03E-74 | 1.10148858 | 0.399 | 0.154 | 3.92E-70 | cDC2.Mgl2 | Srgap3 |
| Nme11 | 2.42E-74 | 0.97170357 | 0.481 | 0.247 | 4.68E-70 | cDC2.Mgl2 | Nme1 |
| Cox5a | 3.07E-74 | 0.47063423 | 0.724 | 0.547 | 5.94E-70 | cDC2.Mgl2 | Cox5a |
| Cd300lf1 | 3.69E-74 | 0.6382197 | 0.597 | 0.334 | 7.13E-70 | cDC2.Mgl2 | Cd300lf |
| Creb51 | 5.55E-74 | 0.910931 | 0.479 | 0.222 | 1.07E-69 | cDC2.Mgl2 | Creb5 |
| Epha21 | 6.61E-74 | 0.46630295 | 0.386 | 0.231 | 1.28E-69 | cDC2.Mgl2 | Epha2 |
| Eif4b | 7.23E-74 | 0.25533261 | 0.744 | 0.589 | 1.40E-69 | cDC2.Mgl2 | Eif4b |
| Gpr171 | 8.26E-74 | 0.27924965 | 0.714 | 0.45 | 1.60E-69 | cDC2.Mgl2 | Gpr171 |
| Cltb1 | 3.25E-73 | 0.66807912 | 0.614 | 0.389 | 6.28E-69 | cDC2.Mgl2 | Cltb |
| Mvk | 1.86E-72 | 0.9123352 | 0.353 | 0.175 | 3.60E-68 | cDC2.Mgl2 | Mvk |
| Lin54 | 7.34E-72 | 0.97791155 | 0.371 | 0.174 | 1.42E-67 | cDC2.Mgl2 | Lin54 |
| Gpr141 | 9.62E-72 | 1.29001024 | 0.269 | 0.087 | 1.86E-67 | cDC2.Mgl2 | Gpr141 |
| Pnpla7 | 1.23E-71 | 1.06784474 | 0.413 | 0.186 | 2.38E-67 | cDC2.Mgl2 | Pnpla7 |
| Dapp11 | 1.43E-71 | 0.28106869 | 0.698 | 0.53 | 2.76E-67 | cDC2.Mgl2 | Dapp1 |
| Slc1a51 | 1.79E-71 | 0.49994766 | 0.614 | 0.399 | 3.47E-67 | cDC2.Mgl2 | Slc1a5 |
| Pold1 | 1.89E-71 | 0.79514387 | 0.273 | 0.156 | 3.65E-67 | cDC2.Mgl2 | Pold1 |
| Arih2 | 5.73E-71 | 0.4259284 | 0.692 | 0.495 | 1.11E-66 | cDC2.Mgl2 | Arih2 |
| Hivep21 | 1.03E-70 | 0.56925559 | 0.562 | 0.339 | 1.99E-66 | cDC2.Mgl2 | Hivep2 |
| Sipa1 | 2.29E-70 | 0.68065493 | 0.587 | 0.348 | 4.43E-66 | cDC2.Mgl2 | Sipa1 |
| Zc3hav11 | 1.39E-69 | 0.31369427 | 0.622 | 0.469 | 2.68E-65 | cDC2.Mgl2 | Zc3hav1 |
| Cox8a | 3.00E-69 | 0.39806314 | 0.805 | 0.635 | 5.81E-65 | cDC2.Mgl2 | Cox8a |
| Slc39a1 | 3.07E-69 | 0.34364224 | 0.783 | 0.629 | 5.93E-65 | cDC2.Mgl2 | Slc39a1 |
| Fundc1 | 4.18E-69 | 0.65407298 | 0.639 | 0.421 | 8.08E-65 | cDC2.Mgl2 | Fundc1 |
| Acsl1 | 1.39E-68 | 0.94242047 | 0.378 | 0.165 | 2.68E-64 | cDC2.Mgl2 | Acsl1 |
| Nsmaf | 1.42E-68 | 0.59079319 | 0.586 | 0.351 | 2.74E-64 | cDC2.Mgl2 | Nsmaf |
| Apbb1ip | 2.85E-68 | 0.29232359 | 0.787 | 0.637 | 5.51E-64 | cDC2.Mgl2 | Apbb1ip |
| Galnt61 | 3.31E-68 | 0.98945784 | 0.423 | 0.196 | 6.41E-64 | cDC2.Mgl2 | Galnt6 |
| Stmn1 | 3.37E-68 | 1.19906012 | 0.263 | 0.113 | 6.52E-64 | cDC2.Mgl2 | Stmn1 |
| Tmem185b1 | 3.49E-68 | 0.71589615 | 0.481 | 0.267 | 6.74E-64 | cDC2.Mgl2 | Tmem185b |
| Hs6st11 | 8.58E-68 | 0.39350899 | 0.566 | 0.393 | 1.66E-63 | cDC2.Mgl2 | Hs6st1 |
| Cbfb | 8.98E-68 | 0.25268924 | 0.808 | 0.653 | 1.74E-63 | cDC2.Mgl2 | Cbfb |
| Ntpcr | 1.02E-67 | 1.19835926 | 0.362 | 0.166 | 1.97E-63 | cDC2.Mgl2 | Ntpcr |
| Btbd1 | 4.53E-67 | 1.12156348 | 0.357 | 0.168 | 8.75E-63 | cDC2.Mgl2 | Btbd1 |
| Dot1l | 4.91E-67 | 0.58057921 | 0.567 | 0.367 | 9.49E-63 | cDC2.Mgl2 | Dot1l |
| Fam162a | 5.54E-67 | 0.85728963 | 0.427 | 0.242 | 1.07E-62 | cDC2.Mgl2 | Fam162a |
| Cass4 | 8.71E-67 | 0.70851187 | 0.472 | 0.268 | 1.68E-62 | cDC2.Mgl2 | Cass4 |
| Nqo2 | 1.45E-66 | 1.31622751 | 0.303 | 0.113 | 2.80E-62 | cDC2.Mgl2 | Nqo2 |
| Ltc4s | 3.15E-66 | 0.89740926 | 0.324 | 0.122 | 6.10E-62 | cDC2.Mgl2 | Ltc4s |
| Gpat4 | 3.47E-66 | 0.54778067 | 0.623 | 0.406 | 6.71E-62 | cDC2.Mgl2 | Gpat4 |
| Ptprj | 3.49E-66 | 0.28055595 | 0.725 | 0.563 | 6.75E-62 | cDC2.Mgl2 | Ptprj |
| Oat | 5.31E-66 | 0.72647114 | 0.53 | 0.315 | 1.03E-61 | cDC2.Mgl2 | Oat |
| Tcp11l2 | 5.53E-66 | 1.16112523 | 0.314 | 0.15 | 1.07E-61 | cDC2.Mgl2 | Tcp11l2 |
| Ppm1g1 | 1.12E-65 | 0.43252727 | 0.628 | 0.448 | 2.17E-61 | cDC2.Mgl2 | Ppm1g |
| Trappc6a | 1.68E-65 | 0.8857931 | 0.49 | 0.268 | 3.24E-61 | cDC2.Mgl2 | Trappc6a |
| Slc35c2 | 2.94E-65 | 0.59404149 | 0.585 | 0.327 | 5.68E-61 | cDC2.Mgl2 | Slc35c2 |
| Pdcl3 | 3.69E-65 | 0.60326451 | 0.588 | 0.382 | 7.13E-61 | cDC2.Mgl2 | Pdcl3 |
| Bloc1s2 | 6.08E-65 | 0.30797562 | 0.721 | 0.539 | 1.17E-60 | cDC2.Mgl2 | Bloc1s2 |
| Rhoh1 | 1.14E-64 | 0.96413329 | 0.409 | 0.195 | 2.21E-60 | cDC2.Mgl2 | Rhoh |
| Gabarapl2 | 1.45E-64 | 0.53729287 | 0.64 | 0.428 | 2.80E-60 | cDC2.Mgl2 | Gabarapl2 |
| Impdh21 | 1.62E-64 | 0.58436305 | 0.526 | 0.36 | 3.14E-60 | cDC2.Mgl2 | Impdh2 |
| Lsm4 | 3.38E-64 | 0.4815482 | 0.644 | 0.433 | 6.54E-60 | cDC2.Mgl2 | Lsm4 |
| Adgre5 | 3.42E-64 | 0.33431187 | 0.671 | 0.453 | 6.62E-60 | cDC2.Mgl2 | Adgre5 |
| Tspan2 | 3.66E-64 | 1.15263272 | 0.305 | 0.122 | 7.08E-60 | cDC2.Mgl2 | Tspan2 |
| Hprt | 4.51E-64 | 0.50105519 | 0.579 | 0.388 | 8.72E-60 | cDC2.Mgl2 | Hprt |
| Pank21 | 1.28E-63 | 0.41538218 | 0.561 | 0.365 | 2.48E-59 | cDC2.Mgl2 | Pank2 |
| Arhgap15 | 1.40E-63 | 0.93862799 | 0.388 | 0.18 | 2.70E-59 | cDC2.Mgl2 | Arhgap15 |
| Tgfbr2 | 1.59E-63 | 0.70805653 | 0.553 | 0.294 | 3.07E-59 | cDC2.Mgl2 | Tgfbr2 |
| Dgat1 | 1.64E-63 | 0.80525988 | 0.379 | 0.2 | 3.18E-59 | cDC2.Mgl2 | Dgat1 |
| Dad11 | 1.88E-63 | 0.26270674 | 0.758 | 0.613 | 3.63E-59 | cDC2.Mgl2 | Dad1 |
| 7-Mar | 2.50E-63 | 0.32000429 | 0.683 | 0.516 | 4.84E-59 | cDC2.Mgl2 | 7-Mar |
| Trim16 | 3.45E-63 | 1.02850325 | 0.29 | 0.119 | 6.67E-59 | cDC2.Mgl2 | Trim16 |
| Ankrd27 | 5.81E-63 | 0.67628258 | 0.461 | 0.256 | 1.12E-58 | cDC2.Mgl2 | Ankrd27 |
| Ccnd11 | 7.46E-63 | 0.27252599 | 0.546 | 0.376 | 1.44E-58 | cDC2.Mgl2 | Ccnd1 |
| Atp5d | 8.57E-63 | 0.27587677 | 0.799 | 0.666 | 1.66E-58 | cDC2.Mgl2 | Atp5d |
| Flot11 | 1.53E-62 | 0.47833965 | 0.616 | 0.427 | 2.95E-58 | cDC2.Mgl2 | Flot1 |
| Cdkn2aipnl | 1.85E-62 | 1.03494339 | 0.33 | 0.142 | 3.58E-58 | cDC2.Mgl2 | Cdkn2aipnl |
| Hspa51 | 2.39E-62 | 0.36295915 | 0.738 | 0.56 | 4.61E-58 | cDC2.Mgl2 | Hspa5 |
| Nfatc21 | 6.58E-62 | 0.51019075 | 0.602 | 0.37 | 1.27E-57 | cDC2.Mgl2 | Nfatc2 |
| Prcp1 | 7.47E-62 | 0.27206782 | 0.705 | 0.563 | 1.44E-57 | cDC2.Mgl2 | Prcp |
| Rhof1 | 8.13E-62 | 0.40552287 | 0.425 | 0.241 | 1.57E-57 | cDC2.Mgl2 | Rhof |
| Sae1 | 8.73E-62 | 0.53108265 | 0.474 | 0.281 | 1.69E-57 | cDC2.Mgl2 | Sae1 |
| Pqlc1 | 1.16E-61 | 0.75776253 | 0.524 | 0.285 | 2.25E-57 | cDC2.Mgl2 | Pqlc1 |
| Srsf1 | 1.52E-61 | 0.31024268 | 0.675 | 0.5 | 2.94E-57 | cDC2.Mgl2 | Srsf1 |
| Ranbp1 | 1.73E-61 | 0.41215236 | 0.557 | 0.416 | 3.34E-57 | cDC2.Mgl2 | Ranbp1 |
| Cnbp | 1.76E-61 | 0.31166642 | 0.724 | 0.552 | 3.41E-57 | cDC2.Mgl2 | Cnbp |
| Comt1 | 2.59E-61 | 0.84051621 | 0.489 | 0.262 | 5.01E-57 | cDC2.Mgl2 | Comt |
| Timm10b1 | 8.05E-61 | 0.63424399 | 0.553 | 0.342 | 1.56E-56 | cDC2.Mgl2 | Timm10b |
| Bri3bp | 1.07E-60 | 0.42179102 | 0.719 | 0.502 | 2.06E-56 | cDC2.Mgl2 | Bri3bp |
| Dnmt1 | 1.79E-60 | 0.55882012 | 0.335 | 0.229 | 3.47E-56 | cDC2.Mgl2 | Dnmt1 |
| Zfp1061 | 2.07E-60 | 0.27831714 | 0.683 | 0.528 | 4.00E-56 | cDC2.Mgl2 | Zfp106 |
| Glrx | 2.12E-60 | 0.69347368 | 0.469 | 0.261 | 4.10E-56 | cDC2.Mgl2 | Glrx |
| Hepacam2 | 2.46E-60 | 0.83761162 | 0.478 | 0.218 | 4.76E-56 | cDC2.Mgl2 | Hepacam2 |
| Nrm | 2.95E-60 | 1.12876744 | 0.319 | 0.143 | 5.70E-56 | cDC2.Mgl2 | Nrm |
| Mbnl1 | 4.71E-60 | 0.38812333 | 0.694 | 0.48 | 9.11E-56 | cDC2.Mgl2 | Mbnl1 |
| Igsf8 | 7.22E-60 | 0.33327599 | 0.592 | 0.401 | 1.40E-55 | cDC2.Mgl2 | Igsf8 |
| Ppp1r18 | 1.03E-59 | 0.26253046 | 0.74 | 0.561 | 1.99E-55 | cDC2.Mgl2 | Ppp1r18 |
| Ndufv3 | 1.11E-59 | 0.48586977 | 0.674 | 0.485 | 2.15E-55 | cDC2.Mgl2 | Ndufv3 |
| Egln1 | 1.15E-59 | 0.53919574 | 0.594 | 0.388 | 2.23E-55 | cDC2.Mgl2 | Egln1 |
| Rinl | 1.46E-59 | 0.87858684 | 0.449 | 0.225 | 2.82E-55 | cDC2.Mgl2 | Rinl |
| Smim31 | 1.93E-59 | 0.63970149 | 0.508 | 0.331 | 3.74E-55 | cDC2.Mgl2 | Smim3 |
| Ppp3r1 | 2.12E-59 | 0.46727604 | 0.648 | 0.466 | 4.09E-55 | cDC2.Mgl2 | Ppp3r1 |
| Gng2 | 7.91E-59 | 0.59506011 | 0.55 | 0.311 | 1.53E-54 | cDC2.Mgl2 | Gng2 |
| Ets2 | 8.46E-59 | 0.4059119 | 0.659 | 0.43 | 1.64E-54 | cDC2.Mgl2 | Ets2 |
| Crlf21 | 1.51E-58 | 0.58710495 | 0.648 | 0.408 | 2.91E-54 | cDC2.Mgl2 | Crlf2 |
| Bzw21 | 2.84E-58 | 0.72583023 | 0.456 | 0.249 | 5.49E-54 | cDC2.Mgl2 | Bzw2 |
| Kpna4 | 3.41E-58 | 0.33309865 | 0.693 | 0.539 | 6.60E-54 | cDC2.Mgl2 | Kpna4 |
| Smarca2 | 3.75E-57 | 0.75757859 | 0.397 | 0.205 | 7.24E-53 | cDC2.Mgl2 | Smarca2 |
| Pgm1 | 4.53E-57 | 0.98872843 | 0.355 | 0.173 | 8.76E-53 | cDC2.Mgl2 | Pgm1 |
| Camk2g1 | 8.56E-57 | 0.61932263 | 0.493 | 0.297 | 1.65E-52 | cDC2.Mgl2 | Camk2g |
| Clic11 | 8.91E-57 | 0.3075042 | 0.772 | 0.607 | 1.72E-52 | cDC2.Mgl2 | Clic1 |
| Trio | 1.11E-56 | 0.30420215 | 0.62 | 0.419 | 2.15E-52 | cDC2.Mgl2 | Trio |
| Snrpf1 | 1.75E-56 | 0.34223951 | 0.679 | 0.511 | 3.38E-52 | cDC2.Mgl2 | Snrpf |
| Ifi27 | 1.98E-56 | 0.90359625 | 0.333 | 0.154 | 3.83E-52 | cDC2.Mgl2 | Ifi27 |
| Tfdp1 | 7.21E-56 | 0.55421188 | 0.409 | 0.264 | 1.39E-51 | cDC2.Mgl2 | Tfdp1 |
| Ptpn4 | 9.98E-56 | 0.4403457 | 0.495 | 0.307 | 1.93E-51 | cDC2.Mgl2 | Ptpn4 |
| Abr1 | 1.18E-55 | 0.40713936 | 0.693 | 0.489 | 2.27E-51 | cDC2.Mgl2 | Abr |
| Cyb5r31 | 2.49E-55 | 0.28091834 | 0.671 | 0.474 | 4.81E-51 | cDC2.Mgl2 | Cyb5r3 |
| Unc119 | 2.73E-55 | 0.89843532 | 0.435 | 0.199 | 5.28E-51 | cDC2.Mgl2 | Unc119 |
| Ralb | 2.76E-55 | 0.50412312 | 0.589 | 0.404 | 5.33E-51 | cDC2.Mgl2 | Ralb |
| Vmp1 | 3.10E-55 | 0.30151654 | 0.709 | 0.532 | 5.98E-51 | cDC2.Mgl2 | Vmp1 |
| Atf61 | 3.35E-55 | 0.52816233 | 0.557 | 0.364 | 6.47E-51 | cDC2.Mgl2 | Atf6 |
| Stt3b | 4.94E-55 | 0.35854749 | 0.599 | 0.402 | 9.55E-51 | cDC2.Mgl2 | Stt3b |
| Mtpn | 7.61E-55 | 0.39396286 | 0.695 | 0.52 | 1.47E-50 | cDC2.Mgl2 | Mtpn |
| Man1a | 8.19E-55 | 0.48052916 | 0.656 | 0.452 | 1.58E-50 | cDC2.Mgl2 | Man1a |
| Gnaq | 9.06E-55 | 0.50190594 | 0.613 | 0.399 | 1.75E-50 | cDC2.Mgl2 | Gnaq |
| Mapk141 | 1.12E-54 | 0.27494459 | 0.655 | 0.479 | 2.16E-50 | cDC2.Mgl2 | Mapk14 |
| Nfam11 | 1.44E-54 | 0.51890042 | 0.514 | 0.3 | 2.78E-50 | cDC2.Mgl2 | Nfam1 |
| Nop56 | 1.45E-54 | 0.6537648 | 0.441 | 0.257 | 2.81E-50 | cDC2.Mgl2 | Nop56 |
| Mfge8 | 1.68E-54 | 0.69366638 | 0.586 | 0.334 | 3.24E-50 | cDC2.Mgl2 | Mfge8 |
| Mri11 | 1.99E-54 | 0.82076468 | 0.37 | 0.189 | 3.85E-50 | cDC2.Mgl2 | Mri1 |
| Pxdc11 | 2.24E-54 | 0.35236862 | 0.494 | 0.332 | 4.32E-50 | cDC2.Mgl2 | Pxdc1 |
| Tmed51 | 2.32E-54 | 0.5627119 | 0.558 | 0.361 | 4.49E-50 | cDC2.Mgl2 | Tmed5 |
| Myl12b | 3.10E-54 | 0.27852101 | 0.755 | 0.609 | 6.00E-50 | cDC2.Mgl2 | Myl12b |
| Pdlim7 | 3.15E-54 | 0.59302259 | 0.453 | 0.254 | 6.09E-50 | cDC2.Mgl2 | Pdlim7 |
| Dhcr241 | 4.04E-54 | 0.561972 | 0.425 | 0.243 | 7.82E-50 | cDC2.Mgl2 | Dhcr24 |
| Rnf1661 | 1.08E-53 | 0.57721843 | 0.518 | 0.315 | 2.10E-49 | cDC2.Mgl2 | Rnf166 |
| Kpna1 | 2.06E-53 | 0.54076845 | 0.487 | 0.328 | 3.98E-49 | cDC2.Mgl2 | Kpna1 |
| Fbxo341 | 2.22E-53 | 0.99712276 | 0.361 | 0.187 | 4.29E-49 | cDC2.Mgl2 | Fbxo34 |
| Zfp6091 | 2.29E-53 | 0.576305 | 0.461 | 0.256 | 4.42E-49 | cDC2.Mgl2 | Zfp609 |
| Atp6ap21 | 3.55E-53 | 0.3237503 | 0.756 | 0.573 | 6.85E-49 | cDC2.Mgl2 | Atp6ap2 |
| Gart1 | 4.21E-53 | 0.87881977 | 0.387 | 0.215 | 8.14E-49 | cDC2.Mgl2 | Gart |
| Baiap2 | 4.26E-53 | 0.72851507 | 0.432 | 0.225 | 8.23E-49 | cDC2.Mgl2 | Baiap2 |
| Nhp2 | 4.62E-53 | 0.52408518 | 0.551 | 0.361 | 8.92E-49 | cDC2.Mgl2 | Nhp2 |
| Csgalnact2 | 5.59E-53 | 0.68064964 | 0.453 | 0.259 | 1.08E-48 | cDC2.Mgl2 | Csgalnact2 |
| Mbnl2 | 9.26E-53 | 0.43633088 | 0.627 | 0.45 | 1.79E-48 | cDC2.Mgl2 | Mbnl2 |
| Pgs11 | 1.70E-52 | 0.45253958 | 0.659 | 0.467 | 3.29E-48 | cDC2.Mgl2 | Pgs1 |
| Traf3ip3 | 6.08E-52 | 0.85732095 | 0.381 | 0.187 | 1.18E-47 | cDC2.Mgl2 | Traf3ip3 |
| Med30 | 8.71E-52 | 0.56697931 | 0.46 | 0.291 | 1.68E-47 | cDC2.Mgl2 | Med30 |
| Topbp1 | 9.64E-52 | 0.71624199 | 0.253 | 0.137 | 1.86E-47 | cDC2.Mgl2 | Topbp1 |
| Mgat21 | 9.84E-52 | 0.48458736 | 0.555 | 0.37 | 1.90E-47 | cDC2.Mgl2 | Mgat2 |
| Lair1 | 1.26E-51 | 0.32591239 | 0.479 | 0.21 | 2.44E-47 | cDC2.Mgl2 | Lair1 |
| Taok3 | 1.57E-51 | 0.29299661 | 0.655 | 0.501 | 3.04E-47 | cDC2.Mgl2 | Taok3 |
| Rapgef1 | 2.01E-51 | 0.38096055 | 0.55 | 0.368 | 3.88E-47 | cDC2.Mgl2 | Rapgef1 |
| Msh2 | 3.49E-51 | 1.1142089 | 0.265 | 0.117 | 6.76E-47 | cDC2.Mgl2 | Msh2 |
| Hnrnpab | 8.09E-51 | 0.39626733 | 0.57 | 0.409 | 1.56E-46 | cDC2.Mgl2 | Hnrnpab |
| Daglb | 1.03E-50 | 0.93682699 | 0.394 | 0.19 | 1.99E-46 | cDC2.Mgl2 | Daglb |
| Tmem170b | 1.98E-50 | 0.55364138 | 0.456 | 0.271 | 3.84E-46 | cDC2.Mgl2 | Tmem170b |
| Elmsan11 | 2.47E-50 | 0.43184729 | 0.514 | 0.335 | 4.77E-46 | cDC2.Mgl2 | Elmsan1 |
| Dusp16 | 3.80E-50 | 0.38077249 | 0.549 | 0.371 | 7.35E-46 | cDC2.Mgl2 | Dusp16 |
| Spred2 | 3.94E-50 | 0.74446996 | 0.355 | 0.186 | 7.63E-46 | cDC2.Mgl2 | Spred2 |
| Afdn1 | 6.49E-50 | 0.48061633 | 0.513 | 0.328 | 1.25E-45 | cDC2.Mgl2 | Afdn |
| Cybb1 | 6.96E-50 | 0.26399003 | 0.616 | 0.291 | 1.35E-45 | cDC2.Mgl2 | Cybb |
| Evl | 1.18E-49 | 0.59278548 | 0.577 | 0.317 | 2.27E-45 | cDC2.Mgl2 | Evl |
| Vps18 | 1.66E-49 | 0.61626184 | 0.466 | 0.296 | 3.20E-45 | cDC2.Mgl2 | Vps18 |
| Ero1l | 1.80E-49 | 0.70161422 | 0.479 | 0.3 | 3.49E-45 | cDC2.Mgl2 | Ero1l |
| Asah1 | 2.06E-49 | 0.35714429 | 0.644 | 0.494 | 3.98E-45 | cDC2.Mgl2 | Asah1 |
| Hsd3b71 | 2.71E-49 | 0.69848131 | 0.396 | 0.239 | 5.24E-45 | cDC2.Mgl2 | Hsd3b7 |
| Rasal3 | 2.97E-49 | 0.73578909 | 0.343 | 0.159 | 5.74E-45 | cDC2.Mgl2 | Rasal3 |
| Ssh21 | 3.05E-49 | 0.51245499 | 0.468 | 0.269 | 5.90E-45 | cDC2.Mgl2 | Ssh2 |
| Tmem881 | 5.63E-49 | 0.74718387 | 0.387 | 0.222 | 1.09E-44 | cDC2.Mgl2 | Tmem88 |
| Tbrg41 | 7.19E-49 | 0.62234611 | 0.474 | 0.277 | 1.39E-44 | cDC2.Mgl2 | Tbrg4 |
| Cdc42se2 | 7.32E-49 | 0.48576634 | 0.59 | 0.404 | 1.41E-44 | cDC2.Mgl2 | Cdc42se2 |
| Nod21 | 8.12E-49 | 0.3222716 | 0.511 | 0.358 | 1.57E-44 | cDC2.Mgl2 | Nod2 |
| Ubr4 | 1.09E-48 | 0.39314616 | 0.616 | 0.451 | 2.10E-44 | cDC2.Mgl2 | Ubr4 |
| Snrpd31 | 1.17E-48 | 0.40652014 | 0.656 | 0.471 | 2.27E-44 | cDC2.Mgl2 | Snrpd3 |
| Set | 1.19E-48 | 0.54647435 | 0.515 | 0.352 | 2.30E-44 | cDC2.Mgl2 | Set |
| Wfdc171 | 2.86E-48 | 0.32642985 | 0.588 | 0.337 | 5.52E-44 | cDC2.Mgl2 | Wfdc17 |
| Cope1 | 7.58E-48 | 0.27176228 | 0.714 | 0.55 | 1.47E-43 | cDC2.Mgl2 | Cope |
| Itpr1 | 8.44E-48 | 0.40883496 | 0.516 | 0.314 | 1.63E-43 | cDC2.Mgl2 | Itpr1 |
| Atp8a1 | 8.57E-48 | 0.4219146 | 0.592 | 0.358 | 1.66E-43 | cDC2.Mgl2 | Atp8a1 |
| P4ha1 | 1.01E-47 | 0.7475081 | 0.435 | 0.252 | 1.95E-43 | cDC2.Mgl2 | P4ha1 |
| Rassf11 | 1.03E-47 | 0.27439368 | 0.549 | 0.395 | 1.98E-43 | cDC2.Mgl2 | Rassf1 |
| Parp11 | 1.58E-47 | 0.51828076 | 0.49 | 0.325 | 3.06E-43 | cDC2.Mgl2 | Parp1 |
| Limk1 | 2.86E-47 | 0.44934421 | 0.459 | 0.279 | 5.52E-43 | cDC2.Mgl2 | Limk1 |
| Pgd | 2.90E-47 | 0.5600893 | 0.642 | 0.409 | 5.60E-43 | cDC2.Mgl2 | Pgd |
| Map4k1 | 5.16E-47 | 0.39612071 | 0.646 | 0.418 | 9.97E-43 | cDC2.Mgl2 | Map4k1 |
| Dip2c | 5.78E-47 | 1.07426169 | 0.307 | 0.137 | 1.12E-42 | cDC2.Mgl2 | Dip2c |
| Stim2 | 6.01E-47 | 0.87400227 | 0.345 | 0.182 | 1.16E-42 | cDC2.Mgl2 | Stim2 |
| Smim14 | 7.84E-47 | 0.36171542 | 0.569 | 0.398 | 1.52E-42 | cDC2.Mgl2 | Smim14 |
| G3bp1 | 9.08E-47 | 0.52930434 | 0.552 | 0.365 | 1.76E-42 | cDC2.Mgl2 | G3bp1 |
| Rasgrp4 | 1.01E-46 | 0.49013283 | 0.627 | 0.398 | 1.96E-42 | cDC2.Mgl2 | Rasgrp4 |
| Atf11 | 1.13E-46 | 0.463814 | 0.612 | 0.423 | 2.18E-42 | cDC2.Mgl2 | Atf1 |
| Snx3 | 1.15E-46 | 0.26008473 | 0.84 | 0.678 | 2.23E-42 | cDC2.Mgl2 | Snx3 |
| Nup214 | 1.47E-46 | 0.63389725 | 0.434 | 0.253 | 2.83E-42 | cDC2.Mgl2 | Nup214 |
| Tmem245 | 1.70E-46 | 0.69284054 | 0.351 | 0.171 | 3.29E-42 | cDC2.Mgl2 | Tmem245 |
| Ptpre1 | 1.91E-46 | 0.406817 | 0.562 | 0.346 | 3.69E-42 | cDC2.Mgl2 | Ptpre |
| Eif2ak4 | 1.99E-46 | 0.67211654 | 0.418 | 0.237 | 3.85E-42 | cDC2.Mgl2 | Eif2ak4 |
| Cptm1l | 2.31E-46 | 0.36086448 | 0.633 | 0.489 | 4.47E-42 | cDC2.Mgl2 | Cptm1l |
| Rtn3 | 2.36E-46 | 0.39706044 | 0.655 | 0.51 | 4.55E-42 | cDC2.Mgl2 | Rtn3 |
| Tpp1 | 2.60E-46 | 0.69356854 | 0.456 | 0.249 | 5.02E-42 | cDC2.Mgl2 | Tpp1 |
| 1-Sep | 2.76E-46 | 0.67741825 | 0.315 | 0.128 | 5.34E-42 | cDC2.Mgl2 | 1-Sep |
| Akr1b31 | 3.82E-46 | 0.65215193 | 0.483 | 0.306 | 7.39E-42 | cDC2.Mgl2 | Akr1b3 |
| Nup1531 | 4.61E-46 | 0.32884331 | 0.588 | 0.434 | 8.91E-42 | cDC2.Mgl2 | Nup153 |
| Hexim1 | 6.19E-46 | 0.3815091 | 0.575 | 0.389 | 1.20E-41 | cDC2.Mgl2 | Hexim1 |
| Rpn21 | 6.59E-46 | 0.49588821 | 0.563 | 0.378 | 1.27E-41 | cDC2.Mgl2 | Rpn2 |
| Fam53c | 6.82E-46 | 0.87792991 | 0.356 | 0.187 | 1.32E-41 | cDC2.Mgl2 | Fam53c |
| Rnaseh2c | 1.05E-45 | 0.81671264 | 0.361 | 0.193 | 2.03E-41 | cDC2.Mgl2 | Rnaseh2c |
| Nfatc11 | 1.28E-45 | 0.54928615 | 0.498 | 0.299 | 2.47E-41 | cDC2.Mgl2 | Nfatc1 |
| Ptpn7 | 1.79E-45 | 1.20861131 | 0.26 | 0.109 | 3.47E-41 | cDC2.Mgl2 | Ptpn7 |
| Tfeb | 2.32E-45 | 0.69889387 | 0.353 | 0.185 | 4.48E-41 | cDC2.Mgl2 | Tfeb |
| Prkd3 | 2.38E-45 | 0.56064753 | 0.469 | 0.286 | 4.60E-41 | cDC2.Mgl2 | Prkd3 |
| Fam3c1 | 2.87E-45 | 0.80576393 | 0.411 | 0.212 | 5.55E-41 | cDC2.Mgl2 | Fam3c |
| Ubtd11 | 6.56E-45 | 0.7395894 | 0.428 | 0.226 | 1.27E-40 | cDC2.Mgl2 | Ubtd1 |
| Golga7 | 7.40E-45 | 0.31153405 | 0.651 | 0.476 | 1.43E-40 | cDC2.Mgl2 | Golga7 |
| Ube2e1 | 1.02E-44 | 0.55906988 | 0.45 | 0.28 | 1.96E-40 | cDC2.Mgl2 | Ube2e1 |
| Prpf19 | 1.76E-44 | 0.51692208 | 0.516 | 0.333 | 3.40E-40 | cDC2.Mgl2 | Prpf19 |
| Mybbp1a1 | 2.25E-44 | 0.45383879 | 0.521 | 0.338 | 4.34E-40 | cDC2.Mgl2 | Mybbp1a |
| Cd84 | 2.72E-44 | 1.34011196 | 0.329 | 0.124 | 5.25E-40 | cDC2.Mgl2 | Cd84 |
| Cryl1 | 4.51E-44 | 1.17973853 | 0.278 | 0.122 | 8.73E-40 | cDC2.Mgl2 | Cryl1 |
| Otub1 | 6.21E-44 | 0.28422203 | 0.653 | 0.521 | 1.20E-39 | cDC2.Mgl2 | Otub1 |
| Fcgr31 | 1.06E-43 | 0.60806563 | 0.611 | 0.329 | 2.05E-39 | cDC2.Mgl2 | Fcgr3 |
| Fdft1 | 1.15E-43 | 0.41876344 | 0.37 | 0.234 | 2.21E-39 | cDC2.Mgl2 | Fdft1 |
| Msmo1 | 1.96E-43 | 0.56585621 | 0.382 | 0.227 | 3.79E-39 | cDC2.Mgl2 | Msmo1 |
| Inpp5b | 2.74E-43 | 0.31289856 | 0.489 | 0.323 | 5.29E-39 | cDC2.Mgl2 | Inpp5b |
| Dock5 | 3.39E-43 | 0.46277314 | 0.441 | 0.263 | 6.55E-39 | cDC2.Mgl2 | Dock5 |
| Ggh | 3.66E-43 | 0.76731453 | 0.324 | 0.163 | 7.08E-39 | cDC2.Mgl2 | Ggh |
| Ap1s1 | 4.79E-43 | 0.39894346 | 0.511 | 0.35 | 9.25E-39 | cDC2.Mgl2 | Ap1s1 |
| Nop101 | 4.90E-43 | 0.53401604 | 0.475 | 0.295 | 9.47E-39 | cDC2.Mgl2 | Nop10 |
| Ssr41 | 5.18E-43 | 0.34182061 | 0.682 | 0.496 | 1.00E-38 | cDC2.Mgl2 | Ssr4 |
| Rnf131 | 5.45E-43 | 0.50687914 | 0.497 | 0.314 | 1.05E-38 | cDC2.Mgl2 | Rnf13 |
| Tspoap1 | 5.51E-43 | 0.52348484 | 0.26 | 0.113 | 1.07E-38 | cDC2.Mgl2 | Tspoap1 |
| Zdhhc14 | 5.66E-43 | 0.80445403 | 0.293 | 0.127 | 1.09E-38 | cDC2.Mgl2 | Zdhhc14 |
| Slc12a61 | 5.83E-43 | 0.71737121 | 0.371 | 0.197 | 1.13E-38 | cDC2.Mgl2 | Slc12a6 |
| Ndufb81 | 7.28E-43 | 0.39271956 | 0.671 | 0.512 | 1.41E-38 | cDC2.Mgl2 | Ndufb8 |
| Ptpa | 7.78E-43 | 0.47728948 | 0.565 | 0.385 | 1.50E-38 | cDC2.Mgl2 | Ptpa |
| Rap2a | 8.22E-43 | 0.28886965 | 0.669 | 0.504 | 1.59E-38 | cDC2.Mgl2 | Rap2a |
| Slc35e11 | 8.72E-43 | 0.42996426 | 0.572 | 0.391 | 1.69E-38 | cDC2.Mgl2 | Slc35e1 |
| Mpp6 | 1.21E-42 | 0.43477164 | 0.51 | 0.339 | 2.34E-38 | cDC2.Mgl2 | Mpp6 |
| Noc2l1 | 1.46E-42 | 0.37105526 | 0.542 | 0.365 | 2.82E-38 | cDC2.Mgl2 | Noc2l |
| Snrpb | 1.58E-42 | 0.37149436 | 0.544 | 0.393 | 3.05E-38 | cDC2.Mgl2 | Snrpb |
| Ahsa1 | 1.78E-42 | 0.28481669 | 0.63 | 0.462 | 3.44E-38 | cDC2.Mgl2 | Ahsa1 |
| Stard4 | 1.80E-42 | 0.56987101 | 0.377 | 0.209 | 3.48E-38 | cDC2.Mgl2 | Stard4 |
| Sptssa | 2.12E-42 | 0.5151336 | 0.482 | 0.294 | 4.10E-38 | cDC2.Mgl2 | Sptssa |
| Lrp5 | 3.19E-42 | 0.57969806 | 0.48 | 0.288 | 6.16E-38 | cDC2.Mgl2 | Lrp5 |
| Srp72 | 3.44E-42 | 0.37654842 | 0.553 | 0.41 | 6.64E-38 | cDC2.Mgl2 | Srp72 |
| Abcg3 | 3.82E-42 | 1.02632239 | 0.268 | 0.098 | 7.38E-38 | cDC2.Mgl2 | Abcg3 |
| Larp4b | 9.02E-42 | 0.31712214 | 0.604 | 0.424 | 1.74E-37 | cDC2.Mgl2 | Larp4b |
| Plk31 | 9.86E-42 | 0.65760696 | 0.368 | 0.205 | 1.91E-37 | cDC2.Mgl2 | Plk3 |
| Ccdc86 | 1.17E-41 | 0.52826197 | 0.506 | 0.327 | 2.25E-37 | cDC2.Mgl2 | Ccdc86 |
| Ncln1 | 1.20E-41 | 0.46103187 | 0.495 | 0.306 | 2.31E-37 | cDC2.Mgl2 | Ncln |
| Timm13 | 1.82E-41 | 0.28399219 | 0.647 | 0.5 | 3.51E-37 | cDC2.Mgl2 | Timm13 |
| Srm | 2.18E-41 | 0.81577988 | 0.265 | 0.137 | 4.21E-37 | cDC2.Mgl2 | Srm |
| Acat11 | 2.20E-41 | 0.48726508 | 0.457 | 0.278 | 4.25E-37 | cDC2.Mgl2 | Acat1 |
| Knop1 | 2.21E-41 | 0.91204319 | 0.295 | 0.135 | 4.27E-37 | cDC2.Mgl2 | Knop1 |
| Pfkl | 2.30E-41 | 0.39483133 | 0.548 | 0.389 | 4.45E-37 | cDC2.Mgl2 | Pfkl |
| Dnajc14 | 2.48E-41 | 0.36146985 | 0.605 | 0.46 | 4.79E-37 | cDC2.Mgl2 | Dnajc14 |
| Rad23b | 2.82E-41 | 0.3200345 | 0.606 | 0.424 | 5.45E-37 | cDC2.Mgl2 | Rad23b |
| Gtf2b | 2.87E-41 | 0.3521881 | 0.51 | 0.366 | 5.55E-37 | cDC2.Mgl2 | Gtf2b |
| Arl5b1 | 3.09E-41 | 0.41138863 | 0.442 | 0.31 | 5.98E-37 | cDC2.Mgl2 | Arl5b |
| Cd300c21 | 3.29E-41 | 0.90572931 | 0.41 | 0.187 | 6.36E-37 | cDC2.Mgl2 | Cd300c2 |
| Eif3b | 3.34E-41 | 0.40767579 | 0.556 | 0.373 | 6.45E-37 | cDC2.Mgl2 | Eif3b |
| Gch11 | 3.43E-41 | 0.44693423 | 0.471 | 0.307 | 6.62E-37 | cDC2.Mgl2 | Gch1 |
| Fuca21 | 4.09E-41 | 0.50918019 | 0.501 | 0.31 | 7.90E-37 | cDC2.Mgl2 | Fuca2 |
| Elf1 | 4.86E-41 | 0.25477719 | 0.636 | 0.49 | 9.39E-37 | cDC2.Mgl2 | Elf1 |
| Anapc11 | 5.38E-41 | 0.30679049 | 0.546 | 0.411 | 1.04E-36 | cDC2.Mgl2 | Anapc11 |
| Ccdc134 | 6.88E-41 | 0.91916696 | 0.289 | 0.13 | 1.33E-36 | cDC2.Mgl2 | Ccdc134 |
| Rheb1 | 7.17E-41 | 0.27734583 | 0.633 | 0.501 | 1.39E-36 | cDC2.Mgl2 | Rheb |
| Jak3 | 1.30E-40 | 0.38595583 | 0.571 | 0.392 | 2.52E-36 | cDC2.Mgl2 | Jak3 |
| Ski1 | 1.67E-40 | 0.41196833 | 0.642 | 0.432 | 3.24E-36 | cDC2.Mgl2 | Ski |
| Dele1 | 1.97E-40 | 0.82858954 | 0.28 | 0.124 | 3.82E-36 | cDC2.Mgl2 | Dele1 |
| Cdk5rap3 | 2.49E-40 | 0.50466645 | 0.47 | 0.294 | 4.81E-36 | cDC2.Mgl2 | Cdk5rap3 |
| Wdfy3 | 2.77E-40 | 0.58412039 | 0.571 | 0.342 | 5.35E-36 | cDC2.Mgl2 | Wdfy3 |
| Srsf2 | 2.96E-40 | 0.30607176 | 0.621 | 0.48 | 5.73E-36 | cDC2.Mgl2 | Srsf2 |
| Dtymk | 3.95E-40 | 0.41417272 | 0.312 | 0.195 | 7.64E-36 | cDC2.Mgl2 | Dtymk |
| Cldn1 | 9.51E-40 | 0.82097952 | 0.273 | 0.143 | 1.84E-35 | cDC2.Mgl2 | Cldn1 |
| Slc48a1 | 9.97E-40 | 0.70172527 | 0.451 | 0.266 | 1.93E-35 | cDC2.Mgl2 | Slc48a1 |
| Napa | 1.25E-39 | 0.35825132 | 0.614 | 0.437 | 2.43E-35 | cDC2.Mgl2 | Napa |
| Fam217b | 1.42E-39 | 0.91452174 | 0.272 | 0.122 | 2.75E-35 | cDC2.Mgl2 | Fam217b |
| Cct2 | 1.77E-39 | 0.43242936 | 0.644 | 0.46 | 3.42E-35 | cDC2.Mgl2 | Cct2 |
| Prkch1 | 3.62E-39 | 0.530781 | 0.401 | 0.225 | 6.99E-35 | cDC2.Mgl2 | Prkch |
| Ttpal | 3.67E-39 | 1.04215081 | 0.288 | 0.133 | 7.10E-35 | cDC2.Mgl2 | Ttpal |
| Fh11 | 3.95E-39 | 0.32464097 | 0.488 | 0.332 | 7.64E-35 | cDC2.Mgl2 | Fh1 |
| Lpcat2 | 4.19E-39 | 0.6756925 | 0.469 | 0.257 | 8.10E-35 | cDC2.Mgl2 | Lpcat2 |
| Tmem160 | 4.26E-39 | 0.4260505 | 0.537 | 0.381 | 8.24E-35 | cDC2.Mgl2 | Tmem160 |
| Abl21 | 4.45E-39 | 0.51365929 | 0.393 | 0.258 | 8.60E-35 | cDC2.Mgl2 | Abl2 |
| Cct7 | 5.32E-39 | 0.42942539 | 0.495 | 0.351 | 1.03E-34 | cDC2.Mgl2 | Cct7 |
| Naa40 | 1.10E-38 | 0.72099391 | 0.36 | 0.203 | 2.14E-34 | cDC2.Mgl2 | Naa40 |
| Rmnd5b | 1.14E-38 | 0.65919164 | 0.361 | 0.194 | 2.21E-34 | cDC2.Mgl2 | Rmnd5b |
| Idh2 | 1.23E-38 | 0.51867624 | 0.391 | 0.229 | 2.39E-34 | cDC2.Mgl2 | Idh2 |
| Prr7 | 1.34E-38 | 0.60077942 | 0.265 | 0.134 | 2.58E-34 | cDC2.Mgl2 | Prr7 |
| Slbp | 1.36E-38 | 0.32998856 | 0.447 | 0.321 | 2.63E-34 | cDC2.Mgl2 | Slbp |
| Trmt1121 | 1.36E-38 | 0.29415505 | 0.622 | 0.464 | 2.63E-34 | cDC2.Mgl2 | Trmt112 |
| Cebpd1 | 1.49E-38 | 0.58781598 | 0.395 | 0.251 | 2.88E-34 | cDC2.Mgl2 | Cebpd |
| Arrb1 | 1.76E-38 | 0.45591209 | 0.561 | 0.369 | 3.40E-34 | cDC2.Mgl2 | Arrb1 |
| Sec13 | 2.68E-38 | 0.34135257 | 0.588 | 0.414 | 5.18E-34 | cDC2.Mgl2 | Sec13 |
| Clint1 | 3.29E-38 | 0.2982856 | 0.621 | 0.45 | 6.37E-34 | cDC2.Mgl2 | Clint1 |
| Nbeal2 | 5.47E-38 | 0.399018 | 0.503 | 0.319 | 1.06E-33 | cDC2.Mgl2 | Nbeal2 |
| Pdxk | 6.05E-38 | 0.60267738 | 0.369 | 0.206 | 1.17E-33 | cDC2.Mgl2 | Pdxk |
| Dhrs1 | 7.39E-38 | 0.5097484 | 0.467 | 0.298 | 1.43E-33 | cDC2.Mgl2 | Dhrs1 |
| Vps26b | 8.93E-38 | 0.46743343 | 0.456 | 0.295 | 1.73E-33 | cDC2.Mgl2 | Vps26b |
| Unc119b | 1.18E-37 | 0.46710521 | 0.477 | 0.236 | 2.29E-33 | cDC2.Mgl2 | Unc119b |
| Lst1 | 1.74E-37 | 0.61673134 | 0.408 | 0.215 | 3.37E-33 | cDC2.Mgl2 | Lst1 |
| Ptges3 | 1.92E-37 | 0.38822498 | 0.521 | 0.359 | 3.70E-33 | cDC2.Mgl2 | Ptges3 |
| Usp381 | 2.05E-37 | 0.4775504 | 0.412 | 0.265 | 3.97E-33 | cDC2.Mgl2 | Usp38 |
| Cdk2 | 3.26E-37 | 0.36148959 | 0.329 | 0.204 | 6.31E-33 | cDC2.Mgl2 | Cdk2 |
| Lrmp1 | 3.85E-37 | 0.65568667 | 0.369 | 0.199 | 7.44E-33 | cDC2.Mgl2 | Lrmp |
| Eif3i | 9.93E-37 | 0.34964971 | 0.55 | 0.385 | 1.92E-32 | cDC2.Mgl2 | Eif3i |

| | | | | | | | |
|---|---|---|---|---|---|---|---|
| Carhsp1 | 1.13E-36 | 0.65438738 | 0.386 | 0.234 | 2.19E-32 | cDC2.Mgl2 | Carhsp1 |
| Asnsd1 | 1.25E-36 | 0.59455082 | 0.403 | 0.247 | 2.42E-32 | cDC2.Mgl2 | Asnsd1 |
| Pgm21 | 1.29E-36 | 0.65528564 | 0.434 | 0.253 | 2.50E-32 | cDC2.Mgl2 | Pgm2 |
| Rnpep1 | 1.42E-36 | 0.3493208 | 0.601 | 0.428 | 2.75E-32 | cDC2.Mgl2 | Rnpep |
| Pgk1 | 1.46E-36 | 0.32293891 | 0.592 | 0.437 | 2.83E-32 | cDC2.Mgl2 | Pgk1 |
| Ptp4a3 | 1.70E-36 | 0.29562794 | 0.383 | 0.214 | 3.29E-32 | cDC2.Mgl2 | Ptp4a3 |
| Idh3a1 | 1.86E-36 | 0.47626041 | 0.401 | 0.262 | 3.60E-32 | cDC2.Mgl2 | Idh3a |
| Adam9 | 2.56E-36 | 0.32042353 | 0.58 | 0.405 | 4.95E-32 | cDC2.Mgl2 | Adam9 |
| Slc29a1 | 2.60E-36 | 0.7536467 | 0.259 | 0.126 | 5.03E-32 | cDC2.Mgl2 | Slc29a1 |
| Arl6ip51 | 3.06E-36 | 0.26734805 | 0.623 | 0.453 | 5.92E-32 | cDC2.Mgl2 | Arl6ip5 |
| Fam234b | 4.85E-36 | 0.76665195 | 0.316 | 0.152 | 9.38E-32 | cDC2.Mgl2 | Fam234b |
| Cetn3 | 4.95E-36 | 0.37205588 | 0.503 | 0.348 | 9.57E-32 | cDC2.Mgl2 | Cetn3 |
| Narf | 4.99E-36 | 0.54631778 | 0.348 | 0.202 | 9.64E-32 | cDC2.Mgl2 | Narf |
| Dbn1 | 1.18E-35 | 0.37774388 | 0.571 | 0.31 | 2.29E-31 | cDC2.Mgl2 | Dbn1 |
| Mars | 1.32E-35 | 0.49181944 | 0.384 | 0.227 | 2.56E-31 | cDC2.Mgl2 | Mars |
| Mafk1 | 1.39E-35 | 0.6818653 | 0.38 | 0.232 | 2.69E-31 | cDC2.Mgl2 | Mafk |
| Dars1 | 1.67E-35 | 0.38077761 | 0.551 | 0.398 | 3.22E-31 | cDC2.Mgl2 | Dars |
| Mfng1 | 1.70E-35 | 0.66184793 | 0.329 | 0.172 | 3.29E-31 | cDC2.Mgl2 | Mfng |
| Jmy | 1.94E-35 | 0.74258384 | 0.291 | 0.167 | 3.75E-31 | cDC2.Mgl2 | Jmy |
| Plxna11 | 2.10E-35 | 0.67196887 | 0.42 | 0.227 | 4.06E-31 | cDC2.Mgl2 | Plxna1 |
| Atp2a21 | 2.50E-35 | 0.29339589 | 0.632 | 0.471 | 4.83E-31 | cDC2.Mgl2 | Atp2a2 |
| Prkacb | 2.89E-35 | 0.46627557 | 0.43 | 0.279 | 5.59E-31 | cDC2.Mgl2 | Prkacb |
| Cct6a | 3.40E-35 | 0.35953562 | 0.492 | 0.361 | 6.57E-31 | cDC2.Mgl2 | Cct6a |
| Retreg1 | 3.90E-35 | 0.69714476 | 0.379 | 0.22 | 7.54E-31 | cDC2.Mgl2 | Retreg1 |
| Cpsf31 | 4.28E-35 | 0.45030838 | 0.413 | 0.265 | 8.27E-31 | cDC2.Mgl2 | Cpsf3 |
| Bicd2 | 5.27E-35 | 0.35319214 | 0.476 | 0.306 | 1.02E-30 | cDC2.Mgl2 | Bicd2 |
| Susd6 | 5.34E-35 | 0.43086432 | 0.575 | 0.403 | 1.03E-30 | cDC2.Mgl2 | Susd6 |
| Chmp2b | 5.86E-35 | 0.44393708 | 0.432 | 0.275 | 1.13E-30 | cDC2.Mgl2 | Chmp2b |
| Mob3b1 | 8.11E-35 | 0.45731683 | 0.386 | 0.242 | 1.57E-30 | cDC2.Mgl2 | Mob3b |
| Hist1h2bc | 8.52E-35 | 0.77783729 | 0.312 | 0.19 | 1.65E-30 | cDC2.Mgl2 | Hist1h2bc |
| Lrrc25 | 1.02E-34 | 0.53342605 | 0.522 | 0.315 | 1.97E-30 | cDC2.Mgl2 | Lrrc25 |
| Emc61 | 1.27E-34 | 0.5152718 | 0.464 | 0.302 | 2.45E-30 | cDC2.Mgl2 | Emc6 |
| Nudt4 | 1.29E-34 | 0.51523249 | 0.333 | 0.19 | 2.49E-30 | cDC2.Mgl2 | Nudt4 |
| Apex11 | 1.29E-34 | 0.56226752 | 0.359 | 0.207 | 2.50E-30 | cDC2.Mgl2 | Apex1 |
| Phb | 1.40E-34 | 0.63109718 | 0.3 | 0.153 | 2.71E-30 | cDC2.Mgl2 | Phb |
| Pxn | 2.22E-34 | 0.29584598 | 0.537 | 0.39 | 4.29E-30 | cDC2.Mgl2 | Pxn |
| Acox1 | 2.27E-34 | 0.59079861 | 0.418 | 0.255 | 4.40E-30 | cDC2.Mgl2 | Acox1 |
| Diaph21 | 2.50E-34 | 0.42092798 | 0.479 | 0.317 | 4.83E-30 | cDC2.Mgl2 | Diaph2 |
| Cpeb2 | 2.61E-34 | 0.76605747 | 0.343 | 0.191 | 5.04E-30 | cDC2.Mgl2 | Cpeb2 |
| Lamtor3 | 2.82E-34 | 0.35136746 | 0.488 | 0.344 | 5.46E-30 | cDC2.Mgl2 | Lamtor3 |
| Ap1s3 | 3.81E-34 | 0.3774421 | 0.495 | 0.305 | 7.37E-30 | cDC2.Mgl2 | Ap1s3 |
| Gngt2 | 4.90E-34 | 0.38625099 | 0.653 | 0.397 | 9.47E-30 | cDC2.Mgl2 | Gngt2 |
| Usp37 | 5.08E-34 | 0.72875248 | 0.272 | 0.139 | 9.82E-30 | cDC2.Mgl2 | Usp37 |
| Bcl21 | 5.24E-34 | 0.61754032 | 0.287 | 0.145 | 1.01E-29 | cDC2.Mgl2 | Bcl2 |
| Atp2c1 | 5.25E-34 | 0.39502143 | 0.535 | 0.369 | 1.02E-29 | cDC2.Mgl2 | Atp2c1 |
| Klf7 | 6.70E-34 | 0.5270264 | 0.34 | 0.189 | 1.30E-29 | cDC2.Mgl2 | Klf7 |
| Lipa | 8.04E-34 | 0.61029998 | 0.462 | 0.258 | 1.55E-29 | cDC2.Mgl2 | Lipa |
| Ripor1 | 9.01E-34 | 0.84616137 | 0.256 | 0.125 | 1.74E-29 | cDC2.Mgl2 | Ripor1 |
| Nagpa1 | 1.03E-33 | 0.69188735 | 0.404 | 0.232 | 1.99E-29 | cDC2.Mgl2 | Nagpa |
| Dusp22 | 1.14E-33 | 0.42032917 | 0.32 | 0.181 | 2.21E-29 | cDC2.Mgl2 | Dusp22 |
| Akt2 | 1.15E-33 | 0.45564978 | 0.461 | 0.294 | 2.22E-29 | cDC2.Mgl2 | Akt2 |
| Cep83 | 1.44E-33 | 0.55898606 | 0.344 | 0.187 | 2.78E-29 | cDC2.Mgl2 | Cep83 |
| Dpy19l1 | 2.25E-33 | 0.30039867 | 0.448 | 0.302 | 4.35E-29 | cDC2.Mgl2 | Dpy19l1 |
| Eif4ebp2 | 2.38E-33 | 0.39203451 | 0.564 | 0.409 | 4.60E-29 | cDC2.Mgl2 | Eif4ebp2 |
| Zfhx3 | 2.44E-33 | 1.06917394 | 0.258 | 0.113 | 4.72E-29 | cDC2.Mgl2 | Zfhx3 |
| Gmfg1 | 2.48E-33 | 0.57409455 | 0.405 | 0.256 | 4.79E-29 | cDC2.Mgl2 | Gmfg |
| Dctpp1 | 2.74E-33 | 0.83532829 | 0.259 | 0.124 | 5.29E-29 | cDC2.Mgl2 | Dctpp1 |
| Arhgap18 | 2.98E-33 | 0.46545569 | 0.545 | 0.326 | 5.76E-29 | cDC2.Mgl2 | Arhgap18 |
| Mgat4a1 | 3.43E-33 | 0.45617171 | 0.424 | 0.258 | 6.64E-29 | cDC2.Mgl2 | Mgat4a |
| Adora2b1 | 6.40E-33 | 0.69555689 | 0.278 | 0.146 | 1.24E-28 | cDC2.Mgl2 | Adora2b |
| Ubap2 | 8.12E-33 | 0.52143069 | 0.364 | 0.217 | 1.57E-28 | cDC2.Mgl2 | Ubap2 |
| Efr3a | 9.06E-33 | 0.47878795 | 0.422 | 0.276 | 1.75E-28 | cDC2.Mgl2 | Efr3a |
| Nolc1 | 9.36E-33 | 0.72609736 | 0.27 | 0.141 | 1.81E-28 | cDC2.Mgl2 | Nolc1 |
| Tbc1d9b1 | 1.04E-32 | 0.37963629 | 0.505 | 0.345 | 2.02E-28 | cDC2.Mgl2 | Tbc1d9b |
| Ech11 | 1.05E-32 | 0.39333629 | 0.505 | 0.345 | 2.02E-28 | cDC2.Mgl2 | Ech1 |
| Mfsd10 | 1.14E-32 | 0.5068904 | 0.342 | 0.204 | 2.20E-28 | cDC2.Mgl2 | Mfsd10 |
| DEPRECATED-I | 1.24E-32 | 0.89629643 | 0.27 | 0.149 | 2.40E-28 | cDC2.Mgl2 | DEPRECATED-ENSMUSG00000039782 |
| Ebi3 | 1.38E-32 | 0.63864516 | 0.369 | 0.183 | 2.66E-28 | cDC2.Mgl2 | Ebi3 |
| Ube2e31 | 1.54E-32 | 0.33871421 | 0.424 | 0.286 | 2.98E-28 | cDC2.Mgl2 | Ube2e3 |
| Ccpg1 | 2.38E-32 | 0.66952791 | 0.376 | 0.21 | 4.60E-28 | cDC2.Mgl2 | Ccpg1 |
| Acot71 | 2.69E-32 | 0.30168597 | 0.684 | 0.522 | 5.20E-28 | cDC2.Mgl2 | Acot7 |
| Nsd21 | 2.99E-32 | 0.34433643 | 0.495 | 0.34 | 5.79E-28 | cDC2.Mgl2 | Nsd2 |
| Lpxn | 3.35E-32 | 0.88582524 | 0.309 | 0.16 | 6.48E-28 | cDC2.Mgl2 | Lpxn |
| Osbpl91 | 6.22E-32 | 0.29868701 | 0.629 | 0.483 | 1.20E-27 | cDC2.Mgl2 | Osbpl9 |
| Tinf2 | 7.06E-32 | 0.48392838 | 0.351 | 0.232 | 1.37E-27 | cDC2.Mgl2 | Tinf2 |
| Gla1 | 8.27E-32 | 0.58480392 | 0.332 | 0.2 | 1.60E-27 | cDC2.Mgl2 | Gla |
| Ndufs7 | 1.37E-31 | 0.33900573 | 0.54 | 0.377 | 2.65E-27 | cDC2.Mgl2 | Ndufs7 |
| Micos10 | 1.38E-31 | 0.31615398 | 0.566 | 0.421 | 2.66E-27 | cDC2.Mgl2 | Micos10 |
| Hltf | 1.67E-31 | 0.77854222 | 0.26 | 0.132 | 3.23E-27 | cDC2.Mgl2 | Hltf |
| Atp5g11 | 1.69E-31 | 0.53657394 | 0.444 | 0.295 | 3.27E-27 | cDC2.Mgl2 | Atp5g1 |
| Ppan | 1.90E-31 | 0.63465657 | 0.274 | 0.135 | 3.68E-27 | cDC2.Mgl2 | Ppan |
| Nars | 1.98E-31 | 0.69993095 | 0.431 | 0.253 | 3.82E-27 | cDC2.Mgl2 | Nars |
| Pigt1 | 2.01E-31 | 0.46181136 | 0.497 | 0.341 | 3.88E-27 | cDC2.Mgl2 | Pigt |
| Osgep1 | 3.09E-31 | 0.38922559 | 0.43 | 0.298 | 5.97E-27 | cDC2.Mgl2 | Osgep |
| Nudt18 | 3.12E-31 | 0.85666499 | 0.293 | 0.163 | 6.03E-27 | cDC2.Mgl2 | Nudt18 |
| Dcaf121 | 3.59E-31 | 0.35008464 | 0.494 | 0.35 | 6.95E-27 | cDC2.Mgl2 | Dcaf12 |
| Coq2 | 3.71E-31 | 0.62191813 | 0.309 | 0.17 | 7.16E-27 | cDC2.Mgl2 | Coq2 |
| Dock41 | 5.95E-31 | 0.67017074 | 0.282 | 0.166 | 1.15E-26 | cDC2.Mgl2 | Dock4 |
| Hectd4 | 5.97E-31 | 0.45390652 | 0.379 | 0.223 | 1.15E-26 | cDC2.Mgl2 | Hectd4 |
| Cgnl1 | 6.21E-31 | 0.70507668 | 0.289 | 0.146 | 1.20E-26 | cDC2.Mgl2 | Cgnl1 |
| Exosc8 | 8.97E-31 | 0.59413096 | 0.358 | 0.214 | 1.73E-26 | cDC2.Mgl2 | Exosc8 |
| Lrba | 9.67E-31 | 0.59797675 | 0.402 | 0.229 | 1.87E-26 | cDC2.Mgl2 | Lrba |
| Tbxas11 | 1.02E-30 | 0.54937442 | 0.51 | 0.296 | 1.98E-26 | cDC2.Mgl2 | Tbxas1 |
| Wbp11 | 1.09E-30 | 0.4004274 | 0.472 | 0.308 | 2.11E-26 | cDC2.Mgl2 | Wbp11 |
| Selenon | 1.14E-30 | 0.8136888 | 0.263 | 0.13 | 2.20E-26 | cDC2.Mgl2 | Selenon |
| Ppp2cb | 1.16E-30 | 0.3343519 | 0.523 | 0.382 | 2.24E-26 | cDC2.Mgl2 | Ppp2cb |
| Ssr2 | 1.17E-30 | 0.2573062 | 0.632 | 0.483 | 2.26E-26 | cDC2.Mgl2 | Ssr2 |
| Sh3pxd2a | 1.43E-30 | 0.41234226 | 0.385 | 0.229 | 2.76E-26 | cDC2.Mgl2 | Sh3pxd2a |
| Sap18b1 | 1.47E-30 | 0.47015208 | 0.408 | 0.275 | 2.84E-26 | cDC2.Mgl2 | Sap18b |
| Pop4 | 1.68E-30 | 0.66419403 | 0.27 | 0.162 | 3.24E-26 | cDC2.Mgl2 | Pop4 |
| Ppfia1 | 1.74E-30 | 0.35663165 | 0.449 | 0.308 | 3.36E-26 | cDC2.Mgl2 | Ppfia1 |
| Slc25a19 | 2.05E-30 | 0.37866523 | 0.454 | 0.303 | 3.97E-26 | cDC2.Mgl2 | Slc25a19 |
| Ckb | 2.43E-30 | 0.32870005 | 0.559 | 0.331 | 4.71E-26 | cDC2.Mgl2 | Ckb |
| Mtr | 2.89E-30 | 0.58405866 | 0.292 | 0.159 | 5.58E-26 | cDC2.Mgl2 | Mtr |
| Paics | 6.34E-30 | 0.53443128 | 0.323 | 0.202 | 1.23E-25 | cDC2.Mgl2 | Paics |
| Pfdn2 | 7.49E-30 | 0.40298873 | 0.465 | 0.305 | 1.45E-25 | cDC2.Mgl2 | Pfdn2 |
| Tmx1 | 7.55E-30 | 0.40441625 | 0.48 | 0.319 | 1.46E-25 | cDC2.Mgl2 | Tmx1 |
| Micos13 | 8.77E-30 | 0.29845925 | 0.568 | 0.405 | 1.70E-25 | cDC2.Mgl2 | Micos13 |
| Pip4k2b1 | 1.17E-29 | 0.27074399 | 0.428 | 0.282 | 2.26E-25 | cDC2.Mgl2 | Pip4k2b |
| Sash3 | 1.24E-29 | 0.31751135 | 0.411 | 0.275 | 2.39E-25 | cDC2.Mgl2 | Sash3 |
| Arhgap39 | 1.35E-29 | 0.72753318 | 0.302 | 0.147 | 2.60E-25 | cDC2.Mgl2 | Arhgap39 |
| Opa3 | 1.58E-29 | 0.43091482 | 0.326 | 0.202 | 3.05E-25 | cDC2.Mgl2 | Opa3 |
| Appl2 | 1.71E-29 | 0.93406933 | 0.255 | 0.118 | 3.31E-25 | cDC2.Mgl2 | Appl2 |
| Lsm71 | 1.77E-29 | 0.29896615 | 0.481 | 0.322 | 3.43E-25 | cDC2.Mgl2 | Lsm7 |
| Med291 | 2.80E-29 | 0.45750135 | 0.369 | 0.241 | 5.42E-25 | cDC2.Mgl2 | Med29 |
| Fkbp5 | 3.63E-29 | 0.59099238 | 0.354 | 0.203 | 7.02E-25 | cDC2.Mgl2 | Fkbp5 |
| Smc1a | 4.07E-29 | 0.37154522 | 0.462 | 0.321 | 7.87E-25 | cDC2.Mgl2 | Smc1a |
| Ide | 4.81E-29 | 0.26848588 | 0.435 | 0.313 | 9.30E-25 | cDC2.Mgl2 | Ide |
| Nomo1 | 5.78E-29 | 0.66742918 | 0.304 | 0.172 | 1.12E-24 | cDC2.Mgl2 | Nomo1 |
| Spryd3 | 1.05E-28 | 0.46808447 | 0.402 | 0.262 | 2.03E-24 | cDC2.Mgl2 | Spryd3 |
| Utp181 | 1.06E-28 | 0.56112553 | 0.359 | 0.226 | 2.06E-24 | cDC2.Mgl2 | Utp18 |
| Mical2 | 1.11E-28 | 0.96608018 | 0.265 | 0.12 | 2.15E-24 | cDC2.Mgl2 | Mical2 |
| Dennd1b | 1.14E-28 | 0.34133757 | 0.525 | 0.36 | 2.20E-24 | cDC2.Mgl2 | Dennd1b |
| Tmem164 | 1.47E-28 | 0.34413023 | 0.427 | 0.281 | 2.84E-24 | cDC2.Mgl2 | Tmem164 |
| Dxo | 1.54E-28 | 0.45033377 | 0.29 | 0.188 | 2.99E-24 | cDC2.Mgl2 | Dxo |
| Ksr1 | 1.57E-28 | 0.6562809 | 0.285 | 0.133 | 3.03E-24 | cDC2.Mgl2 | Ksr1 |
| Mtap | 1.79E-28 | 0.42223729 | 0.292 | 0.172 | 3.46E-24 | cDC2.Mgl2 | Mtap |
| Sulf21 | 2.00E-28 | 0.3560706 | 0.563 | 0.373 | 3.86E-24 | cDC2.Mgl2 | Sulf2 |
| Cuta | 2.83E-28 | 0.27717012 | 0.591 | 0.429 | 5.47E-24 | cDC2.Mgl2 | Cuta |
| Kcnab21 | 3.51E-28 | 0.54896353 | 0.423 | 0.266 | 6.78E-24 | cDC2.Mgl2 | Kcnab2 |
| Rnf24 | 5.16E-28 | 0.67728573 | 0.263 | 0.135 | 9.98E-24 | cDC2.Mgl2 | Rnf24 |
| Nap1l1 | 9.01E-28 | 0.29825523 | 0.44 | 0.318 | 1.74E-23 | cDC2.Mgl2 | Nap1l1 |
| Fuca1 | 9.54E-28 | 0.32755079 | 0.56 | 0.398 | 1.84E-23 | cDC2.Mgl2 | Fuca1 |
| Tomm401 | 9.86E-28 | 0.37897159 | 0.461 | 0.309 | 1.91E-23 | cDC2.Mgl2 | Tomm40 |
| Arhgap25 | 1.01E-27 | 0.51479511 | 0.344 | 0.196 | 1.94E-23 | cDC2.Mgl2 | Arhgap25 |
| Adh5 | 1.14E-27 | 0.27949952 | 0.558 | 0.415 | 2.20E-23 | cDC2.Mgl2 | Adh5 |
| Polr2f | 1.60E-27 | 0.32031979 | 0.415 | 0.292 | 3.09E-23 | cDC2.Mgl2 | Polr2f |
| Snrpd1 | 2.25E-27 | 0.29940236 | 0.457 | 0.342 | 4.36E-23 | cDC2.Mgl2 | Snrpd1 |
| Scp2 | 2.47E-27 | 0.54064924 | 0.45 | 0.3 | 4.77E-23 | cDC2.Mgl2 | Scp2 |
| Atp5md | 2.99E-27 | 0.30867696 | 0.454 | 0.328 | 5.79E-23 | cDC2.Mgl2 | Atp5md |
| Emc101 | 5.24E-27 | 0.26612108 | 0.564 | 0.413 | 1.01E-22 | cDC2.Mgl2 | Emc10 |
| Lpcat4 | 5.59E-27 | 0.56785541 | 0.271 | 0.144 | 1.08E-22 | cDC2.Mgl2 | Lpcat4 |
| Hspd1 | 5.91E-27 | 0.44055448 | 0.394 | 0.258 | 1.14E-22 | cDC2.Mgl2 | Hspd1 |
| Wdr77 | 6.36E-27 | 0.40119736 | 0.29 | 0.164 | 1.23E-22 | cDC2.Mgl2 | Wdr77 |
| Hlx | 7.47E-27 | 0.31155027 | 0.437 | 0.312 | 1.44E-22 | cDC2.Mgl2 | Hlx |
| G6pdx1 | 7.55E-27 | 0.57487882 | 0.357 | 0.218 | 1.46E-22 | cDC2.Mgl2 | G6pdx |
| Ipo9 | 8.15E-27 | 0.51364367 | 0.336 | 0.204 | 1.58E-22 | cDC2.Mgl2 | Ipo9 |
| BC029722 | 8.91E-27 | 0.67362175 | 0.321 | 0.19 | 1.72E-22 | cDC2.Mgl2 | BC029722 |
| Sec24a1 | 9.91E-27 | 0.65818581 | 0.322 | 0.197 | 1.92E-22 | cDC2.Mgl2 | Sec24a |
| Sdhc1 | 1.20E-26 | 0.30544334 | 0.641 | 0.485 | 2.32E-22 | cDC2.Mgl2 | Sdhc |
| Atf4 | 1.20E-26 | 0.38402492 | 0.461 | 0.303 | 2.32E-22 | cDC2.Mgl2 | Atf4 |
| Taf13 | 1.36E-26 | 0.57245885 | 0.279 | 0.162 | 2.64E-22 | cDC2.Mgl2 | Taf13 |
| Rnf180 | 1.37E-26 | 0.29575058 | 0.429 | 0.302 | 2.65E-22 | cDC2.Mgl2 | Rnf180 |
| Snx81 | 1.70E-26 | 0.39377077 | 0.467 | 0.294 | 3.28E-22 | cDC2.Mgl2 | Snx8 |
| Lamtor2 | 1.86E-26 | 0.48486913 | 0.535 | 0.37 | 3.59E-22 | cDC2.Mgl2 | Lamtor2 |
| Selenoh | 2.39E-26 | 0.82411021 | 0.273 | 0.148 | 4.62E-22 | cDC2.Mgl2 | Selenoh |
| Ctns | 2.40E-26 | 0.73479232 | 0.302 | 0.158 | 4.64E-22 | cDC2.Mgl2 | Ctns |
| H2-Ke6 | 2.42E-26 | 0.52397704 | 0.343 | 0.201 | 4.67E-22 | cDC2.Mgl2 | H2-Ke6 |
| Eva1b | 2.61E-26 | 0.26661619 | 0.518 | 0.374 | 5.04E-22 | cDC2.Mgl2 | Eva1b |
| B4galt4 | 2.82E-26 | 0.53151832 | 0.278 | 0.156 | 5.45E-22 | cDC2.Mgl2 | B4galt4 |
| Ppp2r1b | 3.32E-26 | 0.52963942 | 0.328 | 0.201 | 6.42E-22 | cDC2.Mgl2 | Ppp2r1b |
| Ruvbl2 | 3.33E-26 | 0.71312733 | 0.347 | 0.202 | 6.44E-22 | cDC2.Mgl2 | Ruvbl2 |
| Mydgf1 | 3.42E-26 | 0.39573434 | 0.544 | 0.4 | 6.60E-22 | cDC2.Mgl2 | Mydgf |
| Spata13 | 3.43E-26 | 0.60089289 | 0.254 | 0.143 | 6.64E-22 | cDC2.Mgl2 | Spata13 |
| Prelid3b | 4.08E-26 | 0.44042998 | 0.397 | 0.257 | 7.89E-22 | cDC2.Mgl2 | Prelid3b |
| Ifitm5 | 4.68E-26 | 0.8812305 | 0.254 | 0.133 | 9.05E-22 | cDC2.Mgl2 | Ifitm5 |
| DEPRECATED-I | 6.66E-26 | 0.35571096 | 0.487 | 0.339 | 1.29E-21 | cDC2.Mgl2 | DEPRECATED-ENSMUSG00000017221 |
| Tmem214 | 7.74E-26 | 0.50069514 | 0.346 | 0.214 | 1.50E-21 | cDC2.Mgl2 | Tmem214 |
| Aacs | 8.29E-26 | 0.49178886 | 0.328 | 0.186 | 1.60E-21 | cDC2.Mgl2 | Aacs |
| Eif4e3 | 8.39E-26 | 0.46481293 | 0.345 | 0.205 | 1.62E-21 | cDC2.Mgl2 | Eif4e3 |
| Sys1 | 8.92E-26 | 0.35515503 | 0.519 | 0.358 | 1.73E-21 | cDC2.Mgl2 | Sys1 |
| Myg1 | 1.07E-25 | 0.6721721 | 0.26 | 0.131 | 2.07E-21 | cDC2.Mgl2 | Myg1 |
| Pabpc4 | 1.08E-25 | 0.51049859 | 0.356 | 0.217 | 2.09E-21 | cDC2.Mgl2 | Pabpc4 |
| Ascc1 | 1.16E-25 | 0.63157826 | 0.304 | 0.172 | 2.25E-21 | cDC2.Mgl2 | Ascc1 |
| Snx9 | 1.22E-25 | 0.36689227 | 0.606 | 0.408 | 2.37E-21 | cDC2.Mgl2 | Snx9 |
| Bpnt1 | 1.57E-25 | 0.51602422 | 0.358 | 0.22 | 3.04E-21 | cDC2.Mgl2 | Bpnt1 |
| Mthfsl | 1.72E-25 | 0.30714696 | 0.29 | 0.21 | 3.32E-21 | cDC2.Mgl2 | Mthfsl |
| Dgkd1 | 2.21E-25 | 0.36329334 | 0.465 | 0.297 | 4.27E-21 | cDC2.Mgl2 | Dgkd |
| Pcyt2 | 2.31E-25 | 0.45847491 | 0.353 | 0.223 | 4.46E-21 | cDC2.Mgl2 | Pcyt2 |
| Pa2g41 | 2.66E-25 | 0.37282183 | 0.355 | 0.234 | 5.14E-21 | cDC2.Mgl2 | Pa2g4 |
| Ctsl | 3.78E-25 | 0.43171504 | 0.46 | 0.221 | 7.31E-21 | cDC2.Mgl2 | Ctsl |
| Slc30a71 | 4.79E-25 | 0.3695666 | 0.394 | 0.262 | 9.25E-21 | cDC2.Mgl2 | Slc30a7 |
| Susd3 | 4.84E-25 | 1.00204672 | 0.277 | 0.145 | 9.36E-21 | cDC2.Mgl2 | Susd3 |
| Ppp1r11 | 4.88E-25 | 0.30577208 | 0.572 | 0.406 | 9.43E-21 | cDC2.Mgl2 | Ppp1r11 |
| Tsr1 | 6.41E-25 | 0.45706735 | 0.321 | 0.191 | 1.24E-20 | cDC2.Mgl2 | Tsr1 |
| Heatr11 | 7.79E-25 | 0.40638301 | 0.325 | 0.211 | 1.51E-20 | cDC2.Mgl2 | Heatr1 |
| Med14 | 9.04E-25 | 0.49087569 | 0.325 | 0.208 | 1.75E-20 | cDC2.Mgl2 | Med14 |
| Yars | 9.17E-25 | 0.55066949 | 0.298 | 0.165 | 1.77E-20 | cDC2.Mgl2 | Yars |
| Map4k2 | 1.23E-24 | 0.50651449 | 0.37 | 0.216 | 2.37E-20 | cDC2.Mgl2 | Map4k2 |
| Pithd1 | 1.81E-24 | 0.43132353 | 0.323 | 0.192 | 3.50E-20 | cDC2.Mgl2 | Pithd1 |
| Dna21 | 1.88E-24 | 0.43660604 | 0.409 | 0.24 | 3.63E-20 | cDC2.Mgl2 | Dna2 |
| Cmas1 | 1.92E-24 | 0.35360084 | 0.463 | 0.313 | 3.72E-20 | cDC2.Mgl2 | Cmas |
| Usp36 | 2.14E-24 | 0.55299386 | 0.25 | 0.144 | 4.14E-20 | cDC2.Mgl2 | Usp36 |
| Pigs1 | 2.15E-24 | 0.43555534 | 0.425 | 0.291 | 4.15E-20 | cDC2.Mgl2 | Pigs |
| Mogs | 2.92E-24 | 0.68763047 | 0.318 | 0.181 | 5.64E-20 | cDC2.Mgl2 | Mogs |
| Pofut2 | 3.35E-24 | 0.56289664 | 0.309 | 0.184 | 6.48E-20 | cDC2.Mgl2 | Pofut2 |
| Rpe | 3.86E-24 | 0.41969536 | 0.413 | 0.272 | 7.46E-20 | cDC2.Mgl2 | Rpe |
| Sec11a1 | 3.87E-24 | 0.3526499 | 0.544 | 0.38 | 7.48E-20 | cDC2.Mgl2 | Sec11a |
| Mysm11 | 4.04E-24 | 0.37390215 | 0.353 | 0.24 | 7.81E-20 | cDC2.Mgl2 | Mysm1 |
| Pdia41 | 4.24E-24 | 0.29510216 | 0.539 | 0.369 | 8.20E-20 | cDC2.Mgl2 | Pdia4 |
| Ndufa12 | 4.43E-24 | 0.44938761 | 0.38 | 0.251 | 8.57E-20 | cDC2.Mgl2 | Ndufa12 |
| Uqcc2 | 5.07E-24 | 0.35161262 | 0.411 | 0.282 | 9.81E-20 | cDC2.Mgl2 | Uqcc2 |
| Man2c1 | 6.58E-24 | 0.51151183 | 0.42 | 0.263 | 1.27E-19 | cDC2.Mgl2 | Man2c1 |
| Yipf3 | 7.61E-24 | 0.30094954 | 0.467 | 0.325 | 1.47E-19 | cDC2.Mgl2 | Yipf3 |
| Slc14a11 | 8.18E-24 | 0.38510547 | 0.276 | 0.165 | 1.58E-19 | cDC2.Mgl2 | Slc14a1 |
| Aim2 | 1.17E-23 | 0.57128218 | 0.345 | 0.223 | 2.26E-19 | cDC2.Mgl2 | Aim2 |
| Phb2 | 1.27E-23 | 0.26484331 | 0.501 | 0.374 | 2.45E-19 | cDC2.Mgl2 | Phb2 |
| Cers5 | 1.38E-23 | 0.36924439 | 0.441 | 0.295 | 2.68E-19 | cDC2.Mgl2 | Cers5 |
| Cenpx | 1.42E-23 | 0.39379224 | 0.398 | 0.26 | 2.75E-19 | cDC2.Mgl2 | Cenpx |
| Ddx27 | 1.48E-23 | 0.3812812 | 0.345 | 0.223 | 2.86E-19 | cDC2.Mgl2 | Ddx27 |
| Npepl1 | 2.10E-23 | 0.2586064 | 0.469 | 0.339 | 4.05E-19 | cDC2.Mgl2 | Npepl1 |
| Plpbp | 2.23E-23 | 0.32844914 | 0.471 | 0.328 | 4.32E-19 | cDC2.Mgl2 | Plpbp |
| Nelfe1 | 2.26E-23 | 0.29335126 | 0.368 | 0.252 | 4.36E-19 | cDC2.Mgl2 | Nelfe |
| Galk1 | 2.50E-23 | 0.65054001 | 0.262 | 0.147 | 4.83E-19 | cDC2.Mgl2 | Galk1 |
| Hint11 | 2.71E-23 | 0.37964886 | 0.407 | 0.278 | 5.23E-19 | cDC2.Mgl2 | Hint1 |
| Klhdc10 | 3.25E-23 | 0.41649494 | 0.351 | 0.218 | 6.28E-19 | cDC2.Mgl2 | Klhdc10 |
| Txnl1 | 3.42E-23 | 0.27797811 | 0.508 | 0.375 | 6.61E-19 | cDC2.Mgl2 | Txnl1 |
| Dnajc3 | 3.79E-23 | 0.29454282 | 0.577 | 0.412 | 7.32E-19 | cDC2.Mgl2 | Dnajc3 |
| Acsf21 | 4.05E-23 | 0.3081151 | 0.354 | 0.221 | 7.82E-19 | cDC2.Mgl2 | Acsf2 |
| Rwdd11 | 4.23E-23 | 0.46402888 | 0.317 | 0.189 | 8.18E-19 | cDC2.Mgl2 | Rwdd1 |
| Ddx41 | 4.60E-23 | 0.27568284 | 0.36 | 0.224 | 8.88E-19 | cDC2.Mgl2 | Ddx41 |
| Gpn1 | 4.60E-23 | 0.53954841 | 0.292 | 0.173 | 8.89E-19 | cDC2.Mgl2 | Gpn1 |
| Slc25a38 | 6.29E-23 | 0.33328179 | 0.377 | 0.226 | 1.22E-18 | cDC2.Mgl2 | Slc25a38 |
| Gbp7 | 8.85E-23 | 0.87675142 | 0.283 | 0.141 | 1.71E-18 | cDC2.Mgl2 | Gbp7 |
| Batf | 1.03E-22 | 0.57033512 | 0.264 | 0.147 | 1.99E-18 | cDC2.Mgl2 | Batf |
| Aqr | 1.05E-22 | 0.38105447 | 0.354 | 0.231 | 2.03E-18 | cDC2.Mgl2 | Aqr |
| Kbtbd111 | 1.24E-22 | 0.42867316 | 0.334 | 0.222 | 2.40E-18 | cDC2.Mgl2 | Kbtbd11 |
| Gys1 | 1.28E-22 | 0.76290284 | 0.295 | 0.164 | 2.48E-18 | cDC2.Mgl2 | Gys1 |
| Slc35c1 | 1.38E-22 | 0.63565435 | 0.254 | 0.142 | 2.67E-18 | cDC2.Mgl2 | Slc35c1 |
| Rnf126 | 1.56E-22 | 0.42740315 | 0.44 | 0.302 | 3.02E-18 | cDC2.Mgl2 | Rnf126 |
| Tomm6 | 1.60E-22 | 0.39806315 | 0.514 | 0.365 | 3.10E-18 | cDC2.Mgl2 | Tomm6 |
| Tmem250-ps | 2.60E-22 | 0.62358971 | 0.319 | 0.198 | 5.03E-18 | cDC2.Mgl2 | Tmem250-ps |
| Prep1 | 2.76E-22 | 0.39539055 | 0.381 | 0.253 | 5.33E-18 | cDC2.Mgl2 | Prep |
| Pole4 | 2.91E-22 | 0.44599869 | 0.373 | 0.243 | 5.62E-18 | cDC2.Mgl2 | Pole4 |
| Mtln | 3.12E-22 | 0.44789234 | 0.272 | 0.149 | 6.03E-18 | cDC2.Mgl2 | Mtln |
| Micu1 | 3.45E-22 | 0.50661148 | 0.295 | 0.167 | 6.68E-18 | cDC2.Mgl2 | Micu1 |
| Bop11 | 3.52E-22 | 0.44106685 | 0.314 | 0.204 | 6.80E-18 | cDC2.Mgl2 | Bop1 |
| Ino80e | 4.10E-22 | 0.29404514 | 0.366 | 0.254 | 7.92E-18 | cDC2.Mgl2 | Ino80e |
| Dbndd2 | 4.15E-22 | 0.36537014 | 0.336 | 0.204 | 8.03E-18 | cDC2.Mgl2 | Dbndd2 |
| Bscl21 | 4.53E-22 | 0.28027356 | 0.486 | 0.33 | 8.76E-18 | cDC2.Mgl2 | Bscl2 |
| Cox6a1 | 4.67E-22 | 0.27143634 | 0.541 | 0.395 | 9.02E-18 | cDC2.Mgl2 | Cox6a1 |
| Eif2s1 | 4.78E-22 | 0.32581574 | 0.486 | 0.363 | 9.24E-18 | cDC2.Mgl2 | Eif2s1 |
| Fmnl3 | 5.55E-22 | 0.61173271 | 0.269 | 0.142 | 1.07E-17 | cDC2.Mgl2 | Fmnl3 |
| Nipa2 | 6.08E-22 | 0.27888246 | 0.485 | 0.359 | 1.18E-17 | cDC2.Mgl2 | Nipa2 |
| Stx2 | 6.38E-22 | 0.52892478 | 0.34 | 0.203 | 1.23E-17 | cDC2.Mgl2 | Stx2 |
| Rangap1 | 6.74E-22 | 0.35308874 | 0.351 | 0.227 | 1.30E-17 | cDC2.Mgl2 | Rangap1 |
| Nfx1 | 7.66E-22 | 0.26820239 | 0.288 | 0.179 | 1.48E-17 | cDC2.Mgl2 | Nfx1 |
| Rars | 7.94E-22 | 0.56796237 | 0.323 | 0.206 | 1.54E-17 | cDC2.Mgl2 | Rars |
| Atg3 | 8.56E-22 | 0.2834774 | 0.472 | 0.336 | 1.66E-17 | cDC2.Mgl2 | Atg3 |
| Atp5f1 | 8.81E-22 | 0.35904767 | 0.583 | 0.424 | 1.70E-17 | cDC2.Mgl2 | Atp5f1 |
| Phf81 | 1.38E-21 | 0.33238391 | 0.355 | 0.23 | 2.67E-17 | cDC2.Mgl2 | Phf8 |
| Itpr2 | 1.45E-21 | 0.39748043 | 0.299 | 0.184 | 2.80E-17 | cDC2.Mgl2 | Itpr2 |
| Lsm2 | 1.47E-21 | 0.4465149 | 0.318 | 0.208 | 2.85E-17 | cDC2.Mgl2 | Lsm2 |
| Maco1 | 1.71E-21 | 0.40187523 | 0.371 | 0.239 | 3.30E-17 | cDC2.Mgl2 | Maco1 |
| Zbtb111 | 1.90E-21 | 0.49378823 | 0.408 | 0.268 | 3.67E-17 | cDC2.Mgl2 | Zbtb11 |
| Spindoc | 2.18E-21 | 0.59531138 | 0.299 | 0.188 | 4.22E-17 | cDC2.Mgl2 | Spindoc |
| Glb1 | 2.36E-21 | 0.52064083 | 0.313 | 0.184 | 4.56E-17 | cDC2.Mgl2 | Glb1 |
| Bud231 | 2.60E-21 | 0.51109382 | 0.343 | 0.211 | 5.03E-17 | cDC2.Mgl2 | Bud23 |
| Wbp1 | 2.66E-21 | 0.40477452 | 0.344 | 0.215 | 5.14E-17 | cDC2.Mgl2 | Wbp1 |
| Klhl18 | 2.87E-21 | 0.43218658 | 0.352 | 0.211 | 5.55E-17 | cDC2.Mgl2 | Klhl18 |
| Nf2 | 3.16E-21 | 0.28385188 | 0.395 | 0.264 | 6.11E-17 | cDC2.Mgl2 | Nf2 |
| Pno11 | 3.30E-21 | 0.47219435 | 0.317 | 0.213 | 6.38E-17 | cDC2.Mgl2 | Pno1 |
| Slc43a21 | 3.35E-21 | 0.49671721 | 0.391 | 0.209 | 6.47E-17 | cDC2.Mgl2 | Slc43a2 |
| Ptpn22 | 4.43E-21 | 0.71829967 | 0.291 | 0.146 | 8.56E-17 | cDC2.Mgl2 | Ptpn22 |
| Tmco11 | 4.78E-21 | 0.29668332 | 0.406 | 0.268 | 9.23E-17 | cDC2.Mgl2 | Tmco1 |
| Slc30a5 | 4.96E-21 | 0.39554741 | 0.359 | 0.226 | 9.59E-17 | cDC2.Mgl2 | Slc30a5 |
| Ivd | 5.12E-21 | 0.31454576 | 0.288 | 0.175 | 9.91E-17 | cDC2.Mgl2 | Ivd |
| Pycr2 | 5.52E-21 | 0.28608841 | 0.343 | 0.228 | 1.07E-16 | cDC2.Mgl2 | Pycr2 |
| Rere | 6.12E-21 | 0.25889387 | 0.388 | 0.25 | 1.18E-16 | cDC2.Mgl2 | Rere |
| Cnpy2 | 6.18E-21 | 0.45145076 | 0.351 | 0.24 | 1.19E-16 | cDC2.Mgl2 | Cnpy2 |
| Ube2v21 | 6.64E-21 | 0.6720833 | 0.344 | 0.212 | 1.28E-16 | cDC2.Mgl2 | Ube2v2 |
| Vps53 | 7.87E-21 | 0.42085336 | 0.364 | 0.232 | 1.52E-16 | cDC2.Mgl2 | Vps53 |
| Glipr1 | 8.21E-21 | 0.4047924 | 0.326 | 0.21 | 1.59E-16 | cDC2.Mgl2 | Glipr1 |
| Gripap11 | 8.67E-21 | 0.35857735 | 0.377 | 0.25 | 1.68E-16 | cDC2.Mgl2 | Gripap1 |
| Rpp21 | 9.67E-21 | 0.30198635 | 0.341 | 0.222 | 1.87E-16 | cDC2.Mgl2 | Rpp21 |
| Rab3gap2 | 1.00E-20 | 0.27756093 | 0.335 | 0.22 | 1.93E-16 | cDC2.Mgl2 | Rab3gap2 |
| Tsn | 1.08E-20 | 0.31237407 | 0.448 | 0.324 | 2.08E-16 | cDC2.Mgl2 | Tsn |
| Pprc11 | 1.09E-20 | 0.41819593 | 0.313 | 0.195 | 2.11E-16 | cDC2.Mgl2 | Pprc1 |
| Iars | 1.10E-20 | 0.27943481 | 0.368 | 0.245 | 2.13E-16 | cDC2.Mgl2 | Iars |
| Dnajc15 | 1.15E-20 | 0.38363181 | 0.408 | 0.282 | 2.23E-16 | cDC2.Mgl2 | Dnajc15 |
| Cyb5b | 1.37E-20 | 0.36596244 | 0.384 | 0.243 | 2.65E-16 | cDC2.Mgl2 | Cyb5b |
| Hps3 | 1.94E-20 | 0.33786258 | 0.427 | 0.262 | 3.75E-16 | cDC2.Mgl2 | Hps3 |
| Wdfy2 | 2.02E-20 | 0.5747239 | 0.376 | 0.226 | 3.90E-16 | cDC2.Mgl2 | Wdfy2 |
| Abca3 | 2.38E-20 | 0.36256622 | 0.375 | 0.248 | 4.60E-16 | cDC2.Mgl2 | Abca3 |
| Creld2 | 2.44E-20 | 0.48775846 | 0.333 | 0.226 | 4.71E-16 | cDC2.Mgl2 | Creld2 |
| Tmem251 | 2.55E-20 | 0.46397807 | 0.294 | 0.185 | 4.93E-16 | cDC2.Mgl2 | Tmem251 |
| Rnf1411 | 4.26E-20 | 0.42759863 | 0.337 | 0.212 | 8.23E-16 | cDC2.Mgl2 | Rnf141 |
| Ssna1 | 4.53E-20 | 0.2793922 | 0.426 | 0.302 | 8.75E-16 | cDC2.Mgl2 | Ssna1 |
| Abce11 | 4.97E-20 | 0.64005757 | 0.307 | 0.191 | 9.62E-16 | cDC2.Mgl2 | Abce1 |
| Bccip1 | 5.09E-20 | 0.33602763 | 0.389 | 0.281 | 9.84E-16 | cDC2.Mgl2 | Bccip |
| Snapc5 | 5.63E-20 | 0.2697464 | 0.394 | 0.267 | 1.09E-15 | cDC2.Mgl2 | Snapc5 |
| Pex19 | 7.57E-20 | 0.39453943 | 0.328 | 0.213 | 1.46E-15 | cDC2.Mgl2 | Pex19 |
| Ppie | 7.68E-20 | 0.41102045 | 0.343 | 0.226 | 1.49E-15 | cDC2.Mgl2 | Ppie |
| Rbmxl1 | 8.06E-20 | 0.36194532 | 0.282 | 0.181 | 1.56E-15 | cDC2.Mgl2 | Rbmxl1 |
| Sars | 9.60E-20 | 0.29786505 | 0.354 | 0.222 | 1.86E-15 | cDC2.Mgl2 | Sars |
| Chchd1 | 9.68E-20 | 0.49425616 | 0.305 | 0.196 | 1.87E-15 | cDC2.Mgl2 | Chchd1 |
| Edem3 | 1.24E-19 | 0.45416143 | 0.256 | 0.155 | 2.39E-15 | cDC2.Mgl2 | Edem3 |
| Abcf1 | 1.26E-19 | 0.32124514 | 0.414 | 0.299 | 2.43E-15 | cDC2.Mgl2 | Abcf1 |
| Wdr181 | 1.35E-19 | 0.40184309 | 0.327 | 0.218 | 2.62E-15 | cDC2.Mgl2 | Wdr18 |
| Zfp7031 | 1.42E-19 | 0.25279423 | 0.549 | 0.411 | 2.74E-15 | cDC2.Mgl2 | Zfp703 |
| Wdr13 | 1.45E-19 | 0.25437372 | 0.321 | 0.211 | 2.81E-15 | cDC2.Mgl2 | Wdr13 |
| Ddx56 | 1.48E-19 | 0.40499692 | 0.313 | 0.198 | 2.86E-15 | cDC2.Mgl2 | Ddx56 |
| Hk11 | 1.72E-19 | 0.25340478 | 0.538 | 0.39 | 3.33E-15 | cDC2.Mgl2 | Hk1 |
| Plod31 | 1.77E-19 | 0.3994349 | 0.386 | 0.242 | 3.42E-15 | cDC2.Mgl2 | Plod3 |
| Rcor1 | 1.89E-19 | 0.4884658 | 0.401 | 0.27 | 3.65E-15 | cDC2.Mgl2 | Rcor1 |
| Bin3 | 1.89E-19 | 0.27845349 | 0.482 | 0.344 | 3.66E-15 | cDC2.Mgl2 | Bin3 |
| Abl1 | 2.48E-19 | 0.37806587 | 0.424 | 0.301 | 4.79E-15 | cDC2.Mgl2 | Abl1 |
| Nr2c2ap | 2.71E-19 | 0.3088879 | 0.354 | 0.243 | 5.24E-15 | cDC2.Mgl2 | Nr2c2ap |
| Rab20 | 3.76E-19 | 0.42141012 | 0.304 | 0.197 | 7.27E-15 | cDC2.Mgl2 | Rab20 |
| Commd2 | 3.96E-19 | 0.46721203 | 0.277 | 0.17 | 7.67E-15 | cDC2.Mgl2 | Commd2 |
| Cs | 4.14E-19 | 0.28458331 | 0.469 | 0.333 | 8.00E-15 | cDC2.Mgl2 | Cs |
| Slamf8 | 5.00E-19 | 0.38137069 | 0.4 | 0.236 | 9.67E-15 | cDC2.Mgl2 | Slamf8 |
| Ufsp2 | 5.26E-19 | 0.47254695 | 0.351 | 0.23 | 1.02E-14 | cDC2.Mgl2 | Ufsp2 |
| Cd200r1 | 6.93E-19 | 0.55989716 | 0.299 | 0.17 | 1.34E-14 | cDC2.Mgl2 | Cd200r1 |
| Polr1c | 8.59E-19 | 0.35457807 | 0.396 | 0.277 | 1.66E-14 | cDC2.Mgl2 | Polr1c |
| Ppp5c | 9.34E-19 | 0.39373834 | 0.336 | 0.221 | 1.80E-14 | cDC2.Mgl2 | Ppp5c |
| Wdr43 | 9.87E-19 | 0.25705916 | 0.29 | 0.185 | 1.91E-14 | cDC2.Mgl2 | Wdr43 |
| Hes6 | 1.12E-18 | 0.33421824 | 0.289 | 0.2 | 2.17E-14 | cDC2.Mgl2 | Hes6 |
| Reep4 | 1.30E-18 | 0.28701531 | 0.277 | 0.17 | 2.52E-14 | cDC2.Mgl2 | Reep4 |
| Denr | 1.36E-18 | 0.32867009 | 0.407 | 0.275 | 2.64E-14 | cDC2.Mgl2 | Denr |
| Zfyve27 | 1.38E-18 | 0.50425086 | 0.323 | 0.205 | 2.66E-14 | cDC2.Mgl2 | Zfyve27 |
| Ost4 | 1.42E-18 | 0.39877678 | 0.391 | 0.274 | 2.75E-14 | cDC2.Mgl2 | Ost4 |
| Pik3r6 | 1.47E-18 | 0.27147205 | 0.359 | 0.219 | 2.84E-14 | cDC2.Mgl2 | Pik3r6 |
| Tomm40l | 1.51E-18 | 0.38100735 | 0.332 | 0.209 | 2.91E-14 | cDC2.Mgl2 | Tomm40l |
| Gnl31 | 1.53E-18 | 0.39195718 | 0.306 | 0.198 | 2.95E-14 | cDC2.Mgl2 | Gnl3 |
| Dipk1a | 1.64E-18 | 0.49837219 | 0.262 | 0.152 | 3.17E-14 | cDC2.Mgl2 | Dipk1a |
| Itprip | 1.74E-18 | 0.27891775 | 0.362 | 0.242 | 3.36E-14 | cDC2.Mgl2 | Itprip |
| Hspa14 | 2.07E-18 | 0.32403808 | 0.357 | 0.254 | 4.01E-14 | cDC2.Mgl2 | Hspa14 |
| Ndufaf81 | 2.59E-18 | 0.34123797 | 0.349 | 0.235 | 5.01E-14 | cDC2.Mgl2 | Ndufaf8 |
| Timm50 | 2.86E-18 | 0.38152395 | 0.32 | 0.223 | 5.52E-14 | cDC2.Mgl2 | Timm50 |
| Cct5 | 3.60E-18 | 0.26223223 | 0.31 | 0.233 | 6.97E-14 | cDC2.Mgl2 | Cct5 |
| Eif5 | 3.73E-18 | 0.2885347 | 0.347 | 0.237 | 7.20E-14 | cDC2.Mgl2 | Eif5 |
| Lrrc8d | 3.88E-18 | 0.27978504 | 0.337 | 0.232 | 7.50E-14 | cDC2.Mgl2 | Lrrc8d |
| Lyar | 5.18E-18 | 0.4304249 | 0.277 | 0.193 | 1.00E-13 | cDC2.Mgl2 | Lyar |
| Eif4h | 5.50E-18 | 0.26328815 | 0.531 | 0.382 | 1.06E-13 | cDC2.Mgl2 | Eif4h |
| Nudt5 | 6.61E-18 | 0.42519963 | 0.26 | 0.168 | 1.28E-13 | cDC2.Mgl2 | Nudt5 |
| Ctso | 7.09E-18 | 0.41331461 | 0.258 | 0.155 | 1.37E-13 | cDC2.Mgl2 | Ctso |
| Rnaseh2a | 9.12E-18 | 0.46412707 | 0.264 | 0.158 | 1.76E-13 | cDC2.Mgl2 | Rnaseh2a |
| Hspa9 | 9.54E-18 | 0.41810426 | 0.317 | 0.21 | 1.84E-13 | cDC2.Mgl2 | Hspa9 |
| Dram2 | 1.19E-17 | 0.54424401 | 0.25 | 0.146 | 2.30E-13 | cDC2.Mgl2 | Dram2 |
| Usb11 | 1.23E-17 | 0.45577538 | 0.305 | 0.202 | 2.37E-13 | cDC2.Mgl2 | Usb1 |
| Zfp131 | 1.53E-17 | 0.32470681 | 0.408 | 0.279 | 2.95E-13 | cDC2.Mgl2 | Zfp131 |
| Ap1m1 | 1.58E-17 | 0.30981806 | 0.454 | 0.32 | 3.05E-13 | cDC2.Mgl2 | Ap1m1 |
| Ltv1 | 1.64E-17 | 0.46096785 | 0.291 | 0.182 | 3.18E-13 | cDC2.Mgl2 | Ltv1 |
| Suclg1 | 1.65E-17 | 0.39189067 | 0.304 | 0.191 | 3.19E-13 | cDC2.Mgl2 | Suclg1 |
| Ndufb6 | 1.99E-17 | 0.294947 | 0.549 | 0.429 | 3.85E-13 | cDC2.Mgl2 | Ndufb6 |
| Zcchc21 | 2.60E-17 | 0.36965462 | 0.295 | 0.2 | 5.02E-13 | cDC2.Mgl2 | Zcchc2 |
| Gnpat | 3.11E-17 | 0.37857008 | 0.319 | 0.209 | 6.02E-13 | cDC2.Mgl2 | Gnpat |
| Prpf311 | 3.77E-17 | 0.32909963 | 0.421 | 0.305 | 7.29E-13 | cDC2.Mgl2 | Prpf31 |
| Mfsd12 | 4.78E-17 | 0.33974696 | 0.375 | 0.256 | 9.25E-13 | cDC2.Mgl2 | Mfsd12 |
| Taf10 | 6.22E-17 | 0.36416302 | 0.359 | 0.261 | 1.20E-12 | cDC2.Mgl2 | Taf10 |
| Ssr3 | 7.13E-17 | 0.27245705 | 0.526 | 0.398 | 1.38E-12 | cDC2.Mgl2 | Ssr3 |
| C1qbp | 8.04E-17 | 0.52180135 | 0.274 | 0.175 | 1.55E-12 | cDC2.Mgl2 | C1qbp |
| Mnt | 8.58E-17 | 0.3655241 | 0.273 | 0.177 | 1.66E-12 | cDC2.Mgl2 | Mnt |
| Trim28 | 1.20E-16 | 0.26356285 | 0.354 | 0.24 | 2.31E-12 | cDC2.Mgl2 | Trim28 |
| Mki67 | 1.23E-16 | 0.65645617 | 0.259 | 0.111 | 2.39E-12 | cDC2.Mgl2 | Mki67 |
| Nudt8 | 1.31E-16 | 0.34247669 | 0.253 | 0.164 | 2.52E-12 | cDC2.Mgl2 | Nudt8 |
| Gar1 | 1.48E-16 | 0.30196068 | 0.35 | 0.237 | 2.86E-12 | cDC2.Mgl2 | Gar1 |
| Usp14 | 1.81E-16 | 0.38456449 | 0.322 | 0.222 | 3.50E-12 | cDC2.Mgl2 | Usp14 |
| Nubp2 | 2.03E-16 | 0.32857525 | 0.475 | 0.363 | 3.92E-12 | cDC2.Mgl2 | Nubp2 |
| Snrpa1 | 2.13E-16 | 0.41984089 | 0.373 | 0.27 | 4.12E-12 | cDC2.Mgl2 | Snrpa1 |
| Anp32b | 2.16E-16 | 0.43702698 | 0.324 | 0.219 | 4.17E-12 | cDC2.Mgl2 | Anp32b |
| Kri1 | 2.56E-16 | 0.38447775 | 0.351 | 0.247 | 4.95E-12 | cDC2.Mgl2 | Kri1 |
| Dcps | 2.80E-16 | 0.35912626 | 0.261 | 0.161 | 5.41E-12 | cDC2.Mgl2 | Dcps |
| Med27 | 2.90E-16 | 0.25208347 | 0.312 | 0.203 | 5.61E-12 | cDC2.Mgl2 | Med27 |
| Tbkbp11 | 3.35E-16 | 0.53150738 | 0.31 | 0.191 | 6.49E-12 | cDC2.Mgl2 | Tbkbp1 |
| Mcts1 | 4.23E-16 | 0.46185423 | 0.362 | 0.255 | 8.18E-12 | cDC2.Mgl2 | Mcts1 |
| Pxk1 | 4.25E-16 | 0.25958742 | 0.324 | 0.224 | 8.21E-12 | cDC2.Mgl2 | Pxk |
| Tbc1d2b1 | 4.26E-16 | 0.34352801 | 0.34 | 0.23 | 8.24E-12 | cDC2.Mgl2 | Tbc1d2b |
| Acd | 4.49E-16 | 0.34519867 | 0.288 | 0.18 | 8.68E-12 | cDC2.Mgl2 | Acd |
| Prmt1 | 4.77E-16 | 0.26572051 | 0.388 | 0.279 | 9.23E-12 | cDC2.Mgl2 | Prmt1 |
| Fnip1 | 6.13E-16 | 0.25948881 | 0.318 | 0.214 | 1.18E-11 | cDC2.Mgl2 | Fnip1 |
| Jpt2 | 7.78E-16 | 0.27006123 | 0.274 | 0.181 | 1.50E-11 | cDC2.Mgl2 | Jpt2 |
| Prpf38a | 8.66E-16 | 0.42129395 | 0.323 | 0.216 | 1.68E-11 | cDC2.Mgl2 | Prpf38a |
| Morc2a | 1.02E-15 | 0.37553938 | 0.346 | 0.219 | 1.97E-11 | cDC2.Mgl2 | Morc2a |
| Acp5 | 1.03E-15 | 0.44209065 | 0.418 | 0.217 | 1.99E-11 | cDC2.Mgl2 | Acp5 |
| Hdgfl2 | 1.78E-15 | 0.25361327 | 0.383 | 0.278 | 3.43E-11 | cDC2.Mgl2 | Hdgfl2 |
| Mospd3 | 1.89E-15 | 0.36273157 | 0.386 | 0.269 | 3.65E-11 | cDC2.Mgl2 | Mospd3 |
| Zfp706 | 2.28E-15 | 0.48303896 | 0.322 | 0.221 | 4.41E-11 | cDC2.Mgl2 | Zfp706 |
| Enoph1 | 2.47E-15 | 0.30365467 | 0.306 | 0.217 | 4.78E-11 | cDC2.Mgl2 | Enoph1 |
| Eif1b | 2.90E-15 | 0.26436847 | 0.323 | 0.236 | 5.60E-11 | cDC2.Mgl2 | Eif1b |
| Eid3 | 3.75E-15 | 0.33666696 | 0.295 | 0.189 | 7.25E-11 | cDC2.Mgl2 | Eid3 |
| Bckdk | 3.79E-15 | 0.27692989 | 0.309 | 0.218 | 7.32E-11 | cDC2.Mgl2 | Bckdk |
| Aldh16a1 | 4.64E-15 | 0.34702162 | 0.28 | 0.186 | 8.98E-11 | cDC2.Mgl2 | Aldh16a1 |
| Atg71 | 4.76E-15 | 0.25123882 | 0.307 | 0.197 | 9.19E-11 | cDC2.Mgl2 | Atg7 |
| Hspbp1 | 7.59E-15 | 0.43085264 | 0.282 | 0.186 | 1.47E-10 | cDC2.Mgl2 | Hspbp1 |
| Tcf20 | 8.30E-15 | 0.27019053 | 0.4 | 0.286 | 1.60E-10 | cDC2.Mgl2 | Tcf20 |
| Clec4d | 9.89E-15 | 0.58079804 | 0.301 | 0.145 | 1.91E-10 | cDC2.Mgl2 | Clec4d |
| Acer31 | 1.55E-14 | 0.26272322 | 0.328 | 0.215 | 3.00E-10 | cDC2.Mgl2 | Acer3 |
| Gatm | 1.60E-14 | 0.26095583 | 0.468 | 0.295 | 3.10E-10 | cDC2.Mgl2 | Gatm |
| Epop | 1.77E-14 | 0.41437972 | 0.266 | 0.146 | 3.42E-10 | cDC2.Mgl2 | Epop |
| Exoc61 | 2.08E-14 | 0.30528194 | 0.303 | 0.21 | 4.03E-10 | cDC2.Mgl2 | Exoc6 |
| Eif4a3 | 2.19E-14 | 0.41493565 | 0.314 | 0.209 | 4.23E-10 | cDC2.Mgl2 | Eif4a3 |
| Amdhd2 | 2.78E-14 | 0.47280802 | 0.345 | 0.217 | 5.38E-10 | cDC2.Mgl2 | Amdhd2 |
| Bcap29 | 2.92E-14 | 0.36496448 | 0.279 | 0.186 | 5.64E-10 | cDC2.Mgl2 | Bcap29 |
| Sh3glb2 | 3.28E-14 | 0.50385267 | 0.273 | 0.167 | 6.34E-10 | cDC2.Mgl2 | Sh3glb2 |
| Kctd17 | 3.29E-14 | 0.37050128 | 0.271 | 0.184 | 6.36E-10 | cDC2.Mgl2 | Kctd17 |
| Bcl2a1a | 3.40E-14 | 0.3630917 | 0.264 | 0.187 | 6.57E-10 | cDC2.Mgl2 | Bcl2a1a |
| Fam76a | 3.40E-14 | 0.32015195 | 0.298 | 0.201 | 6.57E-10 | cDC2.Mgl2 | Fam76a |
| Chd9 | 3.79E-14 | 0.34645199 | 0.34 | 0.22 | 7.33E-10 | cDC2.Mgl2 | Chd9 |
| Atraid | 5.13E-14 | 0.40461239 | 0.287 | 0.185 | 9.92E-10 | cDC2.Mgl2 | Atraid |
| Foxred1 | 5.32E-14 | 0.32249133 | 0.252 | 0.158 | 1.03E-09 | cDC2.Mgl2 | Foxred1 |
| Zfp652 | 5.41E-14 | 0.38721682 | 0.353 | 0.222 | 1.05E-09 | cDC2.Mgl2 | Zfp652 |
| Nop2 | 6.73E-14 | 0.34407853 | 0.277 | 0.202 | 1.30E-09 | cDC2.Mgl2 | Nop2 |
| Pdrg1 | 7.62E-14 | 0.27129971 | 0.326 | 0.233 | 1.47E-09 | cDC2.Mgl2 | Pdrg1 |
| Foxo3 | 7.84E-14 | 0.35666679 | 0.305 | 0.191 | 1.52E-09 | cDC2.Mgl2 | Foxo3 |
| Adsl | 1.50E-13 | 0.25510191 | 0.282 | 0.186 | 2.90E-09 | cDC2.Mgl2 | Adsl |
| Plekhg3 | 2.07E-13 | 0.3721478 | 0.283 | 0.177 | 4.01E-09 | cDC2.Mgl2 | Plekhg3 |
| Thada | 2.63E-13 | 0.27851142 | 0.263 | 0.169 | 5.09E-09 | cDC2.Mgl2 | Thada |
| Slc25a17 | 3.69E-13 | 0.3143244 | 0.317 | 0.231 | 7.13E-09 | cDC2.Mgl2 | Slc25a17 |
| Hpcal1 | 3.82E-13 | 0.3044895 | 0.307 | 0.21 | 7.39E-09 | cDC2.Mgl2 | Hpcal1 |
| Pdk3 | 3.85E-13 | 0.5103609 | 0.274 | 0.183 | 7.44E-09 | cDC2.Mgl2 | Pdk3 |
| Rasgrp1 | 4.64E-13 | 0.48116563 | 0.256 | 0.148 | 8.98E-09 | cDC2.Mgl2 | Rasgrp1 |
| Shmt2 | 4.69E-13 | 0.37641008 | 0.35 | 0.248 | 9.07E-09 | cDC2.Mgl2 | Shmt2 |
| Ift20 | 7.11E-13 | 0.29651132 | 0.306 | 0.224 | 1.37E-08 | cDC2.Mgl2 | Ift20 |
| Zdhhc8 | 7.38E-13 | 0.30284891 | 0.275 | 0.183 | 1.43E-08 | cDC2.Mgl2 | Zdhhc8 |
| Imp4 | 8.01E-13 | 0.250119 | 0.319 | 0.228 | 1.55E-08 | cDC2.Mgl2 | Imp4 |
| Gtf3a | 8.16E-13 | 0.38958697 | 0.285 | 0.177 | 1.58E-08 | cDC2.Mgl2 | Gtf3a |
| Mtfr1l | 1.04E-12 | 0.45428802 | 0.267 | 0.168 | 2.01E-08 | cDC2.Mgl2 | Mtfr1l |
| Tmem268 | 1.05E-12 | 0.27911071 | 0.268 | 0.173 | 2.02E-08 | cDC2.Mgl2 | Tmem268 |
| Galk2 | 1.07E-12 | 0.27446721 | 0.281 | 0.193 | 2.06E-08 | cDC2.Mgl2 | Galk2 |
| G6pc3 | 1.43E-12 | 0.36617871 | 0.278 | 0.195 | 2.76E-08 | cDC2.Mgl2 | G6pc3 |
| Gnl3l | 2.10E-12 | 0.56039367 | 0.267 | 0.173 | 4.06E-08 | cDC2.Mgl2 | Gnl3l |
| Fam173a | 2.26E-12 | 0.30679201 | 0.33 | 0.225 | 4.38E-08 | cDC2.Mgl2 | Fam173a |
| Nop16 | 2.79E-12 | 0.34579469 | 0.275 | 0.179 | 5.39E-08 | cDC2.Mgl2 | Nop16 |
| Ube2j11 | 2.80E-12 | 0.25453379 | 0.329 | 0.236 | 5.40E-08 | cDC2.Mgl2 | Ube2j1 |
| Rrn3 | 2.95E-12 | 0.25336612 | 0.275 | 0.189 | 5.70E-08 | cDC2.Mgl2 | Rrn3 |
| Dohh | 5.43E-12 | 0.29829507 | 0.267 | 0.191 | 1.05E-07 | cDC2.Mgl2 | Dohh |
| Ccdc115 | 1.08E-11 | 0.3545734 | 0.256 | 0.173 | 2.08E-07 | cDC2.Mgl2 | Ccdc115 |
| Hip11 | 1.21E-11 | 0.32292839 | 0.381 | 0.255 | 2.34E-07 | cDC2.Mgl2 | Hip1 |
| Glrx5 | 1.91E-11 | 0.33576327 | 0.258 | 0.165 | 3.68E-07 | cDC2.Mgl2 | Glrx5 |
| Ten1 | 5.64E-11 | 0.370707 | 0.279 | 0.185 | 1.09E-06 | cDC2.Mgl2 | Ten1 |
| Gdap2 | 1.25E-10 | 0.33673533 | 0.271 | 0.183 | 2.41E-06 | cDC2.Mgl2 | Gdap2 |
| Samd8 | 1.28E-10 | 0.28731656 | 0.267 | 0.178 | 2.47E-06 | cDC2.Mgl2 | Samd8 |
| Ndufa5 | 1.41E-10 | 0.2898812 | 0.263 | 0.177 | 2.72E-06 | cDC2.Mgl2 | Ndufa5 |
| Ipo41 | 1.58E-10 | 0.33652974 | 0.268 | 0.182 | 3.06E-06 | cDC2.Mgl2 | Ipo4 |
| Pold4 | 1.66E-10 | 0.27335466 | 0.471 | 0.366 | 3.21E-06 | cDC2.Mgl2 | Pold4 |
| Papss1 | 2.46E-10 | 0.36307655 | 0.31 | 0.217 | 4.75E-06 | cDC2.Mgl2 | Papss1 |
| Syvn1 | 2.57E-10 | 0.32900069 | 0.281 | 0.202 | 4.97E-06 | cDC2.Mgl2 | Syvn1 |
| Bad | 2.74E-10 | 0.31603646 | 0.281 | 0.19 | 5.30E-06 | cDC2.Mgl2 | Bad |
| Ccl6 | 4.52E-10 | 0.63315167 | 0.321 | 0.148 | 8.74E-06 | cDC2.Mgl2 | Ccl6 |
| Unc45a | 5.35E-10 | 0.31962672 | 0.272 | 0.201 | 1.03E-05 | cDC2.Mgl2 | Unc45a |
| Gde11 | 1.35E-09 | 0.36847734 | 0.287 | 0.189 | 2.62E-05 | cDC2.Mgl2 | Gde1 |
| Wsb2 | 2.03E-09 | 0.31493776 | 0.269 | 0.182 | 3.92E-05 | cDC2.Mgl2 | Wsb2 |
| Apobr | 2.51E-09 | 0.29372723 | 0.281 | 0.188 | 4.85E-05 | cDC2.Mgl2 | Apobr |
| Hectd3 | 3.45E-09 | 0.264601 | 0.312 | 0.224 | 6.68E-05 | cDC2.Mgl2 | Hectd3 |
| Guk1 | 4.06E-09 | 0.25987477 | 0.306 | 0.224 | 7.86E-05 | cDC2.Mgl2 | Guk1 |
| Eml2 | 3.11E-08 | 0.32568665 | 0.252 | 0.179 | 0.00060182 | cDC2.Mgl2 | Eml2 |
| Cisd3 | 1.35E-07 | 0.50714714 | 0.256 | 0.186 | 0.00260332 | cDC2.Mgl2 | Cisd3 |
| Clec9a | 0 | 4.53802834 | 0.831 | 0.061 | 0 | cDC1.Xcr1 | Clec9a |
| Ifi205 | 0 | 3.95431258 | 0.898 | 0.226 | 0 | cDC1.Xcr1 | Ifi205 |
| Rab7b | 0 | 5.14083341 | 0.658 | 0.033 | 0 | cDC1.Xcr1 | Rab7b |
| Tlr3 | 0 | 6.3393755 | 0.635 | 0.032 | 0 | cDC1.Xcr1 | Tlr3 |
| Mpeg1 | 0 | 2.51444638 | 0.953 | 0.358 | 0 | cDC1.Xcr1 | Mpeg1 |
| Agpat3 | 0 | 4.59089647 | 0.632 | 0.044 | 0 | cDC1.Xcr1 | Agpat3 |
| Xcr1 | 0 | 7.63282073 | 0.574 | 0.006 | 0 | cDC1.Xcr1 | Xcr1 |
| Itgae1 | 0 | 2.59842281 | 0.846 | 0.282 | 0 | cDC1.Xcr1 | Itgae |
| Ppt1 | 0 | 3.88725835 | 0.882 | 0.337 | 0 | cDC1.Xcr1 | Ppt1 |
| Gpr141b | 0 | 6.17777249 | 0.55 | 0.014 | 0 | cDC1.Xcr1 | Gpr141b |
| Slc8b1 | 0 | 2.9084631 | 0.771 | 0.267 | 0 | cDC1.Xcr1 | Slc8b1 |
| Btla | 0 | 2.54189963 | 0.761 | 0.272 | 0 | cDC1.Xcr1 | Btla |
| Arsb | 0 | 3.21796147 | 0.656 | 0.174 | 0 | cDC1.Xcr1 | Arsb |
| Sult1a1 | 0 | 4.63109403 | 0.538 | 0.064 | 0 | cDC1.Xcr1 | Sult1a1 |
| Unc119b1 | 0 | 2.62236143 | 0.675 | 0.206 | 0 | cDC1.Xcr1 | Unc119b |
| Naaa | 0 | 3.58841905 | 0.951 | 0.482 | 0 | cDC1.Xcr1 | Naaa |
| Camk1d | 0 | 2.5858465 | 0.782 | 0.324 | 0 | cDC1.Xcr1 | Camk1d |
| Slco3a1 | 0 | 2.51410315 | 0.656 | 0.198 | 0 | cDC1.Xcr1 | Slco3a1 |
| Mycl | 0 | 2.93776073 | 0.609 | 0.151 | 0 | cDC1.Xcr1 | Mycl |
| Ckb1 | 0 | 2.31140878 | 0.755 | 0.301 | 0 | cDC1.Xcr1 | Ckb |
| Asb21 | 0 | 2.85364088 | 0.674 | 0.222 | 0 | cDC1.Xcr1 | Asb2 |
| Dnase1l3 | 0 | 4.14964191 | 0.517 | 0.066 | 0 | cDC1.Xcr1 | Dnase1l3 |
| Txndc15 | 0 | 2.40127406 | 0.697 | 0.247 | 0 | cDC1.Xcr1 | Txndc15 |
| Irf81 | 0 | 2.22411387 | 0.969 | 0.53 | 0 | cDC1.Xcr1 | Irf8 |
| Plet11 | 0 | 1.5607364 | 0.891 | 0.453 | 0 | cDC1.Xcr1 | Plet1 |
| Mctp1 | 0 | 3.1358004 | 0.553 | 0.117 | 0 | cDC1.Xcr1 | Mctp1 |
| Wdfy41 | 0 | 2.2748599 | 0.952 | 0.517 | 0 | cDC1.Xcr1 | Wdfy4 |
| St3gal5 | 0 | 3.47141953 | 0.518 | 0.088 | 0 | cDC1.Xcr1 | St3gal5 |
| Cadm1 | 0 | 4.33621313 | 0.466 | 0.037 | 0 | cDC1.Xcr1 | Cadm1 |
| 3-Sep | 0 | 3.31784076 | 0.518 | 0.092 | 0 | cDC1.Xcr1 | 3-Sep |
| Ttc39a | 0 | 3.22730824 | 0.447 | 0.034 | 0 | cDC1.Xcr1 | Ttc39a |
| Fgd2 | 0 | 1.99539498 | 0.815 | 0.402 | 0 | cDC1.Xcr1 | Fgd2 |
| Fnip2 | 0 | 2.77377956 | 0.532 | 0.12 | 0 | cDC1.Xcr1 | Fnip2 |
| Pianp | 0 | 5.29271801 | 0.428 | 0.016 | 0 | cDC1.Xcr1 | Pianp |
| Dbn11 | 0 | 1.88724055 | 0.703 | 0.294 | 0 | cDC1.Xcr1 | Dbn1 |
| Gatm1 | 0 | 2.01100631 | 0.67 | 0.262 | 0 | cDC1.Xcr1 | Gatm |
| Tlr12 | 0 | 3.80868875 | 0.441 | 0.033 | 0 | cDC1.Xcr1 | Tlr12 |
| Sdad1 | 0 | 2.35190372 | 0.591 | 0.186 | 0 | cDC1.Xcr1 | Sdad1 |
| Acvrl1 | 0 | 2.1134371 | 0.607 | 0.202 | 0 | cDC1.Xcr1 | Acvrl1 |

| | | | | | | | |
|---|---|---|---|---|---|---|---|
| Cyfip2 | 0 | 1.86531008 | 0.675 | 0.277 | 0 | cDC1.Xcr1 | Cyfip2 |
| 6-Sep | 0 | 1.93135524 | 0.725 | 0.332 | 0 | cDC1.Xcr1 | 6-Sep |
| Rab32 | 0 | 2.02845627 | 0.695 | 0.305 | 0 | cDC1.Xcr1 | Rab32 |
| Cyp26b1 | 0 | 7.0336798 | 0.387 | 0.011 | 0 | cDC1.Xcr1 | Cyp26b1 |
| Aif1 | 0 | 1.74430505 | 0.841 | 0.466 | 0 | cDC1.Xcr1 | Aif1 |
| Hepacam21 | 0 | 2.19412216 | 0.571 | 0.209 | 0 | cDC1.Xcr1 | Hepacam2 |
| 1-Mar | 0 | 2.06706521 | 0.599 | 0.239 | 0 | cDC1.Xcr1 | 1-Mar |
| Havcr21 | 0 | 1.57332754 | 0.706 | 0.35 | 0 | cDC1.Xcr1 | Havcr2 |
| Cd36 | 0 | 2.94627614 | 0.404 | 0.049 | 0 | cDC1.Xcr1 | Cd36 |
| Cd24a1 | 0 | 1.15684733 | 0.855 | 0.5 | 0 | cDC1.Xcr1 | Cd24a |
| Jtb | 0 | 1.74585086 | 0.663 | 0.324 | 0 | cDC1.Xcr1 | Jtb |
| Ncoa71 | 0 | 1.33021442 | 0.65 | 0.313 | 0 | cDC1.Xcr1 | Ncoa7 |
| Arhgap181 | 0 | 1.75663233 | 0.645 | 0.315 | 0 | cDC1.Xcr1 | Arhgap18 |
| Anxa61 | 0 | 1.74486188 | 0.849 | 0.529 | 0 | cDC1.Xcr1 | Anxa6 |
| Mlec1 | 0 | 1.65568979 | 0.809 | 0.49 | 0 | cDC1.Xcr1 | Mlec |
| Trim351 | 0 | 1.15370545 | 0.767 | 0.45 | 0 | cDC1.Xcr1 | Trim35 |
| Atp6v0a2 | 0 | 1.69119658 | 0.67 | 0.355 | 0 | cDC1.Xcr1 | Atp6v0a2 |
| Bcl6 | 0 | 1.48898927 | 0.728 | 0.422 | 0 | cDC1.Xcr1 | Bcl6 |
| Naga | 0 | 1.67254168 | 0.881 | 0.586 | 0 | cDC1.Xcr1 | Naga |
| Fchsd2 | 0 | 1.23531735 | 0.708 | 0.421 | 0 | cDC1.Xcr1 | Fchsd2 |
| Gpr1711 | 0 | 1.37617277 | 0.737 | 0.455 | 0 | cDC1.Xcr1 | Gpr171 |
| Dusp4 | 0 | 3.16211908 | 0.343 | 0.062 | 0 | cDC1.Xcr1 | Dusp4 |
| Pik3ap11 | 0 | 1.16439724 | 0.848 | 0.567 | 0 | cDC1.Xcr1 | Pik3ap1 |
| Shtn1 | 0 | 1.30099532 | 0.748 | 0.474 | 0 | cDC1.Xcr1 | Shtn1 |
| Ciita2 | 0 | 1.0617811 | 0.859 | 0.589 | 0 | cDC1.Xcr1 | Ciita |
| Ppm1m1 | 0 | 1.2883093 | 0.841 | 0.573 | 0 | cDC1.Xcr1 | Ppm1m |
| Anpep1 | 0 | 1.23050347 | 0.748 | 0.491 | 0 | cDC1.Xcr1 | Anpep |
| Plekho2 | 0 | 1.29368874 | 0.864 | 0.61 | 0 | cDC1.Xcr1 | Plekho2 |
| Plbd11 | 0 | 1.65939277 | 0.94 | 0.692 | 0 | cDC1.Xcr1 | Plbd1 |
| Man2b11 | 0 | 1.13966518 | 0.841 | 0.599 | 0 | cDC1.Xcr1 | Man2b1 |
| Ece1 | 0 | 1.14589608 | 0.848 | 0.609 | 0 | cDC1.Xcr1 | Ece1 |
| Rgs10 | 0 | 1.23143151 | 0.782 | 0.544 | 0 | cDC1.Xcr1 | Rgs10 |
| Ly6e2 | 0 | 0.48393128 | 0.898 | 0.661 | 0 | cDC1.Xcr1 | Ly6e |
| 9-Sep | 0 | 1.12467485 | 0.824 | 0.592 | 0 | cDC1.Xcr1 | 9-Sep |
| Qpct1 | 0 | 1.50950567 | 0.485 | 0.253 | 0 | cDC1.Xcr1 | Qpct |
| Gm2a1 | 0 | 1.27008443 | 0.989 | 0.775 | 0 | cDC1.Xcr1 | Gm2a |
| Lgals3 | 0 | 1.52657453 | 0.946 | 0.732 | 0 | cDC1.Xcr1 | Lgals3 |
| Odc11 | 0 | 0.90786106 | 0.81 | 0.598 | 0 | cDC1.Xcr1 | Odc1 |
| Hhex | 0 | 1.42451759 | 0.664 | 0.454 | 0 | cDC1.Xcr1 | Hhex |
| Rab43 | 0 | 1.75540395 | 0.939 | 0.737 | 0 | cDC1.Xcr1 | Rab43 |
| Psmb9 | 0 | 1.67175258 | 0.919 | 0.719 | 0 | cDC1.Xcr1 | Psmb9 |
| Psap | 0 | 2.09795925 | 0.962 | 0.771 | 0 | cDC1.Xcr1 | Psap |
| Hspa1b | 0 | 0.665421 | 0.528 | 0.339 | 0 | cDC1.Xcr1 | Hspa1b |
| Tap11 | 0 | 1.1051122 | 0.854 | 0.675 | 0 | cDC1.Xcr1 | Tap1 |
| Tap21 | 0 | 0.97893106 | 0.832 | 0.653 | 0 | cDC1.Xcr1 | Tap2 |
| Inpp5d | 0 | 1.06425428 | 0.835 | 0.657 | 0 | cDC1.Xcr1 | Inpp5d |
| Lims11 | 0 | 1.18391999 | 0.773 | 0.597 | 0 | cDC1.Xcr1 | Lims1 |
| Rtcb | 0 | 1.13190521 | 0.786 | 0.616 | 0 | cDC1.Xcr1 | Rtcb |
| Stmn11 | 0 | 2.0326074 | 0.284 | 0.114 | 0 | cDC1.Xcr1 | Stmn1 |
| H2afx | 0 | 1.66208288 | 0.37 | 0.204 | 0 | cDC1.Xcr1 | H2afx |
| H2-Ab12 | 0 | 0.4783811 | 0.993 | 0.842 | 0 | cDC1.Xcr1 | H2-Ab1 |
| Hist1h1e | 0 | 1.89639741 | 0.281 | 0.134 | 0 | cDC1.Xcr1 | Hist1h1e |
| Fnbp1 | 0 | 1.14227458 | 0.94 | 0.795 | 0 | cDC1.Xcr1 | Fnbp1 |
| Taldo11 | 0 | 1.03772555 | 0.888 | 0.751 | 0 | cDC1.Xcr1 | Taldo1 |
| Jund1 | 0 | 0.76675994 | 0.922 | 0.793 | 0 | cDC1.Xcr1 | Jund |
| Actr3 | 0 | 0.93970665 | 0.781 | 0.657 | 0 | cDC1.Xcr1 | Actr3 |
| Tmsb10 | 0 | 1.40107159 | 0.972 | 0.856 | 0 | cDC1.Xcr1 | Tmsb10 |
| Eef1b2 | 0 | 0.95846598 | 0.866 | 0.754 | 0 | cDC1.Xcr1 | Eef1b2 |
| Kctd12 | 0 | 0.98493938 | 0.876 | 0.764 | 0 | cDC1.Xcr1 | Kctd12 |
| Plekho1 | 0 | 0.84236037 | 0.854 | 0.744 | 0 | cDC1.Xcr1 | Plekho1 |
| Cd521 | 0 | 0.69960825 | 0.947 | 0.844 | 0 | cDC1.Xcr1 | Cd52 |
| Itm2c1 | 0 | 0.64021383 | 0.897 | 0.808 | 0 | cDC1.Xcr1 | Itm2c |
| Zyx | 0 | 0.76588169 | 0.898 | 0.816 | 0 | cDC1.Xcr1 | Zyx |
| Irf5 | 0 | 0.72976819 | 0.926 | 0.848 | 0 | cDC1.Xcr1 | Irf5 |
| Ptms1 | 0 | 0.90844793 | 0.963 | 0.885 | 0 | cDC1.Xcr1 | Ptms |
| Fos1 | 0 | 0.44938421 | 0.749 | 0.678 | 0 | cDC1.Xcr1 | Fos |
| Psme1 | 0 | 0.52693664 | 0.904 | 0.845 | 0 | cDC1.Xcr1 | Psme1 |
| Ucp21 | 0 | 0.57292862 | 0.971 | 0.917 | 0 | cDC1.Xcr1 | Ucp2 |
| S100a11 | 0 | 0.50542664 | 0.963 | 0.912 | 0 | cDC1.Xcr1 | S100a11 |
| Cd742 | 0 | 0.30350011 | 0.998 | 0.967 | 0 | cDC1.Xcr1 | Cd74 |
| Actg11 | 0 | 0.36815083 | 0.957 | 0.936 | 0 | cDC1.Xcr1 | Actg1 |
| Cst31 | 0 | 1.07555431 | 0.999 | 0.984 | 0 | cDC1.Xcr1 | Cst3 |
| Tmsb4x1 | 0 | 0.48380194 | 1 | 0.996 | 0 | cDC1.Xcr1 | Tmsb4x |
| Arap1 | 9.35E-306 | 1.88210853 | 0.608 | 0.222 | 1.81E-301 | cDC1.Xcr1 | Arap1 |
| Zfp3661 | 6.15E-303 | 1.29732605 | 0.74 | 0.432 | 1.19E-298 | cDC1.Xcr1 | Zfp366 |
| Evl1 | 2.70E-298 | 1.93546141 | 0.674 | 0.308 | 5.21E-294 | cDC1.Xcr1 | Evl |
| Gpr651 | 7.67E-295 | 1.26051701 | 0.512 | 0.245 | 1.48E-290 | cDC1.Xcr1 | Gpr65 |
| BC028528 | 2.82E-293 | 2.37517129 | 0.51 | 0.144 | 5.46E-289 | cDC1.Xcr1 | BC028528 |
| Zfp710 | 1.40E-290 | 1.85319895 | 0.645 | 0.304 | 2.70E-286 | cDC1.Xcr1 | Zfp710 |
| Got2 | 1.14E-286 | 1.0648413 | 0.741 | 0.548 | 2.21E-282 | cDC1.Xcr1 | Got2 |
| Sdc3 | 2.21E-286 | 1.15139223 | 0.788 | 0.504 | 4.27E-282 | cDC1.Xcr1 | Sdc3 |
| Dbnl1 | 2.09E-285 | 0.86476833 | 0.88 | 0.752 | 4.04E-281 | cDC1.Xcr1 | Dbnl |
| Sla2 | 1.35E-282 | 1.16028951 | 0.782 | 0.493 | 2.61E-278 | cDC1.Xcr1 | Sla |
| Sdf2l11 | 5.75E-279 | 1.10428559 | 0.686 | 0.456 | 1.11E-274 | cDC1.Xcr1 | Sdf2l1 |
| Dynlt1f | 1.24E-277 | 2.23316308 | 0.553 | 0.209 | 2.40E-273 | cDC1.Xcr1 | Dynlt1f |
| Lcp1 | 1.40E-277 | 0.39638935 | 0.96 | 0.926 | 2.70E-273 | cDC1.Xcr1 | Lcp1 |
| Coro7 | 2.93E-276 | 1.48420963 | 0.689 | 0.357 | 5.66E-272 | cDC1.Xcr1 | Coro7 |
| Hic11 | 1.38E-275 | 1.50050345 | 0.555 | 0.252 | 2.67E-271 | cDC1.Xcr1 | Hic1 |
| Psmb81 | 1.34E-273 | 0.51363815 | 0.876 | 0.832 | 2.60E-269 | cDC1.Xcr1 | Psmb8 |
| Fgl21 | 4.13E-273 | 0.858518 | 0.825 | 0.579 | 7.99E-269 | cDC1.Xcr1 | Fgl2 |
| Hspa8 | 2.17E-268 | 0.44040089 | 0.923 | 0.896 | 4.19E-264 | cDC1.Xcr1 | Hspa8 |
| Gcsam | 1.52E-267 | 5.69294785 | 0.485 | 0.012 | 2.94E-263 | cDC1.Xcr1 | Gcsam |
| Gng102 | 1.74E-266 | 0.93909169 | 0.864 | 0.651 | 3.37E-262 | cDC1.Xcr1 | Gng10 |
| Hspa1a1 | 1.99E-266 | 0.60795227 | 0.507 | 0.334 | 3.84E-262 | cDC1.Xcr1 | Hspa1a |
| BC035044 | 5.05E-263 | 2.25685973 | 0.521 | 0.14 | 9.77E-259 | cDC1.Xcr1 | BC035044 |
| Snx22 | 1.44E-262 | 5.78696139 | 0.515 | 0.013 | 2.79E-258 | cDC1.Xcr1 | Snx22 |
| Tbc1d91 | 6.69E-262 | 1.18363838 | 0.766 | 0.542 | 1.29E-257 | cDC1.Xcr1 | Tbc1d9 |
| Atpif1 | 8.60E-262 | 1.39050442 | 0.725 | 0.444 | 1.66E-257 | cDC1.Xcr1 | Atpif1 |
| Nr4a12 | 2.21E-258 | 0.29309839 | 0.729 | 0.705 | 4.27E-254 | cDC1.Xcr1 | Nr4a1 |
| Tubb51 | 8.24E-258 | 0.44399686 | 0.7 | 0.678 | 1.59E-253 | cDC1.Xcr1 | Tubb5 |
| Unc93b1 | 3.12E-257 | 0.77769923 | 0.855 | 0.719 | 6.04E-253 | cDC1.Xcr1 | Unc93b1 |
| Pttg1ip | 2.87E-256 | 1.53967336 | 0.614 | 0.354 | 5.54E-252 | cDC1.Xcr1 | Pttg1ip |
| Ifitm21 | 5.05E-254 | 0.48564721 | 0.903 | 0.809 | 9.76E-250 | cDC1.Xcr1 | Ifitm2 |
| Sh3bp1 | 1.43E-253 | 0.9775345 | 0.819 | 0.619 | 2.77E-249 | cDC1.Xcr1 | Sh3bp1 |
| Ppp1r111 | 2.43E-251 | 1.36043141 | 0.673 | 0.393 | 4.69E-247 | cDC1.Xcr1 | Ppp1r11 |
| Pomp | 2.08E-247 | 1.04620192 | 0.794 | 0.604 | 4.01E-243 | cDC1.Xcr1 | Pomp |
| Ap1s31 | 4.00E-244 | 1.75468963 | 0.612 | 0.289 | 7.74E-240 | cDC1.Xcr1 | Ap1s3 |
| Actn12 | 4.83E-244 | 0.4683245 | 0.778 | 0.754 | 9.35E-240 | cDC1.Xcr1 | Actn1 |
| Atox1 | 9.81E-244 | 0.59439925 | 0.969 | 0.88 | 1.90E-239 | cDC1.Xcr1 | Atox1 |
| Anxa12 | 4.13E-242 | 1.21657482 | 0.662 | 0.371 | 7.99E-238 | cDC1.Xcr1 | Anxa1 |
| Cdk14 | 2.52E-241 | 2.78748644 | 0.453 | 0.103 | 4.87E-237 | cDC1.Xcr1 | Cdk14 |
| Rab11fip4 | 1.99E-240 | 3.33695747 | 0.438 | 0.057 | 3.85E-236 | cDC1.Xcr1 | Rab11fip4 |
| Itgb22 | 6.18E-237 | 0.62217729 | 0.868 | 0.67 | 1.19E-232 | cDC1.Xcr1 | Itgb2 |
| Pfkp2 | 5.33E-235 | 0.76185071 | 0.86 | 0.684 | 1.03E-230 | cDC1.Xcr1 | Pfkp |
| Tcf19 | 1.50E-233 | 2.72570828 | 0.333 | 0.064 | 2.90E-229 | cDC1.Xcr1 | Tcf19 |
| Klf2 | 4.71E-231 | 1.41487485 | 0.376 | 0.161 | 9.11E-227 | cDC1.Xcr1 | Klf2 |
| Ptpn181 | 1.12E-230 | 1.45173665 | 0.708 | 0.376 | 2.17E-226 | cDC1.Xcr1 | Ptpn18 |
| Il16 | 4.79E-228 | 2.4024497 | 0.419 | 0.105 | 9.27E-224 | cDC1.Xcr1 | Il16 |
| Itgb71 | 7.92E-228 | 0.84583714 | 0.733 | 0.484 | 1.53E-223 | cDC1.Xcr1 | Itgb7 |
| Rasgrp41 | 9.10E-228 | 1.41586191 | 0.706 | 0.391 | 1.76E-223 | cDC1.Xcr1 | Rasgrp4 |
| Grn | 3.59E-227 | 0.55372518 | 0.866 | 0.699 | 6.93E-223 | cDC1.Xcr1 | Grn |
| Coro1a1 | 1.38E-226 | 0.29863285 | 0.941 | 0.909 | 2.67E-222 | cDC1.Xcr1 | Coro1a |
| Slc9a9 | 4.65E-223 | 2.03922132 | 0.493 | 0.171 | 8.98E-219 | cDC1.Xcr1 | Slc9a9 |
| Treml4 | 3.51E-221 | 2.53607157 | 0.332 | 0.073 | 6.79E-217 | cDC1.Xcr1 | Treml4 |
| Ank1 | 1.33E-217 | 5.57828559 | 0.477 | 0.011 | 2.57E-213 | cDC1.Xcr1 | Ank1 |
| Gusb | 2.48E-217 | 1.11051868 | 0.734 | 0.522 | 4.79E-213 | cDC1.Xcr1 | Gusb |
| Slc9a3r2 | 6.63E-214 | 2.84148784 | 0.343 | 0.058 | 1.28E-209 | cDC1.Xcr1 | Slc9a3r2 |
| Tbc1d81 | 2.52E-213 | 0.48210964 | 0.804 | 0.55 | 4.88E-209 | cDC1.Xcr1 | Tbc1d8 |
| Slc25a20 | 5.08E-213 | 1.28021894 | 0.626 | 0.421 | 9.81E-209 | cDC1.Xcr1 | Slc25a20 |
| Ifitm61 | 8.75E-212 | 1.63572075 | 0.46 | 0.216 | 1.69E-207 | cDC1.Xcr1 | Ifitm6 |
| Ldha1 | 1.22E-211 | 0.76325848 | 0.911 | 0.749 | 2.37E-207 | cDC1.Xcr1 | Ldha |
| Fam149a | 3.26E-211 | 3.48501556 | 0.38 | 0.042 | 6.30E-207 | cDC1.Xcr1 | Fam149a |
| Gpcpd1 | 1.82E-205 | 0.85935719 | 0.549 | 0.414 | 3.52E-201 | cDC1.Xcr1 | Gpcpd1 |
| Adam191 | 4.70E-204 | 1.08967401 | 0.57 | 0.228 | 9.08E-200 | cDC1.Xcr1 | Adam19 |
| Rnase6 | 3.34E-202 | 1.56695788 | 0.499 | 0.191 | 6.45E-198 | cDC1.Xcr1 | Rnase6 |
| Ndfip1 | 6.19E-202 | 0.71320439 | 0.793 | 0.694 | 1.20E-197 | cDC1.Xcr1 | Ndfip1 |
| Hfe1 | 8.89E-202 | 1.41372028 | 0.657 | 0.428 | 1.72E-197 | cDC1.Xcr1 | Hfe |
| Cyp27a1 | 7.32E-199 | 1.87249769 | 0.489 | 0.171 | 1.42E-194 | cDC1.Xcr1 | Cyp27a1 |
| Map4k11 | 1.33E-193 | 1.10632115 | 0.676 | 0.42 | 2.58E-189 | cDC1.Xcr1 | Map4k1 |
| Btg21 | 1.63E-191 | 0.50595562 | 0.832 | 0.773 | 3.14E-187 | cDC1.Xcr1 | Btg2 |
| Slamf81 | 1.14E-190 | 1.90649944 | 0.531 | 0.216 | 2.20E-186 | cDC1.Xcr1 | Slamf8 |
| Fmnl21 | 5.05E-187 | 1.16329632 | 0.559 | 0.283 | 9.76E-183 | cDC1.Xcr1 | Fmnl2 |
| Unc1191 | 5.55E-187 | 1.89002221 | 0.504 | 0.194 | 1.07E-182 | cDC1.Xcr1 | Unc119 |
| Ppt2 | 3.38E-186 | 2.42753028 | 0.357 | 0.092 | 6.54E-182 | cDC1.Xcr1 | Ppt2 |
| Wtip | 3.79E-186 | 2.33889687 | 0.388 | 0.084 | 7.33E-182 | cDC1.Xcr1 | Wtip |
| Preb | 1.08E-185 | 1.18137472 | 0.711 | 0.464 | 2.09E-181 | cDC1.Xcr1 | Preb |
| Plpp1 | 9.15E-184 | 4.43463619 | 0.314 | 0.015 | 1.77E-179 | cDC1.Xcr1 | Plpp1 |
| Gng12 | 4.52E-181 | 2.02341316 | 0.403 | 0.103 | 8.74E-177 | cDC1.Xcr1 | Gng12 |
| Rasgrp3 | 5.16E-181 | 3.02854102 | 0.326 | 0.025 | 9.98E-177 | cDC1.Xcr1 | Rasgrp3 |
| Tnfrsf21 | 1.34E-180 | 2.01527553 | 0.474 | 0.159 | 2.60E-176 | cDC1.Xcr1 | Tnfrsf21 |
| Mkrn1 | 1.67E-180 | 1.49512931 | 0.57 | 0.291 | 3.23E-176 | cDC1.Xcr1 | Mkrn1 |
| Commd8 | 7.48E-180 | 1.09047784 | 0.668 | 0.441 | 1.45E-175 | cDC1.Xcr1 | Commd8 |
| Dynll21 | 1.08E-178 | 0.66763892 | 0.691 | 0.5 | 2.08E-174 | cDC1.Xcr1 | Dynll2 |
| Rpn11 | 4.02E-178 | 0.84421458 | 0.658 | 0.456 | 7.78E-174 | cDC1.Xcr1 | Rpn1 |
| Hps31 | 1.63E-177 | 1.78300981 | 0.56 | 0.243 | 3.15E-173 | cDC1.Xcr1 | Hps3 |
| mt-Co21 | 4.16E-177 | 0.26803583 | 0.85 | 0.808 | 8.04E-173 | cDC1.Xcr1 | mt-Co2 |
| Tlr11 | 6.31E-176 | 5.05095742 | 0.336 | 0.012 | 1.22E-171 | cDC1.Xcr1 | Tlr11 |
| H2afy1 | 7.19E-174 | 0.46044169 | 0.909 | 0.855 | 1.39E-169 | cDC1.Xcr1 | H2afy |
| Pdia3 | 1.12E-173 | 0.89321679 | 0.702 | 0.538 | 2.16E-169 | cDC1.Xcr1 | Pdia3 |
| Rbpj | 7.55E-173 | 0.81821318 | 0.697 | 0.539 | 1.46E-168 | cDC1.Xcr1 | Rbpj |
| Ptpn221 | 1.36E-171 | 2.20867006 | 0.412 | 0.128 | 2.63E-167 | cDC1.Xcr1 | Ptpn22 |
| Flt1 | 3.87E-170 | 3.75383519 | 0.301 | 0.027 | 7.49E-166 | cDC1.Xcr1 | Flt1 |
| Mdh1 | 2.78E-169 | 1.04745353 | 0.659 | 0.482 | 5.37E-165 | cDC1.Xcr1 | Mdh1 |
| Zdhhc23 | 5.96E-169 | 2.38365695 | 0.357 | 0.077 | 1.15E-164 | cDC1.Xcr1 | Zdhhc23 |
| Nup2101 | 4.40E-168 | 0.88307171 | 0.728 | 0.468 | 8.51E-164 | cDC1.Xcr1 | Nup210 |
| Esyt2 | 2.07E-167 | 0.89263906 | 0.674 | 0.475 | 4.01E-163 | cDC1.Xcr1 | Esyt2 |
| Eefsec | 1.93E-166 | 2.39978656 | 0.346 | 0.093 | 3.72E-162 | cDC1.Xcr1 | Eefsec |
| Map2k6 | 1.16E-165 | 2.71644975 | 0.325 | 0.069 | 2.25E-161 | cDC1.Xcr1 | Map2k6 |
| H2-Ob1 | 1.47E-165 | 1.41141585 | 0.603 | 0.276 | 2.85E-161 | cDC1.Xcr1 | H2-Ob |
| Myo9a | 3.59E-165 | 1.78439367 | 0.496 | 0.19 | 6.94E-161 | cDC1.Xcr1 | Myo9a |
| Cldnd11 | 1.18E-164 | 0.99721722 | 0.403 | 0.399 | 2.29E-160 | cDC1.Xcr1 | Cldnd1 |
| Pbx1 | 2.29E-162 | 2.62977391 | 0.377 | 0.064 | 4.43E-158 | cDC1.Xcr1 | Pbx1 |
| Slc46a3 | 7.90E-162 | 1.74956891 | 0.438 | 0.143 | 1.53E-157 | cDC1.Xcr1 | Slc46a3 |
| Tbca | 5.14E-160 | 0.92061296 | 0.74 | 0.548 | 9.93E-156 | cDC1.Xcr1 | Tbca |
| Baz1a | 2.43E-155 | 0.76924716 | 0.694 | 0.563 | 4.69E-151 | cDC1.Xcr1 | Baz1a |
| Gpr41 | 3.84E-155 | 2.0679069 | 0.451 | 0.119 | 7.42E-151 | cDC1.Xcr1 | Gpr4 |
| Tmpo | 2.79E-154 | 1.30255929 | 0.398 | 0.203 | 5.38E-150 | cDC1.Xcr1 | Tmpo |
| Batf31 | 2.37E-153 | 0.67128608 | 0.746 | 0.441 | 4.57E-149 | cDC1.Xcr1 | Batf3 |
| Selenos | 2.75E-153 | 1.28628367 | 0.515 | 0.272 | 5.31E-149 | cDC1.Xcr1 | Selenos |
| Ap1b1 | 1.13E-152 | 1.31593029 | 0.678 | 0.426 | 2.18E-148 | cDC1.Xcr1 | Ap1b1 |
| Sema4f | 1.37E-150 | 4.73846433 | 0.311 | 0.012 | 2.65E-146 | cDC1.Xcr1 | Sema4f |
| Rab13 | 4.85E-147 | 2.00521953 | 0.315 | 0.07 | 9.37E-143 | cDC1.Xcr1 | Rab13 |
| Dapk2 | 1.08E-146 | 5.76940302 | 0.306 | 0.007 | 2.09E-142 | cDC1.Xcr1 | Dapk2 |
| Jaml2 | 2.61E-146 | 0.75169652 | 0.778 | 0.546 | 5.05E-142 | cDC1.Xcr1 | Jaml |
| Gpr33 | 1.53E-145 | 5.58762028 | 0.303 | 0.008 | 2.95E-141 | cDC1.Xcr1 | Gpr33 |
| Hcls12 | 2.26E-145 | 0.49764914 | 0.908 | 0.775 | 4.36E-141 | cDC1.Xcr1 | Hcls1 |
| Atp8a11 | 1.41E-144 | 1.36704745 | 0.652 | 0.354 | 2.73E-140 | cDC1.Xcr1 | Atp8a1 |
| Wdr86 | 2.46E-144 | 2.24277181 | 0.338 | 0.078 | 4.76E-140 | cDC1.Xcr1 | Wdr86 |
| Dek | 1.38E-142 | 0.63576011 | 0.65 | 0.535 | 2.67E-138 | cDC1.Xcr1 | Dek |
| Mbnl11 | 8.02E-142 | 0.99027394 | 0.703 | 0.486 | 1.55E-137 | cDC1.Xcr1 | Mbnl1 |
| Stt3b1 | 9.26E-141 | 1.05077215 | 0.559 | 0.416 | 1.79E-136 | cDC1.Xcr1 | Stt3b |
| mt-Co11 | 1.50E-140 | 0.62633274 | 0.746 | 0.62 | 2.90E-136 | cDC1.Xcr1 | mt-Co1 |
| Calr1 | 1.78E-139 | 0.64133336 | 0.724 | 0.59 | 3.44E-135 | cDC1.Xcr1 | Calr |
| Manf | 8.09E-139 | 0.76388335 | 0.643 | 0.504 | 1.56E-134 | cDC1.Xcr1 | Manf |
| Gdpd5 | 1.20E-138 | 2.37855086 | 0.336 | 0.075 | 2.33E-134 | cDC1.Xcr1 | Gdpd5 |
| Rbbp7 | 2.29E-138 | 0.78965016 | 0.61 | 0.463 | 4.42E-134 | cDC1.Xcr1 | Rbbp7 |
| Parvg1 | 4.06E-138 | 1.31806206 | 0.659 | 0.342 | 7.85E-134 | cDC1.Xcr1 | Parvg |
| Zfp367 | 5.31E-138 | 1.96383216 | 0.26 | 0.064 | 1.03E-133 | cDC1.Xcr1 | Zfp367 |
| Spg21 | 5.64E-138 | 1.09075054 | 0.629 | 0.407 | 1.09E-133 | cDC1.Xcr1 | Spg21 |
| Usp10 | 4.45E-136 | 1.60191034 | 0.401 | 0.149 | 8.61E-132 | cDC1.Xcr1 | Usp10 |
| Adgre51 | 7.46E-136 | 0.73164359 | 0.661 | 0.463 | 1.44E-131 | cDC1.Xcr1 | Adgre5 |
| Lrrk2 | 1.24E-134 | 1.47727208 | 0.471 | 0.182 | 2.40E-130 | cDC1.Xcr1 | Lrrk2 |
| Tor1a | 4.00E-134 | 1.07637292 | 0.597 | 0.368 | 7.73E-130 | cDC1.Xcr1 | Tor1a |
| Shmt21 | 1.20E-132 | 1.34601593 | 0.458 | 0.23 | 2.33E-128 | cDC1.Xcr1 | Shmt2 |
| Plekha51 | 1.47E-132 | 1.39529811 | 0.542 | 0.279 | 2.84E-128 | cDC1.Xcr1 | Plekha5 |
| H2afz1 | 1.70E-130 | 0.32111529 | 0.697 | 0.662 | 3.29E-126 | cDC1.Xcr1 | H2afz |
| Flt31 | 2.28E-130 | 0.67501446 | 0.571 | 0.306 | 4.40E-126 | cDC1.Xcr1 | Flt3 |
| Cbfb1 | 3.44E-130 | 0.71780996 | 0.778 | 0.664 | 6.65E-126 | cDC1.Xcr1 | Cbfb |
| Lpcat11 | 4.42E-130 | 0.9624762 | 0.487 | 0.258 | 8.55E-126 | cDC1.Xcr1 | Lpcat1 |
| Cd821 | 2.80E-129 | 0.70537168 | 0.727 | 0.538 | 5.42E-125 | cDC1.Xcr1 | Cd82 |
| Adgra2 | 2.29E-127 | 2.54838063 | 0.292 | 0.047 | 4.42E-123 | cDC1.Xcr1 | Adgra2 |
| Lrba1 | 1.08E-126 | 1.21542521 | 0.451 | 0.226 | 2.08E-122 | cDC1.Xcr1 | Lrba |
| Pi4k2a | 5.11E-126 | 1.18372583 | 0.489 | 0.259 | 9.88E-122 | cDC1.Xcr1 | Pi4k2a |
| Fuca11 | 1.08E-125 | 1.08156907 | 0.588 | 0.398 | 2.09E-121 | cDC1.Xcr1 | Fuca1 |
| Lpar6 | 5.86E-124 | 1.74576911 | 0.338 | 0.137 | 1.13E-119 | cDC1.Xcr1 | Lpar6 |
| Lpgat1 | 9.31E-124 | 1.11210463 | 0.566 | 0.294 | 1.80E-119 | cDC1.Xcr1 | Lpgat1 |
| Eid1 | 3.72E-123 | 0.83797927 | 0.734 | 0.575 | 7.20E-119 | cDC1.Xcr1 | Eid1 |
| Arhgdia | 1.44E-122 | 0.45903094 | 0.891 | 0.83 | 2.78E-118 | cDC1.Xcr1 | Arhgdia |
| F2rl2 | 7.06E-122 | 2.34025428 | 0.266 | 0.074 | 1.37E-117 | cDC1.Xcr1 | F2rl2 |
| Shc1 | 1.62E-121 | 1.13736808 | 0.603 | 0.382 | 3.14E-117 | cDC1.Xcr1 | Shc1 |
| Rnf144b | 3.47E-121 | 2.71814201 | 0.289 | 0.055 | 6.70E-117 | cDC1.Xcr1 | Rnf144b |
| Apobec31 | 8.08E-121 | 0.36516029 | 0.754 | 0.651 | 1.56E-116 | cDC1.Xcr1 | Apobec3 |
| Dock101 | 9.76E-120 | 0.63214778 | 0.772 | 0.632 | 1.89E-115 | cDC1.Xcr1 | Dock10 |
| Mllt3 | 1.49E-119 | 3.12046722 | 0.271 | 0.033 | 2.89E-115 | cDC1.Xcr1 | Mllt3 |
| Clec12a | 4.56E-119 | 1.8703601 | 0.292 | 0.081 | 8.82E-115 | cDC1.Xcr1 | Clec12a |
| Pqlc11 | 5.69E-119 | 1.17541015 | 0.52 | 0.294 | 1.10E-114 | cDC1.Xcr1 | Pqlc1 |
| Ccdc102a | 9.10E-119 | 1.04181796 | 0.63 | 0.388 | 1.76E-114 | cDC1.Xcr1 | Ccdc102a |
| Tm9sf2 | 7.82E-118 | 0.87352222 | 0.596 | 0.423 | 1.51E-113 | cDC1.Xcr1 | Tm9sf2 |
| Pstpip11 | 2.01E-115 | 1.30675092 | 0.559 | 0.278 | 3.89E-111 | cDC1.Xcr1 | Pstpip1 |
| Ndufb11 | 4.95E-114 | 0.93431584 | 0.643 | 0.453 | 9.57E-110 | cDC1.Xcr1 | Ndufb11 |
| Polr1a | 8.37E-114 | 1.67861734 | 0.387 | 0.132 | 1.62E-109 | cDC1.Xcr1 | Polr1a |
| Nsun2 | 1.03E-113 | 1.12888019 | 0.523 | 0.301 | 1.99E-109 | cDC1.Xcr1 | Nsun2 |
| Ccl4 | 1.73E-113 | 0.30392373 | 0.371 | 0.221 | 3.34E-109 | cDC1.Xcr1 | Ccl4 |
| Bri3bp1 | 3.66E-113 | 0.81417979 | 0.705 | 0.512 | 7.08E-109 | cDC1.Xcr1 | Bri3bp |
| Itga3 | 1.72E-112 | 1.77984759 | 0.272 | 0.076 | 3.33E-108 | cDC1.Xcr1 | Itga3 |
| Fbxo4 | 7.80E-109 | 2.13751202 | 0.267 | 0.052 | 1.51E-104 | cDC1.Xcr1 | Fbxo4 |
| Sgms1 | 8.68E-109 | 1.62806125 | 0.316 | 0.103 | 1.68E-104 | cDC1.Xcr1 | Sgms1 |
| Bloc1s21 | 1.37E-108 | 0.67365431 | 0.725 | 0.544 | 2.66E-104 | cDC1.Xcr1 | Bloc1s2 |
| Apobr1 | 7.31E-108 | 1.72769218 | 0.436 | 0.162 | 1.41E-103 | cDC1.Xcr1 | Apobr |
| Ctdsp2 | 5.17E-107 | 0.66918865 | 0.73 | 0.584 | 9.99E-103 | cDC1.Xcr1 | Ctdsp2 |
| Gng21 | 8.83E-107 | 1.26075167 | 0.565 | 0.317 | 1.71E-102 | cDC1.Xcr1 | Gng2 |
| Kctd10 | 1.57E-106 | 0.79616003 | 0.591 | 0.394 | 3.04E-102 | cDC1.Xcr1 | Kctd10 |
| Chuk | 2.35E-106 | 0.9203978 | 0.529 | 0.344 | 4.55E-102 | cDC1.Xcr1 | Chuk |
| Ywhag | 2.68E-106 | 0.53215823 | 0.854 | 0.776 | 5.18E-102 | cDC1.Xcr1 | Ywhag |
| Arf3 | 4.90E-106 | 0.74607738 | 0.817 | 0.684 | 9.47E-102 | cDC1.Xcr1 | Arf3 |
| Znrf2 | 6.28E-106 | 0.76044352 | 0.699 | 0.532 | 1.21E-101 | cDC1.Xcr1 | Znrf2 |
| H2-DMb11 | 2.38E-105 | 0.75139937 | 0.727 | 0.492 | 4.60E-101 | cDC1.Xcr1 | H2-DMb1 |
| Nfatc22 | 2.55E-105 | 1.08226192 | 0.623 | 0.374 | 4.92E-101 | cDC1.Xcr1 | Nfatc2 |
| Khk1 | 2.96E-105 | 1.3538425 | 0.445 | 0.214 | 5.72E-101 | cDC1.Xcr1 | Khk |
| Celf2 | 3.93E-105 | 0.74561996 | 0.672 | 0.44 | 7.60E-101 | cDC1.Xcr1 | Celf2 |
| Ubash3b1 | 5.17E-105 | 0.97928425 | 0.471 | 0.262 | 9.99E-101 | cDC1.Xcr1 | Ubash3b |
| Gna15 | 9.23E-105 | 1.39825818 | 0.413 | 0.148 | 1.78E-100 | cDC1.Xcr1 | Gna15 |
| Tns3 | 1.57E-104 | 1.15186244 | 0.475 | 0.191 | 3.04E-100 | cDC1.Xcr1 | Tns3 |
| Trio1 | 2.12E-104 | 0.82668119 | 0.621 | 0.426 | 4.10E-100 | cDC1.Xcr1 | Trio |
| Gramd2 | 6.33E-104 | 5.61185365 | 0.281 | 0.007 | 1.22E-99 | cDC1.Xcr1 | Gramd2 |
| Fbxl14 | 1.36E-103 | 1.17776526 | 0.451 | 0.225 | 2.63E-99 | cDC1.Xcr1 | Fbxl14 |
| Oxct1 | 2.23E-103 | 1.32938527 | 0.373 | 0.144 | 4.32E-99 | cDC1.Xcr1 | Oxct1 |
| Cmtm61 | 2.51E-103 | 0.55426762 | 0.755 | 0.634 | 4.85E-99 | cDC1.Xcr1 | Cmtm6 |
| Slc7a8 | 1.41E-102 | 2.59800082 | 0.255 | 0.046 | 2.73E-98 | cDC1.Xcr1 | Slc7a8 |
| Cyb5a | 1.62E-102 | 1.06005545 | 0.578 | 0.349 | 3.13E-98 | cDC1.Xcr1 | Cyb5a |
| Arpc2 | 7.68E-102 | 0.50381107 | 0.794 | 0.728 | 1.49E-97 | cDC1.Xcr1 | Arpc2 |
| Morc2a1 | 1.65E-101 | 1.39641622 | 0.433 | 0.206 | 3.18E-97 | cDC1.Xcr1 | Morc2a |
| Laptm4b1 | 7.00E-101 | 0.58507825 | 0.447 | 0.172 | 1.35E-96 | cDC1.Xcr1 | Laptm4b |
| Nudc | 8.09E-101 | 0.81354542 | 0.665 | 0.505 | 1.56E-96 | cDC1.Xcr1 | Nudc |
| Prkar2a1 | 1.70E-99 | 1.05462323 | 0.566 | 0.344 | 3.29E-95 | cDC1.Xcr1 | Prkar2a |
| Cdc14a | 1.84E-99 | 1.28045484 | 0.39 | 0.17 | 3.55E-95 | cDC1.Xcr1 | Cdc14a |
| Ccndbp1 | 8.68E-99 | 1.15776867 | 0.527 | 0.308 | 1.68E-94 | cDC1.Xcr1 | Ccndbp1 |
| Mcm61 | 1.52E-98 | 1.11885647 | 0.282 | 0.189 | 2.94E-94 | cDC1.Xcr1 | Mcm6 |
| Sumo3 | 4.19E-98 | 0.94179614 | 0.581 | 0.378 | 8.10E-94 | cDC1.Xcr1 | Sumo3 |
| Ptma1 | 1.58E-97 | 0.3357783 | 0.697 | 0.676 | 3.06E-93 | cDC1.Xcr1 | Ptma |
| Ptk2 | 3.49E-96 | 2.14328553 | 0.254 | 0.056 | 6.74E-92 | cDC1.Xcr1 | Ptk2 |
| Gsn1 | 1.70E-95 | 0.47399232 | 0.925 | 0.749 | 3.29E-91 | cDC1.Xcr1 | Gsn |
| Cpne3 | 3.51E-95 | 1.02256336 | 0.505 | 0.311 | 6.78E-91 | cDC1.Xcr1 | Cpne3 |
| mt-Atp81 | 3.74E-95 | 0.36213236 | 0.755 | 0.66 | 7.23E-91 | cDC1.Xcr1 | mt-Atp8 |
| Ubfd1 | 1.11E-94 | 1.06434021 | 0.483 | 0.265 | 2.15E-90 | cDC1.Xcr1 | Ubfd1 |
| Skap2 | 2.85E-94 | 0.49127668 | 0.795 | 0.719 | 5.50E-90 | cDC1.Xcr1 | Skap2 |
| Ppib | 5.35E-94 | 0.5152852 | 0.788 | 0.707 | 1.03E-89 | cDC1.Xcr1 | Ppib |
| Cd2ap | 6.05E-93 | 0.67190144 | 0.618 | 0.456 | 1.17E-88 | cDC1.Xcr1 | Cd2ap |
| Pdia61 | 5.45E-92 | 0.57402744 | 0.596 | 0.491 | 1.05E-87 | cDC1.Xcr1 | Pdia6 |
| Lasp1 | 5.50E-92 | 1.02284363 | 0.615 | 0.401 | 1.06E-87 | cDC1.Xcr1 | Lasp1 |
| Sri1 | 6.44E-92 | 0.45471741 | 0.79 | 0.696 | 1.25E-87 | cDC1.Xcr1 | Sri |
| Asap2 | 1.41E-91 | 1.34706071 | 0.339 | 0.117 | 2.73E-87 | cDC1.Xcr1 | Asap2 |
| Cdc42se1 | 1.71E-91 | 0.53449083 | 0.746 | 0.633 | 3.31E-87 | cDC1.Xcr1 | Cdc42se1 |
| Lrp51 | 4.22E-91 | 1.19441411 | 0.507 | 0.289 | 8.17E-87 | cDC1.Xcr1 | Lrp5 |
| Hsp90b1 | 6.31E-91 | 0.68838496 | 0.599 | 0.454 | 1.22E-86 | cDC1.Xcr1 | Hsp90b1 |
| Fam91a1 | 6.62E-91 | 0.7732639 | 0.587 | 0.392 | 1.28E-86 | cDC1.Xcr1 | Fam91a1 |
| Osbpl3 | 1.02E-90 | 0.98547333 | 0.379 | 0.136 | 1.97E-86 | cDC1.Xcr1 | Osbpl3 |
| Ccdc88b1 | 1.09E-90 | 0.58549556 | 0.767 | 0.633 | 2.10E-86 | cDC1.Xcr1 | Ccdc88b |
| Mospd2 | 7.10E-90 | 1.36558127 | 0.393 | 0.201 | 1.37E-85 | cDC1.Xcr1 | Mospd2 |
| Leprotl1 | 1.06E-89 | 1.05407569 | 0.54 | 0.315 | 2.05E-85 | cDC1.Xcr1 | Leprotl1 |
| mt-Atp61 | 2.86E-89 | 0.28676365 | 0.754 | 0.675 | 5.53E-85 | cDC1.Xcr1 | mt-Atp6 |
| Otulinl2 | 7.61E-89 | 0.69279255 | 0.7 | 0.516 | 1.47E-84 | cDC1.Xcr1 | Otulinl |
| Nsmaf1 | 1.41E-87 | 0.94257655 | 0.572 | 0.362 | 2.73E-83 | cDC1.Xcr1 | Nsmaf |
| Itpripl1 | 2.77E-87 | 1.58102348 | 0.32 | 0.11 | 5.36E-83 | cDC1.Xcr1 | Itpripl1 |
| Stx17 | 1.65E-86 | 1.47973445 | 0.357 | 0.152 | 3.18E-82 | cDC1.Xcr1 | Stx17 |
| Abca31 | 1.92E-86 | 1.23696078 | 0.461 | 0.236 | 3.72E-82 | cDC1.Xcr1 | Abca3 |
| Dnmbp | 2.38E-86 | 2.00126407 | 0.289 | 0.081 | 4.61E-82 | cDC1.Xcr1 | Dnmbp |
| Anp32e2 | 3.37E-86 | 0.47923954 | 0.652 | 0.572 | 6.52E-82 | cDC1.Xcr1 | Anp32e |
| Prr5 | 3.78E-86 | 1.69074172 | 0.286 | 0.074 | 7.30E-82 | cDC1.Xcr1 | Prr5 |
| mt-Co31 | 4.07E-86 | 0.42870322 | 0.717 | 0.626 | 7.87E-82 | cDC1.Xcr1 | mt-Co3 |
| Snx21 | 1.44E-85 | 1.702901 | 0.314 | 0.106 | 2.78E-81 | cDC1.Xcr1 | Snx21 |
| Slc35c21 | 2.26E-85 | 1.14193956 | 0.593 | 0.334 | 4.37E-81 | cDC1.Xcr1 | Slc35c2 |
| Acadl1 | 4.43E-85 | 0.76123466 | 0.553 | 0.358 | 8.56E-81 | cDC1.Xcr1 | Acadl |
| Akr1a1 | 1.07E-84 | 0.35470133 | 0.902 | 0.873 | 2.07E-80 | cDC1.Xcr1 | Akr1a1 |
| Rp2 | 3.39E-84 | 0.66245989 | 0.597 | 0.465 | 6.55E-80 | cDC1.Xcr1 | Rp2 |
| Fam117a | 5.18E-84 | 1.13389944 | 0.373 | 0.155 | 1.00E-79 | cDC1.Xcr1 | Fam117a |
| Mfsd14a | 1.20E-83 | 1.30108167 | 0.423 | 0.207 | 2.32E-79 | cDC1.Xcr1 | Mfsd14a |
| Hsph1 | 2.16E-83 | 1.10136136 | 0.277 | 0.15 | 4.18E-79 | cDC1.Xcr1 | Hsph1 |
| Snx31 | 4.42E-83 | 0.52427083 | 0.772 | 0.697 | 8.55E-79 | cDC1.Xcr1 | Snx3 |
| Mical21 | 6.07E-83 | 1.53710742 | 0.317 | 0.115 | 1.17E-78 | cDC1.Xcr1 | Mical2 |
| Gpsm31 | 6.28E-83 | 0.68537316 | 0.798 | 0.688 | 1.21E-78 | cDC1.Xcr1 | Gpsm3 |
| Dpy19l11 | 1.59E-82 | 0.9914287 | 0.507 | 0.296 | 3.06E-78 | cDC1.Xcr1 | Dpy19l1 |
| Plekhj11 | 1.03E-81 | 0.5914783 | 0.619 | 0.474 | 2.00E-77 | cDC1.Xcr1 | Plekhj1 |
| Gpr551 | 1.23E-81 | 0.953847 | 0.473 | 0.289 | 2.39E-77 | cDC1.Xcr1 | Gpr55 |
| Nav1 | 4.89E-81 | 0.63829466 | 0.554 | 0.401 | 9.45E-77 | cDC1.Xcr1 | Nav1 |
| Ctnnbip1 | 4.99E-81 | 1.33822617 | 0.419 | 0.211 | 9.65E-77 | cDC1.Xcr1 | Ctnnbip1 |
| Sh2d3c | 1.84E-80 | 1.59166849 | 0.271 | 0.079 | 3.55E-76 | cDC1.Xcr1 | Sh2d3c |
| Gnb41 | 7.20E-80 | 0.42519391 | 0.499 | 0.285 | 1.39E-75 | cDC1.Xcr1 | Gnb4 |
| mt-Cytb | 7.43E-80 | 0.37202607 | 0.673 | 0.547 | 1.44E-75 | cDC1.Xcr1 | mt-Cytb |
| Ppp1r14a1 | 1.28E-79 | 1.15484992 | 0.488 | 0.213 | 2.48E-75 | cDC1.Xcr1 | Ppp1r14a |
| Stard10 | 6.60E-79 | 0.92053273 | 0.305 | 0.116 | 1.28E-74 | cDC1.Xcr1 | Stard10 |
| Psmc3 | 1.36E-78 | 0.63369898 | 0.675 | 0.554 | 2.63E-74 | cDC1.Xcr1 | Psmc3 |
| Atg31 | 2.02E-78 | 0.94681262 | 0.532 | 0.329 | 3.90E-74 | cDC1.Xcr1 | Atg3 |
| Extl31 | 2.03E-78 | 1.14902424 | 0.462 | 0.25 | 3.92E-74 | cDC1.Xcr1 | Extl3 |
| Dnajc8 | 2.36E-78 | 0.85206169 | 0.583 | 0.415 | 4.57E-74 | cDC1.Xcr1 | Dnajc8 |
| Efhd21 | 3.80E-78 | 0.38301356 | 0.893 | 0.831 | 7.35E-74 | cDC1.Xcr1 | Efhd2 |
| Creld21 | 4.26E-78 | 0.88783303 | 0.384 | 0.22 | 8.23E-74 | cDC1.Xcr1 | Creld2 |
| Abhd17a | 8.14E-78 | 0.89635246 | 0.56 | 0.376 | 1.57E-73 | cDC1.Xcr1 | Abhd17a |
| Supt201 | 1.59E-77 | 0.50133663 | 0.624 | 0.452 | 3.07E-73 | cDC1.Xcr1 | Supt20 |
| Slc33a11 | 1.96E-77 | 0.62495092 | 0.512 | 0.312 | 3.79E-73 | cDC1.Xcr1 | Slc33a1 |
| Tifab | 2.11E-77 | 1.12808733 | 0.422 | 0.175 | 4.08E-73 | cDC1.Xcr1 | Tifab |
| Jag21 | 5.72E-77 | 0.94953359 | 0.491 | 0.246 | 1.11E-72 | cDC1.Xcr1 | Jag2 |
| Amz12 | 6.36E-77 | 0.88991971 | 0.582 | 0.343 | 1.23E-72 | cDC1.Xcr1 | Amz1 |
| Grk2 | 9.32E-77 | 0.40813021 | 0.814 | 0.742 | 1.80E-72 | cDC1.Xcr1 | Grk2 |
| Ccdc107 | 1.26E-76 | 1.16449033 | 0.373 | 0.181 | 2.44E-72 | cDC1.Xcr1 | Ccdc107 |
| Arf5 | 1.62E-76 | 0.50075657 | 0.826 | 0.721 | 3.12E-72 | cDC1.Xcr1 | Arf5 |
| Dctpp11 | 2.05E-76 | 1.58704804 | 0.299 | 0.12 | 3.97E-72 | cDC1.Xcr1 | Dctpp1 |
| Tcea31 | 4.20E-76 | 1.13147881 | 0.439 | 0.202 | 8.12E-72 | cDC1.Xcr1 | Tcea3 |
| Cd81 | 4.22E-76 | 1.05981402 | 0.36 | 0.152 | 8.16E-72 | cDC1.Xcr1 | Cd81 |
| Mical11 | 8.58E-76 | 0.74929774 | 0.648 | 0.453 | 1.66E-71 | cDC1.Xcr1 | Mical1 |
| Slc31a2 | 1.86E-75 | 1.71643739 | 0.336 | 0.121 | 3.59E-71 | cDC1.Xcr1 | Slc31a2 |
| St6gal11 | 2.21E-75 | 0.93900874 | 0.412 | 0.284 | 4.28E-71 | cDC1.Xcr1 | St6gal1 |
| Trib11 | 2.41E-75 | 0.62158775 | 0.594 | 0.46 | 4.67E-71 | cDC1.Xcr1 | Trib1 |
| Ranbp11 | 3.82E-75 | 0.81951354 | 0.56 | 0.42 | 7.38E-71 | cDC1.Xcr1 | Ranbp1 |
| Smpdl3a1 | 4.21E-75 | 0.78038711 | 0.45 | 0.329 | 8.14E-71 | cDC1.Xcr1 | Smpdl3a |
| Dna22 | 5.31E-75 | 1.01474761 | 0.451 | 0.238 | 1.03E-70 | cDC1.Xcr1 | Dna2 |
| Elovl51 | 1.30E-74 | 0.46086043 | 0.624 | 0.521 | 2.51E-70 | cDC1.Xcr1 | Elovl5 |
| Notch21 | 1.41E-74 | 0.55153629 | 0.616 | 0.465 | 2.73E-70 | cDC1.Xcr1 | Notch2 |
| Rab5if | 5.26E-74 | 0.81226613 | 0.621 | 0.454 | 1.02E-69 | cDC1.Xcr1 | Rab5if |
| Snx91 | 1.25E-73 | 0.80372924 | 0.585 | 0.419 | 2.41E-69 | cDC1.Xcr1 | Snx9 |
| Prkd31 | 1.40E-73 | 1.04910062 | 0.462 | 0.294 | 2.70E-69 | cDC1.Xcr1 | Prkd3 |
| Tnfaip8l21 | 1.64E-73 | 1.16641173 | 0.499 | 0.285 | 3.17E-69 | cDC1.Xcr1 | Tnfaip8l2 |
| Esyt11 | 2.13E-73 | 0.42253078 | 0.735 | 0.666 | 4.13E-69 | cDC1.Xcr1 | Esyt1 |
| Iqsec1 | 2.19E-73 | 0.94976585 | 0.456 | 0.24 | 4.24E-69 | cDC1.Xcr1 | Iqsec1 |
| Vars1 | 2.05E-72 | 0.71236061 | 0.571 | 0.381 | 3.97E-68 | cDC1.Xcr1 | Vars |
| Tuba1b | 2.56E-72 | 0.2872069 | 0.555 | 0.56 | 4.96E-68 | cDC1.Xcr1 | Tuba1b |
| Hyou11 | 3.42E-72 | 0.70335976 | 0.521 | 0.363 | 6.60E-68 | cDC1.Xcr1 | Hyou1 |
| Tln12 | 4.92E-72 | 0.39247865 | 0.871 | 0.794 | 9.52E-68 | cDC1.Xcr1 | Tln1 |
| Dlgap41 | 5.94E-72 | 0.46779011 | 0.663 | 0.52 | 1.15E-67 | cDC1.Xcr1 | Dlgap4 |
| Slc25a45 | 6.67E-72 | 1.166974 | 0.444 | 0.239 | 1.29E-67 | cDC1.Xcr1 | Slc25a45 |
| Rnf167 | 1.18E-71 | 1.22166533 | 0.392 | 0.183 | 2.28E-67 | cDC1.Xcr1 | Rnf167 |
| Anks3 | 4.55E-71 | 0.83511771 | 0.428 | 0.252 | 8.79E-67 | cDC1.Xcr1 | Anks3 |
| Cbl | 9.46E-71 | 0.48359549 | 0.714 | 0.6 | 1.83E-66 | cDC1.Xcr1 | Cbl |
| Cd9 | 1.43E-70 | 0.49601623 | 0.797 | 0.65 | 2.76E-66 | cDC1.Xcr1 | Cd9 |
| Adpgk | 2.64E-70 | 0.66380299 | 0.527 | 0.308 | 5.10E-66 | cDC1.Xcr1 | Adpgk |
| Mpc2 | 3.41E-70 | 0.96260198 | 0.482 | 0.313 | 6.59E-66 | cDC1.Xcr1 | Mpc2 |
| mt-Nd1 | 3.88E-70 | 0.42458313 | 0.586 | 0.473 | 7.51E-66 | cDC1.Xcr1 | mt-Nd1 |
| Gnb21 | 1.10E-69 | 0.4482071 | 0.748 | 0.663 | 2.14E-65 | cDC1.Xcr1 | Gnb2 |
| Nadk1 | 1.12E-69 | 0.78271381 | 0.629 | 0.464 | 2.16E-65 | cDC1.Xcr1 | Nadk |
| Txndc5 | 1.17E-69 | 0.7834112 | 0.544 | 0.341 | 2.25E-65 | cDC1.Xcr1 | Txndc5 |
| Cbfa2t3 | 2.21E-69 | 0.29192749 | 0.772 | 0.677 | 4.27E-65 | cDC1.Xcr1 | Cbfa2t3 |
| Fen1 | 5.18E-69 | 0.85669659 | 0.304 | 0.171 | 1.00E-64 | cDC1.Xcr1 | Fen1 |
| Pde1b | 5.28E-69 | 0.79278923 | 0.493 | 0.313 | 1.02E-64 | cDC1.Xcr1 | Pde1b |
| Erlin1 | 9.05E-69 | 0.96609223 | 0.478 | 0.291 | 1.75E-64 | cDC1.Xcr1 | Erlin1 |
| Rasa31 | 1.16E-68 | 0.98254063 | 0.397 | 0.183 | 2.24E-64 | cDC1.Xcr1 | Rasa3 |
| Praf2 | 2.14E-68 | 1.35959408 | 0.286 | 0.114 | 4.14E-64 | cDC1.Xcr1 | Praf2 |
| Ppm1k | 2.61E-68 | 1.54361484 | 0.262 | 0.086 | 5.05E-64 | cDC1.Xcr1 | Ppm1k |
| Arpin1 | 5.09E-68 | 0.91599794 | 0.337 | 0.156 | 9.84E-64 | cDC1.Xcr1 | Arpin |
| Sod1 | 7.99E-68 | 0.44106337 | 0.714 | 0.627 | 1.55E-63 | cDC1.Xcr1 | Sod1 |
| Anxa71 | 1.10E-67 | 0.58756243 | 0.621 | 0.473 | 2.13E-63 | cDC1.Xcr1 | Anxa7 |
| Hipk2 | 1.17E-67 | 0.61686996 | 0.661 | 0.497 | 2.26E-63 | cDC1.Xcr1 | Hipk2 |
| Serf2 | 1.93E-67 | 0.50630654 | 0.838 | 0.741 | 3.73E-63 | cDC1.Xcr1 | Serf2 |
| Dnajc9 | 5.43E-67 | 1.0680782 | 0.267 | 0.149 | 1.05E-62 | cDC1.Xcr1 | Dnajc9 |
| Eif3g | 8.31E-67 | 0.59909056 | 0.652 | 0.516 | 1.61E-62 | cDC1.Xcr1 | Eif3g |
| Pcna1 | 9.38E-67 | 0.50489497 | 0.495 | 0.485 | 1.81E-62 | cDC1.Xcr1 | Pcna |
| Dnajc31 | 1.43E-66 | 0.54458039 | 0.537 | 0.425 | 2.77E-62 | cDC1.Xcr1 | Dnajc3 |
| Kdm7a1 | 2.31E-66 | 0.80429243 | 0.605 | 0.437 | 4.47E-62 | cDC1.Xcr1 | Kdm7a |
| Afg3l1 | 2.94E-66 | 0.85543809 | 0.522 | 0.314 | 5.69E-62 | cDC1.Xcr1 | Afg3l1 |
| 12-Sep | 3.26E-66 | 0.98187458 | 0.338 | 0.13 | 6.31E-62 | cDC1.Xcr1 | 1-Sep |
| Cass41 | 5.42E-66 | 0.78858228 | 0.451 | 0.279 | 1.05E-61 | cDC1.Xcr1 | Cass4 |
| Sigmar11 | 5.69E-66 | 1.10192404 | 0.433 | 0.23 | 1.10E-61 | cDC1.Xcr1 | Sigmar1 |
| Nlk | 8.45E-66 | 1.36566816 | 0.33 | 0.147 | 1.63E-61 | cDC1.Xcr1 | Nlk |
| Wdfy31 | 9.44E-66 | 0.96185969 | 0.531 | 0.358 | 1.83E-61 | cDC1.Xcr1 | Wdfy3 |
| Atp6v0e | 1.71E-65 | 0.47489612 | 0.708 | 0.627 | 3.31E-61 | cDC1.Xcr1 | Atp6v0e |
| Tsta31 | 3.91E-65 | 0.953659 | 0.435 | 0.251 | 7.57E-61 | cDC1.Xcr1 | Tsta3 |
| Itpr11 | 4.85E-65 | 0.75754466 | 0.513 | 0.322 | 9.38E-61 | cDC1.Xcr1 | Itpr1 |
| Stard3nl | 5.76E-65 | 0.9858983 | 0.44 | 0.24 | 1.11E-60 | cDC1.Xcr1 | Stard3nl |
| Rtn41 | 1.09E-64 | 0.34029644 | 0.755 | 0.695 | 2.10E-60 | cDC1.Xcr1 | Rtn4 |
| Slc25a381 | 1.18E-64 | 1.06296325 | 0.424 | 0.222 | 2.28E-60 | cDC1.Xcr1 | Slc25a38 |
| Fam102b | 2.71E-64 | 1.29724115 | 0.358 | 0.171 | 5.25E-60 | cDC1.Xcr1 | Fam102b |
| Jdp21 | 5.33E-64 | 0.58652245 | 0.585 | 0.383 | 1.03E-59 | cDC1.Xcr1 | Jdp2 |
| Mcm21 | 5.50E-64 | 1.21121593 | 0.252 | 0.143 | 1.06E-59 | cDC1.Xcr1 | Mcm2 |
| Cap11 | 7.92E-64 | 0.25044045 | 0.859 | 0.848 | 1.53E-59 | cDC1.Xcr1 | Cap1 |
| Vav11 | 9.45E-64 | 0.61235566 | 0.678 | 0.471 | 1.83E-59 | cDC1.Xcr1 | Vav1 |
| Sh3gl1 | 2.22E-63 | 0.7552681 | 0.55 | 0.335 | 4.29E-59 | cDC1.Xcr1 | Sh3gl1 |
| Map3k4 | 3.16E-63 | 1.97518033 | 0.265 | 0.073 | 6.12E-59 | cDC1.Xcr1 | Map3k4 |
| Slamf7 | 5.16E-63 | 0.34420373 | 0.555 | 0.412 | 9.98E-59 | cDC1.Xcr1 | Slamf7 |
| Snx23 | 6.05E-63 | 0.69347818 | 0.565 | 0.39 | 1.17E-58 | cDC1.Xcr1 | Snx2 |
| Acox31 | 7.88E-63 | 0.75084597 | 0.648 | 0.461 | 1.52E-58 | cDC1.Xcr1 | Acox3 |
| Rala1 | 8.03E-63 | 0.59107049 | 0.547 | 0.394 | 1.55E-58 | cDC1.Xcr1 | Rala |
| Pygo2 | 9.23E-63 | 1.05851883 | 0.373 | 0.205 | 1.78E-58 | cDC1.Xcr1 | Pygo2 |
| Ndufa4 | 1.21E-62 | 1.17263457 | 0.368 | 0.18 | 2.35E-58 | cDC1.Xcr1 | Ndufa4 |
| Ap3d1 | 3.10E-62 | 0.60430289 | 0.625 | 0.482 | 5.99E-58 | cDC1.Xcr1 | Ap3d1 |
| Rab4b | 5.20E-62 | 0.60035884 | 0.654 | 0.521 | 1.01E-57 | cDC1.Xcr1 | Rab4b |
| Mapk142 | 1.67E-61 | 0.66052693 | 0.64 | 0.488 | 3.22E-57 | cDC1.Xcr1 | Mapk14 |
| Trim30a1 | 2.33E-61 | 0.86167669 | 0.451 | 0.252 | 4.50E-57 | cDC1.Xcr1 | Trim30a |
| Usp221 | 2.75E-61 | 0.80612478 | 0.479 | 0.301 | 5.32E-57 | cDC1.Xcr1 | Usp22 |
| Dctn5 | 3.90E-61 | 1.0417451 | 0.468 | 0.266 | 7.54E-57 | cDC1.Xcr1 | Dctn5 |
| Vdac2 | 4.12E-61 | 0.82660912 | 0.509 | 0.361 | 7.96E-57 | cDC1.Xcr1 | Vdac2 |
| Ddost1 | 4.86E-61 | 0.41325606 | 0.697 | 0.605 | 9.41E-57 | cDC1.Xcr1 | Ddost |
| Aff1 | 5.23E-61 | 0.35999727 | 0.681 | 0.587 | 1.01E-56 | cDC1.Xcr1 | Aff1 |
| Palld1 | 9.81E-61 | 0.59115737 | 0.369 | 0.226 | 1.90E-56 | cDC1.Xcr1 | Palld |
| Lmnb12 | 1.21E-60 | 0.34488511 | 0.667 | 0.609 | 2.34E-56 | cDC1.Xcr1 | Lmnb1 |
| Clic12 | 1.22E-60 | 0.48853648 | 0.724 | 0.622 | 2.37E-56 | cDC1.Xcr1 | Clic1 |
| Tomm341 | 1.40E-60 | 0.55587681 | 0.518 | 0.354 | 2.70E-56 | cDC1.Xcr1 | Tomm34 |
| Plin2 | 1.42E-60 | 0.79139947 | 0.597 | 0.394 | 2.75E-56 | cDC1.Xcr1 | Plin2 |
| Ttc71 | 2.36E-60 | 0.47032963 | 0.724 | 0.604 | 4.56E-56 | cDC1.Xcr1 | Ttc7 |
| Atp5b | 2.79E-60 | 0.37468259 | 0.702 | 0.638 | 5.40E-56 | cDC1.Xcr1 | Atp5b |
| C2cd2l | 3.02E-60 | 1.40893584 | 0.339 | 0.146 | 5.84E-56 | cDC1.Xcr1 | C2cd2l |
| Myo1f1 | 7.00E-60 | 0.75343986 | 0.642 | 0.442 | 1.35E-55 | cDC1.Xcr1 | Myo1f |
| Psmb7 | 9.91E-60 | 0.70674752 | 0.531 | 0.402 | 1.92E-55 | cDC1.Xcr1 | Psmb7 |
| Elac2 | 1.28E-59 | 1.52688389 | 0.3 | 0.123 | 2.47E-55 | cDC1.Xcr1 | Elac2 |
| Hspa52 | 2.81E-59 | 0.42588764 | 0.665 | 0.58 | 5.44E-55 | cDC1.Xcr1 | Hspa5 |
| Cyb5r4 | 5.22E-59 | 0.76280595 | 0.605 | 0.434 | 1.01E-54 | cDC1.Xcr1 | Cyb5r4 |
| Ccdc861 | 8.24E-59 | 0.97000092 | 0.54 | 0.327 | 1.59E-54 | cDC1.Xcr1 | Ccdc86 |
| Dennd4b | 1.21E-58 | 0.60603158 | 0.64 | 0.479 | 2.34E-54 | cDC1.Xcr1 | Dennd4b |
| Fam168a | 1.94E-58 | 1.13545448 | 0.376 | 0.202 | 3.75E-54 | cDC1.Xcr1 | Fam168a |
| Arid4a1 | 2.02E-58 | 0.78981483 | 0.451 | 0.267 | 3.90E-54 | cDC1.Xcr1 | Arid4a |
| Rbm471 | 2.12E-58 | 0.84039505 | 0.603 | 0.416 | 4.09E-54 | cDC1.Xcr1 | Rbm47 |
| Sipa1l11 | 2.24E-58 | 0.94842715 | 0.375 | 0.181 | 4.33E-54 | cDC1.Xcr1 | Sipa1l1 |
| Hps5 | 3.53E-58 | 1.15861865 | 0.397 | 0.206 | 6.82E-54 | cDC1.Xcr1 | Hps5 |
| Fdps | 4.91E-58 | 0.44870577 | 0.473 | 0.373 | 9.50E-54 | cDC1.Xcr1 | Fdps |
| Tmem30a | 8.06E-58 | 0.69652882 | 0.478 | 0.328 | 1.56E-53 | cDC1.Xcr1 | Tmem30a |
| Trappc4 | 8.98E-58 | 0.854592 | 0.556 | 0.377 | 1.74E-53 | cDC1.Xcr1 | Trappc4 |
| Mcm31 | 2.56E-57 | 0.67491745 | 0.3 | 0.216 | 4.95E-53 | cDC1.Xcr1 | Mcm3 |
| Hck1 | 4.65E-57 | 0.4757232 | 0.718 | 0.576 | 9.00E-53 | cDC1.Xcr1 | Hck |
| Cox8a1 | 1.03E-56 | 0.47048232 | 0.751 | 0.651 | 1.99E-52 | cDC1.Xcr1 | Cox8a |
| Adgrg52 | 1.52E-56 | 0.62831583 | 0.691 | 0.488 | 2.95E-52 | cDC1.Xcr1 | Adgrg5 |
| Alox5ap2 | 1.89E-56 | 0.36231656 | 0.79 | 0.617 | 3.65E-52 | cDC1.Xcr1 | Alox5ap |
| Ube2s | 2.76E-56 | 0.67127674 | 0.42 | 0.301 | 5.34E-52 | cDC1.Xcr1 | Ube2s |
| Hps4 | 3.35E-56 | 1.48817481 | 0.266 | 0.093 | 6.48E-52 | cDC1.Xcr1 | Hps4 |
| Tram2 | 1.08E-55 | 1.46352149 | 0.26 | 0.088 | 2.08E-51 | cDC1.Xcr1 | Tram2 |
| Scarb11 | 1.12E-55 | 0.82647703 | 0.433 | 0.237 | 2.17E-51 | cDC1.Xcr1 | Scarb1 |
| Gabbr11 | 1.19E-55 | 1.20275461 | 0.332 | 0.157 | 2.30E-51 | cDC1.Xcr1 | Gabbr1 |
| Znrd1 | 1.62E-55 | 0.84780236 | 0.411 | 0.24 | 3.13E-51 | cDC1.Xcr1 | Znrd1 |
| Srrt | 1.63E-55 | 0.70862363 | 0.525 | 0.359 | 3.15E-51 | cDC1.Xcr1 | Srrt |
| Cmtm7 | 2.42E-55 | 0.74545965 | 0.5 | 0.325 | 4.68E-51 | cDC1.Xcr1 | Cmtm7 |
| Sipa11 | 2.62E-55 | 0.71103698 | 0.513 | 0.37 | 5.07E-51 | cDC1.Xcr1 | Sipa1 |
| Chchd3 | 2.64E-55 | 0.80117114 | 0.504 | 0.338 | 5.11E-51 | cDC1.Xcr1 | Chchd3 |
| Naip6 | 2.99E-55 | 1.10052018 | 0.35 | 0.178 | 5.78E-51 | cDC1.Xcr1 | Naip6 |
| Eif4a21 | 4.77E-55 | 0.29841905 | 0.714 | 0.641 | 9.22E-51 | cDC1.Xcr1 | Eif4a2 |
| Gns | 5.76E-55 | 0.52160407 | 0.602 | 0.482 | 1.11E-50 | cDC1.Xcr1 | Gns |
| Tbc1d141 | 8.19E-55 | 0.41373955 | 0.6 | 0.447 | 1.58E-50 | cDC1.Xcr1 | Tbc1d14 |
| Zfp282 | 9.14E-55 | 1.48169654 | 0.279 | 0.103 | 1.77E-50 | cDC1.Xcr1 | Zfp282 |
| Patz1 | 9.66E-55 | 0.85273251 | 0.415 | 0.223 | 1.87E-50 | cDC1.Xcr1 | Patz1 |
| mt-Nd4l1 | 1.29E-54 | 0.44682148 | 0.646 | 0.536 | 2.50E-50 | cDC1.Xcr1 | mt-Nd4l |
| Zfp281 | 1.64E-54 | 1.20083095 | 0.265 | 0.108 | 3.17E-50 | cDC1.Xcr1 | Zfp281 |
| Plp2 | 2.13E-54 | 0.57713693 | 0.65 | 0.491 | 4.12E-50 | cDC1.Xcr1 | Plp2 |
| Ahr1 | 4.29E-54 | 0.43315984 | 0.653 | 0.574 | 8.29E-50 | cDC1.Xcr1 | Ahr |
| Magt1 | 5.97E-54 | 0.49946123 | 0.549 | 0.399 | 1.15E-49 | cDC1.Xcr1 | Magt1 |
| Npm1 | 1.35E-53 | 0.4082777 | 0.746 | 0.662 | 2.61E-49 | cDC1.Xcr1 | Npm1 |
| Lsm41 | 2.00E-53 | 0.54048787 | 0.588 | 0.451 | 3.86E-49 | cDC1.Xcr1 | Lsm4 |
| Ssbp3 | 2.36E-53 | 1.39637332 | 0.271 | 0.107 | 4.56E-49 | cDC1.Xcr1 | Ssbp3 |
| Slc25a39 | 3.16E-53 | 0.74085421 | 0.526 | 0.385 | 6.10E-49 | cDC1.Xcr1 | Slc25a39 |
| Uba2 | 4.85E-53 | 0.6338349 | 0.509 | 0.357 | 9.39E-49 | cDC1.Xcr1 | Uba2 |
| Lsr1 | 6.31E-53 | 0.74464995 | 0.5 | 0.312 | 1.22E-48 | cDC1.Xcr1 | Lsr |
| Ppp1r15b | 8.85E-53 | 0.61130726 | 0.518 | 0.373 | 1.71E-48 | cDC1.Xcr1 | Ppp1r15b |
| Aagab1 | 1.38E-52 | 0.67241348 | 0.478 | 0.314 | 2.67E-48 | cDC1.Xcr1 | Aagab |
| Limk11 | 1.96E-52 | 0.83638415 | 0.444 | 0.288 | 3.79E-48 | cDC1.Xcr1 | Limk1 |
| Hnrnpul2 | 2.63E-52 | 0.33703884 | 0.75 | 0.679 | 5.09E-48 | cDC1.Xcr1 | Hnrnpul2 |
| Casp61 | 2.83E-52 | 0.56884785 | 0.623 | 0.422 | 5.47E-48 | cDC1.Xcr1 | Casp6 |
| Pip4k2a1 | 4.05E-52 | 0.51349977 | 0.727 | 0.625 | 7.84E-48 | cDC1.Xcr1 | Pip4k2a |
| Psma4 | 4.79E-52 | 0.42202433 | 0.674 | 0.58 | 9.26E-48 | cDC1.Xcr1 | Psma4 |
| Gprc5c1 | 4.88E-52 | 0.81930378 | 0.491 | 0.312 | 9.44E-48 | cDC1.Xcr1 | Gprc5c |
| Mbd2 | 5.41E-52 | 0.38392423 | 0.854 | 0.778 | 1.05E-47 | cDC1.Xcr1 | Mbd2 |
| Nono | 7.70E-52 | 0.27243394 | 0.673 | 0.599 | 1.49E-47 | cDC1.Xcr1 | Nono |
| Spcs3 | 1.09E-51 | 0.63186474 | 0.499 | 0.368 | 2.11E-47 | cDC1.Xcr1 | Spcs3 |
| Usp12 | 2.08E-51 | 0.68012282 | 0.566 | 0.424 | 4.02E-47 | cDC1.Xcr1 | Usp12 |

| | | | | | | | |
|---|---|---|---|---|---|---|---|
| Card111 | 2.94E-51 | 0.94147773 | 0.453 | 0.245 | 5.68E-47 | cDC1.Xcr1 | Card11 |
| Tmem229b | 9.23E-51 | 0.6122719 | 0.447 | 0.222 | 1.79E-46 | cDC1.Xcr1 | Tmem229b |
| Oaz1 | 1.07E-50 | 0.58853221 | 0.734 | 0.593 | 2.07E-46 | cDC1.Xcr1 | Oaz1 |
| Rrbp1 | 1.26E-50 | 0.49068245 | 0.749 | 0.635 | 2.43E-46 | cDC1.Xcr1 | Rrbp1 |
| Mpp61 | 1.49E-50 | 0.76578968 | 0.498 | 0.347 | 2.88E-46 | cDC1.Xcr1 | Mpp6 |
| Iars1 | 2.35E-50 | 1.09981356 | 0.407 | 0.241 | 4.54E-46 | cDC1.Xcr1 | Iars |
| Tbkbp12 | 5.79E-50 | 1.40964653 | 0.363 | 0.185 | 1.12E-45 | cDC1.Xcr1 | Tbkbp1 |
| Tbcb | 7.90E-50 | 0.49144772 | 0.631 | 0.51 | 1.53E-45 | cDC1.Xcr1 | Tbcb |
| Ipo5 | 1.04E-49 | 0.6135471 | 0.4 | 0.26 | 2.01E-45 | cDC1.Xcr1 | Ipo5 |
| Rnf4 | 1.38E-49 | 0.5996881 | 0.642 | 0.496 | 2.67E-45 | cDC1.Xcr1 | Rnf4 |
| Tbxas12 | 1.49E-49 | 0.85131444 | 0.498 | 0.306 | 2.88E-45 | cDC1.Xcr1 | Tbxas1 |
| Ruvbl1 | 1.61E-49 | 1.11795666 | 0.292 | 0.122 | 3.11E-45 | cDC1.Xcr1 | Ruvbl1 |
| Dcaf7 | 2.58E-49 | 0.60815875 | 0.615 | 0.485 | 4.99E-45 | cDC1.Xcr1 | Dcaf7 |
| Tox4 | 6.41E-49 | 0.59977642 | 0.547 | 0.397 | 1.24E-44 | cDC1.Xcr1 | Tox4 |
| Orai11 | 8.67E-49 | 0.46269406 | 0.672 | 0.553 | 1.68E-44 | cDC1.Xcr1 | Orai1 |
| Cops91 | 1.08E-48 | 0.43886402 | 0.722 | 0.664 | 2.10E-44 | cDC1.Xcr1 | Cops9 |
| Nlgn2 | 1.31E-48 | 0.91796113 | 0.271 | 0.112 | 2.52E-44 | cDC1.Xcr1 | Nlgn2 |
| Ddx39 | 2.36E-48 | 0.44254932 | 0.445 | 0.355 | 4.57E-44 | cDC1.Xcr1 | Ddx39 |
| H2-DMa2 | 3.84E-48 | 0.39678333 | 0.886 | 0.68 | 7.42E-44 | cDC1.Xcr1 | H2-DMa |
| Prkra | 4.35E-48 | 1.08527147 | 0.257 | 0.107 | 8.42E-44 | cDC1.Xcr1 | Prkra |
| Cdca4 | 4.77E-48 | 0.97148234 | 0.273 | 0.151 | 9.22E-44 | cDC1.Xcr1 | Cdca4 |
| Hnrnpf | 9.02E-48 | 0.30478936 | 0.669 | 0.615 | 1.74E-43 | cDC1.Xcr1 | Hnrnpf |
| Brd81 | 1.11E-47 | 0.62110871 | 0.359 | 0.235 | 2.14E-43 | cDC1.Xcr1 | Brd8 |
| Zfp6521 | 1.63E-47 | 1.02250298 | 0.386 | 0.22 | 3.15E-43 | cDC1.Xcr1 | Zfp652 |
| Ranbp3 | 2.10E-47 | 0.58644788 | 0.496 | 0.35 | 4.07E-43 | cDC1.Xcr1 | Ranbp3 |
| Fli11 | 2.65E-47 | 0.8325243 | 0.485 | 0.295 | 5.12E-43 | cDC1.Xcr1 | Fli1 |
| Tbc1d10c | 3.20E-47 | 0.74378916 | 0.397 | 0.204 | 6.19E-43 | cDC1.Xcr1 | Tbc1d10c |
| Hnrnpm | 3.22E-47 | 0.29678924 | 0.783 | 0.716 | 6.23E-43 | cDC1.Xcr1 | Hnrnpm |
| Erp291 | 3.37E-47 | 0.33571019 | 0.761 | 0.681 | 6.51E-43 | cDC1.Xcr1 | Erp29 |
| Srsf12 | 3.59E-47 | 0.42651211 | 0.608 | 0.519 | 6.94E-43 | cDC1.Xcr1 | Srsf1 |
| Arid2 | 4.34E-47 | 0.662092 | 0.444 | 0.296 | 8.39E-43 | cDC1.Xcr1 | Arid2 |
| Dennd1b1 | 4.54E-47 | 0.65401292 | 0.533 | 0.364 | 8.77E-43 | cDC1.Xcr1 | Dennd1b |
| Mapre11 | 5.14E-47 | 0.27261667 | 0.673 | 0.623 | 9.93E-43 | cDC1.Xcr1 | Mapre1 |
| Ppp1r181 | 6.14E-47 | 0.60892493 | 0.701 | 0.575 | 1.19E-42 | cDC1.Xcr1 | Ppp1r18 |
| Scand1 | 6.25E-47 | 0.40442717 | 0.645 | 0.527 | 1.21E-42 | cDC1.Xcr1 | Scand1 |
| Ppp1r2 | 6.26E-47 | 0.75407057 | 0.478 | 0.301 | 1.21E-42 | cDC1.Xcr1 | Ppp1r2 |
| Erbin | 1.26E-46 | 0.45394244 | 0.564 | 0.467 | 2.43E-42 | cDC1.Xcr1 | Erbin |
| Hectd41 | 1.38E-46 | 0.94964546 | 0.403 | 0.223 | 2.66E-42 | cDC1.Xcr1 | Hectd4 |
| Tlnrd1 | 1.39E-46 | 0.77356551 | 0.439 | 0.261 | 2.69E-42 | cDC1.Xcr1 | Tlnrd1 |
| Myl12b1 | 1.81E-46 | 0.44162996 | 0.717 | 0.622 | 3.50E-42 | cDC1.Xcr1 | Myl12b |
| Synrg | 1.98E-46 | 1.03346149 | 0.415 | 0.231 | 3.83E-42 | cDC1.Xcr1 | Synrg |
| Pum2 | 3.03E-46 | 0.31322717 | 0.683 | 0.616 | 5.86E-42 | cDC1.Xcr1 | Pum2 |
| Cct3 | 4.89E-46 | 0.77366153 | 0.507 | 0.35 | 9.45E-42 | cDC1.Xcr1 | Cct3 |
| Ano6 | 5.42E-46 | 0.92935409 | 0.353 | 0.202 | 1.05E-41 | cDC1.Xcr1 | Ano6 |
| P4hb1 | 7.71E-46 | 0.40690004 | 0.691 | 0.571 | 1.49E-41 | cDC1.Xcr1 | P4hb |
| Map4k5 | 9.44E-46 | 0.79767357 | 0.309 | 0.144 | 1.82E-41 | cDC1.Xcr1 | Map4k5 |
| Rcn2 | 1.05E-45 | 0.99813141 | 0.347 | 0.192 | 2.04E-41 | cDC1.Xcr1 | Rcn2 |
| Cdk2ap21 | 1.20E-45 | 0.4743348 | 0.619 | 0.479 | 2.32E-41 | cDC1.Xcr1 | Cdk2ap2 |
| Txndc16 | 1.59E-45 | 1.23269895 | 0.265 | 0.112 | 3.08E-41 | cDC1.Xcr1 | Txndc16 |
| Slc12a9 | 1.93E-45 | 1.00212269 | 0.401 | 0.21 | 3.73E-41 | cDC1.Xcr1 | Slc12a9 |
| Trit1 | 1.94E-45 | 1.1806122 | 0.309 | 0.133 | 3.76E-41 | cDC1.Xcr1 | Trit1 |
| Eps81 | 2.80E-45 | 0.51201748 | 0.621 | 0.418 | 5.41E-41 | cDC1.Xcr1 | Eps8 |
| Tmem50b1 | 3.11E-45 | 0.68901353 | 0.565 | 0.373 | 6.01E-41 | cDC1.Xcr1 | Tmem50b |
| Gcnt21 | 3.66E-45 | 0.44109292 | 0.442 | 0.478 | 7.07E-41 | cDC1.Xcr1 | Gcnt2 |
| Coa51 | 4.16E-45 | 0.55595731 | 0.574 | 0.424 | 8.05E-41 | cDC1.Xcr1 | Coa5 |
| Leprot | 7.18E-45 | 0.57055381 | 0.504 | 0.391 | 1.39E-40 | cDC1.Xcr1 | Leprot |
| Slc25a24 | 8.83E-45 | 0.86404824 | 0.388 | 0.23 | 1.71E-40 | cDC1.Xcr1 | Slc25a24 |
| Sf3b2 | 2.44E-44 | 0.40741618 | 0.684 | 0.596 | 4.72E-40 | cDC1.Xcr1 | Sf3b2 |
| Plcb22 | 3.11E-44 | 0.80224616 | 0.532 | 0.37 | 6.01E-40 | cDC1.Xcr1 | Plcb2 |
| Osbpl1a | 3.95E-44 | 1.23236077 | 0.252 | 0.112 | 7.63E-40 | cDC1.Xcr1 | Osbpl1a |
| Fbl1 | 3.97E-44 | 0.77631785 | 0.593 | 0.42 | 7.68E-40 | cDC1.Xcr1 | Fbl |
| Klrb1b1 | 4.22E-44 | 0.51585527 | 0.566 | 0.437 | 8.17E-40 | cDC1.Xcr1 | Klrb1b |
| Rinl1 | 9.42E-44 | 1.06465638 | 0.414 | 0.239 | 1.82E-39 | cDC1.Xcr1 | Rinl |
| Selenow | 1.04E-43 | 0.28449403 | 0.691 | 0.592 | 2.01E-39 | cDC1.Xcr1 | Selenow |
| Prpf191 | 1.41E-43 | 0.71568859 | 0.487 | 0.346 | 2.72E-39 | cDC1.Xcr1 | Prpf19 |
| Rnf145 | 1.42E-43 | 0.88402251 | 0.463 | 0.286 | 2.75E-39 | cDC1.Xcr1 | Rnf145 |
| Slc25a101 | 1.51E-43 | 0.47789591 | 0.305 | 0.196 | 2.92E-39 | cDC1.Xcr1 | Slc25a10 |
| Isyna12 | 2.54E-43 | 0.53936613 | 0.555 | 0.411 | 4.92E-39 | cDC1.Xcr1 | Isyna1 |
| Stx21 | 2.67E-43 | 1.17751599 | 0.353 | 0.205 | 5.16E-39 | cDC1.Xcr1 | Stx2 |
| Dpf2 | 5.22E-43 | 0.69391959 | 0.522 | 0.373 | 1.01E-38 | cDC1.Xcr1 | Dpf2 |
| Nudt16 | 5.92E-43 | 1.31449206 | 0.281 | 0.127 | 1.14E-38 | cDC1.Xcr1 | Nudt16 |
| Bin31 | 1.82E-42 | 0.74542165 | 0.502 | 0.345 | 3.52E-38 | cDC1.Xcr1 | Bin3 |
| Stk17b1 | 1.88E-42 | 0.30721479 | 0.748 | 0.642 | 3.63E-38 | cDC1.Xcr1 | Stk17b |
| Pik3cb | 2.02E-42 | 0.75481645 | 0.475 | 0.336 | 3.90E-38 | cDC1.Xcr1 | Pik3cb |
| Rnf1571 | 2.22E-42 | 0.40371678 | 0.306 | 0.134 | 4.28E-38 | cDC1.Xcr1 | Rnf157 |
| Me2 | 2.36E-42 | 0.75181115 | 0.39 | 0.258 | 4.56E-38 | cDC1.Xcr1 | Me2 |
| Dnajb111 | 2.91E-42 | 0.50805925 | 0.569 | 0.454 | 5.62E-38 | cDC1.Xcr1 | Dnajb11 |
| Rbm171 | 4.09E-42 | 0.3129982 | 0.627 | 0.526 | 7.90E-38 | cDC1.Xcr1 | Rbm17 |
| Gmfb | 8.28E-42 | 0.54274283 | 0.531 | 0.389 | 1.60E-37 | cDC1.Xcr1 | Gmfb |
| Fkbp21 | 8.35E-42 | 0.51605142 | 0.493 | 0.362 | 1.61E-37 | cDC1.Xcr1 | Fkbp2 |
| Slbp1 | 9.41E-42 | 0.35458595 | 0.406 | 0.333 | 1.82E-37 | cDC1.Xcr1 | Slbp |
| Rnf187 | 9.89E-42 | 0.32743551 | 0.717 | 0.6 | 1.91E-37 | cDC1.Xcr1 | Rnf187 |
| Lars | 1.02E-41 | 0.772561 | 0.4 | 0.247 | 1.98E-37 | cDC1.Xcr1 | Lars |
| Dpp42 | 1.33E-41 | 0.39827856 | 0.613 | 0.482 | 2.57E-37 | cDC1.Xcr1 | Dpp4 |
| Selenoh1 | 1.40E-41 | 0.90240064 | 0.267 | 0.153 | 2.71E-37 | cDC1.Xcr1 | Selenoh |
| Kmo2 | 2.16E-41 | 0.5174237 | 0.765 | 0.535 | 4.18E-37 | cDC1.Xcr1 | Kmo |
| Dnajb1 | 3.69E-41 | 0.55257228 | 0.39 | 0.318 | 7.14E-37 | cDC1.Xcr1 | Dnajb1 |
| Nde1 | 6.06E-41 | 0.91524788 | 0.276 | 0.148 | 1.17E-36 | cDC1.Xcr1 | Nde1 |
| Bex3 | 7.15E-41 | 0.59376306 | 0.375 | 0.249 | 1.38E-36 | cDC1.Xcr1 | Bex3 |
| Tent2 | 7.63E-41 | 0.48256438 | 0.458 | 0.35 | 1.47E-36 | cDC1.Xcr1 | Tent2 |
| Wbp1l | 8.77E-41 | 0.64045542 | 0.521 | 0.383 | 1.70E-36 | cDC1.Xcr1 | Wbp1l |
| Rab11a | 1.28E-40 | 0.38315966 | 0.522 | 0.39 | 2.47E-36 | cDC1.Xcr1 | Rab11a |
| Tax1bp11 | 1.51E-40 | 0.2586324 | 0.619 | 0.546 | 2.92E-36 | cDC1.Xcr1 | Tax1bp1 |
| Disc1 | 1.73E-40 | 1.4165602 | 0.286 | 0.129 | 3.34E-36 | cDC1.Xcr1 | Disc1 |
| Dnmt11 | 2.66E-40 | 0.59467994 | 0.289 | 0.241 | 5.15E-36 | cDC1.Xcr1 | Dnmt1 |
| Klc2 | 3.51E-40 | 1.11364885 | 0.319 | 0.163 | 6.79E-36 | cDC1.Xcr1 | Klc2 |
| Trim41 | 3.92E-40 | 0.69178949 | 0.408 | 0.243 | 7.59E-36 | cDC1.Xcr1 | Trim41 |
| Clasp11 | 4.20E-40 | 0.31272666 | 0.446 | 0.302 | 8.13E-36 | cDC1.Xcr1 | Clasp1 |
| Iqgap22 | 5.07E-40 | 0.63763714 | 0.498 | 0.324 | 9.81E-36 | cDC1.Xcr1 | Iqgap2 |
| Twsg1 | 5.75E-40 | 0.79304097 | 0.292 | 0.15 | 1.11E-35 | cDC1.Xcr1 | Twsg1 |
| Copg2 | 6.20E-40 | 0.85151413 | 0.385 | 0.238 | 1.20E-35 | cDC1.Xcr1 | Copg2 |
| Psme3 | 6.74E-40 | 0.53043679 | 0.582 | 0.472 | 1.30E-35 | cDC1.Xcr1 | Psme3 |
| Nsmce4a | 7.55E-40 | 0.67219169 | 0.374 | 0.244 | 1.46E-35 | cDC1.Xcr1 | Nsmce4a |
| Aldh21 | 1.14E-39 | 0.45053573 | 0.65 | 0.536 | 2.20E-35 | cDC1.Xcr1 | Aldh2 |
| Pde3b | 2.72E-39 | 0.71588559 | 0.369 | 0.238 | 5.26E-35 | cDC1.Xcr1 | Pde3b |
| Tbrg11 | 4.06E-39 | 0.49616262 | 0.517 | 0.401 | 7.85E-35 | cDC1.Xcr1 | Tbrg1 |
| Itgal | 4.52E-39 | 0.62022937 | 0.32 | 0.164 | 8.73E-35 | cDC1.Xcr1 | Itgal |
| Tmem2141 | 5.47E-39 | 0.89170329 | 0.38 | 0.212 | 1.06E-34 | cDC1.Xcr1 | Tmem214 |
| Ptpn71 | 6.86E-39 | 1.30613285 | 0.259 | 0.114 | 1.33E-34 | cDC1.Xcr1 | Ptpn7 |
| Phyh | 8.03E-39 | 0.66286349 | 0.332 | 0.185 | 1.55E-34 | cDC1.Xcr1 | Phyh |
| Hdac1 | 8.17E-39 | 0.46314299 | 0.531 | 0.416 | 1.58E-34 | cDC1.Xcr1 | Hdac1 |
| Eif3k1 | 8.68E-39 | 0.40606569 | 0.792 | 0.714 | 1.68E-34 | cDC1.Xcr1 | Eif3k |
| Myd882 | 1.39E-38 | 0.44934778 | 0.721 | 0.596 | 2.68E-34 | cDC1.Xcr1 | Myd88 |
| Uqcrfs1 | 1.46E-38 | 0.35077789 | 0.609 | 0.54 | 2.83E-34 | cDC1.Xcr1 | Uqcrfs1 |
| Ifi35 | 1.48E-38 | 0.58651917 | 0.542 | 0.383 | 2.86E-34 | cDC1.Xcr1 | Ifi35 |
| Prrsl1 | 2.34E-38 | 0.45575863 | 0.415 | 0.235 | 4.53E-34 | cDC1.Xcr1 | Prrsl |
| Plekhn11 | 2.47E-38 | 0.62254687 | 0.341 | 0.211 | 4.78E-34 | cDC1.Xcr1 | Plekhn1 |
| Gtf2e2 | 2.49E-38 | 0.89657662 | 0.387 | 0.24 | 4.81E-34 | cDC1.Xcr1 | Gtf2e2 |
| Ccdc69 | 2.67E-38 | 1.31453513 | 0.259 | 0.114 | 5.16E-34 | cDC1.Xcr1 | Ccdc69 |
| Capn15 | 2.81E-38 | 1.18165781 | 0.275 | 0.136 | 5.43E-34 | cDC1.Xcr1 | Capn15 |
| Rubcnl | 6.98E-38 | 0.80932329 | 0.284 | 0.127 | 1.35E-33 | cDC1.Xcr1 | Rubcnl |
| Cisd1 | 7.52E-38 | 0.99043472 | 0.261 | 0.129 | 1.45E-33 | cDC1.Xcr1 | Cisd1 |
| Cnot8 | 9.39E-38 | 0.6567257 | 0.419 | 0.295 | 1.82E-33 | cDC1.Xcr1 | Cnot8 |
| Kdm3b1 | 1.08E-37 | 0.29734831 | 0.632 | 0.522 | 2.09E-33 | cDC1.Xcr1 | Kdm3b |
| Wdfy1 | 1.08E-37 | 1.09568962 | 0.281 | 0.134 | 2.10E-33 | cDC1.Xcr1 | Wdfy1 |
| Chmp1b | 1.33E-37 | 0.65006863 | 0.484 | 0.353 | 2.57E-33 | cDC1.Xcr1 | Chmp1b |
| mt-Nd51 | 1.43E-37 | 0.30887275 | 0.556 | 0.454 | 2.76E-33 | cDC1.Xcr1 | mt-Nd5 |
| Trpc4ap | 1.58E-37 | 0.40083523 | 0.641 | 0.547 | 3.05E-33 | cDC1.Xcr1 | Trpc4ap |
| Ndufa3 | 1.58E-37 | 0.31025693 | 0.741 | 0.657 | 3.06E-33 | cDC1.Xcr1 | Ndufa3 |
| Paxip1 | 1.73E-37 | 0.97429956 | 0.275 | 0.155 | 3.34E-33 | cDC1.Xcr1 | Paxip1 |
| Reep51 | 1.90E-37 | 0.34974051 | 0.695 | 0.605 | 3.68E-33 | cDC1.Xcr1 | Reep5 |
| Vps9d1 | 2.06E-37 | 0.98060851 | 0.338 | 0.183 | 3.97E-33 | cDC1.Xcr1 | Vps9d1 |
| Lrrc251 | 4.66E-37 | 0.70434076 | 0.51 | 0.325 | 9.02E-33 | cDC1.Xcr1 | Lrrc25 |
| Gtf3c6 | 4.78E-37 | 0.7463693 | 0.395 | 0.24 | 9.24E-33 | cDC1.Xcr1 | Gtf3c6 |
| Tut7 | 4.90E-37 | 0.35168057 | 0.623 | 0.551 | 9.47E-33 | cDC1.Xcr1 | Tut7 |
| Stx16 | 5.02E-37 | 0.59484218 | 0.491 | 0.353 | 9.71E-33 | cDC1.Xcr1 | Stx16 |
| Ssb | 5.39E-37 | 0.3428159 | 0.599 | 0.515 | 1.04E-32 | cDC1.Xcr1 | Ssb |
| Atp2c11 | 7.49E-37 | 0.46526018 | 0.5 | 0.382 | 1.45E-32 | cDC1.Xcr1 | Atp2c1 |
| Cltb2 | 7.90E-37 | 0.57565016 | 0.565 | 0.406 | 1.53E-32 | cDC1.Xcr1 | Cltb |
| Npepl11 | 9.25E-37 | 0.57440799 | 0.485 | 0.341 | 1.79E-32 | cDC1.Xcr1 | Npepl1 |
| Etfb1 | 1.30E-36 | 0.58942414 | 0.553 | 0.434 | 2.51E-32 | cDC1.Xcr1 | Etfb |
| Polr2g1 | 1.48E-36 | 0.60666761 | 0.391 | 0.259 | 2.87E-32 | cDC1.Xcr1 | Polr2g |
| Psma5 | 2.52E-36 | 0.45530252 | 0.576 | 0.462 | 4.87E-32 | cDC1.Xcr1 | Psma5 |
| Ndufa2 | 2.81E-36 | 0.30423426 | 0.699 | 0.63 | 5.43E-32 | cDC1.Xcr1 | Ndufa2 |
| Anapc111 | 4.29E-36 | 0.48037542 | 0.513 | 0.422 | 8.30E-32 | cDC1.Xcr1 | Anapc11 |
| Eef2 | 5.06E-36 | 0.26304164 | 0.835 | 0.807 | 9.78E-32 | cDC1.Xcr1 | Eef2 |
| Caprin1 | 5.08E-36 | 0.5905717 | 0.523 | 0.395 | 9.83E-32 | cDC1.Xcr1 | Caprin1 |
| Suox | 6.29E-36 | 0.70471054 | 0.273 | 0.141 | 1.22E-31 | cDC1.Xcr1 | Suox |
| Slfn51 | 7.96E-36 | 0.63439462 | 0.408 | 0.277 | 1.54E-31 | cDC1.Xcr1 | Slfn5 |
| Prkar2b | 8.59E-36 | 0.85764753 | 0.257 | 0.124 | 1.66E-31 | cDC1.Xcr1 | Prkar2b |
| Ahsa11 | 1.02E-35 | 0.35706321 | 0.566 | 0.481 | 1.97E-31 | cDC1.Xcr1 | Ahsa1 |
| Rusc1 | 1.14E-35 | 1.05059184 | 0.276 | 0.153 | 2.20E-31 | cDC1.Xcr1 | Rusc1 |
| Tram11 | 1.26E-35 | 0.28622617 | 0.627 | 0.532 | 2.43E-31 | cDC1.Xcr1 | Tram1 |
| Creg1 | 1.36E-35 | 0.58529373 | 0.534 | 0.4 | 2.62E-31 | cDC1.Xcr1 | Creg1 |
| Phf23 | 1.52E-35 | 0.61981484 | 0.511 | 0.386 | 2.94E-31 | cDC1.Xcr1 | Phf23 |
| Tm9sf3 | 1.55E-35 | 0.25066136 | 0.673 | 0.617 | 2.99E-31 | cDC1.Xcr1 | Tm9sf3 |
| Srsf9 | 2.12E-35 | 0.4262086 | 0.741 | 0.647 | 4.10E-31 | cDC1.Xcr1 | Srsf9 |
| Rab1b1 | 2.70E-35 | 0.55093849 | 0.759 | 0.619 | 5.22E-31 | cDC1.Xcr1 | Rab1b |
| Vps11 | 2.78E-35 | 0.65997137 | 0.429 | 0.279 | 5.37E-31 | cDC1.Xcr1 | Vps11 |
| Psma3 | 3.87E-35 | 0.47390168 | 0.494 | 0.384 | 7.49E-31 | cDC1.Xcr1 | Psma3 |
| Ppp1ca | 4.12E-35 | 0.3258915 | 0.67 | 0.598 | 7.97E-31 | cDC1.Xcr1 | Ppp1ca |
| Dnajc7 | 5.11E-35 | 0.3680135 | 0.523 | 0.424 | 9.88E-31 | cDC1.Xcr1 | Dnajc7 |
| Atp2a22 | 6.34E-35 | 0.35384743 | 0.599 | 0.483 | 1.23E-30 | cDC1.Xcr1 | Atp2a2 |
| Rer1 | 1.33E-34 | 0.55769789 | 0.474 | 0.372 | 2.56E-30 | cDC1.Xcr1 | Rer1 |
| Ccdc121 | 2.77E-34 | 0.38721965 | 0.694 | 0.586 | 5.36E-30 | cDC1.Xcr1 | Ccdc12 |
| Ptpa1 | 2.78E-34 | 0.44455098 | 0.526 | 0.398 | 5.37E-30 | cDC1.Xcr1 | Ptpa |
| Arhgef6 | 3.12E-34 | 0.30412836 | 0.515 | 0.394 | 6.02E-30 | cDC1.Xcr1 | Arhgef6 |
| Klhl181 | 3.65E-34 | 1.01593916 | 0.371 | 0.213 | 7.05E-30 | cDC1.Xcr1 | Klhl18 |
| Adipor2 | 3.83E-34 | 0.57214404 | 0.458 | 0.319 | 7.41E-30 | cDC1.Xcr1 | Adipor2 |
| Cux11 | 4.29E-34 | 0.41443522 | 0.679 | 0.551 | 8.30E-30 | cDC1.Xcr1 | Cux1 |
| Bak1 | 4.50E-34 | 0.27481677 | 0.717 | 0.651 | 8.71E-30 | cDC1.Xcr1 | Bak1 |
| Mtpn1 | 5.92E-34 | 0.45644596 | 0.658 | 0.533 | 1.14E-29 | cDC1.Xcr1 | Mtpn |
| Pcbp11 | 6.74E-34 | 0.25520858 | 0.729 | 0.648 | 1.30E-29 | cDC1.Xcr1 | Pcbp1 |
| Acss1 | 7.30E-34 | 1.06551164 | 0.306 | 0.146 | 1.41E-29 | cDC1.Xcr1 | Acss1 |
| Ppp6r1 | 1.04E-33 | 0.40209747 | 0.667 | 0.564 | 2.01E-29 | cDC1.Xcr1 | Ppp6r1 |
| Gltp | 1.14E-33 | 0.28548679 | 0.665 | 0.584 | 2.21E-29 | cDC1.Xcr1 | Gltp |
| Pafah1b3 | 1.21E-33 | 0.46427028 | 0.464 | 0.314 | 2.35E-29 | cDC1.Xcr1 | Pafah1b3 |
| Fcho1 | 1.25E-33 | 0.48926639 | 0.396 | 0.252 | 2.42E-29 | cDC1.Xcr1 | Fcho1 |
| Dcp2 | 1.38E-33 | 0.57619889 | 0.414 | 0.293 | 2.68E-29 | cDC1.Xcr1 | Dcp2 |
| Eml21 | 1.58E-33 | 1.03333484 | 0.319 | 0.169 | 3.05E-29 | cDC1.Xcr1 | Eml2 |
| H2afv1 | 1.66E-33 | 0.31276427 | 0.382 | 0.341 | 3.21E-29 | cDC1.Xcr1 | H2afv |
| Bub3 | 2.25E-33 | 0.56148816 | 0.414 | 0.297 | 4.35E-29 | cDC1.Xcr1 | Bub3 |
| Pak2 | 2.36E-33 | 0.26129969 | 0.729 | 0.693 | 4.57E-29 | cDC1.Xcr1 | Pak2 |
| Rnf123 | 2.70E-33 | 0.62111428 | 0.363 | 0.217 | 5.23E-29 | cDC1.Xcr1 | Rnf123 |
| Plekhg31 | 3.89E-33 | 0.97506782 | 0.298 | 0.178 | 7.52E-29 | cDC1.Xcr1 | Plekhg3 |
| Prpf8 | 4.10E-33 | 0.3072546 | 0.588 | 0.511 | 7.93E-29 | cDC1.Xcr1 | Prpf8 |
| Dazap1 | 4.13E-33 | 0.2688802 | 0.684 | 0.613 | 7.99E-29 | cDC1.Xcr1 | Dazap1 |
| Sf3b31 | 4.71E-33 | 0.4270692 | 0.526 | 0.408 | 9.10E-29 | cDC1.Xcr1 | Sf3b3 |
| Cuta1 | 4.88E-33 | 0.56613104 | 0.561 | 0.44 | 9.43E-29 | cDC1.Xcr1 | Cuta |
| Snrpa | 5.95E-33 | 0.57800511 | 0.412 | 0.292 | 1.15E-28 | cDC1.Xcr1 | Snrpa |
| Nedd8 | 6.07E-33 | 0.29521961 | 0.781 | 0.713 | 1.17E-28 | cDC1.Xcr1 | Nedd8 |
| Uggt11 | 6.43E-33 | 0.29283492 | 0.598 | 0.494 | 1.24E-28 | cDC1.Xcr1 | Uggt1 |
| Chd3 | 7.71E-33 | 0.67735881 | 0.409 | 0.248 | 1.49E-28 | cDC1.Xcr1 | Chd3 |
| Prpf181 | 1.12E-32 | 0.68223171 | 0.445 | 0.311 | 2.17E-28 | cDC1.Xcr1 | Prpf18 |
| Ptdss1 | 1.32E-32 | 0.60375448 | 0.464 | 0.322 | 2.54E-28 | cDC1.Xcr1 | Ptdss1 |
| Asah11 | 1.58E-32 | 0.40117849 | 0.604 | 0.507 | 3.06E-28 | cDC1.Xcr1 | Asah1 |
| Surf41 | 1.59E-32 | 0.32379503 | 0.587 | 0.532 | 3.08E-28 | cDC1.Xcr1 | Surf4 |
| Derl1 | 1.82E-32 | 0.27564174 | 0.567 | 0.472 | 3.52E-28 | cDC1.Xcr1 | Derl1 |
| Lipe1 | 1.83E-32 | 0.31160268 | 0.581 | 0.45 | 3.54E-28 | cDC1.Xcr1 | Lipe |
| Uchl3 | 2.75E-32 | 0.93524429 | 0.256 | 0.129 | 5.31E-28 | cDC1.Xcr1 | Uchl3 |
| Hspe1 | 3.16E-32 | 0.68333929 | 0.436 | 0.321 | 6.11E-28 | cDC1.Xcr1 | Hspe1 |
| Rae1 | 3.52E-32 | 0.50873685 | 0.392 | 0.257 | 6.80E-28 | cDC1.Xcr1 | Rae1 |
| Rap1gap2 | 4.10E-32 | 0.72720982 | 0.419 | 0.259 | 7.94E-28 | cDC1.Xcr1 | Rap1gap2 |
| Hnrnpd | 4.34E-32 | 0.25910704 | 0.626 | 0.555 | 8.40E-28 | cDC1.Xcr1 | Hnrnpd |
| Hspa141 | 4.48E-32 | 0.64734191 | 0.376 | 0.254 | 8.65E-28 | cDC1.Xcr1 | Hspa14 |
| Nucks1 | 5.22E-32 | 0.77372674 | 0.311 | 0.218 | 1.01E-27 | cDC1.Xcr1 | Nucks1 |
| Cox7c | 6.15E-32 | 0.39712542 | 0.669 | 0.564 | 1.19E-27 | cDC1.Xcr1 | Cox7c |
| Epg5 | 7.76E-32 | 0.93609369 | 0.289 | 0.143 | 1.50E-27 | cDC1.Xcr1 | Epg5 |
| Snrpd11 | 8.31E-32 | 0.46158558 | 0.428 | 0.352 | 1.61E-27 | cDC1.Xcr1 | Snrpd1 |
| Ap1g1 | 8.31E-32 | 0.45460504 | 0.572 | 0.46 | 1.61E-27 | cDC1.Xcr1 | Ap1g1 |
| Ppfibp2 | 9.17E-32 | 1.07064114 | 0.265 | 0.126 | 1.77E-27 | cDC1.Xcr1 | Ppfibp2 |
| Snrpa11 | 1.08E-31 | 0.60184353 | 0.397 | 0.268 | 2.08E-27 | cDC1.Xcr1 | Snrpa1 |
| Rara1 | 1.26E-31 | 0.44067598 | 0.62 | 0.472 | 2.43E-27 | cDC1.Xcr1 | Rara |
| Pak11 | 1.27E-31 | 0.64765563 | 0.576 | 0.414 | 2.46E-27 | cDC1.Xcr1 | Pak1 |
| Acsl51 | 1.41E-31 | 0.34218742 | 0.529 | 0.452 | 2.72E-27 | cDC1.Xcr1 | Acsl5 |
| Snx17 | 1.79E-31 | 0.42879948 | 0.598 | 0.506 | 3.47E-27 | cDC1.Xcr1 | Snx17 |
| Vrk2 | 2.07E-31 | 0.79309572 | 0.305 | 0.169 | 4.00E-27 | cDC1.Xcr1 | Vrk2 |
| Cops5 | 2.17E-31 | 0.57084707 | 0.471 | 0.319 | 4.20E-27 | cDC1.Xcr1 | Cops5 |
| Eif2s2 | 2.21E-31 | 0.84881387 | 0.317 | 0.183 | 4.28E-27 | cDC1.Xcr1 | Eif2s2 |
| Lpcat21 | 2.54E-31 | 0.89048732 | 0.409 | 0.276 | 4.91E-27 | cDC1.Xcr1 | Lpcat2 |
| Fdxr | 2.58E-31 | 1.32863233 | 0.254 | 0.118 | 5.00E-27 | cDC1.Xcr1 | Fdxr |
| Rnps1 | 2.93E-31 | 0.34243797 | 0.521 | 0.415 | 5.67E-27 | cDC1.Xcr1 | Rnps1 |
| Pdia42 | 3.58E-31 | 0.35073669 | 0.475 | 0.387 | 6.92E-27 | cDC1.Xcr1 | Pdia4 |
| Uqcrc2 | 3.76E-31 | 0.40102668 | 0.512 | 0.425 | 7.27E-27 | cDC1.Xcr1 | Uqcrc2 |
| Nup1532 | 4.09E-31 | 0.47034232 | 0.56 | 0.445 | 7.91E-27 | cDC1.Xcr1 | Nup153 |
| Dlg1 | 4.54E-31 | 0.68665811 | 0.46 | 0.311 | 8.77E-27 | cDC1.Xcr1 | Dlg1 |
| Rab2a | 4.55E-31 | 0.40831079 | 0.55 | 0.465 | 8.80E-27 | cDC1.Xcr1 | Rab2a |
| Cdon | 5.00E-31 | 0.64690427 | 0.275 | 0.134 | 9.67E-27 | cDC1.Xcr1 | Cdon |
| Csnk1a1 | 7.10E-31 | 0.37431292 | 0.555 | 0.468 | 1.37E-26 | cDC1.Xcr1 | Csnk1a1 |
| Rab10 | 7.67E-31 | 0.43635359 | 0.51 | 0.378 | 1.48E-26 | cDC1.Xcr1 | Rab10 |
| Reep31 | 8.02E-31 | 0.42537658 | 0.464 | 0.35 | 1.55E-26 | cDC1.Xcr1 | Reep3 |
| Nagk | 9.41E-31 | 0.97592297 | 0.341 | 0.194 | 1.82E-26 | cDC1.Xcr1 | Nagk |
| Arl3 | 9.88E-31 | 0.64400775 | 0.404 | 0.266 | 1.91E-26 | cDC1.Xcr1 | Arl3 |
| Wdfy21 | 1.02E-30 | 1.00176462 | 0.406 | 0.225 | 1.96E-26 | cDC1.Xcr1 | Wdfy2 |
| Lman21 | 1.22E-30 | 0.36288706 | 0.638 | 0.542 | 2.35E-26 | cDC1.Xcr1 | Lman2 |
| Erp44 | 1.38E-30 | 0.5062879 | 0.368 | 0.261 | 2.67E-26 | cDC1.Xcr1 | Erp44 |
| Aldh18a1 | 1.50E-30 | 0.71205419 | 0.339 | 0.198 | 2.91E-26 | cDC1.Xcr1 | Aldh18a1 |
| Mbd3 | 1.51E-30 | 0.36730177 | 0.57 | 0.485 | 2.91E-26 | cDC1.Xcr1 | Mbd3 |
| Vmp11 | 1.70E-30 | 0.32318799 | 0.653 | 0.549 | 3.29E-26 | cDC1.Xcr1 | Vmp1 |
| Gfer | 2.36E-30 | 0.64248964 | 0.35 | 0.202 | 4.56E-26 | cDC1.Xcr1 | Gfer |
| Trpv21 | 2.99E-30 | 0.66373474 | 0.457 | 0.328 | 5.79E-26 | cDC1.Xcr1 | Trpv2 |
| Mat2a | 4.15E-30 | 0.56771784 | 0.598 | 0.521 | 8.01E-26 | cDC1.Xcr1 | Mat2a |
| Xpo1 | 4.38E-30 | 0.27390772 | 0.48 | 0.385 | 8.48E-26 | cDC1.Xcr1 | Xpo1 |
| Lsm21 | 6.67E-30 | 0.64150617 | 0.319 | 0.212 | 1.29E-25 | cDC1.Xcr1 | Lsm2 |
| Lap3 | 6.96E-30 | 1.14863603 | 0.271 | 0.149 | 1.35E-25 | cDC1.Xcr1 | Lap3 |
| Capn11 | 8.04E-30 | 0.35758503 | 0.463 | 0.368 | 1.55E-25 | cDC1.Xcr1 | Capn1 |
| Rbfa | 8.13E-30 | 0.76253131 | 0.319 | 0.187 | 1.57E-25 | cDC1.Xcr1 | Rbfa |
| Dpy30 | 9.04E-30 | 0.49002946 | 0.36 | 0.248 | 1.75E-25 | cDC1.Xcr1 | Dpy30 |
| Ik | 1.02E-29 | 0.29479016 | 0.638 | 0.555 | 1.98E-25 | cDC1.Xcr1 | Ik |
| Nap1l11 | 1.08E-29 | 0.44993408 | 0.425 | 0.325 | 2.08E-25 | cDC1.Xcr1 | Nap1l1 |
| Kit1 | 1.57E-29 | 0.53887101 | 0.553 | 0.393 | 3.03E-25 | cDC1.Xcr1 | Kit |
| Tomm5 | 1.98E-29 | 0.54948687 | 0.425 | 0.307 | 3.82E-25 | cDC1.Xcr1 | Tomm5 |
| Eloa | 2.73E-29 | 0.5030529 | 0.62 | 0.502 | 5.29E-25 | cDC1.Xcr1 | Eloa |
| Cox7b | 2.79E-29 | 0.47411124 | 0.642 | 0.501 | 5.39E-25 | cDC1.Xcr1 | Cox7b |
| Exosc81 | 4.37E-29 | 0.58873011 | 0.308 | 0.228 | 8.46E-25 | cDC1.Xcr1 | Exosc8 |
| Arhgap221 | 4.58E-29 | 0.48809012 | 0.434 | 0.285 | 8.86E-25 | cDC1.Xcr1 | Arhgap22 |
| Grk5 | 4.83E-29 | 1.00294508 | 0.26 | 0.128 | 9.34E-25 | cDC1.Xcr1 | Grk5 |
| Itpkb | 4.88E-29 | 0.45684871 | 0.342 | 0.216 | 9.44E-25 | cDC1.Xcr1 | Itpkb |
| Mtmr41 | 5.89E-29 | 0.27871485 | 0.555 | 0.416 | 1.14E-24 | cDC1.Xcr1 | Mtmr4 |
| Srek1 | 6.14E-29 | 0.46863388 | 0.37 | 0.243 | 1.19E-24 | cDC1.Xcr1 | Srek1 |
| Lpxn1 | 6.27E-29 | 0.96470643 | 0.311 | 0.165 | 1.21E-24 | cDC1.Xcr1 | Lpxn |
| Pttg1 | 1.06E-28 | 0.7002656 | 0.287 | 0.154 | 2.05E-24 | cDC1.Xcr1 | Pttg1 |
| Ap1m11 | 1.47E-28 | 0.59491712 | 0.451 | 0.325 | 2.84E-24 | cDC1.Xcr1 | Ap1m1 |
| Eif5b | 1.62E-28 | 0.66327652 | 0.388 | 0.262 | 3.14E-24 | cDC1.Xcr1 | Eif5b |
| Plpp51 | 1.66E-28 | 0.70008752 | 0.359 | 0.263 | 3.20E-24 | cDC1.Xcr1 | Plpp5 |
| Ssr21 | 3.11E-28 | 0.4575246 | 0.621 | 0.49 | 6.02E-24 | cDC1.Xcr1 | Ssr2 |
| Fam98c1 | 3.41E-28 | 0.42483839 | 0.496 | 0.367 | 6.60E-24 | cDC1.Xcr1 | Fam98c |
| Bin22 | 4.82E-28 | 0.29318852 | 0.776 | 0.633 | 9.32E-24 | cDC1.Xcr1 | Bin2 |
| Dtymk1 | 4.84E-28 | 0.82737858 | 0.297 | 0.202 | 9.36E-24 | cDC1.Xcr1 | Dtymk |
| Sptbn11 | 5.28E-28 | 0.4568547 | 0.368 | 0.235 | 1.02E-23 | cDC1.Xcr1 | Sptbn1 |
| Acaca | 5.44E-28 | 0.5488972 | 0.352 | 0.215 | 1.05E-23 | cDC1.Xcr1 | Acaca |
| Cs1 | 6.31E-28 | 0.47991104 | 0.44 | 0.343 | 1.22E-23 | cDC1.Xcr1 | Cs |
| Anp32a | 9.26E-28 | 0.40971776 | 0.663 | 0.573 | 1.79E-23 | cDC1.Xcr1 | Anp32a |
| Ndufs81 | 1.02E-27 | 0.41342942 | 0.547 | 0.425 | 1.96E-23 | cDC1.Xcr1 | Ndufs8 |
| Ogfr | 1.34E-27 | 0.50324784 | 0.551 | 0.419 | 2.58E-23 | cDC1.Xcr1 | Ogfr |
| Srgap31 | 1.35E-27 | 0.79523526 | 0.314 | 0.178 | 2.61E-23 | cDC1.Xcr1 | Srgap3 |
| Snx29 | 1.36E-27 | 0.7316068 | 0.409 | 0.261 | 2.63E-23 | cDC1.Xcr1 | Snx29 |
| Cystm11 | 1.70E-27 | 0.55250856 | 0.544 | 0.397 | 3.29E-23 | cDC1.Xcr1 | Cystm1 |
| Lemd2 | 2.04E-27 | 0.56286456 | 0.461 | 0.356 | 3.94E-23 | cDC1.Xcr1 | Lemd2 |
| Tma7 | 2.89E-27 | 0.28256004 | 0.605 | 0.535 | 5.59E-23 | cDC1.Xcr1 | Tma7 |
| Arhgap4 | 3.07E-27 | 1.06797171 | 0.293 | 0.151 | 5.94E-23 | cDC1.Xcr1 | Arhgap4 |
| Hip12 | 3.13E-27 | 0.68110986 | 0.396 | 0.256 | 6.05E-23 | cDC1.Xcr1 | Hip1 |
| Inpp4a | 3.22E-27 | 0.54678425 | 0.384 | 0.238 | 6.22E-23 | cDC1.Xcr1 | Inpp4a |
| G3bp2 | 3.60E-27 | 0.50433669 | 0.516 | 0.415 | 6.96E-23 | cDC1.Xcr1 | G3bp2 |
| Rfxap | 4.28E-27 | 0.77774957 | 0.309 | 0.178 | 8.27E-23 | cDC1.Xcr1 | Rfxap |
| Nubp21 | 4.30E-27 | 0.44311026 | 0.488 | 0.364 | 8.32E-23 | cDC1.Xcr1 | Nubp2 |
| Chst111 | 4.82E-27 | 0.2977396 | 0.461 | 0.329 | 9.33E-23 | cDC1.Xcr1 | Chst11 |
| Eif4g32 | 5.48E-27 | 0.30503423 | 0.707 | 0.59 | 1.06E-22 | cDC1.Xcr1 | Eif4g3 |
| Ndufa13 | 7.14E-27 | 0.2581835 | 0.718 | 0.652 | 1.38E-22 | cDC1.Xcr1 | Ndufa13 |
| Pacsin2 | 7.63E-27 | 0.3142083 | 0.684 | 0.598 | 1.47E-22 | cDC1.Xcr1 | Pacsin2 |
| Tars | 7.69E-27 | 0.57421007 | 0.392 | 0.262 | 1.49E-22 | cDC1.Xcr1 | Tars |
| Zfp207 | 8.34E-27 | 0.36399351 | 0.589 | 0.518 | 1.61E-22 | cDC1.Xcr1 | Zfp207 |
| Srp19 | 8.84E-27 | 0.59757968 | 0.409 | 0.285 | 1.71E-22 | cDC1.Xcr1 | Srp19 |
| Tpst2 | 1.18E-26 | 0.57714025 | 0.299 | 0.166 | 2.29E-22 | cDC1.Xcr1 | Tpst2 |
| Ncf41 | 1.20E-26 | 0.66031474 | 0.616 | 0.43 | 2.33E-22 | cDC1.Xcr1 | Ncf4 |
| Ostc1 | 1.28E-26 | 0.46832438 | 0.484 | 0.366 | 2.47E-22 | cDC1.Xcr1 | Ostc |
| Klhdc101 | 1.30E-26 | 0.65340859 | 0.352 | 0.222 | 2.51E-22 | cDC1.Xcr1 | Klhdc10 |
| Eif2s11 | 1.93E-26 | 0.43573157 | 0.493 | 0.366 | 3.74E-22 | cDC1.Xcr1 | Eif2s1 |
| Ppp1r21 | 2.22E-26 | 0.58243324 | 0.36 | 0.231 | 4.30E-22 | cDC1.Xcr1 | Ppp1r21 |
| Jpt21 | 3.05E-26 | 0.74248067 | 0.286 | 0.182 | 5.90E-22 | cDC1.Xcr1 | Jpt2 |
| Sfr1 | 3.12E-26 | 0.42430812 | 0.614 | 0.473 | 6.04E-22 | cDC1.Xcr1 | Sfr1 |
| Arfip1 | 3.37E-26 | 0.68979906 | 0.346 | 0.218 | 6.51E-22 | cDC1.Xcr1 | Arfip1 |
| Ndufs4 | 4.09E-26 | 0.70050797 | 0.369 | 0.258 | 7.92E-22 | cDC1.Xcr1 | Ndufs4 |
| Nup50 | 5.38E-26 | 0.59651751 | 0.294 | 0.202 | 1.04E-21 | cDC1.Xcr1 | Nup50 |
| Shkbp1 | 6.09E-26 | 0.563219 | 0.428 | 0.309 | 1.18E-21 | cDC1.Xcr1 | Shkbp1 |
| Cars | 6.12E-26 | 0.63124075 | 0.26 | 0.149 | 1.18E-21 | cDC1.Xcr1 | Cars |
| Scap | 6.55E-26 | 0.51866206 | 0.359 | 0.236 | 1.27E-21 | cDC1.Xcr1 | Scap |
| Sdhd | 7.15E-26 | 0.35543098 | 0.526 | 0.468 | 1.38E-21 | cDC1.Xcr1 | Sdhd |
| Rab7 | 8.96E-26 | 0.44346784 | 0.612 | 0.519 | 1.73E-21 | cDC1.Xcr1 | Rab7 |
| Tfg1 | 1.02E-25 | 0.25417466 | 0.475 | 0.383 | 1.96E-21 | cDC1.Xcr1 | Tfg |
| Ccdc124 | 1.09E-25 | 0.66575954 | 0.554 | 0.403 | 2.11E-21 | cDC1.Xcr1 | Ccdc124 |
| Usp8 | 1.72E-25 | 0.43750099 | 0.511 | 0.385 | 3.33E-21 | cDC1.Xcr1 | Usp8 |
| Dhx8 | 1.72E-25 | 0.55851858 | 0.358 | 0.229 | 3.33E-21 | cDC1.Xcr1 | Dhx8 |
| Abl11 | 1.81E-25 | 0.51780181 | 0.431 | 0.304 | 3.49E-21 | cDC1.Xcr1 | Abl1 |
| Tor1b | 2.16E-25 | 0.94959066 | 0.263 | 0.144 | 4.17E-21 | cDC1.Xcr1 | Tor1b |
| Stat2 | 2.24E-25 | 0.37241591 | 0.413 | 0.287 | 4.33E-21 | cDC1.Xcr1 | Stat2 |
| Ube2m | 2.38E-25 | 0.28732469 | 0.558 | 0.437 | 4.60E-21 | cDC1.Xcr1 | Ube2m |
| Psmb5 | 2.61E-25 | 0.44100101 | 0.635 | 0.54 | 5.04E-21 | cDC1.Xcr1 | Psmb5 |
| Tardbp | 2.68E-25 | 0.26672699 | 0.565 | 0.461 | 5.19E-21 | cDC1.Xcr1 | Tardbp |
| Arhgef7 | 3.03E-25 | 0.68032668 | 0.341 | 0.225 | 5.86E-21 | cDC1.Xcr1 | Arhgef7 |
| Ttc9c | 3.24E-25 | 0.54827826 | 0.382 | 0.259 | 6.27E-21 | cDC1.Xcr1 | Ttc9c |
| Slc39a7 | 3.40E-25 | 0.34758679 | 0.414 | 0.316 | 6.57E-21 | cDC1.Xcr1 | Slc39a7 |
| Usp33 | 3.72E-25 | 0.75897501 | 0.256 | 0.146 | 7.19E-21 | cDC1.Xcr1 | Usp33 |
| Numb | 3.72E-25 | 0.61577998 | 0.293 | 0.17 | 7.19E-21 | cDC1.Xcr1 | Numb |
| Ppp5c1 | 3.80E-25 | 0.57103538 | 0.352 | 0.222 | 7.35E-21 | cDC1.Xcr1 | Ppp5c |
| Map4k21 | 3.80E-25 | 0.56002349 | 0.352 | 0.225 | 7.35E-21 | cDC1.Xcr1 | Map4k2 |
| Cox5b | 4.40E-25 | 0.3212771 | 0.717 | 0.656 | 8.51E-21 | cDC1.Xcr1 | Cox5b |
| Cliptm1l1 | 5.46E-25 | 0.30762194 | 0.597 | 0.501 | 1.06E-20 | cDC1.Xcr1 | Cliptm1l |
| Ube2k | 6.90E-25 | 0.32449142 | 0.471 | 0.385 | 1.33E-20 | cDC1.Xcr1 | Ube2k |
| Psma1 | 7.15E-25 | 0.28443874 | 0.602 | 0.534 | 1.38E-20 | cDC1.Xcr1 | Psma1 |
| Ppp2r3c | 7.45E-25 | 1.01195704 | 0.283 | 0.161 | 1.44E-20 | cDC1.Xcr1 | Ppp2r3c |
| Nucb11 | 7.80E-25 | 0.39087164 | 0.576 | 0.447 | 1.51E-20 | cDC1.Xcr1 | Nucb1 |
| Derl2 | 7.97E-25 | 0.42026331 | 0.358 | 0.243 | 1.54E-20 | cDC1.Xcr1 | Derl2 |
| Mvb12a | 9.64E-25 | 0.53699807 | 0.447 | 0.315 | 1.86E-20 | cDC1.Xcr1 | Mvb12a |
| Ppan1 | 9.89E-25 | 1.15741952 | 0.268 | 0.141 | 1.91E-20 | cDC1.Xcr1 | Ppan |
| Atf6b | 1.33E-24 | 0.41651949 | 0.357 | 0.251 | 2.56E-20 | cDC1.Xcr1 | Atf6b |
| Naa38 | 1.33E-24 | 0.5405988 | 0.42 | 0.309 | 2.57E-20 | cDC1.Xcr1 | Naa38 |
| Clint11 | 1.42E-24 | 0.40040118 | 0.564 | 0.467 | 2.74E-20 | cDC1.Xcr1 | Clint1 |
| Txnl4a | 1.45E-24 | 0.78384828 | 0.335 | 0.211 | 2.81E-20 | cDC1.Xcr1 | Txnl4a |
| Creb3 | 1.52E-24 | 0.71036056 | 0.326 | 0.207 | 2.95E-20 | cDC1.Xcr1 | Creb3 |
| Dym | 1.76E-24 | 0.76657609 | 0.317 | 0.205 | 3.41E-20 | cDC1.Xcr1 | Dym |
| Sec131 | 2.05E-24 | 0.47318582 | 0.544 | 0.428 | 3.96E-20 | cDC1.Xcr1 | Sec13 |
| Rprd1b | 2.28E-24 | 0.86045225 | 0.25 | 0.141 | 4.41E-20 | cDC1.Xcr1 | Rprd1b |
| Rraga1 | 2.70E-24 | 0.5572018 | 0.354 | 0.243 | 5.23E-20 | cDC1.Xcr1 | Rraga |
| Traf3ip31 | 2.87E-24 | 0.87241092 | 0.325 | 0.205 | 5.55E-20 | cDC1.Xcr1 | Traf3ip3 |
| Kdm3a | 3.04E-24 | 0.42883521 | 0.301 | 0.211 | 5.87E-20 | cDC1.Xcr1 | Kdm3a |
| Cdkn2aipnl1 | 3.72E-24 | 0.71990745 | 0.255 | 0.163 | 7.19E-20 | cDC1.Xcr1 | Cdkn2aipnl |
| Ptpre2 | 5.42E-24 | 0.63586385 | 0.515 | 0.362 | 1.05E-19 | cDC1.Xcr1 | Ptpre |
| Rfx7 | 6.99E-24 | 0.54192171 | 0.337 | 0.206 | 1.35E-19 | cDC1.Xcr1 | Rfx7 |
| Psmd14 | 7.31E-24 | 0.42933255 | 0.475 | 0.379 | 1.41E-19 | cDC1.Xcr1 | Psmd14 |
| Farsa | 7.44E-24 | 0.54187781 | 0.362 | 0.25 | 1.44E-19 | cDC1.Xcr1 | Farsa |
| Vav31 | 8.89E-24 | 0.4916136 | 0.376 | 0.248 | 1.72E-19 | cDC1.Xcr1 | Vav3 |
| Dars2 | 1.16E-23 | 0.44661189 | 0.521 | 0.409 | 2.23E-19 | cDC1.Xcr1 | Dars |
| Hadhb | 1.25E-23 | 0.27050503 | 0.419 | 0.322 | 2.42E-19 | cDC1.Xcr1 | Hadhb |
| Fasn1 | 1.35E-23 | 0.49151431 | 0.474 | 0.334 | 2.62E-19 | cDC1.Xcr1 | Fasn |
| Rabggtb | 1.47E-23 | 0.50462728 | 0.409 | 0.288 | 2.84E-19 | cDC1.Xcr1 | Rabggtb |
| Golph3 | 1.77E-23 | 0.36719722 | 0.585 | 0.474 | 3.43E-19 | cDC1.Xcr1 | Golph3 |
| Fkbp151 | 1.81E-23 | 0.43215934 | 0.529 | 0.405 | 3.51E-19 | cDC1.Xcr1 | Fkbp15 |
| Slc9a7 | 2.06E-23 | 0.80163606 | 0.259 | 0.142 | 3.99E-19 | cDC1.Xcr1 | Slc9a7 |
| Strbp | 2.32E-23 | 0.59052291 | 0.254 | 0.142 | 4.49E-19 | cDC1.Xcr1 | Strbp |
| Rpn22 | 2.53E-23 | 0.30196837 | 0.498 | 0.397 | 4.88E-19 | cDC1.Xcr1 | Rpn2 |
| Trappc1 | 2.53E-23 | 0.54367981 | 0.414 | 0.308 | 4.89E-19 | cDC1.Xcr1 | Trappc1 |
| Tor1aip11 | 2.55E-23 | 0.28206017 | 0.669 | 0.598 | 4.92E-19 | cDC1.Xcr1 | Tor1aip1 |
| Sh2b31 | 2.59E-23 | 0.30995507 | 0.555 | 0.435 | 5.00E-19 | cDC1.Xcr1 | Sh2b3 |
| Psip1 | 2.88E-23 | 0.26916183 | 0.33 | 0.244 | 5.58E-19 | cDC1.Xcr1 | Psip1 |
| Prps1l3 | 3.27E-23 | 0.58033777 | 0.288 | 0.176 | 6.32E-19 | cDC1.Xcr1 | Prps1l3 |
| Nop21 | 3.50E-23 | 0.59846915 | 0.332 | 0.194 | 6.77E-19 | cDC1.Xcr1 | Nop2 |
| Plscr31 | 3.88E-23 | 0.79310816 | 0.387 | 0.254 | 7.51E-19 | cDC1.Xcr1 | Plscr3 |
| Frg1 | 3.98E-23 | 0.42917909 | 0.364 | 0.252 | 7.70E-19 | cDC1.Xcr1 | Frg1 |
| Larp1b | 5.45E-23 | 0.8422294 | 0.26 | 0.149 | 1.05E-18 | cDC1.Xcr1 | Larp1b |
| Rtca | 6.08E-23 | 0.6503345 | 0.354 | 0.223 | 1.17E-18 | cDC1.Xcr1 | Rtca |
| Zswim81 | 6.94E-23 | 0.40842263 | 0.488 | 0.372 | 1.34E-18 | cDC1.Xcr1 | Zswim8 |
| Arl6ip1 | 7.28E-23 | 0.32815537 | 0.344 | 0.267 | 1.41E-18 | cDC1.Xcr1 | Arl6ip1 |
| Gid8 | 8.31E-23 | 0.50569976 | 0.436 | 0.331 | 1.61E-18 | cDC1.Xcr1 | Gid8 |
| Prmt11 | 8.84E-23 | 0.66111847 | 0.404 | 0.28 | 1.71E-18 | cDC1.Xcr1 | Prmt1 |
| Gpi11 | 9.90E-23 | 0.33554544 | 0.722 | 0.658 | 1.91E-18 | cDC1.Xcr1 | Gpi1 |
| Vrk3 | 9.91E-23 | 0.44118326 | 0.463 | 0.336 | 1.92E-18 | cDC1.Xcr1 | Vrk3 |
| Cldn11 | 9.94E-23 | 0.81643854 | 0.262 | 0.15 | 1.92E-18 | cDC1.Xcr1 | Cldn1 |
| Phf10 | 1.01E-22 | 0.6093877 | 0.331 | 0.224 | 1.96E-18 | cDC1.Xcr1 | Phf10 |
| Pura | 1.02E-22 | 0.47958984 | 0.458 | 0.339 | 1.97E-18 | cDC1.Xcr1 | Pura |
| Lamtor21 | 1.07E-22 | 0.33796352 | 0.462 | 0.39 | 2.07E-18 | cDC1.Xcr1 | Lamtor2 |
| Usp7 | 1.07E-22 | 0.27144082 | 0.538 | 0.453 | 2.08E-18 | cDC1.Xcr1 | Usp7 |
| Ddx23 | 1.10E-22 | 0.33128231 | 0.576 | 0.468 | 2.13E-18 | cDC1.Xcr1 | Ddx23 |
| Ncaph2 | 1.15E-22 | 0.43660455 | 0.308 | 0.231 | 2.23E-18 | cDC1.Xcr1 | Ncaph2 |
| Copg1 | 1.23E-22 | 0.34803729 | 0.513 | 0.416 | 2.37E-18 | cDC1.Xcr1 | Copg1 |
| Prdx2 | 1.54E-22 | 0.25536648 | 0.615 | 0.54 | 2.99E-18 | cDC1.Xcr1 | Prdx2 |
| Fkbp51 | 1.55E-22 | 0.40455437 | 0.308 | 0.217 | 3.00E-18 | cDC1.Xcr1 | Fkbp5 |
| Slc25a52 | 1.67E-22 | 0.46655915 | 0.438 | 0.354 | 3.23E-18 | cDC1.Xcr1 | Slc25a5 |
| Cdc42se21 | 1.93E-22 | 0.39978731 | 0.536 | 0.421 | 3.72E-18 | cDC1.Xcr1 | Cdc42se2 |
| Lyst | 2.14E-22 | 0.38180832 | 0.474 | 0.338 | 4.14E-18 | cDC1.Xcr1 | Lyst |
| Ppa1 | 2.17E-22 | 0.71472549 | 0.303 | 0.179 | 4.19E-18 | cDC1.Xcr1 | Ppa1 |
| Anks1 | 2.44E-22 | 0.66992777 | 0.388 | 0.256 | 4.73E-18 | cDC1.Xcr1 | Anks1 |
| Sys11 | 3.23E-22 | 0.4729715 | 0.502 | 0.367 | 6.25E-18 | cDC1.Xcr1 | Sys1 |
| Epn11 | 3.41E-22 | 0.33175337 | 0.779 | 0.693 | 6.59E-18 | cDC1.Xcr1 | Epn1 |
| Aars | 3.58E-22 | 0.38594414 | 0.426 | 0.326 | 6.93E-18 | cDC1.Xcr1 | Aars |
| Naa251 | 3.70E-22 | 0.64307555 | 0.301 | 0.184 | 7.16E-18 | cDC1.Xcr1 | Naa25 |
| Golga2 | 3.76E-22 | 0.45190716 | 0.382 | 0.272 | 7.28E-18 | cDC1.Xcr1 | Golga2 |
| Nvl1 | 3.81E-22 | 0.32705808 | 0.35 | 0.228 | 7.37E-18 | cDC1.Xcr1 | Nvl |
| Trabd | 4.42E-22 | 0.37686765 | 0.386 | 0.288 | 8.55E-18 | cDC1.Xcr1 | Trabd |
| Tm6sf12 | 4.53E-22 | 0.51207431 | 0.621 | 0.429 | 8.76E-18 | cDC1.Xcr1 | Tm6sf1 |
| Nubp1 | 4.71E-22 | 0.36531252 | 0.538 | 0.446 | 9.11E-18 | cDC1.Xcr1 | Nubp1 |
| Ndufa10 | 5.03E-22 | 0.50483965 | 0.506 | 0.396 | 9.73E-18 | cDC1.Xcr1 | Ndufa10 |
| Mef2d1 | 5.18E-22 | 0.28362354 | 0.607 | 0.519 | 1.00E-17 | cDC1.Xcr1 | Mef2d |
| Lacc1 | 5.22E-22 | 0.41342832 | 0.29 | 0.219 | 1.01E-17 | cDC1.Xcr1 | Lacc1 |
| Pts | 5.54E-22 | 0.76684993 | 0.284 | 0.263 | 1.07E-17 | cDC1.Xcr1 | Pts |
| Kpnb1 | 5.60E-22 | 0.41111535 | 0.371 | 0.276 | 1.08E-17 | cDC1.Xcr1 | Kpnb1 |
| Sec61g1 | 5.87E-22 | 0.47922166 | 0.449 | 0.328 | 1.13E-17 | cDC1.Xcr1 | Sec61g |
| Med271 | 6.01E-22 | 0.7723848 | 0.332 | 0.203 | 1.16E-17 | cDC1.Xcr1 | Med27 |
| Tyk2 | 8.08E-22 | 0.293661 | 0.505 | 0.378 | 1.56E-17 | cDC1.Xcr1 | Tyk2 |
| Csk1 | 8.63E-22 | 0.38489655 | 0.668 | 0.55 | 1.67E-17 | cDC1.Xcr1 | Csk |
| Top2b | 9.29E-22 | 0.4424652 | 0.408 | 0.33 | 1.80E-17 | cDC1.Xcr1 | Top2b |
| Micos101 | 9.57E-22 | 0.39523005 | 0.537 | 0.432 | 1.85E-17 | cDC1.Xcr1 | Micos10 |
| Strn4 | 1.08E-21 | 0.44811863 | 0.474 | 0.383 | 2.09E-17 | cDC1.Xcr1 | Strn4 |
| Ube2q1 | 1.09E-21 | 0.40711554 | 0.618 | 0.53 | 2.11E-17 | cDC1.Xcr1 | Ube2q1 |
| Sec31a | 1.23E-21 | 0.39438667 | 0.428 | 0.32 | 2.38E-17 | cDC1.Xcr1 | Sec31a |
| Rtraf1 | 1.38E-21 | 0.43335017 | 0.379 | 0.272 | 2.68E-17 | cDC1.Xcr1 | Rtraf |
| Pip4p1 | 1.48E-21 | 0.42302308 | 0.482 | 0.381 | 2.86E-17 | cDC1.Xcr1 | Pip4p1 |
| Sec61a1 | 1.50E-21 | 0.38271766 | 0.419 | 0.336 | 2.89E-17 | cDC1.Xcr1 | Sec61a1 |
| Tfdp11 | 1.87E-21 | 0.27146447 | 0.339 | 0.283 | 3.61E-17 | cDC1.Xcr1 | Tfdp1 |
| Zmpste24 | 1.88E-21 | 0.63381339 | 0.368 | 0.263 | 3.64E-17 | cDC1.Xcr1 | Zmpste24 |
| Pcbd2 | 1.89E-21 | 0.89848229 | 0.259 | 0.148 | 3.65E-17 | cDC1.Xcr1 | Pcbd2 |
| Cnp | 2.37E-21 | 0.67752872 | 0.36 | 0.229 | 4.59E-17 | cDC1.Xcr1 | Cnp |
| Sf3b5 | 2.53E-21 | 0.62281346 | 0.464 | 0.334 | 4.90E-17 | cDC1.Xcr1 | Sf3b5 |
| Vta1 | 2.60E-21 | 1.05074964 | 0.284 | 0.153 | 5.03E-17 | cDC1.Xcr1 | Vta1 |
| Pdcd2l | 2.62E-21 | 0.66289629 | 0.316 | 0.188 | 5.06E-17 | cDC1.Xcr1 | Pdcd2l |
| Hmox2 | 2.77E-21 | 0.39925711 | 0.513 | 0.41 | 5.35E-17 | cDC1.Xcr1 | Hmox2 |
| Tmem258 | 3.35E-21 | 0.30546532 | 0.54 | 0.473 | 6.47E-17 | cDC1.Xcr1 | Tmem258 |
| Plekhm3 | 3.42E-21 | 0.53847649 | 0.385 | 0.247 | 6.61E-17 | cDC1.Xcr1 | Plekhm3 |
| mt-Nd31 | 3.57E-21 | 0.37867757 | 0.369 | 0.251 | 6.90E-17 | cDC1.Xcr1 | mt-Nd3 |
| Tmem43 | 3.94E-21 | 0.59923521 | 0.273 | 0.171 | 7.62E-17 | cDC1.Xcr1 | Tmem43 |
| Bcap291 | 4.03E-21 | 0.69463311 | 0.299 | 0.185 | 7.80E-17 | cDC1.Xcr1 | Bcap29 |
| Bnip2 | 4.82E-21 | 0.33879481 | 0.548 | 0.466 | 9.32E-17 | cDC1.Xcr1 | Bnip2 |
| Arih21 | 5.23E-21 | 0.28187844 | 0.553 | 0.529 | 1.01E-16 | cDC1.Xcr1 | Arih2 |
| Tsr11 | 5.33E-21 | 0.82715678 | 0.315 | 0.197 | 1.03E-16 | cDC1.Xcr1 | Tsr1 |
| Pole41 | 5.71E-21 | 0.54624322 | 0.347 | 0.252 | 1.10E-16 | cDC1.Xcr1 | Pole4 |
| Hook31 | 5.79E-21 | 0.37689668 | 0.324 | 0.239 | 1.12E-16 | cDC1.Xcr1 | Hook3 |
| Strada | 5.92E-21 | 0.70949087 | 0.263 | 0.161 | 1.14E-16 | cDC1.Xcr1 | Strada |
| Atp5f11 | 6.05E-21 | 0.40978934 | 0.536 | 0.439 | 1.17E-16 | cDC1.Xcr1 | Atp5f1 |
| Atp5e | 6.11E-21 | 0.25452017 | 0.69 | 0.624 | 1.18E-16 | cDC1.Xcr1 | Atp5e |
| Atxn7l3b1 | 6.63E-21 | 0.46339734 | 0.458 | 0.334 | 1.28E-16 | cDC1.Xcr1 | Atxn7l3b |
| Zfp7061 | 6.91E-21 | 0.49013013 | 0.333 | 0.222 | 1.34E-16 | cDC1.Xcr1 | Zfp706 |
| Cdv3 | 7.22E-21 | 0.25515871 | 0.461 | 0.392 | 1.40E-16 | cDC1.Xcr1 | Cdv3 |
| Was2 | 7.56E-21 | 0.39646398 | 0.645 | 0.546 | 1.46E-16 | cDC1.Xcr1 | Was |
| Adsl1 | 7.59E-21 | 0.73050833 | 0.308 | 0.185 | 1.47E-16 | cDC1.Xcr1 | Adsl |
| Vamp31 | 8.27E-21 | 0.47213422 | 0.426 | 0.315 | 1.60E-16 | cDC1.Xcr1 | Vamp3 |
| Gga1 | 8.41E-21 | 0.51386539 | 0.578 | 0.456 | 1.63E-16 | cDC1.Xcr1 | Gga1 |
| Yars1 | 8.93E-21 | 0.82524276 | 0.275 | 0.174 | 1.73E-16 | cDC1.Xcr1 | Yars |
| Mfsd101 | 9.18E-21 | 0.5145746 | 0.33 | 0.211 | 1.77E-16 | cDC1.Xcr1 | Mfsd10 |
| Smarcc1 | 9.85E-21 | 0.30219744 | 0.38 | 0.283 | 1.90E-16 | cDC1.Xcr1 | Smarcc1 |
| Scd21 | 1.11E-20 | 0.58997513 | 0.387 | 0.263 | 2.14E-16 | cDC1.Xcr1 | Scd2 |
| Arrb11 | 1.32E-20 | 0.46771546 | 0.513 | 0.385 | 2.56E-16 | cDC1.Xcr1 | Arrb1 |
| Eif3l1 | 1.56E-20 | 0.25170541 | 0.596 | 0.533 | 3.01E-16 | cDC1.Xcr1 | Eif3l |
| Rgl2 | 1.88E-20 | 0.48745619 | 0.439 | 0.337 | 3.64E-16 | cDC1.Xcr1 | Rgl2 |
| Myo18a1 | 2.02E-20 | 0.27079839 | 0.419 | 0.309 | 3.90E-16 | cDC1.Xcr1 | Myo18a |
| Rpa11 | 2.02E-20 | 0.39077294 | 0.413 | 0.335 | 3.91E-16 | cDC1.Xcr1 | Rpa1 |
| Dpysl22 | 2.07E-20 | 0.26687609 | 0.638 | 0.516 | 3.99E-16 | cDC1.Xcr1 | Dpysl2 |
| Hibadh | 2.11E-20 | 0.38708842 | 0.426 | 0.315 | 4.08E-16 | cDC1.Xcr1 | Hibadh |
| Cct81 | 2.55E-20 | 0.42718016 | 0.469 | 0.36 | 4.93E-16 | cDC1.Xcr1 | Cct8 |
| Alg10b | 2.60E-20 | 0.69060761 | 0.31 | 0.2 | 5.04E-16 | cDC1.Xcr1 | Alg10b |
| Cnot11 | 2.66E-20 | 0.55365022 | 0.287 | 0.181 | 5.15E-16 | cDC1.Xcr1 | Cnot11 |
| Pdcd6ip | 3.02E-20 | 0.27442004 | 0.664 | 0.609 | 5.84E-16 | cDC1.Xcr1 | Pdcd6ip |
| Cln3 | 3.82E-20 | 0.89624479 | 0.279 | 0.152 | 7.38E-16 | cDC1.Xcr1 | Cln3 |
| Otub11 | 4.06E-20 | 0.31952482 | 0.619 | 0.532 | 7.84E-16 | cDC1.Xcr1 | Otub1 |
| Bbx1 | 4.27E-20 | 0.51068841 | 0.326 | 0.224 | 8.25E-16 | cDC1.Xcr1 | Bbx |
| Rnf20 | 4.36E-20 | 0.51429647 | 0.366 | 0.244 | 8.42E-16 | cDC1.Xcr1 | Rnf20 |
| Gatad2a1 | 5.00E-20 | 0.31541332 | 0.551 | 0.476 | 9.67E-16 | cDC1.Xcr1 | Gatad2a |
| Asna1 | 5.81E-20 | 0.58516164 | 0.495 | 0.376 | 1.12E-15 | cDC1.Xcr1 | Asna1 |
| Psmb1 | 5.84E-20 | 0.2662799 | 0.496 | 0.411 | 1.13E-15 | cDC1.Xcr1 | Psmb1 |
| Hmgb1 | 5.91E-20 | 0.25525572 | 0.252 | 0.181 | 1.14E-15 | cDC1.Xcr1 | Hmgb1 |
| Morc31 | 7.26E-20 | 0.3578195 | 0.382 | 0.285 | 1.40E-15 | cDC1.Xcr1 | Morc3 |
| Ints14 | 7.84E-20 | 0.67142054 | 0.272 | 0.176 | 1.52E-15 | cDC1.Xcr1 | Ints14 |
| Anp32b1 | 8.98E-20 | 0.3853575 | 0.304 | 0.226 | 1.74E-15 | cDC1.Xcr1 | Anp32b |
| Qars | 9.08E-20 | 0.39489834 | 0.509 | 0.401 | 1.76E-15 | cDC1.Xcr1 | Qars |
| Zbtb44 | 9.42E-20 | 0.38722048 | 0.301 | 0.221 | 1.82E-15 | cDC1.Xcr1 | Zbtb44 |
| Xxylt1 | 1.07E-19 | 0.92937823 | 0.273 | 0.157 | 2.08E-15 | cDC1.Xcr1 | Xxylt1 |
| Snrpc | 1.47E-19 | 0.30736906 | 0.467 | 0.38 | 2.85E-15 | cDC1.Xcr1 | Snrpc |
| Ric8a | 1.48E-19 | 0.44417318 | 0.491 | 0.382 | 2.86E-15 | cDC1.Xcr1 | Ric8a |
| Ust | 1.59E-19 | 0.54350018 | 0.262 | 0.146 | 3.08E-15 | cDC1.Xcr1 | Ust |
| Dipk1a1 | 1.62E-19 | 0.71902179 | 0.255 | 0.157 | 3.13E-15 | cDC1.Xcr1 | Dipk1a |
| Ppme1 | 1.74E-19 | 0.90087174 | 0.294 | 0.186 | 3.37E-15 | cDC1.Xcr1 | Ppme1 |
| Pum3 | 1.90E-19 | 0.42073227 | 0.398 | 0.282 | 3.68E-15 | cDC1.Xcr1 | Pum3 |
| Tmed91 | 2.12E-19 | 0.27042744 | 0.56 | 0.483 | 4.09E-15 | cDC1.Xcr1 | Tmed9 |
| Pnrc2 | 2.25E-19 | 0.25317572 | 0.402 | 0.31 | 4.35E-15 | cDC1.Xcr1 | Pnrc2 |
| Cyc1 | 2.55E-19 | 0.27606487 | 0.479 | 0.407 | 4.94E-15 | cDC1.Xcr1 | Cyc1 |
| Klf3 | 2.57E-19 | 0.28714528 | 0.469 | 0.374 | 4.97E-15 | cDC1.Xcr1 | Klf3 |
| Mapk1ip1l1 | 3.50E-19 | 0.2929138 | 0.558 | 0.469 | 6.77E-15 | cDC1.Xcr1 | Mapk1ip1l |
| Brpf1 | 3.71E-19 | 0.55312038 | 0.388 | 0.291 | 7.18E-15 | cDC1.Xcr1 | Brpf1 |
| Srsf101 | 3.72E-19 | 0.28140777 | 0.469 | 0.367 | 7.20E-15 | cDC1.Xcr1 | Srsf10 |

| | | | | | | | |
|---|---|---|---|---|---|---|---|
| Timm501 | 3.91E-19 | 0.478327 | 0.331 | 0.224 | 7.55E-15 | cDC1.Xcr1 | Timm50 |
| Sdhb1 | 3.95E-19 | 0.29264831 | 0.523 | 0.449 | 7.63E-15 | cDC1.Xcr1 | Sdhb |
| Twf22 | 4.24E-19 | 0.39375666 | 0.638 | 0.527 | 8.20E-15 | cDC1.Xcr1 | Twf2 |
| Max | 4.90E-19 | 0.37893926 | 0.415 | 0.308 | 9.47E-15 | cDC1.Xcr1 | Max |
| Nelfe2 | 5.56E-19 | 0.38019702 | 0.332 | 0.263 | 1.08E-14 | cDC1.Xcr1 | Nelfe |
| Sem1 | 5.86E-19 | 0.33163707 | 0.706 | 0.635 | 1.13E-14 | cDC1.Xcr1 | Sem1 |
| Psmb21 | 6.03E-19 | 0.29843129 | 0.608 | 0.519 | 1.17E-14 | cDC1.Xcr1 | Psmb2 |
| Pold41 | 7.09E-19 | 0.46921593 | 0.469 | 0.37 | 1.37E-14 | cDC1.Xcr1 | Pold4 |
| Sephs2 | 8.47E-19 | 0.8854979 | 0.252 | 0.148 | 1.64E-14 | cDC1.Xcr1 | Sephs2 |
| Atp5g3 | 8.55E-19 | 0.37599437 | 0.419 | 0.323 | 1.65E-14 | cDC1.Xcr1 | Atp5g3 |
| Zfp8001 | 9.49E-19 | 0.41728543 | 0.386 | 0.276 | 1.83E-14 | cDC1.Xcr1 | Zfp800 |
| Dctn3 | 1.01E-18 | 0.52078892 | 0.451 | 0.337 | 1.96E-14 | cDC1.Xcr1 | Dctn3 |
| Pitpnm11 | 1.03E-18 | 0.36314531 | 0.491 | 0.377 | 1.98E-14 | cDC1.Xcr1 | Pitpnm1 |
| Fgfr1op2 | 1.07E-18 | 0.304726 | 0.577 | 0.476 | 2.07E-14 | cDC1.Xcr1 | Fgfr1op2 |
| Ipo13 | 1.19E-18 | 0.93267581 | 0.267 | 0.15 | 2.30E-14 | cDC1.Xcr1 | Ipo13 |
| Blvra | 1.32E-18 | 0.46022217 | 0.337 | 0.242 | 2.55E-14 | cDC1.Xcr1 | Blvra |
| Usp25 | 1.32E-18 | 0.29172474 | 0.539 | 0.447 | 2.56E-14 | cDC1.Xcr1 | Usp25 |
| Snu13 | 1.39E-18 | 0.33315364 | 0.549 | 0.471 | 2.69E-14 | cDC1.Xcr1 | Snu13 |
| Nudt21 | 1.45E-18 | 0.40906022 | 0.401 | 0.327 | 2.81E-14 | cDC1.Xcr1 | Nudt21 |
| P2ry61 | 1.48E-18 | 0.78892228 | 0.338 | 0.204 | 2.86E-14 | cDC1.Xcr1 | P2ry6 |
| Isca2 | 1.67E-18 | 0.46385998 | 0.25 | 0.161 | 3.23E-14 | cDC1.Xcr1 | Isca2 |
| Trim27 | 1.80E-18 | 0.35659631 | 0.456 | 0.35 | 3.49E-14 | cDC1.Xcr1 | Trim27 |
| Srsf6 | 1.84E-18 | 0.28292293 | 0.681 | 0.62 | 3.55E-14 | cDC1.Xcr1 | Srsf6 |
| Cnot3 | 2.08E-18 | 0.29041035 | 0.407 | 0.324 | 4.02E-14 | cDC1.Xcr1 | Cnot3 |
| Dcaf122 | 2.19E-18 | 0.49034237 | 0.48 | 0.358 | 4.24E-14 | cDC1.Xcr1 | Dcaf12 |
| Atp5j2 | 2.25E-18 | 0.38436176 | 0.667 | 0.581 | 4.34E-14 | cDC1.Xcr1 | Atp5j2 |
| Cbx3 | 2.38E-18 | 0.31280709 | 0.254 | 0.191 | 4.61E-14 | cDC1.Xcr1 | Cbx3 |
| Ccdc93 | 2.40E-18 | 0.63805903 | 0.261 | 0.16 | 4.63E-14 | cDC1.Xcr1 | Ccdc93 |
| Sf3a2 | 2.50E-18 | 0.40366203 | 0.407 | 0.32 | 4.84E-14 | cDC1.Xcr1 | Sf3a2 |
| Agap31 | 2.70E-18 | 0.31343543 | 0.401 | 0.312 | 5.22E-14 | cDC1.Xcr1 | Agap3 |
| Txnip1 | 3.41E-18 | 0.58301735 | 0.408 | 0.281 | 6.58E-14 | cDC1.Xcr1 | Txnip |
| Crtc2 | 3.41E-18 | 0.32163587 | 0.52 | 0.416 | 6.58E-14 | cDC1.Xcr1 | Crtc2 |
| Eif3b1 | 3.43E-18 | 0.35089162 | 0.504 | 0.39 | 6.63E-14 | cDC1.Xcr1 | Eif3b |
| Psen1 | 3.50E-18 | 0.38244735 | 0.395 | 0.312 | 6.77E-14 | cDC1.Xcr1 | Psen1 |
| Lin7c | 3.58E-18 | 0.29527893 | 0.484 | 0.411 | 6.92E-14 | cDC1.Xcr1 | Lin7c |
| Gnai21 | 3.66E-18 | 0.27617192 | 0.907 | 0.862 | 7.07E-14 | cDC1.Xcr1 | Gnai2 |
| Gmpr2 | 4.03E-18 | 0.66103783 | 0.299 | 0.2 | 7.80E-14 | cDC1.Xcr1 | Gmpr2 |
| Tsg101 | 4.46E-18 | 0.52543232 | 0.424 | 0.3 | 8.62E-14 | cDC1.Xcr1 | Tsg101 |
| Med8 | 4.83E-18 | 0.65724379 | 0.33 | 0.229 | 9.34E-14 | cDC1.Xcr1 | Med8 |
| Dock71 | 4.86E-18 | 0.3754973 | 0.385 | 0.29 | 9.39E-14 | cDC1.Xcr1 | Dock7 |
| Tent5c | 5.11E-18 | 0.38276047 | 0.4 | 0.314 | 9.89E-14 | cDC1.Xcr1 | Tent5c |
| Orc41 | 5.35E-18 | 0.29340685 | 0.352 | 0.245 | 1.04E-13 | cDC1.Xcr1 | Orc4 |
| Retreg11 | 6.30E-18 | 0.52497557 | 0.336 | 0.234 | 1.22E-13 | cDC1.Xcr1 | Retreg1 |
| Rai11 | 7.73E-18 | 0.41811729 | 0.341 | 0.232 | 1.49E-13 | cDC1.Xcr1 | Rai1 |
| Pik3cd2 | 8.00E-18 | 0.26095754 | 0.688 | 0.546 | 1.55E-13 | cDC1.Xcr1 | Pik3cd |
| Rbm7 | 8.94E-18 | 0.35971698 | 0.414 | 0.317 | 1.73E-13 | cDC1.Xcr1 | Rbm7 |
| Kif3b | 9.65E-18 | 0.54551008 | 0.411 | 0.324 | 1.87E-13 | cDC1.Xcr1 | Kif3b |
| Gps1 | 9.67E-18 | 0.29601455 | 0.393 | 0.305 | 1.87E-13 | cDC1.Xcr1 | Gps1 |
| Ube2e11 | 1.03E-17 | 0.58173246 | 0.397 | 0.296 | 1.98E-13 | cDC1.Xcr1 | Ube2e1 |
| Slc52a31 | 1.03E-17 | 0.50924452 | 0.25 | 0.144 | 1.99E-13 | cDC1.Xcr1 | Slc52a3 |
| Itpripl21 | 1.06E-17 | 0.26070497 | 0.564 | 0.46 | 2.04E-13 | cDC1.Xcr1 | Itpripl2 |
| Vps361 | 1.14E-17 | 0.69651297 | 0.35 | 0.245 | 2.20E-13 | cDC1.Xcr1 | Vps36 |
| Ebna1bp2 | 1.17E-17 | 0.4444455 | 0.431 | 0.327 | 2.26E-13 | cDC1.Xcr1 | Ebna1bp2 |
| Gar11 | 1.20E-17 | 0.37751391 | 0.338 | 0.243 | 2.33E-13 | cDC1.Xcr1 | Gar1 |
| Agpat11 | 1.31E-17 | 0.39554248 | 0.387 | 0.308 | 2.53E-13 | cDC1.Xcr1 | Agpat1 |
| Arfip2 | 1.38E-17 | 0.58929157 | 0.276 | 0.172 | 2.68E-13 | cDC1.Xcr1 | Arfip2 |
| Eif1ad | 1.47E-17 | 0.45259413 | 0.265 | 0.189 | 2.85E-13 | cDC1.Xcr1 | Eif1ad |
| Sfxn11 | 1.53E-17 | 0.52058478 | 0.365 | 0.267 | 2.95E-13 | cDC1.Xcr1 | Sfxn1 |
| Dync1li2 | 1.80E-17 | 0.44166814 | 0.371 | 0.264 | 3.48E-13 | cDC1.Xcr1 | Dync1li2 |
| Mapk9 | 1.82E-17 | 0.59203294 | 0.303 | 0.209 | 3.52E-13 | cDC1.Xcr1 | Mapk9 |
| Smg5 | 1.95E-17 | 0.42203852 | 0.447 | 0.344 | 3.77E-13 | cDC1.Xcr1 | Smg5 |
| Zbtb7a | 2.01E-17 | 0.32167787 | 0.515 | 0.412 | 3.88E-13 | cDC1.Xcr1 | Zbtb7a |
| Hipk12 | 2.05E-17 | 0.37930046 | 0.733 | 0.603 | 3.96E-13 | cDC1.Xcr1 | Hipk1 |
| Supt16 | 2.06E-17 | 0.34060055 | 0.462 | 0.366 | 3.97E-13 | cDC1.Xcr1 | Supt16 |
| Pcyt21 | 2.11E-17 | 0.49412772 | 0.346 | 0.229 | 4.08E-13 | cDC1.Xcr1 | Pcyt2 |
| Socs11 | 2.55E-17 | 0.36640126 | 0.32 | 0.284 | 4.92E-13 | cDC1.Xcr1 | Socs1 |
| Ptpn111 | 2.59E-17 | 0.40740443 | 0.458 | 0.343 | 5.01E-13 | cDC1.Xcr1 | Ptpn11 |
| Gfm1 | 2.68E-17 | 0.53973326 | 0.272 | 0.174 | 5.18E-13 | cDC1.Xcr1 | Gfm1 |
| Irf91 | 2.73E-17 | 0.43631964 | 0.42 | 0.33 | 5.28E-13 | cDC1.Xcr1 | Irf9 |
| Farsb1 | 2.95E-17 | 0.60375018 | 0.314 | 0.198 | 5.70E-13 | cDC1.Xcr1 | Farsb |
| Ap5b1 | 3.32E-17 | 0.46889338 | 0.251 | 0.156 | 6.43E-13 | cDC1.Xcr1 | Ap5b1 |
| Erlin2 | 3.35E-17 | 0.49514449 | 0.375 | 0.256 | 6.48E-13 | cDC1.Xcr1 | Erlin2 |
| Hcfc1 | 3.55E-17 | 0.28371409 | 0.419 | 0.316 | 6.87E-13 | cDC1.Xcr1 | Hcfc1 |
| Timm17a1 | 4.09E-17 | 0.47906071 | 0.342 | 0.257 | 7.90E-13 | cDC1.Xcr1 | Timm17a |
| Copz1 | 4.29E-17 | 0.2577075 | 0.65 | 0.579 | 8.30E-13 | cDC1.Xcr1 | Copz1 |
| Tpgs1 | 4.52E-17 | 0.48191781 | 0.304 | 0.21 | 8.74E-13 | cDC1.Xcr1 | Tpgs1 |
| Lmbrd1 | 4.54E-17 | 0.62409742 | 0.413 | 0.283 | 8.77E-13 | cDC1.Xcr1 | Lmbrd1 |
| Elob | 4.62E-17 | 0.25940328 | 0.695 | 0.641 | 8.94E-13 | cDC1.Xcr1 | Elob |
| Kat6b | 5.61E-17 | 0.46764483 | 0.261 | 0.161 | 1.08E-12 | cDC1.Xcr1 | Kat6b |
| Hdac41 | 5.80E-17 | 0.75961146 | 0.326 | 0.213 | 1.12E-12 | cDC1.Xcr1 | Hdac4 |
| Psmd13 | 5.98E-17 | 0.28373565 | 0.518 | 0.44 | 1.16E-12 | cDC1.Xcr1 | Psmd13 |
| Larp4 | 6.47E-17 | 0.32992474 | 0.458 | 0.376 | 1.25E-12 | cDC1.Xcr1 | Larp4 |
| Ap2a21 | 6.94E-17 | 0.34170403 | 0.578 | 0.445 | 1.34E-12 | cDC1.Xcr1 | Ap2a2 |
| Dapk3 | 7.14E-17 | 0.55815501 | 0.377 | 0.279 | 1.38E-12 | cDC1.Xcr1 | Dapk3 |
| Lrrc47 | 7.92E-17 | 0.414734 | 0.403 | 0.309 | 1.53E-12 | cDC1.Xcr1 | Lrrc47 |
| Rere1 | 8.22E-17 | 0.43395855 | 0.365 | 0.26 | 1.59E-12 | cDC1.Xcr1 | Rere |
| Golim4 | 8.78E-17 | 0.57084383 | 0.252 | 0.149 | 1.70E-12 | cDC1.Xcr1 | Golim4 |
| Uqcr11 | 9.47E-17 | 0.49322634 | 0.458 | 0.369 | 1.83E-12 | cDC1.Xcr1 | Uqcr11 |
| Ywhaz | 1.30E-16 | 0.60561094 | 0.412 | 0.3 | 2.50E-12 | cDC1.Xcr1 | Ywhaz |
| Cse1l | 1.48E-16 | 0.38789357 | 0.336 | 0.257 | 2.85E-12 | cDC1.Xcr1 | Cse1l |
| Slc25a28 | 1.48E-16 | 0.49163563 | 0.314 | 0.232 | 2.86E-12 | cDC1.Xcr1 | Slc25a28 |
| Dhdds | 1.58E-16 | 0.59746025 | 0.276 | 0.179 | 3.06E-12 | cDC1.Xcr1 | Dhdds |
| Psmd12 | 1.61E-16 | 0.26107876 | 0.434 | 0.339 | 3.10E-12 | cDC1.Xcr1 | Psmd12 |
| Metap1 | 1.64E-16 | 0.4443184 | 0.311 | 0.223 | 3.18E-12 | cDC1.Xcr1 | Metap1 |
| Cops61 | 1.72E-16 | 0.34116576 | 0.548 | 0.46 | 3.33E-12 | cDC1.Xcr1 | Cops6 |
| Lrmp2 | 1.76E-16 | 0.82949593 | 0.317 | 0.215 | 3.40E-12 | cDC1.Xcr1 | Lrmp |
| D17H6S53E | 1.84E-16 | 0.69330502 | 0.251 | 0.144 | 3.56E-12 | cDC1.Xcr1 | D17H6S53E |
| Hint12 | 1.93E-16 | 0.59084492 | 0.385 | 0.287 | 3.72E-12 | cDC1.Xcr1 | Hint1 |
| Cox6a11 | 1.95E-16 | 0.35532755 | 0.489 | 0.41 | 3.77E-12 | cDC1.Xcr1 | Cox6a1 |
| Babam1 | 1.97E-16 | 0.30370266 | 0.491 | 0.412 | 3.80E-12 | cDC1.Xcr1 | Babam1 |
| Kcmf1 | 1.98E-16 | 0.26117457 | 0.509 | 0.445 | 3.82E-12 | cDC1.Xcr1 | Kcmf1 |
| Flnb | 2.26E-16 | 0.29909704 | 0.46 | 0.348 | 4.37E-12 | cDC1.Xcr1 | Flnb |
| Ralbp1 | 2.26E-16 | 0.57961963 | 0.403 | 0.289 | 4.37E-12 | cDC1.Xcr1 | Ralbp1 |
| Rtn11 | 2.43E-16 | 0.43902566 | 0.415 | 0.342 | 4.70E-12 | cDC1.Xcr1 | Rtn1 |
| Tomm70a | 2.57E-16 | 0.36220263 | 0.347 | 0.23 | 4.97E-12 | cDC1.Xcr1 | Tomm70a |
| Psma2 | 4.13E-16 | 0.35641977 | 0.449 | 0.364 | 7.98E-12 | cDC1.Xcr1 | Psma2 |
| Eftud2 | 4.16E-16 | 0.30797262 | 0.478 | 0.399 | 8.04E-12 | cDC1.Xcr1 | Eftud2 |
| Taf4 | 4.38E-16 | 0.47124903 | 0.344 | 0.257 | 8.46E-12 | cDC1.Xcr1 | Taf4 |
| Gdap21 | 5.20E-16 | 0.64380875 | 0.29 | 0.182 | 1.01E-11 | cDC1.Xcr1 | Gdap2 |
| Bcas2 | 5.47E-16 | 0.27071917 | 0.375 | 0.275 | 1.06E-11 | cDC1.Xcr1 | Bcas2 |
| Aarsd1 | 6.04E-16 | 0.36852018 | 0.273 | 0.181 | 1.17E-11 | cDC1.Xcr1 | Aarsd1 |
| Hnrnpab1 | 6.19E-16 | 0.30460345 | 0.493 | 0.43 | 1.20E-11 | cDC1.Xcr1 | Hnrnpab |
| Slc25a41 | 6.25E-16 | 0.34927679 | 0.471 | 0.368 | 1.21E-11 | cDC1.Xcr1 | Slc25a4 |
| Mospd31 | 7.61E-16 | 0.5477926 | 0.381 | 0.274 | 1.47E-11 | cDC1.Xcr1 | Mospd3 |
| Isoc1 | 7.68E-16 | 0.60612252 | 0.272 | 0.176 | 1.49E-11 | cDC1.Xcr1 | Isoc1 |
| Map2k4 | 7.82E-16 | 0.70810232 | 0.257 | 0.154 | 1.51E-11 | cDC1.Xcr1 | Map2k4 |
| Magoh1 | 8.09E-16 | 0.48698125 | 0.435 | 0.355 | 1.56E-11 | cDC1.Xcr1 | Magoh |
| Tbpl1 | 9.23E-16 | 0.67456065 | 0.254 | 0.162 | 1.79E-11 | cDC1.Xcr1 | Tbpl1 |
| Fbxo61 | 9.97E-16 | 0.29364556 | 0.369 | 0.263 | 1.93E-11 | cDC1.Xcr1 | Fbxo6 |
| Raf1 | 1.00E-15 | 0.33326598 | 0.451 | 0.357 | 1.94E-11 | cDC1.Xcr1 | Raf1 |
| Gbf1 | 1.01E-15 | 0.47591059 | 0.358 | 0.249 | 1.95E-11 | cDC1.Xcr1 | Gbf1 |
| Sae11 | 1.26E-15 | 0.54311819 | 0.379 | 0.306 | 2.45E-11 | cDC1.Xcr1 | Sae1 |
| Cdk21 | 1.29E-15 | 0.25718851 | 0.278 | 0.218 | 2.49E-11 | cDC1.Xcr1 | Cdk2 |
| Eed | 1.32E-15 | 0.31747274 | 0.352 | 0.274 | 2.55E-11 | cDC1.Xcr1 | Eed |
| Mgat22 | 1.37E-15 | 0.32103333 | 0.46 | 0.395 | 2.64E-11 | cDC1.Xcr1 | Mgat2 |
| Swi5 | 1.40E-15 | 0.42779725 | 0.431 | 0.336 | 2.71E-11 | cDC1.Xcr1 | Swi5 |
| Set1 | 1.44E-15 | 0.28187222 | 0.447 | 0.371 | 2.78E-11 | cDC1.Xcr1 | Set |
| Uqcrb | 1.48E-15 | 0.48841488 | 0.391 | 0.301 | 2.86E-11 | cDC1.Xcr1 | Uqcrb |
| Ddx212 | 1.52E-15 | 0.2824292 | 0.635 | 0.531 | 2.93E-11 | cDC1.Xcr1 | Ddx21 |
| Ccm2 | 1.82E-15 | 0.39837428 | 0.359 | 0.264 | 3.52E-11 | cDC1.Xcr1 | Ccm2 |
| BC031181 | 1.94E-15 | 0.30144894 | 0.393 | 0.314 | 3.76E-11 | cDC1.Xcr1 | BC031181 |
| Aim21 | 2.06E-15 | 0.30901686 | 0.327 | 0.23 | 3.98E-11 | cDC1.Xcr1 | Aim2 |
| Cnih1 | 2.07E-15 | 0.35965559 | 0.343 | 0.262 | 4.00E-11 | cDC1.Xcr1 | Cnih1 |
| Gadd45gip1 | 2.14E-15 | 0.34252539 | 0.344 | 0.259 | 4.13E-11 | cDC1.Xcr1 | Gadd45gip1 |
| Lypla2 | 2.19E-15 | 0.39705305 | 0.505 | 0.41 | 4.23E-11 | cDC1.Xcr1 | Lypla2 |
| Fam234b1 | 2.42E-15 | 0.61569379 | 0.261 | 0.168 | 4.68E-11 | cDC1.Xcr1 | Fam234b |
| Ndufab1 | 2.54E-15 | 0.64468536 | 0.324 | 0.205 | 4.91E-11 | cDC1.Xcr1 | Ndufab1 |
| Palm1 | 2.55E-15 | 0.31997342 | 0.406 | 0.294 | 4.93E-11 | cDC1.Xcr1 | Palm |
| Nmt2 | 2.63E-15 | 0.52838652 | 0.255 | 0.166 | 5.09E-11 | cDC1.Xcr1 | Nmt2 |
| Ldb1 | 2.82E-15 | 0.28242185 | 0.474 | 0.385 | 5.46E-11 | cDC1.Xcr1 | Ldb1 |
| Fyttd1 | 3.21E-15 | 0.55147458 | 0.335 | 0.244 | 6.20E-11 | cDC1.Xcr1 | Fyttd1 |
| Pip4k2c | 3.33E-15 | 0.68384235 | 0.311 | 0.222 | 6.43E-11 | cDC1.Xcr1 | Pip4k2c |
| Fundc11 | 3.56E-15 | 0.47554563 | 0.531 | 0.449 | 6.89E-11 | cDC1.Xcr1 | Fundc1 |
| Rtf2 | 3.60E-15 | 0.39551852 | 0.548 | 0.486 | 6.97E-11 | cDC1.Xcr1 | Rtf2 |
| Trim26 | 4.31E-15 | 0.38282345 | 0.316 | 0.229 | 8.33E-11 | cDC1.Xcr1 | Trim26 |
| Cct6a1 | 4.85E-15 | 0.30744305 | 0.463 | 0.371 | 9.38E-11 | cDC1.Xcr1 | Cct6a |
| Sgk31 | 4.89E-15 | 0.47653771 | 0.462 | 0.358 | 9.46E-11 | cDC1.Xcr1 | Sgk3 |
| Kars | 5.07E-15 | 0.28894925 | 0.496 | 0.423 | 9.81E-11 | cDC1.Xcr1 | Kars |
| Tomm7 | 5.20E-15 | 0.33196349 | 0.571 | 0.463 | 1.01E-10 | cDC1.Xcr1 | Tomm7 |
| Rnf139 | 6.02E-15 | 0.52144926 | 0.268 | 0.177 | 1.16E-10 | cDC1.Xcr1 | Rnf139 |
| Dtx3 | 6.29E-15 | 0.3663791 | 0.267 | 0.205 | 1.22E-10 | cDC1.Xcr1 | Dtx3 |
| Hpcal11 | 6.92E-15 | 0.51317641 | 0.315 | 0.212 | 1.34E-10 | cDC1.Xcr1 | Hpcal1 |
| Cops7a | 7.10E-15 | 0.55138014 | 0.337 | 0.233 | 1.37E-10 | cDC1.Xcr1 | Cops7a |
| Kat7 | 7.85E-15 | 0.38262027 | 0.314 | 0.224 | 1.52E-10 | cDC1.Xcr1 | Kat7 |
| Fermt32 | 8.16E-15 | 0.40239871 | 0.714 | 0.59 | 1.58E-10 | cDC1.Xcr1 | Fermt3 |
| Usp5 | 8.54E-15 | 0.27926271 | 0.413 | 0.323 | 1.65E-10 | cDC1.Xcr1 | Usp5 |
| Rapgef11 | 9.21E-15 | 0.2923826 | 0.451 | 0.393 | 1.78E-10 | cDC1.Xcr1 | Rapgef1 |
| Lsm72 | 9.73E-15 | 0.40598193 | 0.426 | 0.338 | 1.88E-10 | cDC1.Xcr1 | Lsm7 |
| Ube2n | 9.95E-15 | 0.4061115 | 0.328 | 0.248 | 1.92E-10 | cDC1.Xcr1 | Ube2n |
| Cox7a2 | 1.05E-14 | 0.37689845 | 0.369 | 0.3 | 2.04E-10 | cDC1.Xcr1 | Cox7a2 |
| Arhgap151 | 1.09E-14 | 0.69944566 | 0.297 | 0.205 | 2.10E-10 | cDC1.Xcr1 | Arhgap15 |
| Copb2 | 1.18E-14 | 0.30423191 | 0.473 | 0.395 | 2.28E-10 | cDC1.Xcr1 | Copb2 |
| Camkk11 | 1.18E-14 | 0.5845559 | 0.362 | 0.25 | 2.28E-10 | cDC1.Xcr1 | Camkk1 |
| Cox14 | 1.40E-14 | 0.34467668 | 0.554 | 0.453 | 2.71E-10 | cDC1.Xcr1 | Cox14 |
| Efr3a1 | 1.47E-14 | 0.34030184 | 0.365 | 0.292 | 2.84E-10 | cDC1.Xcr1 | Efr3a |
| Mars1 | 1.54E-14 | 0.32136683 | 0.322 | 0.244 | 2.98E-10 | cDC1.Xcr1 | Mars |
| Trp531 | 1.55E-14 | 0.33643514 | 0.464 | 0.388 | 3.00E-10 | cDC1.Xcr1 | Trp53 |
| Zfp655 | 1.77E-14 | 0.35336221 | 0.283 | 0.193 | 3.43E-10 | cDC1.Xcr1 | Zfp655 |
| Ndufb7 | 1.77E-14 | 0.25262687 | 0.591 | 0.502 | 3.43E-10 | cDC1.Xcr1 | Ndufb7 |
| Eif2ak1 | 1.96E-14 | 0.57203802 | 0.344 | 0.257 | 3.79E-10 | cDC1.Xcr1 | Eif2ak1 |
| Rpia | 2.06E-14 | 0.81741559 | 0.263 | 0.164 | 3.98E-10 | cDC1.Xcr1 | Rpia |
| Pdcl31 | 2.20E-14 | 0.40703012 | 0.504 | 0.406 | 4.26E-10 | cDC1.Xcr1 | Pdcl3 |
| Wdr48 | 2.21E-14 | 0.40714196 | 0.278 | 0.187 | 4.27E-10 | cDC1.Xcr1 | Wdr48 |
| Dnttip1 | 2.24E-14 | 0.63467967 | 0.327 | 0.224 | 4.32E-10 | cDC1.Xcr1 | Dnttip1 |
| Txndc9 | 2.47E-14 | 0.4356648 | 0.451 | 0.356 | 4.78E-10 | cDC1.Xcr1 | Txndc9 |
| Slc35b11 | 2.52E-14 | 0.35430069 | 0.288 | 0.21 | 4.87E-10 | cDC1.Xcr1 | Slc35b1 |
| Phactr2 | 2.61E-14 | 0.42204761 | 0.365 | 0.265 | 5.05E-10 | cDC1.Xcr1 | Phactr2 |
| Pcmt1 | 2.91E-14 | 0.30683944 | 0.359 | 0.296 | 5.62E-10 | cDC1.Xcr1 | Pcmt1 |
| Nfl1 | 2.97E-14 | 0.26970916 | 0.336 | 0.268 | 5.75E-10 | cDC1.Xcr1 | Nfl1 |
| Grcc10 | 3.17E-14 | 0.27277998 | 0.577 | 0.522 | 6.12E-10 | cDC1.Xcr1 | Grcc10 |
| Hexa1 | 3.26E-14 | 0.28739749 | 0.643 | 0.559 | 6.30E-10 | cDC1.Xcr1 | Hexa |
| Ubap2l | 3.28E-14 | 0.25092224 | 0.544 | 0.492 | 6.34E-10 | cDC1.Xcr1 | Ubap2l |
| Hnrnpr | 3.66E-14 | 0.3685045 | 0.363 | 0.279 | 7.08E-10 | cDC1.Xcr1 | Hnrnpr |
| Hypk | 3.91E-14 | 0.40632927 | 0.401 | 0.297 | 7.56E-10 | cDC1.Xcr1 | Hypk |
| Bicd21 | 4.16E-14 | 0.29531082 | 0.406 | 0.325 | 8.04E-10 | cDC1.Xcr1 | Bicd2 |
| Tnpo3 | 4.54E-14 | 0.41834994 | 0.376 | 0.298 | 8.77E-10 | cDC1.Xcr1 | Tnpo3 |
| Adh51 | 4.80E-14 | 0.33077659 | 0.511 | 0.429 | 9.29E-10 | cDC1.Xcr1 | Adh5 |
| Rexo21 | 4.98E-14 | 0.36819701 | 0.509 | 0.416 | 9.63E-10 | cDC1.Xcr1 | Rexo2 |
| Gnl3l1 | 6.07E-14 | 0.42625483 | 0.27 | 0.175 | 1.17E-09 | cDC1.Xcr1 | Gnl3l |
| Hars1 | 6.34E-14 | 0.29054312 | 0.353 | 0.271 | 1.23E-09 | cDC1.Xcr1 | Hars |
| Rnf5 | 6.53E-14 | 0.41180864 | 0.319 | 0.226 | 1.26E-09 | cDC1.Xcr1 | Rnf5 |
| Spcs1 | 6.89E-14 | 0.33734002 | 0.539 | 0.463 | 1.33E-09 | cDC1.Xcr1 | Spcs1 |
| Zfp384 | 7.07E-14 | 0.25929397 | 0.401 | 0.311 | 1.37E-09 | cDC1.Xcr1 | Zfp384 |
| Ndufaf82 | 7.10E-14 | 0.50387982 | 0.339 | 0.241 | 1.37E-09 | cDC1.Xcr1 | Ndufaf8 |
| Polr3k | 7.44E-14 | 0.71780452 | 0.292 | 0.192 | 1.44E-09 | cDC1.Xcr1 | Polr3k |
| Emc7 | 7.55E-14 | 0.40385704 | 0.395 | 0.303 | 1.46E-09 | cDC1.Xcr1 | Emc7 |
| Dtx2 | 8.76E-14 | 0.46793201 | 0.29 | 0.187 | 1.69E-09 | cDC1.Xcr1 | Dtx2 |
| Tomm402 | 1.13E-13 | 0.47702002 | 0.429 | 0.321 | 2.18E-09 | cDC1.Xcr1 | Tomm40 |
| Rbm22 | 1.20E-13 | 0.41316951 | 0.355 | 0.278 | 2.32E-09 | cDC1.Xcr1 | Rbm22 |
| Aldh9a11 | 1.20E-13 | 0.33476848 | 0.544 | 0.472 | 2.32E-09 | cDC1.Xcr1 | Aldh9a1 |
| Cdc5l | 1.23E-13 | 0.32962524 | 0.363 | 0.299 | 2.38E-09 | cDC1.Xcr1 | Cdc5l |
| Rassf52 | 1.32E-13 | 0.26243522 | 0.653 | 0.542 | 2.55E-09 | cDC1.Xcr1 | Rassf5 |
| Idh21 | 1.35E-13 | 0.47662458 | 0.336 | 0.245 | 2.61E-09 | cDC1.Xcr1 | Idh2 |
| Ncbp2 | 1.45E-13 | 0.53522293 | 0.332 | 0.236 | 2.80E-09 | cDC1.Xcr1 | Ncbp2 |
| Abcf2 | 1.55E-13 | 0.75754603 | 0.25 | 0.156 | 3.00E-09 | cDC1.Xcr1 | Abcf2 |
| Vkorc1l1 | 1.56E-13 | 0.25817062 | 0.279 | 0.193 | 3.02E-09 | cDC1.Xcr1 | Vkorc1l1 |
| Drg1 | 1.69E-13 | 0.33154987 | 0.419 | 0.354 | 3.26E-09 | cDC1.Xcr1 | Drg1 |
| Ak31 | 1.80E-13 | 0.33773198 | 0.327 | 0.233 | 3.49E-09 | cDC1.Xcr1 | Ak3 |
| Psmd7 | 1.87E-13 | 0.36937271 | 0.464 | 0.389 | 3.62E-09 | cDC1.Xcr1 | Psmd7 |
| Oaz2 | 1.91E-13 | 0.50736825 | 0.283 | 0.19 | 3.69E-09 | cDC1.Xcr1 | Oaz2 |
| Vhl1 | 1.96E-13 | 0.33139356 | 0.396 | 0.279 | 3.78E-09 | cDC1.Xcr1 | Vhl |
| Fam98b | 2.07E-13 | 0.51690738 | 0.278 | 0.189 | 4.00E-09 | cDC1.Xcr1 | Fam98b |
| Ttc39b | 2.11E-13 | 0.51438672 | 0.29 | 0.192 | 4.07E-09 | cDC1.Xcr1 | Ttc39b |
| Snrpb1 | 2.20E-13 | 0.26001758 | 0.477 | 0.411 | 4.25E-09 | cDC1.Xcr1 | Snrpb |
| Trap1 | 2.22E-13 | 0.45545683 | 0.335 | 0.248 | 4.28E-09 | cDC1.Xcr1 | Trap1 |
| Sec23b1 | 2.31E-13 | 0.27926763 | 0.326 | 0.26 | 4.47E-09 | cDC1.Xcr1 | Sec23b |
| Usf3 | 2.46E-13 | 0.35809782 | 0.312 | 0.22 | 4.76E-09 | cDC1.Xcr1 | Usf3 |
| Dhrs11 | 2.58E-13 | 0.54730702 | 0.415 | 0.314 | 4.99E-09 | cDC1.Xcr1 | Dhrs1 |
| Zfp385a1 | 2.66E-13 | 0.54210607 | 0.482 | 0.391 | 5.14E-09 | cDC1.Xcr1 | Zfp385a |
| Cdk191 | 3.15E-13 | 0.30412592 | 0.304 | 0.233 | 6.09E-09 | cDC1.Xcr1 | Cdk19 |
| Snrpd32 | 3.89E-13 | 0.31778966 | 0.572 | 0.494 | 7.52E-09 | cDC1.Xcr1 | Snrpd3 |
| R3hdm1 | 4.87E-13 | 0.4188909 | 0.317 | 0.233 | 9.42E-09 | cDC1.Xcr1 | R3hdm1 |
| Atxn10 | 5.17E-13 | 0.31507159 | 0.463 | 0.386 | 1.00E-08 | cDC1.Xcr1 | Atxn10 |
| Pcbp21 | 5.31E-13 | 0.26798557 | 0.564 | 0.501 | 1.03E-08 | cDC1.Xcr1 | Pcbp2 |
| Snapc1 | 5.32E-13 | 0.64267275 | 0.25 | 0.175 | 1.03E-08 | cDC1.Xcr1 | Snapc1 |
| Lsm6 | 5.38E-13 | 0.63925295 | 0.272 | 0.178 | 1.04E-08 | cDC1.Xcr1 | Lsm6 |
| Ost41 | 5.46E-13 | 0.50965023 | 0.38 | 0.281 | 1.06E-08 | cDC1.Xcr1 | Ost4 |
| Rpp211 | 5.63E-13 | 0.51204335 | 0.31 | 0.232 | 1.09E-08 | cDC1.Xcr1 | Rpp21 |
| Abcc1 | 6.18E-13 | 0.26466929 | 0.349 | 0.273 | 1.19E-08 | cDC1.Xcr1 | Abcc1 |
| Pygb1 | 6.37E-13 | 0.36680299 | 0.471 | 0.358 | 1.23E-08 | cDC1.Xcr1 | Pygb |
| Gabarapl21 | 6.84E-13 | 0.31021195 | 0.545 | 0.454 | 1.32E-08 | cDC1.Xcr1 | Gabarapl2 |
| Esrra1 | 7.02E-13 | 0.42498134 | 0.456 | 0.351 | 1.36E-08 | cDC1.Xcr1 | Esrra |
| Kdelr21 | 7.10E-13 | 0.38982949 | 0.392 | 0.304 | 1.37E-08 | cDC1.Xcr1 | Kdelr2 |
| Slc25a22 | 7.17E-13 | 0.54181577 | 0.259 | 0.18 | 1.39E-08 | cDC1.Xcr1 | Slc25a22 |
| Cpsf1 | 7.65E-13 | 0.38123544 | 0.314 | 0.226 | 1.48E-08 | cDC1.Xcr1 | Cpsf1 |
| Hspd11 | 7.74E-13 | 0.35204015 | 0.332 | 0.275 | 1.50E-08 | cDC1.Xcr1 | Hspd1 |
| Man1b1 | 7.91E-13 | 0.28357954 | 0.415 | 0.321 | 1.53E-08 | cDC1.Xcr1 | Man1b1 |
| Tbcd1 | 9.39E-13 | 0.25526465 | 0.352 | 0.248 | 1.81E-08 | cDC1.Xcr1 | Tbcd |
| Nacc1 | 9.60E-13 | 0.26074155 | 0.451 | 0.358 | 1.86E-08 | cDC1.Xcr1 | Nacc1 |
| Plekha2 | 9.70E-13 | 0.34246506 | 0.504 | 0.421 | 1.88E-08 | cDC1.Xcr1 | Plekha2 |
| Kat6a | 9.78E-13 | 0.2541831 | 0.452 | 0.368 | 1.89E-08 | cDC1.Xcr1 | Kat6a |
| Psmb3 | 1.03E-12 | 0.45140495 | 0.325 | 0.247 | 2.00E-08 | cDC1.Xcr1 | Psmb3 |
| Naa501 | 1.07E-12 | 0.29873295 | 0.502 | 0.432 | 2.08E-08 | cDC1.Xcr1 | Naa50 |
| Eri1 | 1.12E-12 | 0.32012679 | 0.287 | 0.244 | 2.16E-08 | cDC1.Xcr1 | Eri1 |
| Sestd1 | 1.16E-12 | 0.33459149 | 0.324 | 0.223 | 2.25E-08 | cDC1.Xcr1 | Sestd1 |
| Cox6c | 1.23E-12 | 0.43523755 | 0.391 | 0.301 | 2.38E-08 | cDC1.Xcr1 | Cox6c |
| Eef1g1 | 1.38E-12 | 0.25067249 | 0.585 | 0.53 | 2.67E-08 | cDC1.Xcr1 | Eef1g |
| Eif2ak41 | 1.46E-12 | 0.4448013 | 0.346 | 0.257 | 2.81E-08 | cDC1.Xcr1 | Eif2ak4 |
| Ufm1 | 1.48E-12 | 0.30766444 | 0.431 | 0.348 | 2.86E-08 | cDC1.Xcr1 | Ufm1 |
| Slc35a41 | 1.60E-12 | 0.43302857 | 0.447 | 0.342 | 3.09E-08 | cDC1.Xcr1 | Slc35a4 |
| Pdap1 | 1.67E-12 | 0.25734209 | 0.502 | 0.441 | 3.23E-08 | cDC1.Xcr1 | Pdap1 |
| Csnk2b | 1.70E-12 | 0.30555126 | 0.479 | 0.396 | 3.29E-08 | cDC1.Xcr1 | Csnk2b |
| Nhp21 | 1.99E-12 | 0.32138797 | 0.455 | 0.386 | 3.84E-08 | cDC1.Xcr1 | Nhp2 |
| Cct71 | 2.04E-12 | 0.37196175 | 0.472 | 0.361 | 3.95E-08 | cDC1.Xcr1 | Cct7 |
| Pitpnb | 2.06E-12 | 0.33001723 | 0.314 | 0.241 | 3.98E-08 | cDC1.Xcr1 | Pitpnb |
| Plekhf2 | 2.44E-12 | 0.44104328 | 0.311 | 0.246 | 4.73E-08 | cDC1.Xcr1 | Plekhf2 |
| Mtdh | 2.47E-12 | 0.37487709 | 0.435 | 0.341 | 4.78E-08 | cDC1.Xcr1 | Mtdh |
| Rnf1412 | 2.51E-12 | 0.46704289 | 0.314 | 0.221 | 4.85E-08 | cDC1.Xcr1 | Rnf141 |
| Syvn11 | 2.54E-12 | 0.50187118 | 0.277 | 0.206 | 4.92E-08 | cDC1.Xcr1 | Syvn1 |
| Ercc31 | 2.85E-12 | 0.27439198 | 0.358 | 0.272 | 5.52E-08 | cDC1.Xcr1 | Ercc3 |
| Hspa41 | 2.99E-12 | 0.27137889 | 0.424 | 0.344 | 5.79E-08 | cDC1.Xcr1 | Hspa4 |
| Urm1 | 3.34E-12 | 0.44185375 | 0.284 | 0.211 | 6.45E-08 | cDC1.Xcr1 | Urm1 |
| Tspyl1 | 3.53E-12 | 0.45123643 | 0.42 | 0.323 | 6.82E-08 | cDC1.Xcr1 | Tspyl1 |
| Ccnk | 4.23E-12 | 0.37902327 | 0.363 | 0.28 | 8.17E-08 | cDC1.Xcr1 | Ccnk |
| Tradd | 4.43E-12 | 0.42087111 | 0.293 | 0.209 | 8.57E-08 | cDC1.Xcr1 | Tradd |
| Gars | 4.92E-12 | 0.49845017 | 0.259 | 0.182 | 9.50E-08 | cDC1.Xcr1 | Gars |
| Lman11 | 4.92E-12 | 0.38379468 | 0.333 | 0.261 | 9.52E-08 | cDC1.Xcr1 | Lman1 |
| Tmem219 | 5.07E-12 | 0.49540296 | 0.311 | 0.218 | 9.80E-08 | cDC1.Xcr1 | Tmem219 |
| Fnta | 5.28E-12 | 0.3838453 | 0.314 | 0.231 | 1.02E-07 | cDC1.Xcr1 | Fnta |
| Lgals11 | 5.88E-12 | 0.44458657 | 0.288 | 0.188 | 1.14E-07 | cDC1.Xcr1 | Lgals1 |
| Mindy3 | 6.45E-12 | 0.44380769 | 0.282 | 0.198 | 1.25E-07 | cDC1.Xcr1 | Mindy3 |
| Nfu1 | 6.70E-12 | 0.94564561 | 0.252 | 0.169 | 1.30E-07 | cDC1.Xcr1 | Nfu1 |
| Tmem2561 | 8.28E-12 | 0.2641121 | 0.562 | 0.481 | 1.60E-07 | cDC1.Xcr1 | Tmem256 |
| Ankrd40 | 9.23E-12 | 0.25434345 | 0.445 | 0.358 | 1.79E-07 | cDC1.Xcr1 | Ankrd40 |
| Pmepa1 | 9.48E-12 | 0.27318033 | 0.436 | 0.31 | 1.83E-07 | cDC1.Xcr1 | Pmepa1 |
| Tmem248 | 1.04E-11 | 0.41214517 | 0.324 | 0.252 | 2.00E-07 | cDC1.Xcr1 | Tmem248 |
| Rnf1261 | 1.08E-11 | 0.27267861 | 0.414 | 0.312 | 2.09E-07 | cDC1.Xcr1 | Rnf126 |
| Ubap1 | 1.17E-11 | 0.27571993 | 0.297 | 0.217 | 2.26E-07 | cDC1.Xcr1 | Ubap1 |
| Flot12 | 1.19E-11 | 0.315073 | 0.542 | 0.447 | 2.30E-07 | cDC1.Xcr1 | Flot1 |
| Tmem87a | 1.32E-11 | 0.27601135 | 0.276 | 0.202 | 2.55E-07 | cDC1.Xcr1 | Tmem87a |
| Bckdk1 | 1.67E-11 | 0.37377183 | 0.281 | 0.227 | 3.24E-07 | cDC1.Xcr1 | Bckdk |
| Chchd11 | 1.89E-11 | 0.5550621 | 0.29 | 0.203 | 3.66E-07 | cDC1.Xcr1 | Chchd1 |
| Utp31 | 2.21E-11 | 0.27994844 | 0.305 | 0.229 | 4.27E-07 | cDC1.Xcr1 | Utp3 |
| Gtf2a2 | 2.25E-11 | 0.46867945 | 0.25 | 0.177 | 4.34E-07 | cDC1.Xcr1 | Gtf2a2 |
| Ap1ar | 2.30E-11 | 0.36951218 | 0.35 | 0.246 | 4.45E-07 | cDC1.Xcr1 | Ap1ar |
| Mak16 | 2.43E-11 | 0.65267489 | 0.251 | 0.164 | 4.70E-07 | cDC1.Xcr1 | Mak16 |
| Psmd2 | 2.65E-11 | 0.25527886 | 0.505 | 0.426 | 5.13E-07 | cDC1.Xcr1 | Psmd2 |
| Trappc8 | 2.80E-11 | 0.30641739 | 0.36 | 0.289 | 5.40E-07 | cDC1.Xcr1 | Trappc8 |
| Ndufb10 | 3.07E-11 | 0.37934649 | 0.576 | 0.524 | 5.94E-07 | cDC1.Xcr1 | Ndufb10 |
| Sec23ip | 3.35E-11 | 0.29996487 | 0.284 | 0.208 | 6.48E-07 | cDC1.Xcr1 | Sec23ip |
| Trim281 | 3.36E-11 | 0.34482035 | 0.319 | 0.251 | 6.49E-07 | cDC1.Xcr1 | Trim28 |
| Pik3c3 | 4.57E-11 | 0.32996691 | 0.293 | 0.212 | 8.84E-07 | cDC1.Xcr1 | Pik3c3 |
| Sacm1l | 4.60E-11 | 0.3384692 | 0.366 | 0.282 | 8.89E-07 | cDC1.Xcr1 | Sacm1l |
| Entpd7 | 4.71E-11 | 0.31358744 | 0.283 | 0.206 | 9.11E-07 | cDC1.Xcr1 | Entpd7 |
| Rpe1 | 4.84E-11 | 0.26994043 | 0.349 | 0.289 | 9.35E-07 | cDC1.Xcr1 | Rpe |
| Prpsap1 | 5.23E-11 | 0.44605704 | 0.338 | 0.267 | 1.01E-06 | cDC1.Xcr1 | Prpsap1 |
| Drap11 | 6.20E-11 | 0.29798838 | 0.679 | 0.601 | 1.20E-06 | cDC1.Xcr1 | Drap1 |
| Hsdl2 | 6.61E-11 | 0.45521306 | 0.328 | 0.254 | 1.28E-06 | cDC1.Xcr1 | Hsdl2 |
| Exoc62 | 6.63E-11 | 0.28742501 | 0.294 | 0.215 | 1.28E-06 | cDC1.Xcr1 | Exoc6 |
| Lipa1 | 7.51E-11 | 0.27606683 | 0.369 | 0.283 | 1.45E-06 | cDC1.Xcr1 | Lipa |
| Tial1 | 8.11E-11 | 0.32429348 | 0.423 | 0.373 | 1.57E-06 | cDC1.Xcr1 | Tial1 |
| Hint3 | 9.02E-11 | 0.34329431 | 0.281 | 0.204 | 1.74E-06 | cDC1.Xcr1 | Hint3 |
| C1qbp1 | 9.15E-11 | 0.59064127 | 0.252 | 0.182 | 1.77E-06 | cDC1.Xcr1 | C1qbp |
| Fam104a1 | 1.06E-10 | 0.33664767 | 0.336 | 0.257 | 2.06E-06 | cDC1.Xcr1 | Fam104a |
| Mxd4 | 1.12E-10 | 0.60248115 | 0.277 | 0.208 | 2.16E-06 | cDC1.Xcr1 | Mxd4 |
| Errfi1 | 1.12E-10 | 0.41508698 | 0.316 | 0.236 | 2.17E-06 | cDC1.Xcr1 | Errfi1 |
| Cers51 | 1.16E-10 | 0.2529 | 0.376 | 0.313 | 2.25E-06 | cDC1.Xcr1 | Cers5 |
| Mia3 | 1.17E-10 | 0.2629634 | 0.449 | 0.363 | 2.26E-06 | cDC1.Xcr1 | Mia3 |
| Pycr21 | 1.20E-10 | 0.28972107 | 0.333 | 0.233 | 2.31E-06 | cDC1.Xcr1 | Pycr2 |
| Ccpg11 | 1.24E-10 | 0.31945714 | 0.306 | 0.229 | 2.40E-06 | cDC1.Xcr1 | Ccpg1 |
| Usp39 | 1.28E-10 | 0.50049657 | 0.279 | 0.2 | 2.47E-06 | cDC1.Xcr1 | Usp39 |
| Psmc6 | 1.33E-10 | 0.32363492 | 0.471 | 0.42 | 2.57E-06 | cDC1.Xcr1 | Psmc6 |
| Plpbp1 | 1.41E-10 | 0.32105374 | 0.425 | 0.342 | 2.73E-06 | cDC1.Xcr1 | Plpbp |
| Tsc2 | 1.51E-10 | 0.30971827 | 0.294 | 0.218 | 2.91E-06 | cDC1.Xcr1 | Tsc2 |
| Tufm | 1.97E-10 | 0.4209898 | 0.315 | 0.241 | 3.80E-06 | cDC1.Xcr1 | Tufm |
| Aco2 | 2.21E-10 | 0.445486 | 0.272 | 0.193 | 4.27E-06 | cDC1.Xcr1 | Aco2 |
| Fam207a1 | 2.26E-10 | 0.47997551 | 0.32 | 0.239 | 4.37E-06 | cDC1.Xcr1 | Fam207a |
| Uso1 | 2.36E-10 | 0.3188463 | 0.376 | 0.292 | 4.57E-06 | cDC1.Xcr1 | Uso1 |
| Wdr431 | 2.61E-10 | 0.39310667 | 0.273 | 0.191 | 5.04E-06 | cDC1.Xcr1 | Wdr43 |
| Cript | 2.63E-10 | 0.29990013 | 0.355 | 0.273 | 5.08E-06 | cDC1.Xcr1 | Cript |
| Ppie1 | 2.75E-10 | 0.31581 | 0.317 | 0.235 | 5.32E-06 | cDC1.Xcr1 | Ppie |
| Fgd31 | 2.81E-10 | 0.27747893 | 0.482 | 0.353 | 5.44E-06 | cDC1.Xcr1 | Fgd3 |
| Rbx1 | 2.97E-10 | 0.278576 | 0.338 | 0.275 | 5.74E-06 | cDC1.Xcr1 | Rbx1 |
| Yipf4 | 3.37E-10 | 0.35499851 | 0.29 | 0.213 | 6.51E-06 | cDC1.Xcr1 | Yipf4 |
| Spsb3 | 3.67E-10 | 0.5541429 | 0.27 | 0.206 | 7.10E-06 | cDC1.Xcr1 | Spsb3 |
| Hpf1 | 4.04E-10 | 0.34946438 | 0.265 | 0.196 | 7.81E-06 | cDC1.Xcr1 | Hpf1 |
| Frmd4b | 4.16E-10 | 0.74027998 | 0.267 | 0.173 | 8.04E-06 | cDC1.Xcr1 | Frmd4b |
| Glrx3 | 4.58E-10 | 0.5028166 | 0.327 | 0.229 | 8.85E-06 | cDC1.Xcr1 | Glrx3 |
| Srp14 | 4.69E-10 | 0.47373615 | 0.451 | 0.377 | 9.07E-06 | cDC1.Xcr1 | Srp14 |
| Pik3r2 | 4.71E-10 | 0.4675231 | 0.288 | 0.194 | 9.11E-06 | cDC1.Xcr1 | Pik3r2 |
| Plaa | 4.74E-10 | 0.28886713 | 0.4 | 0.318 | 9.16E-06 | cDC1.Xcr1 | Plaa |
| Syf2 | 5.78E-10 | 0.26292877 | 0.614 | 0.546 | 1.12E-05 | cDC1.Xcr1 | Syf2 |
| Rnf220 | 5.92E-10 | 0.30207831 | 0.491 | 0.424 | 1.14E-05 | cDC1.Xcr1 | Rnf220 |
| Eif4a31 | 6.40E-10 | 0.28443142 | 0.286 | 0.218 | 1.24E-05 | cDC1.Xcr1 | Eif4a3 |
| Rragc | 7.96E-10 | 0.25438468 | 0.385 | 0.307 | 1.54E-05 | cDC1.Xcr1 | Rragc |
| Trnau1ap | 8.18E-10 | 0.34915754 | 0.257 | 0.191 | 1.58E-05 | cDC1.Xcr1 | Trnau1ap |
| Smarcad1 | 8.63E-10 | 0.32505579 | 0.287 | 0.214 | 1.67E-05 | cDC1.Xcr1 | Smarcad1 |
| Ift521 | 1.10E-09 | 0.27599452 | 0.348 | 0.274 | 2.13E-05 | cDC1.Xcr1 | Ift52 |
| Klhdc3 | 1.17E-09 | 0.39634267 | 0.386 | 0.285 | 2.26E-05 | cDC1.Xcr1 | Klhdc3 |
| Nus1 | 1.20E-09 | 0.30449645 | 0.369 | 0.287 | 2.33E-05 | cDC1.Xcr1 | Nus1 |
| Ocstamp1 | 1.25E-09 | 0.47788016 | 0.267 | 0.181 | 2.41E-05 | cDC1.Xcr1 | Ocstamp |
| Arnt1 | 1.32E-09 | 0.27965571 | 0.435 | 0.364 | 2.55E-05 | cDC1.Xcr1 | Arnt |
| Zhx1 | 1.42E-09 | 0.27254011 | 0.27 | 0.202 | 2.75E-05 | cDC1.Xcr1 | Zhx1 |
| Ino80b1 | 1.43E-09 | 0.2600627 | 0.317 | 0.231 | 2.76E-05 | cDC1.Xcr1 | Ino80b |
| Uqcrq | 1.47E-09 | 0.38632483 | 0.321 | 0.259 | 2.83E-05 | cDC1.Xcr1 | Uqcrq |
| Parp121 | 1.50E-09 | 0.5319519 | 0.3 | 0.215 | 2.89E-05 | cDC1.Xcr1 | Parp12 |
| Nars1 | 1.63E-09 | 0.26975501 | 0.346 | 0.276 | 3.16E-05 | cDC1.Xcr1 | Nars |
| Phax | 1.70E-09 | 0.28446006 | 0.287 | 0.218 | 3.29E-05 | cDC1.Xcr1 | Phax |
| Slc35a5 | 1.77E-09 | 0.27909407 | 0.281 | 0.215 | 3.42E-05 | cDC1.Xcr1 | Slc35a5 |
| Meaf61 | 1.83E-09 | 0.2769305 | 0.38 | 0.318 | 3.55E-05 | cDC1.Xcr1 | Meaf6 |
| Eif6 | 1.87E-09 | 0.43407653 | 0.418 | 0.323 | 3.62E-05 | cDC1.Xcr1 | Eif6 |
| Upf3a | 2.03E-09 | 0.39324811 | 0.259 | 0.202 | 3.92E-05 | cDC1.Xcr1 | Upf3a |
| Kdm5a | 2.03E-09 | 0.35884694 | 0.347 | 0.28 | 3.93E-05 | cDC1.Xcr1 | Kdm5a |
| Atp6v1h | 2.10E-09 | 0.39103391 | 0.418 | 0.326 | 4.06E-05 | cDC1.Xcr1 | Atp6v1h |
| Phf14 | 2.15E-09 | 0.38630411 | 0.344 | 0.263 | 4.15E-05 | cDC1.Xcr1 | Phf14 |
| Ndufa121 | 2.19E-09 | 0.29509714 | 0.347 | 0.262 | 4.24E-05 | cDC1.Xcr1 | Ndufa12 |
| Strap | 2.32E-09 | 0.52524494 | 0.273 | 0.208 | 4.49E-05 | cDC1.Xcr1 | Strap |
| Suclg11 | 2.36E-09 | 0.47941186 | 0.275 | 0.201 | 4.57E-05 | cDC1.Xcr1 | Suclg1 |
| Snrpe | 2.66E-09 | 0.289241 | 0.376 | 0.31 | 5.15E-05 | cDC1.Xcr1 | Snrpe |
| Cdk5rap31 | 2.69E-09 | 0.35729448 | 0.401 | 0.314 | 5.20E-05 | cDC1.Xcr1 | Cdk5rap3 |
| Tubgcp3 | 2.80E-09 | 0.34857548 | 0.254 | 0.188 | 5.42E-05 | cDC1.Xcr1 | Tubgcp3 |
| Grpel11 | 2.91E-09 | 0.26828641 | 0.447 | 0.359 | 5.62E-05 | cDC1.Xcr1 | Grpel1 |
| 2-Mar | 3.04E-09 | 0.43447222 | 0.27 | 0.195 | 5.88E-05 | cDC1.Xcr1 | 2-Mar |
| Ddx561 | 3.24E-09 | 0.43890251 | 0.29 | 0.206 | 6.27E-05 | cDC1.Xcr1 | Ddx56 |
| Sharpin | 3.28E-09 | 0.4635433 | 0.331 | 0.258 | 6.34E-05 | cDC1.Xcr1 | Sharpin |
| Atg101 | 3.44E-09 | 0.27538985 | 0.31 | 0.247 | 6.65E-05 | cDC1.Xcr1 | Atg101 |
| Lsm8 | 3.50E-09 | 0.2613008 | 0.273 | 0.226 | 6.77E-05 | cDC1.Xcr1 | Lsm8 |
| Pfdn21 | 3.57E-09 | 0.31265405 | 0.395 | 0.324 | 6.90E-05 | cDC1.Xcr1 | Pfdn2 |
| Cd200r11 | 3.65E-09 | 0.39703064 | 0.257 | 0.182 | 7.05E-05 | cDC1.Xcr1 | Cd200r1 |
| Gabpb2 | 3.67E-09 | 0.27319925 | 0.257 | 0.189 | 7.09E-05 | cDC1.Xcr1 | Gabpb2 |
| Psmd31 | 3.92E-09 | 0.38362664 | 0.413 | 0.339 | 7.57E-05 | cDC1.Xcr1 | Psmd3 |
| Glo1 | 4.25E-09 | 0.68266858 | 0.284 | 0.201 | 8.22E-05 | cDC1.Xcr1 | Glo1 |
| Ptpmt1 | 4.54E-09 | 0.33697371 | 0.27 | 0.198 | 8.78E-05 | cDC1.Xcr1 | Ptpmt1 |
| Ibtk1 | 4.68E-09 | 0.38558618 | 0.275 | 0.213 | 9.05E-05 | cDC1.Xcr1 | Ibtk |
| Idh3a2 | 5.26E-09 | 0.45803431 | 0.359 | 0.275 | 0.00010174 | cDC1.Xcr1 | Idh3a |
| Zdhhc81 | 5.45E-09 | 0.46114564 | 0.254 | 0.19 | 0.00010537 | cDC1.Xcr1 | Zdhhc8 |
| Aurkaip1 | 5.76E-09 | 0.42379771 | 0.434 | 0.399 | 0.00011134 | cDC1.Xcr1 | Aurkaip1 |
| Mcts11 | 5.78E-09 | 0.27367929 | 0.326 | 0.266 | 0.00011182 | cDC1.Xcr1 | Mcts1 |
| BC005624 | 6.18E-09 | 0.44220635 | 0.332 | 0.251 | 0.00011955 | cDC1.Xcr1 | BC005624 |
| Phkg2 | 6.50E-09 | 0.51343655 | 0.26 | 0.175 | 0.00012558 | cDC1.Xcr1 | Phkg2 |
| Ppfibp11 | 6.58E-09 | 0.25956359 | 0.332 | 0.251 | 0.00012718 | cDC1.Xcr1 | Ppfibp1 |
| Map1s | 6.71E-09 | 0.60650792 | 0.288 | 0.206 | 0.00012982 | cDC1.Xcr1 | Map1s |
| Atic1 | 7.26E-09 | 0.27279146 | 0.369 | 0.289 | 0.00014039 | cDC1.Xcr1 | Atic |
| Sar1b1 | 8.07E-09 | 0.50830687 | 0.381 | 0.291 | 0.00015599 | cDC1.Xcr1 | Sar1b |
| Naa401 | 8.42E-09 | 0.38399582 | 0.29 | 0.222 | 0.00016279 | cDC1.Xcr1 | Naa40 |
| Btk1 | 8.85E-09 | 0.42457231 | 0.304 | 0.232 | 0.00017109 | cDC1.Xcr1 | Btk |
| Pacs1 | 9.12E-09 | 0.45424689 | 0.25 | 0.183 | 0.00017634 | cDC1.Xcr1 | Pacs1 |
| Rufy1 | 9.23E-09 | 0.35225941 | 0.336 | 0.27 | 0.00017844 | cDC1.Xcr1 | Rufy1 |
| Exosc3 | 9.43E-09 | 0.34407721 | 0.303 | 0.223 | 0.00018238 | cDC1.Xcr1 | Exosc3 |
| Gipc1 | 9.94E-09 | 0.58587708 | 0.25 | 0.183 | 0.00019224 | cDC1.Xcr1 | Gipc1 |
| Slc30a51 | 1.00E-08 | 0.2621911 | 0.304 | 0.241 | 0.00019369 | cDC1.Xcr1 | Slc30a5 |
| Arhgap251 | 1.12E-08 | 0.52878609 | 0.293 | 0.21 | 0.00021657 | cDC1.Xcr1 | Arhgap25 |
| Ap2s1 | 1.24E-08 | 0.26907918 | 0.561 | 0.471 | 0.00024055 | cDC1.Xcr1 | Ap2s1 |
| Rab6a | 1.35E-08 | 0.33981944 | 0.408 | 0.359 | 0.00026152 | cDC1.Xcr1 | Rab6a |
| Pmpcb | 1.46E-08 | 0.45834039 | 0.331 | 0.246 | 0.00028153 | cDC1.Xcr1 | Pmpcb |
| Phb21 | 1.60E-08 | 0.25144112 | 0.442 | 0.389 | 0.00030956 | cDC1.Xcr1 | Phb2 |
| Ddx271 | 1.65E-08 | 0.25186047 | 0.294 | 0.237 | 0.00031982 | cDC1.Xcr1 | Ddx27 |
| Snrpd2 | 1.90E-08 | 0.44149746 | 0.273 | 0.229 | 0.00036758 | cDC1.Xcr1 | Snrpd2 |
| Wbp12 | 2.05E-08 | 0.31325287 | 0.29 | 0.23 | 0.00039604 | cDC1.Xcr1 | Wbp1 |
| Cnih41 | 2.10E-08 | 0.34085435 | 0.283 | 0.228 | 0.00040632 | cDC1.Xcr1 | Cnih4 |
| Vps41 | 2.50E-08 | 0.44017006 | 0.267 | 0.19 | 0.00048426 | cDC1.Xcr1 | Vps41 |
| Entpd11 | 2.63E-08 | 0.31617611 | 0.321 | 0.276 | 0.00050864 | cDC1.Xcr1 | Entpd1 |
| Prdx1 | 3.07E-08 | 0.3176872 | 0.548 | 0.437 | 0.00059409 | cDC1.Xcr1 | Prdx1 |
| Polr2f1 | 3.47E-08 | 0.28366817 | 0.368 | 0.306 | 0.00067058 | cDC1.Xcr1 | Polr2f |
| Chd91 | 3.56E-08 | 0.27996682 | 0.292 | 0.233 | 0.00068774 | cDC1.Xcr1 | Chd9 |
| Rab31 | 3.56E-08 | 0.52126185 | 0.289 | 0.206 | 0.00068798 | cDC1.Xcr1 | Rab31 |
| Tmem222 | 3.59E-08 | 0.34463397 | 0.271 | 0.205 | 0.00069429 | cDC1.Xcr1 | Tmem222 |
| Lrwd1 | 3.72E-08 | 0.27700372 | 0.29 | 0.233 | 0.00072003 | cDC1.Xcr1 | Lrwd1 |
| Fcf1 | 3.75E-08 | 0.27516536 | 0.35 | 0.288 | 0.00072486 | cDC1.Xcr1 | Fcf1 |
| Gatd3a | 3.80E-08 | 0.62220034 | 0.31 | 0.222 | 0.00073392 | cDC1.Xcr1 | Gatd3a |
| Ufsp21 | 3.85E-08 | 0.284864 | 0.31 | 0.242 | 0.00074431 | cDC1.Xcr1 | Ufsp2 |
| Itga6 | 3.89E-08 | 0.36029184 | 0.26 | 0.182 | 0.00075191 | cDC1.Xcr1 | Itga6 |
| Mob3b2 | 3.90E-08 | 0.45067209 | 0.335 | 0.257 | 0.00075361 | cDC1.Xcr1 | Mob3b |
| Timm10b2 | 4.63E-08 | 0.33427093 | 0.457 | 0.368 | 0.00089523 | cDC1.Xcr1 | Timm10b |
| Acp1 | 5.07E-08 | 0.4207436 | 0.283 | 0.204 | 0.00097962 | cDC1.Xcr1 | Acp1 |
| Ssrd1 | 6.46E-08 | 0.2997096 | 0.567 | 0.491 | 0.00124804 | cDC1.Xcr1 | Ssrd1 |
| Eif4ebp21 | 6.48E-08 | 0.34745076 | 0.484 | 0.429 | 0.00125219 | cDC1.Xcr1 | Eif4ebp2 |
| Txn21 | 7.37E-08 | 0.30211782 | 0.444 | 0.393 | 0.00142468 | cDC1.Xcr1 | Txn2 |
| Snw1 | 8.64E-08 | 0.34770312 | 0.33 | 0.261 | 0.0016698 | cDC1.Xcr1 | Snw1 |
| Dus3l | 9.42E-08 | 0.28014054 | 0.306 | 0.22 | 0.00182136 | cDC1.Xcr1 | Dus3l |
| Ciz11 | 1.09E-07 | 0.34743682 | 0.343 | 0.272 | 0.00210362 | cDC1.Xcr1 | Ciz1 |
| Fbxo42 | 1.16E-07 | 0.34418985 | 0.26 | 0.196 | 0.00225192 | cDC1.Xcr1 | Fbxo42 |
| Ppox | 1.18E-07 | 0.32847188 | 0.263 | 0.203 | 0.00227479 | cDC1.Xcr1 | Ppox |
| Adcy72 | 1.19E-07 | 0.29562367 | 0.512 | 0.397 | 0.00229479 | cDC1.Xcr1 | Adcy7 |
| Cops4 | 1.29E-07 | 0.38061491 | 0.306 | 0.233 | 0.0024875 | cDC1.Xcr1 | Cops4 |
| Pebp1 | 1.34E-07 | 0.30468974 | 0.346 | 0.295 | 0.00260039 | cDC1.Xcr1 | Pebp1 |
| Commd10 | 1.68E-07 | 0.36115311 | 0.303 | 0.219 | 0.00324752 | cDC1.Xcr1 | Commd10 |
| Cd300lf2 | 1.89E-07 | 0.41626763 | 0.479 | 0.366 | 0.00366044 | cDC1.Xcr1 | Cd300lf |
| Tmco12 | 2.25E-07 | 0.31538772 | 0.337 | 0.286 | 0.004351 | cDC1.Xcr1 | Tmco1 |
| Setdb11 | 2.29E-07 | 0.26999011 | 0.309 | 0.235 | 0.0044247 | cDC1.Xcr1 | Setdb1 |
| Exosc51 | 2.44E-07 | 0.26602896 | 0.453 | 0.353 | 0.00472402 | cDC1.Xcr1 | Exosc5 |
| Rexo41 | 2.47E-07 | 0.27046161 | 0.319 | 0.233 | 0.00478423 | cDC1.Xcr1 | Rexo4 |
| Lamtor5 | 2.53E-07 | 0.29083767 | 0.251 | 0.186 | 0.00490082 | cDC1.Xcr1 | Lamtor5 |
| Eif1b1 | 2.56E-07 | 0.44668167 | 0.316 | 0.24 | 0.00495336 | cDC1.Xcr1 | Eif1b |
| Usp191 | 2.66E-07 | 0.30818899 | 0.422 | 0.359 | 0.00513462 | cDC1.Xcr1 | Usp19 |
| Ruvbl21 | 2.73E-07 | 0.26818985 | 0.282 | 0.22 | 0.00527575 | cDC1.Xcr1 | Ruvbl2 |
| Atp5j | 2.82E-07 | 0.43732143 | 0.346 | 0.295 | 0.00545567 | cDC1.Xcr1 | Atp5j |
| S100a1 | 3.05E-07 | 0.56206997 | 0.266 | 0.213 | 0.00588737 | cDC1.Xcr1 | S100a1 |
| Mfsd5 | 3.05E-07 | 0.41807538 | 0.312 | 0.244 | 0.00590072 | cDC1.Xcr1 | Mfsd5 |
| Rex1bd | 3.08E-07 | 0.28854361 | 0.287 | 0.229 | 0.00595291 | cDC1.Xcr1 | Rex1bd |
| Zc3h14 | 3.42E-07 | 0.35981982 | 0.29 | 0.234 | 0.006621 | cDC1.Xcr1 | Zc3h14 |
| Pithd11 | 3.62E-07 | 0.36927685 | 0.266 | 0.207 | 0.00699275 | cDC1.Xcr1 | Pithd1 |
| Ing11 | 3.79E-07 | 0.26904097 | 0.319 | 0.252 | 0.00732218 | cDC1.Xcr1 | Ing1 |
| Uqcr10 | 4.40E-07 | 0.33052721 | 0.315 | 0.262 | 0.00850243 | cDC1.Xcr1 | Uqcr10 |
| Sash31 | 4.62E-07 | 0.29243715 | 0.376 | 0.287 | 0.00893373 | cDC1.Xcr1 | Sash3 |
| Sh2b2 | 4.66E-07 | 0.30060735 | 0.256 | 0.181 | 0.00901729 | cDC1.Xcr1 | Sh2b2 |
| DEPRECATED-I | 5.24E-07 | 0.39591925 | 0.431 | 0.355 | 0.01013153 | cDC1.Xcr1 | DEPRECATED-ENSMUSG00000017221 |
| Lamtor31 | 5.69E-07 | 0.33637228 | 0.447 | 0.356 | 0.01101017 | cDC1.Xcr1 | Lamtor3 |
| Cpsf2 | 5.85E-07 | 0.32722796 | 0.322 | 0.258 | 0.01130746 | cDC1.Xcr1 | Cpsf2 |
| Hspbp11 | 6.38E-07 | 0.33415383 | 0.257 | 0.194 | 0.01233037 | cDC1.Xcr1 | Hspbp1 |
| Hmbox1 | 6.41E-07 | 0.33307806 | 0.261 | 0.185 | 0.01239082 | cDC1.Xcr1 | Hmbox1 |
| Vti1b1 | 6.58E-07 | 0.25303466 | 0.343 | 0.308 | 0.01273084 | cDC1.Xcr1 | Vti1b |
| Psmd10 | 8.97E-07 | 0.34511025 | 0.273 | 0.208 | 0.01734114 | cDC1.Xcr1 | Psmd10 |
| Pgam11 | 1.03E-06 | 0.29648953 | 0.589 | 0.496 | 0.01995052 | cDC1.Xcr1 | Pgam1 |
| Atp5g12 | 1.13E-06 | 0.35633789 | 0.366 | 0.315 | 0.02180714 | cDC1.Xcr1 | Atp5g1 |
| Wdr45b | 1.15E-06 | 0.26128564 | 0.282 | 0.212 | 0.02223817 | cDC1.Xcr1 | Wdr45b |
| Dennd1c | 1.20E-06 | 0.28811189 | 0.339 | 0.278 | 0.0232682 | cDC1.Xcr1 | Dennd1c |
| Vps35 | 1.22E-06 | 0.26273726 | 0.388 | 0.338 | 0.0235265 | cDC1.Xcr1 | Vps35 |
| Twf1 | 1.27E-06 | 0.31118615 | 0.312 | 0.254 | 0.02461903 | cDC1.Xcr1 | Twf1 |
| Narf1 | 1.32E-06 | 0.41403628 | 0.289 | 0.218 | 0.02558209 | cDC1.Xcr1 | Narf |
| Vps29 | 1.48E-06 | 0.37869884 | 0.422 | 0.368 | 0.02856413 | cDC1.Xcr1 | Vps29 |
| Eif2b21 | 1.51E-06 | 0.31894244 | 0.513 | 0.394 | 0.02909873 | cDC1.Xcr1 | Eif2b2 |
| Zc3h15 | 1.65E-06 | 0.30319438 | 0.312 | 0.263 | 0.03183158 | cDC1.Xcr1 | Zc3h15 |
| Akirin1 | 1.98E-06 | 0.33061049 | 0.331 | 0.275 | 0.0383773 | cDC1.Xcr1 | Akirin1 |
| Trappc2l | 2.20E-06 | 0.2525365 | 0.375 | 0.309 | 0.04251244 | cDC1.Xcr1 | Trappc2l |
| Camk2d1 | 2.64E-06 | 0.29536461 | 0.282 | 0.21 | 0.05112492 | cDC1.Xcr1 | Camk2d |
| Ppp1r8 | 2.69E-06 | 0.2952094 | 0.254 | 0.211 | 0.05203569 | cDC1.Xcr1 | Ppp1r8 |
| Rars1 | 2.72E-06 | 0.31731762 | 0.279 | 0.218 | 0.05258515 | cDC1.Xcr1 | Rars |
| Rbmxl11 | 2.84E-06 | 0.36769648 | 0.254 | 0.19 | 0.0549106 | cDC1.Xcr1 | Rbmxl1 |
| Gnl32 | 3.34E-06 | 0.25815246 | 0.26 | 0.211 | 0.06466011 | cDC1.Xcr1 | Gnl3 |
| Cpsf32 | 3.46E-06 | 0.33141366 | 0.35 | 0.282 | 0.06693689 | cDC1.Xcr1 | Cpsf3 |
| Prelid3b1 | 3.57E-06 | 0.29250984 | 0.325 | 0.276 | 0.0689477 | cDC1.Xcr1 | Prelid3b |
| Sucla2 | 3.94E-06 | 0.37929196 | 0.29 | 0.234 | 0.07613724 | cDC1.Xcr1 | Sucla2 |
| Dnajc2 | 4.02E-06 | 0.2560481 | 0.263 | 0.209 | 0.07772744 | cDC1.Xcr1 | Dnajc2 |
| Exosc10 | 6.25E-06 | 0.36234757 | 0.287 | 0.224 | 0.12093391 | cDC1.Xcr1 | Exosc10 |
| Snf8 | 7.10E-06 | 0.26630297 | 0.479 | 0.42 | 0.13733994 | cDC1.Xcr1 | Snf8 |
| Ndufs71 | 7.32E-06 | 0.28338758 | 0.462 | 0.397 | 0.14160463 | cDC1.Xcr1 | Ndufs7 |
| Gnrf1 | 7.72E-06 | 0.28841284 | 0.348 | 0.282 | 0.14935427 | cDC1.Xcr1 | Gnrf1 |
| Arl6ip4 | 9.15E-06 | 0.34744743 | 0.326 | 0.264 | 0.1768109 | cDC1.Xcr1 | Arl6ip4 |
| Sptssa1 | 9.23E-06 | 0.33156612 | 0.368 | 0.323 | 0.17842557 | cDC1.Xcr1 | Sptssa |
| Zbtb22 | 9.87E-06 | 0.30536671 | 0.275 | 0.224 | 0.19076457 | cDC1.Xcr1 | Zbtb22 |
| Lyar1 | 9.93E-06 | 0.27826698 | 0.261 | 0.199 | 0.19191118 | cDC1.Xcr1 | Lyar |
| Fam89b | 1.10E-05 | 0.26503406 | 0.407 | 0.347 | 0.21260705 | cDC1.Xcr1 | Fam89b |
| Nr2c2ap1 | 1.17E-05 | 0.32428316 | 0.32 | 0.253 | 0.22639455 | cDC1.Xcr1 | Nr2c2ap |
| Gmppa | 1.27E-05 | 0.33439481 | 0.265 | 0.21 | 0.24576946 | cDC1.Xcr1 | Gmppa |
| Slc25a11 | 1.30E-05 | 0.31380839 | 0.515 | 0.458 | 0.25075881 | cDC1.Xcr1 | Slc25a11 |
| Sars1 | 1.32E-05 | 0.36674456 | 0.304 | 0.236 | 0.2552493 | cDC1.Xcr1 | Sars |
| Dcaf13 | 1.33E-05 | 0.2595805 | 0.315 | 0.257 | 0.25661814 | cDC1.Xcr1 | Dcaf13 |

| | | | | | | | |
|---|---|---|---|---|---|---|---|
| Dohh1 | 1.73E-05 | 0.38087732 | 0.266 | 0.193 | 0.33398875 | cDC1.Xcr1 | Dohh |
| Washc3 | 1.91E-05 | 0.28710233 | 0.257 | 0.2 | 0.36869392 | cDC1.Xcr1 | Washc3 |
| Baiap21 | 1.94E-05 | 0.36284126 | 0.328 | 0.252 | 0.37472247 | cDC1.Xcr1 | Baiap2 |
| Scp21 | 1.98E-05 | 0.25186081 | 0.376 | 0.319 | 0.38207748 | cDC1.Xcr1 | Scp2 |
| Bbip1 | 2.28E-05 | 0.40817101 | 0.298 | 0.244 | 0.4401503 | cDC1.Xcr1 | Bbip1 |
| Denr1 | 2.52E-05 | 0.51164716 | 0.355 | 0.289 | 0.48728402 | cDC1.Xcr1 | Denr |
| Nme21 | 2.75E-05 | 0.2553838 | 0.415 | 0.371 | 0.53109422 | cDC1.Xcr1 | Nme2 |
| Ubap21 | 3.32E-05 | 0.30804696 | 0.278 | 0.239 | 0.64100139 | cDC1.Xcr1 | Ubap2 |
| Actr10 | 3.38E-05 | 0.33168828 | 0.281 | 0.222 | 0.6540909 | cDC1.Xcr1 | Actr10 |
| Sgk1 | 3.65E-05 | 0.69859267 | 0.362 | 0.173 | 0.70584945 | cDC1.Xcr1 | Sgk1 |
| Celf1 | 3.82E-05 | 0.33490581 | 0.251 | 0.201 | 0.73905975 | cDC1.Xcr1 | Celf1 |
| Spp11 | 4.71E-05 | 0.42829411 | 0.625 | 0.597 | 0.91147124 | cDC1.Xcr1 | Spp1 |
| Pigm | 5.45E-05 | 0.27206096 | 0.255 | 0.194 | 1 | cDC1.Xcr1 | Pigm |
| Nosip | 5.54E-05 | 0.25803936 | 0.268 | 0.213 | 1 | cDC1.Xcr1 | Nosip |
| Gripap12 | 9.48E-05 | 0.31373188 | 0.316 | 0.266 | 1 | cDC1.Xcr1 | Gripap1 |
| Nop102 | 9.54E-05 | 0.27024711 | 0.384 | 0.319 | 1 | cDC1.Xcr1 | Nop10 |
| Hk22 | 0.00014573 | 0.31010684 | 0.564 | 0.465 | 1 | cDC1.Xcr1 | Hk2 |
| Ndufs6 | 0.000164 | 0.3946291 | 0.256 | 0.212 | 1 | cDC1.Xcr1 | Ndufs6 |
| Golga71 | 0.00022572 | 0.26836094 | 0.564 | 0.499 | 1 | cDC1.Xcr1 | Golga7 |
| Fdx2 | 0.00023474 | 0.3318678 | 0.268 | 0.206 | 1 | cDC1.Xcr1 | Fdx2 |
| Mknk1 | 0.00029581 | 0.30200632 | 0.344 | 0.279 | 1 | cDC1.Xcr1 | Mknk1 |
| Nme12 | 0.00031728 | 0.2537233 | 0.347 | 0.281 | 1 | cDC1.Xcr1 | Nme1 |
| Bud232 | 0.00041936 | 0.33844502 | 0.277 | 0.228 | 1 | cDC1.Xcr1 | Bud23 |
| Atg16l1 | 0.00043075 | 0.36999717 | 0.281 | 0.22 | 1 | cDC1.Xcr1 | Atg16l1 |
| Atp5mpl1 | 0.00049226 | 0.27893068 | 0.409 | 0.379 | 1 | cDC1.Xcr1 | Atp5mpl |
| Ntan1 | 0.00051936 | 0.30058844 | 0.272 | 0.219 | 1 | cDC1.Xcr1 | Ntan1 |
| Sod2 | 0.00073406 | 0.2849909 | 0.46 | 0.382 | 1 | cDC1.Xcr1 | Sod2 |
| D5Ertd579e | 0.00079415 | 0.28547379 | 0.288 | 0.239 | 1 | cDC1.Xcr1 | D5Ertd579e |
| Tomm201 | 0.00110101 | 0.30813912 | 0.366 | 0.318 | 1 | cDC1.Xcr1 | Tomm20 |
| Mea11 | 0.00217373 | 0.2564284 | 0.369 | 0.315 | 1 | cDC1.Xcr1 | Mea1 |
| Tmem165 | 0.00246868 | 0.25156473 | 0.278 | 0.23 | 1 | cDC1.Xcr1 | Tmem165 |
| Akt21 | 0.00263459 | 0.44072952 | 0.376 | 0.317 | 1 | cDC1.Xcr1 | Akt2 |
| BC0297221 | 0.00393153 | 0.26793874 | 0.261 | 0.206 | 1 | cDC1.Xcr1 | BC029722 |
| Uqcc3 | 0.0042804 | 0.33798076 | 0.257 | 0.218 | 1 | cDC1.Xcr1 | Uqcc3 |
| Brf1 | 0.0043936 | 0.26767243 | 0.325 | 0.288 | 1 | cDC1.Xcr1 | Brf1 |
| Wdr182 | 0.00614817 | 0.30226449 | 0.278 | 0.231 | 1 | cDC1.Xcr1 | Wdr18 |
| Il1rn | 0 | 3.42792671 | 0.947 | 0.275 | 0 | cDC.inflam. | Il1rn |
| Slc7a11 | 0 | 2.98760874 | 0.908 | 0.246 | 0 | cDC.inflam. | Slc7a11 |
| Tnfrsf91 | 0 | 2.3532431 | 0.893 | 0.252 | 0 | cDC.inflam. | Tnfrsf9 |
| Il4i11 | 0 | 1.93065867 | 0.893 | 0.259 | 0 | cDC.inflam. | Il4i1 |
| Ccr71 | 0 | 1.41309294 | 0.981 | 0.38 | 0 | cDC.inflam. | Ccr7 |
| C3 | 0 | 4.25621728 | 0.675 | 0.076 | 0 | cDC.inflam. | C3 |
| Cd141 | 0 | 2.96320354 | 0.898 | 0.299 | 0 | cDC.inflam. | Cd14 |
| Pla2g71 | 0 | 2.81057372 | 0.796 | 0.219 | 0 | cDC.inflam. | Pla2g7 |
| Cd631 | 0 | 1.8937504 | 0.947 | 0.374 | 0 | cDC.inflam. | Cd63 |
| Ccl221 | 0 | 1.293144 | 0.917 | 0.345 | 0 | cDC.inflam. | Ccl22 |
| Anxa31 | 0 | 0.65487297 | 0.913 | 0.359 | 0 | cDC.inflam. | Anxa3 |
| Arl5c1 | 0 | 1.38691947 | 0.869 | 0.316 | 0 | cDC.inflam. | Arl5c |
| Mmp14 | 0 | 2.46172526 | 0.825 | 0.275 | 0 | cDC.inflam. | Mmp14 |
| Axl1 | 0 | 2.5182944 | 0.767 | 0.219 | 0 | cDC.inflam. | Axl |
| Vegfa | 0 | 2.56685484 | 0.806 | 0.26 | 0 | cDC.inflam. | Vegfa |
| Cxcl21 | 0 | 2.77072266 | 0.859 | 0.343 | 0 | cDC.inflam. | Cxcl2 |
| Ptgs22 | 0 | 2.25283719 | 0.796 | 0.295 | 0 | cDC.inflam. | Ptgs2 |
| Ets21 | 0 | 1.68633224 | 0.942 | 0.454 | 0 | cDC.inflam. | Ets2 |
| Tnfaip21 | 0 | 2.03978332 | 0.893 | 0.406 | 0 | cDC.inflam. | Tnfaip2 |
| Alas1 | 0 | 2.67742777 | 0.665 | 0.192 | 0 | cDC.inflam. | Alas1 |
| Fcgr2b1 | 0 | 1.40390701 | 0.869 | 0.399 | 0 | cDC.inflam. | Fcgr2b |
| Cebpb1 | 0 | 2.31563025 | 0.927 | 0.461 | 0 | cDC.inflam. | Cebpb |
| Olr1 | 0 | 4.11043536 | 0.51 | 0.048 | 0 | cDC.inflam. | Olr1 |
| Nrg1 | 0 | 2.81931273 | 0.558 | 0.097 | 0 | cDC.inflam. | Nrg1 |
| Ddit41 | 0 | 1.75925757 | 0.748 | 0.289 | 0 | cDC.inflam. | Ddit4 |
| Mmp251 | 0 | 0.83451935 | 0.835 | 0.377 | 0 | cDC.inflam. | Mmp25 |
| Acp51 | 0 | 2.77377204 | 0.694 | 0.237 | 0 | cDC.inflam. | Acp5 |
| Tarm1 | 0 | 3.4716073 | 0.515 | 0.062 | 0 | cDC.inflam. | Tarm1 |
| Ccl91 | 0 | 2.48835069 | 0.587 | 0.143 | 0 | cDC.inflam. | Ccl9 |
| F10 | 0 | 3.65047001 | 0.49 | 0.05 | 0 | cDC.inflam. | F10 |
| C1qb | 0 | 2.01382455 | 0.573 | 0.135 | 0 | cDC.inflam. | C1qb |
| Ccrl21 | 0 | 1.7561848 | 0.811 | 0.376 | 0 | cDC.inflam. | Ccrl2 |
| Sgk11 | 0 | 3.10916963 | 0.621 | 0.186 | 0 | cDC.inflam. | Sgk1 |
| Clec4e | 0 | 4.22358201 | 0.49 | 0.059 | 0 | cDC.inflam. | Clec4e |
| Marcksl11 | 0 | 1.0837093 | 0.951 | 0.523 | 0 | cDC.inflam. | Marcksl1 |
| Dcstamp1 | 0 | 2.09131512 | 0.66 | 0.236 | 0 | cDC.inflam. | Dcstamp |
| Clec4n2 | 0 | 1.49031427 | 0.816 | 0.4 | 0 | cDC.inflam. | Clec4n |
| Csf1r1 | 0 | 1.44736712 | 0.791 | 0.389 | 0 | cDC.inflam. | Csf1r |
| C1qa | 0 | 2.10472639 | 0.495 | 0.104 | 0 | cDC.inflam. | C1qa |
| Slc7a2 | 0 | 4.55393652 | 0.422 | 0.031 | 0 | cDC.inflam. | Slc7a2 |
| Ass11 | 0 | 1.39812747 | 0.864 | 0.474 | 0 | cDC.inflam. | Ass1 |
| Pdcd1lg21 | 0 | 1.32076565 | 0.893 | 0.51 | 0 | cDC.inflam. | Pdcd1lg2 |
| Gngt21 | 0 | 1.64918971 | 0.806 | 0.431 | 0 | cDC.inflam. | Gngt2 |
| Tgfbi1 | 0 | 1.35347485 | 0.806 | 0.432 | 0 | cDC.inflam. | Tgfbi |
| Ier31 | 0 | 2.14663723 | 0.806 | 0.443 | 0 | cDC.inflam. | Ier3 |
| Ccl51 | 0 | 0.50138111 | 0.602 | 0.242 | 0 | cDC.inflam. | Ccl5 |
| Cstb1 | 0 | 1.8203208 | 0.917 | 0.558 | 0 | cDC.inflam. | Cstb |
| Ramp31 | 0 | 0.9178087 | 0.927 | 0.577 | 0 | cDC.inflam. | Ramp3 |
| Gadd45b1 | 0 | 1.18088684 | 0.99 | 0.643 | 0 | cDC.inflam. | Gadd45b |
| Map4k41 | 0 | 0.94537823 | 0.961 | 0.615 | 0 | cDC.inflam. | Map4k4 |
| Il1a | 0 | 3.49413153 | 0.408 | 0.065 | 0 | cDC.inflam. | Il1a |
| Fn1 | 0 | 2.54261237 | 0.66 | 0.319 | 0 | cDC.inflam. | Fn1 |
| Rnf19b1 | 0 | 1.33143372 | 0.879 | 0.54 | 0 | cDC.inflam. | Rnf19b |
| Thbs1 | 0 | 2.89022241 | 0.461 | 0.123 | 0 | cDC.inflam. | Thbs1 |
| Ccl172 | 0 | 1.80266541 | 0.82 | 0.489 | 0 | cDC.inflam. | Ccl17 |
| Tnf | 0 | 2.63339473 | 0.519 | 0.189 | 0 | cDC.inflam. | Tnf |
| Slc15a32 | 0 | 1.84211182 | 0.961 | 0.634 | 0 | cDC.inflam. | Slc15a3 |
| Sdc1 | 0 | 2.530031 | 0.456 | 0.133 | 0 | cDC.inflam. | Sdc1 |
| Mmp19 | 0 | 2.68243331 | 0.49 | 0.167 | 0 | cDC.inflam. | Mmp19 |
| Fabp51 | 0 | 0.498529 | 0.854 | 0.532 | 0 | cDC.inflam. | Fabp5 |
| Id21 | 0 | 0.97448444 | 0.961 | 0.642 | 0 | cDC.inflam. | Id2 |
| Procr1 | 0 | 2.12788818 | 0.519 | 0.202 | 0 | cDC.inflam. | Procr |
| AA4671971 | 0 | 0.95141507 | 0.859 | 0.545 | 0 | cDC.inflam. | AA467197 |
| Cd91 | 0 | 1.1881791 | 0.971 | 0.661 | 0 | cDC.inflam. | Cd9 |
| Txnrd11 | 0 | 1.34786008 | 0.932 | 0.628 | 0 | cDC.inflam. | Txnrd1 |
| Lyz21 | 0 | 1.83099444 | 0.583 | 0.282 | 0 | cDC.inflam. | Lyz2 |
| Nr4a31 | 0 | 1.504492 | 0.981 | 0.68 | 0 | cDC.inflam. | Nr4a3 |
| Cxcl1 | 0 | 3.97459037 | 0.306 | 0.029 | 0 | cDC.inflam. | Cxcl1 |
| Cdkn1a1 | 0 | 0.95307192 | 0.995 | 0.724 | 0 | cDC.inflam. | Cdkn1a |
| Nfe2l22 | 0 | 1.27064343 | 0.966 | 0.696 | 0 | cDC.inflam. | Nfe2l2 |
| Nfkbia2 | 0 | 1.41951271 | 0.99 | 0.725 | 0 | cDC.inflam. | Nfkbia |
| Pnrc1 | 0 | 1.0502582 | 0.971 | 0.714 | 0 | cDC.inflam. | Pnrc1 |
| Cxcl161 | 0 | 0.87109045 | 0.971 | 0.718 | 0 | cDC.inflam. | Cxcl16 |
| Clic41 | 0 | 1.07493152 | 0.966 | 0.716 | 0 | cDC.inflam. | Clic4 |
| Tmem1231 | 0 | 0.60977142 | 0.888 | 0.639 | 0 | cDC.inflam. | Tmem123 |
| Dusp51 | 0 | 1.31594408 | 0.981 | 0.733 | 0 | cDC.inflam. | Dusp5 |
| Prdx5 | 0 | 1.4606528 | 0.917 | 0.682 | 0 | cDC.inflam. | Prdx5 |
| H2-M2 | 0 | 3.33735809 | 0.272 | 0.039 | 0 | cDC.inflam. | H2-M2 |
| Mmp121 | 0 | 1.69685244 | 0.5 | 0.271 | 0 | cDC.inflam. | Mmp12 |
| Inhba | 0 | 4.39890106 | 0.267 | 0.043 | 0 | cDC.inflam. | Inhba |
| Dusp11 | 0 | 1.37501079 | 0.951 | 0.73 | 0 | cDC.inflam. | Dusp1 |
| Npc2 | 0 | 0.68150079 | 0.956 | 0.739 | 0 | cDC.inflam. | Npc2 |
| Btg11 | 0 | 0.89629048 | 0.951 | 0.739 | 0 | cDC.inflam. | Btg1 |
| Il1b2 | 0 | 1.08577936 | 0.99 | 0.784 | 0 | cDC.inflam. | Il1b |
| Prkcd1 | 0 | 0.83161069 | 0.971 | 0.781 | 0 | cDC.inflam. | Prkcd |
| Mfge81 | 0 | 1.31339492 | 0.544 | 0.375 | 0 | cDC.inflam. | Mfge8 |
| Rel1 | 0 | 0.76997368 | 0.956 | 0.791 | 0 | cDC.inflam. | Rel |
| Bhlhe401 | 0 | 0.5856722 | 0.966 | 0.854 | 0 | cDC.inflam. | Bhlhe40 |
| Junb2 | 0 | 0.67236787 | 1 | 0.889 | 0 | cDC.inflam. | Junb |
| Ctsz | 0 | 0.91253842 | 0.99 | 0.898 | 0 | cDC.inflam. | Ctsz |
| Zfp361 | 0 | 0.44130925 | 0.99 | 0.907 | 0 | cDC.inflam. | Zfp36 |
| Crip11 | 0 | 0.29639745 | 0.976 | 0.909 | 0 | cDC.inflam. | Crip1 |
| Ucp22 | 0 | 1.09216907 | 0.981 | 0.923 | 0 | cDC.inflam. | Ucp2 |
| Ctss2 | 0 | 0.60870776 | 0.971 | 0.92 | 0 | cDC.inflam. | Ctss |
| Fth11 | 0 | 1.51255726 | 1 | 0.956 | 0 | cDC.inflam. | Fth1 |
| Tmsb4x2 | 0 | 0.34502549 | 1 | 0.996 | 0 | cDC.inflam. | Tmsb4x |
| Anxa41 | 7.34E-308 | 1.19206909 | 0.879 | 0.515 | 1.42E-303 | cDC.inflam. | Anxa4 |
| Abca1 | 5.58E-306 | 3.44663436 | 0.583 | 0.06 | 1.08E-301 | cDC.inflam. | Abca1 |
| Blvrb | 1.24E-302 | 2.97212678 | 0.607 | 0.108 | 2.39E-298 | cDC.inflam. | Blvrb |
| Clec4d1 | 1.03E-298 | 2.97549187 | 0.607 | 0.156 | 1.99E-294 | cDC.inflam. | Clec4d |
| Gda1 | 5.42E-296 | 1.84587454 | 0.83 | 0.309 | 1.05E-291 | cDC.inflam. | Gda |
| Ifrd12 | 4.42E-295 | 0.93444035 | 0.951 | 0.674 | 8.54E-291 | cDC.inflam. | Ifrd1 |
| Prdx11 | 2.95E-294 | 1.46851101 | 0.665 | 0.446 | 5.70E-290 | cDC.inflam. | Prdx1 |
| Tspan32 | 7.21E-294 | 1.33557747 | 0.723 | 0.347 | 1.39E-289 | cDC.inflam. | Tspan3 |
| Bcl2a1b1 | 4.32E-291 | 0.91322587 | 0.976 | 0.673 | 8.36E-287 | cDC.inflam. | Bcl2a1b |
| Nrp21 | 1.75E-288 | 1.60093528 | 0.694 | 0.18 | 3.39E-284 | cDC.inflam. | Nrp2 |
| Cfb | 5.57E-286 | 2.6893059 | 0.427 | 0.077 | 1.08E-281 | cDC.inflam. | Cfb |
| Sod21 | 6.18E-283 | 1.82312291 | 0.733 | 0.38 | 1.19E-278 | cDC.inflam. | Sod2 |
| Nfil31 | 1.57E-279 | 0.97419179 | 0.913 | 0.605 | 3.04E-275 | cDC.inflam. | Nfil3 |
| Pld3 | 1.18E-275 | 2.86876881 | 0.573 | 0.107 | 2.29E-271 | cDC.inflam. | Pld3 |
| Icam11 | 5.59E-275 | 1.19695048 | 0.888 | 0.576 | 1.08E-270 | cDC.inflam. | Icam1 |
| Ppp1r15a1 | 3.99E-274 | 1.08745157 | 0.859 | 0.61 | 7.71E-270 | cDC.inflam. | Ppp1r15a |
| Cybb2 | 3.85E-273 | 1.32044223 | 0.665 | 0.34 | 7.45E-269 | cDC.inflam. | Cybb |
| Rnf1492 | 3.79E-263 | 1.19014324 | 0.947 | 0.621 | 7.32E-259 | cDC.inflam. | Rnf149 |
| Msrb11 | 9.23E-255 | 1.40018906 | 0.917 | 0.525 | 1.78E-250 | cDC.inflam. | Msrb1 |
| Vamp8 | 9.71E-254 | 0.93878052 | 0.937 | 0.728 | 1.88E-249 | cDC.inflam. | Vamp8 |
| Samsn11 | 4.67E-252 | 0.70991946 | 0.922 | 0.512 | 9.03E-248 | cDC.inflam. | Samsn1 |
| Tgif11 | 1.38E-251 | 1.09910284 | 0.874 | 0.561 | 2.66E-247 | cDC.inflam. | Tgif1 |
| Srgn1 | 5.76E-250 | 0.38259931 | 0.971 | 0.906 | 1.11E-245 | cDC.inflam. | Srgn |
| Socs31 | 3.28E-246 | 0.96751682 | 0.903 | 0.596 | 6.34E-242 | cDC.inflam. | Socs3 |
| Plxnd11 | 3.62E-246 | 0.87927749 | 0.908 | 0.605 | 7.00E-242 | cDC.inflam. | Plxnd1 |
| Ccdc88a1 | 1.40E-241 | 1.01100952 | 0.85 | 0.544 | 2.71E-237 | cDC.inflam. | Ccdc88a |
| Ninj1 | 1.39E-236 | 1.96972255 | 0.772 | 0.258 | 2.70E-232 | cDC.inflam. | Ninj1 |
| Sat11 | 8.35E-236 | 0.91447773 | 0.932 | 0.667 | 1.62E-231 | cDC.inflam. | Sat1 |
| Sqstm11 | 2.44E-229 | 1.14253269 | 0.898 | 0.619 | 4.72E-225 | cDC.inflam. | Sqstm1 |
| Sema4a1 | 1.91E-228 | 0.80072876 | 0.879 | 0.589 | 3.69E-224 | cDC.inflam. | Sema4a |
| Psap1 | 1.28E-225 | 0.42443836 | 0.942 | 0.796 | 2.48E-221 | cDC.inflam. | Psap |
| Slfn21 | 7.66E-222 | 1.10182704 | 0.82 | 0.531 | 1.48E-217 | cDC.inflam. | Slfn2 |
| Aldoa1 | 3.83E-221 | 0.92961135 | 0.937 | 0.684 | 7.41E-217 | cDC.inflam. | Aldoa |
| Cd441 | 8.37E-219 | 0.87233687 | 0.859 | 0.586 | 1.62E-214 | cDC.inflam. | Cd44 |
| Tgfbr11 | 5.80E-217 | 1.84816855 | 0.646 | 0.253 | 1.12E-212 | cDC.inflam. | Tgfbr1 |
| Nfkb21 | 3.26E-207 | 0.54396954 | 0.971 | 0.773 | 6.31E-203 | cDC.inflam. | Nfkb2 |
| Zfp36l11 | 5.55E-204 | 0.95571096 | 0.903 | 0.542 | 1.07E-199 | cDC.inflam. | Zfp36l1 |
| Gsr | 9.32E-204 | 1.36490312 | 0.757 | 0.435 | 1.80E-199 | cDC.inflam. | Gsr |
| Cd831 | 1.77E-203 | 0.5008835 | 0.932 | 0.707 | 3.43E-199 | cDC.inflam. | Cd83 |
| Tmem1311 | 5.03E-200 | 0.85558917 | 0.898 | 0.648 | 9.73E-196 | cDC.inflam. | Tmem131 |
| Basp11 | 5.61E-200 | 0.50303482 | 0.956 | 0.819 | 1.08E-195 | cDC.inflam. | Basp1 |
| Slc38a21 | 9.85E-200 | 0.85830516 | 0.971 | 0.706 | 1.90E-195 | cDC.inflam. | Slc38a2 |
| Slc11a1 | 7.46E-198 | 3.3247417 | 0.451 | 0.061 | 1.44E-193 | cDC.inflam. | Slc11a1 |
| Gga21 | 3.27E-194 | 1.50226506 | 0.617 | 0.213 | 6.33E-190 | cDC.inflam. | Gga2 |
| Msr1 | 1.79E-190 | 2.94210006 | 0.466 | 0.05 | 3.46E-186 | cDC.inflam. | Msr1 |
| Tspan331 | 3.03E-190 | 1.38357004 | 0.748 | 0.317 | 5.86E-186 | cDC.inflam. | Tspan33 |
| Rgcc | 1.53E-189 | 2.79395825 | 0.33 | 0.076 | 2.95E-185 | cDC.inflam. | Rgcc |
| Srxn1 | 5.85E-188 | 2.33789989 | 0.558 | 0.13 | 1.13E-183 | cDC.inflam. | Srxn1 |
| Fcrlb | 1.81E-187 | 3.01057729 | 0.471 | 0.065 | 3.51E-183 | cDC.inflam. | Fcrlb |
| Mafb | 3.52E-186 | 2.05777196 | 0.476 | 0.12 | 6.81E-182 | cDC.inflam. | Mafb |
| Osm1 | 1.03E-185 | 1.18522745 | 0.66 | 0.3 | 2.00E-181 | cDC.inflam. | Osm |
| Malt11 | 9.74E-185 | 0.56414084 | 0.917 | 0.703 | 1.88E-180 | cDC.inflam. | Malt1 |
| Fam32a1 | 1.46E-182 | 1.017987 | 0.864 | 0.533 | 2.82E-178 | cDC.inflam. | Fam32a |
| Nfkbiz2 | 5.15E-182 | 1.05139067 | 0.806 | 0.508 | 9.96E-178 | cDC.inflam. | Nfkbiz |
| Csf2rb3 | 7.07E-181 | 0.54025287 | 0.976 | 0.814 | 1.37E-176 | cDC.inflam. | Csf2rb |
| Maff2 | 5.17E-180 | 1.15264535 | 0.874 | 0.503 | 9.99E-176 | cDC.inflam. | Maff |
| Lamp11 | 6.90E-179 | 1.07421461 | 0.811 | 0.449 | 1.33E-174 | cDC.inflam. | Lamp1 |
| Lgals31 | 8.86E-179 | 0.30291748 | 0.898 | 0.761 | 1.71E-174 | cDC.inflam. | Lgals3 |
| Arl5a1 | 1.16E-178 | 1.14530453 | 0.825 | 0.39 | 2.25E-174 | cDC.inflam. | Arl5a |
| Fos2 | 1.13E-175 | 0.48404741 | 0.864 | 0.682 | 2.18E-171 | cDC.inflam. | Fos |
| Hilpda2 | 1.38E-175 | 1.1401612 | 0.665 | 0.349 | 2.68E-171 | cDC.inflam. | Hilpda |
| App2 | 1.09E-174 | 0.57815538 | 0.947 | 0.75 | 2.11E-170 | cDC.inflam. | App |
| Nectin2 | 1.87E-174 | 2.1347664 | 0.592 | 0.145 | 3.62E-170 | cDC.inflam. | Nectin2 |
| Tnfrsf1b1 | 2.89E-174 | 1.05711464 | 0.879 | 0.591 | 5.58E-170 | cDC.inflam. | Tnfrsf1b |
| Tmem176a2 | 5.83E-174 | 0.7835488 | 0.684 | 0.414 | 1.13E-169 | cDC.inflam. | Tmem176a |
| Ehd1 | 8.09E-174 | 1.33649072 | 0.835 | 0.453 | 1.56E-169 | cDC.inflam. | Ehd1 |
| Trf | 8.41E-172 | 1.35361402 | 0.66 | 0.333 | 1.63E-167 | cDC.inflam. | Trf |
| Hvcn11 | 2.79E-169 | 1.91777599 | 0.641 | 0.149 | 5.40E-165 | cDC.inflam. | Hvcn1 |
| Mcl1 | 4.10E-168 | 0.39763748 | 0.985 | 0.903 | 7.93E-164 | cDC.inflam. | Mcl1 |
| H2-Eb2 | 7.87E-167 | 2.11663362 | 0.432 | 0.087 | 1.52E-162 | cDC.inflam. | H2-Eb2 |
| Atp6v0c2 | 1.73E-166 | 0.78927798 | 0.908 | 0.696 | 3.35E-162 | cDC.inflam. | Atp6v0c |
| Zfp7032 | 2.88E-166 | 1.27614195 | 0.782 | 0.422 | 5.56E-162 | cDC.inflam. | Zfp703 |
| Marcks1 | 1.86E-164 | 0.28725496 | 0.976 | 0.786 | 3.60E-160 | cDC.inflam. | Marcks |
| Mxi11 | 2.25E-164 | 1.0750536 | 0.767 | 0.393 | 4.35E-160 | cDC.inflam. | Mxi1 |
| Ccl3 | 9.14E-162 | 2.04017237 | 0.359 | 0.172 | 1.77E-157 | cDC.inflam. | Ccl3 |
| Sgpl1 | 1.10E-161 | 1.13228629 | 0.767 | 0.477 | 2.13E-157 | cDC.inflam. | Sgpl1 |
| Txn11 | 3.44E-160 | 1.01244732 | 0.82 | 0.49 | 6.65E-156 | cDC.inflam. | Txn1 |
| Tmem176b1 | 2.08E-159 | 0.69350155 | 0.733 | 0.474 | 4.01E-155 | cDC.inflam. | Tmem176b |
| Pgd1 | 3.64E-158 | 1.30275162 | 0.752 | 0.441 | 7.03E-154 | cDC.inflam. | Pgd |
| Smox1 | 1.06E-157 | 1.47477297 | 0.728 | 0.333 | 2.06E-153 | cDC.inflam. | Smox |
| Adam81 | 5.05E-157 | 0.75779173 | 0.854 | 0.536 | 9.77E-153 | cDC.inflam. | Adam8 |
| Gna13 | 4.34E-156 | 0.5450566 | 0.966 | 0.807 | 8.40E-152 | cDC.inflam. | Gna13 |
| F11r1 | 8.33E-156 | 1.25198393 | 0.714 | 0.338 | 1.61E-151 | cDC.inflam. | F11r |
| Zeb21 | 8.57E-156 | 1.14673091 | 0.704 | 0.367 | 1.66E-151 | cDC.inflam. | Zeb2 |
| Nfkb12 | 6.21E-154 | 0.52327899 | 0.937 | 0.777 | 1.20E-149 | cDC.inflam. | Nfkb1 |
| Tnfaip31 | 1.07E-153 | 1.03168485 | 0.806 | 0.461 | 2.08E-149 | cDC.inflam. | Tnfaip3 |
| Ikbke | 4.74E-152 | 3.13552048 | 0.447 | 0.047 | 9.17E-148 | cDC.inflam. | Ikbke |
| Sirpa1 | 7.22E-152 | 0.79113599 | 0.879 | 0.587 | 1.40E-147 | cDC.inflam. | Sirpa |
| Lrp11 | 2.01E-151 | 1.3246964 | 0.675 | 0.267 | 3.88E-147 | cDC.inflam. | Lrp1 |
| Ddhd11 | 9.87E-150 | 1.17258379 | 0.68 | 0.352 | 1.91E-145 | cDC.inflam. | Ddhd1 |
| Myo1g1 | 1.76E-149 | 0.26275533 | 0.971 | 0.823 | 3.41E-145 | cDC.inflam. | Myo1g |
| Osgin21 | 6.56E-148 | 1.50931585 | 0.704 | 0.282 | 1.27E-143 | cDC.inflam. | Osgin2 |
| Plaur1 | 1.36E-145 | 1.1057784 | 0.714 | 0.384 | 2.63E-141 | cDC.inflam. | Plaur |
| Grina | 4.42E-145 | 1.28656897 | 0.738 | 0.428 | 8.55E-141 | cDC.inflam. | Grina |
| Trps12 | 3.99E-144 | 0.98549647 | 0.84 | 0.563 | 7.71E-140 | cDC.inflam. | Trps1 |
| Ptafr2 | 3.52E-143 | 1.22239914 | 0.806 | 0.376 | 6.81E-139 | cDC.inflam. | Ptafr |
| Slc3a21 | 1.25E-142 | 0.65962066 | 0.927 | 0.73 | 2.42E-138 | cDC.inflam. | Slc3a2 |
| Arf4 | 2.31E-140 | 0.75232975 | 0.82 | 0.584 | 4.47E-136 | cDC.inflam. | Arf4 |
| Rgs21 | 2.65E-140 | 1.05280666 | 0.733 | 0.443 | 5.13E-136 | cDC.inflam. | Rgs2 |
| Syngr21 | 1.27E-139 | 0.35521432 | 0.961 | 0.842 | 2.46E-135 | cDC.inflam. | Syngr2 |
| Spred11 | 3.20E-138 | 0.79470403 | 0.84 | 0.471 | 6.18E-134 | cDC.inflam. | Spred1 |
| Cd2741 | 2.45E-137 | 0.62669418 | 0.879 | 0.58 | 4.74E-133 | cDC.inflam. | Cd274 |
| Slc16a31 | 9.22E-136 | 1.3018669 | 0.782 | 0.422 | 1.78E-131 | cDC.inflam. | Slc16a3 |
| Pde4b1 | 6.13E-135 | 0.65870585 | 0.947 | 0.682 | 1.19E-130 | cDC.inflam. | Pde4b |
| Etv31 | 1.90E-134 | 0.49477404 | 0.932 | 0.69 | 3.68E-130 | cDC.inflam. | Etv3 |
| Pxdc12 | 8.43E-134 | 1.28996949 | 0.743 | 0.347 | 1.63E-129 | cDC.inflam. | Pxdc1 |
| Cxcr42 | 1.01E-133 | 0.64486407 | 0.869 | 0.532 | 1.95E-129 | cDC.inflam. | Cxcr4 |
| Hivep3 | 6.31E-130 | 2.45846633 | 0.374 | 0.077 | 1.22E-125 | cDC.inflam. | Hivep3 |
| Rab201 | 9.60E-127 | 1.65556413 | 0.621 | 0.2 | 1.86E-122 | cDC.inflam. | Rab20 |
| Atp6v1b21 | 6.32E-126 | 0.79561624 | 0.806 | 0.552 | 1.22E-121 | cDC.inflam. | Atp6v1b2 |
| Capg1 | 2.56E-123 | 1.14962072 | 0.641 | 0.307 | 4.95E-119 | cDC.inflam. | Capg |
| Mllt61 | 6.24E-123 | 0.63219529 | 0.888 | 0.623 | 1.21E-118 | cDC.inflam. | Mllt6 |
| Got1 | 8.19E-123 | 0.60501578 | 0.898 | 0.659 | 1.58E-118 | cDC.inflam. | Got1 |
| Tcirg1 | 9.93E-123 | 0.70088834 | 0.854 | 0.611 | 1.92E-118 | cDC.inflam. | Tcirg1 |
| P2rx41 | 3.64E-122 | 1.07724948 | 0.675 | 0.388 | 7.03E-118 | cDC.inflam. | P2rx4 |
| Ralgds | 4.32E-122 | 2.27377049 | 0.529 | 0.137 | 8.35E-118 | cDC.inflam. | Ralgds |
| Hcar2 | 6.03E-121 | 2.63646591 | 0.354 | 0.11 | 1.17E-116 | cDC.inflam. | Hcar2 |
| Furin1 | 6.17E-121 | 1.05670766 | 0.782 | 0.489 | 1.19E-116 | cDC.inflam. | Furin |
| M6pr | 8.50E-121 | 0.70128058 | 0.883 | 0.692 | 1.64E-116 | cDC.inflam. | M6pr |
| Samhd12 | 3.24E-118 | 0.35453741 | 0.888 | 0.775 | 6.27E-114 | cDC.inflam. | Samhd1 |
| Rai14 | 2.40E-117 | 1.87763014 | 0.553 | 0.154 | 4.65E-113 | cDC.inflam. | Rai14 |
| Ubc | 3.69E-117 | 0.49310096 | 0.951 | 0.825 | 7.13E-113 | cDC.inflam. | Ubc |
| Csrnp11 | 4.76E-116 | 0.67315554 | 0.942 | 0.684 | 9.20E-112 | cDC.inflam. | Csrnp1 |
| H1fx | 5.52E-116 | 2.19191652 | 0.282 | 0.052 | 1.07E-111 | cDC.inflam. | H1fx |
| Lmo4 | 8.39E-116 | 1.46219056 | 0.573 | 0.259 | 1.62E-111 | cDC.inflam. | Lmo4 |
| Tlr21 | 1.02E-115 | 1.313611 | 0.553 | 0.248 | 1.98E-111 | cDC.inflam. | Tlr2 |
| Plek2 | 1.89E-115 | 0.37358055 | 0.947 | 0.852 | 3.66E-111 | cDC.inflam. | Plek |
| Litaf1 | 6.11E-115 | 0.52384868 | 0.942 | 0.77 | 1.18E-110 | cDC.inflam. | Litaf |
| Arg2 | 2.00E-114 | 3.11738678 | 0.345 | 0.045 | 3.87E-110 | cDC.inflam. | Arg2 |
| Cyba | 4.21E-114 | 0.43572426 | 0.966 | 0.811 | 8.15E-110 | cDC.inflam. | Cyba |
| Fem1b | 8.62E-114 | 1.59366685 | 0.544 | 0.221 | 1.67E-109 | cDC.inflam. | Fem1b |
| Pla2g15 | 2.95E-113 | 1.93706038 | 0.427 | 0.134 | 5.71E-109 | cDC.inflam. | Pla2g15 |
| Arhgef10l | 7.86E-112 | 1.83976944 | 0.534 | 0.118 | 1.52E-107 | cDC.inflam. | Arhgef10l |
| Rab11fip11 | 2.87E-111 | 0.77058714 | 0.932 | 0.641 | 5.56E-107 | cDC.inflam. | Rab11fip1 |
| Relb1 | 1.52E-110 | 0.37088966 | 0.888 | 0.531 | 2.93E-106 | cDC.inflam. | Relb |
| Fcer1g1 | 6.98E-109 | 0.31234499 | 0.927 | 0.689 | 1.35E-104 | cDC.inflam. | Fcer1g |
| Ak21 | 3.04E-108 | 0.95517623 | 0.762 | 0.489 | 5.89E-104 | cDC.inflam. | Ak2 |
| Cd401 | 3.03E-107 | 0.59482509 | 0.714 | 0.393 | 5.85E-103 | cDC.inflam. | Cd40 |
| Csf2rb22 | 3.89E-107 | 0.70924848 | 0.835 | 0.504 | 7.52E-103 | cDC.inflam. | Csf2rb2 |
| Tfec | 7.10E-107 | 3.97141336 | 0.301 | 0.018 | 1.37E-102 | cDC.inflam. | Tfec |
| Csf3r | 2.70E-106 | 2.86977833 | 0.252 | 0.065 | 5.22E-102 | cDC.inflam. | Csf3r |
| Src | 2.06E-105 | 1.2709728 | 0.626 | 0.275 | 3.99E-101 | cDC.inflam. | Src |
| Gch12 | 6.28E-103 | 1.21578037 | 0.636 | 0.326 | 1.21E-98 | cDC.inflam. | Gch1 |
| Fam49a1 | 1.91E-102 | 0.57035451 | 0.714 | 0.374 | 3.69E-98 | cDC.inflam. | Fam49a |
| Mt21 | 7.26E-102 | 0.99446471 | 0.451 | 0.197 | 1.40E-97 | cDC.inflam. | Mt2 |
| Cebpa1 | 2.82E-101 | 1.23436141 | 0.636 | 0.3 | 5.45E-97 | cDC.inflam. | Cebpa |
| Sowahc1 | 1.58E-100 | 0.6956443 | 0.811 | 0.496 | 3.05E-96 | cDC.inflam. | Sowahc |
| Nfkbid1 | 1.74E-100 | 0.9644913 | 0.748 | 0.461 | 3.37E-96 | cDC.inflam. | Nfkbid |
| Ncf22 | 2.20E-100 | 0.77023721 | 0.869 | 0.538 | 4.26E-96 | cDC.inflam. | Ncf2 |
| Rhoc1 | 4.22E-100 | 0.92576058 | 0.621 | 0.241 | 8.16E-96 | cDC.inflam. | Rhoc |
| Asap1 | 9.95E-99 | 1.08362149 | 0.718 | 0.4 | 1.92E-94 | cDC.inflam. | Asap1 |
| Pdgfb | 3.79E-96 | 1.88529481 | 0.408 | 0.109 | 7.32E-92 | cDC.inflam. | Pdgfb |
| Rab8b1 | 1.92E-95 | 0.2876717 | 0.937 | 0.75 | 3.72E-91 | cDC.inflam. | Rab8b |
| Pgap21 | 6.53E-94 | 1.19748215 | 0.636 | 0.32 | 1.26E-89 | cDC.inflam. | Pgap2 |
| Nampt1 | 2.85E-93 | 0.86530287 | 0.748 | 0.425 | 5.51E-89 | cDC.inflam. | Nampt |
| Rnf217 | 4.24E-93 | 1.9567128 | 0.447 | 0.124 | 8.19E-89 | cDC.inflam. | Rnf217 |
| Hmox1 | 7.50E-93 | 1.6077729 | 0.374 | 0.116 | 1.45E-88 | cDC.inflam. | Hmox1 |
| Spty2d12 | 1.29E-92 | 0.7510983 | 0.757 | 0.497 | 2.50E-88 | cDC.inflam. | Spty2d1 |
| Pmaip11 | 1.84E-92 | 0.67369555 | 0.835 | 0.541 | 3.55E-88 | cDC.inflam. | Pmaip1 |
| Ripk21 | 4.36E-89 | 1.25486368 | 0.544 | 0.244 | 8.43E-85 | cDC.inflam. | Ripk2 |
| Il21r1 | 6.05E-89 | 0.4908904 | 0.83 | 0.589 | 1.17E-84 | cDC.inflam. | Il21r |
| Gpalpp11 | 9.31E-88 | 1.38317771 | 0.549 | 0.234 | 1.80E-83 | cDC.inflam. | Gpalpp1 |
| Sertad11 | 3.19E-87 | 1.23807116 | 0.607 | 0.278 | 6.17E-83 | cDC.inflam. | Sertad1 |
| Adam17 | 7.59E-87 | 0.98224698 | 0.733 | 0.434 | 1.47E-82 | cDC.inflam. | Adam17 |
| C1qc | 7.14E-85 | 1.72114565 | 0.354 | 0.073 | 1.38E-80 | cDC.inflam. | C1qc |
| Arl4c1 | 8.98E-85 | 0.81567561 | 0.733 | 0.426 | 1.74E-80 | cDC.inflam. | Arl4c |
| Hif1a1 | 1.58E-84 | 0.9699637 | 0.699 | 0.395 | 3.06E-80 | cDC.inflam. | Hif1a |
| Ggta11 | 9.89E-84 | 0.56135913 | 0.859 | 0.595 | 1.91E-79 | cDC.inflam. | Ggta1 |
| Cyth1 | 1.25E-83 | 0.65589169 | 0.864 | 0.58 | 2.43E-79 | cDC.inflam. | Cyth1 |
| C5ar1 | 1.26E-83 | 2.88615926 | 0.316 | 0.056 | 2.44E-79 | cDC.inflam. | C5ar1 |
| Pgk11 | 1.40E-83 | 0.83183531 | 0.723 | 0.455 | 2.70E-79 | cDC.inflam. | Pgk1 |
| Flrt31 | 2.85E-83 | 0.5566276 | 0.583 | 0.344 | 5.52E-79 | cDC.inflam. | Flrt3 |
| Sdc41 | 3.38E-83 | 0.96095148 | 0.67 | 0.296 | 6.53E-79 | cDC.inflam. | Sdc4 |
| Herpud1 | 4.02E-83 | 0.36617874 | 0.83 | 0.631 | 7.78E-79 | cDC.inflam. | Herpud1 |
| Pik3r11 | 9.55E-83 | 0.38663042 | 0.845 | 0.627 | 1.85E-78 | cDC.inflam. | Pik3r1 |
| Skil1 | 7.12E-82 | 0.56917514 | 0.913 | 0.664 | 1.38E-77 | cDC.inflam. | Skil |
| Foxp41 | 8.79E-82 | 0.66712995 | 0.689 | 0.329 | 1.70E-77 | cDC.inflam. | Foxp4 |
| Il7r1 | 5.13E-81 | 0.83786774 | 0.694 | 0.287 | 9.91E-77 | cDC.inflam. | Il7r |
| Myo1c1 | 8.06E-81 | 0.61498254 | 0.806 | 0.586 | 1.56E-76 | cDC.inflam. | Myo1c |
| Itgal1 | 8.29E-81 | 1.15048462 | 0.49 | 0.176 | 1.60E-76 | cDC.inflam. | Itgal |
| Stat11 | 1.46E-80 | 0.51949398 | 0.723 | 0.521 | 2.82E-76 | cDC.inflam. | Stat1 |
| Aldh1a21 | 2.20E-80 | 1.13326618 | 0.267 | 0.166 | 4.25E-76 | cDC.inflam. | Aldh1a2 |
| Smurf11 | 3.94E-80 | 0.90231193 | 0.709 | 0.333 | 7.61E-76 | cDC.inflam. | Smurf1 |
| Dusp161 | 3.09E-79 | 1.20049712 | 0.67 | 0.393 | 5.97E-75 | cDC.inflam. | Dusp16 |
| Dnajc101 | 7.39E-79 | 0.67419336 | 0.646 | 0.398 | 1.43E-74 | cDC.inflam. | Dnajc10 |
| Zbtb182 | 8.42E-79 | 0.51289291 | 0.83 | 0.577 | 1.63E-74 | cDC.inflam. | Zbtb18 |
| Clcf1 | 1.03E-78 | 1.67560599 | 0.5 | 0.214 | 1.99E-74 | cDC.inflam. | Clcf1 |
| Nufip11 | 4.35E-78 | 1.48930065 | 0.544 | 0.211 | 8.41E-74 | cDC.inflam. | Nufip1 |
| Chst112 | 8.65E-78 | 1.11858045 | 0.636 | 0.338 | 1.67E-73 | cDC.inflam. | Chst11 |
| Tpm41 | 1.14E-77 | 0.32574255 | 0.908 | 0.763 | 2.20E-73 | cDC.inflam. | Tpm4 |
| Myo1e1 | 2.27E-77 | 1.63355403 | 0.51 | 0.162 | 4.39E-73 | cDC.inflam. | Myo1e |
| Ier51 | 1.60E-76 | 0.30991245 | 0.927 | 0.76 | 3.10E-72 | cDC.inflam. | Ier5 |
| Ctsc1 | 1.19E-75 | 0.32523071 | 0.791 | 0.677 | 2.29E-71 | cDC.inflam. | Ctsc |
| Zc3h12c1 | 3.43E-75 | 1.12659458 | 0.383 | 0.129 | 6.63E-71 | cDC.inflam. | Zc3h12c |
| Ncoa41 | 3.81E-75 | 0.79999332 | 0.646 | 0.418 | 7.36E-71 | cDC.inflam. | Ncoa4 |
| Phldb11 | 1.88E-74 | 1.2360978 | 0.437 | 0.13 | 3.63E-70 | cDC.inflam. | Phldb1 |
| Man2a1 | 1.95E-74 | 1.27094772 | 0.641 | 0.307 | 3.78E-70 | cDC.inflam. | Man2a1 |
| Zmynd151 | 1.46E-73 | 0.64488079 | 0.539 | 0.199 | 2.82E-69 | cDC.inflam. | Zmynd15 |
| Mfhas11 | 2.27E-73 | 0.76401164 | 0.675 | 0.316 | 4.39E-69 | cDC.inflam. | Mfhas1 |
| Mcemp11 | 2.89E-73 | 1.01850416 | 0.641 | 0.295 | 5.59E-69 | cDC.inflam. | Mcemp1 |
| Epb41l21 | 3.68E-73 | 0.62523816 | 0.743 | 0.454 | 7.11E-69 | cDC.inflam. | Epb41l2 |
| Lipa2 | 9.57E-73 | 1.14353233 | 0.451 | 0.29 | 1.85E-68 | cDC.inflam. | Lipa |
| Zc3h12a2 | 1.69E-72 | 0.80720019 | 0.835 | 0.488 | 3.27E-68 | cDC.inflam. | Zc3h12a |
| Atp1a11 | 2.82E-72 | 0.26020976 | 0.917 | 0.797 | 5.46E-68 | cDC.inflam. | Atp1a1 |
| Nuak21 | 9.32E-72 | 0.349216 | 0.922 | 0.72 | 1.80E-67 | cDC.inflam. | Nuak2 |
| Pik3r51 | 8.59E-71 | 0.34922679 | 0.83 | 0.668 | 1.66E-66 | cDC.inflam. | Pik3r5 |
| Akap51 | 1.11E-70 | 0.94642413 | 0.403 | 0.181 | 2.15E-66 | cDC.inflam. | Akap5 |
| Rap2a1 | 1.31E-70 | 0.65664264 | 0.811 | 0.524 | 2.52E-66 | cDC.inflam. | Rap2a |
| Ctsb | 1.35E-70 | 0.30770102 | 0.893 | 0.723 | 2.60E-66 | cDC.inflam. | Ctsb |
| Vat1 | 5.03E-70 | 1.30449639 | 0.519 | 0.231 | 9.72E-66 | cDC.inflam. | Vat1 |
| Apobec1 | 5.66E-70 | 1.2631428 | 0.35 | 0.119 | 1.10E-65 | cDC.inflam. | Apobec1 |
| Cd861 | 1.15E-69 | 0.40216382 | 0.917 | 0.678 | 2.23E-65 | cDC.inflam. | Cd86 |
| Syk1 | 1.68E-69 | 0.2794006 | 0.932 | 0.789 | 3.24E-65 | cDC.inflam. | Syk |
| Gbp21 | 1.94E-69 | 0.75903056 | 0.51 | 0.246 | 3.74E-65 | cDC.inflam. | Gbp2 |
| Creg11 | 5.58E-69 | 0.77679948 | 0.665 | 0.412 | 1.08E-64 | cDC.inflam. | Creg1 |
| Cd80 | 6.55E-69 | 0.28520162 | 0.869 | 0.663 | 1.27E-64 | cDC.inflam. | Cd80 |
| Sav11 | 1.95E-68 | 0.73710555 | 0.714 | 0.44 | 3.78E-64 | cDC.inflam. | Sav1 |
| Atp6v0b | 1.90E-67 | 0.64858284 | 0.83 | 0.555 | 3.66E-63 | cDC.inflam. | Atp6v0b |
| Adssl11 | 2.25E-67 | 1.17271306 | 0.529 | 0.252 | 4.36E-63 | cDC.inflam. | Adssl1 |
| Zfand2a | 3.18E-67 | 1.31606228 | 0.481 | 0.185 | 6.14E-63 | cDC.inflam. | Zfand2a |
| Smchd1 | 4.23E-67 | 0.76096491 | 0.689 | 0.428 | 8.17E-63 | cDC.inflam. | Smchd1 |
| Plgrkt1 | 9.54E-67 | 0.79647499 | 0.68 | 0.429 | 1.84E-62 | cDC.inflam. | Plgrkt |
| Gpr1321 | 1.07E-66 | 0.25901813 | 0.932 | 0.746 | 2.06E-62 | cDC.inflam. | Gpr132 |
| Gla2 | 1.11E-66 | 1.30111322 | 0.495 | 0.213 | 2.14E-62 | cDC.inflam. | Gla |
| Atp1b31 | 2.67E-66 | 0.82976343 | 0.723 | 0.434 | 5.16E-62 | cDC.inflam. | Atp1b3 |
| Gsto11 | 3.11E-66 | 0.33422812 | 0.796 | 0.603 | 6.02E-62 | cDC.inflam. | Gsto1 |
| Prr131 | 1.29E-65 | 0.74613839 | 0.718 | 0.447 | 2.49E-61 | cDC.inflam. | Prr13 |
| Thap121 | 1.65E-65 | 1.24941404 | 0.568 | 0.231 | 3.19E-61 | cDC.inflam. | Thap12 |
| St3gal11 | 6.51E-65 | 0.65728623 | 0.728 | 0.423 | 1.26E-60 | cDC.inflam. | St3gal1 |
| Tgm22 | 9.07E-65 | 0.50333046 | 0.883 | 0.557 | 1.75E-60 | cDC.inflam. | Tgm2 |
| Slc35b22 | 1.67E-64 | 0.49889686 | 0.825 | 0.571 | 3.22E-60 | cDC.inflam. | Slc35b2 |
| Slc23a2 | 1.07E-63 | 1.55413279 | 0.432 | 0.155 | 2.07E-59 | cDC.inflam. | Slc23a2 |
| Tmem51 | 2.34E-63 | 1.59479354 | 0.422 | 0.125 | 4.53E-59 | cDC.inflam. | Tmem51 |
| Nfat51 | 2.54E-63 | 0.62294094 | 0.704 | 0.393 | 4.92E-59 | cDC.inflam. | Nfat5 |
| Cish1 | 3.59E-62 | 0.72806705 | 0.714 | 0.436 | 6.95E-58 | cDC.inflam. | Cish |
| Lgals3bp1 | 3.65E-62 | 0.70350389 | 0.67 | 0.452 | 7.06E-58 | cDC.inflam. | Lgals3bp |
| Eid31 | 8.11E-62 | 1.62190727 | 0.529 | 0.196 | 1.57E-57 | cDC.inflam. | Eid3 |
| Dusp2 | 2.53E-61 | 1.24382055 | 0.417 | 0.189 | 4.90E-57 | cDC.inflam. | Dusp2 |
| St8sia41 | 2.96E-61 | 0.32499353 | 0.786 | 0.592 | 5.72E-57 | cDC.inflam. | St8sia4 |
| Rasgrp11 | 7.56E-61 | 1.2763209 | 0.451 | 0.157 | 1.46E-56 | cDC.inflam. | Rasgrp1 |
| Ldlr2 | 4.74E-60 | 0.50107068 | 0.752 | 0.498 | 9.16E-56 | cDC.inflam. | Ldlr |
| Fosb2 | 6.33E-60 | 0.28237864 | 0.883 | 0.648 | 1.22E-55 | cDC.inflam. | Fosb |
| Abcg31 | 7.89E-59 | 1.73365383 | 0.325 | 0.122 | 1.53E-54 | cDC.inflam. | Abcg3 |
| Chka1 | 1.20E-58 | 1.1015551 | 0.451 | 0.195 | 2.33E-54 | cDC.inflam. | Chka |
| Plxdc2 | 1.22E-58 | 1.50838364 | 0.282 | 0.101 | 2.35E-54 | cDC.inflam. | Plxdc2 |
| BC005537 | 2.09E-58 | 0.33721445 | 0.888 | 0.734 | 4.04E-54 | cDC.inflam. | BC005537 |
| Colgalt11 | 1.24E-57 | 0.47970547 | 0.845 | 0.61 | 2.39E-53 | cDC.inflam. | Colgalt1 |
| Slc25a25 | 1.32E-57 | 1.28555322 | 0.476 | 0.197 | 2.56E-53 | cDC.inflam. | Slc25a25 |
| Clcn7 | 1.97E-57 | 1.19706503 | 0.5 | 0.189 | 3.81E-53 | cDC.inflam. | Clcn7 |
| Cx3cl11 | 2.65E-57 | 0.87397448 | 0.427 | 0.165 | 5.13E-53 | cDC.inflam. | Cx3cl1 |
| Tnip1 | 3.12E-57 | 0.84451978 | 0.67 | 0.382 | 6.02E-53 | cDC.inflam. | Tnip1 |
| Rhbdf21 | 3.27E-57 | 1.16202935 | 0.636 | 0.27 | 6.33E-53 | cDC.inflam. | Rhbdf2 |
| Phlda1 | 2.24E-56 | 0.99606119 | 0.311 | 0.148 | 4.34E-52 | cDC.inflam. | Phlda1 |
| Adora2a1 | 4.00E-56 | 0.81700645 | 0.539 | 0.177 | 7.72E-52 | cDC.inflam. | Adora2a |
| Clptm11 | 5.10E-56 | 0.56274137 | 0.757 | 0.502 | 9.87E-52 | cDC.inflam. | Clptm1 |
| Bcl2l11 | 6.41E-56 | 0.36138018 | 0.806 | 0.589 | 1.24E-51 | cDC.inflam. | Bcl2l11 |
| DEPRECATED-I | 2.26E-55 | 0.92632023 | 0.524 | 0.225 | 4.36E-51 | cDC.inflam. | DEPRECATED-ENSMUSG00000025025 |
| Gpr351 | 6.38E-55 | 0.7038599 | 0.718 | 0.508 | 1.23E-50 | cDC.inflam. | Gpr35 |
| Dram1 | 2.17E-54 | 1.36205455 | 0.393 | 0.123 | 4.19E-50 | cDC.inflam. | Dram1 |
| Sra1 | 3.55E-54 | 0.88054818 | 0.626 | 0.393 | 6.86E-50 | cDC.inflam. | Sra1 |
| Arrb12 | 4.03E-54 | 0.86712557 | 0.675 | 0.394 | 7.78E-50 | cDC.inflam. | Arrb1 |
| Wnt11 | 4.04E-54 | 1.15865063 | 0.345 | 0.133 | 7.81E-50 | cDC.inflam. | Wnt11 |
| Lrpap11 | 4.60E-54 | 1.0335075 | 0.573 | 0.297 | 8.90E-50 | cDC.inflam. | Lrpap1 |
| Gclm | 1.04E-53 | 1.68679187 | 0.364 | 0.134 | 2.01E-49 | cDC.inflam. | Gclm |
| Rftn11 | 1.88E-53 | 0.31371191 | 0.748 | 0.396 | 3.64E-49 | cDC.inflam. | Rftn1 |
| Ifih11 | 9.59E-53 | 0.77620763 | 0.432 | 0.242 | 1.85E-48 | cDC.inflam. | Ifih1 |
| Slc39a12 | 3.61E-52 | 0.46552409 | 0.85 | 0.65 | 6.97E-48 | cDC.inflam. | Slc39a1 |
| Nfkbie2 | 3.64E-52 | 0.49357915 | 0.786 | 0.557 | 7.04E-48 | cDC.inflam. | Nfkbie |
| Gabarapl11 | 4.66E-52 | 0.57916703 | 0.684 | 0.447 | 9.01E-48 | cDC.inflam. | Gabarapl1 |
| Acp2 | 8.49E-52 | 1.61888383 | 0.432 | 0.125 | 1.64E-47 | cDC.inflam. | Acp2 |
| Jag1 | 8.66E-52 | 1.52216464 | 0.296 | 0.068 | 1.67E-47 | cDC.inflam. | Jag1 |
| Lst11 | 1.30E-51 | 1.20092101 | 0.5 | 0.241 | 2.51E-47 | cDC.inflam. | Lst1 |
| Bri31 | 1.35E-51 | 0.49268257 | 0.796 | 0.548 | 2.61E-47 | cDC.inflam. | Bri3 |
| Egr11 | 1.35E-51 | 0.70106905 | 0.495 | 0.375 | 2.61E-47 | cDC.inflam. | Egr1 |
| Ampd21 | 1.99E-51 | 1.11797145 | 0.563 | 0.238 | 3.85E-47 | cDC.inflam. | Ampd2 |
| Igfbp4 | 3.22E-51 | 0.75241298 | 0.359 | 0.195 | 6.22E-47 | cDC.inflam. | Igfbp4 |
| Bag3 | 6.38E-51 | 0.91584794 | 0.471 | 0.208 | 1.23E-46 | cDC.inflam. | Bag3 |
| Gabarap | 8.42E-51 | 0.34716496 | 0.835 | 0.664 | 1.63E-46 | cDC.inflam. | Gabarap |
| Cyp4f18 | 1.10E-50 | 1.58972157 | 0.34 | 0.118 | 2.13E-46 | cDC.inflam. | Cyp4f18 |
| Slc6a61 | 1.50E-50 | 0.41360353 | 0.825 | 0.619 | 2.90E-46 | cDC.inflam. | Slc6a6 |
| Palld2 | 2.32E-50 | 0.54573436 | 0.461 | 0.24 | 4.48E-46 | cDC.inflam. | Palld |
| Flcn | 3.01E-50 | 1.20426856 | 0.49 | 0.212 | 5.82E-46 | cDC.inflam. | Flcn |
| Pim32 | 3.45E-50 | 0.77658737 | 0.646 | 0.393 | 6.68E-46 | cDC.inflam. | Pim3 |
| Glipr11 | 4.75E-50 | 1.0329421 | 0.485 | 0.221 | 9.18E-46 | cDC.inflam. | Glipr1 |
| Aplp21 | 5.09E-50 | 0.47450268 | 0.752 | 0.464 | 9.83E-46 | cDC.inflam. | Aplp2 |
| Ext1 | 5.16E-50 | 0.74175173 | 0.709 | 0.411 | 9.98E-46 | cDC.inflam. | Ext1 |
| Cited21 | 5.81E-50 | 0.83579385 | 0.451 | 0.24 | 1.12E-45 | cDC.inflam. | Cited2 |
| Cbx41 | 6.38E-50 | 1.11618218 | 0.447 | 0.231 | 1.23E-45 | cDC.inflam. | Cbx4 |
| Mapkapk21 | 1.08E-49 | 0.40721144 | 0.879 | 0.687 | 2.09E-45 | cDC.inflam. | Mapkapk2 |
| St6galnac61 | 1.19E-49 | 1.07073359 | 0.519 | 0.195 | 2.29E-45 | cDC.inflam. | St6galnac6 |
| Kdm6b2 | 2.06E-49 | 0.27840996 | 0.947 | 0.774 | 3.99E-45 | cDC.inflam. | Kdm6b |
| Pvr | 2.12E-49 | 0.77084809 | 0.573 | 0.362 | 4.10E-45 | cDC.inflam. | Pvr |
| Il10rb | 2.23E-49 | 0.83483048 | 0.694 | 0.408 | 4.31E-45 | cDC.inflam. | Il10rb |
| Mitf | 4.13E-49 | 1.90546252 | 0.32 | 0.079 | 7.98E-45 | cDC.inflam. | Mitf |
| Aif11 | 6.71E-49 | 0.41814047 | 0.718 | 0.518 | 1.30E-44 | cDC.inflam. | Aif1 |
| Lfng1 | 1.47E-48 | 0.53831008 | 0.675 | 0.43 | 2.85E-44 | cDC.inflam. | Lfng |
| Atxn7l11 | 2.43E-48 | 0.42606323 | 0.762 | 0.497 | 4.70E-44 | cDC.inflam. | Atxn7l1 |
| Sema6d | 2.63E-48 | 1.20437703 | 0.311 | 0.086 | 5.08E-44 | cDC.inflam. | Sema6d |
| Hexim11 | 4.85E-48 | 0.75684644 | 0.68 | 0.414 | 9.38E-44 | cDC.inflam. | Hexim1 |
| Cdk171 | 6.05E-48 | 0.97985568 | 0.553 | 0.23 | 1.17E-43 | cDC.inflam. | Cdk17 |
| Fosl22 | 6.93E-48 | 0.26699116 | 0.893 | 0.646 | 1.34E-43 | cDC.inflam. | Fosl2 |
| Dpep2 | 1.69E-47 | 1.95868774 | 0.277 | 0.071 | 3.28E-43 | cDC.inflam. | Dpep2 |
| Myof | 1.95E-47 | 0.56864184 | 0.549 | 0.281 | 3.78E-43 | cDC.inflam. | Myof |
| Atf32 | 3.19E-47 | 0.40141577 | 0.723 | 0.474 | 6.17E-43 | cDC.inflam. | Atf3 |
| Car13 | 7.30E-47 | 1.54639713 | 0.345 | 0.076 | 1.41E-42 | cDC.inflam. | Car13 |
| Atp11a1 | 9.82E-47 | 0.33378491 | 0.665 | 0.324 | 1.90E-42 | cDC.inflam. | Atp11a |
| Mdm21 | 3.00E-46 | 0.68044602 | 0.684 | 0.374 | 5.81E-42 | cDC.inflam. | Mdm2 |
| Egr2 | 4.01E-46 | 1.01505242 | 0.388 | 0.16 | 7.75E-42 | cDC.inflam. | Egr2 |
| Nlrp32 | 6.91E-46 | 0.73463142 | 0.757 | 0.433 | 1.34E-41 | cDC.inflam. | Nlrp3 |
| Gsap1 | 1.45E-45 | 0.72836505 | 0.675 | 0.377 | 2.80E-41 | cDC.inflam. | Gsap |
| Insig11 | 1.54E-45 | 0.38210747 | 0.704 | 0.493 | 2.97E-41 | cDC.inflam. | Insig1 |
| Nfe2l11 | 3.75E-45 | 0.41372996 | 0.743 | 0.495 | 7.26E-41 | cDC.inflam. | Nfe2l1 |
| St3gal31 | 9.16E-45 | 0.98118959 | 0.515 | 0.22 | 1.77E-40 | cDC.inflam. | St3gal3 |
| Zswim42 | 9.39E-45 | 0.94279553 | 0.602 | 0.28 | 1.82E-40 | cDC.inflam. | Zswim4 |
| Rassf31 | 1.23E-44 | 0.51354902 | 0.641 | 0.328 | 2.38E-40 | cDC.inflam. | Rassf3 |
| Htra21 | 1.72E-44 | 0.64983359 | 0.558 | 0.275 | 3.32E-40 | cDC.inflam. | Htra2 |
| Rras21 | 2.06E-44 | 0.61464557 | 0.563 | 0.305 | 3.98E-40 | cDC.inflam. | Rras2 |
| Ddx3x | 3.02E-44 | 0.32679431 | 0.879 | 0.692 | 5.83E-40 | cDC.inflam. | Ddx3x |
| Tmem120b1 | 3.66E-44 | 0.97106128 | 0.461 | 0.193 | 7.08E-40 | cDC.inflam. | Tmem120b |
| Flvcr11 | 3.84E-44 | 0.49542309 | 0.485 | 0.243 | 7.42E-40 | cDC.inflam. | Flvcr1 |
| Itgb5 | 1.61E-43 | 1.65240464 | 0.306 | 0.074 | 3.11E-39 | cDC.inflam. | Itgb5 |
| Sqor | 1.77E-43 | 2.26186927 | 0.277 | 0.052 | 3.42E-39 | cDC.inflam. | Sqor |
| Mthfs2 | 2.81E-43 | 1.04131024 | 0.587 | 0.292 | 5.42E-39 | cDC.inflam. | Mthfs |
| Tent5a2 | 5.00E-43 | 0.87261566 | 0.461 | 0.266 | 9.67E-39 | cDC.inflam. | Tent5a |
| Fam117b1 | 8.92E-43 | 0.50826378 | 0.772 | 0.582 | 1.72E-38 | cDC.inflam. | Fam117b |
| Ttc39b1 | 1.24E-42 | 0.81917628 | 0.485 | 0.196 | 2.40E-38 | cDC.inflam. | Ttc39b |
| Ndel12 | 1.40E-42 | 0.42179987 | 0.782 | 0.567 | 2.72E-38 | cDC.inflam. | Ndel1 |
| Ell21 | 1.44E-42 | 0.58848713 | 0.699 | 0.396 | 2.77E-38 | cDC.inflam. | Ell2 |
| Col27a11 | 2.20E-42 | 1.0895193 | 0.291 | 0.102 | 4.26E-38 | cDC.inflam. | Col27a1 |
| Itgax1 | 2.41E-42 | 0.34127105 | 0.83 | 0.613 | 4.65E-38 | cDC.inflam. | Itgax |
| Fbrsl11 | 2.59E-42 | 0.43875984 | 0.631 | 0.333 | 5.00E-38 | cDC.inflam. | Fbrsl1 |
| Cpeb4 | 2.66E-42 | 0.70183823 | 0.568 | 0.309 | 5.15E-38 | cDC.inflam. | Cpeb4 |
| Rab121 | 2.79E-42 | 0.79143619 | 0.505 | 0.235 | 5.40E-38 | cDC.inflam. | Rab12 |
| Tbc1d41 | 4.45E-42 | 0.68594354 | 0.476 | 0.194 | 8.59E-38 | cDC.inflam. | Tbc1d4 |
| Ltc4s1 | 1.72E-41 | 1.13124821 | 0.277 | 0.155 | 3.33E-37 | cDC.inflam. | Ltc4s |
| Sdcbp2 | 4.29E-41 | 0.26830895 | 0.869 | 0.695 | 8.29E-37 | cDC.inflam. | Sdcbp |
| Idh1 | 4.52E-41 | 0.99739727 | 0.422 | 0.16 | 8.74E-37 | cDC.inflam. | Idh1 |
| Gclc | 4.57E-41 | 1.23556605 | 0.369 | 0.15 | 8.84E-37 | cDC.inflam. | Gclc |
| Uhrf2 | 6.15E-41 | 0.96639923 | 0.442 | 0.262 | 1.19E-36 | cDC.inflam. | Uhrf2 |
| Lilrb4a2 | 8.12E-41 | 0.45750161 | 0.85 | 0.56 | 1.57E-36 | cDC.inflam. | Lilrb4a |
| Man1a1 | 1.29E-40 | 0.49940548 | 0.723 | 0.481 | 2.49E-36 | cDC.inflam. | Man1a |
| Glul | 1.68E-40 | 1.0861746 | 0.403 | 0.177 | 3.24E-36 | cDC.inflam. | Glul |
| Elk3 | 1.81E-40 | 0.67191549 | 0.631 | 0.338 | 3.49E-36 | cDC.inflam. | Elk3 |
| Atp6v1e1 | 2.35E-40 | 0.71472533 | 0.709 | 0.475 | 4.55E-36 | cDC.inflam. | Atp6v1e1 |
| Stat5a1 | 2.71E-40 | 0.54416355 | 0.626 | 0.411 | 5.23E-36 | cDC.inflam. | Stat5a |
| Ccl61 | 3.77E-40 | 1.11975896 | 0.354 | 0.173 | 7.29E-36 | cDC.inflam. | Ccl6 |
| Scarb21 | 6.36E-40 | 0.57501833 | 0.583 | 0.364 | 1.23E-35 | cDC.inflam. | Scarb2 |
| Sec61b1 | 8.04E-40 | 0.59002206 | 0.646 | 0.464 | 1.55E-35 | cDC.inflam. | Sec61b |
| Tob22 | 1.49E-39 | 0.48325041 | 0.704 | 0.505 | 2.88E-35 | cDC.inflam. | Tob2 |
| Il2rg1 | 1.94E-39 | 0.32272186 | 0.82 | 0.569 | 3.75E-35 | cDC.inflam. | Il2rg |
| Slc38a6 | 7.13E-39 | 1.33734203 | 0.316 | 0.079 | 1.38E-34 | cDC.inflam. | Slc38a6 |
| Ltbp3 | 1.00E-38 | 1.43696905 | 0.35 | 0.104 | 1.94E-34 | cDC.inflam. | Ltbp3 |
| Plin21 | 2.03E-38 | 0.6032601 | 0.709 | 0.414 | 3.93E-34 | cDC.inflam. | Plin2 |
| Abtb21 | 2.14E-38 | 0.89290318 | 0.471 | 0.183 | 4.14E-34 | cDC.inflam. | Abtb2 |
| Mpc11 | 2.28E-38 | 0.33078654 | 0.738 | 0.558 | 4.41E-34 | cDC.inflam. | Mpc1 |
| Klrb1b2 | 8.11E-38 | 0.51579636 | 0.675 | 0.449 | 1.57E-33 | cDC.inflam. | Klrb1b |

| | | | | | | | |
|---|---|---|---|---|---|---|---|
| Mapkapk31 | 9.82E-38 | 0.61062927 | 0.641 | 0.393 | 1.90E-33 | cDC.inflam. | Mapkapk3 |
| Atp6v0a11 | 2.16E-37 | 0.51800357 | 0.558 | 0.309 | 4.17E-33 | cDC.inflam. | Atp6v0a1 |
| Slc2a12 | 3.65E-37 | 0.69007341 | 0.709 | 0.479 | 7.05E-33 | cDC.inflam. | Slc2a1 |
| S100a12 | 1.17E-36 | 0.94878574 | 0.461 | 0.211 | 2.27E-32 | cDC.inflam. | S100a1 |
| Irak21 | 1.47E-36 | 0.86375224 | 0.699 | 0.361 | 2.83E-32 | cDC.inflam. | Irak2 |
| Dtnbp1 | 2.53E-36 | 0.6449461 | 0.534 | 0.298 | 4.90E-32 | cDC.inflam. | Dtnbp1 |
| Sema4d1 | 3.01E-36 | 0.34650401 | 0.757 | 0.562 | 5.82E-32 | cDC.inflam. | Sema4d |
| Arap11 | 3.15E-36 | 0.56591816 | 0.563 | 0.272 | 6.10E-32 | cDC.inflam. | Arap1 |
| Aco1 | 4.13E-36 | 0.53240862 | 0.485 | 0.281 | 7.99E-32 | cDC.inflam. | Aco1 |
| Ric11 | 6.12E-36 | 0.39302878 | 0.612 | 0.407 | 1.18E-31 | cDC.inflam. | Ric1 |
| Rasgef1b2 | 6.25E-36 | 0.5702312 | 0.505 | 0.354 | 1.21E-31 | cDC.inflam. | Rasgef1b |
| Prdm1 | 6.62E-36 | 1.62232107 | 0.286 | 0.11 | 1.28E-31 | cDC.inflam. | Prdm1 |
| Aldh3b1 | 6.90E-36 | 1.42159412 | 0.296 | 0.103 | 1.33E-31 | cDC.inflam. | Aldh3b1 |
| Sphk1 | 7.71E-36 | 1.22936205 | 0.35 | 0.108 | 1.49E-31 | cDC.inflam. | Sphk1 |
| Asph | 7.85E-36 | 0.68150774 | 0.451 | 0.226 | 1.52E-31 | cDC.inflam. | Asph |
| Adam101 | 9.63E-36 | 0.6230476 | 0.524 | 0.334 | 1.86E-31 | cDC.inflam. | Adam10 |
| Wsb11 | 1.10E-35 | 0.49979345 | 0.786 | 0.554 | 2.13E-31 | cDC.inflam. | Wsb1 |
| Nectin1 | 1.41E-35 | 0.44783676 | 0.675 | 0.476 | 2.73E-31 | cDC.inflam. | Nectin1 |
| Abcc11 | 1.81E-35 | 0.75450012 | 0.524 | 0.275 | 3.51E-31 | cDC.inflam. | Abcc1 |
| Ttc39c | 1.93E-35 | 0.68892316 | 0.364 | 0.132 | 3.72E-31 | cDC.inflam. | Ttc39c |
| Nck1 | 4.23E-35 | 1.19963524 | 0.35 | 0.131 | 8.18E-31 | cDC.inflam. | Nck1 |
| Pcyt1a1 | 6.90E-35 | 0.38662088 | 0.544 | 0.372 | 1.33E-30 | cDC.inflam. | Pcyt1a |
| Igsf6 | 8.01E-35 | 1.38137531 | 0.335 | 0.145 | 1.55E-30 | cDC.inflam. | Igsf6 |
| Mthfd1 | 9.05E-35 | 1.24506234 | 0.461 | 0.215 | 1.75E-30 | cDC.inflam. | Mthfd |
| Itprid21 | 9.27E-35 | 0.37616224 | 0.782 | 0.541 | 1.79E-30 | cDC.inflam. | Itprid2 |
| Dapk11 | 1.05E-34 | 0.58887265 | 0.67 | 0.484 | 2.03E-30 | cDC.inflam. | Dapk1 |
| Plekhb21 | 1.43E-34 | 0.28816612 | 0.743 | 0.534 | 2.77E-30 | cDC.inflam. | Plekhb2 |
| Ifi271 | 2.56E-34 | 1.02625297 | 0.408 | 0.178 | 4.96E-30 | cDC.inflam. | Ifi27 |
| Card191 | 3.32E-34 | 0.65613049 | 0.612 | 0.369 | 6.42E-30 | cDC.inflam. | Card19 |
| Pik3r61 | 6.73E-34 | 0.7063992 | 0.461 | 0.236 | 1.30E-29 | cDC.inflam. | Pik3r6 |
| G6pdx2 | 7.28E-34 | 0.96418297 | 0.456 | 0.236 | 1.41E-29 | cDC.inflam. | G6pdx |
| Sppl2a2 | 7.49E-34 | 0.45102836 | 0.748 | 0.509 | 1.45E-29 | cDC.inflam. | Sppl2a |
| Il6st1 | 8.01E-34 | 0.35036267 | 0.612 | 0.387 | 1.55E-29 | cDC.inflam. | Il6st |
| Mylip | 8.93E-34 | 1.13327669 | 0.35 | 0.158 | 1.73E-29 | cDC.inflam. | Mylip |
| Sulf22 | 1.08E-33 | 0.54543774 | 0.646 | 0.399 | 2.09E-29 | cDC.inflam. | Sulf2 |
| Gls | 1.36E-33 | 0.45619964 | 0.757 | 0.58 | 2.63E-29 | cDC.inflam. | Gls |
| Fhod11 | 1.62E-33 | 0.74229862 | 0.481 | 0.254 | 3.14E-29 | cDC.inflam. | Fhod1 |
| Ankrd33b1 | 1.89E-33 | 0.57996512 | 0.393 | 0.138 | 3.65E-29 | cDC.inflam. | Ankrd33b |
| Cflar1 | 2.68E-33 | 0.57713676 | 0.612 | 0.362 | 5.19E-29 | cDC.inflam. | Cflar |
| Map1lc3b | 4.21E-33 | 0.41521318 | 0.723 | 0.508 | 8.14E-29 | cDC.inflam. | Map1lc3b |
| Pnpla81 | 5.56E-33 | 0.4527198 | 0.485 | 0.227 | 1.08E-28 | cDC.inflam. | Pnpla8 |
| Srgap2 | 6.90E-33 | 1.01406412 | 0.447 | 0.171 | 1.33E-28 | cDC.inflam. | Srgap2 |
| Tnip32 | 8.02E-33 | 0.36893005 | 0.583 | 0.32 | 1.55E-28 | cDC.inflam. | Tnip3 |
| Klf62 | 1.06E-32 | 0.30532577 | 0.675 | 0.486 | 2.05E-28 | cDC.inflam. | Klf6 |
| Mfsd11 | 1.08E-32 | 0.58087965 | 0.553 | 0.365 | 2.08E-28 | cDC.inflam. | Mfsd1 |
| Tnfrsf11a1 | 1.83E-32 | 0.38678149 | 0.602 | 0.346 | 3.54E-28 | cDC.inflam. | Tnfrsf11a |
| Tnfrsf12a1 | 2.93E-32 | 0.59307088 | 0.515 | 0.25 | 5.66E-28 | cDC.inflam. | Tnfrsf12a |
| Tnfsf12 | 3.80E-32 | 1.28374422 | 0.335 | 0.108 | 7.34E-28 | cDC.inflam. | Tnfsf12 |
| Baiap22 | 4.92E-32 | 0.98608256 | 0.485 | 0.255 | 9.51E-28 | cDC.inflam. | Baiap2 |
| Cd811 | 9.24E-32 | 0.46801397 | 0.35 | 0.179 | 1.79E-27 | cDC.inflam. | Cd81 |
| Slc25a37 | 9.87E-32 | 1.44139412 | 0.296 | 0.094 | 1.91E-27 | cDC.inflam. | Slc25a37 |
| Slc26a2 | 1.02E-31 | 1.12567027 | 0.393 | 0.176 | 1.97E-27 | cDC.inflam. | Slc26a2 |
| Inf21 | 2.35E-31 | 0.38858773 | 0.471 | 0.191 | 4.54E-27 | cDC.inflam. | Inf2 |
| Tcn2 | 5.57E-31 | 0.29156746 | 0.757 | 0.583 | 1.08E-26 | cDC.inflam. | Tcn2 |
| Dnmt3a1 | 8.65E-31 | 0.65017476 | 0.553 | 0.339 | 1.67E-26 | cDC.inflam. | Dnmt3a |
| Trem12 | 1.34E-30 | 1.32518204 | 0.34 | 0.178 | 2.59E-26 | cDC.inflam. | Trem1 |
| Tpi11 | 2.64E-30 | 0.41353426 | 0.67 | 0.471 | 5.10E-26 | cDC.inflam. | Tpi1 |
| Ddx46 | 2.68E-30 | 0.58526513 | 0.583 | 0.392 | 5.17E-26 | cDC.inflam. | Ddx46 |
| Lgmn1 | 3.18E-30 | 0.62656562 | 0.558 | 0.297 | 6.16E-26 | cDC.inflam. | Lgmn |
| Iah1 | 3.25E-30 | 0.6526931 | 0.515 | 0.287 | 6.29E-26 | cDC.inflam. | Iah1 |
| Adgre52 | 4.00E-30 | 0.53334088 | 0.728 | 0.485 | 7.73E-26 | cDC.inflam. | Adgre5 |
| Ripk3 | 6.27E-30 | 1.289424 | 0.311 | 0.104 | 1.21E-25 | cDC.inflam. | Ripk3 |
| Apbb1ip1 | 7.04E-30 | 0.36064336 | 0.874 | 0.657 | 1.36E-25 | cDC.inflam. | Apbb1ip |
| Slc5a3 | 7.55E-30 | 0.81513125 | 0.32 | 0.166 | 1.46E-25 | cDC.inflam. | Slc5a3 |
| Vopp11 | 8.89E-30 | 0.27636333 | 0.607 | 0.367 | 1.72E-25 | cDC.inflam. | Vopp1 |
| Isg151 | 1.65E-29 | 0.72695283 | 0.481 | 0.261 | 3.20E-25 | cDC.inflam. | Isg15 |
| Adam91 | 3.11E-29 | 0.30616402 | 0.636 | 0.43 | 6.01E-25 | cDC.inflam. | Adam9 |
| Arrdc41 | 3.14E-29 | 0.41468982 | 0.617 | 0.392 | 6.07E-25 | cDC.inflam. | Arrdc4 |
| Gfpt11 | 4.93E-29 | 0.34721978 | 0.621 | 0.426 | 9.52E-25 | cDC.inflam. | Gfpt1 |
| Pdlim5 | 7.37E-29 | 0.35396609 | 0.689 | 0.456 | 1.42E-24 | cDC.inflam. | Pdlim5 |
| Slc41a11 | 1.20E-28 | 0.65825619 | 0.524 | 0.318 | 2.32E-24 | cDC.inflam. | Slc41a1 |
| Traf5 | 1.37E-28 | 0.77676535 | 0.393 | 0.222 | 2.65E-24 | cDC.inflam. | Traf5 |
| Pkm1 | 1.47E-28 | 0.27168057 | 0.908 | 0.712 | 2.85E-24 | cDC.inflam. | Pkm |
| Fcgr32 | 1.59E-28 | 0.35908333 | 0.592 | 0.374 | 3.07E-24 | cDC.inflam. | Fcgr3 |
| C1qtnf6 | 1.99E-28 | 1.02667733 | 0.35 | 0.116 | 3.84E-24 | cDC.inflam. | C1qtnf6 |
| Atp2b11 | 2.81E-28 | 0.39406245 | 0.675 | 0.477 | 5.43E-24 | cDC.inflam. | Atp2b1 |
| Rapgef21 | 3.58E-28 | 0.68614439 | 0.471 | 0.263 | 6.92E-24 | cDC.inflam. | Rapgef2 |
| Mif1 | 4.66E-28 | 0.5519991 | 0.65 | 0.428 | 9.01E-24 | cDC.inflam. | Mif |
| B4galt52 | 6.05E-28 | 0.40454471 | 0.762 | 0.536 | 1.17E-23 | cDC.inflam. | B4galt5 |
| Tob11 | 6.96E-28 | 0.70860952 | 0.515 | 0.299 | 1.35E-23 | cDC.inflam. | Tob1 |
| Polb1 | 7.44E-28 | 0.47168898 | 0.597 | 0.34 | 1.44E-23 | cDC.inflam. | Polb |
| Tnfsf91 | 1.54E-27 | 0.32533089 | 0.413 | 0.305 | 2.98E-23 | cDC.inflam. | Tnfsf9 |
| Adap2 | 1.66E-27 | 0.9783763 | 0.549 | 0.25 | 3.21E-23 | cDC.inflam. | Adap2 |
| Fut4 | 1.68E-27 | 0.89345138 | 0.35 | 0.138 | 3.25E-23 | cDC.inflam. | Fut4 |
| Pot1b | 2.50E-27 | 0.98727016 | 0.252 | 0.102 | 4.82E-23 | cDC.inflam. | Pot1b |
| Cln5 | 3.26E-27 | 0.51867645 | 0.456 | 0.249 | 6.30E-23 | cDC.inflam. | Cln5 |
| Renbp | 4.18E-27 | 1.03671413 | 0.451 | 0.211 | 8.09E-23 | cDC.inflam. | Renbp |
| Gpr68 | 4.87E-27 | 0.76381161 | 0.422 | 0.208 | 9.41E-23 | cDC.inflam. | Gpr68 |
| Tpra1 | 5.78E-27 | 0.98662908 | 0.306 | 0.115 | 1.12E-22 | cDC.inflam. | Tpra1 |
| Pygl1 | 6.85E-27 | 0.46317494 | 0.68 | 0.366 | 1.32E-22 | cDC.inflam. | Pygl |
| Appl11 | 8.47E-27 | 0.45045315 | 0.587 | 0.409 | 1.64E-22 | cDC.inflam. | Appl1 |
| Zfp1311 | 8.77E-27 | 0.74487858 | 0.549 | 0.293 | 1.69E-22 | cDC.inflam. | Zfp131 |
| Idi1 | 1.18E-26 | 1.0159801 | 0.388 | 0.184 | 2.28E-22 | cDC.inflam. | Idi1 |
| Slc20a11 | 1.43E-26 | 0.59708231 | 0.403 | 0.284 | 2.77E-22 | cDC.inflam. | Slc20a1 |
| Pgam12 | 1.76E-26 | 0.42621505 | 0.66 | 0.505 | 3.41E-22 | cDC.inflam. | Pgam1 |
| Neu1 | 2.77E-26 | 0.97223816 | 0.466 | 0.229 | 5.36E-22 | cDC.inflam. | Neu1 |
| Tbc1d13 | 3.51E-26 | 0.65964798 | 0.505 | 0.292 | 6.79E-22 | cDC.inflam. | Tbc1d13 |
| Atp6v1a1 | 5.64E-26 | 0.41485911 | 0.718 | 0.496 | 1.09E-21 | cDC.inflam. | Atp6v1a |
| Ctsd | 7.43E-26 | 1.03915609 | 0.451 | 0.181 | 1.44E-21 | cDC.inflam. | Ctsd |
| Traf31 | 1.23E-25 | 0.47793535 | 0.49 | 0.253 | 2.39E-21 | cDC.inflam. | Traf3 |
| Ero1l1 | 1.29E-25 | 0.85788649 | 0.534 | 0.326 | 2.50E-21 | cDC.inflam. | Ero1l |
| Itgav1 | 1.38E-25 | 0.66995185 | 0.461 | 0.261 | 2.67E-21 | cDC.inflam. | Itgav |
| Tmbim11 | 1.62E-25 | 0.38656745 | 0.631 | 0.41 | 3.12E-21 | cDC.inflam. | Tmbim1 |
| Fyb | 1.70E-25 | 0.30380993 | 0.471 | 0.26 | 3.28E-21 | cDC.inflam. | Fyb |
| Rbpms | 2.38E-25 | 0.67152966 | 0.359 | 0.178 | 4.60E-21 | cDC.inflam. | Rbpms |
| Alg9 | 3.05E-25 | 0.81456707 | 0.383 | 0.177 | 5.89E-21 | cDC.inflam. | Alg9 |
| Impact1 | 3.22E-25 | 0.65130271 | 0.524 | 0.267 | 6.22E-21 | cDC.inflam. | Impact |
| Pgf2 | 3.49E-25 | 0.49174961 | 0.32 | 0.207 | 6.75E-21 | cDC.inflam. | Pgf |
| Mgat4a2 | 3.84E-25 | 0.55610203 | 0.515 | 0.28 | 7.42E-21 | cDC.inflam. | Mgat4a |
| Ago3 | 4.42E-25 | 0.85687933 | 0.369 | 0.159 | 8.54E-21 | cDC.inflam. | Ago3 |
| Ppt21 | 4.81E-25 | 1.01706922 | 0.306 | 0.128 | 9.30E-21 | cDC.inflam. | Ppt2 |
| Naip2 | 6.07E-25 | 0.95808545 | 0.403 | 0.18 | 1.17E-20 | cDC.inflam. | Naip2 |
| Sbds | 7.00E-25 | 0.26675049 | 0.621 | 0.386 | 1.35E-20 | cDC.inflam. | Sbds |
| P2rx5 | 1.78E-24 | 1.3128162 | 0.34 | 0.115 | 3.45E-20 | cDC.inflam. | P2rx5 |
| Cndp2 | 2.31E-24 | 0.84015145 | 0.456 | 0.278 | 4.46E-20 | cDC.inflam. | Cndp2 |
| Arhgap311 | 2.89E-24 | 0.25984445 | 0.612 | 0.377 | 5.59E-20 | cDC.inflam. | Arhgap31 |
| Mapk31 | 3.76E-24 | 0.34044818 | 0.66 | 0.422 | 7.27E-20 | cDC.inflam. | Mapk3 |
| Fas1 | 5.02E-24 | 0.77830379 | 0.33 | 0.106 | 9.71E-20 | cDC.inflam. | Fas |
| Cd38 | 6.10E-24 | 0.97456067 | 0.306 | 0.14 | 1.18E-19 | cDC.inflam. | Cd38 |
| Slc37a2 | 6.26E-24 | 1.17287336 | 0.257 | 0.1 | 1.21E-19 | cDC.inflam. | Slc37a2 |
| Gpr1831 | 6.84E-24 | 0.33331738 | 0.466 | 0.343 | 1.32E-19 | cDC.inflam. | Gpr183 |
| Pstpip2 | 9.68E-24 | 1.18128693 | 0.282 | 0.144 | 1.87E-19 | cDC.inflam. | Pstpip2 |
| Rreb11 | 9.83E-24 | 0.78171627 | 0.485 | 0.254 | 1.90E-19 | cDC.inflam. | Rreb1 |
| Birc21 | 1.22E-23 | 0.40449655 | 0.49 | 0.249 | 2.36E-19 | cDC.inflam. | Birc2 |
| Acss11 | 1.25E-23 | 1.01812609 | 0.33 | 0.165 | 2.42E-19 | cDC.inflam. | Acss1 |
| Rasa41 | 1.28E-23 | 0.39256037 | 0.583 | 0.334 | 2.48E-19 | cDC.inflam. | Rasa4 |
| Specc1l | 1.30E-23 | 0.44414179 | 0.495 | 0.341 | 2.52E-19 | cDC.inflam. | Specc1l |
| Eea1 | 2.04E-23 | 0.67499501 | 0.417 | 0.194 | 3.94E-19 | cDC.inflam. | Eea1 |
| Cyth3 | 2.42E-23 | 0.73855918 | 0.432 | 0.196 | 4.69E-19 | cDC.inflam. | Cyth3 |
| Sec14l1 | 2.59E-23 | 0.78568559 | 0.413 | 0.203 | 5.01E-19 | cDC.inflam. | Sec14l1 |
| Lacc11 | 2.60E-23 | 0.77887622 | 0.461 | 0.221 | 5.02E-19 | cDC.inflam. | Lacc1 |
| Papss21 | 4.29E-23 | 0.51843926 | 0.316 | 0.107 | 8.29E-19 | cDC.inflam. | Papss2 |
| Tmem106a | 4.62E-23 | 0.4840647 | 0.573 | 0.37 | 8.92E-19 | cDC.inflam. | Tmem106a |
| Dot1l1 | 5.70E-23 | 0.50106285 | 0.66 | 0.394 | 1.10E-18 | cDC.inflam. | Dot1l |
| Plk21 | 5.93E-23 | 0.57422819 | 0.374 | 0.198 | 1.15E-18 | cDC.inflam. | Plk2 |
| Tmem63b1 | 8.12E-23 | 0.98351667 | 0.403 | 0.152 | 1.57E-18 | cDC.inflam. | Tmem63b |
| Rab321 | 8.56E-23 | 0.29322779 | 0.578 | 0.359 | 1.65E-18 | cDC.inflam. | Rab32 |
| Jmy1 | 8.74E-23 | 0.54692003 | 0.388 | 0.183 | 1.69E-18 | cDC.inflam. | Jmy |
| Homer12 | 1.01E-22 | 0.78783355 | 0.539 | 0.317 | 1.95E-18 | cDC.inflam. | Homer1 |
| mt-Nd41 | 1.02E-22 | 0.35139986 | 0.481 | 0.284 | 1.96E-18 | cDC.inflam. | mt-Nd4 |
| Ecm1 | 2.10E-22 | 0.55236738 | 0.35 | 0.174 | 4.05E-18 | cDC.inflam. | Ecm1 |
| Lin541 | 3.52E-22 | 0.84375968 | 0.403 | 0.204 | 6.81E-18 | cDC.inflam. | Lin54 |
| Grk51 | 3.73E-22 | 0.72610387 | 0.35 | 0.141 | 7.22E-18 | cDC.inflam. | Grk5 |
| Nupr1 | 4.01E-22 | 0.76109542 | 0.267 | 0.099 | 7.74E-18 | cDC.inflam. | Nupr1 |
| Tmem651 | 4.15E-22 | 0.48698353 | 0.383 | 0.211 | 8.01E-18 | cDC.inflam. | Tmem65 |
| Mvd | 4.79E-22 | 0.76500055 | 0.296 | 0.139 | 9.26E-18 | cDC.inflam. | Mvd |
| Gpc11 | 4.89E-22 | 0.42478854 | 0.471 | 0.273 | 9.45E-18 | cDC.inflam. | Gpc1 |
| Phlpp11 | 5.44E-22 | 0.44531792 | 0.597 | 0.388 | 1.05E-17 | cDC.inflam. | Phlpp1 |
| Tuba1c1 | 5.74E-22 | 0.26231081 | 0.723 | 0.543 | 1.11E-17 | cDC.inflam. | Tuba1c |
| Abi31 | 6.04E-22 | 0.5013764 | 0.66 | 0.402 | 1.17E-17 | cDC.inflam. | Abi3 |
| Mfsd121 | 6.65E-22 | 0.66797251 | 0.432 | 0.272 | 1.28E-17 | cDC.inflam. | Mfsd12 |
| Gas2l1 | 7.04E-22 | 0.64279385 | 0.325 | 0.161 | 1.36E-17 | cDC.inflam. | Gas2l1 |
| Dynlt3 | 7.31E-22 | 0.55848807 | 0.456 | 0.218 | 1.41E-17 | cDC.inflam. | Dynlt3 |
| Rcbtb2 | 8.07E-22 | 1.24688931 | 0.272 | 0.125 | 1.56E-17 | cDC.inflam. | Rcbtb2 |
| Gstt3 | 1.08E-21 | 0.92172579 | 0.311 | 0.156 | 2.09E-17 | cDC.inflam. | Gstt3 |
| Trib12 | 1.10E-21 | 0.41209083 | 0.675 | 0.473 | 2.13E-17 | cDC.inflam. | Trib1 |
| Tmco3 | 1.18E-21 | 0.98698333 | 0.34 | 0.123 | 2.29E-17 | cDC.inflam. | Tmco3 |
| Itga51 | 1.37E-21 | 0.39445224 | 0.583 | 0.353 | 2.65E-17 | cDC.inflam. | Itga5 |
| Smcr81 | 1.86E-21 | 0.82655309 | 0.505 | 0.279 | 3.59E-17 | cDC.inflam. | Smcr8 |
| Hpse | 2.07E-21 | 0.88408179 | 0.374 | 0.122 | 3.99E-17 | cDC.inflam. | Hpse |
| Rnf1801 | 2.52E-21 | 0.28253582 | 0.515 | 0.318 | 4.87E-17 | cDC.inflam. | Rnf180 |
| Serinc51 | 3.72E-21 | 0.49768194 | 0.345 | 0.155 | 7.19E-17 | cDC.inflam. | Serinc5 |
| Treml41 | 4.59E-21 | 1.30615432 | 0.311 | 0.107 | 8.86E-17 | cDC.inflam. | Treml4 |
| Cnppd11 | 5.08E-21 | 0.37130033 | 0.612 | 0.428 | 9.82E-17 | cDC.inflam. | Cnppd1 |
| Sestd11 | 5.68E-21 | 0.80981994 | 0.471 | 0.229 | 1.10E-16 | cDC.inflam. | Sestd1 |
| Kif13b | 5.82E-21 | 1.22601574 | 0.306 | 0.111 | 1.13E-16 | cDC.inflam. | Kif13b |
| Ccni1 | 5.89E-21 | 0.27906098 | 0.743 | 0.512 | 1.14E-16 | cDC.inflam. | Ccni |
| Dok11 | 6.47E-21 | 0.41026262 | 0.515 | 0.293 | 1.25E-16 | cDC.inflam. | Dok1 |
| Tmem1651 | 7.35E-21 | 0.64074472 | 0.408 | 0.231 | 1.42E-16 | cDC.inflam. | Tmem165 |
| Pfkl1 | 9.60E-21 | 0.47740907 | 0.602 | 0.411 | 1.86E-16 | cDC.inflam. | Pfkl |
| Cers61 | 1.11E-20 | 0.2624484 | 0.752 | 0.529 | 2.14E-16 | cDC.inflam. | Cers6 |
| Crebl21 | 1.17E-20 | 0.73659152 | 0.35 | 0.15 | 2.26E-16 | cDC.inflam. | Crebl2 |
| Ddt1 | 1.48E-20 | 0.37597706 | 0.437 | 0.317 | 2.86E-16 | cDC.inflam. | Ddt |
| Qdpr | 1.54E-20 | 0.48599191 | 0.592 | 0.349 | 2.98E-16 | cDC.inflam. | Qdpr |
| Actr1a1 | 1.61E-20 | 0.28905071 | 0.748 | 0.517 | 3.11E-16 | cDC.inflam. | Actr1a |
| Ghitm | 1.88E-20 | 0.41828111 | 0.573 | 0.414 | 3.63E-16 | cDC.inflam. | Ghitm |
| Ptpn122 | 1.98E-20 | 0.43327811 | 0.646 | 0.454 | 3.83E-16 | cDC.inflam. | Ptpn12 |
| Eif4ebp1 | 2.25E-20 | 1.0661782 | 0.398 | 0.176 | 4.35E-16 | cDC.inflam. | Eif4ebp1 |
| Atp6v0d11 | 3.08E-20 | 0.27677777 | 0.689 | 0.51 | 5.95E-16 | cDC.inflam. | Atp6v0d1 |
| Rit1 | 3.40E-20 | 0.53555789 | 0.354 | 0.179 | 6.58E-16 | cDC.inflam. | Rit1 |
| Vdr2 | 3.61E-20 | 0.25553568 | 0.66 | 0.446 | 6.99E-16 | cDC.inflam. | Vdr |
| Echs1 | 4.36E-20 | 0.72273318 | 0.374 | 0.204 | 8.43E-16 | cDC.inflam. | Echs1 |
| Rab101 | 4.76E-20 | 0.38597858 | 0.578 | 0.392 | 9.20E-16 | cDC.inflam. | Rab10 |
| Rabgef11 | 4.87E-20 | 0.46401705 | 0.636 | 0.399 | 9.43E-16 | cDC.inflam. | Rabgef1 |
| Zfp5981 | 6.00E-20 | 0.32212403 | 0.563 | 0.363 | 1.16E-15 | cDC.inflam. | Zfp598 |
| Abhd12 | 6.15E-20 | 0.48690748 | 0.641 | 0.434 | 1.19E-15 | cDC.inflam. | Abhd12 |
| Casp4 | 6.27E-20 | 0.63389323 | 0.383 | 0.172 | 1.21E-15 | cDC.inflam. | Casp4 |
| Cmtm3 | 9.97E-20 | 0.54320505 | 0.485 | 0.274 | 1.93E-15 | cDC.inflam. | Cmtm3 |
| Tubb61 | 1.11E-19 | 0.4353844 | 0.612 | 0.41 | 2.14E-15 | cDC.inflam. | Tubb6 |
| Cisd2 | 1.13E-19 | 0.31769263 | 0.539 | 0.37 | 2.19E-15 | cDC.inflam. | Cisd2 |
| Sh3kbp1 | 1.17E-19 | 0.81841975 | 0.398 | 0.207 | 2.27E-15 | cDC.inflam. | Sh3kbp1 |
| DEPRECATED-I | 1.18E-19 | 1.16566374 | 0.33 | 0.165 | 2.29E-15 | cDC.inflam. | DEPRECATED-ENSMUSG00000039782 |
| Tgoln11 | 1.44E-19 | 0.53741393 | 0.602 | 0.405 | 2.79E-15 | cDC.inflam. | Tgoln1 |
| Arl5b2 | 3.44E-19 | 0.51328259 | 0.49 | 0.329 | 6.65E-15 | cDC.inflam. | Arl5b |
| Egln3 | 3.82E-19 | 1.06155005 | 0.262 | 0.126 | 7.39E-15 | cDC.inflam. | Egln3 |
| St7 | 5.44E-19 | 0.81619915 | 0.272 | 0.149 | 1.05E-14 | cDC.inflam. | St7 |
| Crlf3 | 6.00E-19 | 0.46820652 | 0.665 | 0.465 | 1.16E-14 | cDC.inflam. | Crlf3 |
| Gramd1a1 | 6.97E-19 | 0.30049989 | 0.573 | 0.37 | 1.35E-14 | cDC.inflam. | Gramd1a |
| Tlnrd11 | 8.49E-19 | 0.53416839 | 0.471 | 0.282 | 1.64E-14 | cDC.inflam. | Tlnrd1 |
| Cpeb21 | 1.77E-18 | 0.79871452 | 0.383 | 0.213 | 3.42E-14 | cDC.inflam. | Cpeb2 |
| Rnf19a | 2.06E-18 | 0.72984367 | 0.33 | 0.165 | 3.98E-14 | cDC.inflam. | Rnf19a |
| Snx14 | 2.25E-18 | 1.15239252 | 0.267 | 0.115 | 4.35E-14 | cDC.inflam. | Snx14 |
| Cep3501 | 3.12E-18 | 0.37908984 | 0.612 | 0.399 | 6.03E-14 | cDC.inflam. | Cep350 |
| Tent5c1 | 3.26E-18 | 0.32253404 | 0.456 | 0.322 | 6.30E-14 | cDC.inflam. | Tent5c |
| Tbc1d10a1 | 3.57E-18 | 0.30144579 | 0.408 | 0.257 | 6.90E-14 | cDC.inflam. | Tbc1d10a |
| Higd1a | 3.83E-18 | 0.40340328 | 0.587 | 0.382 | 7.40E-14 | cDC.inflam. | Higd1a |
| Rhob | 5.62E-18 | 0.50463864 | 0.369 | 0.2 | 1.09E-13 | cDC.inflam. | Rhob |
| Bcl2a1a1 | 6.89E-18 | 0.53914945 | 0.374 | 0.194 | 1.33E-13 | cDC.inflam. | Bcl2a1a |
| Shc11 | 9.01E-18 | 0.38733687 | 0.617 | 0.409 | 1.74E-13 | cDC.inflam. | Shc1 |
| Nus11 | 9.81E-18 | 0.5725971 | 0.485 | 0.292 | 1.90E-13 | cDC.inflam. | Nus1 |
| Dgka | 1.48E-17 | 1.08479025 | 0.277 | 0.101 | 2.86E-13 | cDC.inflam. | Dgka |
| Ckap4 | 1.98E-17 | 0.88811283 | 0.316 | 0.164 | 3.82E-13 | cDC.inflam. | Ckap4 |
| Arpin2 | 2.25E-17 | 0.32907421 | 0.388 | 0.176 | 4.36E-13 | cDC.inflam. | Arpin |
| Nudt91 | 2.49E-17 | 0.32630535 | 0.534 | 0.334 | 4.81E-13 | cDC.inflam. | Nudt9 |
| Atg5 | 3.22E-17 | 0.83552477 | 0.33 | 0.177 | 6.22E-13 | cDC.inflam. | Atg5 |
| Eif4e1 | 3.93E-17 | 0.43162684 | 0.549 | 0.414 | 7.60E-13 | cDC.inflam. | Eif4e |
| Abhd4 | 6.40E-17 | 0.6847163 | 0.398 | 0.208 | 1.24E-12 | cDC.inflam. | Abhd4 |
| Rab71 | 6.67E-17 | 0.27059031 | 0.689 | 0.527 | 1.29E-12 | cDC.inflam. | Rab7 |
| Dgat11 | 7.00E-17 | 0.71838298 | 0.417 | 0.226 | 1.35E-12 | cDC.inflam. | Dgat1 |
| Trip101 | 7.57E-17 | 0.35388959 | 0.442 | 0.248 | 1.46E-12 | cDC.inflam. | Trip10 |
| P4ha11 | 1.04E-16 | 0.77401484 | 0.471 | 0.279 | 2.02E-12 | cDC.inflam. | P4ha1 |
| Med152 | 1.11E-16 | 0.29846763 | 0.68 | 0.444 | 2.15E-12 | cDC.inflam. | Med15 |
| Plk32 | 1.15E-16 | 0.61718138 | 0.374 | 0.23 | 2.21E-12 | cDC.inflam. | Plk3 |
| Nploc41 | 1.21E-16 | 0.32338186 | 0.558 | 0.39 | 2.34E-12 | cDC.inflam. | Nploc4 |
| Nufip2 | 1.21E-16 | 0.38655727 | 0.587 | 0.373 | 2.34E-12 | cDC.inflam. | Nufip2 |
| Lrrc8d1 | 1.54E-16 | 0.80739713 | 0.413 | 0.245 | 2.98E-12 | cDC.inflam. | Lrrc8d |
| Stx61 | 1.60E-16 | 0.30432942 | 0.515 | 0.313 | 3.10E-12 | cDC.inflam. | Stx6 |
| Bcor | 1.64E-16 | 0.54049243 | 0.369 | 0.222 | 3.18E-12 | cDC.inflam. | Bcor |
| Ncor22 | 2.07E-16 | 0.2592025 | 0.505 | 0.324 | 4.00E-12 | cDC.inflam. | Ncor2 |
| Tpp11 | 2.14E-16 | 0.39577199 | 0.471 | 0.28 | 4.14E-12 | cDC.inflam. | Tpp1 |
| Hsd17b11 | 2.15E-16 | 0.38903178 | 0.568 | 0.376 | 4.15E-12 | cDC.inflam. | Hsd17b11 |
| Gpr1071 | 2.15E-16 | 0.40608045 | 0.408 | 0.244 | 4.16E-12 | cDC.inflam. | Gpr107 |
| Tns31 | 3.20E-16 | 0.45878834 | 0.413 | 0.23 | 6.19E-12 | cDC.inflam. | Tns3 |
| Pgrmc1 | 3.23E-16 | 0.4975933 | 0.345 | 0.191 | 6.24E-12 | cDC.inflam. | Pgrmc1 |
| Blcap | 3.32E-16 | 0.90807283 | 0.354 | 0.173 | 6.42E-12 | cDC.inflam. | Blcap |
| Mapk61 | 3.83E-16 | 0.45913527 | 0.471 | 0.281 | 7.40E-12 | cDC.inflam. | Mapk6 |
| Zbtb12 | 4.27E-16 | 0.3623816 | 0.553 | 0.348 | 8.26E-12 | cDC.inflam. | Zbtb1 |
| Sorl12 | 4.96E-16 | 0.68598464 | 0.558 | 0.366 | 9.59E-12 | cDC.inflam. | Sorl1 |
| Ctdnep1 | 5.20E-16 | 0.29756865 | 0.539 | 0.388 | 1.01E-11 | cDC.inflam. | Ctdnep1 |
| Dctn41 | 6.27E-16 | 0.30943227 | 0.592 | 0.397 | 1.21E-11 | cDC.inflam. | Dctn4 |
| Vps37c | 6.30E-16 | 0.33634197 | 0.398 | 0.23 | 1.22E-11 | cDC.inflam. | Vps37c |
| Stimate | 6.48E-16 | 0.72685455 | 0.272 | 0.131 | 1.25E-11 | cDC.inflam. | Stimate |
| Fam214b | 6.55E-16 | 0.64457521 | 0.364 | 0.179 | 1.27E-11 | cDC.inflam. | Fam214b |
| Ctnnd1 | 6.56E-16 | 0.47624579 | 0.413 | 0.187 | 1.27E-11 | cDC.inflam. | Ctnnd1 |
| Ccng21 | 7.17E-16 | 0.25179258 | 0.437 | 0.266 | 1.39E-11 | cDC.inflam. | Ccng2 |
| Afg3l2 | 7.64E-16 | 0.37365795 | 0.403 | 0.264 | 1.48E-11 | cDC.inflam. | Afg3l2 |
| Wsb21 | 7.73E-16 | 0.72279784 | 0.369 | 0.191 | 1.49E-11 | cDC.inflam. | Wsb2 |
| Pon2 | 7.97E-16 | 0.47479717 | 0.408 | 0.22 | 1.54E-11 | cDC.inflam. | Pon2 |
| Atp6v1c1 | 8.23E-16 | 0.45704849 | 0.544 | 0.355 | 1.59E-11 | cDC.inflam. | Atp6v1c1 |
| Rbm38 | 8.82E-16 | 0.35322656 | 0.461 | 0.314 | 1.71E-11 | cDC.inflam. | Rbm38 |
| Plcl1 | 9.75E-16 | 0.97358303 | 0.262 | 0.106 | 1.89E-11 | cDC.inflam. | Plcl1 |
| Trp53inp2 | 9.99E-16 | 0.77313374 | 0.296 | 0.141 | 1.93E-11 | cDC.inflam. | Trp53inp2 |
| Pdlim41 | 1.12E-15 | 0.26358298 | 0.383 | 0.192 | 2.16E-11 | cDC.inflam. | Pdlim4 |
| Ago11 | 1.16E-15 | 0.44331948 | 0.413 | 0.236 | 2.24E-11 | cDC.inflam. | Ago1 |
| Ugcg1 | 1.20E-15 | 0.52420809 | 0.369 | 0.225 | 2.31E-11 | cDC.inflam. | Ugcg |
| Foxo31 | 1.21E-15 | 0.8159453 | 0.374 | 0.206 | 2.35E-11 | cDC.inflam. | Foxo3 |
| Bola2 | 1.28E-15 | 0.2835219 | 0.534 | 0.386 | 2.48E-11 | cDC.inflam. | Bola2 |
| Trim8 | 1.32E-15 | 0.25261735 | 0.578 | 0.379 | 2.55E-11 | cDC.inflam. | Trim8 |
| Klhl21 | 1.49E-15 | 0.80958707 | 0.282 | 0.141 | 2.88E-11 | cDC.inflam. | Klhl21 |
| Ypel3 | 1.69E-15 | 0.50902685 | 0.49 | 0.286 | 3.28E-11 | cDC.inflam. | Ypel3 |
| Cep85l1 | 1.92E-15 | 0.53587027 | 0.427 | 0.259 | 3.72E-11 | cDC.inflam. | Cep85l |
| Apoe | 2.22E-15 | 0.52307411 | 0.573 | 0.335 | 4.29E-11 | cDC.inflam. | Apoe |
| Crk1 | 2.29E-15 | 0.31035057 | 0.587 | 0.411 | 4.42E-11 | cDC.inflam. | Crk |
| Mbnl21 | 2.46E-15 | 0.29798352 | 0.646 | 0.477 | 4.76E-11 | cDC.inflam. | Mbnl2 |
| Galnt121 | 3.39E-15 | 0.36777226 | 0.364 | 0.249 | 6.56E-11 | cDC.inflam. | Galnt12 |
| Psen21 | 4.09E-15 | 0.42471884 | 0.393 | 0.233 | 7.91E-11 | cDC.inflam. | Psen2 |
| Akt1s1 | 5.21E-15 | 0.69713324 | 0.277 | 0.149 | 1.01E-10 | cDC.inflam. | Akt1s1 |
| Tmem63a | 5.25E-15 | 0.39171707 | 0.364 | 0.207 | 1.02E-10 | cDC.inflam. | Tmem63a |
| Golph3l | 6.59E-15 | 0.44586033 | 0.34 | 0.166 | 1.27E-10 | cDC.inflam. | Golph3l |
| St6galnac4 | 7.97E-15 | 0.49387936 | 0.369 | 0.255 | 1.54E-10 | cDC.inflam. | St6galnac4 |
| Cpd | 8.24E-15 | 0.91725578 | 0.257 | 0.126 | 1.59E-10 | cDC.inflam. | Cpd |
| Mocs1 | 1.10E-14 | 1.16146008 | 0.252 | 0.091 | 2.12E-10 | cDC.inflam. | Mocs1 |
| Adipor1 | 1.27E-14 | 0.35183766 | 0.762 | 0.579 | 2.46E-10 | cDC.inflam. | Adipor1 |
| Slc35f6 | 1.42E-14 | 0.63184054 | 0.33 | 0.166 | 2.74E-10 | cDC.inflam. | Slc35f6 |
| Ptpn21 | 1.45E-14 | 0.33498749 | 0.587 | 0.365 | 2.79E-10 | cDC.inflam. | Ptpn2 |
| Gde12 | 1.51E-14 | 0.54906721 | 0.345 | 0.202 | 2.91E-10 | cDC.inflam. | Gde1 |
| Raf11 | 1.65E-14 | 0.30978019 | 0.534 | 0.365 | 3.19E-10 | cDC.inflam. | Raf1 |
| Kdm4a1 | 1.77E-14 | 0.37314969 | 0.505 | 0.3 | 3.42E-10 | cDC.inflam. | Kdm4a |
| Dock1 | 1.80E-14 | 0.2542919 | 0.296 | 0.146 | 3.48E-10 | cDC.inflam. | Dock1 |
| Creb52 | 1.86E-14 | 0.3485401 | 0.437 | 0.264 | 3.59E-10 | cDC.inflam. | Creb5 |
| Camkk2 | 2.30E-14 | 0.65631202 | 0.286 | 0.143 | 4.46E-10 | cDC.inflam. | Camkk2 |
| Zswim6 | 3.89E-14 | 0.95474549 | 0.277 | 0.121 | 7.53E-10 | cDC.inflam. | Zswim6 |
| Scyl2 | 4.48E-14 | 0.32426723 | 0.495 | 0.323 | 8.65E-10 | cDC.inflam. | Scyl2 |
| Galnt7 | 6.22E-14 | 0.46985477 | 0.286 | 0.14 | 1.20E-09 | cDC.inflam. | Galnt7 |
| Acadm | 6.35E-14 | 0.43600542 | 0.369 | 0.216 | 1.23E-09 | cDC.inflam. | Acadm |
| Errfi11 | 6.66E-14 | 0.50926082 | 0.388 | 0.243 | 1.29E-09 | cDC.inflam. | Errfi1 |
| Stx111 | 6.66E-14 | 0.26599094 | 0.437 | 0.273 | 1.29E-09 | cDC.inflam. | Stx11 |
| Inppl1 | 7.78E-14 | 0.65388536 | 0.417 | 0.213 | 1.50E-09 | cDC.inflam. | Inppl1 |
| Twf11 | 1.04E-13 | 0.35092641 | 0.427 | 0.257 | 2.01E-09 | cDC.inflam. | Twf1 |
| Rnf103 | 1.07E-13 | 0.54785418 | 0.364 | 0.212 | 2.07E-09 | cDC.inflam. | Rnf103 |
| Osbpl31 | 1.30E-13 | 0.62135006 | 0.32 | 0.169 | 2.52E-09 | cDC.inflam. | Osbpl3 |
| Lats21 | 1.37E-13 | 0.40896788 | 0.481 | 0.335 | 2.66E-09 | cDC.inflam. | Lats2 |
| Hax1 | 1.40E-13 | 0.35851397 | 0.519 | 0.325 | 2.71E-09 | cDC.inflam. | Hax1 |
| Patl11 | 1.43E-13 | 0.41071321 | 0.485 | 0.309 | 2.77E-09 | cDC.inflam. | Patl1 |
| Rheb2 | 1.59E-13 | 0.25319304 | 0.718 | 0.518 | 3.07E-09 | cDC.inflam. | Rheb |
| Gk | 2.33E-13 | 0.97756531 | 0.252 | 0.126 | 4.51E-09 | cDC.inflam. | Gk |
| Alkbh5 | 2.59E-13 | 0.340491 | 0.626 | 0.43 | 5.01E-09 | cDC.inflam. | Alkbh5 |
| Traf61 | 2.66E-13 | 0.28646608 | 0.578 | 0.369 | 5.15E-09 | cDC.inflam. | Traf6 |
| Tiparp2 | 2.80E-13 | 0.26907643 | 0.602 | 0.427 | 5.42E-09 | cDC.inflam. | Tiparp |
| Susd61 | 2.88E-13 | 0.28642469 | 0.597 | 0.429 | 5.57E-09 | cDC.inflam. | Susd6 |
| Rmnd5a | 3.11E-13 | 0.25467553 | 0.495 | 0.325 | 6.02E-09 | cDC.inflam. | Rmnd5a |
| Slc27a1 | 3.43E-13 | 0.43743794 | 0.311 | 0.177 | 6.63E-09 | cDC.inflam. | Slc27a1 |
| Uap1l1 | 3.49E-13 | 0.64640988 | 0.301 | 0.162 | 6.74E-09 | cDC.inflam. | Uap1l1 |
| Tmem1641 | 3.61E-13 | 0.47951758 | 0.49 | 0.301 | 6.99E-09 | cDC.inflam. | Tmem164 |
| Map2k31 | 3.82E-13 | 0.25498274 | 0.655 | 0.503 | 7.39E-09 | cDC.inflam. | Map2k3 |
| Zfp385a2 | 4.04E-13 | 0.30782014 | 0.592 | 0.398 | 7.81E-09 | cDC.inflam. | Zfp385a |
| Slc41a2 | 4.28E-13 | 0.46323178 | 0.267 | 0.12 | 8.28E-09 | cDC.inflam. | Slc41a2 |
| Slc7a7 | 4.39E-13 | 0.77620963 | 0.291 | 0.149 | 8.50E-09 | cDC.inflam. | Slc7a7 |
| N4bp11 | 5.45E-13 | 0.39295061 | 0.66 | 0.476 | 1.05E-08 | cDC.inflam. | N4bp1 |
| Rab311 | 9.21E-13 | 0.37518567 | 0.413 | 0.212 | 1.78E-08 | cDC.inflam. | Rab31 |
| Npepps | 1.02E-12 | 0.43351422 | 0.437 | 0.253 | 1.96E-08 | cDC.inflam. | Npepps |
| Spire1 | 1.10E-12 | 0.56662082 | 0.374 | 0.201 | 2.13E-08 | cDC.inflam. | Spire1 |
| Thap2 | 1.11E-12 | 0.43506892 | 0.282 | 0.155 | 2.15E-08 | cDC.inflam. | Thap2 |
| Dars21 | 1.20E-12 | 0.97986992 | 0.252 | 0.124 | 2.31E-08 | cDC.inflam. | Dars2 |
| Vps291 | 1.24E-12 | 0.3314533 | 0.544 | 0.369 | 2.40E-08 | cDC.inflam. | Vps29 |
| Mcur11 | 1.49E-12 | 0.41794715 | 0.471 | 0.295 | 2.88E-08 | cDC.inflam. | Mcur1 |
| Pdlim71 | 1.64E-12 | 0.65197608 | 0.456 | 0.285 | 3.17E-08 | cDC.inflam. | Pdlim7 |
| Htatip2 | 1.77E-12 | 0.67427519 | 0.325 | 0.162 | 3.42E-08 | cDC.inflam. | Htatip2 |
| Rufy31 | 1.85E-12 | 0.52537224 | 0.296 | 0.176 | 3.58E-08 | cDC.inflam. | Rufy3 |
| Tmem1271 | 2.05E-12 | 0.51621453 | 0.481 | 0.322 | 3.96E-08 | cDC.inflam. | Tmem127 |
| Mgat4b | 2.31E-12 | 0.49582959 | 0.413 | 0.229 | 4.46E-08 | cDC.inflam. | Mgat4b |
| Nab21 | 2.38E-12 | 0.48131706 | 0.408 | 0.212 | 4.61E-08 | cDC.inflam. | Nab2 |
| Katnbl1 | 2.47E-12 | 0.44698315 | 0.267 | 0.124 | 4.78E-08 | cDC.inflam. | Katnbl1 |
| Pex13 | 2.56E-12 | 0.70582487 | 0.296 | 0.174 | 4.94E-08 | cDC.inflam. | Pex13 |
| Sh3bp41 | 2.56E-12 | 0.35483573 | 0.388 | 0.185 | 4.95E-08 | cDC.inflam. | Sh3bp4 |
| Nsf1 | 2.83E-12 | 0.35265654 | 0.456 | 0.315 | 5.47E-08 | cDC.inflam. | Nsf |
| Ttyh21 | 3.09E-12 | 0.51505116 | 0.32 | 0.162 | 5.97E-08 | cDC.inflam. | Ttyh2 |
| Klf9 | 3.13E-12 | 0.63782841 | 0.252 | 0.113 | 6.05E-08 | cDC.inflam. | Klf9 |
| Cisd31 | 4.36E-12 | 0.46857056 | 0.335 | 0.194 | 8.43E-08 | cDC.inflam. | Cisd3 |
| Eaf1 | 4.64E-12 | 0.65578014 | 0.325 | 0.186 | 8.97E-08 | cDC.inflam. | Eaf1 |
| Ugdh1 | 7.43E-12 | 0.4511667 | 0.49 | 0.3 | 1.44E-07 | cDC.inflam. | Ugdh |
| Slc11a2 | 7.56E-12 | 0.97552447 | 0.262 | 0.124 | 1.46E-07 | cDC.inflam. | Slc11a2 |
| Msl21 | 8.09E-12 | 0.35570028 | 0.621 | 0.431 | 1.56E-07 | cDC.inflam. | Msl2 |
| Slc25a1 | 8.66E-12 | 0.2726691 | 0.529 | 0.349 | 1.68E-07 | cDC.inflam. | Slc25a1 |
| Vps35l1 | 8.88E-12 | 0.29858047 | 0.519 | 0.335 | 1.72E-07 | cDC.inflam. | Vps35l |
| Tcp11l21 | 9.21E-12 | 0.42673421 | 0.286 | 0.177 | 1.78E-07 | cDC.inflam. | Tcp11l2 |
| Lrrc58 | 9.93E-12 | 0.40804733 | 0.282 | 0.167 | 1.92E-07 | cDC.inflam. | Lrrc58 |
| Sc5d | 1.25E-11 | 0.27155053 | 0.262 | 0.149 | 2.42E-07 | cDC.inflam. | Sc5d |
| Rab5b | 1.56E-11 | 0.35203466 | 0.408 | 0.273 | 3.01E-07 | cDC.inflam. | Rab5b |
| Klhl51 | 1.59E-11 | 0.27967345 | 0.505 | 0.328 | 3.08E-07 | cDC.inflam. | Klhl5 |
| Dnajb9 | 2.17E-11 | 0.33507352 | 0.364 | 0.212 | 4.20E-07 | cDC.inflam. | Dnajb9 |
| Slc25a511 | 2.39E-11 | 0.28346031 | 0.437 | 0.31 | 4.63E-07 | cDC.inflam. | Slc25a51 |
| Stard41 | 2.44E-11 | 0.28157897 | 0.369 | 0.236 | 4.71E-07 | cDC.inflam. | Stard4 |
| Prr141 | 2.48E-11 | 0.4720214 | 0.481 | 0.304 | 4.79E-07 | cDC.inflam. | Prr14 |
| DEPRECATED-I | 2.57E-11 | 0.496851 | 0.282 | 0.132 | 4.97E-07 | cDC.inflam. | DEPRECATED-ENSMUSG00000032667 |
| Mdfic1 | 2.69E-11 | 0.33262628 | 0.408 | 0.271 | 5.20E-07 | cDC.inflam. | Mdfic |
| Slc35a3 | 2.72E-11 | 0.50213818 | 0.398 | 0.261 | 5.25E-07 | cDC.inflam. | Slc35a3 |
| Tcf7l2 | 2.84E-11 | 0.79607133 | 0.282 | 0.142 | 5.49E-07 | cDC.inflam. | Tcf7l2 |
| Arap21 | 3.20E-11 | 0.38909136 | 0.437 | 0.317 | 6.19E-07 | cDC.inflam. | Arap2 |
| Mthfr1 | 3.71E-11 | 0.36053472 | 0.553 | 0.319 | 7.17E-07 | cDC.inflam. | Mthfr |
| Bahd11 | 4.08E-11 | 0.29772123 | 0.388 | 0.27 | 7.89E-07 | cDC.inflam. | Bahd1 |
| Sap18b2 | 4.35E-11 | 0.50139039 | 0.403 | 0.296 | 8.40E-07 | cDC.inflam. | Sap18b |
| Slc2a6 | 5.11E-11 | 0.25080059 | 0.32 | 0.199 | 9.89E-07 | cDC.inflam. | Slc2a6 |
| Rab91 | 5.17E-11 | 0.5016381 | 0.316 | 0.174 | 1.00E-06 | cDC.inflam. | Rab9 |
| Ppfibp12 | 5.49E-11 | 0.38470611 | 0.432 | 0.257 | 1.06E-06 | cDC.inflam. | Ppfibp1 |
| Cd722 | 6.07E-11 | 0.2743913 | 0.359 | 0.24 | 1.17E-06 | cDC.inflam. | Cd72 |
| Znrf11 | 6.15E-11 | 0.53924724 | 0.422 | 0.242 | 1.19E-06 | cDC.inflam. | Znrf1 |
| Tmem1891 | 7.37E-11 | 0.28111067 | 0.515 | 0.407 | 1.42E-06 | cDC.inflam. | Tmem189 |
| Pde4d | 8.18E-11 | 0.60179156 | 0.282 | 0.128 | 1.58E-06 | cDC.inflam. | Pde4d |
| Rad54l2 | 9.12E-11 | 0.60502959 | 0.301 | 0.156 | 1.76E-06 | cDC.inflam. | Rad54l2 |
| Mcub1 | 1.03E-10 | 0.50258251 | 0.277 | 0.174 | 1.99E-06 | cDC.inflam. | Mcub |
| Retreg2 | 1.09E-10 | 0.47858529 | 0.413 | 0.247 | 2.11E-06 | cDC.inflam. | Retreg2 |
| Mcfd21 | 1.18E-10 | 0.43443259 | 0.427 | 0.265 | 2.29E-06 | cDC.inflam. | Mcfd2 |
| Prdm2 | 1.23E-10 | 0.30583499 | 0.403 | 0.279 | 2.37E-06 | cDC.inflam. | Prdm2 |
| Tmem198b1 | 1.32E-10 | 0.36090582 | 0.291 | 0.181 | 2.55E-06 | cDC.inflam. | Tmem198b |
| Tns4 | 1.45E-10 | 0.64982366 | 0.272 | 0.111 | 2.79E-06 | cDC.inflam. | Tns4 |
| Atg4c | 1.48E-10 | 0.83826782 | 0.257 | 0.116 | 2.87E-06 | cDC.inflam. | Atg4c |
| Slc9a1 | 1.56E-10 | 0.26910154 | 0.432 | 0.261 | 3.02E-06 | cDC.inflam. | Slc9a1 |
| Mpp5 | 1.72E-10 | 0.71730879 | 0.257 | 0.114 | 3.32E-06 | cDC.inflam. | Mpp5 |
| Pgp | 1.86E-10 | 0.69880244 | 0.272 | 0.144 | 3.60E-06 | cDC.inflam. | Pgp |
| Trim36 | 2.09E-10 | 0.75582335 | 0.286 | 0.145 | 4.04E-06 | cDC.inflam. | Trim36 |
| Serpinb6a | 2.49E-10 | 0.56922789 | 0.272 | 0.12 | 4.81E-06 | cDC.inflam. | Serpinb6a |
| Luzp1 | 2.82E-10 | 0.74828374 | 0.286 | 0.143 | 5.45E-06 | cDC.inflam. | Luzp1 |
| Ppfibp21 | 2.84E-10 | 0.83462523 | 0.291 | 0.142 | 5.50E-06 | cDC.inflam. | Ppfibp2 |
| Smim32 | 3.40E-10 | 0.32942933 | 0.558 | 0.357 | 6.58E-06 | cDC.inflam. | Smim3 |
| Dnajb21 | 3.70E-10 | 0.4026096 | 0.282 | 0.132 | 7.15E-06 | cDC.inflam. | Dnajb2 |
| Acadsb | 4.37E-10 | 0.42084586 | 0.316 | 0.159 | 8.45E-06 | cDC.inflam. | Acadsb |
| Ints6 | 5.61E-10 | 0.53259562 | 0.403 | 0.275 | 1.09E-05 | cDC.inflam. | Ints6 |
| Ift57 | 5.72E-10 | 0.25238795 | 0.296 | 0.171 | 1.11E-05 | cDC.inflam. | Ift57 |
| Chpf2 | 5.74E-10 | 0.35074076 | 0.398 | 0.24 | 1.11E-05 | cDC.inflam. | Chpf2 |
| Sap30l | 6.06E-10 | 0.36149385 | 0.354 | 0.196 | 1.17E-05 | cDC.inflam. | Sap30l |
| Bmpr21 | 6.10E-10 | 0.35186507 | 0.408 | 0.262 | 1.18E-05 | cDC.inflam. | Bmpr2 |
| Rbbp8 | 6.12E-10 | 0.40634218 | 0.456 | 0.268 | 1.18E-05 | cDC.inflam. | Rbbp8 |
| Tesk1 | 6.72E-10 | 0.25543921 | 0.442 | 0.272 | 1.30E-05 | cDC.inflam. | Tesk1 |
| Lpar61 | 6.77E-10 | 0.62457604 | 0.34 | 0.162 | 1.31E-05 | cDC.inflam. | Lpar6 |
| Vdac11 | 7.00E-10 | 0.34525182 | 0.432 | 0.314 | 1.35E-05 | cDC.inflam. | Vdac1 |
| Ahcyl11 | 7.47E-10 | 0.25314906 | 0.573 | 0.416 | 1.44E-05 | cDC.inflam. | Ahcyl1 |
| Ppm1b1 | 7.73E-10 | 0.33801006 | 0.534 | 0.36 | 1.49E-05 | cDC.inflam. | Ppm1b |
| Pkig | 7.95E-10 | 0.34565942 | 0.282 | 0.17 | 1.54E-05 | cDC.inflam. | Pkig |
| Rexf11 | 1.08E-09 | 0.37627235 | 0.33 | 0.219 | 2.09E-05 | cDC.inflam. | Rexf1 |
| Spen | 1.16E-09 | 0.40144042 | 0.422 | 0.258 | 2.23E-05 | cDC.inflam. | Spen |
| Egln2 | 1.80E-09 | 0.32128421 | 0.379 | 0.233 | 3.48E-05 | cDC.inflam. | Egln2 |
| Ipmk | 1.82E-09 | 0.44837117 | 0.32 | 0.177 | 3.51E-05 | cDC.inflam. | Ipmk |
| Pde1b1 | 2.62E-09 | 0.30528554 | 0.5 | 0.336 | 5.07E-05 | cDC.inflam. | Pde1b |
| Zdhhc6 | 2.68E-09 | 0.53564139 | 0.316 | 0.177 | 5.19E-05 | cDC.inflam. | Zdhhc6 |
| Dlat | 3.90E-09 | 0.3860319 | 0.277 | 0.189 | 7.54E-05 | cDC.inflam. | Dlat |
| Rnf181 | 3.97E-09 | 0.3717272 | 0.413 | 0.271 | 7.67E-05 | cDC.inflam. | Rnf181 |
| Suco1 | 4.23E-09 | 0.31137974 | 0.383 | 0.249 | 8.19E-05 | cDC.inflam. | Suco |
| Zfyve1 | 4.42E-09 | 0.42607793 | 0.267 | 0.165 | 8.54E-05 | cDC.inflam. | Zfyve1 |
| Map4k3 | 4.75E-09 | 0.44243116 | 0.257 | 0.143 | 9.18E-05 | cDC.inflam. | Map4k3 |
| Znhit1 | 5.08E-09 | 0.33674475 | 0.451 | 0.338 | 9.82E-05 | cDC.inflam. | Znhit1 |
| Elovl1 | 5.42E-09 | 0.31508179 | 0.437 | 0.294 | 0.00010475 | cDC.inflam. | Elovl1 |
| Ddx1 | 6.97E-09 | 0.49100722 | 0.359 | 0.19 | 0.00013473 | cDC.inflam. | Ddx1 |
| Klf10 | 7.18E-09 | 0.60941405 | 0.282 | 0.158 | 0.00013876 | cDC.inflam. | Klf10 |
| Ubxn2a | 7.52E-09 | 0.26495014 | 0.374 | 0.292 | 0.00014532 | cDC.inflam. | Ubxn2a |
| Washc5 | 9.38E-09 | 0.50697872 | 0.282 | 0.161 | 0.00018134 | cDC.inflam. | Washc5 |
| Mindy2 | 9.65E-09 | 0.3499377 | 0.286 | 0.18 | 0.00018665 | cDC.inflam. | Mindy2 |
| Mtmr121 | 9.68E-09 | 0.28482889 | 0.369 | 0.244 | 0.0001872 | cDC.inflam. | Mtmr12 |
| Selenon1 | 1.15E-08 | 0.63900892 | 0.272 | 0.15 | 0.00022327 | cDC.inflam. | Selenon |
| Bad1 | 1.28E-08 | 0.41625988 | 0.335 | 0.202 | 0.00024751 | cDC.inflam. | Bad |
| Clcn4 | 1.39E-08 | 0.33334038 | 0.32 | 0.224 | 0.00026962 | cDC.inflam. | Clcn4 |
| Samd81 | 1.65E-08 | 0.54174755 | 0.311 | 0.19 | 0.00031997 | cDC.inflam. | Samd8 |
| Faim | 1.92E-08 | 0.34862251 | 0.383 | 0.238 | 0.00037119 | cDC.inflam. | Faim |
| Dock72 | 2.06E-08 | 0.42884812 | 0.471 | 0.298 | 0.00039831 | cDC.inflam. | Dock7 |
| Sdhaf11 | 2.06E-08 | 0.25581165 | 0.32 | 0.181 | 0.0003986 | cDC.inflam. | Sdhaf1 |
| Mospd21 | 2.08E-08 | 0.38562414 | 0.374 | 0.226 | 0.00040143 | cDC.inflam. | Mospd2 |
| Gpn3 | 2.23E-08 | 0.32726891 | 0.267 | 0.151 | 0.00043059 | cDC.inflam. | Gpn3 |
| Arf21 | 2.26E-08 | 0.27130098 | 0.451 | 0.291 | 0.00043784 | cDC.inflam. | Arf2 |
| Tmem9b | 2.31E-08 | 0.31566112 | 0.316 | 0.22 | 0.00044725 | cDC.inflam. | Tmem9b |
| Asap21 | 2.39E-08 | 0.27401303 | 0.257 | 0.148 | 0.00046139 | cDC.inflam. | Asap2 |
| Ssh3 | 2.47E-08 | 0.3426325 | 0.262 | 0.144 | 0.00047799 | cDC.inflam. | Ssh3 |
| Aldoc1 | 2.63E-08 | 0.30801007 | 0.413 | 0.274 | 0.00050854 | cDC.inflam. | Aldoc |
| Ccdc186 | 2.86E-08 | 0.32227303 | 0.267 | 0.159 | 0.0005525 | cDC.inflam. | Ccdc186 |
| Fndc3b1 | 3.86E-08 | 0.31694844 | 0.403 | 0.246 | 0.00074672 | cDC.inflam. | Fndc3b |
| Pgm11 | 4.26E-08 | 0.36261993 | 0.354 | 0.201 | 0.00082399 | cDC.inflam. | Pgm1 |
| Nsmce21 | 4.28E-08 | 0.38306809 | 0.388 | 0.243 | 0.00082772 | cDC.inflam. | Nsmce2 |
| Fam162a1 | 4.28E-08 | 0.28574588 | 0.427 | 0.271 | 0.00082814 | cDC.inflam. | Fam162a |
| C9orf72 | 5.76E-08 | 0.35714369 | 0.282 | 0.145 | 0.0011134 | cDC.inflam. | C9orf72 |
| Blvra1 | 5.82E-08 | 0.32998807 | 0.383 | 0.252 | 0.00112572 | cDC.inflam. | Blvra |
| Acads | 5.97E-08 | 0.29012088 | 0.345 | 0.197 | 0.00115414 | cDC.inflam. | Acads |
| Wipi2 | 7.22E-08 | 0.50667454 | 0.286 | 0.163 | 0.00139623 | cDC.inflam. | Wipi2 |
| Nipsnap3b | 7.37E-08 | 0.35057274 | 0.301 | 0.2 | 0.00142567 | cDC.inflam. | Nipsnap3b |
| Zbtb221 | 7.77E-08 | 0.28457585 | 0.369 | 0.226 | 0.00150183 | cDC.inflam. | Zbtb22 |
| Pik3c2a | 1.02E-07 | 0.50170869 | 0.267 | 0.168 | 0.00196823 | cDC.inflam. | Pik3c2a |
| Tpd52 | 1.07E-07 | 0.58378261 | 0.291 | 0.166 | 0.00206084 | cDC.inflam. | Tpd52 |
| Prdx61 | 1.20E-07 | 0.42012789 | 0.369 | 0.251 | 0.00232659 | cDC.inflam. | Prdx6 |
| Stam2 | 1.32E-07 | 0.61607117 | 0.306 | 0.179 | 0.00254746 | cDC.inflam. | Stam2 |
| Cdc42ep41 | 1.38E-07 | 0.39307286 | 0.306 | 0.193 | 0.00266916 | cDC.inflam. | Cdc42ep4 |
| Tmem882 | 1.41E-07 | 0.39322361 | 0.369 | 0.249 | 0.00272935 | cDC.inflam. | Tmem88 |
| Ncf42 | 1.48E-07 | 0.40744949 | 0.665 | 0.451 | 0.00286729 | cDC.inflam. | Ncf4 |
| Slmap | 1.55E-07 | 0.33996069 | 0.34 | 0.227 | 0.00298867 | cDC.inflam. | Slmap |
| Tollip1 | 1.61E-07 | 0.31701518 | 0.432 | 0.262 | 0.00311332 | cDC.inflam. | Tollip |
| Dtx21 | 2.03E-07 | 0.25716845 | 0.325 | 0.199 | 0.00391599 | cDC.inflam. | Dtx2 |
| Apba3 | 2.16E-07 | 0.49167566 | 0.291 | 0.162 | 0.00417777 | cDC.inflam. | Apba3 |
| Plcb3 | 2.58E-07 | 0.71016688 | 0.296 | 0.145 | 0.00498655 | cDC.inflam. | Plcb3 |
| Tjap1 | 3.20E-07 | 0.27832496 | 0.33 | 0.219 | 0.00618491 | cDC.inflam. | Tjap1 |
| Mob1b | 3.20E-07 | 0.25389016 | 0.345 | 0.223 | 0.00618491 | cDC.inflam. | Mob1b |
| Ggh1 | 3.27E-07 | 0.59660275 | 0.296 | 0.189 | 0.00631874 | cDC.inflam. | Ggh |
| Acox11 | 3.72E-07 | 0.2604981 | 0.398 | 0.281 | 0.00719575 | cDC.inflam. | Acox1 |
| Tmem41b | 4.55E-07 | 0.48318326 | 0.286 | 0.162 | 0.00879129 | cDC.inflam. | Tmem41b |
| Dhx402 | 4.65E-07 | 0.35543147 | 0.354 | 0.238 | 0.00898799 | cDC.inflam. | Dhx40 |
| Dxo1 | 5.46E-07 | 0.31041099 | 0.364 | 0.2 | 0.01054901 | cDC.inflam. | Dxo |
| Hadha | 6.13E-07 | 0.28545837 | 0.485 | 0.339 | 0.01185419 | cDC.inflam. | Hadha |
| Arsa | 6.39E-07 | 0.44628714 | 0.291 | 0.169 | 0.01235562 | cDC.inflam. | Arsa |
| Larp1b1 | 6.71E-07 | 0.35315359 | 0.252 | 0.163 | 0.01297822 | cDC.inflam. | Larp1b |
| Ccr12 | 7.31E-07 | 0.35383314 | 0.422 | 0.277 | 0.0141353 | cDC.inflam. | Ccr1 |
| Nod22 | 8.33E-07 | 0.29030545 | 0.587 | 0.379 | 0.01610085 | cDC.inflam. | Nod2 |
| Foxj2 | 9.64E-07 | 0.40453983 | 0.32 | 0.201 | 0.01863909 | cDC.inflam. | Foxj2 |
| Psmd101 | 1.05E-06 | 0.26117338 | 0.301 | 0.215 | 0.02032285 | cDC.inflam. | Psmd10 |
| Josd1 | 1.21E-06 | 0.37098248 | 0.33 | 0.222 | 0.02331886 | cDC.inflam. | Josd1 |
| Apool | 1.21E-06 | 0.58126603 | 0.257 | 0.133 | 0.02332901 | cDC.inflam. | Apool |
| Ikbkg | 1.37E-06 | 0.29729194 | 0.33 | 0.204 | 0.02657367 | cDC.inflam. | Ikbkg |
| Hook2 | 1.38E-06 | 0.59678377 | 0.272 | 0.154 | 0.0267516 | cDC.inflam. | Hook2 |
| Mroh1 | 1.65E-06 | 0.41246323 | 0.359 | 0.257 | 0.03183913 | cDC.inflam. | Mroh1 |
| Taf1 | 1.75E-06 | 0.28541173 | 0.325 | 0.217 | 0.03387542 | cDC.inflam. | Taf1 |
| Mark41 | 2.19E-06 | 0.28263221 | 0.354 | 0.247 | 0.04236748 | cDC.inflam. | Mark4 |
| Dhps | 2.60E-06 | 0.28249809 | 0.306 | 0.19 | 0.05017544 | cDC.inflam. | Dhps |
| Pop41 | 2.64E-06 | 0.53025966 | 0.291 | 0.178 | 0.05099725 | cDC.inflam. | Pop4 |
| Mast2 | 3.06E-06 | 0.42931799 | 0.267 | 0.136 | 0.05910548 | cDC.inflam. | Mast2 |
| Fnip11 | 3.18E-06 | 0.32782335 | 0.364 | 0.229 | 0.06140962 | cDC.inflam. | Fnip1 |
| Klf71 | 3.27E-06 | 0.25492084 | 0.301 | 0.214 | 0.06318533 | cDC.inflam. | Klf7 |
| Ccnc | 3.55E-06 | 0.27548802 | 0.262 | 0.155 | 0.06855257 | cDC.inflam. | Ccnc |
| Taf12 | 4.01E-06 | 0.46754684 | 0.267 | 0.166 | 0.07757133 | cDC.inflam. | Taf12 |
| Klf21 | 4.98E-06 | 0.573234 | 0.325 | 0.191 | 0.09631169 | cDC.inflam. | Klf2 |
| Mnt1 | 6.10E-06 | 0.27096542 | 0.311 | 0.191 | 0.11793807 | cDC.inflam. | Mnt |
| Mvk1 | 8.77E-06 | 0.34620372 | 0.286 | 0.206 | 0.16961856 | cDC.inflam. | Mvk |
| Sh2b21 | 1.14E-05 | 0.34928234 | 0.252 | 0.19 | 0.22094227 | cDC.inflam. | Sh2b2 |
| Lemd31 | 1.16E-05 | 0.29294192 | 0.393 | 0.299 | 0.22422801 | cDC.inflam. | Lemd3 |
| Entpd71 | 1.16E-05 | 0.4204728 | 0.306 | 0.215 | 0.22501685 | cDC.inflam. | Entpd7 |
| Ccnt1 | 1.46E-05 | 0.3006218 | 0.408 | 0.285 | 0.28208528 | cDC.inflam. | Ccnt1 |
| Zfp52 | 2.41E-05 | 0.68214169 | 0.252 | 0.142 | 0.46523735 | cDC.inflam. | Zfp52 |
| Sipa1l11 | 2.60E-05 | 0.27653142 | 0.325 | 0.208 | 0.50319077 | cDC.inflam. | Sipa1l1 |
| Gse1 | 2.88E-05 | 0.26881361 | 0.33 | 0.204 | 0.55654846 | cDC.inflam. | Gse1 |
| Metap11 | 2.96E-05 | 0.25676212 | 0.325 | 0.233 | 0.57251131 | cDC.inflam. | Metap1 |
| Rad9a | 3.18E-05 | 0.36593992 | 0.257 | 0.167 | 0.61521875 | cDC.inflam. | Rad9a |
| Camk1 | 3.37E-05 | 0.44030078 | 0.277 | 0.147 | 0.65098898 | cDC.inflam. | Camk1 |
| Kdm3a1 | 6.17E-05 | 0.27660965 | 0.369 | 0.219 | 1 | cDC.inflam. | Kdm3a |
| Tmco6 | 6.29E-05 | 0.26322344 | 0.252 | 0.133 | 1 | cDC.inflam. | Tmco6 |

| | | | | | | | |
|---|---|---|---|---|---|---|---|
| Ube2a1 | 0.00011929 | 0.29311386 | 0.408 | 0.282 | 1 | cDC.inflam. | Ube2a |
| Vapa1 | 0.00012727 | 0.25795117 | 0.316 | 0.219 | 1 | cDC.inflam. | Vapa |
| Lpcat3 | 0.00018655 | 0.36624114 | 0.252 | 0.17 | 1 | cDC.inflam. | Lpcat3 |
| Tsc22d4 | 0.00033994 | 0.30939936 | 0.398 | 0.274 | 1 | cDC.inflam. | Tsc22d4 |
| Klhl2 | 0.00037876 | 0.28834727 | 0.252 | 0.167 | 1 | cDC.inflam. | Klhl2 |
| Dop1b | 0.000471 | 0.28203627 | 0.252 | 0.172 | 1 | cDC.inflam. | Dop1b |
| Slc25a331 | 0.00351275 | 0.32221043 | 0.33 | 0.216 | 1 | cDC.inflam. | Slc25a33 |
| Ing12 | 0.00451161 | 0.26277697 | 0.35 | 0.259 | 1 | cDC.inflam. | Ing1 |
| Cln31 | 0.00547408 | 0.2821199 | 0.267 | 0.169 | 1 | cDC.inflam. | Cln3 |
| Siglech | 0 | 8.72717427 | 0.971 | 0.012 | 0 | pDC | Siglech |
| Atp1b1 | 0 | 7.00914477 | 0.964 | 0.031 | 0 | pDC | Atp1b1 |
| Mctp2 | 0 | 8.94191477 | 0.92 | 0.008 | 0 | pDC | Mctp2 |
| Sell1 | 0 | 4.75648779 | 0.956 | 0.112 | 0 | pDC | Sell |
| Ccr9 | 0 | 7.81345674 | 0.839 | 0.02 | 0 | pDC | Ccr9 |
| Spib | 0 | 4.84607672 | 0.861 | 0.07 | 0 | pDC | Spib |
| Smim5 | 0 | 6.23641503 | 0.81 | 0.041 | 0 | pDC | Smim5 |
| Lag3 | 0 | 5.18576381 | 0.854 | 0.087 | 0 | pDC | Lag3 |
| Adam111 | 0 | 3.40593535 | 0.839 | 0.096 | 0 | pDC | Adam11 |
| Ighm | 0 | 5.04801614 | 0.978 | 0.238 | 0 | pDC | Ighm |
| Tex2 | 0 | 5.02026803 | 0.956 | 0.217 | 0 | pDC | Tex2 |
| Sema4b | 0 | 6.04741258 | 0.781 | 0.046 | 0 | pDC | Sema4b |
| Fcrla | 0 | 6.12205564 | 0.766 | 0.042 | 0 | pDC | Fcrla |
| Slco4a1 | 0 | 6.14513446 | 0.752 | 0.034 | 0 | pDC | Slco4a1 |
| Eepd1 | 0 | 4.50632567 | 0.898 | 0.2 | 0 | pDC | Eepd1 |
| Lair11 | 0 | 4.36060752 | 0.912 | 0.243 | 0 | pDC | Lair1 |
| Blnk | 0 | 5.50917208 | 0.715 | 0.054 | 0 | pDC | Blnk |
| Ly6d | 0 | 8.9983437 | 0.65 | 0.004 | 0 | pDC | Ly6d |
| Tcf41 | 0 | 3.78657205 | 0.92 | 0.274 | 0 | pDC | Tcf4 |
| Cd4 | 0 | 7.90603606 | 0.642 | 0.012 | 0 | pDC | Cd4 |
| Tmem229b1 | 0 | 4.21532443 | 0.861 | 0.242 | 0 | pDC | Tmem229b |
| Fam43a | 0 | 5.4387797 | 0.723 | 0.105 | 0 | pDC | Fam43a |
| Fyb1 | 0 | 3.76515994 | 0.869 | 0.252 | 0 | pDC | Fyb |
| Ccl41 | 0 | 4.01922662 | 0.839 | 0.229 | 0 | pDC | Ccl4 |
| Ly6c1 | 0 | 4.65386983 | 0.657 | 0.048 | 0 | pDC | Ly6c1 |
| Cyb561a3 | 0 | 4.16580388 | 0.971 | 0.369 | 0 | pDC | Cyb561a3 |
| Ly6c2 | 0 | 4.19819833 | 0.65 | 0.057 | 0 | pDC | Ly6c2 |
| Runx2 | 0 | 3.44950451 | 0.869 | 0.284 | 0 | pDC | Runx2 |
| Pltp1 | 0 | 2.79045075 | 0.971 | 0.396 | 0 | pDC | Pltp |
| Tsc22d1 | 0 | 3.51668387 | 0.825 | 0.254 | 0 | pDC | Tsc22d1 |
| Mpeg11 | 0 | 3.22343967 | 0.993 | 0.439 | 0 | pDC | Mpeg1 |
| Ly6a1 | 0 | 3.60858416 | 0.701 | 0.147 | 0 | pDC | Ly6a |
| Abhd17b1 | 0 | 3.76889381 | 0.832 | 0.285 | 0 | pDC | Abhd17b |
| Plac81 | 0 | 2.71893921 | 0.956 | 0.472 | 0 | pDC | Plac8 |
| Bst21 | 0 | 4.50151365 | 0.978 | 0.505 | 0 | pDC | Bst2 |
| Slpi | 0 | 5.12961483 | 0.496 | 0.031 | 0 | pDC | Slpi |
| Cd8a | 0 | 5.92006547 | 0.489 | 0.024 | 0 | pDC | Cd8a |
| Slc44a22 | 0 | 2.98890266 | 0.942 | 0.481 | 0 | pDC | Slc44a2 |
| Irf82 | 0 | 2.28489943 | 0.985 | 0.59 | 0 | pDC | Irf8 |
| Pld41 | 0 | 2.7369323 | 0.978 | 0.642 | 0 | pDC | Pld4 |
| Ly6e3 | 0 | 1.99992205 | 0.993 | 0.691 | 0 | pDC | Ly6e |
| Grn1 | 0 | 3.10539864 | 0.993 | 0.719 | 0 | pDC | Grn |
| Serinc3 | 0 | 2.60300773 | 0.934 | 0.667 | 0 | pDC | Serinc3 |
| Ctsb1 | 0 | 2.66207258 | 0.956 | 0.724 | 0 | pDC | Ctsb |
| Bcl11a1 | 7.48E-299 | 3.74639027 | 0.774 | 0.159 | 1.45E-294 | pDC | Bcl11a |
| Ets1 | 7.91E-285 | 5.13845051 | 0.715 | 0.046 | 1.53E-280 | pDC | Ets1 |
| Ctsl1 | 4.15E-284 | 3.76567849 | 0.869 | 0.25 | 8.02E-280 | pDC | Ctsl |
| Tlr7 | 7.95E-281 | 7.52707729 | 0.628 | 0.014 | 1.54E-276 | pDC | Tlr7 |
| Trp53i111 | 9.36E-278 | 3.96383372 | 0.839 | 0.139 | 1.81E-273 | pDC | Trp53i11 |
| Cd71 | 2.19E-266 | 2.8751467 | 0.657 | 0.178 | 4.23E-262 | pDC | Cd7 |
| 13-Sep | 1.45E-264 | 4.13287397 | 0.832 | 0.145 | 2.80E-260 | pDC | 1-Sep |
| Lifr2 | 3.53E-255 | 3.44534785 | 0.869 | 0.262 | 6.82E-251 | pDC | Lifr |
| Ptprcap | 4.19E-253 | 5.53433418 | 0.65 | 0.041 | 8.10E-249 | pDC | Ptprcap |
| Pdzd4 | 2.96E-249 | 6.19516569 | 0.679 | 0.02 | 5.73E-245 | pDC | Pdzd4 |
| Sla21 | 1.79E-241 | 5.72762151 | 0.584 | 0.034 | 3.45E-237 | pDC | Sla2 |
| Npc11 | 5.88E-238 | 3.40359488 | 0.854 | 0.286 | 1.14E-233 | pDC | Npc1 |
| Calcoco1 | 2.50E-230 | 3.80941342 | 0.781 | 0.139 | 4.83E-226 | pDC | Calcoco1 |
| Ripor2 | 5.56E-228 | 4.93894396 | 0.599 | 0.031 | 1.08E-223 | pDC | Ripor2 |
| Slc41a21 | 8.13E-226 | 4.24827695 | 0.774 | 0.108 | 1.57E-221 | pDC | Slc41a2 |
| Pacsin1 | 7.73E-214 | 5.81388917 | 0.606 | 0.025 | 1.49E-209 | pDC | Pacsin1 |
| Cd8b1 | 1.69E-211 | 9.51923757 | 0.54 | 0.003 | 3.26E-207 | pDC | Cd8b1 |
| Cybb3 | 1.70E-203 | 2.87075132 | 0.861 | 0.339 | 3.30E-199 | pDC | Cybb |
| Rell11 | 9.46E-202 | 3.09625284 | 0.869 | 0.329 | 1.83E-197 | pDC | Rell1 |
| Lrp8 | 3.03E-200 | 3.69525809 | 0.555 | 0.088 | 5.87E-196 | pDC | Lrp8 |
| Adpgk1 | 8.06E-199 | 3.00127169 | 0.854 | 0.329 | 1.56E-194 | pDC | Adpgk |
| St8sia42 | 4.87E-194 | 2.22814006 | 0.927 | 0.591 | 9.42E-190 | pDC | St8sia4 |
| Itgal2 | 1.47E-192 | 3.49942924 | 0.803 | 0.172 | 2.84E-188 | pDC | Itgal |
| Mgat1 | 6.20E-191 | 3.0991032 | 0.781 | 0.308 | 1.20E-186 | pDC | Mgat1 |
| Fads3 | 1.91E-188 | 4.3836593 | 0.628 | 0.063 | 3.68E-184 | pDC | Fads3 |
| Snx182 | 9.34E-186 | 2.5933341 | 0.912 | 0.501 | 1.81E-181 | pDC | Snx18 |
| Cxxc5 | 6.14E-184 | 4.64543515 | 0.679 | 0.069 | 1.19E-179 | pDC | Cxxc5 |
| Ptprf | 4.90E-183 | 4.57359818 | 0.474 | 0.032 | 9.48E-179 | pDC | Ptprf |
| Dntt | 4.43E-182 | 9.94117308 | 0.766 | 0.001 | 8.56E-178 | pDC | Dntt |
| Wfs1 | 7.64E-181 | 5.33121841 | 0.547 | 0.034 | 1.48E-176 | pDC | Wfs1 |
| Cox6a2 | 1.44E-180 | 9.36834262 | 0.752 | 0.002 | 2.78E-176 | pDC | Cox6a2 |
| Rnase61 | 1.97E-180 | 3.28019069 | 0.73 | 0.227 | 3.80E-176 | pDC | Rnase6 |
| Rilpl21 | 2.99E-176 | 2.53478943 | 0.92 | 0.485 | 5.79E-172 | pDC | Rilpl2 |
| Atp2b4 | 2.95E-175 | 3.35224721 | 0.686 | 0.138 | 5.71E-171 | pDC | Atp2b4 |
| Rnf1871 | 2.27E-172 | 2.46878474 | 0.898 | 0.611 | 4.40E-168 | pDC | Rnf187 |
| Nucb2 | 2.65E-171 | 3.80756787 | 0.752 | 0.177 | 5.12E-167 | pDC | Nucb2 |
| Rabgap1l1 | 9.03E-171 | 1.92613332 | 0.898 | 0.347 | 1.75E-166 | pDC | Rabgap1l |
| Paqr5 | 9.36E-169 | 6.96789351 | 0.577 | 0.009 | 1.81E-164 | pDC | Paqr5 |
| Chdh | 4.88E-164 | 6.35169304 | 0.569 | 0.008 | 9.44E-160 | pDC | Chdh |
| Dap | 3.10E-163 | 2.6748251 | 0.766 | 0.321 | 5.99E-159 | pDC | Dap |
| Lefty1 | 8.36E-161 | 5.75443398 | 0.453 | 0.012 | 1.62E-156 | pDC | Lefty1 |
| Psap2 | 1.43E-158 | 1.76004036 | 0.985 | 0.797 | 2.77E-154 | pDC | Psap |
| Tifa | 9.97E-157 | 3.58730906 | 0.62 | 0.159 | 1.93E-152 | pDC | Tifa |
| Plxdc1 | 1.03E-155 | 4.61397196 | 0.555 | 0.047 | 2.00E-151 | pDC | Plxdc1 |
| Itgb3 | 7.81E-154 | 4.66034641 | 0.467 | 0.033 | 1.51E-149 | pDC | Itgb3 |
| S1pr4 | 1.56E-153 | 4.93618444 | 0.489 | 0.036 | 3.01E-149 | pDC | S1pr4 |
| Fads2 | 1.13E-148 | 3.9406935 | 0.584 | 0.082 | 2.18E-144 | pDC | Fads2 |
| Igkc | 2.21E-146 | 5.13942404 | 0.292 | 0.013 | 4.26E-142 | pDC | Igkc |
| Atp13a21 | 2.44E-144 | 3.15775271 | 0.737 | 0.196 | 4.72E-140 | pDC | Atp13a2 |
| Rab33b | 1.03E-143 | 3.61216909 | 0.657 | 0.138 | 2.00E-139 | pDC | Rab33b |
| Sik11 | 7.87E-143 | 2.48368117 | 0.934 | 0.574 | 1.52E-138 | pDC | Sik1 |
| Man2a21 | 1.64E-142 | 2.91845105 | 0.81 | 0.271 | 3.17E-138 | pDC | Man2a2 |
| Tmem63a1 | 2.51E-142 | 3.24643471 | 0.723 | 0.199 | 4.86E-138 | pDC | Tmem63a |
| Csf2rb23 | 2.82E-141 | 2.25316315 | 0.898 | 0.507 | 5.46E-137 | pDC | Csf2rb2 |
| Snx5 | 3.83E-140 | 2.92024064 | 0.81 | 0.35 | 7.41E-136 | pDC | Snx5 |
| Ablim1 | 7.16E-140 | 4.46213418 | 0.533 | 0.054 | 1.38E-135 | pDC | Ablim1 |
| Cd37 | 1.03E-139 | 3.07557008 | 0.686 | 0.173 | 2.00E-135 | pDC | Cd37 |
| Snx291 | 4.48E-138 | 3.05114899 | 0.796 | 0.271 | 8.66E-134 | pDC | Snx29 |
| Hs3st1 | 3.55E-137 | 5.82461321 | 0.401 | 0.013 | 6.86E-133 | pDC | Hs3st1 |
| Cd300c | 1.12E-133 | 5.73369484 | 0.387 | 0.014 | 2.17E-129 | pDC | Cd300c |
| Slc49a4 | 8.37E-132 | 2.66765541 | 0.832 | 0.334 | 1.62E-127 | pDC | Slc49a4 |
| Ctsh | 1.26E-131 | 1.67750501 | 0.942 | 0.805 | 2.43E-127 | pDC | Ctsh |
| Ptprs | 3.30E-130 | 2.25939491 | 0.847 | 0.422 | 6.38E-126 | pDC | Ptprs |
| Ldlrad3 | 2.15E-128 | 5.19387198 | 0.474 | 0.032 | 4.16E-124 | pDC | Ldlrad3 |
| Havcr1 | 9.29E-126 | 4.89945386 | 0.474 | 0.021 | 1.80E-121 | pDC | Havcr1 |
| P2ry13 | 1.85E-124 | 4.71955016 | 0.416 | 0.04 | 3.58E-120 | pDC | P2ry13 |
| Pml | 3.53E-124 | 3.80872192 | 0.555 | 0.09 | 6.82E-120 | pDC | Pml |
| Irf2bp22 | 9.53E-124 | 1.77396754 | 0.985 | 0.789 | 1.84E-119 | pDC | Irf2bp2 |
| Selplg1 | 2.54E-123 | 1.88864609 | 0.934 | 0.707 | 4.90E-119 | pDC | Selplg |
| Spn | 1.54E-121 | 3.53877051 | 0.657 | 0.092 | 2.98E-117 | pDC | Spn |
| Atp1a3 | 2.13E-120 | 5.21830818 | 0.387 | 0.028 | 4.13E-116 | pDC | Atp1a3 |
| Rnf122 | 5.71E-119 | 4.89235126 | 0.482 | 0.031 | 1.10E-114 | pDC | Rnf122 |
| Klra17 | 3.32E-118 | 4.97734672 | 0.409 | 0.019 | 6.43E-114 | pDC | Klra17 |
| DEPRECATED-I | 5.26E-116 | 5.43574744 | 0.431 | 0.024 | 1.02E-111 | pDC | DEPRECATED-ENSMUSG00000027636 |
| Epha22 | 3.93E-111 | 2.81560605 | 0.788 | 0.246 | 7.60E-107 | pDC | Epha2 |
| Rac22 | 2.39E-110 | 1.76092801 | 0.985 | 0.73 | 4.62E-106 | pDC | Rac2 |
| D13Ertd608e | 7.42E-109 | 8.32470358 | 0.46 | 0.002 | 1.43E-104 | pDC | D13Ertd608e |
| Bcr | 5.77E-107 | 3.49257952 | 0.628 | 0.138 | 1.12E-102 | pDC | Bcr |
| Pik3ip1 | 1.72E-106 | 4.90834255 | 0.336 | 0.022 | 3.33E-102 | pDC | Pik3ip1 |
| Carmil1 | 1.53E-104 | 3.58414884 | 0.54 | 0.093 | 2.97E-100 | pDC | Carmil1 |
| Cacna1e | 4.01E-101 | 8.57037956 | 0.511 | 0.002 | 7.75E-97 | pDC | Cacna1e |
| Prkca | 1.10E-99 | 4.52618029 | 0.423 | 0.041 | 2.13E-95 | pDC | Prkca |
| Cracr2a | 2.46E-98 | 5.64000702 | 0.431 | 0.024 | 4.75E-94 | pDC | Cracr2a |
| Slc27a11 | 8.47E-98 | 3.36595371 | 0.701 | 0.168 | 1.64E-93 | pDC | Slc27a1 |
| Slc25a12 | 7.33E-96 | 2.9354988 | 0.73 | 0.249 | 1.42E-91 | pDC | Slc25a12 |
| Mzb1 | 3.01E-94 | 8.46996291 | 0.38 | 0.002 | 5.82E-90 | pDC | Mzb1 |
| Gna151 | 8.72E-94 | 3.04944763 | 0.62 | 0.179 | 1.69E-89 | pDC | Gna15 |
| Tubgcp5 | 4.29E-93 | 3.70056261 | 0.511 | 0.087 | 8.30E-89 | pDC | Tubgcp5 |
| Fbxo17 | 1.75E-91 | 5.8957828 | 0.321 | 0.011 | 3.39E-87 | pDC | Fbxo17 |
| Tapt1 | 2.98E-89 | 3.61411231 | 0.496 | 0.104 | 5.76E-85 | pDC | Tapt1 |
| Hpse1 | 3.50E-89 | 3.42363319 | 0.547 | 0.121 | 6.77E-85 | pDC | Hpse |
| Fam214a | 7.03E-89 | 4.41992099 | 0.394 | 0.042 | 1.36E-84 | pDC | Fam214a |
| Tmed31 | 7.46E-89 | 2.49516321 | 0.686 | 0.351 | 1.44E-84 | pDC | Tmed3 |
| Tagap1 | 1.08E-88 | 2.45536668 | 0.591 | 0.225 | 2.08E-84 | pDC | Tagap |
| Traf4 | 3.72E-88 | 3.1757459 | 0.628 | 0.146 | 7.19E-84 | pDC | Traf4 |
| Ifnar21 | 1.04E-85 | 2.17391826 | 0.788 | 0.492 | 2.01E-81 | pDC | Ifnar2 |
| Cdkn2d1 | 5.53E-85 | 2.61628298 | 0.642 | 0.225 | 1.07E-80 | pDC | Cdkn2d |
| Unc93b11 | 1.53E-84 | 1.83168027 | 0.949 | 0.735 | 2.96E-80 | pDC | Unc93b1 |
| Il12a | 1.93E-84 | 6.67843732 | 0.292 | 0.003 | 3.73E-80 | pDC | Il12a |
| Grina1 | 1.74E-83 | 2.22909595 | 0.759 | 0.431 | 3.36E-79 | pDC | Grina |
| Tom1 | 6.32E-83 | 3.99704719 | 0.46 | 0.081 | 1.22E-78 | pDC | Tom1 |
| Lpgat11 | 7.62E-83 | 2.67221794 | 0.693 | 0.328 | 1.47E-78 | pDC | Lpgat1 |
| Cobll1 | 2.22E-82 | 4.98536825 | 0.35 | 0.017 | 4.30E-78 | pDC | Cobll1 |
| Mef2c1 | 4.61E-81 | 2.93261101 | 0.599 | 0.188 | 8.91E-77 | pDC | Mef2c |
| Itsn1 | 4.88E-80 | 3.98814133 | 0.387 | 0.044 | 9.44E-76 | pDC | Itsn1 |
| Fam241a | 2.95E-79 | 2.98026303 | 0.445 | 0.153 | 5.70E-75 | pDC | Fam241a |
| Gas7 | 9.82E-78 | 3.48637092 | 0.518 | 0.099 | 1.90E-73 | pDC | Gas7 |
| Klk1 | 1.65E-76 | 11.061178 | 0.496 | 0.001 | 3.20E-72 | pDC | Klk1 |
| Tfrc | 2.47E-76 | 3.6870072 | 0.496 | 0.088 | 4.78E-72 | pDC | Tfrc |
| Cmah | 2.85E-76 | 6.96716872 | 0.343 | 0.003 | 5.51E-72 | pDC | Cmah |
| Ptprc2 | 4.45E-74 | 1.79756743 | 0.942 | 0.719 | 8.60E-70 | pDC | Ptprc |
| Myb | 8.83E-73 | 3.64982154 | 0.343 | 0.025 | 1.71E-68 | pDC | Myb |
| Glcci1 | 4.55E-72 | 2.82347901 | 0.54 | 0.196 | 8.80E-68 | pDC | Glcci1 |
| Rasgrp2 | 5.97E-71 | 3.60997187 | 0.409 | 0.064 | 1.15E-66 | pDC | Rasgrp2 |
| Stambpl1 | 1.49E-70 | 3.61585202 | 0.438 | 0.07 | 2.89E-66 | pDC | Stambpl1 |
| Gpr1712 | 4.55E-70 | 2.04282155 | 0.752 | 0.493 | 8.80E-66 | pDC | Gpr171 |
| Irf11 | 1.13E-69 | 1.60536725 | 0.781 | 0.453 | 2.19E-65 | pDC | Irf1 |
| Ncf11 | 2.06E-69 | 1.4914915 | 0.898 | 0.743 | 3.98E-65 | pDC | Ncf1 |
| Cxcr3 | 8.17E-69 | 4.15160587 | 0.35 | 0.043 | 1.58E-64 | pDC | Cxcr3 |
| Slc7a52 | 3.01E-68 | 2.01235338 | 0.759 | 0.476 | 5.81E-64 | pDC | Slc7a5 |
| Nceh1 | 6.89E-68 | 3.49323899 | 0.562 | 0.109 | 1.33E-63 | pDC | Nceh1 |
| Ldhb | 1.34E-66 | 5.14753121 | 0.277 | 0.013 | 2.59E-62 | pDC | Ldhb |
| Cd381 | 1.88E-65 | 3.02730804 | 0.518 | 0.137 | 3.63E-61 | pDC | Cd38 |
| Dctn6 | 1.71E-64 | 2.63786197 | 0.672 | 0.23 | 3.31E-60 | pDC | Dctn6 |
| Gria3 | 1.73E-64 | 3.43719253 | 0.277 | 0.027 | 3.34E-60 | pDC | Gria3 |
| Abca5 | 3.19E-64 | 5.10733099 | 0.263 | 0.012 | 6.16E-60 | pDC | Abca5 |
| Plekhm31 | 4.03E-63 | 2.64003183 | 0.693 | 0.257 | 7.78E-59 | pDC | Plekhm3 |
| Txnip2 | 4.65E-63 | 2.14453473 | 0.642 | 0.292 | 8.99E-59 | pDC | Txnip |
| Klhl6 | 5.27E-63 | 3.49031738 | 0.431 | 0.084 | 1.02E-58 | pDC | Klhl6 |
| Klf22 | 2.27E-62 | 2.36424196 | 0.416 | 0.19 | 4.38E-58 | pDC | Klf2 |
| H2-Q71 | 1.11E-61 | 1.52809501 | 0.788 | 0.436 | 2.14E-57 | pDC | H2-Q7 |
| Slc2a3 | 2.70E-61 | 3.40844849 | 0.445 | 0.063 | 5.23E-57 | pDC | Slc2a3 |
| Clec2d1 | 7.56E-61 | 1.40264539 | 0.949 | 0.695 | 1.46E-56 | pDC | Clec2d |
| Fam174a | 1.10E-60 | 2.51406641 | 0.65 | 0.3 | 2.13E-56 | pDC | Fam174a |
| Mn1 | 1.88E-60 | 6.81905098 | 0.314 | 0.002 | 3.63E-56 | pDC | Mn1 |
| Emid1 | 1.15E-59 | 7.54002298 | 0.27 | 0.003 | 2.22E-55 | pDC | Emid1 |
| Iglc3 | 1.65E-59 | 9.7872948 | 0.423 | 0.001 | 3.19E-55 | pDC | Iglc3 |
| Klhl42 | 2.84E-59 | 4.55650494 | 0.263 | 0.021 | 5.49E-55 | pDC | Klhl42 |
| Cnp1 | 3.55E-59 | 2.58225544 | 0.599 | 0.24 | 6.85E-55 | pDC | Cnp |
| Zbtb10 | 6.60E-59 | 2.74849843 | 0.416 | 0.086 | 1.28E-54 | pDC | Zbtb10 |
| Rubcnl1 | 9.24E-59 | 2.95011738 | 0.526 | 0.142 | 1.79E-54 | pDC | Rubcnl |
| Smc6 | 1.07E-57 | 2.5672433 | 0.635 | 0.267 | 2.06E-53 | pDC | Smc6 |
| Ifnar1 | 1.51E-57 | 2.0316348 | 0.73 | 0.448 | 2.92E-53 | pDC | Ifnar1 |
| Neil1 | 2.40E-57 | 3.66064449 | 0.277 | 0.053 | 4.64E-53 | pDC | Neil1 |
| Ly9 | 4.31E-57 | 3.75464513 | 0.401 | 0.056 | 8.33E-53 | pDC | Ly9 |
| Cd180 | 8.95E-57 | 3.7228327 | 0.482 | 0.1 | 1.73E-52 | pDC | Cd180 |
| Dll4 | 1.99E-55 | 2.93306852 | 0.314 | 0.037 | 3.84E-51 | pDC | Dll4 |
| H2-T24 | 2.27E-55 | 3.8457243 | 0.358 | 0.044 | 4.40E-51 | pDC | H2-T24 |
| Trit11 | 4.24E-55 | 2.81256435 | 0.555 | 0.15 | 8.19E-51 | pDC | Trit1 |
| Prr51 | 6.45E-55 | 3.22932397 | 0.489 | 0.098 | 1.25E-50 | pDC | Prr5 |
| Ldlrad4 | 1.12E-54 | 3.34852788 | 0.358 | 0.07 | 2.17E-50 | pDC | Ldlrad4 |
| Dclre1c1 | 5.81E-54 | 2.26027336 | 0.606 | 0.227 | 1.12E-49 | pDC | Dclre1c |
| Slamf6 | 9.95E-54 | 4.38619197 | 0.263 | 0.021 | 1.92E-49 | pDC | Slamf6 |
| Tmc8 | 2.19E-53 | 3.3317483 | 0.409 | 0.065 | 4.23E-49 | pDC | Tmc8 |
| Aff3 | 9.20E-53 | 3.30810849 | 0.409 | 0.069 | 1.78E-48 | pDC | Aff3 |
| H131 | 2.17E-52 | 1.96055656 | 0.803 | 0.456 | 4.19E-48 | pDC | H13 |
| Sh2b22 | 5.48E-52 | 2.89171916 | 0.584 | 0.182 | 1.06E-47 | pDC | Sh2b2 |
| Pik3cd3 | 1.56E-51 | 1.83270832 | 0.839 | 0.561 | 3.02E-47 | pDC | Pik3cd |
| Spns3 | 6.65E-51 | 3.21168779 | 0.394 | 0.079 | 1.29E-46 | pDC | Spns3 |
| Tmem221 | 1.42E-50 | 4.65103185 | 0.27 | 0.018 | 2.75E-46 | pDC | Tmem221 |
| Txndc51 | 7.87E-50 | 2.14383193 | 0.723 | 0.364 | 1.52E-45 | pDC | Txndc5 |
| Tcf12 | 1.99E-49 | 2.50254522 | 0.569 | 0.183 | 3.85E-45 | pDC | Tcf12 |
| Abhd15 | 2.38E-49 | 3.37825434 | 0.387 | 0.066 | 4.60E-45 | pDC | Abhd15 |
| Pecam1 | 2.87E-49 | 2.621314 | 0.474 | 0.131 | 5.55E-45 | pDC | Pecam1 |
| Fam117a1 | 3.35E-49 | 2.65542389 | 0.562 | 0.18 | 6.49E-45 | pDC | Fam117a |
| Atp2a1 | 1.59E-48 | 7.78411406 | 0.277 | 0.002 | 3.08E-44 | pDC | Atp2a1 |
| Gne | 2.86E-48 | 3.19183101 | 0.387 | 0.069 | 5.52E-44 | pDC | Gne |
| Syne2 | 5.78E-48 | 3.18703526 | 0.343 | 0.052 | 1.12E-43 | pDC | Syne2 |
| Tyrobp2 | 9.85E-48 | 1.41423041 | 0.993 | 0.835 | 1.90E-43 | pDC | Tyrobp |
| Als2 | 2.17E-47 | 3.46193607 | 0.336 | 0.064 | 4.20E-43 | pDC | Als2 |
| Inpp4a1 | 3.79E-47 | 2.5410755 | 0.635 | 0.251 | 7.33E-43 | pDC | Inpp4a |
| Pgls | 5.96E-47 | 1.84886062 | 0.759 | 0.515 | 1.15E-42 | pDC | Pgls |
| Npc21 | 7.60E-47 | 1.33318018 | 0.927 | 0.743 | 1.47E-42 | pDC | Npc2 |
| Cd1642 | 8.78E-47 | 1.74539305 | 0.956 | 0.617 | 1.70E-42 | pDC | Cd164 |
| Gtf2i | 2.79E-46 | 2.39102698 | 0.635 | 0.297 | 5.40E-42 | pDC | Gtf2i |
| Sema4d2 | 6.94E-46 | 1.78190573 | 0.905 | 0.561 | 1.34E-41 | pDC | Sema4d |
| Mvb12a1 | 7.12E-46 | 2.18721174 | 0.686 | 0.326 | 1.38E-41 | pDC | Mvb12a |
| Ddr12 | 2.12E-45 | 2.07981285 | 0.708 | 0.356 | 4.10E-41 | pDC | Ddr1 |
| Hvcn12 | 2.35E-45 | 2.39505054 | 0.518 | 0.159 | 4.54E-41 | pDC | Hvcn1 |
| Stk17b2 | 2.71E-45 | 1.67227133 | 0.847 | 0.654 | 5.24E-41 | pDC | Stk17b |
| Them6 | 4.80E-45 | 3.22621753 | 0.263 | 0.053 | 9.28E-41 | pDC | Them6 |
| Cdip1 | 1.26E-44 | 2.7728969 | 0.547 | 0.186 | 2.43E-40 | pDC | Cdip1 |
| Cd471 | 1.67E-44 | 1.26473341 | 0.898 | 0.8 | 3.23E-40 | pDC | Cd47 |
| Slf2 | 2.65E-44 | 2.72253938 | 0.526 | 0.179 | 5.12E-40 | pDC | Slf2 |
| Ivns1abp1 | 3.53E-44 | 1.55340452 | 0.825 | 0.666 | 6.82E-40 | pDC | Ivns1abp |
| Notch1 | 4.45E-44 | 3.04904434 | 0.431 | 0.098 | 8.60E-40 | pDC | Notch1 |
| Cd442 | 5.02E-44 | 1.41105127 | 0.81 | 0.592 | 9.71E-40 | pDC | Cd44 |
| Rhobtb2 | 7.40E-44 | 3.94412194 | 0.358 | 0.06 | 1.43E-39 | pDC | Rhobtb2 |
| B3gnt8 | 5.29E-43 | 3.3825084 | 0.328 | 0.062 | 1.02E-38 | pDC | B3gnt8 |
| Nek61 | 7.30E-43 | 1.96860845 | 0.701 | 0.322 | 1.41E-38 | pDC | Nek6 |
| Inafm2 | 1.42E-41 | 2.78459717 | 0.453 | 0.141 | 2.75E-37 | pDC | Inafm2 |
| Phc1 | 2.65E-41 | 3.15836161 | 0.358 | 0.08 | 5.13E-37 | pDC | Phc1 |
| Rab6b | 9.26E-41 | 3.69280435 | 0.277 | 0.032 | 1.79E-36 | pDC | Rab6b |
| Ptpn62 | 7.23E-40 | 1.60378996 | 0.92 | 0.646 | 1.40E-35 | pDC | Ptpn6 |
| Obscn | 1.46E-39 | 9.17684052 | 0.263 | 0.001 | 2.82E-35 | pDC | Obscn |
| Irf71 | 1.57E-39 | 2.25471804 | 0.599 | 0.162 | 3.03E-35 | pDC | Irf7 |
| Rab27a | 3.81E-39 | 2.09487144 | 0.518 | 0.175 | 7.37E-35 | pDC | Rab27a |
| Trp53inp1 | 5.30E-38 | 2.62733171 | 0.328 | 0.072 | 1.02E-33 | pDC | Trp53inp1 |
| Ints31 | 1.63E-37 | 1.91950763 | 0.54 | 0.277 | 3.16E-33 | pDC | Ints3 |
| Fam3c2 | 1.70E-37 | 2.48730512 | 0.54 | 0.242 | 3.29E-33 | pDC | Fam3c |
| Rnd31 | 5.11E-37 | 1.82788221 | 0.54 | 0.248 | 9.88E-33 | pDC | Rnd3 |
| Rap1a1 | 9.32E-37 | 1.70372313 | 0.737 | 0.552 | 1.80E-32 | pDC | Rap1a |
| L3mbtl3 | 1.31E-36 | 2.83431866 | 0.35 | 0.076 | 2.53E-32 | pDC | L3mbtl3 |
| Cdc14b | 3.42E-36 | 3.99701968 | 0.263 | 0.036 | 6.62E-32 | pDC | Cdc14b |
| Ramp1 | 3.79E-36 | 2.76133908 | 0.401 | 0.077 | 7.33E-32 | pDC | Ramp1 |
| Rftn12 | 4.96E-36 | 1.62639447 | 0.759 | 0.4 | 9.59E-32 | pDC | Rftn1 |
| Msi21 | 7.02E-36 | 1.99084232 | 0.672 | 0.231 | 1.36E-31 | pDC | Msi2 |
| Dhcr7 | 7.60E-36 | 2.59661198 | 0.409 | 0.125 | 1.47E-31 | pDC | Dhcr7 |
| Jade1 | 8.43E-36 | 2.82366509 | 0.409 | 0.087 | 1.63E-31 | pDC | Jade1 |
| Tspan131 | 1.29E-35 | 1.4883984 | 0.942 | 0.726 | 2.50E-31 | pDC | Tspan13 |
| Ogt | 1.60E-35 | 1.83560098 | 0.679 | 0.369 | 3.09E-31 | pDC | Ogt |
| Rgl1 | 2.82E-35 | 3.12986872 | 0.365 | 0.06 | 5.46E-31 | pDC | Rgl1 |
| Fgfr1op21 | 7.46E-35 | 1.67558848 | 0.781 | 0.484 | 1.44E-30 | pDC | Fgfr1op2 |
| Sp31 | 1.59E-34 | 1.99107233 | 0.526 | 0.29 | 3.08E-30 | pDC | Sp3 |
| Ikzf11 | 1.53E-33 | 1.60433688 | 0.825 | 0.576 | 2.97E-29 | pDC | Ikzf1 |
| Rasa32 | 6.86E-33 | 2.24934223 | 0.54 | 0.209 | 1.33E-28 | pDC | Rasa3 |
| Stoml1 | 1.05E-32 | 2.35272526 | 0.35 | 0.097 | 2.03E-28 | pDC | Stoml1 |
| Apc1 | 1.81E-32 | 2.27455115 | 0.518 | 0.218 | 3.50E-28 | pDC | Apc |
| Ptp4a31 | 2.82E-32 | 2.35993977 | 0.533 | 0.238 | 5.45E-28 | pDC | Ptp4a3 |
| Orai3 | 3.89E-32 | 2.21234001 | 0.504 | 0.221 | 7.52E-28 | pDC | Orai3 |
| Slc15a41 | 4.06E-32 | 1.94957638 | 0.635 | 0.364 | 7.85E-28 | pDC | Slc15a4 |
| Klf131 | 9.89E-31 | 1.90121831 | 0.62 | 0.377 | 1.91E-26 | pDC | Klf13 |
| Emp31 | 1.35E-30 | 1.5443233 | 0.854 | 0.454 | 2.62E-26 | pDC | Emp3 |
| Adk | 3.35E-30 | 3.62392915 | 0.277 | 0.043 | 6.48E-26 | pDC | Adk |
| Pdcd4 | 3.64E-30 | 2.14696443 | 0.569 | 0.218 | 7.03E-26 | pDC | Pdcd4 |
| Il7r2 | 4.63E-30 | 1.73839276 | 0.62 | 0.295 | 8.95E-26 | pDC | Il7r |
| Gna11 | 6.99E-30 | 3.15295517 | 0.343 | 0.077 | 1.35E-25 | pDC | Gna11 |
| Sec11c1 | 1.46E-29 | 1.53716992 | 0.766 | 0.504 | 2.83E-25 | pDC | Sec11c |
| Grm8 | 1.75E-28 | 12.3896326 | 0.54 | 0 | 3.39E-24 | pDC | Grm8 |
| Slc44a1 | 4.91E-28 | 2.25517955 | 0.431 | 0.132 | 9.49E-24 | pDC | Slc44a1 |
| Tlr9 | 1.09E-27 | 2.46434254 | 0.438 | 0.148 | 2.10E-23 | pDC | Tlr9 |
| Ifi47 | 1.35E-27 | 2.49757777 | 0.482 | 0.115 | 2.61E-23 | pDC | Ifi47 |
| Hivep31 | 1.40E-27 | 3.10884768 | 0.299 | 0.083 | 2.71E-23 | pDC | Hivep3 |
| Sort1 | 1.86E-27 | 2.00001089 | 0.358 | 0.111 | 3.59E-23 | pDC | Sort1 |
| Tifab1 | 2.95E-27 | 2.04617864 | 0.474 | 0.208 | 5.71E-23 | pDC | Tifab |
| Capn5 | 3.41E-27 | 2.6617379 | 0.263 | 0.049 | 6.58E-23 | pDC | Capn5 |
| Zcchc24 | 4.64E-27 | 2.56238862 | 0.431 | 0.151 | 8.97E-23 | pDC | Zcchc24 |
| Snx30 | 7.96E-27 | 2.88493785 | 0.38 | 0.106 | 1.54E-22 | pDC | Snx30 |
| St8sia1 | 1.05E-26 | 2.26056665 | 0.277 | 0.075 | 2.04E-22 | pDC | St8sia1 |
| Creb3l2 | 2.22E-26 | 2.68944712 | 0.255 | 0.073 | 4.29E-22 | pDC | Creb3l2 |
| Cd209d | 3.50E-26 | 2.29831105 | 0.314 | 0.078 | 6.77E-22 | pDC | Cd209d |
| Ppm1h | 3.99E-26 | 2.08202669 | 0.438 | 0.203 | 7.71E-22 | pDC | Ppm1h |
| Cd331 | 4.40E-26 | 1.9996235 | 0.423 | 0.258 | 8.51E-22 | pDC | Cd33 |
| Tbc1d82 | 6.75E-26 | 1.02851959 | 0.847 | 0.584 | 1.30E-21 | pDC | Tbc1d8 |
| Slc29a31 | 1.19E-25 | 2.06033702 | 0.635 | 0.286 | 2.29E-21 | pDC | Slc29a3 |
| Edem2 | 2.55E-25 | 1.88050734 | 0.62 | 0.416 | 4.94E-21 | pDC | Edem2 |
| Arhgap17 | 3.29E-25 | 2.22595569 | 0.584 | 0.303 | 6.35E-21 | pDC | Arhgap17 |
| Tram12 | 4.86E-25 | 1.82047788 | 0.847 | 0.539 | 9.40E-21 | pDC | Tram1 |
| Pafah1b31 | 5.77E-25 | 1.79999362 | 0.628 | 0.331 | 1.12E-20 | pDC | Pafah1b3 |
| Zfp363 | 1.32E-24 | 0.27583898 | 0.964 | 0.909 | 2.56E-20 | pDC | Zfp36 |
| Pycr22 | 2.51E-24 | 2.16761789 | 0.518 | 0.242 | 4.86E-20 | pDC | Pycr2 |
| Hnrnpul1 | 3.25E-23 | 1.42647172 | 0.905 | 0.745 | 6.28E-19 | pDC | Hnrnpul1 |
| B3gnt2 | 5.89E-23 | 2.21402719 | 0.387 | 0.119 | 1.14E-18 | pDC | B3gnt2 |
| Serpinh1 | 6.17E-23 | 2.27116549 | 0.277 | 0.078 | 1.19E-18 | pDC | Serpinh1 |
| Ccpg12 | 7.78E-23 | 2.09264197 | 0.562 | 0.233 | 1.51E-18 | pDC | Ccpg1 |
| Pde7a | 1.15E-22 | 2.55658149 | 0.416 | 0.13 | 2.23E-18 | pDC | Pde7a |
| Rhbdf22 | 1.34E-22 | 2.07155121 | 0.577 | 0.277 | 2.60E-18 | pDC | Rhbdf2 |
| Cyfip21 | 1.50E-22 | 1.85010768 | 0.686 | 0.332 | 2.89E-18 | pDC | Cyfip2 |
| Tsc22d3 | 2.97E-22 | 2.05473109 | 0.482 | 0.247 | 5.74E-18 | pDC | Tsc22d3 |
| Rasal31 | 3.31E-22 | 2.28920869 | 0.453 | 0.187 | 6.39E-18 | pDC | Rasal3 |
| Scap1 | 3.56E-22 | 2.28432578 | 0.54 | 0.248 | 6.88E-18 | pDC | Scap |
| Ctso1 | 5.06E-22 | 2.63087531 | 0.445 | 0.167 | 9.78E-18 | pDC | Ctso |
| 1-Jun | 8.71E-22 | 1.14679096 | 0.788 | 0.674 | 1.68E-17 | pDC | Jun |
| Padi21 | 8.77E-22 | 2.10051562 | 0.438 | 0.155 | 1.70E-17 | pDC | Padi2 |
| Pkig1 | 1.32E-21 | 2.29016025 | 0.453 | 0.166 | 2.56E-17 | pDC | Pkig |
| Tom1l2 | 1.73E-21 | 2.56181541 | 0.307 | 0.087 | 3.34E-17 | pDC | Tom1l2 |
| Zdhhc141 | 3.75E-21 | 2.39083324 | 0.416 | 0.152 | 7.25E-17 | pDC | Zdhhc14 |
| Ugcg2 | 4.36E-21 | 1.82214779 | 0.445 | 0.225 | 8.42E-17 | pDC | Ugcg |
| Ppfia4 | 8.82E-21 | 2.47550205 | 0.365 | 0.141 | 1.70E-16 | pDC | Ppfia4 |
| Hdac51 | 1.14E-20 | 2.12239189 | 0.518 | 0.216 | 2.21E-16 | pDC | Hdac5 |
| Emc3 | 1.19E-20 | 2.16423304 | 0.431 | 0.224 | 2.29E-16 | pDC | Emc3 |
| Serp11 | 1.20E-20 | 2.16686431 | 0.496 | 0.226 | 2.32E-16 | pDC | Serp1 |
| Tbcel | 1.85E-20 | 2.40039835 | 0.314 | 0.096 | 3.59E-16 | pDC | Tbcel |
| Man1c1 | 3.74E-20 | 2.55963131 | 0.35 | 0.12 | 7.24E-16 | pDC | Man1c1 |
| Atf6b1 | 5.81E-20 | 2.06465574 | 0.518 | 0.261 | 1.12E-15 | pDC | Atf6b |
| Pdpk1 | 9.47E-20 | 1.89379459 | 0.533 | 0.304 | 1.83E-15 | pDC | Pdpk1 |
| Card112 | 4.20E-19 | 1.75226189 | 0.606 | 0.269 | 8.11E-15 | pDC | Card11 |
| Zc3h12c2 | 7.33E-19 | 1.9983545 | 0.343 | 0.133 | 1.42E-14 | pDC | Zc3h12c |
| Panx1 | 8.93E-19 | 2.56154363 | 0.27 | 0.088 | 1.73E-14 | pDC | Panx1 |
| Chst15 | 1.82E-18 | 1.96598145 | 0.35 | 0.156 | 3.51E-14 | pDC | Chst15 |
| Cmtm71 | 2.36E-18 | 1.77423319 | 0.642 | 0.345 | 4.56E-14 | pDC | Cmtm7 |
| C1galt1 | 2.67E-18 | 2.46760297 | 0.314 | 0.114 | 5.17E-14 | pDC | C1galt1 |
| Ap1ar1 | 2.90E-18 | 1.98116136 | 0.423 | 0.258 | 5.62E-14 | pDC | Ap1ar |
| Serinc52 | 4.14E-18 | 1.76492873 | 0.453 | 0.155 | 8.01E-14 | pDC | Serinc5 |
| Galk11 | 4.74E-18 | 2.45294932 | 0.423 | 0.162 | 9.17E-14 | pDC | Galk1 |
| Ptar1 | 1.03E-17 | 2.2190205 | 0.416 | 0.164 | 1.98E-13 | pDC | Ptar1 |
| Itpr12 | 1.23E-17 | 1.72075739 | 0.664 | 0.344 | 2.38E-13 | pDC | Itpr1 |
| Sil1 | 1.24E-17 | 2.22128092 | 0.314 | 0.121 | 2.40E-13 | pDC | Sil1 |
| Bloc1s22 | 2.22E-17 | 1.4969648 | 0.781 | 0.568 | 4.30E-13 | pDC | Bloc1s2 |
| Cd691 | 2.44E-17 | 1.88247159 | 0.358 | 0.187 | 4.71E-13 | pDC | Cd69 |
| Gns1 | 2.48E-17 | 1.68813577 | 0.737 | 0.495 | 4.79E-13 | pDC | Gns |
| Nxf11 | 2.85E-17 | 1.20009965 | 0.693 | 0.495 | 5.50E-13 | pDC | Nxf1 |
| Sh2b1 | 2.91E-17 | 2.11837343 | 0.445 | 0.176 | 5.62E-13 | pDC | Sh2b1 |
| Zdhhc13 | 5.57E-17 | 2.47361862 | 0.358 | 0.125 | 1.08E-12 | pDC | Zdhhc13 |
| Lbr2 | 6.63E-17 | 1.69004007 | 0.65 | 0.356 | 1.28E-12 | pDC | Lbr |
| Trib13 | 8.33E-17 | 1.64357 | 0.715 | 0.475 | 1.61E-12 | pDC | Trib1 |
| Plaur2 | 8.46E-17 | 1.37866795 | 0.606 | 0.391 | 1.64E-12 | pDC | Plaur |
| N4bp3 | 1.10E-16 | 2.49843823 | 0.277 | 0.106 | 2.13E-12 | pDC | N4bp3 |
| Vps8 | 2.88E-16 | 2.03302785 | 0.401 | 0.175 | 5.57E-12 | pDC | Vps8 |
| Rgp1 | 2.92E-16 | 2.07303782 | 0.358 | 0.148 | 5.64E-12 | pDC | Rgp1 |
| Slc39a141 | 3.03E-16 | 1.98438068 | 0.416 | 0.184 | 5.86E-12 | pDC | Slc39a14 |
| Fam20b | 3.45E-16 | 2.3731645 | 0.438 | 0.19 | 6.66E-12 | pDC | Fam20b |
| Evi2a1 | 4.69E-16 | 1.51980205 | 0.715 | 0.47 | 9.06E-12 | pDC | Evi2a |
| Dnajc13 | 5.18E-16 | 1.68551291 | 0.562 | 0.337 | 1.00E-11 | pDC | Dnajc13 |
| Atp6v1d | 5.42E-16 | 1.56055844 | 0.657 | 0.45 | 1.05E-11 | pDC | Atp6v1d |
| Smim141 | 6.95E-16 | 1.66375127 | 0.65 | 0.425 | 1.34E-11 | pDC | Smim14 |
| Btg3 | 7.46E-16 | 2.31049695 | 0.35 | 0.131 | 1.44E-11 | pDC | Btg3 |
| Adipor21 | 9.13E-16 | 1.61328449 | 0.504 | 0.337 | 1.77E-11 | pDC | Adipor2 |
| Cds2 | 9.27E-16 | 2.11225338 | 0.423 | 0.195 | 1.79E-11 | pDC | Cds2 |
| Clec12a1 | 1.22E-15 | 2.49496131 | 0.285 | 0.11 | 2.37E-11 | pDC | Clec12a |
| Xbp12 | 1.49E-15 | 1.19947641 | 0.891 | 0.706 | 2.88E-11 | pDC | Xbp1 |
| Zfp810 | 1.55E-15 | 2.32470726 | 0.255 | 0.07 | 3.00E-11 | pDC | Zfp810 |
| Calhm2 | 1.67E-15 | 2.18610792 | 0.35 | 0.121 | 3.23E-11 | pDC | Calhm2 |
| Nptn | 1.78E-15 | 1.52696012 | 0.708 | 0.537 | 3.45E-11 | pDC | Nptn |
| Slc9a91 | 1.82E-15 | 2.06385601 | 0.504 | 0.215 | 3.52E-11 | pDC | Slc9a9 |
| Ppp1r16b | 2.13E-15 | 1.60991633 | 0.518 | 0.292 | 4.11E-11 | pDC | Ppp1r16b |
| Kctd121 | 2.38E-15 | 1.15098771 | 0.964 | 0.777 | 4.60E-11 | pDC | Kctd12 |
| Saraf | 2.93E-15 | 1.84168235 | 0.672 | 0.397 | 5.67E-11 | pDC | Saraf |
| Gpr18 | 3.71E-15 | 2.63307956 | 0.27 | 0.094 | 7.17E-11 | pDC | Gpr18 |
| Manf1 | 3.92E-15 | 1.49769774 | 0.818 | 0.518 | 7.58E-11 | pDC | Manf |
| Elf1 | 4.90E-15 | 1.37098364 | 0.65 | 0.443 | 9.47E-11 | pDC | Elf1 |
| Man1a2 | 4.96E-15 | 1.99701736 | 0.445 | 0.216 | 9.60E-11 | pDC | Man1a2 |
| Vps37a | 5.82E-15 | 1.90153569 | 0.423 | 0.216 | 1.13E-10 | pDC | Vps37a |
| Hsp90b11 | 5.93E-15 | 1.55040896 | 0.708 | 0.471 | 1.15E-10 | pDC | Hsp90b1 |
| Dgke | 7.03E-15 | 2.43711676 | 0.263 | 0.108 | 1.36E-10 | pDC | Dgke |
| Ppif | 8.35E-15 | 2.25915108 | 0.277 | 0.115 | 1.61E-10 | pDC | Ppif |
| Fgfr11 | 1.13E-14 | 1.65330423 | 0.482 | 0.22 | 2.19E-10 | pDC | Fgfr1 |
| Rai12 | 1.71E-14 | 1.85783795 | 0.438 | 0.244 | 3.30E-10 | pDC | Rai1 |
| Acap11 | 1.77E-14 | 1.63095025 | 0.65 | 0.315 | 3.42E-10 | pDC | Acap1 |
| Klhl24 | 3.74E-14 | 2.03015857 | 0.38 | 0.164 | 7.23E-10 | pDC | Klhl24 |
| Mroh11 | 4.17E-14 | 1.78492658 | 0.482 | 0.255 | 8.07E-10 | pDC | Mroh1 |
| Tm7sf2 | 5.02E-14 | 2.23023508 | 0.27 | 0.09 | 9.70E-10 | pDC | Tm7sf2 |
| Selenot1 | 6.80E-14 | 1.58729091 | 0.555 | 0.362 | 1.32E-09 | pDC | Selenot |
| Stard5 | 1.21E-13 | 1.92218806 | 0.255 | 0.113 | 2.35E-09 | pDC | Stard5 |
| Eif2ak3 | 1.35E-13 | 1.57685418 | 0.365 | 0.207 | 2.61E-09 | pDC | Eif2ak3 |
| Hgh1 | 2.07E-13 | 2.46964116 | 0.277 | 0.081 | 4.01E-09 | pDC | Hgh1 |
| Ceacam11 | 2.45E-13 | 1.43813097 | 0.394 | 0.139 | 4.74E-09 | pDC | Ceacam1 |
| Sub11 | 2.58E-13 | 1.30700876 | 0.708 | 0.565 | 4.99E-09 | pDC | Sub1 |
| Pqlc3 | 2.85E-13 | 1.95408093 | 0.394 | 0.188 | 5.51E-09 | pDC | Pqlc3 |
| Fos3 | 3.21E-13 | 0.5095253 | 0.752 | 0.687 | 6.21E-09 | pDC | Fos |
| Arhgef61 | 3.41E-13 | 1.3785297 | 0.613 | 0.408 | 6.60E-09 | pDC | Arhgef6 |
| Nnt | 3.64E-13 | 2.30776343 | 0.365 | 0.161 | 7.04E-09 | pDC | Nnt |
| Grk61 | 5.61E-13 | 1.50719809 | 0.708 | 0.435 | 1.09E-08 | pDC | Grk6 |
| Fuca22 | 6.48E-13 | 1.5583887 | 0.526 | 0.341 | 1.25E-08 | pDC | Fuca2 |
| Fam160b1 | 6.92E-13 | 1.97649614 | 0.394 | 0.198 | 1.34E-08 | pDC | Fam160b1 |
| Tram21 | 7.31E-13 | 2.20609068 | 0.255 | 0.112 | 1.41E-08 | pDC | Tram2 |
| Prxl2c | 7.87E-13 | 2.09421008 | 0.27 | 0.116 | 1.52E-08 | pDC | Prxl2c |
| Pink1 | 9.36E-13 | 1.7379506 | 0.438 | 0.239 | 1.81E-08 | pDC | Pink1 |
| Il161 | 1.23E-12 | 1.97895214 | 0.38 | 0.149 | 2.37E-08 | pDC | Il16 |
| Nprl3 | 1.68E-12 | 2.06572672 | 0.27 | 0.113 | 3.24E-08 | pDC | Nprl3 |
| Snn | 1.71E-12 | 1.83241458 | 0.255 | 0.076 | 3.31E-08 | pDC | Snn |
| Mfsd122 | 4.28E-12 | 1.77541829 | 0.489 | 0.273 | 8.28E-08 | pDC | Mfsd12 |
| Sept111 | 4.65E-12 | 1.79887532 | 0.438 | 0.238 | 8.99E-08 | pDC | 11-Sep |
| Cep192 | 4.74E-12 | 1.93196125 | 0.394 | 0.162 | 9.16E-08 | pDC | Cep192 |
| Zeb22 | 5.12E-12 | 1.53566411 | 0.708 | 0.371 | 9.90E-08 | pDC | Zeb2 |
| Acbd5 | 5.27E-12 | 1.47449569 | 0.445 | 0.267 | 1.02E-07 | pDC | Acbd5 |
| Clec10a1 | 5.32E-12 | 1.25582504 | 0.672 | 0.296 | 1.03E-07 | pDC | Clec10a |
| Mxd41 | 6.93E-12 | 1.79524419 | 0.416 | 0.213 | 1.34E-07 | pDC | Mxd4 |
| Arfgef2 | 1.31E-11 | 2.22383748 | 0.263 | 0.107 | 2.53E-07 | pDC | Arfgef2 |
| Xrcc6 | 1.37E-11 | 2.26725278 | 0.255 | 0.097 | 2.65E-07 | pDC | Xrcc6 |
| Slc7a6os | 1.60E-11 | 1.49551614 | 0.416 | 0.234 | 3.09E-07 | pDC | Slc7a6os |
| Ubxn4 | 1.61E-11 | 1.61780339 | 0.701 | 0.472 | 3.11E-07 | pDC | Ubxn4 |
| AU040320 | 1.95E-11 | 1.64771363 | 0.401 | 0.221 | 3.78E-07 | pDC | AU040320 |
| Slamf92 | 2.19E-11 | 1.91865418 | 0.445 | 0.22 | 4.23E-07 | pDC | Slamf9 |
| Cds1 | 2.24E-11 | 2.13874621 | 0.336 | 0.132 | 4.33E-07 | pDC | Cds1 |
| Tm7sf3 | 2.69E-11 | 2.39328838 | 0.292 | 0.129 | 5.20E-07 | pDC | Tm7sf3 |
| Dgcr6 | 2.86E-11 | 1.95420403 | 0.27 | 0.111 | 5.52E-07 | pDC | Dgcr6 |
| Satb1 | 3.28E-11 | 1.58006465 | 0.255 | 0.083 | 6.33E-07 | pDC | Satb1 |
| Vav12 | 3.67E-11 | 1.36065514 | 0.708 | 0.499 | 7.09E-07 | pDC | Vav1 |
| Amfr | 3.98E-11 | 1.62644923 | 0.62 | 0.375 | 7.69E-07 | pDC | Amfr |
| Ube2q2 | 4.24E-11 | 2.00997816 | 0.328 | 0.165 | 8.20E-07 | pDC | Ube2q2 |
| H2-K11 | 4.33E-11 | 0.80213634 | 0.985 | 0.905 | 8.36E-07 | pDC | H2-K1 |
| Ech12 | 4.89E-11 | 1.50692844 | 0.518 | 0.372 | 9.45E-07 | pDC | Ech1 |
| Ero1lb | 5.41E-11 | 1.56262004 | 0.511 | 0.35 | 1.05E-06 | pDC | Ero1lb |
| Spin11 | 6.41E-11 | 1.68360018 | 0.453 | 0.24 | 1.24E-06 | pDC | Spin1 |
| Uggt12 | 7.89E-11 | 1.32261547 | 0.672 | 0.506 | 1.52E-06 | pDC | Uggt1 |
| Fndc3a | 8.18E-11 | 1.46205727 | 0.555 | 0.361 | 1.58E-06 | pDC | Fndc3a |
| Gapt | 1.02E-10 | 2.02600781 | 0.299 | 0.152 | 1.98E-06 | pDC | Gapt |
| Snx92 | 1.04E-10 | 1.61251062 | 0.635 | 0.44 | 2.02E-06 | pDC | Snx9 |
| Gng121 | 1.25E-10 | 1.79358597 | 0.372 | 0.145 | 2.41E-06 | pDC | Gng12 |
| Edem31 | 1.27E-10 | 2.00809154 | 0.307 | 0.17 | 2.46E-06 | pDC | Edem3 |
| Bst1 | 1.30E-10 | 2.15808633 | 0.27 | 0.112 | 2.52E-06 | pDC | Bst1 |
| Rnf1231 | 1.50E-10 | 1.61904972 | 0.431 | 0.235 | 2.90E-06 | pDC | Rnf123 |
| Trim11 | 1.89E-10 | 2.27863721 | 0.27 | 0.118 | 3.66E-06 | pDC | Trim11 |
| Tmem50a1 | 2.85E-10 | 1.3593122 | 0.686 | 0.493 | 5.52E-06 | pDC | Tmem50a |
| Slfn8 | 3.19E-10 | 1.9205457 | 0.277 | 0.093 | 6.16E-06 | pDC | Slfn8 |
| Ern1 | 3.61E-10 | 1.53565205 | 0.467 | 0.277 | 6.99E-06 | pDC | Ern1 |
| Tmco61 | 3.93E-10 | 2.10579821 | 0.307 | 0.133 | 7.59E-06 | pDC | Tmco6 |
| Mapkapk22 | 5.87E-10 | 1.08649059 | 0.803 | 0.692 | 1.14E-05 | pDC | Mapkapk2 |
| R3hdm4 | 5.99E-10 | 1.51260692 | 0.708 | 0.522 | 1.16E-05 | pDC | R3hdm4 |
| Cnot81 | 6.08E-10 | 1.57149569 | 0.533 | 0.309 | 1.18E-05 | pDC | Cnot8 |
| Tmed8 | 6.30E-10 | 1.83107252 | 0.314 | 0.132 | 1.22E-05 | pDC | Tmed8 |
| Nr3c1 | 9.05E-10 | 1.65742289 | 0.46 | 0.245 | 1.75E-05 | pDC | Nr3c1 |
| Vars2 | 1.17E-09 | 1.42212398 | 0.679 | 0.404 | 2.26E-05 | pDC | Vars |
| Cirbp1 | 1.37E-09 | 1.17250223 | 0.577 | 0.367 | 2.64E-05 | pDC | Cirbp |
| Sf11 | 1.37E-09 | 1.21581506 | 0.423 | 0.232 | 2.65E-05 | pDC | Sf1 |
| Trmt2a | 1.47E-09 | 1.59840284 | 0.401 | 0.232 | 2.85E-05 | pDC | Trmt2a |
| Clec2i | 1.48E-09 | 1.49078616 | 0.285 | 0.146 | 2.86E-05 | pDC | Clec2i |
| Macf12 | 1.48E-09 | 1.35448668 | 0.584 | 0.419 | 2.86E-05 | pDC | Macf1 |
| Hadhb1 | 1.67E-09 | 1.70504514 | 0.511 | 0.333 | 3.22E-05 | pDC | Hadhb |
| Rubcn1 | 1.97E-09 | 1.45202944 | 0.328 | 0.179 | 3.81E-05 | pDC | Rubcn |
| Vps13a | 2.03E-09 | 1.80485221 | 0.343 | 0.144 | 3.92E-05 | pDC | Vps13a |
| Nudcd31 | 2.09E-09 | 1.19490166 | 0.526 | 0.335 | 4.04E-05 | pDC | Nudcd3 |
| Tlr121 | 2.54E-09 | 1.98166132 | 0.307 | 0.093 | 4.91E-05 | pDC | Tlr12 |
| Gnas1 | 2.92E-09 | 0.97965135 | 0.818 | 0.73 | 5.64E-05 | pDC | Gnas |
| Itm2b | 3.25E-09 | 1.0607062 | 0.927 | 0.839 | 6.28E-05 | pDC | Itm2b |
| Foxp11 | 3.54E-09 | 1.24182706 | 0.606 | 0.412 | 6.84E-05 | pDC | Foxp1 |
| Med101 | 3.86E-09 | 1.49202548 | 0.423 | 0.292 | 7.47E-05 | pDC | Med10 |
| Ier52 | 3.89E-09 | 0.96769966 | 0.949 | 0.762 | 7.52E-05 | pDC | Ier5 |
| Prr132 | 4.22E-09 | 1.18802522 | 0.635 | 0.453 | 8.17E-05 | pDC | Prr13 |
| Stat21 | 4.30E-09 | 1.6073855 | 0.489 | 0.302 | 8.31E-05 | pDC | Stat2 |
| Dedd2 | 4.36E-09 | 2.27739235 | 0.328 | 0.148 | 8.42E-05 | pDC | Dedd2 |
| Chst121 | 4.37E-09 | 1.76990252 | 0.445 | 0.224 | 8.45E-05 | pDC | Chst12 |
| Nr2c2ap2 | 4.67E-09 | 1.68195018 | 0.445 | 0.259 | 9.02E-05 | pDC | Nr2c2ap |
| E2f4 | 9.12E-09 | 1.29036413 | 0.591 | 0.485 | 0.00017627 | pDC | E2f4 |
| Map3k8 | 9.20E-09 | 1.33032459 | 0.431 | 0.248 | 0.00017795 | pDC | Map3k8 |
| Trim25 | 9.90E-09 | 1.13067429 | 0.759 | 0.588 | 0.00019132 | pDC | Trim25 |
| Dmac2 | 1.34E-08 | 1.71841537 | 0.292 | 0.152 | 0.0002595 | pDC | Dmac2 |
| Clcn5 | 1.35E-08 | 2.29655815 | 0.299 | 0.127 | 0.0002601 | pDC | Clcn5 |
| Slc9a11 | 1.39E-08 | 1.65857089 | 0.467 | 0.262 | 0.00026846 | pDC | Slc9a1 |
| Rab3gap1 | 1.50E-08 | 1.61495786 | 0.387 | 0.235 | 0.00029014 | pDC | Rab3gap1 |
| Cd822 | 1.67E-08 | 1.14341302 | 0.715 | 0.564 | 0.00032367 | pDC | Cd82 |
| Sh2d3c1 | 1.74E-08 | 1.94542338 | 0.263 | 0.106 | 0.00033686 | pDC | Sh2d3c |
| Arhgap271 | 1.80E-08 | 1.46416302 | 0.496 | 0.285 | 0.00034758 | pDC | Arhgap27 |
| Surf42 | 1.82E-08 | 1.26771435 | 0.708 | 0.536 | 0.00035236 | pDC | Surf4 |
| Dock21 | 1.99E-08 | 0.99738322 | 0.752 | 0.609 | 0.00038424 | pDC | Dock2 |
| Emc102 | 2.02E-08 | 1.42744039 | 0.606 | 0.437 | 0.00039062 | pDC | Emc10 |
| Crebrf | 2.27E-08 | 1.75748575 | 0.35 | 0.191 | 0.00043899 | pDC | Crebrf |
| Fam122a | 2.28E-08 | 1.97589549 | 0.27 | 0.12 | 0.00044048 | pDC | Fam122a |
| Scpep11 | 2.35E-08 | 1.18956927 | 0.81 | 0.62 | 0.00045519 | pDC | Scpep1 |
| Ccdc97 | 2.68E-08 | 1.36158907 | 0.46 | 0.279 | 0.0005187 | pDC | Ccdc97 |
| Mndal | 3.19E-08 | 1.59194231 | 0.285 | 0.148 | 0.00061634 | pDC | Mndal |
| Srgap32 | 3.41E-08 | 1.9179683 | 0.38 | 0.195 | 0.00065847 | pDC | Srgap3 |
| Pip5k1c1 | 3.56E-08 | 1.07173294 | 0.81 | 0.723 | 0.00068923 | pDC | Pip5k1c |
| H2afy2 | 4.66E-08 | 0.30686885 | 0.898 | 0.863 | 0.00090077 | pDC | H2afy |

| | | | | | | | |
|---|---|---|---|---|---|---|---|
| Ptms2 | 4.85E-08 | 0.32269615 | 0.964 | 0.896 | 0.00093688 | pDC | Ptms |
| Adss | 5.62E-08 | 1.43508883 | 0.496 | 0.336 | 0.00108618 | pDC | Adss |
| Jund2 | 7.49E-08 | 0.39598667 | 0.876 | 0.812 | 0.00144823 | pDC | Jund |
| Bscl22 | 8.19E-08 | 1.62571339 | 0.489 | 0.356 | 0.00158396 | pDC | Bscl2 |
| Slc50a1 | 8.52E-08 | 1.61324273 | 0.35 | 0.177 | 0.00164644 | pDC | Slc50a1 |
| Parp81 | 1.01E-07 | 1.24729848 | 0.504 | 0.301 | 0.00194377 | pDC | Parp8 |
| Ctcf1 | 1.03E-07 | 1.45744795 | 0.38 | 0.271 | 0.00198872 | pDC | Ctcf |
| Zfp280d | 1.10E-07 | 1.76965693 | 0.299 | 0.143 | 0.00211709 | pDC | Zfp280d |
| Cd300lf3 | 1.13E-07 | 1.25817078 | 0.547 | 0.38 | 0.00218447 | pDC | Cd300lf |
| Aacs1 | 1.21E-07 | 1.58396043 | 0.372 | 0.209 | 0.00233263 | pDC | Aacs |
| Znrf12 | 1.30E-07 | 1.32664632 | 0.409 | 0.245 | 0.00250595 | pDC | Znrf1 |
| Reep52 | 1.55E-07 | 1.02828227 | 0.737 | 0.616 | 0.00299086 | pDC | Reep5 |
| Abhd17a1 | 1.69E-07 | 1.66479079 | 0.65 | 0.399 | 0.00327323 | pDC | Abhd17a |
| Sqle | 1.79E-07 | 1.34865936 | 0.416 | 0.26 | 0.00345495 | pDC | Sqle |
| Sdc42 | 1.94E-07 | 1.74195911 | 0.526 | 0.305 | 0.00376009 | pDC | Sdc4 |
| Fam234b2 | 1.96E-07 | 1.73697974 | 0.372 | 0.178 | 0.00379391 | pDC | Fam234b |
| Sh3bp42 | 2.26E-07 | 1.13260578 | 0.343 | 0.19 | 0.00436166 | pDC | Sh3bp4 |
| Ostm1 | 2.49E-07 | 1.66111627 | 0.372 | 0.223 | 0.00481214 | pDC | Ostm1 |
| Mef2d2 | 2.62E-07 | 1.33106938 | 0.686 | 0.528 | 0.00506992 | pDC | Mef2d |
| Sash32 | 2.66E-07 | 1.68080612 | 0.511 | 0.295 | 0.00513708 | pDC | Sash3 |
| Prps2 | 2.70E-07 | 2.11099262 | 0.255 | 0.116 | 0.00522056 | pDC | Prps2 |
| Rufy11 | 2.77E-07 | 1.73816655 | 0.489 | 0.275 | 0.00534956 | pDC | Rufy1 |
| Abi32 | 2.88E-07 | 1.45974191 | 0.642 | 0.407 | 0.00556557 | pDC | Abi3 |
| Rmnd5b1 | 3.28E-07 | 1.72023708 | 0.409 | 0.221 | 0.00634437 | pDC | Rmnd5b |
| Wdr81 | 3.57E-07 | 1.62572942 | 0.314 | 0.188 | 0.00690972 | pDC | Wdr81 |
| Coro71 | 4.31E-07 | 1.18540669 | 0.613 | 0.405 | 0.00833651 | pDC | Coro7 |
| P4hb2 | 4.57E-07 | 0.26274846 | 0.54 | 0.592 | 0.00884493 | pDC | P4hb |
| AW549877 | 4.65E-07 | 1.67228335 | 0.285 | 0.131 | 0.00899146 | pDC | AW549877 |
| Sap30bp | 5.36E-07 | 1.53865145 | 0.328 | 0.19 | 0.01037266 | pDC | Sap30bp |
| Slc25a281 | 5.87E-07 | 1.41343882 | 0.387 | 0.241 | 0.01135658 | pDC | Slc25a28 |
| Abr2 | 7.21E-07 | 1.27155072 | 0.679 | 0.523 | 0.01393771 | pDC | Abr |
| Cerk1 | 7.38E-07 | 1.76055844 | 0.38 | 0.206 | 0.01426748 | pDC | Cerk |
| Gltp1 | 8.44E-07 | 1.15302432 | 0.796 | 0.592 | 0.01631383 | pDC | Gltp |
| Dcaf71 | 8.68E-07 | 1.20318797 | 0.65 | 0.502 | 0.01678488 | pDC | Dcaf7 |
| Me21 | 8.98E-07 | 1.45648851 | 0.453 | 0.274 | 0.01737138 | pDC | Me2 |
| H2-M31 | 9.08E-07 | 1.12314126 | 0.54 | 0.412 | 0.01756209 | pDC | H2-M3 |
| Trim241 | 1.02E-06 | 1.20096474 | 0.328 | 0.187 | 0.01970525 | pDC | Trim24 |
| Zxdb | 1.09E-06 | 1.50062324 | 0.255 | 0.133 | 0.02109264 | pDC | Zxdb |
| Fdft11 | 1.12E-06 | 1.39494468 | 0.409 | 0.255 | 0.02167185 | pDC | Fdft1 |
| AI467606 | 1.32E-06 | 2.02382045 | 0.255 | 0.12 | 0.02546341 | pDC | AI467606 |
| Dnajc32 | 1.35E-06 | 1.48819666 | 0.657 | 0.437 | 0.02610523 | pDC | Dnajc3 |
| Calr2 | 1.49E-06 | 0.32885107 | 0.642 | 0.611 | 0.02884524 | pDC | Calr |
| Hhex1 | 1.55E-06 | 0.32062997 | 0.518 | 0.487 | 0.02987458 | pDC | Hhex |
| Srsf51 | 1.55E-06 | 1.10199709 | 0.766 | 0.638 | 0.02995688 | pDC | Srsf5 |
| Irgm11 | 1.78E-06 | 1.6372537 | 0.343 | 0.177 | 0.03433174 | pDC | Irgm1 |
| Dpm31 | 1.82E-06 | 1.67971437 | 0.394 | 0.234 | 0.03523343 | pDC | Dpm3 |
| Ptbp11 | 1.87E-06 | 0.32764747 | 0.693 | 0.684 | 0.03621129 | pDC | Ptbp1 |
| Selenos1 | 2.07E-06 | 1.46460443 | 0.482 | 0.306 | 0.04010487 | pDC | Selenos |
| Mumo11 | 2.08E-06 | 1.45744597 | 0.416 | 0.252 | 0.04030626 | pDC | Mumo1 |
| Prkag2 | 2.57E-06 | 1.56291855 | 0.423 | 0.201 | 0.04966494 | pDC | Prkag2 |
| Rbmx | 2.62E-06 | 1.52659807 | 0.285 | 0.131 | 0.05073747 | pDC | Rbmx |
| Pdia43 | 2.66E-06 | 1.39761408 | 0.613 | 0.396 | 0.05152052 | pDC | Pdia4 |
| Tagln22 | 2.81E-06 | 0.44362029 | 0.971 | 0.892 | 0.05425082 | pDC | Tagln2 |
| Fdps1 | 2.85E-06 | 1.23124994 | 0.511 | 0.386 | 0.05516098 | pDC | Fdps |
| Txndc11 | 3.21E-06 | 1.71472597 | 0.584 | 0.347 | 0.06201818 | pDC | Txndc11 |
| Atf7ip1 | 3.54E-06 | 1.15293319 | 0.46 | 0.328 | 0.06845635 | pDC | Atf7ip |
| Ilf3 | 3.66E-06 | 1.23407196 | 0.423 | 0.274 | 0.07080042 | pDC | Ilf3 |
| Ppp6r11 | 3.92E-06 | 1.17198949 | 0.701 | 0.577 | 0.07581847 | pDC | Ppp6r1 |
| Cdk5rap32 | 4.60E-06 | 1.5939903 | 0.489 | 0.323 | 0.08899619 | pDC | Cdk5rap3 |
| Rexo22 | 4.80E-06 | 1.20738231 | 0.577 | 0.427 | 0.09274372 | pDC | Rexo2 |
| Rmnd5a1 | 4.81E-06 | 1.18582443 | 0.467 | 0.328 | 0.09290393 | pDC | Rmnd5a |
| Dtx22 | 4.83E-06 | 1.56487514 | 0.365 | 0.2 | 0.09336821 | pDC | Dtx2 |
| Alox5ap3 | 5.18E-06 | 0.42861467 | 0.861 | 0.639 | 0.10013379 | pDC | Alox5ap |
| Cox4i1 | 5.96E-06 | 0.34373821 | 0.547 | 0.567 | 0.11516084 | pDC | Cox4i1 |
| Cbx42 | 6.02E-06 | 1.46169048 | 0.409 | 0.235 | 0.11635675 | pDC | Cbx4 |
| Stx72 | 6.71E-06 | 1.09272771 | 0.788 | 0.679 | 0.12978417 | pDC | Stx7 |
| Prkcb1 | 6.72E-06 | 1.39578093 | 0.416 | 0.225 | 0.13000602 | pDC | Prkcb |
| Sp1001 | 7.83E-06 | 1.14994662 | 0.511 | 0.315 | 0.15137209 | pDC | Sp100 |
| Tgfbr21 | 9.29E-06 | 1.35493686 | 0.54 | 0.338 | 0.17957106 | pDC | Tgfbr2 |
| Pmepa11 | 1.15E-05 | 1.50620455 | 0.533 | 0.324 | 0.22329155 | pDC | Pmepa1 |
| Mlec2 | 1.21E-05 | 0.42319831 | 0.599 | 0.54 | 0.23471976 | pDC | Mlec |
| Retreg3 | 1.23E-05 | 1.44897862 | 0.416 | 0.295 | 0.23730497 | pDC | Retreg3 |
| Xxylt11 | 1.36E-05 | 1.65327595 | 0.299 | 0.173 | 0.26261138 | pDC | Xxylt1 |
| Lmf1 | 1.38E-05 | 1.45388304 | 0.292 | 0.19 | 0.26595816 | pDC | Lmf1 |
| Uso11 | 1.46E-05 | 1.24691091 | 0.423 | 0.302 | 0.28319242 | pDC | Uso1 |
| Snrpf2 | 1.51E-05 | 0.26923083 | 0.445 | 0.546 | 0.29199613 | pDC | Snrpf |
| Gatad11 | 1.60E-05 | 1.35561781 | 0.416 | 0.274 | 0.30859046 | pDC | Gatad1 |
| Frmd4a | 1.65E-05 | 1.17739946 | 0.409 | 0.274 | 0.31869739 | pDC | Frmd4a |
| Armcx3 | 1.69E-05 | 1.54052043 | 0.255 | 0.143 | 0.32746167 | pDC | Armcx3 |
| Gpr1461 | 1.72E-05 | 1.51227067 | 0.343 | 0.184 | 0.33242702 | pDC | Gpr146 |
| Trappc51 | 1.72E-05 | 1.23312075 | 0.409 | 0.28 | 0.33275693 | pDC | Trappc5 |
| Rmc1 | 1.83E-05 | 1.80577443 | 0.307 | 0.166 | 0.35448522 | pDC | Rmc1 |
| Smim15 | 1.89E-05 | 1.52530222 | 0.277 | 0.172 | 0.36585631 | pDC | Smim15 |
| Eef21 | 2.00E-05 | 0.43687706 | 0.832 | 0.811 | 0.38633407 | pDC | Eef2 |
| Kat6b1 | 2.16E-05 | 1.58374867 | 0.299 | 0.174 | 0.41697236 | pDC | Kat6b |
| Paip1 | 2.26E-05 | 1.2897586 | 0.401 | 0.291 | 0.43759318 | pDC | Paip1 |
| Krtcap2 | 2.31E-05 | 1.39830881 | 0.285 | 0.183 | 0.44600237 | pDC | Krtcap2 |
| Inpp5k | 2.36E-05 | 1.53090026 | 0.277 | 0.175 | 0.4558745 | pDC | Inpp5k |
| Alg1 | 2.60E-05 | 1.78346921 | 0.277 | 0.137 | 0.50336023 | pDC | Alg1 |
| Eef2k1 | 2.65E-05 | 1.69798812 | 0.343 | 0.177 | 0.5130006 | pDC | Eef2k |
| Ipo91 | 2.88E-05 | 1.60817953 | 0.358 | 0.226 | 0.55604347 | pDC | Ipo9 |
| Itgb23 | 2.92E-05 | 0.50850381 | 0.891 | 0.697 | 0.5653194 | pDC | Itgb2 |
| Dbnl2 | 3.06E-05 | 0.31905289 | 0.825 | 0.771 | 0.59095747 | pDC | Dbnl |
| Axl2 | 3.09E-05 | 1.2404036 | 0.387 | 0.242 | 0.59671767 | pDC | Axl2 |
| Trim30a2 | 3.12E-05 | 1.25548651 | 0.416 | 0.281 | 0.60396359 | pDC | Trim30a |
| Gpcpd11 | 3.23E-05 | 1.1335077 | 0.642 | 0.43 | 0.62356417 | pDC | Gpcpd1 |
| Inpp5d1 | 3.29E-05 | 0.29770679 | 0.672 | 0.686 | 0.63667372 | pDC | Inpp5d |
| Slc30a52 | 3.34E-05 | 1.52249457 | 0.409 | 0.247 | 0.6453084 | pDC | Slc30a5 |
| Spata131 | 3.39E-05 | 1.56109192 | 0.292 | 0.16 | 0.65525878 | pDC | Spata13 |
| Atp1a12 | 3.51E-05 | 0.98432231 | 0.898 | 0.799 | 0.67902079 | pDC | Atp1a1 |
| Epn12 | 3.68E-05 | 0.3712047 | 0.774 | 0.705 | 0.71172503 | pDC | Epn1 |
| Exoc41 | 3.81E-05 | 1.02504166 | 0.489 | 0.346 | 0.73616254 | pDC | Exoc4 |
| Ubl7 | 3.88E-05 | 1.15673728 | 0.431 | 0.344 | 0.74997556 | pDC | Ubl7 |
| Rnf132 | 4.01E-05 | 1.25097091 | 0.511 | 0.345 | 0.77470228 | pDC | Rnf13 |
| Wrn | 4.09E-05 | 1.22097407 | 0.307 | 0.196 | 0.79160556 | pDC | Wrn |
| Ubqln1 | 4.23E-05 | 0.40632671 | 0.438 | 0.508 | 0.8174213 | pDC | Ubqln1 |
| Mbnl12 | 4.36E-05 | 1.29142296 | 0.693 | 0.516 | 0.84225686 | pDC | Mbnl1 |
| Mettl23 | 4.51E-05 | 1.70141312 | 0.27 | 0.158 | 0.87262166 | pDC | Mettl23 |
| Tspyl11 | 5.07E-05 | 1.23904975 | 0.467 | 0.335 | 0.98055764 | pDC | Tspyl1 |
| Mat2b | 5.32E-05 | 1.64830246 | 0.307 | 0.194 | 1 | pDC | Mat2b |
| Jaml3 | 5.72E-05 | 1.11736548 | 0.781 | 0.578 | 1 | pDC | Jaml |
| Ube2g21 | 5.76E-05 | 0.25919321 | 0.299 | 0.402 | 1 | pDC | Ube2g2 |
| Pkp3 | 6.04E-05 | 1.07127841 | 0.336 | 0.217 | 1 | pDC | Pkp3 |
| Fryl | 6.42E-05 | 1.00822498 | 0.555 | 0.37 | 1 | pDC | Fryl |
| Ypel31 | 6.65E-05 | 1.16559019 | 0.423 | 0.291 | 1 | pDC | Ypel3 |
| Mpnd | 6.93E-05 | 1.15296161 | 0.496 | 0.337 | 1 | pDC | Mpnd |
| Slc3a22 | 8.27E-05 | 0.52755454 | 0.781 | 0.736 | 1 | pDC | Slc3a2 |
| Etnk11 | 8.91E-05 | 0.73196112 | 0.701 | 0.534 | 1 | pDC | Etnk1 |
| Lrrc471 | 9.55E-05 | 1.27608429 | 0.46 | 0.32 | 1 | pDC | Lrrc47 |
| Susd62 | 9.66E-05 | 1.17084763 | 0.547 | 0.432 | 1 | pDC | Susd6 |
| Tbc1d10c1 | 0.00010036 | 0.99275702 | 0.328 | 0.232 | 1 | pDC | Tbc1d10c |
| Apobec32 | 0.00010048 | 0.90238415 | 0.803 | 0.664 | 1 | pDC | Apobec3 |
| Snrpd12 | 0.0001005 | 0.29771454 | 0.292 | 0.366 | 1 | pDC | Snrpd1 |
| Ssr12 | 0.00010556 | 1.11506618 | 0.752 | 0.611 | 1 | pDC | Ssr1 |
| D8Ertd738e1 | 0.00010606 | 0.25983867 | 0.511 | 0.56 | 1 | pDC | D8Ertd738e |
| Dnajc91 | 0.00010893 | 1.39987155 | 0.343 | 0.163 | 1 | pDC | Dnajc9 |
| Tgfbr12 | 0.0001191 | 1.37862526 | 0.401 | 0.265 | 1 | pDC | Tgfbr1 |
| Hmgcr1 | 0.00012646 | 1.1288751 | 0.708 | 0.543 | 1 | pDC | Hmgcr |
| Cnpy31 | 0.00012819 | 1.05264516 | 0.672 | 0.487 | 1 | pDC | Cnpy3 |
| Psip11 | 0.00012898 | 0.32137238 | 0.146 | 0.261 | 1 | pDC | Psip1 |
| Wapl | 0.00014529 | 0.25482604 | 0.263 | 0.361 | 1 | pDC | Wapl |
| Ppm1m2 | 0.00015153 | 0.39344276 | 0.635 | 0.616 | 1 | pDC | Ppm1m |
| Ahnak1 | 0.00016195 | 0.78422721 | 0.788 | 0.485 | 1 | pDC | Ahnak |
| Fam91a11 | 0.00016355 | 0.31170397 | 0.292 | 0.427 | 1 | pDC | Fam91a1 |
| H2-D1 | 0.00016414 | 0.98150221 | 0.73 | 0.602 | 1 | pDC | H2-D1 |
| Dazap11 | 0.00017358 | 0.30793219 | 0.599 | 0.625 | 1 | pDC | Dazap1 |
| Arhgef101 | 0.00017724 | 1.30718779 | 0.35 | 0.197 | 1 | pDC | Arhgef10 |
| Atp6v1h1 | 0.00018294 | 1.15173821 | 0.467 | 0.337 | 1 | pDC | Atp6v1h |
| Lgals12 | 0.00019389 | 1.6584832 | 0.372 | 0.2 | 1 | pDC | Lgals1 |
| Chic2 | 0.00020502 | 1.40808217 | 0.321 | 0.19 | 1 | pDC | Chic2 |
| Srsf13 | 0.00021007 | 0.3994001 | 0.46 | 0.535 | 1 | pDC | Srsf1 |
| Map4k51 | 0.00022826 | 1.7573091 | 0.299 | 0.167 | 1 | pDC | Map4k5 |
| Nfkbid2 | 0.00023154 | 0.9972768 | 0.591 | 0.469 | 1 | pDC | Nfkbid |
| Ecd | 0.00023738 | 1.3311695 | 0.292 | 0.185 | 1 | pDC | Ecd |
| Atp2a31 | 0.0002402 | 1.22417419 | 0.416 | 0.272 | 1 | pDC | Atp2a3 |
| Slc7a6 | 0.0002429 | 1.49908863 | 0.255 | 0.151 | 1 | pDC | Slc7a6 |
| Usf2 | 0.00025883 | 0.32718075 | 0.547 | 0.603 | 1 | pDC | Usf2 |
| N4bp2l21 | 0.00025921 | 1.32356322 | 0.394 | 0.286 | 1 | pDC | N4bp2l2 |
| Pik3ap12 | 0.0002646 | 1.13932202 | 0.898 | 0.604 | 1 | pDC | Pik3ap1 |
| Slc25a42 | 0.00026739 | 0.28369557 | 0.292 | 0.387 | 1 | pDC | Slc25a4 |
| Scimp2 | 0.00027569 | 1.48673411 | 0.511 | 0.298 | 1 | pDC | Scimp |
| Fcer1g2 | 0.0002805 | 0.54422724 | 0.92 | 0.693 | 1 | pDC | Fcer1g |
| Rad54l21 | 0.00029467 | 1.36458266 | 0.263 | 0.159 | 1 | pDC | Rad54l2 |
| Mdh11 | 0.00031138 | 0.36216174 | 0.474 | 0.512 | 1 | pDC | Mdh1 |
| Il10rb1 | 0.00032345 | 1.09308298 | 0.533 | 0.417 | 1 | pDC | Il10rb |
| Zbtb381 | 0.00032693 | 1.14837349 | 0.27 | 0.177 | 1 | pDC | Zbtb38 |
| Map4k22 | 0.00033707 | 1.38145233 | 0.358 | 0.242 | 1 | pDC | Map4k2 |
| Gnptab1 | 0.0003392 | 1.09357555 | 0.511 | 0.341 | 1 | pDC | Gnptab |
| H2-Q41 | 0.00037062 | 0.96918272 | 0.35 | 0.191 | 1 | pDC | H2-Q4 |
| Dmxl1 | 0.00037428 | 1.56725672 | 0.336 | 0.202 | 1 | pDC | Dmxl1 |
| Setd21 | 0.00038144 | 1.07145564 | 0.445 | 0.365 | 1 | pDC | Setd2 |
| Asah12 | 0.00038608 | 0.3785103 | 0.445 | 0.525 | 1 | pDC | Asah1 |
| Lrch11 | 0.00042411 | 1.17881437 | 0.474 | 0.364 | 1 | pDC | Lrch1 |
| Cdh12 | 0.00044097 | 0.54817777 | 0.708 | 0.449 | 1 | pDC | Cdh1 |
| Ly862 | 0.00049359 | 1.11706889 | 0.664 | 0.529 | 1 | pDC | Ly86 |
| Zc3h12d | 0.00051724 | 1.3405762 | 0.292 | 0.158 | 1 | pDC | Zc3h12d |
| Ndufv31 | 0.00052556 | 0.25772337 | 0.453 | 0.523 | 1 | pDC | Ndufv3 |
| Trabd1 | 0.00055626 | 1.39595758 | 0.431 | 0.3 | 1 | pDC | Trabd |
| Kdelr1 | 0.00059131 | 1.0521198 | 0.547 | 0.417 | 1 | pDC | Kdelr1 |
| Bri3bp2 | 0.00060149 | 0.27036132 | 0.526 | 0.544 | 1 | pDC | Bri3bp |
| Dgkd2 | 0.00060636 | 1.10341403 | 0.46 | 0.325 | 1 | pDC | Dgkd |
| Qk | 0.00066259 | 0.33430658 | 0.65 | 0.674 | 1 | pDC | Qk |
| Dpp8 | 0.00068515 | 0.35851343 | 0.234 | 0.354 | 1 | pDC | Dpp8 |
| Mtss11 | 0.00070572 | 0.42249096 | 0.467 | 0.472 | 1 | pDC | Mtss1 |
| Pdia31 | 0.0007179 | 0.35218168 | 0.599 | 0.563 | 1 | pDC | Pdia3 |
| Atp1b32 | 0.00076577 | 1.09340262 | 0.511 | 0.444 | 1 | pDC | Atp1b3 |
| Arap12 | 0.00083752 | 1.17904501 | 0.518 | 0.278 | 1 | pDC | Arap1 |
| Strn31 | 0.00088087 | 1.29536108 | 0.401 | 0.279 | 1 | pDC | Strn3 |
| Leprotl11 | 0.00090983 | 1.26890238 | 0.46 | 0.348 | 1 | pDC | Leprotl1 |
| Ostc2 | 0.00092585 | 0.29789374 | 0.307 | 0.388 | 1 | pDC | Ostc |
| Erg28 | 0.0009491 | 1.36319936 | 0.314 | 0.18 | 1 | pDC | Erg28 |
| Lbh1 | 0.00095192 | 1.42944902 | 0.489 | 0.34 | 1 | pDC | Lbh |
| Tmem106b | 0.00098384 | 1.15949513 | 0.285 | 0.182 | 1 | pDC | Tmem106b |
| Atl2 | 0.00098444 | 1.39998926 | 0.255 | 0.15 | 1 | pDC | Atl2 |
| BC0055371 | 0.00102821 | 0.36544142 | 0.752 | 0.74 | 1 | pDC | BC005537 |
| Ankle21 | 0.00104621 | 1.20815443 | 0.343 | 0.238 | 1 | pDC | Ankle2 |
| Gsr1 | 0.00105211 | 0.69401199 | 0.518 | 0.446 | 1 | pDC | Gsr |
| Clcn3 | 0.00106511 | 1.63175883 | 0.277 | 0.168 | 1 | pDC | Clcn3 |
| Plxdc21 | 0.00113966 | 1.73325531 | 0.285 | 0.104 | 1 | pDC | Plxdc2 |
| Fam193b | 0.00119875 | 0.89007934 | 0.328 | 0.229 | 1 | pDC | Fam193b |
| Anks31 | 0.00126188 | 1.45045545 | 0.409 | 0.277 | 1 | pDC | Anks3 |
| U2af2 | 0.00126278 | 0.26154798 | 0.613 | 0.645 | 1 | pDC | U2af2 |
| Fcho11 | 0.0012966 | 1.27596004 | 0.365 | 0.273 | 1 | pDC | Fcho1 |
| Tmed21 | 0.00130841 | 1.14050317 | 0.577 | 0.42 | 1 | pDC | Tmed2 |
| Etfb2 | 0.00133374 | 0.29234822 | 0.401 | 0.454 | 1 | pDC | Etfb |
| Mesd | 0.00135915 | 1.55233342 | 0.255 | 0.152 | 1 | pDC | Mesd |
| Fbl2 | 0.00139379 | 0.30392699 | 0.394 | 0.449 | 1 | pDC | Fbl |
| Pnisr | 0.00143929 | 0.68093671 | 0.416 | 0.329 | 1 | pDC | Pnisr |
| Creb31 | 0.00145745 | 1.30209843 | 0.328 | 0.223 | 1 | pDC | Creb3 |
| Mgrn1 | 0.0014599 | 0.29950946 | 0.328 | 0.421 | 1 | pDC | Mgrn1 |
| Coq21 | 0.0016028 | 1.39110716 | 0.27 | 0.195 | 1 | pDC | Coq2 |
| Ctnnd22 | 0.00160973 | 0.71206686 | 0.409 | 0.331 | 1 | pDC | Ctnnd2 |
| Fndc3b2 | 0.00163865 | 1.07452535 | 0.35 | 0.25 | 1 | pDC | Fndc3b |
| Cyth42 | 0.00168913 | 1.04011814 | 0.869 | 0.629 | 1 | pDC | Cyth4 |
| Gaa1 | 0.00171318 | 1.36411564 | 0.409 | 0.281 | 1 | pDC | Gaa |
| Itpkb1 | 0.00173727 | 0.96127445 | 0.328 | 0.234 | 1 | pDC | Itpkb |
| Rapgef12 | 0.00176907 | 1.3033798 | 0.562 | 0.398 | 1 | pDC | Rapgef1 |
| Rab102 | 0.00179749 | 0.29636636 | 0.321 | 0.402 | 1 | pDC | Rab10 |
| Haao | 0.00183497 | 1.1464032 | 0.358 | 0.222 | 1 | pDC | Haao |
| Asph1 | 0.00185041 | 1.32912758 | 0.409 | 0.23 | 1 | pDC | Asph |
| Rnf1671 | 0.00187689 | 1.16157038 | 0.321 | 0.214 | 1 | pDC | Rnf167 |
| Atf62 | 0.00187972 | 0.99611913 | 0.511 | 0.397 | 1 | pDC | Atf6 |
| Copz11 | 0.00188728 | 0.29008718 | 0.555 | 0.592 | 1 | pDC | Copz1 |
| Npepps1 | 0.00200389 | 0.27083437 | 0.161 | 0.264 | 1 | pDC | Npepps |
| Rassf53 | 0.00202177 | 0.33939873 | 0.569 | 0.56 | 1 | pDC | Rassf5 |
| Prpf38b1 | 0.00202392 | 0.32133702 | 0.416 | 0.484 | 1 | pDC | Prpf38b |
| Ddx17 | 0.00202897 | 0.62087177 | 0.803 | 0.762 | 1 | pDC | Ddx17 |
| Slc39a71 | 0.00203867 | 1.10586832 | 0.423 | 0.329 | 1 | pDC | Slc39a7 |
| Ttc141 | 0.00208703 | 1.11017081 | 0.328 | 0.235 | 1 | pDC | Ttc14 |
| Ehmt2 | 0.00211881 | 0.31725326 | 0.54 | 0.585 | 1 | pDC | Ehmt2 |
| Unc13d | 0.00223625 | 1.32661446 | 0.38 | 0.298 | 1 | pDC | Unc13d |
| Snu131 | 0.00228759 | 0.43418418 | 0.453 | 0.485 | 1 | pDC | Snu13 |
| Hp1bp31 | 0.00230573 | 1.00499571 | 0.584 | 0.449 | 1 | pDC | Hp1bp3 |
| Trp532 | 0.00234637 | 0.28313767 | 0.336 | 0.402 | 1 | pDC | Trp53 |
| Bmyc | 0.00236094 | 0.9712755 | 0.409 | 0.289 | 1 | pDC | Bmyc |
| Hibadh1 | 0.00241031 | 1.06972026 | 0.467 | 0.33 | 1 | pDC | Hibadh |
| Rassf41 | 0.00245095 | 0.98182668 | 0.606 | 0.424 | 1 | pDC | Rassf4 |
| Fermt33 | 0.00245583 | 0.40113322 | 0.599 | 0.61 | 1 | pDC | Fermt3 |
| Flt32 | 0.0024597 | 0.74781062 | 0.474 | 0.345 | 1 | pDC | Flt3 |
| Phactr21 | 0.00259417 | 1.33030959 | 0.416 | 0.277 | 1 | pDC | Phactr2 |
| Phf2 | 0.00260379 | 0.9882665 | 0.343 | 0.263 | 1 | pDC | Phf2 |
| Atxn2l | 0.00262111 | 0.54632654 | 0.606 | 0.611 | 1 | pDC | Atxn2l |
| Mycl1 | 0.00263462 | 1.21987387 | 0.343 | 0.222 | 1 | pDC | Mycl |
| Slc12a62 | 0.00265808 | 1.42559118 | 0.343 | 0.227 | 1 | pDC | Slc12a6 |
| Acsl11 | 0.00267483 | 1.15034259 | 0.285 | 0.204 | 1 | pDC | Acsl1 |
| Tspyl2 | 0.00267717 | 1.36945857 | 0.255 | 0.144 | 1 | pDC | Tspyl2 |
| Ndufb9 | 0.0027413 | 0.29816449 | 0.431 | 0.482 | 1 | pDC | Ndufb9 |
| Nasp1 | 0.00274377 | 0.25299949 | 0.197 | 0.26 | 1 | pDC | Nasp |
| Adssl12 | 0.00276927 | 1.25096633 | 0.401 | 0.26 | 1 | pDC | Adssl1 |
| Cib1 | 0.00279254 | 1.49368731 | 0.328 | 0.212 | 1 | pDC | Cib1 |
| Rrbp11 | 0.00285569 | 0.95370757 | 0.781 | 0.65 | 1 | pDC | Rrbp1 |
| Baz2a | 0.00296563 | 1.15796175 | 0.54 | 0.414 | 1 | pDC | Baz2a |
| Jtb1 | 0.00298395 | 0.34655058 | 0.387 | 0.378 | 1 | pDC | Jtb |
| Brd4 | 0.0030277 | 0.27436561 | 0.591 | 0.67 | 1 | pDC | Brd4 |
| Mtdh1 | 0.00313338 | 1.15059819 | 0.489 | 0.352 | 1 | pDC | Mtdh |
| Sep-91 | 0.00323607 | 0.8910399 | 0.759 | 0.626 | 1 | pDC | 9-Sep |
| Fam107b1 | 0.00325693 | 1.06904437 | 0.569 | 0.432 | 1 | pDC | Fam107b |
| Hmgcs12 | 0.00356092 | 0.75709737 | 0.489 | 0.443 | 1 | pDC | Hmgcs1 |
| Eif5b1 | 0.00372335 | 0.50833994 | 0.204 | 0.284 | 1 | pDC | Eif5b |
| C2cd5 | 0.00379268 | 1.53386566 | 0.27 | 0.162 | 1 | pDC | C2cd5 |
| Zfp36l12 | 0.00396398 | 0.8675359 | 0.693 | 0.553 | 1 | pDC | Zfp36l1 |
| Golga21 | 0.00397808 | 1.49006464 | 0.372 | 0.288 | 1 | pDC | Golga2 |
| Eps151 | 0.00408461 | 0.88876346 | 0.584 | 0.454 | 1 | pDC | Eps15 |
| Zfp292 | 0.00408567 | 1.26301334 | 0.394 | 0.281 | 1 | pDC | Zfp292 |
| Mfsd51 | 0.00415515 | 0.30410827 | 0.175 | 0.257 | 1 | pDC | Mfsd5 |
| Dhx37 | 0.00461373 | 1.3299684 | 0.255 | 0.165 | 1 | pDC | Dhx37 |
| Osgep2 | 0.00465998 | 0.36085998 | 0.255 | 0.325 | 1 | pDC | Osgep |
| Gbp81 | 0.00468386 | 1.06829412 | 0.292 | 0.17 | 1 | pDC | Gbp8 |
| Ints1 | 0.00468494 | 1.31471614 | 0.401 | 0.279 | 1 | pDC | Ints1 |
| Pcnp | 0.00470355 | 0.28731035 | 0.27 | 0.324 | 1 | pDC | Pcnp |
| Nab12 | 0.00470843 | 0.30839475 | 0.504 | 0.538 | 1 | pDC | Nab1 |
| Mindy21 | 0.00478864 | 1.36956365 | 0.277 | 0.182 | 1 | pDC | Mindy2 |
| Tomm51 | 0.00479888 | 0.38850649 | 0.263 | 0.328 | 1 | pDC | Tomm5 |
| Rer11 | 0.00483185 | 0.40833439 | 0.336 | 0.39 | 1 | pDC | Rer1 |
| Arcn11 | 0.00487119 | 0.38542297 | 0.518 | 0.542 | 1 | pDC | Arcn1 |
| Adam192 | 0.00493806 | 0.92578277 | 0.445 | 0.278 | 1 | pDC | Adam19 |
| Pex5 | 0.00498285 | 1.55105741 | 0.255 | 0.152 | 1 | pDC | Pex5 |
| Sdhb2 | 0.00500819 | 0.25076627 | 0.387 | 0.463 | 1 | pDC | Sdhb |
| Ell22 | 0.00516054 | 1.06294069 | 0.533 | 0.405 | 1 | pDC | Ell2 |
| Pttg11 | 0.00518463 | 1.18812274 | 0.255 | 0.174 | 1 | pDC | Pttg1 |
| Nrd11 | 0.00529382 | 0.36216157 | 0.489 | 0.49 | 1 | pDC | Nrd1 |
| Smc1a1 | 0.00532254 | 0.42582781 | 0.307 | 0.349 | 1 | pDC | Smc1a |
| Zfp1312 | 0.00543743 | 1.03593326 | 0.387 | 0.301 | 1 | pDC | Zfp131 |
| Tmem2581 | 0.00552602 | 0.29483193 | 0.467 | 0.484 | 1 | pDC | Tmem258 |
| Vsir1 | 0.00565741 | 0.98600559 | 0.496 | 0.372 | 1 | pDC | Vsir |
| Galnt2 | 0.00574415 | 1.03847487 | 0.445 | 0.322 | 1 | pDC | Galnt2 |
| Dennd4b1 | 0.00575915 | 0.41719129 | 0.445 | 0.507 | 1 | pDC | Dennd4b |
| H2-Ke61 | 0.00588273 | 1.10470556 | 0.285 | 0.227 | 1 | pDC | H2-Ke6 |
| Kynu | 0.0060728 | 0.90982708 | 0.496 | 0.341 | 1 | pDC | Kynu |
| Serbp1 | 0.0061345 | 0.28347324 | 0.489 | 0.536 | 1 | pDC | Serbp1 |
| Eid11 | 0.00617361 | 0.36556023 | 0.599 | 0.601 | 1 | pDC | Eid1 |
| Psmb101 | 0.00631524 | 0.39470509 | 0.474 | 0.502 | 1 | pDC | Psmb10 |
| Dhx9 | 0.00640232 | 1.12024258 | 0.664 | 0.516 | 1 | pDC | Dhx9 |
| Tapbpl1 | 0.00659543 | 0.92061053 | 0.387 | 0.292 | 1 | pDC | Tapbpl |
| Cnot21 | 0.00662777 | 1.01259925 | 0.46 | 0.377 | 1 | pDC | Cnot2 |
| Lnpep | 0.00670407 | 0.41234894 | 0.606 | 0.651 | 1 | pDC | Lnpep |
| Rab351 | 0.00677972 | 0.2853419 | 0.372 | 0.449 | 1 | pDC | Rab35 |
| Sf3b41 | 0.0067892 | 0.47946967 | 0.182 | 0.289 | 1 | pDC | Sf3b4 |
| Anp32e3 | 0.00680985 | 0.3739179 | 0.577 | 0.585 | 1 | pDC | Anp32e |
| Ctsa2 | 0.00682947 | 0.98173943 | 0.854 | 0.657 | 1 | pDC | Ctsa |
| Lyn1 | 0.00684312 | 0.45943138 | 0.715 | 0.681 | 1 | pDC | Lyn |
| B2m1 | 0.00689264 | 0.77408716 | 0.92 | 0.818 | 1 | pDC | B2m |
| Ifi2031 | 0.00704719 | 1.2493385 | 0.321 | 0.179 | 1 | pDC | Ifi203 |
| Plbd2 | 0.00736389 | 0.29747469 | 0.321 | 0.397 | 1 | pDC | Plbd2 |
| Prcp2 | 0.00740108 | 0.55108419 | 0.686 | 0.588 | 1 | pDC | Prcp |
| Tmed92 | 0.00741073 | 1.07312822 | 0.599 | 0.493 | 1 | pDC | Tmed9 |
| Hspa53 | 0.00752973 | 0.34764574 | 0.65 | 0.592 | 1 | pDC | Hspa5 |
| Mthfd14a1 | 0.00753598 | 1.28348103 | 0.35 | 0.239 | 1 | pDC | Mthfd14a |
| Kbtbd2 | 0.00756617 | 0.29641439 | 0.197 | 0.26 | 1 | pDC | Kbtbd2 |
| Tle5 | 0.00758426 | 0.28526237 | 0.511 | 0.576 | 1 | pDC | Tle5 |
| Prpf81 | 0.00776944 | 0.27113418 | 0.511 | 0.524 | 1 | pDC | Prpf8 |
| Tmem179b1 | 0.00780468 | 0.40246557 | 0.219 | 0.296 | 1 | pDC | Tmem179b |
| Tet31 | 0.00782044 | 0.42540413 | 0.358 | 0.407 | 1 | pDC | Tet3 |
| Atp2a23 | 0.00786962 | 0.43735432 | 0.467 | 0.502 | 1 | pDC | Atp2a2 |
| Tollip2 | 0.00795371 | 0.36907348 | 0.197 | 0.271 | 1 | pDC | Tollip |
| Usp4 | 0.00809436 | 0.37448776 | 0.423 | 0.433 | 1 | pDC | Usp4 |
| Hpf11 | 0.00823585 | 1.41080731 | 0.292 | 0.205 | 1 | pDC | Hpf1 |
| Adam171 | 0.00846316 | 0.25755493 | 0.372 | 0.448 | 1 | pDC | Adam17 |
| Ddx5 | 0.0088018 | 0.4077836 | 0.672 | 0.659 | 1 | pDC | Ddx5 |
| Ndufc1 | 0.00900637 | 0.2561167 | 0.292 | 0.332 | 1 | pDC | Ndufc1 |
| Cul3 | 0.00912514 | 0.47819656 | 0.474 | 0.548 | 1 | pDC | Cul3 |
| Slc25a191 | 0.0094184 | 1.16246827 | 0.416 | 0.329 | 1 | pDC | Slc25a19 |
| Rab5a | 0.00946396 | 0.27288403 | 0.409 | 0.466 | 1 | pDC | Rab5a |
| Snx61 | 0.00947141 | 0.38381644 | 0.204 | 0.262 | 1 | pDC | Snx6 |
| Smarcd2 | 0.00949935 | 0.27235308 | 0.358 | 0.399 | 1 | pDC | Smarcd2 |
| Sipa12 | 0.00977543 | 0.50140765 | 0.365 | 0.394 | 1 | pDC | Sipa1 |
| Ppt22 | 0.00989662 | 1.56132349 | 0.255 | 0.132 | 1 | pDC | Ppt2 |
| C1qc1 | 0 | 5.58432126 | 0.99 | 0.067 | 0 | TAM.contam. | C1qc |
| Maf | 0 | 6.46790716 | 0.939 | 0.028 | 0 | TAM.contam. | Maf |
| C1qa1 | 0 | 5.48285384 | 0.99 | 0.103 | 0 | TAM.contam. | C1qa |
| Pf4 | 0 | 6.7136268 | 0.919 | 0.032 | 0 | TAM.contam. | Pf4 |
| Fcrls | 0 | 7.40809427 | 0.899 | 0.018 | 0 | TAM.contam. | Fcrls |
| C3ar1 | 0 | 5.02293096 | 0.929 | 0.05 | 0 | TAM.contam. | C3ar1 |
| C1qb1 | 0 | 5.0006276 | 1 | 0.135 | 0 | TAM.contam. | C1qb |
| Mafb1 | 0 | 4.36680666 | 0.98 | 0.117 | 0 | TAM.contam. | Mafb |
| Abca11 | 0 | 4.45584664 | 0.919 | 0.064 | 0 | TAM.contam. | Abca1 |
| C5ar11 | 0 | 4.64568321 | 0.889 | 0.05 | 0 | TAM.contam. | C5ar1 |
| Mrc11 | 0 | 3.71446784 | 0.99 | 0.16 | 0 | TAM.contam. | Mrc1 |
| Selenop1 | 0 | 4.6780664 | 1 | 0.196 | 0 | TAM.contam. | Selenop |
| Ms4a7 | 0 | 7.01954693 | 0.818 | 0.015 | 0 | TAM.contam. | Ms4a7 |
| Msr11 | 0 | 4.38047974 | 0.848 | 0.052 | 0 | TAM.contam. | Msr1 |
| Hmox11 | 0 | 5.01042698 | 0.899 | 0.111 | 0 | TAM.contam. | Hmox1 |
| Timp2 | 0 | 3.47238914 | 0.879 | 0.105 | 0 | TAM.contam. | Timp2 |
| Gas6 | 0 | 6.93544713 | 0.778 | 0.007 | 0 | TAM.contam. | Gas6 |
| Adgre1 | 0 | 4.09104285 | 0.838 | 0.07 | 0 | TAM.contam. | Adgre1 |
| Stab1 | 0 | 5.14448478 | 0.818 | 0.051 | 0 | TAM.contam. | Stab1 |
| Pld31 | 0 | 3.06529646 | 0.869 | 0.111 | 0 | TAM.contam. | Pld3 |
| Ms4a4a1 | 0 | 3.18933407 | 0.909 | 0.157 | 0 | TAM.contam. | Ms4a4a |
| Aoah | 0 | 5.34073215 | 0.768 | 0.021 | 0 | TAM.contam. | Aoah |
| Mertk | 0 | 3.56250076 | 0.798 | 0.055 | 0 | TAM.contam. | Mertk |
| Rhob1 | 0 | 2.94868589 | 0.929 | 0.192 | 0 | TAM.contam. | Rhob |
| Ctsd1 | 0 | 3.96041502 | 0.909 | 0.177 | 0 | TAM.contam. | Ctsd |
| Pla2g151 | 0 | 2.98823187 | 0.859 | 0.132 | 0 | TAM.contam. | Pla2g15 |
| Ecm11 | 0 | 2.20399983 | 0.889 | 0.167 | 0 | TAM.contam. | Ecm1 |
| Lyz22 | 0 | 4.00434022 | 1 | 0.28 | 0 | TAM.contam. | Lyz2 |
| Rab11fip51 | 0 | 3.05371003 | 0.869 | 0.153 | 0 | TAM.contam. | Rab11fip5 |
| Lgmn2 | 0 | 3.96667715 | 1 | 0.294 | 0 | TAM.contam. | Lgmn |
| F13a1 | 0 | 5.53725989 | 0.737 | 0.031 | 0 | TAM.contam. | F13a1 |
| Itgb51 | 0 | 3.88415178 | 0.768 | 0.07 | 0 | TAM.contam. | Itgb5 |
| Ctsl2 | 0 | 2.53253117 | 0.949 | 0.253 | 0 | TAM.contam. | Ctsl |
| Slc11a11 | 0 | 3.5242355 | 0.758 | 0.063 | 0 | TAM.contam. | Slc11a1 |
| Rnase4 | 0 | 5.25733016 | 0.717 | 0.026 | 0 | TAM.contam. | Rnase4 |
| Dusp6 | 0 | 2.5616072 | 0.828 | 0.142 | 0 | TAM.contam. | Dusp6 |
| Rab3il11 | 0 | 2.45563745 | 0.919 | 0.239 | 0 | TAM.contam. | Rab3il1 |
| Sash11 | 0 | 2.44040666 | 0.838 | 0.16 | 0 | TAM.contam. | Sash1 |
| Cd812 | 0 | 2.12638333 | 0.848 | 0.173 | 0 | TAM.contam. | Cd81 |
| Atp13a22 | 0 | 2.02787412 | 0.869 | 0.198 | 0 | TAM.contam. | Atp13a2 |
| Pla2g72 | 0 | 2.08546289 | 0.899 | 0.229 | 0 | TAM.contam. | Pla2g7 |
| Apoe1 | 0 | 6.88093975 | 1 | 0.332 | 0 | TAM.contam. | Apoe |
| Dhrs31 | 0 | 2.39709768 | 0.889 | 0.24 | 0 | TAM.contam. | Dhrs3 |
| Ccl92 | 0 | 2.07776905 | 0.798 | 0.149 | 0 | TAM.contam. | Ccl9 |
| Lrp13 | 0 | 1.8994059 | 0.919 | 0.271 | 0 | TAM.contam. | Lrp1 |
| Cebpa2 | 0 | 2.12040583 | 0.949 | 0.301 | 0 | TAM.contam. | Cebpa |
| Ninj11 | 0 | 2.29795109 | 0.899 | 0.267 | 0 | TAM.contam. | Ninj1 |
| Cd93 | 0 | 3.99342401 | 0.667 | 0.037 | 0 | TAM.contam. | Cd93 |
| Trf1 | 0 | 3.08007689 | 0.949 | 0.334 | 0 | TAM.contam. | Trf |
| Glul1 | 0 | 2.36772039 | 0.788 | 0.175 | 0 | TAM.contam. | Glul |
| Ccl62 | 0 | 3.08421077 | 0.778 | 0.169 | 0 | TAM.contam. | Ccl6 |
| Igfbp41 | 0 | 1.65073991 | 0.798 | 0.19 | 0 | TAM.contam. | Igfbp4 |
| Fcgr33 | 0 | 2.23839716 | 0.97 | 0.371 | 0 | TAM.contam. | Fcgr3 |
| Cd300ld5 | 0 | 5.65094641 | 0.616 | 0.017 | 0 | TAM.contam. | Cd300ld5 |
| Csf1r2 | 0 | 2.14912616 | 0.99 | 0.394 | 0 | TAM.contam. | Csf1r |
| Serpinb6a1 | 0 | 2.58466761 | 0.707 | 0.115 | 0 | TAM.contam. | Serpinb6a |
| Zeb23 | 0 | 1.80517201 | 0.96 | 0.369 | 0 | TAM.contam. | Zeb2 |
| Siglec1 | 0 | 5.39963729 | 0.606 | 0.015 | 0 | TAM.contam. | Siglec1 |
| Fcgr2b2 | 0 | 2.25846747 | 0.99 | 0.406 | 0 | TAM.contam. | Fcgr2b |
| Cd632 | 0 | 1.52736553 | 0.97 | 0.386 | 0 | TAM.contam. | Cd63 |
| Tmem86a | 0 | 2.97732438 | 0.677 | 0.095 | 0 | TAM.contam. | Tmem86a |
| Thbs11 | 0 | 3.13808547 | 0.707 | 0.125 | 0 | TAM.contam. | Thbs1 |
| Fam20c | 0 | 3.97581784 | 0.606 | 0.029 | 0 | TAM.contam. | Fam20c |
| Itgam2 | 0 | 1.52378009 | 0.929 | 0.353 | 0 | TAM.contam. | Itgam |
| Slc43a22 | 0 | 2.15218993 | 0.808 | 0.233 | 0 | TAM.contam. | Slc43a2 |
| Mmp141 | 0 | 1.5351047 | 0.859 | 0.286 | 0 | TAM.contam. | Mmp14 |
| Ccl31 | 0 | 2.13337778 | 0.727 | 0.169 | 0 | TAM.contam. | Ccl3 |
| Vsir2 | 0 | 1.903299 | 0.919 | 0.365 | 0 | TAM.contam. | Vsir |
| Ccl7 | 0 | 6.53130478 | 0.566 | 0.014 | 0 | TAM.contam. | Ccl7 |
| Plau1 | 0 | 2.3804893 | 0.697 | 0.151 | 0 | TAM.contam. | Plau |
| Pltp2 | 0 | 1.41240502 | 0.939 | 0.401 | 0 | TAM.contam. | Pltp |
| Tlr22 | 0 | 1.64908271 | 0.778 | 0.25 | 0 | TAM.contam. | Tlr2 |
| Cebpb2 | 0 | 1.72554429 | 0.99 | 0.47 | 0 | TAM.contam. | Cebpb |
| Lamp12 | 0 | 1.54820002 | 0.97 | 0.454 | 0 | TAM.contam. | Lamp1 |
| Cd142 | 0 | 1.39014444 | 0.828 | 0.313 | 0 | TAM.contam. | Cd14 |
| Plin22 | 0 | 2.05206857 | 0.929 | 0.416 | 0 | TAM.contam. | Plin2 |
| Tgfbi2 | 0 | 1.17994489 | 0.949 | 0.437 | 0 | TAM.contam. | Tgfbi |
| Ptgs1 | 0 | 4.99973746 | 0.525 | 0.02 | 0 | TAM.contam. | Ptgs1 |
| Ms4a6d1 | 0 | 1.28644508 | 0.859 | 0.363 | 0 | TAM.contam. | Ms4a6d |
| Ccl8 | 0 | 7.3005451 | 0.505 | 0.013 | 0 | TAM.contam. | Ccl8 |
| Dab21 | 0 | 1.63735427 | 0.97 | 0.482 | 0 | TAM.contam. | Dab2 |
| Acp52 | 0 | 2.19088469 | 0.727 | 0.246 | 0 | TAM.contam. | Acp5 |
| Fn11 | 0 | 2.58888613 | 0.798 | 0.323 | 0 | TAM.contam. | Fn1 |
| C4b | 0 | 6.72611617 | 0.485 | 0.011 | 0 | TAM.contam. | C4b |
| Ccl2 | 0 | 6.15606023 | 0.485 | 0.015 | 0 | TAM.contam. | Ccl2 |
| Arg1 | 0 | 4.20775155 | 0.525 | 0.062 | 0 | TAM.contam. | Arg1 |
| Ier32 | 0 | 1.91136419 | 0.909 | 0.448 | 0 | TAM.contam. | Ier3 |
| Lgals3bp2 | 0 | 1.53019919 | 0.909 | 0.452 | 0 | TAM.contam. | Lgals3bp |
| Egr12 | 0 | 1.85566628 | 0.828 | 0.371 | 0 | TAM.contam. | Egr1 |
| Fcgrt1 | 0 | 1.6346308 | 0.939 | 0.488 | 0 | TAM.contam. | Fcgrt |
| Il1rn1 | 0 | 1.24252814 | 0.737 | 0.293 | 0 | TAM.contam. | Il1rn |
| Cxcl22 | 0 | 1.5162863 | 0.788 | 0.356 | 0 | TAM.contam. | Cxcl2 |
| Vcam1 | 0 | 3.70417599 | 0.455 | 0.041 | 0 | TAM.contam. | Vcam1 |
| Creg12 | 0 | 1.43213638 | 0.828 | 0.414 | 0 | TAM.contam. | Creg1 |
| Aif12 | 0 | 0.99010364 | 0.929 | 0.519 | 0 | TAM.contam. | Aif1 |
| Fabp52 | 0 | 0.55161824 | 0.939 | 0.537 | 0 | TAM.contam. | Fabp5 |
| Cd682 | 0 | 1.31962976 | 0.96 | 0.558 | 0 | TAM.contam. | Cd68 |
| Sirpa2 | 0 | 1.27098716 | 0.99 | 0.592 | 0 | TAM.contam. | Sirpa |
| Marcksl12 | 0 | 0.49385274 | 0.919 | 0.533 | 0 | TAM.contam. | Marcksl1 |
| Cstb2 | 0 | 1.12292005 | 0.949 | 0.565 | 0 | TAM.contam. | Cstb |
| Gdf15 | 0 | 5.27604011 | 0.394 | 0.019 | 0 | TAM.contam. | Gdf15 |
| Cxcr43 | 0 | 0.69960368 | 0.909 | 0.538 | 0 | TAM.contam. | Cxcr4 |
| Atf33 | 0 | 0.80663465 | 0.848 | 0.477 | 0 | TAM.contam. | Atf3 |
| Rnf1493 | 0 | 0.90983579 | 0.99 | 0.627 | 0 | TAM.contam. | Rnf149 |
| Tcn21 | 0 | 0.97262655 | 0.939 | 0.583 | 0 | TAM.contam. | Tcn2 |
| Ccl42 | 0 | 0.75277095 | 0.566 | 0.239 | 0 | TAM.contam. | Ccl4 |
| Ctsa3 | 0 | 1.11571418 | 0.98 | 0.656 | 0 | TAM.contam. | Ctsa |
| Ctsc2 | 0 | 1.63305752 | 0.99 | 0.676 | 0 | TAM.contam. | Ctsc |
| Serinc31 | 0 | 1.20872961 | 0.98 | 0.668 | 0 | TAM.contam. | Serinc3 |
| 2-Jun | 0 | 1.310993 | 0.98 | 0.671 | 0 | TAM.contam. | Jun |
| Sat12 | 0 | 1.0497167 | 0.97 | 0.672 | 0 | TAM.contam. | Sat1 |
| Sdcbp3 | 0 | 0.98319226 | 0.99 | 0.697 | 0 | TAM.contam. | Sdcbp |
| Fcer1g3 | 0 | 0.9928945 | 0.98 | 0.693 | 0 | TAM.contam. | Fcer1g |
| Fos4 | 0 | 1.10832405 | 0.97 | 0.683 | 0 | TAM.contam. | Fos |
| Grn2 | 0 | 1.46429159 | 1 | 0.721 | 0 | TAM.contam. | Grn |
| Ctsb2 | 0 | 2.94033839 | 1 | 0.725 | 0 | TAM.contam. | Ctsb |
| Atp6v0c3 | 0 | 0.79574012 | 0.97 | 0.699 | 0 | TAM.contam. | Atp6v0c |
| Dusp13 | 0 | 0.8156407 | 1 | 0.734 | 0 | TAM.contam. | Dusp1 |
| Spp12 | 0 | 0.49989246 | 0.859 | 0.596 | 0 | TAM.contam. | Spp1 |
| Nfkbia3 | 0 | 0.55246992 | 0.99 | 0.731 | 0 | TAM.contam. | Nfkbia |
| Npc22 | 0 | 0.92931284 | 1 | 0.743 | 0 | TAM.contam. | Npc2 |
| Klf61 | 0 | 0.29764717 | 0.97 | 0.721 | 0 | TAM.contam. | Klf6 |
| Cd163 | 0 | 5.71456566 | 0.253 | 0.01 | 0 | TAM.contam. | Cd163 |
| Cxcl162 | 0 | 0.25611952 | 0.96 | 0.724 | 0 | TAM.contam. | Cxcl16 |
| App3 | 0 | 0.84803288 | 0.99 | 0.754 | 0 | TAM.contam. | App |
| Il4ra1 | 0 | 0.72029054 | 0.99 | 0.764 | 0 | TAM.contam. | Il4ra |
| Kctd122 | 0 | 0.4690021 | 1 | 0.777 | 0 | TAM.contam. | Kctd12 |
| Psap3 | 0 | 0.50279927 | 1 | 0.798 | 0 | TAM.contam. | Psap |
| Marcks2 | 0 | 0.36554839 | 0.99 | 0.79 | 0 | TAM.contam. | Marcks |
| Ctsh1 | 0 | 0.3742924 | 0.98 | 0.805 | 0 | TAM.contam. | Ctsh |
| Itm2b1 | 0 | 1.78018096 | 1 | 0.839 | 0 | TAM.contam. | Itm2b |
| B2m2 | 0 | 0.37606996 | 0.96 | 0.818 | 0 | TAM.contam. | B2m |
| Akr1a11 | 0 | 0.37819007 | 1 | 0.875 | 0 | TAM.contam. | Akr1a1 |
| Ctsz1 | 0 | 0.51014165 | 1 | 0.899 | 0 | TAM.contam. | Ctsz |
| Mcl11 | 0 | 0.31853545 | 1 | 0.904 | 0 | TAM.contam. | Mcl1 |
| Clta | 0 | 0.47272522 | 1 | 0.904 | 0 | TAM.contam. | Clta |
| H2-K12 | 0 | 0.32208934 | 1 | 0.906 | 0 | TAM.contam. | H2-K1 |
| Zfp364 | 0 | 0.25239603 | 1 | 0.909 | 0 | TAM.contam. | Zfp36 |
| Ctss3 | 0 | 0.93873319 | 1 | 0.92 | 0 | TAM.contam. | Ctss |
| Fth12 | 0 | 0.65357505 | 1 | 0.957 | 0 | TAM.contam. | Fth1 |
| Cmklr1 | 2.27E-305 | 3.93448732 | 0.667 | 0.037 | 4.39E-301 | TAM.contam. | Cmklr1 |
| Ralgds1 | 2.42E-305 | 2.50676203 | 0.747 | 0.141 | 4.68E-301 | TAM.contam. | Ralgds |
| Hexa2 | 5.37E-301 | 1.15133788 | 0.96 | 0.565 | 1.04E-296 | TAM.contam. | Hexa |
| Zfp7033 | 1.46E-300 | 1.30743753 | 0.949 | 0.427 | 2.82E-296 | TAM.contam. | Zfp703 |
| Fcna | 1.93E-300 | 5.74286492 | 0.434 | 0.011 | 3.73E-296 | TAM.contam. | Fcna |
| Mpeg12 | 2.71E-299 | 0.39740716 | 0.919 | 0.445 | 5.24E-295 | TAM.contam. | Mpeg1 |
| Nxpe5 | 1.19E-296 | 3.62742235 | 0.707 | 0.064 | 2.31E-292 | TAM.contam. | Nxpe5 |
| Stard8 | 2.12E-296 | 4.05196788 | 0.505 | 0.041 | 4.10E-292 | TAM.contam. | Stard8 |
| Snx24 | 1.46E-294 | 1.60826249 | 0.899 | 0.409 | 2.82E-290 | TAM.contam. | Snx2 |
| Ubc1 | 3.41E-291 | 0.35124533 | 0.99 | 0.827 | 6.59E-287 | TAM.contam. | Ubc |
| Mmp13 | 3.56E-288 | 2.68543438 | 0.354 | 0.058 | 6.88E-284 | TAM.contam. | Mmp13 |
| Lrp6 | 3.66E-288 | 1.70935987 | 0.909 | 0.35 | 7.08E-284 | TAM.contam. | Lrp6 |
| Sdc31 | 1.65E-283 | 0.85939598 | 0.949 | 0.542 | 3.19E-279 | TAM.contam. | Sdc3 |
| Abcc3 | 3.21E-281 | 4.25567636 | 0.616 | 0.033 | 6.21E-277 | TAM.contam. | Abcc3 |
| Emp11 | 1.10E-280 | 1.43865375 | 0.778 | 0.258 | 2.13E-276 | TAM.contam. | Emp1 |
| Txnip3 | 2.33E-280 | 1.29101615 | 0.818 | 0.292 | 4.50E-276 | TAM.contam. | Txnip |
| Unc93b12 | 1.82E-276 | 0.56204653 | 1 | 0.736 | 3.52E-272 | TAM.contam. | Unc93b1 |
| Prdx51 | 7.09E-273 | 0.61821802 | 0.97 | 0.686 | 1.37E-268 | TAM.contam. | Prdx5 |
| Folr2 | 1.69E-272 | 7.04495903 | 0.616 | 0.007 | 3.27E-268 | TAM.contam. | Folr2 |
| Pid11 | 6.99E-271 | 1.74184824 | 0.889 | 0.242 | 1.35E-266 | TAM.contam. | Pid1 |
| Rab202 | 2.30E-270 | 1.83295145 | 0.808 | 0.205 | 4.44E-266 | TAM.contam. | Rab20 |
| Qk1 | 2.28E-269 | 0.84135148 | 0.96 | 0.668 | 4.40E-265 | TAM.contam. | Qk |
| P2ry62 | 2.88E-268 | 2.0705596 | 0.869 | 0.213 | 5.56E-264 | TAM.contam. | P2ry6 |
| Gas71 | 1.93E-267 | 2.59111564 | 0.747 | 0.097 | 3.73E-263 | TAM.contam. | Gas7 |
| Cyba1 | 4.51E-264 | 0.38274874 | 0.98 | 0.814 | 8.72E-260 | TAM.contam. | Cyba |
| Ccl12 | 6.04E-260 | 7.58686658 | 0.576 | 0.005 | 1.17E-255 | TAM.contam. | Ccl12 |
| Tpp12 | 4.69E-259 | 1.86335017 | 0.859 | 0.277 | 9.07E-255 | TAM.contam. | Tpp1 |
| Tnf1 | 2.39E-256 | 1.24746714 | 0.707 | 0.192 | 4.61E-252 | TAM.contam. | Tnf |
| Npl | 3.22E-255 | 5.81235662 | 0.606 | 0.012 | 6.23E-251 | TAM.contam. | Npl |
| Camk11 | 3.87E-255 | 2.538402 | 0.788 | 0.139 | 7.49E-251 | TAM.contam. | Camk1 |
| Litaf2 | 5.35E-253 | 0.46257031 | 0.97 | 0.773 | 1.03E-248 | TAM.contam. | Litaf |

| | | | | | | | |
|---|---|---|---|---|---|---|---|
| Lmna1 | 1.85E-251 | 1.20649345 | 0.798 | 0.25 | 3.58E-247 | TAM.contam. | Lmna |
| Hspa1a2 | 1.05E-249 | 0.33756219 | 0.687 | 0.356 | 2.03E-245 | TAM.contam. | Hspa1a |
| Rgl11 | 3.68E-249 | 2.80652419 | 0.737 | 0.055 | 7.11E-245 | TAM.contam. | Rgl1 |
| Cd841 | 1.70E-245 | 2.35955945 | 0.798 | 0.15 | 3.28E-241 | TAM.contam. | Cd84 |
| Zfp36l13 | 1.14E-244 | 0.59443946 | 0.929 | 0.549 | 2.19E-240 | TAM.contam. | Zfp36l1 |
| Ccrl22 | 2.56E-243 | 0.99971813 | 0.808 | 0.385 | 4.95E-239 | TAM.contam. | Ccrl2 |
| Sdc43 | 5.62E-242 | 1.35283896 | 0.869 | 0.3 | 1.09E-237 | TAM.contam. | Sdc4 |
| Phyhd1 | 2.88E-241 | 3.94956437 | 0.566 | 0.029 | 5.56E-237 | TAM.contam. | Phyhd1 |
| Pmp22 | 6.83E-240 | 5.87118279 | 0.434 | 0.008 | 1.32E-235 | TAM.contam. | Pmp22 |
| Tgfbr22 | 1.15E-237 | 1.59483305 | 0.859 | 0.333 | 2.22E-233 | TAM.contam. | Tgfbr2 |
| Fcgr11 | 1.03E-233 | 2.12308598 | 0.727 | 0.16 | 1.99E-229 | TAM.contam. | Fcgr1 |
| Tmem37 | 1.14E-231 | 5.23526091 | 0.586 | 0.011 | 2.20E-227 | TAM.contam. | Tmem37 |
| Ifrd13 | 5.86E-228 | 0.41150346 | 0.96 | 0.68 | 1.13E-223 | TAM.contam. | Ifrd1 |
| Gna12 | 3.75E-227 | 1.57969792 | 0.808 | 0.317 | 7.24E-223 | TAM.contam. | Gna12 |
| C31 | 1.43E-226 | 2.43812868 | 0.636 | 0.09 | 2.76E-222 | TAM.contam. | C3 |
| Sgk12 | 1.66E-226 | 1.50720903 | 0.646 | 0.195 | 3.21E-222 | TAM.contam. | Sgk1 |
| Vegfa1 | 3.76E-224 | 1.2559524 | 0.646 | 0.275 | 7.28E-220 | TAM.contam. | Vegfa |
| Itga9 | 3.69E-223 | 6.46368106 | 0.687 | 0.007 | 7.14E-219 | TAM.contam. | Itga9 |
| Axl2 | 5.81E-222 | 1.28316955 | 0.677 | 0.233 | 1.12E-217 | TAM.contam. | Axl |
| Clec4d2 | 2.60E-221 | 1.95495082 | 0.626 | 0.165 | 5.02E-217 | TAM.contam. | Clec4d |
| Tnfrsf92 | 6.35E-220 | 0.47577534 | 0.626 | 0.271 | 1.23E-215 | TAM.contam. | Tnfrsf9 |
| Blvrb1 | 9.54E-219 | 2.31968815 | 0.768 | 0.115 | 1.84E-214 | TAM.contam. | Blvrb |
| Slc7a81 | 2.63E-218 | 3.03859605 | 0.707 | 0.067 | 5.08E-214 | TAM.contam. | Slc7a8 |
| Gda2 | 2.99E-218 | 1.29695527 | 0.808 | 0.321 | 5.78E-214 | TAM.contam. | Gda |
| Slc15a33 | 1.37E-216 | 0.37113375 | 0.97 | 0.641 | 2.65E-212 | TAM.contam. | Slc15a3 |
| Apobec11 | 5.32E-216 | 2.11968567 | 0.697 | 0.117 | 1.03E-211 | TAM.contam. | Apobec1 |
| Arap13 | 8.58E-216 | 1.37304822 | 0.859 | 0.273 | 1.66E-211 | TAM.contam. | Arap1 |
| Vat11 | 1.96E-214 | 1.65348718 | 0.788 | 0.232 | 3.79E-210 | TAM.contam. | Vat1 |
| Trem2 | 2.74E-213 | 4.48526714 | 0.576 | 0.018 | 5.30E-209 | TAM.contam. | Trem2 |
| Rab1a1 | 4.98E-213 | 0.83315505 | 0.939 | 0.618 | 9.63E-209 | TAM.contam. | Rab1a |
| Rnf19b2 | 1.02E-209 | 0.56561087 | 0.919 | 0.547 | 1.97E-205 | TAM.contam. | Rnf19b |
| Adipor11 | 1.08E-207 | 0.81820022 | 0.939 | 0.58 | 2.09E-203 | TAM.contam. | Adipor1 |
| Adap21 | 5.36E-205 | 1.70478031 | 0.859 | 0.251 | 1.04E-200 | TAM.contam. | Adap2 |
| Apoc2 | 2.00E-203 | 4.51389959 | 0.485 | 0.014 | 3.86E-199 | TAM.contam. | Apoc2 |
| Cxcl11 | 7.98E-203 | 2.454814 | 0.253 | 0.036 | 1.54E-198 | TAM.contam. | Cxcl1 |
| Lhfpl2 | 4.57E-202 | 4.08931857 | 0.414 | 0.028 | 8.83E-198 | TAM.contam. | Lhfpl2 |
| Ehd11 | 1.08E-200 | 0.9611024 | 0.889 | 0.46 | 2.09E-196 | TAM.contam. | Ehd1 |
| Mt11 | 5.63E-198 | 0.28672889 | 0.949 | 0.599 | 1.09E-193 | TAM.contam. | Mt1 |
| Xdh | 3.27E-197 | 2.67048032 | 0.616 | 0.078 | 6.33E-193 | TAM.contam. | Xdh |
| Ets22 | 1.98E-196 | 0.8018267 | 0.859 | 0.466 | 3.83E-192 | TAM.contam. | Ets2 |
| Snx62 | 1.30E-195 | 1.63145096 | 0.758 | 0.25 | 2.52E-191 | TAM.contam. | Snx6 |
| Adam92 | 1.34E-195 | 1.13104066 | 0.889 | 0.43 | 2.60E-191 | TAM.contam. | Adam9 |
| Prxl2b | 1.20E-194 | 3.07402225 | 0.667 | 0.068 | 2.31E-190 | TAM.contam. | Prxl2b |
| Pdgfc | 2.23E-193 | 4.10571293 | 0.576 | 0.021 | 4.31E-189 | TAM.contam. | Pdgfc |
| Itsn11 | 8.23E-193 | 3.41187598 | 0.636 | 0.042 | 1.59E-188 | TAM.contam. | Itsn1 |
| Il10rb2 | 9.87E-193 | 1.16371425 | 0.889 | 0.41 | 1.91E-188 | TAM.contam. | Il10rb |
| Cyth11 | 5.25E-191 | 0.63888711 | 0.949 | 0.584 | 1.01E-186 | TAM.contam. | Cyth1 |
| Fosb3 | 1.66E-190 | 0.35240021 | 0.919 | 0.652 | 3.21E-186 | TAM.contam. | Fosb |
| mt-Nd6 | 5.48E-188 | 0.44987667 | 0.929 | 0.516 | 1.06E-183 | TAM.contam. | mt-Nd6 |
| Wfdc172 | 1.46E-187 | 1.04415051 | 0.758 | 0.377 | 2.83E-183 | TAM.contam. | Wfdc17 |
| Mfsd13 | 1.50E-186 | 1.35491739 | 0.859 | 0.363 | 2.91E-182 | TAM.contam. | Mfsd1 |
| Wwp1 | 9.91E-185 | 2.41500215 | 0.646 | 0.122 | 1.92E-180 | TAM.contam. | Wwp1 |
| Ms4a6b1 | 1.08E-184 | 1.6918197 | 0.697 | 0.185 | 2.09E-180 | TAM.contam. | Ms4a6b |
| Nrp1 | 3.66E-184 | 2.59079998 | 0.667 | 0.087 | 7.08E-180 | TAM.contam. | Nrp1 |
| Fcgr4 | 5.64E-183 | 2.86181822 | 0.606 | 0.076 | 1.09E-178 | TAM.contam. | Fcgr4 |
| Pnrc11 | 2.16E-182 | 0.26962452 | 0.97 | 0.72 | 4.17E-178 | TAM.contam. | Pnrc1 |
| Atp6v1b22 | 7.86E-182 | 0.74628936 | 0.96 | 0.554 | 1.52E-177 | TAM.contam. | Atp6v1b2 |
| Cfb1 | 1.36E-181 | 2.27123631 | 0.586 | 0.082 | 2.63E-177 | TAM.contam. | Cfb |
| Eid12 | 4.60E-180 | 0.67900281 | 0.96 | 0.593 | 8.90E-176 | TAM.contam. | Eid1 |
| Sqstm12 | 3.46E-179 | 0.44494586 | 0.949 | 0.624 | 6.68E-175 | TAM.contam. | Sqstm1 |
| Ehd41 | 4.90E-179 | 1.28432196 | 0.879 | 0.364 | 9.48E-175 | TAM.contam. | Ehd4 |
| Lst12 | 9.20E-178 | 1.66495306 | 0.798 | 0.241 | 1.78E-173 | TAM.contam. | Lst1 |
| Abhd121 | 2.79E-176 | 1.17366359 | 0.889 | 0.434 | 5.40E-172 | TAM.contam. | Abhd12 |
| Tmem176b2 | 3.44E-176 | 0.50219636 | 0.909 | 0.476 | 6.66E-172 | TAM.contam. | Tmem176b |
| Grina2 | 6.63E-176 | 0.9994694 | 0.909 | 0.431 | 1.28E-171 | TAM.contam. | Grina |
| Tspan34 | 6.09E-174 | 0.66264039 | 0.778 | 0.354 | 1.18E-169 | TAM.contam. | Tspan3 |
| Wdfy32 | 2.68E-172 | 1.26135164 | 0.919 | 0.375 | 5.18E-168 | TAM.contam. | Wdfy3 |
| Gns2 | 7.58E-172 | 0.93116035 | 0.929 | 0.493 | 1.46E-167 | TAM.contam. | Gns |
| Hpgds | 1.53E-171 | 5.20680351 | 0.586 | 0.016 | 2.95E-167 | TAM.contam. | Hpgds |
| Bri32 | 1.70E-171 | 0.66698702 | 0.929 | 0.551 | 3.28E-167 | TAM.contam. | Bri3 |
| Tmem106a1 | 1.97E-170 | 1.26736867 | 0.899 | 0.368 | 3.81E-166 | TAM.contam. | Tmem106a |
| Slfn22 | 5.36E-170 | 0.5088493 | 0.909 | 0.535 | 1.04E-165 | TAM.contam. | Slfn2 |
| Plxnd12 | 8.01E-170 | 0.34011169 | 0.929 | 0.611 | 1.55E-165 | TAM.contam. | Plxnd1 |
| Lrrc252 | 3.05E-169 | 1.40404731 | 0.899 | 0.344 | 5.89E-165 | TAM.contam. | Lrrc25 |
| Ap1b11 | 2.37E-168 | 1.02749036 | 0.889 | 0.458 | 4.59E-164 | TAM.contam. | Ap1b1 |
| Acp21 | 7.15E-167 | 2.28643695 | 0.707 | 0.126 | 1.38E-162 | TAM.contam. | Acp2 |
| Abca9 | 7.97E-167 | 3.18128751 | 0.515 | 0.032 | 1.54E-162 | TAM.contam. | Abca9 |
| Klf63 | 2.57E-165 | 0.76358752 | 0.859 | 0.486 | 4.97E-161 | TAM.contam. | Klf6 |
| Adam151 | 4.44E-165 | 1.59834782 | 0.859 | 0.264 | 8.59E-161 | TAM.contam. | Adam15 |
| Ifi2071 | 6.20E-165 | 1.38257373 | 0.747 | 0.224 | 1.20E-160 | TAM.contam. | Ifi207 |
| Il1a1 | 1.74E-163 | 1.76932974 | 0.364 | 0.073 | 3.37E-159 | TAM.contam. | Il1a |
| Clec4n3 | 9.85E-163 | 0.60743473 | 0.828 | 0.409 | 1.90E-158 | TAM.contam. | Clec4n |
| Sec14l11 | 6.25E-162 | 1.60453596 | 0.747 | 0.201 | 1.21E-157 | TAM.contam. | Sec14l1 |
| Cbr2 | 2.38E-161 | 8.19553235 | 0.424 | 0.003 | 4.60E-157 | TAM.contam. | Cbr2 |
| BC0055372 | 6.66E-161 | 0.36154095 | 0.949 | 0.736 | 1.29E-156 | TAM.contam. | BC005537 |
| Rab5c | 7.58E-161 | 0.44344218 | 0.949 | 0.7 | 1.47E-156 | TAM.contam. | Rab5c |
| Tnfsf13 | 1.32E-160 | 3.00590978 | 0.515 | 0.053 | 2.55E-156 | TAM.contam. | Tnfsf13 |
| Rcn3 | 1.00E-159 | 2.06067451 | 0.566 | 0.123 | 1.94E-155 | TAM.contam. | Rcn3 |
| Slc3a23 | 1.11E-159 | 0.30940195 | 0.929 | 0.734 | 2.14E-155 | TAM.contam. | Slc3a2 |
| Tgfbr13 | 1.74E-158 | 1.32934835 | 0.697 | 0.26 | 3.37E-154 | TAM.contam. | Tgfbr1 |
| Hebp1 | 2.06E-158 | 3.52969413 | 0.495 | 0.032 | 3.98E-154 | TAM.contam. | Hebp1 |
| Cd382 | 1.57E-156 | 1.96174674 | 0.646 | 0.137 | 3.03E-152 | TAM.contam. | Cd38 |
| 8-Sep | 1.44E-154 | 2.18856552 | 0.596 | 0.096 | 2.78E-150 | TAM.contam. | 8-Sep |
| Igf1 | 9.82E-152 | 4.70899955 | 0.414 | 0.008 | 1.90E-147 | TAM.contam. | Igf1 |
| Cndp21 | 3.34E-151 | 1.53807182 | 0.798 | 0.275 | 6.45E-147 | TAM.contam. | Cndp2 |
| Pmepa12 | 1.46E-150 | 1.04767411 | 0.727 | 0.322 | 2.83E-146 | TAM.contam. | Pmepa1 |
| Clec4a1 | 1.18E-149 | 2.57892767 | 0.515 | 0.066 | 2.29E-145 | TAM.contam. | Clec4a1 |
| Ppp1r15a2 | 1.77E-148 | 0.2936288 | 0.899 | 0.615 | 3.42E-144 | TAM.contam. | Ppp1r15a |
| Cd300ld | 7.56E-148 | 2.46061584 | 0.596 | 0.106 | 1.46E-143 | TAM.contam. | Cd300ld |
| Hacd4 | 1.23E-147 | 3.28873886 | 0.535 | 0.044 | 2.37E-143 | TAM.contam. | Hacd4 |
| Ski2 | 1.57E-147 | 0.93544341 | 0.949 | 0.462 | 3.03E-143 | TAM.contam. | Ski |
| Sgpl11 | 1.91E-147 | 0.7936791 | 0.848 | 0.481 | 3.70E-143 | TAM.contam. | Sgpl1 |
| Aldoa2 | 1.42E-146 | 0.34712107 | 0.99 | 0.689 | 2.75E-142 | TAM.contam. | Aldoa |
| Cited22 | 8.45E-144 | 1.1463644 | 0.677 | 0.24 | 1.63E-139 | TAM.contam. | Cited2 |
| Atp6ap11 | 3.18E-143 | 0.67035994 | 0.899 | 0.554 | 6.14E-139 | TAM.contam. | Atp6ap1 |
| Plk22 | 4.83E-143 | 1.29874712 | 0.677 | 0.196 | 9.33E-139 | TAM.contam. | Plk2 |
| Fyb2 | 4.98E-143 | 1.06764885 | 0.838 | 0.257 | 9.62E-139 | TAM.contam. | Fyb |
| Atp2b12 | 3.04E-142 | 0.77943827 | 0.909 | 0.477 | 5.88E-138 | TAM.contam. | Atp2b1 |
| Man1a3 | 4.62E-142 | 0.75982143 | 0.909 | 0.482 | 8.92E-138 | TAM.contam. | Man1a |
| Nrp22 | 2.72E-141 | 0.87885437 | 0.758 | 0.19 | 5.26E-137 | TAM.contam. | Nrp2 |
| Lamp21 | 3.64E-141 | 0.70612418 | 0.939 | 0.551 | 7.04E-137 | TAM.contam. | Lamp2 |
| Scpep12 | 9.71E-141 | 0.41567208 | 0.949 | 0.619 | 1.88E-136 | TAM.contam. | Scpep1 |
| H3f3b1 | 7.48E-139 | 0.29808049 | 0.939 | 0.721 | 1.45E-134 | TAM.contam. | H3f3b |
| Rab312 | 1.69E-138 | 1.427076 | 0.768 | 0.209 | 3.27E-134 | TAM.contam. | Rab31 |
| Rnpep2 | 1.81E-138 | 1.03334313 | 0.899 | 0.453 | 3.49E-134 | TAM.contam. | Rnpep |
| Mgst1 | 2.46E-138 | 4.82435333 | 0.333 | 0.01 | 4.76E-134 | TAM.contam. | Mgst1 |
| Csf3r1 | 3.16E-136 | 1.80591707 | 0.525 | 0.064 | 6.10E-132 | TAM.contam. | Csf3r |
| Scamp5 | 6.69E-136 | 5.89924979 | 0.505 | 0.006 | 1.29E-131 | TAM.contam. | Scamp5 |
| Tnfsf121 | 2.67E-135 | 2.37878337 | 0.677 | 0.106 | 5.15E-131 | TAM.contam. | Tnfsf12 |
| B4galt11 | 3.89E-135 | 0.69595923 | 0.899 | 0.488 | 7.52E-131 | TAM.contam. | B4galt1 |
| Rnf1302 | 4.96E-134 | 0.84914652 | 0.939 | 0.492 | 9.60E-130 | TAM.contam. | Rnf130 |
| Tnfaip32 | 9.86E-133 | 0.44422219 | 0.788 | 0.469 | 1.91E-128 | TAM.contam. | Tnfaip3 |
| Mcfd22 | 1.12E-131 | 1.43907045 | 0.778 | 0.261 | 2.16E-127 | TAM.contam. | Mcfd2 |
| Myo7a | 1.38E-131 | 2.82443221 | 0.505 | 0.055 | 2.66E-127 | TAM.contam. | Myo7a |
| Entpd12 | 3.24E-131 | 1.27404347 | 0.727 | 0.274 | 6.26E-127 | TAM.contam. | Entpd1 |
| Igsf61 | 5.76E-130 | 2.03050498 | 0.687 | 0.142 | 1.11E-125 | TAM.contam. | Igsf6 |
| Lrg1 | 4.21E-128 | 3.60185551 | 0.273 | 0.018 | 8.14E-124 | TAM.contam. | Lrg1 |
| Cacna1d | 6.22E-128 | 4.79048917 | 0.434 | 0.011 | 1.20E-123 | TAM.contam. | Cacna1d |
| Slc48a11 | 3.39E-127 | 1.22320194 | 0.667 | 0.294 | 6.55E-123 | TAM.contam. | Slc48a1 |
| Tfec1 | 7.61E-127 | 3.79906409 | 0.455 | 0.021 | 1.47E-122 | TAM.contam. | Tfec |
| Clec4a3 | 8.87E-127 | 2.07822016 | 0.657 | 0.082 | 1.72E-122 | TAM.contam. | Clec4a3 |
| Pdpn | 1.37E-126 | 3.14074722 | 0.293 | 0.026 | 2.64E-122 | TAM.contam. | Pdpn |
| Spred12 | 2.64E-126 | 0.58884253 | 0.929 | 0.477 | 5.11E-122 | TAM.contam. | Spred1 |
| Nisch1 | 2.87E-126 | 0.43501936 | 0.939 | 0.564 | 5.55E-122 | TAM.contam. | Nisch |
| Metrnl1 | 3.91E-126 | 1.19763896 | 0.818 | 0.33 | 7.55E-122 | TAM.contam. | Metrnl |
| Slc6a8 | 5.37E-126 | 3.08061725 | 0.374 | 0.023 | 1.04E-121 | TAM.contam. | Slc6a8 |
| Lipa3 | 8.70E-126 | 1.30597592 | 0.737 | 0.288 | 1.68E-121 | TAM.contam. | Lipa |
| Lair12 | 1.08E-125 | 0.89288043 | 0.667 | 0.253 | 2.09E-121 | TAM.contam. | Lair1 |
| Lilrb4a3 | 2.03E-125 | 0.325436 | 0.939 | 0.565 | 3.93E-121 | TAM.contam. | Lilrb4a |
| Mmp122 | 4.00E-125 | 0.59148278 | 0.444 | 0.277 | 7.73E-121 | TAM.contam. | Mmp12 |
| Prune2 | 1.05E-124 | 3.8876641 | 0.465 | 0.025 | 2.04E-120 | TAM.contam. | Prune2 |
| Rin2 | 1.07E-124 | 2.20104843 | 0.596 | 0.096 | 2.07E-120 | TAM.contam. | Rin2 |
| Tbxas13 | 2.17E-124 | 1.10557862 | 0.848 | 0.326 | 4.20E-120 | TAM.contam. | Tbxas1 |
| Scn1b | 3.29E-124 | 2.78043672 | 0.515 | 0.054 | 6.37E-120 | TAM.contam. | Scn1b |
| Ap2m1 | 6.13E-124 | 0.46494161 | 0.939 | 0.635 | 1.19E-119 | TAM.contam. | Ap2m1 |
| Ccr51 | 5.60E-123 | 0.57071161 | 0.939 | 0.44 | 1.08E-118 | TAM.contam. | Ccr5 |
| Mdfic2 | 1.13E-122 | 1.30674405 | 0.798 | 0.266 | 2.18E-118 | TAM.contam. | Mdfic |
| Msrb12 | 2.01E-122 | 0.45966567 | 0.869 | 0.534 | 3.88E-118 | TAM.contam. | Msrb1 |
| Aplp22 | 2.64E-122 | 0.68999004 | 0.919 | 0.467 | 5.11E-118 | TAM.contam. | Aplp2 |
| Sh3glb11 | 3.27E-122 | 0.30049652 | 0.879 | 0.662 | 6.32E-118 | TAM.contam. | Sh3glb1 |
| Npc12 | 1.18E-121 | 1.12491775 | 0.788 | 0.292 | 2.27E-117 | TAM.contam. | Npc1 |
| Wdr261 | 1.61E-121 | 0.33415335 | 0.96 | 0.68 | 3.12E-117 | TAM.contam. | Wdr26 |
| Slc6a62 | 1.70E-120 | 0.41936821 | 0.909 | 0.621 | 3.29E-116 | TAM.contam. | Slc6a6 |
| Tmem50a2 | 9.33E-120 | 0.80807268 | 0.889 | 0.491 | 1.80E-115 | TAM.contam. | Tmem50a |
| Dgkz1 | 2.26E-119 | 0.99246789 | 0.848 | 0.353 | 4.38E-115 | TAM.contam. | Dgkz |
| Camk2d2 | 1.45E-117 | 1.43952784 | 0.768 | 0.21 | 2.81E-113 | TAM.contam. | Camk2d |
| Cd300c22 | 2.10E-117 | 1.44222644 | 0.707 | 0.22 | 4.05E-113 | TAM.contam. | Cd300c2 |
| Furin2 | 3.91E-117 | 0.55731576 | 0.889 | 0.494 | 7.57E-113 | TAM.contam. | Furin |
| Rap2b1 | 1.49E-116 | 0.38046717 | 0.828 | 0.381 | 2.89E-112 | TAM.contam. | Rap2b |
| Aldh22 | 1.93E-116 | 0.50893642 | 0.919 | 0.547 | 3.72E-112 | TAM.contam. | Aldh2 |
| Rcbtb21 | 7.83E-116 | 2.03615803 | 0.626 | 0.122 | 1.51E-111 | TAM.contam. | Rcbtb2 |
| Tgif12 | 1.24E-115 | 0.40309755 | 0.879 | 0.567 | 2.40E-111 | TAM.contam. | Tgif1 |
| Mt22 | 3.52E-114 | 0.7886634 | 0.556 | 0.201 | 6.80E-110 | TAM.contam. | Mt2 |
| Atp6v1a2 | 4.19E-114 | 0.75368139 | 0.909 | 0.497 | 8.10E-110 | TAM.contam. | Atp6v1a |
| Slc27a12 | 9.24E-114 | 1.41761251 | 0.697 | 0.172 | 1.79E-109 | TAM.contam. | Slc27a1 |
| Tcirg11 | 1.16E-113 | 0.3066609 | 0.96 | 0.615 | 2.24E-109 | TAM.contam. | Tcirg1 |
| Rnasek | 7.91E-113 | 0.47556461 | 0.909 | 0.618 | 1.53E-108 | TAM.contam. | Rnasek |
| Ctnnd11 | 1.45E-112 | 1.20732084 | 0.737 | 0.185 | 2.80E-108 | TAM.contam. | Ctnnd1 |
| Dusp7 | 1.87E-112 | 2.00264448 | 0.566 | 0.105 | 3.62E-108 | TAM.contam. | Dusp7 |
| Frmd4b1 | 1.02E-111 | 1.7233001 | 0.707 | 0.178 | 1.98E-107 | TAM.contam. | Frmd4b |
| Plod1 | 1.05E-111 | 2.60096996 | 0.566 | 0.078 | 2.04E-107 | TAM.contam. | Plod1 |
| Lfng2 | 1.21E-111 | 0.50862685 | 0.838 | 0.432 | 2.33E-107 | TAM.contam. | Lfng |
| Psd3 | 1.58E-111 | 2.41406187 | 0.475 | 0.04 | 3.06E-107 | TAM.contam. | Psd3 |
| Slc36a1 | 2.69E-110 | 2.47697217 | 0.576 | 0.067 | 5.19E-106 | TAM.contam. | Slc36a1 |
| Tmem119 | 4.50E-110 | 3.00857683 | 0.374 | 0.02 | 8.70E-106 | TAM.contam. | Tmem119 |
| Ctla2b | 7.88E-110 | 3.43517541 | 0.343 | 0.034 | 1.52E-105 | TAM.contam. | Ctla2b |
| Renbp1 | 3.39E-109 | 1.33164698 | 0.808 | 0.209 | 6.56E-105 | TAM.contam. | Renbp |
| Nucb12 | 6.59E-109 | 0.64512382 | 0.838 | 0.46 | 1.27E-104 | TAM.contam. | Nucb1 |
| Sirt21 | 8.60E-109 | 0.65517418 | 0.818 | 0.403 | 1.66E-104 | TAM.contam. | Sirt2 |
| Uap1l11 | 4.49E-108 | 1.66052606 | 0.606 | 0.159 | 8.68E-104 | TAM.contam. | Uap1l1 |
| Ncf23 | 5.94E-108 | 0.39759506 | 0.889 | 0.545 | 1.15E-103 | TAM.contam. | Ncf2 |
| Aldh7a1 | 1.33E-107 | 2.7550001 | 0.465 | 0.057 | 2.57E-103 | TAM.contam. | Aldh7a1 |
| P2rx42 | 1.36E-107 | 0.72648344 | 0.788 | 0.392 | 2.63E-103 | TAM.contam. | P2rx4 |
| Adam172 | 7.39E-107 | 0.74595657 | 0.818 | 0.439 | 1.43E-102 | TAM.contam. | Adam17 |
| Cd300ld3 | 1.65E-106 | 4.07170994 | 0.404 | 0.016 | 3.19E-102 | TAM.contam. | Cd300ld3 |
| Nrg11 | 8.59E-106 | 1.53485348 | 0.525 | 0.108 | 1.66E-101 | TAM.contam. | Nrg1 |
| Ccr13 | 9.18E-106 | 1.20508584 | 0.788 | 0.273 | 1.78E-101 | TAM.contam. | Ccr1 |
| Tmem1892 | 1.16E-105 | 0.8929368 | 0.818 | 0.404 | 2.25E-101 | TAM.contam. | Tmem189 |
| Serpine1 | 1.40E-105 | 1.18142524 | 0.293 | 0.125 | 2.71E-101 | TAM.contam. | Serpine1 |
| Slc38a61 | 1.66E-105 | 2.18364298 | 0.545 | 0.08 | 3.20E-101 | TAM.contam. | Slc38a6 |
| Tmbim6 | 2.36E-105 | 0.32799192 | 0.909 | 0.674 | 4.57E-101 | TAM.contam. | Tmbim6 |
| Dse | 4.04E-105 | 1.91318784 | 0.606 | 0.114 | 7.80E-101 | TAM.contam. | Dse |
| Hsbp1 | 9.09E-105 | 0.51913823 | 0.848 | 0.491 | 1.76E-100 | TAM.contam. | Hsbp1 |
| Rreb12 | 1.14E-104 | 1.25623204 | 0.778 | 0.253 | 2.20E-100 | TAM.contam. | Rreb1 |
| Sox4 | 3.84E-104 | 1.36040329 | 0.364 | 0.082 | 7.42E-100 | TAM.contam. | Sox4 |
| Tnfsf14 | 3.87E-104 | 2.79042787 | 0.364 | 0.033 | 7.48E-100 | TAM.contam. | Tnfsf14 |
| Myo5a | 6.67E-104 | 0.66354971 | 0.828 | 0.45 | 1.29E-99 | TAM.contam. | Myo5a |
| Ap2a22 | 8.60E-104 | 0.6437315 | 0.788 | 0.46 | 1.66E-99 | TAM.contam. | Ap2a2 |
| Tmem176a3 | 2.38E-103 | 0.38902718 | 0.838 | 0.416 | 4.60E-99 | TAM.contam. | Tmem176a |
| Atp6ap22 | 1.40E-102 | 0.48474709 | 0.929 | 0.602 | 2.71E-98 | TAM.contam. | Atp6ap2 |
| Tlr4 | 1.65E-102 | 3.15373779 | 0.515 | 0.04 | 3.19E-98 | TAM.contam. | Tlr4 |
| Zfand52 | 4.30E-102 | 0.27889727 | 0.919 | 0.644 | 8.30E-98 | TAM.contam. | Zfand5 |
| Klhl9 | 4.87E-102 | 0.94728156 | 0.758 | 0.321 | 9.42E-98 | TAM.contam. | Klhl9 |
| Irs21 | 3.35E-101 | 0.51160074 | 0.838 | 0.458 | 6.49E-97 | TAM.contam. | Irs2 |
| Zc3h12a3 | 1.02E-100 | 0.49475426 | 0.808 | 0.496 | 1.97E-96 | TAM.contam. | Zc3h12a |
| Sh3bp21 | 1.66E-100 | 1.30710744 | 0.818 | 0.244 | 3.21E-96 | TAM.contam. | Sh3bp2 |
| Glmp | 1.81E-100 | 0.53214852 | 0.848 | 0.501 | 3.50E-96 | TAM.contam. | Glmp |
| C5ar2 | 2.07E-100 | 3.73912169 | 0.455 | 0.019 | 4.00E-96 | TAM.contam. | C5ar2 |
| Phlda11 | 4.67E-100 | 1.28968896 | 0.505 | 0.148 | 9.03E-96 | TAM.contam. | Phlda1 |
| Flcn1 | 8.93E-100 | 1.25996591 | 0.727 | 0.214 | 1.73E-95 | TAM.contam. | Flcn |
| Atxn1 | 1.77E-99 | 1.30049197 | 0.545 | 0.084 | 3.42E-95 | TAM.contam. | Atxn1 |
| Man2a11 | 2.07E-99 | 1.04765174 | 0.778 | 0.311 | 4.00E-95 | TAM.contam. | Man2a1 |
| Man1c11 | 2.14E-99 | 1.90186474 | 0.596 | 0.117 | 4.13E-95 | TAM.contam. | Man1c1 |
| Mef2a1 | 4.39E-99 | 0.81854614 | 0.879 | 0.394 | 8.48E-95 | TAM.contam. | Mef2a |
| Eps152 | 6.35E-99 | 0.46261443 | 0.828 | 0.45 | 1.23E-94 | TAM.contam. | Eps15 |
| Lyl1 | 7.54E-99 | 2.54634732 | 0.495 | 0.065 | 1.46E-94 | TAM.contam. | Lyl1 |
| Cmtm31 | 1.90E-98 | 1.00011098 | 0.758 | 0.274 | 3.68E-94 | TAM.contam. | Cmtm3 |
| Gab1 | 5.36E-98 | 3.28580825 | 0.404 | 0.027 | 1.04E-93 | TAM.contam. | Gab1 |
| Mapkapk23 | 1.46E-97 | 0.25162485 | 0.97 | 0.689 | 2.83E-93 | TAM.contam. | Mapkapk2 |
| Ap2s11 | 2.02E-97 | 0.68441001 | 0.848 | 0.478 | 3.91E-93 | TAM.contam. | Ap2s1 |
| Slco2b1 | 2.23E-97 | 6.94211364 | 0.444 | 0.003 | 4.31E-93 | TAM.contam. | Slco2b1 |
| Myof1 | 1.49E-96 | 0.84406544 | 0.758 | 0.282 | 2.88E-92 | TAM.contam. | Myof |
| P4ha12 | 1.98E-96 | 1.14870137 | 0.707 | 0.279 | 3.83E-92 | TAM.contam. | P4ha1 |
| Wsb12 | 4.76E-96 | 0.37313281 | 0.879 | 0.557 | 9.20E-92 | TAM.contam. | Wsb1 |
| P2ry12 | 5.24E-96 | 3.0525496 | 0.384 | 0.034 | 1.01E-91 | TAM.contam. | P2ry12 |
| Ncoa42 | 6.88E-96 | 0.64574164 | 0.828 | 0.419 | 1.33E-91 | TAM.contam. | Ncoa4 |
| Scarb22 | 1.81E-95 | 0.8234847 | 0.818 | 0.364 | 3.50E-91 | TAM.contam. | Scarb2 |
| Atp6v0a12 | 1.95E-95 | 0.69699998 | 0.808 | 0.309 | 3.77E-91 | TAM.contam. | Atp6v0a1 |
| Galnt21 | 9.12E-95 | 1.0287964 | 0.798 | 0.316 | 1.76E-90 | TAM.contam. | Galnt2 |
| Appl21 | 9.98E-95 | 1.76916455 | 0.586 | 0.135 | 1.93E-90 | TAM.contam. | Appl2 |
| Ms4a6c1 | 1.47E-94 | 0.82763786 | 0.707 | 0.271 | 2.83E-90 | TAM.contam. | Ms4a6c |
| Atp6v0b1 | 2.09E-94 | 0.46199822 | 0.909 | 0.559 | 4.03E-90 | TAM.contam. | Atp6v0b |
| Tnfaip22 | 2.81E-94 | 0.3903111 | 0.818 | 0.418 | 5.43E-90 | TAM.contam. | Tnfaip2 |
| C4a | 4.03E-94 | 5.07850412 | 0.263 | 0.004 | 7.79E-90 | TAM.contam. | C4a |
| P2rx7 | 1.05E-93 | 2.8268205 | 0.515 | 0.054 | 2.02E-89 | TAM.contam. | P2rx7 |
| Arrdc42 | 1.58E-93 | 0.71291152 | 0.838 | 0.392 | 3.06E-89 | TAM.contam. | Arrdc4 |
| Arhgef10l1 | 5.76E-93 | 1.27596306 | 0.626 | 0.125 | 1.11E-88 | TAM.contam. | Arhgef10l |
| Cd332 | 8.97E-93 | 0.95965728 | 0.707 | 0.253 | 1.73E-88 | TAM.contam. | Cd33 |
| Rabac11 | 1.02E-92 | 0.35572871 | 0.889 | 0.536 | 1.96E-88 | TAM.contam. | Rabac1 |
| Adrb2 | 1.07E-92 | 1.10790853 | 0.424 | 0.113 | 2.07E-88 | TAM.contam. | Adrb2 |
| Lrp121 | 2.43E-92 | 1.73059818 | 0.616 | 0.157 | 4.69E-88 | TAM.contam. | Lrp12 |
| Tnfrsf11a2 | 9.28E-92 | 0.45703174 | 0.788 | 0.348 | 1.79E-87 | TAM.contam. | Tnfrsf11a |
| Osm2 | 9.38E-91 | 0.55587399 | 0.687 | 0.308 | 1.81E-86 | TAM.contam. | Osm |
| Fnip21 | 1.14E-90 | 1.19398084 | 0.596 | 0.178 | 2.20E-86 | TAM.contam. | Fnip2 |
| Gpr652 | 1.27E-90 | 0.57491427 | 0.707 | 0.28 | 2.46E-86 | TAM.contam. | Gpr65 |
| Snx82 | 1.60E-90 | 0.94609267 | 0.828 | 0.317 | 3.10E-86 | TAM.contam. | Snx8 |
| Itga61 | 1.86E-90 | 1.14539774 | 0.596 | 0.187 | 3.59E-86 | TAM.contam. | Itga6 |
| Cfh | 2.28E-90 | 1.82061468 | 0.444 | 0.109 | 4.40E-86 | TAM.contam. | Cfh |
| Osbpl81 | 2.39E-90 | 0.40754817 | 0.838 | 0.439 | 4.62E-86 | TAM.contam. | Osbpl8 |
| Sertad12 | 8.98E-90 | 0.81088761 | 0.677 | 0.284 | 1.74E-85 | TAM.contam. | Sertad1 |
| Usf21 | 1.72E-89 | 0.28507626 | 0.909 | 0.596 | 3.32E-85 | TAM.contam. | Usf2 |
| Pcyox1 | 3.47E-89 | 1.70945444 | 0.525 | 0.117 | 6.72E-85 | TAM.contam. | Pcyox1 |
| Capg2 | 4.48E-89 | 0.63175878 | 0.798 | 0.311 | 8.66E-85 | TAM.contam. | Capg |
| Pdia44 | 4.61E-89 | 0.58204275 | 0.788 | 0.394 | 8.92E-85 | TAM.contam. | Pdia4 |
| Fblim1 | 1.51E-88 | 3.08454794 | 0.394 | 0.025 | 2.91E-84 | TAM.contam. | Fblim1 |
| Dtx41 | 1.61E-88 | 1.43196809 | 0.545 | 0.136 | 3.11E-84 | TAM.contam. | Dtx4 |
| Cybb4 | 6.10E-88 | 0.50965468 | 0.727 | 0.345 | 1.18E-83 | TAM.contam. | Cybb |
| Tmem87b | 6.36E-88 | 1.58026703 | 0.596 | 0.121 | 1.23E-83 | TAM.contam. | Tmem87b |
| Mmp191 | 7.88E-88 | 1.25652305 | 0.606 | 0.172 | 1.52E-83 | TAM.contam. | Mmp19 |
| Snx292 | 2.82E-87 | 0.85637906 | 0.818 | 0.275 | 5.44E-83 | TAM.contam. | Snx29 |
| Clptm12 | 3.33E-87 | 0.42657529 | 0.838 | 0.506 | 6.43E-83 | TAM.contam. | Clptm1 |
| Tmem104 | 1.43E-86 | 1.17449109 | 0.646 | 0.186 | 2.77E-82 | TAM.contam. | Tmem104 |
| Tspan4 | 3.54E-86 | 2.22098577 | 0.455 | 0.065 | 6.85E-82 | TAM.contam. | Tspan4 |
| Cflar2 | 3.75E-86 | 0.54708737 | 0.828 | 0.363 | 7.25E-82 | TAM.contam. | Cflar |
| Mxi12 | 5.58E-86 | 0.30242211 | 0.778 | 0.4 | 1.08E-81 | TAM.contam. | Mxi1 |
| Sulf23 | 9.84E-86 | 0.59925941 | 0.818 | 0.401 | 1.90E-81 | TAM.contam. | Sulf2 |
| Ypel32 | 1.01E-85 | 0.9907549 | 0.747 | 0.286 | 1.94E-81 | TAM.contam. | Ypel3 |
| Cltc1 | 2.45E-85 | 0.51537688 | 0.838 | 0.525 | 4.73E-81 | TAM.contam. | Cltc |
| Sesn1 | 2.74E-85 | 2.76341174 | 0.434 | 0.046 | 5.30E-81 | TAM.contam. | Sesn1 |
| Sod3 | 2.90E-85 | 0.83267628 | 0.293 | 0.063 | 5.60E-81 | TAM.contam. | Sod3 |
| Comt2 | 3.36E-85 | 1.02144107 | 0.747 | 0.296 | 6.50E-81 | TAM.contam. | Comt |
| Ddhd12 | 2.25E-84 | 0.67193506 | 0.788 | 0.357 | 4.36E-80 | TAM.contam. | Ddhd1 |
| Dstn1 | 1.19E-83 | 0.25930369 | 0.869 | 0.476 | 2.31E-79 | TAM.contam. | Dstn |
| Cyfip11 | 1.49E-83 | 0.33816115 | 0.848 | 0.559 | 2.89E-79 | TAM.contam. | Cyfip1 |
| Amdhd21 | 1.61E-83 | 1.24120414 | 0.657 | 0.233 | 3.11E-79 | TAM.contam. | Amdhd2 |
| Cenpb | 2.12E-83 | 0.67287666 | 0.899 | 0.402 | 4.11E-79 | TAM.contam. | Cenpb |
| Hes1 | 3.77E-83 | 1.87579242 | 0.273 | 0.049 | 7.28E-79 | TAM.contam. | Hes1 |
| Hist1h2bc1 | 6.48E-83 | 0.86512388 | 0.636 | 0.205 | 1.25E-78 | TAM.contam. | Hist1h2bc |
| Slc13a3 | 8.67E-83 | 2.58421312 | 0.333 | 0.047 | 1.68E-78 | TAM.contam. | Slc13a3 |
| Lonrf3 | 4.29E-82 | 3.63713279 | 0.354 | 0.017 | 8.30E-78 | TAM.contam. | Lonrf3 |
| DEPRECATED-I | 5.01E-82 | 2.58986878 | 0.434 | 0.054 | 9.69E-78 | TAM.contam. | DEPRECATED-ENSMUSG00000002103 |
| Snx51 | 1.08E-81 | 0.8027072 | 0.788 | 0.354 | 2.09E-77 | TAM.contam. | Snx5 |
| Tsku | 1.85E-81 | 3.86552255 | 0.333 | 0.015 | 3.58E-77 | TAM.contam. | Tsku |
| Fkbp8 | 1.97E-81 | 0.2636893 | 0.97 | 0.661 | 3.81E-77 | TAM.contam. | Fkbp8 |
| Plbd21 | 2.19E-81 | 0.71728875 | 0.758 | 0.388 | 4.24E-77 | TAM.contam. | Plbd2 |
| Tbc1d22a | 2.33E-81 | 0.73268176 | 0.798 | 0.372 | 4.50E-77 | TAM.contam. | Tbc1d22a |
| Rxra1 | 2.68E-81 | 1.32679615 | 0.596 | 0.171 | 5.18E-77 | TAM.contam. | Rxra |
| Fam234a | 6.17E-81 | 1.41830551 | 0.626 | 0.177 | 1.19E-76 | TAM.contam. | Fam234a |
| Lrpap12 | 1.05E-80 | 0.98206426 | 0.707 | 0.3 | 2.02E-76 | TAM.contam. | Lrpap1 |
| Nupr11 | 1.87E-80 | 1.5067187 | 0.535 | 0.097 | 3.62E-76 | TAM.contam. | Nupr1 |
| Ifnar22 | 5.15E-80 | 0.47310198 | 0.869 | 0.493 | 9.96E-76 | TAM.contam. | Ifnar2 |
| Gpr34 | 6.00E-80 | 3.25008117 | 0.303 | 0.022 | 1.16E-75 | TAM.contam. | Gpr34 |
| Lmo22 | 9.18E-80 | 0.73800416 | 0.788 | 0.375 | 1.78E-75 | TAM.contam. | Lmo2 |
| Mdm22 | 9.20E-80 | 0.5964503 | 0.687 | 0.381 | 1.78E-75 | TAM.contam. | Mdm2 |
| Gpx3 | 1.93E-79 | 1.96828503 | 0.323 | 0.045 | 3.73E-75 | TAM.contam. | Gpx3 |
| S1pr1 | 7.25E-79 | 2.04959246 | 0.343 | 0.021 | 1.40E-74 | TAM.contam. | S1pr1 |
| Atp1b33 | 1.12E-78 | 0.72307411 | 0.879 | 0.437 | 2.17E-74 | TAM.contam. | Atp1b3 |
| Rab322 | 1.15E-78 | 0.64720814 | 0.788 | 0.359 | 2.22E-74 | TAM.contam. | Rab32 |
| Ptafr3 | 2.22E-78 | 0.40071681 | 0.818 | 0.385 | 4.29E-74 | TAM.contam. | Ptafr |
| Kank2 | 2.90E-78 | 2.77002525 | 0.323 | 0.016 | 5.60E-74 | TAM.contam. | Kank2 |
| Plxnb21 | 3.14E-78 | 0.64135908 | 0.848 | 0.388 | 6.07E-74 | TAM.contam. | Plxnb2 |
| Rbpj1 | 6.25E-78 | 0.39869159 | 0.929 | 0.557 | 1.21E-73 | TAM.contam. | Rbpj |
| Cerk2 | 1.45E-77 | 1.33185053 | 0.707 | 0.2 | 2.81E-73 | TAM.contam. | Cerk |
| Galc | 6.21E-77 | 1.50862656 | 0.485 | 0.069 | 1.20E-72 | TAM.contam. | Galc |
| Vmp12 | 6.56E-77 | 0.30481149 | 0.879 | 0.559 | 1.27E-72 | TAM.contam. | Vmp1 |
| Klf23 | 9.37E-77 | 0.83725932 | 0.697 | 0.186 | 1.81E-72 | TAM.contam. | Klf2 |
| Arl8a1 | 1.43E-76 | 0.39191619 | 0.869 | 0.5 | 2.77E-72 | TAM.contam. | Arl8a |
| Adcy3 | 1.46E-76 | 5.19556767 | 0.263 | 0.004 | 2.83E-72 | TAM.contam. | Adcy3 |
| Tsc22d31 | 3.27E-76 | 0.71516921 | 0.687 | 0.244 | 6.33E-72 | TAM.contam. | Tsc22d3 |
| Rasgef1b3 | 6.67E-76 | 0.41773083 | 0.768 | 0.352 | 1.29E-71 | TAM.contam. | Rasgef1b |
| Cemip2 | 7.03E-76 | 2.0933983 | 0.414 | 0.063 | 1.36E-71 | TAM.contam. | Cemip2 |
| Nek62 | 7.62E-76 | 0.46686644 | 0.768 | 0.324 | 1.47E-71 | TAM.contam. | Nek6 |
| Tceal9 | 9.32E-76 | 1.4869202 | 0.646 | 0.154 | 1.80E-71 | TAM.contam. | Tceal9 |
| Mpp1 | 9.42E-76 | 1.79616976 | 0.495 | 0.096 | 1.82E-71 | TAM.contam. | Mpp1 |
| Ppp1r12c | 1.46E-75 | 0.50890516 | 0.919 | 0.498 | 2.82E-71 | TAM.contam. | Ppp1r12c |
| Cracr2b | 2.98E-75 | 3.86148298 | 0.323 | 0.015 | 5.77E-71 | TAM.contam. | Cracr2b |
| Smox2 | 4.68E-75 | 0.57767816 | 0.768 | 0.341 | 9.05E-71 | TAM.contam. | Smox |
| Pdgfa | 8.04E-75 | 2.38599644 | 0.374 | 0.036 | 1.55E-70 | TAM.contam. | Pdgfa |
| Etv51 | 1.43E-74 | 1.24343666 | 0.636 | 0.21 | 2.76E-70 | TAM.contam. | Etv5 |
| Cpd1 | 2.13E-74 | 1.55505111 | 0.485 | 0.125 | 4.12E-70 | TAM.contam. | Cpd |
| Tmem81 | 3.67E-74 | 0.40906289 | 0.717 | 0.375 | 7.10E-70 | TAM.contam. | Tmem8 |
| Lgals92 | 4.03E-74 | 0.90420017 | 0.747 | 0.279 | 7.79E-70 | TAM.contam. | Lgals9 |
| Bach11 | 1.68E-73 | 0.2830619 | 0.869 | 0.525 | 3.25E-69 | TAM.contam. | Bach1 |
| Mtus1 | 2.77E-73 | 2.12674846 | 0.424 | 0.054 | 5.35E-69 | TAM.contam. | Mtus1 |
| Sypl1 | 3.67E-73 | 1.0800049 | 0.717 | 0.248 | 7.10E-69 | TAM.contam. | Sypl |
| Zfp6221 | 3.83E-73 | 0.69376933 | 0.667 | 0.33 | 7.41E-69 | TAM.contam. | Zfp622 |
| Tspan321 | 4.50E-73 | 3.83493839 | 0.293 | 0.008 | 8.69E-69 | TAM.contam. | Tspan32 |
| Idh11 | 4.81E-73 | 0.97846071 | 0.535 | 0.163 | 9.30E-69 | TAM.contam. | Idh1 |
| Ophn1 | 1.12E-72 | 4.91226761 | 0.283 | 0.006 | 2.17E-68 | TAM.contam. | Ophn1 |
| Mtss12 | 1.93E-72 | 0.40546427 | 0.788 | 0.465 | 3.73E-68 | TAM.contam. | Mtss1 |
| Cebpg | 4.06E-72 | 0.59760089 | 0.667 | 0.288 | 7.85E-68 | TAM.contam. | Cebpg |
| Epb41l22 | 4.37E-72 | 0.37492646 | 0.768 | 0.459 | 8.46E-68 | TAM.contam. | Epb41l2 |
| Cebpd2 | 8.76E-72 | 0.71754641 | 0.727 | 0.27 | 1.69E-67 | TAM.contam. | Cebpd |
| Cela1 | 1.16E-71 | 6.28009078 | 0.303 | 0.003 | 2.23E-67 | TAM.contam. | Cela1 |
| Dusp31 | 1.22E-71 | 0.28783284 | 0.869 | 0.523 | 2.36E-67 | TAM.contam. | Dusp3 |
| Plod32 | 1.27E-71 | 1.01035508 | 0.687 | 0.261 | 2.46E-67 | TAM.contam. | Plod3 |
| Clcn71 | 1.33E-71 | 1.07635671 | 0.576 | 0.194 | 2.57E-67 | TAM.contam. | Clcn7 |
| Tpi12 | 1.35E-71 | 0.32955247 | 0.798 | 0.472 | 2.61E-67 | TAM.contam. | Tpi1 |
| Itgal3 | 1.57E-71 | 0.63377149 | 0.515 | 0.182 | 3.04E-67 | TAM.contam. | Itgal |
| Ddit42 | 1.87E-71 | 0.39870967 | 0.616 | 0.301 | 3.62E-67 | TAM.contam. | Ddit4 |
| Bcl2l12 | 2.94E-71 | 0.37778098 | 0.707 | 0.295 | 5.69E-67 | TAM.contam. | Bcl2l1 |
| Nceh11 | 2.95E-71 | 1.53283075 | 0.525 | 0.114 | 5.70E-67 | TAM.contam. | Nceh1 |
| Src1 | 4.58E-71 | 0.57180645 | 0.677 | 0.282 | 8.85E-67 | TAM.contam. | Src |
| P2ry131 | 7.67E-71 | 2.02435249 | 0.354 | 0.045 | 1.48E-66 | TAM.contam. | P2ry13 |
| Gla3 | 1.49E-70 | 1.03616994 | 0.697 | 0.216 | 2.87E-66 | TAM.contam. | Gla |
| Ly863 | 5.99E-70 | 0.3063532 | 0.879 | 0.526 | 1.16E-65 | TAM.contam. | Ly86 |
| Slc35f61 | 7.09E-70 | 1.39901777 | 0.596 | 0.165 | 1.37E-65 | TAM.contam. | Slc35f6 |
| Pon21 | 1.19E-69 | 0.93323276 | 0.596 | 0.22 | 2.31E-65 | TAM.contam. | Pon2 |
| F101 | 1.33E-69 | 1.56046506 | 0.263 | 0.064 | 2.56E-65 | TAM.contam. | F10 |
| Id3 | 2.55E-69 | 1.4110803 | 0.374 | 0.096 | 4.93E-65 | TAM.contam. | Id3 |
| Ikbke1 | 2.85E-69 | 1.84835411 | 0.485 | 0.055 | 5.50E-65 | TAM.contam. | Ikbke |
| Daglb1 | 3.49E-69 | 1.21588896 | 0.727 | 0.219 | 6.74E-65 | TAM.contam. | Daglb |
| Gadd45g2 | 6.87E-69 | 0.61317404 | 0.576 | 0.236 | 1.33E-64 | TAM.contam. | Gadd45g |
| Emilin21 | 7.30E-69 | 0.35635728 | 0.808 | 0.398 | 1.41E-64 | TAM.contam. | Emilin2 |
| Slc37a21 | 7.61E-69 | 1.54641659 | 0.525 | 0.098 | 1.47E-64 | TAM.contam. | Slc37a2 |
| Arl4c2 | 9.13E-69 | 0.40723276 | 0.879 | 0.429 | 1.77E-64 | TAM.contam. | Arl4c |
| Tlr71 | 1.21E-68 | 1.9385032 | 0.384 | 0.024 | 2.35E-64 | TAM.contam. | Tlr7 |
| Arrdc3 | 3.69E-68 | 1.84559731 | 0.455 | 0.076 | 7.13E-64 | TAM.contam. | Arrdc3 |
| Tnfaip8l22 | 4.01E-68 | 0.75159144 | 0.677 | 0.312 | 7.75E-64 | TAM.contam. | Tnfaip8l2 |
| Inpp4b | 4.12E-68 | 2.41575896 | 0.333 | 0.022 | 7.96E-64 | TAM.contam. | Inpp4b |
| Hfe2 | 5.86E-68 | 0.36166028 | 0.869 | 0.457 | 1.13E-63 | TAM.contam. | Hfe |
| Cln8 | 5.99E-68 | 1.82617234 | 0.596 | 0.129 | 1.16E-63 | TAM.contam. | Cln8 |
| Itgav2 | 7.52E-68 | 0.70092187 | 0.636 | 0.262 | 1.45E-63 | TAM.contam. | Itgav |
| Asah13 | 1.08E-67 | 0.38037605 | 0.879 | 0.516 | 2.08E-63 | TAM.contam. | Asah1 |
| Trim81 | 1.14E-67 | 0.59171721 | 0.828 | 0.379 | 2.20E-63 | TAM.contam. | Trim8 |
| Gas2l3 | 2.27E-67 | 3.05955992 | 0.404 | 0.042 | 4.38E-63 | TAM.contam. | Gas2l3 |
| Lyz1 | 4.80E-67 | 3.16394018 | 0.343 | 0.019 | 9.28E-63 | TAM.contam. | Lyz1 |
| Cdk9 | 1.25E-66 | 0.32414387 | 0.879 | 0.554 | 2.41E-62 | TAM.contam. | Cdk9 |
| Ccni2 | 1.34E-66 | 0.25156999 | 0.859 | 0.515 | 2.58E-62 | TAM.contam. | Ccni |
| Fgd4 | 3.30E-66 | 3.21981074 | 0.323 | 0.016 | 6.39E-62 | TAM.contam. | Fgd4 |
| Sema4b1 | 3.47E-66 | 1.2011754 | 0.414 | 0.059 | 6.71E-62 | TAM.contam. | Sema4b |
| Nampt2 | 7.34E-66 | 0.35869734 | 0.778 | 0.432 | 1.42E-61 | TAM.contam. | Nampt |
| Lgals8 | 8.53E-66 | 1.23748453 | 0.606 | 0.146 | 1.65E-61 | TAM.contam. | Lgals8 |
| Etv1 | 9.81E-66 | 5.98899841 | 0.273 | 0.003 | 1.90E-61 | TAM.contam. | Etv1 |
| Sh2b23 | 1.04E-65 | 1.03753401 | 0.556 | 0.186 | 2.00E-61 | TAM.contam. | Sh2b2 |
| Fcrlb1 | 1.50E-65 | 1.71982721 | 0.384 | 0.076 | 2.90E-61 | TAM.contam. | Fcrlb |
| Tpst21 | 2.73E-65 | 1.03075596 | 0.646 | 0.179 | 5.27E-61 | TAM.contam. | Tpst2 |
| Tlr8 | 6.91E-65 | 3.46072438 | 0.343 | 0.02 | 1.34E-60 | TAM.contam. | Tlr8 |
| Pip4p11 | 7.75E-65 | 0.53218103 | 0.788 | 0.389 | 1.50E-60 | TAM.contam. | Pip4p1 |
| Zswim61 | 2.20E-64 | 1.56093714 | 0.535 | 0.119 | 4.26E-60 | TAM.contam. | Zswim6 |
| Tceal8 | 2.24E-64 | 1.57324767 | 0.465 | 0.094 | 4.33E-60 | TAM.contam. | Tceal8 |
| Cln32 | 2.67E-64 | 1.13713332 | 0.586 | 0.165 | 5.17E-60 | TAM.contam. | Cln3 |
| Ang | 4.30E-64 | 5.71886749 | 0.263 | 0.003 | 8.32E-60 | TAM.contam. | Ang |
| Zfand2a1 | 4.45E-64 | 0.95626659 | 0.616 | 0.189 | 8.60E-60 | TAM.contam. | Zfand2a |
| Mapk32 | 4.59E-64 | 0.3307801 | 0.788 | 0.425 | 8.88E-60 | TAM.contam. | Mapk3 |
| Bscl23 | 8.98E-64 | 0.5166595 | 0.727 | 0.352 | 1.74E-59 | TAM.contam. | Bscl2 |
| Tex264 | 1.12E-63 | 1.34097667 | 0.465 | 0.125 | 2.16E-59 | TAM.contam. | Tex264 |
| Eif4ebp11 | 1.75E-63 | 1.18587877 | 0.566 | 0.177 | 3.38E-59 | TAM.contam. | Eif4ebp1 |
| Twf12 | 2.34E-63 | 0.87109003 | 0.717 | 0.254 | 4.52E-59 | TAM.contam. | Twf1 |
| Cab39 | 5.58E-63 | 0.33928499 | 0.869 | 0.528 | 1.08E-58 | TAM.contam. | Cab39 |
| Clec4a2 | 5.98E-63 | 2.0960385 | 0.475 | 0.077 | 1.16E-58 | TAM.contam. | Clec4a2 |
| Gaa2 | 6.53E-63 | 0.74119698 | 0.707 | 0.276 | 1.26E-58 | TAM.contam. | Gaa |
| Dmxl2 | 9.38E-63 | 2.56496593 | 0.384 | 0.037 | 1.81E-58 | TAM.contam. | Dmxl2 |
| Fez2 | 1.07E-62 | 1.26171075 | 0.566 | 0.136 | 2.06E-58 | TAM.contam. | Fez2 |
| Abcc5 | 1.84E-62 | 1.55544022 | 0.394 | 0.053 | 3.55E-58 | TAM.contam. | Abcc5 |
| Mindy1 | 2.01E-62 | 1.68498187 | 0.414 | 0.07 | 3.88E-58 | TAM.contam. | Mindy1 |
| Gng122 | 2.57E-62 | 1.21606126 | 0.596 | 0.143 | 4.96E-58 | TAM.contam. | Gng12 |
| Washc2 | 2.61E-62 | 0.34070231 | 0.859 | 0.43 | 5.05E-58 | TAM.contam. | Washc2 |
| Atp6v1c11 | 5.34E-62 | 0.67854177 | 0.758 | 0.355 | 1.03E-57 | TAM.contam. | Atp6v1c1 |
| Soat11 | 5.92E-62 | 0.50496124 | 0.778 | 0.405 | 1.15E-57 | TAM.contam. | Soat1 |
| Pgd2 | 7.15E-62 | 0.49501786 | 0.838 | 0.446 | 1.38E-57 | TAM.contam. | Pgd |
| Pitpnc1 | 8.16E-62 | 1.76433126 | 0.485 | 0.079 | 1.58E-57 | TAM.contam. | Pitpnc1 |
| Ncstn1 | 1.15E-61 | 0.4112485 | 0.798 | 0.439 | 2.22E-57 | TAM.contam. | Ncstn |
| Tmem652 | 1.59E-61 | 1.03020291 | 0.646 | 0.209 | 3.07E-57 | TAM.contam. | Tmem65 |
| Atp2a24 | 3.02E-61 | 0.32937599 | 0.818 | 0.495 | 5.84E-57 | TAM.contam. | Atp2a2 |
| Kif13b1 | 3.48E-61 | 1.29480948 | 0.455 | 0.112 | 6.73E-57 | TAM.contam. | Kif13b |
| Rragc1 | 3.49E-61 | 0.79106404 | 0.717 | 0.312 | 6.76E-57 | TAM.contam. | Rragc |
| Atp6v1e11 | 6.62E-61 | 0.43986562 | 0.879 | 0.477 | 1.28E-56 | TAM.contam. | Atp6v1e1 |
| Ulk2 | 9.32E-61 | 1.25595006 | 0.475 | 0.14 | 1.80E-56 | TAM.contam. | Ulk2 |
| Ip6k11 | 1.02E-60 | 0.43312688 | 0.778 | 0.389 | 1.98E-56 | TAM.contam. | Ip6k1 |
| Tut71 | 1.09E-60 | 0.25771217 | 0.889 | 0.556 | 2.11E-56 | TAM.contam. | Tut7 |
| Neu11 | 1.19E-60 | 0.95073164 | 0.626 | 0.231 | 2.30E-56 | TAM.contam. | Neu1 |
| Socs21 | 1.19E-60 | 0.25670127 | 0.727 | 0.281 | 2.31E-56 | TAM.contam. | Socs2 |
| Ubtd12 | 4.24E-60 | 0.91686082 | 0.646 | 0.257 | 8.21E-56 | TAM.contam. | Ubtd1 |
| Slc9a92 | 4.40E-60 | 0.99976868 | 0.596 | 0.215 | 8.51E-56 | TAM.contam. | Slc9a9 |
| Arg21 | 5.38E-60 | 1.89265426 | 0.354 | 0.051 | 1.04E-55 | TAM.contam. | Arg2 |
| Ldlrap1 | 1.32E-59 | 1.6062892 | 0.444 | 0.072 | 2.55E-55 | TAM.contam. | Ldlrap1 |
| Fli12 | 1.34E-59 | 0.6418357 | 0.768 | 0.317 | 2.59E-55 | TAM.contam. | Fli1 |
| Sod22 | 2.12E-59 | 0.59549853 | 0.778 | 0.386 | 4.10E-55 | TAM.contam. | Sod2 |
| Xylt2 | 5.21E-59 | 2.1726725 | 0.434 | 0.06 | 1.01E-54 | TAM.contam. | Xylt2 |
| Ctdsp1 | 5.23E-59 | 0.46216226 | 0.818 | 0.426 | 1.01E-54 | TAM.contam. | Ctdsp1 |
| Slc12a71 | 6.18E-59 | 0.89208539 | 0.636 | 0.216 | 1.20E-54 | TAM.contam. | Slc12a7 |
| Man2b2 | 7.31E-59 | 0.56668469 | 0.717 | 0.328 | 1.41E-54 | TAM.contam. | Man2b2 |
| Tbc1d142 | 7.45E-59 | 0.29533889 | 0.818 | 0.465 | 1.44E-54 | TAM.contam. | Tbc1d14 |
| Fchsd21 | 1.04E-58 | 0.35506023 | 0.869 | 0.459 | 2.00E-54 | TAM.contam. | Fchsd2 |
| Hexim12 | 1.25E-58 | 0.36946255 | 0.828 | 0.416 | 2.42E-54 | TAM.contam. | Hexim1 |
| Stom1 | 1.63E-58 | 0.88508198 | 0.636 | 0.197 | 3.15E-54 | TAM.contam. | Stom |
| Ephx1 | 2.42E-58 | 2.55738446 | 0.253 | 0.022 | 4.67E-54 | TAM.contam. | Ephx1 |
| Mid1ip1 | 2.65E-58 | 1.21856867 | 0.485 | 0.108 | 5.13E-54 | TAM.contam. | Mid1ip1 |
| Pnpla21 | 2.81E-58 | 0.47219801 | 0.848 | 0.429 | 5.43E-54 | TAM.contam. | Pnpla2 |
| Nfxl1 | 2.86E-58 | 1.55450421 | 0.525 | 0.106 | 5.53E-54 | TAM.contam. | Nfxl1 |
| Fndc3a1 | 3.85E-58 | 0.568149 | 0.707 | 0.36 | 7.45E-54 | TAM.contam. | Fndc3a |
| Os9 | 4.75E-58 | 0.35478333 | 0.879 | 0.466 | 9.18E-54 | TAM.contam. | Os9 |
| Tor4a | 6.28E-58 | 1.20956989 | 0.596 | 0.142 | 1.21E-53 | TAM.contam. | Tor4a |
| Nfic | 6.45E-58 | 1.1139159 | 0.525 | 0.142 | 1.25E-53 | TAM.contam. | Nfic |
| Tob12 | 5.77E-57 | 0.60177962 | 0.556 | 0.303 | 1.12E-52 | TAM.contam. | Tob1 |
| Cd300lb | 6.67E-57 | 2.25628253 | 0.374 | 0.051 | 1.29E-52 | TAM.contam. | Cd300lb |
| Adam102 | 7.24E-57 | 0.61724309 | 0.758 | 0.334 | 1.40E-52 | TAM.contam. | Adam10 |
| Laptm4a | 8.61E-57 | 0.74900803 | 0.657 | 0.269 | 1.66E-52 | TAM.contam. | Laptm4a |
| Saraf1 | 8.76E-57 | 0.58320398 | 0.818 | 0.396 | 1.69E-52 | TAM.contam. | Saraf |
| Mitf1 | 9.25E-57 | 1.93082622 | 0.424 | 0.082 | 1.79E-52 | TAM.contam. | Mitf |
| Klra2 | 1.47E-56 | 2.80128171 | 0.293 | 0.025 | 2.83E-52 | TAM.contam. | Klra2 |
| Tpcn2 | 1.66E-56 | 1.6256747 | 0.434 | 0.098 | 3.21E-52 | TAM.contam. | Tpcn2 |
| Isg152 | 2.30E-56 | 0.50444995 | 0.586 | 0.264 | 4.44E-52 | TAM.contam. | Isg15 |
| Vps26a1 | 2.81E-56 | 0.36863458 | 0.778 | 0.403 | 5.44E-52 | TAM.contam. | Vps26a |
| Cd371 | 4.10E-56 | 0.63087525 | 0.586 | 0.179 | 7.94E-52 | TAM.contam. | Cd37 |
| Fndc3b3 | 4.80E-56 | 0.68342598 | 0.667 | 0.244 | 9.28E-52 | TAM.contam. | Fndc3b |
| Ifi272 | 8.25E-56 | 1.0322218 | 0.586 | 0.18 | 1.59E-51 | TAM.contam. | Ifi27 |
| Mafg1 | 1.18E-55 | 0.75589894 | 0.646 | 0.23 | 2.28E-51 | TAM.contam. | Mafg |
| Wbp21 | 1.27E-55 | 0.70531244 | 0.727 | 0.325 | 2.46E-51 | TAM.contam. | Wbp2 |
| Tnfrsf211 | 1.60E-55 | 0.77309998 | 0.626 | 0.202 | 3.09E-51 | TAM.contam. | Tnfrsf21 |
| Sdc11 | 1.64E-55 | 0.5818064 | 0.273 | 0.144 | 3.17E-51 | TAM.contam. | Sdc1 |
| Arc1 | 2.33E-55 | 0.39144603 | 0.354 | 0.096 | 4.50E-51 | TAM.contam. | Arc |
| Tcp11l22 | 2.36E-55 | 1.03047695 | 0.566 | 0.174 | 4.57E-51 | TAM.contam. | Tcp11l2 |
| Pea15a1 | 2.46E-55 | 0.55560298 | 0.808 | 0.377 | 4.75E-51 | TAM.contam. | Pea15a |
| Lcp22 | 3.30E-55 | 0.59073834 | 0.788 | 0.354 | 6.37E-51 | TAM.contam. | Lcp2 |
| Gpalpp12 | 4.91E-55 | 0.7265161 | 0.636 | 0.239 | 9.49E-51 | TAM.contam. | Gpalpp1 |
| Ms4a14 | 5.41E-55 | 6.91464015 | 0.283 | 0.002 | 1.05E-50 | TAM.contam. | Ms4a14 |
| St6galnac41 | 8.19E-55 | 0.71541604 | 0.626 | 0.252 | 1.58E-50 | TAM.contam. | St6galnac4 |
| Sorbs3 | 1.14E-54 | 6.13975324 | 0.293 | 0.003 | 2.21E-50 | TAM.contam. | Sorbs3 |
| Card192 | 1.27E-54 | 0.7107445 | 0.798 | 0.37 | 2.45E-50 | TAM.contam. | Card19 |
| Rai141 | 1.43E-54 | 0.9343457 | 0.515 | 0.164 | 2.76E-50 | TAM.contam. | Rai14 |
| Alox5 | 1.43E-54 | 2.46649502 | 0.263 | 0.019 | 2.77E-50 | TAM.contam. | Alox5 |
| Ero1l2 | 2.41E-54 | 0.5009475 | 0.606 | 0.329 | 4.66E-50 | TAM.contam. | Ero1l |
| Jmy2 | 2.51E-54 | 0.91273105 | 0.556 | 0.184 | 4.85E-50 | TAM.contam. | Jmy |
| Rnf133 | 3.80E-54 | 0.64622265 | 0.717 | 0.342 | 7.34E-50 | TAM.contam. | Rnf13 |
| Pon3 | 4.15E-54 | 2.95956942 | 0.293 | 0.027 | 8.02E-50 | TAM.contam. | Pon3 |
| Cnppd12 | 4.19E-54 | 0.32367552 | 0.788 | 0.429 | 8.10E-50 | TAM.contam. | Cnppd1 |
| Slc38a7 | 4.77E-54 | 2.08679731 | 0.404 | 0.048 | 9.22E-50 | TAM.contam. | Slc38a7 |
| Pacc1 | 6.31E-54 | 1.36190638 | 0.485 | 0.133 | 1.22E-49 | TAM.contam. | Pacc1 |
| Mrfap1 | 1.14E-53 | 0.27297304 | 0.788 | 0.497 | 2.20E-49 | TAM.contam. | Mrfap1 |
| Sqor1 | 1.15E-53 | 2.25518894 | 0.394 | 0.054 | 2.23E-49 | TAM.contam. | Sqor |
| Alkbh51 | 1.73E-53 | 0.42554219 | 0.788 | 0.431 | 3.35E-49 | TAM.contam. | Alkbh5 |
| Akap7 | 2.11E-53 | 2.27196028 | 0.263 | 0.041 | 4.08E-49 | TAM.contam. | Akap7 |
| Rbfa1 | 2.29E-53 | 0.99110757 | 0.636 | 0.2 | 4.43E-49 | TAM.contam. | Rbfa |
| Hvcn13 | 3.32E-53 | 0.707526 | 0.525 | 0.162 | 6.41E-49 | TAM.contam. | Hvcn1 |
| Pfkl2 | 5.72E-53 | 0.34190855 | 0.808 | 0.411 | 1.11E-48 | TAM.contam. | Pfkl |
| Mylip1 | 6.01E-53 | 0.92333347 | 0.525 | 0.159 | 1.16E-48 | TAM.contam. | Mylip |
| Slc5a31 | 6.71E-53 | 0.92892047 | 0.525 | 0.166 | 1.30E-48 | TAM.contam. | Slc5a3 |
| Dnajc131 | 6.74E-53 | 0.53840682 | 0.727 | 0.336 | 1.30E-48 | TAM.contam. | Dnajc13 |
| Wdr911 | 9.79E-53 | 0.77368764 | 0.636 | 0.229 | 1.89E-48 | TAM.contam. | Wdr91 |
| Hbp11 | 1.05E-52 | 0.39932262 | 0.707 | 0.312 | 2.03E-48 | TAM.contam. | Hbp1 |
| Ntpcr1 | 1.17E-52 | 1.08801199 | 0.545 | 0.196 | 2.26E-48 | TAM.contam. | Ntpcr |
| Vegfb | 1.38E-52 | 1.34601161 | 0.465 | 0.094 | 2.67E-48 | TAM.contam. | Vegfb |
| Ttyh22 | 1.43E-52 | 0.72223454 | 0.586 | 0.16 | 2.77E-48 | TAM.contam. | Ttyh2 |
| Rusc2 | 1.84E-52 | 2.8993873 | 0.253 | 0.025 | 3.56E-48 | TAM.contam. | Rusc2 |
| Mfsd111 | 1.84E-52 | 1.52773949 | 0.455 | 0.095 | 3.56E-48 | TAM.contam. | Mfsd11 |
| Gbp71 | 3.53E-52 | 1.20458984 | 0.465 | 0.162 | 6.82E-48 | TAM.contam. | Gbp7 |

| | | | | | | | |
|---|---|---|---|---|---|---|---|
| Rassf42 | 5.16E-52 | 0.26918314 | 0.838 | 0.421 | 9.98E-48 | TAM.contam. | Rassf4 |
| Osbpl11 | 5.40E-52 | 1.20428057 | 0.485 | 0.133 | 1.04E-47 | TAM.contam. | Osbpl11 |
| Ndufc21 | 6.06E-52 | 0.32315375 | 0.828 | 0.517 | 1.17E-47 | TAM.contam. | Ndufc2 |
| Idua | 6.13E-52 | 1.36188693 | 0.495 | 0.118 | 1.18E-47 | TAM.contam. | Idua |
| Slc11a21 | 6.42E-52 | 1.52760169 | 0.485 | 0.122 | 1.24E-47 | TAM.contam. | Slc11a2 |
| Dennd1a | 1.05E-51 | 1.31077009 | 0.525 | 0.125 | 2.03E-47 | TAM.contam. | Dennd1a |
| Sipa1l2 | 1.68E-51 | 1.99496181 | 0.404 | 0.058 | 3.26E-47 | TAM.contam. | Sipa1l2 |
| Pdlim42 | 2.02E-51 | 0.35720815 | 0.576 | 0.192 | 3.91E-47 | TAM.contam. | Pdlim4 |
| Snx93 | 3.43E-51 | 0.33683921 | 0.808 | 0.438 | 6.62E-47 | TAM.contam. | Snx9 |
| Pde2a | 4.21E-51 | 2.09925059 | 0.343 | 0.048 | 8.15E-47 | TAM.contam. | Pde2a |
| Rasgrp12 | 1.85E-50 | 0.88272755 | 0.515 | 0.162 | 3.57E-46 | TAM.contam. | Rasgrp1 |
| Susd63 | 1.98E-50 | 0.34458267 | 0.768 | 0.429 | 3.82E-46 | TAM.contam. | Susd6 |
| Idh22 | 2.28E-50 | 0.7824602 | 0.697 | 0.251 | 4.40E-46 | TAM.contam. | Idh2 |
| Pdgfb1 | 2.57E-50 | 1.16971885 | 0.444 | 0.115 | 4.97E-46 | TAM.contam. | Pdgfb |
| Tmem140 | 2.98E-50 | 1.44904352 | 0.384 | 0.055 | 5.75E-46 | TAM.contam. | Tmem140 |
| Plxna12 | 5.68E-50 | 0.92687278 | 0.636 | 0.256 | 1.10E-45 | TAM.contam. | Plxna1 |
| Irf72 | 6.84E-50 | 0.74505594 | 0.414 | 0.169 | 1.32E-45 | TAM.contam. | Irf7 |
| Ptpn182 | 7.50E-50 | 0.40469656 | 0.859 | 0.421 | 1.45E-45 | TAM.contam. | Ptpn18 |
| Cln51 | 8.91E-50 | 0.84855751 | 0.636 | 0.25 | 1.72E-45 | TAM.contam. | Cln5 |
| Inppl11 | 9.13E-50 | 0.83231426 | 0.657 | 0.213 | 1.77E-45 | TAM.contam. | Inppl1 |
| Rgmb1 | 9.52E-50 | 0.6655641 | 0.586 | 0.161 | 1.84E-45 | TAM.contam. | Rgmb |
| Jag11 | 1.15E-49 | 1.10433309 | 0.283 | 0.073 | 2.22E-45 | TAM.contam. | Jag1 |
| Gmfb1 | 1.30E-49 | 0.38435151 | 0.707 | 0.406 | 2.51E-45 | TAM.contam. | Gmfb |
| Rcan1 | 2.17E-49 | 1.96781386 | 0.343 | 0.075 | 4.19E-45 | TAM.contam. | Rcan1 |
| Alas11 | 2.76E-49 | 0.84050237 | 0.586 | 0.204 | 5.34E-45 | TAM.contam. | Alas1 |
| Gpt2 | 2.87E-49 | 2.9353472 | 0.263 | 0.02 | 5.54E-45 | TAM.contam. | Gpt2 |
| Map112 | 3.97E-49 | 0.6763758 | 0.667 | 0.258 | 7.68E-45 | TAM.contam. | Map11 |
| Dennd4c | 5.80E-49 | 1.75280571 | 0.434 | 0.075 | 1.12E-44 | TAM.contam. | Dennd4c |
| Elk31 | 8.30E-49 | 0.29460349 | 0.697 | 0.343 | 1.61E-44 | TAM.contam. | Elk3 |
| Ethe1 | 1.11E-48 | 1.3352212 | 0.455 | 0.096 | 2.15E-44 | TAM.contam. | Ethe1 |
| Ssh31 | 1.24E-48 | 1.0813628 | 0.566 | 0.141 | 2.40E-44 | TAM.contam. | Ssh3 |
| Man2a22 | 1.93E-48 | 0.52652136 | 0.687 | 0.278 | 3.72E-44 | TAM.contam. | Man2a2 |
| Adssl13 | 2.64E-48 | 0.60464215 | 0.606 | 0.257 | 5.11E-44 | TAM.contam. | Adssl1 |
| Reep32 | 3.37E-48 | 0.40930405 | 0.778 | 0.36 | 6.52E-44 | TAM.contam. | Reep3 |
| Ifi27l2a1 | 3.81E-48 | 0.53905697 | 0.475 | 0.214 | 7.36E-44 | TAM.contam. | Ifi27l2a |
| Palld3 | 5.45E-48 | 0.28316067 | 0.545 | 0.243 | 1.05E-43 | TAM.contam. | Palld |
| Chp2 | 7.45E-48 | 7.08213357 | 0.273 | 0.001 | 1.44E-43 | TAM.contam. | Chp2 |
| Mfsd123 | 1.08E-47 | 0.61088618 | 0.657 | 0.271 | 2.09E-43 | TAM.contam. | Mfsd12 |
| Zcchc241 | 1.56E-47 | 0.96527862 | 0.545 | 0.15 | 3.02E-43 | TAM.contam. | Zcchc24 |
| Sh3gl11 | 1.64E-47 | 0.32872927 | 0.677 | 0.355 | 3.17E-43 | TAM.contam. | Sh3gl1 |
| Slc16a10 | 1.81E-47 | 2.21262899 | 0.333 | 0.039 | 3.50E-43 | TAM.contam. | Slc16a10 |
| Cd302 | 2.36E-47 | 1.84867254 | 0.364 | 0.069 | 4.56E-43 | TAM.contam. | Cd302 |
| Nacc2 | 2.68E-47 | 1.67260234 | 0.343 | 0.05 | 5.18E-43 | TAM.contam. | Nacc2 |
| Mtr1 | 3.06E-47 | 1.26515802 | 0.576 | 0.176 | 5.92E-43 | TAM.contam. | Mtr |
| Eaf11 | 3.95E-47 | 0.90344839 | 0.606 | 0.184 | 7.64E-43 | TAM.contam. | Eaf1 |
| Arhgap10 | 4.44E-47 | 2.61164267 | 0.263 | 0.025 | 8.57E-43 | TAM.contam. | Arhgap10 |
| Hs1bp3 | 4.51E-47 | 1.99387518 | 0.394 | 0.05 | 8.71E-43 | TAM.contam. | Hs1bp3 |
| Gng22 | 4.99E-47 | 0.37156695 | 0.778 | 0.348 | 9.66E-43 | TAM.contam. | Gng2 |
| Cyb5r1 | 6.69E-47 | 1.67900397 | 0.465 | 0.095 | 1.29E-42 | TAM.contam. | Cyb5r1 |
| Kdelr11 | 8.27E-47 | 0.28684938 | 0.747 | 0.414 | 1.60E-42 | TAM.contam. | Kdelr1 |
| Snta1 | 1.03E-46 | 2.37144194 | 0.303 | 0.029 | 2.00E-42 | TAM.contam. | Snta1 |
| Ccng22 | 1.09E-46 | 0.27305276 | 0.626 | 0.266 | 2.12E-42 | TAM.contam. | Ccng2 |
| Mgat11 | 1.38E-46 | 0.48288783 | 0.717 | 0.313 | 2.66E-42 | TAM.contam. | Mgat1 |
| Ncor23 | 1.64E-46 | 0.37030847 | 0.707 | 0.324 | 3.18E-42 | TAM.contam. | Ncor2 |
| Ltc4s2 | 1.81E-46 | 0.79139753 | 0.475 | 0.154 | 3.50E-42 | TAM.contam. | Ltc4s |
| Secisbp2l | 2.59E-46 | 0.47082534 | 0.747 | 0.343 | 5.00E-42 | TAM.contam. | Secisbp2l |
| Tlr1 | 3.30E-46 | 1.66033315 | 0.434 | 0.081 | 6.37E-42 | TAM.contam. | Tlr1 |
| Ltbr1 | 4.47E-46 | 0.55643023 | 0.717 | 0.339 | 8.65E-42 | TAM.contam. | Ltbr |
| Gas2l11 | 4.97E-46 | 1.0895179 | 0.556 | 0.16 | 9.61E-42 | TAM.contam. | Gas2l1 |
| Nt5dc2 | 5.47E-46 | 2.19803938 | 0.273 | 0.05 | 1.06E-41 | TAM.contam. | Nt5dc2 |
| Sephs21 | 6.79E-46 | 1.27626362 | 0.545 | 0.158 | 1.31E-41 | TAM.contam. | Sephs2 |
| Hk3 | 7.84E-46 | 1.94135961 | 0.364 | 0.061 | 1.52E-41 | TAM.contam. | Hk3 |
| Arhgef71 | 1.21E-45 | 0.68948492 | 0.657 | 0.235 | 2.34E-41 | TAM.contam. | Arhgef7 |
| Ypel51 | 1.47E-45 | 0.42444644 | 0.697 | 0.323 | 2.85E-41 | TAM.contam. | Ypel5 |
| Atp6v0d12 | 1.83E-45 | 0.28873946 | 0.869 | 0.511 | 3.53E-41 | TAM.contam. | Atp6v0d1 |
| Dock11 | 2.64E-45 | 0.67834632 | 0.515 | 0.145 | 5.11E-41 | TAM.contam. | Dock1 |
| Slc7a21 | 2.66E-45 | 1.62229184 | 0.343 | 0.041 | 5.14E-41 | TAM.contam. | Slc7a2 |
| Rgcc1 | 3.38E-45 | 1.36299619 | 0.303 | 0.082 | 6.53E-41 | TAM.contam. | Rgcc |
| Dip2c1 | 3.85E-45 | 1.10578272 | 0.515 | 0.162 | 7.44E-41 | TAM.contam. | Dip2c |
| Slc25a512 | 4.79E-45 | 0.59846633 | 0.657 | 0.308 | 9.26E-41 | TAM.contam. | Slc25a51 |
| Kif1c | 5.67E-45 | 0.54973597 | 0.747 | 0.317 | 1.10E-40 | TAM.contam. | Kif1c |
| Dclre1c2 | 5.89E-45 | 0.6568293 | 0.677 | 0.228 | 1.14E-40 | TAM.contam. | Dclre1c |
| Adamtsl4 | 6.22E-45 | 2.20172502 | 0.283 | 0.044 | 1.20E-40 | TAM.contam. | Adamtsl4 |
| Prdx12 | 9.62E-45 | 0.3073305 | 0.737 | 0.449 | 1.86E-40 | TAM.contam. | Prdx1 |
| Slc25a371 | 1.47E-44 | 1.39924011 | 0.414 | 0.096 | 2.84E-40 | TAM.contam. | Slc25a37 |
| Arhgef3 | 1.60E-44 | 1.08290392 | 0.475 | 0.143 | 3.09E-40 | TAM.contam. | Arhgef3 |
| Aldh3b11 | 1.67E-44 | 1.32764014 | 0.475 | 0.104 | 3.23E-40 | TAM.contam. | Aldh3b1 |
| Wls | 2.10E-44 | 1.21983013 | 0.485 | 0.112 | 4.06E-40 | TAM.contam. | Wls |
| Naglu | 2.50E-44 | 1.08749198 | 0.455 | 0.142 | 4.84E-40 | TAM.contam. | Naglu |
| Fam53b | 3.14E-44 | 0.74626577 | 0.545 | 0.124 | 6.06E-40 | TAM.contam. | Fam53b |
| Pgm12 | 3.32E-44 | 0.75963448 | 0.545 | 0.201 | 6.42E-40 | TAM.contam. | Pgm1 |
| Serinc11 | 5.16E-44 | 0.55415541 | 0.788 | 0.394 | 9.98E-40 | TAM.contam. | Serinc1 |
| Dnase1l1 | 6.58E-44 | 0.64278198 | 0.596 | 0.232 | 1.27E-39 | TAM.contam. | Dnase1l1 |
| Clcn51 | 6.85E-44 | 1.28399301 | 0.515 | 0.124 | 1.33E-39 | TAM.contam. | Clcn5 |
| Cant1 | 8.44E-44 | 0.58725598 | 0.717 | 0.276 | 1.63E-39 | TAM.contam. | Cant1 |
| Manbal | 8.51E-44 | 0.75801257 | 0.566 | 0.217 | 1.64E-39 | TAM.contam. | Manbal |
| Phf231 | 1.01E-43 | 0.42894064 | 0.778 | 0.399 | 1.95E-39 | TAM.contam. | Phf23 |
| Lrrc8a | 1.45E-43 | 1.18526099 | 0.434 | 0.095 | 2.80E-39 | TAM.contam. | Lrrc8a |
| Dok31 | 1.69E-43 | 1.18679746 | 0.626 | 0.149 | 3.27E-39 | TAM.contam. | Dok3 |
| Blnk1 | 1.83E-43 | 1.02995788 | 0.424 | 0.065 | 3.53E-39 | TAM.contam. | Blnk |
| Abcg32 | 2.20E-43 | 1.20774682 | 0.444 | 0.124 | 4.25E-39 | TAM.contam. | Abcg3 |
| Rhbdf23 | 2.30E-43 | 0.43231595 | 0.646 | 0.278 | 4.44E-39 | TAM.contam. | Rhbdf2 |
| Ifi204 | 2.48E-43 | 1.69763387 | 0.374 | 0.064 | 4.80E-39 | TAM.contam. | Ifi204 |
| Mocs11 | 2.63E-43 | 1.14705436 | 0.404 | 0.091 | 5.08E-39 | TAM.contam. | Mocs1 |
| Smcr82 | 4.12E-43 | 0.5566239 | 0.636 | 0.281 | 7.97E-39 | TAM.contam. | Smcr8 |
| Ap1s11 | 4.33E-43 | 0.37765125 | 0.737 | 0.373 | 8.38E-39 | TAM.contam. | Ap1s1 |
| Prkacb1 | 4.36E-43 | 0.4738027 | 0.636 | 0.301 | 8.43E-39 | TAM.contam. | Prkacb |
| Mvb12b1 | 4.72E-43 | 0.79092403 | 0.535 | 0.214 | 9.12E-39 | TAM.contam. | Mvb12b |
| Usp81 | 5.66E-43 | 0.39207102 | 0.778 | 0.398 | 1.09E-38 | TAM.contam. | Usp8 |
| Dok12 | 7.47E-43 | 0.41893364 | 0.707 | 0.294 | 1.44E-38 | TAM.contam. | Dok1 |
| Bsg | 7.85E-43 | 0.49275839 | 0.687 | 0.339 | 1.52E-38 | TAM.contam. | Bsg |
| Tmem511 | 8.49E-43 | 0.93928139 | 0.505 | 0.13 | 1.64E-38 | TAM.contam. | Tmem51 |
| Fam219a | 9.21E-43 | 1.27920615 | 0.404 | 0.097 | 1.78E-38 | TAM.contam. | Fam219a |
| Ern11 | 1.12E-42 | 0.49902355 | 0.596 | 0.276 | 2.16E-38 | TAM.contam. | Ern1 |
| Tmx11 | 2.01E-42 | 0.4669229 | 0.727 | 0.342 | 3.90E-38 | TAM.contam. | Tmx1 |
| Slc29a32 | 2.05E-42 | 0.34387712 | 0.636 | 0.289 | 3.97E-38 | TAM.contam. | Slc29a3 |
| Trappc3 | 2.52E-42 | 0.51546512 | 0.667 | 0.299 | 4.87E-38 | TAM.contam. | Trappc3 |
| Pik33 | 3.23E-42 | 0.71886426 | 0.616 | 0.228 | 6.25E-38 | TAM.contam. | Pik3 |
| Ppp3ca1 | 3.66E-42 | 0.55290303 | 0.667 | 0.277 | 7.08E-38 | TAM.contam. | Ppp3ca |
| Pnp1 | 3.90E-42 | 0.29326221 | 0.758 | 0.464 | 7.55E-38 | TAM.contam. | Pnp |
| Tex261 | 5.25E-42 | 0.68172314 | 0.667 | 0.288 | 1.02E-37 | TAM.contam. | Tex261 |
| Tifab2 | 5.61E-42 | 0.50907568 | 0.505 | 0.209 | 1.09E-37 | TAM.contam. | Tifab |
| Scyl21 | 7.36E-42 | 0.52772917 | 0.707 | 0.323 | 1.42E-37 | TAM.contam. | Scyl2 |
| Igf1r | 1.05E-41 | 1.03996816 | 0.434 | 0.1 | 2.04E-37 | TAM.contam. | Igf1r |
| Naa80 | 1.17E-41 | 1.12641422 | 0.374 | 0.101 | 2.27E-37 | TAM.contam. | Naa80 |
| Rnf1031 | 1.64E-41 | 0.66818452 | 0.596 | 0.211 | 3.17E-37 | TAM.contam. | Rnf103 |
| Lilr4b2 | 1.82E-41 | 0.25338077 | 0.828 | 0.437 | 3.52E-37 | TAM.contam. | Lilr4b |
| S100a13 | 2.06E-41 | 0.58418216 | 0.535 | 0.215 | 3.99E-37 | TAM.contam. | S100a1 |
| Srgap21 | 2.10E-41 | 1.01378682 | 0.525 | 0.175 | 4.07E-37 | TAM.contam. | Srgap2 |
| Aco11 | 2.13E-41 | 0.49497017 | 0.636 | 0.283 | 4.11E-37 | TAM.contam. | Aco1 |
| Pdxk1 | 2.49E-41 | 0.74826514 | 0.586 | 0.23 | 4.81E-37 | TAM.contam. | Pdxk |
| Plagl21 | 2.60E-41 | 0.51843985 | 0.545 | 0.256 | 5.03E-37 | TAM.contam. | Plagl2 |
| Mgat5 | 3.05E-41 | 1.01121803 | 0.566 | 0.14 | 5.90E-37 | TAM.contam. | Mgat5 |
| Pttg1ip1 | 3.76E-41 | 0.3487555 | 0.808 | 0.388 | 7.27E-37 | TAM.contam. | Pttg1ip |
| Tlr5 | 5.51E-41 | 2.06424404 | 0.384 | 0.06 | 1.07E-36 | TAM.contam. | Tlr5 |
| Tmem591 | 6.37E-41 | 0.25763979 | 0.737 | 0.404 | 1.23E-36 | TAM.contam. | Tmem59 |
| Serinc53 | 6.77E-41 | 0.33938865 | 0.465 | 0.157 | 1.31E-36 | TAM.contam. | Serinc5 |
| Mocs2 | 7.19E-41 | 0.95461462 | 0.475 | 0.15 | 1.39E-36 | TAM.contam. | Mocs2 |
| Baiap23 | 8.30E-41 | 0.70590957 | 0.636 | 0.257 | 1.60E-36 | TAM.contam. | Baiap2 |
| Sema4c1 | 8.85E-41 | 0.44497074 | 0.505 | 0.18 | 1.71E-36 | TAM.contam. | Sema4c |
| Cmtm72 | 1.04E-40 | 0.26856114 | 0.727 | 0.346 | 2.01E-36 | TAM.contam. | Cmtm7 |
| Foxo32 | 1.08E-40 | 0.68371802 | 0.566 | 0.206 | 2.09E-36 | TAM.contam. | Foxo3 |
| Dnajb91 | 1.20E-40 | 0.82814891 | 0.596 | 0.211 | 2.31E-36 | TAM.contam. | Dnajb9 |
| Ppm1a1 | 2.83E-40 | 0.30772547 | 0.737 | 0.386 | 5.47E-36 | TAM.contam. | Ppm1a |
| Trim47 | 3.48E-40 | 2.78231319 | 0.273 | 0.025 | 6.72E-36 | TAM.contam. | Trim47 |
| Tbc1d9b2 | 9.56E-40 | 0.33705775 | 0.707 | 0.369 | 1.85E-35 | TAM.contam. | Tbc1d9b |
| Pstpip21 | 1.36E-39 | 1.20176407 | 0.495 | 0.143 | 2.62E-35 | TAM.contam. | Pstpip2 |
| Sra11 | 1.48E-39 | 0.31690828 | 0.707 | 0.396 | 2.86E-35 | TAM.contam. | Sra1 |
| Hivep32 | 3.37E-39 | 1.27252108 | 0.414 | 0.083 | 6.52E-35 | TAM.contam. | Hivep3 |
| Slc45a4 | 3.54E-39 | 1.40783128 | 0.444 | 0.115 | 6.84E-35 | TAM.contam. | Slc45a4 |
| Atp9b1 | 3.64E-39 | 0.51210058 | 0.737 | 0.326 | 7.04E-35 | TAM.contam. | Atp9b |
| Chka2 | 4.84E-39 | 0.33265583 | 0.505 | 0.2 | 9.36E-35 | TAM.contam. | Chka |
| Fkbp22 | 6.80E-39 | 0.43646797 | 0.758 | 0.375 | 1.31E-34 | TAM.contam. | Fkbp2 |
| Tubb2a1 | 1.32E-38 | 0.26117753 | 0.646 | 0.288 | 2.56E-34 | TAM.contam. | Tubb2a |
| Retreg21 | 1.60E-38 | 0.59915682 | 0.636 | 0.246 | 3.09E-34 | TAM.contam. | Retreg2 |
| Atg2a | 1.70E-38 | 0.36027214 | 0.747 | 0.336 | 3.29E-34 | TAM.contam. | Atg2a |
| Cyp4f181 | 1.90E-38 | 1.08448049 | 0.465 | 0.12 | 3.67E-34 | TAM.contam. | Cyp4f18 |
| Chpf21 | 2.20E-38 | 0.73900565 | 0.606 | 0.24 | 4.24E-34 | TAM.contam. | Chpf2 |
| Map3k81 | 2.55E-38 | 0.26407846 | 0.576 | 0.247 | 4.93E-34 | TAM.contam. | Map3k8 |
| Sumf1 | 2.72E-38 | 0.30183367 | 0.707 | 0.391 | 5.26E-34 | TAM.contam. | Sumf1 |
| Rnasel | 2.81E-38 | 1.07489268 | 0.455 | 0.12 | 5.44E-34 | TAM.contam. | Rnasel |
| Gpr1072 | 2.82E-38 | 0.50759806 | 0.586 | 0.244 | 5.45E-34 | TAM.contam. | Gpr107 |
| Gclm1 | 3.45E-38 | 0.83640684 | 0.333 | 0.14 | 6.66E-34 | TAM.contam. | Gclm |
| Lmbrd11 | 3.56E-38 | 0.48861947 | 0.707 | 0.296 | 6.88E-34 | TAM.contam. | Lmbrd1 |
| Ifih12 | 4.03E-38 | 0.67937215 | 0.606 | 0.242 | 7.80E-34 | TAM.contam. | Ifih1 |
| Kcnab22 | 4.49E-38 | 0.52038659 | 0.646 | 0.289 | 8.68E-34 | TAM.contam. | Kcnab2 |
| Snx301 | 4.84E-38 | 1.0219679 | 0.414 | 0.107 | 9.36E-34 | TAM.contam. | Snx30 |
| Slc20a12 | 4.84E-38 | 0.395481 | 0.626 | 0.282 | 9.36E-34 | TAM.contam. | Slc20a1 |
| Ddrgk1 | 6.18E-38 | 0.28774374 | 0.677 | 0.333 | 1.20E-33 | TAM.contam. | Ddrgk1 |
| Ndufa1 | 9.95E-38 | 0.2549443 | 0.808 | 0.522 | 1.92E-33 | TAM.contam. | Ndufa1 |
| C2cd2l1 | 1.06E-37 | 0.87835391 | 0.495 | 0.171 | 2.05E-33 | TAM.contam. | C2cd2l |
| Lacc12 | 1.15E-37 | 0.52999273 | 0.616 | 0.223 | 2.21E-33 | TAM.contam. | Lacc1 |
| Abhd41 | 1.27E-37 | 0.66815707 | 0.586 | 0.208 | 2.46E-33 | TAM.contam. | Abhd4 |
| Il18bp | 1.31E-37 | 0.92524287 | 0.293 | 0.076 | 2.53E-33 | TAM.contam. | Il18bp |
| Arl11 | 1.41E-37 | 1.34983523 | 0.364 | 0.073 | 2.72E-33 | TAM.contam. | Arl11 |
| Vps13c | 1.62E-37 | 1.15337343 | 0.404 | 0.13 | 3.13E-33 | TAM.contam. | Vps13c |
| Nufip21 | 2.15E-37 | 0.31406388 | 0.687 | 0.375 | 4.15E-33 | TAM.contam. | Nufip2 |
| Tmem63a2 | 2.39E-37 | 0.38272879 | 0.535 | 0.207 | 4.61E-33 | TAM.contam. | Tmem63a |
| Abcc12 | 2.78E-37 | 0.48692384 | 0.576 | 0.279 | 5.37E-33 | TAM.contam. | Abcc1 |
| Tmem1652 | 3.02E-37 | 0.55024192 | 0.576 | 0.231 | 5.84E-33 | TAM.contam. | Tmem165 |
| Emilin1 | 3.91E-37 | 1.62244227 | 0.343 | 0.055 | 7.56E-33 | TAM.contam. | Emilin1 |
| Gde13 | 4.21E-37 | 0.73970056 | 0.525 | 0.201 | 8.14E-33 | TAM.contam. | Gde1 |
| Wdfy22 | 4.54E-37 | 0.59506934 | 0.596 | 0.247 | 8.78E-33 | TAM.contam. | Wdfy2 |
| Sfxn31 | 4.60E-37 | 0.25930044 | 0.626 | 0.312 | 8.89E-33 | TAM.contam. | Sfxn3 |
| Sec62 | 5.60E-37 | 0.62696876 | 0.768 | 0.339 | 1.08E-32 | TAM.contam. | Sec62 |
| Itpkb2 | 5.95E-37 | 0.35625062 | 0.586 | 0.23 | 1.15E-32 | TAM.contam. | Itpkb |
| Nus12 | 7.06E-37 | 0.65607609 | 0.677 | 0.292 | 1.36E-32 | TAM.contam. | Nus1 |
| Lpcat22 | 7.26E-37 | 0.52129587 | 0.566 | 0.292 | 1.40E-32 | TAM.contam. | Lpcat2 |
| Eea11 | 7.91E-37 | 0.55190435 | 0.485 | 0.197 | 1.53E-32 | TAM.contam. | Eea1 |
| Akt1s11 | 8.28E-37 | 0.62075839 | 0.475 | 0.148 | 1.60E-32 | TAM.contam. | Akt1s1 |
| Agtrap | 1.06E-36 | 1.63105423 | 0.343 | 0.064 | 2.05E-32 | TAM.contam. | Agtrap |
| Bcl101 | 1.27E-36 | 0.37441413 | 0.707 | 0.319 | 2.45E-32 | TAM.contam. | Bcl10 |
| Slc31a21 | 1.33E-36 | 0.61788226 | 0.465 | 0.149 | 2.57E-32 | TAM.contam. | Slc31a2 |
| Hgsnat | 1.34E-36 | 1.02880713 | 0.384 | 0.122 | 2.58E-32 | TAM.contam. | Hgsnat |
| Ago31 | 1.36E-36 | 0.77279872 | 0.515 | 0.161 | 2.63E-32 | TAM.contam. | Ago3 |
| Mapk62 | 1.64E-36 | 0.55731461 | 0.646 | 0.281 | 3.18E-32 | TAM.contam. | Mapk6 |
| Cnpy21 | 1.70E-36 | 0.56950507 | 0.606 | 0.254 | 3.29E-32 | TAM.contam. | Cnpy2 |
| Cbx43 | 2.06E-36 | 0.6884768 | 0.626 | 0.232 | 3.99E-32 | TAM.contam. | Cbx4 |
| Rassf8 | 2.79E-36 | 1.57700306 | 0.354 | 0.063 | 5.40E-32 | TAM.contam. | Rassf8 |
| Insr | 3.31E-36 | 0.92702388 | 0.434 | 0.111 | 6.40E-32 | TAM.contam. | Insr |
| Sgpp11 | 3.37E-36 | 0.50671172 | 0.636 | 0.353 | 6.51E-32 | TAM.contam. | Sgpp1 |
| Ubn1 | 5.42E-36 | 0.26989718 | 0.657 | 0.315 | 1.05E-31 | TAM.contam. | Ubn1 |
| Ly96 | 5.68E-36 | 1.15115319 | 0.384 | 0.059 | 1.10E-31 | TAM.contam. | Ly96 |
| Ralbp11 | 5.72E-36 | 0.37226309 | 0.646 | 0.301 | 1.11E-31 | TAM.contam. | Ralbp1 |
| Gbp9 | 6.07E-36 | 1.2368194 | 0.414 | 0.079 | 1.17E-31 | TAM.contam. | Gbp9 |
| Cables1 | 7.02E-36 | 1.78471161 | 0.273 | 0.032 | 1.36E-31 | TAM.contam. | Cables1 |
| Syngr1 | 8.38E-36 | 6.11348858 | 0.253 | 0.001 | 1.62E-31 | TAM.contam. | Syngr1 |
| Rnf150 | 8.41E-36 | 1.21894544 | 0.374 | 0.121 | 1.63E-31 | TAM.contam. | Rnf150 |
| Selenon2 | 8.86E-36 | 0.98930038 | 0.485 | 0.149 | 1.71E-31 | TAM.contam. | Selenon |
| Iffo21 | 1.26E-35 | 0.33757081 | 0.535 | 0.186 | 2.43E-31 | TAM.contam. | Iffo2 |
| Fam89b1 | 1.32E-35 | 0.30610323 | 0.646 | 0.351 | 2.56E-31 | TAM.contam. | Fam89b |
| Trp53inp21 | 1.41E-35 | 0.88355195 | 0.495 | 0.141 | 2.73E-31 | TAM.contam. | Trp53inp2 |
| Leprot1 | 1.62E-35 | 0.26408693 | 0.778 | 0.402 | 3.14E-31 | TAM.contam. | Leprot |
| Mcoln2 | 1.65E-35 | 1.96969723 | 0.293 | 0.047 | 3.19E-31 | TAM.contam. | Mcoln2 |
| Fam174a1 | 1.88E-35 | 0.29798199 | 0.636 | 0.303 | 3.64E-31 | TAM.contam. | Fam174a |
| Dram21 | 1.91E-35 | 1.01942341 | 0.475 | 0.16 | 3.69E-31 | TAM.contam. | Dram2 |
| Dpp7 | 2.71E-35 | 1.51783213 | 0.364 | 0.084 | 5.25E-31 | TAM.contam. | Dpp7 |
| Skp1a | 2.93E-35 | 0.4212185 | 0.717 | 0.325 | 5.67E-31 | TAM.contam. | Skp1a |
| Snx27 | 3.05E-35 | 0.81271038 | 0.444 | 0.133 | 5.90E-31 | TAM.contam. | Snx27 |
| Pnpla82 | 3.92E-35 | 0.42154293 | 0.556 | 0.231 | 7.58E-31 | TAM.contam. | Pnpla8 |
| Stk25 | 5.06E-35 | 0.42988437 | 0.667 | 0.259 | 9.79E-31 | TAM.contam. | Stk25 |
| Hagh | 5.90E-35 | 0.99366887 | 0.495 | 0.148 | 1.14E-30 | TAM.contam. | Hagh |
| Hlx1 | 6.50E-35 | 0.28202242 | 0.697 | 0.329 | 1.26E-30 | TAM.contam. | Hlx |
| Ccdc931 | 6.57E-35 | 0.49857045 | 0.475 | 0.171 | 1.27E-30 | TAM.contam. | Ccdc93 |
| Exoc5 | 9.18E-35 | 0.37959522 | 0.747 | 0.376 | 1.78E-30 | TAM.contam. | Exoc5 |
| Zfhx31 | 1.04E-34 | 0.97182156 | 0.485 | 0.133 | 2.01E-30 | TAM.contam. | Zfhx3 |
| Tns32 | 1.09E-34 | 0.51116639 | 0.586 | 0.23 | 2.11E-30 | TAM.contam. | Tns3 |
| Ulk1 | 1.13E-34 | 1.0510873 | 0.414 | 0.125 | 2.18E-30 | TAM.contam. | Ulk1 |
| Trim14 | 1.36E-34 | 0.3363113 | 0.586 | 0.244 | 2.63E-30 | TAM.contam. | Trim14 |
| Epb41l1 | 1.43E-34 | 1.54323572 | 0.313 | 0.058 | 2.76E-30 | TAM.contam. | Epb41l1 |
| Rab122 | 1.66E-34 | 0.31337453 | 0.525 | 0.241 | 3.22E-30 | TAM.contam. | Rab12 |
| B3gnt7 | 2.10E-34 | 1.82528629 | 0.253 | 0.054 | 4.05E-30 | TAM.contam. | B3gnt7 |
| Srxn11 | 2.75E-34 | 0.65370569 | 0.434 | 0.141 | 5.31E-30 | TAM.contam. | Srxn1 |
| Tlr13 | 2.87E-34 | 1.20738579 | 0.455 | 0.124 | 5.56E-30 | TAM.contam. | Tlr13 |
| Anapc2 | 4.00E-34 | 0.27819303 | 0.717 | 0.38 | 7.74E-30 | TAM.contam. | Anapc2 |
| Sptlc21 | 4.91E-34 | 0.72987484 | 0.717 | 0.29 | 9.48E-30 | TAM.contam. | Sptlc2 |
| Egln21 | 5.76E-34 | 0.49304689 | 0.606 | 0.231 | 1.11E-29 | TAM.contam. | Egln2 |
| Cd151 | 7.55E-34 | 1.0203115 | 0.374 | 0.082 | 1.46E-29 | TAM.contam. | Cd151 |
| Topors | 7.64E-34 | 0.82892187 | 0.485 | 0.15 | 1.48E-29 | TAM.contam. | Topors |
| DEPRECATED-I | 8.30E-34 | 1.00849782 | 0.444 | 0.132 | 1.60E-29 | TAM.contam. | DEPRECATED-ENSMUSG00000032667 |
| Sptssa2 | 9.89E-34 | 0.33497604 | 0.657 | 0.324 | 1.91E-29 | TAM.contam. | Sptssa |
| Tbc1d131 | 1.06E-33 | 0.43909736 | 0.687 | 0.293 | 2.06E-29 | TAM.contam. | Tbc1d13 |
| Ugcg3 | 1.23E-33 | 0.74906027 | 0.636 | 0.223 | 2.38E-29 | TAM.contam. | Ugcg |
| Sbf21 | 1.26E-33 | 0.52503506 | 0.495 | 0.15 | 2.43E-29 | TAM.contam. | Sbf2 |
| Klf101 | 1.70E-33 | 0.85862278 | 0.515 | 0.156 | 3.29E-29 | TAM.contam. | Klf10 |
| Arid5b | 1.71E-33 | 1.3276379 | 0.384 | 0.084 | 3.30E-29 | TAM.contam. | Arid5b |
| Blvra2 | 1.80E-33 | 0.97827865 | 0.616 | 0.25 | 3.47E-29 | TAM.contam. | Blvra |
| H6pd | 2.20E-33 | 1.02990741 | 0.374 | 0.128 | 4.25E-29 | TAM.contam. | H6pd |
| Gtf3c2 | 2.27E-33 | 0.30237341 | 0.717 | 0.369 | 4.39E-29 | TAM.contam. | Gtf3c2 |
| Npepps2 | 2.53E-33 | 0.49356808 | 0.566 | 0.255 | 4.90E-29 | TAM.contam. | Npepps |
| Tlnrd12 | 2.63E-33 | 0.34976303 | 0.586 | 0.284 | 5.09E-29 | TAM.contam. | Tlnrd1 |
| Crat | 3.15E-33 | 1.63138507 | 0.404 | 0.093 | 6.09E-29 | TAM.contam. | Crat |
| Slc7a71 | 3.25E-33 | 0.9769497 | 0.444 | 0.149 | 6.28E-29 | TAM.contam. | Slc7a7 |
| Gpr1462 | 3.30E-33 | 0.69141492 | 0.515 | 0.182 | 6.39E-29 | TAM.contam. | Gpr146 |
| Mob3c | 3.67E-33 | 1.04164325 | 0.394 | 0.093 | 7.09E-29 | TAM.contam. | Mob3c |
| Dtnbp11 | 4.68E-33 | 0.30276297 | 0.707 | 0.3 | 9.05E-29 | TAM.contam. | Dtnbp1 |
| Mef2c2 | 5.15E-33 | 0.52391245 | 0.495 | 0.193 | 9.95E-29 | TAM.contam. | Mef2c |
| Calm2 | 6.32E-33 | 0.26987862 | 0.707 | 0.405 | 1.22E-28 | TAM.contam. | Calm2 |
| Zfp467 | 7.60E-33 | 1.20152775 | 0.364 | 0.06 | 1.47E-28 | TAM.contam. | Zfp467 |
| 8-Mar | 8.13E-33 | 2.85257412 | 0.253 | 0.024 | 1.57E-28 | TAM.contam. | 8-Mar |
| Fmnl31 | 1.00E-32 | 0.7887451 | 0.525 | 0.159 | 1.94E-28 | TAM.contam. | Fmnl3 |
| Slc30a1 | 1.07E-32 | 1.19815594 | 0.495 | 0.124 | 2.06E-28 | TAM.contam. | Slc30a1 |
| Pnkd1 | 1.22E-32 | 0.38751267 | 0.414 | 0.164 | 2.36E-28 | TAM.contam. | Pnkd |
| Dbi1 | 1.48E-32 | 0.40193776 | 0.667 | 0.307 | 2.86E-28 | TAM.contam. | Dbi |
| Hcar21 | 1.51E-32 | 0.60052753 | 0.354 | 0.116 | 2.92E-28 | TAM.contam. | Hcar2 |
| Sh3kbp11 | 1.68E-32 | 0.5963069 | 0.525 | 0.209 | 3.24E-28 | TAM.contam. | Sh3kbp1 |
| Il162 | 1.73E-32 | 0.57160685 | 0.434 | 0.15 | 3.35E-28 | TAM.contam. | Il16 |
| Tmem1642 | 1.91E-32 | 0.54873665 | 0.657 | 0.302 | 3.70E-28 | TAM.contam. | Tmem164 |
| Mboat71 | 2.23E-32 | 0.3198048 | 0.687 | 0.341 | 4.31E-28 | TAM.contam. | Mboat7 |
| Abi33 | 2.26E-32 | 0.30122191 | 0.697 | 0.407 | 4.37E-28 | TAM.contam. | Abi3 |
| Wdr811 | 2.39E-32 | 0.68138409 | 0.515 | 0.185 | 4.62E-28 | TAM.contam. | Wdr81 |
| Mindy22 | 2.48E-32 | 0.6207496 | 0.485 | 0.178 | 4.79E-28 | TAM.contam. | Mindy2 |
| Agap1 | 2.74E-32 | 0.60211796 | 0.414 | 0.11 | 5.31E-28 | TAM.contam. | Agap1 |
| Tmem431 | 2.86E-32 | 0.62745516 | 0.485 | 0.182 | 5.53E-28 | TAM.contam. | Tmem43 |
| Ly91 | 3.17E-32 | 1.11027597 | 0.253 | 0.061 | 6.14E-28 | TAM.contam. | Ly9 |
| Slc25a102 | 3.27E-32 | 0.52045695 | 0.535 | 0.207 | 6.33E-28 | TAM.contam. | Slc25a10 |
| Ptgr1 | 5.74E-32 | 1.36842714 | 0.343 | 0.067 | 1.11E-27 | TAM.contam. | Ptgr1 |
| Myo1e2 | 5.98E-32 | 0.73527065 | 0.485 | 0.17 | 1.16E-27 | TAM.contam. | Myo1e |
| Fam193b1 | 6.28E-32 | 0.25824616 | 0.576 | 0.225 | 1.21E-27 | TAM.contam. | Fam193b |
| Atp6v1h2 | 6.86E-32 | 0.33206018 | 0.677 | 0.334 | 1.33E-27 | TAM.contam. | Atp6v1h |
| Tmf1 | 1.01E-31 | 0.37435083 | 0.677 | 0.322 | 1.95E-27 | TAM.contam. | Tmf1 |
| Hadh | 1.09E-31 | 0.85294163 | 0.424 | 0.113 | 2.11E-27 | TAM.contam. | Hadh |
| Rufy32 | 1.20E-31 | 0.41410078 | 0.495 | 0.174 | 2.31E-27 | TAM.contam. | Rufy3 |
| Mroh12 | 1.43E-31 | 0.28300755 | 0.606 | 0.254 | 2.77E-27 | TAM.contam. | Mroh1 |
| Rb1 | 1.52E-31 | 0.5805962 | 0.586 | 0.223 | 2.95E-27 | TAM.contam. | Rb1 |
| Dag1 | 1.84E-31 | 1.01749004 | 0.495 | 0.137 | 3.55E-27 | TAM.contam. | Dag1 |
| Tesk11 | 1.94E-31 | 0.46200644 | 0.626 | 0.272 | 3.76E-27 | TAM.contam. | Tesk1 |
| Tbc1d201 | 2.00E-31 | 0.38206777 | 0.677 | 0.311 | 3.87E-27 | TAM.contam. | Tbc1d20 |
| Zfp3841 | 2.01E-31 | 0.42243856 | 0.677 | 0.319 | 3.88E-27 | TAM.contam. | Zfp384 |
| Tom11 | 2.04E-31 | 1.01007862 | 0.394 | 0.085 | 3.95E-27 | TAM.contam. | Tom1 |
| Slc16a61 | 2.07E-31 | 0.73282667 | 0.566 | 0.215 | 4.01E-27 | TAM.contam. | Slc16a6 |
| Tmem159 | 2.20E-31 | 1.69848352 | 0.374 | 0.065 | 4.25E-27 | TAM.contam. | Tmem159 |
| Nlrp1b | 2.61E-31 | 1.77496854 | 0.303 | 0.057 | 5.05E-27 | TAM.contam. | Nlrp1b |
| Ocrl | 2.62E-31 | 1.15929458 | 0.354 | 0.072 | 5.06E-27 | TAM.contam. | Ocrl |
| Armcx31 | 2.75E-31 | 0.9045173 | 0.475 | 0.14 | 5.31E-27 | TAM.contam. | Armcx3 |
| Bag31 | 2.75E-31 | 0.4132351 | 0.556 | 0.212 | 5.32E-27 | TAM.contam. | Bag3 |
| Stat5b1 | 2.80E-31 | 0.26287268 | 0.677 | 0.351 | 5.41E-27 | TAM.contam. | Stat5b |
| Gnpda1 | 2.99E-31 | 0.49248834 | 0.556 | 0.231 | 5.78E-27 | TAM.contam. | Gnpda1 |
| Arl2 | 3.22E-31 | 0.81493843 | 0.505 | 0.158 | 6.23E-27 | TAM.contam. | Arl2 |
| Notch11 | 3.27E-31 | 1.09460381 | 0.374 | 0.102 | 6.32E-27 | TAM.contam. | Notch1 |
| Dnajb121 | 4.14E-31 | 0.2599996 | 0.596 | 0.314 | 8.01E-27 | TAM.contam. | Dnajb12 |
| Tec | 4.74E-31 | 0.44103821 | 0.495 | 0.184 | 9.16E-27 | TAM.contam. | Tec |
| Pnpla71 | 5.14E-31 | 0.55200612 | 0.475 | 0.224 | 9.94E-27 | TAM.contam. | Pnpla7 |
| Akr1b10 | 5.18E-31 | 0.91880638 | 0.465 | 0.132 | 1.00E-26 | TAM.contam. | Akr1b10 |
| Ipmk1 | 6.56E-31 | 0.45906272 | 0.475 | 0.177 | 1.27E-26 | TAM.contam. | Ipmk |
| Vipas39 | 7.05E-31 | 0.51407165 | 0.525 | 0.204 | 1.36E-26 | TAM.contam. | Vipas39 |
| DEPRECATED-I | 9.29E-31 | 0.59878956 | 0.475 | 0.166 | 1.80E-26 | TAM.contam. | DEPRECATED-ENSMUSG00000039782 |
| Numa11 | 9.53E-31 | 0.26205133 | 0.667 | 0.336 | 1.84E-26 | TAM.contam. | Numa1 |
| Atn1 | 1.01E-30 | 0.29607981 | 0.626 | 0.262 | 1.95E-26 | TAM.contam. | Atn1 |
| Bst11 | 1.11E-30 | 1.157168 | 0.354 | 0.112 | 2.14E-26 | TAM.contam. | Bst1 |
| Zfp6522 | 1.13E-30 | 0.66452137 | 0.586 | 0.24 | 2.18E-26 | TAM.contam. | Zfp652 |
| Fbh1 | 1.41E-30 | 0.64115368 | 0.576 | 0.237 | 2.73E-26 | TAM.contam. | Fbh1 |
| Fnip12 | 1.66E-30 | 0.6236734 | 0.556 | 0.228 | 3.21E-26 | TAM.contam. | Fnip1 |
| Nkiras2 | 1.71E-30 | 0.55441226 | 0.545 | 0.265 | 3.31E-26 | TAM.contam. | Nkiras2 |
| Ppp1r211 | 1.82E-30 | 0.34053468 | 0.596 | 0.245 | 3.52E-26 | TAM.contam. | Ppp1r21 |
| Mgat4a3 | 2.62E-30 | 0.34705914 | 0.636 | 0.283 | 5.06E-26 | TAM.contam. | Mgat4a |
| Stxbp1 | 2.71E-30 | 0.97376018 | 0.424 | 0.091 | 5.23E-26 | TAM.contam. | Stxbp1 |
| Cd723 | 3.59E-30 | 0.43837139 | 0.566 | 0.238 | 6.94E-26 | TAM.contam. | Cd72 |
| Patl12 | 3.70E-30 | 0.37236937 | 0.657 | 0.309 | 7.16E-26 | TAM.contam. | Patl1 |
| Lmbrd2 | 3.78E-30 | 1.20248149 | 0.404 | 0.111 | 7.30E-26 | TAM.contam. | Lmbrd2 |
| Trrap1 | 4.00E-30 | 0.51504538 | 0.545 | 0.24 | 7.73E-26 | TAM.contam. | Trrap |
| Fam102b1 | 4.26E-30 | 0.69112333 | 0.535 | 0.195 | 8.23E-26 | TAM.contam. | Fam102b |
| Retreg31 | 5.49E-30 | 0.43588641 | 0.545 | 0.294 | 1.06E-25 | TAM.contam. | Retreg3 |
| Dmxl11 | 5.94E-30 | 0.56435951 | 0.495 | 0.2 | 1.15E-25 | TAM.contam. | Dmxl1 |
| Ppt23 | 6.44E-30 | 0.76512694 | 0.465 | 0.128 | 1.24E-25 | TAM.contam. | Ppt2 |
| Agap32 | 6.52E-30 | 0.50501272 | 0.677 | 0.32 | 1.26E-25 | TAM.contam. | Agap3 |
| Brox | 6.87E-30 | 0.57068263 | 0.606 | 0.214 | 1.33E-25 | TAM.contam. | Brox |
| Dock73 | 7.57E-30 | 0.25528211 | 0.667 | 0.298 | 1.46E-25 | TAM.contam. | Dock7 |
| Rapgef22 | 7.96E-30 | 0.392047 | 0.626 | 0.265 | 1.54E-25 | TAM.contam. | Rapgef2 |
| Fmn1 | 9.56E-30 | 0.9533657 | 0.263 | 0.053 | 1.85E-25 | TAM.contam. | Fmn1 |
| Tspan142 | 1.22E-29 | 0.25261042 | 0.636 | 0.304 | 2.36E-25 | TAM.contam. | Tspan14 |
| Slc44a11 | 1.32E-29 | 0.42369694 | 0.444 | 0.134 | 2.56E-25 | TAM.contam. | Slc44a1 |
| Oxr1 | 1.34E-29 | 0.87583389 | 0.374 | 0.126 | 2.59E-25 | TAM.contam. | Oxr1 |
| Hdac52 | 1.71E-29 | 0.34775937 | 0.596 | 0.217 | 3.32E-25 | TAM.contam. | Hdac5 |
| Ltbp31 | 1.78E-29 | 0.78276607 | 0.404 | 0.109 | 3.44E-25 | TAM.contam. | Ltbp3 |
| B3gnt21 | 1.78E-29 | 0.89231344 | 0.424 | 0.12 | 3.44E-25 | TAM.contam. | B3gnt2 |
| Tmem250-ps1 | 1.84E-29 | 0.47442028 | 0.556 | 0.215 | 3.56E-25 | TAM.contam. | Tmem250-ps |
| Dpp9 | 1.86E-29 | 0.31899203 | 0.687 | 0.348 | 3.60E-25 | TAM.contam. | Dpp9 |
| Bmf1 | 1.87E-29 | 0.44402442 | 0.505 | 0.2 | 3.62E-25 | TAM.contam. | Bmf |
| Rfk1 | 1.94E-29 | 0.30134751 | 0.606 | 0.277 | 3.76E-25 | TAM.contam. | Rfk |
| Specc1l1 | 1.99E-29 | 0.25756223 | 0.677 | 0.341 | 3.84E-25 | TAM.contam. | Specc1l |
| Cpeb22 | 2.10E-29 | 0.51428982 | 0.545 | 0.214 | 4.06E-25 | TAM.contam. | Cpeb2 |
| Atraid1 | 2.29E-29 | 0.66111984 | 0.475 | 0.199 | 4.42E-25 | TAM.contam. | Atraid |
| Plin3 | 2.31E-29 | 0.34080574 | 0.687 | 0.332 | 4.47E-25 | TAM.contam. | Plin3 |
| Afg3l21 | 2.36E-29 | 0.70162601 | 0.616 | 0.263 | 4.55E-25 | TAM.contam. | Afg3l2 |
| Tfe31 | 2.87E-29 | 0.32953041 | 0.707 | 0.309 | 5.55E-25 | TAM.contam. | Tfe3 |
| Sord | 3.04E-29 | 1.5967456 | 0.303 | 0.06 | 5.88E-25 | TAM.contam. | Sord |
| Lpxn2 | 3.13E-29 | 0.88580597 | 0.515 | 0.182 | 6.04E-25 | TAM.contam. | Lpxn |
| Snx12 | 3.58E-29 | 0.35144567 | 0.636 | 0.326 | 6.92E-25 | TAM.contam. | Snx12 |
| Zdhhc51 | 3.73E-29 | 0.25920986 | 0.677 | 0.351 | 7.22E-25 | TAM.contam. | Zdhhc5 |
| Dpep21 | 3.79E-29 | 1.21144098 | 0.333 | 0.074 | 7.34E-25 | TAM.contam. | Dpep2 |
| Impact2 | 4.75E-29 | 0.47440122 | 0.626 | 0.27 | 9.18E-25 | TAM.contam. | Impact |
| Fam214b1 | 5.92E-29 | 0.4993394 | 0.485 | 0.181 | 1.15E-24 | TAM.contam. | Fam214b |
| Mesd1 | 6.18E-29 | 0.91204654 | 0.505 | 0.148 | 1.20E-24 | TAM.contam. | Mesd |
| Ctso2 | 6.43E-29 | 0.73801064 | 0.495 | 0.168 | 1.24E-24 | TAM.contam. | Ctso |
| Zfp90 | 7.47E-29 | 1.17244295 | 0.343 | 0.079 | 1.44E-24 | TAM.contam. | Zfp90 |
| Ube2l61 | 7.87E-29 | 0.25735756 | 0.465 | 0.19 | 1.52E-24 | TAM.contam. | Ube2l6 |
| Ube2a2 | 9.54E-29 | 0.36486899 | 0.606 | 0.28 | 1.84E-24 | TAM.contam. | Ube2a |
| Rora | 9.54E-29 | 1.14674794 | 0.253 | 0.04 | 1.84E-24 | TAM.contam. | Rora |
| Smad3 | 1.00E-28 | 1.16583153 | 0.343 | 0.064 | 1.94E-24 | TAM.contam. | Smad3 |
| Wdr41 | 1.03E-28 | 0.72116679 | 0.455 | 0.171 | 2.00E-24 | TAM.contam. | Wdr41 |
| Pald1 | 1.17E-28 | 1.165361 | 0.313 | 0.089 | 2.26E-24 | TAM.contam. | Pald1 |
| Cul4a | 1.20E-28 | 0.39516263 | 0.525 | 0.218 | 2.32E-24 | TAM.contam. | Cul4a |
| Trpv4 | 1.22E-28 | 1.49487107 | 0.303 | 0.043 | 2.35E-24 | TAM.contam. | Trpv4 |
| Ggnbp21 | 1.36E-28 | 0.50093658 | 0.636 | 0.26 | 2.64E-24 | TAM.contam. | Ggnbp2 |
| Rexf12 | 1.42E-28 | 0.48393132 | 0.505 | 0.218 | 2.74E-24 | TAM.contam. | Rexf1 |
| Hist3h2a | 1.57E-28 | 0.73525143 | 0.404 | 0.119 | 3.04E-24 | TAM.contam. | Hist3h2a |
| Pgpep1 | 1.67E-28 | 1.25473445 | 0.323 | 0.058 | 3.22E-24 | TAM.contam. | Pgpep1 |
| Plcb31 | 1.70E-28 | 0.59699513 | 0.374 | 0.146 | 3.28E-24 | TAM.contam. | Plcb3 |
| Psen22 | 1.73E-28 | 0.36222098 | 0.576 | 0.232 | 3.34E-24 | TAM.contam. | Psen2 |
| Cdk172 | 1.73E-28 | 0.35039576 | 0.525 | 0.237 | 3.34E-24 | TAM.contam. | Cdk17 |
| Ptgr2 | 1.91E-28 | 0.61462206 | 0.444 | 0.173 | 3.70E-24 | TAM.contam. | Ptgr2 |
| Glb11 | 2.10E-28 | 0.4629426 | 0.556 | 0.202 | 4.06E-24 | TAM.contam. | Glb1 |
| Kif3a | 2.35E-28 | 1.74724491 | 0.283 | 0.043 | 4.55E-24 | TAM.contam. | Kif3a |
| Vapb | 2.94E-28 | 0.42302986 | 0.636 | 0.315 | 5.68E-24 | TAM.contam. | Vapb |
| Car131 | 3.10E-28 | 0.88306068 | 0.343 | 0.082 | 5.99E-24 | TAM.contam. | Car13 |
| Cebpzos | 3.31E-28 | 0.69684356 | 0.394 | 0.13 | 6.40E-24 | TAM.contam. | Cebpzos |
| Ankrd401 | 3.41E-28 | 0.25874444 | 0.677 | 0.366 | 6.59E-24 | TAM.contam. | Ankrd40 |
| Smim20 | 3.63E-28 | 0.50303571 | 0.525 | 0.204 | 7.03E-24 | TAM.contam. | Smim20 |
| Tgfbrap1 | 4.96E-28 | 1.1377425 | 0.424 | 0.13 | 9.58E-24 | TAM.contam. | Tgfbrap1 |
| Mon2 | 6.64E-28 | 0.70032088 | 0.596 | 0.238 | 1.28E-23 | TAM.contam. | Mon2 |
| Ptgfrn | 9.48E-28 | 0.91717719 | 0.283 | 0.069 | 1.83E-23 | TAM.contam. | Ptgfrn |
| Prdx4 | 1.11E-27 | 0.99009011 | 0.424 | 0.133 | 2.14E-23 | TAM.contam. | Prdx4 |
| H2afv2 | 1.24E-27 | 0.30240056 | 0.727 | 0.34 | 2.41E-23 | TAM.contam. | H2afv |
| Snx11 | 1.28E-27 | 0.45761291 | 0.515 | 0.156 | 2.47E-23 | TAM.contam. | Snx11 |
| Abl12 | 1.30E-27 | 0.45051101 | 0.697 | 0.317 | 2.51E-23 | TAM.contam. | Abl1 |
| Ets11 | 1.38E-27 | 0.57595896 | 0.253 | 0.061 | 2.68E-23 | TAM.contam. | Ets1 |
| Slc23a21 | 1.46E-27 | 0.72849723 | 0.495 | 0.16 | 2.82E-23 | TAM.contam. | Slc23a2 |
| Tmem168 | 1.71E-27 | 0.60695936 | 0.556 | 0.218 | 3.31E-23 | TAM.contam. | Tmem168 |
| Grsf11 | 1.73E-27 | 0.28983544 | 0.606 | 0.286 | 3.34E-23 | TAM.contam. | Grsf1 |
| Zfp704 | 1.80E-27 | 1.25551851 | 0.293 | 0.061 | 3.49E-23 | TAM.contam. | Zfp704 |
| Manba | 2.08E-27 | 0.7128372 | 0.414 | 0.158 | 4.03E-23 | TAM.contam. | Manba |
| Zdhhc142 | 2.10E-27 | 0.74014169 | 0.475 | 0.153 | 4.07E-23 | TAM.contam. | Zdhhc14 |
| Klhl241 | 2.46E-27 | 0.70839174 | 0.485 | 0.163 | 4.75E-23 | TAM.contam. | Klhl24 |
| Btbd2 | 3.15E-27 | 0.95648843 | 0.465 | 0.142 | 6.09E-23 | TAM.contam. | Btbd2 |
| Pttg12 | 3.20E-27 | 0.56929259 | 0.465 | 0.17 | 6.19E-23 | TAM.contam. | Pttg1 |
| Lamtor1 | 3.25E-27 | 0.4001521 | 0.545 | 0.218 | 6.28E-23 | TAM.contam. | Lamtor1 |
| Tpra11 | 3.37E-27 | 0.9207454 | 0.384 | 0.117 | 6.52E-23 | TAM.contam. | Tpra1 |
| Pign | 3.53E-27 | 0.70393579 | 0.485 | 0.15 | 6.83E-23 | TAM.contam. | Pign |
| Etfrf1 | 3.87E-27 | 0.68343112 | 0.313 | 0.09 | 7.49E-23 | TAM.contam. | Etfrf1 |
| Ppp1cc | 4.90E-27 | 0.36020542 | 0.636 | 0.282 | 9.48E-23 | TAM.contam. | Ppp1cc |
| Dok21 | 6.03E-27 | 0.42097078 | 0.424 | 0.146 | 1.16E-22 | TAM.contam. | Dok2 |
| Ndufs72 | 6.05E-27 | 0.25167952 | 0.768 | 0.401 | 1.17E-22 | TAM.contam. | Ndufs7 |
| Znrf13 | 6.12E-27 | 0.31631217 | 0.566 | 0.243 | 1.18E-22 | TAM.contam. | Znrf1 |
| Qdpr1 | 7.16E-27 | 0.28566178 | 0.657 | 0.353 | 1.38E-22 | TAM.contam. | Qdpr |
| Ogt1 | 8.18E-27 | 0.25046703 | 0.667 | 0.372 | 1.58E-22 | TAM.contam. | Ogt |
| Casp41 | 8.98E-27 | 0.49523202 | 0.505 | 0.174 | 1.74E-22 | TAM.contam. | Casp4 |
| Nagk1 | 9.00E-27 | 0.44858076 | 0.556 | 0.211 | 1.74E-22 | TAM.contam. | Nagk |
| Spg20 | 9.79E-27 | 1.17943979 | 0.293 | 0.052 | 1.89E-22 | TAM.contam. | Spg20 |
| Rnf1662 | 1.02E-26 | 0.33705125 | 0.707 | 0.347 | 1.97E-22 | TAM.contam. | Rnf166 |
| Vps292 | 1.07E-26 | 0.29262808 | 0.747 | 0.369 | 2.08E-22 | TAM.contam. | Vps29 |
| Cept11 | 1.49E-26 | 0.41912153 | 0.636 | 0.278 | 2.88E-22 | TAM.contam. | Cept1 |
| Rubcn2 | 1.50E-26 | 0.40342025 | 0.495 | 0.177 | 2.90E-22 | TAM.contam. | Rubcn |
| Scamp1 | 1.52E-26 | 0.89973493 | 0.384 | 0.118 | 2.95E-22 | TAM.contam. | Scamp1 |
| Tmem191 | 1.79E-26 | 0.28506702 | 0.636 | 0.267 | 3.46E-22 | TAM.contam. | Tmem19 |
| Bmpr22 | 1.84E-26 | 0.27342135 | 0.566 | 0.262 | 3.56E-22 | TAM.contam. | Bmpr2 |
| Rhoh2 | 1.86E-26 | 0.30270341 | 0.515 | 0.23 | 3.60E-22 | TAM.contam. | Rhoh |
| Prkcb2 | 1.95E-26 | 0.39801376 | 0.545 | 0.224 | 3.78E-22 | TAM.contam. | Prkcb |
| Ckap41 | 1.98E-26 | 0.71080184 | 0.535 | 0.163 | 3.83E-22 | TAM.contam. | Ckap4 |
| Nfix | 2.03E-26 | 0.7741906 | 0.424 | 0.132 | 3.93E-22 | TAM.contam. | Nfix |
| Hpse2 | 2.13E-26 | 0.59348555 | 0.384 | 0.128 | 4.13E-22 | TAM.contam. | Hpse |
| Crtap | 2.13E-26 | 0.71496389 | 0.404 | 0.135 | 4.13E-22 | TAM.contam. | Crtap |
| Klc4 | 2.36E-26 | 0.82353083 | 0.374 | 0.095 | 4.55E-22 | TAM.contam. | Klc4 |
| Ccdc22 | 2.37E-26 | 0.82234336 | 0.455 | 0.13 | 4.57E-22 | TAM.contam. | Ccdc22 |
| Parp141 | 2.45E-26 | 0.2795359 | 0.626 | 0.284 | 4.73E-22 | TAM.contam. | Parp14 |
| Rnf1451 | 2.50E-26 | 0.32313602 | 0.646 | 0.308 | 4.83E-22 | TAM.contam. | Rnf145 |
| Micu11 | 2.53E-26 | 0.52393939 | 0.455 | 0.186 | 4.90E-22 | TAM.contam. | Micu1 |
| Hbegf | 2.55E-26 | 0.55118984 | 0.313 | 0.112 | 4.93E-22 | TAM.contam. | Hbegf |
| Washc51 | 2.65E-26 | 0.72792465 | 0.434 | 0.161 | 5.12E-22 | TAM.contam. | Washc5 |
| Map2k71 | 2.88E-26 | 0.35591606 | 0.576 | 0.239 | 5.57E-22 | TAM.contam. | Map2k7 |
| Usp21 | 3.07E-26 | 0.6665781 | 0.465 | 0.159 | 5.93E-22 | TAM.contam. | Usp21 |
| Apc2 | 3.10E-26 | 0.339276 | 0.535 | 0.22 | 5.99E-22 | TAM.contam. | Apc |
| St13 | 3.50E-26 | 0.42281979 | 0.616 | 0.234 | 6.77E-22 | TAM.contam. | St13 |
| Phf20 | 4.16E-26 | 0.57045959 | 0.465 | 0.174 | 8.05E-22 | TAM.contam. | Phf20 |
| 11-Mar | 6.70E-26 | 0.44609267 | 0.646 | 0.291 | 1.30E-21 | TAM.contam. | 1-Mar |
| Ppp1r14b | 6.91E-26 | 0.36103862 | 0.505 | 0.243 | 1.34E-21 | TAM.contam. | Ppp1r14b |
| Atf71 | 8.42E-26 | 0.299551 | 0.535 | 0.23 | 1.63E-21 | TAM.contam. | Atf7 |
| Zfyve21 | 8.81E-26 | 1.39433776 | 0.253 | 0.047 | 1.70E-21 | TAM.contam. | Zfyve21 |
| Nnt1 | 8.88E-26 | 0.42249218 | 0.444 | 0.161 | 1.72E-21 | TAM.contam. | Nnt |
| Fbxo30 | 1.18E-25 | 1.2136453 | 0.343 | 0.081 | 2.28E-21 | TAM.contam. | Fbxo30 |
| Etfdh | 1.29E-25 | 0.58041719 | 0.394 | 0.144 | 2.50E-21 | TAM.contam. | Etfdh |
| Vti1b2 | 1.85E-25 | 0.30129353 | 0.646 | 0.307 | 3.59E-21 | TAM.contam. | Vti1b |
| Arhgap12 | 1.94E-25 | 1.07589473 | 0.323 | 0.108 | 3.74E-21 | TAM.contam. | Arhgap12 |
| Thbd | 3.08E-25 | 1.16454818 | 0.253 | 0.064 | 5.95E-21 | TAM.contam. | Thbd |
| Ascc31 | 3.65E-25 | 0.37194255 | 0.646 | 0.319 | 7.06E-21 | TAM.contam. | Ascc3 |
| Peak1 | 3.91E-25 | 0.63082322 | 0.384 | 0.114 | 7.57E-21 | TAM.contam. | Peak1 |
| Ash2l | 4.36E-25 | 0.39643187 | 0.535 | 0.231 | 8.42E-21 | TAM.contam. | Ash2l |
| Ier5l | 4.48E-25 | 0.88776269 | 0.354 | 0.133 | 8.67E-21 | TAM.contam. | Ier5l |
| Grk52 | 4.78E-25 | 0.5597409 | 0.414 | 0.144 | 9.25E-21 | TAM.contam. | Grk5 |
| Rnf111 | 5.02E-25 | 0.33530123 | 0.707 | 0.371 | 9.71E-21 | TAM.contam. | Rnf111 |
| Vps351 | 5.58E-25 | 0.3034772 | 0.717 | 0.339 | 1.08E-20 | TAM.contam. | Vps35 |
| Acads1 | 5.72E-25 | 0.44850144 | 0.515 | 0.197 | 1.11E-20 | TAM.contam. | Acads |
| Pank3 | 5.74E-25 | 0.29289989 | 0.576 | 0.277 | 1.11E-20 | TAM.contam. | Pank3 |
| Tmem9b1 | 6.53E-25 | 0.57493423 | 0.515 | 0.218 | 1.26E-20 | TAM.contam. | Tmem9b |
| Mepce1 | 7.22E-25 | 0.27157707 | 0.576 | 0.233 | 1.40E-20 | TAM.contam. | Mepce |
| Txndc12 | 7.81E-25 | 0.83801453 | 0.384 | 0.136 | 1.51E-20 | TAM.contam. | Txndc12 |
| Tbc1d2b2 | 7.84E-25 | 0.67112327 | 0.545 | 0.245 | 1.52E-20 | TAM.contam. | Tbc1d2b |
| Kansl1l | 8.78E-25 | 0.91844587 | 0.293 | 0.06 | 1.70E-20 | TAM.contam. | Kansl1l |
| Lpin1 | 9.10E-25 | 1.0319877 | 0.384 | 0.104 | 1.76E-20 | TAM.contam. | Lpin1 |
| Dbndd21 | 9.31E-25 | 0.41352958 | 0.525 | 0.223 | 1.80E-20 | TAM.contam. | Dbndd2 |
| Trpm7 | 9.94E-25 | 0.42019064 | 0.566 | 0.22 | 1.92E-20 | TAM.contam. | Trpm7 |
| Mecp2 | 1.12E-24 | 0.34456818 | 0.444 | 0.167 | 2.16E-20 | TAM.contam. | Mecp2 |
| Map3k2 | 1.15E-24 | 0.32418461 | 0.586 | 0.271 | 2.23E-20 | TAM.contam. | Map3k2 |
| Zzz3 | 1.24E-24 | 0.3080288 | 0.475 | 0.243 | 2.41E-20 | TAM.contam. | Zzz3 |
| Ripk1 | 1.33E-24 | 0.39219116 | 0.687 | 0.381 | 2.57E-20 | TAM.contam. | Ripk1 |
| Sec24a2 | 1.67E-24 | 0.56258665 | 0.535 | 0.214 | 3.24E-20 | TAM.contam. | Sec24a |
| Oasl21 | 1.74E-24 | 0.30024321 | 0.465 | 0.205 | 3.37E-20 | TAM.contam. | Oasl2 |
| Vps411 | 1.83E-24 | 0.41605669 | 0.444 | 0.198 | 3.53E-20 | TAM.contam. | Vps41 |
| Mga | 1.83E-24 | 0.56184058 | 0.444 | 0.171 | 3.54E-20 | TAM.contam. | Mga |
| Gbp3 | 2.25E-24 | 0.72028777 | 0.475 | 0.152 | 4.34E-20 | TAM.contam. | Gbp3 |
| C1qtnf61 | 2.25E-24 | 0.31116971 | 0.333 | 0.122 | 4.35E-20 | TAM.contam. | C1qtnf6 |
| Mxd42 | 2.34E-24 | 0.42458372 | 0.545 | 0.212 | 4.52E-20 | TAM.contam. | Mxd4 |
| Akirin2 | 2.35E-24 | 0.26903796 | 0.556 | 0.272 | 4.55E-20 | TAM.contam. | Akirin2 |
| Cgrrf1 | 2.49E-24 | 1.11684066 | 0.313 | 0.069 | 4.82E-20 | TAM.contam. | Cgrrf1 |
| Sfmbt1 | 2.70E-24 | 0.46552584 | 0.475 | 0.171 | 5.21E-20 | TAM.contam. | Sfmbt1 |
| Fads1 | 3.19E-24 | 1.05476711 | 0.343 | 0.103 | 6.17E-20 | TAM.contam. | Fads1 |
| Tmco13 | 3.35E-24 | 0.40726021 | 0.586 | 0.288 | 6.48E-20 | TAM.contam. | Tmco1 |
| Ube2g1 | 3.62E-24 | 0.34791056 | 0.485 | 0.234 | 7.00E-20 | TAM.contam. | Ube2g1 |
| Slc25a36 | 3.66E-24 | 0.33673972 | 0.545 | 0.209 | 7.08E-20 | TAM.contam. | Slc25a36 |
| Pi4k2a1 | 4.57E-24 | 0.27658724 | 0.677 | 0.288 | 8.84E-20 | TAM.contam. | Pi4k2a |
| Acox12 | 4.94E-24 | 0.46540195 | 0.677 | 0.278 | 9.54E-20 | TAM.contam. | Acox1 |
| Svbp | 5.21E-24 | 1.11054125 | 0.273 | 0.07 | 1.01E-19 | TAM.contam. | Svbp |
| Rbm381 | 5.31E-24 | 0.29383802 | 0.566 | 0.315 | 1.03E-19 | TAM.contam. | Rbm38 |
| Spryd31 | 6.04E-24 | 0.2886929 | 0.586 | 0.283 | 1.17E-19 | TAM.contam. | Spryd3 |
| Smim10l1 | 6.57E-24 | 0.30542042 | 0.475 | 0.2 | 1.27E-19 | TAM.contam. | Smim10l1 |
| Erf | 7.26E-24 | 0.47492135 | 0.545 | 0.258 | 1.40E-19 | TAM.contam. | Erf |
| Mast3 | 8.07E-24 | 0.3118034 | 0.596 | 0.283 | 1.56E-19 | TAM.contam. | Mast3 |
| Crebzf1 | 8.40E-24 | 0.26360372 | 0.616 | 0.278 | 1.62E-19 | TAM.contam. | Crebzf |
| Tmem2681 | 8.97E-24 | 0.57219774 | 0.485 | 0.185 | 1.73E-19 | TAM.contam. | Tmem268 |
| Vps81 | 1.01E-23 | 0.64583707 | 0.495 | 0.175 | 1.96E-19 | TAM.contam. | Vps8 |
| Papss22 | 1.23E-23 | 0.37288943 | 0.434 | 0.109 | 2.39E-19 | TAM.contam. | Papss2 |
| Ctps2 | 1.46E-23 | 0.5793824 | 0.424 | 0.122 | 2.83E-19 | TAM.contam. | Ctps2 |
| Htatip21 | 1.58E-23 | 0.430708 | 0.384 | 0.164 | 3.06E-19 | TAM.contam. | Htatip2 |
| Cyhr1 | 1.64E-23 | 0.66331137 | 0.444 | 0.144 | 3.18E-19 | TAM.contam. | Cyhr1 |
| Agpat31 | 1.87E-23 | 0.25109326 | 0.495 | 0.132 | 3.62E-19 | TAM.contam. | Agpat3 |
| Txlna | 2.58E-23 | 0.36797614 | 0.616 | 0.279 | 4.98E-19 | TAM.contam. | Txlna |
| Tm9sf1 | 2.72E-23 | 0.32822216 | 0.586 | 0.28 | 5.27E-19 | TAM.contam. | Tm9sf1 |
| Tmem883 | 2.82E-23 | 0.48587628 | 0.586 | 0.247 | 5.46E-19 | TAM.contam. | Tmem88 |

| | | | | | | | |
|---|---|---|---|---|---|---|---|
| Il18 | 2.94E-23 | 1.05591017 | 0.293 | 0.074 | 5.68E-19 | TAM.contam. | Il18 |
| Fadd | 2.95E-23 | 0.32612036 | 0.465 | 0.168 | 5.69E-19 | TAM.contam. | Fadd |
| Dyrk1b | 3.36E-23 | 0.98844625 | 0.303 | 0.072 | 6.49E-19 | TAM.contam. | Dyrk1b |
| Zbp1 | 4.08E-23 | 0.86111155 | 0.414 | 0.149 | 7.88E-19 | TAM.contam. | Zbp1 |
| Zfp6551 | 4.15E-23 | 0.63373587 | 0.505 | 0.202 | 8.03E-19 | TAM.contam. | Zfp655 |
| Serp12 | 4.38E-23 | 0.46524506 | 0.556 | 0.227 | 8.47E-19 | TAM.contam. | Serp1 |
| Zfp991 | 5.02E-23 | 1.75187732 | 0.273 | 0.051 | 9.71E-19 | TAM.contam. | Zfp991 |
| Sirpb1c | 7.27E-23 | 1.45219927 | 0.283 | 0.045 | 1.41E-18 | TAM.contam. | Sirpb1c |
| Smim71 | 7.44E-23 | 0.48881824 | 0.545 | 0.261 | 1.44E-18 | TAM.contam. | Smim7 |
| Parp13 | 7.51E-23 | 0.37327477 | 0.727 | 0.349 | 1.45E-18 | TAM.contam. | Parp1 |
| Tbc1d23 | 7.82E-23 | 0.8203081 | 0.374 | 0.118 | 1.51E-18 | TAM.contam. | Tbc1d23 |
| Sin3a | 8.24E-23 | 0.39225679 | 0.657 | 0.292 | 1.59E-18 | TAM.contam. | Sin3a |
| Lat21 | 9.06E-23 | 0.28837275 | 0.616 | 0.277 | 1.75E-18 | TAM.contam. | Lat2 |
| Fzd7 | 9.49E-23 | 0.83204068 | 0.333 | 0.129 | 1.83E-18 | TAM.contam. | Fzd7 |
| Gnptg | 1.08E-22 | 0.6360364 | 0.384 | 0.128 | 2.09E-18 | TAM.contam. | Gnptg |
| Bahd12 | 1.10E-22 | 0.29043153 | 0.586 | 0.268 | 2.13E-18 | TAM.contam. | Bahd1 |
| Rasa1 | 1.17E-22 | 0.74755608 | 0.394 | 0.156 | 2.26E-18 | TAM.contam. | Rasa1 |
| Arhgap252 | 1.21E-22 | 0.38051735 | 0.586 | 0.217 | 2.34E-18 | TAM.contam. | Arhgap25 |
| Lamc1 | 1.24E-22 | 1.07076305 | 0.263 | 0.049 | 2.39E-18 | TAM.contam. | Lamc1 |
| Emc11 | 1.31E-22 | 0.28710385 | 0.515 | 0.246 | 2.54E-18 | TAM.contam. | Emc1 |
| Cryl11 | 2.11E-22 | 0.77009909 | 0.424 | 0.146 | 4.08E-18 | TAM.contam. | Cryl1 |
| Strn | 2.18E-22 | 0.39327409 | 0.535 | 0.193 | 4.22E-18 | TAM.contam. | Strn |
| Pcmtd1 | 2.21E-22 | 0.81589393 | 0.364 | 0.11 | 4.26E-18 | TAM.contam. | Pcmtd1 |
| Pik3r21 | 2.69E-22 | 0.71906272 | 0.505 | 0.204 | 5.20E-18 | TAM.contam. | Pik3r2 |
| Cpt1a1 | 2.87E-22 | 0.43938943 | 0.485 | 0.227 | 5.55E-18 | TAM.contam. | Cpt1a |
| Dennd6a | 2.88E-22 | 0.60709245 | 0.455 | 0.144 | 5.56E-18 | TAM.contam. | Dennd6a |
| Ilvbl | 3.69E-22 | 0.84029763 | 0.424 | 0.125 | 7.14E-18 | TAM.contam. | Ilvbl |
| Klhl211 | 3.88E-22 | 0.47133144 | 0.384 | 0.142 | 7.50E-18 | TAM.contam. | Klhl21 |
| Katnbl11 | 5.36E-22 | 0.83712498 | 0.364 | 0.125 | 1.04E-17 | TAM.contam. | Katnbl1 |
| Alg11 | 5.45E-22 | 0.70880466 | 0.414 | 0.135 | 1.05E-17 | TAM.contam. | Alg1 |
| Fam3c3 | 5.73E-22 | 0.2511526 | 0.545 | 0.244 | 1.11E-17 | TAM.contam. | Fam3c |
| Sppl3 | 5.89E-22 | 0.34476346 | 0.657 | 0.302 | 1.14E-17 | TAM.contam. | Sppl3 |
| Prmt2 | 5.93E-22 | 1.18330898 | 0.253 | 0.051 | 1.15E-17 | TAM.contam. | Prmt2 |
| Arhgef12 | 5.95E-22 | 1.60027926 | 0.253 | 0.042 | 1.15E-17 | TAM.contam. | Arhgef12 |
| Tm9sf4 | 6.53E-22 | 0.55811922 | 0.455 | 0.184 | 1.26E-17 | TAM.contam. | Tm9sf4 |
| Pxk2 | 7.80E-22 | 0.40131604 | 0.556 | 0.237 | 1.51E-17 | TAM.contam. | Pxk |
| Slc29a11 | 8.49E-22 | 0.81279818 | 0.424 | 0.146 | 1.64E-17 | TAM.contam. | Slc29a1 |
| Nufip12 | 9.58E-22 | 0.39613149 | 0.475 | 0.22 | 1.85E-17 | TAM.contam. | Nufip1 |
| Fbxo421 | 1.00E-21 | 0.61410926 | 0.515 | 0.2 | 1.94E-17 | TAM.contam. | Fbxo42 |
| Tom1l21 | 1.09E-21 | 0.87832218 | 0.333 | 0.089 | 2.11E-17 | TAM.contam. | Tom1l2 |
| Pwwp2b | 1.24E-21 | 1.72122211 | 0.263 | 0.043 | 2.39E-17 | TAM.contam. | Pwwp2b |
| Calhm21 | 1.34E-21 | 0.46439539 | 0.414 | 0.122 | 2.59E-17 | TAM.contam. | Calhm2 |
| Carhsp11 | 1.39E-21 | 0.30962811 | 0.556 | 0.257 | 2.69E-17 | TAM.contam. | Carhsp1 |
| Cyp4v3 | 1.67E-21 | 1.14002104 | 0.354 | 0.101 | 3.23E-17 | TAM.contam. | Cyp4v3 |
| Slc30a53 | 1.70E-21 | 0.5353812 | 0.576 | 0.245 | 3.29E-17 | TAM.contam. | Slc30a5 |
| Pias4 | 1.85E-21 | 0.44462131 | 0.343 | 0.102 | 3.57E-17 | TAM.contam. | Pias4 |
| Nck11 | 1.85E-21 | 0.63479735 | 0.374 | 0.135 | 3.58E-17 | TAM.contam. | Nck1 |
| Kremen1 | 1.85E-21 | 0.29069367 | 0.333 | 0.118 | 3.59E-17 | TAM.contam. | Kremen1 |
| Nagpa2 | 1.94E-21 | 0.43534086 | 0.525 | 0.259 | 3.75E-17 | TAM.contam. | Nagpa |
| Xaf1 | 1.99E-21 | 0.94675878 | 0.343 | 0.113 | 3.85E-17 | TAM.contam. | Xaf1 |
| Gadd45a | 2.06E-21 | 0.88933046 | 0.303 | 0.076 | 3.98E-17 | TAM.contam. | Gadd45a |
| Bad2 | 2.22E-21 | 0.34870344 | 0.495 | 0.202 | 4.30E-17 | TAM.contam. | Bad |
| Rit11 | 2.26E-21 | 0.3440626 | 0.455 | 0.181 | 4.37E-17 | TAM.contam. | Rit1 |
| Tmem106b1 | 3.19E-21 | 0.42396959 | 0.444 | 0.18 | 6.17E-17 | TAM.contam. | Tmem106b |
| Frrs1 | 3.60E-21 | 0.79704225 | 0.364 | 0.111 | 6.95E-17 | TAM.contam. | Frrs1 |
| Map1lc3a | 3.86E-21 | 0.50742342 | 0.404 | 0.148 | 7.47E-17 | TAM.contam. | Map1lc3a |
| Oas2 | 4.10E-21 | 1.06496754 | 0.333 | 0.087 | 7.93E-17 | TAM.contam. | Oas2 |
| Cav2 | 4.20E-21 | 1.09736058 | 0.313 | 0.07 | 8.12E-17 | TAM.contam. | Cav2 |
| Susd31 | 4.41E-21 | 0.51022831 | 0.354 | 0.166 | 8.52E-17 | TAM.contam. | Susd3 |
| Spast | 4.52E-21 | 0.3822294 | 0.525 | 0.218 | 8.75E-17 | TAM.contam. | Spast |
| Rnft1 | 4.71E-21 | 0.75550373 | 0.424 | 0.136 | 9.10E-17 | TAM.contam. | Rnft1 |
| Sft2d1 | 4.95E-21 | 0.6866604 | 0.424 | 0.158 | 9.57E-17 | TAM.contam. | Sft2d1 |
| Sphk11 | 6.14E-21 | 0.36532429 | 0.313 | 0.114 | 1.19E-16 | TAM.contam. | Sphk1 |
| Nbr1 | 7.06E-21 | 0.91465591 | 0.374 | 0.128 | 1.36E-16 | TAM.contam. | Nbr1 |
| Rbl21 | 7.68E-21 | 0.30109747 | 0.525 | 0.214 | 1.49E-16 | TAM.contam. | Rbl2 |
| Phc3 | 8.57E-21 | 0.42019988 | 0.475 | 0.2 | 1.66E-16 | TAM.contam. | Phc3 |
| Herc1 | 8.97E-21 | 0.32696271 | 0.596 | 0.257 | 1.73E-16 | TAM.contam. | Herc1 |
| Pdk11 | 9.04E-21 | 0.57319394 | 0.505 | 0.199 | 1.75E-16 | TAM.contam. | Pdk1 |
| Dhdh | 9.09E-21 | 0.94957276 | 0.414 | 0.105 | 1.76E-16 | TAM.contam. | Dhdh |
| Ugp2 | 9.56E-21 | 0.43839742 | 0.424 | 0.175 | 1.85E-16 | TAM.contam. | Ugp2 |
| Trim65 | 1.01E-20 | 0.57392444 | 0.293 | 0.085 | 1.96E-16 | TAM.contam. | Trim65 |
| Gng5 | 1.09E-20 | 0.58530744 | 0.444 | 0.179 | 2.12E-16 | TAM.contam. | Gng5 |
| Dram11 | 1.11E-20 | 0.76503969 | 0.455 | 0.128 | 2.15E-16 | TAM.contam. | Dram1 |
| Tpd521 | 1.25E-20 | 0.4741779 | 0.434 | 0.166 | 2.42E-16 | TAM.contam. | Tpd52 |
| Wsb22 | 1.40E-20 | 0.33073503 | 0.495 | 0.192 | 2.71E-16 | TAM.contam. | Wsb2 |
| Ptpro1 | 1.55E-20 | 0.25974541 | 0.444 | 0.156 | 3.00E-16 | TAM.contam. | Ptpro |
| Heatr5a1 | 1.57E-20 | 0.3311358 | 0.515 | 0.243 | 3.04E-16 | TAM.contam. | Heatr5a |
| Evi5 | 1.83E-20 | 1.21338331 | 0.273 | 0.06 | 3.54E-16 | TAM.contam. | Evi5 |
| Senp5 | 1.84E-20 | 0.28186078 | 0.515 | 0.203 | 3.56E-16 | TAM.contam. | Senp5 |
| Egln31 | 2.46E-20 | 0.55810306 | 0.354 | 0.127 | 4.75E-16 | TAM.contam. | Egln3 |
| Arnt2 | 2.78E-20 | 0.25466569 | 0.727 | 0.368 | 5.37E-16 | TAM.contam. | Arnt |
| Ppp1r16a1 | 3.18E-20 | 0.31713708 | 0.455 | 0.18 | 6.15E-16 | TAM.contam. | Ppp1r16a |
| Hgf | 3.35E-20 | 0.65878876 | 0.253 | 0.044 | 6.48E-16 | TAM.contam. | Hgf |
| Zwint | 3.44E-20 | 0.34385008 | 0.495 | 0.223 | 6.64E-16 | TAM.contam. | Zwint |
| Echs11 | 3.53E-20 | 0.52471277 | 0.505 | 0.205 | 6.82E-16 | TAM.contam. | Echs1 |
| Clec5a | 3.79E-20 | 0.68731104 | 0.343 | 0.134 | 7.33E-16 | TAM.contam. | Clec5a |
| Tle4 | 3.86E-20 | 0.54861298 | 0.424 | 0.152 | 7.46E-16 | TAM.contam. | Tle4 |
| Cacul1 | 4.11E-20 | 0.29668978 | 0.646 | 0.286 | 7.95E-16 | TAM.contam. | Cacul1 |
| Cpq | 4.62E-20 | 1.06164209 | 0.313 | 0.084 | 8.94E-16 | TAM.contam. | Cpq |
| Tcf7l21 | 5.29E-20 | 0.55245004 | 0.394 | 0.143 | 1.02E-15 | TAM.contam. | Tcf7l2 |
| Deptor | 5.92E-20 | 1.14129711 | 0.293 | 0.067 | 1.15E-15 | TAM.contam. | Deptor |
| Akirin11 | 6.25E-20 | 0.28950784 | 0.515 | 0.28 | 1.21E-15 | TAM.contam. | Akirin1 |
| Ankrd50 | 6.41E-20 | 0.78681415 | 0.394 | 0.116 | 1.24E-15 | TAM.contam. | Ankrd50 |
| Zdhhc9 | 7.26E-20 | 0.75333158 | 0.424 | 0.129 | 1.40E-15 | TAM.contam. | Zdhhc9 |
| Gpaa1 | 7.44E-20 | 0.4034888 | 0.323 | 0.136 | 1.44E-15 | TAM.contam. | Gpaa1 |
| Eef2k2 | 1.03E-19 | 0.58722666 | 0.495 | 0.175 | 2.00E-15 | TAM.contam. | Eef2k |
| Nop561 | 1.20E-19 | 0.29845013 | 0.586 | 0.286 | 2.32E-15 | TAM.contam. | Nop56 |
| Oasl1 | 1.21E-19 | 0.45091455 | 0.293 | 0.077 | 2.35E-15 | TAM.contam. | Oasl1 |
| Fam126a | 1.23E-19 | 0.43672054 | 0.354 | 0.141 | 2.37E-15 | TAM.contam. | Fam126a |
| Stard9 | 1.28E-19 | 1.02040197 | 0.354 | 0.093 | 2.47E-15 | TAM.contam. | Stard9 |
| Peli2 | 1.30E-19 | 0.77851565 | 0.384 | 0.1 | 2.51E-15 | TAM.contam. | Peli2 |
| Tk2 | 1.44E-19 | 0.53905377 | 0.354 | 0.124 | 2.79E-15 | TAM.contam. | Tk2 |
| Samd82 | 1.57E-19 | 0.58260363 | 0.444 | 0.19 | 3.04E-15 | TAM.contam. | Samd8 |
| Galns | 1.74E-19 | 0.99446153 | 0.323 | 0.099 | 3.37E-15 | TAM.contam. | Galns |
| Lap31 | 1.81E-19 | 0.48160439 | 0.404 | 0.164 | 3.51E-15 | TAM.contam. | Lap3 |
| Ell | 1.85E-19 | 0.67718627 | 0.434 | 0.122 | 3.57E-15 | TAM.contam. | Ell |
| Slc31a1 | 1.95E-19 | 0.67085503 | 0.354 | 0.142 | 3.76E-15 | TAM.contam. | Slc31a1 |
| Necap1 | 1.96E-19 | 0.80639699 | 0.293 | 0.094 | 3.79E-15 | TAM.contam. | Necap1 |
| Dclre1a | 1.99E-19 | 1.08647979 | 0.283 | 0.056 | 3.85E-15 | TAM.contam. | Dclre1a |
| Dicer1 | 2.13E-19 | 0.53269502 | 0.444 | 0.172 | 4.12E-15 | TAM.contam. | Dicer1 |
| Ube2q21 | 2.27E-19 | 0.53477305 | 0.455 | 0.164 | 4.39E-15 | TAM.contam. | Ube2q2 |
| Mapre21 | 2.38E-19 | 0.2799205 | 0.495 | 0.234 | 4.61E-15 | TAM.contam. | Mapre2 |
| Tbc1d24 | 2.72E-19 | 1.35664889 | 0.273 | 0.056 | 5.26E-15 | TAM.contam. | Tbc1d24 |
| Cln6 | 2.94E-19 | 0.56215127 | 0.374 | 0.129 | 5.68E-15 | TAM.contam. | Cln6 |
| Prkab1 | 3.10E-19 | 0.91808916 | 0.263 | 0.075 | 5.99E-15 | TAM.contam. | Prkab1 |
| Daam1 | 3.37E-19 | 0.41929485 | 0.354 | 0.121 | 6.51E-15 | TAM.contam. | Daam1 |
| Egr21 | 3.48E-19 | 0.3691947 | 0.465 | 0.164 | 6.72E-15 | TAM.contam. | Egr2 |
| Ino80c | 3.56E-19 | 0.25728761 | 0.495 | 0.212 | 6.88E-15 | TAM.contam. | Ino80c |
| Tssc4 | 3.65E-19 | 0.68693536 | 0.323 | 0.099 | 7.05E-15 | TAM.contam. | Tssc4 |
| Kbtbd21 | 4.07E-19 | 0.32173559 | 0.515 | 0.253 | 7.87E-15 | TAM.contam. | Kbtbd2 |
| Naip21 | 4.53E-19 | 0.48163883 | 0.455 | 0.183 | 8.75E-15 | TAM.contam. | Naip2 |
| Tmx4 | 5.11E-19 | 0.63029226 | 0.404 | 0.122 | 9.88E-15 | TAM.contam. | Tmx4 |
| Pip5k1a | 5.19E-19 | 0.39220263 | 0.404 | 0.162 | 1.00E-14 | TAM.contam. | Pip5k1a |
| Oxct11 | 5.21E-19 | 0.61230115 | 0.505 | 0.174 | 1.01E-14 | TAM.contam. | Oxct1 |
| Ano61 | 5.30E-19 | 0.29019308 | 0.586 | 0.219 | 1.02E-14 | TAM.contam. | Ano6 |
| Pef1 | 5.58E-19 | 0.29497827 | 0.404 | 0.164 | 1.08E-14 | TAM.contam. | Pef1 |
| Atg51 | 5.89E-19 | 0.52106044 | 0.455 | 0.178 | 1.14E-14 | TAM.contam. | Atg5 |
| Ube4a1 | 5.91E-19 | 0.35446645 | 0.495 | 0.244 | 1.14E-14 | TAM.contam. | Ube4a |
| Atg4c1 | 6.14E-19 | 0.50116593 | 0.333 | 0.118 | 1.19E-14 | TAM.contam. | Atg4c |
| Pkig2 | 6.63E-19 | 0.46477381 | 0.444 | 0.169 | 1.28E-14 | TAM.contam. | Pkig |
| Ice1 | 7.82E-19 | 0.48646536 | 0.485 | 0.162 | 1.51E-14 | TAM.contam. | Ice1 |
| Pigs2 | 8.52E-19 | 0.29457067 | 0.636 | 0.31 | 1.65E-14 | TAM.contam. | Pigs |
| Tlk1 | 9.23E-19 | 0.30127738 | 0.535 | 0.215 | 1.79E-14 | TAM.contam. | Tlk1 |
| Aida | 9.50E-19 | 0.46549522 | 0.404 | 0.142 | 1.84E-14 | TAM.contam. | Aida |
| Zfand2b | 1.03E-18 | 0.38041435 | 0.394 | 0.163 | 1.99E-14 | TAM.contam. | Zfand2b |
| As3mt | 1.08E-18 | 0.83320579 | 0.333 | 0.101 | 2.08E-14 | TAM.contam. | As3mt |
| AW5498771 | 1.52E-18 | 0.29151629 | 0.374 | 0.13 | 2.95E-14 | TAM.contam. | AW549877 |
| Ap5s1 | 1.58E-18 | 1.2917735 | 0.273 | 0.066 | 3.06E-14 | TAM.contam. | Ap5s1 |
| Anks11 | 1.61E-18 | 0.33574086 | 0.636 | 0.271 | 3.11E-14 | TAM.contam. | Anks1 |
| Slc36a4 | 1.90E-18 | 0.81733441 | 0.323 | 0.1 | 3.68E-14 | TAM.contam. | Slc36a4 |
| Bcl2l2 | 2.26E-18 | 0.36075223 | 0.354 | 0.131 | 4.36E-14 | TAM.contam. | Bcl2l2 |
| Azin11 | 2.35E-18 | 0.27590209 | 0.566 | 0.3 | 4.54E-14 | TAM.contam. | Azin1 |
| Adi1 | 2.46E-18 | 0.83312642 | 0.455 | 0.139 | 4.75E-14 | TAM.contam. | Adi1 |
| Zfp787 | 2.50E-18 | 0.78489633 | 0.313 | 0.089 | 4.83E-14 | TAM.contam. | Zfp787 |
| B4gat1 | 3.09E-18 | 0.9690421 | 0.303 | 0.085 | 5.98E-14 | TAM.contam. | B4gat1 |
| Atxn2 | 3.27E-18 | 0.52874559 | 0.414 | 0.167 | 6.33E-14 | TAM.contam. | Atxn2 |
| Acad8 | 3.38E-18 | 0.95111588 | 0.303 | 0.09 | 6.53E-14 | TAM.contam. | Acad8 |
| Snap29 | 3.47E-18 | 0.27414602 | 0.566 | 0.267 | 6.71E-14 | TAM.contam. | Snap29 |
| Lpar62 | 3.78E-18 | 0.429126 | 0.485 | 0.163 | 7.31E-14 | TAM.contam. | Lpar6 |
| Snapc4 | 3.82E-18 | 0.92329188 | 0.333 | 0.088 | 7.38E-14 | TAM.contam. | Snapc4 |
| Comtd1 | 4.14E-18 | 0.7947739 | 0.323 | 0.09 | 8.01E-14 | TAM.contam. | Comtd1 |
| Vezt | 4.16E-18 | 1.10328651 | 0.273 | 0.064 | 8.04E-14 | TAM.contam. | Vezt |
| Washc4 | 4.25E-18 | 0.35294635 | 0.465 | 0.192 | 8.22E-14 | TAM.contam. | Washc4 |
| Bbip11 | 5.09E-18 | 0.27545004 | 0.535 | 0.247 | 9.84E-14 | TAM.contam. | Bbip1 |
| Pgm2l1 | 5.40E-18 | 1.19907072 | 0.333 | 0.092 | 1.04E-13 | TAM.contam. | Pgm2l1 |
| Lonp1 | 6.20E-18 | 0.3015326 | 0.576 | 0.248 | 1.20E-13 | TAM.contam. | Lonp1 |
| Kifc3 | 6.77E-18 | 1.01959487 | 0.273 | 0.07 | 1.31E-13 | TAM.contam. | Kifc3 |
| Dexi2 | 7.07E-18 | 0.78404469 | 0.364 | 0.127 | 1.37E-13 | TAM.contam. | Dexi2 |
| Zdhhc7 | 7.39E-18 | 0.29327528 | 0.444 | 0.189 | 1.43E-13 | TAM.contam. | Zdhhc7 |
| Chst122 | 8.04E-18 | 0.32042061 | 0.495 | 0.224 | 1.55E-13 | TAM.contam. | Chst12 |
| Pkd2 | 8.35E-18 | 0.92259918 | 0.253 | 0.05 | 1.61E-13 | TAM.contam. | Pkd2 |
| D17H6S53E1 | 9.28E-18 | 0.36527934 | 0.394 | 0.157 | 1.79E-13 | TAM.contam. | D17H6S53E |
| Prdm11 | 1.00E-17 | 0.53248576 | 0.374 | 0.112 | 1.93E-13 | TAM.contam. | Prdm1 |
| Nop22 | 1.14E-17 | 0.39637452 | 0.465 | 0.211 | 2.20E-13 | TAM.contam. | Nop2 |
| Mcoln1 | 1.39E-17 | 0.2899255 | 0.455 | 0.179 | 2.69E-13 | TAM.contam. | Mcoln1 |
| Naa20 | 1.41E-17 | 0.29997863 | 0.444 | 0.201 | 2.72E-13 | TAM.contam. | Naa20 |
| Gpr137 | 1.53E-17 | 0.38623575 | 0.414 | 0.152 | 2.95E-13 | TAM.contam. | Gpr137 |
| Hccs | 1.79E-17 | 0.34479852 | 0.465 | 0.168 | 3.47E-13 | TAM.contam. | Hccs |
| Ripk31 | 2.04E-17 | 0.62489517 | 0.273 | 0.109 | 3.95E-13 | TAM.contam. | Ripk3 |
| Bet1 | 2.06E-17 | 0.74539702 | 0.414 | 0.158 | 3.97E-13 | TAM.contam. | Bet1 |
| Slu7 | 2.13E-17 | 0.29892231 | 0.475 | 0.183 | 4.11E-13 | TAM.contam. | Slu7 |
| Apba31 | 2.19E-17 | 0.3312604 | 0.394 | 0.162 | 4.24E-13 | TAM.contam. | Apba3 |
| Dhx34 | 2.46E-17 | 0.93264643 | 0.293 | 0.077 | 4.76E-13 | TAM.contam. | Dhx34 |
| Wdr7 | 2.46E-17 | 0.53523611 | 0.404 | 0.149 | 4.76E-13 | TAM.contam. | Wdr7 |
| Mapk91 | 2.82E-17 | 0.39331125 | 0.465 | 0.219 | 5.45E-13 | TAM.contam. | Mapk9 |
| Rbms2 | 3.75E-17 | 0.49768404 | 0.384 | 0.135 | 7.26E-13 | TAM.contam. | Rbms2 |
| Scaf8 | 3.88E-17 | 0.27694825 | 0.505 | 0.203 | 7.51E-13 | TAM.contam. | Scaf8 |
| Mettl9 | 4.06E-17 | 0.25467387 | 0.465 | 0.231 | 7.86E-13 | TAM.contam. | Mettl9 |
| Tmem242 | 4.18E-17 | 0.30016612 | 0.515 | 0.226 | 8.09E-13 | TAM.contam. | Tmem242 |
| Mfap3 | 4.28E-17 | 0.45122473 | 0.424 | 0.169 | 8.27E-13 | TAM.contam. | Mfap3 |
| Rybp1 | 5.42E-17 | 0.36703656 | 0.414 | 0.201 | 1.05E-12 | TAM.contam. | Rybp |
| Cpd4 | 5.56E-17 | 0.37019748 | 0.465 | 0.214 | 1.07E-12 | TAM.contam. | Cpd4 |
| Fam13b | 5.88E-17 | 0.29288521 | 0.525 | 0.205 | 1.14E-12 | TAM.contam. | Fam13b |
| Xylt1 | 5.93E-17 | 0.67268955 | 0.263 | 0.071 | 1.15E-12 | TAM.contam. | Xylt1 |
| Spire11 | 6.04E-17 | 0.25348641 | 0.455 | 0.203 | 1.17E-12 | TAM.contam. | Spire1 |
| Spidr | 6.14E-17 | 0.73149419 | 0.323 | 0.106 | 1.19E-12 | TAM.contam. | Spidr |
| Ipo131 | 7.00E-17 | 0.27137244 | 0.404 | 0.164 | 1.35E-12 | TAM.contam. | Ipo13 |
| Depdc5 | 7.61E-17 | 0.49832333 | 0.414 | 0.151 | 1.47E-12 | TAM.contam. | Depdc5 |
| Fam45a | 7.67E-17 | 0.57410925 | 0.343 | 0.121 | 1.48E-12 | TAM.contam. | Fam45a |
| Slc39a9 | 7.91E-17 | 0.30147575 | 0.404 | 0.172 | 1.53E-12 | TAM.contam. | Slc39a9 |
| Ap5b11 | 7.96E-17 | 0.41097861 | 0.434 | 0.166 | 1.54E-12 | TAM.contam. | Ap5b1 |
| Mms19 | 8.12E-17 | 0.33624907 | 0.394 | 0.161 | 1.57E-12 | TAM.contam. | Mms19 |
| Fzd5 | 8.26E-17 | 0.72105554 | 0.343 | 0.109 | 1.60E-12 | TAM.contam. | Fzd5 |
| Prxl2c1 | 8.78E-17 | 0.43696078 | 0.374 | 0.116 | 1.70E-12 | TAM.contam. | Prxl2c |
| Mpv17 | 8.78E-17 | 0.59398946 | 0.364 | 0.118 | 1.70E-12 | TAM.contam. | Mpv17 |
| Polg2 | 9.11E-17 | 0.34059216 | 0.293 | 0.135 | 1.76E-12 | TAM.contam. | Polg2 |
| Sh3glb21 | 9.43E-17 | 0.42368309 | 0.414 | 0.183 | 1.82E-12 | TAM.contam. | Sh3glb2 |
| Ndfip2 | 9.51E-17 | 0.70523381 | 0.364 | 0.122 | 1.84E-12 | TAM.contam. | Ndfip2 |
| Ngrn | 9.53E-17 | 0.78580812 | 0.263 | 0.069 | 1.84E-12 | TAM.contam. | Ngrn |
| Fam111a | 9.69E-17 | 0.84437018 | 0.313 | 0.106 | 1.87E-12 | TAM.contam. | Fam111a |
| Zfp992 | 9.69E-17 | 0.50366843 | 0.354 | 0.119 | 1.87E-12 | TAM.contam. | Zfp992 |
| Smarcal1 | 1.03E-16 | 0.26442275 | 0.354 | 0.138 | 2.00E-12 | TAM.contam. | Smarcal1 |
| Surf2 | 1.04E-16 | 0.51892723 | 0.323 | 0.093 | 2.00E-12 | TAM.contam. | Surf2 |
| Dido1 | 1.05E-16 | 0.35686401 | 0.424 | 0.165 | 2.04E-12 | TAM.contam. | Dido1 |
| Helz | 1.06E-16 | 0.25164215 | 0.414 | 0.202 | 2.04E-12 | TAM.contam. | Helz |
| Crbn | 1.37E-16 | 0.30034346 | 0.374 | 0.137 | 2.66E-12 | TAM.contam. | Crbn |
| Slc39a13 | 1.51E-16 | 0.69603546 | 0.273 | 0.093 | 2.92E-12 | TAM.contam. | Slc39a13 |
| Zfp326 | 1.58E-16 | 0.55447051 | 0.333 | 0.147 | 3.06E-12 | TAM.contam. | Zfp326 |
| Hmg20a | 2.02E-16 | 0.46886733 | 0.414 | 0.161 | 3.90E-12 | TAM.contam. | Hmg20a |
| Pdcl | 2.09E-16 | 0.56843793 | 0.394 | 0.155 | 4.05E-12 | TAM.contam. | Pdcl |
| Cdkal1 | 2.31E-16 | 0.37208093 | 0.414 | 0.152 | 4.47E-12 | TAM.contam. | Cdkal1 |
| Exoc3 | 2.32E-16 | 0.33506262 | 0.455 | 0.187 | 4.48E-12 | TAM.contam. | Exoc3 |
| Siae | 2.44E-16 | 1.00233406 | 0.333 | 0.089 | 4.72E-12 | TAM.contam. | Siae |
| Kctd6 | 2.56E-16 | 1.23785786 | 0.263 | 0.052 | 4.94E-12 | TAM.contam. | Kctd6 |
| Rab28 | 2.69E-16 | 0.44001977 | 0.414 | 0.18 | 5.20E-12 | TAM.contam. | Rab28 |
| Cryzl1 | 2.83E-16 | 0.72440538 | 0.343 | 0.108 | 5.47E-12 | TAM.contam. | Cryzl1 |
| Dhx57 | 3.10E-16 | 0.40919485 | 0.424 | 0.145 | 5.99E-12 | TAM.contam. | Dhx57 |
| Golga3 | 3.54E-16 | 0.30754302 | 0.374 | 0.15 | 6.84E-12 | TAM.contam. | Golga3 |
| Pros1 | 3.55E-16 | 0.5501044 | 0.333 | 0.111 | 6.86E-12 | TAM.contam. | Pros1 |
| Chst14 | 3.56E-16 | 0.75225901 | 0.313 | 0.082 | 6.88E-12 | TAM.contam. | Chst14 |
| Pex14 | 3.66E-16 | 0.42448254 | 0.495 | 0.216 | 7.08E-12 | TAM.contam. | Pex14 |
| Dhrs7 | 3.86E-16 | 0.81052772 | 0.323 | 0.115 | 7.45E-12 | TAM.contam. | Dhrs7 |
| Pogk | 3.92E-16 | 0.48433512 | 0.293 | 0.085 | 7.58E-12 | TAM.contam. | Pogk |
| Dedd | 4.08E-16 | 0.30563553 | 0.414 | 0.177 | 7.89E-12 | TAM.contam. | Dedd |
| Chd6 | 5.41E-16 | 0.2530163 | 0.465 | 0.166 | 1.05E-11 | TAM.contam. | Chd6 |
| Abcf21 | 5.63E-16 | 0.53398413 | 0.434 | 0.166 | 1.09E-11 | TAM.contam. | Abcf2 |
| Tmem38b | 5.64E-16 | 1.44297368 | 0.253 | 0.062 | 1.09E-11 | TAM.contam. | Tmem38b |
| Fech | 5.68E-16 | 0.62828781 | 0.384 | 0.117 | 1.10E-11 | TAM.contam. | Fech |
| Tifa1 | 5.69E-16 | 0.25353937 | 0.455 | 0.166 | 1.10E-11 | TAM.contam. | Tifa |
| Heatr12 | 6.27E-16 | 0.28897266 | 0.485 | 0.228 | 1.21E-11 | TAM.contam. | Heatr1 |
| Uba5 | 6.34E-16 | 0.26306401 | 0.455 | 0.168 | 1.23E-11 | TAM.contam. | Uba5 |
| Svil1 | 6.39E-16 | 0.58230147 | 0.515 | 0.188 | 1.24E-11 | TAM.contam. | Svil |
| Slc27a4 | 6.57E-16 | 0.45403625 | 0.323 | 0.115 | 1.27E-11 | TAM.contam. | Slc27a4 |
| Zfp653 | 6.72E-16 | 0.43811707 | 0.333 | 0.098 | 1.30E-11 | TAM.contam. | Zfp653 |
| Irgm2 | 6.75E-16 | 1.03149633 | 0.273 | 0.06 | 1.31E-11 | TAM.contam. | Irgm2 |
| Arl14ep | 6.95E-16 | 1.00661788 | 0.293 | 0.088 | 1.34E-11 | TAM.contam. | Arl14ep |
| Clpx | 7.20E-16 | 0.70596666 | 0.323 | 0.109 | 1.39E-11 | TAM.contam. | Clpx |
| St71 | 7.41E-16 | 0.70136331 | 0.414 | 0.149 | 1.43E-11 | TAM.contam. | St7 |
| Kif2a | 8.26E-16 | 0.40822585 | 0.434 | 0.152 | 1.60E-11 | TAM.contam. | Kif2a |
| Ubxn6 | 9.02E-16 | 0.26209688 | 0.424 | 0.201 | 1.74E-11 | TAM.contam. | Ubxn6 |
| Ado | 9.16E-16 | 0.39235006 | 0.374 | 0.15 | 1.77E-11 | TAM.contam. | Ado |
| Asb8 | 9.59E-16 | 0.48227351 | 0.384 | 0.134 | 1.85E-11 | TAM.contam. | Asb8 |
| Pias2 | 1.08E-15 | 0.25511027 | 0.414 | 0.175 | 2.08E-11 | TAM.contam. | Pias2 |
| Pccb | 1.11E-15 | 0.83330396 | 0.323 | 0.089 | 2.15E-11 | TAM.contam. | Pccb |
| Agpat2 | 1.12E-15 | 0.89698749 | 0.263 | 0.067 | 2.16E-11 | TAM.contam. | Agpat2 |
| Srd5a3 | 1.27E-15 | 0.61802115 | 0.333 | 0.106 | 2.46E-11 | TAM.contam. | Srd5a3 |
| Sp1 | 1.34E-15 | 0.28857018 | 0.475 | 0.21 | 2.59E-11 | TAM.contam. | Sp1 |
| Ankhd1 | 1.63E-15 | 0.4218204 | 0.444 | 0.191 | 3.15E-11 | TAM.contam. | Ankhd1 |
| Scrib | 1.66E-15 | 0.32343468 | 0.374 | 0.134 | 3.22E-11 | TAM.contam. | Scrib |
| Fam114a1 | 1.79E-15 | 0.90432303 | 0.253 | 0.056 | 3.46E-11 | TAM.contam. | Fam114a1 |
| Slc12a63 | 1.86E-15 | 0.4370386 | 0.455 | 0.225 | 3.59E-11 | TAM.contam. | Slc12a6 |
| Golph3l1 | 1.92E-15 | 0.45310014 | 0.414 | 0.168 | 3.72E-11 | TAM.contam. | Golph3l |
| Hs2st1 | 2.26E-15 | 0.52783025 | 0.384 | 0.148 | 4.37E-11 | TAM.contam. | Hs2st1 |
| Btbd7 | 2.42E-15 | 0.35519397 | 0.444 | 0.183 | 4.68E-11 | TAM.contam. | Btbd7 |
| Opa31 | 2.53E-15 | 0.29867663 | 0.455 | 0.221 | 4.90E-11 | TAM.contam. | Opa3 |
| Gzf1 | 2.85E-15 | 0.64230509 | 0.293 | 0.08 | 5.52E-11 | TAM.contam. | Gzf1 |
| Kdm4b | 3.30E-15 | 0.36595403 | 0.404 | 0.164 | 6.38E-11 | TAM.contam. | Kdm4b |
| Entpd72 | 3.35E-15 | 0.32134374 | 0.465 | 0.213 | 6.48E-11 | TAM.contam. | Entpd7 |
| Lix1l | 3.65E-15 | 0.60306423 | 0.253 | 0.065 | 7.05E-11 | TAM.contam. | Lix1l |
| Ndst2 | 3.69E-15 | 0.65350958 | 0.404 | 0.124 | 7.14E-11 | TAM.contam. | Ndst2 |
| BC003965 | 4.30E-15 | 0.5440551 | 0.323 | 0.102 | 8.31E-11 | TAM.contam. | BC003965 |
| Usp361 | 4.42E-15 | 0.43011549 | 0.414 | 0.159 | 8.54E-11 | TAM.contam. | Usp36 |
| Fbxw17 | 4.56E-15 | 0.55055833 | 0.323 | 0.084 | 8.82E-11 | TAM.contam. | Fbxw17 |
| Ddx11 | 5.01E-15 | 0.34683741 | 0.465 | 0.192 | 9.68E-11 | TAM.contam. | Ddx1 |
| Utp15 | 5.44E-15 | 0.3595332 | 0.404 | 0.151 | 1.05E-10 | TAM.contam. | Utp15 |
| Ddx19a | 5.60E-15 | 0.26485556 | 0.515 | 0.24 | 1.08E-10 | TAM.contam. | Ddx19a |
| Nhlrc3 | 5.63E-15 | 0.82405878 | 0.273 | 0.08 | 1.09E-10 | TAM.contam. | Nhlrc3 |
| Lpcat31 | 5.94E-15 | 0.31934317 | 0.394 | 0.169 | 1.15E-10 | TAM.contam. | Lpcat3 |
| Hacd3 | 6.19E-15 | 0.32849686 | 0.303 | 0.124 | 1.20E-10 | TAM.contam. | Hacd3 |
| Tmub2 | 6.20E-15 | 0.34324971 | 0.424 | 0.184 | 1.20E-10 | TAM.contam. | Tmub2 |
| Ostm11 | 6.27E-15 | 0.42030883 | 0.465 | 0.222 | 1.21E-10 | TAM.contam. | Ostm1 |
| Stim1 | 6.43E-15 | 0.294299 | 0.455 | 0.19 | 1.24E-10 | TAM.contam. | Stim1 |
| Chid1 | 6.50E-15 | 0.61403069 | 0.394 | 0.152 | 1.26E-10 | TAM.contam. | Chid1 |
| Pla2g4a1 | 6.99E-15 | 0.65292733 | 0.394 | 0.154 | 1.35E-10 | TAM.contam. | Pla2g4a |
| Fut8 | 7.36E-15 | 0.32383316 | 0.384 | 0.144 | 1.42E-10 | TAM.contam. | Fut8 |
| Nars2 | 7.64E-15 | 0.25795984 | 0.556 | 0.282 | 1.48E-10 | TAM.contam. | Nars |
| Tcp11l1 | 8.05E-15 | 0.47303663 | 0.253 | 0.09 | 1.56E-10 | TAM.contam. | Tcp11l1 |
| Trp53inp11 | 8.30E-15 | 0.27817629 | 0.283 | 0.075 | 1.60E-10 | TAM.contam. | Trp53inp1 |
| Akap10 | 8.72E-15 | 0.26330633 | 0.444 | 0.194 | 1.69E-10 | TAM.contam. | Akap10 |
| Zfp2631 | 8.89E-15 | 0.2680883 | 0.495 | 0.278 | 1.72E-10 | TAM.contam. | Zfp263 |
| Phf1 | 9.01E-15 | 0.56726906 | 0.374 | 0.114 | 1.74E-10 | TAM.contam. | Phf1 |
| Irgq | 9.12E-15 | 0.52338334 | 0.293 | 0.079 | 1.76E-10 | TAM.contam. | Irgq |
| Emsy | 9.49E-15 | 0.28341719 | 0.394 | 0.145 | 1.83E-10 | TAM.contam. | Emsy |
| Acsl3 | 9.62E-15 | 0.65056557 | 0.293 | 0.095 | 1.86E-10 | TAM.contam. | Acsl3 |
| Fbxl3 | 9.72E-15 | 0.37224992 | 0.394 | 0.115 | 1.88E-10 | TAM.contam. | Fbxl3 |
| Arid3a | 1.01E-14 | 0.37926038 | 0.404 | 0.145 | 1.95E-10 | TAM.contam. | Arid3a |
| Nr2f6 | 1.04E-14 | 0.35621692 | 0.354 | 0.129 | 2.00E-10 | TAM.contam. | Nr2f6 |
| Ifi211 | 1.24E-14 | 0.32784034 | 0.364 | 0.128 | 2.40E-10 | TAM.contam. | Ifi211 |
| Rcn21 | 1.49E-14 | 0.27422982 | 0.475 | 0.212 | 2.88E-10 | TAM.contam. | Rcn2 |
| Terf2 | 1.53E-14 | 0.31558747 | 0.384 | 0.154 | 2.96E-10 | TAM.contam. | Terf2 |
| Mex3d | 1.64E-14 | 0.62213341 | 0.364 | 0.11 | 3.17E-10 | TAM.contam. | Mex3d |
| Map4k31 | 1.89E-14 | 0.25729484 | 0.364 | 0.143 | 3.65E-10 | TAM.contam. | Map4k3 |
| Mtm1 | 1.89E-14 | 1.22309746 | 0.323 | 0.094 | 3.66E-10 | TAM.contam. | Mtm1 |
| Exoc8 | 2.12E-14 | 0.36032206 | 0.394 | 0.133 | 4.10E-10 | TAM.contam. | Exoc8 |
| Gnl33 | 2.17E-14 | 0.27456641 | 0.444 | 0.215 | 4.19E-10 | TAM.contam. | Gnl3 |
| Xpr1 | 2.41E-14 | 0.43199051 | 0.404 | 0.164 | 4.65E-10 | TAM.contam. | Xpr1 |
| Gxylt1 | 2.47E-14 | 0.30918974 | 0.444 | 0.202 | 4.78E-10 | TAM.contam. | Gxylt1 |
| Ipo92 | 3.54E-14 | 0.27872471 | 0.505 | 0.224 | 6.83E-10 | TAM.contam. | Ipo9 |
| Arl15 | 3.56E-14 | 1.03488145 | 0.273 | 0.066 | 6.88E-10 | TAM.contam. | Arl15 |
| Zfp319 | 4.10E-14 | 0.5885001 | 0.333 | 0.089 | 7.92E-10 | TAM.contam. | Zfp319 |
| Zmym5 | 4.41E-14 | 0.35986025 | 0.323 | 0.115 | 8.52E-10 | TAM.contam. | Zmym5 |
| Rnf7 | 4.46E-14 | 0.46176319 | 0.424 | 0.168 | 8.61E-10 | TAM.contam. | Rnf7 |
| Ccdc84 | 4.62E-14 | 0.69182129 | 0.263 | 0.08 | 8.94E-10 | TAM.contam. | Ccdc84 |
| Galnt71 | 4.94E-14 | 0.2523029 | 0.384 | 0.141 | 9.55E-10 | TAM.contam. | Galnt7 |
| Rbx11 | 5.06E-14 | 0.29991495 | 0.586 | 0.279 | 9.78E-10 | TAM.contam. | Rbx1 |
| Zfp715 | 5.63E-14 | 0.5067812 | 0.394 | 0.115 | 1.09E-09 | TAM.contam. | Zfp715 |
| Tmem11 | 5.72E-14 | 0.29295014 | 0.444 | 0.203 | 1.11E-09 | TAM.contam. | Tmem11 |
| Zfp687 | 6.21E-14 | 0.3280237 | 0.404 | 0.157 | 1.20E-09 | TAM.contam. | Zfp687 |
| Acvr1b | 6.32E-14 | 0.52300672 | 0.384 | 0.15 | 1.22E-09 | TAM.contam. | Acvr1b |
| Iqsec2 | 6.73E-14 | 0.72447547 | 0.253 | 0.064 | 1.30E-09 | TAM.contam. | Iqsec2 |
| Pecam11 | 6.93E-14 | 0.25201458 | 0.354 | 0.136 | 1.34E-09 | TAM.contam. | Pecam1 |
| Wdr11 | 7.37E-14 | 0.74085428 | 0.343 | 0.125 | 1.43E-09 | TAM.contam. | Wdr11 |
| Creld1 | 7.83E-14 | 0.31925007 | 0.323 | 0.131 | 1.51E-09 | TAM.contam. | Creld1 |
| Slfn81 | 8.19E-14 | 0.67400363 | 0.333 | 0.094 | 1.58E-09 | TAM.contam. | Slfn8 |
| Fbxo7 | 8.49E-14 | 0.30702377 | 0.343 | 0.147 | 1.64E-09 | TAM.contam. | Fbxo7 |
| Als21 | 8.80E-14 | 0.5364479 | 0.253 | 0.068 | 1.70E-09 | TAM.contam. | Als2 |
| Sntb2 | 8.85E-14 | 0.55891958 | 0.283 | 0.086 | 1.71E-09 | TAM.contam. | Sntb2 |
| Coq8b | 9.72E-14 | 0.60876519 | 0.293 | 0.098 | 1.88E-09 | TAM.contam. | Coq8b |
| Tecpr1 | 9.72E-14 | 0.50863999 | 0.293 | 0.097 | 1.88E-09 | TAM.contam. | Tecpr1 |
| Isca21 | 9.91E-14 | 0.26518633 | 0.424 | 0.17 | 1.92E-09 | TAM.contam. | Isca2 |
| Tmem265 | 1.06E-13 | 0.34343836 | 0.343 | 0.13 | 2.05E-09 | TAM.contam. | Tmem265 |
| Tstd2 | 1.25E-13 | 0.26263829 | 0.364 | 0.132 | 2.42E-09 | TAM.contam. | Tstd2 |
| Nkapd1 | 1.30E-13 | 0.31731649 | 0.323 | 0.118 | 2.52E-09 | TAM.contam. | Nkapd1 |
| Ampd3 | 1.33E-13 | 0.63447938 | 0.273 | 0.094 | 2.58E-09 | TAM.contam. | Ampd3 |
| Umps | 1.33E-13 | 0.81402192 | 0.293 | 0.094 | 2.58E-09 | TAM.contam. | Umps |
| Aktip | 1.36E-13 | 0.68967708 | 0.303 | 0.087 | 2.64E-09 | TAM.contam. | Aktip |
| Plekhm1 | 1.38E-13 | 0.57805479 | 0.354 | 0.135 | 2.67E-09 | TAM.contam. | Plekhm1 |
| Ift46 | 1.38E-13 | 0.32489452 | 0.354 | 0.138 | 2.67E-09 | TAM.contam. | Ift46 |
| Ubap22 | 1.52E-13 | 0.32285102 | 0.515 | 0.24 | 2.95E-09 | TAM.contam. | Ubap2 |
| Nod1 | 1.68E-13 | 0.50209169 | 0.333 | 0.113 | 3.25E-09 | TAM.contam. | Nod1 |
| Mbd6 | 1.72E-13 | 0.52732343 | 0.333 | 0.12 | 3.32E-09 | TAM.contam. | Mbd6 |
| Abcb7 | 1.81E-13 | 0.37411062 | 0.333 | 0.154 | 3.50E-09 | TAM.contam. | Abcb7 |
| Pik3c2a1 | 1.86E-13 | 0.37781414 | 0.414 | 0.167 | 3.60E-09 | TAM.contam. | Pik3c2a |
| Ccs | 1.94E-13 | 0.74241864 | 0.313 | 0.098 | 3.75E-09 | TAM.contam. | Ccs |
| Dmac1 | 2.03E-13 | 0.44893613 | 0.394 | 0.175 | 3.92E-09 | TAM.contam. | Dmac1 |
| Txnrd2 | 2.14E-13 | 0.30264056 | 0.333 | 0.129 | 4.13E-09 | TAM.contam. | Txnrd2 |
| Pde8a | 2.19E-13 | 0.39947836 | 0.303 | 0.118 | 4.23E-09 | TAM.contam. | Pde8a |
| Lancl1 | 2.75E-13 | 0.36602183 | 0.354 | 0.127 | 5.31E-09 | TAM.contam. | Lancl1 |
| Luzp11 | 2.98E-13 | 0.5358371 | 0.354 | 0.145 | 5.77E-09 | TAM.contam. | Luzp1 |
| Mphosph8 | 3.15E-13 | 0.64040533 | 0.293 | 0.107 | 6.10E-09 | TAM.contam. | Mphosph8 |
| Slc30a9 | 3.24E-13 | 0.27982732 | 0.444 | 0.197 | 6.27E-09 | TAM.contam. | Slc30a9 |
| Caap1 | 3.49E-13 | 0.25923972 | 0.364 | 0.14 | 6.74E-09 | TAM.contam. | Caap1 |
| Opa1 | 3.80E-13 | 0.33865931 | 0.404 | 0.19 | 7.35E-09 | TAM.contam. | Opa1 |
| Trp53rka | 4.14E-13 | 0.87770512 | 0.253 | 0.069 | 8.01E-09 | TAM.contam. | Trp53rka |
| Tbcel1 | 4.53E-13 | 0.49943973 | 0.293 | 0.098 | 8.77E-09 | TAM.contam. | Tbcel |
| Golga5 | 4.68E-13 | 0.28264112 | 0.374 | 0.173 | 9.04E-09 | TAM.contam. | Golga5 |
| Prr14l | 4.70E-13 | 0.27368855 | 0.424 | 0.22 | 9.09E-09 | TAM.contam. | Prr14l |
| Glo11 | 4.70E-13 | 0.27572005 | 0.444 | 0.21 | 9.09E-09 | TAM.contam. | Glo1 |
| Sprtn | 4.92E-13 | 0.86413136 | 0.313 | 0.094 | 9.51E-09 | TAM.contam. | Sprtn |
| Mat2b1 | 5.20E-13 | 0.35105699 | 0.404 | 0.193 | 1.00E-08 | TAM.contam. | Mat2b |
| Ccnh | 5.37E-13 | 0.68268241 | 0.273 | 0.081 | 1.04E-08 | TAM.contam. | Ccnh |
| Clock | 5.57E-13 | 0.66801886 | 0.354 | 0.107 | 1.08E-08 | TAM.contam. | Clock |
| Rab11fip2 | 5.86E-13 | 0.62494983 | 0.253 | 0.082 | 1.13E-08 | TAM.contam. | Rab11fip2 |
| Wdr45 | 6.56E-13 | 0.33589941 | 0.343 | 0.13 | 1.27E-08 | TAM.contam. | Wdr45 |
| Snapin | 6.73E-13 | 0.40213817 | 0.354 | 0.142 | 1.30E-08 | TAM.contam. | Snapin |
| Ifi471 | 6.98E-13 | 0.34911044 | 0.323 | 0.121 | 1.35E-08 | TAM.contam. | Ifi47 |
| Irgm12 | 7.04E-13 | 0.30033596 | 0.444 | 0.176 | 1.36E-08 | TAM.contam. | Irgm1 |
| Eif2ak2 | 7.44E-13 | 0.25163382 | 0.404 | 0.172 | 1.44E-08 | TAM.contam. | Eif2ak2 |
| Vill | 7.98E-13 | 0.29448491 | 0.313 | 0.16 | 1.54E-08 | TAM.contam. | Vill |
| Slc10a3 | 8.19E-13 | 0.52025707 | 0.283 | 0.103 | 1.58E-08 | TAM.contam. | Slc10a3 |
| Enc1 | 8.59E-13 | 0.30219743 | 0.384 | 0.143 | 1.66E-08 | TAM.contam. | Enc1 |
| Sap130 | 8.72E-13 | 0.31499426 | 0.505 | 0.21 | 1.69E-08 | TAM.contam. | Sap130 |
| Fmr11 | 9.82E-13 | 0.3332541 | 0.505 | 0.255 | 1.90E-08 | TAM.contam. | Fmr1 |
| Tmem2191 | 1.02E-12 | 0.31369803 | 0.495 | 0.228 | 1.96E-08 | TAM.contam. | Tmem219 |
| Rft1 | 1.04E-12 | 0.64456306 | 0.303 | 0.107 | 2.01E-08 | TAM.contam. | Rft1 |
| Parvb | 1.07E-12 | 0.83281981 | 0.283 | 0.072 | 2.06E-08 | TAM.contam. | Parvb |
| Abraxas2 | 1.14E-12 | 0.28558396 | 0.374 | 0.152 | 2.20E-08 | TAM.contam. | Abraxas2 |
| Dvl2 | 1.15E-12 | 0.41919443 | 0.323 | 0.123 | 2.22E-08 | TAM.contam. | Dvl2 |
| Smpd1 | 1.16E-12 | 0.95368441 | 0.303 | 0.089 | 2.23E-08 | TAM.contam. | Smpd1 |
| Ctdp1 | 1.23E-12 | 0.5397973 | 0.333 | 0.129 | 2.37E-08 | TAM.contam. | Ctdp1 |
| Jkamp | 1.28E-12 | 0.56692198 | 0.313 | 0.109 | 2.48E-08 | TAM.contam. | Jkamp |
| Arsa1 | 1.35E-12 | 0.26805056 | 0.414 | 0.169 | 2.61E-08 | TAM.contam. | Arsa |
| Prdm4 | 1.42E-12 | 0.92714775 | 0.354 | 0.102 | 2.74E-08 | TAM.contam. | Prdm4 |
| Czib | 1.53E-12 | 0.36430175 | 0.374 | 0.149 | 2.95E-08 | TAM.contam. | Czib |
| Elof1 | 1.53E-12 | 0.6568482 | 0.404 | 0.156 | 2.97E-08 | TAM.contam. | Elof1 |
| Pikfyve | 1.57E-12 | 0.35357127 | 0.313 | 0.097 | 3.03E-08 | TAM.contam. | Pikfyve |
| Cdk7 | 1.57E-12 | 0.54098455 | 0.374 | 0.134 | 3.03E-08 | TAM.contam. | Cdk7 |
| Pml1 | 1.57E-12 | 0.32104104 | 0.333 | 0.098 | 3.04E-08 | TAM.contam. | Pml |
| Mob4 | 1.83E-12 | 0.50154651 | 0.485 | 0.193 | 3.54E-08 | TAM.contam. | Mob4 |
| Zfp574 | 1.91E-12 | 0.33797957 | 0.303 | 0.111 | 3.70E-08 | TAM.contam. | Zfp574 |
| Cnnm3 | 2.23E-12 | 0.30709911 | 0.303 | 0.105 | 4.31E-08 | TAM.contam. | Cnnm3 |
| Rnf185 | 2.30E-12 | 0.62869832 | 0.283 | 0.081 | 4.45E-08 | TAM.contam. | Rnf185 |
| Ndufs5 | 2.36E-12 | 0.27037879 | 0.424 | 0.201 | 4.56E-08 | TAM.contam. | Ndufs5 |
| Ufl1 | 2.48E-12 | 0.34463669 | 0.394 | 0.148 | 4.79E-08 | TAM.contam. | Ufl1 |
| Smad5 | 2.49E-12 | 0.34838846 | 0.323 | 0.111 | 4.82E-08 | TAM.contam. | Smad5 |
| Atrn | 2.55E-12 | 0.40569227 | 0.293 | 0.109 | 4.93E-08 | TAM.contam. | Atrn |
| Stam | 2.59E-12 | 0.27864564 | 0.323 | 0.126 | 5.01E-08 | TAM.contam. | Stam |
| Commd9 | 2.72E-12 | 0.55868989 | 0.263 | 0.082 | 5.27E-08 | TAM.contam. | Commd9 |
| Cyb561d2 | 2.95E-12 | 0.78319599 | 0.283 | 0.105 | 5.70E-08 | TAM.contam. | Cyb561d2 |
| Gfm2 | 3.16E-12 | 0.26458215 | 0.273 | 0.104 | 6.11E-08 | TAM.contam. | Gfm2 |
| Lcorl | 3.88E-12 | 0.41935835 | 0.323 | 0.105 | 7.51E-08 | TAM.contam. | Lcorl |
| Pdzd11 | 3.95E-12 | 0.29223105 | 0.364 | 0.144 | 7.63E-08 | TAM.contam. | Pdzd11 |
| Hsdl1 | 4.21E-12 | 0.46804607 | 0.354 | 0.117 | 8.14E-08 | TAM.contam. | Hsdl1 |
| Ttf1 | 4.30E-12 | 0.27360785 | 0.273 | 0.091 | 8.30E-08 | TAM.contam. | Ttf1 |
| Pilrb11 | 4.92E-12 | 0.59952304 | 0.495 | 0.204 | 9.51E-08 | TAM.contam. | Pilrb1 |
| Umad1 | 5.17E-12 | 0.49636377 | 0.343 | 0.127 | 9.99E-08 | TAM.contam. | Umad1 |
| Noct | 5.27E-12 | 0.35326556 | 0.354 | 0.136 | 1.02E-07 | TAM.contam. | Noct |
| Dtx23 | 5.34E-12 | 0.26616655 | 0.404 | 0.2 | 1.03E-07 | TAM.contam. | Dtx2 |
| Tceanc2 | 5.65E-12 | 0.4036079 | 0.414 | 0.174 | 1.09E-07 | TAM.contam. | Tceanc2 |
| Xpot | 5.84E-12 | 0.57762467 | 0.313 | 0.107 | 1.13E-07 | TAM.contam. | Xpot |
| Irf2bpl | 5.85E-12 | 0.37475133 | 0.333 | 0.093 | 1.13E-07 | TAM.contam. | Irf2bpl |
| Zbtb4 | 6.01E-12 | 0.35480991 | 0.374 | 0.128 | 1.16E-07 | TAM.contam. | Zbtb4 |
| Mad1l1 | 6.11E-12 | 0.53601158 | 0.333 | 0.129 | 1.18E-07 | TAM.contam. | Mad1l1 |
| Ppp4r1 | 6.22E-12 | 0.41668521 | 0.303 | 0.122 | 1.20E-07 | TAM.contam. | Ppp4r1 |
| Ints6l | 6.62E-12 | 0.36534153 | 0.303 | 0.118 | 1.28E-07 | TAM.contam. | Ints6l |
| Triap1 | 6.95E-12 | 0.71228901 | 0.364 | 0.126 | 1.34E-07 | TAM.contam. | Triap1 |
| Zfp84 | 7.63E-12 | 0.37355916 | 0.273 | 0.103 | 1.48E-07 | TAM.contam. | Zfp84 |
| Hspa13 | 7.96E-12 | 0.30001602 | 0.323 | 0.119 | 1.54E-07 | TAM.contam. | Hspa13 |
| Cnep1r1 | 8.74E-12 | 0.32168981 | 0.354 | 0.153 | 1.69E-07 | TAM.contam. | Cnep1r1 |
| Ddhd2 | 8.98E-12 | 0.27462053 | 0.394 | 0.179 | 1.74E-07 | TAM.contam. | Ddhd2 |
| Bap1 | 9.20E-12 | 0.27885069 | 0.404 | 0.16 | 1.78E-07 | TAM.contam. | Bap1 |
| Xrcc4 | 9.59E-12 | 0.63555985 | 0.273 | 0.103 | 1.85E-07 | TAM.contam. | Xrcc4 |
| Slc25a44 | 9.84E-12 | 0.60409799 | 0.273 | 0.09 | 1.90E-07 | TAM.contam. | Slc25a44 |
| Pik3ca | 1.07E-11 | 0.36490243 | 0.394 | 0.161 | 2.07E-07 | TAM.contam. | Pik3ca |
| Spata132 | 1.14E-11 | 0.26280063 | 0.333 | 0.16 | 2.20E-07 | TAM.contam. | Spata13 |
| Parp10 | 1.15E-11 | 0.61802988 | 0.343 | 0.13 | 2.23E-07 | TAM.contam. | Parp10 |
| Sec22a | 1.39E-11 | 0.82272161 | 0.253 | 0.076 | 2.69E-07 | TAM.contam. | Sec22a |
| Gtf2h1 | 1.45E-11 | 0.32523683 | 0.364 | 0.153 | 2.81E-07 | TAM.contam. | Gtf2h1 |
| Irf2bp1 | 1.51E-11 | 0.40279006 | 0.364 | 0.14 | 2.92E-07 | TAM.contam. | Irf2bp1 |
| Riok2 | 1.52E-11 | 0.2929842 | 0.404 | 0.166 | 2.94E-07 | TAM.contam. | Riok2 |
| Mbtps2 | 1.56E-11 | 0.67811141 | 0.313 | 0.103 | 3.01E-07 | TAM.contam. | Mbtps2 |
| AB124611 | 2.03E-11 | 0.27266084 | 0.283 | 0.133 | 3.93E-07 | TAM.contam. | AB124611 |
| Gabpb1 | 2.13E-11 | 0.33716475 | 0.374 | 0.169 | 4.12E-07 | TAM.contam. | Gabpb1 |
| Tcf121 | 2.16E-11 | 0.40705985 | 0.394 | 0.189 | 4.18E-07 | TAM.contam. | Tcf12 |
| Mcat | 2.35E-11 | 0.60541319 | 0.273 | 0.075 | 4.54E-07 | TAM.contam. | Mcat |
| Foxo4 | 2.35E-11 | 0.48998404 | 0.253 | 0.094 | 4.55E-07 | TAM.contam. | Foxo4 |
| Cwc27 | 2.37E-11 | 0.61909337 | 0.283 | 0.099 | 4.57E-07 | TAM.contam. | Cwc27 |
| Pkdcc | 2.50E-11 | 0.53098558 | 0.273 | 0.094 | 4.83E-07 | TAM.contam. | Pkdcc |
| Dnajb4 | 2.56E-11 | 0.85054207 | 0.263 | 0.063 | 4.94E-07 | TAM.contam. | Dnajb4 |
| Fbxo9 | 2.72E-11 | 0.41813095 | 0.283 | 0.1 | 5.25E-07 | TAM.contam. | Fbxo9 |
| Lztr1 | 2.83E-11 | 0.60654727 | 0.333 | 0.129 | 5.48E-07 | TAM.contam. | Lztr1 |
| Fam3a | 2.91E-11 | 0.65874402 | 0.303 | 0.1 | 5.62E-07 | TAM.contam. | Fam3a |
| Raph1 | 2.94E-11 | 0.59959299 | 0.303 | 0.103 | 5.68E-07 | TAM.contam. | Raph1 |
| Phka2 | 3.07E-11 | 0.5593571 | 0.293 | 0.117 | 5.94E-07 | TAM.contam. | Phka2 |
| B3gat3 | 3.20E-11 | 0.30139427 | 0.394 | 0.162 | 6.20E-07 | TAM.contam. | B3gat3 |
| Tada3 | 3.34E-11 | 0.36346344 | 0.333 | 0.133 | 6.46E-07 | TAM.contam. | Tada3 |
| Hscb | 3.97E-11 | 0.31917356 | 0.273 | 0.101 | 7.67E-07 | TAM.contam. | Hscb |
| DEPRECATED-I | 4.14E-11 | 0.32167884 | 0.485 | 0.23 | 8.00E-07 | TAM.contam. | DEPRECATED-ENSMUSG00000022521 |
| Rpain | 4.30E-11 | 0.33003935 | 0.333 | 0.141 | 8.32E-07 | TAM.contam. | Rpain |
| Erlec1 | 4.31E-11 | 0.32188962 | 0.293 | 0.105 | 8.33E-07 | TAM.contam. | Erlec1 |
| Stk11ip | 5.87E-11 | 0.25362051 | 0.343 | 0.129 | 1.13E-06 | TAM.contam. | Stk11ip |
| Fdx1 | 6.57E-11 | 0.25242029 | 0.273 | 0.111 | 1.27E-06 | TAM.contam. | Fdx1 |
| Mzt1 | 6.65E-11 | 0.29544348 | 0.374 | 0.165 | 1.28E-06 | TAM.contam. | Mzt1 |
| Arel1 | 6.93E-11 | 0.37638106 | 0.364 | 0.173 | 1.34E-06 | TAM.contam. | Arel1 |
| Stard51 | 6.96E-11 | 0.70197252 | 0.333 | 0.112 | 1.35E-06 | TAM.contam. | Stard5 |
| Chchd7 | 7.64E-11 | 0.34006144 | 0.263 | 0.093 | 1.48E-06 | TAM.contam. | Chchd7 |
| Klhl7 | 7.72E-11 | 0.44775069 | 0.273 | 0.091 | 1.49E-06 | TAM.contam. | Klhl7 |
| Gtf3c3 | 8.15E-11 | 0.36357444 | 0.293 | 0.121 | 1.58E-06 | TAM.contam. | Gtf3c3 |
| Klf11 | 8.21E-11 | 0.69623512 | 0.253 | 0.086 | 1.59E-06 | TAM.contam. | Klf11 |
| Coq5 | 8.59E-11 | 0.38953871 | 0.273 | 0.105 | 1.66E-06 | TAM.contam. | Coq5 |
| Dnajc24 | 9.25E-11 | 0.50584827 | 0.303 | 0.125 | 1.79E-06 | TAM.contam. | Dnajc24 |
| Sh3rf1 | 9.47E-11 | 0.34355284 | 0.253 | 0.088 | 1.83E-06 | TAM.contam. | Sh3rf1 |
| Fbxo22 | 9.96E-11 | 0.39682463 | 0.354 | 0.145 | 1.92E-06 | TAM.contam. | Fbxo22 |
| Rnf2 | 1.07E-10 | 0.45960484 | 0.374 | 0.134 | 2.06E-06 | TAM.contam. | Rnf2 |
| Brd9 | 1.21E-10 | 0.25847121 | 0.323 | 0.137 | 2.34E-06 | TAM.contam. | Brd9 |
| Spns1 | 1.24E-10 | 0.62340073 | 0.283 | 0.088 | 2.39E-06 | TAM.contam. | Spns1 |
| Cacfd1 | 1.25E-10 | 0.33702924 | 0.333 | 0.133 | 2.42E-06 | TAM.contam. | Cacfd1 |
| Txndc161 | 1.28E-10 | 0.37605473 | 0.303 | 0.133 | 2.47E-06 | TAM.contam. | Txndc16 |
| Clp1 | 1.35E-10 | 0.33483877 | 0.343 | 0.139 | 2.61E-06 | TAM.contam. | Clp1 |
| Dock42 | 1.48E-10 | 0.26149863 | 0.424 | 0.183 | 2.86E-06 | TAM.contam. | Dock4 |
| Dip2a | 1.61E-10 | 0.26626754 | 0.333 | 0.147 | 3.11E-06 | TAM.contam. | Dip2a |
| Pex2 | 1.65E-10 | 0.30520693 | 0.343 | 0.139 | 3.20E-06 | TAM.contam. | Pex2 |
| Lman2l | 1.90E-10 | 0.35445325 | 0.313 | 0.14 | 3.67E-06 | TAM.contam. | Lman2l |
| Zbtb34 | 1.92E-10 | 0.35272105 | 0.273 | 0.1 | 3.72E-06 | TAM.contam. | Zbtb34 |
| Slc35a1 | 2.87E-10 | 0.56474315 | 0.263 | 0.089 | 5.54E-06 | TAM.contam. | Slc35a1 |
| Farsa1 | 3.22E-10 | 0.29170924 | 0.495 | 0.264 | 6.22E-06 | TAM.contam. | Farsa |
| Agpat5 | 3.43E-10 | 0.3982885 | 0.343 | 0.144 | 6.63E-06 | TAM.contam. | Agpat5 |
| Snx141 | 3.74E-10 | 0.55955562 | 0.313 | 0.118 | 7.23E-06 | TAM.contam. | Snx14 |
| Vps50 | 3.78E-10 | 0.41678758 | 0.263 | 0.113 | 7.31E-06 | TAM.contam. | Vps50 |
| Mcf2l | 3.94E-10 | 0.53307768 | 0.273 | 0.096 | 7.63E-06 | TAM.contam. | Mcf2l |
| Coq10a | 4.00E-10 | 0.71898905 | 0.253 | 0.115 | 7.74E-06 | TAM.contam. | Coq10a |
| Hps41 | 4.23E-10 | 0.26732697 | 0.293 | 0.117 | 8.18E-06 | TAM.contam. | Hps4 |
| Bloc1s4 | 4.43E-10 | 0.37942986 | 0.313 | 0.111 | 8.57E-06 | TAM.contam. | Bloc1s4 |
| Azi2 | 4.70E-10 | 0.32683575 | 0.313 | 0.114 | 9.09E-06 | TAM.contam. | Azi2 |
| Pag1 | 5.85E-10 | 0.40817372 | 0.293 | 0.123 | 1.13E-05 | TAM.contam. | Pag1 |
| Bet1l | 5.93E-10 | 0.28199666 | 0.293 | 0.11 | 1.15E-05 | TAM.contam. | Bet1l |
| Ldah | 6.14E-10 | 0.64279448 | 0.333 | 0.138 | 1.19E-05 | TAM.contam. | Ldah |
| Wdr83 | 6.75E-10 | 0.28982776 | 0.283 | 0.095 | 1.31E-05 | TAM.contam. | Wdr83 |
| Nbeal1 | 6.93E-10 | 0.35390529 | 0.313 | 0.12 | 1.34E-05 | TAM.contam. | Nbeal1 |
| Tm7sf31 | 7.12E-10 | 0.47434765 | 0.313 | 0.129 | 1.38E-05 | TAM.contam. | Tm7sf3 |
| Tor3a | 7.79E-10 | 0.58126639 | 0.333 | 0.141 | 1.51E-05 | TAM.contam. | Tor3a |
| Ebag9 | 8.15E-10 | 0.27653439 | 0.303 | 0.122 | 1.58E-05 | TAM.contam. | Ebag9 |
| Abtb1 | 8.27E-10 | 0.25126291 | 0.263 | 0.108 | 1.60E-05 | TAM.contam. | Abtb1 |
| Dusp12 | 8.67E-10 | 0.34905197 | 0.253 | 0.079 | 1.68E-05 | TAM.contam. | Dusp12 |
| Upf3b | 8.77E-10 | 0.41646371 | 0.273 | 0.108 | 1.70E-05 | TAM.contam. | Upf3b |
| Dcun1d3 | 8.84E-10 | 0.32935285 | 0.293 | 0.105 | 1.71E-05 | TAM.contam. | Dcun1d3 |
| Cyp20a1 | 9.09E-10 | 0.71382449 | 0.273 | 0.098 | 1.76E-05 | TAM.contam. | Cyp20a1 |
| Vkorc1 | 9.29E-10 | 0.27730346 | 0.313 | 0.124 | 1.80E-05 | TAM.contam. | Vkorc1 |

| | | | | | | | |
|---|---|---|---|---|---|---|---|
| Zfp639 | 1.04E-09 | 0.76666834 | 0.283 | 0.107 | 2.02E-05 | TAM.contam. | Zfp639 |
| Rab40c | 1.08E-09 | 0.28837471 | 0.333 | 0.126 | 2.09E-05 | TAM.contam. | Rab40c |
| Lsm1 | 1.23E-09 | 0.25358519 | 0.343 | 0.141 | 2.38E-05 | TAM.contam. | Lsm1 |
| Aak1 | 1.54E-09 | 0.29577959 | 0.283 | 0.117 | 2.98E-05 | TAM.contam. | Aak1 |
| Slc41a3 | 1.65E-09 | 0.64639051 | 0.263 | 0.084 | 3.19E-05 | TAM.contam. | Slc41a3 |
| Zxdc | 1.74E-09 | 0.52177367 | 0.263 | 0.097 | 3.36E-05 | TAM.contam. | Zxdc |
| Tbcc | 1.74E-09 | 0.2507452 | 0.333 | 0.133 | 3.37E-05 | TAM.contam. | Tbcc |
| Bnip3l | 1.80E-09 | 0.30714671 | 0.303 | 0.121 | 3.47E-05 | TAM.contam. | Bnip3l |
| Coq9 | 1.92E-09 | 0.63944162 | 0.323 | 0.113 | 3.71E-05 | TAM.contam. | Coq9 |
| Lnpk | 1.97E-09 | 0.30472938 | 0.273 | 0.118 | 3.81E-05 | TAM.contam. | Lnpk |
| Pou2f1 | 2.06E-09 | 0.39533538 | 0.354 | 0.129 | 3.99E-05 | TAM.contam. | Pou2f1 |
| Wdyhv1 | 2.23E-09 | 0.30574288 | 0.263 | 0.088 | 4.30E-05 | TAM.contam. | Wdyhv1 |
| Mtif2 | 2.65E-09 | 0.33231962 | 0.263 | 0.095 | 5.13E-05 | TAM.contam. | Mtif2 |
| Sgms11 | 2.71E-09 | 0.42917549 | 0.364 | 0.133 | 5.25E-05 | TAM.contam. | Sgms1 |
| Rfng | 3.37E-09 | 0.33121471 | 0.263 | 0.1 | 6.52E-05 | TAM.contam. | Rfng |
| Zscan29 | 3.48E-09 | 0.61806564 | 0.263 | 0.092 | 6.73E-05 | TAM.contam. | Zscan29 |
| Heatr5b | 3.49E-09 | 0.36790739 | 0.313 | 0.122 | 6.74E-05 | TAM.contam. | Heatr5b |
| Abhd17c | 3.84E-09 | 0.34355304 | 0.283 | 0.115 | 7.42E-05 | TAM.contam. | Abhd17c |
| Alg8 | 4.55E-09 | 0.53617676 | 0.263 | 0.099 | 8.80E-05 | TAM.contam. | Alg8 |
| Megf8 | 4.73E-09 | 0.37246473 | 0.323 | 0.126 | 9.14E-05 | TAM.contam. | Megf8 |
| Pctp | 4.91E-09 | 0.28730692 | 0.283 | 0.109 | 9.49E-05 | TAM.contam. | Pctp |
| Zfyve19 | 4.91E-09 | 0.29574714 | 0.303 | 0.129 | 9.50E-05 | TAM.contam. | Zfyve19 |
| Ctbs | 5.17E-09 | 0.57172673 | 0.293 | 0.107 | 0.00010005 | TAM.contam. | Ctbs |
| Zcchc7 | 5.56E-09 | 0.42032866 | 0.333 | 0.163 | 0.00010759 | TAM.contam. | Zcchc7 |
| Brms1l | 6.39E-09 | 0.29128695 | 0.273 | 0.099 | 0.00012357 | TAM.contam. | Brms1l |
| Pgp1 | 6.85E-09 | 0.25659972 | 0.343 | 0.145 | 0.00013237 | TAM.contam. | Pgp |
| Polr2h | 7.42E-09 | 0.63895308 | 0.343 | 0.153 | 0.00014352 | TAM.contam. | Polr2h |
| Ston2 | 8.17E-09 | 0.50126679 | 0.253 | 0.092 | 0.00015805 | TAM.contam. | Ston2 |
| Zcchc17 | 8.22E-09 | 0.41390418 | 0.313 | 0.147 | 0.00015895 | TAM.contam. | Zcchc17 |
| Zfp654 | 9.19E-09 | 0.34379323 | 0.303 | 0.129 | 0.00017772 | TAM.contam. | Zfp654 |
| Rundc1 | 9.66E-09 | 0.2589979 | 0.313 | 0.112 | 0.0001868 | TAM.contam. | Rundc1 |
| Yaf2 | 1.00E-08 | 0.77232339 | 0.263 | 0.099 | 0.0001941 | TAM.contam. | Yaf2 |
| Gpatch11 | 1.11E-08 | 0.2964175 | 0.293 | 0.139 | 0.00021415 | TAM.contam. | Gpatch11 |
| Mul1 | 1.11E-08 | 0.45192065 | 0.253 | 0.096 | 0.00021487 | TAM.contam. | Mul1 |
| Urb2 | 1.11E-08 | 0.61453155 | 0.263 | 0.093 | 0.00021487 | TAM.contam. | Urb2 |
| Epc2 | 1.17E-08 | 0.26649437 | 0.293 | 0.139 | 0.00022556 | TAM.contam. | Epc2 |
| Lcmt1 | 1.20E-08 | 0.68032755 | 0.313 | 0.123 | 0.00023253 | TAM.contam. | Lcmt1 |
| Abhd5 | 1.26E-08 | 0.69620236 | 0.273 | 0.093 | 0.00024408 | TAM.contam. | Abhd5 |
| Apmap | 1.27E-08 | 0.41007587 | 0.354 | 0.177 | 0.00024543 | TAM.contam. | Apmap |
| Pwp1 | 1.29E-08 | 0.40993407 | 0.293 | 0.106 | 0.00024973 | TAM.contam. | Pwp1 |
| Tsfm | 1.53E-08 | 0.60215039 | 0.303 | 0.134 | 0.00029599 | TAM.contam. | Tsfm |
| Gpatch2 | 1.76E-08 | 0.41923941 | 0.283 | 0.131 | 0.00034093 | TAM.contam. | Gpatch2 |
| Hdac7 | 2.27E-08 | 0.36807836 | 0.283 | 0.1 | 0.00043907 | TAM.contam. | Hdac7 |
| Osbpl1a1 | 2.46E-08 | 0.27872386 | 0.303 | 0.131 | 0.00047483 | TAM.contam. | Osbpl1a |
| Ap5m1 | 3.03E-08 | 0.3089392 | 0.283 | 0.131 | 0.00058508 | TAM.contam. | Ap5m1 |
| Ppp2r2d | 3.10E-08 | 0.28598655 | 0.313 | 0.134 | 0.00059907 | TAM.contam. | Ppp2r2d |
| Pelp1 | 3.15E-08 | 0.29216376 | 0.374 | 0.156 | 0.00060942 | TAM.contam. | Pelp1 |
| Med22 | 3.71E-08 | 0.35903102 | 0.323 | 0.141 | 0.00071784 | TAM.contam. | Med22 |
| Tusc2 | 5.19E-08 | 0.59716455 | 0.263 | 0.101 | 0.00100435 | TAM.contam. | Tusc2 |
| Acp6 | 5.68E-08 | 0.76244231 | 0.253 | 0.098 | 0.00109826 | TAM.contam. | Acp6 |
| Ppp1r13b | 5.82E-08 | 0.59203844 | 0.313 | 0.123 | 0.00112619 | TAM.contam. | Ppp1r13b |
| Cd1801 | 6.63E-08 | 0.32951547 | 0.253 | 0.107 | 0.00128143 | TAM.contam. | Cd180 |
| Kdm5b | 7.24E-08 | 0.36795458 | 0.253 | 0.09 | 0.00139928 | TAM.contam. | Kdm5b |
| Fam149b | 7.84E-08 | 0.27180421 | 0.253 | 0.103 | 0.00151609 | TAM.contam. | Fam149b |
| Cdkn2aipnl2 | 8.00E-08 | 0.25187403 | 0.354 | 0.174 | 0.00154744 | TAM.contam. | Cdkn2aipnl |
| Shprh | 8.42E-08 | 0.32811061 | 0.283 | 0.117 | 0.00162702 | TAM.contam. | Shprh |
| Eef1aknmt | 9.44E-08 | 0.71307186 | 0.263 | 0.091 | 0.00182523 | TAM.contam. | Eef1aknmt |
| Acap3 | 9.60E-08 | 0.28394767 | 0.253 | 0.101 | 0.00185541 | TAM.contam. | Acap3 |
| Dpy19l4 | 9.75E-08 | 0.31103583 | 0.253 | 0.087 | 0.00188464 | TAM.contam. | Dpy19l4 |
| Flywch1 | 1.15E-07 | 0.3505047 | 0.283 | 0.108 | 0.00221524 | TAM.contam. | Flywch1 |
| Cysltr12 | 1.33E-07 | 0.33917076 | 0.434 | 0.208 | 0.00257253 | TAM.contam. | Cysltr1 |
| Cog4 | 1.37E-07 | 0.36491084 | 0.263 | 0.087 | 0.00264922 | TAM.contam. | Cog4 |
| Zfp646 | 1.93E-07 | 0.43386964 | 0.253 | 0.084 | 0.00373448 | TAM.contam. | Zfp646 |
| Nsmf | 2.02E-07 | 0.27151767 | 0.253 | 0.087 | 0.00390387 | TAM.contam. | Nsmf |
| Mavs | 3.34E-07 | 0.34433855 | 0.303 | 0.119 | 0.00644818 | TAM.contam. | Mavs |
| C1galt1c1 | 3.62E-07 | 0.25699704 | 0.384 | 0.164 | 0.00700481 | TAM.contam. | C1galt1c1 |
| Fkbp4 | 3.70E-07 | 0.32314707 | 0.333 | 0.149 | 0.00715315 | TAM.contam. | Fkbp4 |
| Deaf1 | 4.64E-07 | 0.35088176 | 0.263 | 0.104 | 0.00896208 | TAM.contam. | Deaf1 |
| Zfp24 | 6.61E-07 | 0.31425053 | 0.253 | 0.111 | 0.01277301 | TAM.contam. | Zfp24 |
| Fyco1 | 6.62E-07 | 0.26701545 | 0.263 | 0.125 | 0.01280196 | TAM.contam. | Fyco1 |
| Abcb1b | 1.62E-06 | 0.37014717 | 0.253 | 0.098 | 0.03131481 | TAM.contam. | Abcb1b |
| Klf16 | 1.93E-06 | 0.29001887 | 0.343 | 0.167 | 0.0372731 | TAM.contam. | Klf16 |
| Mtln1 | 2.00E-06 | 0.29412522 | 0.333 | 0.169 | 0.03873022 | TAM.contam. | Mtln |
| Gtf3c4 | 2.48E-06 | 0.36868126 | 0.263 | 0.118 | 0.04796232 | TAM.contam. | Gtf3c4 |
| Zbtb17 | 3.55E-06 | 0.31348961 | 0.313 | 0.14 | 0.06865737 | TAM.contam. | Zbtb17 |
| Adck1 | 6.39E-06 | 0.25973986 | 0.263 | 0.118 | 0.12358029 | TAM.contam. | Adck1 |
| Grhpr | 1.37E-05 | 0.39854735 | 0.273 | 0.118 | 0.26498174 | TAM.contam. | Grhpr |
| Heatr3 | 1.55E-05 | 0.27908123 | 0.263 | 0.121 | 0.29940313 | TAM.contam. | Heatr3 |
| Skiv2l | 2.53E-05 | 0.30182453 | 0.293 | 0.142 | 0.49002508 | TAM.contam. | Skiv2l |
| Elk1 | 2.85E-05 | 0.35347929 | 0.253 | 0.112 | 0.55031664 | TAM.contam. | Elk1 |

| | | | | | | | | |
|---|---|---|---|---|---|---|---|---|
| Sft2d2 | 2.29E-36 | 0.51164379 | 0.259 | 0.196 | 4.43E-32 | 1 | Sft2d2 | neutrophil.2 |
| Pitpna | 1.58E-35 | 0.34823804 | 0.359 | 0.282 | 3.05E-31 | 1 | Pitpna | neutrophil.2 |
| Rnasek | 3.49E-35 | 0.47658531 | 0.254 | 0.194 | 6.75E-31 | 1 | Rnasek | neutrophil.2 |
| Csf2rb | 4.18E-35 | 0.26189384 | 0.639 | 0.537 | 8.08E-31 | 1 | Csf2rb | neutrophil.2 |
| Brd4 | 4.74E-35 | 0.39933484 | 0.336 | 0.269 | 9.16E-31 | 1 | Brd4 | neutrophil.2 |
| Clec2d | 1.40E-34 | 0.35547708 | 0.408 | 0.335 | 2.72E-30 | 1 | Clec2d | neutrophil.2 |
| Ankrd13a | 3.56E-34 | 0.344683 | 0.294 | 0.23 | 6.89E-30 | 1 | Ankrd13a | neutrophil.2 |
| Pstpip1 | 4.73E-34 | 0.39886687 | 0.312 | 0.245 | 9.14E-30 | 1 | Pstpip1 | neutrophil.2 |
| Atp6v0b | 2.27E-33 | 0.35324149 | 0.354 | 0.284 | 4.39E-29 | 1 | Atp6v0b | neutrophil.2 |
| Aff1 | 6.36E-33 | 0.33361174 | 0.546 | 0.45 | 1.23E-28 | 1 | Aff1 | neutrophil.2 |
| Rab8a | 8.14E-33 | 0.36504496 | 0.359 | 0.287 | 1.57E-28 | 1 | Rab8a | neutrophil.2 |
| Pnpla2 | 8.82E-33 | 0.38847841 | 0.278 | 0.206 | 1.70E-28 | 1 | Pnpla2 | neutrophil.2 |
| Cd52 | 1.07E-32 | 0.35234554 | 0.655 | 0.553 | 2.07E-28 | 1 | Cd52 | neutrophil.2 |
| N4bp1 | 3.53E-32 | 0.33958419 | 0.501 | 0.403 | 6.83E-28 | 1 | N4bp1 | neutrophil.2 |
| Fuca2 | 4.83E-32 | 0.4879683 | 0.277 | 0.208 | 9.33E-28 | 1 | Fuca2 | neutrophil.2 |
| Cab39 | 1.27E-31 | 0.44740034 | 0.277 | 0.214 | 2.46E-27 | 1 | Cab39 | neutrophil.2 |
| Eif4g2 | 1.36E-31 | 0.31497126 | 0.369 | 0.29 | 2.62E-27 | 1 | Eif4g2 | neutrophil.2 |
| Lcp2 | 1.48E-31 | 0.38562698 | 0.496 | 0.403 | 2.87E-27 | 1 | Lcp2 | neutrophil.2 |
| Cytip | 8.92E-31 | 0.37655613 | 0.427 | 0.345 | 1.72E-26 | 1 | Cytip | neutrophil.2 |
| Ptpn1 | 8.94E-31 | 0.32242118 | 0.496 | 0.403 | 1.73E-26 | 1 | Ptpn1 | neutrophil.2 |
| Ezr | 7.52E-30 | 0.28540862 | 0.492 | 0.364 | 1.45E-25 | 1 | Ezr | neutrophil.2 |
| Adipor1 | 1.08E-29 | 0.28887238 | 0.663 | 0.57 | 2.09E-25 | 1 | Adipor1 | neutrophil.2 |
| Lnpep | 1.76E-29 | 0.36286581 | 0.366 | 0.294 | 3.41E-25 | 1 | Lnpep | neutrophil.2 |
| Calm1 | 2.13E-29 | 0.32055187 | 0.413 | 0.337 | 4.12E-25 | 1 | Calm1 | neutrophil.2 |
| Gpsm3 | 8.59E-29 | 0.29825464 | 0.417 | 0.352 | 1.66E-24 | 1 | Gpsm3 | neutrophil.2 |
| Efhd2 | 1.82E-28 | 0.29967408 | 0.548 | 0.455 | 3.52E-24 | 1 | Efhd2 | neutrophil.2 |
| Nadk | 6.51E-28 | 0.27666795 | 0.462 | 0.387 | 1.26E-23 | 1 | Nadk | neutrophil.2 |
| Tm9sf3 | 1.13E-27 | 0.38434657 | 0.263 | 0.2 | 2.19E-23 | 1 | Tm9sf3 | neutrophil.2 |
| Itgam | 2.27E-27 | 0.2947114 | 0.678 | 0.581 | 4.39E-23 | 1 | Itgam | neutrophil.2 |
| Prrc2c | 2.43E-27 | 0.40160153 | 0.257 | 0.193 | 4.69E-23 | 1 | Prrc2c | neutrophil.2 |
| Atxn2l | 2.78E-26 | 0.40236041 | 0.251 | 0.196 | 5.38E-22 | 1 | Atxn2l | neutrophil.2 |
| Ifngr1 | 2.19E-25 | 0.36610888 | 0.276 | 0.214 | 4.23E-21 | 1 | Ifngr1 | neutrophil.2 |
| Selplg | 4.66E-25 | 0.41350587 | 0.465 | 0.387 | 9.00E-21 | 1 | Selplg | neutrophil.2 |
| Pxn | 6.18E-25 | 0.41567084 | 0.291 | 0.228 | 1.19E-20 | 1 | Pxn | neutrophil.2 |
| Gcnt2 | 6.62E-25 | 0.41191849 | 0.343 | 0.277 | 1.28E-20 | 1 | Gcnt2 | neutrophil.2 |
| Arhgdia | 1.08E-24 | 0.2973652 | 0.329 | 0.263 | 2.08E-20 | 1 | Arhgdia | neutrophil.2 |
| Wipf1 | 2.00E-24 | 0.38586196 | 0.402 | 0.327 | 3.86E-20 | 1 | Wipf1 | neutrophil.2 |
| Fam32a | 2.85E-23 | 0.3835227 | 0.288 | 0.228 | 5.51E-19 | 1 | Fam32a | neutrophil.2 |
| Atf7ip | 3.72E-23 | 0.41111122 | 0.261 | 0.204 | 7.19E-19 | 1 | Atf7ip | neutrophil.2 |
| Pnp | 1.33E-22 | 0.3499025 | 0.286 | 0.219 | 2.57E-18 | 1 | Pnp | neutrophil.2 |
| Abi1 | 1.58E-22 | 0.31592339 | 0.38 | 0.31 | 3.06E-18 | 1 | Abi1 | neutrophil.2 |
| Mdm2 | 2.26E-22 | 0.30087841 | 0.267 | 0.211 | 4.36E-18 | 1 | Mdm2 | neutrophil.2 |
| Myh9 | 2.69E-22 | 0.25190653 | 0.314 | 0.255 | 5.20E-18 | 1 | Myh9 | neutrophil.2 |
| Unc93b1 | 5.01E-22 | 0.48619474 | 0.26 | 0.206 | 9.69E-18 | 1 | Unc93b1 | neutrophil.2 |
| Hp | 7.44E-22 | 0.33217019 | 0.525 | 0.464 | 1.44E-17 | 1 | Hp | neutrophil.2 |
| Pum2 | 2.15E-21 | 0.34554142 | 0.337 | 0.274 | 4.16E-17 | 1 | Pum2 | neutrophil.2 |
| Tnfaip2 | 5.92E-21 | 0.25949127 | 0.845 | 0.722 | 1.14E-16 | 1 | Tnfaip2 | neutrophil.2 |
| Cd37 | 1.32E-20 | 0.32677569 | 0.251 | 0.204 | 2.55E-16 | 1 | Cd37 | neutrophil.2 |
| Ip6k1 | 1.75E-20 | 0.33790657 | 0.322 | 0.272 | 3.38E-16 | 1 | Ip6k1 | neutrophil.2 |
| M6pr | 6.83E-20 | 0.35652197 | 0.27 | 0.219 | 1.32E-15 | 1 | M6pr | neutrophil.2 |
| Inpp5d | 8.79E-20 | 0.31082164 | 0.402 | 0.337 | 1.70E-15 | 1 | Inpp5d | neutrophil.2 |
| Rab1a | 2.09E-18 | 0.26764648 | 0.357 | 0.295 | 4.04E-14 | 1 | Rab1a | neutrophil.2 |
| Ankrd11 | 3.98E-18 | 0.3185832 | 0.265 | 0.214 | 7.70E-14 | 1 | Ankrd11 | neutrophil.2 |
| Cap1 | 6.88E-18 | 0.26588798 | 0.674 | 0.586 | 1.33E-13 | 1 | Cap1 | neutrophil.2 |
| Dnm2 | 1.05E-17 | 0.33044464 | 0.278 | 0.227 | 2.02E-13 | 1 | Dnm2 | neutrophil.2 |
| Bcl3 | 1.46E-17 | 0.25854754 | 0.428 | 0.372 | 2.83E-13 | 1 | Bcl3 | neutrophil.2 |
| Cd33 | 1.57E-17 | 0.29134789 | 0.61 | 0.518 | 3.03E-13 | 1 | Cd33 | neutrophil.2 |
| Lyn | 1.66E-17 | 0.28659408 | 0.469 | 0.409 | 3.21E-13 | 1 | Lyn | neutrophil.2 |
| Bst1 | 1.95E-17 | 0.35128163 | 0.282 | 0.232 | 3.76E-13 | 1 | Bst1 | neutrophil.2 |
| Rbm47 | 3.12E-17 | 0.38086278 | 0.266 | 0.217 | 6.04E-13 | 1 | Rbm47 | neutrophil.2 |
| Capzb | 5.86E-16 | 0.33328554 | 0.423 | 0.362 | 1.13E-11 | 1 | Capzb | neutrophil.2 |
| Nuak2 | 1.19E-15 | 0.34547496 | 0.335 | 0.27 | 2.30E-11 | 1 | Nuak2 | neutrophil.2 |
| Tarm1 | 2.18E-15 | 0.41814978 | 0.267 | 0.246 | 4.21E-11 | 1 | Tarm1 | neutrophil.2 |
| Nckap1l | 1.17E-14 | 0.25806515 | 0.361 | 0.311 | 2.26E-10 | 1 | Nckap1l | neutrophil.2 |
| Mef2d | 2.35E-14 | 0.29398339 | 0.253 | 0.208 | 4.54E-10 | 1 | Mef2d | neutrophil.2 |
| Lbr | 3.44E-14 | 0.39373783 | 0.251 | 0.203 | 6.66E-10 | 1 | Lbr | neutrophil.2 |
| Arpc2 | 4.71E-14 | 0.34231909 | 0.279 | 0.236 | 9.10E-10 | 1 | Arpc2 | neutrophil.2 |
| Atp6v0e | 2.52E-13 | 0.30024932 | 0.254 | 0.212 | 4.88E-09 | 1 | Atp6v0e | neutrophil.2 |
| Rab32 | 5.07E-13 | 0.33459539 | 0.279 | 0.233 | 9.80E-09 | 1 | Rab32 | neutrophil.2 |
| Cox17 | 7.57E-13 | 0.26224243 | 0.428 | 0.369 | 1.46E-08 | 1 | Cox17 | neutrophil.2 |
| Dazap2 | 8.81E-13 | 0.37668605 | 0.278 | 0.236 | 1.70E-08 | 1 | Dazap2 | neutrophil.2 |
| Rara | 9.26E-13 | 0.33385125 | 0.368 | 0.313 | 1.79E-08 | 1 | Rara | neutrophil.2 |
| Uba1 | 1.15E-11 | 0.27543387 | 0.283 | 0.229 | 2.23E-07 | 1 | Uba1 | neutrophil.2 |
| Ppp2ca | 1.12E-10 | 0.29370847 | 0.265 | 0.224 | 2.16E-06 | 1 | Ppp2ca | neutrophil.2 |
| Gabarap | 1.38E-10 | 0.29484467 | 0.445 | 0.387 | 2.68E-06 | 1 | Gabarap | neutrophil.2 |
| Hbegf | 1.59E-10 | 0.29528322 | 0.252 | 0.186 | 3.08E-06 | 1 | Hbegf | neutrophil.2 |
| Sema4a | 1.73E-09 | 0.2547872 | 0.438 | 0.376 | 3.35E-05 | 1 | Sema4a | neutrophil.2 |
| Atp1b3 | 2.96E-09 | 0.28534162 | 0.254 | 0.251 | 5.72E-05 | 1 | Atp1b3 | neutrophil.2 |
| Atp6ap2 | 3.81E-08 | 0.28981735 | 0.352 | 0.295 | 7.37E-04 | 1 | Atp6ap2 | neutrophil.2 |
| Arrdc4 | 4.79E-08 | 0.2507135 | 0.292 | 0.256 | 9.26E-04 | 1 | Arrdc4 | neutrophil.2 |
| Myl12b | 8.37E-08 | 0.27456243 | 0.251 | 0.215 | 0.00161871 | 1 | Myl12b | neutrophil.2 |
| Zfp36l1 | 1.49E-06 | 0.27749775 | 0.34 | 0.298 | 0.02871276 | 1 | Zfp36l1 | neutrophil.2 |
| Pten | 2.92E-05 | 0.26898123 | 0.269 | 0.235 | 0.56483913 | 1 | Pten | neutrophil.2 |
| Il1f9 | 0 | 2.29532097 | 0.886 | 0.335 | 0 | 2 | Il1f9 | neutrophil.3 |
| Slpi | 0 | 2.51359552 | 0.879 | 0.4 | 0 | 2 | Slpi | neutrophil.3 |
| Sqstm1 | 0 | 1.92128812 | 0.857 | 0.417 | 0 | 2 | Sqstm1 | neutrophil.3 |
| C3 | 0 | 1.45224759 | 0.86 | 0.42 | 0 | 2 | C3 | neutrophil.3 |
| Mpeg1 | 0 | 1.89454529 | 0.767 | 0.332 | 0 | 2 | Mpeg1 | neutrophil.3 |
| Sema4d | 0 | 1.79525104 | 0.917 | 0.503 | 0 | 2 | Sema4d | neutrophil.3 |
| Marcks | 0 | 1.37427204 | 0.907 | 0.514 | 0 | 2 | Marcks | neutrophil.3 |
| Raf1 | 0 | 1.79240345 | 0.599 | 0.209 | 0 | 2 | Raf1 | neutrophil.3 |
| Rilpl2 | 0 | 2.02478074 | 0.53 | 0.142 | 0 | 2 | Rilpl2 | neutrophil.3 |
| Ehd1 | 0 | 1.26530647 | 0.787 | 0.408 | 0 | 2 | Ehd1 | neutrophil.3 |
| Ctsz | 0 | 1.43870076 | 0.76 | 0.384 | 0 | 2 | Ctsz | neutrophil.3 |
| Ctsb | 0 | 1.73037436 | 0.881 | 0.508 | 0 | 2 | Ctsb | neutrophil.3 |
| Hmox1 | 0 | 2.05786953 | 0.561 | 0.197 | 0 | 2 | Hmox1 | neutrophil.3 |
| Ninj1 | 0 | 1.16956592 | 0.857 | 0.493 | 0 | 2 | Ninj1 | neutrophil.3 |
| Ctsd | 0 | 1.50797199 | 0.65 | 0.294 | 0 | 2 | Ctsd | neutrophil.3 |
| Tnip1 | 0 | 1.63606851 | 0.541 | 0.185 | 0 | 2 | Tnip1 | neutrophil.3 |
| Ptafr | 0 | 1.27026783 | 0.891 | 0.538 | 0 | 2 | Ptafr | neutrophil.3 |
| Chil1 | 0 | 2.45488658 | 0.47 | 0.125 | 0 | 2 | Chil1 | neutrophil.3 |
| Cstb | 0 | 1.52507423 | 0.738 | 0.394 | 0 | 2 | Cstb | neutrophil.3 |
| Sod2 | 0 | 1.50217609 | 0.6 | 0.258 | 0 | 2 | Sod2 | neutrophil.3 |
| Pgd | 0 | 1.69135526 | 0.627 | 0.289 | 0 | 2 | Pgd | neutrophil.3 |
| Nampt | 0 | 1.96061557 | 0.529 | 0.191 | 0 | 2 | Nampt | neutrophil.3 |
| Taldo1 | 0 | 1.42789902 | 0.707 | 0.369 | 0 | 2 | Taldo1 | neutrophil.3 |
| Ubap1 | 0 | 2.10650148 | 0.467 | 0.13 | 0 | 2 | Ubap1 | neutrophil.3 |
| Ikbke | 0 | 1.31163935 | 0.56 | 0.224 | 0 | 2 | Ikbke | neutrophil.3 |
| Zeb2 | 0 | 1.40991405 | 0.556 | 0.221 | 0 | 2 | Zeb2 | neutrophil.3 |
| Abcc1 | 0 | 2.81238451 | 0.395 | 0.06 | 0 | 2 | Abcc1 | neutrophil.3 |
| Treml4 | 0 | 1.39554011 | 0.594 | 0.259 | 0 | 2 | Treml4 | neutrophil.3 |
| Creg1 | 0 | 2.26298283 | 0.493 | 0.159 | 0 | 2 | Creg1 | neutrophil.3 |
| Cd274 | 0 | 1.12469259 | 0.727 | 0.398 | 0 | 2 | Cd274 | neutrophil.3 |
| Ezr | 0 | 1.45484281 | 0.656 | 0.331 | 0 | 2 | Ezr | neutrophil.3 |
| Zdhhc18 | 0 | 1.61395896 | 0.574 | 0.254 | 0 | 2 | Zdhhc18 | neutrophil.3 |
| Slc7a11 | 0 | 1.11579431 | 0.925 | 0.609 | 0 | 2 | Slc7a11 | neutrophil.3 |
| Apbb1ip | 0 | 1.13679555 | 0.752 | 0.436 | 0 | 2 | Apbb1ip | neutrophil.3 |
| Acod1 | 0 | 1.05439842 | 0.89 | 0.576 | 0 | 2 | Acod1 | neutrophil.3 |
| Gclm | 0 | 2.98504391 | 0.373 | 0.061 | 0 | 2 | Gclm | neutrophil.3 |
| Klf2 | 0 | 1.1043211 | 0.786 | 0.477 | 0 | 2 | Klf2 | neutrophil.3 |
| Abr | 0 | 1.15801881 | 0.673 | 0.366 | 0 | 2 | Abr | neutrophil.3 |
| Lfng | 0 | 1.12474138 | 0.645 | 0.339 | 0 | 2 | Lfng | neutrophil.3 |
| Bcl2l11 | 0 | 1.02381877 | 0.685 | 0.38 | 0 | 2 | Bcl2l11 | neutrophil.3 |
| Bri3 | 0 | 1.42081185 | 0.519 | 0.214 | 0 | 2 | Bri3 | neutrophil.3 |
| Sema4a | 0 | 1.11128362 | 0.639 | 0.334 | 0 | 2 | Sema4a | neutrophil.3 |
| Card19 | 0 | 1.02022982 | 0.645 | 0.342 | 0 | 2 | Card19 | neutrophil.3 |
| Traf1 | 0 | 0.99836424 | 0.557 | 0.255 | 0 | 2 | Traf1 | neutrophil.3 |
| Klf6 | 0 | 0.99024252 | 0.723 | 0.421 | 0 | 2 | Klf6 | neutrophil.3 |
| Osgin1 | 0 | 1.58710662 | 0.497 | 0.196 | 0 | 2 | Osgin1 | neutrophil.3 |
| Lgals3 | 0 | 1.64334891 | 0.514 | 0.218 | 0 | 2 | Lgals3 | neutrophil.3 |
| Ptpn1 | 0 | 1.15798143 | 0.664 | 0.369 | 0 | 2 | Ptpn1 | neutrophil.3 |
| Lipa | 0 | 2.63946346 | 0.378 | 0.083 | 0 | 2 | Lipa | neutrophil.3 |
| Sgms2 | 0 | 1.62671654 | 0.456 | 0.162 | 0 | 2 | Sgms2 | neutrophil.3 |
| Rnf19b | 0 | 1.00275664 | 0.693 | 0.399 | 0 | 2 | Rnf19b | neutrophil.3 |
| Prdx5 | 0 | 1.49019036 | 0.842 | 0.551 | 0 | 2 | Prdx5 | neutrophil.3 |
| Sgpl1 | 0 | 1.50014415 | 0.488 | 0.199 | 0 | 2 | Sgpl1 | neutrophil.3 |
| Prr13 | 0 | 1.56910992 | 0.473 | 0.185 | 0 | 2 | Prr13 | neutrophil.3 |
| Ccl3 | 0 | 0.90014192 | 0.845 | 0.558 | 0 | 2 | Ccl3 | neutrophil.3 |
| Mllt6 | 0 | 1.2444671 | 0.484 | 0.2 | 0 | 2 | Mllt6 | neutrophil.3 |
| Slc11a1 | 0 | 0.79858604 | 0.818 | 0.534 | 0 | 2 | Slc11a1 | neutrophil.3 |
| Icam1 | 0 | 1.01485557 | 0.868 | 0.585 | 0 | 2 | Icam1 | neutrophil.3 |
| Adgre5 | 0 | 0.99665144 | 0.897 | 0.617 | 0 | 2 | Adgre5 | neutrophil.3 |
| Txn1 | 0 | 1.21041694 | 0.538 | 0.259 | 0 | 2 | Txn1 | neutrophil.3 |
| Nod1 | 0 | 1.70086635 | 0.407 | 0.128 | 0 | 2 | Nod1 | neutrophil.3 |
| Csf1 | 0 | 1.33854333 | 0.553 | 0.275 | 0 | 2 | Csf1 | neutrophil.3 |
| Fmnl1 | 0 | 0.94215581 | 0.683 | 0.406 | 0 | 2 | Fmnl1 | neutrophil.3 |
| Ltb | 0 | 1.32956066 | 0.502 | 0.228 | 0 | 2 | Ltb | neutrophil.3 |
| Nadk | 0 | 0.91008944 | 0.625 | 0.354 | 0 | 2 | Nadk | neutrophil.3 |
| Slc8b1 | 0 | 2.00566215 | 0.365 | 0.094 | 0 | 2 | Slc8b1 | neutrophil.3 |
| N4bp1 | 0 | 0.91856324 | 0.643 | 0.375 | 0 | 2 | N4bp1 | neutrophil.3 |
| Mreg | 0 | 1.96048384 | 0.358 | 0.092 | 0 | 2 | Mreg | neutrophil.3 |
| Adam17 | 0 | 1.26731816 | 0.468 | 0.203 | 0 | 2 | Adam17 | neutrophil.3 |
| Ptprj | 0 | 1.61511189 | 0.412 | 0.15 | 0 | 2 | Ptprj | neutrophil.3 |
| Pmaip1 | 0 | 0.89215087 | 0.539 | 0.277 | 0 | 2 | Pmaip1 | neutrophil.3 |
| Cd14 | 0 | 0.27481377 | 0.922 | 0.662 | 0 | 2 | Cd14 | neutrophil.3 |
| Csf1r | 0 | 1.22117175 | 0.48 | 0.223 | 0 | 2 | Csf1r | neutrophil.3 |
| Ccdc88b | 0 | 0.84906586 | 0.589 | 0.332 | 0 | 2 | Ccdc88b | neutrophil.3 |
| Fgd3 | 0 | 1.03256107 | 0.559 | 0.303 | 0 | 2 | Fgd3 | neutrophil.3 |
| Msrb1 | 0 | 0.80413933 | 0.854 | 0.603 | 0 | 2 | Msrb1 | neutrophil.3 |
| Mafg | 0 | 1.82383282 | 0.336 | 0.086 | 0 | 2 | Mafg | neutrophil.3 |
| Atp6v0c | 0 | 0.81197993 | 0.826 | 0.576 | 0 | 2 | Atp6v0c | neutrophil.3 |
| Ccl4 | 0 | 0.72142773 | 0.555 | 0.307 | 0 | 2 | Ccl4 | neutrophil.3 |
| S100a8 | 0 | 0.75413305 | 0.822 | 0.574 | 0 | 2 | S100a8 | neutrophil.3 |
| Fas | 0 | 1.2320299 | 0.41 | 0.162 | 0 | 2 | Fas | neutrophil.3 |
| Gsap | 0 | 1.04746034 | 0.468 | 0.221 | 0 | 2 | Gsap | neutrophil.3 |
| Ncoa3 | 0 | 1.48214826 | 0.405 | 0.16 | 0 | 2 | Ncoa3 | neutrophil.3 |
| C5ar1 | 0 | 0.70168433 | 0.844 | 0.602 | 0 | 2 | C5ar1 | neutrophil.3 |
| Slc2a6 | 0 | 1.58435745 | 0.385 | 0.144 | 0 | 2 | Slc2a6 | neutrophil.3 |
| Metrnl | 0 | 1.70297692 | 0.381 | 0.142 | 0 | 2 | Metrnl | neutrophil.3 |
| Clec4d | 0 | 0.65511817 | 0.941 | 0.702 | 0 | 2 | Clec4d | neutrophil.3 |
| Stx11 | 0 | 0.97218546 | 0.426 | 0.188 | 0 | 2 | Stx11 | neutrophil.3 |
| Rab20 | 0 | 0.62261642 | 0.723 | 0.486 | 0 | 2 | Rab20 | neutrophil.3 |
| Tlr2 | 0 | 0.9755826 | 0.466 | 0.231 | 0 | 2 | Tlr2 | neutrophil.3 |
| F10 | 0 | 1.45438393 | 0.388 | 0.153 | 0 | 2 | F10 | neutrophil.3 |
| Rhog | 0 | 0.76173289 | 0.716 | 0.481 | 0 | 2 | Rhog | neutrophil.3 |
| Spn | 0 | 2.778872 | 0.274 | 0.041 | 0 | 2 | Spn | neutrophil.3 |
| Resf1 | 0 | 0.92352211 | 0.584 | 0.351 | 0 | 2 | Resf1 | neutrophil.3 |
| Irak3 | 0 | 1.63832918 | 0.339 | 0.108 | 0 | 2 | Irak3 | neutrophil.3 |
| Kpna4 | 0 | 0.91721273 | 0.502 | 0.271 | 0 | 2 | Kpna4 | neutrophil.3 |
| Prkcd | 0 | 0.77417707 | 0.581 | 0.35 | 0 | 2 | Prkcd | neutrophil.3 |
| Arid5b | 0 | 1.6642926 | 0.337 | 0.107 | 0 | 2 | Arid5b | neutrophil.3 |
| Aldoa | 0 | 0.71791639 | 0.583 | 0.354 | 0 | 2 | Aldoa | neutrophil.3 |
| S100a6 | 0 | 0.61132594 | 0.691 | 0.462 | 0 | 2 | S100a6 | neutrophil.3 |
| Tnf | 0 | 0.90922837 | 0.778 | 0.55 | 0 | 2 | Tnf | neutrophil.3 |
| Cd9 | 0 | 1.41827022 | 0.979 | 0.752 | 0 | 2 | Cd9 | neutrophil.3 |
| Itgb2 | 0 | 0.90297581 | 0.821 | 0.596 | 0 | 2 | Itgb2 | neutrophil.3 |
| Pnrc1 | 0 | 0.74582388 | 0.865 | 0.642 | 0 | 2 | Pnrc1 | neutrophil.3 |
| Cflar | 0 | 1.11763101 | 0.401 | 0.18 | 0 | 2 | Cflar | neutrophil.3 |
| Mefv | 0 | 1.15374288 | 0.411 | 0.19 | 0 | 2 | Mefv | neutrophil.3 |
| Ywhag | 0 | 0.87575225 | 0.508 | 0.288 | 0 | 2 | Ywhag | neutrophil.3 |
| Clec4e | 0 | 0.68113009 | 0.797 | 0.579 | 0 | 2 | Clec4e | neutrophil.3 |
| Flna | 0 | 0.87010656 | 0.498 | 0.28 | 0 | 2 | Flna | neutrophil.3 |
| Igsf6 | 0 | 0.72692831 | 0.552 | 0.335 | 0 | 2 | Igsf6 | neutrophil.3 |
| Fpr2 | 0 | 0.96115763 | 0.522 | 0.305 | 0 | 2 | Fpr2 | neutrophil.3 |
| H2-D1 | 0 | 1.03433942 | 0.46 | 0.243 | 0 | 2 | H2-D1 | neutrophil.3 |
| Abi1 | 0 | 0.87152598 | 0.502 | 0.286 | 0 | 2 | Abi1 | neutrophil.3 |
| Stk10 | 0 | 1.84376868 | 0.306 | 0.09 | 0 | 2 | Stk10 | neutrophil.3 |
| Cd82 | 0 | 0.82659762 | 0.663 | 0.448 | 0 | 2 | Cd82 | neutrophil.3 |
| Diaph1 | 0 | 0.84803505 | 0.541 | 0.326 | 0 | 2 | Diaph1 | neutrophil.3 |
| Bach1 | 0 | 1.15684875 | 0.407 | 0.192 | 0 | 2 | Bach1 | neutrophil.3 |
| Sat1 | 0 | 0.66788282 | 0.682 | 0.468 | 0 | 2 | Sat1 | neutrophil.3 |
| Cyba | 0 | 0.71282708 | 0.612 | 0.398 | 0 | 2 | Cyba | neutrophil.3 |
| Psap | 0 | 1.34650326 | 0.515 | 0.302 | 0 | 2 | Psap | neutrophil.3 |
| Asprv1 | 0 | 1.46299114 | 0.42 | 0.207 | 0 | 2 | Asprv1 | neutrophil.3 |
| Efhd2 | 0 | 0.63301754 | 0.648 | 0.436 | 0 | 2 | Efhd2 | neutrophil.3 |
| Irak2 | 0 | 0.64430472 | 0.665 | 0.453 | 0 | 2 | Irak2 | neutrophil.3 |
| B2m | 0 | 0.62199478 | 0.753 | 0.542 | 0 | 2 | B2m | neutrophil.3 |
| Map7d1 | 0 | 0.79485882 | 0.468 | 0.258 | 0 | 2 | Map7d1 | neutrophil.3 |
| Iqsec1 | 0 | 1.14176114 | 0.454 | 0.244 | 0 | 2 | Iqsec1 | neutrophil.3 |
| Sh2b2 | 0 | 0.83871078 | 0.456 | 0.247 | 0 | 2 | Sh2b2 | neutrophil.3 |
| Emp3 | 0 | 1.80386248 | 0.315 | 0.106 | 0 | 2 | Emp3 | neutrophil.3 |
| Riok3 | 0 | 1.01713762 | 0.468 | 0.259 | 0 | 2 | Riok3 | neutrophil.3 |
| Hbp1 | 0 | 1.19911369 | 0.362 | 0.153 | 0 | 2 | Hbp1 | neutrophil.3 |
| Relb | 0 | 1.28764681 | 0.324 | 0.116 | 0 | 2 | Relb | neutrophil.3 |
| Srxn1 | 0 | 1.8204168 | 0.297 | 0.09 | 0 | 2 | Srxn1 | neutrophil.3 |
| Hck | 0 | 0.78988398 | 0.472 | 0.266 | 0 | 2 | Hck | neutrophil.3 |
| Lasp1 | 0 | 0.94581085 | 0.423 | 0.218 | 0 | 2 | Lasp1 | neutrophil.3 |
| Mapkapk2 | 0 | 0.59505964 | 0.855 | 0.65 | 0 | 2 | Mapkapk2 | neutrophil.3 |
| Cd300a | 0 | 0.45936362 | 0.65 | 0.446 | 0 | 2 | Cd300a | neutrophil.3 |
| Cd44 | 0 | 0.5993995 | 0.901 | 0.698 | 0 | 2 | Cd44 | neutrophil.3 |
| Rab11fip1 | 0 | 0.5893141 | 0.68 | 0.477 | 0 | 2 | Rab11fip1 | neutrophil.3 |
| Arhgef3 | 0 | 1.57808239 | 0.3 | 0.097 | 0 | 2 | Arhgef3 | neutrophil.3 |
| Il2rg | 0 | 1.46595023 | 0.307 | 0.105 | 0 | 2 | Il2rg | neutrophil.3 |
| Surf4 | 0 | 1.14710345 | 0.378 | 0.178 | 0 | 2 | Surf4 | neutrophil.3 |
| Agpat4 | 0 | 1.78282747 | 0.268 | 0.068 | 0 | 2 | Agpat4 | neutrophil.3 |
| Plgrkt | 0 | 1.34220536 | 0.325 | 0.125 | 0 | 2 | Plgrkt | neutrophil.3 |
| Zswim4 | 0 | 0.82416396 | 0.476 | 0.276 | 0 | 2 | Zswim4 | neutrophil.3 |
| Basp1 | 0 | 0.45274338 | 0.698 | 0.498 | 0 | 2 | Basp1 | neutrophil.3 |
| Rab32 | 0 | 1.00976483 | 0.406 | 0.207 | 0 | 2 | Rab32 | neutrophil.3 |
| Arf2 | 0 | 1.14667786 | 0.395 | 0.196 | 0 | 2 | Arf2 | neutrophil.3 |
| Nfkb2 | 0 | 0.36187526 | 0.659 | 0.46 | 0 | 2 | Nfkb2 | neutrophil.3 |
| Smox | 0 | 0.43488365 | 0.648 | 0.452 | 0 | 2 | Smox | neutrophil.3 |
| Slfn2 | 0 | 0.42657286 | 0.716 | 0.52 | 0 | 2 | Slfn2 | neutrophil.3 |
| Plekho2 | 0 | 0.4876017 | 0.727 | 0.532 | 0 | 2 | Plekho2 | neutrophil.3 |
| Mapk6 | 0 | 1.38506482 | 0.305 | 0.112 | 0 | 2 | Mapk6 | neutrophil.3 |
| Trem3 | 0 | 0.56205632 | 0.556 | 0.365 | 0 | 2 | Trem3 | neutrophil.3 |
| Cd53 | 0 | 0.82654185 | 0.85 | 0.66 | 0 | 2 | Cd53 | neutrophil.3 |
| Txnrd1 | 0 | 0.68049908 | 0.742 | 0.552 | 0 | 2 | Txnrd1 | neutrophil.3 |
| Anxa11 | 0 | 0.84732074 | 0.469 | 0.279 | 0 | 2 | Anxa11 | neutrophil.3 |
| Capg | 0 | 0.89089264 | 0.376 | 0.187 | 0 | 2 | Capg | neutrophil.3 |
| Tnfrsf1b | 0 | 0.85520618 | 0.739 | 0.55 | 0 | 2 | Tnfrsf1b | neutrophil.3 |
| Atf7ip | 0 | 1.11687687 | 0.37 | 0.182 | 0 | 2 | Atf7ip | neutrophil.3 |
| Marcksl1 | 0 | 0.94711081 | 0.964 | 0.776 | 0 | 2 | Marcksl1 | neutrophil.3 |
| St3gal1 | 0 | 0.83281948 | 0.425 | 0.238 | 0 | 2 | St3gal1 | neutrophil.3 |
| Sh3bp2 | 0 | 1.36947419 | 0.297 | 0.111 | 0 | 2 | Sh3bp2 | neutrophil.3 |
| Zfand3 | 0 | 0.72774517 | 0.465 | 0.28 | 0 | 2 | Zfand3 | neutrophil.3 |
| Nfe2l2 | 0 | 0.49384406 | 0.75 | 0.566 | 0 | 2 | Nfe2l2 | neutrophil.3 |
| Iqgap1 | 0 | 0.50560057 | 0.784 | 0.6 | 0 | 2 | Iqgap1 | neutrophil.3 |
| Znrf1 | 0 | 1.40692325 | 0.287 | 0.103 | 0 | 2 | Znrf1 | neutrophil.3 |
| Itgal | 0 | 0.6990007 | 0.58 | 0.396 | 0 | 2 | Itgal | neutrophil.3 |
| Slc9a3r1 | 0 | 0.53428545 | 0.546 | 0.363 | 0 | 2 | Slc9a3r1 | neutrophil.3 |
| Sh2d3c | 0 | 1.13133372 | 0.308 | 0.125 | 0 | 2 | Sh2d3c | neutrophil.3 |
| Mbnl2 | 0 | 1.08262601 | 0.385 | 0.203 | 0 | 2 | Mbnl2 | neutrophil.3 |
| Pla2g7 | 0 | 0.71812599 | 0.613 | 0.431 | 0 | 2 | Pla2g7 | neutrophil.3 |
| Hspa5 | 0 | 0.62917846 | 0.532 | 0.352 | 0 | 2 | Hspa5 | neutrophil.3 |
| Gna13 | 0 | 0.50964039 | 0.532 | 0.352 | 0 | 2 | Gna13 | neutrophil.3 |
| Cln5 | 0 | 1.32770729 | 0.28 | 0.1 | 0 | 2 | Cln5 | neutrophil.3 |
| Ncf4 | 0 | 0.54683374 | 0.518 | 0.338 | 0 | 2 | Ncf4 | neutrophil.3 |
| S100a9 | 0 | 0.26734403 | 0.756 | 0.577 | 0 | 2 | S100a9 | neutrophil.3 |
| Map4k4 | 0 | 1.10657043 | 0.331 | 0.152 | 0 | 2 | Map4k4 | neutrophil.3 |
| Rhbdf2 | 0 | 1.01771843 | 0.323 | 0.144 | 0 | 2 | Rhbdf2 | neutrophil.3 |
| Ptbp3 | 0 | 0.4930153 | 0.566 | 0.388 | 0 | 2 | Ptbp3 | neutrophil.3 |
| Ankrd44 | 0 | 1.33715734 | 0.288 | 0.111 | 0 | 2 | Ankrd44 | neutrophil.3 |
| Ralgds | 0 | 0.91487665 | 0.358 | 0.181 | 0 | 2 | Ralgds | neutrophil.3 |
| Alas1 | 0 | 0.66552104 | 0.418 | 0.242 | 0 | 2 | Alas1 | neutrophil.3 |
| Stat6 | 0 | 0.74462034 | 0.413 | 0.237 | 0 | 2 | Stat6 | neutrophil.3 |
| Oga | 0 | 0.81472686 | 0.407 | 0.231 | 0 | 2 | Oga | neutrophil.3 |
| Erbin | 0 | 0.84507662 | 0.405 | 0.23 | 0 | 2 | Erbin | neutrophil.3 |
| Tuba1c | 0 | 0.58942196 | 0.517 | 0.342 | 0 | 2 | Tuba1c | neutrophil.3 |
| Cdc42ep2 | 0 | 1.49872546 | 0.275 | 0.101 | 0 | 2 | Cdc42ep2 | neutrophil.3 |
| Dedd2 | 0 | 0.64251625 | 0.408 | 0.235 | 0 | 2 | Dedd2 | neutrophil.3 |
| Cxcr4 | 0 | 0.46233566 | 0.719 | 0.546 | 0 | 2 | Cxcr4 | neutrophil.3 |
| Fpr1 | 0 | 1.50145422 | 0.272 | 0.101 | 0 | 2 | Fpr1 | neutrophil.3 |
| Ifnar1 | 0 | 1.330113 | 0.272 | 0.103 | 0 | 2 | Ifnar1 | neutrophil.3 |
| Igf2r | 0 | 1.26740317 | 0.283 | 0.116 | 0 | 2 | Igf2r | neutrophil.3 |
| Siglece | 0 | 0.70608693 | 0.573 | 0.406 | 0 | 2 | Siglece | neutrophil.3 |
| Msn | 0 | 0.47923121 | 0.905 | 0.74 | 0 | 2 | Msn | neutrophil.3 |
| Spen | 0 | 0.9788735 | 0.313 | 0.149 | 0 | 2 | Spen | neutrophil.3 |
| Rhob | 0 | 0.44363621 | 0.496 | 0.332 | 0 | 2 | Rhob | neutrophil.3 |
| Atp6ap2 | 0 | 0.79971248 | 0.44 | 0.277 | 0 | 2 | Atp6ap2 | neutrophil.3 |
| Ptpn23 | 0 | 1.24149714 | 0.268 | 0.107 | 0 | 2 | Ptpn23 | neutrophil.3 |
| Map3k8 | 0 | 0.83375405 | 0.329 | 0.168 | 0 | 2 | Map3k8 | neutrophil.3 |
| Foxn2 | 0 | 0.98813722 | 0.298 | 0.138 | 0 | 2 | Foxn2 | neutrophil.3 |
| Snx10 | 0 | 1.38611999 | 0.252 | 0.093 | 0 | 2 | Snx10 | neutrophil.3 |
| Bst1 | 0 | 0.89233579 | 0.372 | 0.214 | 0 | 2 | Bst1 | neutrophil.3 |
| Tlr13 | 0 | 1.22832309 | 0.285 | 0.127 | 0 | 2 | Tlr13 | neutrophil.3 |
| Bhlhe40 | 0 | 0.31357567 | 0.687 | 0.531 | 0 | 2 | Bhlhe40 | neutrophil.3 |
| Xylt1 | 0 | 0.85042023 | 0.374 | 0.218 | 0 | 2 | Xylt1 | neutrophil.3 |
| Arl8a | 0 | 1.06001526 | 0.285 | 0.13 | 0 | 2 | Arl8a | neutrophil.3 |
| Rapgef1 | 0 | 0.98678712 | 0.313 | 0.16 | 0 | 2 | Rapgef1 | neutrophil.3 |
| Psmb5 | 0 | 1.06767833 | 0.269 | 0.12 | 0 | 2 | Psmb5 | neutrophil.3 |
| Plin2 | 0 | 0.54247692 | 0.475 | 0.326 | 0 | 2 | Plin2 | neutrophil.3 |
| Rap1b | 0 | 0.38491369 | 0.699 | 0.55 | 0 | 2 | Rap1b | neutrophil.3 |
| Cd68 | 0 | 0.95692742 | 0.36 | 0.212 | 0 | 2 | Cd68 | neutrophil.3 |
| Hivep1 | 0 | 1.08571919 | 0.274 | 0.127 | 0 | 2 | Hivep1 | neutrophil.3 |
| Cyfip2 | 0 | 1.24806477 | 0.254 | 0.108 | 0 | 2 | Cyfip2 | neutrophil.3 |
| Fyb | 0 | 0.76751153 | 0.31 | 0.164 | 0 | 2 | Fyb | neutrophil.3 |
| Tbc1d2b | 0 | 1.12727352 | 0.272 | 0.127 | 0 | 2 | Tbc1d2b | neutrophil.3 |
| Pnpla2 | 0 | 0.79097427 | 0.337 | 0.195 | 0 | 2 | Pnpla2 | neutrophil.3 |
| Ubc | 0 | 0.53130003 | 0.915 | 0.777 | 0 | 2 | Ubc | neutrophil.3 |
| Dennd3 | 0 | 0.94963147 | 0.277 | 0.142 | 0 | 2 | Dennd3 | neutrophil.3 |
| Il1rn | 0 | 0.5303443 | 0.954 | 0.819 | 0 | 2 | Il1rn | neutrophil.3 |
| Casp4 | 0 | 0.92732033 | 0.297 | 0.162 | 0 | 2 | Casp4 | neutrophil.3 |
| Clec4n | 0 | 0.43070472 | 0.69 | 0.56 | 0 | 2 | Clec4n | neutrophil.3 |
| Dhx40 | 0 | 0.40819173 | 0.446 | 0.329 | 0 | 2 | Dhx40 | neutrophil.3 |
| Anxa2 | 0 | 0.47736649 | 0.675 | 0.559 | 0 | 2 | Anxa2 | neutrophil.3 |
| Vasp | 0 | 0.27282231 | 0.915 | 0.802 | 0 | 2 | Vasp | neutrophil.3 |
| Chka | 0 | 0.79972823 | 0.251 | 0.145 | 0 | 2 | Chka | neutrophil.3 |
| Nfkbia | 0 | 0.28694698 | 0.957 | 0.867 | 0 | 2 | Nfkbia | neutrophil.3 |
| Fth1 | 0 | 0.92013851 | 0.994 | 0.924 | 0 | 2 | Fth1 | neutrophil.3 |
| Atp6v1a | 3.83E-307 | 0.82808002 | 0.33 | 0.199 | 7.41E-303 | 2 | Atp6v1a | neutrophil.3 |
| Asap1 | 4.41E-301 | 0.94872854 | 0.255 | 0.118 | 8.53E-297 | 2 | Asap1 | neutrophil.3 |
| Atp6v1b2 | 6.28E-296 | 0.71587168 | 0.409 | 0.258 | 1.22E-291 | 2 | Atp6v1b2 | neutrophil.3 |
| Fcgr2b | 3.84E-294 | 0.6092518 | 0.353 | 0.226 | 7.42E-290 | 2 | Fcgr2b | neutrophil.3 |
| Wsb1 | 6.31E-294 | 0.32496431 | 0.676 | 0.524 | 1.22E-289 | 2 | Wsb1 | neutrophil.3 |
| Smcr8 | 4.37E-293 | 0.83306339 | 0.326 | 0.172 | 8.46E-289 | 2 | Smcr8 | neutrophil.3 |
| BC005537 | 3.74E-289 | 0.33499234 | 0.592 | 0.425 | 7.22E-285 | 2 | BC005537 | neutrophil.3 |
| App | 1.32E-286 | 0.52007861 | 0.615 | 0.44 | 2.55E-282 | 2 | App | neutrophil.3 |
| Birc3 | 3.50E-284 | 0.3713131 | 0.589 | 0.413 | 6.77E-280 | 2 | Birc3 | neutrophil.3 |
| Ubtd1 | 4.52E-283 | 0.95118192 | 0.29 | 0.152 | 8.75E-279 | 2 | Ubtd1 | neutrophil.3 |
| Dock2 | 6.66E-281 | 0.68490177 | 0.397 | 0.242 | 1.29E-276 | 2 | Dock2 | neutrophil.3 |
| Slc3a2 | 8.69E-280 | 0.55950551 | 0.531 | 0.359 | 1.68E-275 | 2 | Slc3a2 | neutrophil.3 |
| Olr1 | 1.36E-275 | 0.47143465 | 0.324 | 0.179 | 2.62E-271 | 2 | Olr1 | neutrophil.3 |
| Ube2f | 5.12E-275 | 0.93328317 | 0.266 | 0.122 | 9.90E-271 | 2 | Ube2f | neutrophil.3 |
| Alcam | 7.75E-270 | 0.88184526 | 0.312 | 0.166 | 1.50E-265 | 2 | Alcam | neutrophil.3 |
| Atf3 | 9.33E-270 | 0.35185622 | 0.538 | 0.361 | 1.80E-265 | 2 | Atf3 | neutrophil.3 |
| Stk38 | 2.46E-269 | 0.79002959 | 0.303 | 0.164 | 4.76E-265 | 2 | Stk38 | neutrophil.3 |
| Ripor2 | 1.38E-268 | 0.66615442 | 0.328 | 0.194 | 2.66E-264 | 2 | Ripor2 | neutrophil.3 |
| Ostf1 | 3.90E-268 | 0.57404682 | 0.481 | 0.332 | 7.54E-264 | 2 | Ostf1 | neutrophil.3 |
| Pkm | 2.30E-266 | 0.41761682 | 0.64 | 0.475 | 4.45E-262 | 2 | Pkm | neutrophil.3 |
| Midn | 1.39E-263 | 0.42654587 | 0.546 | 0.398 | 2.68E-259 | 2 | Midn | neutrophil.3 |
| Atp1b3 | 9.21E-261 | 0.71249017 | 0.404 | 0.219 | 1.78E-256 | 2 | Atp1b3 | neutrophil.3 |
| M6pr | 2.86E-259 | 0.75190601 | 0.347 | 0.204 | 5.53E-255 | 2 | M6pr | neutrophil.3 |
| Atp2b1 | 1.53E-257 | 0.66467223 | 0.382 | 0.247 | 2.95E-253 | 2 | Atp2b1 | neutrophil.3 |
| Blcap | 2.92E-257 | 0.71648984 | 0.366 | 0.214 | 5.65E-253 | 2 | Blcap | neutrophil.3 |
| Plec | 6.04E-257 | 0.60618263 | 0.318 | 0.178 | 1.17E-252 | 2 | Plec | neutrophil.3 |
| Nedd9 | 8.66E-257 | 0.94916734 | 0.253 | 0.128 | 1.67E-252 | 2 | Nedd9 | neutrophil.3 |
| Bcl3 | 1.24E-256 | 0.43944014 | 0.501 | 0.358 | 2.41E-252 | 2 | Bcl3 | neutrophil.3 |
| Atp6v1e1 | 3.13E-255 | 0.51234448 | 0.449 | 0.312 | 6.06E-251 | 2 | Atp6v1e1 | neutrophil.3 |
| Pten | 7.06E-255 | 0.7615549 | 0.358 | 0.217 | 1.37E-250 | 2 | Pten | neutrophil.3 |
| Lrrc25 | 7.49E-249 | 0.7195909 | 0.37 | 0.228 | 1.45E-244 | 2 | Lrrc25 | neutrophil.3 |
| Tatdn2 | 1.86E-247 | 0.62365632 | 0.352 | 0.221 | 3.61E-243 | 2 | Tatdn2 | neutrophil.3 |
| Tbc1d10b | 1.30E-245 | 0.87627967 | 0.257 | 0.128 | 2.51E-241 | 2 | Tbc1d10b | neutrophil.3 |
| Mbp | 1.06E-240 | 0.76848812 | 0.29 | 0.156 | 2.06E-236 | 2 | Mbp | neutrophil.3 |
| Havcr2 | 4.31E-240 | 0.767469 | 0.266 | 0.143 | 8.33E-236 | 2 | Havcr2 | neutrophil.3 |
| Chd4 | 2.90E-235 | 0.37215829 | 0.411 | 0.266 | 5.60E-231 | 2 | Chd4 | neutrophil.3 |
| Ehbp1l1 | 1.12E-232 | 0.46128915 | 0.347 | 0.207 | 2.16E-228 | 2 | Ehbp1l1 | neutrophil.3 |
| Klf3 | 1.74E-229 | 0.68604654 | 0.339 | 0.198 | 3.36E-225 | 2 | Klf3 | neutrophil.3 |
| Ptpn6 | 1.17E-227 | 0.3922709 | 0.574 | 0.437 | 2.25E-223 | 2 | Ptpn6 | neutrophil.3 |
| Cd63 | 1.42E-227 | 0.35727131 | 0.494 | 0.36 | 2.75E-223 | 2 | Cd63 | neutrophil.3 |
| Rela | 6.21E-227 | 0.64736755 | 0.389 | 0.242 | 1.20E-222 | 2 | Rela | neutrophil.3 |
| Rhoa | 1.63E-224 | 0.57231523 | 0.436 | 0.281 | 3.16E-220 | 2 | Rhoa | neutrophil.3 |
| Notch2 | 5.65E-223 | 0.55047174 | 0.423 | 0.276 | 1.09E-218 | 2 | Notch2 | neutrophil.3 |
| Pts | 2.40E-221 | 0.82202087 | 0.257 | 0.151 | 4.65E-217 | 2 | Pts | neutrophil.3 |
| Lcp2 | 2.82E-221 | 0.40714267 | 0.526 | 0.398 | 5.46E-217 | 2 | Lcp2 | neutrophil.3 |
| Rffl | 6.38E-219 | 0.75379401 | 0.269 | 0.139 | 1.23E-214 | 2 | Rffl | neutrophil.3 |
| Myo1f | 8.84E-219 | 0.44253815 | 0.468 | 0.325 | 1.71E-214 | 2 | Myo1f | neutrophil.3 |
| Itgb1 | 1.74E-218 | 0.5867845 | 0.373 | 0.226 | 3.36E-214 | 2 | Itgb1 | neutrophil.3 |
| Fbrs | 1.48E-215 | 0.2665807 | 0.519 | 0.371 | 2.87E-211 | 2 | Fbrs | neutrophil.3 |
| Nfkbib | 1.13E-204 | 0.7136902 | 0.26 | 0.146 | 2.18E-200 | 2 | Nfkbib | neutrophil.3 |
| Tax1bp1 | 6.19E-202 | 0.52934061 | 0.436 | 0.29 | 1.20E-197 | 2 | Tax1bp1 | neutrophil.3 |
| Lcor | 1.45E-199 | 0.72929187 | 0.257 | 0.149 | 2.80E-195 | 2 | Lcor | neutrophil.3 |
| Ubald2 | 2.38E-199 | 0.62648147 | 0.367 | 0.234 | 4.60E-195 | 2 | Ubald2 | neutrophil.3 |
| Fuca2 | 3.15E-199 | 0.75712543 | 0.321 | 0.2 | 6.09E-195 | 2 | Fuca2 | neutrophil.3 |
| Itga5 | 3.73E-198 | 0.70515467 | 0.256 | 0.139 | 7.21E-194 | 2 | Itga5 | neutrophil.3 |
| Trps1 | 3.52E-191 | 0.76367463 | 0.264 | 0.147 | 6.81E-187 | 2 | Trps1 | neutrophil.3 |
| Il10rb | 4.14E-191 | 0.36323766 | 0.535 | 0.397 | 8.01E-187 | 2 | Il10rb | neutrophil.3 |
| Gabarap | 1.20E-189 | 0.49759762 | 0.52 | 0.372 | 2.32E-185 | 2 | Gabarap | neutrophil.3 |
| Gpr132 | 3.58E-186 | 0.38842073 | 0.431 | 0.297 | 6.92E-182 | 2 | Gpr132 | neutrophil.3 |
| Dgkz | 4.96E-186 | 0.56647203 | 0.311 | 0.192 | 9.59E-182 | 2 | Dgkz | neutrophil.3 |
| Fndc3b | 3.19E-182 | 0.61132185 | 0.355 | 0.218 | 6.16E-178 | 2 | Fndc3b | neutrophil.3 |
| Sell | 2.15E-181 | 0.43080618 | 0.427 | 0.31 | 4.15E-177 | 2 | Sell | neutrophil.3 |
| Wwp2 | 2.74E-181 | 0.73161843 | 0.269 | 0.149 | 5.30E-177 | 2 | Wwp2 | neutrophil.3 |
| Pdlim7 | 1.45E-179 | 0.32014351 | 0.44 | 0.304 | 2.81E-175 | 2 | Pdlim7 | neutrophil.3 |
| Psd4 | 2.83E-179 | 0.42020632 | 0.368 | 0.235 | 5.46E-175 | 2 | Psd4 | neutrophil.3 |
| Cnppd1 | 1.22E-177 | 0.50655064 | 0.364 | 0.236 | 2.36E-173 | 2 | Cnppd1 | neutrophil.3 |
| Atp1a1 | 3.76E-176 | 0.37098883 | 0.442 | 0.326 | 7.26E-172 | 2 | Atp1a1 | neutrophil.3 |
| Kras | 7.76E-172 | 0.52522737 | 0.349 | 0.218 | 1.50E-167 | 2 | Kras | neutrophil.3 |
| Pglyrp1 | 1.39E-170 | 0.53421618 | 0.316 | 0.214 | 2.69E-166 | 2 | Pglyrp1 | neutrophil.3 |
| Gnai3 | 7.05E-168 | 0.49270857 | 0.402 | 0.267 | 1.36E-163 | 2 | Gnai3 | neutrophil.3 |
| Bcl10 | 5.09E-166 | 0.72550685 | 0.252 | 0.138 | 9.84E-162 | 2 | Bcl10 | neutrophil.3 |
| Rab8a | 1.05E-165 | 0.4515483 | 0.402 | 0.279 | 2.03E-161 | 2 | Rab8a | neutrophil.3 |
| Retreg1 | 1.65E-165 | 0.65246754 | 0.351 | 0.226 | 3.20E-161 | 2 | Retreg1 | neutrophil.3 |
| Mtmr6 | 3.53E-163 | 0.51435309 | 0.307 | 0.193 | 6.82E-159 | 2 | Mtmr6 | neutrophil.3 |
| Pitpna | 1.30E-162 | 0.44525839 | 0.399 | 0.275 | 2.51E-158 | 2 | Pitpna | neutrophil.3 |
| Hectd1 | 3.57E-161 | 0.66587563 | 0.262 | 0.143 | 6.89E-157 | 2 | Hectd1 | neutrophil.3 |
| Tmem164 | 1.13E-156 | 0.48366768 | 0.35 | 0.236 | 2.18E-152 | 2 | Tmem164 | neutrophil.3 |
| Ccnl2 | 8.53E-156 | 0.40102792 | 0.374 | 0.251 | 1.65E-151 | 2 | Ccnl2 | neutrophil.3 |
| Map1lc3b | 1.01E-149 | 0.39494703 | 0.473 | 0.339 | 1.96E-145 | 2 | Map1lc3b | neutrophil.3 |
| Tmbim6 | 1.36E-149 | 0.40704309 | 0.549 | 0.407 | 2.62E-145 | 2 | Tmbim6 | neutrophil.3 |
| Susd6 | 6.33E-148 | 0.29203484 | 0.513 | 0.386 | 1.22E-143 | 2 | Susd6 | neutrophil.3 |
| Brd2 | 1.92E-147 | 0.3536489 | 0.41 | 0.276 | 3.72E-143 | 2 | Brd2 | neutrophil.3 |
| Atp6v0b | 3.61E-146 | 0.4537622 | 0.399 | 0.276 | 6.98E-142 | 2 | Atp6v0b | neutrophil.3 |
| Cass4 | 1.02E-145 | 0.28375507 | 0.503 | 0.377 | 1.98E-141 | 2 | Cass4 | neutrophil.3 |
| Fbxl5 | 2.01E-143 | 0.63336457 | 0.278 | 0.178 | 3.89E-139 | 2 | Fbxl5 | neutrophil.3 |
| Fus | 5.09E-141 | 0.25451892 | 0.527 | 0.403 | 9.83E-137 | 2 | Fus | neutrophil.3 |
| Rbpj | 2.63E-139 | 0.47440811 | 0.335 | 0.224 | 5.09E-135 | 2 | Rbpj | neutrophil.3 |
| Lrp10 | 7.72E-135 | 0.37181428 | 0.43 | 0.32 | 1.49E-130 | 2 | Lrp10 | neutrophil.3 |
| Clta | 3.80E-133 | 0.28600604 | 0.461 | 0.336 | 7.34E-129 | 2 | Clta | neutrophil.3 |
| Atox1 | 8.60E-132 | 0.35899015 | 0.409 | 0.303 | 1.66E-127 | 2 | Atox1 | neutrophil.3 |
| Irf5 | 1.73E-131 | 0.25495804 | 0.331 | 0.229 | 3.35E-127 | 2 | Irf5 | neutrophil.3 |
| Clec2d | 1.73E-131 | 0.29068243 | 0.453 | 0.327 | 3.35E-127 | 2 | Clec2d | neutrophil.3 |
| H3f3a | 3.54E-131 | 0.30011373 | 0.522 | 0.401 | 6.85E-127 | 2 | H3f3a | neutrophil.3 |
| Myo1c | 1.04E-129 | 0.44335818 | 0.296 | 0.189 | 2.00E-125 | 2 | Myo1c | neutrophil.3 |
| Tubb4b | 5.55E-129 | 0.40445534 | 0.268 | 0.176 | 1.07E-124 | 2 | Tubb4b | neutrophil.3 |
| Sin3b | 3.13E-126 | 0.43872164 | 0.276 | 0.193 | 6.05E-122 | 2 | Sin3b | neutrophil.3 |
| Elmo1 | 6.98E-125 | 0.40364916 | 0.266 | 0.174 | 1.35E-120 | 2 | Elmo1 | neutrophil.3 |
| Pirb | 1.47E-123 | 0.31536297 | 0.456 | 0.341 | 2.83E-119 | 2 | Pirb | neutrophil.3 |
| Cast | 1.90E-123 | 0.58489136 | 0.253 | 0.168 | 3.68E-119 | 2 | Cast | neutrophil.3 |
| Tgoln1 | 2.70E-122 | 0.35731836 | 0.458 | 0.358 | 5.22E-118 | 2 | Tgoln1 | neutrophil.3 |
| Lpcat2 | 1.85E-121 | 0.52794627 | 0.285 | 0.181 | 3.58E-117 | 2 | Lpcat2 | neutrophil.3 |
| Ypel5 | 2.50E-121 | 0.55100417 | 0.259 | 0.16 | 4.83E-117 | 2 | Ypel5 | neutrophil.3 |
| Clk1 | 2.75E-120 | 0.25423956 | 0.48 | 0.373 | 5.32E-116 | 2 | Clk1 | neutrophil.3 |
| Mvp | 1.73E-118 | 0.42534938 | 0.308 | 0.205 | 3.35E-114 | 2 | Mvp | neutrophil.3 |
| Rara | 5.76E-118 | 0.39013718 | 0.398 | 0.308 | 1.11E-113 | 2 | Rara | neutrophil.3 |
| Pfkp | 1.20E-114 | 0.40014494 | 0.277 | 0.186 | 2.32E-110 | 2 | Pfkp | neutrophil.3 |
| Tmem189 | 2.86E-108 | 0.40093286 | 0.318 | 0.211 | 5.53E-104 | 2 | Tmem189 | neutrophil.3 |
| Tapbp | 3.11E-108 | 0.30514219 | 0.454 | 0.341 | 6.02E-104 | 2 | Tapbp | neutrophil.3 |
| Ppp1r10 | 3.33E-103 | 0.25187576 | 0.343 | 0.261 | 6.45E-99 | 2 | Ppp1r10 | neutrophil.3 |
| AA467197 | 6.93E-103 | 0.29866233 | 0.4 | 0.297 | 1.34E-98 | 2 | AA467197 | neutrophil.3 |
| Rab1a | 9.95E-102 | 0.29939493 | 0.392 | 0.289 | 1.92E-97 | 2 | Rab1a | neutrophil.3 |
| Cltc | 1.17E-99 | 0.37065277 | 0.292 | 0.199 | 2.26E-95 | 2 | Cltc | neutrophil.3 |
| Hnrnph2 | 2.03E-99 | 0.47789763 | 0.297 | 0.203 | 3.92E-95 | 2 | Hnrnph2 | neutrophil.3 |
| Nfat5 | 4.05E-99 | 0.37125039 | 0.279 | 0.187 | 7.83E-95 | 2 | Nfat5 | neutrophil.3 |
| Ccni | 5.01E-98 | 0.47678928 | 0.272 | 0.178 | 9.69E-94 | 2 | Ccni | neutrophil.3 |
| Sde2 | 3.96E-97 | 0.39632916 | 0.31 | 0.213 | 7.67E-93 | 2 | Sde2 | neutrophil.3 |
| Jak1 | 5.31E-97 | 0.29418152 | 0.408 | 0.309 | 1.03E-92 | 2 | Jak1 | neutrophil.3 |
| Sft2d2 | 7.88E-94 | 0.4505914 | 0.28 | 0.192 | 1.52E-89 | 2 | Sft2d2 | neutrophil.3 |
| Carhsp1 | 1.39E-87 | 0.45256817 | 0.299 | 0.223 | 2.70E-83 | 2 | Carhsp1 | neutrophil.3 |
| Sbno1 | 5.77E-86 | 0.37273123 | 0.27 | 0.186 | 1.12E-81 | 2 | Sbno1 | neutrophil.3 |
| Rcsd1 | 2.54E-84 | 0.32973453 | 0.259 | 0.187 | 4.90E-80 | 2 | Rcsd1 | neutrophil.3 |
| Shisa5 | 1.53E-83 | 0.35917231 | 0.346 | 0.26 | 2.95E-79 | 2 | Shisa5 | neutrophil.3 |
| Gla | 1.66E-81 | 0.3554682 | 0.325 | 0.232 | 3.22E-77 | 2 | Gla | neutrophil.3 |
| Bax | 2.62E-80 | 0.35517742 | 0.25 | 0.168 | 5.07E-76 | 2 | Bax | neutrophil.3 |
| Brd4 | 3.21E-78 | 0.26211667 | 0.361 | 0.265 | 6.21E-74 | 2 | Brd4 | neutrophil.3 |
| Clic1 | 5.68E-77 | 0.40946949 | 0.274 | 0.192 | 1.10E-72 | 2 | Clic1 | neutrophil.3 |
| Heca | 6.37E-75 | 0.26945736 | 0.288 | 0.203 | 1.23E-70 | 2 | Heca | neutrophil.3 |
| Wfdc17 | 6.12E-72 | 0.38239079 | 0.273 | 0.188 | 1.18E-67 | 2 | Wfdc17 | neutrophil.3 |
| Csf2rb2 | 3.33E-71 | 0.31952866 | 0.255 | 0.184 | 6.43E-67 | 2 | Csf2rb2 | neutrophil.3 |
| Gdi1 | 1.60E-70 | 0.30625626 | 0.256 | 0.181 | 3.09E-66 | 2 | Gdi1 | neutrophil.3 |
| Ip6k1 | 4.81E-70 | 0.28815535 | 0.357 | 0.265 | 9.30E-66 | 2 | Ip6k1 | neutrophil.3 |
| Nptn | 6.42E-70 | 0.38187635 | 0.251 | 0.169 | 1.24E-65 | 2 | Nptn | neutrophil.3 |
| Cbl | 1.55E-68 | 0.32968168 | 0.259 | 0.186 | 3.00E-64 | 2 | Cbl | neutrophil.3 |
| Tpm4 | 2.46E-65 | 0.31637617 | 0.372 | 0.274 | 4.76E-61 | 2 | Tpm4 | neutrophil.3 |
| Slc6a6 | 1.46E-64 | 0.30726373 | 0.297 | 0.224 | 2.81E-60 | 2 | Slc6a6 | neutrophil.3 |
| Usp32 | 4.69E-64 | 0.25268031 | 0.254 | 0.174 | 9.06E-60 | 2 | Usp32 | neutrophil.3 |
| Tnfrsf23 | 2.63E-61 | 0.28086544 | 0.288 | 0.214 | 5.09E-57 | 2 | Tnfrsf23 | neutrophil.3 |
| Tgfbr1 | 4.34E-61 | 0.30816508 | 0.28 | 0.195 | 8.40E-57 | 2 | Tgfbr1 | neutrophil.3 |
| Gmfg | 1.21E-59 | 0.29866155 | 0.268 | 0.198 | 2.33E-55 | 2 | Gmfg | neutrophil.3 |
| Myl12b | 4.00E-55 | 0.27774992 | 0.282 | 0.209 | 7.73E-51 | 2 | Myl12b | neutrophil.3 |
| Uba1 | 7.82E-54 | 0.31477454 | 0.305 | 0.225 | 1.51E-49 | 2 | Uba1 | neutrophil.3 |
| Tle3 | 2.86E-52 | 0.25768081 | 0.321 | 0.261 | 5.52E-48 | 2 | Tle3 | neutrophil.3 |
| Pxn | 1.29E-51 | 0.25366275 | 0.298 | 0.228 | 2.49E-47 | 2 | Pxn | neutrophil.3 |
| Atp6v0d1 | 1.80E-51 | 0.28345916 | 0.269 | 0.206 | 3.47E-47 | 2 | Atp6v0d1 | neutrophil.3 |
| Gsk3a | 3.01E-51 | 0.29272954 | 0.276 | 0.206 | 5.81E-47 | 2 | Gsk3a | neutrophil.3 |
| Ppp1r2 | 1.44E-49 | 0.28883277 | 0.259 | 0.19 | 2.78E-45 | 2 | Ppp1r2 | neutrophil.3 |
| Atp6v0e | 3.62E-40 | 0.26359294 | 0.268 | 0.21 | 6.99E-36 | 2 | Atp6v0e | neutrophil.3 |
| Siglecf | 0 | 1.84748413 | 0.611 | 0.154 | 0 | 3 | Siglecf | neutrophil.4 |
| Isy1 | 0 | 1.68273792 | 0.585 | 0.191 | 0 | 3 | Isy1 | neutrophil.4 |
| Sik1 | 0 | 1.26026132 | 0.65 | 0.273 | 0 | 3 | Sik1 | neutrophil.4 |
| Emd | 0 | 1.62862518 | 0.581 | 0.225 | 0 | 3 | Emd | neutrophil.4 |
| Fos | 0 | 1.40926813 | 0.867 | 0.523 | 0 | 3 | Fos | neutrophil.4 |
| Cdk11b | 0 | 2.03353868 | 0.485 | 0.142 | 0 | 3 | Cdk11b | neutrophil.4 |
| Nr4a3 | 0 | 1.16392874 | 0.733 | 0.392 | 0 | 3 | Nr4a3 | neutrophil.4 |
| Cpeb2 | 0 | 1.0017609 | 0.715 | 0.378 | 0 | 3 | Cpeb2 | neutrophil.4 |
| Nuak2 | 0 | 1.27943337 | 0.577 | 0.242 | 0 | 3 | Nuak2 | neutrophil.4 |
| Cwc25 | 0 | 2.28730253 | 0.473 | 0.144 | 0 | 3 | Cwc25 | neutrophil.4 |
| Cst3 | 0 | 0.94441588 | 0.88 | 0.555 | 0 | 3 | Cst3 | neutrophil.4 |
| Hexb | 0 | 1.87883404 | 0.426 | 0.101 | 0 | 3 | Hexb | neutrophil.4 |
| DEPRECATED-l | 0 | 0.99741788 | 0.657 | 0.338 | 0 | 3 | DEPRECATED-l | neutrophil.4 |
| Dennd4a | 0 | 0.77416157 | 0.702 | 0.383 | 0 | 3 | Dennd4a | neutrophil.4 |
| Antxr2 | 0 | 0.93896656 | 0.828 | 0.511 | 0 | 3 | Antxr2 | neutrophil.4 |
| Cdkn1a | 0 | 1.09237703 | 0.894 | 0.589 | 0 | 3 | Cdkn1a | neutrophil.4 |
| Wnk1 | 0 | 0.94415463 | 0.641 | 0.339 | 0 | 3 | Wnk1 | neutrophil.4 |
| Man1b1 | 0 | 2.02192878 | 0.401 | 0.102 | 0 | 3 | Man1b1 | neutrophil.4 |
| Ifrd1 | 0 | 0.85970382 | 0.807 | 0.508 | 0 | 3 | Ifrd1 | neutrophil.4 |
| Entpd3 | 0 | 1.51789056 | 0.438 | 0.143 | 0 | 3 | Entpd3 | neutrophil.4 |
| Bcl2l1 | 0 | 0.92979269 | 0.578 | 0.283 | 0 | 3 | Bcl2l1 | neutrophil.4 |
| Scimp | 0 | 1.41328697 | 0.406 | 0.12 | 0 | 3 | Scimp | neutrophil.4 |
| Rab43 | 0 | 1.27338718 | 0.477 | 0.191 | 0 | 3 | Rab43 | neutrophil.4 |
| Uck2 | 0 | 1.42222336 | 0.435 | 0.151 | 0 | 3 | Uck2 | neutrophil.4 |
| Ndel1 | 0 | 0.86609283 | 0.753 | 0.469 | 0 | 3 | Ndel1 | neutrophil.4 |
| Itgax | 0 | 0.75884019 | 0.575 | 0.298 | 0 | 3 | Itgax | neutrophil.4 |
| Fosl2 | 0 | 0.91794466 | 0.931 | 0.655 | 0 | 3 | Fosl2 | neutrophil.4 |
| Gsn | 0 | 0.61493796 | 0.758 | 0.484 | 0 | 3 | Gsn | neutrophil.4 |
| Ski | 0 | 0.49816176 | 0.659 | 0.385 | 0 | 3 | Ski | neutrophil.4 |
| Crem | 0 | 1.98681199 | 0.456 | 0.182 | 0 | 3 | Crem | neutrophil.4 |
| Hnrnpa2b1 | 0 | 0.75341687 | 0.563 | 0.293 | 0 | 3 | Hnrnpa2b1 | neutrophil.4 |

| | | | | | | | | |
|---|---|---|---|---|---|---|---|---|
| Tiparp | 0 | 1.13786875 | 0.5 | 0.234 | 0 | 3 | Tiparp | neutrophil.4 |
| Alox5 | 0 | 0.9344144 | 0.434 | 0.17 | 0 | 3 | Alox5 | neutrophil.4 |
| Il1r2 | 0 | 1.13782347 | 0.978 | 0.716 | 0 | 3 | Il1r2 | neutrophil.4 |
| Csrnp1 | 0 | 0.48973632 | 0.935 | 0.674 | 0 | 3 | Csrnp1 | neutrophil.4 |
| H2afy | 0 | 1.2931316 | 0.393 | 0.133 | 0 | 3 | H2afy | neutrophil.4 |
| Lmnb1 | 0 | 0.38992616 | 0.703 | 0.444 | 0 | 3 | Lmnb1 | neutrophil.4 |
| Stk17b | 0 | 0.77464706 | 0.556 | 0.298 | 0 | 3 | Stk17b | neutrophil.4 |
| Cmtm6 | 0 | 0.59029865 | 0.651 | 0.393 | 0 | 3 | Cmtm6 | neutrophil.4 |
| Tinf2 | 0 | 1.32736252 | 0.406 | 0.15 | 0 | 3 | Tinf2 | neutrophil.4 |
| Fam49b | 0 | 0.55787194 | 0.626 | 0.37 | 0 | 3 | Fam49b | neutrophil.4 |
| Picalm | 0 | 0.60781076 | 0.683 | 0.428 | 0 | 3 | Picalm | neutrophil.4 |
| Vsir | 0 | 0.51813648 | 0.677 | 0.422 | 0 | 3 | Vsir | neutrophil.4 |
| Rabgef1 | 0 | 0.63793425 | 0.662 | 0.408 | 0 | 3 | Rabgef1 | neutrophil.4 |
| H3f3b | 0 | 0.65519756 | 0.811 | 0.559 | 0 | 3 | H3f3b | neutrophil.4 |
| Nr4a2 | 0 | 3.189252 | 0.298 | 0.047 | 0 | 3 | Nr4a2 | neutrophil.4 |
| Cnbp | 0 | 1.46644381 | 0.347 | 0.096 | 0 | 3 | Cnbp | neutrophil.4 |
| Ly6e | 0 | 0.5706767 | 0.591 | 0.341 | 0 | 3 | Ly6e | neutrophil.4 |
| Jdp2 | 0 | 0.66052475 | 0.658 | 0.411 | 0 | 3 | Jdp2 | neutrophil.4 |
| Nup98 | 0 | 1.06841368 | 0.445 | 0.203 | 0 | 3 | Nup98 | neutrophil.4 |
| Abi3 | 0 | 1.03211928 | 0.375 | 0.133 | 0 | 3 | Abi3 | neutrophil.4 |
| Nabp1 | 0 | 0.58179543 | 0.612 | 0.371 | 0 | 3 | Nabp1 | neutrophil.4 |
| Myadm | 0 | 0.59034604 | 0.541 | 0.302 | 0 | 3 | Myadm | neutrophil.4 |
| Per1 | 0 | 0.53533117 | 0.623 | 0.385 | 0 | 3 | Per1 | neutrophil.4 |
| Arrb2 | 0 | 0.49790447 | 0.59 | 0.352 | 0 | 3 | Arrb2 | neutrophil.4 |
| Skil | 0 | 0.48775203 | 0.782 | 0.545 | 0 | 3 | Skil | neutrophil.4 |
| Ptk2b | 0 | 0.4265415 | 0.635 | 0.398 | 0 | 3 | Ptk2b | neutrophil.4 |
| Plxnc1 | 0 | 1.50974104 | 0.321 | 0.085 | 0 | 3 | Plxnc1 | neutrophil.4 |
| Arih1 | 0 | 0.77562409 | 0.539 | 0.306 | 0 | 3 | Arih1 | neutrophil.4 |
| Cd80 | 0 | 0.48869151 | 0.754 | 0.523 | 0 | 3 | Cd80 | neutrophil.4 |
| Ms4a6d | 0 | 2.0690786 | 0.287 | 0.057 | 0 | 3 | Ms4a6d | neutrophil.4 |
| Dot1l | 0 | 1.06713388 | 0.399 | 0.17 | 0 | 3 | Dot1l | neutrophil.4 |
| Coq10b | 0 | 0.92049726 | 0.42 | 0.191 | 0 | 3 | Coq10b | neutrophil.4 |
| Il4ra | 0 | 0.56903013 | 0.702 | 0.473 | 0 | 3 | Il4ra | neutrophil.4 |
| Stat3 | 0 | 0.57189933 | 0.585 | 0.356 | 0 | 3 | Stat3 | neutrophil.4 |
| Camkk1 | 0 | 0.9082573 | 0.393 | 0.167 | 0 | 3 | Camkk1 | neutrophil.4 |
| Arih2 | 0 | 0.50119368 | 0.596 | 0.371 | 0 | 3 | Arih2 | neutrophil.4 |
| Ptgs2 | 0 | 0.9790571 | 0.831 | 0.607 | 0 | 3 | Ptgs2 | neutrophil.4 |
| Clec4n | 0 | 0.45552551 | 0.78 | 0.556 | 0 | 3 | Clec4n | neutrophil.4 |
| Rin3 | 0 | 0.66598633 | 0.456 | 0.233 | 0 | 3 | Rin3 | neutrophil.4 |
| Rac2 | 0 | 0.37152723 | 0.907 | 0.684 | 0 | 3 | Rac2 | neutrophil.4 |
| Homer1 | 0 | 0.47986354 | 0.55 | 0.328 | 0 | 3 | Homer1 | neutrophil.4 |
| Bcl2a1b | 0 | 0.40016975 | 0.755 | 0.533 | 0 | 3 | Bcl2a1b | neutrophil.4 |
| Laptm5 | 0 | 0.46934526 | 0.898 | 0.677 | 0 | 3 | Laptm5 | neutrophil.4 |
| Ccrl2 | 0 | 0.7832602 | 0.836 | 0.615 | 0 | 3 | Ccrl2 | neutrophil.4 |
| Spty2d1 | 0 | 1.28694312 | 0.416 | 0.197 | 0 | 3 | Spty2d1 | neutrophil.4 |
| Tcirg1 | 0 | 0.63041056 | 0.411 | 0.192 | 0 | 3 | Tcirg1 | neutrophil.4 |
| Cotl1 | 0 | 0.30747828 | 0.811 | 0.593 | 0 | 3 | Cotl1 | neutrophil.4 |
| Ptgs1 | 0 | 1.58318579 | 0.293 | 0.075 | 0 | 3 | Ptgs1 | neutrophil.4 |
| B4galnt1 | 0 | 1.2382184 | 0.331 | 0.114 | 0 | 3 | B4galnt1 | neutrophil.4 |
| Tarm1 | 0 | 0.80671442 | 0.441 | 0.224 | 0 | 3 | Tarm1 | neutrophil.4 |
| Igfbp6 | 0 | 1.53445012 | 0.316 | 0.099 | 0 | 3 | Igfbp6 | neutrophil.4 |
| Tnfrsf1a | 0 | 0.4244807 | 0.744 | 0.527 | 0 | 3 | Tnfrsf1a | neutrophil.4 |
| Pde4b | 0 | 0.51248465 | 0.641 | 0.425 | 0 | 3 | Pde4b | neutrophil.4 |
| Coro2a | 0 | 0.86760128 | 0.37 | 0.155 | 0 | 3 | Coro2a | neutrophil.4 |
| Ltc4s | 0 | 1.54105517 | 0.285 | 0.07 | 0 | 3 | Ltc4s | neutrophil.4 |
| Tut7 | 0 | 0.47643261 | 0.643 | 0.429 | 0 | 3 | Tut7 | neutrophil.4 |
| Spata13 | 0 | 0.80026765 | 0.47 | 0.257 | 0 | 3 | Spata13 | neutrophil.4 |
| Nr4a1 | 0 | 0.82080264 | 0.926 | 0.715 | 0 | 3 | Nr4a1 | neutrophil.4 |
| Ttyh3 | 0 | 0.43621845 | 0.532 | 0.321 | 0 | 3 | Ttyh3 | neutrophil.4 |
| Nop53 | 0 | 1.65128118 | 0.34 | 0.129 | 0 | 3 | Nop53 | neutrophil.4 |
| Prpf8 | 0 | 1.27413691 | 0.322 | 0.111 | 0 | 3 | Prpf8 | neutrophil.4 |
| Ctsa | 0 | 0.68096028 | 0.424 | 0.214 | 0 | 3 | Ctsa | neutrophil.4 |
| Vegfa | 0 | 1.04418228 | 0.422 | 0.213 | 0 | 3 | Vegfa | neutrophil.4 |
| Gcnt2 | 0 | 0.6745267 | 0.474 | 0.265 | 0 | 3 | Gcnt2 | neutrophil.4 |
| Arhgdib | 0 | 0.55975012 | 0.501 | 0.293 | 0 | 3 | Arhgdib | neutrophil.4 |
| Pfn1 | 0 | 0.29620868 | 0.733 | 0.525 | 0 | 3 | Pfn1 | neutrophil.4 |
| Irf2bp2 | 0 | 0.32458997 | 0.65 | 0.443 | 0 | 3 | Irf2bp2 | neutrophil.4 |
| Hspa8 | 0 | 0.35432017 | 0.604 | 0.397 | 0 | 3 | Hspa8 | neutrophil.4 |
| Egr1 | 0 | 0.54574321 | 0.933 | 0.728 | 0 | 3 | Egr1 | neutrophil.4 |
| Sf3b1 | 0 | 0.41116673 | 0.669 | 0.464 | 0 | 3 | Sf3b1 | neutrophil.4 |
| Hipk1 | 0 | 0.76576245 | 0.415 | 0.21 | 0 | 3 | Hipk1 | neutrophil.4 |
| Hnrnpul1 | 0 | 0.52500626 | 0.478 | 0.273 | 0 | 3 | Hnrnpul1 | neutrophil.4 |
| Rab5c | 0 | 0.58151527 | 0.476 | 0.271 | 0 | 3 | Rab5c | neutrophil.4 |
| Nfil3 | 0 | 0.69448148 | 0.53 | 0.327 | 0 | 3 | Nfil3 | neutrophil.4 |
| Atf1 | 0 | 0.52596908 | 0.438 | 0.235 | 0 | 3 | Atf1 | neutrophil.4 |
| P2ry13 | 0 | 0.9676537 | 0.354 | 0.151 | 0 | 3 | P2ry13 | neutrophil.4 |
| Slc3a2 | 0 | 0.59600009 | 0.568 | 0.365 | 0 | 3 | Slc3a2 | neutrophil.4 |
| Gm2a | 0 | 0.36616798 | 0.628 | 0.426 | 0 | 3 | Gm2a | neutrophil.4 |
| Cytip | 0 | 0.4870466 | 0.537 | 0.336 | 0 | 3 | Cytip | neutrophil.4 |
| Atp6v0a1 | 0 | 0.59654483 | 0.44 | 0.239 | 0 | 3 | Atp6v0a1 | neutrophil.4 |
| Ucp2 | 0 | 0.3116982 | 0.673 | 0.473 | 0 | 3 | Ucp2 | neutrophil.4 |
| Mt1 | 0 | 1.94018188 | 0.284 | 0.084 | 0 | 3 | Mt1 | neutrophil.4 |
| Mmp9 | 0 | 0.58423087 | 0.837 | 0.638 | 0 | 3 | Mmp9 | neutrophil.4 |
| Rheb | 0 | 1.13759419 | 0.326 | 0.129 | 0 | 3 | Rheb | neutrophil.4 |
| Slc35e1 | 0 | 1.20346351 | 0.312 | 0.115 | 0 | 3 | Slc35e1 | neutrophil.4 |
| Neurl3 | 0 | 0.3933128 | 0.766 | 0.57 | 0 | 3 | Neurl3 | neutrophil.4 |
| Stk11 | 0 | 0.63012219 | 0.413 | 0.218 | 0 | 3 | Stk11 | neutrophil.4 |
| Coro1a | 0 | 0.25234809 | 0.784 | 0.591 | 0 | 3 | Coro1a | neutrophil.4 |
| Aff1 | 0 | 0.50740689 | 0.638 | 0.445 | 0 | 3 | Aff1 | neutrophil.4 |
| Csk | 0 | 0.80460294 | 0.354 | 0.161 | 0 | 3 | Csk | neutrophil.4 |
| Slc38a1 | 0 | 1.47337543 | 0.279 | 0.087 | 0 | 3 | Slc38a1 | neutrophil.4 |
| Clec5a | 0 | 0.7814189 | 0.422 | 0.23 | 0 | 3 | Clec5a | neutrophil.4 |
| Hk2 | 0 | 0.61360043 | 0.361 | 0.169 | 0 | 3 | Hk2 | neutrophil.4 |
| Enah | 0 | 1.33346654 | 0.254 | 0.063 | 0 | 3 | Enah | neutrophil.4 |
| D8Ertd738e | 0 | 0.81164334 | 0.371 | 0.18 | 0 | 3 | D8Ertd738e | neutrophil.4 |
| Pim3 | 0 | 1.70940633 | 0.258 | 0.067 | 0 | 3 | Pim3 | neutrophil.4 |
| Trio | 0 | 1.38556831 | 0.272 | 0.082 | 0 | 3 | Trio | neutrophil.4 |
| Cd52 | 0 | 0.36780519 | 0.74 | 0.55 | 0 | 3 | Cd52 | neutrophil.4 |
| Prrc2c | 0 | 0.73915499 | 0.371 | 0.183 | 0 | 3 | Prrc2c | neutrophil.4 |
| Fosb | 0 | 0.99618084 | 0.456 | 0.268 | 0 | 3 | Fosb | neutrophil.4 |
| Ccnl1 | 0 | 0.27215198 | 0.748 | 0.56 | 0 | 3 | Ccnl1 | neutrophil.4 |
| Naaa | 0 | 0.81606016 | 0.344 | 0.158 | 0 | 3 | Naaa | neutrophil.4 |
| Spag9 | 0 | 0.44408704 | 0.573 | 0.387 | 0 | 3 | Spag9 | neutrophil.4 |
| Tmem14c | 0 | 1.10093788 | 0.294 | 0.109 | 0 | 3 | Tmem14c | neutrophil.4 |
| Arf4 | 0 | 0.86571118 | 0.372 | 0.188 | 0 | 3 | Arf4 | neutrophil.4 |
| Ctss | 0 | 0.51077552 | 0.44 | 0.257 | 0 | 3 | Ctss | neutrophil.4 |
| AA467197 | 0 | 0.4507779 | 0.475 | 0.293 | 0 | 3 | AA467197 | neutrophil.4 |
| Vps37b | 0 | 0.4530911 | 0.604 | 0.422 | 0 | 3 | Vps37b | neutrophil.4 |
| Otulinl | 0 | 0.98424854 | 0.307 | 0.126 | 0 | 3 | Otulinl | neutrophil.4 |
| Actg1 | 0 | 0.25847796 | 0.857 | 0.676 | 0 | 3 | Actg1 | neutrophil.4 |
| Ddx5 | 0 | 0.38767377 | 0.602 | 0.422 | 0 | 3 | Ddx5 | neutrophil.4 |
| Runx1 | 0 | 0.76558104 | 0.348 | 0.168 | 0 | 3 | Runx1 | neutrophil.4 |
| Reep5 | 0 | 0.8583126 | 0.321 | 0.142 | 0 | 3 | Reep5 | neutrophil.4 |
| mt-Atp6 | 0 | 0.50484388 | 0.378 | 0.199 | 0 | 3 | mt-Atp6 | neutrophil.4 |
| Ptprc | 0 | 0.27987945 | 0.688 | 0.51 | 0 | 3 | Ptprc | neutrophil.4 |
| Ier2 | 0 | 0.34191446 | 0.718 | 0.54 | 0 | 3 | Ier2 | neutrophil.4 |
| Csf2rb | 0 | 0.40361182 | 0.713 | 0.535 | 0 | 3 | Csf2rb | neutrophil.4 |
| B4galt1 | 0 | 0.55215523 | 0.409 | 0.232 | 0 | 3 | B4galt1 | neutrophil.4 |
| Ncf2 | 0 | 0.2548892 | 0.752 | 0.576 | 0 | 3 | Ncf2 | neutrophil.4 |
| Cep83 | 0 | 0.95193374 | 0.28 | 0.107 | 0 | 3 | Cep83 | neutrophil.4 |
| Atp2a3 | 0 | 0.92618294 | 0.275 | 0.103 | 0 | 3 | Atp2a3 | neutrophil.4 |
| Rgs2 | 0 | 0.73278979 | 0.352 | 0.18 | 0 | 3 | Rgs2 | neutrophil.4 |
| Man2b1 | 0 | 0.56174806 | 0.344 | 0.172 | 0 | 3 | Man2b1 | neutrophil.4 |
| Atp13a3 | 0 | 0.9550818 | 0.281 | 0.111 | 0 | 3 | Atp13a3 | neutrophil.4 |
| Btg2 | 0 | 0.86966774 | 0.99 | 0.821 | 0 | 3 | Btg2 | neutrophil.4 |
| Dad1 | 0 | 1.13100695 | 0.255 | 0.091 | 0 | 3 | Dad1 | neutrophil.4 |
| Irs2 | 0 | 0.72329402 | 0.345 | 0.183 | 0 | 3 | Irs2 | neutrophil.4 |
| Dusp5 | 0 | 0.36901401 | 0.796 | 0.64 | 0 | 3 | Dusp5 | neutrophil.4 |
| Ppp1r15a | 0 | 0.32853424 | 0.817 | 0.664 | 0 | 3 | Ppp1r15a | neutrophil.4 |
| Nlrp12 | 0 | 1.09883317 | 0.267 | 0.115 | 0 | 3 | Nlrp12 | neutrophil.4 |
| Gadd45b | 0 | 0.42792677 | 0.916 | 0.764 | 0 | 3 | Gadd45b | neutrophil.4 |
| Cebpd | 0 | 0.57547177 | 0.439 | 0.287 | 0 | 3 | Cebpd | neutrophil.4 |
| Rhoq | 0 | 0.96613535 | 0.314 | 0.163 | 0 | 3 | Rhoq | neutrophil.4 |
| Tfip11 | 0 | 1.08170796 | 0.271 | 0.122 | 0 | 3 | Tfip11 | neutrophil.4 |
| Malt1 | 0 | 0.80973236 | 0.323 | 0.178 | 0 | 3 | Malt1 | neutrophil.4 |
| Phlda1 | 0 | 0.81661147 | 0.305 | 0.16 | 0 | 3 | Phlda1 | neutrophil.4 |
| G0s2 | 0 | 0.46999711 | 0.708 | 0.571 | 0 | 3 | G0s2 | neutrophil.4 |
| Fam110a | 0 | 0.80454967 | 0.291 | 0.155 | 0 | 3 | Fam110a | neutrophil.4 |
| Dusp2 | 0 | 0.3373142 | 0.5 | 0.366 | 0 | 3 | Dusp2 | neutrophil.4 |
| Plk3 | 0 | 0.259599 | 0.591 | 0.463 | 0 | 3 | Plk3 | neutrophil.4 |
| Pim1 | 0 | 0.4248376 | 0.988 | 0.876 | 0 | 3 | Pim1 | neutrophil.4 |
| Osbpl9 | 0 | 0.50290338 | 0.37 | 0.259 | 0 | 3 | Osbpl9 | neutrophil.4 |
| Il1b | 0 | 0.61370147 | 0.998 | 0.921 | 0 | 3 | Il1b | neutrophil.4 |
| Junb | 0 | 0.55622104 | 0.997 | 0.928 | 0 | 3 | Junb | neutrophil.4 |
| Hbegf | 2.21E-302 | 0.62387557 | 0.317 | 0.183 | 4.27E-298 | 3 | Hbegf | neutrophil.4 |
| Sfpq | 2.05E-292 | 0.39348916 | 0.396 | 0.208 | 3.96E-288 | 3 | Sfpq | neutrophil.4 |
| Fmnl3 | 1.48E-290 | 0.84037207 | 0.32 | 0.146 | 2.87E-286 | 3 | Fmnl3 | neutrophil.4 |
| Gngt2 | 9.06E-289 | 0.59332299 | 0.368 | 0.181 | 1.75E-284 | 3 | Gngt2 | neutrophil.4 |
| Ndufb1-ps | 1.28E-287 | 0.90622693 | 0.277 | 0.114 | 2.47E-283 | 3 | Ndufb1-ps | neutrophil.4 |
| Eif4g2 | 1.91E-286 | 0.4036901 | 0.473 | 0.282 | 3.69E-282 | 3 | Eif4g2 | neutrophil.4 |
| Slc25a3 | 1.21E-284 | 0.63812038 | 0.402 | 0.221 | 2.34E-280 | 3 | Slc25a3 | neutrophil.4 |
| Sirpa | 2.00E-284 | 0.26050857 | 0.624 | 0.442 | 3.87E-280 | 3 | Sirpa | neutrophil.4 |
| Actr2 | 7.38E-277 | 0.55773527 | 0.431 | 0.248 | 1.43E-272 | 3 | Actr2 | neutrophil.4 |
| Nfkbie | 1.20E-275 | 0.43234344 | 0.439 | 0.294 | 2.31E-271 | 3 | Nfkbie | neutrophil.4 |
| Mob1a | 1.54E-268 | 0.5633964 | 0.423 | 0.233 | 2.99E-264 | 3 | Mob1a | neutrophil.4 |
| Srsf6 | 1.61E-268 | 0.48277364 | 0.342 | 0.168 | 3.11E-264 | 3 | Srsf6 | neutrophil.4 |
| Dhx15 | 1.05E-267 | 0.75388285 | 0.271 | 0.112 | 2.03E-263 | 3 | Dhx15 | neutrophil.4 |
| Trim25 | 7.81E-266 | 0.45827908 | 0.303 | 0.178 | 1.51E-261 | 3 | Trim25 | neutrophil.4 |
| Atrnl1 | 3.39E-265 | 0.63634414 | 0.404 | 0.226 | 6.56E-261 | 3 | Atrnl1 | neutrophil.4 |
| Jak2 | 4.33E-262 | 0.27066192 | 0.645 | 0.467 | 8.38E-258 | 3 | Jak2 | neutrophil.4 |
| Sipa1 | 6.95E-262 | 0.85379598 | 0.293 | 0.124 | 1.34E-257 | 3 | Sipa1 | neutrophil.4 |
| Ptbp1 | 1.08E-259 | 0.6642485 | 0.291 | 0.133 | 2.09E-255 | 3 | Ptbp1 | neutrophil.4 |
| Ell2 | 1.25E-259 | 0.30703183 | 0.499 | 0.332 | 2.42E-255 | 3 | Ell2 | neutrophil.4 |
| Eif5a | 2.73E-259 | 0.49651621 | 0.441 | 0.287 | 5.29E-255 | 3 | Eif5a | neutrophil.4 |
| Nab1 | 1.80E-257 | 0.2896422 | 0.504 | 0.348 | 3.48E-253 | 3 | Nab1 | neutrophil.4 |
| Pum2 | 2.86E-253 | 0.50261452 | 0.436 | 0.265 | 5.53E-249 | 3 | Pum2 | neutrophil.4 |
| Hist1h2bc | 3.67E-252 | 0.70546082 | 0.314 | 0.171 | 7.10E-248 | 3 | Hist1h2bc | neutrophil.4 |
| Cers6 | 3.00E-251 | 0.40143212 | 0.457 | 0.269 | 5.80E-247 | 3 | Cers6 | neutrophil.4 |
| Plcg2 | 3.95E-249 | 0.68713325 | 0.284 | 0.126 | 7.63E-245 | 3 | Plcg2 | neutrophil.4 |
| Sh3bgrl3 | 4.47E-248 | 0.2660251 | 0.562 | 0.388 | 8.65E-244 | 3 | Sh3bgrl3 | neutrophil.4 |
| mt-Nd1 | 2.90E-247 | 0.60105504 | 0.281 | 0.13 | 5.61E-243 | 3 | mt-Nd1 | neutrophil.4 |
| Adprh | 1.04E-244 | 0.30115246 | 0.559 | 0.384 | 2.02E-240 | 3 | Adprh | neutrophil.4 |
| Cd244a | 5.20E-240 | 0.53000468 | 0.342 | 0.187 | 1.01E-235 | 3 | Cd244a | neutrophil.4 |
| Ndst1 | 4.70E-239 | 0.51899505 | 0.359 | 0.191 | 9.09E-235 | 3 | Ndst1 | neutrophil.4 |
| Cd177 | 3.37E-238 | 0.6762088 | 0.257 | 0.143 | 6.51E-234 | 3 | Cd177 | neutrophil.4 |
| Cd300c2 | 1.93E-237 | 0.83981902 | 0.262 | 0.117 | 3.73E-233 | 3 | Cd300c2 | neutrophil.4 |
| Sbno2 | 9.60E-232 | 0.30495744 | 0.553 | 0.386 | 1.86E-227 | 3 | Sbno2 | neutrophil.4 |
| Was | 6.04E-231 | 0.71486466 | 0.326 | 0.162 | 1.17E-226 | 3 | Was | neutrophil.4 |
| Sh3bp1 | 2.09E-230 | 0.52476096 | 0.334 | 0.164 | 4.04E-226 | 3 | Sh3bp1 | neutrophil.4 |
| Mdm2 | 2.69E-229 | 0.51234712 | 0.327 | 0.207 | 5.21E-225 | 3 | Mdm2 | neutrophil.4 |
| Vamp8 | 2.89E-229 | 0.37059636 | 0.456 | 0.285 | 5.60E-225 | 3 | Vamp8 | neutrophil.4 |
| Hsd17b11 | 5.72E-229 | 0.61831361 | 0.279 | 0.138 | 1.11E-224 | 3 | Hsd17b11 | neutrophil.4 |
| Slco3a1 | 8.05E-229 | 0.78173352 | 0.289 | 0.135 | 1.56E-224 | 3 | Slco3a1 | neutrophil.4 |
| Prkcb | 5.44E-227 | 0.44470273 | 0.29 | 0.144 | 1.05E-222 | 3 | Prkcb | neutrophil.4 |
| U2af2 | 1.17E-226 | 0.60110607 | 0.305 | 0.148 | 2.27E-222 | 3 | U2af2 | neutrophil.4 |
| Inpp5d | 4.21E-226 | 0.35381744 | 0.505 | 0.328 | 8.15E-222 | 3 | Inpp5d | neutrophil.4 |
| Ndufa13 | 6.11E-226 | 0.65919685 | 0.29 | 0.137 | 1.18E-221 | 3 | Ndufa13 | neutrophil.4 |
| Actn1 | 9.17E-226 | 0.4120346 | 0.467 | 0.288 | 1.77E-221 | 3 | Actn1 | neutrophil.4 |
| 7-Mar | 1.24E-225 | 0.38975548 | 0.498 | 0.337 | 2.40E-221 | 3 | 7-Mar | neutrophil.4 |
| Cd300ld | 3.03E-224 | 0.36856653 | 0.511 | 0.328 | 5.85E-220 | 3 | Cd300ld | neutrophil.4 |
| Nfatc1 | 1.42E-223 | 0.72872852 | 0.272 | 0.128 | 2.74E-219 | 3 | Nfatc1 | neutrophil.4 |
| Grn | 6.01E-220 | 0.44833093 | 0.323 | 0.172 | 1.16E-215 | 3 | Grn | neutrophil.4 |
| Nedd8 | 5.84E-216 | 0.63126577 | 0.308 | 0.154 | 1.13E-211 | 3 | Nedd8 | neutrophil.4 |
| Tra2a | 1.78E-214 | 0.30203572 | 0.473 | 0.3 | 3.43E-210 | 3 | Tra2a | neutrophil.4 |
| Ppp1r3b | 2.92E-212 | 0.42560997 | 0.431 | 0.267 | 5.65E-208 | 3 | Ppp1r3b | neutrophil.4 |
| Arl5b | 7.71E-212 | 0.62602554 | 0.29 | 0.168 | 1.49E-207 | 3 | Arl5b | neutrophil.4 |
| Gcnt1 | 8.88E-209 | 0.70023246 | 0.291 | 0.145 | 1.72E-204 | 3 | Gcnt1 | neutrophil.4 |
| mt-Co3 | 9.05E-209 | 0.38831858 | 0.322 | 0.177 | 1.75E-204 | 3 | mt-Co3 | neutrophil.4 |
| mt-Atp8 | 6.30E-208 | 0.29906132 | 0.335 | 0.181 | 1.22E-203 | 3 | mt-Atp8 | neutrophil.4 |
| Chtop | 8.79E-208 | 0.66579112 | 0.263 | 0.118 | 1.70E-203 | 3 | Chtop | neutrophil.4 |
| Nudt4 | 5.16E-207 | 0.47505611 | 0.369 | 0.233 | 9.97E-203 | 3 | Nudt4 | neutrophil.4 |
| Cdkn1b | 8.19E-203 | 0.42568214 | 0.326 | 0.207 | 1.58E-198 | 3 | Cdkn1b | neutrophil.4 |
| Zcchc2 | 1.76E-201 | 0.64747445 | 0.293 | 0.16 | 3.41E-197 | 3 | Zcchc2 | neutrophil.4 |
| Cfp | 3.28E-200 | 0.40290246 | 0.407 | 0.257 | 6.33E-196 | 3 | Cfp | neutrophil.4 |
| Sla | 4.44E-200 | 0.51882889 | 0.303 | 0.152 | 8.59E-196 | 3 | Sla | neutrophil.4 |
| Khdc4 | 3.67E-198 | 0.64975958 | 0.253 | 0.115 | 7.10E-194 | 3 | Khdc4 | neutrophil.4 |
| Dnajb6 | 4.31E-198 | 0.62256673 | 0.265 | 0.136 | 8.34E-194 | 3 | Dnajb6 | neutrophil.4 |
| Prex1 | 7.82E-198 | 0.35053223 | 0.444 | 0.268 | 1.51E-193 | 3 | Prex1 | neutrophil.4 |
| Ankrd13a | 1.36E-191 | 0.34214278 | 0.379 | 0.224 | 2.62E-187 | 3 | Ankrd13a | neutrophil.4 |
| Fam168b | 1.97E-191 | 0.5669244 | 0.307 | 0.161 | 3.81E-187 | 3 | Fam168b | neutrophil.4 |
| Mdh2 | 3.62E-191 | 0.67999145 | 0.255 | 0.124 | 6.99E-187 | 3 | Mdh2 | neutrophil.4 |
| Cyth4 | 8.38E-190 | 0.29099257 | 0.413 | 0.277 | 1.62E-185 | 3 | Cyth4 | neutrophil.4 |
| Rab1b | 9.85E-190 | 0.68895898 | 0.26 | 0.122 | 1.90E-185 | 3 | Rab1b | neutrophil.4 |
| Hnrnpm | 1.06E-188 | 0.31423831 | 0.347 | 0.197 | 2.05E-184 | 3 | Hnrnpm | neutrophil.4 |
| Rbm3 | 6.25E-187 | 0.51465674 | 0.272 | 0.129 | 1.21E-182 | 3 | Rbm3 | neutrophil.4 |
| Tgfbr2 | 4.77E-186 | 0.52588698 | 0.282 | 0.14 | 9.22E-182 | 3 | Tgfbr2 | neutrophil.4 |
| Wdr1 | 4.53E-183 | 0.41428435 | 0.377 | 0.225 | 8.75E-179 | 3 | Wdr1 | neutrophil.4 |
| Srrm2 | 2.13E-180 | 0.39262285 | 0.361 | 0.208 | 4.11E-176 | 3 | Srrm2 | neutrophil.4 |
| R3hdm4 | 4.21E-180 | 0.59316229 | 0.308 | 0.153 | 8.13E-176 | 3 | R3hdm4 | neutrophil.4 |
| Rbm39 | 3.07E-178 | 0.42435073 | 0.315 | 0.169 | 5.94E-174 | 3 | Rbm39 | neutrophil.4 |
| Syncrip | 2.75E-176 | 0.46428781 | 0.263 | 0.126 | 5.32E-172 | 3 | Syncrip | neutrophil.4 |
| Calm2 | 5.08E-176 | 0.35854632 | 0.362 | 0.222 | 9.82E-172 | 3 | Calm2 | neutrophil.4 |
| Eif4a2 | 5.78E-176 | 0.25977109 | 0.375 | 0.237 | 1.12E-171 | 3 | Eif4a2 | neutrophil.4 |
| Ctdsp1 | 1.41E-175 | 0.40119118 | 0.301 | 0.166 | 2.73E-171 | 3 | Ctdsp1 | neutrophil.4 |
| Papola | 4.39E-175 | 0.43905539 | 0.269 | 0.14 | 8.49E-171 | 3 | Papola | neutrophil.4 |
| Acly | 9.82E-175 | 0.45104887 | 0.288 | 0.155 | 1.90E-170 | 3 | Acly | neutrophil.4 |
| Ubl3 | 2.83E-174 | 0.45570251 | 0.308 | 0.18 | 5.48E-170 | 3 | Ubl3 | neutrophil.4 |
| Bzw1 | 1.59E-171 | 0.59454468 | 0.266 | 0.137 | 3.08E-167 | 3 | Bzw1 | neutrophil.4 |
| Tcn2 | 1.89E-171 | 0.3368704 | 0.401 | 0.259 | 3.66E-167 | 3 | Tcn2 | neutrophil.4 |
| Ctdsp2 | 8.18E-170 | 0.37186645 | 0.326 | 0.192 | 1.58E-165 | 3 | Ctdsp2 | neutrophil.4 |
| Ankrd11 | 5.54E-168 | 0.44074401 | 0.352 | 0.207 | 1.07E-163 | 3 | Ankrd11 | neutrophil.4 |
| Pstpip1 | 2.33E-165 | 0.33512729 | 0.388 | 0.24 | 4.50E-161 | 3 | Pstpip1 | neutrophil.4 |
| Dennd5a | 7.76E-164 | 0.28121426 | 0.43 | 0.29 | 1.50E-159 | 3 | Dennd5a | neutrophil.4 |
| Ago2 | 9.88E-164 | 0.29543416 | 0.376 | 0.233 | 1.91E-159 | 3 | Ago2 | neutrophil.4 |
| mt-Co1 | 1.72E-163 | 0.35793313 | 0.27 | 0.151 | 3.32E-159 | 3 | mt-Co1 | neutrophil.4 |
| Hnrnpk | 1.28E-162 | 0.46953139 | 0.291 | 0.155 | 2.47E-158 | 3 | Hnrnpk | neutrophil.4 |
| Arpc3 | 8.02E-161 | 0.42040149 | 0.396 | 0.249 | 1.55E-156 | 3 | Arpc3 | neutrophil.4 |
| Purb | 1.48E-160 | 0.26694297 | 0.318 | 0.185 | 2.87E-156 | 3 | Purb | neutrophil.4 |
| Nfam1 | 1.61E-160 | 0.32773007 | 0.451 | 0.292 | 3.11E-156 | 3 | Nfam1 | neutrophil.4 |
| Tgfbr1 | 7.62E-160 | 0.35333402 | 0.317 | 0.196 | 1.47E-155 | 3 | Tgfbr1 | neutrophil.4 |
| mt-Cytb | 1.89E-158 | 0.33969347 | 0.337 | 0.18 | 3.65E-154 | 3 | mt-Cytb | neutrophil.4 |
| Rnf130 | 4.49E-156 | 0.5569039 | 0.259 | 0.138 | 8.69E-152 | 3 | Rnf130 | neutrophil.4 |
| Hnrnpf | 6.85E-156 | 0.55449231 | 0.289 | 0.156 | 1.32E-151 | 3 | Hnrnpf | neutrophil.4 |
| Ppp2ca | 1.03E-154 | 0.38865075 | 0.351 | 0.216 | 1.99E-150 | 3 | Ppp2ca | neutrophil.4 |
| Btbd9 | 2.89E-152 | 0.44033521 | 0.302 | 0.177 | 5.59E-148 | 3 | Btbd9 | neutrophil.4 |
| Arid5a | 3.38E-151 | 0.46303922 | 0.296 | 0.168 | 6.54E-147 | 3 | Arid5a | neutrophil.4 |
| Jmjd1c | 6.44E-151 | 0.3690803 | 0.362 | 0.234 | 1.25E-146 | 3 | Jmjd1c | neutrophil.4 |
| Ppp2cb | 3.15E-149 | 0.42787942 | 0.297 | 0.166 | 6.09E-145 | 3 | Ppp2cb | neutrophil.4 |
| Elob | 8.99E-149 | 0.44539231 | 0.305 | 0.173 | 1.74E-144 | 3 | Elob | neutrophil.4 |
| Ythdf3 | 9.34E-148 | 0.3112661 | 0.359 | 0.226 | 1.81E-143 | 3 | Ythdf3 | neutrophil.4 |
| Ifngr1 | 7.70E-146 | 0.30332663 | 0.323 | 0.212 | 1.49E-141 | 3 | Ifngr1 | neutrophil.4 |
| Calm3 | 1.22E-144 | 0.34361407 | 0.362 | 0.222 | 2.36E-140 | 3 | Calm3 | neutrophil.4 |
| Stk24 | 1.65E-144 | 0.41536002 | 0.33 | 0.196 | 3.19E-140 | 3 | Stk24 | neutrophil.4 |
| Eif4a1 | 3.83E-139 | 0.36530118 | 0.332 | 0.201 | 7.40E-135 | 3 | Eif4a1 | neutrophil.4 |
| Cox8a | 9.11E-139 | 0.36914791 | 0.341 | 0.202 | 1.76E-134 | 3 | Cox8a | neutrophil.4 |
| Arrdc4 | 2.94E-138 | 0.28381099 | 0.367 | 0.249 | 5.69E-134 | 3 | Arrdc4 | neutrophil.4 |
| Lbr | 3.77E-138 | 0.43899904 | 0.334 | 0.196 | 7.29E-134 | 3 | Lbr | neutrophil.4 |
| Arf3 | 6.45E-138 | 0.31941849 | 0.327 | 0.194 | 1.25E-133 | 3 | Arf3 | neutrophil.4 |
| Ifitm5 | 5.02E-137 | 0.46374226 | 0.274 | 0.166 | 9.71E-133 | 3 | Ifitm5 | neutrophil.4 |
| Cbfb | 1.18E-136 | 0.30177638 | 0.289 | 0.162 | 2.29E-132 | 3 | Cbfb | neutrophil.4 |
| Tpst2 | 5.43E-136 | 0.56526657 | 0.256 | 0.138 | 1.05E-131 | 3 | Tpst2 | neutrophil.4 |
| Ranbp2 | 1.08E-135 | 0.43143424 | 0.251 | 0.136 | 2.09E-131 | 3 | Ranbp2 | neutrophil.4 |
| Gramd1a | 1.34E-134 | 0.36550788 | 0.263 | 0.147 | 2.60E-130 | 3 | Gramd1a | neutrophil.4 |
| Ralb | 4.62E-134 | 0.48328712 | 0.264 | 0.144 | 8.94E-130 | 3 | Ralb | neutrophil.4 |
| Nrros | 8.08E-133 | 0.43619156 | 0.296 | 0.174 | 1.56E-128 | 3 | Nrros | neutrophil.4 |
| Rbm25 | 1.96E-129 | 0.32113149 | 0.289 | 0.167 | 3.78E-125 | 3 | Rbm25 | neutrophil.4 |
| Chp1 | 3.03E-126 | 0.3983616 | 0.33 | 0.209 | 5.85E-122 | 3 | Chp1 | neutrophil.4 |
| Cdc34 | 2.57E-125 | 0.41135811 | 0.256 | 0.139 | 4.97E-121 | 3 | Cdc34 | neutrophil.4 |
| Pcbp1 | 1.14E-124 | 0.42820697 | 0.286 | 0.163 | 2.21E-120 | 3 | Pcbp1 | neutrophil.4 |
| Abhd2 | 1.43E-124 | 0.46988194 | 0.265 | 0.148 | 2.76E-120 | 3 | Abhd2 | neutrophil.4 |
| Csde1 | 5.22E-122 | 0.50083573 | 0.254 | 0.134 | 1.01E-117 | 3 | Csde1 | neutrophil.4 |
| Tspan14 | 1.20E-121 | 0.51031252 | 0.251 | 0.142 | 2.32E-117 | 3 | Tspan14 | neutrophil.4 |
| Kdm7a | 1.96E-120 | 0.37844267 | 0.299 | 0.191 | 3.79E-116 | 3 | Kdm7a | neutrophil.4 |
| Dgat1 | 5.20E-120 | 0.37837484 | 0.314 | 0.177 | 1.01E-115 | 3 | Dgat1 | neutrophil.4 |
| Zmiz1 | 7.15E-118 | 0.43191202 | 0.282 | 0.166 | 1.38E-113 | 3 | Zmiz1 | neutrophil.4 |
| Rrbp1 | 7.18E-117 | 0.25684899 | 0.261 | 0.152 | 1.39E-112 | 3 | Rrbp1 | neutrophil.4 |
| Sf3b2 | 7.60E-117 | 0.37380747 | 0.281 | 0.163 | 1.47E-112 | 3 | Sf3b2 | neutrophil.4 |
| Setd5 | 9.14E-112 | 0.34007617 | 0.261 | 0.163 | 1.77E-107 | 3 | Setd5 | neutrophil.4 |
| B4galt3 | 9.56E-111 | 0.50004002 | 0.256 | 0.145 | 1.85E-106 | 3 | B4galt3 | neutrophil.4 |
| Arpc4 | 4.88E-110 | 0.34186363 | 0.303 | 0.187 | 9.43E-106 | 3 | Arpc4 | neutrophil.4 |
| Pdcd6ip | 5.56E-106 | 0.27279462 | 0.275 | 0.168 | 1.08E-101 | 3 | Pdcd6ip | neutrophil.4 |
| Unc93b1 | 1.70E-105 | 0.34225115 | 0.317 | 0.202 | 3.29E-101 | 3 | Unc93b1 | neutrophil.4 |
| Arf6 | 2.05E-105 | 0.27606707 | 0.327 | 0.217 | 3.97E-101 | 3 | Arf6 | neutrophil.4 |
| Ikzf1 | 2.40E-104 | 0.37877884 | 0.272 | 0.158 | 4.63E-100 | 3 | Ikzf1 | neutrophil.4 |
| Lpp | 1.91E-102 | 0.41732089 | 0.26 | 0.152 | 3.69E-98 | 3 | Lpp | neutrophil.4 |
| Atp6v1f | 4.17E-99 | 0.28264327 | 0.275 | 0.17 | 8.06E-95 | 3 | Atp6v1f | neutrophil.4 |
| Wbp1l | 1.58E-96 | 0.36125218 | 0.254 | 0.152 | 3.05E-92 | 3 | Wbp1l | neutrophil.4 |
| Sdf4 | 1.17E-95 | 0.36046138 | 0.261 | 0.159 | 2.26E-91 | 3 | Sdf4 | neutrophil.4 |
| R3hdm2 | 4.73E-92 | 0.33983383 | 0.264 | 0.164 | 9.15E-88 | 3 | R3hdm2 | neutrophil.4 |
| mt-Nd4l | 6.72E-92 | 0.28990396 | 0.253 | 0.136 | 1.30E-87 | 3 | mt-Nd4l | neutrophil.4 |
| Itprip | 1.65E-86 | 0.25552989 | 0.293 | 0.191 | 3.19E-82 | 3 | Itprip | neutrophil.4 |
| Tbl1xr1 | 1.02E-85 | 0.2774685 | 0.273 | 0.171 | 1.96E-81 | 3 | Tbl1xr1 | neutrophil.4 |
| Ube2d3 | 6.19E-84 | 0.25717142 | 0.265 | 0.167 | 1.20E-79 | 3 | Ube2d3 | neutrophil.4 |
| Arhgap9 | 5.03E-79 | 0.25683591 | 0.289 | 0.194 | 9.73E-75 | 3 | Arhgap9 | neutrophil.4 |
| Fgr | 0 | 0.29986996 | 0.349 | 0.614 | 0 | 4 | Fgr | neutrophil.5 |
| Ier2 | 0 | 0.25638118 | 0.333 | 0.579 | 0 | 4 | Ier2 | neutrophil.5 |
| Mapkapk2 | 0 | 0.40133994 | 0.464 | 0.703 | 0 | 4 | Mapkapk2 | neutrophil.5 |
| Neurl3 | 0 | 0.26951517 | 0.38 | 0.609 | 0 | 4 | Neurl3 | neutrophil.5 |
| Bcl2a1b | 0 | 0.30417621 | 0.348 | 0.576 | 0 | 4 | Bcl2a1b | neutrophil.5 |
| Fcer1g | 0 | 0.25758565 | 0.61 | 0.837 | 0 | 4 | Fcer1g | neutrophil.5 |
| Jund | 0 | 0.38933313 | 0.519 | 0.746 | 0 | 4 | Jund | neutrophil.5 |
| Ninj1 | 0 | 0.25829888 | 0.351 | 0.572 | 0 | 4 | Ninj1 | neutrophil.5 |
| Cst3 | 0 | 0.3278783 | 0.389 | 0.609 | 0 | 4 | Cst3 | neutrophil.5 |
| Rab11fip1 | 0 | 0.34218622 | 0.312 | 0.528 | 0 | 4 | Rab11fip1 | neutrophil.5 |
| Prdx5 | 0 | 0.53272122 | 0.406 | 0.617 | 0 | 4 | Prdx5 | neutrophil.5 |
| Kdm6b | 0 | 0.44117593 | 0.594 | 0.8 | 0 | 4 | Kdm6b | neutrophil.5 |
| Maff | 0 | 0.51307571 | 0.39 | 0.59 | 0 | 4 | Maff | neutrophil.5 |
| Clec4e | 0 | 0.33846459 | 0.433 | 0.631 | 0 | 4 | Clec4e | neutrophil.5 |
| Ncf1 | 0 | 0.46802566 | 0.447 | 0.644 | 0 | 4 | Ncf1 | neutrophil.5 |
| Tyrobp | 0 | 0.42926763 | 0.679 | 0.871 | 0 | 4 | Tyrobp | neutrophil.5 |
| Sema4d | 0 | 0.43571298 | 0.403 | 0.589 | 0 | 4 | Sema4d | neutrophil.5 |
| Cd274 | 0 | 0.32696472 | 0.29 | 0.468 | 0 | 4 | Cd274 | neutrophil.5 |
| Sqstm1 | 0 | 0.39355126 | 0.332 | 0.506 | 0 | 4 | Sqstm1 | neutrophil.5 |
| Adgre5 | 0 | 0.44367873 | 0.511 | 0.678 | 0 | 4 | Adgre5 | neutrophil.5 |
| Ier5 | 0 | 0.72871065 | 0.597 | 0.762 | 0 | 4 | Ier5 | neutrophil.5 |
| Basp1 | 0 | 0.54892054 | 0.387 | 0.544 | 0 | 4 | Basp1 | neutrophil.5 |
| Mcl1 | 0 | 0.40037142 | 0.857 | 0.944 | 0 | 4 | Mcl1 | neutrophil.5 |
| Tnf | 0 | 0.52846614 | 0.518 | 0.595 | 0 | 4 | Tnf | neutrophil.5 |
| Il1a | 0 | 0.27905439 | 0.488 | 0.557 | 0 | 4 | Il1a | neutrophil.5 |
| Nfkbia | 0 | 0.34096297 | 0.821 | 0.887 | 0 | 4 | Nfkbia | neutrophil.5 |
| Cebpb | 0 | 0.68877513 | 0.887 | 0.935 | 0 | 4 | Cebpb | neutrophil.5 |
| C3 | 2.94E-305 | 0.60297109 | 0.381 | 0.505 | 5.69E-301 | 4 | C3 | neutrophil.5 |
| Plekho2 | 6.07E-303 | 0.30602775 | 0.335 | 0.583 | 1.17E-298 | 4 | Plekho2 | neutrophil.5 |
| Bcl2l11 | 1.23E-274 | 0.45237208 | 0.267 | 0.446 | 2.38E-270 | 4 | Bcl2l11 | neutrophil.5 |
| Gsn | 1.59E-273 | 0.50504346 | 0.344 | 0.529 | 3.08E-269 | 4 | Gsn | neutrophil.5 |
| Cfl1 | 9.95E-257 | 0.32557597 | 0.322 | 0.596 | 1.92E-252 | 4 | Cfl1 | neutrophil.5 |
| Marcks | 3.19E-255 | 0.82374872 | 0.46 | 0.591 | 6.17E-251 | 4 | Marcks | neutrophil.5 |
| Ctsz | 5.83E-255 | 0.26981572 | 0.262 | 0.464 | 1.13E-250 | 4 | Ctsz | neutrophil.5 |
| Gpx1 | 3.85E-253 | 0.28548596 | 0.289 | 0.495 | 7.44E-249 | 4 | Gpx1 | neutrophil.5 |
| Pfn1 | 4.80E-249 | 0.36065666 | 0.316 | 0.568 | 9.28E-245 | 4 | Pfn1 | neutrophil.5 |
| Irf1 | 2.17E-247 | 0.3373056 | 0.224 | 0.356 | 4.20E-243 | 4 | Irf1 | neutrophil.5 |
| Klf2 | 1.37E-246 | 0.71478763 | 0.404 | 0.541 | 2.65E-242 | 4 | Klf2 | neutrophil.5 |
| Dusp16 | 3.08E-244 | 0.3659853 | 0.272 | 0.487 | 5.95E-240 | 4 | Dusp16 | neutrophil.5 |
| Apbb1ip | 5.48E-236 | 0.3511229 | 0.283 | 0.507 | 1.06E-231 | 4 | Apbb1ip | neutrophil.5 |
| Ehd1 | 1.55E-231 | 0.54907215 | 0.31 | 0.487 | 3.00E-227 | 4 | Ehd1 | neutrophil.5 |
| Csf1 | 1.58E-208 | 0.44413446 | 0.217 | 0.332 | 3.05E-204 | 4 | Csf1 | neutrophil.5 |
| Irf2bp2 | 2.34E-208 | 0.31729804 | 0.276 | 0.482 | 4.53E-204 | 4 | Irf2bp2 | neutrophil.5 |
| Arpc1b | 3.72E-207 | 0.44501099 | 0.306 | 0.544 | 7.19E-203 | 4 | Arpc1b | neutrophil.5 |
| Cd300a | 4.34E-200 | 0.44478305 | 0.308 | 0.495 | 8.40E-196 | 4 | Cd300a | neutrophil.5 |
| Lst1 | 2.86E-190 | 0.34729052 | 0.24 | 0.448 | 5.54E-186 | 4 | Lst1 | neutrophil.5 |
| Irak2 | 4.13E-183 | 0.54798078 | 0.301 | 0.504 | 7.99E-179 | 4 | Irak2 | neutrophil.5 |
| Igsf6 | 1.07E-182 | 0.33120242 | 0.201 | 0.386 | 2.08E-178 | 4 | Igsf6 | neutrophil.5 |
| Myd88 | 5.44E-179 | 0.29442763 | 0.212 | 0.432 | 1.05E-174 | 4 | Myd88 | neutrophil.5 |
| Oasl2 | 2.62E-178 | 0.52247949 | 0.221 | 0.387 | 5.07E-174 | 4 | Oasl2 | neutrophil.5 |
| Nfkb2 | 2.33E-175 | 0.53334178 | 0.314 | 0.509 | 4.50E-171 | 4 | Nfkb2 | neutrophil.5 |
| Plin2 | 1.18E-174 | 0.3069828 | 0.203 | 0.363 | 2.27E-170 | 4 | Plin2 | neutrophil.5 |
| Ski | 4.53E-168 | 0.26639414 | 0.228 | 0.432 | 8.76E-164 | 4 | Ski | neutrophil.5 |
| Ezr | 2.51E-167 | 0.45107706 | 0.231 | 0.4 | 4.85E-163 | 4 | Ezr | neutrophil.5 |
| Adprh | 1.05E-164 | 0.26750557 | 0.199 | 0.42 | 2.03E-160 | 4 | Adprh | neutrophil.5 |
| H3f3a | 2.60E-164 | 0.29484023 | 0.208 | 0.439 | 5.03E-160 | 4 | H3f3a | neutrophil.5 |
| Arrb2 | 3.01E-163 | 0.26910268 | 0.196 | 0.394 | 5.83E-159 | 4 | Arrb2 | neutrophil.5 |
| Clec2d | 1.12E-159 | 0.48291929 | 0.201 | 0.36 | 2.16E-155 | 4 | Clec2d | neutrophil.5 |
| Vmp1 | 1.47E-152 | 0.37860201 | 0.215 | 0.413 | 2.83E-148 | 4 | Vmp1 | neutrophil.5 |
| AA467197 | 3.81E-151 | 0.51934398 | 0.181 | 0.325 | 7.36E-147 | 4 | AA467197 | neutrophil.5 |
| Card19 | 2.52E-149 | 0.47614679 | 0.226 | 0.408 | 4.87E-145 | 4 | Card19 | neutrophil.5 |
| Susd6 | 2.03E-148 | 0.42590831 | 0.223 | 0.422 | 3.93E-144 | 4 | Susd6 | neutrophil.5 |
| Srsf5 | 2.49E-148 | 0.31888934 | 0.226 | 0.446 | 4.82E-144 | 4 | Srsf5 | neutrophil.5 |
| Csf2ra | 7.70E-148 | 0.61257021 | 0.307 | 0.504 | 1.49E-143 | 4 | Csf2ra | neutrophil.5 |
| Treml4 | 6.89E-147 | 0.27934942 | 0.176 | 0.328 | 1.33E-142 | 4 | Treml4 | neutrophil.5 |
| Ncf4 | 1.48E-143 | 0.34348849 | 0.187 | 0.384 | 2.87E-139 | 4 | Ncf4 | neutrophil.5 |
| Vav1 | 8.91E-143 | 0.27273988 | 0.176 | 0.374 | 1.72E-138 | 4 | Vav1 | neutrophil.5 |
| Pcf11 | 1.22E-142 | 0.37429882 | 0.226 | 0.431 | 2.36E-138 | 4 | Pcf11 | neutrophil.5 |
| Cd63 | 5.67E-142 | 0.49359551 | 0.228 | 0.396 | 1.10E-137 | 4 | Cd63 | neutrophil.5 |
| Gpr84 | 9.75E-142 | 0.45806388 | 0.177 | 0.277 | 1.89E-137 | 4 | Gpr84 | neutrophil.5 |
| Skap2 | 2.10E-141 | 0.28359873 | 0.184 | 0.388 | 4.05E-137 | 4 | Skap2 | neutrophil.5 |
| Cd74 | 8.24E-141 | 0.50832631 | 0.25 | 0.387 | 1.59E-136 | 4 | Cd74 | neutrophil.5 |
| Fgd3 | 1.50E-139 | 0.3852327 | 0.186 | 0.36 | 2.90E-135 | 4 | Fgd3 | neutrophil.5 |
| Slc9a3r1 | 1.63E-136 | 0.49398606 | 0.224 | 0.408 | 3.15E-132 | 4 | Slc9a3r1 | neutrophil.5 |
| Tgfb1 | 1.18E-134 | 0.49000447 | 0.245 | 0.415 | 2.27E-130 | 4 | Tgfb1 | neutrophil.5 |
| Zdhhc18 | 4.16E-134 | 0.31312465 | 0.154 | 0.322 | 8.05E-130 | 4 | Zdhhc18 | neutrophil.5 |
| Pmaip1 | 1.49E-130 | 0.50936784 | 0.203 | 0.332 | 2.88E-126 | 4 | Pmaip1 | neutrophil.5 |
| Rgs3 | 2.75E-130 | 0.28369664 | 0.154 | 0.31 | 5.31E-126 | 4 | Rgs3 | neutrophil.5 |
| Gpsm3 | 7.36E-127 | 0.3875212 | 0.189 | 0.378 | 1.42E-122 | 4 | Gpsm3 | neutrophil.5 |
| Atox1 | 1.88E-124 | 0.38986979 | 0.164 | 0.334 | 3.64E-120 | 4 | Atox1 | neutrophil.5 |
| Tapbp | 1.69E-122 | 0.53685352 | 0.2 | 0.373 | 3.27E-118 | 4 | Tapbp | neutrophil.5 |
| Atp1a1 | 1.75E-120 | 0.44153196 | 0.205 | 0.357 | 3.38E-116 | 4 | Atp1a1 | neutrophil.5 |
| Mbd2 | 2.14E-120 | 0.48233166 | 0.193 | 0.364 | 4.13E-116 | 4 | Mbd2 | neutrophil.5 |
| Slc35b2 | 4.23E-119 | 0.76632878 | 0.291 | 0.406 | 8.17E-115 | 4 | Slc35b2 | neutrophil.5 |
| Ctss | 1.63E-118 | 0.26125453 | 0.138 | 0.29 | 3.14E-114 | 4 | Ctss | neutrophil.5 |
| Ffar2 | 2.14E-117 | 0.36241835 | 0.158 | 0.263 | 4.14E-113 | 4 | Ffar2 | neutrophil.5 |
| Pilra | 2.82E-117 | 0.25909044 | 0.148 | 0.314 | 5.45E-113 | 4 | Pilra | neutrophil.5 |
| Fbrs | 3.64E-117 | 0.4488021 | 0.229 | 0.41 | 7.04E-113 | 4 | Fbrs | neutrophil.5 |
| Ankrd33b | 1.18E-116 | 0.28041344 | 0.158 | 0.317 | 2.28E-112 | 4 | Ankrd33b | neutrophil.5 |
| Traf1 | 2.59E-116 | 0.88218881 | 0.252 | 0.311 | 5.00E-112 | 4 | Traf1 | neutrophil.5 |
| Cdc42 | 6.98E-115 | 0.26958228 | 0.142 | 0.331 | 1.35E-110 | 4 | Cdc42 | neutrophil.5 |
| Osgin1 | 1.51E-113 | 0.34405848 | 0.137 | 0.257 | 2.92E-109 | 4 | Osgin1 | neutrophil.5 |
| Ltb | 5.12E-113 | 0.45320073 | 0.155 | 0.285 | 9.91E-109 | 4 | Ltb | neutrophil.5 |
| Atp6v1b2 | 1.57E-110 | 0.28543783 | 0.137 | 0.296 | 3.04E-106 | 4 | Atp6v1b2 | neutrophil.5 |
| Rel | 2.37E-110 | 0.29171516 | 0.15 | 0.301 | 4.58E-106 | 4 | Rel | neutrophil.5 |
| Gna13 | 5.83E-110 | 0.69315286 | 0.25 | 0.393 | 1.13E-105 | 4 | Gna13 | neutrophil.5 |
| Vamp8 | 1.08E-109 | 0.31995118 | 0.148 | 0.317 | 2.08E-105 | 4 | Vamp8 | neutrophil.5 |
| Pitpna | 3.14E-109 | 0.30548088 | 0.133 | 0.309 | 6.07E-105 | 4 | Pitpna | neutrophil.5 |
| Clta | 4.71E-107 | 0.5213428 | 0.201 | 0.37 | 9.10E-103 | 4 | Clta | neutrophil.5 |
| Tnip1 | 2.06E-105 | 0.35126523 | 0.13 | 0.256 | 3.98E-101 | 4 | Tnip1 | neutrophil.5 |
| Gsap | 4.90E-105 | 0.27462924 | 0.119 | 0.275 | 9.47E-101 | 4 | Gsap | neutrophil.5 |
| Arrdc4 | 7.85E-105 | 0.43019334 | 0.134 | 0.273 | 1.52E-100 | 4 | Arrdc4 | neutrophil.5 |
| Arap1 | 1.27E-104 | 0.32118701 | 0.172 | 0.344 | 2.45E-100 | 4 | Arap1 | neutrophil.5 |
| Rabac1 | 3.63E-104 | 0.40396817 | 0.153 | 0.328 | 7.02E-100 | 4 | Rabac1 | neutrophil.5 |
| Ttyh3 | 2.26E-102 | 0.54471955 | 0.199 | 0.357 | 4.38E-98 | 4 | Ttyh3 | neutrophil.5 |
| Atp6v0a1 | 1.67E-100 | 0.43413475 | 0.138 | 0.273 | 3.22E-96 | 4 | Atp6v0a1 | neutrophil.5 |
| Dusp2 | 4.31E-97 | 1.00111604 | 0.312 | 0.387 | 8.34E-93 | 4 | Dusp2 | neutrophil.5 |
| Rhoa | 4.93E-94 | 0.45162821 | 0.153 | 0.32 | 9.54E-90 | 4 | Rhoa | neutrophil.5 |
| Hnrnpul1 | 8.77E-94 | 0.46913376 | 0.155 | 0.309 | 1.70E-89 | 4 | Hnrnpul1 | neutrophil.5 |
| Lrp10 | 1.28E-93 | 0.59917185 | 0.191 | 0.351 | 2.48E-89 | 4 | Lrp10 | neutrophil.5 |
| Arf1 | 6.76E-93 | 0.29560923 | 0.135 | 0.304 | 1.31E-88 | 4 | Arf1 | neutrophil.5 |
| Bri3 | 1.39E-92 | 0.3795736 | 0.142 | 0.277 | 2.69E-88 | 4 | Bri3 | neutrophil.5 |
| Chd4 | 1.34E-90 | 0.48117138 | 0.158 | 0.302 | 2.59E-86 | 4 | Chd4 | neutrophil.5 |
| Lasp1 | 1.87E-89 | 0.3012596 | 0.119 | 0.264 | 3.61E-85 | 4 | Lasp1 | neutrophil.5 |
| Brk1 | 2.73E-89 | 0.34321011 | 0.127 | 0.281 | 5.28E-85 | 4 | Brk1 | neutrophil.5 |
| Itgb1 | 6.99E-86 | 0.34125165 | 0.125 | 0.261 | 1.35E-81 | 4 | Itgb1 | neutrophil.5 |
| Cnppd1 | 3.68E-85 | 0.29918245 | 0.128 | 0.268 | 7.12E-81 | 4 | Cnppd1 | neutrophil.5 |
| Psd4 | 3.88E-83 | 0.3616945 | 0.133 | 0.268 | 7.50E-79 | 4 | Psd4 | neutrophil.5 |
| Rin3 | 1.62E-82 | 0.4631606 | 0.133 | 0.269 | 3.12E-78 | 4 | Rin3 | neutrophil.5 |
| Adam17 | 4.34E-81 | 0.45432778 | 0.129 | 0.258 | 8.40E-77 | 4 | Adam17 | neutrophil.5 |
| Fes | 1.02E-80 | 0.31815542 | 0.126 | 0.268 | 1.98E-76 | 4 | Fes | neutrophil.5 |
| Ccnl2 | 4.69E-80 | 0.36428995 | 0.135 | 0.283 | 9.06E-76 | 4 | Ccnl2 | neutrophil.5 |
| Cdc42se1 | 9.62E-80 | 0.32240554 | 0.116 | 0.259 | 1.86E-75 | 4 | Cdc42se1 | neutrophil.5 |
| Ywhag | 1.67E-79 | 0.76363639 | 0.213 | 0.335 | 3.23E-75 | 4 | Ywhag | neutrophil.5 |
| Map7d1 | 1.92E-79 | 0.61017132 | 0.166 | 0.304 | 3.71E-75 | 4 | Map7d1 | neutrophil.5 |
| Sh2b2 | 6.37E-79 | 0.7637037 | 0.179 | 0.291 | 1.23E-74 | 4 | Sh2b2 | neutrophil.5 |
| Ddx17 | 7.26E-78 | 0.4207115 | 0.158 | 0.304 | 1.40E-73 | 4 | Ddx17 | neutrophil.5 |
| Pdlim7 | 1.23E-77 | 0.69561348 | 0.202 | 0.338 | 2.37E-73 | 4 | Pdlim7 | neutrophil.5 |
| Chchd2 | 1.69E-75 | 0.75683225 | 0.187 | 0.32 | 3.27E-71 | 4 | Chchd2 | neutrophil.5 |
| Med13l | 8.05E-72 | 0.4287467 | 0.13 | 0.272 | 1.56E-67 | 4 | Med13l | neutrophil.5 |
| Pip5k1c | 4.18E-70 | 0.50370246 | 0.13 | 0.258 | 8.09E-66 | 4 | Pip5k1c | neutrophil.5 |
| Gnai2 | 9.91E-70 | 0.98238192 | 0.314 | 0.446 | 1.92E-65 | 4 | Gnai2 | neutrophil.5 |
| Mllt6 | 1.80E-69 | 0.63445202 | 0.158 | 0.256 | 3.49E-65 | 4 | Mllt6 | neutrophil.5 |
| Ppp1r9b | 9.41E-67 | 0.63943543 | 0.163 | 0.283 | 1.82E-62 | 4 | Ppp1r9b | neutrophil.5 |
| Gnb1 | 2.49E-66 | 0.51490494 | 0.132 | 0.26 | 4.81E-62 | 4 | Gnb1 | neutrophil.5 |
| Trir | 3.04E-66 | 0.62200704 | 0.156 | 0.292 | 5.89E-62 | 4 | Trir | neutrophil.5 |
| Dedd2 | 1.18E-64 | 0.75558044 | 0.176 | 0.272 | 2.28E-60 | 4 | Dedd2 | neutrophil.5 |
| Anp32a | 1.93E-64 | 0.52693141 | 0.16 | 0.288 | 3.74E-60 | 4 | Anp32a | neutrophil.5 |
| Slc39a1 | 1.11E-63 | 0.63180174 | 0.148 | 0.263 | 2.14E-59 | 4 | Slc39a1 | neutrophil.5 |
| Atf1 | 2.98E-57 | 0.80849747 | 0.162 | 0.267 | 5.76E-53 | 4 | Atf1 | neutrophil.5 |
| Col1a1 | 4.03E-54 | 0.65533565 | 0.255 | 0.37 | 7.79E-50 | 4 | Col1a1 | neutrophil.5 |
| Ikbke | 2.79E-37 | 1.25678616 | 0.264 | 0.284 | 5.39E-33 | 4 | Ikbke | neutrophil.5 |
| Col3a1 | 4.95E-34 | 0.69958438 | 0.181 | 0.271 | 9.56E-30 | 4 | Col3a1 | neutrophil.5 |
| Syne1 | 0 | 6.36996501 | 0.43 | 0.031 | 0 | 5 | Syne1 | neutrophil.6 |
| Jund | 0 | 0.34296909 | 0.387 | 0.753 | 0 | 5 | Jund | neutrophil.6 |
| Tpt1 | 0 | 0.29727523 | 0.243 | 0.6 | 0 | 5 | Tpt1 | neutrophil.6 |
| Rap1b | 0 | 0.27276792 | 0.246 | 0.599 | 0 | 5 | Rap1b | neutrophil.6 |
| Iqgap1 | 0 | 0.42884421 | 0.303 | 0.655 | 0 | 5 | Iqgap1 | neutrophil.6 |
| Gsr | 0 | 0.57827843 | 0.511 | 0.833 | 0 | 5 | Gsr | neutrophil.6 |
| Cd80 | 0 | 0.28047286 | 0.253 | 0.571 | 0 | 5 | Cd80 | neutrophil.6 |
| Vasp | 0 | 0.50930046 | 0.526 | 0.843 | 0 | 5 | Vasp | neutrophil.6 |
| Themis2 | 0 | 0.27932459 | 0.212 | 0.524 | 0 | 5 | Themis2 | neutrophil.6 |
| Csf2rb | 0 | 0.5768147 | 0.264 | 0.576 | 0 | 5 | Csf2rb | neutrophil.6 |
| Cd24a | 0 | 0.54708549 | 0.376 | 0.663 | 0 | 5 | Cd24a | neutrophil.6 |
| Skil | 0 | 0.71664349 | 0.311 | 0.591 | 0 | 5 | Skil | neutrophil.6 |
| Cotl1 | 0 | 0.8921201 | 0.368 | 0.637 | 0 | 5 | Cotl1 | neutrophil.6 |
| Eef1a1 | 0 | 0.77892328 | 0.333 | 0.587 | 0 | 5 | Eef1a1 | neutrophil.6 |
| Man1a | 0 | 5.41320525 | 0.292 | 0.039 | 0 | 5 | Man1a | neutrophil.6 |
| Basp1 | 0 | 0.27898373 | 0.305 | 0.548 | 0 | 5 | Basp1 | neutrophil.6 |
| Tgm2 | 0 | 0.78007705 | 0.508 | 0.745 | 0 | 5 | Tgm2 | neutrophil.6 |
| S100a6 | 0 | 0.73973668 | 0.308 | 0.515 | 0 | 5 | S100a6 | neutrophil.6 |
| Tmsb4x | 0 | 0.88149859 | 0.807 | 0.932 | 0 | 5 | Tmsb4x | neutrophil.6 |
| Thbs1 | 0 | 0.51437099 | 0.31 | 0.298 | 0 | 5 | Thbs1 | neutrophil.6 |
| Cfl1 | 1.90E-299 | 0.37154358 | 0.251 | 0.599 | 3.68E-295 | 5 | Cfl1 | neutrophil.6 |
| Ncf1 | 1.40E-273 | 1.17352373 | 0.444 | 0.643 | 2.70E-269 | 5 | Ncf1 | neutrophil.6 |
| Nabp1 | 2.94E-253 | 0.53693847 | 0.194 | 0.414 | 5.68E-249 | 5 | Nabp1 | neutrophil.6 |
| Cmip | 5.40E-252 | 0.30510659 | 0.203 | 0.527 | 1.04E-247 | 5 | Cmip | neutrophil.6 |
| Clu | 1.33E-247 | 1.77364924 | 0.578 | 0.566 | 2.57E-243 | 5 | Clu | neutrophil.6 |
| Ncf2 | 8.63E-239 | 1.11182478 | 0.419 | 0.609 | 1.67E-234 | 5 | Ncf2 | neutrophil.6 |
| Tut7 | 3.30E-238 | 0.4537235 | 0.194 | 0.472 | 6.38E-234 | 5 | Tut7 | neutrophil.6 |
| mt-Co2 | 2.29E-235 | 0.99784847 | 0.271 | 0.351 | 4.43E-231 | 5 | mt-Co2 | neutrophil.6 |
| Cd74 | 1.98E-230 | 0.41531312 | 0.18 | 0.391 | 3.84E-226 | 5 | Cd74 | neutrophil.6 |
| Tes | 3.35E-229 | 0.35298314 | 0.167 | 0.394 | 6.47E-225 | 5 | Tes | neutrophil.6 |
| Dhx40 | 2.95E-223 | 0.80825383 | 0.196 | 0.36 | 5.70E-219 | 5 | Dhx40 | neutrophil.6 |
| Rsrp1 | 3.39E-213 | 0.65339571 | 0.26 | 0.497 | 6.55E-209 | 5 | Rsrp1 | neutrophil.6 |
| Homer1 | 6.01E-205 | 0.69962696 | 0.203 | 0.365 | 1.16E-200 | 5 | Homer1 | neutrophil.6 |
| Antxr2 | 2.04E-203 | 1.3906379 | 0.461 | 0.555 | 3.94E-199 | 5 | Antxr2 | neutrophil.6 |
| BC005537 | 6.14E-203 | 0.7013357 | 0.235 | 0.469 | 1.19E-198 | 5 | BC005537 | neutrophil.6 |
| Rab8b | 1.90E-198 | 0.6972706 | 0.229 | 0.458 | 3.67E-194 | 5 | Rab8b | neutrophil.6 |
| Csf2ra | 2.53E-195 | 0.76169556 | 0.258 | 0.506 | 4.90E-191 | 5 | Csf2ra | neutrophil.6 |
| Mknk2 | 5.68E-189 | 0.48092632 | 0.187 | 0.446 | 1.10E-184 | 5 | Mknk2 | neutrophil.6 |
| Fosb | 2.48E-185 | 0.59594118 | 0.157 | 0.3 | 4.79E-181 | 5 | Fosb | neutrophil.6 |
| Ptbp3 | 2.56E-182 | 0.49875048 | 0.175 | 0.436 | 4.95E-178 | 5 | Ptbp3 | neutrophil.6 |
| Gnai2 | 2.05E-179 | 0.36446622 | 0.179 | 0.454 | 3.97E-175 | 5 | Gnai2 | neutrophil.6 |
| Lyst | 6.54E-178 | 0.40050442 | 0.141 | 0.368 | 1.27E-173 | 5 | Lyst | neutrophil.6 |
| Nsd3 | 1.54E-175 | 0.97631809 | 0.269 | 0.443 | 2.98E-171 | 5 | Nsd3 | neutrophil.6 |
| Clk1 | 2.76E-175 | 0.36069358 | 0.151 | 0.409 | 5.34E-171 | 5 | Clk1 | neutrophil.6 |
| Fus | 6.90E-175 | 0.44079967 | 0.172 | 0.442 | 1.33E-170 | 5 | Fus | neutrophil.6 |
| Tgfb1 | 2.45E-173 | 0.60155188 | 0.194 | 0.417 | 4.73E-169 | 5 | Tgfb1 | neutrophil.6 |
| Cd63 | 9.08E-170 | 0.63676409 | 0.195 | 0.397 | 1.76E-165 | 5 | Cd63 | neutrophil.6 |
| Pla2g7 | 2.79E-169 | 1.15239521 | 0.329 | 0.472 | 5.40E-165 | 5 | Pla2g7 | neutrophil.6 |
| Hspa8 | 2.82E-167 | 0.707292 | 0.227 | 0.435 | 5.45E-163 | 5 | Hspa8 | neutrophil.6 |
| Lamp2 | 2.77E-166 | 0.45486106 | 0.141 | 0.374 | 5.36E-162 | 5 | Lamp2 | neutrophil.6 |
| Son | 2.04E-165 | 0.44486035 | 0.164 | 0.417 | 3.94E-161 | 5 | Son | neutrophil.6 |

| | | | | | | | | |
|---|---|---|---|---|---|---|---|---|
| Nab1 | 2.30E-165 | 0.4874046 | 0.15 | 0.381 | 4.44E-161 | 5 | Nab1 | neutrophil.6 |
| Itm2b | 5.26E-165 | 0.69946869 | 0.211 | 0.444 | 1.02E-160 | 5 | Itm2b | neutrophil.6 |
| Nfkb2 | 5.96E-158 | 1.09273215 | 0.316 | 0.507 | 1.15E-153 | 5 | Nfkb2 | neutrophil.6 |
| Tln1 | 6.04E-157 | 0.47852239 | 0.159 | 0.4 | 1.17E-152 | 5 | Tln1 | neutrophil.6 |
| Gpr132 | 1.96E-154 | 0.37034617 | 0.146 | 0.333 | 3.79E-150 | 5 | Gpr132 | neutrophil.6 |
| H2-K1 | 4.75E-154 | 1.15164322 | 0.356 | 0.539 | 9.18E-150 | 5 | H2-K1 | neutrophil.6 |
| Vmp1 | 3.84E-152 | 0.76198259 | 0.211 | 0.412 | 7.43E-148 | 5 | Vmp1 | neutrophil.6 |
| Pcf11 | 2.50E-149 | 0.69515173 | 0.199 | 0.432 | 4.84E-145 | 5 | Pcf11 | neutrophil.6 |
| Pygl | 7.81E-149 | 0.63278784 | 0.18 | 0.395 | 1.51E-144 | 5 | Pygl | neutrophil.6 |
| Mboat7 | 3.20E-147 | 0.32894834 | 0.115 | 0.345 | 6.18E-143 | 5 | Mboat7 | neutrophil.6 |
| Lrrfip1 | 1.28E-145 | 0.55203549 | 0.148 | 0.37 | 2.48E-141 | 5 | Lrrfip1 | neutrophil.6 |
| Ell2 | 1.28E-145 | 0.85154694 | 0.187 | 0.363 | 2.48E-141 | 5 | Ell2 | neutrophil.6 |
| Arpc5 | 1.27E-144 | 0.56379398 | 0.156 | 0.377 | 2.46E-140 | 5 | Arpc5 | neutrophil.6 |
| Nfkbie | 2.27E-144 | 0.40063148 | 0.122 | 0.324 | 4.40E-140 | 5 | Nfkbie | neutrophil.6 |
| Rab44 | 4.80E-142 | 1.38743079 | 0.422 | 0.516 | 9.28E-138 | 5 | Rab44 | neutrophil.6 |
| Mbd2 | 1.64E-140 | 0.58023461 | 0.153 | 0.366 | 3.17E-136 | 5 | Mbd2 | neutrophil.6 |
| Spp1 | 2.32E-138 | 1.79000005 | 0.506 | 0.518 | 4.49E-134 | 5 | Spp1 | neutrophil.6 |
| Tra2a | 5.53E-138 | 0.57393892 | 0.143 | 0.333 | 1.07E-133 | 5 | Tra2a | neutrophil.6 |
| Irf2bp2 | 2.88E-137 | 0.88311331 | 0.273 | 0.481 | 5.58E-133 | 5 | Irf2bp2 | neutrophil.6 |
| Map2k3 | 1.66E-136 | 0.57774283 | 0.154 | 0.366 | 3.21E-132 | 5 | Map2k3 | neutrophil.6 |
| Srsf5 | 2.28E-134 | 0.80153521 | 0.222 | 0.445 | 4.40E-130 | 5 | Srsf5 | neutrophil.6 |
| Pdlim7 | 2.31E-133 | 0.58284046 | 0.137 | 0.341 | 4.46E-129 | 5 | Pdlim7 | neutrophil.6 |
| Vim | 1.00E-132 | 1.55794146 | 0.408 | 0.443 | 1.93E-128 | 5 | Vim | neutrophil.6 |
| Ldha | 1.15E-131 | 0.51244437 | 0.141 | 0.308 | 2.22E-127 | 5 | Ldha | neutrophil.6 |
| Ccdc88b | 4.33E-131 | 1.0463402 | 0.205 | 0.389 | 8.37E-127 | 5 | Ccdc88b | neutrophil.6 |
| Ppp1r10 | 5.74E-131 | 0.34784255 | 0.119 | 0.286 | 1.11E-126 | 5 | Ppp1r10 | neutrophil.6 |
| Gls | 2.56E-129 | 0.84051977 | 0.199 | 0.378 | 4.95E-125 | 5 | Gls | neutrophil.6 |
| Zeb2 | 2.55E-128 | 0.44199355 | 0.135 | 0.289 | 4.93E-124 | 5 | Zeb2 | neutrophil.6 |
| Cdc42 | 4.03E-128 | 0.38565838 | 0.108 | 0.332 | 7.79E-124 | 5 | Cdc42 | neutrophil.6 |
| Serinc3 | 4.80E-128 | 0.94125075 | 0.244 | 0.459 | 9.27E-124 | 5 | Serinc3 | neutrophil.6 |
| Dennd5a | 8.76E-128 | 0.28887506 | 0.104 | 0.32 | 1.69E-123 | 5 | Dennd5a | neutrophil.6 |
| Ski | 3.22E-126 | 1.0353334 | 0.244 | 0.429 | 6.23E-122 | 5 | Ski | neutrophil.6 |
| Adrb2 | 4.08E-126 | 0.90472645 | 0.213 | 0.371 | 7.88E-122 | 5 | Adrb2 | neutrophil.6 |
| Akt1 | 6.72E-125 | 0.54749025 | 0.128 | 0.332 | 1.30E-120 | 5 | Akt1 | neutrophil.6 |
| Prex1 | 1.61E-122 | 0.29596113 | 0.1 | 0.302 | 3.11E-118 | 5 | Prex1 | neutrophil.6 |
| Flna | 3.58E-122 | 0.87488063 | 0.17 | 0.328 | 6.93E-118 | 5 | Flna | neutrophil.6 |
| Id1 | 4.79E-121 | 0.32243773 | 0.102 | 0.255 | 9.26E-117 | 5 | Id1 | neutrophil.6 |
| Raf1 | 2.40E-119 | 0.53611542 | 0.131 | 0.287 | 4.63E-115 | 5 | Raf1 | neutrophil.6 |
| Bin2 | 4.44E-117 | 0.40663533 | 0.118 | 0.295 | 8.58E-113 | 5 | Bin2 | neutrophil.6 |
| Gsap | 3.86E-116 | 0.4497737 | 0.094 | 0.276 | 7.47E-112 | 5 | Gsap | neutrophil.6 |
| Furin | 1.03E-113 | 0.82982183 | 0.182 | 0.366 | 2.00E-109 | 5 | Furin | neutrophil.6 |
| Rlim | 1.31E-113 | 0.39877766 | 0.109 | 0.307 | 2.52E-109 | 5 | Rlim | neutrophil.6 |
| Hnrnpa2b1 | 6.80E-113 | 0.86381478 | 0.158 | 0.337 | 1.32E-108 | 5 | Hnrnpa2b1 | neutrophil.6 |
| Chd2 | 3.74E-109 | 0.70085978 | 0.136 | 0.324 | 7.23E-105 | 5 | Chd2 | neutrophil.6 |
| Cers6 | 7.96E-109 | 0.67702903 | 0.127 | 0.302 | 1.54E-104 | 5 | Cers6 | neutrophil.6 |
| Slc35b2 | 5.32E-108 | 1.15745415 | 0.261 | 0.407 | 1.03E-103 | 5 | Slc35b2 | neutrophil.6 |
| Ccnl2 | 1.07E-107 | 0.30223767 | 0.089 | 0.285 | 2.08E-103 | 5 | Ccnl2 | neutrophil.6 |
| Chchd2 | 3.36E-107 | 0.62219436 | 0.142 | 0.322 | 6.50E-103 | 5 | Chchd2 | neutrophil.6 |
| Itgb1 | 1.05E-106 | 0.26059434 | 0.087 | 0.263 | 2.03E-102 | 5 | Itgb1 | neutrophil.6 |
| Pkn1 | 1.51E-106 | 0.58523099 | 0.127 | 0.315 | 2.92E-102 | 5 | Pkn1 | neutrophil.6 |
| Rtn3 | 1.55E-105 | 0.29192858 | 0.08 | 0.266 | 3.00E-101 | 5 | Rtn3 | neutrophil.6 |
| Ago2 | 1.29E-104 | 0.27368603 | 0.086 | 0.261 | 2.49E-100 | 5 | Ago2 | neutrophil.6 |
| Gadd45a | 1.42E-104 | 0.89672611 | 0.192 | 0.288 | 2.75E-100 | 5 | Gadd45a | neutrophil.6 |
| Fbrs | 2.51E-104 | 1.07475847 | 0.224 | 0.409 | 4.85E-100 | 5 | Fbrs | neutrophil.6 |
| Eif4g2 | 3.14E-103 | 0.72858594 | 0.133 | 0.317 | 6.08E-99 | 5 | Eif4g2 | neutrophil.6 |
| Cux1 | 5.99E-103 | 0.52531013 | 0.091 | 0.263 | 1.16E-98 | 5 | Cux1 | neutrophil.6 |
| Ctnnb1 | 9.82E-100 | 0.99252149 | 0.176 | 0.339 | 1.90E-95 | 5 | Ctnnb1 | neutrophil.6 |
| Arhgdia | 1.07E-99 | 0.50485134 | 0.103 | 0.287 | 2.07E-95 | 5 | Arhgdia | neutrophil.6 |
| Rtn4 | 4.85E-99 | 0.4028346 | 0.078 | 0.254 | 9.38E-95 | 5 | Rtn4 | neutrophil.6 |
| Dock2 | 9.43E-99 | 0.50657827 | 0.107 | 0.281 | 1.82E-94 | 5 | Dock2 | neutrophil.6 |
| Calm2 | 5.57E-97 | 0.26509547 | 0.083 | 0.25 | 1.08E-92 | 5 | Calm2 | neutrophil.6 |
| Trir | 1.67E-96 | 0.58806147 | 0.115 | 0.294 | 3.23E-92 | 5 | Trir | neutrophil.6 |
| Wdr1 | 6.79E-96 | 0.45095627 | 0.083 | 0.254 | 1.31E-91 | 5 | Wdr1 | neutrophil.6 |
| Alas1 | 2.82E-94 | 1.09541633 | 0.14 | 0.282 | 5.44E-90 | 5 | Alas1 | neutrophil.6 |
| Chd4 | 1.10E-93 | 0.9129653 | 0.163 | 0.301 | 2.14E-89 | 5 | Chd4 | neutrophil.6 |
| Gnb1 | 3.60E-92 | 0.45186731 | 0.089 | 0.262 | 6.97E-88 | 5 | Gnb1 | neutrophil.6 |
| Psd4 | 3.74E-92 | 0.62172513 | 0.115 | 0.269 | 7.22E-88 | 5 | Psd4 | neutrophil.6 |
| Dennd4a | 5.03E-91 | 1.42173096 | 0.312 | 0.428 | 9.73E-87 | 5 | Dennd4a | neutrophil.6 |
| Slc25a3 | 9.14E-91 | 0.52195861 | 0.092 | 0.253 | 1.77E-86 | 5 | Slc25a3 | neutrophil.6 |
| Wdr26 | 5.78E-89 | 0.99315601 | 0.158 | 0.317 | 1.12E-84 | 5 | Wdr26 | neutrophil.6 |
| Ythdf3 | 1.45E-87 | 0.50239206 | 0.09 | 0.252 | 2.81E-83 | 5 | Ythdf3 | neutrophil.6 |
| Hck | 2.54E-87 | 1.01094937 | 0.173 | 0.311 | 4.90E-83 | 5 | Hck | neutrophil.6 |
| Rel | 3.90E-86 | 1.08127756 | 0.166 | 0.299 | 7.54E-82 | 5 | Rel | neutrophil.6 |
| Stk40 | 1.51E-84 | 0.73221442 | 0.098 | 0.26 | 2.92E-80 | 5 | Stk40 | neutrophil.6 |
| Tmod3 | 1.55E-84 | 0.71777928 | 0.113 | 0.272 | 2.99E-80 | 5 | Tmod3 | neutrophil.6 |
| Osbpl9 | 2.48E-81 | 1.05321156 | 0.152 | 0.28 | 4.80E-77 | 5 | Osbpl9 | neutrophil.6 |
| Atg2a | 5.75E-80 | 0.87340181 | 0.121 | 0.257 | 1.11E-75 | 5 | Atg2a | neutrophil.6 |
| Arap1 | 7.96E-80 | 1.05943185 | 0.189 | 0.342 | 1.54E-75 | 5 | Arap1 | neutrophil.6 |
| Kras | 2.65E-77 | 0.75755683 | 0.108 | 0.25 | 5.12E-73 | 5 | Kras | neutrophil.6 |
| Cdk9 | 1.17E-76 | 0.74293544 | 0.117 | 0.253 | 2.26E-72 | 5 | Cdk9 | neutrophil.6 |
| Eif4a2 | 8.71E-72 | 0.89315481 | 0.135 | 0.262 | 1.68E-67 | 5 | Eif4a2 | neutrophil.6 |
| Fes | 3.50E-71 | 0.83638218 | 0.127 | 0.267 | 6.77E-67 | 5 | Fes | neutrophil.6 |
| Irf5 | 9.40E-71 | 1.07549962 | 0.137 | 0.254 | 1.82E-66 | 5 | Irf5 | neutrophil.6 |
| Ankrd13a | 3.29E-69 | 0.92806318 | 0.114 | 0.251 | 6.36E-65 | 5 | Ankrd13a | neutrophil.6 |
| Neurl3 | 2.94E-67 | 1.69456707 | 0.536 | 0.597 | 5.68E-63 | 5 | Neurl3 | neutrophil.6 |
| Med13 | 4.49E-65 | 0.90168776 | 0.116 | 0.251 | 8.67E-61 | 5 | Med13 | neutrophil.6 |
| Kdm2a | 3.83E-63 | 1.4165337 | 0.24 | 0.369 | 7.40E-59 | 5 | Kdm2a | neutrophil.6 |
| Myh9 | 7.71E-62 | 1.11904767 | 0.145 | 0.275 | 1.49E-57 | 5 | Myh9 | neutrophil.6 |
| Srsf7 | 1.25E-60 | 1.02677368 | 0.154 | 0.261 | 2.42E-56 | 5 | Srsf7 | neutrophil.6 |
| Ddx17 | 6.23E-57 | 1.28368926 | 0.177 | 0.301 | 1.20E-52 | 5 | Ddx17 | neutrophil.6 |
| Ptpre | 3.17E-42 | 1.42019241 | 0.212 | 0.306 | 6.13E-38 | 5 | Ptpre | neutrophil.6 |
| Arhgap45 | 6.05E-33 | 1.45909646 | 0.175 | 0.259 | 1.17E-28 | 5 | Arhgap45 | neutrophil.6 |
| Cyp4f18 | 7.43E-32 | 1.7069937 | 0.296 | 0.303 | 1.44E-27 | 5 | Cyp4f18 | neutrophil.6 |
| Ankrd33b | 1.98E-28 | 1.60725033 | 0.252 | 0.309 | 3.82E-24 | 5 | Ankrd33b | neutrophil.6 |
| Anp32a | 1.70E-22 | 1.612735 | 0.217 | 0.283 | 3.28E-18 | 5 | Anp32a | neutrophil.6 |
| Ccl6 | 5.94E-07 | 1.90864413 | 0.266 | 0.203 | 0.01148006 | 5 | Ccl6 | neutrophil.6 |
| Col1a1 | 3.41E-05 | 2.0519922 | 0.357 | 0.362 | 0.6595955 | 5 | Col1a1 | neutrophil.6 |
| Adam8 | 2.52E-04 | 2.115737 | 0.35 | 0.288 | 1 | 5 | Adam8 | neutrophil.6 |
| Isg15 | 0 | 2.78995218 | 0.796 | 0.231 | 0 | 6 | Isg15 | neutrophil.7 |
| Gbp2 | 0 | 4.78905164 | 0.598 | 0.057 | 0 | 6 | Gbp2 | neutrophil.7 |
| Stat1 | 0 | 3.0965967 | 0.608 | 0.116 | 0 | 6 | Stat1 | neutrophil.7 |
| Irf7 | 0 | 2.73636409 | 0.517 | 0.107 | 0 | 6 | Irf7 | neutrophil.7 |
| Parp14 | 0 | 2.99878679 | 0.485 | 0.085 | 0 | 6 | Parp14 | neutrophil.7 |
| Igtp | 0 | 4.32315951 | 0.416 | 0.03 | 0 | 6 | Igtp | neutrophil.7 |
| Tap1 | 0 | 2.57386106 | 0.469 | 0.09 | 0 | 6 | Tap1 | neutrophil.7 |
| Rtp4 | 0 | 3.00091148 | 0.425 | 0.058 | 0 | 6 | Rtp4 | neutrophil.7 |
| Oasl2 | 0 | 1.00175643 | 0.718 | 0.356 | 0 | 6 | Oasl2 | neutrophil.7 |
| Oas3 | 0 | 3.27439608 | 0.414 | 0.052 | 0 | 6 | Oas3 | neutrophil.7 |
| Samhd1 | 0 | 1.38188688 | 0.821 | 0.46 | 0 | 6 | Samhd1 | neutrophil.7 |
| Irgm1 | 0 | 4.3330379 | 0.385 | 0.028 | 0 | 6 | Irgm1 | neutrophil.7 |
| Rsad2 | 0 | 4.53419286 | 0.383 | 0.032 | 0 | 6 | Rsad2 | neutrophil.7 |
| Ifi47 | 0 | 5.41416183 | 0.355 | 0.011 | 0 | 6 | Ifi47 | neutrophil.7 |
| Slfn5 | 0 | 3.76956091 | 0.381 | 0.04 | 0 | 6 | Slfn5 | neutrophil.7 |
| Trafd1 | 0 | 2.52518728 | 0.428 | 0.087 | 0 | 6 | Trafd1 | neutrophil.7 |
| Gbp5 | 0 | 5.17184591 | 0.358 | 0.018 | 0 | 6 | Gbp5 | neutrophil.7 |
| Ifitm3 | 0 | 2.30953472 | 0.461 | 0.123 | 0 | 6 | Ifitm3 | neutrophil.7 |
| Slfn4 | 0 | 2.21951572 | 0.695 | 0.362 | 0 | 6 | Slfn4 | neutrophil.7 |
| Rnf213 | 0 | 2.73967382 | 0.386 | 0.064 | 0 | 6 | Rnf213 | neutrophil.7 |
| Irf1 | 0 | 1.81629403 | 0.637 | 0.33 | 0 | 6 | Irf1 | neutrophil.7 |
| Isg20 | 0 | 2.51872956 | 0.387 | 0.085 | 0 | 6 | Isg20 | neutrophil.7 |
| B2m | 0 | 0.90155855 | 0.843 | 0.564 | 0 | 6 | B2m | neutrophil.7 |
| Ctss | 0 | 1.07953923 | 0.54 | 0.264 | 0 | 6 | Ctss | neutrophil.7 |
| Tapbp | 0 | 0.981981 | 0.62 | 0.346 | 0 | 6 | Tapbp | neutrophil.7 |
| Acod1 | 0 | 0.93917132 | 0.889 | 0.616 | 0 | 6 | Acod1 | neutrophil.7 |
| Ly6e | 0 | 1.09638223 | 0.629 | 0.356 | 0 | 6 | Ly6e | neutrophil.7 |
| Zbp1 | 0 | 3.33756888 | 0.304 | 0.032 | 0 | 6 | Zbp1 | neutrophil.7 |
| Nlrc5 | 0 | 2.63392232 | 0.322 | 0.052 | 0 | 6 | Nlrc5 | neutrophil.7 |
| Ifih1 | 0 | 2.73843378 | 0.315 | 0.048 | 0 | 6 | Ifih1 | neutrophil.7 |
| Cd274 | 0 | 0.88921172 | 0.704 | 0.441 | 0 | 6 | Cd274 | neutrophil.7 |
| Fgl2 | 0 | 0.70558898 | 0.736 | 0.475 | 0 | 6 | Fgl2 | neutrophil.7 |
| Helz2 | 0 | 2.05794258 | 0.345 | 0.084 | 0 | 6 | Helz2 | neutrophil.7 |
| Trim30a | 0 | 1.63359289 | 0.387 | 0.131 | 0 | 6 | Trim30a | neutrophil.7 |
| Psmb8 | 0 | 1.77799898 | 0.355 | 0.107 | 0 | 6 | Psmb8 | neutrophil.7 |
| Oas2 | 0 | 2.58827589 | 0.294 | 0.049 | 0 | 6 | Oas2 | neutrophil.7 |
| Tap2 | 0 | 1.43012066 | 0.412 | 0.167 | 0 | 6 | Tap2 | neutrophil.7 |
| H2-T24 | 0 | 2.88034518 | 0.284 | 0.039 | 0 | 6 | H2-T24 | neutrophil.7 |
| Gbp7 | 0 | 3.16501735 | 0.278 | 0.034 | 0 | 6 | Gbp7 | neutrophil.7 |
| Irgm2 | 0 | 3.25277646 | 0.27 | 0.03 | 0 | 6 | Irgm2 | neutrophil.7 |
| Nampt | 0 | 0.98617953 | 0.477 | 0.237 | 0 | 6 | Nampt | neutrophil.7 |
| Dtx3l | 0 | 2.8168443 | 0.272 | 0.038 | 0 | 6 | Dtx3l | neutrophil.7 |
| Clec2d | 0 | 0.81626534 | 0.568 | 0.336 | 0 | 6 | Clec2d | neutrophil.7 |
| Sell | 0 | 0.83779418 | 0.547 | 0.318 | 0 | 6 | Sell | neutrophil.7 |
| Plac8 | 0 | 1.11931878 | 0.445 | 0.219 | 0 | 6 | Plac8 | neutrophil.7 |
| Sp100 | 0 | 1.97400676 | 0.299 | 0.075 | 0 | 6 | Sp100 | neutrophil.7 |
| Slc15a3 | 0 | 0.56679679 | 0.872 | 0.649 | 0 | 6 | Slc15a3 | neutrophil.7 |
| Fcgr4 | 0 | 1.48974567 | 0.35 | 0.128 | 0 | 6 | Fcgr4 | neutrophil.7 |
| Psmb9 | 0 | 2.11342178 | 0.272 | 0.062 | 0 | 6 | Psmb9 | neutrophil.7 |
| Tlr2 | 0 | 0.86888149 | 0.469 | 0.261 | 0 | 6 | Tlr2 | neutrophil.7 |
| H2-K1 | 0 | 0.40657827 | 0.724 | 0.516 | 0 | 6 | H2-K1 | neutrophil.7 |
| Tnfrsf1a | 0 | 0.64145886 | 0.748 | 0.542 | 0 | 6 | Tnfrsf1a | neutrophil.7 |
| Hdc | 0 | 0.45864723 | 0.823 | 0.618 | 0 | 6 | Hdc | neutrophil.7 |
| Parp10 | 0 | 1.77438648 | 0.274 | 0.07 | 0 | 6 | Parp10 | neutrophil.7 |
| Slfn2 | 0 | 0.43251254 | 0.741 | 0.544 | 0 | 6 | Slfn2 | neutrophil.7 |
| Smox | 0 | 0.50159193 | 0.671 | 0.476 | 0 | 6 | Smox | neutrophil.7 |
| Pde4b | 0 | 0.5268239 | 0.631 | 0.44 | 0 | 6 | Pde4b | neutrophil.7 |
| Grina | 0 | 0.43355463 | 0.862 | 0.672 | 0 | 6 | Grina | neutrophil.7 |
| Ifitm1 | 0 | 0.42847661 | 0.807 | 0.624 | 0 | 6 | Ifitm1 | neutrophil.7 |
| Csrnp1 | 0 | 0.31316327 | 0.868 | 0.696 | 0 | 6 | Csrnp1 | neutrophil.7 |
| Tnfaip2 | 0 | 0.75440006 | 0.904 | 0.736 | 0 | 6 | Tnfaip2 | neutrophil.7 |
| Socs3 | 0 | 0.44890376 | 0.715 | 0.549 | 0 | 6 | Socs3 | neutrophil.7 |
| Hcar2 | 0 | 0.49135213 | 0.823 | 0.66 | 0 | 6 | Hcar2 | neutrophil.7 |
| Ptgs2 | 0 | 0.44975782 | 0.787 | 0.625 | 0 | 6 | Ptgs2 | neutrophil.7 |
| Mxd1 | 0 | 0.49015418 | 0.909 | 0.748 | 0 | 6 | Mxd1 | neutrophil.7 |
| Ccrl2 | 0 | 0.5143299 | 0.79 | 0.633 | 0 | 6 | Ccrl2 | neutrophil.7 |
| Osm | 0 | 0.6040247 | 0.468 | 0.312 | 0 | 6 | Osm | neutrophil.7 |
| S100a11 | 0 | 0.31919254 | 0.829 | 0.676 | 0 | 6 | S100a11 | neutrophil.7 |
| Ifitm2 | 0 | 0.29631664 | 0.836 | 0.684 | 0 | 6 | Ifitm2 | neutrophil.7 |
| Clec4d | 0 | 0.29857445 | 0.875 | 0.736 | 0 | 6 | Clec4d | neutrophil.7 |
| Nfkbiz | 0 | 0.28458838 | 0.813 | 0.674 | 0 | 6 | Nfkbiz | neutrophil.7 |
| Ets2 | 0 | 0.30526006 | 0.93 | 0.795 | 0 | 6 | Ets2 | neutrophil.7 |
| Ppp1r15a | 0 | 0.25257958 | 0.809 | 0.675 | 0 | 6 | Ppp1r15a | neutrophil.7 |
| Egr1 | 0 | 0.27147393 | 0.876 | 0.745 | 0 | 6 | Egr1 | neutrophil.7 |
| Marcksl1 | 0 | 0.25642434 | 0.929 | 0.802 | 0 | 6 | Marcksl1 | neutrophil.7 |
| Cd14 | 0 | 0.29094234 | 0.822 | 0.701 | 0 | 6 | Cd14 | neutrophil.7 |
| Gadd45b | 0 | 0.25795461 | 0.892 | 0.775 | 0 | 6 | Gadd45b | neutrophil.7 |
| S100a9 | 0 | 0.27123183 | 0.709 | 0.603 | 0 | 6 | S100a9 | neutrophil.7 |
| Icam1 | 0 | 0.31896485 | 0.729 | 0.629 | 0 | 6 | Icam1 | neutrophil.7 |
| Ccl4 | 0 | 0.31427737 | 0.442 | 0.345 | 0 | 6 | Ccl4 | neutrophil.7 |
| Dusp1 | 0 | 0.30831726 | 0.933 | 0.845 | 0 | 6 | Dusp1 | neutrophil.7 |
| Zfp36 | 0 | 0.34385982 | 0.957 | 0.877 | 0 | 6 | Zfp36 | neutrophil.7 |
| Il1b | 0 | 0.36493015 | 0.995 | 0.926 | 0 | 6 | Il1b | neutrophil.7 |
| Srgn | 0 | 0.28185876 | 0.997 | 0.952 | 0 | 6 | Srgn | neutrophil.7 |
| Cd52 | 7.18E-296 | 0.41793902 | 0.756 | 0.562 | 1.39E-291 | 6 | Cd52 | neutrophil.7 |
| Entpd1 | 9.02E-295 | 0.42333876 | 0.682 | 0.487 | 1.74E-290 | 6 | Entpd1 | neutrophil.7 |
| Btg1 | 3.41E-294 | 0.28980287 | 0.837 | 0.673 | 6.59E-290 | 6 | Btg1 | neutrophil.7 |
| Socs1 | 7.35E-291 | 1.02496631 | 0.301 | 0.14 | 1.42E-286 | 6 | Socs1 | neutrophil.7 |
| Nr4a3 | 5.06E-289 | 0.303858 | 0.559 | 0.425 | 9.78E-285 | 6 | Nr4a3 | neutrophil.7 |
| Ccr1 | 2.48E-282 | 0.47168407 | 0.67 | 0.48 | 4.79E-278 | 6 | Ccr1 | neutrophil.7 |
| Slfn1 | 5.59E-274 | 1.25562913 | 0.352 | 0.15 | 1.08E-269 | 6 | Slfn1 | neutrophil.7 |
| Ptafr | 2.60E-267 | 0.35219945 | 0.765 | 0.59 | 5.03E-263 | 6 | Ptafr | neutrophil.7 |
| Trem1 | 5.27E-264 | 0.36300002 | 0.742 | 0.577 | 1.02E-259 | 6 | Trem1 | neutrophil.7 |
| Themis2 | 9.40E-257 | 0.39068968 | 0.703 | 0.493 | 1.82E-252 | 6 | Themis2 | neutrophil.7 |
| DEPRECATED-I | 9.35E-247 | 0.65411677 | 0.556 | 0.342 | 1.81E-242 | 6 | DEPRECATED-I | neutrophil.7 |
| Samsn1 | 3.33E-245 | 0.35739128 | 0.702 | 0.539 | 6.45E-241 | 6 | Samsn1 | neutrophil.7 |
| Atf3 | 3.14E-243 | 0.30519944 | 0.528 | 0.384 | 6.07E-239 | 6 | Atf3 | neutrophil.7 |
| Sp140 | 4.34E-218 | 0.98668132 | 0.348 | 0.157 | 8.38E-214 | 6 | Sp140 | neutrophil.7 |
| Cxcr2 | 1.09E-214 | 0.3901914 | 0.665 | 0.485 | 2.11E-210 | 6 | Cxcr2 | neutrophil.7 |
| Lmnb1 | 1.85E-214 | 0.37965508 | 0.649 | 0.464 | 3.58E-210 | 6 | Lmnb1 | neutrophil.7 |
| Trim30b | 3.74E-208 | 0.82314427 | 0.42 | 0.221 | 7.23E-204 | 6 | Trim30b | neutrophil.7 |
| Sdcbp | 1.72E-199 | 0.27137272 | 0.786 | 0.614 | 3.33E-195 | 6 | Sdcbp | neutrophil.7 |
| Vps37b | 3.20E-197 | 0.28326293 | 0.56 | 0.437 | 6.18E-193 | 6 | Vps37b | neutrophil.7 |
| Birc3 | 1.11E-196 | 0.35533088 | 0.597 | 0.435 | 2.15E-192 | 6 | Birc3 | neutrophil.7 |
| Rnf19b | 1.63E-195 | 0.37802424 | 0.613 | 0.441 | 3.15E-191 | 6 | Rnf19b | neutrophil.7 |
| Homer1 | 8.70E-195 | 0.33508154 | 0.486 | 0.347 | 1.68E-190 | 6 | Homer1 | neutrophil.7 |
| Casp4 | 1.61E-193 | 0.91765397 | 0.361 | 0.176 | 3.12E-189 | 6 | Casp4 | neutrophil.7 |
| Snx20 | 1.67E-189 | 0.281302 | 0.711 | 0.549 | 3.23E-185 | 6 | Snx20 | neutrophil.7 |
| H2-D1 | 1.53E-187 | 0.69142512 | 0.487 | 0.269 | 2.97E-183 | 6 | H2-D1 | neutrophil.7 |
| Tor1aip1 | 1.00E-182 | 0.93701453 | 0.339 | 0.17 | 1.94E-178 | 6 | Tor1aip1 | neutrophil.7 |
| Trib1 | 1.10E-181 | 0.28566808 | 0.685 | 0.529 | 2.12E-177 | 6 | Trib1 | neutrophil.7 |
| Rab7 | 2.12E-174 | 0.41946104 | 0.604 | 0.432 | 4.10E-170 | 6 | Rab7 | neutrophil.7 |
| Lsp1 | 2.21E-174 | 0.66825301 | 0.461 | 0.28 | 4.28E-170 | 6 | Lsp1 | neutrophil.7 |
| Psme1 | 2.20E-172 | 1.07563453 | 0.279 | 0.121 | 4.25E-168 | 6 | Psme1 | neutrophil.7 |
| Aff1 | 2.36E-172 | 0.33860769 | 0.629 | 0.459 | 4.56E-168 | 6 | Aff1 | neutrophil.7 |
| Ndel1 | 3.84E-168 | 0.2515306 | 0.643 | 0.494 | 7.42E-164 | 6 | Ndel1 | neutrophil.7 |
| Lcp2 | 3.30E-166 | 0.44709851 | 0.593 | 0.41 | 6.38E-162 | 6 | Lcp2 | neutrophil.7 |
| Zc3hav1 | 1.97E-157 | 0.64406829 | 0.395 | 0.215 | 3.80E-153 | 6 | Zc3hav1 | neutrophil.7 |
| Trim56 | 1.83E-154 | 0.98411074 | 0.273 | 0.115 | 3.54E-150 | 6 | Trim56 | neutrophil.7 |
| Ccdc88b | 5.11E-149 | 0.3176142 | 0.534 | 0.368 | 9.88E-145 | 6 | Ccdc88b | neutrophil.7 |
| Tmbim6 | 1.93E-145 | 0.42436808 | 0.603 | 0.422 | 3.74E-141 | 6 | Tmbim6 | neutrophil.7 |
| Fbxl5 | 1.49E-143 | 0.64345805 | 0.313 | 0.189 | 2.87E-139 | 6 | Fbxl5 | neutrophil.7 |
| Smchd1 | 1.29E-142 | 0.99468671 | 0.26 | 0.115 | 2.49E-138 | 6 | Smchd1 | neutrophil.7 |
| Gda | 7.97E-137 | 0.51327282 | 0.415 | 0.263 | 1.54E-132 | 6 | Gda | neutrophil.7 |
| Rabgef1 | 1.69E-133 | 0.27656124 | 0.579 | 0.43 | 3.28E-129 | 6 | Rabgef1 | neutrophil.7 |
| Usp25 | 2.79E-131 | 0.82618636 | 0.283 | 0.135 | 5.40E-127 | 6 | Usp25 | neutrophil.7 |
| Shisa5 | 8.06E-126 | 0.54724762 | 0.43 | 0.267 | 1.56E-121 | 6 | Shisa5 | neutrophil.7 |
| Trim30d | 2.87E-123 | 0.96825916 | 0.275 | 0.125 | 5.56E-119 | 6 | Trim30d | neutrophil.7 |
| Lrg1 | 2.52E-119 | 0.30316301 | 0.417 | 0.295 | 4.87E-115 | 6 | Lrg1 | neutrophil.7 |
| Cers6 | 1.14E-117 | 0.39601497 | 0.413 | 0.284 | 2.21E-113 | 6 | Cers6 | neutrophil.7 |
| Gsap | 3.42E-114 | 0.44482674 | 0.418 | 0.255 | 6.61E-110 | 6 | Gsap | neutrophil.7 |
| Sod2 | 1.60E-111 | 0.3378152 | 0.403 | 0.312 | 3.09E-107 | 6 | Sod2 | neutrophil.7 |
| Lamp2 | 2.06E-110 | 0.27581835 | 0.505 | 0.351 | 3.99E-106 | 6 | Lamp2 | neutrophil.7 |
| Fpr2 | 1.73E-107 | 0.35447829 | 0.455 | 0.336 | 3.34E-103 | 6 | Fpr2 | neutrophil.7 |
| Gk | 1.03E-104 | 0.66213352 | 0.272 | 0.147 | 1.99E-100 | 6 | Gk | neutrophil.7 |
| Egr3 | 1.06E-104 | 0.36149773 | 0.298 | 0.198 | 2.04E-100 | 6 | Egr3 | neutrophil.7 |
| Nod2 | 7.53E-103 | 0.54502693 | 0.325 | 0.189 | 1.46E-98 | 6 | Nod2 | neutrophil.7 |
| Tgoln1 | 8.63E-103 | 0.29537862 | 0.513 | 0.368 | 1.67E-98 | 6 | Tgoln1 | neutrophil.7 |
| Stat3 | 2.23E-102 | 0.27247532 | 0.535 | 0.374 | 4.31E-98 | 6 | Stat3 | neutrophil.7 |
| Myd88 | 1.31E-101 | 0.28900079 | 0.585 | 0.407 | 2.53E-97 | 6 | Myd88 | neutrophil.7 |
| Unc93b1 | 1.91E-100 | 0.59569054 | 0.357 | 0.208 | 3.69E-96 | 6 | Unc93b1 | neutrophil.7 |
| Asprv1 | 7.12E-98 | 0.29549076 | 0.321 | 0.24 | 1.38E-93 | 6 | Asprv1 | neutrophil.7 |
| Cd300ld | 1.19E-92 | 0.30976458 | 0.458 | 0.343 | 2.30E-88 | 6 | Cd300ld | neutrophil.7 |
| Cmtr1 | 1.17E-89 | 0.63144541 | 0.281 | 0.141 | 2.26E-85 | 6 | Cmtr1 | neutrophil.7 |
| Sbno2 | 2.25E-88 | 0.25557477 | 0.531 | 0.399 | 4.36E-84 | 6 | Sbno2 | neutrophil.7 |
| Peli1 | 3.96E-87 | 0.51569143 | 0.294 | 0.177 | 7.65E-83 | 6 | Peli1 | neutrophil.7 |
| Il1rap | 4.39E-85 | 0.52252443 | 0.358 | 0.218 | 8.49E-81 | 6 | Il1rap | neutrophil.7 |
| Crem | 2.35E-82 | 0.25239607 | 0.286 | 0.21 | 4.54E-78 | 6 | Crem | neutrophil.7 |
| Upp1 | 1.09E-80 | 0.27332245 | 0.371 | 0.26 | 2.10E-76 | 6 | Upp1 | neutrophil.7 |
| Pnp | 4.42E-80 | 0.41381744 | 0.352 | 0.225 | 8.55E-76 | 6 | Pnp | neutrophil.7 |
| Ccnyl1 | 3.76E-78 | 0.54292034 | 0.295 | 0.169 | 7.27E-74 | 6 | Ccnyl1 | neutrophil.7 |
| Wfdc17 | 5.78E-78 | 0.37464641 | 0.31 | 0.197 | 1.12E-73 | 6 | Wfdc17 | neutrophil.7 |
| Chmp4b | 4.86E-71 | 0.50549106 | 0.339 | 0.212 | 9.39E-67 | 6 | Chmp4b | neutrophil.7 |
| Rasip1 | 1.31E-67 | 0.48669312 | 0.272 | 0.165 | 2.53E-63 | 6 | Rasip1 | neutrophil.7 |
| Cd47 | 3.39E-67 | 0.25827604 | 0.411 | 0.283 | 6.56E-63 | 6 | Cd47 | neutrophil.7 |
| Bst1 | 6.77E-67 | 0.35465078 | 0.35 | 0.235 | 1.31E-62 | 6 | Bst1 | neutrophil.7 |
| Il13ra1 | 3.37E-66 | 0.32418603 | 0.404 | 0.283 | 6.51E-62 | 6 | Il13ra1 | neutrophil.7 |
| Rhoh | 2.15E-64 | 0.33326408 | 0.349 | 0.235 | 4.15E-60 | 6 | Rhoh | neutrophil.7 |
| Tspo | 1.35E-62 | 0.41406144 | 0.282 | 0.176 | 2.62E-58 | 6 | Tspo | neutrophil.7 |
| Glrx | 2.28E-62 | 0.45865007 | 0.269 | 0.171 | 4.41E-58 | 6 | Glrx | neutrophil.7 |
| Il18rap | 1.82E-60 | 0.39163249 | 0.331 | 0.211 | 3.52E-56 | 6 | Il18rap | neutrophil.7 |
| Coq10b | 2.61E-57 | 0.28663525 | 0.31 | 0.213 | 5.05E-53 | 6 | Coq10b | neutrophil.7 |
| Rab5c | 8.17E-55 | 0.25623306 | 0.407 | 0.289 | 1.58E-50 | 6 | Rab5c | neutrophil.7 |
| Selenon | 4.94E-54 | 0.32578359 | 0.307 | 0.21 | 9.54E-50 | 6 | Selenon | neutrophil.7 |
| Trim25 | 2.68E-53 | 0.26480919 | 0.296 | 0.186 | 5.17E-49 | 6 | Trim25 | neutrophil.7 |
| Emb | 1.24E-52 | 0.34461496 | 0.296 | 0.189 | 2.39E-48 | 6 | Emb | neutrophil.7 |
| Ilrun | 1.02E-49 | 0.4523226 | 0.264 | 0.16 | 1.97E-45 | 6 | Ilrun | neutrophil.7 |
| Jaml | 2.90E-48 | 0.3327711 | 0.254 | 0.18 | 5.60E-44 | 6 | Jaml | neutrophil.7 |
| Nod1 | 3.16E-48 | 0.37190231 | 0.265 | 0.171 | 6.12E-44 | 6 | Nod1 | neutrophil.7 |
| Zcchc2 | 5.01E-45 | 0.36495398 | 0.27 | 0.17 | 9.69E-41 | 6 | Zcchc2 | neutrophil.7 |
| Mcemp1 | 1.83E-43 | 0.36814942 | 0.271 | 0.19 | 3.55E-39 | 6 | Mcemp1 | neutrophil.7 |
| Lpcat2 | 1.74E-39 | 0.30674315 | 0.29 | 0.194 | 3.37E-35 | 6 | Lpcat2 | neutrophil.7 |
| Snap23 | 2.01E-36 | 0.33797644 | 0.282 | 0.183 | 3.89E-32 | 6 | Snap23 | neutrophil.7 |
| Tob1 | 0 | 2.30530282 | 0.781 | 0.345 | 0 | 7 | Tob1 | neutrophil.8 |
| Jun | 0 | 2.20192708 | 0.893 | 0.487 | 0 | 7 | Jun | neutrophil.8 |
| Dnajb9 | 0 | 2.1870835 | 0.587 | 0.206 | 0 | 7 | Dnajb9 | neutrophil.8 |
| Ints6 | 0 | 2.33951915 | 0.55 | 0.175 | 0 | 7 | Ints6 | neutrophil.8 |
| Hexim1 | 0 | 2.07417264 | 0.794 | 0.432 | 0 | 7 | Hexim1 | neutrophil.8 |
| Astl | 0 | 2.61561958 | 0.513 | 0.156 | 0 | 7 | Astl | neutrophil.8 |
| Ppp1r10 | 0 | 1.97801275 | 0.607 | 0.257 | 0 | 7 | Ppp1r10 | neutrophil.8 |
| Cited2 | 0 | 1.98228727 | 0.637 | 0.293 | 0 | 7 | Cited2 | neutrophil.8 |
| Plekhg2 | 0 | 2.90653241 | 0.423 | 0.085 | 0 | 7 | Plekhg2 | neutrophil.8 |
| Iqcn | 0 | 2.97709627 | 0.373 | 0.053 | 0 | 7 | Iqcn | neutrophil.8 |
| Id2 | 0 | 2.37391566 | 0.451 | 0.15 | 0 | 7 | Id2 | neutrophil.8 |
| Srsf7 | 0 | 1.79935977 | 0.532 | 0.239 | 0 | 7 | Srsf7 | neutrophil.8 |
| Nabp1 | 0 | 1.49942955 | 0.675 | 0.384 | 0 | 7 | Nabp1 | neutrophil.8 |
| Pcna | 0 | 1.706669 | 0.52 | 0.242 | 0 | 7 | Pcna | neutrophil.8 |
| Adrb2 | 0 | 1.58239097 | 0.624 | 0.346 | 0 | 7 | Adrb2 | neutrophil.8 |
| Tubb4b | 0 | 2.00608805 | 0.453 | 0.178 | 0 | 7 | Tubb4b | neutrophil.8 |
| Adra2b | 0 | 2.45962497 | 0.346 | 0.085 | 0 | 7 | Adra2b | neutrophil.8 |
| Dhx40 | 0 | 1.50963407 | 0.587 | 0.336 | 0 | 7 | Dhx40 | neutrophil.8 |
| Pnp | 0 | 1.65604128 | 0.468 | 0.219 | 0 | 7 | Pnp | neutrophil.8 |
| Kctd12 | 0 | 1.35935572 | 0.867 | 0.623 | 0 | 7 | Kctd12 | neutrophil.8 |
| BC005537 | 0 | 1.32027105 | 0.683 | 0.441 | 0 | 7 | BC005537 | neutrophil.8 |
| Rsrp1 | 0 | 1.24644037 | 0.708 | 0.469 | 0 | 7 | Rsrp1 | neutrophil.8 |
| Brd2 | 0 | 1.48415572 | 0.524 | 0.287 | 0 | 7 | Brd2 | neutrophil.8 |
| Gadd45g | 0 | 2.2516661 | 0.342 | 0.109 | 0 | 7 | Gadd45g | neutrophil.8 |
| Hbegf | 0 | 1.86469972 | 0.417 | 0.186 | 0 | 7 | Hbegf | neutrophil.8 |
| Rhob | 0 | 1.35558052 | 0.577 | 0.349 | 0 | 7 | Rhob | neutrophil.8 |
| Id1 | 0 | 1.50896105 | 0.458 | 0.233 | 0 | 7 | Id1 | neutrophil.8 |
| Egr1 | 0 | 1.63029108 | 0.965 | 0.74 | 0 | 7 | Egr1 | neutrophil.8 |
| Gstp3 | 0 | 2.00303432 | 0.346 | 0.128 | 0 | 7 | Gstp3 | neutrophil.8 |
| Polr2a | 0 | 1.45150224 | 0.525 | 0.311 | 0 | 7 | Polr2a | neutrophil.8 |
| Cbx4 | 0 | 1.90443522 | 0.356 | 0.146 | 0 | 7 | Cbx4 | neutrophil.8 |
| Tmem230 | 0 | 2.68054104 | 0.266 | 0.063 | 0 | 7 | Tmem230 | neutrophil.8 |
| Slc20a1 | 0 | 1.48367106 | 0.453 | 0.251 | 0 | 7 | Slc20a1 | neutrophil.8 |
| Polg2 | 0 | 2.3650508 | 0.253 | 0.061 | 0 | 7 | Polg2 | neutrophil.8 |
| Gadd45b | 0 | 0.78259439 | 0.942 | 0.773 | 0 | 7 | Gadd45b | neutrophil.8 |
| Ier3 | 0 | 1.03112178 | 0.953 | 0.828 | 0 | 7 | Ier3 | neutrophil.8 |
| Dusp1 | 0 | 0.66134808 | 0.955 | 0.844 | 0 | 7 | Dusp1 | neutrophil.8 |
| Zfp36 | 0 | 0.69096371 | 0.975 | 0.876 | 0 | 7 | Zfp36 | neutrophil.8 |
| Nfkbia | 0 | 0.63987229 | 0.959 | 0.878 | 0 | 7 | Nfkbia | neutrophil.8 |
| Txnip | 4.96E-291 | 0.97227888 | 0.672 | 0.464 | 9.60E-287 | 7 | Txnip | neutrophil.8 |
| Rnd1 | 7.57E-284 | 1.64443184 | 0.351 | 0.162 | 1.46E-279 | 7 | Rnd1 | neutrophil.8 |
| Sgk1 | 9.75E-284 | 1.79468385 | 0.317 | 0.134 | 1.88E-279 | 7 | Sgk1 | neutrophil.8 |
| Cebpd | 1.16E-274 | 1.29229004 | 0.525 | 0.292 | 2.24E-270 | 7 | Cebpd | neutrophil.8 |
| Srsf2 | 1.27E-272 | 1.74192096 | 0.365 | 0.151 | 2.46E-268 | 7 | Srsf2 | neutrophil.8 |
| Jund | 1.37E-268 | 0.83760755 | 0.85 | 0.723 | 2.64E-264 | 7 | Jund | neutrophil.8 |
| Gls | 5.72E-261 | 1.09958709 | 0.555 | 0.355 | 1.10E-256 | 7 | Gls | neutrophil.8 |
| Socs1 | 1.37E-259 | 1.72887576 | 0.336 | 0.138 | 2.66E-255 | 7 | Socs1 | neutrophil.8 |
| Icam4 | 3.26E-259 | 1.67221582 | 0.272 | 0.115 | 6.30E-255 | 7 | Icam4 | neutrophil.8 |
| Fos | 4.41E-226 | 0.76084224 | 0.758 | 0.553 | 8.52E-222 | 7 | Fos | neutrophil.8 |
| Ier2 | 5.26E-209 | 0.91002711 | 0.711 | 0.553 | 1.02E-204 | 7 | Ier2 | neutrophil.8 |
| Egr2 | 3.15E-203 | 1.56702262 | 0.309 | 0.153 | 6.09E-199 | 7 | Egr2 | neutrophil.8 |
| Sertad1 | 1.01E-188 | 1.02648132 | 0.641 | 0.462 | 1.94E-184 | 7 | Sertad1 | neutrophil.8 |
| Hsp90aa1 | 8.20E-182 | 1.34970905 | 0.385 | 0.194 | 1.59E-177 | 7 | Hsp90aa1 | neutrophil.8 |
| Mylip | 4.66E-181 | 1.58109337 | 0.317 | 0.147 | 9.00E-177 | 7 | Mylip | neutrophil.8 |
| H3f3b | 5.22E-180 | 0.90035288 | 0.731 | 0.581 | 1.01E-175 | 7 | H3f3b | neutrophil.8 |
| H2afz | 3.69E-179 | 1.69271539 | 0.294 | 0.123 | 7.13E-175 | 7 | H2afz | neutrophil.8 |
| Tnf | 1.40E-178 | 0.50930566 | 0.759 | 0.581 | 2.70E-174 | 7 | Tnf | neutrophil.8 |
| Calm2 | 1.57E-176 | 1.31799362 | 0.422 | 0.229 | 3.04E-172 | 7 | Calm2 | neutrophil.8 |
| Eif4a2 | 6.59E-171 | 1.22660081 | 0.413 | 0.245 | 1.28E-166 | 7 | Eif4a2 | neutrophil.8 |
| Dusp2 | 2.02E-170 | 0.96196299 | 0.571 | 0.371 | 3.91E-166 | 7 | Dusp2 | neutrophil.8 |
| Slc7a5 | 3.97E-162 | 1.69251289 | 0.255 | 0.113 | 7.68E-158 | 7 | Slc7a5 | neutrophil.8 |
| Irs2 | 2.00E-159 | 1.38014535 | 0.379 | 0.192 | 3.87E-155 | 7 | Irs2 | neutrophil.8 |
| Phlda1 | 1.33E-158 | 1.35755498 | 0.271 | 0.172 | 2.57E-154 | 7 | Phlda1 | neutrophil.8 |
| Btg2 | 2.26E-157 | 0.59727643 | 0.96 | 0.835 | 4.36E-153 | 7 | Btg2 | neutrophil.8 |
| Ier5 | 1.77E-151 | 0.62858343 | 0.848 | 0.745 | 3.43E-147 | 7 | Ier5 | neutrophil.8 |
| Egln3 | 3.72E-150 | 0.84436591 | 0.538 | 0.392 | 7.19E-146 | 7 | Egln3 | neutrophil.8 |
| Fosb | 3.77E-142 | 1.04335598 | 0.417 | 0.283 | 7.28E-138 | 7 | Fosb | neutrophil.8 |
| Dnaja1 | 4.15E-136 | 1.28566452 | 0.329 | 0.183 | 8.02E-132 | 7 | Dnaja1 | neutrophil.8 |
| Dynll1 | 7.75E-136 | 1.43178892 | 0.274 | 0.139 | 1.50E-131 | 7 | Dynll1 | neutrophil.8 |
| Rgs2 | 1.55E-128 | 1.2163387 | 0.337 | 0.193 | 3.00E-124 | 7 | Rgs2 | neutrophil.8 |
| Egr3 | 6.54E-115 | 1.1116848 | 0.302 | 0.198 | 1.26E-110 | 7 | Egr3 | neutrophil.8 |
| Chka | 2.76E-109 | 1.13203163 | 0.273 | 0.157 | 5.34E-105 | 7 | Chka | neutrophil.8 |
| Syf2 | 6.04E-108 | 1.17337844 | 0.342 | 0.216 | 1.17E-103 | 7 | Syf2 | neutrophil.8 |
| Sap18b | 7.87E-97 | 1.41264915 | 0.289 | 0.131 | 1.52E-92 | 7 | Sap18b | neutrophil.8 |
| Ddit3 | 1.82E-91 | 1.14603358 | 0.349 | 0.211 | 3.53E-87 | 7 | Ddit3 | neutrophil.8 |
| Jpt1 | 2.89E-90 | 0.88879495 | 0.358 | 0.224 | 5.58E-86 | 7 | Jpt1 | neutrophil.8 |
| Fus | 2.98E-87 | 0.76962084 | 0.544 | 0.418 | 5.77E-83 | 7 | Fus | neutrophil.8 |
| Atf3 | 1.41E-74 | 0.71150978 | 0.478 | 0.387 | 2.72E-70 | 7 | Atf3 | neutrophil.8 |
| Gpx1 | 6.17E-74 | 0.64079374 | 0.567 | 0.476 | 1.19E-69 | 7 | Gpx1 | neutrophil.8 |
| Tmem88 | 3.78E-70 | 0.91633121 | 0.381 | 0.257 | 7.32E-66 | 7 | Tmem88 | neutrophil.8 |
| Irf1 | 5.06E-69 | 0.76840215 | 0.5 | 0.338 | 9.79E-65 | 7 | Irf1 | neutrophil.8 |
| Klf10 | 3.00E-63 | 1.07104139 | 0.264 | 0.161 | 5.80E-59 | 7 | Klf10 | neutrophil.8 |
| Rbm38 | 9.86E-63 | 0.87032706 | 0.414 | 0.296 | 1.91E-58 | 7 | Rbm38 | neutrophil.8 |
| Mat2a | 1.69E-62 | 0.71845531 | 0.295 | 0.206 | 3.26E-58 | 7 | Mat2a | neutrophil.8 |
| Sra1 | 6.04E-62 | 0.90694748 | 0.352 | 0.261 | 1.17E-57 | 7 | Sra1 | neutrophil.8 |
| Ifitm5 | 1.83E-61 | 0.95540364 | 0.267 | 0.174 | 3.54E-57 | 7 | Ifitm5 | neutrophil.8 |
| Icam1 | 6.54E-58 | 0.38617242 | 0.733 | 0.629 | 1.26E-53 | 7 | Icam1 | neutrophil.8 |
| Nufip2 | 9.87E-54 | 0.78490668 | 0.31 | 0.216 | 1.91E-49 | 7 | Nufip2 | neutrophil.8 |
| Ppp1r15a | 2.83E-51 | 0.53645008 | 0.76 | 0.677 | 5.47E-47 | 7 | Ppp1r15a | neutrophil.8 |
| Chd4 | 3.19E-51 | 0.67866978 | 0.376 | 0.287 | 6.17E-47 | 7 | Chd4 | neutrophil.8 |
| Zfp36l2 | 4.78E-49 | 0.62932739 | 0.439 | 0.336 | 9.25E-45 | 7 | Zfp36l2 | neutrophil.8 |
| Ubc | 3.06E-48 | 0.46638824 | 0.884 | 0.796 | 5.92E-44 | 7 | Ubc | neutrophil.8 |
| Calr | 1.02E-45 | 0.99157817 | 0.263 | 0.168 | 1.97E-41 | 7 | Calr | neutrophil.8 |
| Mmp9 | 1.91E-42 | 0.36765677 | 0.747 | 0.657 | 3.69E-38 | 7 | Mmp9 | neutrophil.8 |
| Srsf5 | 7.77E-39 | 0.58844564 | 0.511 | 0.426 | 1.50E-34 | 7 | Srsf5 | neutrophil.8 |
| Mknk2 | 2.86E-35 | 0.56030226 | 0.505 | 0.425 | 5.53E-31 | 7 | Mknk2 | neutrophil.8 |
| Ffar2 | 1.71E-34 | 0.60626859 | 0.355 | 0.25 | 3.31E-30 | 7 | Ffar2 | neutrophil.8 |
| Wsb1 | 1.81E-34 | 0.5339604 | 0.637 | 0.546 | 3.50E-30 | 7 | Wsb1 | neutrophil.8 |
| Junb | 5.02E-34 | 0.33857769 | 0.982 | 0.934 | 9.71E-30 | 7 | Junb | neutrophil.8 |
| Gla | 3.08E-32 | 0.77600077 | 0.339 | 0.243 | 5.95E-28 | 7 | Gla | neutrophil.8 |
| Lfng | 3.17E-31 | 0.56577167 | 0.496 | 0.386 | 6.13E-27 | 7 | Lfng | neutrophil.8 |
| Midn | 3.56E-31 | 0.54059155 | 0.516 | 0.419 | 6.88E-27 | 7 | Midn | neutrophil.8 |
| Rgs3 | 6.90E-31 | 0.59774522 | 0.378 | 0.294 | 1.33E-26 | 7 | Rgs3 | neutrophil.8 |
| Marcksl1 | 1.38E-30 | 0.39545004 | 0.913 | 0.803 | 2.66E-26 | 7 | Marcksl1 | neutrophil.8 |
| Chchd2 | 4.57E-29 | 0.56937186 | 0.364 | 0.308 | 8.83E-25 | 7 | Chchd2 | neutrophil.8 |
| Cd24a | 4.68E-29 | 0.34755466 | 0.701 | 0.641 | 9.04E-25 | 7 | Cd24a | neutrophil.8 |
| Sat1 | 1.29E-28 | 0.46000428 | 0.538 | 0.503 | 2.49E-24 | 7 | Sat1 | neutrophil.8 |
| Pmaip1 | 1.64E-24 | 0.46970808 | 0.381 | 0.319 | 3.18E-20 | 7 | Pmaip1 | neutrophil.8 |
| Hspa8 | 3.87E-24 | 0.48697117 | 0.517 | 0.416 | 7.48E-20 | 7 | Hspa8 | neutrophil.8 |
| Dedd2 | 9.36E-24 | 0.61121388 | 0.34 | 0.261 | 1.81E-19 | 7 | Dedd2 | neutrophil.8 |
| Ctnnb1 | 6.38E-23 | 0.51806586 | 0.404 | 0.324 | 1.23E-18 | 7 | Ctnnb1 | neutrophil.8 |
| Sptan1 | 4.04E-22 | 0.58883873 | 0.295 | 0.231 | 7.82E-18 | 7 | Sptan1 | neutrophil.8 |
| Smcr8 | 2.24E-21 | 0.72491575 | 0.267 | 0.195 | 4.33E-17 | 7 | Smcr8 | neutrophil.8 |
| Cnppd1 | 2.25E-21 | 0.60828188 | 0.317 | 0.255 | 4.35E-17 | 7 | Cnppd1 | neutrophil.8 |
| mt-Co2 | 3.23E-21 | 0.52336246 | 0.48 | 0.338 | 6.25E-17 | 7 | mt-Co2 | neutrophil.8 |
| Calm1 | 6.54E-20 | 0.542437 | 0.421 | 0.347 | 1.26E-15 | 7 | Calm1 | neutrophil.8 |
| Hspa5 | 2.17E-18 | 0.56183634 | 0.469 | 0.379 | 4.19E-14 | 7 | Hspa5 | neutrophil.8 |
| H2-K1 | 8.58E-17 | 0.34171907 | 0.583 | 0.523 | 1.66E-12 | 7 | H2-K1 | neutrophil.8 |
| Oasl2 | 1.42E-16 | 0.51348934 | 0.384 | 0.375 | 2.74E-12 | 7 | Oasl2 | neutrophil.8 |
| Camkk1 | 1.13E-14 | 0.51854506 | 0.258 | 0.191 | 2.18E-10 | 7 | Camkk1 | neutrophil.8 |
| Vsir | 1.38E-14 | 0.37210025 | 0.516 | 0.448 | 2.66E-10 | 7 | Vsir | neutrophil.8 |
| Slc38a2 | 5.88E-14 | 0.42129801 | 0.533 | 0.466 | 1.14E-09 | 7 | Slc38a2 | neutrophil.8 |
| mt-Cytb | 9.29E-14 | 0.57763476 | 0.298 | 0.193 | 1.80E-09 | 7 | mt-Cytb | neutrophil.8 |
| Ddx5 | 9.29E-13 | 0.45225156 | 0.492 | 0.44 | 1.80E-08 | 7 | Ddx5 | neutrophil.8 |
| Irf2bp2 | 9.71E-13 | 0.31379334 | 0.548 | 0.463 | 1.88E-08 | 7 | Irf2bp2 | neutrophil.8 |
| Atp6v0a1 | 1.10E-12 | 0.54048827 | 0.349 | 0.258 | 2.14E-08 | 7 | Atp6v0a1 | neutrophil.8 |
| mt-Co3 | 4.56E-12 | 0.56612993 | 0.296 | 0.189 | 8.82E-08 | 7 | mt-Co3 | neutrophil.8 |
| Lrg1 | 9.36E-12 | 0.4819885 | 0.383 | 0.297 | 1.81E-07 | 7 | Lrg1 | neutrophil.8 |
| Cdk2ap2 | 2.77E-11 | 0.38477292 | 0.513 | 0.448 | 5.36E-07 | 7 | Cdk2ap2 | neutrophil.8 |
| Eif4a1 | 1.02E-10 | 0.57166943 | 0.263 | 0.214 | 1.97E-06 | 7 | Eif4a1 | neutrophil.8 |
| P4hb | 1.25E-10 | 0.47725679 | 0.357 | 0.302 | 2.43E-06 | 7 | P4hb | neutrophil.8 |
| Lst1 | 1.79E-10 | 0.39520671 | 0.446 | 0.432 | 3.46E-06 | 7 | Lst1 | neutrophil.8 |
| Pstpip1 | 2.29E-10 | 0.43906126 | 0.309 | 0.255 | 4.42E-06 | 7 | Pstpip1 | neutrophil.8 |
| Arhgap45 | 2.47E-10 | 0.32726261 | 0.303 | 0.25 | 4.78E-06 | 7 | Arhgap45 | neutrophil.8 |
| Klf2 | 2.51E-10 | 0.40977519 | 0.596 | 0.527 | 4.86E-06 | 7 | Klf2 | neutrophil.8 |
| Fes | 2.57E-10 | 0.43874734 | 0.313 | 0.255 | 4.96E-06 | 7 | Fes | neutrophil.8 |
| Homer1 | 3.43E-10 | 0.37526521 | 0.389 | 0.352 | 6.62E-06 | 7 | Homer1 | neutrophil.8 |
| Ucp2 | 5.30E-10 | 0.40751236 | 0.563 | 0.493 | 1.02E-05 | 7 | Ucp2 | neutrophil.8 |
| Rin3 | 8.56E-10 | 0.43997961 | 0.311 | 0.256 | 1.65E-05 | 7 | Rin3 | neutrophil.8 |
| mt-Atp6 | 1.43E-09 | 0.4378445 | 0.299 | 0.216 | 2.76E-05 | 7 | mt-Atp6 | neutrophil.8 |
| Rara | 6.27E-09 | 0.44785537 | 0.375 | 0.321 | 1.21E-04 | 7 | Rara | neutrophil.8 |
| Zfand5 | 1.78E-08 | 0.40890576 | 0.35 | 0.3 | 3.45E-04 | 7 | Zfand5 | neutrophil.8 |
| Cdk9 | 8.82E-08 | 0.37310469 | 0.278 | 0.242 | 0.00170601 | 7 | Cdk9 | neutrophil.8 |
| Ppp1r9b | 1.76E-07 | 0.36913217 | 0.321 | 0.271 | 0.00339566 | 7 | Ppp1r9b | neutrophil.8 |
| Il4ra | 5.87E-07 | 0.31274924 | 0.558 | 0.497 | 0.01134394 | 7 | Il4ra | neutrophil.8 |
| Hnrnpa2b1 | 3.24E-06 | 0.36967473 | 0.354 | 0.323 | 0.06271128 | 7 | Hnrnpa2b1 | neutrophil.8 |
| Col4a3bp | 4.77E-06 | 0.38909249 | 0.259 | 0.214 | 0.09225521 | 7 | Col4a3bp | neutrophil.8 |
| mt-Atp8 | 6.51E-06 | 0.44031795 | 0.269 | 0.195 | 0.12588265 | 7 | mt-Atp8 | neutrophil.8 |
| Tgfbi | 3.48E-05 | 0.34171523 | 0.407 | 0.357 | 0.67235482 | 7 | Tgfbi | neutrophil.8 |
| Fgl2 | 7.28E-05 | 0.32862598 | 0.558 | 0.485 | 1 | 7 | Fgl2 | neutrophil.8 |
| Cmtm6 | 1.20E-04 | 0.33565744 | 0.468 | 0.421 | 1 | 7 | Cmtm6 | neutrophil.8 |
| Il17ra | 2.24E-04 | 0.39072534 | 0.284 | 0.247 | 1 | 7 | Il17ra | neutrophil.8 |
| Cdkn1a | 4.22E-04 | 0.27719274 | 0.707 | 0.62 | 1 | 7 | Cdkn1a | neutrophil.8 |
| Vegfa | 6.13E-04 | 0.31818018 | 0.321 | 0.233 | 1 | 7 | Vegfa | neutrophil.8 |
| Cfp | 0.00104634 | 0.33188185 | 0.308 | 0.273 | 1 | 7 | Cfp | neutrophil.8 |
| Cd164 | 0.00180739 | 0.28052929 | 0.333 | 0.306 | 1 | 7 | Cd164 | neutrophil.8 |
| Tnfrsf23 | 0.00236392 | 0.3908414 | 0.255 | 0.225 | 1 | 7 | Tnfrsf23 | neutrophil.8 |
| Wipf1 | 0.00377412 | 0.29315108 | 0.369 | 0.339 | 1 | 7 | Wipf1 | neutrophil.8 |
| Cd68 | 0.00753104 | 0.39678764 | 0.269 | 0.235 | 1 | 7 | Cd68 | neutrophil.8 |
| Ero1l | 0 | 3.57978758 | 0.861 | 0.222 | 0 | 8 | Ero1l | neutrophil.9 |
| Hilpda | 0 | 3.34255243 | 0.795 | 0.177 | 0 | 8 | Hilpda | neutrophil.9 |
| Ldha | 0 | 2.51457669 | 0.803 | 0.276 | 0 | 8 | Ldha | neutrophil.9 |
| Hk2 | 0 | 3.03126762 | 0.679 | 0.171 | 0 | 8 | Hk2 | neutrophil.9 |
| Vegfa | 0 | 2.45012278 | 0.691 | 0.219 | 0 | 8 | Vegfa | neutrophil.9 |
| Tpi1 | 0 | 2.81375118 | 0.59 | 0.13 | 0 | 8 | Tpi1 | neutrophil.9 |
| Egln3 | 0 | 1.81445927 | 0.818 | 0.383 | 0 | 8 | Egln3 | neutrophil.9 |
| Gpi1 | 0 | 2.88821422 | 0.559 | 0.132 | 0 | 8 | Gpi1 | neutrophil.9 |
| Mif | 0 | 2.62845272 | 0.496 | 0.109 | 0 | 8 | Mif | neutrophil.9 |
| Higd1a | 0 | 3.33739131 | 0.431 | 0.065 | 0 | 8 | Higd1a | neutrophil.9 |
| Egln1 | 0 | 2.91875369 | 0.441 | 0.082 | 0 | 8 | Egln1 | neutrophil.9 |
| Basp1 | 0 | 1.28439114 | 0.875 | 0.518 | 0 | 8 | Basp1 | neutrophil.9 |
| P4ha1 | 0 | 2.557972 | 0.435 | 0.103 | 0 | 8 | P4ha1 | neutrophil.9 |
| Pgam1 | 0 | 1.76744109 | 0.502 | 0.173 | 0 | 8 | Pgam1 | neutrophil.9 |
| Fnip2 | 0 | 2.78888003 | 0.433 | 0.105 | 0 | 8 | Fnip2 | neutrophil.9 |
| Pfkl | 0 | 2.3318712 | 0.432 | 0.114 | 0 | 8 | Pfkl | neutrophil.9 |
| Aldoa | 0 | 1.39526956 | 0.691 | 0.381 | 0 | 8 | Aldoa | neutrophil.9 |
| Alkbh5 | 0 | 2.00555934 | 0.47 | 0.164 | 0 | 8 | Alkbh5 | neutrophil.9 |
| Slc2a1 | 0 | 1.41721076 | 0.772 | 0.472 | 0 | 8 | Slc2a1 | neutrophil.9 |
| Pgk1 | 0 | 1.89805369 | 0.442 | 0.144 | 0 | 8 | Pgk1 | neutrophil.9 |
| Pfkp | 0 | 1.78339517 | 0.487 | 0.191 | 0 | 8 | Pfkp | neutrophil.9 |
| Tmem189 | 0 | 1.97841622 | 0.51 | 0.218 | 0 | 8 | Tmem189 | neutrophil.9 |
| Hmox1 | 0 | 1.79730831 | 0.537 | 0.248 | 0 | 8 | Hmox1 | neutrophil.9 |
| P4hb | 0 | 1.41468381 | 0.58 | 0.294 | 0 | 8 | P4hb | neutrophil.9 |
| Hsp90aa1 | 0 | 2.31128327 | 0.477 | 0.193 | 0 | 8 | Hsp90aa1 | neutrophil.9 |
| Npepps | 0 | 2.02094791 | 0.405 | 0.123 | 0 | 8 | Npepps | neutrophil.9 |
| Ifrd1 | 0 | 1.20807839 | 0.807 | 0.532 | 0 | 8 | Ifrd1 | neutrophil.9 |
| F10 | 0 | 1.48796528 | 0.456 | 0.183 | 0 | 8 | F10 | neutrophil.9 |
| Atp6v1a | 0 | 1.68890486 | 0.484 | 0.211 | 0 | 8 | Atp6v1a | neutrophil.9 |
| Gna13 | 0 | 1.25441028 | 0.639 | 0.372 | 0 | 8 | Gna13 | neutrophil.9 |
| Jun | 0 | 1.64365715 | 0.763 | 0.498 | 0 | 8 | Jun | neutrophil.9 |
| Pkm | 0 | 1.12874922 | 0.753 | 0.493 | 0 | 8 | Pkm | neutrophil.9 |
| Bsg | 0 | 2.14278583 | 0.351 | 0.099 | 0 | 8 | Bsg | neutrophil.9 |
| Gsn | 0 | 1.23418624 | 0.751 | 0.506 | 0 | 8 | Gsn | neutrophil.9 |
| Fam162a | 0 | 2.85969536 | 0.292 | 0.048 | 0 | 8 | Fam162a | neutrophil.9 |
| Fndc3a | 0 | 2.2572827 | 0.347 | 0.104 | 0 | 8 | Fndc3a | neutrophil.9 |
| Dedd2 | 0 | 1.45377596 | 0.496 | 0.256 | 0 | 8 | Dedd2 | neutrophil.9 |
| Dnajb9 | 0 | 1.50680609 | 0.45 | 0.216 | 0 | 8 | Dnajb9 | neutrophil.9 |
| Hcar2 | 0 | 0.78354426 | 0.885 | 0.659 | 0 | 8 | Hcar2 | neutrophil.9 |
| Atp6v1c1 | 0 | 2.01479282 | 0.314 | 0.101 | 0 | 8 | Atp6v1c1 | neutrophil.9 |
| Rhob | 0 | 1.20964776 | 0.564 | 0.352 | 0 | 8 | Rhob | neutrophil.9 |
| Gys1 | 0 | 2.56428038 | 0.259 | 0.048 | 0 | 8 | Gys1 | neutrophil.9 |
| Atf3 | 0 | 1.24933777 | 0.591 | 0.384 | 0 | 8 | Atf3 | neutrophil.9 |
| Ccrl2 | 0 | 0.83245341 | 0.837 | 0.633 | 0 | 8 | Ccrl2 | neutrophil.9 |
| Zfp36l2 | 0 | 1.41997045 | 0.537 | 0.334 | 0 | 8 | Zfp36l2 | neutrophil.9 |
| Cited2 | 0 | 1.3915485 | 0.503 | 0.303 | 0 | 8 | Cited2 | neutrophil.9 |
| Pfkfb3 | 0 | 2.03418715 | 0.291 | 0.093 | 0 | 8 | Pfkfb3 | neutrophil.9 |
| Ndrg1 | 0 | 2.02062759 | 0.31 | 0.113 | 0 | 8 | Ndrg1 | neutrophil.9 |
| Ccl3 | 0 | 1.16020029 | 0.792 | 0.6 | 0 | 8 | Ccl3 | neutrophil.9 |
| Ddit3 | 0 | 1.51521443 | 0.401 | 0.211 | 0 | 8 | Ddit3 | neutrophil.9 |
| Sdcbp | 0 | 0.93252355 | 0.8 | 0.616 | 0 | 8 | Sdcbp | neutrophil.9 |
| Dnaja1 | 0 | 1.6576566 | 0.367 | 0.183 | 0 | 8 | Dnaja1 | neutrophil.9 |
| Ddit4 | 0 | 2.26404296 | 0.271 | 0.091 | 0 | 8 | Ddit4 | neutrophil.9 |
| Egr1 | 0 | 0.87884927 | 0.904 | 0.746 | 0 | 8 | Egr1 | neutrophil.9 |
| Dusp5 | 0 | 0.72332208 | 0.801 | 0.652 | 0 | 8 | Dusp5 | neutrophil.9 |
| Cxcl2 | 0 | 0.29081267 | 0.916 | 0.887 | 0 | 8 | Cxcl2 | neutrophil.9 |
| Nfkbia | 0 | 0.32399977 | 0.884 | 0.882 | 0 | 8 | Nfkbia | neutrophil.9 |
| Soat1 | 7.43E-301 | 2.0002062 | 0.334 | 0.114 | 1.44E-296 | 8 | Soat1 | neutrophil.9 |

| | | | | | | | | |
|---|---|---|---|---|---|---|---|---|
| Mbnl2 | 1.17E-300 | 1.48360947 | 0.479 | 0.224 | 2.27E-296 | 8 | Mbnl2 | neutrophil.9 |
| Tob1 | 8.33E-299 | 1.02188794 | 0.539 | 0.361 | 1.61E-294 | 8 | Tob1 | neutrophil.9 |
| Cstb | 8.74E-299 | 1.08745441 | 0.66 | 0.445 | 1.69E-294 | 8 | Cstb | neutrophil.9 |
| Mreg | 6.81E-283 | 1.71231912 | 0.333 | 0.13 | 1.32E-278 | 8 | Mreg | neutrophil.9 |
| Tiparp | 1.20E-274 | 1.29675887 | 0.464 | 0.258 | 2.32E-270 | 8 | Tiparp | neutrophil.9 |
| Lgals3 | 7.85E-269 | 1.2874874 | 0.481 | 0.26 | 1.52E-264 | 8 | Lgals3 | neutrophil.9 |
| Cdkn1a | 1.52E-261 | 0.51137489 | 0.82 | 0.617 | 2.95E-257 | 8 | Cdkn1a | neutrophil.9 |
| Osbpl8 | 2.32E-254 | 2.00203876 | 0.264 | 0.094 | 4.50E-250 | 8 | Osbpl8 | neutrophil.9 |
| Rgcc | 8.47E-246 | 1.12610794 | 0.541 | 0.289 | 1.64E-241 | 8 | Rgcc | neutrophil.9 |
| Phlda1 | 1.96E-243 | 1.3730332 | 0.331 | 0.171 | 3.80E-239 | 8 | Phlda1 | neutrophil.9 |
| Tarm1 | 1.64E-242 | 1.11809952 | 0.423 | 0.243 | 3.17E-238 | 8 | Tarm1 | neutrophil.9 |
| Nabp1 | 3.21E-240 | 0.84507531 | 0.59 | 0.391 | 6.21E-236 | 8 | Nabp1 | neutrophil.9 |
| Kdm3a | 1.37E-239 | 1.68419262 | 0.354 | 0.13 | 2.65E-235 | 8 | Kdm3a | neutrophil.9 |
| Gla | 1.80E-238 | 1.3913438 | 0.459 | 0.239 | 3.48E-234 | 8 | Gla | neutrophil.9 |
| Fau | 4.03E-238 | 0.93672112 | 0.763 | 0.537 | 7.79E-234 | 8 | Fau | neutrophil.9 |
| Id2 | 1.82E-237 | 1.33260374 | 0.302 | 0.16 | 3.52E-233 | 8 | Id2 | neutrophil.9 |
| Polr2a | 3.51E-229 | 1.17900869 | 0.494 | 0.315 | 6.78E-225 | 8 | Polr2a | neutrophil.9 |
| Hexim1 | 1.43E-227 | 0.86286779 | 0.595 | 0.445 | 2.77E-223 | 8 | Hexim1 | neutrophil.9 |
| Maff | 4.59E-225 | 0.76482898 | 0.773 | 0.567 | 8.88E-221 | 8 | Maff | neutrophil.9 |
| Hspa8 | 1.16E-223 | 0.95880378 | 0.57 | 0.416 | 2.24E-219 | 8 | Hspa8 | neutrophil.9 |
| Hbegf | 2.15E-220 | 1.18745408 | 0.354 | 0.192 | 4.16E-216 | 8 | Hbegf | neutrophil.9 |
| Bhlhe40 | 2.63E-207 | 0.76483952 | 0.723 | 0.551 | 5.09E-203 | 8 | Bhlhe40 | neutrophil.9 |
| Irs2 | 4.95E-205 | 1.18271621 | 0.388 | 0.195 | 9.57E-201 | 8 | Irs2 | neutrophil.9 |
| Suco | 9.56E-198 | 1.5516184 | 0.332 | 0.149 | 1.85E-193 | 8 | Suco | neutrophil.9 |
| Fth1 | 7.13E-192 | 0.39465476 | 0.966 | 0.934 | 1.38E-187 | 8 | Fth1 | neutrophil.9 |
| Ier3 | 1.27E-186 | 0.36195837 | 0.894 | 0.832 | 2.46E-182 | 8 | Ier3 | neutrophil.9 |
| H3f3b | 2.18E-185 | 0.72775109 | 0.733 | 0.582 | 4.21E-181 | 8 | H3f3b | neutrophil.9 |
| Ctsl | 8.47E-180 | 1.5411834 | 0.284 | 0.132 | 1.64E-175 | 8 | Ctsl | neutrophil.9 |
| Ccl4 | 1.51E-178 | 0.57569114 | 0.433 | 0.346 | 2.91E-174 | 8 | Ccl4 | neutrophil.9 |
| Tbc1d15 | 3.61E-173 | 1.59565923 | 0.334 | 0.155 | 6.99E-169 | 8 | Tbc1d15 | neutrophil.9 |
| Srsf7 | 1.04E-167 | 0.97773525 | 0.432 | 0.247 | 2.01E-163 | 8 | Srsf7 | neutrophil.9 |
| Zfp292 | 3.62E-165 | 1.58757881 | 0.344 | 0.139 | 7.00E-161 | 8 | Zfp292 | neutrophil.9 |
| Ralgds | 4.48E-161 | 1.11930878 | 0.361 | 0.206 | 8.65E-157 | 8 | Ralgds | neutrophil.9 |
| Kctd12 | 2.90E-160 | 0.45800376 | 0.665 | 0.635 | 5.61E-156 | 8 | Kctd12 | neutrophil.9 |
| Ints6 | 4.15E-157 | 0.95391477 | 0.334 | 0.188 | 8.03E-153 | 8 | Ints6 | neutrophil.9 |
| Arrb1 | 3.89E-155 | 0.9935057 | 0.336 | 0.163 | 7.52E-151 | 8 | Arrb1 | neutrophil.9 |
| Tatdn2 | 6.19E-154 | 1.07412097 | 0.41 | 0.237 | 1.20E-149 | 8 | Tatdn2 | neutrophil.9 |
| Hk1 | 1.41E-153 | 1.30671439 | 0.331 | 0.152 | 2.73E-149 | 8 | Hk1 | neutrophil.9 |
| BC005537 | 7.32E-153 | 0.65065725 | 0.572 | 0.449 | 1.42E-148 | 8 | BC005537 | neutrophil.9 |
| B4galt1 | 3.29E-152 | 1.05805333 | 0.446 | 0.245 | 6.35E-148 | 8 | B4galt1 | neutrophil.9 |
| Fos | 1.09E-151 | 0.39918611 | 0.696 | 0.558 | 2.11E-147 | 8 | Fos | neutrophil.9 |
| Furin | 1.75E-144 | 0.84954458 | 0.512 | 0.347 | 3.38E-140 | 8 | Furin | neutrophil.9 |
| Pcna | 8.28E-141 | 0.9606123 | 0.43 | 0.249 | 1.60E-136 | 8 | Pcna | neutrophil.9 |
| Hspa5 | 4.55E-139 | 0.88133725 | 0.506 | 0.378 | 8.79E-135 | 8 | Hspa5 | neutrophil.9 |
| Ell | 2.17E-137 | 1.45673284 | 0.289 | 0.121 | 4.20E-133 | 8 | Ell | neutrophil.9 |
| Gns | 1.09E-131 | 1.40576295 | 0.255 | 0.105 | 2.10E-127 | 8 | Gns | neutrophil.9 |
| Tcirg1 | 1.35E-130 | 1.09159646 | 0.39 | 0.21 | 2.62E-126 | 8 | Tcirg1 | neutrophil.9 |
| Wsb1 | 1.31E-129 | 0.70972232 | 0.69 | 0.545 | 2.54E-125 | 8 | Wsb1 | neutrophil.9 |
| Slc3a2 | 3.50E-128 | 0.72658974 | 0.573 | 0.382 | 6.78E-124 | 8 | Slc3a2 | neutrophil.9 |
| Rnh1 | 1.13E-126 | 1.16825803 | 0.311 | 0.15 | 2.18E-122 | 8 | Rnh1 | neutrophil.9 |
| Ppp1r3b | 2.59E-126 | 1.03859527 | 0.524 | 0.277 | 5.00E-122 | 8 | Ppp1r3b | neutrophil.9 |
| Pik3ap1 | 1.80E-125 | 0.49778324 | 0.785 | 0.659 | 3.48E-121 | 8 | Pik3ap1 | neutrophil.9 |
| Tubb6 | 4.59E-125 | 1.43267934 | 0.253 | 0.093 | 8.88E-121 | 8 | Tubb6 | neutrophil.9 |
| Mafk | 1.55E-124 | 1.2586787 | 0.299 | 0.149 | 2.99E-120 | 8 | Mafk | neutrophil.9 |
| Nfkbiz | 1.41E-121 | 0.38726846 | 0.763 | 0.678 | 2.73E-117 | 8 | Nfkbiz | neutrophil.9 |
| Atp6v0c | 4.64E-121 | 0.60426772 | 0.77 | 0.614 | 8.97E-117 | 8 | Atp6v0c | neutrophil.9 |
| Jund | 4.35E-120 | 0.43074407 | 0.824 | 0.725 | 8.41E-116 | 8 | Jund | neutrophil.9 |
| Sh2d3c | 1.09E-119 | 1.13426708 | 0.263 | 0.153 | 2.11E-115 | 8 | Sh2d3c | neutrophil.9 |
| Il1rn | 8.87E-119 | 0.26200439 | 0.897 | 0.84 | 1.72E-114 | 8 | Il1rn | neutrophil.9 |
| Acly | 2.62E-117 | 0.9535641 | 0.271 | 0.166 | 5.07E-113 | 8 | Acly | neutrophil.9 |
| Ptgs2 | 1.72E-116 | 0.28424107 | 0.71 | 0.63 | 3.33E-112 | 8 | Ptgs2 | neutrophil.9 |
| Crem | 7.15E-113 | 0.53350642 | 0.281 | 0.211 | 1.38E-108 | 8 | Crem | neutrophil.9 |
| Upp1 | 3.82E-112 | 1.02642656 | 0.407 | 0.26 | 7.38E-108 | 8 | Upp1 | neutrophil.9 |
| Nek6 | 1.47E-105 | 1.06468719 | 0.352 | 0.188 | 2.85E-101 | 8 | Nek6 | neutrophil.9 |
| Sap18b | 1.10E-103 | 1.28288378 | 0.287 | 0.134 | 2.13E-99 | 8 | Sap18b | neutrophil.9 |
| Usf2 | 3.59E-103 | 1.00517121 | 0.358 | 0.195 | 6.94E-99 | 8 | Usf2 | neutrophil.9 |
| Fnip1 | 3.72E-103 | 1.33465109 | 0.274 | 0.124 | 7.19E-99 | 8 | Fnip1 | neutrophil.9 |
| Zfand5 | 1.05E-100 | 0.91967334 | 0.43 | 0.298 | 2.04E-96 | 8 | Zfand5 | neutrophil.9 |
| Sh2b2 | 3.80E-97 | 0.85704408 | 0.426 | 0.277 | 7.34E-93 | 8 | Sh2b2 | neutrophil.9 |
| Thbs1 | 6.73E-96 | 0.32159973 | 0.36 | 0.296 | 1.30E-91 | 8 | Thbs1 | neutrophil.9 |
| Mknk2 | 2.68E-93 | 0.57218018 | 0.575 | 0.423 | 5.18E-89 | 8 | Mknk2 | neutrophil.9 |
| Zeb2 | 7.27E-89 | 0.85615046 | 0.402 | 0.274 | 1.41E-84 | 8 | Zeb2 | neutrophil.9 |
| Trim25 | 1.25E-84 | 0.80065899 | 0.311 | 0.187 | 2.41E-80 | 8 | Trim25 | neutrophil.9 |
| Mdm2 | 1.88E-84 | 0.92578758 | 0.324 | 0.217 | 3.64E-80 | 8 | Mdm2 | neutrophil.9 |
| Dnajc5 | 5.68E-83 | 1.04195352 | 0.285 | 0.147 | 1.10E-78 | 8 | Dnajc5 | neutrophil.9 |
| Pnpla7 | 3.31E-81 | 1.03910355 | 0.271 | 0.143 | 6.41E-77 | 8 | Pnpla7 | neutrophil.9 |
| M6pr | 3.59E-80 | 0.81640538 | 0.372 | 0.223 | 6.93E-76 | 8 | M6pr | neutrophil.9 |
| Atg2a | 1.30E-79 | 0.93595845 | 0.329 | 0.245 | 2.51E-75 | 8 | Atg2a | neutrophil.9 |
| Bcl2l1 | 5.99E-77 | 0.72448802 | 0.477 | 0.311 | 1.16E-72 | 8 | Bcl2l1 | neutrophil.9 |
| Ago2 | 5.07E-75 | 0.81571837 | 0.353 | 0.245 | 9.80E-71 | 8 | Ago2 | neutrophil.9 |
| Tnfrsf23 | 1.39E-74 | 0.97316254 | 0.357 | 0.221 | 2.69E-70 | 8 | Tnfrsf23 | neutrophil.9 |
| Sav1 | 1.39E-74 | 1.0145488 | 0.256 | 0.132 | 2.69E-70 | 8 | Sav1 | neutrophil.9 |
| Ubc | 9.71E-74 | 0.35360667 | 0.875 | 0.798 | 1.88E-69 | 8 | Ubc | neutrophil.9 |
| Plin2 | 7.19E-73 | 0.65247856 | 0.495 | 0.346 | 1.39E-68 | 8 | Plin2 | neutrophil.9 |
| Ctsb | 1.37E-72 | 0.51866027 | 0.719 | 0.567 | 2.65E-68 | 8 | Ctsb | neutrophil.9 |
| Calr | 2.14E-72 | 1.02876758 | 0.297 | 0.168 | 4.13E-68 | 8 | Calr | neutrophil.9 |
| Tgif1 | 3.13E-70 | 0.47796209 | 0.62 | 0.498 | 6.06E-66 | 8 | Tgif1 | neutrophil.9 |
| Cbx4 | 4.15E-70 | 0.94878093 | 0.26 | 0.153 | 8.03E-66 | 8 | Cbx4 | neutrophil.9 |
| Osbpl9 | 8.89E-68 | 0.48694796 | 0.334 | 0.269 | 1.72E-63 | 8 | Osbpl9 | neutrophil.9 |
| Tra2a | 1.10E-67 | 0.66244239 | 0.45 | 0.315 | 2.13E-63 | 8 | Tra2a | neutrophil.9 |
| Gadd45b | 1.30E-67 | 0.28263307 | 0.885 | 0.777 | 2.52E-63 | 8 | Gadd45b | neutrophil.9 |
| Hdlbp | 1.86E-65 | 0.99751147 | 0.256 | 0.137 | 3.59E-61 | 8 | Hdlbp | neutrophil.9 |
| Rbm38 | 2.73E-65 | 0.65030983 | 0.406 | 0.298 | 5.27E-61 | 8 | Rbm38 | neutrophil.9 |
| Ccnl1 | 4.64E-65 | 0.4391629 | 0.687 | 0.578 | 8.97E-61 | 8 | Ccnl1 | neutrophil.9 |
| Rsrc2 | 2.05E-64 | 0.96316008 | 0.292 | 0.174 | 3.97E-60 | 8 | Rsrc2 | neutrophil.9 |
| Irak2 | 2.33E-64 | 0.60081003 | 0.633 | 0.484 | 4.51E-60 | 8 | Irak2 | neutrophil.9 |
| Slbp | 1.55E-63 | 0.83003176 | 0.256 | 0.142 | 3.01E-59 | 8 | Slbp | neutrophil.9 |
| Vav1 | 3.39E-63 | 0.70625493 | 0.488 | 0.354 | 6.56E-59 | 8 | Vav1 | neutrophil.9 |
| Ranbp2 | 1.08E-62 | 0.96955755 | 0.265 | 0.145 | 2.08E-58 | 8 | Ranbp2 | neutrophil.9 |
| Eef1a1 | 2.94E-62 | 0.48189705 | 0.738 | 0.564 | 5.68E-58 | 8 | Eef1a1 | neutrophil.9 |
| Ostf1 | 5.85E-61 | 0.59774121 | 0.477 | 0.353 | 1.13E-56 | 8 | Ostf1 | neutrophil.9 |
| Eif4a2 | 6.89E-61 | 0.60816225 | 0.374 | 0.249 | 1.33E-56 | 8 | Eif4a2 | neutrophil.9 |
| Rbpj | 1.01E-60 | 0.78146957 | 0.359 | 0.238 | 1.95E-56 | 8 | Rbpj | neutrophil.9 |
| Cpeb4 | 1.75E-59 | 0.99167546 | 0.262 | 0.157 | 3.38E-55 | 8 | Cpeb4 | neutrophil.9 |
| Optn | 1.77E-59 | 0.92943588 | 0.265 | 0.147 | 3.43E-55 | 8 | Optn | neutrophil.9 |
| Ncoa4 | 2.12E-59 | 0.87350888 | 0.27 | 0.168 | 4.10E-55 | 8 | Ncoa4 | neutrophil.9 |
| Fcgr3 | 3.16E-58 | 0.47376031 | 0.599 | 0.483 | 6.11E-54 | 8 | Fcgr3 | neutrophil.9 |
| Rsrp1 | 3.51E-58 | 0.39792201 | 0.598 | 0.477 | 6.78E-54 | 8 | Rsrp1 | neutrophil.9 |
| Tuba1c | 2.53E-57 | 0.46760021 | 0.485 | 0.367 | 4.89E-53 | 8 | Tuba1c | neutrophil.9 |
| Tpt1 | 9.14E-57 | 0.49582049 | 0.718 | 0.571 | 1.77E-52 | 8 | Tpt1 | neutrophil.9 |
| Atp6v1d | 9.02E-56 | 0.91190205 | 0.257 | 0.146 | 1.74E-51 | 8 | Atp6v1d | neutrophil.9 |
| Srsf2 | 1.18E-55 | 0.71384122 | 0.263 | 0.158 | 2.29E-51 | 8 | Srsf2 | neutrophil.9 |
| Sra1 | 1.79E-55 | 0.62943263 | 0.397 | 0.261 | 3.45E-51 | 8 | Sra1 | neutrophil.9 |
| AA467197 | 2.18E-55 | 0.57918206 | 0.437 | 0.31 | 4.22E-51 | 8 | AA467197 | neutrophil.9 |
| Ppp1r15a | 2.92E-54 | 0.37848121 | 0.798 | 0.677 | 5.65E-50 | 8 | Ppp1r15a | neutrophil.9 |
| Cd63 | 1.05E-51 | 0.45522743 | 0.502 | 0.379 | 2.02E-47 | 8 | Cd63 | neutrophil.9 |
| Malt1 | 7.81E-48 | 0.70681372 | 0.281 | 0.191 | 1.51E-43 | 8 | Malt1 | neutrophil.9 |
| Bax | 1.12E-47 | 0.80314797 | 0.299 | 0.177 | 2.16E-43 | 8 | Bax | neutrophil.9 |
| Chchd2 | 6.33E-47 | 0.50032174 | 0.419 | 0.306 | 1.22E-42 | 8 | Chchd2 | neutrophil.9 |
| Calm2 | 1.03E-46 | 0.60496838 | 0.342 | 0.234 | 1.99E-42 | 8 | Calm2 | neutrophil.9 |
| Vsir | 5.33E-46 | 0.45248283 | 0.551 | 0.448 | 1.03E-41 | 8 | Vsir | neutrophil.9 |
| Brd2 | 5.45E-46 | 0.55319719 | 0.393 | 0.295 | 1.05E-41 | 8 | Brd2 | neutrophil.9 |
| Map3k8 | 6.78E-45 | 0.83539407 | 0.282 | 0.192 | 1.31E-40 | 8 | Map3k8 | neutrophil.9 |
| Gadd45a | 1.15E-43 | 0.50130395 | 0.347 | 0.279 | 2.23E-39 | 8 | Gadd45a | neutrophil.9 |
| Sertad1 | 3.51E-42 | 0.43184632 | 0.557 | 0.468 | 6.78E-38 | 8 | Sertad1 | neutrophil.9 |
| Plbd1 | 5.83E-42 | 0.76744521 | 0.284 | 0.221 | 1.13E-37 | 8 | Plbd1 | neutrophil.9 |
| Klf6 | 9.14E-42 | 0.40961965 | 0.55 | 0.47 | 1.77E-37 | 8 | Klf6 | neutrophil.9 |
| Mef2d | 3.44E-41 | 0.67002364 | 0.317 | 0.212 | 6.66E-37 | 8 | Mef2d | neutrophil.9 |
| Slc20a1 | 3.82E-41 | 0.51363789 | 0.335 | 0.259 | 7.39E-37 | 8 | Slc20a1 | neutrophil.9 |
| Chka | 4.94E-41 | 0.77562709 | 0.252 | 0.16 | 9.56E-37 | 8 | Chka | neutrophil.9 |
| Zmiz1 | 7.09E-40 | 0.77193322 | 0.279 | 0.176 | 1.37E-35 | 8 | Zmiz1 | neutrophil.9 |
| Safb2 | 4.88E-39 | 0.58162008 | 0.304 | 0.193 | 9.44E-35 | 8 | Safb2 | neutrophil.9 |
| Sat1 | 5.05E-39 | 0.4011113 | 0.549 | 0.503 | 9.77E-35 | 8 | Sat1 | neutrophil.9 |
| Atf4 | 2.75E-38 | 0.71747611 | 0.251 | 0.159 | 5.31E-34 | 8 | Atf4 | neutrophil.9 |
| Flot1 | 7.41E-38 | 0.55445039 | 0.295 | 0.194 | 1.43E-33 | 8 | Flot1 | neutrophil.9 |
| Gls | 4.74E-36 | 0.2827859 | 0.436 | 0.363 | 9.16E-32 | 8 | Gls | neutrophil.9 |
| Tcf25 | 7.99E-36 | 0.42563592 | 0.289 | 0.195 | 1.54E-31 | 8 | Tcf25 | neutrophil.9 |
| Ctsz | 3.41E-35 | 0.36747702 | 0.539 | 0.446 | 6.60E-31 | 8 | Ctsz | neutrophil.9 |
| Pnrc1 | 5.21E-34 | 0.36324691 | 0.788 | 0.676 | 1.01E-29 | 8 | Pnrc1 | neutrophil.9 |
| Psd4 | 3.83E-32 | 0.40261985 | 0.352 | 0.255 | 7.40E-28 | 8 | Psd4 | neutrophil.9 |
| Runx1 | 5.59E-32 | 0.60006541 | 0.273 | 0.186 | 1.08E-27 | 8 | Runx1 | neutrophil.9 |
| Ppp1r10 | 7.35E-31 | 0.33965209 | 0.349 | 0.272 | 1.42E-26 | 8 | Ppp1r10 | neutrophil.9 |
| Emilin2 | 4.10E-30 | 0.31123896 | 0.669 | 0.598 | 7.93E-26 | 8 | Emilin2 | neutrophil.9 |
| Tubb4b | 4.74E-30 | 0.48697149 | 0.266 | 0.189 | 9.16E-26 | 8 | Tubb4b | neutrophil.9 |
| Hnrnpf | 5.31E-30 | 0.56321448 | 0.251 | 0.168 | 1.03E-25 | 8 | Hnrnpf | neutrophil.9 |
| Ctnnb1 | 3.28E-29 | 0.38095242 | 0.438 | 0.324 | 6.33E-25 | 8 | Ctnnb1 | neutrophil.9 |
| Cd274 | 1.31E-28 | 0.46390829 | 0.542 | 0.452 | 2.53E-24 | 8 | Cd274 | neutrophil.9 |
| Dot1l | 1.63E-28 | 0.4662146 | 0.265 | 0.194 | 3.14E-24 | 8 | Dot1l | neutrophil.9 |
| Snx18 | 1.99E-28 | 0.50891996 | 0.396 | 0.294 | 3.85E-24 | 8 | Snx18 | neutrophil.9 |
| Fam49b | 2.03E-28 | 0.28438672 | 0.446 | 0.398 | 3.92E-24 | 8 | Fam49b | neutrophil.9 |
| Hnrnpa2b1 | 3.67E-28 | 0.30939784 | 0.41 | 0.322 | 7.10E-24 | 8 | Hnrnpa2b1 | neutrophil.9 |
| Jmjd1c | 1.46E-27 | 0.56381079 | 0.338 | 0.245 | 2.83E-23 | 8 | Jmjd1c | neutrophil.9 |
| Eif4g2 | 1.75E-27 | 0.43535493 | 0.416 | 0.3 | 3.39E-23 | 8 | Eif4g2 | neutrophil.9 |
| Il23a | 9.25E-27 | 0.25184112 | 0.261 | 0.225 | 1.79E-22 | 8 | Il23a | neutrophil.9 |
| Gsk3a | 1.43E-26 | 0.59715108 | 0.309 | 0.214 | 2.77E-22 | 8 | Gsk3a | neutrophil.9 |
| Siglecf | 4.23E-26 | 0.2890646 | 0.285 | 0.204 | 8.17E-22 | 8 | Siglecf | neutrophil.9 |
| Cd53 | 5.88E-26 | 0.33726759 | 0.756 | 0.691 | 1.14E-21 | 8 | Cd53 | neutrophil.9 |
| Slc39a1 | 6.74E-26 | 0.33833489 | 0.314 | 0.252 | 1.30E-21 | 8 | Slc39a1 | neutrophil.9 |
| Arf4 | 1.85E-25 | 0.44688281 | 0.275 | 0.207 | 3.58E-21 | 8 | Arf4 | neutrophil.9 |
| Psap | 1.93E-25 | 0.41638189 | 0.391 | 0.336 | 3.73E-21 | 8 | Psap | neutrophil.9 |
| Tax1bp1 | 3.28E-25 | 0.44471381 | 0.412 | 0.311 | 6.34E-21 | 8 | Tax1bp1 | neutrophil.9 |
| Atp6v1e1 | 4.23E-25 | 0.41692341 | 0.406 | 0.333 | 8.18E-21 | 8 | Atp6v1e1 | neutrophil.9 |
| Hnrnpu | 6.44E-25 | 0.42552386 | 0.281 | 0.195 | 1.25E-20 | 8 | Hnrnpu | neutrophil.9 |
| Dgkz | 1.32E-24 | 0.58203531 | 0.294 | 0.209 | 2.56E-20 | 8 | Dgkz | neutrophil.9 |
| Eif4a1 | 2.95E-24 | 0.44897203 | 0.308 | 0.213 | 5.70E-20 | 8 | Eif4a1 | neutrophil.9 |
| Srsf6 | 3.37E-24 | 0.26901671 | 0.265 | 0.185 | 6.52E-20 | 8 | Srsf6 | neutrophil.9 |
| Card19 | 5.24E-24 | 0.44701516 | 0.463 | 0.392 | 1.01E-19 | 8 | Card19 | neutrophil.9 |
| Map4k4 | 2.63E-23 | 0.51026817 | 0.257 | 0.18 | 5.08E-19 | 8 | Map4k4 | neutrophil.9 |
| Xylt1 | 3.27E-23 | 0.51613253 | 0.325 | 0.242 | 6.32E-19 | 8 | Xylt1 | neutrophil.9 |
| Spty2d1 | 4.63E-23 | 0.52228206 | 0.311 | 0.219 | 8.95E-19 | 8 | Spty2d1 | neutrophil.9 |
| Sin3b | 1.40E-22 | 0.5302398 | 0.274 | 0.204 | 2.72E-18 | 8 | Sin3b | neutrophil.9 |
| Rnasek | 3.58E-22 | 0.61072038 | 0.259 | 0.203 | 6.92E-18 | 8 | Rnasek | neutrophil.9 |
| Tgoln1 | 1.67E-21 | 0.47735461 | 0.474 | 0.371 | 3.22E-17 | 8 | Tgoln1 | neutrophil.9 |
| Arrdc4 | 4.22E-21 | 0.37952063 | 0.317 | 0.261 | 8.16E-17 | 8 | Arrdc4 | neutrophil.9 |
| Ctsa | 4.90E-20 | 0.49247117 | 0.304 | 0.236 | 9.48E-16 | 8 | Ctsa | neutrophil.9 |
| Clk1 | 3.76E-19 | 0.30674852 | 0.466 | 0.389 | 7.27E-15 | 8 | Clk1 | neutrophil.9 |
| Rabgef1 | 4.48E-19 | 0.25222561 | 0.517 | 0.435 | 8.65E-15 | 8 | Rabgef1 | neutrophil.9 |
| Ptbp3 | 1.94E-18 | 0.31325239 | 0.499 | 0.416 | 3.76E-14 | 8 | Ptbp3 | neutrophil.9 |
| Atxn2l | 9.13E-18 | 0.39246734 | 0.27 | 0.203 | 1.76E-13 | 8 | Atxn2l | neutrophil.9 |
| Adipor1 | 1.77E-17 | 0.27384489 | 0.659 | 0.584 | 3.42E-13 | 8 | Adipor1 | neutrophil.9 |
| Smcr8 | 5.88E-17 | 0.38276914 | 0.266 | 0.196 | 1.14E-12 | 8 | Smcr8 | neutrophil.9 |
| Cab39 | 6.02E-17 | 0.42373815 | 0.291 | 0.223 | 1.16E-12 | 8 | Cab39 | neutrophil.9 |
| Syf2 | 7.03E-17 | 0.2807518 | 0.287 | 0.22 | 1.36E-12 | 8 | Syf2 | neutrophil.9 |
| Per1 | 2.66E-16 | 0.26340292 | 0.467 | 0.411 | 5.15E-12 | 8 | Per1 | neutrophil.9 |
| Csf2rb2 | 2.74E-16 | 0.46115845 | 0.261 | 0.193 | 5.29E-12 | 8 | Csf2rb2 | neutrophil.9 |
| Atp6ap2 | 1.02E-15 | 0.36619289 | 0.389 | 0.302 | 1.97E-11 | 8 | Atp6ap2 | neutrophil.9 |
| Cnppd1 | 1.14E-15 | 0.35718234 | 0.307 | 0.256 | 2.20E-11 | 8 | Cnppd1 | neutrophil.9 |
| Edem1 | 3.75E-15 | 0.40388112 | 0.261 | 0.197 | 7.25E-11 | 8 | Edem1 | neutrophil.9 |
| Entpd3 | 6.33E-15 | 0.46766575 | 0.258 | 0.174 | 1.22E-10 | 8 | Entpd3 | neutrophil.9 |
| Lnpep | 6.88E-15 | 0.30154754 | 0.374 | 0.305 | 1.33E-10 | 8 | Lnpep | neutrophil.9 |
| Ddx5 | 7.25E-15 | 0.27560056 | 0.521 | 0.44 | 1.40E-10 | 8 | Ddx5 | neutrophil.9 |
| Pum2 | 1.60E-14 | 0.31471891 | 0.345 | 0.283 | 3.09E-10 | 8 | Pum2 | neutrophil.9 |
| Jak1 | 3.04E-14 | 0.30929637 | 0.382 | 0.324 | 5.88E-10 | 8 | Jak1 | neutrophil.9 |
| Actn4 | 1.22E-13 | 0.39251195 | 0.282 | 0.212 | 2.36E-09 | 8 | Actn4 | neutrophil.9 |
| Lasp1 | 2.03E-13 | 0.33539875 | 0.317 | 0.251 | 3.92E-09 | 8 | Lasp1 | neutrophil.9 |
| Cd68 | 5.35E-13 | 0.43584459 | 0.285 | 0.235 | 1.03E-08 | 8 | Cd68 | neutrophil.9 |
| Kpna4 | 6.31E-13 | 0.32622483 | 0.373 | 0.309 | 1.22E-08 | 8 | Kpna4 | neutrophil.9 |
| Atp6v0e | 6.52E-13 | 0.47058377 | 0.285 | 0.217 | 1.26E-08 | 8 | Atp6v0e | neutrophil.9 |
| Id1 | 8.06E-13 | 0.34164696 | 0.29 | 0.243 | 1.56E-08 | 8 | Id1 | neutrophil.9 |
| Pnp | 1.48E-12 | 0.41202415 | 0.289 | 0.23 | 2.86E-08 | 8 | Pnp | neutrophil.9 |
| Stat6 | 1.94E-12 | 0.38924513 | 0.326 | 0.266 | 3.75E-08 | 8 | Stat6 | neutrophil.9 |
| Nisch | 3.90E-12 | 0.26699986 | 0.251 | 0.19 | 7.53E-08 | 8 | Nisch | neutrophil.9 |
| Sgpl1 | 8.58E-12 | 0.40816175 | 0.311 | 0.247 | 1.66E-07 | 8 | Sgpl1 | neutrophil.9 |
| Retreg1 | 1.42E-11 | 0.36403444 | 0.31 | 0.246 | 2.74E-07 | 8 | Retreg1 | neutrophil.9 |
| Arhgdia | 1.66E-11 | 0.28414737 | 0.343 | 0.272 | 3.21E-07 | 8 | Arhgdia | neutrophil.9 |
| Snx3 | 2.27E-11 | 0.36394566 | 0.306 | 0.241 | 4.39E-07 | 8 | Snx3 | neutrophil.9 |
| Atp6v1b2 | 1.16E-10 | 0.30211071 | 0.332 | 0.283 | 2.25E-06 | 8 | Atp6v1b2 | neutrophil.9 |
| Raly | 1.37E-10 | 0.26961044 | 0.293 | 0.226 | 2.65E-06 | 8 | Raly | neutrophil.9 |
| Sun2 | 1.57E-10 | 0.31259269 | 0.295 | 0.234 | 3.04E-06 | 8 | Sun2 | neutrophil.9 |
| Hnrnph2 | 1.86E-10 | 0.36071056 | 0.28 | 0.217 | 3.59E-06 | 8 | Hnrnph2 | neutrophil.9 |
| Srrm2 | 1.92E-10 | 0.25467669 | 0.271 | 0.224 | 3.71E-06 | 8 | Srrm2 | neutrophil.9 |
| Sde2 | 1.26E-09 | 0.35201971 | 0.283 | 0.228 | 2.43E-05 | 8 | Sde2 | neutrophil.9 |
| Erbin | 1.27E-09 | 0.36996579 | 0.323 | 0.258 | 2.46E-05 | 8 | Erbin | neutrophil.9 |
| Bri3 | 2.49E-09 | 0.31446323 | 0.331 | 0.265 | 4.81E-05 | 8 | Bri3 | neutrophil.9 |
| Surf4 | 4.14E-09 | 0.36780464 | 0.267 | 0.21 | 8.00E-05 | 8 | Surf4 | neutrophil.9 |
| Atp6v1g1 | 2.42E-08 | 0.28132519 | 0.402 | 0.347 | 4.68E-04 | 8 | Atp6v1g1 | neutrophil.9 |
| Cdkn1b | 9.95E-08 | 0.2873413 | 0.268 | 0.219 | 0.00192419 | 8 | Cdkn1b | neutrophil.9 |
| Ubald2 | 1.72E-07 | 0.32610703 | 0.316 | 0.255 | 0.00331766 | 8 | Ubald2 | neutrophil.9 |
| Sh3glb1 | 2.48E-07 | 0.3172918 | 0.263 | 0.217 | 0.00479063 | 8 | Sh3glb1 | neutrophil.9 |
| Atp6v0d1 | 6.93E-07 | 0.28179371 | 0.25 | 0.216 | 0.01339105 | 8 | Atp6v0d1 | neutrophil.9 |
| Igfbp4 | 0 | 3.6518375 | 0.692 | 0.092 | 0 | 9 | Igfbp4 | neutrophil.10 |
| Tmsb10 | 0 | 3.12580025 | 0.687 | 0.102 | 0 | 9 | Tmsb10 | neutrophil.10 |
| Krt8 | 0 | 3.13785442 | 0.71 | 0.131 | 0 | 9 | Krt8 | neutrophil.10 |
| Anxa5 | 0 | 3.38401367 | 0.685 | 0.107 | 0 | 9 | Anxa5 | neutrophil.10 |
| Lmna | 0 | 2.61351694 | 0.664 | 0.109 | 0 | 9 | Lmna | neutrophil.10 |
| Krt18 | 0 | 3.06106196 | 0.628 | 0.083 | 0 | 9 | Krt18 | neutrophil.10 |
| Cyb5r3 | 0 | 4.79819306 | 0.602 | 0.062 | 0 | 9 | Cyb5r3 | neutrophil.10 |
| Fn1 | 0 | 3.57043801 | 0.722 | 0.188 | 0 | 9 | Fn1 | neutrophil.10 |
| Eef1b2 | 0 | 3.19374458 | 0.644 | 0.111 | 0 | 9 | Eef1b2 | neutrophil.10 |
| Vim | 0 | 2.94884562 | 0.961 | 0.431 | 0 | 9 | Vim | neutrophil.10 |
| Myof | 0 | 3.68933787 | 0.562 | 0.043 | 0 | 9 | Myof | neutrophil.10 |
| Ccnd1 | 0 | 4.63526444 | 0.544 | 0.026 | 0 | 9 | Ccnd1 | neutrophil.10 |
| Ecm1 | 0 | 3.62256123 | 0.611 | 0.094 | 0 | 9 | Ecm1 | neutrophil.10 |
| Aldh2 | 0 | 2.78714098 | 0.669 | 0.154 | 0 | 9 | Aldh2 | neutrophil.10 |
| S100a10 | 0 | 2.90953848 | 0.596 | 0.097 | 0 | 9 | S100a10 | neutrophil.10 |
| Hsp90ab1 | 0 | 3.09332928 | 0.596 | 0.098 | 0 | 9 | Hsp90ab1 | neutrophil.10 |
| Npm1 | 0 | 3.18743928 | 0.563 | 0.065 | 0 | 9 | Npm1 | neutrophil.10 |
| Rack1 | 0 | 3.02050968 | 0.599 | 0.113 | 0 | 9 | Rack1 | neutrophil.10 |
| Mprip | 0 | 2.82328014 | 0.574 | 0.094 | 0 | 9 | Mprip | neutrophil.10 |
| Tpm1 | 0 | 3.12001063 | 0.529 | 0.056 | 0 | 9 | Tpm1 | neutrophil.10 |
| Eef2 | 0 | 2.35798657 | 0.625 | 0.162 | 0 | 9 | Eef2 | neutrophil.10 |
| Gnas | 0 | 2.27807062 | 0.614 | 0.152 | 0 | 9 | Gnas | neutrophil.10 |
| Igfbp7 | 0 | 3.67103664 | 0.548 | 0.091 | 0 | 9 | Igfbp7 | neutrophil.10 |
| Cystm1 | 0 | 2.98987761 | 0.53 | 0.074 | 0 | 9 | Cystm1 | neutrophil.10 |
| Fbln2 | 0 | 3.89960578 | 0.507 | 0.055 | 0 | 9 | Fbln2 | neutrophil.10 |
| Ptms | 0 | 1.93262479 | 0.61 | 0.16 | 0 | 9 | Ptms | neutrophil.10 |
| Anxa4 | 0 | 3.19363842 | 0.493 | 0.05 | 0 | 9 | Anxa4 | neutrophil.10 |
| Net1 | 0 | 5.26469644 | 0.474 | 0.035 | 0 | 9 | Net1 | neutrophil.10 |
| Cdkn2a | 0 | 3.758 | 0.469 | 0.035 | 0 | 9 | Cdkn2a | neutrophil.10 |
| Ybx1 | 0 | 2.39241867 | 0.558 | 0.125 | 0 | 9 | Ybx1 | neutrophil.10 |
| Tagln2 | 0 | 1.65236792 | 0.665 | 0.233 | 0 | 9 | Tagln2 | neutrophil.10 |
| Col18a1 | 0 | 4.03604 | 0.473 | 0.041 | 0 | 9 | Col18a1 | neutrophil.10 |
| mt-Co2 | 0 | 2.72199897 | 0.768 | 0.338 | 0 | 9 | mt-Co2 | neutrophil.10 |
| Cavin1 | 0 | 3.45603074 | 0.463 | 0.034 | 0 | 9 | Cavin1 | neutrophil.10 |
| Cpe | 0 | 4.53462236 | 0.458 | 0.029 | 0 | 9 | Cpe | neutrophil.10 |
| S100a6 | 0 | 2.67759756 | 0.918 | 0.494 | 0 | 9 | S100a6 | neutrophil.10 |
| Sptbn1 | 0 | 3.64084368 | 0.456 | 0.033 | 0 | 9 | Sptbn1 | neutrophil.10 |
| Plec | 0 | 1.87257237 | 0.616 | 0.195 | 0 | 9 | Plec | neutrophil.10 |
| Tm4sf1 | 0 | 1.93718284 | 0.592 | 0.171 | 0 | 9 | Tm4sf1 | neutrophil.10 |
| Dst | 0 | 4.51784345 | 0.43 | 0.017 | 0 | 9 | Dst | neutrophil.10 |
| mt-Co3 | 0 | 2.71378021 | 0.599 | 0.187 | 0 | 9 | mt-Co3 | neutrophil.10 |
| Prdx2 | 0 | 3.40163832 | 0.445 | 0.037 | 0 | 9 | Prdx2 | neutrophil.10 |
| Nedd4 | 0 | 3.62097165 | 0.433 | 0.026 | 0 | 9 | Nedd4 | neutrophil.10 |
| Ctnna1 | 0 | 2.12331474 | 0.499 | 0.095 | 0 | 9 | Ctnna1 | neutrophil.10 |
| Cttn | 0 | 3.43639075 | 0.432 | 0.029 | 0 | 9 | Cttn | neutrophil.10 |
| Kif1c | 0 | 3.83780189 | 0.471 | 0.069 | 0 | 9 | Kif1c | neutrophil.10 |
| Spp1 | 0 | 2.75751348 | 0.911 | 0.51 | 0 | 9 | Spp1 | neutrophil.10 |
| Cldn18 | 0 | 3.10189803 | 0.454 | 0.054 | 0 | 9 | Cldn18 | neutrophil.10 |
| mt-Co1 | 0 | 2.54920042 | 0.559 | 0.159 | 0 | 9 | mt-Co1 | neutrophil.10 |
| Plac8 | 0 | 1.46419787 | 0.62 | 0.224 | 0 | 9 | Plac8 | neutrophil.10 |
| Selenow | 0 | 3.21714765 | 0.43 | 0.034 | 0 | 9 | Selenow | neutrophil.10 |
| Clu | 0 | 4.21468743 | 0.956 | 0.56 | 0 | 9 | Clu | neutrophil.10 |
| F2r | 0 | 3.45534388 | 0.438 | 0.047 | 0 | 9 | F2r | neutrophil.10 |
| Flna | 0 | 1.65063979 | 0.701 | 0.311 | 0 | 9 | Flna | neutrophil.10 |
| Actn4 | 0 | 1.55677393 | 0.596 | 0.208 | 0 | 9 | Actn4 | neutrophil.10 |
| mt-Atp6 | 0 | 2.43380578 | 0.599 | 0.213 | 0 | 9 | mt-Atp6 | neutrophil.10 |
| Axl | 0 | 3.113503 | 0.425 | 0.041 | 0 | 9 | Axl | neutrophil.10 |
| mt-Atp8 | 0 | 2.48124819 | 0.576 | 0.193 | 0 | 9 | mt-Atp8 | neutrophil.10 |
| Tspo | 0 | 1.60309511 | 0.555 | 0.175 | 0 | 9 | Tspo | neutrophil.10 |
| Tpm4 | 0 | 1.33305953 | 0.661 | 0.284 | 0 | 9 | Tpm4 | neutrophil.10 |
| Timp2 | 0 | 2.7049536 | 0.452 | 0.079 | 0 | 9 | Timp2 | neutrophil.10 |
| Eef1a1 | 0 | 2.75677385 | 0.935 | 0.564 | 0 | 9 | Eef1a1 | neutrophil.10 |
| Ltbp3 | 0 | 3.79316484 | 0.393 | 0.022 | 0 | 9 | Ltbp3 | neutrophil.10 |
| Ahnak | 0 | 2.14836209 | 0.45 | 0.08 | 0 | 9 | Ahnak | neutrophil.10 |
| Eif3k | 0 | 2.1135687 | 0.477 | 0.109 | 0 | 9 | Eif3k | neutrophil.10 |
| Anxa3 | 0 | 1.86507079 | 0.497 | 0.131 | 0 | 9 | Anxa3 | neutrophil.10 |
| Kctd10 | 0 | 3.56102906 | 0.44 | 0.075 | 0 | 9 | Kctd10 | neutrophil.10 |
| Eef1g | 0 | 2.97591074 | 0.406 | 0.044 | 0 | 9 | Eef1g | neutrophil.10 |
| Adam9 | 0 | 2.15088585 | 0.445 | 0.085 | 0 | 9 | Adam9 | neutrophil.10 |
| Serpinb6a | 0 | 2.9912265 | 0.4 | 0.042 | 0 | 9 | Serpinb6a | neutrophil.10 |
| Akr1a1 | 0 | 1.49909459 | 0.547 | 0.192 | 0 | 9 | Akr1a1 | neutrophil.10 |
| mt-Cytb | 0 | 2.72193268 | 0.547 | 0.192 | 0 | 9 | mt-Cytb | neutrophil.10 |
| Flnb | 0 | 2.94860979 | 0.402 | 0.049 | 0 | 9 | Flnb | neutrophil.10 |
| Hspa8 | 0 | 0.92999548 | 0.768 | 0.416 | 0 | 9 | Hspa8 | neutrophil.10 |
| Pabpc1 | 0 | 2.05918864 | 0.485 | 0.133 | 0 | 9 | Pabpc1 | neutrophil.10 |
| Plp2 | 0 | 1.31456588 | 0.566 | 0.214 | 0 | 9 | Plp2 | neutrophil.10 |
| Ncl | 0 | 2.2457422 | 0.462 | 0.112 | 0 | 9 | Ncl | neutrophil.10 |
| Las1l | 0 | 3.21635377 | 0.388 | 0.041 | 0 | 9 | Las1l | neutrophil.10 |
| Sh3pxd2a | 0 | 5.11059882 | 0.36 | 0.014 | 0 | 9 | Sh3pxd2a | neutrophil.10 |
| Hnrnpa1 | 0 | 2.80562881 | 0.401 | 0.056 | 0 | 9 | Hnrnpa1 | neutrophil.10 |
| Eif4b | 0 | 2.24635005 | 0.412 | 0.067 | 0 | 9 | Eif4b | neutrophil.10 |
| Map4 | 0 | 2.31265182 | 0.441 | 0.102 | 0 | 9 | Map4 | neutrophil.10 |
| Fam129b | 0 | 3.14532374 | 0.377 | 0.039 | 0 | 9 | Fam129b | neutrophil.10 |
| Fau | 0 | 1.50831955 | 0.878 | 0.54 | 0 | 9 | Fau | neutrophil.10 |
| Actn1 | 0 | 1.11362763 | 0.64 | 0.304 | 0 | 9 | Actn1 | neutrophil.10 |
| Dstn | 0 | 1.68069646 | 0.454 | 0.119 | 0 | 9 | Dstn | neutrophil.10 |
| Krt7 | 0 | 3.12582531 | 0.386 | 0.051 | 0 | 9 | Krt7 | neutrophil.10 |
| Cenpb | 0 | 3.41761834 | 0.393 | 0.062 | 0 | 9 | Cenpb | neutrophil.10 |
| Uba1 | 0 | 1.14746394 | 0.564 | 0.233 | 0 | 9 | Uba1 | neutrophil.10 |
| Ppib | 0 | 2.06476164 | 0.416 | 0.089 | 0 | 9 | Ppib | neutrophil.10 |
| Cox7a2l | 0 | 2.68013449 | 0.386 | 0.06 | 0 | 9 | Cox7a2l | neutrophil.10 |
| Csde1 | 0 | 1.73438811 | 0.467 | 0.142 | 0 | 9 | Csde1 | neutrophil.10 |
| mt-Nd5 | 0 | 2.83804031 | 0.436 | 0.112 | 0 | 9 | mt-Nd5 | neutrophil.10 |
| Tnnt2 | 0 | 3.65525796 | 0.339 | 0.016 | 0 | 9 | Tnnt2 | neutrophil.10 |
| mt-Nd1 | 0 | 2.80751762 | 0.462 | 0.142 | 0 | 9 | mt-Nd1 | neutrophil.10 |
| Ctse | 0 | 3.34430131 | 0.367 | 0.047 | 0 | 9 | Ctse | neutrophil.10 |
| Ehmt2 | 0 | 3.00414492 | 0.353 | 0.033 | 0 | 9 | Ehmt2 | neutrophil.10 |
| Msln | 0 | 3.66207879 | 0.349 | 0.031 | 0 | 9 | Msln | neutrophil.10 |
| Sh3glb1 | 0 | 1.14647242 | 0.529 | 0.213 | 0 | 9 | Sh3glb1 | neutrophil.10 |
| Tpt1 | 0 | 2.00614485 | 0.887 | 0.571 | 0 | 9 | Tpt1 | neutrophil.10 |
| Gsto1 | 0 | 1.92977921 | 0.405 | 0.091 | 0 | 9 | Gsto1 | neutrophil.10 |
| mt-Nd4l | 0 | 2.62393889 | 0.457 | 0.144 | 0 | 9 | mt-Nd4l | neutrophil.10 |
| Ppp1r14b | 0 | 3.67908428 | 0.331 | 0.019 | 0 | 9 | Ppp1r14b | neutrophil.10 |
| Cgnl1 | 0 | 5.76264112 | 0.317 | 0.006 | 0 | 9 | Cgnl1 | neutrophil.10 |
| Ctnnb1 | 0 | 0.91741457 | 0.633 | 0.323 | 0 | 9 | Ctnnb1 | neutrophil.10 |
| Itgb1 | 0 | 1.18992931 | 0.554 | 0.246 | 0 | 9 | Itgb1 | neutrophil.10 |
| Dap | 0 | 2.1865984 | 0.378 | 0.07 | 0 | 9 | Dap | neutrophil.10 |
| Capn2 | 0 | 2.74384613 | 0.337 | 0.03 | 0 | 9 | Capn2 | neutrophil.10 |
| Ctnnd1 | 0 | 2.57675362 | 0.344 | 0.04 | 0 | 9 | Ctnnd1 | neutrophil.10 |
| Col4a1 | 0 | 2.57528314 | 0.343 | 0.04 | 0 | 9 | Col4a1 | neutrophil.10 |
| Cdc37 | 0 | 1.72828885 | 0.4 | 0.097 | 0 | 9 | Cdc37 | neutrophil.10 |
| Atp5b | 0 | 1.56434161 | 0.425 | 0.122 | 0 | 9 | Atp5b | neutrophil.10 |
| Anxa2 | 0 | 1.57314054 | 0.875 | 0.574 | 0 | 9 | Anxa2 | neutrophil.10 |
| Myl12a | 0 | 1.30793248 | 0.523 | 0.222 | 0 | 9 | Myl12a | neutrophil.10 |
| Npc2 | 0 | 1.36963125 | 0.465 | 0.165 | 0 | 9 | Npc2 | neutrophil.10 |
| Rhoc | 0 | 2.58330423 | 0.344 | 0.044 | 0 | 9 | Rhoc | neutrophil.10 |
| Ppia | 0 | 1.85052309 | 0.375 | 0.077 | 0 | 9 | Ppia | neutrophil.10 |
| Wfdc2 | 0 | 3.24361455 | 0.329 | 0.034 | 0 | 9 | Wfdc2 | neutrophil.10 |
| Sdc1 | 0 | 1.93181354 | 0.367 | 0.073 | 0 | 9 | Sdc1 | neutrophil.10 |
| Cox4i1 | 0 | 1.85369821 | 0.375 | 0.083 | 0 | 9 | Cox4i1 | neutrophil.10 |
| Itga3 | 0 | 2.71264869 | 0.32 | 0.03 | 0 | 9 | Itga3 | neutrophil.10 |
| Cdh1 | 0 | 2.09834869 | 0.339 | 0.052 | 0 | 9 | Cdh1 | neutrophil.10 |
| Wwtr1 | 0 | 3.45827535 | 0.307 | 0.02 | 0 | 9 | Wwtr1 | neutrophil.10 |
| Ubxn1 | 0 | 1.40047992 | 0.459 | 0.175 | 0 | 9 | Ubxn1 | neutrophil.10 |
| Ptgfrn | 0 | 2.92737243 | 0.312 | 0.03 | 0 | 9 | Ptgfrn | neutrophil.10 |
| Bsg | 0 | 1.56108479 | 0.386 | 0.104 | 0 | 9 | Bsg | neutrophil.10 |
| Pfdn1 | 0 | 3.21146662 | 0.304 | 0.022 | 0 | 9 | Pfdn1 | neutrophil.10 |
| Anxa1 | 0 | 1.33860299 | 0.416 | 0.134 | 0 | 9 | Anxa1 | neutrophil.10 |
| Krt19 | 0 | 2.61247001 | 0.336 | 0.055 | 0 | 9 | Krt19 | neutrophil.10 |
| Ctsl | 0 | 1.33836667 | 0.414 | 0.133 | 0 | 9 | Ctsl | neutrophil.10 |
| Itm2b | 0 | 0.76769747 | 0.704 | 0.423 | 0 | 9 | Itm2b | neutrophil.10 |
| Myef2 | 0 | 4.13866066 | 0.29 | 0.01 | 0 | 9 | Myef2 | neutrophil.10 |
| Naca | 0 | 1.98916129 | 0.343 | 0.064 | 0 | 9 | Naca | neutrophil.10 |
| Elob | 0 | 1.13129457 | 0.463 | 0.184 | 0 | 9 | Elob | neutrophil.10 |
| Map1b | 0 | 5.02830936 | 0.287 | 0.008 | 0 | 9 | Map1b | neutrophil.10 |
| Ptprs | 0 | 3.60121707 | 0.296 | 0.018 | 0 | 9 | Ptprs | neutrophil.10 |
| Fkbp1a | 0 | 2.41938828 | 0.316 | 0.039 | 0 | 9 | Fkbp1a | neutrophil.10 |
| Huwe1 | 0 | 2.04709778 | 0.361 | 0.084 | 0 | 9 | Huwe1 | neutrophil.10 |
| Avil | 0 | 3.5666809 | 0.29 | 0.016 | 0 | 9 | Avil | neutrophil.10 |
| Eif3f | 0 | 2.24465317 | 0.323 | 0.05 | 0 | 9 | Eif3f | neutrophil.10 |
| Mtch1 | 0 | 1.61139863 | 0.357 | 0.085 | 0 | 9 | Mtch1 | neutrophil.10 |
| Pgk1 | 0 | 1.36656251 | 0.421 | 0.151 | 0 | 9 | Pgk1 | neutrophil.10 |
| App | 0 | 0.78321749 | 0.736 | 0.466 | 0 | 9 | App | neutrophil.10 |
| Crip1 | 0 | 2.45281203 | 0.308 | 0.04 | 0 | 9 | Crip1 | neutrophil.10 |
| Dhx9 | 0 | 1.74836066 | 0.335 | 0.069 | 0 | 9 | Dhx9 | neutrophil.10 |
| Sod1 | 0 | 1.66164553 | 0.353 | 0.087 | 0 | 9 | Sod1 | neutrophil.10 |
| Sh3d19 | 0 | 3.50134448 | 0.276 | 0.013 | 0 | 9 | Sh3d19 | neutrophil.10 |
| Eif3l | 0 | 1.96381016 | 0.323 | 0.06 | 0 | 9 | Eif3l | neutrophil.10 |
| Ano1 | 0 | 2.97637406 | 0.283 | 0.021 | 0 | 9 | Ano1 | neutrophil.10 |
| Sh3bgrl | 0 | 2.35922466 | 0.303 | 0.042 | 0 | 9 | Sh3bgrl | neutrophil.10 |
| Ybx3 | 0 | 3.35626593 | 0.28 | 0.02 | 0 | 9 | Ybx3 | neutrophil.10 |
| Eif3h | 0 | 2.68502158 | 0.295 | 0.035 | 0 | 9 | Eif3h | neutrophil.10 |
| Lgals1 | 0 | 3.5348045 | 0.276 | 0.016 | 0 | 9 | Lgals1 | neutrophil.10 |
| Tubb5 | 0 | 1.82634779 | 0.335 | 0.077 | 0 | 9 | Tubb5 | neutrophil.10 |
| S100a1 | 0 | 3.10532801 | 0.28 | 0.023 | 0 | 9 | S100a1 | neutrophil.10 |
| Chd3 | 0 | 3.59519191 | 0.272 | 0.015 | 0 | 9 | Chd3 | neutrophil.10 |
| mt-Nd6 | 0 | 1.64802209 | 0.39 | 0.134 | 0 | 9 | mt-Nd6 | neutrophil.10 |
| Pls3 | 0 | 3.18616878 | 0.274 | 0.019 | 0 | 9 | Pls3 | neutrophil.10 |
| Nme2 | 0 | 2.90686678 | 0.28 | 0.027 | 0 | 9 | Nme2 | neutrophil.10 |
| Dync1li2 | 0 | 2.44158362 | 0.304 | 0.052 | 0 | 9 | Dync1li2 | neutrophil.10 |
| Prdx1 | 0 | 1.69551039 | 0.312 | 0.063 | 0 | 9 | Prdx1 | neutrophil.10 |
| Agrn | 0 | 2.17840824 | 0.293 | 0.044 | 0 | 9 | Agrn | neutrophil.10 |
| Pkm | 0 | 0.5550607 | 0.748 | 0.499 | 0 | 9 | Pkm | neutrophil.10 |
| Itga6 | 0 | 2.2835451 | 0.283 | 0.034 | 0 | 9 | Itga6 | neutrophil.10 |
| Capns1 | 0 | 1.61420886 | 0.331 | 0.084 | 0 | 9 | Capns1 | neutrophil.10 |
| Ipo5 | 0 | 2.60632449 | 0.274 | 0.028 | 0 | 9 | Ipo5 | neutrophil.10 |
| Lamb1 | 0 | 3.15477513 | 0.263 | 0.018 | 0 | 9 | Lamb1 | neutrophil.10 |
| Tcp1 | 0 | 2.42315367 | 0.28 | 0.035 | 0 | 9 | Tcp1 | neutrophil.10 |
| Slc25a4 | 0 | 3.36682969 | 0.255 | 0.012 | 0 | 9 | Slc25a4 | neutrophil.10 |
| Palld | 0 | 3.56052389 | 0.26 | 0.017 | 0 | 9 | Palld | neutrophil.10 |
| Stard10 | 0 | 2.83948968 | 0.266 | 0.024 | 0 | 9 | Stard10 | neutrophil.10 |
| Npnt | 0 | 3.02950157 | 0.264 | 0.023 | 0 | 9 | Npnt | neutrophil.10 |
| Selenop | 0 | 2.02176039 | 0.301 | 0.06 | 0 | 9 | Selenop | neutrophil.10 |
| mt-Nd4 | 0 | 2.62612256 | 0.304 | 0.063 | 0 | 9 | mt-Nd4 | neutrophil.10 |
| Kars | 0 | 2.20160472 | 0.283 | 0.042 | 0 | 9 | Kars | neutrophil.10 |
| Cd63 | 0 | 0.56181273 | 0.619 | 0.379 | 0 | 9 | Cd63 | neutrophil.10 |
| Camk2n1 | 0 | 3.230298 | 0.254 | 0.015 | 0 | 9 | Camk2n1 | neutrophil.10 |
| S100a4 | 0 | 2.54970826 | 0.291 | 0.053 | 0 | 9 | S100a4 | neutrophil.10 |
| Cfl1 | 0 | 0.64123977 | 0.809 | 0.572 | 0 | 9 | Cfl1 | neutrophil.10 |
| Itgb4 | 0 | 3.07514414 | 0.254 | 0.019 | 0 | 9 | Itgb4 | neutrophil.10 |
| Cct2 | 0 | 2.17308357 | 0.27 | 0.036 | 0 | 9 | Cct2 | neutrophil.10 |
| S100a11 | 0 | 1.27835408 | 0.914 | 0.68 | 0 | 9 | S100a11 | neutrophil.10 |
| Cdc42bpg | 0 | 3.57616467 | 0.254 | 0.023 | 0 | 9 | Cdc42bpg | neutrophil.10 |
| Pdia3 | 0 | 1.73154706 | 0.3 | 0.071 | 0 | 9 | Pdia3 | neutrophil.10 |
| Pura | 0 | 2.43056459 | 0.278 | 0.051 | 0 | 9 | Pura | neutrophil.10 |
| Stmn1 | 0 | 2.64585444 | 0.251 | 0.024 | 0 | 9 | Stmn1 | neutrophil.10 |
| Col6a1 | 0 | 2.12597708 | 0.272 | 0.046 | 0 | 9 | Col6a1 | neutrophil.10 |
| Sprr1a | 0 | 3.18187982 | 0.258 | 0.034 | 0 | 9 | Sprr1a | neutrophil.10 |
| Igfbp3 | 0 | 2.12604588 | 0.284 | 0.062 | 0 | 9 | Igfbp3 | neutrophil.10 |
| Dynll2 | 0 | 3.14536909 | 0.251 | 0.032 | 0 | 9 | Dynll2 | neutrophil.10 |
| mt-Nd2 | 0 | 2.54650075 | 0.267 | 0.058 | 0 | 9 | mt-Nd2 | neutrophil.10 |
| Actg1 | 0 | 0.32765488 | 0.89 | 0.694 | 0 | 9 | Actg1 | neutrophil.10 |
| mt-Nd3 | 0 | 2.30805832 | 0.251 | 0.066 | 0 | 9 | mt-Nd3 | neutrophil.10 |
| Msn | 0 | 0.77334057 | 0.942 | 0.766 | 0 | 9 | Msn | neutrophil.10 |
| Tmsb4x | 0 | 0.46240596 | 0.992 | 0.922 | 0 | 9 | Tmsb4x | neutrophil.10 |
| Ddost | 4.66E-305 | 1.49883057 | 0.311 | 0.081 | 9.00E-301 | 9 | Ddost | neutrophil.10 |
| Atp5j2 | 1.20E-301 | 1.69430082 | 0.348 | 0.09 | 2.31E-297 | 9 | Atp5j2 | neutrophil.10 |
| Slc25a39 | 2.85E-301 | 2.10370674 | 0.28 | 0.048 | 5.51E-297 | 9 | Slc25a39 | neutrophil.10 |
| Golim4 | 7.66E-301 | 2.40720378 | 0.288 | 0.04 | 1.48E-296 | 9 | Golim4 | neutrophil.10 |
| Tecr | 1.82E-299 | 1.56530093 | 0.301 | 0.071 | 3.52E-295 | 9 | Tecr | neutrophil.10 |
| Txndc17 | 4.69E-297 | 1.91918224 | 0.279 | 0.046 | 9.06E-293 | 9 | Txndc17 | neutrophil.10 |
| Btf3 | 1.81E-293 | 1.49198251 | 0.364 | 0.106 | 3.50E-289 | 9 | Btf3 | neutrophil.10 |
| Tm9sf3 | 1.08E-289 | 1.05278856 | 0.475 | 0.207 | 2.08E-285 | 9 | Tm9sf3 | neutrophil.10 |
| Fxr1 | 4.83E-289 | 1.59778549 | 0.378 | 0.09 | 9.34E-285 | 9 | Fxr1 | neutrophil.10 |
| Galnt1 | 1.16E-281 | 1.61530968 | 0.312 | 0.073 | 2.24E-277 | 9 | Galnt1 | neutrophil.10 |
| Cox7c | 5.70E-279 | 1.82446721 | 0.297 | 0.054 | 1.10E-274 | 9 | Cox7c | neutrophil.10 |
| Col6a2 | 6.10E-279 | 1.76144574 | 0.252 | 0.049 | 1.18E-274 | 9 | Col6a2 | neutrophil.10 |
| Arpc1b | 4.99E-278 | 0.40687822 | 0.756 | 0.523 | 9.65E-274 | 9 | Arpc1b | neutrophil.10 |
| Hsbp1 | 3.41E-276 | 1.34547254 | 0.337 | 0.102 | 6.59E-272 | 9 | Hsbp1 | neutrophil.10 |
| Hdlbp | 1.80E-272 | 1.07196326 | 0.422 | 0.137 | 3.49E-268 | 9 | Hdlbp | neutrophil.10 |
| Nono | 2.57E-272 | 1.55984126 | 0.319 | 0.074 | 4.97E-268 | 9 | Nono | neutrophil.10 |
| Cox6b1 | 3.82E-270 | 1.38946374 | 0.371 | 0.111 | 7.38E-266 | 9 | Cox6b1 | neutrophil.10 |
| Mlec | 2.31E-267 | 1.46904007 | 0.3 | 0.075 | 4.47E-263 | 9 | Mlec | neutrophil.10 |
| Cops9 | 2.71E-265 | 1.51708504 | 0.327 | 0.09 | 5.23E-261 | 9 | Cops9 | neutrophil.10 |
| Ganab | 1.61E-262 | 1.67341369 | 0.274 | 0.061 | 3.10E-258 | 9 | Ganab | neutrophil.10 |
| Calm3 | 3.75E-262 | 0.94115947 | 0.505 | 0.234 | 7.25E-258 | 9 | Calm3 | neutrophil.10 |
| Dynlrb1 | 2.64E-261 | 1.45682468 | 0.363 | 0.105 | 5.09E-257 | 9 | Dynlrb1 | neutrophil.10 |
| Tle5 | 1.23E-260 | 1.56113108 | 0.305 | 0.078 | 2.38E-256 | 9 | Tle5 | neutrophil.10 |
| Snrpf | 6.05E-259 | 2.09268007 | 0.254 | 0.042 | 1.17E-254 | 9 | Snrpf | neutrophil.10 |
| Metap2 | 5.69E-258 | 2.07194563 | 0.255 | 0.035 | 1.10E-253 | 9 | Metap2 | neutrophil.10 |
| Tkt | 1.17E-256 | 1.14482151 | 0.408 | 0.141 | 2.26E-252 | 9 | Tkt | neutrophil.10 |
| Idh3b | 1.38E-256 | 1.79543276 | 0.26 | 0.048 | 2.67E-252 | 9 | Idh3b | neutrophil.10 |
| Ndufa13 | 4.53E-254 | 1.10900265 | 0.422 | 0.15 | 8.77E-250 | 9 | Ndufa13 | neutrophil.10 |
| Ndufa2 | 4.92E-254 | 1.19804738 | 0.42 | 0.135 | 9.52E-250 | 9 | Ndufa2 | neutrophil.10 |
| Akt1 | 6.38E-254 | 0.71972491 | 0.6 | 0.314 | 1.23E-249 | 9 | Akt1 | neutrophil.10 |
| Serpine1 | 9.15E-253 | 0.46966008 | 0.264 | 0.081 | 1.77E-248 | 9 | Serpine1 | neutrophil.10 |
| Hipk2 | 9.74E-249 | 2.33064426 | 0.263 | 0.039 | 1.88E-244 | 9 | Hipk2 | neutrophil.10 |
| Calr | 2.74E-248 | 1.18517609 | 0.417 | 0.169 | 5.29E-244 | 9 | Calr | neutrophil.10 |
| Myh9 | 2.39E-245 | 0.88013584 | 0.541 | 0.261 | 4.63E-241 | 9 | Myh9 | neutrophil.10 |
| Canx | 3.85E-245 | 1.46194191 | 0.352 | 0.099 | 7.45E-241 | 9 | Canx | neutrophil.10 |
| Pdia6 | 7.56E-244 | 1.26301065 | 0.29 | 0.082 | 1.46E-239 | 9 | Pdia6 | neutrophil.10 |
| Atpif1 | 6.58E-243 | 2.04436287 | 0.252 | 0.039 | 1.27E-238 | 9 | Atpif1 | neutrophil.10 |
| Fkbp8 | 1.29E-242 | 1.08542637 | 0.446 | 0.158 | 2.49E-238 | 9 | Fkbp8 | neutrophil.10 |
| Ndufs2 | 6.00E-237 | 1.3576945 | 0.312 | 0.081 | 1.16E-232 | 9 | Ndufs2 | neutrophil.10 |
| Dync1h1 | 1.89E-234 | 1.40912638 | 0.333 | 0.104 | 3.65E-230 | 9 | Dync1h1 | neutrophil.10 |
| Rtn4 | 5.62E-229 | 0.70832746 | 0.463 | 0.239 | 1.09E-224 | 9 | Rtn4 | neutrophil.10 |
| Hdgf | 1.33E-226 | 1.93405862 | 0.279 | 0.044 | 2.57E-222 | 9 | Hdgf | neutrophil.10 |
| Ptma | 1.62E-226 | 1.35255109 | 0.336 | 0.088 | 3.13E-222 | 9 | Ptma | neutrophil.10 |
| Serf2 | 1.01E-223 | 0.98875622 | 0.436 | 0.173 | 1.96E-219 | 9 | Serf2 | neutrophil.10 |
| Rtcb | 2.72E-222 | 2.10790959 | 0.252 | 0.039 | 5.27E-218 | 9 | Rtcb | neutrophil.10 |
| Spint2 | 4.56E-221 | 1.7615556 | 0.251 | 0.051 | 8.81E-217 | 9 | Spint2 | neutrophil.10 |
| Drap1 | 9.30E-215 | 0.9166866 | 0.432 | 0.181 | 1.80E-210 | 9 | Drap1 | neutrophil.10 |
| Psmd8 | 2.64E-214 | 1.69590325 | 0.258 | 0.055 | 5.11E-210 | 9 | Psmd8 | neutrophil.10 |
| S100a13 | 1.42E-213 | 1.46503331 | 0.259 | 0.059 | 2.75E-209 | 9 | S100a13 | neutrophil.10 |
| Sfr1 | 2.04E-213 | 1.48020166 | 0.304 | 0.073 | 3.95E-209 | 9 | Sfr1 | neutrophil.10 |
| Sptan1 | 1.41E-212 | 0.79941453 | 0.491 | 0.23 | 2.73E-208 | 9 | Sptan1 | neutrophil.10 |
| Ssr2 | 1.69E-212 | 1.42437431 | 0.256 | 0.066 | 3.26E-208 | 9 | Ssr2 | neutrophil.10 |
| Ldha | 6.42E-211 | 0.48443561 | 0.554 | 0.293 | 1.24E-206 | 9 | Ldha | neutrophil.10 |
| Vcp | 6.63E-206 | 0.93605555 | 0.396 | 0.144 | 1.28E-201 | 9 | Vcp | neutrophil.10 |
| Ugdh | 8.58E-206 | 1.41660411 | 0.271 | 0.066 | 1.66E-201 | 9 | Ugdh | neutrophil.10 |
| Golm1 | 1.24E-204 | 1.42002647 | 0.315 | 0.082 | 2.39E-200 | 9 | Golm1 | neutrophil.10 |
| Slc4a2 | 8.85E-203 | 1.43494789 | 0.292 | 0.069 | 1.71E-198 | 9 | Slc4a2 | neutrophil.10 |
| Slc25a3 | 3.58E-199 | 0.74017228 | 0.495 | 0.238 | 6.91E-195 | 9 | Slc25a3 | neutrophil.10 |
| Uqcrc1 | 7.98E-199 | 1.21731272 | 0.355 | 0.114 | 1.54E-194 | 9 | Uqcrc1 | neutrophil.10 |
| 2-Sep | 4.71E-194 | 1.23194884 | 0.278 | 0.079 | 9.10E-190 | 9 | 2-Sep | neutrophil.10 |
| Park7 | 8.91E-194 | 1.43947302 | 0.297 | 0.077 | 1.72E-189 | 9 | Park7 | neutrophil.10 |
| Atxn10 | 8.94E-194 | 1.25769764 | 0.317 | 0.084 | 1.73E-189 | 9 | Atxn10 | neutrophil.10 |
| Calm1 | 1.22E-193 | 0.45382254 | 0.591 | 0.347 | 2.35E-189 | 9 | Calm1 | neutrophil.10 |
| Cnn2 | 1.06E-191 | 0.96466753 | 0.325 | 0.136 | 2.05E-187 | 9 | Cnn2 | neutrophil.10 |
| Ndufa3 | 3.47E-191 | 1.05342219 | 0.365 | 0.135 | 6.70E-187 | 9 | Ndufa3 | neutrophil.10 |
| Sri | 8.58E-191 | 1.10009726 | 0.357 | 0.116 | 1.66E-186 | 9 | Sri | neutrophil.10 |
| Smarca4 | 1.06E-189 | 1.28679005 | 0.295 | 0.073 | 2.06E-185 | 9 | Smarca4 | neutrophil.10 |
| Bag1 | 2.30E-189 | 1.23756449 | 0.317 | 0.095 | 4.44E-185 | 9 | Bag1 | neutrophil.10 |
| Lgals3 | 1.83E-188 | 0.42741312 | 0.506 | 0.265 | 3.53E-184 | 9 | Lgals3 | neutrophil.10 |
| Ssr4 | 7.84E-187 | 1.40584587 | 0.255 | 0.068 | 1.52E-182 | 9 | Ssr4 | neutrophil.10 |
| Dnajc10 | 1.63E-185 | 1.30468304 | 0.278 | 0.074 | 3.15E-181 | 9 | Dnajc10 | neutrophil.10 |
| Rock2 | 1.21E-179 | 1.45488401 | 0.297 | 0.074 | 2.33E-175 | 9 | Rock2 | neutrophil.10 |
| Mif | 1.42E-178 | 1.01263904 | 0.344 | 0.12 | 2.74E-174 | 9 | Mif | neutrophil.10 |

| | | | | | | | | |
|---|---|---|---|---|---|---|---|---|
| Anp32a | 3.34E-178 | 0.62974126 | 0.515 | 0.274 | 6.46E-174 | 9 | Anp32a | neutrophil.10 |
| Setd3 | 2.19E-174 | 1.70988429 | 0.27 | 0.052 | 4.24E-170 | 9 | Setd3 | neutrophil.10 |
| Atp1a1 | 1.16E-172 | 0.31080332 | 0.579 | 0.342 | 2.25E-168 | 9 | Atp1a1 | neutrophil.10 |
| Smarcc2 | 1.08E-169 | 1.25279626 | 0.335 | 0.11 | 2.09E-165 | 9 | Smarcc2 | neutrophil.10 |
| Pdap1 | 2.66E-169 | 1.23553621 | 0.272 | 0.077 | 5.13E-165 | 9 | Pdap1 | neutrophil.10 |
| Myo18a | 2.85E-169 | 1.30073601 | 0.304 | 0.096 | 5.50E-165 | 9 | Myo18a | neutrophil.10 |
| Atp5d | 2.47E-167 | 0.94650536 | 0.352 | 0.118 | 4.78E-163 | 9 | Atp5d | neutrophil.10 |
| Lamp1 | 1.94E-162 | 0.94722028 | 0.336 | 0.127 | 3.75E-158 | 9 | Lamp1 | neutrophil.10 |
| Hsp90aa1 | 4.77E-162 | 0.64450586 | 0.438 | 0.2 | 9.22E-158 | 9 | Hsp90aa1 | neutrophil.10 |
| Ndfip1 | 2.07E-160 | 0.66868546 | 0.393 | 0.189 | 4.01E-156 | 9 | Ndfip1 | neutrophil.10 |
| Ndufa6 | 2.36E-157 | 1.24617317 | 0.288 | 0.078 | 4.57E-153 | 9 | Ndufa6 | neutrophil.10 |
| Pgam1 | 1.32E-155 | 0.79435704 | 0.425 | 0.182 | 2.56E-151 | 9 | Pgam1 | neutrophil.10 |
| Txn1 | 1.83E-154 | 0.41631622 | 0.522 | 0.304 | 3.53E-150 | 9 | Txn1 | neutrophil.10 |
| 9-Sep | 5.15E-154 | 1.21607588 | 0.279 | 0.075 | 9.95E-150 | 9 | 9-Sep | neutrophil.10 |
| Chchd2 | 3.62E-152 | 0.41143047 | 0.506 | 0.307 | 6.99E-148 | 9 | Chchd2 | neutrophil.10 |
| Psmc3 | 2.36E-150 | 1.15611416 | 0.266 | 0.082 | 4.57E-146 | 9 | Psmc3 | neutrophil.10 |
| Clptm1 | 3.81E-150 | 0.90035972 | 0.305 | 0.123 | 7.37E-146 | 9 | Clptm1 | neutrophil.10 |
| Nisch | 8.44E-150 | 0.69823509 | 0.42 | 0.189 | 1.63E-145 | 9 | Nisch | neutrophil.10 |
| Bcap31 | 7.95E-149 | 0.92405864 | 0.256 | 0.09 | 1.54E-144 | 9 | Bcap31 | neutrophil.10 |
| Atp5e | 1.82E-145 | 1.14027121 | 0.295 | 0.1 | 3.53E-141 | 9 | Atp5e | neutrophil.10 |
| Arhgdia | 2.79E-145 | 0.33578096 | 0.493 | 0.271 | 5.40E-141 | 9 | Arhgdia | neutrophil.10 |
| Snx9 | 6.08E-142 | 1.35296181 | 0.256 | 0.064 | 1.18E-137 | 9 | Snx9 | neutrophil.10 |
| Capg | 8.20E-142 | 0.51740405 | 0.433 | 0.216 | 1.58E-137 | 9 | Capg | neutrophil.10 |
| Ube2m | 1.36E-141 | 1.31726388 | 0.264 | 0.075 | 2.63E-137 | 9 | Ube2m | neutrophil.10 |
| Gpi1 | 7.84E-141 | 0.73093015 | 0.367 | 0.145 | 1.52E-136 | 9 | Gpi1 | neutrophil.10 |
| Map4k4 | 1.44E-138 | 0.89989479 | 0.409 | 0.179 | 2.78E-134 | 9 | Map4k4 | neutrophil.10 |
| Ywhag | 4.19E-137 | 0.30808039 | 0.538 | 0.323 | 8.11E-133 | 9 | Ywhag | neutrophil.10 |
| Rbm3 | 7.20E-137 | 0.73324689 | 0.363 | 0.142 | 1.39E-132 | 9 | Rbm3 | neutrophil.10 |
| Tuba1a | 1.84E-135 | 0.84038737 | 0.293 | 0.121 | 3.55E-131 | 9 | Tuba1a | neutrophil.10 |
| Ssr3 | 7.27E-131 | 1.03596162 | 0.279 | 0.08 | 1.41E-126 | 9 | Ssr3 | neutrophil.10 |
| Myo1c | 1.07E-128 | 0.53722557 | 0.428 | 0.204 | 2.07E-124 | 9 | Myo1c | neutrophil.10 |
| Trim56 | 1.57E-128 | 1.01875964 | 0.321 | 0.12 | 3.04E-124 | 9 | Trim56 | neutrophil.10 |
| Epn1 | 3.47E-124 | 0.73190142 | 0.441 | 0.192 | 6.71E-120 | 9 | Epn1 | neutrophil.10 |
| Pfdn5 | 7.91E-124 | 1.0275058 | 0.312 | 0.109 | 1.53E-119 | 9 | Pfdn5 | neutrophil.10 |
| Fis1 | 4.40E-123 | 0.82994237 | 0.371 | 0.143 | 8.50E-119 | 9 | Fis1 | neutrophil.10 |
| Ywhae | 6.22E-123 | 0.8776102 | 0.328 | 0.123 | 1.20E-118 | 9 | Ywhae | neutrophil.10 |
| Mmp14 | 7.19E-123 | 0.77316973 | 0.282 | 0.092 | 1.39E-118 | 9 | Mmp14 | neutrophil.10 |
| Ndufa7 | 9.22E-123 | 1.00653362 | 0.267 | 0.089 | 1.78E-118 | 9 | Ndufa7 | neutrophil.10 |
| Glud1 | 1.97E-122 | 0.60807571 | 0.365 | 0.158 | 3.81E-118 | 9 | Glud1 | neutrophil.10 |
| Cox8a | 3.12E-118 | 0.51746865 | 0.416 | 0.215 | 6.04E-114 | 9 | Cox8a | neutrophil.10 |
| Tcf20 | 2.80E-116 | 1.35866493 | 0.255 | 0.069 | 5.41E-112 | 9 | Tcf20 | neutrophil.10 |
| Glmp | 4.99E-116 | 0.86881681 | 0.262 | 0.096 | 9.64E-112 | 9 | Glmp | neutrophil.10 |
| Clta | 8.70E-116 | 0.29917056 | 0.604 | 0.353 | 1.68E-111 | 9 | Clta | neutrophil.10 |
| Got2 | 2.39E-115 | 1.03635476 | 0.25 | 0.088 | 4.63E-111 | 9 | Got2 | neutrophil.10 |
| Ube2r2 | 4.46E-115 | 0.95637652 | 0.286 | 0.101 | 8.62E-111 | 9 | Ube2r2 | neutrophil.10 |
| Copa | 6.55E-115 | 0.65766457 | 0.34 | 0.144 | 1.27E-110 | 9 | Copa | neutrophil.10 |
| P4hb | 3.17E-114 | 0.3282626 | 0.531 | 0.301 | 6.13E-110 | 9 | P4hb | neutrophil.10 |
| Cers2 | 1.00E-113 | 0.70162179 | 0.323 | 0.131 | 1.94E-109 | 9 | Cers2 | neutrophil.10 |
| Acly | 7.35E-112 | 0.61111595 | 0.38 | 0.167 | 1.42E-107 | 9 | Acly | neutrophil.10 |
| Snd1 | 1.17E-110 | 0.82087162 | 0.307 | 0.105 | 2.27E-106 | 9 | Snd1 | neutrophil.10 |
| Reep5 | 1.33E-110 | 0.61634982 | 0.349 | 0.16 | 2.57E-106 | 9 | Reep5 | neutrophil.10 |
| Rrbp1 | 2.05E-109 | 0.5671123 | 0.349 | 0.161 | 3.96E-105 | 9 | Rrbp1 | neutrophil.10 |
| Kif5b | 1.07E-107 | 0.78971684 | 0.312 | 0.126 | 2.07E-103 | 9 | Kif5b | neutrophil.10 |
| Nucb1 | 3.62E-106 | 0.81897036 | 0.315 | 0.131 | 7.01E-102 | 9 | Nucb1 | neutrophil.10 |
| Khsrp | 7.07E-106 | 0.96453376 | 0.282 | 0.088 | 1.37E-101 | 9 | Khsrp | neutrophil.10 |
| H13 | 1.64E-103 | 0.73324634 | 0.328 | 0.146 | 3.16E-99 | 9 | H13 | neutrophil.10 |
| Gnb1 | 2.86E-103 | 0.48086478 | 0.466 | 0.247 | 5.54E-99 | 9 | Gnb1 | neutrophil.10 |
| Sf3b2 | 2.41E-102 | 0.4878645 | 0.364 | 0.173 | 4.67E-98 | 9 | Sf3b2 | neutrophil.10 |
| Atp2a2 | 1.41E-101 | 0.96366018 | 0.252 | 0.081 | 2.72E-97 | 9 | Atp2a2 | neutrophil.10 |
| Bzw1 | 2.24E-100 | 0.51516872 | 0.309 | 0.149 | 4.33E-96 | 9 | Bzw1 | neutrophil.10 |
| 7-Sep | 2.48E-98 | 0.5122841 | 0.34 | 0.17 | 4.79E-94 | 9 | 7-Sep | neutrophil.10 |
| Cope | 9.96E-97 | 0.76609832 | 0.26 | 0.095 | 1.93E-92 | 9 | Cope | neutrophil.10 |
| Itpripl2 | 6.97E-96 | 0.78591841 | 0.321 | 0.134 | 1.35E-91 | 9 | Itpripl2 | neutrophil.10 |
| Mib1 | 9.35E-96 | 0.93191999 | 0.307 | 0.111 | 1.81E-91 | 9 | Mib1 | neutrophil.10 |
| Pfkl | 2.27E-94 | 0.56539062 | 0.287 | 0.124 | 4.39E-90 | 9 | Pfkl | neutrophil.10 |
| Paip2 | 2.45E-94 | 0.57109079 | 0.315 | 0.138 | 4.75E-90 | 9 | Paip2 | neutrophil.10 |
| Ccnd2 | 1.61E-93 | 0.81348011 | 0.262 | 0.109 | 3.12E-89 | 9 | Ccnd2 | neutrophil.10 |
| Ubr4 | 2.08E-93 | 0.83759068 | 0.308 | 0.128 | 4.02E-89 | 9 | Ubr4 | neutrophil.10 |
| Mlf2 | 1.06E-92 | 0.65287644 | 0.324 | 0.138 | 2.05E-88 | 9 | Mlf2 | neutrophil.10 |
| Syncrip | 1.96E-92 | 0.54453182 | 0.332 | 0.138 | 3.78E-88 | 9 | Syncrip | neutrophil.10 |
| Tpi1 | 2.47E-92 | 0.45513455 | 0.337 | 0.145 | 4.78E-88 | 9 | Tpi1 | neutrophil.10 |
| Rnf10 | 3.69E-92 | 0.46803805 | 0.401 | 0.196 | 7.13E-88 | 9 | Rnf10 | neutrophil.10 |
| Cyth2 | 1.17E-91 | 0.91852333 | 0.264 | 0.083 | 2.25E-87 | 9 | Cyth2 | neutrophil.10 |
| Hk1 | 4.31E-91 | 0.62737337 | 0.359 | 0.156 | 8.34E-87 | 9 | Hk1 | neutrophil.10 |
| Ogdh | 6.73E-90 | 0.8088782 | 0.268 | 0.097 | 1.30E-85 | 9 | Ogdh | neutrophil.10 |
| Nedd8 | 1.07E-89 | 0.42972823 | 0.341 | 0.169 | 2.06E-85 | 9 | Nedd8 | neutrophil.10 |
| Dctn2 | 2.21E-89 | 0.76742172 | 0.278 | 0.109 | 4.27E-85 | 9 | Dctn2 | neutrophil.10 |
| Vcl | 2.72E-89 | 0.54689069 | 0.267 | 0.11 | 5.25E-85 | 9 | Vcl | neutrophil.10 |
| Itm2c | 3.26E-89 | 0.56731395 | 0.296 | 0.134 | 6.31E-85 | 9 | Itm2c | neutrophil.10 |
| Csnk1g2 | 4.94E-89 | 0.95527336 | 0.287 | 0.095 | 9.55E-85 | 9 | Csnk1g2 | neutrophil.10 |
| Ahcyl1 | 5.12E-89 | 0.7417275 | 0.255 | 0.095 | 9.89E-85 | 9 | Ahcyl1 | neutrophil.10 |
| Esyt2 | 6.88E-88 | 0.56814275 | 0.291 | 0.122 | 1.33E-83 | 9 | Esyt2 | neutrophil.10 |
| Hist1h2bc | 9.66E-88 | 0.40194123 | 0.317 | 0.186 | 1.87E-83 | 9 | Hist1h2bc | neutrophil.10 |
| Ywhab | 9.88E-88 | 0.5712987 | 0.299 | 0.122 | 1.91E-83 | 9 | Ywhab | neutrophil.10 |
| Dctn1 | 1.12E-86 | 0.92274956 | 0.283 | 0.099 | 2.16E-82 | 9 | Dctn1 | neutrophil.10 |
| Sin3b | 5.47E-86 | 0.36685991 | 0.39 | 0.204 | 1.06E-81 | 9 | Sin3b | neutrophil.10 |
| Ndufb1-ps | 6.67E-86 | 0.57132103 | 0.295 | 0.13 | 1.29E-81 | 9 | Ndufb1-ps | neutrophil.10 |
| Ube2v1 | 1.31E-85 | 0.6398853 | 0.279 | 0.119 | 2.53E-81 | 9 | Ube2v1 | neutrophil.10 |
| Sdha | 1.72E-85 | 0.61498489 | 0.272 | 0.116 | 3.32E-81 | 9 | Sdha | neutrophil.10 |
| Emp3 | 3.70E-84 | 0.29459891 | 0.312 | 0.139 | 7.16E-80 | 9 | Emp3 | neutrophil.10 |
| Prelid1 | 2.33E-83 | 0.46181385 | 0.352 | 0.164 | 4.51E-79 | 9 | Prelid1 | neutrophil.10 |
| Odc1 | 4.82E-83 | 0.62790833 | 0.256 | 0.095 | 9.31E-79 | 9 | Odc1 | neutrophil.10 |
| Ctsh | 2.34E-82 | 0.35591214 | 0.258 | 0.119 | 4.52E-78 | 9 | Ctsh | neutrophil.10 |
| Rhoa | 2.53E-82 | 0.27654292 | 0.525 | 0.304 | 4.89E-78 | 9 | Rhoa | neutrophil.10 |
| Acsl5 | 1.59E-81 | 0.60304598 | 0.29 | 0.119 | 3.07E-77 | 9 | Acsl5 | neutrophil.10 |
| Kmt5a | 2.20E-81 | 0.5776246 | 0.313 | 0.136 | 4.24E-77 | 9 | Kmt5a | neutrophil.10 |
| Flii | 1.90E-80 | 0.50897234 | 0.312 | 0.135 | 3.68E-76 | 9 | Flii | neutrophil.10 |
| Dazap1 | 4.01E-80 | 0.64383372 | 0.271 | 0.097 | 7.76E-76 | 9 | Dazap1 | neutrophil.10 |
| Samd4b | 1.75E-78 | 0.91185874 | 0.278 | 0.096 | 3.39E-74 | 9 | Samd4b | neutrophil.10 |
| Clic1 | 1.64E-77 | 0.44287795 | 0.388 | 0.203 | 3.18E-73 | 9 | Clic1 | neutrophil.10 |
| Chmp5 | 1.19E-75 | 0.71204155 | 0.28 | 0.118 | 2.30E-71 | 9 | Chmp5 | neutrophil.10 |
| Tmem59 | 1.93E-75 | 0.52631577 | 0.264 | 0.122 | 3.73E-71 | 9 | Tmem59 | neutrophil.10 |
| Add1 | 4.60E-75 | 0.53737271 | 0.323 | 0.148 | 8.90E-71 | 9 | Add1 | neutrophil.10 |
| Zfp106 | 1.65E-74 | 0.52268318 | 0.384 | 0.195 | 3.19E-70 | 9 | Zfp106 | neutrophil.10 |
| H2afy | 2.52E-73 | 0.38203341 | 0.349 | 0.161 | 4.87E-69 | 9 | H2afy | neutrophil.10 |
| Mapk3 | 8.89E-73 | 0.63647492 | 0.256 | 0.107 | 1.72E-68 | 9 | Mapk3 | neutrophil.10 |
| Myl12b | 1.36E-71 | 0.35442824 | 0.382 | 0.219 | 2.63E-67 | 9 | Myl12b | neutrophil.10 |
| Mapre1 | 1.46E-71 | 0.33181092 | 0.364 | 0.195 | 2.83E-67 | 9 | Mapre1 | neutrophil.10 |
| Mtpn | 1.85E-71 | 0.72099113 | 0.251 | 0.1 | 3.59E-67 | 9 | Mtpn | neutrophil.10 |
| Hnrnpl | 8.92E-70 | 0.58616091 | 0.259 | 0.096 | 1.72E-65 | 9 | Hnrnpl | neutrophil.10 |
| Mt1 | 7.31E-69 | 0.6626873 | 0.254 | 0.105 | 1.41E-64 | 9 | Mt1 | neutrophil.10 |
| Ppp2r1a | 6.65E-68 | 0.40296631 | 0.28 | 0.136 | 1.29E-63 | 9 | Ppp2r1a | neutrophil.10 |
| Ap2m1 | 5.61E-63 | 0.32248016 | 0.336 | 0.169 | 1.09E-58 | 9 | Ap2m1 | neutrophil.10 |
| Srp9 | 6.89E-63 | 0.56221517 | 0.255 | 0.114 | 1.33E-58 | 9 | Srp9 | neutrophil.10 |
| Qsox1 | 1.49E-62 | 0.45334829 | 0.297 | 0.149 | 2.88E-58 | 9 | Qsox1 | neutrophil.10 |
| Ccni | 3.21E-60 | 0.3769601 | 0.364 | 0.192 | 6.20E-56 | 9 | Ccni | neutrophil.10 |
| Hnrnpul2 | 2.29E-59 | 0.39899016 | 0.317 | 0.151 | 4.43E-55 | 9 | Hnrnpul2 | neutrophil.10 |
| Polr1d | 1.59E-57 | 0.52856078 | 0.251 | 0.096 | 3.07E-53 | 9 | Polr1d | neutrophil.10 |
| Ptbp1 | 5.27E-57 | 0.26857205 | 0.296 | 0.149 | 1.02E-52 | 9 | Ptbp1 | neutrophil.10 |
| Bag6 | 6.49E-57 | 0.45194723 | 0.256 | 0.107 | 1.26E-52 | 9 | Bag6 | neutrophil.10 |
| Luc7l2 | 1.73E-56 | 0.34486004 | 0.34 | 0.188 | 3.34E-52 | 9 | Luc7l2 | neutrophil.10 |
| Colgalt1 | 8.98E-56 | 0.30744293 | 0.252 | 0.122 | 1.74E-51 | 9 | Colgalt1 | neutrophil.10 |
| Grcc10 | 5.26E-55 | 0.43725065 | 0.278 | 0.129 | 1.02E-50 | 9 | Grcc10 | neutrophil.10 |
| Usp9x | 1.89E-54 | 0.45182813 | 0.255 | 0.111 | 3.65E-50 | 9 | Usp9x | neutrophil.10 |
| Tram1 | 1.10E-53 | 0.62069776 | 0.252 | 0.106 | 2.13E-49 | 9 | Tram1 | neutrophil.10 |
| Pcbp2 | 2.13E-53 | 0.50891302 | 0.263 | 0.126 | 4.11E-49 | 9 | Pcbp2 | neutrophil.10 |
| Nrd1 | 1.48E-51 | 0.29373974 | 0.291 | 0.159 | 2.86E-47 | 9 | Nrd1 | neutrophil.10 |
| Trip12 | 8.48E-51 | 0.36983945 | 0.324 | 0.17 | 1.64E-46 | 9 | Trip12 | neutrophil.10 |
| Cox5b | 2.01E-49 | 0.44357528 | 0.264 | 0.126 | 3.88E-45 | 9 | Cox5b | neutrophil.10 |
| Psme1 | 3.64E-48 | 0.43288522 | 0.272 | 0.127 | 7.03E-44 | 9 | Psme1 | neutrophil.10 |
| Ssr1 | 1.60E-47 | 0.48435206 | 0.25 | 0.124 | 3.09E-43 | 9 | Ssr1 | neutrophil.10 |
| Aamp | 3.47E-46 | 0.25667883 | 0.251 | 0.123 | 6.72E-42 | 9 | Aamp | neutrophil.10 |
| Aff4 | 1.01E-39 | 0.28802072 | 0.26 | 0.139 | 1.95E-35 | 9 | Aff4 | neutrophil.10 |
| Cast | 2.35E-39 | 0.26325147 | 0.316 | 0.18 | 4.55E-35 | 9 | Cast | neutrophil.10 |
| Ik | 1.72E-38 | 0.28357443 | 0.345 | 0.187 | 3.33E-34 | 9 | Ik | neutrophil.10 |
| Lin7c | 2.67E-37 | 0.48368881 | 0.256 | 0.117 | 5.17E-33 | 9 | Lin7c | neutrophil.10 |
| Spop | 1.48E-35 | 0.34507542 | 0.262 | 0.136 | 2.85E-31 | 9 | Spop | neutrophil.10 |
| Prpf8 | 1.48E-33 | 0.25216596 | 0.251 | 0.134 | 2.85E-29 | 9 | Prpf8 | neutrophil.10 |
| Mafb | 0 | 6.18463854 | 0.573 | 0.019 | 0 | 10 | Mafb | neutrophil.11 |
| Apoe | 0 | 3.92301361 | 0.683 | 0.231 | 0 | 10 | Apoe | neutrophil.11 |
| Fn1 | 0 | 3.26209466 | 0.633 | 0.195 | 0 | 10 | Fn1 | neutrophil.11 |
| H2-Ab1 | 0 | 4.53754506 | 0.518 | 0.087 | 0 | 10 | H2-Ab1 | neutrophil.11 |
| Cd74 | 0 | 3.06277725 | 0.754 | 0.375 | 0 | 10 | Cd74 | neutrophil.11 |
| H2-Eb1 | 0 | 4.51895094 | 0.467 | 0.092 | 0 | 10 | H2-Eb1 | neutrophil.11 |
| Lyz2 | 0 | 3.26898103 | 0.533 | 0.163 | 0 | 10 | Lyz2 | neutrophil.11 |
| H2-Aa | 0 | 3.4044882 | 0.412 | 0.079 | 0 | 10 | H2-Aa | neutrophil.11 |
| Lrp1 | 6.98E-306 | 4.50998871 | 0.503 | 0.048 | 1.35E-301 | 10 | Lrp1 | neutrophil.11 |
| C1qb | 2.10E-275 | 3.873342 | 0.322 | 0.044 | 4.06E-271 | 10 | C1qb | neutrophil.11 |
| Pld4 | 7.30E-264 | 5.13051502 | 0.266 | 0.008 | 1.41E-259 | 10 | Pld4 | neutrophil.11 |
| Vim | 4.71E-218 | 2.14956445 | 0.814 | 0.439 | 9.11E-214 | 10 | Vim | neutrophil.11 |
| Ctss | 5.19E-217 | 2.13079651 | 0.598 | 0.277 | 1.00E-212 | 10 | Ctss | neutrophil.11 |
| S100a4 | 4.24E-141 | 2.88457176 | 0.327 | 0.056 | 8.21E-137 | 10 | S100a4 | neutrophil.11 |
| Ctsc | 1.87E-127 | 3.0798725 | 0.487 | 0.094 | 3.61E-123 | 10 | Ctsc | neutrophil.11 |
| Ifi30 | 1.33E-118 | 3.25141716 | 0.387 | 0.077 | 2.58E-114 | 10 | Ifi30 | neutrophil.11 |
| Klf4 | 2.87E-118 | 3.56366402 | 0.271 | 0.033 | 5.55E-114 | 10 | Klf4 | neutrophil.11 |
| Ahnak | 1.06E-112 | 2.93621695 | 0.387 | 0.085 | 2.05E-108 | 10 | Ahnak | neutrophil.11 |
| Hspa1a | 4.35E-106 | 3.48701656 | 0.271 | 0.042 | 8.41E-102 | 10 | Hspa1a | neutrophil.11 |
| Cebpa | 1.38E-101 | 3.62940357 | 0.332 | 0.054 | 2.66E-97 | 10 | Cebpa | neutrophil.11 |
| Sowahc | 3.20E-79 | 3.75397303 | 0.312 | 0.047 | 6.18E-75 | 10 | Sowahc | neutrophil.11 |
| Crip1 | 2.08E-76 | 2.91262502 | 0.266 | 0.044 | 4.02E-72 | 10 | Crip1 | neutrophil.11 |
| mt-Co2 | 1.26E-69 | 1.26449217 | 0.543 | 0.345 | 2.43E-65 | 10 | mt-Co2 | neutrophil.11 |
| Tmsb10 | 4.34E-59 | 2.15718698 | 0.337 | 0.111 | 8.40E-55 | 10 | Tmsb10 | neutrophil.11 |
| mt-Nd6 | 3.56E-55 | 2.11868676 | 0.392 | 0.137 | 6.87E-51 | 10 | mt-Nd6 | neutrophil.11 |
| Tgfbi | 9.99E-47 | 1.33755851 | 0.523 | 0.359 | 1.93E-42 | 10 | Tgfbi | neutrophil.11 |
| Myo1g | 1.00E-46 | 2.31885478 | 0.276 | 0.08 | 1.94E-42 | 10 | Myo1g | neutrophil.11 |
| Ucp2 | 9.38E-43 | 1.18947916 | 0.603 | 0.496 | 1.81E-38 | 10 | Ucp2 | neutrophil.11 |
| Zfp703 | 2.25E-40 | 2.12939207 | 0.387 | 0.124 | 4.35E-36 | 10 | Zfp703 | neutrophil.11 |
| Jun | 5.49E-39 | 0.85128123 | 0.749 | 0.507 | 1.06E-34 | 10 | Jun | neutrophil.11 |
| Fos | 7.49E-30 | 0.46611808 | 0.678 | 0.563 | 1.45E-25 | 10 | Fos | neutrophil.11 |
| Trps1 | 8.96E-30 | 1.55907049 | 0.352 | 0.166 | 1.73E-25 | 10 | Trps1 | neutrophil.11 |
| Plac8 | 3.72E-29 | 0.75826182 | 0.302 | 0.23 | 7.19E-25 | 10 | Plac8 | neutrophil.11 |
| Lgals3 | 3.96E-29 | 1.29044719 | 0.472 | 0.268 | 7.66E-25 | 10 | Lgals3 | neutrophil.11 |
| Ptms | 3.86E-28 | 1.44549919 | 0.362 | 0.166 | 7.46E-24 | 10 | Ptms | neutrophil.11 |
| Runx3 | 9.08E-28 | 1.89839934 | 0.251 | 0.095 | 1.76E-23 | 10 | Runx3 | neutrophil.11 |
| mt-Co1 | 1.22E-27 | 1.44738306 | 0.382 | 0.164 | 2.35E-23 | 10 | mt-Co1 | neutrophil.11 |
| Csf1r | 4.34E-27 | 1.52359162 | 0.503 | 0.266 | 8.38E-23 | 10 | Csf1r | neutrophil.11 |
| mt-Atp8 | 8.35E-27 | 1.09669268 | 0.412 | 0.198 | 1.62E-22 | 10 | mt-Atp8 | neutrophil.11 |
| mt-Co3 | 2.56E-24 | 1.35444658 | 0.367 | 0.193 | 4.94E-20 | 10 | mt-Co3 | neutrophil.11 |
| mt-Atp6 | 5.39E-23 | 1.05745079 | 0.342 | 0.219 | 1.04E-18 | 10 | mt-Atp6 | neutrophil.11 |
| Lmna | 1.25E-22 | 1.5557202 | 0.296 | 0.117 | 2.41E-18 | 10 | Lmna | neutrophil.11 |
| Jund | 1.13E-21 | 0.88010475 | 0.824 | 0.729 | 2.19E-17 | 10 | Jund | neutrophil.11 |
| mt-Nd4l | 8.21E-21 | 1.03478599 | 0.352 | 0.149 | 1.59E-16 | 10 | mt-Nd4l | neutrophil.11 |
| Psap | 2.74E-20 | 1.12754979 | 0.533 | 0.338 | 5.29E-16 | 10 | Psap | neutrophil.11 |
| Tagln2 | 1.93E-19 | 1.31731407 | 0.437 | 0.24 | 3.74E-15 | 10 | Tagln2 | neutrophil.11 |
| Clta | 2.01E-19 | 0.98526798 | 0.533 | 0.357 | 3.88E-15 | 10 | Clta | neutrophil.11 |
| Npc2 | 2.99E-19 | 1.54102697 | 0.357 | 0.169 | 5.78E-15 | 10 | Npc2 | neutrophil.11 |
| Gpx1 | 7.89E-19 | 0.54159054 | 0.558 | 0.48 | 1.53E-14 | 10 | Gpx1 | neutrophil.11 |
| Gm2a | 3.42E-17 | 0.96314568 | 0.573 | 0.449 | 6.61E-13 | 10 | Gm2a | neutrophil.11 |
| Tgfb1 | 1.37E-16 | 0.93405892 | 0.503 | 0.402 | 2.65E-12 | 10 | Tgfb1 | neutrophil.11 |
| Cyth4 | 1.83E-16 | 1.1475181 | 0.432 | 0.293 | 3.53E-12 | 10 | Cyth4 | neutrophil.11 |
| Ly6e | 1.25E-15 | 0.63188234 | 0.392 | 0.371 | 2.42E-11 | 10 | Ly6e | neutrophil.11 |
| mt-Nd5 | 1.52E-15 | 1.1178856 | 0.261 | 0.117 | 2.93E-11 | 10 | mt-Nd5 | neutrophil.11 |
| Malt1 | 2.80E-15 | 1.61460346 | 0.327 | 0.194 | 5.42E-11 | 10 | Malt1 | neutrophil.11 |
| Id2 | 2.86E-15 | 1.65066355 | 0.367 | 0.164 | 5.53E-11 | 10 | Id2 | neutrophil.11 |
| Zfp36l2 | 7.91E-15 | 0.84882436 | 0.513 | 0.341 | 1.53E-10 | 10 | Zfp36l2 | neutrophil.11 |
| Sgk1 | 1.73E-14 | 1.43926418 | 0.261 | 0.143 | 3.34E-10 | 10 | Sgk1 | neutrophil.11 |
| Fosb | 3.23E-14 | 0.86366607 | 0.457 | 0.289 | 6.25E-10 | 10 | Fosb | neutrophil.11 |
| Irf2bp2 | 3.74E-14 | 0.86381743 | 0.598 | 0.467 | 7.23E-10 | 10 | Irf2bp2 | neutrophil.11 |
| Rrbp1 | 7.50E-14 | 1.46490085 | 0.312 | 0.164 | 1.45E-09 | 10 | Rrbp1 | neutrophil.11 |
| Tcirg1 | 1.24E-13 | 1.54246694 | 0.397 | 0.217 | 2.41E-09 | 10 | Tcirg1 | neutrophil.11 |
| Atf3 | 3.87E-13 | 0.50655366 | 0.492 | 0.391 | 7.49E-09 | 10 | Atf3 | neutrophil.11 |
| Prkcd | 4.06E-13 | 0.94120066 | 0.513 | 0.39 | 7.85E-09 | 10 | Prkcd | neutrophil.11 |
| Ctsh | 8.91E-13 | 1.03416419 | 0.261 | 0.121 | 1.72E-08 | 10 | Ctsh | neutrophil.11 |
| Fcgr2b | 8.97E-13 | 1.00861693 | 0.412 | 0.248 | 1.73E-08 | 10 | Fcgr2b | neutrophil.11 |
| Akr1a1 | 1.09E-12 | 1.13452534 | 0.367 | 0.198 | 2.10E-08 | 10 | Akr1a1 | neutrophil.11 |
| Plec | 1.61E-12 | 1.03080596 | 0.327 | 0.202 | 3.11E-08 | 10 | Plec | neutrophil.11 |
| Epn1 | 3.63E-12 | 1.21783833 | 0.317 | 0.196 | 7.02E-08 | 10 | Epn1 | neutrophil.11 |
| Gnai2 | 4.08E-12 | 0.92356851 | 0.558 | 0.435 | 7.90E-08 | 10 | Gnai2 | neutrophil.11 |
| Irf5 | 1.06E-11 | 1.25549582 | 0.397 | 0.246 | 2.05E-07 | 10 | Irf5 | neutrophil.11 |
| Eif4g3 | 2.34E-11 | 1.04705001 | 0.291 | 0.177 | 4.53E-07 | 10 | Eif4g3 | neutrophil.11 |
| Srsf6 | 4.80E-11 | 1.35300018 | 0.327 | 0.188 | 9.28E-07 | 10 | Srsf6 | neutrophil.11 |
| Dusp6 | 1.23E-10 | 1.35206675 | 0.302 | 0.151 | 2.38E-06 | 10 | Dusp6 | neutrophil.11 |
| H2afy | 1.38E-10 | 1.05202159 | 0.342 | 0.163 | 2.66E-06 | 10 | H2afy | neutrophil.11 |
| Sem1 | 3.10E-10 | 1.3896003 | 0.367 | 0.209 | 5.99E-06 | 10 | Sem1 | neutrophil.11 |
| Srsf2 | 3.47E-10 | 1.2508222 | 0.286 | 0.162 | 6.71E-06 | 10 | Srsf2 | neutrophil.11 |
| Eef2 | 3.52E-10 | 1.72021932 | 0.312 | 0.169 | 6.80E-06 | 10 | Eef2 | neutrophil.11 |
| Rhob | 3.89E-10 | 0.74184958 | 0.472 | 0.36 | 7.52E-06 | 10 | Rhob | neutrophil.11 |
| mt-Cytb | 5.87E-10 | 0.62332165 | 0.352 | 0.198 | 1.14E-05 | 10 | mt-Cytb | neutrophil.11 |
| Grn | 6.20E-10 | 0.93361934 | 0.302 | 0.189 | 1.20E-05 | 10 | Grn | neutrophil.11 |
| Sat1 | 6.64E-10 | 0.37932316 | 0.563 | 0.505 | 1.28E-05 | 10 | Sat1 | neutrophil.11 |
| Bin2 | 7.71E-10 | 1.02961558 | 0.437 | 0.282 | 1.49E-05 | 10 | Bin2 | neutrophil.11 |
| Ifngr1 | 9.27E-10 | 0.92133773 | 0.307 | 0.225 | 1.79E-05 | 10 | Ifngr1 | neutrophil.11 |
| Uck2 | 1.07E-09 | 1.1111311 | 0.337 | 0.184 | 2.08E-05 | 10 | Uck2 | neutrophil.11 |
| Itm2b | 3.72E-09 | 0.77059113 | 0.508 | 0.428 | 7.20E-05 | 10 | Itm2b | neutrophil.11 |
| Gna13 | 6.98E-09 | 1.0177866 | 0.508 | 0.382 | 1.35E-04 | 10 | Gna13 | neutrophil.11 |
| Vcp | 7.22E-09 | 1.27924647 | 0.271 | 0.148 | 1.40E-04 | 10 | Vcp | neutrophil.11 |
| Tcf25 | 1.19E-08 | 1.00111739 | 0.357 | 0.198 | 2.30E-04 | 10 | Tcf25 | neutrophil.11 |
| Fau | 1.19E-08 | 0.50692182 | 0.603 | 0.546 | 2.31E-04 | 10 | Fau | neutrophil.11 |
| Eef1a1 | 1.41E-08 | 0.40533745 | 0.628 | 0.57 | 2.73E-04 | 10 | Eef1a1 | neutrophil.11 |
| Cyth1 | 1.52E-08 | 1.13326365 | 0.296 | 0.18 | 2.95E-04 | 10 | Cyth1 | neutrophil.11 |
| Cebpb | 2.75E-08 | 0.25812733 | 0.92 | 0.932 | 5.33E-04 | 10 | Cebpb | neutrophil.11 |
| Golph3 | 2.82E-08 | 0.86739405 | 0.317 | 0.197 | 5.45E-04 | 10 | Golph3 | neutrophil.11 |
| Zeb2 | 2.94E-08 | 0.85271505 | 0.457 | 0.278 | 5.69E-04 | 10 | Zeb2 | neutrophil.11 |
| Rel | 5.94E-08 | 0.84014485 | 0.392 | 0.29 | 0.0011493 | 10 | Rel | neutrophil.11 |
| Serf2 | 9.27E-08 | 1.1814458 | 0.291 | 0.177 | 0.00179284 | 10 | Serf2 | neutrophil.11 |
| Gnas | 9.86E-08 | 0.93876355 | 0.261 | 0.16 | 0.00190683 | 10 | Gnas | neutrophil.11 |
| Oaz1 | 1.06E-07 | 1.39590442 | 0.266 | 0.144 | 0.0020529 | 10 | Oaz1 | neutrophil.11 |
| Yy1 | 1.12E-07 | 0.88487379 | 0.342 | 0.219 | 0.00216061 | 10 | Yy1 | neutrophil.11 |
| Celf2 | 1.20E-07 | 1.15631354 | 0.271 | 0.148 | 0.00232246 | 10 | Celf2 | neutrophil.11 |
| Nfkb1 | 1.29E-07 | 0.79821215 | 0.312 | 0.214 | 0.00248703 | 10 | Nfkb1 | neutrophil.11 |
| Glud1 | 1.96E-07 | 1.37157086 | 0.276 | 0.161 | 0.00379283 | 10 | Glud1 | neutrophil.11 |
| Plin2 | 3.49E-07 | 0.45333574 | 0.442 | 0.351 | 0.00675656 | 10 | Plin2 | neutrophil.11 |
| Bach1 | 3.70E-07 | 0.70945454 | 0.296 | 0.229 | 0.00715144 | 10 | Bach1 | neutrophil.11 |
| H3f3a | 4.15E-07 | 0.78147318 | 0.487 | 0.422 | 0.00801407 | 10 | H3f3a | neutrophil.11 |
| Mat2a | 4.71E-07 | 0.71536503 | 0.342 | 0.21 | 0.00910444 | 10 | Mat2a | neutrophil.11 |
| Ppp1r12c | 4.86E-07 | 1.15529138 | 0.251 | 0.148 | 0.00939978 | 10 | Ppp1r12c | neutrophil.11 |
| Phlda1 | 5.75E-07 | 1.1477025 | 0.312 | 0.177 | 0.01111625 | 10 | Phlda1 | neutrophil.11 |
| Cytip | 7.71E-07 | 0.39232594 | 0.402 | 0.36 | 0.01491263 | 10 | Cytip | neutrophil.11 |
| F10 | 8.17E-07 | 0.38525623 | 0.332 | 0.193 | 0.01579789 | 10 | F10 | neutrophil.11 |
| Fes | 8.28E-07 | 0.45658657 | 0.357 | 0.257 | 0.01600872 | 10 | Fes | neutrophil.11 |
| Ybx1 | 8.36E-07 | 0.95472866 | 0.251 | 0.132 | 0.01615432 | 10 | Ybx1 | neutrophil.11 |
| Cst3 | 1.13E-06 | 0.34666568 | 0.648 | 0.593 | 0.02192196 | 10 | Cst3 | neutrophil.11 |
| Nisch | 1.22E-06 | 0.77684513 | 0.302 | 0.192 | 0.02361031 | 10 | Nisch | neutrophil.11 |
| H2-K1 | 1.44E-06 | 0.48882079 | 0.583 | 0.526 | 0.02793266 | 10 | H2-K1 | neutrophil.11 |
| Zfp131 | 1.58E-06 | 0.37512768 | 0.286 | 0.195 | 0.0304639 | 10 | Zfp131 | neutrophil.11 |
| Ctsa | 1.65E-06 | 0.67048068 | 0.337 | 0.238 | 0.03184251 | 10 | Ctsa | neutrophil.11 |
| Pitpna | 1.83E-06 | 0.82569565 | 0.372 | 0.296 | 0.03546932 | 10 | Pitpna | neutrophil.11 |
| Qk | 4.83E-06 | 0.86514399 | 0.322 | 0.217 | 0.093292 | 10 | Qk | neutrophil.11 |
| Pfn1 | 5.45E-06 | 0.60270548 | 0.608 | 0.549 | 0.10534146 | 10 | Pfn1 | neutrophil.11 |
| Arl8a | 9.48E-06 | 0.79019777 | 0.261 | 0.156 | 0.18330111 | 10 | Arl8a | neutrophil.11 |
| Kctd12 | 1.02E-05 | 0.5112719 | 0.739 | 0.636 | 0.19700941 | 10 | Kctd12 | neutrophil.11 |
| Thbs1 | 1.33E-05 | 0.46724525 | 0.482 | 0.298 | 0.25752831 | 10 | Thbs1 | neutrophil.11 |
| Atox1 | 1.59E-05 | 0.76722189 | 0.442 | 0.321 | 0.30646438 | 10 | Atox1 | neutrophil.11 |
| Hck | 1.90E-05 | 0.36531064 | 0.342 | 0.301 | 0.36709011 | 10 | Hck | neutrophil.11 |
| Irs2 | 1.94E-05 | 0.88434615 | 0.312 | 0.202 | 0.37433036 | 10 | Irs2 | neutrophil.11 |
| Arhgef1 | 2.16E-05 | 0.5326922 | 0.332 | 0.246 | 0.41744283 | 10 | Arhgef1 | neutrophil.11 |
| Id1 | 2.34E-05 | 0.96512334 | 0.317 | 0.244 | 0.45332039 | 10 | Id1 | neutrophil.11 |
| Csf2ra | 2.63E-05 | 0.44790396 | 0.553 | 0.489 | 0.50822073 | 10 | Csf2ra | neutrophil.11 |
| Acly | 3.68E-05 | 0.6479889 | 0.251 | 0.17 | 0.71245316 | 10 | Acly | neutrophil.11 |
| Runx1 | 4.05E-05 | 0.71526823 | 0.276 | 0.189 | 0.78259204 | 10 | Runx1 | neutrophil.11 |
| Snx18 | 4.20E-05 | 0.43331338 | 0.352 | 0.297 | 0.81292221 | 10 | Snx18 | neutrophil.11 |
| Cbfb | 5.03E-05 | 0.7360291 | 0.276 | 0.176 | 0.97321573 | 10 | Cbfb | neutrophil.11 |
| Vegfa | 7.86E-05 | 0.78763625 | 0.337 | 0.237 | 1 | 10 | Vegfa | neutrophil.11 |
| Edem1 | 1.00E-04 | 0.70751006 | 0.261 | 0.2 | 1 | 10 | Edem1 | neutrophil.11 |
| Cfp | 1.34E-04 | 0.57220588 | 0.387 | 0.274 | 1 | 10 | Cfp | neutrophil.11 |
| Cux1 | 1.93E-04 | 0.89906763 | 0.312 | 0.251 | 1 | 10 | Cux1 | neutrophil.11 |
| Arhgap45 | 2.21E-04 | 0.67196729 | 0.342 | 0.253 | 1 | 10 | Arhgap45 | neutrophil.11 |
| Stk38 | 2.53E-04 | 0.75577295 | 0.266 | 0.188 | 1 | 10 | Stk38 | neutrophil.11 |
| Ehbp1l1 | 4.88E-04 | 0.3768002 | 0.291 | 0.231 | 1 | 10 | Ehbp1l1 | neutrophil.11 |
| Ctsz | 4.94E-04 | 0.37803312 | 0.497 | 0.449 | 1 | 10 | Ctsz | neutrophil.11 |
| Lrrfip1 | 5.59E-04 | 0.5285259 | 0.397 | 0.355 | 1 | 10 | Lrrfip1 | neutrophil.11 |
| Purb | 6.53E-04 | 0.6323238 | 0.291 | 0.2 | 1 | 10 | Purb | neutrophil.11 |
| Arhgdia | 0.00108134 | 0.58662601 | 0.332 | 0.275 | 1 | 10 | Arhgdia | neutrophil.11 |
| Dgkz | 0.0012562 | 0.75561571 | 0.286 | 0.212 | 1 | 10 | Dgkz | neutrophil.11 |
| App | 0.00128393 | 0.40419219 | 0.533 | 0.47 | 1 | 10 | App | neutrophil.11 |
| Tnrc18 | 0.00162689 | 0.46316461 | 0.296 | 0.226 | 1 | 10 | Tnrc18 | neutrophil.11 |
| Tmem189 | 0.00180668 | 0.71397513 | 0.342 | 0.229 | 1 | 10 | Tmem189 | neutrophil.11 |
| Plbd1 | 0.00182719 | 0.27705018 | 0.256 | 0.223 | 1 | 10 | Plbd1 | neutrophil.11 |
| Cebpd | 0.00239643 | 0.3423854 | 0.337 | 0.304 | 1 | 10 | Cebpd | neutrophil.11 |
| Akap13 | 0.00239752 | 0.69393411 | 0.261 | 0.195 | 1 | 10 | Akap13 | neutrophil.11 |
| Myl12a | 0.00323751 | 0.52919688 | 0.296 | 0.227 | 1 | 10 | Myl12a | neutrophil.11 |
| Rara | 0.00339749 | 0.44945893 | 0.357 | 0.323 | 1 | 10 | Rara | neutrophil.11 |
| Hspa8 | 0.0035445 | 0.42519445 | 0.462 | 0.422 | 1 | 10 | Hspa8 | neutrophil.11 |
| Ywhag | 0.00426361 | 0.53480551 | 0.387 | 0.326 | 1 | 10 | Ywhag | neutrophil.11 |
| Sfpq | 0.00468871 | 0.47736664 | 0.286 | 0.23 | 1 | 10 | Sfpq | neutrophil.11 |
| Ier2 | 0.00511242 | 0.25978499 | 0.608 | 0.561 | 1 | 10 | Ier2 | neutrophil.11 |
| Actn4 | 0.00595272 | 0.65720471 | 0.307 | 0.214 | 1 | 10 | Actn4 | neutrophil.11 |
| D8Ertd738e | 0.00667391 | 0.94331686 | 0.261 | 0.202 | 1 | 10 | D8Ertd738e | neutrophil.11 |
| Pcf11 | 0.00693422 | 0.39803552 | 0.472 | 0.416 | 1 | 10 | Pcf11 | neutrophil.11 |
| Kdm7a | 0.00724896 | 0.74634881 | 0.276 | 0.203 | 1 | 10 | Kdm7a | neutrophil.11 |
| Crk | 0.00732401 | 0.54000142 | 0.276 | 0.213 | 1 | 10 | Crk | neutrophil.11 |
| Anp32a | 0.00794733 | 0.29854832 | 0.332 | 0.278 | 1 | 10 | Anp32a | neutrophil.11 |
| Zfand5 | 0.00806871 | 0.283116 | 0.332 | 0.303 | 1 | 10 | Zfand5 | neutrophil.11 |
| Gda | 0.00898416 | 0.2924374 | 0.327 | 0.271 | 1 | 10 | Gda | neutrophil.11 |
| Bax | 0.00974269 | 0.55890142 | 0.266 | 0.182 | 1 | 10 | Bax | neutrophil.11 |

| | p_val | avg_log2FC | pct.1 | pct.2 | p_val_adj | cluster | gene |
|---|---|---|---|---|---|---|---|
| Foxp3 | 0 | 6.28000981 | 0.804 | 0.015 | 0 | regulatory Tc | Foxp3 |
| Tnfrsf4 | 0 | 3.46301248 | 0.763 | 0.153 | 0 | regulatory Tc | Tnfrsf4 |
| Il2ra | 0 | 3.26180762 | 0.791 | 0.208 | 0 | regulatory Tc | Il2ra |
| Klrg1 | 0 | 4.00222945 | 0.631 | 0.072 | 0 | regulatory Tc | Klrg1 |
| Ikzf2 | 0 | 2.71776959 | 0.769 | 0.228 | 0 | regulatory Tc | Ikzf2 |
| Tnfrsf9 | 0 | 2.72355917 | 0.803 | 0.298 | 0 | regulatory Tc | Tnfrsf9 |
| Dusp4 | 0 | 2.53645962 | 0.663 | 0.182 | 0 | regulatory Tc | Dusp4 |
| Itgav | 0 | 2.59879006 | 0.726 | 0.253 | 0 | regulatory Tc | Itgav |
| Arl5a | 0 | 2.91117338 | 0.602 | 0.153 | 0 | regulatory Tc | Arl5a |
| Gpr55 | 0 | 3.02472583 | 0.524 | 0.086 | 0 | regulatory Tc | Gpr55 |
| Ctla4 | 0 | 2.34380107 | 0.818 | 0.382 | 0 | regulatory Tc | Ctla4 |
| Cd4 | 0 | 0.92100408 | 0.659 | 0.228 | 0 | regulatory Tc | Cd4 |
| Cd81 | 0 | 3.24375912 | 0.491 | 0.062 | 0 | regulatory Tc | Cd81 |
| Slc25a19 | 0 | 2.4895076 | 0.603 | 0.175 | 0 | regulatory Tc | Slc25a19 |
| Il1rl1 | 0 | 2.91529036 | 0.47 | 0.054 | 0 | regulatory Tc | Il1rl1 |
| Slc15a3 | 0 | 2.86602897 | 0.508 | 0.094 | 0 | regulatory Tc | Slc15a3 |
| Itgae | 0 | 1.91124602 | 0.572 | 0.172 | 0 | regulatory Tc | Itgae |
| Maf | 0 | 1.64687621 | 0.701 | 0.307 | 0 | regulatory Tc | Maf |
| Areg | 0 | 3.38279119 | 0.45 | 0.057 | 0 | regulatory Tc | Areg |
| Ass1 | 0 | 1.85810743 | 0.569 | 0.18 | 0 | regulatory Tc | Ass1 |
| Ccr8 | 0 | 2.32523878 | 0.519 | 0.143 | 0 | regulatory Tc | Ccr8 |
| Tnfrsf18 | 0 | 1.77162689 | 0.79 | 0.422 | 0 | regulatory Tc | Tnfrsf18 |
| Cd5 | 0 | 1.29343979 | 0.724 | 0.365 | 0 | regulatory Tc | Cd5 |
| Gadd45b | 0 | 1.02846023 | 0.77 | 0.425 | 0 | regulatory Tc | Gadd45b |
| Snx18 | 0 | 1.50398149 | 0.682 | 0.343 | 0 | regulatory Tc | Snx18 |
| Itm2c | 0 | 1.70768474 | 0.782 | 0.445 | 0 | regulatory Tc | Itm2c |
| Ighm | 0 | 2.12390428 | 0.493 | 0.157 | 0 | regulatory Tc | Ighm |
| Ly6a | 0 | 1.49349319 | 0.62 | 0.291 | 0 | regulatory Tc | Ly6a |
| Hopx | 0 | 2.20483854 | 0.722 | 0.394 | 0 | regulatory Tc | Hopx |
| Ddit4 | 0 | 1.9001555 | 0.567 | 0.24 | 0 | regulatory Tc | Ddit4 |
| Gm2a | 0 | 1.66038578 | 0.657 | 0.331 | 0 | regulatory Tc | Gm2a |
| Glrx | 0 | 1.83816324 | 0.582 | 0.256 | 0 | regulatory Tc | Glrx |
| Odc1 | 0 | 1.65534168 | 0.875 | 0.566 | 0 | regulatory Tc | Odc1 |
| Mxd1 | 0 | 1.66986611 | 0.695 | 0.387 | 0 | regulatory Tc | Mxd1 |
| Samsn1 | 0 | 1.24505727 | 0.675 | 0.374 | 0 | regulatory Tc | Samsn1 |
| Capg | 0 | 1.20373829 | 0.67 | 0.371 | 0 | regulatory Tc | Capg |
| Rel | 0 | 1.6449875 | 0.606 | 0.312 | 0 | regulatory Tc | Rel |
| Cd74 | 0 | 1.44606065 | 0.571 | 0.283 | 0 | regulatory Tc | Cd74 |
| Cd2 | 0 | 1.19274466 | 0.809 | 0.529 | 0 | regulatory Tc | Cd2 |
| Zfp36l1 | 0 | 1.07837919 | 0.706 | 0.428 | 0 | regulatory Tc | Zfp36l1 |
| Icos | 0 | 0.82385321 | 0.757 | 0.482 | 0 | regulatory Tc | Icos |
| Traf1 | 0 | 1.25198729 | 0.739 | 0.468 | 0 | regulatory Tc | Traf1 |
| Tnfrsf1b | 0 | 1.33128499 | 0.752 | 0.489 | 0 | regulatory Tc | Tnfrsf1b |
| Gzmb | 0 | 1.10635084 | 0.554 | 0.316 | 0 | regulatory Tc | Gzmb |
| Pim1 | 0 | 1.05835216 | 0.874 | 0.655 | 0 | regulatory Tc | Pim1 |
| S100a4 | 0 | 0.54186874 | 0.79 | 0.576 | 0 | regulatory Tc | S100a4 |
| Penk | 0 | 2.20884539 | 0.252 | 0.041 | 0 | regulatory Tc | Penk |
| Wnk1 | 0 | 1.00821597 | 0.745 | 0.556 | 0 | regulatory Tc | Wnk1 |
| Nfkbia | 0 | 0.69538958 | 0.895 | 0.707 | 0 | regulatory Tc | Nfkbia |
| Nr4a1 | 0 | 0.62656295 | 0.792 | 0.621 | 0 | regulatory Tc | Nr4a1 |
| Itgb7 | 0 | 0.59782622 | 0.797 | 0.63 | 0 | regulatory Tc | Itgb7 |
| Serinc3 | 0 | 0.89320601 | 0.849 | 0.704 | 0 | regulatory Tc | Serinc3 |
| Jund | 0 | 0.88605515 | 0.901 | 0.794 | 0 | regulatory Tc | Jund |
| Vim | 0 | 1.17344505 | 0.914 | 0.819 | 0 | regulatory Tc | Vim |
| Ly6e | 0 | 0.76917328 | 0.894 | 0.801 | 0 | regulatory Tc | Ly6e |
| Srgn | 0 | 0.72313635 | 0.915 | 0.877 | 0 | regulatory Tc | Srgn |
| S100a10 | 0 | 0.29319303 | 0.79 | 0.771 | 0 | regulatory Tc | S100a10 |
| Rrad | 1.18E-305 | 1.87337737 | 0.457 | 0.186 | 2.27E-301 | regulatory Tc | Rrad |
| Fam110a | 4.95E-296 | 2.00075778 | 0.569 | 0.195 | 9.57E-292 | regulatory Tc | Fam110a |
| Bcl2l1 | 2.10E-295 | 1.77731408 | 0.506 | 0.208 | 4.06E-291 | regulatory Tc | Bcl2l1 |
| Stx11 | 5.66E-294 | 1.91992522 | 0.516 | 0.19 | 1.09E-289 | regulatory Tc | Stx11 |
| Trac | 3.15E-286 | 0.43397929 | 0.819 | 0.62 | 6.09E-282 | regulatory Tc | Trac |
| Cd83 | 3.82E-267 | 3.23977559 | 0.323 | 0.054 | 7.38E-263 | regulatory Tc | Cd83 |
| Rora | 8.52E-264 | 0.89378991 | 0.652 | 0.376 | 1.65E-259 | regulatory Tc | Rora |
| Lmna | 8.24E-257 | 1.05501107 | 0.489 | 0.27 | 1.59E-252 | regulatory Tc | Lmna |
| Tent5a | 6.62E-253 | 2.06997169 | 0.422 | 0.141 | 1.28E-248 | regulatory Tc | Tent5a |
| Il2rg | 5.96E-251 | 0.54861622 | 0.841 | 0.767 | 1.15E-246 | regulatory Tc | Il2rg |
| Stat1 | 5.61E-249 | 0.92259874 | 0.573 | 0.378 | 1.08E-244 | regulatory Tc | Stat1 |
| Nfkbiz | 2.31E-247 | 0.96742559 | 0.654 | 0.388 | 4.46E-243 | regulatory Tc | Nfkbiz |
| H2-K1 | 1.02E-244 | 0.36421913 | 0.958 | 0.941 | 1.97E-240 | regulatory Tc | H2-K1 |
| Rab11fip1 | 1.52E-242 | 1.43373585 | 0.568 | 0.261 | 2.94E-238 | regulatory Tc | Rab11fip1 |
| Crem | 9.70E-242 | 0.5843648 | 0.808 | 0.627 | 1.88E-237 | regulatory Tc | Crem |
| N4bp1 | 3.66E-241 | 1.86159862 | 0.559 | 0.223 | 7.08E-237 | regulatory Tc | N4bp1 |
| Hcls1 | 4.83E-238 | 0.69823565 | 0.737 | 0.593 | 9.34E-234 | regulatory Tc | Hcls1 |
| Tspan13 | 1.87E-237 | 1.5310406 | 0.531 | 0.233 | 3.61E-233 | regulatory Tc | Tspan13 |
| Pkm | 7.40E-236 | 0.91210141 | 0.591 | 0.4 | 1.43E-231 | regulatory Tc | Pkm |
| Lamc1 | 5.98E-233 | 2.59799895 | 0.48 | 0.097 | 1.16E-228 | regulatory Tc | Lamc1 |
| Ets2 | 1.46E-230 | 2.40055114 | 0.46 | 0.116 | 2.83E-226 | regulatory Tc | Ets2 |
| Ctsz | 1.29E-227 | 1.87859621 | 0.417 | 0.152 | 2.49E-223 | regulatory Tc | Ctsz |
| Ctss | 1.02E-219 | 1.20609246 | 0.62 | 0.357 | 1.98E-215 | regulatory Tc | Ctss |
| Med15 | 6.74E-218 | 1.50693991 | 0.55 | 0.269 | 1.30E-213 | regulatory Tc | Med15 |
| Picalm | 6.57E-216 | 1.21137776 | 0.561 | 0.299 | 1.27E-211 | regulatory Tc | Picalm |
| Trp53i11 | 4.29E-215 | 1.42499698 | 0.518 | 0.213 | 8.29E-211 | regulatory Tc | Trp53i11 |
| Bmyc | 6.20E-215 | 3.86418883 | 0.398 | 0.043 | 1.20E-210 | regulatory Tc | Bmyc |
| Nrip1 | 1.48E-214 | 1.71819579 | 0.485 | 0.186 | 2.85E-210 | regulatory Tc | Nrip1 |
| Shisa5 | 1.11E-213 | 0.39871963 | 0.904 | 0.857 | 2.14E-209 | regulatory Tc | Shisa5 |
| Fth1 | 6.77E-213 | 0.43439535 | 0.85 | 0.787 | 1.31E-208 | regulatory Tc | Fth1 |
| Nfkb2 | 8.86E-213 | 0.86088203 | 0.71 | 0.471 | 1.71E-208 | regulatory Tc | Nfkb2 |
| Got1 | 8.17E-208 | 1.79716626 | 0.421 | 0.156 | 1.58E-203 | regulatory Tc | Got1 |
| Psen2 | 9.90E-208 | 1.75946955 | 0.484 | 0.17 | 1.91E-203 | regulatory Tc | Psen2 |
| Lmnb1 | 4.63E-207 | 0.85309777 | 0.488 | 0.314 | 8.95E-203 | regulatory Tc | Lmnb1 |
| Bcl3 | 3.64E-206 | 1.16933966 | 0.562 | 0.273 | 7.03E-202 | regulatory Tc | Bcl3 |
| Lrrc32 | 3.54E-204 | 5.00278172 | 0.401 | 0.016 | 6.85E-200 | regulatory Tc | Lrrc32 |
| Gata3 | 5.77E-202 | 0.9420001 | 0.526 | 0.302 | 1.12E-197 | regulatory Tc | Gata3 |
| Hacd3 | 3.51E-199 | 1.76568712 | 0.455 | 0.165 | 6.78E-195 | regulatory Tc | Hacd3 |
| Ttc39c | 3.57E-199 | 2.02249044 | 0.47 | 0.147 | 6.90E-195 | regulatory Tc | Ttc39c |
| Nr4a3 | 1.75E-194 | 0.7852358 | 0.615 | 0.43 | 3.39E-190 | regulatory Tc | Nr4a3 |
| Cmtm7 | 2.15E-194 | 1.07340255 | 0.488 | 0.263 | 4.16E-190 | regulatory Tc | Cmtm7 |
| M6pr | 2.61E-192 | 0.8962131 | 0.505 | 0.331 | 5.04E-188 | regulatory Tc | M6pr |
| Coro2a | 5.65E-188 | 1.44506809 | 0.509 | 0.229 | 1.09E-183 | regulatory Tc | Coro2a |
| Sdf4 | 4.34E-187 | 0.85030056 | 0.698 | 0.507 | 8.38E-183 | regulatory Tc | Sdf4 |
| Mapkapk2 | 2.08E-186 | 1.07004389 | 0.579 | 0.352 | 4.02E-182 | regulatory Tc | Mapkapk2 |
| Jak2 | 3.66E-186 | 0.91967249 | 0.602 | 0.376 | 7.09E-182 | regulatory Tc | Jak2 |
| Sla | 1.36E-185 | 0.83665809 | 0.656 | 0.433 | 2.63E-181 | regulatory Tc | Sla |
| Sik1 | 1.23E-183 | 0.76834181 | 0.704 | 0.501 | 2.37E-179 | regulatory Tc | Sik1 |
| Dusp1 | 5.28E-183 | 0.42126946 | 0.721 | 0.593 | 1.02E-178 | regulatory Tc | Dusp1 |
| Sdc4 | 8.05E-183 | 2.96325904 | 0.401 | 0.055 | 1.56E-178 | regulatory Tc | Sdc4 |
| Ifrd1 | 2.88E-182 | 0.82721334 | 0.631 | 0.443 | 5.57E-178 | regulatory Tc | Ifrd1 |
| Plxnd1 | 3.40E-181 | 1.54731471 | 0.441 | 0.161 | 6.58E-177 | regulatory Tc | Plxnd1 |
| Crip1 | 6.26E-181 | 0.33163548 | 0.656 | 0.58 | 1.21E-176 | regulatory Tc | Crip1 |
| Ndfip1 | 1.72E-179 | 0.54799264 | 0.801 | 0.688 | 3.32E-175 | regulatory Tc | Ndfip1 |
| Anp32a | 3.78E-177 | 0.49220195 | 0.724 | 0.606 | 7.31E-173 | regulatory Tc | Anp32a |
| Swap70 | 1.38E-175 | 2.44326763 | 0.372 | 0.09 | 2.67E-171 | regulatory Tc | Swap70 |
| H2afz | 1.55E-174 | 0.89004807 | 0.468 | 0.315 | 2.99E-170 | regulatory Tc | H2afz |
| Flnb | 1.83E-174 | 1.82462168 | 0.378 | 0.136 | 3.53E-170 | regulatory Tc | Flnb |
| Pcyt1a | 3.39E-173 | 3.17735126 | 0.329 | 0.057 | 6.55E-169 | regulatory Tc | Pcyt1a |
| H1f0 | 5.28E-172 | 1.20377708 | 0.331 | 0.146 | 1.02E-167 | regulatory Tc | H1f0 |
| Samhd1 | 6.39E-172 | 0.76440995 | 0.581 | 0.385 | 1.24E-167 | regulatory Tc | Samhd1 |
| Id2 | 1.22E-168 | 0.48753228 | 0.694 | 0.575 | 2.37E-164 | regulatory Tc | Id2 |
| Ccr2 | 1.47E-167 | 0.90912365 | 0.477 | 0.263 | 2.83E-163 | regulatory Tc | Ccr2 |
| Egln3 | 2.78E-167 | 2.3721072 | 0.323 | 0.07 | 5.37E-163 | regulatory Tc | Egln3 |
| Ifi27l2a | 7.70E-167 | 1.36474377 | 0.393 | 0.158 | 1.49E-162 | regulatory Tc | Ifi27l2a |
| Icam1 | 1.65E-161 | 0.80277246 | 0.557 | 0.358 | 3.18E-157 | regulatory Tc | Icam1 |
| Mfge8 | 1.06E-158 | 1.21276497 | 0.537 | 0.272 | 2.05E-154 | regulatory Tc | Mfge8 |
| Ltb | 1.78E-157 | 0.51795704 | 0.749 | 0.626 | 3.44E-153 | regulatory Tc | Ltb |
| Rilpl2 | 9.57E-157 | 1.24386739 | 0.439 | 0.231 | 1.85E-152 | regulatory Tc | Rilpl2 |
| Ikzf4 | 5.29E-154 | 3.22492055 | 0.354 | 0.055 | 1.02E-149 | regulatory Tc | Ikzf4 |
| Slc25a51 | 1.76E-153 | 1.5566344 | 0.399 | 0.168 | 3.39E-149 | regulatory Tc | Slc25a51 |
| Cish | 3.65E-152 | 0.91404357 | 0.59 | 0.327 | 7.06E-148 | regulatory Tc | Cish |
| Adap1 | 1.22E-151 | 2.13049753 | 0.381 | 0.113 | 2.36E-147 | regulatory Tc | Adap1 |
| Ccnd2 | 6.16E-151 | 0.49801582 | 0.61 | 0.484 | 1.19E-146 | regulatory Tc | Ccnd2 |
| Sema4d | 1.59E-150 | 0.44950947 | 0.788 | 0.683 | 3.08E-146 | regulatory Tc | Sema4d |
| Zc3h12d | 6.04E-150 | 1.89375849 | 0.381 | 0.111 | 1.17E-145 | regulatory Tc | Zc3h12d |
| Ldha | 3.49E-149 | 0.41737123 | 0.609 | 0.513 | 6.75E-145 | regulatory Tc | Ldha |
| Myo1g | 9.19E-149 | 0.75189882 | 0.595 | 0.433 | 1.78E-144 | regulatory Tc | Myo1g |
| Ell2 | 3.32E-147 | 1.60245523 | 0.411 | 0.154 | 6.42E-143 | regulatory Tc | Ell2 |
| Sema4b | 5.06E-144 | 1.47247749 | 0.396 | 0.136 | 9.79E-140 | regulatory Tc | Sema4b |
| Cebpb | 7.59E-144 | 1.07134069 | 0.495 | 0.272 | 1.47E-139 | regulatory Tc | Cebpb |
| Pcsk1 | 8.88E-143 | 4.93918355 | 0.252 | 0.011 | 1.72E-138 | regulatory Tc | Pcsk1 |
| Uap1 | 1.04E-142 | 1.37361409 | 0.365 | 0.167 | 2.01E-138 | regulatory Tc | Uap1 |
| Ndrg1 | 6.46E-142 | 2.02035587 | 0.3 | 0.078 | 1.25E-137 | regulatory Tc | Ndrg1 |
| Tank | 7.78E-141 | 1.62694529 | 0.467 | 0.191 | 1.50E-136 | regulatory Tc | Tank |
| Neb | 9.83E-137 | 3.97647518 | 0.348 | 0.034 | 1.90E-132 | regulatory Tc | Neb |
| Csrnp1 | 1.85E-136 | 0.81440394 | 0.567 | 0.34 | 3.58E-132 | regulatory Tc | Csrnp1 |
| Syngr2 | 2.88E-135 | 1.51152818 | 0.4 | 0.169 | 5.58E-131 | regulatory Tc | Syngr2 |
| Izumo1r | 6.30E-134 | 2.84149989 | 0.313 | 0.044 | 1.22E-129 | regulatory Tc | Izumo1r |
| Il1r2 | 1.01E-133 | 1.31607308 | 0.295 | 0.067 | 1.95E-129 | regulatory Tc | Il1r2 |
| Vrk1 | 1.26E-133 | 1.46220373 | 0.34 | 0.12 | 2.44E-129 | regulatory Tc | Vrk1 |
| Cd6 | 3.63E-133 | 0.85990052 | 0.516 | 0.286 | 7.02E-129 | regulatory Tc | Cd6 |
| Snx20 | 1.87E-130 | 1.22270232 | 0.458 | 0.212 | 3.61E-126 | regulatory Tc | Snx20 |
| Slc9a3r1 | 2.65E-130 | 0.75232837 | 0.605 | 0.44 | 5.12E-126 | regulatory Tc | Slc9a3r1 |
| Cdkn1a | 3.17E-130 | 0.84915348 | 0.391 | 0.193 | 6.13E-126 | regulatory Tc | Cdkn1a |
| Sdcbp2 | 3.49E-129 | 1.98354019 | 0.35 | 0.103 | 6.75E-125 | regulatory Tc | Sdcbp2 |
| Ppp1r15a | 2.26E-127 | 0.57887631 | 0.601 | 0.449 | 4.36E-123 | regulatory Tc | Ppp1r15a |
| Rhof | 4.90E-127 | 0.85642246 | 0.552 | 0.353 | 9.48E-123 | regulatory Tc | Rhof |
| Snx9 | 4.01E-126 | 2.02327975 | 0.376 | 0.11 | 7.76E-122 | regulatory Tc | Snx9 |
| Mknk2 | 1.19E-124 | 0.49981362 | 0.704 | 0.573 | 2.30E-120 | regulatory Tc | Mknk2 |
| Cep85l | 1.44E-124 | 3.17773215 | 0.305 | 0.043 | 2.79E-120 | regulatory Tc | Cep85l |
| Plec | 4.79E-123 | 0.57458169 | 0.656 | 0.502 | 9.27E-119 | regulatory Tc | Plec |
| Map2k3 | 2.26E-120 | 0.84596001 | 0.488 | 0.296 | 4.37E-116 | regulatory Tc | Map2k3 |
| Igtp | 9.57E-120 | 1.07712069 | 0.321 | 0.141 | 1.85E-115 | regulatory Tc | Igtp |
| Il4ra | 3.61E-119 | 0.84678017 | 0.522 | 0.309 | 6.97E-115 | regulatory Tc | Il4ra |
| Itgb8 | 2.97E-118 | 5.07975387 | 0.296 | 0.013 | 5.74E-114 | regulatory Tc | Itgb8 |
| Hilpda | 8.25E-118 | 0.95784259 | 0.394 | 0.208 | 1.60E-113 | regulatory Tc | Hilpda |
| Dgat2 | 1.03E-117 | 5.41431231 | 0.318 | 0.01 | 2.00E-113 | regulatory Tc | Dgat2 |
| Vmp1 | 3.70E-117 | 0.98811324 | 0.524 | 0.308 | 7.15E-113 | regulatory Tc | Vmp1 |
| Tmem123 | 2.15E-116 | 0.97344156 | 0.515 | 0.327 | 4.16E-112 | regulatory Tc | Tmem123 |
| Gem | 2.46E-116 | 1.11017181 | 0.391 | 0.224 | 4.75E-112 | regulatory Tc | Gem |
| Chd3 | 9.83E-116 | 0.81653669 | 0.534 | 0.316 | 1.90E-111 | regulatory Tc | Chd3 |
| Pqlc1 | 1.19E-115 | 2.5028735 | 0.284 | 0.05 | 2.30E-111 | regulatory Tc | Pqlc1 |
| Ptprcap | 1.69E-115 | 0.6291613 | 0.681 | 0.52 | 3.26E-111 | regulatory Tc | Ptprcap |
| Jdp2 | 3.82E-115 | 1.77193647 | 0.286 | 0.116 | 7.38E-111 | regulatory Tc | Jdp2 |
| Dgat1 | 9.23E-115 | 0.76976807 | 0.446 | 0.27 | 1.78E-110 | regulatory Tc | Dgat1 |
| Slc16a3 | 7.50E-113 | 1.78521518 | 0.276 | 0.094 | 1.45E-108 | regulatory Tc | Slc16a3 |
| Nfkbid | 1.48E-109 | 0.54823306 | 0.574 | 0.395 | 2.86E-105 | regulatory Tc | Nfkbid |
| Mdh1 | 1.99E-109 | 0.84132077 | 0.46 | 0.297 | 3.85E-105 | regulatory Tc | Mdh1 |
| Sh3rf1 | 1.13E-108 | 2.95717738 | 0.282 | 0.045 | 2.18E-104 | regulatory Tc | Sh3rf1 |
| Batf | 3.59E-108 | 0.92931214 | 0.392 | 0.223 | 6.95E-104 | regulatory Tc | Batf |
| Rgs10 | 4.50E-108 | 0.92085991 | 0.407 | 0.205 | 8.71E-104 | regulatory Tc | Rgs10 |
| Lpcat1 | 1.14E-107 | 2.11250883 | 0.291 | 0.077 | 2.21E-103 | regulatory Tc | Lpcat1 |
| Phlda1 | 1.66E-107 | 1.20375447 | 0.295 | 0.13 | 3.21E-103 | regulatory Tc | Phlda1 |
| Spty2d1 | 2.58E-107 | 0.98294294 | 0.419 | 0.236 | 4.99E-103 | regulatory Tc | Spty2d1 |
| Apaf1 | 8.62E-107 | 1.15585971 | 0.423 | 0.223 | 1.67E-102 | regulatory Tc | Apaf1 |
| Cd44 | 2.06E-106 | 0.34808261 | 0.581 | 0.491 | 3.99E-102 | regulatory Tc | Cd44 |
| Tspan32 | 6.80E-106 | 1.62871653 | 0.322 | 0.08 | 1.31E-101 | regulatory Tc | Tspan32 |
| Fam129a | 4.62E-104 | 0.77625899 | 0.512 | 0.337 | 8.93E-100 | regulatory Tc | Fam129a |
| Kdm6b | 7.12E-104 | 0.59218543 | 0.623 | 0.435 | 1.38E-99 | regulatory Tc | Kdm6b |
| Mxi1 | 1.27E-103 | 1.34904618 | 0.409 | 0.184 | 2.45E-99 | regulatory Tc | Mxi1 |
| Rbm38 | 1.23E-102 | 0.82230964 | 0.507 | 0.319 | 2.39E-98 | regulatory Tc | Rbm38 |
| Nfkb1 | 8.56E-102 | 0.56755298 | 0.58 | 0.441 | 1.66E-97 | regulatory Tc | Nfkb1 |
| Ncoa3 | 2.17E-101 | 0.92453403 | 0.458 | 0.272 | 4.20E-97 | regulatory Tc | Ncoa3 |
| Il6st | 2.92E-101 | 1.28589738 | 0.321 | 0.122 | 5.65E-97 | regulatory Tc | Il6st |
| Ctsb | 6.13E-101 | 0.65111684 | 0.526 | 0.364 | 1.19E-96 | regulatory Tc | Ctsb |
| Jak3 | 8.52E-101 | 0.65863964 | 0.609 | 0.429 | 1.65E-96 | regulatory Tc | Jak3 |
| Ubald2 | 2.30E-100 | 0.58421661 | 0.681 | 0.523 | 4.46E-96 | regulatory Tc | Ubald2 |
| Hif1a | 1.73E-99 | 0.95046736 | 0.463 | 0.261 | 3.34E-95 | regulatory Tc | Hif1a |
| Tob2 | 6.64E-99 | 0.80406051 | 0.433 | 0.263 | 1.28E-94 | regulatory Tc | Tob2 |
| Anxa11 | 8.50E-99 | 0.70779883 | 0.485 | 0.325 | 1.64E-94 | regulatory Tc | Anxa11 |
| Ptma | 2.43E-98 | 0.28737875 | 0.511 | 0.432 | 4.70E-94 | regulatory Tc | Ptma |
| Mbd2 | 8.97E-98 | 0.38391194 | 0.783 | 0.709 | 1.73E-93 | regulatory Tc | Mbd2 |
| Tiam1 | 4.64E-97 | 1.11274398 | 0.34 | 0.128 | 8.97E-93 | regulatory Tc | Tiam1 |
| Ubl3 | 1.05E-96 | 0.52792841 | 0.574 | 0.423 | 2.03E-92 | regulatory Tc | Ubl3 |
| Tent5c | 1.09E-96 | 1.25165937 | 0.358 | 0.143 | 2.11E-92 | regulatory Tc | Tent5c |
| Arl5c | 1.39E-96 | 1.33368436 | 0.272 | 0.099 | 2.70E-92 | regulatory Tc | Arl5c |
| Bclaf3 | 3.81E-96 | 1.72658939 | 0.274 | 0.092 | 7.36E-92 | regulatory Tc | Bclaf3 |
| Hmgcr | 3.40E-95 | 1.07496477 | 0.364 | 0.201 | 6.58E-91 | regulatory Tc | Hmgcr |
| Ncmap | 3.55E-95 | 4.37664315 | 0.262 | 0.016 | 6.86E-91 | regulatory Tc | Ncmap |
| Ecm1 | 5.28E-93 | 1.27091698 | 0.324 | 0.117 | 1.02E-88 | regulatory Tc | Ecm1 |
| Wls | 1.21E-92 | 1.59931612 | 0.319 | 0.117 | 2.33E-88 | regulatory Tc | Wls |
| Skil | 2.86E-92 | 0.51933521 | 0.602 | 0.456 | 5.53E-88 | regulatory Tc | Skil |
| Tgif1 | 2.92E-92 | 0.39734772 | 0.648 | 0.495 | 5.65E-88 | regulatory Tc | Tgif1 |
| Adam9 | 5.22E-92 | 1.47552211 | 0.328 | 0.125 | 1.01E-87 | regulatory Tc | Adam9 |
| Zc3h12a | 2.55E-91 | 0.61707023 | 0.576 | 0.403 | 4.93E-87 | regulatory Tc | Zc3h12a |
| Mllt6 | 1.81E-90 | 0.60146018 | 0.601 | 0.465 | 3.50E-86 | regulatory Tc | Mllt6 |
| Cpd | 1.88E-89 | 1.22652448 | 0.271 | 0.097 | 3.63E-85 | regulatory Tc | Cpd |
| Sypl | 2.72E-88 | 1.18293849 | 0.3 | 0.126 | 5.26E-84 | regulatory Tc | Sypl |
| Cpm | 5.71E-88 | 1.45291356 | 0.267 | 0.081 | 1.10E-83 | regulatory Tc | Cpm |
| Gbp3 | 1.11E-87 | 1.8856935 | 0.263 | 0.081 | 2.14E-83 | regulatory Tc | Gbp3 |
| Dot1l | 1.31E-87 | 0.99252127 | 0.377 | 0.199 | 2.54E-83 | regulatory Tc | Dot1l |
| Aebp2 | 4.84E-87 | 0.49623619 | 0.619 | 0.473 | 9.36E-83 | regulatory Tc | Aebp2 |
| Myo1e | 1.19E-86 | 1.29731187 | 0.317 | 0.133 | 2.29E-82 | regulatory Tc | Myo1e |
| Bmp2k | 1.40E-86 | 1.43039386 | 0.331 | 0.126 | 2.72E-82 | regulatory Tc | Bmp2k |
| Peli1 | 5.45E-85 | 0.59984836 | 0.628 | 0.457 | 1.05E-80 | regulatory Tc | Peli1 |
| Tnip1 | 7.94E-85 | 0.75197644 | 0.498 | 0.317 | 1.53E-80 | regulatory Tc | Tnip1 |
| Inf2 | 1.61E-84 | 1.59332566 | 0.285 | 0.096 | 3.12E-80 | regulatory Tc | Inf2 |
| Traf3 | 2.47E-84 | 1.41508022 | 0.323 | 0.118 | 4.77E-80 | regulatory Tc | Traf3 |
| Smad7 | 3.78E-83 | 0.42268189 | 0.565 | 0.43 | 7.30E-79 | regulatory Tc | Smad7 |
| Tnfrsf8 | 6.30E-83 | 6.46685481 | 0.275 | 0.006 | 1.22E-78 | regulatory Tc | Tnfrsf8 |
| Lrba | 1.22E-82 | 1.17287182 | 0.31 | 0.142 | 2.36E-78 | regulatory Tc | Lrba |
| Gpx1 | 1.28E-82 | 0.29215424 | 0.467 | 0.344 | 2.47E-78 | regulatory Tc | Gpx1 |
| Slc35b2 | 1.31E-82 | 1.05971677 | 0.38 | 0.214 | 2.53E-78 | regulatory Tc | Slc35b2 |
| Grb2 | 3.05E-82 | 0.83999667 | 0.437 | 0.257 | 5.89E-78 | regulatory Tc | Grb2 |
| Lrp10 | 2.47E-81 | 0.36854228 | 0.629 | 0.517 | 4.77E-77 | regulatory Tc | Lrp10 |
| Pmepa1 | 4.63E-81 | 0.97799551 | 0.38 | 0.205 | 8.96E-77 | regulatory Tc | Pmepa1 |
| Ap3b1 | 4.99E-81 | 0.81071762 | 0.432 | 0.29 | 9.65E-77 | regulatory Tc | Ap3b1 |
| Cyba | 5.28E-81 | 0.42708933 | 0.593 | 0.461 | 1.02E-76 | regulatory Tc | Cyba |
| Ppm1b | 1.18E-80 | 0.72293244 | 0.523 | 0.353 | 2.28E-76 | regulatory Tc | Ppm1b |
| Mapkapk3 | 1.08E-78 | 0.84355443 | 0.435 | 0.257 | 2.09E-74 | regulatory Tc | Mapkapk3 |
| Mmd | 2.07E-78 | 1.07079798 | 0.355 | 0.206 | 4.00E-74 | regulatory Tc | Mmd |
| Ppp1r16b | 1.22E-75 | 0.48741019 | 0.676 | 0.542 | 2.37E-71 | regulatory Tc | Ppp1r16b |
| Spin1 | 1.53E-75 | 0.85944136 | 0.371 | 0.194 | 2.95E-71 | regulatory Tc | Spin1 |
| Rhog | 3.32E-75 | 0.49086966 | 0.537 | 0.392 | 6.42E-71 | regulatory Tc | Rhog |
| Ccrl2 | 5.32E-74 | 1.43758283 | 0.264 | 0.101 | 1.03E-69 | regulatory Tc | Ccrl2 |
| Vgll4 | 6.20E-74 | 0.51502748 | 0.554 | 0.45 | 1.20E-69 | regulatory Tc | Vgll4 |
| Birc3 | 2.01E-73 | 0.50213706 | 0.446 | 0.315 | 3.89E-69 | regulatory Tc | Birc3 |
| Slc5a3 | 5.20E-73 | 1.22019879 | 0.298 | 0.136 | 1.01E-68 | regulatory Tc | Slc5a3 |
| Prdm1 | 4.85E-72 | 1.27125446 | 0.252 | 0.091 | 9.38E-68 | regulatory Tc | Prdm1 |
| Il3ra | 5.31E-72 | 1.4228373 | 0.321 | 0.134 | 1.03E-67 | regulatory Tc | Il3ra |
| Isy1 | 2.83E-70 | 0.59070041 | 0.595 | 0.465 | 5.48E-66 | regulatory Tc | Isy1 |
| Il21r | 2.08E-69 | 0.59707355 | 0.593 | 0.386 | 4.02E-65 | regulatory Tc | Il21r |
| Gpr65 | 2.40E-69 | 0.88132397 | 0.354 | 0.216 | 4.64E-65 | regulatory Tc | Gpr65 |
| Pkp3 | 4.63E-67 | 0.85492103 | 0.356 | 0.199 | 8.95E-63 | regulatory Tc | Pkp3 |
| Sbno2 | 7.71E-67 | 0.68414227 | 0.371 | 0.216 | 1.49E-62 | regulatory Tc | Sbno2 |
| Slc26a2 | 1.06E-66 | 1.35003059 | 0.26 | 0.099 | 2.05E-62 | regulatory Tc | Slc26a2 |
| Cyb5a | 2.94E-66 | 0.85840674 | 0.338 | 0.179 | 5.68E-62 | regulatory Tc | Cyb5a |
| Camk4 | 4.08E-65 | 0.70413337 | 0.555 | 0.366 | 7.89E-61 | regulatory Tc | Camk4 |
| Ier5 | 3.35E-64 | 0.56034706 | 0.588 | 0.43 | 6.48E-60 | regulatory Tc | Ier5 |
| Csrp1 | 5.59E-64 | 1.04598555 | 0.324 | 0.17 | 1.08E-59 | regulatory Tc | Csrp1 |
| Crlf2 | 1.88E-63 | 1.04644534 | 0.386 | 0.213 | 3.64E-59 | regulatory Tc | Crlf2 |
| Nfil3 | 8.75E-63 | 0.77614417 | 0.397 | 0.246 | 1.69E-58 | regulatory Tc | Nfil3 |
| Tap2 | 9.60E-63 | 0.33745356 | 0.621 | 0.512 | 1.86E-58 | regulatory Tc | Tap2 |
| Ccr4 | 3.00E-62 | 0.65193154 | 0.271 | 0.12 | 5.80E-58 | regulatory Tc | Ccr4 |
| Grn | 4.61E-62 | 1.08723491 | 0.26 | 0.112 | 8.92E-58 | regulatory Tc | Grn |
| Helz2 | 1.33E-61 | 0.79647344 | 0.405 | 0.253 | 2.57E-57 | regulatory Tc | Helz2 |
| Ptbp1 | 1.72E-61 | 0.35803593 | 0.602 | 0.512 | 3.32E-57 | regulatory Tc | Ptbp1 |
| Slc41a1 | 1.92E-61 | 1.08904634 | 0.266 | 0.12 | 3.70E-57 | regulatory Tc | Slc41a1 |
| H2-Q6 | 6.68E-61 | 0.47618697 | 0.83 | 0.716 | 1.29E-56 | regulatory Tc | H2-Q6 |
| Ehd1 | 2.96E-60 | 1.06219375 | 0.313 | 0.158 | 5.72E-56 | regulatory Tc | Ehd1 |
| Capn2 | 5.45E-60 | 0.31955373 | 0.478 | 0.381 | 1.05E-55 | regulatory Tc | Capn2 |
| Vcp | 5.96E-60 | 0.38980765 | 0.558 | 0.455 | 1.15E-55 | regulatory Tc | Vcp |
| Vps54 | 2.13E-59 | 0.69319059 | 0.501 | 0.343 | 4.11E-55 | regulatory Tc | Vps54 |
| Atxn7l1 | 3.43E-59 | 0.77720654 | 0.401 | 0.246 | 6.63E-55 | regulatory Tc | Atxn7l1 |
| Jpt1 | 8.60E-59 | 0.32943045 | 0.614 | 0.507 | 1.66E-54 | regulatory Tc | Jpt1 |
| Cyfip1 | 1.70E-58 | 1.26396049 | 0.25 | 0.1 | 3.29E-54 | regulatory Tc | Cyfip1 |
| Elovl5 | 3.38E-58 | 0.66051433 | 0.402 | 0.248 | 6.54E-54 | regulatory Tc | Elovl5 |
| Rab43 | 1.38E-56 | 0.57522843 | 0.457 | 0.323 | 2.66E-52 | regulatory Tc | Rab43 |
| Npc2 | 1.63E-56 | 0.29046997 | 0.548 | 0.445 | 3.16E-52 | regulatory Tc | Npc2 |
| Mier1 | 5.00E-56 | 0.44419156 | 0.474 | 0.354 | 9.67E-52 | regulatory Tc | Mier1 |
| Fyb | 7.39E-56 | 0.32091299 | 0.614 | 0.5 | 1.43E-51 | regulatory Tc | Fyb |
| Tap1 | 9.34E-56 | 0.41782241 | 0.552 | 0.456 | 1.81E-51 | regulatory Tc | Tap1 |
| Pim3 | 9.90E-56 | 0.91671143 | 0.297 | 0.158 | 1.91E-51 | regulatory Tc | Pim3 |
| Aldoa | 1.74E-55 | 0.3234374 | 0.529 | 0.434 | 3.37E-51 | regulatory Tc | Aldoa |
| Dad1 | 4.01E-55 | 0.33425055 | 0.583 | 0.467 | 7.75E-51 | regulatory Tc | Dad1 |
| Xbp1 | 6.55E-55 | 0.58166188 | 0.323 | 0.215 | 1.27E-50 | regulatory Tc | Xbp1 |
| Zbtb7b | 2.78E-54 | 1.08100623 | 0.263 | 0.11 | 5.38E-50 | regulatory Tc | Zbtb7b |
| Lbh | 4.99E-54 | 0.40436132 | 0.552 | 0.437 | 9.64E-50 | regulatory Tc | Lbh |
| Sntb1 | 7.13E-54 | 0.83268181 | 0.328 | 0.163 | 1.38E-49 | regulatory Tc | Sntb1 |
| Havcr2 | 1.68E-53 | 1.23667264 | 0.304 | 0.126 | 3.26E-49 | regulatory Tc | Havcr2 |
| Insig1 | 1.87E-53 | 0.54440577 | 0.306 | 0.197 | 3.62E-49 | regulatory Tc | Insig1 |
| Edem1 | 1.94E-53 | 0.46411889 | 0.518 | 0.389 | 3.75E-49 | regulatory Tc | Edem1 |
| Ehbp1l1 | 2.76E-53 | 0.51555062 | 0.415 | 0.276 | 5.33E-49 | regulatory Tc | Ehbp1l1 |
| Birc2 | 2.91E-53 | 1.04661371 | 0.318 | 0.163 | 5.63E-49 | regulatory Tc | Birc2 |
| Zbtb1 | 8.05E-53 | 0.47944412 | 0.49 | 0.367 | 1.56E-48 | regulatory Tc | Zbtb1 |
| Pmaip1 | 2.12E-52 | 0.96158731 | 0.271 | 0.122 | 4.09E-48 | regulatory Tc | Pmaip1 |
| Isg20 | 2.54E-52 | 0.99961583 | 0.262 | 0.127 | 4.91E-48 | regulatory Tc | Isg20 |
| Grap | 2.92E-52 | 0.42932842 | 0.46 | 0.348 | 5.65E-48 | regulatory Tc | Grap |
| Igflr1 | 3.08E-52 | 1.41665498 | 0.264 | 0.114 | 5.96E-48 | regulatory Tc | Igflr1 |
| Pak2 | 5.19E-52 | 0.29128204 | 0.581 | 0.492 | 1.00E-47 | regulatory Tc | Pak2 |
| Nfkbie | 5.40E-51 | 0.96393146 | 0.306 | 0.159 | 1.04E-46 | regulatory Tc | Nfkbie |
| Snx2 | 6.21E-51 | 0.48650678 | 0.311 | 0.206 | 1.20E-46 | regulatory Tc | Snx2 |
| Stat5a | 7.58E-51 | 1.01591702 | 0.301 | 0.167 | 1.47E-46 | regulatory Tc | Stat5a |
| Rara | 1.36E-50 | 0.57803565 | 0.406 | 0.257 | 2.63E-46 | regulatory Tc | Rara |
| Rab8b | 3.74E-50 | 0.35893362 | 0.633 | 0.525 | 7.24E-46 | regulatory Tc | Rab8b |
| Syt11 | 5.72E-50 | 1.02021352 | 0.25 | 0.114 | 1.11E-45 | regulatory Tc | Syt11 |
| Slc39a1 | 1.72E-49 | 0.69246549 | 0.331 | 0.197 | 3.34E-45 | regulatory Tc | Slc39a1 |
| H3f3a | 1.85E-49 | 0.2504688 | 0.536 | 0.445 | 3.59E-45 | regulatory Tc | H3f3a |
| Glcci1 | 1.88E-49 | 0.86062273 | 0.319 | 0.181 | 3.63E-45 | regulatory Tc | Glcci1 |
| Arf4 | 2.86E-49 | 0.50293568 | 0.407 | 0.295 | 5.53E-45 | regulatory Tc | Arf4 |
| Itpkb | 3.31E-49 | 0.35659922 | 0.543 | 0.402 | 6.39E-45 | regulatory Tc | Itpkb |
| Fbxo11 | 7.43E-49 | 0.67454147 | 0.345 | 0.213 | 1.44E-44 | regulatory Tc | Fbxo11 |
| Edf1 | 1.04E-48 | 0.46608572 | 0.404 | 0.287 | 2.01E-44 | regulatory Tc | Edf1 |
| Cd27 | 1.30E-48 | 1.03971898 | 0.311 | 0.168 | 2.52E-44 | regulatory Tc | Cd27 |
| Tapbp | 1.36E-48 | 0.3344683 | 0.56 | 0.468 | 2.62E-44 | regulatory Tc | Tapbp |
| Rhoh | 1.40E-48 | 0.41278736 | 0.434 | 0.302 | 2.70E-44 | regulatory Tc | Rhoh |
| H2-D1 | 2.27E-48 | 0.40139791 | 0.582 | 0.469 | 4.39E-44 | regulatory Tc | H2-D1 |
| Axxl1 | 2.84E-48 | 0.58135427 | 0.331 | 0.208 | 5.50E-44 | regulatory Tc | Axxl1 |
| Tmem154 | 8.79E-48 | 0.7732239 | 0.251 | 0.119 | 1.70E-43 | regulatory Tc | Tmem154 |
| Fam3c | 1.01E-47 | 0.73451455 | 0.324 | 0.181 | 1.96E-43 | regulatory Tc | Fam3c |
| Reps1 | 1.48E-47 | 0.64980411 | 0.348 | 0.213 | 2.86E-43 | regulatory Tc | Reps1 |
| Man1a | 5.10E-47 | 0.66630899 | 0.413 | 0.27 | 9.86E-43 | regulatory Tc | Man1a |
| Relb | 6.24E-47 | 0.88586273 | 0.35 | 0.194 | 1.21E-42 | regulatory Tc | Relb |
| Traf4 | 1.06E-46 | 0.97645055 | 0.278 | 0.156 | 2.06E-42 | regulatory Tc | Traf4 |
| Slc25a3 | 2.58E-46 | 0.25503182 | 0.622 | 0.542 | 4.98E-42 | regulatory Tc | Slc25a3 |
| Gna15 | 1.24E-45 | 0.57065685 | 0.31 | 0.186 | 2.39E-41 | regulatory Tc | Gna15 |
| Sfr1 | 1.35E-45 | 0.34807927 | 0.397 | 0.291 | 2.61E-41 | regulatory Tc | Sfr1 |
| Ifnar1 | 2.22E-45 | 0.67961783 | 0.38 | 0.259 | 4.30E-41 | regulatory Tc | Ifnar1 |
| Rab1a | 3.79E-45 | 0.39695239 | 0.423 | 0.314 | 7.33E-41 | regulatory Tc | Rab1a |
| Nt5e | 5.34E-45 | 0.60044339 | 0.308 | 0.193 | 1.03E-40 | regulatory Tc | Nt5e |
| Mdfic | 1.11E-44 | 0.94745773 | 0.286 | 0.161 | 2.15E-40 | regulatory Tc | Mdfic |
| Sytl2 | 1.81E-44 | 0.65973588 | 0.36 | 0.217 | 3.50E-40 | regulatory Tc | Sytl2 |
| Atl3 | 1.09E-43 | 0.4711039 | 0.437 | 0.321 | 2.10E-39 | regulatory Tc | Atl3 |
| Fosb | 4.90E-43 | 0.30772744 | 0.372 | 0.274 | 9.47E-39 | regulatory Tc | Fosb |
| Adprh | 5.65E-43 | 0.51790051 | 0.362 | 0.245 | 1.09E-38 | regulatory Tc | Adprh |
| Cst3 | 6.20E-43 | 0.4713235 | 0.451 | 0.338 | 1.20E-38 | regulatory Tc | Cst3 |
| Hivep1 | 1.08E-42 | 0.66305427 | 0.287 | 0.165 | 2.09E-38 | regulatory Tc | Hivep1 |
| Lilrb4a | 1.83E-42 | 1.12153521 | 0.272 | 0.156 | 3.53E-38 | regulatory Tc | Lilrb4a |
| Arf1 | 5.52E-42 | 0.33561053 | 0.498 | 0.381 | 1.07E-37 | regulatory Tc | Arf1 |
| Ciapin1 | 1.17E-41 | 0.98364826 | 0.278 | 0.161 | 2.25E-37 | regulatory Tc | Ciapin1 |
| Cast | 2.40E-41 | 0.35656994 | 0.446 | 0.328 | 4.64E-37 | regulatory Tc | Cast |
| Cdc42se2 | 3.37E-41 | 0.37826706 | 0.43 | 0.317 | 6.51E-37 | regulatory Tc | Cdc42se2 |
| Arhgap31 | 3.95E-41 | 0.70046118 | 0.256 | 0.136 | 7.64E-37 | regulatory Tc | Arhgap31 |
| Ssbp3 | 6.76E-41 | 0.66705584 | 0.295 | 0.178 | 1.31E-36 | regulatory Tc | Ssbp3 |
| Ywhae | 8.42E-41 | 0.41619416 | 0.381 | 0.274 | 1.63E-36 | regulatory Tc | Ywhae |
| Cyth1 | 8.78E-41 | 0.27009779 | 0.498 | 0.362 | 1.70E-36 | regulatory Tc | Cyth1 |
| Wsb1 | 1.24E-40 | 0.31590937 | 0.526 | 0.399 | 2.40E-36 | regulatory Tc | Wsb1 |
| Arf6 | 1.83E-40 | 0.27748544 | 0.568 | 0.482 | 3.55E-36 | regulatory Tc | Arf6 |
| Snx3 | 2.56E-40 | 0.36912726 | 0.441 | 0.329 | 4.94E-36 | regulatory Tc | Snx3 |
| Gbp7 | 3.66E-40 | 0.80441635 | 0.286 | 0.167 | 7.08E-36 | regulatory Tc | Gbp7 |
| Kdm2b | 4.46E-40 | 0.74418929 | 0.307 | 0.195 | 8.62E-36 | regulatory Tc | Kdm2b |
| Ubash3b | 6.76E-40 | 0.80892459 | 0.314 | 0.186 | 1.31E-35 | regulatory Tc | Ubash3b |
| Scyl2 | 1.67E-39 | 0.63154449 | 0.315 | 0.194 | 3.22E-35 | regulatory Tc | Scyl2 |
| Cd274 | 3.43E-39 | 0.82414081 | 0.288 | 0.165 | 6.63E-35 | regulatory Tc | Cd274 |
| Xpo6 | 3.69E-39 | 0.50808264 | 0.381 | 0.272 | 7.14E-35 | regulatory Tc | Xpo6 |
| Gnai2 | 8.62E-39 | 0.29479425 | 0.582 | 0.499 | 1.67E-34 | regulatory Tc | Gnai2 |
| Plagl2 | 2.68E-38 | 0.79765902 | 0.26 | 0.139 | 5.19E-34 | regulatory Tc | Plagl2 |
| Eed | 3.93E-38 | 0.60804951 | 0.288 | 0.174 | 7.60E-34 | regulatory Tc | Eed |
| Zmiz2 | 1.34E-37 | 0.67985439 | 0.372 | 0.242 | 2.59E-33 | regulatory Tc | Zmiz2 |
| Spint2 | 1.41E-37 | 0.60059877 | 0.275 | 0.158 | 2.73E-33 | regulatory Tc | Spint2 |
| Ywhah | 2.24E-37 | 0.27434418 | 0.391 | 0.319 | 4.33E-33 | regulatory Tc | Ywhah |
| Tgfb1 | 1.16E-36 | 0.29415729 | 0.569 | 0.427 | 2.25E-32 | regulatory Tc | Tgfb1 |
| Prkch | 1.59E-36 | 0.33540725 | 0.568 | 0.425 | 3.08E-32 | regulatory Tc | Prkch |
| Adam17 | 2.90E-36 | 0.3925273 | 0.314 | 0.2 | 5.61E-32 | regulatory Tc | Adam17 |
| Spcs1 | 4.35E-36 | 0.38269998 | 0.358 | 0.269 | 8.40E-32 | regulatory Tc | Spcs1 |
| Msi2 | 4.81E-36 | 0.52672794 | 0.433 | 0.318 | 9.30E-32 | regulatory Tc | Msi2 |
| Arl6ip1 | 1.15E-35 | 0.4314372 | 0.317 | 0.224 | 2.23E-31 | regulatory Tc | Arl6ip1 |
| Faim | 1.37E-35 | 0.94420433 | 0.253 | 0.134 | 2.65E-31 | regulatory Tc | Faim |
| Cldnd1 | 4.82E-35 | 0.36747234 | 0.283 | 0.197 | 9.33E-31 | regulatory Tc | Cldnd1 |
| Kbtbd11 | 7.32E-35 | 0.44187458 | 0.388 | 0.289 | 1.42E-30 | regulatory Tc | Kbtbd11 |
| Hk1 | 1.02E-34 | 0.2765104 | 0.439 | 0.348 | 1.97E-30 | regulatory Tc | Hk1 |
| Sdhb | 1.20E-34 | 0.37296514 | 0.337 | 0.253 | 2.31E-30 | regulatory Tc | Sdhb |
| Srebf2 | 2.27E-34 | 0.33963439 | 0.42 | 0.325 | 4.39E-30 | regulatory Tc | Srebf2 |
| Malt1 | 7.71E-34 | 0.43080801 | 0.436 | 0.338 | 1.49E-29 | regulatory Tc | Malt1 |
| Rap1gds1 | 8.70E-34 | 0.29797769 | 0.364 | 0.266 | 1.68E-29 | regulatory Tc | Rap1gds1 |
| Cd200r1 | 1.24E-33 | 1.18781078 | 0.257 | 0.122 | 2.39E-29 | regulatory Tc | Cd200r1 |
| Ptpn7 | 1.65E-33 | 0.36555832 | 0.328 | 0.241 | 3.19E-29 | regulatory Tc | Ptpn7 |
| Agfg1 | 2.03E-33 | 0.44115363 | 0.447 | 0.319 | 3.92E-29 | regulatory Tc | Agfg1 |
| Srf | 2.48E-33 | 0.35505535 | 0.31 | 0.229 | 4.80E-29 | regulatory Tc | Srf |
| Pfkl | 2.57E-33 | 0.56506621 | 0.27 | 0.165 | 4.98E-29 | regulatory Tc | Pfkl |
| Trp53inp1 | 2.98E-33 | 0.35459258 | 0.336 | 0.246 | 5.77E-29 | regulatory Tc | Trp53inp1 |
| Sae1 | 5.38E-33 | 0.25792976 | 0.266 | 0.205 | 1.04E-28 | regulatory Tc | Sae1 |
| Tnik | 9.09E-33 | 0.63738753 | 0.259 | 0.151 | 1.76E-28 | regulatory Tc | Tnik |
| Mxd4 | 1.50E-32 | 0.43381178 | 0.356 | 0.248 | 2.90E-28 | regulatory Tc | Mxd4 |
| Ncf4 | 3.15E-32 | 0.52022314 | 0.353 | 0.225 | 6.08E-28 | regulatory Tc | Ncf4 |
| Rab21 | 5.06E-32 | 0.34244509 | 0.535 | 0.448 | 9.78E-28 | regulatory Tc | Rab21 |
| Plaat3 | 9.92E-32 | 0.34028107 | 0.475 | 0.372 | 1.92E-27 | regulatory Tc | Plaat3 |
| Dennd4a | 5.83E-31 | 0.48677678 | 0.522 | 0.391 | 1.13E-26 | regulatory Tc | Dennd4a |
| Rnpep | 9.09E-31 | 0.47667883 | 0.263 | 0.165 | 1.76E-26 | regulatory Tc | Rnpep |
| Rabgap1l | 1.45E-30 | 0.4068831 | 0.519 | 0.42 | 2.80E-26 | regulatory Tc | Rabgap1l |
| Zc3h15 | 3.02E-30 | 0.6059222 | 0.288 | 0.186 | 5.83E-26 | regulatory Tc | Zc3h15 |
| Epb41l2 | 4.02E-30 | 0.56988656 | 0.258 | 0.163 | 7.78E-26 | regulatory Tc | Epb41l2 |
| 7-Sep | 5.20E-30 | 0.27065958 | 0.466 | 0.381 | 1.01E-25 | regulatory Tc | 7-Sep |
| Cd69 | 8.41E-30 | 0.30484624 | 0.394 | 0.312 | 1.63E-25 | regulatory Tc | Cd69 |
| Camk2d | 9.13E-30 | 0.54927856 | 0.268 | 0.162 | 1.76E-25 | regulatory Tc | Camk2d |
| Galnt2 | 1.40E-29 | 0.34333508 | 0.448 | 0.342 | 2.70E-25 | regulatory Tc | Galnt2 |
| Rab27a | 1.93E-29 | 0.51919464 | 0.35 | 0.239 | 3.72E-25 | regulatory Tc | Rab27a |
| Zbtb38 | 2.29E-29 | 0.54423655 | 0.253 | 0.15 | 4.43E-25 | regulatory Tc | Zbtb38 |
| Slc2a3 | 2.39E-29 | 0.71342217 | 0.318 | 0.202 | 4.62E-25 | regulatory Tc | Slc2a3 |
| Il12rb1 | 2.44E-29 | 0.35468833 | 0.263 | 0.184 | 4.72E-25 | regulatory Tc | Il12rb1 |
| Slc2a1 | 3.69E-29 | 0.54241541 | 0.276 | 0.19 | 7.14E-25 | regulatory Tc | Slc2a1 |
| Osbpl3 | 3.75E-29 | 0.51382659 | 0.376 | 0.252 | 7.25E-25 | regulatory Tc | Osbpl3 |
| Ptp4a2 | 5.13E-29 | 0.46560047 | 0.366 | 0.262 | 9.91E-25 | regulatory Tc | Ptp4a2 |
| Sec11c | 5.24E-29 | 0.26092292 | 0.418 | 0.342 | 1.01E-24 | regulatory Tc | Sec11c |
| Sqstm1 | 1.53E-28 | 0.34024318 | 0.512 | 0.398 | 2.95E-24 | regulatory Tc | Sqstm1 |
| Fam169b | 1.72E-28 | 0.65991895 | 0.279 | 0.146 | 3.33E-24 | regulatory Tc | Fam169b |
| Ube2b | 2.13E-28 | 0.28639007 | 0.505 | 0.411 | 4.11E-24 | regulatory Tc | Ube2b |
| Ube4b | 4.19E-28 | 0.74144867 | 0.267 | 0.165 | 8.11E-24 | regulatory Tc | Ube4b |
| Atxn1 | 4.89E-28 | 0.55642305 | 0.362 | 0.235 | 9.45E-24 | regulatory Tc | Atxn1 |
| Vamp8 | 1.51E-27 | 0.38475541 | 0.446 | 0.332 | 2.92E-23 | regulatory Tc | Vamp8 |
| Ccr5 | 1.86E-27 | 0.39098332 | 0.398 | 0.265 | 3.60E-23 | regulatory Tc | Ccr5 |
| Pitpnm1 | 1.94E-27 | 0.38464308 | 0.416 | 0.316 | 3.75E-23 | regulatory Tc | Pitpnm1 |
| Cmtm6 | 2.36E-27 | 0.29972884 | 0.35 | 0.26 | 4.56E-23 | regulatory Tc | Cmtm6 |
| Phc3 | 2.90E-27 | 0.37771236 | 0.364 | 0.263 | 5.61E-23 | regulatory Tc | Phc3 |
| Fbxw11 | 5.36E-27 | 0.51775247 | 0.283 | 0.187 | 1.04E-22 | regulatory Tc | Fbxw11 |
| Zfp703 | 8.96E-27 | 0.62339912 | 0.263 | 0.163 | 1.73E-22 | regulatory Tc | Zfp703 |
| Tsc22d4 | 1.29E-26 | 0.33718674 | 0.347 | 0.252 | 2.49E-22 | regulatory Tc | Tsc22d4 |
| H13 | 1.39E-26 | 0.28583642 | 0.388 | 0.292 | 2.68E-22 | regulatory Tc | H13 |
| Slamf1 | 1.90E-26 | 0.47944014 | 0.278 | 0.174 | 3.67E-22 | regulatory Tc | Slamf1 |
| Gstp3 | 9.85E-26 | 0.37471899 | 0.29 | 0.192 | 1.90E-21 | regulatory Tc | Gstp3 |
| Ythdc1 | 1.02E-25 | 0.41084503 | 0.328 | 0.227 | 1.97E-21 | regulatory Tc | Ythdc1 |
| Ccng1 | 2.52E-25 | 0.31344786 | 0.323 | 0.24 | 4.87E-21 | regulatory Tc | Ccng1 |
| Galnt6 | 2.77E-25 | 0.46701122 | 0.312 | 0.214 | 5.35E-21 | regulatory Tc | Galnt6 |
| Mtmr3 | 2.83E-25 | 0.36229294 | 0.252 | 0.175 | 5.47E-21 | regulatory Tc | Mtmr3 |
| Abracl | 2.83E-25 | 0.26619498 | 0.365 | 0.274 | 5.47E-21 | regulatory Tc | Abracl |
| Parp14 | 3.72E-25 | 0.41327323 | 0.301 | 0.232 | 7.19E-21 | regulatory Tc | Parp14 |
| Cisd2 | 9.28E-25 | 0.50707961 | 0.297 | 0.2 | 1.79E-20 | regulatory Tc | Cisd2 |
| Kcnn4 | 1.56E-24 | 0.34383322 | 0.323 | 0.209 | 3.01E-20 | regulatory Tc | Kcnn4 |
| Tnk2 | 1.65E-24 | 0.30497193 | 0.347 | 0.242 | 3.20E-20 | regulatory Tc | Tnk2 |
| Serpinb6a | 2.43E-24 | 0.44654298 | 0.25 | 0.169 | 4.71E-20 | regulatory Tc | Serpinb6a |
| Rac1 | 2.49E-24 | 0.47033373 | 0.259 | 0.18 | 4.81E-20 | regulatory Tc | Rac1 |
| Arrdc1 | 2.91E-24 | 0.60806411 | 0.259 | 0.165 | 5.63E-20 | regulatory Tc | Arrdc1 |
| Ccdc6 | 4.37E-24 | 0.52152651 | 0.303 | 0.209 | 8.45E-20 | regulatory Tc | Ccdc6 |
| Glmp | 4.94E-24 | 0.33682666 | 0.386 | 0.283 | 9.56E-20 | regulatory Tc | Glmp |
| Mex3c | 6.30E-24 | 0.34507928 | 0.313 | 0.222 | 1.22E-19 | regulatory Tc | Mex3c |
| Mtdh | 6.90E-24 | 0.35402551 | 0.282 | 0.197 | 1.33E-19 | regulatory Tc | Mtdh |
| Retreg1 | 7.64E-24 | 0.29835064 | 0.252 | 0.157 | 1.48E-19 | regulatory Tc | Retreg1 |
| Pgk1 | 8.16E-24 | 0.35729875 | 0.256 | 0.193 | 1.58E-19 | regulatory Tc | Pgk1 |
| Fkbp1a | 9.90E-24 | 0.37737831 | 0.298 | 0.215 | 1.91E-19 | regulatory Tc | Fkbp1a |
| Qk | 1.15E-23 | 0.38932422 | 0.302 | 0.209 | 2.23E-19 | regulatory Tc | Qk |
| Sat1 | 1.17E-23 | 0.29867909 | 0.43 | 0.34 | 2.26E-19 | regulatory Tc | Sat1 |
| Traf2 | 3.27E-23 | 0.42298858 | 0.315 | 0.229 | 6.33E-19 | regulatory Tc | Traf2 |
| Itpk1 | 4.09E-23 | 0.36520283 | 0.381 | 0.288 | 7.91E-19 | regulatory Tc | Itpk1 |
| Necap2 | 4.16E-23 | 0.29285324 | 0.26 | 0.175 | 8.05E-19 | regulatory Tc | Necap2 |
| Adora2a | 1.06E-22 | 0.58490409 | 0.27 | 0.175 | 2.06E-18 | regulatory Tc | Adora2a |
| Ssh1 | 1.47E-22 | 0.46430636 | 0.304 | 0.202 | 2.84E-18 | regulatory Tc | Ssh1 |
| Srsf1 | 1.78E-22 | 0.31402222 | 0.318 | 0.252 | 3.43E-18 | regulatory Tc | Srsf1 |
| Map4k2 | 2.09E-22 | 0.37269374 | 0.425 | 0.313 | 4.04E-18 | regulatory Tc | Map4k2 |
| Plk3 | 2.44E-22 | 0.41221276 | 0.303 | 0.201 | 4.72E-18 | regulatory Tc | Plk3 |
| Ubxn7 | 2.84E-22 | 0.42168319 | 0.301 | 0.214 | 5.49E-18 | regulatory Tc | Ubxn7 |
| Dennd5a | 3.20E-22 | 0.35197756 | 0.311 | 0.205 | 6.20E-18 | regulatory Tc | Dennd5a |
| Txndc17 | 3.52E-22 | 0.37979018 | 0.313 | 0.221 | 6.81E-18 | regulatory Tc | Txndc17 |
| Atp2b4 | 4.08E-22 | 0.26003798 | 0.404 | 0.304 | 7.88E-18 | regulatory Tc | Atp2b4 |
| Psmb5 | 4.23E-22 | 0.26009335 | 0.296 | 0.231 | 8.18E-18 | regulatory Tc | Psmb5 |
| Ctnnb1 | 5.52E-22 | 0.27047856 | 0.412 | 0.329 | 1.07E-17 | regulatory Tc | Ctnnb1 |
| Tmc8 | 7.33E-22 | 0.47822842 | 0.267 | 0.178 | 1.42E-17 | regulatory Tc | Tmc8 |
| Serinc1 | 9.93E-22 | 0.39604869 | 0.313 | 0.221 | 1.92E-17 | regulatory Tc | Serinc1 |
| Prdx2 | 1.75E-21 | 0.30359526 | 0.272 | 0.206 | 3.39E-17 | regulatory Tc | Prdx2 |
| Cmip | 2.18E-21 | 0.2709455 | 0.544 | 0.444 | 4.21E-17 | regulatory Tc | Cmip |
| Aff1 | 3.66E-21 | 0.2674725 | 0.4 | 0.305 | 7.08E-17 | regulatory Tc | Aff1 |
| Metap2 | 4.59E-21 | 0.28699256 | 0.259 | 0.185 | 8.88E-17 | regulatory Tc | Metap2 |
| Sp3 | 4.79E-21 | 0.27610356 | 0.346 | 0.261 | 9.27E-17 | regulatory Tc | Sp3 |
| Csnk1g3 | 6.96E-21 | 0.38804173 | 0.275 | 0.193 | 1.35E-16 | regulatory Tc | Csnk1g3 |
| Derl1 | 7.27E-21 | 0.30294779 | 0.34 | 0.255 | 1.41E-16 | regulatory Tc | Derl1 |
| Stat3 | 1.06E-20 | 0.29657439 | 0.605 | 0.489 | 2.05E-16 | regulatory Tc | Stat3 |
| Nab2 | 2.07E-20 | 0.50658784 | 0.257 | 0.168 | 4.00E-16 | regulatory Tc | Nab2 |
| Tmem173 | 2.08E-20 | 0.37407678 | 0.314 | 0.218 | 4.02E-16 | regulatory Tc | Tmem173 |
| Rab37 | 2.57E-20 | 0.42464347 | 0.25 | 0.166 | 4.96E-16 | regulatory Tc | Rab37 |
| Cflar | 7.05E-20 | 0.32149106 | 0.26 | 0.184 | 1.36E-15 | regulatory Tc | Cflar |
| Zfp362 | 7.35E-20 | 0.36889353 | 0.255 | 0.166 | 1.42E-15 | regulatory Tc | Zfp362 |
| Ypel3 | 9.69E-20 | 0.32768832 | 0.487 | 0.392 | 1.87E-15 | regulatory Tc | Ypel3 |
| Baz1b | 1.34E-19 | 0.277596 | 0.284 | 0.211 | 2.59E-15 | regulatory Tc | Baz1b |
| Nadk | 1.54E-19 | 0.27722484 | 0.255 | 0.183 | 2.98E-15 | regulatory Tc | Nadk |
| Ikbke | 2.13E-19 | 0.56225105 | 0.253 | 0.157 | 4.13E-15 | regulatory Tc | Ikbke |
| Nfkbib | 3.98E-19 | 0.33250559 | 0.317 | 0.237 | 7.69E-15 | regulatory Tc | Nfkbib |
| Tmem134 | 1.61E-18 | 0.31711685 | 0.329 | 0.25 | 3.12E-14 | regulatory Tc | Tmem134 |
| Lsm8 | 2.08E-18 | 0.50617996 | 0.28 | 0.2 | 4.03E-14 | regulatory Tc | Lsm8 |
| Ubxn4 | 2.10E-18 | 0.30244862 | 0.315 | 0.238 | 4.05E-14 | regulatory Tc | Ubxn4 |
| Adipor2 | 2.17E-18 | 0.28136265 | 0.274 | 0.202 | 4.20E-14 | regulatory Tc | Adipor2 |
| Eif4e | 5.11E-18 | 0.25006056 | 0.257 | 0.194 | 9.89E-14 | regulatory Tc | Eif4e |
| Itgb3 | 7.64E-18 | 0.34045941 | 0.293 | 0.205 | 1.48E-13 | regulatory Tc | Itgb3 |
| Col4a3bp | 8.96E-18 | 0.48456716 | 0.286 | 0.2 | 1.73E-13 | regulatory Tc | Col4a3bp |
| Slc38a10 | 1.05E-17 | 0.2585525 | 0.284 | 0.208 | 2.02E-13 | regulatory Tc | Slc38a10 |
| Pear1 | 9.78E-17 | 0.32090515 | 0.301 | 0.224 | 1.89E-12 | regulatory Tc | Pear1 |
| Pygb | 2.89E-16 | 0.31747997 | 0.288 | 0.214 | 5.58E-12 | regulatory Tc | Pygb |
| Ap2s1 | 3.02E-16 | 0.28737117 | 0.258 | 0.191 | 5.83E-12 | regulatory Tc | Ap2s1 |
| Arl5b | 4.95E-16 | 0.30923146 | 0.296 | 0.212 | 9.56E-12 | regulatory Tc | Arl5b |
| Smarce1 | 8.38E-16 | 0.29151511 | 0.261 | 0.2 | 1.62E-11 | regulatory Tc | Smarce1 |
| Ints6 | 9.00E-16 | 0.28167072 | 0.287 | 0.212 | 1.74E-11 | regulatory Tc | Ints6 |

| | | | | | | | |
|---|---|---|---|---|---|---|---|
| Amfr | 2.42E-15 | 0.33536225 | 0.258 | 0.186 | 4.68E-11 | regulatory Tc | Amfr |
| Fgfr1op2 | 7.02E-15 | 0.25689124 | 0.367 | 0.29 | 1.36E-10 | regulatory Tc | Fgfr1op2 |
| Cdk6 | 8.99E-15 | 0.31114182 | 0.336 | 0.213 | 1.74E-10 | regulatory Tc | Cdk6 |
| Socs1 | 9.04E-15 | 0.28326562 | 0.293 | 0.257 | 1.75E-10 | regulatory Tc | Socs1 |
| Pde4d | 1.25E-14 | 0.30107966 | 0.395 | 0.304 | 2.42E-10 | regulatory Tc | Pde4d |
| Il10ra | 1.99E-14 | 0.2975332 | 0.255 | 0.18 | 3.85E-10 | regulatory Tc | Il10ra |
| Sik3 | 3.14E-14 | 0.29650151 | 0.261 | 0.192 | 6.06E-10 | regulatory Tc | Sik3 |
| Cbx7 | 4.06E-14 | 0.26455275 | 0.277 | 0.201 | 7.85E-10 | regulatory Tc | Cbx7 |
| Ppp1cb | 7.92E-14 | 0.28499239 | 0.34 | 0.268 | 1.53E-09 | regulatory Tc | Ppp1cb |
| Atp9b | 7.95E-14 | 0.31494588 | 0.287 | 0.212 | 1.54E-09 | regulatory Tc | Atp9b |
| Ccng2 | 1.72E-13 | 0.29900214 | 0.309 | 0.248 | 3.32E-09 | regulatory Tc | Ccng2 |
| Ccnl2 | 4.41E-13 | 0.29681773 | 0.433 | 0.349 | 8.52E-09 | regulatory Tc | Ccnl2 |
| Itpr2 | 5.42E-13 | 0.25097461 | 0.27 | 0.201 | 1.05E-08 | regulatory Tc | Itpr2 |
| Crebzf | 7.15E-13 | 0.47381139 | 0.278 | 0.214 | 1.38E-08 | regulatory Tc | Crebzf |
| Anxa5 | 7.24E-13 | 0.25385685 | 0.354 | 0.274 | 1.40E-08 | regulatory Tc | Anxa5 |
| Nfat5 | 3.08E-12 | 0.43100206 | 0.308 | 0.226 | 5.95E-08 | regulatory Tc | Nfat5 |
| Orai1 | 3.47E-12 | 0.29803749 | 0.319 | 0.249 | 6.70E-08 | regulatory Tc | Orai1 |
| Ptms | 5.22E-12 | 0.41699778 | 0.461 | 0.349 | 1.01E-07 | regulatory Tc | Ptms |
| Rab11b | 8.16E-12 | 0.2639246 | 0.298 | 0.229 | 1.58E-07 | regulatory Tc | Rab11b |
| Max | 8.89E-12 | 0.27635151 | 0.278 | 0.222 | 1.72E-07 | regulatory Tc | Max |
| Ppfia1 | 1.08E-11 | 0.35901152 | 0.261 | 0.193 | 2.09E-07 | regulatory Tc | Ppfia1 |
| Gimap7 | 2.44E-11 | 0.32594864 | 0.278 | 0.197 | 4.71E-07 | regulatory Tc | Gimap7 |
| Aplp2 | 3.78E-11 | 0.2832483 | 0.252 | 0.194 | 7.31E-07 | regulatory Tc | Aplp2 |
| Riok3 | 1.28E-10 | 0.26315717 | 0.308 | 0.242 | 2.47E-06 | regulatory Tc | Riok3 |
| Tbcb | 1.80E-10 | 0.26238517 | 0.269 | 0.211 | 3.49E-06 | regulatory Tc | Tbcb |
| Rsbn1l | 5.90E-10 | 0.25266497 | 0.295 | 0.233 | 1.14E-05 | regulatory Tc | Rsbn1l |
| Sh2b1 | 2.89E-09 | 0.26556961 | 0.294 | 0.239 | 5.60E-05 | regulatory Tc | Sh2b1 |
| Rgs16 | 0.00065169 | 0.46036257 | 0.322 | 0.196 | 1 | regulatory Tc | Rgs16 |
| S1pr1 | 2.48E-232 | 2.80601558 | 0.549 | 0.148 | 4.80E-228 | CD4 T cells | S1pr1 |
| Eef1a1 | 3.96E-191 | 0.41189907 | 0.964 | 0.942 | 7.65E-187 | CD4 T cells | Eef1a1 |
| Ramp3 | 2.05E-187 | 2.37036615 | 0.428 | 0.173 | 3.97E-183 | CD4 T cells | Ramp3 |
| Tcf7 | 2.39E-174 | 2.64628346 | 0.526 | 0.135 | 4.63E-170 | CD4 T cells | Tcf7 |
| Cd40lg | 1.00E-75 | 2.46321207 | 0.275 | 0.068 | 1.93E-71 | CD4 T cells | Cd40lg |
| Dusp10 | 2.38E-75 | 1.82781239 | 0.449 | 0.309 | 4.60E-71 | CD4 T cells | Dusp10 |
| Zfp36l2 | 5.95E-65 | 1.07104841 | 0.816 | 0.684 | 1.15E-60 | CD4 T cells | Zfp36l2 |
| Klf2 | 2.79E-53 | 1.41541688 | 0.469 | 0.245 | 5.40E-49 | CD4 T cells | Klf2 |
| Hspa8 | 1.17E-49 | 0.32809989 | 0.718 | 0.766 | 2.27E-45 | CD4 T cells | Hspa8 |
| Hspa1b | 1.50E-41 | 1.30762622 | 0.31 | 0.206 | 2.90E-37 | CD4 T cells | Hspa1b |
| Dgka | 1.65E-34 | 1.4574218 | 0.688 | 0.507 | 3.18E-30 | CD4 T cells | Dgka |
| Cd53 | 1.39E-31 | 0.28930824 | 0.695 | 0.74 | 2.69E-27 | CD4 T cells | Cd53 |
| Rack1 | 1.87E-31 | 0.5150223 | 0.697 | 0.693 | 3.62E-27 | CD4 T cells | Rack1 |
| Tnfaip3 | 1.15E-30 | 1.00867958 | 0.74 | 0.749 | 2.22E-26 | CD4 T cells | Tnfaip3 |
| Fau | 1.27E-29 | 0.53241892 | 0.871 | 0.857 | 2.45E-25 | CD4 T cells | Fau |
| Npm1 | 7.17E-29 | 0.33095486 | 0.375 | 0.432 | 1.39E-24 | CD4 T cells | Npm1 |
| Klf3 | 8.40E-29 | 1.72197334 | 0.27 | 0.119 | 1.62E-24 | CD4 T cells | Klf3 |
| Gramd3 | 1.89E-27 | 1.29508356 | 0.516 | 0.374 | 3.66E-23 | CD4 T cells | Gramd3 |
| Arhgef1 | 3.02E-25 | 0.30271978 | 0.741 | 0.803 | 5.84E-21 | CD4 T cells | Arhgef1 |
| Junb | 6.32E-24 | 0.54147023 | 0.92 | 0.893 | 1.22E-19 | CD4 T cells | Junb |
| Lef1 | 1.82E-23 | 1.90415657 | 0.317 | 0.191 | 3.53E-19 | CD4 T cells | Lef1 |
| mt-Atp6 | 8.80E-23 | 0.29076069 | 0.514 | 0.56 | 1.70E-18 | CD4 T cells | mt-Atp6 |
| mt-Co3 | 2.79E-22 | 0.26381454 | 0.423 | 0.48 | 5.39E-18 | CD4 T cells | mt-Co3 |
| Pnrc1 | 2.85E-21 | 0.30829905 | 0.758 | 0.765 | 5.51E-17 | CD4 T cells | Pnrc1 |
| Lat | 2.88E-21 | 0.37269782 | 0.662 | 0.645 | 5.56E-17 | CD4 T cells | Lat |
| Myh9 | 2.94E-21 | 0.26362259 | 0.443 | 0.54 | 5.69E-17 | CD4 T cells | Myh9 |
| Sf3b1 | 4.55E-21 | 0.33971197 | 0.585 | 0.636 | 8.79E-17 | CD4 T cells | Sf3b1 |
| Arid5a | 4.73E-21 | 1.53026842 | 0.31 | 0.173 | 9.14E-17 | CD4 T cells | Arid5a |
| Jak1 | 2.30E-20 | 0.32134631 | 0.531 | 0.629 | 4.45E-16 | CD4 T cells | Jak1 |
| Eef1b2 | 2.44E-20 | 0.43817042 | 0.512 | 0.548 | 4.72E-16 | CD4 T cells | Eef1b2 |
| Tpt1 | 3.93E-20 | 0.70946289 | 0.955 | 0.897 | 7.60E-16 | CD4 T cells | Tpt1 |
| Trbc2 | 4.95E-20 | 0.4665914 | 0.551 | 0.556 | 9.56E-16 | CD4 T cells | Trbc2 |
| Nup210 | 1.71E-19 | 0.28038337 | 0.37 | 0.441 | 3.30E-15 | CD4 T cells | Nup210 |
| Lbr | 2.05E-18 | 0.27261004 | 0.347 | 0.429 | 3.97E-14 | CD4 T cells | Lbr |
| Ifngr1 | 8.88E-18 | 0.40018457 | 0.823 | 0.834 | 1.72E-13 | CD4 T cells | Ifngr1 |
| Dazap1 | 1.33E-17 | 0.31542062 | 0.342 | 0.432 | 2.57E-13 | CD4 T cells | Dazap1 |
| Chtop | 3.62E-17 | 0.27505366 | 0.29 | 0.403 | 7.01E-13 | CD4 T cells | Chtop |
| mt-Cytb | 6.60E-17 | 0.29575635 | 0.383 | 0.408 | 1.28E-12 | CD4 T cells | mt-Cytb |
| mt-Co1 | 1.18E-16 | 0.30746501 | 0.443 | 0.485 | 2.29E-12 | CD4 T cells | mt-Co1 |
| Morf4l1 | 1.39E-16 | 0.25362166 | 0.37 | 0.465 | 2.68E-12 | CD4 T cells | Morf4l1 |
| Btg2 | 1.65E-16 | 0.96963998 | 0.735 | 0.607 | 3.20E-12 | CD4 T cells | Btg2 |
| Fam107b | 3.18E-16 | 0.40216608 | 0.564 | 0.641 | 6.16E-12 | CD4 T cells | Fam107b |
| Rgs1 | 4.07E-16 | 0.28339645 | 0.478 | 0.583 | 7.87E-12 | CD4 T cells | Rgs1 |
| Trip12 | 5.36E-16 | 0.25694207 | 0.362 | 0.479 | 1.04E-11 | CD4 T cells | Trip12 |
| Son | 5.48E-16 | 0.25577444 | 0.486 | 0.598 | 1.06E-11 | CD4 T cells | Son |
| Ripor2 | 1.37E-15 | 1.27741129 | 0.395 | 0.269 | 2.64E-11 | CD4 T cells | Ripor2 |
| Smap2 | 2.25E-15 | 0.26934735 | 0.365 | 0.472 | 4.36E-11 | CD4 T cells | Smap2 |
| Wdr26 | 3.20E-15 | 0.29540987 | 0.441 | 0.531 | 6.18E-11 | CD4 T cells | Wdr26 |
| Mcl1 | 1.72E-14 | 0.37044908 | 0.615 | 0.672 | 3.32E-10 | CD4 T cells | Mcl1 |
| Pabpc1 | 1.91E-14 | 0.35219512 | 0.333 | 0.414 | 3.69E-10 | CD4 T cells | Pabpc1 |
| Nop53 | 5.29E-14 | 0.27215005 | 0.436 | 0.513 | 1.02E-09 | CD4 T cells | Nop53 |
| Tob1 | 6.10E-14 | 1.24642701 | 0.338 | 0.263 | 1.18E-09 | CD4 T cells | Tob1 |
| Ldb1 | 1.70E-13 | 0.2773041 | 0.428 | 0.503 | 3.30E-09 | CD4 T cells | Ldb1 |
| Rtcb | 2.39E-13 | 0.26130917 | 0.201 | 0.296 | 4.61E-09 | CD4 T cells | Rtcb |
| Fus | 5.62E-13 | 0.32967375 | 0.395 | 0.454 | 1.09E-08 | CD4 T cells | Fus |
| Eif4g3 | 5.74E-13 | 0.27402786 | 0.307 | 0.411 | 1.11E-08 | CD4 T cells | Eif4g3 |
| Per1 | 1.01E-12 | 0.27895756 | 0.461 | 0.534 | 1.96E-08 | CD4 T cells | Per1 |
| B4galnt1 | 1.04E-12 | 0.26355434 | 0.32 | 0.363 | 2.00E-08 | CD4 T cells | B4galnt1 |
| mt-Nd4l | 1.15E-12 | 0.26330521 | 0.292 | 0.365 | 2.22E-08 | CD4 T cells | mt-Nd4l |
| Clec2d | 1.17E-12 | 0.28485923 | 0.436 | 0.498 | 2.27E-08 | CD4 T cells | Clec2d |
| Lnpep | 2.31E-12 | 0.29724203 | 0.433 | 0.52 | 4.48E-08 | CD4 T cells | Lnpep |
| Huwe1 | 3.55E-12 | 0.27920123 | 0.32 | 0.41 | 6.86E-08 | CD4 T cells | Huwe1 |
| Ets1 | 4.83E-12 | 0.7266142 | 0.949 | 0.906 | 9.34E-08 | CD4 T cells | Ets1 |
| Sell | 1.25E-11 | 0.58192991 | 0.209 | 0.31 | 2.41E-07 | CD4 T cells | Sell |
| Psip1 | 3.98E-11 | 0.31438684 | 0.255 | 0.328 | 7.69E-07 | CD4 T cells | Psip1 |
| Brd4 | 5.07E-11 | 0.40860074 | 0.468 | 0.527 | 9.80E-07 | CD4 T cells | Brd4 |
| Ubc | 5.47E-11 | 0.37765164 | 0.58 | 0.632 | 1.06E-06 | CD4 T cells | Ubc |
| Def6 | 5.55E-11 | 0.25861255 | 0.277 | 0.366 | 1.07E-06 | CD4 T cells | Def6 |
| Tpp2 | 9.31E-11 | 0.25612273 | 0.25 | 0.337 | 1.80E-06 | CD4 T cells | Tpp2 |
| Ppp1r12a | 1.35E-10 | 0.27712748 | 0.468 | 0.531 | 2.62E-06 | CD4 T cells | Ppp1r12a |
| Dock2 | 1.84E-10 | 0.44801204 | 0.635 | 0.689 | 3.57E-06 | CD4 T cells | Dock2 |
| Supt4a | 3.13E-10 | 0.28287334 | 0.332 | 0.41 | 6.06E-06 | CD4 T cells | Supt4a |
| Emg1 | 5.51E-10 | 0.40297845 | 0.262 | 0.311 | 1.07E-05 | CD4 T cells | Emg1 |
| Tgfbr2 | 6.04E-10 | 1.176106 | 0.609 | 0.476 | 1.17E-05 | CD4 T cells | Tgfbr2 |
| Ndufa6 | 7.31E-10 | 0.30862778 | 0.267 | 0.338 | 1.41E-05 | CD4 T cells | Ndufa6 |
| Ubap2l | 1.01E-09 | 0.25872746 | 0.206 | 0.294 | 1.95E-05 | CD4 T cells | Ubap2l |
| Rasal3 | 1.03E-09 | 0.36881438 | 0.469 | 0.513 | 1.99E-05 | CD4 T cells | Rasal3 |
| Foxo1 | 1.21E-09 | 0.35362937 | 0.332 | 0.403 | 2.34E-05 | CD4 T cells | Foxo1 |
| Ube2d3 | 1.34E-09 | 0.32315743 | 0.249 | 0.327 | 2.60E-05 | CD4 T cells | Ube2d3 |
| Socs3 | 1.41E-09 | 1.3740858 | 0.287 | 0.209 | 2.72E-05 | CD4 T cells | Socs3 |
| Fam102a | 1.76E-09 | 1.18016139 | 0.493 | 0.393 | 3.40E-05 | CD4 T cells | Fam102a |
| Sbno21 | 1.90E-09 | 0.29087993 | 0.226 | 0.307 | 3.68E-05 | CD4 T cells | Sbno2 |
| Prpf4b | 2.04E-09 | 0.26641168 | 0.224 | 0.301 | 3.94E-05 | CD4 T cells | Prpf4b |
| Cytip | 2.09E-09 | 0.59634053 | 0.71 | 0.715 | 4.05E-05 | CD4 T cells | Cytip |
| Satb1 | 2.56E-09 | 1.44326513 | 0.383 | 0.312 | 4.95E-05 | CD4 T cells | Satb1 |
| Nek7 | 3.37E-09 | 0.25646177 | 0.192 | 0.274 | 6.52E-05 | CD4 T cells | Nek7 |
| Raf1 | 3.79E-09 | 0.25105521 | 0.232 | 0.323 | 7.33E-05 | CD4 T cells | Raf1 |
| Ppp1r15a1 | 3.83E-09 | 0.32121853 | 0.511 | 0.531 | 7.41E-05 | CD4 T cells | Ppp1r15a |
| Hsp90aa1 | 3.93E-09 | 0.60174848 | 0.348 | 0.378 | 7.60E-05 | CD4 T cells | Hsp90aa1 |
| Akap13 | 4.18E-09 | 0.3758921 | 0.509 | 0.573 | 8.08E-05 | CD4 T cells | Akap13 |
| mt-Atp8 | 4.28E-09 | 0.5038628 | 0.522 | 0.524 | 8.27E-05 | CD4 T cells | mt-Atp8 |
| Cnot1 | 4.46E-09 | 0.29176937 | 0.211 | 0.291 | 8.62E-05 | CD4 T cells | Cnot1 |
| Acin1 | 4.92E-09 | 0.43881824 | 0.322 | 0.413 | 9.52E-05 | CD4 T cells | Acin1 |
| Gmfg | 5.57E-09 | 0.3002484 | 0.257 | 0.338 | 0.00010778 | CD4 T cells | Gmfg |
| Eif3h | 7.44E-09 | 0.35752791 | 0.224 | 0.284 | 0.00014385 | CD4 T cells | Eif3h |
| Grk6 | 7.48E-09 | 0.37078462 | 0.327 | 0.388 | 0.00014469 | CD4 T cells | Grk6 |
| Hexim1 | 7.69E-09 | 0.30802771 | 0.313 | 0.403 | 0.00014859 | CD4 T cells | Hexim1 |
| Hipk3 | 8.33E-09 | 0.31489233 | 0.191 | 0.267 | 0.00016113 | CD4 T cells | Hipk3 |
| Ctla2a | 9.12E-09 | 0.61179966 | 0.378 | 0.426 | 0.00017628 | CD4 T cells | Ctla2a |
| Gata31 | 9.29E-09 | 0.42101315 | 0.362 | 0.427 | 0.00017962 | CD4 T cells | Gata3 |
| Kcmf1 | 1.26E-08 | 0.30305928 | 0.171 | 0.257 | 0.00024376 | CD4 T cells | Kcmf1 |
| Eif4ebp2 | 1.40E-08 | 0.28272001 | 0.235 | 0.304 | 0.00027121 | CD4 T cells | Eif4ebp2 |
| Srsf7 | 1.67E-08 | 0.49782024 | 0.367 | 0.399 | 0.00032376 | CD4 T cells | Srsf7 |
| Wdr82 | 1.82E-08 | 0.34160814 | 0.235 | 0.321 | 0.00035257 | CD4 T cells | Wdr82 |
| Tmem64 | 2.01E-08 | 1.04297817 | 0.287 | 0.215 | 0.00038923 | CD4 T cells | Tmem64 |
| Ablim1 | 2.13E-08 | 0.34248999 | 0.378 | 0.483 | 0.00041122 | CD4 T cells | Ablim1 |
| Kansl1 | 2.20E-08 | 0.2970321 | 0.315 | 0.401 | 0.00042478 | CD4 T cells | Kansl1 |
| Dync1h1 | 2.21E-08 | 0.38011121 | 0.294 | 0.373 | 0.00042684 | CD4 T cells | Dync1h1 |
| Srrm1 | 2.33E-08 | 0.33886754 | 0.249 | 0.321 | 0.00045069 | CD4 T cells | Srrm1 |
| Erbin | 2.33E-08 | 0.31931587 | 0.292 | 0.378 | 0.0004512 | CD4 T cells | Erbin |
| Brd2 | 2.43E-08 | 0.3082773 | 0.299 | 0.378 | 0.00047077 | CD4 T cells | Brd2 |
| Creb1 | 2.46E-08 | 0.29359277 | 0.275 | 0.349 | 0.00047539 | CD4 T cells | Creb1 |
| Prkacb | 2.75E-08 | 0.26503139 | 0.235 | 0.313 | 0.0005308 | CD4 T cells | Prkacb |
| Cggbp1 | 2.80E-08 | 0.35895123 | 0.282 | 0.366 | 0.00054072 | CD4 T cells | Cggbp1 |
| Saraf | 3.56E-08 | 0.48733332 | 0.595 | 0.595 | 0.00068912 | CD4 T cells | Saraf |
| Gtpbp1 | 3.57E-08 | 0.32843252 | 0.221 | 0.296 | 0.00068953 | CD4 T cells | Gtpbp1 |
| Pds5a | 3.76E-08 | 0.37484048 | 0.232 | 0.31 | 0.00072689 | CD4 T cells | Pds5a |
| Eef1g | 3.96E-08 | 0.55587642 | 0.255 | 0.291 | 0.0007649 | CD4 T cells | Eef1g |
| G3bp2 | 3.98E-08 | 0.31149846 | 0.216 | 0.299 | 0.00076947 | CD4 T cells | G3bp2 |
| Fyb1 | 4.28E-08 | 0.55544004 | 0.552 | 0.561 | 0.00082772 | CD4 T cells | Fyb |
| Sh3kbp1 | 4.48E-08 | 0.30149742 | 0.25 | 0.315 | 0.00086695 | CD4 T cells | Sh3kbp1 |
| Kcnn41 | 5.06E-08 | 0.2657665 | 0.201 | 0.278 | 0.00097778 | CD4 T cells | Kcnn4 |
| Sec24c | 6.39E-08 | 0.25439873 | 0.221 | 0.304 | 0.00123482 | CD4 T cells | Sec24c |
| Ythdf3 | 6.74E-08 | 0.2914874 | 0.318 | 0.391 | 0.00130342 | CD4 T cells | Ythdf3 |
| Kif21b | 6.76E-08 | 0.25131681 | 0.391 | 0.465 | 0.00130641 | CD4 T cells | Kif21b |
| Hnrnpll | 7.13E-08 | 0.27998374 | 0.206 | 0.268 | 0.00137857 | CD4 T cells | Hnrnpll |
| Srsf5 | 7.61E-08 | 0.26018229 | 0.367 | 0.443 | 0.00147216 | CD4 T cells | Srsf5 |
| Csde1 | 8.19E-08 | 0.34699372 | 0.237 | 0.301 | 0.0015843 | CD4 T cells | Csde1 |
| Pfdn5 | 8.39E-08 | 0.31137257 | 0.287 | 0.37 | 0.00162218 | CD4 T cells | Pfdn5 |
| Dnaja1 | 8.67E-08 | 0.38613739 | 0.299 | 0.359 | 0.00167539 | CD4 T cells | Dnaja1 |
| Pde4d1 | 8.69E-08 | 0.34959217 | 0.3 | 0.358 | 0.00168077 | CD4 T cells | Pde4d |
| Gak | 8.73E-08 | 0.25184012 | 0.189 | 0.266 | 0.00168704 | CD4 T cells | Gak |
| Smc4 | 8.79E-08 | 0.7306417 | 0.226 | 0.266 | 0.00170026 | CD4 T cells | Smc4 |
| Ttc39b | 9.01E-08 | 0.26732034 | 0.327 | 0.42 | 0.00174226 | CD4 T cells | Ttc39b |
| Rexo2 | 9.09E-08 | 0.40501297 | 0.197 | 0.25 | 0.00175826 | CD4 T cells | Rexo2 |
| Gpsm3 | 1.21E-07 | 0.39693147 | 0.441 | 0.49 | 0.00233815 | CD4 T cells | Gpsm3 |
| Lrch4 | 1.49E-07 | 0.39993996 | 0.353 | 0.42 | 0.0028797 | CD4 T cells | Lrch4 |
| Ccnl1 | 1.56E-07 | 0.33821498 | 0.403 | 0.475 | 0.00301473 | CD4 T cells | Ccnl1 |
| Ssh2 | 1.64E-07 | 1.11541225 | 0.299 | 0.232 | 0.00316218 | CD4 T cells | Ssh2 |
| Rnf166 | 1.77E-07 | 0.2648596 | 0.259 | 0.315 | 0.00342182 | CD4 T cells | Rnf166 |
| Add1 | 1.85E-07 | 0.46352588 | 0.3 | 0.353 | 0.0035734 | CD4 T cells | Add1 |
| Arid1a | 2.04E-07 | 0.27538806 | 0.244 | 0.327 | 0.00393826 | CD4 T cells | Arid1a |
| Map3k14 | 2.21E-07 | 1.14546362 | 0.279 | 0.19 | 0.00428068 | CD4 T cells | Map3k14 |
| Faah | 2.29E-07 | 1.26965143 | 0.322 | 0.238 | 0.00442851 | CD4 T cells | Faah |
| Sorl1 | 2.40E-07 | 1.04577741 | 0.352 | 0.263 | 0.00464166 | CD4 T cells | Sorl1 |
| Cyld | 2.47E-07 | 0.36898701 | 0.325 | 0.405 | 0.00478341 | CD4 T cells | Cyld |
| Phf12 | 2.67E-07 | 0.28955967 | 0.245 | 0.313 | 0.00515443 | CD4 T cells | Phf12 |
| Clk1 | 2.78E-07 | 0.47711011 | 0.647 | 0.676 | 0.00536571 | CD4 T cells | Clk1 |
| Rock1 | 2.86E-07 | 0.38434604 | 0.318 | 0.389 | 0.00553112 | CD4 T cells | Rock1 |
| Tle3 | 2.95E-07 | 0.39434226 | 0.348 | 0.409 | 0.00569505 | CD4 T cells | Tle3 |
| Csnk1g2 | 3.36E-07 | 0.28438036 | 0.265 | 0.331 | 0.0064997 | CD4 T cells | Csnk1g2 |
| Setd5 | 4.07E-07 | 0.29315159 | 0.227 | 0.309 | 0.00786147 | CD4 T cells | Setd5 |
| Slc44a2 | 4.17E-07 | 0.28926547 | 0.26 | 0.322 | 0.00806979 | CD4 T cells | Slc44a2 |
| Kpna4 | 4.23E-07 | 0.27022949 | 0.211 | 0.291 | 0.00818175 | CD4 T cells | Kpna4 |
| Clk4 | 4.27E-07 | 0.26941988 | 0.184 | 0.267 | 0.00825622 | CD4 T cells | Clk4 |
| Arfgef1 | 4.43E-07 | 0.3099979 | 0.199 | 0.275 | 0.0085613 | CD4 T cells | Arfgef1 |
| Ddx5 | 7.21E-07 | 0.53314915 | 0.516 | 0.561 | 0.01393469 | CD4 T cells | Ddx5 |
| Clu | 7.70E-07 | 0.29352653 | 0.544 | 0.574 | 0.01488404 | CD4 T cells | Clu |
| Stat6 | 7.90E-07 | 0.36111051 | 0.287 | 0.345 | 0.01527934 | CD4 T cells | Stat6 |
| Ube3a | 8.05E-07 | 0.28706914 | 0.191 | 0.262 | 0.01556382 | CD4 T cells | Ube3a |
| Ppm1a | 8.30E-07 | 0.2644517 | 0.219 | 0.304 | 0.0160452 | CD4 T cells | Ppm1a |
| Ppp1r9b | 8.33E-07 | 0.34183151 | 0.418 | 0.489 | 0.01610313 | CD4 T cells | Ppp1r9b |
| Kmt2e | 8.70E-07 | 0.33383379 | 0.282 | 0.349 | 0.01681179 | CD4 T cells | Kmt2e |
| Ube2h | 9.15E-07 | 0.37413336 | 0.3 | 0.365 | 0.01769566 | CD4 T cells | Ube2h |
| Bclaf1 | 9.54E-07 | 0.31113083 | 0.242 | 0.31 | 0.01844857 | CD4 T cells | Bclaf1 |
| Tmem234 | 9.87E-07 | 0.33592922 | 0.265 | 0.334 | 0.01908813 | CD4 T cells | Tmem234 |
| Man2a1 | 1.12E-06 | 0.27438332 | 0.217 | 0.284 | 0.02156785 | CD4 T cells | Man2a1 |
| Slfn2 | 1.12E-06 | 1.12260808 | 0.595 | 0.447 | 0.02174037 | CD4 T cells | Slfn2 |
| Nfatc3 | 1.13E-06 | 0.39077892 | 0.305 | 0.374 | 0.02191249 | CD4 T cells | Nfatc3 |
| Nfatc2 | 1.28E-06 | 0.34554304 | 0.308 | 0.395 | 0.02480193 | CD4 T cells | Nfatc2 |
| Dyrk1a | 1.53E-06 | 0.36054826 | 0.204 | 0.262 | 0.02961721 | CD4 T cells | Dyrk1a |
| Ppp1cc | 1.58E-06 | 0.49755765 | 0.297 | 0.347 | 0.0305499 | CD4 T cells | Ppp1cc |
| Atp2b1 | 1.83E-06 | 0.2728334 | 0.294 | 0.364 | 0.03533039 | CD4 T cells | Atp2b1 |
| 6-Sep | 1.85E-06 | 0.36196823 | 0.294 | 0.334 | 0.0356935 | CD4 T cells | 6-Sep |
| Dennd1b | 1.87E-06 | 0.3353006 | 0.216 | 0.296 | 0.03622969 | CD4 T cells | Dennd1b |
| Rc3h1 | 2.00E-06 | 0.26643776 | 0.191 | 0.262 | 0.03860639 | CD4 T cells | Rc3h1 |
| Galnt61 | 2.07E-06 | 0.31184512 | 0.222 | 0.271 | 0.04004245 | CD4 T cells | Galnt6 |
| Ptpn22 | 2.22E-06 | 0.48902401 | 0.559 | 0.564 | 0.04292077 | CD4 T cells | Ptpn22 |
| Frmd8 | 2.26E-06 | 0.44761553 | 0.237 | 0.292 | 0.04379088 | CD4 T cells | Frmd8 |
| Atp13a3 | 2.39E-06 | 0.43959167 | 0.211 | 0.267 | 0.04626481 | CD4 T cells | Atp13a3 |
| Mprip | 2.78E-06 | 0.30961975 | 0.192 | 0.255 | 0.05366031 | CD4 T cells | Mprip |
| Cdkn1b | 2.92E-06 | 0.54222843 | 0.561 | 0.551 | 0.05643373 | CD4 T cells | Cdkn1b |
| Birc6 | 3.06E-06 | 0.5885776 | 0.353 | 0.395 | 0.05911365 | CD4 T cells | Birc6 |
| Usp38 | 3.20E-06 | 0.30644514 | 0.191 | 0.253 | 0.06190517 | CD4 T cells | Usp38 |
| Xrn2 | 3.83E-06 | 0.38197768 | 0.274 | 0.326 | 0.07413034 | CD4 T cells | Xrn2 |
| Ifi47 | 3.96E-06 | 0.42254825 | 0.194 | 0.258 | 0.07655413 | CD4 T cells | Ifi47 |
| Srcap | 4.04E-06 | 0.42785693 | 0.282 | 0.348 | 0.07808065 | CD4 T cells | Srcap |
| Nufip2 | 4.40E-06 | 0.38243623 | 0.216 | 0.29 | 0.08516245 | CD4 T cells | Nufip2 |
| Mat2a | 4.51E-06 | 0.38505631 | 0.265 | 0.329 | 0.08719146 | CD4 T cells | Mat2a |
| Inpp4b | 4.85E-06 | 0.57417459 | 0.4 | 0.402 | 0.09374904 | CD4 T cells | Inpp4b |
| Ep400 | 5.29E-06 | 0.45262652 | 0.259 | 0.321 | 0.10225129 | CD4 T cells | Ep400 |
| Gna13 | 5.33E-06 | 0.52256063 | 0.653 | 0.71 | 0.10300391 | CD4 T cells | Gna13 |
| Pcbp2 | 5.39E-06 | 0.5490894 | 0.254 | 0.307 | 0.1041304 | CD4 T cells | Pcbp2 |
| Ncoa2 | 6.72E-06 | 0.30586707 | 0.222 | 0.282 | 0.12993185 | CD4 T cells | Ncoa2 |
| Smad4 | 6.86E-06 | 0.51965153 | 0.289 | 0.34 | 0.13263583 | CD4 T cells | Smad4 |
| Msl2 | 6.93E-06 | 0.42524284 | 0.284 | 0.359 | 0.13405308 | CD4 T cells | Msl2 |
| Vps37b | 8.48E-06 | 0.75250367 | 0.796 | 0.726 | 0.16398125 | CD4 T cells | Vps37b |
| Dusp11 | 8.60E-06 | 0.37025019 | 0.269 | 0.32 | 0.16632022 | CD4 T cells | Dusp11 |
| Dnajb6 | 8.77E-06 | 0.29498647 | 0.245 | 0.333 | 0.16961294 | CD4 T cells | Dnajb6 |
| Qrich1 | 9.69E-06 | 0.28782116 | 0.189 | 0.256 | 0.18729841 | CD4 T cells | Qrich1 |
| Smchd1 | 1.71E-05 | 0.45524938 | 0.235 | 0.271 | 0.3313852 | CD4 T cells | Smchd1 |
| Usp7 | 2.42E-05 | 0.45877821 | 0.229 | 0.284 | 0.46814725 | CD4 T cells | Usp7 |
| Setd1b | 2.82E-05 | 0.35595183 | 0.192 | 0.25 | 0.54432301 | CD4 T cells | Setd1b |
| Sirt7 | 2.95E-05 | 0.26318786 | 0.186 | 0.252 | 0.56953552 | CD4 T cells | Sirt7 |
| Exoc4 | 3.17E-05 | 0.3473435 | 0.226 | 0.275 | 0.61241115 | CD4 T cells | Exoc4 |
| Rbbp6 | 3.32E-05 | 0.28010047 | 0.227 | 0.287 | 0.64181948 | CD4 T cells | Rbbp6 |
| Sub1 | 3.58E-05 | 0.44525297 | 0.227 | 0.275 | 0.69229954 | CD4 T cells | Sub1 |
| Nsd1 | 3.65E-05 | 0.49757876 | 0.386 | 0.442 | 0.70650902 | CD4 T cells | Nsd1 |
| Rsrc2 | 3.67E-05 | 0.43939354 | 0.337 | 0.378 | 0.70865827 | CD4 T cells | Rsrc2 |
| Kras | 4.24E-05 | 0.39976791 | 0.197 | 0.253 | 0.81996576 | CD4 T cells | Kras |
| Dnajc7 | 4.36E-05 | 0.4735593 | 0.211 | 0.257 | 0.84310645 | CD4 T cells | Dnajc7 |
| Mrtfa | 4.67E-05 | 0.38825859 | 0.206 | 0.259 | 0.90296213 | CD4 T cells | Mrtfa |
| Ythdc11 | 5.05E-05 | 0.38254007 | 0.226 | 0.287 | 0.97642285 | CD4 T cells | Ythdc1 |
| Gabarapl2 | 5.61E-05 | 0.46803652 | 0.262 | 0.311 | 1 | CD4 T cells | Gabarapl2 |
| Mepce | 5.64E-05 | 0.28835649 | 0.214 | 0.281 | 1 | CD4 T cells | Mepce |
| Sec11a | 5.78E-05 | 0.52941175 | 0.254 | 0.296 | 1 | CD4 T cells | Sec11a |
| Rasa3 | 6.09E-05 | 0.95299227 | 0.323 | 0.262 | 1 | CD4 T cells | Rasa3 |
| Cirbp | 6.19E-05 | 0.35001627 | 0.249 | 0.294 | 1 | CD4 T cells | Cirbp |
| Apbb1ip | 6.31E-05 | 0.49616332 | 0.441 | 0.478 | 1 | CD4 T cells | Apbb1ip |
| Polr2a | 7.27E-05 | 0.48123441 | 0.335 | 0.372 | 1 | CD4 T cells | Polr2a |
| Fosl2 | 7.60E-05 | 0.60688644 | 0.615 | 0.652 | 1 | CD4 T cells | Fosl2 |
| Cox7a2l | 8.82E-05 | 0.63819415 | 0.3 | 0.334 | 1 | CD4 T cells | Cox7a2l |
| Ascc3 | 8.87E-05 | 0.27448908 | 0.194 | 0.252 | 1 | CD4 T cells | Ascc3 |
| Cnot6l | 8.92E-05 | 0.52346556 | 0.368 | 0.424 | 1 | CD4 T cells | Cnot6l |
| Prpf38b | 9.12E-05 | 0.56630667 | 0.254 | 0.3 | 1 | CD4 T cells | Prpf38b |
| Itgb31 | 9.53E-05 | 0.44977379 | 0.224 | 0.255 | 1 | CD4 T cells | Itgb3 |
| Epc1 | 0.00011192 | 0.43724423 | 0.29 | 0.36 | 1 | CD4 T cells | Epc1 |
| Nipbl | 0.00011574 | 0.37373307 | 0.232 | 0.289 | 1 | CD4 T cells | Nipbl |
| P2ry10 | 0.00011897 | 0.57821976 | 0.489 | 0.516 | 1 | CD4 T cells | P2ry10 |
| Stk38 | 0.00012674 | 1.13380093 | 0.343 | 0.291 | 1 | CD4 T cells | Stk38 |
| Rbm39 | 0.00013204 | 0.41167461 | 0.32 | 0.375 | 1 | CD4 T cells | Rbm39 |
| Tnfaip8 | 0.00013327 | 1.13094088 | 0.352 | 0.306 | 1 | CD4 T cells | Tnfaip8 |
| Znrf1 | 0.00014177 | 0.95929914 | 0.468 | 0.39 | 1 | CD4 T cells | Znrf1 |
| Arhgap45 | 0.00014702 | 0.57325571 | 0.74 | 0.713 | 1 | CD4 T cells | Arhgap45 |
| Rbm5 | 0.00016521 | 0.44778876 | 0.289 | 0.354 | 1 | CD4 T cells | Rbm5 |
| N4bp2l2 | 0.00018657 | 0.40920587 | 0.222 | 0.264 | 1 | CD4 T cells | N4bp2l2 |
| Stim1 | 0.00022235 | 0.38908292 | 0.212 | 0.266 | 1 | CD4 T cells | Stim1 |
| Itpkb1 | 0.00024523 | 0.45633289 | 0.434 | 0.482 | 1 | CD4 T cells | Itpkb |
| Arid4b | 0.00024998 | 0.31776058 | 0.207 | 0.275 | 1 | CD4 T cells | Arid4b |
| Gpr132 | 0.00027332 | 0.56510696 | 0.587 | 0.601 | 1 | CD4 T cells | Gpr132 |
| Furin | 0.0003123 | 0.64420234 | 0.401 | 0.462 | 1 | CD4 T cells | Furin |
| Crlf3 | 0.00031643 | 0.63159147 | 0.383 | 0.425 | 1 | CD4 T cells | Crlf3 |
| Chd7 | 0.00034659 | 0.33108597 | 0.267 | 0.315 | 1 | CD4 T cells | Chd7 |
| Zgpat | 0.0003709 | 0.38430909 | 0.202 | 0.261 | 1 | CD4 T cells | Zgpat |
| Ccm2 | 0.00038181 | 0.5109879 | 0.269 | 0.31 | 1 | CD4 T cells | Ccm2 |
| Rbm4 | 0.00038424 | 0.38843062 | 0.214 | 0.268 | 1 | CD4 T cells | Rbm4 |
| Esyt2 | 0.00038731 | 0.40087307 | 0.363 | 0.421 | 1 | CD4 T cells | Esyt2 |
| Coq10b | 0.00041903 | 0.40844355 | 0.262 | 0.302 | 1 | CD4 T cells | Coq10b |
| Gsk3b | 0.0004678 | 0.48531778 | 0.262 | 0.29 | 1 | CD4 T cells | Gsk3b |
| 7-Mar | 0.00048591 | 0.50657197 | 0.264 | 0.315 | 1 | CD4 T cells | 7-Mar |
| Pik3cd | 0.0005331 | 0.49481571 | 0.441 | 0.452 | 1 | CD4 T cells | Pik3cd |
| Nt5e1 | 0.00055647 | 0.40756208 | 0.222 | 0.257 | 1 | CD4 T cells | Nt5e |
| Epb41 | 0.00055686 | 1.0379779 | 0.297 | 0.234 | 1 | CD4 T cells | Epb41 |
| Herc1 | 0.00056601 | 0.46560217 | 0.224 | 0.262 | 1 | CD4 T cells | Herc1 |
| Zc3hav1 | 0.00058114 | 0.56193682 | 0.594 | 0.612 | 1 | CD4 T cells | Zc3hav1 |
| Top2b | 0.00058937 | 0.48031861 | 0.259 | 0.286 | 1 | CD4 T cells | Top2b |
| Chd1 | 0.00059335 | 0.47819193 | 0.217 | 0.278 | 1 | CD4 T cells | Chd1 |
| Lpxn | 0.00067 | 0.45111279 | 0.312 | 0.339 | 1 | CD4 T cells | Lpxn |
| Zfp292 | 0.00069554 | 0.39959335 | 0.297 | 0.344 | 1 | CD4 T cells | Zfp292 |
| Fosb1 | 0.00086058 | 0.32949152 | 0.269 | 0.333 | 1 | CD4 T cells | Fosb |
| Tsc22d3 | 0.00101305 | 0.98034353 | 0.33 | 0.3 | 1 | CD4 T cells | Tsc22d3 |
| Prkcq | 0.00113566 | 1.12859215 | 0.299 | 0.24 | 1 | CD4 T cells | Prkcq |
| Ifi203 | 0.00126287 | 0.37728732 | 0.24 | 0.271 | 1 | CD4 T cells | Ifi203 |
| Gimap9 | 0.00141284 | 0.49239239 | 0.229 | 0.263 | 1 | CD4 T cells | Gimap9 |
| Add3 | 0.00149238 | 0.64472985 | 0.277 | 0.305 | 1 | CD4 T cells | Add3 |
| Use1 | 0.0015848 | 0.5429699 | 0.287 | 0.316 | 1 | CD4 T cells | Use1 |
| Ankrd44 | 0.00166604 | 0.4124872 | 0.265 | 0.302 | 1 | CD4 T cells | Ankrd44 |
| Rbm33 | 0.00171245 | 0.44197003 | 0.264 | 0.312 | 1 | CD4 T cells | Rbm33 |
| B4galt1 | 0.00178912 | 0.50707751 | 0.325 | 0.367 | 1 | CD4 T cells | B4galt1 |
| Nisch | 0.00179038 | 0.49031636 | 0.303 | 0.353 | 1 | CD4 T cells | Nisch |
| Arhgef18 | 0.00183524 | 0.89996912 | 0.446 | 0.372 | 1 | CD4 T cells | Arhgef18 |
| Gimap6 | 0.00183934 | 0.54060502 | 0.436 | 0.468 | 1 | CD4 T cells | Gimap6 |
| Aff11 | 0.00195375 | 0.44633661 | 0.327 | 0.359 | 1 | CD4 T cells | Aff1 |
| Chd2 | 0.00212116 | 0.36622863 | 0.343 | 0.395 | 1 | CD4 T cells | Chd2 |
| Irf2bp2 | 0.00213592 | 0.81549262 | 0.511 | 0.487 | 1 | CD4 T cells | Irf2bp2 |
| Tet3 | 0.00215332 | 0.49283923 | 0.224 | 0.265 | 1 | CD4 T cells | Tet3 |
| Brd1 | 0.00218009 | 0.46788609 | 0.226 | 0.264 | 1 | CD4 T cells | Brd1 |
| Il7r | 0.00222108 | 0.82794096 | 0.449 | 0.382 | 1 | CD4 T cells | Il7r |
| Stk4 | 0.00233314 | 0.52701985 | 0.257 | 0.301 | 1 | CD4 T cells | Stk4 |
| Gatad2b | 0.00233484 | 0.51188184 | 0.257 | 0.286 | 1 | CD4 T cells | Gatad2b |
| Arhgef3 | 0.00241671 | 0.46063888 | 0.257 | 0.298 | 1 | CD4 T cells | Arhgef3 |
| Gvin1 | 0.00261887 | 0.27058391 | 0.265 | 0.295 | 1 | CD4 T cells | Gvin1 |
| Elk4 | 0.00274143 | 0.29587705 | 0.249 | 0.289 | 1 | CD4 T cells | Elk4 |
| Scaf11 | 0.00304176 | 0.62404997 | 0.249 | 0.278 | 1 | CD4 T cells | Scaf11 |
| Klf6 | 0.00314367 | 0.56522926 | 0.502 | 0.529 | 1 | CD4 T cells | Klf6 |
| Gramd1a | 0.00335831 | 0.61770826 | 0.395 | 0.414 | 1 | CD4 T cells | Gramd1a |
| Prkd2 | 0.00344096 | 0.46053556 | 0.257 | 0.288 | 1 | CD4 T cells | Prkd2 |
| Mycbp2 | 0.00381898 | 0.53093328 | 0.289 | 0.329 | 1 | CD4 T cells | Mycbp2 |
| Birc31 | 0.00388623 | 0.57726458 | 0.365 | 0.386 | 1 | CD4 T cells | Birc3 |
| Zfp36 | 0.00418959 | 0.70665416 | 0.59 | 0.628 | 1 | CD4 T cells | Zfp36 |
| Dnm2 | 0.00424218 | 0.59086648 | 0.303 | 0.331 | 1 | CD4 T cells | Dnm2 |
| Znrf2 | 0.0044341 | 0.53743582 | 0.343 | 0.376 | 1 | CD4 T cells | Znrf2 |
| Cd28 | 0.00530986 | 0.98144961 | 0.425 | 0.35 | 1 | CD4 T cells | Cd28 |
| Smg1 | 0.00532931 | 0.54209261 | 0.302 | 0.331 | 1 | CD4 T cells | Smg1 |
| Susd6 | 0.00551399 | 0.53374045 | 0.284 | 0.319 | 1 | CD4 T cells | Susd6 |
| Ubash3a | 0.00597044 | 0.62512033 | 0.282 | 0.31 | 1 | CD4 T cells | Ubash3a |
| Ubn1 | 0.00675117 | 0.45685876 | 0.235 | 0.271 | 1 | CD4 T cells | Ubn1 |
| Tagap | 0.00722666 | 0.44765689 | 0.294 | 0.3 | 1 | CD4 T cells | Tagap |
| Ier2 | 0.00727062 | 0.81348287 | 0.554 | 0.528 | 1 | CD4 T cells | Ier2 |
| H2-Q4 | 0.00747656 | 0.34587419 | 0.247 | 0.29 | 1 | CD4 T cells | H2-Q4 |
| Itk | 0.00861317 | 0.82928827 | 0.788 | 0.703 | 1 | CD4 T cells | Itk |
| Cdk11b | 0.00879338 | 0.40566169 | 0.269 | 0.287 | 1 | CD4 T cells | Cdk11b |
| Hipk1 | 0.00938884 | 0.71923618 | 0.423 | 0.419 | 1 | CD4 T cells | Hipk1 |
| Nkg7 | 0 | 2.73389816 | 0.837 | 0.17 | 0 | CD8 T cells | Nkg7 |
| Cd8a | 0 | 5.03811061 | 0.664 | 0.022 | 0 | CD8 T cells | Cd8a |
| Cd8b1 | 0 | 4.20525964 | 0.66 | 0.058 | 0 | CD8 T cells | Cd8b1 |
| Ctsw | 0 | 1.86054106 | 0.739 | 0.221 | 0 | CD8 T cells | Ctsw |
| Ms4a4b | 0 | 1.96473497 | 0.729 | 0.224 | 0 | CD8 T cells | Ms4a4b |
| Ccl5 | 0 | 2.7903148 | 0.761 | 0.288 | 0 | CD8 T cells | Ccl5 |
| Fasl | 0 | 2.32482042 | 0.539 | 0.095 | 0 | CD8 T cells | Fasl |
| Themis | 0 | 2.31295765 | 0.536 | 0.11 | 0 | CD8 T cells | Themis |
| Klrc1 | 0 | 2.66831915 | 0.46 | 0.071 | 0 | CD8 T cells | Klrc1 |
| Lag3 | 0 | 2.08882433 | 0.449 | 0.135 | 0 | CD8 T cells | Lag3 |
| Thy1 | 0 | 1.22692107 | 0.912 | 0.624 | 0 | CD8 T cells | Thy1 |
| Cd7 | 0 | 1.99634029 | 0.401 | 0.121 | 0 | CD8 T cells | Cd7 |
| Nr4a2 | 0 | 1.56581957 | 0.482 | 0.214 | 0 | CD8 T cells | Nr4a2 |
| Cd3g | 0 | 1.14487067 | 0.779 | 0.534 | 0 | CD8 T cells | Cd3g |
| Trbc21 | 0 | 1.06139072 | 0.741 | 0.509 | 0 | CD8 T cells | Trbc2 |
| Trbv31 | 0 | 4.2079443 | 0.254 | 0.029 | 0 | CD8 T cells | Trbv31 |
| Pdcd1 | 0 | 1.10650102 | 0.587 | 0.364 | 0 | CD8 T cells | Pdcd1 |
| Trbc1 | 0 | 1.21379075 | 0.568 | 0.359 | 0 | CD8 T cells | Trbc1 |
| Ccl4 | 0 | 2.36305241 | 0.272 | 0.065 | 0 | CD8 T cells | Ccl4 |
| Cd3e | 0 | 0.90647548 | 0.855 | 0.675 | 0 | CD8 T cells | Cd3e |
| Rgs161 | 0 | 1.12136211 | 0.392 | 0.23 | 0 | CD8 T cells | Rgs16 |
| Eef1a11 | 0 | 0.45085134 | 0.987 | 0.934 | 0 | CD8 T cells | Eef1a1 |
| Tmsb4x | 0 | 0.44242924 | 0.994 | 0.976 | 0 | CD8 T cells | Tmsb4x |
| Ptpn221 | 8.24E-283 | 0.99297199 | 0.792 | 0.506 | 1.59E-278 | CD8 T cells | Ptpn22 |
| Ifng | 4.15E-281 | 2.64833186 | 0.312 | 0.05 | 8.03E-277 | CD8 T cells | Ifng |
| Cd52 | 9.25E-260 | 0.55335416 | 0.913 | 0.812 | 1.79E-255 | CD8 T cells | Cd52 |
| B4galnt11 | 3.45E-253 | 1.32679188 | 0.607 | 0.296 | 6.67E-249 | CD8 T cells | B4galnt1 |
| Gzmk | 3.59E-244 | 5.81339119 | 0.306 | 0.008 | 6.94E-240 | CD8 T cells | Gzmk |
| Sh2d2a | 2.04E-224 | 0.71779298 | 0.744 | 0.61 | 3.94E-220 | CD8 T cells | Sh2d2a |
| Klrd1 | 2.84E-224 | 1.97042972 | 0.384 | 0.071 | 5.48E-220 | CD8 T cells | Klrd1 |
| Sema4a | 1.82E-216 | 1.58982158 | 0.54 | 0.175 | 3.51E-212 | CD8 T cells | Sema4a |
| Zyx | 5.97E-209 | 1.12333653 | 0.672 | 0.408 | 1.15E-204 | CD8 T cells | Zyx |
| Lgals3 | 9.85E-206 | 1.46010669 | 0.398 | 0.172 | 1.90E-201 | CD8 T cells | Lgals3 |
| Itgal | 3.76E-201 | 0.7429402 | 0.849 | 0.662 | 7.27E-197 | CD8 T cells | Itgal |
| Laptm5 | 3.60E-199 | 0.40208147 | 0.94 | 0.88 | 6.96E-195 | CD8 T cells | Laptm5 |
| Cd160 | 1.62E-196 | 2.96501256 | 0.25 | 0.033 | 3.13E-192 | CD8 T cells | Cd160 |
| Cxcr6 | 9.17E-193 | 1.07120213 | 0.539 | 0.24 | 1.77E-188 | CD8 T cells | Cxcr6 |
| Gramd31 | 1.49E-182 | 0.95367264 | 0.596 | 0.339 | 2.88E-178 | CD8 T cells | Gramd3 |
| Ugcg | 1.85E-175 | 1.04946956 | 0.458 | 0.19 | 3.58E-171 | CD8 T cells | Ugcg |
| Itgb2 | 8.59E-173 | 0.72767769 | 0.844 | 0.651 | 1.66E-168 | CD8 T cells | Itgb2 |
| Tox | 9.27E-166 | 1.21781983 | 0.422 | 0.273 | 1.79E-161 | CD8 T cells | Tox |
| Ostf1 | 1.05E-161 | 1.02868986 | 0.597 | 0.361 | 2.03E-157 | CD8 T cells | Ostf1 |
| Tbx21 | 3.42E-161 | 1.5959865 | 0.482 | 0.133 | 6.61E-157 | CD8 T cells | Tbx21 |
| Anxa2 | 1.24E-158 | 0.57805739 | 0.636 | 0.446 | 2.40E-154 | CD8 T cells | Anxa2 |
| Wipf1 | 2.19E-156 | 0.94842243 | 0.695 | 0.457 | 4.23E-152 | CD8 T cells | Wipf1 |
| Sorl11 | 2.08E-153 | 1.2384207 | 0.521 | 0.211 | 4.02E-149 | CD8 T cells | Sorl1 |
| Cd3d | 2.89E-148 | 0.56545579 | 0.755 | 0.6 | 5.59E-144 | CD8 T cells | Cd3d |
| Ikzf1 | 4.89E-145 | 0.76875535 | 0.731 | 0.484 | 9.45E-141 | CD8 T cells | Ikzf1 |
| Chsy1 | 3.39E-144 | 1.3404515 | 0.47 | 0.223 | 6.56E-140 | CD8 T cells | Chsy1 |
| Ms4a6b | 4.19E-142 | 1.10984469 | 0.528 | 0.262 | 8.10E-138 | CD8 T cells | Ms4a6b |
| Slamf7 | 9.75E-135 | 2.74534448 | 0.286 | 0.04 | 1.88E-130 | CD8 T cells | Slamf7 |
| Rac2 | 1.71E-129 | 0.38274748 | 0.958 | 0.884 | 3.30E-125 | CD8 T cells | Rac2 |
| Pde7a | 7.65E-127 | 1.58615511 | 0.413 | 0.129 | 1.48E-122 | CD8 T cells | Pde7a |
| Ctsd | 2.22E-126 | 1.364873 | 0.427 | 0.183 | 4.28E-122 | CD8 T cells | Ctsd |
| Gtf2i | 5.43E-126 | 1.21693592 | 0.443 | 0.19 | 1.05E-121 | CD8 T cells | Gtf2i |
| Fyn | 1.41E-120 | 0.89713115 | 0.656 | 0.453 | 2.72E-116 | CD8 T cells | Fyn |
| Ybx3 | 6.63E-120 | 1.37086665 | 0.435 | 0.203 | 1.28E-115 | CD8 T cells | Ybx3 |
| Bcl2 | 2.65E-119 | 1.2325296 | 0.372 | 0.131 | 5.12E-115 | CD8 T cells | Bcl2 |
| Lck | 5.91E-117 | 0.76038028 | 0.806 | 0.567 | 1.14E-112 | CD8 T cells | Lck |
| Bcl2l11 | 1.62E-116 | 0.85062901 | 0.454 | 0.247 | 3.13E-112 | CD8 T cells | Bcl2l11 |
| B4galt5 | 2.82E-116 | 1.21329974 | 0.38 | 0.163 | 5.46E-112 | CD8 T cells | B4galt5 |
| Serpinb9 | 8.49E-116 | 1.27403076 | 0.352 | 0.13 | 1.64E-111 | CD8 T cells | Serpinb9 |
| Bhlhe40 | 1.42E-114 | 0.34316614 | 0.774 | 0.677 | 2.74E-110 | CD8 T cells | Bhlhe40 |
| Pik3cd1 | 9.56E-111 | 0.71987041 | 0.629 | 0.406 | 1.85E-106 | CD8 T cells | Pik3cd |
| Pde3b | 1.22E-109 | 1.2519504 | 0.406 | 0.137 | 2.36E-105 | CD8 T cells | Pde3b |
| Cnn2 | 5.76E-107 | 0.54281708 | 0.773 | 0.626 | 1.11E-102 | CD8 T cells | Cnn2 |
| Lcp1 | 3.39E-102 | 0.45099758 | 0.862 | 0.744 | 6.55E-98 | CD8 T cells | Lcp1 |
| Arsb | 4.40E-102 | 1.29905119 | 0.295 | 0.114 | 8.50E-98 | CD8 T cells | Arsb |
| Nedd9 | 4.73E-101 | 1.00175848 | 0.479 | 0.268 | 9.14E-97 | CD8 T cells | Nedd9 |
| Sh2d1a | 1.30E-100 | 1.40153285 | 0.307 | 0.116 | 2.52E-96 | CD8 T cells | Sh2d1a |
| Rgs3 | 3.59E-100 | 0.8627376 | 0.667 | 0.42 | 6.94E-96 | CD8 T cells | Rgs3 |
| Gng2 | 9.25E-100 | 0.92676854 | 0.521 | 0.29 | 1.79E-95 | CD8 T cells | Gng2 |
| Prf1 | 3.39E-99 | 0.91981207 | 0.317 | 0.088 | 6.55E-95 | CD8 T cells | Prf1 |
| Nsmaf | 1.25E-96 | 1.03047258 | 0.426 | 0.231 | 2.41E-92 | CD8 T cells | Nsmaf |
| Pstpip1 | 4.81E-94 | 0.91223735 | 0.521 | 0.309 | 9.30E-90 | CD8 T cells | Pstpip1 |
| Cd226 | 1.02E-93 | 1.60905295 | 0.292 | 0.092 | 1.98E-89 | CD8 T cells | Cd226 |
| Fau1 | 3.15E-93 | 0.34302444 | 0.932 | 0.841 | 6.08E-89 | CD8 T cells | Fau |
| Plek | 1.15E-92 | 0.89744307 | 0.337 | 0.145 | 2.22E-88 | CD8 T cells | Plek |
| Itgb1 | 4.14E-91 | 0.81969315 | 0.547 | 0.327 | 8.00E-87 | CD8 T cells | Itgb1 |
| Itk1 | 2.90E-90 | 0.39317436 | 0.842 | 0.681 | 5.60E-86 | CD8 T cells | Itk |
| Rnf1661 | 6.51E-90 | 0.99583877 | 0.492 | 0.263 | 1.26E-85 | CD8 T cells | Rnf166 |
| Fam78a | 5.42E-89 | 1.28988416 | 0.341 | 0.13 | 1.05E-84 | CD8 T cells | Fam78a |
| Il21r1 | 8.42E-88 | 0.49694523 | 0.625 | 0.463 | 1.63E-83 | CD8 T cells | Il21r |
| Serpina3g | 4.83E-85 | 0.95704352 | 0.359 | 0.186 | 9.35E-81 | CD8 T cells | Serpina3g |
| Litaf | 2.25E-83 | 0.85969986 | 0.486 | 0.293 | 4.35E-79 | CD8 T cells | Litaf |
| Cd82 | 9.73E-83 | 0.33118503 | 0.825 | 0.742 | 1.88E-78 | CD8 T cells | Cd82 |
| Zap70 | 1.67E-82 | 0.6154026 | 0.64 | 0.462 | 3.23E-78 | CD8 T cells | Zap70 |
| Stk39 | 5.09E-82 | 1.38777895 | 0.269 | 0.093 | 9.85E-78 | CD8 T cells | Stk39 |
| Lyst | 1.34E-80 | 1.41913718 | 0.301 | 0.121 | 2.59E-76 | CD8 T cells | Lyst |
| Adgre5 | 3.82E-80 | 0.70418386 | 0.583 | 0.383 | 7.38E-76 | CD8 T cells | Adgre5 |
| Nlrc5 | 5.83E-80 | 0.61881848 | 0.622 | 0.398 | 1.13E-75 | CD8 T cells | Nlrc5 |
| Cst7 | 6.60E-80 | 1.04567547 | 0.347 | 0.181 | 1.28E-75 | CD8 T cells | Cst7 |
| Prex1 | 8.84E-79 | 0.58048721 | 0.701 | 0.536 | 1.71E-74 | CD8 T cells | Prex1 |
| Rpa2 | 5.60E-78 | 1.98094587 | 0.281 | 0.084 | 1.08E-73 | CD8 T cells | Rpa2 |
| Lax1 | 3.15E-76 | 1.11099258 | 0.377 | 0.168 | 6.10E-72 | CD8 T cells | Lax1 |
| Stk17b | 2.96E-75 | 0.4562055 | 0.797 | 0.666 | 5.73E-71 | CD8 T cells | Stk17b |
| Tmsb10 | 7.05E-75 | 0.32447092 | 0.876 | 0.792 | 1.36E-70 | CD8 T cells | Tmsb10 |
| Trac1 | 6.73E-74 | 0.36167822 | 0.876 | 0.687 | 1.30E-69 | CD8 T cells | Trac |
| Cotl1 | 7.22E-74 | 0.33582459 | 0.87 | 0.783 | 1.40E-69 | CD8 T cells | Cotl1 |
| Chst12 | 3.71E-72 | 1.24533362 | 0.261 | 0.108 | 7.17E-68 | CD8 T cells | Chst12 |
| Cd96 | 6.34E-72 | 1.07811045 | 0.312 | 0.117 | 1.23E-67 | CD8 T cells | Cd96 |
| Il2rb | 7.41E-72 | 0.25594549 | 0.843 | 0.76 | 1.43E-67 | CD8 T cells | Il2rb |
| Plekha2 | 9.73E-72 | 0.69663168 | 0.496 | 0.308 | 1.88E-67 | CD8 T cells | Plekha2 |
| Pdcd4 | 2.11E-71 | 0.52939742 | 0.69 | 0.502 | 4.07E-67 | CD8 T cells | Pdcd4 |
| Zfp36l21 | 4.79E-71 | 0.41511333 | 0.782 | 0.678 | 9.26E-67 | CD8 T cells | Zfp36l2 |
| Lfng | 9.59E-71 | 0.6620831 | 0.595 | 0.387 | 1.85E-66 | CD8 T cells | Lfng |
| Cblb | 1.01E-68 | 0.53513509 | 0.563 | 0.415 | 1.95E-64 | CD8 T cells | Cblb |
| Adgrg5 | 2.24E-68 | 1.24578764 | 0.257 | 0.105 | 4.33E-64 | CD8 T cells | Adgrg5 |
| Tgfbr21 | 1.21E-67 | 0.50180558 | 0.636 | 0.456 | 2.34E-63 | CD8 T cells | Tgfbr2 |
| Evl | 1.73E-67 | 0.87280747 | 0.467 | 0.188 | 3.34E-63 | CD8 T cells | Evl |
| Clta | 3.86E-66 | 0.58870875 | 0.711 | 0.548 | 7.46E-62 | CD8 T cells | Clta |
| Rbpj | 6.22E-65 | 0.60306253 | 0.429 | 0.302 | 1.20E-60 | CD8 T cells | Rbpj |
| Nfatc1 | 5.53E-63 | 0.70943825 | 0.448 | 0.289 | 1.07E-58 | CD8 T cells | Nfatc1 |
| Cxcr3 | 1.80E-62 | 1.5169653 | 0.337 | 0.145 | 3.48E-58 | CD8 T cells | Cxcr3 |
| Lsp1 | 2.91E-62 | 0.47755735 | 0.703 | 0.554 | 5.62E-58 | CD8 T cells | Lsp1 |
| Ptms1 | 2.36E-61 | 0.48926832 | 0.521 | 0.38 | 4.56E-57 | CD8 T cells | Ptms |
| H2-Q7 | 4.35E-60 | 0.34640061 | 0.859 | 0.728 | 8.40E-56 | CD8 T cells | H2-Q7 |
| Psmb9 | 5.81E-59 | 0.73455742 | 0.56 | 0.372 | 1.12E-54 | CD8 T cells | Psmb9 |
| Plac8 | 4.94E-58 | 0.73895874 | 0.255 | 0.174 | 9.56E-54 | CD8 T cells | Plac8 |
| Epsti1 | 6.15E-58 | 0.79815539 | 0.433 | 0.245 | 1.19E-53 | CD8 T cells | Epsti1 |
| Ptprc | 7.34E-58 | 0.36829769 | 0.823 | 0.713 | 1.42E-53 | CD8 T cells | Ptprc |
| Hcst | 5.75E-56 | 1.13104094 | 0.354 | 0.174 | 1.11E-51 | CD8 T cells | Hcst |
| Fam102a1 | 9.41E-56 | 0.46842635 | 0.532 | 0.373 | 1.82E-51 | CD8 T cells | Fam102a |
| Efhd2 | 6.41E-55 | 0.88011969 | 0.395 | 0.226 | 1.24E-50 | CD8 T cells | Efhd2 |
| Cd48 | 9.69E-55 | 0.80877596 | 0.473 | 0.298 | 1.87E-50 | CD8 T cells | Cd48 |
| Hip1r | 1.56E-54 | 0.7149062 | 0.413 | 0.258 | 3.02E-50 | CD8 T cells | Hip1r |
| Ccnd3 | 4.43E-54 | 0.78057813 | 0.506 | 0.303 | 8.57E-50 | CD8 T cells | Ccnd3 |
| Arap2 | 7.46E-54 | 0.79426538 | 0.296 | 0.151 | 1.44E-49 | CD8 T cells | Arap2 |
| Esyt1 | 2.93E-53 | 0.38087115 | 0.74 | 0.609 | 5.67E-49 | CD8 T cells | Esyt1 |
| Klrk1 | 4.17E-53 | 0.99205088 | 0.281 | 0.102 | 8.07E-49 | CD8 T cells | Klrk1 |
| Gpr68 | 2.70E-52 | 0.75904509 | 0.461 | 0.294 | 5.22E-48 | CD8 T cells | Gpr68 |
| Cox17 | 5.47E-52 | 0.61172773 | 0.424 | 0.308 | 1.06E-47 | CD8 T cells | Cox17 |
| Psmb8 | 1.14E-51 | 0.56543318 | 0.659 | 0.506 | 2.21E-47 | CD8 T cells | Psmb8 |
| Rack11 | 2.62E-51 | 0.39789073 | 0.815 | 0.663 | 5.07E-47 | CD8 T cells | Rack1 |
| Sub11 | 3.97E-50 | 0.76419559 | 0.407 | 0.236 | 7.67E-46 | CD8 T cells | Sub1 |
| Anxa6 | 1.74E-49 | 0.41360408 | 0.776 | 0.634 | 3.37E-45 | CD8 T cells | Anxa6 |
| Pglyrp1 | 1.25E-48 | 0.45160328 | 0.35 | 0.266 | 2.42E-44 | CD8 T cells | Pglyrp1 |
| Iqsec1 | 1.53E-48 | 0.96598607 | 0.292 | 0.141 | 2.95E-44 | CD8 T cells | Iqsec1 |
| Rassf5 | 6.63E-48 | 0.64071141 | 0.636 | 0.452 | 1.28E-43 | CD8 T cells | Rassf5 |
| Arhgap9 | 9.34E-48 | 0.55720721 | 0.614 | 0.472 | 1.81E-43 | CD8 T cells | Arhgap9 |
| Runx3 | 5.56E-47 | 0.5112732 | 0.624 | 0.457 | 1.07E-42 | CD8 T cells | Runx3 |
| Gdi2 | 1.39E-46 | 0.86332488 | 0.377 | 0.234 | 2.68E-42 | CD8 T cells | Gdi2 |
| Ctsc | 2.66E-46 | 0.79061488 | 0.326 | 0.184 | 5.14E-42 | CD8 T cells | Ctsc |
| Sh2d3c | 7.27E-46 | 1.21799564 | 0.288 | 0.126 | 1.41E-41 | CD8 T cells | Sh2d3c |
| Grk2 | 8.12E-46 | 0.45192365 | 0.646 | 0.517 | 1.57E-41 | CD8 T cells | Grk2 |
| Btg21 | 8.85E-46 | 0.26706899 | 0.727 | 0.596 | 1.71E-41 | CD8 T cells | Btg2 |
| Dnajc9 | 3.46E-45 | 0.56832172 | 0.393 | 0.264 | 6.69E-41 | CD8 T cells | Dnajc9 |
| Casp3 | 3.61E-45 | 0.73687966 | 0.348 | 0.219 | 6.99E-41 | CD8 T cells | Casp3 |
| Dgkz | 6.50E-45 | 0.44696799 | 0.571 | 0.459 | 1.26E-40 | CD8 T cells | Dgkz |
| Spn | 8.68E-45 | 0.30050831 | 0.774 | 0.663 | 1.68E-40 | CD8 T cells | Spn |
| Eef1b21 | 1.24E-44 | 0.45320682 | 0.659 | 0.515 | 2.40E-40 | CD8 T cells | Eef1b2 |
| Slfn21 | 1.35E-44 | 0.45749234 | 0.614 | 0.427 | 2.60E-40 | CD8 T cells | Slfn2 |
| Adam19 | 8.69E-44 | 0.72727313 | 0.432 | 0.246 | 1.68E-39 | CD8 T cells | Adam19 |
| Padi2 | 1.42E-43 | 0.95776468 | 0.329 | 0.174 | 2.75E-39 | CD8 T cells | Padi2 |
| Atp10a | 1.89E-43 | 0.95202311 | 0.265 | 0.125 | 3.66E-39 | CD8 T cells | Atp10a |
| Gimap4 | 6.88E-43 | 0.29007785 | 0.698 | 0.55 | 1.33E-38 | CD8 T cells | Gimap4 |
| Grk61 | 3.35E-42 | 0.46409898 | 0.495 | 0.353 | 6.48E-38 | CD8 T cells | Grk6 |

| | | | | | | | |
|---|---|---|---|---|---|---|---|
| Prkcq1 | 7.66E-42 | 0.78728673 | 0.389 | 0.212 | 1.48E-37 | CD8 T cells | Prkcq |
| Sep-61 | 5.13E-41 | 0.52633961 | 0.442 | 0.301 | 9.91E-37 | CD8 T cells | 6-Sep |
| Rab11fip4 | 6.36E-41 | 0.9836703 | 0.254 | 0.116 | 1.23E-36 | CD8 T cells | Rab11fip4 |
| Dnajc15 | 1.38E-39 | 1.00416038 | 0.302 | 0.155 | 2.66E-35 | CD8 T cells | Dnajc15 |
| Dusp2 | 3.74E-39 | 0.75811364 | 0.318 | 0.186 | 7.23E-35 | CD8 T cells | Dusp2 |
| Rnf19a | 7.01E-39 | 0.57715312 | 0.285 | 0.17 | 1.35E-34 | CD8 T cells | Rnf19a |
| Lpxn1 | 1.27E-38 | 0.58466023 | 0.471 | 0.303 | 2.45E-34 | CD8 T cells | Lpxn |
| Ccni | 1.59E-38 | 0.43177461 | 0.495 | 0.364 | 3.07E-34 | CD8 T cells | Ccni |
| Hspa81 | 4.27E-38 | 0.28429421 | 0.847 | 0.739 | 8.26E-34 | CD8 T cells | Hspa8 |
| Eif3f | 4.48E-38 | 0.52024751 | 0.57 | 0.422 | 8.67E-34 | CD8 T cells | Eif3f |
| Mink1 | 2.44E-37 | 0.84205377 | 0.302 | 0.158 | 4.72E-33 | CD8 T cells | Mink1 |
| Stat4 | 6.74E-37 | 0.44728454 | 0.49 | 0.348 | 1.30E-32 | CD8 T cells | Stat4 |
| Epb411 | 1.00E-36 | 0.74341995 | 0.379 | 0.208 | 1.93E-32 | CD8 T cells | Epb41 |
| B4galt11 | 1.68E-36 | 0.57192539 | 0.507 | 0.327 | 3.24E-32 | CD8 T cells | B4galt1 |
| Rapgef1 | 1.78E-36 | 0.6014593 | 0.439 | 0.295 | 3.44E-32 | CD8 T cells | Rapgef1 |
| Prkch1 | 3.05E-36 | 0.40649715 | 0.601 | 0.475 | 5.89E-32 | CD8 T cells | Prkch |
| Prkacb1 | 3.18E-36 | 0.68494067 | 0.443 | 0.27 | 6.15E-32 | CD8 T cells | Prkacb |
| Rnf138 | 3.52E-36 | 0.74221443 | 0.396 | 0.246 | 6.81E-32 | CD8 T cells | Rnf138 |
| Gramd1b | 2.25E-35 | 0.62982659 | 0.351 | 0.223 | 4.34E-31 | CD8 T cells | Gramd1b |
| Inpp4b1 | 6.89E-35 | 0.46031687 | 0.51 | 0.374 | 1.33E-30 | CD8 T cells | Inpp4b |
| Nol7 | 1.94E-34 | 0.79000806 | 0.332 | 0.204 | 3.75E-30 | CD8 T cells | Nol7 |
| Myl12a | 2.70E-34 | 0.37087944 | 0.618 | 0.498 | 5.21E-30 | CD8 T cells | Myl12a |
| Sp100 | 5.18E-34 | 0.60512334 | 0.487 | 0.344 | 1.00E-29 | CD8 T cells | Sp100 |
| Gch1 | 7.94E-34 | 0.59194415 | 0.304 | 0.178 | 1.53E-29 | CD8 T cells | Gch1 |
| Selplg | 1.09E-33 | 0.31939513 | 0.724 | 0.628 | 2.12E-29 | CD8 T cells | Selplg |
| Dock10 | 1.62E-33 | 0.48052798 | 0.494 | 0.368 | 3.14E-29 | CD8 T cells | Dock10 |
| Cd281 | 2.36E-33 | 0.62890833 | 0.485 | 0.327 | 4.57E-29 | CD8 T cells | Cd28 |
| Tagap1 | 4.48E-33 | 0.61424244 | 0.362 | 0.283 | 8.66E-29 | CD8 T cells | Tagap |
| Znrf21 | 1.41E-32 | 0.60376289 | 0.507 | 0.339 | 2.72E-28 | CD8 T cells | Znrf2 |
| Hip1 | 1.61E-32 | 0.67511069 | 0.291 | 0.171 | 3.11E-28 | CD8 T cells | Hip1 |
| Slc20a1 | 2.82E-32 | 0.76151909 | 0.333 | 0.195 | 5.46E-28 | CD8 T cells | Slc20a1 |
| Rsrp1 | 3.22E-32 | 0.32301231 | 0.627 | 0.493 | 6.22E-28 | CD8 T cells | Rsrp1 |
| Eif3k | 3.26E-32 | 0.51855753 | 0.57 | 0.413 | 6.30E-28 | CD8 T cells | Eif3k |
| Hnrnpa1 | 7.46E-32 | 0.56070916 | 0.443 | 0.316 | 1.44E-27 | CD8 T cells | Hnrnpa1 |
| Ptger4 | 7.88E-32 | 0.64226467 | 0.313 | 0.192 | 1.52E-27 | CD8 T cells | Ptger4 |
| Lnpep1 | 1.23E-30 | 0.39645816 | 0.624 | 0.481 | 2.38E-26 | CD8 T cells | Lnpep |
| Akna | 2.94E-30 | 0.37536565 | 0.589 | 0.475 | 5.69E-26 | CD8 T cells | Akna |
| Pld3 | 3.55E-30 | 0.35567535 | 0.489 | 0.369 | 6.86E-26 | CD8 T cells | Pld3 |
| Arpc5 | 4.75E-30 | 0.35083992 | 0.625 | 0.502 | 9.19E-26 | CD8 T cells | Arpc5 |
| Grap2 | 6.05E-30 | 0.49921762 | 0.525 | 0.388 | 1.17E-25 | CD8 T cells | Grap2 |
| Rab8a | 1.31E-29 | 0.40347916 | 0.53 | 0.401 | 2.54E-25 | CD8 T cells | Rab8a |
| Tubb5 | 3.42E-29 | 0.27994385 | 0.503 | 0.427 | 6.62E-25 | CD8 T cells | Tubb5 |
| Ech1 | 7.06E-29 | 0.63483824 | 0.367 | 0.247 | 1.37E-24 | CD8 T cells | Ech1 |
| Gnptg | 7.74E-29 | 0.71427201 | 0.29 | 0.177 | 1.50E-24 | CD8 T cells | Gnptg |
| Arid5a1 | 9.03E-29 | 0.56765478 | 0.303 | 0.16 | 1.75E-24 | CD8 T cells | Arid5a |
| Nbeal2 | 1.03E-28 | 0.65553166 | 0.474 | 0.317 | 1.99E-24 | CD8 T cells | Nbeal2 |
| Pced1b | 2.92E-28 | 0.63429925 | 0.36 | 0.227 | 5.65E-24 | CD8 T cells | Pced1b |
| Sh3bgrl3 | 6.47E-28 | 0.28922371 | 0.792 | 0.695 | 1.25E-23 | CD8 T cells | Sh3bgrl3 |
| Sh3kbp11 | 1.62E-27 | 0.48180555 | 0.416 | 0.281 | 3.13E-23 | CD8 T cells | Sh3kbp1 |
| Ubac2 | 1.69E-27 | 0.70566968 | 0.332 | 0.207 | 3.26E-23 | CD8 T cells | Ubac2 |
| Man2b2 | 2.01E-27 | 0.70393057 | 0.302 | 0.19 | 3.89E-23 | CD8 T cells | Man2b2 |
| Sesn3 | 2.14E-27 | 0.48824777 | 0.35 | 0.255 | 4.14E-23 | CD8 T cells | Sesn3 |
| Rbm3 | 7.08E-27 | 0.48420524 | 0.494 | 0.359 | 1.37E-22 | CD8 T cells | Rbm3 |
| Capzb | 8.26E-27 | 0.34956796 | 0.652 | 0.549 | 1.60E-22 | CD8 T cells | Capzb |
| Mbnl1 | 7.71E-26 | 0.27676802 | 0.628 | 0.53 | 1.49E-21 | CD8 T cells | Mbnl1 |
| Dock8 | 8.47E-26 | 0.29561225 | 0.485 | 0.397 | 1.64E-21 | CD8 T cells | Dock8 |
| Agfg11 | 1.17E-25 | 0.30682915 | 0.464 | 0.367 | 2.27E-21 | CD8 T cells | Agfg1 |
| Ubash3a1 | 1.19E-25 | 0.45521856 | 0.415 | 0.279 | 2.31E-21 | CD8 T cells | Ubash3a |
| Tma7 | 1.19E-25 | 0.5243449 | 0.412 | 0.287 | 2.31E-21 | CD8 T cells | Tma7 |
| Btg3 | 1.49E-25 | 0.66391558 | 0.264 | 0.159 | 2.89E-21 | CD8 T cells | Btg3 |
| Hipk11 | 1.90E-25 | 0.48538756 | 0.535 | 0.391 | 3.67E-21 | CD8 T cells | Hipk1 |
| Sipa1 | 2.28E-25 | 0.53398653 | 0.425 | 0.302 | 4.41E-21 | CD8 T cells | Sipa1 |
| Runx2 | 3.87E-25 | 0.5130729 | 0.42 | 0.293 | 7.47E-21 | CD8 T cells | Runx2 |
| Slc44a21 | 4.37E-25 | 0.37235397 | 0.415 | 0.29 | 8.45E-21 | CD8 T cells | Slc44a2 |
| Ywhag | 5.21E-25 | 0.44921244 | 0.408 | 0.288 | 1.01E-20 | CD8 T cells | Ywhag |
| Cdk61 | 7.39E-25 | 0.55028861 | 0.335 | 0.263 | 1.43E-20 | CD8 T cells | Cdk6 |
| Myh91 | 9.58E-25 | 0.32794907 | 0.626 | 0.504 | 1.85E-20 | CD8 T cells | Myh9 |
| Abr | 1.09E-24 | 0.49616511 | 0.372 | 0.264 | 2.10E-20 | CD8 T cells | Abr |
| Naca | 2.07E-24 | 0.5789055 | 0.478 | 0.336 | 4.00E-20 | CD8 T cells | Naca |
| Snrpg | 6.53E-24 | 0.63861676 | 0.314 | 0.195 | 1.26E-19 | CD8 T cells | Snrpg |
| Cnot6l1 | 1.21E-23 | 0.33154814 | 0.508 | 0.395 | 2.34E-19 | CD8 T cells | Cnot6l |
| Usp48 | 1.47E-23 | 0.5687393 | 0.294 | 0.185 | 2.84E-19 | CD8 T cells | Usp48 |
| Mbp | 1.61E-23 | 0.4709507 | 0.369 | 0.253 | 3.11E-19 | CD8 T cells | Mbp |
| Cdc42se1 | 1.70E-23 | 0.33431679 | 0.48 | 0.368 | 3.28E-19 | CD8 T cells | Cdc42se1 |
| Ifi2031 | 2.24E-23 | 0.45454517 | 0.355 | 0.246 | 4.32E-19 | CD8 T cells | Ifi203 |
| Abcb1a | 5.75E-23 | 0.49287269 | 0.289 | 0.179 | 1.11E-18 | CD8 T cells | Abcb1a |
| Emg11 | 8.31E-23 | 0.50779907 | 0.394 | 0.283 | 1.61E-18 | CD8 T cells | Emg1 |
| Ndufa10 | 9.82E-23 | 0.75874997 | 0.314 | 0.193 | 1.90E-18 | CD8 T cells | Ndufa10 |
| Mta3 | 1.93E-22 | 0.46144083 | 0.411 | 0.304 | 3.73E-18 | CD8 T cells | Mta3 |
| Rinl | 2.40E-22 | 0.35412238 | 0.515 | 0.378 | 4.65E-18 | CD8 T cells | Rinl |
| Csnk2b | 2.48E-22 | 0.67099773 | 0.341 | 0.219 | 4.80E-18 | CD8 T cells | Csnk2b |
| Mcm6 | 2.79E-22 | 0.64543462 | 0.318 | 0.21 | 5.40E-18 | CD8 T cells | Mcm6 |
| Arrb2 | 3.95E-22 | 0.35565343 | 0.527 | 0.422 | 7.64E-18 | CD8 T cells | Arrb2 |
| Bcl2a1b | 1.04E-21 | 0.42144286 | 0.419 | 0.316 | 2.01E-17 | CD8 T cells | Bcl2a1b |
| Gpr171 | 1.14E-21 | 0.65672084 | 0.277 | 0.165 | 2.20E-17 | CD8 T cells | Gpr171 |
| Ndufb10 | 1.31E-21 | 0.60756648 | 0.351 | 0.237 | 2.53E-17 | CD8 T cells | Ndufb10 |
| Cox8a | 1.37E-21 | 0.38828726 | 0.441 | 0.327 | 2.65E-17 | CD8 T cells | Cox8a |
| Supt4a1 | 1.57E-21 | 0.29518574 | 0.487 | 0.38 | 3.03E-17 | CD8 T cells | Supt4a |
| Arhgdib | 1.83E-21 | 0.28358222 | 0.811 | 0.681 | 3.53E-17 | CD8 T cells | Arhgdib |
| Vsir | 2.27E-21 | 0.26698432 | 0.525 | 0.401 | 4.40E-17 | CD8 T cells | Vsir |
| Slc28a2 | 2.41E-21 | 0.52090032 | 0.305 | 0.193 | 4.66E-17 | CD8 T cells | Slc28a2 |
| Rasal31 | 3.17E-21 | 0.38114863 | 0.608 | 0.483 | 6.13E-17 | CD8 T cells | Rasal3 |
| Rassf2 | 3.41E-21 | 0.61580059 | 0.299 | 0.176 | 6.59E-17 | CD8 T cells | Rassf2 |
| Kif13b | 4.47E-21 | 0.67469351 | 0.26 | 0.163 | 8.64E-17 | CD8 T cells | Kif13b |
| Psme1 | 4.76E-21 | 0.41693161 | 0.638 | 0.495 | 9.21E-17 | CD8 T cells | Psme1 |
| Ankrd441 | 6.26E-21 | 0.36847873 | 0.39 | 0.275 | 1.21E-16 | CD8 T cells | Ankrd44 |
| Ndufa13 | 1.02E-20 | 0.28952891 | 0.53 | 0.431 | 1.97E-16 | CD8 T cells | Ndufa13 |
| Chd71 | 1.07E-20 | 0.31971587 | 0.387 | 0.29 | 2.07E-16 | CD8 T cells | Chd7 |
| Tle31 | 1.19E-20 | 0.32732832 | 0.486 | 0.382 | 2.30E-16 | CD8 T cells | Tle3 |
| Adcy7 | 1.42E-20 | 0.39929252 | 0.414 | 0.298 | 2.75E-16 | CD8 T cells | Adcy7 |
| Cox5a | 1.69E-20 | 0.59597311 | 0.359 | 0.239 | 3.27E-16 | CD8 T cells | Cox5a |
| Gatad2a | 2.28E-20 | 0.41881135 | 0.446 | 0.347 | 4.41E-16 | CD8 T cells | Gatad2a |
| H2afv | 2.72E-20 | 0.39189805 | 0.361 | 0.286 | 5.26E-16 | CD8 T cells | H2afv |
| Ablim11 | 6.69E-20 | 0.34607753 | 0.566 | 0.447 | 1.29E-15 | CD8 T cells | Ablim1 |
| Bin2 | 8.86E-20 | 0.2982054 | 0.718 | 0.598 | 1.71E-15 | CD8 T cells | Bin2 |
| Pfdn51 | 1.94E-19 | 0.40975828 | 0.454 | 0.338 | 3.75E-15 | CD8 T cells | Pfdn5 |
| Atxn11 | 2.07E-19 | 0.38241386 | 0.383 | 0.281 | 4.00E-15 | CD8 T cells | Atxn1 |
| Plcg1 | 2.23E-19 | 0.34886505 | 0.446 | 0.33 | 4.31E-15 | CD8 T cells | Plcg1 |
| Ndufa3 | 2.42E-19 | 0.39519285 | 0.478 | 0.351 | 4.67E-15 | CD8 T cells | Ndufa3 |
| Etfb | 2.91E-19 | 0.56029438 | 0.301 | 0.196 | 5.62E-15 | CD8 T cells | Etfb |
| Bola2 | 4.78E-19 | 0.54947014 | 0.317 | 0.213 | 9.23E-15 | CD8 T cells | Bola2 |
| Eif3h1 | 5.21E-19 | 0.52707843 | 0.366 | 0.255 | 1.01E-14 | CD8 T cells | Eif3h |
| Txnip | 5.43E-19 | 0.38326807 | 0.47 | 0.338 | 1.05E-14 | CD8 T cells | Txnip |
| Txn1 | 5.87E-19 | 0.54493094 | 0.388 | 0.265 | 1.14E-14 | CD8 T cells | Txn1 |
| Gosr2 | 7.71E-19 | 0.56859075 | 0.301 | 0.193 | 1.49E-14 | CD8 T cells | Gosr2 |
| Ndufa61 | 9.91E-19 | 0.45723953 | 0.432 | 0.305 | 1.92E-14 | CD8 T cells | Ndufa6 |
| Atp8b2 | 1.64E-18 | 0.61051352 | 0.307 | 0.202 | 3.18E-14 | CD8 T cells | Atp8b2 |
| Flt3l | 2.09E-18 | 0.69426035 | 0.303 | 0.193 | 4.05E-14 | CD8 T cells | Flt3l |
| Inpp1 | 2.42E-18 | 0.64286345 | 0.287 | 0.168 | 4.69E-14 | CD8 T cells | Inpp1 |
| Hp1bp3 | 2.44E-18 | 0.3479538 | 0.46 | 0.357 | 4.71E-14 | CD8 T cells | Hp1bp3 |
| Btf3 | 2.56E-18 | 0.44949945 | 0.498 | 0.371 | 4.96E-14 | CD8 T cells | Btf3 |
| Tbc1d10c | 3.50E-18 | 0.28096688 | 0.615 | 0.525 | 6.77E-14 | CD8 T cells | Tbc1d10c |
| E2f4 | 3.80E-18 | 0.32784712 | 0.432 | 0.327 | 7.34E-14 | CD8 T cells | E2f4 |
| Tbc1d2b | 4.28E-18 | 0.5933202 | 0.287 | 0.185 | 8.27E-14 | CD8 T cells | Tbc1d2b |
| Psmb10 | 8.44E-18 | 0.49417061 | 0.312 | 0.214 | 1.63E-13 | CD8 T cells | Psmb10 |
| Glipr2 | 9.03E-18 | 0.55140934 | 0.305 | 0.208 | 1.75E-13 | CD8 T cells | Glipr2 |
| Ppp1ca | 9.90E-18 | 0.43280553 | 0.473 | 0.366 | 1.91E-13 | CD8 T cells | Ppp1ca |
| Tnfaip81 | 1.15E-17 | 0.37274331 | 0.402 | 0.289 | 2.22E-13 | CD8 T cells | Tnfaip8 |
| Acadl | 1.43E-17 | 0.53199068 | 0.271 | 0.175 | 2.77E-13 | CD8 T cells | Acadl |
| Map4k1 | 1.72E-17 | 0.52487715 | 0.322 | 0.219 | 3.32E-13 | CD8 T cells | Map4k1 |
| Cox7a2l1 | 3.09E-17 | 0.31616231 | 0.404 | 0.312 | 5.96E-13 | CD8 T cells | Cox7a2l |
| Ppia | 3.36E-17 | 0.28221053 | 0.473 | 0.364 | 6.50E-13 | CD8 T cells | Ppia |
| Snrpf | 3.41E-17 | 0.33276552 | 0.402 | 0.299 | 6.60E-13 | CD8 T cells | Snrpf |
| Actr1b | 3.92E-17 | 0.42683906 | 0.283 | 0.195 | 7.59E-13 | CD8 T cells | Actr1b |
| Myo1f | 4.00E-17 | 0.41531983 | 0.406 | 0.32 | 7.74E-13 | CD8 T cells | Myo1f |
| H2-M3 | 4.18E-17 | 0.51336057 | 0.27 | 0.172 | 8.07E-13 | CD8 T cells | H2-M3 |
| Baz1a | 4.25E-17 | 0.41754325 | 0.417 | 0.303 | 8.23E-13 | CD8 T cells | Baz1a |
| Unc119b | 4.40E-17 | 0.50507951 | 0.281 | 0.177 | 8.51E-13 | CD8 T cells | Unc119b |
| Idh3a | 4.60E-17 | 0.53777257 | 0.257 | 0.166 | 8.90E-13 | CD8 T cells | Idh3a |
| Aak1 | 4.71E-17 | 0.37523067 | 0.329 | 0.223 | 9.10E-13 | CD8 T cells | Aak1 |
| Arhgdia | 6.21E-17 | 0.34288788 | 0.609 | 0.485 | 1.20E-12 | CD8 T cells | Arhgdia |
| Atp6v0a2 | 6.51E-17 | 0.55832843 | 0.305 | 0.205 | 1.26E-12 | CD8 T cells | Atp6v0a2 |
| Plcl2 | 7.57E-17 | 0.456248 | 0.283 | 0.187 | 1.46E-12 | CD8 T cells | Plcl2 |
| Hspa1b1 | 9.02E-17 | 0.44691827 | 0.253 | 0.21 | 1.74E-12 | CD8 T cells | Hspa1b |
| Atf7ip | 1.06E-16 | 0.42181227 | 0.396 | 0.294 | 2.05E-12 | CD8 T cells | Atf7ip |
| Otulin | 1.22E-16 | 0.47581926 | 0.318 | 0.226 | 2.36E-12 | CD8 T cells | Otulin |
| Egr1 | 1.27E-16 | 0.54606073 | 0.334 | 0.269 | 2.46E-12 | CD8 T cells | Egr1 |
| Acsl5 | 1.43E-16 | 0.60789377 | 0.315 | 0.212 | 2.76E-12 | CD8 T cells | Acsl5 |
| Tpst2 | 3.25E-16 | 0.40138816 | 0.42 | 0.315 | 6.28E-12 | CD8 T cells | Tpst2 |
| Dynlrb1 | 4.75E-16 | 0.34794214 | 0.443 | 0.354 | 9.19E-12 | CD8 T cells | Dynlrb1 |
| Eef1g1 | 7.56E-16 | 0.53574804 | 0.379 | 0.264 | 1.46E-11 | CD8 T cells | Eef1g |
| Mif2 | 7.85E-16 | 0.34156907 | 0.39 | 0.293 | 1.52E-11 | CD8 T cells | Mif2 |
| Arhgef31 | 8.52E-16 | 0.47718612 | 0.377 | 0.273 | 1.65E-11 | CD8 T cells | Arhgef3 |
| Usp25 | 9.14E-16 | 0.27039425 | 0.444 | 0.35 | 1.77E-11 | CD8 T cells | Usp25 |
| Gltp | 1.22E-15 | 0.46785762 | 0.308 | 0.221 | 2.35E-11 | CD8 T cells | Gltp |
| Dap | 1.23E-15 | 0.31983848 | 0.262 | 0.199 | 2.38E-11 | CD8 T cells | Dap |
| Ptp4a3 | 1.31E-15 | 0.28984582 | 0.285 | 0.213 | 2.54E-11 | CD8 T cells | Ptp4a3 |
| Acot7 | 1.45E-15 | 0.31439175 | 0.434 | 0.363 | 2.80E-11 | CD8 T cells | Acot7 |
| Lasp1 | 2.32E-15 | 0.34958955 | 0.337 | 0.248 | 4.48E-11 | CD8 T cells | Lasp1 |
| Havcr21 | 2.41E-15 | 0.39458887 | 0.264 | 0.209 | 4.66E-11 | CD8 T cells | Havcr2 |
| Dpysl2 | 3.97E-15 | 0.42606524 | 0.34 | 0.245 | 7.67E-11 | CD8 T cells | Dpysl2 |
| Xrn21 | 5.54E-15 | 0.36837012 | 0.414 | 0.296 | 1.07E-10 | CD8 T cells | Xrn2 |
| Spata13 | 8.98E-15 | 0.3660871 | 0.512 | 0.399 | 1.74E-10 | CD8 T cells | Spata13 |
| Cwc15 | 1.36E-14 | 0.33328446 | 0.425 | 0.34 | 2.63E-10 | CD8 T cells | Cwc15 |
| Serbp1 | 1.49E-14 | 0.32140649 | 0.485 | 0.369 | 2.88E-10 | CD8 T cells | Serbp1 |
| Frmd81 | 2.10E-14 | 0.3207251 | 0.368 | 0.266 | 4.05E-10 | CD8 T cells | Frmd8 |
| Hivep2 | 2.71E-14 | 0.42258975 | 0.277 | 0.192 | 5.24E-10 | CD8 T cells | Hivep2 |
| Rassf1 | 3.68E-14 | 0.33713228 | 0.257 | 0.187 | 7.11E-10 | CD8 T cells | Rassf1 |
| Serf2 | 3.70E-14 | 0.34913114 | 0.493 | 0.384 | 7.15E-10 | CD8 T cells | Serf2 |
| Ppp3r1 | 3.83E-14 | 0.50801233 | 0.271 | 0.183 | 7.41E-10 | CD8 T cells | Ppp3r1 |
| Kars | 4.45E-14 | 0.5599451 | 0.348 | 0.231 | 8.60E-10 | CD8 T cells | Kars |
| Anapc11 | 5.22E-14 | 0.43204828 | 0.282 | 0.197 | 1.01E-09 | CD8 T cells | Anapc11 |
| Smg7 | 6.64E-14 | 0.49718919 | 0.297 | 0.211 | 1.28E-09 | CD8 T cells | Smg7 |
| Dnajc3 | 8.11E-14 | 0.49562851 | 0.341 | 0.242 | 1.57E-09 | CD8 T cells | Dnajc3 |
| Cdkn2aipnl | 8.41E-14 | 0.53094747 | 0.25 | 0.171 | 1.63E-09 | CD8 T cells | Cdkn2aipnl |
| Inpp5b | 9.16E-14 | 0.51972921 | 0.278 | 0.184 | 1.77E-09 | CD8 T cells | Inpp5b |
| Tomm7 | 1.01E-13 | 0.333416 | 0.329 | 0.234 | 1.96E-09 | CD8 T cells | Tomm7 |
| Med14 | 1.36E-13 | 0.41915438 | 0.279 | 0.208 | 2.63E-09 | CD8 T cells | Med14 |
| Nudt21 | 1.49E-13 | 0.58488865 | 0.276 | 0.184 | 2.88E-09 | CD8 T cells | Nudt21 |
| Ubxn1 | 2.25E-13 | 0.34909343 | 0.422 | 0.315 | 4.35E-09 | CD8 T cells | Ubxn1 |
| Npm11 | 2.26E-13 | 0.31637425 | 0.517 | 0.403 | 4.38E-09 | CD8 T cells | Npm1 |
| Ogdh | 2.30E-13 | 0.26452572 | 0.461 | 0.39 | 4.45E-09 | CD8 T cells | Ogdh |
| Cct7 | 2.39E-13 | 0.55462824 | 0.294 | 0.194 | 4.61E-09 | CD8 T cells | Cct7 |
| Bin1 | 2.70E-13 | 0.28132373 | 0.387 | 0.301 | 5.22E-09 | CD8 T cells | Bin1 |
| Gng10 | 2.74E-13 | 0.57647918 | 0.257 | 0.175 | 5.30E-09 | CD8 T cells | Gng10 |
| Dazap11 | 2.74E-13 | 0.27744149 | 0.497 | 0.403 | 5.31E-09 | CD8 T cells | Dazap1 |
| Jarid2 | 2.80E-13 | 0.44242269 | 0.302 | 0.204 | 5.41E-09 | CD8 T cells | Jarid2 |
| Uggt1 | 3.11E-13 | 0.39562704 | 0.3 | 0.219 | 6.02E-09 | CD8 T cells | Uggt1 |
| Fundc2 | 4.11E-13 | 0.43962827 | 0.268 | 0.183 | 7.95E-09 | CD8 T cells | Fundc2 |
| Efr3a | 5.22E-13 | 0.54020773 | 0.26 | 0.179 | 1.01E-08 | CD8 T cells | Efr3a |
| Ubap2l1 | 5.43E-13 | 0.44991712 | 0.351 | 0.267 | 1.05E-08 | CD8 T cells | Ubap2l |
| Cyld1 | 5.75E-13 | 0.38326269 | 0.484 | 0.374 | 1.11E-08 | CD8 T cells | Cyld |
| Eef1d | 5.82E-13 | 0.60715396 | 0.253 | 0.157 | 1.13E-08 | CD8 T cells | Eef1d |
| Cd2bp2 | 6.09E-13 | 0.53481911 | 0.348 | 0.244 | 1.18E-08 | CD8 T cells | Cd2bp2 |
| Nptn | 6.38E-13 | 0.30217942 | 0.317 | 0.234 | 1.23E-08 | CD8 T cells | Nptn |
| Cox5b | 6.68E-13 | 0.38206318 | 0.397 | 0.305 | 1.29E-08 | CD8 T cells | Cox5b |
| Nop10 | 7.86E-13 | 0.39218083 | 0.257 | 0.175 | 1.52E-08 | CD8 T cells | Nop10 |
| Rtcb1 | 8.00E-13 | 0.40674344 | 0.364 | 0.266 | 1.55E-08 | CD8 T cells | Rtcb |
| Snrpe | 9.30E-13 | 0.44896444 | 0.276 | 0.187 | 1.80E-08 | CD8 T cells | Snrpe |
| Tcirg1 | 9.40E-13 | 0.49320565 | 0.256 | 0.183 | 1.82E-08 | CD8 T cells | Tcirg1 |
| Tmed5 | 1.06E-12 | 0.46242997 | 0.267 | 0.185 | 2.04E-08 | CD8 T cells | Tmed5 |
| Arpc4 | 1.08E-12 | 0.2687434 | 0.435 | 0.342 | 2.09E-08 | CD8 T cells | Arpc4 |
| Was | 1.17E-12 | 0.25193557 | 0.467 | 0.368 | 2.27E-08 | CD8 T cells | Was |
| Cox7c | 1.82E-12 | 0.27228531 | 0.379 | 0.303 | 3.51E-08 | CD8 T cells | Cox7c |
| Ptpn11 | 1.98E-12 | 0.47923518 | 0.265 | 0.183 | 3.83E-08 | CD8 T cells | Ptpn11 |
| Hdac1 | 2.00E-12 | 0.3477643 | 0.367 | 0.275 | 3.86E-08 | CD8 T cells | Hdac1 |
| Nsd2 | 2.04E-12 | 0.57715755 | 0.293 | 0.189 | 3.95E-08 | CD8 T cells | Nsd2 |
| Lims1 | 2.43E-12 | 0.37751486 | 0.314 | 0.227 | 4.70E-08 | CD8 T cells | Lims1 |
| Lsm4 | 2.66E-12 | 0.27497752 | 0.323 | 0.242 | 5.15E-08 | CD8 T cells | Lsm4 |
| Atp5o | 2.85E-12 | 0.39418868 | 0.293 | 0.209 | 5.50E-08 | CD8 T cells | Atp5o |
| Ifi27 | 3.16E-12 | 0.30194853 | 0.347 | 0.266 | 6.10E-08 | CD8 T cells | Ifi27 |
| Med28 | 4.05E-12 | 0.43009661 | 0.311 | 0.231 | 7.83E-08 | CD8 T cells | Med28 |
| R3hdm4 | 4.17E-12 | 0.32718158 | 0.36 | 0.278 | 8.06E-08 | CD8 T cells | R3hdm4 |
| Suco | 4.39E-12 | 0.34324297 | 0.253 | 0.17 | 8.49E-08 | CD8 T cells | Suco |
| Tes | 4.58E-12 | 0.27595352 | 0.499 | 0.407 | 8.86E-08 | CD8 T cells | Tes |
| Pcna | 5.40E-12 | 0.29180347 | 0.406 | 0.316 | 1.04E-07 | CD8 T cells | Pcna |
| Arpc2 | 7.42E-12 | 0.28956673 | 0.489 | 0.418 | 1.44E-07 | CD8 T cells | Arpc2 |
| Zgpat1 | 8.04E-12 | 0.3121722 | 0.329 | 0.236 | 1.55E-07 | CD8 T cells | Zgpat |
| Nlrc3 | 8.68E-12 | 0.37277575 | 0.362 | 0.271 | 1.68E-07 | CD8 T cells | Nlrc3 |
| Gimap71 | 8.74E-12 | 0.41500748 | 0.298 | 0.225 | 1.69E-07 | CD8 T cells | Gimap7 |
| Plekhj1 | 9.38E-12 | 0.39216105 | 0.356 | 0.26 | 1.81E-07 | CD8 T cells | Plekhj1 |
| Dynll1 | 1.05E-11 | 0.44877597 | 0.268 | 0.192 | 2.02E-07 | CD8 T cells | Dynll1 |
| Cbx4 | 1.05E-11 | 0.34984541 | 0.438 | 0.336 | 2.04E-07 | CD8 T cells | Cbx4 |
| Tmem167 | 1.15E-11 | 0.44105408 | 0.261 | 0.184 | 2.22E-07 | CD8 T cells | Tmem167 |
| Herc11 | 1.35E-11 | 0.31945302 | 0.329 | 0.24 | 2.60E-07 | CD8 T cells | Herc1 |
| Itga4 | 1.42E-11 | 0.29634441 | 0.331 | 0.266 | 2.75E-07 | CD8 T cells | Itga4 |
| Esyt21 | 1.82E-11 | 0.25250782 | 0.495 | 0.395 | 3.52E-07 | CD8 T cells | Esyt2 |
| Cmc1 | 2.17E-11 | 0.35458895 | 0.255 | 0.181 | 4.20E-07 | CD8 T cells | Cmc1 |
| Micos13 | 2.46E-11 | 0.37289115 | 0.265 | 0.199 | 4.76E-07 | CD8 T cells | Micos13 |
| Nek71 | 2.88E-11 | 0.3232874 | 0.317 | 0.251 | 5.57E-07 | CD8 T cells | Nek7 |
| Tiprl | 2.89E-11 | 0.27282941 | 0.299 | 0.222 | 5.58E-07 | CD8 T cells | Tiprl |
| Eif4b | 2.95E-11 | 0.27126708 | 0.441 | 0.33 | 5.70E-07 | CD8 T cells | Eif4b |
| Akr1b3 | 3.82E-11 | 0.36139462 | 0.302 | 0.222 | 7.39E-07 | CD8 T cells | Akr1b3 |
| Ptpn6 | 4.66E-11 | 0.3902884 | 0.266 | 0.219 | 9.02E-07 | CD8 T cells | Ptpn6 |
| Bscl2 | 4.75E-11 | 0.39629363 | 0.311 | 0.221 | 9.18E-07 | CD8 T cells | Bscl2 |
| Ndufb7 | 4.89E-11 | 0.3241703 | 0.369 | 0.294 | 9.46E-07 | CD8 T cells | Ndufb7 |
| Higd1a | 5.07E-11 | 0.33727598 | 0.343 | 0.252 | 9.80E-07 | CD8 T cells | Higd1a |
| Cnot3 | 5.33E-11 | 0.49872855 | 0.291 | 0.196 | 1.03E-06 | CD8 T cells | Cnot3 |
| Kmt2e1 | 5.42E-11 | 0.29480552 | 0.408 | 0.324 | 1.05E-06 | CD8 T cells | Kmt2e |
| Cbx3 | 6.54E-11 | 0.43549112 | 0.255 | 0.174 | 1.26E-06 | CD8 T cells | Cbx3 |
| Dync1h11 | 6.57E-11 | 0.28565653 | 0.436 | 0.346 | 1.27E-06 | CD8 T cells | Dync1h1 |
| Xpo1 | 6.69E-11 | 0.54469927 | 0.274 | 0.193 | 1.29E-06 | CD8 T cells | Xpo1 |
| Igbp1 | 8.94E-11 | 0.37444919 | 0.281 | 0.207 | 1.73E-06 | CD8 T cells | Igbp1 |
| Ociad1 | 9.68E-11 | 0.29033536 | 0.28 | 0.209 | 1.87E-06 | CD8 T cells | Ociad1 |
| Ttc39b1 | 1.16E-10 | 0.27958456 | 0.49 | 0.389 | 2.25E-06 | CD8 T cells | Ttc39b |
| Tapbpl | 1.18E-10 | 0.42884315 | 0.332 | 0.24 | 2.28E-06 | CD8 T cells | Tapbpl |
| Parvg | 1.23E-10 | 0.48932003 | 0.289 | 0.208 | 2.39E-06 | CD8 T cells | Parvg |
| Mfng | 1.25E-10 | 0.40228441 | 0.271 | 0.199 | 2.41E-06 | CD8 T cells | Mfng |
| Ppp3cb | 1.94E-10 | 0.32359089 | 0.357 | 0.282 | 3.76E-06 | CD8 T cells | Ppp3cb |
| Zfp598 | 2.49E-10 | 0.27888721 | 0.25 | 0.188 | 4.82E-06 | CD8 T cells | Zfp598 |
| Copb1 | 2.74E-10 | 0.34447499 | 0.311 | 0.24 | 5.31E-06 | CD8 T cells | Copb1 |
| Sms | 2.75E-10 | 0.2767699 | 0.304 | 0.234 | 5.33E-06 | CD8 T cells | Sms |
| Psmb1 | 3.02E-10 | 0.39475865 | 0.288 | 0.203 | 5.83E-06 | CD8 T cells | Psmb1 |
| Eif3l | 3.10E-10 | 0.4227922 | 0.333 | 0.234 | 5.99E-06 | CD8 T cells | Eif3l |
| Arhgef6 | 3.47E-10 | 0.33538567 | 0.287 | 0.211 | 6.72E-06 | CD8 T cells | Arhgef6 |
| Med30 | 3.82E-10 | 0.42811724 | 0.252 | 0.18 | 7.39E-06 | CD8 T cells | Med30 |
| Ndufa1 | 3.93E-10 | 0.2891022 | 0.38 | 0.289 | 7.59E-06 | CD8 T cells | Ndufa1 |
| Pitpnc1 | 4.20E-10 | 0.29566605 | 0.298 | 0.223 | 8.11E-06 | CD8 T cells | Pitpnc1 |
| Gsk3a | 4.53E-10 | 0.36143536 | 0.364 | 0.264 | 8.76E-06 | CD8 T cells | Gsk3a |
| Clec2d1 | 4.59E-10 | 0.27910675 | 0.584 | 0.467 | 8.87E-06 | CD8 T cells | Clec2d |
| Ndufb1-ps | 4.73E-10 | 0.41463604 | 0.324 | 0.238 | 9.15E-06 | CD8 T cells | Ndufb1-ps |
| Eif3g | 4.81E-10 | 0.28841349 | 0.324 | 0.247 | 9.29E-06 | CD8 T cells | Eif3g |
| Ilk | 4.89E-10 | 0.31195328 | 0.315 | 0.251 | 9.45E-06 | CD8 T cells | Ilk |
| Dars | 5.07E-10 | 0.43070375 | 0.3 | 0.216 | 9.79E-06 | CD8 T cells | Dars |
| Rbck1 | 5.23E-10 | 0.35913811 | 0.314 | 0.243 | 1.01E-05 | CD8 T cells | Rbck1 |
| Scamp3 | 5.55E-10 | 0.39996815 | 0.292 | 0.21 | 1.07E-05 | CD8 T cells | Scamp3 |
| Suz12 | 7.43E-10 | 0.29213717 | 0.273 | 0.214 | 1.44E-05 | CD8 T cells | Suz12 |
| Gga1 | 9.17E-10 | 0.30930195 | 0.269 | 0.191 | 1.77E-05 | CD8 T cells | Gga1 |
| Ide | 9.75E-10 | 0.25623321 | 0.378 | 0.316 | 1.88E-05 | CD8 T cells | Ide |
| Slc35e1 | 1.14E-09 | 0.39882168 | 0.292 | 0.209 | 2.20E-05 | CD8 T cells | Slc35e1 |
| Mndal | 1.16E-09 | 0.32430964 | 0.274 | 0.194 | 2.24E-05 | CD8 T cells | Mndal |
| Zfp91 | 1.43E-09 | 0.26415022 | 0.362 | 0.28 | 2.77E-05 | CD8 T cells | Zfp91 |
| Crybg1 | 1.99E-09 | 0.33923657 | 0.25 | 0.179 | 3.84E-05 | CD8 T cells | Crybg1 |
| Nras | 2.52E-09 | 0.37835852 | 0.258 | 0.189 | 4.87E-05 | CD8 T cells | Nras |
| Mbd3 | 3.32E-09 | 0.29577491 | 0.298 | 0.232 | 6.43E-05 | CD8 T cells | Mbd3 |
| Hdhd5 | 3.80E-09 | 0.36379619 | 0.266 | 0.185 | 7.35E-05 | CD8 T cells | Hdhd5 |
| Myl6 | 5.83E-09 | 0.3146363 | 0.385 | 0.29 | 0.00011274 | CD8 T cells | Myl6 |
| Rdx | 5.99E-09 | 0.25495356 | 0.255 | 0.186 | 0.00011577 | CD8 T cells | Rdx |
| Becn1 | 6.17E-09 | 0.2840848 | 0.382 | 0.306 | 0.00011934 | CD8 T cells | Becn1 |
| Atp6v0e | 6.24E-09 | 0.33053791 | 0.337 | 0.26 | 0.00012061 | CD8 T cells | Atp6v0e |
| Tkt | 6.86E-09 | 0.32126449 | 0.379 | 0.295 | 0.00013261 | CD8 T cells | Tkt |
| Sec11a1 | 7.17E-09 | 0.33227691 | 0.374 | 0.27 | 0.00013856 | CD8 T cells | Sec11a |
| Ppp2r5c | 7.99E-09 | 0.27837908 | 0.416 | 0.331 | 0.00015446 | CD8 T cells | Ppp2r5c |
| Sugt1 | 8.09E-09 | 0.34185375 | 0.288 | 0.222 | 0.00015649 | CD8 T cells | Sugt1 |
| Sdhc | 8.65E-09 | 0.30023418 | 0.3 | 0.213 | 0.00016731 | CD8 T cells | Sdhc |
| Tomm22 | 1.01E-08 | 0.42631182 | 0.291 | 0.206 | 0.00019508 | CD8 T cells | Tomm22 |
| Kdm7a | 1.27E-08 | 0.34753256 | 0.253 | 0.202 | 0.00024604 | CD8 T cells | Kdm7a |
| Adam10 | 1.51E-08 | 0.30168009 | 0.263 | 0.2 | 0.00029162 | CD8 T cells | Adam10 |
| Nin | 1.51E-08 | 0.47747177 | 0.263 | 0.186 | 0.00029166 | CD8 T cells | Nin |
| Hmgb1 | 2.16E-08 | 0.32037162 | 0.267 | 0.193 | 0.000418 | CD8 T cells | Hmgb1 |
| Snrpd3 | 2.21E-08 | 0.3378974 | 0.294 | 0.218 | 0.00042757 | CD8 T cells | Snrpd3 |
| Cox6b1 | 2.32E-08 | 0.2694256 | 0.367 | 0.285 | 0.00044893 | CD8 T cells | Cox6b1 |
| Pabpc11 | 2.37E-08 | 0.28587866 | 0.474 | 0.388 | 0.00045761 | CD8 T cells | Pabpc1 |
| Med10 | 2.60E-08 | 0.34325581 | 0.264 | 0.193 | 0.00050198 | CD8 T cells | Med10 |
| Trmt112 | 2.70E-08 | 0.33129495 | 0.322 | 0.24 | 0.00052119 | CD8 T cells | Trmt112 |
| Rbbp4 | 2.88E-08 | 0.29106939 | 0.391 | 0.323 | 0.00055723 | CD8 T cells | Rbbp4 |
| Stt3b | 3.70E-08 | 0.30435201 | 0.263 | 0.211 | 0.00071478 | CD8 T cells | Stt3b |
| Larp1 | 3.93E-08 | 0.34488891 | 0.322 | 0.237 | 0.00076038 | CD8 T cells | Larp1 |
| Psma1 | 4.70E-08 | 0.29250583 | 0.29 | 0.222 | 0.00090816 | CD8 T cells | Psma1 |
| Actr3 | 5.15E-08 | 0.25199818 | 0.503 | 0.4 | 0.00099507 | CD8 T cells | Actr3 |
| Mast3 | 5.75E-08 | 0.28622753 | 0.363 | 0.283 | 0.00111089 | CD8 T cells | Mast3 |
| Tbc1d10b | 5.86E-08 | 0.29625071 | 0.268 | 0.193 | 0.00113294 | CD8 T cells | Tbc1d10b |
| Anxa7 | 5.89E-08 | 0.25875285 | 0.25 | 0.194 | 0.00113825 | CD8 T cells | Anxa7 |
| Hcfc1 | 7.73E-08 | 0.46434633 | 0.275 | 0.193 | 0.00149511 | CD8 T cells | Hcfc1 |
| Bex3 | 8.31E-08 | 0.28748068 | 0.253 | 0.196 | 0.00160708 | CD8 T cells | Bex3 |
| Rer1 | 8.34E-08 | 0.2566538 | 0.299 | 0.234 | 0.00161264 | CD8 T cells | Rer1 |
| Otub1 | 8.70E-08 | 0.2898821 | 0.281 | 0.207 | 0.00168175 | CD8 T cells | Otub1 |
| Cct2 | 9.43E-08 | 0.35921568 | 0.318 | 0.235 | 0.00182348 | CD8 T cells | Cct2 |
| Sumo2 | 9.72E-08 | 0.37687575 | 0.269 | 0.201 | 0.0018792 | CD8 T cells | Sumo2 |
| Pdpk1 | 1.02E-07 | 0.25225708 | 0.281 | 0.208 | 0.00196242 | CD8 T cells | Pdpk1 |
| Chchd3 | 1.15E-07 | 0.35506837 | 0.279 | 0.2 | 0.00222751 | CD8 T cells | Chchd3 |
| Ubl5 | 1.17E-07 | 0.31625645 | 0.355 | 0.29 | 0.00225719 | CD8 T cells | Ubl5 |
| Snrpb | 1.22E-07 | 0.34679993 | 0.268 | 0.203 | 0.00236264 | CD8 T cells | Snrpb |
| Dnmt1 | 1.31E-07 | 0.34054806 | 0.286 | 0.214 | 0.00253464 | CD8 T cells | Dnmt1 |
| Ttc7 | 1.36E-07 | 0.26325407 | 0.324 | 0.267 | 0.00262366 | CD8 T cells | Ttc7 |
| Gtpbp11 | 1.46E-07 | 0.27929633 | 0.346 | 0.273 | 0.00281791 | CD8 T cells | Gtpbp1 |
| Twf2 | 1.48E-07 | 0.4219524 | 0.262 | 0.2 | 0.00285328 | CD8 T cells | Twf2 |
| Rp9 | 1.77E-07 | 0.30808089 | 0.276 | 0.225 | 0.00343135 | CD8 T cells | Rp9 |
| Gapvd1 | 2.30E-07 | 0.26068705 | 0.261 | 0.199 | 0.0044454 | CD8 T cells | Gapvd1 |
| Gmfg1 | 3.95E-07 | 0.26328737 | 0.391 | 0.313 | 0.00764081 | CD8 T cells | Gmfg |
| Kcnab2 | 4.13E-07 | 0.25187877 | 0.403 | 0.322 | 0.00798565 | CD8 T cells | Kcnab2 |
| Anp32e | 4.22E-07 | 0.26005035 | 0.407 | 0.321 | 0.00816234 | CD8 T cells | Anp32e |
| Zbtb7a | 5.64E-07 | 0.25668159 | 0.348 | 0.279 | 0.01091214 | CD8 T cells | Zbtb7a |
| Atp5l | 6.93E-07 | 0.30435536 | 0.273 | 0.205 | 0.01339544 | CD8 T cells | Atp5l |
| Atf1 | 6.99E-07 | 0.30337259 | 0.335 | 0.253 | 0.01351648 | CD8 T cells | Atf1 |
| Wdr821 | 7.67E-07 | 0.25154525 | 0.373 | 0.296 | 0.01482874 | CD8 T cells | Wdr82 |
| Psmd13 | 8.44E-07 | 0.32719034 | 0.265 | 0.186 | 0.01632664 | CD8 T cells | Psmd13 |
| Capza1 | 9.80E-07 | 0.28610739 | 0.303 | 0.243 | 0.01893824 | CD8 T cells | Capza1 |
| Map4k4 | 1.28E-06 | 0.30749222 | 0.296 | 0.235 | 0.0246749 | CD8 T cells | Map4k4 |
| Fip1l1 | 1.91E-06 | 0.37565533 | 0.256 | 0.191 | 0.03690044 | CD8 T cells | Fip1l1 |
| Dnajc14 | 1.96E-06 | 0.40328251 | 0.298 | 0.234 | 0.03787134 | CD8 T cells | Dnajc14 |
| Chd8 | 2.83E-06 | 0.3739384 | 0.25 | 0.186 | 0.05463709 | CD8 T cells | Chd8 |
| Glyr1 | 3.13E-06 | 0.25358111 | 0.288 | 0.227 | 0.06051539 | CD8 T cells | Glyr1 |
| Etfa | 4.50E-06 | 0.34240254 | 0.258 | 0.199 | 0.08704552 | CD8 T cells | Etfa |
| Mrtfa1 | 5.24E-06 | 0.25576877 | 0.312 | 0.238 | 0.10137019 | CD8 T cells | Mrtfa |
| Iqgap2 | 6.71E-06 | 0.36349637 | 0.267 | 0.213 | 0.12970805 | CD8 T cells | Iqgap2 |
| Lin7c | 6.71E-06 | 0.30450838 | 0.27 | 0.215 | 0.129809 | CD8 T cells | Lin7c |
| Nipal3 | 6.83E-06 | 0.31725376 | 0.256 | 0.192 | 0.13205885 | CD8 T cells | Nipal3 |
| Psma5 | 7.93E-06 | 0.41283638 | 0.257 | 0.185 | 0.15327794 | CD8 T cells | Psma5 |
| Ndufa8 | 8.90E-06 | 0.25782657 | 0.344 | 0.283 | 0.17201813 | CD8 T cells | Ndufa8 |
| Uqcrc2 | 1.03E-05 | 0.26366889 | 0.268 | 0.221 | 0.19847664 | CD8 T cells | Uqcrc2 |
| Hnrnpf | 1.92E-05 | 0.26105669 | 0.46 | 0.382 | 0.37113411 | CD8 T cells | Hnrnpf |
| Eif3i | 2.28E-05 | 0.30991482 | 0.254 | 0.191 | 0.44052558 | CD8 T cells | Eif3i |
| Foxn2 | 3.60E-05 | 0.30162878 | 0.261 | 0.196 | 0.69601173 | CD8 T cells | Foxn2 |
| Mta1 | 5.44E-05 | 0.26932368 | 0.303 | 0.235 | 1 | CD8 T cells | Mta1 |
| Prdm2 | 8.79E-05 | 0.27362113 | 0.261 | 0.208 | 1 | CD8 T cells | Prdm2 |
| Abracl1 | 0.00010212 | 0.2677658 | 0.378 | 0.308 | 1 | CD8 T cells | Abracl |
| Pafah1b2 | 0.00014014 | 0.28870075 | 0.278 | 0.217 | 1 | CD8 T cells | Pafah1b2 |
| Snx17 | 0.00040763 | 0.29896047 | 0.277 | 0.234 | 1 | CD8 T cells | Snx17 |
| Mtpn | 0.00041968 | 0.26268461 | 0.254 | 0.198 | 1 | CD8 T cells | Mtpn |
| Cdc34 | 0.00049311 | 0.26089702 | 0.255 | 0.197 | 1 | CD8 T cells | Cdc34 |
| Csnk2a1 | 0.00050335 | 0.30337072 | 0.258 | 0.212 | 1 | CD8 T cells | Csnk2a1 |
| R3hdm2 | 0.00096413 | 0.26562176 | 0.259 | 0.214 | 1 | CD8 T cells | R3hdm2 |
| Hdac7 | 0.00142587 | 0.27200711 | 0.28 | 0.219 | 1 | CD8 T cells | Hdac7 |
| Tcof1 | 0.00202186 | 0.27852354 | 0.252 | 0.198 | 1 | CD8 T cells | Tcof1 |
| Abhd8 | 0.00569643 | 0.24834491 | 0.25 | 0.193 | 1 | CD8 T cells | Abhd8 |
| Cd163l1 | 0 | 7.86467913 | 0.911 | 0.01 | 0 | gamma-delta T | Cd163l1 |
| Trdc | 0 | 6.15946739 | 0.911 | 0.028 | 0 | gamma-delta T | Trdc |
| Blk | 0 | 6.54436204 | 0.787 | 0.016 | 0 | gamma-delta T | Blk |
| Tmem176a | 0 | 4.08927628 | 0.795 | 0.08 | 0 | gamma-delta T | Tmem176a |
| Zbtb16 | 0 | 3.91373696 | 0.756 | 0.049 | 0 | gamma-delta T | Zbtb16 |
| Tmem176b | 0 | 3.86897294 | 0.767 | 0.084 | 0 | gamma-delta T | Tmem176b |
| Rorc | 0 | 4.06594952 | 0.729 | 0.088 | 0 | gamma-delta T | Rorc |
| Tcrg-C1 | 0 | 5.89501543 | 0.655 | 0.016 | 0 | gamma-delta T | Tcrg-C1 |
| Selenop | 0 | 2.75087328 | 0.775 | 0.278 | 0 | gamma-delta T | Selenop |
| Il17a | 0 | 4.99999624 | 0.326 | 0.02 | 0 | gamma-delta T | Il17a |
| Il1r1 | 2.44E-272 | 4.54296323 | 0.574 | 0.04 | 4.72E-268 | gamma-delta T | Il1r1 |
| Aqp3 | 2.74E-240 | 4.28887801 | 0.558 | 0.044 | 5.30E-236 | gamma-delta T | Aqp3 |
| Mmp25 | 2.31E-237 | 3.56409154 | 0.616 | 0.068 | 4.46E-233 | gamma-delta T | Mmp25 |
| Serpinb1a | 2.07E-232 | 3.69526464 | 0.547 | 0.053 | 4.01E-228 | gamma-delta T | Serpinb1a |
| Il23r | 3.33E-214 | 4.2908809 | 0.5 | 0.046 | 6.44E-210 | gamma-delta T | Il23r |
| F2r | 3.04E-205 | 2.64790347 | 0.771 | 0.262 | 5.87E-201 | gamma-delta T | F2r |
| Tcrg-V6 | 8.92E-184 | 8.688697 | 0.682 | 0.004 | 1.72E-179 | gamma-delta T | Tcrg-V6 |
| Pik3ap1 | 1.58E-174 | 2.93180336 | 0.647 | 0.142 | 3.05E-170 | gamma-delta T | Pik3ap1 |
| Cxcr61 | 3.07E-156 | 2.16633636 | 0.752 | 0.276 | 5.93E-152 | gamma-delta T | Cxcr6 |
| Smox | 2.58E-137 | 2.58660371 | 0.62 | 0.183 | 4.99E-133 | gamma-delta T | Smox |
| Tmem641 | 2.66E-128 | 2.30100669 | 0.628 | 0.202 | 5.14E-124 | gamma-delta T | Tmem64 |
| Sox13 | 3.01E-123 | 7.31013464 | 0.496 | 0.004 | 5.82E-119 | gamma-delta T | Sox13 |
| Tcrg-C4 | 4.28E-122 | 3.87742118 | 0.376 | 0.024 | 8.28E-118 | gamma-delta T | Tcrg-C4 |
| Tns4 | 1.15E-111 | 3.7179733 | 0.345 | 0.031 | 2.23E-107 | gamma-delta T | Tns4 |
| Abi3bp | 6.63E-109 | 5.11054364 | 0.306 | 0.011 | 1.28E-104 | gamma-delta T | Abi3bp |
| Adgrg51 | 1.03E-104 | 2.60262316 | 0.531 | 0.115 | 1.99E-100 | gamma-delta T | Adgrg5 |
| Pde5a | 3.72E-101 | 5.04880992 | 0.283 | 0.011 | 7.20E-97 | gamma-delta T | Pde5a |
| Ly6g5b | 4.40E-100 | 3.35545054 | 0.388 | 0.056 | 8.50E-96 | gamma-delta T | Ly6g5b |
| Clnk | 1.09E-96 | 3.99795909 | 0.322 | 0.02 | 2.11E-92 | gamma-delta T | Clnk |
| Il17re | 4.97E-89 | 3.96555202 | 0.298 | 0.021 | 9.60E-85 | gamma-delta T | Il17re |
| Jaml | 2.16E-88 | 2.50037311 | 0.45 | 0.085 | 4.18E-84 | gamma-delta T | Jaml |
| Kcnk1 | 8.83E-83 | 6.31899756 | 0.271 | 0.004 | 1.71E-78 | gamma-delta T | Kcnk1 |
| Nr1d1 | 1.13E-79 | 3.77392141 | 0.267 | 0.023 | 2.18E-75 | gamma-delta T | Nr1d1 |
| Il18r1 | 3.32E-77 | 1.4505643 | 0.771 | 0.383 | 6.42E-73 | gamma-delta T | Il18r1 |
| Arrb1 | 3.28E-76 | 2.26486273 | 0.523 | 0.167 | 6.34E-72 | gamma-delta T | Arrb1 |
| Scpep1 | 5.79E-76 | 3.00271472 | 0.337 | 0.085 | 1.12E-71 | gamma-delta T | Scpep1 |
| Ikzf3 | 1.75E-75 | 1.46399742 | 0.837 | 0.535 | 3.38E-71 | gamma-delta T | Ikzf3 |
| Igf1r | 1.99E-73 | 2.86741145 | 0.353 | 0.064 | 3.86E-69 | gamma-delta T | Igf1r |
| Znrf11 | 1.16E-70 | 1.60138911 | 0.729 | 0.382 | 2.24E-66 | gamma-delta T | Znrf1 |
| Lgals3bp | 9.06E-59 | 1.85026561 | 0.554 | 0.229 | 1.75E-54 | gamma-delta T | Lgals3bp |
| 8-Sep | 2.40E-57 | 2.60684547 | 0.353 | 0.075 | 4.64E-53 | gamma-delta T | 8-Sep |
| Ramp1 | 5.43E-55 | 2.10173433 | 0.519 | 0.181 | 1.05E-50 | gamma-delta T | Ramp1 |
| Atrnl1 | 2.06E-51 | 2.70374146 | 0.306 | 0.07 | 3.99E-47 | gamma-delta T | Atrnl1 |
| Mgat5 | 2.60E-51 | 1.91060762 | 0.523 | 0.216 | 5.02E-47 | gamma-delta T | Mgat5 |
| Lax11 | 9.24E-43 | 1.74731787 | 0.465 | 0.196 | 1.79E-38 | gamma-delta T | Lax1 |
| Fgl2 | 3.47E-42 | 1.28557464 | 0.802 | 0.46 | 6.71E-38 | gamma-delta T | Fgl2 |
| Cd247 | 7.37E-41 | 1.28841919 | 0.752 | 0.495 | 1.43E-36 | gamma-delta T | Cd247 |
| Emb | 9.44E-39 | 0.95390093 | 0.779 | 0.579 | 1.83E-34 | gamma-delta T | Emb |
| St6galnac3 | 3.83E-38 | 2.14206336 | 0.372 | 0.108 | 7.40E-34 | gamma-delta T | St6galnac3 |
| Pi4k2a | 1.62E-37 | 2.17932203 | 0.333 | 0.115 | 3.13E-33 | gamma-delta T | Pi4k2a |
| Gabbr1 | 3.86E-37 | 1.53206978 | 0.504 | 0.237 | 7.46E-33 | gamma-delta T | Gabbr1 |
| Rexo21 | 1.33E-33 | 1.73727217 | 0.519 | 0.23 | 2.57E-29 | gamma-delta T | Rexo2 |
| Cd164 | 5.09E-33 | 1.2532883 | 0.787 | 0.552 | 9.83E-29 | gamma-delta T | Cd164 |
| Sdc1 | 3.95E-29 | 1.82369453 | 0.283 | 0.091 | 7.64E-25 | gamma-delta T | Sdc1 |
| Psap | 3.72E-26 | 1.35195393 | 0.519 | 0.303 | 7.20E-22 | gamma-delta T | Psap |
| Il7r1 | 5.14E-26 | 1.09444084 | 0.62 | 0.378 | 9.94E-22 | gamma-delta T | Il7r |
| Itpr3 | 2.04E-25 | 1.72821172 | 0.298 | 0.124 | 3.94E-21 | gamma-delta T | Itpr3 |
| Irak2 | 1.53E-24 | 1.54608124 | 0.453 | 0.229 | 2.95E-20 | gamma-delta T | Irak2 |
| Irs2 | 1.99E-23 | 1.32275084 | 0.391 | 0.182 | 3.84E-19 | gamma-delta T | Irs2 |
| Dap1 | 3.01E-23 | 1.74507334 | 0.419 | 0.201 | 5.81E-19 | gamma-delta T | Dap |
| Dock21 | 4.42E-23 | 0.86869593 | 0.802 | 0.676 | 8.54E-19 | gamma-delta T | Dock2 |
| Lmo4 | 8.49E-23 | 1.63511952 | 0.314 | 0.086 | 1.64E-18 | gamma-delta T | Lmo4 |
| Ccr21 | 1.14E-22 | 1.13339063 | 0.674 | 0.361 | 2.20E-18 | gamma-delta T | Ccr2 |
| Arhgap26 | 1.24E-22 | 1.53172968 | 0.388 | 0.161 | 2.39E-18 | gamma-delta T | Arhgap26 |
| Hspa1b2 | 2.58E-22 | 0.9068113 | 0.353 | 0.211 | 4.99E-18 | gamma-delta T | Hspa1b |
| Marveld1 | 3.01E-22 | 2.0235297 | 0.256 | 0.087 | 5.81E-18 | gamma-delta T | Marveld1 |
| Prkx | 4.97E-22 | 1.56354774 | 0.364 | 0.18 | 9.61E-18 | gamma-delta T | Prkx |
| Tnfsf14 | 3.12E-21 | 1.95031713 | 0.252 | 0.071 | 6.03E-17 | gamma-delta T | Tnfsf14 |
| Atp8a1 | 1.24E-20 | 1.66599013 | 0.271 | 0.104 | 2.39E-16 | gamma-delta T | Atp8a1 |
| Rasgrp2 | 1.43E-20 | 1.34452188 | 0.302 | 0.132 | 2.77E-16 | gamma-delta T | Rasgrp2 |
| Arnt2 | 2.01E-20 | 1.8145877 | 0.349 | 0.144 | 3.89E-16 | gamma-delta T | Arnt2 |
| Rcbtb2 | 2.40E-20 | 1.84954511 | 0.271 | 0.139 | 4.65E-16 | gamma-delta T | Rcbtb2 |
| Smpdl3a | 2.91E-20 | 1.42480141 | 0.341 | 0.18 | 5.63E-16 | gamma-delta T | Smpdl3a |
| S100a6 | 7.06E-20 | 1.0438063 | 0.798 | 0.588 | 1.37E-15 | gamma-delta T | S100a6 |
| Zfp652 | 9.50E-19 | 1.43138213 | 0.364 | 0.149 | 1.84E-14 | gamma-delta T | Zfp652 |
| Crmp1 | 9.63E-19 | 1.8179302 | 0.337 | 0.147 | 1.86E-14 | gamma-delta T | Crmp1 |
| Man2a2 | 1.00E-18 | 1.28482792 | 0.461 | 0.241 | 1.94E-14 | gamma-delta T | Man2a2 |
| Rasa31 | 5.15E-18 | 1.00596918 | 0.465 | 0.259 | 9.96E-14 | gamma-delta T | Rasa3 |
| Slc16a6 | 5.33E-18 | 1.70393226 | 0.318 | 0.157 | 1.03E-13 | gamma-delta T | Slc16a6 |
| Slc12a7 | 7.05E-18 | 1.41110959 | 0.337 | 0.15 | 1.36E-13 | gamma-delta T | Slc12a7 |
| Itm2b | 3.86E-17 | 0.85209673 | 0.911 | 0.761 | 7.47E-13 | gamma-delta T | Itm2b |
| Ncor1 | 7.45E-17 | 1.06328508 | 0.636 | 0.45 | 1.44E-12 | gamma-delta T | Ncor1 |
| Jun | 5.27E-16 | 0.89870204 | 0.457 | 0.339 | 1.02E-11 | gamma-delta T | Jun |
| Gpr183 | 7.36E-16 | 1.26356849 | 0.504 | 0.287 | 1.42E-11 | gamma-delta T | Gpr183 |
| Top2b1 | 1.09E-15 | 1.08154917 | 0.442 | 0.275 | 2.10E-11 | gamma-delta T | Top2b |
| Txnip1 | 1.40E-15 | 0.73416128 | 0.473 | 0.359 | 2.71E-11 | gamma-delta T | Txnip |
| Clint1 | 2.67E-15 | 1.13981533 | 0.612 | 0.406 | 5.17E-11 | gamma-delta T | Clint1 |
| Ltb4r1 | 5.31E-15 | 1.66448359 | 0.295 | 0.125 | 1.03E-10 | gamma-delta T | Ltb4r1 |
| Furin1 | 6.60E-15 | 0.87774604 | 0.647 | 0.445 | 1.28E-10 | gamma-delta T | Furin |
| Myadm | 1.86E-14 | 1.48731794 | 0.302 | 0.142 | 3.60E-10 | gamma-delta T | Myadm |
| Klf21 | 3.42E-14 | 0.63599188 | 0.388 | 0.265 | 6.60E-10 | gamma-delta T | Klf2 |
| Tnf | 3.53E-14 | 1.32179277 | 0.271 | 0.127 | 6.82E-10 | gamma-delta T | Tnf |
| Klhl6 | 2.14E-13 | 1.21903097 | 0.271 | 0.136 | 4.14E-09 | gamma-delta T | Klhl6 |
| Vps37b1 | 1.41E-12 | 0.5252795 | 0.806 | 0.73 | 2.72E-08 | gamma-delta T | Vps37b |
| Abi3 | 1.53E-12 | 1.45614329 | 0.283 | 0.132 | 2.95E-08 | gamma-delta T | Abi3 |
| Nipal31 | 2.02E-12 | 1.3410135 | 0.345 | 0.198 | 3.91E-08 | gamma-delta T | Nipal3 |
| S100a41 | 2.31E-12 | 0.88799021 | 0.919 | 0.677 | 4.46E-08 | gamma-delta T | S100a4 |
| Lats2 | 7.96E-12 | 1.11356144 | 0.473 | 0.277 | 1.54E-07 | gamma-delta T | Lats2 |
| Notch2 | 1.44E-11 | 1.45383833 | 0.326 | 0.165 | 2.77E-07 | gamma-delta T | Notch2 |
| Adam8 | 1.69E-11 | 1.4975448 | 0.326 | 0.173 | 3.26E-07 | gamma-delta T | Adam8 |
| S1pr11 | 2.17E-11 | 0.76216252 | 0.36 | 0.185 | 4.20E-07 | gamma-delta T | S1pr1 |
| Dctn1 | 3.02E-11 | 1.42559417 | 0.345 | 0.192 | 5.84E-07 | gamma-delta T | Dctn1 |
| S100a11 | 3.19E-11 | 0.99187892 | 0.86 | 0.657 | 6.16E-07 | gamma-delta T | S100a11 |
| Cdk2ap2 | 4.61E-11 | 1.12829609 | 0.539 | 0.366 | 8.91E-07 | gamma-delta T | Cdk2ap2 |
| Rasgrp1 | 6.94E-11 | 0.92437503 | 0.516 | 0.371 | 1.34E-06 | gamma-delta T | Rasgrp1 |
| Gpx11 | 1.54E-10 | 1.1941946 | 0.593 | 0.399 | 2.98E-06 | gamma-delta T | Gpx1 |
| Lsp11 | 3.68E-10 | 0.93762203 | 0.736 | 0.576 | 7.11E-06 | gamma-delta T | Lsp1 |
| 11-Sep | 4.72E-10 | 1.29249228 | 0.477 | 0.286 | 9.13E-06 | gamma-delta T | 11-Sep |
| Cd2261 | 5.94E-10 | 1.2775537 | 0.283 | 0.124 | 1.15E-05 | gamma-delta T | Cd226 |
| Tspo | 6.40E-10 | 0.95061809 | 0.694 | 0.487 | 1.24E-05 | gamma-delta T | Tspo |
| Golm1 | 7.80E-10 | 1.20814204 | 0.298 | 0.165 | 1.51E-05 | gamma-delta T | Golm1 |
| Rarg | 1.06E-09 | 1.24068371 | 0.306 | 0.175 | 2.06E-05 | gamma-delta T | Rarg |
| Fam102a2 | 1.17E-09 | 0.81539755 | 0.585 | 0.395 | 2.26E-05 | gamma-delta T | Fam102a |
| Agpat4 | 2.28E-09 | 1.44689617 | 0.271 | 0.148 | 4.41E-05 | gamma-delta T | Agpat4 |
| Eif4e3 | 2.61E-09 | 1.25112136 | 0.275 | 0.148 | 5.05E-05 | gamma-delta T | Eif4e3 |
| Acsbg1 | 5.23E-09 | 1.22721377 | 0.287 | 0.183 | 0.00010118 | gamma-delta T | Acsbg1 |
| Btg22 | 5.93E-09 | 0.51100106 | 0.748 | 0.615 | 0.0001146 | gamma-delta T | Btg2 |
| Tatdn2 | 6.17E-09 | 0.91774041 | 0.547 | 0.419 | 0.00011925 | gamma-delta T | Tatdn2 |
| Arhgdib1 | 7.57E-09 | 0.28729349 | 0.791 | 0.703 | 0.00014637 | gamma-delta T | Arhgdib |
| Itgb11 | 1.25E-08 | 0.74553252 | 0.484 | 0.365 | 0.000242 | gamma-delta T | Itgb1 |
| Map3k141 | 3.30E-08 | 1.068222 | 0.349 | 0.193 | 0.00063722 | gamma-delta T | Map3k14 |
| Cd3g1 | 3.32E-08 | 0.83527416 | 0.74 | 0.575 | 0.00064208 | gamma-delta T | Cd3g |
| Stk24 | 4.01E-08 | 0.96271591 | 0.578 | 0.45 | 0.00077623 | gamma-delta T | Stk24 |
| Macf1 | 5.45E-08 | 0.6892338 | 0.647 | 0.535 | 0.00105319 | gamma-delta T | Macf1 |
| Senp6 | 1.09E-07 | 1.06552175 | 0.372 | 0.249 | 0.00210265 | gamma-delta T | Senp6 |
| Bcl2a1b1 | 1.57E-07 | 1.0069821 | 0.481 | 0.329 | 0.00303441 | gamma-delta T | Bcl2a1b |
| Cnot6 | 1.99E-07 | 1.27270539 | 0.333 | 0.211 | 0.00384348 | gamma-delta T | Cnot6 |
| Atf6 | 2.68E-07 | 1.2133418 | 0.283 | 0.162 | 0.00518997 | gamma-delta T | Atf6 |
| Tnfrsf25 | 3.96E-07 | 1.27515177 | 0.256 | 0.127 | 0.00766473 | gamma-delta T | Tnfrsf25 |
| Laptm51 | 5.32E-07 | 0.621475 | 0.961 | 0.889 | 0.01028959 | gamma-delta T | Laptm5 |
| Jmy | 5.55E-07 | 0.66574266 | 0.388 | 0.285 | 0.01072766 | gamma-delta T | Jmy |
| Uhrf2 | 5.79E-07 | 0.94987903 | 0.446 | 0.291 | 0.01119945 | gamma-delta T | Uhrf2 |

| | | | | | | | |
|---|---|---|---|---|---|---|---|
| Selplg1 | 6.66E-07 | 0.73632508 | 0.783 | 0.64 | 0.01288043 | gamma-delta T | Selplg |
| Atp2a3 | 6.91E-07 | 0.87856914 | 0.504 | 0.36 | 0.01336324 | gamma-delta T | Atp2a3 |
| mt-Nd6 | 7.77E-07 | 0.60294324 | 0.562 | 0.419 | 0.01502658 | gamma-delta T | mt-Nd6 |
| Slc37a3 | 1.21E-06 | 0.96292286 | 0.26 | 0.156 | 0.02339018 | gamma-delta T | Slc37a3 |
| Parp1 | 1.42E-06 | 1.39940842 | 0.302 | 0.17 | 0.02737482 | gamma-delta T | Parp1 |
| Prdx6 | 1.68E-06 | 1.24824658 | 0.256 | 0.153 | 0.03241888 | gamma-delta T | Prdx6 |
| mt-Co11 | 1.71E-06 | 0.77083655 | 0.62 | 0.473 | 0.0331518 | gamma-delta T | mt-Co1 |
| Il12rb11 | 2.44E-06 | 1.17950316 | 0.372 | 0.218 | 0.04714934 | gamma-delta T | Il12rb1 |
| Hdhd51 | 3.15E-06 | 0.90218865 | 0.326 | 0.195 | 0.06082174 | gamma-delta T | Hdhd5 |
| Ppp1r9b1 | 4.06E-06 | 0.72755638 | 0.616 | 0.473 | 0.07855385 | gamma-delta T | Ppp1r9b |
| Ero1l | 5.04E-06 | 0.9514144 | 0.446 | 0.276 | 0.09742873 | gamma-delta T | Ero1l |
| Rnf1662 | 5.81E-06 | 0.88974265 | 0.461 | 0.301 | 0.11232007 | gamma-delta T | Rnf166 |
| Hipk12 | 9.58E-06 | 0.81792586 | 0.527 | 0.414 | 0.18529929 | gamma-delta T | Hipk1 |
| Smad3 | 1.11E-05 | 0.82829163 | 0.388 | 0.26 | 0.21464543 | gamma-delta T | Smad3 |
| Ripor21 | 1.57E-05 | 0.47018067 | 0.36 | 0.279 | 0.30352215 | gamma-delta T | Ripor2 |
| Vsir1 | 1.90E-05 | 0.85246129 | 0.55 | 0.419 | 0.3679472 | gamma-delta T | Vsir |
| Mapk6 | 1.95E-05 | 1.11661152 | 0.376 | 0.248 | 0.37734244 | gamma-delta T | Mapk6 |
| Ptpra | 2.19E-05 | 1.06026128 | 0.333 | 0.215 | 0.42288897 | gamma-delta T | Ptpra |
| Plxnd11 | 2.79E-05 | 1.02128092 | 0.516 | 0.297 | 0.53962557 | gamma-delta T | Plxnd1 |
| Arhgap25 | 3.07E-05 | 0.83359898 | 0.353 | 0.245 | 0.59270658 | gamma-delta T | Arhgap25 |
| Flna | 3.82E-05 | 0.79069324 | 0.686 | 0.549 | 0.73835063 | gamma-delta T | Flna |
| Rnf7 | 5.14E-05 | 1.13578299 | 0.267 | 0.166 | 0.99388062 | gamma-delta T | Rnf7 |
| Ahcyl2 | 5.16E-05 | 1.00491829 | 0.267 | 0.169 | 0.99761322 | gamma-delta T | Ahcyl2 |
| Pde3b1 | 6.15E-05 | 0.88955288 | 0.291 | 0.185 | 1 | gamma-delta T | Pde3b |
| Tcf71 | 9.09E-05 | 0.43363993 | 0.31 | 0.173 | 1 | gamma-delta T | Tcf7 |
| Pdcd11 | 0.00010021 | 0.78987163 | 0.612 | 0.397 | 1 | gamma-delta T | Pdcd1 |
| mt-Atp81 | 0.0001042 | 0.57147565 | 0.647 | 0.517 | 1 | gamma-delta T | mt-Atp8 |
| Antxr2 | 0.00010755 | 0.87179075 | 0.329 | 0.229 | 1 | gamma-delta T | Antxr2 |
| Pik3cd2 | 0.00011077 | 0.73063996 | 0.628 | 0.441 | 1 | gamma-delta T | Pik3cd |
| Tagln2 | 0.00011247 | 0.44312895 | 0.841 | 0.73 | 1 | gamma-delta T | Tagln2 |
| Cerk | 0.00013783 | 0.93405289 | 0.271 | 0.169 | 1 | gamma-delta T | Cerk |
| Atp1b3 | 0.00014256 | 0.76488483 | 0.566 | 0.446 | 1 | gamma-delta T | Atp1b3 |
| Ier21 | 0.00014959 | 0.53944625 | 0.612 | 0.527 | 1 | gamma-delta T | Ier2 |
| Zap701 | 0.00016023 | 0.78384067 | 0.616 | 0.491 | 1 | gamma-delta T | Zap70 |
| mt-Co31 | 0.00019778 | 0.73393019 | 0.628 | 0.466 | 1 | gamma-delta T | mt-Co3 |
| Fchsd2 | 0.00020151 | 1.03645781 | 0.298 | 0.185 | 1 | gamma-delta T | Fchsd2 |
| Rgs11 | 0.00021904 | 0.43690717 | 0.632 | 0.567 | 1 | gamma-delta T | Rgs1 |
| Pdia4 | 0.00026955 | 1.13713421 | 0.364 | 0.266 | 1 | gamma-delta T | Pdia4 |
| Itgb71 | 0.00031667 | 0.66662321 | 0.868 | 0.71 | 1 | gamma-delta T | Itgb7 |
| mt-Atp61 | 0.00037451 | 0.55116183 | 0.671 | 0.549 | 1 | gamma-delta T | mt-Atp6 |
| Rinl1 | 0.00038986 | 0.78925505 | 0.527 | 0.399 | 1 | gamma-delta T | Rinl |
| Stim11 | 0.00042241 | 0.82510345 | 0.357 | 0.255 | 1 | gamma-delta T | Stim1 |
| Tor1aip1 | 0.00062133 | 0.82489273 | 0.496 | 0.395 | 1 | gamma-delta T | Tor1aip1 |
| Ywhab | 0.00062862 | 0.33797298 | 0.388 | 0.373 | 1 | gamma-delta T | Ywhab |
| Atp2a2 | 0.0006791 | 0.27776151 | 0.403 | 0.39 | 1 | gamma-delta T | Atp2a2 |
| Myo1f1 | 0.00094039 | 0.97658952 | 0.446 | 0.331 | 1 | gamma-delta T | Myo1f |
| Ahnak | 0.00096368 | 0.4202903 | 0.826 | 0.714 | 1 | gamma-delta T | Ahnak |
| Ppp1ca1 | 0.00101852 | 0.2779062 | 0.384 | 0.387 | 1 | gamma-delta T | Ppp1ca |
| Sema4a1 | 0.00112518 | 0.75951581 | 0.372 | 0.241 | 1 | gamma-delta T | Sema4a |
| S1pr4 | 0.00118923 | 0.47887443 | 0.446 | 0.385 | 1 | gamma-delta T | S1pr4 |
| mt-Nd1 | 0.00142599 | 0.80910685 | 0.461 | 0.308 | 1 | gamma-delta T | mt-Nd1 |
| Park7 | 0.00145882 | 0.26198438 | 0.38 | 0.359 | 1 | gamma-delta T | Park7 |
| Sema4d1 | 0.00186055 | 0.28333883 | 0.76 | 0.737 | 1 | gamma-delta T | Sema4d |
| Pde4b | 0.00247643 | 0.76623859 | 0.388 | 0.296 | 1 | gamma-delta T | Pde4b |
| S100a101 | 0.00255504 | 0.82253493 | 0.946 | 0.772 | 1 | gamma-delta T | S100a10 |
| Pdia3 | 0.00265612 | 0.30033224 | 0.368 | 0.36 | 1 | gamma-delta T | Pdia3 |
| Galnt1 | 0.00272731 | 0.83634288 | 0.496 | 0.351 | 1 | gamma-delta T | Galnt1 |
| Slc25a24 | 0.0028107 | 0.90303805 | 0.271 | 0.18 | 1 | gamma-delta T | Slc25a24 |
| Akt1 | 0.00292507 | 0.33853661 | 0.388 | 0.391 | 1 | gamma-delta T | Akt1 |
| Selenot | 0.00296249 | 0.8605133 | 0.326 | 0.243 | 1 | gamma-delta T | Selenot |
| Kcnab21 | 0.00338129 | 0.87835856 | 0.465 | 0.332 | 1 | gamma-delta T | Kcnab2 |
| Atp2b11 | 0.00339631 | 0.65549446 | 0.442 | 0.351 | 1 | gamma-delta T | Atp2b1 |
| Tspan14 | 0.00381784 | 0.91773536 | 0.275 | 0.196 | 1 | gamma-delta T | Tspan14 |
| Tecpr1 | 0.00423431 | 0.55601228 | 0.411 | 0.327 | 1 | gamma-delta T | Tecpr1 |
| Tle5 | 0.00519189 | 0.28242439 | 0.43 | 0.462 | 1 | gamma-delta T | Tle5 |
| Rassf21 | 0.00526519 | 0.72276488 | 0.267 | 0.197 | 1 | gamma-delta T | Rassf2 |
| Itgal1 | 0.00588058 | 0.59693217 | 0.806 | 0.693 | 1 | gamma-delta T | Itgal |
| Rapgef2 | 0.00652542 | 0.75233524 | 0.283 | 0.198 | 1 | gamma-delta T | Rapgef2 |
| Atox1 | 0.00665577 | 0.25001841 | 0.225 | 0.264 | 1 | gamma-delta T | Atox1 |
| Hprt | 0.00684557 | 0.29558163 | 0.306 | 0.287 | 1 | gamma-delta T | Hprt |
| Aagab | 0.00776696 | 0.71300835 | 0.275 | 0.213 | 1 | gamma-delta T | Aagab |
| Pacs1 | 0.00792741 | 0.7489818 | 0.298 | 0.24 | 1 | gamma-delta T | Pacs1 |
| Runx1 | 0.00834078 | 0.75969954 | 0.279 | 0.206 | 1 | gamma-delta T | Runx1 |
| Tab2 | 0.00862069 | 0.6548475 | 0.477 | 0.414 | 1 | gamma-delta T | Tab2 |
| Ric1 | 0.0086946 | 0.68871761 | 0.291 | 0.208 | 1 | gamma-delta T | Ric1 |
| Anxa61 | 0.00881313 | 0.36040046 | 0.756 | 0.657 | 1 | gamma-delta T | Anxa6 |
| Fli1 | 0.00891011 | 0.65978883 | 0.55 | 0.442 | 1 | gamma-delta T | Fli1 |
| Slco3a1 | 0.0090918 | 0.72210095 | 0.298 | 0.214 | 1 | gamma-delta T | Slco3a1 |
| Fcer1g | 0 | 5.9502618 | 0.897 | 0.048 | 0 | NK cells | Fcer1g |
| Tyrobp | 0 | 6.02379302 | 0.873 | 0.044 | 0 | NK cells | Tyrobp |
| Nkg71 | 0 | 2.15521107 | 0.919 | 0.25 | 0 | NK cells | Nkg7 |
| Spry2 | 0 | 5.096226 | 0.667 | 0.037 | 0 | NK cells | Spry2 |
| Gzma | 0 | 3.52958606 | 0.682 | 0.066 | 0 | NK cells | Gzma |
| Ugcg1 | 0 | 3.05165988 | 0.797 | 0.196 | 0 | NK cells | Ugcg |
| Klrk11 | 0 | 3.42735602 | 0.687 | 0.091 | 0 | NK cells | Klrk1 |
| Klre1 | 0 | 4.66858886 | 0.616 | 0.038 | 0 | NK cells | Klre1 |
| Prf11 | 0 | 4.15262542 | 0.648 | 0.09 | 0 | NK cells | Prf1 |
| Ctsw1 | 0 | 2.03008421 | 0.839 | 0.281 | 0 | NK cells | Ctsw |
| Irf8 | 0 | 4.34869809 | 0.677 | 0.127 | 0 | NK cells | Irf8 |
| Klrb1b | 0 | 5.85404745 | 0.518 | 0.015 | 0 | NK cells | Klrb1b |
| Klrd11 | 0 | 3.01889848 | 0.589 | 0.095 | 0 | NK cells | Klrd1 |
| Car2 | 0 | 4.08215978 | 0.54 | 0.049 | 0 | NK cells | Car2 |
| Xcl1 | 0 | 3.93121623 | 0.496 | 0.028 | 0 | NK cells | Xcl1 |
| Tbx211 | 0 | 2.57769115 | 0.633 | 0.166 | 0 | NK cells | Tbx21 |
| Ccl51 | 0 | 1.9535704 | 0.809 | 0.346 | 0 | NK cells | Ccl5 |
| Evl1 | 0 | 2.13348936 | 0.643 | 0.209 | 0 | NK cells | Evl |
| Anxa21 | 0 | 1.74738816 | 0.834 | 0.454 | 0 | NK cells | Anxa2 |
| Il2rb1 | 0 | 1.25328455 | 0.939 | 0.763 | 0 | NK cells | Il2rb |
| Sell1 | 1.72E-304 | 2.14656136 | 0.636 | 0.27 | 3.32E-300 | NK cells | Sell |
| Cd71 | 5.54E-297 | 2.24618826 | 0.601 | 0.141 | 1.07E-292 | NK cells | Cd7 |
| Atp1b1 | 7.79E-279 | 4.97892471 | 0.526 | 0.019 | 1.51E-274 | NK cells | Atp1b1 |
| Eomes | 9.27E-273 | 4.02999101 | 0.455 | 0.046 | 1.79E-268 | NK cells | Eomes |
| Adamts14 | 3.09E-268 | 5.75102396 | 0.531 | 0.018 | 5.97E-264 | NK cells | Adamts14 |
| Plek1 | 3.74E-265 | 2.6333533 | 0.577 | 0.15 | 7.24E-261 | NK cells | Plek |
| Txk | 1.11E-245 | 2.46245071 | 0.66 | 0.18 | 2.14E-241 | NK cells | Txk |
| Serpinb91 | 6.40E-241 | 2.64152066 | 0.548 | 0.142 | 1.24E-236 | NK cells | Serpinb9 |
| Bcl2l111 | 1.89E-239 | 1.8416894 | 0.621 | 0.26 | 3.66E-235 | NK cells | Bcl2l11 |
| Cd244a | 5.15E-234 | 3.7689421 | 0.484 | 0.049 | 9.97E-230 | NK cells | Cd244a |
| Gpc1 | 2.58E-233 | 3.25882923 | 0.435 | 0.068 | 4.99E-229 | NK cells | Gpc1 |
| Bcl21 | 2.69E-216 | 2.26835479 | 0.553 | 0.147 | 5.19E-212 | NK cells | Bcl2 |
| Klrc11 | 3.91E-210 | 2.17344847 | 0.45 | 0.123 | 7.56E-206 | NK cells | Klrc1 |
| Chsy11 | 2.59E-200 | 2.25020024 | 0.658 | 0.239 | 5.00E-196 | NK cells | Chsy1 |
| Itgax | 8.66E-193 | 4.98602609 | 0.372 | 0.018 | 1.67E-188 | NK cells | Itgax |
| Samd3 | 6.78E-188 | 4.3501594 | 0.386 | 0.022 | 1.31E-183 | NK cells | Samd3 |
| Itgb12 | 1.37E-184 | 1.4976035 | 0.731 | 0.34 | 2.64E-180 | NK cells | Itgb1 |
| Arsb1 | 1.61E-179 | 2.90349564 | 0.496 | 0.12 | 3.10E-175 | NK cells | Arsb |
| Sulf2 | 1.12E-176 | 4.51136381 | 0.377 | 0.021 | 2.17E-172 | NK cells | Sulf2 |
| Klri2 | 8.76E-171 | 6.43928875 | 0.352 | 0.009 | 1.69E-166 | NK cells | Klri2 |
| Ms4a4b1 | 6.65E-167 | 1.57727945 | 0.709 | 0.292 | 1.29E-162 | NK cells | Ms4a4b |
| Gas7 | 2.51E-158 | 4.23221375 | 0.337 | 0.017 | 4.85E-154 | NK cells | Gas7 |
| Syk | 2.61E-156 | 4.34078976 | 0.367 | 0.02 | 5.05E-152 | NK cells | Syk |
| Arl4d | 2.68E-154 | 4.21393684 | 0.308 | 0.021 | 5.18E-150 | NK cells | Arl4d |
| Itgam | 3.12E-147 | 4.45443667 | 0.313 | 0.029 | 6.04E-143 | NK cells | Itgam |
| Vps37b2 | 1.14E-146 | 0.55185427 | 0.841 | 0.725 | 2.20E-142 | NK cells | Vps37b |
| Styk1 | 1.53E-145 | 4.86236297 | 0.298 | 0.009 | 2.96E-141 | NK cells | Styk1 |
| Ccl41 | 2.94E-137 | 2.19833549 | 0.318 | 0.088 | 5.69E-133 | NK cells | Ccl4 |
| Tmsb4x1 | 8.25E-137 | 0.35923959 | 0.995 | 0.978 | 1.60E-132 | NK cells | Tmsb4x |
| Fasl1 | 6.79E-134 | 2.02146983 | 0.509 | 0.156 | 1.31E-129 | NK cells | Fasl |
| Rin3 | 1.13E-132 | 2.29135133 | 0.506 | 0.142 | 2.18E-128 | NK cells | Rin3 |
| Plac81 | 1.06E-131 | 1.11193643 | 0.337 | 0.177 | 2.06E-127 | NK cells | Plac8 |
| Il12rb2 | 2.37E-130 | 3.22906465 | 0.374 | 0.043 | 4.58E-126 | NK cells | Il12rb2 |
| Lat2 | 1.67E-116 | 2.72938369 | 0.364 | 0.083 | 3.24E-112 | NK cells | Lat2 |
| Serpinb6b | 1.62E-114 | 2.76793889 | 0.345 | 0.072 | 3.14E-110 | NK cells | Serpinb6b |
| Klrb1f | 4.92E-114 | 4.24941571 | 0.264 | 0.015 | 9.51E-110 | NK cells | Klrb1f |
| Tm6sf1 | 2.83E-113 | 2.78748699 | 0.386 | 0.068 | 5.47E-109 | NK cells | Tm6sf1 |
| Sema4a2 | 2.66E-112 | 1.59468347 | 0.567 | 0.22 | 5.14E-108 | NK cells | Sema4a |
| Hhex | 3.53E-112 | 4.41717098 | 0.279 | 0.018 | 6.82E-108 | NK cells | Hhex |
| Pik3ap11 | 1.03E-107 | 2.06419248 | 0.487 | 0.14 | 1.99E-103 | NK cells | Pik3ap1 |
| Cd9 | 1.52E-106 | 2.21747934 | 0.438 | 0.112 | 2.93E-102 | NK cells | Cd9 |
| Pik3r1 | 5.01E-106 | 1.70240532 | 0.567 | 0.26 | 9.68E-102 | NK cells | Pik3r1 |
| Emid1 | 5.08E-104 | 4.78997265 | 0.254 | 0.01 | 9.82E-100 | NK cells | Emid1 |
| Klrb1c | 2.86E-103 | 7.80908119 | 0.472 | 0.003 | 5.52E-99 | NK cells | Klrb1c |
| Tmsb101 | 1.07E-100 | 0.5437371 | 0.885 | 0.802 | 2.06E-96 | NK cells | Tmsb10 |
| Plcg2 | 4.49E-99 | 4.46530334 | 0.267 | 0.015 | 8.68E-95 | NK cells | Plcg2 |
| Rnf130 | 7.13E-99 | 2.73396535 | 0.364 | 0.067 | 1.38E-94 | NK cells | Rnf130 |
| Eef1a12 | 5.89E-98 | 0.34880595 | 0.971 | 0.942 | 1.14E-93 | NK cells | Eef1a1 |
| Gimap61 | 2.47E-96 | 1.06824427 | 0.694 | 0.445 | 4.77E-92 | NK cells | Gimap6 |
| Cxxc5 | 6.04E-96 | 4.2059317 | 0.264 | 0.019 | 1.17E-91 | NK cells | Cxxc5 |
| Ifitm10 | 1.30E-95 | 2.72496736 | 0.301 | 0.044 | 2.50E-91 | NK cells | Ifitm10 |
| Ptprc1 | 4.58E-89 | 0.93831181 | 0.89 | 0.721 | 8.86E-85 | NK cells | Ptprc |
| Lgals3bp1 | 5.00E-89 | 1.20147701 | 0.46 | 0.227 | 9.67E-85 | NK cells | Lgals3bp |
| Gzmb1 | 4.05E-88 | 0.63836227 | 0.572 | 0.43 | 7.82E-84 | NK cells | Gzmb |
| Zyx1 | 1.26E-87 | 1.26720926 | 0.731 | 0.438 | 2.43E-83 | NK cells | Zyx |
| Slamf71 | 2.16E-86 | 2.41118881 | 0.303 | 0.07 | 4.17E-82 | NK cells | Slamf7 |
| Gimap41 | 1.68E-85 | 0.9405037 | 0.822 | 0.559 | 3.25E-81 | NK cells | Gimap4 |
| Serpina3g1 | 1.39E-83 | 1.56626164 | 0.438 | 0.202 | 2.69E-79 | NK cells | Serpina3g |
| Spp1 | 8.86E-83 | 0.46512956 | 0.511 | 0.494 | 1.71E-78 | NK cells | Spp1 |
| Ccr51 | 2.71E-82 | 1.40544931 | 0.579 | 0.314 | 5.24E-78 | NK cells | Ccr5 |
| Pecam1 | 7.81E-81 | 1.50696674 | 0.315 | 0.126 | 1.51E-76 | NK cells | Pecam1 |
| Aoah | 6.62E-80 | 6.96552333 | 0.32 | 0.003 | 1.28E-75 | NK cells | Aoah |
| Rhoq | 2.75E-78 | 1.95645805 | 0.291 | 0.088 | 5.31E-74 | NK cells | Rhoq |
| Epstl11 | 1.44E-77 | 1.53296647 | 0.528 | 0.261 | 2.79E-73 | NK cells | Epstl1 |
| Atp8b4 | 9.94E-76 | 2.0183054 | 0.35 | 0.088 | 1.92E-71 | NK cells | Atp8b4 |
| Pde2a | 7.77E-73 | 2.00305943 | 0.391 | 0.127 | 1.50E-68 | NK cells | Pde2a |
| B4galt51 | 1.40E-71 | 1.72688174 | 0.443 | 0.186 | 2.71E-67 | NK cells | B4galt5 |
| Il18rap | 1.30E-70 | 1.55796949 | 0.577 | 0.282 | 2.52E-66 | NK cells | Il18rap |
| Anxa1 | 1.57E-69 | 2.05479276 | 0.286 | 0.073 | 3.04E-65 | NK cells | Anxa1 |
| Xdh | 1.19E-68 | 2.70266354 | 0.284 | 0.064 | 2.30E-64 | NK cells | Xdh |
| Lrrk1 | 1.25E-68 | 2.97128397 | 0.286 | 0.05 | 2.41E-64 | NK cells | Lrrk1 |
| Cd1601 | 1.50E-66 | 2.15578915 | 0.269 | 0.06 | 2.90E-62 | NK cells | Cd160 |
| Mctp2 | 6.58E-66 | 2.61788689 | 0.279 | 0.047 | 1.27E-61 | NK cells | Mctp2 |
| Gsap | 1.01E-64 | 2.46487452 | 0.328 | 0.087 | 1.95E-60 | NK cells | Gsap |
| Itga2 | 1.52E-62 | 2.63927036 | 0.284 | 0.062 | 2.94E-58 | NK cells | Itga2 |
| Ifng1 | 7.83E-62 | 1.70800077 | 0.306 | 0.085 | 1.51E-57 | NK cells | Ifng |
| Gpr1711 | 3.18E-61 | 1.53959705 | 0.396 | 0.17 | 6.15E-57 | NK cells | Gpr171 |
| Padi21 | 1.61E-58 | 1.67471168 | 0.418 | 0.186 | 3.12E-54 | NK cells | Padi2 |
| Klra4 | 3.43E-57 | 9.83388729 | 0.269 | 0.001 | 6.63E-53 | NK cells | Klra4 |
| Trbc11 | 5.99E-54 | 0.75804321 | 0.667 | 0.378 | 1.16E-49 | NK cells | Trbc1 |
| Pik3r5 | 1.44E-53 | 1.42307414 | 0.355 | 0.144 | 2.79E-49 | NK cells | Pik3r5 |
| Cd47 | 8.84E-53 | 0.82342697 | 0.689 | 0.542 | 1.71E-48 | NK cells | Cd47 |
| Adgre51 | 1.21E-52 | 1.24720243 | 0.611 | 0.407 | 2.34E-48 | NK cells | Adgre5 |
| Sh2d3c1 | 1.76E-51 | 1.71870888 | 0.381 | 0.139 | 3.40E-47 | NK cells | Sh2d3c |
| B4galnt12 | 4.37E-51 | 0.99588646 | 0.565 | 0.341 | 8.45E-47 | NK cells | B4galnt1 |
| Rgs31 | 1.66E-50 | 1.05285444 | 0.714 | 0.448 | 3.22E-46 | NK cells | Rgs3 |
| Itgal2 | 6.58E-50 | 0.82748862 | 0.836 | 0.687 | 1.27E-45 | NK cells | Itgal |
| Efhd21 | 3.72E-49 | 1.37585302 | 0.479 | 0.241 | 7.19E-45 | NK cells | Efhd2 |
| Ptpn61 | 1.90E-48 | 1.34007893 | 0.406 | 0.214 | 3.68E-44 | NK cells | Ptpn6 |
| Myc | 7.42E-46 | 1.50771925 | 0.264 | 0.13 | 1.43E-41 | NK cells | Myc |
| Ncr1 | 2.96E-45 | 10.5521996 | 0.672 | 0.001 | 5.73E-41 | NK cells | Ncr1 |
| Ctsd1 | 5.25E-45 | 1.51244891 | 0.472 | 0.211 | 1.02E-40 | NK cells | Ctsd |
| Jak11 | 9.51E-45 | 0.85493929 | 0.756 | 0.606 | 1.84E-40 | NK cells | Jak1 |
| Rgs12 | 1.77E-44 | 0.31442727 | 0.579 | 0.57 | 3.43E-40 | NK cells | Rgs1 |
| Gramd4 | 5.18E-44 | 1.45027094 | 0.357 | 0.148 | 1.00E-39 | NK cells | Gramd4 |
| Zfp36l22 | 1.35E-42 | 0.38382501 | 0.817 | 0.689 | 2.61E-38 | NK cells | Zfp36l2 |
| Ctla2b | 2.77E-42 | 1.47647584 | 0.364 | 0.146 | 5.35E-38 | NK cells | Ctla2b |
| Dnajc151 | 4.43E-41 | 1.17863027 | 0.328 | 0.172 | 8.56E-37 | NK cells | Dnajc15 |
| Ptpn222 | 2.90E-40 | 0.59672408 | 0.724 | 0.549 | 5.61E-36 | NK cells | Ptpn22 |
| Chst121 | 3.33E-40 | 1.5138567 | 0.306 | 0.124 | 6.45E-36 | NK cells | Chst12 |
| Itgb21 | 2.76E-39 | 0.92949812 | 0.807 | 0.679 | 5.34E-35 | NK cells | Itgb2 |
| Prkcb | 3.88E-39 | 1.77388674 | 0.379 | 0.158 | 7.50E-35 | NK cells | Prkcb |
| Txnip2 | 2.69E-35 | 0.73981668 | 0.54 | 0.349 | 5.19E-31 | NK cells | Txnip |
| Gem1 | 6.12E-34 | 0.71078924 | 0.474 | 0.298 | 1.18E-29 | NK cells | Gem |
| Btg23 | 2.11E-33 | 0.35368306 | 0.709 | 0.615 | 4.08E-29 | NK cells | Btg2 |
| Pdcd41 | 4.50E-33 | 0.64694029 | 0.682 | 0.527 | 8.69E-29 | NK cells | Pdcd4 |
| Sh2d2a1 | 2.08E-32 | 0.47647682 | 0.726 | 0.629 | 4.02E-28 | NK cells | Sh2d2a |
| Sh3bgrl31 | 2.22E-32 | 0.46810431 | 0.802 | 0.707 | 4.28E-28 | NK cells | Sh3bgrl3 |
| Tbc1d2b1 | 8.03E-32 | 1.35231682 | 0.379 | 0.19 | 1.55E-27 | NK cells | Tbc1d2b |
| Klra8 | 1.73E-31 | 7.79755679 | 0.252 | 0.001 | 3.34E-27 | NK cells | Klra8 |
| Ptpn12 | 2.23E-31 | 1.73876857 | 0.289 | 0.112 | 4.31E-27 | NK cells | Ptpn12 |
| mt-Co2 | 2.16E-30 | 0.36199061 | 0.743 | 0.681 | 4.17E-26 | NK cells | mt-Co2 |
| Klf22 | 5.14E-30 | 1.00788464 | 0.438 | 0.257 | 9.93E-26 | NK cells | Klf2 |
| Gvin11 | 9.02E-30 | 1.03460322 | 0.501 | 0.274 | 1.74E-25 | NK cells | Gvin1 |
| Maff | 3.56E-29 | 1.16350567 | 0.262 | 0.115 | 6.88E-25 | NK cells | Maff |
| Cd521 | 1.82E-28 | 0.35172563 | 0.912 | 0.825 | 3.51E-24 | NK cells | Cd52 |
| Stat41 | 2.23E-28 | 0.82958089 | 0.553 | 0.361 | 4.30E-24 | NK cells | Stat4 |
| Prex11 | 9.04E-28 | 0.83129154 | 0.731 | 0.555 | 1.75E-23 | NK cells | Prex1 |
| Ikzf11 | 3.63E-27 | 0.75869849 | 0.719 | 0.517 | 7.01E-23 | NK cells | Ikzf1 |
| Grcc10 | 9.80E-27 | 0.63036585 | 0.528 | 0.414 | 1.89E-22 | NK cells | Grcc10 |
| Nr4a21 | 1.11E-26 | 0.80654095 | 0.401 | 0.256 | 2.15E-22 | NK cells | Nr4a2 |
| Pde3b2 | 1.03E-25 | 0.8400974 | 0.362 | 0.176 | 1.98E-21 | NK cells | Pde3b |
| Dennd4a1 | 1.19E-25 | 0.53345661 | 0.565 | 0.451 | 2.30E-21 | NK cells | Dennd4a |
| Ern1 | 3.50E-25 | 0.63603753 | 0.391 | 0.262 | 6.76E-21 | NK cells | Ern1 |
| Lta4h | 3.77E-25 | 1.17805385 | 0.435 | 0.265 | 7.29E-21 | NK cells | Lta4h |
| Rad54l2 | 5.29E-24 | 1.41238205 | 0.303 | 0.138 | 1.02E-19 | NK cells | Rad54l2 |
| Sephs2 | 6.08E-24 | 1.22485717 | 0.269 | 0.137 | 1.18E-19 | NK cells | Sephs2 |
| Ctla2a1 | 8.30E-24 | 0.40323767 | 0.543 | 0.41 | 1.60E-19 | NK cells | Ctla2a |
| Itga41 | 9.66E-24 | 1.11673233 | 0.477 | 0.262 | 1.87E-19 | NK cells | Itga4 |
| Iqsec11 | 1.11E-23 | 1.45716009 | 0.33 | 0.158 | 2.15E-19 | NK cells | Iqsec1 |
| Pglyrp11 | 1.52E-23 | 0.52788142 | 0.369 | 0.275 | 2.95E-19 | NK cells | Pglyrp1 |
| Rras2 | 2.45E-23 | 1.58361413 | 0.271 | 0.109 | 4.73E-19 | NK cells | Rras2 |
| Dock81 | 8.05E-23 | 0.45643835 | 0.494 | 0.407 | 1.56E-18 | NK cells | Dock8 |
| Gimap91 | 9.44E-23 | 1.10014678 | 0.43 | 0.244 | 1.82E-18 | NK cells | Gimap9 |
| Sep-62 | 3.31E-22 | 0.70555522 | 0.455 | 0.318 | 6.39E-18 | NK cells | 6-Sep |
| Lnpep2 | 5.54E-22 | 0.69752765 | 0.65 | 0.498 | 1.07E-17 | NK cells | Lnpep |
| Litaf1 | 7.10E-22 | 0.67220074 | 0.44 | 0.322 | 1.37E-17 | NK cells | Litaf |
| Grina | 1.43E-21 | 0.72635092 | 0.477 | 0.334 | 2.76E-17 | NK cells | Grina |
| Esyt11 | 1.68E-21 | 0.65055088 | 0.768 | 0.624 | 3.26E-17 | NK cells | Esyt1 |
| Usp481 | 2.79E-21 | 0.98985221 | 0.35 | 0.195 | 5.39E-17 | NK cells | Usp48 |
| Prkacb2 | 9.28E-21 | 0.89301342 | 0.455 | 0.291 | 1.79E-16 | NK cells | Prkacb |
| Ripor22 | 1.60E-20 | 0.78239202 | 0.447 | 0.269 | 3.10E-16 | NK cells | Ripor2 |
| Lfng1 | 5.45E-20 | 0.82683321 | 0.575 | 0.416 | 1.05E-15 | NK cells | Lfng |
| Dusp21 | 7.71E-20 | 0.81973389 | 0.347 | 0.201 | 1.49E-15 | NK cells | Dusp2 |
| Gse1 | 9.57E-20 | 1.27239127 | 0.271 | 0.115 | 1.85E-15 | NK cells | Gse1 |
| Metrnl | 2.37E-19 | 1.31607248 | 0.281 | 0.142 | 4.58E-15 | NK cells | Metrnl |
| Ccnd31 | 3.89E-19 | 0.87852505 | 0.467 | 0.333 | 7.52E-15 | NK cells | Ccnd3 |
| Etfb1 | 6.53E-19 | 0.82228964 | 0.303 | 0.21 | 1.26E-14 | NK cells | Etfb |
| Sept111 | 1.77E-18 | 1.01043298 | 0.455 | 0.282 | 3.43E-14 | NK cells | 11-Sep |
| Lpp | 2.27E-18 | 1.10495402 | 0.279 | 0.142 | 4.39E-14 | NK cells | Lpp |
| Gramd1a1 | 3.50E-18 | 0.48444697 | 0.45 | 0.409 | 6.78E-14 | NK cells | Gramd1a |
| mt-Co32 | 6.55E-18 | 0.32576464 | 0.474 | 0.474 | 1.27E-13 | NK cells | mt-Co3 |
| Hsd11b1 | 6.79E-18 | 1.64932176 | 0.291 | 0.149 | 1.31E-13 | NK cells | Hsd11b1 |
| Runx31 | 8.90E-18 | 0.54738536 | 0.594 | 0.482 | 1.72E-13 | NK cells | Runx3 |
| Pik3cd3 | 2.28E-17 | 0.51141985 | 0.587 | 0.439 | 4.41E-13 | NK cells | Pik3cd |
| Rpn1 | 2.64E-17 | 0.62482915 | 0.279 | 0.211 | 5.11E-13 | NK cells | Rpn1 |
| Pdia6 | 3.38E-17 | 0.54862491 | 0.264 | 0.211 | 6.54E-13 | NK cells | Pdia6 |
| Tcf72 | 3.51E-17 | 0.40774674 | 0.318 | 0.168 | 6.78E-13 | NK cells | Tcf7 |
| Ilk1 | 3.56E-17 | 0.94369568 | 0.399 | 0.252 | 6.88E-13 | NK cells | Ilk |
| Spata131 | 7.14E-17 | 0.89078092 | 0.582 | 0.408 | 1.38E-12 | NK cells | Spata13 |
| Os9 | 9.86E-17 | 0.6116953 | 0.381 | 0.271 | 1.91E-12 | NK cells | Os9 |
| Tln1 | 1.23E-16 | 0.54693224 | 0.714 | 0.624 | 2.38E-12 | NK cells | Tln1 |
| Nup2101 | 1.74E-16 | 0.57048675 | 0.54 | 0.424 | 3.36E-12 | NK cells | Nup210 |
| H2afy | 1.88E-16 | 0.56588115 | 0.687 | 0.556 | 3.63E-12 | NK cells | H2afy |
| Eif5a | 1.98E-16 | 0.6673916 | 0.443 | 0.327 | 3.83E-12 | NK cells | Eif5a |
| Gimap5 | 2.00E-16 | 0.97693072 | 0.357 | 0.21 | 3.87E-12 | NK cells | Gimap5 |
| Bin21 | 3.89E-16 | 0.630287 | 0.768 | 0.61 | 7.52E-12 | NK cells | Bin2 |
| Arap21 | 6.52E-16 | 1.05626931 | 0.308 | 0.169 | 1.26E-11 | NK cells | Arap2 |
| mt-Nd11 | 8.20E-16 | 0.56160203 | 0.374 | 0.311 | 1.58E-11 | NK cells | mt-Nd1 |
| B4galt12 | 1.03E-15 | 0.72930867 | 0.496 | 0.351 | 1.99E-11 | NK cells | B4galt1 |
| Otulin1 | 2.04E-15 | 0.85549729 | 0.35 | 0.236 | 3.95E-11 | NK cells | Otulin |
| Clcf1 | 2.09E-15 | 0.88663372 | 0.276 | 0.154 | 4.04E-11 | NK cells | Clcf1 |
| Atp2a31 | 2.19E-15 | 0.80201448 | 0.462 | 0.359 | 4.23E-11 | NK cells | Atp2a3 |
| Arap1 | 2.40E-15 | 0.899227 | 0.337 | 0.209 | 4.64E-11 | NK cells | Arap1 |
| Serf21 | 2.43E-15 | 0.69622736 | 0.516 | 0.396 | 4.70E-11 | NK cells | Serf2 |
| Ifngr11 | 3.49E-15 | 0.26698817 | 0.885 | 0.828 | 6.75E-11 | NK cells | Ifngr1 |
| mt-Atp62 | 3.51E-15 | 0.25020671 | 0.56 | 0.554 | 6.79E-11 | NK cells | mt-Atp6 |
| Lasp11 | 5.04E-15 | 0.94780773 | 0.394 | 0.255 | 9.74E-11 | NK cells | Lasp1 |
| Npm12 | 1.64E-14 | 0.27876268 | 0.465 | 0.422 | 3.18E-10 | NK cells | Npm1 |
| Dok2 | 1.65E-14 | 1.16958701 | 0.394 | 0.259 | 3.19E-10 | NK cells | Dok2 |
| Nabp1 | 1.83E-14 | 0.58048379 | 0.472 | 0.38 | 3.54E-10 | NK cells | Nabp1 |
| Ssr2 | 1.89E-14 | 0.92078895 | 0.313 | 0.221 | 3.66E-10 | NK cells | Ssr2 |
| Ankrd13a | 2.27E-14 | 0.87176331 | 0.462 | 0.325 | 4.38E-10 | NK cells | Ankrd13a |
| Preb | 3.80E-14 | 0.87259496 | 0.318 | 0.202 | 7.34E-10 | NK cells | Preb |
| Ptp4a31 | 3.97E-14 | 0.84390898 | 0.306 | 0.221 | 7.68E-10 | NK cells | Ptp4a3 |
| Map7d1 | 4.11E-14 | 0.94598228 | 0.357 | 0.233 | 7.95E-10 | NK cells | Map7d1 |
| Nfatc11 | 4.96E-14 | 0.83671797 | 0.467 | 0.309 | 9.59E-10 | NK cells | Nfatc1 |
| Eef1g2 | 6.68E-14 | 0.33053528 | 0.328 | 0.283 | 1.29E-09 | NK cells | Eef1g |
| Kcnab22 | 8.39E-14 | 0.91549894 | 0.457 | 0.328 | 1.62E-09 | NK cells | Kcnab2 |
| Klhl61 | 9.58E-14 | 0.85775583 | 0.252 | 0.133 | 1.85E-09 | NK cells | Klhl6 |
| Slc12a71 | 1.08E-13 | 0.85824764 | 0.279 | 0.15 | 2.09E-09 | NK cells | Slc12a7 |
| Gatad2a1 | 1.30E-13 | 0.63847939 | 0.491 | 0.356 | 2.52E-09 | NK cells | Gatad2a |
| Ubac21 | 1.52E-13 | 0.66451264 | 0.355 | 0.222 | 2.94E-09 | NK cells | Ubac2 |
| Selplg2 | 2.05E-13 | 0.53663625 | 0.738 | 0.639 | 3.96E-09 | NK cells | Selplg |
| Cdyl | 2.08E-13 | 1.07205806 | 0.306 | 0.171 | 4.01E-09 | NK cells | Cdyl |
| Il10rb | 2.49E-13 | 1.06289984 | 0.389 | 0.243 | 4.81E-09 | NK cells | Il10rb |
| Ddx10 | 3.72E-13 | 0.61519667 | 0.262 | 0.167 | 7.19E-09 | NK cells | Ddx10 |
| Rasa32 | 5.10E-13 | 0.72839412 | 0.406 | 0.258 | 9.86E-09 | NK cells | Rasa3 |
| Satb11 | 8.16E-13 | 0.60958786 | 0.469 | 0.307 | 1.58E-08 | NK cells | Satb1 |
| Cst71 | 9.76E-13 | 0.81458962 | 0.303 | 0.206 | 1.89E-08 | NK cells | Cst7 |
| Wipf11 | 1.06E-12 | 0.54346822 | 0.623 | 0.495 | 2.05E-08 | NK cells | Wipf1 |
| Sub12 | 1.29E-12 | 0.53208123 | 0.362 | 0.262 | 2.49E-08 | NK cells | Sub1 |
| Me2 | 1.55E-12 | 1.07365271 | 0.271 | 0.152 | 2.99E-08 | NK cells | Me2 |
| Eef1b22 | 1.74E-12 | 0.27997771 | 0.567 | 0.542 | 3.36E-08 | NK cells | Eef1b2 |
| Ssr1 | 2.51E-12 | 0.49119345 | 0.369 | 0.3 | 4.85E-08 | NK cells | Ssr1 |
| Ppm1g | 3.20E-12 | 0.89098854 | 0.43 | 0.3 | 6.19E-08 | NK cells | Ppm1g |
| Plekha21 | 3.58E-12 | 0.5701247 | 0.455 | 0.336 | 6.92E-08 | NK cells | Plekha2 |
| Arhgef181 | 3.74E-12 | 0.64802305 | 0.489 | 0.371 | 7.24E-08 | NK cells | Arhgef18 |
| P4hb | 4.23E-12 | 0.43892293 | 0.301 | 0.257 | 8.19E-08 | NK cells | P4hb |
| Phb2 | 4.69E-12 | 0.67555394 | 0.259 | 0.177 | 9.07E-08 | NK cells | Phb2 |
| Lax12 | 4.71E-12 | 0.90455697 | 0.355 | 0.197 | 9.11E-08 | NK cells | Lax1 |
| Sdcbp | 5.06E-12 | 0.67199081 | 0.513 | 0.399 | 9.78E-08 | NK cells | Sdcbp |
| Cdk2ap1 | 8.08E-12 | 0.95919323 | 0.252 | 0.155 | 1.56E-07 | NK cells | Cdk2ap1 |
| Arhgap261 | 1.08E-11 | 0.83255796 | 0.274 | 0.163 | 2.09E-07 | NK cells | Arhgap26 |
| Tab21 | 1.36E-11 | 0.63386078 | 0.504 | 0.41 | 2.62E-07 | NK cells | Tab2 |
| Iqgap1 | 2.34E-11 | 0.449424 | 0.707 | 0.615 | 4.52E-07 | NK cells | Iqgap1 |
| Sptbn1 | 2.77E-11 | 0.67865806 | 0.445 | 0.342 | 5.35E-07 | NK cells | Sptbn1 |
| Mgat1 | 3.09E-11 | 0.97894178 | 0.286 | 0.167 | 5.98E-07 | NK cells | Mgat1 |
| Mlxip | 4.85E-11 | 0.56094244 | 0.33 | 0.232 | 9.37E-07 | NK cells | Mlxip |
| Elk41 | 6.31E-11 | 0.69052684 | 0.396 | 0.274 | 1.22E-06 | NK cells | Elk4 |
| Atp8b21 | 6.45E-11 | 0.80765845 | 0.313 | 0.215 | 1.25E-06 | NK cells | Atp8b2 |
| Gyg | 7.00E-11 | 1.07588857 | 0.262 | 0.152 | 1.35E-06 | NK cells | Gyg |
| Sipa11 | 7.20E-11 | 0.64306453 | 0.445 | 0.316 | 1.39E-06 | NK cells | Sipa1 |
| Arhgef2 | 7.33E-11 | 0.82342057 | 0.44 | 0.309 | 1.42E-06 | NK cells | Arhgef2 |
| mt-Cytb1 | 1.12E-10 | 0.27217056 | 0.425 | 0.404 | 2.16E-06 | NK cells | mt-Cytb |
| Myo1f2 | 2.60E-10 | 0.72129817 | 0.467 | 0.326 | 5.03E-06 | NK cells | Myo1f |
| Trappc8 | 2.62E-10 | 0.88936239 | 0.254 | 0.152 | 5.06E-06 | NK cells | Trappc8 |
| Lcp11 | 3.02E-10 | 0.28249554 | 0.812 | 0.764 | 5.83E-06 | NK cells | Lcp1 |
| Ier22 | 3.40E-10 | 0.38346084 | 0.601 | 0.525 | 6.58E-06 | NK cells | Ier2 |
| Sun2 | 3.62E-10 | 0.5822519 | 0.511 | 0.398 | 6.99E-06 | NK cells | Sun2 |
| Ube2g2 | 4.54E-10 | 1.018348 | 0.269 | 0.157 | 8.78E-06 | NK cells | Ube2g2 |
| Sp1001 | 4.93E-10 | 0.59673708 | 0.487 | 0.362 | 9.52E-06 | NK cells | Sp100 |
| F2r1 | 6.65E-10 | 0.34891187 | 0.369 | 0.281 | 1.29E-05 | NK cells | F2r |
| Kri1 | 7.83E-10 | 0.83599689 | 0.259 | 0.153 | 1.51E-05 | NK cells | Kri1 |
| Kctd12 | 9.69E-10 | 0.54693978 | 0.416 | 0.274 | 1.87E-05 | NK cells | Kctd12 |
| Cuta | 1.21E-09 | 0.63649272 | 0.333 | 0.257 | 2.34E-05 | NK cells | Cuta |
| Irak21 | 1.26E-09 | 0.70231178 | 0.345 | 0.231 | 2.44E-05 | NK cells | Irak2 |
| Stt3b1 | 1.63E-09 | 0.35299288 | 0.262 | 0.218 | 3.14E-05 | NK cells | Stt3b |
| Gnas | 3.50E-09 | 0.41983192 | 0.643 | 0.57 | 6.77E-05 | NK cells | Gnas |
| Spcs3 | 3.78E-09 | 0.69231942 | 0.264 | 0.202 | 7.31E-05 | NK cells | Spcs3 |
| Vars | 3.92E-09 | 0.5016203 | 0.399 | 0.315 | 7.58E-05 | NK cells | Vars |
| Eif4b1 | 4.87E-09 | 0.40887772 | 0.425 | 0.346 | 9.42E-05 | NK cells | Eif4b |
| Mndal1 | 6.42E-09 | 0.58808754 | 0.274 | 0.205 | 0.00012406 | NK cells | Mndal |
| Hdhd52 | 6.59E-09 | 0.68558708 | 0.293 | 0.193 | 0.0001274 | NK cells | Hdhd5 |
| Sytl3 | 1.35E-08 | 0.84560943 | 0.286 | 0.183 | 0.00026173 | NK cells | Sytl3 |
| Lgals9 | 1.45E-08 | 0.76871631 | 0.32 | 0.223 | 0.00027965 | NK cells | Lgals9 |
| Prdx5 | 1.59E-08 | 0.32025895 | 0.374 | 0.358 | 0.000307 | NK cells | Prdx5 |
| Xrn22 | 1.59E-08 | 0.58730799 | 0.421 | 0.311 | 0.00030772 | NK cells | Xrn2 |
| Abr1 | 1.81E-08 | 0.43172874 | 0.35 | 0.28 | 0.00034906 | NK cells | Abr |
| Tnfrsf1a | 1.85E-08 | 0.95378136 | 0.254 | 0.145 | 0.00035819 | NK cells | Tnfrsf1a |
| Apobec3 | 2.17E-08 | 0.43175199 | 0.567 | 0.495 | 0.00041964 | NK cells | Apobec3 |
| mt-Nd5 | 2.50E-08 | 0.25396257 | 0.267 | 0.247 | 0.00048401 | NK cells | mt-Nd5 |
| Psmb101 | 2.96E-08 | 0.79685503 | 0.328 | 0.225 | 0.00057206 | NK cells | Psmb10 |
| Ptk2b | 4.54E-08 | 0.36145806 | 0.609 | 0.529 | 0.00087798 | NK cells | Ptk2b |
| Cmc11 | 5.85E-08 | 1.01365521 | 0.271 | 0.189 | 0.00113148 | NK cells | Cmc1 |
| mt-Nd4l1 | 7.22E-08 | 0.35034522 | 0.406 | 0.352 | 0.00139669 | NK cells | mt-Nd4l |
| Gimap8 | 9.48E-08 | 0.43370666 | 0.592 | 0.504 | 0.00183351 | NK cells | Gimap8 |
| Nptn1 | 1.18E-07 | 0.76503354 | 0.357 | 0.242 | 0.00227845 | NK cells | Nptn |
| Cblb1 | 1.66E-07 | 0.26753638 | 0.511 | 0.439 | 0.00321247 | NK cells | Cblb |
| Znrf22 | 2.21E-07 | 0.43630719 | 0.465 | 0.365 | 0.00426508 | NK cells | Znrf2 |
| Pitpnc11 | 3.21E-07 | 0.5718974 | 0.33 | 0.23 | 0.006197 | NK cells | Pitpnc1 |
| Il18r11 | 3.55E-07 | 0.35793444 | 0.538 | 0.39 | 0.0068596 | NK cells | Il18r1 |
| Supt4a2 | 3.56E-07 | 0.33438902 | 0.46 | 0.396 | 0.00687368 | NK cells | Supt4a |
| Usp3 | 3.61E-07 | 0.54312796 | 0.281 | 0.203 | 0.006988 | NK cells | Usp3 |
| Dnajc31 | 3.61E-07 | 0.42704351 | 0.315 | 0.257 | 0.00698919 | NK cells | Dnajc3 |
| Mapk14 | 3.89E-07 | 0.58759131 | 0.323 | 0.219 | 0.00752512 | NK cells | Mapk14 |
| Snu13 | 6.55E-07 | 0.52425447 | 0.328 | 0.259 | 0.01266585 | NK cells | Snu13 |
| Pdcd6ip | 7.57E-07 | 0.39122419 | 0.509 | 0.431 | 0.01464089 | NK cells | Pdcd6ip |
| Fam49b | 9.22E-07 | 0.40817557 | 0.526 | 0.467 | 0.01782864 | NK cells | Fam49b |
| Il16 | 1.17E-06 | 0.55415717 | 0.418 | 0.323 | 0.02271246 | NK cells | Il16 |
| Upf1 | 1.28E-06 | 0.69603609 | 0.403 | 0.308 | 0.02475816 | NK cells | Upf1 |
| Ndufa101 | 1.28E-06 | 0.50755538 | 0.276 | 0.212 | 0.02479371 | NK cells | Ndufa10 |
| Zfp5981 | 1.29E-06 | 0.60777239 | 0.259 | 0.195 | 0.02490215 | NK cells | Zfp598 |
| Serbp11 | 1.31E-06 | 0.38428888 | 0.462 | 0.386 | 0.02532338 | NK cells | Serbp1 |
| Otulinl | 1.32E-06 | 0.72294325 | 0.35 | 0.239 | 0.02555116 | NK cells | Otulinl |
| Ahnak1 | 1.33E-06 | 0.37352963 | 0.792 | 0.713 | 0.02580954 | NK cells | Ahnak |
| Foxp1 | 1.47E-06 | 0.49357932 | 0.306 | 0.225 | 0.02839826 | NK cells | Foxp1 |
| Capzb1 | 1.58E-06 | 0.39828798 | 0.672 | 0.561 | 0.0306144 | NK cells | Capzb |
| Khdc4 | 1.85E-06 | 0.52600369 | 0.342 | 0.254 | 0.03568001 | NK cells | Khdc4 |
| Ctsc1 | 1.86E-06 | 0.82598845 | 0.293 | 0.206 | 0.03596674 | NK cells | Ctsc |
| Trp53 | 2.41E-06 | 0.6460825 | 0.386 | 0.292 | 0.04655156 | NK cells | Trp53 |
| Fxyd5 | 2.44E-06 | 0.26604009 | 0.685 | 0.602 | 0.04708882 | NK cells | Fxyd5 |
| Kidins220 | 2.50E-06 | 0.5211149 | 0.259 | 0.191 | 0.04841338 | NK cells | Kidins220 |
| Clec2d2 | 3.12E-06 | 0.37208693 | 0.599 | 0.481 | 0.06023115 | NK cells | Clec2d |
| Ybx31 | 3.28E-06 | 0.41218402 | 0.323 | 0.243 | 0.06336932 | NK cells | Ybx3 |
| Unc119b1 | 3.43E-06 | 0.37985562 | 0.252 | 0.193 | 0.06625918 | NK cells | Unc119b |
| Dennd1b1 | 3.90E-06 | 0.549982 | 0.377 | 0.279 | 0.07538982 | NK cells | Dennd1b |
| Ankrd442 | 4.12E-06 | 0.59129513 | 0.379 | 0.291 | 0.079623 | NK cells | Ankrd44 |
| Cux1 | 4.29E-06 | 0.55020792 | 0.274 | 0.21 | 0.08302111 | NK cells | Cux1 |
| Ly9 | 4.84E-06 | 0.8763156 | 0.284 | 0.192 | 0.09349859 | NK cells | Ly9 |
| Gramd1b1 | 6.14E-06 | 0.4719613 | 0.296 | 0.245 | 0.11879391 | NK cells | Gramd1b |
| Hnrnpa11 | 7.01E-06 | 0.35894261 | 0.374 | 0.338 | 0.13555184 | NK cells | Hnrnpa1 |
| Camk2g | 7.42E-06 | 0.71818239 | 0.259 | 0.17 | 0.14337024 | NK cells | Camk2g |
| Spag9 | 7.74E-06 | 0.46064338 | 0.274 | 0.215 | 0.14961457 | NK cells | Spag9 |
| Gmfb | 8.10E-06 | 0.38756619 | 0.291 | 0.223 | 0.15665025 | NK cells | Gmfb |
| Tmed51 | 9.93E-06 | 0.74931885 | 0.262 | 0.196 | 0.19205392 | NK cells | Tmed5 |
| Pnpla2 | 1.04E-05 | 0.63651831 | 0.323 | 0.235 | 0.20023503 | NK cells | Pnpla2 |
| Lman2 | 1.08E-05 | 0.30130251 | 0.35 | 0.324 | 0.20789045 | NK cells | Lman2 |
| Rabep1 | 1.17E-05 | 0.51008716 | 0.257 | 0.183 | 0.22661459 | NK cells | Rabep1 |
| Stat61 | 1.23E-05 | 0.47106908 | 0.408 | 0.332 | 0.23849126 | NK cells | Stat6 |
| Eif3i1 | 1.36E-05 | 0.29756067 | 0.252 | 0.2 | 0.26305589 | NK cells | Eif3i |
| Atp11b | 1.51E-05 | 0.30497994 | 0.284 | 0.204 | 0.29287302 | NK cells | Atp11b |
| Tes1 | 1.76E-05 | 0.32424151 | 0.494 | 0.42 | 0.3395821 | NK cells | Tes |
| Ostf11 | 1.80E-05 | 0.28876036 | 0.447 | 0.405 | 0.34773091 | NK cells | Ostf1 |
| Pfn1 | 1.95E-05 | 0.32264346 | 0.875 | 0.79 | 0.37794199 | NK cells | Pfn1 |
| Arpc1b | 2.09E-05 | 0.31035752 | 0.626 | 0.584 | 0.40359691 | NK cells | Arpc1b |
| Cyfip2 | 2.23E-05 | 0.50810784 | 0.601 | 0.485 | 0.43070431 | NK cells | Cyfip2 |
| Tmed9 | 2.31E-05 | 0.29882631 | 0.328 | 0.281 | 0.44728772 | NK cells | Tmed9 |
| Cope | 2.70E-05 | 0.51292767 | 0.328 | 0.271 | 0.52127838 | NK cells | Cope |
| Pabpc12 | 2.91E-05 | 0.32598996 | 0.462 | 0.4 | 0.56241798 | NK cells | Pabpc1 |
| Tbc1d10c1 | 3.06E-05 | 0.39577221 | 0.633 | 0.535 | 0.5919479 | NK cells | Tbc1d10c |
| 1-Sep | 3.21E-05 | 0.37762312 | 0.775 | 0.687 | 0.62105714 | NK cells | 1-Sep |
| Eml4 | 3.52E-05 | 0.27450775 | 0.301 | 0.228 | 0.68103362 | NK cells | Eml4 |
| Ssr4 | 3.62E-05 | 0.32527322 | 0.291 | 0.253 | 0.70051561 | NK cells | Ssr4 |
| Drap1 | 3.64E-05 | 0.25811915 | 0.416 | 0.374 | 0.70470383 | NK cells | Drap1 |
| Ndufs2 | 3.68E-05 | 0.39035327 | 0.333 | 0.277 | 0.71137924 | NK cells | Ndufs2 |
| Atp6v1b2 | 3.69E-05 | 0.81224973 | 0.279 | 0.194 | 0.7133037 | NK cells | Atp6v1b2 |
| Pde7a1 | 3.77E-05 | 0.30882097 | 0.271 | 0.179 | 0.72947245 | NK cells | Pde7a |
| Ifnar2 | 4.69E-05 | 0.3556188 | 0.269 | 0.204 | 0.90764424 | NK cells | Ifnar2 |
| Tiprl1 | 5.18E-05 | 0.41928312 | 0.281 | 0.233 | 1 | NK cells | Tiprl |
| Tnrc18 | 5.29E-05 | 0.57429409 | 0.279 | 0.211 | 1 | NK cells | Tnrc18 |
| Stxbp2 | 6.15E-05 | 0.46733745 | 0.291 | 0.219 | 1 | NK cells | Stxbp2 |
| Xpo11 | 6.59E-05 | 0.63373129 | 0.274 | 0.203 | 1 | NK cells | Xpo1 |
| Arl6ip5 | 7.32E-05 | 0.39509522 | 0.567 | 0.47 | 1 | NK cells | Arl6ip5 |
| Ahdc1 | 7.60E-05 | 0.43133244 | 0.257 | 0.199 | 1 | NK cells | Ahdc1 |
| Grk62 | 8.04E-05 | 0.37449959 | 0.452 | 0.375 | 1 | NK cells | Grk6 |
| Tardbp | 8.25E-05 | 0.50328186 | 0.347 | 0.272 | 1 | NK cells | Tardbp |
| Pear11 | 8.46E-05 | 0.53524032 | 0.357 | 0.256 | 1 | NK cells | Pear1 |
| Rfc2 | 8.81E-05 | 0.92231329 | 0.259 | 0.173 | 1 | NK cells | Rfc2 |
| Slc44a22 | 8.90E-05 | 0.3081031 | 0.347 | 0.312 | 1 | NK cells | Slc44a2 |
| Eid1 | 8.90E-05 | 0.33261955 | 0.279 | 0.225 | 1 | NK cells | Eid1 |
| Slc35e11 | 0.00010092 | 0.38689262 | 0.271 | 0.222 | 1 | NK cells | Slc35e1 |
| Hadhb | 0.00010117 | 0.62915555 | 0.333 | 0.266 | 1 | NK cells | Hadhb |
| Acly | 0.00011428 | 0.29640719 | 0.455 | 0.387 | 1 | NK cells | Acly |
| Psme3 | 0.00011785 | 0.43026624 | 0.313 | 0.275 | 1 | NK cells | Psme3 |
| Ddx21 | 0.00012038 | 0.32748563 | 0.44 | 0.396 | 1 | NK cells | Ddx21 |
| Glyr11 | 0.00012039 | 0.58593255 | 0.301 | 0.234 | 1 | NK cells | Glyr1 |
| Reep5 | 0.00013953 | 0.41991517 | 0.462 | 0.381 | 1 | NK cells | Reep5 |
| Herc3 | 0.00014874 | 0.36422532 | 0.254 | 0.192 | 1 | NK cells | Herc3 |
| Mrtfa2 | 0.00015271 | 0.35236609 | 0.291 | 0.25 | 1 | NK cells | Mrtfa |
| Babam1 | 0.00015374 | 0.50379658 | 0.262 | 0.206 | 1 | NK cells | Babam1 |
| Tmem189 | 0.00016016 | 0.71206155 | 0.281 | 0.22 | 1 | NK cells | Tmem189 |
| Zgpat2 | 0.00016849 | 0.46572152 | 0.315 | 0.249 | 1 | NK cells | Zgpat |
| Hip1r1 | 0.00017265 | 0.32870394 | 0.34 | 0.284 | 1 | NK cells | Hip1r |
| Ppp2cb | 0.00021911 | 0.32955132 | 0.311 | 0.25 | 1 | NK cells | Ppp2cb |
| Snrpf1 | 0.00022121 | 0.28254011 | 0.364 | 0.315 | 1 | NK cells | Snrpf |
| Bag1 | 0.00023154 | 0.45668168 | 0.374 | 0.309 | 1 | NK cells | Bag1 |
| Rnh1 | 0.00023281 | 0.55909079 | 0.276 | 0.218 | 1 | NK cells | Rnh1 |
| Rp91 | 0.00023295 | 0.35729526 | 0.286 | 0.231 | 1 | NK cells | Rp9 |
| Ddost | 0.00025114 | 0.49349593 | 0.406 | 0.337 | 1 | NK cells | Ddost |
| Canx | 0.00026645 | 0.39482046 | 0.308 | 0.252 | 1 | NK cells | Canx |
| Arhgef32 | 0.00028436 | 0.48295428 | 0.35 | 0.289 | 1 | NK cells | Arhgef3 |
| Adamts10 | 0.00029325 | 0.32942933 | 0.289 | 0.239 | 1 | NK cells | Adamts10 |
| Rapgef11 | 0.00031075 | 0.52924062 | 0.389 | 0.318 | 1 | NK cells | Rapgef1 |
| Tatdn21 | 0.00031297 | 0.28870622 | 0.489 | 0.42 | 1 | NK cells | Tatdn2 |
| Eif3d | 0.00031476 | 0.4065617 | 0.318 | 0.274 | 1 | NK cells | Eif3d |
| Rtcb2 | 0.00035347 | 0.39041924 | 0.333 | 0.281 | 1 | NK cells | Rtcb |
| Cox5b1 | 0.00039273 | 0.26403912 | 0.345 | 0.322 | 1 | NK cells | Cox5b |
| Abhd2 | 0.00039338 | 0.58769917 | 0.406 | 0.306 | 1 | NK cells | Abhd2 |
| Sppl2a | 0.00040967 | 0.49045887 | 0.262 | 0.207 | 1 | NK cells | Sppl2a |
| Add11 | 0.00042628 | 0.37102086 | 0.399 | 0.342 | 1 | NK cells | Add1 |
| Gimap72 | 0.0004683 | 0.491728 | 0.301 | 0.235 | 1 | NK cells | Gimap7 |
| Ndufs8 | 0.00050051 | 0.46871707 | 0.254 | 0.19 | 1 | NK cells | Ndufs8 |
| Ganab | 0.0005042 | 0.36529925 | 0.311 | 0.255 | 1 | NK cells | Ganab |
| Nsmaf1 | 0.00062151 | 0.52957752 | 0.347 | 0.263 | 1 | NK cells | Nsmaf |
| Rab8a1 | 0.00063526 | 0.32642122 | 0.496 | 0.421 | 1 | NK cells | Rab8a |
| Mien1 | 0.00064527 | 0.3979783 | 0.267 | 0.215 | 1 | NK cells | Mien1 |
| Nbeal21 | 0.00064535 | 0.33857035 | 0.406 | 0.344 | 1 | NK cells | Nbeal2 |
| Tsc22d31 | 0.00065821 | 0.39741279 | 0.357 | 0.299 | 1 | NK cells | Tsc22d3 |
| Fgd3 | 0.00072039 | 0.46345786 | 0.267 | 0.212 | 1 | NK cells | Fgd3 |
| Polr2e | 0.00072236 | 0.46900857 | 0.308 | 0.248 | 1 | NK cells | Polr2e |
| Snrpd31 | 0.00075814 | 0.35904394 | 0.262 | 0.231 | 1 | NK cells | Snrpd3 |
| Midn | 0.0007976 | 0.30081115 | 0.479 | 0.41 | 1 | NK cells | Midn |
| Herpud1 | 0.0008166 | 0.58570605 | 0.379 | 0.282 | 1 | NK cells | Herpud1 |
| Prpf4b1 | 0.00084789 | 0.36396161 | 0.355 | 0.287 | 1 | NK cells | Prpf4b |
| Irak1 | 0.00087021 | 0.35247321 | 0.279 | 0.236 | 1 | NK cells | Irak1 |
| Ide1 | 0.0008751 | 0.28804041 | 0.386 | 0.323 | 1 | NK cells | Ide |
| Rnf4 | 0.00088127 | 0.39518319 | 0.374 | 0.311 | 1 | NK cells | Rnf4 |
| Plekhj11 | 0.0009555 | 0.37224668 | 0.325 | 0.275 | 1 | NK cells | Plekhj1 |
| Plp2 | 0.00100873 | 0.38980883 | 0.352 | 0.32 | 1 | NK cells | Plp2 |
| Gtpbp12 | 0.00104888 | 0.33728111 | 0.347 | 0.282 | 1 | NK cells | Gtpbp1 |
| Brk1 | 0.00112252 | 0.46374747 | 0.44 | 0.349 | 1 | NK cells | Brk1 |
| Rrbp1 | 0.00120603 | 0.36259733 | 0.311 | 0.25 | 1 | NK cells | Rrbp1 |
| Pten | 0.00129235 | 0.36269436 | 0.35 | 0.296 | 1 | NK cells | Pten |
| Cox8a1 | 0.00145594 | 0.29999699 | 0.394 | 0.346 | 1 | NK cells | Cox8a |
| Mob3a | 0.00145841 | 0.45677168 | 0.347 | 0.271 | 1 | NK cells | Mob3a |
| Efr3a1 | 0.00149349 | 0.51994159 | 0.257 | 0.19 | 1 | NK cells | Efr3a |
| Nelfb | 0.00150229 | 0.54585853 | 0.252 | 0.185 | 1 | NK cells | Nelfb |
| Nrbp1 | 0.00150675 | 0.4300598 | 0.276 | 0.212 | 1 | NK cells | Nrbp1 |
| Fryl | 0.00154075 | 0.39121782 | 0.425 | 0.355 | 1 | NK cells | Fryl |

| | | | | | | | |
|---|---|---|---|---|---|---|---|
| Rbm26 | 0.00154849 | 0.38231511 | 0.267 | 0.212 | 1 | NK cells | Rbm26 |
| Cox7b | 0.00165582 | 0.36482971 | 0.284 | 0.256 | 1 | NK cells | Cox7b |
| Maea | 0.00173739 | 0.44894284 | 0.279 | 0.232 | 1 | NK cells | Maea |
| Ist1 | 0.00180758 | 0.4102322 | 0.298 | 0.239 | 1 | NK cells | Ist1 |
| Ndel1 | 0.0018669 | 0.38046247 | 0.252 | 0.19 | 1 | NK cells | Ndel1 |
| Tmem258 | 0.00198114 | 0.42248089 | 0.293 | 0.25 | 1 | NK cells | Tmem258 |
| Tram1 | 0.00215346 | 0.30346953 | 0.54 | 0.454 | 1 | NK cells | Tram1 |
| Calm2 | 0.0024765 | 0.42902676 | 0.306 | 0.245 | 1 | NK cells | Calm2 |
| Cnn21 | 0.00250429 | 0.29925791 | 0.697 | 0.651 | 1 | NK cells | Cnn2 |
| Stag2 | 0.00276133 | 0.45039899 | 0.271 | 0.222 | 1 | NK cells | Stag2 |
| Sf3a1 | 0.00319417 | 0.54434898 | 0.252 | 0.199 | 1 | NK cells | Sf3a1 |
| Bcap31 | 0.00320115 | 0.5021411 | 0.359 | 0.289 | 1 | NK cells | Bcap31 |
| Tapbpl1 | 0.00333215 | 0.33907595 | 0.32 | 0.253 | 1 | NK cells | Tapbpl |
| Heca | 0.00342975 | 0.30622198 | 0.531 | 0.448 | 1 | NK cells | Heca |
| Ptbp3 | 0.00358802 | 0.32983845 | 0.606 | 0.54 | 1 | NK cells | Ptbp3 |
| Rasa1 | 0.00379468 | 0.37228688 | 0.271 | 0.217 | 1 | NK cells | Rasa1 |
| Rbbp41 | 0.00388788 | 0.27445999 | 0.372 | 0.334 | 1 | NK cells | Rbbp4 |
| Cd84 | 0.00400701 | 0.3324359 | 0.306 | 0.257 | 1 | NK cells | Cd84 |
| Atp5j2 | 0.004097 | 0.33728151 | 0.374 | 0.312 | 1 | NK cells | Atp5j2 |
| Idh3g | 0.00429123 | 0.28300602 | 0.279 | 0.234 | 1 | NK cells | Idh3g |
| Smad31 | 0.00435895 | 0.46973635 | 0.313 | 0.262 | 1 | NK cells | Smad3 |
| Ptpn2 | 0.00446165 | 0.52901035 | 0.262 | 0.203 | 1 | NK cells | Ptpn2 |
| Map4k41 | 0.00468725 | 0.2712953 | 0.306 | 0.242 | 1 | NK cells | Map4k4 |
| Acsl51 | 0.00481022 | 0.45040611 | 0.281 | 0.228 | 1 | NK cells | Acsl5 |
| Psma4 | 0.00482781 | 0.30456108 | 0.323 | 0.294 | 1 | NK cells | Psma4 |
| Hsp90b1 | 0.00486435 | 0.28820013 | 0.335 | 0.288 | 1 | NK cells | Hsp90b1 |
| Zmat2 | 0.00522357 | 0.33945566 | 0.262 | 0.215 | 1 | NK cells | Zmat2 |
| Tpm4 | 0.00553803 | 0.37085957 | 0.619 | 0.522 | 1 | NK cells | Tpm4 |
| Inpp5b1 | 0.00572523 | 0.48335984 | 0.267 | 0.198 | 1 | NK cells | Inpp5b |
| Ndufb11 | 0.00610947 | 0.34210205 | 0.337 | 0.296 | 1 | NK cells | Ndufb11 |
| Nin1 | 0.00618946 | 0.28391615 | 0.252 | 0.197 | 1 | NK cells | Nin |
| Idh3b | 0.00623889 | 0.31886182 | 0.296 | 0.265 | 1 | NK cells | Idh3b |
| Vav1 | 0.00638959 | 0.44221528 | 0.389 | 0.333 | 1 | NK cells | Vav1 |
| Tnks2 | 0.00681709 | 0.34325995 | 0.318 | 0.252 | 1 | NK cells | Tnks2 |
| Bin11 | 0.00689373 | 0.33911695 | 0.352 | 0.315 | 1 | NK cells | Bin1 |
| Ccar2 | 0.00689843 | 0.37111319 | 0.254 | 0.193 | 1 | NK cells | Ccar2 |
| Psenen | 0.00694671 | 0.3179075 | 0.438 | 0.392 | 1 | NK cells | Psenen |
| Cap1 | 0.00830471 | 0.38778607 | 0.8 | 0.72 | 1 | NK cells | Cap1 |
| Flii | 0.00856782 | 0.49672623 | 0.298 | 0.245 | 1 | NK cells | Flii |
| Tmem1731 | 0.00887331 | 0.26156716 | 0.303 | 0.265 | 1 | NK cells | Tmem173 |
| Tmod3 | 0.00941193 | 0.27274845 | 0.43 | 0.363 | 1 | NK cells | Tmod3 |
| St8sia4 | 0.00964606 | 0.38911292 | 0.347 | 0.295 | 1 | NK cells | St8sia4 |
| Git2 | 0.00969949 | 0.32914761 | 0.394 | 0.321 | 1 | NK cells | Git2 |
| Isyna1 | 0.00981137 | 0.37063318 | 0.254 | 0.212 | 1 | NK cells | Isyna1 |
| Zbtb161 | 0 | 4.4516483 | 0.912 | 0.058 | 0 | innate lymphc | Zbtb16 |
| Lmo41 | 0 | 4.68228324 | 0.824 | 0.074 | 0 | innate lymphc | Lmo4 |
| Pxdc1 | 1.59E-280 | 4.595939 | 0.585 | 0.042 | 3.07E-276 | innate lymphc | Pxdc1 |
| Ramp31 | 5.40E-255 | 2.35325445 | 0.742 | 0.185 | 1.04E-250 | innate lymphc | Ramp3 |
| Tmem176a1 | 7.97E-239 | 3.15056982 | 0.748 | 0.095 | 1.54E-234 | innate lymphc | Tmem176a |
| Sdc11 | 2.83E-233 | 3.51035618 | 0.553 | 0.086 | 5.47E-229 | innate lymphc | Sdc1 |
| Tmem176b1 | 1.12E-209 | 2.95116919 | 0.667 | 0.1 | 2.17E-205 | innate lymphc | Tmem176b |
| Vps37b3 | 2.81E-202 | 1.17743394 | 0.925 | 0.728 | 5.43E-198 | innate lymphc | Vps37b |
| Cd40lg1 | 2.55E-170 | 2.89443243 | 0.503 | 0.079 | 4.94E-166 | innate lymphc | Cd40lg |
| Rorc1 | 1.02E-151 | 2.68492307 | 0.616 | 0.104 | 1.98E-147 | innate lymphc | Rorc |
| Lgals31 | 2.66E-149 | 2.27429848 | 0.535 | 0.207 | 5.15E-145 | innate lymphc | Lgals3 |
| Ramp11 | 5.57E-130 | 2.55151948 | 0.66 | 0.183 | 1.08E-125 | innate lymphc | Ramp1 |
| Il17rb | 5.46E-127 | 4.42760703 | 0.346 | 0.03 | 1.06E-122 | innate lymphc | Il17rb |
| Il23r1 | 2.73E-122 | 3.45348642 | 0.503 | 0.054 | 5.29E-118 | innate lymphc | Il23r |
| Il17re1 | 5.57E-121 | 4.1620829 | 0.377 | 0.024 | 1.08E-116 | innate lymphc | Il17re |
| Il18r12 | 2.55E-118 | 1.60050137 | 0.836 | 0.388 | 4.94E-114 | innate lymphc | Il18r1 |
| Serpinb1a1 | 3.55E-115 | 2.84248651 | 0.459 | 0.065 | 6.86E-111 | innate lymphc | Serpinb1a |
| Igf1r1 | 2.60E-107 | 3.0879963 | 0.497 | 0.066 | 5.02E-103 | innate lymphc | Igf1r |
| Mmp251 | 1.68E-104 | 2.68189022 | 0.547 | 0.081 | 3.25E-100 | innate lymphc | Mmp25 |
| Il7r2 | 1.03E-100 | 1.58793051 | 0.818 | 0.377 | 1.99E-96 | innate lymphc | Il7r |
| Ret | 1.64E-100 | 4.58046262 | 0.296 | 0.018 | 3.17E-96 | innate lymphc | Ret |
| Sptssa | 1.65E-99 | 2.2802698 | 0.591 | 0.212 | 3.18E-95 | innate lymphc | Sptssa |
| Selenop1 | 3.62E-84 | 1.74550011 | 0.679 | 0.291 | 6.99E-80 | innate lymphc | Selenop |
| Tmem642 | 6.16E-84 | 2.05319076 | 0.654 | 0.21 | 1.19E-79 | innate lymphc | Tmem64 |
| Cxcr62 | 4.45E-80 | 1.78057317 | 0.742 | 0.286 | 8.60E-76 | innate lymphc | Cxcr6 |
| Il1r11 | 1.92E-79 | 2.76542547 | 0.447 | 0.055 | 3.71E-75 | innate lymphc | Il1r1 |
| Rgcc | 7.99E-76 | 1.94705552 | 0.472 | 0.195 | 1.54E-71 | innate lymphc | Rgcc |
| Lingo4 | 3.78E-75 | 4.24040047 | 0.27 | 0.013 | 7.30E-71 | innate lymphc | Lingo4 |
| Rnf144a | 1.24E-74 | 3.27046217 | 0.352 | 0.041 | 2.40E-70 | innate lymphc | Rnf144a |
| Aqp31 | 5.94E-72 | 2.65997605 | 0.447 | 0.058 | 1.15E-67 | innate lymphc | Aqp3 |
| Smox1 | 9.85E-66 | 1.93993515 | 0.585 | 0.193 | 1.90E-61 | innate lymphc | Smox |
| Blk1 | 3.28E-64 | 2.41233686 | 0.39 | 0.044 | 6.35E-60 | innate lymphc | Blk |
| Ly6g5b1 | 5.84E-64 | 2.86187505 | 0.346 | 0.064 | 1.13E-59 | innate lymphc | Ly6g5b |
| Emb1 | 3.68E-63 | 1.04321169 | 0.899 | 0.579 | 7.12E-59 | innate lymphc | Emb |
| Adgrg52 | 8.36E-63 | 2.24068791 | 0.566 | 0.122 | 1.62E-58 | innate lymphc | Adgrg5 |
| F2r2 | 4.80E-56 | 1.68838263 | 0.635 | 0.277 | 9.28E-52 | innate lymphc | F2r |
| Btg24 | 1.89E-52 | 0.86921584 | 0.824 | 0.616 | 3.66E-48 | innate lymphc | Btg2 |
| Klf4 | 1.79E-51 | 2.22115969 | 0.302 | 0.104 | 3.46E-47 | innate lymphc | Klf4 |
| Cwc25 | 1.56E-50 | 2.13825526 | 0.39 | 0.149 | 3.02E-46 | innate lymphc | Cwc25 |
| Znrf12 | 1.36E-48 | 1.25002241 | 0.736 | 0.388 | 2.62E-44 | innate lymphc | Znrf1 |
| Jmy1 | 4.93E-48 | 1.42443443 | 0.547 | 0.282 | 9.53E-44 | innate lymphc | Jmy |
| Gclc | 6.09E-48 | 2.32042491 | 0.396 | 0.114 | 1.18E-43 | innate lymphc | Gclc |
| mt-Atp63 | 1.22E-47 | 1.05809371 | 0.824 | 0.546 | 2.36E-43 | innate lymphc | mt-Atp6 |
| Furin2 | 1.45E-46 | 1.17968278 | 0.73 | 0.446 | 2.81E-42 | innate lymphc | Furin |
| Qpct | 1.22E-44 | 3.42787635 | 0.264 | 0.027 | 2.35E-40 | innate lymphc | Qpct |
| Icos1 | 4.88E-44 | 0.95436192 | 0.855 | 0.619 | 9.44E-40 | innate lymphc | Icos |
| Mgat51 | 1.32E-42 | 1.68246291 | 0.597 | 0.22 | 2.54E-38 | innate lymphc | Mgat5 |
| Prkd3 | 8.04E-42 | 2.1252052 | 0.371 | 0.105 | 1.55E-37 | innate lymphc | Prkd3 |
| Cdk11b1 | 1.30E-41 | 1.41433309 | 0.484 | 0.279 | 2.52E-37 | innate lymphc | Cdk11b |
| Fosl21 | 2.33E-40 | 0.75385202 | 0.78 | 0.644 | 4.50E-36 | innate lymphc | Fosl2 |
| Tcf73 | 2.89E-40 | 1.3508758 | 0.509 | 0.17 | 5.59E-36 | innate lymphc | Tcf7 |
| Sep-81 | 3.71E-40 | 2.45790709 | 0.34 | 0.08 | 7.16E-36 | innate lymphc | 8-Sep |
| mt-Atp82 | 1.96E-39 | 1.02152598 | 0.792 | 0.515 | 3.79E-35 | innate lymphc | mt-Atp8 |
| Fos | 2.99E-38 | 0.95055931 | 0.478 | 0.33 | 5.78E-34 | innate lymphc | Fos |
| Cd1641 | 1.28E-36 | 1.18060315 | 0.792 | 0.556 | 2.48E-32 | innate lymphc | Cd164 |
| Gabbr11 | 2.75E-34 | 1.22222593 | 0.522 | 0.242 | 5.32E-30 | innate lymphc | Gabbr1 |
| mt-Co21 | 9.36E-34 | 0.76251527 | 0.899 | 0.679 | 1.81E-29 | innate lymphc | mt-Co2 |
| Tex2 | 1.43E-32 | 1.16783982 | 0.742 | 0.43 | 2.77E-28 | innate lymphc | Tex2 |
| Prrt1 | 6.19E-32 | 2.64960445 | 0.258 | 0.043 | 1.20E-27 | innate lymphc | Prrt1 |
| Kctd121 | 1.52E-31 | 0.97519365 | 0.503 | 0.278 | 2.94E-27 | innate lymphc | Kctd12 |
| Maff1 | 2.03E-31 | 1.74274874 | 0.302 | 0.121 | 3.93E-27 | innate lymphc | Maff |
| Evl2 | 4.62E-31 | 1.1925077 | 0.654 | 0.23 | 8.93E-27 | innate lymphc | Evl |
| Mtss1 | 1.90E-30 | 2.26234216 | 0.258 | 0.045 | 3.67E-26 | innate lymphc | Mtss1 |
| Crem1 | 6.63E-29 | 0.62805085 | 0.78 | 0.72 | 1.28E-24 | innate lymphc | Crem |
| St3gal6 | 1.67E-27 | 2.35903857 | 0.296 | 0.055 | 3.23E-23 | innate lymphc | St3gal6 |
| Pde3b3 | 7.29E-27 | 1.42107431 | 0.484 | 0.181 | 1.41E-22 | innate lymphc | Pde3b |
| Map3k142 | 9.94E-27 | 1.25855601 | 0.409 | 0.194 | 1.92E-22 | innate lymphc | Map3k14 |
| Ikzf31 | 2.00E-25 | 0.96166984 | 0.843 | 0.541 | 3.87E-21 | innate lymphc | Ikzf3 |
| Cxcr4 | 2.59E-24 | 0.61947241 | 0.748 | 0.558 | 5.01E-20 | innate lymphc | Cxcr4 |
| Bcl2a1b2 | 5.31E-24 | 1.19816813 | 0.572 | 0.329 | 1.03E-19 | innate lymphc | Bcl2a1b |
| mt-Cytb2 | 8.09E-23 | 0.96371022 | 0.66 | 0.397 | 1.56E-18 | innate lymphc | mt-Cytb |
| Rab4a | 1.30E-22 | 1.89447026 | 0.283 | 0.079 | 2.51E-18 | innate lymphc | Rab4a |
| Tnfaip31 | 2.19E-22 | 0.46368357 | 0.767 | 0.747 | 4.24E-18 | innate lymphc | Tnfaip3 |
| Clint11 | 2.22E-22 | 1.03968652 | 0.66 | 0.408 | 4.29E-18 | innate lymphc | Clint1 |
| Irak22 | 4.41E-22 | 1.30672879 | 0.44 | 0.234 | 8.52E-18 | innate lymphc | Irak2 |
| Irf2bp21 | 4.93E-22 | 0.81086039 | 0.686 | 0.484 | 9.53E-18 | innate lymphc | Irf2bp2 |
| mt-Nd4l2 | 5.03E-22 | 1.08061031 | 0.597 | 0.349 | 9.73E-18 | innate lymphc | mt-Nd4l |
| Pi4k2a1 | 5.32E-22 | 1.83061682 | 0.358 | 0.118 | 1.03E-17 | innate lymphc | Pi4k2a |
| Tbl1x | 6.38E-22 | 2.12553744 | 0.365 | 0.112 | 1.23E-17 | innate lymphc | Tbl1x |
| Tiparp | 1.24E-20 | 1.53758667 | 0.308 | 0.137 | 2.40E-16 | innate lymphc | Tiparp |
| Abi31 | 1.74E-20 | 1.66903605 | 0.346 | 0.133 | 3.36E-16 | innate lymphc | Abi3 |
| Man2a21 | 7.56E-20 | 1.31392889 | 0.497 | 0.244 | 1.46E-15 | innate lymphc | Man2a2 |
| Notch21 | 1.18E-19 | 1.46850857 | 0.377 | 0.167 | 2.28E-15 | innate lymphc | Notch2 |
| Prdx61 | 1.88E-19 | 1.63456866 | 0.39 | 0.151 | 3.63E-15 | innate lymphc | Prdx6 |
| Slc16a61 | 2.40E-19 | 1.67586051 | 0.34 | 0.159 | 4.64E-15 | innate lymphc | Slc16a6 |
| Itk2 | 2.78E-19 | 0.53026787 | 0.843 | 0.709 | 5.38E-15 | innate lymphc | Itk |
| mt-Co12 | 3.90E-19 | 0.81733868 | 0.717 | 0.473 | 7.53E-15 | innate lymphc | mt-Co1 |
| Ero1l1 | 6.53E-19 | 1.2039727 | 0.516 | 0.277 | 1.26E-14 | innate lymphc | Ero1l |
| Thy11 | 2.24E-18 | 0.77560724 | 0.912 | 0.674 | 4.32E-14 | innate lymphc | Thy1 |
| mt-Co33 | 2.37E-18 | 0.87405889 | 0.673 | 0.467 | 4.58E-14 | innate lymphc | mt-Co3 |
| Prkx1 | 2.55E-17 | 1.42692952 | 0.39 | 0.183 | 4.92E-13 | innate lymphc | Prkx |
| Irs21 | 3.63E-17 | 1.11657439 | 0.377 | 0.186 | 7.01E-13 | innate lymphc | Irs2 |
| S1pr12 | 3.97E-17 | 0.79260239 | 0.377 | 0.188 | 7.67E-13 | innate lymphc | S1pr1 |
| Rexo22 | 5.03E-17 | 1.26772715 | 0.459 | 0.237 | 9.72E-13 | innate lymphc | Rexo2 |
| Zfp6521 | 6.97E-17 | 1.24814788 | 0.333 | 0.154 | 1.35E-12 | innate lymphc | Zfp652 |
| Acat1 | 1.26E-16 | 2.1297037 | 0.333 | 0.104 | 2.44E-12 | innate lymphc | Acat1 |
| Elmsan1 | 1.32E-16 | 1.18549988 | 0.321 | 0.162 | 2.55E-12 | innate lymphc | Elmsan1 |
| Rarg1 | 1.57E-16 | 1.31103007 | 0.384 | 0.175 | 3.04E-12 | innate lymphc | Rarg |
| Psap1 | 2.16E-15 | 1.19496913 | 0.547 | 0.306 | 4.18E-11 | innate lymphc | Psap |
| Atp8b41 | 2.58E-15 | 1.51128043 | 0.296 | 0.103 | 4.99E-11 | innate lymphc | Atp8b4 |
| Jaml1 | 4.10E-15 | 1.44320228 | 0.296 | 0.097 | 7.94E-11 | innate lymphc | Jaml |
| Galnt12 | 6.75E-15 | 1.60252544 | 0.365 | 0.13 | 1.31E-10 | innate lymphc | Galnt12 |
| Dennd4c | 1.03E-14 | 1.48433473 | 0.302 | 0.122 | 1.99E-10 | innate lymphc | Dennd4c |
| Lgals3bp2 | 1.17E-14 | 1.10789227 | 0.497 | 0.237 | 2.27E-10 | innate lymphc | Lgals3bp |
| Tatdn22 | 1.45E-14 | 0.99621657 | 0.648 | 0.418 | 2.81E-10 | innate lymphc | Tatdn2 |
| Slco3a11 | 1.77E-14 | 1.23972102 | 0.428 | 0.211 | 3.42E-10 | innate lymphc | Slco3a1 |
| mt-Nd61 | 3.44E-14 | 0.76155213 | 0.604 | 0.421 | 6.65E-10 | innate lymphc | mt-Nd6 |
| Pgm2l1 | 3.60E-14 | 1.30752233 | 0.277 | 0.117 | 6.96E-10 | innate lymphc | Pgm2l1 |
| Arrb11 | 9.57E-14 | 1.3613768 | 0.384 | 0.179 | 1.85E-09 | innate lymphc | Arrb1 |
| Epas1 | 1.15E-13 | 1.32489137 | 0.352 | 0.15 | 2.22E-09 | innate lymphc | Epas1 |
| St6galnac31 | 1.51E-13 | 1.41289596 | 0.314 | 0.115 | 2.92E-09 | innate lymphc | St6galnac3 |
| Slc12a72 | 1.58E-13 | 1.19730155 | 0.314 | 0.155 | 3.06E-09 | innate lymphc | Slc12a7 |
| Gpr1321 | 1.59E-13 | 0.70190748 | 0.736 | 0.595 | 3.08E-09 | innate lymphc | Gpr132 |
| Elf4e31 | 2.16E-13 | 1.40765652 | 0.314 | 0.15 | 4.17E-09 | innate lymphc | Elf4e3 |
| D16Ertd472e | 2.53E-13 | 0.90437573 | 0.604 | 0.422 | 4.90E-09 | innate lymphc | D16Ertd472e |
| Cd961 | 3.85E-13 | 1.35359839 | 0.358 | 0.149 | 7.45E-09 | innate lymphc | Cd96 |
| Emp1 | 4.97E-13 | 1.47008014 | 0.277 | 0.099 | 9.62E-09 | innate lymphc | Emp1 |
| Ahcyl21 | 1.19E-12 | 1.36339005 | 0.314 | 0.17 | 2.30E-08 | innate lymphc | Ahcyl2 |
| Usp36 | 2.72E-12 | 1.33656757 | 0.277 | 0.123 | 5.26E-08 | innate lymphc | Usp36 |
| Ncor11 | 4.85E-12 | 0.82762689 | 0.642 | 0.454 | 9.37E-08 | innate lymphc | Ncor1 |
| mt-Nd2 | 1.40E-11 | 1.21187458 | 0.277 | 0.13 | 2.71E-07 | innate lymphc | mt-Nd2 |
| 1-Jun | 1.75E-11 | 0.62733372 | 0.465 | 0.341 | 3.39E-07 | innate lymphc | Jun |
| Klf31 | 1.77E-11 | 0.77594688 | 0.289 | 0.132 | 3.43E-07 | innate lymphc | Klf3 |
| Galnt11 | 1.78E-11 | 1.01050372 | 0.604 | 0.351 | 3.44E-07 | innate lymphc | Galnt1 |
| Smpdl3a1 | 1.84E-11 | 1.20981889 | 0.371 | 0.182 | 3.57E-07 | innate lymphc | Smpdl3a |
| Ppp1r3b | 1.95E-11 | 1.53756588 | 0.289 | 0.121 | 3.76E-07 | innate lymphc | Ppp1r3b |
| Rnf71 | 2.59E-11 | 1.16425872 | 0.314 | 0.166 | 5.01E-07 | innate lymphc | Rnf7 |
| Tnfrsf251 | 2.60E-11 | 1.24251749 | 0.314 | 0.128 | 5.02E-07 | innate lymphc | Tnfrsf25 |
| Skap1 | 3.68E-11 | 0.69086713 | 0.497 | 0.342 | 7.12E-07 | innate lymphc | Skap1 |
| Pde4b1 | 3.70E-11 | 0.95276471 | 0.459 | 0.295 | 7.16E-07 | innate lymphc | Pde4b |
| Gpx12 | 6.73E-11 | 0.96305407 | 0.579 | 0.404 | 1.30E-06 | innate lymphc | Gpx1 |
| Junb1 | 9.02E-11 | 0.26491364 | 0.931 | 0.895 | 1.74E-06 | innate lymphc | Junb |
| Dctn11 | 1.72E-10 | 1.32448584 | 0.377 | 0.194 | 3.32E-06 | innate lymphc | Dctn1 |
| Fam102a3 | 2.41E-10 | 0.75953095 | 0.61 | 0.398 | 4.65E-06 | innate lymphc | Fam102a |
| Gpr1831 | 2.58E-10 | 1.04703583 | 0.541 | 0.29 | 4.99E-06 | innate lymphc | Gpr183 |
| Rasa33 | 2.85E-10 | 0.6624796 | 0.478 | 0.263 | 5.52E-06 | innate lymphc | Rasa3 |
| Prr7 | 2.87E-10 | 1.00192773 | 0.308 | 0.162 | 5.55E-06 | innate lymphc | Prr7 |
| Cd2262 | 3.54E-10 | 1.13265613 | 0.302 | 0.126 | 6.84E-06 | innate lymphc | Cd226 |
| Hspa1b3 | 5.02E-10 | 0.55387819 | 0.333 | 0.215 | 9.70E-06 | innate lymphc | Hspa1b |
| Camk1d | 5.43E-10 | 1.24348249 | 0.346 | 0.162 | 1.05E-05 | innate lymphc | Camk1d |
| Dgat11 | 6.57E-10 | 0.90514925 | 0.535 | 0.357 | 1.27E-05 | innate lymphc | Dgat1 |
| Top2b2 | 7.77E-10 | 0.95714077 | 0.428 | 0.279 | 1.50E-05 | innate lymphc | Top2b |
| Fosb2 | 1.11E-09 | 0.5945525 | 0.403 | 0.323 | 2.15E-05 | innate lymphc | Fosb |
| Slc37a31 | 1.59E-09 | 1.00653082 | 0.302 | 0.157 | 3.08E-05 | innate lymphc | Slc37a3 |
| Cited2 | 5.87E-09 | 0.94243665 | 0.333 | 0.254 | 0.00011341 | innate lymphc | Cited2 |
| Satb12 | 6.14E-09 | 0.69542919 | 0.478 | 0.315 | 0.00011865 | innate lymphc | Satb1 |
| Slc9a9 | 6.43E-09 | 1.08627636 | 0.314 | 0.157 | 0.00012432 | innate lymphc | Slc9a9 |
| Acsl4 | 9.78E-09 | 1.07916509 | 0.434 | 0.245 | 0.00018914 | innate lymphc | Acsl4 |
| Epb412 | 1.06E-08 | 0.78570089 | 0.365 | 0.238 | 0.00020463 | innate lymphc | Epb41 |
| Aak11 | 1.27E-08 | 0.83637276 | 0.403 | 0.239 | 0.00024602 | innate lymphc | Aak1 |
| Sept112 | 1.41E-08 | 0.87853434 | 0.44 | 0.291 | 0.00027334 | innate lymphc | 11-Sep |
| mt-Nd3 | 2.14E-08 | 1.33353541 | 0.296 | 0.138 | 0.00041381 | innate lymphc | mt-Nd3 |
| Mapk141 | 4.58E-08 | 0.95135155 | 0.409 | 0.221 | 0.00088498 | innate lymphc | Mapk14 |
| mt-Nd4 | 5.22E-08 | 0.84741176 | 0.302 | 0.167 | 0.00101008 | innate lymphc | mt-Nd4 |
| Eprs | 5.54E-08 | 1.14650802 | 0.346 | 0.196 | 0.00107033 | innate lymphc | Eprs |
| Ltb4r11 | 6.20E-08 | 1.3256798 | 0.302 | 0.128 | 0.00119798 | innate lymphc | Ltb4r1 |
| Ctdsp2 | 9.26E-08 | 0.75574467 | 0.484 | 0.343 | 0.00179115 | innate lymphc | Ctdsp2 |
| Gch11 | 9.69E-08 | 0.85764569 | 0.314 | 0.2 | 0.00187261 | innate lymphc | Gch1 |
| Smap21 | 9.79E-08 | 0.79058841 | 0.654 | 0.454 | 0.00189275 | innate lymphc | Smap2 |
| mt-Nd12 | 1.07E-07 | 0.7707419 | 0.503 | 0.31 | 0.00207014 | innate lymphc | mt-Nd1 |
| Nt5e2 | 1.19E-07 | 0.7861076 | 0.39 | 0.249 | 0.00229998 | innate lymphc | Nt5e |
| Lax13 | 1.34E-07 | 0.77225274 | 0.358 | 0.205 | 0.00258577 | innate lymphc | Lax1 |
| Cd841 | 1.80E-07 | 0.78711961 | 0.409 | 0.256 | 0.00348611 | innate lymphc | Cd84 |
| Tspan141 | 2.73E-07 | 0.9978352 | 0.358 | 0.194 | 0.00528737 | innate lymphc | Tspan14 |
| Itgal3 | 3.19E-07 | 0.50877372 | 0.849 | 0.694 | 0.00616569 | innate lymphc | Itgal |
| Bzw2 | 3.72E-07 | 1.09794218 | 0.352 | 0.198 | 0.00719252 | innate lymphc | Bzw2 |
| Ly6e1 | 4.33E-07 | 0.57622398 | 0.975 | 0.846 | 0.00838022 | innate lymphc | Ly6e |
| Arhgef182 | 4.77E-07 | 0.45072258 | 0.509 | 0.377 | 0.00922142 | innate lymphc | Arhgef18 |
| Kmt2d | 4.83E-07 | 0.52878878 | 0.667 | 0.497 | 0.00933674 | innate lymphc | Kmt2d |
| Tor1aip11 | 4.95E-07 | 0.76008617 | 0.56 | 0.395 | 0.00956688 | innate lymphc | Tor1aip1 |
| Srpk1 | 5.95E-07 | 0.64311154 | 0.585 | 0.443 | 0.01150296 | innate lymphc | Srpk1 |
| Atn1 | 7.63E-07 | 0.72751062 | 0.415 | 0.267 | 0.01475671 | innate lymphc | Atn1 |
| Sun21 | 8.50E-07 | 0.6338513 | 0.616 | 0.4 | 0.01643213 | innate lymphc | Sun2 |
| mt-Nd51 | 9.41E-07 | 0.75446735 | 0.365 | 0.245 | 0.0181967 | innate lymphc | mt-Nd5 |
| Arl5b1 | 1.17E-06 | 0.81879606 | 0.371 | 0.253 | 0.02259885 | innate lymphc | Arl5b |
| Sep-63 | 1.20E-06 | 0.83786649 | 0.535 | 0.323 | 0.02326112 | innate lymphc | 6-Sep |
| Thada | 1.40E-06 | 0.93626406 | 0.258 | 0.136 | 0.02698919 | innate lymphc | Thada |
| Csrnp11 | 1.41E-06 | 0.45419968 | 0.535 | 0.457 | 0.02718858 | innate lymphc | Csrnp1 |
| Dock22 | 1.42E-06 | 0.46991308 | 0.792 | 0.679 | 0.02745172 | innate lymphc | Dock2 |
| Arpc3 | 1.49E-06 | 0.86017524 | 0.572 | 0.393 | 0.02887343 | innate lymphc | Arpc3 |
| Wbp1l | 1.67E-06 | 0.61941765 | 0.371 | 0.261 | 0.03223864 | innate lymphc | Wbp1l |
| Pfkp | 1.70E-06 | 0.73065305 | 0.535 | 0.356 | 0.03293447 | innate lymphc | Pfkp |
| Parp11 | 1.76E-06 | 1.32484632 | 0.314 | 0.172 | 0.03397678 | innate lymphc | Parp1 |
| Per11 | 2.78E-06 | 0.49789534 | 0.654 | 0.522 | 0.05377752 | innate lymphc | Per1 |
| Laptm52 | 3.57E-06 | 0.4144917 | 0.975 | 0.89 | 0.06907989 | innate lymphc | Laptm5 |
| Sema4a3 | 4.77E-06 | 0.55602549 | 0.409 | 0.243 | 0.09226287 | innate lymphc | Sema4a |
| Med13l | 4.92E-06 | 0.66570226 | 0.358 | 0.231 | 0.09520153 | innate lymphc | Med13l |
| Smad32 | 4.97E-06 | 0.59328967 | 0.403 | 0.262 | 0.0961773 | innate lymphc | Smad3 |
| Arhgap262 | 6.16E-06 | 0.71302783 | 0.321 | 0.167 | 0.11915276 | innate lymphc | Arhgap26 |
| Fryl1 | 6.47E-06 | 0.51470396 | 0.491 | 0.356 | 0.12499568 | innate lymphc | Fryl |
| Lsp12 | 6.99E-06 | 0.61691788 | 0.748 | 0.579 | 0.13510608 | innate lymphc | Lsp1 |
| Cept1 | 7.64E-06 | 1.03027099 | 0.258 | 0.162 | 0.14767911 | innate lymphc | Cept1 |
| Hdhd53 | 7.74E-06 | 0.8134764 | 0.314 | 0.198 | 0.14959574 | innate lymphc | Hdhd5 |
| Mppe1 | 9.00E-06 | 0.99551671 | 0.252 | 0.132 | 0.1739743 | innate lymphc | Mppe1 |
| Atf61 | 9.92E-06 | 0.96020278 | 0.314 | 0.164 | 0.19178436 | innate lymphc | Atf6 |
| Slc38a1 | 1.16E-05 | 0.87219785 | 0.465 | 0.292 | 0.22390408 | innate lymphc | Slc38a1 |
| Itpr21 | 1.17E-05 | 0.64517543 | 0.346 | 0.234 | 0.22643537 | innate lymphc | Itpr2 |
| Cnot61 | 1.20E-05 | 0.95042505 | 0.358 | 0.212 | 0.23156935 | innate lymphc | Cnot6 |
| Gna131 | 1.26E-05 | 0.36465351 | 0.78 | 0.701 | 0.24434127 | innate lymphc | Gna13 |
| Pi4ka | 1.36E-05 | 0.80685978 | 0.327 | 0.218 | 0.26300366 | innate lymphc | Pi4ka |
| Atp2b12 | 1.62E-05 | 0.56670643 | 0.491 | 0.351 | 0.31385597 | innate lymphc | Atp2b1 |
| Cstb | 2.80E-05 | 0.92178249 | 0.314 | 0.231 | 0.54157934 | innate lymphc | Cstb |
| Fnbp1 | 2.94E-05 | 0.43057167 | 0.792 | 0.668 | 0.56800728 | innate lymphc | Fnbp1 |
| Mib1 | 3.13E-05 | 0.8749655 | 0.302 | 0.191 | 0.60437833 | innate lymphc | Mib1 |
| Stk17b1 | 3.17E-05 | 0.33520904 | 0.774 | 0.69 | 0.61307759 | innate lymphc | Stk17b |
| Fchsd21 | 3.21E-05 | 0.79808982 | 0.283 | 0.187 | 0.61990303 | innate lymphc | Fchsd2 |
| Rassf11 | 3.83E-05 | 0.99831282 | 0.314 | 0.197 | 0.74097553 | innate lymphc | Rassf1 |
| Fam129a1 | 4.36E-05 | 0.65890167 | 0.61 | 0.424 | 0.84392669 | innate lymphc | Fam129a |
| Il12rb12 | 4.38E-05 | 1.02130829 | 0.346 | 0.222 | 0.84617734 | innate lymphc | Il12rb1 |
| Maml1 | 5.25E-05 | 0.92584304 | 0.321 | 0.2 | 1 | innate lymphc | Maml1 |
| Nop531 | 6.77E-05 | 0.49386154 | 0.522 | 0.504 | 1 | innate lymphc | Nop53 |
| Rnf1663 | 7.01E-05 | 0.70934804 | 0.447 | 0.304 | 1 | innate lymphc | Rnf166 |
| Orai2 | 7.49E-05 | 0.72805248 | 0.277 | 0.166 | 1 | innate lymphc | Orai2 |
| Dusp12 | 8.70E-05 | 0.33849936 | 0.755 | 0.657 | 1 | innate lymphc | Dusp1 |
| Diaph1 | 9.69E-05 | 0.59262821 | 0.572 | 0.433 | 1 | innate lymphc | Diaph1 |
| Uhrf1bp1l | 9.75E-05 | 0.81461686 | 0.283 | 0.16 | 1 | innate lymphc | Uhrf1bp1l |
| Gtf2i1 | 0.00010111 | 0.80603256 | 0.384 | 0.236 | 1 | innate lymphc | Gtf2i |
| Ndel11 | 0.00010188 | 0.79400595 | 0.264 | 0.193 | 1 | innate lymphc | Ndel1 |
| Xlr4c | 0.00010316 | 0.75126541 | 0.264 | 0.153 | 1 | innate lymphc | Xlr4c |
| Aff4 | 0.00011984 | 0.53022168 | 0.403 | 0.262 | 1 | innate lymphc | Aff4 |
| Inpp5b2 | 0.00012256 | 0.78397726 | 0.283 | 0.2 | 1 | innate lymphc | Inpp5b |
| Sytl1 | 0.00017621 | 0.94174868 | 0.258 | 0.154 | 1 | innate lymphc | Sytl1 |
| Fam168b | 0.00017843 | 0.611606 | 0.509 | 0.38 | 1 | innate lymphc | Fam168b |
| Prrc2c | 0.00017979 | 0.55355727 | 0.66 | 0.539 | 1 | innate lymphc | Prrc2c |
| Elf1 | 0.00019205 | 0.58670556 | 0.686 | 0.516 | 1 | innate lymphc | Elf1 |
| Wac | 0.00023044 | 0.89882683 | 0.314 | 0.226 | 1 | innate lymphc | Wac |
| Rab19 | 0.00023202 | 0.92202028 | 0.27 | 0.143 | 1 | innate lymphc | Rab19 |
| Zfp445 | 0.00026253 | 0.85247945 | 0.358 | 0.202 | 1 | innate lymphc | Zfp445 |
| Macf11 | 0.00027302 | 0.30898553 | 0.635 | 0.538 | 1 | innate lymphc | Macf1 |
| 12-Sep | 0.00027895 | 0.36416935 | 0.799 | 0.691 | 1 | innate lymphc | 1-Sep |
| Chmp1b | 0.00028003 | 0.81973101 | 0.415 | 0.297 | 1 | innate lymphc | Chmp1b |
| Ahr | 0.00030049 | 1.06392734 | 0.258 | 0.128 | 1 | innate lymphc | Ahr |
| Rb1 | 0.00030472 | 0.7727586 | 0.27 | 0.163 | 1 | innate lymphc | Rb1 |
| Dnajb61 | 0.0003746 | 0.58511557 | 0.428 | 0.32 | 1 | innate lymphc | Dnajb6 |
| Rai1 | 0.0003771 | 0.53809053 | 0.321 | 0.213 | 1 | innate lymphc | Rai1 |
| Tle4 | 0.00039272 | 0.98488705 | 0.264 | 0.149 | 1 | innate lymphc | Tle4 |
| Tob21 | 0.00041097 | 0.41561484 | 0.415 | 0.351 | 1 | innate lymphc | Tob2 |
| Eef1a13 | 0.00043285 | 0.40951102 | 1 | 0.943 | 1 | innate lymphc | Eef1a1 |
| Mycbp21 | 0.00044039 | 0.5440698 | 0.453 | 0.32 | 1 | innate lymphc | Mycbp2 |
| Pik3cd4 | 0.00046437 | 0.47809968 | 0.673 | 0.443 | 1 | innate lymphc | Pik3cd |
| Rnf145 | 0.00047038 | 0.66926593 | 0.371 | 0.248 | 1 | innate lymphc | Rnf145 |
| Rin31 | 0.00049254 | 0.74085973 | 0.27 | 0.167 | 1 | innate lymphc | Rin3 |
| Runx11 | 0.00052557 | 0.76188211 | 0.314 | 0.207 | 1 | innate lymphc | Runx1 |
| Aff12 | 0.00060111 | 0.31297858 | 0.377 | 0.354 | 1 | innate lymphc | Aff1 |
| Coq10b1 | 0.00063847 | 0.6555622 | 0.434 | 0.293 | 1 | innate lymphc | Coq10b |
| Utrn | 0.00067391 | 0.39853478 | 0.34 | 0.253 | 1 | innate lymphc | Utrn |
| Selplg3 | 0.00067804 | 0.48471424 | 0.818 | 0.642 | 1 | innate lymphc | Selplg |
| Tfip11 | 0.0007249 | 0.77144804 | 0.258 | 0.172 | 1 | innate lymphc | Tfip11 |
| Arid5a2 | 0.00077439 | 0.4305111 | 0.283 | 0.186 | 1 | innate lymphc | Arid5a |
| Grk63 | 0.00077482 | 0.5816935 | 0.553 | 0.376 | 1 | innate lymphc | Grk6 |
| Pkn1 | 0.00080221 | 0.44402689 | 0.648 | 0.506 | 1 | innate lymphc | Pkn1 |
| Slc25a241 | 0.00086018 | 0.78373249 | 0.277 | 0.182 | 1 | innate lymphc | Slc25a24 |
| Cpm1 | 0.00091908 | 0.6151224 | 0.277 | 0.176 | 1 | innate lymphc | Cpm |
| Rora1 | 0.00092892 | 0.47196129 | 0.736 | 0.514 | 1 | innate lymphc | Rora |
| Lats21 | 0.00095814 | 0.5625245 | 0.39 | 0.284 | 1 | innate lymphc | Lats2 |
| Rnf19b | 0.00109455 | 0.58151247 | 0.421 | 0.306 | 1 | innate lymphc | Rnf19b |
| Vsir2 | 0.00119649 | 0.52691411 | 0.579 | 0.421 | 1 | innate lymphc | Vsir |
| Gramd1a2 | 0.00122431 | 0.4341803 | 0.547 | 0.408 | 1 | innate lymphc | Gramd1a |
| Gpcpd1 | 0.0012903 | 0.72528633 | 0.277 | 0.185 | 1 | innate lymphc | Gpcpd1 |
| Sp4 | 0.00135036 | 0.73317729 | 0.252 | 0.153 | 1 | innate lymphc | Sp4 |
| Klk8 | 0.00139164 | 1.03783073 | 0.277 | 0.154 | 1 | innate lymphc | Klk8 |
| Tpt11 | 0.00178629 | 0.26837487 | 0.975 | 0.901 | 1 | innate lymphc | Tpt1 |
| Tecpr11 | 0.0019212 | 0.27630025 | 0.396 | 0.329 | 1 | innate lymphc | Tecpr1 |
| Nlk | 0.00193971 | 0.63802735 | 0.252 | 0.16 | 1 | innate lymphc | Nlk |
| Golm11 | 0.001957 | 0.58886843 | 0.258 | 0.169 | 1 | innate lymphc | Golm1 |
| Nudt3 | 0.00216435 | 0.54424769 | 0.308 | 0.211 | 1 | innate lymphc | Nudt3 |
| Fli11 | 0.00222471 | 0.48761118 | 0.553 | 0.444 | 1 | innate lymphc | Fli1 |
| Hipk13 | 0.00224707 | 0.51915573 | 0.541 | 0.416 | 1 | innate lymphc | Hipk1 |
| Ppp1r11 | 0.00226642 | 0.88390275 | 0.365 | 0.233 | 1 | innate lymphc | Ppp1r11 |
| Ogt | 0.00229583 | 0.49308067 | 0.541 | 0.393 | 1 | innate lymphc | Ogt |
| Eef1b23 | 0.00254251 | 0.56323163 | 0.711 | 0.539 | 1 | innate lymphc | Eef1b2 |
| Cabin1 | 0.00266723 | 0.79573918 | 0.258 | 0.175 | 1 | innate lymphc | Cabin1 |
| Rnf40 | 0.00272211 | 0.71924352 | 0.314 | 0.203 | 1 | innate lymphc | Rnf40 |
| Rbm51 | 0.0033009 | 0.61694046 | 0.44 | 0.343 | 1 | innate lymphc | Rbm5 |
| Mbp1 | 0.00338392 | 0.43559806 | 0.415 | 0.271 | 1 | innate lymphc | Mbp |
| Faah1 | 0.00342496 | 0.43107508 | 0.327 | 0.246 | 1 | innate lymphc | Faah |
| Rasgrp11 | 0.00358204 | 0.45558538 | 0.491 | 0.375 | 1 | innate lymphc | Rasgrp1 |
| Ankrd40 | 0.00370859 | 0.81863632 | 0.321 | 0.225 | 1 | innate lymphc | Ankrd40 |
| Ankrd11 | 0.00396405 | 0.3728644 | 0.528 | 0.427 | 1 | innate lymphc | Ankrd11 |
| Gpsm31 | 0.00399677 | 0.41107814 | 0.61 | 0.481 | 1 | innate lymphc | Gpsm3 |
| Sik11 | 0.00433118 | 0.45842778 | 0.767 | 0.603 | 1 | innate lymphc | Sik1 |
| Rbm331 | 0.00440769 | 0.40781954 | 0.415 | 0.303 | 1 | innate lymphc | Rbm33 |
| Paxbp1 | 0.0052776 | 0.38980257 | 0.314 | 0.213 | 1 | innate lymphc | Paxbp1 |
| Pde7a2 | 0.00528281 | 0.40147772 | 0.289 | 0.183 | 1 | innate lymphc | Pde7a |
| Rack12 | 0.00533634 | 0.46567825 | 0.836 | 0.689 | 1 | innate lymphc | Rack1 |
| Kdm6a | 0.00614201 | 0.70442088 | 0.333 | 0.223 | 1 | innate lymphc | Kdm6a |
| Ppp1r15a2 | 0.00614535 | 0.2573077 | 0.585 | 0.527 | 1 | innate lymphc | Ppp1r15a |
| Aebp21 | 0.00618424 | 0.47417045 | 0.667 | 0.546 | 1 | innate lymphc | Aebp2 |
| Ssh21 | 0.00637751 | 0.33671138 | 0.346 | 0.236 | 1 | innate lymphc | Ssh2 |
| Ubr4 | 0.00676393 | 0.51385112 | 0.34 | 0.289 | 1 | innate lymphc | Ubr4 |
| Selenot1 | 0.00689033 | 0.67050814 | 0.346 | 0.244 | 1 | innate lymphc | Selenot |
| Crebbp | 0.00704853 | 0.50551188 | 0.377 | 0.29 | 1 | innate lymphc | Crebbp |
| Tprgl | 0.00788279 | 0.81925302 | 0.283 | 0.193 | 1 | innate lymphc | Tprgl |
| Il27ra | 0.00853867 | 0.41719641 | 0.421 | 0.331 | 1 | innate lymphc | Il27ra |
| Eml41 | 0.00976458 | 0.60822257 | 0.289 | 0.232 | 1 | innate lymphc | Eml4 |
