## Supplementary Table 3 for "Discovery of a Linked Constellation of Gene Expression Revealed by Local Editing of Fibroblasts in Tumors"

Differentially expressed genes (DEGs) of AAV-gRNA groups using pairwise pseudobulk differential gene expression analysis comparing control (=AAV-gTrac) to experimental groups (AAV-gOsmr, AAV-gTgfbr2, or AAV-gIl1r1)

Sheet1: AAV-gTrac vs AAV-gOsmr
Sheet2: AAV-gTrac vs AAV-gTgfbr2
Sheet3: AAV-gTrac vs AAV-gIl1r1
DEGs filtered by fdr < 0.05

AAV-gTrac vs AAV-gOsmr
DEGs filtered by fdr < 0.05

| | primerid | Pr(>Chisq) | coef | ci.hi | ci.lo | fdr | direction |
|---|---|---|---|---|---|---|---|
| 1 | Klf5 | 6.90E-07 | -0.1648658 | -0.1269732 | -0.2027584 | 0.003773 | down |
| 2 | Lmo4 | 1.48E-06 | 0.31266108 | 0.39310957 | 0.23221259 | 0.003773 | up |
| 3 | Rarb | 4.31E-07 | -0.1709226 | -0.1406224 | -0.2012227 | 0.003773 | down |
| 4 | Stard13 | 1.57E-06 | -0.2166671 | -0.1510206 | -0.2823136 | 0.003773 | down |
| 5 | mt-Nd6 | 1.47E-06 | -0.462599 | -0.3450188 | -0.5801791 | 0.003773 | down |
| 6 | Bmpr2 | 4.05E-06 | 0.32513457 | 0.41033271 | 0.23993642 | 0.00635782 | up |
| 7 | Pex6 | 3.45E-06 | -0.1189857 | -0.0797985 | -0.158173 | 0.00635782 | down |
| 8 | Sema3f | 4.24E-06 | 0.16957427 | 0.20736403 | 0.13178451 | 0.00635782 | up |
| 9 | Krt7 | 7.44E-06 | -0.1035622 | -0.0764229 | -0.1307016 | 0.00743439 | down |
| 10 | Mn1 | 6.27E-06 | 0.14274333 | 0.17679607 | 0.10869058 | 0.00743439 | up |
| 11 | Slc6a6 | 6.86E-06 | 0.2332126 | 0.30494016 | 0.16148503 | 0.00743439 | up |
| 12 | Tnfrsf11b | 6.40E-06 | 0.31931519 | 0.39310067 | 0.2455297 | 0.00743439 | up |
| 13 | Pfkl | 9.44E-06 | -0.2475072 | -0.1862851 | -0.3087293 | 0.00871064 | down |
| 14 | Plcb1 | 1.02E-05 | 0.11917823 | 0.14860383 | 0.08975263 | 0.00877401 | up |
| 15 | Luzp1 | 1.15E-05 | 0.13261868 | 0.17968595 | 0.08555142 | 0.00916821 | up |
| 16 | Ice1 | 1.49E-05 | 0.17313144 | 0.22552489 | 0.12073799 | 0.01074749 | up |
| 17 | Ifitm1 | 1.52E-05 | 0.39759117 | 0.51099368 | 0.28418866 | 0.01074749 | up |
| 18 | Tbc1d8 | 1.67E-05 | -0.0411589 | -0.0265139 | -0.0558039 | 0.01115717 | down |
| 19 | Msln | 1.80E-05 | -0.058235 | -0.0398457 | -0.0766243 | 0.01137143 | down |
| 20 | Antxr1 | 2.76E-05 | 0.43302509 | 0.67098781 | 0.19506238 | 0.01332858 | up |
| 21 | Cdkn1b | 2.63E-05 | -0.1761028 | -0.127581 | -0.2246246 | 0.01332858 | down |
| 22 | Hilpda | 2.62E-05 | -0.2492495 | -0.1666647 | -0.3318342 | 0.01332858 | down |
| 23 | Pla2g4a | 2.78E-05 | -0.1781525 | -0.125876 | -0.230429 | 0.01332858 | down |
| 24 | Sobp | 2.55E-05 | 0.07511963 | 0.10438948 | 0.04584978 | 0.01332858 | up |
| 25 | Zfp395 | 2.31E-05 | -0.193107 | -0.1370762 | -0.2491379 | 0.01332858 | down |
| 26 | Gpi1 | 3.17E-05 | -0.2805835 | -0.201106 | -0.3600611 | 0.01461653 | down |
| 27 | Gpr84 | 3.41E-05 | 0.06625329 | 0.08429631 | 0.04821027 | 0.01515851 | up |
| 28 | Wdr91 | 3.55E-05 | -0.1295961 | -0.0729724 | -0.1862198 | 0.01520001 | down |
| 29 | Bid | 4.51E-05 | 0.05331036 | 0.06951949 | 0.03710124 | 0.01693046 | up |
| 30 | Ccdc80 | 4.94E-05 | -0.361223 | -0.2698233 | -0.4526227 | 0.01693046 | down |
| 31 | Fscn1 | 4.37E-05 | -0.1415972 | -0.0941055 | -0.1890889 | 0.01693046 | down |
| 32 | Il1a | 4.73E-05 | 0.07862046 | 0.10462211 | 0.05261881 | 0.01693046 | up |
| 33 | Porcn | 4.86E-05 | 0.22487159 | 0.29581295 | 0.15393024 | 0.01693046 | up |
| 34 | Sipa1 | 4.38E-05 | -0.179954 | -0.1377785 | -0.2221294 | 0.01693046 | down |
| 35 | Soga1 | 4.81E-05 | -0.1423321 | -0.0930646 | -0.1915995 | 0.01693046 | down |
| 36 | Myo7a | 5.26E-05 | 0.41265573 | 0.54187171 | 0.28343975 | 0.01753337 | up |
| 37 | Chd7 | 5.71E-05 | 0.06878855 | 0.08923232 | 0.04834477 | 0.01776915 | up |
| 38 | Ddit4l | 5.97E-05 | -0.1174242 | -0.0735553 | -0.1612931 | 0.01776915 | down |
| 39 | Gucy1b1 | 5.52E-05 | -0.0965766 | -0.0663064 | -0.1268468 | 0.01776915 | down |
| 40 | Ptges | 5.84E-05 | 0.18842124 | 0.25397125 | 0.12287123 | 0.01776915 | up |
| 41 | Smox | 6.13E-05 | -0.132915 | -0.0724379 | -0.193392 | 0.01776915 | down |
| 42 | Ube2o | 6.22E-05 | -0.1415768 | -0.099431 | -0.1837225 | 0.01776915 | down |
| 43 | Anxa10 | 6.51E-05 | -0.0473693 | -0.0342616 | -0.0604769 | 0.01814398 | down |
| 44 | Olfml2b | 6.66E-05 | 0.22161584 | 0.30183087 | 0.14140081 | 0.01814398 | up |
| 45 | Hmga2 | 7.24E-05 | 0.10840323 | 0.14107758 | 0.07572887 | 0.01886817 | up |
| 46 | Tbx15 | 7.19E-05 | 0.28657321 | 0.38792684 | 0.18521958 | 0.01886817 | up |
| 47 | Etfbkmt | 8.61E-05 | -0.1309149 | -0.0843602 | -0.1774697 | 0.0198618 | down |
| 48 | Fam3c | 8.14E-05 | 0.15426393 | 0.20714077 | 0.10138709 | 0.0198618 | up |
| 49 | Gde1 | 8.55E-05 | -0.1320996 | -0.0747877 | -0.1894114 | 0.0198618 | down |
| 50 | Nectin2 | 8.59E-05 | 0.22793198 | 0.30722355 | 0.14864041 | 0.0198618 | up |
| 51 | Pea15a | 8.04E-05 | -0.1732948 | -0.1235846 | -0.223005 | 0.0198618 | down |
| 52 | Ppp1r3b | 8.21E-05 | -0.1203164 | -0.0662289 | -0.1744039 | 0.0198618 | down |
| 53 | Tnn | 9.45E-05 | 0.817367 | 1.23369417 | 0.40103983 | 0.02136785 | up |
| 54 | Mif | 9.64E-05 | -0.2320046 | -0.1688953 | -0.2951138 | 0.02140336 | down |
| 55 | Metap1 | 0.00010814 | 0.09872637 | 0.1397714 | 0.05768133 | 0.02321485 | up |
| 56 | Ttyh2 | 0.00010843 | -0.2942204 | -0.2022411 | -0.3861997 | 0.02321485 | down |
| 57 | Ccng2 | 0.00013348 | -0.1641561 | -0.0956407 | -0.2326715 | 0.02378113 | down |
| 58 | Crispld2 | 0.00013329 | 0.88066147 | 1.1780086 | 0.58331434 | 0.02378113 | up |
| 59 | Dlc1 | 0.00011898 | -0.2543921 | -0.1671941 | -0.3415902 | 0.02378113 | down |
| 60 | Gabarap | 0.00014659 | 0.16597945 | 0.24534766 | 0.08661124 | 0.02378113 | up |
| 61 | Gys1 | 0.00014128 | -0.177731 | -0.1098037 | -0.2456583 | 0.02378113 | down |
| 62 | Hhipl1 | 0.00014327 | 0.26624641 | 0.37132115 | 0.16117168 | 0.02378113 | up |
| 63 | Hic1 | 0.0001481 | -0.2456266 | -0.1423776 | -0.3488755 | 0.02378113 | down |
| 64 | Krt19 | 0.00014221 | -0.2663621 | -0.176213 | -0.3565111 | 0.02378113 | down |
| 65 | Lgals4 | 0.00011413 | -0.0883115 | -0.0605628 | -0.1160603 | 0.02378113 | down |
| 66 | Nr2f1 | 0.00013503 | -0.1475635 | -0.0903561 | -0.2047708 | 0.02378113 | down |
| 67 | P4ha1 | 0.0001309 | -0.3030727 | -0.1678114 | -0.438334 | 0.02378113 | down |
| 68 | Pdk1 | 0.0001445 | -0.1948315 | -0.1314488 | -0.2582143 | 0.02378113 | down |
| 69 | Pgp | 0.00014694 | -0.1711069 | -0.1114126 | -0.2308013 | 0.02378113 | down |
| 70 | Ralgds | 0.00012434 | 0.13151846 | 0.1747047 | 0.08833223 | 0.02378113 | up |
| 71 | Smg6 | 0.00013406 | -0.2167807 | -0.1418798 | -0.2916816 | 0.02378113 | down |
| 72 | Smtnl2 | 0.00013328 | -0.0704979 | -0.0514998 | -0.0894959 | 0.02378113 | down |
| 73 | Snx33 | 0.00014876 | 0.1060734 | 0.14813292 | 0.06401388 | 0.02378113 | up |
| 74 | Tec | 0.0001204 | -0.0598027 | -0.0374189 | -0.0821866 | 0.02378113 | down |
| 75 | Tmem9 | 0.00014722 | 0.13160828 | 0.17883817 | 0.08437839 | 0.02378113 | up |
| 76 | Mamdc2 | 0.00015457 | 0.44227877 | 0.67799151 | 0.20656602 | 0.02436757 | up |
| 77 | Tpsb2 | 0.00015649 | -0.0312359 | -0.018748 | -0.0437237 | 0.02436757 | down |
| 78 | Mpp2 | 0.00016009 | -0.1530459 | -0.112821 | -0.1932708 | 0.02460851 | down |
| 79 | Gsta4 | 0.00016687 | -0.069757 | -0.0538741 | -0.08564 | 0.02532644 | down |
| 80 | Slc2a1 | 0.00017449 | -0.2952412 | -0.1974293 | -0.3930531 | 0.02615155 | down |
| 81 | Draxin | 0.000182 | 0.28326935 | 0.37487767 | 0.19166102 | 0.0263209 | up |
| 82 | Pcca | 0.0001822 | -0.1344496 | -0.0777023 | -0.1911969 | 0.0263209 | down |
| 83 | Ubxn6 | 0.00017902 | 0.12396442 | 0.20884731 | 0.03908153 | 0.0263209 | up |
| 84 | Tbx18 | 0.00018505 | 0.13514937 | 0.21237332 | 0.05792542 | 0.02641374 | up |
| 85 | Cdh1 | 0.00019109 | -0.0483942 | -0.0331103 | -0.0636782 | 0.0267303 | down |
| 86 | Slc16a3 | 0.00019173 | -0.2835817 | -0.1806579 | -0.3865055 | 0.0267303 | down |
| 87 | Clip1 | 0.00019785 | 0.27362511 | 0.36517158 | 0.18207865 | 0.02695641 | up |
| 88 | Cxcl5 | 0.00019603 | 0.50964674 | 0.72327829 | 0.29601519 | 0.02695641 | up |
| 89 | Lrrn2 | 0.00021129 | 0.11681809 | 0.16253359 | 0.07110259 | 0.02827017 | up |
| 90 | Osmr | 0.0002122 | -0.34769 | -0.1622881 | -0.5330918 | 0.02827017 | down |
| 91 | Abca9 | 0.00022946 | -0.1641362 | -0.0702564 | -0.258016 | 0.02961296 | down |
| 92 | Cxcl16 | 0.00022969 | 0.29051722 | 0.39586936 | 0.18516507 | 0.02961296 | up |
| 93 | Ly75 | 0.00022932 | 0.15176136 | 0.20979166 | 0.09373106 | 0.02961296 | up |
| 94 | Stk25 | 0.0002386 | -0.1054846 | -0.0630886 | -0.1478806 | 0.03043382 | down |
| 95 | Ubald2 | 0.00024375 | -0.1703943 | -0.1062458 | -0.2345427 | 0.03076349 | down |
| 96 | Abca8a | 0.00024928 | -0.158464 | -0.1096468 | -0.2072813 | 0.03076775 | down |
| 97 | Glmp | 0.00025148 | 0.10568681 | 0.14571145 | 0.06566218 | 0.03076775 | up |
| 98 | Nek6 | 0.0002499 | 0.30074255 | 0.40685303 | 0.19463207 | 0.03076775 | up |
| 99 | Ciart | 0.0002636 | -0.0336589 | -0.021859 | -0.0454588 | 0.03175189 | down |
| 100 | Gcn1 | 0.00027134 | -0.1186601 | -0.0734709 | -0.1638493 | 0.03175189 | down |
| 101 | Mycbp2 | 0.00026849 | 0.20866293 | 0.26779933 | 0.14952653 | 0.03175189 | up |
| 102 | Rasgrf1 | 0.00027276 | -0.0898573 | -0.0423045 | -0.1374102 | 0.03175189 | down |
| 103 | Slc7a8 | 0.00027172 | -0.1586669 | -0.0951389 | -0.2221949 | 0.03175189 | down |
| 104 | Krt8 | 0.00027603 | -0.2973929 | -0.1873903 | -0.4073955 | 0.03182356 | down |
| 105 | Tjap1 | 0.00028237 | 0.15781722 | 0.21506473 | 0.10056971 | 0.03224419 | up |
| 106 | Arhgap8 | 0.00029646 | 0.02591873 | 0.03577878 | 0.01605868 | 0.03324875 | up |
| 107 | Cdh3 | 0.00029906 | 0.07034997 | 0.09827422 | 0.04242572 | 0.03324875 | up |
| 108 | Mettl21a | 0.00029949 | -0.0922635 | -0.0539707 | -0.1305564 | 0.03324875 | down |
| 109 | Acsl4 | 0.00030458 | -0.191312 | -0.1281557 | -0.2544682 | 0.03350345 | down |
| 110 | Yy1 | 0.00031455 | -0.1391312 | -0.0815883 | -0.1966741 | 0.03428562 | down |
| 111 | Tnfsf11 | 0.00032189 | 0.29815142 | 0.42173977 | 0.17456308 | 0.03476984 | up |
| 112 | Rgmb | 0.00034586 | -0.2597409 | -0.1619918 | -0.35749 | 0.03702511 | down |
| 113 | Galnt3 | 0.00035812 | -0.0229085 | -0.0137845 | -0.0320325 | 0.03772718 | down |
| 114 | Ids | 0.00036353 | -0.1211775 | -0.078648 | -0.163707 | 0.03772718 | down |
| 115 | Rin2 | 0.00036249 | 0.1710642 | 0.25772466 | 0.08440373 | 0.03772718 | up |
| 116 | Sertad3 | 0.000365 | -0.0812846 | -0.0483953 | -0.1141738 | 0.03772718 | down |
| 117 | Gcnt2 | 0.00037566 | 0.17194599 | 0.25950217 | 0.08438981 | 0.03785027 | up |
| 118 | Tgtp2 | 0.00037268 | 0.02873585 | 0.04100488 | 0.01646683 | 0.03785027 | up |
| 119 | Tmem63a | 0.00037555 | 0.11383441 | 0.15951098 | 0.06815783 | 0.03785027 | up |
| 120 | Vstm4 | 0.00038406 | -0.1475858 | -0.0636423 | -0.2315294 | 0.03837449 | down |
| 121 | Dennd2a | 0.0003895 | -0.3080512 | -0.1896522 | -0.4264501 | 0.03859627 | down |
| 122 | Phlda3 | 0.00039946 | -0.1030759 | -0.0527079 | -0.1534438 | 0.03925816 | down |
| 123 | Hmgn3 | 0.00040691 | -0.1112639 | -0.0635514 | -0.1589764 | 0.03935955 | down |
| 124 | Spint2 | 0.00040705 | -0.0524251 | -0.0374556 | -0.0673946 | 0.03935955 | down |
| 125 | Cend1 | 0.00041077 | 0.01465982 | 0.02332416 | 0.00599548 | 0.03940142 | up |
| 126 | Fam160a1 | 0.00042992 | 0.05920693 | 0.07984386 | 0.03857 | 0.03995912 | up |
| 127 | Lyn | 0.00042143 | 0.19383278 | 0.26502708 | 0.12263847 | 0.03995912 | up |
| 128 | Megf11 | 0.00042354 | -0.0661665 | -0.0401905 | -0.0921426 | 0.03995912 | down |
| 129 | Vdr | 0.00042704 | 0.09912317 | 0.13508737 | 0.06315898 | 0.03995912 | up |
| 130 | Igtp | 0.00044093 | 0.11682533 | 0.19210049 | 0.04155017 | 0.04052359 | up |
| 131 | Krt18 | 0.00044275 | -0.1438449 | -0.0960897 | -0.1916001 | 0.04052359 | down |
| 132 | Acat3 | 0.0004583 | 0.09128608 | 0.11898295 | 0.06358921 | 0.04162896 | up |
| 133 | Adcyap1r1 | 0.00046618 | -0.1003543 | -0.0619787 | -0.1387298 | 0.04202617 | down |
| 134 | Gpd2 | 0.0004707 | 0.05283376 | 0.07754028 | 0.02812724 | 0.0421169 | up |
| 135 | Itgb4 | 0.00048289 | -0.054353 | -0.0378465 | -0.0708595 | 0.04288741 | down |
| 136 | C1qtnf1 | 0.00049185 | 0.3119769 | 0.43715566 | 0.18679813 | 0.04336239 | up |
| 137 | Iigp1 | 0.00050156 | 0.20845401 | 0.39336867 | 0.02353935 | 0.04389544 | up |
| 138 | Irx3 | 0.00051321 | -0.1994575 | -0.1023276 | -0.2965873 | 0.04428441 | down |
| 139 | Scarb2 | 0.00051339 | 0.18819986 | 0.27515549 | 0.10124423 | 0.04428441 | up |
| 140 | Akr1b10 | 0.00053727 | 0.09665675 | 0.13198125 | 0.06133226 | 0.04443401 | up |
| 141 | Cdk14 | 0.00052335 | 0.17725033 | 0.25095275 | 0.10354792 | 0.04443401 | up |
| 142 | Col23a1 | 0.00054106 | 0.66989693 | 1.00732833 | 0.33246553 | 0.04443401 | up |
| 143 | Itga7 | 0.00052319 | -0.050669 | -0.0246639 | -0.0766742 | 0.04443401 | down |
| 144 | Ldhb | 0.00053948 | 0.06698533 | 0.09109687 | 0.04287379 | 0.04443401 | up |
| 145 | Plcl1 | 0.00053339 | -0.0794141 | -0.0414428 | -0.1173853 | 0.04443401 | down |
| 146 | Socs5 | 0.00052874 | 0.10532773 | 0.16555018 | 0.04510527 | 0.04443401 | up |
| 147 | Fam162a | 0.00055091 | -0.2554808 | -0.1468648 | -0.3640969 | 0.04493508 | down |
| 148 | Zfp516 | 0.00056014 | -0.2015911 | -0.1187207 | -0.2844614 | 0.0453791 | down |
| 149 | Cystm1 | 0.00057232 | -0.1300532 | -0.0863508 | -0.1737557 | 0.04574731 | down |
| 150 | Zc3h12a | 0.00057224 | 0.14840069 | 0.20663209 | 0.09016929 | 0.04574731 | up |
| 151 | Atp2b4 | 0.00058811 | 0.1432397 | 0.18775458 | 0.09872483 | 0.04608801 | up |
| 152 | Etv5 | 0.00058778 | -0.1477319 | -0.0707449 | -0.2247189 | 0.04608801 | down |
| 153 | Fbxl14 | 0.00058453 | -0.1052594 | -0.0472175 | -0.1633013 | 0.04608801 | down |
| 154 | Il27ra | 0.00059488 | -0.0496299 | -0.0333466 | -0.0659132 | 0.046316 | down |
| 155 | Epas1 | 0.00061286 | 0.42877256 | 0.59380046 | 0.26374466 | 0.04680349 | up |
| 156 | Fam117b | 0.00061224 | -0.1434239 | -0.0875324 | -0.1993154 | 0.04680349 | down |
| 157 | Nhsl1 | 0.00060994 | -0.1090279 | -0.059808 | -0.1582479 | 0.04680349 | down |
| 158 | Cpe | 0.00061767 | -0.2762977 | -0.1751035 | -0.3774918 | 0.04687247 | down |
| 159 | Bhlhe41 | 0.00063304 | -0.2234041 | -0.1413037 | -0.3055045 | 0.04761622 | down |
| 160 | Cadm4 | 0.00065571 | -0.1618076 | -0.0620685 | -0.2615467 | 0.04761622 | down |
| 161 | Clcn3 | 0.00065124 | -0.0604741 | -0.0376916 | -0.0832566 | 0.04761622 | down |
| 162 | Fah | 0.00064703 | 0.23444779 | 0.33038425 | 0.13851134 | 0.04761622 | up |
| 163 | Irf1 | 0.00065195 | 0.30461618 | 0.42597542 | 0.18325694 | 0.04761622 | up |
| 164 | Mcu | 0.00063565 | -0.0943093 | -0.0553903 | -0.1332282 | 0.04761622 | down |
| 165 | Sod2 | 0.00064793 | 0.4527837 | 0.64911234 | 0.25645506 | 0.04761622 | up |
| 166 | Tmem170b | 0.00065924 | -0.0676017 | -0.0403064 | -0.0948971 | 0.04761622 | down |
| 167 | Zc3h12c | 0.00067229 | 0.04927839 | 0.07601416 | 0.02254262 | 0.04826791 | up |
| 168 | Lonrf1 | 0.00068893 | -0.0404917 | -0.0216025 | -0.0593809 | 0.04882008 | down |
| 169 | Lratd1 | 0.0006882 | -0.0662618 | -0.0452613 | -0.0872622 | 0.04882008 | down |
| 170 | Plcd1 | 0.00069219 | 0.12524462 | 0.17042046 | 0.08006878 | 0.04882008 | up |
| 171 | Ltbp2 | 0.00069852 | 0.75292832 | 1.04275228 | 0.46310437 | 0.04897838 | up |
| 172 | P2rx4 | 0.00070442 | 0.18333139 | 0.24938443 | 0.11727835 | 0.0491045 | up |

AAV-gTrac vs AAV-gTgfbr2
DEGs filtered by fdr < 0.05

| | primerid | Pr(>Chisq) | coef | ci.hi | ci.lo | fdr | direction |
|---|---|---|---|---|---|---|---|
| 1 | Sav1 | 2.39E-10 | 0.46156751 | 0.51406857 | 0.40906645 | 2.87E-06 | up |
| 2 | P2rx4 | 6.38E-09 | 0.38296038 | 0.43394034 | 0.33198042 | 3.83E-05 | up |
| 3 | Cxcl5 | 1.23E-08 | 0.85779148 | 1.05389815 | 0.66168482 | 4.90E-05 | up |
| 4 | Dhrs3 | 1.67E-08 | -1.069294 | -0.8827532 | -1.2558347 | 5.01E-05 | down |
| 5 | Hgf | 4.69E-08 | 0.38201653 | 0.4535975 | 0.31043556 | 0.00011262 | up |
| 6 | Pea15a | 5.78E-08 | -0.330367 | -0.2702989 | -0.3904351 | 0.00011572 | down |
| 7 | Plcb1 | 1.01E-07 | 0.25031671 | 0.31413367 | 0.18649976 | 0.00017257 | up |
| 8 | Ptprj | 1.23E-07 | 0.20773677 | 0.24944716 | 0.16602639 | 0.00018459 | up |
| 9 | Fbn2 | 1.83E-07 | -0.5586636 | -0.4495674 | -0.6677598 | 0.00021983 | down |
| 10 | Ubxn6 | 1.80E-07 | 0.30434696 | 0.38805114 | 0.22064279 | 0.00021983 | up |
| 11 | Sipa1l1 | 2.95E-07 | 0.32061463 | 0.4001281 | 0.24110116 | 0.0003216 | up |
| 12 | Rasa3 | 3.33E-07 | -0.2665692 | -0.1973735 | -0.3357648 | 0.00033362 | down |
| 13 | Ak3 | 3.90E-07 | 0.37476868 | 0.45735197 | 0.29218538 | 0.00036025 | up |
| 14 | Ercc1 | 4.21E-07 | -0.1746842 | -0.1266185 | -0.2227498 | 0.00036119 | down |
| 15 | Pi15 | 4.58E-07 | 0.33043349 | 0.42974856 | 0.23111842 | 0.00036656 | up |
| 16 | Dlg2 | 5.42E-07 | 0.17784977 | 0.23712821 | 0.11857133 | 0.00039721 | up |
| 17 | Loxl3 | 5.62E-07 | -0.6149766 | -0.4609976 | -0.7689556 | 0.00039721 | down |
| 18 | Ehbp1l1 | 8.10E-07 | -0.277347 | -0.2026458 | -0.3520482 | 0.00054006 | down |
| 19 | Il1r1 | 1.32E-06 | 0.52345098 | 0.6228671 | 0.42403485 | 0.00083477 | up |
| 20 | Lrig1 | 1.43E-06 | 0.25485552 | 0.30740427 | 0.20230677 | 0.00085642 | up |
| 21 | Acot7 | 1.60E-06 | -0.3086892 | -0.2290652 | -0.3883132 | 0.00091699 | down |
| 22 | Ccdc80 | 1.90E-06 | -0.4684035 | -0.2667041 | -0.6701028 | 0.00099095 | down |
| 23 | Chst2 | 1.89E-06 | -0.4720548 | -0.3410138 | -0.6030958 | 0.00099095 | down |
| 24 | Endod1 | 2.33E-06 | -0.1903425 | -0.1433662 | -0.2373189 | 0.00113532 | down |
| 25 | Tmem63a | 2.36E-06 | 0.17626092 | 0.2257576 | 0.12676425 | 0.00113532 | up |
| 26 | Tra2b | 2.47E-06 | 0.15497695 | 0.20345652 | 0.10649738 | 0.00113946 | up |
| 27 | Xdh | 2.82E-06 | 0.72547641 | 0.91649723 | 0.53445559 | 0.00125305 | up |
| 28 | Capn1 | 3.08E-06 | 0.33102596 | 0.41933153 | 0.2427204 | 0.00129935 | up |
| 29 | Pycr1 | 3.14E-06 | -0.3003328 | -0.2158765 | -0.384789 | 0.00129935 | down |
| 30 | Timm8b | 3.52E-06 | 0.1598102 | 0.20694588 | 0.11267452 | 0.00140664 | up |
| 31 | Cspg4 | 3.98E-06 | -0.4686495 | -0.2647952 | -0.6725038 | 0.00154193 | down |
| 32 | Scarb2 | 4.39E-06 | 0.30925179 | 0.41594091 | 0.20256267 | 0.00164616 | up |
| 33 | Tmem38b | 5.35E-06 | 0.2396059 | 0.30185984 | 0.17735196 | 0.00194728 | up |
| 34 | Herc4 | 5.67E-06 | 0.14336079 | 0.18303993 | 0.10368165 | 0.00200225 | up |
| 35 | Lama4 | 5.96E-06 | 0.4518098 | 0.59147335 | 0.31214625 | 0.00201473 | up |
| 36 | Tmem100 | 6.04E-06 | 0.28072186 | 0.37453913 | 0.18690459 | 0.00201473 | up |
| 37 | Ralgds | 6.21E-06 | 0.16547525 | 0.21194182 | 0.11900868 | 0.00201548 | up |
| 38 | Prkcsh | 6.49E-06 | -0.2868694 | -0.2190233 | -0.3547155 | 0.00205056 | down |
| 39 | Il1rap | 7.14E-06 | 0.55658145 | 0.69726714 | 0.41589576 | 0.00219861 | up |
| 40 | Cav1 | 8.53E-06 | 0.84124716 | 1.07233346 | 0.61016086 | 0.002327 | up |
| 41 | Coa3 | 8.24E-06 | -0.1457245 | -0.1000914 | -0.1913576 | 0.002327 | down |
| 42 | Megf11 | 7.79E-06 | -0.1089555 | -0.0879612 | -0.1299498 | 0.002327 | down |
| 43 | Prkd1 | 8.48E-06 | -0.1624374 | -0.1036046 | -0.2212703 | 0.002327 | down |
| 44 | Stom | 8.34E-06 | 0.28053127 | 0.3695796 | 0.19148293 | 0.002327 | up |
| 45 | Ankrd54 | 9.46E-06 | -0.1218577 | -0.0793597 | -0.1643557 | 0.00252284 | down |
| 46 | Srpr | 1.02E-05 | -0.1995421 | -0.147287 | -0.2517972 | 0.00263977 | down |
| 47 | Tgfb3 | 1.03E-05 | -0.6510207 | -0.4279258 | -0.8741157 | 0.00263977 | down |
| 48 | Col8a1 | 1.19E-05 | -1.3309401 | -1.0160533 | -1.6458269 | 0.00296827 | down |
| 49 | Dcn | 1.24E-05 | 1.40724959 | 1.84019911 | 0.97430006 | 0.00304949 | up |
| 50 | Actn4 | 1.56E-05 | -0.4162342 | -0.2963162 | -0.5361522 | 0.00314039 | down |
| 51 | Arhgap21 | 1.36E-05 | 0.14728102 | 0.19696421 | 0.09759783 | 0.00314039 | up |
| 52 | Bcl2 | 1.67E-05 | -0.1364473 | -0.096516 | -0.1763786 | 0.00314039 | down |
| 53 | Chsy1 | 1.59E-05 | -0.2513918 | -0.1698756 | -0.3329079 | 0.00314039 | down |
| 54 | Emilin1 | 1.64E-05 | -0.3472824 | -0.2374141 | -0.4571508 | 0.00314039 | down |
| 55 | Eps8 | 1.33E-05 | 0.33052683 | 0.42188168 | 0.23917197 | 0.00314039 | up |
| 56 | Fetub | 1.58E-05 | 0.14231098 | 0.19259779 | 0.09202416 | 0.00314039 | up |
| 57 | Jak2 | 1.59E-05 | 0.31327622 | 0.41104988 | 0.21550256 | 0.00314039 | up |
| 58 | Piezo2 | 1.42E-05 | -0.4818237 | -0.1996255 | -0.7640219 | 0.00314039 | down |
| 59 | Pstpip1 | 1.60E-05 | -0.2293637 | -0.1588858 | -0.2998417 | 0.00314039 | down |
| 60 | Rfx1 | 1.66E-05 | -0.083494 | -0.0466627 | -0.1203254 | 0.00314039 | down |
| 61 | Rspo4 | 1.34E-05 | 0.18648098 | 0.35336832 | 0.01959363 | 0.00314039 | up |
| 62 | Tmem98 | 1.57E-05 | -0.263368 | -0.1955297 | -0.3312062 | 0.00314039 | down |
| 63 | Tprkb | 1.41E-05 | 0.13179143 | 0.16999612 | 0.09358674 | 0.00314039 | up |
| 64 | Ubald2 | 1.48E-05 | -0.1885396 | -0.1326816 | -0.2443975 | 0.00314039 | down |
| 65 | Fzd2 | 1.75E-05 | -0.3491024 | -0.2184063 | -0.4797985 | 0.00322292 | down |
| 66 | Pdxk | 1.78E-05 | 0.16056076 | 0.20520517 | 0.11591634 | 0.00324181 | up |
| 67 | Spry1 | 1.81E-05 | -0.5387573 | -0.3849268 | -0.6925878 | 0.00324548 | down |
| 68 | Dad1 | 1.91E-05 | -0.333284 | -0.2072618 | -0.4593062 | 0.00337702 | down |
| 69 | Hic1 | 2.01E-05 | -0.3683099 | -0.2039908 | -0.532629 | 0.00344182 | down |
| 70 | Tbx15 | 1.99E-05 | 0.33488626 | 0.43500519 | 0.23476732 | 0.00344182 | up |
| 71 | Mprip | 2.04E-05 | -0.401297 | -0.3105844 | -0.4920097 | 0.00345251 | down |
| 72 | Acap2 | 2.48E-05 | 0.18638925 | 0.24688806 | 0.12589043 | 0.00413909 | up |
| 73 | Ppp4r2 | 2.62E-05 | 0.14021415 | 0.18490874 | 0.09551956 | 0.00431246 | up |
| 74 | Ddit4l | 2.75E-05 | -0.150949 | -0.0939565 | -0.2079414 | 0.00431594 | down |
| 75 | Efemp2 | 2.69E-05 | 0.36997471 | 0.55448264 | 0.18546678 | 0.00431594 | up |
| 76 | Pmpcb | 2.71E-05 | -0.2234589 | -0.1522909 | -0.294627 | 0.00431594 | down |
| 77 | Prkx | 2.77E-05 | 0.18190949 | 0.24228043 | 0.12153855 | 0.00431594 | up |
| 78 | Nfe2l2 | 2.84E-05 | 0.41664 | 0.58706277 | 0.24621723 | 0.0043196 | up |
| 79 | Schip1 | 2.81E-05 | -0.3648348 | -0.2507984 | -0.4788712 | 0.0043196 | down |
| 80 | Xylt1 | 3.06E-05 | 0.18783748 | 0.23740791 | 0.13826704 | 0.00459741 | up |
| 81 | Nfil3 | 3.17E-05 | 0.37511126 | 0.48422767 | 0.26599485 | 0.00469363 | up |
| 82 | Mindy1 | 3.25E-05 | 0.12475514 | 0.17038995 | 0.07912033 | 0.0047554 | up |
| 83 | Coq8b | 3.50E-05 | -0.1711497 | -0.104462 | -0.2378373 | 0.00505518 | down |
| 84 | Adam19 | 3.76E-05 | -0.5933526 | -0.4246703 | -0.7620349 | 0.00507124 | down |
| 85 | Fscn1 | 3.71E-05 | -0.4596438 | -0.325748 | -0.5935396 | 0.00507124 | down |
| 86 | Il13ra1 | 3.75E-05 | 0.49871779 | 0.67534841 | 0.32208717 | 0.00507124 | up |
| 87 | Mtch1 | 3.75E-05 | -0.5294164 | -0.3106539 | -0.7481789 | 0.00507124 | down |
| 88 | Phldb2 | 3.59E-05 | -0.530722 | -0.3722825 | -0.6891615 | 0.00507124 | down |
| 89 | Selenos | 3.71E-05 | -0.1677313 | -0.108347 | -0.2271156 | 0.00507124 | down |
| 90 | Rbp1 | 3.81E-05 | -0.7998906 | -0.5432227 | -1.0565586 | 0.00508253 | down |
| 91 | Lgals3 | 3.89E-05 | 0.80489033 | 1.08849306 | 0.5212876 | 0.0051108 | up |
| 92 | Sod2 | 3.92E-05 | 0.51853264 | 0.66157504 | 0.37549023 | 0.0051108 | up |
| 93 | Ahdc1 | 4.16E-05 | -0.2225062 | -0.1550111 | -0.2900013 | 0.00532783 | down |
| 94 | Drd1 | 4.18E-05 | 0.08690419 | 0.10827503 | 0.06553335 | 0.00532783 | up |
| 95 | Glrx | 4.22E-05 | 0.33597272 | 0.44761517 | 0.22433027 | 0.00532783 | up |
| 96 | C2cd2l | 4.27E-05 | -0.1121929 | -0.0731243 | -0.1512615 | 0.00534324 | down |
| 97 | Cntfr | 4.45E-05 | -0.1192574 | -0.0900401 | -0.1484747 | 0.00550135 | down |
| 98 | Stat6 | 4.51E-05 | 0.16110048 | 0.21371593 | 0.10848502 | 0.00553062 | up |
| 99 | Cyba | 4.60E-05 | -0.1290111 | -0.0815171 | -0.176505 | 0.00558412 | down |
| 100 | Osmr | 4.66E-05 | 0.57904404 | 0.81297315 | 0.34511492 | 0.00559311 | up |
| 101 | Spef1 | 4.74E-05 | -0.1560379 | -0.1068193 | -0.2052565 | 0.00563201 | down |
| 102 | Lnx2 | 4.90E-05 | -0.1387607 | -0.0975778 | -0.1799436 | 0.00577018 | down |
| 103 | Tmem59l | 5.02E-05 | -0.0805054 | -0.0393437 | -0.1216671 | 0.00585237 | down |
| 104 | Hmga2 | 5.14E-05 | 0.16117307 | 0.25575984 | 0.06658631 | 0.00591563 | up |
| 105 | S100a4 | 5.17E-05 | 0.89985587 | 1.18082073 | 0.61889101 | 0.00591563 | up |
| 106 | Lrig3 | 5.25E-05 | -0.1011824 | -0.0691552 | -0.1332096 | 0.00594833 | down |
| 107 | Btbd3 | 5.40E-05 | 0.15232805 | 0.20956559 | 0.09509051 | 0.00596703 | up |
| 108 | Kctd5 | 5.47E-05 | -0.1993879 | -0.1401186 | -0.2586571 | 0.00596703 | down |
| 109 | Lefty1 | 5.42E-05 | 0.07890017 | 0.10808754 | 0.04971281 | 0.00596703 | up |
| 110 | Wdr36 | 5.44E-05 | 0.12041979 | 0.16390913 | 0.07693046 | 0.00596703 | up |
| 111 | Bicc1 | 5.58E-05 | -0.2171672 | -0.1361923 | -0.2981422 | 0.00603136 | down |
| 112 | Eci2 | 5.85E-05 | 0.19238912 | 0.25105534 | 0.1337229 | 0.006247 | up |
| 113 | Krt7 | 5.88E-05 | -0.1125433 | -0.0846303 | -0.1404563 | 0.006247 | down |
| 114 | Cdc42ep2 | 6.10E-05 | 0.24759573 | 0.33097039 | 0.16422107 | 0.00629991 | up |
| 115 | Lamp2 | 6.19E-05 | 0.60667149 | 1.01562935 | 0.19771362 | 0.00629991 | up |
| 116 | Lrig2 | 6.06E-05 | 0.14479084 | 0.19314836 | 0.09643332 | 0.00629991 | up |
| 117 | Lum | 6.19E-05 | 1.0029031 | 1.30238201 | 0.70342418 | 0.00629991 | up |
| 118 | Skil | 6.16E-05 | -0.4080978 | -0.277289 | -0.5389065 | 0.00629991 | down |
| 119 | Mxra7 | 6.30E-05 | -0.386146 | -0.2665967 | -0.5056954 | 0.00630091 | down |
| 120 | Sema4b | 6.30E-05 | 0.116744 | 0.16168393 | 0.07180407 | 0.00630091 | up |
| 121 | Tmem128 | 6.43E-05 | 0.17516371 | 0.22655683 | 0.12377058 | 0.0063768 | up |
| 122 | Tbc1d1 | 6.61E-05 | -0.2243046 | -0.1520276 | -0.2965816 | 0.00650682 | down |
| 123 | Cdh13 | 6.82E-05 | 0.22053595 | 0.28989351 | 0.15117839 | 0.0066397 | up |
| 124 | Trpm7 | 6.86E-05 | 0.1244575 | 0.17082433 | 0.07809067 | 0.0066397 | up |
| 125 | Rab7b | 7.00E-05 | 0.12363564 | 0.17750439 | 0.06976688 | 0.0067275 | up |
| 126 | Ap3s1 | 7.21E-05 | 0.14794561 | 0.2234978 | 0.07239342 | 0.00682226 | up |
| 127 | Ldb1 | 7.22E-05 | 0.13838601 | 0.18468787 | 0.09208416 | 0.00682226 | up |
| 128 | Atp2c1 | 7.31E-05 | -0.1396794 | -0.084294 | -0.1950649 | 0.00685235 | down |
| 129 | Actn1 | 7.54E-05 | -0.6454738 | -0.4477172 | -0.8432304 | 0.00701646 | down |
| 130 | Srpx2 | 7.66E-05 | -0.863556 | -0.5772946 | -1.1498175 | 0.00707615 | down |
| 131 | Mef2c | 7.75E-05 | 0.1344088 | 0.21477381 | 0.05404378 | 0.00710449 | up |
| 132 | Acsbg1 | 8.01E-05 | -0.0773615 | -0.028331 | -0.1263921 | 0.00728402 | down |
| 133 | Dock11 | 8.17E-05 | 0.13261008 | 0.17882501 | 0.08639515 | 0.00737604 | up |
| 134 | Gpnbp2 | 8.30E-05 | 0.24388819 | 0.3223955 | 0.16538089 | 0.00738403 | up |
| 135 | Mast2 | 8.30E-05 | -0.2236961 | -0.1455544 | -0.3018377 | 0.00738403 | down |
| 136 | Acta2 | 8.56E-05 | -1.340333 | -0.9131665 | -1.7674995 | 0.00740149 | down |
| 137 | Inppl1 | 8.47E-05 | -0.139893 | -0.0925312 | -0.1872547 | 0.00740149 | down |
| 138 | Ngly1 | 8.56E-05 | 0.13162935 | 0.17562133 | 0.08763737 | 0.00740149 | up |
| 139 | Tubb3 | 8.57E-05 | -0.1218016 | -0.0852278 | -0.1583753 | 0.00740149 | down |
| 140 | Dsip1l | 9.01E-05 | -0.2117122 | -0.1281678 | -0.2952566 | 0.0075654 | down |
| 141 | Leprotl1 | 8.98E-05 | -0.1934133 | -0.126219 | -0.2606076 | 0.0075654 | down |
| 142 | Tdp1 | 8.85E-05 | -0.0975614 | -0.0656191 | -0.1295037 | 0.0075654 | down |
| 143 | Tgfb1i1 | 9.01E-05 | -0.2949925 | -0.1957067 | -0.3942784 | 0.0075654 | down |
| 144 | Gdpd5 | 9.17E-05 | -0.1707155 | -0.1082166 | -0.2332145 | 0.00764582 | down |
| 145 | Camk4 | 9.30E-05 | -0.0281403 | -0.0123958 | -0.0438848 | 0.00769004 | down |
| 146 | Fgd5 | 9.41E-05 | -0.1555616 | -0.0690134 | -0.2421098 | 0.00769004 | down |
| 147 | Gask1b | 9.42E-05 | -0.8995576 | -0.6128832 | -1.186232 | 0.00769004 | down |
| 148 | Dcps | 9.61E-05 | -0.1085947 | -0.0527491 | -0.1644402 | 0.00779285 | down |
| 149 | Cpq | 9.83E-05 | 0.51102603 | 0.69215722 | 0.32989483 | 0.00792188 | up |
| 150 | Bcr | 9.98E-05 | -0.1326808 | -0.0747576 | -0.190604 | 0.00793581 | down |
| 151 | Lratd1 | 9.95E-05 | 0.36095246 | 0.52875194 | 0.19315298 | 0.00793581 | up |
| 152 | Zmpste24 | 0.00010068 | -0.1621661 | -0.1008877 | -0.2234444 | 0.00795178 | down |
| 153 | Slc39a10 | 0.00010315 | 0.17874581 | 0.26797136 | 0.08952025 | 0.0080937 | up |
| 154 | Gabarap | 0.00010487 | 0.16530141 | 0.23831519 | 0.09228762 | 0.00814147 | up |
| 155 | Ttyh3 | 0.00010512 | -0.3376334 | -0.2243776 | -0.4508893 | 0.00814147 | down |
| 156 | Ipmk | 0.00010625 | 0.25395664 | 0.33800758 | 0.1699057 | 0.00817667 | up |
| 157 | Cd302 | 0.00011031 | 0.67938242 | 0.92510293 | 0.43366191 | 0.00836014 | up |
| 158 | Micos10 | 0.00011073 | -0.142577 | -0.093456 | -0.191698 | 0.00836014 | down |
| 159 | Psmb1 | 0.00011039 | 0.16090125 | 0.21379176 | 0.10801075 | 0.00836014 | up |
| 160 | Hoxc8 | 0.00011326 | -0.1332295 | -0.0789434 | -0.1875156 | 0.00849794 | down |
| 161 | Srprb | 0.00011459 | -0.2868309 | -0.1766915 | -0.3969703 | 0.00854432 | down |
| 162 | Pkig | 0.00011535 | -0.3479209 | -0.2284192 | -0.4674226 | 0.00854805 | down |
| 163 | Gnai1 | 0.00012004 | 0.16454111 | 0.25689858 | 0.07218363 | 0.00884112 | up |
| 164 | Lrch3 | 0.00012206 | 0.13152839 | 0.18041979 | 0.082637 | 0.00888546 | up |
| 165 | Tspan5 | 0.00012212 | 0.34820197 | 0.45936633 | 0.2370376 | 0.00888546 | up |
| 166 | Sf3b5 | 0.00012504 | -0.3117099 | -0.2030713 | -0.4203485 | 0.00904304 | down |
| 167 | Thra | 0.00012736 | 0.16680874 | 0.22057942 | 0.11303806 | 0.0091551 | up |
| 168 | Rnf40 | 0.00012814 | -0.0931879 | -0.0458784 | -0.1404974 | 0.00915692 | down |
| 169 | Rnaseh2c | 0.00013133 | -0.2260559 | -0.1429197 | -0.309192 | 0.00932289 | down |
| 170 | Serinc2 | 0.00013202 | -0.3993534 | -0.2379864 | -0.5607203 | 0.00932289 | down |
| 171 | Ephb4 | 0.00013781 | -0.1121348 | -0.0669501 | -0.1573194 | 0.00950156 | down |
| 172 | Itpkb | 0.00013851 | -0.1302784 | -0.0912528 | -0.169304 | 0.00950156 | down |
| 173 | Lpin3 | 0.00013769 | -0.175503 | -0.1213876 | -0.2296183 | 0.00950156 | down |
| 174 | Maml1 | 0.00013746 | -0.1354525 | -0.0846367 | -0.1862684 | 0.00950156 | down |
| 175 | Mocs1 | 0.00013765 | 0.13652676 | 0.18090731 | 0.0921462 | 0.00950156 | up |
| 176 | Phlda3 | 0.00014044 | -0.1650155 | -0.1137025 | -0.2163285 | 0.00957924 | down |
| 177 | Ptgir | 0.00014162 | 0.36034743 | 0.47614018 | 0.24455468 | 0.00960515 | up |
| 178 | Dnal4 | 0.00014527 | -0.0927395 | -0.0580573 | -0.1274217 | 0.00979744 | down |
| 179 | Ccrl2 | 0.00014882 | -0.0658002 | -0.0414553 | -0.0901451 | 0.00981631 | down |
| 180 | Krt8 | 0.00014796 | -0.3401874 | -0.2068627 | -0.4735121 | 0.00981631 | down |
| 181 | Ncam1 | 0.00015127 | -0.8270384 | -0.5400514 | -1.1140254 | 0.00981631 | down |
| 182 | Ncbp2 | 0.00014741 | -0.1767005 | -0.1043348 | -0.2490662 | 0.00981631 | down |
| 183 | Odc1 | 0.00015063 | -0.5392594 | -0.3474822 | -0.7310366 | 0.00981631 | down |
| 184 | Phc1 | 0.00014993 | -0.0722902 | -0.042135 | -0.1024454 | 0.00981631 | down |
| 185 | Tgfbr2 | 0.00015077 | -0.4074651 | -0.1716446 | -0.6432855 | 0.00981631 | down |
| 186 | Eif3i | 0.00015352 | 0.12040164 | 0.16731364 | 0.07348964 | 0.00986993 | up |
| 187 | Fuca1 | 0.00015374 | 0.1823112 | 0.24948644 | 0.11513596 | 0.00986993 | up |
| 188 | Bace1 | 0.00015796 | -0.5035451 | -0.3327591 | -0.6743311 | 0.00987586 | down |
| 189 | Clptm1l | 0.00015886 | -0.1390146 | -0.092982 | -0.1850472 | 0.00987586 | down |
| 190 | Ltbp1 | 0.00015959 | 0.79817812 | 1.07641176 | 0.51994447 | 0.00987586 | up |
| 191 | Nr3c1 | 0.00015957 | 0.34177344 | 0.46800082 | 0.21554606 | 0.00987586 | up |
| 192 | Ovr1 | 0.000159 | 0.21436791 | 0.27990618 | 0.14882965 | 0.00987586 | up |
| 193 | Prrc2a | 0.00015482 | -0.1200856 | -0.076901 | -0.1632703 | 0.00987586 | down |
| 194 | Srsf9 | 0.00015744 | -0.2639883 | -0.1671757 | -0.3608009 | 0.00987586 | down |
| 195 | Cpxm1 | 0.00016372 | 0.68794446 | 1.01130916 | 0.36457975 | 0.01006983 | up |
| 196 | Map3k1 | 0.00016441 | 0.13304627 | 0.17916417 | 0.08692836 | 0.01006983 | up |
| 197 | Nckap1 | 0.00016675 | 0.3292232 | 0.44654155 | 0.21190485 | 0.01008896 | up |
| 198 | Riok3 | 0.00016616 | 0.17397029 | 0.23673096 | 0.11120962 | 0.01008896 | up |
| 199 | Txnip | 0.00016724 | 0.32472652 | 0.45133042 | 0.19812262 | 0.01008896 | up |
| 200 | Ctsl | 0.00017776 | 0.52619831 | 0.72551125 | 0.32688538 | 0.01064231 | up |
| 201 | Inava | 0.00017871 | -0.2426353 | -0.1257828 | -0.3594878 | 0.01064231 | down |
| 202 | Ivns1abp | 0.00017907 | 0.16222737 | 0.21880389 | 0.10565085 | 0.01064231 | up |
| 203 | Alg1 | 0.00018908 | 0.1767166 | 0.24862453 | 0.10480866 | 0.01094967 | up |
| 204 | Ak1 | 0.00018809 | -0.4606457 | -0.3000194 | -0.6212719 | 0.01094967 | down |
| 205 | Hoxa4 | 0.00018972 | -0.0530382 | -0.033755 | -0.0723215 | 0.01094967 | down |
| 206 | Kmt2c | 0.00018869 | 0.13766771 | 0.19271461 | 0.08262081 | 0.01094967 | up |
| 207 | Ndufc1 | 0.00018865 | -0.159639 | -0.1037633 | -0.2155147 | 0.01094967 | down |
| 208 | Slc39a6 | 0.00018865 | 0.18892968 | 0.26746954 | 0.11038983 | 0.01094967 | up |
| 209 | Arhgap42 | 0.00019363 | 0.24321537 | 0.33319524 | 0.1532355 | 0.01108457 | up |
| 210 | Galm | 0.0001939 | 0.09540741 | 0.13013232 | 0.0606825 | 0.01108457 | up |
| 211 | Osbpl9 | 0.00019719 | 0.3093063 | 0.41772332 | 0.20088928 | 0.01111387 | up |
| 212 | Scp2 | 0.00019683 | 0.10865323 | 0.15473871 | 0.06256775 | 0.01111387 | up |
| 213 | Socs5 | 0.00019578 | 0.16404492 | 0.23143311 | 0.09665673 | 0.01111387 | up |
| 214 | Aldh7a1 | 0.00019907 | -0.2188627 | -0.1363404 | -0.3013849 | 0.01114595 | down |
| 215 | Ifngr2 | 0.00019962 | 0.20816678 | 0.30348659 | 0.11284697 | 0.01114595 | up |
| 216 | Jup | 0.00020239 | -0.2558953 | -0.1716515 | -0.3401392 | 0.01124836 | down |
| 217 | Cotl1 | 0.00020409 | -0.4728004 | -0.2631357 | -0.682465 | 0.01129093 | down |
| 218 | Rab28 | 0.00020679 | 0.19128325 | 0.26388999 | 0.11867651 | 0.01138765 | up |
| 219 | Stat3 | 0.0002079 | 0.26639193 | 0.38341484 | 0.14936902 | 0.01139639 | up |
| 220 | Ajuba | 0.00020943 | -0.1498777 | -0.0916798 | -0.2080757 | 0.01142806 | down |
| 221 | Ccdc6 | 0.00021261 | -0.3302602 | -0.2109614 | -0.4495589 | 0.01154947 | down |
| 222 | Dnajb11 | 0.00021831 | -0.2372839 | -0.153311 | -0.3212567 | 0.01180542 | down |
| 223 | Ndufs4 | 0.00021971 | 0.16673172 | 0.22117145 | 0.11229199 | 0.01182808 | up |
| 224 | Slk | 0.0002207 | 0.37327691 | 0.50702425 | 0.23952957 | 0.01182832 | up |
| 225 | BC034090 | 0.00022402 | -0.1264132 | -0.0774058 | -0.1754206 | 0.01185411 | down |
| 226 | Tank | 0.00022359 | 0.14976812 | 0.19847963 | 0.1010566 | 0.01185411 | up |
| 227 | Tspan3 | 0.00022415 | -0.2449754 | -0.1007084 | -0.3892423 | 0.01185411 | down |
| 228 | Tuba8 | 0.00022904 | 0.05650589 | 0.08132871 | 0.03168307 | 0.01205995 | up |
| 229 | Efnb1 | 0.0002313 | -0.3050764 | -0.186651 | -0.4235018 | 0.01207295 | down |
| 230 | Snap47 | 0.00023071 | -0.1874602 | -0.1177577 | -0.2571627 | 0.01207295 | down |
| 231 | Elf1 | 0.00023317 | 0.19226275 | 0.26078231 | 0.12374319 | 0.01208934 | up |
| 232 | Slc25a5 | 0.00023363 | 0.35994623 | 0.49637211 | 0.22352034 | 0.01208934 | up |
| 233 | Ube2m | 0.00023623 | -0.1607049 | -0.095614 | -0.2257958 | 0.01217145 | down |
| 234 | Anp32a | 0.00023877 | -0.3969944 | -0.2448895 | -0.5490993 | 0.01220217 | down |
| 235 | Golm1 | 0.00024089 | -0.251863 | -0.1424418 | -0.3612843 | 0.01220217 | down |
| 236 | Pml | 0.00024076 | -0.27432 | -0.1664599 | -0.38218 | 0.01220217 | down |
| 237 | Rab1a | 0.00023909 | 0.13703998 | 0.21535256 | 0.05872739 | 0.01220217 | up |
| 238 | Cmtm7 | 0.00024416 | -0.1352778 | -0.0612166 | -0.209339 | 0.01231579 | down |
| 239 | Alg1 | 0.00024703 | -0.1295934 | -0.0765785 | -0.1826083 | 0.01240827 | down |
| 240 | Gipc1 | 0.0002496 | -0.1656989 | -0.0954053 | -0.2359924 | 0.01243572 | down |
| 241 | Med6 | 0.00025044 | 0.13740715 | 0.19766088 | 0.07715343 | 0.01243572 | up |
| 242 | Plpp5 | 0.00025068 | -0.2953688 | -0.1902658 | -0.4004718 | 0.01243572 | down |
| 243 | Ifnar1 | 0.00025818 | 0.26337733 | 0.36411447 | 0.16264019 | 0.01270277 | up |
| 244 | Ugp2 | 0.00025759 | 0.44731315 | 0.61297809 | 0.2816482 | 0.01270277 | up |
| 245 | Fam110c | 0.0002681 | 0.17312686 | 0.23485054 | 0.11140318 | 0.01285252 | up |
| 246 | Ifitm2 | 0.00026694 | 0.29563321 | 0.38509305 | 0.20617336 | 0.01285252 | up |
| 247 | Klf3 | 0.00026601 | 0.16591984 | 0.23391802 | 0.09792166 | 0.01285252 | up |
| 248 | Ndufb2 | 0.0002675 | -0.1394348 | -0.063133 | -0.2157365 | 0.01285252 | down |
| 249 | Rap1a | 0.00026622 | 0.18191554 | 0.25094612 | 0.11288496 | 0.01285252 | up |
| 250 | Tmem51 | 0.00026411 | 0.09148042 | 0.14970782 | 0.03325302 | 0.01285252 | up |
| 251 | Tro | 0.00026872 | -0.0453984 | -0.0274394 | -0.0633575 | 0.01285252 | down |
| 252 | Gngt2 | 0.00027062 | 0.18622206 | 0.26786342 | 0.10458071 | 0.01289217 | up |
| 253 | Ccdc74a | 0.00027354 | 0.03436909 | 0.04757419 | 0.02116399 | 0.01292865 | up |
| 254 | Tfr2 | 0.00027333 | -0.1503043 | -0.090494 | -0.2101146 | 0.01292865 | down |
| 255 | Arhgef2 | 0.00027666 | -0.2217341 | -0.1421473 | -0.3013209 | 0.01302476 | down |
| 256 | Chst11 | 0.00028112 | -0.5351335 | -0.3351994 | -0.7350675 | 0.0130719 | down |
| 257 | Eml4 | 0.00028272 | -0.2018341 | -0.1136322 | -0.290036 | 0.0130719 | down |
| 258 | Mob1b | 0.00028311 | 0.12203149 | 0.17424093 | 0.06982205 | 0.0130719 | up |
| 259 | Mrgprf | 0.00028069 | 0.24101193 | 0.32608254 | 0.15594132 | 0.0130719 | up |
| 260 | Rnf41 | 0.0002793 | -0.1376249 | -0.0683498 | -0.2068999 | 0.0130719 | down |
| 261 | Acp2 | 0.00028465 | 0.17457903 | 0.23926882 | 0.10988924 | 0.01309289 | up |
| 262 | Arfip2 | 0.00028944 | -0.1601013 | -0.1057187 | -0.2144839 | 0.01325341 | down |
| 263 | Ldoc1 | 0.00029035 | 0.03142937 | 0.04146643 | 0.02139231 | 0.01325341 | up |
| 264 | Ndel1 | 0.00029219 | 0.18969226 | 0.26345115 | 0.11593338 | 0.01328706 | up |
| 265 | Dlc1 | 0.00029708 | -0.3773858 | -0.2581127 | -0.4966588 | 0.01345835 | down |
| 266 | Csnk1g1 | 0.00030241 | 0.09022553 | 0.13687546 | 0.04357561 | 0.01354653 | up |
| 267 | Hcfc1 | 0.00030191 | -0.1028587 | -0.0541557 | -0.1515617 | 0.01354653 | down |
| 268 | Hfe | 0.00030067 | 0.13351814 | 0.20014609 | 0.06689019 | 0.01354653 | up |
| 269 | Shb | 0.0003047 | -0.2051826 | -0.1348096 | -0.2755556 | 0.01359827 | down |
| 270 | Cenpb | 0.00030839 | -0.2372554 | -0.1402705 | -0.3342402 | 0.0137118 | down |
| 271 | Abtb2 | 0.00031912 | -0.1230347 | -0.0680904 | -0.1779789 | 0.01409482 | down |
| 272 | Cacfd1 | 0.0003217 | -0.2157905 | -0.1201519 | -0.3114291 | 0.01409482 | down |
| 273 | Mycbp2 | 0.00032152 | 0.24488417 | 0.33621067 | 0.15355766 | 0.01409482 | up |
| 274 | Xpnpep1 | 0.00032095 | -0.1721279 | -0.1065872 | -0.2376687 | 0.01409482 | down |
| 275 | Yipf6 | 0.00032508 | 0.08814854 | 0.12464413 | 0.05165295 | 0.01419129 | up |
| 276 | Tmem132a | 0.0003274 | -0.1716552 | -0.102599 | -0.2407114 | 0.01424072 | down |
| 277 | Irgq | 0.00033288 | -0.12746 | -0.086805 | -0.1681151 | 0.01437472 | down |
| 278 | Zfp622 | 0.00033187 | -0.0991189 | -0.0561896 | -0.1420482 | 0.01437472 | down |
| 279 | Nt5e | 0.0003354 | 0.11486214 | 0.21737405 | 0.01235022 | 0.01443202 | up |
| 280 | Maea | 0.00033897 | -0.1139223 | -0.0683821 | -0.1594625 | 0.01453317 | down |
| 281 | Cd47 | 0.00034268 | 0.51954299 | 0.72199192 | 0.31709405 | 0.01453677 | up |
| 282 | Pla1a | 0.00034197 | 0.49707121 | 0.75420802 | 0.23993441 | 0.01453677 | up |
| 283 | Serpini1 | 0.00034241 | 0.04613749 | 0.07244869 | 0.01982629 | 0.01453677 | up |
| 284 | Axl1 | 0.00035727 | 0.14638711 | 0.21270648 | 0.08006774 | 0.01464703 | up |
| 285 | Bnip3l | 0.0003586 | 0.20582232 | 0.31871159 | 0.09293305 | 0.01464703 | up |
| 286 | Dok1 | 0.00035783 | -0.4783747 | -0.2904359 | -0.6663135 | 0.01464703 | down |
| 287 | Ece1 | 0.00036236 | 0.24715839 | 0.34033763 | 0.15397915 | 0.01464703 | up |
| 288 | Fam110a | 0.00034827 | -0.1816379 | -0.0789199 | -0.2843559 | 0.01464703 | down |
| 289 | Flot2 | 0.00034878 | 0.21847302 | 0.30227186 | 0.13467419 | 0.01464703 | up |
| 290 | Gpc2 | 0.00036159 | -0.0354258 | -0.0228739 | -0.0479777 | 0.01464703 | down |
| 291 | Gtf3c1 | 0.00034898 | -0.0833022 | -0.0437978 | -0.1228066 | 0.01464703 | down |
| 292 | Itpkc | 0.00035754 | -0.0843653 | -0.045814 | -0.1229167 | 0.01464703 | down |
| 293 | Klc2 | 0.00036222 | -0.0989025 | -0.062749 | -0.1350559 | 0.01464703 | down |
| 294 | Lgalsl | 0.00035447 | -0.3488235 | -0.2172717 | -0.4803754 | 0.01464703 | down |
| 295 | Pard6a | 0.00035774 | -0.0560352 | -0.0355971 | -0.0764732 | 0.01464703 | down |
| 296 | Sat1 | 0.00036081 | 0.41592064 | 0.56840223 | 0.26343906 | 0.01464703 | up |
| 297 | Zdhhc16 | 0.00035929 | -0.2751694 | -0.1596116 | -0.3907272 | 0.01464703 | down |
| 298 | Syap1 | 0.00036546 | 0.17235177 | 0.24343834 | 0.1012652 | 0.01472273 | up |
| 299 | Sox13 | 0.00036831 | 0.06147719 | 0.08432673 | 0.03862766 | 0.01478765 | up |
| 300 | Filip1 | 0.00037105 | 0.05582744 | 0.09092514 | 0.02072974 | 0.01484815 | up |
| 301 | Abracl | 0.00037649 | -0.2395324 | -0.152239 | -0.3268259 | 0.01496879 | down |
| 302 | Wnt11 | 0.00037656 | -0.3914502 | -0.2206488 | -0.5622515 | 0.01496879 | down |
| 303 | Tnfaip2 | 0.00038099 | 0.34670921 | 0.55808403 | 0.13533439 | 0.01509503 | up |
| 304 | Ash1l | 0.00038912 | 0.13096819 | 0.18254274 | 0.07939363 | 0.01536632 | up |
| 305 | Psmd13 | 0.00039465 | -0.1471705 | -0.0765544 | -0.2177865 | 0.01553354 | down |
| 306 | Entpd7 | 0.00039719 | -0.3285509 | -0.2034039 | -0.4536979 | 0.01558269 | down |
| 307 | Naa15 | 0.00039899 | 0.17445714 | 0.2485378 | 0.10037648 | 0.01560204 | up |
| 308 | Ppp1r13l | 0.00040124 | -0.1472179 | -0.0897105 | -0.2047252 | 0.01560881 | down |
| 309 | Vangl2 | 0.00040176 | -0.1338113 | -0.0795314 | -0.1880912 | 0.01560881 | down |
| 310 | Tmcc2 | 0.00040306 | -0.1006464 | -0.051789 | -0.1495037 | 0.01560902 | down |
| 311 | Mpzl2 | 0.00040788 | -0.0383096 | -0.0229172 | -0.0537019 | 0.01574454 | down |
| 312 | Myl6 | 0.00040992 | -0.3154773 | -0.2012814 | -0.4296731 | 0.01577274 | down |
| 313 | Oaz2 | 0.00041314 | -0.1992574 | -0.1115435 | -0.2869712 | 0.01584565 | down |
| 314 | Pnpla8 | 0.00041567 | 0.23630779 | 0.33135987 | 0.14125571 | 0.01587123 | up |
| 315 | Tagln | 0.00041645 | -1.0673137 | -0.6351008 | -1.4995265 | 0.01587123 | down |
| 316 | Dusp6 | 0.00042005 | -0.3919242 | -0.2275508 | -0.5562977 | 0.01595803 | down |
| 317 | Por | 0.00042629 | -0.2077707 | -0.1381403 | -0.277401 | 0.01614381 | down |
| 318 | Foxk1 | 0.00043377 | -0.1263081 | -0.0830688 | -0.1695475 | 0.01633105 | down |
| 319 | Ppp1r11 | 0.00043531 | -0.1088236 | -0.0609815 | -0.1566657 | 0.01633105 | down |
| 320 | Prelid3b | 0.00043408 | 0.10945177 | 0.1689377 | 0.04996584 | 0.01633105 | up |
| 321 | Crlf1 | 0.00043811 | -0.4327528 | -0.1846758 | -0.6808298 | 0.01638492 | down |
| 322 | Arg1 | 0.00045121 | -0.2996467 | -0.163791 | -0.4355024 | 0.01641445 | down |
| 323 | Cttnbp2nl | 0.00044989 | -0.11137 | -0.0646628 | -0.1580772 | 0.01641445 | down |
| 324 | Eif2b2 | 0.00044167 | -0.1286277 | -0.0601954 | -0.1970599 | 0.01641445 | down |
| 325 | Grik2 | 0.00044887 | 0.14543317 | 0.22571689 | 0.06514945 | 0.01641445 | up |
| 326 | Nmd3 | 0.00044542 | 0.13602644 | 0.17966894 | 0.09238394 | 0.01641445 | up |
| 327 | Nrip1 | 0.00044408 | -0.2737811 | -0.1541171 | -0.3934452 | 0.01641445 | down |
| 328 | Pgrmc2 | 0.00044964 | -0.2174238 | -0.1344141 | -0.3004336 | 0.01641445 | down |
| 329 | Relch | 0.00044978 | 0.11156859 | 0.17867927 | 0.04445792 | 0.01641445 | up |
| 330 | Tbc1d16 | 0.00044851 | -0.0862742 | -0.0537449 | -0.1188036 | 0.01641445 | down |
| 331 | Vegfb | 0.00045316 | 0.07619305 | 0.11883731 | 0.03354879 | 0.01643565 | up |
| 332 | Gltp | 0.00045731 | 0.12707943 | 0.17562066 | 0.0785382 | 0.01647776 | up |
| 333 | Mical3 | 0.00045814 | -0.123336 | -0.0660615 | -0.1806104 | 0.01647776 | down |
| 334 | Zhx1 | 0.00045844 | 0.14557067 | 0.21109484 | 0.08004651 | 0.01647776 | up |
| 335 | Sub1 | 0.00045986 | 0.18863353 | 0.26316047 | 0.11410659 | 0.01647933 | up |
| 336 | Slc44a1 | 0.00046158 | 0.3323017 | 0.47305716 | 0.19154624 | 0.01649192 | up |
| 337 | Tnfsf11 | 0.00046405 | 0.19643622 | 0.3397199 | 0.05315254 | 0.01653107 | up |
| 338 | Pfk3 | 0.00046727 | -0.1739119 | -0.1107387 | -0.2370851 | 0.0165475 | down |
| 339 | Tdrd7 | 0.00046616 | 0.10895219 | 0.15882628 | 0.05907809 | 0.0165475 | up |
| 340 | Tmem65 | 0.00046891 | 0.31516189 | 0.43995342 | 0.19037036 | 0.01655681 | up |
| 341 | Sphk2 | 0.00047217 | -0.0679498 | -0.0323605 | -0.103539 | 0.01662278 | down |
| 342 | Fosl1 | 0.00047768 | 0.36754031 | 0.49773039 | 0.23735023 | 0.01666384 | up |
| 343 | Papss2 | 0.00047889 | -0.1770346 | -0.1100866 | -0.2439825 | 0.01666384 | down |
| 344 | Stmn4 | 0.00047537 | -0.1107549 | -0.0640278 | -0.157482 | 0.01666384 | down |
| 345 | Zrsr2 | 0.00047747 | -0.1167445 | -0.0567839 | -0.1767052 | 0.01666384 | down |
| 346 | DEPRECATED-[illegible] | 0.00048194 | 0.08808763 | 0.12619162 | 0.04998365 | 0.01672168 | up |
| 347 | Lrrk1 | 0.0004861 | -0.1841573 | -0.0713749 | -0.2969396 | 0.01681729 | down |
| 348 | Arrdc3 | 0.00048786 | 0.31628411 | 0.46867193 | 0.16389628 | 0.01682089 | up |
| 349 | Ncstn | 0.000489 | 0.14882995 | 0.19777225 | 0.09988764 | 0.01682089 | up |
| 350 | Lrrc58 | 0.00049096 | -0.117278 | -0.0598309 | -0.1747251 | 0.01683977 | down |
| 351 | Wnt9a | 0.0004967 | -0.2151781 | -0.1262877 | -0.3040684 | 0.01698818 | down |
| 352 | Rspo3 | 0.00049987 | 0.09201618 | 0.13899728 | 0.04503508 | 0.0170481 | up |
| 353 | Adamts2 | 0.00050283 | 0.34163621 | 0.53092227 | 0.15235015 | 0.01710036 | up |
| 354 | Lzts2 | 0.00050692 | -0.2817952 | -0.1669435 | -0.3966468 | 0.01715688 | down |
| 355 | Myh9 | 0.00050878 | -0.4301412 | -0.2731959 | -0.5870866 | 0.01715688 | down |
| 356 | Tcn2 | 0.00050741 | 0.19714843 | 0.28818718 | 0.10610968 | 0.01715688 | up |
| 357 | Fam220a.1 | 0.00051471 | 0.1218388 | 0.18365052 | 0.06002708 | 0.01719092 | up |
| 358 | Kctd11 | 0.0005153 | -0.3248414 | -0.1949103 | -0.4547725 | 0.01719092 | down |
| 359 | Thap11 | 0.00051272 | -0.1114616 | -0.0610384 | -0.1618848 | 0.01719092 | down |
| 360 | Ypel5 | 0.00051551 | 0.18709025 | 0.26738446 | 0.10679605 | 0.01719092 | up |
| 361 | Col18a1 | 0.00051794 | 1.0262393 | 1.51596994 | 0.53650865 | 0.01722388 | up |
| 362 | Cdkn1c | 0.00052039 | 0.13057162 | 0.19143403 | 0.06970922 | 0.01725766 | up |
| 363 | Ptprt | 0.0005228 | 0.05569582 | 0.09734874 | 0.01404291 | 0.01728974 | up |
| 364 | En1 | 0.00052519 | -0.2890749 | -0.1524205 | -0.4257292 | 0.01732115 | down |
| 365 | Csnk1a1 | 0.0005357 | 0.13348132 | 0.18139665 | 0.08556598 | 0.01761936 | up |
| 366 | Pgd | 0.00054317 | 0.20807278 | 0.29401671 | 0.12212886 | 0.01781624 | up |
| 367 | Gpalpp1 | 0.00054851 | 0.11128822 | 0.1566667 | 0.06590975 | 0.01784529 | up |
| 368 | Nit2 | 0.00054704 | 0.0820625 | 0.11665758 | 0.04746742 | 0.01784529 | up |
| 369 | Sp6 | 0.00054755 | -0.0675631 | -0.0438906 | -0.0912357 | 0.01784529 | down |
| 370 | Rasgrp2 | 0.00055328 | 0.12889526 | 0.21221241 | 0.04557812 | 0.01795154 | up |
| 371 | Ptges3l | 0.00055939 | -0.0822144 | -0.0497219 | -0.1147069 | 0.01800393 | down |
| 372 | Rock2 | 0.00055879 | 0.28829291 | 0.40298654 | 0.17359929 | 0.01800393 | up |
| 373 | Zeb2 | 0.00055866 | 0.21696595 | 0.32144771 | 0.11248419 | 0.01800393 | up |
| 374 | Gnpda2 | 0.00056758 | 0.14421659 | 0.22720123 | 0.06123196 | 0.01807368 | up |
| 375 | Hsbp1 | 0.00056681 | -0.1187328 | -0.0667413 | -0.1707242 | 0.01807368 | down |
| 376 | Rrm2b | 0.00056721 | 0.07820385 | 0.12707707 | 0.02933063 | 0.01807368 | up |
| 377 | Vps26b | 0.00056574 | -0.1395317 | -0.0877602 | -0.1913031 | 0.01807368 | down |
| 378 | Wipi1 | 0.00057223 | 0.08897634 | 0.1977519 | -0.0197992 | 0.01817375 | up |
| 379 | Cd3eap | 0.0005771 | -0.060346 | -0.0356448 | -0.0850473 | 0.01823189 | down |
| 380 | Fdx1 | 0.00057637 | 0.14438595 | 0.20163047 | 0.08714143 | 0.01823189 | up |
| 381 | Ggcx | 0.00057991 | -0.2166834 | -0.128145 | -0.3052218 | 0.01827255 | down |
| 382 | Foxo3 | 0.00058321 | -0.1644867 | -0.0958744 | -0.233099 | 0.01832825 | down |
| 383 | Tbca | 0.00058608 | 0.19276935 | 0.26939119 | 0.11614751 | 0.01837046 | up |
| 384 | Cdk20 | 0.00058881 | 0.13196698 | 0.18184745 | 0.08208651 | 0.01840813 | up |
| 385 | Ly96 | 0.00059602 | 0.12225619 | 0.1843144 | 0.06019799 | 0.01853676 | up |
| 386 | Tmsb4x | 0.00059515 | -0.5711326 | -0.2503221 | -0.8919431 | 0.01853676 | down |
| 387 | Atp6v1c1 | 0.00059969 | 0.20026789 | 0.27960009 | 0.12093568 | 0.01860288 | up |
| 388 | Grcc10 | 0.00060161 | -0.130433 | -0.0769804 | -0.1838856 | 0.0186141 | down |
| 389 | Il19 | 0.00060865 | 0.08536645 | 0.13144537 | 0.03928753 | 0.01878369 | up |
| 390 | Jag1 | 0.00061458 | -0.2441668 | -0.1431967 | -0.345137 | 0.01886968 | down |
| 391 | Ramp3 | 0.00061343 | 0.04264605 | 0.0603783 | 0.02491381 | 0.01886968 | up |
| 392 | Naa20 | 0.00061819 | 0.14114132 | 0.19332685 | 0.0889558 | 0.01888404 | up |
| 393 | Synj2 | 0.0006171 | -0.1765858 | -0.1120406 | -0.2411311 | 0.01888404 | down |
| 394 | Rhoq | 0.00062025 | 0.27457684 | 0.43081728 | 0.1183364 | 0.01889861 | up |
| 395 | Prkcb | 0.00062332 | 0.08834228 | 0.15099723 | 0.02568733 | 0.01894408 | up |
| 396 | Adprm | 0.00062587 | 0.06297576 | 0.08836378 | 0.03758775 | 0.01897364 | up |
| 397 | Bcl2l11 | 0.00063191 | -0.1283178 | -0.0803338 | -0.1763017 | 0.01908769 | down |
| 398 | Ccdc82 | 0.00063576 | 0.10753525 | 0.14777311 | 0.06729739 | 0.01908769 | up |
| 399 | Fdxr | 0.00063599 | -0.2357917 | -0.1412519 | -0.3303316 | 0.01908769 | down |
| 400 | Podnl1 | 0.00063595 | -0.421904 | -0.0752955 | -0.7685125 | 0.01908769 | down |
| 401 | Draxin | 0.00064163 | 0.31738487 | 0.44963695 | 0.18513278 | 0.01914742 | up |
| 402 | Ubtd2 | 0.00064257 | 0.16071112 | 0.22680353 | 0.09461872 | 0.01914742 | up |
| 403 | Zbtb7b | 0.00064277 | -0.1159103 | -0.0735146 | -0.158306 | 0.01914742 | down |
| 404 | Nt5dc2 | 0.00064673 | -0.5143561 | -0.3061665 | -0.7225458 | 0.01921788 | down |
| 405 | Dnm1 | 0.00064991 | 0.10523792 | 0.15624428 | 0.05423156 | 0.0192647 | up |
| 406 | Ppp1r35 | 0.00065299 | -0.0679995 | -0.0367314 | -0.0992675 | 0.01930814 | down |
| 407 | Crk | 0.00065511 | -0.1953427 | -0.1146308 | -0.2760546 | 0.0193233 | down |
| 408 | St6galnac5 | 0.0006609 | 0.07015167 | 0.13083446 | 0.00946887 | 0.01944621 | up |
| 409 | Creb3 | 0.00066713 | -0.1181824 | -0.0695119 | -0.1668529 | 0.01958158 | down |
| 410 | Got2 | 0.00067653 | -0.2714781 | -0.1512042 | -0.391752 | 0.01980914 | down |
| 411 | Ereg | 0.00068847 | 0.29190451 | 0.48603893 | 0.09777008 | 0.02006102 | up |
| 412 | Sipa1l3 | 0.00068729 | -0.1172505 | -0.0669341 | -0.1675669 | 0.02006102 | down |
| 413 | Wdfy2 | 0.00069264 | 0.09343134 | 0.13662896 | 0.05023372 | 0.02013362 | up |
| 414 | Slc52a2 | 0.0006964 | -0.065099 | -0.0268801 | -0.103318 | 0.02014536 | down |
| 415 | Timm50 | 0.00069587 | -0.11367 | -0.0631807 | -0.1641593 | 0.02014536 | down |
| 416 | Adcy7 | 0.00069971 | -0.3254633 | -0.1412832 | -0.5096434 | 0.02019229 | down |
| 417 | Gna13 | 0.00070596 | -0.1134675 | -0.0615293 | -0.1654057 | 0.02032374 | down |
| 418 | Abraxas2 | 0.00071615 | 0.07940539 | 0.11397516 | 0.04483561 | 0.02037601 | up |
| 419 | Aldh18a1 | 0.00071234 | -0.2866178 | -0.1666943 | -0.4065413 | 0.02037601 | down |
| 420 | Nsa2 | 0.00071626 | 0.33861477 | 0.49199137 | 0.18523818 | 0.02037601 | up |
| 421 | Tgfbr3 | 0.00071401 | 0.643118 | 0.98794525 | 0.29829075 | 0.02037601 | up |
| 422 | Vrk3 | 0.00071587 | -0.1312325 | -0.0545343 | -0.2079307 | 0.02037601 | down |
| 423 | Relt | 0.00071883 | -0.0665839 | -0.0423357 | -0.0908322 | 0.02040072 | down |
| 424 | Arrb1 | 0.00073353 | 0.22284351 | 0.32573059 | 0.11995643 | 0.02047913 | up |
| 425 | Atxn3 | 0.00072731 | 0.10677693 | 0.15739552 | 0.05615834 | 0.02047913 | up |
| 426 | Cpz | 0.00072558 | 0.3388456 | 0.53008987 | 0.14760132 | 0.02047913 | up |
| 427 | Dazap2 | 0.00072881 | 0.12122967 | 0.18603462 | 0.05642472 | 0.02047913 | up |
| 428 | H2-Q10 | 0.00073125 | 0.0343251 | 0.04770965 | 0.02094055 | 0.02047913 | up |
| 429 | Prep | 0.00073294 | -0.1056655 | -0.0436531 | -0.1676779 | 0.02047913 | down |
| 430 | Psap | 0.00072754 | 0.38487043 | 0.5462638 | 0.22347705 | 0.02047913 | up |
| 431 | Pmepa1 | 0.00074036 | -0.9918139 | -0.5716306 | -1.4119972 | 0.02057254 | down |
| 432 | Prrx1 | 0.00074202 | 0.37425008 | 0.49548533 | 0.25301482 | 0.02057254 | up |
| 433 | Trmt11 | 0.00074003 | 0.06322422 | 0.09222494 | 0.0342235 | 0.02057254 | up |
| 434 | Naxe | 0.00074447 | 0.12734806 | 0.19398996 | 0.06070616 | 0.02059314 | up |
| 435 | Cnpy3 | 0.00075678 | -0.1924942 | -0.1121792 | -0.2728091 | 0.0207026 | down |
| 436 | Epha2 | 0.00075486 | -0.0799758 | -0.0587865 | -0.1011651 | 0.0207026 | down |
| 437 | Hp1bp3 | 0.00075656 | 0.09393797 | 0.13697794 | 0.050898 | 0.0207026 | up |
| 438 | Sybu | 0.00075405 | 0.01208006 | 0.018355 | 0.00580511 | 0.0207026 | up |
| 439 | Tln2 | 0.00075705 | -0.3717303 | -0.2236391 | -0.5198214 | 0.0207026 | down |
| 440 | Arrdc4 | 0.0007766 | 0.16945975 | 0.26740143 | 0.07151807 | 0.02071787 | up |
| 441 | Baiap2l1 | 0.00077374 | -0.0618268 | -0.0341735 | -0.0894801 | 0.02071787 | down |
| 442 | Bambi | 0.00076761 | -0.0806576 | -0.0531905 | -0.1081248 | 0.02071787 | down |
| 443 | Bzw1 | 0.00076447 | 0.29671906 | 0.49741737 | 0.09602075 | 0.02071787 | up |
| 444 | Crlf2 | 0.00077145 | -0.1040593 | -0.0626552 | -0.1454635 | 0.02071787 | down |
| 445 | Fgf10 | 0.00077572 | 0.20101312 | 0.27961923 | 0.12240701 | 0.02071787 | up |
| 446 | Lipa | 0.0007692 | 0.15651177 | 0.22987827 | 0.08314527 | 0.02071787 | up |
| 447 | Mdfi | 0.00077123 | -0.2389301 | -0.1465129 | -0.3313473 | 0.02071787 | down |
| 448 | Pros1 | 0.00077225 | 0.34313033 | 0.53187133 | 0.15438932 | 0.02071787 | up |
| 449 | Srp68 | 0.00077064 | -0.1659851 | -0.0949807 | -0.2369895 | 0.02071787 | down |
| 450 | Stc1 | 0.00076587 | 0.22014593 | 0.31670938 | 0.12358249 | 0.02071787 | up |
| 451 | Rapgef2 | 0.00078478 | -0.1031409 | -0.0568366 | -0.1494453 | 0.02075171 | down |
| 452 | Snw1 | 0.00078235 | 0.18546562 | 0.25738564 | 0.1135456 | 0.02075171 | up |
| 453 | Stt3b | 0.00078427 | 0.12629752 | 0.20525343 | 0.04734161 | 0.02075171 | up |
| 454 | Unc5b | 0.00078235 | -0.3589525 | 0.02941151 | -0.7473164 | 0.02075171 | down |
| 455 | Mtrex | 0.00078956 | 0.08799078 | 0.12752593 | 0.04845563 | 0.02078661 | up |
| 456 | Usp24 | 0.00078956 | 0.14160548 | 0.20385059 | 0.07936037 | 0.02078661 | up |
| 457 | Cbfb | 0.00080271 | -0.3143823 | -0.1828365 | -0.4459282 | 0.02082391 | down |
| 458 | Cyth1 | 0.00080065 | -0.1029244 | -0.0565612 | -0.1492877 | 0.02082391 | down |
| 459 | Dgkd | 0.00080116 | -0.102204 | -0.065497 | -0.1389111 | 0.02082391 | down |
| 460 | Grb10 | 0.00079284 | 0.34270325 | 0.4996936 | 0.18571289 | 0.02082391 | up |
| 461 | Hoxb2 | 0.00079891 | -0.2013111 | -0.115252 | -0.2873702 | 0.02082391 | down |
| 462 | Mospd2 | 0.00079716 | 0.07000156 | 0.10584656 | 0.03415656 | 0.02082391 | up |
| 463 | Pdp1 | 0.00080312 | 0.12732986 | 0.16912784 | 0.08553188 | 0.02082391 | up |
| 464 | Nadk | 0.00080844 | 0.2568109 | 0.36322312 | 0.15039867 | 0.02091652 | up |
| 465 | Zbtb1 | 0.00081071 | 0.0947396 | 0.14133651 | 0.0481427 | 0.02093021 | up |
| 466 | Arl1 | 0.00081906 | 0.29723482 | 0.45477678 | 0.13969285 | 0.02093162 | up |
| 467 | Cavin4 | 0.00081606 | -0.0286001 | -0.0149952 | -0.0422051 | 0.02093162 | down |
| 468 | Cnnm4 | 0.00081674 | -0.1858048 | -0.1103357 | -0.2612739 | 0.02093162 | down |
| 469 | Pgf | 0.00081948 | -0.1856029 | -0.0986406 | -0.2725651 | 0.02093162 | down |
| 470 | Steap2 | 0.00081729 | 0.21078223 | 0.33000804 | 0.09155643 | 0.02093162 | up |
| 471 | Agpat1 | 0.00082994 | -0.1000184 | -0.0533831 | -0.1466536 | 0.02093409 | down |
| 472 | Arhgef40 | 0.00083527 | -0.2688087 | -0.1540106 | -0.3836068 | 0.02093409 | down |
| 473 | Atp2b1 | 0.00083072 | 0.19002315 | 0.25797963 | 0.12206667 | 0.02093409 | up |
| 474 | Gcn1 | 0.00082806 | -0.1873426 | -0.1076682 | -0.2670171 | 0.02093409 | down |
| 475 | Il1rl2 | 0.00083034 | 0.25855358 | 0.38061405 | 0.13649311 | 0.02093409 | up |
| 476 | Ptpmt1 | 0.00082416 | 0.16478085 | 0.23273283 | 0.09682886 | 0.02093409 | up |
| 477 | Rfx7 | 0.00083423 | 0.08135368 | 0.1228543 | 0.03985306 | 0.02093409 | up |
| 478 | Sorbs1 | 0.0008326 | -0.0678762 | -0.0452401 | -0.0905124 | 0.02093409 | down |
| 479 | Tmem243 | 0.00082324 | 0.08349879 | 0.11541816 | 0.05157942 | 0.02093409 | up |
| 480 | Pwwp2b | 0.00084324 | 0.03861096 | 0.05789383 | 0.01932808 | 0.02108973 | up |
| 481 | Kdm5c | 0.00084526 | -0.1149716 | -0.053721 | -0.1762222 | 0.02109643 | down |
| 482 | Elf3e | 0.00085061 | 0.37516569 | 0.54045723 | 0.20987416 | 0.02115419 | up |
| 483 | Gxylt2 | 0.0008511 | -0.3856353 | -0.2354319 | -0.5358387 | 0.02115419 | down |
| 484 | Fez1 | 0.00086094 | 0.20895463 | 0.30985836 | 0.10805091 | 0.02117958 | up |

| | | | | | | | |
|---|---|---|---|---|---|---|---|
| 485 | Jade1 | 0.00085503 | -0.2611085 | -0.1475053 | -0.3747118 | 0.02117958 | down |
| 486 | Lrrc15 | 0.00085972 | -1.7315871 | -1.0623361 | -2.4008382 | 0.02117958 | down |
| 487 | Scpep1 | 0.00085736 | 0.27388391 | 0.44723949 | 0.10052834 | 0.02117958 | up |
| 488 | Xpa | 0.00085793 | 0.14450912 | 0.20907006 | 0.07994817 | 0.02117958 | up |
| 489 | Stk24 | 0.00086311 | 0.21274658 | 0.29977536 | 0.1257178 | 0.02118935 | up |
| 490 | Tma7 | 0.00086539 | 0.2228533 | 0.31419422 | 0.13151238 | 0.02120211 | up |
| 491 | Ap4s1 | 0.00088563 | 0.10201935 | 0.14646624 | 0.05757247 | 0.02164844 | up |
| 492 | Ccdc122 | 0.00089429 | 0.12237765 | 0.18676788 | 0.05798742 | 0.02164844 | up |
| 493 | Hpse | 0.00089358 | 0.29059594 | 0.43205033 | 0.14914154 | 0.02164844 | up |
| 494 | Mdk | 0.00088747 | -0.3243381 | -0.1608419 | -0.4878343 | 0.02164844 | down |
| 495 | Peg3 | 0.0008916 | -0.2651395 | -0.1644534 | -0.3658257 | 0.02164844 | down |
| 496 | Prkca | 0.00089443 | 0.08508454 | 0.1218797 | 0.04828938 | 0.02164844 | up |
| 497 | Ndufa8 | 0.00089626 | -0.1831677 | -0.0894604 | -0.276875 | 0.02164915 | down |
| 498 | Cds2 | 0.00089816 | 0.2092312 | 0.31356533 | 0.10489708 | 0.02165138 | up |
| 499 | Safb2 | 0.0009016 | -0.1431631 | -0.075312 | -0.2110141 | 0.0216908 | down |
| 500 | Ier5l | 0.00090433 | -0.2633906 | -0.1549108 | -0.3718703 | 0.02171289 | down |
| 501 | Gjb4 | 0.00090816 | -0.0202481 | -0.010161 | -0.0303353 | 0.02176147 | down |
| 502 | Gpr39 | 0.00091199 | -0.1712076 | -0.1103096 | -0.2321056 | 0.02180957 | down |
| 503 | Impdh1 | 0.00091412 | 0.10216487 | 0.15101719 | 0.05331255 | 0.02181701 | up |
| 504 | Bicd2 | 0.00092848 | -0.1120455 | -0.062063 | -0.162028 | 0.02210072 | down |
| 505 | Ctps2 | 0.00093588 | 0.23073544 | 0.32597854 | 0.13549234 | 0.02210072 | up |
| 506 | Igfbp7 | 0.00093876 | -0.4625186 | -0.3186832 | -0.6063541 | 0.02210072 | down |
| 507 | Kcnmb1 | 0.00093098 | -0.0912942 | -0.0425965 | -0.139992 | 0.02210072 | down |
| 508 | Rrm1 | 0.00093296 | -0.2608045 | -0.1664432 | -0.3551657 | 0.02210072 | down |
| 509 | Snx14 | 0.00093889 | 0.14426564 | 0.22229684 | 0.06623444 | 0.02210072 | up |
| 510 | Taf9b | 0.00093672 | 0.08134171 | 0.11973788 | 0.04294553 | 0.02210072 | up |
| 511 | Ttc39c | 0.00094469 | 0.07114689 | 0.10483156 | 0.03746222 | 0.02219363 | up |
| 512 | Snx30 | 0.00095079 | -0.1159446 | -0.0667996 | -0.1650897 | 0.02229341 | down |
| 513 | Dusp7 | 0.00095851 | 0.18634873 | 0.2582368 | 0.11446065 | 0.02234572 | up |
| 514 | Lonp2 | 0.0009586 | 0.12809919 | 0.17849025 | 0.07770813 | 0.02234572 | up |
| 515 | Wrn | 0.00095831 | 0.0722315 | 0.10439527 | 0.04005102 | 0.02234572 | up |
| 516 | Cmklr1 | 0.0009653 | -0.2106638 | -0.1100022 | -0.3113254 | 0.02241258 | down |
| 517 | Irf1 | 0.00096624 | -0.1400547 | -0.0688268 | -0.2112826 | 0.02241258 | down |
| 518 | Zfp52 | 0.00096707 | 0.05543344 | 0.08466672 | 0.02620016 | 0.02241258 | up |
| 519 | Ier5 | 0.00096998 | -0.3053991 | -0.1819071 | -0.4288911 | 0.02241466 | down |
| 520 | Mmp19 | 0.0009709 | 0.25164904 | 0.35042135 | 0.15287673 | 0.02241466 | up |
| 521 | Avpi1 | 0.00097406 | -0.0867698 | -0.0508702 | -0.1226694 | 0.02242337 | down |
| 522 | Rnft1 | 0.00097501 | 0.32832343 | 0.46531879 | 0.19132808 | 0.02242337 | up |
| 523 | Lrat | 0.00097721 | -0.0758001 | -0.0391437 | -0.1124565 | 0.02243104 | down |
| 524 | Itb | 0.00098804 | 0.11524078 | 0.17124864 | 0.05923292 | 0.02263637 | up |
| 525 | Onecut2 | 0.0009917 | -0.0582717 | -0.037404 | -0.0791394 | 0.02267681 | down |
| 526 | Cyb5a | 0.0009965 | 0.46321386 | 0.6572473 | 0.26918043 | 0.02274016 | up |
| 527 | Tmx4 | 0.00099826 | 0.08460386 | 0.12934185 | 0.03986586 | 0.02274016 | up |
| 528 | Kctd9 | 0.00100227 | 0.17377176 | 0.24432802 | 0.1032155 | 0.02278826 | up |
| 529 | Arhgap26 | 0.00100614 | 0.03560649 | 0.05008783 | 0.02112516 | 0.02283308 | up |
| 530 | Naf1 | 0.00102331 | 0.12153534 | 0.18552439 | 0.05754628 | 0.02300527 | up |
| 531 | Pbp2 | 0.00102211 | -0.0567441 | -0.0302773 | -0.0832109 | 0.02300527 | down |
| 532 | Rnf115 | 0.00102192 | 0.15577959 | 0.23569064 | 0.07586854 | 0.02300527 | up |
| 533 | Smtn | 0.00102201 | -0.2778488 | -0.1566226 | -0.3990751 | 0.02300527 | down |
| 534 | Tnrc18 | 0.00101769 | -0.1992715 | -0.115026 | -0.2835171 | 0.02300527 | down |
| 535 | Ralbp1 | 0.0010297 | 0.15068125 | 0.21534963 | 0.08601286 | 0.02310577 | up |
| 536 | Timmdc1 | 0.00104111 | -0.1013924 | -0.0504633 | -0.1523216 | 0.02331824 | down |
| 537 | Zim1 | 0.00104579 | -0.019913 | -0.0082283 | -0.0315978 | 0.02337946 | down |
| 538 | Lrrfip1 | 0.00104932 | -0.162994 | -0.0893555 | -0.2366325 | 0.02341458 | down |
| 539 | Pla2r1 | 0.00105602 | 0.2046597 | 0.28633995 | 0.12297944 | 0.02352055 | up |
| 540 | Mospd1 | 0.0010663 | 0.10843988 | 0.15382116 | 0.06305861 | 0.02366171 | up |
| 541 | Tmem119 | 0.00106454 | -0.6040856 | -0.3269463 | -0.8812249 | 0.02366171 | down |
| 542 | Lpcat1 | 0.00107664 | -0.176191 | -0.1089614 | -0.2434207 | 0.02384698 | down |
| 543 | Cdk11b | 0.00108267 | -0.0973441 | -0.0500429 | -0.1446453 | 0.02384856 | down |
| 544 | Clybl | 0.00108195 | 0.09773568 | 0.13957881 | 0.05589256 | 0.02384856 | up |
| 545 | Itgb3 | 0.00108151 | -0.0593786 | 0.10188136 | -0.2206385 | 0.02384856 | down |
| 546 | Hprt | 0.00108959 | 0.15464034 | 0.2135896 | 0.09569109 | 0.02391317 | up |
| 547 | Nras | 0.00108848 | 0.15331438 | 0.22011649 | 0.08651227 | 0.02391317 | up |
| 548 | Stab1 | 0.00109613 | -0.0539135 | -0.0381744 | -0.0696527 | 0.02401284 | down |
| 549 | Cit | 0.00110029 | -0.1136753 | -0.071438 | -0.1559127 | 0.02405882 | down |
| 550 | Pot1a | 0.00110224 | 0.05586652 | 0.08412654 | 0.02760649 | 0.02405882 | up |
| 551 | Slc9b2 | 0.001108 | 0.06314589 | 0.10251279 | 0.023779 | 0.02414062 | up |
| 552 | Arhgap32 | 0.00111625 | 0.11653122 | 0.18003867 | 0.05302376 | 0.02427644 | up |
| 553 | Amigo3 | 0.00112398 | -0.1288124 | -0.0580351 | -0.1995896 | 0.02431247 | down |
| 554 | Chmp2b | 0.00112055 | 0.17967865 | 0.2577558 | 0.10160151 | 0.02431247 | up |
| 555 | Spindoc | 0.00112302 | -0.1375002 | -0.0647644 | -0.210236 | 0.02431247 | down |
| 556 | Mtmr3 | 0.00112801 | 0.10159875 | 0.15278955 | 0.05040795 | 0.02435564 | up |
| 557 | Alb | 0.00113532 | 0.04553306 | 0.06205544 | 0.02901068 | 0.02439019 | up |
| 558 | Retreg3 | 0.0011357 | -0.1094652 | -0.0596635 | -0.159267 | 0.02439019 | down |
| 559 | Txn1 | 0.00113506 | 0.20447368 | 0.30349828 | 0.10544907 | 0.02439019 | up |
| 560 | Crebzf | 0.00114407 | -0.1183273 | -0.0638022 | -0.1728523 | 0.0244659 | down |
| 561 | Nlgn2 | 0.00114534 | 0.28885159 | 0.43498063 | 0.14272255 | 0.0244659 | up |
| 562 | Stxbp1 | 0.00114515 | 0.11115278 | 0.17056309 | 0.05174246 | 0.0244659 | up |
| 563 | Cdadc1 | 0.00115419 | 0.14477786 | 0.20663763 | 0.08291808 | 0.02453808 | up |
| 564 | Hint1 | 0.00115343 | 0.14751236 | 0.20938605 | 0.08563867 | 0.02453808 | up |
| 565 | Luc7l3 | 0.0011569 | -0.1007052 | -0.0577842 | -0.1436262 | 0.02453808 | down |
| 566 | Rhobtb3 | 0.00115603 | 0.23313272 | 0.33409191 | 0.13217352 | 0.02453808 | up |
| 567 | Fgd3 | 0.00116098 | -0.0351045 | -0.0219815 | -0.0482276 | 0.02458133 | down |
| 568 | Ak2 | 0.0011705 | -0.3747923 | -0.2098058 | -0.5397787 | 0.0247105 | down |
| 569 | C1qtnf1 | 0.00117385 | 0.23175753 | 0.34207221 | 0.12144286 | 0.0247105 | up |
| 570 | Ctf1 | 0.00117265 | -0.0389235 | -0.0234484 | -0.0543986 | 0.0247105 | down |
| 571 | Eng | 0.00117532 | -0.1882482 | -0.1044435 | -0.272053 | 0.0247105 | down |
| 572 | BC016579 | 0.001185 | -0.0116508 | -0.0053341 | -0.0179674 | 0.02481574 | down |
| 573 | Mtx1 | 0.00118652 | -0.0762016 | -0.0353051 | -0.117098 | 0.02481574 | down |
| 574 | Zfp280c | 0.00118377 | 0.06886637 | 0.10659059 | 0.03114214 | 0.02481574 | up |
| 575 | Cyth3 | 0.00119376 | 0.32036495 | 0.44086264 | 0.19986725 | 0.02488044 | up |
| 576 | Spcs2 | 0.00119201 | 0.07381473 | 0.10161774 | 0.04601171 | 0.02488044 | up |
| 577 | Fblim1 | 0.00119816 | -0.3128276 | -0.1281553 | -0.4974999 | 0.02488565 | down |
| 578 | Glul | 0.00119781 | 0.45881215 | 0.65630091 | 0.26132338 | 0.02488565 | up |
| 579 | Camkk1 | 0.00120944 | -0.0151767 | 0.00954567 | -0.0398991 | 0.02507651 | down |
| 580 | Cox8a | 0.00121359 | -0.1664324 | -0.0804713 | -0.2523934 | 0.02511914 | down |
| 581 | Etf1 | 0.00121987 | 0.19276465 | 0.27924514 | 0.10628416 | 0.02516243 | up |
| 582 | Naaladl2 | 0.00121926 | 0.06999389 | 0.09477246 | 0.04521531 | 0.02516243 | up |
| 583 | Sh3bp4 | 0.0012307 | -0.0992554 | -0.0636391 | -0.1348717 | 0.0253423 | down |
| 584 | Plbd1 | 0.00124562 | -0.0136665 | -0.0035954 | -0.0237375 | 0.02548292 | down |
| 585 | Vamp7 | 0.00124282 | 0.22167593 | 0.32568083 | 0.11767104 | 0.02548292 | up |
| 586 | Zc3h7a | 0.00124602 | 0.09886131 | 0.1466243 | 0.05109832 | 0.02548292 | up |
| 587 | Zfp512b | 0.00124477 | 0.12780816 | 0.18576701 | 0.06984931 | 0.02548292 | up |
| 588 | Adam12 | 0.00127116 | -0.5477636 | -0.3234728 | -0.7720544 | 0.02574433 | down |
| 589 | Cblb | 0.00126329 | 0.29248895 | 0.42503782 | 0.15994007 | 0.02574433 | up |
| 590 | Fbxo27 | 0.0012812 | 0.02760413 | 0.04780796 | 0.0074003 | 0.02574433 | up |
| 591 | Gsr | 0.00128239 | 0.20824372 | 0.29871648 | 0.11777096 | 0.02574433 | up |
| 592 | Mcoln2 | 0.00126791 | 0.22888857 | 0.34873533 | 0.10904181 | 0.02574433 | up |
| 593 | Mettl9 | 0.00127506 | 0.25376873 | 0.39771175 | 0.10982572 | 0.02574433 | up |
| 594 | Neo1 | 0.00127706 | -0.1667541 | -0.097089 | -0.2364192 | 0.02574433 | down |
| 595 | Ntrk3 | 0.00127171 | -0.0144082 | -0.0073907 | -0.0214258 | 0.02574433 | down |
| 596 | Scaf11 | 0.00127707 | 0.10527961 | 0.15603463 | 0.05452459 | 0.02574433 | up |
| 597 | Tgfb1 | 0.00127415 | -0.5364402 | -0.28353 | -0.7893504 | 0.02574433 | down |
| 598 | Zranb2 | 0.00128237 | -0.0838629 | -0.0389392 | -0.1287865 | 0.02574433 | down |
| 599 | Hoxc6 | 0.0012877 | -0.1951776 | -0.1058671 | -0.2844881 | 0.0257647 | down |
| 600 | Vwa5a | 0.00128606 | 0.32583045 | 0.46877786 | 0.18288304 | 0.0257647 | up |
| 601 | Epas1 | 0.00129205 | 0.53006977 | 0.7616643 | 0.29847524 | 0.02580881 | up |
| 602 | Ctsa | 0.00129871 | 0.20137392 | 0.28544084 | 0.11730701 | 0.02589874 | up |
| 603 | Dip2a | 0.00130195 | -0.180617 | -0.1023699 | -0.2588642 | 0.02592015 | down |
| 604 | Dpysl3 | 0.00130946 | -0.4558328 | -0.2745379 | -0.6371277 | 0.02593895 | down |
| 605 | Lig1 | 0.00130918 | -0.2570873 | -0.1462367 | -0.3679379 | 0.02593895 | down |
| 606 | Plekha3 | 0.00130752 | -0.1143148 | -0.053732 | -0.1748975 | 0.02593895 | down |
| 607 | Ube2f | 0.00131153 | -0.1879943 | -0.1042056 | -0.2717829 | 0.02593895 | down |
| 608 | Lnpep | 0.00132941 | 0.18940038 | 0.27226413 | 0.10653664 | 0.02624929 | up |
| 609 | Szrd1 | 0.00133462 | -0.1634204 | -0.0992073 | -0.2276334 | 0.02630891 | down |
| 610 | Zfp800 | 0.00133846 | 0.06010432 | 0.09386015 | 0.02634848 | 0.02634139 | up |
| 611 | Ganab | 0.00134989 | -0.1080082 | -0.0592218 | -0.1567947 | 0.02652275 | down |
| 612 | Dld | 0.00135425 | 0.20329003 | 0.2926135 | 0.11396656 | 0.0265649 | up |
| 613 | Ftsj1 | 0.00135911 | -0.1035766 | -0.0471174 | -0.1600357 | 0.02661686 | down |
| 614 | Galnt1 | 0.00136627 | 0.20050358 | 0.27628809 | 0.12471908 | 0.02671347 | up |
| 615 | Tmem263 | 0.00137548 | -0.0765396 | -0.0359476 | -0.1171317 | 0.02684978 | down |
| 616 | Itprip | 0.00138461 | -0.1270979 | -0.0642041 | -0.1899917 | 0.02698415 | down |
| 617 | Clcn4 | 0.00139122 | -0.1078173 | -0.0603789 | -0.1552556 | 0.02702529 | down |
| 618 | Dennd1a | 0.00138952 | 0.1428977 | 0.21812489 | 0.06767051 | 0.02702529 | up |
| 619 | Smyd3 | 0.00139366 | -0.0443276 | 0.00046823 | -0.0891234 | 0.02702894 | down |
| 620 | Aida | 0.00140078 | 0.51922561 | 0.74513195 | 0.29331926 | 0.02704438 | up |
| 621 | Ddx50 | 0.00140347 | 0.20367571 | 0.29441807 | 0.11293336 | 0.02704438 | up |
| 622 | Gtdc1 | 0.00140231 | 0.05421202 | 0.081232 | 0.02719204 | 0.02704438 | up |
| 623 | Ust | 0.00140198 | -0.1930393 | -0.0988082 | -0.2872705 | 0.02704438 | down |
| 624 | Nfic | 0.00140638 | -0.2588768 | -0.1473358 | -0.3704177 | 0.02705708 | down |
| 625 | Mmp13 | 0.00141196 | -1.544634 | -0.8136974 | -2.2755707 | 0.02707754 | down |
| 626 | Pdgfrl | 0.00141128 | -0.2656081 | -0.1562927 | -0.3749235 | 0.02707754 | down |
| 627 | Cda | 0.00142278 | 0.06099011 | 0.0999172 | 0.02206303 | 0.02711183 | up |
| 628 | Phf5a | 0.00141996 | -0.1273806 | -0.0436158 | -0.2111453 | 0.02711183 | down |
| 629 | Tbc1d12 | 0.00141754 | 0.10687594 | 0.17266264 | 0.04108924 | 0.02711183 | up |
| 630 | Tnk2 | 0.0014209 | -0.1680175 | -0.0814555 | -0.2545796 | 0.02711183 | down |
| 631 | Itgb8 | 0.00143454 | 0.10690842 | 0.18304977 | 0.03076708 | 0.02724949 | up |
| 632 | Tpi1 | 0.00143396 | 0.21293481 | 0.35512651 | 0.07074311 | 0.02724949 | up |
| 633 | Ctsc | 0.00145536 | -0.7339866 | -0.1736096 | -1.2943635 | 0.02731249 | down |
| 634 | Epb41l3 | 0.00146288 | 0.34168648 | 0.49304882 | 0.19032414 | 0.02731249 | up |
| 635 | Galnt11 | 0.00146213 | 0.09503089 | 0.13686659 | 0.05319519 | 0.02731249 | up |
| 636 | Gtf2a2 | 0.00144576 | 0.17570893 | 0.25682209 | 0.09459577 | 0.02731249 | up |
| 637 | Hdgfl2 | 0.00145934 | -0.0997071 | -0.0495328 | -0.1498814 | 0.02731249 | down |
| 638 | Oga | 0.00145813 | 0.19051096 | 0.28893743 | 0.0920845 | 0.02731249 | up |
| 639 | Ppm1g | 0.00144756 | -0.1495564 | -0.0706321 | -0.2284807 | 0.02731249 | down |
| 640 | Ppm1m | 0.00145358 | -0.1688258 | -0.0672534 | -0.2703982 | 0.02731249 | down |
| 641 | Ralgps2 | 0.00144372 | -0.0344743 | -0.0180777 | -0.050871 | 0.02731249 | down |
| 642 | Sprr1a | 0.00145927 | -0.1115934 | -0.0627086 | -0.1604782 | 0.02731249 | down |
| 643 | Tfpi2 | 0.00145916 | -0.6653455 | -0.3645231 | -0.9661678 | 0.02731249 | down |
| 644 | Il6ra | 0.0014669 | 0.12278032 | 0.1949821 | 0.05057854 | 0.0273449 | up |
| 645 | Slc25a32 | 0.00147048 | 0.1451559 | 0.23262934 | 0.05768247 | 0.0273692 | up |
| 646 | S100a6 | 0.00147715 | 0.48553559 | 0.64772221 | 0.32334896 | 0.02745084 | up |
| 647 | Nmnat2 | 0.00148634 | 0.03324686 | 0.05479174 | 0.01170198 | 0.02753629 | up |
| 648 | Zfp948 | 0.00148562 | 0.07679628 | 0.12668724 | 0.02690532 | 0.02753629 | up |
| 649 | Myo18a | 0.00149184 | -0.0575416 | -0.0304964 | -0.0845868 | 0.02755305 | down |
| 650 | Spint2 | 0.00148981 | -0.0564724 | -0.0306981 | -0.0822467 | 0.02755305 | down |
| 651 | Anapc1 | 0.00150311 | -0.1056327 | -0.0545331 | -0.1567322 | 0.02771872 | down |
| 652 | Kcnn2 | 0.00150611 | 0.03146939 | 0.0415322 | 0.02140657 | 0.02773133 | up |
| 653 | Mthfsd | 0.00150969 | -0.1139171 | -0.0671105 | -0.1607237 | 0.02775469 | down |
| 654 | Nrd1 | 0.00151315 | 0.13569193 | 0.19669977 | 0.07468408 | 0.02777587 | up |
| 655 | Bid | 0.00153072 | 0.0345998 | 0.04970389 | 0.01949571 | 0.0280027 | up |
| 656 | Tpm2 | 0.00153171 | -0.7765715 | -0.3679726 | -1.1851704 | 0.0280027 | down |
| 657 | Tpsb2 | 0.00153251 | -0.0286354 | -0.0155495 | -0.0417214 | 0.0280027 | down |
| 658 | Pigx | 0.00153625 | 0.115627 | 0.19215541 | 0.0390986 | 0.02802844 | up |
| 659 | Wdr19 | 0.00154299 | -0.0627462 | 0.0090918 | -0.1345916 | 0.02810858 | down |
| 660 | Calcrl | 0.00154947 | 0.36708957 | 0.55995568 | 0.17422346 | 0.02813936 | up |
| 661 | Mlec | 0.00155405 | -0.2417615 | -0.1451395 | -0.3383834 | 0.02813936 | down |
| 662 | Pign | 0.00155241 | 0.14643164 | 0.21469673 | 0.07816655 | 0.02813936 | up |
| 663 | Ss18 | 0.00155282 | -0.1452779 | -0.0835086 | -0.2070472 | 0.02813936 | down |
| 664 | Coro7 | 0.00156435 | -0.0581104 | -0.0246067 | -0.091614 | 0.02819826 | down |
| 665 | Obsl1 | 0.00156265 | -0.2813271 | -0.1535642 | -0.4090901 | 0.02819826 | down |
| 666 | Ugdh | 0.00156104 | 0.26664327 | 0.45057173 | 0.08271481 | 0.02819826 | up |
| 667 | Fam129a | 0.00157615 | 0.20674701 | 0.31465864 | 0.09883537 | 0.028233 | up |
| 668 | Kctd13 | 0.00158363 | -0.0743309 | -0.04394 | -0.1047218 | 0.028233 | down |
| 669 | Nudt9 | 0.00158334 | -0.1843831 | -0.0979413 | -0.2708248 | 0.028233 | down |
| 670 | Ptcd2 | 0.00158064 | 0.08309972 | 0.13056455 | 0.0356349 | 0.028233 | up |
| 671 | Spg7 | 0.0015789 | -0.144427 | -0.0756175 | -0.2132365 | 0.028233 | down |
| 672 | Fndc3a | 0.00159587 | 0.2803429 | 0.40514871 | 0.15553708 | 0.02850958 | up |
| 673 | Snx33 | 0.00159973 | 0.11433447 | 0.16179726 | 0.06687167 | 0.02853602 | up |
| 674 | Dse | 0.00160795 | 0.17445351 | 0.25468116 | 0.09422585 | 0.0286402 | up |
| 675 | Fam234b | 0.00162096 | -0.075437 | -0.0472555 | -0.1036186 | 0.02882905 | down |
| 676 | Smim41 | 0.00163039 | 0.14328891 | 0.22458481 | 0.06199302 | 0.02895386 | up |
| 677 | Plaat3 | 0.00163884 | -0.4120799 | -0.2383659 | -0.5857938 | 0.02898529 | down |
| 678 | Rpe | 0.00163767 | 0.10529851 | 0.1539968 | 0.05660022 | 0.02898529 | up |
| 679 | Zfp157 | 0.0016394 | 0.02803775 | 0.05201918 | 0.00405632 | 0.02898529 | up |
| 680 | Slc25a46 | 0.00165019 | 0.09566831 | 0.14401785 | 0.04731877 | 0.0291331 | up |
| 681 | Tmem165 | 0.00165929 | 0.12644156 | 0.18293454 | 0.06994858 | 0.0292508 | up |
| 682 | Eef1a1 | 0.00166461 | 0.26905856 | 0.37740363 | 0.1607135 | 0.02927168 | up |
| 683 | Pank1 | 0.00166535 | -0.0360187 | -0.0211991 | -0.0508383 | 0.02927168 | down |
| 684 | Zfyve27 | 0.00166834 | -0.0909717 | -0.037823 | -0.1441204 | 0.02928132 | down |
| 685 | Acadsb | 0.00167441 | 0.27218037 | 0.39691313 | 0.14744761 | 0.02931703 | up |
| 686 | Fam173a | 0.00167526 | -0.0944277 | -0.0534749 | -0.1353805 | 0.02931703 | down |
| 687 | Pdgfa | 0.00168663 | -0.2121752 | -0.0985542 | -0.3257961 | 0.02947303 | down |
| 688 | Jmjd1c | 0.00169626 | 0.12993917 | 0.18838204 | 0.0714963 | 0.02959817 | up |
| 689 | Peli2 | 0.00169957 | 0.12786388 | 0.1721225 | 0.08360526 | 0.02961291 | up |
| 690 | Cops4 | 0.00170718 | 0.09868864 | 0.14562209 | 0.0517552 | 0.02965948 | up |
| 691 | Kcnq3 | 0.00170682 | -0.0605231 | -0.030562 | -0.0904841 | 0.02965948 | down |
| 692 | Mtx2 | 0.00171563 | 0.13510299 | 0.19205134 | 0.07815463 | 0.02972031 | up |
| 693 | Tspyl1 | 0.00171414 | -0.1283393 | -0.0621808 | -0.1944977 | 0.02972031 | down |
| 694 | Mios | 0.00172782 | 0.14153926 | 0.19617429 | 0.08690424 | 0.02975973 | up |
| 695 | Mtpap | 0.00172728 | 0.10403981 | 0.14829566 | 0.05978396 | 0.02975973 | up |
| 696 | Slc24a3 | 0.0017277 | -0.0460399 | -0.0278806 | -0.0641993 | 0.02975973 | down |
| 697 | Slc38a4 | 0.00172761 | -0.118341 | -0.0402195 | -0.1964626 | 0.02975973 | down |
| 698 | Cyth2 | 0.00173279 | -0.1691651 | -0.0790798 | -0.2592503 | 0.02980242 | down |
| 699 | Tulp4 | 0.0017396 | -0.1195842 | -0.0610564 | -0.1781121 | 0.02987683 | down |
| 700 | Mtm1 | 0.00174471 | 0.05942177 | 0.08188775 | 0.03695579 | 0.02992049 | up |
| 701 | Pcdh7 | 0.00174713 | 0.29851512 | 0.43376706 | 0.16326318 | 0.02992049 | up |
| 702 | Mmp17 | 0.00175699 | -0.0632184 | -0.0335516 | -0.0928852 | 0.0300291 | down |
| 703 | Ndrg1 | 0.0017643 | 0.42453716 | 0.65748479 | 0.19158954 | 0.0300291 | up |
| 704 | Pde1a | 0.00176545 | 0.14495416 | 0.20409257 | 0.08581574 | 0.0300291 | up |
| 705 | Ptpn1 | 0.00176367 | 0.27779927 | 0.39788924 | 0.1577093 | 0.0300291 | up |
| 706 | Snap23 | 0.00176598 | 0.08068925 | 0.12289451 | 0.03848399 | 0.0300291 | up |
| 707 | Camta2 | 0.00177278 | -0.068449 | -0.029069 | -0.107829 | 0.03006485 | down |
| 708 | Ebpl | 0.00177931 | -0.1178489 | -0.0736953 | -0.1620025 | 0.03006485 | down |
| 709 | Pdcd10 | 0.00177913 | 0.17980887 | 0.25868698 | 0.10093077 | 0.03006485 | up |
| 710 | Pfdn2 | 0.0017806 | -0.1187835 | -0.0630265 | -0.1745404 | 0.03006485 | down |
| 711 | Zfp639 | 0.0017806 | 0.15399753 | 0.21466535 | 0.0933297 | 0.03006485 | up |
| 712 | Anxa10 | 0.00178614 | -0.0451538 | -0.0308008 | -0.0595068 | 0.0300738 | down |
| 713 | Frg2f1 | 0.0017842 | -0.0220949 | -0.0103698 | -0.0338199 | 0.0300738 | down |
| 714 | Pwp1 | 0.00179178 | 0.05660287 | 0.09185728 | 0.02134846 | 0.03012655 | up |
| 715 | Amacr | 0.00179871 | -0.0958926 | -0.0635135 | -0.1282716 | 0.03020064 | down |
| 716 | Akt1s1 | 0.001814 | -0.2469112 | -0.1323831 | -0.3614392 | 0.03041482 | down |
| 717 | Pofut2 | 0.0018243 | -0.2848987 | -0.1339671 | -0.4358303 | 0.03054498 | down |
| 718 | B3gnt8 | 0.00183749 | -0.0469949 | -0.027052 | -0.0669377 | 0.03068021 | down |
| 719 | Tns4 | 0.00183526 | -0.0438871 | -0.0281207 | -0.0596535 | 0.03068021 | down |
| 720 | Arl10 | 0.0018441 | -0.0609678 | -0.036974 | -0.0849616 | 0.0307478 | down |
| 721 | Fjx1 | 0.00184703 | -0.1941308 | -0.0674947 | -0.3207669 | 0.03075397 | down |
| 722 | Ahsa1 | 0.00186101 | -0.1426264 | -0.0706837 | -0.214569 | 0.03078962 | down |
| 723 | Casp3 | 0.001862 | -0.1154126 | -0.0510208 | -0.1798044 | 0.03078962 | down |
| 724 | Fam8a1 | 0.00185523 | 0.15198071 | 0.22785878 | 0.07610265 | 0.03078962 | up |
| 725 | Rflnb | 0.00186041 | -0.2690584 | -0.0691386 | -0.4689781 | 0.03078962 | down |
| 726 | Tm2d3 | 0.00186 | 0.08692093 | 0.12792192 | 0.04591995 | 0.03078962 | up |
| 727 | Zbtb44 | 0.00187038 | 0.15084068 | 0.21869268 | 0.08298869 | 0.03088574 | up |
| 728 | Dennd3 | 0.00188183 | -0.0662059 | -0.0408397 | -0.0915721 | 0.03094716 | down |
| 729 | Mettl26 | 0.00188022 | -0.0344062 | -0.014518 | -0.0542944 | 0.03094716 | down |
| 730 | Trim46 | 0.00187898 | -0.0583496 | -0.0242321 | -0.0924671 | 0.03094716 | down |
| 731 | Arhgap35 | 0.0018881 | 0.33232308 | 0.47956328 | 0.18508287 | 0.03100764 | up |
| 732 | Apold1 | 0.00189784 | 0.02805729 | 0.04547839 | 0.01063619 | 0.03112504 | up |
| 733 | Col25a1 | 0.00191041 | 0.04848171 | 0.09282836 | 0.00413505 | 0.03124645 | up |
| 734 | Pdlim7 | 0.00191044 | -0.3098954 | -0.1751277 | -0.4446632 | 0.03124645 | down |
| 735 | DEPRECATED- | 0.00192373 | -0.3149461 | -0.1193928 | -0.5104994 | 0.03131425 | down |
| 736 | Dnlz | 0.00191845 | -0.0875969 | -0.0218114 | -0.1533824 | 0.03131425 | down |
| 737 | Igf2bp2 | 0.00192723 | 0.1709634 | 0.24958883 | 0.09233798 | 0.03131425 | up |
| 738 | Slc38a6 | 0.00192763 | -0.0901052 | -0.0509156 | -0.1292948 | 0.03131425 | down |
| 739 | Tmem268 | 0.00192458 | -0.0624053 | -0.0294212 | -0.0953894 | 0.03131425 | down |
| 740 | Vps26a | 0.00193619 | 0.12358647 | 0.18145857 | 0.06571437 | 0.0314108 | up |
| 741 | Hs3st3b1 | 0.00194043 | -0.0790801 | -0.0376309 | -0.1205292 | 0.03142409 | down |
| 742 | Rgl2 | 0.00194225 | -0.0966681 | -0.0573371 | -0.1359992 | 0.03142409 | down |
| 743 | Avpr1a | 0.00195896 | -0.3554855 | -0.2074856 | -0.5034854 | 0.03148229 | down |
| 744 | Klc4 | 0.00195866 | -0.068114 | -0.0361039 | -0.1001241 | 0.03148229 | down |
| 745 | Lrrc8a | 0.00195229 | -0.100629 | -0.0440481 | -0.1572099 | 0.03148229 | down |
| 746 | Smtd1 | 0.00195856 | 0.10020046 | 0.14528548 | 0.05511544 | 0.03148229 | up |
| 747 | Slc16a1 | 0.00195801 | 0.48951432 | 0.77464674 | 0.2043819 | 0.03148229 | up |
| 748 | Nipbl | 0.00196474 | 0.32439459 | 0.47760754 | 0.17118164 | 0.03149085 | up |
| 749 | 2-Sep | 0.00196447 | 0.16235863 | 0.26543469 | 0.05928257 | 0.03149085 | up |
| 750 | Asap1 | 0.00196898 | 0.10747497 | 0.1642546 | 0.05069534 | 0.03151688 | up |
| 751 | Tubb6 | 0.0019722 | -0.5072793 | -0.2698819 | -0.7446766 | 0.03152632 | down |
| 752 | Fam149a | 0.00197995 | 0.08406198 | 0.12860355 | 0.03952041 | 0.03160818 | up |
| 753 | Slc22a17 | 0.00198355 | -0.135533 | -0.0671936 | -0.2038724 | 0.0316236 | down |
| 754 | Dennd2a | 0.00199066 | -0.192074 | -0.0986373 | -0.2855108 | 0.03169261 | down |
| 755 | Pgk1 | 0.00199316 | 0.35412953 | 0.57896803 | 0.12929103 | 0.03169261 | up |
| 756 | Mapk1ip1l | 0.00200433 | -0.1625159 | -0.084882 | -0.2401498 | 0.03178952 | down |
| 757 | Pgm5 | 0.00200455 | -0.0164961 | -0.0091355 | -0.0238567 | 0.03178952 | down |
| 758 | Sorcs2 | 0.00201133 | -0.6643519 | -0.3869033 | -0.9418004 | 0.0318549 | down |
| 759 | Csnk1e | 0.00202387 | -0.0999465 | -0.0505003 | -0.1493927 | 0.03192003 | down |
| 760 | Erlec1 | 0.00203408 | 0.29699366 | 0.43012522 | 0.16386209 | 0.03192003 | up |
| 761 | Foxg1 | 0.00203387 | -0.0372025 | -0.0180977 | -0.0563072 | 0.03192003 | down |
| 762 | Mcoln1 | 0.00203292 | 0.11053024 | 0.16408905 | 0.05697143 | 0.03192003 | up |
| 763 | Mterf3 | 0.00203937 | 0.0756269 | 0.11642154 | 0.03483226 | 0.03192003 | up |
| 764 | Sipa1 | 0.0020368 | -0.1088829 | -0.0324484 | -0.1853173 | 0.03192003 | down |
| 765 | Tle5 | 0.00201906 | -0.0986833 | -0.0503901 | -0.1469765 | 0.03192003 | down |
| 766 | Zbtb38 | 0.00202904 | -0.1250208 | -0.0412113 | -0.2088303 | 0.03192003 | down |
| 767 | Zfp277 | 0.00203314 | 0.11298297 | 0.17945745 | 0.04650849 | 0.03192003 | up |
| 768 | Anks1 | 0.00205184 | 0.08146899 | 0.12038451 | 0.04255347 | 0.03207341 | up |
| 769 | Wdr59 | 0.00205844 | 0.09561383 | 0.15660271 | 0.03462494 | 0.03213474 | up |
| 770 | Cdo1 | 0.00207364 | 0.14112996 | 0.22447041 | 0.05778951 | 0.03228795 | up |
| 771 | Zfp362 | 0.00207207 | -0.1497739 | -0.0882258 | -0.211322 | 0.03228795 | down |
| 772 | Capza2 | 0.00210836 | 0.29961235 | 0.45194402 | 0.14728068 | 0.03251963 | up |
| 773 | Commd6 | 0.00209567 | 0.12731366 | 0.19358209 | 0.06104522 | 0.03251963 | up |
| 774 | Dohh | 0.00210149 | -0.1053348 | -0.0585256 | -0.152144 | 0.03251963 | down |
| 775 | Krt18 | 0.00209169 | -0.1512764 | -0.0770874 | -0.2254653 | 0.03251963 | down |
| 776 | Nynrin | 0.00210842 | -0.1536615 | -0.0616127 | -0.2457103 | 0.03251963 | down |
| 777 | Prpf40a | 0.0021129 | 0.08558346 | 0.13989257 | 0.03127434 | 0.03251963 | up |
| 778 | Rap2c | 0.00211056 | 0.285687 | 0.43441696 | 0.13695703 | 0.03251963 | up |
| 779 | Rxrb | 0.00210284 | -0.1128856 | -0.0632568 | -0.1625144 | 0.03251963 | down |
| 780 | Slc2a9 | 0.00210585 | 0.02670462 | 0.03954862 | 0.01386063 | 0.03251963 | up |
| 781 | Rap1gap2 | 0.00212272 | -0.0488633 | -0.022654 | -0.0750725 | 0.03262903 | down |
| 782 | Glis2 | 0.00212572 | -0.2652322 | -0.1469117 | -0.3835527 | 0.03263337 | down |
| 783 | Epha3 | 0.00212849 | -0.2428448 | -0.1123576 | -0.3733321 | 0.03263413 | down |
| 784 | Abhd13 | 0.00213367 | -0.2109387 | -0.1117849 | -0.3100925 | 0.03267185 | down |
| 785 | Kif5c | 0.00213887 | -0.0286229 | -0.0143779 | -0.042868 | 0.0327098 | down |
| 786 | Impact | 0.00214575 | 0.14244358 | 0.20460655 | 0.08028062 | 0.03277324 | up |
| 787 | Cox17 | 0.00215117 | -0.1578731 | -0.0741544 | -0.2415919 | 0.03281417 | down |
| 788 | Atxn7l3b | 0.00216875 | -0.1111162 | -0.0611952 | -0.1610372 | 0.03297913 | down |
| 789 | Eif2ak4 | 0.00217022 | -0.1127265 | -0.0550958 | -0.1703572 | 0.03297913 | down |
| 790 | Pld3 | 0.00216847 | 0.12152409 | 0.17517313 | 0.06787505 | 0.03297913 | up |
| 791 | Cul5 | 0.00217351 | 0.13374076 | 0.19873858 | 0.06874294 | 0.03298729 | up |
| 792 | Trnt1 | 0.00218997 | 0.07221899 | 0.10873985 | 0.03569814 | 0.03319523 | up |
| 793 | Hoxa3 | 0.00219564 | -0.1561153 | -0.0650706 | -0.2471599 | 0.03322391 | down |
| 794 | Sh3rf1 | 0.0021974 | -0.1582062 | -0.0625641 | -0.2538484 | 0.03322391 | down |
| 795 | Zfp90 | 0.00220747 | 0.03677619 | 0.05832762 | 0.01522475 | 0.03333418 | up |
| 796 | Cep164 | 0.00223376 | -0.0752028 | -0.0382639 | -0.1121418 | 0.03367071 | down |
| 797 | Usp28 | 0.00223536 | 0.05656004 | 0.08129094 | 0.03182913 | 0.03367071 | up |
| 798 | Acsl4 | 0.00224518 | -0.1972293 | -0.1108891 | -0.2835695 | 0.03377617 | down |
| 799 | Casp4 | 0.00225879 | 0.20810714 | 0.32523865 | 0.09097563 | 0.03382986 | up |
| 800 | Gpr137c | 0.00225272 | -0.0139879 | -0.0020581 | -0.0259176 | 0.03382986 | down |
| 801 | Jak1 | 0.00226002 | 0.32395836 | 0.4978165 | 0.15010021 | 0.03382986 | up |
| 802 | Kif7 | 0.00225496 | -0.0502323 | -0.0256056 | -0.0748589 | 0.03382986 | down |
| 803 | Foxp4 | 0.00227451 | -0.131229 | -0.0704438 | -0.1920141 | 0.03400433 | down |
| 804 | Baz1b | 0.00227848 | -0.1268841 | -0.0592048 | -0.1945634 | 0.03402135 | down |
| 805 | Ccdc107 | 0.00228879 | -0.1078017 | -0.0650894 | -0.150514 | 0.03402754 | down |
| 806 | Dyrk2 | 0.00229307 | -0.1561144 | -0.056169 | -0.2560597 | 0.03402754 | down |
| 807 | Igsf10 | 0.00228237 | 0.27943491 | 0.41618681 | 0.142683 | 0.03402754 | up |
| 808 | Kif26b | 0.00228644 | -0.5951937 | -0.2305873 | -0.9598001 | 0.03402754 | down |
| 809 | Sox4 | 0.00229305 | -0.208874 | -0.1138908 | -0.3038572 | 0.03402754 | down |
| 810 | Haus7 | 0.00230672 | -0.125172 | -0.0603299 | -0.1900142 | 0.03415523 | down |
| 811 | Tbc1d10b | 0.00231021 | -0.1150122 | -0.0507354 | -0.179289 | 0.03415523 | down |
| 812 | Zfp68 | 0.00231011 | 0.11019576 | 0.18801407 | 0.03237746 | 0.03415523 | up |
| 813 | Cd2bp2 | 0.00232058 | -0.1536324 | -0.0707949 | -0.2364699 | 0.03422431 | down |
| 814 | Nop16 | 0.00231916 | -0.0845228 | -0.0348769 | -0.1341686 | 0.03422431 | down |
| 815 | Nectin3 | 0.00233215 | 0.1196343 | 0.17686586 | 0.06240274 | 0.03435275 | up |
| 816 | Anxa7 | 0.00233666 | 0.38447364 | 0.59357598 | 0.17537129 | 0.03437703 | up |
| 817 | Arhgef19 | 0.00234441 | -0.1687011 | -0.0925724 | -0.2448297 | 0.03440672 | down |
| 818 | Spock3 | 0.00234272 | -0.2678334 | -0.0603539 | -0.4753129 | 0.03440672 | down |
| 819 | Insc | 0.00235326 | -0.0332947 | -0.0106035 | -0.0559858 | 0.03441102 | down |
| 820 | Plekho1 | 0.00235331 | -0.1999677 | -0.1049992 | -0.2949362 | 0.03441102 | down |
| 821 | St6galnac4 | 0.00235116 | -0.1425553 | -0.0796235 | -0.2054871 | 0.03441102 | down |
| 822 | Ptcd3 | 0.00235624 | 0.07271974 | 0.11174391 | 0.03369557 | 0.03441197 | up |
| 823 | Nptn | 0.00236174 | 0.36731532 | 0.55064823 | 0.18398241 | 0.03445038 | up |
| 824 | a | 0.00237462 | 0.31052853 | 0.47926996 | 0.14178711 | 0.03459626 | up |
| 825 | Gsf1 | 0.00239318 | 0.05439088 | 0.08399674 | 0.02478502 | 0.03474024 | up |
| 826 | Nfkb2 | 0.00239318 | -0.1402673 | -0.0574173 | -0.2231172 | 0.03474024 | down |
| 827 | Tmem204 | 0.00239229 | -0.2800682 | 0.0050621 | -0.5651986 | 0.03474024 | down |
| 828 | Apol9b | 0.00240816 | -0.0589866 | -0.0269054 | -0.0910679 | 0.03486566 | down |
| 829 | Bok | 0.00241104 | -0.1188008 | -0.0350214 | -0.2025802 | 0.03486566 | down |
| 830 | Ggta1 | 0.00242674 | 0.25338888 | 0.36745716 | 0.13932061 | 0.03486566 | up |
| 831 | Mbd1 | 0.00242482 | -0.1027051 | -0.0507525 | -0.1546577 | 0.03486566 | down |
| 832 | Mex3a | 0.00242913 | -0.106813 | -0.0476185 | -0.1660075 | 0.03486566 | down |
| 833 | Ngef | 0.00241178 | -0.0318 | -0.0177201 | -0.04588 | 0.03486566 | down |
| 834 | Snx2 | 0.00241766 | 0.13293501 | 0.20021123 | 0.06565878 | 0.03486566 | up |
| 835 | Tmem223 | 0.00242757 | -0.1043204 | -0.049542 | -0.1590988 | 0.03486566 | down |
| 836 | Ubn1 | 0.00243087 | -0.0811211 | -0.0350388 | -0.1272033 | 0.03486566 | down |
| 837 | Yipf4 | 0.00241615 | 0.21666391 | 0.31908437 | 0.11424345 | 0.03486566 | up |
| 838 | Rad11b | 0.00243762 | -0.2832407 | -0.1599899 | -0.4064915 | 0.03492084 | down |
| 839 | Atrx | 0.00245646 | 0.18220673 | 0.26611521 | 0.09829826 | 0.03510754 | up |
| 840 | Ccdc102a | 0.00247603 | -0.2493339 | -0.1282454 | -0.3704225 | 0.03510754 | down |
| 841 | Cct4 | 0.00247962 | 0.17700336 | 0.28602901 | 0.06797771 | 0.03510754 | up |
| 842 | Itga5 | 0.00247443 | -0.3858665 | -0.2240086 | -0.5477244 | 0.03510754 | down |
| 843 | Khdrbs3 | 0.002459 | -0.0923418 | -0.0271338 | -0.1575498 | 0.03510754 | down |
| 844 | Rnf130 | 0.00247253 | 0.30573024 | 0.44532607 | 0.1661344 | 0.03510754 | up |
| 845 | Slc20a1 | 0.0024799 | -0.3536804 | -0.1508886 | -0.5564721 | 0.03510754 | down |
| 846 | Snrnp70 | 0.00246035 | -0.1177909 | -0.0607801 | -0.1748016 | 0.03510754 | down |
| 847 | Tacc1 | 0.0024665 | 0.23460324 | 0.34706309 | 0.12214339 | 0.03510754 | up |
| 848 | Tysnd1 | 0.00247095 | -0.1337968 | -0.0649671 | -0.2026265 | 0.03510754 | down |
| 849 | Rftn1 | 0.00248552 | -0.1751756 | -0.0977792 | -0.2525721 | 0.03514562 | down |
| 850 | Neurl2 | 0.00249453 | -0.0232769 | -0.0092917 | -0.0372621 | 0.03519021 | down |
| 851 | Sertad4 | 0.00249224 | -0.2257916 | -0.130084 | -0.3214992 | 0.03519021 | down |
| 852 | Gprasp1 | 0.0025005 | -0.0523891 | -0.0259308 | -0.0788473 | 0.03523295 | down |
| 853 | Ankib1 | 0.00251022 | 0.12043585 | 0.19856158 | 0.04231012 | 0.03524579 | up |
| 854 | Fyttd1 | 0.00250568 | 0.17361422 | 0.24544847 | 0.10177997 | 0.03524579 | up |
| 855 | Tnfrsf11b | 0.00250948 | 0.23380017 | 0.34496093 | 0.1226394 | 0.03524579 | up |
| 856 | Grem2 | 0.00252099 | -0.571831 | -0.3038142 | -0.8398478 | 0.03535569 | down |
| 857 | Cfap74 | 0.00252602 | -0.0482378 | -0.0281862 | -0.0682894 | 0.03536102 | down |
| 858 | Fcf1 | 0.00252726 | 0.13678566 | 0.19906641 | 0.0745049 | 0.03536102 | up |
| 859 | Nat14 | 0.0025321 | -0.0381945 | -0.0184835 | -0.0579056 | 0.03538753 | down |
| 860 | Ntrk2 | 0.00253623 | 0.4392881 | 0.69761957 | 0.18095663 | 0.03540398 | up |
| 861 | Ctdsp1 | 0.00254204 | -0.0917368 | -0.0391613 | -0.1443122 | 0.03544388 | down |
| 862 | Slit3 | 0.00254738 | 0.25899175 | 0.47947479 | 0.03850871 | 0.03547708 | up |
| 863 | Ccn1 | 0.00255225 | -0.4957069 | -0.2697823 | -0.7216316 | 0.03550374 | down |
| 864 | Acer3 | 0.00257656 | 0.07437757 | 0.11569702 | 0.03305812 | 0.03570413 | up |
| 865 | Fabp5 | 0.00257751 | -0.2655393 | -0.1759181 | -0.3551605 | 0.03570413 | down |
| 866 | Fam78b | 0.0025838 | 0.2351421 | 0.35388104 | 0.11640316 | 0.03570413 | up |
| 867 | Fzd5 | 0.00258361 | -0.0889531 | -0.0487384 | -0.1291677 | 0.03570413 | down |
| 868 | Habp4 | 0.00258631 | -0.0651927 | -0.017996 | -0.1123895 | 0.03570413 | down |
| 869 | Mrtfa | 0.00258534 | -0.1452651 | -0.0652575 | -0.2252726 | 0.03570413 | down |
| 870 | Mt3 | 0.00259045 | 0.07309252 | 0.1387665 | 0.00741853 | 0.03570413 | up |
| 871 | Rnasek | 0.00258952 | -0.1950256 | -0.107758 | -0.2822933 | 0.03570413 | down |
| 872 | Zfp654 | 0.00259999 | 0.17087028 | 0.26005299 | 0.08168757 | 0.03579454 | up |
| 873 | Ldlrad3 | 0.00261672 | -0.0797264 | -0.0393055 | -0.1201473 | 0.03598359 | down |
| 874 | Afg3l1 | 0.00262829 | -0.1913095 | -0.0989509 | -0.2836681 | 0.03610143 | down |
| 875 | Atic | 0.00264221 | -0.1811141 | -0.0908524 | -0.2713757 | 0.03613404 | down |
| 876 | Dnm3 | 0.00263704 | -0.2544031 | -0.1396675 | -0.3691387 | 0.03613404 | down |
| 877 | Kank2 | 0.00264623 | -0.0976753 | -0.0508389 | -0.1445117 | 0.03613404 | down |
| 878 | Ldha | 0.00264712 | 0.17396519 | 0.2721295 | 0.07580088 | 0.03613404 | up |
| 879 | Mcam | 0.00265174 | -0.1563223 | -0.0713303 | -0.2413143 | 0.03613404 | down |
| 880 | Sdcbp | 0.00265069 | 0.22864432 | 0.37938957 | 0.07789908 | 0.03613404 | up |
| 881 | Sdk1 | 0.00264777 | 0.29068045 | 0.42161443 | 0.15974648 | 0.03613404 | up |
| 882 | Cars | 0.00266049 | -0.1483173 | -0.0793519 | -0.2172827 | 0.03621229 | down |
| 883 | Yy1 | 0.00266576 | -0.122658 | -0.0635902 | -0.1817257 | 0.03624284 | down |
| 884 | Cdc73 | 0.00267637 | 0.08980483 | 0.13346007 | 0.04614959 | 0.03626393 | up |
| 885 | Med25 | 0.00267513 | -0.1056656 | -0.0520275 | -0.1593037 | 0.03626393 | down |
| 886 | Rnf185 | 0.00267141 | -0.0889961 | -0.0530592 | -0.1249329 | 0.03626393 | down |
| 887 | Gnl2 | 0.00268336 | 0.11381194 | 0.16489264 | 0.06273124 | 0.03631766 | up |
| 888 | Qpct | 0.00268974 | 0.32929064 | 0.48767188 | 0.1709094 | 0.03635184 | up |
| 889 | Slc30a5 | 0.00269194 | -0.1059829 | -0.0491398 | -0.1628261 | 0.03635184 | down |
| 890 | Cdh24 | 0.0027087 | -0.0511513 | -0.0177591 | -0.0845435 | 0.03640687 | down |
| 891 | Eif5b | 0.0027046 | 0.14502874 | 0.21406725 | 0.07599022 | 0.03640687 | up |
| 892 | Mmp9 | 0.00271118 | -0.532848 | -0.2654971 | -0.8001989 | 0.03640687 | down |
| 893 | Rint1 | 0.00270698 | -0.1019932 | -0.0236823 | -0.1803042 | 0.03640687 | down |
| 894 | Spopl | 0.0027037 | 0.08342868 | 0.1276984 | 0.03915896 | 0.03640687 | up |
| 895 | Ranbp2 | 0.00273194 | 0.1800332 | 0.2658594 | 0.094207 | 0.03664467 | up |
| 896 | Apobec3 | 0.00273538 | -0.0891945 | -0.0484659 | -0.129923 | 0.03664983 | down |
| 897 | Tmem131 | 0.00274929 | 0.19731418 | 0.2959158 | 0.09871257 | 0.03679515 | up |
| 898 | Ifi207 | 0.00275418 | 0.49561974 | 0.72755759 | 0.2636819 | 0.03681955 | up |
| 899 | B9d1 | 0.00276856 | -0.0608215 | -0.0308631 | -0.0907798 | 0.0369061 | down |
| 900 | Hspb7 | 0.00276988 | -0.047178 | -0.0186633 | -0.0756926 | 0.0369061 | down |
| 901 | Stxbp6 | 0.00276927 | 0.0905291 | 0.16421618 | 0.01684203 | 0.0369061 | up |
| 902 | Mien1 | 0.00277309 | -0.1430881 | -0.0624252 | -0.2237511 | 0.03690786 | down |
| 903 | Bet1 | 0.0027789 | -0.1215739 | -0.0526251 | -0.1905226 | 0.03692414 | down |
| 904 | Dcun1d1 | 0.00278828 | 0.03185202 | 0.06802069 | -0.0043166 | 0.03692414 | up |
| 905 | Fam214a | 0.0027815 | -0.141915 | -0.0423341 | -0.2414958 | 0.03692414 | down |
| 906 | Rnf20 | 0.00278701 | -0.0913691 | -0.0462301 | -0.1365081 | 0.03692414 | down |
| 907 | St3gal2 | 0.00278969 | -0.1240567 | -0.0691913 | -0.178922 | 0.03692414 | down |
| 908 | Camk2d | 0.00279411 | 0.23115464 | 0.34261763 | 0.11969164 | 0.03694199 | up |
| 909 | Rnf26 | 0.00280688 | -0.0597161 | -0.0253303 | -0.0941018 | 0.03706999 | down |
| 910 | Ebp | 0.00281243 | -0.2621184 | -0.1389654 | -0.3852714 | 0.03710245 | down |
| 911 | Znhit1 | 0.00282948 | -0.088529 | -0.0404622 | -0.1365958 | 0.03728641 | down |
| 912 | Fgf16 | 0.00283638 | 0.02145418 | 0.03494111 | 0.00796726 | 0.03733631 | up |
| 913 | Ipo9 | 0.00284542 | -0.1363316 | -0.0645156 | -0.2081476 | 0.03733684 | down |
| 914 | Pdzd4 | 0.0028438 | -0.0643672 | -0.0299627 | -0.0987717 | 0.03733684 | down |
| 915 | Prkag1 | 0.00284575 | 0.08888501 | 0.13748346 | 0.04028656 | 0.03733684 | up |
| 916 | Ap3m1 | 0.00285789 | 0.07479733 | 0.11101603 | 0.03857862 | 0.03741442 | up |
| 917 | Zfp740 | 0.00285751 | 0.18200504 | 0.29075554 | 0.07325453 | 0.03741442 | up |
| 918 | Uqcr11 | 0.00286336 | -0.1612337 | -0.0783429 | -0.2441245 | 0.03744511 | down |
| 919 | Ier3ip1 | 0.00288404 | -0.115908 | -0.0562648 | -0.1755513 | 0.03767459 | down |
| 920 | Plin2 | 0.00289465 | -0.2332132 | -0.1198761 | -0.3465504 | 0.03773102 | down |
| 921 | Sectm1b | 0.00289169 | 0.05447653 | 0.08863038 | 0.02032268 | 0.03773102 | up |
| 922 | Zcchc2 | 0.00290023 | 0.06179745 | 0.09591231 | 0.02768259 | 0.03776272 | up |
| 923 | Wnt7b | 0.00291055 | -0.0216761 | -0.0114371 | -0.0319151 | 0.03781504 | down |
| 924 | Zfp655 | 0.0029084 | 0.08096161 | 0.12363305 | 0.03829017 | 0.03781504 | up |
| 925 | Hook3 | 0.00291371 | 0.16033498 | 0.24006211 | 0.08060785 | 0.03781525 | up |
| 926 | Calr | 0.00293849 | -0.1163267 | -0.061656 | -0.1709973 | 0.03804177 | down |
| 927 | Xrcc1 | 0.00294067 | -0.0956318 | -0.0485885 | -0.1426751 | 0.03804177 | down |
| 928 | Yif1a | 0.00293811 | 0.14703571 | 0.22290104 | 0.07117038 | 0.03804177 | up |
| 929 | Enho | 0.00294581 | -0.118743 | -0.0281861 | -0.2092998 | 0.03806717 | down |
| 930 | Jdp2 | 0.00295842 | -0.2654344 | -0.1403059 | -0.3905628 | 0.038189 | down |
| 931 | Echs1 | 0.00298585 | 0.11732498 | 0.17372424 | 0.06092571 | 0.03850174 | up |
| 932 | Defb48 | 0.00299005 | -0.0330366 | -0.0171868 | -0.0488864 | 0.0385146 | down |
| 933 | Clu | 0.00300292 | -0.8196043 | -0.3234855 | -1.3157231 | 0.03860413 | down |
| 934 | Lrp3 | 0.00300665 | 0.13870351 | 0.20146417 | 0.07594286 | 0.03860413 | up |
| 935 | Rxra | 0.00300665 | 0.11801361 | 0.16696595 | 0.06906128 | 0.03860413 | up |
| 936 | Hdac11 | 0.00306135 | 0.06676674 | 0.0960353 | 0.03749817 | 0.03926443 | up |
| 937 | Larp6 | 0.00307054 | 0.06476136 | 0.11768671 | 0.01183602 | 0.03934028 | up |
| 938 | Bnc2 | 0.00308987 | -0.2352008 | -0.1090302 | -0.3613714 | 0.03937045 | down |
| 939 | Eps15 | 0.00309209 | 0.17366596 | 0.25325761 | 0.0940743 | 0.03937045 | up |
| 940 | Gk | 0.00308845 | 0.16993842 | 0.28438098 | 0.05549586 | 0.03937045 | up |
| 941 | Mthfr | 0.00308158 | -0.0553321 | -0.0182461 | -0.0924181 | 0.03937045 | down |
| 942 | Sntb1 | 0.00309907 | 0.07842477 | 0.12542246 | 0.03142708 | 0.03937045 | up |
| 943 | Syne3 | 0.00309913 | 0.10087673 | 0.14487842 | 0.05687504 | 0.03937045 | up |
| 944 | Top2b | 0.00309633 | 0.19173699 | 0.2825763 | 0.10089768 | 0.03937045 | up |
| 945 | Zbtb8os | 0.00309151 | 0.07273061 | 0.10764585 | 0.03781536 | 0.03937045 | up |
| 946 | Stxbp5 | 0.00311538 | 0.15552083 | 0.23555233 | 0.07548934 | 0.03953497 | up |
| 947 | Sim1 | 0.00313581 | -0.0171235 | -0.0085268 | -0.0257201 | 0.03975227 | down |
| 948 | Zcchc24 | 0.0031402 | -0.2872165 | -0.1319798 | -0.4424533 | 0.03976593 | down |
| 949 | Matr3 | 0.00314686 | 0.15641571 | 0.23501154 | 0.07781988 | 0.03979011 | up |
| 950 | Nbea | 0.00314874 | 0.17854396 | 0.29765113 | 0.05943679 | 0.03979011 | up |
| 951 | Creb3l2 | 0.0031593 | -0.2722286 | -0.1307094 | -0.4137478 | 0.03988159 | down |
| 952 | Arfgef1 | 0.00318011 | 0.16239794 | 0.25662716 | 0.06816872 | 0.04007217 | up |
| 953 | Cyp20a1 | 0.00318107 | 0.23552328 | 0.3505787 | 0.12046785 | 0.04007217 | up |
| 954 | Camk2n2 | 0.00319144 | -0.0605245 | -0.0309139 | -0.0901351 | 0.0400766 | down |
| 955 | Csrnp1 | 0.0031859 | -0.2059117 | -0.1064149 | -0.3054084 | 0.0400766 | down |
| 956 | Tmem151b | 0.0031885 | -0.0321962 | -0.0173378 | -0.0470545 | 0.0400766 | down |
| 957 | Agrn | 0.00320454 | -0.1566992 | -0.0735152 | -0.2398832 | 0.04008325 | down |
| 958 | Irak2 | 0.00320532 | -0.0401936 | -0.0238228 | -0.0565644 | 0.04008325 | down |
| 959 | Med30 | 0.00319888 | 0.08806336 | 0.13776465 | 0.03836207 | 0.04008325 | up |
| 960 | Txlna | 0.00319652 | -0.1898368 | -0.0972428 | -0.2824308 | 0.04008325 | down |
| 961 | Atl3 | 0.0032169 | 0.12364301 | 0.19083397 | 0.05645206 | 0.04014443 | up |
| 962 | Cd55 | 0.0032161 | 0.26047591 | 0.49293037 | 0.02802144 | 0.04014443 | up |
| 963 | Gng4 | 0.00322459 | 0.02399165 | 0.03864023 | 0.00934308 | 0.04019851 | up |
| 964 | Tmem248 | 0.00324399 | -0.1871338 | -0.0989829 | -0.2752847 | 0.04039848 | down |
| 965 | Eif3f | 0.00327428 | 0.20869341 | 0.34624941 | 0.07113741 | 0.04064921 | up |
| 966 | Tnrc6c | 0.0032682 | -0.0969605 | -0.0492942 | -0.1446267 | 0.04064921 | down |
| 967 | Tyw5 | 0.00327355 | 0.05500153 | 0.0865378 | 0.02346526 | 0.04064921 | up |
| 968 | Cdon | 0.00328847 | -0.1599025 | -0.0835546 | -0.2362505 | 0.0407831 | down |
| 969 | Npl | 0.00329759 | -0.0327803 | -0.0166157 | -0.048945 | 0.04085402 | down |
| 970 | Ccdc90b | 0.00332272 | 0.17953149 | 0.26970049 | 0.08936249 | 0.04112292 | up |
| 971 | Adgra3 | 0.00336345 | -0.1018586 | -0.0517322 | -0.151985 | 0.04137752 | down |
| 972 | Aff1 | 0.00334725 | -0.1411525 | -0.0703881 | -0.211917 | 0.04137752 | down |

| | | | | | | | |
|---|---|---|---|---|---|---|---|
| 973 | Elmo2 | 0.00336397 | 0.14782166 | 0.22121689 | 0.07442643 | 0.04137752 | up |
| 974 | Renbp | 0.00335945 | 0.09986384 | 0.14742757 | 0.05230011 | 0.04137752 | up |
| 975 | Wtap | 0.00336202 | 0.14761122 | 0.22705644 | 0.06816599 | 0.04137752 | up |
| 976 | Zfpm1 | 0.00335239 | -0.1202791 | -0.0595612 | -0.1809969 | 0.04137752 | down |
| 977 | Adgre1 | 0.00339545 | -0.0080189 | -3.80E-05 | -0.0159999 | 0.04172195 | down |
| 978 | Fto | 0.00340132 | -0.1227043 | -0.0602412 | -0.1851674 | 0.04175141 | down |
| 979 | Ackr4 | 0.00340548 | 0.04858001 | 0.07662097 | 0.02053906 | 0.0417597 | up |
| 980 | Ppp1r9b | 0.00341552 | -0.1346849 | -0.0693056 | -0.2000641 | 0.04184015 | down |
| 981 | Dpysl2 | 0.00341911 | 0.35794449 | 0.53453841 | 0.18135056 | 0.04184139 | up |
| 982 | Ppp3cb | 0.00343181 | 0.11173233 | 0.17554244 | 0.04792222 | 0.04195402 | up |
| 983 | Pla2g2d | 0.00343909 | 0.07371064 | 0.11612017 | 0.03130112 | 0.04200023 | up |
| 984 | Pcyox1 | 0.00346372 | -0.0936993 | -0.0464666 | -0.140932 | 0.04225811 | down |
| 985 | Nom1 | 0.0034691 | 0.06154247 | 0.09915723 | 0.02392771 | 0.04228076 | up |
| 986 | Polr2k | 0.00347316 | 0.07583519 | 0.1101644 | 0.04150598 | 0.04228731 | up |
| 987 | Taok1 | 0.00348918 | 0.26443648 | 0.39691376 | 0.13195919 | 0.04239632 | up |
| 988 | Tnnt1 | 0.00348878 | 0.03405867 | 0.04893995 | 0.01917739 | 0.04239632 | up |
| 989 | Oxca2 | 0.00349941 | -0.0288733 | -0.0131628 | -0.0445837 | 0.04247768 | down |
| 990 | Tpra1 | 0.00350815 | -0.0390563 | -0.0071575 | -0.0709551 | 0.04254078 | down |
| 991 | Marveld1 | 0.00351429 | -0.2048706 | -0.100543 | -0.3091983 | 0.04257217 | down |
| 992 | Ncln | 0.0035183 | -0.0758741 | -0.031657 | -0.1200912 | 0.04257783 | down |
| 993 | H1fx | 0.00355376 | -0.2297803 | -0.0872781 | -0.3722826 | 0.04292042 | down |
| 994 | Iqsec1 | 0.0035504 | 0.25662326 | 0.38451844 | 0.12872808 | 0.04292042 | up |
| 995 | Macrod2 | 0.00356133 | -0.0378901 | -0.019227 | -0.0565533 | 0.04296862 | down |
| 996 | Pde4dip | 0.00357055 | -0.1605833 | -0.095533 | -0.2256336 | 0.0430366 | down |
| 997 | Msn | 0.00357576 | -0.2764627 | -0.1515685 | -0.4013569 | 0.04305619 | down |
| 998 | Amot | 0.00359049 | -0.0934596 | -0.0443764 | -0.1425428 | 0.04319017 | down |
| 999 | Ppp1r12c | 0.0036029 | -0.1638911 | -0.0830868 | -0.2446954 | 0.04325279 | down |
| 1000 | Shroom4 | 0.00360089 | -0.0587024 | -0.0361952 | -0.0812096 | 0.04325279 | down |
| 1001 | Gcnt4 | 0.0036124 | 0.37513788 | 0.84883295 | -0.0985572 | 0.0432803 | up |
| 1002 | Spn | 0.00360997 | -0.1222312 | -0.0644905 | -0.1799718 | 0.0432803 | down |
| 1003 | Pla2g4a | 0.00363287 | -0.118462 | -0.0603094 | -0.1766147 | 0.04347721 | down |
| 1004 | Tcirg1 | 0.00363608 | -0.2360692 | -0.1228104 | -0.349328 | 0.04347721 | down |
| 1005 | Sh2b2 | 0.00364322 | -0.0513636 | -0.0184598 | -0.0842674 | 0.04351929 | down |
| 1006 | Zfp830 | 0.00365318 | -0.0652753 | -0.0347422 | -0.0958084 | 0.04359486 | down |
| 1007 | Col4a3bp | 0.00365902 | -0.1078945 | -0.0443315 | -0.1714576 | 0.04362117 | down |
| 1008 | Cox5a | 0.00367512 | -0.1780642 | -0.087124 | -0.2690045 | 0.04372626 | down |
| 1009 | Fkbp5 | 0.00367269 | -0.1919366 | -0.0968084 | -0.2870648 | 0.04372626 | down |
| 1010 | Mocs2 | 0.00368293 | 0.16679346 | 0.2481017 | 0.08548521 | 0.04377587 | up |
| 1011 | Prxl2c | 0.00369731 | 0.16346456 | 0.24691661 | 0.0800125 | 0.04390326 | up |
| 1012 | Il16 | 0.00371692 | 0.25655182 | 0.37636652 | 0.13673711 | 0.04409253 | up |
| 1013 | C1d | 0.00372304 | 0.17336485 | 0.25554415 | 0.09118555 | 0.0441215 | up |
| 1014 | Snx1 | 0.00372693 | 0.19423037 | 0.29217849 | 0.09628225 | 0.0441241 | up |
| 1015 | Klhl33 | 0.00373387 | -0.072074 | -0.0370061 | -0.107142 | 0.04416265 | down |
| 1016 | Ndufb1-ps | 0.00373979 | 0.11062488 | 0.1684018 | 0.05284797 | 0.04418912 | up |
| 1017 | Csrp1 | 0.00375714 | -0.8068462 | -0.4118288 | -1.2018637 | 0.04435049 | down |
| 1018 | Poglut3 | 0.00376107 | 0.24682974 | 0.3703292 | 0.12333028 | 0.0443533 | up |
| 1019 | Abi2 | 0.00377216 | 0.12687893 | 0.19279092 | 0.06096694 | 0.0444404 | up |
| 1020 | Arb3 | 0.00379104 | 0.05858696 | 0.09008752 | 0.02708641 | 0.04448824 | up |
| 1021 | Ccdc117 | 0.00378724 | -0.0646033 | -0.0305424 | -0.0986642 | 0.04448824 | down |
| 1022 | Marcks | 0.00379061 | -0.3032732 | -0.1431929 | -0.4633535 | 0.04448824 | down |
| 1023 | Whrn | 0.00378374 | -0.0959264 | -0.0554091 | -0.1364438 | 0.04448824 | down |
| 1024 | Rsrc1 | 0.00379957 | 0.0836422 | 0.12076771 | 0.04651669 | 0.04454473 | up |
| 1025 | Ahr | 0.00381279 | 0.37785595 | 0.57266757 | 0.18304433 | 0.0446157 | up |
| 1026 | Ankrd13c | 0.00381752 | 0.12337945 | 0.17732351 | 0.06943538 | 0.0446157 | up |
| 1027 | Snrnp200 | 0.00382049 | 0.08630918 | 0.13316842 | 0.03944994 | 0.0446157 | up |
| 1028 | Sobp | 0.00381944 | 0.08074897 | 0.15804411 | 0.00345383 | 0.0446157 | up |
| 1029 | Cyp4f16 | 0.00382535 | -0.0871793 | -0.0455283 | -0.1288303 | 0.04462907 | down |
| 1030 | Irf2 | 0.00383478 | 0.1261411 | 0.18602848 | 0.06625371 | 0.04465237 | up |
| 1031 | Ubtf | 0.00383262 | 0.14504877 | 0.21995401 | 0.07014353 | 0.04465237 | up |
| 1032 | Nup50 | 0.00384085 | -0.0867574 | -0.0379385 | -0.1355764 | 0.04467962 | down |
| 1033 | Nol3 | 0.00384704 | 0.09294968 | 0.14263015 | 0.04326922 | 0.0447083 | up |
| 1034 | F2r | 0.00385601 | -0.4060872 | -0.2085451 | -0.6036293 | 0.04476919 | down |
| 1035 | Serpinh1 | 0.00386303 | -0.4121886 | -0.2275566 | -0.5968206 | 0.04480743 | down |
| 1036 | Lig3 | 0.00387887 | -0.0852057 | 0.00804541 | -0.1784567 | 0.04494774 | down |
| 1037 | Bivm | 0.0038827 | 0.12425602 | 0.19776106 | 0.05075098 | 0.04494874 | up |
| 1038 | Snapc1 | 0.00388903 | 0.08426222 | 0.13336299 | 0.03516144 | 0.04497862 | up |
| 1039 | Glod4 | 0.00389949 | 0.12169258 | 0.18252415 | 0.06086101 | 0.04505614 | up |
| 1040 | Leprot | 0.00391344 | 0.1287243 | 0.21476174 | 0.04268686 | 0.04513055 | up |
| 1041 | Prima1 | 0.00391205 | -0.0152454 | -0.0080287 | -0.0224622 | 0.04513055 | down |
| 1042 | Actr1b | 0.00392679 | -0.2108279 | -0.0940931 | -0.3275628 | 0.04519765 | down |
| 1043 | Zfp608 | 0.00392622 | 0.15842311 | 0.23231529 | 0.08453094 | 0.04519765 | up |
| 1044 | Eea1 | 0.00393681 | 0.13004502 | 0.19469094 | 0.0653991 | 0.04522626 | up |
| 1045 | Rsrp1 | 0.00393447 | -0.2130586 | -0.1056641 | -0.320453 | 0.04522626 | down |
| 1046 | Fam189a2 | 0.00395566 | -0.0265632 | -0.0129745 | -0.0401519 | 0.04530143 | down |
| 1047 | Frmd6 | 0.00395431 | 0.31585939 | 0.47899694 | 0.15272183 | 0.04530143 | up |
| 1048 | Gnai2 | 0.00396222 | -0.2175287 | -0.072196 | -0.3628614 | 0.04530143 | down |
| 1049 | P2ry2 | 0.00396211 | 0.20732227 | 0.33512988 | 0.07951466 | 0.04530143 | up |
| 1050 | Sptlc2 | 0.00395785 | 0.10804272 | 0.16259655 | 0.05348889 | 0.04530143 | up |
| 1051 | Cd59a | 0.00397917 | -0.1459094 | -0.0510979 | -0.2407208 | 0.04545184 | down |
| 1052 | Cyb561a3 | 0.00399475 | -0.1004258 | -0.0598639 | -0.1409878 | 0.04554317 | down |
| 1053 | Ocrl | 0.00399393 | 0.09755608 | 0.14455163 | 0.05056052 | 0.04554317 | up |
| 1054 | Fut11 | 0.00401877 | 0.15483392 | 0.23103048 | 0.07863736 | 0.04577358 | up |
| 1055 | C1qc | 0.0040235 | -0.0481861 | -0.0241594 | -0.0722128 | 0.04578238 | down |
| 1056 | Fam210a | 0.00402717 | 0.12707197 | 0.18611203 | 0.06803191 | 0.04578238 | up |
| 1057 | Epha7 | 0.00404631 | 0.03924551 | 0.07747163 | 0.00101939 | 0.0459254 | up |
| 1058 | Gbx2 | 0.00405299 | -0.0658844 | -0.0313915 | -0.1003773 | 0.0459254 | down |
| 1059 | Pgghg | 0.00404751 | -0.0657135 | 0.01043908 | -0.1418661 | 0.0459254 | down |
| 1060 | Stambp | 0.00405505 | 0.08491551 | 0.12617768 | 0.04365333 | 0.0459254 | up |
| 1061 | Myl12b | 0.00407174 | 0.18678142 | 0.27590005 | 0.09766279 | 0.04607094 | up |
| 1062 | Stub1 | 0.00408439 | -0.1934755 | -0.0732184 | -0.3137326 | 0.04617055 | down |
| 1063 | Rpp25 | 0.00410485 | -0.0216243 | -0.0129473 | -0.0303013 | 0.04635819 | down |
| 1064 | Ipp | 0.00412195 | -0.0385512 | -0.0167882 | -0.0603142 | 0.04646384 | down |
| 1065 | Tmem167 | 0.00412193 | 0.23354723 | 0.40476527 | 0.0623292 | 0.04646384 | up |
| 1066 | Gmppb | 0.00413062 | -0.1645824 | -0.0413557 | -0.2878092 | 0.04647436 | down |
| 1067 | Nfe2l1 | 0.00412676 | -0.0949615 | -0.0505616 | -0.1393613 | 0.04647436 | down |
| 1068 | Apip | 0.00414026 | 0.07130346 | 0.11049486 | 0.03211207 | 0.04653918 | up |
| 1069 | Tha1 | 0.00415128 | 0.05887718 | 0.08188608 | 0.03586828 | 0.04657578 | up |
| 1070 | Traf4 | 0.00414854 | -0.13315 | -0.0431379 | -0.223162 | 0.04657578 | down |
| 1071 | Lamb2 | 0.00419676 | 0.2839683 | 0.41930247 | 0.14863414 | 0.04704206 | up |
| 1072 | Ccn2 | 0.00420778 | -0.7583979 | -0.3888537 | -1.1279421 | 0.04709047 | down |
| 1073 | Sh3glb2 | 0.00420892 | -0.0893361 | -0.0401246 | -0.1385476 | 0.04709047 | down |
| 1074 | Atg4d | 0.00421534 | -0.055976 | -0.0205897 | -0.0913624 | 0.04710734 | down |
| 1075 | Fkbp11 | 0.00421827 | -0.3000027 | -0.1482146 | -0.4517909 | 0.04710734 | down |
| 1076 | Snai2 | 0.00422561 | 0.15405352 | 0.27554981 | 0.03255723 | 0.04714543 | up |
| 1077 | Evi5 | 0.0042674 | 0.08751053 | 0.13487644 | 0.04014462 | 0.04747923 | up |
| 1078 | Pex6 | 0.00426677 | -0.0905938 | -0.0353521 | -0.1458354 | 0.04747923 | down |
| 1079 | Ttbk2 | 0.00426256 | 0.05547168 | 0.08588875 | 0.02505461 | 0.04747923 | up |
| 1080 | Tppp3 | 0.00427196 | 0.15919685 | 0.44159939 | -0.1232057 | 0.04748602 | up |
| 1081 | Megf8 | 0.00428154 | -0.068607 | -0.0147478 | -0.1224662 | 0.04754842 | down |
| 1082 | Dsel | 0.00430046 | 0.23120995 | 0.34949735 | 0.11292255 | 0.04771442 | up |
| 1083 | Elmo1 | 0.00431202 | 0.09840831 | 0.16144701 | 0.03536961 | 0.04777653 | up |
| 1084 | Ndufa4l2 | 0.00431451 | 0.60510087 | 0.89642155 | 0.31378019 | 0.04777653 | up |
| 1085 | Slitrk2 | 0.00431943 | 0.02608495 | 0.03757971 | 0.01459018 | 0.04777653 | up |
| 1086 | Stard3nl | 0.00432198 | 0.1349061 | 0.20317398 | 0.06663821 | 0.04777653 | up |
| 1087 | Rarb | 0.0043306 | -0.0950741 | -0.06143 | -0.1287182 | 0.0478278 | down |
| 1088 | Tfip11 | 0.00433867 | -0.0826146 | -0.0412857 | -0.1239435 | 0.04787292 | down |
| 1089 | P3h3 | 0.004349 | -0.1090269 | -0.0488033 | -0.1692505 | 0.04794285 | down |
| 1090 | Ccnh | 0.00436241 | 0.06878388 | 0.10883776 | 0.02873 | 0.04804656 | up |
| 1091 | Naca | 0.00437072 | 0.28159949 | 0.4577251 | 0.10547389 | 0.04809393 | up |
| 1092 | N4bp1 | 0.00438361 | 0.09610096 | 0.14945363 | 0.0427483 | 0.04819166 | up |
| 1093 | Arhgap12 | 0.00439093 | 0.07179516 | 0.12097139 | 0.02261892 | 0.04822793 | up |
| 1094 | Bhlhe41 | 0.00440606 | -0.2530317 | -0.1050003 | -0.4010631 | 0.04830575 | down |
| 1095 | Rbm4 | 0.00440507 | -0.1773263 | -0.0866434 | -0.2680092 | 0.04830575 | down |
| 1096 | Nbn | 0.00442723 | 0.05204545 | 0.08224045 | 0.02185045 | 0.04849354 | up |
| 1097 | Pde1b | 0.00443154 | -0.0613812 | -0.0360226 | -0.0867399 | 0.04849648 | down |
| 1098 | Xpo4 | 0.00443895 | 0.0525721 | 0.08489399 | 0.02025022 | 0.04853333 | up |
| 1099 | Nipa1 | 0.0044456 | 0.05857322 | 0.0923848 | 0.02476165 | 0.04856181 | up |
| 1100 | Asns | 0.00445587 | 0.11644675 | 0.19522493 | 0.03766857 | 0.04858657 | up |
| 1101 | Pcdh1 | 0.00445596 | 0.12806512 | 0.21751017 | 0.03862006 | 0.04858657 | up |
| 1102 | Lpcat4 | 0.00447383 | -0.0803002 | -0.0391456 | -0.1214548 | 0.04869486 | down |
| 1103 | Wdr26 | 0.00447401 | 0.11815863 | 0.19389617 | 0.04242109 | 0.04869486 | up |
| 1104 | Zfp638 | 0.00447933 | 0.10260306 | 0.161512 | 0.04369412 | 0.04870866 | up |
| 1105 | Exd2 | 0.00449712 | -0.0545191 | -0.0240241 | -0.085014 | 0.0487696 | down |
| 1106 | Ptprb | 0.00449649 | -0.039828 | -0.0192702 | -0.0603858 | 0.0487696 | down |
| 1107 | Rbm15b | 0.00449191 | -0.0852227 | -0.0400408 | -0.1304047 | 0.0487696 | down |
| 1108 | Cog8 | 0.0045155 | -0.0432944 | -0.0147292 | -0.0718596 | 0.04892466 | down |
| 1109 | Dcaf6 | 0.00452502 | 0.08882091 | 0.13664802 | 0.0409938 | 0.04894466 | up |
| 1110 | Eef1akmt4 | 0.00452957 | -0.158286 | -0.0765035 | -0.2400685 | 0.04894466 | down |
| 1111 | Katnbl1 | 0.00452811 | 0.08109004 | 0.12533348 | 0.03684661 | 0.04894466 | up |
| 1112 | Dram1 | 0.00454179 | 0.31430003 | 0.47839067 | 0.1502094 | 0.04898219 | up |
| 1113 | Heatr5a | 0.00454082 | -0.1315426 | -0.061763 | -0.2013221 | 0.04898219 | down |
| 1114 | Map1b | 0.00454529 | -0.1236713 | -0.0645048 | -0.1828378 | 0.04898219 | down |
| 1115 | Lss | 0.00456625 | 0.07943838 | 0.12722622 | 0.03165054 | 0.04916394 | up |
| 1116 | Nptxr | 0.0045823 | -0.0835823 | -0.0439029 | -0.1232617 | 0.04929257 | down |
| 1117 | Rnf5 | 0.00461005 | -0.0792441 | -0.0239418 | -0.1345463 | 0.04950174 | down |
| 1118 | Soga1 | 0.00461009 | -0.0885494 | -0.038127 | -0.1389719 | 0.04950174 | down |
| 1119 | Zbed3 | 0.00461411 | -0.1070962 | -0.0466752 | -0.1675173 | 0.04950174 | down |
| 1120 | Thnsl2 | 0.00462087 | 0.05183955 | 0.08087108 | 0.02280803 | 0.0495299 | up |
| 1121 | Mme | 0.00463796 | 0.15210814 | 0.25600918 | 0.0482071 | 0.04964042 | up |
| 1122 | Rgl1 | 0.00463945 | -0.1820873 | -0.0928951 | -0.2712796 | 0.04964042 | down |
| 1123 | Gpr27 | 0.00465032 | 0.11741233 | 0.22726641 | 0.00755825 | 0.04971241 | up |
| 1124 | Acaca | 0.00465594 | 0.07817948 | 0.12533108 | 0.03102787 | 0.04972824 | up |
| 1125 | Bcl9l | 0.00466704 | -0.1243213 | -0.0663653 | -0.1822772 | 0.04975831 | down |
| 1126 | Crybg3 | 0.00466296 | -0.0515221 | -0.0217707 | -0.0812736 | 0.04975831 | down |
| 1127 | Gda | 0.00467903 | 0.70232604 | 1.05863935 | 0.34601272 | 0.04984183 | up |
| 1128 | Rbm7 | 0.00469052 | 0.10311027 | 0.16262701 | 0.04359352 | 0.04991993 | up |

AAV-gTrac vs AAV-gIl1r1
DEGs filtered by fdr < 0.05

| | primerid | Pr(>Chisq) | coef | ci.hi | ci.lo | fdr | direction |
|---|---|---|---|---|---|---|---|
| 1 | Gramd1b | 3.87E-06 | -0.0785554 | -0.054239 | -0.1028717 | 0.02273716 | down |
| 2 | Utp11 | 2.10E-06 | 0.18040265 | 0.23338914 | 0.12741616 | 0.02273716 | up |
| 3 | Slc7a6 | 6.86E-06 | 0.17566055 | 0.23056255 | 0.12075855 | 0.02684691 | up |
| 4 | Aggf1 | 5.55E-05 | 0.12111751 | 0.17297732 | 0.06925771 | 0.04701699 | up |
| 5 | Chsy1 | 4.04E-05 | 0.18823889 | 0.26589027 | 0.11058751 | 0.04701699 | up |
| 6 | Clic4 | 2.19E-05 | 0.22420355 | 0.30401456 | 0.14439255 | 0.04701699 | up |
| 7 | Creb3 | 5.14E-05 | 0.17529381 | 0.23170528 | 0.11888234 | 0.04701699 | up |
| 8 | Exoc5 | 4.55E-05 | 0.16389603 | 0.2290093 | 0.09878276 | 0.04701699 | up |
| 9 | Ier3ip1 | 2.15E-05 | 0.2789655 | 0.35482831 | 0.20310268 | 0.04701699 | up |
| 10 | Krt7 | 3.01E-05 | -0.1031892 | -0.0749236 | -0.1314547 | 0.04701699 | down |
| 11 | Mcpt2 | 5.61E-05 | -0.1237551 | -0.0903876 | -0.1571226 | 0.04701699 | down |
| 12 | Psmd14 | 3.11E-05 | 0.22868398 | 0.28641638 | 0.17095158 | 0.04701699 | up |
| 13 | Ptprj | 3.52E-05 | 0.11673019 | 0.1574061 | 0.07605428 | 0.04701699 | up |
| 14 | Thra | 4.81E-05 | -0.1547134 | -0.1072617 | -0.202165 | 0.04701699 | down |
| 15 | Tspan8 | 6.81E-05 | -0.0608505 | -0.0443245 | -0.0773766 | 0.04701699 | down |
| 16 | Ubxn6 | 6.16E-05 | 0.11555936 | 0.19003797 | 0.04108075 | 0.04701699 | up |
| 17 | Zfp395 | 6.75E-05 | -0.1771617 | -0.1162267 | -0.2380968 | 0.04701699 | down |
