## Supplementary Table Reagents for "Discovery of a Linked Constellation of Gene Expression Revealed by Local Editing of Fibroblasts in Tumors"

| **Reagent or Resource** | **Source** | **Identifier** |
|---|---|---|
| antibodies | | |
| anti-mouse CD16/32 (clone 2.4G2) | Tonbo Bioscie | 70-0161-U500 |
| anti-CD206-BV605 (clone C068C2) | BioLegend | |
| anti-CD11b BUV737 (clone M1/70) | BD Biosciences | |
| anti-CD31 PerCP-Cy5.5 (clone 390) | BioLegend | |
| anti-I-A/I-E BV650 (clone M5/114.15.2) | BioLegend | |
| anti-mouse B220 (clone RA3-6B2) BV785 | BioLegend | |
| anti-mouse CD24 (M1/69) BV711 | BD Biosciences | |
| anti-mouse CD45 (clone 30-F11) BV650 | BioLegend | |
| anti-mouse CD45 (clone 30-F11) BUV395 | BD Biosciences | |
| anti-mouse CD64 (clone X54-5/7.1) PE-Cy7 | BioLegend | |
| anti-mouse F4/80 (clone BM8) BV510 | BioLegend | |
| anti-Ly6G (clone 1A8) BV785 | BioLegend | |
| anti-CD90.2 (clone 30-H12) BV785 | BioLegend | |
| anti-CD90.2 (clone 30-H12) AF647 | BioLegend | |
| anti-NK1.1 (clone PK136) BV785 | BioLegend | |
| anti-SiglecF (clone E50-2440) BV785 | BioLegend | |
| anti-CD9 (clone MZ3) APC-Fire750 | BioLegend | |
| anti-TREM2 (clone 237920) APC | Novus Biologicals | |
| anti-Ly6C (clone HK1.4) AF700 | BioLegend | |
| anti-Ly6C (clone HK1.4) BV605 | BioLegend | |
| anti-Ly6C (clone HK1.4) APC-Cy7 | BioLegend | |
| anti-E-Cadherin (CD324) (clone DECMA-1) PE-Cy7 | BioLegend | |
| anti-NCAM1 (CD56) PE | Novus Biologicals | |
| anti-PDPN (clone 8.1.1) APC-Fire750 | BioLegend | |
| anti-PDPN (clone 8.1.1) APC | BioLegend | |
| anti-SCA-1 (Ly6A/E) BV510 | BioLegend | |
| anti-PDGFRa (CD140a) SB702 | ThermoFisher | |
| anti-MCAM (CD146) BUV737 | BD Biosciences | |
| anti-OSMR | Novus Biologicals | |
| anti-IL1R1 (CD121a) (clone JAMA-147) PE | BioLegend | |
| anti-IL1R1 (CD121a) (clone 35F5) AF647 | BD Biosciences | |
| anti-TGFBR2 APC | Novus Biologicals | |
| Donkey anti-Goat PE | Jackson ImmunoResearch | |
| Donkey anti-Goat AF647 | Jackson ImmunoResearch | |
| anti-human/mouse phospho-STAT3 (Y705) AF647 | BD Bioscience | RRID: AB_647144 |
| anti-human/mouse phospho-SMAD2/3 AF647 | BD Bioscience | RRID: AB_2716578 |
| **Oligonucleotides** | | |
| 18s_fwd | GGATGCGTGCATTTATCAGA | |
| 18s_rev | GGCGACTACCATCGAAAGTT | |
| Dpt_fwd | CAGAGTACCCAAGCCACTATGG | |
| Dpt_rev | GGCACATGATGAACTTCCACTGG | |
| Pi16_fwd | AACTGGCACGAGGAGCATGAGT | |
| Pi16_rev | GCCAATTCTCTCAGTCTTGCTCC | |
| Acta2_fwd | TGCTGACAGAGGCACCACTGAA | |
| Acta2_rev | CAGTTGTACGTCCAGAGGCATAG | |
| Postn_fwd | CAGCAAACCACTTTCACCGACC | |
| Postn_rev | AGAAGGCGTTGGTCCATGCTCA | |
| titer_U6_Forward | ggactatcatatgcttaccgt | |
| titer_U6_Reverse | ggtgtttcgtcctttccaca | |
| **Commercial plasmids** | | |
| 7m8 (RepCap plasmid) | Addgene | 64839 |
| pAAV-DJ (RepCap plasmid) | Cell Biolabs | VPK-420-DJ |
| pAAV2/1 (RepCap plasmid) | Addgene | 112862 |
| pHelper (helper plasmid) | Takara | 6234 |
| pL-CRISPR.EFS.GFP | Addgene | 57818 |
| pscAAV-CAG-EGFP | Addgene | 83279 |
| pAAV-U6-gRNA-CBh-mCherry | Addgene | 91947 |
| **Other plasmids** | | |
| pscAAV-hU6-gRNA-CBh-mCherry | this study | |
| pscAAV-hU6-gSCR-CBh-mCherry | this study | |
| pscAAV-hU6-gTrac-CBh-mCherry | this study | |
| pscAAV-hU6-gThy1-CBh-mCherry | this study | |
| pscAAV-hU6-gPdpn-CBh-mCherry | this study | |
| pscAAV-hU6-gTgfbr2-CBh-mCherry | this study | |
| pscAAV-hU6-gOsmr-CBh-mCherry | this study | |
| pscAAV-hU6-gIl1r1-CBh-mCherry | this study | |
| pscAAV-hU6-gThy1-mU6-gPdpn-EFS-mCherry | this study | |
| pscAAV-hU6-gTgfbr2-mU6-gOsmr-EFS-mCherry | this study | |
| pscAAV-hU6-gTgfbr2-mU6-gIl1r1-EFS-mCherry | this study | |
| pscAAV-hU6-gOsmr-mU6-gIl1r1-EFS-mCherry | this study | |
| pscAAV-KIKO-U6-gThy1-p2a-mCherry-STOP-400 | this study | |
| pscAAV-KIKO-U6-gThy1-p2a-mCherry-STOPpA-400 | this study | |
| pscAAV-KIKO-U6-gThy1-p2a-mCherry-T2A-400 | this study | |
| pscAAV-KIKO-U6-gTgfbr2-p2a-mCherry-STOPpA-400 | this study | |
| pscAAV-KIKO-U6-gOsmr-p2a-mCherry-STOPpA-400 | this study | |
| pscAAV-KIKO-U6-gIl1r1-p2a-mCherry-STOPpA-400 | this study | |
| **Experimental models: Cell lines** | | |
| YUMM5.2 (mouse melanoma cell line) | ATCC | CRL-3367 |
| HY19636 (mouse pancreatic cancer cell line) | H. Ying (MD Anderson) | |
| NIH/3T3 (immortalized mouse fibroblast cell line) | ATCC | CRL-1658 |
| NIH/3T3.Cas9-EGFP | this paper | |
| HEK293T | ATCC | CRL-3216 |
| **Experimental models: Organisms/strains** | | |
| Mouse: wild-type C57BL/6 | The Jackson Laboratory | |
| Mouse: Pdgfra-Cre | The Jackson Laboratory | |
| Mouse: Pdgfra-CreERT2 | The Jackson Laboratory | |
| Mouse: Rosa26-LSL-Cas9-EGFP | The Jackson Laboratory | |
| Mouse: Cthrc1-CreERT2 | T. Tsukui and D. Sheppard (UCSF) | |
| Mouse: Ai14 | The Jackson Laboratory | |
| Mouse: Col1a2-CreERT2 | Bin Zhou | PMID: 28650345 |
| Mouse: Tgfbr2-Exon2-fl/fl | Harold Moses | PMID: 11857781 |
| **Commercial Kits** | | |
| AAVpro® Purification Kit Midi (All Serotypes) | Takara | 6675 |
| SsoAdvanced Universal SYBR Green Supermix | Bio-Rad | 1725270 |
| RNeasy Micro Kit | Qiagen | 74004 |
| iScript Reverse Transcription Supermix | Bio-Rad | 1708840 |
| **Bacterial strains** | | |
| DH5alpha | NEB | C2987 |
| NEBstable | NEB | C3040 |
| **Chemicals, peptides, and recombinant proteins** | | |
| EDTA | Teknova | E0306 |
| 16% paraformaldehyde | Electron Micr | 15710 |
| DNase I | Millipore Sign | 10104159001 |
| Collagenase IX | Millipore Sign | C7657 |
| Hyaluronidase | Worthington | LS005477 |
| Collagenase D | Millipore Sign | 11088858001 |
| Methanol | Millipore Sign | 179337 |
| recombinant mouse oncostatin M | Biolegend | 762802 |
| recombinant mouse TGF-b1 | Biolegend | 763102 |
| **Reagents** | | |
| DMEM | Gibco | 11995065 |
| beta-mercaptoethanol | Gibco | 21985-023 |
| fetal bovine serum | Gemini Bio | 900-308 |
| Penicillin-Streptomycin-Glutamine (100X) | Gibco | 10378016 |
| polyethylenimine, linear, MW 25000 | Polysciences, | 23966 |
| DNase | NEB | B0303S |
| Proteinase K | Qiagen | 1114886 |
| Green tattoo paste | Fine Science T | 2420101 |
| OCT | Sakura | 4583 |
| Lab-Tek Chamber Slide System, 8-well | Lab-Tek | 177445 |
| bovine serum albumin | Sigma-Aldrich | A4503 |
| mouse serum | Jackson Immu | 015-000-120 |
| Triton X-100 | Millipore Sign | T8787 |
| **Deposited data** | | |
| Mouse spatiotemporal scRNAseq wound data set | Hu & Kuhn et | GEO: GSE204777 |
| Mouse pancreatic cancer fibroblast scRNAseq data set | Krishnamurty | ArrayExpress: E-MTAB-12028 |
| **Software and algorithms** | | |
| R environment | R Developmer | https://cran.r-project.org |
| Flow Jo | v10.10.0 | flowjo.com |
| Imaris 9.2.1 | Bitplane | https://imaris.oxinst.com/products/imaris-for-cell-biologists?gad=1&amp%3Bgclid=Cj0KCQjw3a2iBhCFARIsAD4jQB2RkztA4vyhmDUQ3_eo2WwjysDt-NhFrH-evt8PxW5s5oKpw_WYJhcaAuhZEALw_wcB |
| Ggpubr (R package) | https://cran.r-project.org/web/packages/ggpubr/index.html | |
| Seurat (R Package) | v5.0.1 | satijalab.org/seurat |
| Biorender | https://www.biorender.com | |
