## Supplementary Table gRNA sequences for "Discovery of a Linked Constellation of Gene Expression Revealed by Local Editing of Fibroblasts in Tumors"

| gRNA | target sequence |
|---|---|
| gCtnnb1 | TGACCTGATGGAGTTGGACA |
| gCxcl5 | ATGGCGAGATGGAACCGCTG |
| gIl1r1 | GAACATATCCAACACCATCT |
| gOsmr | GAACATATCCAACACCATCT |
| gPdpn | GAAGATGATATTGTGACCCC |
| gSCR | GCTTAGTTACGCGTGGACGA |
| gTgfbr2 | GTCCACAGGACGATATGCAG |
| gThy1 | GCTGGTCACCTTCTGCCCTC |
| gTnfrsf1a | GTATCCCCATCAGCAGAGCC |
| gTrac | TATGGATTCCAAGAGCAATG |
